# An organism-wide atlas of hormonal signaling based on the mouse lemur single-cell transcriptome

**DOI:** 10.1101/2021.12.13.472243

**Authors:** Shixuan Liu, Camille Ezran, Michael F. Z. Wang, Zhengda Li, Kyle Awayan, The Tabula Microcebus Consortium, Jonathon Z. Long, Iwijn De Vlaminck, Sheng Wang, Jacques Epelbaum, Christin Kuo, Jeremy Terrien, Mark A. Krasnow, James E. Ferrell

## Abstract

Hormones mediate long-range cell communication in multicellular organisms and play vital roles in normal physiology, metabolism, and health. Using the newly-completed organism-wide single cell transcriptional atlas of a non-human primate, the mouse lemur (*Microcebus murinus*), we have systematically identified hormone-producing and -target cells for 84 classes of hormones, and have created a browsable atlas for hormone signaling that reveals previously unreported sites of hormone regulation and species-specific rewiring. Hormone ligands and receptors exhibited cell-type-dependent, stereotypical expression patterns, and their transcriptional profiles faithfully classified the molecular cell type identities, despite their comprising less than 1% of the transcriptome. Cells of similar cell types further display stage, subtype or organ-dependent specification of hormonal signaling, reflecting the precise control of global hormonal regulation. By linking ligand-expressing cells to the cells expressing the corresponding receptor, we constructed an organism-wide map of the hormonal cell communication network. This network was remarkably densely and robustly connected and included a myriad of feedback circuits. Although it includes classical hierarchical circuits (e.g. pituitary → peripheral endocrine gland → diverse cell types), the hormonal network is overall highly distributed without obvious network hubs or axes. Cross-species comparisons among humans, lemurs, and mice suggest that the mouse lemur better models human hormonal signaling, than does the mouse. Hormonal genes show a higher evolutionary conservation between human and lemur vs. human and mouse at both the genomic level (orthology-mapping and sequence identity) and the transcriptional level (cell type expression patterns). This primate hormone atlas provides a powerful resource to facilitate discovery of regulation on an organism-wide scale and at single-cell resolution, complementing the single-site-focused strategy of classical endocrine studies. The network nature of hormone regulation and the principles discovered here further emphasize the importance of a systems approach to understanding hormone regulation.

## Introduction

Hormones are chemical messengers that circulate through the bloodstream and control and coordinate functions of both nearby and distant cells. Since the discovery of the first hormone, secretin over a hundred years ago^1^, about 100 mammalian hormones have been identified^2^. Over the decades, these hormones have been purified, sequenced, and synthesized, their receptors have been identified, and the intracellular signaling pathways they regulate have been explored^3^. Hormones regulate diverse biological processes ranging from growth and development, reproduction, metabolism, and immune responses, to mood and behavior. Many hormones are also therapeutically significant. For example, the discovery of insulin^4^ changed type I diabetes from an inevitably-fatal disease to a manageable one. Likewise, corticosteroids, a class of small molecule adrenal hormones, are used to treat diverse pathological conditions, including the inflammation attendant to COVID-19, where corticosteroids reduce the death rate in severe cases by ∼30%^5,6^.

The classical endocrine hormones are produced by secretory cells residing in one of the nine endocrine glands. However, it is now clear that other cell types, like fat cells and leukocytes, secrete global regulators into the circulatory system^7^. In addition, some endocrine functions, like menstrual cycles in human females, depend upon interplay and feedback between multiple hormones and cell types^8^. For these reasons, a complete understanding of endocrine physiology requires comprehensive, global approaches. Variations in hormone regulation and function in different species further complicate the situation. Although comparative studies have shown that many hormones are highly conserved across vertebrates^3^, the hormone-producing and -receiving cells and their physiological functions are sometimes different^9^.

The grey mouse lemur (*Microcebus murinus*) is a small non-human primate with a lifespan of 5-13 years. Because of its small size, rapid reproduction, and close genetic proximity to humans, the mouse lemur is an appealing model organism for studies of primate biology, behavior, aging, and disease^10–14^. As described in detail elsewhere^15,16^, we have recently created, using single-cell RNA-sequencing (scRNAseq), a molecular cell atlas of the mouse lemur by profiling mRNA sequences from ∼225,000 cells from 27 mouse lemur organs and tissues from 4 individuals (https://tabula-microcebus.ds.czbiohub.org/). Given the ethical concerns of primate studies, tissue samples used in the study were collected opportunistically from aged lemurs. Histopathology was assessed for all organs^17^; many looked histologically normal, although age- or pathology-related changes were detected in certain tissues such as spontaneous tumors in the uteri of two animals and metastasis in the lung of one^16,17^. Presumably, the overall hormone network is still operative in these lemurs, but the age of the animals is both a limitation and an opportunity to study disease-associated changes in hormone profiles. Nevertheless, this extensive, organism-wide dataset provides an unprecedented opportunity to study global hormone regulation at single cell resolution in a non-human primate.

In this study, we present a comprehensive analysis of primate hormone signaling based on the information in the mouse lemur molecular cell atlas^15^. We systematically mapped hormone-producing and target cells for 84 hormone classes, and created a browsable atlas that depicts gene expression of hormone ligands, modulators, and receptors. These systematic analyses detected both canonical and unreported sites of hormone secretion and action. We also analyzed the global expression patterns of hormone ligands and receptors, constructed and characterized an organism-wide cell communication network for hormone signaling, and systematically searched the network for feedback circuits. This analysis revealed a few remarkable principles of hormone signaling and a long list of potentially novel feedback mechanisms. Lastly, we compared hormonal gene expression across humans, lemurs, and mice in selected tissues and cell types, and examined their conservation in expression patterns. This revealed both hormonal genes with a highly conserved expression pattern and ones that show a primate or species-specific expression pattern.

The hormone atlas provides a useful resource and a general approach to study hormone regulation and evolution on an organism-wide scale and at single-cell resolution, complementing the more reductionistic strategies typical in classical endocrine studies. The network nature of hormone regulation and principles discovered here further emphasize the importance of a systems approach in understanding hormone regulation.

## Results

### A comprehensive table of genes involved in the biosynthesis and sensing of human and mouse lemur hormones

Although many hormones have been studied for decades, there is not a well-annotated database of hormone ligands, synthases, processing enzymes, and receptors. Therefore, we integrated multiple sources to compile a comprehensive gene table for human hormone signaling (**Table 1**). For the hormone list, we mainly referenced the *Handbook of Hormones*^2^, the Wikipedia *<u>List of</u> <u>human hormones</u>*, and the *Handbook of Biologically Active Peptides*^18^, and added newly discovered hormones including asprosin^19^ and the endocrine fibroblast growth factors FGF21, FGF19, and FGF23^20^. To collect the genes involved in hormone production, we assembled the ligand genes for peptide and protein hormones and the hormone synthesis genes for small molecule hormones. Maturation or activation of many hormones requires specific enzymes such as angiotensin-converting enzyme (ACE) for angiotensin, the deiodinases DIO1/2 for thyroid hormones, and prohormone processing enzymes (e.g., PCSK1, PCSK2, CPE) for many peptide hormones^18^. In addition, hormone function may involve endocrine modulators such as IGF binding proteins (IGFBPs) for IGF1 and specific plasma binding proteins for most steroid hormones^21,22^. Accordingly, we added these processing enzymes and modulators for the corresponding hormones. Finally, we curated the hormone receptor genes that are responsible for signaling in the target cells. Most hormones have characterized ligands and receptors, except for endomorphins (tetrapeptides), the ligand gene of which remains unknown. In total, we collected information for 84 groups of hormones and 350 genes, including 102 ligand genes, 41 synthases, 33 processing enzymes/modulators, and 174 receptor genes.

When a hormone included multiple ligand, receptor, synthase, or processing enzyme genes, we incorporated AND/OR logic gates among the genes to determine if a cell type produces or is targeted by a hormone (**Table 1**). When two or more of the genes were required for signaling, genes were connected by AND (&) gates. For example, amylin receptors are heterodimeric complexes of CALCR and RAMP proteins, and both subunits are required for function; thus, CALCR and RAMP were connected by an AND symbol. Similarly, AND gates were applied among hormone synthases that catalyze different steps, and between hormone ligands and the corresponding processing enzymes. On the other hand, OR (|) gates were applied when only one of the genes is required for function. For example, there are nine adrenergic receptor genes, all of which function independently and therefore were linked by OR gates.

We next looked up the mouse lemur orthologs of the human hormonal genes in the most up-to-date mouse lemur genome assembly (Mmur_3.0) with both NCBI and Ensembl ortholog databases (**Table 2**). Almost all (98.9%, 346/350) of the human hormone-related genes were annotated in the mouse lemur genome, and most (98.0%, 339/346) were one-to-one mappings; 7 were many-to-one. The four genes without lemur orthologs (LHB, GALP, INSL4, and DRD5) and 7 many-to-one mappings could have resulted from incomplete annotation of the mouse lemur genome. For one lemur-unannotated gene LHB (luteinizing hormone beta subunit), we identified by sequence homology^23^ a candidate LHB gene region that has high sequence similarity and the conserved neighboring gene (RUVBL2) as human LHB^24^ and was selectively expressed in the expected cells (pituitary gonadotrophs) (**Fig. S1**-luteinizing hormone). We concluded that the identified region is likely the mouse lemur LHB gene and included it in follow-up analyses. Thus, at least 347/350 (99.1%), and possibly all, of the human hormonal genes are present in the mouse lemur. For comparison, the mouse genome lacks orthologs for six genes (98.3% present, 344/350), five of which are known pseudogene or have been lost in the mouse genome (GNRH2, MCHR2, MLN, MLNR, NPBWR2 (Gm14489)) (**Table 2**). In addition, 14 human-mouse orthologs are one-to-many (6), many-to-one (4), or many-to-many (4) mappings, suggesting gene family expansions or contractions in mice or humans. The percentages of human hormonal orthologs present in lemur or mouse genomes were higher than the percentage of all protein-coding genes (PCGs) (>98.9% vs. 89.5% in the mouse lemur genome, and 98.3% vs. 89.8% in the mouse genome). Most (85.8%) human hormone-related proteins were more similar in sequence to their lemur orthologs than to their mouse homologs (**Fig. S2a**). These findings underscore the close similarity between human and lemur hormonal genes.

### Organism-wide identification of hormone-producing and target cells

We systematically identified the source and target cells for each hormone by examining the expression of the hormone ligand, synthase, and receptor genes across 739 lemur cell types from 27 different tissues profiled by 10x scRNAseq^15^. Source cells were defined as cell types that expressed above-threshold levels of the ligand (for protein/peptide hormones) or synthase(s) (for small molecule hormones) plus any necessary modulator genes. Target cells were those that expressed hormone receptor genes. We generated a browsable set of figures and a web portal (https://tabula-microcebus.ds.czbiohub.org/hormone-atlas) that depict the global expression profiles of the hormonal genes (**Fig. S1**, **Fig. S3**, **Table 3**). **Fig. 1a-g** shows the expression of four representative examples of hormones (in red) and their receptor(s) (in blue) in the identified cell types across the seven, color-coded cell compartments (epithelial, endothelial, stromal, neural, germ, lymphoid and myeloid cells). This analysis identified many canonical sites of hormone secretion and targets, affirming the validity of the single-cell transcriptomic measurements and analysis approach. The analysis also revealed many less-recognized or unreported hormone source/target cells and interesting global patterns of hormone signaling. Below, we highlight a few examples of well-known, lesser-known, and previously unknown sources and target cells. We also showcase examples where hormonal expression is unique in the mouse lemur versus mice or humans.

**Fig. 1.**
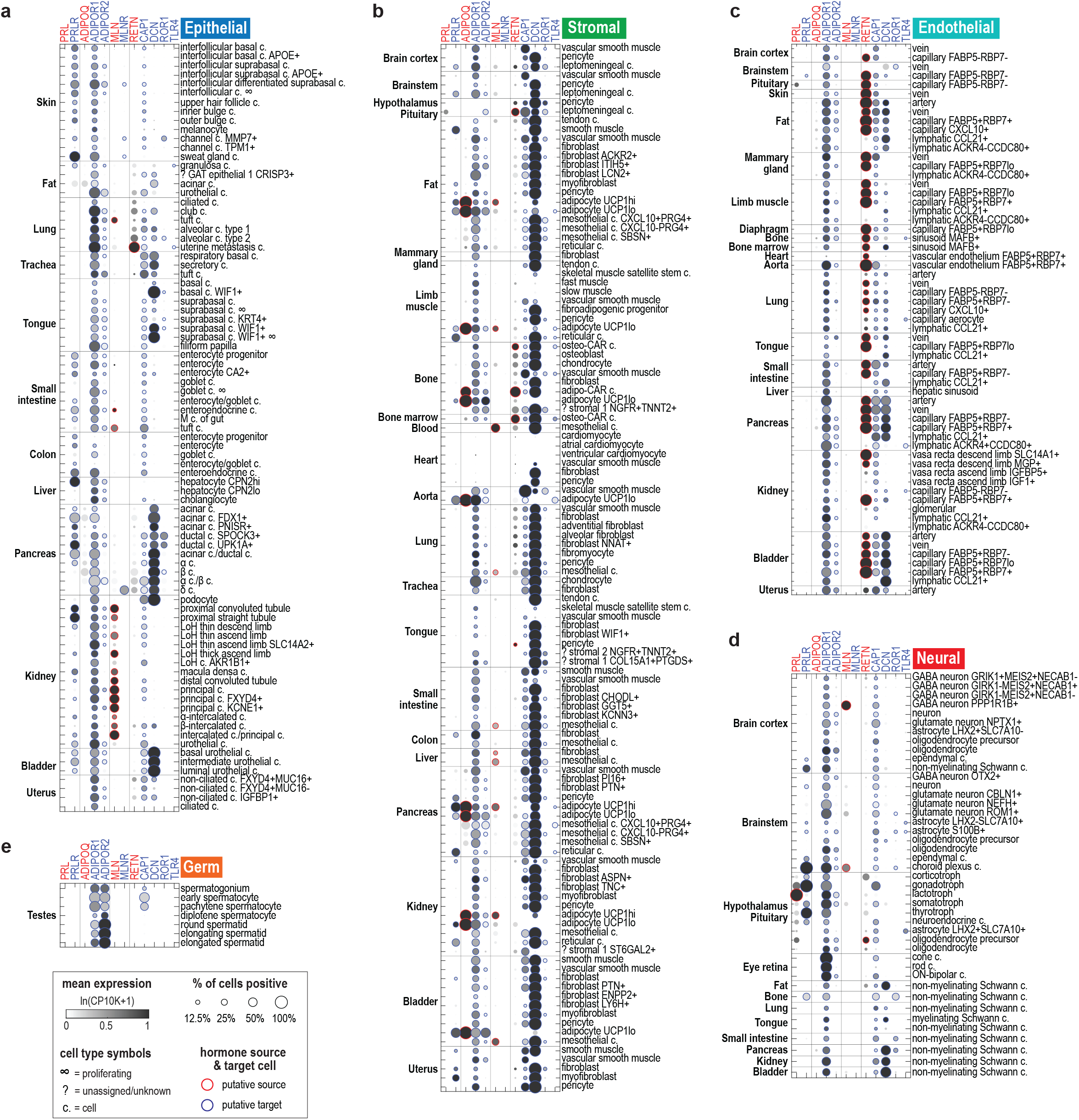
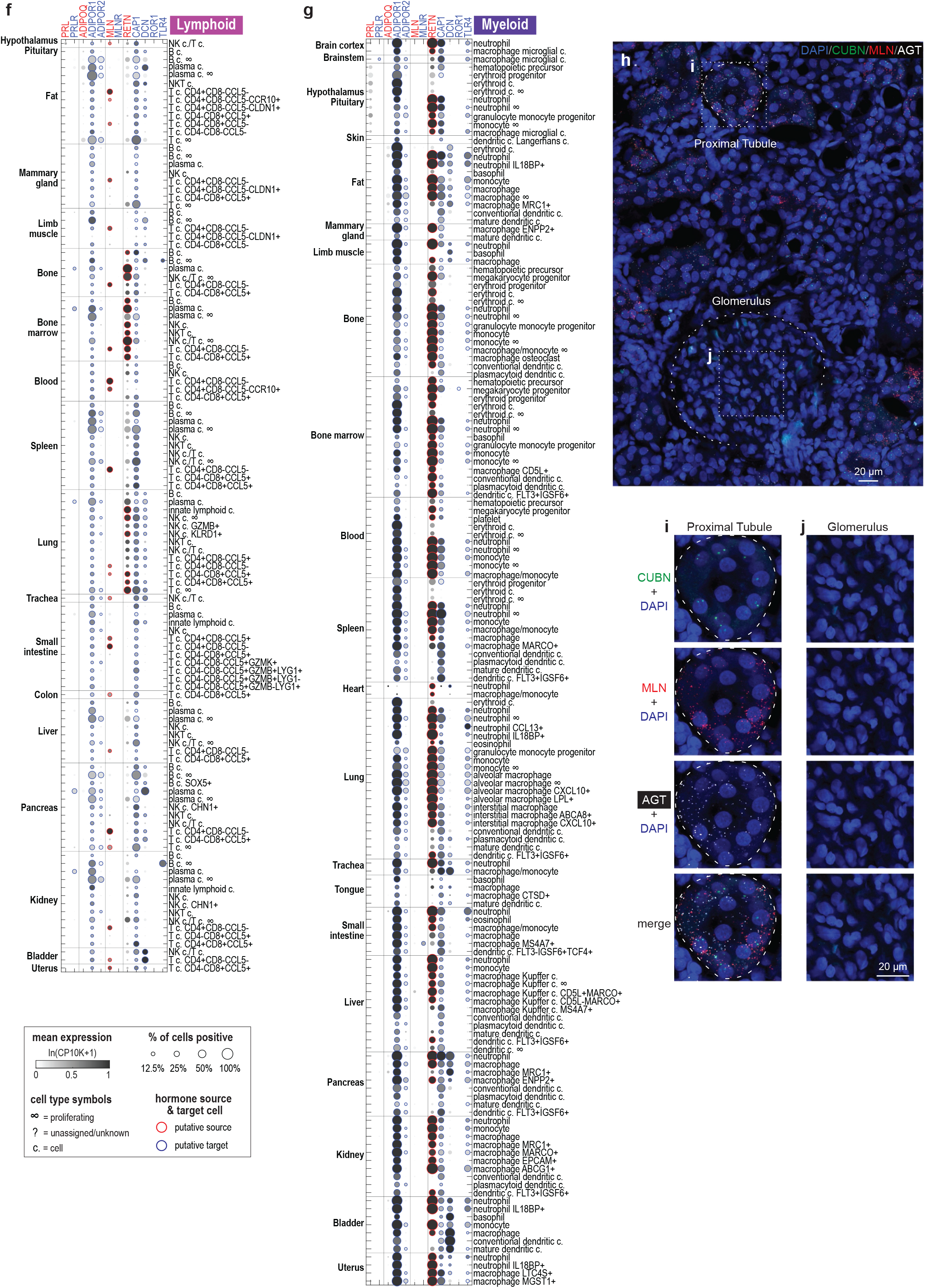
Expression of selected hormone ligands and receptors across mouse lemur cell types. **a-g**. Dot plot showing expression of four exemplary sets of hormone ligands and receptors across the ∼700 cell types from the mouse lemur cell atlas, including: prolactin (ligand: PRL, receptor: PRLR), adiponectin (ligand: ADIPOQ, receptors: ADIPOR1, ADIPOR2), motilin (ligand: MLN, receptor: MLNR), and resistin (ligand: RETN, receptors: CAP1, DCN, ROR1, TLR4). Cell types are arranged by compartment: epithelial cells (**a**), stromal cells (**b**), endothelial cells (**c**), neural cells (**d**), germ cells (**e**), lymphoid cells (**f**), and myeloid cells (**g**). Within each panel, rows are cell types ordered by tissue of origin. Cell type names are labeled on the right, organs and tissues on the left, and gene symbols on the top (red for ligand genes, blue for receptor genes). Dots represent average gene expression (circle color) and percent of cells positive for the gene (circle size). The red/blue circle edges indicate expression was above threshold (see **Methods**) for the ligand or receptor. Cell types that are potentially low quality (e.g., low cell count) are excluded from this plot. Note that granulosa cells and acinar cells, which presumably are derived from the ovary and pancreas, respectively, were also found in perigonadal and mesenteric fat samples. This was likely caused by accidentally including small pieces of adjacent tissues during tissue procurement. A complete set of figures that display ligand and receptor gene expression for all 84 hormone classes can be found in **Fig. S1** and cross-species expression patterns in **Fig. S12**. **h-j**. Single plane confocal images of mouse lemur kidney assayed by RNAscope for two hormones, motilin (MLN) and angiotensin (AGT), and the proximal tubule marker cubilin (CUBN). Cells were also stained with DAPI to visualize nuclei. Note the MLN and AGT expression in the CUBN+ proximal tubule cells and not in the glomerulus, which is mainly composed of podocytes and glomerular capillary cells. Panel **h** shows a global view (merged 20× tile scans) with a single glomerulus in the view and many proximal tubules. The glomerulus and a representative proximal tubule are outlined by dashed curves. Panels **i** and **j** are magnified views of the boxed areas in **h** showing a proximal tubule and glomerulus respectively. See also Fig. 3b for scRNAseq data on MLN and AGT expression across kidney nephron epithelium.

#### Prolactin (PRL)

Prolactin is a classical endocrine hormone^25^ secreted by pituitary lactotrophs. In addition to its milk-promoting effect in females, PRL also plays pleiotropic roles in both sexes^2^. These roles include regulation of water and electrolyte balance in the kidney, and control of skin cell proliferation and hair/feather growth and shedding^26^. Prolactin also plays a major role in energy homeostasis by regulating cell growth and metabolism in the liver, adipose tissues, and pancreas in both sexes^27^, and may be involved in the immune response^28^. These sex-non-specific functions are conserved, not just among milk-producing mammals, but across all vertebrates.

As expected, prolactin transcripts were selectively expressed in the lactotrophs of the lemur pituitary gland, and were not detected in other tissues (**Fig. 1a-g, Fig. S1**-prolactin). The prolactin receptor (PRLR) was found in both male and female lemurs in the expected cell types, including kidney nephron tubule cells, skin epithelial cells, pancreatic ductal and acinar cells, hepatocytes, intestinal epithelial cells, adipocytes, reticular cells, plasma cells, choroid plexus cells, and the neighboring TSH-secreting thyrotrophs and FSH/LH-secreting gonadotrophs in the pituitary gland (**Fig. 1a-g, Fig. S1**-prolactin). Thus, scRNAseq data confirm that prolactin signaling is similar in mouse lemurs and other animals.

#### Adiponectin (ADIPOQ) and leptin (LEP)

Mouse lemurs undergo dramatic seasonal changes in body weight and fat reserves^29–31^ and therefore may experience seasonally-varying levels of adipokines. Adiponectin and leptin are two important metabolic hormones secreted by adipocytes^32,33^. Although adipocytes were originally regarded only as nutrient storage cells, it is now clear that they also serve an endocrine role^34^. Surprisingly, leptin transcripts were almost undetectable in the adipocytes captured in the atlas (see discussion in the accompanying manuscript^16^) (**Fig. S1**-leptin).

However, adiponectin was abundantly and specifically expressed in mouse lemur adipocytes captured from multiple tissues, including four adipose depots (subcutaneous, mesenteric, interscapular brown, and perigonadal), as well as kidney, limb muscle, pancreas, and bone (**Fig. 1a-g**, **Fig. S1**-adiponectin). Both UCP1-low white-like and UCP1-high brown-like adipocytes highly expressed ADIPOQ.

Previous research has characterized tissue-specific functions of adiponectin in the liver, pancreas, skeletal muscle, endothelial cells, macrophages, and adipocytes^35^. While we indeed observed expression of the adiponectin receptor ADIPOR1 in these tissues in the mouse lemur, we also found ADIPOR1 to be abundantly expressed in almost all lemur cell types (**Fig. 1a-g**). Similar ubiquitous expression of ADIPOR1 has been found in humans and mice (**Fig. S12**). These results suggest that adiponectin exerts an organism-wide effect through ADIPOR1.

In comparison, ADIPOR2, the other adiponectin receptor, was expressed more selectively, most notably in the male germ cells and adipocytes (**Fig. 1a-g**). Interestingly, the lemur expression pattern of ADIPOR2 differed from that of the human and mouse in which ADIPOR2 was highly expressed in the liver and only lowly expressed in the testis^36^ (**Fig. S12**). Alveolar macrophages from the mice, but not humans or lemurs, also showed high ADIPOR2 expression (**Fig. S12**), suggesting differential ADIPOR2 effects across the three species.

#### Motilin (MLN)

Motilin is canonically considered a gastrointestinal hormone that is secreted by enteroendocrine cells and regulates gastrointestinal smooth muscle movement^18^. However, rodents have lost both the motilin and the motilin receptor genes in evolution^37^ (**Table 2**). The lack of motilin signaling in rodent models has hampered progress on further characterizing motilin function as well as the development of motilin agonists for the treatment of gastroparesis^38^. Consistent with the canonical view, we observed motilin expression in lemur enteroendocrine cells in the small intestine but not colon (**Fig. 1a-g**). However, we also observed a broad range of other motilin-expressing cell types, including kidney nephron tubule cells, chemosensory tuft cells from both the airway (lung and trachea) and small intestine, certain brain neurons, adipocytes (higher in UCP1-high brown-like adipocytes than in white-like adipocytes), mesothelial cells, and the majority of the CD4+CD8-CCL5- helper T cells (**Fig. 1a-g**, **Fig. S1**- motilin). The receptor was found to be most highly expressed in pancreatic delta cells and several epithelial cell types of the skin (e.g., sweat gland cells). Together, this expression pattern suggests that motilin is involved in a diverse array of mouse lemur physiological functions in addition to the regulation of gut motility.

Motilin had been detected in the human brain^18^, as is the case in the lemur (**Fig. 1d**), but was not previously known to be expressed in the kidney nephron tubule cells. Therefore, we performed RNAscope, a single molecule mRNA fluorescent in situ hybridization (smFISH) assay to further test whether motilin is expressed in the mouse lemur kidney. This revealed abundant motilin transcripts in the epithelial cells of the proximal tubules but not in the glomerulus, which are mainly composed of podocytes and glomerular capillary cells (**Fig. 1h-j**). This supports the validity of the scRNAseq results (**Fig. 1a-g**). It will be interesting to examine, in future studies, motilin expression in other uncanonical cell types outside the intestine, explore what functions motilin plays in these cells, and whether their expression is conserved in humans.

#### Resistin (RETN)

Resistin is a phylogenetically recent hormone, present only in mammals^2^. It is expressed mainly by adipocytes in mice but by macrophages, monocytes, and neutrophils in humans^39,40^. In the mouse lemur, we found that resistin is highly expressed in macrophages, monocytes, and neutrophils, and nearly absent from adipocytes (**Fig. 1a-g, Fig. S1**-resistin), as it is in humans (**Fig. S12**). Interestingly, resistin is also highly expressed in many of the vascular endothelial cell types and in metastatic uterine tumor cells, suggesting additional sources for this hormone in mouse lemurs (**Fig. S12**).

#### Prohormone processing enzymes

Prohormone processing enzymes are required for the maturation of many peptide hormones, including insulin, adrenocorticotropic hormone (ACTH), gastric inhibitory polypeptide (GIP), and somatostatin^18^. The required processing enzymes have been identified for some but not all hormones^41,42^. In this study, we considered both the ligand and the known or suggested processing enzymes when determining if a cell type produces a hormone. Selective expression of the processing enzymes can help distinguish hormones that share the same precursor gene. For example, proglucagon (GCG) is the precursor for both glucagon and glucagon-like peptide-1 (GLP-1). GCG was expressed in both pancreatic α cells and intestinal enteroendocrine cells, but the two cell types expressed different processing enzymes: α cells expressed only PCSK2 which is required for glucagon synthesis, and enteroendocrine cells expressed only PCSK1 which is required for GLP-1 synthesis (**Fig. S1**- glucagon). Thus, as is the case in mice and humans^43^, α cells are sources of glucagon and enteroendocrine cells are sources of GLP-1 in the mouse lemur.

Interestingly, we noticed that some cell types in the mouse lemur transcribed prohormone genes without expressing the respective processing enzymes. For example, proopiomelanocortin (POMC), which is the precursor of ACTH, melanocyte-stimulating hormone (MSH), β- endorphin, and lipotropin, is first processed by PCSK1^44^. As expected, ACTH-secreting corticotrophs expressed both POMC and the necessary processing enzymes, but lemur male germ cells expressed abundant POMC without detectable PCSK1 transcripts (**Fig. S1**- proopiomelanocortin-derived hormones). Similarly, human and mouse male germ cells expressed POMC but no PCSK1 (**Fig. S12**). In addition, lemur male germ cells expressed GIP, which also requires PCSK1 processing; the intestinal and airway tuft cells expressed renin (REN) without the necessary PCSK1 (**Fig. S1**). These results suggest several possible hypotheses: different peptidases are involved, the prohormone transcripts are not translated, the prohormone proteins may function without proteolysis, or the processing enzymes may be expressed below detection levels in the sampled cells.

#### Plasma binding proteins

Plasma binding proteins are not only passive hormone transporters, but can affect hormone functions by modulating hormone half-life and levels of free, active hormones accessible to target cells. Other plasma binding proteins have been reported to potentiate the binding of hormones to their targets^45^.

Steroid and thyroid hormones are transported in the blood by plasma binding proteins of the albumin family. While serum albumin (ALB) binds steroids with low specificity and affinity, four other albumins bind these hormones with high specificity and nanomolar affinity, including sex hormone binding globulin (SHBG) for androgens and estrogens, vitamin D binding protein (GC) for vitamin D, SERPINA6 for cortisol and progesterone, and SERPINA7 for thyroid hormones^21,22^ (**Table 1**). Consistent with the notion that these hormone-binding albumins are produced in the liver^21,22^, we identified specific expressions of SHBG, SERPINA6, and SERPINA7 in the lemur hepatocytes, and interestingly higher levels in the CPN2hi versus the CPN2lo hepatocyte subtypes (**Fig. 3f, Fig. S1**). GC was also expressed in the lemur hepatocytes with a higher level in the CPN2hi subtype; however, we also detected notable expression in pancreatic endocrine, ductal, enteroendocrine cells, and cholangiocytes. These novel sources may contribute to the circulating pool of GC or may have local or exocrine effects, such as regulating intestinal vitamin-D absorption^46^.

In addition, peptide hormones IGF1 and activin are known to be bound by IGF binding proteins (IGFBPs) and FST respectively. In contrast to albumins, IGFBPs and FST were expressed by multiple distinct cell types and showed high cell type variability (**Fig. S1**).

In summary, we have depicted the organism-wide, single-cell expression patterns for hormones, synthases, hormone modulators, and receptors from 84 classes of hormones, and have systematically identified their sources and targets (**Fig. S1**). This global analysis detected canonical cell types underlying hormone secretion and target functions while also revealing previously unreported source and target cells. This organism-wide single-cell hormone atlas should facilitate the systematic study of hormone regulation in an unbiased, global manner, paving the way for the discovery of novel regulation and evolution of hormone signaling.

### Hormonal gene expression is highly cell type specific

Next, we clustered mouse lemur cell types based on the transcriptional profiles of the hormonal genes (**Fig. 2a-b**). We identified 74 clusters and summarized the hormones and hormone receptors that were expressed by each cluster (**Table 4**). We noted that cells of the same type almost always clustered closely irrespective of their tissue of origin, or of the sex or identity of the sampled individual, whereas different cell types were well-separated (**Fig. 2b**, **Fig. S4**, **Table 4**). For example, all lymphatic endothelial cells clustered closely and showed characteristic IGF1 expression despite their coming from nine different tissues and four individuals (cluster 45, **Fig. 3b**, **Fig. S1**, **Table 4**). Mesothelial cells sampled from eight visceral organs formed a cluster distant from other stromal cells and showed distinctive expression of NPR1, CFB, PTGIS, ITGB3, and PNMT (cluster 35, **Fig. 2b**, **Fig. S1**, **Table 4**). Although there are only ∼300 hormonal genes, ∼1% of all detected genes, their expression pattern proved to be indicative of cell type identity regardless of tissue source or location.

**Fig. 2.**
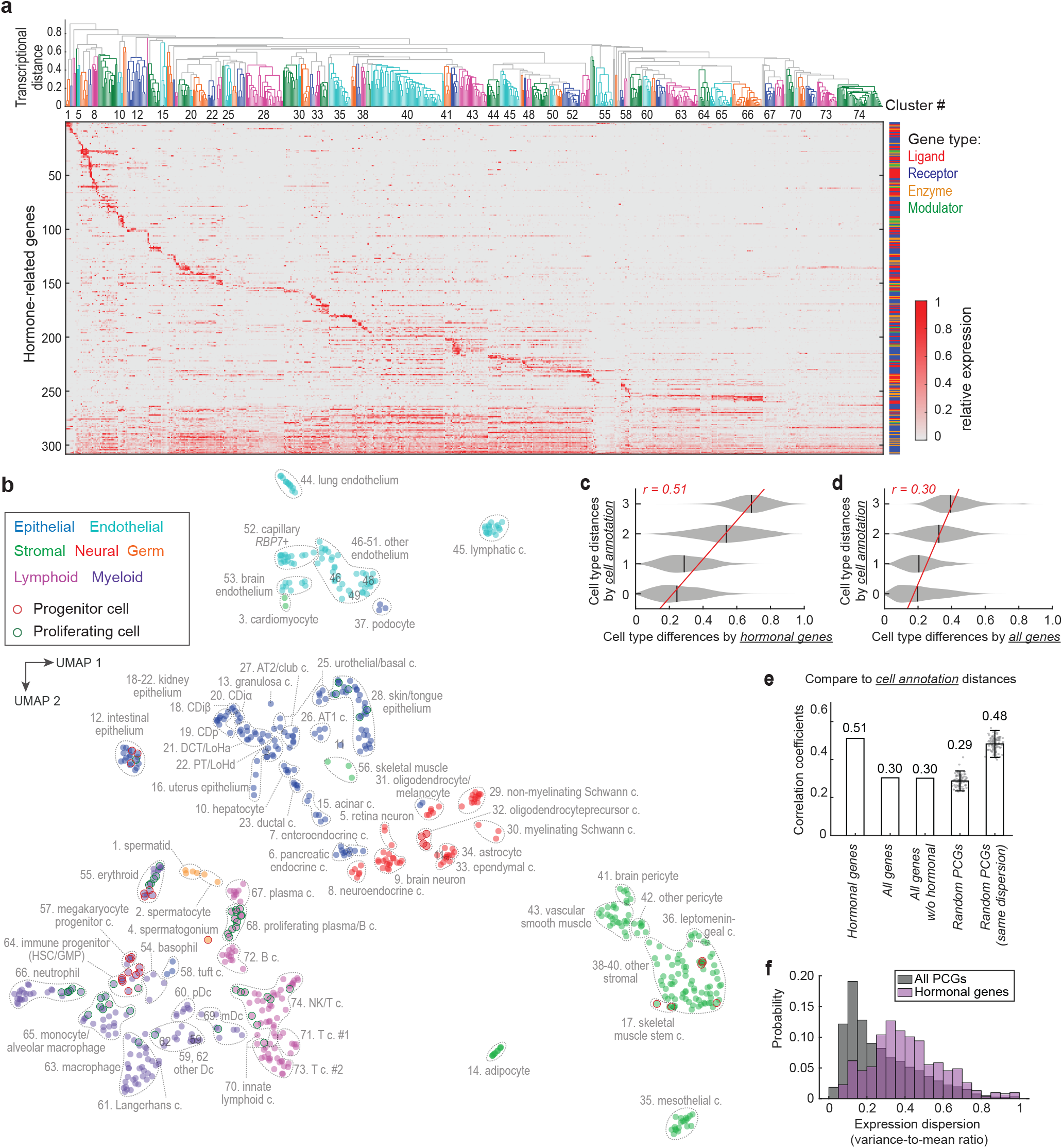
Hormonal gene expression is cell type specific. **a**. Clustering of mouse lemur cell types by hormonal gene expression. At the top is a dendrogram of the hierarchical clustering. Clusters of cell types are color-coded, and cluster numbers are labeled as space allows. The cell types included in each cluster are listed in **Table 4**. The lower part of the panel shows a heatmap of relative gene expression levels. Cluster-specific genes are ordered based on the cluster with the highest expression level, with the genes preferentially expressed in the left-hand cell type clusters on the top. Broadly expressed genes (expressed in more than 35% of cell types) are placed in the bottom and ordered from least to most broadly expressed. **b**. UMAP visualization of mouse lemur cell types based on the expression of hormonal genes. Circles are cell types (unique combination of annotation name and tissue of origin) and color-coded by cell type compartment types (epithelial, neural, germ, stromal, endothelial, lymphoid, and myeloid). Dashed lines circumscribe the cell type clusters as in panel **a**, and cluster IDs and names were labeled nearby. One extremely distant cluster (# 35. mesothelial cells) was shifted up in UMAP-2 towards the center of the figure for display purposes. **c**. Hormonal (**c**) or full transcriptome (**d**) - based cell type pairwise distances compared to the benchmark cell annotation-based distance. *r* represents Pearson’s correlation coefficient. Red lines show linear fitting of the data. **e**. Correlation coefficients of cell type transcriptional distances based on different sets of genes with the benchmark cell annotation-based distance. **f**. Distributions of expression variability (dispersion) of the hormonal genes or all protein-coding genes (PCGs).

To further test this conclusion, we compared cell type clustering by either the hormonal genes (∼300 genes) or the full transcriptome (>30,000 genes). To visualize the results, we projected the high-dimensional transcriptional data into two-dimensional cell type maps by uniform manifold approximation and projection (UMAP), which illustrate cell type similarity based on the selected genes. The cell type UMAPs by either the hormonal genes (∼300 genes) (**Fig. 2b**) or the full transcriptome (**Fig. S5a**) were remarkably similar (see **Fig. S7c** for comparison). The cell type pairwise distances defined by hormonal genes versus the full transcriptome were also positively correlated, with a Pearson correlation coefficient (*r*) of 0.70 (**Fig. S5b**). There were nevertheless minor differences between the two UMAPs. For example, airway and intestinal tuft cells are both chemosensory but are involved in different functions^47^. In the space of hormonal genes, tuft cells from different tissues formed one cluster (cluster 58) with distinctive expression of renin (REN), cyclooxygenase 1 (PTGS1), arachidonate 5-lipoxygenase-activating protein (ALOX5AP, required for leukotriene production), and motilin (MLN) (**Fig. 2b**, **Fig. S1**, **Table 4**). However, in the space of all genes, the lung, trachea, and intestinal tuft cells were separated from each other (labeled as 58A*, 58B*, and 58C* in **Fig. S3**). Another notable exception was skin melanocytes, which clustered with oligodendrocytes (cluster 31) by only the hormone-related genes (**Fig. 2b**) but clustered more closely with skin epithelial cell types in the space of all genes (**Fig. S5a**, oligodendrocytes and melanocytes labeled as 31A and 31B, respectively). Thus, cell types were overall similarly clustered by the hormonal genes or the full transcriptomes, the exceptions showcase rare examples of converged hormonal signaling in distant cell types.

We then assessed how well hormonal gene expression discriminates among different cell types compared to all genes or random sets of PCGs (**Fig. 2c-e, Fig. S5b-e**). As a gold standard, we used the cell annotations described in the accompanying manuscript^15^, which are based on expression of canonical cell markers and iterative clustering with highly variable genes. We then scored how well the cell type pairwise transcriptional distances for these different sets of genes correlated with a cell annotation-based distance (**Fig. S6c**). This annotation-based distance is defined as an integer between 0 and 3: 0 when cell types were annotated with identical names but from different tissues, 1 when the cell annotation names were different but classical histological assignments (Cell Ontology^48^) were identical (i.e., different molecular subtypes), 2 when classical Cell Ontologies were different but cell types were from the same compartment (e.g., both were epithelial), and 3 if cells were different cell types and from different compartments (e.g., endothelial vs. stromal). The pairwise transcriptional distances based on the hormonal genes alone correlated with this annotation-based distance better than that based on the full transcriptome (*r* = 0.51 vs. *r* = 0.30) (**Fig. 2c-e**). Likewise, the hormonal-based distances correlated better than distances from random sets of PCGs (*r* = 0.51 vs. *r* = 0.29 ± 0.03, mean ± S.D., *n* = 100) (**Fig. 2e**). This appeared to be because hormonal genes were more variably expressed across different cell types than average PCGs (**Fig. 2f**); although the full transcriptome contains more information, the discriminative power is diluted by the large number of low-specificity genes. In support of this idea, when sets of randomly selected PCGs with similar expression variability (dispersion) as hormonal genes were used, the correlation coefficients were similar to that of the hormonal genes (**Fig. 2e**).

As a further test, we used a continuous Cell Ontology^48^-based metric as the benchmark (**Fig. S5e**). This metric was calculated as the mean of the Cell Ontology embedding-based distances from the Cell Ontology graph and the text-based distances from the text descriptions of the Cell Ontology terms^49^. The hormonal-based cell type pairwise distances correlated better with this Cell Ontology-based distance than those based on all genes or random sets of PCGs (*r* = 0.45 vs. *r* = 0.23, 0.22 ± 0.04) (**Fig. S5e**), as was the case with the discrete annotation-based distance. Thus, by both measures, hormonal genes do an unusually good job of predicting cell identity.

Finally, we examined if the expression of hormone ligands or receptors alone could distinguish cell types. Interestingly, for most cell type clusters, hormone ligands or receptors alone classified cell types similarly (**Fig. S7a-c**). This suggests that most cell types have stereotypical expression of genes involved in both incoming and outgoing hormonal signaling. A small fraction of clusters were mainly defined by only the ligands or receptors. For example, clusters of pituitary neuroendocrine cells (cluster 7) and tuft cells (cluster 21) were distinguished mainly by hormone ligands, whereas clusters of the skeletal muscle stem cells (cluster 17) and lung endothelial cells (cluster 51) were mostly based on differences in hormone receptor expression (**Fig. S7c**).

In summary, we found that cells generally show stereotypical and cell-type-specific expression of hormone ligands and receptors. Thus, hormone secretion and sensing essentially labels the identity of the cells.

### Specific hormonal gene expression in cell subtypes

Hormonal expression could sometimes distinguish cell subtypes that are at different developmental stages, from different locations within an organ, from different organs, or from normal vs. pathological tissue. Below we highlight several such examples (**Fig. 3**).

**Fig. 3.**
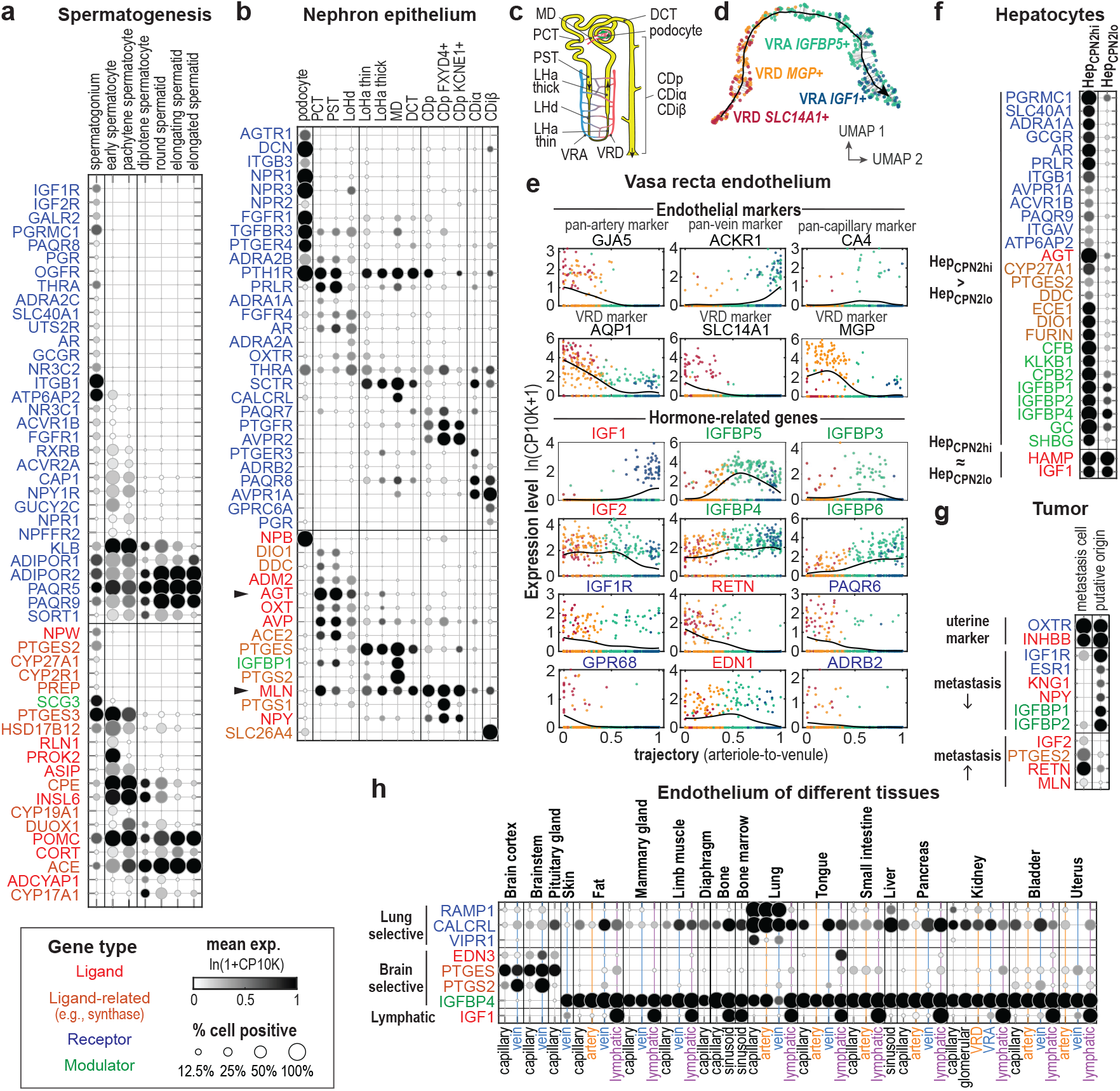
Specific hormonal regulation in related cell types. Expression of hormone ligand, synthase, modulator, and receptor genes that are differentially expressed in (**a**) male germ cells across different stages of spermatogenesis, (**b**) epithelial cells along the spatial axis of the kidney nephron tubule, (**e**) endothelial cells along the arteriole-to-venule spatial axis of the kidney vasa recta, (**f**) the CPN2hi vs CPN2lo subtypes of hepatocytes, (**g**) between metastatic tumor cells in the lung of lemur L2 and the putative cell type of metastatic origin (MUC16+ non-ciliated uterus epithelial cells) sampled from L3, and (**h**) endothelial cells of the lung and brain compared to other organs. Panel **c** shows a diagram of the nephron and vasa recta that illustrates the spatial distribution of the respective cell types, related to panels **b**, **d** and **e**. Panel **d** shows a UMAP visualization of the vasa recta cells based on expression of all genes. The black line shows the detected arteriole-to-venule trajectory with the arrow pointing in the direction of blood flow. Cells are color-coded according to the cell type annotation names in the atlas. Grey lines show the alignment of the individual cells to the trajectory. Panels **a-b, f-h** are dot plots showing average gene expression and percent of positive cells of the indicated genes and cell types, merging data from different lemur individuals. In **a**, cell types are arranged along the course of the spermatogenesis. In **b**, cell types are arranged according to their spatial distribution along the kidney nephron tubule. Solid arrowheads point to AGT and MLN; their kidney expression was confirmed by RNAscope (Fig. 1h**-j**). In **h**, endothelial cells are arranged by tissue of origin. In **e**, scatter plots show gene expression levels along the arteriole-to-venule trajectory for the endothelial markers and hormonal genes that are differentially expressed. Shown here is vasa recta data from one animal. Similar results were obtained from two other lemurs. **PCT**: proximal convoluted tubule. **PT**: proximal tubule. **PST**: proximal straight tubule (also called loop of Henle thick descending limb). **LHd**: loop of Henle thin descending limb. **LHa thin**: loop of Henle thin ascending limb. **LHa thick**: loop of Henle thick ascending limb. **DCT**: distal convoluted tubule. **MD**: macula densa. **CDp**: collecting duct principal cell. **CDiα**: collecting duct α intercalated cell. **CDiβ**: collecting duct β intercalated cell. **D** shows the fitted arteriole-to-venule trajectory of the kidney vasa recta capillary network (upper panel) and single-cell gene expression levels along the trajectory for the endothelial markers and differentially expressed hormone-related genes. Shown here are representative results from one of three lemur vasa recta samples. **VRD**: vasa recta descending section. **VRA**: vasa recta ascending section.

#### Spermatogenesis

Spermatogenesis occurs continuously and asynchronously in the male testis^50^. Thus, sampling the testis at one fixed time point can capture snapshots of the entire process. The atlas includes seven sperm and sperm progenitor cell types that are at different spermatogenesis stages as indicated by the expression of known stage markers^15^. The seven cell types fall into three major stages: the undifferentiated spermatogonia, the spermatocytes (which undergo meiosis), and the haploid spermatids that elongate and form mature sperms. Interestingly, these cells exhibited stage-dependent hormone signaling, with over 50 differentially expressed hormone ligands, enzymes, and receptors (**Fig. 3a**). Previous studies on spermatogenesis have mainly focused on testosterone and on the supporting cells rather than sperms and sperm progenitors^51,52^. The present data suggest that sperms and sperm progenitor cells are not only differentially regulated by a variety of hormones but may also secrete a changing array of signaling molecules as they mature. The data also clarify some incompletely understood aspects of sperm regulation. For example, progesterone was known to activate a sperm-specific calcium channel prior fertilization to induce Ca^2+^ influx, but the receptor(s) responsible for this effect were unidentified^53^. The present data show that two non-classical progesterone membrane receptors (PAQR5/9) are highly expressed in the lemur spermatids and therefore may mediate the process (**Fig. 3a**). Interestingly, two additional membrane progesterone receptors (PAQR8, PGRMC1) were specifically expressed in the undifferentiated spermatogonia (**Fig. 3a**), suggesting that progesterone may also regulate early spermatogenesis^54^. In addition, insulin-like factor 6 (INSL6), a member of the secretory relaxin family, was selectively expressed in the lemur spermatocytes (**Fig. 3a**). Supporting a function in these cells, knockout of INSL6 arrests spermatogenesis at late meiotic prophase in mice^55^. This analysis suggests that sperms and sperm progenitor cells may directly communicate with other cells via hormones that regulate spermatogenesis and sperm functions. Cross-species comparison among human, lemur, and mouse germ cells identified many of the spermatogonia hormonal genes that exhibited conserved dynamics among all three species, and others showing primate/species-specific expression dynamics (**Fig. 7e-g**, **Fig. S13a-b**, and more discussion below).

#### Kidney nephron and vasa recta

Although discrete cell types were common in the mouse lemur cell atlas, we noted several cell types that displayed a continuum of gene expression, including the epithelial cells of the nephron and the endothelium of the vasa recta, the blood vessels that supply the renal medulla^15^ (**Fig. 3d**). Interestingly, these cells also displayed gradients of hormone signaling along their spatial axes (**Fig. 3b-e**).

Nephrons are single-cell layered, one-directional tubules that serve as the structural and functional units of the kidney. We identified a large number of hormonal genes specific to different sections of the nephron epithelium (**Fig. 3b-c**). Some of these were receptors for hormones known to target specific nephron sections to regulate urine formation. For example, the antidiuretic vasopressin specifically promotes water reabsorption in the collecting duct principal cells^2^, and indeed, the vasopressin receptor AVPR2 was expressed exclusively in the lemur collecting duct principal cells. Interestingly, the collecting duct intercalated cells expressed another vasopressin receptor, AVPR1A (**Fig. 3b**), which has been recently reported to mediate the regulation of acid homeostasis by vasopressin^56^. Although the kidney epithelium is not commonly considered to be an endocrine tissue, many hormone ligands/synthases were expressed in different nephron sections, including angiotensin (AGT), adrenomedullin 2 (ADM2), motilin (MLN), and neuropeptide Y (NPY). Consistent with the scRNAseq results, smFISH by RNAscope confirmed expression of MLN and AGT in the nephron proximal tubule cells but not in the podocytes and capillary cells of the glomerulus (**Fig. 1h-j**). Taken together, these data suggest that nephron epithelium may play important roles in hormone production in addition to urine formation.

The vasa recta aligns parallel to the nephron tubules in the kidney medulla (**Fig. 3c**) and functions collaboratively with the nephrons in urine formation^57^. The vasa recta cells also exhibited a gradual shift in hormonal signaling along its arteriole-to-venule spatial gradient. (**Fig. 3d**). **Fig. 3e** shows the hormonal genes that are significantly differentially expressed along the vasa recta axis. Notably, several genes involved in IGF signaling were differentially expressed: IGF1, which was highly expressed in the lemur lymphatic endothelial cells but almost absent from other vascular endothelial types (**Fig. S1**, **Fig. 3h**), was highly expressed at the lymphatic- like^58^ venule end of the vasa recta (**Fig. 3e**); in contrast, IGF2 and IGF1R were more highly expressed in the arteriole end; and four IGFBPs (IGFBP3-6), which modulate IGF signaling, also exhibited different spatial patterns (**Fig. 3e**). This spatial organization of IGF signaling may be evolutionarily conserved, as a recent study on mouse kidney vasculature development shows a similar pattern of IGF1 expression in the mouse vasa recta^59^.

#### Hepatocyte subtypes

The mouse lemur cell atlas identified multiple novel molecular subtypes of known cell types. For example, the liver hepatocytes of the mouse lemur clustered into two discrete subtypes, the CPN2 high and CPN2 low hepatocytes. They differed significantly by total transcript count, and may correspond to the small diploid and large polyploid hepatocytes^15^.

These subtypes were found in both females and males (**Fig. S6**), and seems to be independent of the known zonal heterogeneity^15^. Interestingly, a number of hormone-related genes were differentially expressed (**Fig. 3f**). Hormones including glucagon, androgens, and prolactin appear to selectively target the CPN2hi hepatocytes as indicated by differential expression of the corresponding receptors (GCGR, AR, PRLR). In addition, angiotensin (AGT), a canonical hepatic hormone, was expressed at a much higher level in the CPN2hi hepatocytes, whereas no subtype difference was detected in the expression of hepcidin (HAMP) or IGF1, two other hormones abundantly produced by the liver.

We also identified similar hepatocytes in humans and mice and many of the subtype-specific hormonal expressions were conserved (**Fig. S13c-d**). The hepatocyte subtypes and the subtype specific hormonal expression were found in both male and female lemurs studied (**Fig. S8**), so are not attributable to sex-specific differences in hormone regulation. However, there may exist other sex-dependent differences in hormone regulation of hepatocytes as observed in other species.

The liver is a central organ of metabolism and plays important roles in hormonal regulation. The detected heterogeneity in hormone signaling suggests potentially important functional divergence between the two subtypes. A deeper understanding of the mechanisms and regulation of hepatocyte hormone heterogeneity would be particularly relevant to the study of mouse lemur-specific physiology, like hibernation/torpor and seasonal body weight changes^30^.

#### Metastatic vs. primary tumor cells

The atlas also contains diseased cells, including potential primary endometrial tumor cells in the uterus (MUC16+ non-ciliated uterus epithelial cells) and lung metastatic cells of endometrial cancer origin, as assessed by tumor gene signatures and histology (see accompanying manuscripts^15,16^). Both tumor cell types clustered with the non-tumor epithelial cells of the uterus^16^ and shared characteristic expression of the oxytocin receptor (OXTR) and the inhibin beta B subunit (INHBB) (**Fig. 3g**, **Fig. S1**, **Table 4**), suggesting that the lung metastatic tumor originated from cells resembling the primary MUC16+ uterine tumor cells. Certain aspects of hormone signaling seem to have been altered in the metastatic tumor (**Fig. 3g**). Notably, the tumor cells showed a loss of estrogen receptors (ESR1), a change associated with high grade and advanced stage endometrial cancers in humans^60^ and more often reported in high mortality type 2 human endometrial cancers^61^.

Interestingly, the metastatic tumor cells also acquired new abilities in hormone production. For example, elevated expression of prostaglandin E2 synthase (PTGES2) was present in metastatic cells (**Fig. 3g**). PGE2 has been suggested to be a tumorigenic factor in many cancers^62^. A recent study identified that the PGE2 synthase PTGES2 was elevated in human endometrial cancer tissues and found that PGE2 promotes cell proliferation and invasion in *in vitro* cell assays^63^.

Resistin was also found to be highly expressed in the metastatic cells (**Fig. 3g**). Resistin has been reported to promote invasiveness of breast cancer cells *in vitro*^64^. Thus, endometrial tumors may utilize autocrine resistin signaling to promote metastasis. The shift in hormone signaling identified in the lemur metastatic endometrial tumor is similar overall to that reported in the human. This suggests that the shifts in hormone signaling may be an important step in tumor development and metastasis. Mouse lemurs may serve as a useful animal model to study human cancers that are not well modeled in mice such as endometrial cancers^16^.

#### Vascular endothelium

While hormonal gene expression is generally determined more by cell type than by organ of origin, lung and brain vascular endothelial cells from arteries, veins, and capillaries all exhibited unique organ-specific hormonal signatures (**Fig. 3h**).

Lung endothelial cells expressed higher levels of the hormone receptors VIPR1, CALCRL, and RAMP1 (**Fig. 3h**) than did endothelial cells from other organs. VIPR1 is a receptor for vasoactive intestinal peptide (VIP), a strong bronchodilator and one of the most abundant signaling peptides in the lung^65^. Lung -specific VIPR1 expression was also observed in human but not mouse endothelial cells (**Fig. S13e**, see more discussion below). RAMP1 forms heterodimeric receptors with CALCR or CALCRL that are regulated by several hormone ligands including CGRP, adrenomedullin, amylin, and calcitonin, all of which have been reported to have vasodilatory effects^2^. Interestingly, RAMP1 is expressed in the lung endothelium of the mouse lemur but not in mice or humans, whereas RAMP2 is expressed in all three species and RAMP3 is only highly expressed in humans (**Fig. S13e**, see more discussion below). These results suggest that lung vascular endothelium is a unique hormonal target in comparison to vasculature elsewhere, possibly because of the distinctive nature of the pulmonary circulation (e.g., high volume, low pressure) compared to the systemic circulation.

In contrast, brain vascular endothelium was found to express higher levels of a few hormone ligands/synthases. Endothelin-3 (ET-3), a member of the vasoconstricting endothelin family, was selectively expressed in the brain vascular endothelium (**Fig. 3h**). ET-3 peptide has been detected at various brain locations in rodents, but it remained unsettled what cells secreted the peptide and what physiological role it played^66^. Our results suggest that ET-3 is synthesized by the endothelial cells in the brain, which could target the nearby EDNRB-expressing SLC7A10+ astrocytes and the EDNRA-expressing brain vascular smooth muscle cells and pericytes (**Fig. S1**-endothelin). Additionally, the brain endothelium also expressed high levels of PTGES and PTGS2/COX-2 (**Fig. 3h**), which are the synthases of prostaglandin E2 (PGE2). PGE2 has been suggested to regulate inflammation-induced fever in mice^67,68^. Interestingly, brain endothelial cells also selectively lacked expression of IGFBP4, which was abundantly expressed in the endothelium of all other tissues in lemurs (**Fig. 3h**), as in humans and mice (**Fig. S12**).

These organ-specific hormone signatures were expressed at higher levels in endothelial cells than other cell types from the same organ (**Fig. S1**). Thus, it is unlikely that they resulted from cross-contamination (signal spreading). Altogether, lung and brain endothelial cells showcase rare examples of organ-specific (and cell type-independent) hormonal signaling, which may underlie organ-specific control of the blood flow.

In summary, we have highlighted five examples where closely related cells show detailed specification in their hormone signaling. These examples demonstrate the precise and specialized control of hormonal regulation across different cell types and/or organs.

### The hormonal cell communication network is densely connected

Long-range hormonal signaling connects both nearby and distant cells through the bloodstream and enables cross-organ communication. By linking the source cells that produce a hormone ligand to the target cells that express the corresponding receptor, we constructed a global cell communication network for mouse lemur hormone signaling (**Fig. 4a**). Nodes of the network are cell types and edges are directed from hormone source to target cells.

**Fig. 4.**
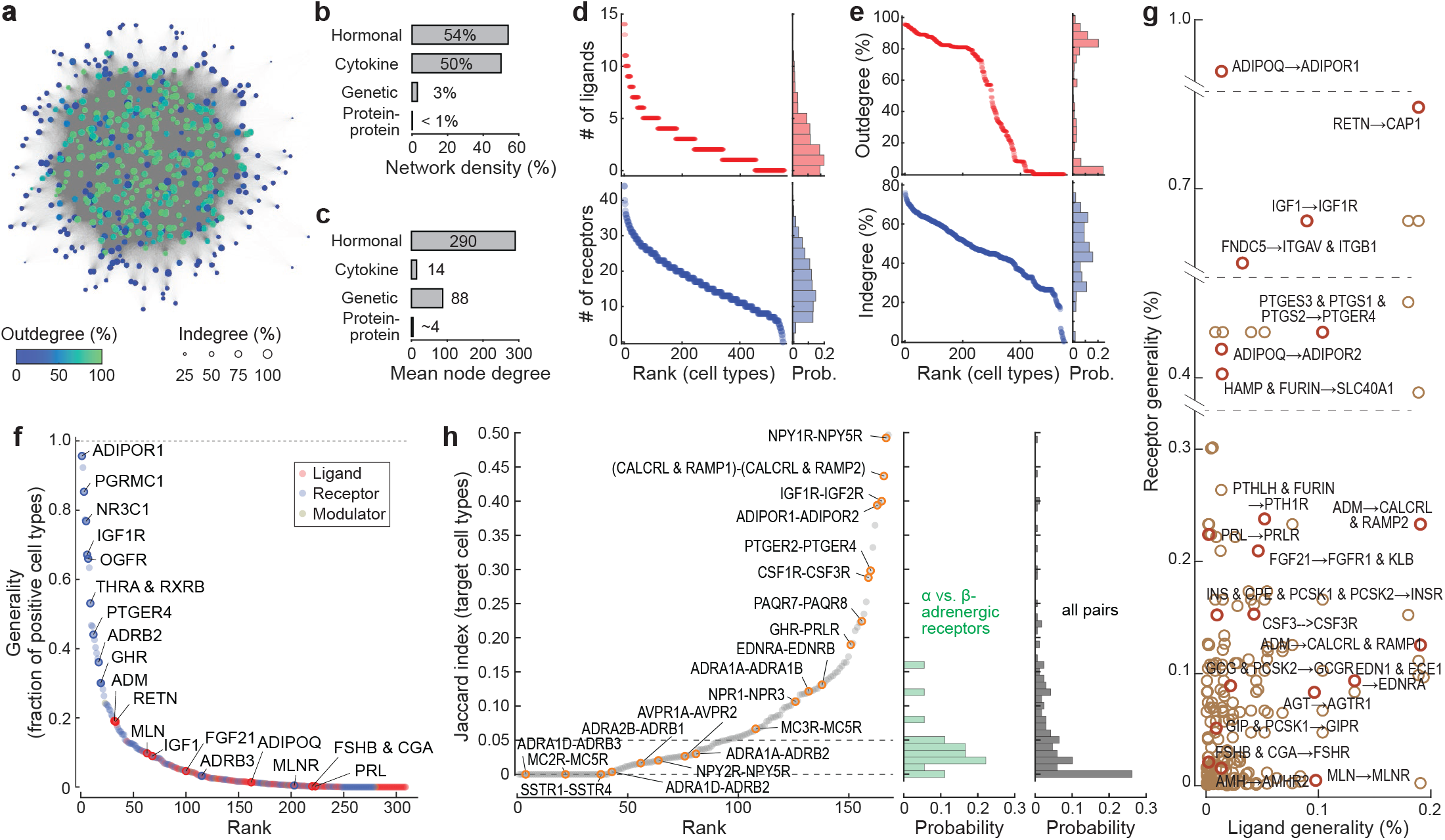
The hormonal cell-cell communication network. **a.** A representation of the network as constructed by force-directed graph drawing. Nodes (cell types) are color-coded according to the node outdegree (% of cell types connected to the current node by outgoing edges) and the node circle size shows the node indegree (% of cell types connected to the current node by incoming edges). Network edges are directed and connect nodes expressing a hormone ligand to all nodes expressing the corresponding hormone receptor. **b-c**. Bar plot of network density (**b**) and average node degree (i.e., the number of connected nodes per node) (**c**) for different biological networks. **d.** Number of ligand and receptor types expressed by the network nodes (cell types), ranked from most to fewest. **e.** Indegrees and outdegrees of the network nodes (cell types), ranked from high to low values. Node degree is normalized to the total number of nodes and cluster size and measures the percentage of cell types connected from or to a network node. **f.** Generality score of all hormone ligands and receptors ranked from the most generally expressed to most selectively expressed. Generality is defined as the percentage of nodes (cluster size normalized) positively expressing the gene(s) involved in ligand synthesis or receptor binding. Also see **Fig. S11a** for generality score defined as the number of cell type clusters positively expressing the gene(s). **g.** Scatter plot showing generality scores of the hormone ligands and corresponding receptors. Each circle is a unique ligand-receptor pair. **h.** Ranking (left) and distribution (right) of Jaccard indices of receptor pairs binding to the same ligand across all hormone receptors.

The hormone network was remarkably densely connected; 54% of all possible directed edges were found to be present. In fact, in the graphical depiction of the network (**Fig. 4a**), individual edges are hardly distinguishable because the edges are so dense. The density of the hormone network is comparable to that of the cytokine network^69^, and much higher than the densities of other characterized biological networks (**Fig. 4b**). For example, the network densities of the yeast, fly, and human protein-protein interaction networks are all estimated to be less than 1%^70–72^, and the yeast genetic interaction network density is ∼3%^73^. The average degree of a node in the hormonal network was also exceptionally high (**Fig. 4c**), emphasizing that the average cell communicates with a large number of different cell types. The dense connections between cell types by hormonal regulation are also highly robust: 72% of all existing edges are connected by more than one ligand-receptor pair, and the network maintains high density connection when randomly removing 10% of the ligand-receptor pairs (**Fig. S9a**).

The hormone network has a high density because most cell types express multiple hormone ligands and receptors (**Fig. 4d**). On average, cell types expressed 18.3 ± 8.2 (mean ± S.D.) receptors or receptor complexes. In comparison, fewer hormone ligands were expressed (2.8 ± 2.6). Only a few cell types expressed 10 or more hormone ligands, and 37.6% of cell types expressed a single ligand or no ligand. This indicates that cells in general respond to more hormones than they produce.

Classical endocrine hormone-producing cells such as enteroendocrine cells, pancreatic endocrine cells, hepatocytes, and neurons were among cells secreting the largest numbers of hormone ligands (**Fig. S10**). Unexpectedly, many stromal cells such as fibroblasts also secreted a large number of ligands, including IGF1, adrenomedullin, angiotensin, and asprosin (**Fig. S10**, **Fig. S1**). Stromal cells were also particularly rich in the expression of hormone receptors, making them among the most connected cell types in the hormone network (as indicated by their high “hubs” and “betweenness” scores, **Fig. S10**). Given the abundance of stromal cells across various tissues, these results suggest stromal cells must play vital roles in endocrine regulation.

Canonically, hormones are believed to be regulated in a hierarchical fashion. Among the most well-known examples are the neuroendocrine cell types of the anterior pituitary—corticotrophs, thyrotrophs, gonadotrophs, lactotrophs, and somatotrophs. These cells are components of regulatory cascades, receiving regulatory signals from one or a few hormones produced in the hypothalamus and in turn producing one or a few hormones to regulate a peripheral endocrine gland. Yet these cells were found to express numerous receptors, ranging from 18 (corticotrophs) to 30 (somatotrophs) (**Table 5**). These cells appeared to be regulated by growth factors, neuropeptides, neurotransmitters (e.g., dopamine), classical hypothalamic hormones. Likewise, they were found to produce hormones in addition to the classical hormones (**Table 5**). For example, corticotrophs expressed galanin (GAL), gastrin (GAS), and cortistatin (CORT), as well as the ACTH precursor (POMC) and its processing enzymes PCSK1, PCSK2, and CPE.

Taken together, this analysis suggests that virtually all cell types across the body are extensively engaged in cell communication via hormonal signaling and that the connections are likely evolutionarily robust to random mutations that disturb hormone regulation. The dense connection of the network results in a large number of highly connected nodes and no obvious network cores (**Fig. 4a**, **Fig. S10**). This differs from the commonly studied scale-free networks, which are dominated by a small number of highly-connected hub nodes^74^. Studies on hormone regulation usually focus on one specific site of secretion and/or target; our analysis emphasizes the importance of a systems view in understanding hormone regulation.

### A small number of hormones are global regulators

We next examined properties of the hormone network edges. In contrast to the protein-protein interaction network and the genetic interaction network, the edges of the hormone communication network are directional and can be distinguished into different types according to the specific ligand-receptor pairing. The hormone network included 350 different edge types that are selected pairings of 129 ligands and 175 receptors.

Some ligands and receptors were expressed in highly specific cell types while others were more broadly expressed. We therefore quantified for each ligand or receptor a generality score to evaluate the range of its expression across different cell types in the atlas (**Fig. 4f-g, Fig. S11a, Table 6**). Strikingly, a small number of hormone receptors were globally expressed. As mentioned above, adiponectin receptor 1 (ADIPOR1), was expressed in 95.7% of the cell types. Other broadly expressed hormone receptors included the renin receptor ATP6AP2 (92.3%), the membrane-associated progesterone receptor PGRMC1 (85.3%), the cortisol receptor NR3C1 (76.8%), a putative resistin receptor CAP1 (77.3%), the IGF1 receptor IGF1R (67.1%), the opioid growth factor receptor OGFR (66.0%), and the thyroid hormone receptor THRA in complex with RXRB (53.1%). While it has been recognized that some of these hormones exert organism-wide effects, our results suggest that they target not only a broad range of tissues but likely regulate a large portion or perhaps majority of the cells of the body.

However, the majority of hormone ligands and receptors were expressed in specific cell types; ∼85% of the hormone ligands and ∼56% of the hormone receptors were detected in less than 5% of the cell types. Certain hormones like follicle stimulating hormone (FSHB and CGA) were only expressed by a single cell type. Only a few hormone ligands were relatively broadly expressed, including adrenomedullin (ADM, 19.1%), acylation stimulating protein (C3, 19.4%), IGF2 (18.0%), and resistin (RETN, 18.9%). The generality score followed a power-law distribution toward the tail where a small number of globally expressed receptors dominated the distribution (**Fig. S11b-c**). This resulted in a bimodal distribution of the network node outdegree (abundance of outgoing edges/downstream regulation) (**Fig. 4a, e**, **Fig****. S10**). Cells expressing hormone ligand(s) to the high-generality receptors (e.g., adipocytes which secrete adiponectin) sent globally connected outgoing edges, and other cells that only expressed ligand(s) to specific receptors sent a much smaller number of outgoing edges. In contrast, the nodes’ indegree (abundance of incoming edges/upstream regulation) was more homogeneously distributed because of the higher specificity of hormone ligand expression and large number of hormone receptors expressed per cell type (**Fig. 4d-e**, **Fig. S10**).

### Alternative receptors for the same hormone tend to be expressed in mutually exclusive cell types

Many hormones can bind to multiple receptors, which may mediate different functions (**Table 1**). We therefore examined if receptors for the same ligand tend to be co-expressed or if their expression is mutually exclusive. To quantitatively compare the target cell type profiles, we calculated a Jaccard index (JI, intersection over union) for each qualifying receptor pair, namely the ratio of cell types that co-expressed both receptors to the cell types that expressed either of the receptors. This analysis revealed that the majority of receptor pairs had extremely low JIs: 24% (41/168) of the receptor pairs had a JI equaling zero, meaning no overlap in their target cell types, and 56% (94/161) had a JI of less than 5% (**Fig. 4h**). A telling case is the adrenoceptors; although epinephrine (adrenaline) targets many cell types, different adrenoceptor subtypes were selectively employed in different cell types (**Fig. 4h, Fig. S1**-epinephrine/norepinephrine). For example, adrenoceptors ADRA1D and ADRB3 shared no target cell types; ADRA1D was expressed broadly in vascular smooth muscle cells and pericytes of multiple organs, certain glial cells, and leptomeningeal cells, whereas ADRB3 mainly targeted adipocytes. Vasopressin receptors AVPR1A and AVPR2 also had less than 3% overlap in their target cell types; AVPR2 was predominantly expressed in the kidney collecting duct principal cells, while AVPR1A was highly expressed in the kidney collecting duct intercalated cells and broadly expressed in many stromal cells (**Fig. S1**-vasopressin).

A few receptor pairs showed a tendency toward co-expression, suggesting possible interactions in cell signaling. Supporting this possibility, a top ranked pair was the IGF receptors IGF1R and IGF2R (JI=0.40), which are known to interact in IGF2 signaling. IGF2R internalizes after IGF2 binding and clears IGF2 from the cell surface, and therefore functions as a competitive inhibitor of IGF2-IGF1R signaling^75,76^. Two other top ranked pairs were NPY1R/NPY5R (JI=0.49), receptors for neuropeptide Y, peptide YY and pancreatic polypeptide^77^, and ADIPOR1/ADIPOR2 (0.39), receptors for adiponectin. Coexpression of NPY1R and NPY5R has been identified in the rodent brain, and recent studies indicate that the two GPCRs may form heterodimers, resulting in modulated agonist binding properties^78,79^. Similarly, biochemical studies on ADIPOR1 and ADIPOR2 in cell culture systems suggest that the two receptors can form heterodimers promoting membrane expression of ADIPOR2^80,81^.

Taken together, this analysis showed that the expression of distinct receptors for the same hormone tends to be mutually exclusive, reflecting functional divergence and specification among hormone receptor subtypes. The rare cases of co-expressed receptors appear to involve functional interaction or cross-regulation between the receptor pairs.

### Feedback circuits are widespread in the hormonal cell communication network

Hormone regulation often involves feedback control. We therefore searched the mouse lemur hormonal cell-cell communication network to systematically identify potential feedback regulation. For simplicity, we focused on two-node feedback circuits, namely cell type pairs that are connected by edges in both directions (**Fig. 5a**). This systematic search revealed a vast number of feedback circuits; 32% of all node pairs formed feedback circuits, showing that feedback control is widespread in the hormone network (**Table 7**). We then tested if feedback circuits are enriched in the hormone network, i.e. more prevalent than would be expected by chance. We applied degree-preserving random rewiring^82^ to randomly permute edge connections without changing node degree (i.e., network edge number and distribution). **Fig. 5b** shows the distribution of the feedback circuit counts from such randomly permuted networks (*n* = 1000).

**Fig. 5.**
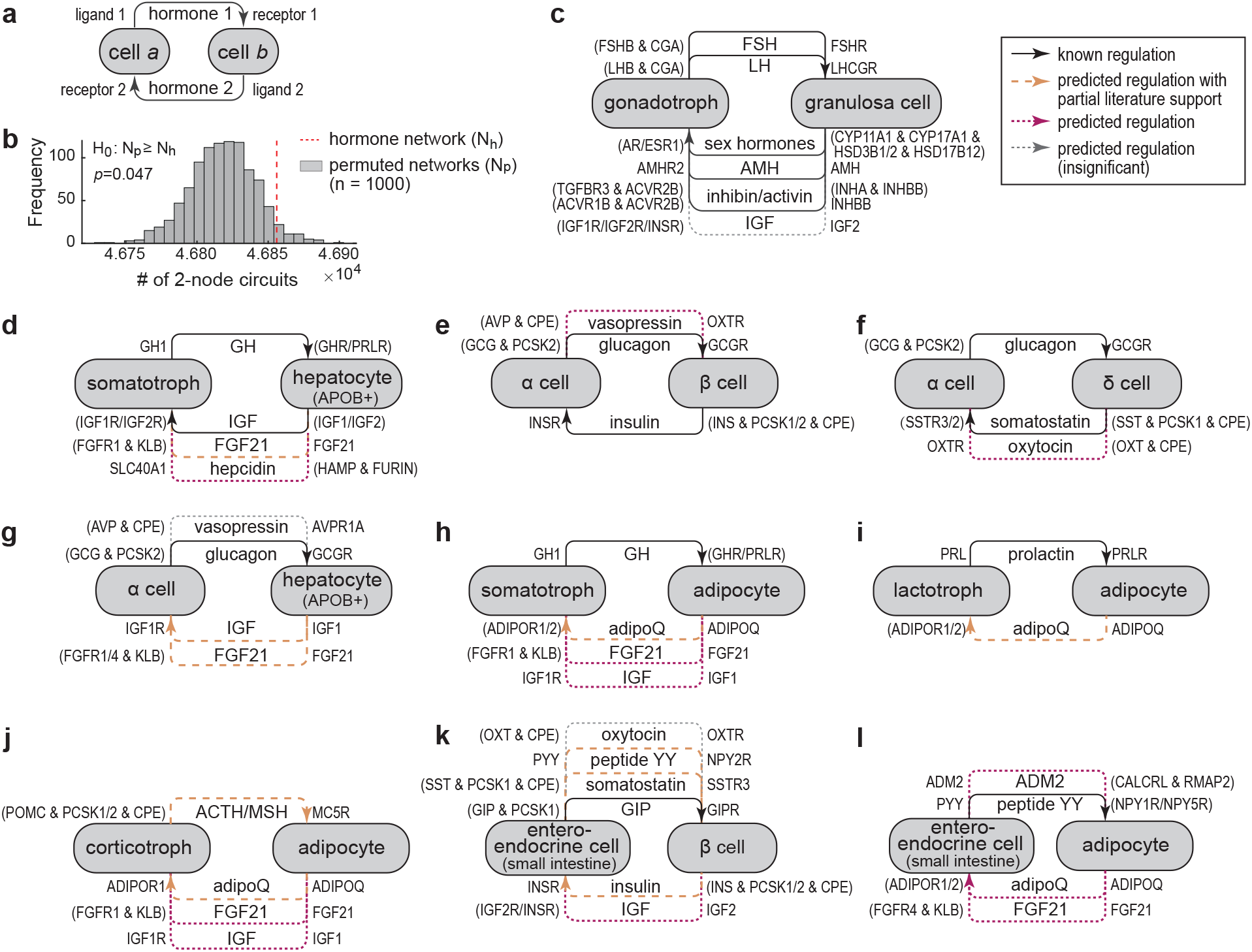
Feedback circuits in the hormonal cell-cell communication network. **a.** Definition of two-node feedback circuits. **b**. Comparison of the number of 2-node feedback circuits identified in the hormonal cell-cell communication network (red dashed line) and that of permuted networks (*n* = 1000). Permutation preserved node outdegree and indegree. **c-h**. Examples of two- node feedback circuits identified in the network, focusing on the endocrine cell types. Solid black arrows indicate known regulation, dashed orange arrows indicate predicted regulation with partial literature support, and dotted arrows indicate predicted regulation without earlier knowledge that are potentially biologically relevant (red) or insignificant (grey). A leg is considered biologically insignificant if the connected hormone-producing cell type is unlikely to be the major source of the hormone (cells had lower expression levels than the canonical major source cells) and if the signaling is not local. Parentheses are used to group multiple genes that are expressed in the relevant cell type that function either together (&) or independently (/).

Results suggest that feedback circuits are enriched in the hormone network compared to random permutations (*p*=0.047). However, the difference was small (<1% on average). Additionally, we randomly removed 10% of ligand-receptor pairs to test how perturbations to ligand-receptor binding affect feedback distribution of the network. This analysis found that the network feedback circuits remained abundant despite edge removal (**Fig. S9b**) as cell-cell connections usually involved multiple ligand-receptor pairs (**Fig. 4d-e**). Taken together, this analysis suggests that feedback circuits are abundantly and robustly distributed as a result of the dense connection of the hormone network. Evolution may have further enriched hormonal feedback circuits as suggested by the random rewiring analysis.

We next reviewed selected feedback circuits between endocrine cell types (**Fig. 5c-l**). The analysis successfully detected well-known feedback regulation. **Fig. 5c** shows the detected feedback loop between pituitary gonadotrophs and the estrogen-producing granulosa cells of the ovary (fortuitously obtained from female perigonadal fat), a classical example of hormonal feedback control that is evolutionarily conserved across vertebrates^83^. Consistent with humans, lemur gonadotrophs expressed two highly-conserved gonadotropins, follicle stimulating hormone (FSH, a dimer of FSHB and CGA) and luteinizing hormone (LH, LHB/CGA dimer), which target granulosa cells through the corresponding receptors FSHR and LHCGR. Granulosa cells, in turn, expressed key synthases required for the production of sex hormones, which feed back to gonadotrophs through androgen and estrogen receptors (AR, ESR1). Additional feedback regulation from granulosa cells to gonadotrophs was also mediated through anti-Müllerian hormone (AMH) and inhibin/activin signaling. We also detected the well-established feedback loop between pituitary somatotrophs and hepatocytes in the liver, which was mediated through growth hormone (GH), IGF1^2,84^, and potentially FGF21^85^ and hepcidin signaling (**Fig. 5d**). Additionally, feedback circuits were detected between pancreatic α and β cells through glucagon and insulin signaling and between α and δ cells through glucagon and somatostatin signaling^86^ (**Fig. 5e-f**).

The analysis also revealed a long list of potential endocrine feedback circuits that have not been reported, or only sparsely reported (**Fig. 5g-l**). For example, we detected a feedback loop between pancreatic α cells and hepatocytes mediated by glucagon, IGF1 and FGF21 (**Fig. 5g**). Regulation of hepatocyte glycogenolysis and gluconeogenesis by glucagon is well-established^87^. Less is known about how glucagon affects hepatocyte hormone secretion, and whether the two cell types were involved in feedback regulation. Interestingly, two previous studies have separately shown that injection of glucagon increases plasma FGF21 in humans^88^ and injection of FGF21 reduces glucagon in mice^89^. A recent study also observed that plasma IGF1 and glucagon levels are negatively correlated in non-diabetic humans and found that IGF1 negatively modulates glucagon secretion in an α cell culture model^90^. Considering that the liver is a major source of circulating IGF1 and FGF21, these data suggest that α cells likely form a negative feedback loop with hepatocytes in maintaining metabolic homeostasis.

Additionally, adipocytes, through adiponectin, formed possible feedback loops with pituitary neuroendocrine cells, including the growth hormone secreting somatotrophs, the prolactin secreting lactotrophs, and the ACTH secreting corticotrophs (**Fig. 5h-j**). It has been reported that growth hormone, prolactin, and ACTH reduce adiponectin secretion in adipocytes^91–93^, but much less is known about the effect of adiponectin on pituitary neuroendocrine cells. Interestingly, a recent study using cultured primary pituitary cells from two non-human primate species found that adiponectin decreases growth hormone and ACTH release and increases prolactin release^94^. Altogether, it suggests that adipocytes form a negative feedback loop with lactotrophs which possibly stabilizes prolactin and adiponectin levels, and form double-negative feedback loops with somatotrophs and corticotrophs, possibly functioning as bistable switches^95^. It remains to be characterized how adiponectin affects pituitary hormone secretion, what are the physiological functions of the pituitary-adipocyte feedback control, and whether this regulation is specific to primates.

Some of the individual legs of the feedback loops shown in **Fig. 5** may not be biologically significant. For example, oxytocin and vasopressin are part of the feedback circuits among pancreatic endocrine cells (**Fig. 5e-f**), between pancreatic α cells and hepatocytes (**Fig. 5g**), and between enteroendocrine cells and pancreatic β cells (**Fig. 5k**). The intra-pancreatic feedback regulation (**Fig. 5e-f**) may be plausible local circuits, but the vasopressin and oxytocin mediated cross-organ communications (**Fig. 5g, k**) seem less plausible, because most of the oxytocin and vasopressin in the bulk circulation is derived from specific hypothalamic neurons and secreted in the posterior pituitary^2^. The lemur pancreatic endocrine and enteroendocrine cells expressed lower levels of oxytocin and vasopressin than many of the neuronal cell types (**Fig. S1**-oxytocin, vasopressin), thus are unlikely major sources of these hormones in circulation despite their potential role in local communications.

Several others of the regulatory circuits identified here may not fit well with the conventional view, but could actually be biologically significant, such as IGF1 and FGF21 mediated communication from adipocytes to somatotrophs, corticotrophs, and enteroendocrine cells (**Fig. 5h, j, l**). Canonically, circulating IGF1 and FGF21 are believed to be predominantly produced by hepatocytes^20,96,97^. However, we identified high IGF1 expression in the mouse lemur stromal compartment, in particular adipocytes and fibroblasts (**Fig. S1**-IGF1). FGF21 expression levels were also comparable between adipocytes and hepatocytes (**Fig. S1**-FGF21). Given that adipocytes are abundant in the mouse lemur and their weight varies by season, it is plausible that adipocytes contribute significantly to the production and temporal variations of the circulating IGF1 and FGF21.

Taken together, this analysis identified an extensive list of potential feedback circuits in the global hormonal cell communication network (**Table 7**), providing a rich resource for generating hypotheses on endocrine regulations and to compare their regulatory mechanisms in different species.

### Exploring hormonal regulation of the circannual rhythms that control organismal biology

The mouse lemur hormone atlas may bring new insights to the study of mouse lemur-specific physiology, such as their circannual rhythms^31,98^, social interactions^99^, and aging^12–14,100^. As an adaptation to seasonal environmental changes, mouse lemurs have evolved striking seasonal rhythms in metabolism, behavior, and reproduction that persist throughout their 5–10-year lifespan. Mouse lemurs build fat reserves in preparation for the winter and enter daily torpor, a hypometabolic state, to preserve energy, while in the summer the animals maintain a lean body and regrow their reproductive organs for mating. The global nature of these phenotypic changes suggests that they may be hormonally mediated. To date, only 12 circulating hormones have been measured in mouse lemurs, but all showed dramatic seasonal changes (**Fig. 6a**, **Table 8**). This suggests that hormones are likely the key messengers coordinating and driving various seasonal changes among different organs. The hormone atlas suggests that these seasonally changing hormones target a broad array of cells and tissues; 95% of the cell types possessed receptors for at least one of these 12 hormones (**Fig. 6b-l**). Several of the seasonal hormones, such as cortisol, thyroxine, and IGF1 are global regulators, targeting over 50% of the cell types (**Fig. 6e-g**). Others, such as GLP and GIP, are highly specific, targeting 0.2% and 5% of the cell types, respectively (**Fig. 6h, k**). Estrogen, androgen, insulin, PYY/PPY target many cell types, but their receptor expression was restricted to only certain cell type compartments (e.g., stromal, epithelial) (**Fig. 6c-d, i-j**). For melatonin, receptor expression was not detected (**Fig. 6l**), possibly due to incomplete sampling of melatonin-expressing brain regions. Comparing across cell types, stromal and epithelial cell types tend to be targeted by most seasonal hormones (**Fig. 6c-l**). These data suggest that seasonal rhythms engage not only tissues that demonstrate obvious morphological changes (e.g., adipose tissues and gonads), but involve global shifts in cellular metabolism and physiology. While it is now clear that ligands of at least 12 hormones are produced in a seasonal manner in the mouse lemur, it remains to be explored if more hormones are seasonal and if hormone receptors are seasonally expressed to control mouse lemur seasonal rhythms.

**Fig. 6.**
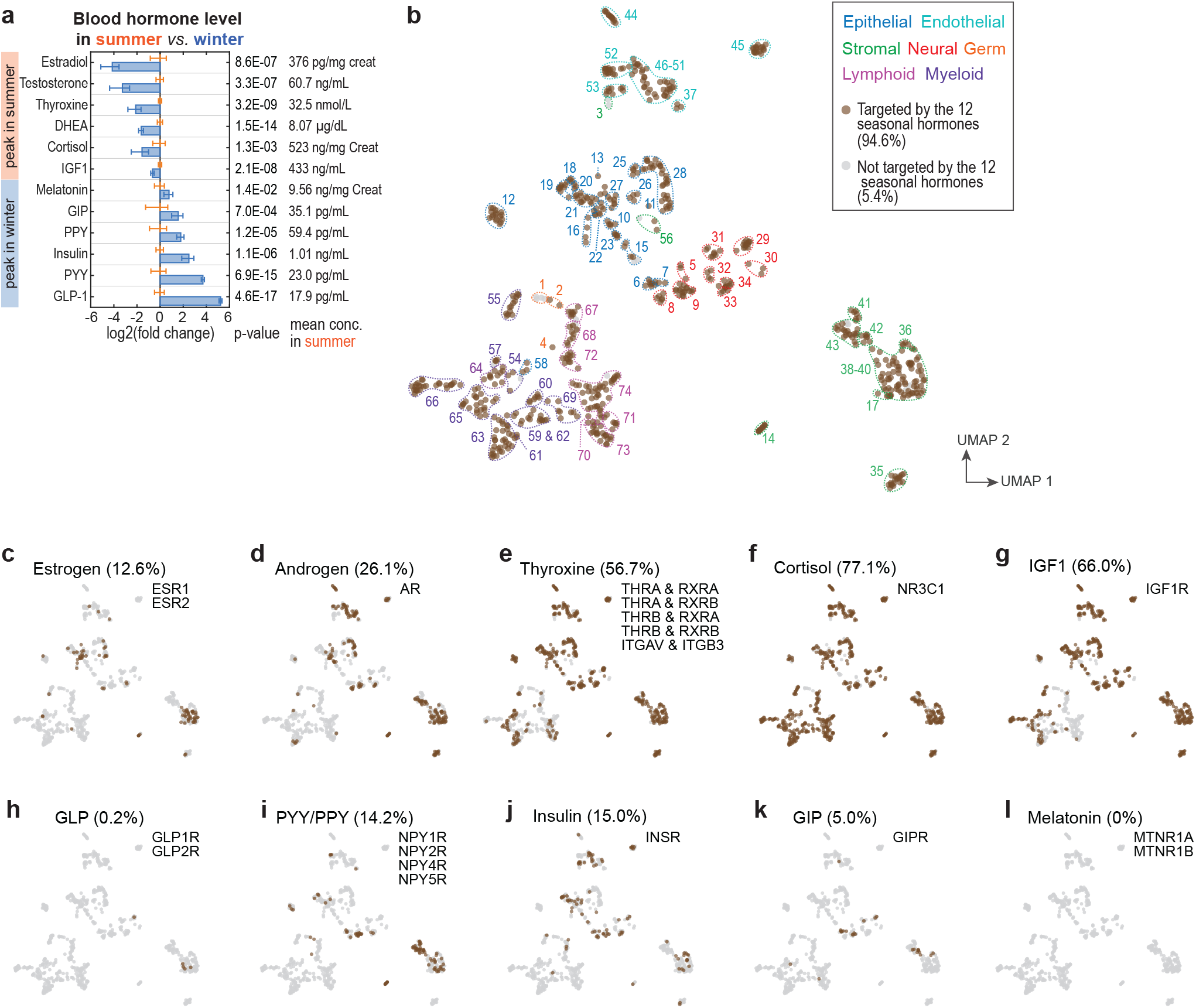
Seasonality of hormone concentrations in the mouse lemur affects almost all cells and tissues. **a**. Concentrations of twelve hormones measured under long-photoperiod (daily 14:10 h light:dark, summer-like) and short-photoperiod (10:14 h light:dark, winter-like) conditions in captive mouse lemurs. Animals were kept indoors under constant temperature (24°) and with photoperiods alternated between long and short photoperiods every 6 months. Bars represent fold changes in hormone concentration compared to average summer levels. Error bars represent 95% confidence intervals. *p*-values were calculated by 2-sample *t*-tests. Data are listed in **Table 8**, collected from literature as follows and supplemented with additional new measurements: estradiol^137,138^, testosterone^139–141^, thyroxine^137,142^ (and additional data in **Table 8**), DHEA^138^, cortisol^143^ (and additional data in **Table 8**), IGF1^144^, melatonin^145^, and gut hormones (GIP, PPY, insulin, PYY, and GLP-1)^146,147^. **b.** Mouse lemur cell types targeted by any of these 12 seasonal hormones. Cell types are displayed in the same UMAP plot as in Fig. 2b. Circles are cell types and are color coded according to whether they express receptors for any of the 12 hormones. Dashed lines circumscribe the cell type clusters and are colored according to cell type compartments. **c-l**. Mouse lemur cell types targeted by each of these 12 seasonal hormones displayed in the same UMAP format as in **b**.

### Evolution of hormonal gene expression in the human, lemur, and mouse

Lastly, to study evolution of hormone signaling, we compiled comparable scRNAseq datasets of human and mouse and performed cross-species comparison of hormonal gene expression across human, mouse lemur, and mouse. Six tissues were studied, including lung, skeletal muscle, liver (epithelium only), testis (germ cells only), and two immune tissues, bone marrow and spleen (immune cells only). For consistency, cells from the human and mouse datasets were reannotated through the same pipeline and with the same cell type markers as for the mouse lemur cell atlas. Data from different species were further integrated and cell annotations unified to ensure that cell types are comparable across species^15^. This resulted in 62 orthologous cell types with enough cells for comparisons in all three species. We next identified the hormonal genes with one-to- one-to-one orthology mapping across the three species and examined if the gene exists in all the analyzed scRNAseq datasets (**Table 2**). A total of 295 genes were identified and their cross-species expression patterns plotted in **Fig. S12**. By comparing the correlation of the overall hormonal gene expression, we found that human and lemur showed a higher (*p-*value = 0.006) correlation coefficient (*r* = 0.50) compared to that between human and mouse (*r* = 0.48) (**Fig. 7a**), suggesting that overall hormonal gene expression is more similar between humans and lemurs compared to mice. For comparison, the mean sequence identity between these 295 human and lemur one-to-one orthologs was 0.86, and for human and mouse orthologs 0.82 (**Fig. S2b**). Thus, humans are more similar to lemurs in both gene sequences and expression patterns, compared to mice.

Next, we examined individual hormonal genes and compared the conservation of their expression patterns between the orthologous cell types of human and lemur vs. that between human and mouse. For each gene, we calculated the correlation coefficient for expression in the 62 orthologous cell types for human vs. lemur (i.e., *r*(human, lemur)), human vs. mouse (*r*(human, mouse)), and lemur vs. mouse (*r*(lemur, mouse)), and we used the correlation coefficients as a gauge of expression pattern similarity (**Fig. 7b-c**). Many of the genes showed a conserved expression among all three species, such as endothelium-specific RAMP2, liver-specific GC, and fibroblast- and pericyte-specific EDNRA (**Fig. 7b-d**, **Fig. S12**).

Interestingly, four genes (RETN, VIPR1, PROK2, and PTGDR) were highly correlated between human and lemur but poorly correlated between human and mouse (**Fig. 7b-c**), suggesting that their expression patterns are conserved in primates and/or diverged in mice. As discussed above, VIPR1 (vasoactive intestinal peptide receptor 1) was selectively expressed in the lung endothelial cells of humans and lemurs but not mice (**Fig. 7d**, **Fig. 13a**). RETN (resistin) was most highly expressed in the neutrophils, monocytes, and macrophages in humans and lemurs but not mice (**Fig. 7d**). Similarly, PTGDR (prostaglandin D2 receptor) was expressed in the natural killer (NK) and NK T cells in humans and lemurs but not mice (**Fig. 7d**). PROK2 (prokineticin 2) was selectively expressed in neutrophils and at lower levels in other myeloid cells in humans and lemurs, but was not expressed in these cell types in mice; instead it was expressed in spermatocytes (**Fig. 7d**).

**Fig. 7.**
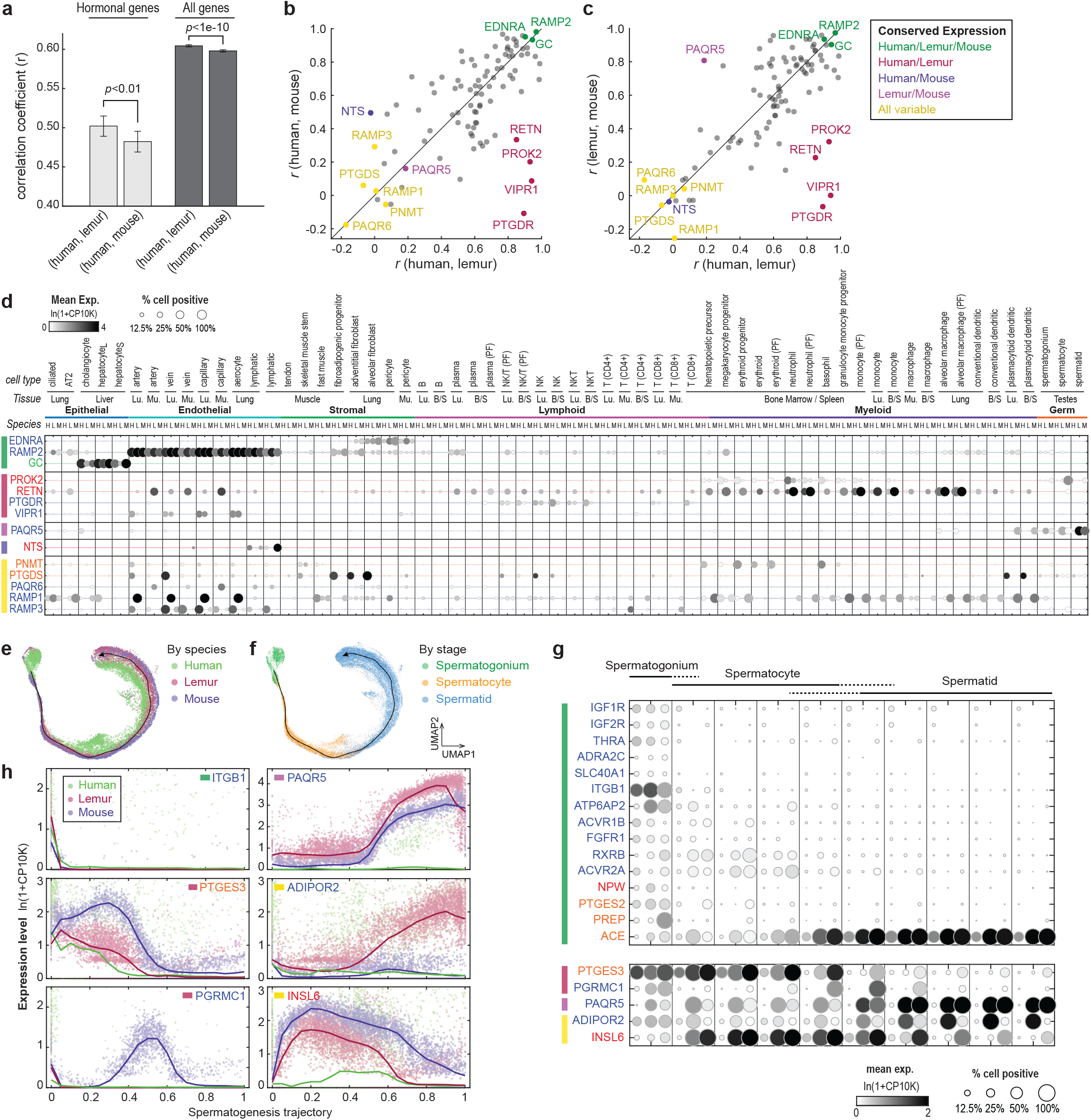
Cross-species comparisons of hormonal gene expression across humans, mouse lemurs, and mice. **a**. Correlation coefficients of overall expression patterns for all the hormonal one-to-one orthologs (left) or all one-to-one orthologs (right) between human and lemur vs. between human and mouse. *p*-value indicates the significance of a higher correlation between human and lemur vs. between human and mouse, calculated by comparing the two correlation coefficients using Fisher’s Z-transformation. **b**. Comparison of correlation coefficients between the expression patterns of human and lemur (*x*-axis) versus that between human and mouse (*y*- axis) for individual hormonal genes. **c**. Comparison of correlation coefficients between the expression patterns of human and lemur (*x*-axis) versus between lemur and mouse (*y*-axis) for individual hormonal genes. Black lines in **b** and **c** indicate 1-1 relationship between the two coefficients. **d**. Dot plot showing expression patterns of highly species-conserved and species- variable example genes in the 62 orthologous cell types across all three species. Rows are orthologous genes, indicated with the respective human gene symbols. Columns are cell types, displayed as triplets of the respective expression in humans, lemurs, and mice. Vertical grey lines segment different cell type triplets. Cell types are ordered first by compartments, then by tissue, and finally by species. **e-f**. Species-integrated UMAPs of the male germ cells with cells color coded by species (**e**) or spermatogenesis stage (**f**). Black lines indicate the identified spermatogenesis trajectory with arrows pointing to the direction of maturation. **g**. Dot plot showing cross-species expression patterns of spermatogenesis stage-specific hormonal genes in germ cells grouped and ordered by the integrated spermatogenesis trajectory. Cell grouping in the UMAP format is shown in **Fig. S13a**. **e**. Scatter plots showing the species-conserved or species-variable expression patterns of example genes in **g**. Points are single cells color coded by species. Solid curves show a moving average of the expression level along the spermatogenesis trajectory. In panels **d**, **g**, and **h**, colored bars next to the genes indicate the species conservation patterns as in panels **b**-**c**.

In contrast, only one gene, neurotensin (NTS) was found to be more similar between human and mouse vs. that of human and lemur, when applying the same threshold (**Fig. 7b-c**). Among the cell types examined in all three species, NTS was selectively expressed in the lymphatic cells in human and mouse but not lemur (**Fig. 7d**). Across all mouse lemur atlas cell types, NTS was most highly expressed in the small intestine enteroendocrine cells (**Fig. S1**- neurotensin/neuromedin N), which is also the canonical site of NTS secretion in humans^18^.

However, we currently lack cross-species scRNAseq data of the enteroendocrine cells for comparison. Presumably, the major site of NTS secretion is conserved across the three species (in enteroendocrine cells), whereas the uncanonical lymphatic NTS expression is specific in humans and mice but not lemurs. It will be interesting to compare, in human and mouse, levels of NTS expression in the lymphatic cells to that of enteroendocrine cells and explore the function of this uncanonical source of NTS.

We also identified one gene (PAQR5, a membrane progesterone receptor) whose expression pattern is more similar between lemur and mouse than between either of the species and human (**Fig. 7c**). Both lemur and mouse spermatids show the highest PAQR5 expression, whereas human cholangiocytes show the highest expression among the cell types examined (**Fig. 7d**).

In addition, a number of genes showed low conservation in their expression patterns among all three species, such as RAMP1, RAMP3, PTGDS, PNMT, PAQR6 (**Fig. 7b-d**). For example, RAMP1, as previously discussed, was most highly expressed in the lung vascular endothelial cells in the lemur. However, mouse myeloid cells, such as monocytes, macrophages, and dendritic cells show the highest RAMP1 expression. In humans, RAMP1 was moderately expressed in the lung ciliated epithelium, hepatocytes (CPN2hi subtype), and skeletal muscle cells, two of which also displayed moderate RAMP1 expression in the lemur and mouse.

Lastly, the cross-species dataset included male germ cells which form a continuous gradient of expression patterns (**Fig. 7e-f**). Spermatogenesis is known to have undergone extensive molecular evolution^101^. Thus, we analyzed the expression dynamics of the stage-specific hormonal genes identified in **Fig. 3a** and examined their conservation across the three species (**Fig. 7g-h**, **Fig. S13a-b**). Although many spermatogonia-specific hormonal genes showed a conserved expression across the three species, some genes displayed species differences. Some involve small shifts or modifications in the expression dynamics. For example, consistent across all three species, PTGES expressed in spermatogonia and spermatocytes and not in spermatids (**Fig. 7g-h**). However, in humans and lemurs, PTGES expression was gradually reduced in the course of sperm differentiation, whereas in mice expression was maintained at a high level through mid to late spermatocytes (**Fig. 7g-h**). Other species-specific expression involved lack of expression, or substantially reduced expression. Examples include the previously discussed PAQR5, which was highly expressed in spermatids in lemurs and mice, but only slightly expressed in human spermatids. In addition, PGRMC1 was expressed only in spermatogonia in humans and lemurs, and was highly expressed during the spermatocyte-spermatid transition in mice (**Fig, 7g-h**).

Taken together, cross-species comparison revealed an overall more similar pattern of hormonal gene expression between human and lemur vs. human and mouse. Analysis of individual hormonal genes identified interesting examples that are highly conserved or display primate/species specific expression. This analysis revealed interesting candidates for future studies to understand the evolution of hormone signaling.

## Discussion

Hormonal signaling is vital to multicellular life, allowing cells to communicate with both surrounding cells and remote cells in other tissues. In this study, we systematically mapped hormone-producing and -target cells across the 739 cell types from 27 different tissues from the mouse lemur molecular cell atlas. We generated a browsable atlas that depicts gene expression of ligands, modulators, and receptors for 84 hormone classes (**Fig. S1**, https://tabula-microcebus.ds.czbiohub.org/hormone-atlas). This dataset complements the scale and resolution of classical endocrine studies and brings a cellular and molecular foundation for studying primate hormone regulation.

The hormone atlas also provides an opportunity to extract global principles of hormone signaling. The hormone-related genes exhibited cell-type-dependent, stereotypical expression patterns, and their transcriptional profiles faithfully define cell type identities, despite their constituting less than 1% of the transcriptome (**Fig. 2**). Ligands or receptors alone also classified cell types well, with certain clusters mostly defined by their ligands and others by their receptors (**Fig. S7**). Cell type clusters defined by the hormonal genes often show further cell type, subtype, or stage-dependent specification within clusters, such as the extensive shifts in hormonal signaling during spermatogenesis, across the spatial gradient of the kidney nephron epithelium and vasa recta endothelial, between subtypes of hepatocytes, and during tumor metastasis (**Fig. 3a-g**). There is also an interesting organ-dependent clustering of the lung and brain vascular endothelial cells, showcasing a rare example of organ-specific rather than cell type-specific hormone signatures (**Fig. 3h**). Altogether, it is remarkable how well a cell’s intercellular communication repertoire encapsulates the cell’s identity, indicating precise and highly specialized hormonal control in different cells across the body.

To study the properties of global hormonal regulation, we constructed an organism-wide cell communication network for mouse lemur hormone signaling, by connecting the hormone-producing cells to the corresponding target cells (**Fig. 4**). This hormone network was exceptionally dense (**Fig. 4a-b**), which mainly resulted from two network features. First, most cells in the network expressed multiple hormone receptors and/or ligands (**Fig. 4d**). Second, although most hormone ligands and receptors were expressed in specific cell types, a small fraction of hormone receptors were nearly ubiquitously expressed, resulting in almost global connection of the relevant hormone-producing cells (**Fig. 4e-g**). Thanks to its dense connection, the hormone network is highly robust to random perturbations that delete individual edges (**Fig. S9**). Although the regulatory circuits of individual hormones may be hierarchical, as classically viewed, overall the network is decentralized, which de-emphasizes the contribution of any one hormone-receptor pair. Many of the hormone-secreting cells are diffusely distributed throughout the organism, in line with the concept of a diffuse neuroendocrine system espoused by Feyrter^102^. It is worth noting that, although changes to a single hormone-receptor pair may only minimally affect the overall hormone network structure, the impact to animal physiology and metabolism could be significant. Information flow in the hormone network depends not only on whether an edge exists but also the identity of the connecting edge(s), i.e., the hormone-receptor pair(s). Interestingly, despite the density of the hormone signaling network, subtypes of hormone receptors that bind to the same ligand were generally expressed in a mutually exclusive fashion, supporting the notion that different hormone receptor subtypes usually mediate different physiological functions through separate target cells (**Fig. 4h**).

Feedback regulation was widespread in hormonal signaling, and it was robust to network perturbations (**Fig. 5b**, **Fig. S9B**). The analysis identified well-known feedback circuits among several endocrine cell pairs and detected additional, previously unreported, hormones involved in feedback control (**Fig. 5c-f**). We also identified many novel feedback circuits (**Fig. 5g-l**, **Table 7**) which may be involved in adaptation or homeostasis (negative feedback circuits) or in generating switch-like responses (positive and double-negative feedback circuits). It is worth emphasizing that not all detected feedback circuits will necessarily be biologically significant. In particular, for hormones that travel and mix in the bulk circulation for long-distance cell communication, signals from a particular hormone-producing cell type could be overwhelmed by other cell types that produce larger amounts of the same hormone. Future analysis may include information on cell numbers of individual cell types and genes responsible for rate-limiting steps to more systematically compare total hormone production in each cell type and discriminate major and minor hormone sources. Nevertheless, this analysis represents a first step to systematically identify endocrine feedback circuits for future functional validation and characterization. Our findings may also inform studies of the systems biology and network topology of hormone signaling (see for example Karin et al.^103^).

Mouse lemurs display dramatic seasonal changes in body weight, hibernation propensity, and reproductive activities^31,98,104^. Such seasonal rhythms are likely regulated, at least in part, by seasonally-varying long-range hormones that coordinate the changes across different organs of the body (**Fig. 6a**). The hormone atlas shows that almost all (95%) of the mouse lemur cell types are regulated by seasonally varying hormones (**Fig. 6b**). This suggests that most, if not all, mouse lemur cells across all organs of the body experience seasonal changes in metabolism, growth, and function. By systematic mapping of hormone secreting and target cells, the atlas also points to candidate cell types and signaling to explore the upstream and downstream regulation for each of the seasonally varying hormones. Given that all 12 hormones measured in the mouse lemur showed significant seasonal changes, we suspect that more hormones will prove to be seasonal. Future work should systematically characterize seasonal patterns of hormones with higher temporal resolution, and to sample animals in different seasons to detect seasonal expression changes in hormone receptors, ligands, and synthases, which could indicate seasonal restructuring of the hormone network. Circannual rhythms are widespread in nature^105,106^ and humans display seasonally varying hormone levels and cellular transcriptomes^107–109^. The findings of the present study may apply to species beyond the mouse lemur, and may bring new insights for season-associated human diseases^110–112^.

Mouse lemurs are genetically closer to humans than mice are. Likewise, the lemur better models human hormonal signaling than does the mouse. At the genomic level, there were fewer hormonal genes that were lost (or became pseudogenes) in lemurs (3) than mice (6). More human genes had one-to-one mapping in lemur (338, if counting LHB) than mouse (330). Further, more human genes (85.8%) had a higher sequence identity with their lemur ortholog than their mouse ortholog (**Fig. S2**). Given that mouse lemur genome and gene annotations remain lower in quality than those of humans and mice, some of the lemur genes may be unannotated or incorrectly annotated, like LHB. Thus, the above comparisons may underestimate human-lemur similarity. Lastly, by comparing the expression patterns of the one-to-one orthologs across the three species, we also found mouse lemurs had an overall higher similarity with humans compared to mice (**Fig. 7a**). For hormonal genes that show a highly species-specific expression pattern, more (4) were conserved between humans and lemurs and unique in mice compared to the opposite (only 1) (**Fig. 7b**).

Single-cell RNA sequencing is a powerful tool for assaying cellular function and activity, but there are limitations. Although the technique has improved rapidly in sensitivity, accuracy, and efficiency, scRNAseq still faces low capture efficiency and high transcript dropout^113^. With the large number of cells assayed in the mouse lemur molecular cell atlas, we averaged gene expression across cells of the same cell type to smooth the noise in individual cells from dropout. In the analysis of hormone regulation, genes involved in hormone production, such as ligands and modulators, tended to be abundantly expressed; but certain hormone receptor proteins may be low in copy number and slow in turnover, and therefore may result in false-negative errors. In addition, there can be false-positive errors caused by contamination from other cells in the same sample (i.e., signal spreading). This is most notable for abundantly expressed ligand genes and tissues affected by endogenous digestive enzymes (e.g., pancreas). In this study, we scored positive expression of a hormone ligand only when both the ligand gene and the necessary modulators (such as prohormone processing enzymes) were present. This approach mitigated false-positive detection of highly abundant ligands. Lastly, scRNAseq has low capture efficiency for certain cells such as neurons and large sized adipocytes. Particularly relevant for the study, hypothalamic neurons were not captured in the current lemur dataset. Nevertheless, the mouse lemur cell atlas included several neuronal subtypes from the brain cortex, brainstem, and retina, as well as both UCP1-high and UCP1-low adipocytes from multiple tissues^15^. Although not yet a complete nor ideal sampling, the mouse lemur cell atlas is one of the most comprehensive and accurately-annotated organism-wide scRNAseq datasets reported to date. Other organism-wide scRNAseq atlases have been created or are being created for other species, including mouse^114–116^, ascidian^117^, macaques^118,119^, and human^120^, bringing exciting resources to the study of the evolution of hormone signaling.

Finally, it is worth emphasizing that the mouse lemur scRNAseq atlas used in the study was constructed with four aged and diseased donors. For example, both females had uterine tumors with metastasis, and all four lemurs exhibited chronic kidney disease which may have affected the transcriptional profiles of the kidney cells. Nonetheless, in the cross-species analysis, we have shown that hormone profiles of individual cell types are overall conserved across human, lemur, and mouse. Human and lemur actually correlated better in their overall transcriptional and hormonal profiles than between human and mouse, despite the fact that mouse data were derived from younger, healthier animals^114^ and that human data was possibly affected by different pathologies^120^. While previous studies reported that concentrations of hormones may change over age^121^, we suspect that the global principles of hormone signaling—e.g. a densely connected, decentralized network with abundant feedback regulation —learned through the mouse lemur atlas may also be pertain to younger lemurs and likely apply to other species. In addition, we have taken advantage of the disease conditions of the mouse lemurs, such as the two females with primary and metastatic uterine adenocarcinoma, which revealed interesting metastasis associated changes in hormone profiles.

Traditionally, biologists focus on one particular cell type and/or organ system. However, hormone signaling by its very nature connects different cell types and organs. The emerging organism-wide atlases open a new door in understanding the cross-organ communication systems that allow organisms to function as a reliable robust whole.

## Methods

### Animals husbandry, tissue procurement and processing

Details on animal husbandry and sample collection and processing have been described in accompanying manuscripts^15,17^. In brief, mouse lemurs were maintained for noninvasive phenotyping and genetic research as approved by the Stanford University Administrative Panel on Laboratory Animal Care (APLAC #27439) and in accordance with the *Guide for the Care and Use of Laboratory Animals*^122^. Animals were housed indoors in an AAALAC-accredited facility in a temperature (24°) and light-controlled environment (daily 14:10 h and 10:14 h light:dark alternating every 6 months to stimulate seasonal breeding behavior and metabolic changes). Animals in declining health that did not respond to standard therapy were euthanized by pentobarbital overdose under isoflurane and tissue samples were collected for scRNAseq and histopathology analysis. Details of animal histopathology can be found in Casey et al.^17^ and animal age and tissue sampling/processing pipelines has been described in the accompanying manuscript^15^. Pathologies common to all four animals were fibrous osteodystrophy, chronic renal disease, uterine tumors (females only), cataracts, and osteoarthritis. Tissues not described in Casey et al.^17^ were histologically normal.

### Mapping and counting transcript reads

The single-cell gene expression matrix of the mouse lemur cell atlas was obtained from the accompanying manuscript^15^ where methods and parameters to extract the transcript counts can be found. In this study, we used only the 10x droplet-based sequencing derived data for all analysis. In brief, the *Microcebus murinus* genome assembly (Mmur 3.0, NCBI) was used to create a genome FASTA file. Sequencing reads were processed with Cell Ranger (version 2.2, 10x Genomics) to generate cell barcodes and the gene expression matrix. We additionally counted reads for three genes (MC1R, SCT, LHB) that are unannotated in Mmur 3.0, NCBI. For MC1R and SCT, we used their annotation in the Ensembl *Microcebus murinus* genome. For LHB, we used the uTAR-scRNA-seq pipeline described by Wang et al.^23^ to first predict its chromosome location in the mouse lemur genome. In brief, the pipeline employs the approach of groHMM^123^ to detect transcriptionally active regions (TARs) from aligned sequencing data and annotates these TARs as unannotated (uTARs) or annotated (aTARs) based on their overlap with the existing annotation. For the LHB_uTAR counts, we uncovered uTARs from the mouse lemur pituitary and hypothalamus dataset and used BLASTn^124,125^ to find nucleotide sequence homology of all uTARs against human LHB. We found one uTAR at position NC_033681.1:19143249-19145949 with high sequence homology to human and macaque LHB. The LHB_uTAR is highly differentially expressed in the expected cell type (gonadotroph). The LHB_uTAR is located next to gene RUVBL2, in conservation with the co-location pattern of the genes in humans and mice^24^. We therefore conclude that the identified LHB_uTAR is the likely coding region of the mouse lemur LHB gene. We then used Drop-seq tools^126^ to extract transcript counts for the three genes (MC1R, SCT, LHB_uTAR).

### Orthology and protein sequence analysis

Gene orthology mapping across human, mouse lemur, and mouse was based on the combination of both the NCBI and Ensembl ortholog databases. Details to combine the two databases were described in the accompanying manuscript^15^. To compare sequence similarity between orthologs of human and mouse or lemur, protein sequences were retrieved from NCBI according to protein RefSeq using the Matlab built-in function getgenpept. Conversion between gene ID and protein Refseq identifier was performed using bioDBnet (https://biodbnet.abcc.ncifcrf.gov/db/db2db.php). We aligned ortholog protein sequences and used the maximal scores in the cases of multiple transcript sequences and non one-to-one ortholog mapping. Global alignment was carried out using the Needleman-Wunsch algorithm and default parameters of the function nwalign.

### Identification of hormone-producing and target cells

The functional concentration is different for different proteins, so we applied adaptive thresholding to determine if a gene is positively expressed in a cell type, where cell type is defined as a unique combination of the cells’ annotation and tissue of origin, as described in the accompanying Tabula Microcebus manuscript^15^. For example, vein cells of the lung and brain were considered two different cell types. Cell types that were of low quality (low gene count or transcript count) or annotated as technical doublets were removed from the analysis. For each gene, two cell type parameters were used, mean expression level and percentage of nonzero (positive) cells. Single-cell expression levels were first normalized by total transcript counts, multiplied by 10,000, averaged across all cells in the cell type, then natural log transformed (i.e., log(1+Average_UMI_count_Per_10K_Transcripts); note that 1 is added to the average so that genes with zero transcripts will yield zero rather than negative infinity). To determine positive expression at the cell type level, we applied adaptive thresholding that compares the expression to the maximally expressing cell types. The algorithm involves 4 parameters which were selected to capture a list of well-studied cell types and hormonal genes. We first applied a low-level absolute threshold on average expression level (0.037) and percent of nonzero cells (5%) below either of which we considered the expression to be negative. Second, we calculated a “maximal” expression level of the gene by searching the high-expression levels across the atlas.

Specifically, we considered the 99th percentile of the cells that passed the first thresholding to be the maximum to increase the robustness of the analysis. Lastly, we applied relative thresholding of the “maximal” level (for circulating proteins such as ligand genes at 20% and other gene types at 5%). Cell types that passed all thresholding criteria were considered to have functional expression of the gene. We used a higher relative threshold for the ligand genes because 1) ligands are secreted into the circulation so a highly expressing source outshadows a low expressing source, whereas non-ligand products of other genes (e.g., receptors, synthases, processing enzymes) function inside individual cells; 2) we noted that ligand genes were usually expressed at higher levels and were more easily affected by cross-contamination (signal spreading) inside a tissue. For hormone ligands and receptors that included multiple genes, we applied AND or OR logic gates as indicated in **Table 1** to determine if a cell type expressed the ligand or receptor. For an AND logic gate, a functional cell type should express all its components above the functional level; while for an OR logic gate, expression of any of its components would suffice.

### Cell type distances, clustering, and visualization

We performed hierarchical clustering of the cell types included in the mouse lemur atlas based on the transcriptional profiles of hormone ligand, synthase, modulator, and receptor genes. Analysis was carried out at the cell type level and relative average gene expression levels were used to calculate cell-type pairwise distances. To test consistency of hormone-related gene expression among mouse lemur individuals, we separated cells from different lemurs. Because circulating immune cells sampled from different tissues were already kept as different cell types, immune cell types were not separated for different lemurs to avoid overcrowding of the data. We found that this simplification only slightly influenced the clustering results of individual cell types and did not affect the overall clustering pattern (**Fig. S4**). Next, cell types were filtered to remove potentially low quality cell types that were: 1) low in cell number, 2) technical doublets, or 3) low in transcript and/or gene count. We also manually corrected for notable cross-contamination in certain cell types. For example, almost all non-endocrine cell types of the pancreas showed high expression of insulin (INS) and proglucagon (GCG), probably caused by cross-contamination due to autolysis in the pancreas during tissue processing. We therefore removed INS and GCG expression from all non-endocrine pancreatic cell types. Additionally, we corrected contamination of GUCA2A, GUCA2B, GIP, NTS, and SST from stromal, endothelial, and immune cells of the intestine (small intestine and colon). Matlab built-in functions were used in the hierarchical clustering analysis. We used cosine distances to calculate cell type pairwise distances and then applied average-linkage to construct the dendrogram (cophenetic correlation coefficient = 0.81). Cell type clusters were then divided at a threshold of 0.49 with minor manual inputs to further divide certain large clusters or merge an apparently close neighboring leaf. We obtained 74 total clusters and numbered them according to their sequence in the dendrogram shown in **Fig. 2a**. We also named the clusters according to the shared characteristics of its cell types. Cell types included in each cluster are listed in **Table 4** with the list of hormones, hormone receptors, and hormone-related genes that were expressed in the majority (≥ 50%) of the cell types for each cluster.

In addition to visualizing the cell type clustering via dendrogram, we also applied uniform manifold approximation and projection (UMAP)^127^ to embed the high dimensional expression data to 2D using a Matlab UMAP package^128^. Cell type pairwise cosine distances were used as input when generating the UMAPs.

For comparative analysis, we additionally calculated cell type pairwise distances by the transcriptomes of all genes, the non-hormonal transcriptome, 100 random sets of PCGs (same number of genes as the hormonal genes), 100 random sets of PCGs controlling for expression variability (dispersion). These transcriptional distances were calculated similarly as the hormone-based distances were, with relative expression levels of genes used to calculate pairwise cosine distances. To select for gene sets with similar distribution of expression specificity, we first binned genes based on their cell type expression dispersion (i.e., variance-to-mean ratio) to 21 uniform bins and then randomly selected the same number of genes in each bin as the hormonal genes.

These transcriptional distances were compared to a benchmark cell annotation-based distance by Pearson correlation. Cell annotations, as defined in the accompanying manuscript^15^, was based on expression of canonical cell markers and iterative clustering with highly variable genes. This annotation-based distance was defined as an integer between 0 and 3 based on the following criteria: distance is 0 when cell types were annotated with identical names but from different tissues, 1 when our cell annotation names were different but classical histological assignments (Cell Ontology^48^) were identical (i.e., different molecular subtypes), 2 when classical Cell Ontologies were different but cell types were from the same compartment (e.g., both were epithelial), and 3 if cells were different cell types and from different compartments (e.g., endothelial vs. stromal).

We additionally calculated a Cell Ontology-based distance. Each cell type was assigned a Cell Ontology term (i.e., its classical histological cell type category) during cell type annotation^15^. Cell Ontology distances were calculated as the mean of Cell Ontology embedding-based distances and text-based distances^49^. Cell Ontology embedding-based distances represent distances between cell types in the Cell Ontology graph. Text-based distances were calculated using an adapted version of Sentence-BERT^49,129^.

To estimate the quality of the clustering results, we calculated silhouette values and compared cluster average silhouette values based on only the hormone ligand genes, hormone receptor genes, or the full transcriptome. Silhouette values for a cell type *i* is from the equation *s*(*i*) = 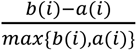, for cell types belonging to a cluster with more than one member. *a*(*i*) is the mean in-cluster distance, namely the mean distance between cell type *i* and all other cell types in the same cluster. *b*(*i*) is the minimal out-cluster distance, namely the minimal mean distance between cell type *i* and members of other clusters.

### Trajectory analysis

Analysis was performed independently for kidney samples from three mouse lemur individuals and for the cross-species spermatogenesis dataset using an in-house program written in Matlab. To detect the vasa recta arteriole-to-venule trajectory, data were first pre-processed following the standard procedure as in Seurat^130^. In brief, single cell transcript counts for all sequenced genes were normalized to total transcript counts, log transformed, and linearly scaled so that each gene has a mean expression of 0 and standard deviation of 1. Then genes with high dispersion (variance-to-mean ratio) compared to other genes with the same mean expression were selected as highly variable genes for principal component analysis (PCA).

Following PCA, the top 20 principal components that were not driven by extreme outlier data points or immediate early genes were used to generate a 2D UMAP using cell-cell Euclidean distances as input^127^. For the cross-species spermatogenesis, we used species-integrated UMAP of the testis germ cells^15^. To quantify the trajectories, we detected the probability density ridge of the data points in the UMAP using automated image processing. The direction of the trajectory was manually assigned according to marker gene expression. Individual cells were aligned to the trajectory by finding the shortest connecting point to the trajectory. Data points that were too distant from the trajectory were deemed outliers and removed from the following analysis. To detect the vasa recta hormone-related genes that follow a trajectory dependent expression pattern, we calculated the Spearman correlation coefficient between the gene expression level and 20 unimodal patterns. These preassigned patterns have their data smoothly changing along the trajectory and have a single peak. We let the 20 patterns have their peaks uniformly distributed from 0 to 1. This analysis tested, for each gene, whether its distribution along the trajectory follows one or more of the 20 unimodal patterns. All *p* values were then multi-testing corrected to detect significant genes and their associated patterns. Lastly, we compared results from the three kidney samples and kept the genes that showed consistent patterns in all three samples.

### Identification of differentially expressed genes

We used the Wilcoxon rank-sum test to detect differentially expressed hormone genes among cell populations. Genes that were expressed above functional level in any of the analyzed cell populations were included in the analysis. For hormone-related genes differentially expressed during spermatogenesis, we tested whether each gene is differentially expressed in one of the three sperm cell clusters compared to the other two clusters. For the analysis of endothelial cells, we tested whether each gene is differentially expressed in the lung or brain endothelial cells compared to all other endothelial cells. For the kidney nephron analysis, we tested whether each gene is differentially expressed in one of the nephron clusters compared to the rest of the nephron epithelial cells. For the hepatocyte analysis, we tested whether each gene is differentially expressed in one subtype compared to the other in both the mouse lemur and the mouse. The mouse hepatocyte data was obtained from the Tabula Muris Senis^114^ smartseq2 plate-based sequencing of mouse hepatocytes. For the tumor cell analysis, we tested whether each gene is differentially expressed in the metastatic cells versus the MUC16+ non-ciliated uterus epithelial cells. All *p* values were multi-testing corrected, and then all genes with corrected *p* values less than 0.05 were examined to confirm consistent expression of the gene in all relevant cell types and mouse lemur individuals.

### Constructing and analyzing the hormonal cell communication network

Cells with the same cell type annotation and tissue of origin were grouped as a cell type/node. Data from different lemur individuals were merged in the analysis. Cell types that were likely low quality were removed. Edge connections were determined independently for all possible cell type pairs. For example, if cell type A expressed a hormone ligand and cell type B expressed the corresponding hormone receptor(s), a directed edge from A to B was drawn. Note that this allows self-loops on any cell type that expresses both the ligand and corresponding receptor. When multiple directed edges connected two cell types, they were merged for the purpose of edge counting.

Network density is defined as the number of edges divided by the number of possible edges in the network. Because the hormone and cytokine networks are directed, its network density is calculated as 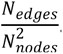. For comparison, the density of an undirected network is calculated as 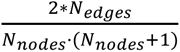 if it allows self-loops (e.g., protein interaction network), or 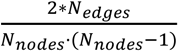 if it does not allow self-loops (e.g., genetic-interaction network).

Outdegree and indegree are node features and defined as the relative number of outgoing edges (node expressing a ligand) and incoming edges (node expressing a receptor). Edge counts were normalized by cell type abundance so that node degrees range from 0-100% and represent the fraction of the cell types that the node connects with by outgoing or incoming edges. To normalize the node degrees, we used a weighted cell type abundance determined by the clustering so that each cluster contributes the same weight. This is because certain cell types like circulating immune cells are likely over-represented in the network as they appear in multiple tissues and would be categorized as different cell types by our definition. We did not merge circulating cell types from different tissues because it is challenging for some cell types like T cells to determine whether they are circulating or potentially tissue specific or tissue-resident.

We also tried simply normalizing the node degree by the total number of nodes; similar conclusions were reached using either method.

### Calculating generality scores for hormone ligands and receptors

To calculate generality scores for the hormone ligands and receptors, we counted the cell types that express the ligand or receptor. Cell types that were likely low quality were removed from the analysis. Cell type counts were normalized by cell type abundances similarly as was the node degree, so that the generality score was not dominated by over-represented cell types. In comparison, we also counted the cell type clusters that express the hormone ligands and receptors. Here we defined that a gene was expressed in a cell type cluster if more than 50% of the cell types in the cluster positively expressed the gene.

To analyze the distribution of generality scores, we tested if the distribution followed normal, exponential, lognormal, or gamma distributions. We first performed parametric distribution fitting and then performed Kolmogorov–Smirnov tests comparing the fitted versus data cumulative distribution functions. This analysis used Matlab built-in functions fitdist and kstest. Additionally, we tested if the data followed a power law distribution using plfit and plpva functions^131^.

### Calculating target cell Jaccard indices for different receptors that bind to the same ligand

We calculated Jaccard indices for pairs of receptors that 1) can bind to the same ligand as described in **Table 1, and 2**) were both expressed in at least one atlas cell type. Jaccard index here is calculated as the number of cell types that express both receptors divided by the number of cell types that express either of the receptors.

### Network perturbation analysis

We performed two types of network perturbations: degree-preserving random rewiring and random edge removal. For degree-preserving random rewiring, we applied the algorithm developed by Maslov and Sneppen^82^. The algorithm starts with the original network and performs repeated rewiring steps. For each rewiring, it randomly identifies two edges (e.g., A→B and C→D) and swaps the connections (i.e., A→D and C→B). This allows constructing a random network while preserving the node indegree and outdegree of the original network. For the perturbation by edge removal, we randomly selected 10% of the edge types (ligand-receptor pairs) and removed these edges from the original network. Note for this analysis, multiple directed edges connecting the same cell type pairs were not merged. Therefore, the connection between two cell types would be removed only if all of its redundant edges (of different ligand-receptor pairs) were selected to be removed.

### Hormone concentration measurements (related to Fig. 6)

Cortisol concentrations (ng/ml) were measured in duplicates from urine samples by Cortisol Urine ELISA (LDN®, catalog no. MS E-5100). Urine creatinine (mg/ml) was measured using MicrovueTM Creatinine Elisa kit (Quidel R Corporation, catalog no. 8009). Creatinine concentration was used to normalize urine measurements as an indicator of renal filtration activity. Cortisol measurements are thus expressed in ng/mg creatine. Thyroxine (T4, nmol/L) was assayed in plasma samples using T4 ELISA kit (LDN®, catalog no. TF E-2400) and the 6-point standard curve was adapted from 0 to 100 nmol/L.

### RNAscope experiments

Mouse lemur specific RNAscope® probes for MLN, AGT, and CUBN were designed and produced by Advanced Cell Diagnostics, Inc. (ACD) (cat #. 1260231, 1263051, 1263061). The kidney samples analyzed were collected from the same lemurs profiled by scRNAseq^15^ and analyzed in this study. Samples were procured, fixed, dehydrated, OCT-embedded, and biobanked at -80°. Cryosections (15 µm thick coronal sections) of the kidney were prepared using a Leica Cm3050s cryostat, stored overnight at -80°, and subsequently processed using the standard RNAscope multiplex fluorescent v2 protocol for fixed-frozen samples (UM 323100 Rev B), with the RNAscope® Multiplex Fluorescent Reagent Kit v2 (cat#. 323270). In brief, sections were first pre-treated during which tissue slides were PBS washed, baked, post-fixed, dehydrated by ethanol, treated with hydrogen peroxide, submerged in RNAscope 1X Target Retrieval Reagent (99°) for 5 min, and finally treated with RNAscope Protease III. Next, sections were hybridized with all three probes for 2h at 40°. To visualize the probes, sections were incubated with each of the three RNAscope Multiplex FL v2 AMP for 30 min at 40°, and then with RNAscope Multiplex FL v2 HRP for 15 min at 40°, and the respective TSA Vivid Dye for 30 min at 40°, and finally the RNAscope Multiplex FL v2 HRP blocker for 15 min at 40°. Lastly, tissue slides were mounted with Prolong Gold Antifade Mounting containing DPAI and stored in 4° before imaging. Images were taken using a Zeiss LSM 880 Confocal Laser Scanning Microscope under the Airyscan mode. The MLN, AGT, and CUBN channels were stained with TSA Vivid Fluorophore 570, 650, and 520, and imaged using the Cy3, Cy5, and FITC filters, respectively.

### Cross-species comparisons of hormonal gene expression

We compared cells of the lung, skeletal muscle, liver, testis, bone marrow, and spleen. All lemur data was from the 10x data of the accompanying Tabula Microcebus manuscript^15^. Human data were from the 10x data of the Tabula Sapiens^120^, except for the lung which we used the 10x data from the Human Lung Cell Atlas^132^ and the testis which we used drop-seq data from Shami et al.^133^. Mouse data were from 10x data of the Tabula Muris Senis^114^ except for the testis which we used 10x data from Ernst et al.^134^. For lung and muscle, all cell types were included in the analysis, whereas we only analyzed the epithelial cells for the liver, immune cells for the bone marrow and spleen, and germ cells for the testis. We used one-to-one orthologs across all three species. This resulted in ∼15,000 genes (324 hormonal genes), ∼13,000 of which were included in all scRNAseq datasets (295 hormonal genes).

To unify cell type annotations, human and mouse datasets were first separately re-annotated according to the same pipeline and marker genes as for the lemur data. Next, to ensure cell types are comparable across species, we applied Portal^135^ to integrate data (one-to-one orthologs only) from different species. Portal projects data into a space that minimizes species differences, from which an integrated UMAP is generated to visualize cell clustering from different species. Integration was performed separately for each tissue, except for bone marrow and spleen which were integrated jointly. We manually inspected each integration UMAP and ensured that cells of the same type showed reasonable cross-species co-clustering and separation from other cell types. We also made small modifications to the cell annotations/clustering during this process to unify annotations across species. For example, proliferating cells might co-cluster with the main non-proliferating population in the original dataset if the number of cells are too few, but cluster separately with the proliferating populations of the other species in the integrated UMAP. In such a scenario, we would separately annotate these cells as a proliferating subtype. Additional details to compile the cross-species scRNAseq datasets were described in the accompanying manuscript^15^.

Cell types with more than 15 cells in all three species were then used for the cross-species comparisons, resulting in a total of 62 orthologous cell types (62*3=186 cell type entries in all three species). Cell type average expressions were calculated separately for each species. Note that cellular expression levels in the species-integrated dataset were normalized by the sum of only the one-to-one orthologs, rather than all the genes, thus at high levels compared to the non-integrated data. Cross-species similarity was calculated separately for individual genes. We first identified genes that are at least moderately expressed in the analyzed cell types (≥3 cell type entries with mean expression >1), which resulted in a total of 90 genes for follow-up analysis.

Expression levels were then normalized by the maximally expressed cell type in each species and cross-species correlation coefficients between human and lemur (*r*(human, lemur)) and between human and mouse (*r*(human, mouse)) were calculated. Δ*r* = *r*(human, lemur)-*r*(human, mouse) was calculated. Lastly, we identified genes that are more conserved between humans and lemurs compared to mice (Δ*r*>0.5), and in contrast genes that are more conserved between humans and mice compared to lemurs (Δ*r*<-0.5). To compare overall similarity of transcriptional profiles of the hormonal genes or all genes, we concatenated cell type expression profiles for individual genes and calculated correlation coefficients between human and lemur and between human and mouse. *p*-values of the difference in correlation coefficients were calculated using Fisher’s Z-transformation^136^.

## Supporting information

Fig. S1a. Ligand and receptor gene expression for 84 hormone classes across the mouse lemur cell atlas, with cell types ordered by cell type.

Fig. S1b. Ligand and receptor gene expression for 84 hormone classes across the mouse lemur cell atlas, with cell types ordered by tissue.

Table 1. Genes involved in the biosynthesis and sensing of 84 classes of hormones.

Table 2. Lemur and mouse orthologs of the human hormonal genes in Table 1.

Table 3. Cell types with positive expression of each of the hormones and receptors.

Table 4. Clusters of mouse lemur cell types according to the expression of hormonal genes.

Table 5. Hormones that regulate or are produced by the anterior pituitary neuroendocrine cells.

Table 6. Generality scores of the hormone ligands, modulators and receptors.

Table 7. Two node feedback circuits detected from the mouse lemur hormone cell-cell communication network.

Table 8. Measurements of concentrations of mouse lemur seasonally-changing hormones.

## Acknowledgments

We are grateful to Sandra Schmid, Uri Alon, Chaitan Khosla, Mathieu Lemaire, the Krasnow lab, and the Ferrell lab for helpful discussions and comments on the manuscript. This work is supported by NIH grant R35 GM131792 to J. E. Ferrell, funding from Wall Center to M. A. Krasnow, and a fellowship from the Wu Tsai Neurosciences Institute to S. Liu. M. A. Krasnow is an investigator of the Howard Hughes Medical Institute.

## Author Contributions

S. Liu, M.A. Krasnow, and J.E. Ferrell conceived the project. S. Liu designed and performed all the analyses with input from C. Ezran, Z. Li, C. Kuo, J. Epelbaum, J. Terrien, M.A. Krasnow, and J.E. Ferrell. S. Liu, C. Kuo, and J. Long assembled the hormone list. M.F.Z. Wang and I.D. Vlaminck performed uTAR analysis on the candidate lemur LHB gene. S. Wang calculated the cell ontology distances. S. Liu, J. Epelbaum, and J. Terrien analyzed the seasonally oscillating hormones. S. Liu and C. Kuo performed the RNAscope experiments. K. Awayan constructed the interactive web portal. The Tabula Microcebus Consortium constructed the mouse lemur molecular cell atlas. S. Liu and J.E. Ferrell wrote the manuscript with input from all authors.

## Supplemental Tables

**Table 1. Genes involved in the biosynthesis and sensing of 84 classes of hormones**. Rows show hormones and columns show the hormone class name, hormone name(s), type of hormones, symbol of the human genes for the respective hormone ligands, synthases and other enzymes, receptors, and plasma binding proteins, whether the hormone synthesis or maturation requires coordination of multiple tissues, years discovered, classical sites of secretion and targets, approximate plasma concentrations in humans, and references.

**Table 2. Lemur and mouse orthologs of the human hormonal genes in Table 1**. Rows show genes and columns show gene type, NCBI and Ensembl gene IDs, NCBI and Ensembl gene symbols of human, lemur, and mouse genes respectively, as well as the ortholog type for the human-lemur and human-mouse ortholog mapping. Each row shows a unique human gene. Entries in columns “Symbol_Lemur” and “Symbol_Lemur” may include multiple genes (separated by commas) for one-to-many mappings. The “all orthology” tab includes the orthology mapping for all hormonal genes. The “non one-to-one orthology” tab shows only the genes with non one-to-one mapping in either lemur or mouse. The “cross-species missing” tabs show the genes with one-to-one orthology mapping but were not included in the cross-species analysis because they were missing from one or more of the analyzed datasets.

**Table 3. Cell types with positive expression of each of the hormones and receptors.** Rows are hormone ligands or receptors and columns show the respective hormone class, hormone name(s), hormone type, entry type (ligand or receptor), related genes, and list of positive cell types.

**Table 4. Clusters of mouse lemur cell types according to the expression of hormonal genes.** Rows show clusters and columns show cluster ID, name, cell types included in the cluster, and the hormones, receptors, and hormone-related genes that were positively expressed in the majority (≥50%) of the cluster cell types.

**Table 5. Hormones that regulate or are produced by the anterior pituitary neuroendocrine cells.** Rows show the hormones and hormone receptors expressed in individual pituitary neuroendocrine cell types. Columns show the corresponding genes, hormone names, and upstream/downstream cell types.

**Table 6. Generality scores of the hormone ligands, modulators and receptors.** Rows show hormone ligands, modulators, or receptors, ordered by descending generality. Columns show entry types, symbols of human and mouse lemur genes, whether the entry includes a single or multiple genes, and generality scores calculated as the percentage of positive cell types and the number of positive clusters.

**Table 7. Two node feedback circuits detected from the mouse lemur hormone cell-cell communication network.** Rows show the detected circuits and columns show the two cell types and hormone signaling that connects the cell types.

**Table 8. Measurements of concentrations of mouse lemur seasonally-changing hormones.** Rows show individual measurements and columns show the season of measurements, concentration, hormone names and concentration unit, as well as the samples used in the measurements.

---

**Fig. S1.** Dot plot of hormone ligand, receptor, and modulator gene expression for 84 classes of hormones across the mouse lemur cell atlas. Each dot plot shows one of the 84 hormone groups and one of the five cell compartment groups (1. epithelial/neural/germ cells (combined for space saving); 2. endothelial cells; 3. stromal cells; 4. lymphoid cells; and 5. myeloid cells). The hormone group name and cell compartment type is indicated on the top right of the figure. The dot plots are arranged in a polar coordinate style. Radial lines represent individual cell types, with cell type names labeled inside the inner circle and the tissue source labeled outside the outer circle. Cell types are arranged, by either their tissue of origin or by their cell type name. Circular arcs represent the related genes for this hormone group (red for ligands, light red for synthases and processing enzymes, green for modulators, and blue for receptor genes). The dark colored dots on the intersections of the radial lines and circular arcs depict mean gene expression levels (by dot shade) and percent of positive cells (by dot size) for the respective gene and cell type. The outermost red or blue dots indicate positive expression of one of the hormone(s) (red) or hormone receptor(s) (blue), or both (purple) for the hormone group by the respective cell type (see **Methods**). The outermost grey dots indicate potentially low-quality cell types (e.g., low cell number (<10) and pancreatic cells that show notable signal contamination) with outlines indicating whether or not the cell type expresses one the hormone ligand(s) (red outline), receptor(s) (blue outline), both (purple outline), or neither (no outline). These figures can be assessed interactively at https://tabula-microcebus.ds.czbiohub.org/hormone-atlas.

**Fig. S2.**
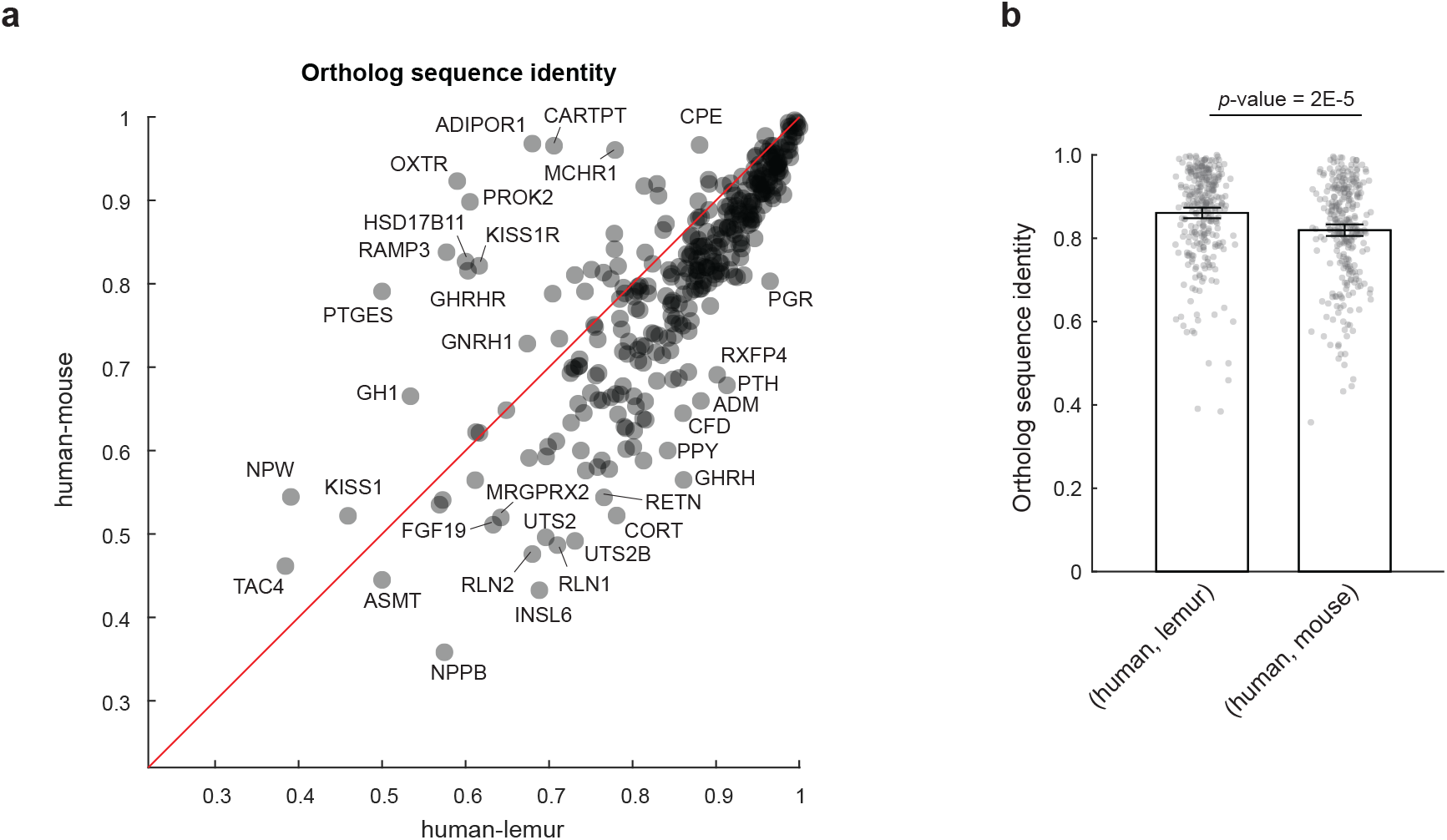
Protein sequences of hormonal genes are generally more similar between human and mouse lemur than human and mouse. **a**. Scatter plot comparing pairwise gene sequence identity between human and lemur vs. human and mouse. Each circle is a hormone-related protein. Sequences were aligned by the Needleman-Wunsch algorithm and sequence identity was calculated as the percentage of identical amino acids. Points below the red line are proteins more similar between human and mouse lemur than between human and mouse. **b**. Bar plot showing mean and 95% confidence interval of the average sequence identity between human and lemur vs. human and mouse. Grey dots show individual genes.

**Fig. S3.**
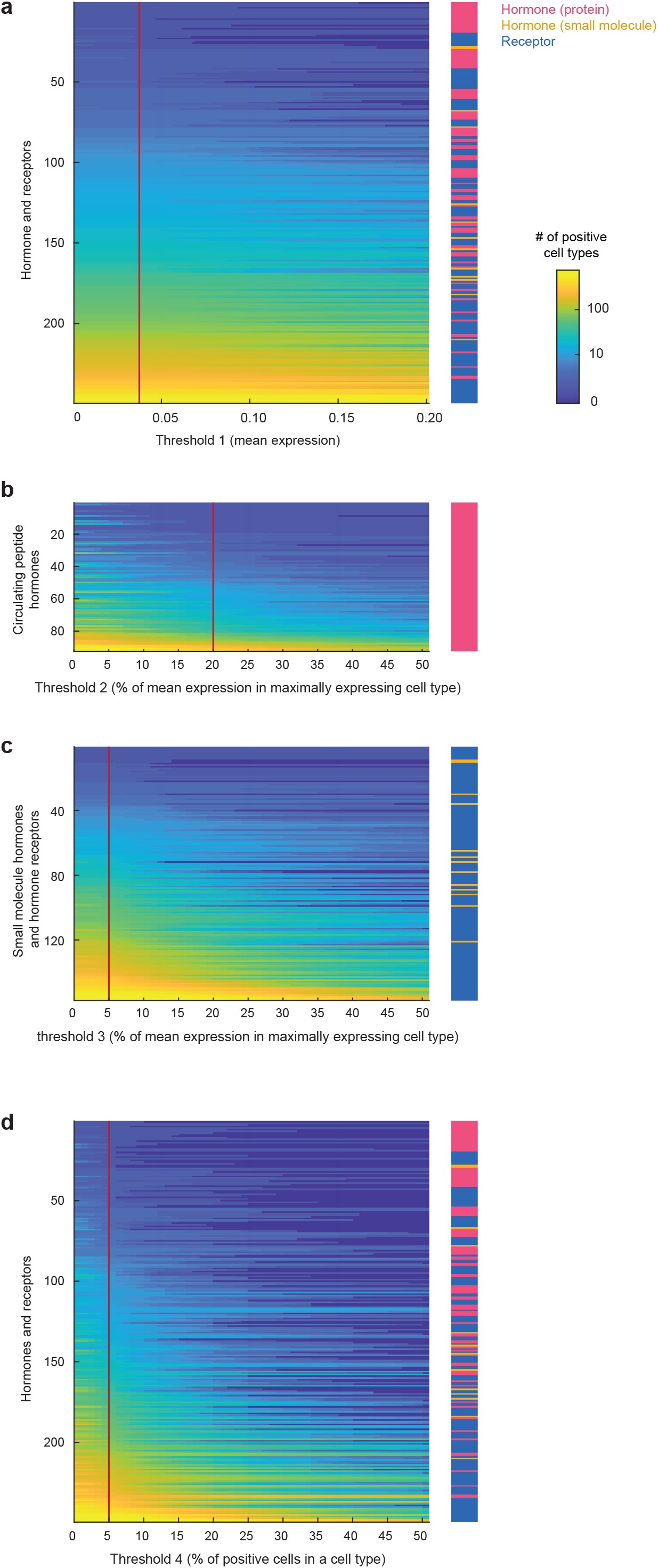
Number of positive cell types for each hormone ligand and receptor when changing the parameters for adaptive thresholding. Parameters include cell type mean expression (**a**), percent of mean expression relative to the maximally expressing cell types separately for circulating peptide hormones (ligand genes) (**b**) or small molecule hormones (synthase genes) and hormone receptors (**c**), and the percent of positive cells in a cell type (**d**). Red vertical lines represent the parameters used in the follow-up analysis. Each row represents one hormone or receptor, ordered by the number of positive cell types at the selected parameter.

**Fig. S4.**
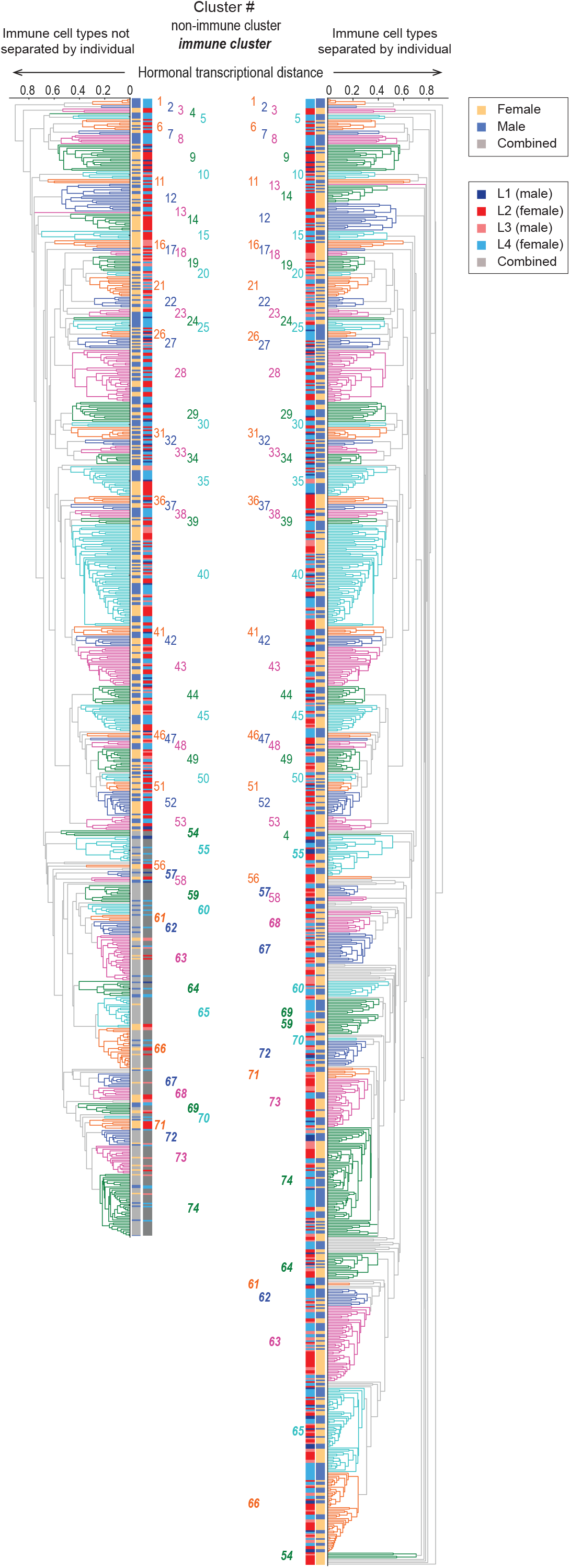
Cell types clustered similarly in both sexes and across sampled mouse lemurs. Shown here are cell type clustering dendrograms with cluster IDs labeled nearby and two color bars indicating the lemur individual and sex the cell type sampled from. The two dendrograms compare cell type clustering patterns when separating (right) or not separating (left) immune populations from different lemur individuals.

**Fig. S5.**
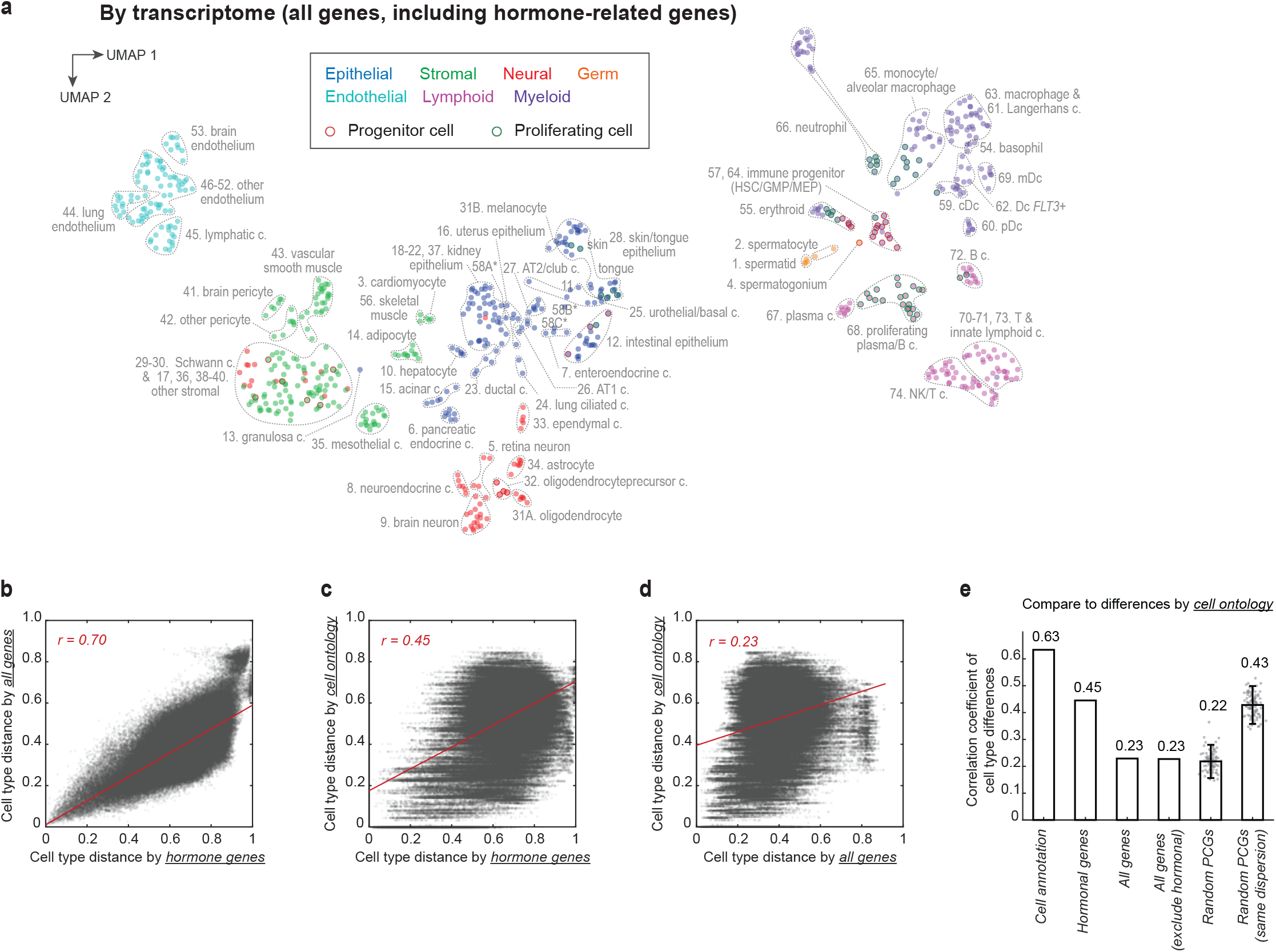
Comparison of cell type clustering by the hormonal genes with that by the full transcriptome. **a.** UMAP visualization of mouse lemur cell types based on the full transcriptome. Circles (cell types) are color-coded by cell compartment types (epithelial, neural, germ, stromal, endothelial, lymphoid, and myeloid). Dashed lines circumscribe cell type clusters as in Fig. 2a**-b** and cluster IDs and names are labeled next to the cell type clusters. Cell types that were clustered by the hormonal genes but not the full transcriptome are labeled separately (i.e., 31B and 31A; 58A, 58B and 58C). Note some cell types that formed discrete clusters based on hormonal gene expression intermingled with other clusters when mapped based on the non-hormonal transcriptome (e.g., cluster 46-52), and therefore were annotated together. **b-d**. Scatter plots and correlation of cell type pairwise distances based on hormonal genes vs. that of all genes (**b**), hormonal genes vs. Cell Ontology (**c**), and all genes vs. Cell Ontology (**d**). **e**. Bar plot of correlation coefficients comparing cell type pairwise distances of different metrics compared to a benchmark Cell Ontology-based distance. Also see Fig. 2e for comparisons with the benchmark of cell annotation-based distances.

**Fig. S6.**
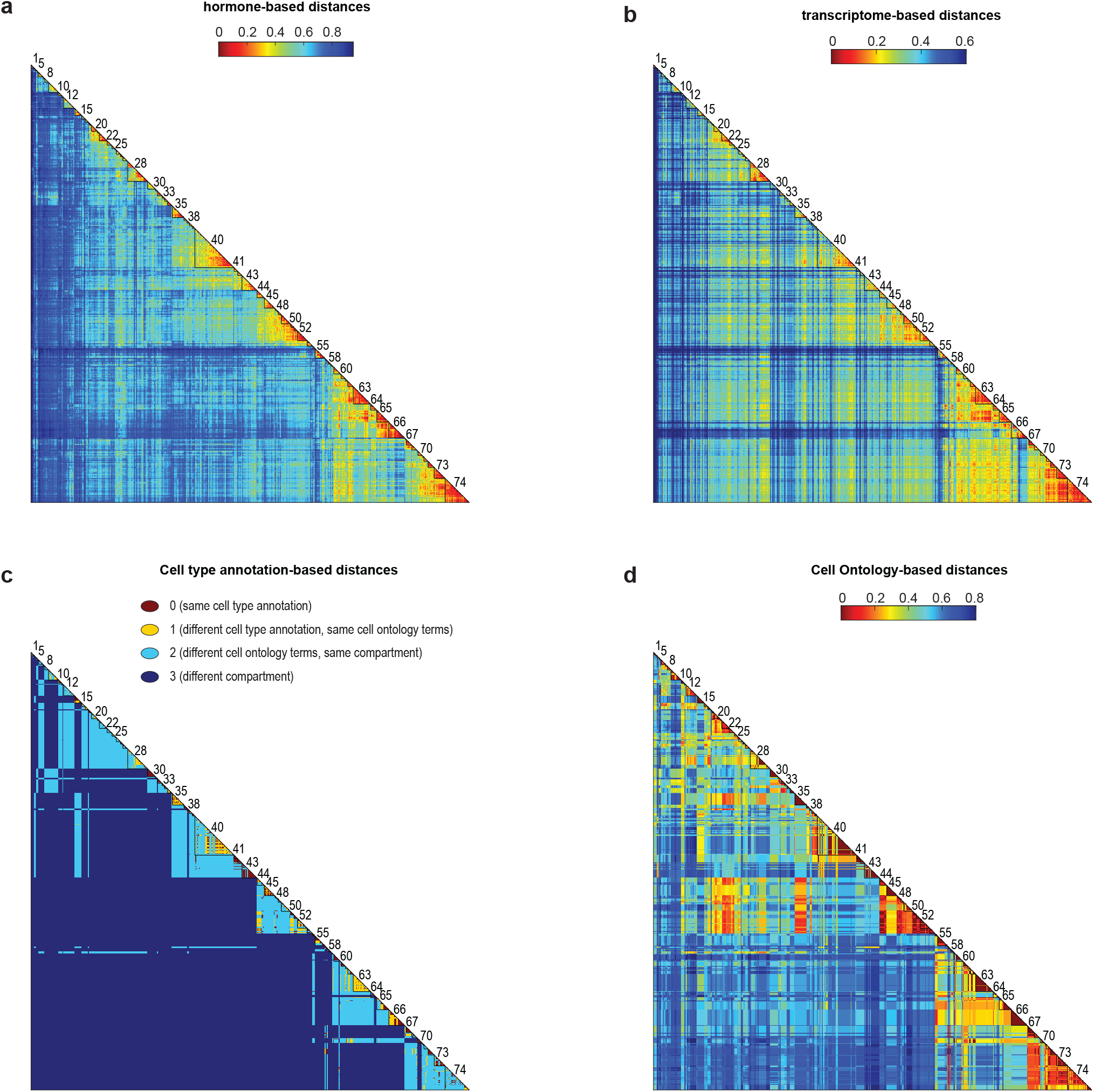
Heatmap representation of cell type pairwise distances based on the hormonal genes (a), the full transcriptome (b), the cell type annotations (c), and Cell Ontology (d). Cell types were ordered by cluster as in Fig. 2a, and cluster IDs were labeled along the diagonal for large clusters.

**Fig. S7.**
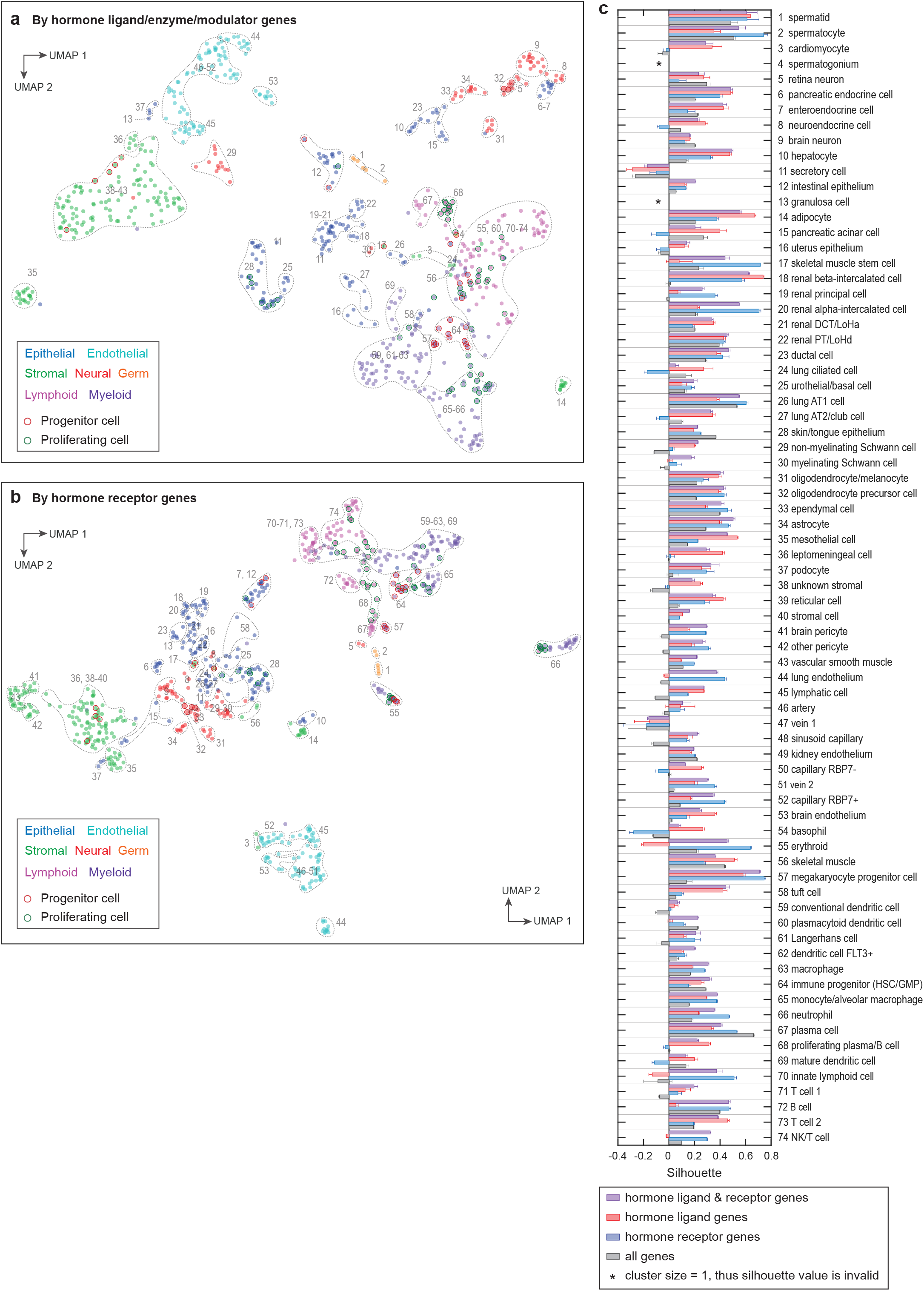
Classification of mouse lemur cell types using only the hormone ligand or the receptor genes. **a-b**, UMAP visualization of cell types based on the transcriptional distances of only the hormone ligands/modulators (**a**) or the hormone receptors (**b**). Circles (cell types) are color-coded by the cell compartment types (epithelial, neural, germ, stromal, endothelial, lymphoid, and myeloid). Dashed lines circumscribe cell type clusters as in Fig. 2a**-b** and cluster IDs are annotated nearby. As UMAPs qualitatively display relationships among data points, the extremely distant clusters in panel **a** (i.e., cluster 35. mesothelial cells by hormone ligand genes) were shifted towards the center of the figure for display purposes. **c.** Cluster averaged silhouette values of the 74 cell type clusters by both hormone ligand and receptor genes (black), only the hormone ligand genes (red), only the hormone receptor genes (blue), or the non-hormonal transcriptome (blank). The silhouette values are a measure of how well each cell type cluster was classified (see **Methods**). Asterisks (*) indicate clusters with only a single cell type, thus silhouette values cannot be calculated.

**Fig. S8.**
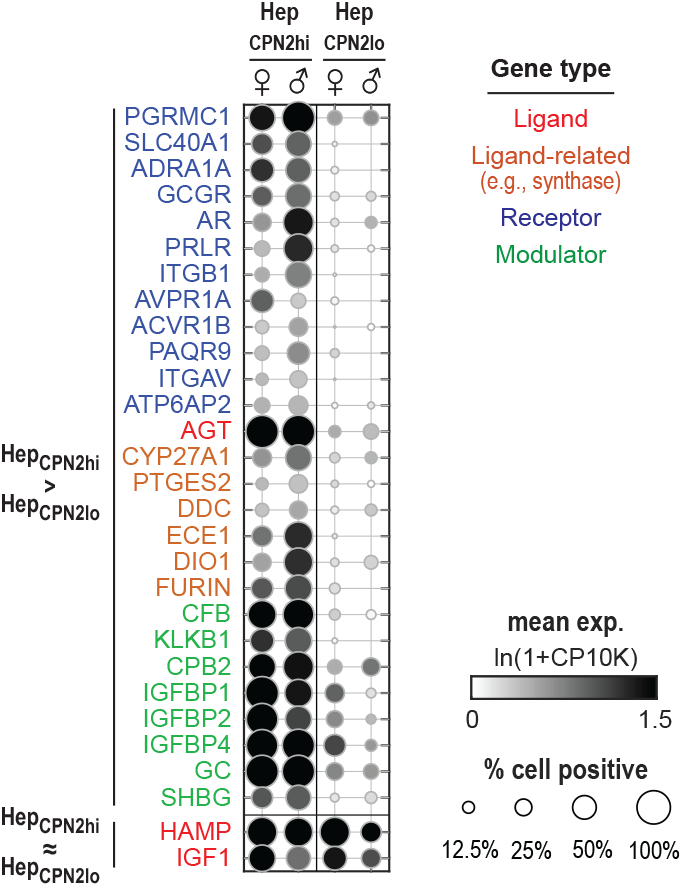
Hormonal gene expression in two hepatocyte subtypes plotted separately for male and female animals. (in comparison to Fig. 3f).

**Fig. S9.**
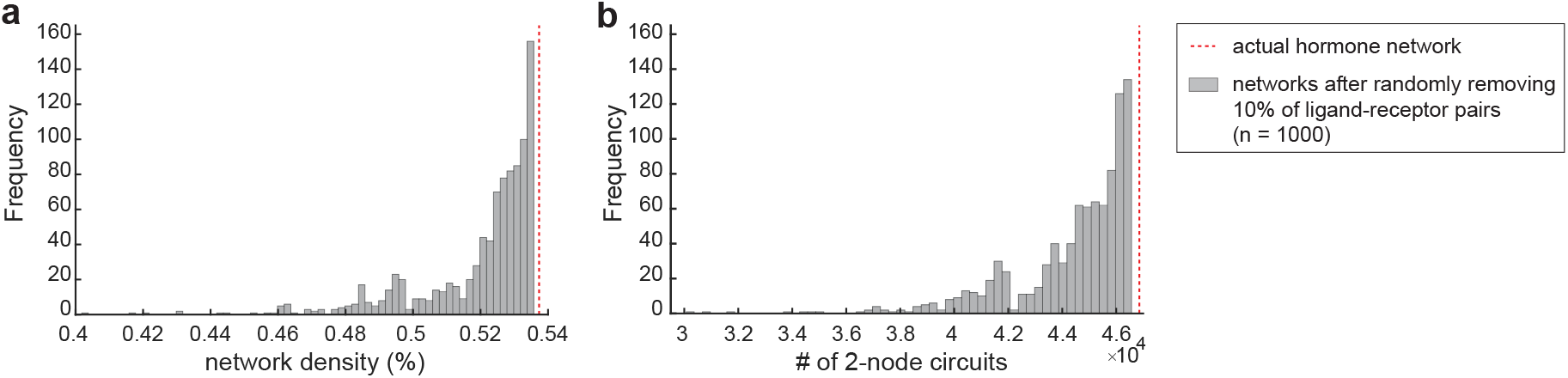
The hormonal cell-cell communication network is robust to edge removal. Distribution of network density (**a**) and number of 2-node feedback circuits (**b**) in networks where 10% of ligand-receptor connections were randomly removed.

**Fig. S10.**
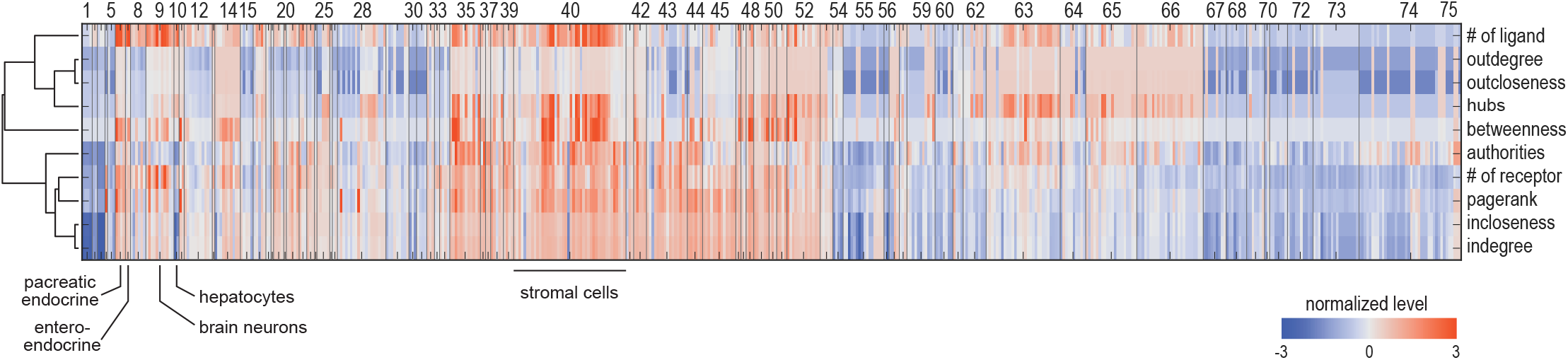
Node properties of the hormonal cell-cell communication network. Heatmap of indegree, outdegree, number of hormone ligands and receptors expressed, and six node centrality measurements (hubs, authorities, incloseness, outcloseness, pagerank, and betweenness) for all nodes (cell types) of the hormonal cell-cell communication network. Because the scales of the node properties are different, shown here are normalized values linearly scaled by median and median-absolute-deviation. Cell types (columns) are ordered by the clustering of the hormonal genes (as in Fig. 2a) and cluster numbers are labeled as space allowed. Node properties (rows) are ordered by hierarchical clustering of the normalized values.

**Fig. S11.**
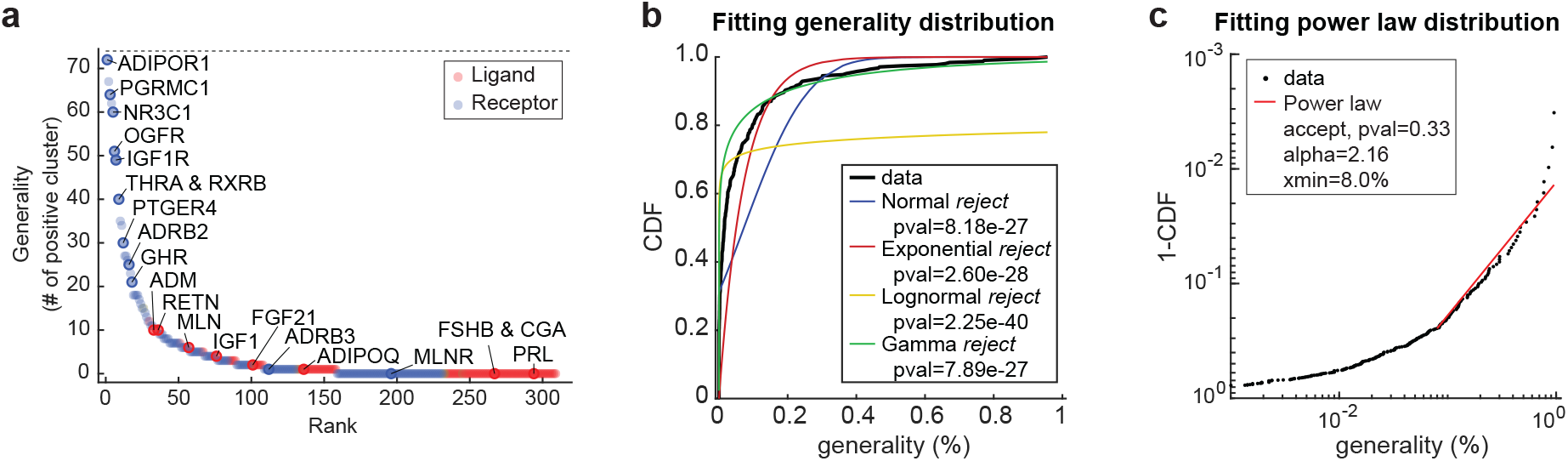
Generality scores for the hormone ligands and receptors. **a.** Generality score, defined as the number of clusters with positive expression of the ligand or receptor (see Fig. 4f for comparison), ranked from most generally expressed to most selectively expressed. **b-c**. Fitting the distribution of the ligand and receptor generality scores. Normal, exponential, log-normal, and gamma distributions failed to fit the distribution (**b**). The tail of the distribution can be approximated by a power law distribution *prob*(x) ∝ *x^-α^* for *x* ≥ *x_min_* (**c**).

**Fig. S12.**
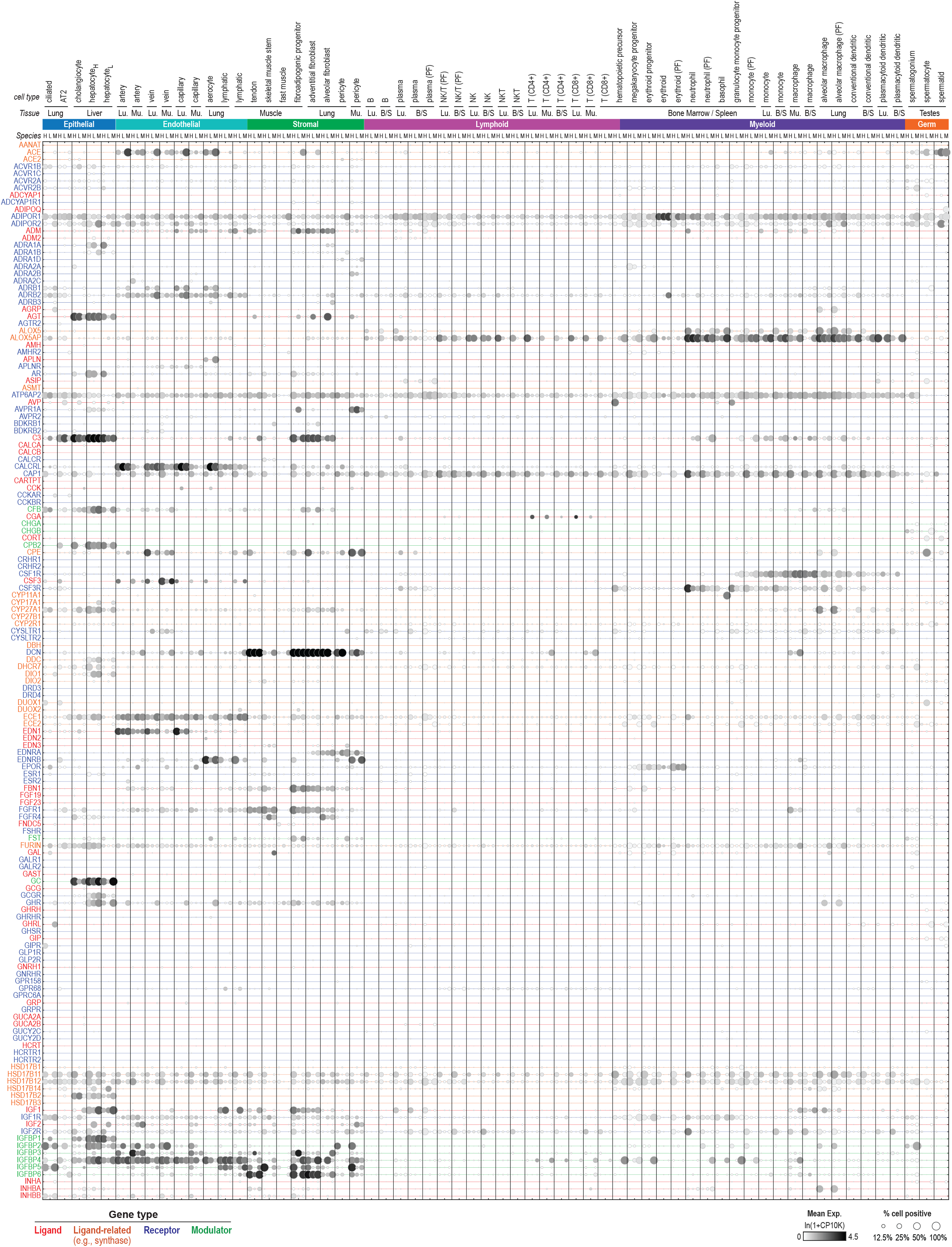
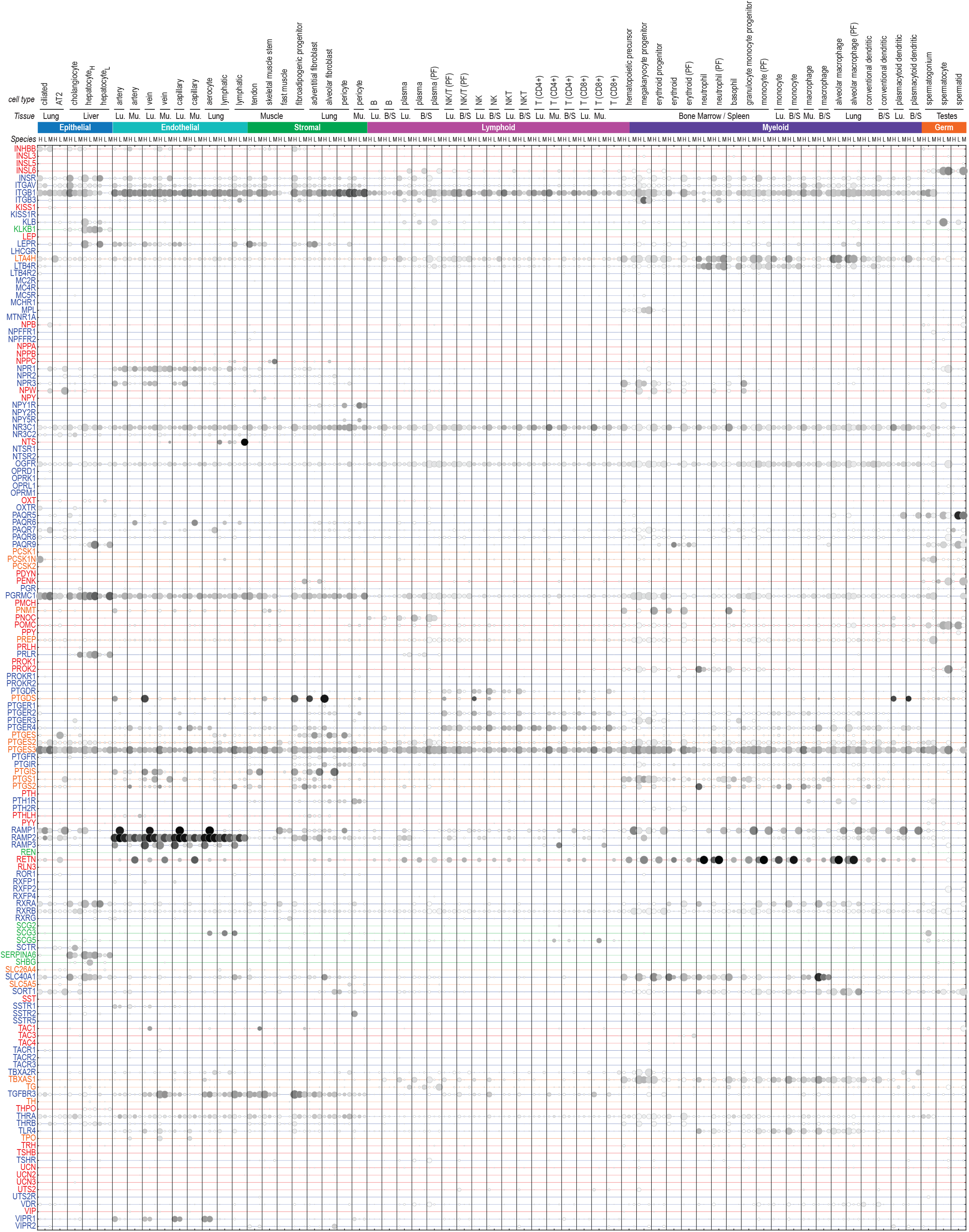
Expression of the hormonal genes across human, lemur, and mouse cell types. Expression is visualized by a dot plot with dot color indicating the cell type average expression and dot size indicating the percent of positive cells in the cell type. Rows are 295 one-to-one-to-one orthologs of the hormonal genes, labeled with the respective human symbols, and ordered alphabetically. Columns are arranged as triplets showing respective expressions in humans, lemurs, and mice. Triplets of different cell types are segregated by vertical gray lines. The orthologous cell type triplets are ordered first by compartments (e.g., epithelial, stromal), then by tissue, and lastly by the cell type annotation.

**Fig. S13.**
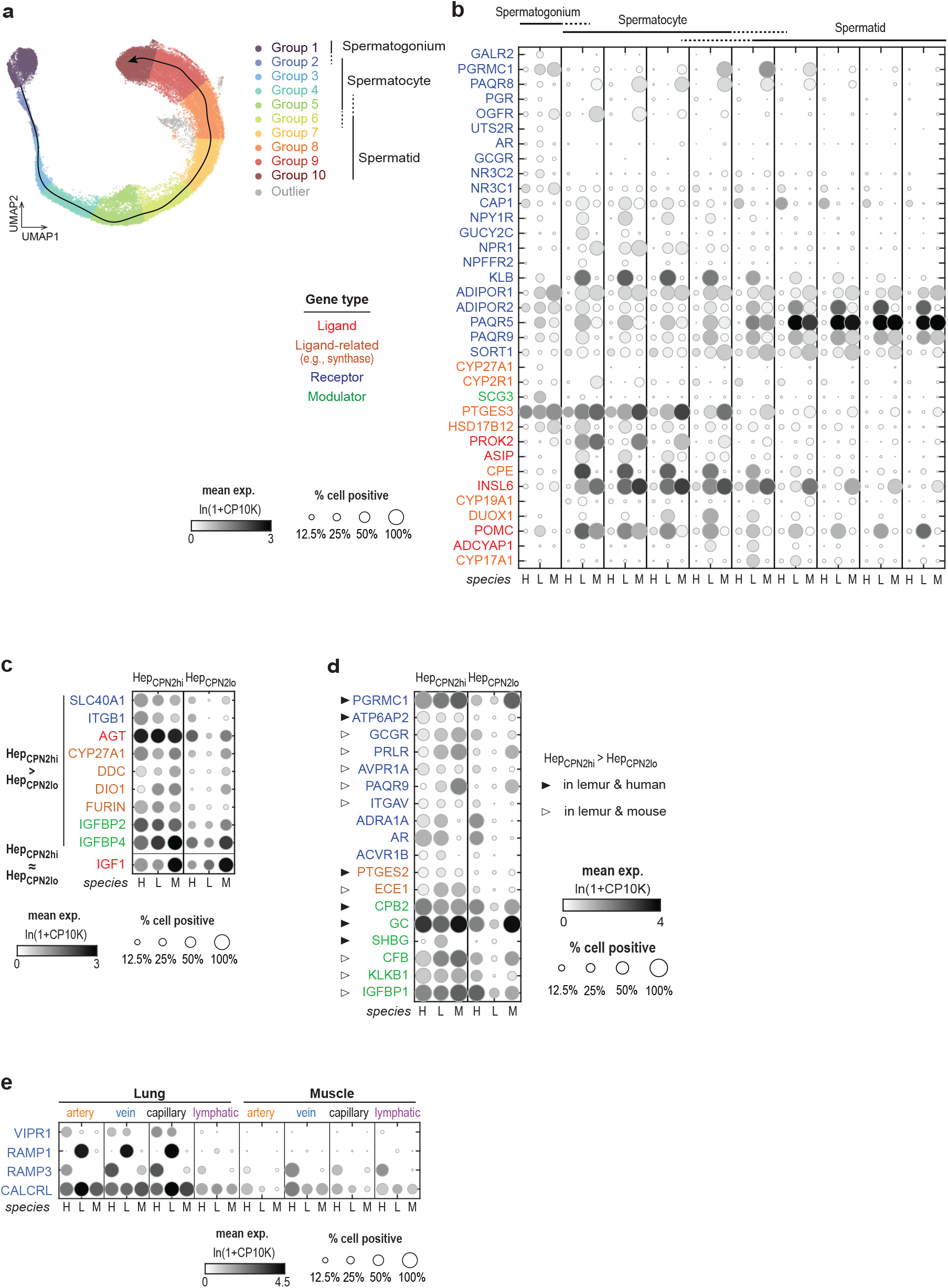
Cross-species comparison of hormonal gene expression related to examples in Fig. 3. **a**. Group of male germ cells by the species-integrated trajectory, related to Fig. 7g. **b**. Expression of the spermatogenesis stage-specific hormonal genes identified in Fig. 3a across human, lemur, and mouse. Shown here are examples with variable expression patterns across the three species. Also see Fig. 7g**-h** for additional examples with either conserved or variable expression patterns across the three species. **c-d**. Expression of the hepatocyte subtype related hormonal genes in the two hepatocyte subtypes across human, lemur, and mouse. The two panels separately show the genes with conserved (**c**) or variable (**d**) expression patterns across the three species. **a**. Expression of the lung-specific hormone signatures in the lung and muscle endothelial cells across human, lemur, and mouse.

## The Tabula Microcebus Consortium

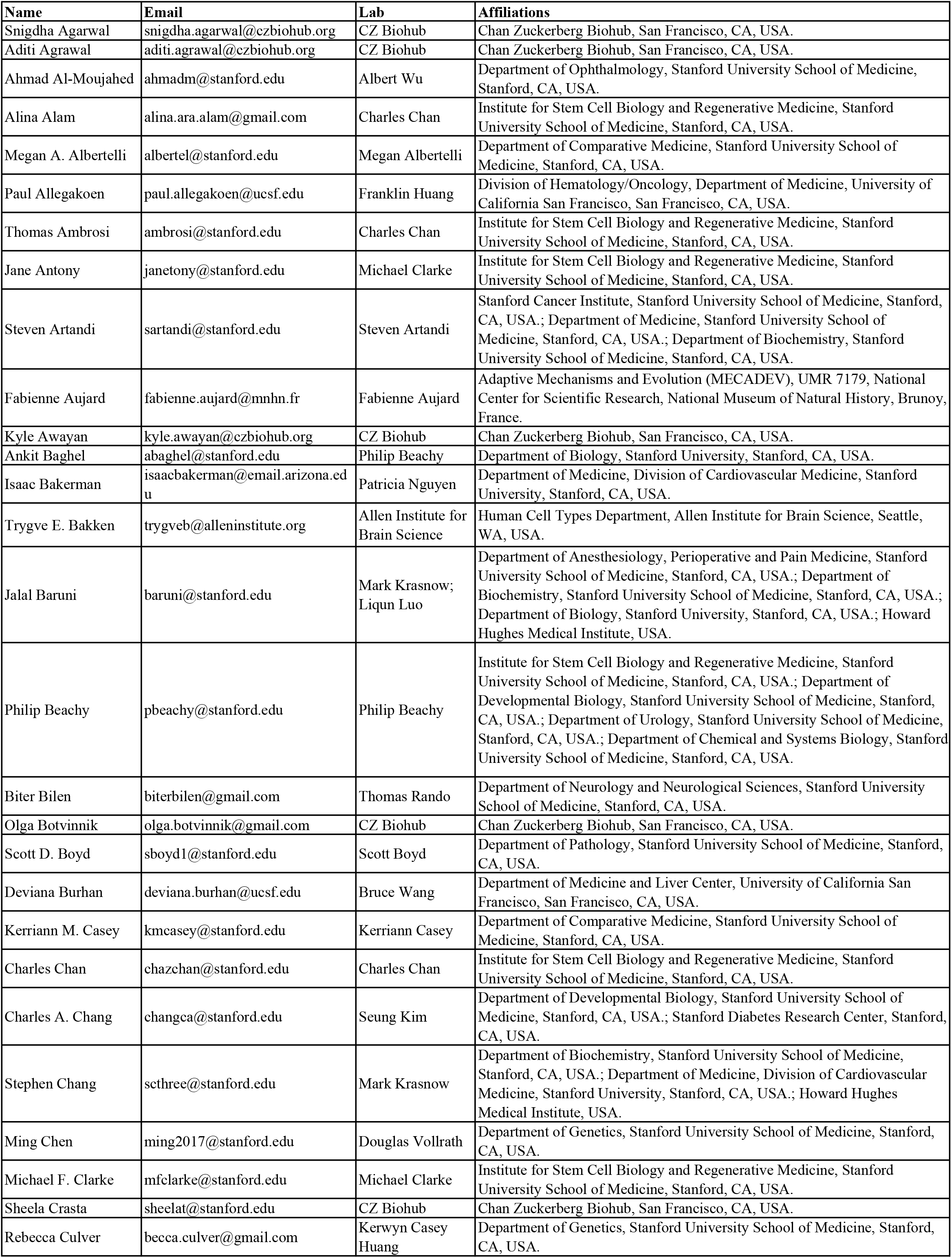
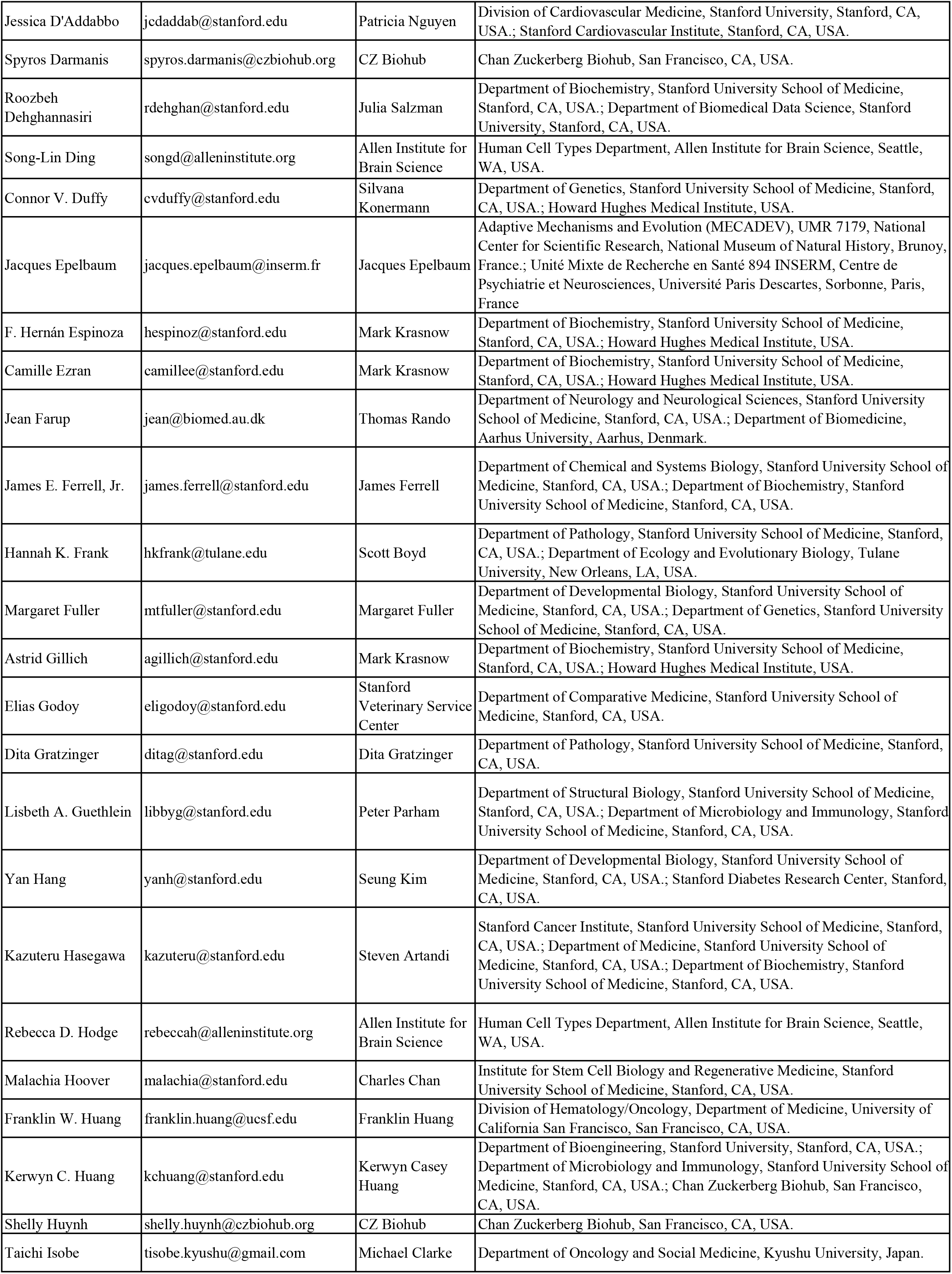
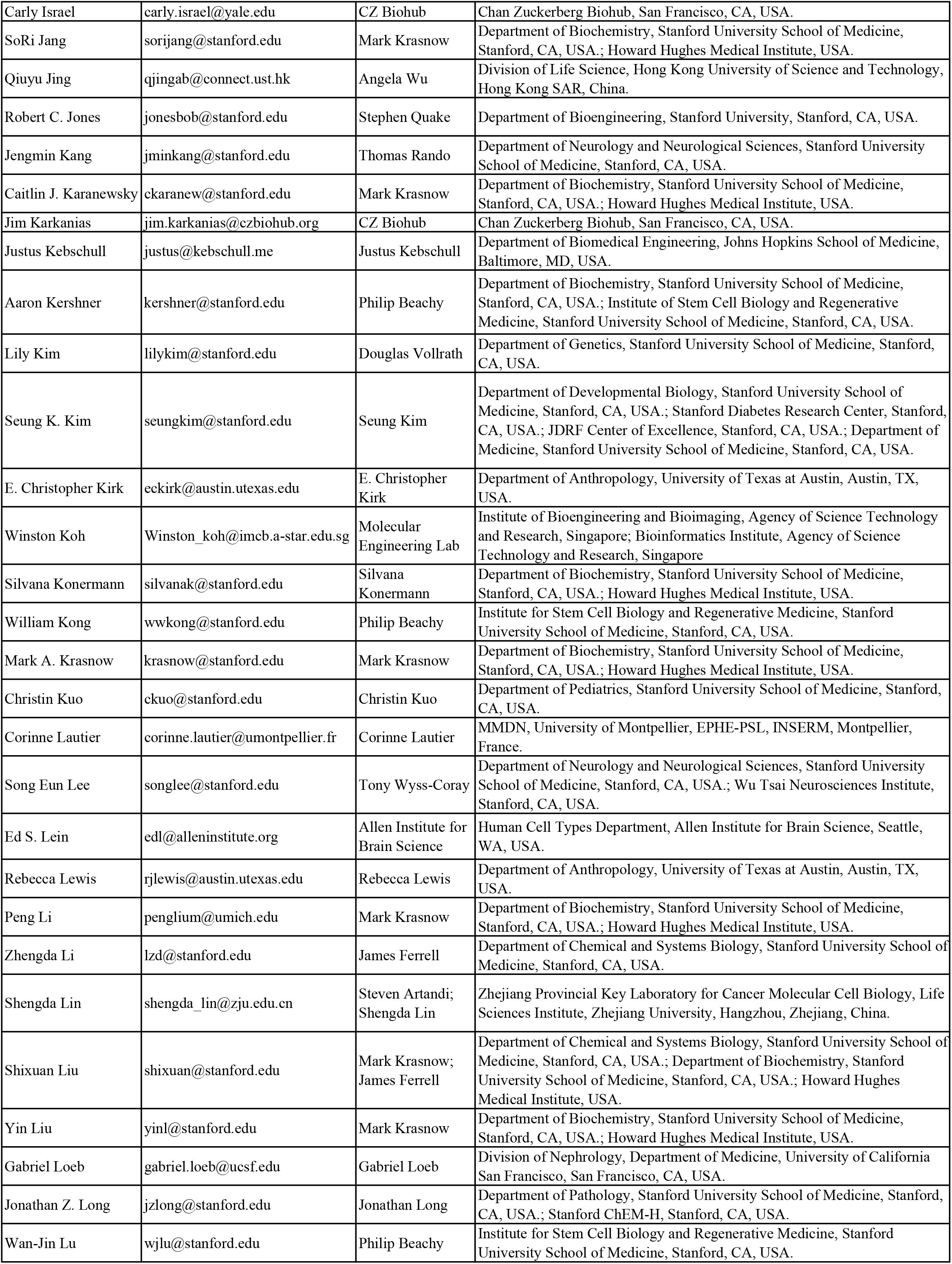
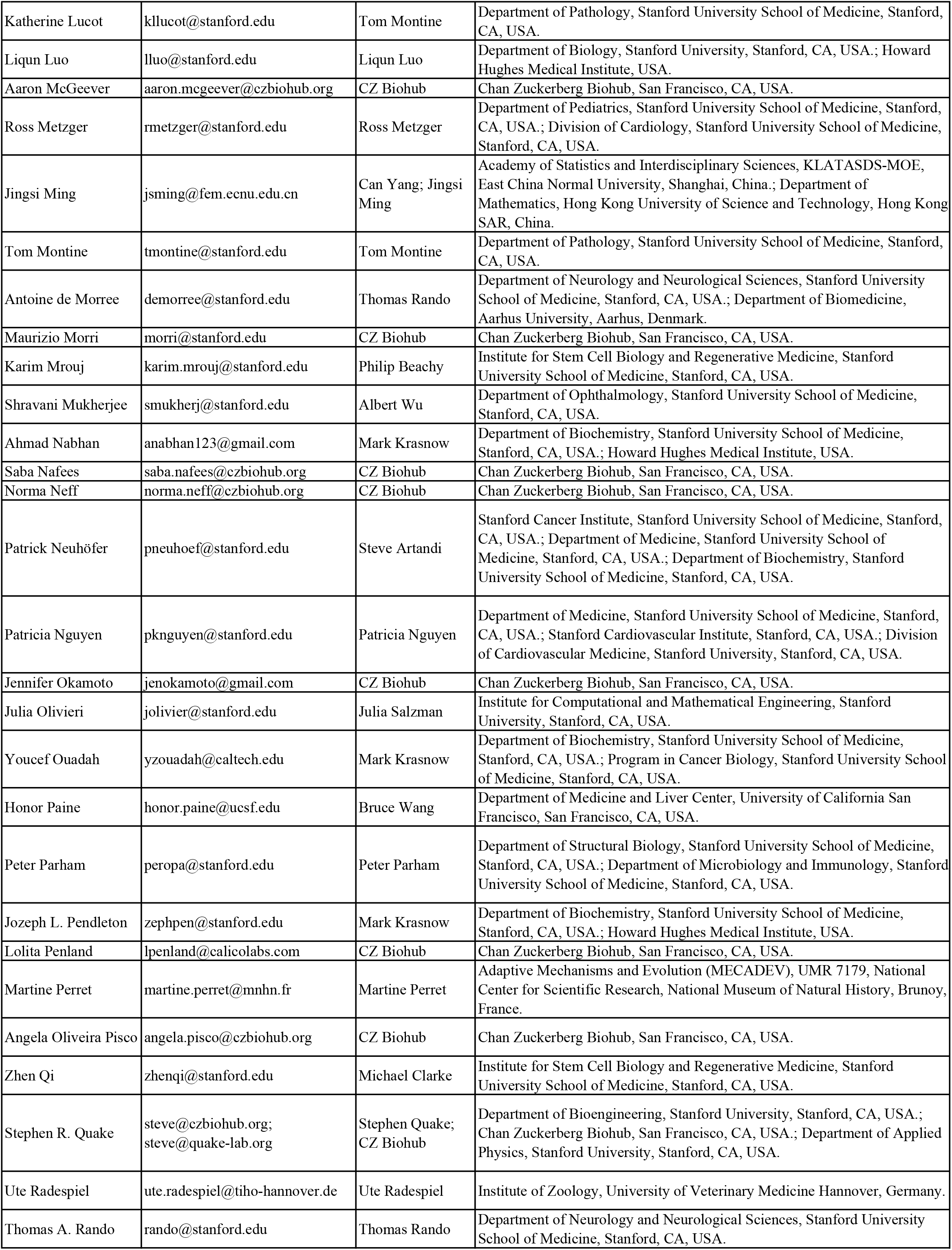
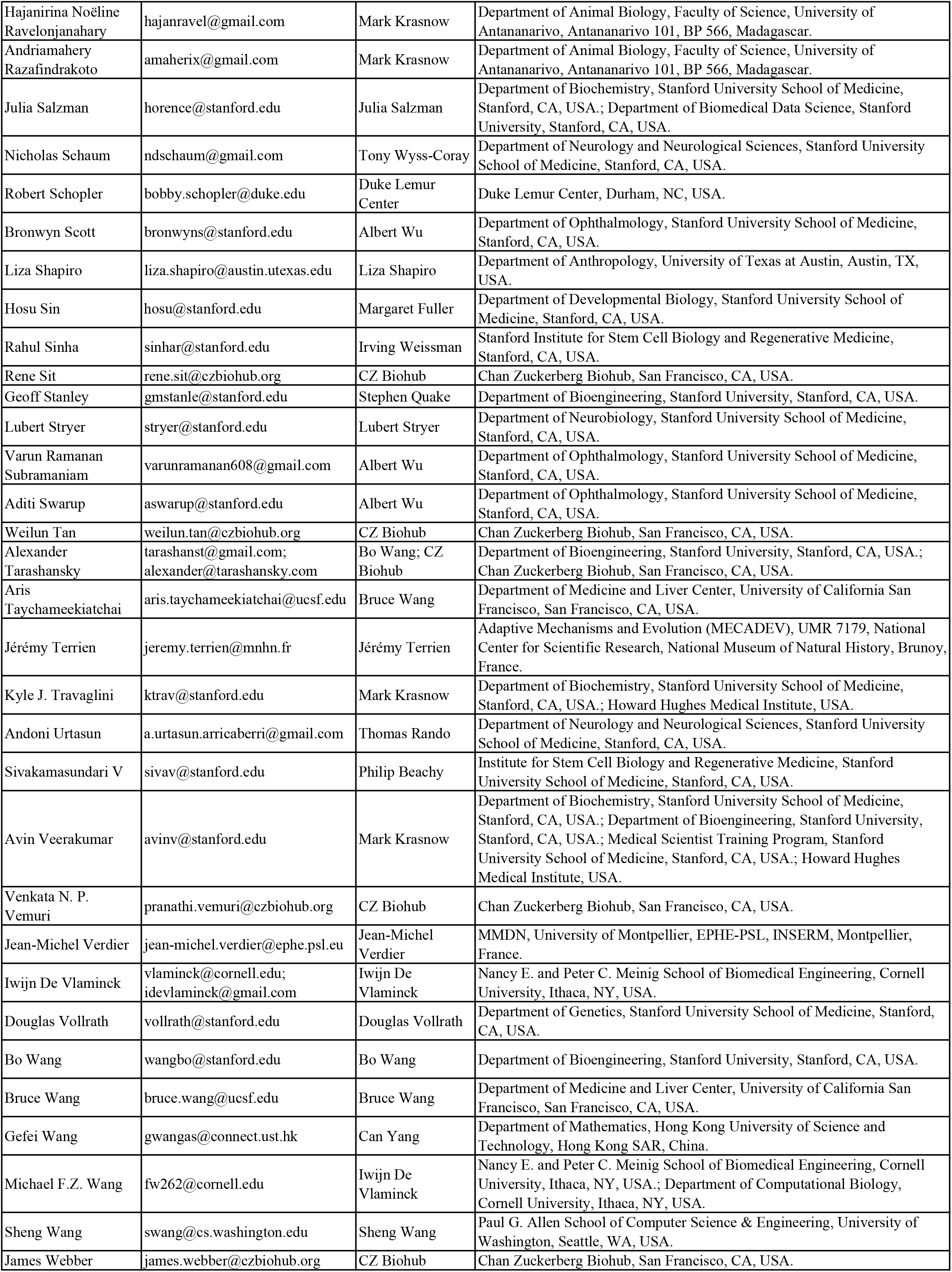
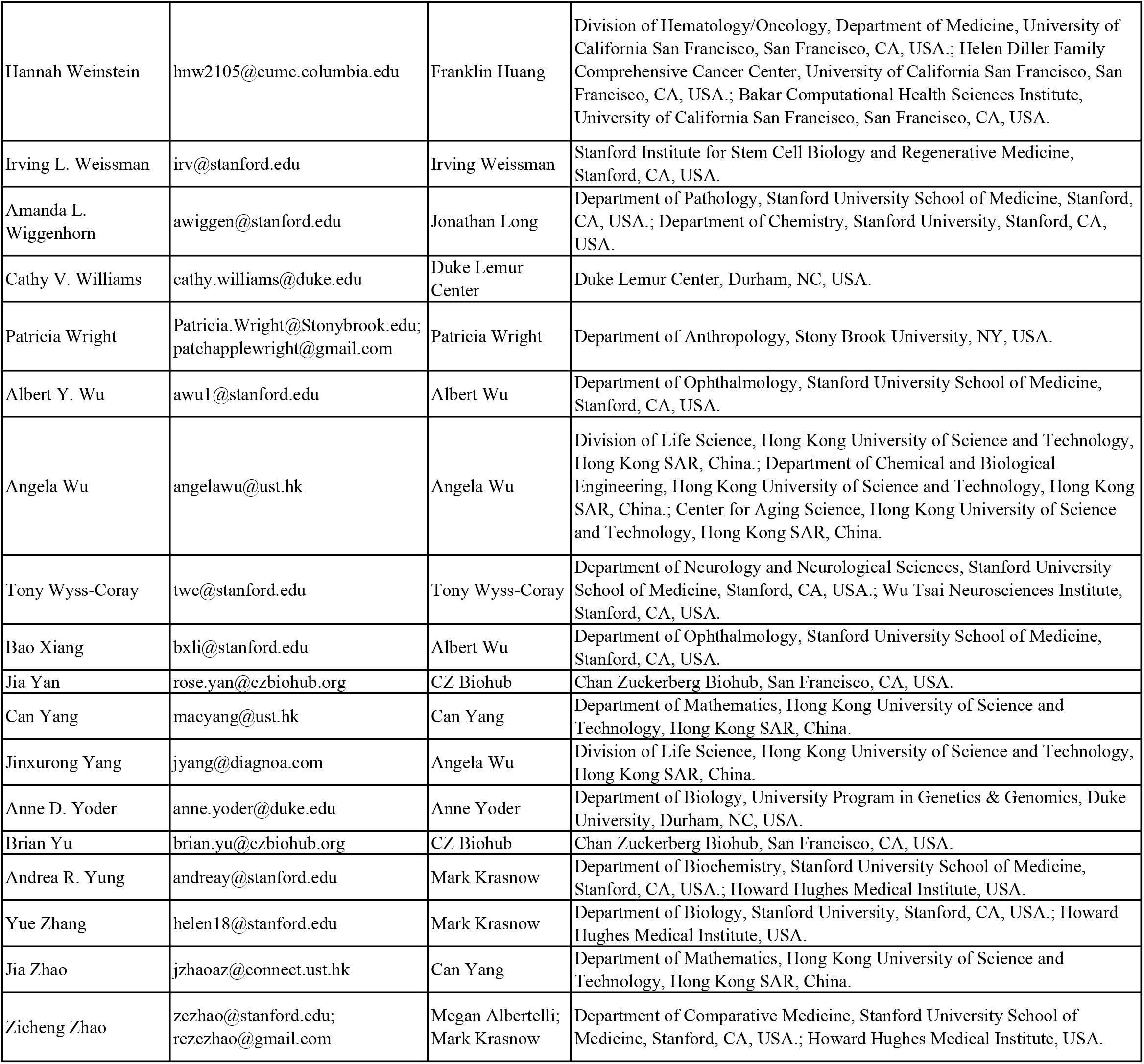

