## Supplementary material for "An organism-wide atlas of hormonal signaling based on the mouse lemur single-cell transcriptome": Fig. S1a. Ligand and receptor gene expression for 84 hormone classes across the mouse lemur cell atlas, with cell types ordered by cell type.

**Acylation stimulating protein (ASP)**
- epithelial/neural/germ

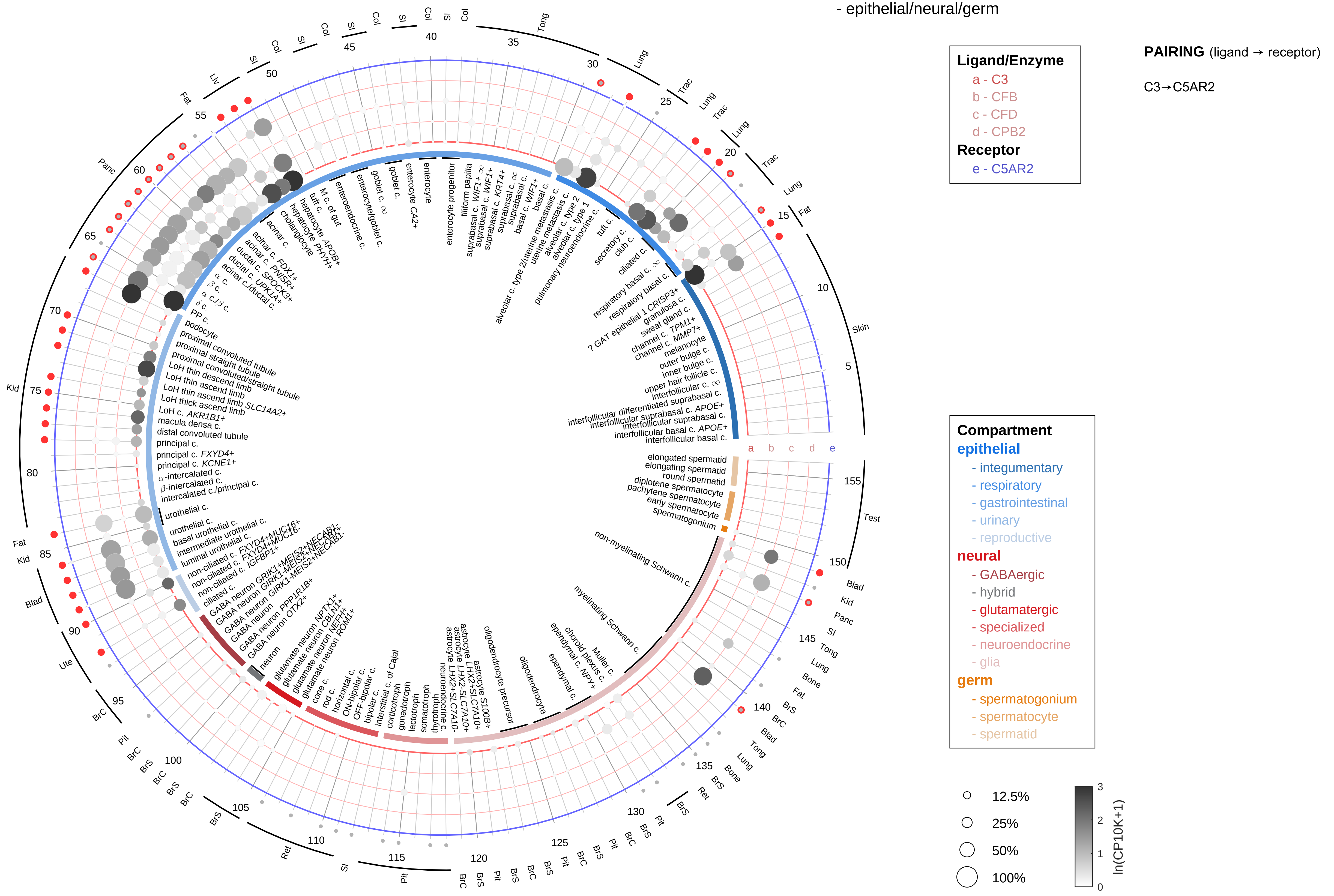

| Abbrev. | Tissue |
|---|---|
| Aort | Aorta |
| Blad | Bladder |
| Blo | Blood |
| Bone | Bone |
| BM | Bone marrow |
| BrC | Brain cortex |
| BrS | Brainstem |
| Col | Colon |
| Dia | Diaphragm |
| Ret | Eye retina |
| Fat | Fat |
| Hrt | Heart |
| Pit | Hypothalamus/Pituitary |
| Kid | Kidney |
| LM | Limb muscle |
| Liv | Liver |
| Lung | Lung |
| MG | Mammary gland |
| Panc | Pancreas |
| Skin | Skin |
| SI | Small intestine |
| Spl | Spleen |
| Test | Testes |
| Thym | Thymus |
| Tong | Tongue |
| Trac | Trachea |
| Ute | Uterus |

#### Acylation stimulating protein (ASP)
- endothelial

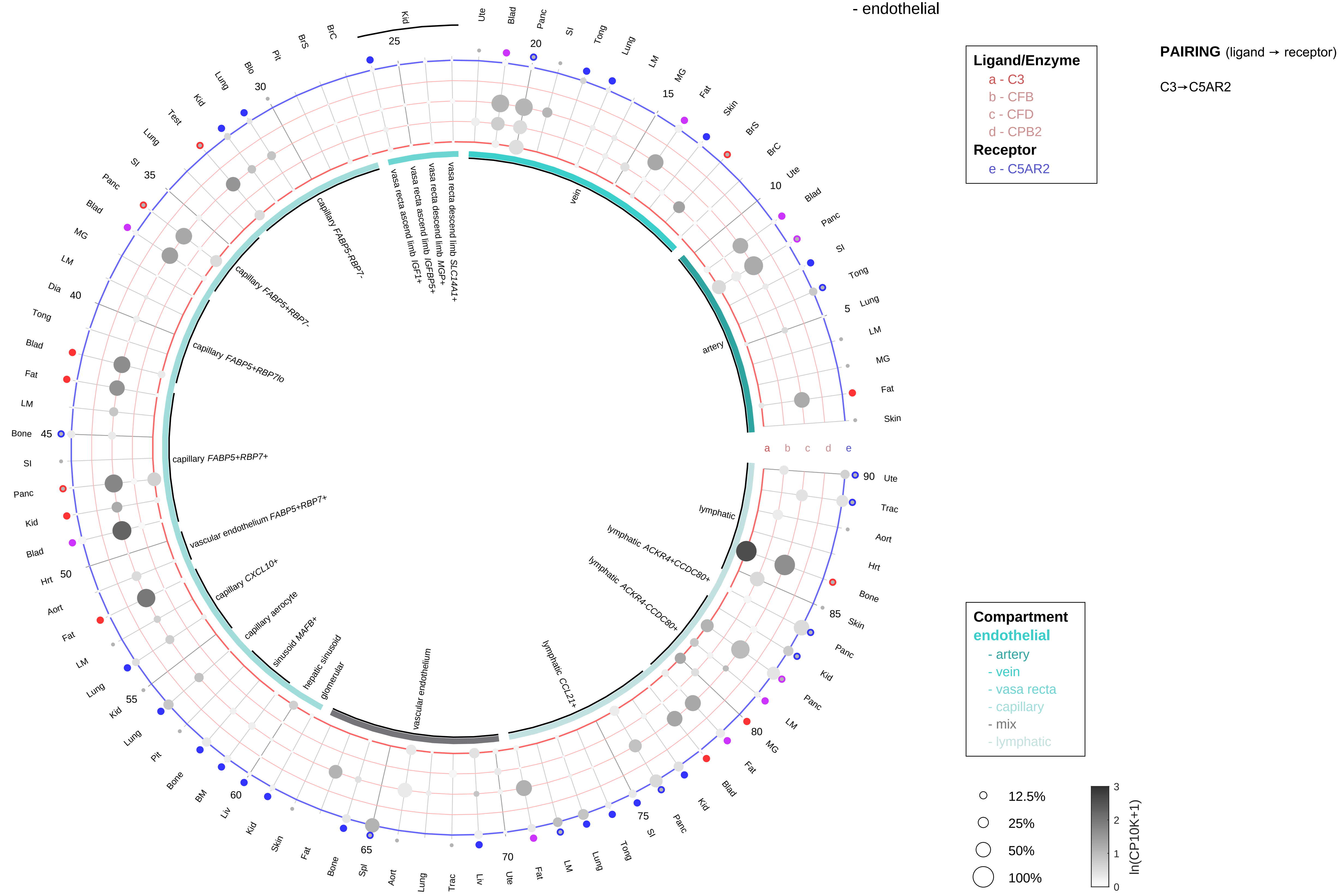

| Abbrev. | Tissue |
|---|---|
| Aort | Aorta |
| Blad | Bladder |
| Blo | Blood |
| Bone | Bone |
| BM | Bone marrow |
| BrC | Brain cortex |
| BrS | Brainstem |
| Col | Colon |
| Dia | Diaphragm |
| Ret | Eye retina |
| Fat | Fat |
| Hrt | Heart |
| Pit | Hypothalamus/Pituitary |
| Kid | Kidney |
| LM | Limb muscle |
| Liv | Liver |
| Lung | Lung |
| MG | Mammary gland |
| Panc | Pancreas |
| Skin | Skin |
| SI | Small intestine |
| Spl | Spleen |
| Test | Testes |
| Thym | Thymus |
| Tong | Tongue |
| Trac | Trachea |
| Ute | Uterus |

#### Acylation stimulating protein (ASP)
- stromal

**Ligand/Enzyme**
- a - C3
- b - CFB
- c - CFD
- d - CPB2

**Receptor**
- e - C5AR2

**PAIRING** (ligand → receptor)

C3→C5AR2

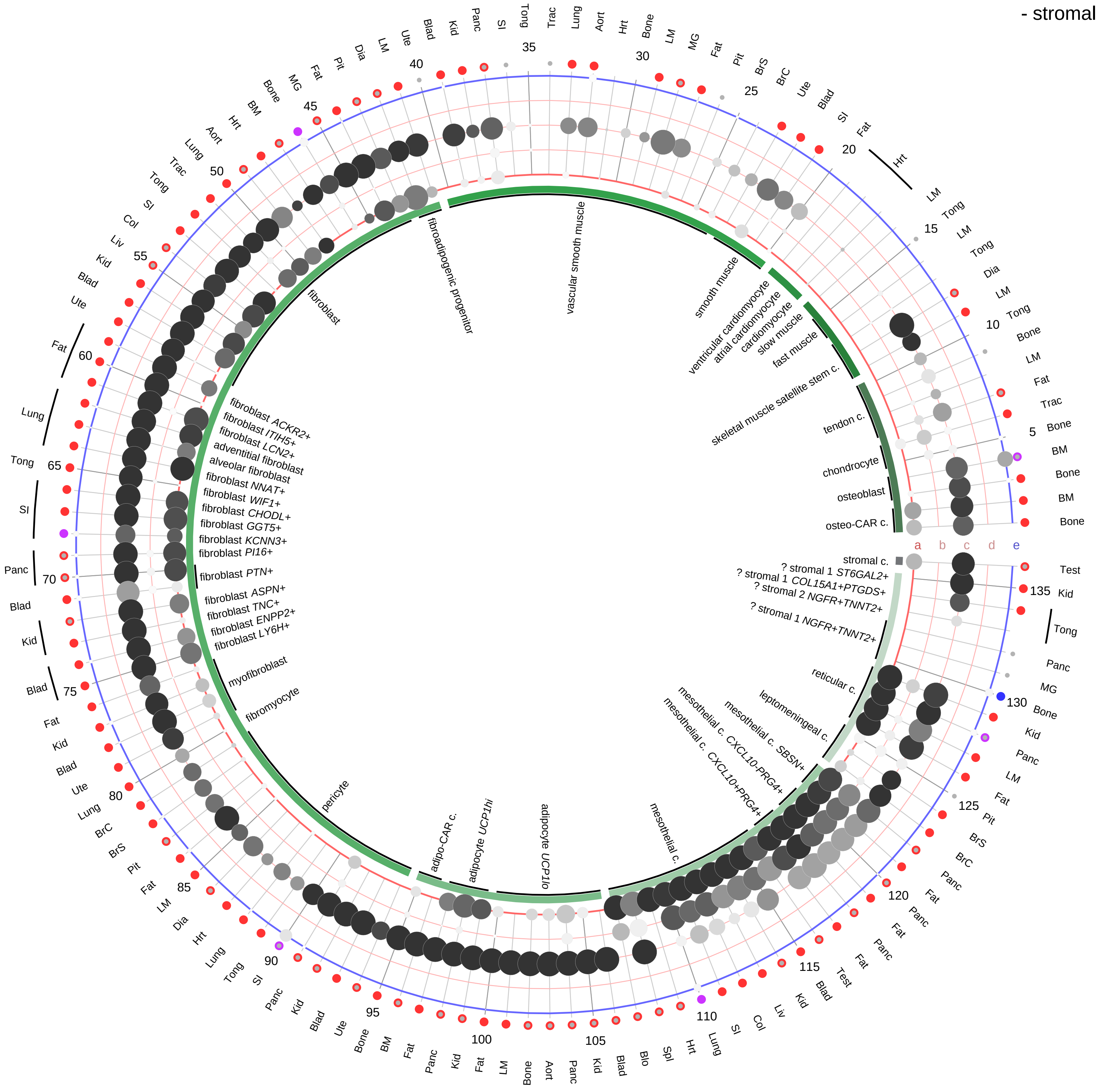

| Abbrev. | Tissue |
|---|---|
| Aort | Aorta |
| Blad | Bladder |
| Blo | Blood |
| Bone | Bone |
| BM | Bone marrow |
| BrC | Brain cortex |
| BrS | Brainstem |
| Col | Colon |
| Dia | Diaphragm |
| Ret | Eye retina |
| Fat | Fat |
| Hrt | Heart |
| Pit | Hypothalamus/Pituitary |
| Kid | Kidney |
| LM | Limb muscle |
| Liv | Liver |
| Lung | Lung |
| MG | Mammary gland |
| Panc | Pancreas |
| Skin | Skin |
| SI | Small intestine |
| Spl | Spleen |
| Test | Testes |
| Thym | Thymus |
| Tong | Tongue |
| Trac | Trachea |
| Ute | Uterus |

**Compartment**

**stromal**
- bone
- skeletal muscle
- cardiac muscle
- smooth muscle
- fibroblast
- adipocyte
- mesothelial
- other
- mix

12.5%
25%
50%
100%

ln(CP10K+1): 0, 1, 2, 3

**Acylation stimulating protein (ASP)**
- lymphoid

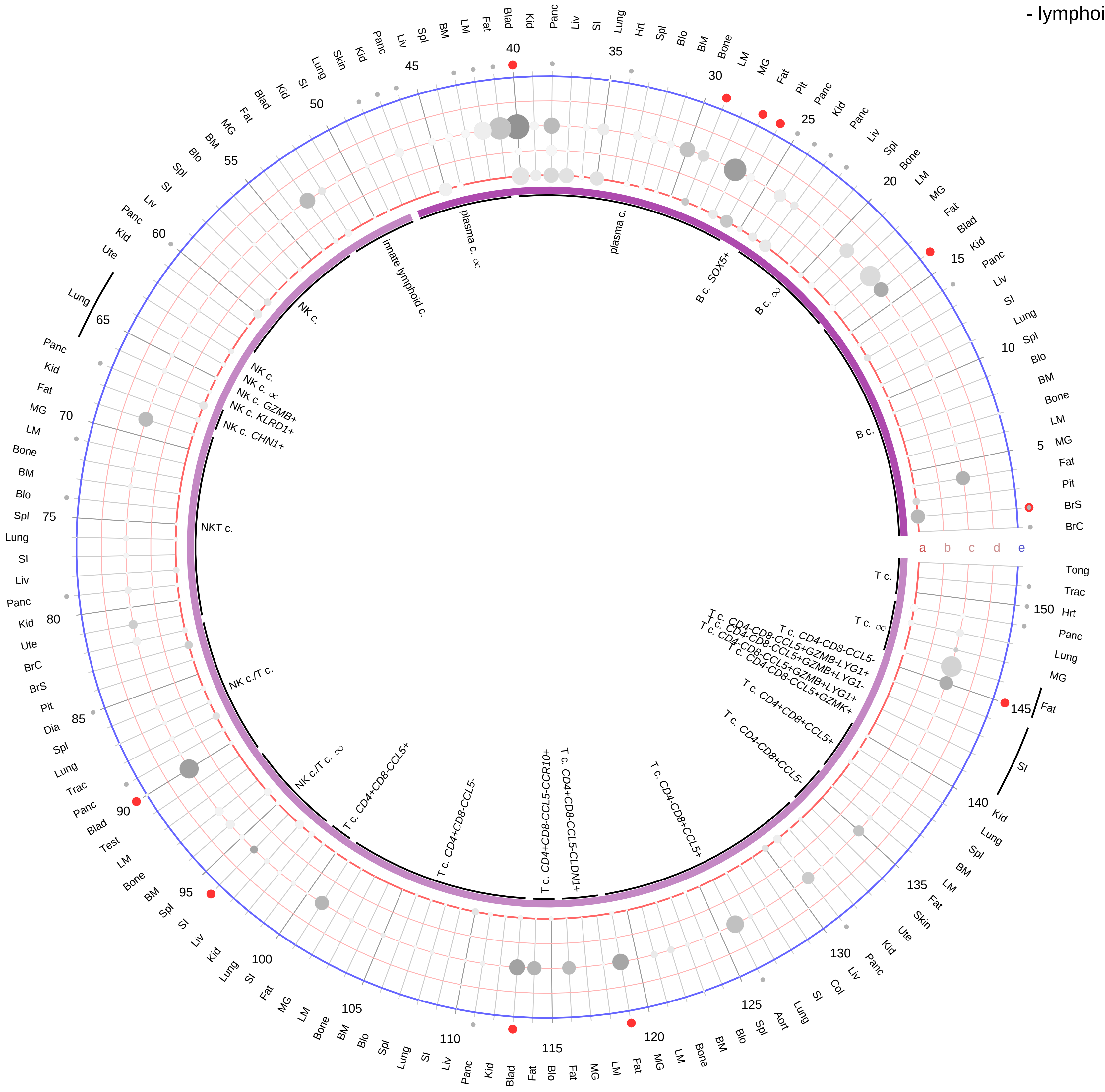

| Abbrev. | Tissue |
|---|---|
| Aort | Aorta |
| Blad | Bladder |
| Blo | Blood |
| Bone | Bone |
| BM | Bone marrow |
| BrC | Brain cortex |
| BrS | Brainstem |
| Col | Colon |
| Dia | Diaphragm |
| Ret | Eye retina |
| Fat | Fat |
| Hrt | Heart |
| Pit | Hypothalamus/Pituitary |
| Kid | Kidney |
| LM | Limb muscle |
| Liv | Liver |
| Lung | Lung |
| MG | Mammary gland |
| Panc | Pancreas |
| Skin | Skin |
| SI | Small intestine |
| Spl | Spleen |
| Test | Testes |
| Thym | Thymus |
| Tong | Tongue |
| Trac | Trachea |
| Ute | Uterus |

**Ligand/Enzyme**
- a - C3
- b - CFB
- c - CFD
- d - CPB2

**Receptor**
- e - C5AR2

**PAIRING** (ligand → receptor)

C3→C5AR2

**Compartment**

**lymphoid**
- B cell
- NK/T cell

12.5%
25%
50%
100%

ln(CP10K+1): 0, 1, 2, 3

**Acylation stimulating protein (ASP)**
- myeloid

**PAIRING** (ligand → receptor)

C3→C5AR2

**Ligand/Enzyme**
- a - C3
- b - CFB
- c - CFD
- d - CPB2

**Receptor**
- e - C5AR2

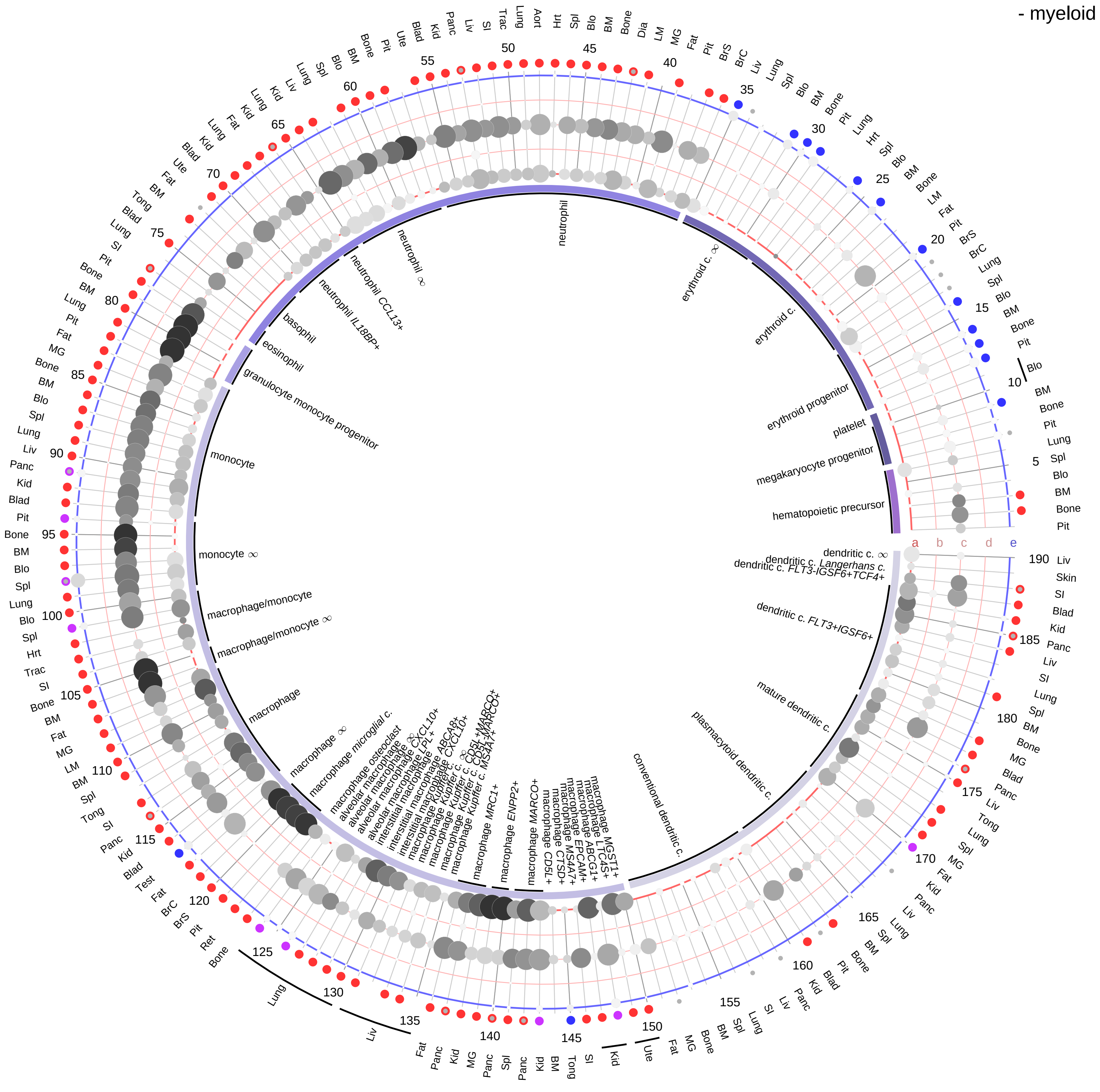

**Compartment**
**hematopoietic**
- hematopoietic precursor

**megakaryocyte-erythroid**
- megakaryoid
- erythroid

**myeloid**
- granulocyte
- granulocyte monocyte progenitor
- monocyte/macrophage
- dendritic

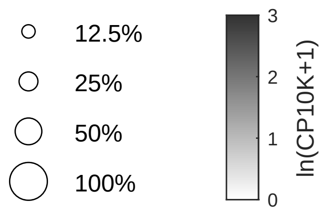

| Abbrev. | Tissue |
|---|---|
| Aort | Aorta |
| Blad | Bladder |
| Blo | Blood |
| Bone | Bone |
| BM | Bone marrow |
| BrC | Brain cortex |
| BrS | Brainstem |
| Col | Colon |
| Dia | Diaphragm |
| Ret | Eye retina |
| Fat | Fat |
| Hrt | Heart |
| Pit | Hypothalamus/Pituitary |
| Kid | Kidney |
| LM | Limb muscle |
| Liv | Liver |
| Lung | Lung |
| MG | Mammary gland |
| Panc | Pancreas |
| Skin | Skin |
| SI | Small intestine |
| Spl | Spleen |
| Test | Testes |
| Thym | Thymus |
| Tong | Tongue |
| Trac | Trachea |
| Ute | Uterus |

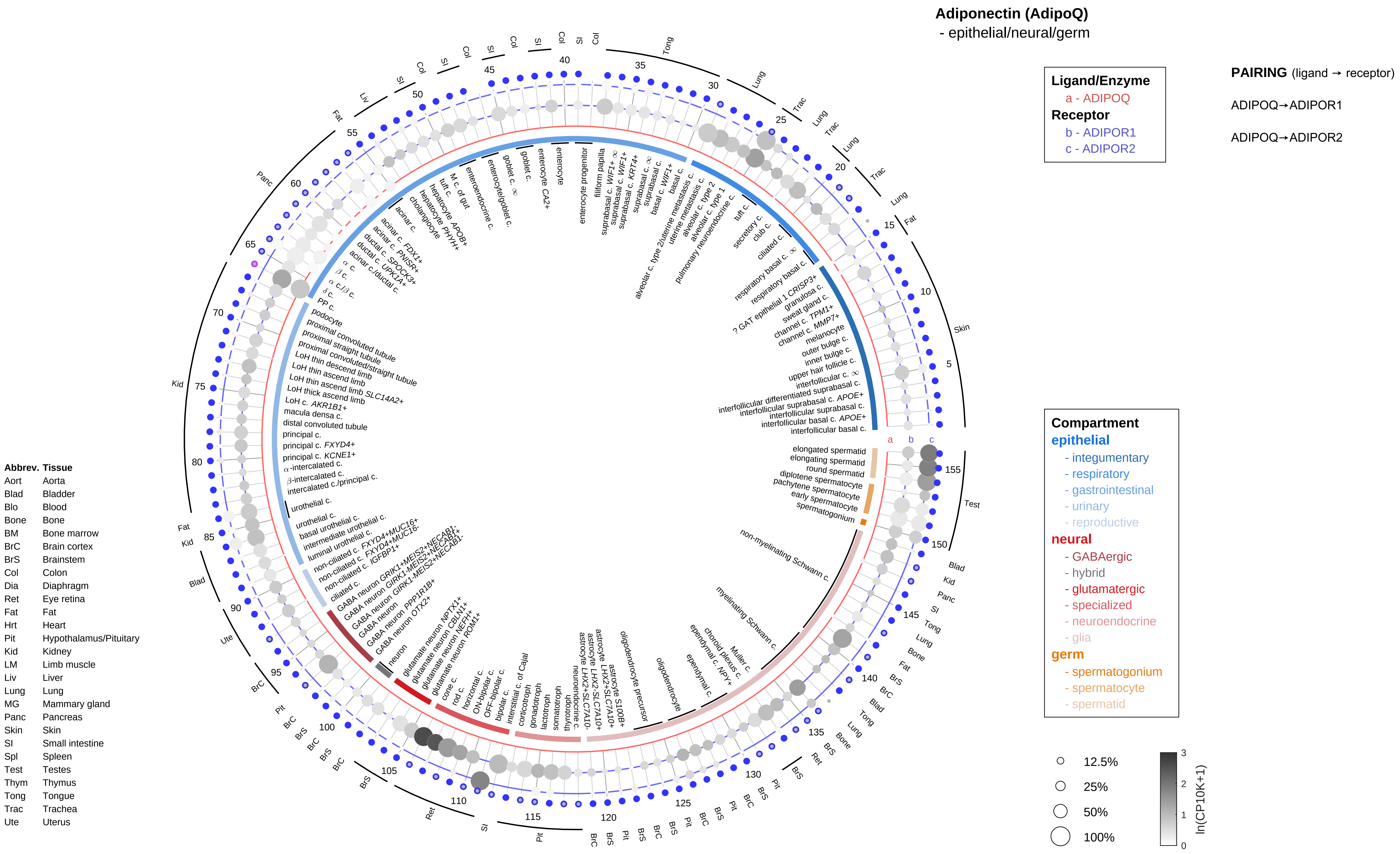

Adiponectin (AdipoQ)
- epithelial/neural/germ
Ligand/Enzyme
a - ADIPOQ
Receptor
b - ADIPOR1
c - ADIPOR2
PAIRING (ligand → receptor)
ADIPOQ→ADIPOR1
ADIPOQ→ADIPOR2
Compartment
epithelial
- integumentary
- respiratory
- gastrointestinal
- urinary
- reproductive
neural
- GABAergic
- hybrid
- glutamatergic
- specialized
- neuroendocrine
- glia
germ
- spermatogonium
- spermatocyte
- spermatid
12.5%
25%
50%
100%
ln(CP10K+1)
Abbrev. Tissue
Aort Aorta
Blad Bladder
Blo Blood
Bone Bone
BM Bone marrow
BrC Brain cortex
BrS Brainstem
Col Colon
Dia Diaphragm
Ret Eye retina
Fat Fat
Hrt Heart
Pit Hypothalamus/Pituitary
Kid Kidney
LM Limb muscle
Liv Liver
Lung Lung
MG Mammary gland
Panc Pancreas
Skin Skin
SI Small intestine
Spl Spleen
Test Testes
Thym Thymus
Tong Tongue
Trac Trachea
Ute Uterus

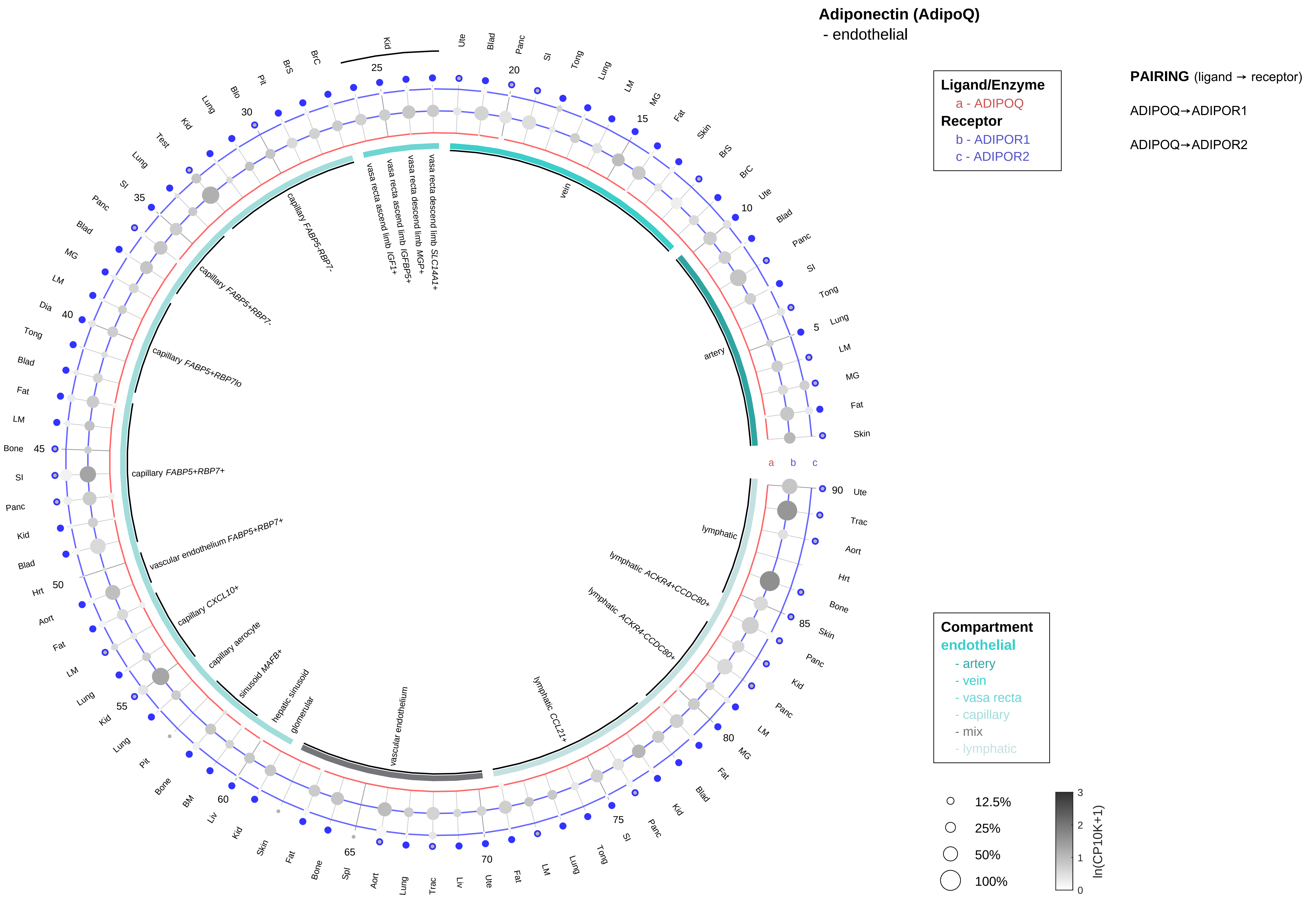

| Abbrev. | Tissue |
|---|---|
| Aort | Aorta |
| Blad | Bladder |
| Blo | Blood |
| Bone | Bone |
| BM | Bone marrow |
| BrC | Brain cortex |
| BrS | Brainstem |
| Col | Colon |
| Dia | Diaphragm |
| Ret | Eye retina |
| Fat | Fat |
| Hrt | Heart |
| Pit | Hypothalamus/Pituitary |
| Kid | Kidney |
| LM | Limb muscle |
| Liv | Liver |
| Lung | Lung |
| MG | Mammary gland |
| Panc | Pancreas |
| Skin | Skin |
| SI | Small intestine |
| Spl | Spleen |
| Test | Testes |
| Thym | Thymus |
| Tong | Tongue |
| Trac | Trachea |
| Ute | Uterus |

#### Adiponectin (AdipoQ)

- stromal

**Ligand/Enzyme**
a - ADIPOQ
**Receptor**
b - ADIPOR1
c - ADIPOR2

**PAIRING** (ligand → receptor)

ADIPOQ→ADIPOR1

ADIPOQ→ADIPOR2

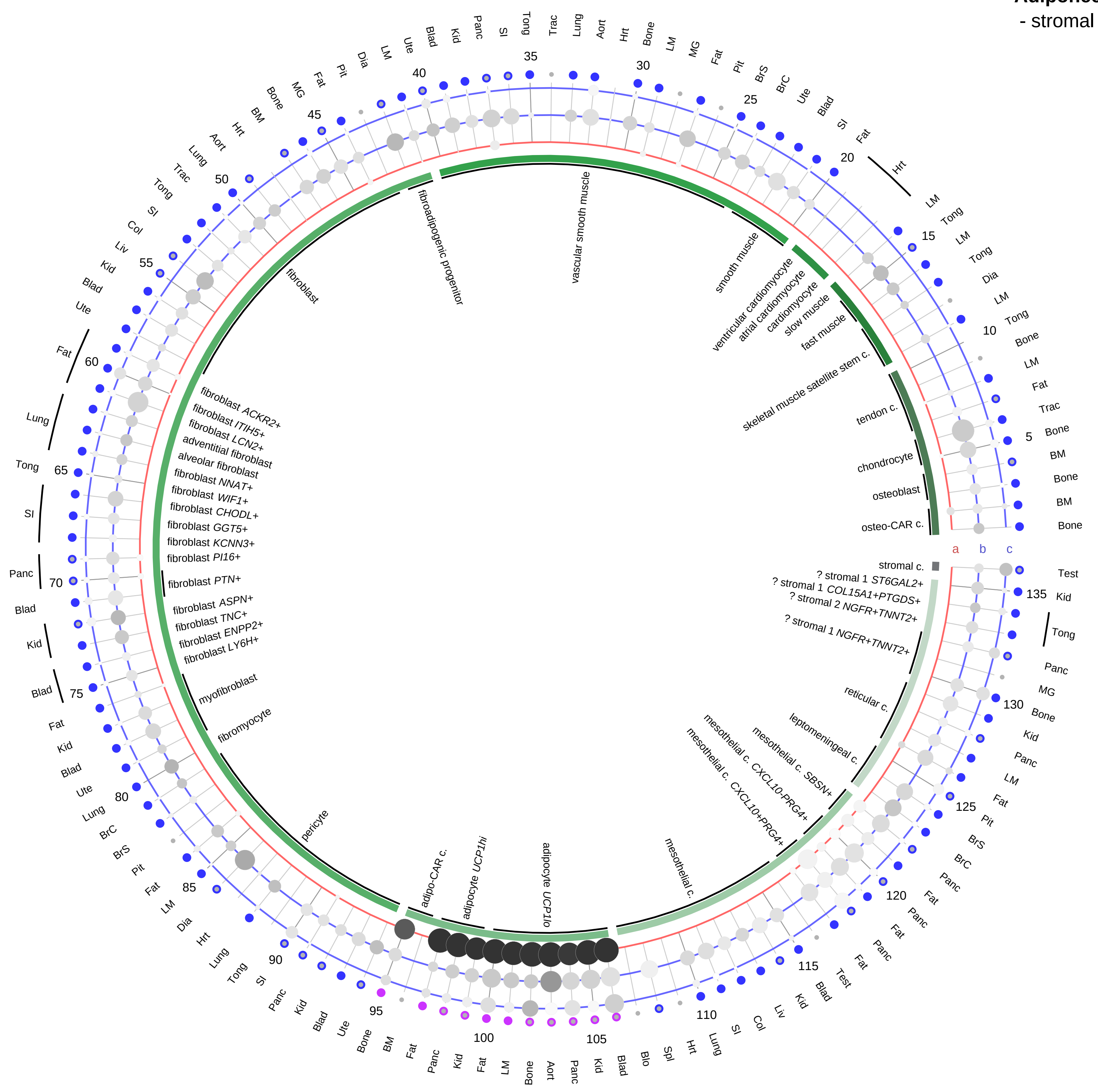

**Compartment**
**stromal**
- bone
- skeletal muscle
- cardiac muscle
- smooth muscle
- fibroblast
- adipocyte
- mesothelial
- other
- mix

| Abbrev. | Tissue |
|---|---|
| Aort | Aorta |
| Blad | Bladder |
| Blo | Blood |
| Bone | Bone |
| BM | Bone marrow |
| BrC | Brain cortex |
| BrS | Brainstem |
| Col | Colon |
| Dia | Diaphragm |
| Ret | Eye retina |
| Fat | Fat |
| Hrt | Heart |
| Pit | Hypothalamus/Pituitary |
| Kid | Kidney |
| LM | Limb muscle |
| Liv | Liver |
| Lung | Lung |
| MG | Mammary gland |
| Panc | Pancreas |
| Skin | Skin |
| SI | Small intestine |
| Spl | Spleen |
| Test | Testes |
| Thym | Thymus |
| Tong | Tongue |
| Trac | Trachea |
| Ute | Uterus |

**Adiponectin (AdipoQ)**
- lymphoid

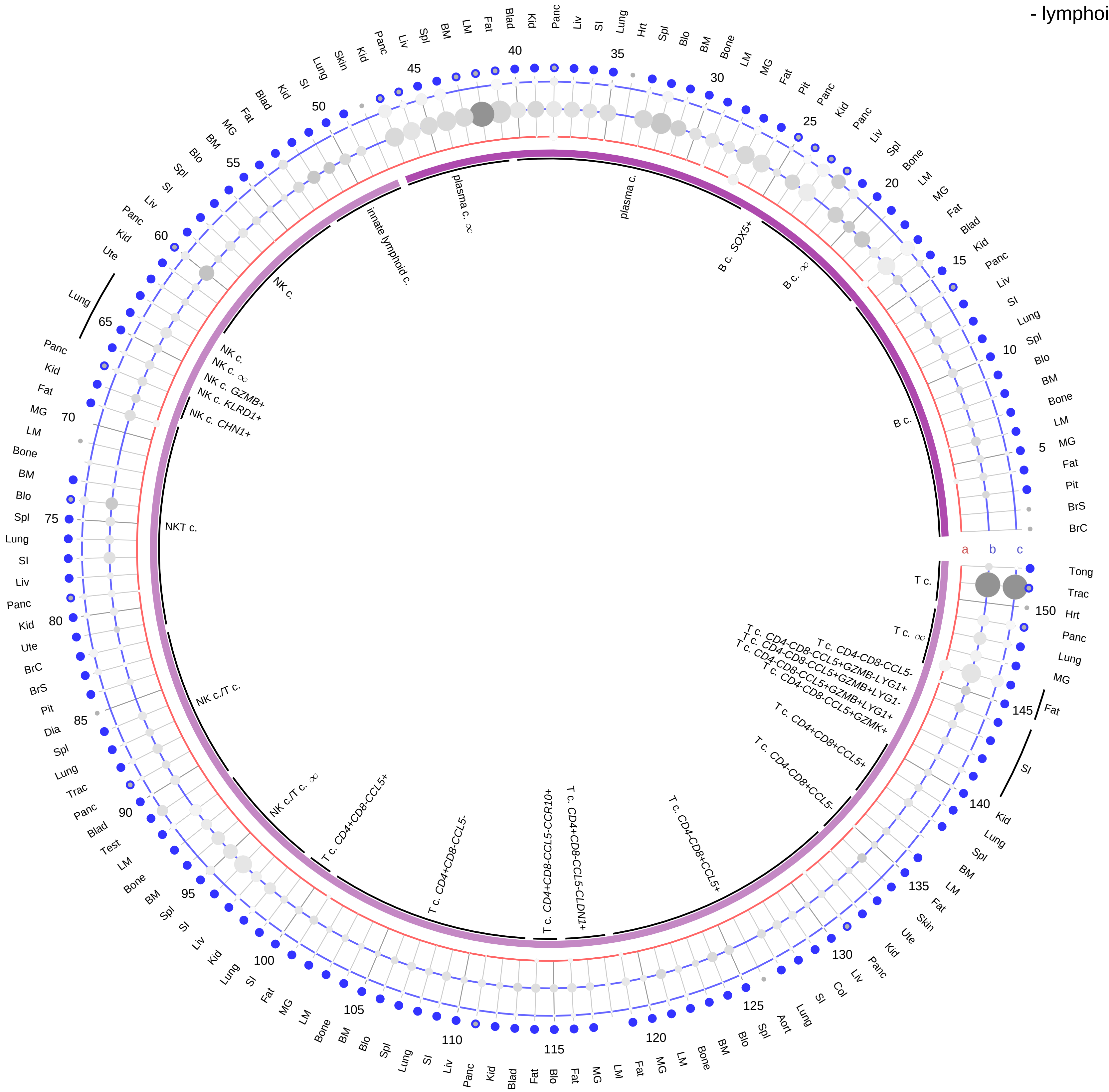

**Ligand/Enzyme**
a - ADIPOQ
**Receptor**
b - ADIPOR1
c - ADIPOR2

**PAIRING** (ligand → receptor)

ADIPOQ→ADIPOR1

ADIPOQ→ADIPOR2

| Abbrev. | Tissue |
|---|---|
| Aort | Aorta |
| Blad | Bladder |
| Blo | Blood |
| Bone | Bone |
| BM | Bone marrow |
| BrC | Brain cortex |
| BrS | Brainstem |
| Col | Colon |
| Dia | Diaphragm |
| Ret | Eye retina |
| Fat | Fat |
| Hrt | Heart |
| Pit | Hypothalamus/Pituitary |
| Kid | Kidney |
| LM | Limb muscle |
| Liv | Liver |
| Lung | Lung |
| MG | Mammary gland |
| Panc | Pancreas |
| Skin | Skin |
| SI | Small intestine |
| Spl | Spleen |
| Test | Testes |
| Thym | Thymus |
| Tong | Tongue |
| Trac | Trachea |
| Ute | Uterus |

**Compartment**
**lymphoid**
- B cell
- NK/T cell

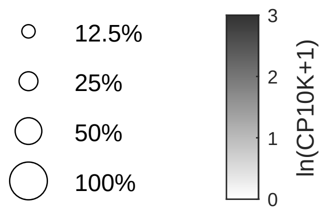

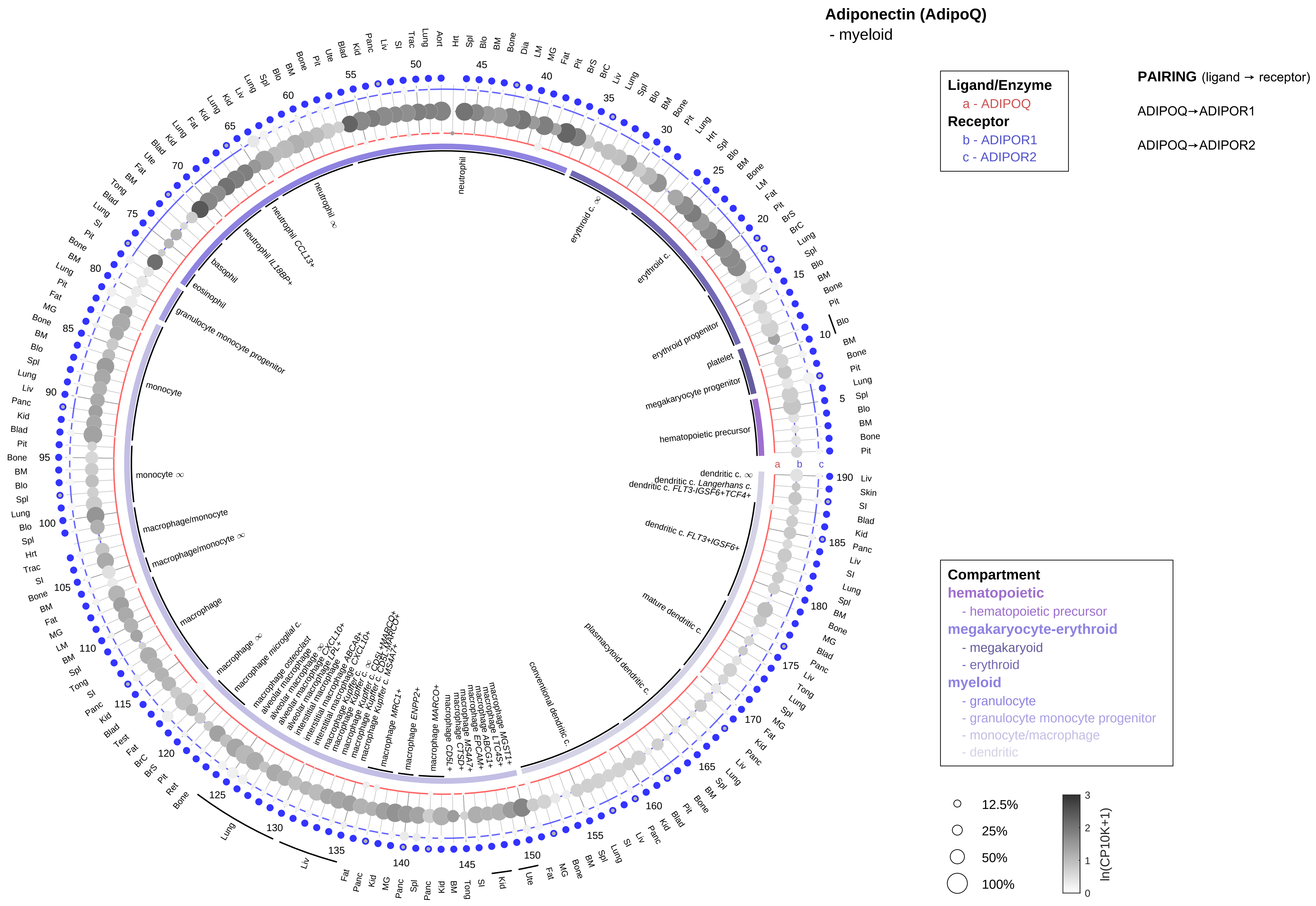

| Abbrev. | Tissue |
|---|---|
| Aort | Aorta |
| Blad | Bladder |
| Blo | Blood |
| Bone | Bone |
| BM | Bone marrow |
| BrC | Brain cortex |
| BrS | Brainstem |
| Col | Colon |
| Dia | Diaphragm |
| Ret | Eye retina |
| Fat | Fat |
| Hrt | Heart |
| Pit | Hypothalamus/Pituitary |
| Kid | Kidney |
| LM | Limb muscle |
| Liv | Liver |
| Lung | Lung |
| MG | Mammary gland |
| Panc | Pancreas |
| Skin | Skin |
| SI | Small intestine |
| Spl | Spleen |
| Test | Testes |
| Thym | Thymus |
| Tong | Tongue |
| Trac | Trachea |
| Ute | Uterus |

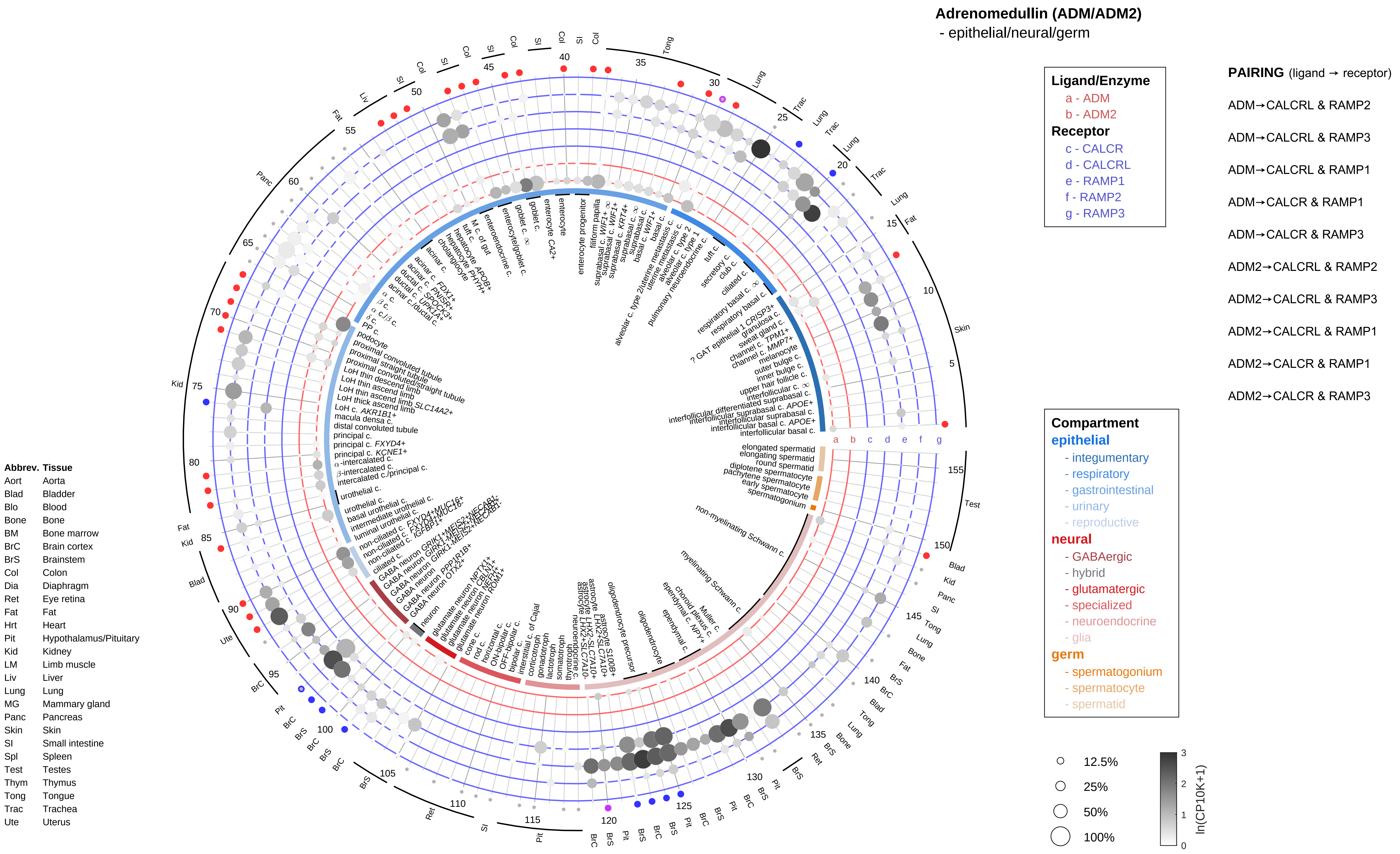
Adrenomedullin (ADM/ADM2)
- epithelial/neural/germ
Ligand/Enzyme
a - ADM
b - ADM2
Receptor
c - CALCR
d - CALCRL
e - RAMP1
f - RAMP2
g - RAMP3
PAIRING (ligand → receptor)
ADM→CALCRL & RAMP2
ADM→CALCRL & RAMP3
ADM→CALCRL & RAMP1
ADM→CALCR & RAMP1
ADM→CALCR & RAMP3
ADM2→CALCRL & RAMP2
ADM2→CALCRL & RAMP3
ADM2→CALCRL & RAMP1
ADM2→CALCR & RAMP1
ADM2→CALCR & RAMP3
Compartment
epithelial
- integumentary
- respiratory
- gastrointestinal
- urinary
- reproductive
neural
- GABAergic
- hybrid
- glutamatergic
- specialized
- neuroendocrine
- glia
germ
- spermatogonium
- spermatocyte
- spermatid
12.5%
25%
50%
100%
ln(CP10K+1)
Abbrev. Tissue
Aort Aorta
Blad Bladder
Blo Blood
Bone Bone
BM Bone marrow
BrC Brain cortex
BrS Brainstem
Col Colon
Dia Diaphragm
Ret Eye retina
Fat Fat
Hrt Heart
Pit Hypothalamus/Pituitary
Kid Kidney
LM Limb muscle
Liv Liver
Lung Lung
MG Mammary gland
Panc Pancreas
Skin Skin
SI Small intestine
Spl Spleen
Test Testes
Thym Thymus
Tong Tongue
Trac Trachea
Ute Uterus

**Adrenomedullin (ADM/ADM2)**
- endothelial

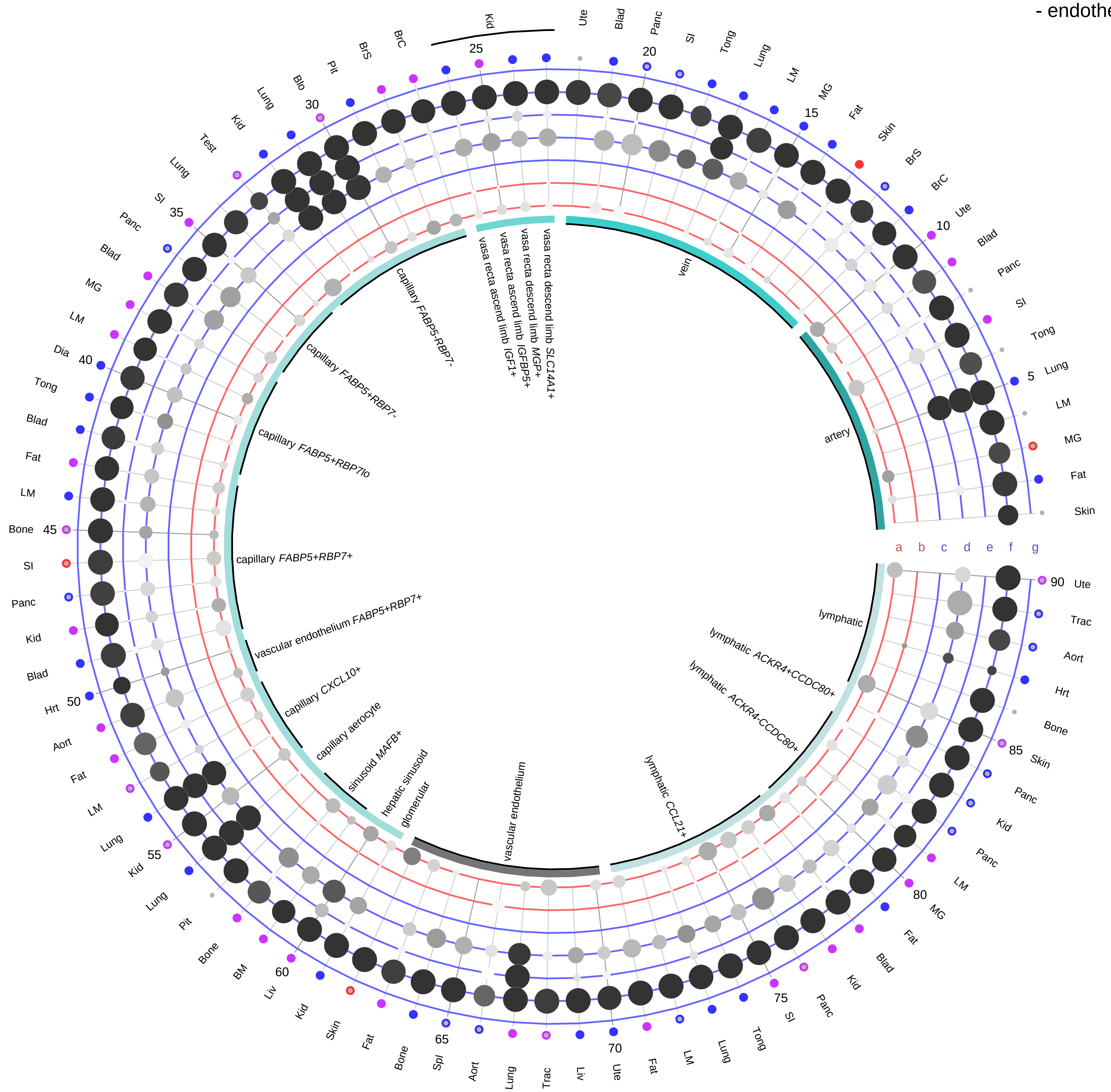

| Ligand/Enzyme |
|---|
| a - ADM |
| b - ADM2 |
| **Receptor** |
| c - CALCR |
| d - CALCRL |
| e - RAMP1 |
| f - RAMP2 |
| g - RAMP3 |

**PAIRING** (ligand → receptor)

ADM→CALCRL & RAMP2

ADM→CALCRL & RAMP3

ADM→CALCRL & RAMP1

ADM→CALCR & RAMP1

ADM→CALCR & RAMP3

ADM2→CALCRL & RAMP2

ADM2→CALCRL & RAMP3

ADM2→CALCRL & RAMP1

ADM2→CALCR & RAMP1

ADM2→CALCR & RAMP3

**Compartment**
**endothelial**
- artery
- vein
- vasa recta
- capillary
- mix
- lymphatic

12.5%
25%
50%
100%

ln(CP10K+1): 0, 1, 2, 3

| Abbrev. | Tissue |
|---|---|
| Aort | Aorta |
| Blad | Bladder |
| Blo | Blood |
| Bone | Bone |
| BM | Bone marrow |
| BrC | Brain cortex |
| BrS | Brainstem |
| Col | Colon |
| Dia | Diaphragm |
| Ret | Eye retina |
| Fat | Fat |
| Hrt | Heart |
| Pit | Hypothalamus/Pituitary |
| Kid | Kidney |
| LM | Limb muscle |
| Liv | Liver |
| Lung | Lung |
| MG | Mammary gland |
| Panc | Pancreas |
| Skin | Skin |
| SI | Small intestine |
| Spl | Spleen |
| Test | Testes |
| Thym | Thymus |
| Tong | Tongue |
| Trac | Trachea |
| Ute | Uterus |

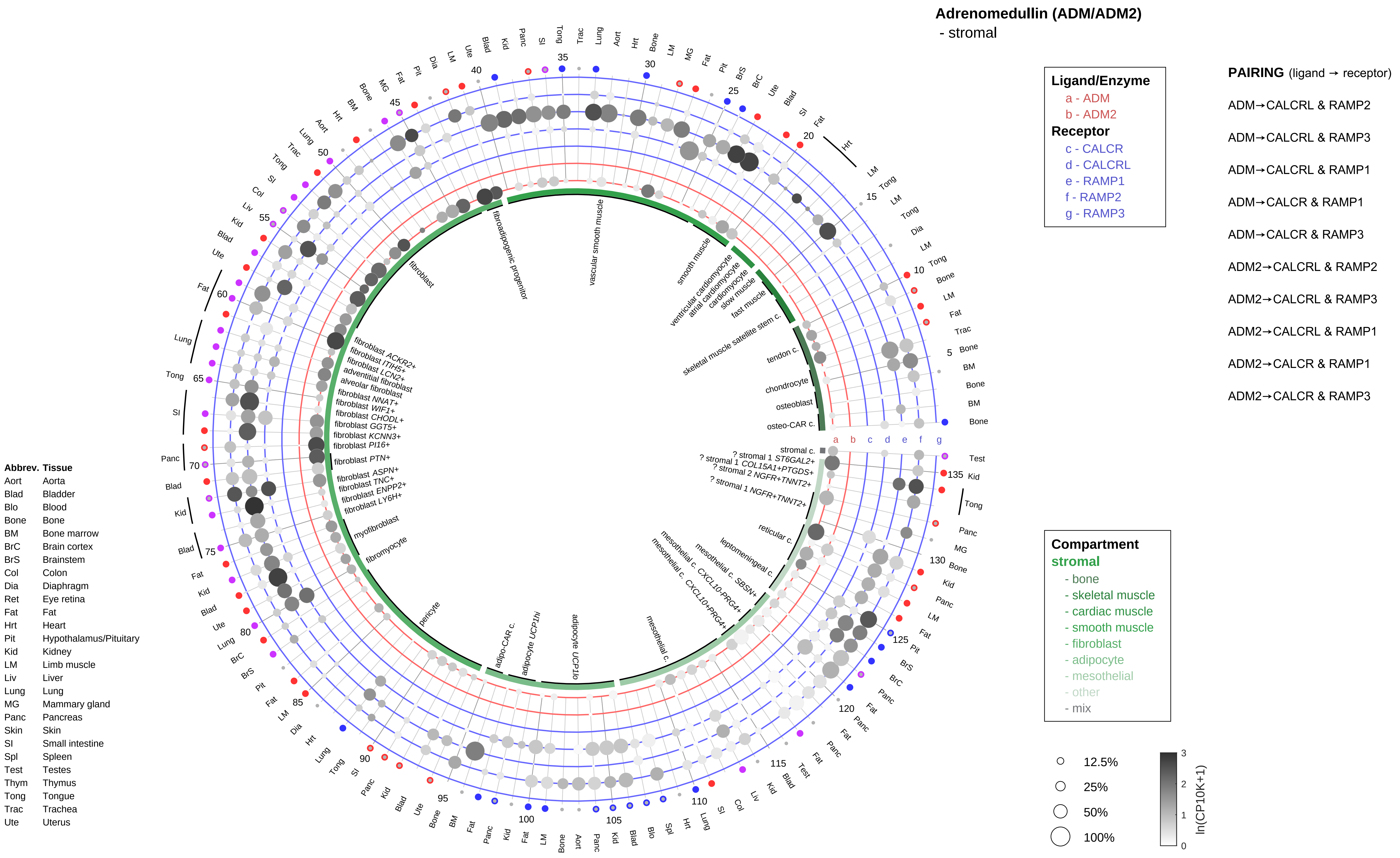
Adrenomedullin (ADM/ADM2)
- stromal
Ligand/Enzyme
a - ADM
b - ADM2
Receptor
c - CALCR
d - CALCRL
e - RAMP1
f - RAMP2
g - RAMP3
PAIRING (ligand → receptor)
ADM→CALCRL & RAMP2
ADM→CALCRL & RAMP3
ADM→CALCRL & RAMP1
ADM→CALCR & RAMP1
ADM→CALCR & RAMP3
ADM2→CALCRL & RAMP2
ADM2→CALCRL & RAMP3
ADM2→CALCRL & RAMP1
ADM2→CALCR & RAMP1
ADM2→CALCR & RAMP3
Compartment
stromal
- bone
- skeletal muscle
- cardiac muscle
- smooth muscle
- fibroblast
- adipocyte
- mesothelial
- other
- mix
12.5%
25%
50%
100%
ln(CP10K+1)
Abbrev. Tissue
Aort Aorta
Blad Bladder
Blo Blood
Bone Bone
BM Bone marrow
BrC Brain cortex
BrS Brainstem
Col Colon
Dia Diaphragm
Ret Eye retina
Fat Fat
Hrt Heart
Pit Hypothalamus/Pituitary
Kid Kidney
LM Limb muscle
Liv Liver
Lung Lung
MG Mammary gland
Panc Pancreas
Skin Skin
SI Small intestine
Spl Spleen
Test Testes
Thym Thymus
Tong Tongue
Trac Trachea
Ute Uterus

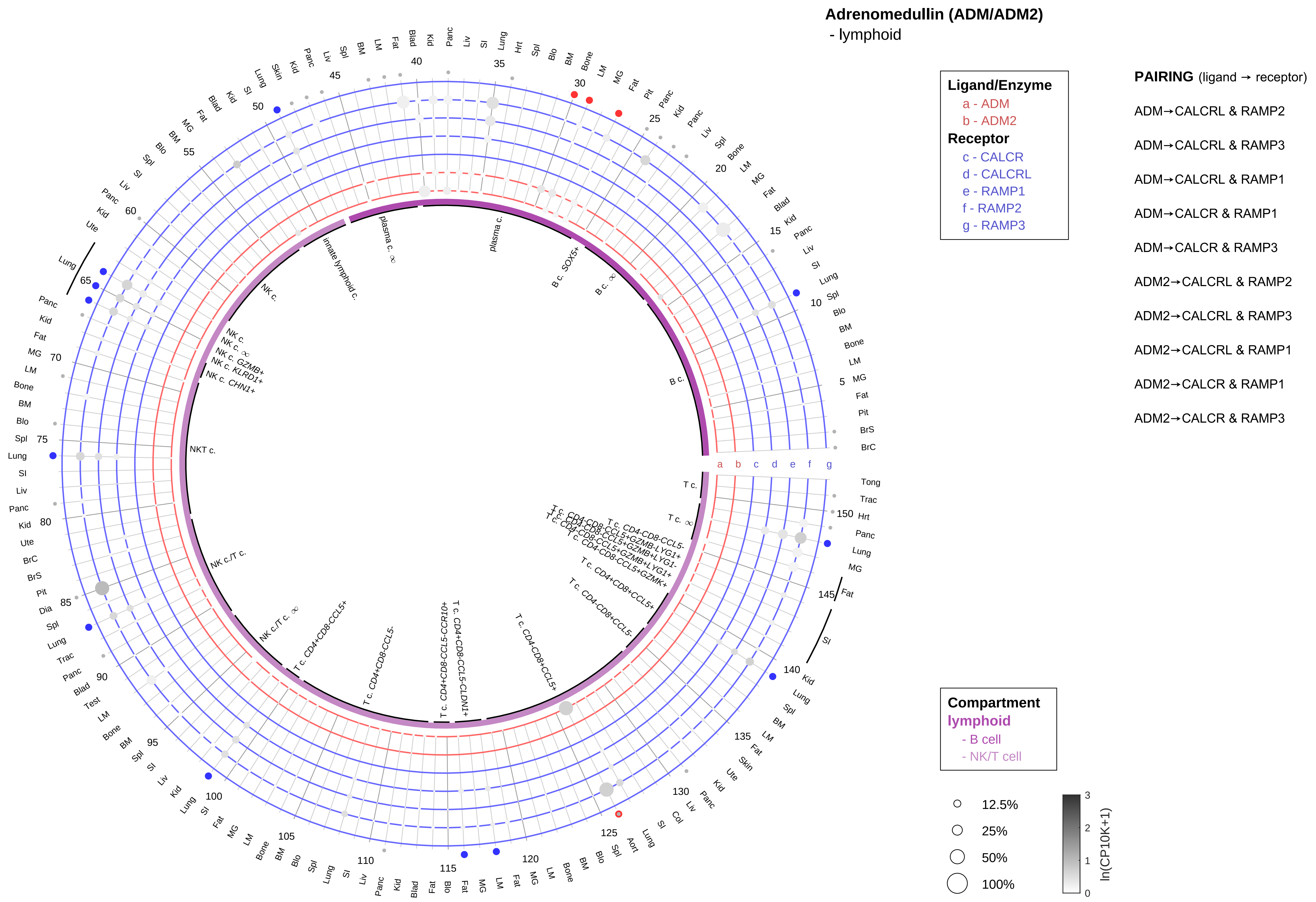

| Abbrev. | Tissue |
|---|---|
| Aort | Aorta |
| Blad | Bladder |
| Blo | Blood |
| Bone | Bone |
| BM | Bone marrow |
| BrC | Brain cortex |
| BrS | Brainstem |
| Col | Colon |
| Dia | Diaphragm |
| Ret | Eye retina |
| Fat | Fat |
| Hrt | Heart |
| Pit | Hypothalamus/Pituitary |
| Kid | Kidney |
| LM | Limb muscle |
| Liv | Liver |
| Lung | Lung |
| MG | Mammary gland |
| Panc | Pancreas |
| Skin | Skin |
| SI | Small intestine |
| Spl | Spleen |
| Test | Testes |
| Thym | Thymus |
| Tong | Tongue |
| Trac | Trachea |
| Ute | Uterus |

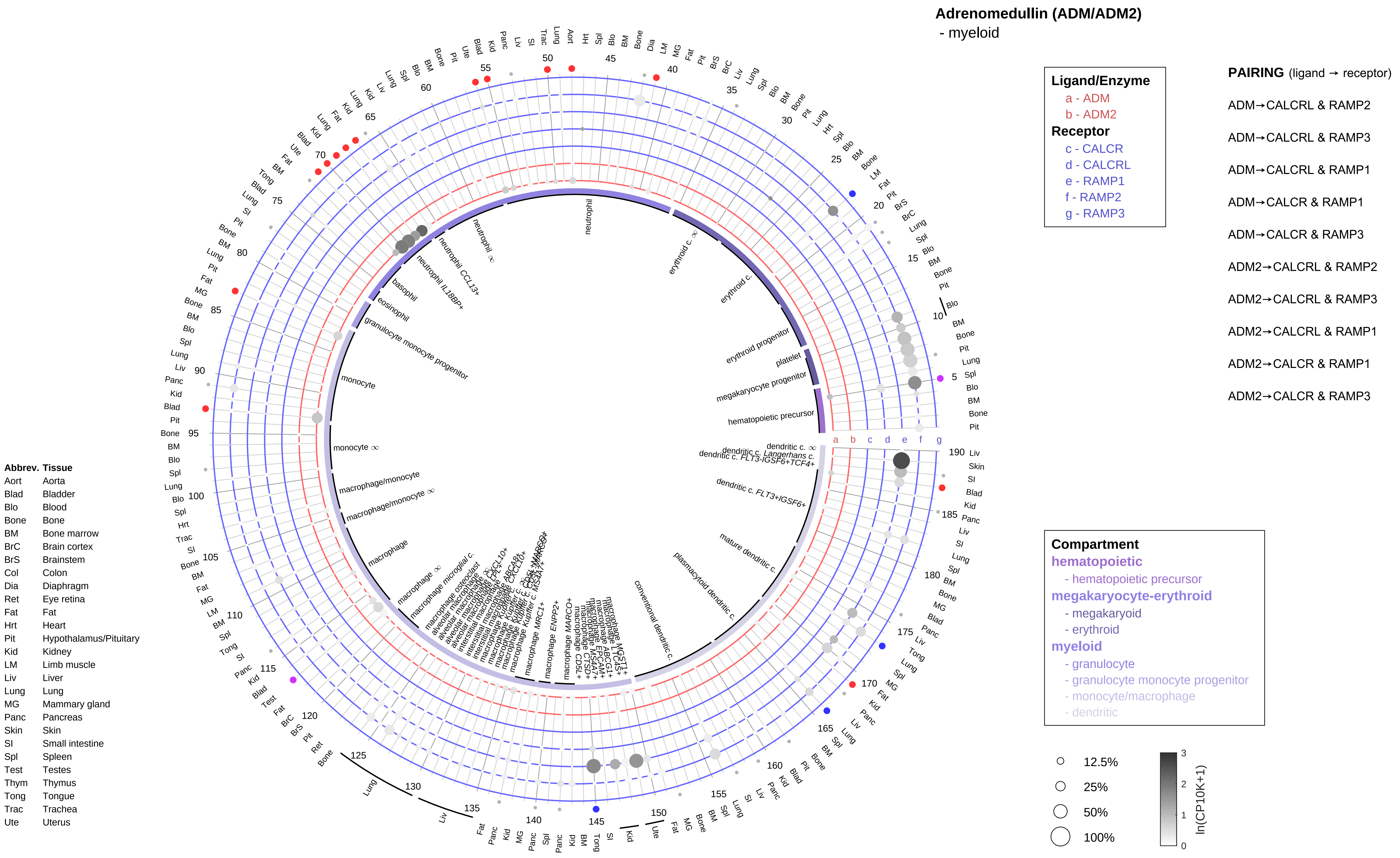
Adrenomedullin (ADM/ADM2)
- myeloid
Ligand/Enzyme
a - ADM
b - ADM2
Receptor
c - CALCR
d - CALCRL
e - RAMP1
f - RAMP2
g - RAMP3
PAIRING (ligand → receptor)
ADM→CALCRL & RAMP2
ADM→CALCRL & RAMP3
ADM→CALCRL & RAMP1
ADM→CALCR & RAMP1
ADM→CALCR & RAMP3
ADM2→CALCRL & RAMP2
ADM2→CALCRL & RAMP3
ADM2→CALCRL & RAMP1
ADM2→CALCR & RAMP1
ADM2→CALCR & RAMP3
Compartment
hematopoietic
- hematopoietic precursor
megakaryocyte-erythroid
- megakaryoid
- erythroid
myeloid
- granulocyte
- granulocyte monocyte progenitor
- monocyte/macrophage
- dendritic
12.5%
25%
50%
100%
ln(CP10K+1)
Abbrev. Tissue
Aort Aorta
Blad Bladder
Blo Blood
Bone Bone
BM Bone marrow
BrC Brain cortex
BrS Brainstem
Col Colon
Dia Diaphragm
Ret Eye retina
Fat Fat
Hrt Heart
Pit Hypothalamus/Pituitary
Kid Kidney
LM Limb muscle
Liv Liver
Lung Lung
MG Mammary gland
Panc Pancreas
Skin Skin
SI Small intestine
Spl Spleen
Test Testes
Thym Thymus
Tong Tongue
Trac Trachea
Ute Uterus

**Agouti related neuropeptide (AGRP)**
- epithelial/neural/germ

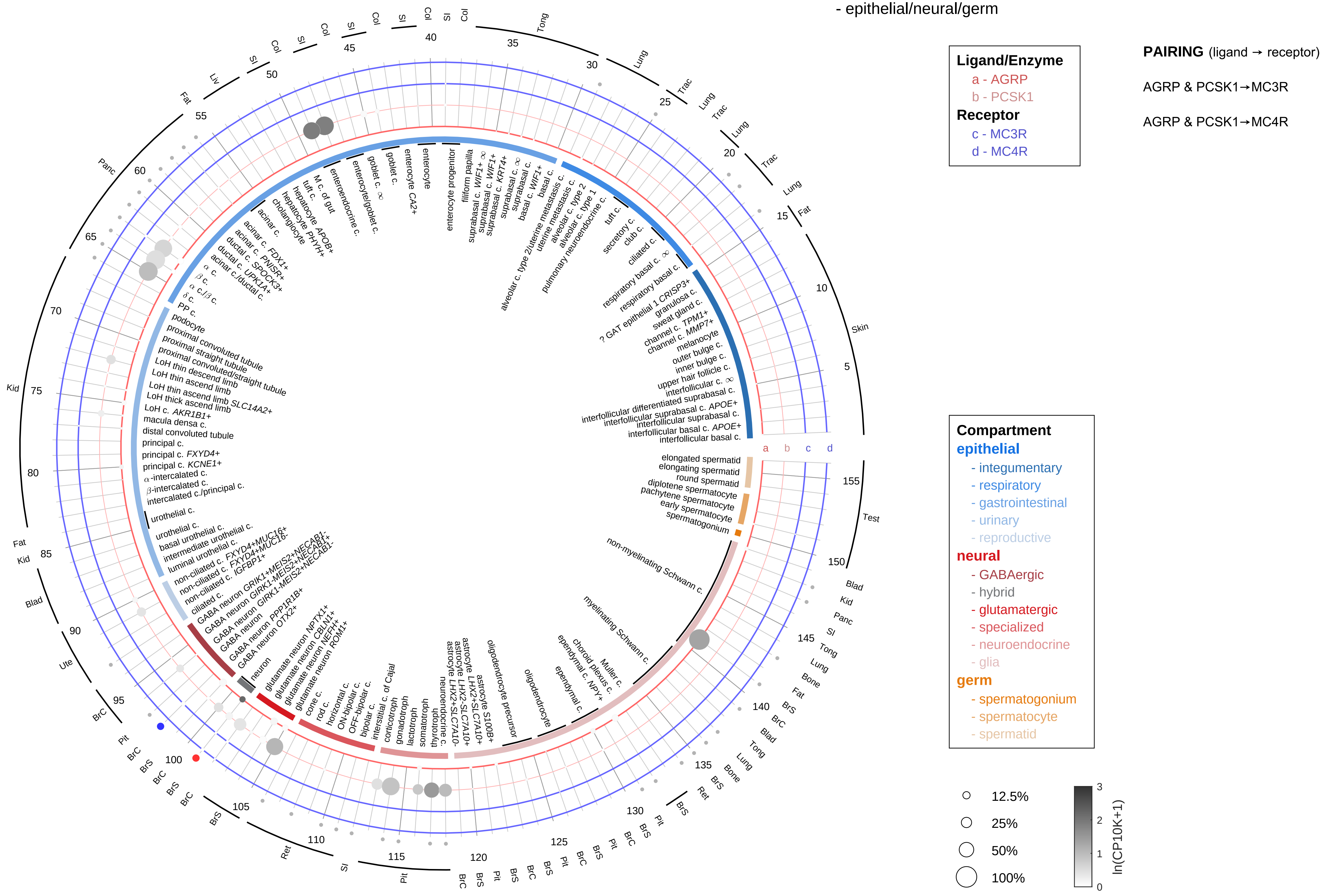

| Abbrev. | Tissue |
|---|---|
| Aort | Aorta |
| Blad | Bladder |
| Blo | Blood |
| Bone | Bone |
| BM | Bone marrow |
| BrC | Brain cortex |
| BrS | Brainstem |
| Col | Colon |
| Dia | Diaphragm |
| Ret | Eye retina |
| Fat | Fat |
| Hrt | Heart |
| Pit | Hypothalamus/Pituitary |
| Kid | Kidney |
| LM | Limb muscle |
| Liv | Liver |
| Lung | Lung |
| MG | Mammary gland |
| Panc | Pancreas |
| Skin | Skin |
| SI | Small intestine |
| Spl | Spleen |
| Test | Testes |
| Thym | Thymus |
| Tong | Tongue |
| Trac | Trachea |
| Ute | Uterus |

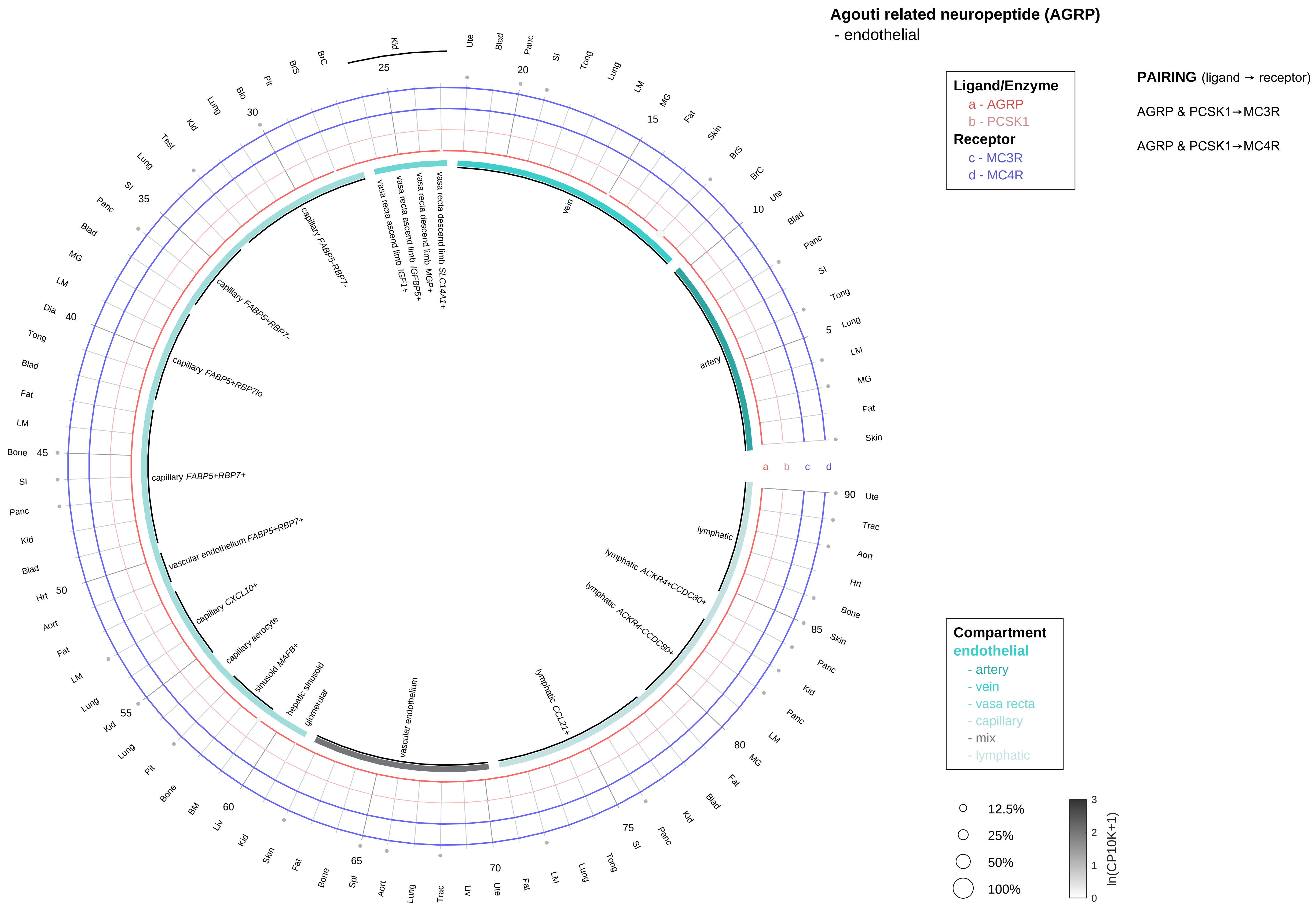

| Abbrev. | Tissue |
|---|---|
| Aort | Aorta |
| Blad | Bladder |
| Blo | Blood |
| Bone | Bone |
| BM | Bone marrow |
| BrC | Brain cortex |
| BrS | Brainstem |
| Col | Colon |
| Dia | Diaphragm |
| Ret | Eye retina |
| Fat | Fat |
| Hrt | Heart |
| Pit | Hypothalamus/Pituitary |
| Kid | Kidney |
| LM | Limb muscle |
| Liv | Liver |
| Lung | Lung |
| MG | Mammary gland |
| Panc | Pancreas |
| Skin | Skin |
| SI | Small intestine |
| Spl | Spleen |
| Test | Testes |
| Thym | Thymus |
| Tong | Tongue |
| Trac | Trachea |
| Ute | Uterus |

#### Agouti related neuropeptide (AGRP)
- stromal

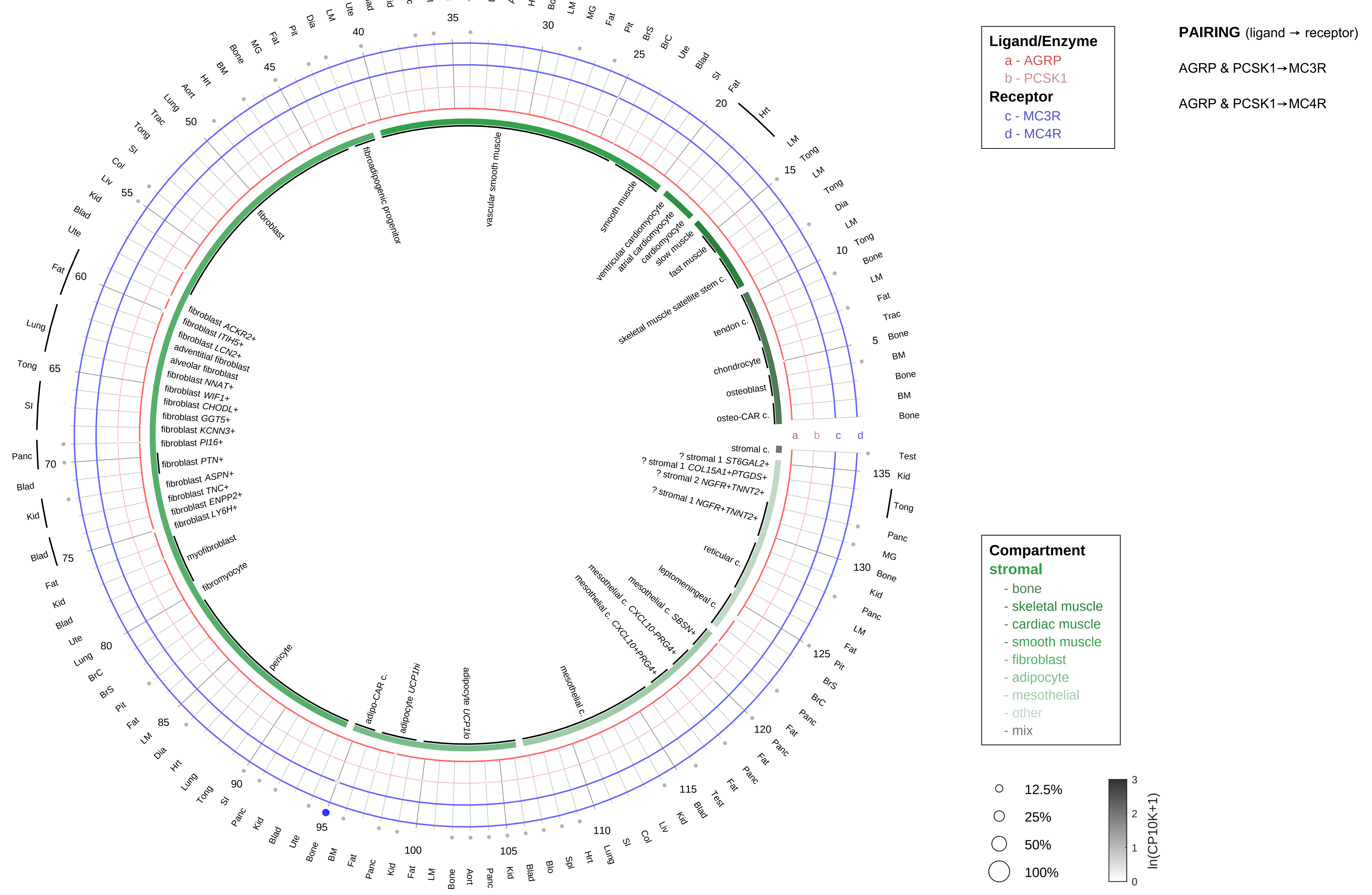

| Abbrev. | Tissue |
|---|---|
| Aort | Aorta |
| Blad | Bladder |
| Blo | Blood |
| Bone | Bone |
| BM | Bone marrow |
| BrC | Brain cortex |
| BrS | Brainstem |
| Col | Colon |
| Dia | Diaphragm |
| Ret | Eye retina |
| Fat | Fat |
| Hrt | Heart |
| Pit | Hypothalamus/Pituitary |
| Kid | Kidney |
| LM | Limb muscle |
| Liv | Liver |
| Lung | Lung |
| MG | Mammary gland |
| Panc | Pancreas |
| Skin | Skin |
| SI | Small intestine |
| Spl | Spleen |
| Test | Testes |
| Thym | Thymus |
| Tong | Tongue |
| Trac | Trachea |
| Ute | Uterus |

**Agouti related neuropeptide (AGRP)**
- lymphoid

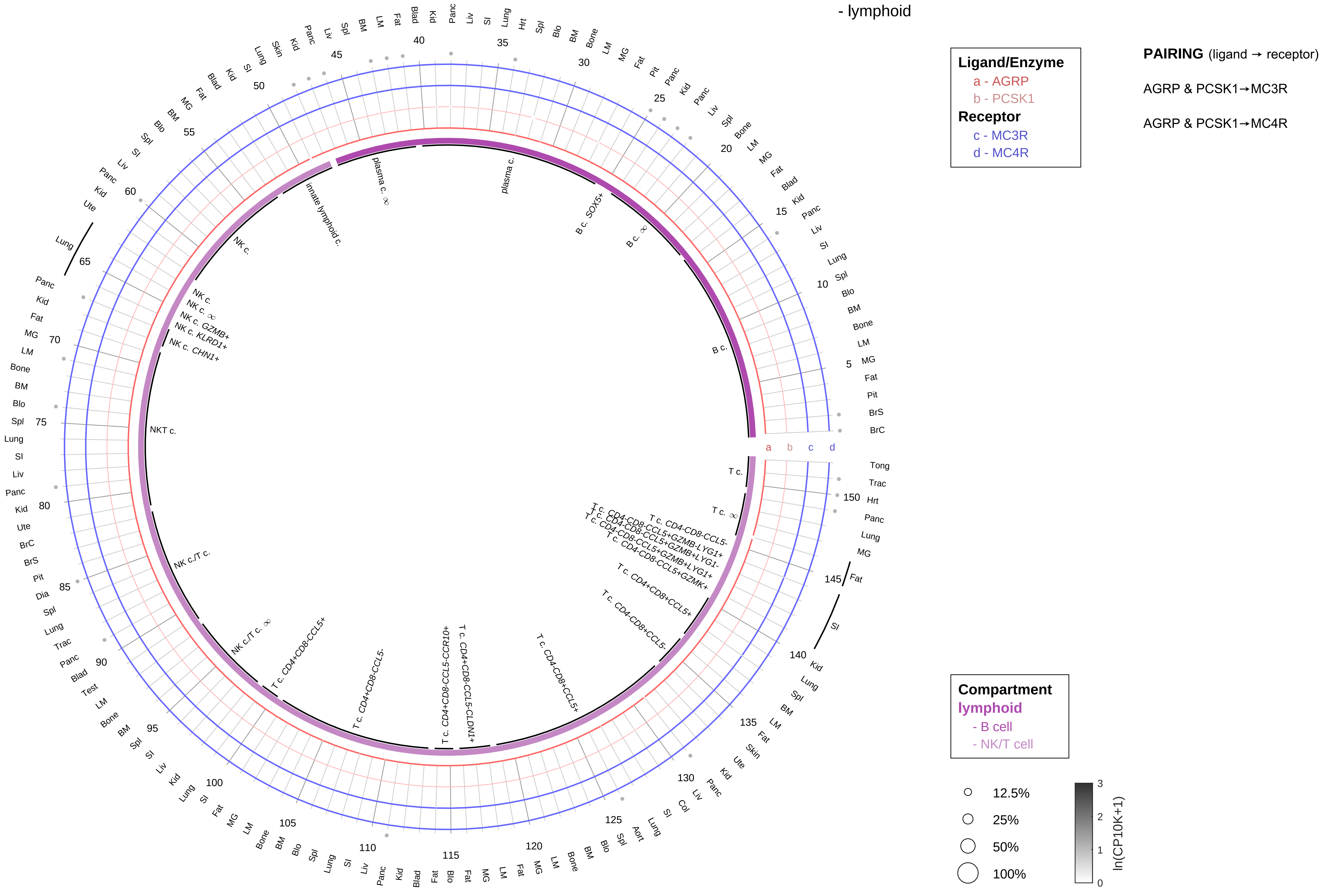

| Abbrev. | Tissue |
|---|---|
| Aort | Aorta |
| Blad | Bladder |
| Blo | Blood |
| Bone | Bone |
| BM | Bone marrow |
| BrC | Brain cortex |
| BrS | Brainstem |
| Col | Colon |
| Dia | Diaphragm |
| Ret | Eye retina |
| Fat | Fat |
| Hrt | Heart |
| Pit | Hypothalamus/Pituitary |
| Kid | Kidney |
| LM | Limb muscle |
| Liv | Liver |
| Lung | Lung |
| MG | Mammary gland |
| Panc | Pancreas |
| Skin | Skin |
| SI | Small intestine |
| Spl | Spleen |
| Test | Testes |
| Thym | Thymus |
| Tong | Tongue |
| Trac | Trachea |
| Ute | Uterus |

**Agouti related neuropeptide (AGRP)**
- myeloid

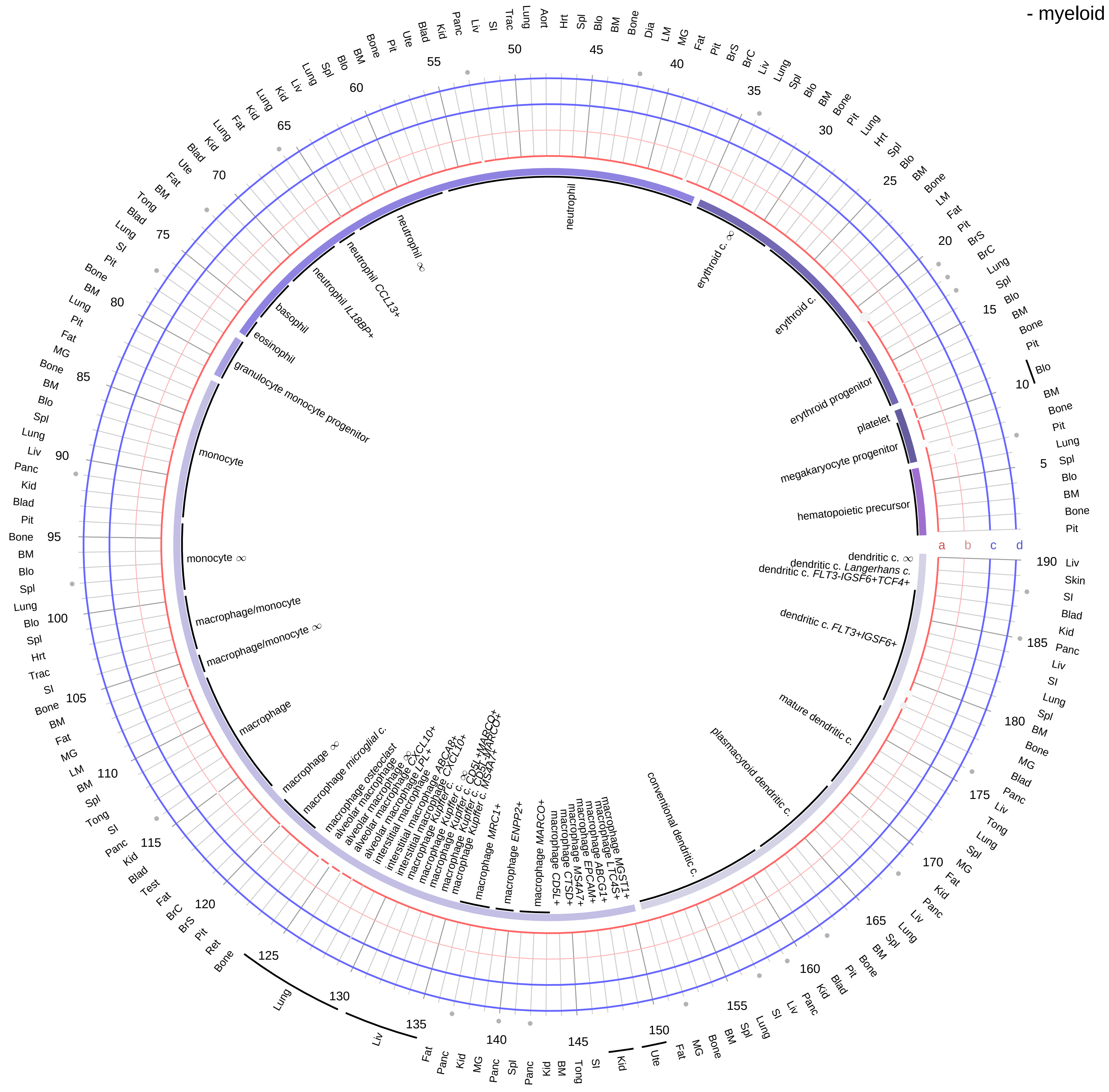

**Ligand/Enzyme**
- a - AGRP
- b - PCSK1

**Receptor**
- c - MC3R
- d - MC4R

**PAIRING** (ligand → receptor)

AGRP & PCSK1→MC3R

AGRP & PCSK1→MC4R

**Compartment**

**hematopoietic**
- hematopoietic precursor

**megakaryocyte-erythroid**
- megakaryoid
- erythroid

**myeloid**
- granulocyte
- granulocyte monocyte progenitor
- monocyte/macrophage
- dendritic

12.5%
25%
50%
100%

ln(CP10K+1): 0, 1, 2, 3

| Abbrev. | Tissue |
|---|---|
| Aort | Aorta |
| Blad | Bladder |
| Blo | Blood |
| Bone | Bone |
| BM | Bone marrow |
| BrC | Brain cortex |
| BrS | Brainstem |
| Col | Colon |
| Dia | Diaphragm |
| Ret | Eye retina |
| Fat | Fat |
| Hrt | Heart |
| Pit | Hypothalamus/Pituitary |
| Kid | Kidney |
| LM | Limb muscle |
| Liv | Liver |
| Lung | Lung |
| MG | Mammary gland |
| Panc | Pancreas |
| Skin | Skin |
| SI | Small intestine |
| Spl | Spleen |
| Test | Testes |
| Thym | Thymus |
| Tong | Tongue |
| Trac | Trachea |
| Ute | Uterus |

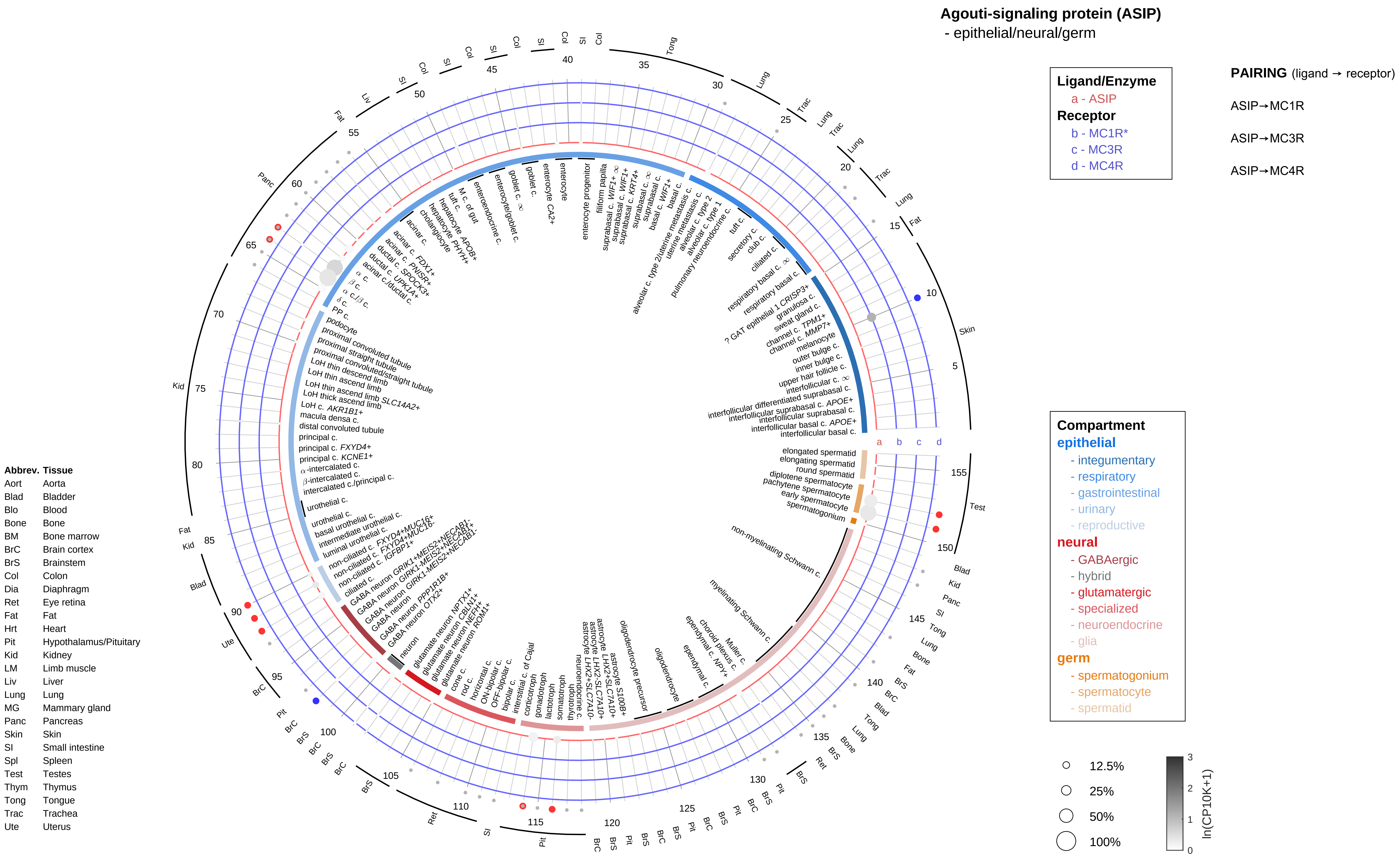

Agouti-signaling protein (ASIP)
- epithelial/neural/germ
Ligand/Enzyme
a - ASIP
Receptor
b - MC1R*
c - MC3R
d - MC4R
PAIRING (ligand → receptor)
ASIP→MC1R
ASIP→MC3R
ASIP→MC4R
Compartment
epithelial
- integumentary
- respiratory
- gastrointestinal
- urinary
- reproductive
neural
- GABAergic
- hybrid
- glutamatergic
- specialized
- neuroendocrine
- glia
germ
- spermatogonium
- spermatocyte
- spermatid
12.5%
25%
50%
100%
ln(CP10K+1)
Abbrev. Tissue
Aort Aorta
Blad Bladder
Blo Blood
Bone Bone
BM Bone marrow
BrC Brain cortex
BrS Brainstem
Col Colon
Dia Diaphragm
Ret Eye retina
Fat Fat
Hrt Heart
Pit Hypothalamus/Pituitary
Kid Kidney
LM Limb muscle
Liv Liver
Lung Lung
MG Mammary gland
Panc Pancreas
Skin Skin
SI Small intestine
Spl Spleen
Test Testes
Thym Thymus
Tong Tongue
Trac Trachea
Ute Uterus

**Agouti-signaling protein (ASIP)**
- endothelial

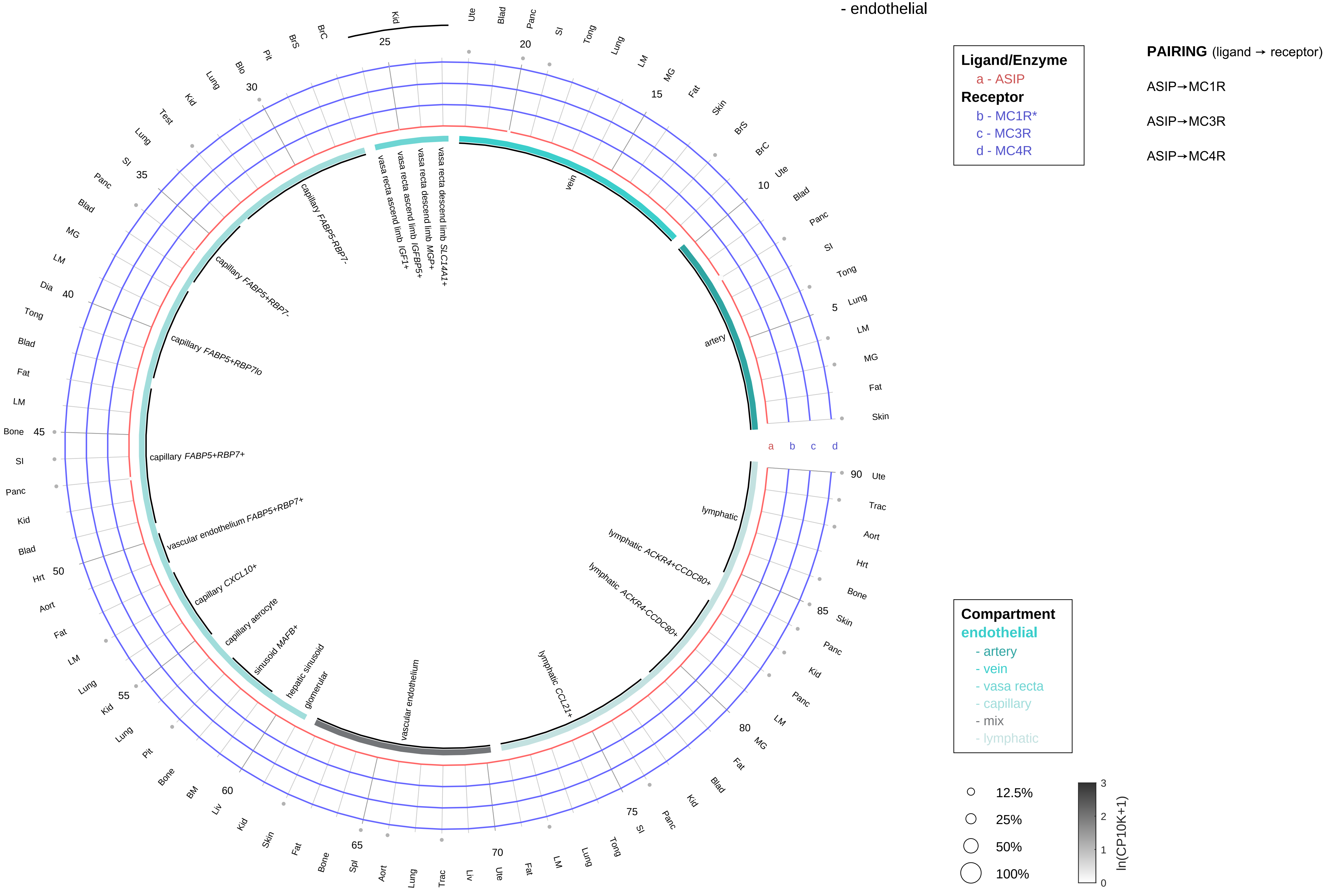

| Abbrev. | Tissue |
|---|---|
| Aort | Aorta |
| Blad | Bladder |
| Blo | Blood |
| Bone | Bone |
| BM | Bone marrow |
| BrC | Brain cortex |
| BrS | Brainstem |
| Col | Colon |
| Dia | Diaphragm |
| Ret | Eye retina |
| Fat | Fat |
| Hrt | Heart |
| Pit | Hypothalamus/Pituitary |
| Kid | Kidney |
| LM | Limb muscle |
| Liv | Liver |
| Lung | Lung |
| MG | Mammary gland |
| Panc | Pancreas |
| Skin | Skin |
| SI | Small intestine |
| Spl | Spleen |
| Test | Testes |
| Thym | Thymus |
| Tong | Tongue |
| Trac | Trachea |
| Ute | Uterus |

**Agouti-signaling protein (ASIP)**
- stromal

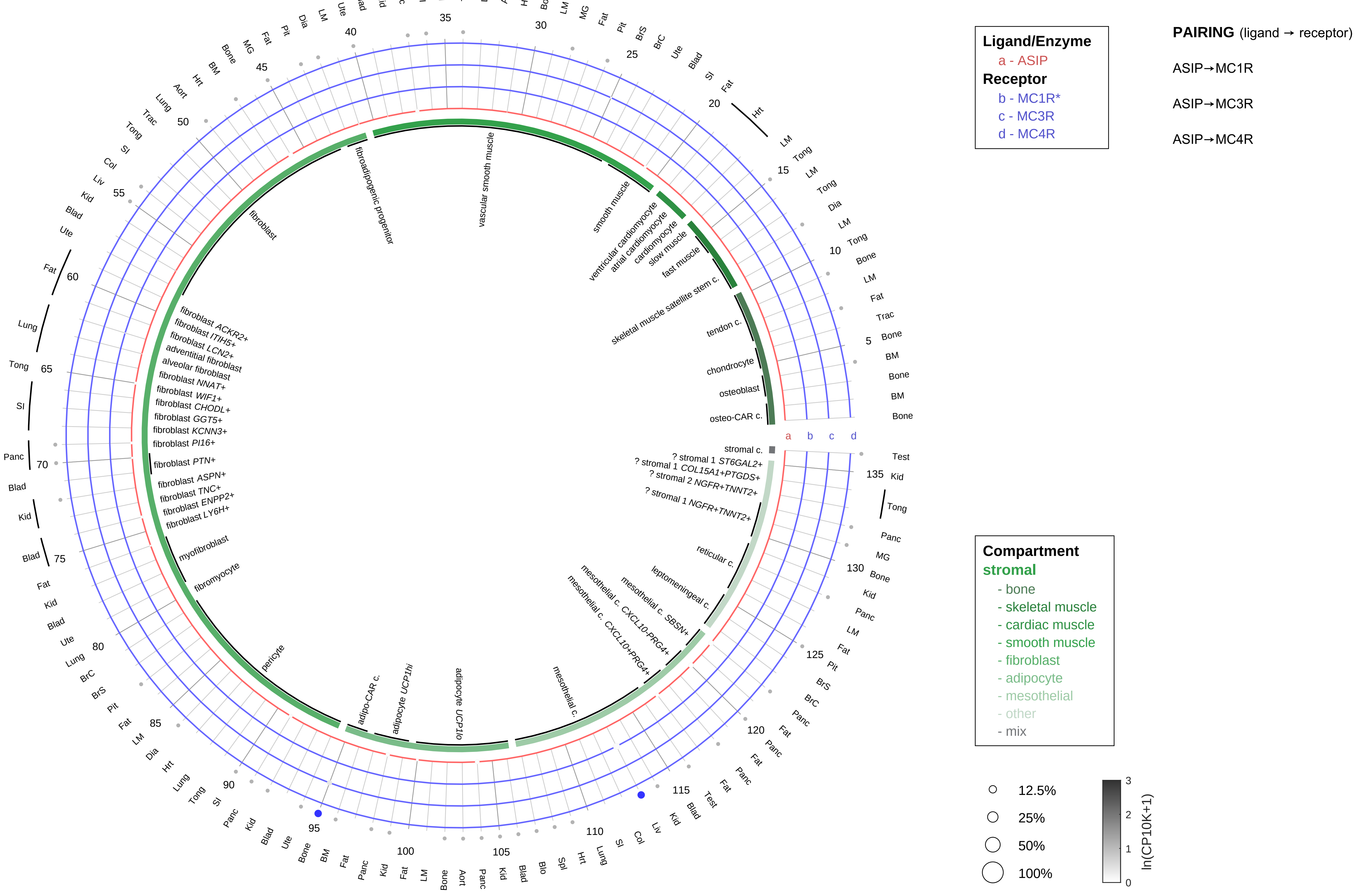

| Abbrev. | Tissue |
|---|---|
| Aort | Aorta |
| Blad | Bladder |
| Blo | Blood |
| Bone | Bone |
| BM | Bone marrow |
| BrC | Brain cortex |
| BrS | Brainstem |
| Col | Colon |
| Dia | Diaphragm |
| Ret | Eye retina |
| Fat | Fat |
| Hrt | Heart |
| Pit | Hypothalamus/Pituitary |
| Kid | Kidney |
| LM | Limb muscle |
| Liv | Liver |
| Lung | Lung |
| MG | Mammary gland |
| Panc | Pancreas |
| Skin | Skin |
| SI | Small intestine |
| Spl | Spleen |
| Test | Testes |
| Thym | Thymus |
| Tong | Tongue |
| Trac | Trachea |
| Ute | Uterus |

**Agouti-signaling protein (ASIP)**
- lymphoid

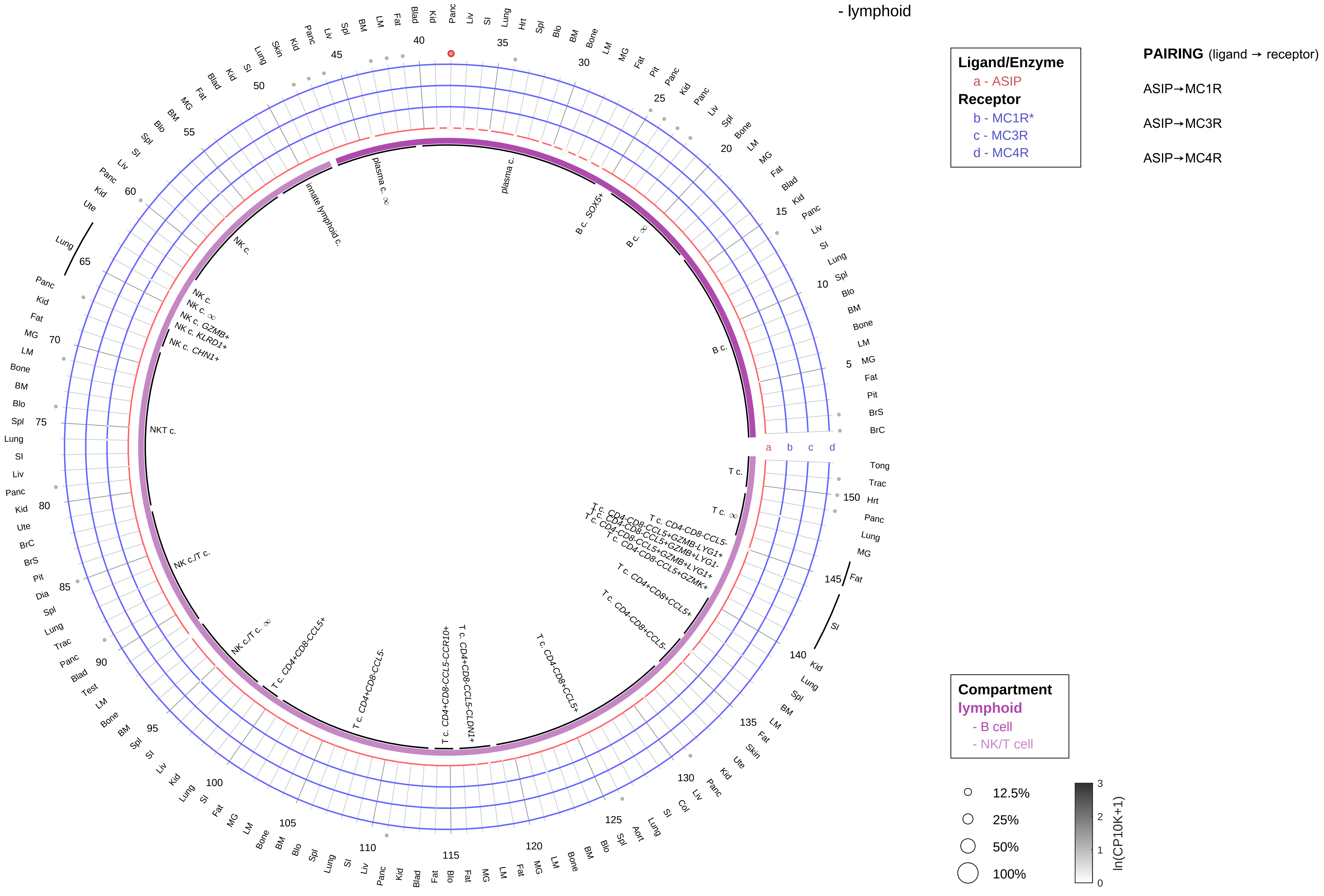

| Abbrev. | Tissue |
|---|---|
| Aort | Aorta |
| Blad | Bladder |
| Blo | Blood |
| Bone | Bone |
| BM | Bone marrow |
| BrC | Brain cortex |
| BrS | Brainstem |
| Col | Colon |
| Dia | Diaphragm |
| Ret | Eye retina |
| Fat | Fat |
| Hrt | Heart |
| Pit | Hypothalamus/Pituitary |
| Kid | Kidney |
| LM | Limb muscle |
| Liv | Liver |
| Lung | Lung |
| MG | Mammary gland |
| Panc | Pancreas |
| Skin | Skin |
| SI | Small intestine |
| Spl | Spleen |
| Test | Testes |
| Thym | Thymus |
| Tong | Tongue |
| Trac | Trachea |
| Ute | Uterus |

**Agouti-signaling protein (ASIP)**
- myeloid

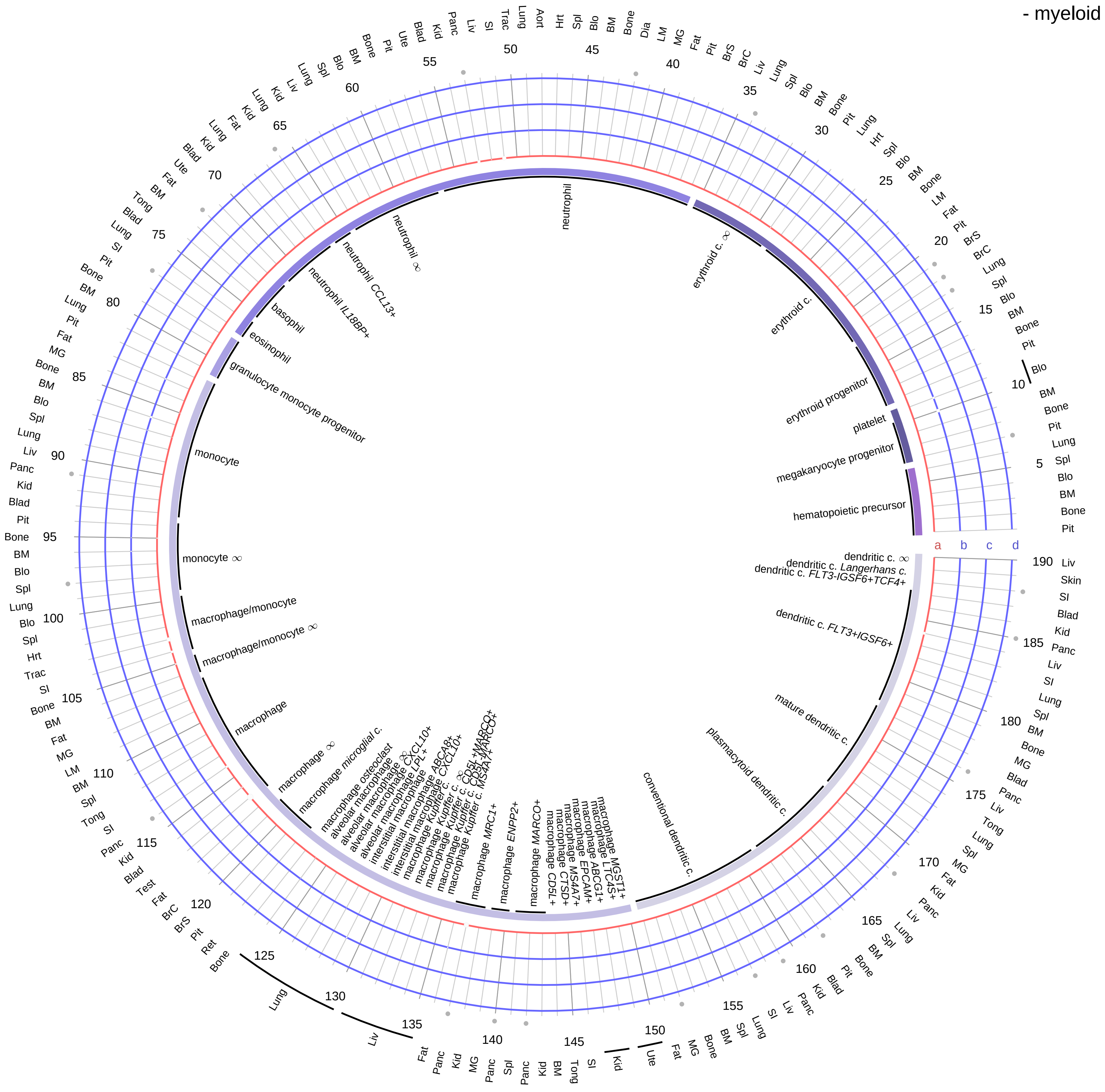

**Ligand/Enzyme**
a - ASIP
**Receptor**
b - MC1R*
c - MC3R
d - MC4R

**PAIRING** (ligand → receptor)

ASIP→MC1R

ASIP→MC3R

ASIP→MC4R

| Abbrev. | Tissue |
|---|---|
| Aort | Aorta |
| Blad | Bladder |
| Blo | Blood |
| Bone | Bone |
| BM | Bone marrow |
| BrC | Brain cortex |
| BrS | Brainstem |
| Col | Colon |
| Dia | Diaphragm |
| Ret | Eye retina |
| Fat | Fat |
| Hrt | Heart |
| Pit | Hypothalamus/Pituitary |
| Kid | Kidney |
| LM | Limb muscle |
| Liv | Liver |
| Lung | Lung |
| MG | Mammary gland |
| Panc | Pancreas |
| Skin | Skin |
| SI | Small intestine |
| Spl | Spleen |
| Test | Testes |
| Thym | Thymus |
| Tong | Tongue |
| Trac | Trachea |
| Ute | Uterus |

**Compartment**
**hematopoietic**
- hematopoietic precursor
**megakaryocyte-erythroid**
- megakaryoid
- erythroid
**myeloid**
- granulocyte
- granulocyte monocyte progenitor
- monocyte/macrophage
- dendritic

12.5%
25%
50%
100%

ln(CP10K+1): 0, 1, 2, 3

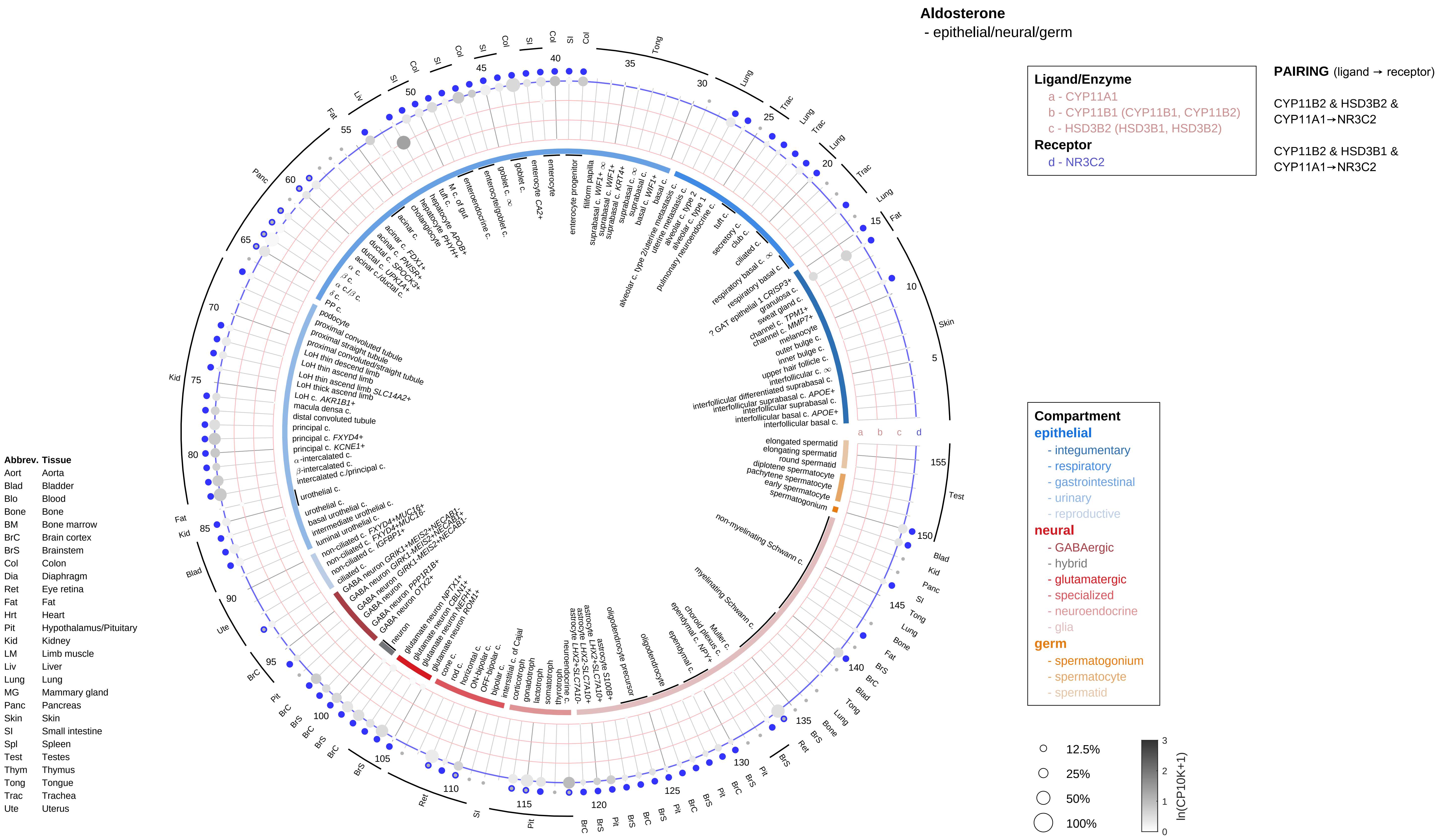

Aldosterone
- epithelial/neural/germ
Ligand/Enzyme
a - CYP11A1
b - CYP11B1 (CYP11B1, CYP11B2)
c - HSD3B2 (HSD3B1, HSD3B2)
Receptor
d - NR3C2
PAIRING (ligand → receptor)
CYP11B2 & HSD3B2 & CYP11A1→NR3C2
CYP11B2 & HSD3B1 & CYP11A1→NR3C2
Compartment
epithelial
- integumentary
- respiratory
- gastrointestinal
- urinary
- reproductive
neural
- GABAergic
- hybrid
- glutamatergic
- specialized
- neuroendocrine
- glia
germ
- spermatogonium
- spermatocyte
- spermatid
12.5%
25%
50%
100%
ln(CP10K+1)
0
1
2
3
Abbrev. Tissue
Aort Aorta
Blad Bladder
Blo Blood
Bone Bone
BM Bone marrow
BrC Brain cortex
BrS Brainstem
Col Colon
Dia Diaphragm
Ret Eye retina
Fat Fat
Hrt Heart
Pit Hypothalamus/Pituitary
Kid Kidney
LM Limb muscle
Liv Liver
Lung Lung
MG Mammary gland
Panc Pancreas
Skin Skin
SI Small intestine
Spl Spleen
Test Testes
Thym Thymus
Tong Tongue
Trac Trachea
Ute Uterus

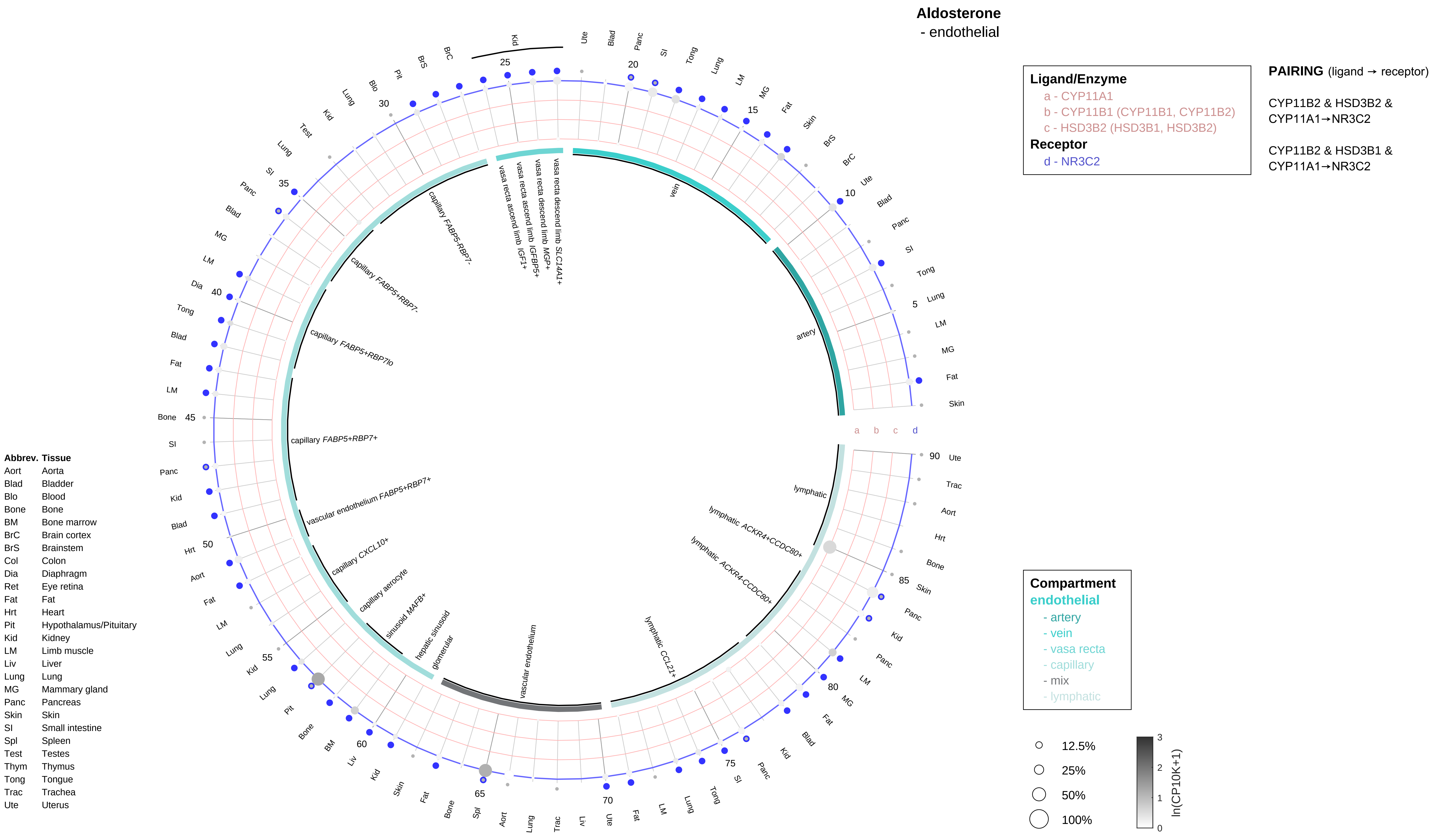

Aldosterone
- endothelial
Ligand/Enzyme
a - CYP11A1
b - CYP11B1 (CYP11B1, CYP11B2)
c - HSD3B2 (HSD3B1, HSD3B2)
Receptor
d - NR3C2
PAIRING (ligand → receptor)
CYP11B2 & HSD3B2 & CYP11A1→NR3C2
CYP11B2 & HSD3B1 & CYP11A1→NR3C2
Compartment
endothelial
- artery
- vein
- vasa recta
- capillary
- mix
- lymphatic
12.5%
25%
50%
100%
ln(CP10K+1)
Abbrev. Tissue
Aort Aorta
Blad Bladder
Blo Blood
Bone Bone
BM Bone marrow
BrC Brain cortex
BrS Brainstem
Col Colon
Dia Diaphragm
Ret Eye retina
Fat Fat
Hrt Heart
Pit Hypothalamus/Pituitary
Kid Kidney
LM Limb muscle
Liv Liver
Lung Lung
MG Mammary gland
Panc Pancreas
Skin Skin
SI Small intestine
Spl Spleen
Test Testes
Thym Thymus
Tong Tongue
Trac Trachea
Ute Uterus
vein
artery
lymphatic
lymphatic ACKR4+CCDC80+
lymphatic ACKR4-CCDC80+
lymphatic CCL21+
vascular endothelium
glomerular
hepatic sinusoid
sinusoid MAFB+
capillary aerocyte
capillary CXCL10+
vascular endothelium FABP5+RBP7+
capillary FABP5+RBP7+
capillary FABP5+RBP7lo
capillary FABP5+RBP7-
capillary FABP5-RBP7-
vasa recta ascend limb IGF1+
vasa recta ascend limb IGFBP5+
vasa recta descend limb MGP+
vasa recta descend limb SLC14A1+

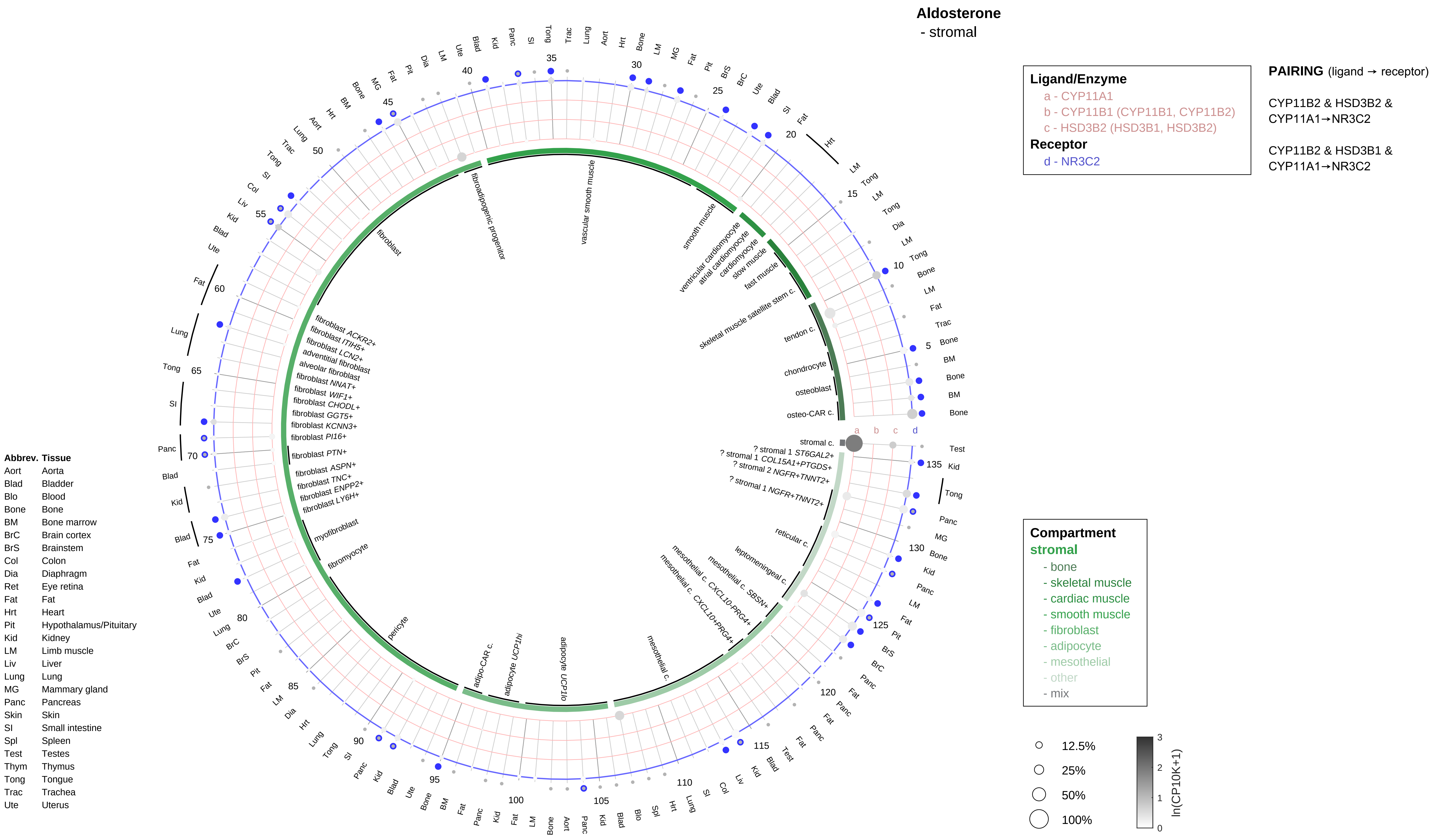

Aldosterone
- stromal
Ligand/Enzyme
a - CYP11A1
b - CYP11B1 (CYP11B1, CYP11B2)
c - HSD3B2 (HSD3B1, HSD3B2)
Receptor
d - NR3C2
PAIRING (ligand → receptor)
CYP11B2 & HSD3B2 & CYP11A1→NR3C2
CYP11B2 & HSD3B1 & CYP11A1→NR3C2
Compartment
stromal
- bone
- skeletal muscle
- cardiac muscle
- smooth muscle
- fibroblast
- adipocyte
- mesothelial
- other
- mix
12.5%
25%
50%
100%
ln(CP10K+1)
Abbrev. Tissue
Aort Aorta
Blad Bladder
Blo Blood
Bone Bone
BM Bone marrow
BrC Brain cortex
BrS Brainstem
Col Colon
Dia Diaphragm
Ret Eye retina
Fat Fat
Hrt Heart
Pit Hypothalamus/Pituitary
Kid Kidney
LM Limb muscle
Liv Liver
Lung Lung
MG Mammary gland
Panc Pancreas
Skin Skin
SI Small intestine
Spl Spleen
Test Testes
Thym Thymus
Tong Tongue
Trac Trachea
Ute Uterus
stromal c.
? stromal 1 ST6GAL2+
? stromal 1 COL15A1+PTGDS+
? stromal 2 NGFR+TNNT2+
? stromal 1 NGFR+TNNT2+
reticular c.
leptomeningeal c.
mesothelial c. SBSN+
mesothelial c. CXCL10-PRG4+
mesothelial c. CXCL10+PRG4+
mesothelial c.
adipocyte UCP1lo
adipocyte UCP1hi
adipo-CAR c.
pericyte
fibromyocyte
myofibroblast
fibroblast LY6H+
fibroblast ENPP2+
fibroblast TNC+
fibroblast ASPN+
fibroblast PTN+
fibroblast PI16+
fibroblast KCNN3+
fibroblast GGT5+
fibroblast CHODL+
fibroblast WIF1+
fibroblast NNAT+
alveolar fibroblast
adventitial fibroblast
fibroblast LCN2+
fibroblast ITIH5+
fibroblast ACKR2+
fibroblast
fibroadipogenic progenitor
vascular smooth muscle
smooth muscle
ventricular cardiomyocyte
atrial cardiomyocyte
cardiomyocyte
slow muscle
fast muscle
skeletal muscle satellite stem c.
tendon c.
chondrocyte
osteoblast
osteo-CAR c.

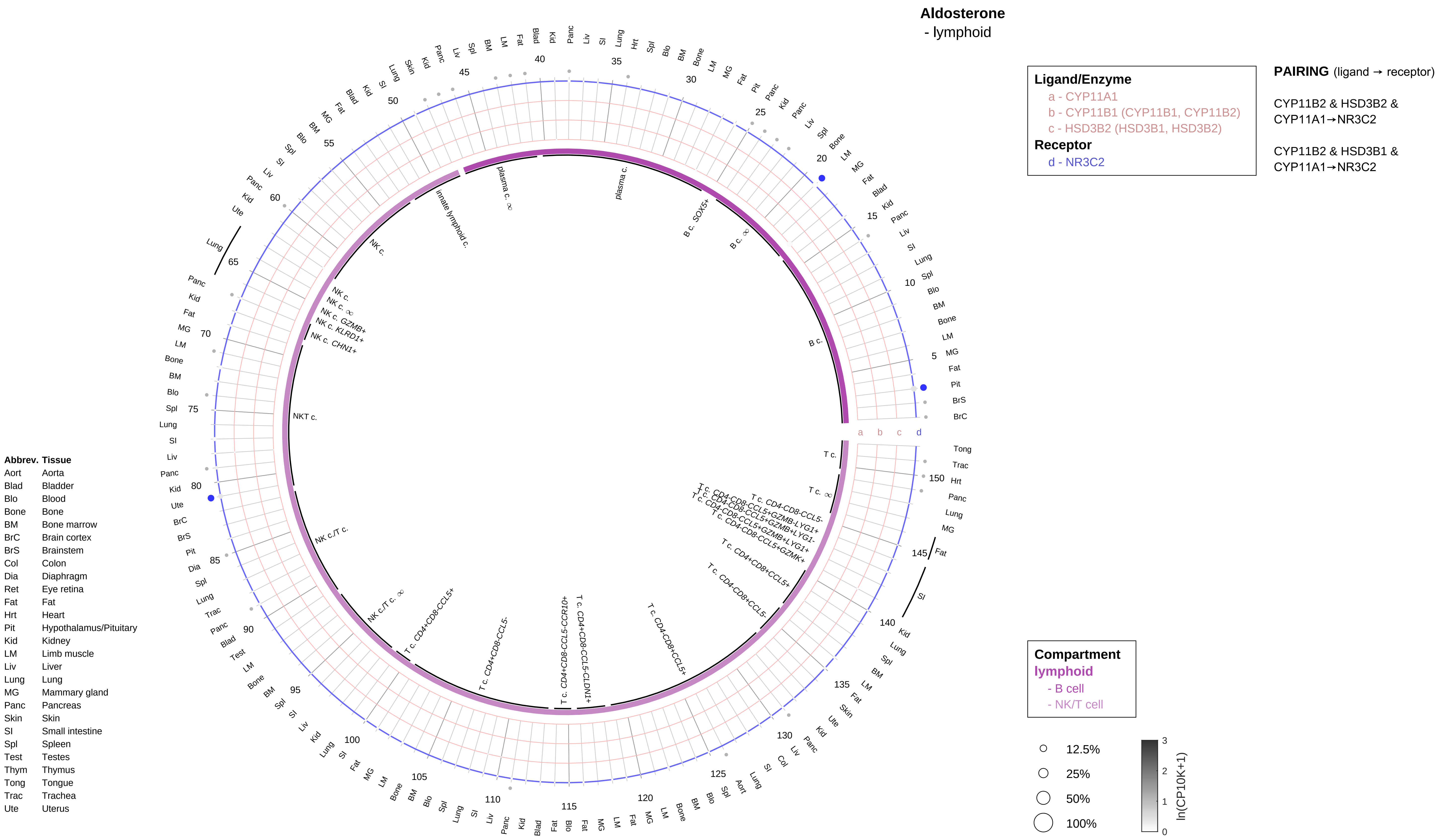
Aldosterone
- lymphoid
Ligand/Enzyme
a - CYP11A1
b - CYP11B1 (CYP11B1, CYP11B2)
c - HSD3B2 (HSD3B1, HSD3B2)
Receptor
d - NR3C2
PAIRING (ligand → receptor)
CYP11B2 & HSD3B2 & CYP11A1→NR3C2
CYP11B2 & HSD3B1 & CYP11A1→NR3C2
Compartment
lymphoid
- B cell
- NK/T cell
12.5%
25%
50%
100%
ln(CP10K+1)
Abbrev. Tissue
Aort Aorta
Blad Bladder
Blo Blood
Bone Bone
BM Bone marrow
BrC Brain cortex
BrS Brainstem
Col Colon
Dia Diaphragm
Ret Eye retina
Fat Fat
Hrt Heart
Pit Hypothalamus/Pituitary
Kid Kidney
LM Limb muscle
Liv Liver
Lung Lung
MG Mammary gland
Panc Pancreas
Skin Skin
SI Small intestine
Spl Spleen
Test Testes
Thym Thymus
Tong Tongue
Trac Trachea
Ute Uterus
plasma c.
plasma c. ∞
B c. SOX5+
B c. ∞
B c.
innate lymphoid c.
NK c.
NK c. ∞
NK c. GZMB+
NK c. KLRD1+
NK c. CHN1+
NKT c.
NK c./T c.
NK c./T c. ∞
T c. CD4+CD8-CCL5+
T c. CD4+CD8-CCL5-
T c. CD4+CD8-CCL5-CCR10+
T c. CD4+CD8-CCL5-CLDN1+
T c. CD4-CD8+CCL5+
T c. CD4-CD8+CCL5-
T c.
T c. ∞
T c. CD4-CD8-CCL5-
T c. CD4-CD8-CCL5+GZMB-LYG1+
T c. CD4-CD8-CCL5+GZMB+LYG1-
T c. CD4-CD8-CCL5+GZMB+LYG1+
T c. CD4-CD8-CCL5+GZMK+

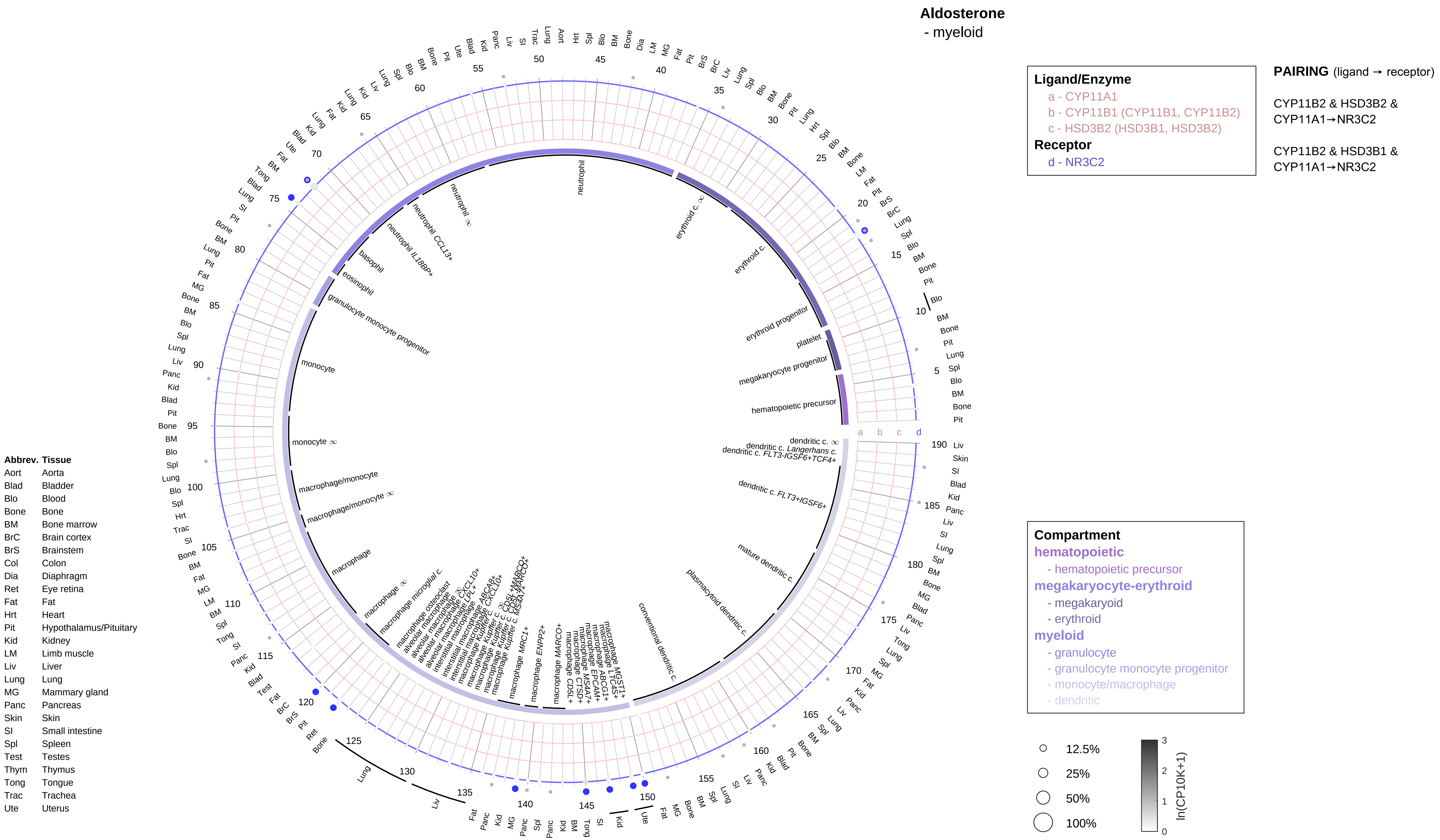

Aldosterone
- myeloid
Ligand/Enzyme
a - CYP11A1
b - CYP11B1 (CYP11B1, CYP11B2)
c - HSD3B2 (HSD3B1, HSD3B2)
Receptor
d - NR3C2
PAIRING (ligand → receptor)
CYP11B2 & HSD3B2 & CYP11A1→NR3C2
CYP11B2 & HSD3B1 & CYP11A1→NR3C2
Compartment
hematopoietic
- hematopoietic precursor
megakaryocyte-erythroid
- megakaryoid
- erythroid
myeloid
- granulocyte
- granulocyte monocyte progenitor
- monocyte/macrophage
- dendritic
12.5%
25%
50%
100%
ln(CP10K+1)
Abbrev. Tissue
Aort Aorta
Blad Bladder
Blo Blood
Bone Bone
BM Bone marrow
BrC Brain cortex
BrS Brainstem
Col Colon
Dia Diaphragm
Ret Eye retina
Fat Fat
Hrt Heart
Pit Hypothalamus/Pituitary
Kid Kidney
LM Limb muscle
Liv Liver
Lung Lung
MG Mammary gland
Panc Pancreas
Skin Skin
SI Small intestine
Spl Spleen
Test Testes
Thym Thymus
Tong Tongue
Trac Trachea
Ute Uterus

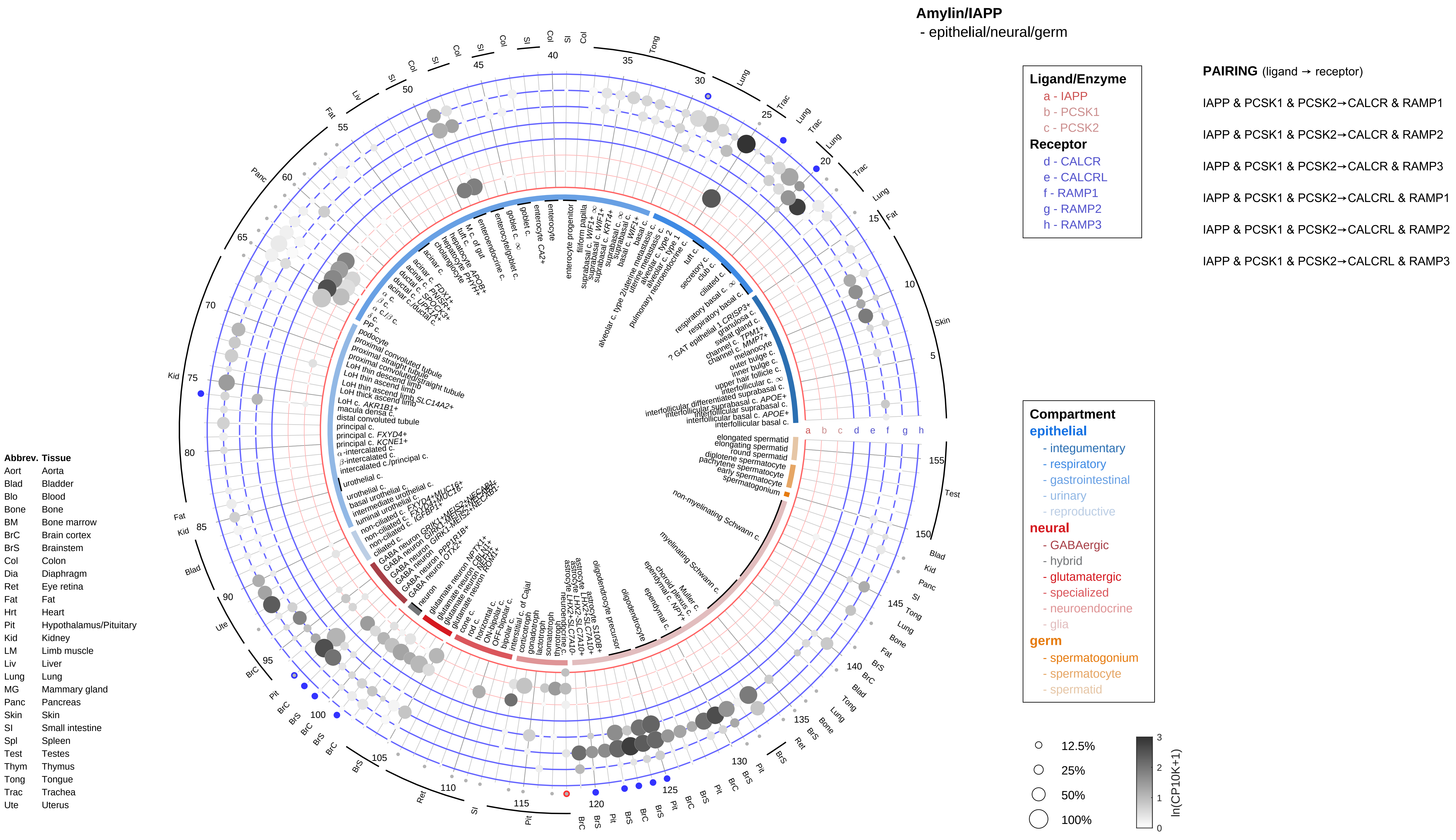
Amylin/IAPP
- epithelial/neural/germ
Ligand/Enzyme
a - IAPP
b - PCSK1
c - PCSK2
Receptor
d - CALCR
e - CALCRL
f - RAMP1
g - RAMP2
h - RAMP3
PAIRING (ligand → receptor)
IAPP & PCSK1 & PCSK2→CALCR & RAMP1
IAPP & PCSK1 & PCSK2→CALCR & RAMP2
IAPP & PCSK1 & PCSK2→CALCR & RAMP3
IAPP & PCSK1 & PCSK2→CALCRL & RAMP1
IAPP & PCSK1 & PCSK2→CALCRL & RAMP2
IAPP & PCSK1 & PCSK2→CALCRL & RAMP3
Compartment
epithelial
- integumentary
- respiratory
- gastrointestinal
- urinary
- reproductive
neural
- GABAergic
- hybrid
- glutamatergic
- specialized
- neuroendocrine
- glia
germ
- spermatogonium
- spermatocyte
- spermatid
12.5%
25%
50%
100%
ln(CP10K+1)
Abbrev. Tissue
Aort Aorta
Blad Bladder
Blo Blood
Bone Bone
BM Bone marrow
BrC Brain cortex
BrS Brainstem
Col Colon
Dia Diaphragm
Ret Eye retina
Fat Fat
Hrt Heart
Pit Hypothalamus/Pituitary
Kid Kidney
LM Limb muscle
Liv Liver
Lung Lung
MG Mammary gland
Panc Pancreas
Skin Skin
SI Small intestine
Spl Spleen
Test Testes
Thym Thymus
Tong Tongue
Trac Trachea
Ute Uterus

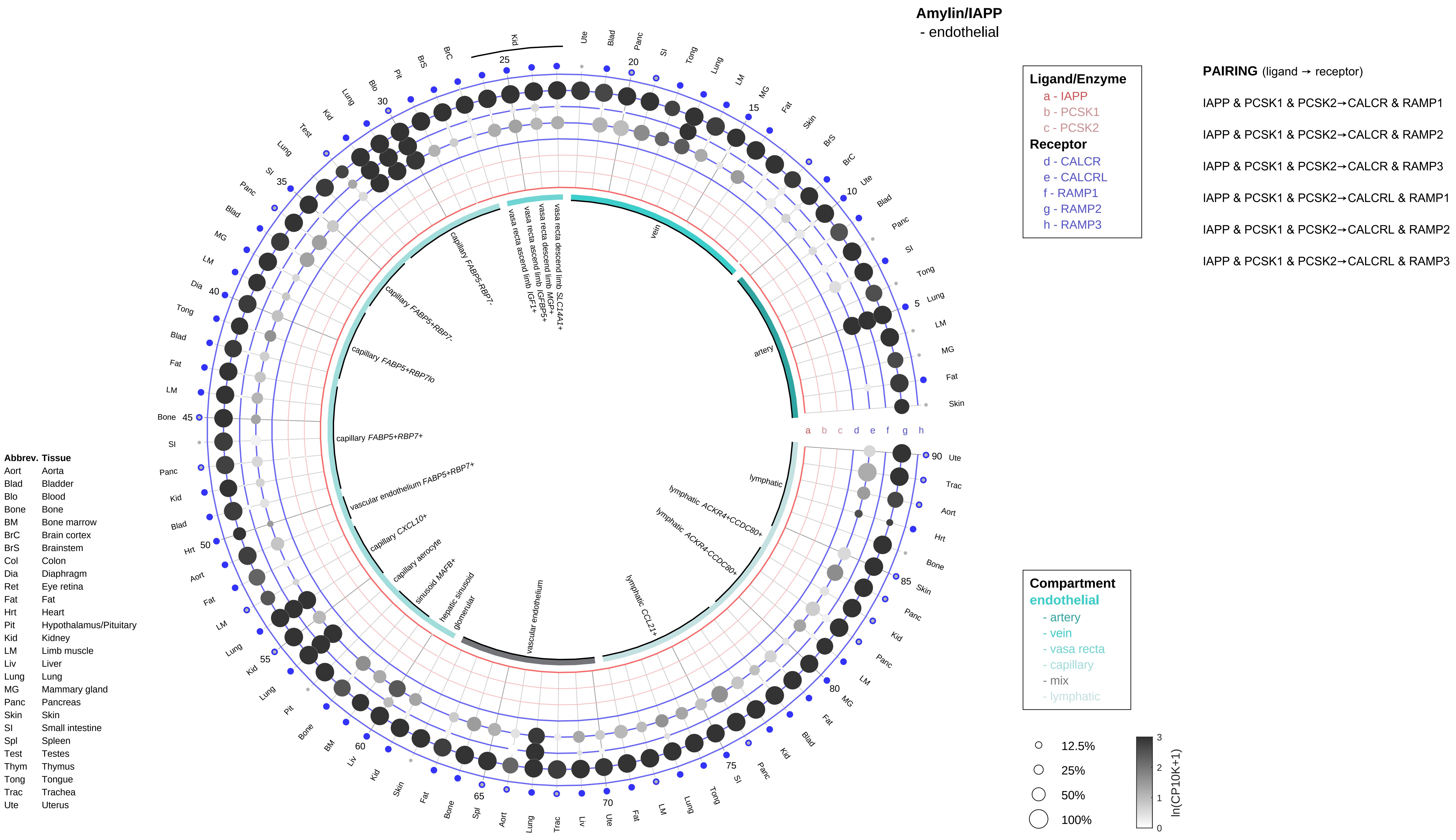

Amylin/IAPP
- endothelial
Ligand/Enzyme
a - IAPP
b - PCSK1
c - PCSK2
Receptor
d - CALCR
e - CALCRL
f - RAMP1
g - RAMP2
h - RAMP3
PAIRING (ligand → receptor)
IAPP & PCSK1 & PCSK2→CALCR & RAMP1
IAPP & PCSK1 & PCSK2→CALCR & RAMP2
IAPP & PCSK1 & PCSK2→CALCR & RAMP3
IAPP & PCSK1 & PCSK2→CALCRL & RAMP1
IAPP & PCSK1 & PCSK2→CALCRL & RAMP2
IAPP & PCSK1 & PCSK2→CALCRL & RAMP3
Compartment
endothelial
- artery
- vein
- vasa recta
- capillary
- mix
- lymphatic
12.5%
25%
50%
100%
ln(CP10K+1)
Abbrev. Tissue
Aort Aorta
Blad Bladder
Blo Blood
Bone Bone
BM Bone marrow
BrC Brain cortex
BrS Brainstem
Col Colon
Dia Diaphragm
Ret Eye retina
Fat Fat
Hrt Heart
Pit Hypothalamus/Pituitary
Kid Kidney
LM Limb muscle
Liv Liver
Lung Lung
MG Mammary gland
Panc Pancreas
Skin Skin
SI Small intestine
Spl Spleen
Test Testes
Thym Thymus
Tong Tongue
Trac Trachea
Ute Uterus
artery
vein
vasa recta descend limb SLC14A1+
vasa recta descend limb MGP+
vasa recta ascend limb IGFBP5+
vasa recta ascend limb IGF1+
capillary FABP5-RBP7-
capillary FABP5+RBP7-
capillary FABP5+RBP7lo
capillary FABP5+RBP7+
vascular endothelium FABP5+RBP7+
capillary CXCL10+
capillary aerocyte
sinusoid MAFB+
hepatic sinusoid
glomerular
vascular endothelium
lymphatic CCL21+
lymphatic ACKR4-CCDC80+
lymphatic ACKR4+CCDC80+
lymphatic

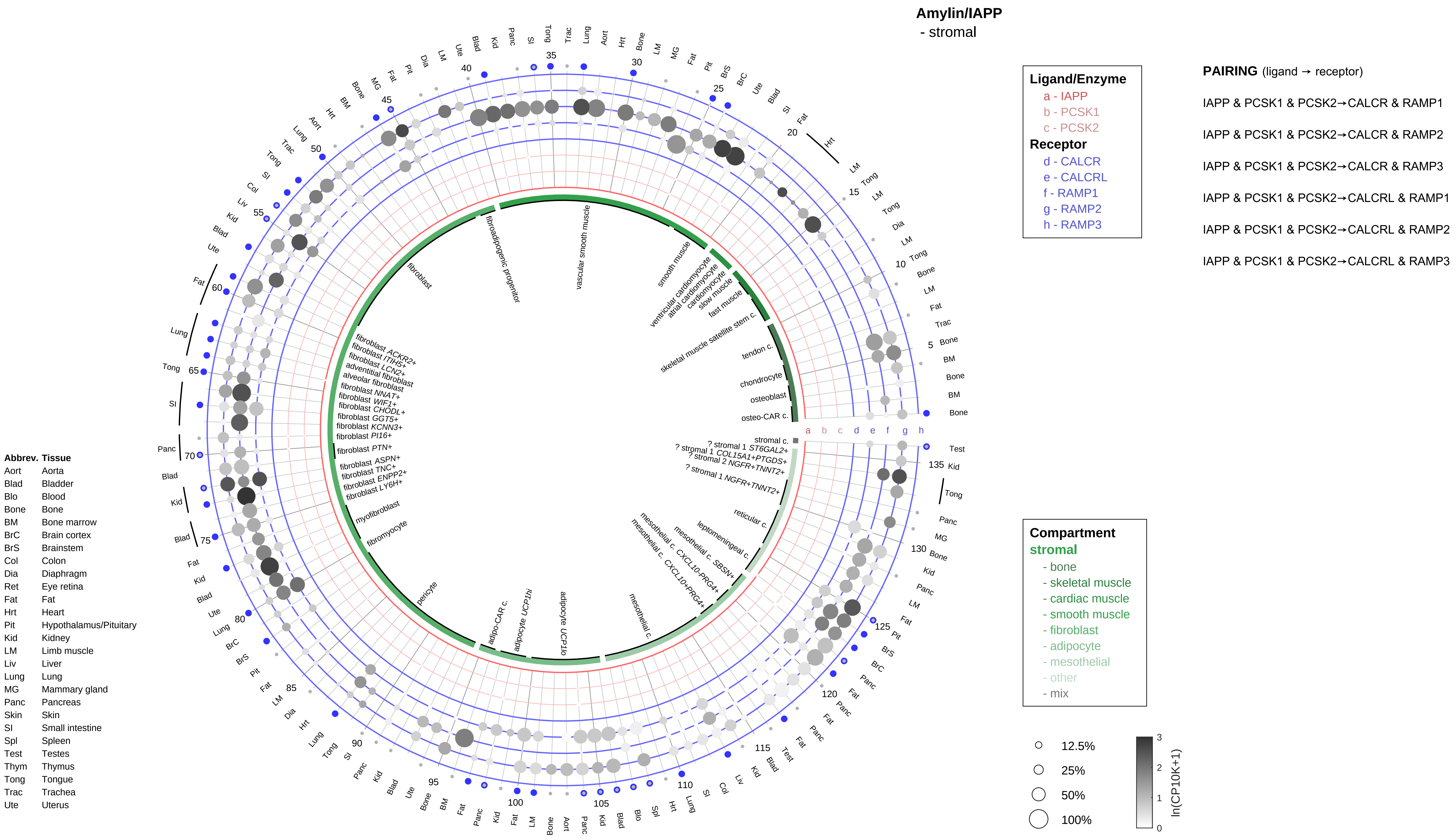

Amylin/IAPP
- stromal
Ligand/Enzyme
a - IAPP
b - PCSK1
c - PCSK2
Receptor
d - CALCR
e - CALCRL
f - RAMP1
g - RAMP2
h - RAMP3
PAIRING (ligand → receptor)
IAPP & PCSK1 & PCSK2→CALCR & RAMP1
IAPP & PCSK1 & PCSK2→CALCR & RAMP2
IAPP & PCSK1 & PCSK2→CALCR & RAMP3
IAPP & PCSK1 & PCSK2→CALCRL & RAMP1
IAPP & PCSK1 & PCSK2→CALCRL & RAMP2
IAPP & PCSK1 & PCSK2→CALCRL & RAMP3
Compartment
stromal
- bone
- skeletal muscle
- cardiac muscle
- smooth muscle
- fibroblast
- adipocyte
- mesothelial
- other
- mix
12.5%
25%
50%
100%
ln(CP10K+1)
Abbrev. Tissue
Aort Aorta
Blad Bladder
Blo Blood
Bone Bone
BM Bone marrow
BrC Brain cortex
BrS Brainstem
Col Colon
Dia Diaphragm
Ret Eye retina
Fat Fat
Hrt Heart
Pit Hypothalamus/Pituitary
Kid Kidney
LM Limb muscle
Liv Liver
Lung Lung
MG Mammary gland
Panc Pancreas
Skin Skin
SI Small intestine
Spl Spleen
Test Testes
Thym Thymus
Tong Tongue
Trac Trachea
Ute Uterus
vascular smooth muscle
smooth muscle
ventricular cardiomyocyte
atrial cardiomyocyte
cardiomyocyte
slow muscle
fast muscle
skeletal muscle satellite stem c.
tendon c.
chondrocyte
osteoblast
osteo-CAR c.
stromal c.
? stromal 1 ST6GAL2+
? stromal 1 COL15A1+PTGDS+
? stromal 2 NGFR+TNNT2+
? stromal 1 NGFR+TNNT2+
reticular c.
leptomeningeal c.
mesothelial c. SBSN+
mesothelial c. CXCL10-PRG4+
mesothelial c. CXCL10+PRG4+
mesothelial c.
adipocyte UCP1lo
adipocyte UCP1hi
adipo-CAR c.
pericyte
fibromyocyte
myofibroblast
fibroblast LY6H+
fibroblast ENPP2+
fibroblast TNC+
fibroblast ASPN+
fibroblast PTN+
fibroblast PI16+
fibroblast KCNN3+
fibroblast GGT5+
fibroblast CHODL+
fibroblast WIF1+
fibroblast NNAT+
alveolar fibroblast
adventitial fibroblast
fibroblast LCN2+
fibroblast ITIH5+
fibroblast ACKR2+
fibroblast
fibroadipogenic progenitor

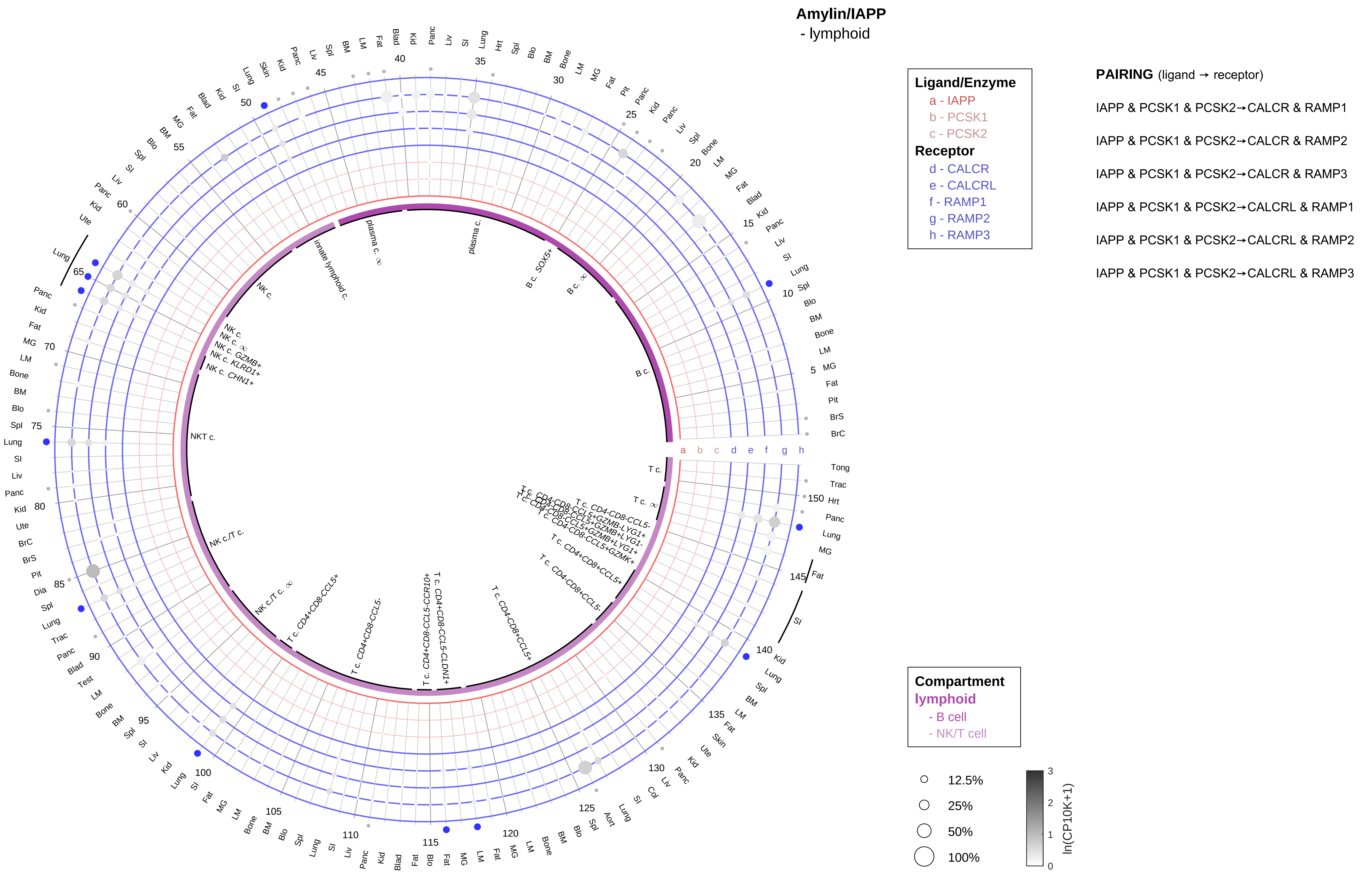

| Abbrev. | Tissue |
|---|---|
| Aort | Aorta |
| Blad | Bladder |
| Blo | Blood |
| Bone | Bone |
| BM | Bone marrow |
| BrC | Brain cortex |
| BrS | Brainstem |
| Col | Colon |
| Dia | Diaphragm |
| Ret | Eye retina |
| Fat | Fat |
| Hrt | Heart |
| Pit | Hypothalamus/Pituitary |
| Kid | Kidney |
| LM | Limb muscle |
| Liv | Liver |
| Lung | Lung |
| MG | Mammary gland |
| Panc | Pancreas |
| Skin | Skin |
| SI | Small intestine |
| Spl | Spleen |
| Test | Testes |
| Thym | Thymus |
| Tong | Tongue |
| Trac | Trachea |
| Ute | Uterus |

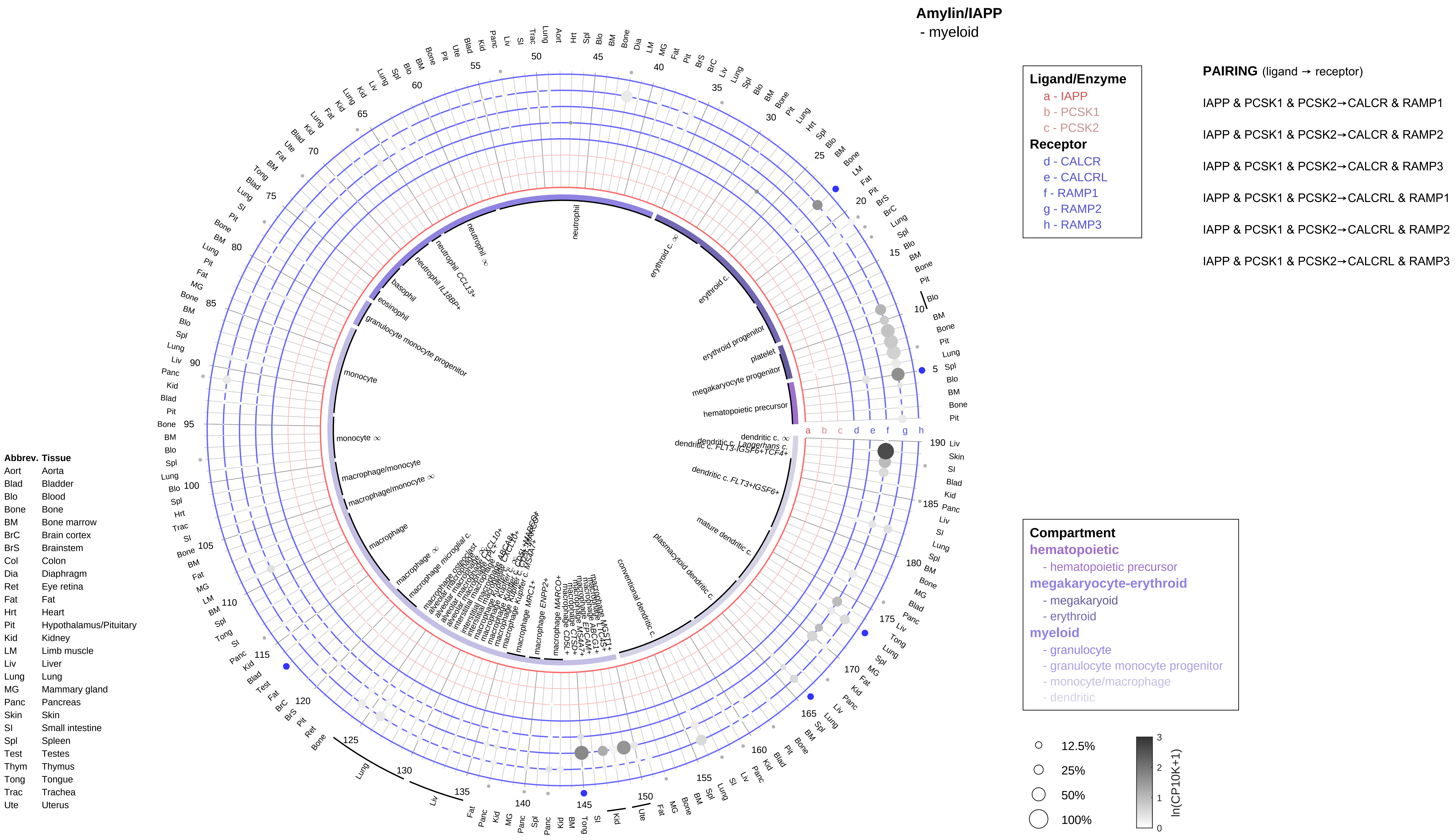

Amylin/IAPP
- myeloid
Ligand/Enzyme
a - IAPP
b - PCSK1
c - PCSK2
Receptor
d - CALCR
e - CALCRL
f - RAMP1
g - RAMP2
h - RAMP3
PAIRING (ligand → receptor)
IAPP & PCSK1 & PCSK2→CALCR & RAMP1
IAPP & PCSK1 & PCSK2→CALCR & RAMP2
IAPP & PCSK1 & PCSK2→CALCR & RAMP3
IAPP & PCSK1 & PCSK2→CALCRL & RAMP1
IAPP & PCSK1 & PCSK2→CALCRL & RAMP2
IAPP & PCSK1 & PCSK2→CALCRL & RAMP3
Compartment
hematopoietic
- hematopoietic precursor
megakaryocyte-erythroid
- megakaryoid
- erythroid
myeloid
- granulocyte
- granulocyte monocyte progenitor
- monocyte/macrophage
- dendritic
12.5%
25%
50%
100%
ln(CP10K+1)
Abbrev. Tissue
Aort Aorta
Blad Bladder
Blo Blood
Bone Bone
BM Bone marrow
BrC Brain cortex
BrS Brainstem
Col Colon
Dia Diaphragm
Ret Eye retina
Fat Fat
Hrt Heart
Pit Hypothalamus/Pituitary
Kid Kidney
LM Limb muscle
Liv Liver
Lung Lung
MG Mammary gland
Panc Pancreas
Skin Skin
SI Small intestine
Spl Spleen
Test Testes
Thym Thymus
Tong Tongue
Trac Trachea
Ute Uterus

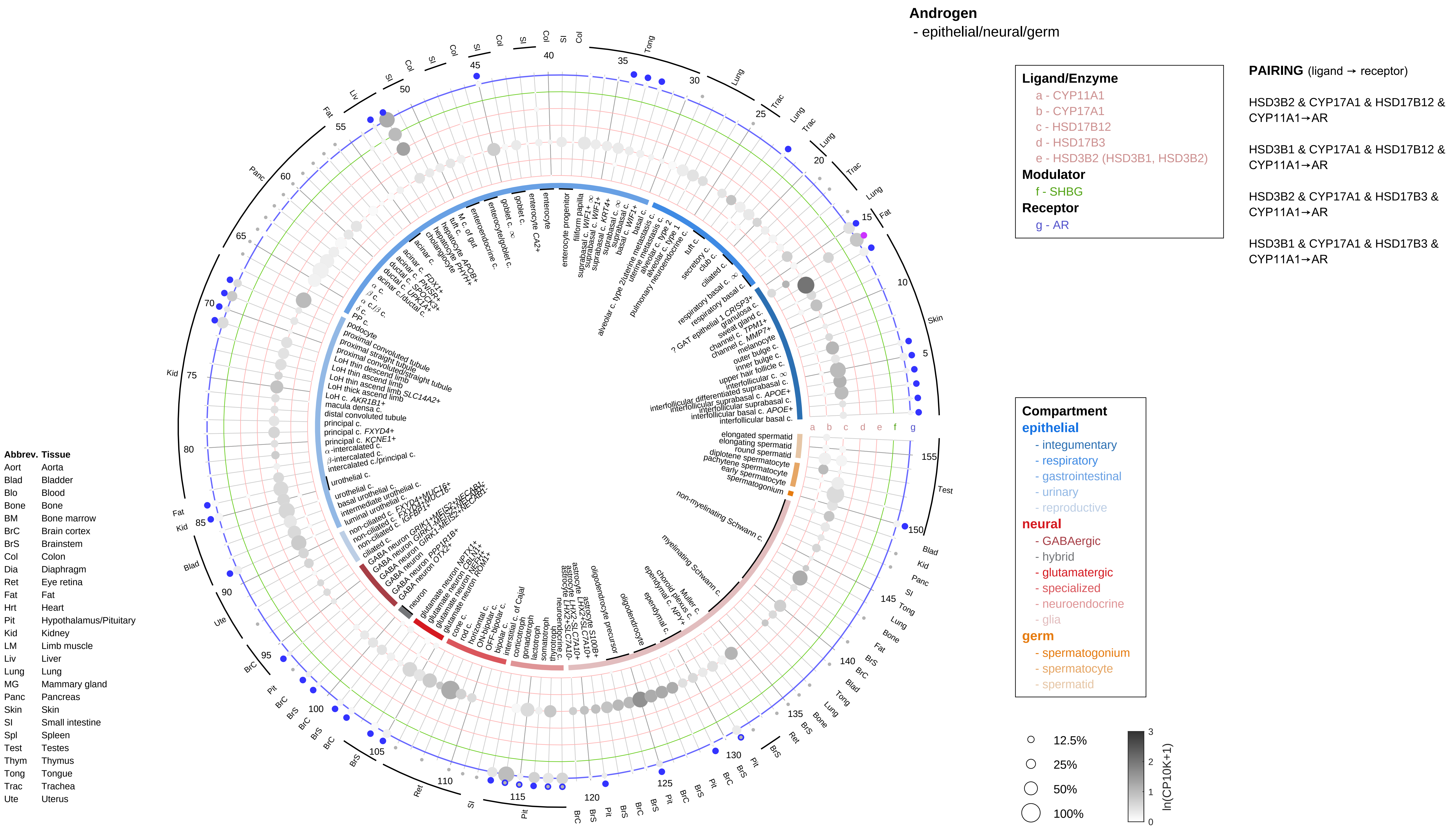
Androgen
- epithelial/neural/germ
Ligand/Enzyme
a - CYP11A1
b - CYP17A1
c - HSD17B12
d - HSD17B3
e - HSD3B2 (HSD3B1, HSD3B2)
Modulator
f - SHBG
Receptor
g - AR
PAIRING (ligand → receptor)
HSD3B2 & CYP17A1 & HSD17B12 & CYP11A1→AR
HSD3B1 & CYP17A1 & HSD17B12 & CYP11A1→AR
HSD3B2 & CYP17A1 & HSD17B3 & CYP11A1→AR
HSD3B1 & CYP17A1 & HSD17B3 & CYP11A1→AR
Compartment
epithelial
- integumentary
- respiratory
- gastrointestinal
- urinary
- reproductive
neural
- GABAergic
- hybrid
- glutamatergic
- specialized
- neuroendocrine
- glia
germ
- spermatogonium
- spermatocyte
- spermatid
12.5%
25%
50%
100%
ln(CP10K+1)
Abbrev. Tissue
Aort Aorta
Blad Bladder
Blo Blood
Bone Bone
BM Bone marrow
BrC Brain cortex
BrS Brainstem
Col Colon
Dia Diaphragm
Ret Eye retina
Fat Fat
Hrt Heart
Pit Hypothalamus/Pituitary
Kid Kidney
LM Limb muscle
Liv Liver
Lung Lung
MG Mammary gland
Panc Pancreas
Skin Skin
SI Small intestine
Spl Spleen
Test Testes
Thym Thymus
Tong Tongue
Trac Trachea
Ute Uterus

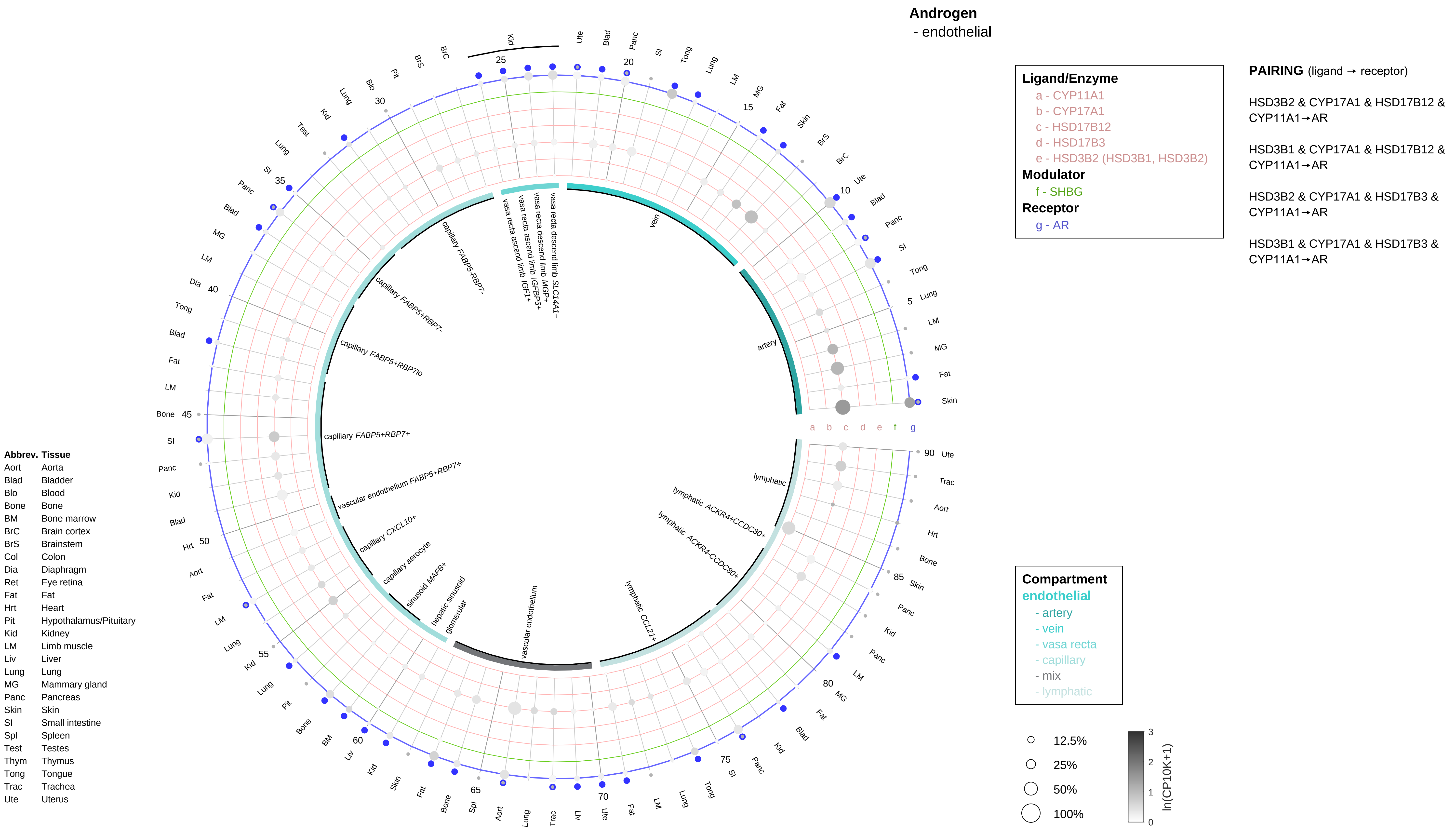
Androgen
- endothelial
Ligand/Enzyme
a - CYP11A1
b - CYP17A1
c - HSD17B12
d - HSD17B3
e - HSD3B2 (HSD3B1, HSD3B2)
Modulator
f - SHBG
Receptor
g - AR
PAIRING (ligand → receptor)
HSD3B2 & CYP17A1 & HSD17B12 & CYP11A1→AR
HSD3B1 & CYP17A1 & HSD17B12 & CYP11A1→AR
HSD3B2 & CYP17A1 & HSD17B3 & CYP11A1→AR
HSD3B1 & CYP17A1 & HSD17B3 & CYP11A1→AR
Compartment
endothelial
- artery
- vein
- vasa recta
- capillary
- mix
- lymphatic
12.5%
25%
50%
100%
ln(CP10K+1)
a b c d e f g
artery
vein
vasa recta descend limb SLC14A1+
vasa recta descend limb MGP+
vasa recta ascend limb IGFBP5+
vasa recta ascend limb IGF1+
capillary FABP5-RBP7-
capillary FABP5+RBP7-
capillary FABP5+RBP7lo
capillary FABP5+RBP7+
vascular endothelium FABP5+RBP7+
capillary CXCL10+
capillary aerocyte
sinusoid MAFB+
hepatic sinusoid
glomerular
vascular endothelium
lymphatic CCL21+
lymphatic ACKR4-CCDC80+
lymphatic ACKR4+CCDC80+
lymphatic
Abbrev. Tissue
Aort Aorta
Blad Bladder
Blo Blood
Bone Bone
BM Bone marrow
BrC Brain cortex
BrS Brainstem
Col Colon
Dia Diaphragm
Ret Eye retina
Fat Fat
Hrt Heart
Pit Hypothalamus/Pituitary
Kid Kidney
LM Limb muscle
Liv Liver
Lung Lung
MG Mammary gland
Panc Pancreas
Skin Skin
SI Small intestine
Spl Spleen
Test Testes
Thym Thymus
Tong Tongue
Trac Trachea
Ute Uterus

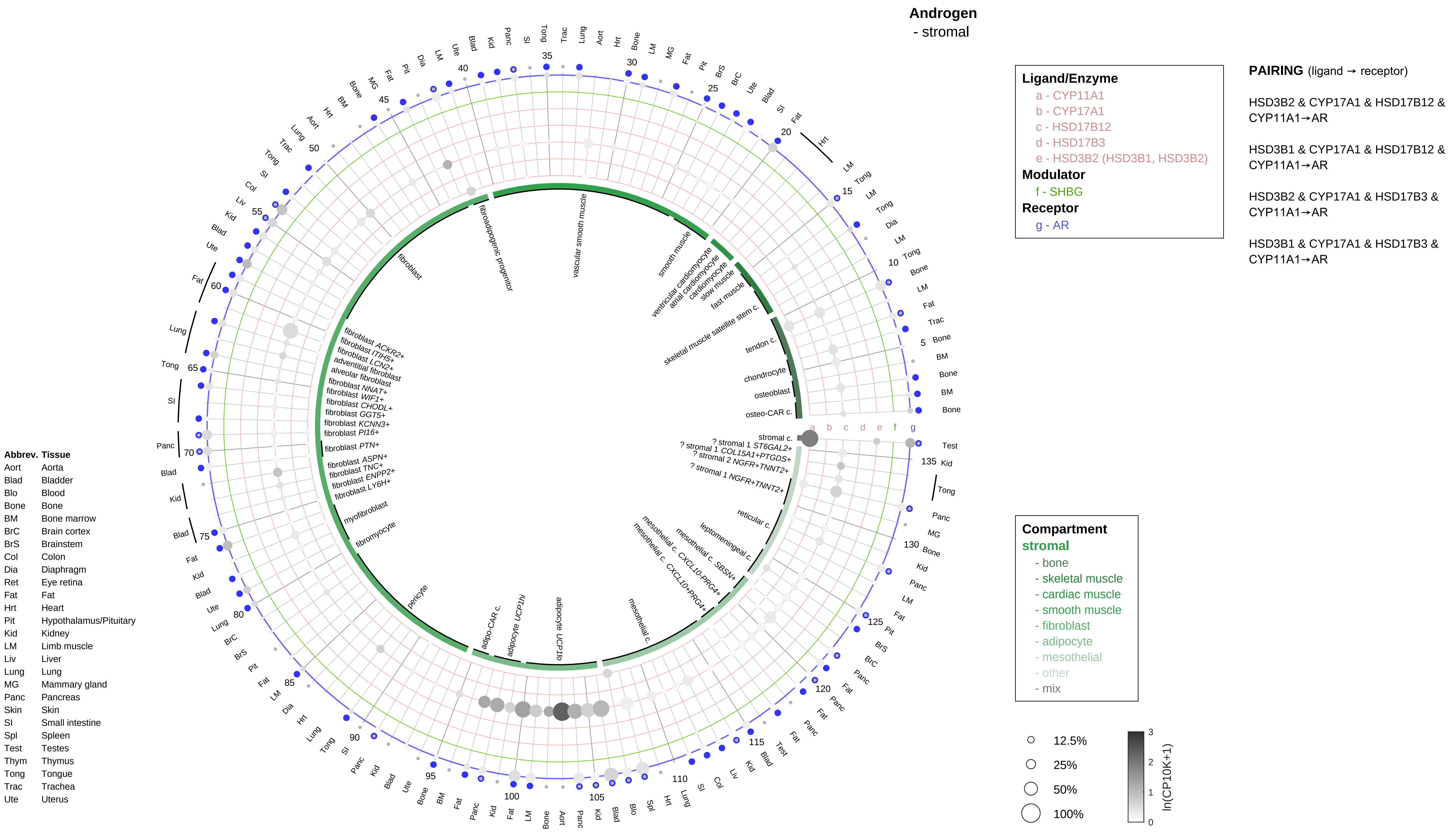

Androgen
- stromal
Ligand/Enzyme
a - CYP11A1
b - CYP17A1
c - HSD17B12
d - HSD17B3
e - HSD3B2 (HSD3B1, HSD3B2)
Modulator
f - SHBG
Receptor
g - AR
PAIRING (ligand → receptor)
HSD3B2 & CYP17A1 & HSD17B12 & CYP11A1→AR
HSD3B1 & CYP17A1 & HSD17B12 & CYP11A1→AR
HSD3B2 & CYP17A1 & HSD17B3 & CYP11A1→AR
HSD3B1 & CYP17A1 & HSD17B3 & CYP11A1→AR
Compartment
stromal
- bone
- skeletal muscle
- cardiac muscle
- smooth muscle
- fibroblast
- adipocyte
- mesothelial
- other
- mix
12.5%
25%
50%
100%
ln(CP10K+1)
Abbrev. Tissue
Aort Aorta
Blad Bladder
Blo Blood
Bone Bone
BM Bone marrow
BrC Brain cortex
BrS Brainstem
Col Colon
Dia Diaphragm
Ret Eye retina
Fat Fat
Hrt Heart
Pit Hypothalamus/Pituitary
Kid Kidney
LM Limb muscle
Liv Liver
Lung Lung
MG Mammary gland
Panc Pancreas
Skin Skin
SI Small intestine
Spl Spleen
Test Testes
Thym Thymus
Tong Tongue
Trac Trachea
Ute Uterus

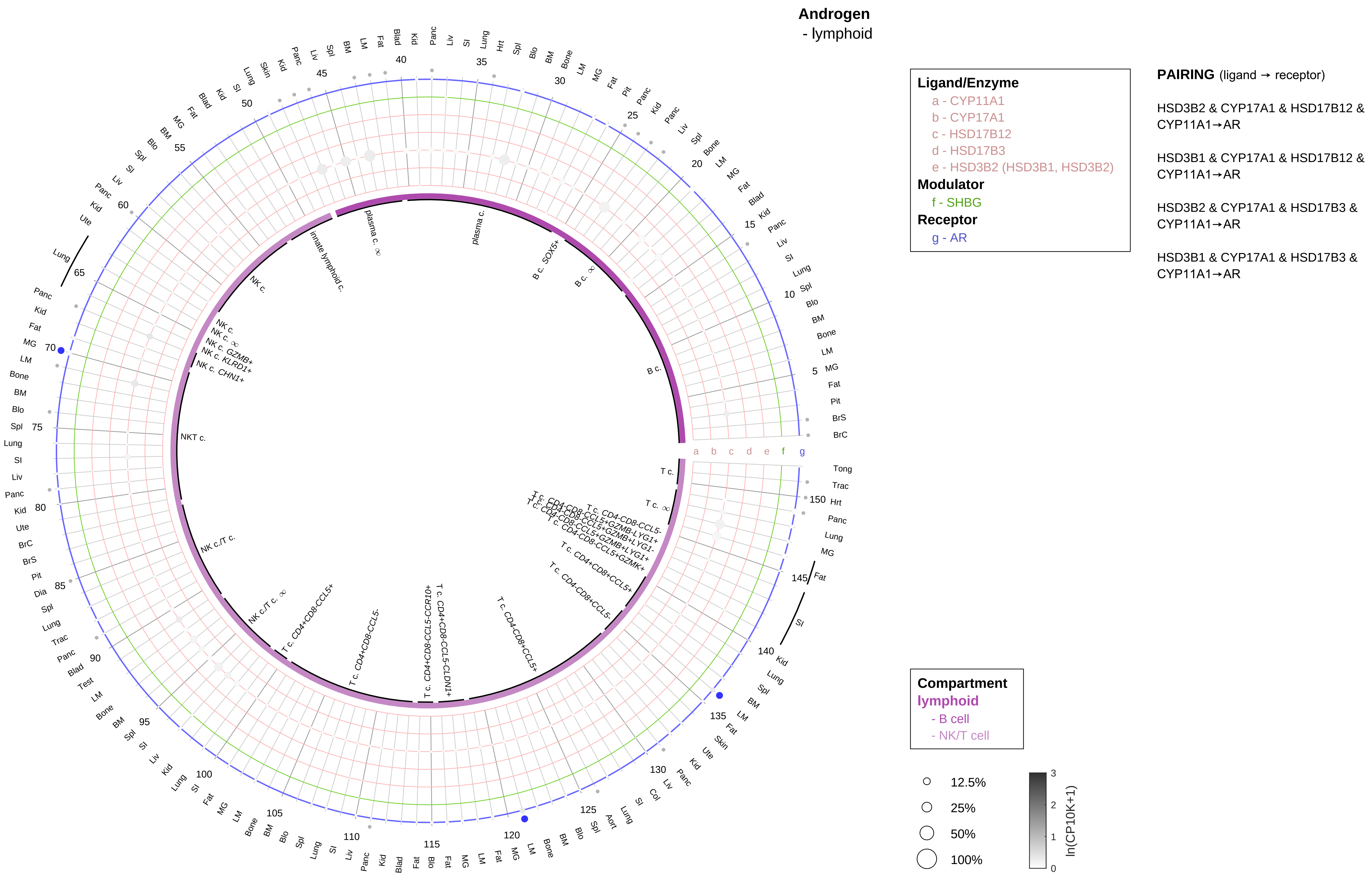

| Abbrev. | Tissue |
|---|---|
| Aort | Aorta |
| Blad | Bladder |
| Blo | Blood |
| Bone | Bone |
| BM | Bone marrow |
| BrC | Brain cortex |
| BrS | Brainstem |
| Col | Colon |
| Dia | Diaphragm |
| Ret | Eye retina |
| Fat | Fat |
| Hrt | Heart |
| Pit | Hypothalamus/Pituitary |
| Kid | Kidney |
| LM | Limb muscle |
| Liv | Liver |
| Lung | Lung |
| MG | Mammary gland |
| Panc | Pancreas |
| Skin | Skin |
| SI | Small intestine |
| Spl | Spleen |
| Test | Testes |
| Thym | Thymus |
| Tong | Tongue |
| Trac | Trachea |
| Ute | Uterus |

| Abbrev. | Tissue |
|---|---|
| Aort | Aorta |
| Blad | Bladder |
| Blo | Blood |
| Bone | Bone |
| BM | Bone marrow |
| BrC | Brain cortex |
| BrS | Brainstem |
| Col | Colon |
| Dia | Diaphragm |
| Ret | Eye retina |
| Fat | Fat |
| Hrt | Heart |
| Pit | Hypothalamus/Pituitary |
| Kid | Kidney |
| LM | Limb muscle |
| Liv | Liver |
| Lung | Lung |
| MG | Mammary gland |
| Panc | Pancreas |
| Skin | Skin |
| SI | Small intestine |
| Spl | Spleen |
| Test | Testes |
| Thym | Thymus |
| Tong | Tongue |
| Trac | Trachea |
| Ute | Uterus |

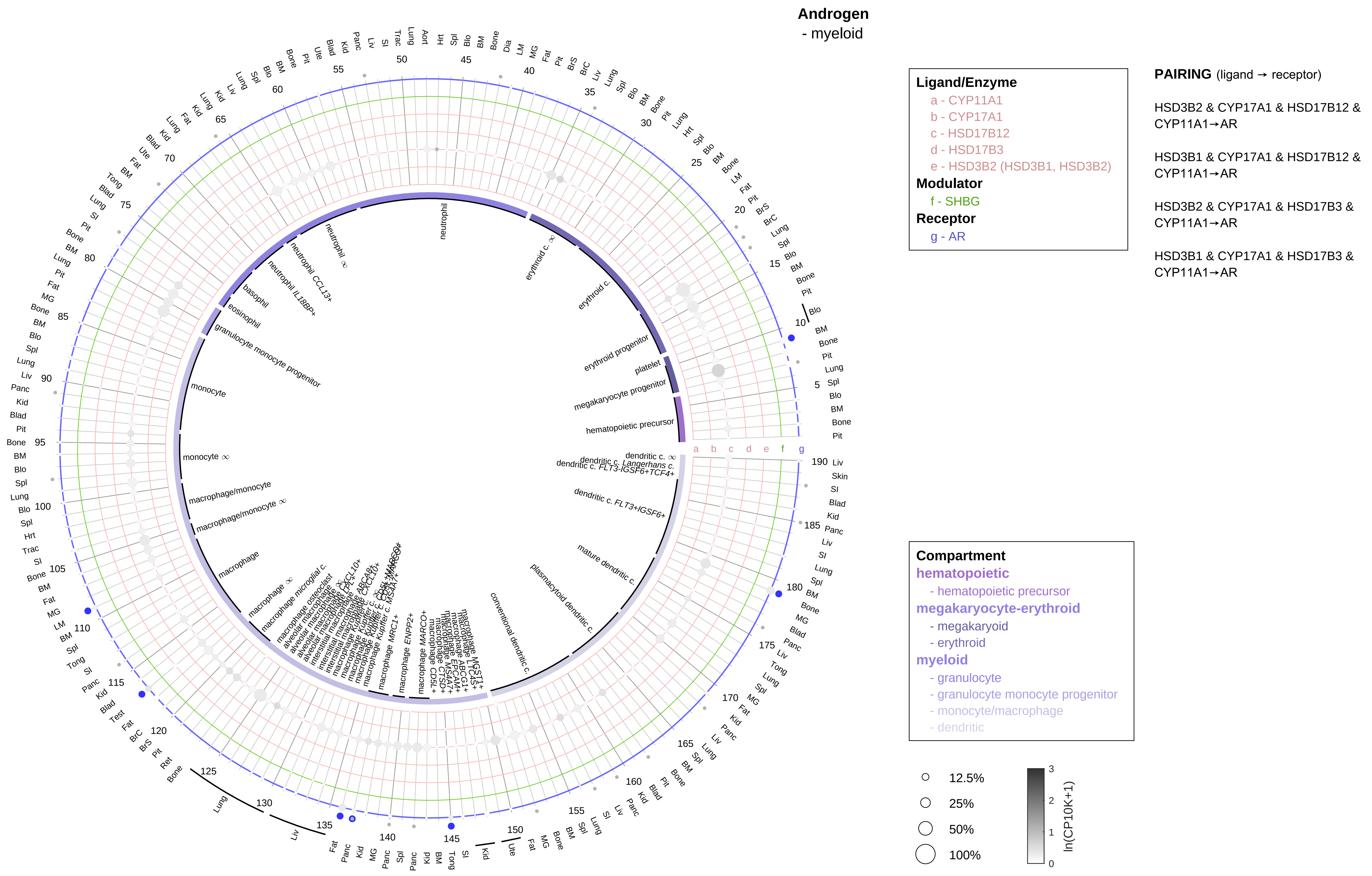

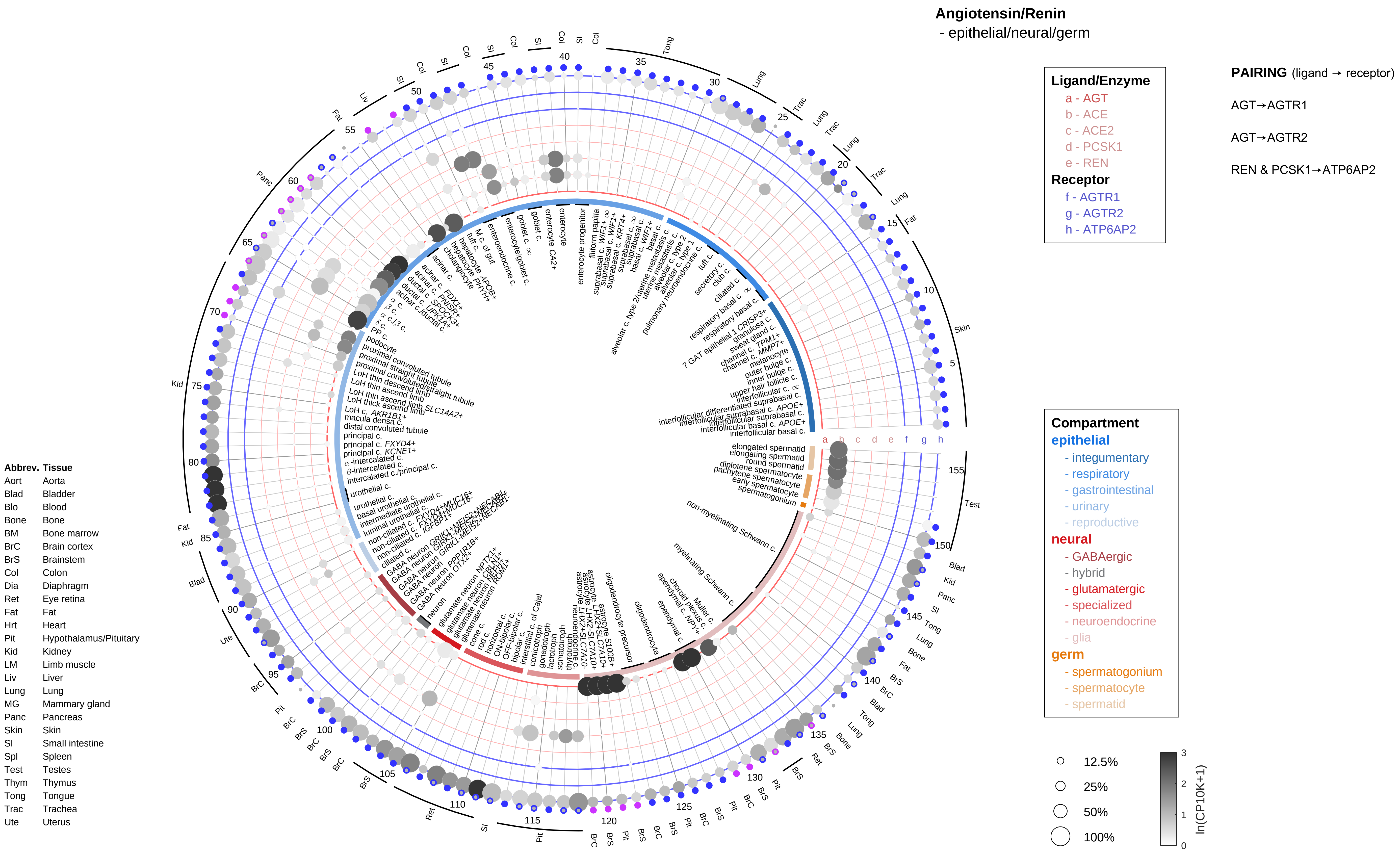

Angiotensin/Renin
- epithelial/neural/germ
Ligand/Enzyme
a - AGT
b - ACE
c - ACE2
d - PCSK1
e - REN
Receptor
f - AGTR1
g - AGTR2
h - ATP6AP2
PAIRING (ligand → receptor)
AGT→AGTR1
AGT→AGTR2
REN & PCSK1→ATP6AP2
Compartment
epithelial
- integumentary
- respiratory
- gastrointestinal
- urinary
- reproductive
neural
- GABAergic
- hybrid
- glutamatergic
- specialized
- neuroendocrine
- glia
germ
- spermatogonium
- spermatocyte
- spermatid
12.5%
25%
50%
100%
ln(CP10K+1)
Abbrev. Tissue
Aort Aorta
Blad Bladder
Blo Blood
Bone Bone
BM Bone marrow
BrC Brain cortex
BrS Brainstem
Col Colon
Dia Diaphragm
Ret Eye retina
Fat Fat
Hrt Heart
Pit Hypothalamus/Pituitary
Kid Kidney
LM Limb muscle
Liv Liver
Lung Lung
MG Mammary gland
Panc Pancreas
Skin Skin
SI Small intestine
Spl Spleen
Test Testes
Thym Thymus
Tong Tongue
Trac Trachea
Ute Uterus

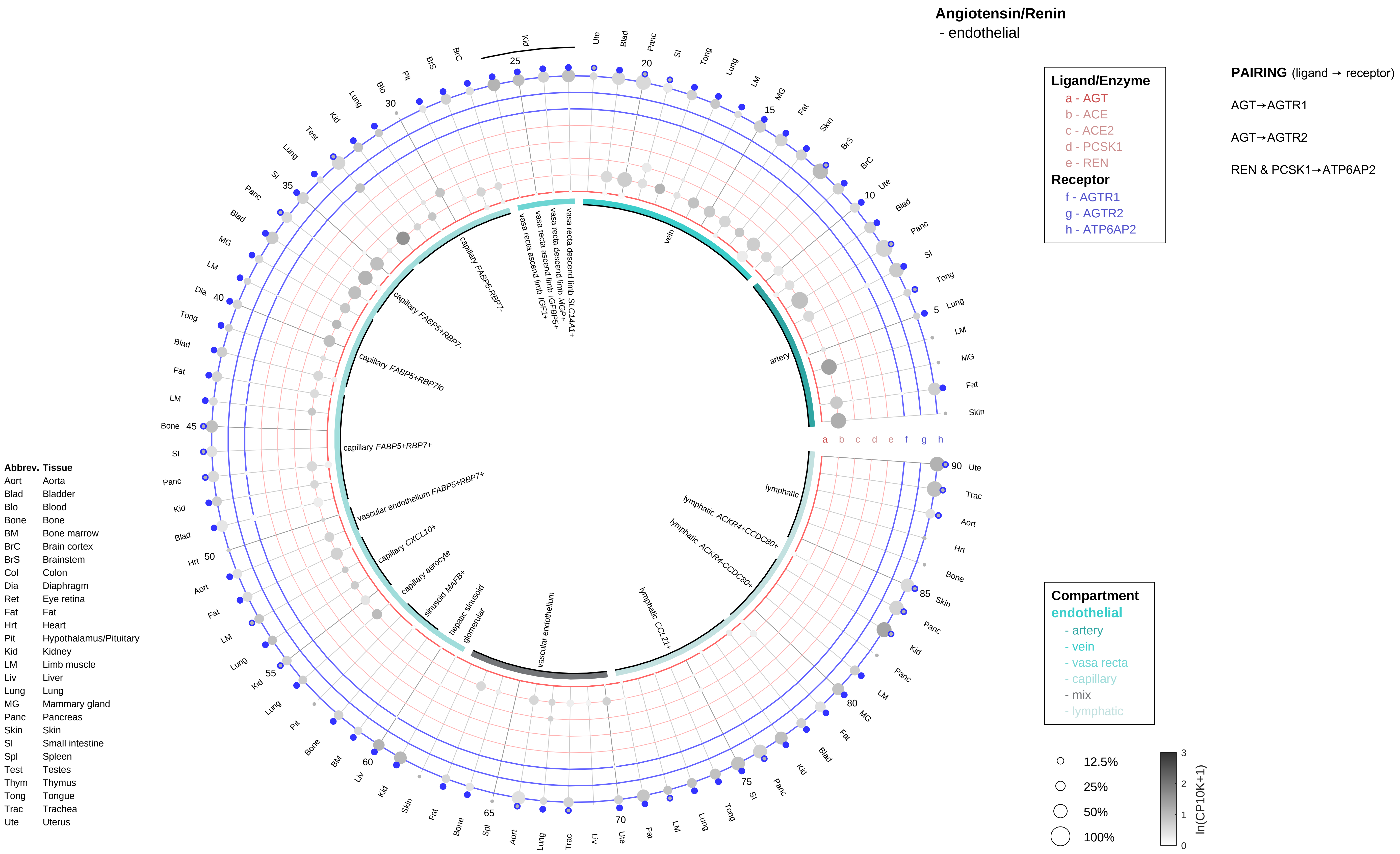
Angiotensin/Renin
- endothelial
Ligand/Enzyme
a - AGT
b - ACE
c - ACE2
d - PCSK1
e - REN
Receptor
f - AGTR1
g - AGTR2
h - ATP6AP2
PAIRING (ligand → receptor)
AGT→AGTR1
AGT→AGTR2
REN & PCSK1→ATP6AP2
Compartment
endothelial
- artery
- vein
- vasa recta
- capillary
- mix
- lymphatic
12.5%
25%
50%
100%
ln(CP10K+1)
Abbrev. Tissue
Aort Aorta
Blad Bladder
Blo Blood
Bone Bone
BM Bone marrow
BrC Brain cortex
BrS Brainstem
Col Colon
Dia Diaphragm
Ret Eye retina
Fat Fat
Hrt Heart
Pit Hypothalamus/Pituitary
Kid Kidney
LM Limb muscle
Liv Liver
Lung Lung
MG Mammary gland
Panc Pancreas
Skin Skin
SI Small intestine
Spl Spleen
Test Testes
Thym Thymus
Tong Tongue
Trac Trachea
Ute Uterus
vein
artery
vasa recta descend limb SLC14A1+
vasa recta descend limb MGP+
vasa recta ascend limb IGFBP5+
vasa recta ascend limb IGF1+
capillary FABP5-RBP7-
capillary FABP5+RBP7-
capillary FABP5+RBP7lo
capillary FABP5+RBP7+
vascular endothelium FABP5+RBP7+
capillary CXCL10+
capillary aerocyte
sinusoid MAFB+
hepatic sinusoid
glomerular
vascular endothelium
lymphatic CCL21+
lymphatic ACKR4-CCDC80+
lymphatic ACKR4+CCDC80+
lymphatic

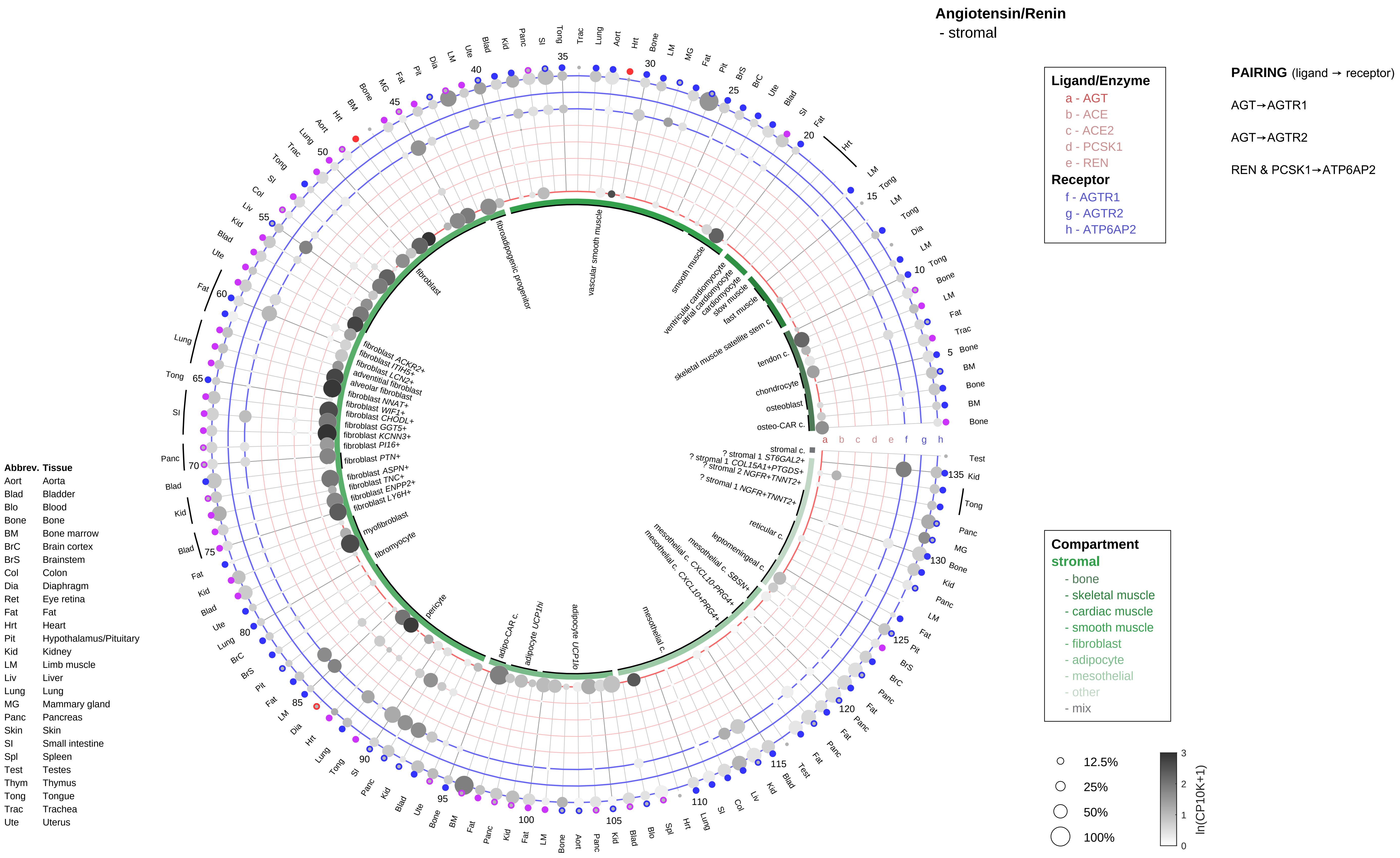

Angiotensin/Renin
- stromal
Ligand/Enzyme
a - AGT
b - ACE
c - ACE2
d - PCSK1
e - REN
Receptor
f - AGTR1
g - AGTR2
h - ATP6AP2
PAIRING (ligand → receptor)
AGT→AGTR1
AGT→AGTR2
REN & PCSK1→ATP6AP2
Compartment
stromal
- bone
- skeletal muscle
- cardiac muscle
- smooth muscle
- fibroblast
- adipocyte
- mesothelial
- other
- mix
12.5%
25%
50%
100%
ln(CP10K+1)
Abbrev. Tissue
Aort Aorta
Blad Bladder
Blo Blood
Bone Bone
BM Bone marrow
BrC Brain cortex
BrS Brainstem
Col Colon
Dia Diaphragm
Ret Eye retina
Fat Fat
Hrt Heart
Pit Hypothalamus/Pituitary
Kid Kidney
LM Limb muscle
Liv Liver
Lung Lung
MG Mammary gland
Panc Pancreas
Skin Skin
SI Small intestine
Spl Spleen
Test Testes
Thym Thymus
Tong Tongue
Trac Trachea
Ute Uterus

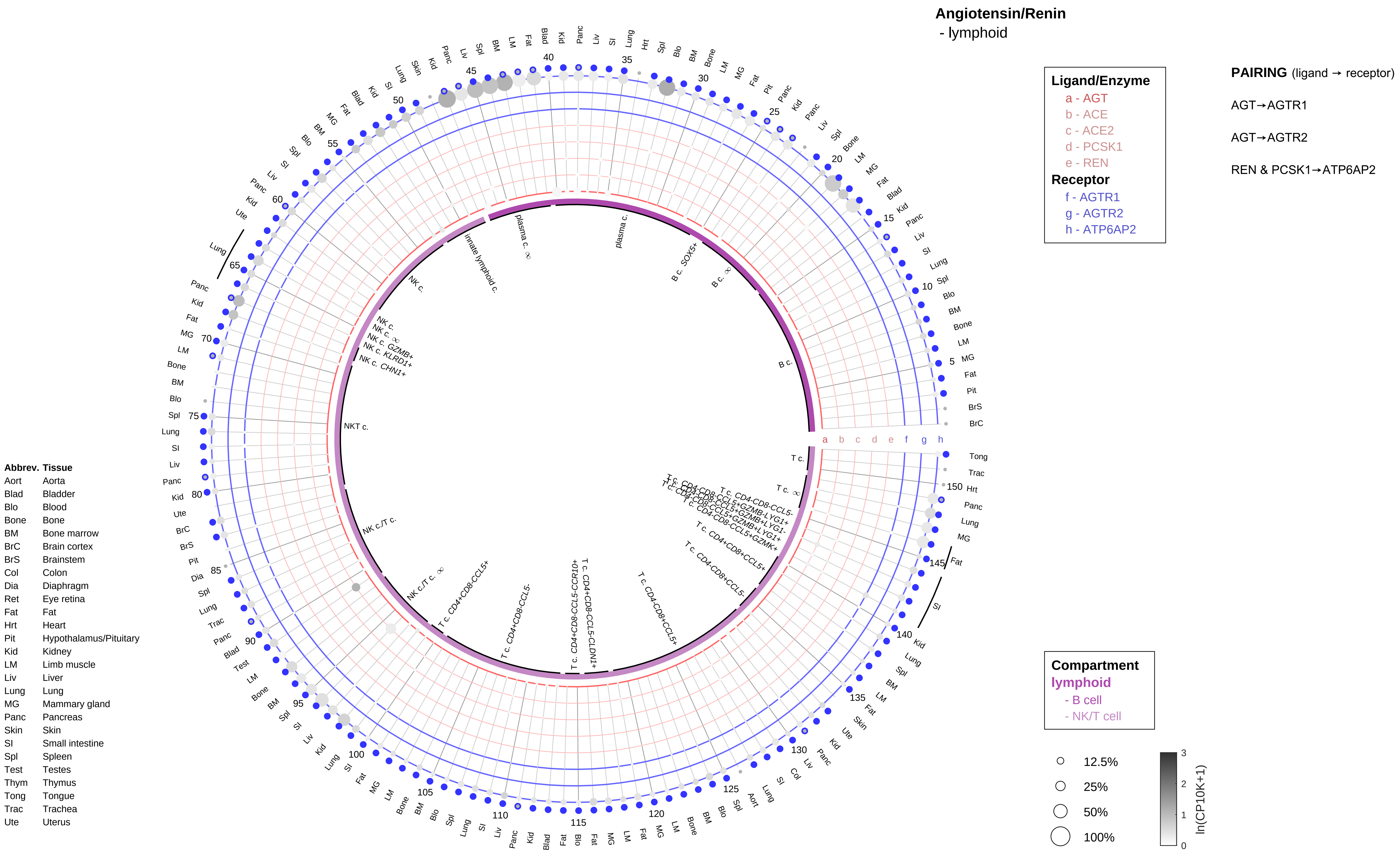

Angiotensin/Renin
- lymphoid
Ligand/Enzyme
a - AGT
b - ACE
c - ACE2
d - PCSK1
e - REN
Receptor
f - AGTR1
g - AGTR2
h - ATP6AP2
PAIRING (ligand → receptor)
AGT→AGTR1
AGT→AGTR2
REN & PCSK1→ATP6AP2
Compartment
lymphoid
- B cell
- NK/T cell
12.5%
25%
50%
100%
ln(CP10K+1)
Abbrev. Tissue
Aort Aorta
Blad Bladder
Blo Blood
Bone Bone
BM Bone marrow
BrC Brain cortex
BrS Brainstem
Col Colon
Dia Diaphragm
Ret Eye retina
Fat Fat
Hrt Heart
Pit Hypothalamus/Pituitary
Kid Kidney
LM Limb muscle
Liv Liver
Lung Lung
MG Mammary gland
Panc Pancreas
Skin Skin
SI Small intestine
Spl Spleen
Test Testes
Thym Thymus
Tong Tongue
Trac Trachea
Ute Uterus

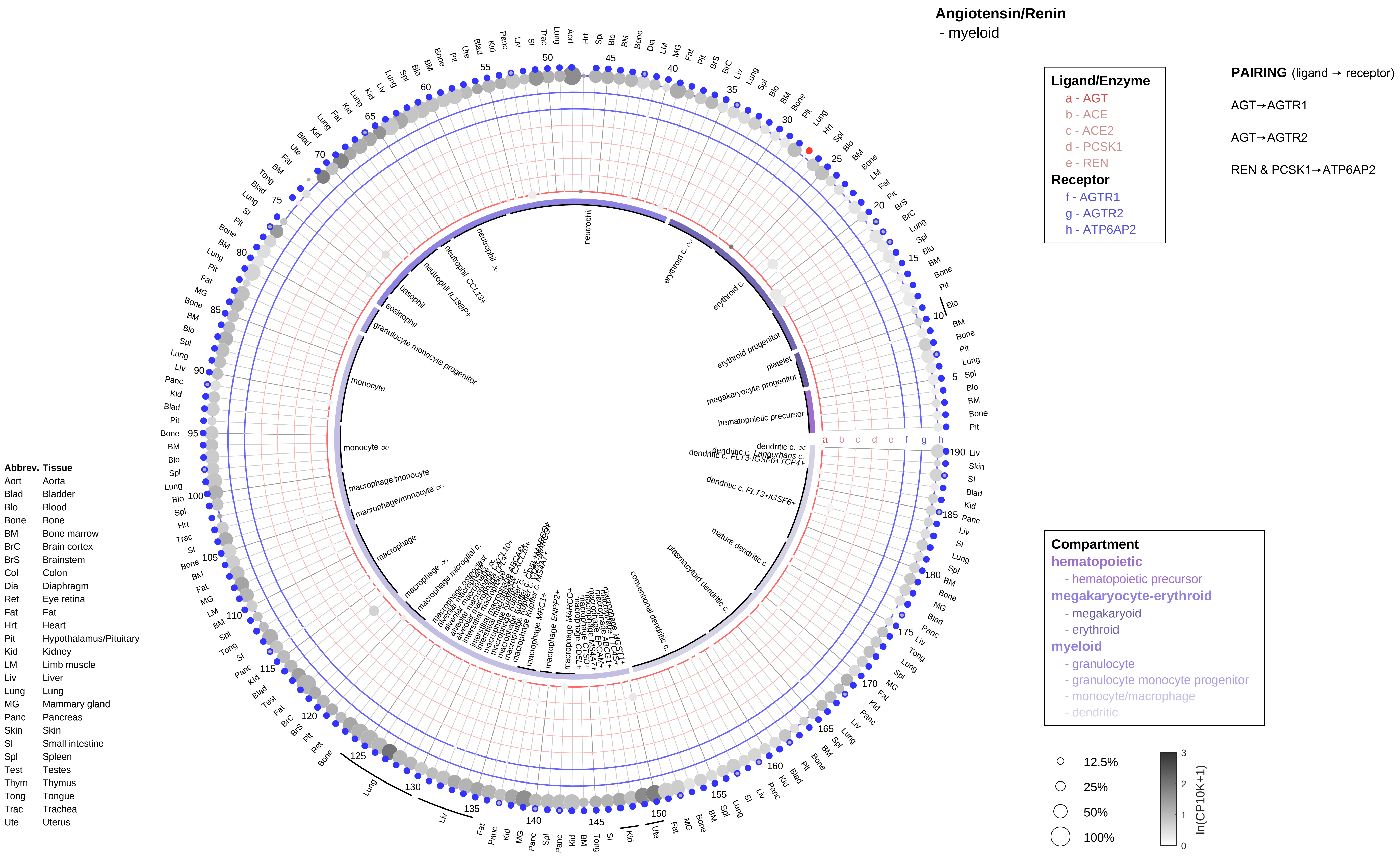
Angiotensin/Renin
- myeloid
Ligand/Enzyme
a - AGT
b - ACE
c - ACE2
d - PCSK1
e - REN
Receptor
f - AGTR1
g - AGTR2
h - ATP6AP2
PAIRING (ligand → receptor)
AGT→AGTR1
AGT→AGTR2
REN & PCSK1→ATP6AP2
Compartment
hematopoietic
- hematopoietic precursor
megakaryocyte-erythroid
- megakaryoid
- erythroid
myeloid
- granulocyte
- granulocyte monocyte progenitor
- monocyte/macrophage
- dendritic
12.5%
25%
50%
100%
ln(CP10K+1)
Abbrev. Tissue
Aort Aorta
Blad Bladder
Blo Blood
Bone Bone
BM Bone marrow
BrC Brain cortex
BrS Brainstem
Col Colon
Dia Diaphragm
Ret Eye retina
Fat Fat
Hrt Heart
Pit Hypothalamus/Pituitary
Kid Kidney
LM Limb muscle
Liv Liver
Lung Lung
MG Mammary gland
Panc Pancreas
Skin Skin
SI Small intestine
Spl Spleen
Test Testes
Thym Thymus
Tong Tongue
Trac Trachea
Ute Uterus
neutrophil
erythroid c.
erythroid progenitor
platelet
megakaryocyte progenitor
hematopoietic precursor
basophil
eosinophil
granulocyte monocyte progenitor
monocyte
macrophage/monocyte
macrophage
conventional dendritic c.
plasmacytoid dendritic c.
mature dendritic c.
dendritic c. FLT3+IGSF6+

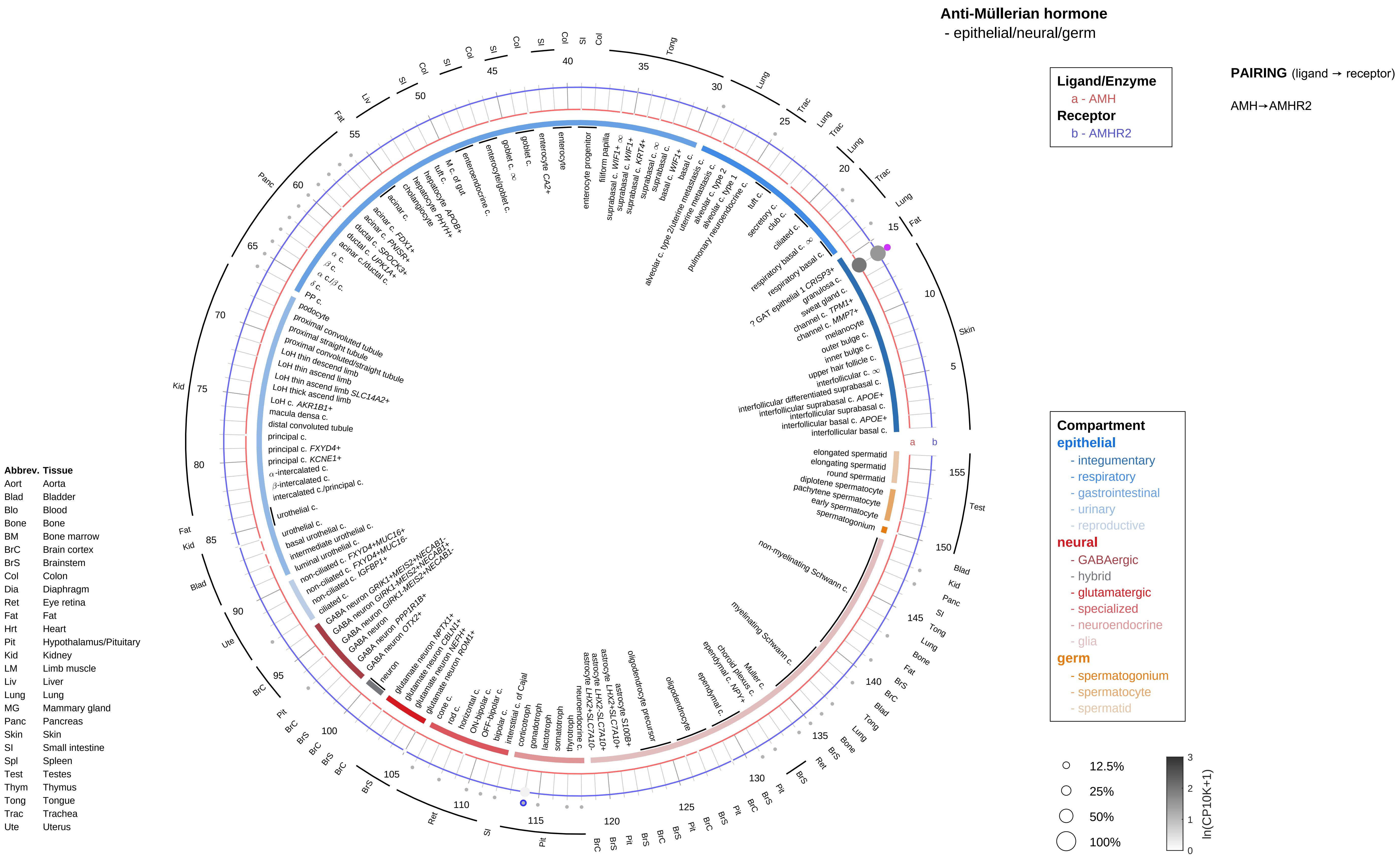
Anti-Müllerian hormone
- epithelial/neural/germ
Ligand/Enzyme
a - AMH
Receptor
b - AMHR2
PAIRING (ligand → receptor)
AMH→AMHR2
Compartment
epithelial
- integumentary
- respiratory
- gastrointestinal
- urinary
- reproductive
neural
- GABAergic
- hybrid
- glutamatergic
- specialized
- neuroendocrine
- glia
germ
- spermatogonium
- spermatocyte
- spermatid
12.5%
25%
50%
100%
ln(CP10K+1)
Abbrev. Tissue
Aort Aorta
Blad Bladder
Blo Blood
Bone Bone
BM Bone marrow
BrC Brain cortex
BrS Brainstem
Col Colon
Dia Diaphragm
Ret Eye retina
Fat Fat
Hrt Heart
Pit Hypothalamus/Pituitary
Kid Kidney
LM Limb muscle
Liv Liver
Lung Lung
MG Mammary gland
Panc Pancreas
Skin Skin
SI Small intestine
Spl Spleen
Test Testes
Thym Thymus
Tong Tongue
Trac Trachea
Ute Uterus

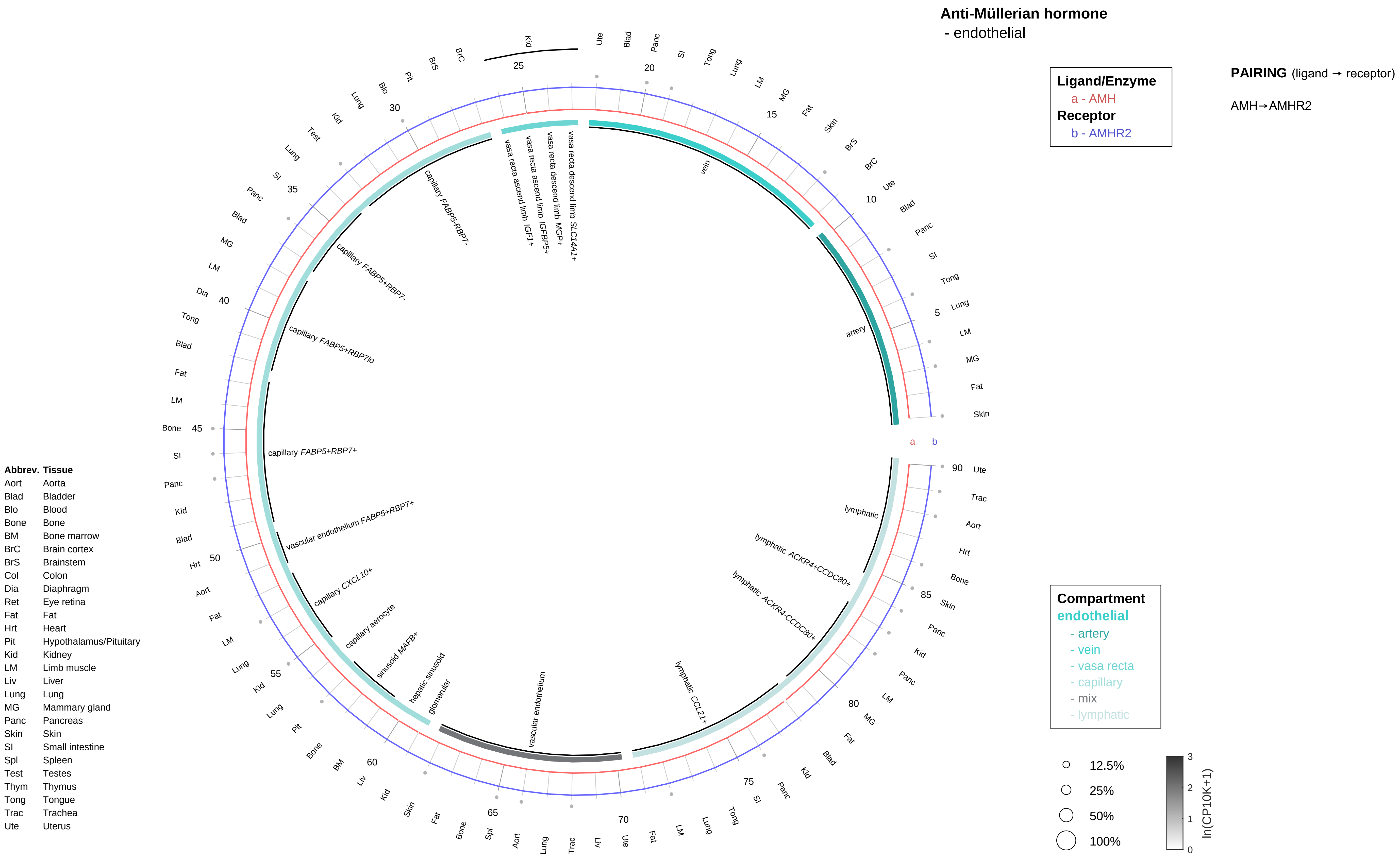
Anti-Müllerian hormone
- endothelial
Ligand/Enzyme
a - AMH
Receptor
b - AMHR2
PAIRING (ligand → receptor)
AMH→AMHR2
Compartment
endothelial
- artery
- vein
- vasa recta
- capillary
- mix
- lymphatic
12.5%
25%
50%
100%
ln(CP10K+1)
0
1
2
3
a
b
artery
vein
vasa recta descend limb SLC14A1+
vasa recta descend limb MGP+
vasa recta ascend limb IGFBP5+
vasa recta ascend limb IGF1+
capillary FABP5-RBP7-
capillary FABP5+RBP7-
capillary FABP5+RBP7lo
capillary FABP5+RBP7+
vascular endothelium FABP5+RBP7+
capillary CXCL10+
capillary aerocyte
sinusoid MAFB+
hepatic sinusoid
glomerular
vascular endothelium
lymphatic CCL21+
lymphatic ACKR4-CCDC80+
lymphatic ACKR4+CCDC80+
lymphatic
Abbrev. Tissue
Aort Aorta
Blad Bladder
Blo Blood
Bone Bone
BM Bone marrow
BrC Brain cortex
BrS Brainstem
Col Colon
Dia Diaphragm
Ret Eye retina
Fat Fat
Hrt Heart
Pit Hypothalamus/Pituitary
Kid Kidney
LM Limb muscle
Liv Liver
Lung Lung
MG Mammary gland
Panc Pancreas
Skin Skin
SI Small intestine
Spl Spleen
Test Testes
Thym Thymus
Tong Tongue
Trac Trachea
Ute Uterus

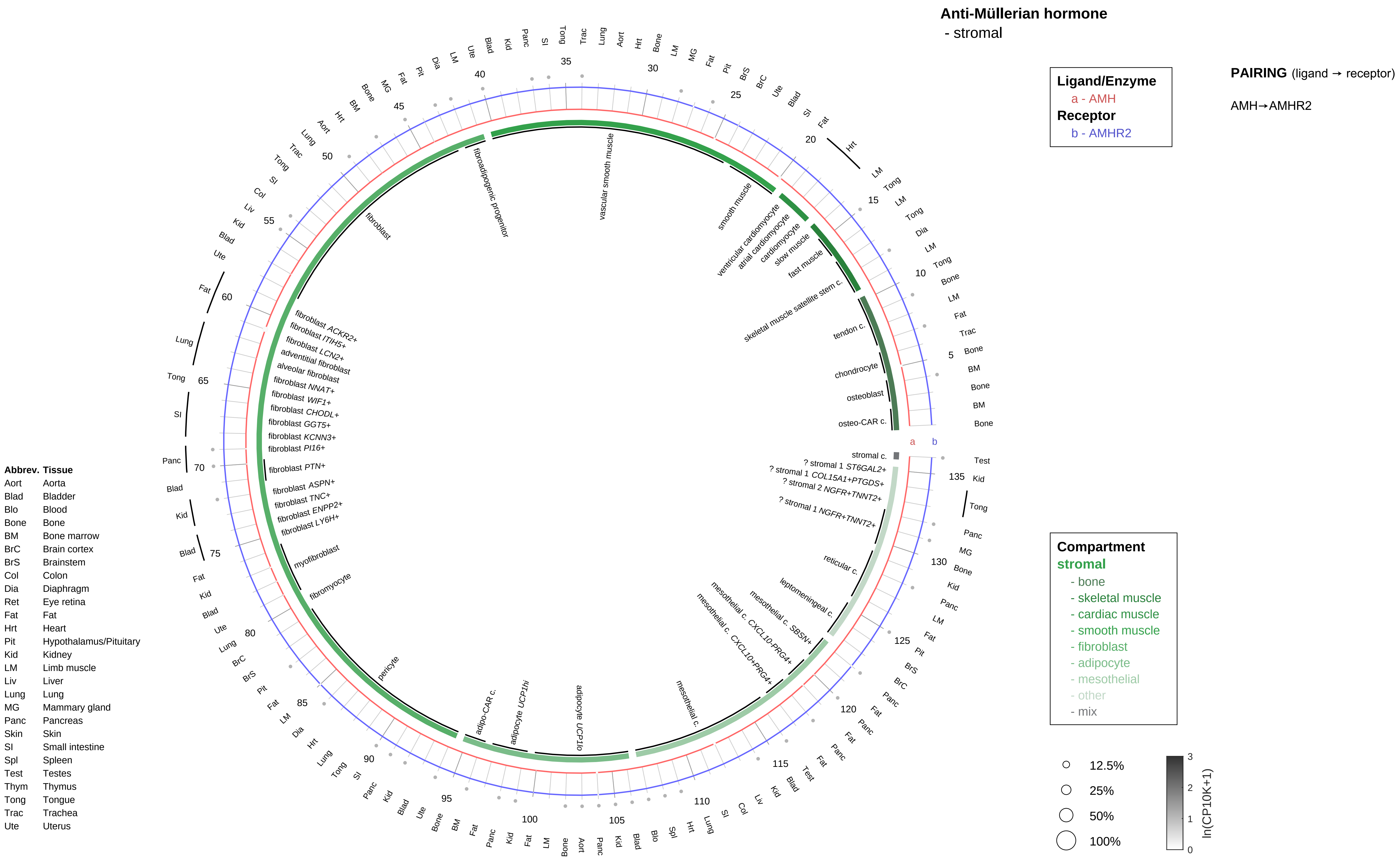

Anti-Müllerian hormone
- stromal
Ligand/Enzyme
a - AMH
Receptor
b - AMHR2
PAIRING (ligand → receptor)
AMH→AMHR2
Compartment
stromal
- bone
- skeletal muscle
- cardiac muscle
- smooth muscle
- fibroblast
- adipocyte
- mesothelial
- other
- mix
12.5%
25%
50%
100%
ln(CP10K+1)
Abbrev. Tissue
Aort Aorta
Blad Bladder
Blo Blood
Bone Bone
BM Bone marrow
BrC Brain cortex
BrS Brainstem
Col Colon
Dia Diaphragm
Ret Eye retina
Fat Fat
Hrt Heart
Pit Hypothalamus/Pituitary
Kid Kidney
LM Limb muscle
Liv Liver
Lung Lung
MG Mammary gland
Panc Pancreas
Skin Skin
SI Small intestine
Spl Spleen
Test Testes
Thym Thymus
Tong Tongue
Trac Trachea
Ute Uterus
vascular smooth muscle
smooth muscle
ventricular cardiomyocyte
atrial cardiomyocyte
cardiomyocyte
slow muscle
fast muscle
skeletal muscle satellite stem c.
tendon c.
chondrocyte
osteoblast
osteo-CAR c.
stromal c.
? stromal 1 ST6GAL2+
? stromal 1 COL15A1+PTGDS+
? stromal 2 NGFR+TNNT2+
? stromal 1 NGFR+TNNT2+
reticular c.
leptomeningeal c.
mesothelial c. SBSN+
mesothelial c. CXCL10-PRG4+
mesothelial c. CXCL10+PRG4+
mesothelial c.
adipocyte UCP1lo
adipocyte UCP1hi
adipo-CAR c.
pericyte
fibromyocyte
myofibroblast
fibroblast LY6H+
fibroblast ENPP2+
fibroblast TNC+
fibroblast ASPN+
fibroblast PTN+
fibroblast PI16+
fibroblast KCNN3+
fibroblast GGT5+
fibroblast CHODL+
fibroblast WIF1+
fibroblast NNAT+
alveolar fibroblast
adventitial fibroblast
fibroblast LCN2+
fibroblast ITIH5+
fibroblast ACKR2+
fibroblast
fibroadipogenic progenitor

**Anti-Müllerian hormone**
- lymphoid

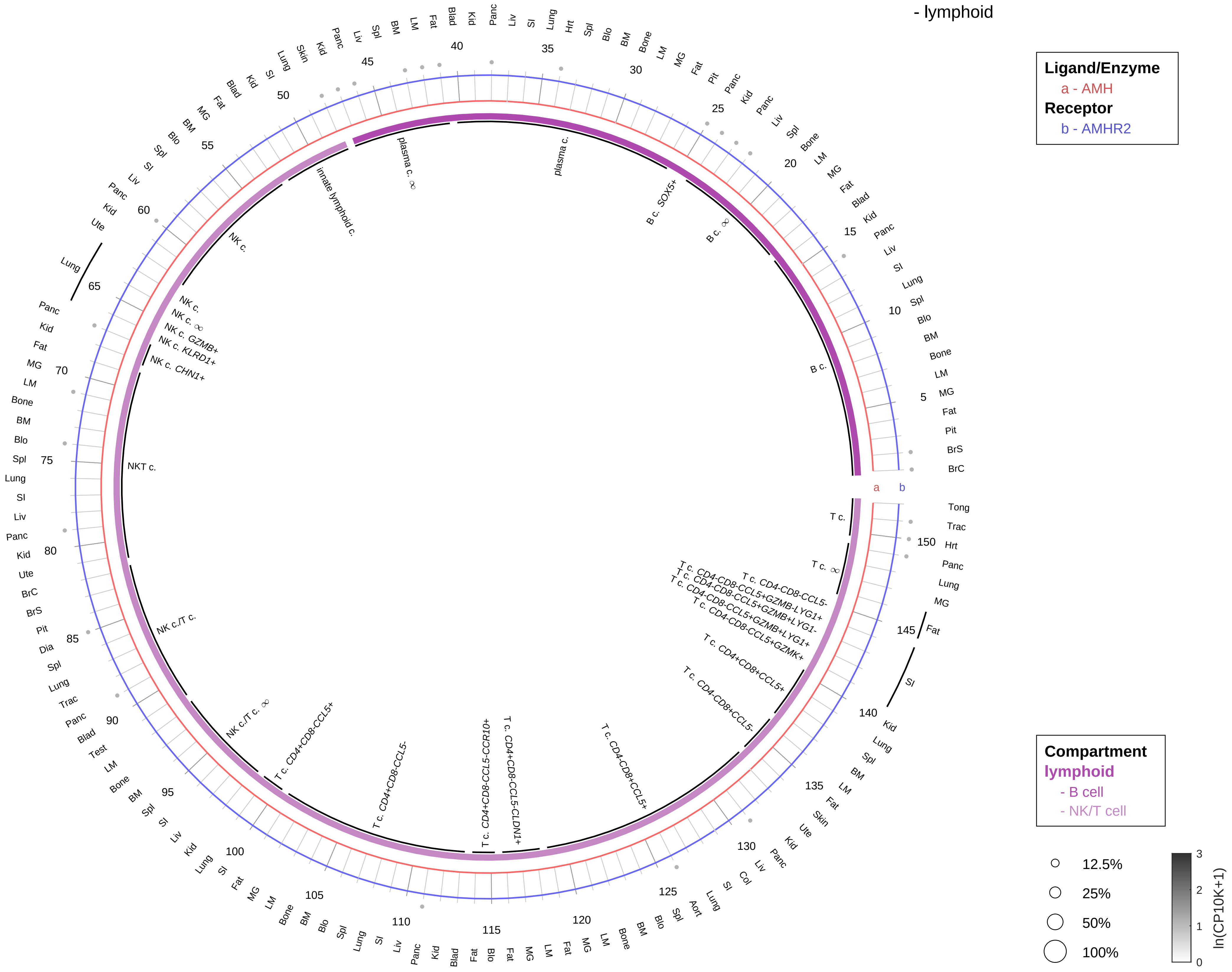

**PAIRING** (ligand → receptor)

AMH→AMHR2

| Abbrev. | Tissue |
|---|---|
| Aort | Aorta |
| Blad | Bladder |
| Blo | Blood |
| Bone | Bone |
| BM | Bone marrow |
| BrC | Brain cortex |
| BrS | Brainstem |
| Col | Colon |
| Dia | Diaphragm |
| Ret | Eye retina |
| Fat | Fat |
| Hrt | Heart |
| Pit | Hypothalamus/Pituitary |
| Kid | Kidney |
| LM | Limb muscle |
| Liv | Liver |
| Lung | Lung |
| MG | Mammary gland |
| Panc | Pancreas |
| Skin | Skin |
| SI | Small intestine |
| Spl | Spleen |
| Test | Testes |
| Thym | Thymus |
| Tong | Tongue |
| Trac | Trachea |
| Ute | Uterus |

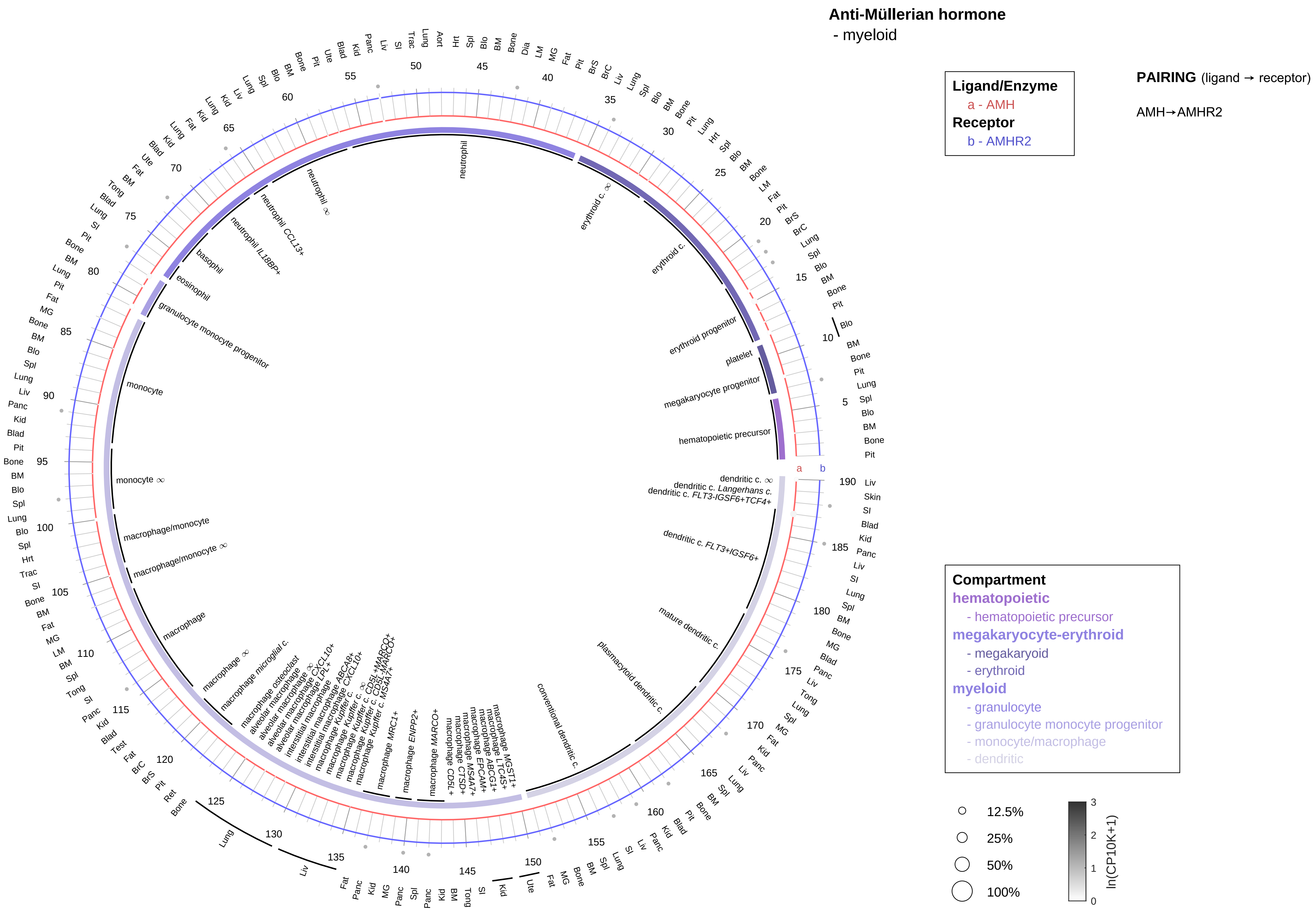

| Abbrev. | Tissue |
|---|---|
| Aort | Aorta |
| Blad | Bladder |
| Blo | Blood |
| Bone | Bone |
| BM | Bone marrow |
| BrC | Brain cortex |
| BrS | Brainstem |
| Col | Colon |
| Dia | Diaphragm |
| Ret | Eye retina |
| Fat | Fat |
| Hrt | Heart |
| Pit | Hypothalamus/Pituitary |
| Kid | Kidney |
| LM | Limb muscle |
| Liv | Liver |
| Lung | Lung |
| MG | Mammary gland |
| Panc | Pancreas |
| Skin | Skin |
| SI | Small intestine |
| Spl | Spleen |
| Test | Testes |
| Thym | Thymus |
| Tong | Tongue |
| Trac | Trachea |
| Ute | Uterus |

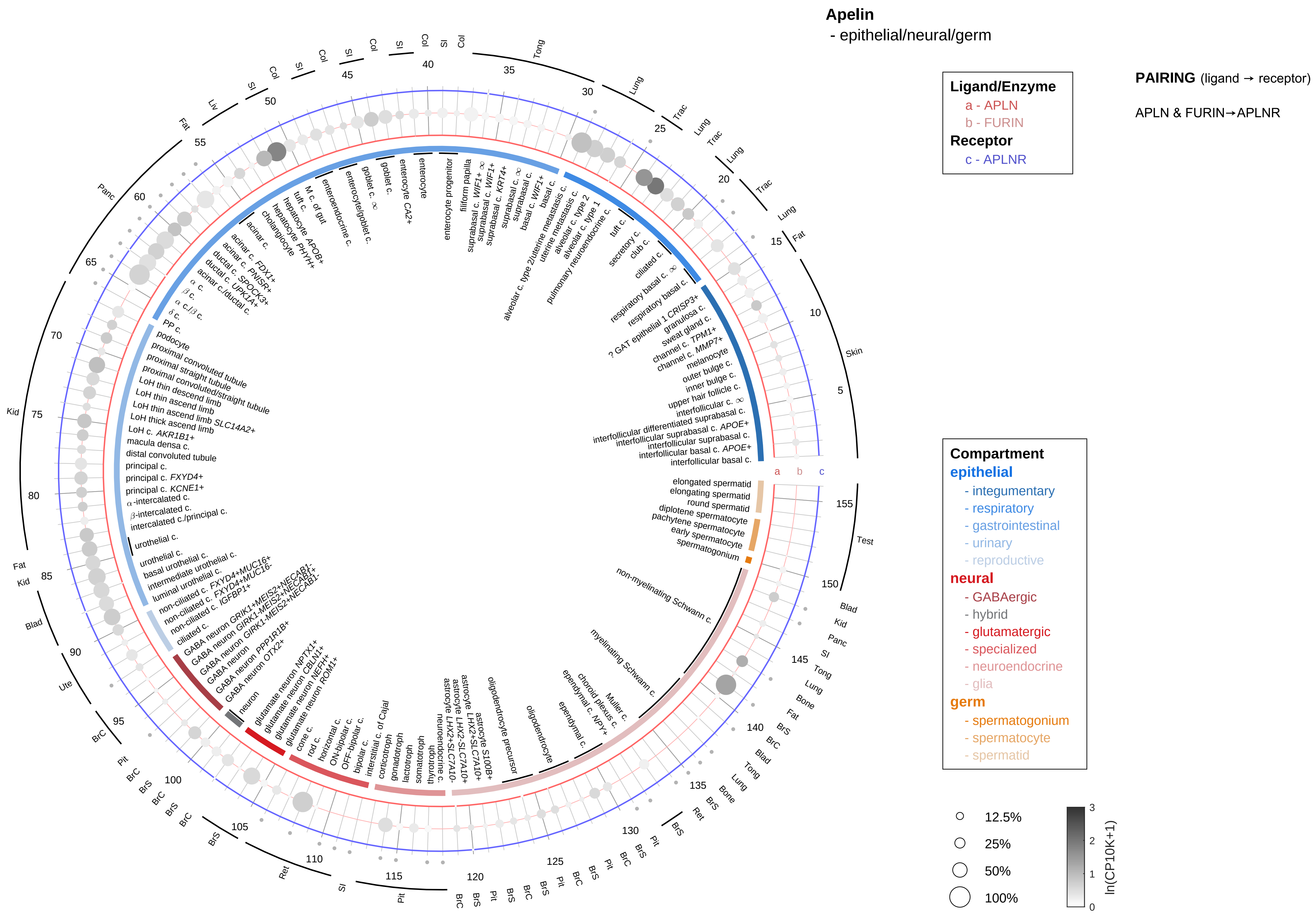

| Abbrev. | Tissue |
|---|---|
| Aort | Aorta |
| Blad | Bladder |
| Blo | Blood |
| Bone | Bone |
| BM | Bone marrow |
| BrC | Brain cortex |
| BrS | Brainstem |
| Col | Colon |
| Dia | Diaphragm |
| Ret | Eye retina |
| Fat | Fat |
| Hrt | Heart |
| Pit | Hypothalamus/Pituitary |
| Kid | Kidney |
| LM | Limb muscle |
| Liv | Liver |
| Lung | Lung |
| MG | Mammary gland |
| Panc | Pancreas |
| Skin | Skin |
| SI | Small intestine |
| Spl | Spleen |
| Test | Testes |
| Thym | Thymus |
| Tong | Tongue |
| Trac | Trachea |
| Ute | Uterus |

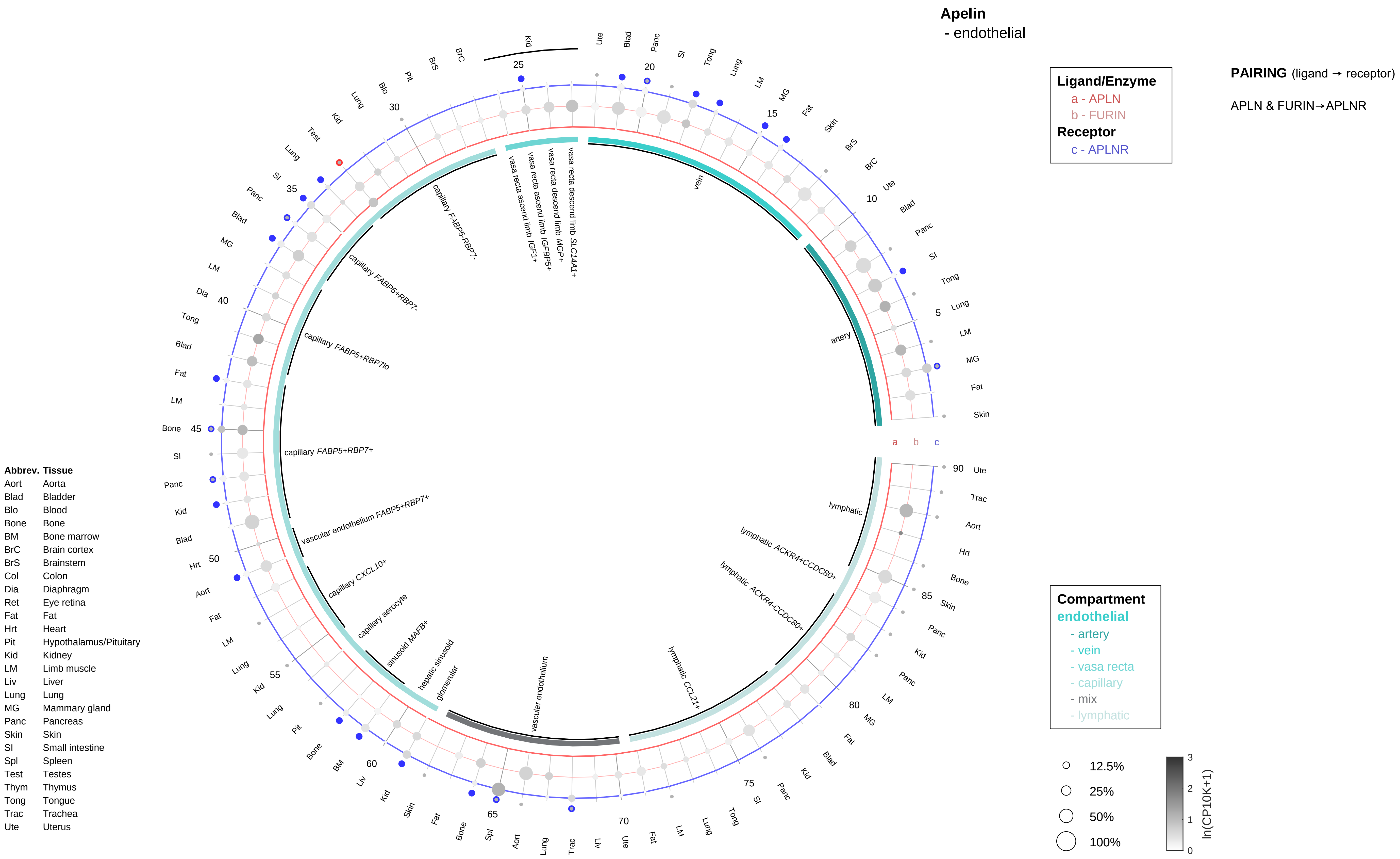
Apelin
- endothelial
Ligand/Enzyme
a - APLN
b - FURIN
Receptor
c - APLNR
PAIRING (ligand → receptor)
APLN & FURIN→APLNR
Compartment
endothelial
- artery
- vein
- vasa recta
- capillary
- mix
- lymphatic
12.5%
25%
50%
100%
ln(CP10K+1)
Abbrev. Tissue
Aort Aorta
Blad Bladder
Blo Blood
Bone Bone
BM Bone marrow
BrC Brain cortex
BrS Brainstem
Col Colon
Dia Diaphragm
Ret Eye retina
Fat Fat
Hrt Heart
Pit Hypothalamus/Pituitary
Kid Kidney
LM Limb muscle
Liv Liver
Lung Lung
MG Mammary gland
Panc Pancreas
Skin Skin
SI Small intestine
Spl Spleen
Test Testes
Thym Thymus
Tong Tongue
Trac Trachea
Ute Uterus
vein
artery
lymphatic
lymphatic ACKR4+CCDC80+
lymphatic ACKR4-CCDC80+
lymphatic CCL21+
vascular endothelium
glomerular
hepatic sinusoid
sinusoid MAFB+
capillary aerocyte
capillary CXCL10+
vascular endothelium FABP5+RBP7+
capillary FABP5+RBP7+
capillary FABP5+RBP7lo
capillary FABP5+RBP7-
capillary FABP5-RBP7-
vasa recta ascend limb IGF1+
vasa recta ascend limb IGFBP5+
vasa recta descend limb MGP+
vasa recta descend limb SLC14A1+

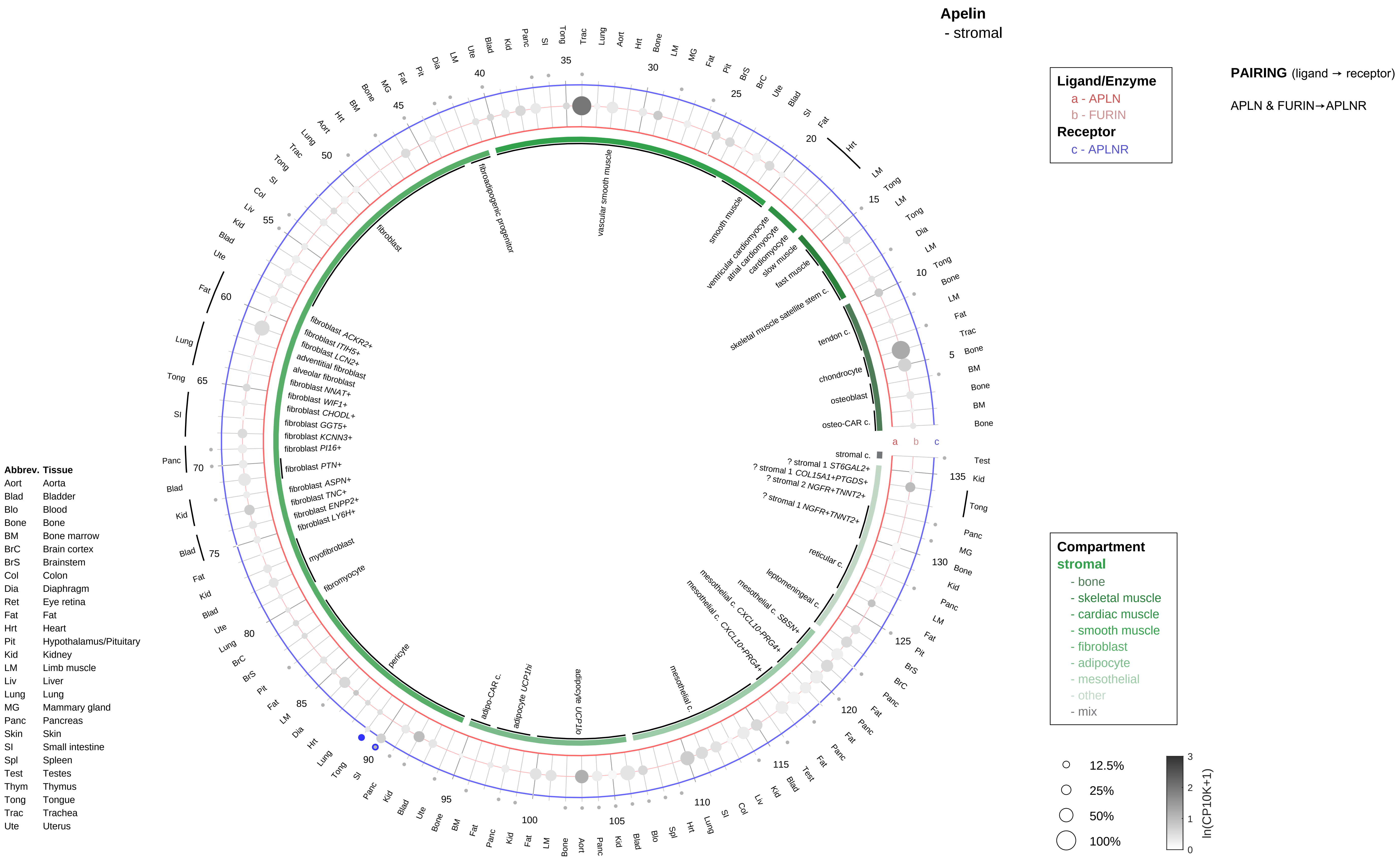

Apelin
- stromal
Ligand/Enzyme
a - APLN
b - FURIN
Receptor
c - APLNR
PAIRING (ligand → receptor)
APLN & FURIN→APLNR
Compartment
stromal
- bone
- skeletal muscle
- cardiac muscle
- smooth muscle
- fibroblast
- adipocyte
- mesothelial
- other
- mix
12.5%
25%
50%
100%
ln(CP10K+1)
Abbrev. Tissue
Aort Aorta
Blad Bladder
Blo Blood
Bone Bone
BM Bone marrow
BrC Brain cortex
BrS Brainstem
Col Colon
Dia Diaphragm
Ret Eye retina
Fat Fat
Hrt Heart
Pit Hypothalamus/Pituitary
Kid Kidney
LM Limb muscle
Liv Liver
Lung Lung
MG Mammary gland
Panc Pancreas
Skin Skin
SI Small intestine
Spl Spleen
Test Testes
Thym Thymus
Tong Tongue
Trac Trachea
Ute Uterus
vascular smooth muscle
fibroadipogenic progenitor
fibroblast
smooth muscle
ventricular cardiomyocyte
atrial cardiomyocyte
cardiomyocyte
slow muscle
fast muscle
skeletal muscle satellite stem c.
tendon c.
chondrocyte
osteoblast
osteo-CAR c.
stromal c.
? stromal 1 ST6GAL2+
? stromal 1 COL15A1+PTGDS+
? stromal 2 NGFR+TNNT2+
? stromal 1 NGFR+TNNT2+
reticular c.
leptomeningeal c.
mesothelial c. SBSN+
mesothelial c. CXCL10-PRG4+
mesothelial c. CXCL10+PRG4+
mesothelial c.
adipocyte UCP1lo
adipocyte UCP1hi
adipo-CAR c.
pericyte
fibromyocyte
myofibroblast
fibroblast LY6H+
fibroblast ENPP2+
fibroblast TNC+
fibroblast ASPN+
fibroblast PTN+
fibroblast PI16+
fibroblast KCNN3+
fibroblast GGT5+
fibroblast CHODL+
fibroblast WIF1+
fibroblast NNAT+
alveolar fibroblast
adventitial fibroblast
fibroblast LCN2+
fibroblast ITIH5+
fibroblast ACKR2+

**Apelin**
- lymphoid

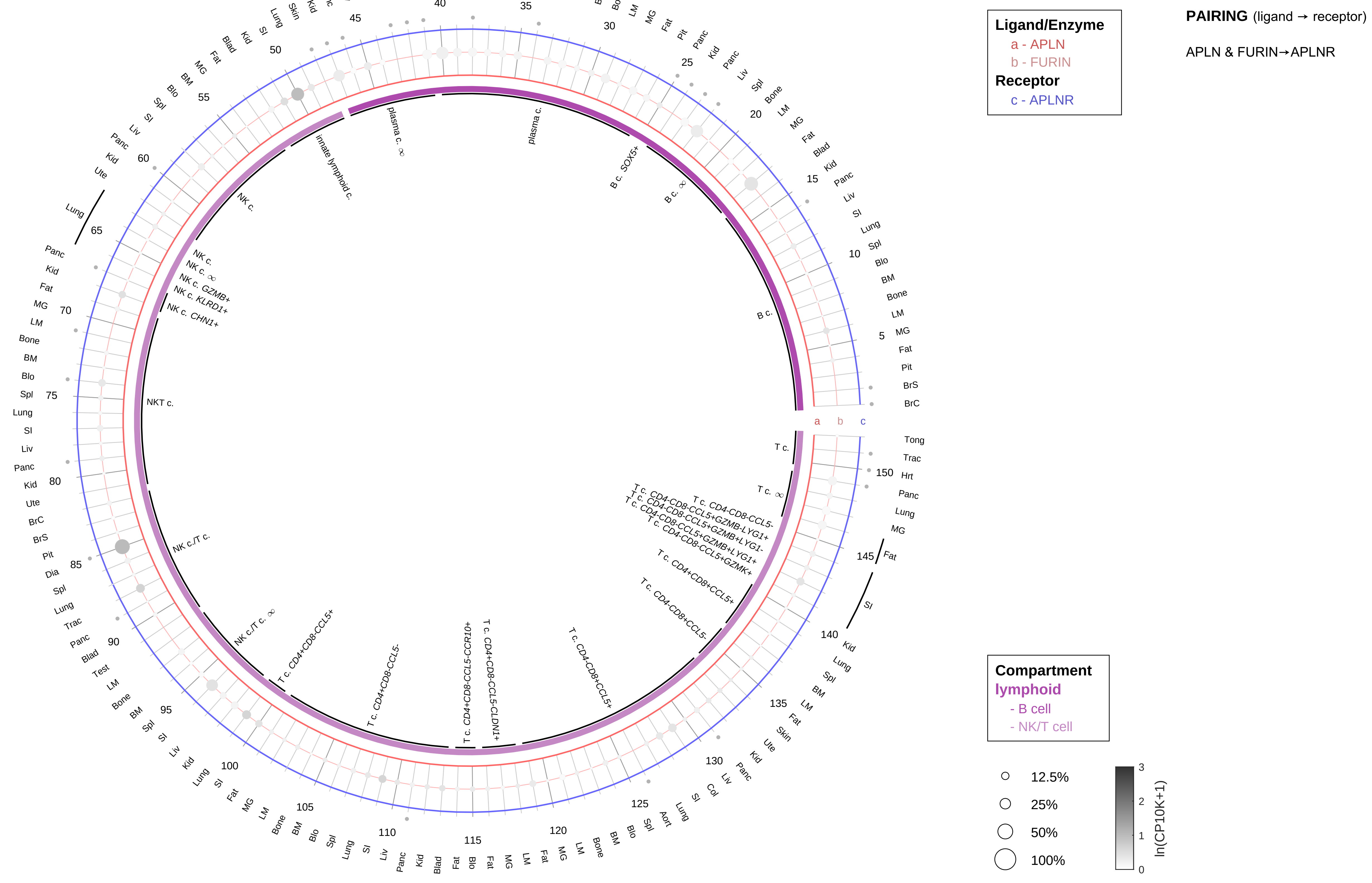

| Abbrev. | Tissue |
|---|---|
| Aort | Aorta |
| Blad | Bladder |
| Blo | Blood |
| Bone | Bone |
| BM | Bone marrow |
| BrC | Brain cortex |
| BrS | Brainstem |
| Col | Colon |
| Dia | Diaphragm |
| Ret | Eye retina |
| Fat | Fat |
| Hrt | Heart |
| Pit | Hypothalamus/Pituitary |
| Kid | Kidney |
| LM | Limb muscle |
| Liv | Liver |
| Lung | Lung |
| MG | Mammary gland |
| Panc | Pancreas |
| Skin | Skin |
| SI | Small intestine |
| Spl | Spleen |
| Test | Testes |
| Thym | Thymus |
| Tong | Tongue |
| Trac | Trachea |
| Ute | Uterus |

#### Apelin
- myeloid

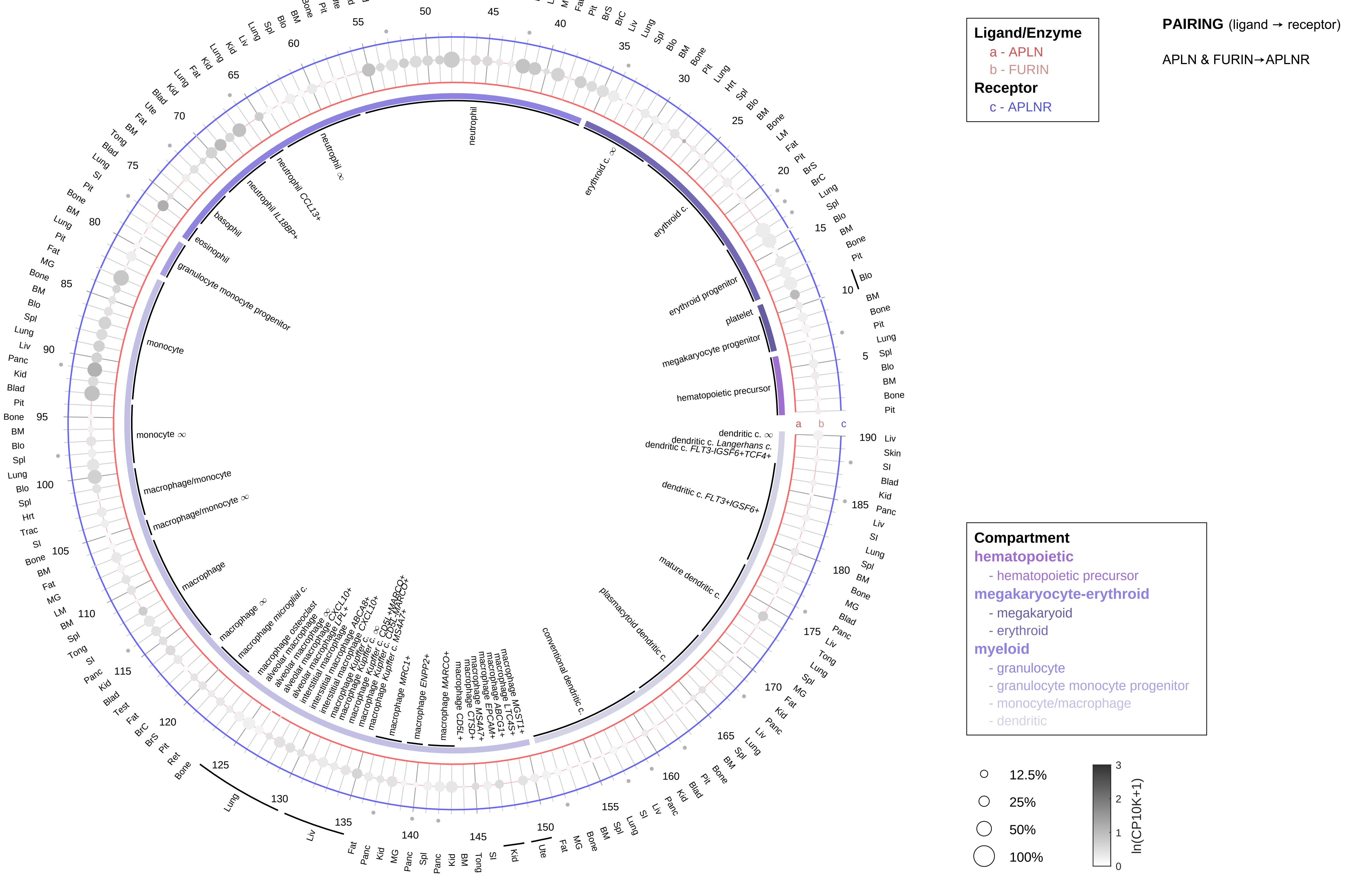

| Abbrev. | Tissue |
|---|---|
| Aort | Aorta |
| Blad | Bladder |
| Blo | Blood |
| Bone | Bone |
| BM | Bone marrow |
| BrC | Brain cortex |
| BrS | Brainstem |
| Col | Colon |
| Dia | Diaphragm |
| Ret | Eye retina |
| Fat | Fat |
| Hrt | Heart |
| Pit | Hypothalamus/Pituitary |
| Kid | Kidney |
| LM | Limb muscle |
| Liv | Liver |
| Lung | Lung |
| MG | Mammary gland |
| Panc | Pancreas |
| Skin | Skin |
| SI | Small intestine |
| Spl | Spleen |
| Test | Testes |
| Thym | Thymus |
| Tong | Tongue |
| Trac | Trachea |
| Ute | Uterus |

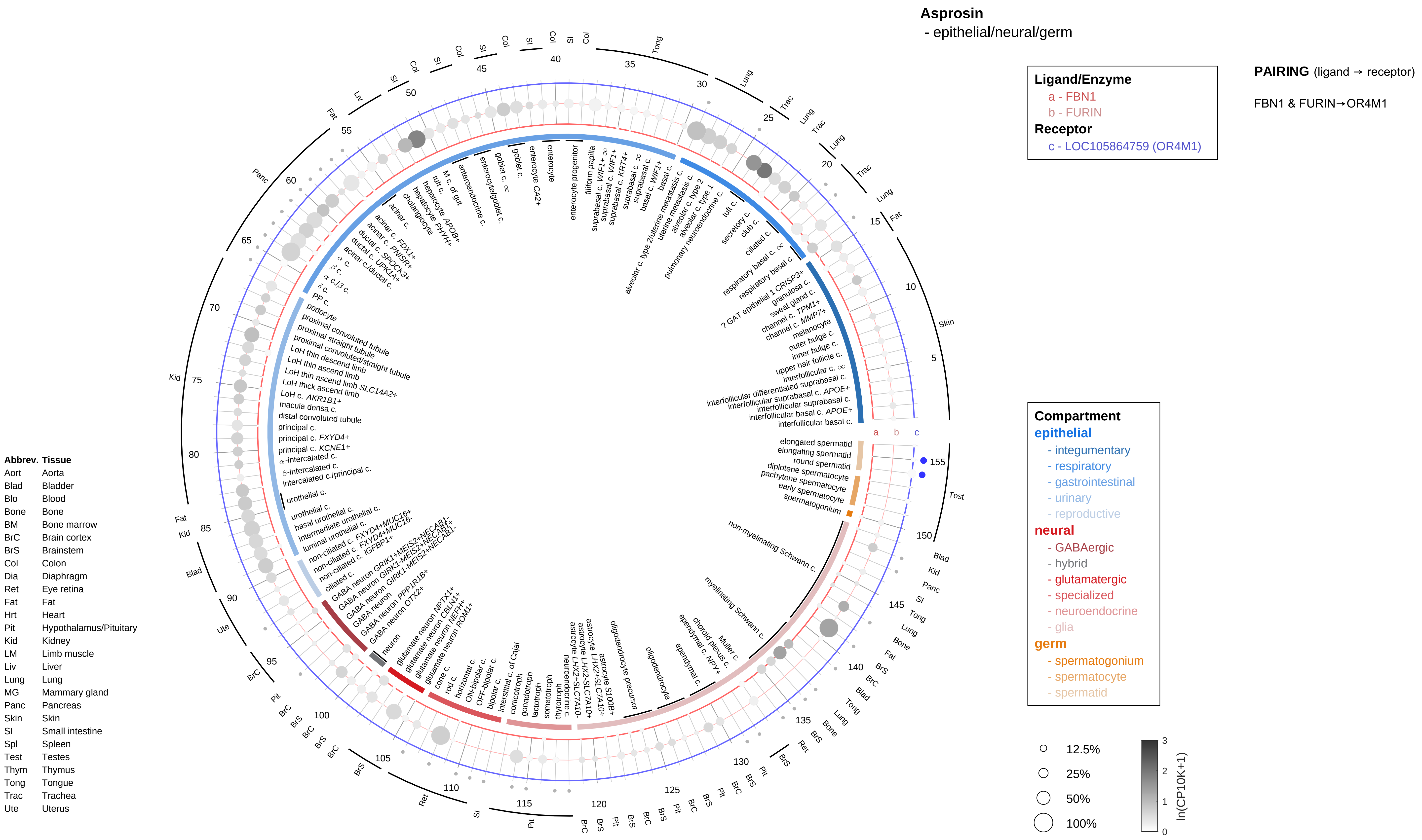

Asprosin
- epithelial/neural/germ
Ligand/Enzyme
a - FBN1
b - FURIN
Receptor
c - LOC105864759 (OR4M1)
PAIRING (ligand → receptor)
FBN1 & FURIN→OR4M1
Compartment
epithelial
- integumentary
- respiratory
- gastrointestinal
- urinary
- reproductive
neural
- GABAergic
- hybrid
- glutamatergic
- specialized
- neuroendocrine
- glia
germ
- spermatogonium
- spermatocyte
- spermatid
12.5%
25%
50%
100%
ln(CP10K+1)
Abbrev. Tissue
Aort Aorta
Blad Bladder
Blo Blood
Bone Bone
BM Bone marrow
BrC Brain cortex
BrS Brainstem
Col Colon
Dia Diaphragm
Ret Eye retina
Fat Fat
Hrt Heart
Pit Hypothalamus/Pituitary
Kid Kidney
LM Limb muscle
Liv Liver
Lung Lung
MG Mammary gland
Panc Pancreas
Skin Skin
SI Small intestine
Spl Spleen
Test Testes
Thym Thymus
Tong Tongue
Trac Trachea
Ute Uterus

**Asprosin**
- endothelial

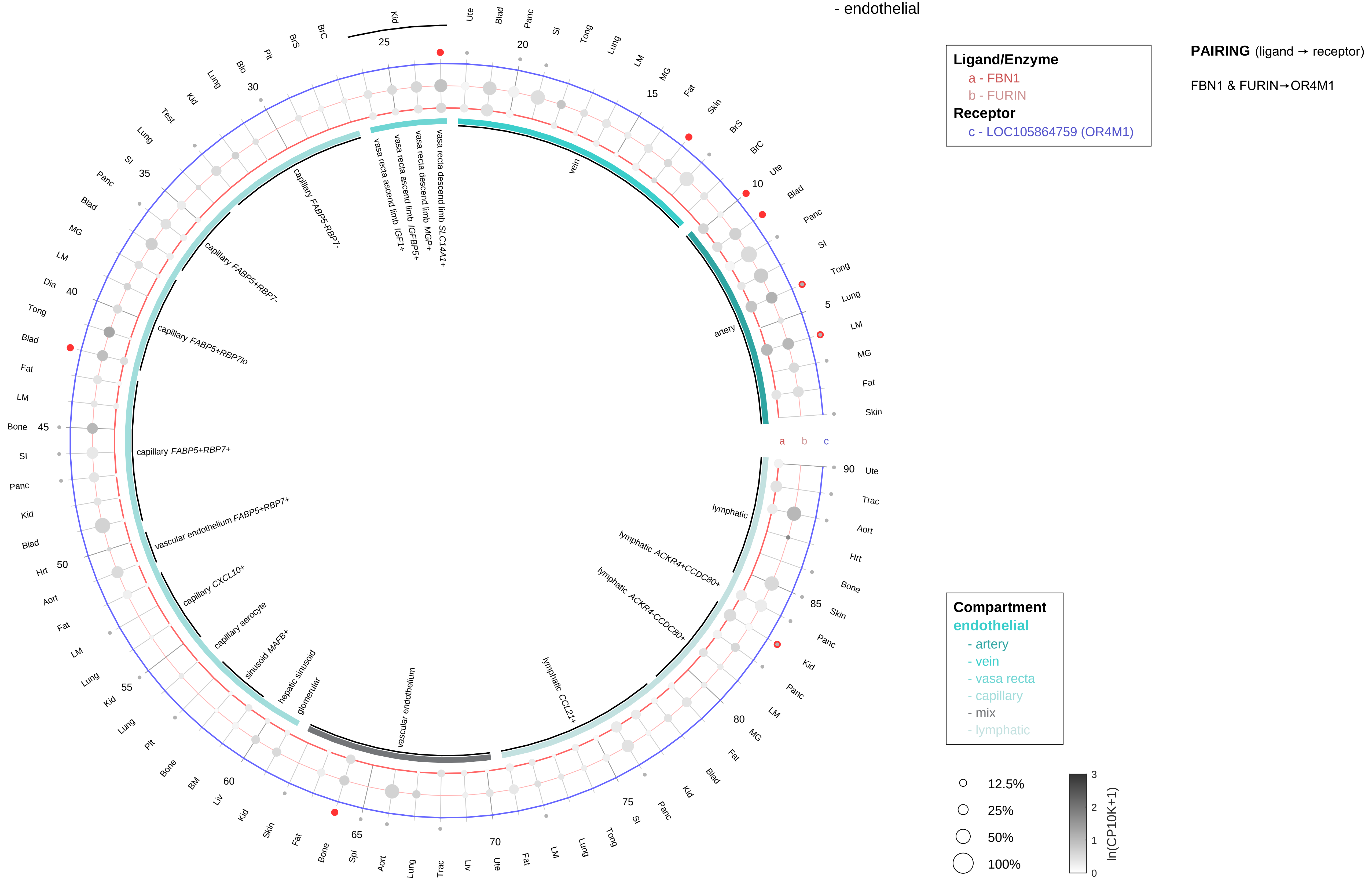

| Abbrev. | Tissue |
|---|---|
| Aort | Aorta |
| Blad | Bladder |
| Blo | Blood |
| Bone | Bone |
| BM | Bone marrow |
| BrC | Brain cortex |
| BrS | Brainstem |
| Col | Colon |
| Dia | Diaphragm |
| Ret | Eye retina |
| Fat | Fat |
| Hrt | Heart |
| Pit | Hypothalamus/Pituitary |
| Kid | Kidney |
| LM | Limb muscle |
| Liv | Liver |
| Lung | Lung |
| MG | Mammary gland |
| Panc | Pancreas |
| Skin | Skin |
| SI | Small intestine |
| Spl | Spleen |
| Test | Testes |
| Thym | Thymus |
| Tong | Tongue |
| Trac | Trachea |
| Ute | Uterus |

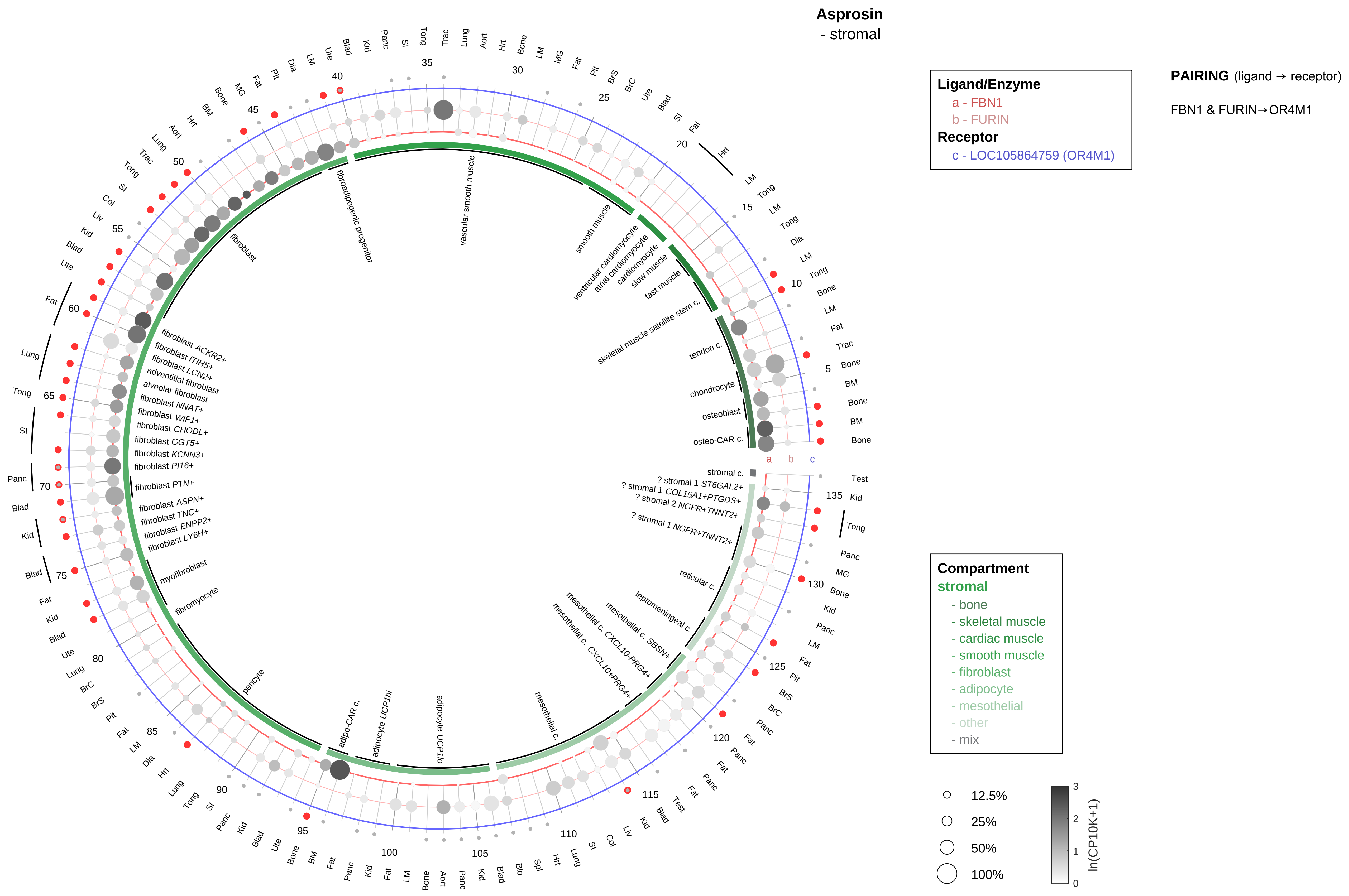

| Abbrev. | Tissue |
|---|---|
| Aort | Aorta |
| Blad | Bladder |
| Blo | Blood |
| Bone | Bone |
| BM | Bone marrow |
| BrC | Brain cortex |
| BrS | Brainstem |
| Col | Colon |
| Dia | Diaphragm |
| Ret | Eye retina |
| Fat | Fat |
| Hrt | Heart |
| Pit | Hypothalamus/Pituitary |
| Kid | Kidney |
| LM | Limb muscle |
| Liv | Liver |
| Lung | Lung |
| MG | Mammary gland |
| Panc | Pancreas |
| Skin | Skin |
| SI | Small intestine |
| Spl | Spleen |
| Test | Testes |
| Thym | Thymus |
| Tong | Tongue |
| Trac | Trachea |
| Ute | Uterus |

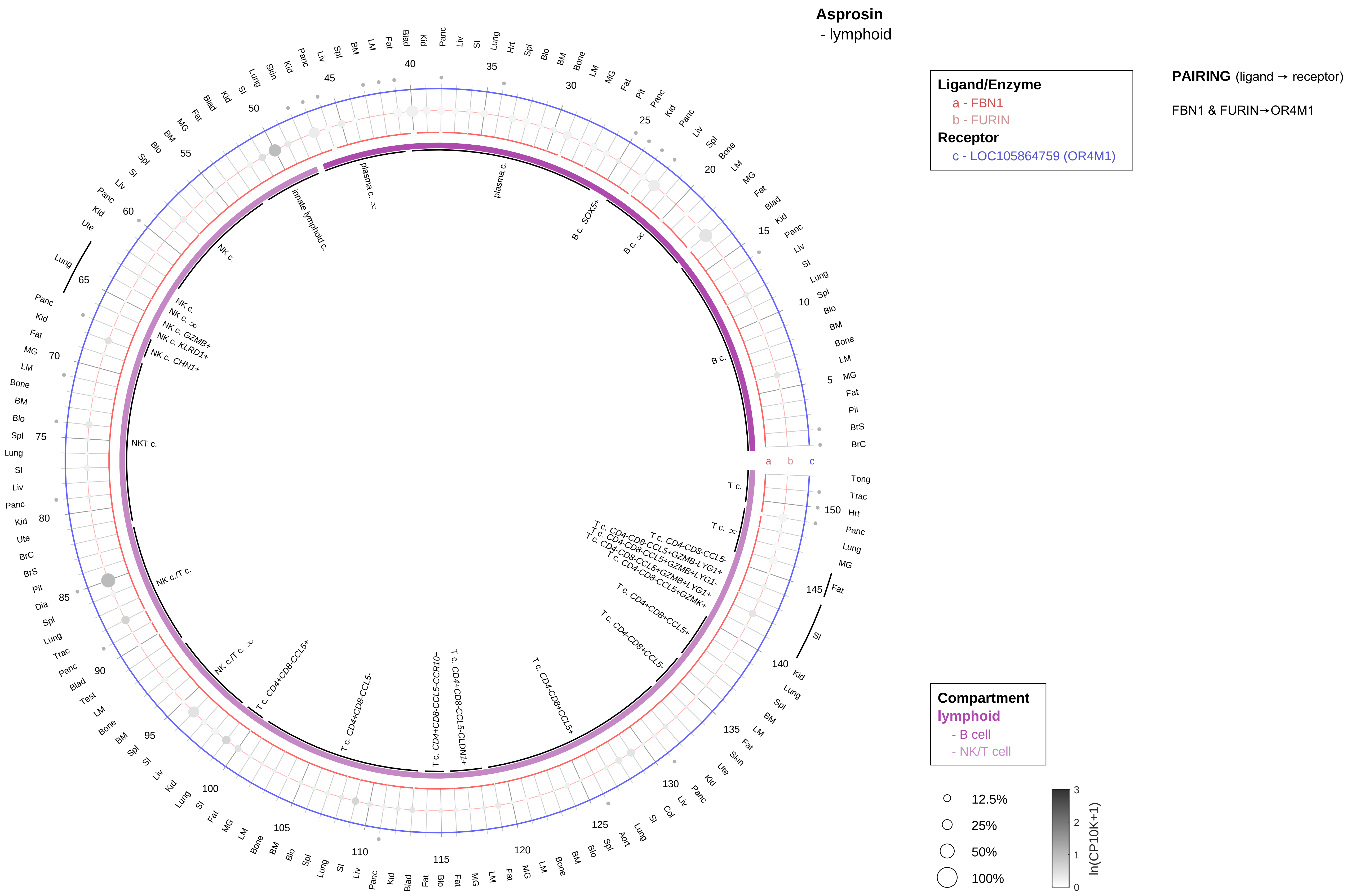

| Abbrev. | Tissue |
|---|---|
| Aort | Aorta |
| Blad | Bladder |
| Blo | Blood |
| Bone | Bone |
| BM | Bone marrow |
| BrC | Brain cortex |
| BrS | Brainstem |
| Col | Colon |
| Dia | Diaphragm |
| Ret | Eye retina |
| Fat | Fat |
| Hrt | Heart |
| Pit | Hypothalamus/Pituitary |
| Kid | Kidney |
| LM | Limb muscle |
| Liv | Liver |
| Lung | Lung |
| MG | Mammary gland |
| Panc | Pancreas |
| Skin | Skin |
| SI | Small intestine |
| Spl | Spleen |
| Test | Testes |
| Thym | Thymus |
| Tong | Tongue |
| Trac | Trachea |
| Ute | Uterus |

**Asprosin**
- myeloid

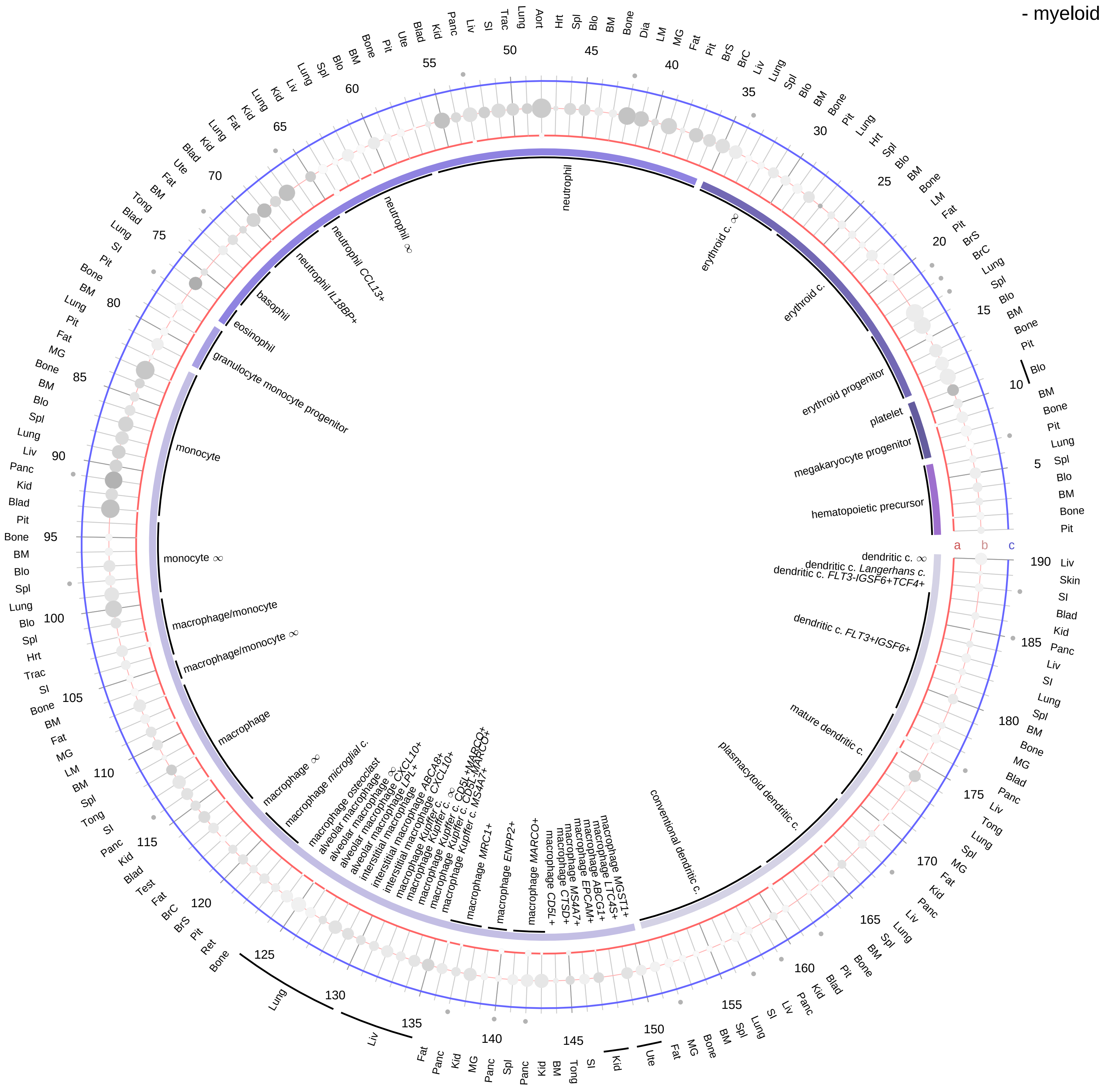

**Ligand/Enzyme**
a - FBN1
b - FURIN
**Receptor**
c - LOC105864759 (OR4M1)

**PAIRING** (ligand → receptor)

FBN1 & FURIN→OR4M1

**Compartment**
**hematopoietic**
- hematopoietic precursor
**megakaryocyte-erythroid**
- megakaryoid
- erythroid
**myeloid**
- granulocyte
- granulocyte monocyte progenitor
- monocyte/macrophage
- dendritic

12.5%
25%
50%
100%

ln(CP10K+1)
0
1
2
3

| Abbrev. | Tissue |
|---|---|
| Aort | Aorta |
| Blad | Bladder |
| Blo | Blood |
| Bone | Bone |
| BM | Bone marrow |
| BrC | Brain cortex |
| BrS | Brainstem |
| Col | Colon |
| Dia | Diaphragm |
| Ret | Eye retina |
| Fat | Fat |
| Hrt | Heart |
| Pit | Hypothalamus/Pituitary |
| Kid | Kidney |
| LM | Limb muscle |
| Liv | Liver |
| Lung | Lung |
| MG | Mammary gland |
| Panc | Pancreas |
| Skin | Skin |
| SI | Small intestine |
| Spl | Spleen |
| Test | Testes |
| Thym | Thymus |
| Tong | Tongue |
| Trac | Trachea |
| Ute | Uterus |

**Calcitonin/CGRP**
- epithelial/neural/germ

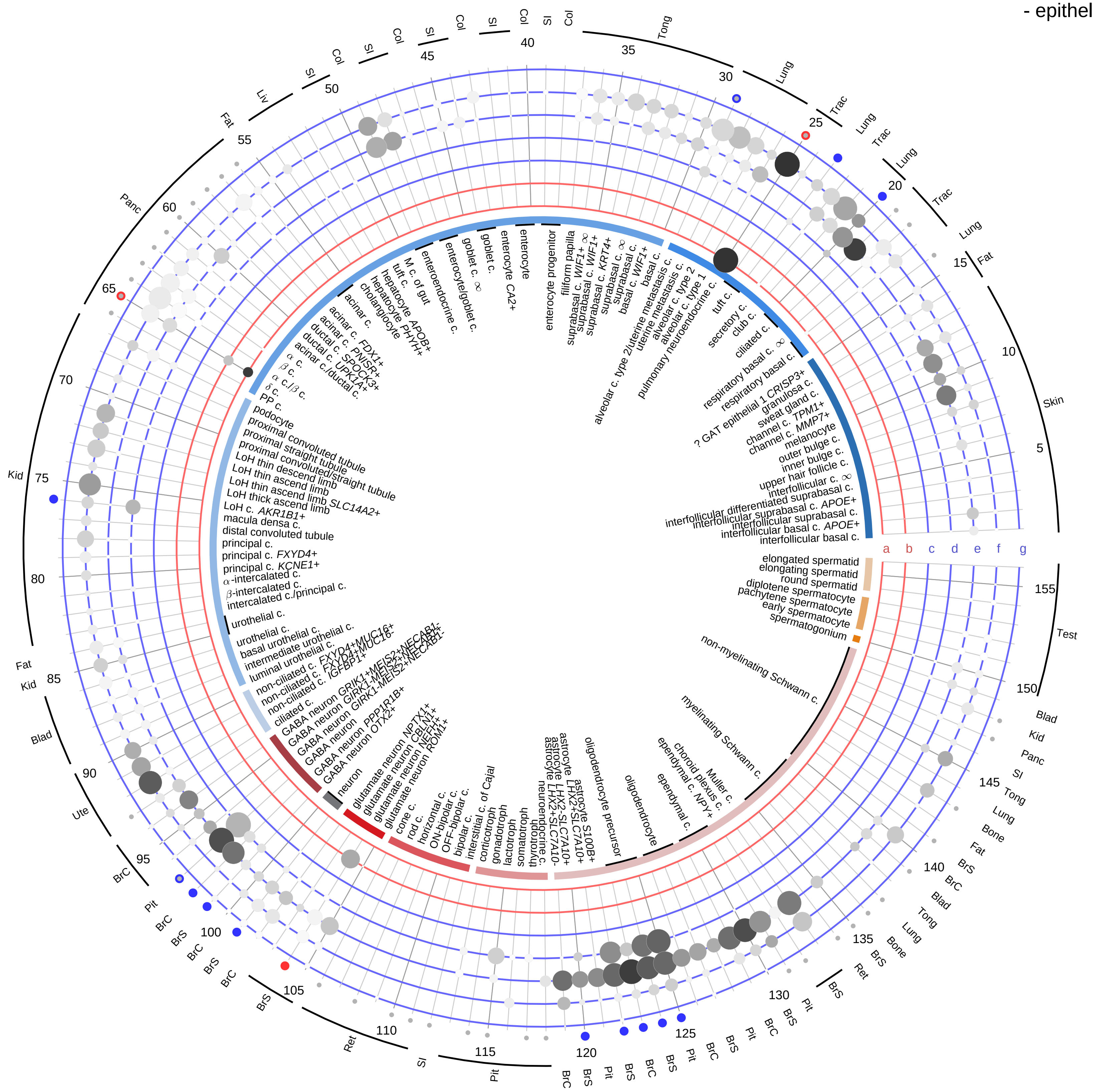

| Ligand/Enzyme |
|---|
| a - CALCA |
| b - CALCB |
| **Receptor** |
| c - CALCR |
| d - CALCRL |
| e - RAMP1 |
| f - RAMP2 |
| g - RAMP3 |

**PAIRING** (ligand → receptor)

CALCA→CALCR

CALCA→CALCRL & RAMP1

CALCA→CALCRL & RAMP2

CALCA→CALCRL & RAMP3

CALCA→CALCR & RAMP1

CALCA→CALCR & RAMP3

CALCB→CALCR

CALCB→CALCRL & RAMP1

CALCB→CALCRL & RAMP2

CALCB→CALCRL & RAMP3

CALCB→CALCR & RAMP1

CALCB→CALCR & RAMP3

**Compartment**
**epithelial**
- integumentary
- respiratory
- gastrointestinal
- urinary
- reproductive

**neural**
- GABAergic
- hybrid
- glutamatergic
- specialized
- neuroendocrine
- glia

**germ**
- spermatogonium
- spermatocyte
- spermatid

12.5%
25%
50%
100%

ln(CP10K+1): 0, 1, 2, 3

| Abbrev. | Tissue |
|---|---|
| Aort | Aorta |
| Blad | Bladder |
| Blo | Blood |
| Bone | Bone |
| BM | Bone marrow |
| BrC | Brain cortex |
| BrS | Brainstem |
| Col | Colon |
| Dia | Diaphragm |
| Ret | Eye retina |
| Fat | Fat |
| Hrt | Heart |
| Pit | Hypothalamus/Pituitary |
| Kid | Kidney |
| LM | Limb muscle |
| Liv | Liver |
| Lung | Lung |
| MG | Mammary gland |
| Panc | Pancreas |
| Skin | Skin |
| SI | Small intestine |
| Spl | Spleen |
| Test | Testes |
| Thym | Thymus |
| Tong | Tongue |
| Trac | Trachea |
| Ute | Uterus |

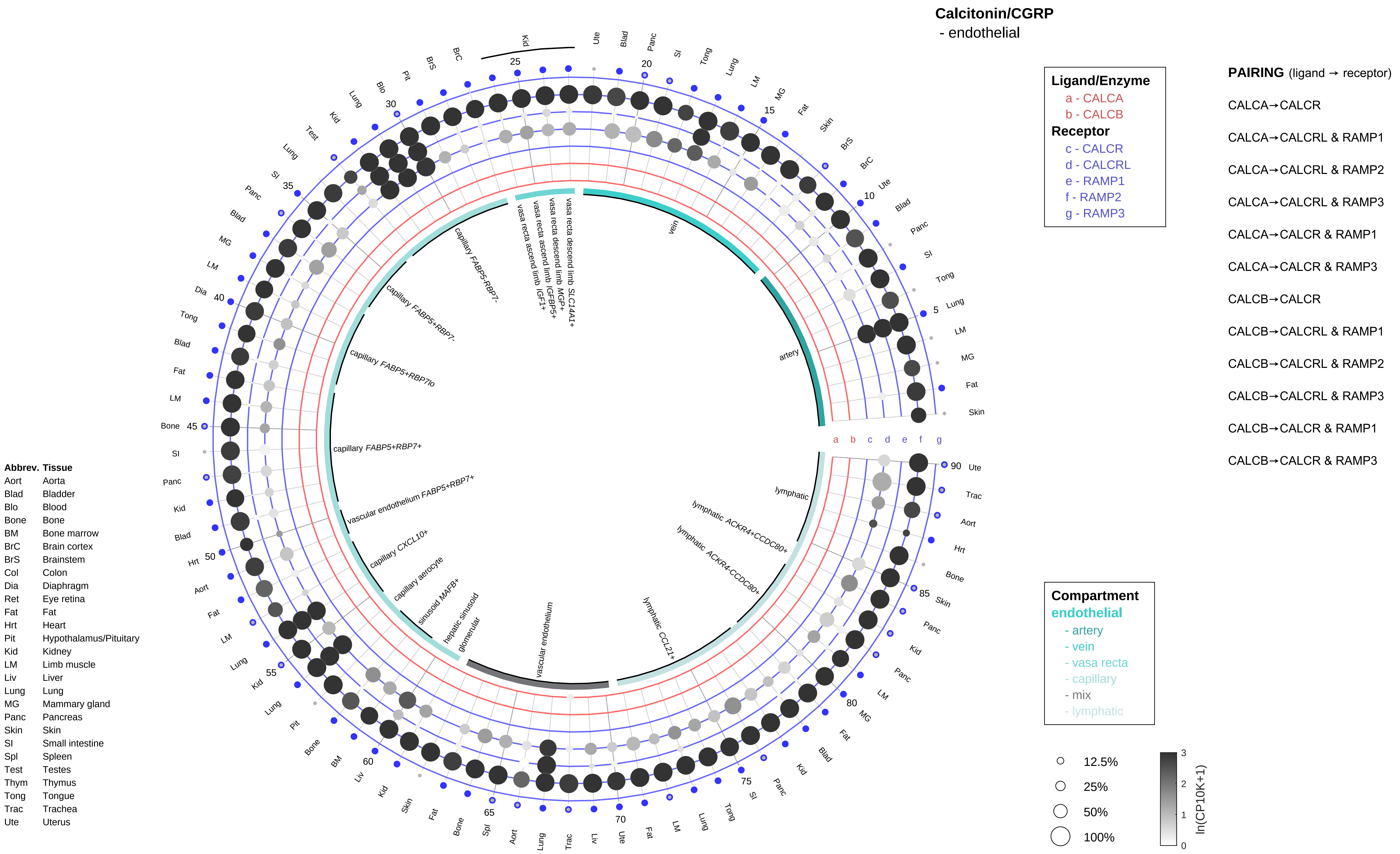

Calcitonin/CGRP
- endothelial
Ligand/Enzyme
a - CALCA
b - CALCB
Receptor
c - CALCR
d - CALCRL
e - RAMP1
f - RAMP2
g - RAMP3
PAIRING (ligand → receptor)
CALCA→CALCR
CALCA→CALCRL & RAMP1
CALCA→CALCRL & RAMP2
CALCA→CALCRL & RAMP3
CALCA→CALCR & RAMP1
CALCA→CALCR & RAMP3
CALCB→CALCR
CALCB→CALCRL & RAMP1
CALCB→CALCRL & RAMP2
CALCB→CALCRL & RAMP3
CALCB→CALCR & RAMP1
CALCB→CALCR & RAMP3
Compartment
endothelial
- artery
- vein
- vasa recta
- capillary
- mix
- lymphatic
12.5%
25%
50%
100%
ln(CP10K+1)
Abbrev. Tissue
Aort Aorta
Blad Bladder
Blo Blood
Bone Bone
BM Bone marrow
BrC Brain cortex
BrS Brainstem
Col Colon
Dia Diaphragm
Ret Eye retina
Fat Fat
Hrt Heart
Pit Hypothalamus/Pituitary
Kid Kidney
LM Limb muscle
Liv Liver
Lung Lung
MG Mammary gland
Panc Pancreas
Skin Skin
SI Small intestine
Spl Spleen
Test Testes
Thym Thymus
Tong Tongue
Trac Trachea
Ute Uterus

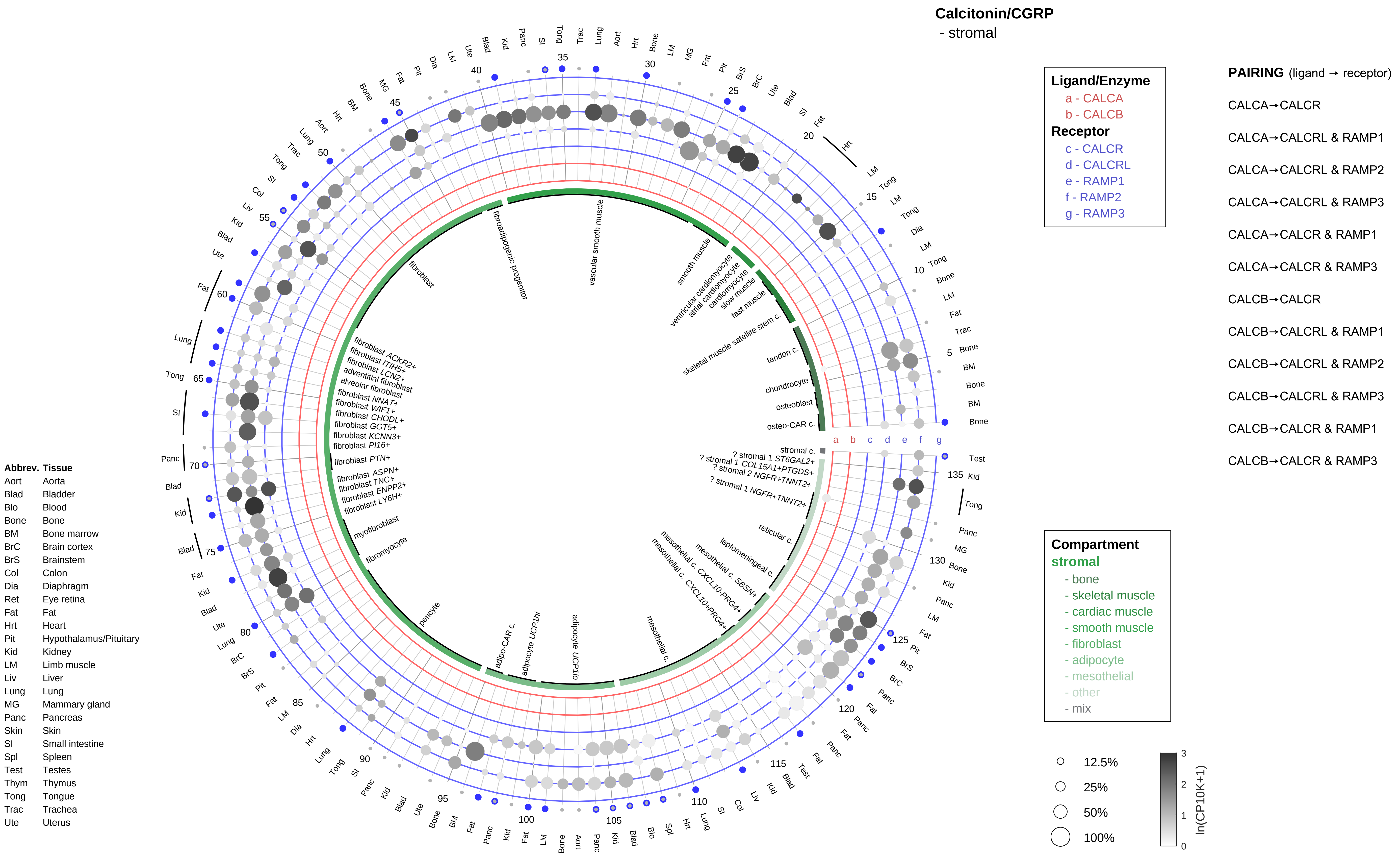
Calcitonin/CGRP
- stromal
Ligand/Enzyme
a - CALCA
b - CALCB
Receptor
c - CALCR
d - CALCRL
e - RAMP1
f - RAMP2
g - RAMP3
PAIRING (ligand → receptor)
CALCA→CALCR
CALCA→CALCRL & RAMP1
CALCA→CALCRL & RAMP2
CALCA→CALCRL & RAMP3
CALCA→CALCR & RAMP1
CALCA→CALCR & RAMP3
CALCB→CALCR
CALCB→CALCRL & RAMP1
CALCB→CALCRL & RAMP2
CALCB→CALCRL & RAMP3
CALCB→CALCR & RAMP1
CALCB→CALCR & RAMP3
Compartment
stromal
- bone
- skeletal muscle
- cardiac muscle
- smooth muscle
- fibroblast
- adipocyte
- mesothelial
- other
- mix
12.5%
25%
50%
100%
ln(CP10K+1)
Abbrev. Tissue
Aort Aorta
Blad Bladder
Blo Blood
Bone Bone
BM Bone marrow
BrC Brain cortex
BrS Brainstem
Col Colon
Dia Diaphragm
Ret Eye retina
Fat Fat
Hrt Heart
Pit Hypothalamus/Pituitary
Kid Kidney
LM Limb muscle
Liv Liver
Lung Lung
MG Mammary gland
Panc Pancreas
Skin Skin
SI Small intestine
Spl Spleen
Test Testes
Thym Thymus
Tong Tongue
Trac Trachea
Ute Uterus

**Calcitonin/CGRP**
- lymphoid

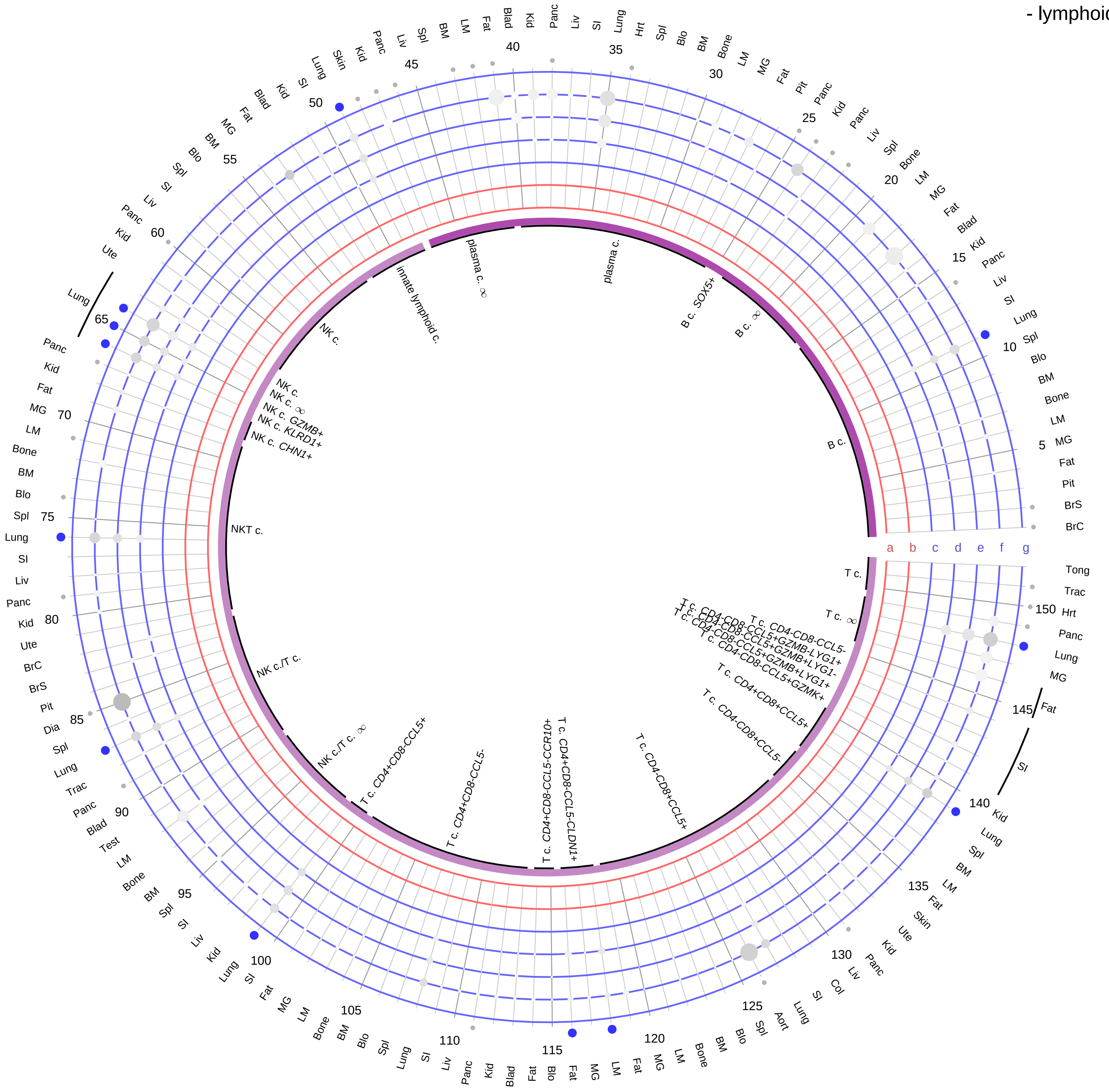

**Ligand/Enzyme**
- a - CALCA
- b - CALCB

**Receptor**
- c - CALCR
- d - CALCRL
- e - RAMP1
- f - RAMP2
- g - RAMP3

**PAIRING** (ligand → receptor)

CALCA→CALCR

CALCA→CALCRL & RAMP1

CALCA→CALCRL & RAMP2

CALCA→CALCRL & RAMP3

CALCA→CALCR & RAMP1

CALCA→CALCR & RAMP3

CALCB→CALCR

CALCB→CALCRL & RAMP1

CALCB→CALCRL & RAMP2

CALCB→CALCRL & RAMP3

CALCB→CALCR & RAMP1

CALCB→CALCR & RAMP3

| Abbrev. | Tissue |
|---|---|
| Aort | Aorta |
| Blad | Bladder |
| Blo | Blood |
| Bone | Bone |
| BM | Bone marrow |
| BrC | Brain cortex |
| BrS | Brainstem |
| Col | Colon |
| Dia | Diaphragm |
| Ret | Eye retina |
| Fat | Fat |
| Hrt | Heart |
| Pit | Hypothalamus/Pituitary |
| Kid | Kidney |
| LM | Limb muscle |
| Liv | Liver |
| Lung | Lung |
| MG | Mammary gland |
| Panc | Pancreas |
| Skin | Skin |
| SI | Small intestine |
| Spl | Spleen |
| Test | Testes |
| Thym | Thymus |
| Tong | Tongue |
| Trac | Trachea |
| Ute | Uterus |

**Compartment**

**lymphoid**
- B cell
- NK/T cell

12.5%
25%
50%
100%

ln(CP10K+1): 0 1 2 3

**Calcitonin/CGRP**
- myeloid

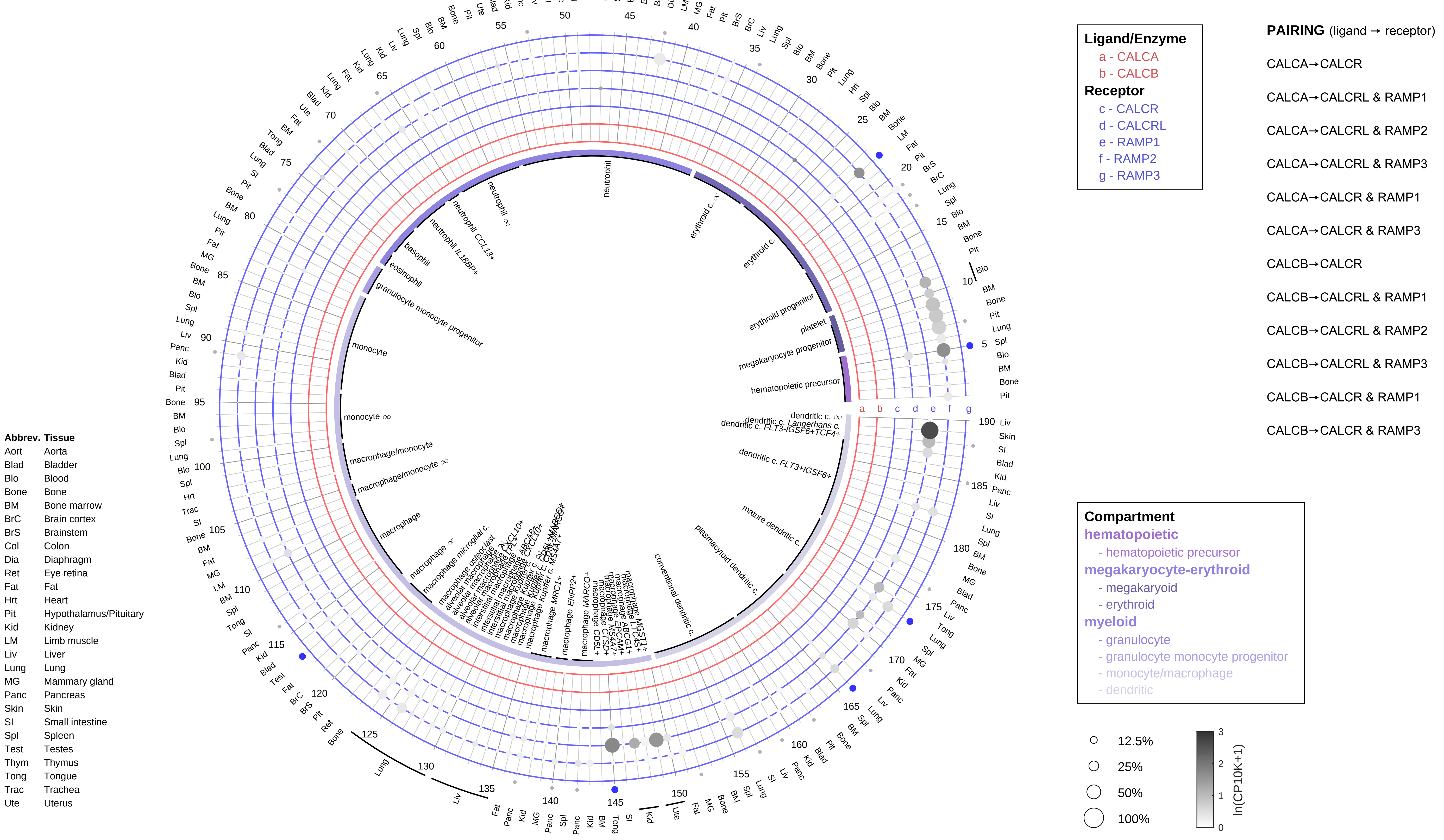

#### Cholecalciferol/Calcidiol/Calcitriol
- epithelial/neural/germ

**PAIRING** (ligand → receptor)

CYP27B1→VDR & RXRA

CYP27B1→VDR & RXRB

CYP27B1→VDR & RXRG

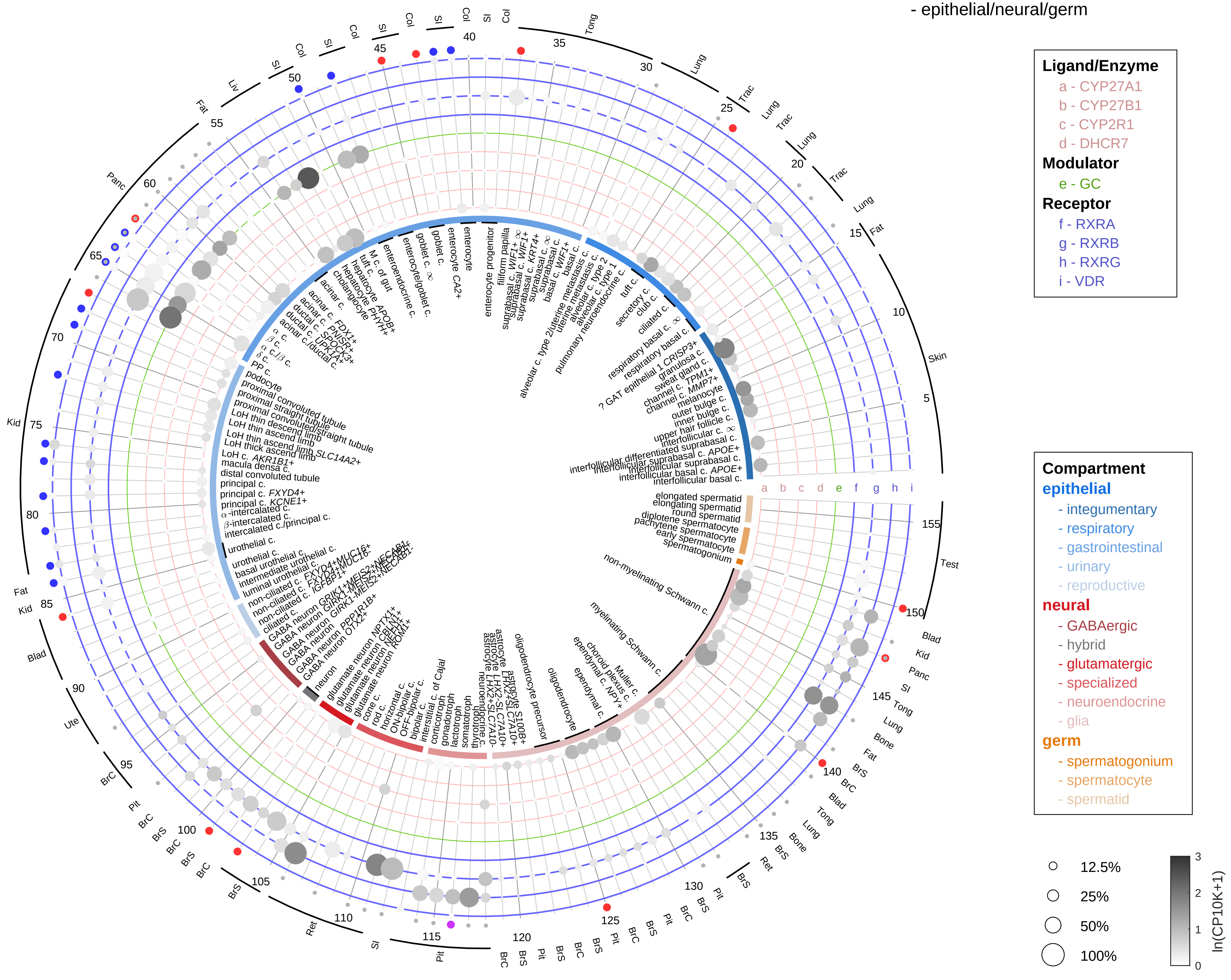

| Abbrev. | Tissue |
|---|---|
| Aort | Aorta |
| Blad | Bladder |
| Blo | Blood |
| Bone | Bone |
| BM | Bone marrow |
| BrC | Brain cortex |
| BrS | Brainstem |
| Col | Colon |
| Dia | Diaphragm |
| Ret | Eye retina |
| Fat | Fat |
| Hrt | Heart |
| Pit | Hypothalamus/Pituitary |
| Kid | Kidney |
| LM | Limb muscle |
| Liv | Liver |
| Lung | Lung |
| MG | Mammary gland |
| Panc | Pancreas |
| Skin | Skin |
| SI | Small intestine |
| Spl | Spleen |
| Test | Testes |
| Thym | Thymus |
| Tong | Tongue |
| Trac | Trachea |
| Ute | Uterus |

**Cholecalciferol/Calcidiol/Calcitriol**
- endothelial

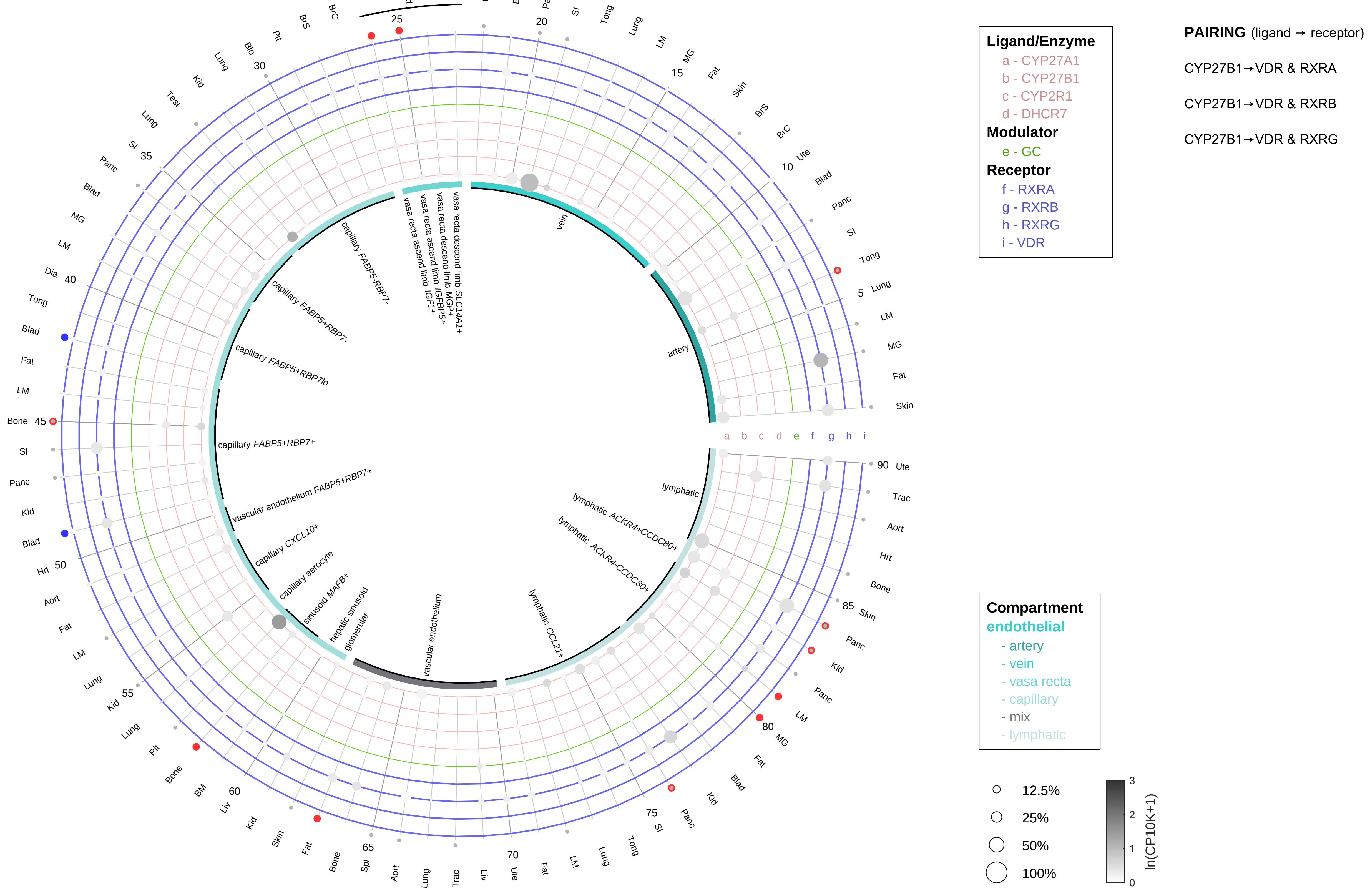

| Abbrev. | Tissue |
|---|---|
| Aort | Aorta |
| Blad | Bladder |
| Blo | Blood |
| Bone | Bone |
| BM | Bone marrow |
| BrC | Brain cortex |
| BrS | Brainstem |
| Col | Colon |
| Dia | Diaphragm |
| Ret | Eye retina |
| Fat | Fat |
| Hrt | Heart |
| Pit | Hypothalamus/Pituitary |
| Kid | Kidney |
| LM | Limb muscle |
| Liv | Liver |
| Lung | Lung |
| MG | Mammary gland |
| Panc | Pancreas |
| Skin | Skin |
| SI | Small intestine |
| Spl | Spleen |
| Test | Testes |
| Thym | Thymus |
| Tong | Tongue |
| Trac | Trachea |
| Ute | Uterus |

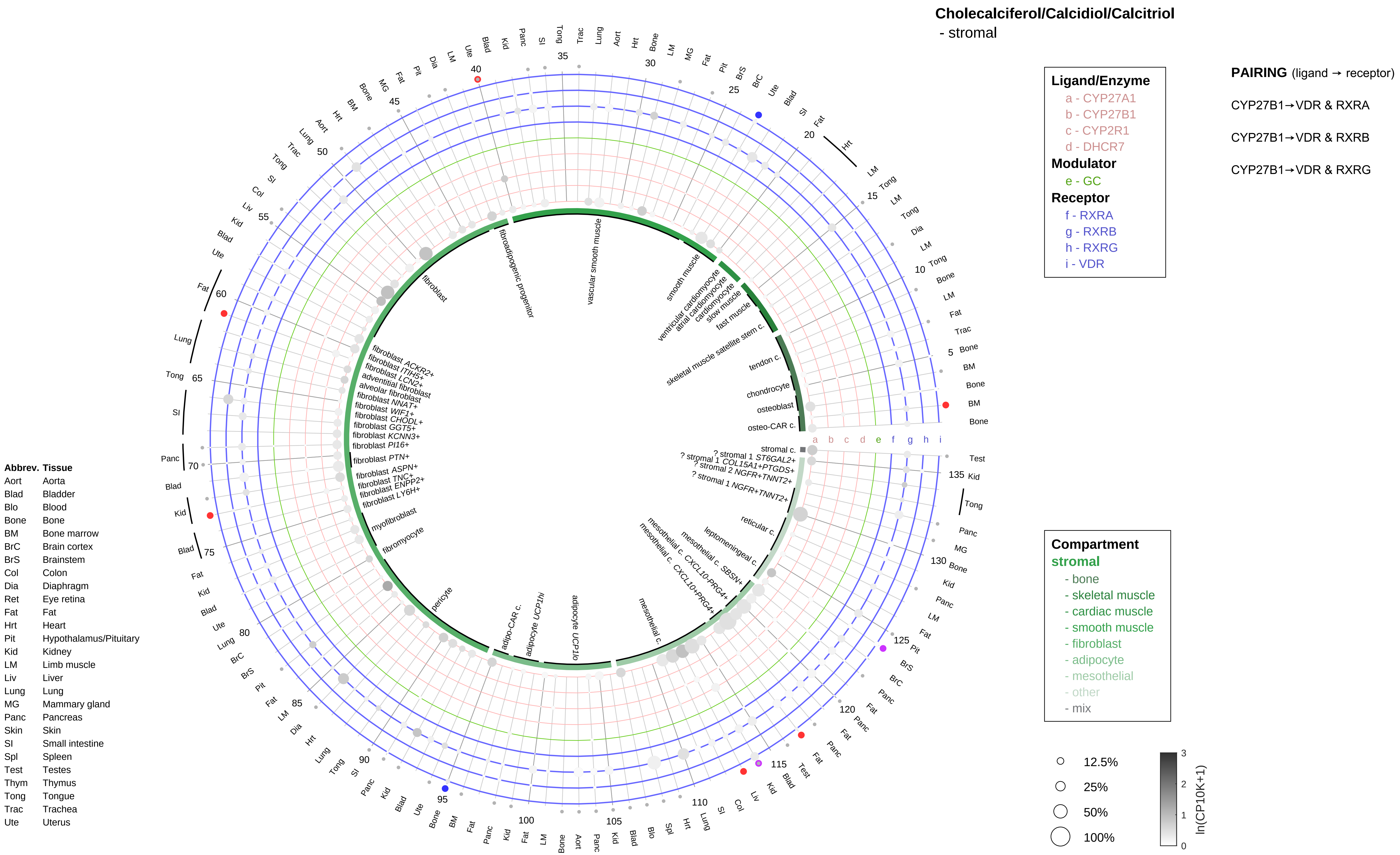
Cholecalciferol/Calcidiol/Calcitriol
- stromal
Ligand/Enzyme
a - CYP27A1
b - CYP27B1
c - CYP2R1
d - DHCR7
Modulator
e - GC
Receptor
f - RXRA
g - RXRB
h - RXRG
i - VDR
PAIRING (ligand → receptor)
CYP27B1→VDR & RXRA
CYP27B1→VDR & RXRB
CYP27B1→VDR & RXRG
Compartment
stromal
- bone
- skeletal muscle
- cardiac muscle
- smooth muscle
- fibroblast
- adipocyte
- mesothelial
- other
- mix
12.5%
25%
50%
100%
ln(CP10K+1)
Abbrev. Tissue
Aort Aorta
Blad Bladder
Blo Blood
Bone Bone
BM Bone marrow
BrC Brain cortex
BrS Brainstem
Col Colon
Dia Diaphragm
Ret Eye retina
Fat Fat
Hrt Heart
Pit Hypothalamus/Pituitary
Kid Kidney
LM Limb muscle
Liv Liver
Lung Lung
MG Mammary gland
Panc Pancreas
Skin Skin
SI Small intestine
Spl Spleen
Test Testes
Thym Thymus
Tong Tongue
Trac Trachea
Ute Uterus
fibroblast
fibroadipogenic progenitor
vascular smooth muscle
smooth muscle
ventricular cardiomyocyte
atrial cardiomyocyte
cardiomyocyte
slow muscle
fast muscle
skeletal muscle satellite stem c.
tendon c.
chondrocyte
osteoblast
osteo-CAR c.
stromal c.
? stromal 1 ST6GAL2+
? stromal 1 COL15A1+PTGDS+
? stromal 2 NGFR+TNNT2+
? stromal 1 NGFR+TNNT2+
reticular c.
leptomeningeal c.
mesothelial c. SBSN+
mesothelial c. CXCL10-PRG4+
mesothelial c. CXCL10+PRG4+
mesothelial c.
adipocyte UCP1lo
adipocyte UCP1hi
adipo-CAR c.
pericyte
fibromyocyte
myofibroblast
fibroblast LY6H+
fibroblast ENPP2+
fibroblast TNC+
fibroblast ASPN+
fibroblast PTN+
fibroblast PI16+
fibroblast KCNN3+
fibroblast GGT5+
fibroblast CHODL+
fibroblast WIF1+
fibroblast NNAT+
alveolar fibroblast
adventitial fibroblast
fibroblast LCN2+
fibroblast ITIH5+
fibroblast ACKR2+

**Cholecalciferol/Calcidiol/Calcitriol**
- lymphoid

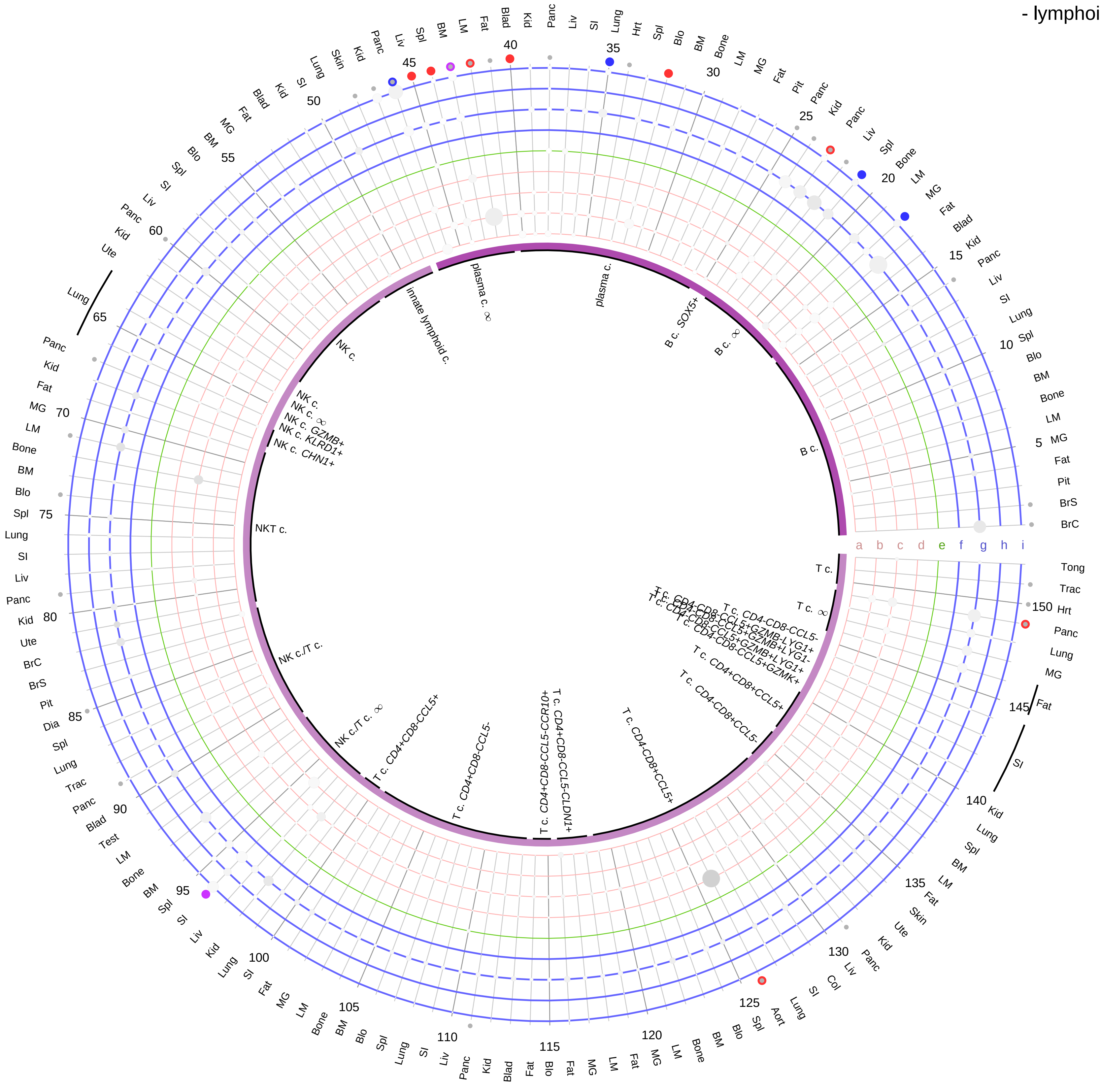

**PAIRING** (ligand → receptor)

CYP27B1→VDR & RXRA

CYP27B1→VDR & RXRB

CYP27B1→VDR & RXRG

| Abbrev. | Tissue |
|---|---|
| Aort | Aorta |
| Blad | Bladder |
| Blo | Blood |
| Bone | Bone |
| BM | Bone marrow |
| BrC | Brain cortex |
| BrS | Brainstem |
| Col | Colon |
| Dia | Diaphragm |
| Ret | Eye retina |
| Fat | Fat |
| Hrt | Heart |
| Pit | Hypothalamus/Pituitary |
| Kid | Kidney |
| LM | Limb muscle |
| Liv | Liver |
| Lung | Lung |
| MG | Mammary gland |
| Panc | Pancreas |
| Skin | Skin |
| SI | Small intestine |
| Spl | Spleen |
| Test | Testes |
| Thym | Thymus |
| Tong | Tongue |
| Trac | Trachea |
| Ute | Uterus |

**Cholecalciferol/Calcidiol/Calcitriol**
- myeloid

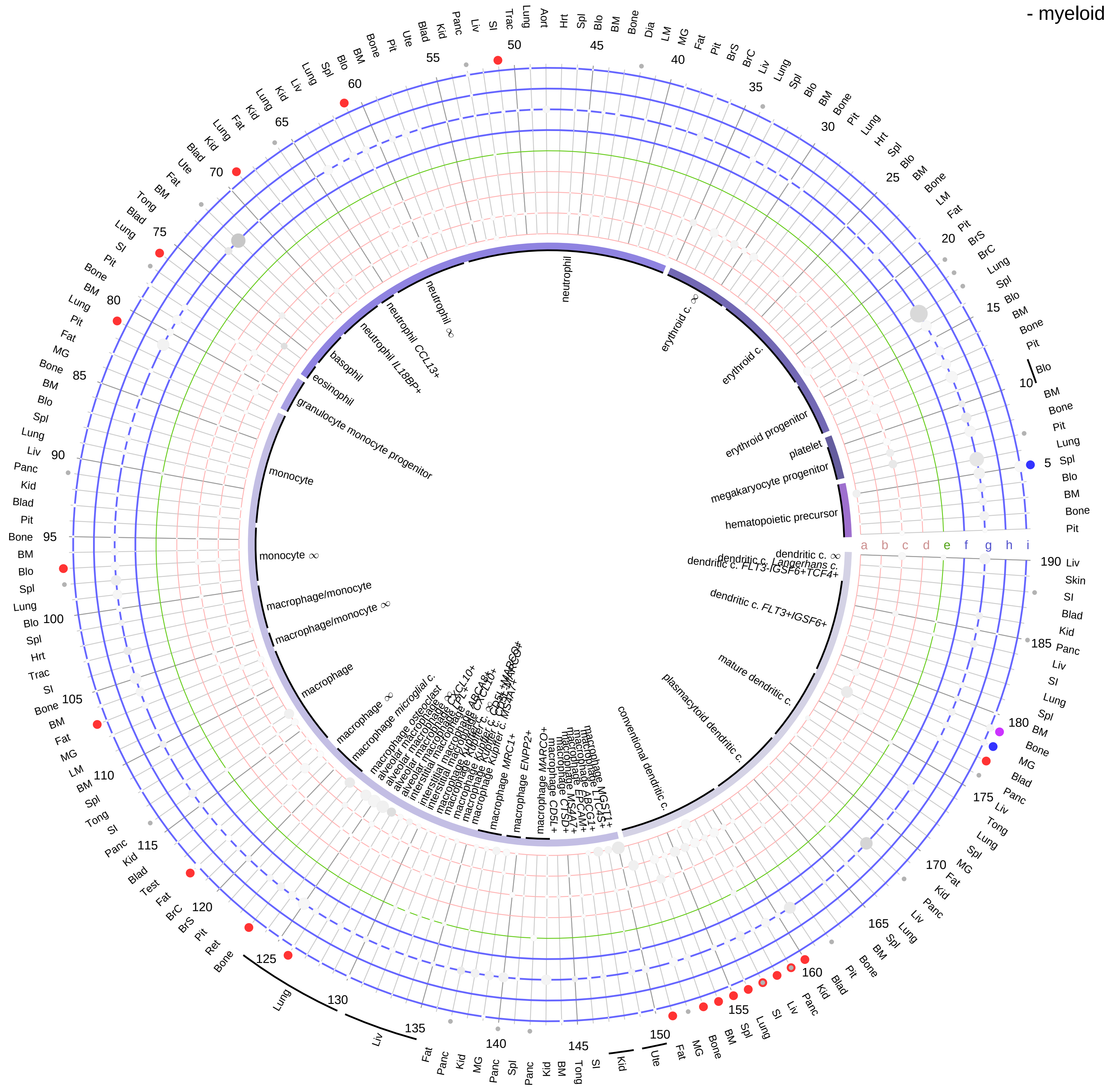

| Abbrev. | Tissue |
|---|---|
| Aort | Aorta |
| Blad | Bladder |
| Blo | Blood |
| Bone | Bone |
| BM | Bone marrow |
| BrC | Brain cortex |
| BrS | Brainstem |
| Col | Colon |
| Dia | Diaphragm |
| Ret | Eye retina |
| Fat | Fat |
| Hrt | Heart |
| Pit | Hypothalamus/Pituitary |
| Kid | Kidney |
| LM | Limb muscle |
| Liv | Liver |
| Lung | Lung |
| MG | Mammary gland |
| Panc | Pancreas |
| Skin | Skin |
| SI | Small intestine |
| Spl | Spleen |
| Test | Testes |
| Thym | Thymus |
| Tong | Tongue |
| Trac | Trachea |
| Ute | Uterus |

**Ligand/Enzyme**
- a - CYP27A1
- b - CYP27B1
- c - CYP2R1
- d - DHCR7

**Modulator**
- e - GC

**Receptor**
- f - RXRA
- g - RXRB
- h - RXRG
- i - VDR

**PAIRING** (ligand → receptor)

CYP27B1→VDR & RXRA

CYP27B1→VDR & RXRB

CYP27B1→VDR & RXRG

**Compartment**

**hematopoietic**
- hematopoietic precursor

**megakaryocyte-erythroid**
- megakaryoid
- erythroid

**myeloid**
- granulocyte
- granulocyte monocyte progenitor
- monocyte/macrophage
- dendritic

12.5%
25%
50%
100%

ln(CP10K+1): 0, 1, 2, 3

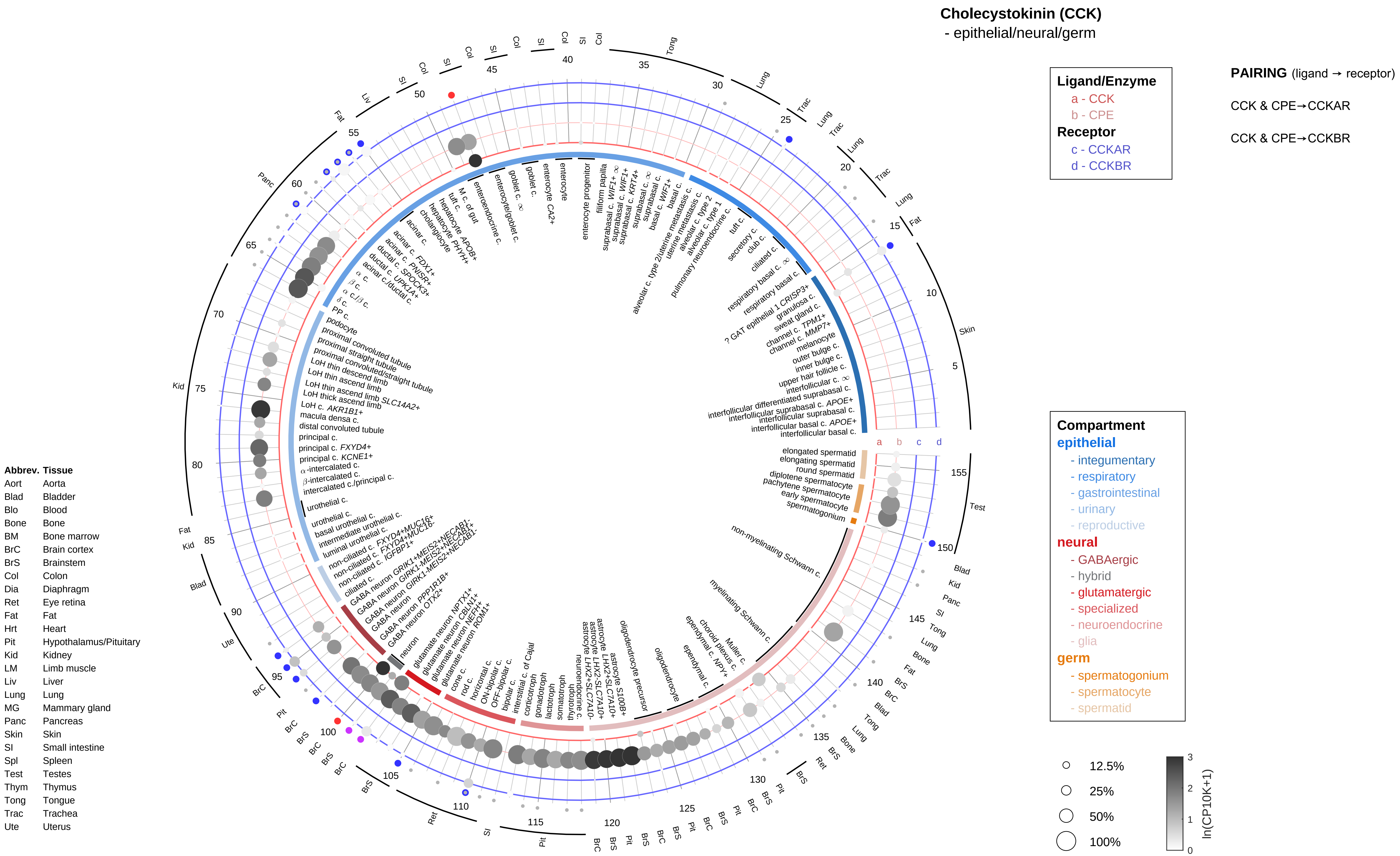

Cholecystokinin (CCK)
- epithelial/neural/germ
Ligand/Enzyme
a - CCK
b - CPE
Receptor
c - CCKAR
d - CCKBR
PAIRING (ligand → receptor)
CCK & CPE→CCKAR
CCK & CPE→CCKBR
Compartment
epithelial
- integumentary
- respiratory
- gastrointestinal
- urinary
- reproductive
neural
- GABAergic
- hybrid
- glutamatergic
- specialized
- neuroendocrine
- glia
germ
- spermatogonium
- spermatocyte
- spermatid
12.5%
25%
50%
100%
ln(CP10K+1)
Abbrev. Tissue
Aort Aorta
Blad Bladder
Blo Blood
Bone Bone
BM Bone marrow
BrC Brain cortex
BrS Brainstem
Col Colon
Dia Diaphragm
Ret Eye retina
Fat Fat
Hrt Heart
Pit Hypothalamus/Pituitary
Kid Kidney
LM Limb muscle
Liv Liver
Lung Lung
MG Mammary gland
Panc Pancreas
Skin Skin
SI Small intestine
Spl Spleen
Test Testes
Thym Thymus
Tong Tongue
Trac Trachea
Ute Uterus

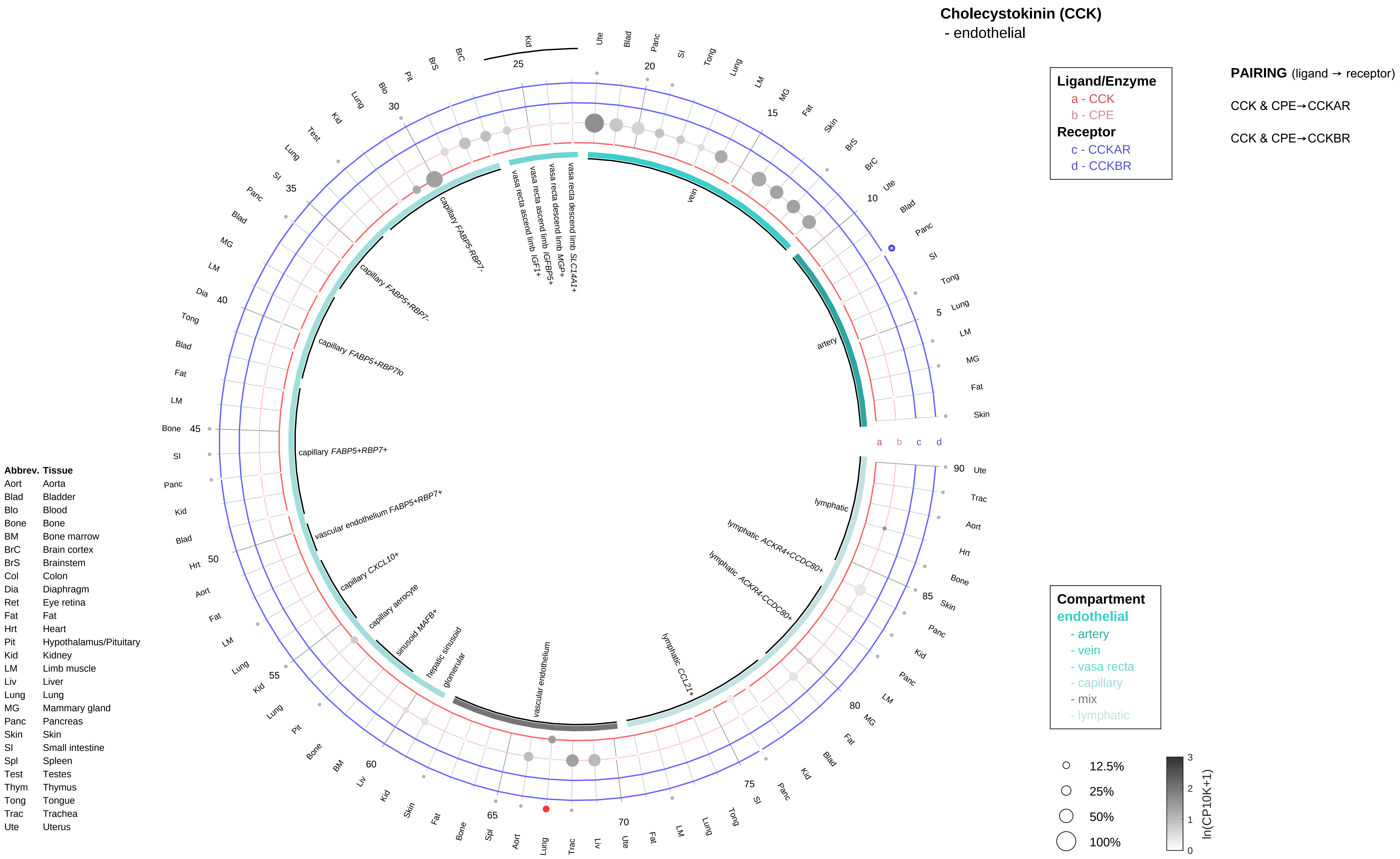

Cholecystokinin (CCK)
- endothelial
Ligand/Enzyme
a - CCK
b - CPE
Receptor
c - CCKAR
d - CCKBR
PAIRING (ligand → receptor)
CCK & CPE→CCKAR
CCK & CPE→CCKBR
Compartment
endothelial
- artery
- vein
- vasa recta
- capillary
- mix
- lymphatic
12.5%
25%
50%
100%
ln(CP10K+1)
artery
vein
vasa recta descend limb SLC14A1+
vasa recta descend limb MGP+
vasa recta ascend limb IGFBP5+
vasa recta ascend limb IGF1+
capillary FABP5-RBP7-
capillary FABP5+RBP7-
capillary FABP5+RBP7lo
capillary FABP5+RBP7+
vascular endothelium FABP5+RBP7+
capillary CXCL10+
capillary aerocyte
sinusoid MAFB+
hepatic sinusoid
glomerular
vascular endothelium
lymphatic CCL21+
lymphatic ACKR4-CCDC80+
lymphatic ACKR4+CCDC80+
lymphatic
Abbrev. Tissue
Aort Aorta
Blad Bladder
Blo Blood
Bone Bone
BM Bone marrow
BrC Brain cortex
BrS Brainstem
Col Colon
Dia Diaphragm
Ret Eye retina
Fat Fat
Hrt Heart
Pit Hypothalamus/Pituitary
Kid Kidney
LM Limb muscle
Liv Liver
Lung Lung
MG Mammary gland
Panc Pancreas
Skin Skin
SI Small intestine
Spl Spleen
Test Testes
Thym Thymus
Tong Tongue
Trac Trachea
Ute Uterus

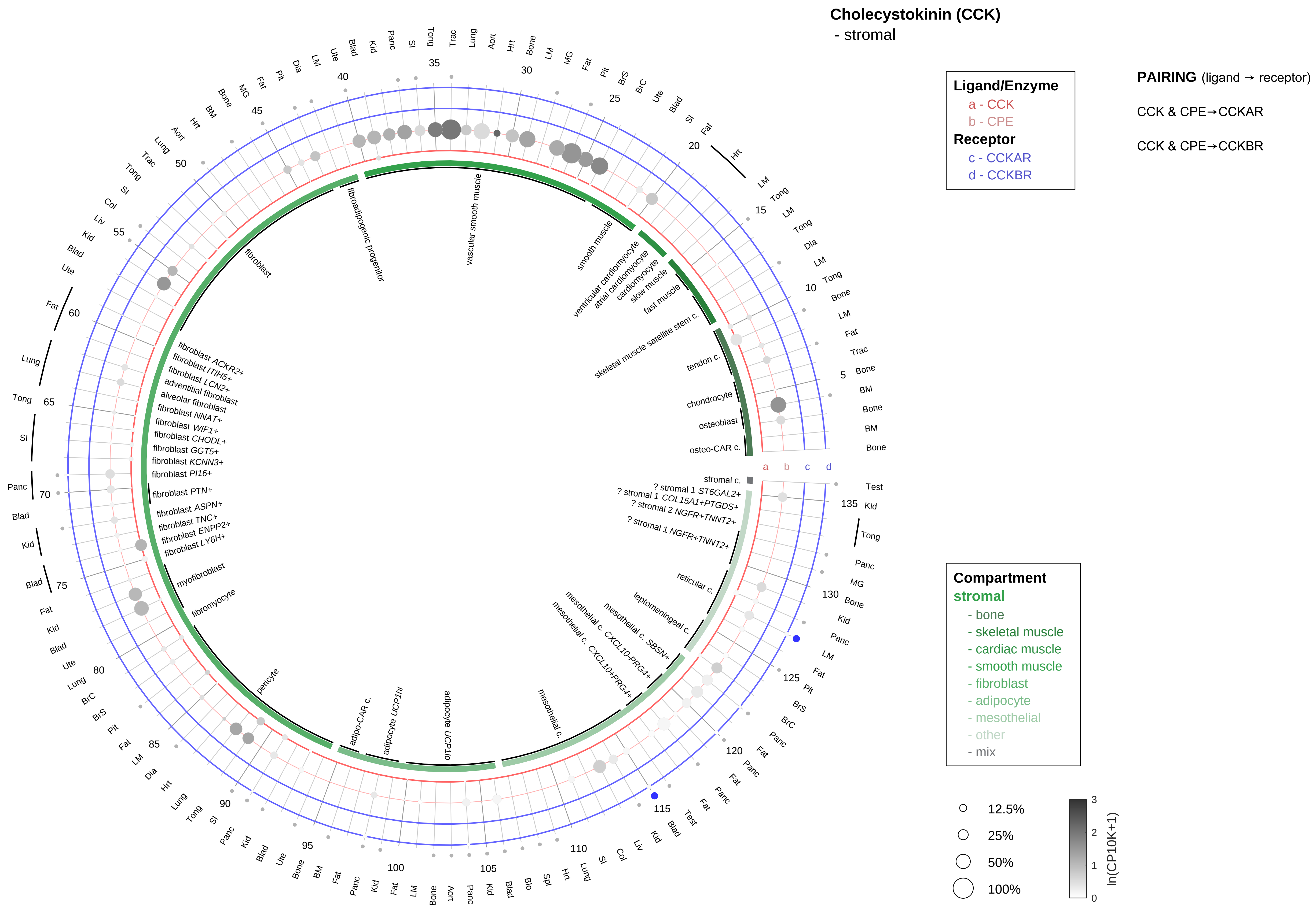

| Abbrev. | Tissue |
|---|---|
| Aort | Aorta |
| Blad | Bladder |
| Blo | Blood |
| Bone | Bone |
| BM | Bone marrow |
| BrC | Brain cortex |
| BrS | Brainstem |
| Col | Colon |
| Dia | Diaphragm |
| Ret | Eye retina |
| Fat | Fat |
| Hrt | Heart |
| Pit | Hypothalamus/Pituitary |
| Kid | Kidney |
| LM | Limb muscle |
| Liv | Liver |
| Lung | Lung |
| MG | Mammary gland |
| Panc | Pancreas |
| Skin | Skin |
| SI | Small intestine |
| Spl | Spleen |
| Test | Testes |
| Thym | Thymus |
| Tong | Tongue |
| Trac | Trachea |
| Ute | Uterus |

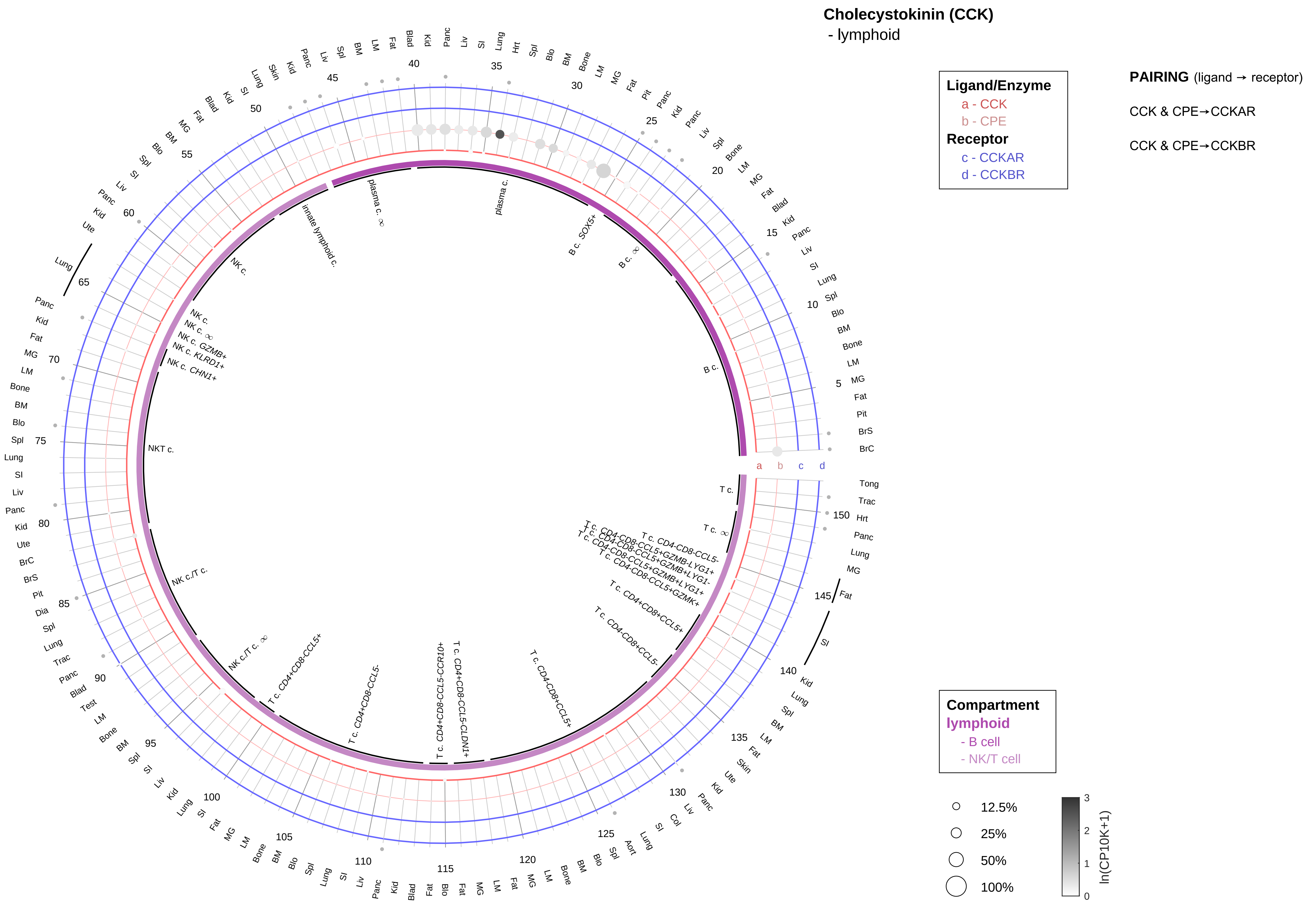

| Abbrev. | Tissue |
|---|---|
| Aort | Aorta |
| Blad | Bladder |
| Blo | Blood |
| Bone | Bone |
| BM | Bone marrow |
| BrC | Brain cortex |
| BrS | Brainstem |
| Col | Colon |
| Dia | Diaphragm |
| Ret | Eye retina |
| Fat | Fat |
| Hrt | Heart |
| Pit | Hypothalamus/Pituitary |
| Kid | Kidney |
| LM | Limb muscle |
| Liv | Liver |
| Lung | Lung |
| MG | Mammary gland |
| Panc | Pancreas |
| Skin | Skin |
| SI | Small intestine |
| Spl | Spleen |
| Test | Testes |
| Thym | Thymus |
| Tong | Tongue |
| Trac | Trachea |
| Ute | Uterus |

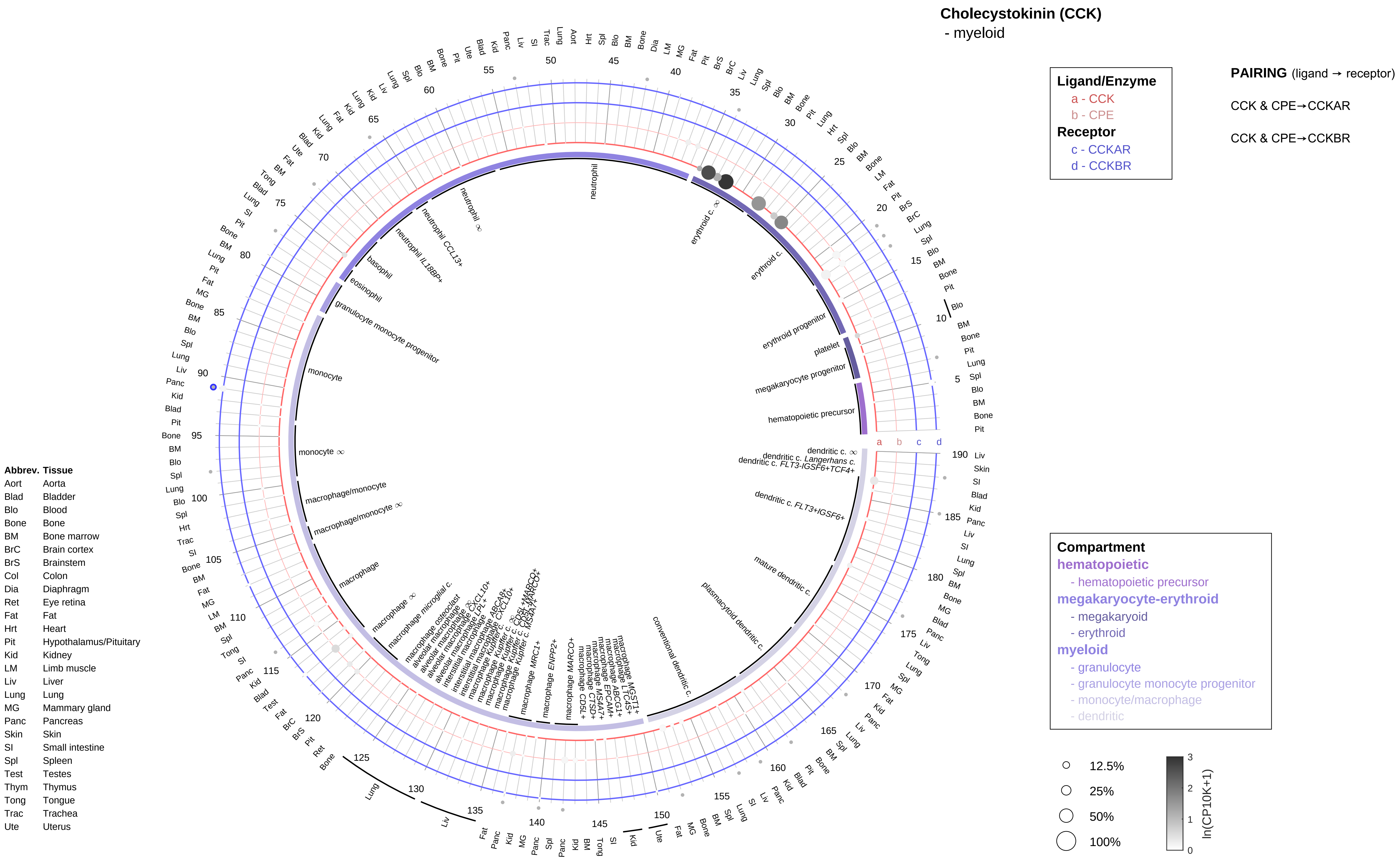
Cholecystokinin (CCK)
- myeloid
Ligand/Enzyme
a - CCK
b - CPE
Receptor
c - CCKAR
d - CCKBR
PAIRING (ligand → receptor)
CCK & CPE→CCKAR
CCK & CPE→CCKBR
Compartment
hematopoietic
- hematopoietic precursor
megakaryocyte-erythroid
- megakaryoid
- erythroid
myeloid
- granulocyte
- granulocyte monocyte progenitor
- monocyte/macrophage
- dendritic
12.5%
25%
50%
100%
ln(CP10K+1)
Abbrev. Tissue
Aort Aorta
Blad Bladder
Blo Blood
Bone Bone
BM Bone marrow
BrC Brain cortex
BrS Brainstem
Col Colon
Dia Diaphragm
Ret Eye retina
Fat Fat
Hrt Heart
Pit Hypothalamus/Pituitary
Kid Kidney
LM Limb muscle
Liv Liver
Lung Lung
MG Mammary gland
Panc Pancreas
Skin Skin
SI Small intestine
Spl Spleen
Test Testes
Thym Thymus
Tong Tongue
Trac Trachea
Ute Uterus

#### Cocaine- and amphetamine-regulated transcript (CART)
- epithelial/neural/germ

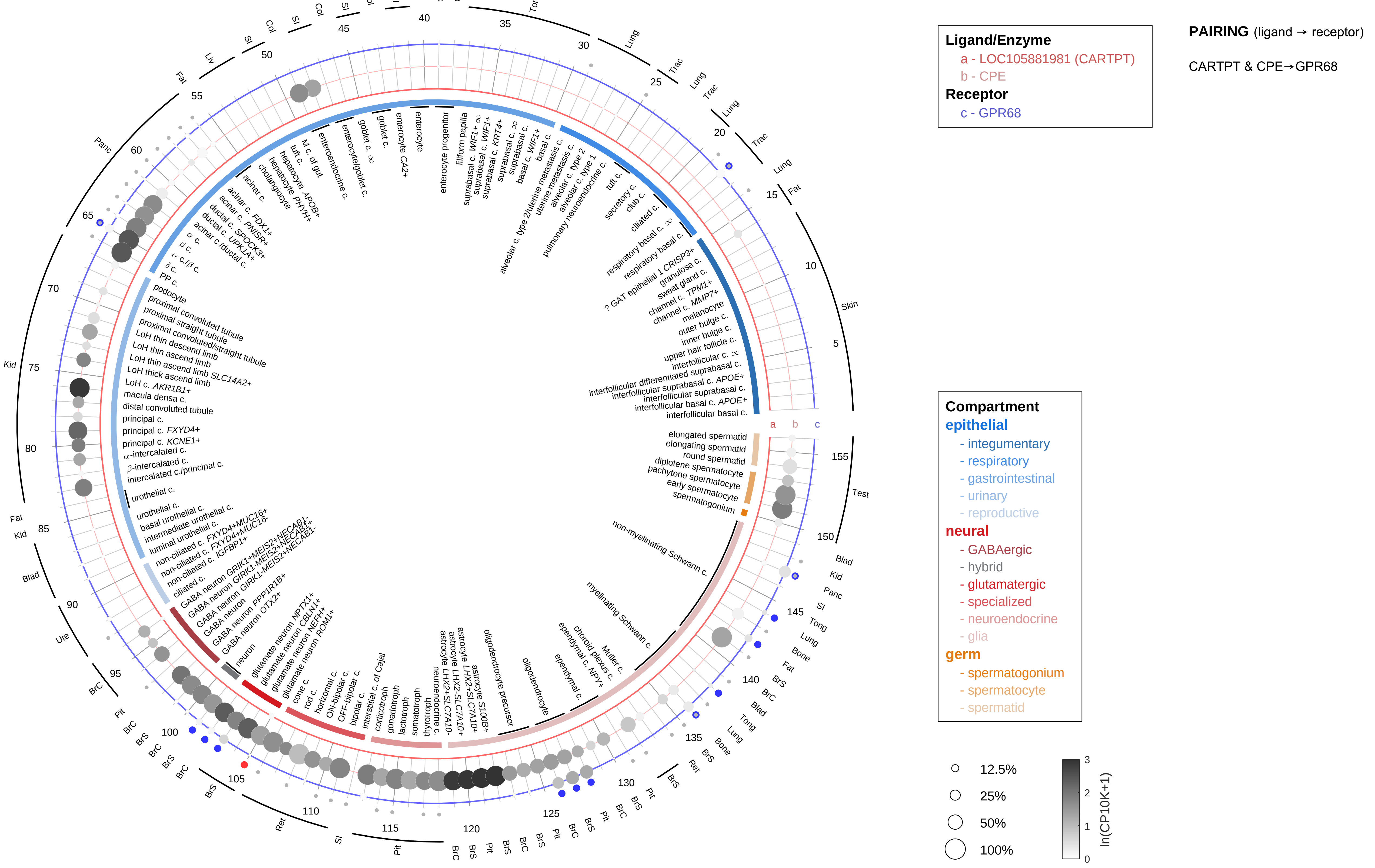

| Abbrev. | Tissue |
|---|---|
| Aort | Aorta |
| Blad | Bladder |
| Blo | Blood |
| Bone | Bone |
| BM | Bone marrow |
| BrC | Brain cortex |
| BrS | Brainstem |
| Col | Colon |
| Dia | Diaphragm |
| Ret | Eye retina |
| Fat | Fat |
| Hrt | Heart |
| Pit | Hypothalamus/Pituitary |
| Kid | Kidney |
| LM | Limb muscle |
| Liv | Liver |
| Lung | Lung |
| MG | Mammary gland |
| Panc | Pancreas |
| Skin | Skin |
| SI | Small intestine |
| Spl | Spleen |
| Test | Testes |
| Thym | Thymus |
| Tong | Tongue |
| Trac | Trachea |
| Ute | Uterus |

#### Cocaine- and amphetamine-regulated transcript (CART)
- endothelial

**Ligand/Enzyme**
a - LOC105881981 (CARTPT)
b - CPE
**Receptor**
c - GPR68

**PAIRING** (ligand → receptor)

CARTPT & CPE→GPR68

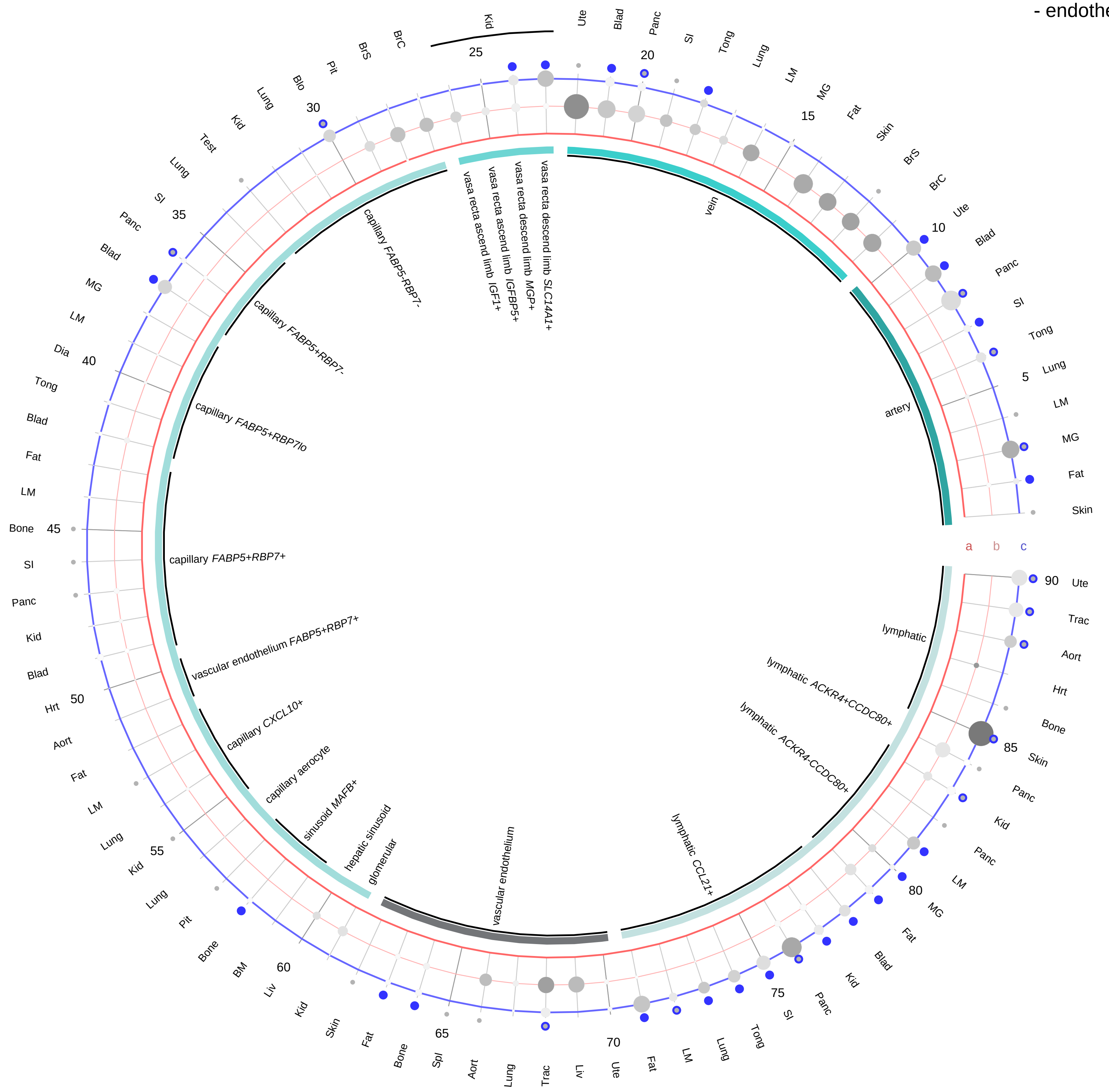

| Abbrev. | Tissue |
|---|---|
| Aort | Aorta |
| Blad | Bladder |
| Blo | Blood |
| Bone | Bone |
| BM | Bone marrow |
| BrC | Brain cortex |
| BrS | Brainstem |
| Col | Colon |
| Dia | Diaphragm |
| Ret | Eye retina |
| Fat | Fat |
| Hrt | Heart |
| Pit | Hypothalamus/Pituitary |
| Kid | Kidney |
| LM | Limb muscle |
| Liv | Liver |
| Lung | Lung |
| MG | Mammary gland |
| Panc | Pancreas |
| Skin | Skin |
| SI | Small intestine |
| Spl | Spleen |
| Test | Testes |
| Thym | Thymus |
| Tong | Tongue |
| Trac | Trachea |
| Ute | Uterus |

**Compartment**
**endothelial**
- artery
- vein
- vasa recta
- capillary
- mix
- lymphatic

12.5%
25%
50%
100%

ln(CP10K+1): 0, 1, 2, 3

**Cocaine- and amphetamine-regulated transcript (CART)**
- stromal

**Ligand/Enzyme**
a - LOC105881981 (CARTPT)
b - CPE
**Receptor**
c - GPR68

**PAIRING** (ligand → receptor)

CARTPT & CPE→GPR68

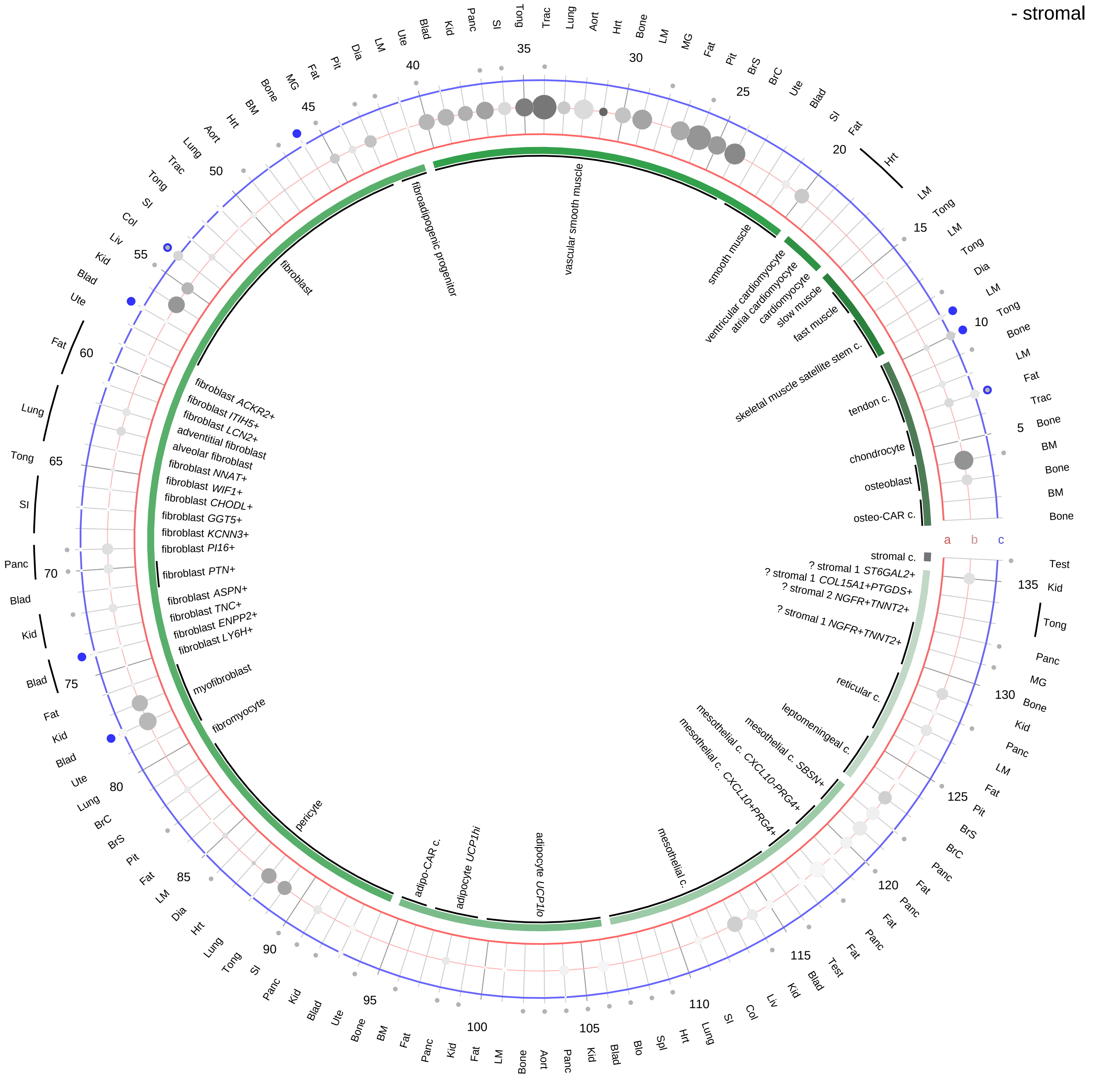

| Abbrev. | Tissue |
|---|---|
| Aort | Aorta |
| Blad | Bladder |
| Blo | Blood |
| Bone | Bone |
| BM | Bone marrow |
| BrC | Brain cortex |
| BrS | Brainstem |
| Col | Colon |
| Dia | Diaphragm |
| Ret | Eye retina |
| Fat | Fat |
| Hrt | Heart |
| Pit | Hypothalamus/Pituitary |
| Kid | Kidney |
| LM | Limb muscle |
| Liv | Liver |
| Lung | Lung |
| MG | Mammary gland |
| Panc | Pancreas |
| Skin | Skin |
| SI | Small intestine |
| Spl | Spleen |
| Test | Testes |
| Thym | Thymus |
| Tong | Tongue |
| Trac | Trachea |
| Ute | Uterus |

**Compartment**
**stromal**
- bone
- skeletal muscle
- cardiac muscle
- smooth muscle
- fibroblast
- adipocyte
- mesothelial
- other
- mix

12.5%
25%
50%
100%

ln(CP10K+1): 0, 1, 2, 3

**Cocaine- and amphetamine-regulated transcript (CART)**
- lymphoid

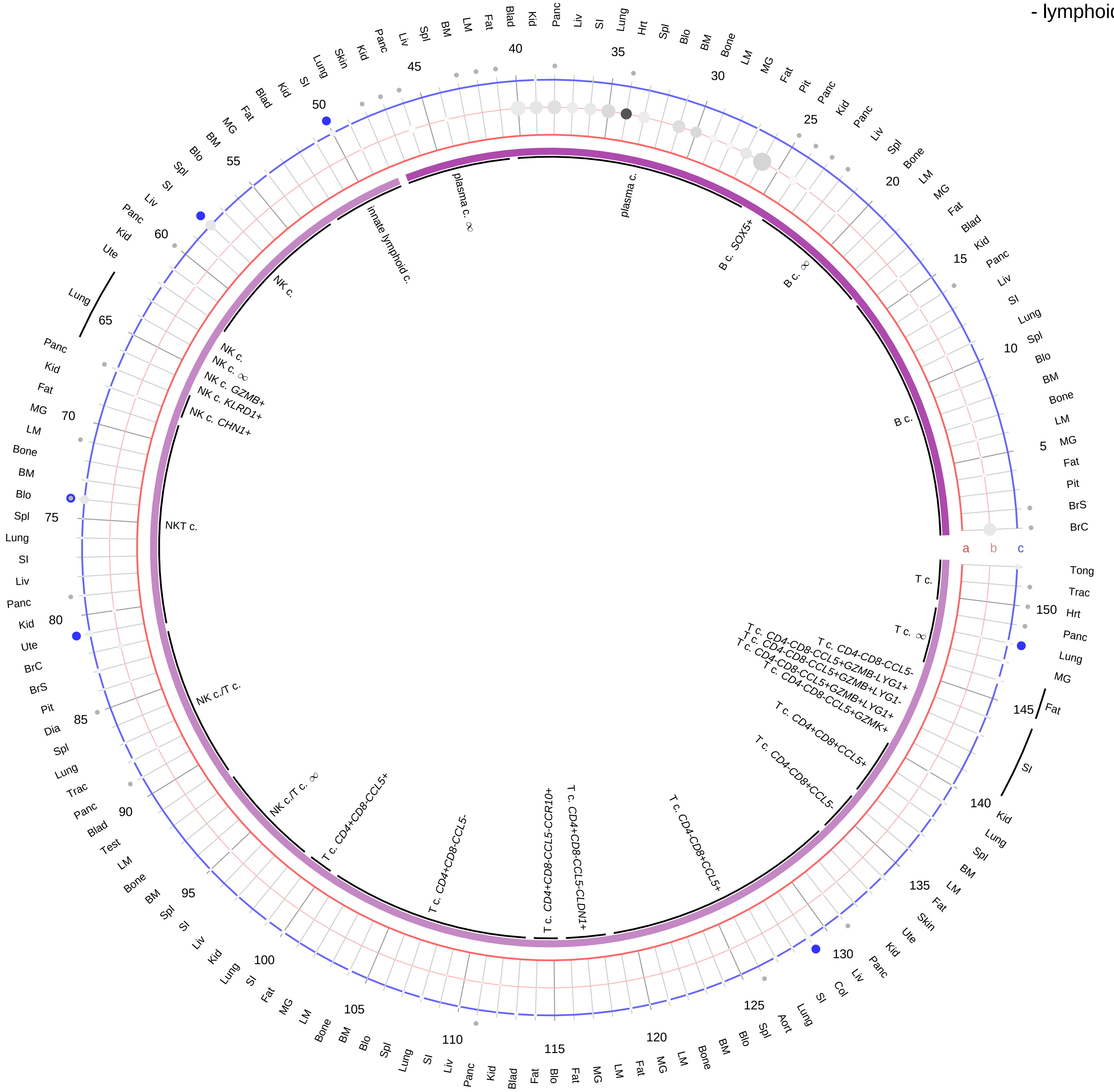

| Abbrev. | Tissue |
|---|---|
| Aort | Aorta |
| Blad | Bladder |
| Blo | Blood |
| Bone | Bone |
| BM | Bone marrow |
| BrC | Brain cortex |
| BrS | Brainstem |
| Col | Colon |
| Dia | Diaphragm |
| Ret | Eye retina |
| Fat | Fat |
| Hrt | Heart |
| Pit | Hypothalamus/Pituitary |
| Kid | Kidney |
| LM | Limb muscle |
| Liv | Liver |
| Lung | Lung |
| MG | Mammary gland |
| Panc | Pancreas |
| Skin | Skin |
| SI | Small intestine |
| Spl | Spleen |
| Test | Testes |
| Thym | Thymus |
| Tong | Tongue |
| Trac | Trachea |
| Ute | Uterus |

**Ligand/Enzyme**
- a - LOC105881981 (CARTPT)
- b - CPE

**Receptor**
- c - GPR68

**PAIRING** (ligand → receptor)

CARTPT & CPE→GPR68

**Compartment**
**lymphoid**
- B cell
- NK/T cell

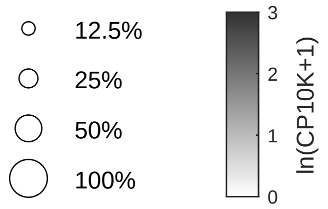

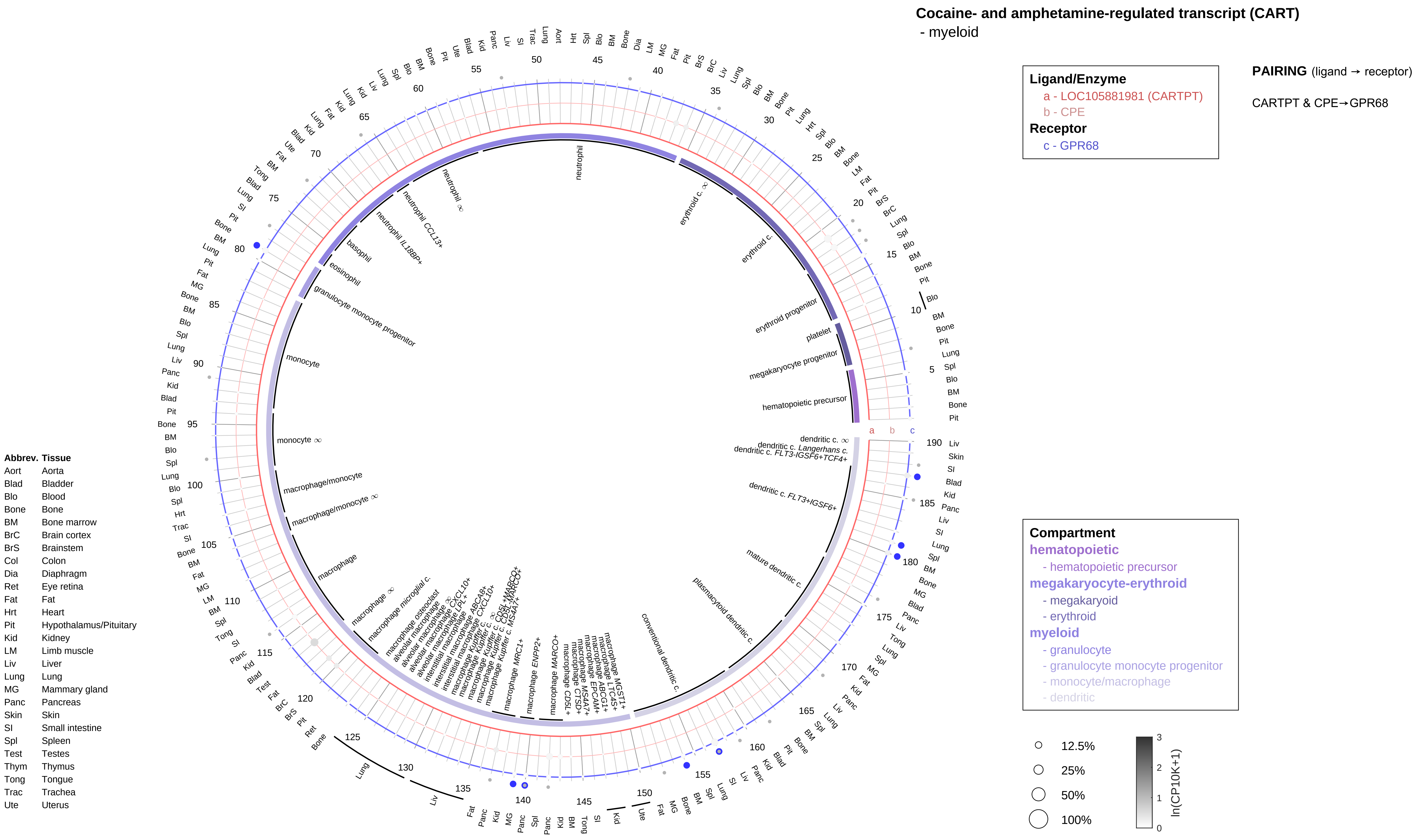

Cocaine- and amphetamine-regulated transcript (CART)
- myeloid
Ligand/Enzyme
a - LOC105881981 (CARTPT)
b - CPE
Receptor
c - GPR68
PAIRING (ligand → receptor)
CARTPT & CPE→GPR68
Compartment
hematopoietic
- hematopoietic precursor
megakaryocyte-erythroid
- megakaryoid
- erythroid
myeloid
- granulocyte
- granulocyte monocyte progenitor
- monocyte/macrophage
- dendritic
12.5%
25%
50%
100%
ln(CP10K+1)
Abbrev. Tissue
Aort Aorta
Blad Bladder
Blo Blood
Bone Bone
BM Bone marrow
BrC Brain cortex
BrS Brainstem
Col Colon
Dia Diaphragm
Ret Eye retina
Fat Fat
Hrt Heart
Pit Hypothalamus/Pituitary
Kid Kidney
LM Limb muscle
Liv Liver
Lung Lung
MG Mammary gland
Panc Pancreas
Skin Skin
SI Small intestine
Spl Spleen
Test Testes
Thym Thymus
Tong Tongue
Trac Trachea
Ute Uterus
neutrophil
erythroid c. ∞
erythroid c.
erythroid progenitor
platelet
megakaryocyte progenitor
hematopoietic precursor
dendritic c. ∞
dendritic c. Langerhans c.
dendritic c. FLT3-IGSF6+TCF4+
dendritic c. FLT3+IGSF6+
mature dendritic c.
plasmacytoid dendritic c.
conventional dendritic c.
neutrophil ∞
neutrophil CCL13+
neutrophil IL18BP+
basophil
eosinophil
granulocyte monocyte progenitor
monocyte
monocyte ∞
macrophage/monocyte
macrophage/monocyte ∞
macrophage
macrophage ∞
macrophage microglial c.
macrophage osteoclast
macrophage MRC1+
macrophage ENPP2+
macrophage MARCO+
macrophage CD5L+

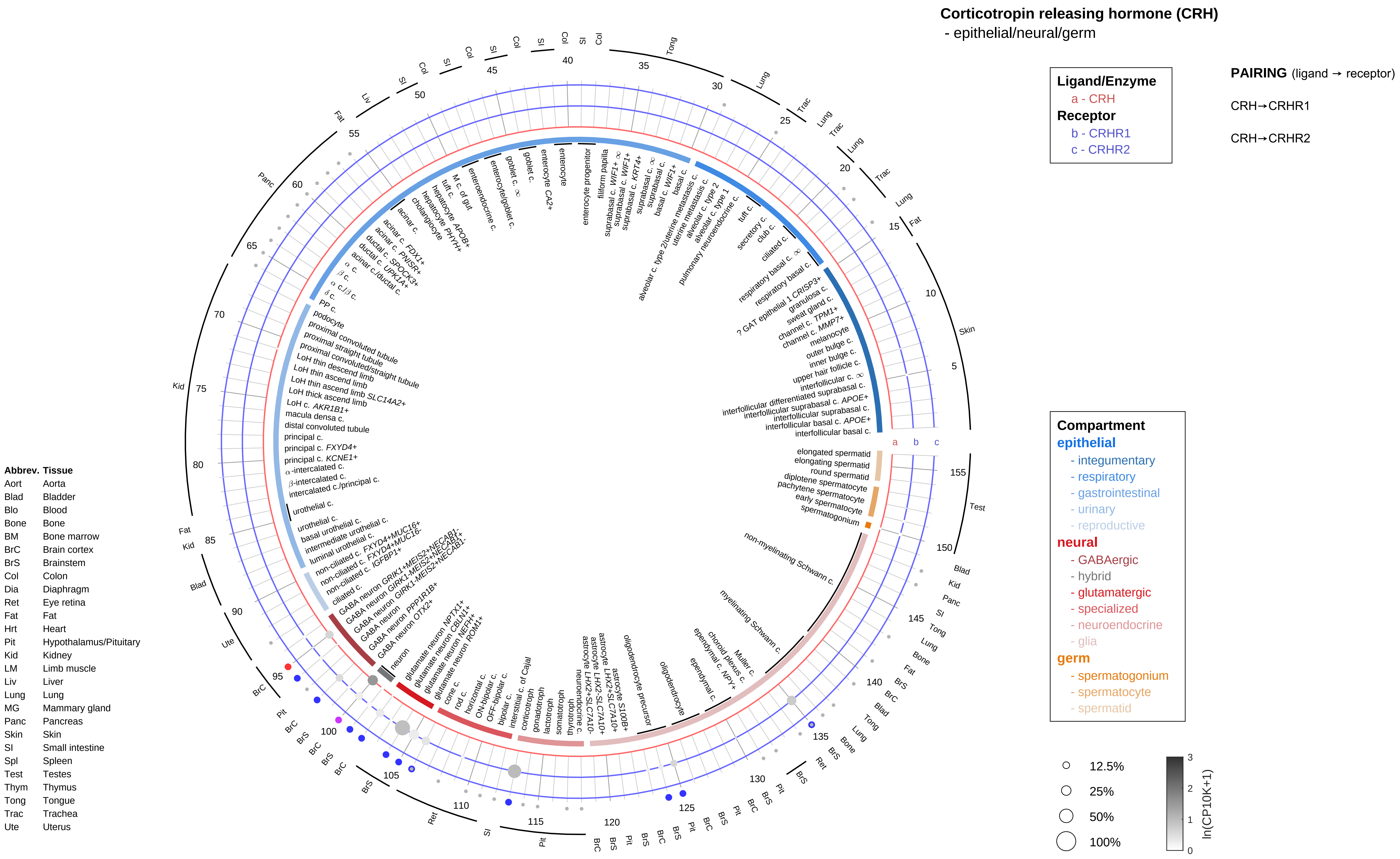

Corticotropin releasing hormone (CRH)
- epithelial/neural/germ
Ligand/Enzyme
a - CRH
Receptor
b - CRHR1
c - CRHR2
PAIRING (ligand → receptor)
CRH→CRHR1
CRH→CRHR2
Compartment
epithelial
- integumentary
- respiratory
- gastrointestinal
- urinary
- reproductive
neural
- GABAergic
- hybrid
- glutamatergic
- specialized
- neuroendocrine
- glia
germ
- spermatogonium
- spermatocyte
- spermatid
12.5%
25%
50%
100%
ln(CP10K+1)
Abbrev. Tissue
Aort Aorta
Blad Bladder
Blo Blood
Bone Bone
BM Bone marrow
BrC Brain cortex
BrS Brainstem
Col Colon
Dia Diaphragm
Ret Eye retina
Fat Fat
Hrt Heart
Pit Hypothalamus/Pituitary
Kid Kidney
LM Limb muscle
Liv Liver
Lung Lung
MG Mammary gland
Panc Pancreas
Skin Skin
SI Small intestine
Spl Spleen
Test Testes
Thym Thymus
Tong Tongue
Trac Trachea
Ute Uterus

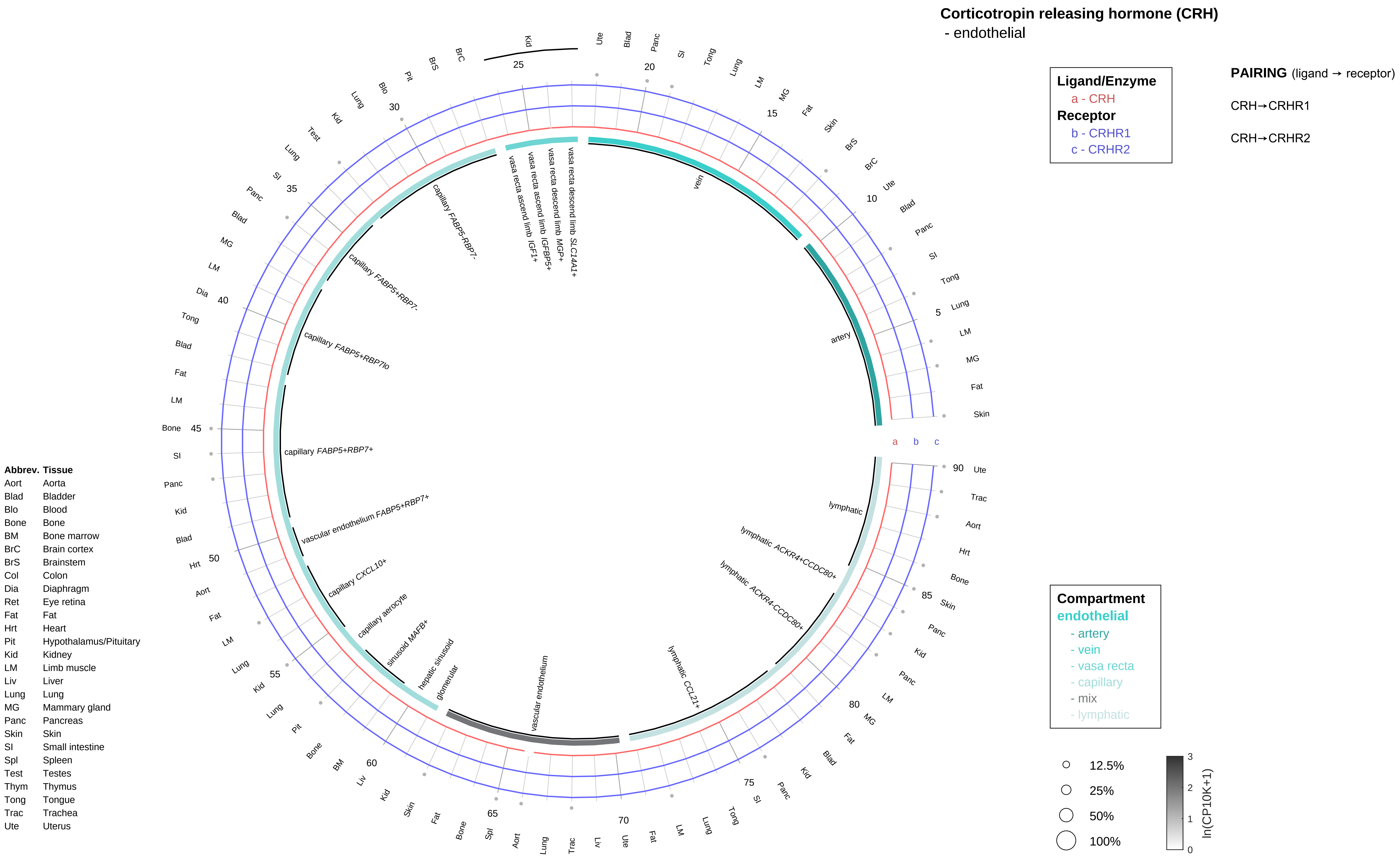

Corticotropin releasing hormone (CRH)
- endothelial
Ligand/Enzyme
a - CRH
Receptor
b - CRHR1
c - CRHR2
PAIRING (ligand → receptor)
CRH→CRHR1
CRH→CRHR2
Compartment
endothelial
- artery
- vein
- vasa recta
- capillary
- mix
- lymphatic
12.5%
25%
50%
100%
ln(CP10K+1)
vein
artery
vasa recta descend limb SLC14A1+
vasa recta descend limb MGP+
vasa recta ascend limb IGFBP5+
vasa recta ascend limb IGF1+
capillary FABP5-RBP7-
capillary FABP5+RBP7-
capillary FABP5+RBP7lo
capillary FABP5+RBP7+
vascular endothelium FABP5+RBP7+
capillary CXCL10+
capillary aerocyte
sinusoid MAFB+
hepatic sinusoid
glomerular
vascular endothelium
lymphatic CCL21+
lymphatic ACKR4-CCDC80+
lymphatic ACKR4+CCDC80+
lymphatic
Abbrev. Tissue
Aort Aorta
Blad Bladder
Blo Blood
Bone Bone
BM Bone marrow
BrC Brain cortex
BrS Brainstem
Col Colon
Dia Diaphragm
Ret Eye retina
Fat Fat
Hrt Heart
Pit Hypothalamus/Pituitary
Kid Kidney
LM Limb muscle
Liv Liver
Lung Lung
MG Mammary gland
Panc Pancreas
Skin Skin
SI Small intestine
Spl Spleen
Test Testes
Thym Thymus
Tong Tongue
Trac Trachea
Ute Uterus

**Corticotropin releasing hormone (CRH)**
- stromal

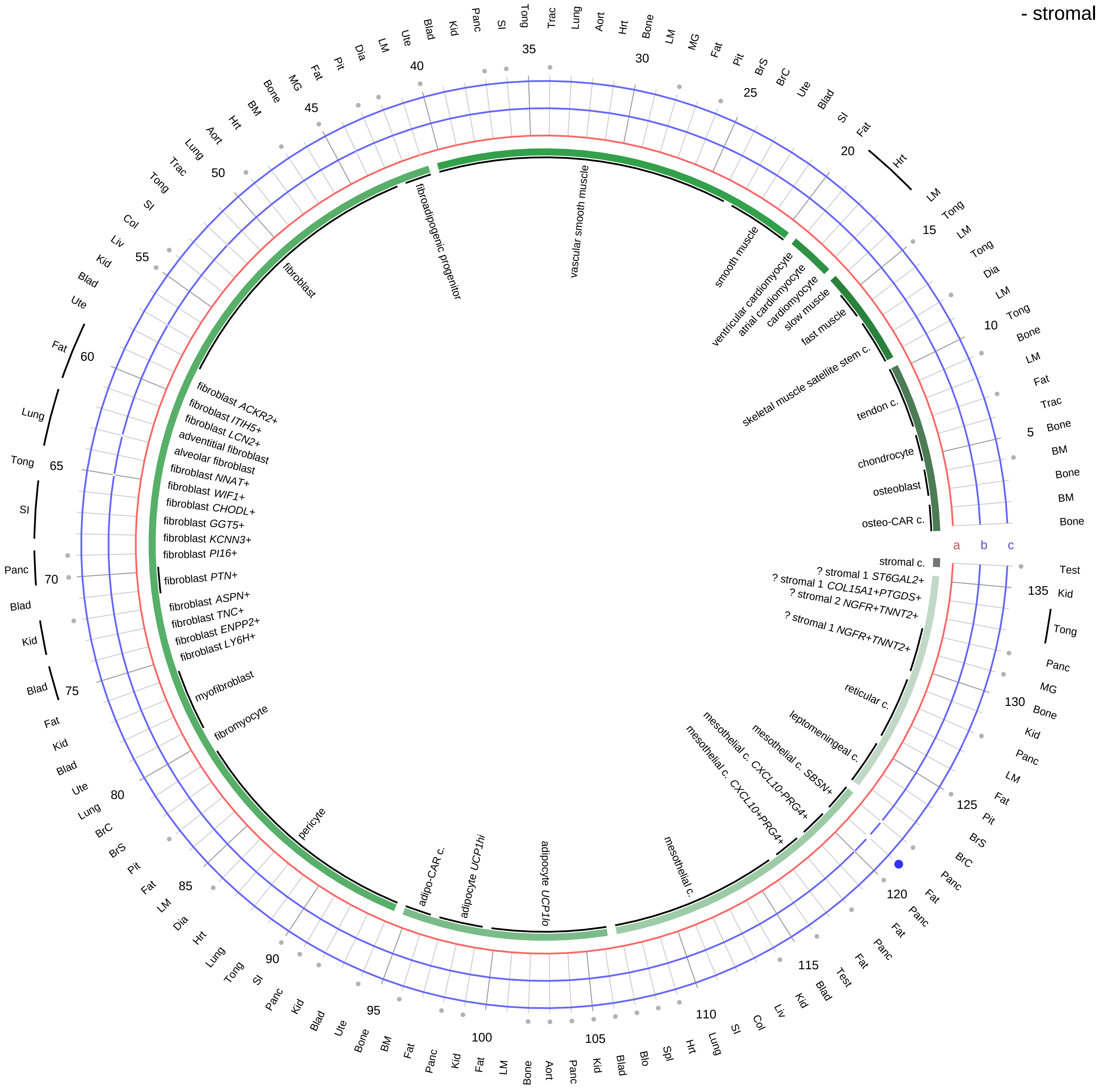

| **Ligand/Enzyme** |
|---|
| a - CRH |
| **Receptor** |
| b - CRHR1 |
| c - CRHR2 |

**PAIRING** (ligand → receptor)

CRH→CRHR1

CRH→CRHR2

| **Compartment** |
|---|
| **stromal** |
| - bone |
| - skeletal muscle |
| - cardiac muscle |
| - smooth muscle |
| - fibroblast |
| - adipocyte |
| - mesothelial |
| - other |
| - mix |

12.5%
25%
50%
100%

ln(CP10K+1): 0, 1, 2, 3

| Abbrev. | Tissue |
|---|---|
| Aort | Aorta |
| Blad | Bladder |
| Blo | Blood |
| Bone | Bone |
| BM | Bone marrow |
| BrC | Brain cortex |
| BrS | Brainstem |
| Col | Colon |
| Dia | Diaphragm |
| Ret | Eye retina |
| Fat | Fat |
| Hrt | Heart |
| Pit | Hypothalamus/Pituitary |
| Kid | Kidney |
| LM | Limb muscle |
| Liv | Liver |
| Lung | Lung |
| MG | Mammary gland |
| Panc | Pancreas |
| Skin | Skin |
| SI | Small intestine |
| Spl | Spleen |
| Test | Testes |
| Thym | Thymus |
| Tong | Tongue |
| Trac | Trachea |
| Ute | Uterus |

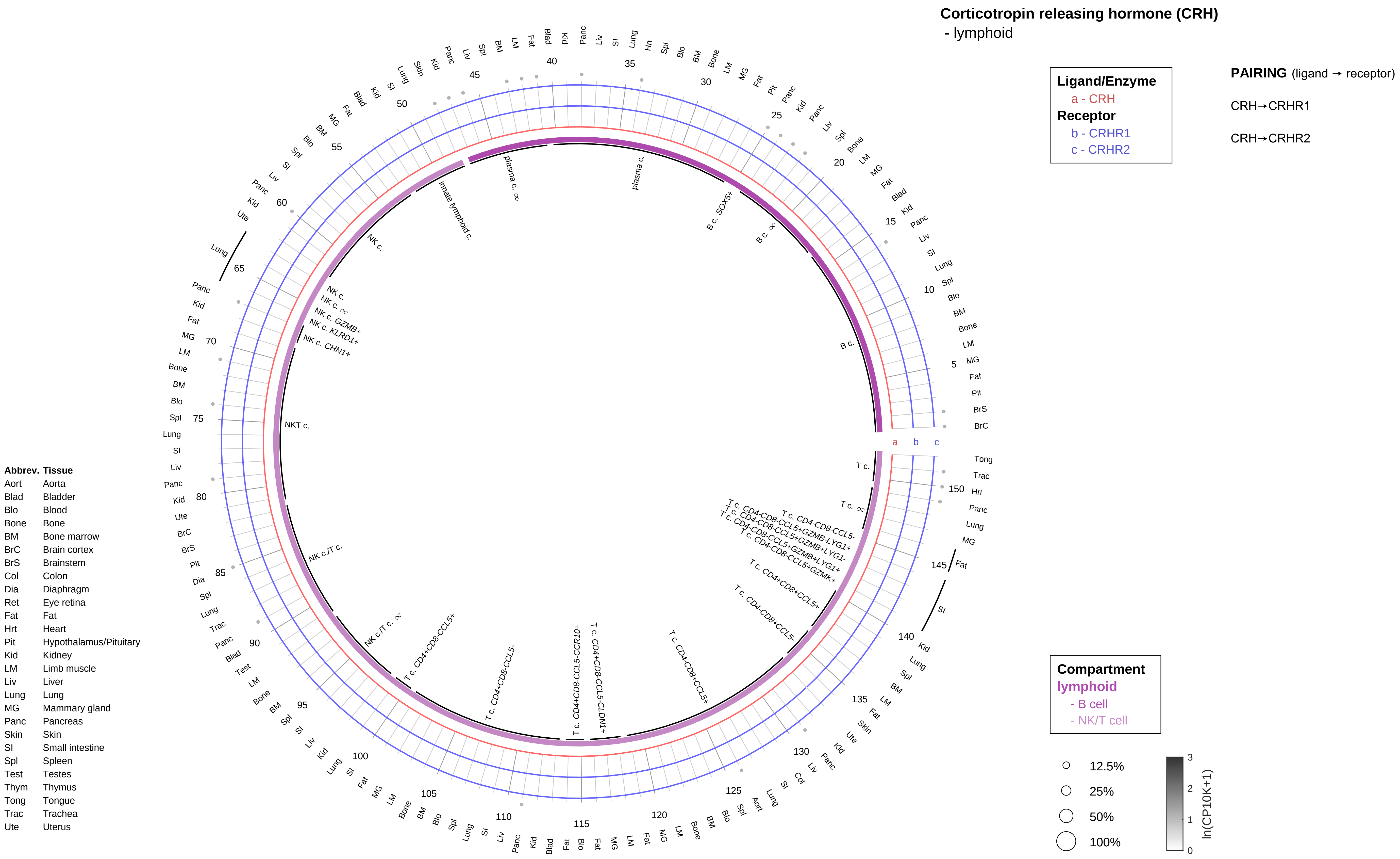
Corticotropin releasing hormone (CRH)
- lymphoid
Ligand/Enzyme
a - CRH
Receptor
b - CRHR1
c - CRHR2
PAIRING (ligand → receptor)
CRH→CRHR1
CRH→CRHR2
Compartment
lymphoid
- B cell
- NK/T cell
12.5%
25%
50%
100%
ln(CP10K+1)
Abbrev. Tissue
Aort Aorta
Blad Bladder
Blo Blood
Bone Bone
BM Bone marrow
BrC Brain cortex
BrS Brainstem
Col Colon
Dia Diaphragm
Ret Eye retina
Fat Fat
Hrt Heart
Pit Hypothalamus/Pituitary
Kid Kidney
LM Limb muscle
Liv Liver
Lung Lung
MG Mammary gland
Panc Pancreas
Skin Skin
SI Small intestine
Spl Spleen
Test Testes
Thym Thymus
Tong Tongue
Trac Trachea
Ute Uterus

**Corticotropin releasing hormone (CRH)**
- myeloid

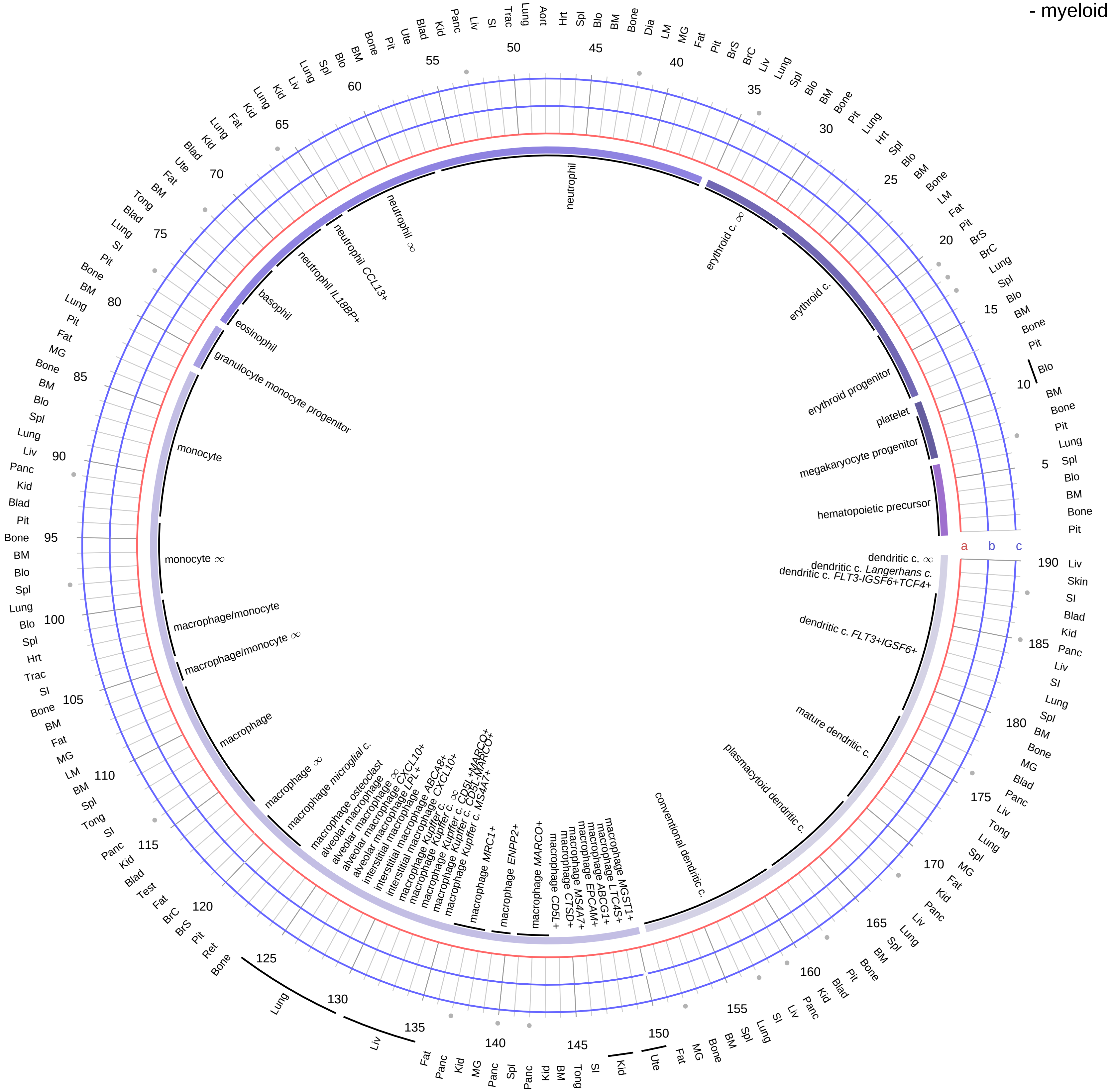

| Ligand/Enzyme |
|---|
| a - CRH |
| **Receptor** |
| b - CRHR1 |
| c - CRHR2 |

**PAIRING** (ligand → receptor)

CRH→CRHR1

CRH→CRHR2

| Abbrev. | Tissue |
|---|---|
| Aort | Aorta |
| Blad | Bladder |
| Blo | Blood |
| Bone | Bone |
| BM | Bone marrow |
| BrC | Brain cortex |
| BrS | Brainstem |
| Col | Colon |
| Dia | Diaphragm |
| Ret | Eye retina |
| Fat | Fat |
| Hrt | Heart |
| Pit | Hypothalamus/Pituitary |
| Kid | Kidney |
| LM | Limb muscle |
| Liv | Liver |
| Lung | Lung |
| MG | Mammary gland |
| Panc | Pancreas |
| Skin | Skin |
| SI | Small intestine |
| Spl | Spleen |
| Test | Testes |
| Thym | Thymus |
| Tong | Tongue |
| Trac | Trachea |
| Ute | Uterus |

**Compartment**
**hematopoietic**
- hematopoietic precursor

**megakaryocyte-erythroid**
- megakaryocyte
- erythroid

**myeloid**
- granulocyte
- granulocyte monocyte progenitor
- monocyte/macrophage
- dendritic

12.5%
25%
50%
100%

ln(CP10K+1): 0, 1, 2, 3

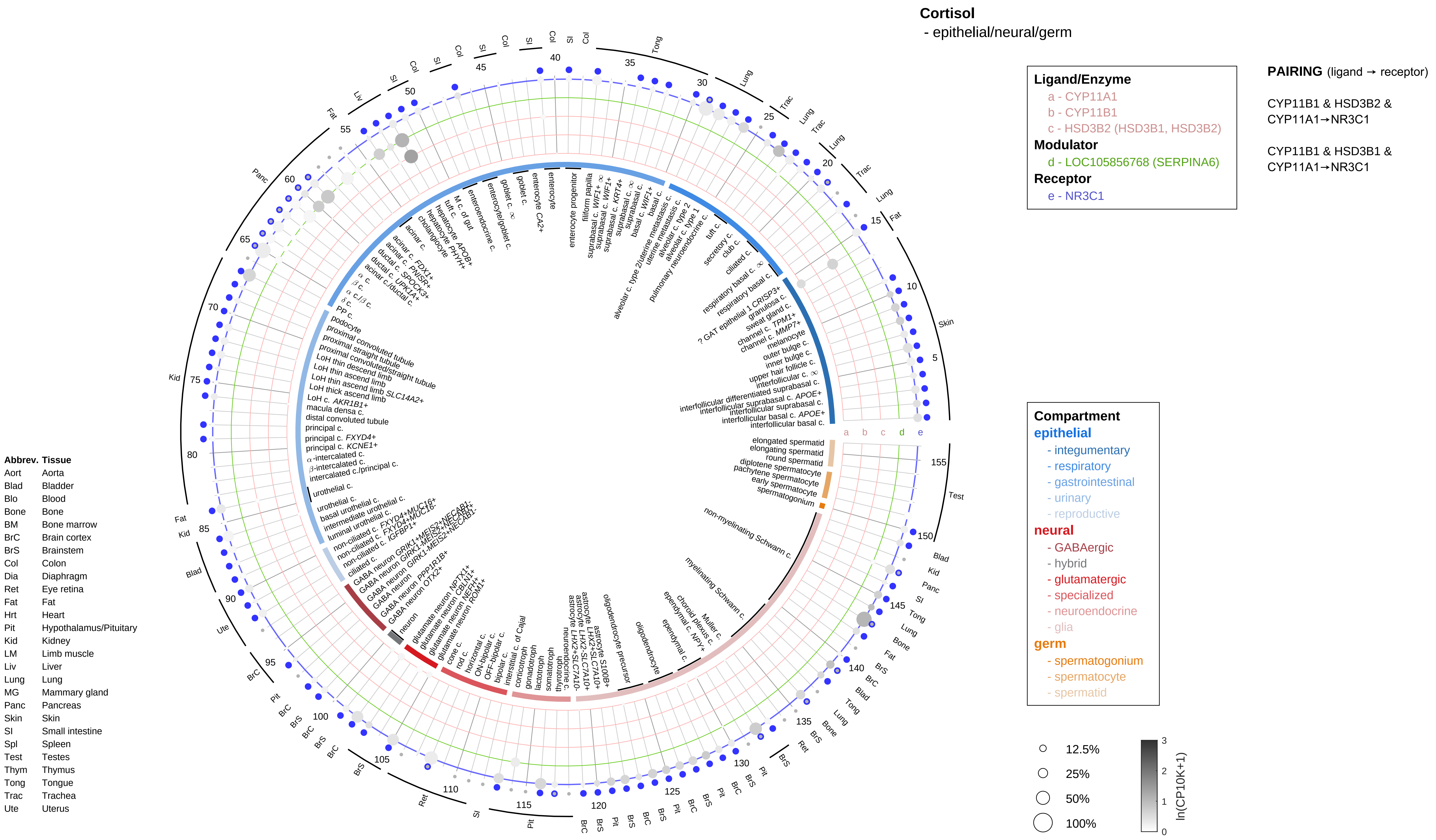
Cortisol
- epithelial/neural/germ
Ligand/Enzyme
a - CYP11A1
b - CYP11B1
c - HSD3B2 (HSD3B1, HSD3B2)
Modulator
d - LOC105856768 (SERPINA6)
Receptor
e - NR3C1
PAIRING (ligand → receptor)
CYP11B1 & HSD3B2 & CYP11A1→NR3C1
CYP11B1 & HSD3B1 & CYP11A1→NR3C1
Compartment
epithelial
- integumentary
- respiratory
- gastrointestinal
- urinary
- reproductive
neural
- GABAergic
- hybrid
- glutamatergic
- specialized
- neuroendocrine
- glia
germ
- spermatogonium
- spermatocyte
- spermatid
12.5%
25%
50%
100%
ln(CP10K+1)
Abbrev. Tissue
Aort Aorta
Blad Bladder
Blo Blood
Bone Bone
BM Bone marrow
BrC Brain cortex
BrS Brainstem
Col Colon
Dia Diaphragm
Ret Eye retina
Fat Fat
Hrt Heart
Pit Hypothalamus/Pituitary
Kid Kidney
LM Limb muscle
Liv Liver
Lung Lung
MG Mammary gland
Panc Pancreas
Skin Skin
SI Small intestine
Spl Spleen
Test Testes
Thym Thymus
Tong Tongue
Trac Trachea
Ute Uterus

**Cortisol**
- endothelial

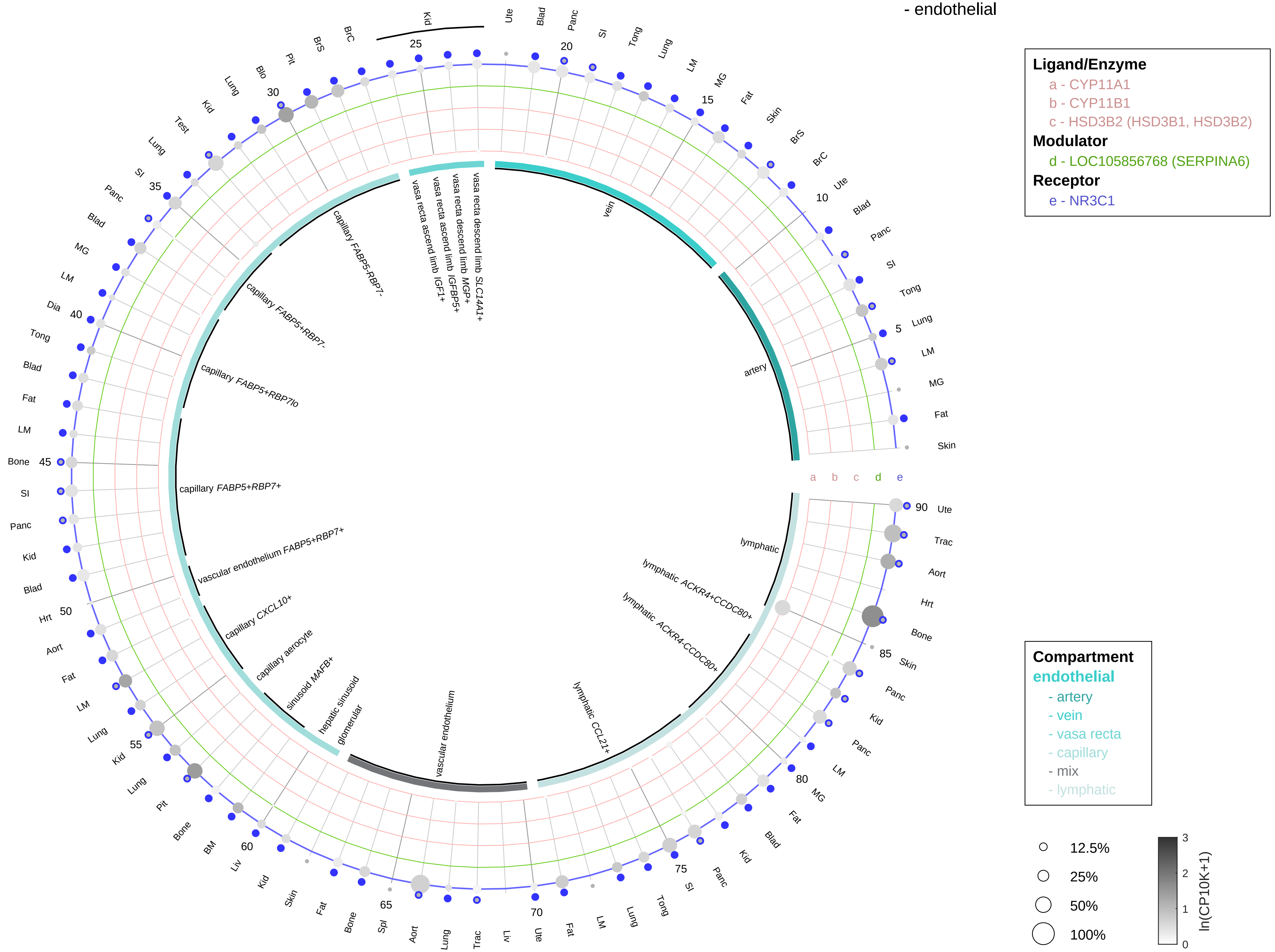

**PAIRING** (ligand → receptor)

CYP11B1 & HSD3B2 & CYP11A1→NR3C1

CYP11B1 & HSD3B1 & CYP11A1→NR3C1

| Abbrev. | Tissue |
|---|---|
| Aort | Aorta |
| Blad | Bladder |
| Blo | Blood |
| Bone | Bone |
| BM | Bone marrow |
| BrC | Brain cortex |
| BrS | Brainstem |
| Col | Colon |
| Dia | Diaphragm |
| Ret | Eye retina |
| Fat | Fat |
| Hrt | Heart |
| Pit | Hypothalamus/Pituitary |
| Kid | Kidney |
| LM | Limb muscle |
| Liv | Liver |
| Lung | Lung |
| MG | Mammary gland |
| Panc | Pancreas |
| Skin | Skin |
| SI | Small intestine |
| Spl | Spleen |
| Test | Testes |
| Thym | Thymus |
| Tong | Tongue |
| Trac | Trachea |
| Ute | Uterus |

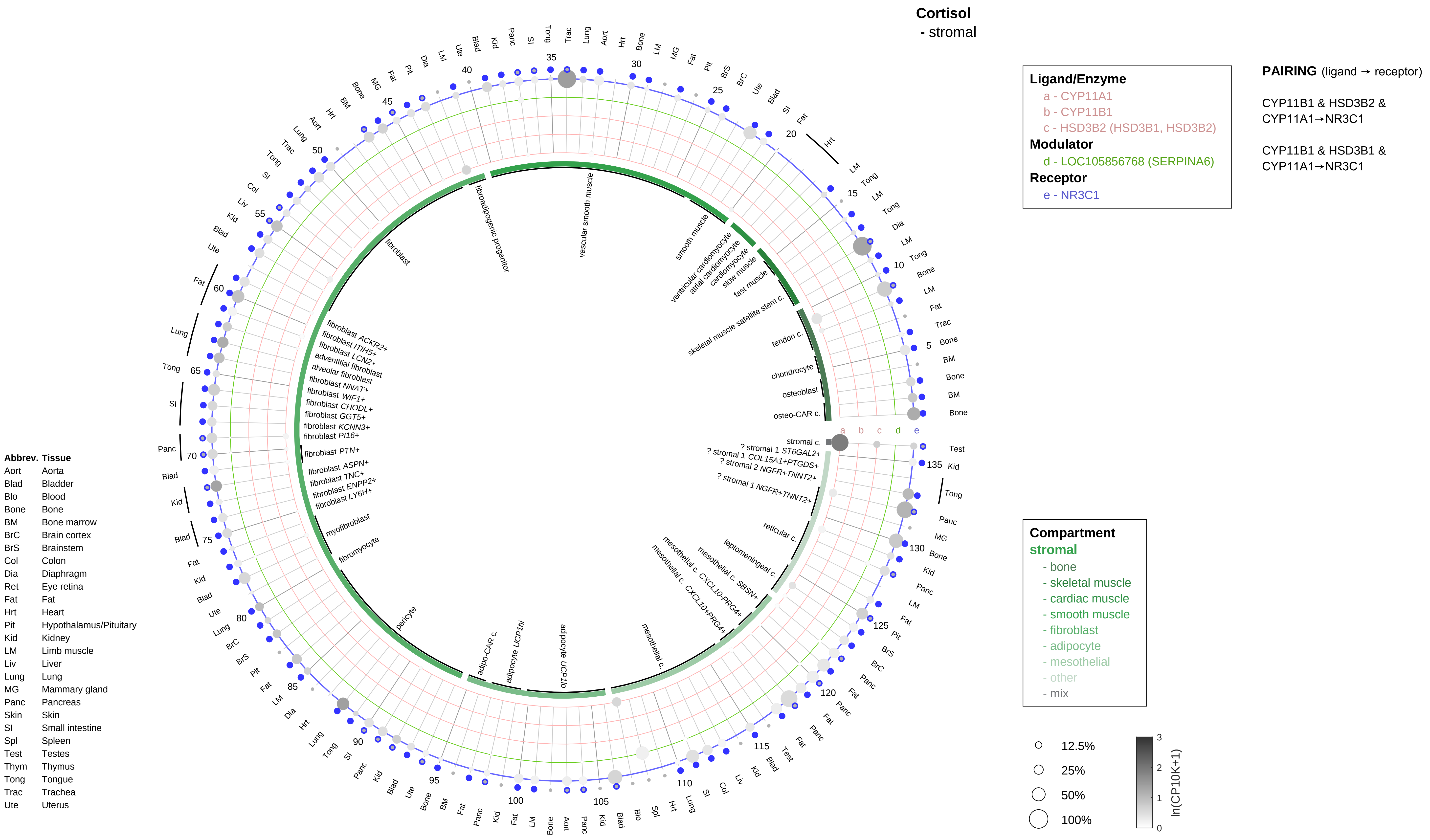
Cortisol
- stromal
Ligand/Enzyme
a - CYP11A1
b - CYP11B1
c - HSD3B2 (HSD3B1, HSD3B2)
Modulator
d - LOC105856768 (SERPINA6)
Receptor
e - NR3C1
PAIRING (ligand → receptor)
CYP11B1 & HSD3B2 & CYP11A1→NR3C1
CYP11B1 & HSD3B1 & CYP11A1→NR3C1
Compartment
stromal
- bone
- skeletal muscle
- cardiac muscle
- smooth muscle
- fibroblast
- adipocyte
- mesothelial
- other
- mix
12.5%
25%
50%
100%
ln(CP10K+1)
Abbrev. Tissue
Aort Aorta
Blad Bladder
Blo Blood
Bone Bone
BM Bone marrow
BrC Brain cortex
BrS Brainstem
Col Colon
Dia Diaphragm
Ret Eye retina
Fat Fat
Hrt Heart
Pit Hypothalamus/Pituitary
Kid Kidney
LM Limb muscle
Liv Liver
Lung Lung
MG Mammary gland
Panc Pancreas
Skin Skin
SI Small intestine
Spl Spleen
Test Testes
Thym Thymus
Tong Tongue
Trac Trachea
Ute Uterus
osteo-CAR c.
osteoblast
chondrocyte
tendon c.
skeletal muscle satellite stem c.
fast muscle
slow muscle
cardiomyocyte
atrial cardiomyocyte
ventricular cardiomyocyte
smooth muscle
vascular smooth muscle
fibroadipogenic progenitor
fibroblast
fibroblast ACKR2+
fibroblast ITIH5+
fibroblast LCN2+
adventitial fibroblast
alveolar fibroblast
fibroblast NNAT+
fibroblast WIF1+
fibroblast CHODL+
fibroblast GGT5+
fibroblast KCNN3+
fibroblast PI16+
fibroblast PTN+
fibroblast ASPN+
fibroblast TNC+
fibroblast ENPP2+
fibroblast LY6H+
myofibroblast
fibromyocyte
pericyte
adipo-CAR c.
adipocyte UCP1hi
adipocyte UCP1lo
mesothelial c.
mesothelial c. CXCL10-PRG4+
mesothelial c. CXCL10-PRG4+
mesothelial c. SBSN+
leptomeningeal c.
reticular c.
? stromal 1 NGFR+TNNT2+
? stromal 2 NGFR+TNNT2+
? stromal 1 COL15A1+PTGDS+
? stromal 1 ST6GAL2+
stromal c.

#### Cortisol
- lymphoid

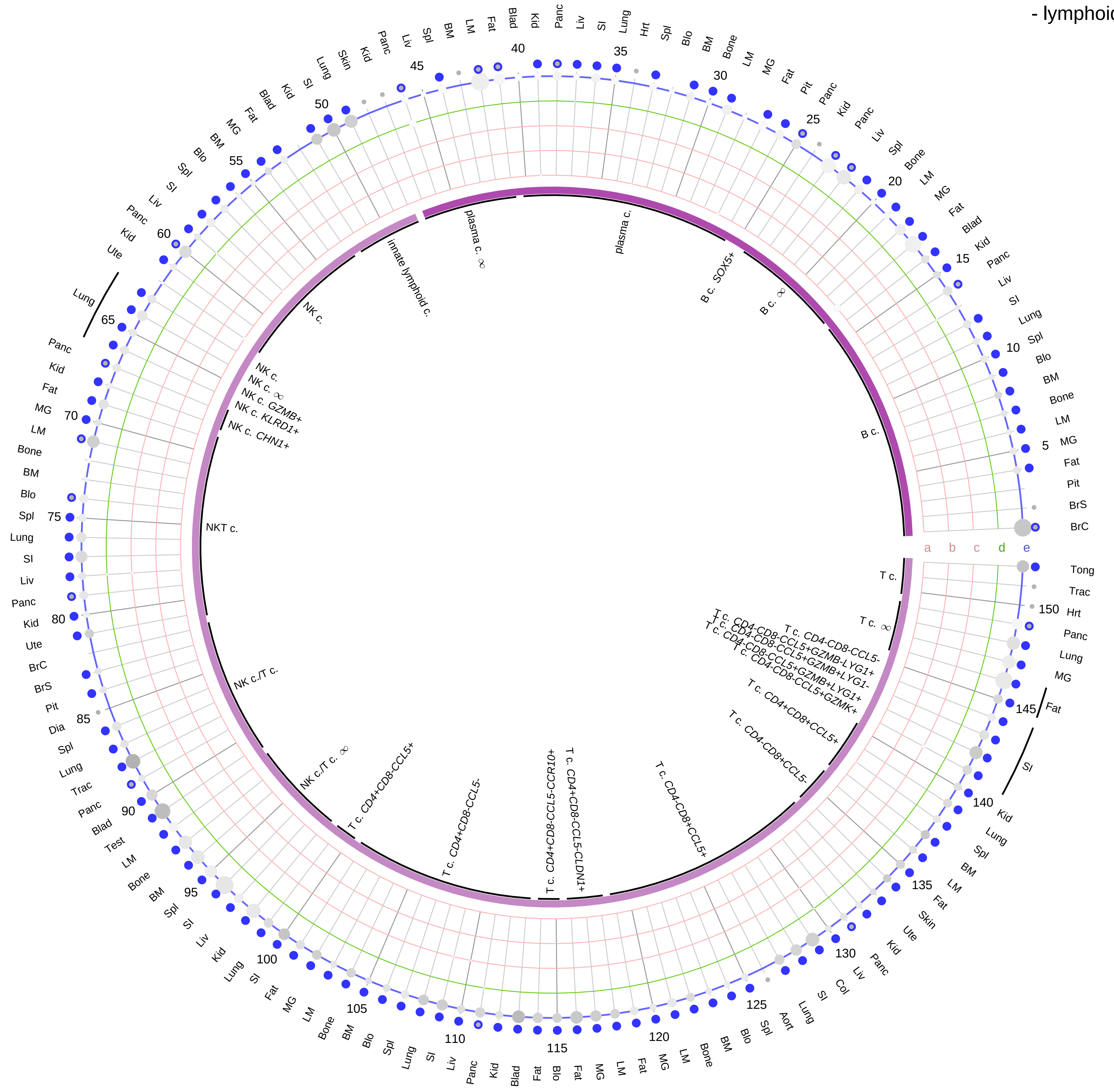

**Ligand/Enzyme**
a - CYP11A1
b - CYP11B1
c - HSD3B2 (HSD3B1, HSD3B2)
**Modulator**
d - LOC105856768 (SERPINA6)
**Receptor**
e - NR3C1

**PAIRING** (ligand → receptor)

CYP11B1 & HSD3B2 & CYP11A1→NR3C1

CYP11B1 & HSD3B1 & CYP11A1→NR3C1

**Compartment**
**lymphoid**
- B cell
- NK/T cell

12.5%
25%
50%
100%

ln(CP10K+1): 0, 1, 2, 3

| Abbrev. | Tissue |
|---|---|
| Aort | Aorta |
| Blad | Bladder |
| Blo | Blood |
| Bone | Bone |
| BM | Bone marrow |
| BrC | Brain cortex |
| BrS | Brainstem |
| Col | Colon |
| Dia | Diaphragm |
| Ret | Eye retina |
| Fat | Fat |
| Hrt | Heart |
| Pit | Hypothalamus/Pituitary |
| Kid | Kidney |
| LM | Limb muscle |
| Liv | Liver |
| Lung | Lung |
| MG | Mammary gland |
| Panc | Pancreas |
| Skin | Skin |
| SI | Small intestine |
| Spl | Spleen |
| Test | Testes |
| Thym | Thymus |
| Tong | Tongue |
| Trac | Trachea |
| Ute | Uterus |

**Cortisol**
- myeloid

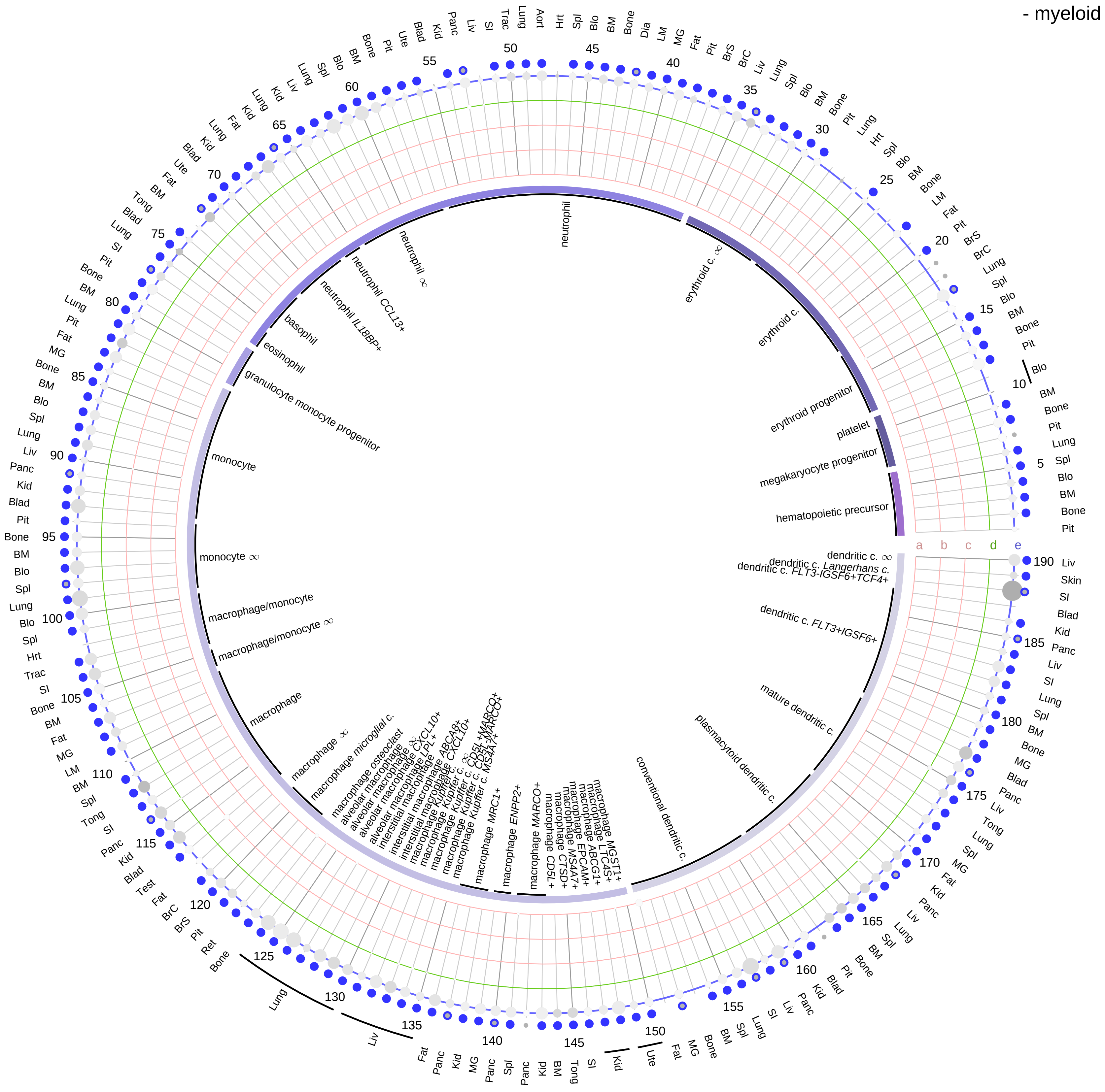

| Ligand/Enzyme |
|---|
| a - CYP11A1 |
| b - CYP11B1 |
| c - HSD3B2 (HSD3B1, HSD3B2) |
| **Modulator** |
| d - LOC105856768 (SERPINA6) |
| **Receptor** |
| e - NR3C1 |

**PAIRING** (ligand → receptor)

CYP11B1 & HSD3B2 & CYP11A1→NR3C1

CYP11B1 & HSD3B1 & CYP11A1→NR3C1

| Abbrev. | Tissue |
|---|---|
| Aort | Aorta |
| Blad | Bladder |
| Blo | Blood |
| Bone | Bone |
| BM | Bone marrow |
| BrC | Brain cortex |
| BrS | Brainstem |
| Col | Colon |
| Dia | Diaphragm |
| Ret | Eye retina |
| Fat | Fat |
| Hrt | Heart |
| Pit | Hypothalamus/Pituitary |
| Kid | Kidney |
| LM | Limb muscle |
| Liv | Liver |
| Lung | Lung |
| MG | Mammary gland |
| Panc | Pancreas |
| Skin | Skin |
| SI | Small intestine |
| Spl | Spleen |
| Test | Testes |
| Thym | Thymus |
| Tong | Tongue |
| Trac | Trachea |
| Ute | Uterus |

**Compartment**
**hematopoietic**
- hematopoietic precursor

**megakaryocyte-erythroid**
- megakaryoid
- erythroid

**myeloid**
- granulocyte
- granulocyte monocyte progenitor
- monocyte/macrophage
- dendritic

12.5%
25%
50%
100%

ln(CP10K+1): 0, 1, 2, 3

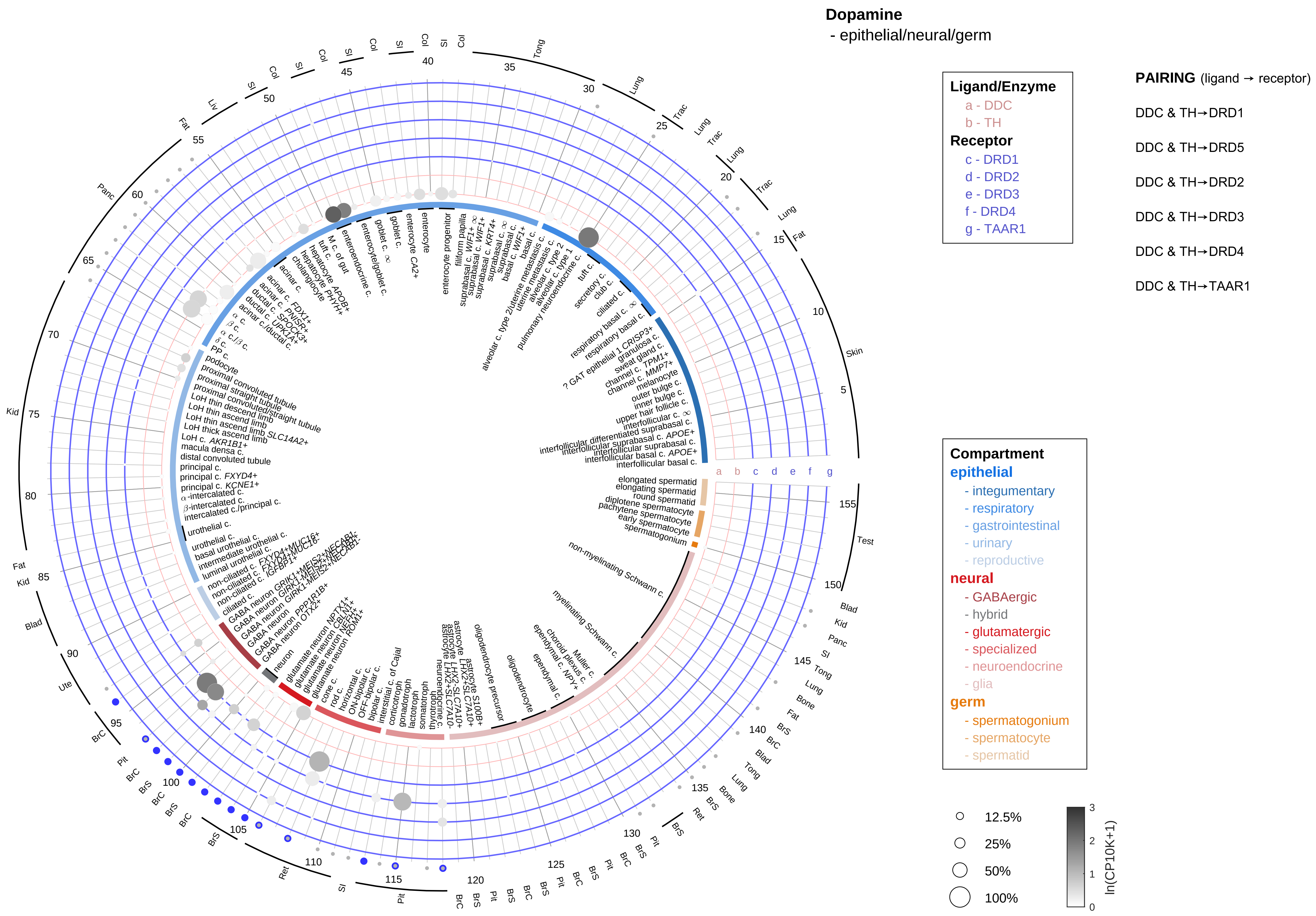

| Abbrev. | Tissue |
|---|---|
| Aort | Aorta |
| Blad | Bladder |
| Blo | Blood |
| Bone | Bone |
| BM | Bone marrow |
| BrC | Brain cortex |
| BrS | Brainstem |
| Col | Colon |
| Dia | Diaphragm |
| Ret | Eye retina |
| Fat | Fat |
| Hrt | Heart |
| Pit | Hypothalamus/Pituitary |
| Kid | Kidney |
| LM | Limb muscle |
| Liv | Liver |
| Lung | Lung |
| MG | Mammary gland |
| Panc | Pancreas |
| Skin | Skin |
| SI | Small intestine |
| Spl | Spleen |
| Test | Testes |
| Thym | Thymus |
| Tong | Tongue |
| Trac | Trachea |
| Ute | Uterus |

**Dopamine**
- endothelial

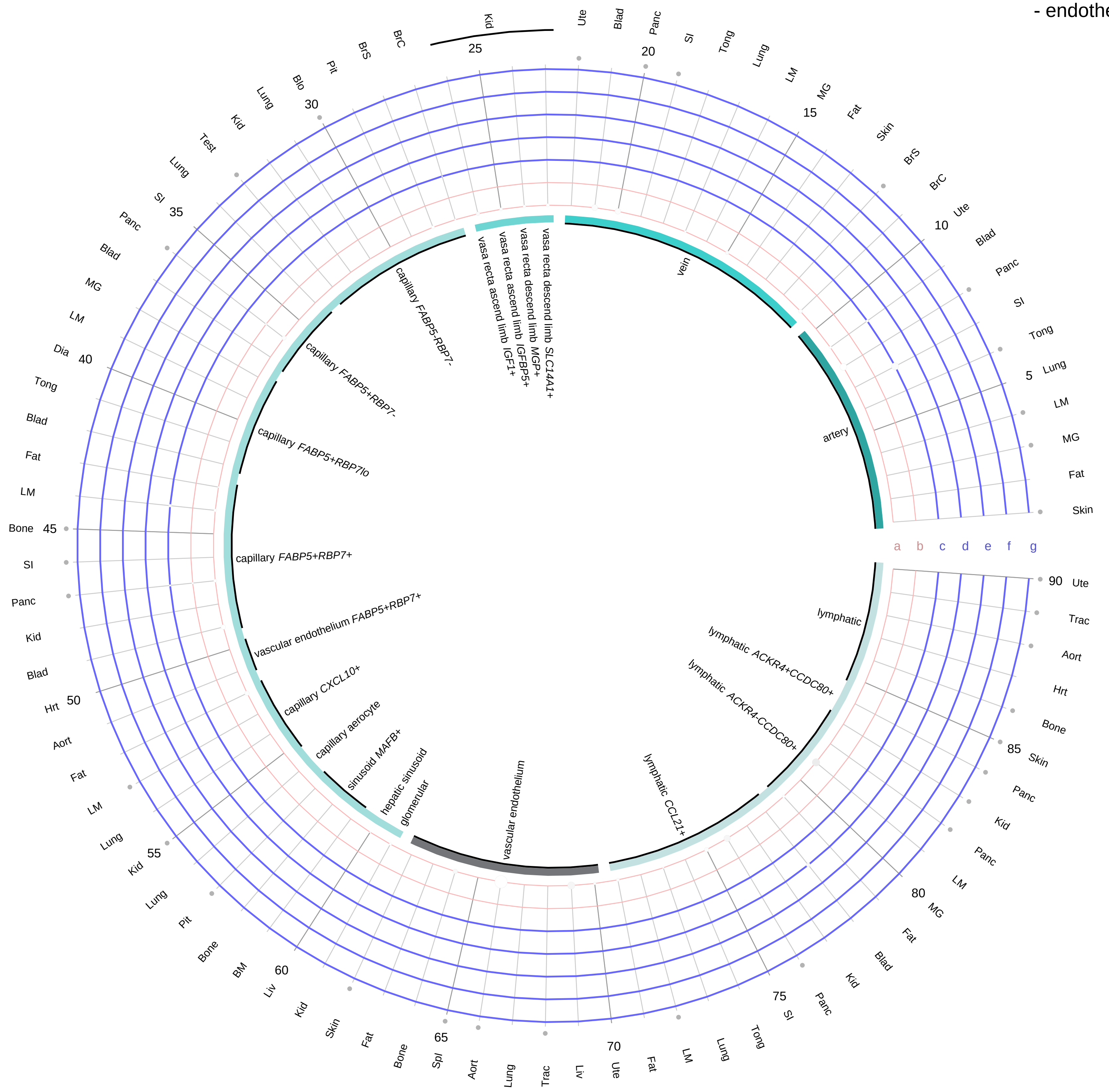

**Ligand/Enzyme**
- a - DDC
- b - TH

**Receptor**
- c - DRD1
- d - DRD2
- e - DRD3
- f - DRD4
- g - TAAR1

**PAIRING** (ligand → receptor)

DDC & TH→DRD1

DDC & TH→DRD5

DDC & TH→DRD2

DDC & TH→DRD3

DDC & TH→DRD4

DDC & TH→TAAR1

**Compartment**
**endothelial**
- artery
- vein
- vasa recta
- capillary
- mix
- lymphatic

12.5%
25%
50%
100%

ln(CP10K+1): 0 1 2 3

| Abbrev. | Tissue |
|---|---|
| Aort | Aorta |
| Blad | Bladder |
| Blo | Blood |
| Bone | Bone |
| BM | Bone marrow |
| BrC | Brain cortex |
| BrS | Brainstem |
| Col | Colon |
| Dia | Diaphragm |
| Ret | Eye retina |
| Fat | Fat |
| Hrt | Heart |
| Pit | Hypothalamus/Pituitary |
| Kid | Kidney |
| LM | Limb muscle |
| Liv | Liver |
| Lung | Lung |
| MG | Mammary gland |
| Panc | Pancreas |
| Skin | Skin |
| SI | Small intestine |
| Spl | Spleen |
| Test | Testes |
| Thym | Thymus |
| Tong | Tongue |
| Trac | Trachea |
| Ute | Uterus |

**Dopamine**
- stromal

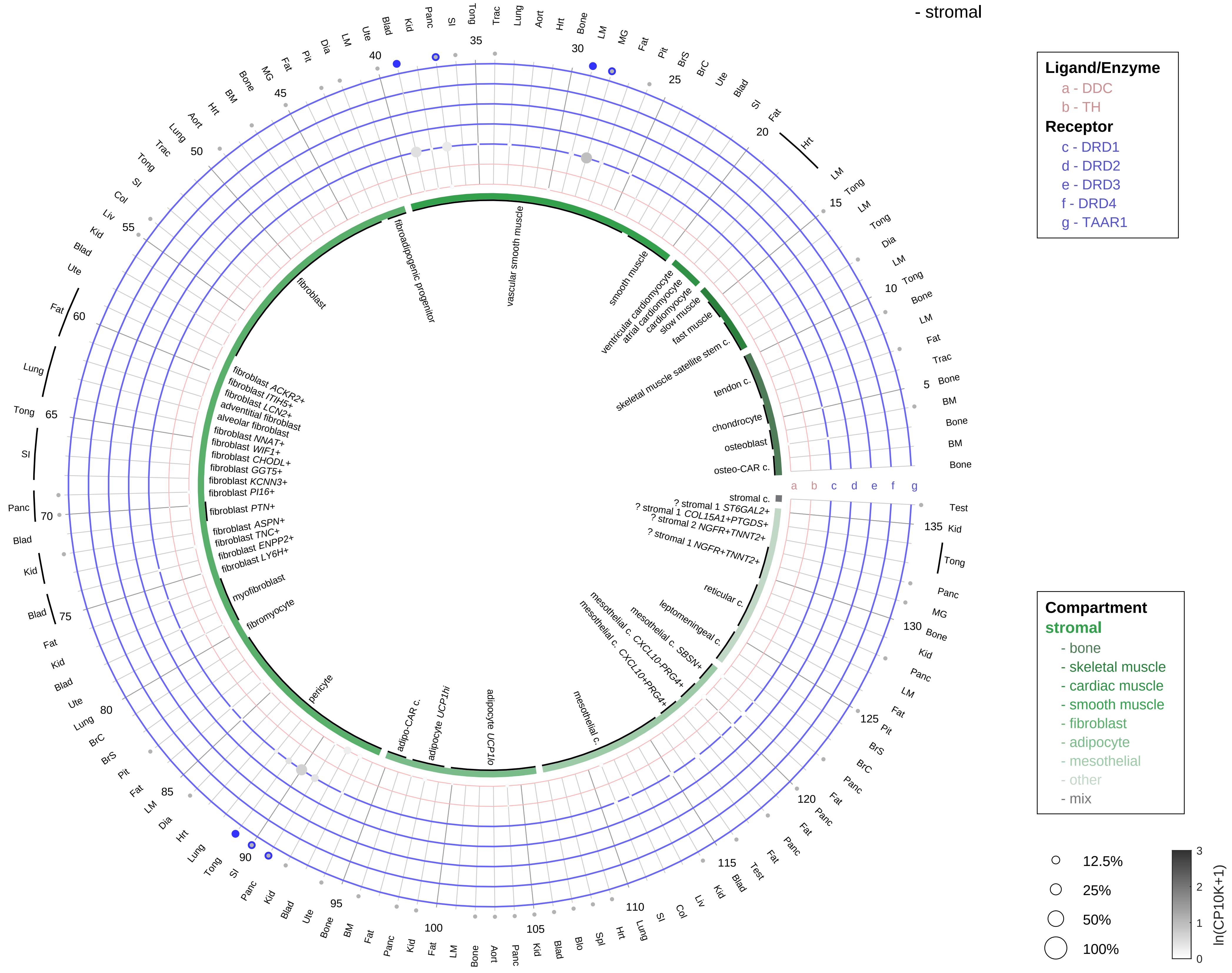

**PAIRING** (ligand → receptor)

DDC & TH→DRD1

DDC & TH→DRD5

DDC & TH→DRD2

DDC & TH→DRD3

DDC & TH→DRD4

DDC & TH→TAAR1

| Abbrev. | Tissue |
|---|---|
| Aort | Aorta |
| Blad | Bladder |
| Blo | Blood |
| Bone | Bone |
| BM | Bone marrow |
| BrC | Brain cortex |
| BrS | Brainstem |
| Col | Colon |
| Dia | Diaphragm |
| Ret | Eye retina |
| Fat | Fat |
| Hrt | Heart |
| Pit | Hypothalamus/Pituitary |
| Kid | Kidney |
| LM | Limb muscle |
| Liv | Liver |
| Lung | Lung |
| MG | Mammary gland |
| Panc | Pancreas |
| Skin | Skin |
| SI | Small intestine |
| Spl | Spleen |
| Test | Testes |
| Thym | Thymus |
| Tong | Tongue |
| Trac | Trachea |
| Ute | Uterus |

**Dopamine**
- lymphoid

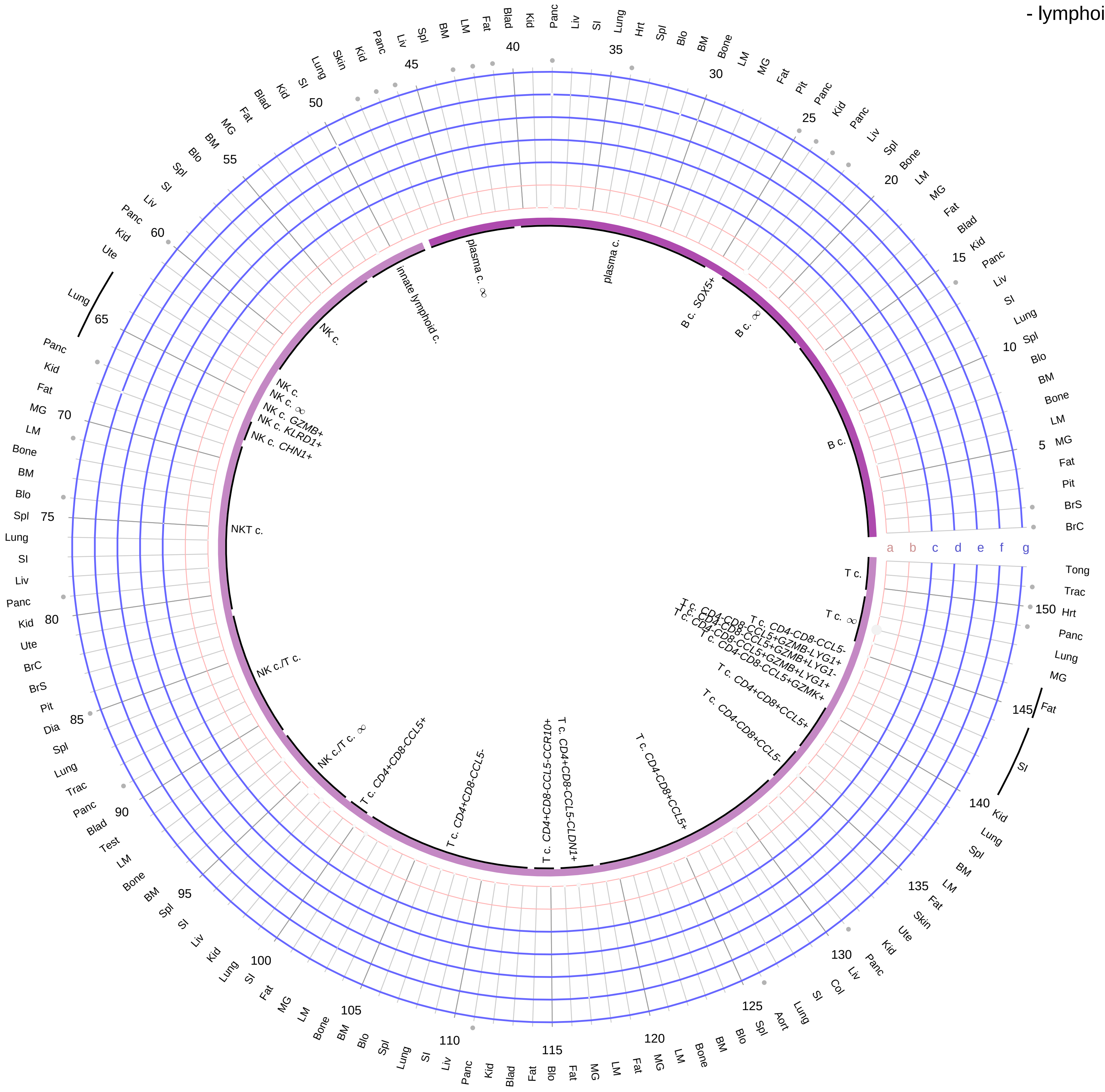

**Ligand/Enzyme**
a - DDC
b - TH

**Receptor**
c - DRD1
d - DRD2
e - DRD3
f - DRD4
g - TAAR1

**PAIRING** (ligand → receptor)

DDC & TH→DRD1

DDC & TH→DRD5

DDC & TH→DRD2

DDC & TH→DRD3

DDC & TH→DRD4

DDC & TH→TAAR1

**Compartment**
**lymphoid**
- B cell
- NK/T cell

12.5%
25%
50%
100%

ln(CP10K+1): 0 1 2 3

| Abbrev. | Tissue |
|---|---|
| Aort | Aorta |
| Blad | Bladder |
| Blo | Blood |
| Bone | Bone |
| BM | Bone marrow |
| BrC | Brain cortex |
| BrS | Brainstem |
| Col | Colon |
| Dia | Diaphragm |
| Ret | Eye retina |
| Fat | Fat |
| Hrt | Heart |
| Pit | Hypothalamus/Pituitary |
| Kid | Kidney |
| LM | Limb muscle |
| Liv | Liver |
| Lung | Lung |
| MG | Mammary gland |
| Panc | Pancreas |
| Skin | Skin |
| SI | Small intestine |
| Spl | Spleen |
| Test | Testes |
| Thym | Thymus |
| Tong | Tongue |
| Trac | Trachea |
| Ute | Uterus |

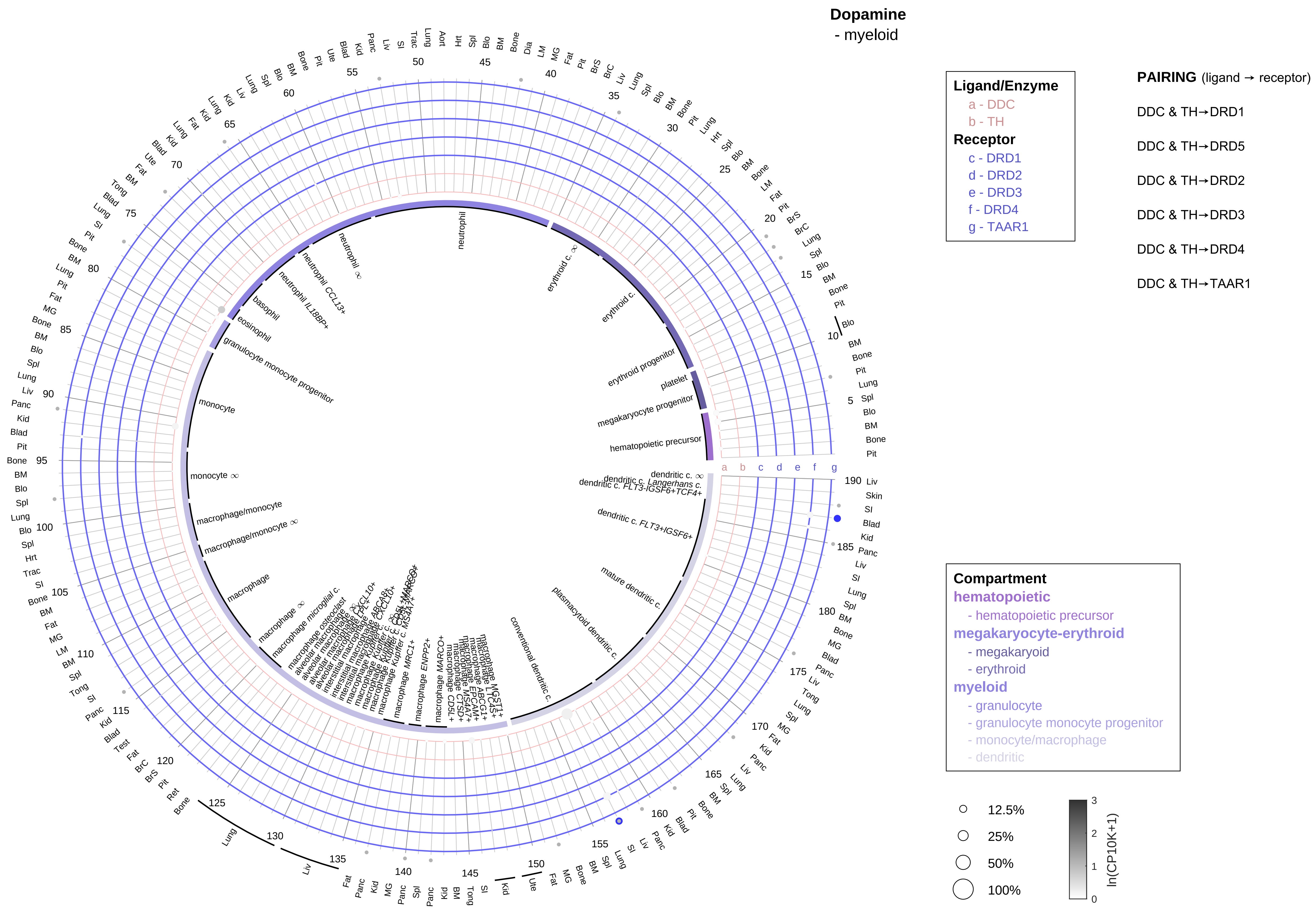

| Abbrev. | Tissue |
|---|---|
| Aort | Aorta |
| Blad | Bladder |
| Blo | Blood |
| Bone | Bone |
| BM | Bone marrow |
| BrC | Brain cortex |
| BrS | Brainstem |
| Col | Colon |
| Dia | Diaphragm |
| Ret | Eye retina |
| Fat | Fat |
| Hrt | Heart |
| Pit | Hypothalamus/Pituitary |
| Kid | Kidney |
| LM | Limb muscle |
| Liv | Liver |
| Lung | Lung |
| MG | Mammary gland |
| Panc | Pancreas |
| Skin | Skin |
| SI | Small intestine |
| Spl | Spleen |
| Test | Testes |
| Thym | Thymus |
| Tong | Tongue |
| Trac | Trachea |
| Ute | Uterus |

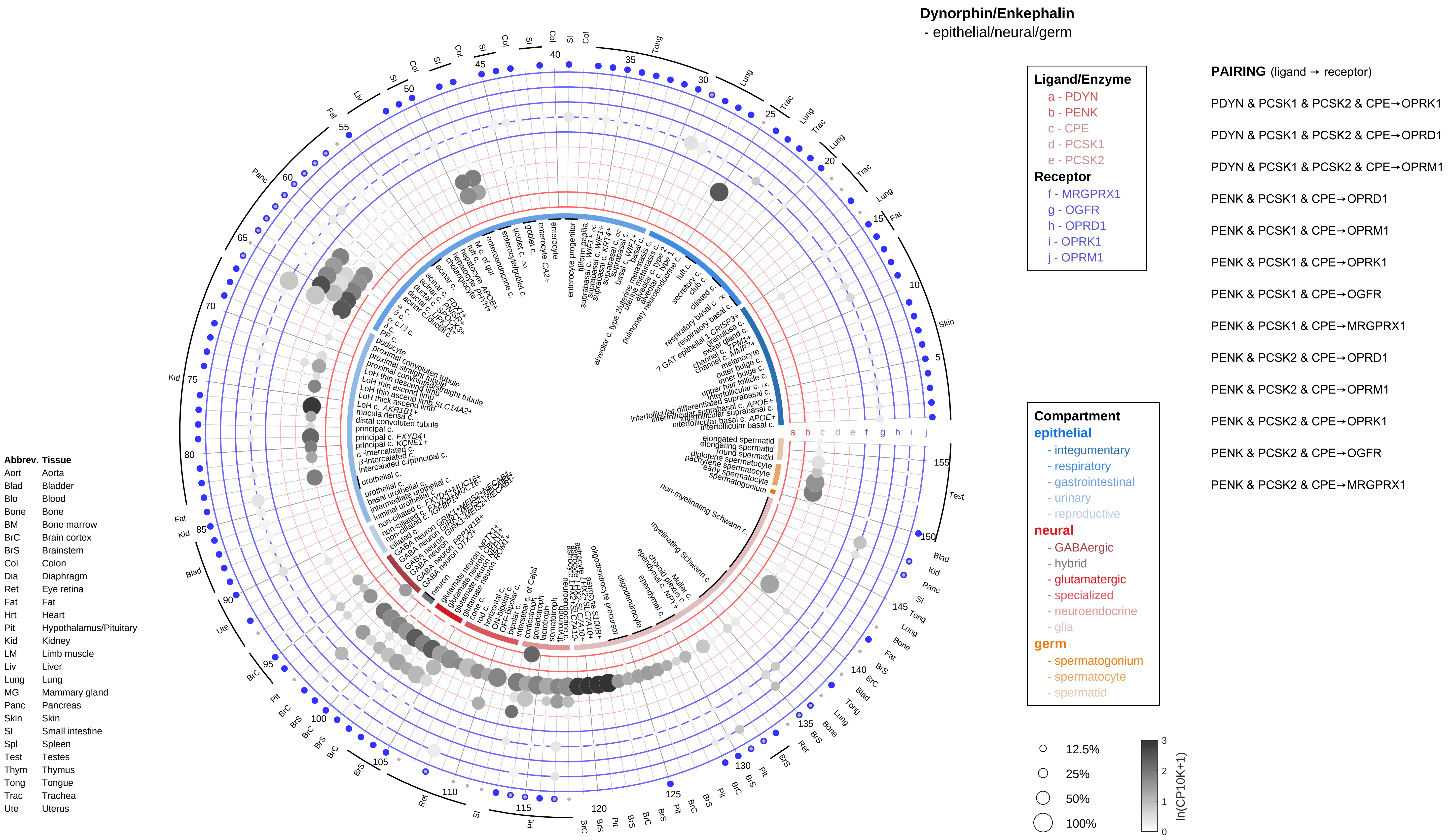

Dynorphin/Enkephalin
- epithelial/neural/germ
Ligand/Enzyme
a - PDYN
b - PENK
c - CPE
d - PCSK1
e - PCSK2
Receptor
f - MRGPRX1
g - OGFR
h - OPRD1
i - OPRK1
j - OPRM1
PAIRING (ligand → receptor)
PDYN & PCSK1 & PCSK2 & CPE→OPRK1
PDYN & PCSK1 & PCSK2 & CPE→OPRD1
PDYN & PCSK1 & PCSK2 & CPE→OPRM1
PENK & PCSK1 & CPE→OPRD1
PENK & PCSK1 & CPE→OPRM1
PENK & PCSK1 & CPE→OPRK1
PENK & PCSK1 & CPE→OGFR
PENK & PCSK1 & CPE→MRGPRX1
PENK & PCSK2 & CPE→OPRD1
PENK & PCSK2 & CPE→OPRM1
PENK & PCSK2 & CPE→OPRK1
PENK & PCSK2 & CPE→OGFR
PENK & PCSK2 & CPE→MRGPRX1
Compartment
epithelial
- integumentary
- respiratory
- gastrointestinal
- urinary
- reproductive
neural
- GABAergic
- hybrid
- glutamatergic
- specialized
- neuroendocrine
- glia
germ
- spermatogonium
- spermatocyte
- spermatid
12.5%
25%
50%
100%
ln(CP10K+1)
Abbrev. Tissue
Aort Aorta
Blad Bladder
Blo Blood
Bone Bone
BM Bone marrow
BrC Brain cortex
BrS Brainstem
Col Colon
Dia Diaphragm
Ret Eye retina
Fat Fat
Hrt Heart
Pit Hypothalamus/Pituitary
Kid Kidney
LM Limb muscle
Liv Liver
Lung Lung
MG Mammary gland
Panc Pancreas
Skin Skin
SI Small intestine
Spl Spleen
Test Testes
Thym Thymus
Tong Tongue
Trac Trachea
Ute Uterus

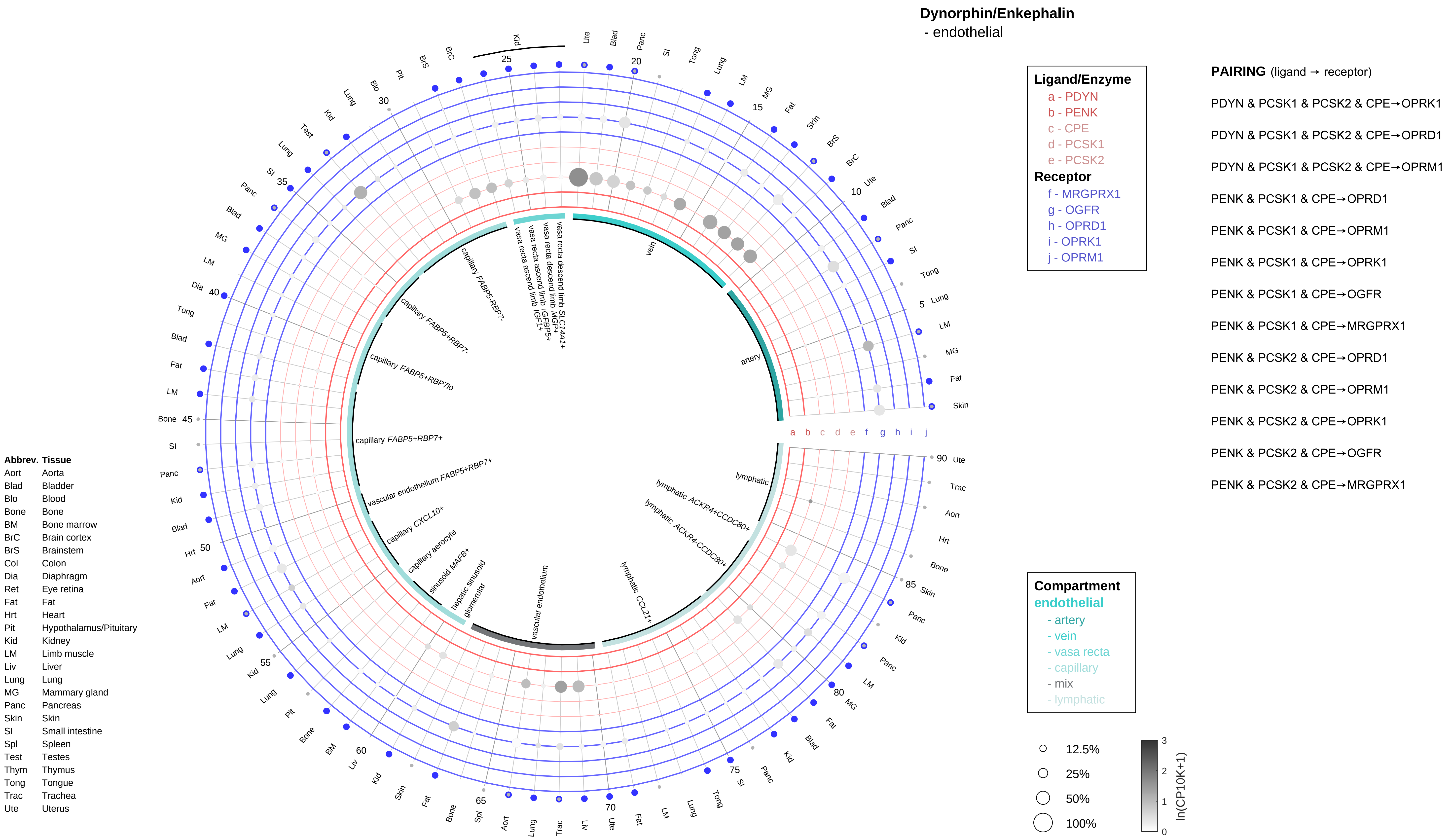

Dynorphin/Enkephalin
- endothelial
Ligand/Enzyme
a - PDYN
b - PENK
c - CPE
d - PCSK1
e - PCSK2
Receptor
f - MRGPRX1
g - OGFR
h - OPRD1
i - OPRK1
j - OPRM1
PAIRING (ligand → receptor)
PDYN & PCSK1 & PCSK2 & CPE→OPRK1
PDYN & PCSK1 & PCSK2 & CPE→OPRD1
PDYN & PCSK1 & PCSK2 & CPE→OPRM1
PENK & PCSK1 & CPE→OPRD1
PENK & PCSK1 & CPE→OPRM1
PENK & PCSK1 & CPE→OPRK1
PENK & PCSK1 & CPE→OGFR
PENK & PCSK1 & CPE→MRGPRX1
PENK & PCSK2 & CPE→OPRD1
PENK & PCSK2 & CPE→OPRM1
PENK & PCSK2 & CPE→OPRK1
PENK & PCSK2 & CPE→OGFR
PENK & PCSK2 & CPE→MRGPRX1
Compartment
endothelial
- artery
- vein
- vasa recta
- capillary
- mix
- lymphatic
12.5%
25%
50%
100%
ln(CP10K+1)
vein
artery
lymphatic
lymphatic ACKR4+CCDC80+
lymphatic ACKR4-CCDC80+
lymphatic CCL21+
vascular endothelium
glomerular
hepatic sinusoid
sinusoid MAFB+
capillary aerocyte
capillary CXCL10+
vascular endothelium FABP5+RBP7+
capillary FABP5+RBP7+
capillary FABP5+RBP7lo
capillary FABP5+RBP7-
capillary FABP5-RBP7-
vasa recta ascend limb IGF1+
vasa recta ascend limb IGFBP5+
vasa recta descend limb MGP+
vasa recta descend limb SLC14A1+
Abbrev. Tissue
Aort Aorta
Blad Bladder
Blo Blood
Bone Bone
BM Bone marrow
BrC Brain cortex
BrS Brainstem
Col Colon
Dia Diaphragm
Ret Eye retina
Fat Fat
Hrt Heart
Pit Hypothalamus/Pituitary
Kid Kidney
LM Limb muscle
Liv Liver
Lung Lung
MG Mammary gland
Panc Pancreas
Skin Skin
SI Small intestine
Spl Spleen
Test Testes
Thym Thymus
Tong Tongue
Trac Trachea
Ute Uterus

**Dynorphin/Enkephalin**
- stromal

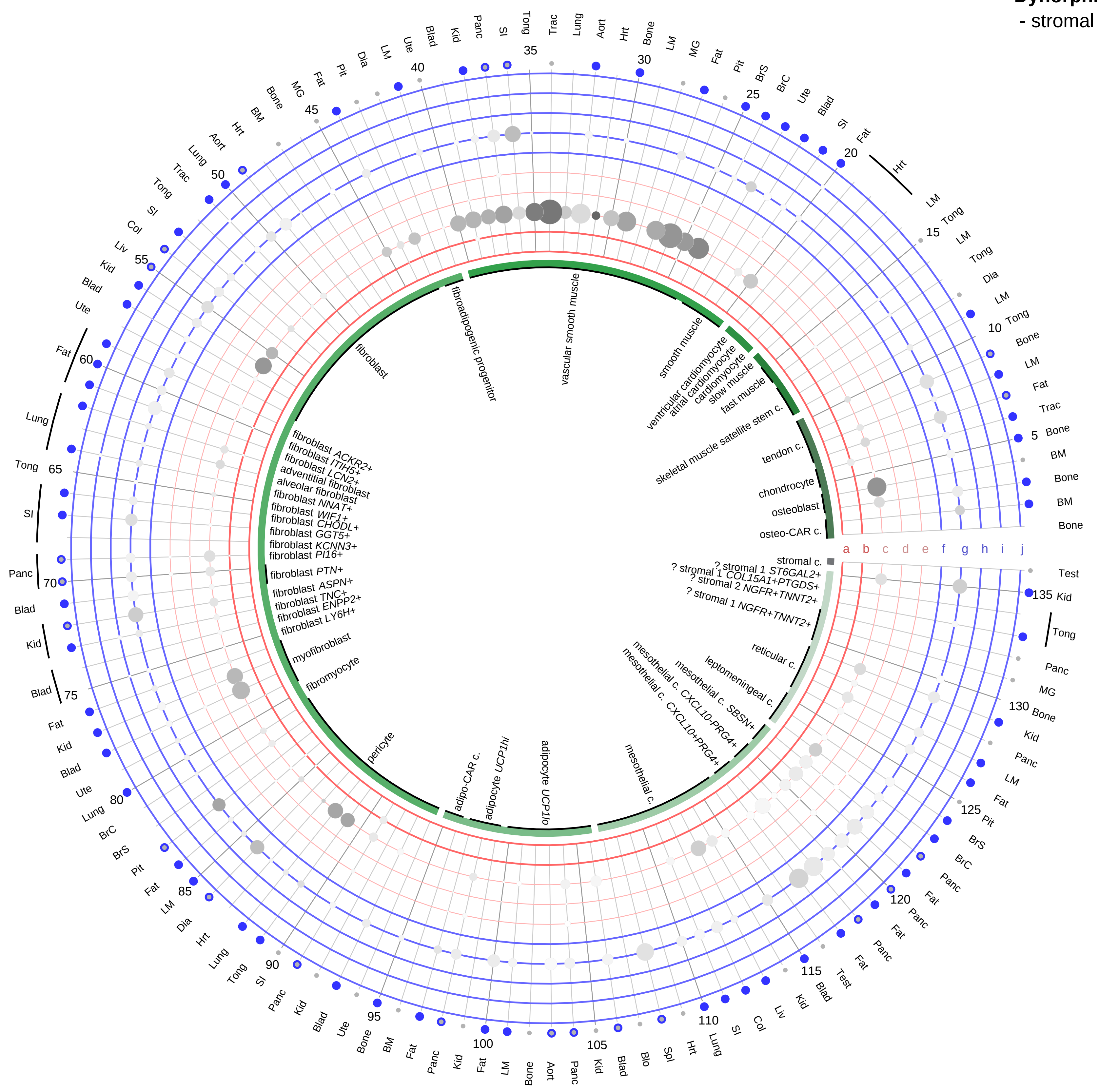

**Ligand/Enzyme**
- a - PDYN
- b - PENK
- c - CPE
- d - PCSK1
- e - PCSK2

**Receptor**
- f - MRGPRX1
- g - OGFR
- h - OPRD1
- i - OPRK1
- j - OPRM1

**PAIRING** (ligand → receptor)

PDYN & PCSK1 & PCSK2 & CPE→OPRK1

PDYN & PCSK1 & PCSK2 & CPE→OPRD1

PDYN & PCSK1 & PCSK2 & CPE→OPRM1

PENK & PCSK1 & CPE→OPRD1

PENK & PCSK1 & CPE→OPRM1

PENK & PCSK1 & CPE→OPRK1

PENK & PCSK1 & CPE→OGFR

PENK & PCSK1 & CPE→MRGPRX1

PENK & PCSK2 & CPE→OPRD1

PENK & PCSK2 & CPE→OPRM1

PENK & PCSK2 & CPE→OPRK1

PENK & PCSK2 & CPE→OGFR

PENK & PCSK2 & CPE→MRGPRX1

**Compartment**

**stromal**
- bone
- skeletal muscle
- cardiac muscle
- smooth muscle
- fibroblast
- adipocyte
- mesothelial
- other
- mix

12.5%
25%
50%
100%

ln(CP10K+1): 0, 1, 2, 3

| Abbrev. | Tissue |
|---|---|
| Aort | Aorta |
| Blad | Bladder |
| Blo | Blood |
| Bone | Bone |
| BM | Bone marrow |
| BrC | Brain cortex |
| BrS | Brainstem |
| Col | Colon |
| Dia | Diaphragm |
| Ret | Eye retina |
| Fat | Fat |
| Hrt | Heart |
| Pit | Hypothalamus/Pituitary |
| Kid | Kidney |
| LM | Limb muscle |
| Liv | Liver |
| Lung | Lung |
| MG | Mammary gland |
| Panc | Pancreas |
| Skin | Skin |
| SI | Small intestine |
| Spl | Spleen |
| Test | Testes |
| Thym | Thymus |
| Tong | Tongue |
| Trac | Trachea |
| Ute | Uterus |

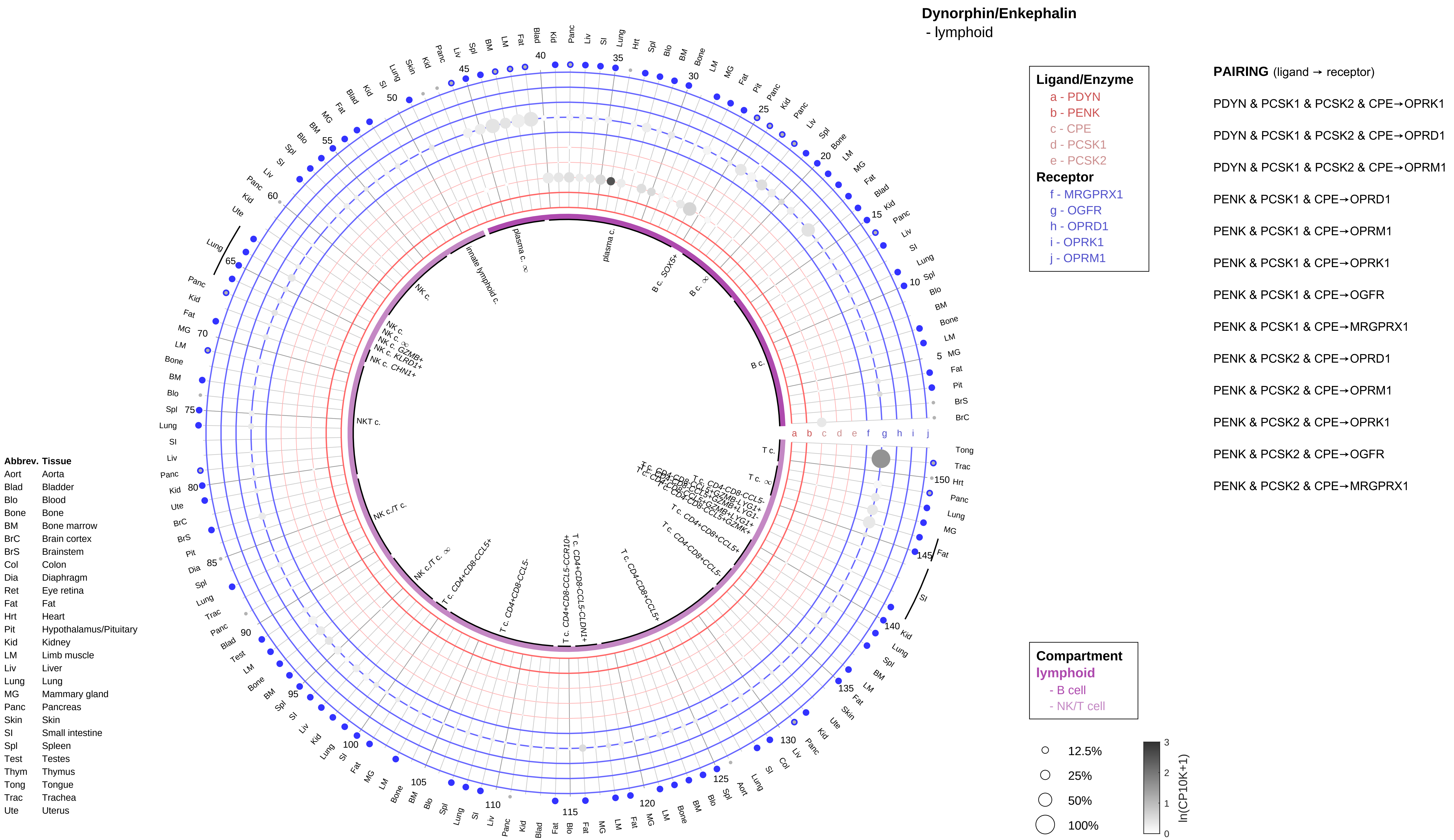
Dynorphin/Enkephalin
- lymphoid
Ligand/Enzyme
a - PDYN
b - PENK
c - CPE
d - PCSK1
e - PCSK2
Receptor
f - MRGPRX1
g - OGFR
h - OPRD1
i - OPRK1
j - OPRM1
PAIRING (ligand → receptor)
PDYN & PCSK1 & PCSK2 & CPE→OPRK1
PDYN & PCSK1 & PCSK2 & CPE→OPRD1
PDYN & PCSK1 & PCSK2 & CPE→OPRM1
PENK & PCSK1 & CPE→OPRD1
PENK & PCSK1 & CPE→OPRM1
PENK & PCSK1 & CPE→OPRK1
PENK & PCSK1 & CPE→OGFR
PENK & PCSK1 & CPE→MRGPRX1
PENK & PCSK2 & CPE→OPRD1
PENK & PCSK2 & CPE→OPRM1
PENK & PCSK2 & CPE→OPRK1
PENK & PCSK2 & CPE→OGFR
PENK & PCSK2 & CPE→MRGPRX1
Compartment
lymphoid
- B cell
- NK/T cell
12.5%
25%
50%
100%
ln(CP10K+1)
Abbrev. Tissue
Aort Aorta
Blad Bladder
Blo Blood
Bone Bone
BM Bone marrow
BrC Brain cortex
BrS Brainstem
Col Colon
Dia Diaphragm
Ret Eye retina
Fat Fat
Hrt Heart
Pit Hypothalamus/Pituitary
Kid Kidney
LM Limb muscle
Liv Liver
Lung Lung
MG Mammary gland
Panc Pancreas
Skin Skin
SI Small intestine
Spl Spleen
Test Testes
Thym Thymus
Tong Tongue
Trac Trachea
Ute Uterus

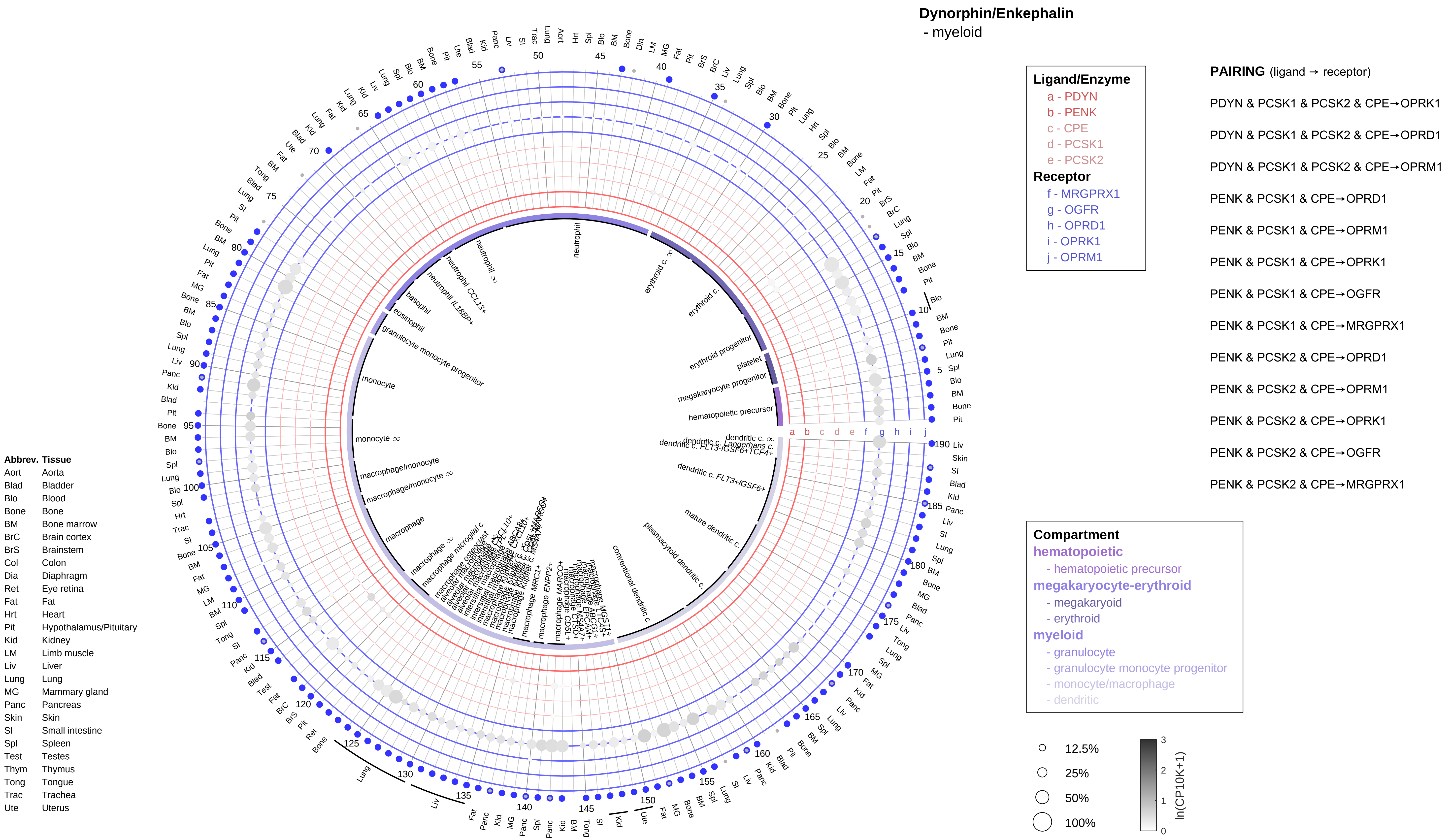
Dynorphin/Enkephalin
- myeloid
Ligand/Enzyme
a - PDYN
b - PENK
c - CPE
d - PCSK1
e - PCSK2
Receptor
f - MRGPRX1
g - OGFR
h - OPRD1
i - OPRK1
j - OPRM1
PAIRING (ligand → receptor)
PDYN & PCSK1 & PCSK2 & CPE→OPRK1
PDYN & PCSK1 & PCSK2 & CPE→OPRD1
PDYN & PCSK1 & PCSK2 & CPE→OPRM1
PENK & PCSK1 & CPE→OPRD1
PENK & PCSK1 & CPE→OPRM1
PENK & PCSK1 & CPE→OPRK1
PENK & PCSK1 & CPE→OGFR
PENK & PCSK1 & CPE→MRGPRX1
PENK & PCSK2 & CPE→OPRD1
PENK & PCSK2 & CPE→OPRM1
PENK & PCSK2 & CPE→OPRK1
PENK & PCSK2 & CPE→OGFR
PENK & PCSK2 & CPE→MRGPRX1
Compartment
hematopoietic
- hematopoietic precursor
megakaryocyte-erythroid
- megakaryoid
- erythroid
myeloid
- granulocyte
- granulocyte monocyte progenitor
- monocyte/macrophage
- dendritic
12.5%
25%
50%
100%
ln(CP10K+1)
Abbrev. Tissue
Aort Aorta
Blad Bladder
Blo Blood
Bone Bone
BM Bone marrow
BrC Brain cortex
BrS Brainstem
Col Colon
Dia Diaphragm
Ret Eye retina
Fat Fat
Hrt Heart
Pit Hypothalamus/Pituitary
Kid Kidney
LM Limb muscle
Liv Liver
Lung Lung
MG Mammary gland
Panc Pancreas
Skin Skin
SI Small intestine
Spl Spleen
Test Testes
Thym Thymus
Tong Tongue
Trac Trachea
Ute Uterus
neutrophil
erythroid c.
erythroid progenitor
platelet
megakaryocyte progenitor
hematopoietic precursor
dendritic c. FLT3+IGSF6+
mature dendritic c.
plasmacytoid dendritic c.
conventional dendritic c.
macrophage
monocyte
basophil
eosinophil
granulocyte monocyte progenitor

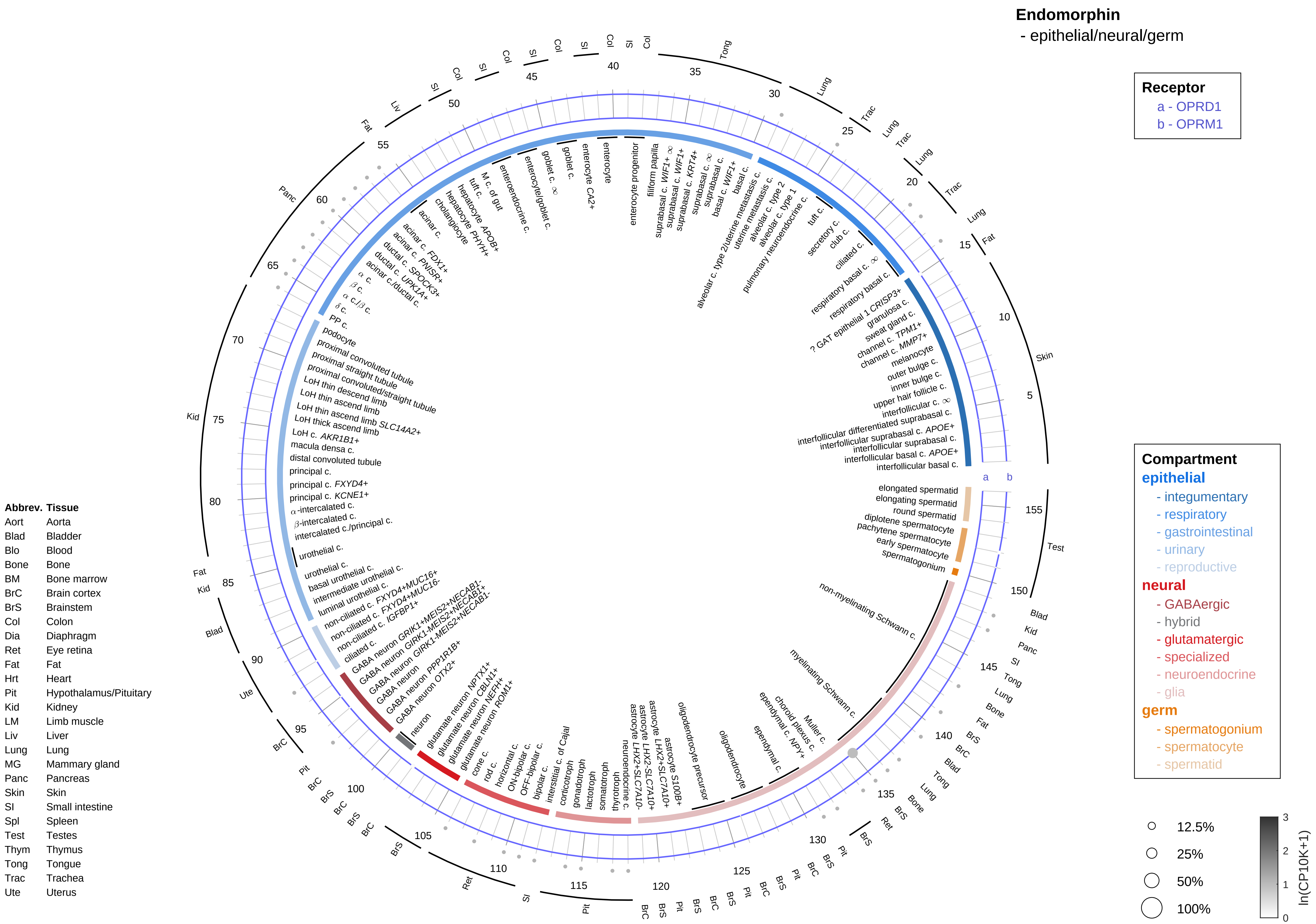
Endomorphin
- epithelial/neural/germ
Receptor
a - OPRD1
b - OPRM1
Compartment
epithelial
- integumentary
- respiratory
- gastrointestinal
- urinary
- reproductive
neural
- GABAergic
- hybrid
- glutamatergic
- specialized
- neuroendocrine
- glia
germ
- spermatogonium
- spermatocyte
- spermatid
12.5%
25%
50%
100%
ln(CP10K+1)
Abbrev. Tissue
Aort Aorta
Blad Bladder
Blo Blood
Bone Bone
BM Bone marrow
BrC Brain cortex
BrS Brainstem
Col Colon
Dia Diaphragm
Ret Eye retina
Fat Fat
Hrt Heart
Pit Hypothalamus/Pituitary
Kid Kidney
LM Limb muscle
Liv Liver
Lung Lung
MG Mammary gland
Panc Pancreas
Skin Skin
SI Small intestine
Spl Spleen
Test Testes
Thym Thymus
Tong Tongue
Trac Trachea
Ute Uterus

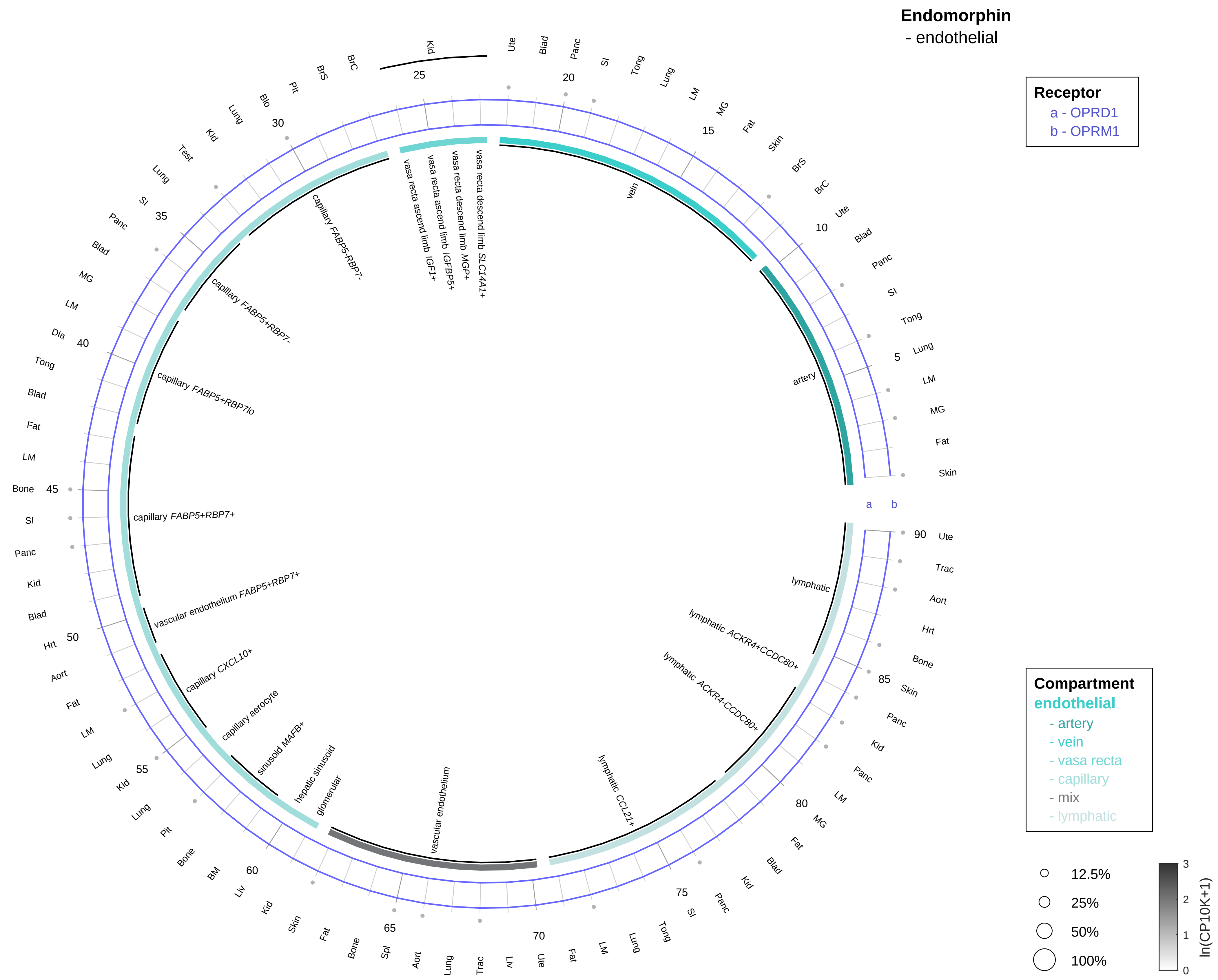

| Abbrev. | Tissue |
|---|---|
| Aort | Aorta |
| Blad | Bladder |
| Blo | Blood |
| Bone | Bone |
| BM | Bone marrow |
| BrC | Brain cortex |
| BrS | Brainstem |
| Col | Colon |
| Dia | Diaphragm |
| Ret | Eye retina |
| Fat | Fat |
| Hrt | Heart |
| Pit | Hypothalamus/Pituitary |
| Kid | Kidney |
| LM | Limb muscle |
| Liv | Liver |
| Lung | Lung |
| MG | Mammary gland |
| Panc | Pancreas |
| Skin | Skin |
| SI | Small intestine |
| Spl | Spleen |
| Test | Testes |
| Thym | Thymus |
| Tong | Tongue |
| Trac | Trachea |
| Ute | Uterus |

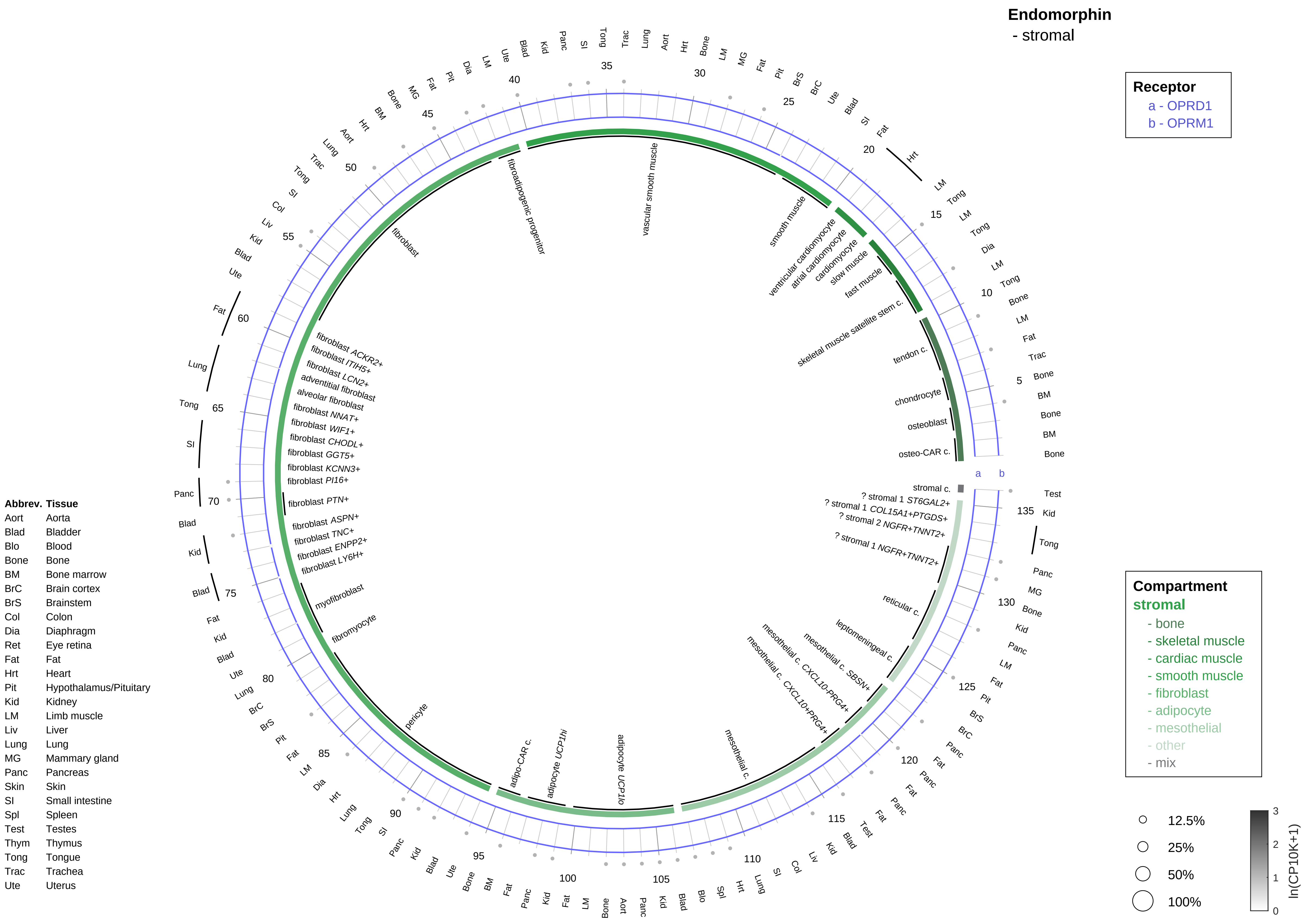
Endomorphin
- stromal
Receptor
a - OPRD1
b - OPRM1
Compartment
stromal
- bone
- skeletal muscle
- cardiac muscle
- smooth muscle
- fibroblast
- adipocyte
- mesothelial
- other
- mix
12.5%
25%
50%
100%
ln(CP10K+1)
Abbrev. Tissue
Aort Aorta
Blad Bladder
Blo Blood
Bone Bone
BM Bone marrow
BrC Brain cortex
BrS Brainstem
Col Colon
Dia Diaphragm
Ret Eye retina
Fat Fat
Hrt Heart
Pit Hypothalamus/Pituitary
Kid Kidney
LM Limb muscle
Liv Liver
Lung Lung
MG Mammary gland
Panc Pancreas
Skin Skin
SI Small intestine
Spl Spleen
Test Testes
Thym Thymus
Tong Tongue
Trac Trachea
Ute Uterus
vascular smooth muscle
smooth muscle
ventricular cardiomyocyte
atrial cardiomyocyte
cardiomyocyte
slow muscle
fast muscle
skeletal muscle satellite stem c.
tendon c.
chondrocyte
osteoblast
osteo-CAR c.
stromal c.
? stromal 1 ST6GAL2+
? stromal 1 COL15A1+PTGDS+
? stromal 2 NGFR+TNNT2+
? stromal 1 NGFR+TNNT2+
reticular c.
leptomeningeal c.
mesothelial c. SBSN+
mesothelial c. CXCL10-PRG4+
mesothelial c. CXCL10+PRG4+
mesothelial c.
adipocyte UCP1lo
adipocyte UCP1hi
adipo-CAR c.
pericyte
fibromyocyte
myofibroblast
fibroblast LY6H+
fibroblast ENPP2+
fibroblast TNC+
fibroblast ASPN+
fibroblast PTN+
fibroblast PI16+
fibroblast KCNN3+
fibroblast GGT5+
fibroblast CHODL+
fibroblast WIF1+
fibroblast NNAT+
alveolar fibroblast
adventitial fibroblast
fibroblast LCN2+
fibroblast ITIH5+
fibroblast ACKR2+
fibroblast
fibroadipogenic progenitor

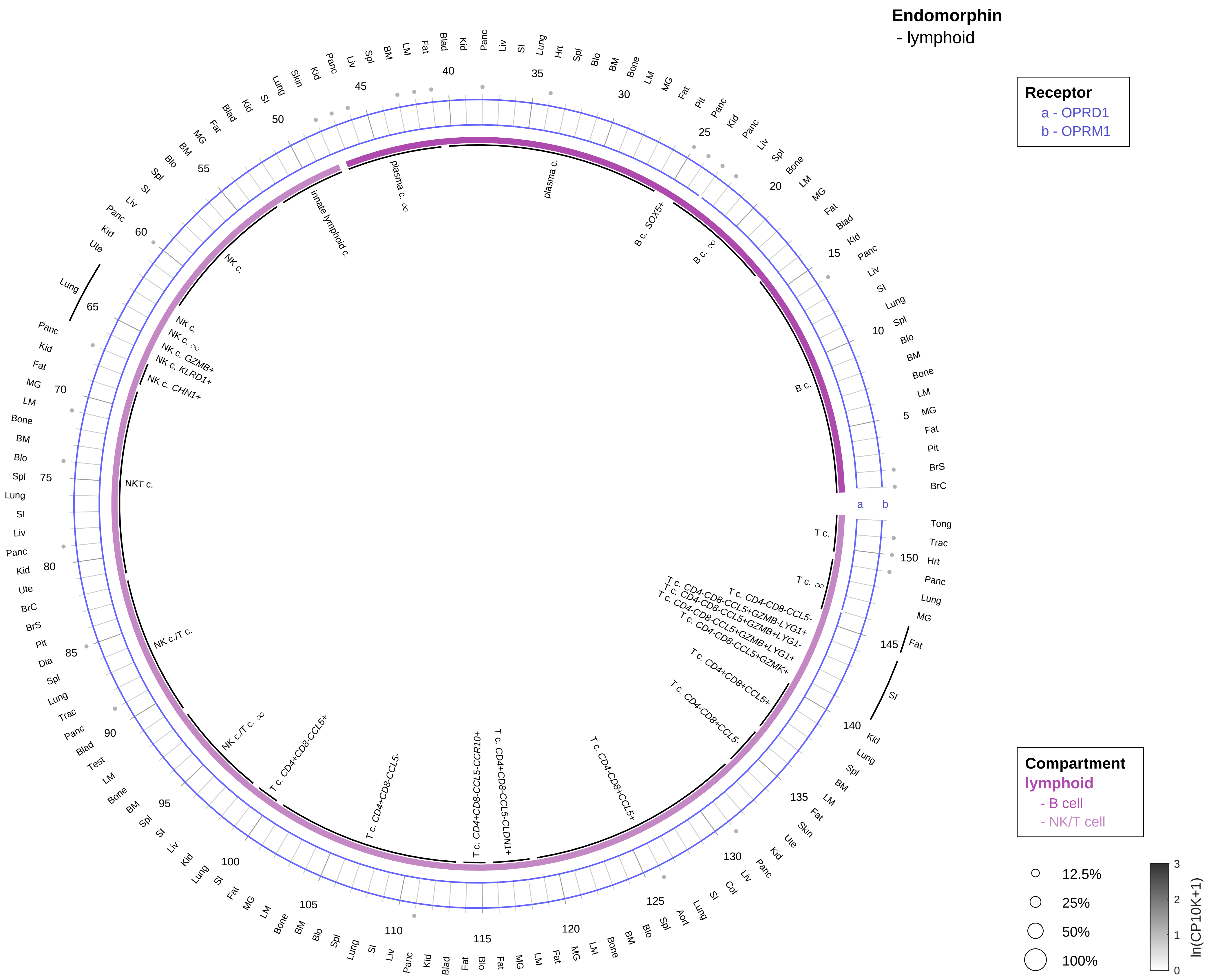

| Abbrev. | Tissue |
|---|---|
| Aort | Aorta |
| Blad | Bladder |
| Blo | Blood |
| Bone | Bone |
| BM | Bone marrow |
| BrC | Brain cortex |
| BrS | Brainstem |
| Col | Colon |
| Dia | Diaphragm |
| Ret | Eye retina |
| Fat | Fat |
| Hrt | Heart |
| Pit | Hypothalamus/Pituitary |
| Kid | Kidney |
| LM | Limb muscle |
| Liv | Liver |
| Lung | Lung |
| MG | Mammary gland |
| Panc | Pancreas |
| Skin | Skin |
| SI | Small intestine |
| Spl | Spleen |
| Test | Testes |
| Thym | Thymus |
| Tong | Tongue |
| Trac | Trachea |
| Ute | Uterus |

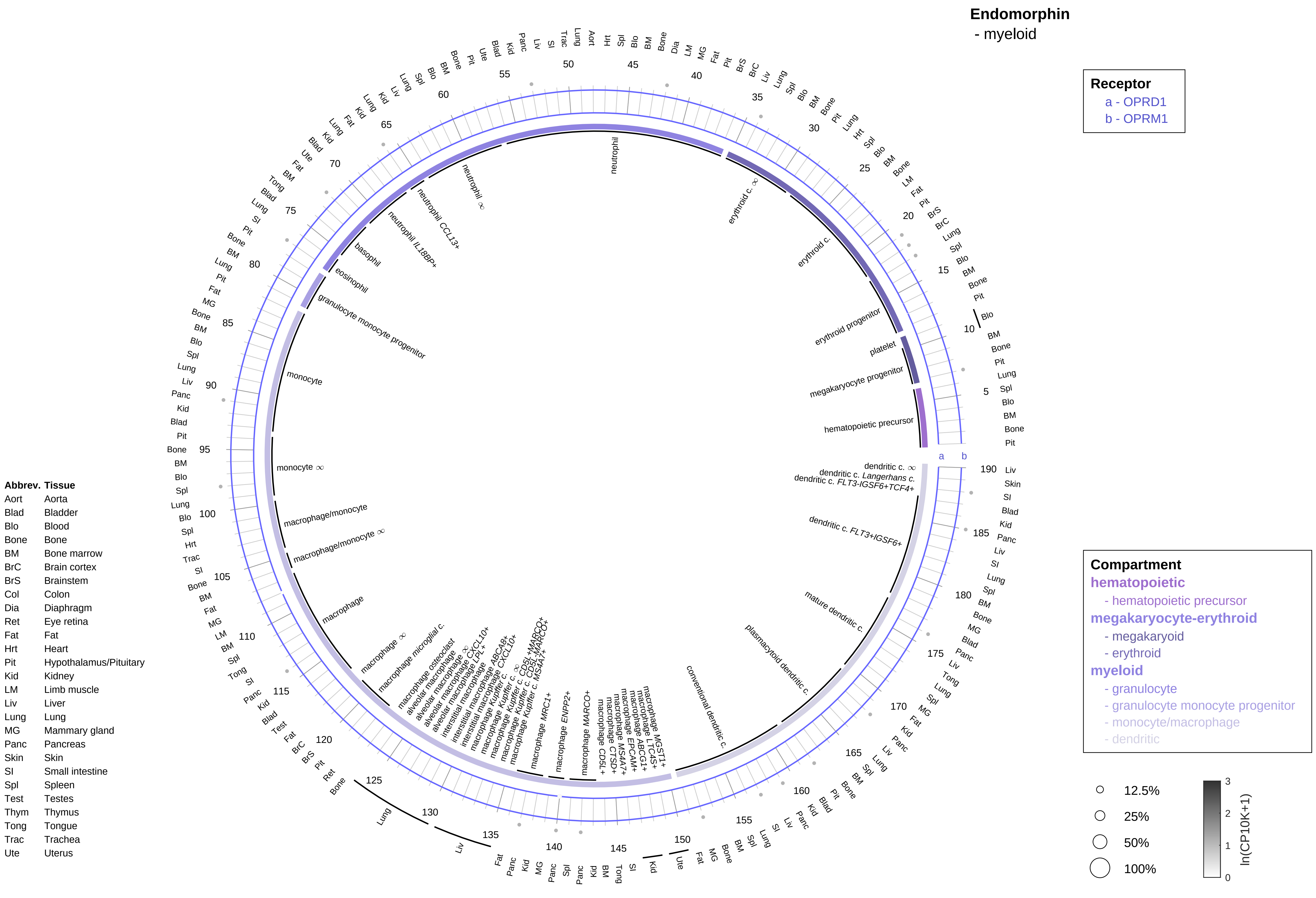

Endomorphin
- myeloid
Receptor
a - OPRD1
b - OPRM1
Compartment
hematopoietic
- hematopoietic precursor
megakaryocyte-erythroid
- megakaryoid
- erythroid
myeloid
- granulocyte
- granulocyte monocyte progenitor
- monocyte/macrophage
- dendritic
12.5%
25%
50%
100%
ln(CP10K+1)
0
1
2
3
Abbrev. Tissue
Aort Aorta
Blad Bladder
Blo Blood
Bone Bone
BM Bone marrow
BrC Brain cortex
BrS Brainstem
Col Colon
Dia Diaphragm
Ret Eye retina
Fat Fat
Hrt Heart
Pit Hypothalamus/Pituitary
Kid Kidney
LM Limb muscle
Liv Liver
Lung Lung
MG Mammary gland
Panc Pancreas
Skin Skin
SI Small intestine
Spl Spleen
Test Testes
Thym Thymus
Tong Tongue
Trac Trachea
Ute Uterus
hematopoietic precursor
megakaryocyte progenitor
platelet
erythroid progenitor
erythroid c.
erythroid c. ∞
neutrophil
neutrophil ∞
neutrophil CCL13+
neutrophil IL18BP+
basophil
eosinophil
granulocyte monocyte progenitor
monocyte
monocyte ∞
macrophage/monocyte
macrophage/monocyte ∞
macrophage
macrophage ∞
macrophage microglial c.
macrophage osteoclast
alveolar macrophage
macrophage MRC1+
macrophage ENPP2+
macrophage MARCO+
macrophage CD5L+
macrophage CTSD+
macrophage MS4A7+
macrophage EPCAM+
macrophage ABCG1+
macrophage LTC4S+
macrophage MGST1+
conventional dendritic c.
plasmacytoid dendritic c.
mature dendritic c.
dendritic c. FLT3+IGSF6+
dendritic c. FLT3-IGSF6+TCF4+
dendritic c. Langerhans c.
dendritic c. ∞

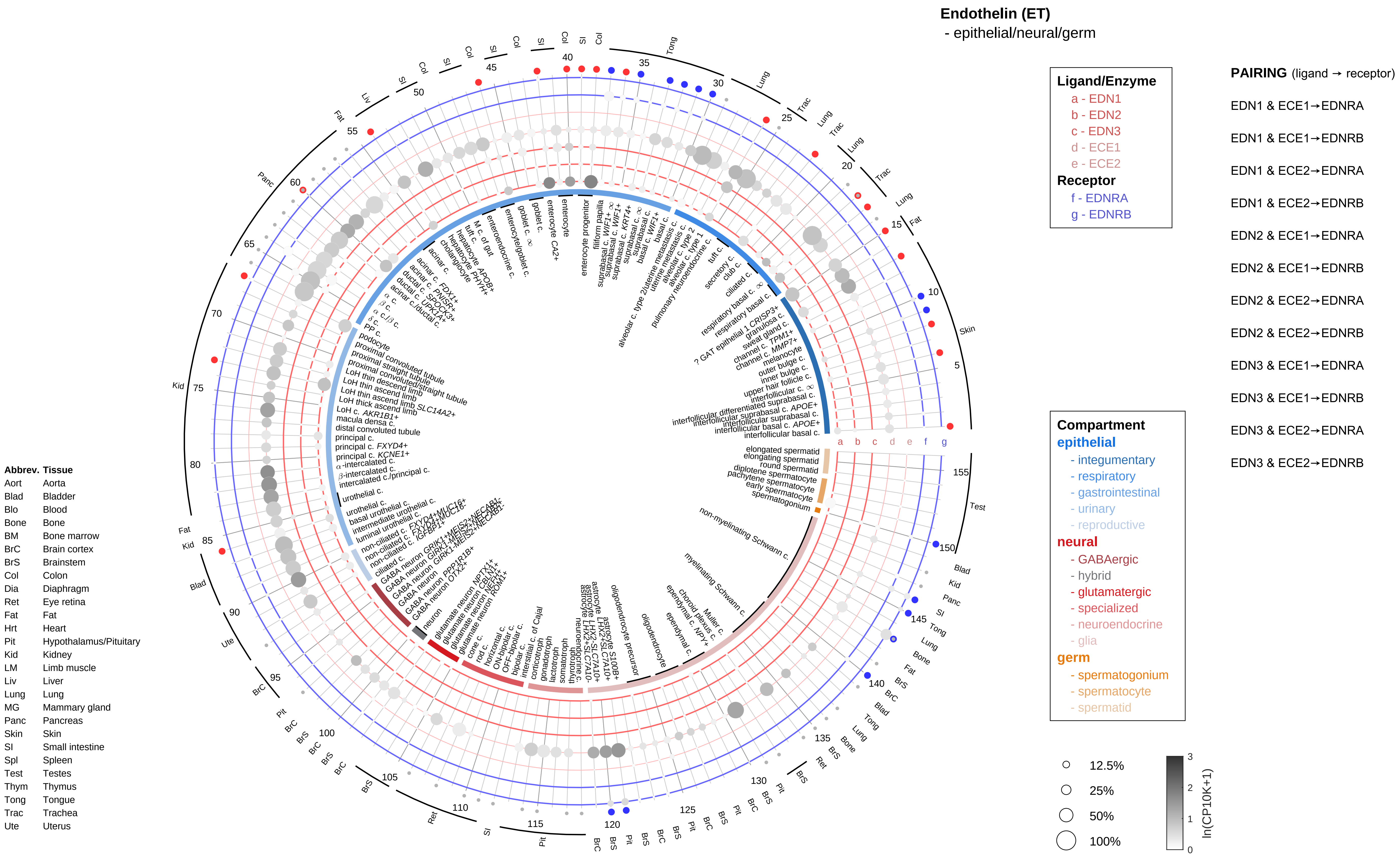
Endothelin (ET)
- epithelial/neural/germ
Ligand/Enzyme
a - EDN1
b - EDN2
c - EDN3
d - ECE1
e - ECE2
Receptor
f - EDNRA
g - EDNRB
PAIRING (ligand → receptor)
EDN1 & ECE1→EDNRA
EDN1 & ECE1→EDNRB
EDN1 & ECE2→EDNRA
EDN1 & ECE2→EDNRB
EDN2 & ECE1→EDNRA
EDN2 & ECE1→EDNRB
EDN2 & ECE2→EDNRA
EDN2 & ECE2→EDNRB
EDN3 & ECE1→EDNRA
EDN3 & ECE1→EDNRB
EDN3 & ECE2→EDNRA
EDN3 & ECE2→EDNRB
Compartment
epithelial
- integumentary
- respiratory
- gastrointestinal
- urinary
- reproductive
neural
- GABAergic
- hybrid
- glutamatergic
- specialized
- neuroendocrine
- glia
germ
- spermatogonium
- spermatocyte
- spermatid
12.5%
25%
50%
100%
ln(CP10K+1)
Abbrev. Tissue
Aort Aorta
Blad Bladder
Blo Blood
Bone Bone
BM Bone marrow
BrC Brain cortex
BrS Brainstem
Col Colon
Dia Diaphragm
Ret Eye retina
Fat Fat
Hrt Heart
Pit Hypothalamus/Pituitary
Kid Kidney
LM Limb muscle
Liv Liver
Lung Lung
MG Mammary gland
Panc Pancreas
Skin Skin
SI Small intestine
Spl Spleen
Test Testes
Thym Thymus
Tong Tongue
Trac Trachea
Ute Uterus

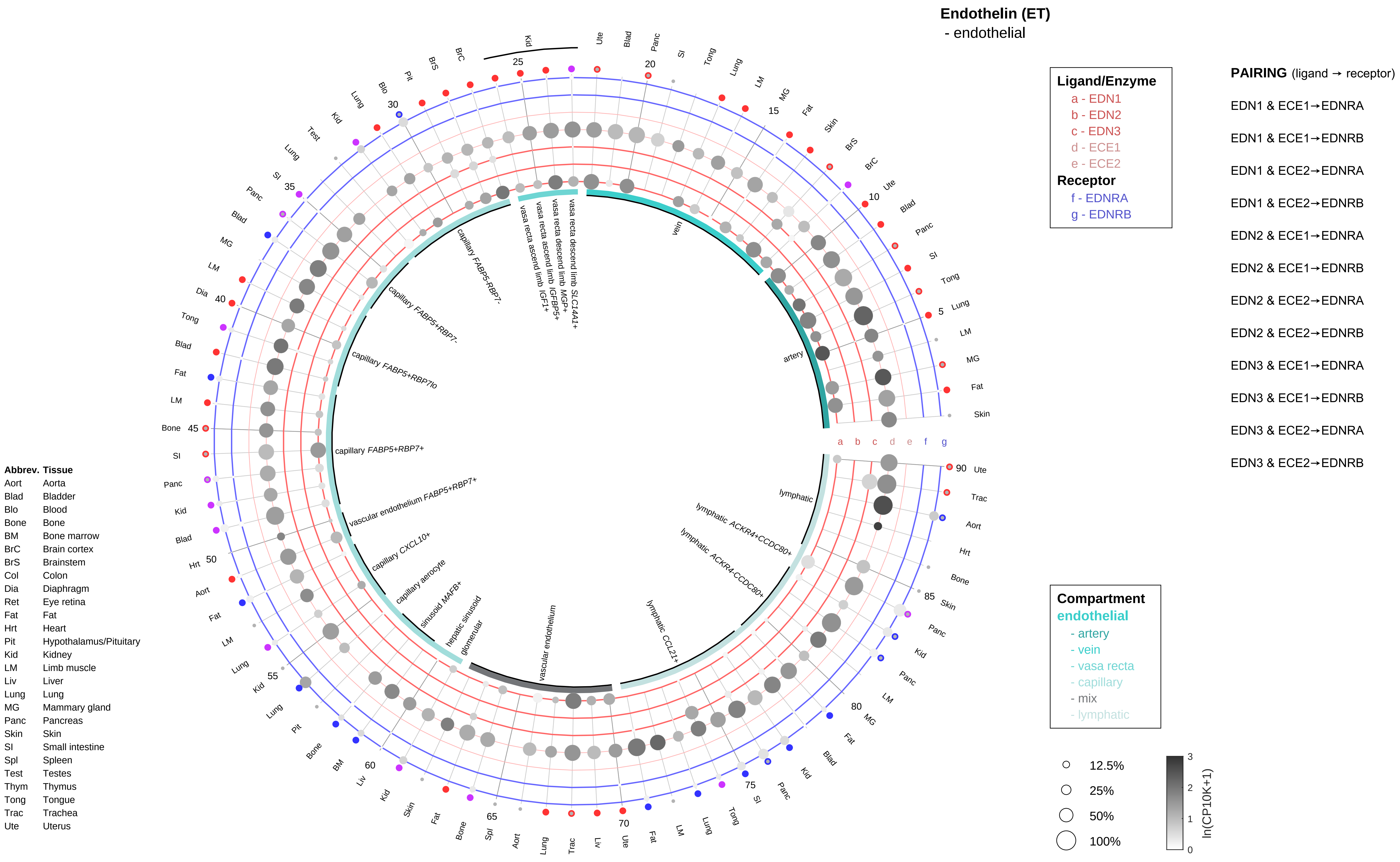

Endothelin (ET)
- endothelial
Ligand/Enzyme
a - EDN1
b - EDN2
c - EDN3
d - ECE1
e - ECE2
Receptor
f - EDNRA
g - EDNRB
PAIRING (ligand → receptor)
EDN1 & ECE1→EDNRA
EDN1 & ECE1→EDNRB
EDN1 & ECE2→EDNRA
EDN1 & ECE2→EDNRB
EDN2 & ECE1→EDNRA
EDN2 & ECE1→EDNRB
EDN2 & ECE2→EDNRA
EDN2 & ECE2→EDNRB
EDN3 & ECE1→EDNRA
EDN3 & ECE1→EDNRB
EDN3 & ECE2→EDNRA
EDN3 & ECE2→EDNRB
Compartment
endothelial
- artery
- vein
- vasa recta
- capillary
- mix
- lymphatic
12.5%
25%
50%
100%
ln(CP10K+1)
Abbrev. Tissue
Aort Aorta
Blad Bladder
Blo Blood
Bone Bone
BM Bone marrow
BrC Brain cortex
BrS Brainstem
Col Colon
Dia Diaphragm
Ret Eye retina
Fat Fat
Hrt Heart
Pit Hypothalamus/Pituitary
Kid Kidney
LM Limb muscle
Liv Liver
Lung Lung
MG Mammary gland
Panc Pancreas
Skin Skin
SI Small intestine
Spl Spleen
Test Testes
Thym Thymus
Tong Tongue
Trac Trachea
Ute Uterus
artery
vein
vasa recta descend limb SLC14A1+
vasa recta descend limb MGP+
vasa recta ascend limb IGFBP5+
vasa recta ascend limb IGF1+
capillary FABP5-RBP7-
capillary FABP5+RBP7-
capillary FABP5+RBP7lo
capillary FABP5+RBP7+
vascular endothelium FABP5+RBP7+
capillary CXCL10+
capillary aerocyte
sinusoid MAFB+
hepatic sinusoid
glomerular
vascular endothelium
lymphatic CCL21+
lymphatic ACKR4-CCDC80+
lymphatic ACKR4+CCDC80+
lymphatic

#### Endothelin (ET)
- stromal

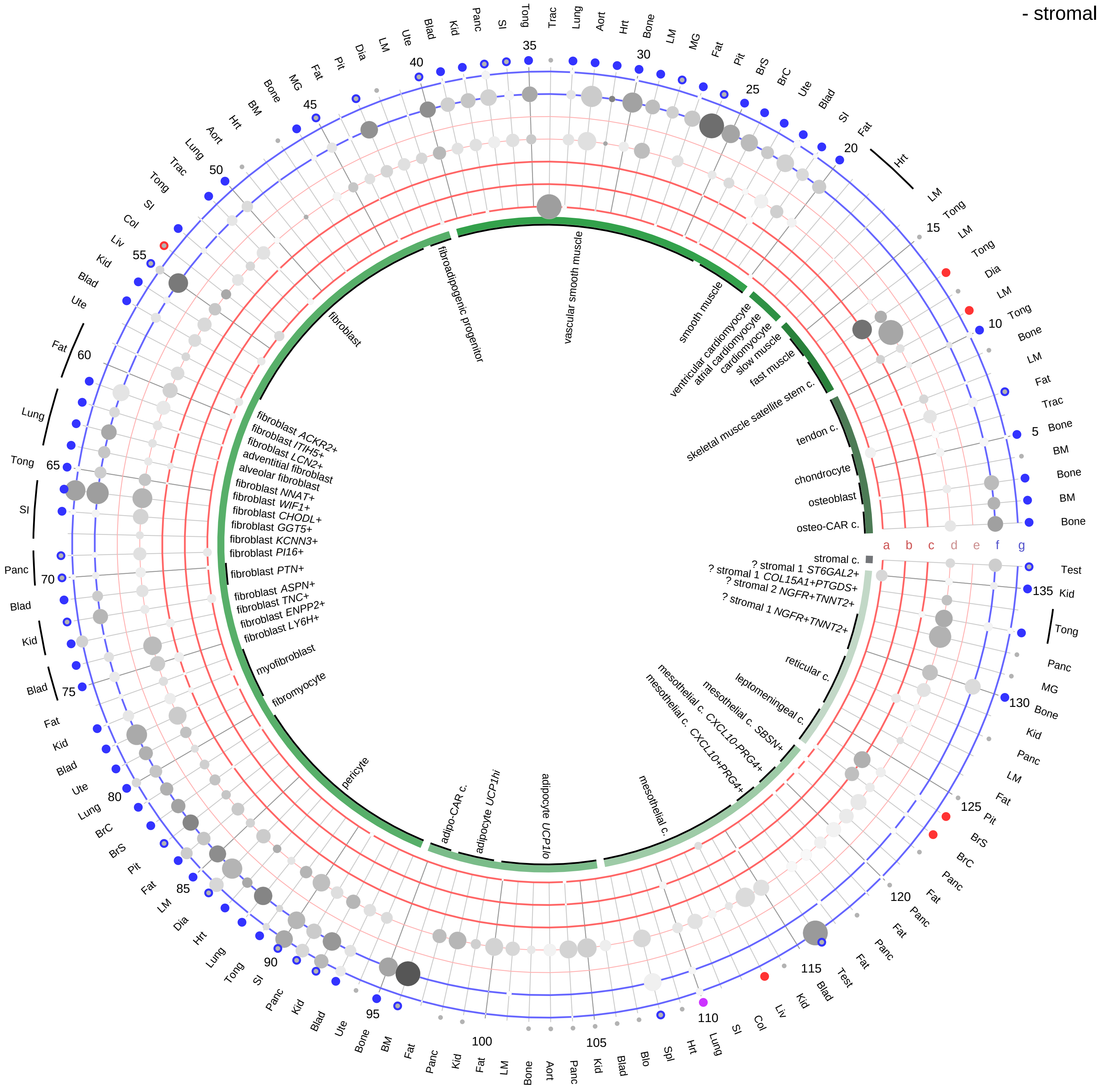

**Ligand/Enzyme**
- a - EDN1
- b - EDN2
- c - EDN3
- d - ECE1
- e - ECE2

**Receptor**
- f - EDNRA
- g - EDNRB

**PAIRING** (ligand → receptor)

EDN1 & ECE1→EDNRA
EDN1 & ECE1→EDNRB
EDN1 & ECE2→EDNRA
EDN1 & ECE2→EDNRB
EDN2 & ECE1→EDNRA
EDN2 & ECE1→EDNRB
EDN2 & ECE2→EDNRA
EDN2 & ECE2→EDNRB
EDN3 & ECE1→EDNRA
EDN3 & ECE1→EDNRB
EDN3 & ECE2→EDNRA
EDN3 & ECE2→EDNRB

**Compartment**
**stromal**
- bone
- skeletal muscle
- cardiac muscle
- smooth muscle
- fibroblast
- adipocyte
- mesothelial
- other
- mix

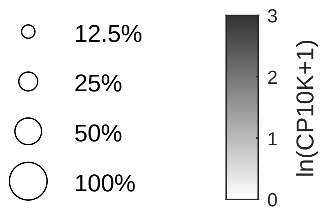

| Abbrev. | Tissue |
|---|---|
| Aort | Aorta |
| Blad | Bladder |
| Blo | Blood |
| Bone | Bone |
| BM | Bone marrow |
| BrC | Brain cortex |
| BrS | Brainstem |
| Col | Colon |
| Dia | Diaphragm |
| Ret | Eye retina |
| Fat | Fat |
| Hrt | Heart |
| Pit | Hypothalamus/Pituitary |
| Kid | Kidney |
| LM | Limb muscle |
| Liv | Liver |
| Lung | Lung |
| MG | Mammary gland |
| Panc | Pancreas |
| Skin | Skin |
| SI | Small intestine |
| Spl | Spleen |
| Test | Testes |
| Thym | Thymus |
| Tong | Tongue |
| Trac | Trachea |
| Ute | Uterus |

| Abbrev. | Tissue |
|---|---|
| Aort | Aorta |
| Blad | Bladder |
| Blo | Blood |
| Bone | Bone |
| BM | Bone marrow |
| BrC | Brain cortex |
| BrS | Brainstem |
| Col | Colon |
| Dia | Diaphragm |
| Ret | Eye retina |
| Fat | Fat |
| Hrt | Heart |
| Pit | Hypothalamus/Pituitary |
| Kid | Kidney |
| LM | Limb muscle |
| Liv | Liver |
| Lung | Lung |
| MG | Mammary gland |
| Panc | Pancreas |
| Skin | Skin |
| SI | Small intestine |
| Spl | Spleen |
| Test | Testes |
| Thym | Thymus |
| Tong | Tongue |
| Trac | Trachea |
| Ute | Uterus |

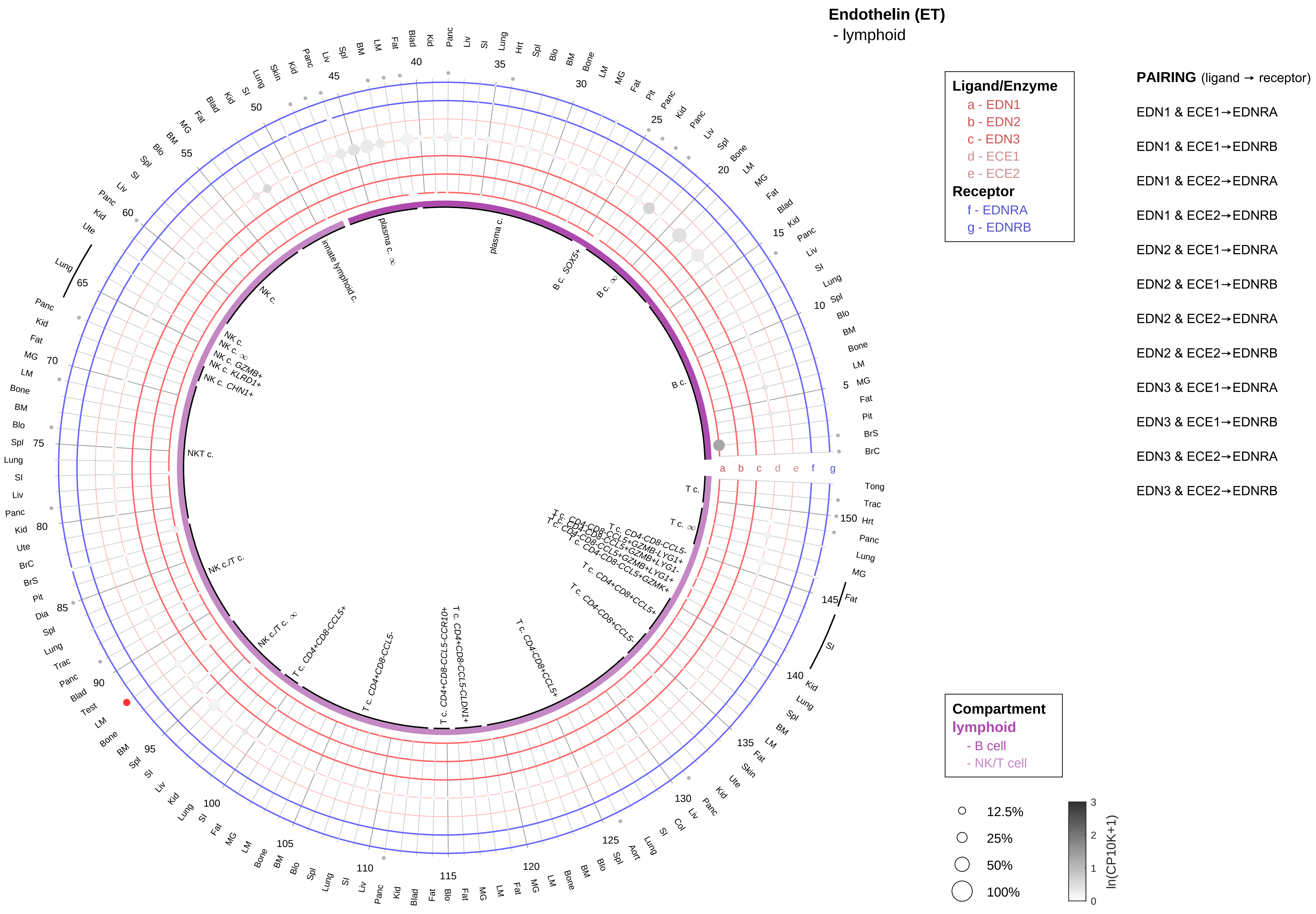

**Endothelin (ET)**
- myeloid

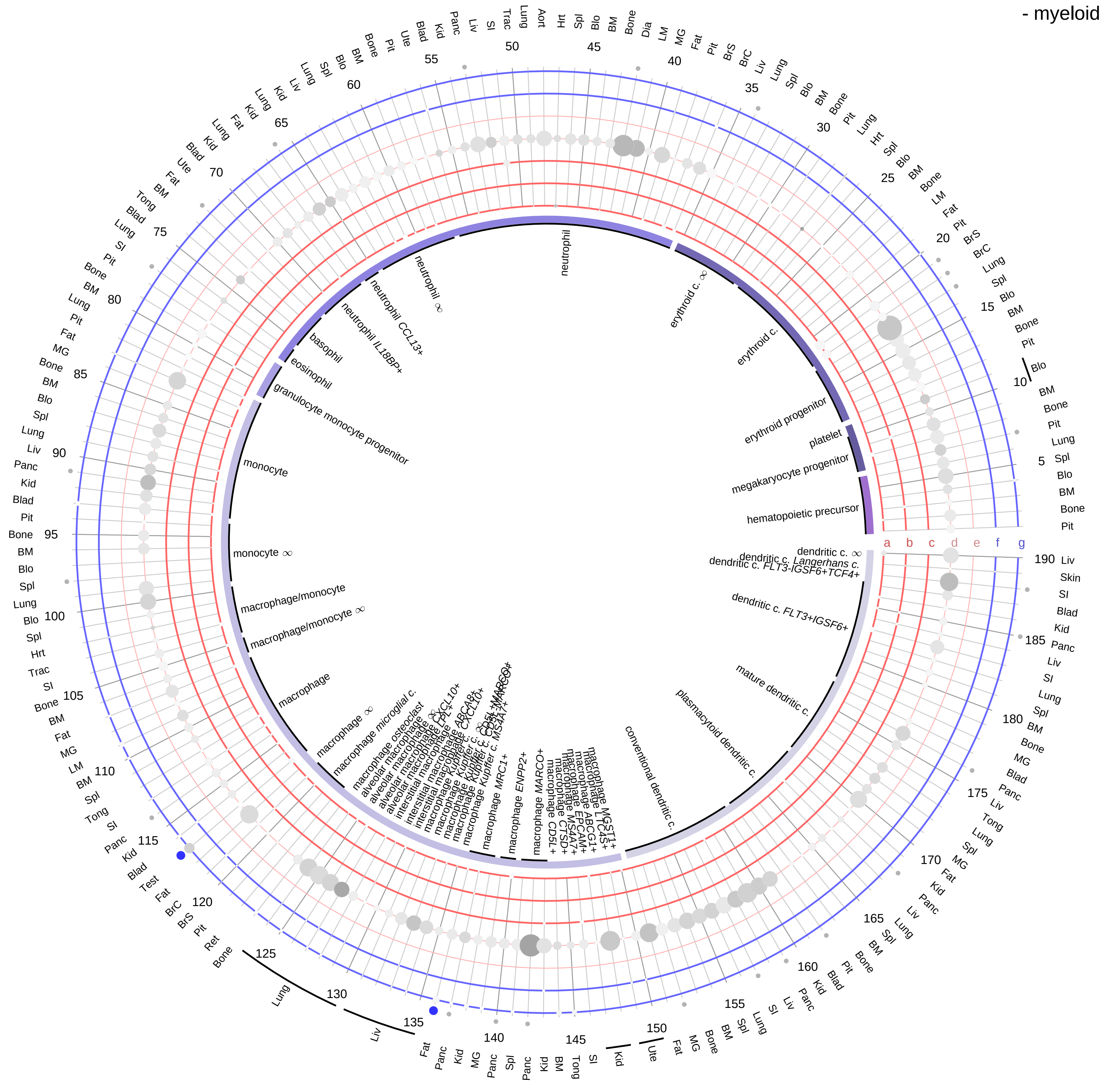

**Ligand/Enzyme**
a - EDN1
b - EDN2
c - EDN3
d - ECE1
e - ECE2
**Receptor**
f - EDNRA
g - EDNRB

**PAIRING** (ligand → receptor)

EDN1 & ECE1→EDNRA
EDN1 & ECE1→EDNRB
EDN1 & ECE2→EDNRA
EDN1 & ECE2→EDNRB
EDN2 & ECE1→EDNRA
EDN2 & ECE1→EDNRB
EDN2 & ECE2→EDNRA
EDN2 & ECE2→EDNRB
EDN3 & ECE1→EDNRA
EDN3 & ECE1→EDNRB
EDN3 & ECE2→EDNRA
EDN3 & ECE2→EDNRB

**Compartment**
**hematopoietic**
- hematopoietic precursor
**megakaryocyte-erythroid**
- megakaryoid
- erythroid
**myeloid**
- granulocyte
- granulocyte monocyte progenitor
- monocyte/macrophage
- dendritic

12.5%
25%
50%
100%

ln(CP10K+1): 0, 1, 2, 3

| Abbrev. | Tissue |
|---|---|
| Aort | Aorta |
| Blad | Bladder |
| Blo | Blood |
| Bone | Bone |
| BM | Bone marrow |
| BrC | Brain cortex |
| BrS | Brainstem |
| Col | Colon |
| Dia | Diaphragm |
| Ret | Eye retina |
| Fat | Fat |
| Hrt | Heart |
| Pit | Hypothalamus/Pituitary |
| Kid | Kidney |
| LM | Limb muscle |
| Liv | Liver |
| Lung | Lung |
| MG | Mammary gland |
| Panc | Pancreas |
| Skin | Skin |
| SI | Small intestine |
| Spl | Spleen |
| Test | Testes |
| Thym | Thymus |
| Tong | Tongue |
| Trac | Trachea |
| Ute | Uterus |

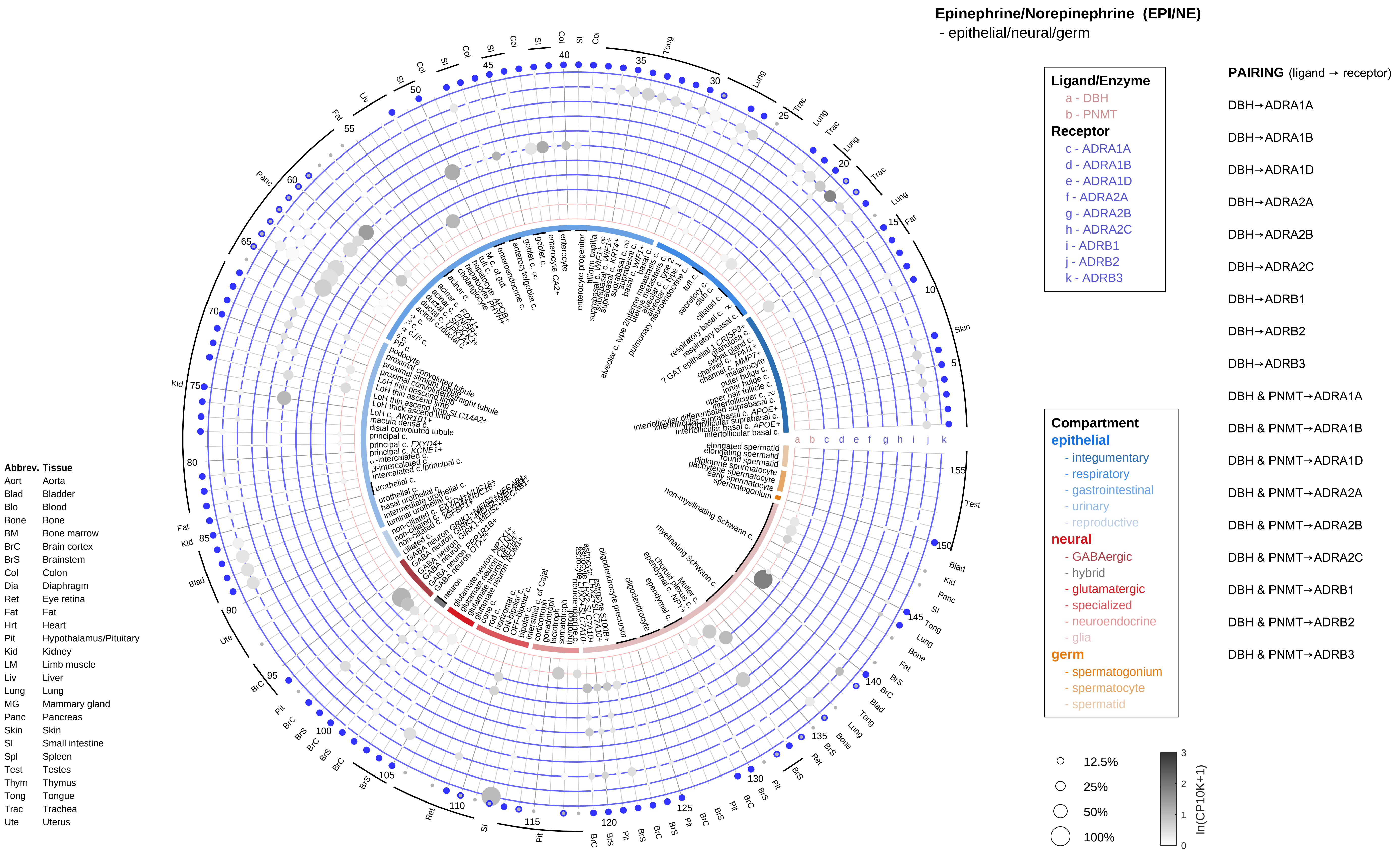
Epinephrine/Norepinephrine (EPI/NE)
- epithelial/neural/germ
Ligand/Enzyme
a - DBH
b - PNMT
Receptor
c - ADRA1A
d - ADRA1B
e - ADRA1D
f - ADRA2A
g - ADRA2B
h - ADRA2C
i - ADRB1
j - ADRB2
k - ADRB3
PAIRING (ligand → receptor)
DBH→ADRA1A
DBH→ADRA1B
DBH→ADRA1D
DBH→ADRA2A
DBH→ADRA2B
DBH→ADRA2C
DBH→ADRB1
DBH→ADRB2
DBH→ADRB3
DBH & PNMT→ADRA1A
DBH & PNMT→ADRA1B
DBH & PNMT→ADRA1D
DBH & PNMT→ADRA2A
DBH & PNMT→ADRA2B
DBH & PNMT→ADRA2C
DBH & PNMT→ADRB1
DBH & PNMT→ADRB2
DBH & PNMT→ADRB3
Compartment
epithelial
- integumentary
- respiratory
- gastrointestinal
- urinary
- reproductive
neural
- GABAergic
- hybrid
- glutamatergic
- specialized
- neuroendocrine
- glia
germ
- spermatogonium
- spermatocyte
- spermatid
12.5%
25%
50%
100%
ln(CP10K+1)
Abbrev. Tissue
Aort Aorta
Blad Bladder
Blo Blood
Bone Bone
BM Bone marrow
BrC Brain cortex
BrS Brainstem
Col Colon
Dia Diaphragm
Ret Eye retina
Fat Fat
Hrt Heart
Pit Hypothalamus/Pituitary
Kid Kidney
LM Limb muscle
Liv Liver
Lung Lung
MG Mammary gland
Panc Pancreas
Skin Skin
SI Small intestine
Spl Spleen
Test Testes
Thym Thymus
Tong Tongue
Trac Trachea
Ute Uterus

#### Epinephrine/Norepinephrine (EPI/NE)
- endothelial

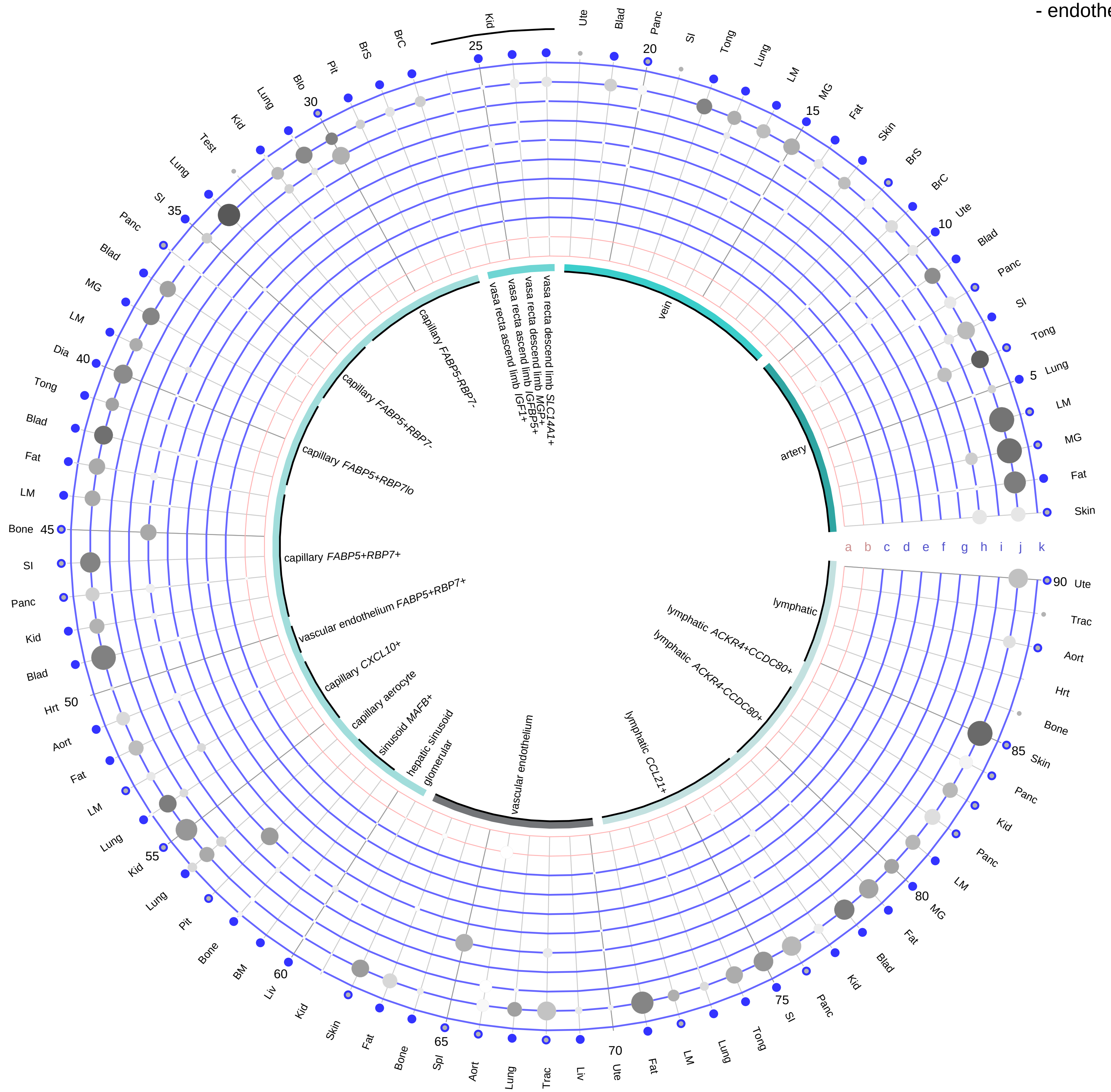

**Ligand/Enzyme**
a - DBH
b - PNMT

**Receptor**
c - ADRA1A
d - ADRA1B
e - ADRA1D
f - ADRA2A
g - ADRA2B
h - ADRA2C
i - ADRB1
j - ADRB2
k - ADRB3

**PAIRING** (ligand → receptor)

DBH→ADRA1A
DBH→ADRA1B
DBH→ADRA1D
DBH→ADRA2A
DBH→ADRA2B
DBH→ADRA2C
DBH→ADRB1
DBH→ADRB2
DBH→ADRB3
DBH & PNMT→ADRA1A
DBH & PNMT→ADRA1B
DBH & PNMT→ADRA1D
DBH & PNMT→ADRA2A
DBH & PNMT→ADRA2B
DBH & PNMT→ADRA2C
DBH & PNMT→ADRB1
DBH & PNMT→ADRB2
DBH & PNMT→ADRB3

**Compartment**
**endothelial**
- artery
- vein
- vasa recta
- capillary
- mix
- lymphatic

12.5%
25%
50%
100%

ln(CP10K+1): 0, 1, 2, 3

| Abbrev. | Tissue |
|---|---|
| Aort | Aorta |
| Blad | Bladder |
| Blo | Blood |
| Bone | Bone |
| BM | Bone marrow |
| BrC | Brain cortex |
| BrS | Brainstem |
| Col | Colon |
| Dia | Diaphragm |
| Ret | Eye retina |
| Fat | Fat |
| Hrt | Heart |
| Pit | Hypothalamus/Pituitary |
| Kid | Kidney |
| LM | Limb muscle |
| Liv | Liver |
| Lung | Lung |
| MG | Mammary gland |
| Panc | Pancreas |
| Skin | Skin |
| SI | Small intestine |
| Spl | Spleen |
| Test | Testes |
| Thym | Thymus |
| Tong | Tongue |
| Trac | Trachea |
| Ute | Uterus |

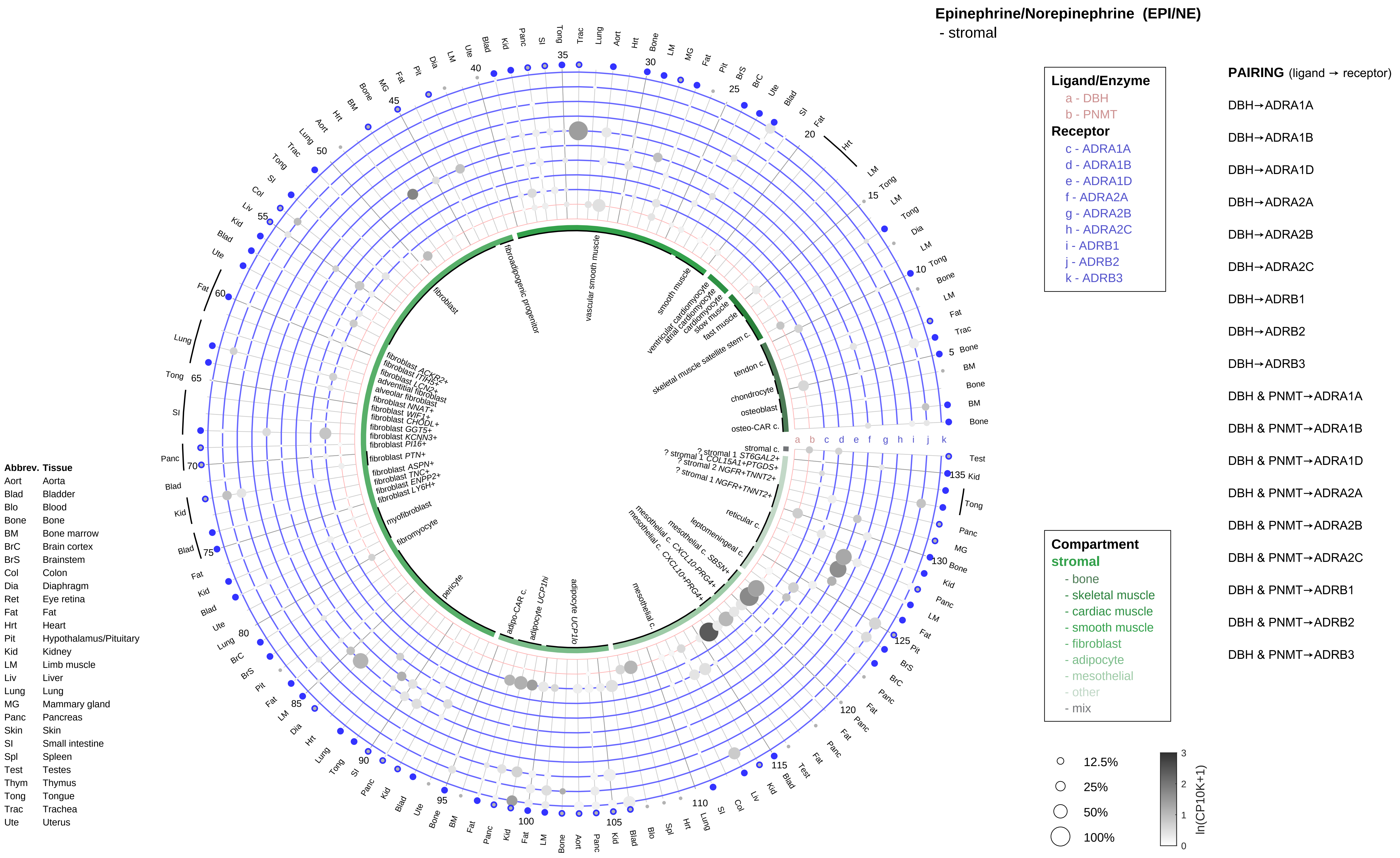
Epinephrine/Norepinephrine (EPI/NE)
- stromal
Ligand/Enzyme
a - DBH
b - PNMT
Receptor
c - ADRA1A
d - ADRA1B
e - ADRA1D
f - ADRA2A
g - ADRA2B
h - ADRA2C
i - ADRB1
j - ADRB2
k - ADRB3
PAIRING (ligand → receptor)
DBH→ADRA1A
DBH→ADRA1B
DBH→ADRA1D
DBH→ADRA2A
DBH→ADRA2B
DBH→ADRA2C
DBH→ADRB1
DBH→ADRB2
DBH→ADRB3
DBH & PNMT→ADRA1A
DBH & PNMT→ADRA1B
DBH & PNMT→ADRA1D
DBH & PNMT→ADRA2A
DBH & PNMT→ADRA2B
DBH & PNMT→ADRA2C
DBH & PNMT→ADRB1
DBH & PNMT→ADRB2
DBH & PNMT→ADRB3
Compartment
stromal
- bone
- skeletal muscle
- cardiac muscle
- smooth muscle
- fibroblast
- adipocyte
- mesothelial
- other
- mix
12.5%
25%
50%
100%
ln(CP10K+1)
Abbrev. Tissue
Aort Aorta
Blad Bladder
Blo Blood
Bone Bone
BM Bone marrow
BrC Brain cortex
BrS Brainstem
Col Colon
Dia Diaphragm
Ret Eye retina
Fat Fat
Hrt Heart
Pit Hypothalamus/Pituitary
Kid Kidney
LM Limb muscle
Liv Liver
Lung Lung
MG Mammary gland
Panc Pancreas
Skin Skin
SI Small intestine
Spl Spleen
Test Testes
Thym Thymus
Tong Tongue
Trac Trachea
Ute Uterus
fibroblast
fibroadipogenic progenitor
vascular smooth muscle
smooth muscle
ventricular cardiomyocyte
atrial cardiomyocyte
cardiomyocyte
slow muscle
fast muscle
skeletal muscle satellite stem c.
tendon c.
chondrocyte
osteoblast
osteo-CAR c.
stromal c.
myofibroblast
fibromyocyte
pericyte
adipo-CAR c.
adipocyte UCP1hi
adipocyte UCP1lo
mesothelial c.
reticular c.
leptomeningeal c.

**Epinephrine/Norepinephrine (EPI/NE)**
- lymphoid

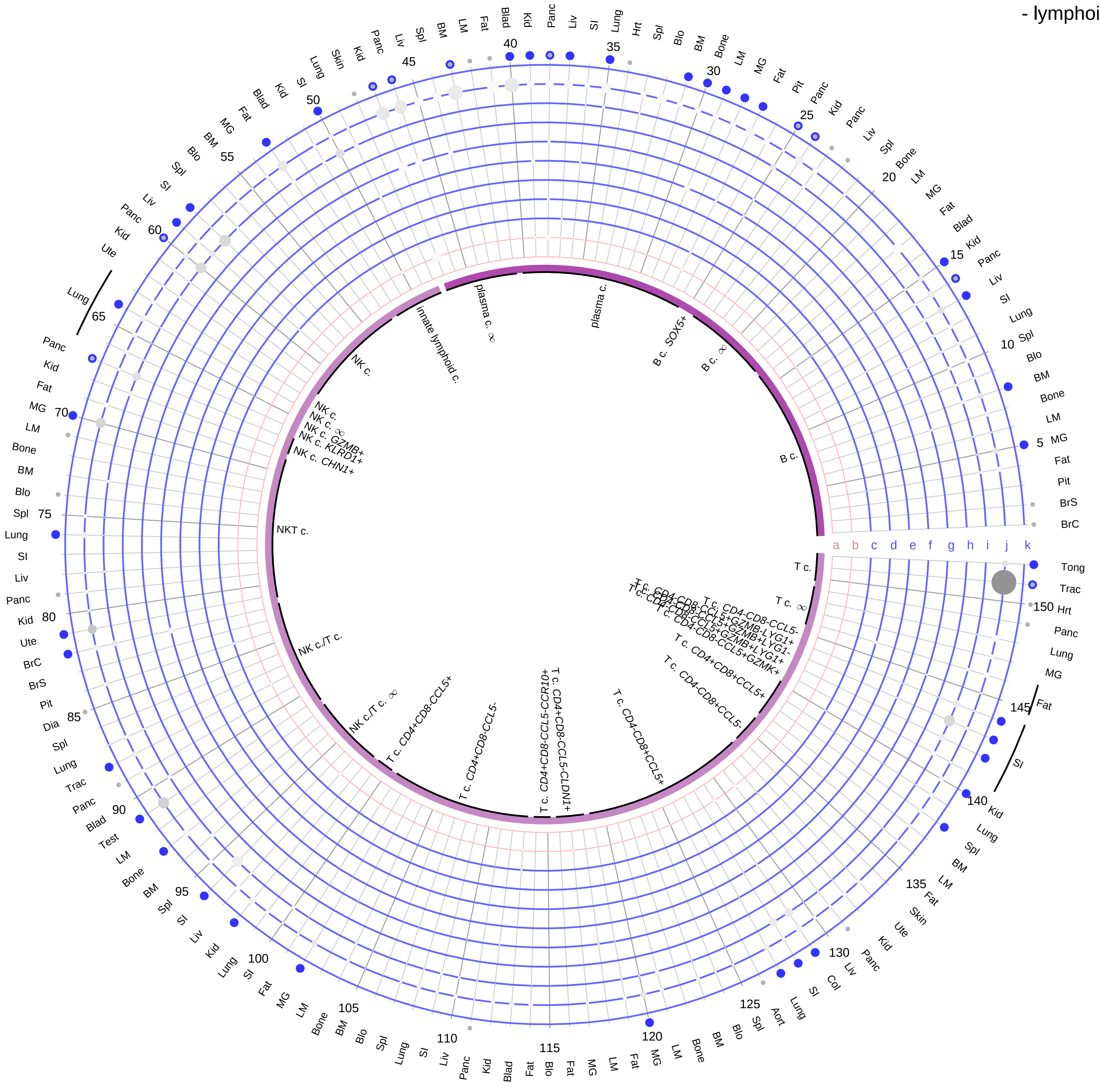

**Ligand/Enzyme**
- a - DBH
- b - PNMT

**Receptor**
- c - ADRA1A
- d - ADRA1B
- e - ADRA1D
- f - ADRA2A
- g - ADRA2B
- h - ADRA2C
- i - ADRB1
- j - ADRB2
- k - ADRB3

**PAIRING** (ligand → receptor)

DBH→ADRA1A
DBH→ADRA1B
DBH→ADRA1D
DBH→ADRA2A
DBH→ADRA2B
DBH→ADRA2C
DBH→ADRB1
DBH→ADRB2
DBH→ADRB3
DBH & PNMT→ADRA1A
DBH & PNMT→ADRA1B
DBH & PNMT→ADRA1D
DBH & PNMT→ADRA2A
DBH & PNMT→ADRA2B
DBH & PNMT→ADRA2C
DBH & PNMT→ADRB1
DBH & PNMT→ADRB2
DBH & PNMT→ADRB3

**Compartment**
**lymphoid**
- B cell
- NK/T cell

12.5%
25%
50%
100%

ln(CP10K+1): 0, 1, 2, 3

| Abbrev. | Tissue |
|---|---|
| Aort | Aorta |
| Blad | Bladder |
| Blo | Blood |
| Bone | Bone |
| BM | Bone marrow |
| BrC | Brain cortex |
| BrS | Brainstem |
| Col | Colon |
| Dia | Diaphragm |
| Ret | Eye retina |
| Fat | Fat |
| Hrt | Heart |
| Pit | Hypothalamus/Pituitary |
| Kid | Kidney |
| LM | Limb muscle |
| Liv | Liver |
| Lung | Lung |
| MG | Mammary gland |
| Panc | Pancreas |
| Skin | Skin |
| SI | Small intestine |
| Spl | Spleen |
| Test | Testes |
| Thym | Thymus |
| Tong | Tongue |
| Trac | Trachea |
| Ute | Uterus |

**Epinephrine/Norepinephrine (EPI/NE)**
- myeloid

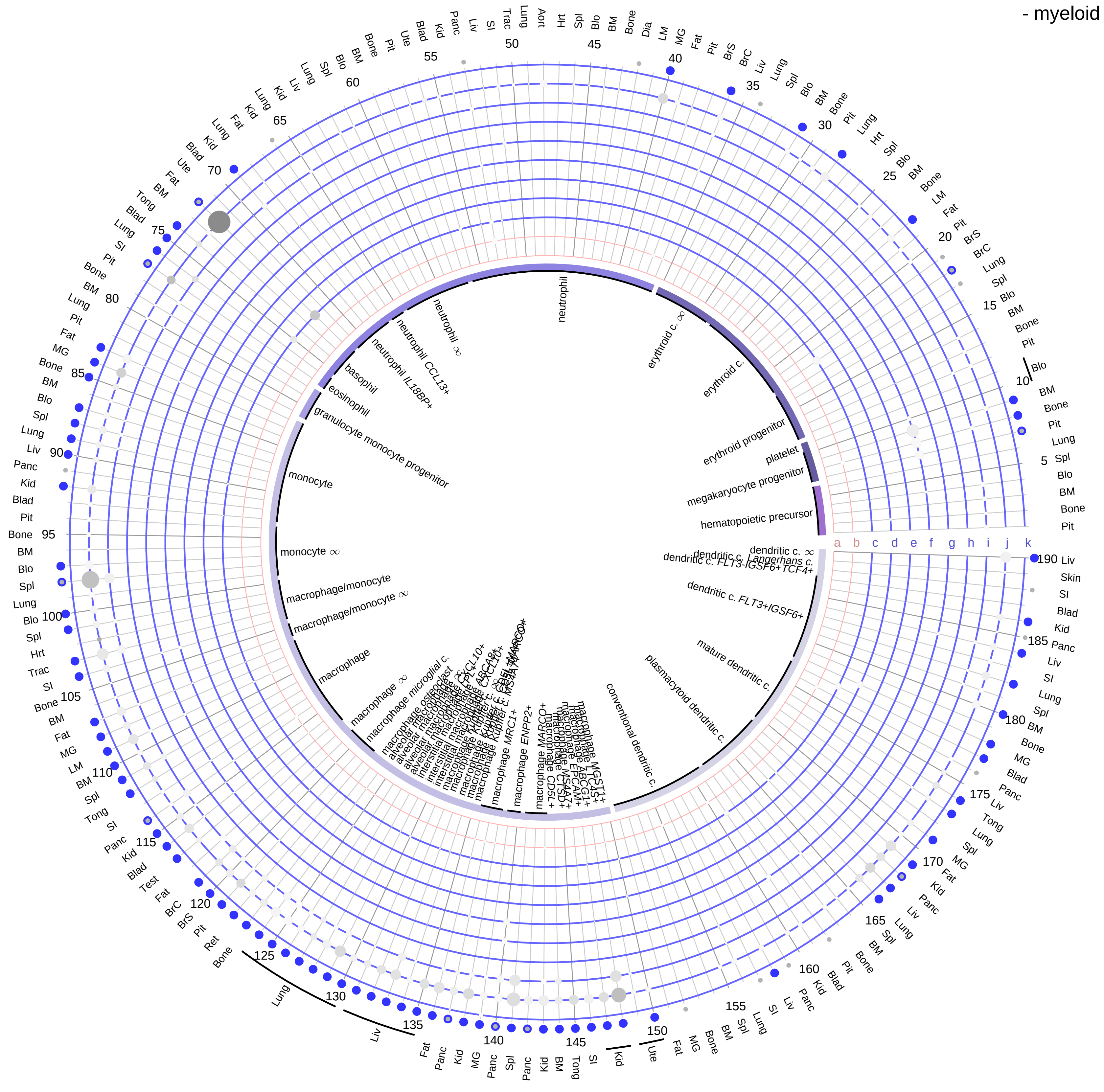

**Ligand/Enzyme**
- a - DBH
- b - PNMT

**Receptor**
- c - ADRA1A
- d - ADRA1B
- e - ADRA1D
- f - ADRA2A
- g - ADRA2B
- h - ADRA2C
- i - ADRB1
- j - ADRB2
- k - ADRB3

**PAIRING** (ligand → receptor)

DBH→ADRA1A
DBH→ADRA1B
DBH→ADRA1D
DBH→ADRA2A
DBH→ADRA2B
DBH→ADRA2C
DBH→ADRB1
DBH→ADRB2
DBH→ADRB3
DBH & PNMT→ADRA1A
DBH & PNMT→ADRA1B
DBH & PNMT→ADRA1D
DBH & PNMT→ADRA2A
DBH & PNMT→ADRA2B
DBH & PNMT→ADRA2C
DBH & PNMT→ADRB1
DBH & PNMT→ADRB2
DBH & PNMT→ADRB3

**Compartment**
**hematopoietic**
- hematopoietic precursor

**megakaryocyte-erythroid**
- megakaryoid
- erythroid

**myeloid**
- granulocyte
- granulocyte monocyte progenitor
- monocyte/macrophage
- dendritic

12.5%
25%
50%
100%

ln(CP10K+1): 0 1 2 3

| Abbrev. | Tissue |
|---|---|
| Aort | Aorta |
| Blad | Bladder |
| Blo | Blood |
| Bone | Bone |
| BM | Bone marrow |
| BrC | Brain cortex |
| BrS | Brainstem |
| Col | Colon |
| Dia | Diaphragm |
| Ret | Eye retina |
| Fat | Fat |
| Hrt | Heart |
| Pit | Hypothalamus/Pituitary |
| Kid | Kidney |
| LM | Limb muscle |
| Liv | Liver |
| Lung | Lung |
| MG | Mammary gland |
| Panc | Pancreas |
| Skin | Skin |
| SI | Small intestine |
| Spl | Spleen |
| Test | Testes |
| Thym | Thymus |
| Tong | Tongue |
| Trac | Trachea |
| Ute | Uterus |

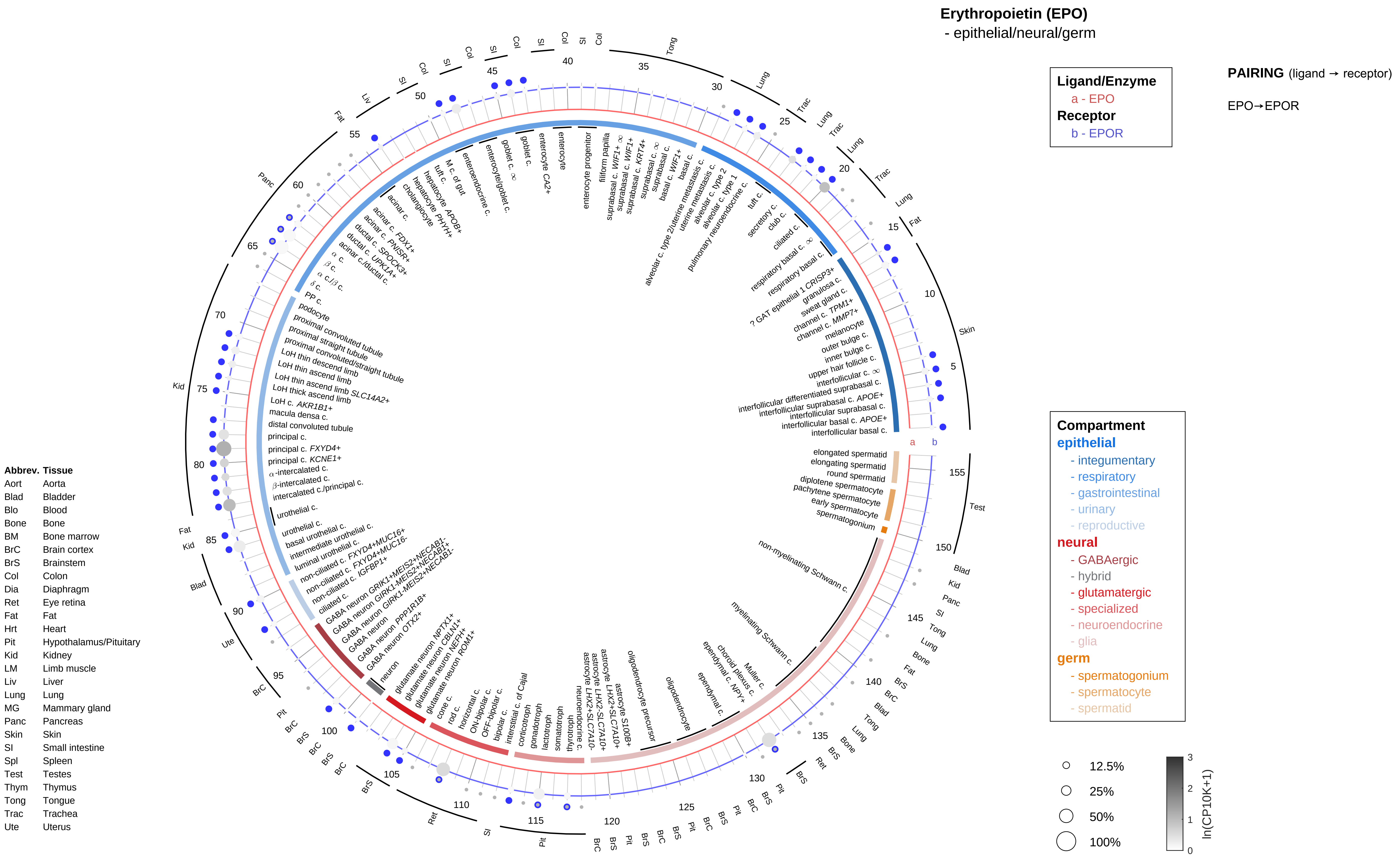

Erythropoietin (EPO)
- epithelial/neural/germ
Ligand/Enzyme
a - EPO
Receptor
b - EPOR
PAIRING (ligand → receptor)
EPO→EPOR
Compartment
epithelial
- integumentary
- respiratory
- gastrointestinal
- urinary
- reproductive
neural
- GABAergic
- hybrid
- glutamatergic
- specialized
- neuroendocrine
- glia
germ
- spermatogonium
- spermatocyte
- spermatid
12.5%
25%
50%
100%
ln(CP10K+1)
Abbrev. Tissue
Aort Aorta
Blad Bladder
Blo Blood
Bone Bone
BM Bone marrow
BrC Brain cortex
BrS Brainstem
Col Colon
Dia Diaphragm
Ret Eye retina
Fat Fat
Hrt Heart
Pit Hypothalamus/Pituitary
Kid Kidney
LM Limb muscle
Liv Liver
Lung Lung
MG Mammary gland
Panc Pancreas
Skin Skin
SI Small intestine
Spl Spleen
Test Testes
Thym Thymus
Tong Tongue
Trac Trachea
Ute Uterus

**Erythropoietin (EPO)**
- endothelial

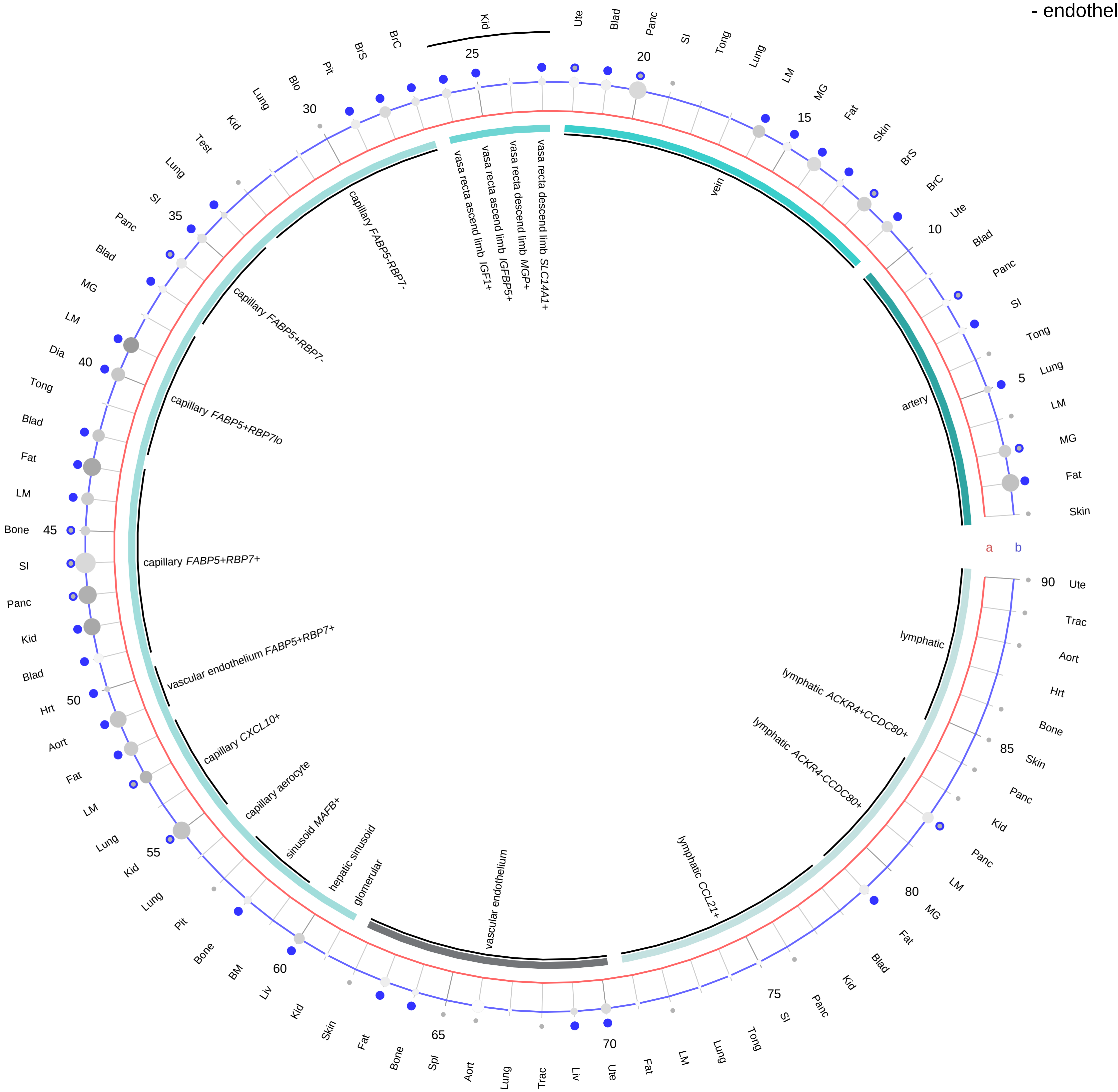

**Ligand/Enzyme**
a - EPO
**Receptor**
b - EPOR

**PAIRING** (ligand → receptor)

EPO→EPOR

**Compartment**
**endothelial**
- artery
- vein
- vasa recta
- capillary
- mix
- lymphatic

12.5%
25%
50%
100%

ln(CP10K+1): 0, 1, 2, 3

| Abbrev. | Tissue |
|---|---|
| Aort | Aorta |
| Blad | Bladder |
| Blo | Blood |
| Bone | Bone |
| BM | Bone marrow |
| BrC | Brain cortex |
| BrS | Brainstem |
| Col | Colon |
| Dia | Diaphragm |
| Ret | Eye retina |
| Fat | Fat |
| Hrt | Heart |
| Pit | Hypothalamus/Pituitary |
| Kid | Kidney |
| LM | Limb muscle |
| Liv | Liver |
| Lung | Lung |
| MG | Mammary gland |
| Panc | Pancreas |
| Skin | Skin |
| SI | Small intestine |
| Spl | Spleen |
| Test | Testes |
| Thym | Thymus |
| Tong | Tongue |
| Trac | Trachea |
| Ute | Uterus |

**Erythropoietin (EPO)**
- stromal

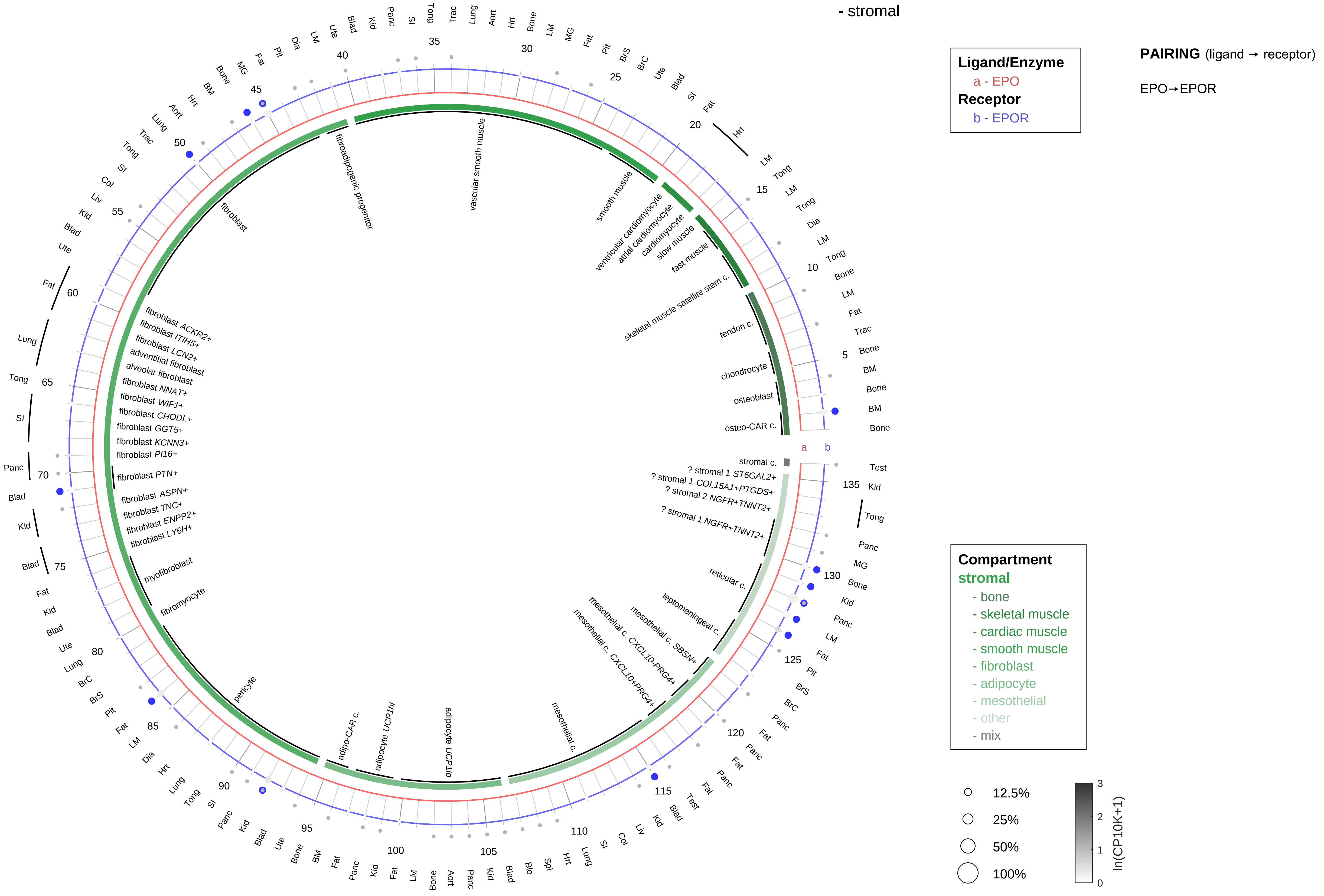

| Abbrev. | Tissue |
|---|---|
| Aort | Aorta |
| Blad | Bladder |
| Blo | Blood |
| Bone | Bone |
| BM | Bone marrow |
| BrC | Brain cortex |
| BrS | Brainstem |
| Col | Colon |
| Dia | Diaphragm |
| Ret | Eye retina |
| Fat | Fat |
| Hrt | Heart |
| Pit | Hypothalamus/Pituitary |
| Kid | Kidney |
| LM | Limb muscle |
| Liv | Liver |
| Lung | Lung |
| MG | Mammary gland |
| Panc | Pancreas |
| Skin | Skin |
| SI | Small intestine |
| Spl | Spleen |
| Test | Testes |
| Thym | Thymus |
| Tong | Tongue |
| Trac | Trachea |
| Ute | Uterus |

**Erythropoietin (EPO)**
- lymphoid

**Ligand/Enzyme**
a - EPO
**Receptor**
b - EPOR

**PAIRING** (ligand → receptor)

EPO→EPOR

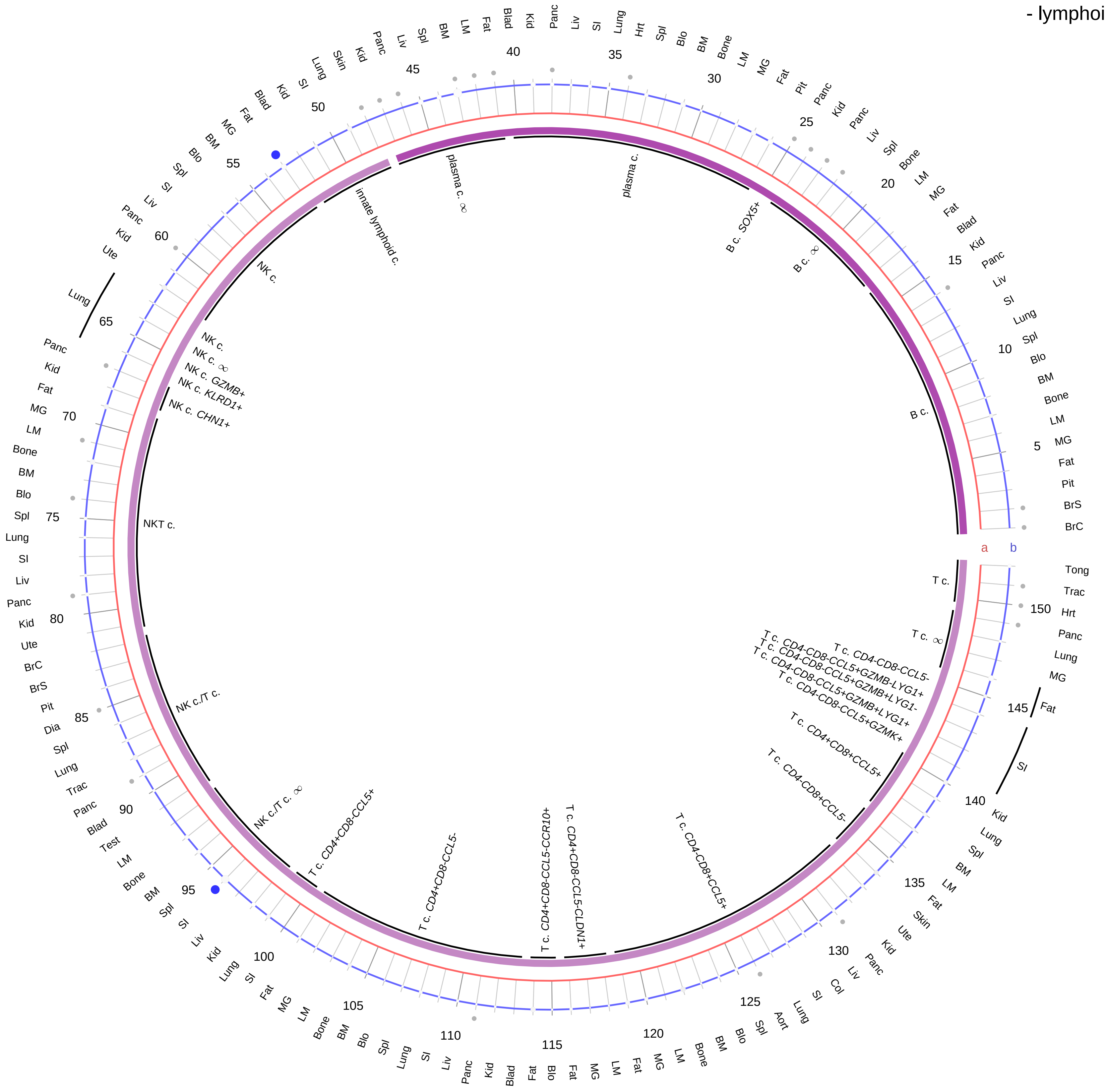

| Abbrev. | Tissue |
|---|---|
| Aort | Aorta |
| Blad | Bladder |
| Blo | Blood |
| Bone | Bone |
| BM | Bone marrow |
| BrC | Brain cortex |
| BrS | Brainstem |
| Col | Colon |
| Dia | Diaphragm |
| Ret | Eye retina |
| Fat | Fat |
| Hrt | Heart |
| Pit | Hypothalamus/Pituitary |
| Kid | Kidney |
| LM | Limb muscle |
| Liv | Liver |
| Lung | Lung |
| MG | Mammary gland |
| Panc | Pancreas |
| Skin | Skin |
| SI | Small intestine |
| Spl | Spleen |
| Test | Testes |
| Thym | Thymus |
| Tong | Tongue |
| Trac | Trachea |
| Ute | Uterus |

**Compartment**
**lymphoid**
- B cell
- NK/T cell

12.5%
25%
50%
100%

ln(CP10K+1): 0, 1, 2, 3

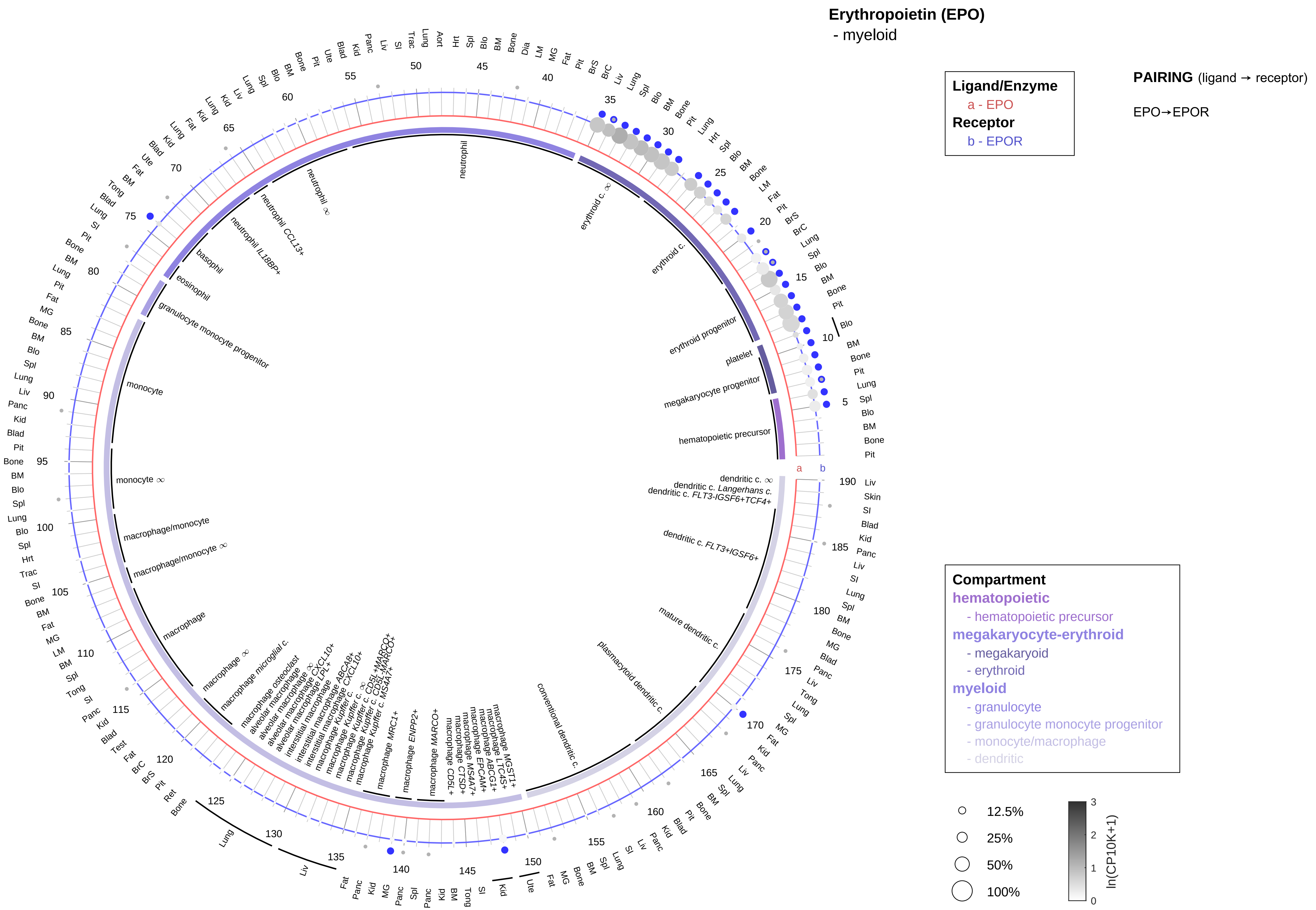

| Abbrev. | Tissue |
|---|---|
| Aort | Aorta |
| Blad | Bladder |
| Blo | Blood |
| Bone | Bone |
| BM | Bone marrow |
| BrC | Brain cortex |
| BrS | Brainstem |
| Col | Colon |
| Dia | Diaphragm |
| Ret | Eye retina |
| Fat | Fat |
| Hrt | Heart |
| Pit | Hypothalamus/Pituitary |
| Kid | Kidney |
| LM | Limb muscle |
| Liv | Liver |
| Lung | Lung |
| MG | Mammary gland |
| Panc | Pancreas |
| Skin | Skin |
| SI | Small intestine |
| Spl | Spleen |
| Test | Testes |
| Thym | Thymus |
| Tong | Tongue |
| Trac | Trachea |
| Ute | Uterus |

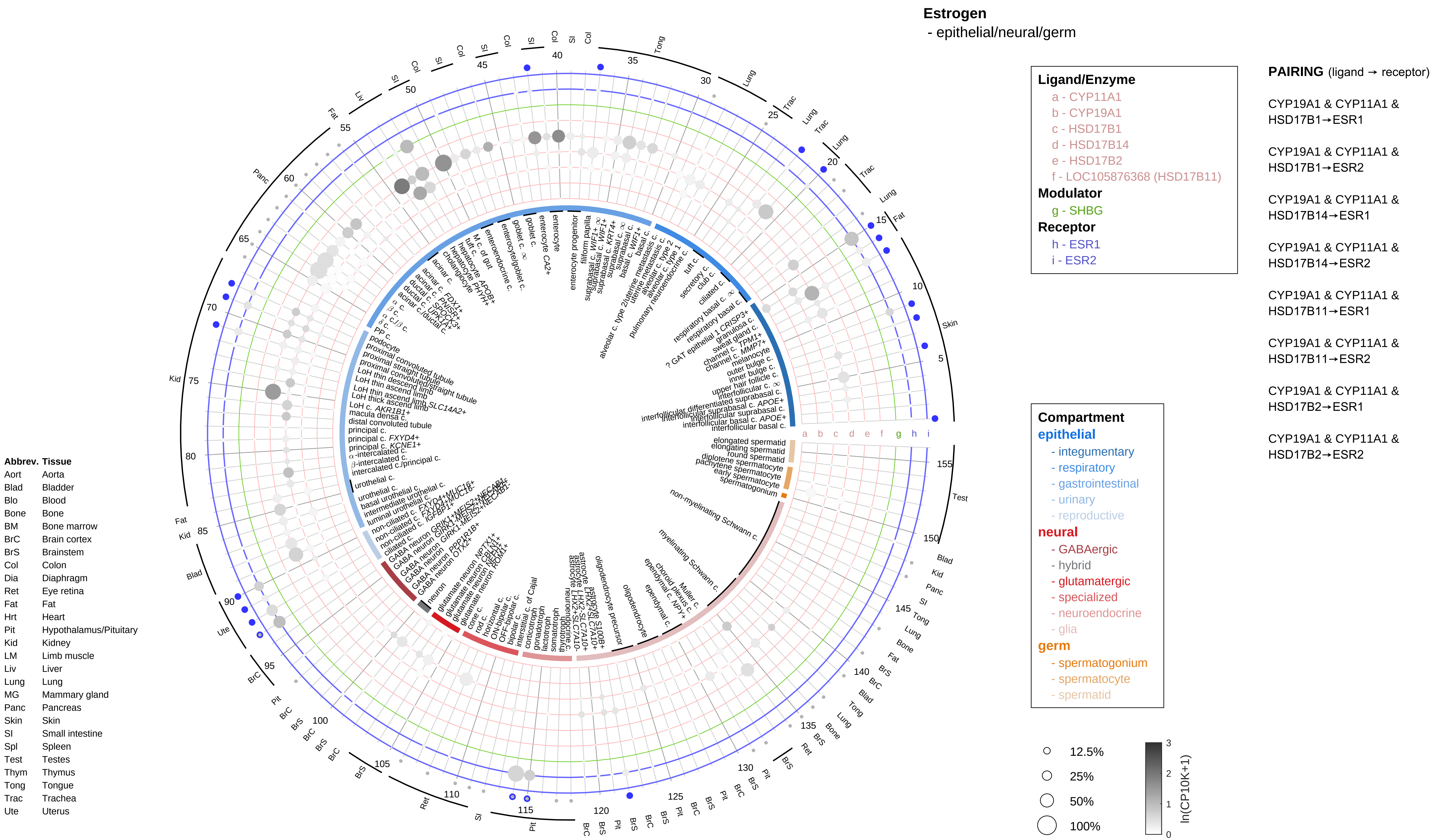
Estrogen
- epithelial/neural/germ
Ligand/Enzyme
a - CYP11A1
b - CYP19A1
c - HSD17B1
d - HSD17B14
e - HSD17B2
f - LOC105876368 (HSD17B11)
Modulator
g - SHBG
Receptor
h - ESR1
i - ESR2
PAIRING (ligand → receptor)
CYP19A1 & CYP11A1 & HSD17B1→ESR1
CYP19A1 & CYP11A1 & HSD17B1→ESR2
CYP19A1 & CYP11A1 & HSD17B14→ESR1
CYP19A1 & CYP11A1 & HSD17B14→ESR2
CYP19A1 & CYP11A1 & HSD17B11→ESR1
CYP19A1 & CYP11A1 & HSD17B11→ESR2
CYP19A1 & CYP11A1 & HSD17B2→ESR1
CYP19A1 & CYP11A1 & HSD17B2→ESR2
Compartment
epithelial
- integumentary
- respiratory
- gastrointestinal
- urinary
- reproductive
neural
- GABAergic
- hybrid
- glutamatergic
- specialized
- neuroendocrine
- glia
germ
- spermatogonium
- spermatocyte
- spermatid
12.5%
25%
50%
100%
ln(CP10K+1)
0
1
2
3
Abbrev. Tissue
Aort Aorta
Blad Bladder
Blo Blood
Bone Bone
BM Bone marrow
BrC Brain cortex
BrS Brainstem
Col Colon
Dia Diaphragm
Ret Eye retina
Fat Fat
Hrt Heart
Pit Hypothalamus/Pituitary
Kid Kidney
LM Limb muscle
Liv Liver
Lung Lung
MG Mammary gland
Panc Pancreas
Skin Skin
SI Small intestine
Spl Spleen
Test Testes
Thym Thymus
Tong Tongue
Trac Trachea
Ute Uterus

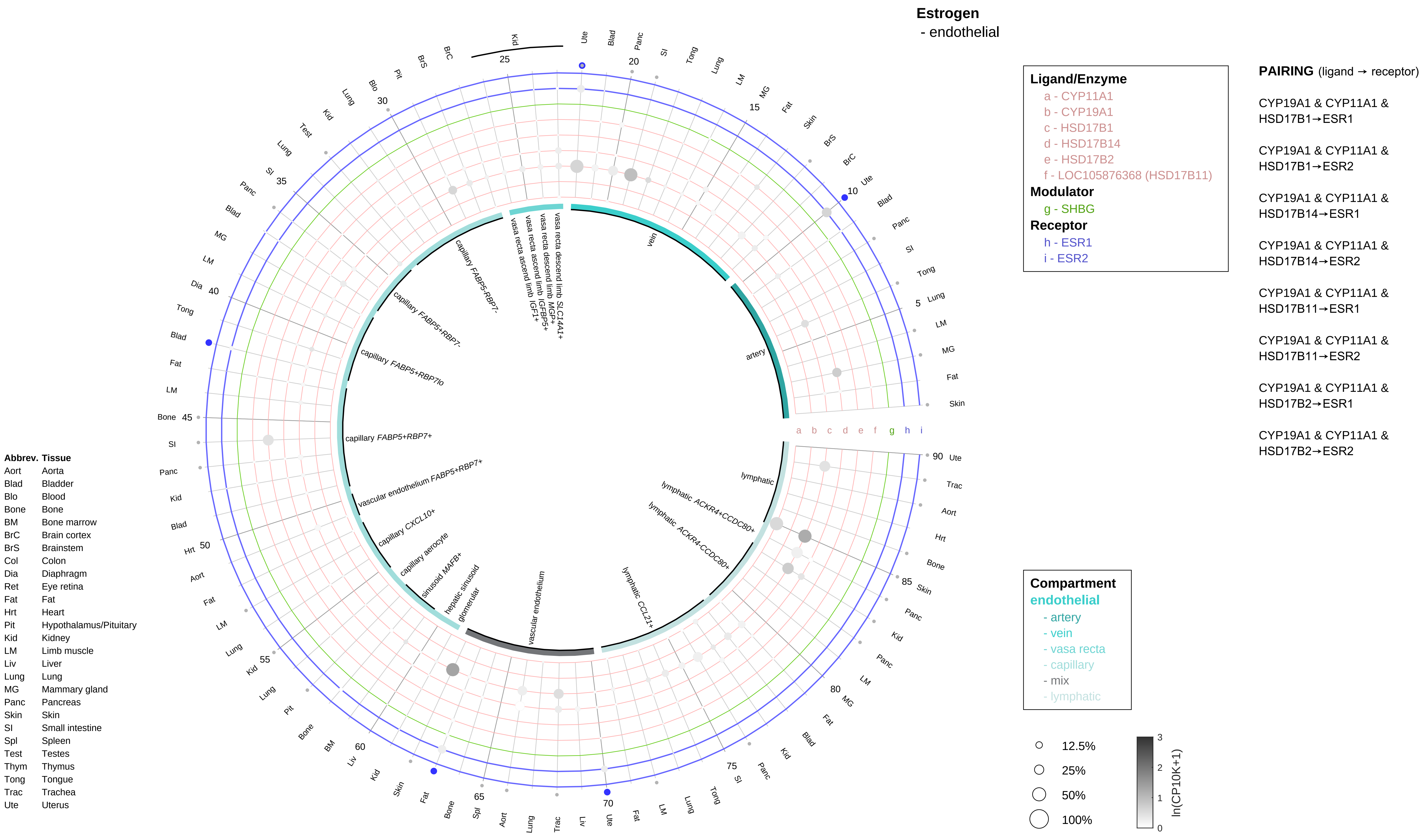

Estrogen
- endothelial
Ligand/Enzyme
a - CYP11A1
b - CYP19A1
c - HSD17B1
d - HSD17B14
e - HSD17B2
f - LOC105876368 (HSD17B11)
Modulator
g - SHBG
Receptor
h - ESR1
i - ESR2
PAIRING (ligand → receptor)
CYP19A1 & CYP11A1 & HSD17B1→ESR1
CYP19A1 & CYP11A1 & HSD17B1→ESR2
CYP19A1 & CYP11A1 & HSD17B14→ESR1
CYP19A1 & CYP11A1 & HSD17B14→ESR2
CYP19A1 & CYP11A1 & HSD17B11→ESR1
CYP19A1 & CYP11A1 & HSD17B11→ESR2
CYP19A1 & CYP11A1 & HSD17B2→ESR1
CYP19A1 & CYP11A1 & HSD17B2→ESR2
vein
artery
vasa recta descend limb SLC14A1+
vasa recta descend limb MGP+
vasa recta ascend limb IGFBP5+
vasa recta ascend limb IGF1+
capillary FABP5-RBP7-
capillary FABP5+RBP7-
capillary FABP5+RBP7lo
capillary FABP5+RBP7+
vascular endothelium FABP5+RBP7+
capillary CXCL10+
capillary aerocyte
sinusoid MAFB+
hepatic sinusoid
glomerular
vascular endothelium
lymphatic CCL21+
lymphatic ACKR4-CCDC80+
lymphatic ACKR4+CCDC80+
lymphatic
a b c d e f g h i
Abbrev. Tissue
Aort Aorta
Blad Bladder
Blo Blood
Bone Bone
BM Bone marrow
BrC Brain cortex
BrS Brainstem
Col Colon
Dia Diaphragm
Ret Eye retina
Fat Fat
Hrt Heart
Pit Hypothalamus/Pituitary
Kid Kidney
LM Limb muscle
Liv Liver
Lung Lung
MG Mammary gland
Panc Pancreas
Skin Skin
SI Small intestine
Spl Spleen
Test Testes
Thym Thymus
Tong Tongue
Trac Trachea
Ute Uterus
Compartment
endothelial
- artery
- vein
- vasa recta
- capillary
- mix
- lymphatic
12.5%
25%
50%
100%
ln(CP10K+1)
0
1
2
3

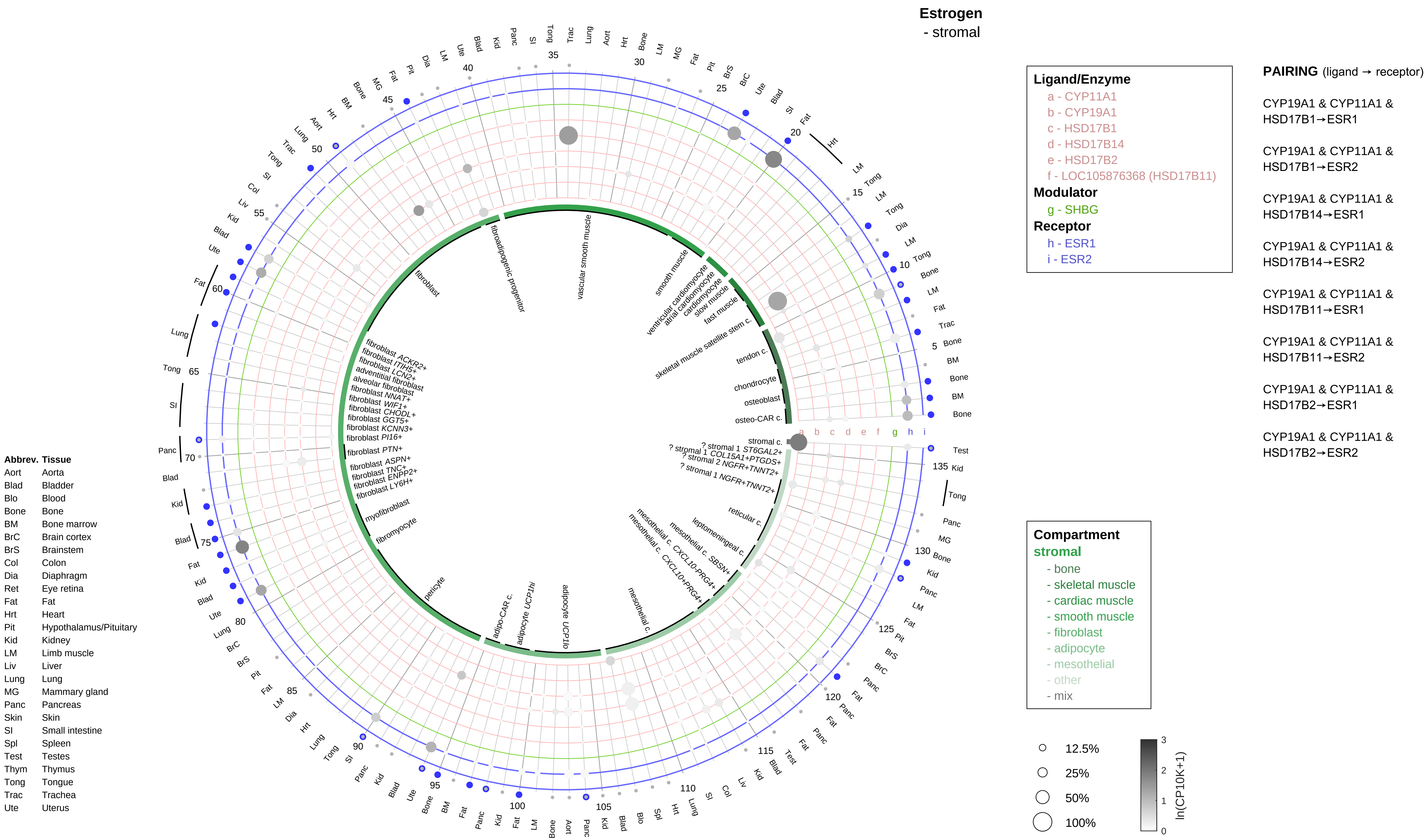

Estrogen
- stromal
Ligand/Enzyme
a - CYP11A1
b - CYP19A1
c - HSD17B1
d - HSD17B14
e - HSD17B2
f - LOC105876368 (HSD17B11)
Modulator
g - SHBG
Receptor
h - ESR1
i - ESR2
PAIRING (ligand → receptor)
CYP19A1 & CYP11A1 & HSD17B1→ESR1
CYP19A1 & CYP11A1 & HSD17B1→ESR2
CYP19A1 & CYP11A1 & HSD17B14→ESR1
CYP19A1 & CYP11A1 & HSD17B14→ESR2
CYP19A1 & CYP11A1 & HSD17B11→ESR1
CYP19A1 & CYP11A1 & HSD17B11→ESR2
CYP19A1 & CYP11A1 & HSD17B2→ESR1
CYP19A1 & CYP11A1 & HSD17B2→ESR2
Compartment
stromal
- bone
- skeletal muscle
- cardiac muscle
- smooth muscle
- fibroblast
- adipocyte
- mesothelial
- other
- mix
12.5%
25%
50%
100%
ln(CP10K+1)
Abbrev. Tissue
Aort Aorta
Blad Bladder
Blo Blood
Bone Bone
BM Bone marrow
BrC Brain cortex
BrS Brainstem
Col Colon
Dia Diaphragm
Ret Eye retina
Fat Fat
Hrt Heart
Pit Hypothalamus/Pituitary
Kid Kidney
LM Limb muscle
Liv Liver
Lung Lung
MG Mammary gland
Panc Pancreas
Skin Skin
SI Small intestine
Spl Spleen
Test Testes
Thym Thymus
Tong Tongue
Trac Trachea
Ute Uterus
vascular smooth muscle
fibroadipogenic progenitor
fibroblast
smooth muscle
ventricular cardiomyocyte
atrial cardiomyocyte
cardiomyocyte
slow muscle
fast muscle
skeletal muscle satellite stem c.
tendon c.
chondrocyte
osteoblast
osteo-CAR c.
stromal c.
? stromal 1 ST6GAL2+
? stromal 1 COL15A1+PTGDS+
? stromal 2 NGFR+TNNT2+
? stromal 1 NGFR+TNNT2+
reticular c.
leptomeningeal c.
mesothelial c. SBSN+
mesothelial c. CXCL10+PRG4+
mesothelial c. CXCL10+PRG4+
mesothelial c.
adipocyte UCP1lo
adipocyte UCP1hi
adipo-CAR c.
pericyte
fibromyocyte
myofibroblast
fibroblast LY6H+
fibroblast ENPP2+
fibroblast TNC+
fibroblast ASPN+
fibroblast PTN+
fibroblast PI16+
fibroblast KCNN3+
fibroblast GGT5+
fibroblast CHODL+
fibroblast WIF1+
fibroblast NNAT+
alveolar fibroblast
adventitial fibroblast
fibroblast LCN2+
fibroblast ITIH5+
fibroblast ACKR2+

**Estrogen**
- lymphoid

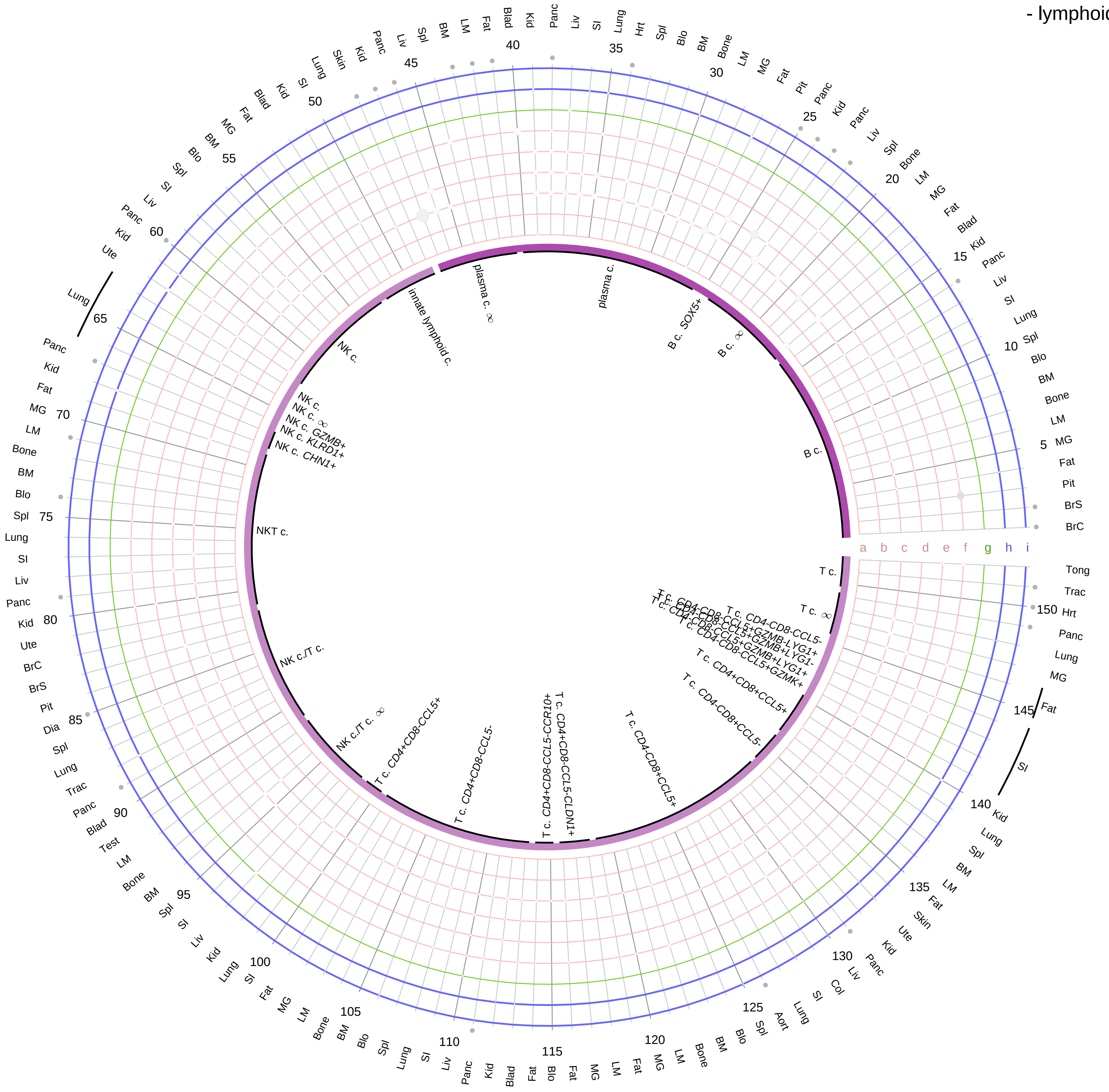

**Ligand/Enzyme**
- a - CYP11A1
- b - CYP19A1
- c - HSD17B1
- d - HSD17B14
- e - HSD17B2
- f - LOC105876368 (HSD17B11)

**Modulator**
- g - SHBG

**Receptor**
- h - ESR1
- i - ESR2

**PAIRING** (ligand → receptor)

CYP19A1 & CYP11A1 & HSD17B1→ESR1

CYP19A1 & CYP11A1 & HSD17B1→ESR2

CYP19A1 & CYP11A1 & HSD17B14→ESR1

CYP19A1 & CYP11A1 & HSD17B14→ESR2

CYP19A1 & CYP11A1 & HSD17B11→ESR1

CYP19A1 & CYP11A1 & HSD17B11→ESR2

CYP19A1 & CYP11A1 & HSD17B2→ESR1

CYP19A1 & CYP11A1 & HSD17B2→ESR2

**Compartment**
**lymphoid**
- B cell
- NK/T cell

12.5%
25%
50%
100%

ln(CP10K+1) 0 1 2 3

| Abbrev. | Tissue |
|---|---|
| Aort | Aorta |
| Blad | Bladder |
| Blo | Blood |
| Bone | Bone |
| BM | Bone marrow |
| BrC | Brain cortex |
| BrS | Brainstem |
| Col | Colon |
| Dia | Diaphragm |
| Ret | Eye retina |
| Fat | Fat |
| Hrt | Heart |
| Pit | Hypothalamus/Pituitary |
| Kid | Kidney |
| LM | Limb muscle |
| Liv | Liver |
| Lung | Lung |
| MG | Mammary gland |
| Panc | Pancreas |
| Skin | Skin |
| SI | Small intestine |
| Spl | Spleen |
| Test | Testes |
| Thym | Thymus |
| Tong | Tongue |
| Trac | Trachea |
| Ute | Uterus |

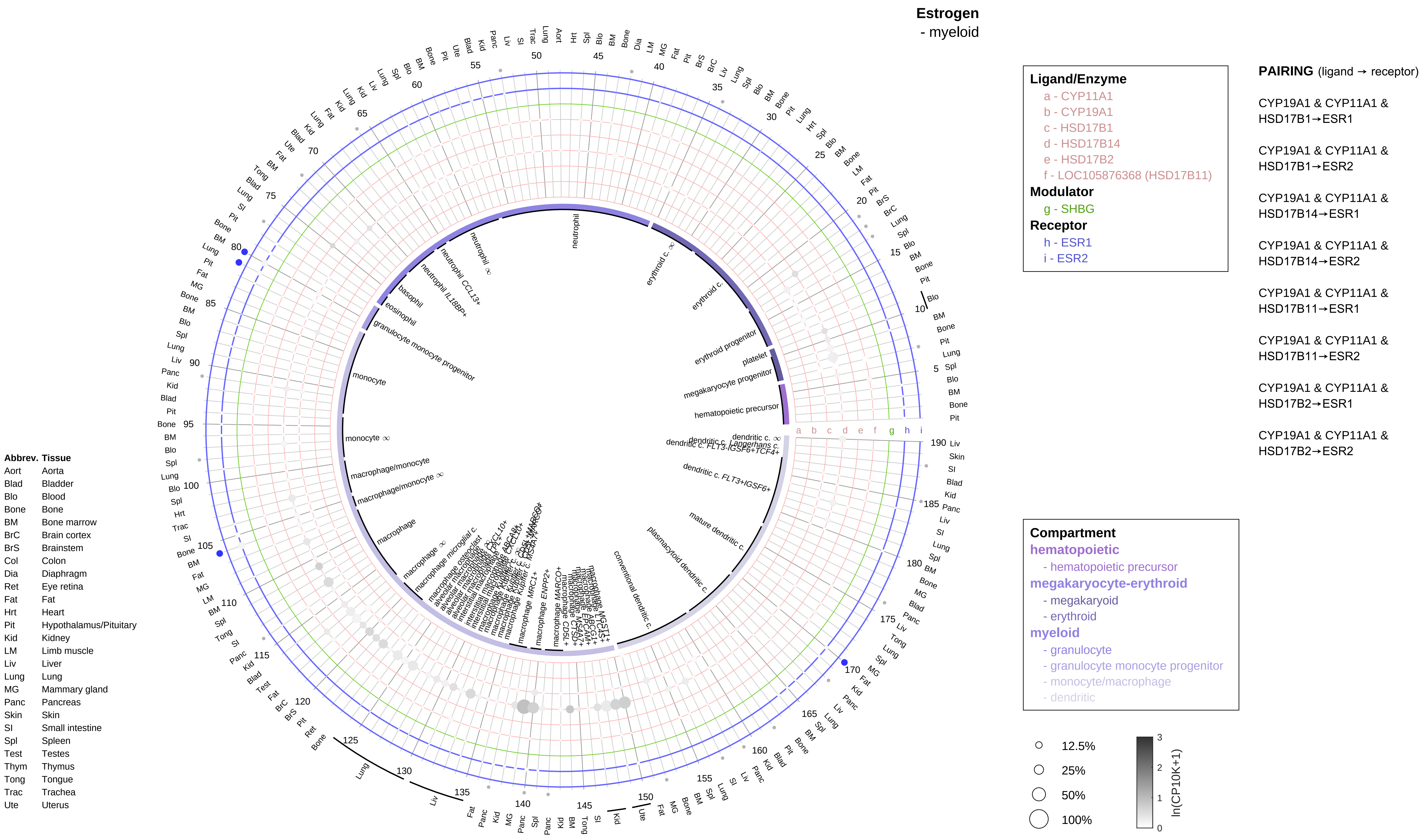
Estrogen
- myeloid
Ligand/Enzyme
a - CYP11A1
b - CYP19A1
c - HSD17B1
d - HSD17B14
e - HSD17B2
f - LOC105876368 (HSD17B11)
Modulator
g - SHBG
Receptor
h - ESR1
i - ESR2
PAIRING (ligand → receptor)
CYP19A1 & CYP11A1 & HSD17B1→ESR1
CYP19A1 & CYP11A1 & HSD17B1→ESR2
CYP19A1 & CYP11A1 & HSD17B14→ESR1
CYP19A1 & CYP11A1 & HSD17B14→ESR2
CYP19A1 & CYP11A1 & HSD17B11→ESR1
CYP19A1 & CYP11A1 & HSD17B11→ESR2
CYP19A1 & CYP11A1 & HSD17B2→ESR1
CYP19A1 & CYP11A1 & HSD17B2→ESR2
Compartment
hematopoietic
- hematopoietic precursor
megakaryocyte-erythroid
- megakaryoid
- erythroid
myeloid
- granulocyte
- granulocyte monocyte progenitor
- monocyte/macrophage
- dendritic
12.5%
25%
50%
100%
ln(CP10K+1)
Abbrev. Tissue
Aort Aorta
Blad Bladder
Blo Blood
Bone Bone
BM Bone marrow
BrC Brain cortex
BrS Brainstem
Col Colon
Dia Diaphragm
Ret Eye retina
Fat Fat
Hrt Heart
Pit Hypothalamus/Pituitary
Kid Kidney
LM Limb muscle
Liv Liver
Lung Lung
MG Mammary gland
Panc Pancreas
Skin Skin
SI Small intestine
Spl Spleen
Test Testes
Thym Thymus
Tong Tongue
Trac Trachea
Ute Uterus
neutrophil
erythroid c.
erythroid progenitor
platelet
megakaryocyte progenitor
hematopoietic precursor
dendritic c. FLT3+IGSF6+
mature dendritic c.
plasmacytoid dendritic c.
conventional dendritic c.
monocyte
macrophage/monocyte
macrophage
basophil
eosinophil
granulocyte monocyte progenitor

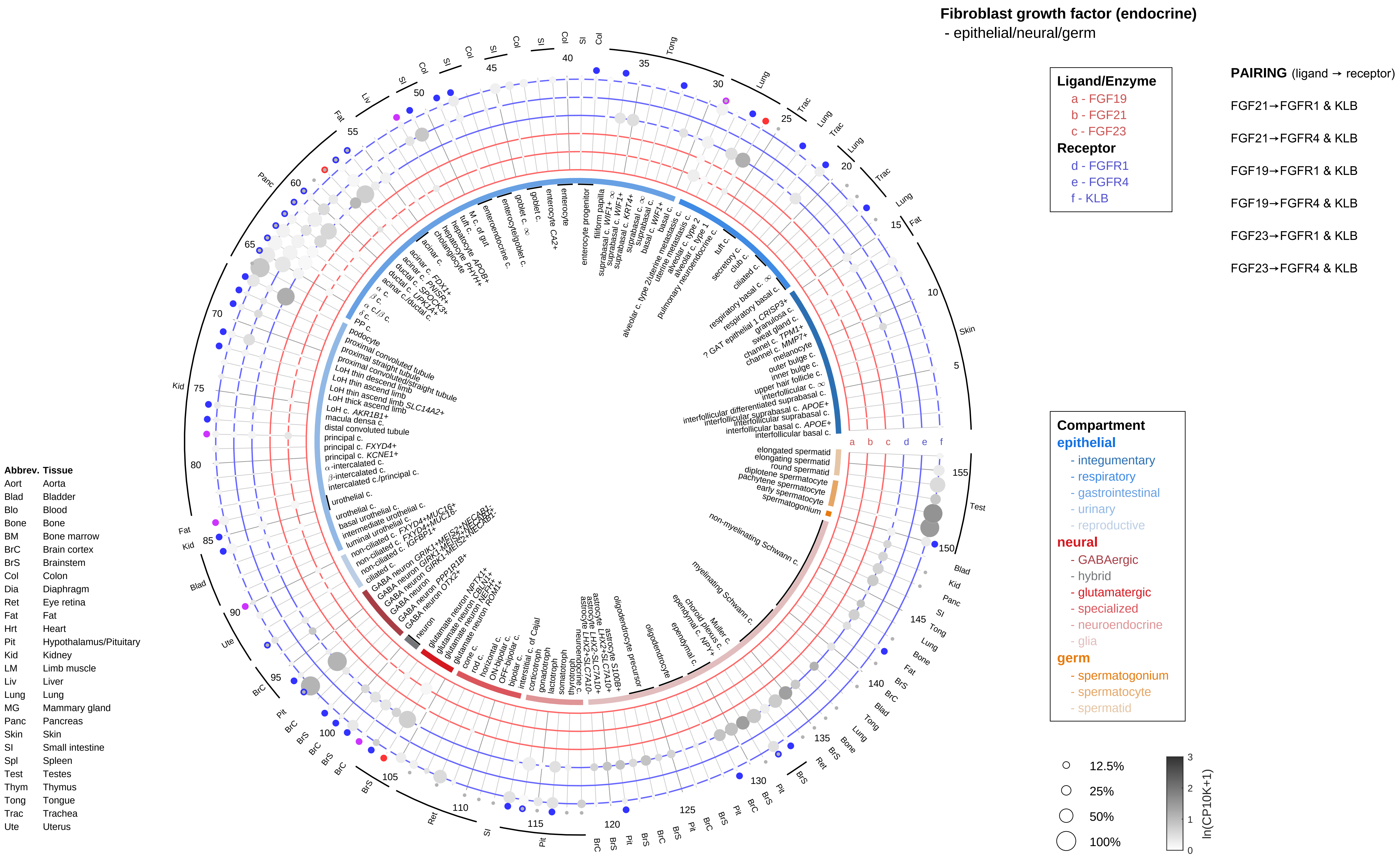

Fibroblast growth factor (endocrine)
- epithelial/neural/germ
Ligand/Enzyme
a - FGF19
b - FGF21
c - FGF23
Receptor
d - FGFR1
e - FGFR4
f - KLB
PAIRING (ligand → receptor)
FGF21→FGFR1 & KLB
FGF21→FGFR4 & KLB
FGF19→FGFR1 & KLB
FGF19→FGFR4 & KLB
FGF23→FGFR1 & KLB
FGF23→FGFR4 & KLB
Compartment
epithelial
- integumentary
- respiratory
- gastrointestinal
- urinary
- reproductive
neural
- GABAergic
- hybrid
- glutamatergic
- specialized
- neuroendocrine
- glia
germ
- spermatogonium
- spermatocyte
- spermatid
12.5%
25%
50%
100%
ln(CP10K+1)
Abbrev. Tissue
Aort Aorta
Blad Bladder
Blo Blood
Bone Bone
BM Bone marrow
BrC Brain cortex
BrS Brainstem
Col Colon
Dia Diaphragm
Ret Eye retina
Fat Fat
Hrt Heart
Pit Hypothalamus/Pituitary
Kid Kidney
LM Limb muscle
Liv Liver
Lung Lung
MG Mammary gland
Panc Pancreas
Skin Skin
SI Small intestine
Spl Spleen
Test Testes
Thym Thymus
Tong Tongue
Trac Trachea
Ute Uterus

**Fibroblast growth factor (endocrine)**
- endothelial

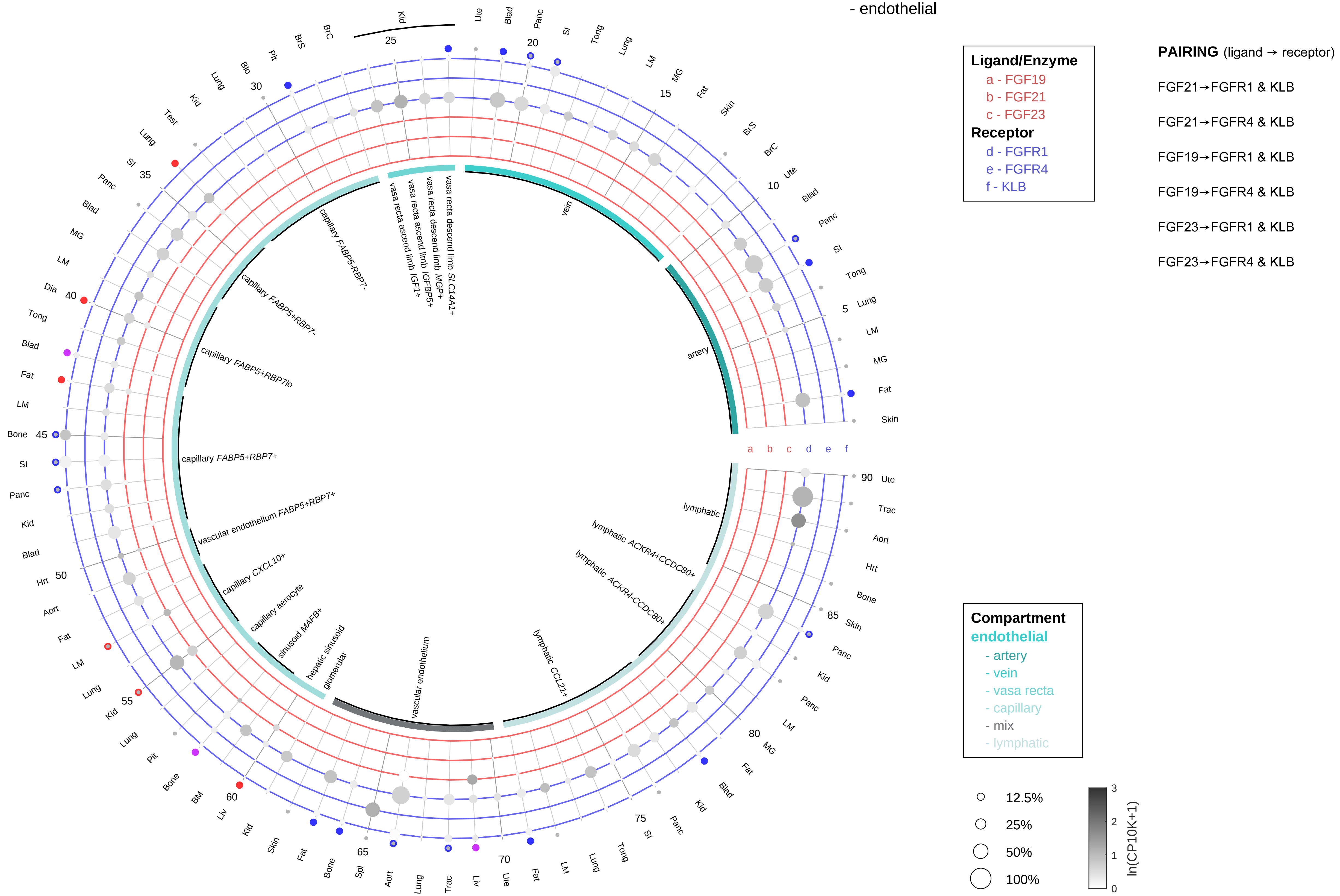

| Abbrev. | Tissue |
|---|---|
| Aort | Aorta |
| Blad | Bladder |
| Blo | Blood |
| Bone | Bone |
| BM | Bone marrow |
| BrC | Brain cortex |
| BrS | Brainstem |
| Col | Colon |
| Dia | Diaphragm |
| Ret | Eye retina |
| Fat | Fat |
| Hrt | Heart |
| Pit | Hypothalamus/Pituitary |
| Kid | Kidney |
| LM | Limb muscle |
| Liv | Liver |
| Lung | Lung |
| MG | Mammary gland |
| Panc | Pancreas |
| Skin | Skin |
| SI | Small intestine |
| Spl | Spleen |
| Test | Testes |
| Thym | Thymus |
| Tong | Tongue |
| Trac | Trachea |
| Ute | Uterus |

**Fibroblast growth factor (endocrine)**
- stromal

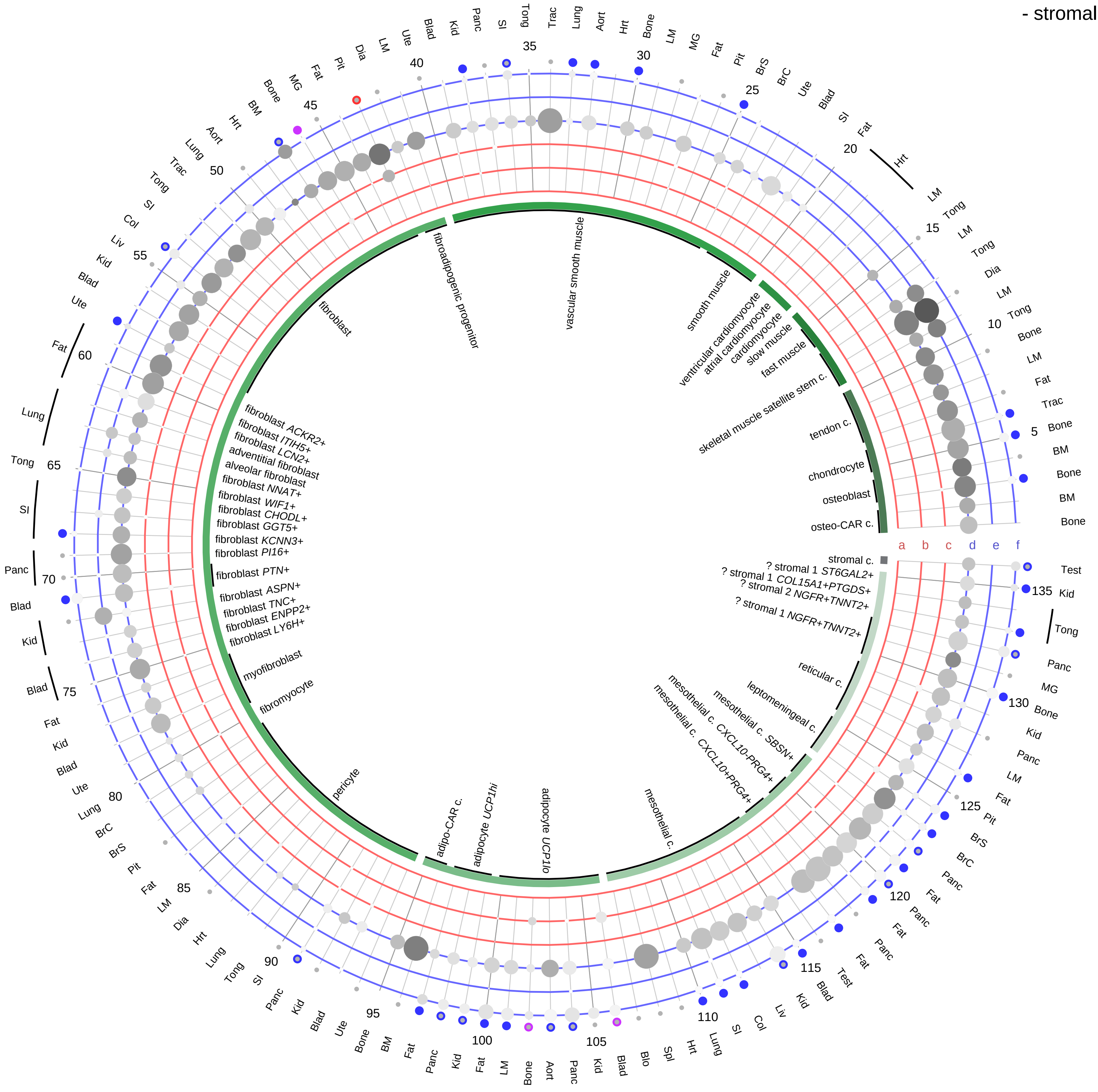

**Ligand/Enzyme**
- a - FGF19
- b - FGF21
- c - FGF23

**Receptor**
- d - FGFR1
- e - FGFR4
- f - KLB

**PAIRING** (ligand → receptor)

FGF21→FGFR1 & KLB

FGF21→FGFR4 & KLB

FGF19→FGFR1 & KLB

FGF19→FGFR4 & KLB

FGF23→FGFR1 & KLB

FGF23→FGFR4 & KLB

**Compartment**
**stromal**
- bone
- skeletal muscle
- cardiac muscle
- smooth muscle
- fibroblast
- adipocyte
- mesothelial
- other
- mix

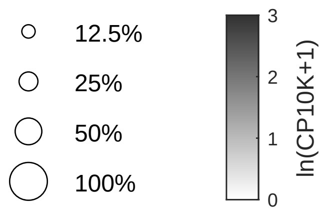

| Abbrev. | Tissue |
|---|---|
| Aort | Aorta |
| Blad | Bladder |
| Blo | Blood |
| Bone | Bone |
| BM | Bone marrow |
| BrC | Brain cortex |
| BrS | Brainstem |
| Col | Colon |
| Dia | Diaphragm |
| Ret | Eye retina |
| Fat | Fat |
| Hrt | Heart |
| Pit | Hypothalamus/Pituitary |
| Kid | Kidney |
| LM | Limb muscle |
| Liv | Liver |
| Lung | Lung |
| MG | Mammary gland |
| Panc | Pancreas |
| Skin | Skin |
| SI | Small intestine |
| Spl | Spleen |
| Test | Testes |
| Thym | Thymus |
| Tong | Tongue |
| Trac | Trachea |
| Ute | Uterus |

**Fibroblast growth factor (endocrine)**
- lymphoid

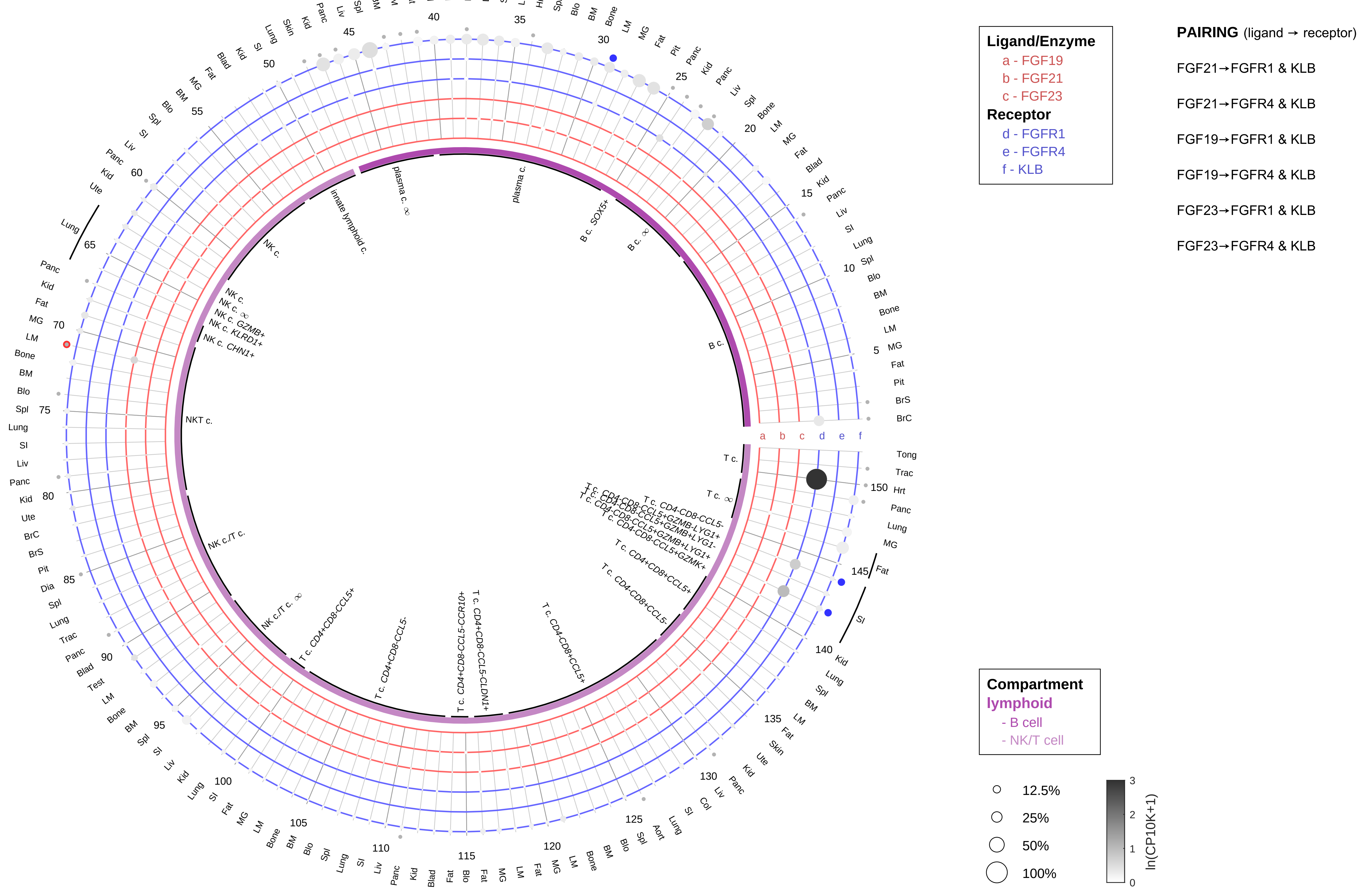

| Abbrev. | Tissue |
|---|---|
| Aort | Aorta |
| Blad | Bladder |
| Blo | Blood |
| Bone | Bone |
| BM | Bone marrow |
| BrC | Brain cortex |
| BrS | Brainstem |
| Col | Colon |
| Dia | Diaphragm |
| Ret | Eye retina |
| Fat | Fat |
| Hrt | Heart |
| Pit | Hypothalamus/Pituitary |
| Kid | Kidney |
| LM | Limb muscle |
| Liv | Liver |
| Lung | Lung |
| MG | Mammary gland |
| Panc | Pancreas |
| Skin | Skin |
| SI | Small intestine |
| Spl | Spleen |
| Test | Testes |
| Thym | Thymus |
| Tong | Tongue |
| Trac | Trachea |
| Ute | Uterus |

**Fibroblast growth factor (endocrine)**
- myeloid

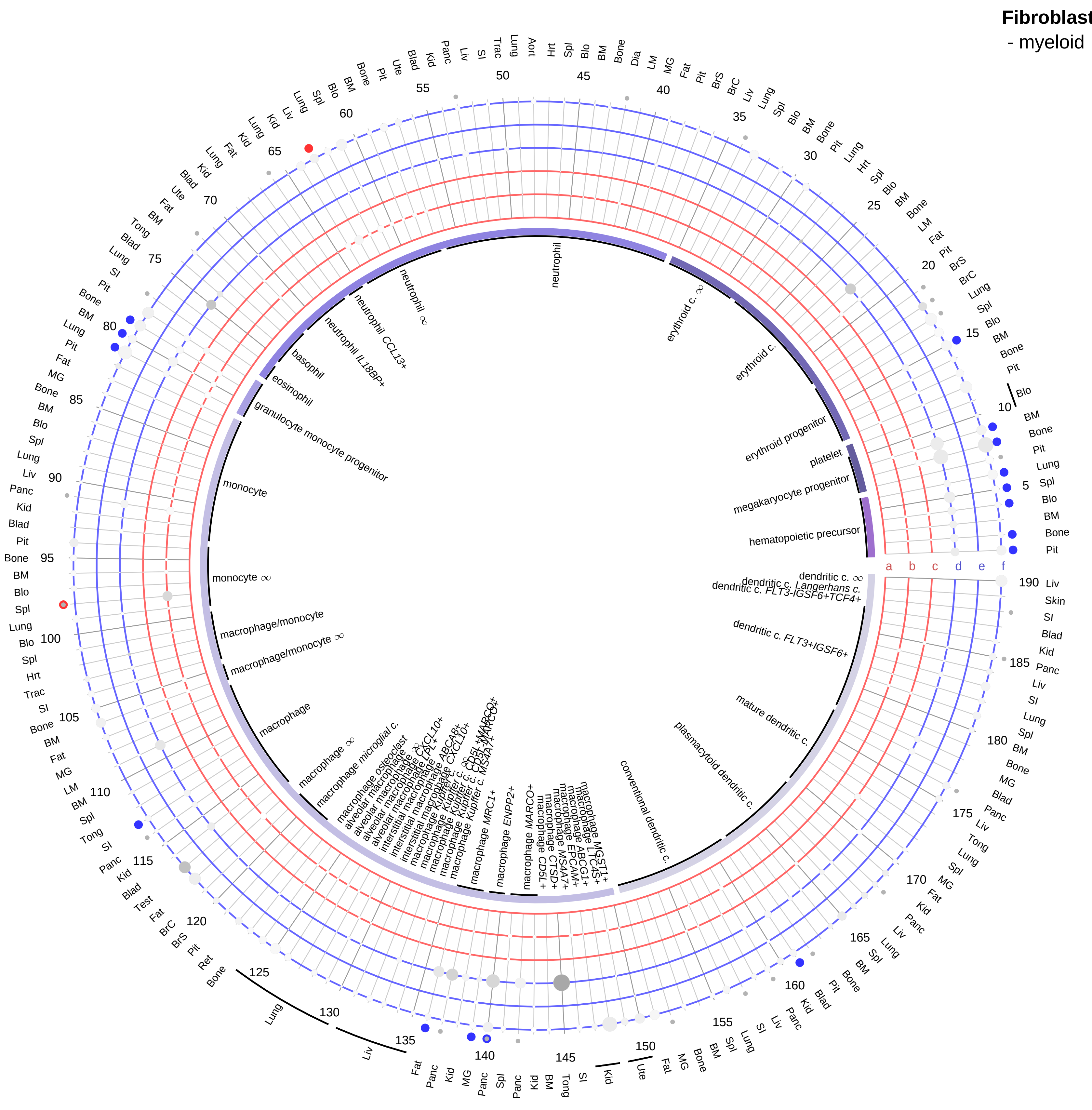

| Ligand/Enzyme |
|---|
| a - FGF19 |
| b - FGF21 |
| c - FGF23 |
| **Receptor** |
| d - FGFR1 |
| e - FGFR4 |
| f - KLB |

**PAIRING** (ligand → receptor)

FGF21→FGFR1 & KLB

FGF21→FGFR4 & KLB

FGF19→FGFR1 & KLB

FGF19→FGFR4 & KLB

FGF23→FGFR1 & KLB

FGF23→FGFR4 & KLB

**Compartment**
**hematopoietic**
- hematopoietic precursor
**megakaryocyte-erythroid**
- megakaryoid
- erythroid
**myeloid**
- granulocyte
- granulocyte monocyte progenitor
- monocyte/macrophage
- dendritic

12.5%
25%
50%
100%

ln(CP10K+1): 0, 1, 2, 3

| Abbrev. | Tissue |
|---|---|
| Aort | Aorta |
| Blad | Bladder |
| Blo | Blood |
| Bone | Bone |
| BM | Bone marrow |
| BrC | Brain cortex |
| BrS | Brainstem |
| Col | Colon |
| Dia | Diaphragm |
| Ret | Eye retina |
| Fat | Fat |
| Hrt | Heart |
| Pit | Hypothalamus/Pituitary |
| Kid | Kidney |
| LM | Limb muscle |
| Liv | Liver |
| Lung | Lung |
| MG | Mammary gland |
| Panc | Pancreas |
| Skin | Skin |
| SI | Small intestine |
| Spl | Spleen |
| Test | Testes |
| Thym | Thymus |
| Tong | Tongue |
| Trac | Trachea |
| Ute | Uterus |

**Follicle stimulating hormone (FSH)**
- epithelial/neural/germ

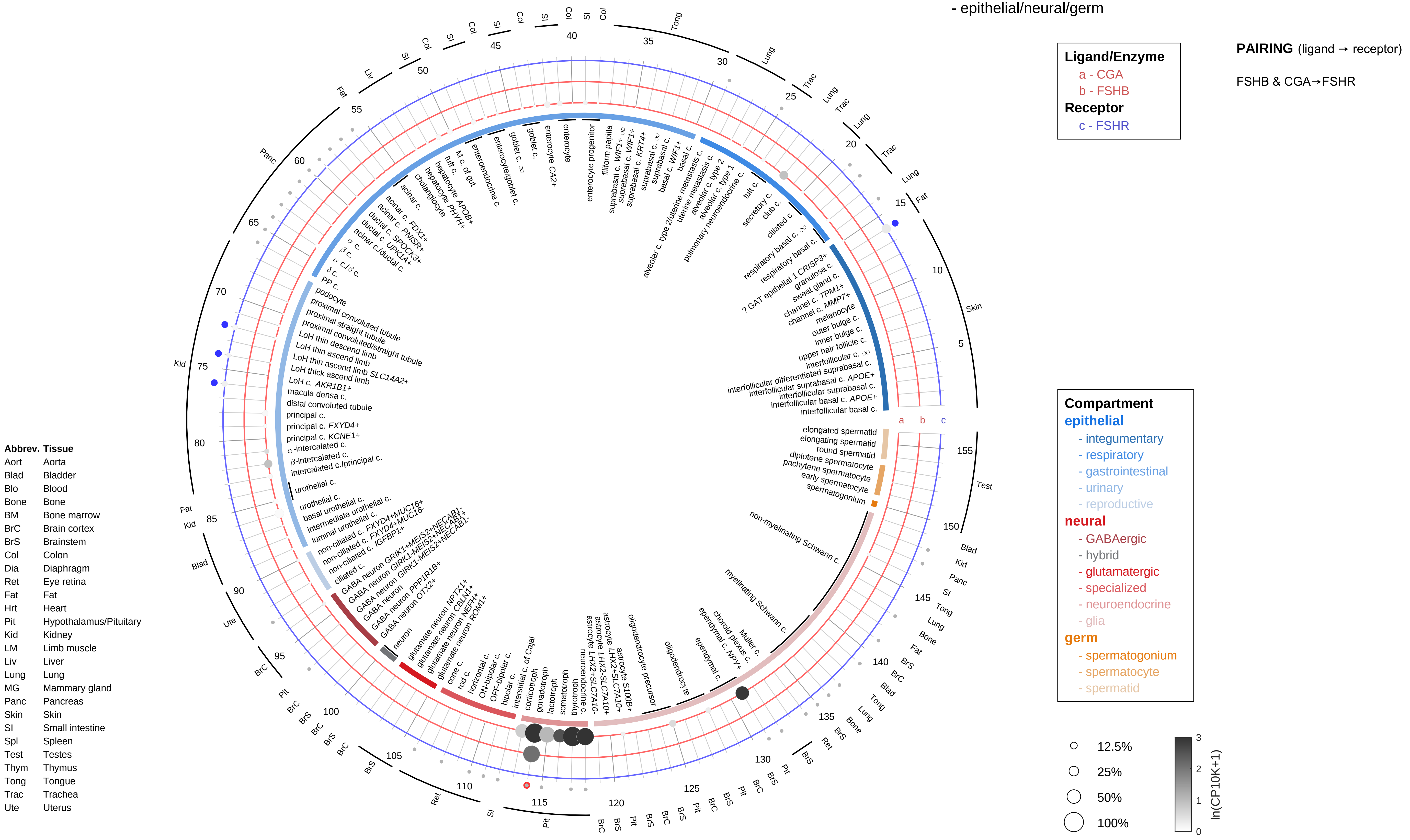

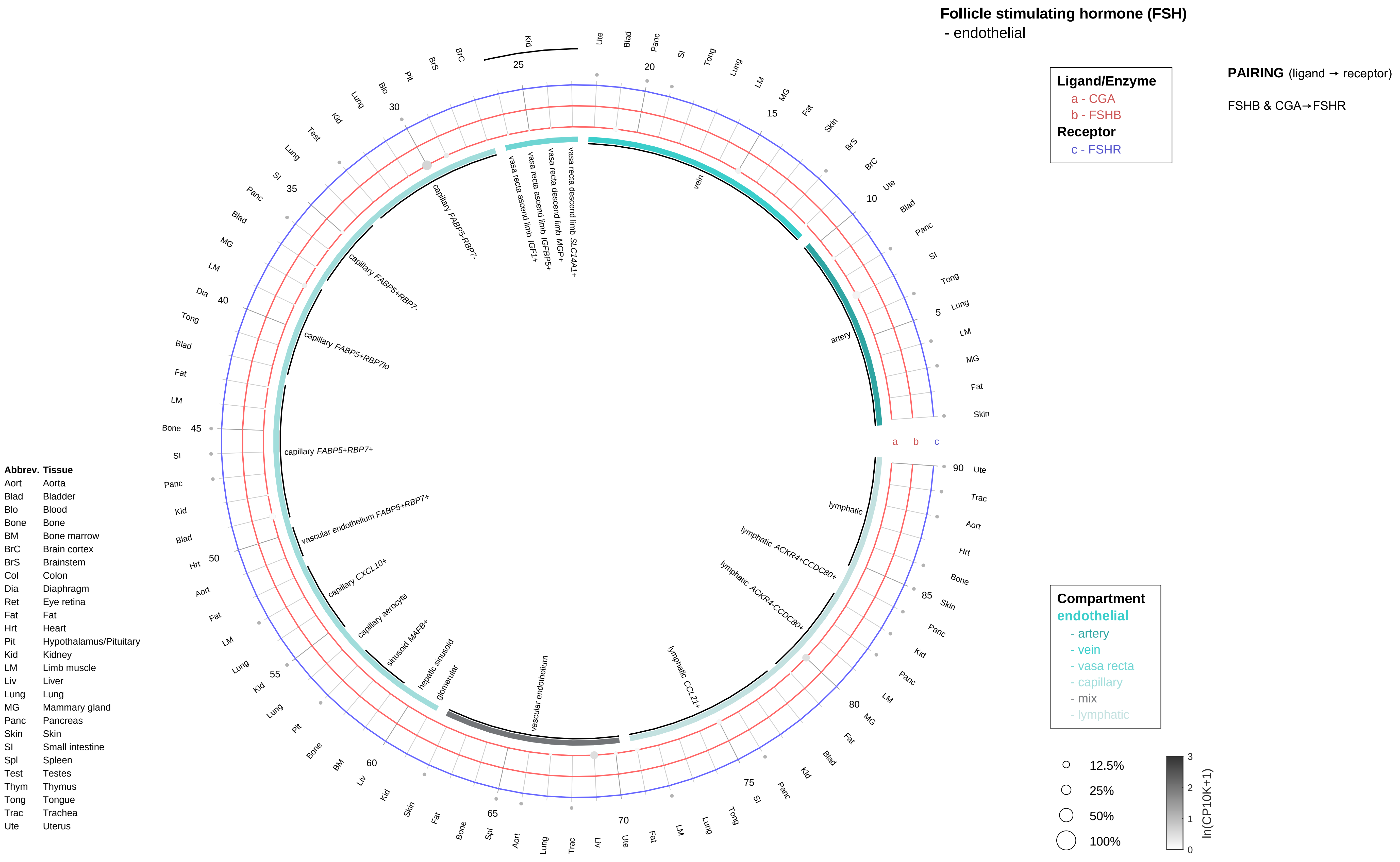

Follicle stimulating hormone (FSH)
- endothelial
Ligand/Enzyme
a - CGA
b - FSHB
Receptor
c - FSHR
PAIRING (ligand → receptor)
FSHB & CGA→FSHR
Compartment
endothelial
- artery
- vein
- vasa recta
- capillary
- mix
- lymphatic
12.5%
25%
50%
100%
ln(CP10K+1)
Abbrev. Tissue
Aort Aorta
Blad Bladder
Blo Blood
Bone Bone
BM Bone marrow
BrC Brain cortex
BrS Brainstem
Col Colon
Dia Diaphragm
Ret Eye retina
Fat Fat
Hrt Heart
Pit Hypothalamus/Pituitary
Kid Kidney
LM Limb muscle
Liv Liver
Lung Lung
MG Mammary gland
Panc Pancreas
Skin Skin
SI Small intestine
Spl Spleen
Test Testes
Thym Thymus
Tong Tongue
Trac Trachea
Ute Uterus
artery
vein
vasa recta descend limb SLC14A1+
vasa recta descend limb MGP+
vasa recta ascend limb IGFBP5+
vasa recta ascend limb IGF1+
capillary FABP5-RBP7-
capillary FABP5+RBP7-
capillary FABP5+RBP7lo
capillary FABP5+RBP7+
vascular endothelium FABP5+RBP7+
capillary CXCL10+
capillary aerocyte
sinusoid MAFB+
hepatic sinusoid
glomerular
vascular endothelium
lymphatic CCL21+
lymphatic ACKR4-CCDC80+
lymphatic ACKR4+CCDC80+
lymphatic

**Follicle stimulating hormone (FSH)**
- stromal

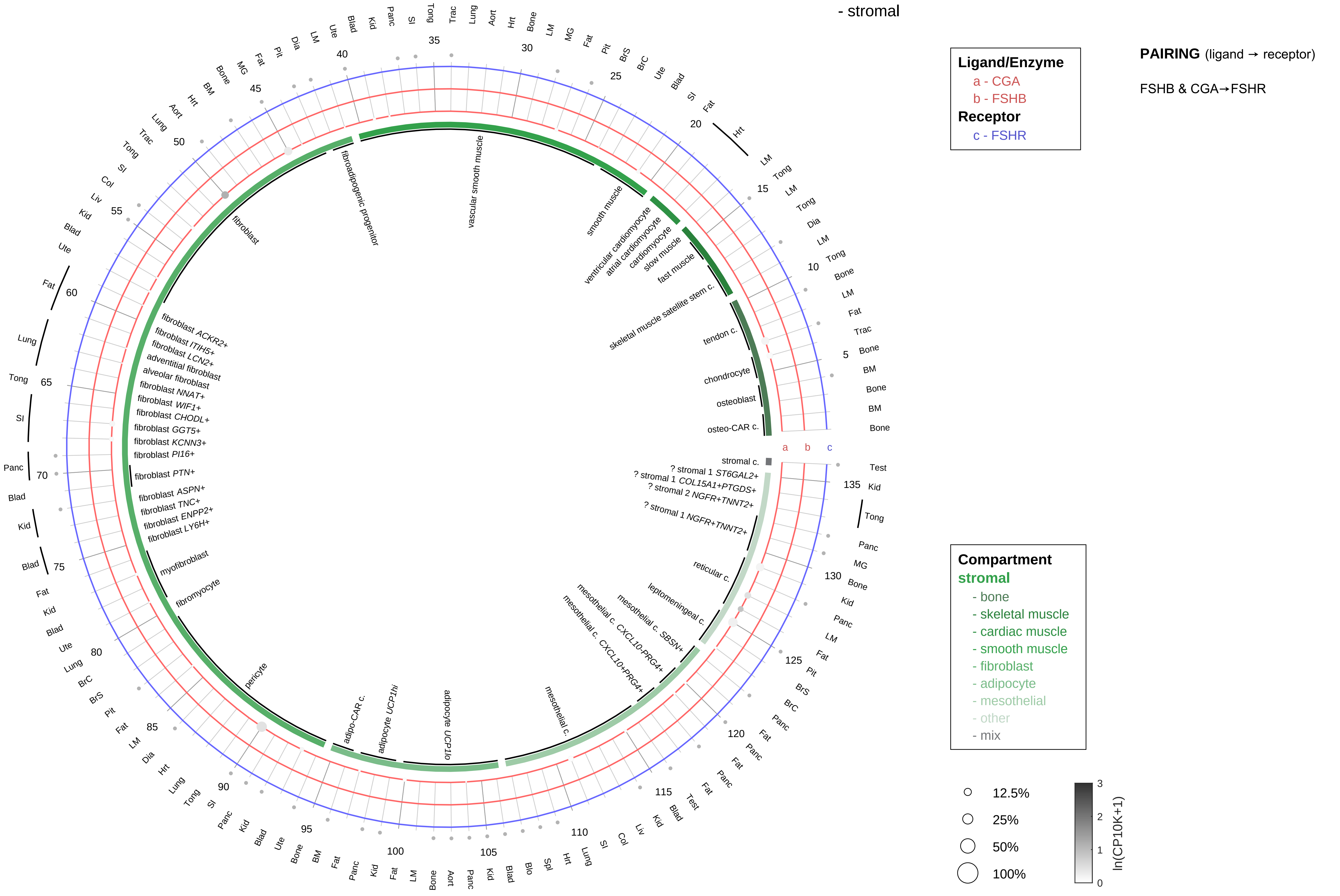

| Abbrev. | Tissue |
|---|---|
| Aort | Aorta |
| Blad | Bladder |
| Blo | Blood |
| Bone | Bone |
| BM | Bone marrow |
| BrC | Brain cortex |
| BrS | Brainstem |
| Col | Colon |
| Dia | Diaphragm |
| Ret | Eye retina |
| Fat | Fat |
| Hrt | Heart |
| Pit | Hypothalamus/Pituitary |
| Kid | Kidney |
| LM | Limb muscle |
| Liv | Liver |
| Lung | Lung |
| MG | Mammary gland |
| Panc | Pancreas |
| Skin | Skin |
| SI | Small intestine |
| Spl | Spleen |
| Test | Testes |
| Thym | Thymus |
| Tong | Tongue |
| Trac | Trachea |
| Ute | Uterus |

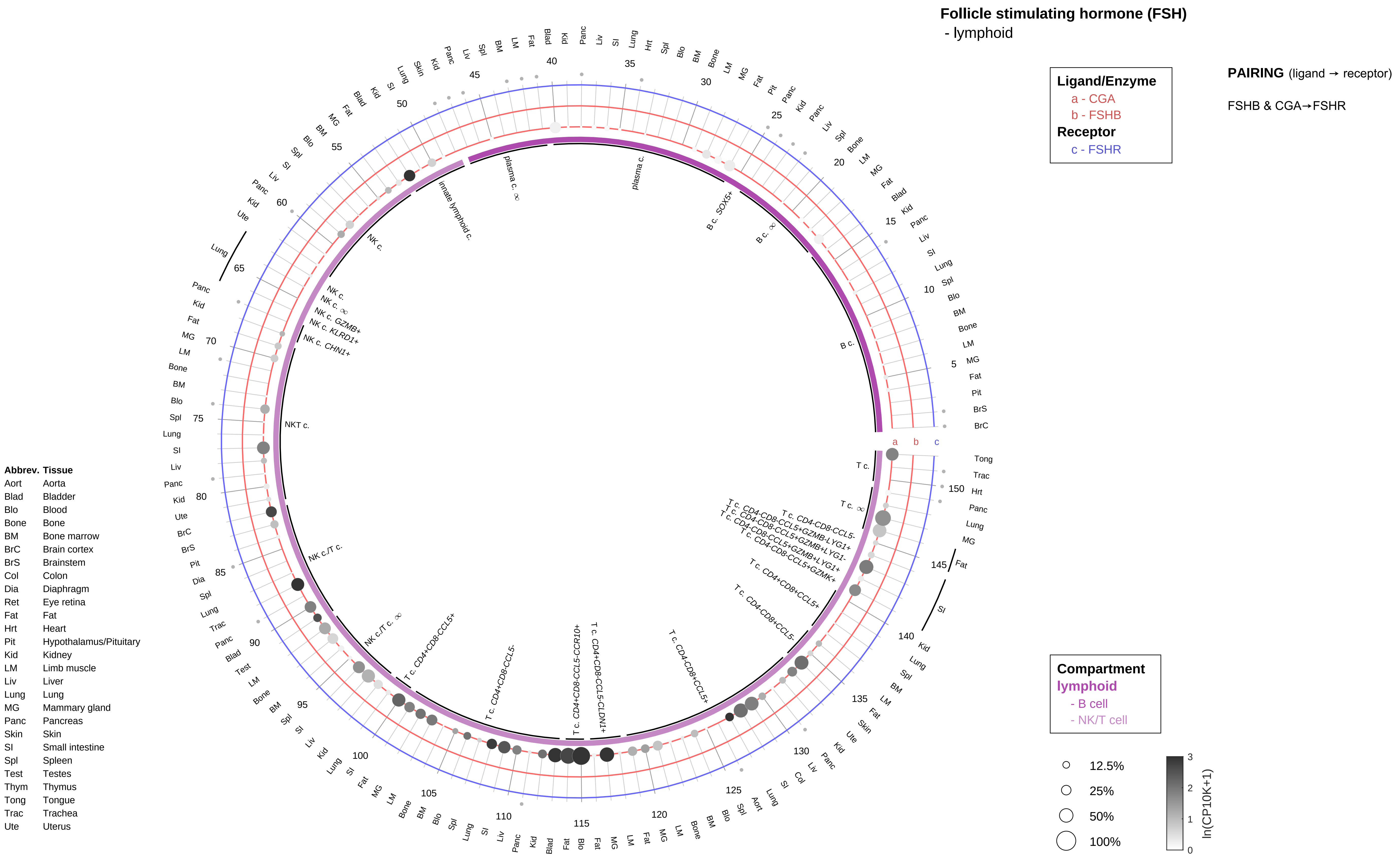
Follicle stimulating hormone (FSH)
- lymphoid
Ligand/Enzyme
a - CGA
b - FSHB
Receptor
c - FSHR
PAIRING (ligand → receptor)
FSHB & CGA→FSHR
Compartment
lymphoid
- B cell
- NK/T cell
12.5%
25%
50%
100%
ln(CP10K+1)
Abbrev. Tissue
Aort Aorta
Blad Bladder
Blo Blood
Bone Bone
BM Bone marrow
BrC Brain cortex
BrS Brainstem
Col Colon
Dia Diaphragm
Ret Eye retina
Fat Fat
Hrt Heart
Pit Hypothalamus/Pituitary
Kid Kidney
LM Limb muscle
Liv Liver
Lung Lung
MG Mammary gland
Panc Pancreas
Skin Skin
SI Small intestine
Spl Spleen
Test Testes
Thym Thymus
Tong Tongue
Trac Trachea
Ute Uterus

#### Follicle stimulating hormone (FSH)
- myeloid

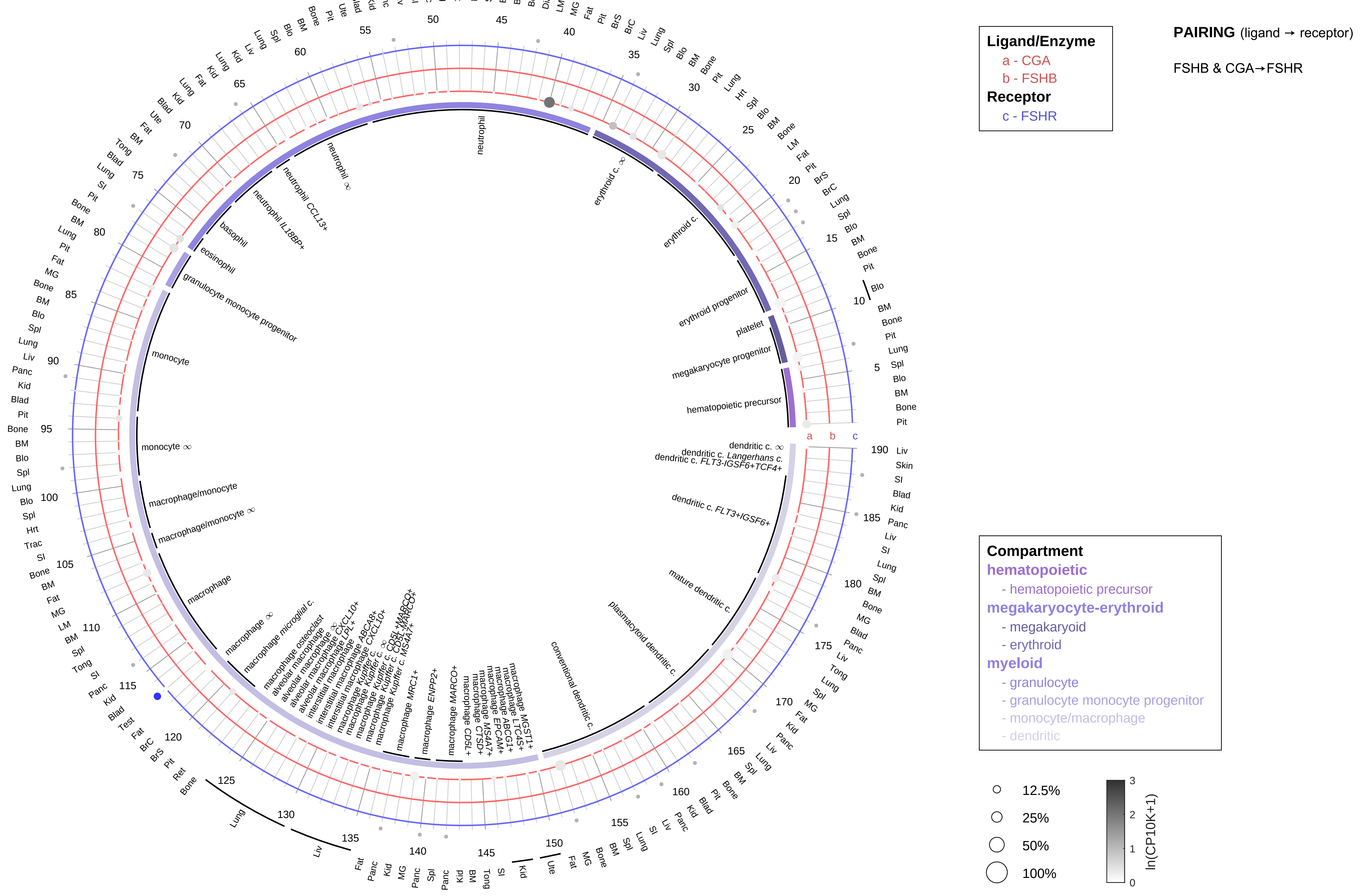

| Abbrev. | Tissue |
|---|---|
| Aort | Aorta |
| Blad | Bladder |
| Blo | Blood |
| Bone | Bone |
| BM | Bone marrow |
| BrC | Brain cortex |
| BrS | Brainstem |
| Col | Colon |
| Dia | Diaphragm |
| Ret | Eye retina |
| Fat | Fat |
| Hrt | Heart |
| Pit | Hypothalamus/Pituitary |
| Kid | Kidney |
| LM | Limb muscle |
| Liv | Liver |
| Lung | Lung |
| MG | Mammary gland |
| Panc | Pancreas |
| Skin | Skin |
| SI | Small intestine |
| Spl | Spleen |
| Test | Testes |
| Thym | Thymus |
| Tong | Tongue |
| Trac | Trachea |
| Ute | Uterus |

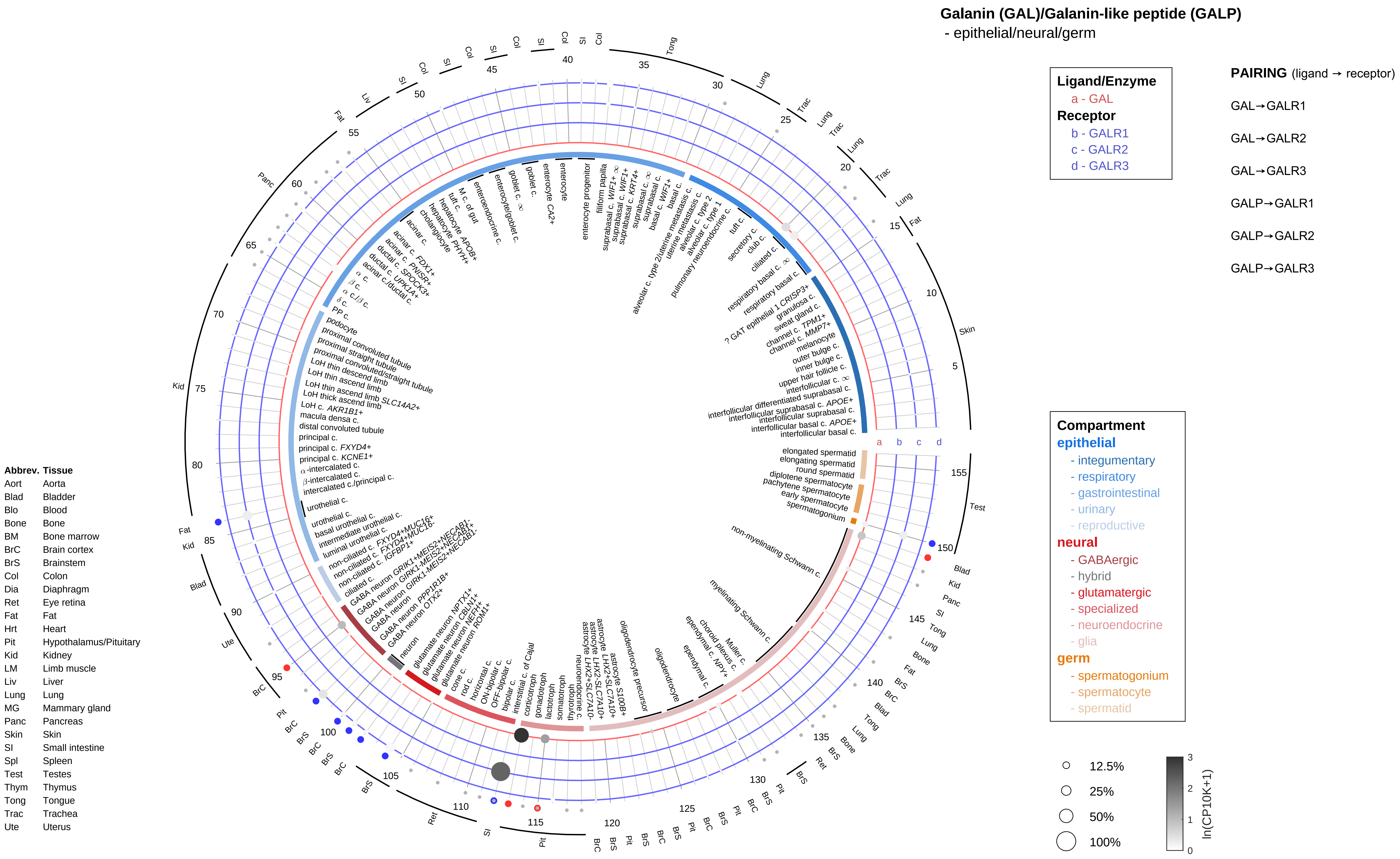

Galanin (GAL)/Galanin-like peptide (GALP)
- epithelial/neural/germ
Ligand/Enzyme
a - GAL
Receptor
b - GALR1
c - GALR2
d - GALR3
PAIRING (ligand → receptor)
GAL→GALR1
GAL→GALR2
GAL→GALR3
GALP→GALR1
GALP→GALR2
GALP→GALR3
Compartment
epithelial
- integumentary
- respiratory
- gastrointestinal
- urinary
- reproductive
neural
- GABAergic
- hybrid
- glutamatergic
- specialized
- neuroendocrine
- glia
germ
- spermatogonium
- spermatocyte
- spermatid
12.5%
25%
50%
100%
ln(CP10K+1)
Abbrev. Tissue
Aort Aorta
Blad Bladder
Blo Blood
Bone Bone
BM Bone marrow
BrC Brain cortex
BrS Brainstem
Col Colon
Dia Diaphragm
Ret Eye retina
Fat Fat
Hrt Heart
Pit Hypothalamus/Pituitary
Kid Kidney
LM Limb muscle
Liv Liver
Lung Lung
MG Mammary gland
Panc Pancreas
Skin Skin
SI Small intestine
Spl Spleen
Test Testes
Thym Thymus
Tong Tongue
Trac Trachea
Ute Uterus

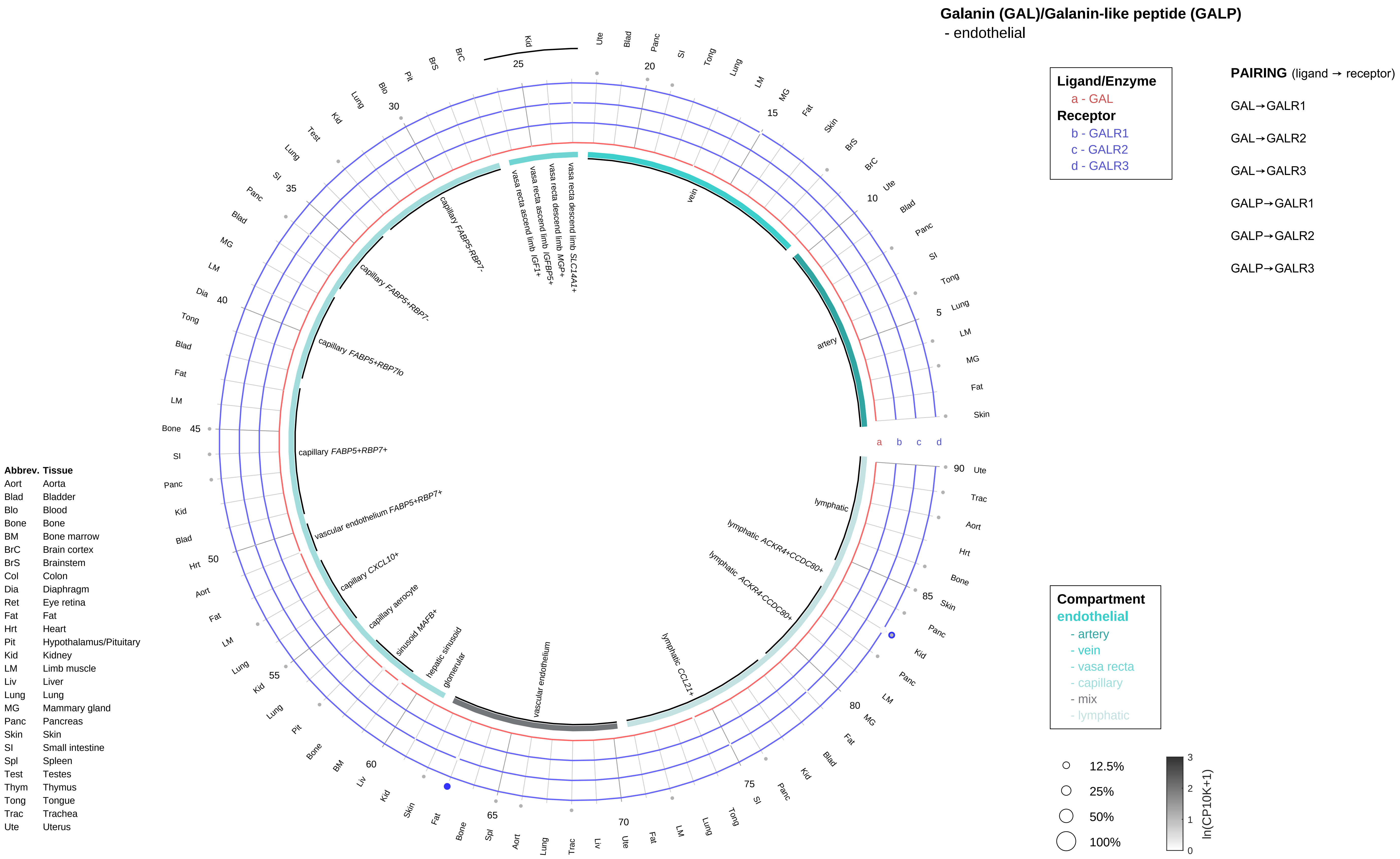
Galanin (GAL)/Galanin-like peptide (GALP)
- endothelial
Ligand/Enzyme
a - GAL
Receptor
b - GALR1
c - GALR2
d - GALR3
PAIRING (ligand → receptor)
GAL→GALR1
GAL→GALR2
GAL→GALR3
GALP→GALR1
GALP→GALR2
GALP→GALR3
Compartment
endothelial
- artery
- vein
- vasa recta
- capillary
- mix
- lymphatic
12.5%
25%
50%
100%
ln(CP10K+1)
Abbrev. Tissue
Aort Aorta
Blad Bladder
Blo Blood
Bone Bone
BM Bone marrow
BrC Brain cortex
BrS Brainstem
Col Colon
Dia Diaphragm
Ret Eye retina
Fat Fat
Hrt Heart
Pit Hypothalamus/Pituitary
Kid Kidney
LM Limb muscle
Liv Liver
Lung Lung
MG Mammary gland
Panc Pancreas
Skin Skin
SI Small intestine
Spl Spleen
Test Testes
Thym Thymus
Tong Tongue
Trac Trachea
Ute Uterus
vein
artery
lymphatic
lymphatic ACKR4+CCDC80+
lymphatic ACKR4-CCDC80+
lymphatic CCL21+
vascular endothelium
glomerular
hepatic sinusoid
sinusoid MAFB+
capillary aerocyte
capillary CXCL10+
vascular endothelium FABP5+RBP7+
capillary FABP5+RBP7+
capillary FABP5+RBP7lo
capillary FABP5+RBP7-
capillary FABP5-RBP7-
vasa recta ascend limb IGF1+
vasa recta ascend limb IGFBP5+
vasa recta descend limb MGP+
vasa recta descend limb SLC14A1+

#### Galanin (GAL)/Galanin-like peptide (GALP)
- stromal

**Ligand/Enzyme**
a - GAL
**Receptor**
b - GALR1
c - GALR2
d - GALR3

**PAIRING** (ligand → receptor)

GAL→GALR1

GAL→GALR2

GAL→GALR3

GALP→GALR1

GALP→GALR2

GALP→GALR3

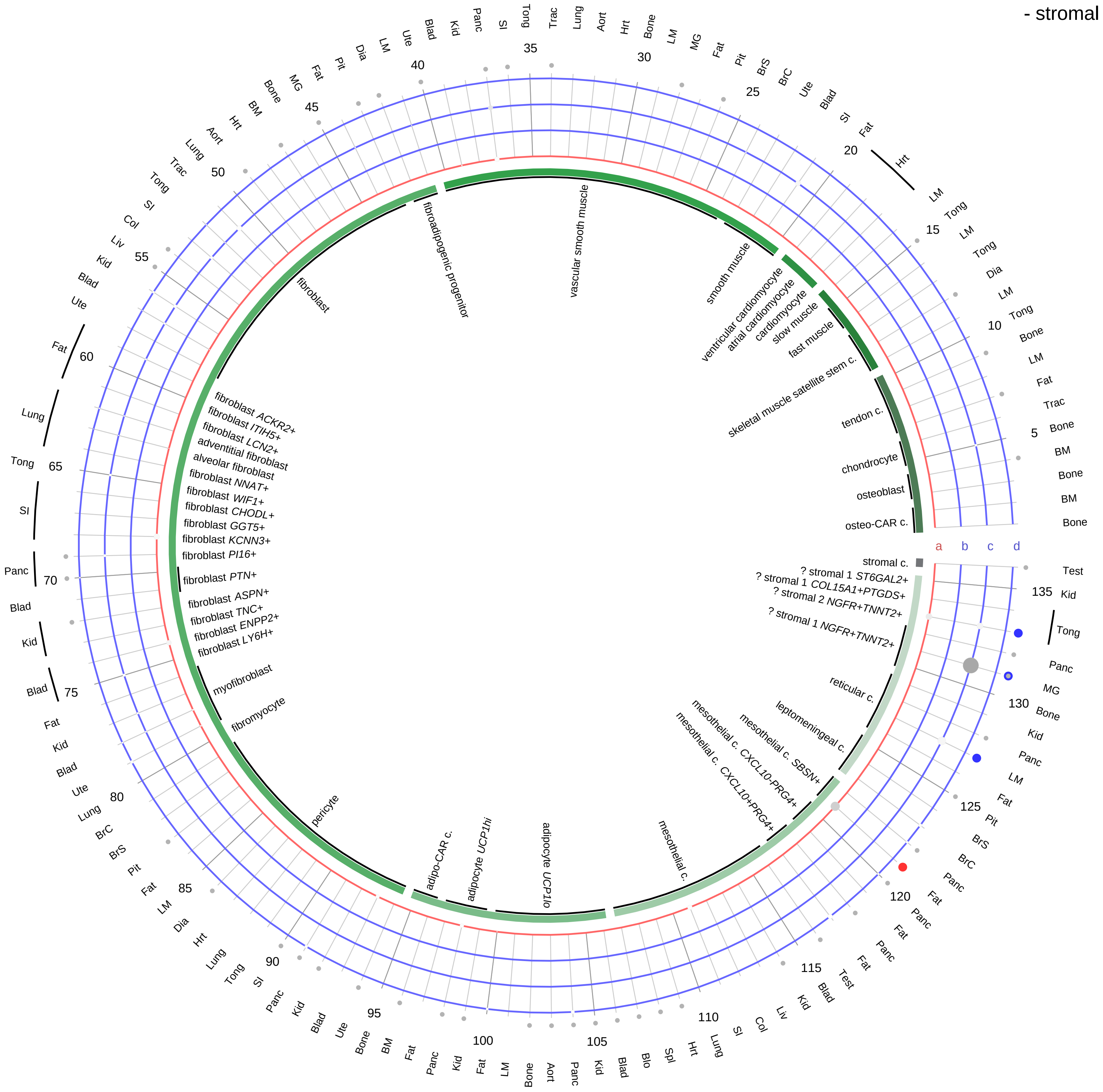

| Abbrev. | Tissue |
|---|---|
| Aort | Aorta |
| Blad | Bladder |
| Blo | Blood |
| Bone | Bone |
| BM | Bone marrow |
| BrC | Brain cortex |
| BrS | Brainstem |
| Col | Colon |
| Dia | Diaphragm |
| Ret | Eye retina |
| Fat | Fat |
| Hrt | Heart |
| Pit | Hypothalamus/Pituitary |
| Kid | Kidney |
| LM | Limb muscle |
| Liv | Liver |
| Lung | Lung |
| MG | Mammary gland |
| Panc | Pancreas |
| Skin | Skin |
| SI | Small intestine |
| Spl | Spleen |
| Test | Testes |
| Thym | Thymus |
| Tong | Tongue |
| Trac | Trachea |
| Ute | Uterus |

**Compartment**
**stromal**
- bone
- skeletal muscle
- cardiac muscle
- smooth muscle
- fibroblast
- adipocyte
- mesothelial
- other
- mix

12.5%
25%
50%
100%

ln(CP10K+1): 0, 1, 2, 3

**Galanin (GAL)/Galanin-like peptide (GALP)**
- lymphoid

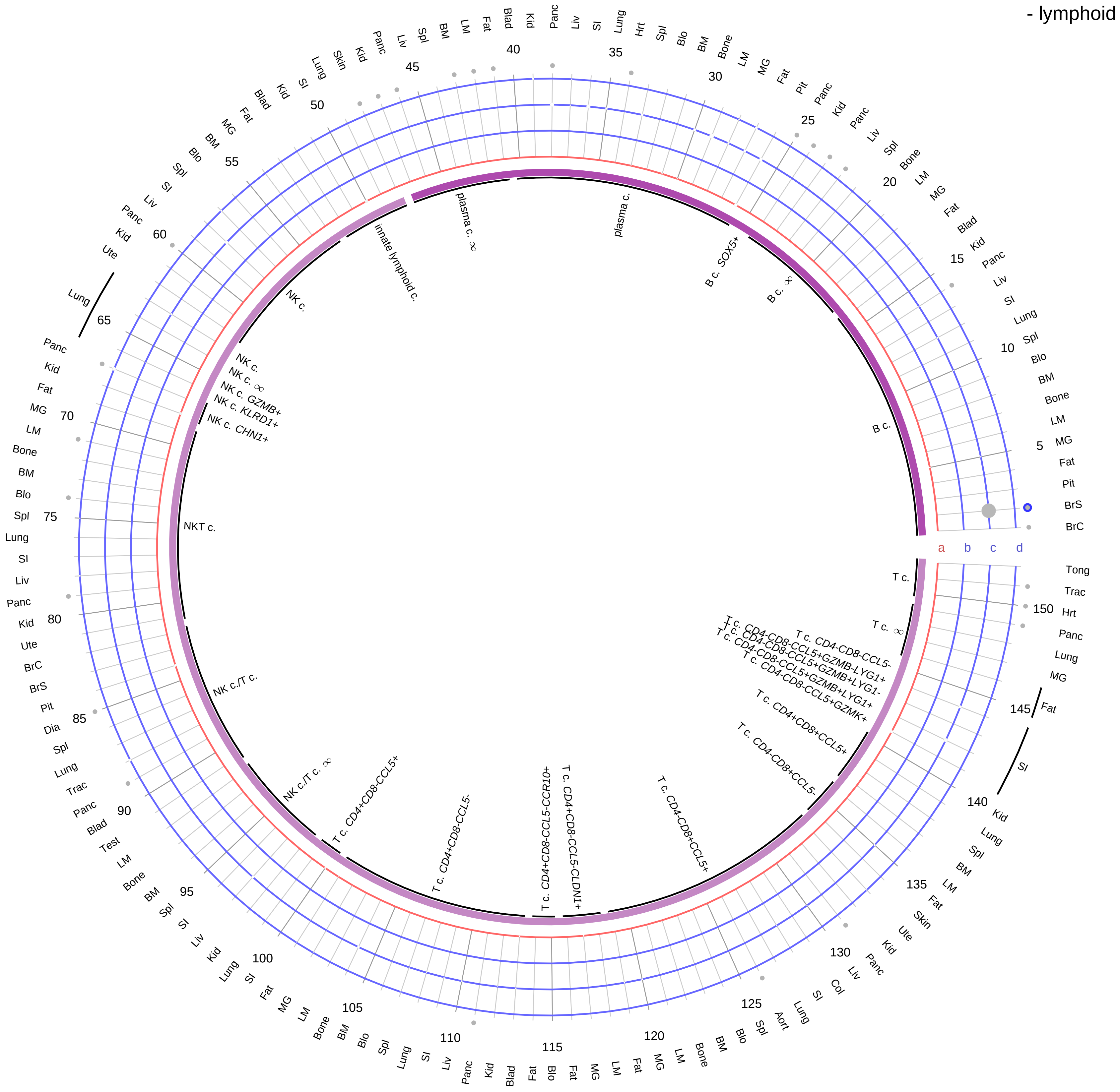

**Ligand/Enzyme**
- a - GAL

**Receptor**
- b - GALR1
- c - GALR2
- d - GALR3

**PAIRING** (ligand → receptor)

GAL→GALR1

GAL→GALR2

GAL→GALR3

GALP→GALR1

GALP→GALR2

GALP→GALR3

**Compartment**

**lymphoid**
- B cell
- NK/T cell

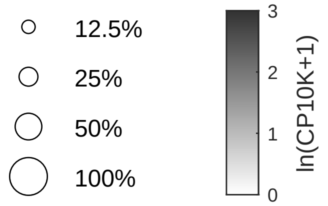

| Abbrev. | Tissue |
|---|---|
| Aort | Aorta |
| Blad | Bladder |
| Blo | Blood |
| Bone | Bone |
| BM | Bone marrow |
| BrC | Brain cortex |
| BrS | Brainstem |
| Col | Colon |
| Dia | Diaphragm |
| Ret | Eye retina |
| Fat | Fat |
| Hrt | Heart |
| Pit | Hypothalamus/Pituitary |
| Kid | Kidney |
| LM | Limb muscle |
| Liv | Liver |
| Lung | Lung |
| MG | Mammary gland |
| Panc | Pancreas |
| Skin | Skin |
| SI | Small intestine |
| Spl | Spleen |
| Test | Testes |
| Thym | Thymus |
| Tong | Tongue |
| Trac | Trachea |
| Ute | Uterus |

**Galanin (GAL)/Galanin-like peptide (GALP)**
- myeloid

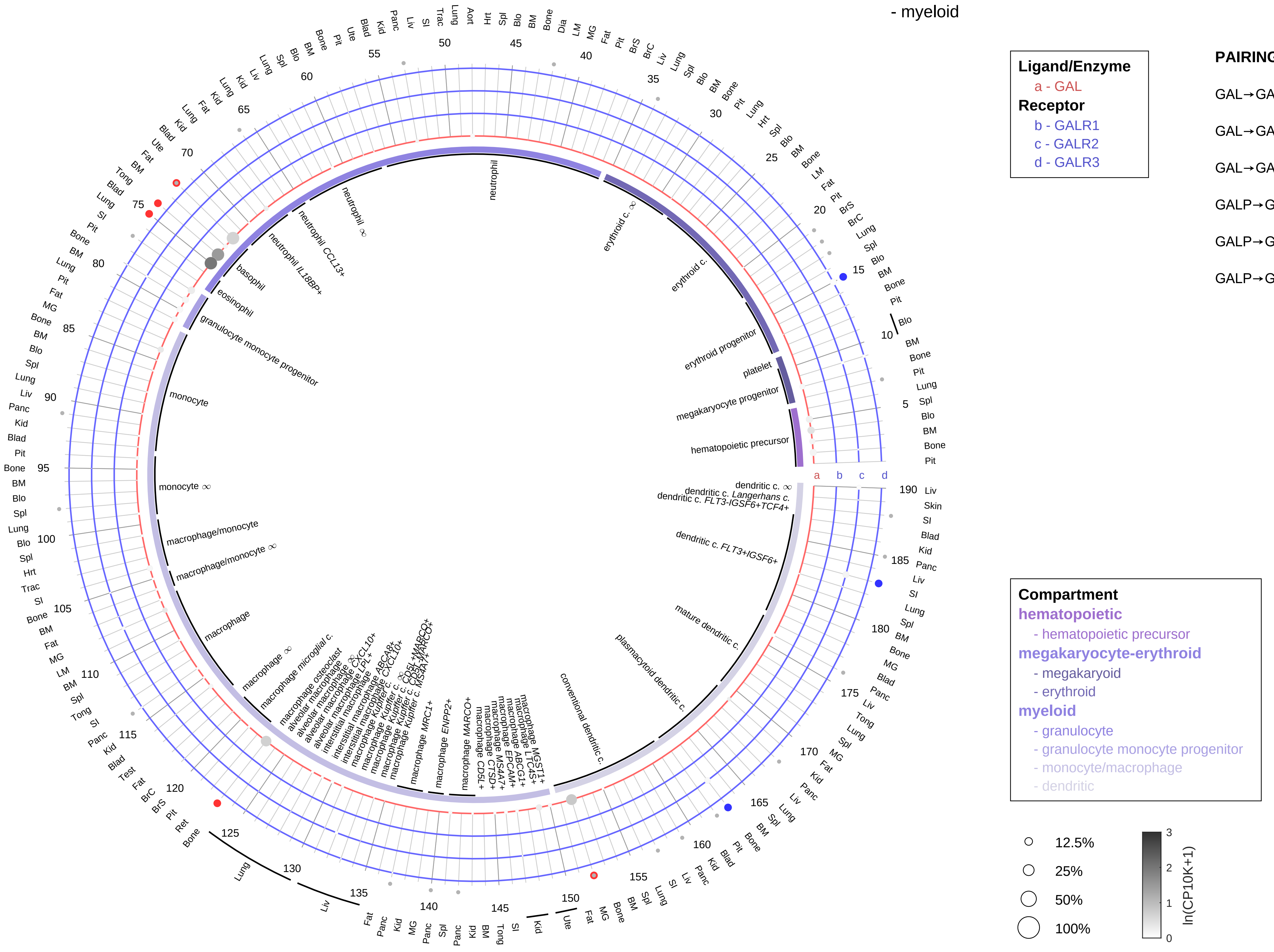

| Abbrev. | Tissue |
|---|---|
| Aort | Aorta |
| Blad | Bladder |
| Blo | Blood |
| Bone | Bone |
| BM | Bone marrow |
| BrC | Brain cortex |
| BrS | Brainstem |
| Col | Colon |
| Dia | Diaphragm |
| Ret | Eye retina |
| Fat | Fat |
| Hrt | Heart |
| Pit | Hypothalamus/Pituitary |
| Kid | Kidney |
| LM | Limb muscle |
| Liv | Liver |
| Lung | Lung |
| MG | Mammary gland |
| Panc | Pancreas |
| Skin | Skin |
| SI | Small intestine |
| Spl | Spleen |
| Test | Testes |
| Thym | Thymus |
| Tong | Tongue |
| Trac | Trachea |
| Ute | Uterus |

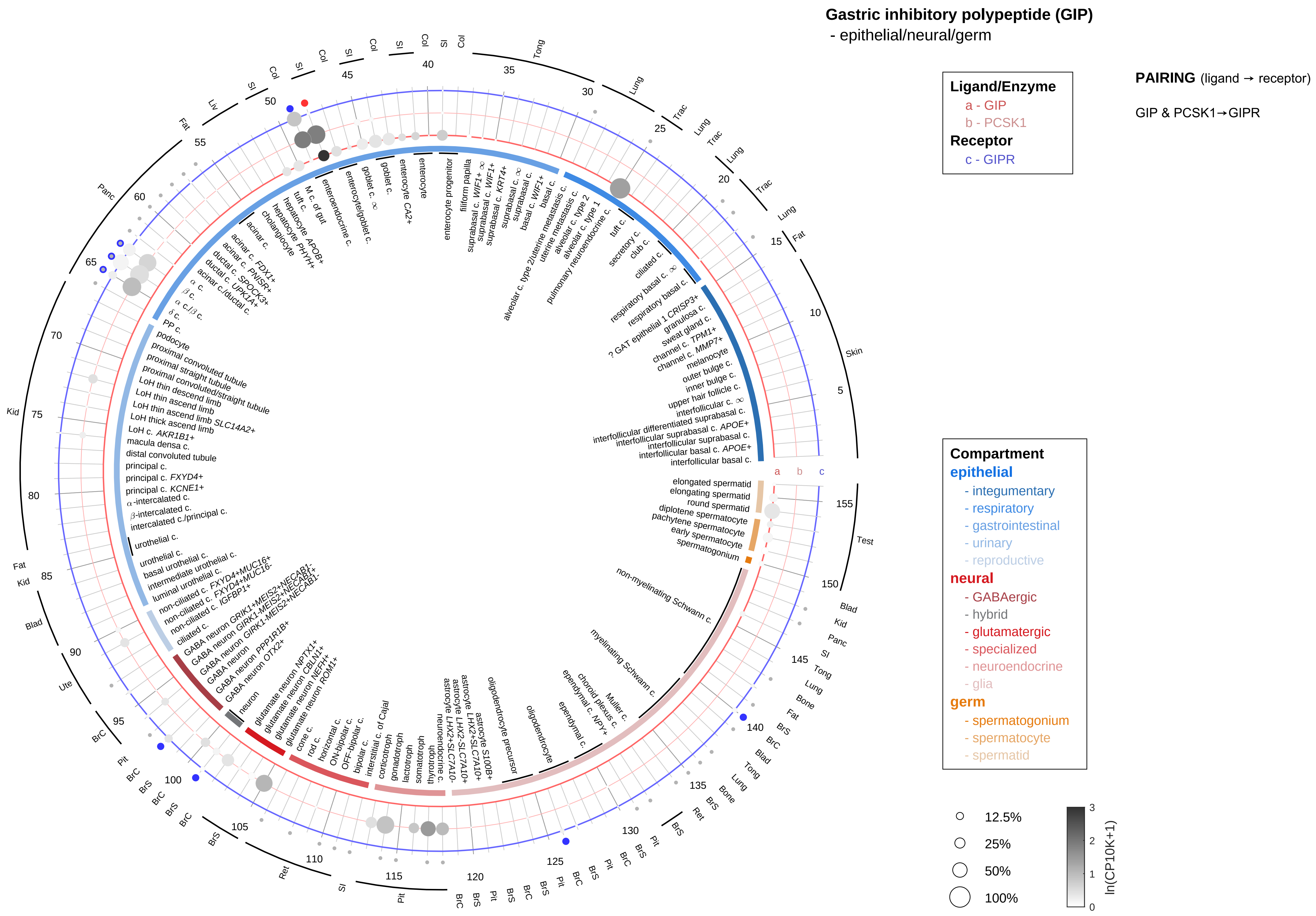

| Abbrev. | Tissue |
|---|---|
| Aort | Aorta |
| Blad | Bladder |
| Blo | Blood |
| Bone | Bone |
| BM | Bone marrow |
| BrC | Brain cortex |
| BrS | Brainstem |
| Col | Colon |
| Dia | Diaphragm |
| Ret | Eye retina |
| Fat | Fat |
| Hrt | Heart |
| Pit | Hypothalamus/Pituitary |
| Kid | Kidney |
| LM | Limb muscle |
| Liv | Liver |
| Lung | Lung |
| MG | Mammary gland |
| Panc | Pancreas |
| Skin | Skin |
| SI | Small intestine |
| Spl | Spleen |
| Test | Testes |
| Thym | Thymus |
| Tong | Tongue |
| Trac | Trachea |
| Ute | Uterus |

**Gastric inhibitory polypeptide (GIP)**
- endothelial

**PAIRING** (ligand → receptor)

GIP & PCSK1→GIPR

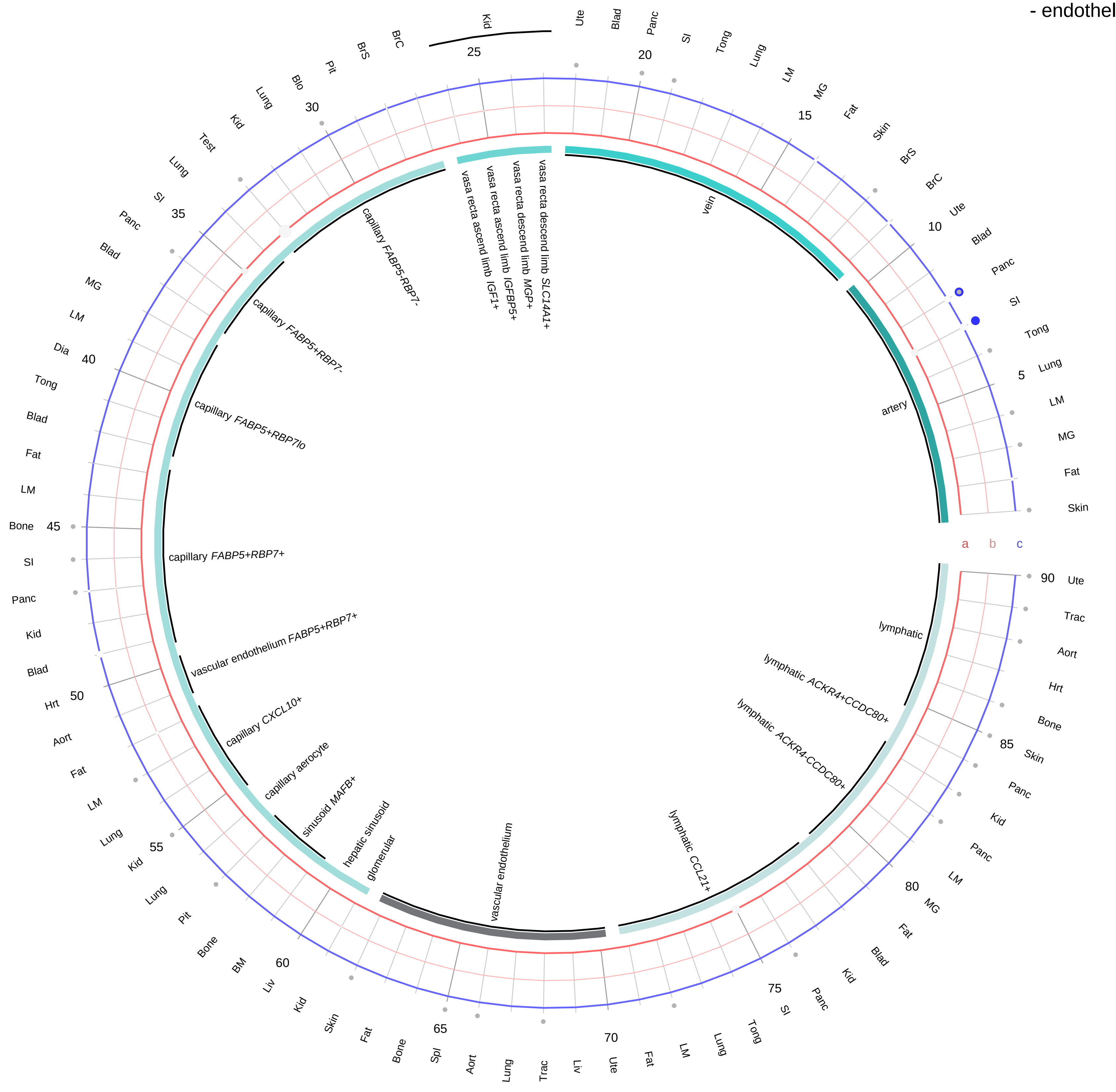

**Ligand/Enzyme**
- a - GIP
- b - PCSK1

**Receptor**
- c - GIPR

**Compartment**
**endothelial**
- artery
- vein
- vasa recta
- capillary
- mix
- lymphatic

12.5%
25%
50%
100%

ln(CP10K+1): 0, 1, 2, 3

| Abbrev. | Tissue |
|---|---|
| Aort | Aorta |
| Blad | Bladder |
| Blo | Blood |
| Bone | Bone |
| BM | Bone marrow |
| BrC | Brain cortex |
| BrS | Brainstem |
| Col | Colon |
| Dia | Diaphragm |
| Ret | Eye retina |
| Fat | Fat |
| Hrt | Heart |
| Pit | Hypothalamus/Pituitary |
| Kid | Kidney |
| LM | Limb muscle |
| Liv | Liver |
| Lung | Lung |
| MG | Mammary gland |
| Panc | Pancreas |
| Skin | Skin |
| SI | Small intestine |
| Spl | Spleen |
| Test | Testes |
| Thym | Thymus |
| Tong | Tongue |
| Trac | Trachea |
| Ute | Uterus |

#### Gastric inhibitory polypeptide (GIP)
- stromal

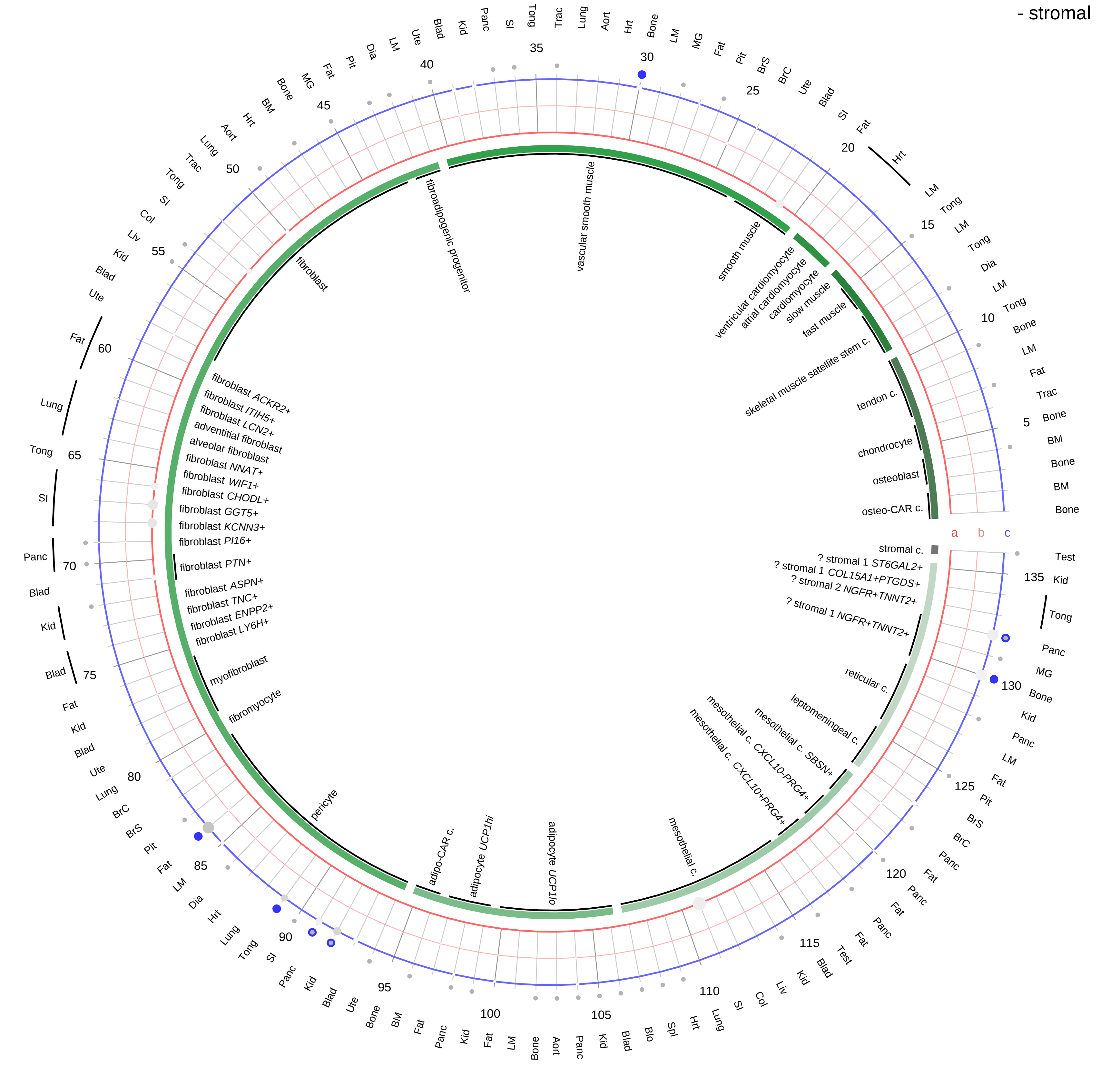

**Ligand/Enzyme**
- a - GIP
- b - PCSK1

**Receptor**
- c - GIPR

**PAIRING** (ligand → receptor)

GIP & PCSK1→GIPR

**Compartment**
**stromal**
- bone
- skeletal muscle
- cardiac muscle
- smooth muscle
- fibroblast
- adipocyte
- mesothelial
- other
- mix

12.5%
25%
50%
100%

ln(CP10K+1): 0, 1, 2, 3

| Abbrev. | Tissue |
|---|---|
| Aort | Aorta |
| Blad | Bladder |
| Blo | Blood |
| Bone | Bone |
| BM | Bone marrow |
| BrC | Brain cortex |
| BrS | Brainstem |
| Col | Colon |
| Dia | Diaphragm |
| Ret | Eye retina |
| Fat | Fat |
| Hrt | Heart |
| Pit | Hypothalamus/Pituitary |
| Kid | Kidney |
| LM | Limb muscle |
| Liv | Liver |
| Lung | Lung |
| MG | Mammary gland |
| Panc | Pancreas |
| Skin | Skin |
| SI | Small intestine |
| Spl | Spleen |
| Test | Testes |
| Thym | Thymus |
| Tong | Tongue |
| Trac | Trachea |
| Ute | Uterus |

**Gastric inhibitory polypeptide (GIP)**
- lymphoid

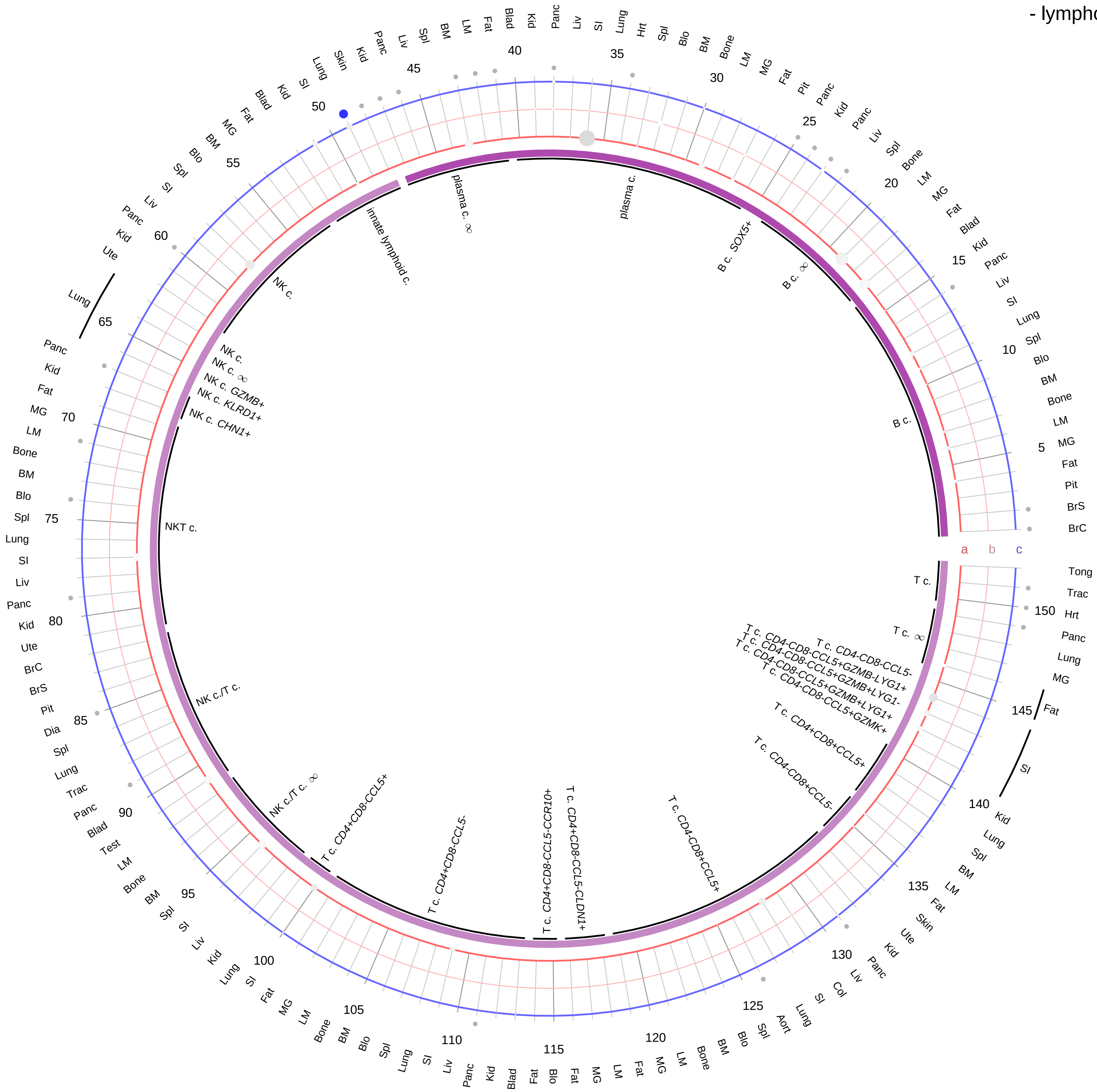

**Ligand/Enzyme**
a - GIP
b - PCSK1
**Receptor**
c - GIPR

**PAIRING** (ligand → receptor)

GIP & PCSK1→GIPR

**Compartment**
**lymphoid**
- B cell
- NK/T cell

12.5%
25%
50%
100%

ln(CP10K+1): 0, 1, 2, 3

| Abbrev. | Tissue |
|---|---|
| Aort | Aorta |
| Blad | Bladder |
| Blo | Blood |
| Bone | Bone |
| BM | Bone marrow |
| BrC | Brain cortex |
| BrS | Brainstem |
| Col | Colon |
| Dia | Diaphragm |
| Ret | Eye retina |
| Fat | Fat |
| Hrt | Heart |
| Pit | Hypothalamus/Pituitary |
| Kid | Kidney |
| LM | Limb muscle |
| Liv | Liver |
| Lung | Lung |
| MG | Mammary gland |
| Panc | Pancreas |
| Skin | Skin |
| SI | Small intestine |
| Spl | Spleen |
| Test | Testes |
| Thym | Thymus |
| Tong | Tongue |
| Trac | Trachea |
| Ute | Uterus |

**Gastric inhibitory polypeptide (GIP)**
- myeloid

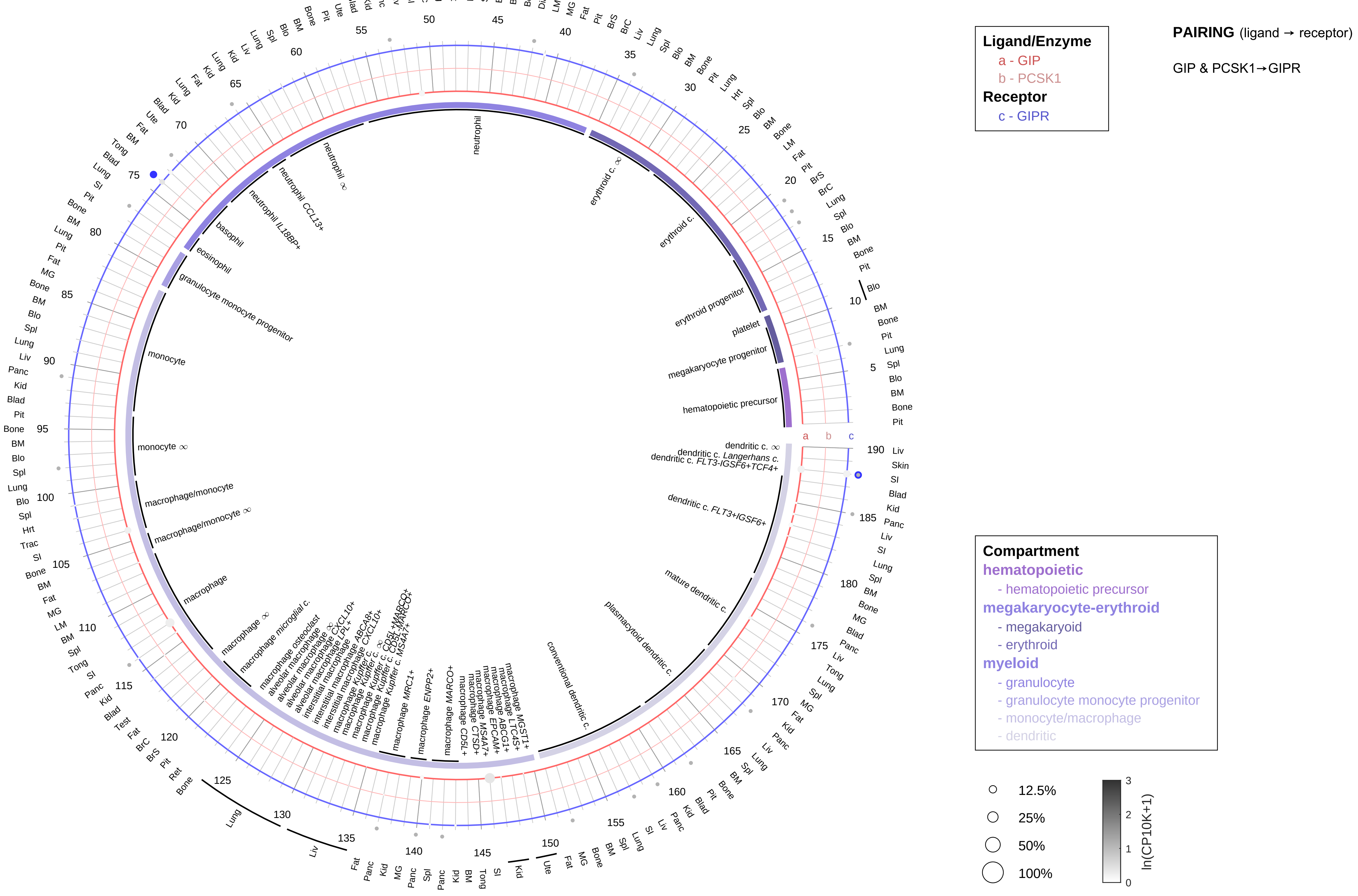

| Abbrev. | Tissue |
|---|---|
| Aort | Aorta |
| Blad | Bladder |
| Blo | Blood |
| Bone | Bone |
| BM | Bone marrow |
| BrC | Brain cortex |
| BrS | Brainstem |
| Col | Colon |
| Dia | Diaphragm |
| Ret | Eye retina |
| Fat | Fat |
| Hrt | Heart |
| Pit | Hypothalamus/Pituitary |
| Kid | Kidney |
| LM | Limb muscle |
| Liv | Liver |
| Lung | Lung |
| MG | Mammary gland |
| Panc | Pancreas |
| Skin | Skin |
| SI | Small intestine |
| Spl | Spleen |
| Test | Testes |
| Thym | Thymus |
| Tong | Tongue |
| Trac | Trachea |
| Ute | Uterus |

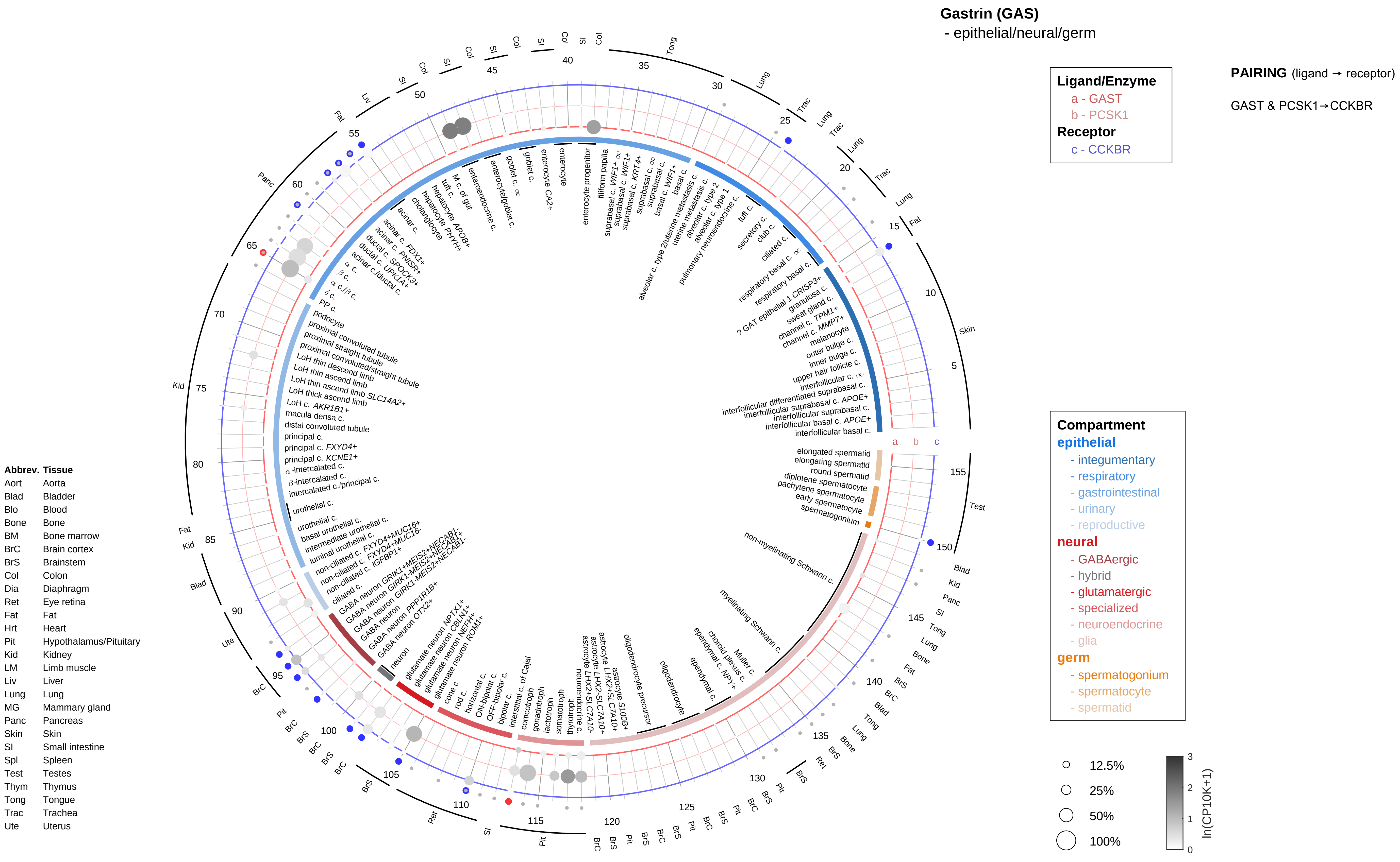
Gastrin (GAS)
- epithelial/neural/germ
Ligand/Enzyme
a - GAST
b - PCSK1
Receptor
c - CCKBR
PAIRING (ligand → receptor)
GAST & PCSK1→CCKBR
Compartment
epithelial
- integumentary
- respiratory
- gastrointestinal
- urinary
- reproductive
neural
- GABAergic
- hybrid
- glutamatergic
- specialized
- neuroendocrine
- glia
germ
- spermatogonium
- spermatocyte
- spermatid
12.5%
25%
50%
100%
ln(CP10K+1)
Abbrev. Tissue
Aort Aorta
Blad Bladder
Blo Blood
Bone Bone
BM Bone marrow
BrC Brain cortex
BrS Brainstem
Col Colon
Dia Diaphragm
Ret Eye retina
Fat Fat
Hrt Heart
Pit Hypothalamus/Pituitary
Kid Kidney
LM Limb muscle
Liv Liver
Lung Lung
MG Mammary gland
Panc Pancreas
Skin Skin
SI Small intestine
Spl Spleen
Test Testes
Thym Thymus
Tong Tongue
Trac Trachea
Ute Uterus

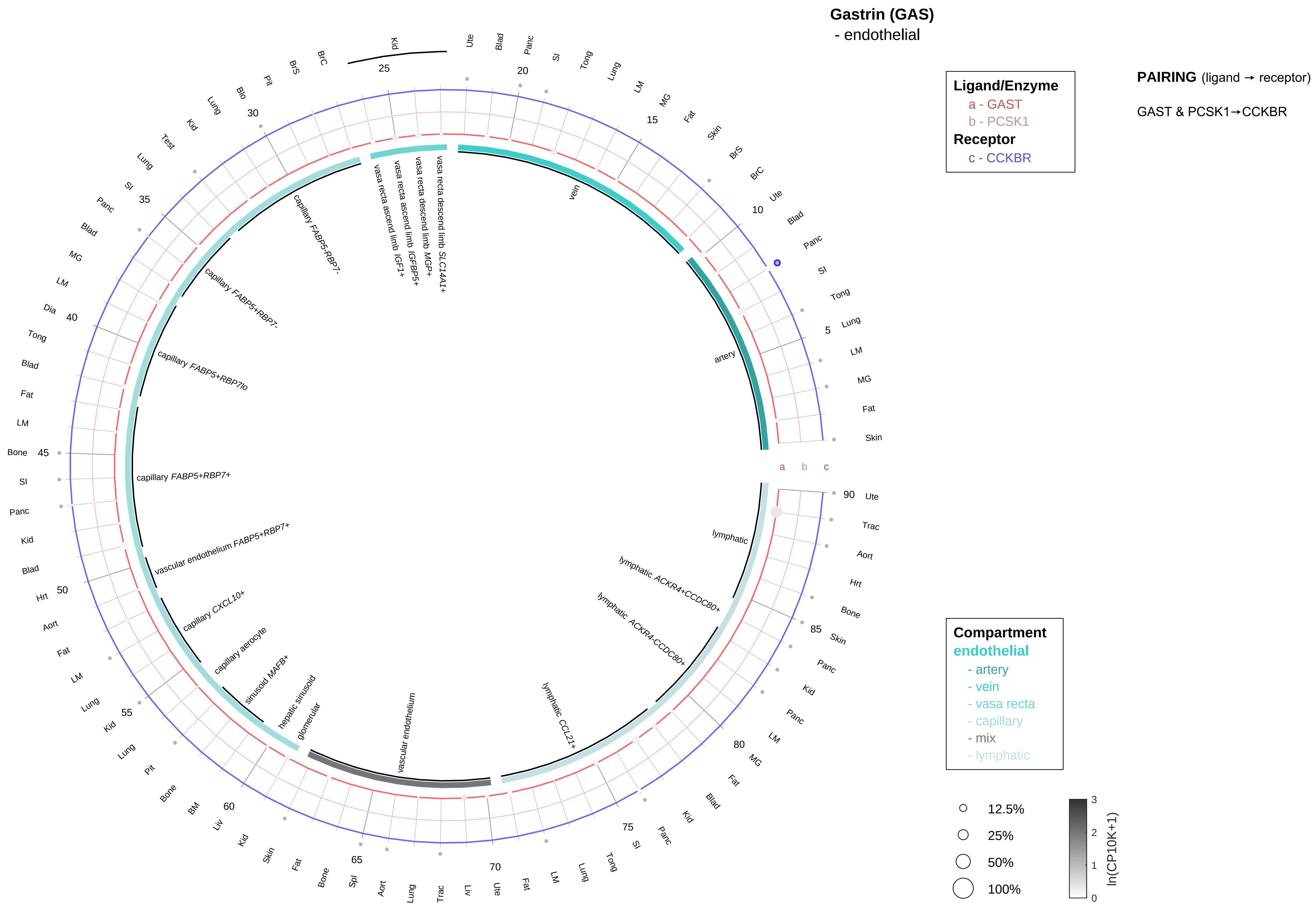

| Abbrev. | Tissue |
|---|---|
| Aort | Aorta |
| Blad | Bladder |
| Blo | Blood |
| Bone | Bone |
| BM | Bone marrow |
| BrC | Brain cortex |
| BrS | Brainstem |
| Col | Colon |
| Dia | Diaphragm |
| Ret | Eye retina |
| Fat | Fat |
| Hrt | Heart |
| Pit | Hypothalamus/Pituitary |
| Kid | Kidney |
| LM | Limb muscle |
| Liv | Liver |
| Lung | Lung |
| MG | Mammary gland |
| Panc | Pancreas |
| Skin | Skin |
| SI | Small intestine |
| Spl | Spleen |
| Test | Testes |
| Thym | Thymus |
| Tong | Tongue |
| Trac | Trachea |
| Ute | Uterus |

**Gastrin (GAS)**
- stromal

**PAIRING** (ligand → receptor)

GAST & PCSK1→CCKBR

**Ligand/Enzyme**
a - GAST
b - PCSK1
**Receptor**
c - CCKBR

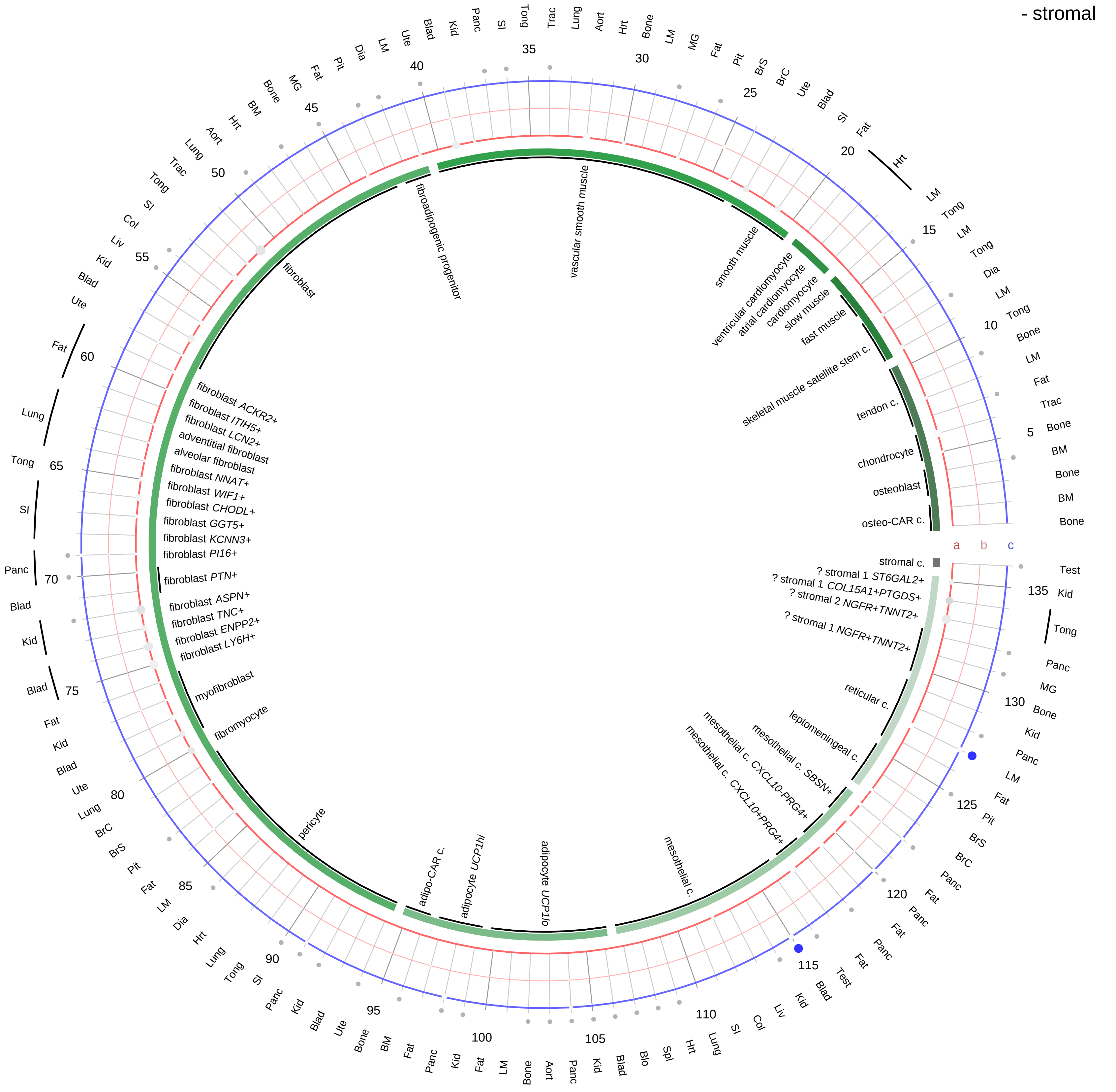

| Abbrev. | Tissue |
|---|---|
| Aort | Aorta |
| Blad | Bladder |
| Blo | Blood |
| Bone | Bone |
| BM | Bone marrow |
| BrC | Brain cortex |
| BrS | Brainstem |
| Col | Colon |
| Dia | Diaphragm |
| Ret | Eye retina |
| Fat | Fat |
| Hrt | Heart |
| Pit | Hypothalamus/Pituitary |
| Kid | Kidney |
| LM | Limb muscle |
| Liv | Liver |
| Lung | Lung |
| MG | Mammary gland |
| Panc | Pancreas |
| Skin | Skin |
| SI | Small intestine |
| Spl | Spleen |
| Test | Testes |
| Thym | Thymus |
| Tong | Tongue |
| Trac | Trachea |
| Ute | Uterus |

**Compartment**
**stromal**
- bone
- skeletal muscle
- cardiac muscle
- smooth muscle
- fibroblast
- adipocyte
- mesothelial
- other
- mix

12.5%
25%
50%
100%

ln(CP10K+1): 0, 1, 2, 3

**Gastrin (GAS)**
- lymphoid

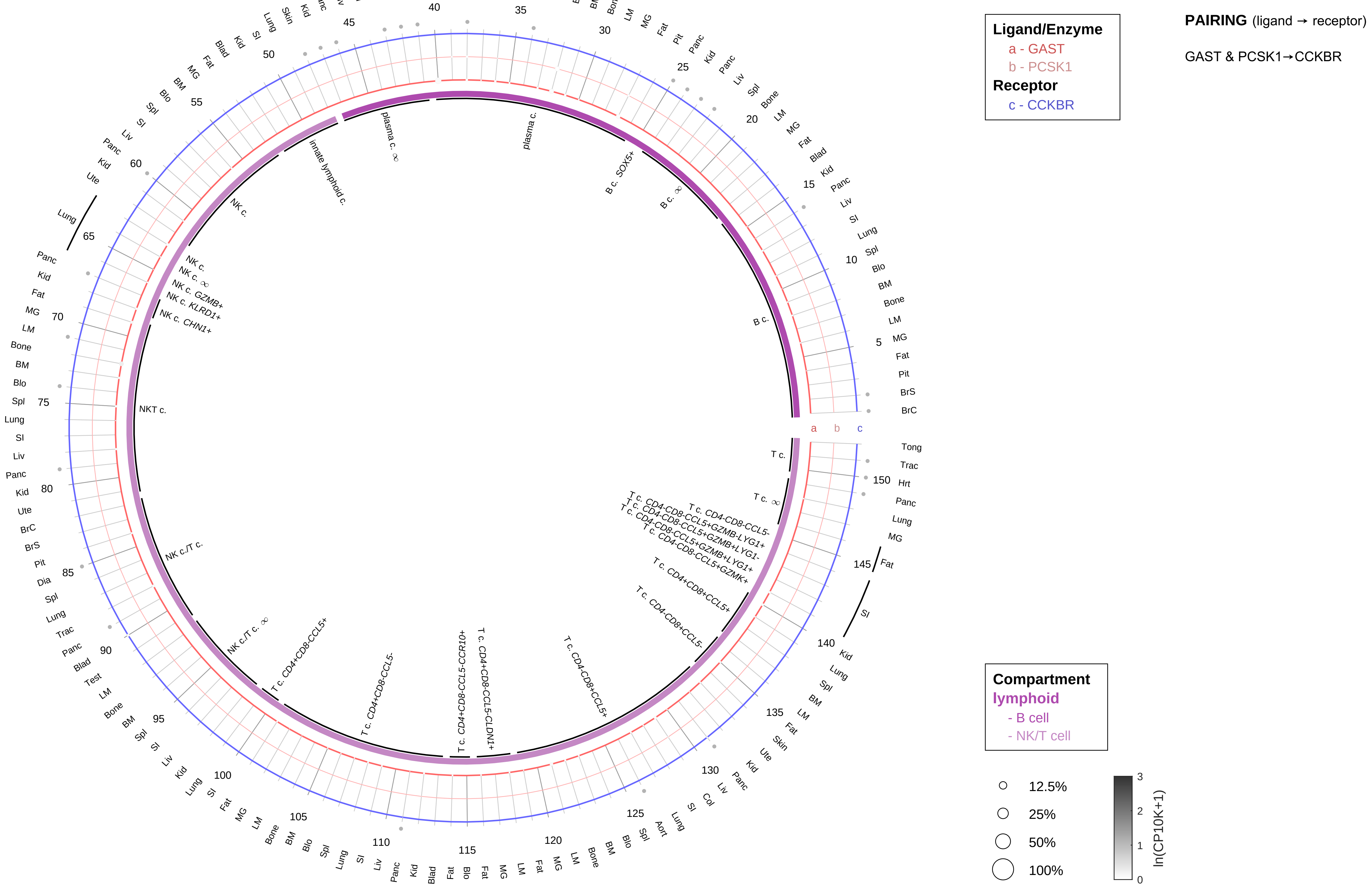

| Abbrev. | Tissue |
|---|---|
| Aort | Aorta |
| Blad | Bladder |
| Blo | Blood |
| Bone | Bone |
| BM | Bone marrow |
| BrC | Brain cortex |
| BrS | Brainstem |
| Col | Colon |
| Dia | Diaphragm |
| Ret | Eye retina |
| Fat | Fat |
| Hrt | Heart |
| Pit | Hypothalamus/Pituitary |
| Kid | Kidney |
| LM | Limb muscle |
| Liv | Liver |
| Lung | Lung |
| MG | Mammary gland |
| Panc | Pancreas |
| Skin | Skin |
| SI | Small intestine |
| Spl | Spleen |
| Test | Testes |
| Thym | Thymus |
| Tong | Tongue |
| Trac | Trachea |
| Ute | Uterus |

#### Gastrin (GAS)

- myeloid

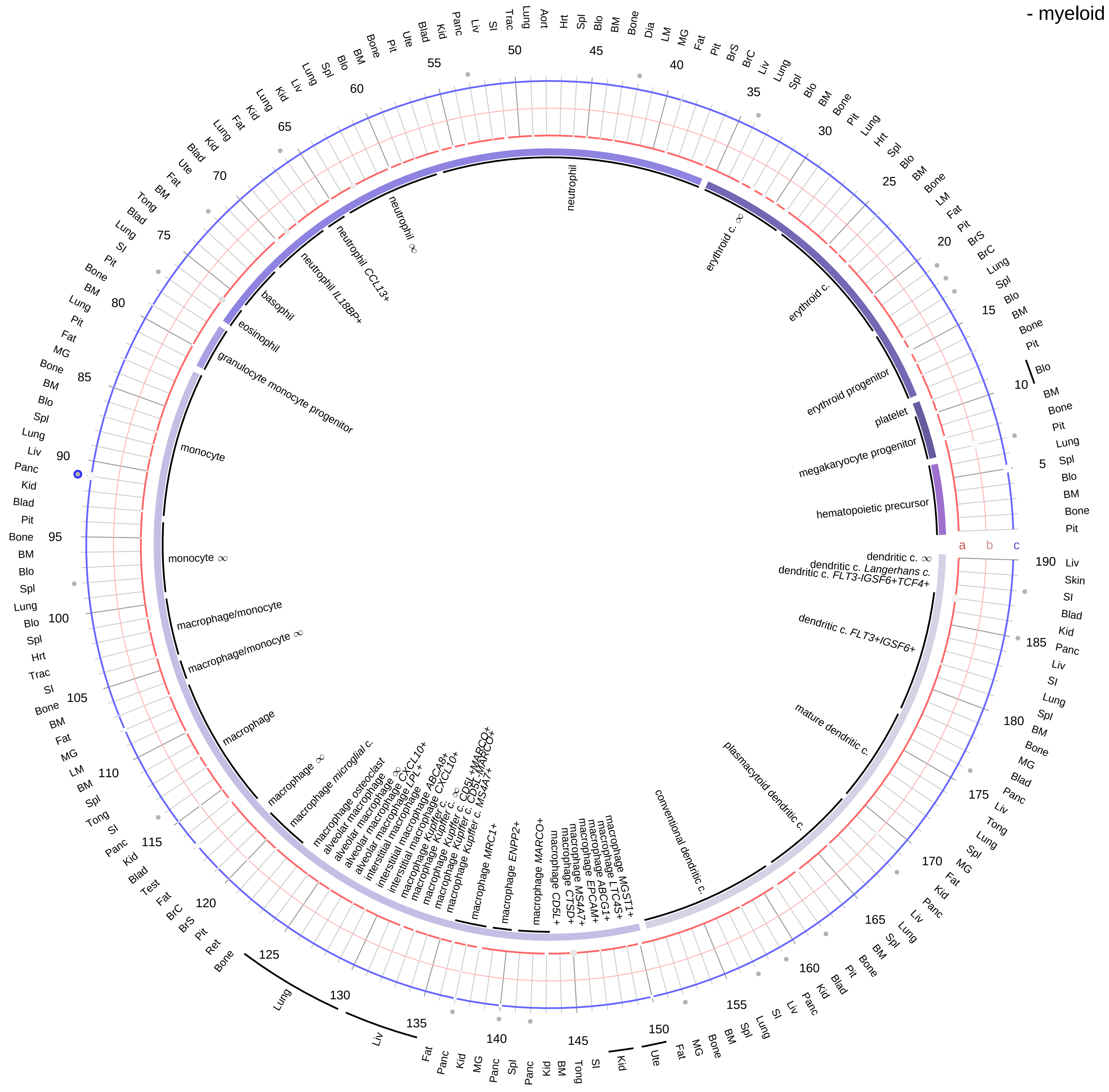

**Ligand/Enzyme**
- a - GAST
- b - PCSK1

**Receptor**
- c - CCKBR

**PAIRING** (ligand → receptor)

GAST & PCSK1→CCKBR

**Compartment**

**hematopoietic**
- hematopoietic precursor

**megakaryocyte-erythroid**
- megakaryoid
- erythroid

**myeloid**
- granulocyte
- granulocyte monocyte progenitor
- monocyte/macrophage
- dendritic

12.5%
25%
50%
100%

ln(CP10K+1): 0, 1, 2, 3

| Abbrev. | Tissue |
|---|---|
| Aort | Aorta |
| Blad | Bladder |
| Blo | Blood |
| Bone | Bone |
| BM | Bone marrow |
| BrC | Brain cortex |
| BrS | Brainstem |
| Col | Colon |
| Dia | Diaphragm |
| Ret | Eye retina |
| Fat | Fat |
| Hrt | Heart |
| Pit | Hypothalamus/Pituitary |
| Kid | Kidney |
| LM | Limb muscle |
| Liv | Liver |
| Lung | Lung |
| MG | Mammary gland |
| Panc | Pancreas |
| Skin | Skin |
| SI | Small intestine |
| Spl | Spleen |
| Test | Testes |
| Thym | Thymus |
| Tong | Tongue |
| Trac | Trachea |
| Ute | Uterus |

**Gastrin-releasing peptide (GRP)**
- epithelial/neural/germ

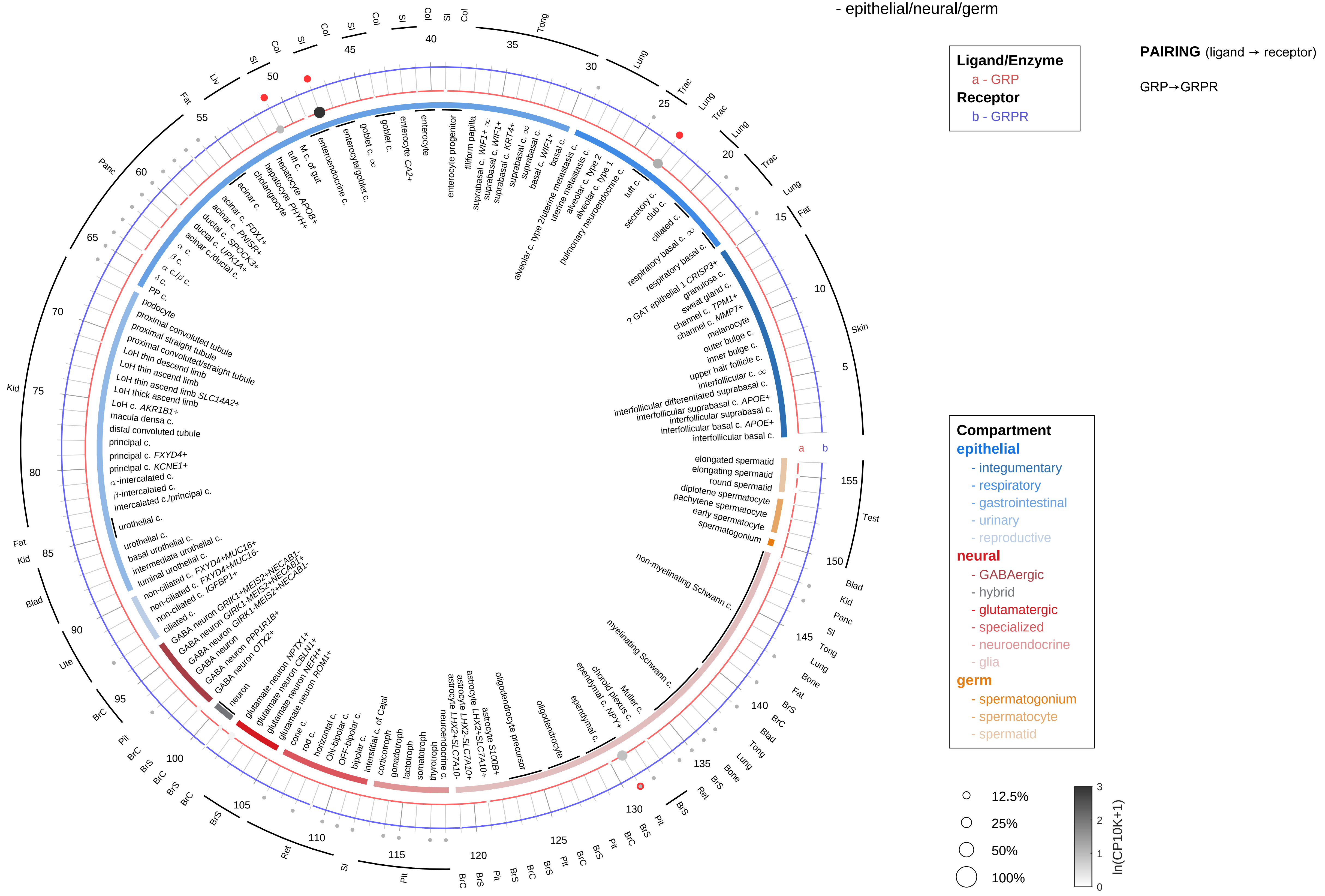

| Abbrev. | Tissue |
|---|---|
| Aort | Aorta |
| Blad | Bladder |
| Blo | Blood |
| Bone | Bone |
| BM | Bone marrow |
| BrC | Brain cortex |
| BrS | Brainstem |
| Col | Colon |
| Dia | Diaphragm |
| Ret | Eye retina |
| Fat | Fat |
| Hrt | Heart |
| Pit | Hypothalamus/Pituitary |
| Kid | Kidney |
| LM | Limb muscle |
| Liv | Liver |
| Lung | Lung |
| MG | Mammary gland |
| Panc | Pancreas |
| Skin | Skin |
| SI | Small intestine |
| Spl | Spleen |
| Test | Testes |
| Thym | Thymus |
| Tong | Tongue |
| Trac | Trachea |
| Ute | Uterus |

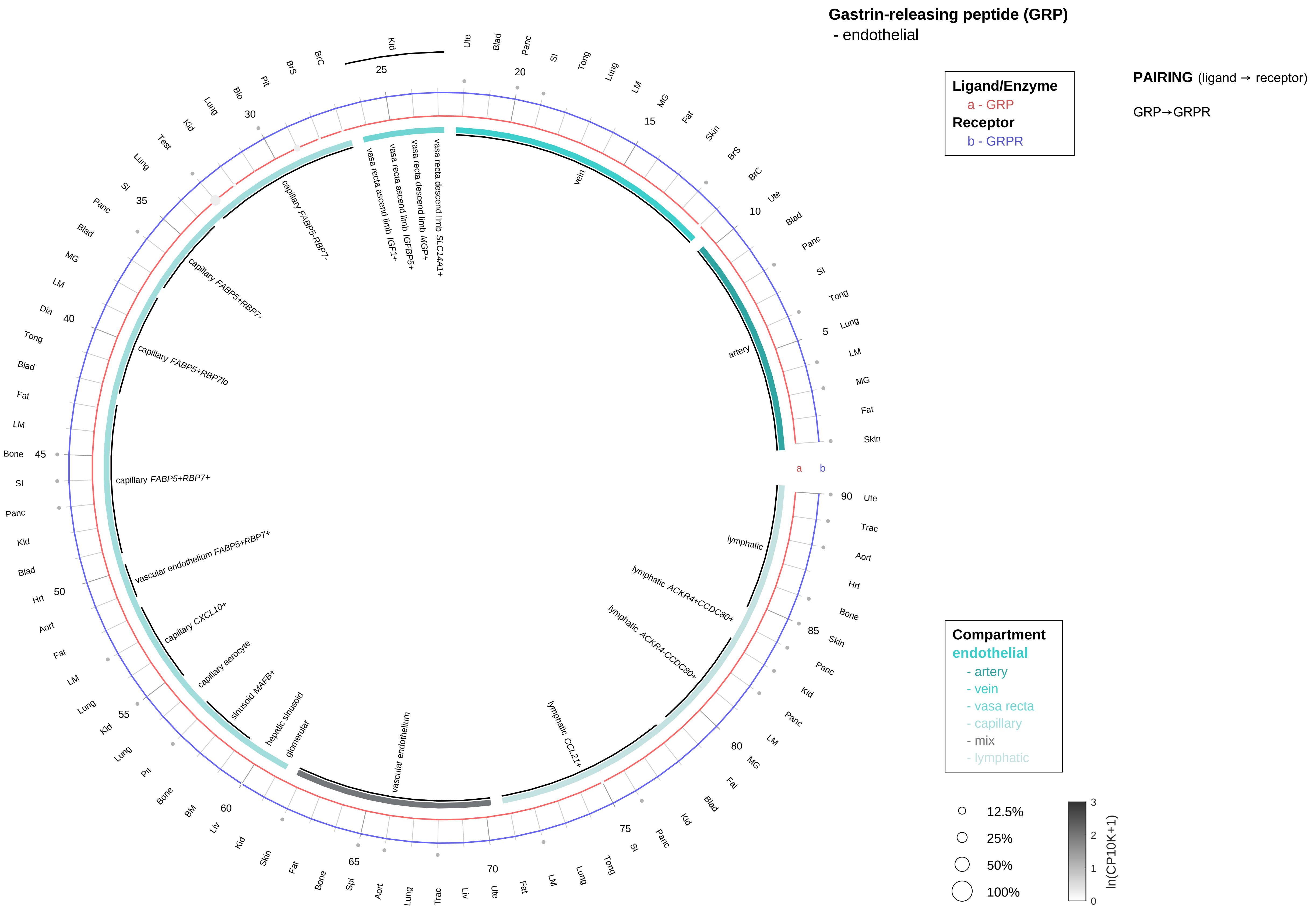

| Abbrev. | Tissue |
|---|---|
| Aort | Aorta |
| Blad | Bladder |
| Blo | Blood |
| Bone | Bone |
| BM | Bone marrow |
| BrC | Brain cortex |
| BrS | Brainstem |
| Col | Colon |
| Dia | Diaphragm |
| Ret | Eye retina |
| Fat | Fat |
| Hrt | Heart |
| Pit | Hypothalamus/Pituitary |
| Kid | Kidney |
| LM | Limb muscle |
| Liv | Liver |
| Lung | Lung |
| MG | Mammary gland |
| Panc | Pancreas |
| Skin | Skin |
| SI | Small intestine |
| Spl | Spleen |
| Test | Testes |
| Thym | Thymus |
| Tong | Tongue |
| Trac | Trachea |
| Ute | Uterus |

#### Gastrin-releasing peptide (GRP)
- stromal

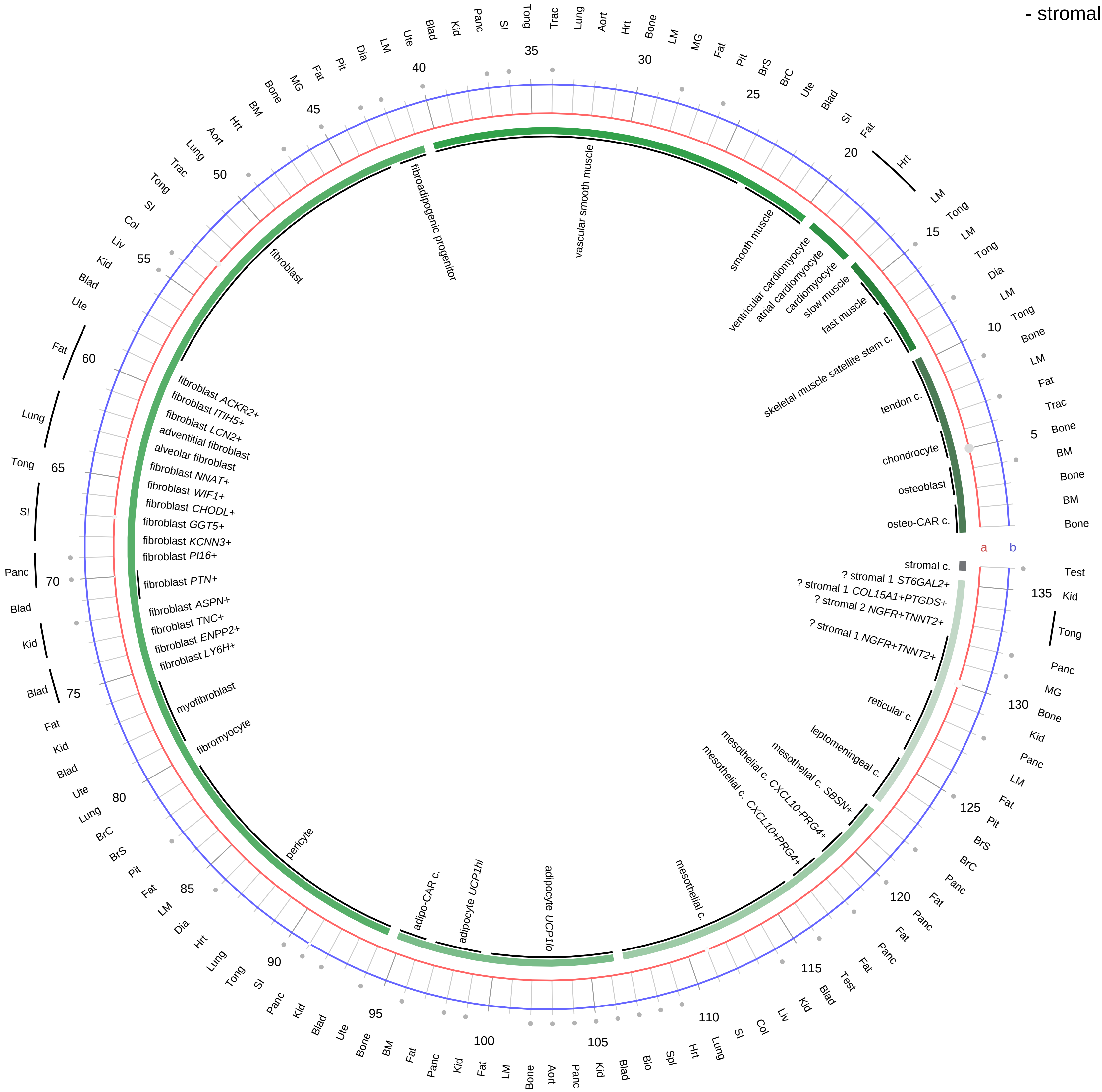

**Ligand/Enzyme**
a - GRP
**Receptor**
b - GRPR

**PAIRING** (ligand → receptor)

GRP→GRPR

| Abbrev. | Tissue |
|---|---|
| Aort | Aorta |
| Blad | Bladder |
| Blo | Blood |
| Bone | Bone |
| BM | Bone marrow |
| BrC | Brain cortex |
| BrS | Brainstem |
| Col | Colon |
| Dia | Diaphragm |
| Ret | Eye retina |
| Fat | Fat |
| Hrt | Heart |
| Pit | Hypothalamus/Pituitary |
| Kid | Kidney |
| LM | Limb muscle |
| Liv | Liver |
| Lung | Lung |
| MG | Mammary gland |
| Panc | Pancreas |
| Skin | Skin |
| SI | Small intestine |
| Spl | Spleen |
| Test | Testes |
| Thym | Thymus |
| Tong | Tongue |
| Trac | Trachea |
| Ute | Uterus |

**Compartment**
**stromal**
- bone
- skeletal muscle
- cardiac muscle
- smooth muscle
- fibroblast
- adipocyte
- mesothelial
- other
- mix

12.5%
25%
50%
100%

ln(CP10K+1): 0, 1, 2, 3

**Gastrin-releasing peptide (GRP)**
- lymphoid

**PAIRING** (ligand → receptor)

GRP→GRPR

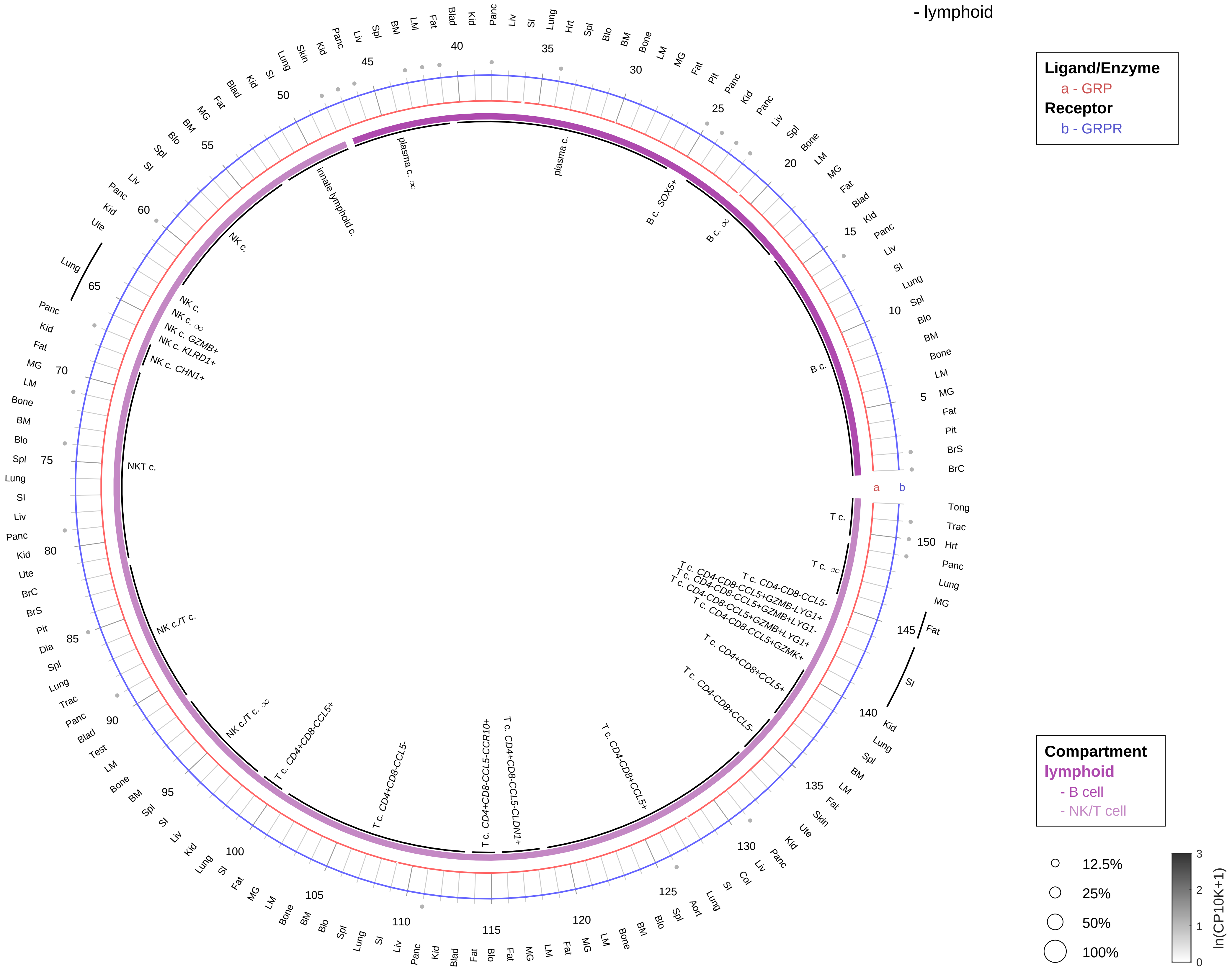

| Abbrev. | Tissue |
|---|---|
| Aort | Aorta |
| Blad | Bladder |
| Blo | Blood |
| Bone | Bone |
| BM | Bone marrow |
| BrC | Brain cortex |
| BrS | Brainstem |
| Col | Colon |
| Dia | Diaphragm |
| Ret | Eye retina |
| Fat | Fat |
| Hrt | Heart |
| Pit | Hypothalamus/Pituitary |
| Kid | Kidney |
| LM | Limb muscle |
| Liv | Liver |
| Lung | Lung |
| MG | Mammary gland |
| Panc | Pancreas |
| Skin | Skin |
| SI | Small intestine |
| Spl | Spleen |
| Test | Testes |
| Thym | Thymus |
| Tong | Tongue |
| Trac | Trachea |
| Ute | Uterus |

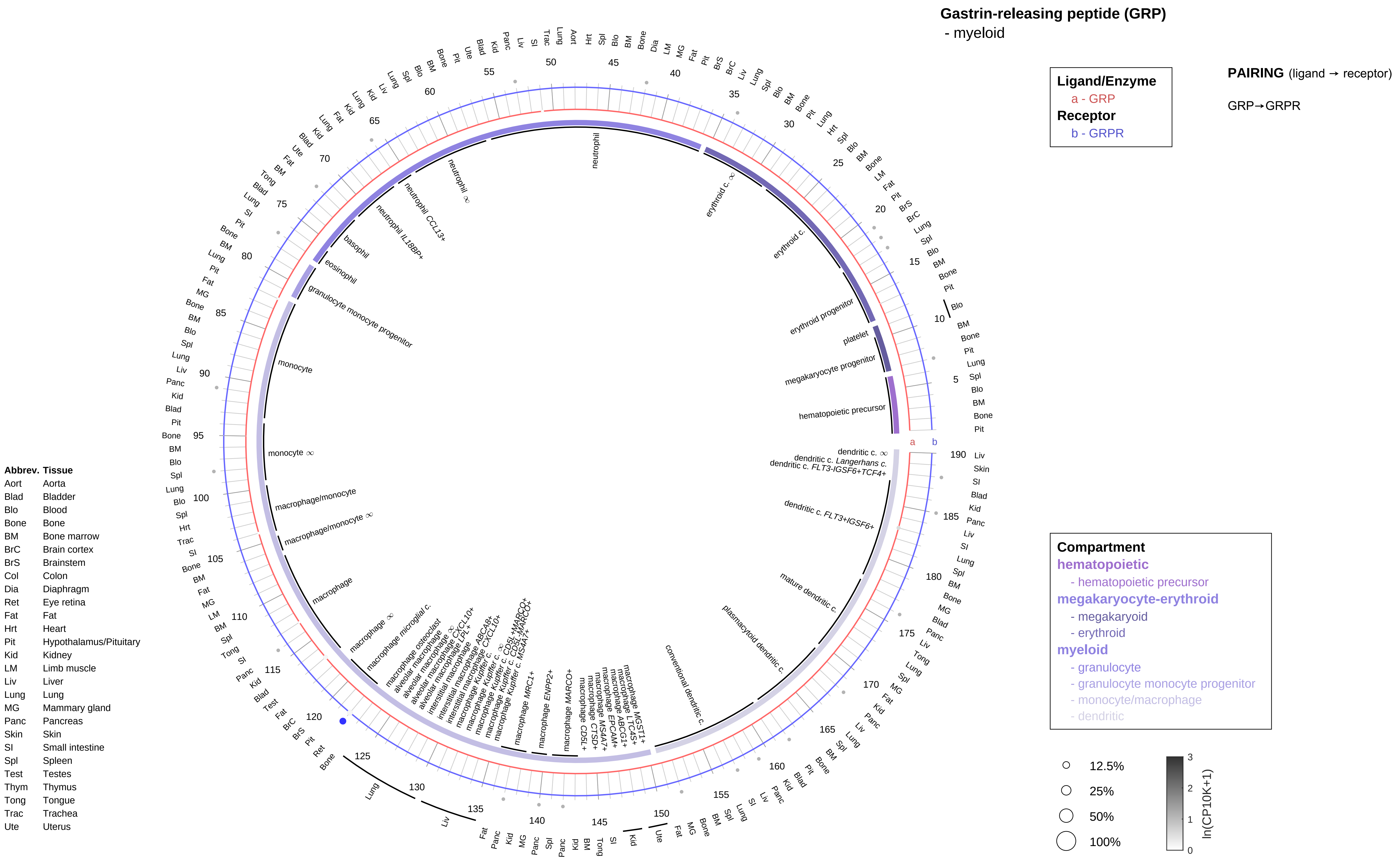

Gastrin-releasing peptide (GRP)
- myeloid
Ligand/Enzyme
a - GRP
Receptor
b - GRPR
PAIRING (ligand → receptor)
GRP→GRPR
Compartment
hematopoietic
- hematopoietic precursor
megakaryocyte-erythroid
- megakaryoid
- erythroid
myeloid
- granulocyte
- granulocyte monocyte progenitor
- monocyte/macrophage
- dendritic
12.5%
25%
50%
100%
ln(CP10K+1)
Abbrev. Tissue
Aort Aorta
Blad Bladder
Blo Blood
Bone Bone
BM Bone marrow
BrC Brain cortex
BrS Brainstem
Col Colon
Dia Diaphragm
Ret Eye retina
Fat Fat
Hrt Heart
Pit Hypothalamus/Pituitary
Kid Kidney
LM Limb muscle
Liv Liver
Lung Lung
MG Mammary gland
Panc Pancreas
Skin Skin
SI Small intestine
Spl Spleen
Test Testes
Thym Thymus
Tong Tongue
Trac Trachea
Ute Uterus
neutrophil
erythroid c. ∞
erythroid c.
erythroid progenitor
platelet
megakaryocyte progenitor
hematopoietic precursor
dendritic c. ∞
dendritic c. Langerhans c.
dendritic c. FLT3-IGSF6+TCF4+
dendritic c. FLT3+IGSF6+
mature dendritic c.
plasmacytoid dendritic c.
conventional dendritic c.
neutrophil ∞
neutrophil CCL13+
neutrophil IL18BP+
basophil
eosinophil
granulocyte monocyte progenitor
monocyte
monocyte ∞
macrophage/monocyte
macrophage/monocyte ∞
macrophage
macrophage ∞
macrophage microglial c.
macrophage osteoclast
macrophage MRC1+
macrophage ENPP2+
macrophage MARCO+

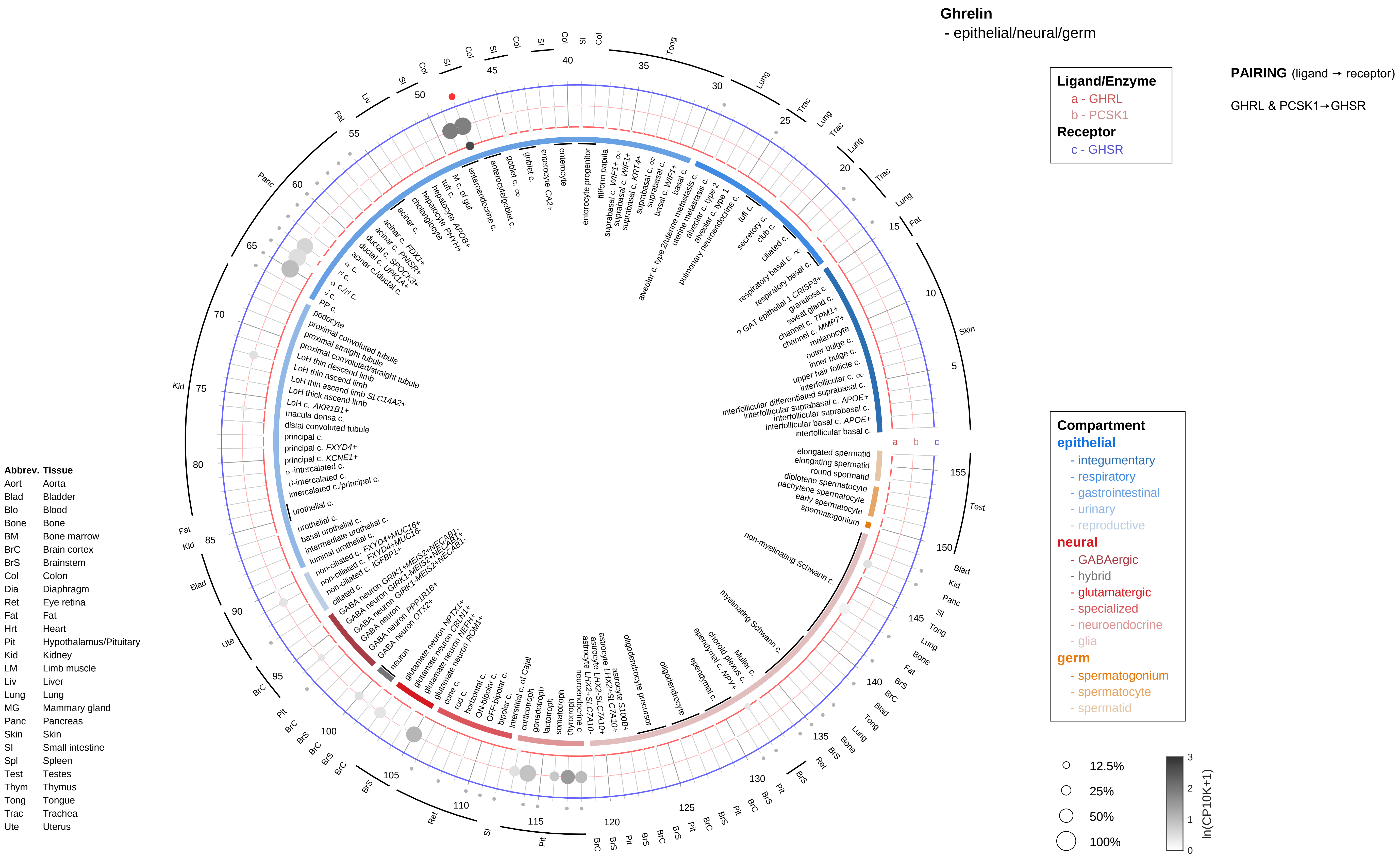
Ghrelin
- epithelial/neural/germ
Ligand/Enzyme
a - GHRL
b - PCSK1
Receptor
c - GHSR
PAIRING (ligand → receptor)
GHRL & PCSK1→GHSR
Compartment
epithelial
- integumentary
- respiratory
- gastrointestinal
- urinary
- reproductive
neural
- GABAergic
- hybrid
- glutamatergic
- specialized
- neuroendocrine
- glia
germ
- spermatogonium
- spermatocyte
- spermatid
12.5%
25%
50%
100%
ln(CP10K+1)
Abbrev. Tissue
Aort Aorta
Blad Bladder
Blo Blood
Bone Bone
BM Bone marrow
BrC Brain cortex
BrS Brainstem
Col Colon
Dia Diaphragm
Ret Eye retina
Fat Fat
Hrt Heart
Pit Hypothalamus/Pituitary
Kid Kidney
LM Limb muscle
Liv Liver
Lung Lung
MG Mammary gland
Panc Pancreas
Skin Skin
SI Small intestine
Spl Spleen
Test Testes
Thym Thymus
Tong Tongue
Trac Trachea
Ute Uterus

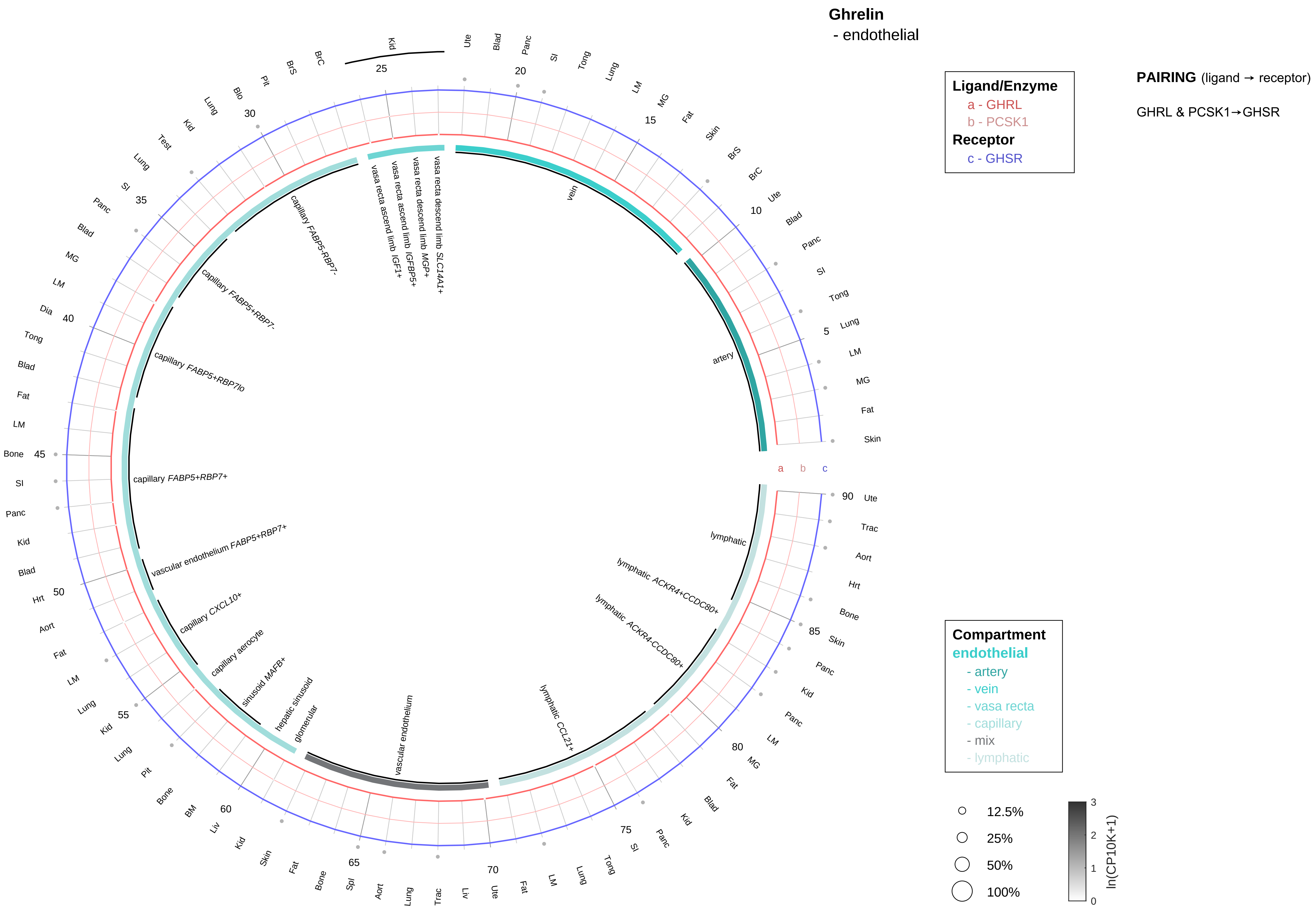

| Abbrev. | Tissue |
|---|---|
| Aort | Aorta |
| Blad | Bladder |
| Blo | Blood |
| Bone | Bone |
| BM | Bone marrow |
| BrC | Brain cortex |
| BrS | Brainstem |
| Col | Colon |
| Dia | Diaphragm |
| Ret | Eye retina |
| Fat | Fat |
| Hrt | Heart |
| Pit | Hypothalamus/Pituitary |
| Kid | Kidney |
| LM | Limb muscle |
| Liv | Liver |
| Lung | Lung |
| MG | Mammary gland |
| Panc | Pancreas |
| Skin | Skin |
| SI | Small intestine |
| Spl | Spleen |
| Test | Testes |
| Thym | Thymus |
| Tong | Tongue |
| Trac | Trachea |
| Ute | Uterus |

**Ghrelin**
- stromal

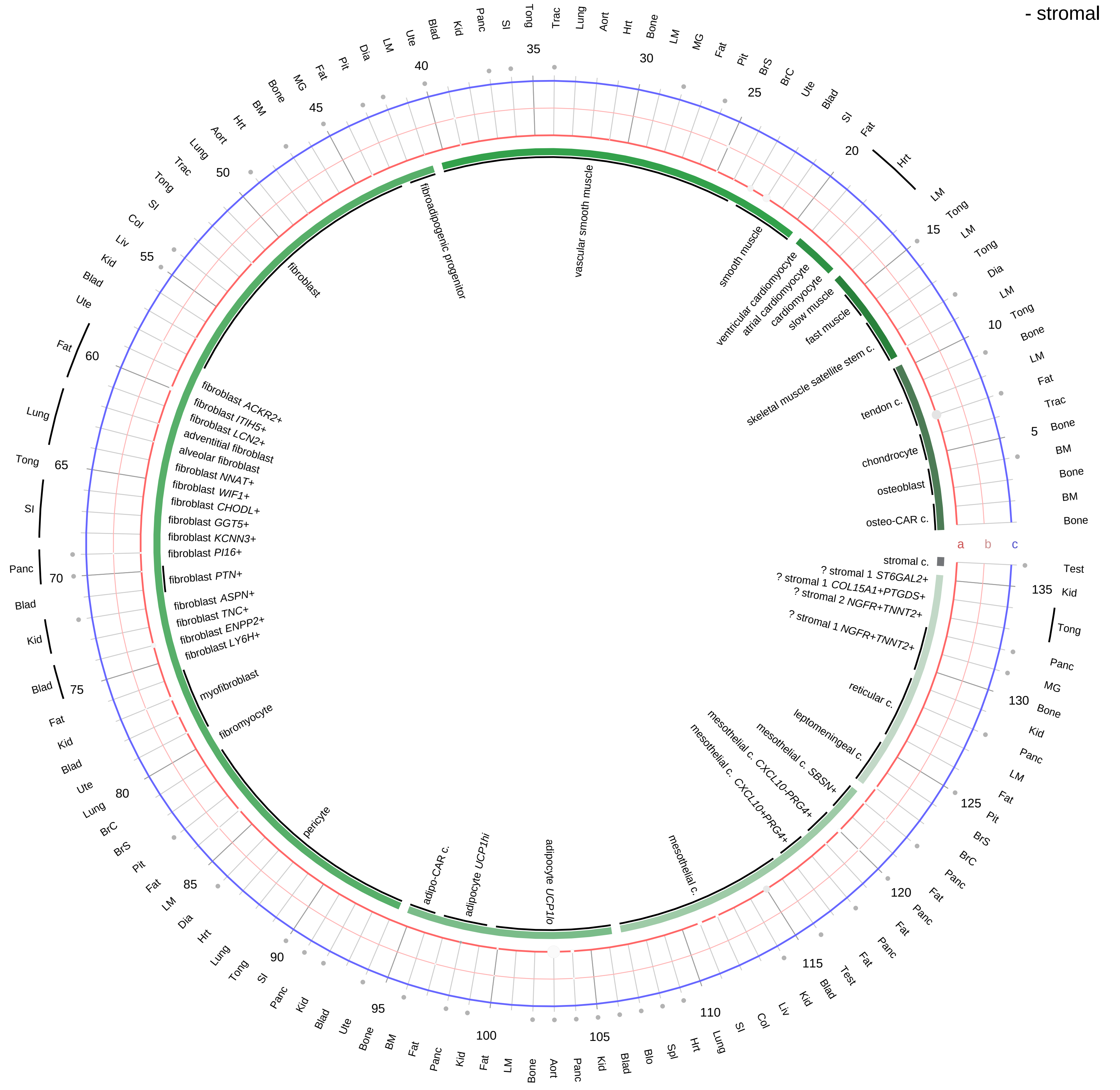

**Ligand/Enzyme**
- a - GHRL
- b - PCSK1

**Receptor**
- c - GHSR

**PAIRING** (ligand → receptor)

GHRL & PCSK1→GHSR

**Compartment**

**stromal**
- bone
- skeletal muscle
- cardiac muscle
- smooth muscle
- fibroblast
- adipocyte
- mesothelial
- other
- mix

12.5%
25%
50%
100%

ln(CP10K+1): 0, 1, 2, 3

| Abbrev. | Tissue |
|---|---|
| Aort | Aorta |
| Blad | Bladder |
| Blo | Blood |
| Bone | Bone |
| BM | Bone marrow |
| BrC | Brain cortex |
| BrS | Brainstem |
| Col | Colon |
| Dia | Diaphragm |
| Ret | Eye retina |
| Fat | Fat |
| Hrt | Heart |
| Pit | Hypothalamus/Pituitary |
| Kid | Kidney |
| LM | Limb muscle |
| Liv | Liver |
| Lung | Lung |
| MG | Mammary gland |
| Panc | Pancreas |
| Skin | Skin |
| SI | Small intestine |
| Spl | Spleen |
| Test | Testes |
| Thym | Thymus |
| Tong | Tongue |
| Trac | Trachea |
| Ute | Uterus |

**Ghrelin**
- lymphoid

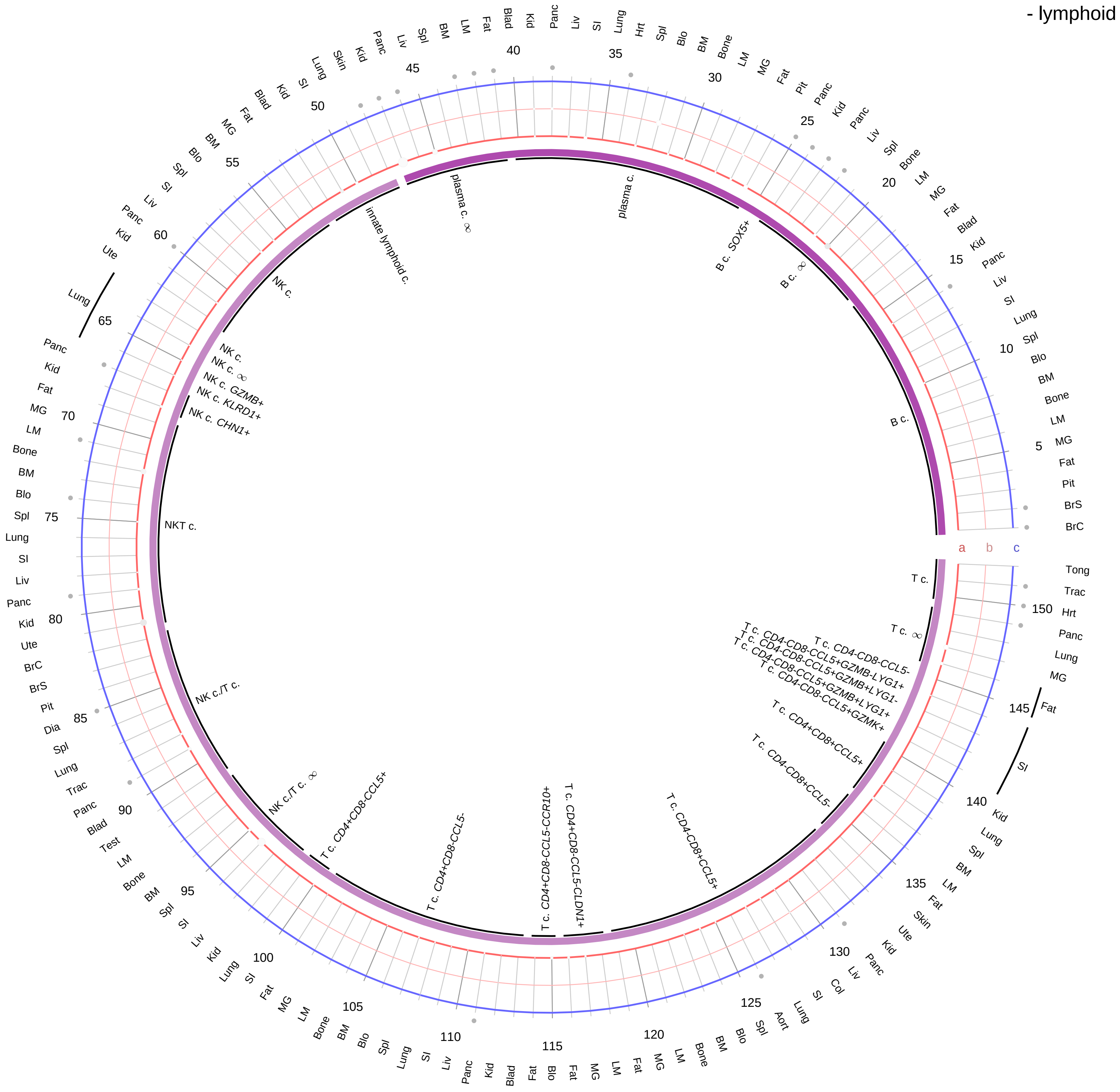

**Ligand/Enzyme**
a - GHRL
b - PCSK1
**Receptor**
c - GHSR

**PAIRING** (ligand → receptor)

GHRL & PCSK1→GHSR

**Compartment**
**lymphoid**
- B cell
- NK/T cell

12.5%
25%
50%
100%

ln(CP10K+1): 0, 1, 2, 3

| Abbrev. | Tissue |
|---|---|
| Aort | Aorta |
| Blad | Bladder |
| Blo | Blood |
| Bone | Bone |
| BM | Bone marrow |
| BrC | Brain cortex |
| BrS | Brainstem |
| Col | Colon |
| Dia | Diaphragm |
| Ret | Eye retina |
| Fat | Fat |
| Hrt | Heart |
| Pit | Hypothalamus/Pituitary |
| Kid | Kidney |
| LM | Limb muscle |
| Liv | Liver |
| Lung | Lung |
| MG | Mammary gland |
| Panc | Pancreas |
| Skin | Skin |
| SI | Small intestine |
| Spl | Spleen |
| Test | Testes |
| Thym | Thymus |
| Tong | Tongue |
| Trac | Trachea |
| Ute | Uterus |

**Ghrelin**
- myeloid

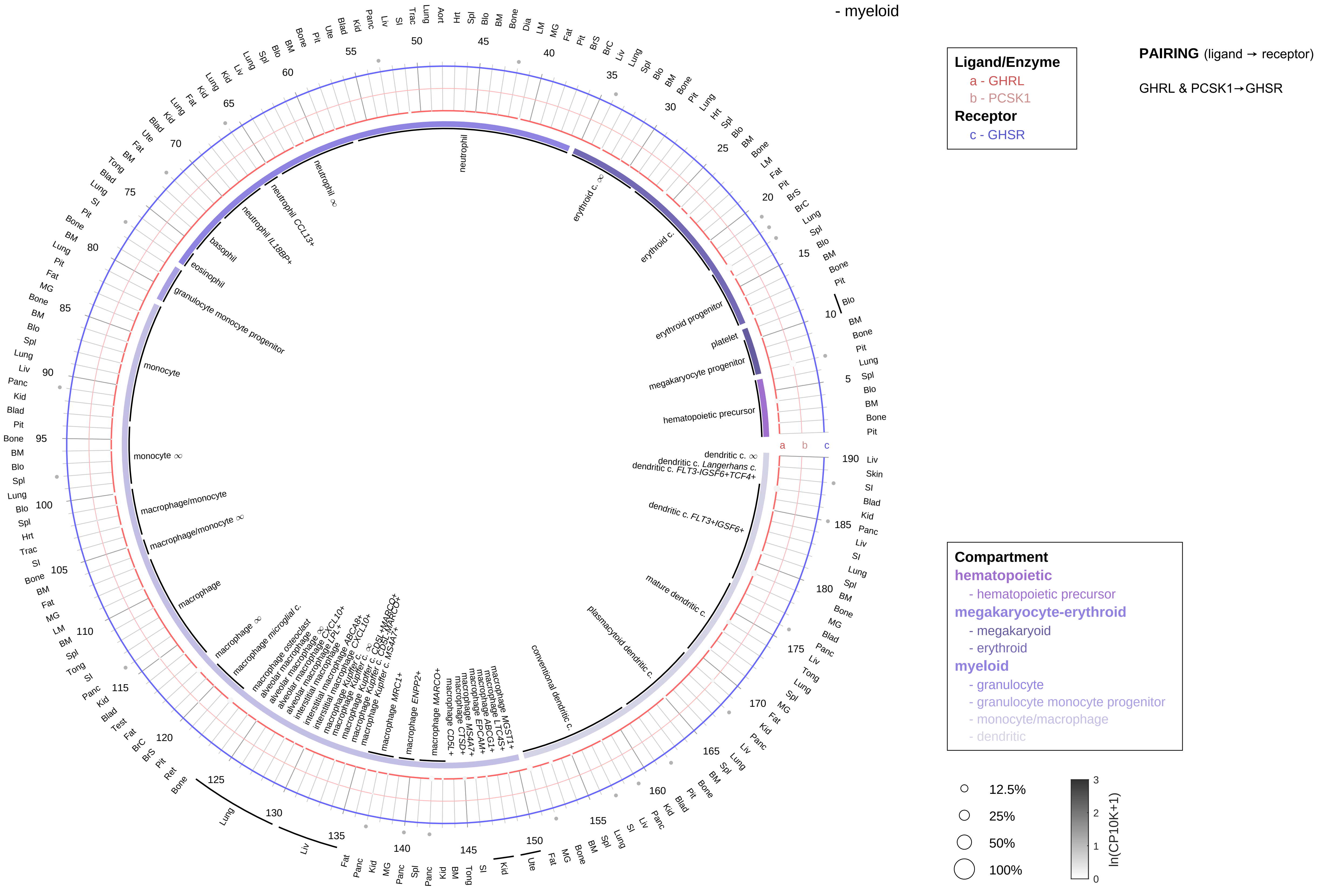

| Abbrev. | Tissue |
|---|---|
| Aort | Aorta |
| Blad | Bladder |
| Blo | Blood |
| Bone | Bone |
| BM | Bone marrow |
| BrC | Brain cortex |
| BrS | Brainstem |
| Col | Colon |
| Dia | Diaphragm |
| Ret | Eye retina |
| Fat | Fat |
| Hrt | Heart |
| Pit | Hypothalamus/Pituitary |
| Kid | Kidney |
| LM | Limb muscle |
| Liv | Liver |
| Lung | Lung |
| MG | Mammary gland |
| Panc | Pancreas |
| Skin | Skin |
| SI | Small intestine |
| Spl | Spleen |
| Test | Testes |
| Thym | Thymus |
| Tong | Tongue |
| Trac | Trachea |
| Ute | Uterus |

**Glucagon/GLP**
- epithelial/neural/germ

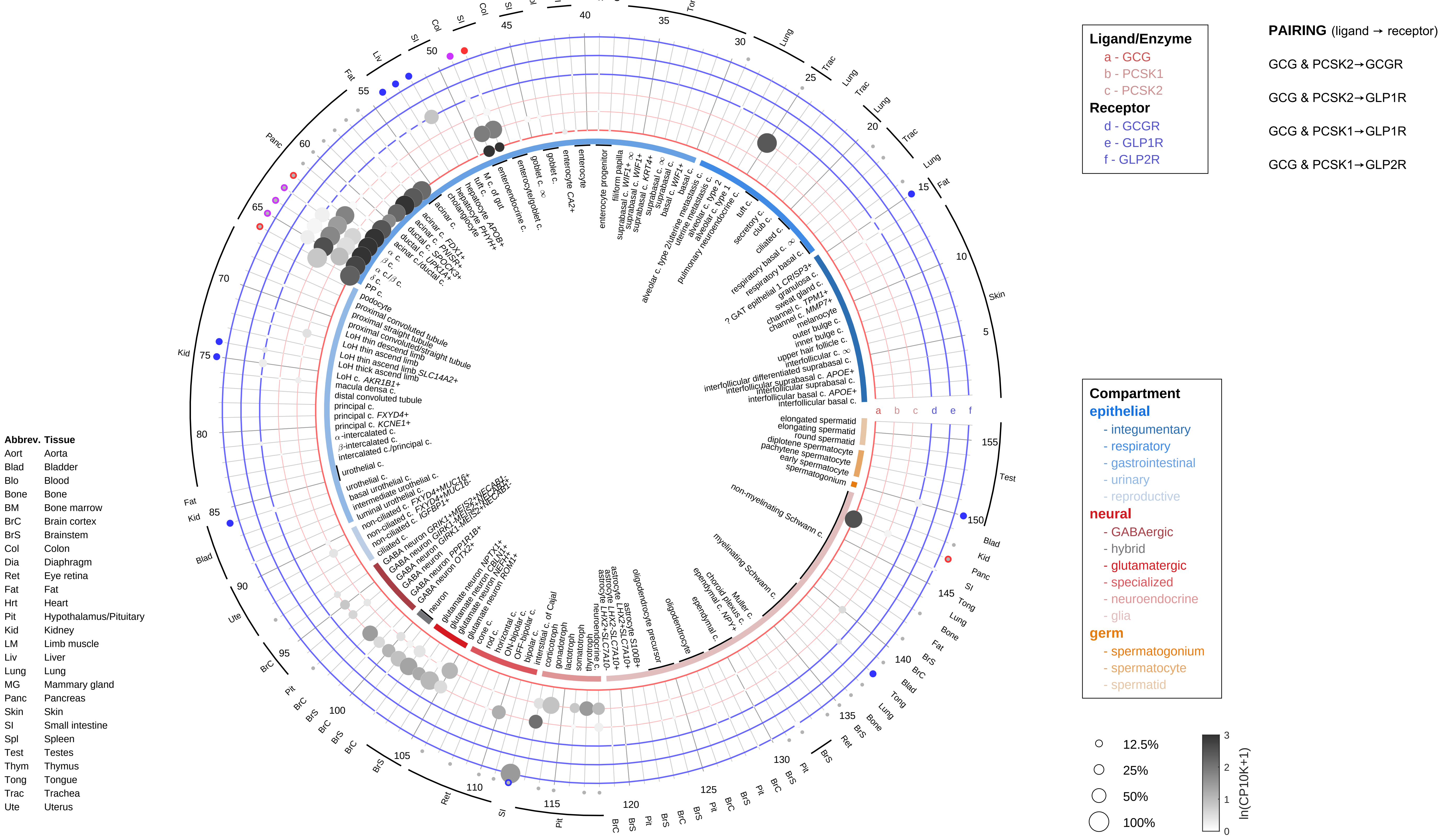

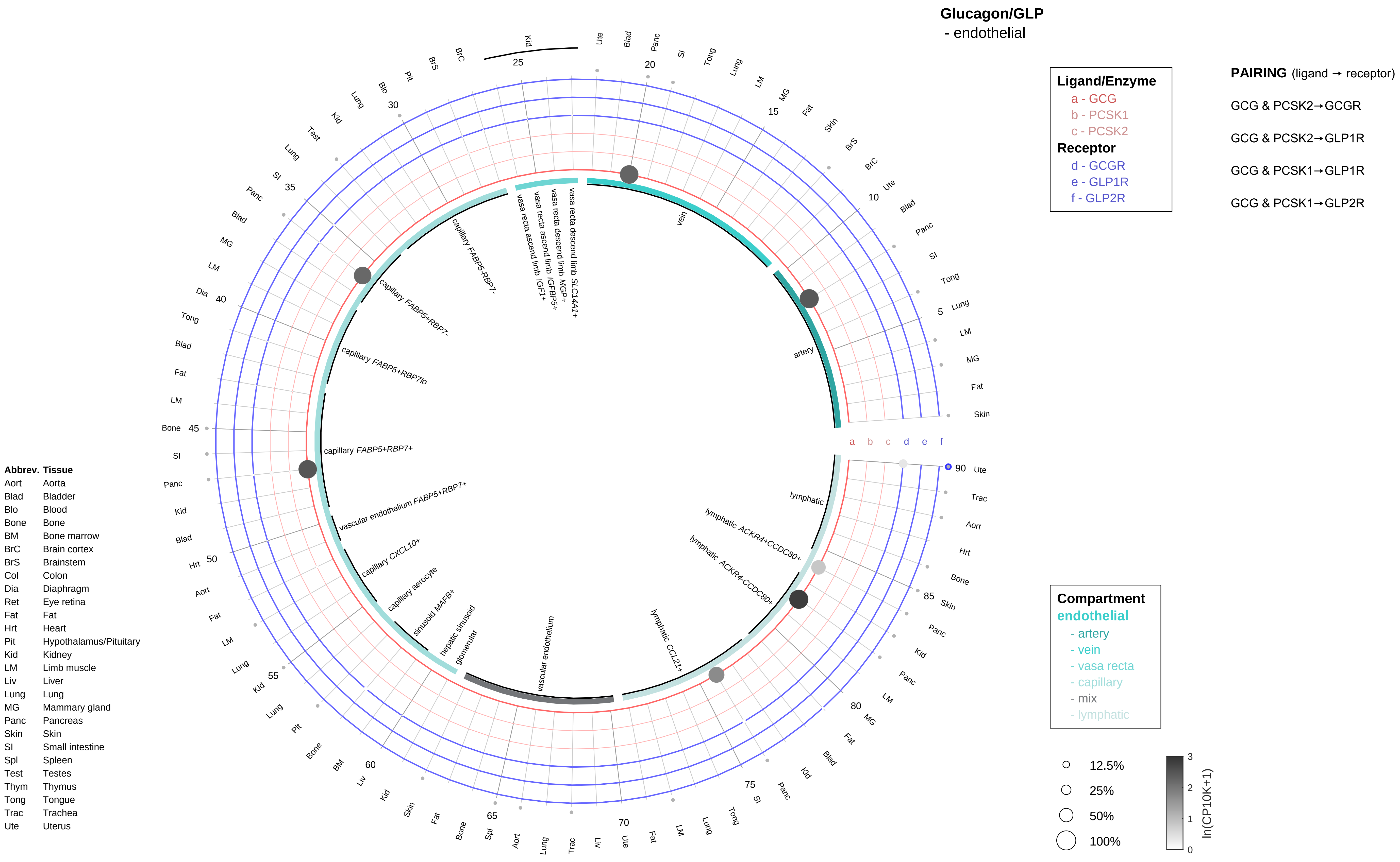

Glucagon/GLP
- endothelial
Ligand/Enzyme
a - GCG
b - PCSK1
c - PCSK2
Receptor
d - GCGR
e - GLP1R
f - GLP2R
PAIRING (ligand → receptor)
GCG & PCSK2→GCGR
GCG & PCSK2→GLP1R
GCG & PCSK1→GLP1R
GCG & PCSK1→GLP2R
vein
artery
vasa recta descend limb SLC14A1+
vasa recta descend limb MGP+
vasa recta ascend limb IGFBP5+
vasa recta ascend limb IGF1+
capillary FABP5-RBP7-
capillary FABP5+RBP7-
capillary FABP5+RBP7lo
capillary FABP5+RBP7+
vascular endothelium FABP5+RBP7+
capillary CXCL10+
capillary aerocyte
sinusoid MAFB+
hepatic sinusoid
glomerular
vascular endothelium
lymphatic CCL21+
lymphatic ACKR4-CCDC80+
lymphatic ACKR4+CCDC80+
lymphatic
Compartment
endothelial
- artery
- vein
- vasa recta
- capillary
- mix
- lymphatic
12.5%
25%
50%
100%
ln(CP10K+1)
Abbrev. Tissue
Aort Aorta
Blad Bladder
Blo Blood
Bone Bone
BM Bone marrow
BrC Brain cortex
BrS Brainstem
Col Colon
Dia Diaphragm
Ret Eye retina
Fat Fat
Hrt Heart
Pit Hypothalamus/Pituitary
Kid Kidney
LM Limb muscle
Liv Liver
Lung Lung
MG Mammary gland
Panc Pancreas
Skin Skin
SI Small intestine
Spl Spleen
Test Testes
Thym Thymus
Tong Tongue
Trac Trachea
Ute Uterus

**Glucagon/GLP**
- stromal

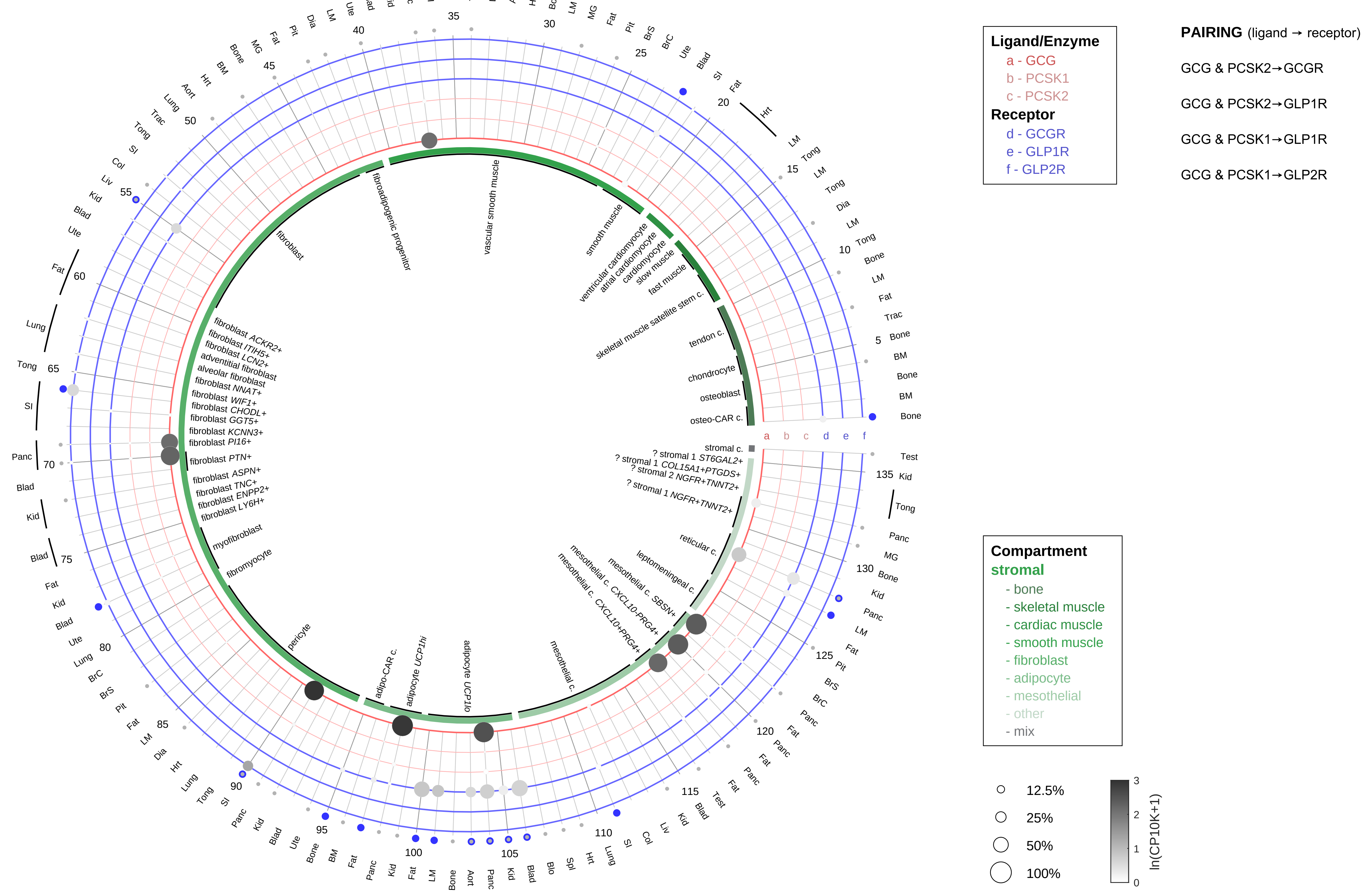

| Abbrev. | Tissue |
|---|---|
| Aort | Aorta |
| Blad | Bladder |
| Blo | Blood |
| Bone | Bone |
| BM | Bone marrow |
| BrC | Brain cortex |
| BrS | Brainstem |
| Col | Colon |
| Dia | Diaphragm |
| Ret | Eye retina |
| Fat | Fat |
| Hrt | Heart |
| Pit | Hypothalamus/Pituitary |
| Kid | Kidney |
| LM | Limb muscle |
| Liv | Liver |
| Lung | Lung |
| MG | Mammary gland |
| Panc | Pancreas |
| Skin | Skin |
| SI | Small intestine |
| Spl | Spleen |
| Test | Testes |
| Thym | Thymus |
| Tong | Tongue |
| Trac | Trachea |
| Ute | Uterus |

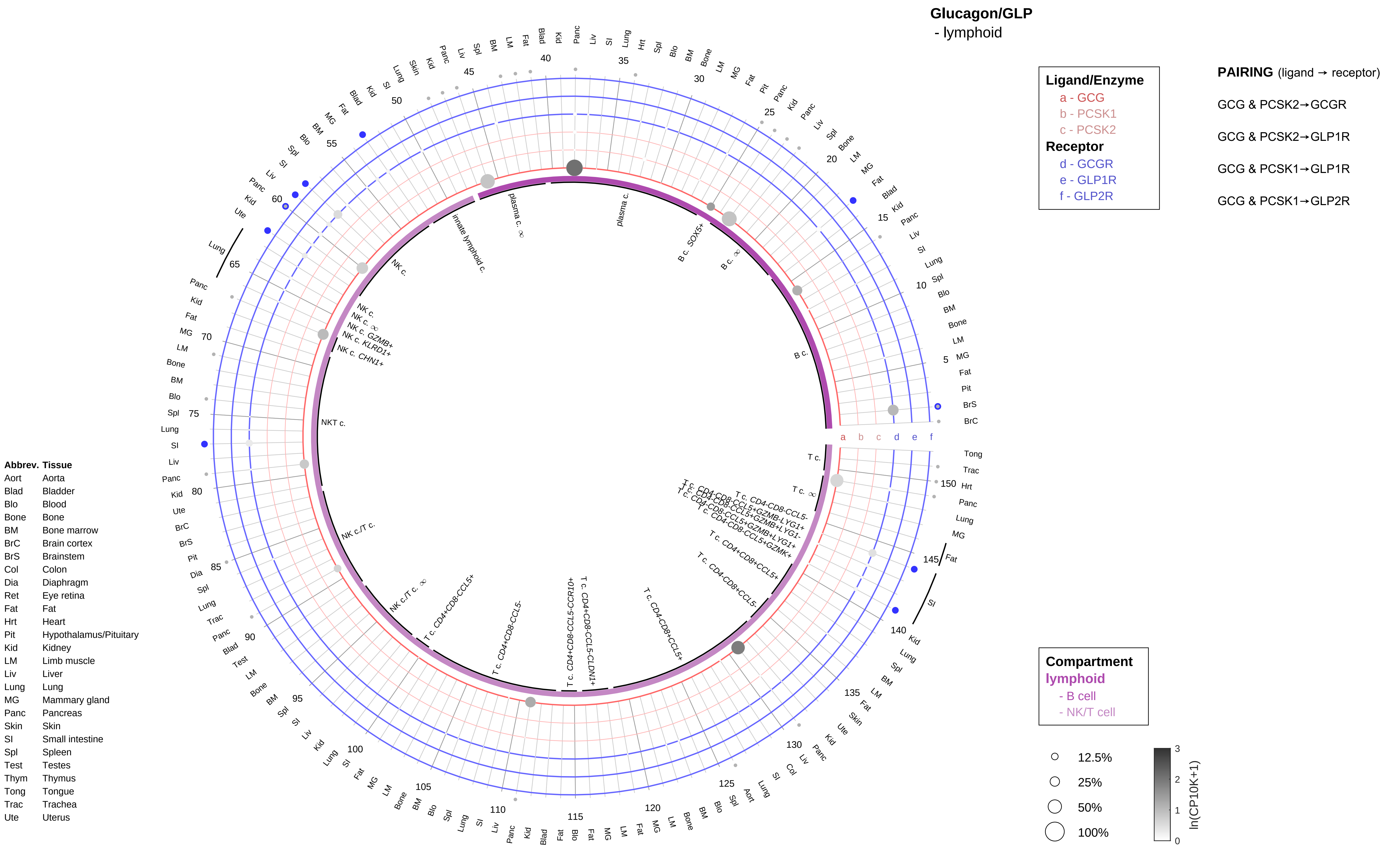

Glucagon/GLP
- lymphoid
Ligand/Enzyme
a - GCG
b - PCSK1
c - PCSK2
Receptor
d - GCGR
e - GLP1R
f - GLP2R
PAIRING (ligand → receptor)
GCG & PCSK2→GCGR
GCG & PCSK2→GLP1R
GCG & PCSK1→GLP1R
GCG & PCSK1→GLP2R
Compartment
lymphoid
- B cell
- NK/T cell
12.5%
25%
50%
100%
ln(CP10K+1)
Abbrev. Tissue
Aort Aorta
Blad Bladder
Blo Blood
Bone Bone
BM Bone marrow
BrC Brain cortex
BrS Brainstem
Col Colon
Dia Diaphragm
Ret Eye retina
Fat Fat
Hrt Heart
Pit Hypothalamus/Pituitary
Kid Kidney
LM Limb muscle
Liv Liver
Lung Lung
MG Mammary gland
Panc Pancreas
Skin Skin
SI Small intestine
Spl Spleen
Test Testes
Thym Thymus
Tong Tongue
Trac Trachea
Ute Uterus
plasma c.
B c. SOX5+
B c.
NKT c.
NK c./T c.
T c.
innate lymphoid c.
NK c.

**Glucagon/GLP**
- myeloid

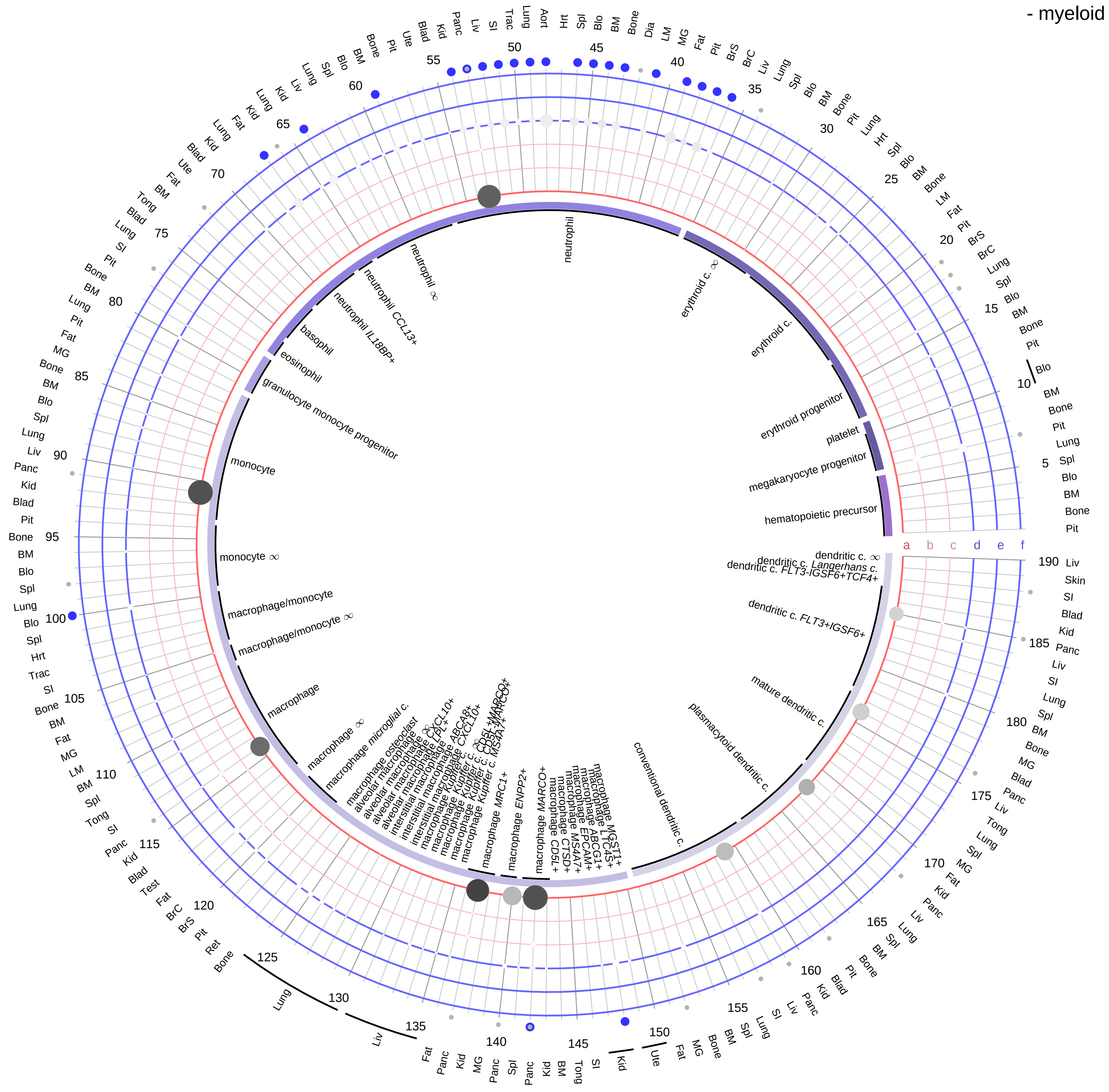

| Ligand/Enzyme |
|---|
| a - GCG |
| b - PCSK1 |
| c - PCSK2 |
| **Receptor** |
| d - GCGR |
| e - GLP1R |
| f - GLP2R |

**PAIRING** (ligand → receptor)

GCG & PCSK2→GCGR

GCG & PCSK2→GLP1R

GCG & PCSK1→GLP1R

GCG & PCSK1→GLP2R

| Abbrev. | Tissue |
|---|---|
| Aort | Aorta |
| Blad | Bladder |
| Blo | Blood |
| Bone | Bone |
| BM | Bone marrow |
| BrC | Brain cortex |
| BrS | Brainstem |
| Col | Colon |
| Dia | Diaphragm |
| Ret | Eye retina |
| Fat | Fat |
| Hrt | Heart |
| Pit | Hypothalamus/Pituitary |
| Kid | Kidney |
| LM | Limb muscle |
| Liv | Liver |
| Lung | Lung |
| MG | Mammary gland |
| Panc | Pancreas |
| Skin | Skin |
| SI | Small intestine |
| Spl | Spleen |
| Test | Testes |
| Thym | Thymus |
| Tong | Tongue |
| Trac | Trachea |
| Ute | Uterus |

**Compartment**
**hematopoietic**
- hematopoietic precursor
**megakaryocyte-erythroid**
- megakaryoid
- erythroid
**myeloid**
- granulocyte
- granulocyte monocyte progenitor
- monocyte/macrophage
- dendritic

12.5%
25%
50%
100%

ln(CP10K+1): 0, 1, 2, 3

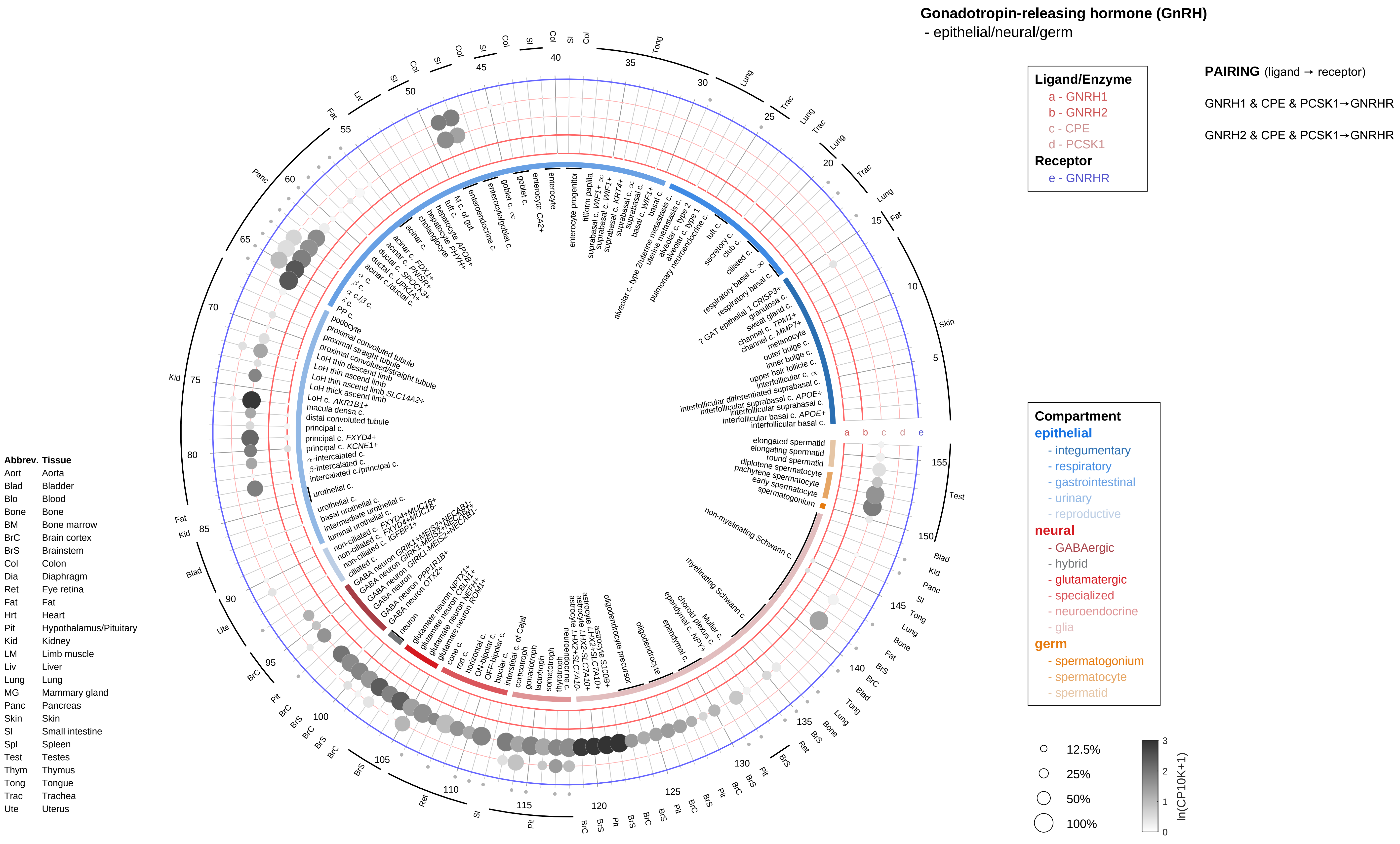

Gonadotropin-releasing hormone (GnRH)
- epithelial/neural/germ
Ligand/Enzyme
a - GNRH1
b - GNRH2
c - CPE
d - PCSK1
Receptor
e - GNRHR
PAIRING (ligand → receptor)
GNRH1 & CPE & PCSK1→GNRHR
GNRH2 & CPE & PCSK1→GNRHR
Compartment
epithelial
- integumentary
- respiratory
- gastrointestinal
- urinary
- reproductive
neural
- GABAergic
- hybrid
- glutamatergic
- specialized
- neuroendocrine
- glia
germ
- spermatogonium
- spermatocyte
- spermatid
12.5%
25%
50%
100%
ln(CP10K+1)
Abbrev. Tissue
Aort Aorta
Blad Bladder
Blo Blood
Bone Bone
BM Bone marrow
BrC Brain cortex
BrS Brainstem
Col Colon
Dia Diaphragm
Ret Eye retina
Fat Fat
Hrt Heart
Pit Hypothalamus/Pituitary
Kid Kidney
LM Limb muscle
Liv Liver
Lung Lung
MG Mammary gland
Panc Pancreas
Skin Skin
SI Small intestine
Spl Spleen
Test Testes
Thym Thymus
Tong Tongue
Trac Trachea
Ute Uterus

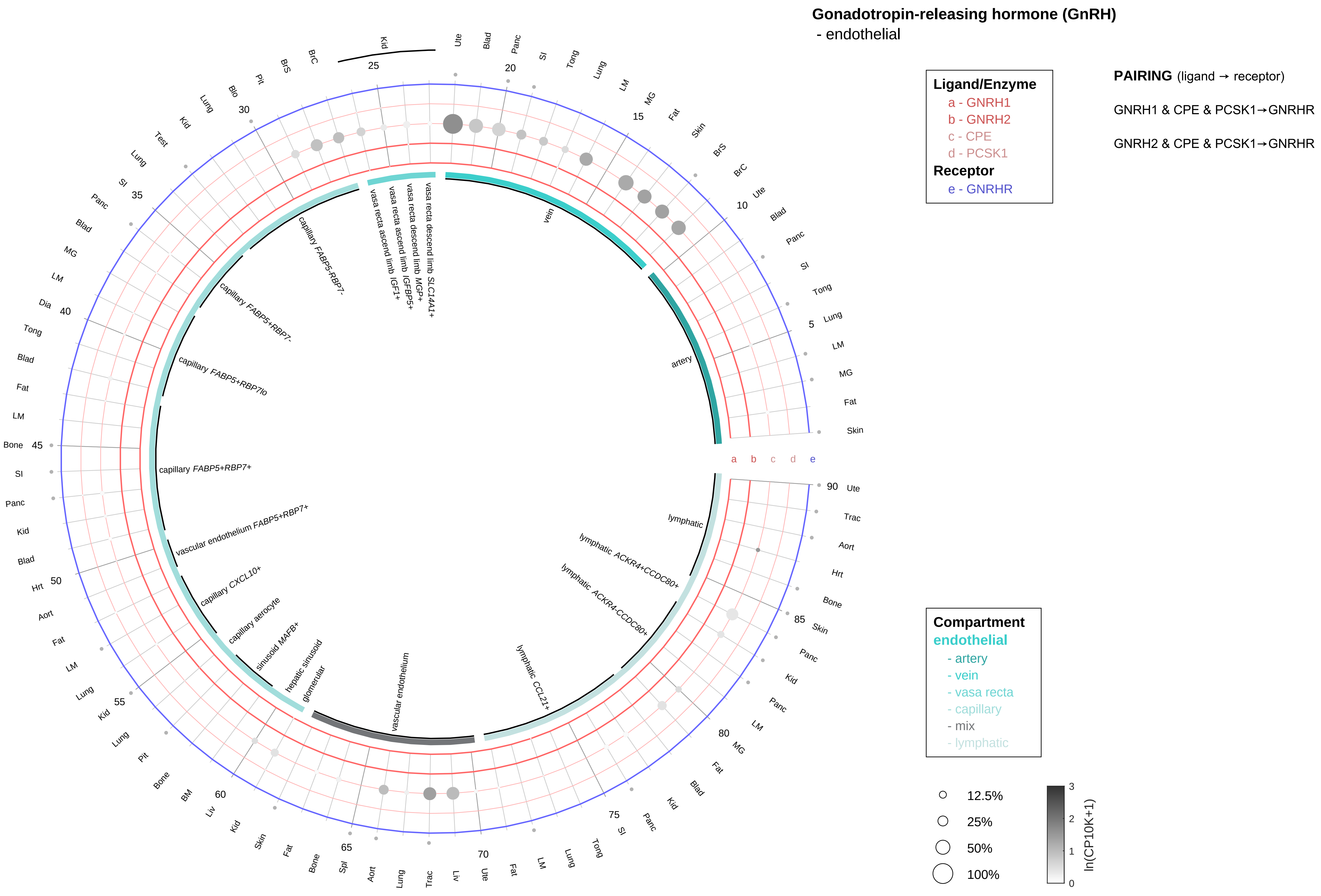

| Abbrev. | Tissue |
|---|---|
| Aort | Aorta |
| Blad | Bladder |
| Blo | Blood |
| Bone | Bone |
| BM | Bone marrow |
| BrC | Brain cortex |
| BrS | Brainstem |
| Col | Colon |
| Dia | Diaphragm |
| Ret | Eye retina |
| Fat | Fat |
| Hrt | Heart |
| Pit | Hypothalamus/Pituitary |
| Kid | Kidney |
| LM | Limb muscle |
| Liv | Liver |
| Lung | Lung |
| MG | Mammary gland |
| Panc | Pancreas |
| Skin | Skin |
| SI | Small intestine |
| Spl | Spleen |
| Test | Testes |
| Thym | Thymus |
| Tong | Tongue |
| Trac | Trachea |
| Ute | Uterus |

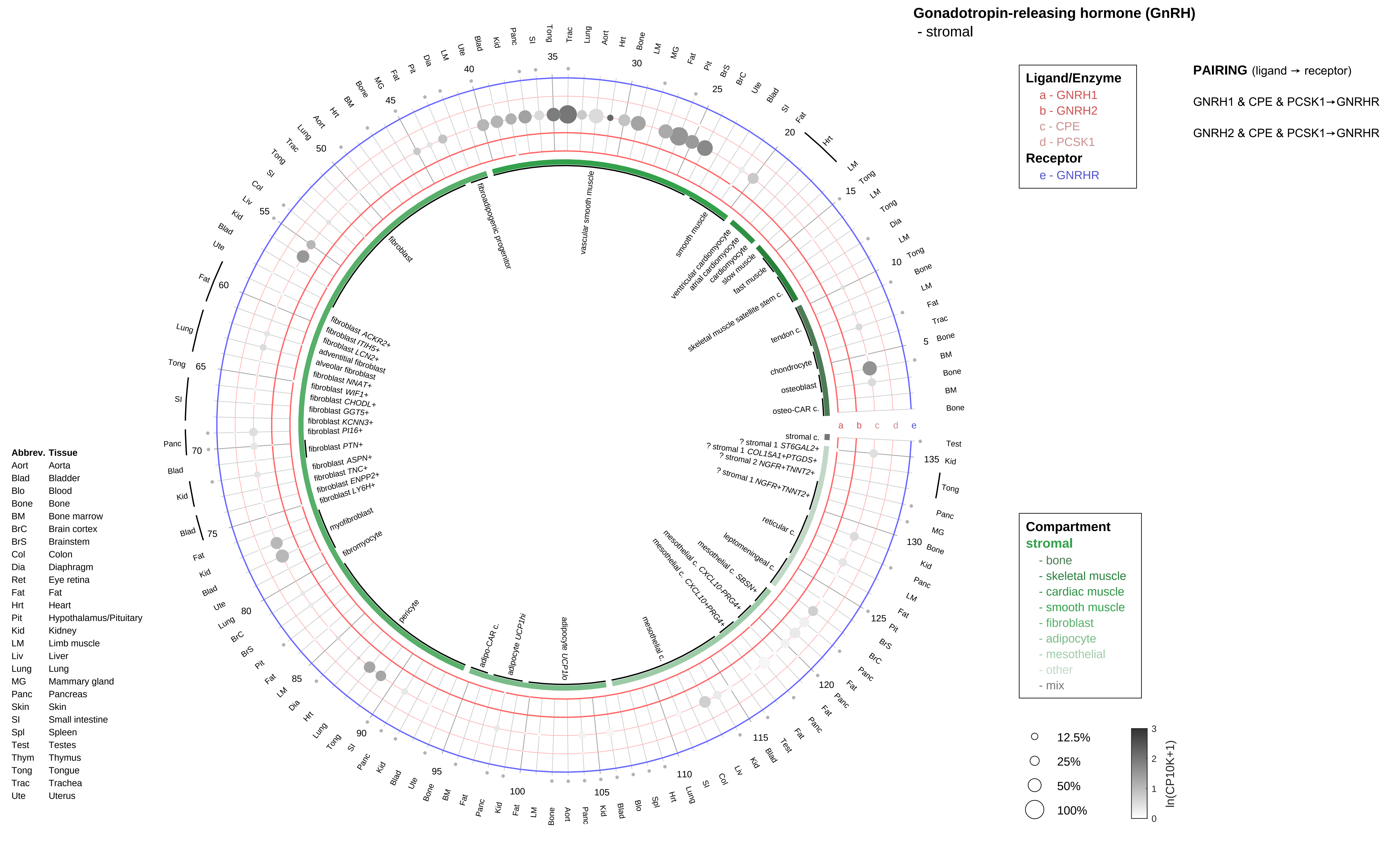
Gonadotropin-releasing hormone (GnRH)
- stromal
Ligand/Enzyme
a - GNRH1
b - GNRH2
c - CPE
d - PCSK1
Receptor
e - GNRHR
PAIRING (ligand → receptor)
GNRH1 & CPE & PCSK1→GNRHR
GNRH2 & CPE & PCSK1→GNRHR
Compartment
stromal
- bone
- skeletal muscle
- cardiac muscle
- smooth muscle
- fibroblast
- adipocyte
- mesothelial
- other
- mix
12.5%
25%
50%
100%
ln(CP10K+1)
Abbrev. Tissue
Aort Aorta
Blad Bladder
Blo Blood
Bone Bone
BM Bone marrow
BrC Brain cortex
BrS Brainstem
Col Colon
Dia Diaphragm
Ret Eye retina
Fat Fat
Hrt Heart
Pit Hypothalamus/Pituitary
Kid Kidney
LM Limb muscle
Liv Liver
Lung Lung
MG Mammary gland
Panc Pancreas
Skin Skin
SI Small intestine
Spl Spleen
Test Testes
Thym Thymus
Tong Tongue
Trac Trachea
Ute Uterus
fibroblast
fibroadipogenic progenitor
vascular smooth muscle
smooth muscle
ventricular cardiomyocyte
atrial cardiomyocyte
cardiomyocyte
slow muscle
fast muscle
skeletal muscle satellite stem c.
tendon c.
chondrocyte
osteoblast
osteo-CAR c.
stromal c.
? stromal 1 ST6GAL2+
? stromal 1 COL15A1+PTGDS+
? stromal 2 NGFR+TNNT2+
? stromal 1 NGFR+TNNT2+
reticular c.
leptomeningeal c.
mesothelial c. SBSN+
mesothelial c. CXCL10-PRG4+
mesothelial c. CXCL10+PRG4+
mesothelial c.
adipocyte UCP1lo
adipocyte UCP1hi
adipo-CAR c.
pericyte
fibromyocyte
myofibroblast
fibroblast LY6H+
fibroblast ENPP2+
fibroblast TNC+
fibroblast ASPN+
fibroblast PTN+
fibroblast PI16+
fibroblast KCNN3+
fibroblast GGT5+
fibroblast CHODL+
fibroblast WIF1+
fibroblast NNAT+
alveolar fibroblast
adventitial fibroblast
fibroblast LCN2+
fibroblast ITIH5+
fibroblast ACKR2+

**Gonadotropin-releasing hormone (GnRH)**
- lymphoid

**Ligand/Enzyme**
a - GNRH1
b - GNRH2
c - CPE
d - PCSK1
**Receptor**
e - GNRHR

**PAIRING** (ligand → receptor)

GNRH1 & CPE & PCSK1→GNRHR

GNRH2 & CPE & PCSK1→GNRHR

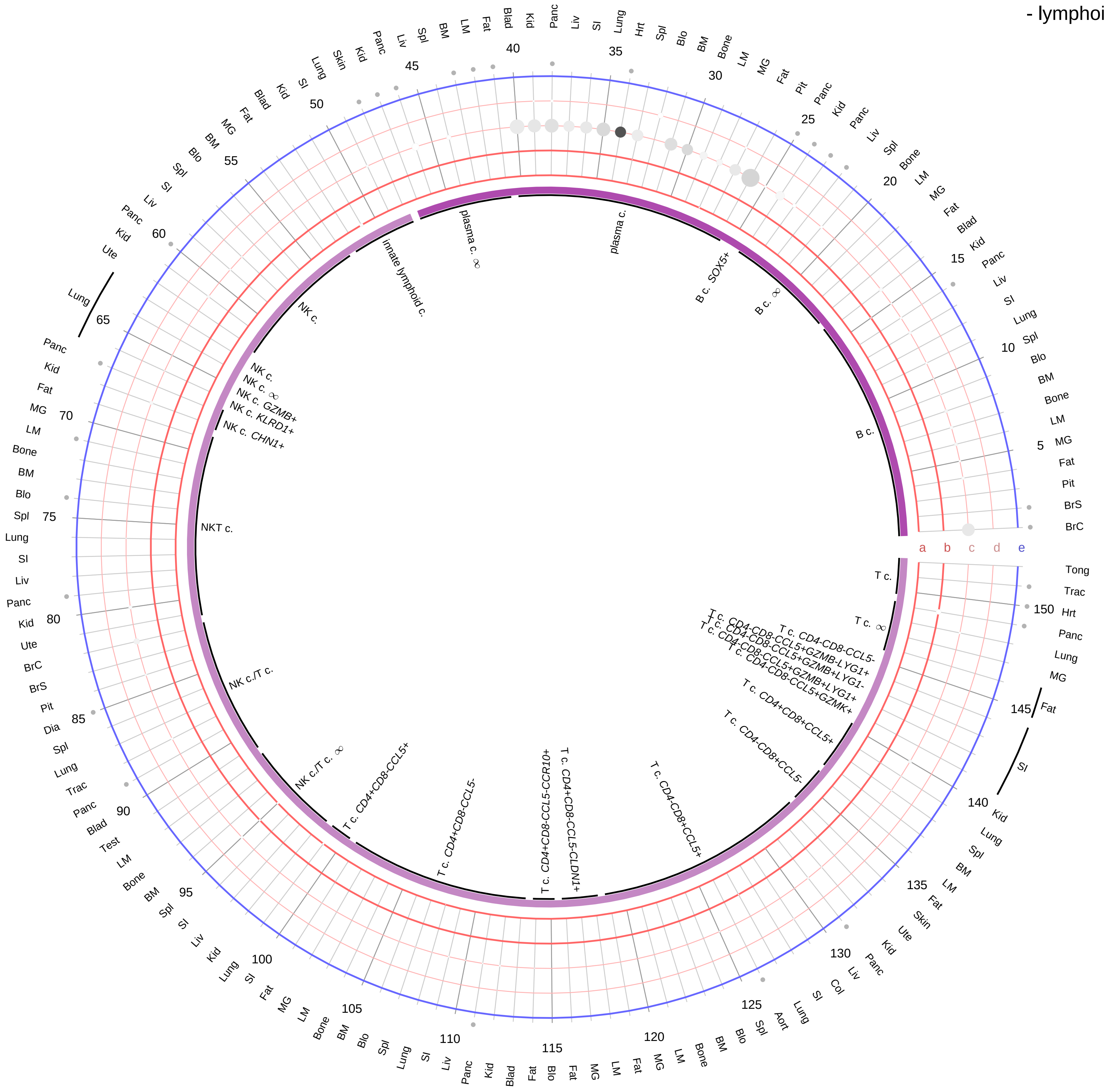

| Abbrev. | Tissue |
|---|---|
| Aort | Aorta |
| Blad | Bladder |
| Blo | Blood |
| Bone | Bone |
| BM | Bone marrow |
| BrC | Brain cortex |
| BrS | Brainstem |
| Col | Colon |
| Dia | Diaphragm |
| Ret | Eye retina |
| Fat | Fat |
| Hrt | Heart |
| Pit | Hypothalamus/Pituitary |
| Kid | Kidney |
| LM | Limb muscle |
| Liv | Liver |
| Lung | Lung |
| MG | Mammary gland |
| Panc | Pancreas |
| Skin | Skin |
| SI | Small intestine |
| Spl | Spleen |
| Test | Testes |
| Thym | Thymus |
| Tong | Tongue |
| Trac | Trachea |
| Ute | Uterus |

**Compartment**
**lymphoid**
- B cell
- NK/T cell

12.5%
25%
50%
100%

ln(CP10K+1)
0 1 2 3

**Gonadotropin-releasing hormone (GnRH)**
- myeloid

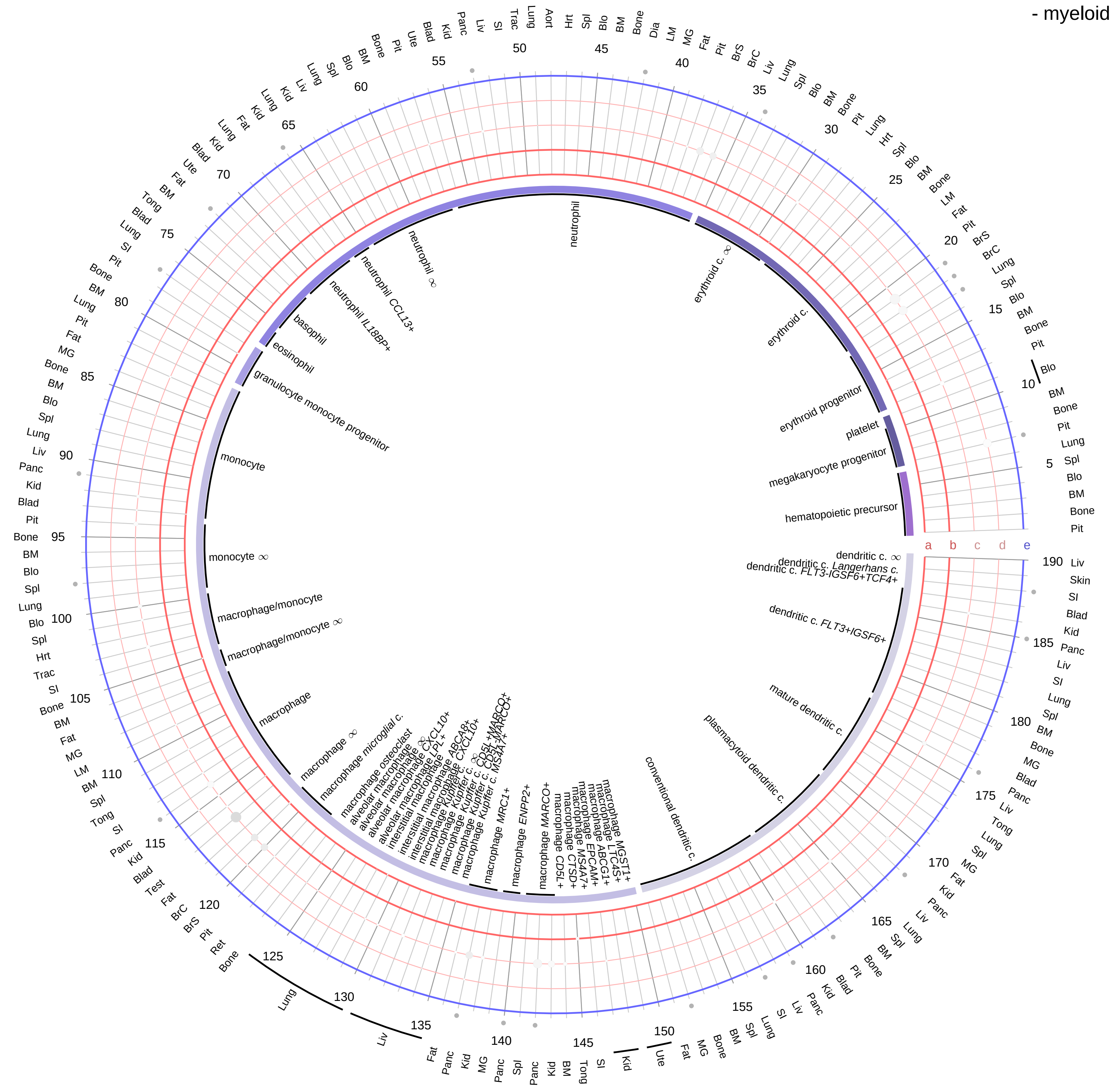

**Ligand/Enzyme**
- a - GNRH1
- b - GNRH2
- c - CPE
- d - PCSK1

**Receptor**
- e - GNRHR

**PAIRING** (ligand → receptor)

GNRH1 & CPE & PCSK1→GNRHR

GNRH2 & CPE & PCSK1→GNRHR

| Abbrev. | Tissue |
|---|---|
| Aort | Aorta |
| Blad | Bladder |
| Blo | Blood |
| Bone | Bone |
| BM | Bone marrow |
| BrC | Brain cortex |
| BrS | Brainstem |
| Col | Colon |
| Dia | Diaphragm |
| Ret | Eye retina |
| Fat | Fat |
| Hrt | Heart |
| Pit | Hypothalamus/Pituitary |
| Kid | Kidney |
| LM | Limb muscle |
| Liv | Liver |
| Lung | Lung |
| MG | Mammary gland |
| Panc | Pancreas |
| Skin | Skin |
| SI | Small intestine |
| Spl | Spleen |
| Test | Testes |
| Thym | Thymus |
| Tong | Tongue |
| Trac | Trachea |
| Ute | Uterus |

**Compartment**

**hematopoietic**
- hematopoietic precursor

**megakaryocyte-erythroid**
- megakaryoid
- erythroid

**myeloid**
- granulocyte
- granulocyte monocyte progenitor
- monocyte/macrophage
- dendritic

12.5%
25%
50%
100%

ln(CP10K+1): 0, 1, 2, 3

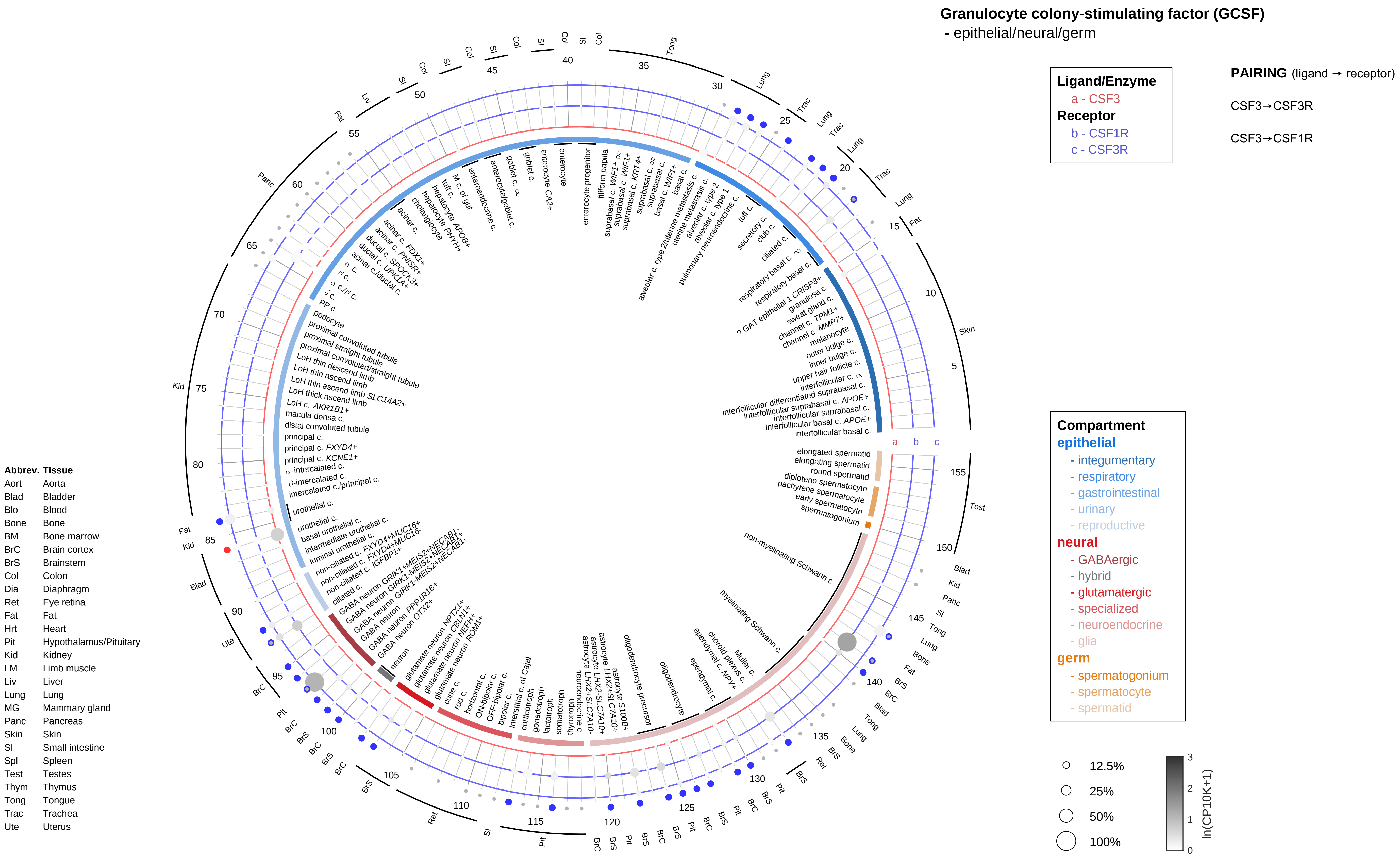

Granulocyte colony-stimulating factor (GCSF)
- epithelial/neural/germ
Ligand/Enzyme
a - CSF3
Receptor
b - CSF1R
c - CSF3R
PAIRING (ligand → receptor)
CSF3→CSF3R
CSF3→CSF1R
Compartment
epithelial
- integumentary
- respiratory
- gastrointestinal
- urinary
- reproductive
neural
- GABAergic
- hybrid
- glutamatergic
- specialized
- neuroendocrine
- glia
germ
- spermatogonium
- spermatocyte
- spermatid
12.5%
25%
50%
100%
ln(CP10K+1)
Abbrev. Tissue
Aort Aorta
Blad Bladder
Blo Blood
Bone Bone
BM Bone marrow
BrC Brain cortex
BrS Brainstem
Col Colon
Dia Diaphragm
Ret Eye retina
Fat Fat
Hrt Heart
Pit Hypothalamus/Pituitary
Kid Kidney
LM Limb muscle
Liv Liver
Lung Lung
MG Mammary gland
Panc Pancreas
Skin Skin
SI Small intestine
Spl Spleen
Test Testes
Thym Thymus
Tong Tongue
Trac Trachea
Ute Uterus

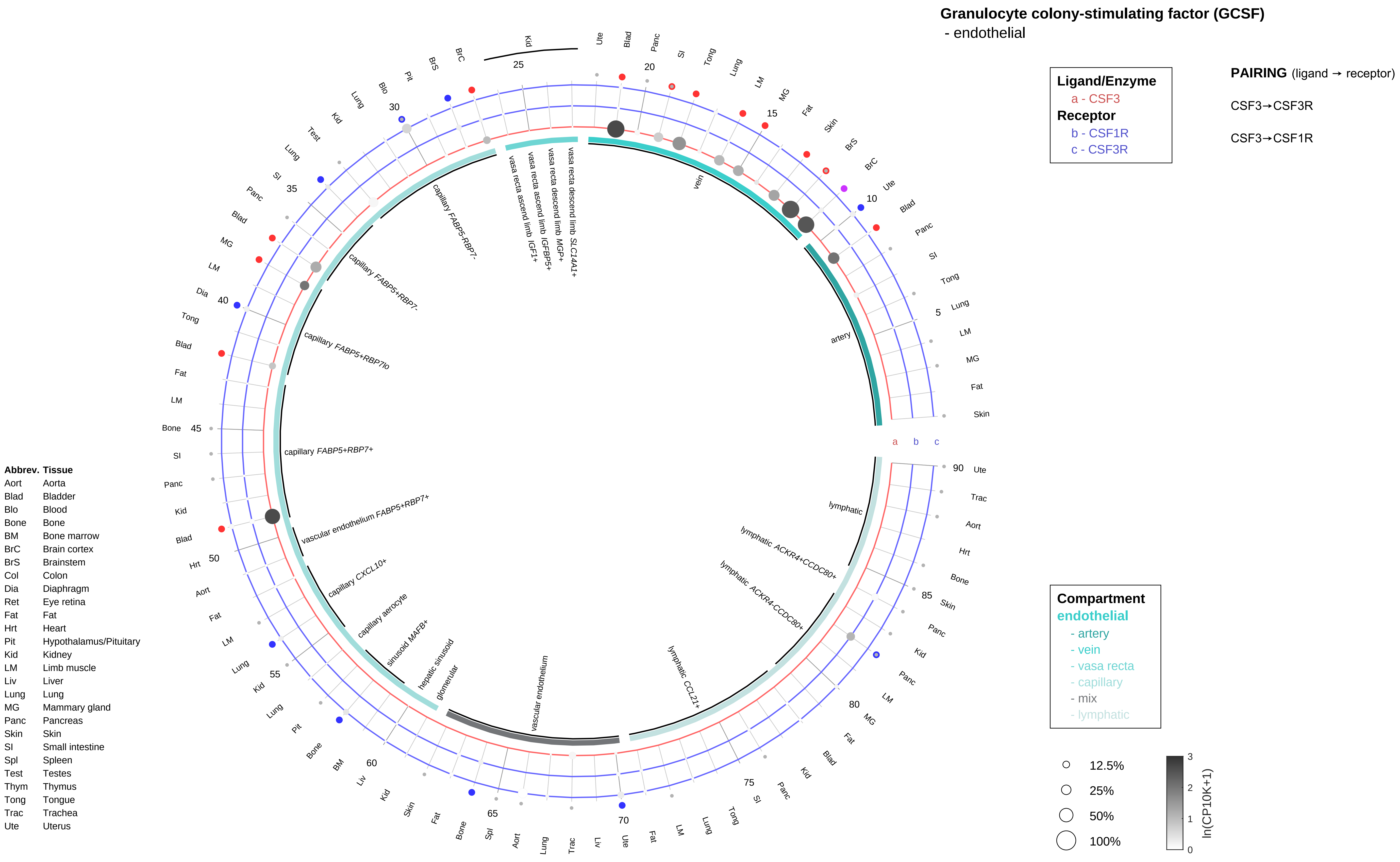

Granulocyte colony-stimulating factor (GCSF)
- endothelial
Ligand/Enzyme
a - CSF3
Receptor
b - CSF1R
c - CSF3R
PAIRING (ligand → receptor)
CSF3→CSF3R
CSF3→CSF1R
Compartment
endothelial
- artery
- vein
- vasa recta
- capillary
- mix
- lymphatic
12.5%
25%
50%
100%
ln(CP10K+1)
artery
vein
vasa recta descend limb SLC14A1+
vasa recta descend limb MGP+
vasa recta ascend limb IGFBP5+
vasa recta ascend limb IGF1+
capillary FABP5-RBP7-
capillary FABP5+RBP7-
capillary FABP5+RBP7lo
capillary FABP5+RBP7+
vascular endothelium FABP5+RBP7+
capillary CXCL10+
capillary aerocyte
sinusoid MAFB+
hepatic sinusoid
glomerular
vascular endothelium
lymphatic CCL21+
lymphatic ACKR4-CCDC80+
lymphatic ACKR4+CCDC80+
lymphatic
Abbrev. Tissue
Aort Aorta
Blad Bladder
Blo Blood
Bone Bone
BM Bone marrow
BrC Brain cortex
BrS Brainstem
Col Colon
Dia Diaphragm
Ret Eye retina
Fat Fat
Hrt Heart
Pit Hypothalamus/Pituitary
Kid Kidney
LM Limb muscle
Liv Liver
Lung Lung
MG Mammary gland
Panc Pancreas
Skin Skin
SI Small intestine
Spl Spleen
Test Testes
Thym Thymus
Tong Tongue
Trac Trachea
Ute Uterus

#### Granulocyte colony-stimulating factor (GCSF)

- stromal

**Ligand/Enzyme**
a - CSF3
**Receptor**
b - CSF1R
c - CSF3R

**PAIRING** (ligand → receptor)

CSF3→CSF3R

CSF3→CSF1R

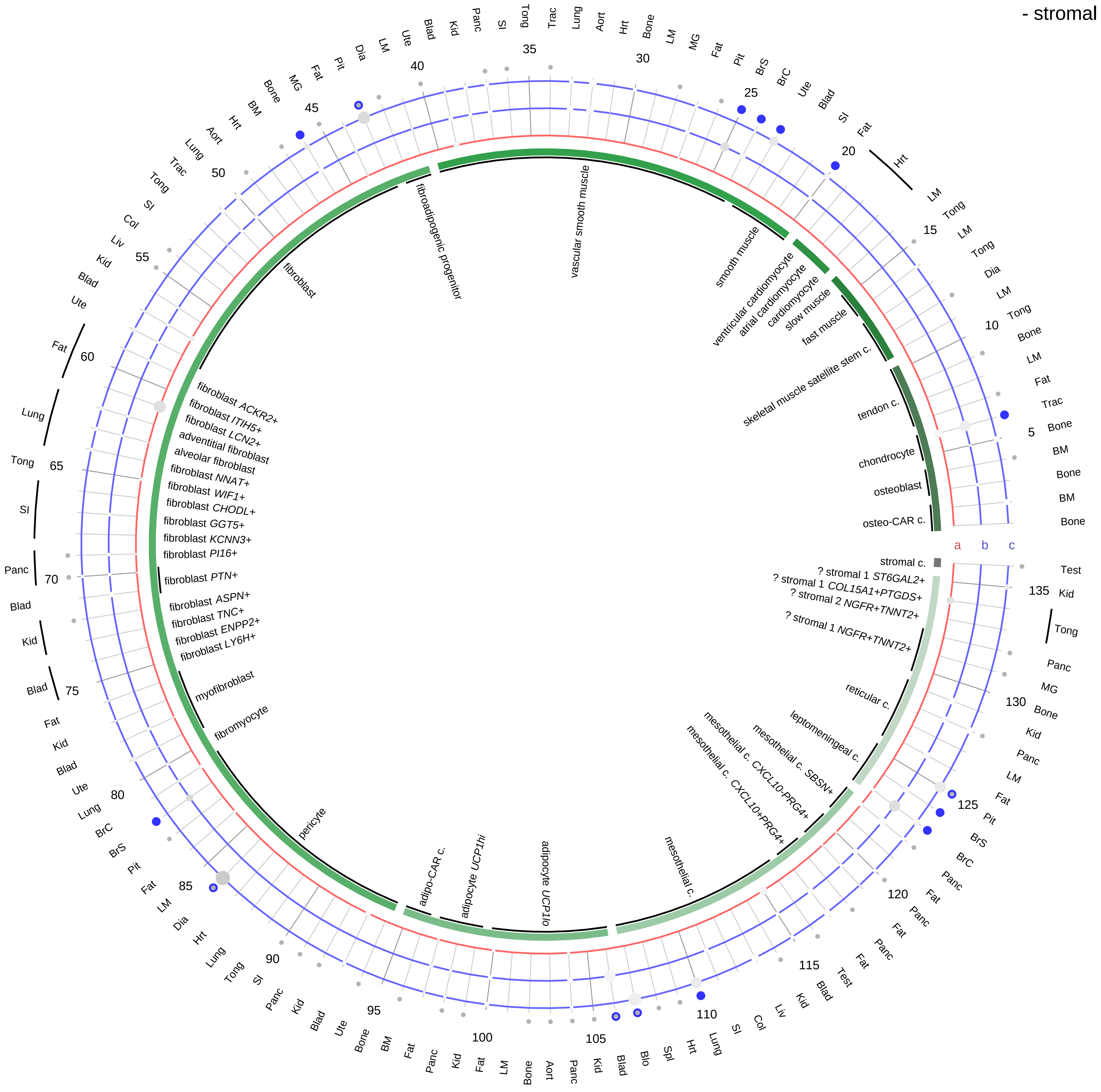

**Compartment**
**stromal**
- bone
- skeletal muscle
- cardiac muscle
- smooth muscle
- fibroblast
- adipocyte
- mesothelial
- other
- mix

12.5%
25%
50%
100%

ln(CP10K+1): 0, 1, 2, 3

| Abbrev. | Tissue |
|---|---|
| Aort | Aorta |
| Blad | Bladder |
| Blo | Blood |
| Bone | Bone |
| BM | Bone marrow |
| BrC | Brain cortex |
| BrS | Brainstem |
| Col | Colon |
| Dia | Diaphragm |
| Ret | Eye retina |
| Fat | Fat |
| Hrt | Heart |
| Pit | Hypothalamus/Pituitary |
| Kid | Kidney |
| LM | Limb muscle |
| Liv | Liver |
| Lung | Lung |
| MG | Mammary gland |
| Panc | Pancreas |
| Skin | Skin |
| SI | Small intestine |
| Spl | Spleen |
| Test | Testes |
| Thym | Thymus |
| Tong | Tongue |
| Trac | Trachea |
| Ute | Uterus |

**Granulocyte colony-stimulating factor (GCSF)**
- lymphoid

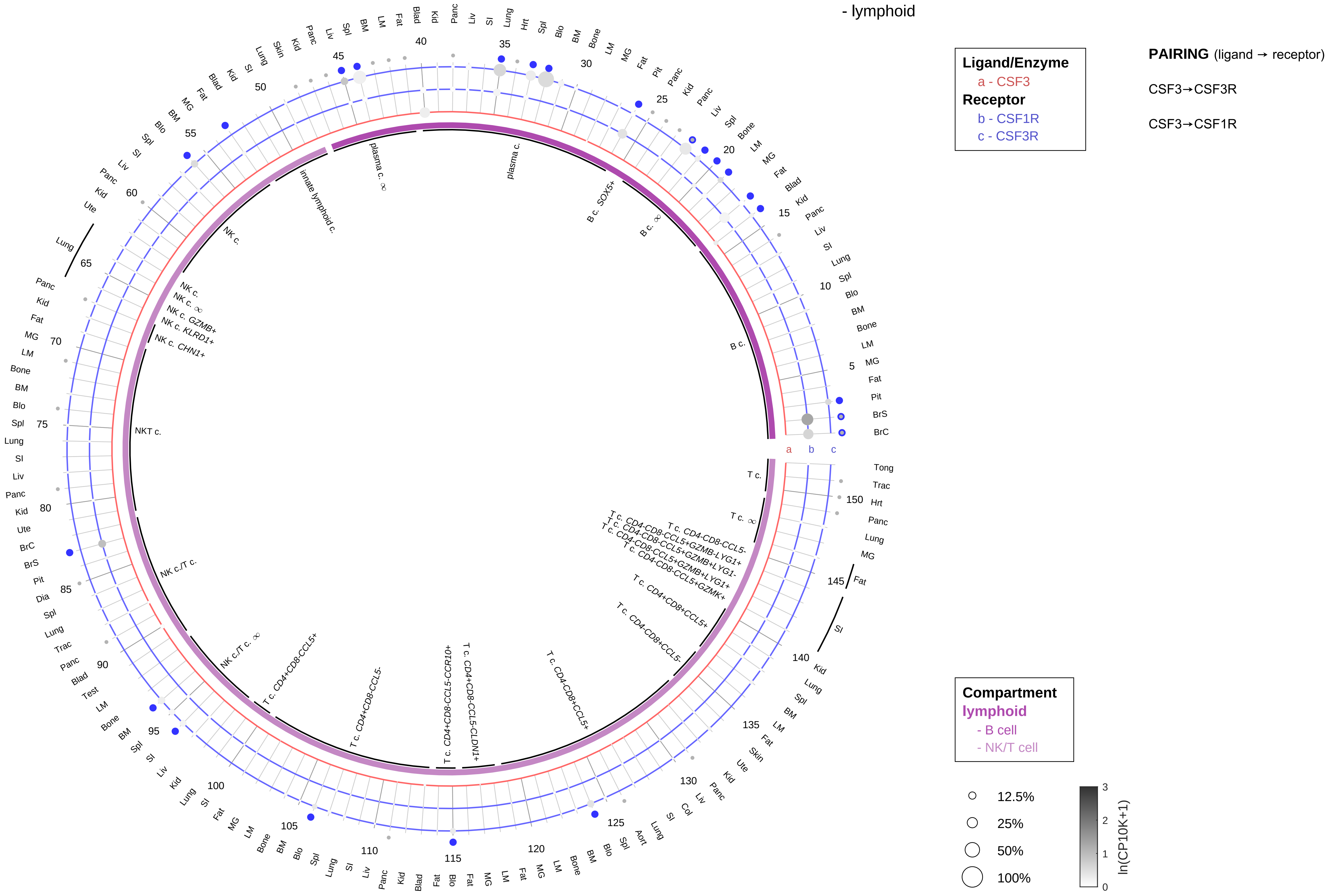

| Abbrev. | Tissue |
|---|---|
| Aort | Aorta |
| Blad | Bladder |
| Blo | Blood |
| Bone | Bone |
| BM | Bone marrow |
| BrC | Brain cortex |
| BrS | Brainstem |
| Col | Colon |
| Dia | Diaphragm |
| Ret | Eye retina |
| Fat | Fat |
| Hrt | Heart |
| Pit | Hypothalamus/Pituitary |
| Kid | Kidney |
| LM | Limb muscle |
| Liv | Liver |
| Lung | Lung |
| MG | Mammary gland |
| Panc | Pancreas |
| Skin | Skin |
| SI | Small intestine |
| Spl | Spleen |
| Test | Testes |
| Thym | Thymus |
| Tong | Tongue |
| Trac | Trachea |
| Ute | Uterus |

**Granulocyte colony-stimulating factor (GCSF)**
- myeloid

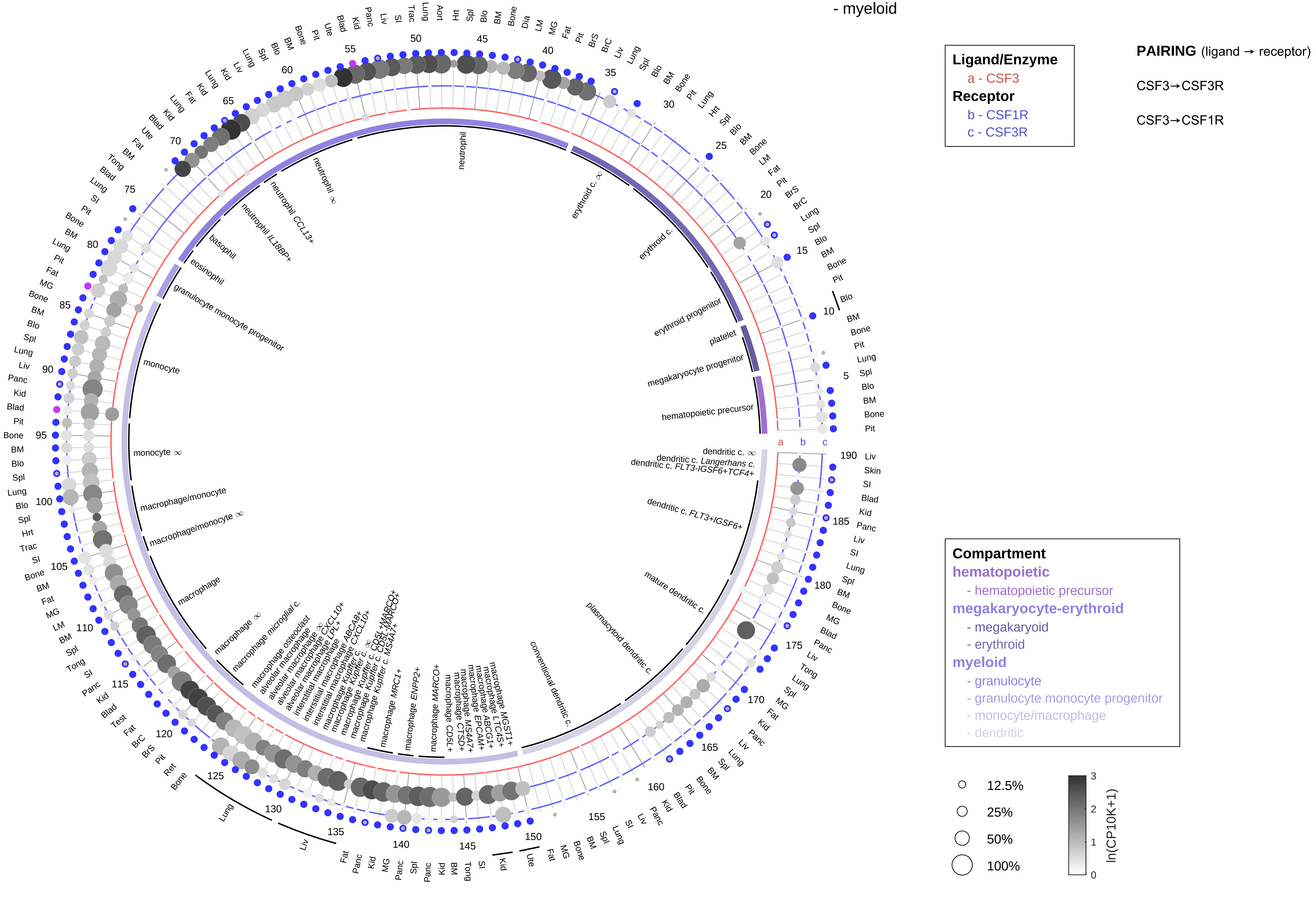

| Abbrev. | Tissue |
|---|---|
| Aort | Aorta |
| Blad | Bladder |
| Blo | Blood |
| Bone | Bone |
| BM | Bone marrow |
| BrC | Brain cortex |
| BrS | Brainstem |
| Col | Colon |
| Dia | Diaphragm |
| Ret | Eye retina |
| Fat | Fat |
| Hrt | Heart |
| Pit | Hypothalamus/Pituitary |
| Kid | Kidney |
| LM | Limb muscle |
| Liv | Liver |
| Lung | Lung |
| MG | Mammary gland |
| Panc | Pancreas |
| Skin | Skin |
| SI | Small intestine |
| Spl | Spleen |
| Test | Testes |
| Thym | Thymus |
| Tong | Tongue |
| Trac | Trachea |
| Ute | Uterus |

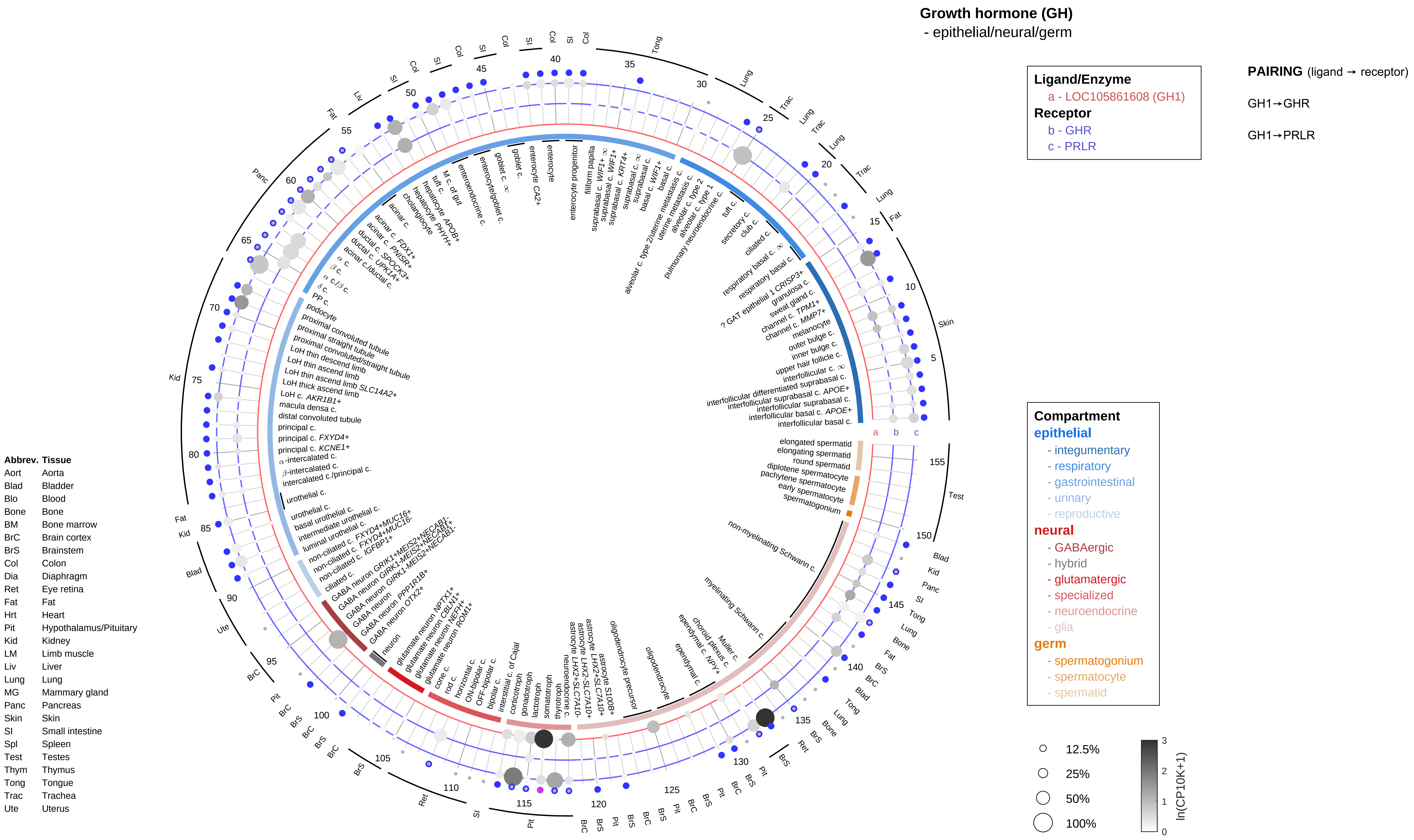
Growth hormone (GH)
- epithelial/neural/germ
Ligand/Enzyme
a - LOC105861608 (GH1)
Receptor
b - GHR
c - PRLR
PAIRING (ligand → receptor)
GH1→GHR
GH1→PRLR
Compartment
epithelial
- integumentary
- respiratory
- gastrointestinal
- urinary
- reproductive
neural
- GABAergic
- hybrid
- glutamatergic
- specialized
- neuroendocrine
- glia
germ
- spermatogonium
- spermatocyte
- spermatid
12.5%
25%
50%
100%
ln(CP10K+1)
Abbrev. Tissue
Aort Aorta
Blad Bladder
Blo Blood
Bone Bone
BM Bone marrow
BrC Brain cortex
BrS Brainstem
Col Colon
Dia Diaphragm
Ret Eye retina
Fat Fat
Hrt Heart
Pit Hypothalamus/Pituitary
Kid Kidney
LM Limb muscle
Liv Liver
Lung Lung
MG Mammary gland
Panc Pancreas
Skin Skin
SI Small intestine
Spl Spleen
Test Testes
Thym Thymus
Tong Tongue
Trac Trachea
Ute Uterus

#### Growth hormone (GH)
- endothelial

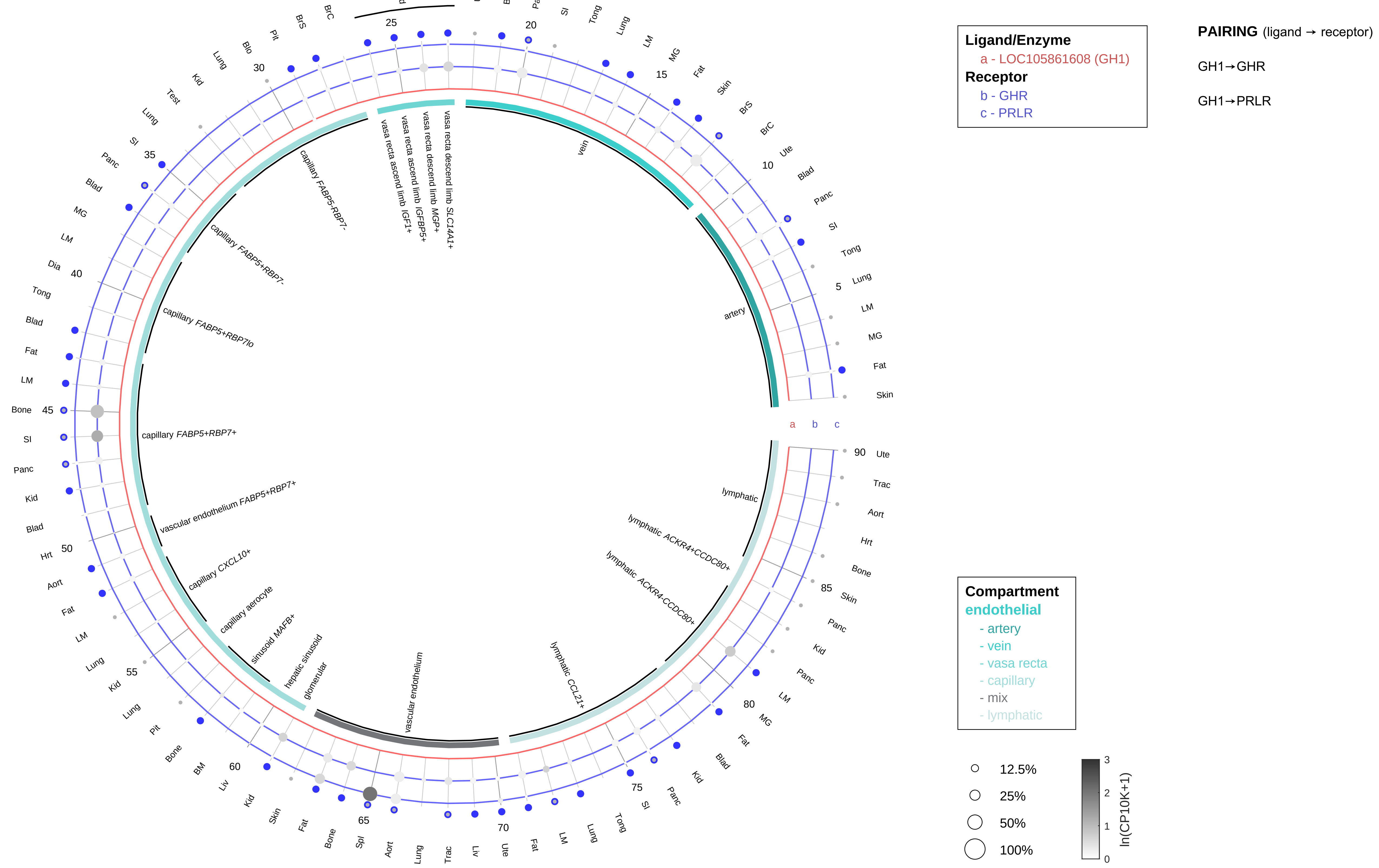

| Abbrev. | Tissue |
|---|---|
| Aort | Aorta |
| Blad | Bladder |
| Blo | Blood |
| Bone | Bone |
| BM | Bone marrow |
| BrC | Brain cortex |
| BrS | Brainstem |
| Col | Colon |
| Dia | Diaphragm |
| Ret | Eye retina |
| Fat | Fat |
| Hrt | Heart |
| Pit | Hypothalamus/Pituitary |
| Kid | Kidney |
| LM | Limb muscle |
| Liv | Liver |
| Lung | Lung |
| MG | Mammary gland |
| Panc | Pancreas |
| Skin | Skin |
| SI | Small intestine |
| Spl | Spleen |
| Test | Testes |
| Thym | Thymus |
| Tong | Tongue |
| Trac | Trachea |
| Ute | Uterus |

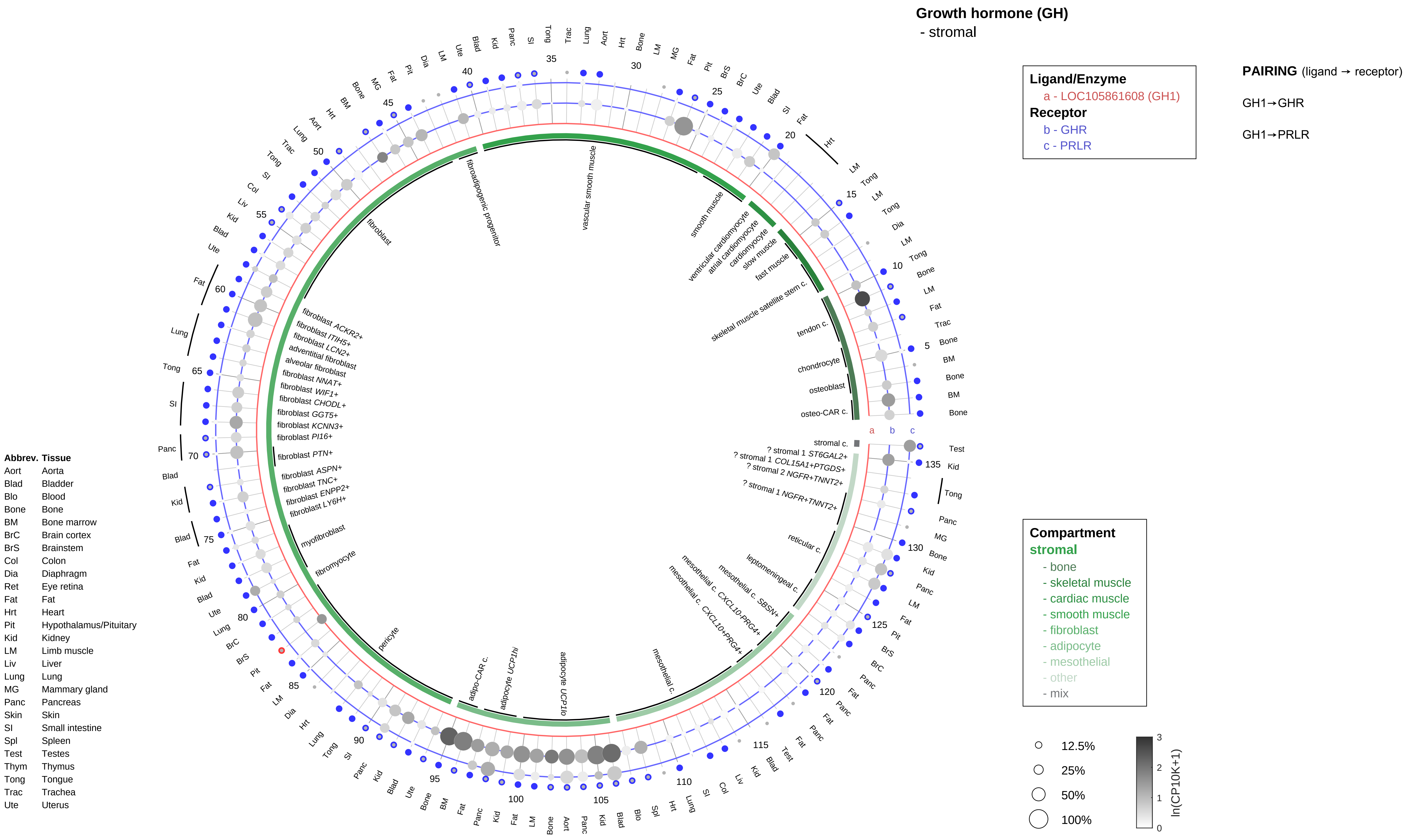

Growth hormone (GH)
- stromal
Ligand/Enzyme
a - LOC105861608 (GH1)
Receptor
b - GHR
c - PRLR
PAIRING (ligand → receptor)
GH1→GHR
GH1→PRLR
Compartment
stromal
- bone
- skeletal muscle
- cardiac muscle
- smooth muscle
- fibroblast
- adipocyte
- mesothelial
- other
- mix
12.5%
25%
50%
100%
ln(CP10K+1)
Abbrev. Tissue
Aort Aorta
Blad Bladder
Blo Blood
Bone Bone
BM Bone marrow
BrC Brain cortex
BrS Brainstem
Col Colon
Dia Diaphragm
Ret Eye retina
Fat Fat
Hrt Heart
Pit Hypothalamus/Pituitary
Kid Kidney
LM Limb muscle
Liv Liver
Lung Lung
MG Mammary gland
Panc Pancreas
Skin Skin
SI Small intestine
Spl Spleen
Test Testes
Thym Thymus
Tong Tongue
Trac Trachea
Ute Uterus

**Growth hormone (GH)**
- lymphoid

**Ligand/Enzyme**
a - LOC105861608 (GH1)
**Receptor**
b - GHR
c - PRLR

**PAIRING** (ligand → receptor)

GH1→GHR

GH1→PRLR

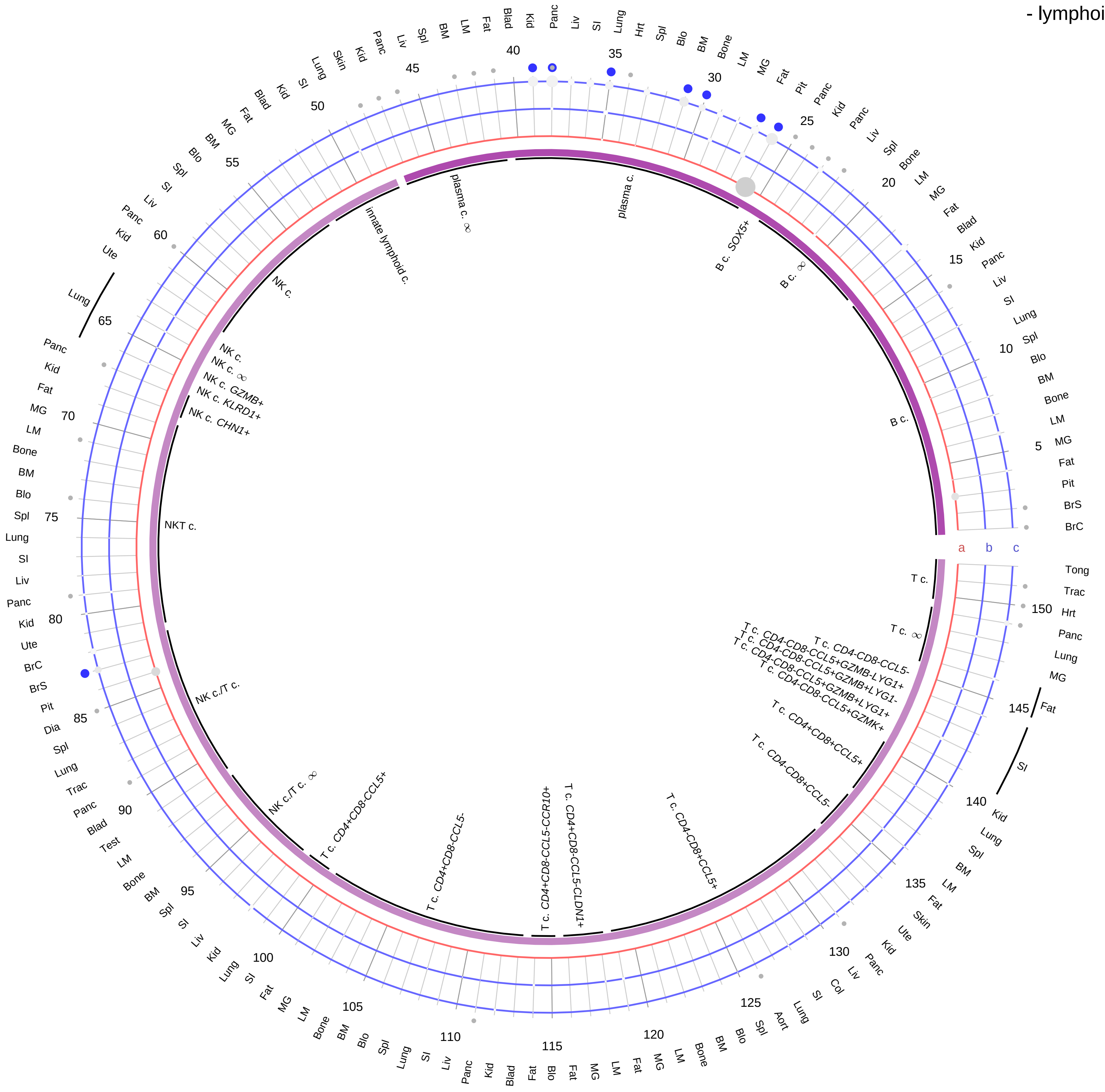

| Abbrev. | Tissue |
|---|---|
| Aort | Aorta |
| Blad | Bladder |
| Blo | Blood |
| Bone | Bone |
| BM | Bone marrow |
| BrC | Brain cortex |
| BrS | Brainstem |
| Col | Colon |
| Dia | Diaphragm |
| Ret | Eye retina |
| Fat | Fat |
| Hrt | Heart |
| Pit | Hypothalamus/Pituitary |
| Kid | Kidney |
| LM | Limb muscle |
| Liv | Liver |
| Lung | Lung |
| MG | Mammary gland |
| Panc | Pancreas |
| Skin | Skin |
| SI | Small intestine |
| Spl | Spleen |
| Test | Testes |
| Thym | Thymus |
| Tong | Tongue |
| Trac | Trachea |
| Ute | Uterus |

**Compartment**
**lymphoid**
- B cell
- NK/T cell

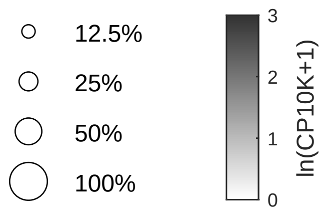

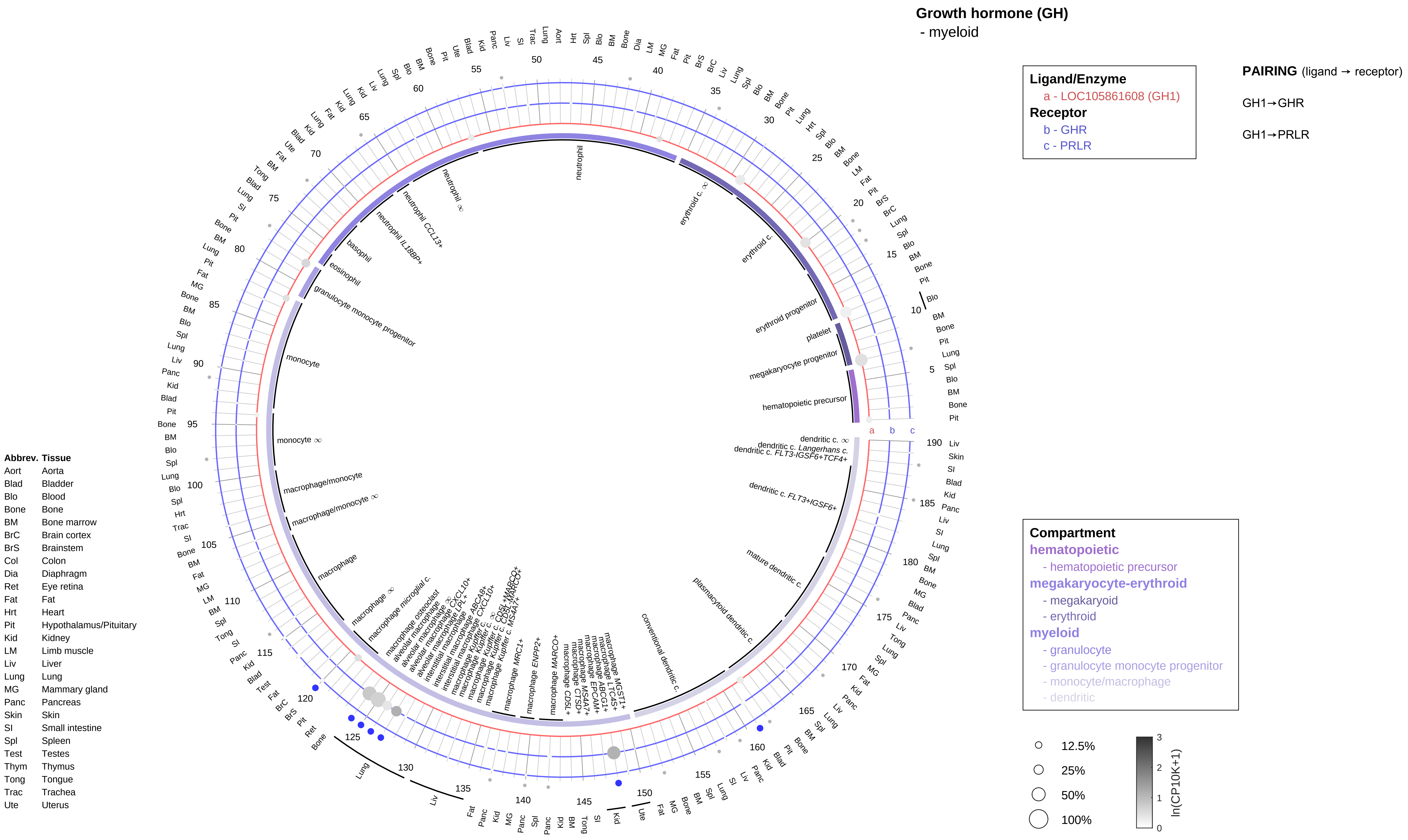

Growth hormone (GH)
- myeloid
Ligand/Enzyme
a - LOC105861608 (GH1)
Receptor
b - GHR
c - PRLR
PAIRING (ligand → receptor)
GH1→GHR
GH1→PRLR
Compartment
hematopoietic
- hematopoietic precursor
megakaryocyte-erythroid
- megakaryoid
- erythroid
myeloid
- granulocyte
- granulocyte monocyte progenitor
- monocyte/macrophage
- dendritic
12.5%
25%
50%
100%
ln(CP10K+1)
Abbrev. Tissue
Aort Aorta
Blad Bladder
Blo Blood
Bone Bone
BM Bone marrow
BrC Brain cortex
BrS Brainstem
Col Colon
Dia Diaphragm
Ret Eye retina
Fat Fat
Hrt Heart
Pit Hypothalamus/Pituitary
Kid Kidney
LM Limb muscle
Liv Liver
Lung Lung
MG Mammary gland
Panc Pancreas
Skin Skin
SI Small intestine
Spl Spleen
Test Testes
Thym Thymus
Tong Tongue
Trac Trachea
Ute Uterus
neutrophil
erythroid c. ∞
erythroid c.
erythroid progenitor
platelet
megakaryocyte progenitor
hematopoietic precursor
dendritic c. ∞
dendritic c. Langerhans c.
dendritic c. FLT3-IGSF6+TCF4+
dendritic c. FLT3+IGSF6+
mature dendritic c.
plasmacytoid dendritic c.
conventional dendritic c.
neutrophil ∞
neutrophil CCL13+
neutrophil IL18BP+
basophil
eosinophil
granulocyte monocyte progenitor
monocyte
monocyte ∞
macrophage/monocyte
macrophage/monocyte ∞
macrophage
macrophage ∞
macrophage microglial c.
macrophage osteoclast
macrophage MRC1+
macrophage ENPP2+
macrophage MARCO+

#### Growth hormone releasing hormone (GHRH)
- epithelial/neural/germ

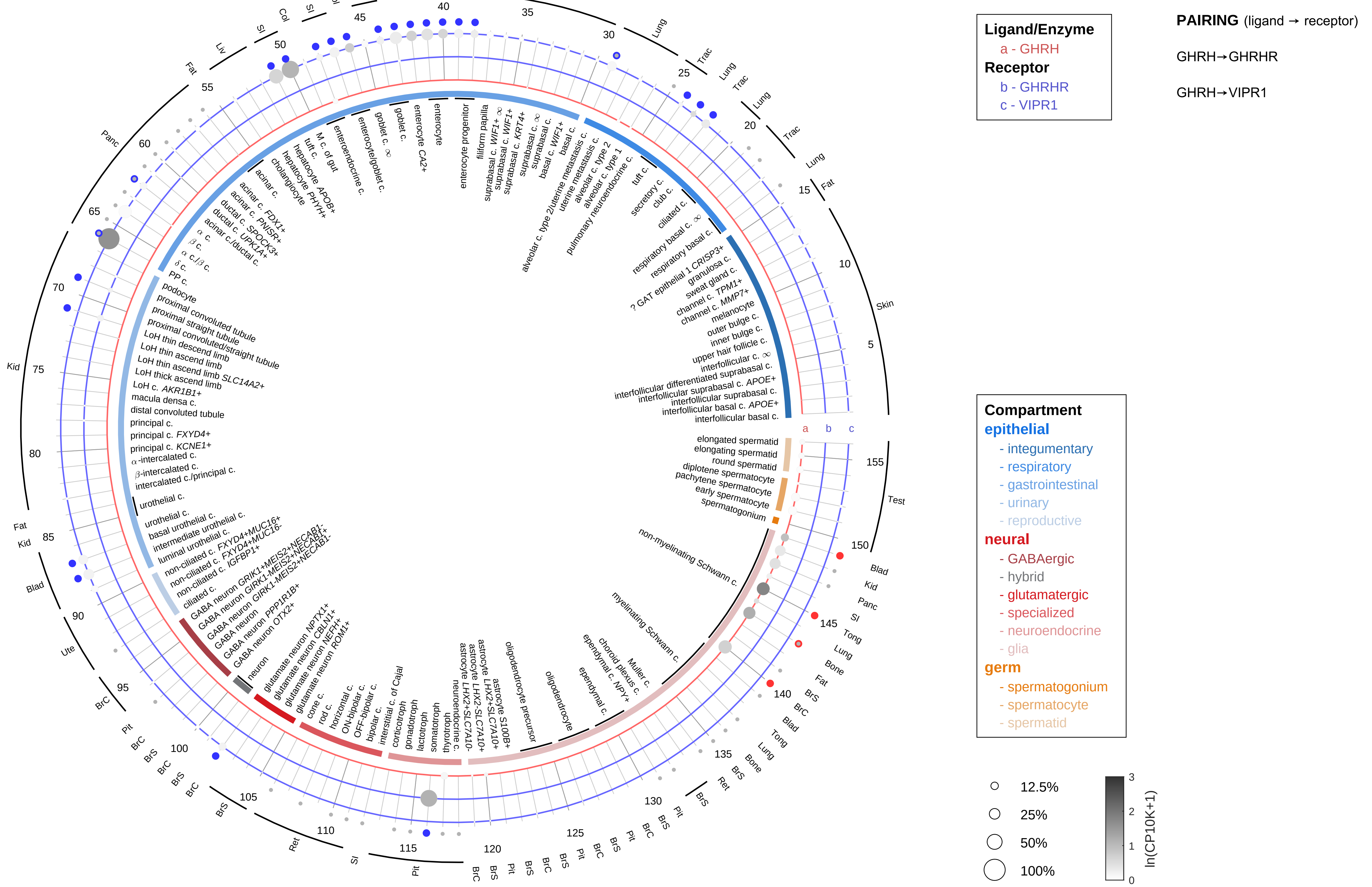

| Abbrev. | Tissue |
|---|---|
| Aort | Aorta |
| Blad | Bladder |
| Blo | Blood |
| Bone | Bone |
| BM | Bone marrow |
| BrC | Brain cortex |
| BrS | Brainstem |
| Col | Colon |
| Dia | Diaphragm |
| Ret | Eye retina |
| Fat | Fat |
| Hrt | Heart |
| Pit | Hypothalamus/Pituitary |
| Kid | Kidney |
| LM | Limb muscle |
| Liv | Liver |
| Lung | Lung |
| MG | Mammary gland |
| Panc | Pancreas |
| Skin | Skin |
| SI | Small intestine |
| Spl | Spleen |
| Test | Testes |
| Thym | Thymus |
| Tong | Tongue |
| Trac | Trachea |
| Ute | Uterus |

**Growth hormone releasing hormone (GHRH)**
- endothelial

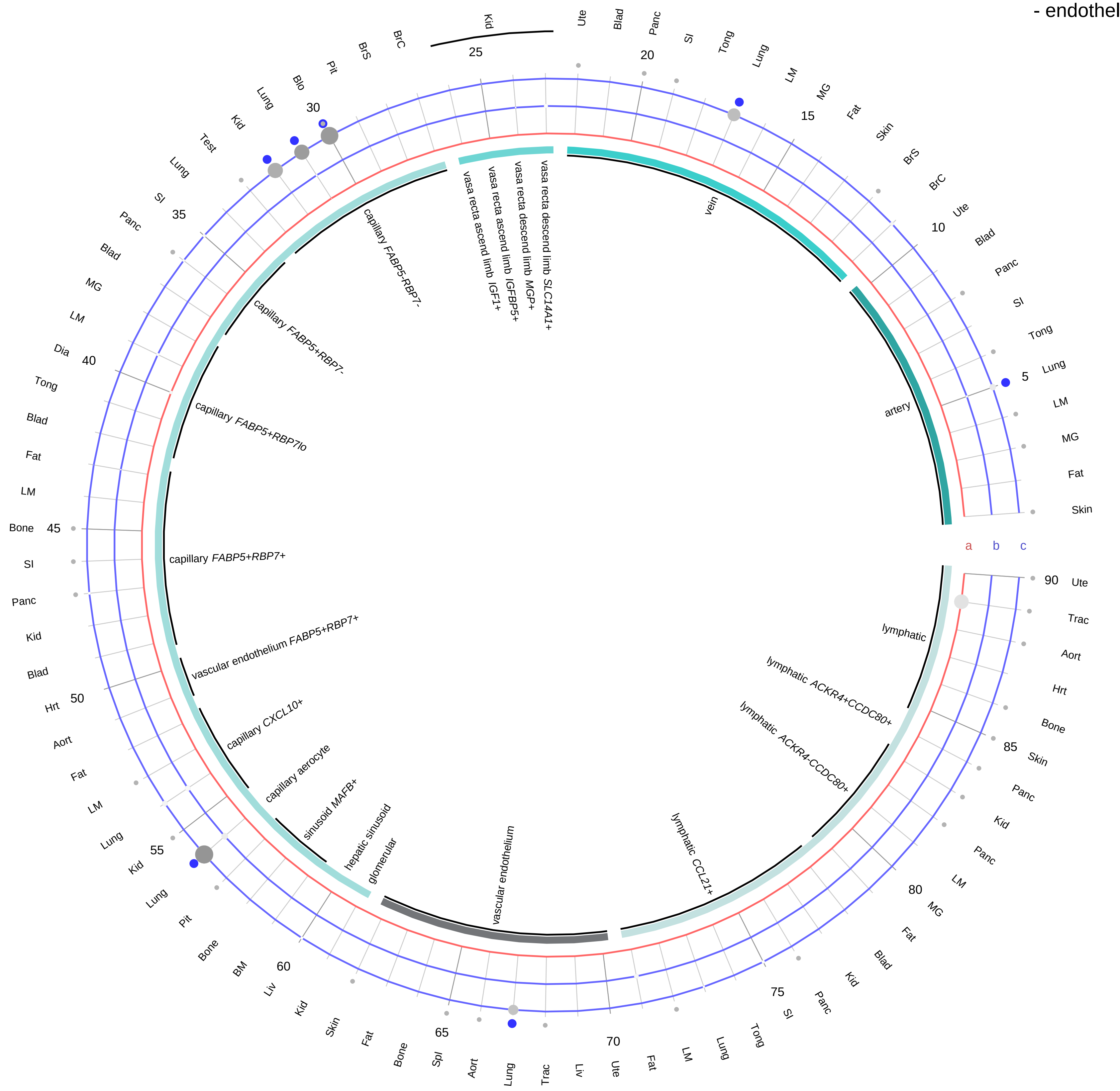

**Ligand/Enzyme**
- a - GHRH

**Receptor**
- b - GHRHR
- c - VIPR1

**PAIRING** (ligand → receptor)

GHRH→GHRHR

GHRH→VIPR1

**Compartment**
**endothelial**
- artery
- vein
- vasa recta
- capillary
- mix
- lymphatic

12.5%
25%
50%
100%

ln(CP10K+1): 0, 1, 2, 3

| Abbrev. | Tissue |
|---|---|
| Aort | Aorta |
| Blad | Bladder |
| Blo | Blood |
| Bone | Bone |
| BM | Bone marrow |
| BrC | Brain cortex |
| BrS | Brainstem |
| Col | Colon |
| Dia | Diaphragm |
| Ret | Eye retina |
| Fat | Fat |
| Hrt | Heart |
| Pit | Hypothalamus/Pituitary |
| Kid | Kidney |
| LM | Limb muscle |
| Liv | Liver |
| Lung | Lung |
| MG | Mammary gland |
| Panc | Pancreas |
| Skin | Skin |
| SI | Small intestine |
| Spl | Spleen |
| Test | Testes |
| Thym | Thymus |
| Tong | Tongue |
| Trac | Trachea |
| Ute | Uterus |

**Growth hormone releasing hormone (GHRH)**
- stromal

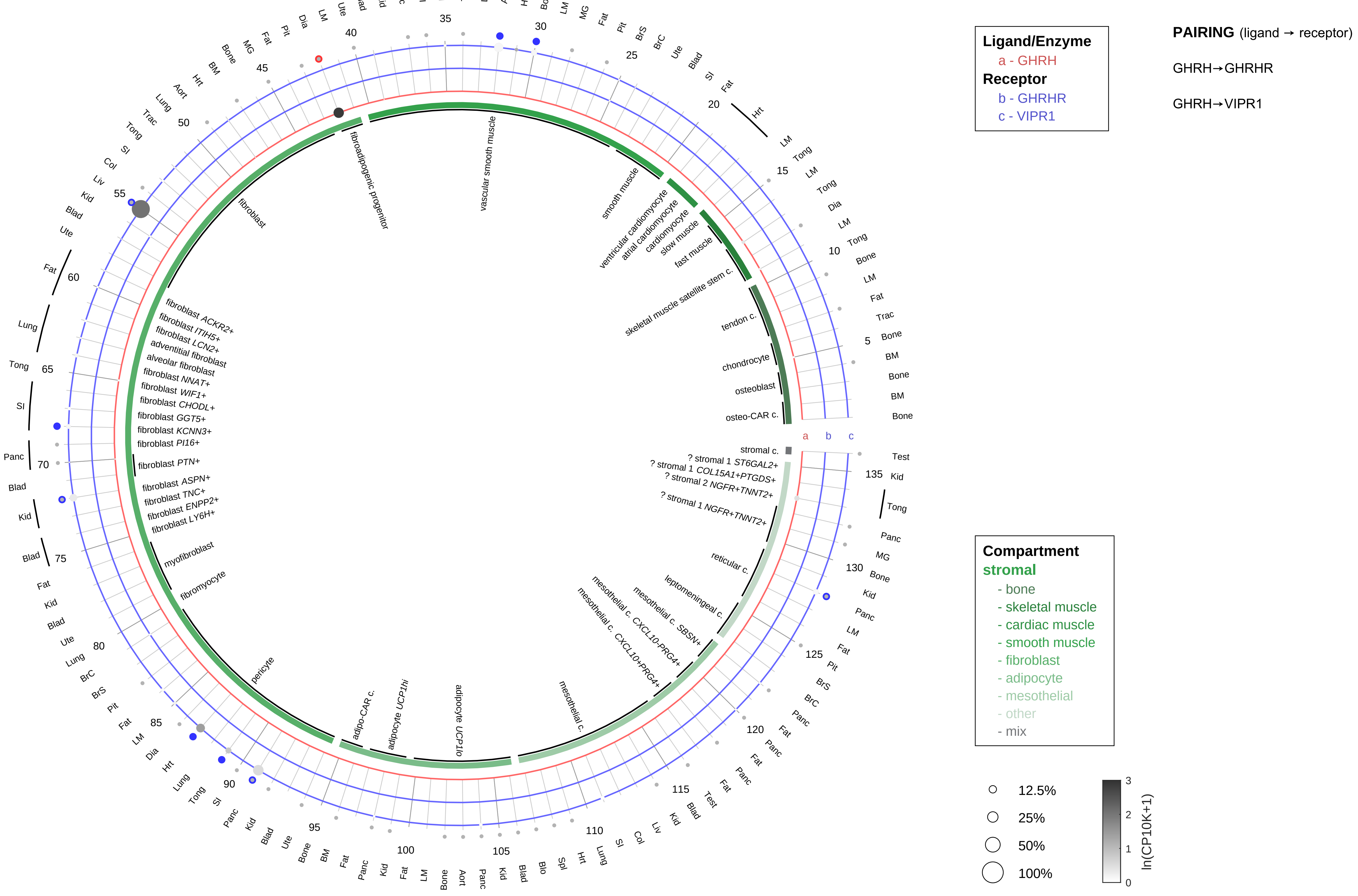

| Abbrev. | Tissue |
|---|---|
| Aort | Aorta |
| Blad | Bladder |
| Blo | Blood |
| Bone | Bone |
| BM | Bone marrow |
| BrC | Brain cortex |
| BrS | Brainstem |
| Col | Colon |
| Dia | Diaphragm |
| Ret | Eye retina |
| Fat | Fat |
| Hrt | Heart |
| Pit | Hypothalamus/Pituitary |
| Kid | Kidney |
| LM | Limb muscle |
| Liv | Liver |
| Lung | Lung |
| MG | Mammary gland |
| Panc | Pancreas |
| Skin | Skin |
| SI | Small intestine |
| Spl | Spleen |
| Test | Testes |
| Thym | Thymus |
| Tong | Tongue |
| Trac | Trachea |
| Ute | Uterus |

**Growth hormone releasing hormone (GHRH)**
- lymphoid

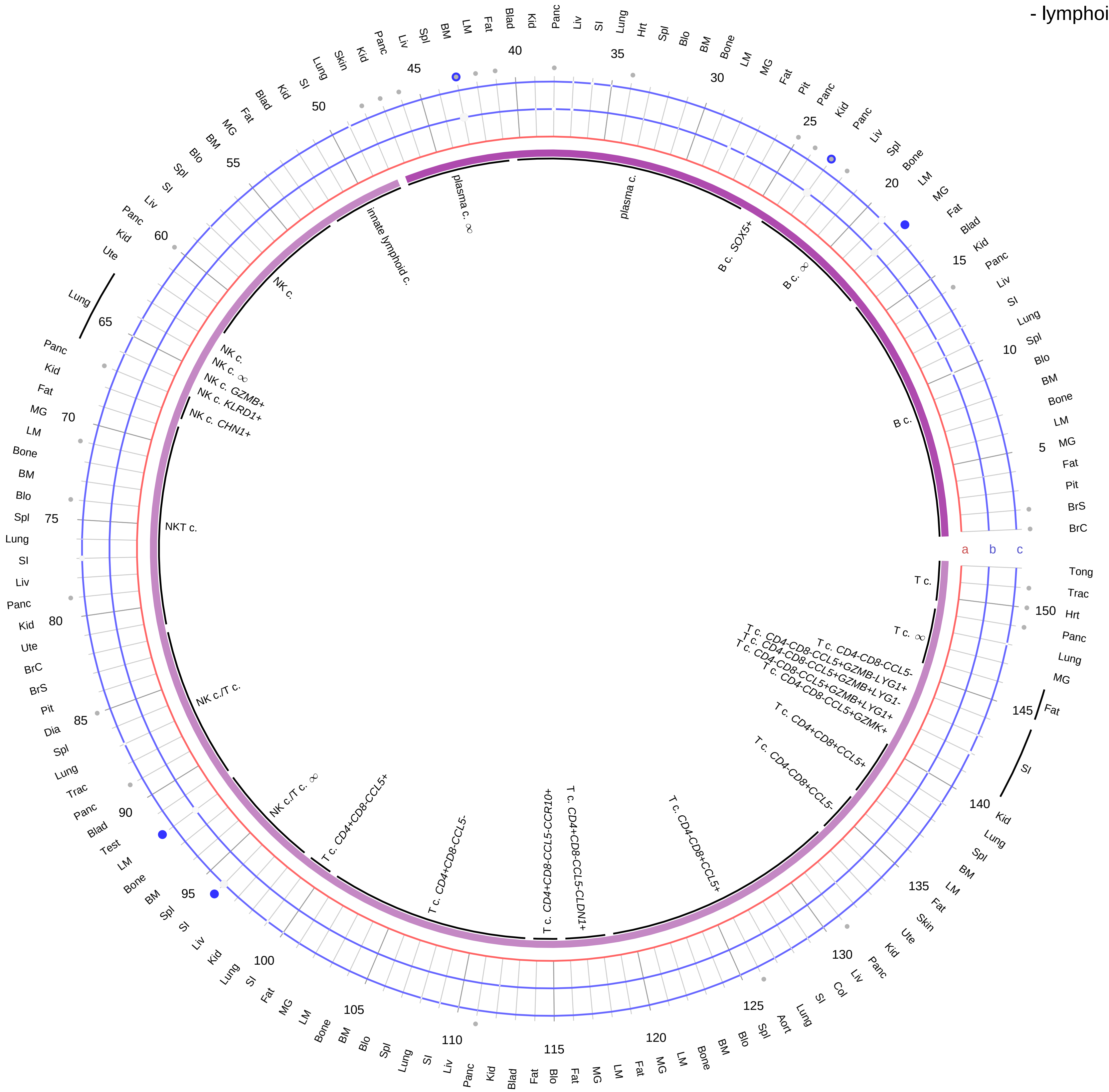

| Abbrev. | Tissue |
|---|---|
| Aort | Aorta |
| Blad | Bladder |
| Blo | Blood |
| Bone | Bone |
| BM | Bone marrow |
| BrC | Brain cortex |
| BrS | Brainstem |
| Col | Colon |
| Dia | Diaphragm |
| Ret | Eye retina |
| Fat | Fat |
| Hrt | Heart |
| Pit | Hypothalamus/Pituitary |
| Kid | Kidney |
| LM | Limb muscle |
| Liv | Liver |
| Lung | Lung |
| MG | Mammary gland |
| Panc | Pancreas |
| Skin | Skin |
| SI | Small intestine |
| Spl | Spleen |
| Test | Testes |
| Thym | Thymus |
| Tong | Tongue |
| Trac | Trachea |
| Ute | Uterus |

**Ligand/Enzyme**
a - GHRH
**Receptor**
b - GHRHR
c - VIPR1

**PAIRING** (ligand → receptor)

GHRH→GHRHR

GHRH→VIPR1

**Compartment**
**lymphoid**
- B cell
- NK/T cell

12.5%
25%
50%
100%

ln(CP10K+1): 0, 1, 2, 3

**Growth hormone releasing hormone (GHRH)**
- myeloid

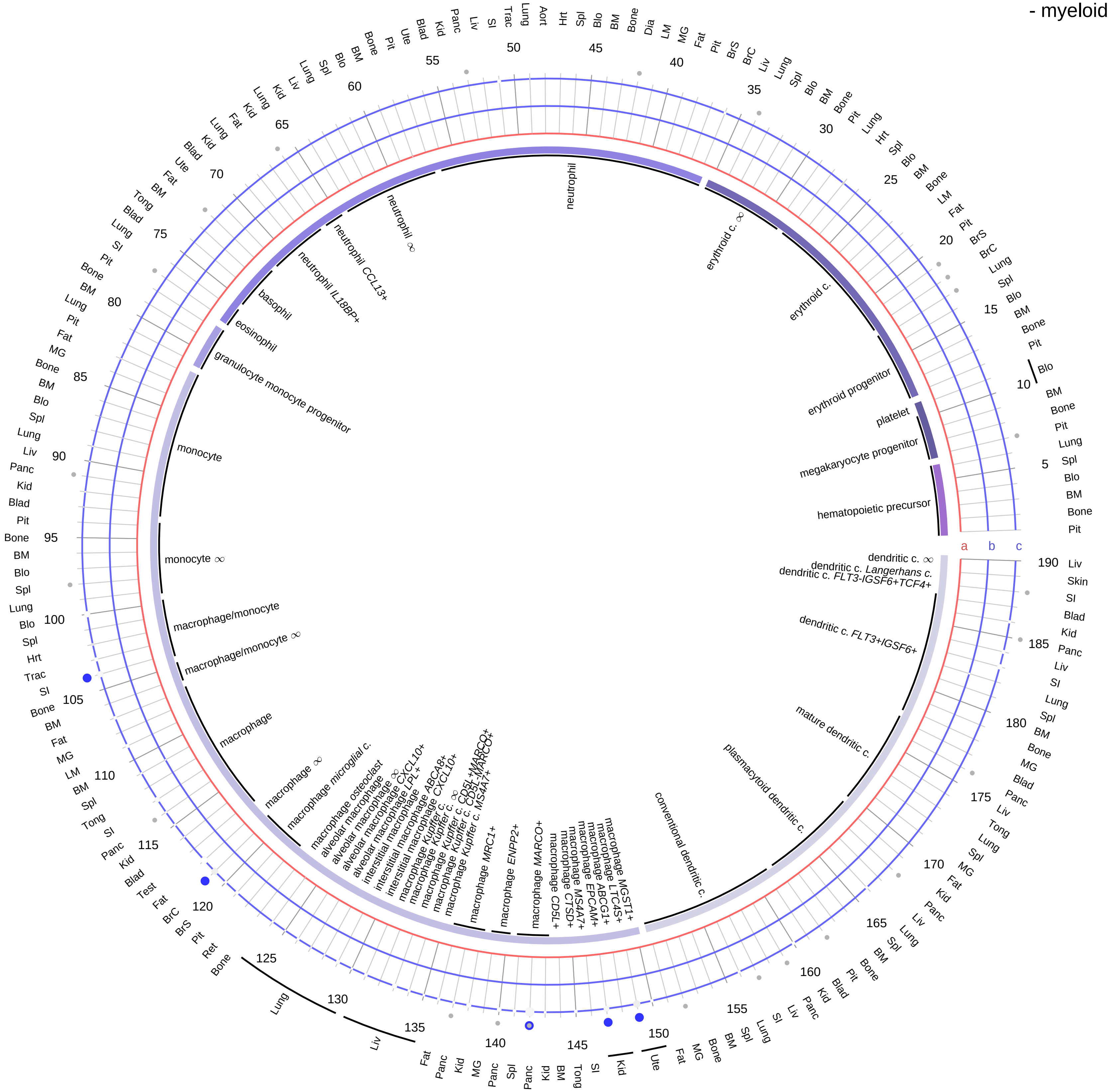

**Ligand/Enzyme**
a - GHRH
**Receptor**
b - GHRHR
c - VIPR1

**PAIRING** (ligand → receptor)

GHRH→GHRHR

GHRH→VIPR1

**Compartment**
**hematopoietic**
- hematopoietic precursor
**megakaryocyte-erythroid**
- megakaryoid
- erythroid
**myeloid**
- granulocyte
- granulocyte monocyte progenitor
- monocyte/macrophage
- dendritic

12.5%
25%
50%
100%

ln(CP10K+1)
0
1
2
3

| Abbrev. | Tissue |
|---|---|
| Aort | Aorta |
| Blad | Bladder |
| Blo | Blood |
| Bone | Bone |
| BM | Bone marrow |
| BrC | Brain cortex |
| BrS | Brainstem |
| Col | Colon |
| Dia | Diaphragm |
| Ret | Eye retina |
| Fat | Fat |
| Hrt | Heart |
| Pit | Hypothalamus/Pituitary |
| Kid | Kidney |
| LM | Limb muscle |
| Liv | Liver |
| Lung | Lung |
| MG | Mammary gland |
| Panc | Pancreas |
| Skin | Skin |
| SI | Small intestine |
| Spl | Spleen |
| Test | Testes |
| Thym | Thymus |
| Tong | Tongue |
| Trac | Trachea |
| Ute | Uterus |

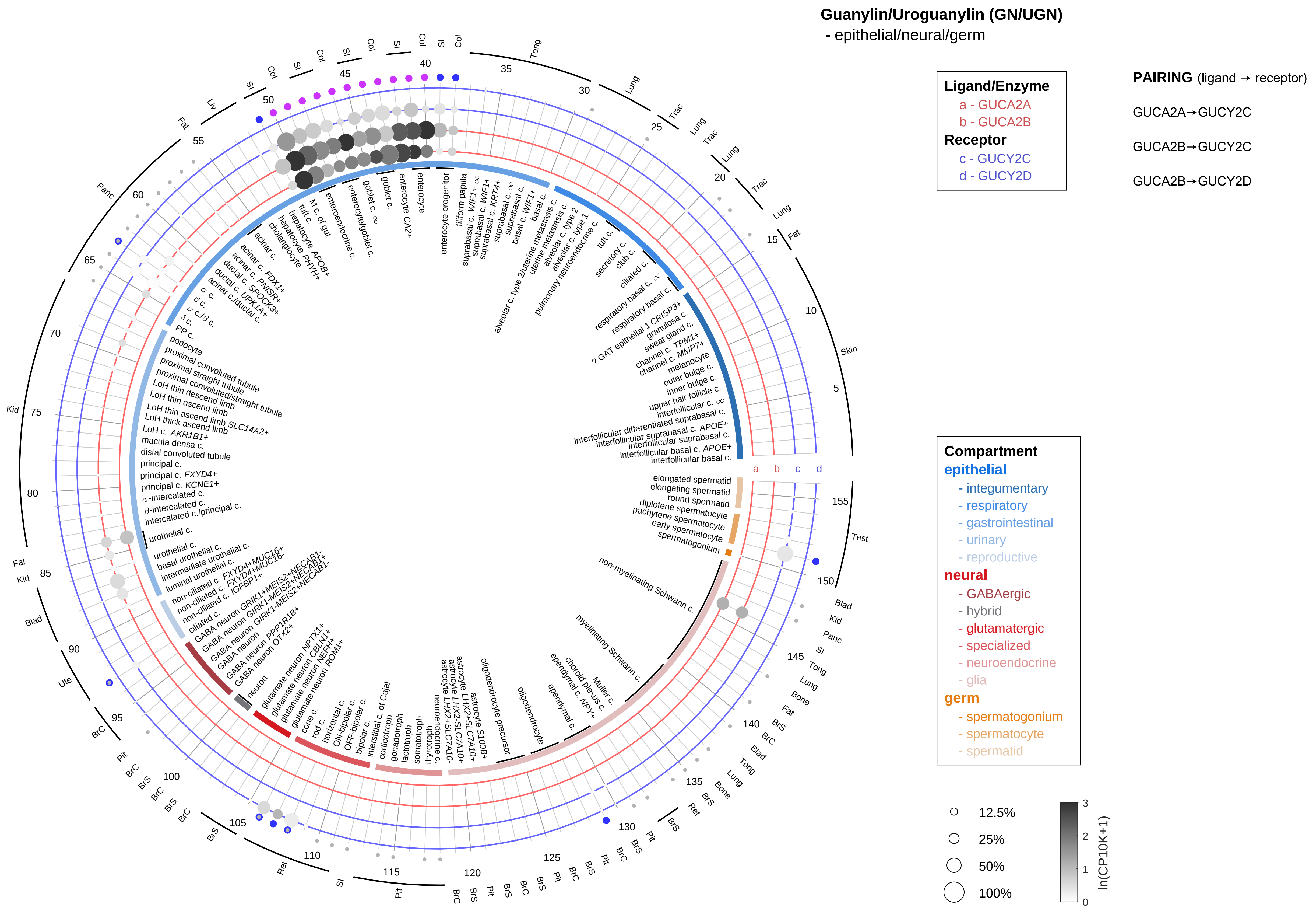

| Abbrev. | Tissue |
|---|---|
| Aort | Aorta |
| Blad | Bladder |
| Blo | Blood |
| Bone | Bone |
| BM | Bone marrow |
| BrC | Brain cortex |
| BrS | Brainstem |
| Col | Colon |
| Dia | Diaphragm |
| Ret | Eye retina |
| Fat | Fat |
| Hrt | Heart |
| Pit | Hypothalamus/Pituitary |
| Kid | Kidney |
| LM | Limb muscle |
| Liv | Liver |
| Lung | Lung |
| MG | Mammary gland |
| Panc | Pancreas |
| Skin | Skin |
| SI | Small intestine |
| Spl | Spleen |
| Test | Testes |
| Thym | Thymus |
| Tong | Tongue |
| Trac | Trachea |
| Ute | Uterus |

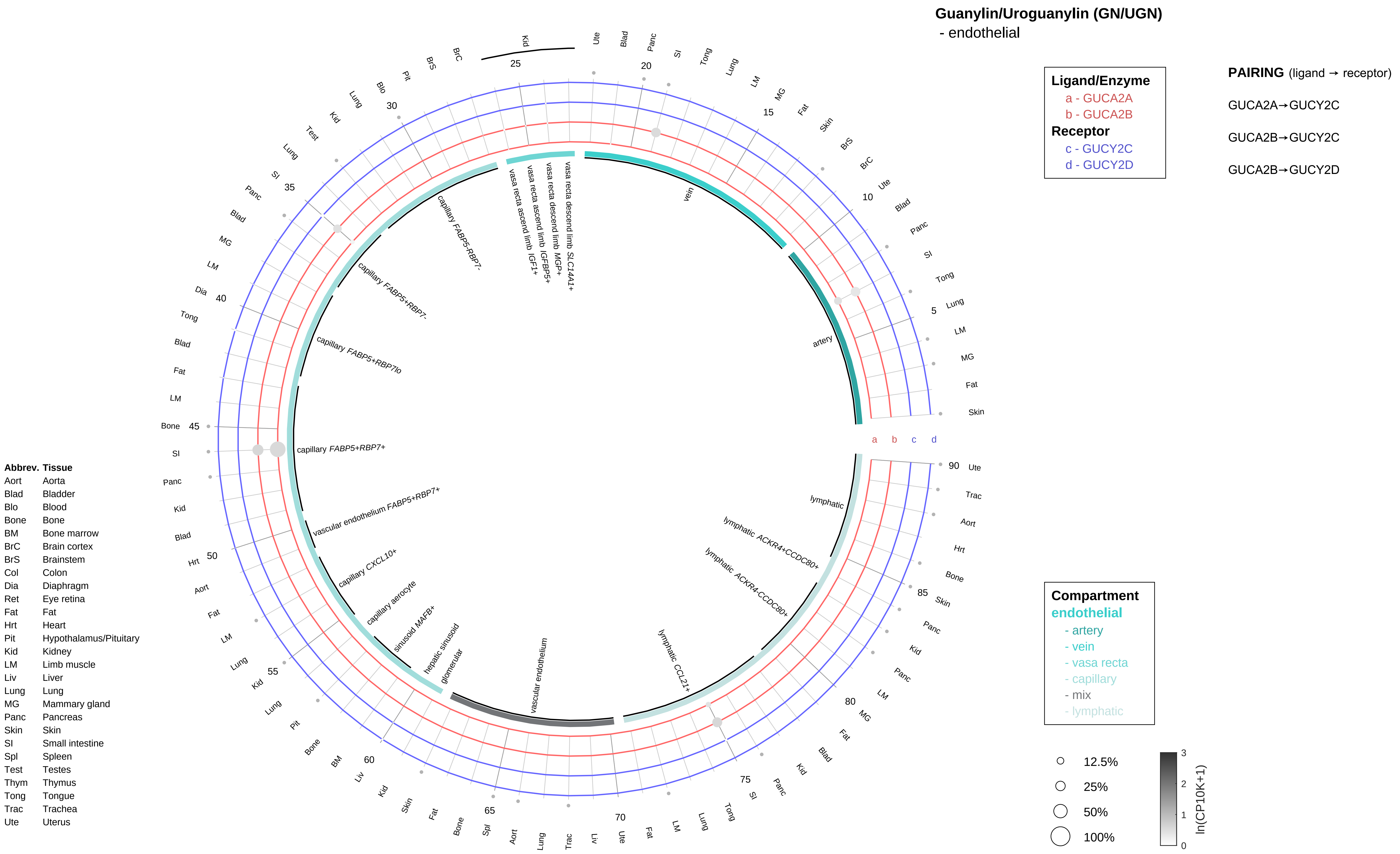

Guanylin/Uroguanylin (GN/UGN)
- endothelial
Ligand/Enzyme
a - GUCA2A
b - GUCA2B
Receptor
c - GUCY2C
d - GUCY2D
PAIRING (ligand → receptor)
GUCA2A→GUCY2C
GUCA2B→GUCY2C
GUCA2B→GUCY2D
Compartment
endothelial
- artery
- vein
- vasa recta
- capillary
- mix
- lymphatic
12.5%
25%
50%
100%
ln(CP10K+1)
artery
vein
vasa recta descend limb SLC14A1+
vasa recta descend limb MGP+
vasa recta ascend limb IGFBP5+
vasa recta ascend limb IGF1+
capillary FABP5-RBP7-
capillary FABP5+RBP7-
capillary FABP5+RBP7lo
capillary FABP5+RBP7+
vascular endothelium FABP5+RBP7+
capillary CXCL10+
capillary aerocyte
sinusoid MAFB+
hepatic sinusoid
glomerular
vascular endothelium
lymphatic CCL21+
lymphatic ACKR4-CCDC80+
lymphatic ACKR4+CCDC80+
lymphatic
Abbrev. Tissue
Aort Aorta
Blad Bladder
Blo Blood
Bone Bone
BM Bone marrow
BrC Brain cortex
BrS Brainstem
Col Colon
Dia Diaphragm
Ret Eye retina
Fat Fat
Hrt Heart
Pit Hypothalamus/Pituitary
Kid Kidney
LM Limb muscle
Liv Liver
Lung Lung
MG Mammary gland
Panc Pancreas
Skin Skin
SI Small intestine
Spl Spleen
Test Testes
Thym Thymus
Tong Tongue
Trac Trachea
Ute Uterus

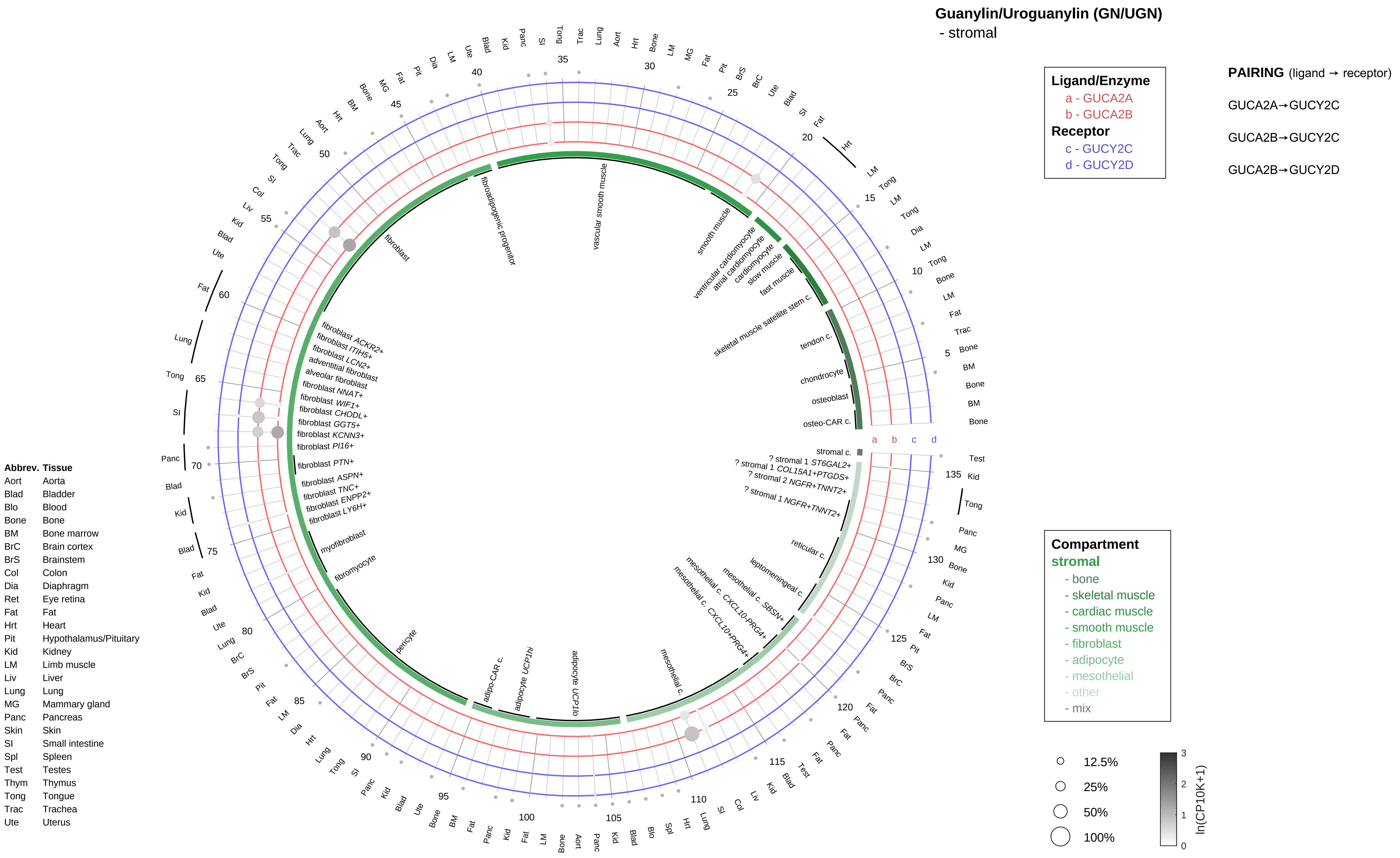

Guanylin/Uroguanylin (GN/UGN)
- stromal
Ligand/Enzyme
a - GUCA2A
b - GUCA2B
Receptor
c - GUCY2C
d - GUCY2D
PAIRING (ligand → receptor)
GUCA2A→GUCY2C
GUCA2B→GUCY2C
GUCA2B→GUCY2D
Compartment
stromal
- bone
- skeletal muscle
- cardiac muscle
- smooth muscle
- fibroblast
- adipocyte
- mesothelial
- other
- mix
12.5%
25%
50%
100%
ln(CP10K+1)
fibroadipogenic progenitor
vascular smooth muscle
smooth muscle
ventricular cardiomyocyte
atrial cardiomyocyte
cardiomyocyte
slow muscle
fast muscle
skeletal muscle satellite stem c.
tendon c.
chondrocyte
osteoblast
osteo-CAR c.
stromal c.
? stromal 1 ST6GAL2+
? stromal 1 COL15A1+PTGDS+
? stromal 2 NGFR+TNNT2+
? stromal 1 NGFR+TNNT2+
reticular c.
leptomeningeal c.
mesothelial c. SBSN+
mesothelial c. CXCL10-PRG4+
mesothelial c. CXCL10+PRG4+
mesothelial c.
adipocyte UCP1lo
adipocyte UCP1hi
adipo-CAR c.
pericyte
fibromyocyte
myofibroblast
fibroblast LY6H+
fibroblast ENPP2+
fibroblast TNC+
fibroblast ASPN+
fibroblast PTN+
fibroblast PI16+
fibroblast KCNN3+
fibroblast GGT5+
fibroblast CHODL+
fibroblast WIF1+
fibroblast NNAT+
alveolar fibroblast
adventitial fibroblast
fibroblast LCN2+
fibroblast ITIH5+
fibroblast ACKR2+
fibroblast
Abbrev. Tissue
Aort Aorta
Blad Bladder
Blo Blood
Bone Bone
BM Bone marrow
BrC Brain cortex
BrS Brainstem
Col Colon
Dia Diaphragm
Ret Eye retina
Fat Fat
Hrt Heart
Pit Hypothalamus/Pituitary
Kid Kidney
LM Limb muscle
Liv Liver
Lung Lung
MG Mammary gland
Panc Pancreas
Skin Skin
SI Small intestine
Spl Spleen
Test Testes
Thym Thymus
Tong Tongue
Trac Trachea
Ute Uterus

**Guanylin/Uroguanylin (GN/UGN)**
- lymphoid

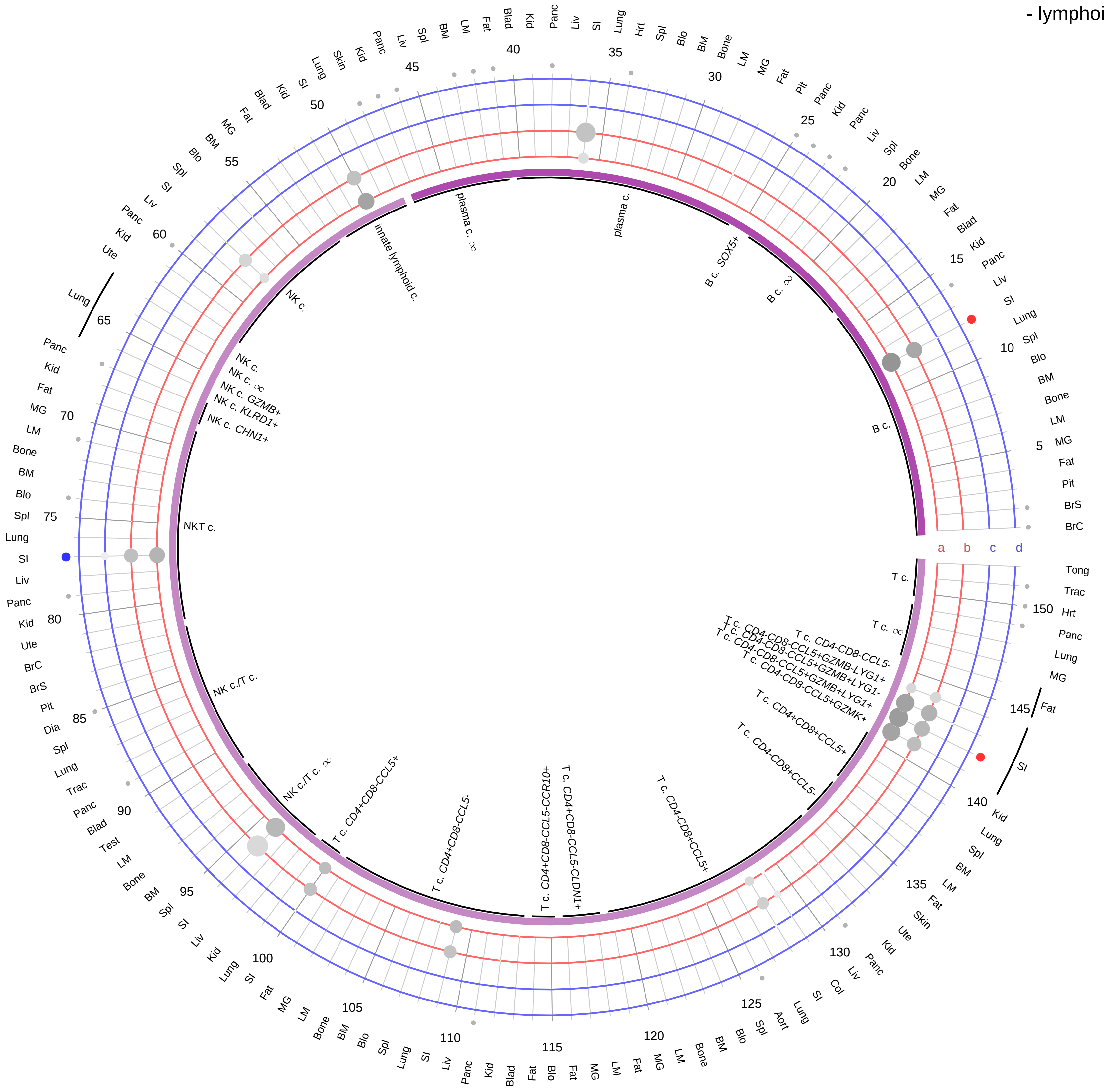

**Ligand/Enzyme**
a - GUCA2A
b - GUCA2B
**Receptor**
c - GUCY2C
d - GUCY2D

**PAIRING** (ligand → receptor)

GUCA2A→GUCY2C

GUCA2B→GUCY2C

GUCA2B→GUCY2D

**Compartment**
**lymphoid**
- B cell
- NK/T cell

12.5%
25%
50%
100%

ln(CP10K+1): 0, 1, 2, 3

| Abbrev. | Tissue |
|---|---|
| Aort | Aorta |
| Blad | Bladder |
| Blo | Blood |
| Bone | Bone |
| BM | Bone marrow |
| BrC | Brain cortex |
| BrS | Brainstem |
| Col | Colon |
| Dia | Diaphragm |
| Ret | Eye retina |
| Fat | Fat |
| Hrt | Heart |
| Pit | Hypothalamus/Pituitary |
| Kid | Kidney |
| LM | Limb muscle |
| Liv | Liver |
| Lung | Lung |
| MG | Mammary gland |
| Panc | Pancreas |
| Skin | Skin |
| SI | Small intestine |
| Spl | Spleen |
| Test | Testes |
| Thym | Thymus |
| Tong | Tongue |
| Trac | Trachea |
| Ute | Uterus |

#### Guanylin/Uroguanylin (GN/UGN)
- myeloid

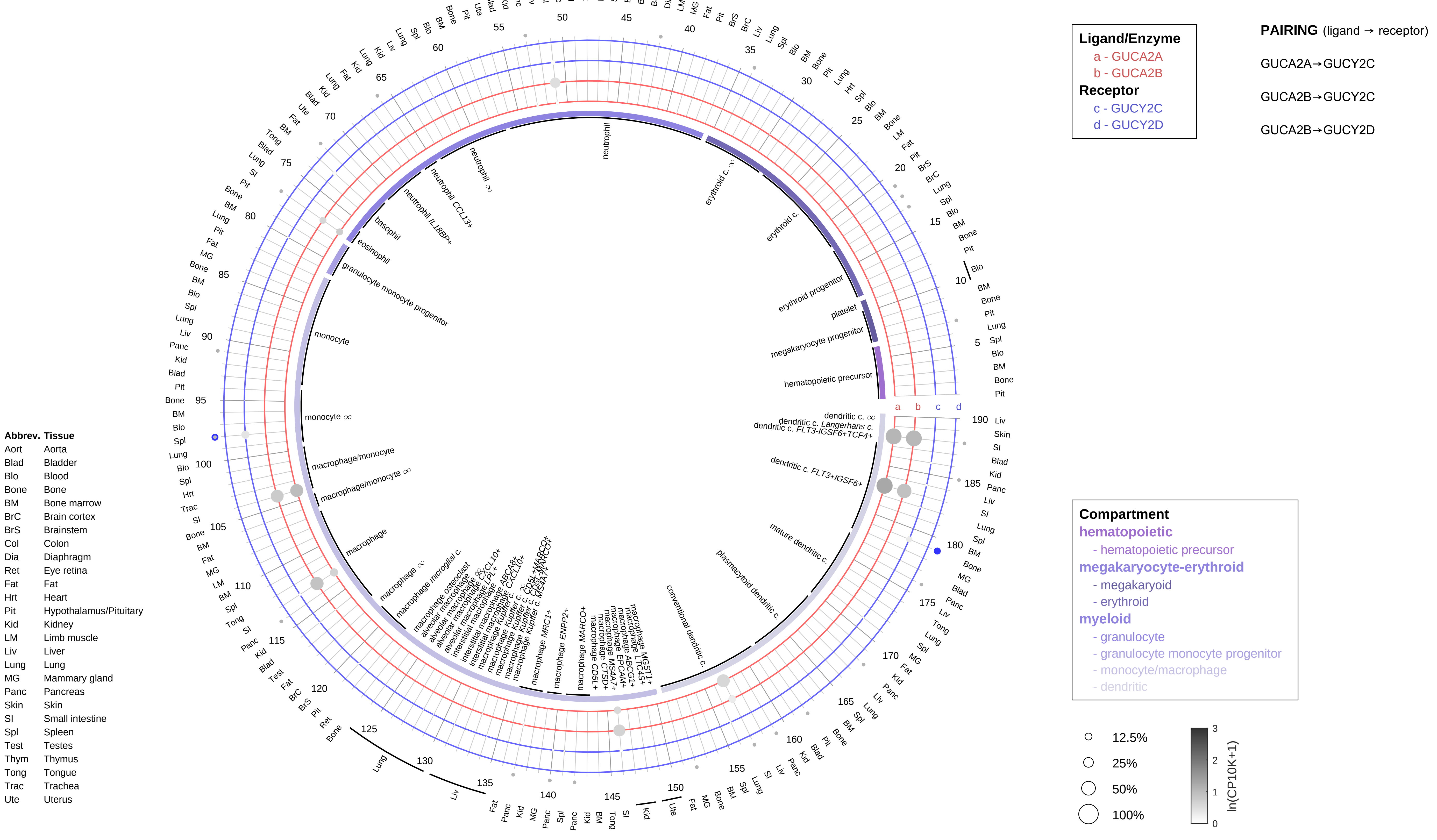

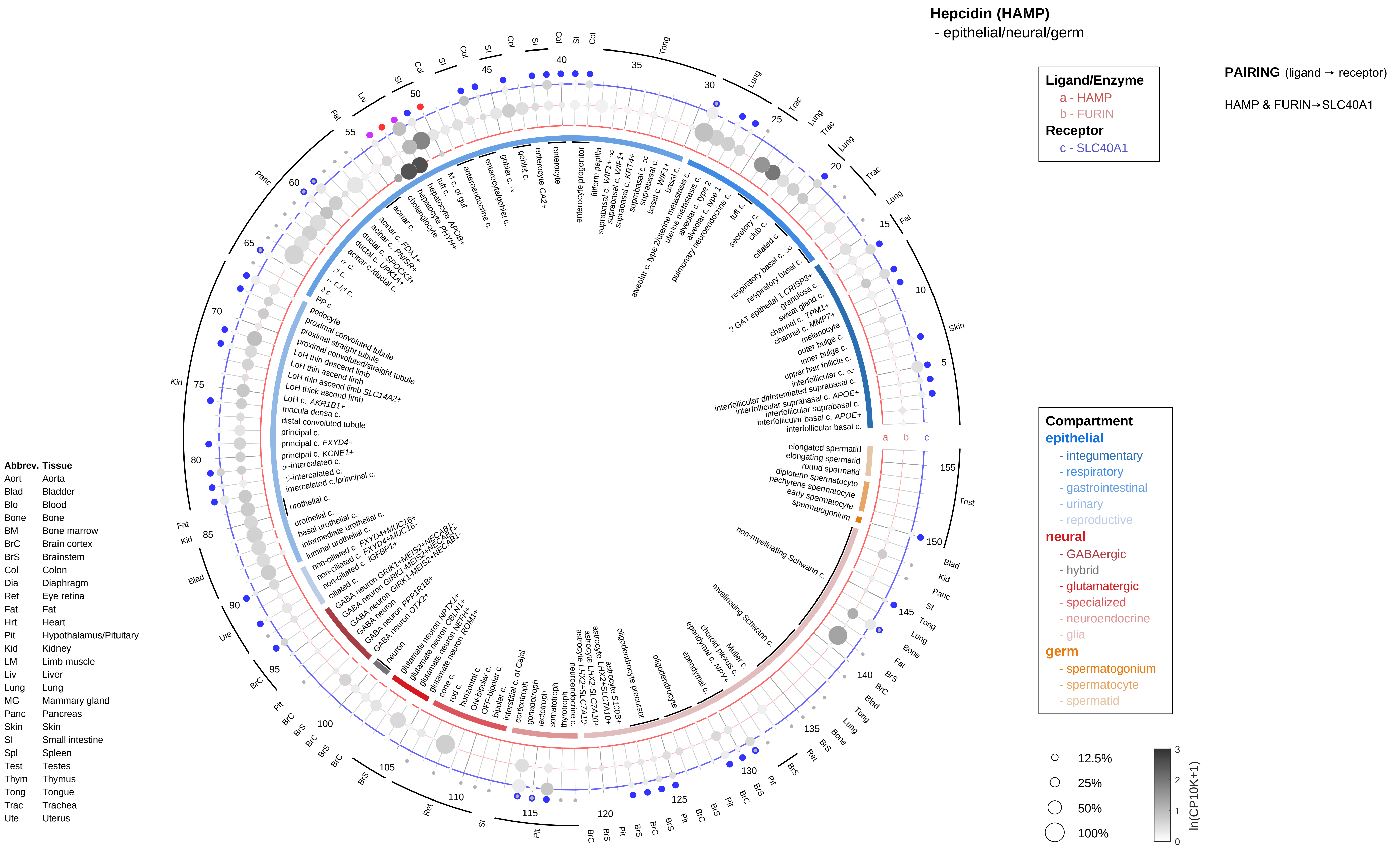

Hepcidin (HAMP)
- epithelial/neural/germ
Ligand/Enzyme
a - HAMP
b - FURIN
Receptor
c - SLC40A1
PAIRING (ligand → receptor)
HAMP & FURIN→SLC40A1
Compartment
epithelial
- integumentary
- respiratory
- gastrointestinal
- urinary
- reproductive
neural
- GABAergic
- hybrid
- glutamatergic
- specialized
- neuroendocrine
- glia
germ
- spermatogonium
- spermatocyte
- spermatid
12.5%
25%
50%
100%
ln(CP10K+1)
Abbrev. Tissue
Aort Aorta
Blad Bladder
Blo Blood
Bone Bone
BM Bone marrow
BrC Brain cortex
BrS Brainstem
Col Colon
Dia Diaphragm
Ret Eye retina
Fat Fat
Hrt Heart
Pit Hypothalamus/Pituitary
Kid Kidney
LM Limb muscle
Liv Liver
Lung Lung
MG Mammary gland
Panc Pancreas
Skin Skin
SI Small intestine
Spl Spleen
Test Testes
Thym Thymus
Tong Tongue
Trac Trachea
Ute Uterus

**Hepcidin (HAMP)**
- endothelial

**PAIRING** (ligand → receptor)

HAMP & FURIN→SLC40A1

**Ligand/Enzyme**
- a - HAMP
- b - FURIN

**Receptor**
- c - SLC40A1

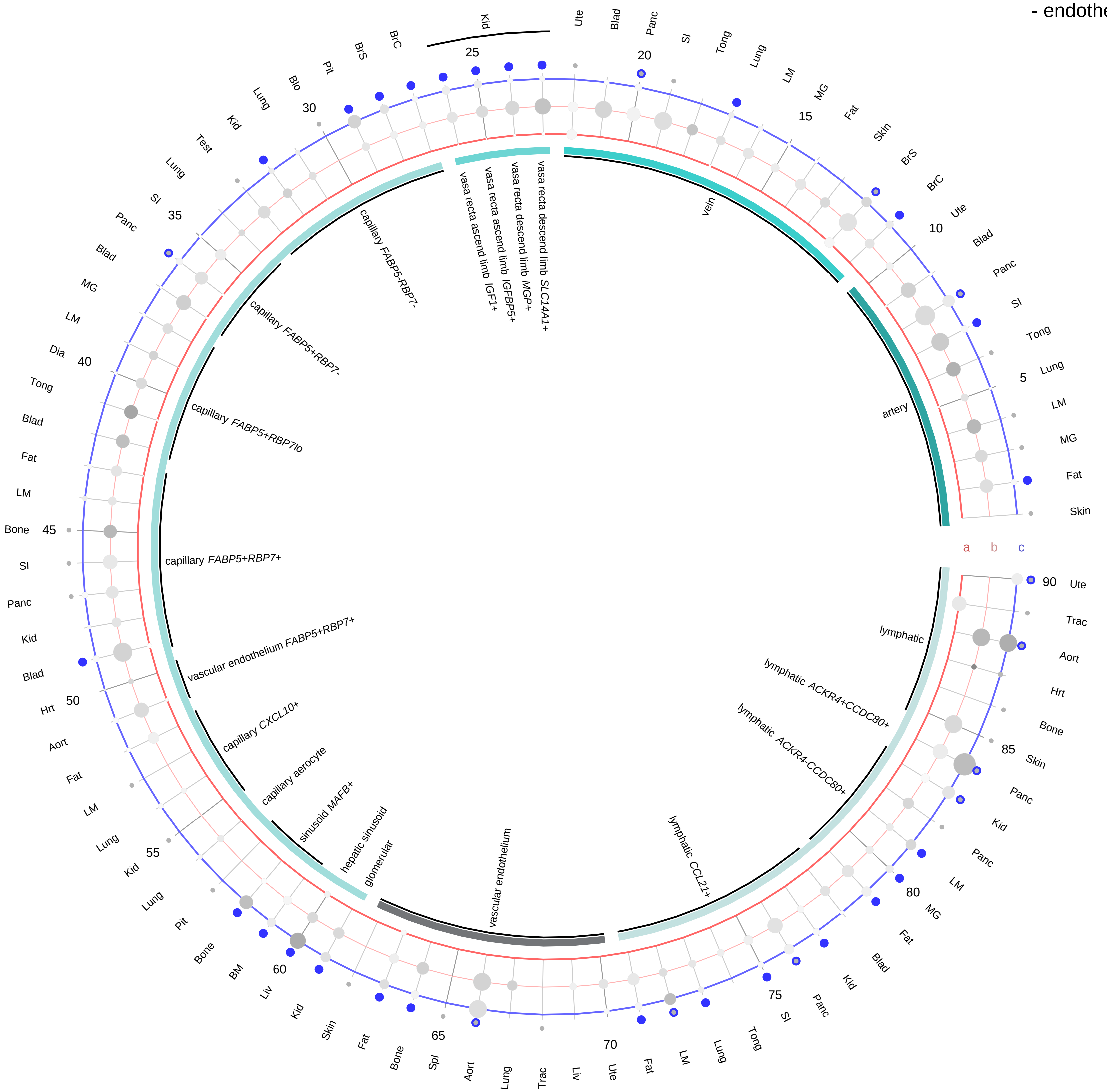

**Compartment**
**endothelial**
- artery
- vein
- vasa recta
- capillary
- mix
- lymphatic

12.5%
25%
50%
100%

ln(CP10K+1): 0, 1, 2, 3

| Abbrev. | Tissue |
|---|---|
| Aort | Aorta |
| Blad | Bladder |
| Blo | Blood |
| Bone | Bone |
| BM | Bone marrow |
| BrC | Brain cortex |
| BrS | Brainstem |
| Col | Colon |
| Dia | Diaphragm |
| Ret | Eye retina |
| Fat | Fat |
| Hrt | Heart |
| Pit | Hypothalamus/Pituitary |
| Kid | Kidney |
| LM | Limb muscle |
| Liv | Liver |
| Lung | Lung |
| MG | Mammary gland |
| Panc | Pancreas |
| Skin | Skin |
| SI | Small intestine |
| Spl | Spleen |
| Test | Testes |
| Thym | Thymus |
| Tong | Tongue |
| Trac | Trachea |
| Ute | Uterus |

**Hepcidin (HAMP)**
- stromal

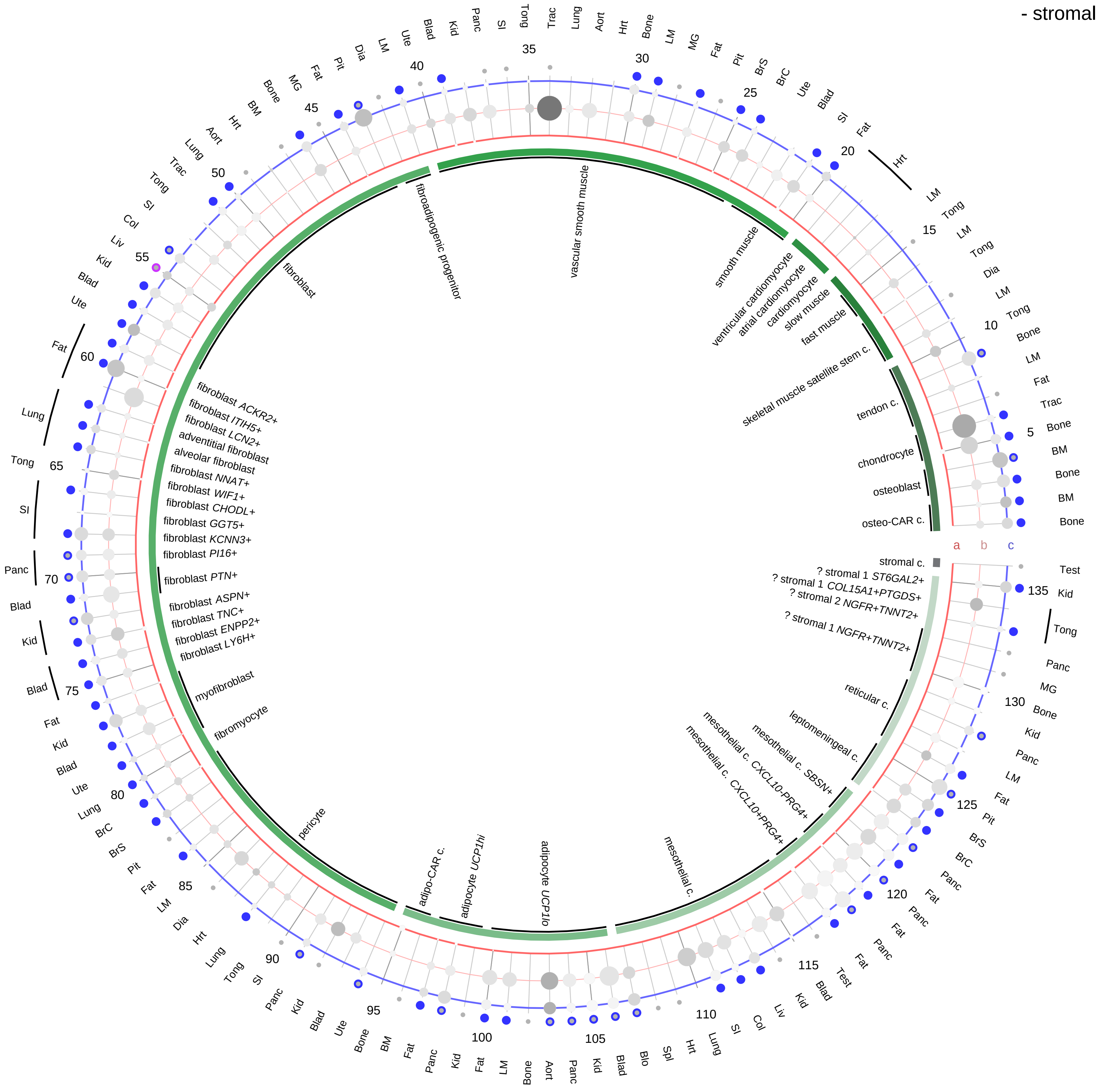

**Ligand/Enzyme**
a - HAMP
b - FURIN
**Receptor**
c - SLC40A1

**PAIRING** (ligand → receptor)

HAMP & FURIN→SLC40A1

**Compartment**
**stromal**
- bone
- skeletal muscle
- cardiac muscle
- smooth muscle
- fibroblast
- adipocyte
- mesothelial
- other
- mix

12.5%
25%
50%
100%

ln(CP10K+1)

| Abbrev. | Tissue |
|---|---|
| Aort | Aorta |
| Blad | Bladder |
| Blo | Blood |
| Bone | Bone |
| BM | Bone marrow |
| BrC | Brain cortex |
| BrS | Brainstem |
| Col | Colon |
| Dia | Diaphragm |
| Ret | Eye retina |
| Fat | Fat |
| Hrt | Heart |
| Pit | Hypothalamus/Pituitary |
| Kid | Kidney |
| LM | Limb muscle |
| Liv | Liver |
| Lung | Lung |
| MG | Mammary gland |
| Panc | Pancreas |
| Skin | Skin |
| SI | Small intestine |
| Spl | Spleen |
| Test | Testes |
| Thym | Thymus |
| Tong | Tongue |
| Trac | Trachea |
| Ute | Uterus |

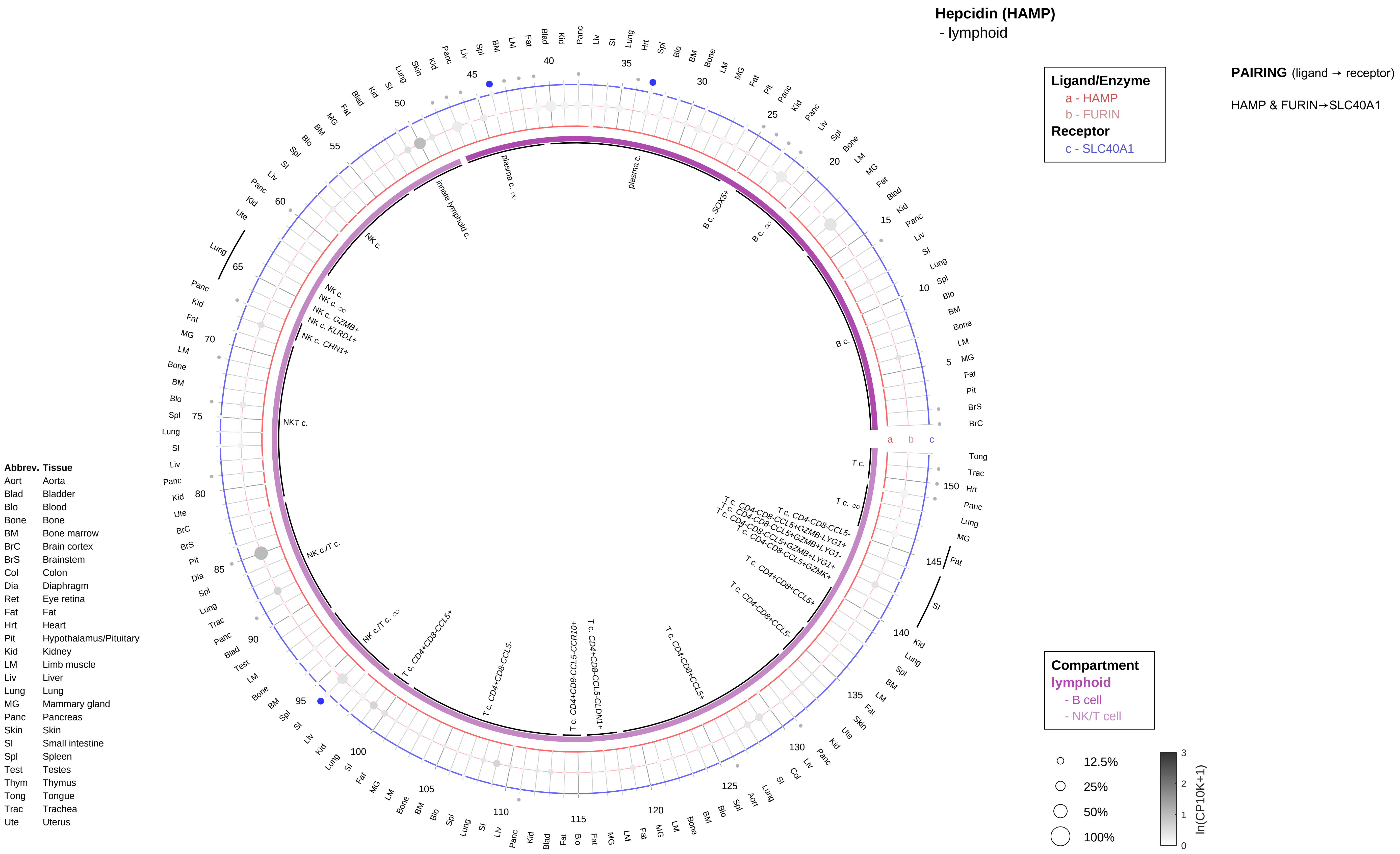
Hepcidin (HAMP)
- lymphoid
Ligand/Enzyme
a - HAMP
b - FURIN
Receptor
c - SLC40A1
PAIRING (ligand → receptor)
HAMP & FURIN→SLC40A1
Compartment
lymphoid
- B cell
- NK/T cell
12.5%
25%
50%
100%
ln(CP10K+1)
Abbrev. Tissue
Aort Aorta
Blad Bladder
Blo Blood
Bone Bone
BM Bone marrow
BrC Brain cortex
BrS Brainstem
Col Colon
Dia Diaphragm
Ret Eye retina
Fat Fat
Hrt Heart
Pit Hypothalamus/Pituitary
Kid Kidney
LM Limb muscle
Liv Liver
Lung Lung
MG Mammary gland
Panc Pancreas
Skin Skin
SI Small intestine
Spl Spleen
Test Testes
Thym Thymus
Tong Tongue
Trac Trachea
Ute Uterus
plasma c.
innate lymphoid c.
B c. SOX5+
B c.
NK c.
NK c. GZMB+
NK c. KLRD1+
NK c. CHN1+
NKT c.
NK c./T c.
T c.
T c. CD4-CD8-CCL5-
T c. CD4-CD8-CCL5+GZMB-LYG1+
T c. CD4-CD8-CCL5+GZMB+LYG1-
T c. CD4-CD8-CCL5+GZMB+LYG1+
T c. CD4-CD8-CCL5+GZMK+
T c. CD4+CD8+CCL5+
T c. CD4-CD8+CCL5-
T c. CD4-CD8+CCL5+
T c. CD4+CD8-CCL5-CLDN1+
T c. CD4+CD8-CCL5-CCR10+
T c. CD4+CD8-CCL5-
T c. CD4+CD8-CCL5+

**Hepcidin (HAMP)**
- myeloid

| |
|---|
| **Ligand/Enzyme** |
| a - HAMP |
| b - FURIN |
| **Receptor** |
| c - SLC40A1 |

**PAIRING** (ligand → receptor)

HAMP & FURIN→SLC40A1

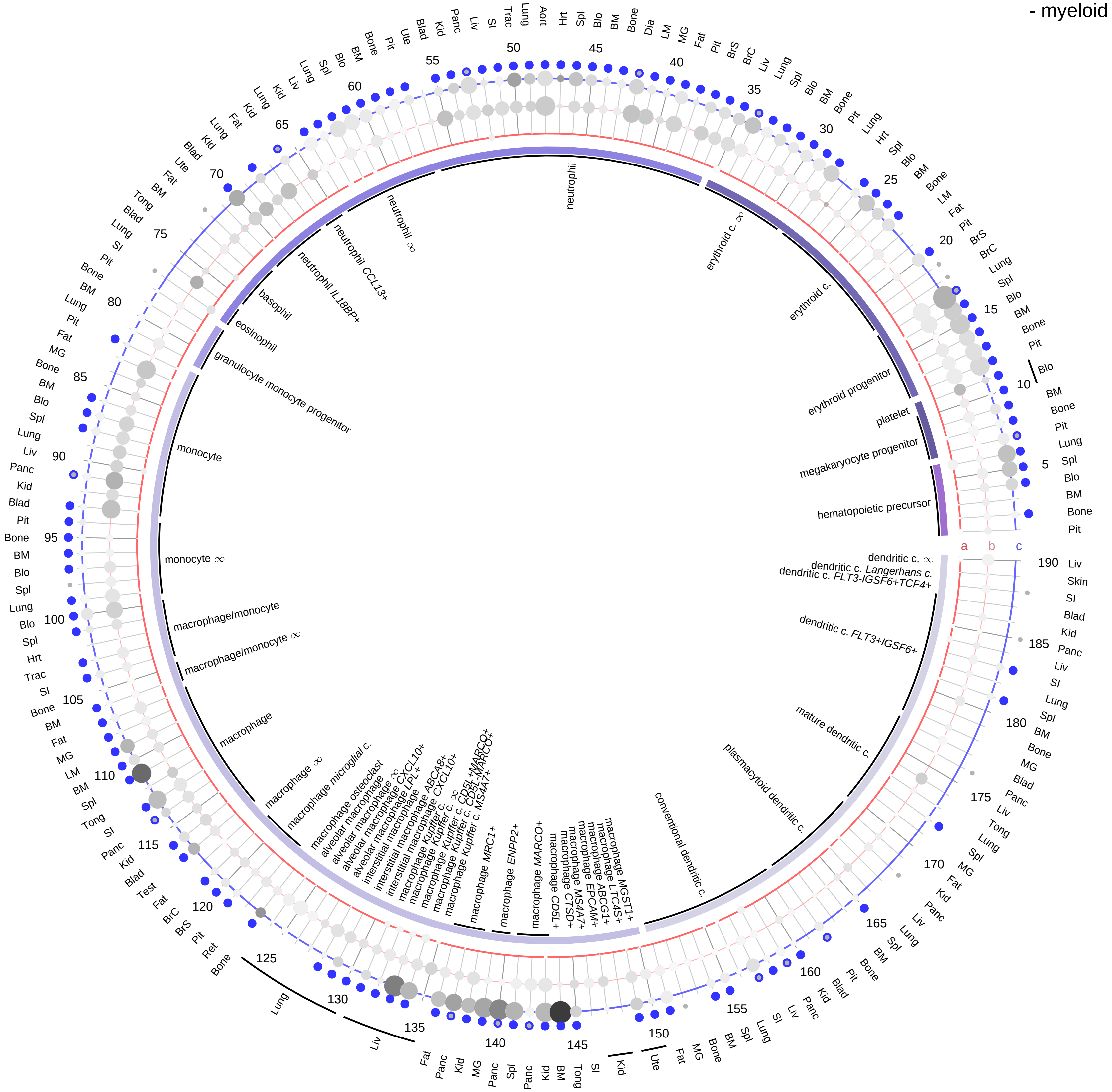

| Abbrev. | Tissue |
|---|---|
| Aort | Aorta |
| Blad | Bladder |
| Blo | Blood |
| Bone | Bone |
| BM | Bone marrow |
| BrC | Brain cortex |
| BrS | Brainstem |
| Col | Colon |
| Dia | Diaphragm |
| Ret | Eye retina |
| Fat | Fat |
| Hrt | Heart |
| Pit | Hypothalamus/Pituitary |
| Kid | Kidney |
| LM | Limb muscle |
| Liv | Liver |
| Lung | Lung |
| MG | Mammary gland |
| Panc | Pancreas |
| Skin | Skin |
| SI | Small intestine |
| Spl | Spleen |
| Test | Testes |
| Thym | Thymus |
| Tong | Tongue |
| Trac | Trachea |
| Ute | Uterus |

**Compartment**

**hematopoietic**
- hematopoietic precursor

**megakaryocyte-erythroid**
- megakaryoid
- erythroid

**myeloid**
- granulocyte
- granulocyte monocyte progenitor
- monocyte/macrophage
- dendritic

12.5%
25%
50%
100%

ln(CP10K+1): 0, 1, 2, 3

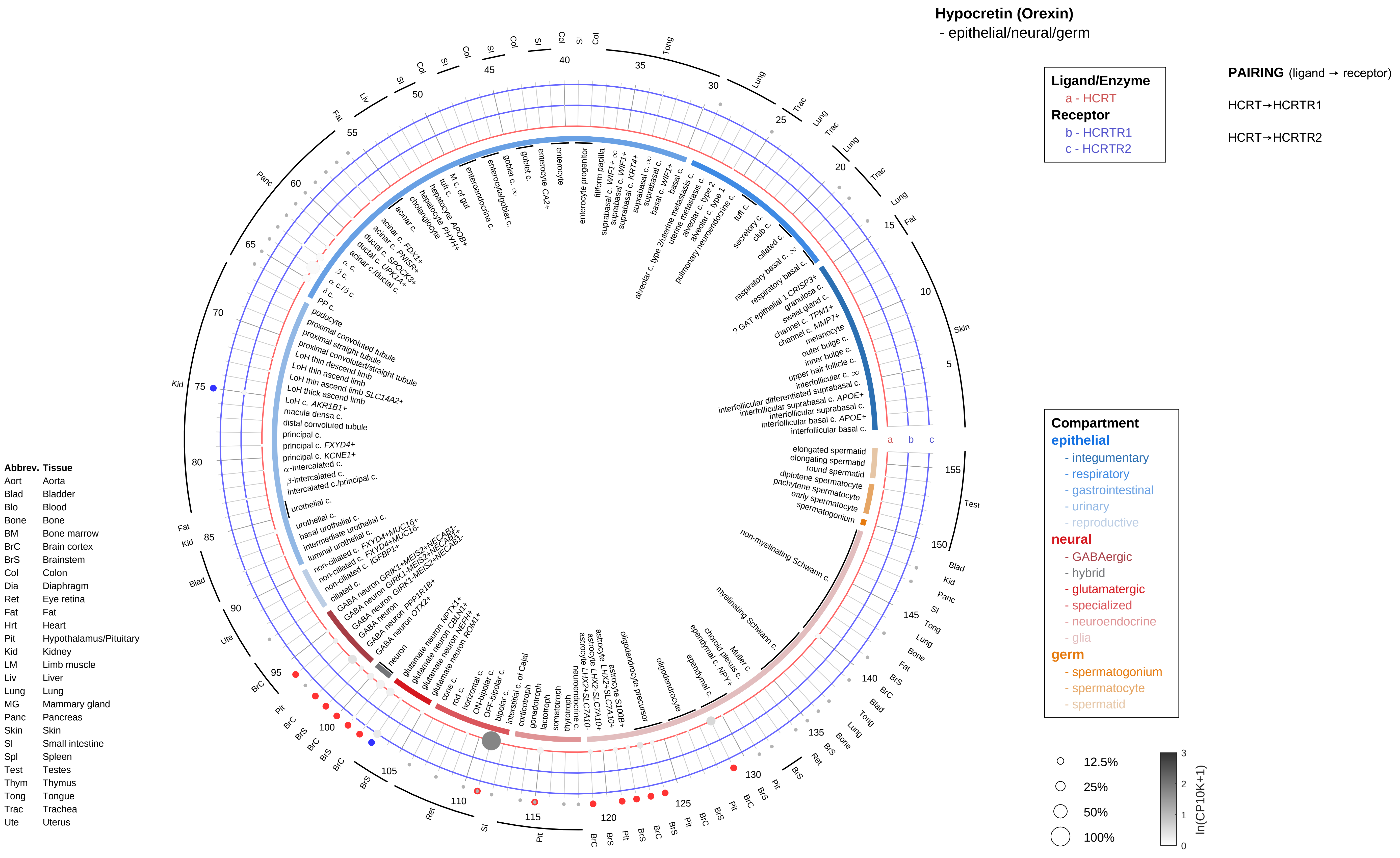
Hypocretin (Orexin)
- epithelial/neural/germ
Ligand/Enzyme
a - HCRT
Receptor
b - HCRTR1
c - HCRTR2
PAIRING (ligand → receptor)
HCRT→HCRTR1
HCRT→HCRTR2
Compartment
epithelial
- integumentary
- respiratory
- gastrointestinal
- urinary
- reproductive
neural
- GABAergic
- hybrid
- glutamatergic
- specialized
- neuroendocrine
- glia
germ
- spermatogonium
- spermatocyte
- spermatid
12.5%
25%
50%
100%
ln(CP10K+1)
Abbrev. Tissue
Aort Aorta
Blad Bladder
Blo Blood
Bone Bone
BM Bone marrow
BrC Brain cortex
BrS Brainstem
Col Colon
Dia Diaphragm
Ret Eye retina
Fat Fat
Hrt Heart
Pit Hypothalamus/Pituitary
Kid Kidney
LM Limb muscle
Liv Liver
Lung Lung
MG Mammary gland
Panc Pancreas
Skin Skin
SI Small intestine
Spl Spleen
Test Testes
Thym Thymus
Tong Tongue
Trac Trachea
Ute Uterus

**Hypocretin (Orexin)**
- endothelial

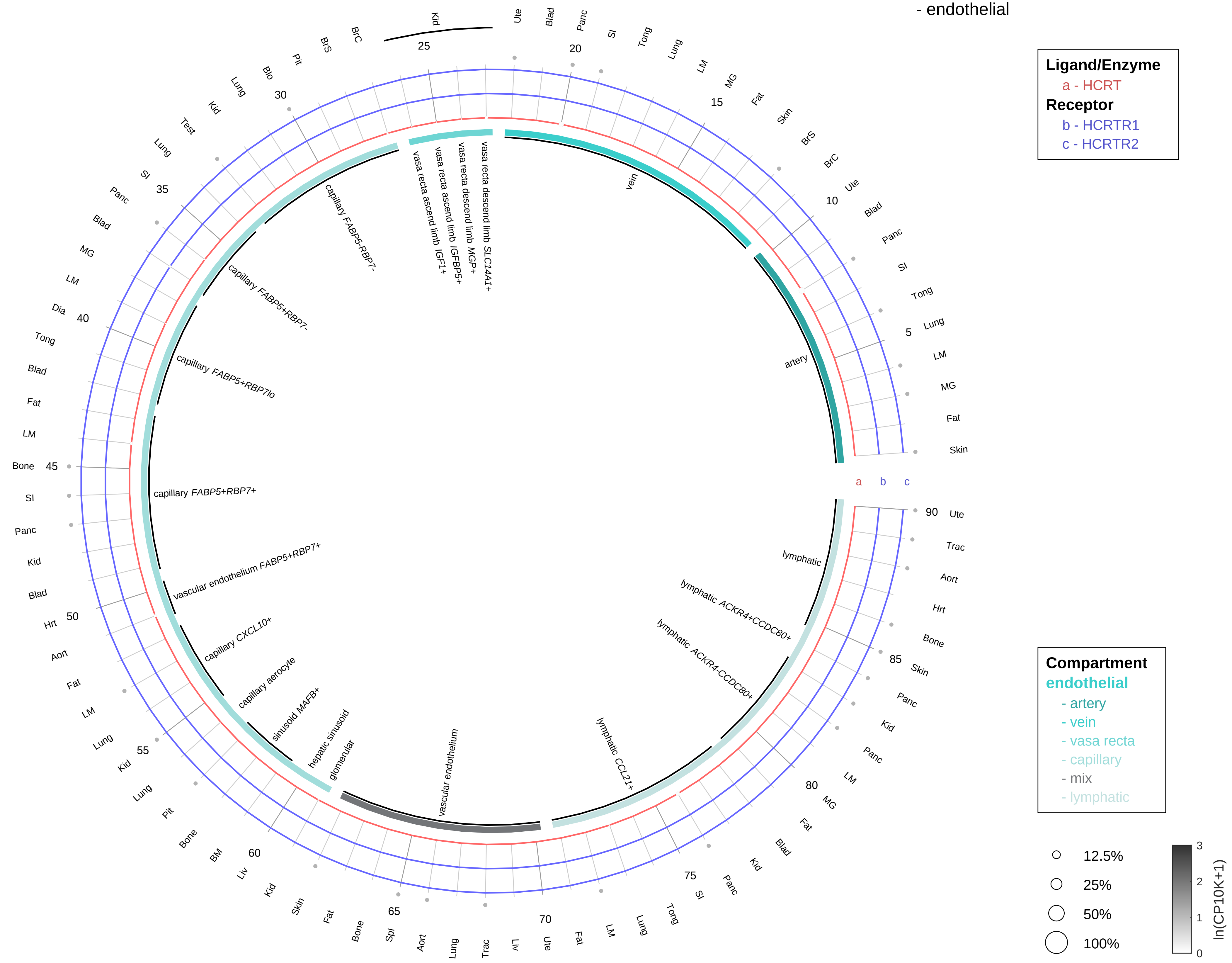

**PAIRING** (ligand → receptor)

HCRT→HCRTR1

HCRT→HCRTR2

| Abbrev. | Tissue |
|---|---|
| Aort | Aorta |
| Blad | Bladder |
| Blo | Blood |
| Bone | Bone |
| BM | Bone marrow |
| BrC | Brain cortex |
| BrS | Brainstem |
| Col | Colon |
| Dia | Diaphragm |
| Ret | Eye retina |
| Fat | Fat |
| Hrt | Heart |
| Pit | Hypothalamus/Pituitary |
| Kid | Kidney |
| LM | Limb muscle |
| Liv | Liver |
| Lung | Lung |
| MG | Mammary gland |
| Panc | Pancreas |
| Skin | Skin |
| SI | Small intestine |
| Spl | Spleen |
| Test | Testes |
| Thym | Thymus |
| Tong | Tongue |
| Trac | Trachea |
| Ute | Uterus |

#### Hypocretin (Orexin)
- stromal

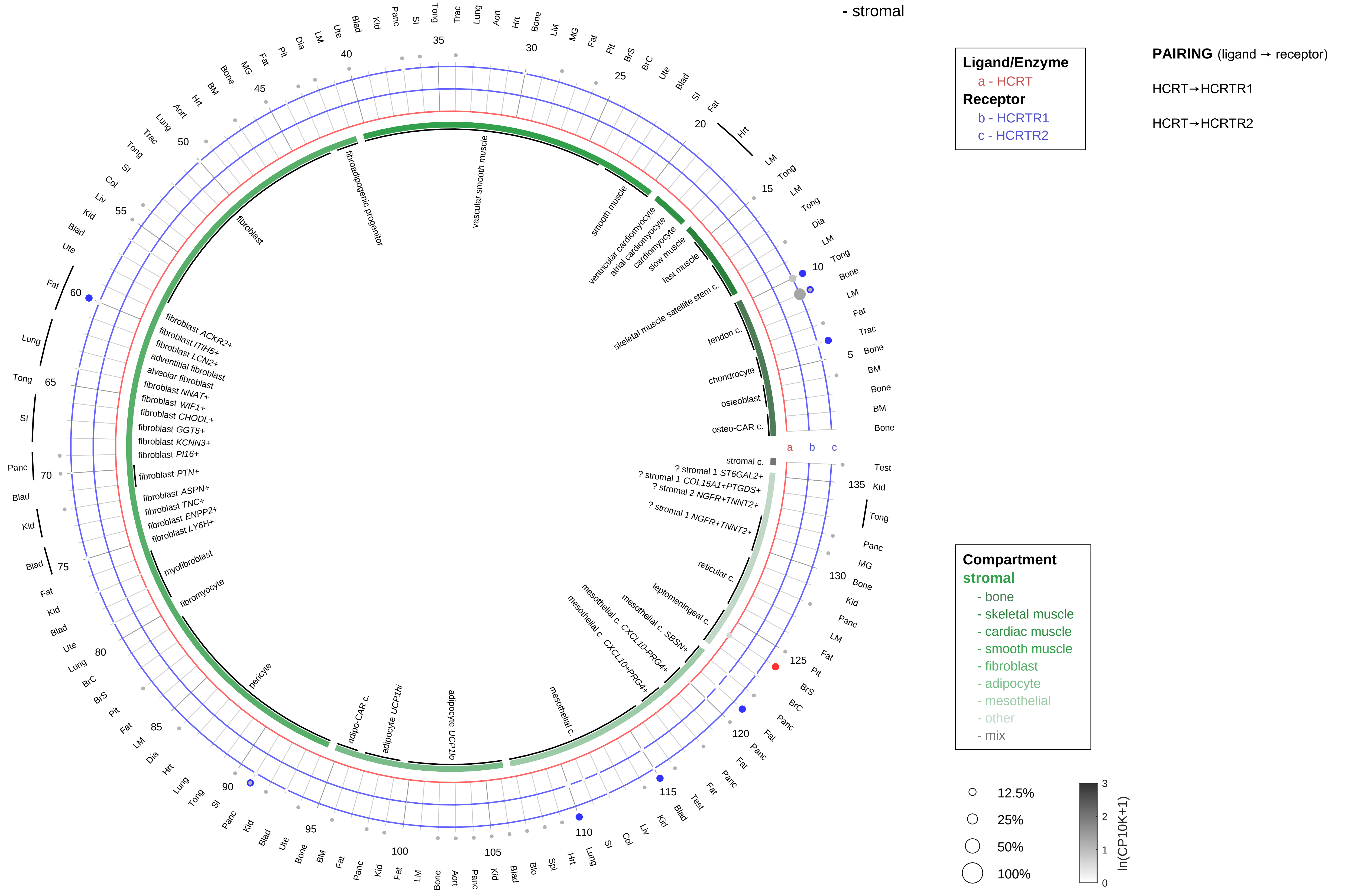

| Abbrev. | Tissue |
|---|---|
| Aort | Aorta |
| Blad | Bladder |
| Blo | Blood |
| Bone | Bone |
| BM | Bone marrow |
| BrC | Brain cortex |
| BrS | Brainstem |
| Col | Colon |
| Dia | Diaphragm |
| Ret | Eye retina |
| Fat | Fat |
| Hrt | Heart |
| Pit | Hypothalamus/Pituitary |
| Kid | Kidney |
| LM | Limb muscle |
| Liv | Liver |
| Lung | Lung |
| MG | Mammary gland |
| Panc | Pancreas |
| Skin | Skin |
| SI | Small intestine |
| Spl | Spleen |
| Test | Testes |
| Thym | Thymus |
| Tong | Tongue |
| Trac | Trachea |
| Ute | Uterus |

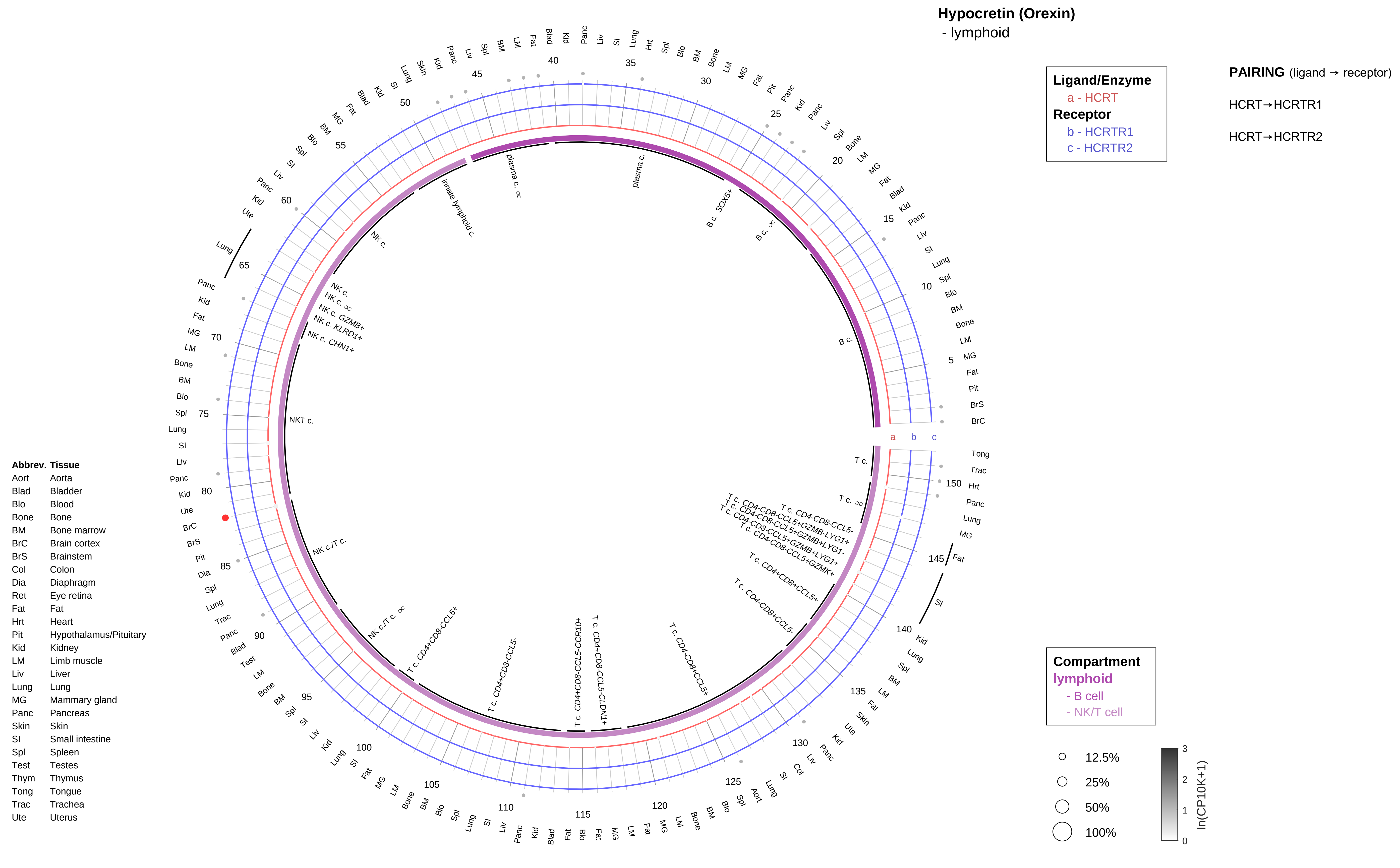
Hypocretin (Orexin)
- lymphoid
Ligand/Enzyme
a - HCRT
Receptor
b - HCRTR1
c - HCRTR2
PAIRING (ligand → receptor)
HCRT→HCRTR1
HCRT→HCRTR2
Compartment
lymphoid
- B cell
- NK/T cell
12.5%
25%
50%
100%
ln(CP10K+1)
Abbrev. Tissue
Aort Aorta
Blad Bladder
Blo Blood
Bone Bone
BM Bone marrow
BrC Brain cortex
BrS Brainstem
Col Colon
Dia Diaphragm
Ret Eye retina
Fat Fat
Hrt Heart
Pit Hypothalamus/Pituitary
Kid Kidney
LM Limb muscle
Liv Liver
Lung Lung
MG Mammary gland
Panc Pancreas
Skin Skin
SI Small intestine
Spl Spleen
Test Testes
Thym Thymus
Tong Tongue
Trac Trachea
Ute Uterus

**Hypocretin (Orexin)**
- myeloid

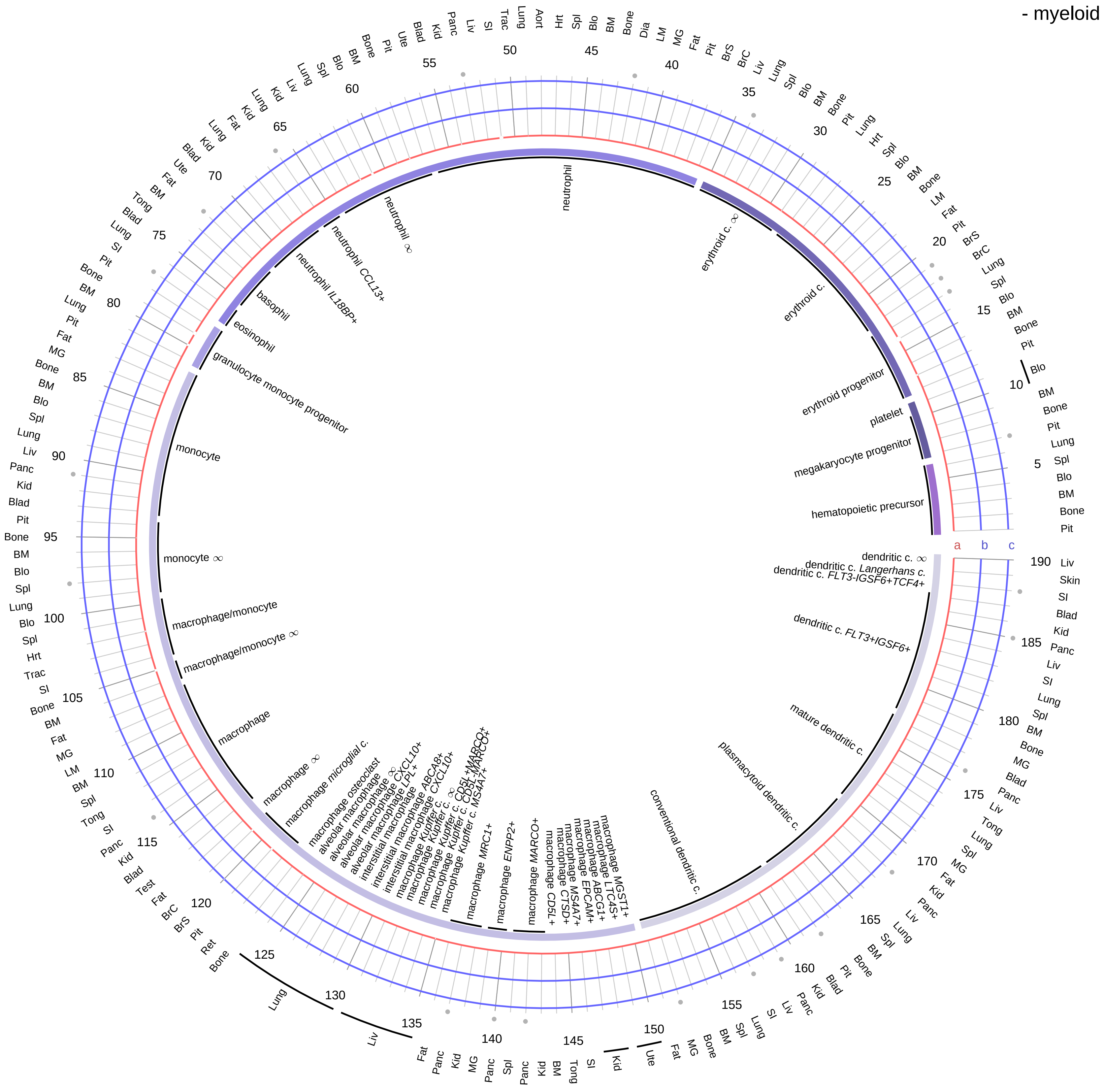

| Ligand/Enzyme |
|---|
| a - HCRT |
| **Receptor** |
| b - HCRTR1 |
| c - HCRTR2 |

**PAIRING** (ligand → receptor)

HCRT→HCRTR1

HCRT→HCRTR2

| Abbrev. | Tissue |
|---|---|
| Aort | Aorta |
| Blad | Bladder |
| Blo | Blood |
| Bone | Bone |
| BM | Bone marrow |
| BrC | Brain cortex |
| BrS | Brainstem |
| Col | Colon |
| Dia | Diaphragm |
| Ret | Eye retina |
| Fat | Fat |
| Hrt | Heart |
| Pit | Hypothalamus/Pituitary |
| Kid | Kidney |
| LM | Limb muscle |
| Liv | Liver |
| Lung | Lung |
| MG | Mammary gland |
| Panc | Pancreas |
| Skin | Skin |
| SI | Small intestine |
| Spl | Spleen |
| Test | Testes |
| Thym | Thymus |
| Tong | Tongue |
| Trac | Trachea |
| Ute | Uterus |

**Compartment**

**hematopoietic**
- hematopoietic precursor

**megakaryocyte-erythroid**
- megakaryoid
- erythroid

**myeloid**
- granulocyte
- granulocyte monocyte progenitor
- monocyte/macrophage
- dendritic

12.5%
25%
50%
100%

ln(CP10K+1): 0, 1, 2, 3

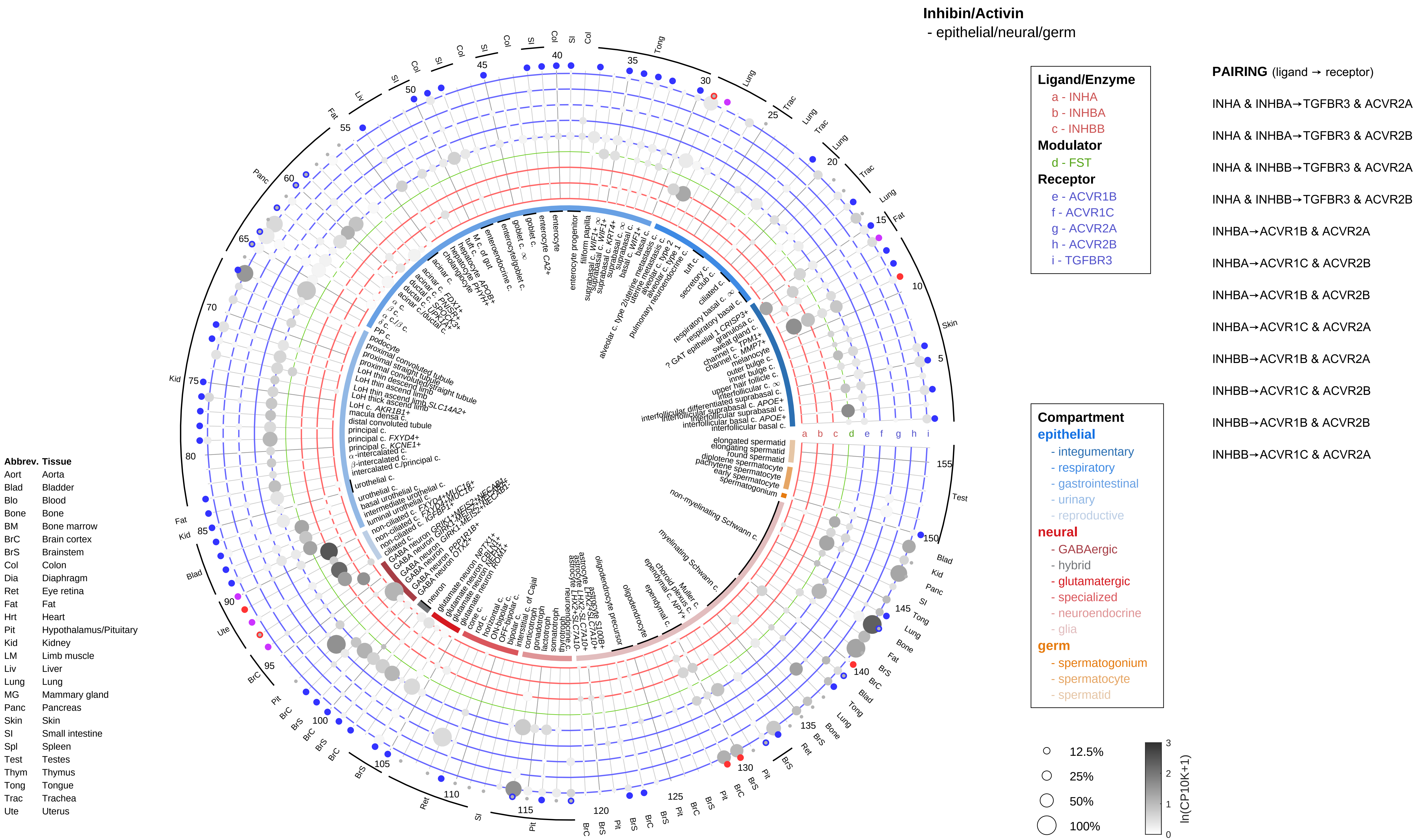

Inhibin/Activin
- epithelial/neural/germ
Ligand/Enzyme
a - INHA
b - INHBA
c - INHBB
Modulator
d - FST
Receptor
e - ACVR1B
f - ACVR1C
g - ACVR2A
h - ACVR2B
i - TGFBR3
PAIRING (ligand → receptor)
INHA & INHBA→TGFBR3 & ACVR2A
INHA & INHBA→TGFBR3 & ACVR2B
INHA & INHBB→TGFBR3 & ACVR2A
INHA & INHBB→TGFBR3 & ACVR2B
INHBA→ACVR1B & ACVR2A
INHBA→ACVR1C & ACVR2B
INHBA→ACVR1B & ACVR2B
INHBA→ACVR1C & ACVR2A
INHBB→ACVR1B & ACVR2A
INHBB→ACVR1C & ACVR2B
INHBB→ACVR1B & ACVR2B
INHBB→ACVR1C & ACVR2A
Compartment
epithelial
- integumentary
- respiratory
- gastrointestinal
- urinary
- reproductive
neural
- GABAergic
- hybrid
- glutamatergic
- specialized
- neuroendocrine
- glia
germ
- spermatogonium
- spermatocyte
- spermatid
12.5%
25%
50%
100%
ln(CP10K+1)
Abbrev. Tissue
Aort Aorta
Blad Bladder
Blo Blood
Bone Bone
BM Bone marrow
BrC Brain cortex
BrS Brainstem
Col Colon
Dia Diaphragm
Ret Eye retina
Fat Fat
Hrt Heart
Pit Hypothalamus/Pituitary
Kid Kidney
LM Limb muscle
Liv Liver
Lung Lung
MG Mammary gland
Panc Pancreas
Skin Skin
SI Small intestine
Spl Spleen
Test Testes
Thym Thymus
Tong Tongue
Trac Trachea
Ute Uterus

**Inhibin/Activin**
- endothelial

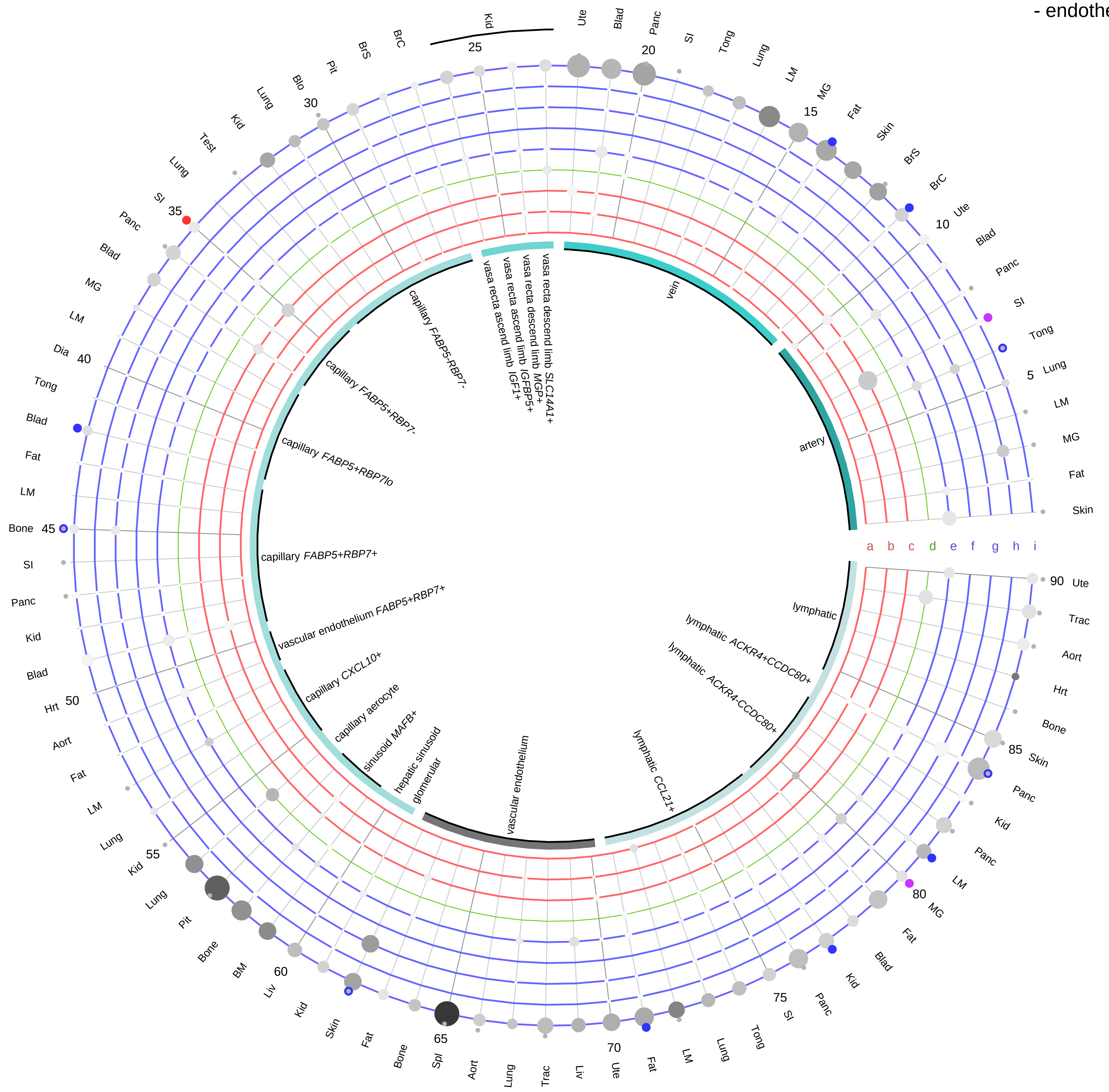

**Ligand/Enzyme**
- a - INHA
- b - INHBA
- c - INHBB

**Modulator**
- d - FST

**Receptor**
- e - ACVR1B
- f - ACVR1C
- g - ACVR2A
- h - ACVR2B
- i - TGFBR3

**PAIRING** (ligand → receptor)

INHA & INHBA→TGFBR3 & ACVR2A
INHA & INHBA→TGFBR3 & ACVR2B
INHA & INHBB→TGFBR3 & ACVR2A
INHA & INHBB→TGFBR3 & ACVR2B
INHBA→ACVR1B & ACVR2A
INHBA→ACVR1C & ACVR2B
INHBA→ACVR1B & ACVR2B
INHBA→ACVR1C & ACVR2A
INHBB→ACVR1B & ACVR2A
INHBB→ACVR1C & ACVR2B
INHBB→ACVR1B & ACVR2B
INHBB→ACVR1C & ACVR2A

**Compartment**
**endothelial**
- artery
- vein
- vasa recta
- capillary
- mix
- lymphatic

12.5%
25%
50%
100%

ln(CP10K+1): 0, 1, 2, 3

| Abbrev. | Tissue |
|---|---|
| Aort | Aorta |
| Blad | Bladder |
| Blo | Blood |
| Bone | Bone |
| BM | Bone marrow |
| BrC | Brain cortex |
| BrS | Brainstem |
| Col | Colon |
| Dia | Diaphragm |
| Ret | Eye retina |
| Fat | Fat |
| Hrt | Heart |
| Pit | Hypothalamus/Pituitary |
| Kid | Kidney |
| LM | Limb muscle |
| Liv | Liver |
| Lung | Lung |
| MG | Mammary gland |
| Panc | Pancreas |
| Skin | Skin |
| SI | Small intestine |
| Spl | Spleen |
| Test | Testes |
| Thym | Thymus |
| Tong | Tongue |
| Trac | Trachea |
| Ute | Uterus |

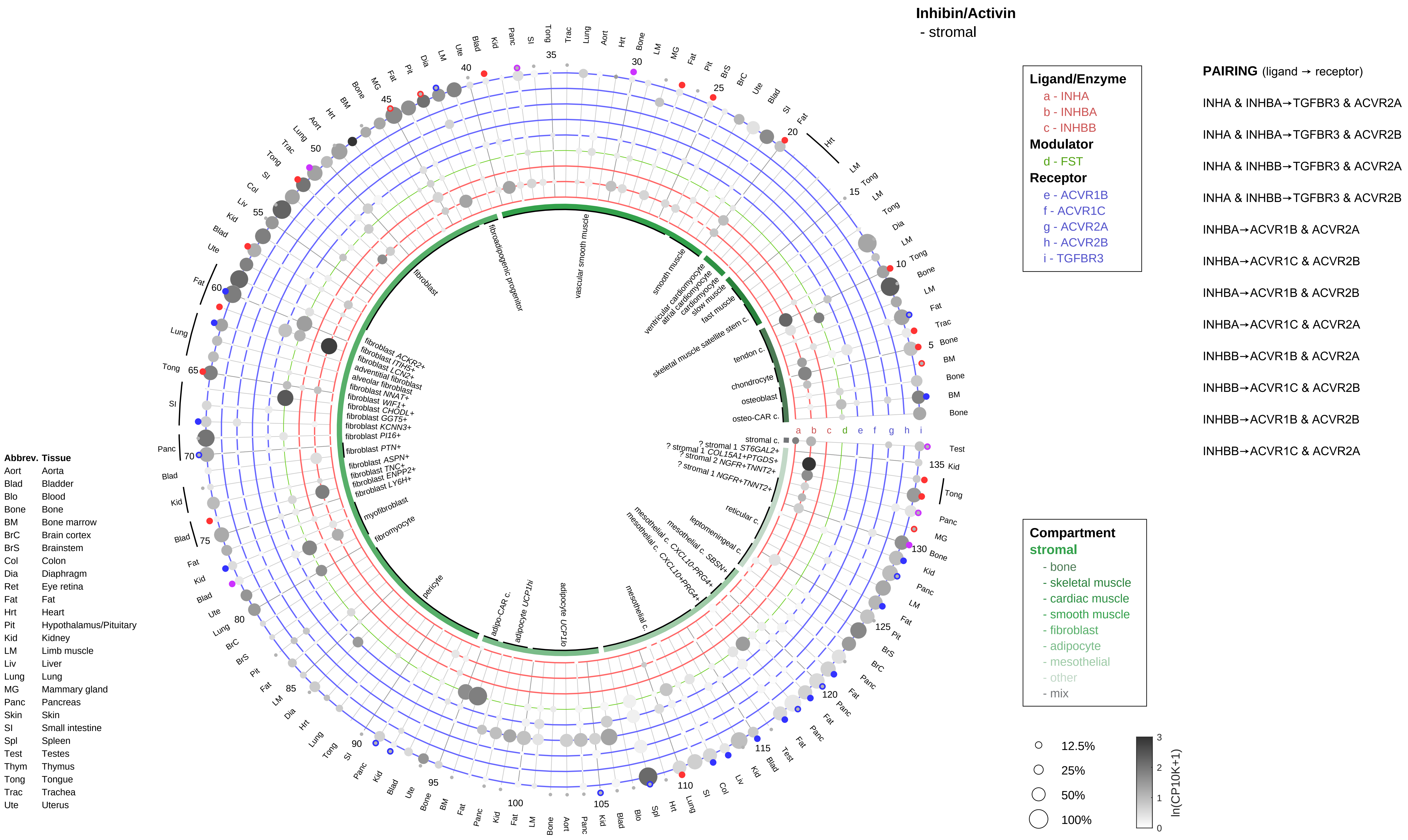
Inhibin/Activin
- stromal
Ligand/Enzyme
a - INHA
b - INHBA
c - INHBB
Modulator
d - FST
Receptor
e - ACVR1B
f - ACVR1C
g - ACVR2A
h - ACVR2B
i - TGFBR3
PAIRING (ligand → receptor)
INHA & INHBA→TGFBR3 & ACVR2A
INHA & INHBA→TGFBR3 & ACVR2B
INHA & INHBB→TGFBR3 & ACVR2A
INHA & INHBB→TGFBR3 & ACVR2B
INHBA→ACVR1B & ACVR2A
INHBA→ACVR1C & ACVR2B
INHBA→ACVR1B & ACVR2B
INHBA→ACVR1C & ACVR2A
INHBB→ACVR1B & ACVR2A
INHBB→ACVR1C & ACVR2B
INHBB→ACVR1B & ACVR2B
INHBB→ACVR1C & ACVR2A
Compartment
stromal
- bone
- skeletal muscle
- cardiac muscle
- smooth muscle
- fibroblast
- adipocyte
- mesothelial
- other
- mix
12.5%
25%
50%
100%
ln(CP10K+1)
Abbrev. Tissue
Aort Aorta
Blad Bladder
Blo Blood
Bone Bone
BM Bone marrow
BrC Brain cortex
BrS Brainstem
Col Colon
Dia Diaphragm
Ret Eye retina
Fat Fat
Hrt Heart
Pit Hypothalamus/Pituitary
Kid Kidney
LM Limb muscle
Liv Liver
Lung Lung
MG Mammary gland
Panc Pancreas
Skin Skin
SI Small intestine
Spl Spleen
Test Testes
Thym Thymus
Tong Tongue
Trac Trachea
Ute Uterus
fibroadipogenic progenitor
vascular smooth muscle
fibroblast
smooth muscle
ventricular cardiomyocyte
atrial cardiomyocyte
cardiomyocyte
slow muscle
fast muscle
skeletal muscle satellite stem c.
tendon c.
chondrocyte
osteoblast
osteo-CAR c.
stromal c.
? stromal 1 ST6GAL2+
? stromal 1 COL15A1+PTGDS+
? stromal 2 NGFR+TNNT2+
? stromal 1 NGFR+TNNT2+
reticular c.
leptomeningeal c.
mesothelial c. SBSN+
mesothelial c. CXCL10-PRG4+
mesothelial c. CXCL10+PRG4+
mesothelial c.
adipocyte UCP1lo
adipocyte UCP1hi
adipo-CAR c.
pericyte
fibromyocyte
myofibroblast
fibroblast LY6H+
fibroblast ENPP2+
fibroblast TNC+
fibroblast ASPN+
fibroblast PTN+
fibroblast PI16+
fibroblast KCNN3+
fibroblast GGT5+
fibroblast CHODL+
fibroblast WIF1+
fibroblast NNAT+
alveolar fibroblast
adventitial fibroblast
fibroblast LCN2+
fibroblast ITIH5+
fibroblast ACKR2+

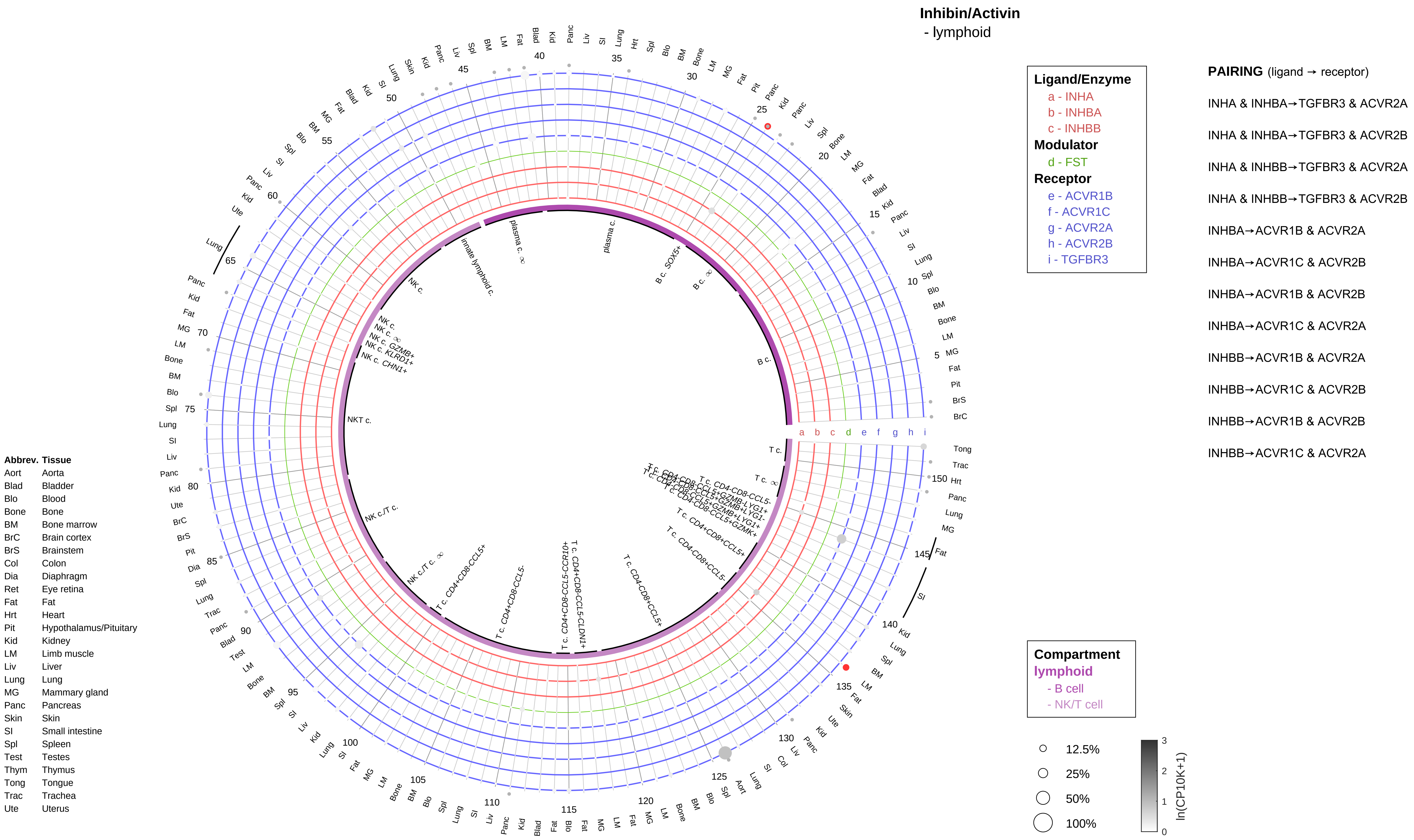

Inhibin/Activin
- lymphoid
Ligand/Enzyme
a - INHA
b - INHBA
c - INHBB
Modulator
d - FST
Receptor
e - ACVR1B
f - ACVR1C
g - ACVR2A
h - ACVR2B
i - TGFBR3
PAIRING (ligand → receptor)
INHA & INHBA→TGFBR3 & ACVR2A
INHA & INHBA→TGFBR3 & ACVR2B
INHA & INHBB→TGFBR3 & ACVR2A
INHA & INHBB→TGFBR3 & ACVR2B
INHBA→ACVR1B & ACVR2A
INHBA→ACVR1C & ACVR2B
INHBA→ACVR1B & ACVR2B
INHBA→ACVR1C & ACVR2A
INHBB→ACVR1B & ACVR2A
INHBB→ACVR1C & ACVR2B
INHBB→ACVR1B & ACVR2B
INHBB→ACVR1C & ACVR2A
Compartment
lymphoid
- B cell
- NK/T cell
12.5%
25%
50%
100%
ln(CP10K+1)
Abbrev. Tissue
Aort Aorta
Blad Bladder
Blo Blood
Bone Bone
BM Bone marrow
BrC Brain cortex
BrS Brainstem
Col Colon
Dia Diaphragm
Ret Eye retina
Fat Fat
Hrt Heart
Pit Hypothalamus/Pituitary
Kid Kidney
LM Limb muscle
Liv Liver
Lung Lung
MG Mammary gland
Panc Pancreas
Skin Skin
SI Small intestine
Spl Spleen
Test Testes
Thym Thymus
Tong Tongue
Trac Trachea
Ute Uterus

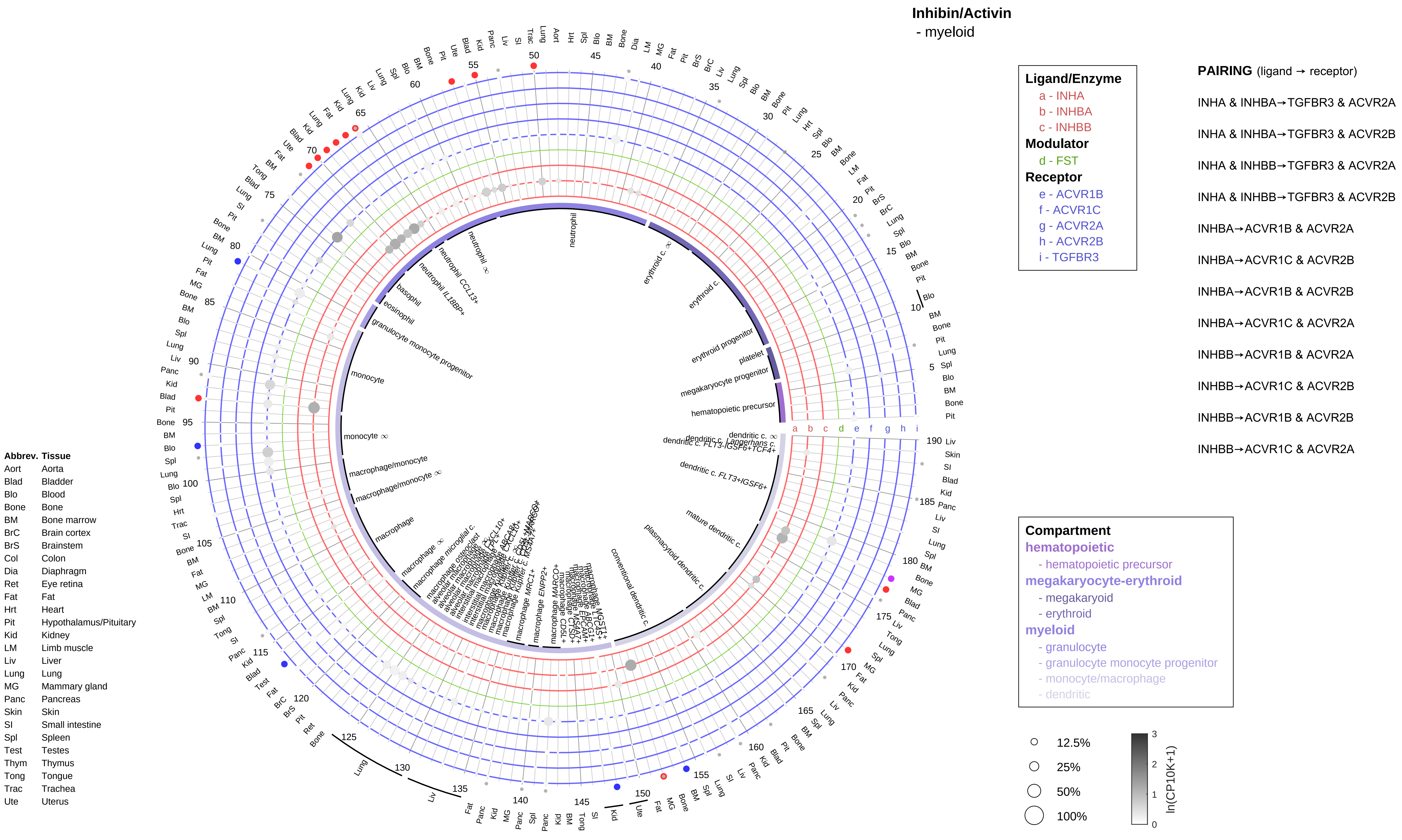

Inhibin/Activin
- myeloid
Ligand/Enzyme
a - INHA
b - INHBA
c - INHBB
Modulator
d - FST
Receptor
e - ACVR1B
f - ACVR1C
g - ACVR2A
h - ACVR2B
i - TGFBR3
PAIRING (ligand → receptor)
INHA & INHBA→TGFBR3 & ACVR2A
INHA & INHBA→TGFBR3 & ACVR2B
INHA & INHBB→TGFBR3 & ACVR2A
INHA & INHBB→TGFBR3 & ACVR2B
INHBA→ACVR1B & ACVR2A
INHBA→ACVR1C & ACVR2B
INHBA→ACVR1B & ACVR2B
INHBA→ACVR1C & ACVR2A
INHBB→ACVR1B & ACVR2A
INHBB→ACVR1C & ACVR2B
INHBB→ACVR1B & ACVR2B
INHBB→ACVR1C & ACVR2A
Compartment
hematopoietic
- hematopoietic precursor
megakaryocyte-erythroid
- megakaryoid
- erythroid
myeloid
- granulocyte
- granulocyte monocyte progenitor
- monocyte/macrophage
- dendritic
12.5%
25%
50%
100%
ln(CP10K+1)
Abbrev. Tissue
Aort Aorta
Blad Bladder
Blo Blood
Bone Bone
BM Bone marrow
BrC Brain cortex
BrS Brainstem
Col Colon
Dia Diaphragm
Ret Eye retina
Fat Fat
Hrt Heart
Pit Hypothalamus/Pituitary
Kid Kidney
LM Limb muscle
Liv Liver
Lung Lung
MG Mammary gland
Panc Pancreas
Skin Skin
SI Small intestine
Spl Spleen
Test Testes
Thym Thymus
Tong Tongue
Trac Trachea
Ute Uterus

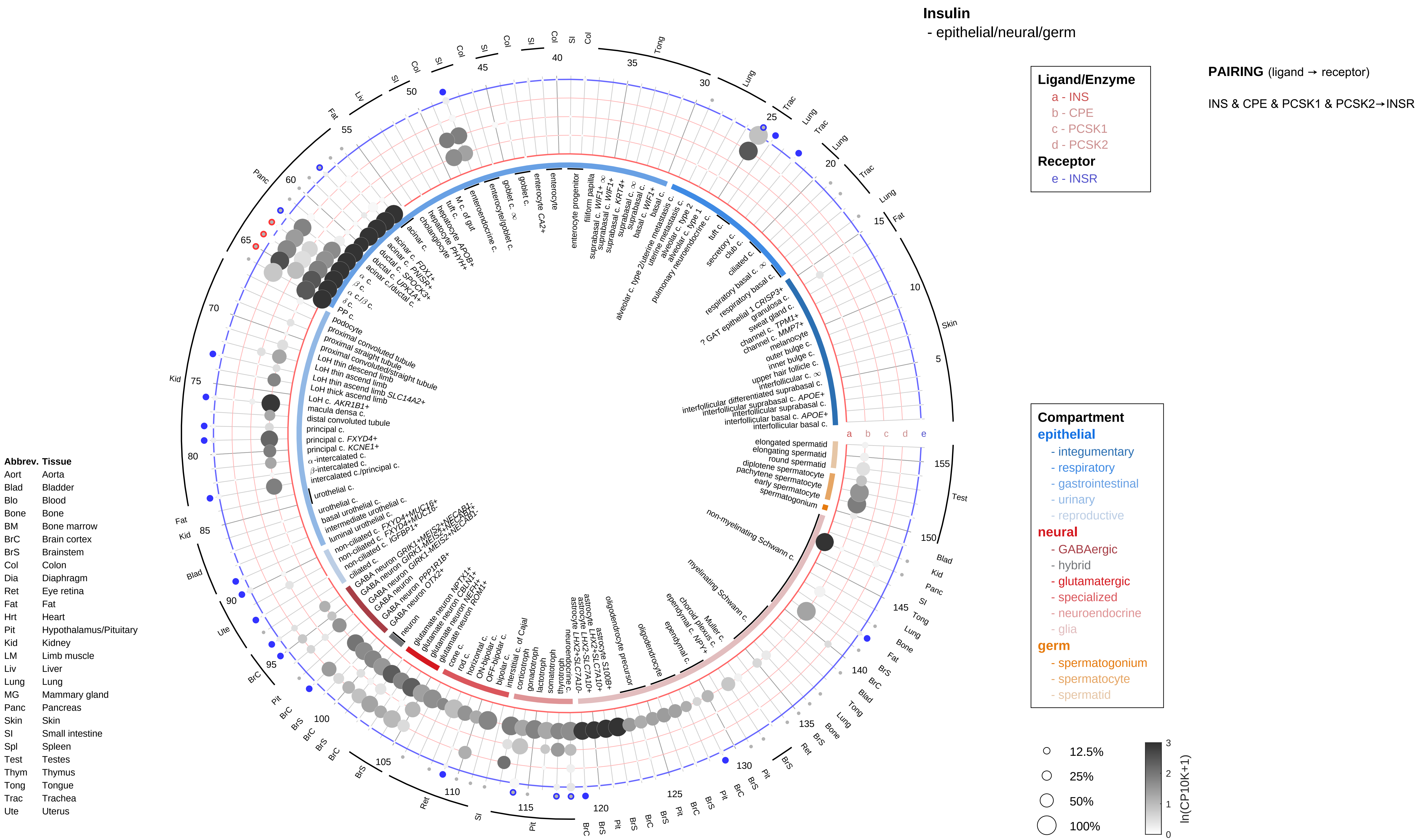
Insulin
- epithelial/neural/germ
Ligand/Enzyme
a - INS
b - CPE
c - PCSK1
d - PCSK2
Receptor
e - INSR
PAIRING (ligand → receptor)
INS & CPE & PCSK1 & PCSK2→INSR
Compartment
epithelial
- integumentary
- respiratory
- gastrointestinal
- urinary
- reproductive
neural
- GABAergic
- hybrid
- glutamatergic
- specialized
- neuroendocrine
- glia
germ
- spermatogonium
- spermatocyte
- spermatid
12.5%
25%
50%
100%
ln(CP10K+1)
Abbrev. Tissue
Aort Aorta
Blad Bladder
Blo Blood
Bone Bone
BM Bone marrow
BrC Brain cortex
BrS Brainstem
Col Colon
Dia Diaphragm
Ret Eye retina
Fat Fat
Hrt Heart
Pit Hypothalamus/Pituitary
Kid Kidney
LM Limb muscle
Liv Liver
Lung Lung
MG Mammary gland
Panc Pancreas
Skin Skin
SI Small intestine
Spl Spleen
Test Testes
Thym Thymus
Tong Tongue
Trac Trachea
Ute Uterus

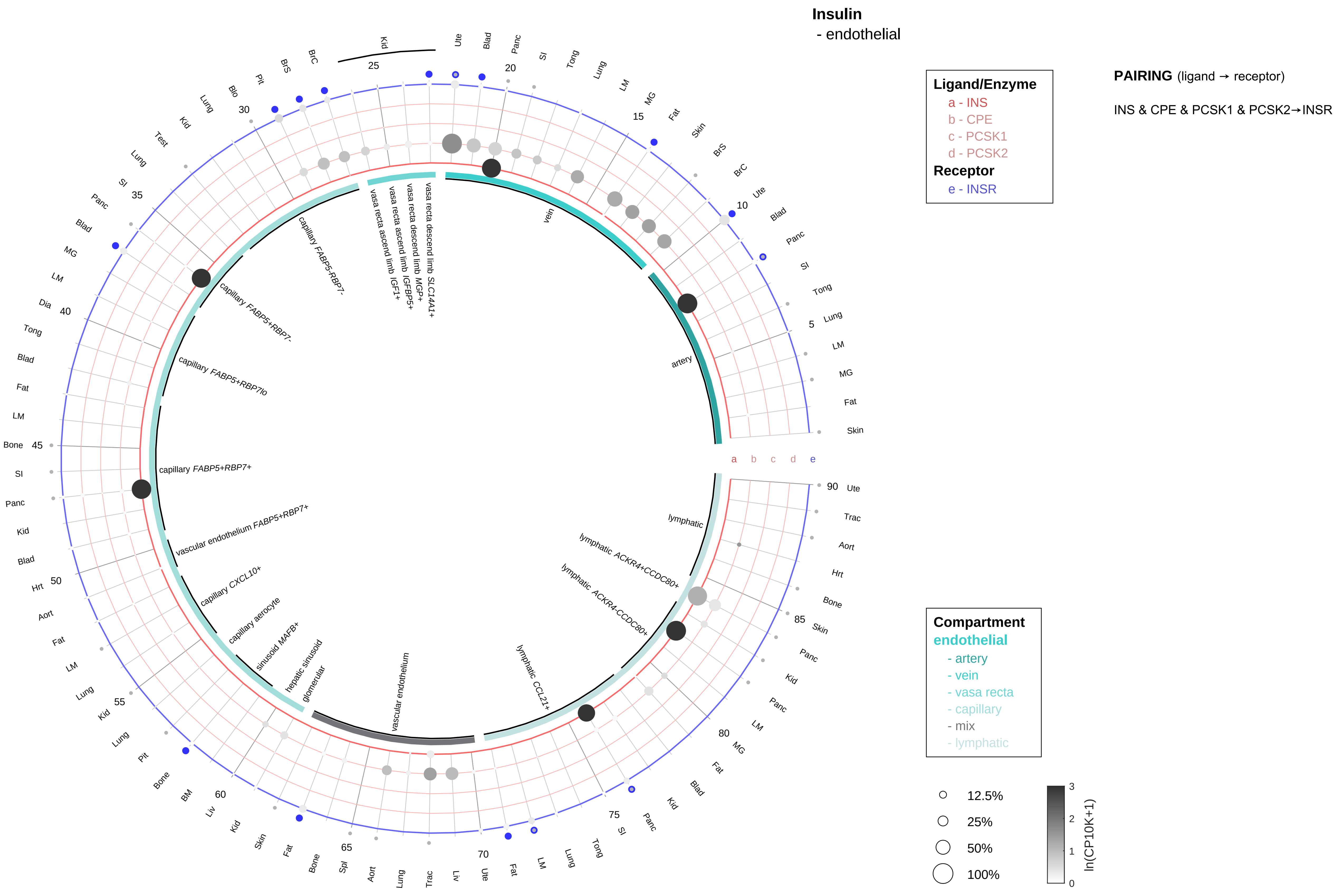

| Abbrev. | Tissue |
|---|---|
| Aort | Aorta |
| Blad | Bladder |
| Blo | Blood |
| Bone | Bone |
| BM | Bone marrow |
| BrC | Brain cortex |
| BrS | Brainstem |
| Col | Colon |
| Dia | Diaphragm |
| Ret | Eye retina |
| Fat | Fat |
| Hrt | Heart |
| Pit | Hypothalamus/Pituitary |
| Kid | Kidney |
| LM | Limb muscle |
| Liv | Liver |
| Lung | Lung |
| MG | Mammary gland |
| Panc | Pancreas |
| Skin | Skin |
| SI | Small intestine |
| Spl | Spleen |
| Test | Testes |
| Thym | Thymus |
| Tong | Tongue |
| Trac | Trachea |
| Ute | Uterus |

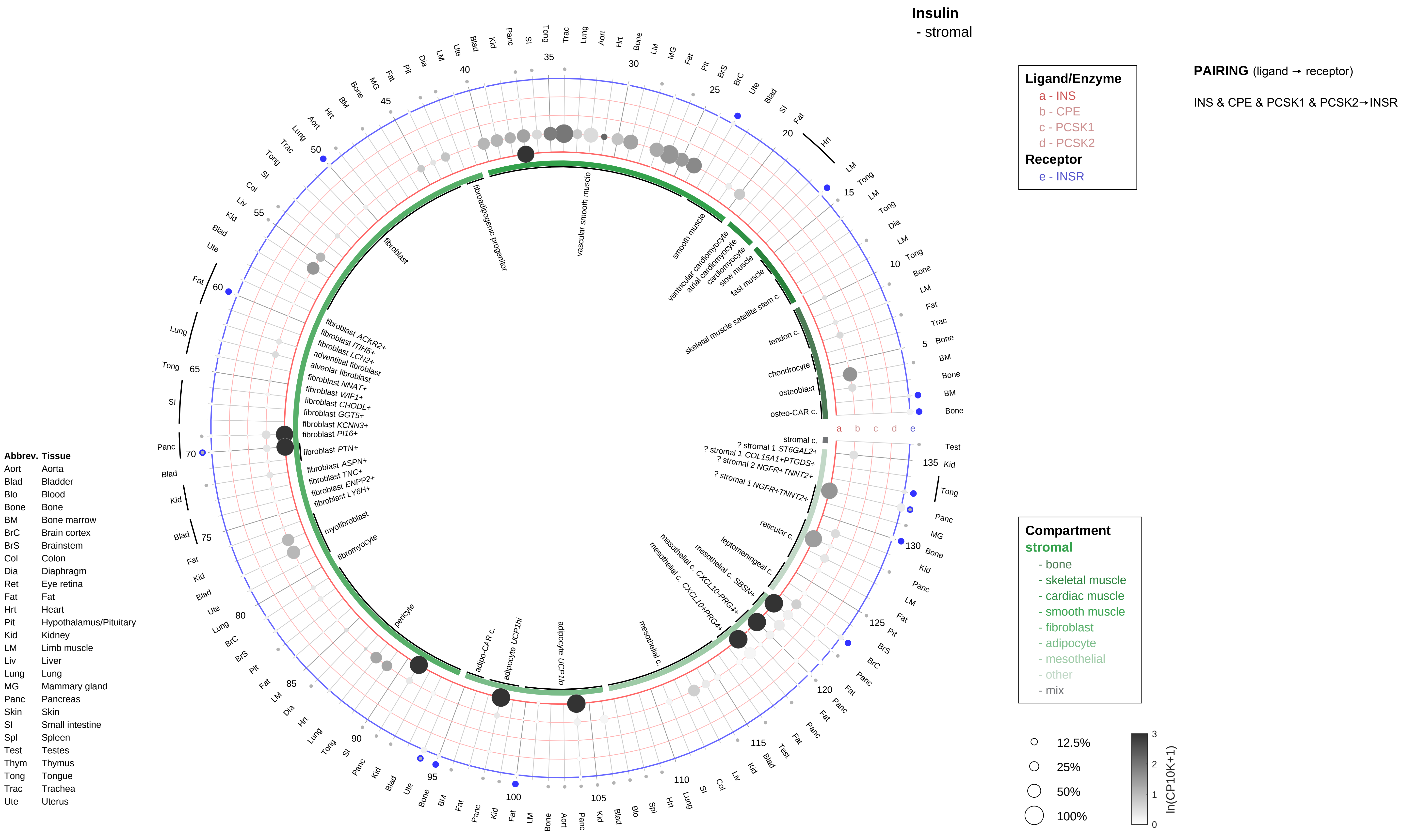

Insulin
- stromal
Ligand/Enzyme
a - INS
b - CPE
c - PCSK1
d - PCSK2
Receptor
e - INSR
PAIRING (ligand → receptor)
INS & CPE & PCSK1 & PCSK2→INSR
Compartment
stromal
- bone
- skeletal muscle
- cardiac muscle
- smooth muscle
- fibroblast
- adipocyte
- mesothelial
- other
- mix
12.5%
25%
50%
100%
ln(CP10K+1)
Abbrev. Tissue
Aort Aorta
Blad Bladder
Blo Blood
Bone Bone
BM Bone marrow
BrC Brain cortex
BrS Brainstem
Col Colon
Dia Diaphragm
Ret Eye retina
Fat Fat
Hrt Heart
Pit Hypothalamus/Pituitary
Kid Kidney
LM Limb muscle
Liv Liver
Lung Lung
MG Mammary gland
Panc Pancreas
Skin Skin
SI Small intestine
Spl Spleen
Test Testes
Thym Thymus
Tong Tongue
Trac Trachea
Ute Uterus

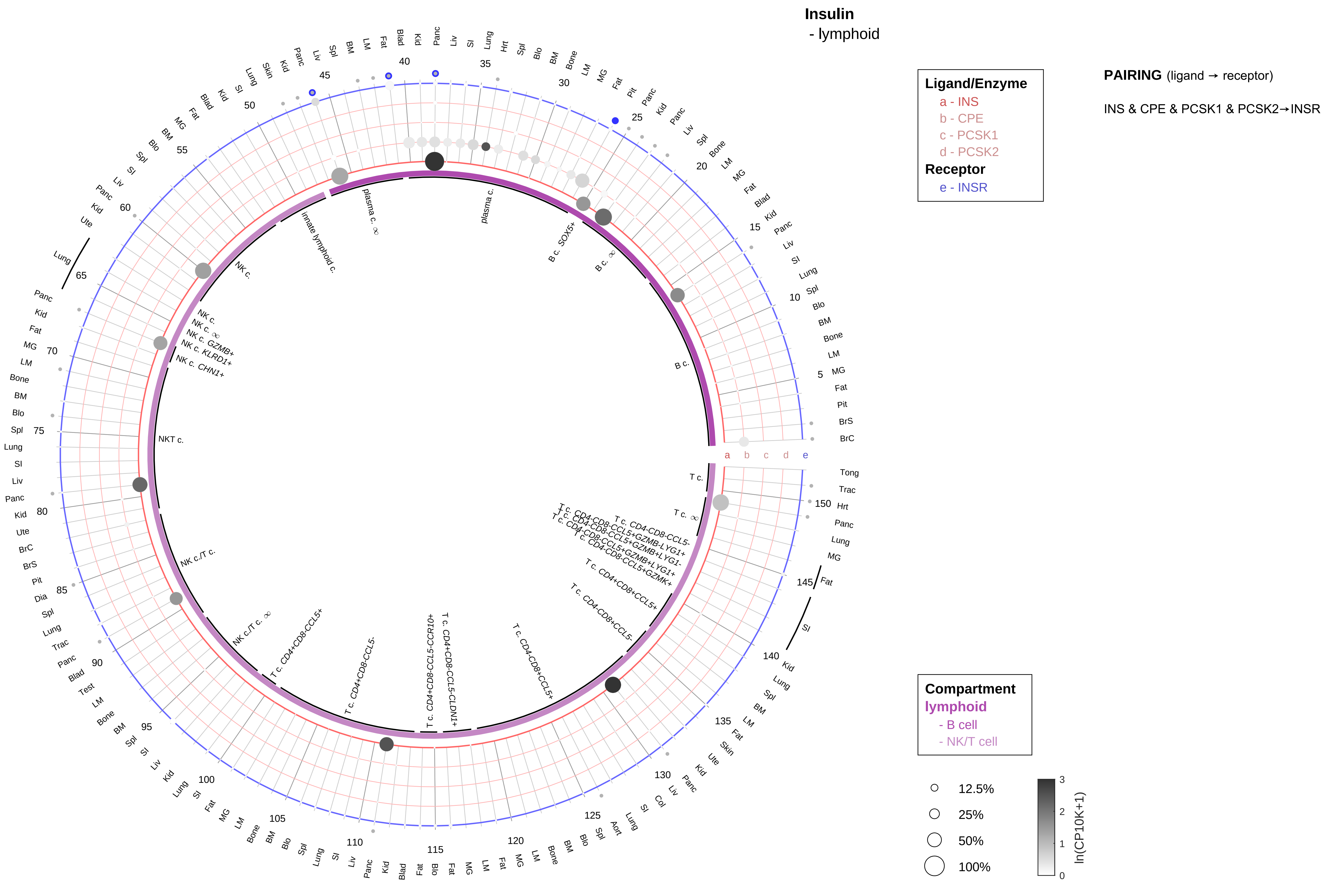

| Abbrev. | Tissue |
|---|---|
| Aort | Aorta |
| Blad | Bladder |
| Blo | Blood |
| Bone | Bone |
| BM | Bone marrow |
| BrC | Brain cortex |
| BrS | Brainstem |
| Col | Colon |
| Dia | Diaphragm |
| Ret | Eye retina |
| Fat | Fat |
| Hrt | Heart |
| Pit | Hypothalamus/Pituitary |
| Kid | Kidney |
| LM | Limb muscle |
| Liv | Liver |
| Lung | Lung |
| MG | Mammary gland |
| Panc | Pancreas |
| Skin | Skin |
| SI | Small intestine |
| Spl | Spleen |
| Test | Testes |
| Thym | Thymus |
| Tong | Tongue |
| Trac | Trachea |
| Ute | Uterus |

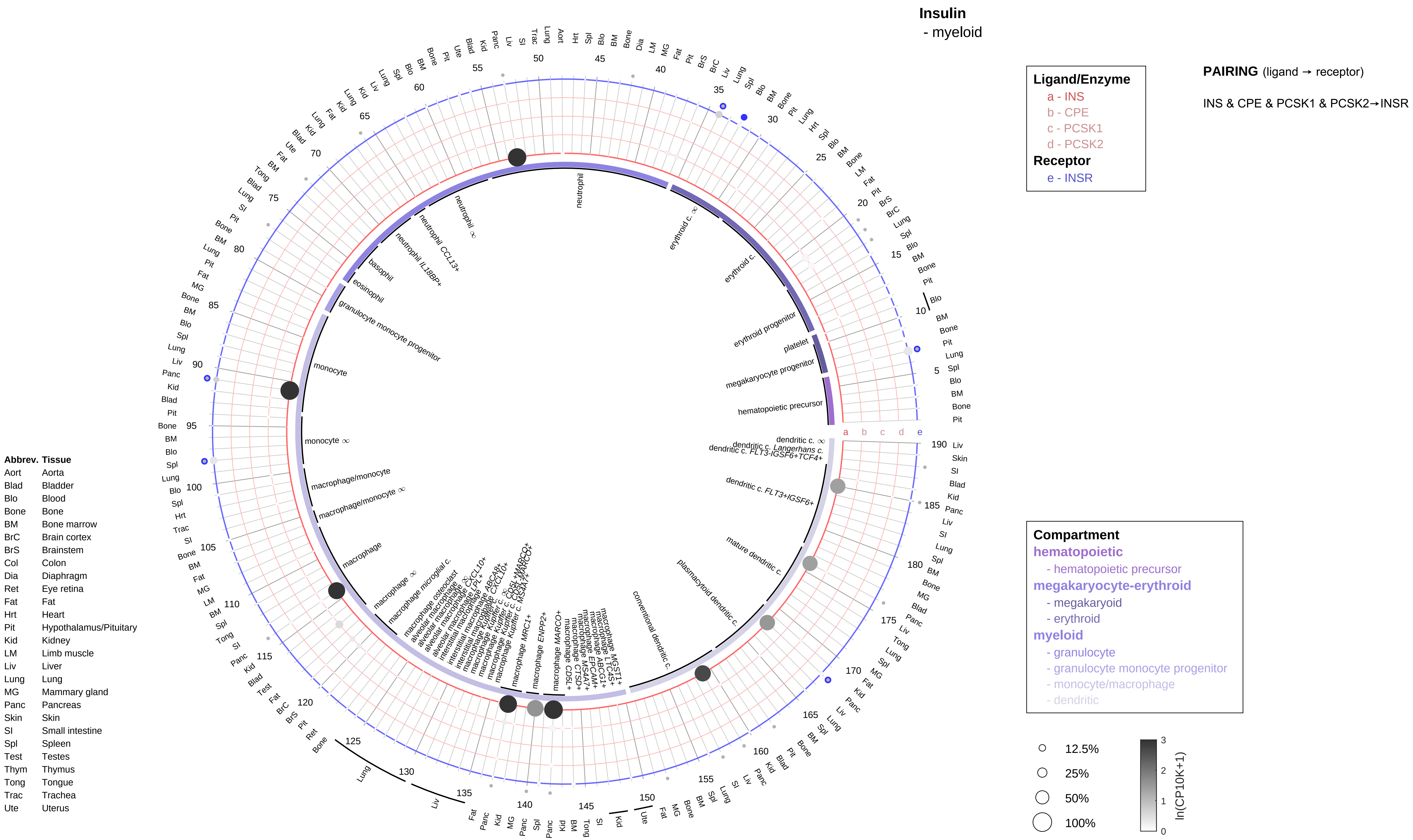
Insulin
- myeloid
Ligand/Enzyme
a - INS
b - CPE
c - PCSK1
d - PCSK2
Receptor
e - INSR
PAIRING (ligand → receptor)
INS & CPE & PCSK1 & PCSK2→INSR
Compartment
hematopoietic
- hematopoietic precursor
megakaryocyte-erythroid
- megakaryoid
- erythroid
myeloid
- granulocyte
- granulocyte monocyte progenitor
- monocyte/macrophage
- dendritic
12.5%
25%
50%
100%
ln(CP10K+1)
Abbrev. Tissue
Aort Aorta
Blad Bladder
Blo Blood
Bone Bone
BM Bone marrow
BrC Brain cortex
BrS Brainstem
Col Colon
Dia Diaphragm
Ret Eye retina
Fat Fat
Hrt Heart
Pit Hypothalamus/Pituitary
Kid Kidney
LM Limb muscle
Liv Liver
Lung Lung
MG Mammary gland
Panc Pancreas
Skin Skin
SI Small intestine
Spl Spleen
Test Testes
Thym Thymus
Tong Tongue
Trac Trachea
Ute Uterus

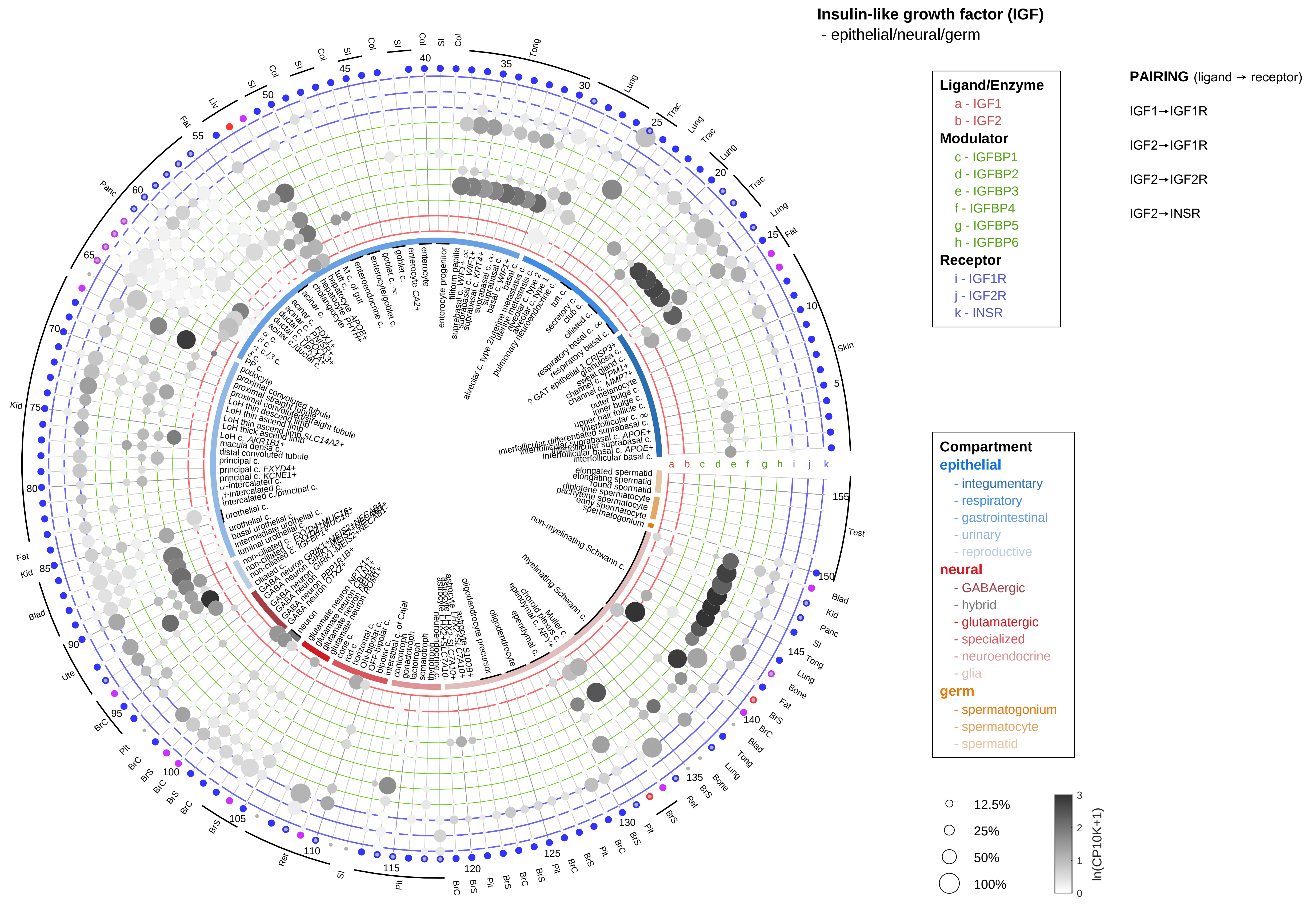

| Abbrev. | Tissue |
|---|---|
| Aort | Aorta |
| Blad | Bladder |
| Blo | Blood |
| Bone | Bone |
| BM | Bone marrow |
| BrC | Brain cortex |
| BrS | Brainstem |
| Col | Colon |
| Dia | Diaphragm |
| Ret | Eye retina |
| Fat | Fat |
| Hrt | Heart |
| Pit | Hypothalamus/Pituitary |
| Kid | Kidney |
| LM | Limb muscle |
| Liv | Liver |
| Lung | Lung |
| MG | Mammary gland |
| Panc | Pancreas |
| Skin | Skin |
| SI | Small intestine |
| Spl | Spleen |
| Test | Testes |
| Thym | Thymus |
| Tong | Tongue |
| Trac | Trachea |
| Ute | Uterus |

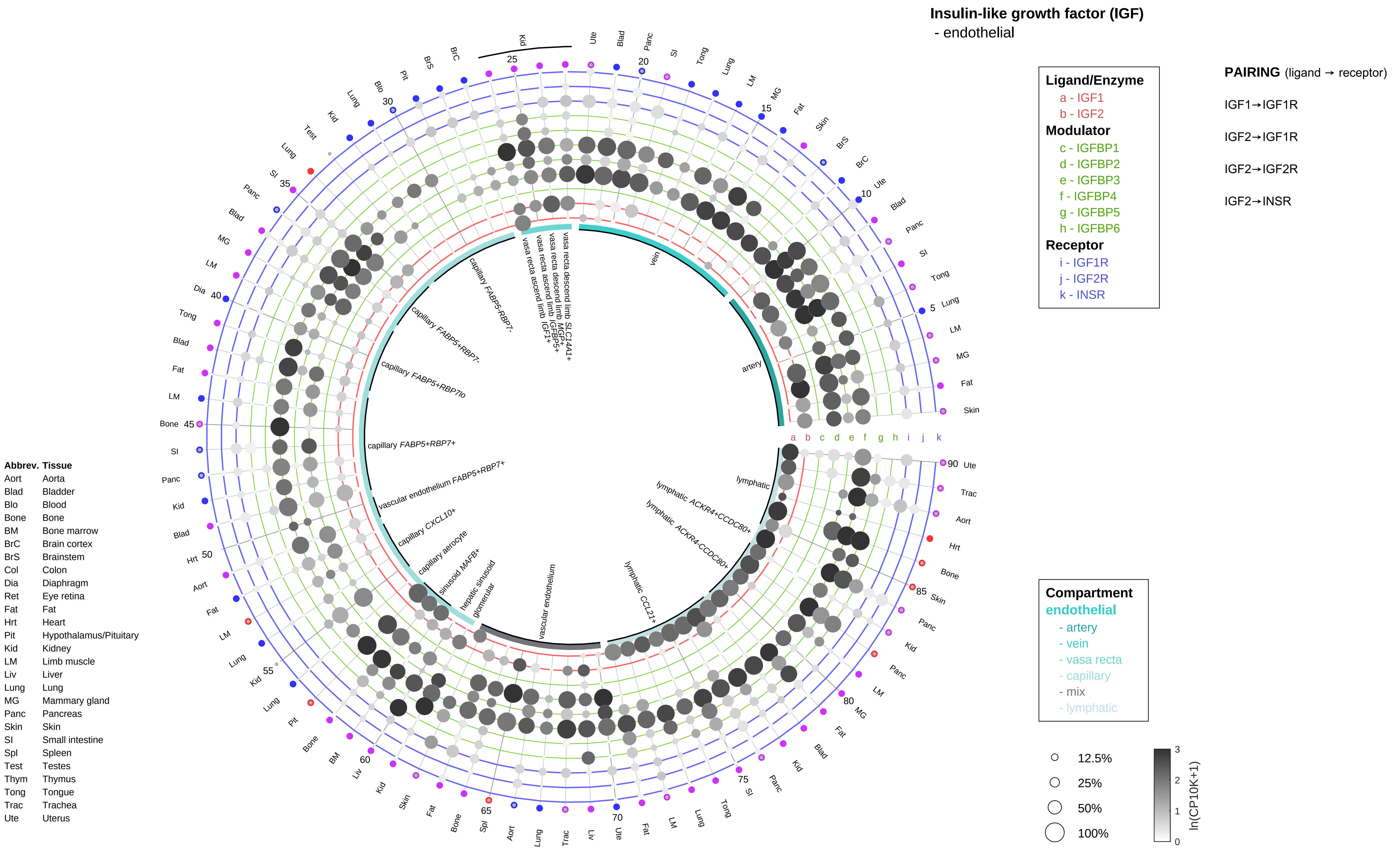
Insulin-like growth factor (IGF)
- endothelial
Ligand/Enzyme
a - IGF1
b - IGF2
Modulator
c - IGFBP1
d - IGFBP2
e - IGFBP3
f - IGFBP4
g - IGFBP5
h - IGFBP6
Receptor
i - IGF1R
j - IGF2R
k - INSR
PAIRING (ligand → receptor)
IGF1→IGF1R
IGF2→IGF1R
IGF2→IGF2R
IGF2→INSR
Compartment
endothelial
- artery
- vein
- vasa recta
- capillary
- mix
- lymphatic
12.5%
25%
50%
100%
ln(CP10K+1)
Abbrev. Tissue
Aort Aorta
Blad Bladder
Blo Blood
Bone Bone
BM Bone marrow
BrC Brain cortex
BrS Brainstem
Col Colon
Dia Diaphragm
Ret Eye retina
Fat Fat
Hrt Heart
Pit Hypothalamus/Pituitary
Kid Kidney
LM Limb muscle
Liv Liver
Lung Lung
MG Mammary gland
Panc Pancreas
Skin Skin
SI Small intestine
Spl Spleen
Test Testes
Thym Thymus
Tong Tongue
Trac Trachea
Ute Uterus
vein
artery
lymphatic
lymphatic ACKR4+CCDC80+
lymphatic ACKR4-CCDC80-
lymphatic CCL21+
vascular endothelium
glomerular
hepatic sinusoid
sinusoid MAFB+
capillary aerocyte
capillary CXCL10+
vascular endothelium FABP5+RBP7+
capillary FABP5+RBP7+
capillary FABP5+RBP7lo
capillary FABP5+RBP7-
capillary FABP5-RBP7-
vasa recta ascend limb IGF1+
vasa recta ascend limb IGFBP5+
vasa recta descend limb MGP+
vasa recta descend limb SLC14A1+

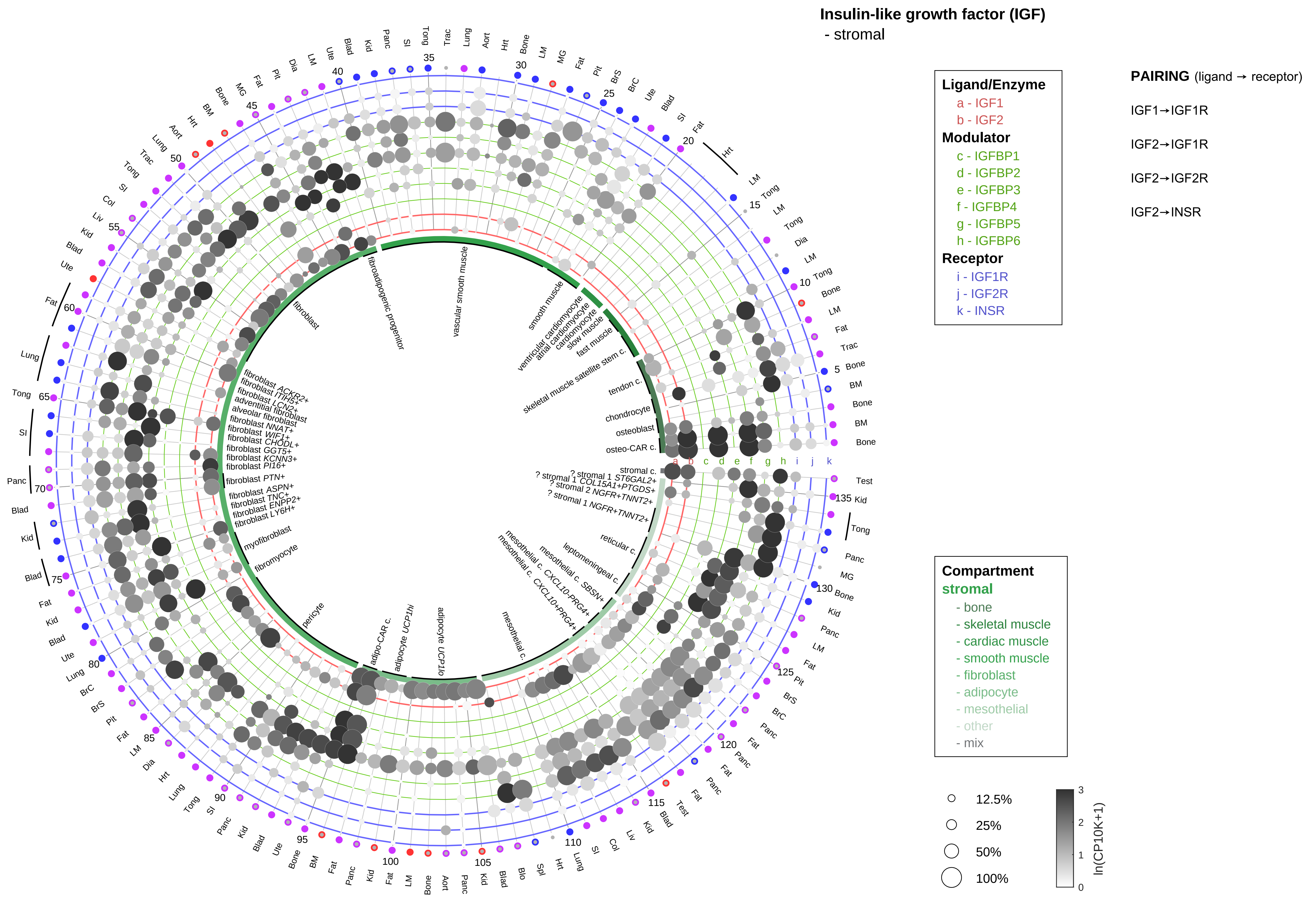

| Abbrev. | Tissue |
|---|---|
| Aort | Aorta |
| Blad | Bladder |
| Blo | Blood |
| Bone | Bone |
| BM | Bone marrow |
| BrC | Brain cortex |
| BrS | Brainstem |
| Col | Colon |
| Dia | Diaphragm |
| Ret | Eye retina |
| Fat | Fat |
| Hrt | Heart |
| Pit | Hypothalamus/Pituitary |
| Kid | Kidney |
| LM | Limb muscle |
| Liv | Liver |
| Lung | Lung |
| MG | Mammary gland |
| Panc | Pancreas |
| Skin | Skin |
| SI | Small intestine |
| Spl | Spleen |
| Test | Testes |
| Thym | Thymus |
| Tong | Tongue |
| Trac | Trachea |
| Ute | Uterus |

#### Insulin-like growth factor (IGF)
- lymphoid

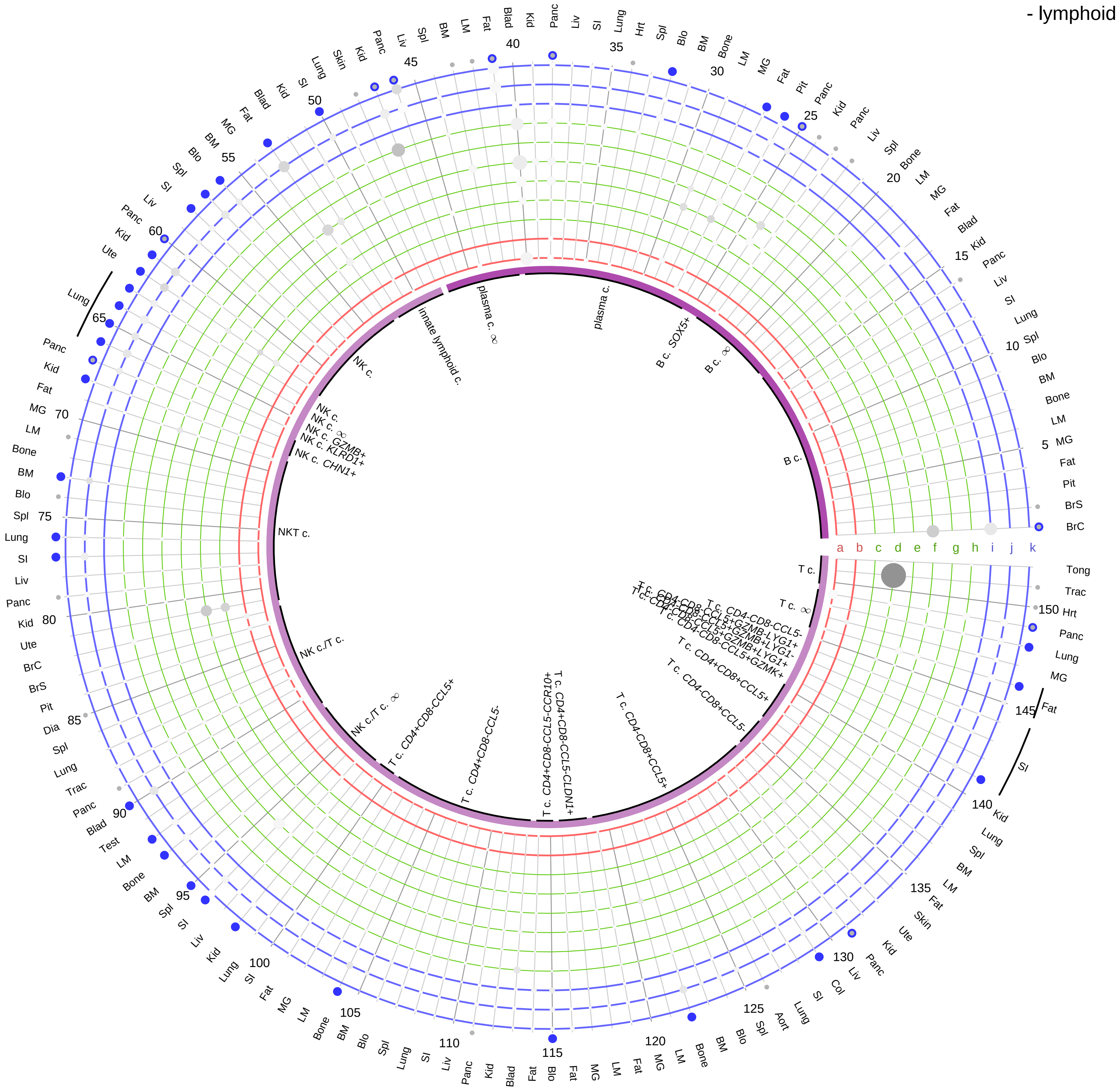

**Ligand/Enzyme**
- a - IGF1
- b - IGF2

**Modulator**
- c - IGFBP1
- d - IGFBP2
- e - IGFBP3
- f - IGFBP4
- g - IGFBP5
- h - IGFBP6

**Receptor**
- i - IGF1R
- j - IGF2R
- k - INSR

**PAIRING** (ligand → receptor)

IGF1→IGF1R

IGF2→IGF1R

IGF2→IGF2R

IGF2→INSR

| Abbrev. | Tissue |
|---|---|
| Aort | Aorta |
| Blad | Bladder |
| Blo | Blood |
| Bone | Bone |
| BM | Bone marrow |
| BrC | Brain cortex |
| BrS | Brainstem |
| Col | Colon |
| Dia | Diaphragm |
| Ret | Eye retina |
| Fat | Fat |
| Hrt | Heart |
| Pit | Hypothalamus/Pituitary |
| Kid | Kidney |
| LM | Limb muscle |
| Liv | Liver |
| Lung | Lung |
| MG | Mammary gland |
| Panc | Pancreas |
| Skin | Skin |
| SI | Small intestine |
| Spl | Spleen |
| Test | Testes |
| Thym | Thymus |
| Tong | Tongue |
| Trac | Trachea |
| Ute | Uterus |

**Compartment**
**lymphoid**
- B cell
- NK/T cell

12.5%
25%
50%
100%

ln(CP10K+1) 0 1 2 3

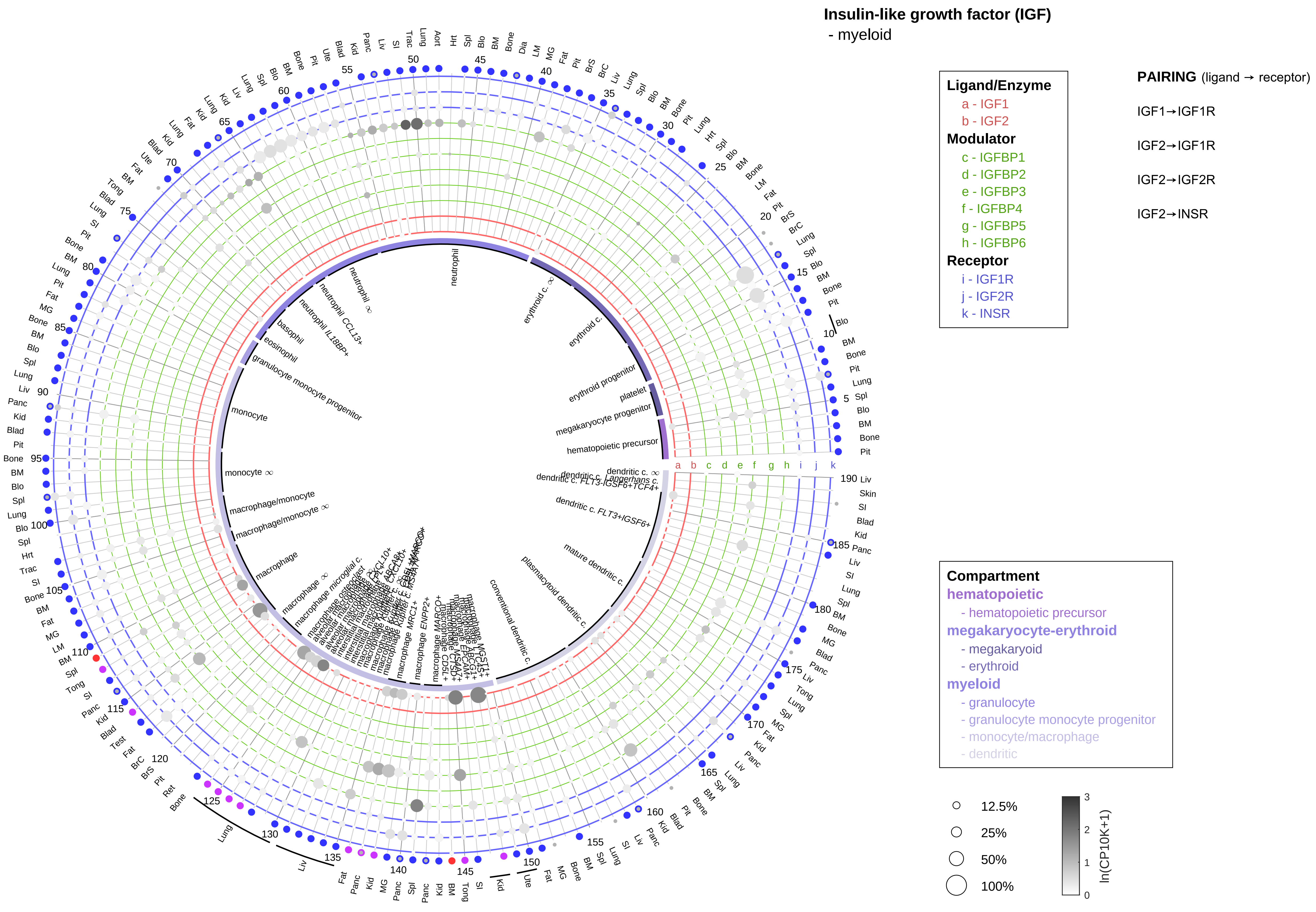

| Abbrev. | Tissue |
|---|---|
| Aort | Aorta |
| Blad | Bladder |
| Blo | Blood |
| Bone | Bone |
| BM | Bone marrow |
| BrC | Brain cortex |
| BrS | Brainstem |
| Col | Colon |
| Dia | Diaphragm |
| Ret | Eye retina |
| Fat | Fat |
| Hrt | Heart |
| Pit | Hypothalamus/Pituitary |
| Kid | Kidney |
| LM | Limb muscle |
| Liv | Liver |
| Lung | Lung |
| MG | Mammary gland |
| Panc | Pancreas |
| Skin | Skin |
| SI | Small intestine |
| Spl | Spleen |
| Test | Testes |
| Thym | Thymus |
| Tong | Tongue |
| Trac | Trachea |
| Ute | Uterus |

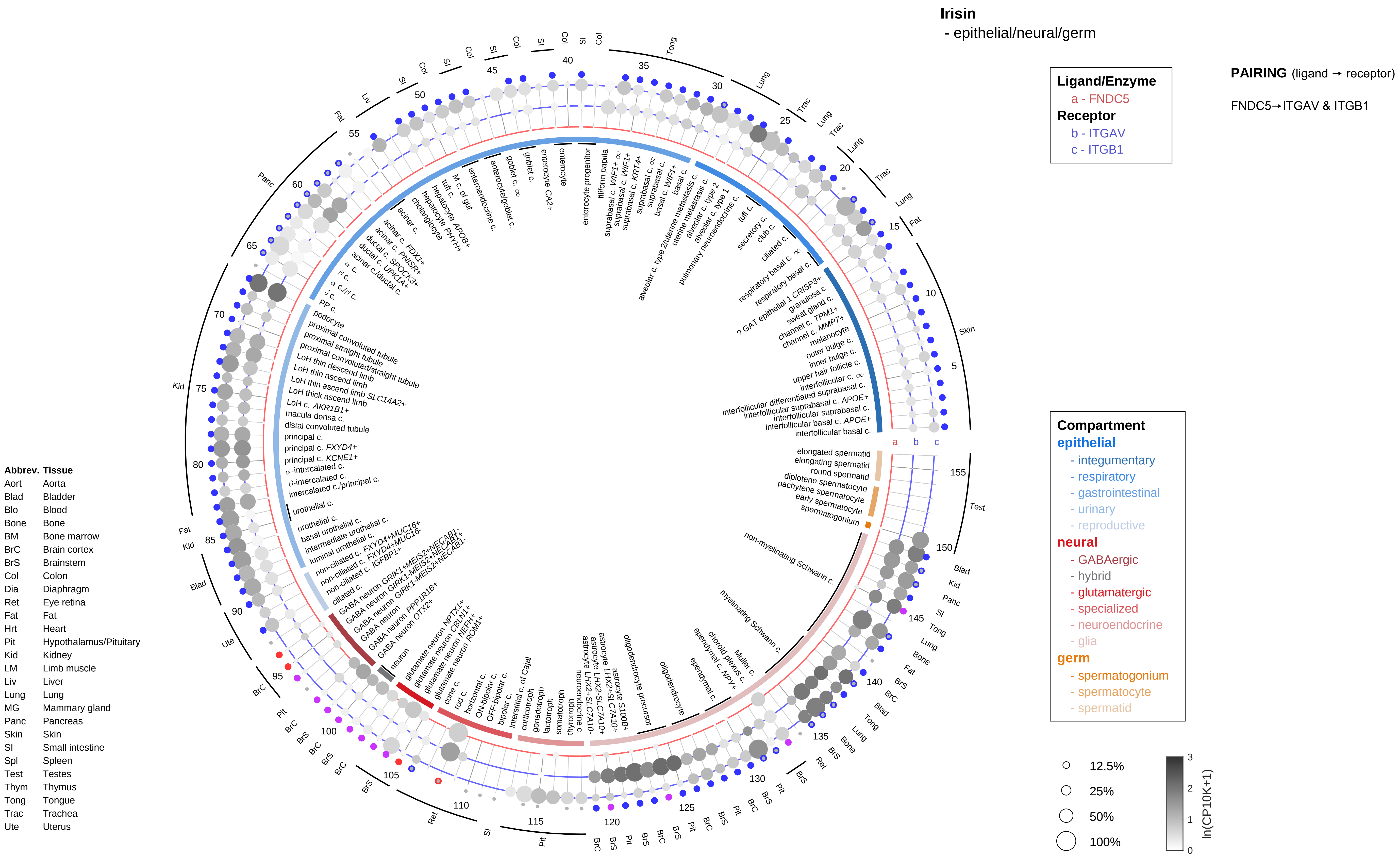

Irisin
- epithelial/neural/germ
Ligand/Enzyme
a - FNDC5
Receptor
b - ITGAV
c - ITGB1
PAIRING (ligand → receptor)
FNDC5→ITGAV & ITGB1
Compartment
epithelial
- integumentary
- respiratory
- gastrointestinal
- urinary
- reproductive
neural
- GABAergic
- hybrid
- glutamatergic
- specialized
- neuroendocrine
- glia
germ
- spermatogonium
- spermatocyte
- spermatid
12.5%
25%
50%
100%
ln(CP10K+1)
Abbrev. Tissue
Aort Aorta
Blad Bladder
Blo Blood
Bone Bone
BM Bone marrow
BrC Brain cortex
BrS Brainstem
Col Colon
Dia Diaphragm
Ret Eye retina
Fat Fat
Hrt Heart
Pit Hypothalamus/Pituitary
Kid Kidney
LM Limb muscle
Liv Liver
Lung Lung
MG Mammary gland
Panc Pancreas
Skin Skin
SI Small intestine
Spl Spleen
Test Testes
Thym Thymus
Tong Tongue
Trac Trachea
Ute Uterus
interfollicular basal c.
interfollicular basal c. APOE+
interfollicular suprabasal c.
interfollicular suprabasal c. APOE+
interfollicular differentiated suprabasal c.
interfollicular c. ∞
upper hair follicle c.
inner bulge c.
outer bulge c.
melanocyte
channel c. MMP7+
channel c. TPM1+
sweat gland c.
granulosa c.
? GAT epithelial 1 CRISP3+
respiratory basal c.
respiratory basal c. ∞
ciliated c.
club c.
secretory c.
tuft c.
pulmonary neuroendocrine c.
alveolar c. type 1
alveolar c. type 2
uterine metastasis c.
alveolar c. type 2/uterine metastasis c.
basal c.
basal c. WIF1+
suprabasal c.
suprabasal c. ∞
suprabasal c. KRT4+
suprabasal c. WIF1+
suprabasal c. WIF1+ ∞
filiform papilla
enterocyte progenitor
enterocyte
enterocyte CA2+
goblet c.
goblet c. ∞
enterocyte/goblet c.
enteroendocrine c.
M c. of gut
tuft c.
hepatocyte APOB+
hepatocyte PHYH+
cholangiocyte
acinar c.
acinar c. FDX1+
acinar c. PNISR+
ductal c. SPOCK3+
ductal c. UPK1A+
acinar c./ductal c.
α c.
β c.
α c./β c.
δ c.
PP c.
podocyte
proximal convoluted tubule
proximal straight tubule
proximal convoluted/straight tubule
LoH thin descend limb
LoH thin ascend limb
LoH thin ascend limb SLC14A2+
LoH thick ascend limb
LoH c. AKR1B1+
macula densa c.
distal convoluted tubule
principal c.
principal c. FXYD4+
principal c. KCNE1+
α-intercalated c.
β-intercalated c.
intercalated c./principal c.
urothelial c.
urothelial c.
basal urothelial c.
intermediate urothelial c.
luminal urothelial c.
non-ciliated c. FXYD4+MUC16+
non-ciliated c. FXYD4+MUC16-
non-ciliated c. IGFBP1+
ciliated c.
GABA neuron GRIK1+MEIS2+NECAB1-
GABA neuron GRIK1-MEIS2+NECAB1+
GABA neuron GRIK1-MEIS2+NECAB1-
GABA neuron PPP1R1B+
GABA neuron OTX2+
GABA neuron
neuron
glutamate neuron NPTX1+
glutamate neuron CBLN1+
glutamate neuron NEFH+
glutamate neuron ROM1+
cone c.
rod c.
horizontal c.
ON-bipolar c.
OFF-bipolar c.
bipolar c.
interstitial c. of Cajal
corticotroph
gonadotroph
lactotroph
somatotroph
thyrotroph
neuroendocrine c.
astrocyte LHX2-SLC7A10+
astrocyte LHX2+SLC7A10-
astrocyte LHX2+SLC7A10+
astrocyte S100B+
oligodendrocyte precursor
oligodendrocyte
ependymal c.
ependymal c. NPY+
choroid plexus c.
Muller c.
myelinating Schwann c.
non-myelinating Schwann c.
spermatogonium
early spermatocyte
pachytene spermatocyte
diplotene spermatocyte
round spermatid
elongating spermatid
elongated spermatid

**Irisin**
- endothelial

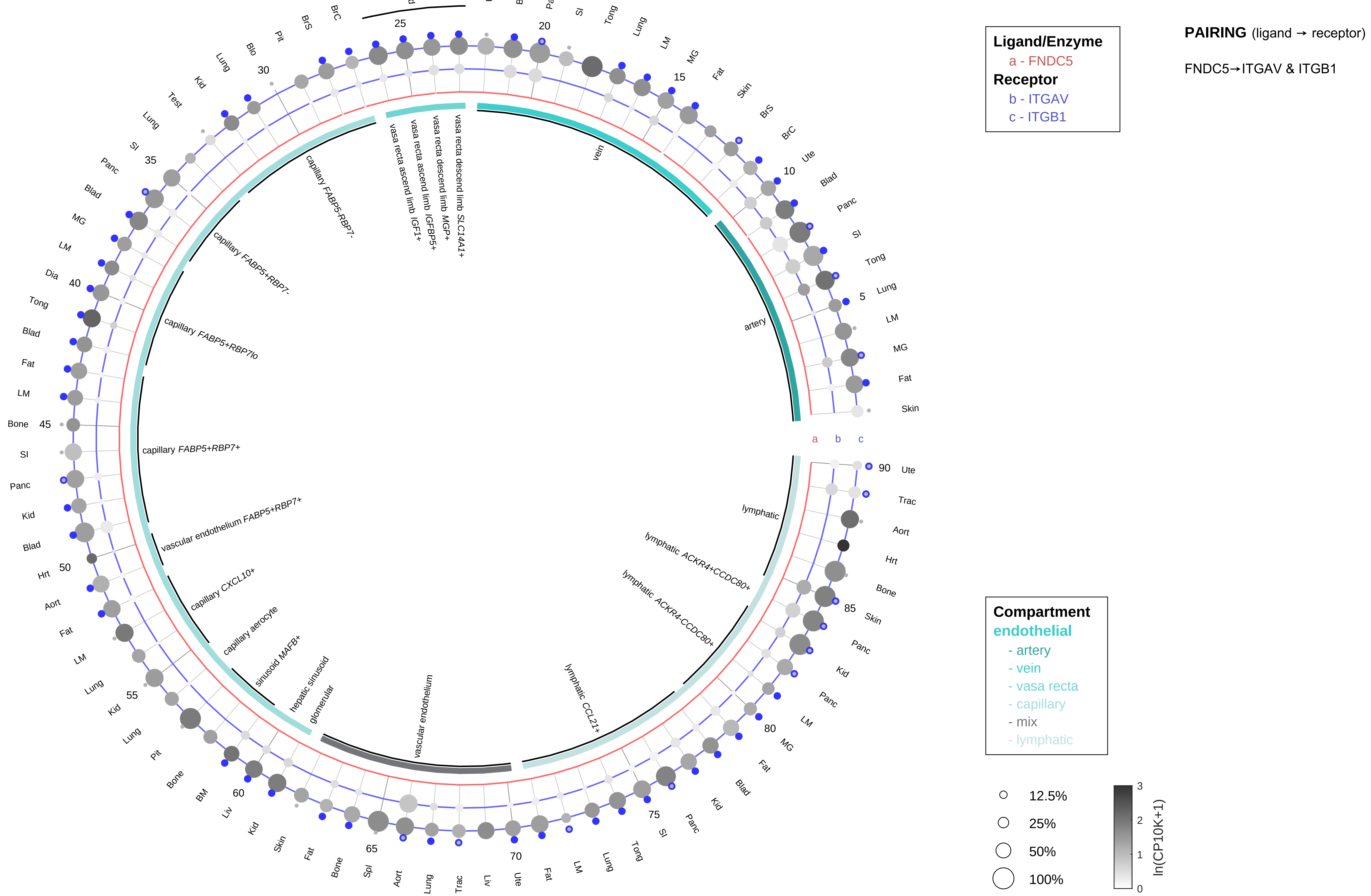

| Abbrev. | Tissue |
|---|---|
| Aort | Aorta |
| Blad | Bladder |
| Blo | Blood |
| Bone | Bone |
| BM | Bone marrow |
| BrC | Brain cortex |
| BrS | Brainstem |
| Col | Colon |
| Dia | Diaphragm |
| Ret | Eye retina |
| Fat | Fat |
| Hrt | Heart |
| Pit | Hypothalamus/Pituitary |
| Kid | Kidney |
| LM | Limb muscle |
| Liv | Liver |
| Lung | Lung |
| MG | Mammary gland |
| Panc | Pancreas |
| Skin | Skin |
| SI | Small intestine |
| Spl | Spleen |
| Test | Testes |
| Thym | Thymus |
| Tong | Tongue |
| Trac | Trachea |
| Ute | Uterus |

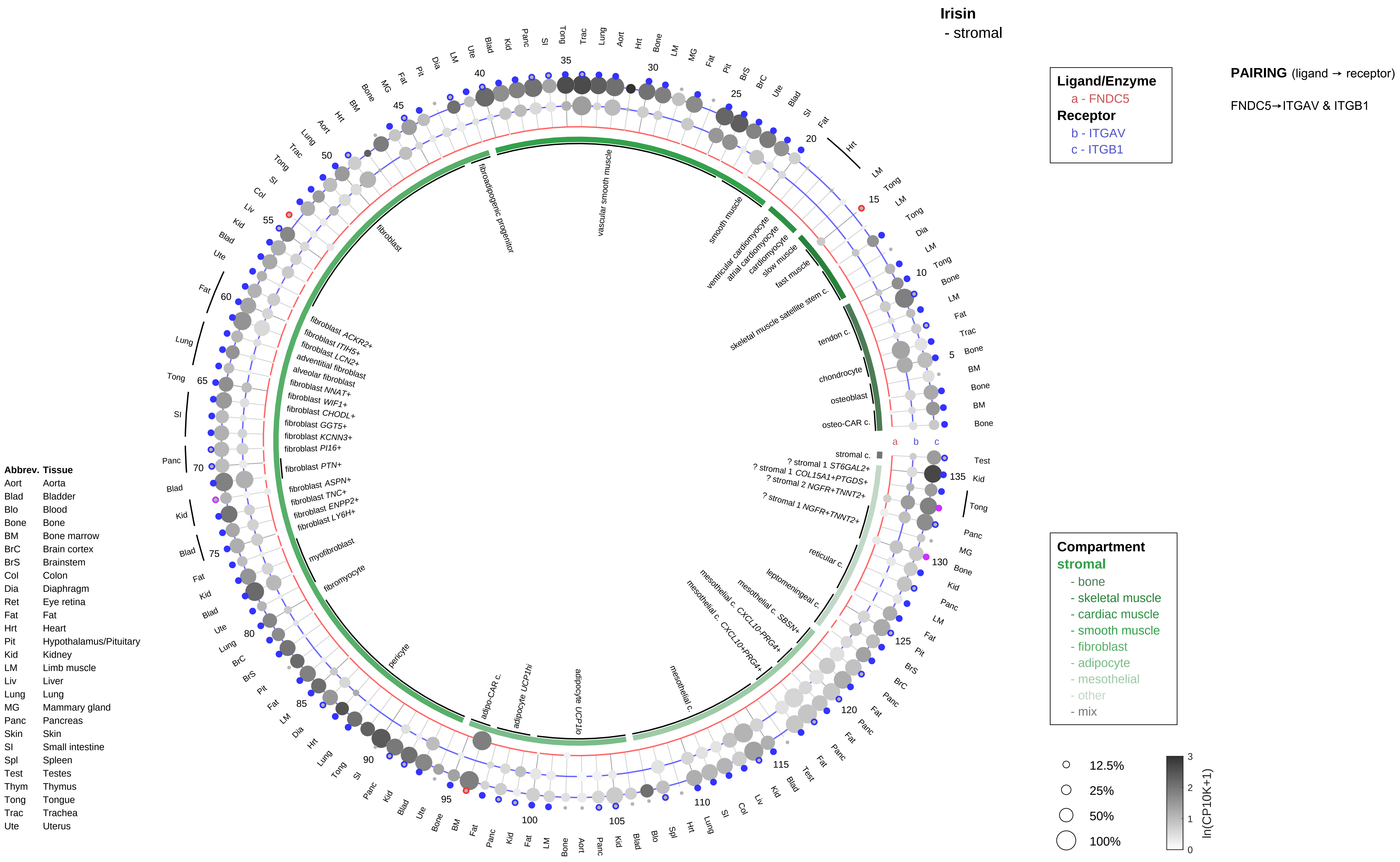

Irisin
- stromal
Ligand/Enzyme
a - FNDC5
Receptor
b - ITGAV
c - ITGB1
PAIRING (ligand → receptor)
FNDC5→ITGAV & ITGB1
Compartment
stromal
- bone
- skeletal muscle
- cardiac muscle
- smooth muscle
- fibroblast
- adipocyte
- mesothelial
- other
- mix
12.5%
25%
50%
100%
ln(CP10K+1)
a
b
c
fibroadipogenic progenitor
vascular smooth muscle
smooth muscle
ventricular cardiomyocyte
atrial cardiomyocyte
cardiomyocyte
slow muscle
fast muscle
skeletal muscle satellite stem c.
tendon c.
chondrocyte
osteoblast
osteo-CAR c.
stromal c.
? stromal 1 ST6GAL2+
? stromal 1 COL15A1+PTGDS+
? stromal 2 NGFR+TNNT2+
? stromal 1 NGFR+TNNT2+
reticular c.
leptomeningeal c.
mesothelial c. SBSN+
mesothelial c. CXCL10-PRG4+
mesothelial c. CXCL10+PRG4+
mesothelial c.
adipocyte UCP1lo
adipocyte UCP1hi
adipo-CAR c.
pericyte
fibromyocyte
myofibroblast
fibroblast LY6H+
fibroblast ENPP2+
fibroblast TNC+
fibroblast ASPN+
fibroblast PTN+
fibroblast PI16+
fibroblast KCNN3+
fibroblast GGT5+
fibroblast CHODL+
fibroblast WIF1+
fibroblast NNAT+
alveolar fibroblast
adventitial fibroblast
fibroblast LCN2+
fibroblast ITIH5+
fibroblast ACKR2+
fibroblast
Abbrev. Tissue
Aort Aorta
Blad Bladder
Blo Blood
Bone Bone
BM Bone marrow
BrC Brain cortex
BrS Brainstem
Col Colon
Dia Diaphragm
Ret Eye retina
Fat Fat
Hrt Heart
Pit Hypothalamus/Pituitary
Kid Kidney
LM Limb muscle
Liv Liver
Lung Lung
MG Mammary gland
Panc Pancreas
Skin Skin
SI Small intestine
Spl Spleen
Test Testes
Thym Thymus
Tong Tongue
Trac Trachea
Ute Uterus

**Irisin**
- lymphoid

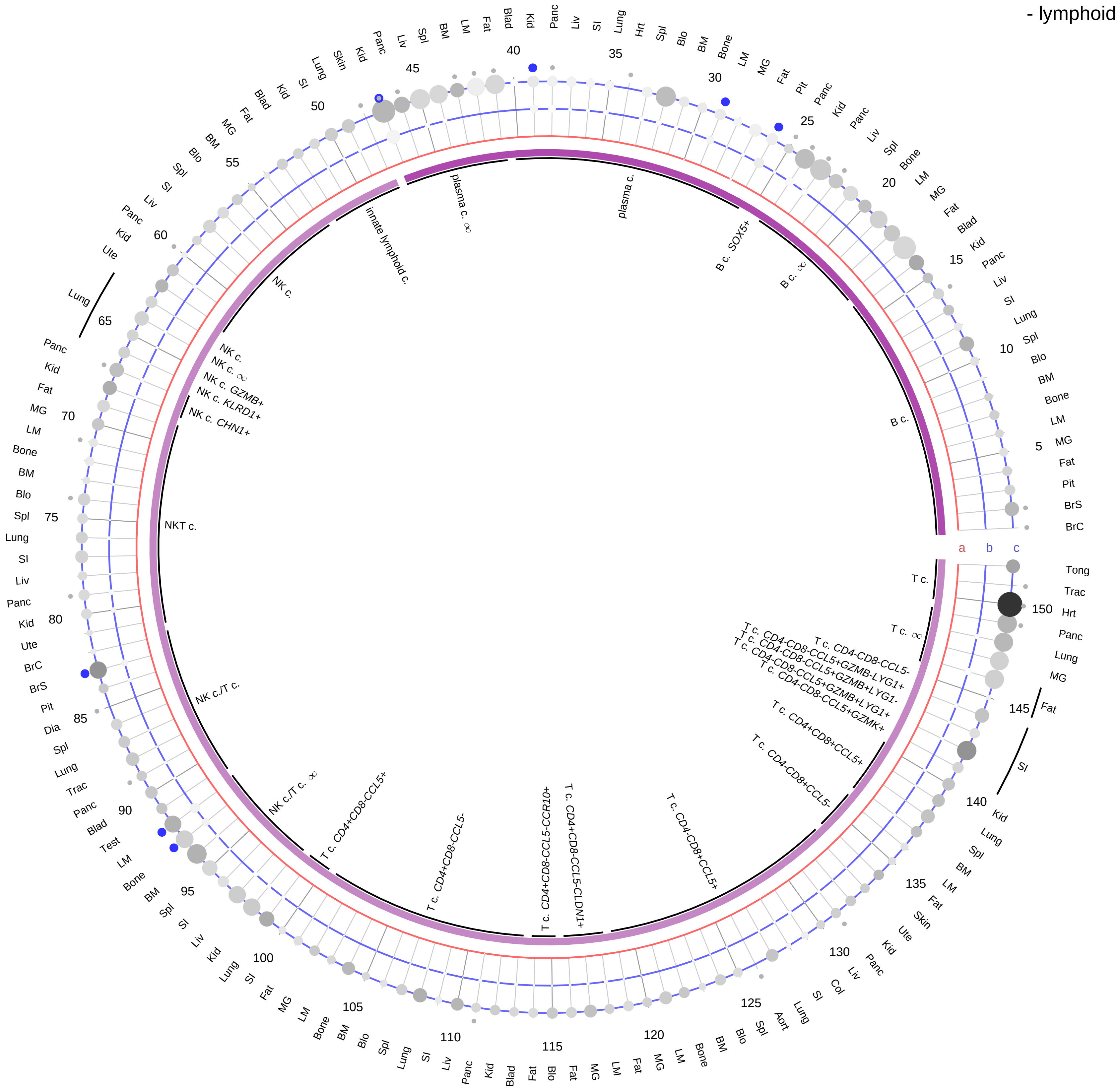

| Ligand/Enzyme |
|---|
| a - FNDC5 |
| **Receptor** |
| b - ITGAV |
| c - ITGB1 |

**PAIRING** (ligand → receptor)

FNDC5→ITGAV & ITGB1

| Abbrev. | Tissue |
|---|---|
| Aort | Aorta |
| Blad | Bladder |
| Blo | Blood |
| Bone | Bone |
| BM | Bone marrow |
| BrC | Brain cortex |
| BrS | Brainstem |
| Col | Colon |
| Dia | Diaphragm |
| Ret | Eye retina |
| Fat | Fat |
| Hrt | Heart |
| Pit | Hypothalamus/Pituitary |
| Kid | Kidney |
| LM | Limb muscle |
| Liv | Liver |
| Lung | Lung |
| MG | Mammary gland |
| Panc | Pancreas |
| Skin | Skin |
| SI | Small intestine |
| Spl | Spleen |
| Test | Testes |
| Thym | Thymus |
| Tong | Tongue |
| Trac | Trachea |
| Ute | Uterus |

**Compartment**
**lymphoid**
- B cell
- NK/T cell

12.5%
25%
50%
100%

ln(CP10K+1): 0, 1, 2, 3

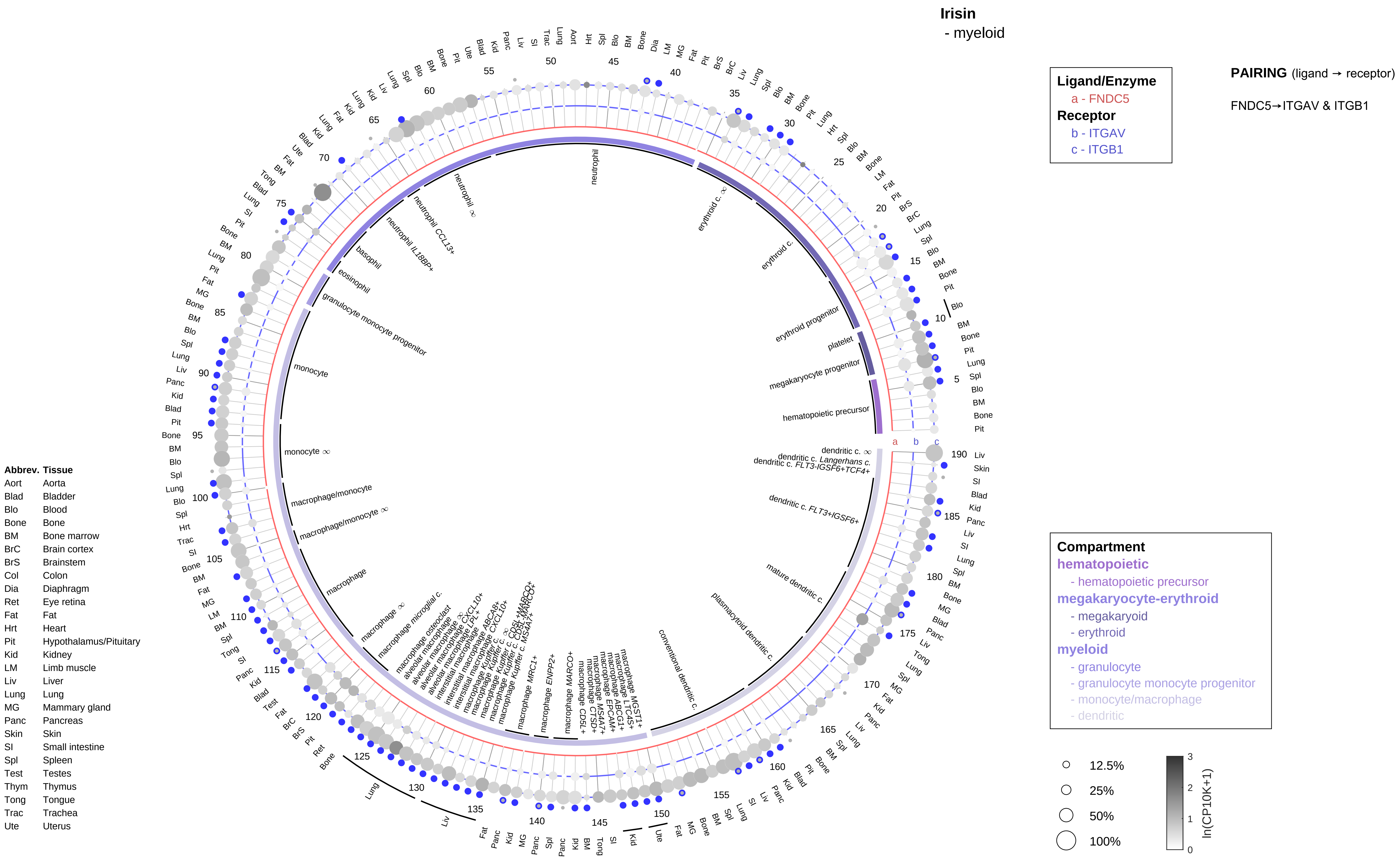
Irisin
- myeloid
Ligand/Enzyme
a - FNDC5
Receptor
b - ITGAV
c - ITGB1
PAIRING (ligand → receptor)
FNDC5→ITGAV & ITGB1
Compartment
hematopoietic
- hematopoietic precursor
megakaryocyte-erythroid
- megakaryoid
- erythroid
myeloid
- granulocyte
- granulocyte monocyte progenitor
- monocyte/macrophage
- dendritic
12.5%
25%
50%
100%
ln(CP10K+1)
Abbrev. Tissue
Aort Aorta
Blad Bladder
Blo Blood
Bone Bone
BM Bone marrow
BrC Brain cortex
BrS Brainstem
Col Colon
Dia Diaphragm
Ret Eye retina
Fat Fat
Hrt Heart
Pit Hypothalamus/Pituitary
Kid Kidney
LM Limb muscle
Liv Liver
Lung Lung
MG Mammary gland
Panc Pancreas
Skin Skin
SI Small intestine
Spl Spleen
Test Testes
Thym Thymus
Tong Tongue
Trac Trachea
Ute Uterus

#### Kininogen/Bradykinin/Kallikrein
- epithelial/neural/germ

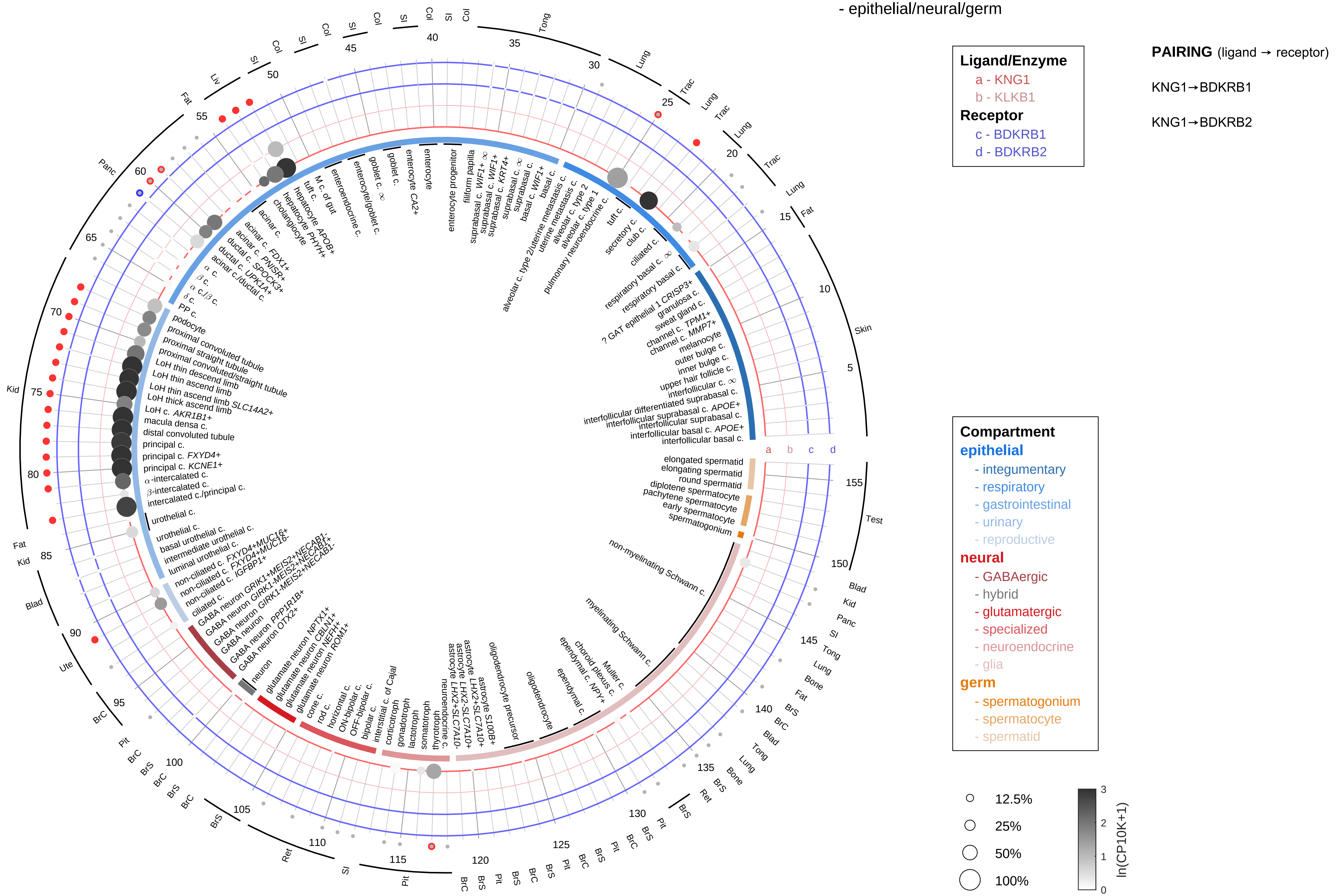

| Abbrev. | Tissue |
|---|---|
| Aort | Aorta |
| Blad | Bladder |
| Blo | Blood |
| Bone | Bone |
| BM | Bone marrow |
| BrC | Brain cortex |
| BrS | Brainstem |
| Col | Colon |
| Dia | Diaphragm |
| Ret | Eye retina |
| Fat | Fat |
| Hrt | Heart |
| Pit | Hypothalamus/Pituitary |
| Kid | Kidney |
| LM | Limb muscle |
| Liv | Liver |
| Lung | Lung |
| MG | Mammary gland |
| Panc | Pancreas |
| Skin | Skin |
| SI | Small intestine |
| Spl | Spleen |
| Test | Testes |
| Thym | Thymus |
| Tong | Tongue |
| Trac | Trachea |
| Ute | Uterus |

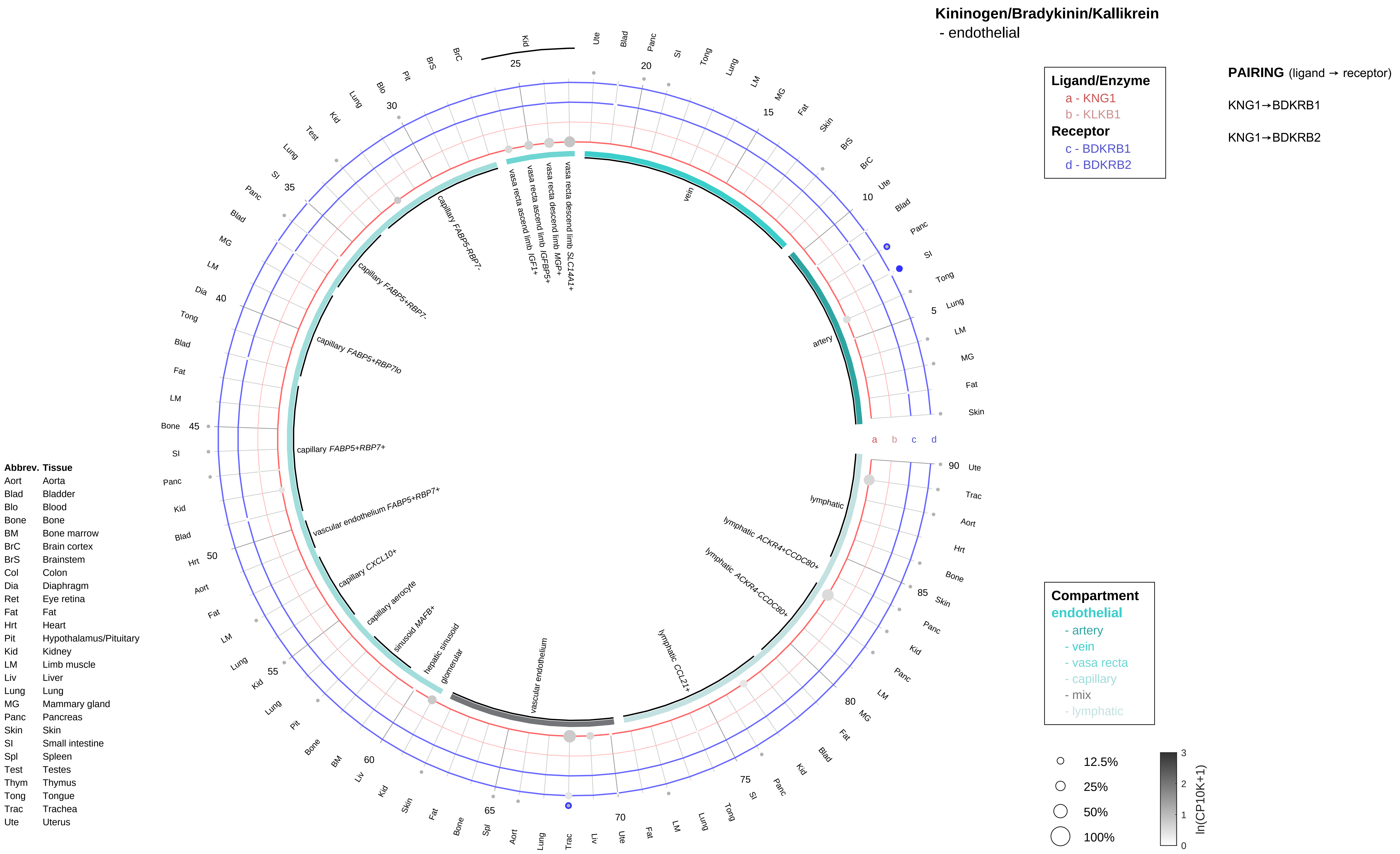

Kininogen/Bradykinin/Kallikrein
- endothelial
Ligand/Enzyme
a - KNG1
b - KLKB1
Receptor
c - BDKRB1
d - BDKRB2
PAIRING (ligand → receptor)
KNG1→BDKRB1
KNG1→BDKRB2
Compartment
endothelial
- artery
- vein
- vasa recta
- capillary
- mix
- lymphatic
12.5%
25%
50%
100%
ln(CP10K+1)
vein
artery
lymphatic
lymphatic ACKR4+CCDC80+
lymphatic ACKR4-CCDC80+
lymphatic CCL21+
vascular endothelium
glomerular
hepatic sinusoid
sinusoid MAFB+
capillary aerocyte
capillary CXCL10+
vascular endothelium FABP5+RBP7+
capillary FABP5+RBP7+
capillary FABP5+RBP7lo
capillary FABP5+RBP7-
capillary FABP5-RBP7-
vasa recta ascend limb IGF1+
vasa recta ascend limb IGFBP5+
vasa recta descend limb MGP+
vasa recta descend limb SLC14A1+
Abbrev. Tissue
Aort Aorta
Blad Bladder
Blo Blood
Bone Bone
BM Bone marrow
BrC Brain cortex
BrS Brainstem
Col Colon
Dia Diaphragm
Ret Eye retina
Fat Fat
Hrt Heart
Pit Hypothalamus/Pituitary
Kid Kidney
LM Limb muscle
Liv Liver
Lung Lung
MG Mammary gland
Panc Pancreas
Skin Skin
SI Small intestine
Spl Spleen
Test Testes
Thym Thymus
Tong Tongue
Trac Trachea
Ute Uterus

#### Kininogen/Bradykinin/Kallikrein
- stromal

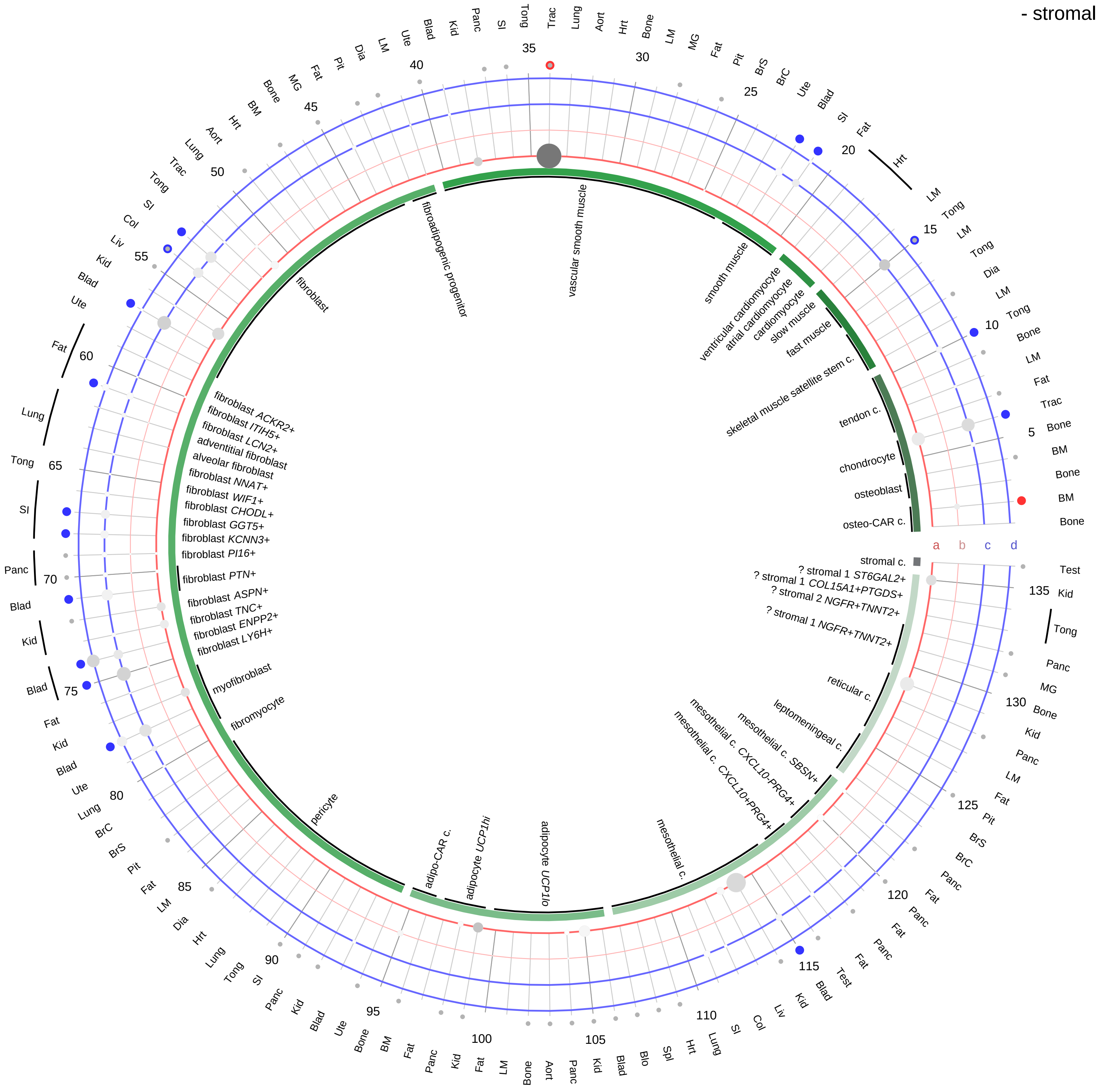

**Ligand/Enzyme**
- a - KNG1
- b - KLKB1

**Receptor**
- c - BDKRB1
- d - BDKRB2

**PAIRING** (ligand → receptor)

KNG1→BDKRB1

KNG1→BDKRB2

| Abbrev. | Tissue |
|---|---|
| Aort | Aorta |
| Blad | Bladder |
| Blo | Blood |
| Bone | Bone |
| BM | Bone marrow |
| BrC | Brain cortex |
| BrS | Brainstem |
| Col | Colon |
| Dia | Diaphragm |
| Ret | Eye retina |
| Fat | Fat |
| Hrt | Heart |
| Pit | Hypothalamus/Pituitary |
| Kid | Kidney |
| LM | Limb muscle |
| Liv | Liver |
| Lung | Lung |
| MG | Mammary gland |
| Panc | Pancreas |
| Skin | Skin |
| SI | Small intestine |
| Spl | Spleen |
| Test | Testes |
| Thym | Thymus |
| Tong | Tongue |
| Trac | Trachea |
| Ute | Uterus |

**Compartment**
**stromal**
- bone
- skeletal muscle
- cardiac muscle
- smooth muscle
- fibroblast
- adipocyte
- mesothelial
- other
- mix

12.5%
25%
50%
100%

ln(CP10K+1) 0 1 2 3

**Kininogen/Bradykinin/Kallikrein**
- lymphoid

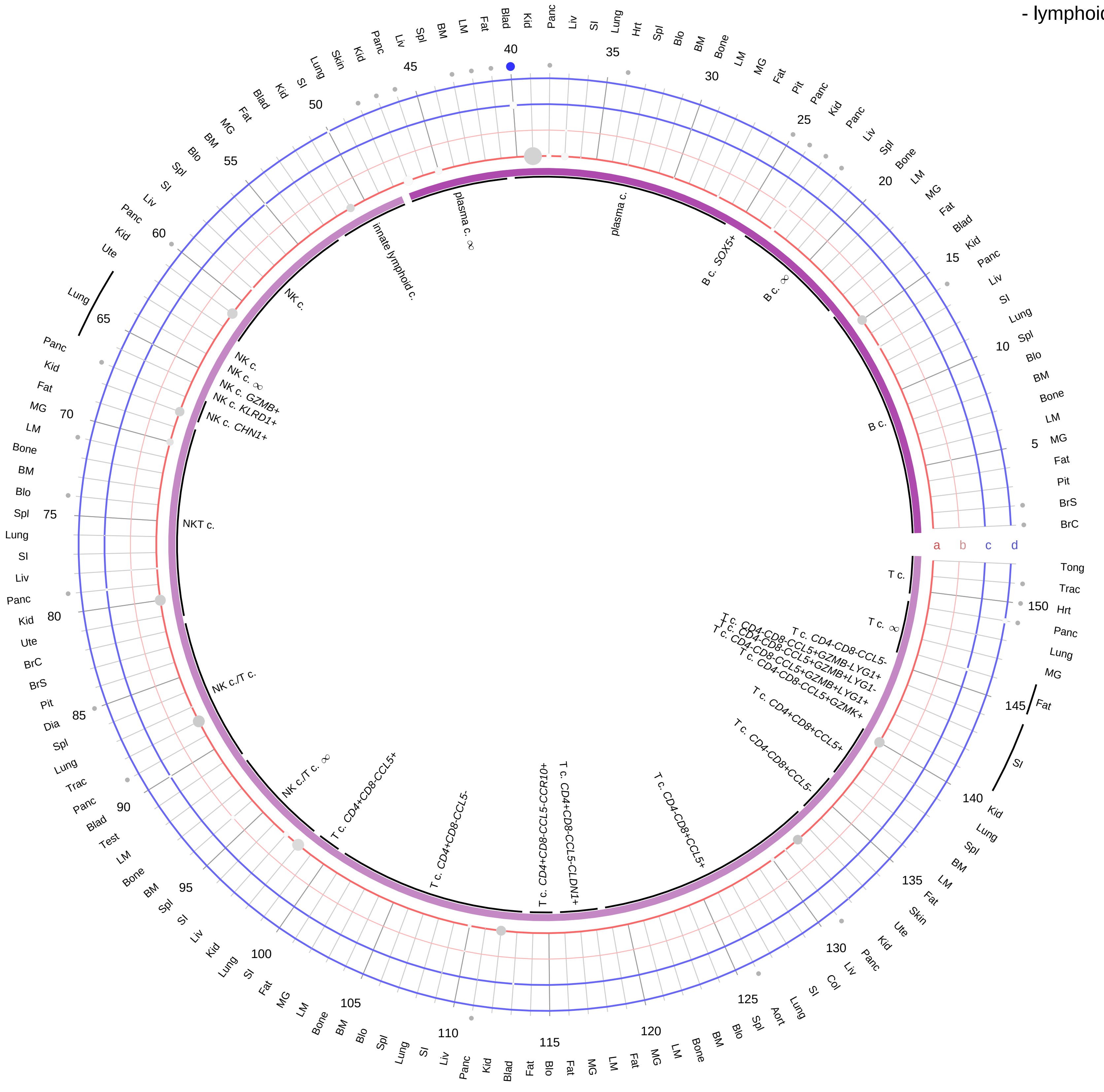

**Ligand/Enzyme**
a - KNG1
b - KLKB1
**Receptor**
c - BDKRB1
d - BDKRB2

**PAIRING** (ligand → receptor)

KNG1→BDKRB1

KNG1→BDKRB2

**Compartment**
**lymphoid**
- B cell
- NK/T cell

12.5%
25%
50%
100%

ln(CP10K+1): 0, 1, 2, 3

| Abbrev. | Tissue |
|---|---|
| Aort | Aorta |
| Blad | Bladder |
| Blo | Blood |
| Bone | Bone |
| BM | Bone marrow |
| BrC | Brain cortex |
| BrS | Brainstem |
| Col | Colon |
| Dia | Diaphragm |
| Ret | Eye retina |
| Fat | Fat |
| Hrt | Heart |
| Pit | Hypothalamus/Pituitary |
| Kid | Kidney |
| LM | Limb muscle |
| Liv | Liver |
| Lung | Lung |
| MG | Mammary gland |
| Panc | Pancreas |
| Skin | Skin |
| SI | Small intestine |
| Spl | Spleen |
| Test | Testes |
| Thym | Thymus |
| Tong | Tongue |
| Trac | Trachea |
| Ute | Uterus |

**Kininogen/Bradykinin/Kallikrein**
- myeloid

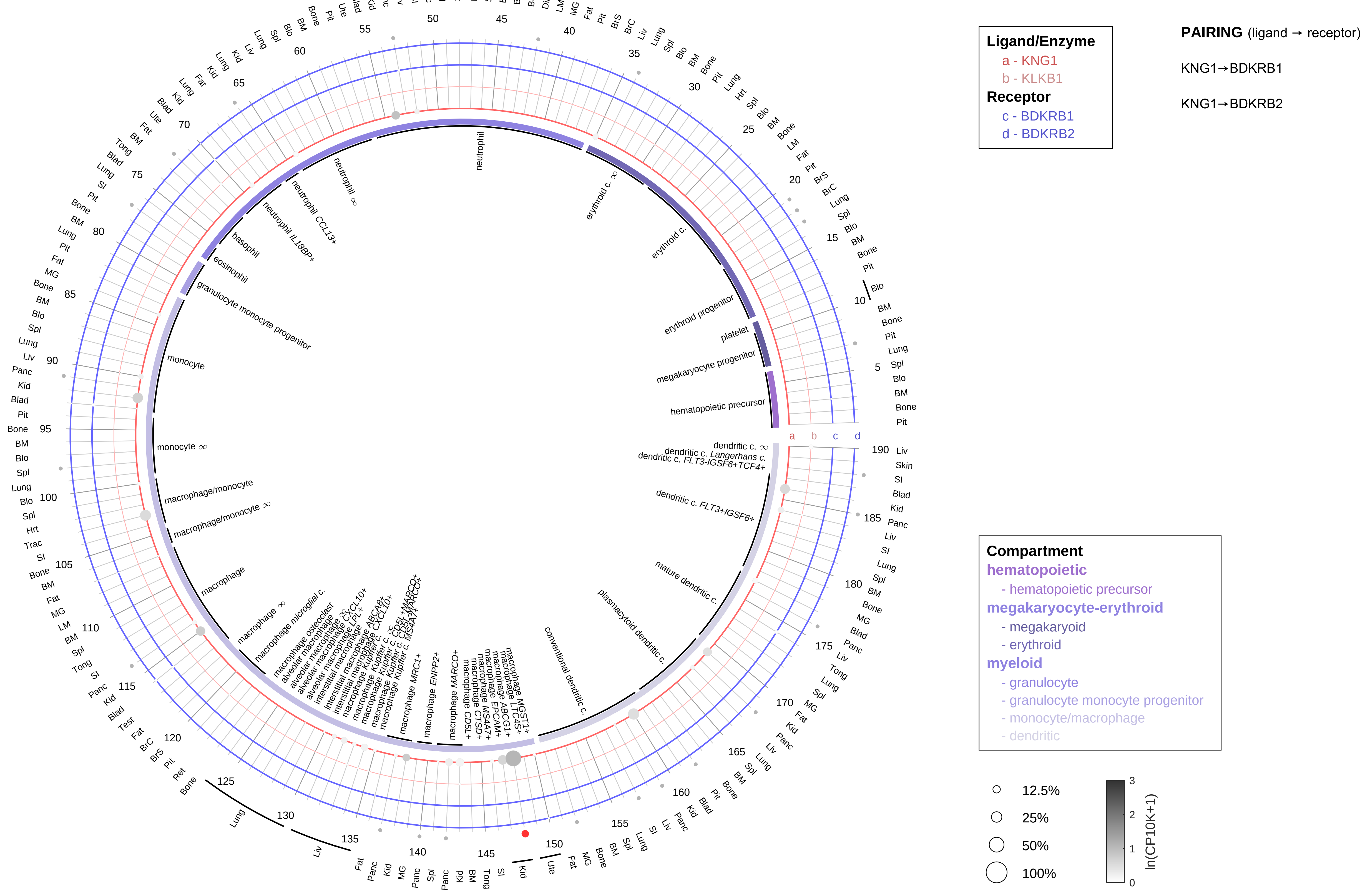

| Abbrev. | Tissue |
|---|---|
| Aort | Aorta |
| Blad | Bladder |
| Blo | Blood |
| Bone | Bone |
| BM | Bone marrow |
| BrC | Brain cortex |
| BrS | Brainstem |
| Col | Colon |
| Dia | Diaphragm |
| Ret | Eye retina |
| Fat | Fat |
| Hrt | Heart |
| Pit | Hypothalamus/Pituitary |
| Kid | Kidney |
| LM | Limb muscle |
| Liv | Liver |
| Lung | Lung |
| MG | Mammary gland |
| Panc | Pancreas |
| Skin | Skin |
| SI | Small intestine |
| Spl | Spleen |
| Test | Testes |
| Thym | Thymus |
| Tong | Tongue |
| Trac | Trachea |
| Ute | Uterus |

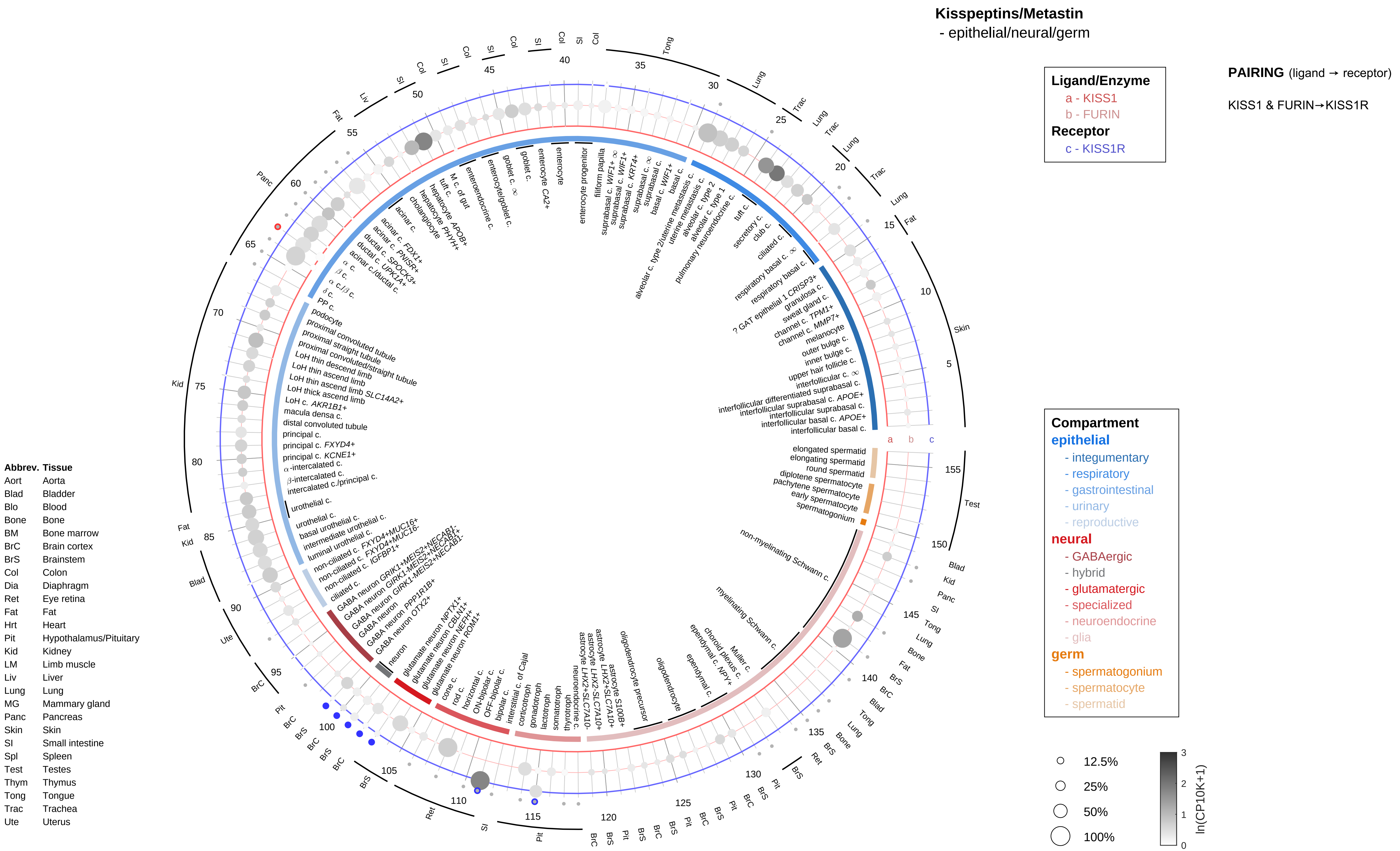

Kisspeptins/Metastin
- epithelial/neural/germ
Ligand/Enzyme
a - KISS1
b - FURIN
Receptor
c - KISS1R
PAIRING (ligand → receptor)
KISS1 & FURIN→KISS1R
Compartment
epithelial
- integumentary
- respiratory
- gastrointestinal
- urinary
- reproductive
neural
- GABAergic
- hybrid
- glutamatergic
- specialized
- neuroendocrine
- glia
germ
- spermatogonium
- spermatocyte
- spermatid
12.5%
25%
50%
100%
ln(CP10K+1)
Abbrev. Tissue
Aort Aorta
Blad Bladder
Blo Blood
Bone Bone
BM Bone marrow
BrC Brain cortex
BrS Brainstem
Col Colon
Dia Diaphragm
Ret Eye retina
Fat Fat
Hrt Heart
Pit Hypothalamus/Pituitary
Kid Kidney
LM Limb muscle
Liv Liver
Lung Lung
MG Mammary gland
Panc Pancreas
Skin Skin
SI Small intestine
Spl Spleen
Test Testes
Thym Thymus
Tong Tongue
Trac Trachea
Ute Uterus

**Kisspeptins/Metastin**
- endothelial

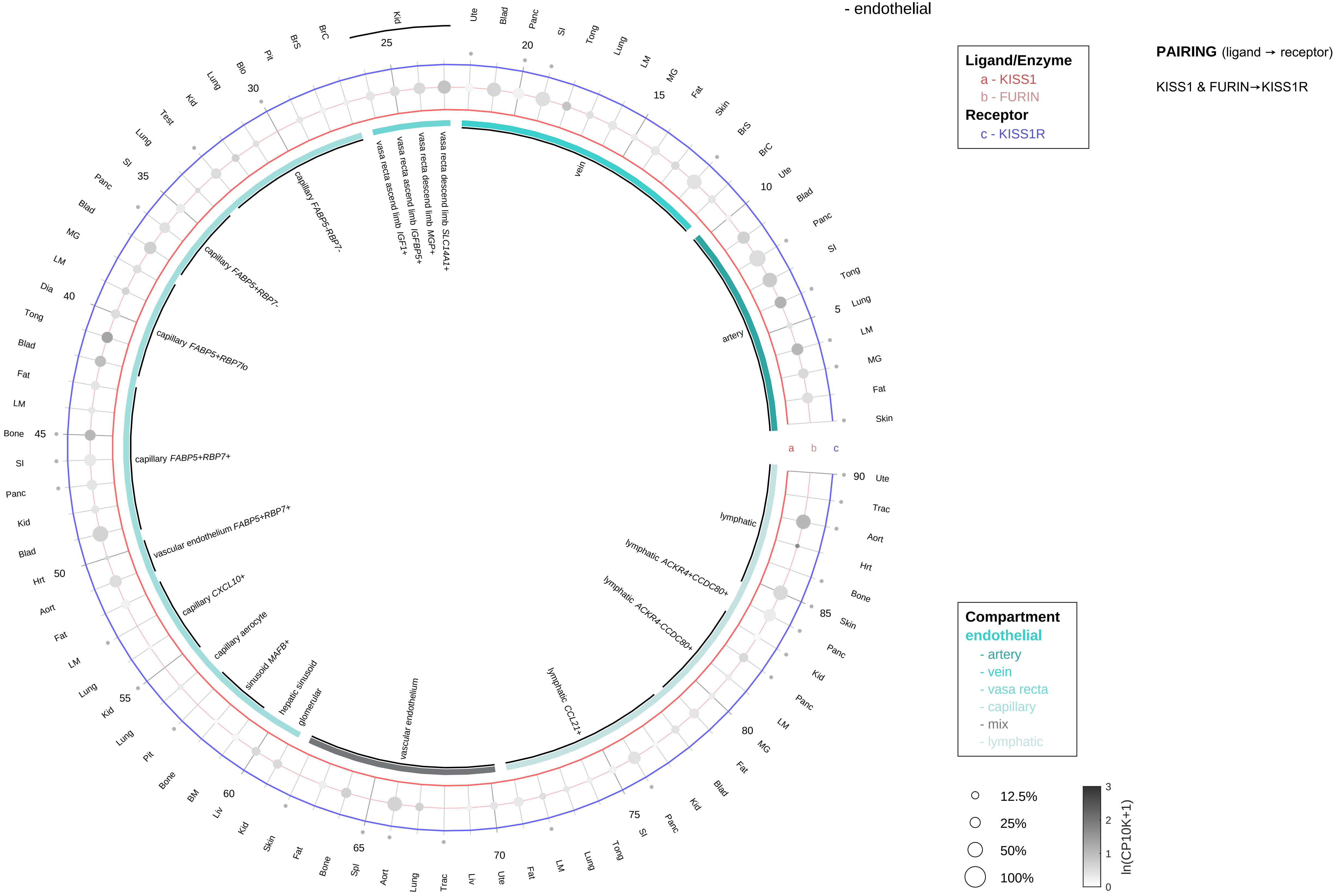

| Abbrev. | Tissue |
|---|---|
| Aort | Aorta |
| Blad | Bladder |
| Blo | Blood |
| Bone | Bone |
| BM | Bone marrow |
| BrC | Brain cortex |
| BrS | Brainstem |
| Col | Colon |
| Dia | Diaphragm |
| Ret | Eye retina |
| Fat | Fat |
| Hrt | Heart |
| Pit | Hypothalamus/Pituitary |
| Kid | Kidney |
| LM | Limb muscle |
| Liv | Liver |
| Lung | Lung |
| MG | Mammary gland |
| Panc | Pancreas |
| Skin | Skin |
| SI | Small intestine |
| Spl | Spleen |
| Test | Testes |
| Thym | Thymus |
| Tong | Tongue |
| Trac | Trachea |
| Ute | Uterus |

**Kisspeptins/Metastin**
- stromal

**Ligand/Enzyme**
- a - KISS1
- b - FURIN

**Receptor**
- c - KISS1R

**PAIRING** (ligand → receptor)

KISS1 & FURIN→KISS1R

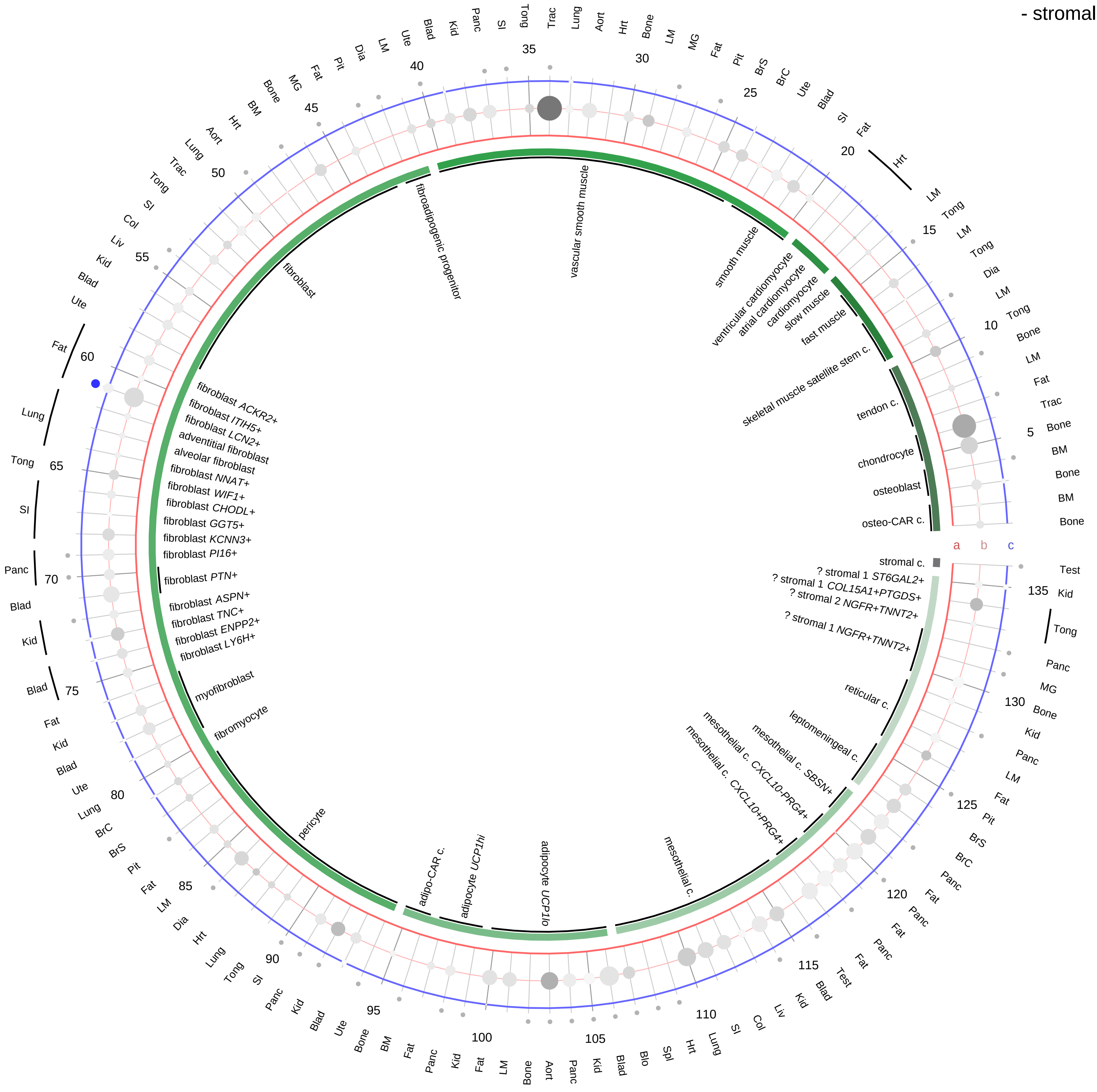

| Abbrev. | Tissue |
|---|---|
| Aort | Aorta |
| Blad | Bladder |
| Blo | Blood |
| Bone | Bone |
| BM | Bone marrow |
| BrC | Brain cortex |
| BrS | Brainstem |
| Col | Colon |
| Dia | Diaphragm |
| Ret | Eye retina |
| Fat | Fat |
| Hrt | Heart |
| Pit | Hypothalamus/Pituitary |
| Kid | Kidney |
| LM | Limb muscle |
| Liv | Liver |
| Lung | Lung |
| MG | Mammary gland |
| Panc | Pancreas |
| Skin | Skin |
| SI | Small intestine |
| Spl | Spleen |
| Test | Testes |
| Thym | Thymus |
| Tong | Tongue |
| Trac | Trachea |
| Ute | Uterus |

**Compartment**
**stromal**
- bone
- skeletal muscle
- cardiac muscle
- smooth muscle
- fibroblast
- adipocyte
- mesothelial
- other
- mix

12.5%
25%
50%
100%

ln(CP10K+1): 0, 1, 2, 3

Kisspeptins/Metastin
- lymphoid

PAIRING (ligand → receptor)

KISS1 & FURIN→KISS1R

**Ligand/Enzyme**
a - KISS1
b - FURIN
**Receptor**
c - KISS1R

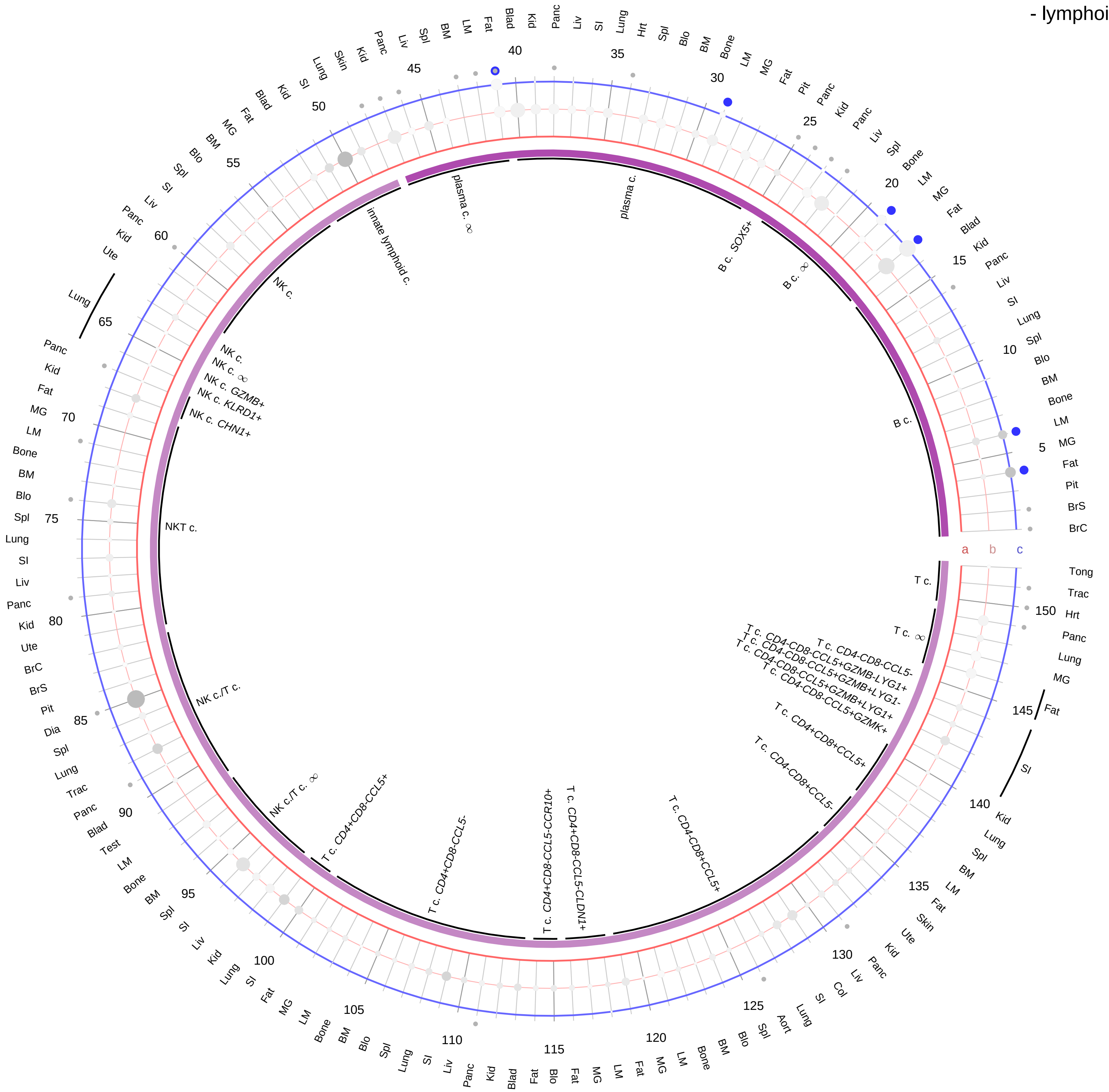

| Abbrev. | Tissue |
|---|---|
| Aort | Aorta |
| Blad | Bladder |
| Blo | Blood |
| Bone | Bone |
| BM | Bone marrow |
| BrC | Brain cortex |
| BrS | Brainstem |
| Col | Colon |
| Dia | Diaphragm |
| Ret | Eye retina |
| Fat | Fat |
| Hrt | Heart |
| Pit | Hypothalamus/Pituitary |
| Kid | Kidney |
| LM | Limb muscle |
| Liv | Liver |
| Lung | Lung |
| MG | Mammary gland |
| Panc | Pancreas |
| Skin | Skin |
| SI | Small intestine |
| Spl | Spleen |
| Test | Testes |
| Thym | Thymus |
| Tong | Tongue |
| Trac | Trachea |
| Ute | Uterus |

**Compartment**
**lymphoid**
- B cell
- NK/T cell

12.5%
25%
50%
100%

ln(CP10K+1)
0
1
2
3

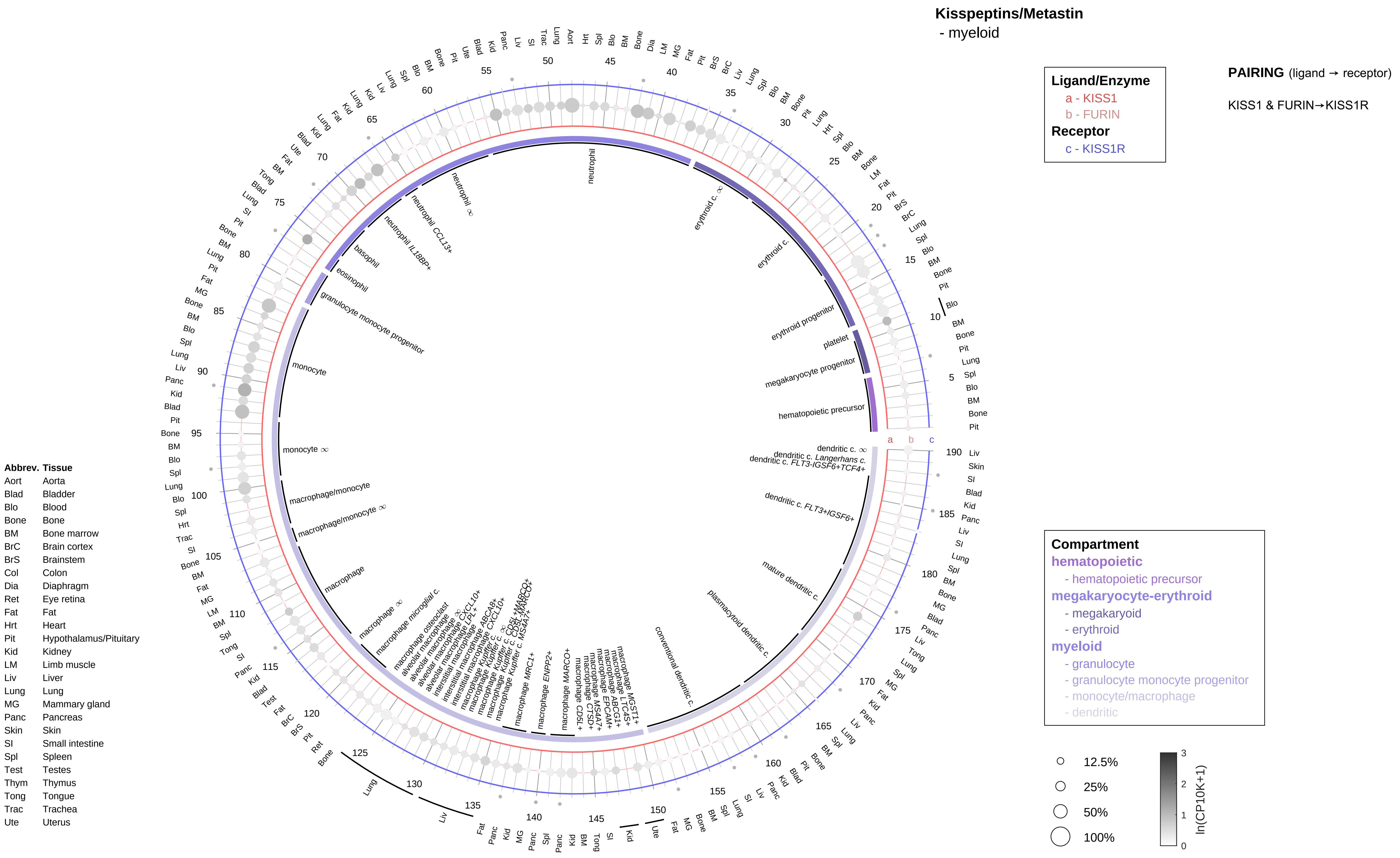
Kisspeptins/Metastin
- myeloid
Ligand/Enzyme
a - KISS1
b - FURIN
Receptor
c - KISS1R
PAIRING (ligand → receptor)
KISS1 & FURIN→KISS1R
Compartment
hematopoietic
- hematopoietic precursor
megakaryocyte-erythroid
- megakaryoid
- erythroid
myeloid
- granulocyte
- granulocyte monocyte progenitor
- monocyte/macrophage
- dendritic
12.5%
25%
50%
100%
ln(CP10K+1)
Abbrev. Tissue
Aort Aorta
Blad Bladder
Blo Blood
Bone Bone
BM Bone marrow
BrC Brain cortex
BrS Brainstem
Col Colon
Dia Diaphragm
Ret Eye retina
Fat Fat
Hrt Heart
Pit Hypothalamus/Pituitary
Kid Kidney
LM Limb muscle
Liv Liver
Lung Lung
MG Mammary gland
Panc Pancreas
Skin Skin
SI Small intestine
Spl Spleen
Test Testes
Thym Thymus
Tong Tongue
Trac Trachea
Ute Uterus

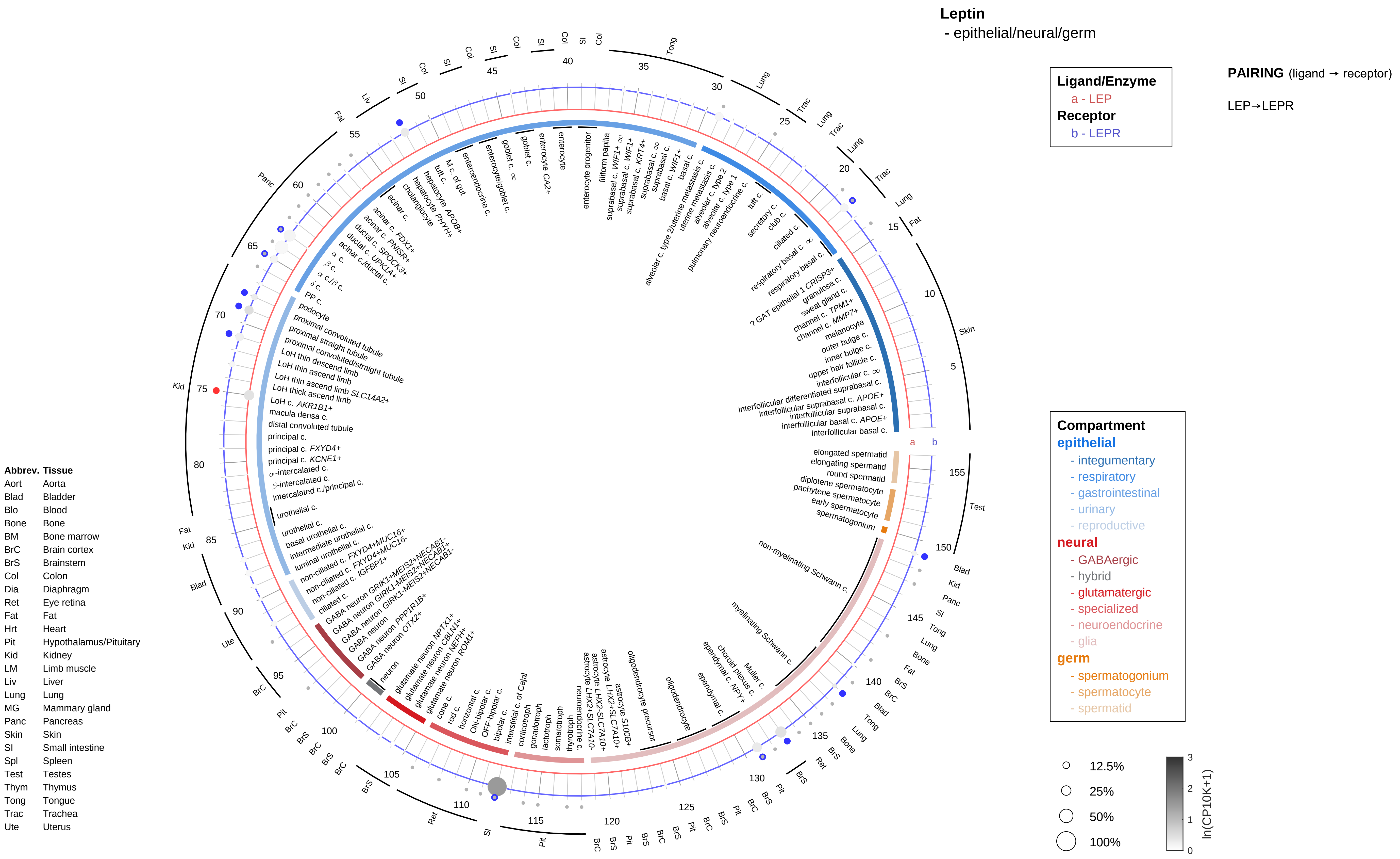

Leptin
- epithelial/neural/germ
Ligand/Enzyme
a - LEP
Receptor
b - LEPR
PAIRING (ligand → receptor)
LEP→LEPR
Compartment
epithelial
- integumentary
- respiratory
- gastrointestinal
- urinary
- reproductive
neural
- GABAergic
- hybrid
- glutamatergic
- specialized
- neuroendocrine
- glia
germ
- spermatogonium
- spermatocyte
- spermatid
12.5%
25%
50%
100%
ln(CP10K+1)
Abbrev. Tissue
Aort Aorta
Blad Bladder
Blo Blood
Bone Bone
BM Bone marrow
BrC Brain cortex
BrS Brainstem
Col Colon
Dia Diaphragm
Ret Eye retina
Fat Fat
Hrt Heart
Pit Hypothalamus/Pituitary
Kid Kidney
LM Limb muscle
Liv Liver
Lung Lung
MG Mammary gland
Panc Pancreas
Skin Skin
SI Small intestine
Spl Spleen
Test Testes
Thym Thymus
Tong Tongue
Trac Trachea
Ute Uterus

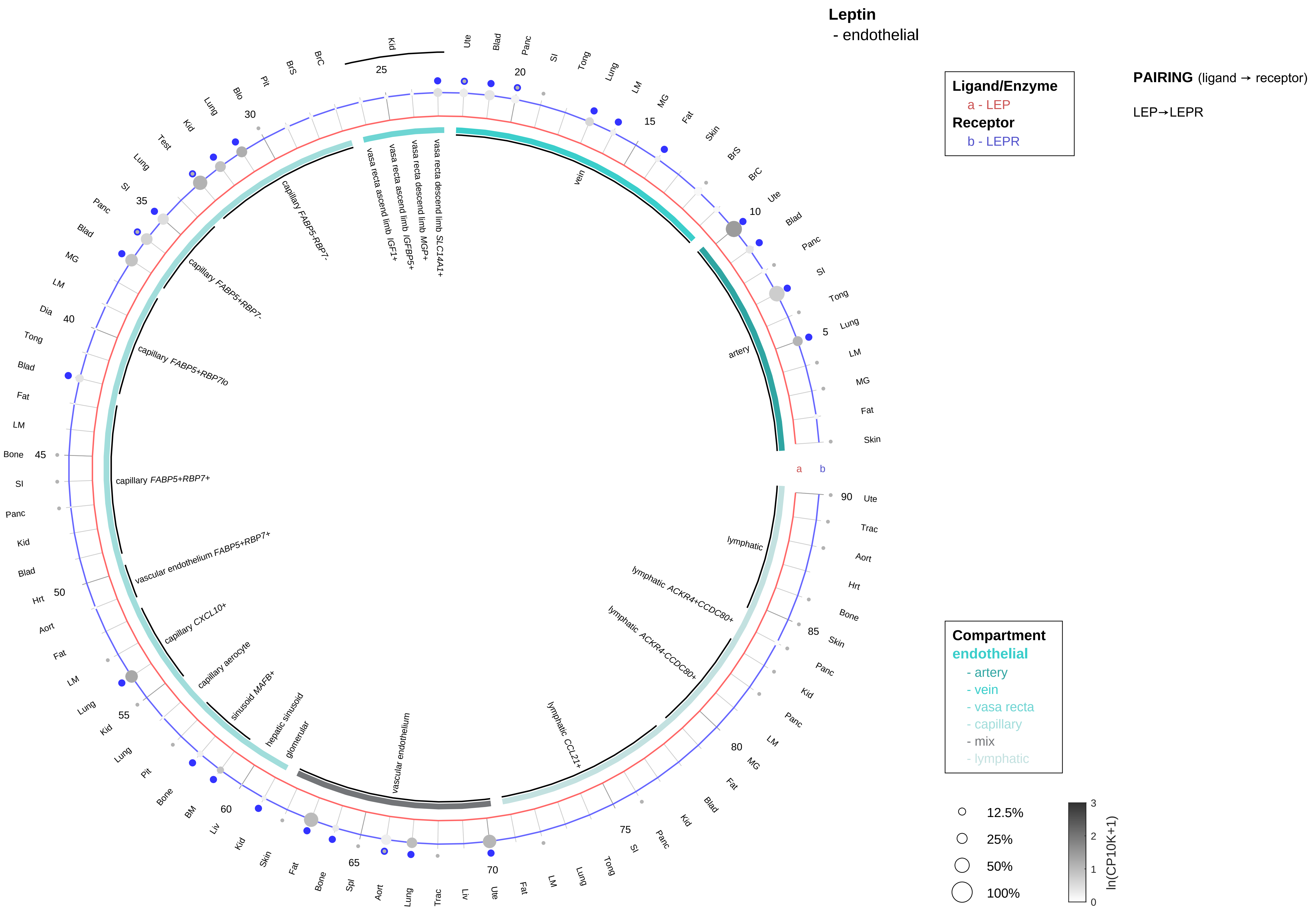

| Abbrev. | Tissue |
|---|---|
| Aort | Aorta |
| Blad | Bladder |
| Blo | Blood |
| Bone | Bone |
| BM | Bone marrow |
| BrC | Brain cortex |
| BrS | Brainstem |
| Col | Colon |
| Dia | Diaphragm |
| Ret | Eye retina |
| Fat | Fat |
| Hrt | Heart |
| Pit | Hypothalamus/Pituitary |
| Kid | Kidney |
| LM | Limb muscle |
| Liv | Liver |
| Lung | Lung |
| MG | Mammary gland |
| Panc | Pancreas |
| Skin | Skin |
| SI | Small intestine |
| Spl | Spleen |
| Test | Testes |
| Thym | Thymus |
| Tong | Tongue |
| Trac | Trachea |
| Ute | Uterus |

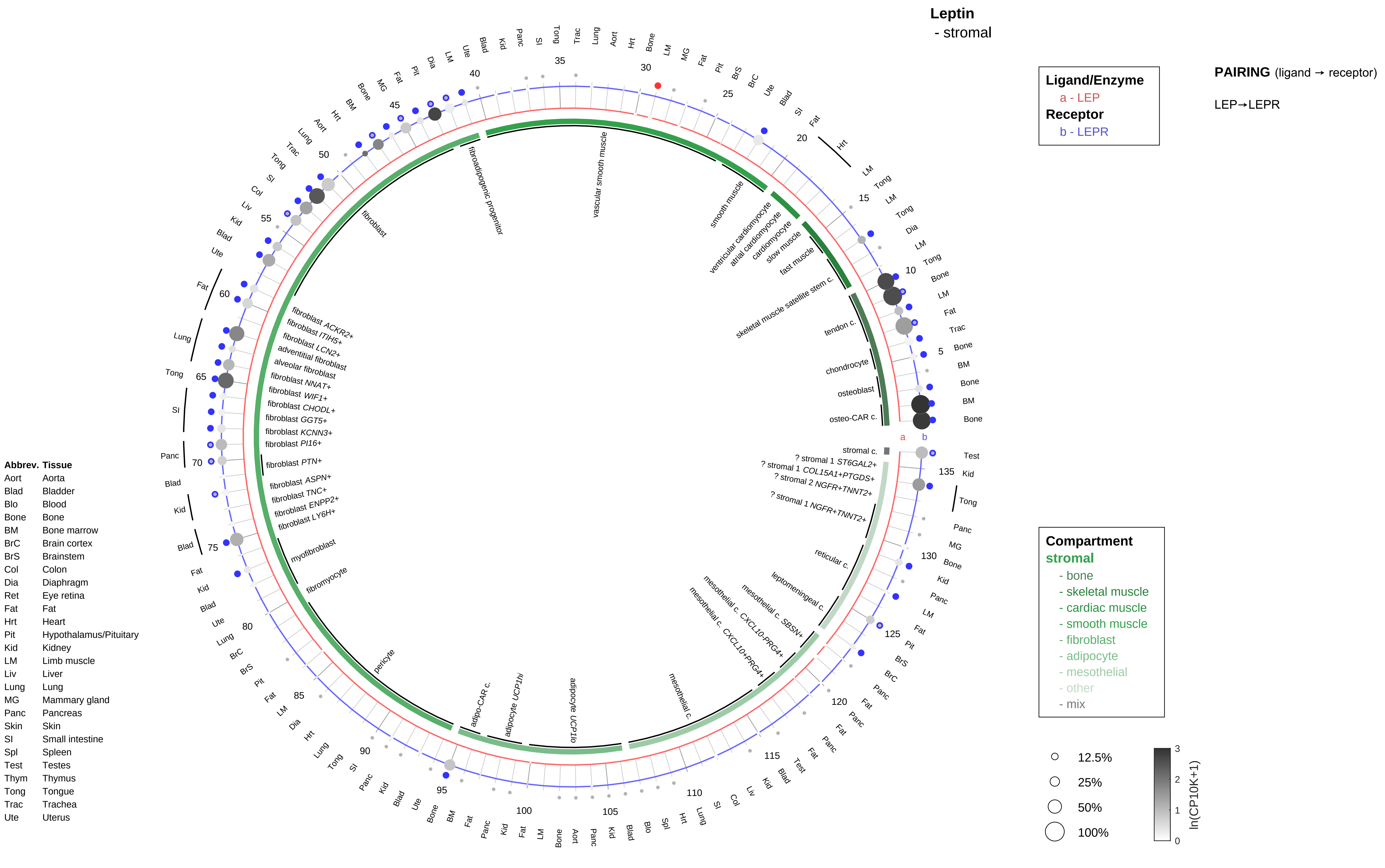

Leptin
- stromal
Ligand/Enzyme
a - LEP
Receptor
b - LEPR
PAIRING (ligand → receptor)
LEP→LEPR
Compartment
stromal
- bone
- skeletal muscle
- cardiac muscle
- smooth muscle
- fibroblast
- adipocyte
- mesothelial
- other
- mix
12.5%
25%
50%
100%
ln(CP10K+1)
Abbrev. Tissue
Aort Aorta
Blad Bladder
Blo Blood
Bone Bone
BM Bone marrow
BrC Brain cortex
BrS Brainstem
Col Colon
Dia Diaphragm
Ret Eye retina
Fat Fat
Hrt Heart
Pit Hypothalamus/Pituitary
Kid Kidney
LM Limb muscle
Liv Liver
Lung Lung
MG Mammary gland
Panc Pancreas
Skin Skin
SI Small intestine
Spl Spleen
Test Testes
Thym Thymus
Tong Tongue
Trac Trachea
Ute Uterus

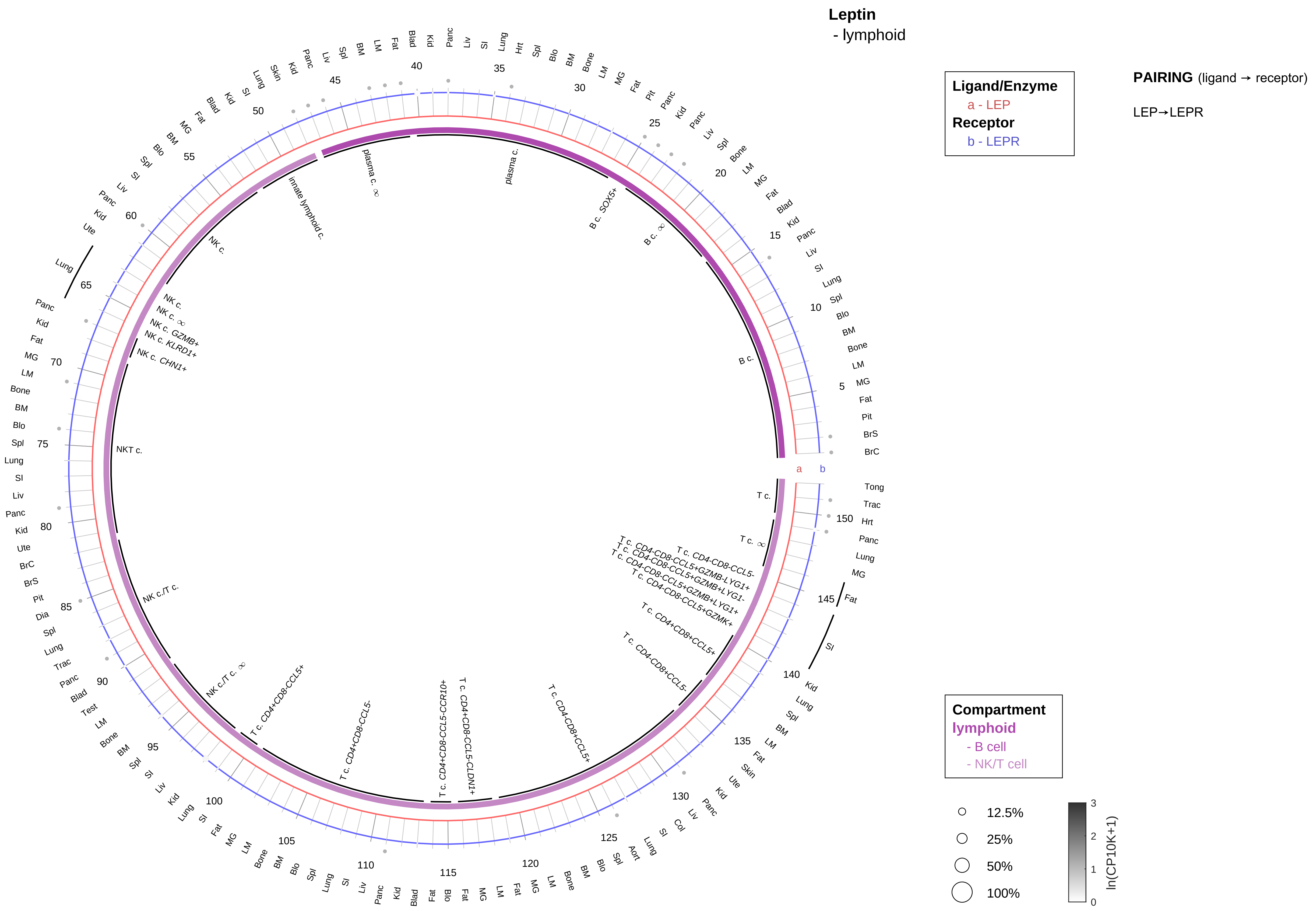

| Abbrev. | Tissue |
|---|---|
| Aort | Aorta |
| Blad | Bladder |
| Blo | Blood |
| Bone | Bone |
| BM | Bone marrow |
| BrC | Brain cortex |
| BrS | Brainstem |
| Col | Colon |
| Dia | Diaphragm |
| Ret | Eye retina |
| Fat | Fat |
| Hrt | Heart |
| Pit | Hypothalamus/Pituitary |
| Kid | Kidney |
| LM | Limb muscle |
| Liv | Liver |
| Lung | Lung |
| MG | Mammary gland |
| Panc | Pancreas |
| Skin | Skin |
| SI | Small intestine |
| Spl | Spleen |
| Test | Testes |
| Thym | Thymus |
| Tong | Tongue |
| Trac | Trachea |
| Ute | Uterus |

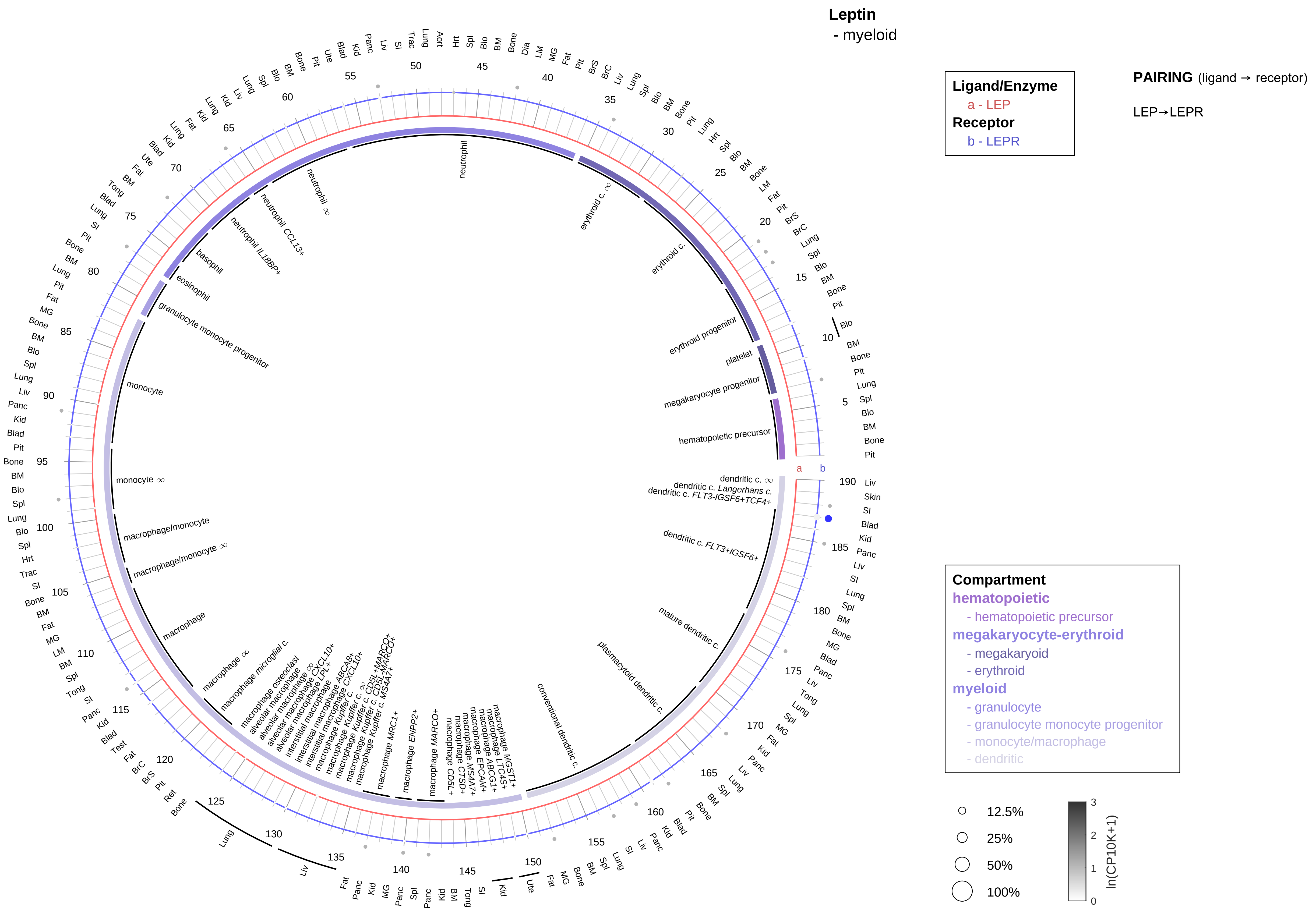

| Abbrev. | Tissue |
|---|---|
| Aort | Aorta |
| Blad | Bladder |
| Blo | Blood |
| Bone | Bone |
| BM | Bone marrow |
| BrC | Brain cortex |
| BrS | Brainstem |
| Col | Colon |
| Dia | Diaphragm |
| Ret | Eye retina |
| Fat | Fat |
| Hrt | Heart |
| Pit | Hypothalamus/Pituitary |
| Kid | Kidney |
| LM | Limb muscle |
| Liv | Liver |
| Lung | Lung |
| MG | Mammary gland |
| Panc | Pancreas |
| Skin | Skin |
| SI | Small intestine |
| Spl | Spleen |
| Test | Testes |
| Thym | Thymus |
| Tong | Tongue |
| Trac | Trachea |
| Ute | Uterus |

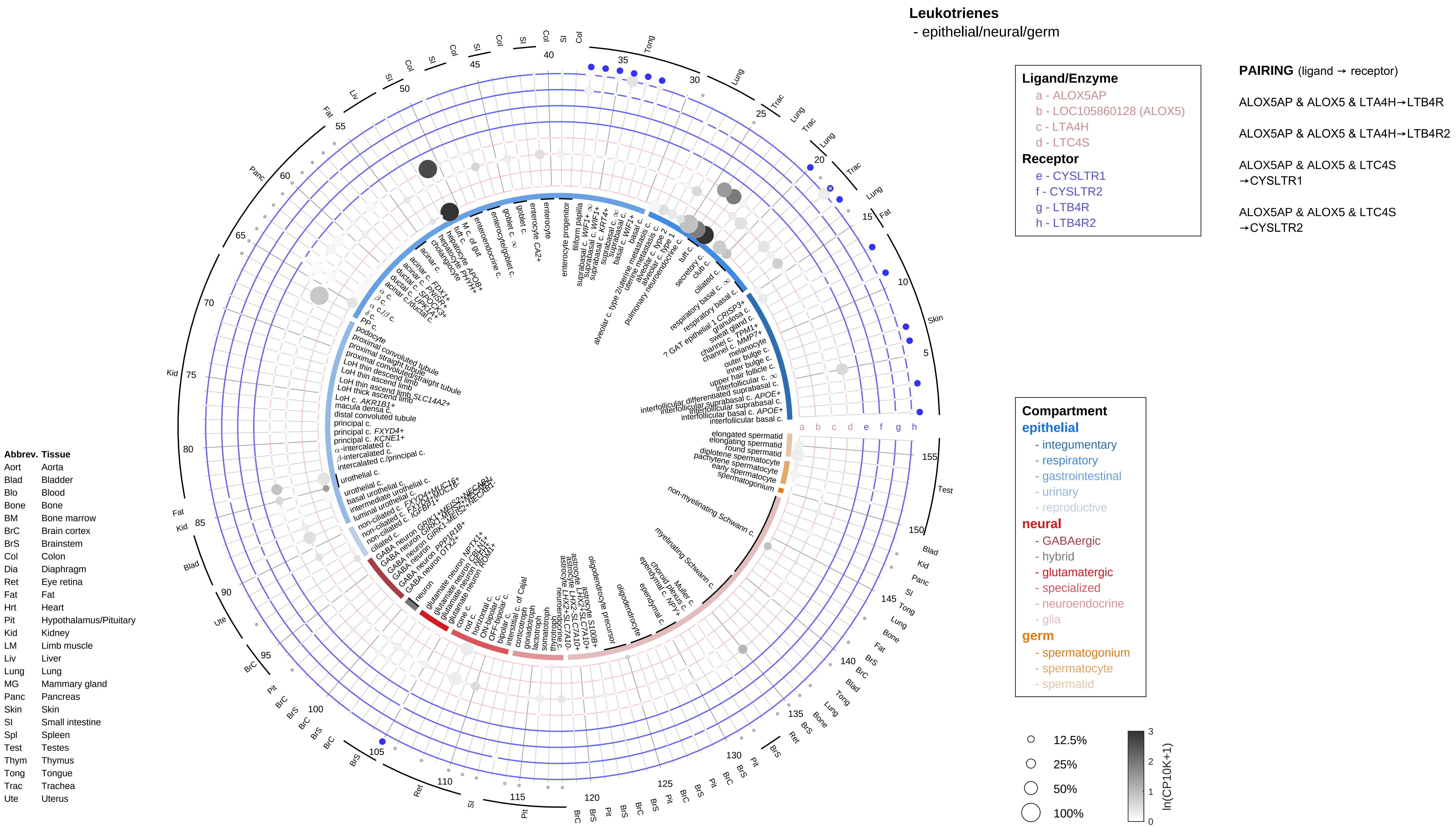
Leukotrienes
- epithelial/neural/germ
Ligand/Enzyme
a - ALOX5AP
b - LOC105860128 (ALOX5)
c - LTA4H
d - LTC4S
Receptor
e - CYSLTR1
f - CYSLTR2
g - LTB4R
h - LTB4R2
PAIRING (ligand → receptor)
ALOX5AP & ALOX5 & LTA4H→LTB4R
ALOX5AP & ALOX5 & LTA4H→LTB4R2
ALOX5AP & ALOX5 & LTC4S →CYSLTR1
ALOX5AP & ALOX5 & LTC4S →CYSLTR2
Compartment
epithelial
- integumentary
- respiratory
- gastrointestinal
- urinary
- reproductive
neural
- GABAergic
- hybrid
- glutamatergic
- specialized
- neuroendocrine
- glia
germ
- spermatogonium
- spermatocyte
- spermatid
12.5%
25%
50%
100%
ln(CP10K+1)
Abbrev. Tissue
Aort Aorta
Blad Bladder
Blo Blood
Bone Bone
BM Bone marrow
BrC Brain cortex
BrS Brainstem
Col Colon
Dia Diaphragm
Ret Eye retina
Fat Fat
Hrt Heart
Pit Hypothalamus/Pituitary
Kid Kidney
LM Limb muscle
Liv Liver
Lung Lung
MG Mammary gland
Panc Pancreas
Skin Skin
SI Small intestine
Spl Spleen
Test Testes
Thym Thymus
Tong Tongue
Trac Trachea
Ute Uterus

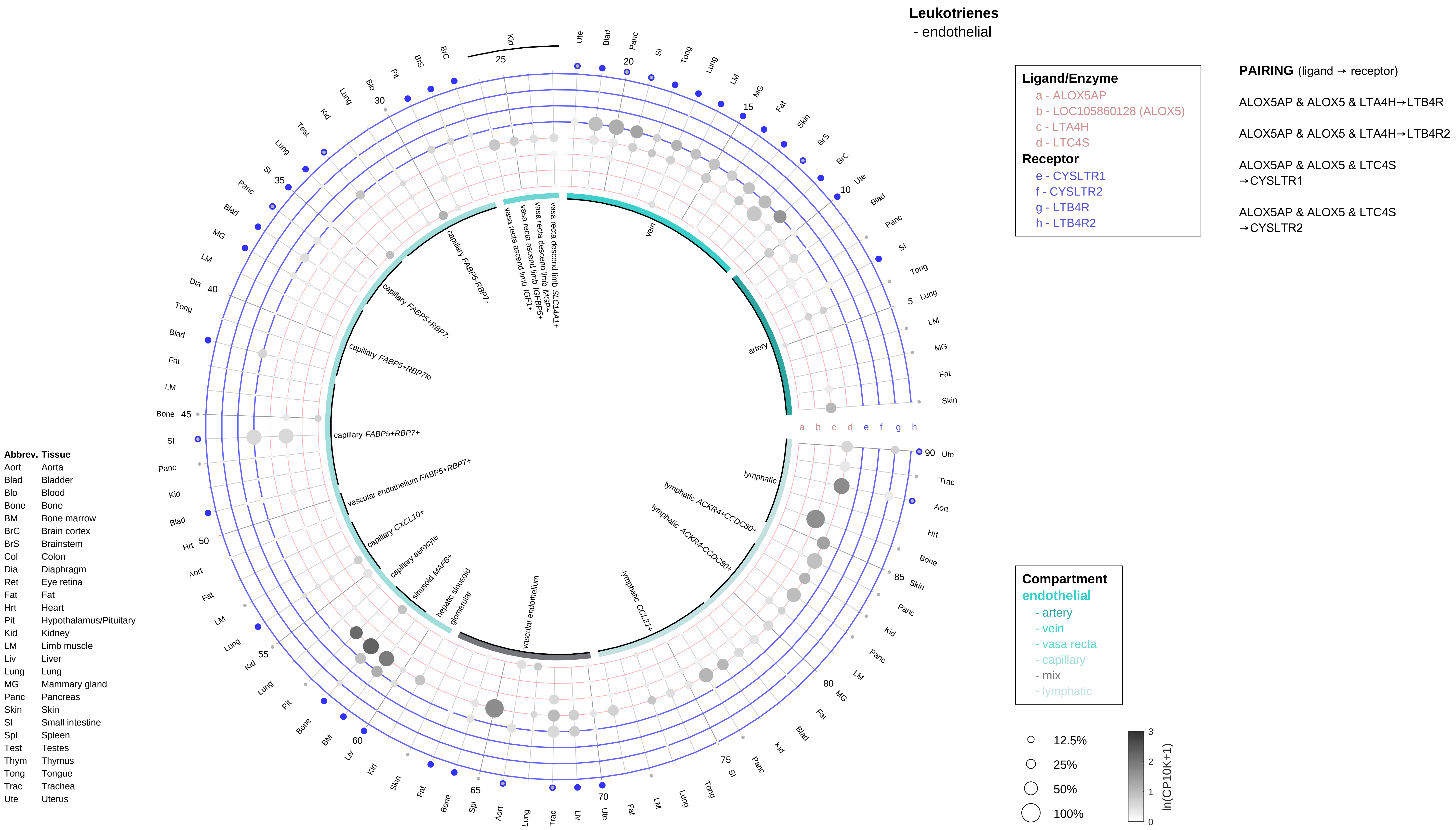

Leukotrienes
- endothelial
Ligand/Enzyme
a - ALOX5AP
b - LOC105860128 (ALOX5)
c - LTA4H
d - LTC4S
Receptor
e - CYSLTR1
f - CYSLTR2
g - LTB4R
h - LTB4R2
PAIRING (ligand → receptor)
ALOX5AP & ALOX5 & LTA4H→LTB4R
ALOX5AP & ALOX5 & LTA4H→LTB4R2
ALOX5AP & ALOX5 & LTC4S →CYSLTR1
ALOX5AP & ALOX5 & LTC4S →CYSLTR2
Abbrev. Tissue
Aort Aorta
Blad Bladder
Blo Blood
Bone Bone
BM Bone marrow
BrC Brain cortex
BrS Brainstem
Col Colon
Dia Diaphragm
Ret Eye retina
Fat Fat
Hrt Heart
Pit Hypothalamus/Pituitary
Kid Kidney
LM Limb muscle
Liv Liver
Lung Lung
MG Mammary gland
Panc Pancreas
Skin Skin
SI Small intestine
Spl Spleen
Test Testes
Thym Thymus
Tong Tongue
Trac Trachea
Ute Uterus
Compartment
endothelial
- artery
- vein
- vasa recta
- capillary
- mix
- lymphatic
12.5%
25%
50%
100%
ln(CP10K+1)

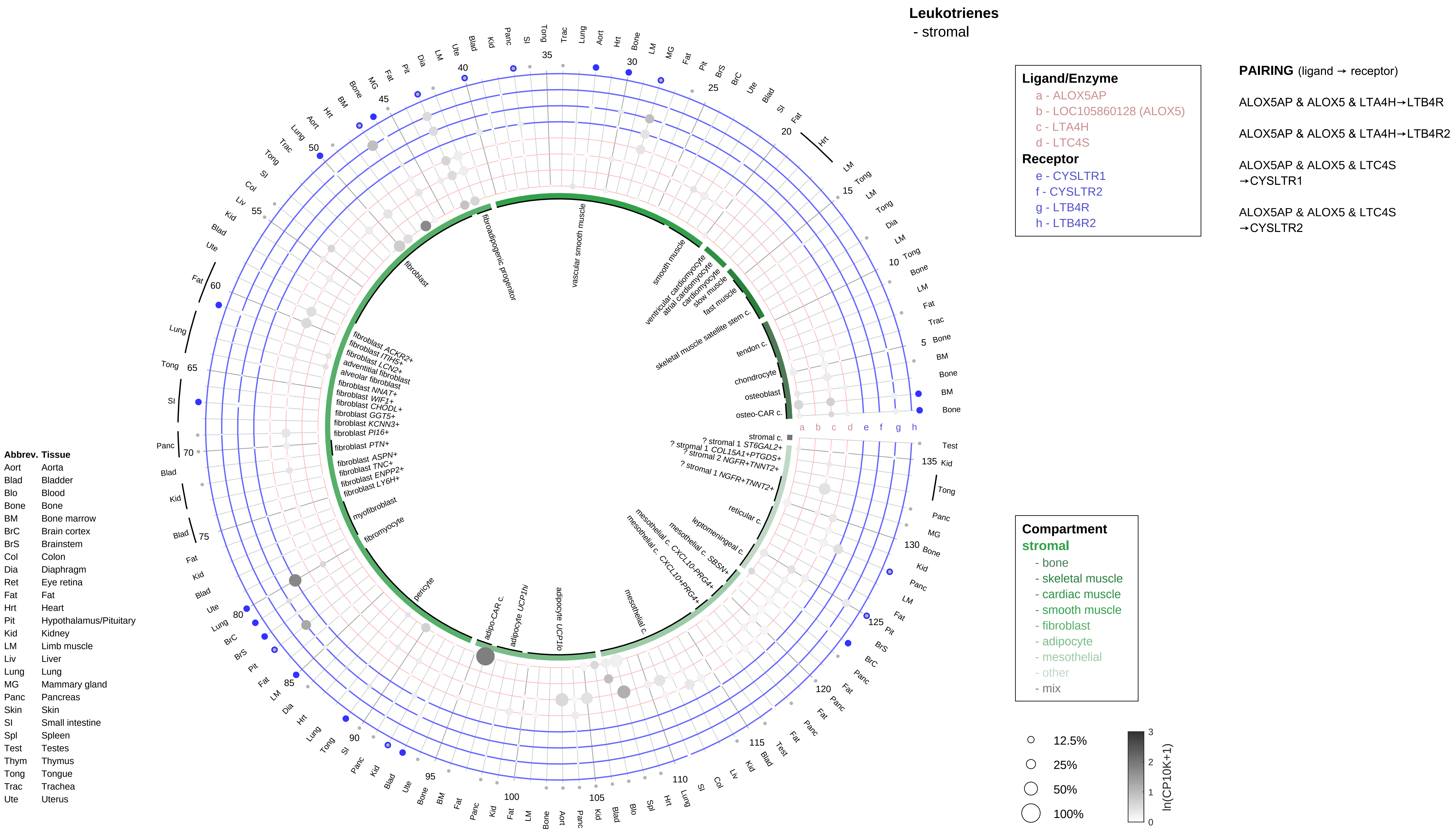

Leukotrienes
- stromal
Ligand/Enzyme
a - ALOX5AP
b - LOC105860128 (ALOX5)
c - LTA4H
d - LTC4S
Receptor
e - CYSLTR1
f - CYSLTR2
g - LTB4R
h - LTB4R2
PAIRING (ligand → receptor)
ALOX5AP & ALOX5 & LTA4H→LTB4R
ALOX5AP & ALOX5 & LTA4H→LTB4R2
ALOX5AP & ALOX5 & LTC4S →CYSLTR1
ALOX5AP & ALOX5 & LTC4S →CYSLTR2
Compartment
stromal
- bone
- skeletal muscle
- cardiac muscle
- smooth muscle
- fibroblast
- adipocyte
- mesothelial
- other
- mix
12.5%
25%
50%
100%
ln(CP10K+1)
Abbrev. Tissue
Aort Aorta
Blad Bladder
Blo Blood
Bone Bone
BM Bone marrow
BrC Brain cortex
BrS Brainstem
Col Colon
Dia Diaphragm
Ret Eye retina
Fat Fat
Hrt Heart
Pit Hypothalamus/Pituitary
Kid Kidney
LM Limb muscle
Liv Liver
Lung Lung
MG Mammary gland
Panc Pancreas
Skin Skin
SI Small intestine
Spl Spleen
Test Testes
Thym Thymus
Tong Tongue
Trac Trachea
Ute Uterus
fibroblast
fibroadipogenic progenitor
vascular smooth muscle
smooth muscle
ventricular cardiomyocyte
atrial cardiomyocyte
cardiomyocyte
slow muscle
fast muscle
skeletal muscle satellite stem c.
tendon c.
chondrocyte
osteoblast
osteo-CAR c.
stromal c.
? stromal 1 ST6GAL2+
? stromal 1 COL15A1+PTGDS+
? stromal 2 NGFR+TNNT2+
? stromal 1 NGFR+TNNT2+
reticular c.
leptomeningeal c.
mesothelial c. SBSN+
mesothelial c. CXCL10-PRG4+
mesothelial c. CXCL10+PRG4+
mesothelial c.
adipocyte UCP1lo
adipocyte UCP1hi
adipo-CAR c.
pericyte
fibromyocyte
myofibroblast
fibroblast LY6H+
fibroblast ENPP2+
fibroblast TNC+
fibroblast ASPN+
fibroblast PTN+
fibroblast PI16+
fibroblast KCNN3+
fibroblast GGT5+
fibroblast CHODL+
fibroblast WIF1+
fibroblast NNAT+
alveolar fibroblast
adventitial fibroblast
fibroblast LCN2+
fibroblast ITIH5+
fibroblast ACKR2+

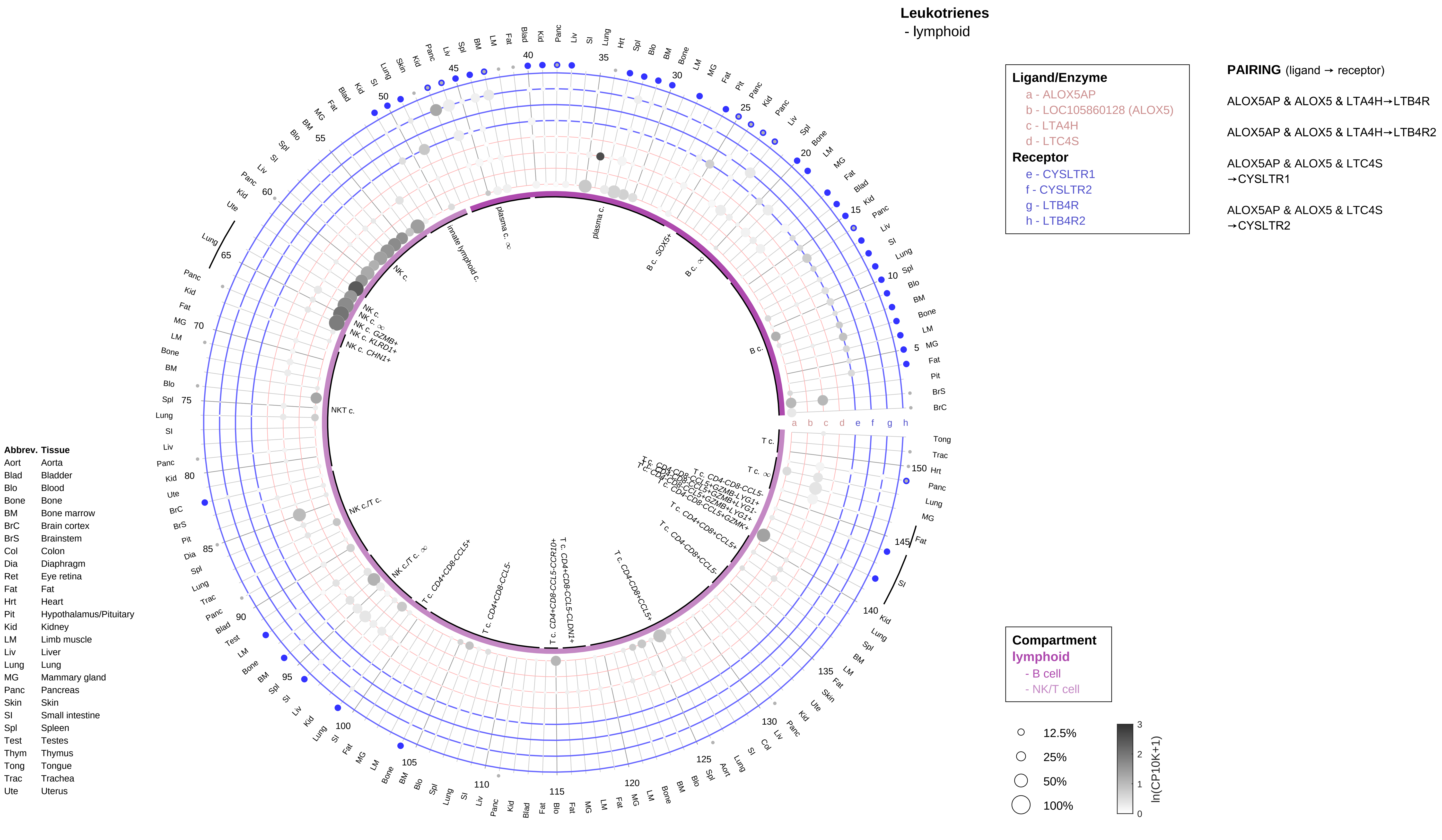
Leukotrienes
- lymphoid
Ligand/Enzyme
a - ALOX5AP
b - LOC105860128 (ALOX5)
c - LTA4H
d - LTC4S
Receptor
e - CYSLTR1
f - CYSLTR2
g - LTB4R
h - LTB4R2
PAIRING (ligand → receptor)
ALOX5AP & ALOX5 & LTA4H→LTB4R
ALOX5AP & ALOX5 & LTA4H→LTB4R2
ALOX5AP & ALOX5 & LTC4S →CYSLTR1
ALOX5AP & ALOX5 & LTC4S →CYSLTR2
Compartment
lymphoid
- B cell
- NK/T cell
12.5%
25%
50%
100%
ln(CP10K+1)
Abbrev. Tissue
Aort Aorta
Blad Bladder
Blo Blood
Bone Bone
BM Bone marrow
BrC Brain cortex
BrS Brainstem
Col Colon
Dia Diaphragm
Ret Eye retina
Fat Fat
Hrt Heart
Pit Hypothalamus/Pituitary
Kid Kidney
LM Limb muscle
Liv Liver
Lung Lung
MG Mammary gland
Panc Pancreas
Skin Skin
SI Small intestine
Spl Spleen
Test Testes
Thym Thymus
Tong Tongue
Trac Trachea
Ute Uterus

**Leukotrienes**
- myeloid

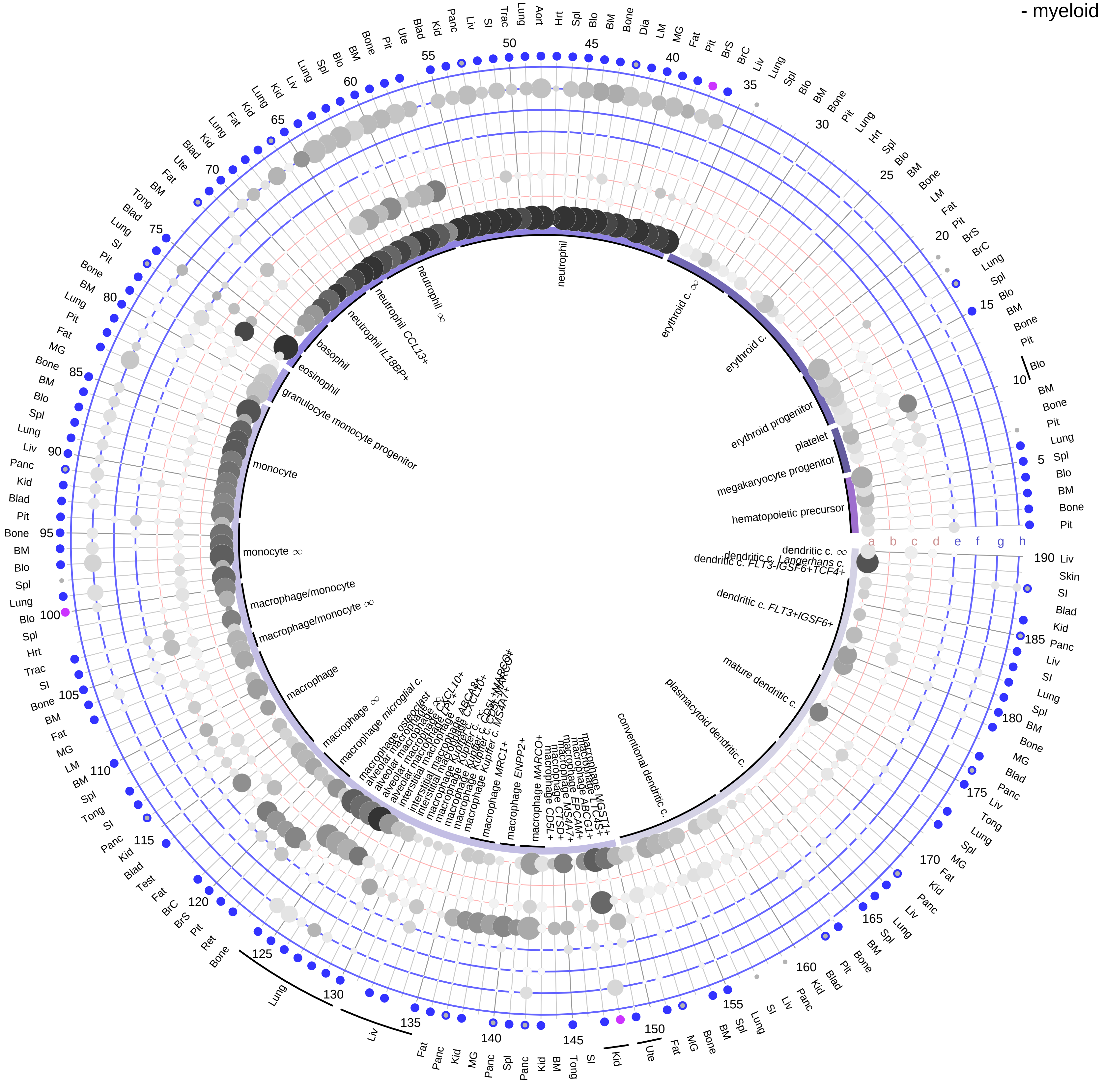

**Ligand/Enzyme**
- a - ALOX5AP
- b - LOC105860128 (ALOX5)
- c - LTA4H
- d - LTC4S

**Receptor**
- e - CYSLTR1
- f - CYSLTR2
- g - LTB4R
- h - LTB4R2

**PAIRING** (ligand → receptor)

ALOX5AP & ALOX5 & LTA4H→LTB4R

ALOX5AP & ALOX5 & LTA4H→LTB4R2

ALOX5AP & ALOX5 & LTC4S →CYSLTR1

ALOX5AP & ALOX5 & LTC4S →CYSLTR2

| Abbrev. | Tissue |
|---|---|
| Aort | Aorta |
| Blad | Bladder |
| Blo | Blood |
| Bone | Bone |
| BM | Bone marrow |
| BrC | Brain cortex |
| BrS | Brainstem |
| Col | Colon |
| Dia | Diaphragm |
| Ret | Eye retina |
| Fat | Fat |
| Hrt | Heart |
| Pit | Hypothalamus/Pituitary |
| Kid | Kidney |
| LM | Limb muscle |
| Liv | Liver |
| Lung | Lung |
| MG | Mammary gland |
| Panc | Pancreas |
| Skin | Skin |
| SI | Small intestine |
| Spl | Spleen |
| Test | Testes |
| Thym | Thymus |
| Tong | Tongue |
| Trac | Trachea |
| Ute | Uterus |

**Compartment**
- **hematopoietic**
  - hematopoietic precursor
- **megakaryocyte-erythroid**
  - megakaryoid
  - erythroid
- **myeloid**
  - granulocyte
  - granulocyte monocyte progenitor
  - monocyte/macrophage
  - dendritic

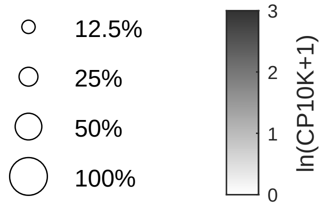

**Luteinizing hormone (LH)**
- epithelial/neural/germ

**Ligand/Enzyme**
- a - CGA
- b - LHB_uTAR

**Receptor**
- c - LHCGR

**PAIRING** (ligand → receptor)

LHB & CGA→LHCGR

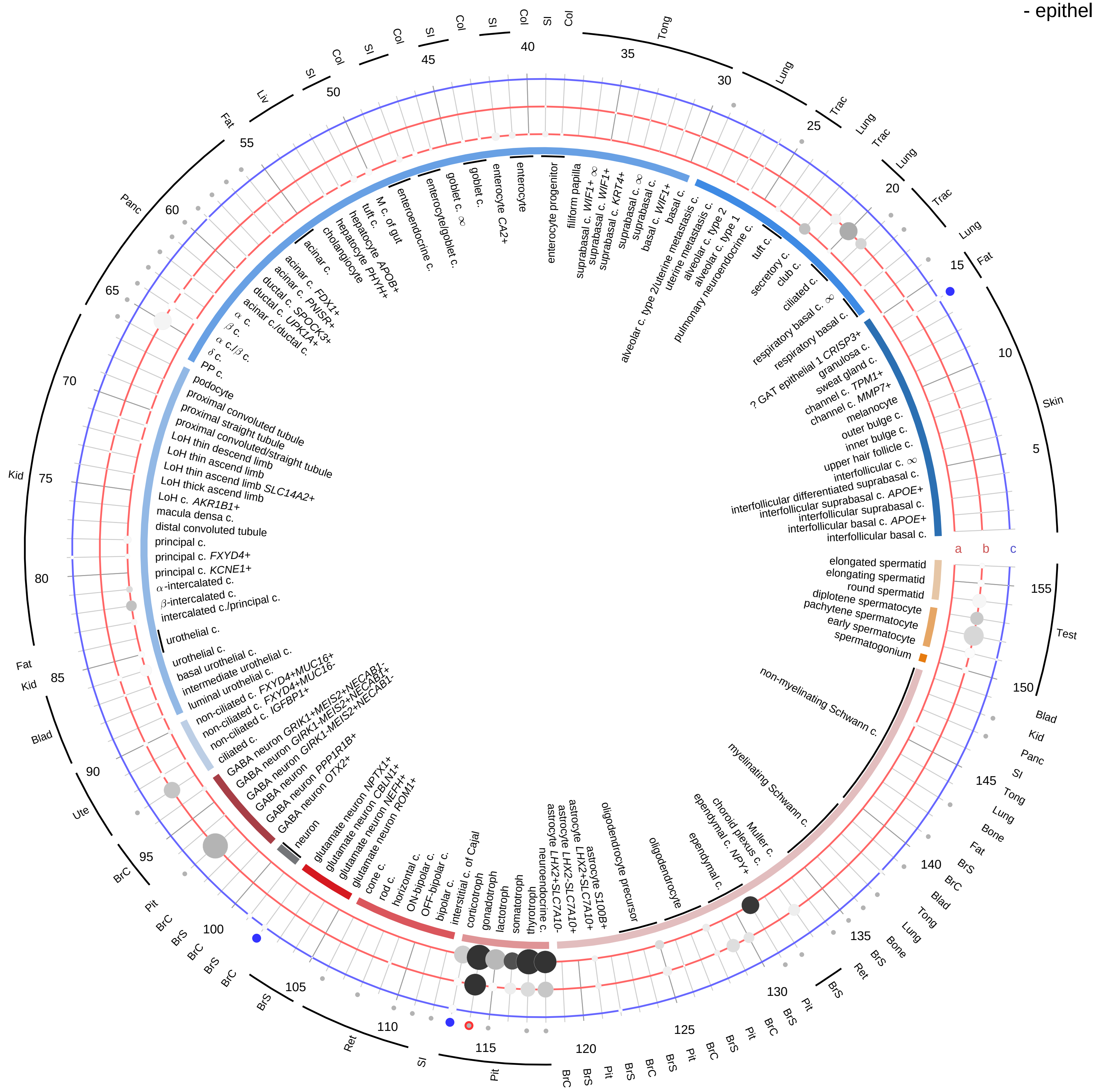

**Compartment**

**epithelial**
- integumentary
- respiratory
- gastrointestinal
- urinary
- reproductive

**neural**
- GABAergic
- hybrid
- glutamatergic
- specialized
- neuroendocrine
- glia

**germ**
- spermatogonium
- spermatocyte
- spermatid

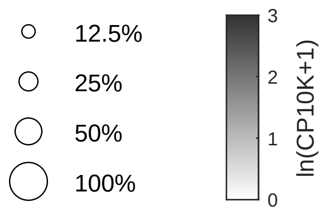

| Abbrev. | Tissue |
|---|---|
| Aort | Aorta |
| Blad | Bladder |
| Blo | Blood |
| Bone | Bone |
| BM | Bone marrow |
| BrC | Brain cortex |
| BrS | Brainstem |
| Col | Colon |
| Dia | Diaphragm |
| Ret | Eye retina |
| Fat | Fat |
| Hrt | Heart |
| Pit | Hypothalamus/Pituitary |
| Kid | Kidney |
| LM | Limb muscle |
| Liv | Liver |
| Lung | Lung |
| MG | Mammary gland |
| Panc | Pancreas |
| Skin | Skin |
| SI | Small intestine |
| Spl | Spleen |
| Test | Testes |
| Thym | Thymus |
| Tong | Tongue |
| Trac | Trachea |
| Ute | Uterus |

**Luteinizing hormone (LH)**
- endothelial

**PAIRING** (ligand → receptor)

LHB & CGA→LHCGR

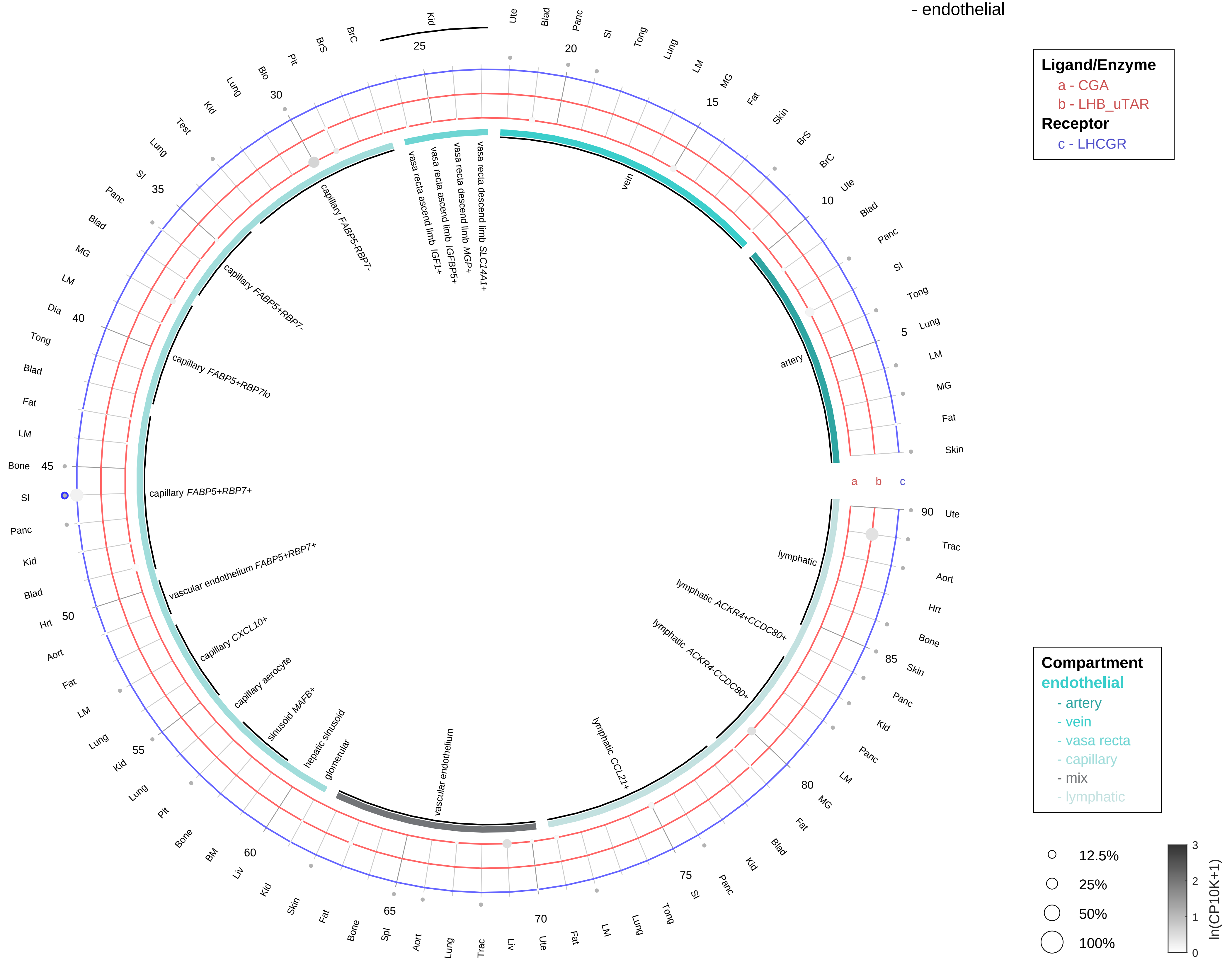

| Abbrev. | Tissue |
|---|---|
| Aort | Aorta |
| Blad | Bladder |
| Blo | Blood |
| Bone | Bone |
| BM | Bone marrow |
| BrC | Brain cortex |
| BrS | Brainstem |
| Col | Colon |
| Dia | Diaphragm |
| Ret | Eye retina |
| Fat | Fat |
| Hrt | Heart |
| Pit | Hypothalamus/Pituitary |
| Kid | Kidney |
| LM | Limb muscle |
| Liv | Liver |
| Lung | Lung |
| MG | Mammary gland |
| Panc | Pancreas |
| Skin | Skin |
| SI | Small intestine |
| Spl | Spleen |
| Test | Testes |
| Thym | Thymus |
| Tong | Tongue |
| Trac | Trachea |
| Ute | Uterus |

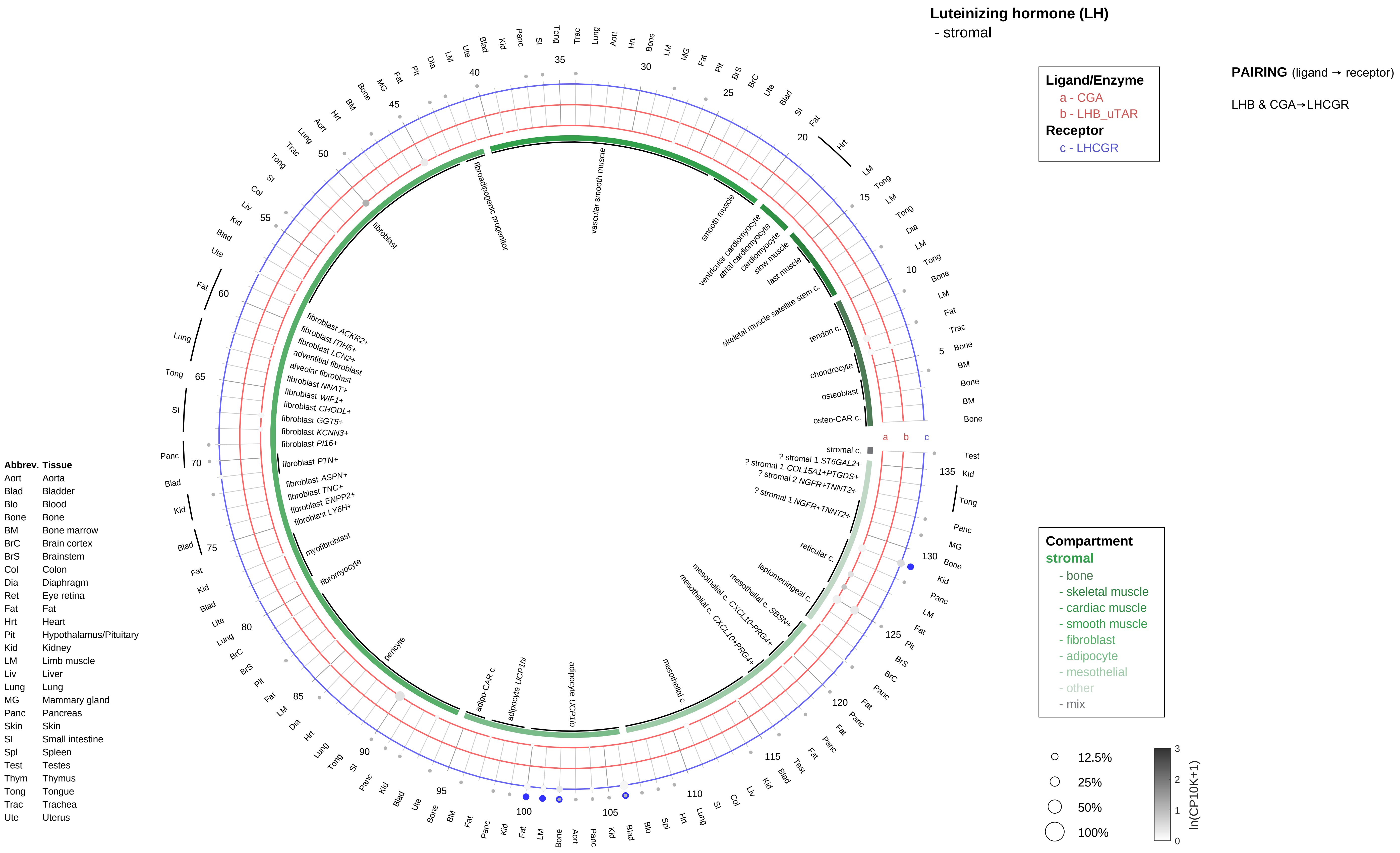

Luteinizing hormone (LH)
- stromal
Ligand/Enzyme
a - CGA
b - LHB_uTAR
Receptor
c - LHCGR
PAIRING (ligand → receptor)
LHB & CGA→LHCGR
Compartment
stromal
- bone
- skeletal muscle
- cardiac muscle
- smooth muscle
- fibroblast
- adipocyte
- mesothelial
- other
- mix
12.5%
25%
50%
100%
ln(CP10K+1)
Abbrev. Tissue
Aort Aorta
Blad Bladder
Blo Blood
Bone Bone
BM Bone marrow
BrC Brain cortex
BrS Brainstem
Col Colon
Dia Diaphragm
Ret Eye retina
Fat Fat
Hrt Heart
Pit Hypothalamus/Pituitary
Kid Kidney
LM Limb muscle
Liv Liver
Lung Lung
MG Mammary gland
Panc Pancreas
Skin Skin
SI Small intestine
Spl Spleen
Test Testes
Thym Thymus
Tong Tongue
Trac Trachea
Ute Uterus
fibroadipogenic progenitor
vascular smooth muscle
smooth muscle
ventricular cardiomyocyte
atrial cardiomyocyte
cardiomyocyte
slow muscle
fast muscle
skeletal muscle satellite stem c.
tendon c.
chondrocyte
osteoblast
osteo-CAR c.
stromal c.
? stromal 1 ST6GAL2+
? stromal 1 COL15A1+PTGDS+
? stromal 2 NGFR+TNNT2+
? stromal 1 NGFR+TNNT2+
reticular c.
leptomeningeal c.
mesothelial c. SBSN+
mesothelial c. CXCL10-PRG4+
mesothelial c. CXCL10+PRG4+
mesothelial c.
adipocyte UCP1lo
adipocyte UCP1hi
adipo-CAR c.
pericyte
fibromyocyte
myofibroblast
fibroblast LY6H+
fibroblast ENPP2+
fibroblast TNC+
fibroblast ASPN+
fibroblast PTN+
fibroblast PI16+
fibroblast KCNN3+
fibroblast GGT5+
fibroblast CHODL+
fibroblast WIF1+
fibroblast NNAT+
alveolar fibroblast
adventitial fibroblast
fibroblast LCN2+
fibroblast ITIH5+
fibroblast ACKR2+
fibroblast

#### Luteinizing hormone (LH)
- lymphoid

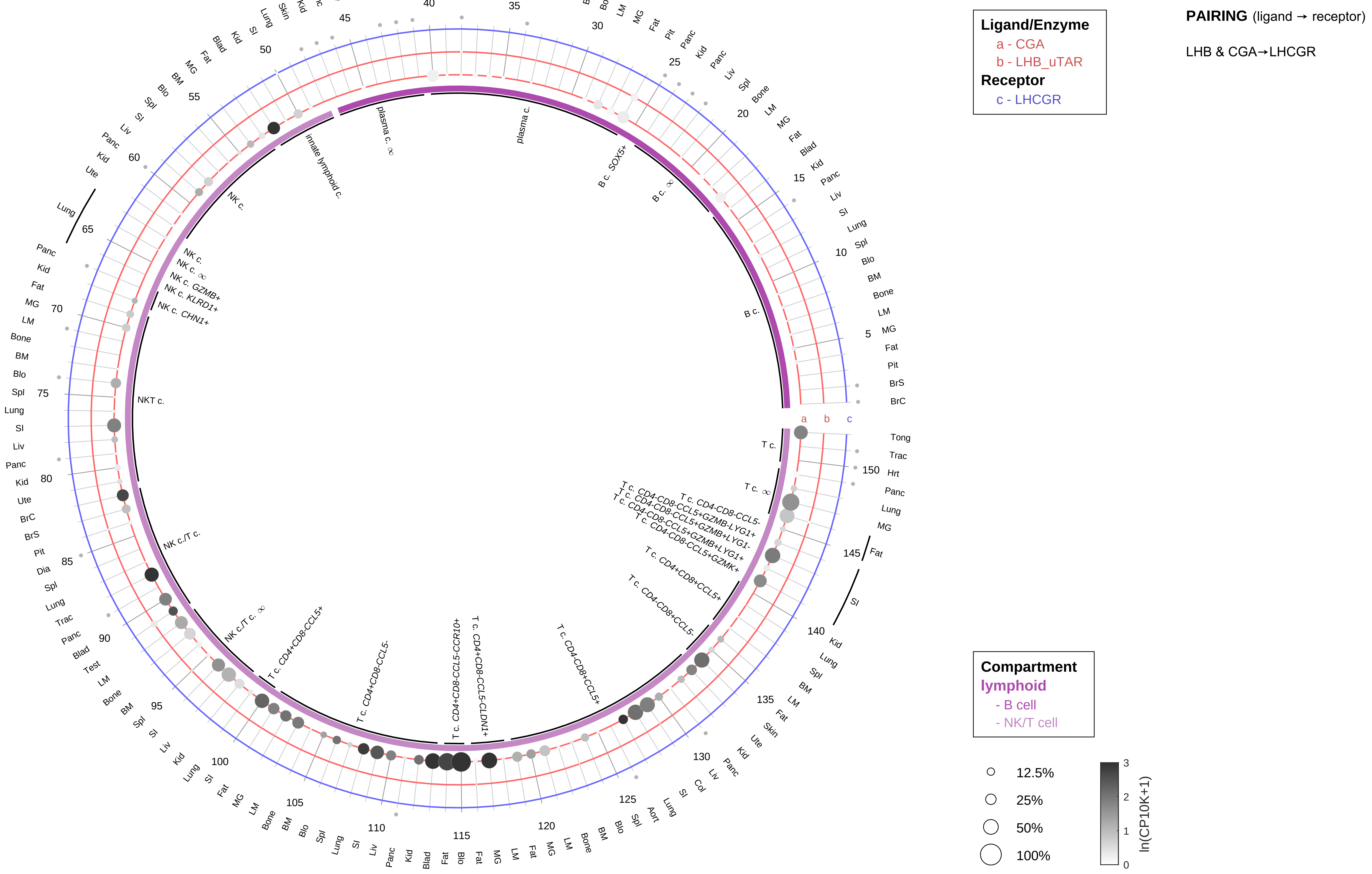

| Abbrev. | Tissue |
|---|---|
| Aort | Aorta |
| Blad | Bladder |
| Blo | Blood |
| Bone | Bone |
| BM | Bone marrow |
| BrC | Brain cortex |
| BrS | Brainstem |
| Col | Colon |
| Dia | Diaphragm |
| Ret | Eye retina |
| Fat | Fat |
| Hrt | Heart |
| Pit | Hypothalamus/Pituitary |
| Kid | Kidney |
| LM | Limb muscle |
| Liv | Liver |
| Lung | Lung |
| MG | Mammary gland |
| Panc | Pancreas |
| Skin | Skin |
| SI | Small intestine |
| Spl | Spleen |
| Test | Testes |
| Thym | Thymus |
| Tong | Tongue |
| Trac | Trachea |
| Ute | Uterus |

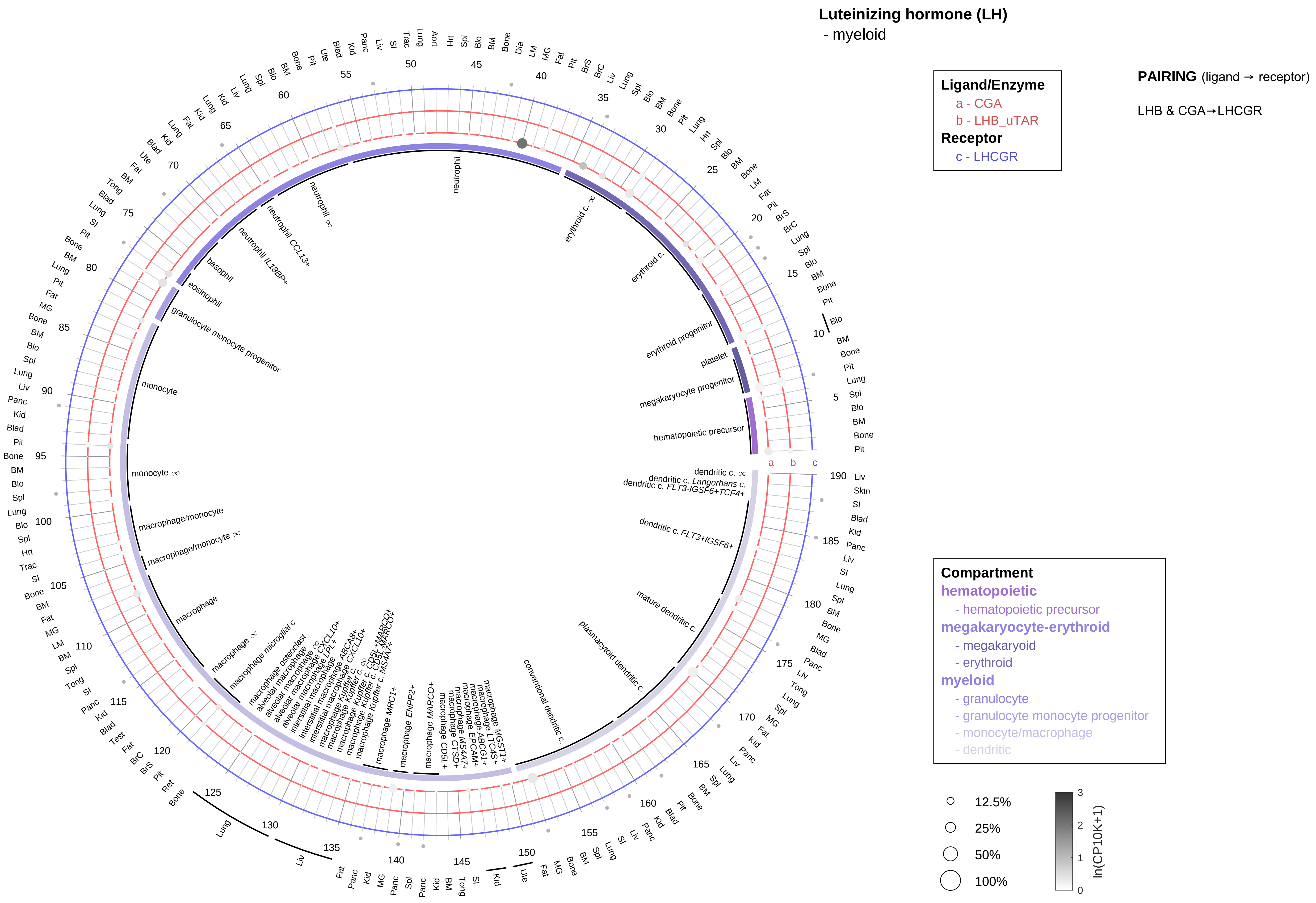

| Abbrev. | Tissue |
|---|---|
| Aort | Aorta |
| Blad | Bladder |
| Blo | Blood |
| Bone | Bone |
| BM | Bone marrow |
| BrC | Brain cortex |
| BrS | Brainstem |
| Col | Colon |
| Dia | Diaphragm |
| Ret | Eye retina |
| Fat | Fat |
| Hrt | Heart |
| Pit | Hypothalamus/Pituitary |
| Kid | Kidney |
| LM | Limb muscle |
| Liv | Liver |
| Lung | Lung |
| MG | Mammary gland |
| Panc | Pancreas |
| Skin | Skin |
| SI | Small intestine |
| Spl | Spleen |
| Test | Testes |
| Thym | Thymus |
| Tong | Tongue |
| Trac | Trachea |
| Ute | Uterus |

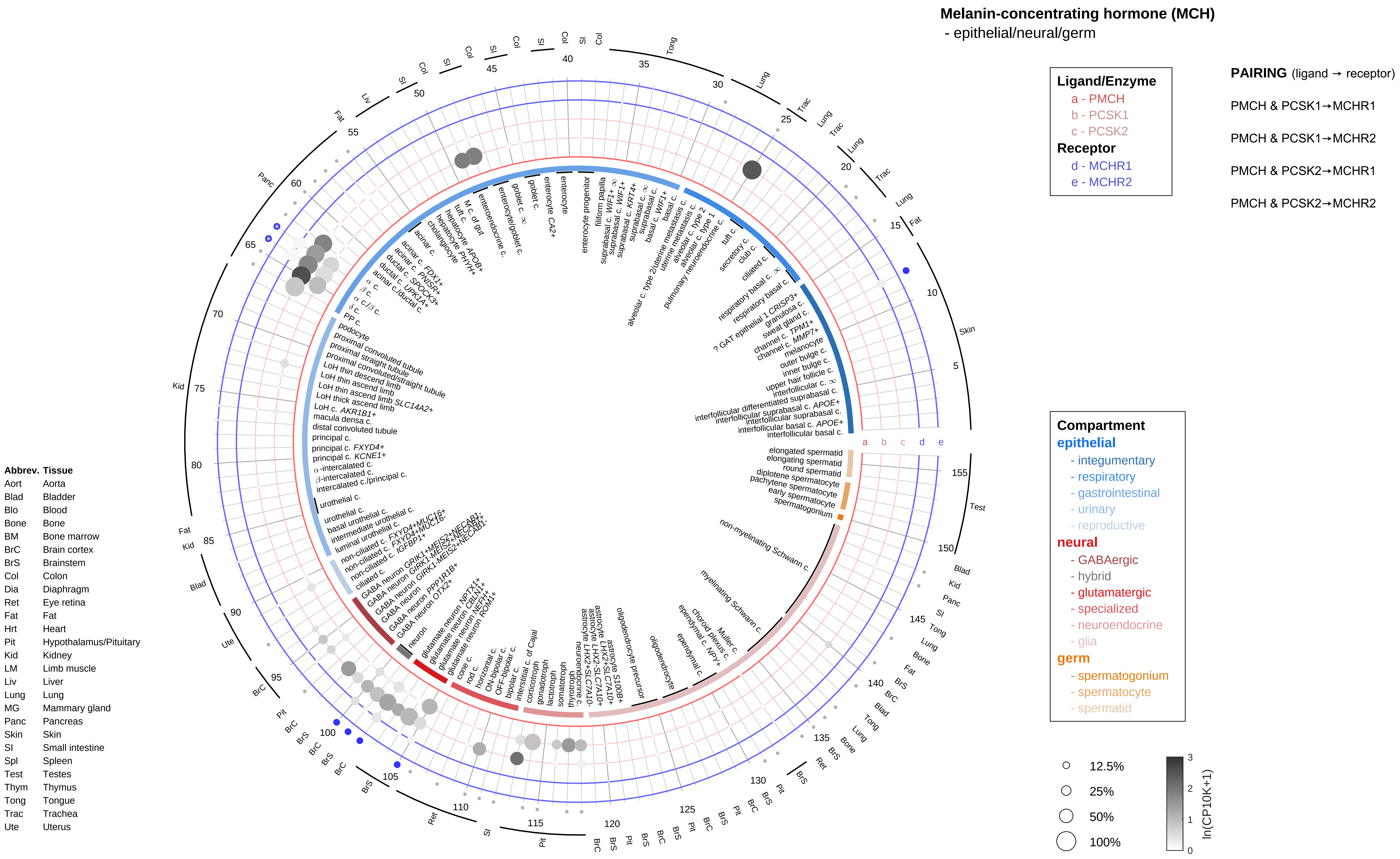
Melanin-concentrating hormone (MCH)
- epithelial/neural/germ
Ligand/Enzyme
a - PMCH
b - PCSK1
c - PCSK2
Receptor
d - MCHR1
e - MCHR2
PAIRING (ligand → receptor)
PMCH & PCSK1→MCHR1
PMCH & PCSK1→MCHR2
PMCH & PCSK2→MCHR1
PMCH & PCSK2→MCHR2
Compartment
epithelial
- integumentary
- respiratory
- gastrointestinal
- urinary
- reproductive
neural
- GABAergic
- hybrid
- glutamatergic
- specialized
- neuroendocrine
- glia
germ
- spermatogonium
- spermatocyte
- spermatid
12.5%
25%
50%
100%
ln(CP10K+1)
Abbrev. Tissue
Aort Aorta
Blad Bladder
Blo Blood
Bone Bone
BM Bone marrow
BrC Brain cortex
BrS Brainstem
Col Colon
Dia Diaphragm
Ret Eye retina
Fat Fat
Hrt Heart
Pit Hypothalamus/Pituitary
Kid Kidney
LM Limb muscle
Liv Liver
Lung Lung
MG Mammary gland
Panc Pancreas
Skin Skin
SI Small intestine
Spl Spleen
Test Testes
Thym Thymus
Tong Tongue
Trac Trachea
Ute Uterus

**Melanin-concentrating hormone (MCH)**
- endothelial

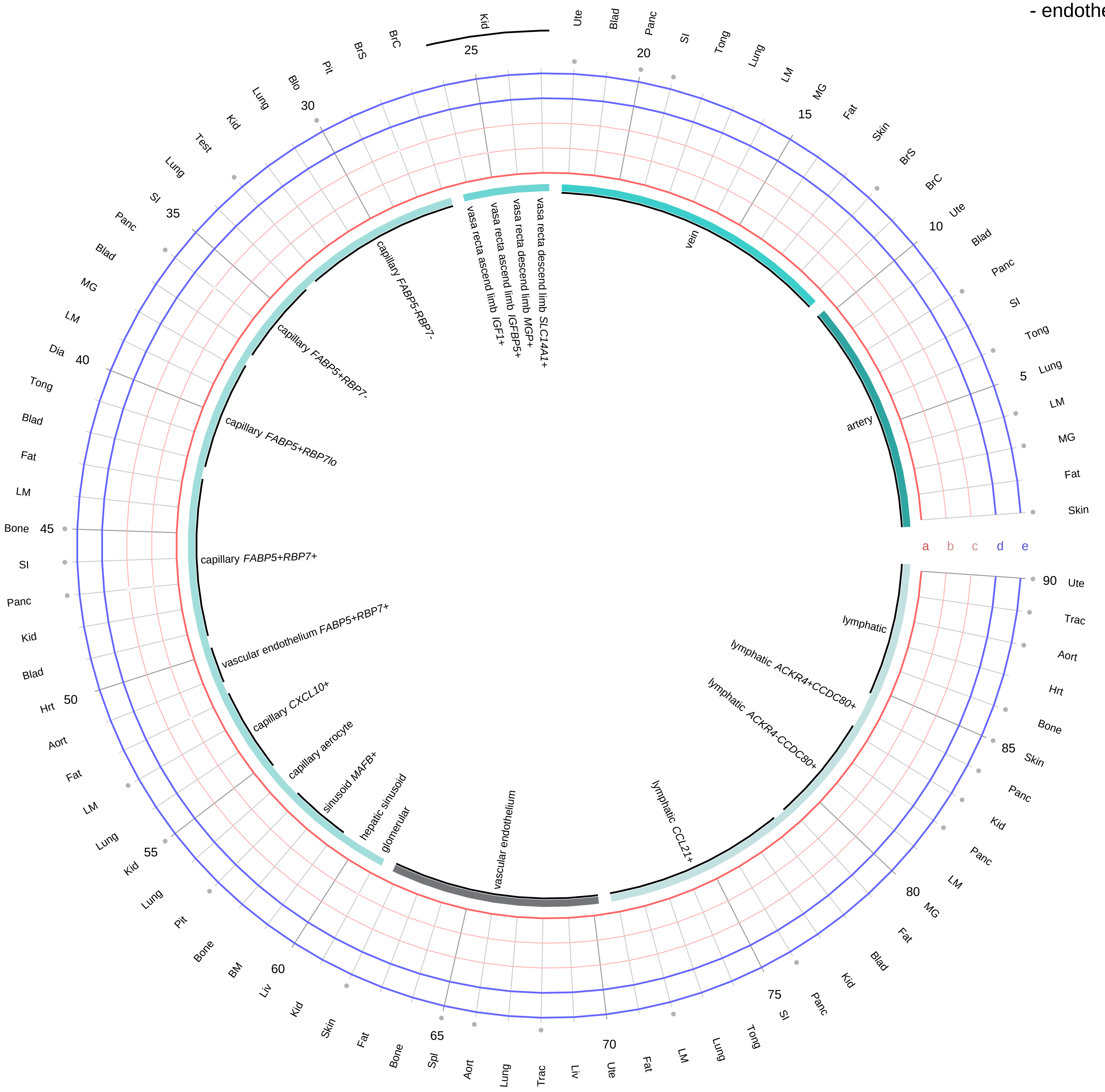

**Ligand/Enzyme**
- a - PMCH
- b - PCSK1
- c - PCSK2

**Receptor**
- d - MCHR1
- e - MCHR2

**PAIRING** (ligand → receptor)

PMCH & PCSK1→MCHR1

PMCH & PCSK1→MCHR2

PMCH & PCSK2→MCHR1

PMCH & PCSK2→MCHR2

| Abbrev. | Tissue |
|---|---|
| Aort | Aorta |
| Blad | Bladder |
| Blo | Blood |
| Bone | Bone |
| BM | Bone marrow |
| BrC | Brain cortex |
| BrS | Brainstem |
| Col | Colon |
| Dia | Diaphragm |
| Ret | Eye retina |
| Fat | Fat |
| Hrt | Heart |
| Pit | Hypothalamus/Pituitary |
| Kid | Kidney |
| LM | Limb muscle |
| Liv | Liver |
| Lung | Lung |
| MG | Mammary gland |
| Panc | Pancreas |
| Skin | Skin |
| SI | Small intestine |
| Spl | Spleen |
| Test | Testes |
| Thym | Thymus |
| Tong | Tongue |
| Trac | Trachea |
| Ute | Uterus |

**Compartment**
**endothelial**
- artery
- vein
- vasa recta
- capillary
- mix
- lymphatic

12.5%
25%
50%
100%

ln(CP10K+1): 0, 1, 2, 3

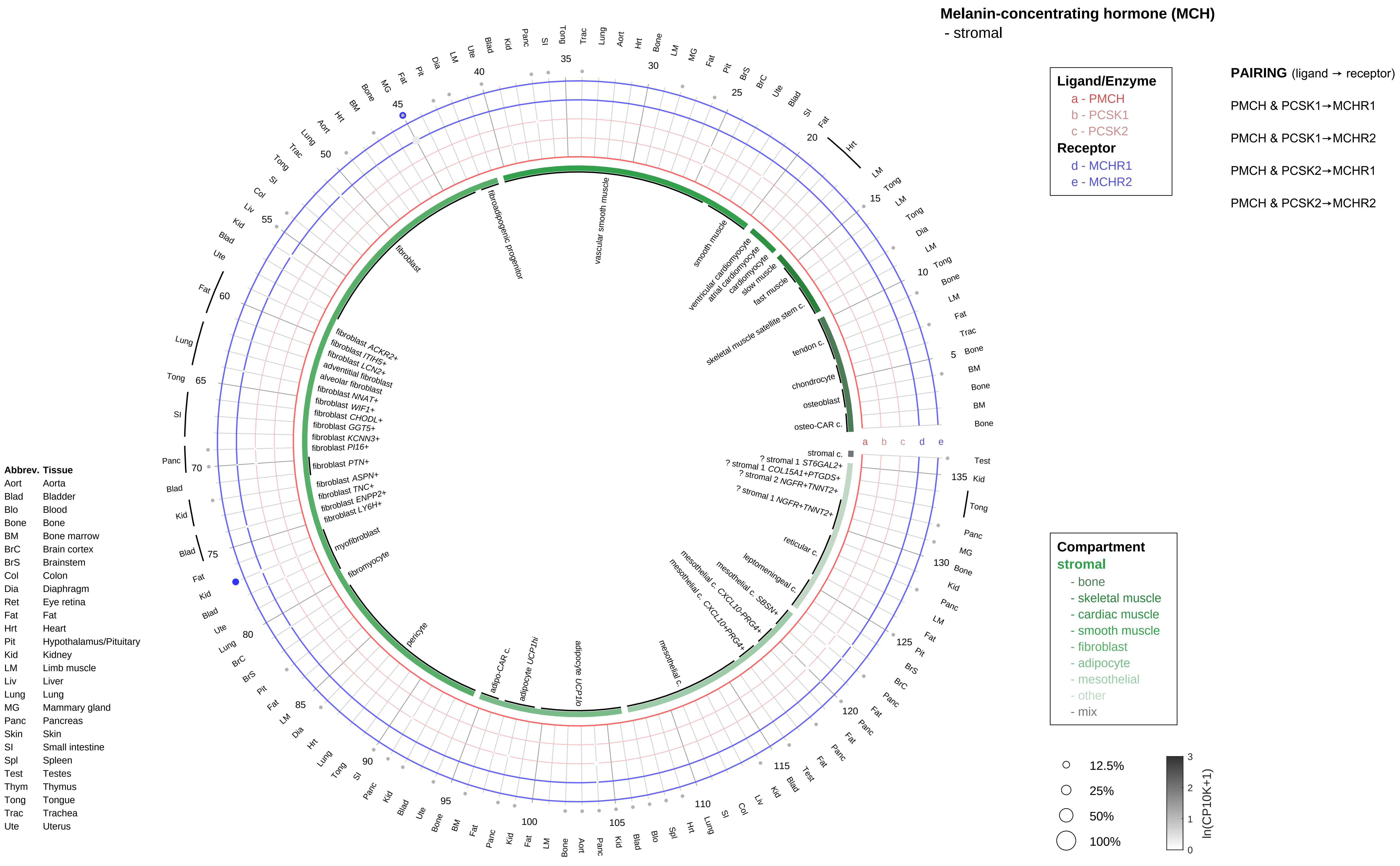
Melanin-concentrating hormone (MCH)
- stromal
Ligand/Enzyme
a - PMCH
b - PCSK1
c - PCSK2
Receptor
d - MCHR1
e - MCHR2
PAIRING (ligand → receptor)
PMCH & PCSK1→MCHR1
PMCH & PCSK1→MCHR2
PMCH & PCSK2→MCHR1
PMCH & PCSK2→MCHR2
Compartment
stromal
- bone
- skeletal muscle
- cardiac muscle
- smooth muscle
- fibroblast
- adipocyte
- mesothelial
- other
- mix
12.5%
25%
50%
100%
ln(CP10K+1)
Abbrev. Tissue
Aort Aorta
Blad Bladder
Blo Blood
Bone Bone
BM Bone marrow
BrC Brain cortex
BrS Brainstem
Col Colon
Dia Diaphragm
Ret Eye retina
Fat Fat
Hrt Heart
Pit Hypothalamus/Pituitary
Kid Kidney
LM Limb muscle
Liv Liver
Lung Lung
MG Mammary gland
Panc Pancreas
Skin Skin
SI Small intestine
Spl Spleen
Test Testes
Thym Thymus
Tong Tongue
Trac Trachea
Ute Uterus
vascular smooth muscle
smooth muscle
ventricular cardiomyocyte
atrial cardiomyocyte
cardiomyocyte
slow muscle
fast muscle
skeletal muscle satellite stem c.
tendon c.
chondrocyte
osteoblast
osteo-CAR c.
stromal c.
? stromal 1 ST6GAL2+
? stromal 1 COL15A1+PTGDS+
? stromal 2 NGFR+TNNT2+
? stromal 1 NGFR+TNNT2+
reticular c.
leptomeningeal c.
mesothelial c. SBSN+
mesothelial c. CXCL10-PRG4+
mesothelial c. CXCL10+PRG4+
mesothelial c.
adipocyte UCP1lo
adipocyte UCP1hi
adipo-CAR c.
pericyte
fibromyocyte
myofibroblast
fibroblast LY6H+
fibroblast ENPP2+
fibroblast TNC+
fibroblast ASPN+
fibroblast PTN+
fibroblast PI16+
fibroblast KCNN3+
fibroblast GGT5+
fibroblast CHODL+
fibroblast WIF1+
fibroblast NNAT+
alveolar fibroblast
adventitial fibroblast
fibroblast LCN2+
fibroblast ITIH5+
fibroblast ACKR2+
fibroblast
fibroadipogenic progenitor

**Melanin-concentrating hormone (MCH)**
- lymphoid

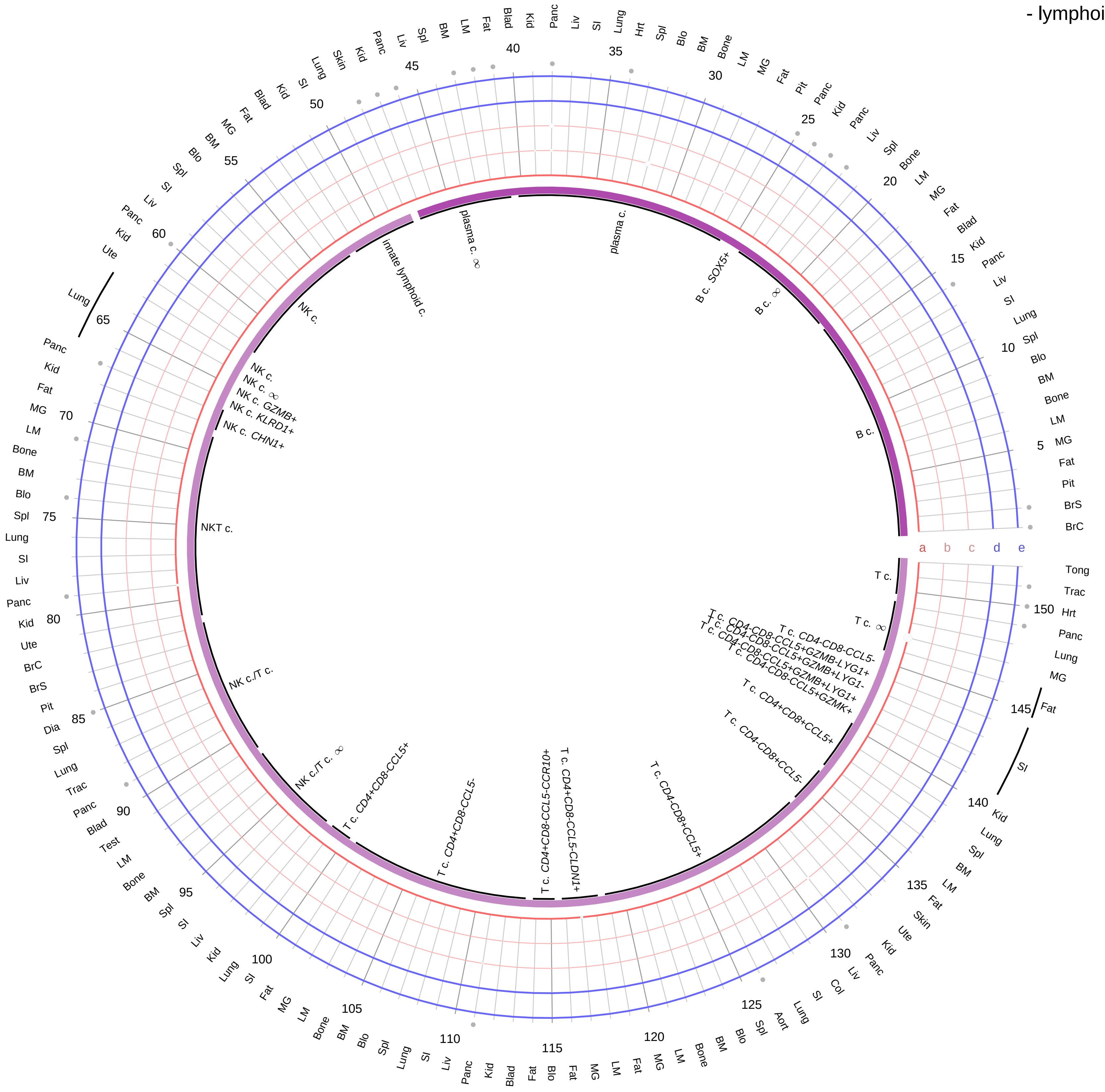

**Ligand/Enzyme**
- a - PMCH
- b - PCSK1
- c - PCSK2

**Receptor**
- d - MCHR1
- e - MCHR2

**PAIRING** (ligand → receptor)

PMCH & PCSK1→MCHR1

PMCH & PCSK1→MCHR2

PMCH & PCSK2→MCHR1

PMCH & PCSK2→MCHR2

**Compartment**
**lymphoid**
- B cell
- NK/T cell

12.5%
25%
50%
100%

ln(CP10K+1): 0, 1, 2, 3

| Abbrev. | Tissue |
|---|---|
| Aort | Aorta |
| Blad | Bladder |
| Blo | Blood |
| Bone | Bone |
| BM | Bone marrow |
| BrC | Brain cortex |
| BrS | Brainstem |
| Col | Colon |
| Dia | Diaphragm |
| Ret | Eye retina |
| Fat | Fat |
| Hrt | Heart |
| Pit | Hypothalamus/Pituitary |
| Kid | Kidney |
| LM | Limb muscle |
| Liv | Liver |
| Lung | Lung |
| MG | Mammary gland |
| Panc | Pancreas |
| Skin | Skin |
| SI | Small intestine |
| Spl | Spleen |
| Test | Testes |
| Thym | Thymus |
| Tong | Tongue |
| Trac | Trachea |
| Ute | Uterus |

**Melanin-concentrating hormone (MCH)**
- myeloid

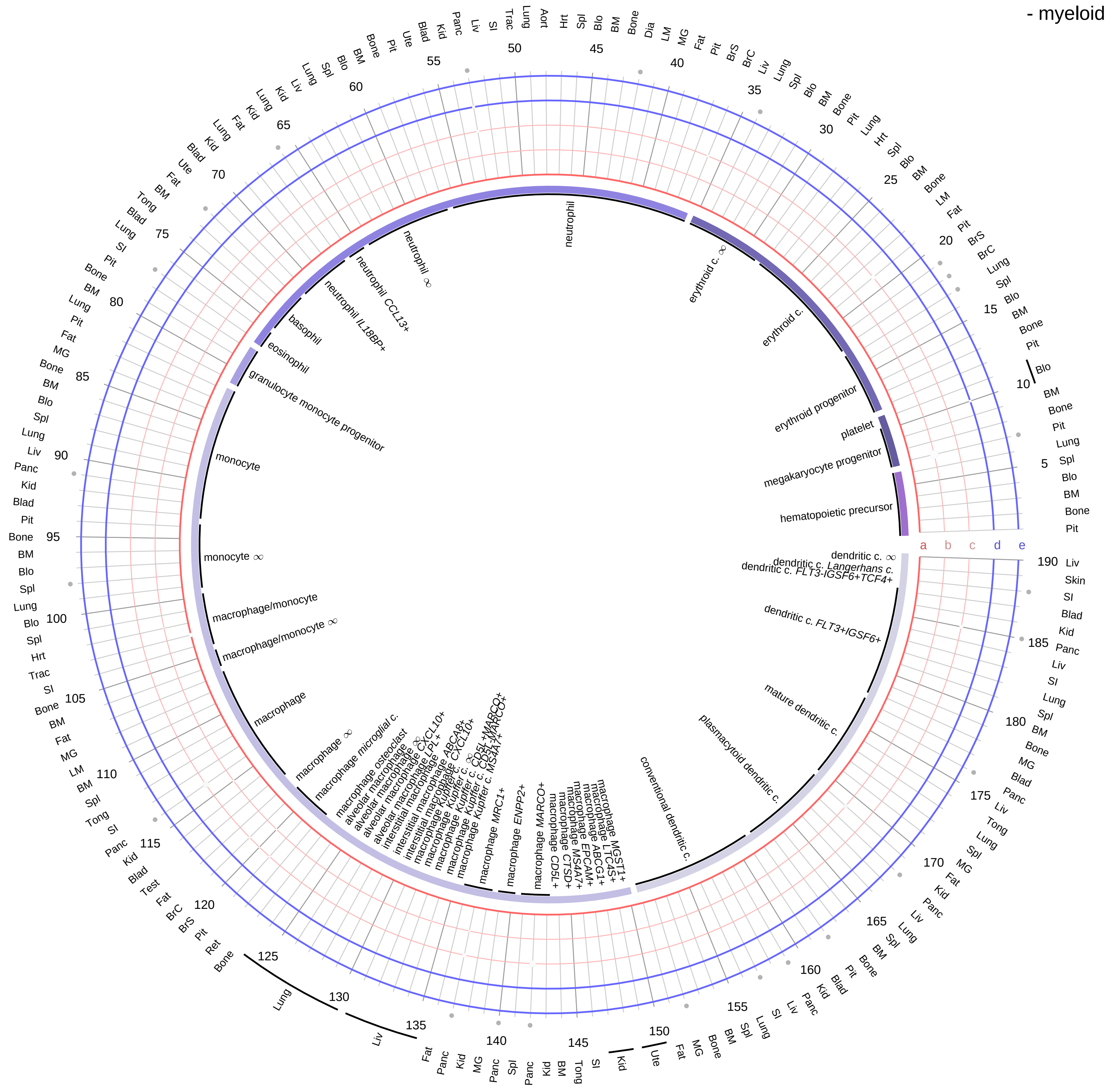

| Ligand/Enzyme |
|---|
| a - PMCH |
| b - PCSK1 |
| c - PCSK2 |
| **Receptor** |
| d - MCHR1 |
| e - MCHR2 |

**PAIRING** (ligand → receptor)

PMCH & PCSK1→MCHR1

PMCH & PCSK1→MCHR2

PMCH & PCSK2→MCHR1

PMCH & PCSK2→MCHR2

**Compartment**
**hematopoietic**
- hematopoietic precursor
**megakaryocyte-erythroid**
- megakaryoid
- erythroid
**myeloid**
- granulocyte
- granulocyte monocyte progenitor
- monocyte/macrophage
- dendritic

12.5%
25%
50%
100%

ln(CP10K+1): 0, 1, 2, 3

| Abbrev. | Tissue |
|---|---|
| Aort | Aorta |
| Blad | Bladder |
| Blo | Blood |
| Bone | Bone |
| BM | Bone marrow |
| BrC | Brain cortex |
| BrS | Brainstem |
| Col | Colon |
| Dia | Diaphragm |
| Ret | Eye retina |
| Fat | Fat |
| Hrt | Heart |
| Pit | Hypothalamus/Pituitary |
| Kid | Kidney |
| LM | Limb muscle |
| Liv | Liver |
| Lung | Lung |
| MG | Mammary gland |
| Panc | Pancreas |
| Skin | Skin |
| SI | Small intestine |
| Spl | Spleen |
| Test | Testes |
| Thym | Thymus |
| Tong | Tongue |
| Trac | Trachea |
| Ute | Uterus |

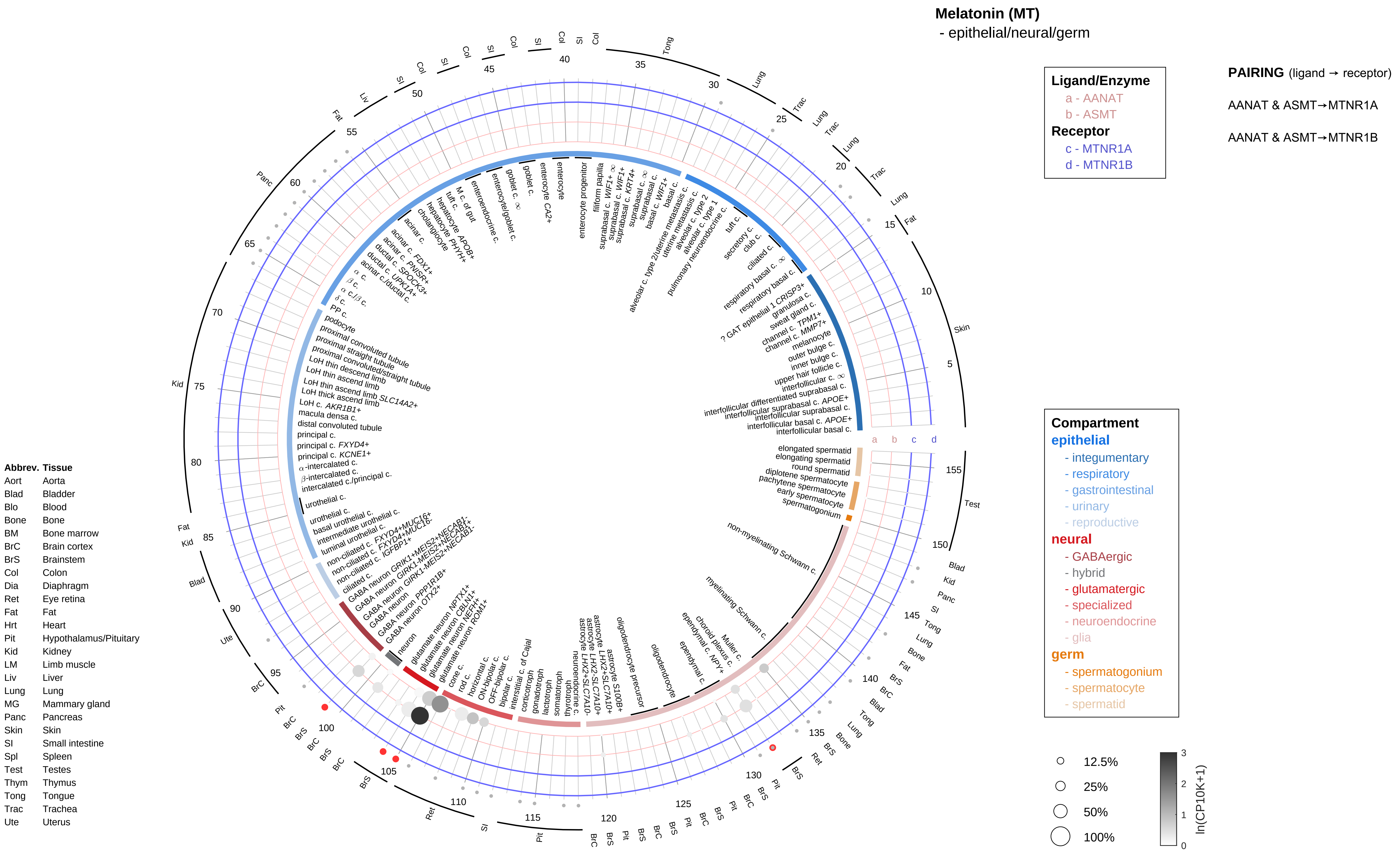

Melatonin (MT)
- epithelial/neural/germ
Ligand/Enzyme
a - AANAT
b - ASMT
Receptor
c - MTNR1A
d - MTNR1B
PAIRING (ligand → receptor)
AANAT & ASMT→MTNR1A
AANAT & ASMT→MTNR1B
Compartment
epithelial
- integumentary
- respiratory
- gastrointestinal
- urinary
- reproductive
neural
- GABAergic
- hybrid
- glutamatergic
- specialized
- neuroendocrine
- glia
germ
- spermatogonium
- spermatocyte
- spermatid
12.5%
25%
50%
100%
ln(CP10K+1)
Abbrev. Tissue
Aort Aorta
Blad Bladder
Blo Blood
Bone Bone
BM Bone marrow
BrC Brain cortex
BrS Brainstem
Col Colon
Dia Diaphragm
Ret Eye retina
Fat Fat
Hrt Heart
Pit Hypothalamus/Pituitary
Kid Kidney
LM Limb muscle
Liv Liver
Lung Lung
MG Mammary gland
Panc Pancreas
Skin Skin
SI Small intestine
Spl Spleen
Test Testes
Thym Thymus
Tong Tongue
Trac Trachea
Ute Uterus

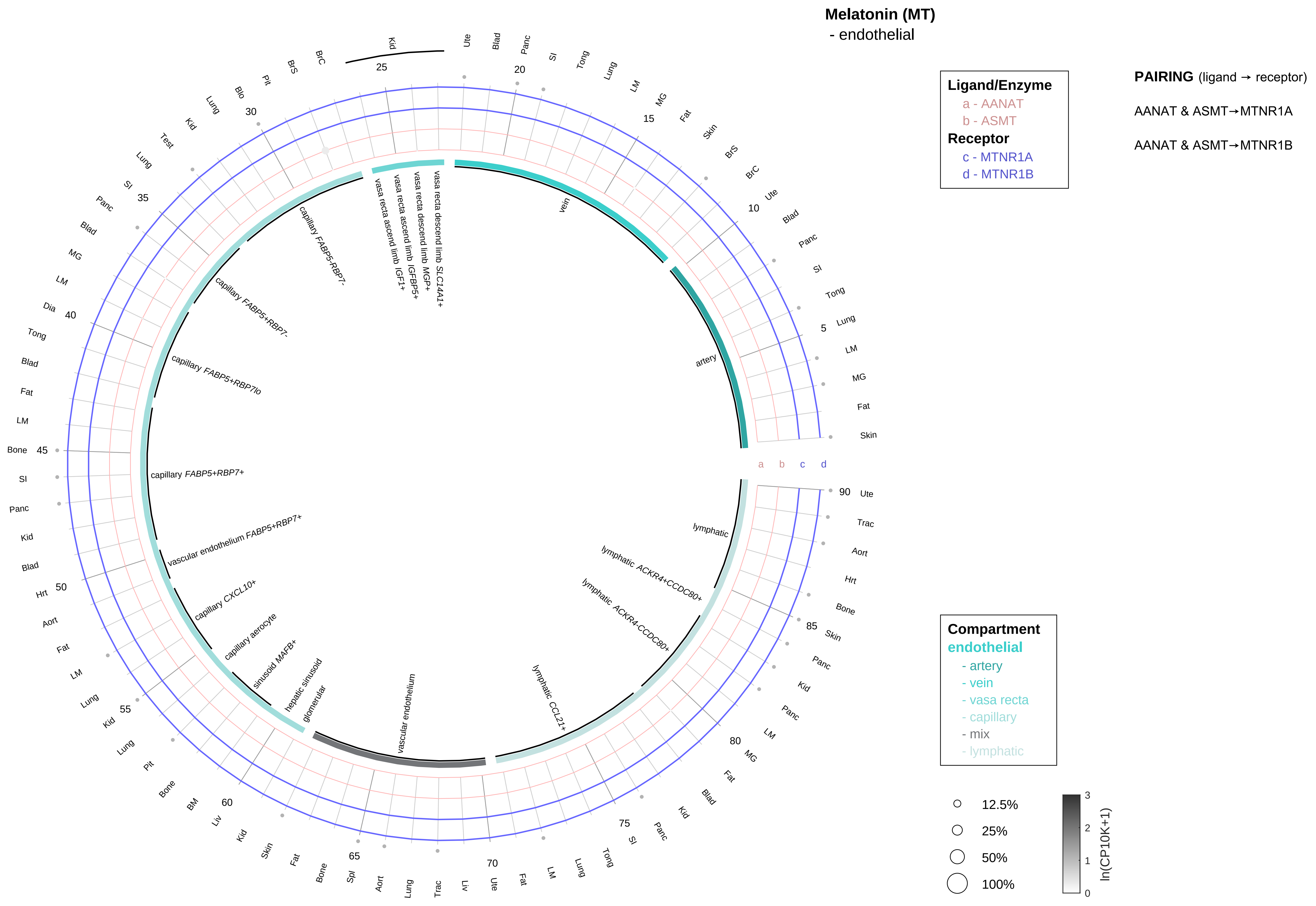

| Abbrev. | Tissue |
|---|---|
| Aort | Aorta |
| Blad | Bladder |
| Blo | Blood |
| Bone | Bone |
| BM | Bone marrow |
| BrC | Brain cortex |
| BrS | Brainstem |
| Col | Colon |
| Dia | Diaphragm |
| Ret | Eye retina |
| Fat | Fat |
| Hrt | Heart |
| Pit | Hypothalamus/Pituitary |
| Kid | Kidney |
| LM | Limb muscle |
| Liv | Liver |
| Lung | Lung |
| MG | Mammary gland |
| Panc | Pancreas |
| Skin | Skin |
| SI | Small intestine |
| Spl | Spleen |
| Test | Testes |
| Thym | Thymus |
| Tong | Tongue |
| Trac | Trachea |
| Ute | Uterus |

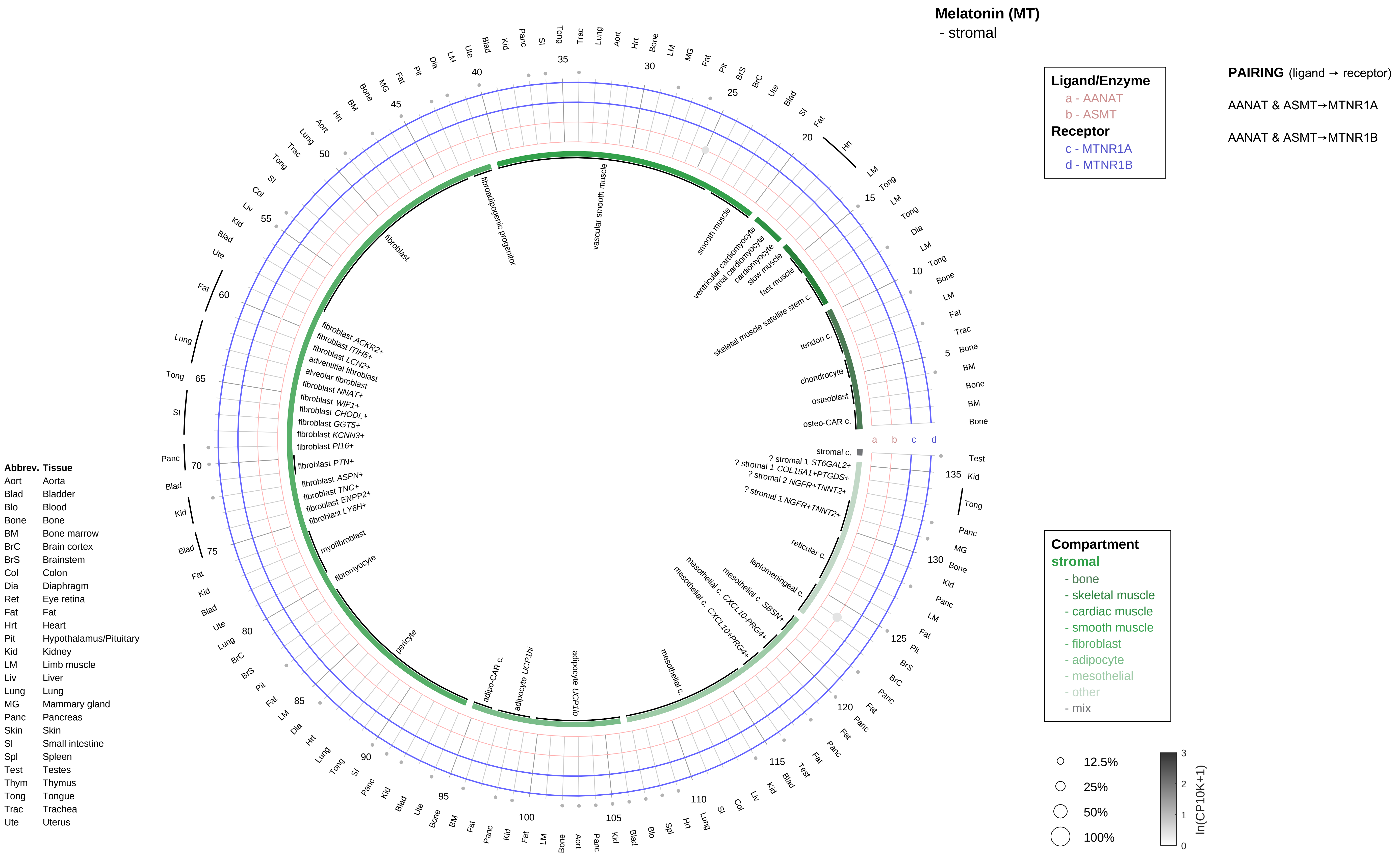

Melatonin (MT)
- stromal
Ligand/Enzyme
a - AANAT
b - ASMT
Receptor
c - MTNR1A
d - MTNR1B
PAIRING (ligand → receptor)
AANAT & ASMT→MTNR1A
AANAT & ASMT→MTNR1B
Compartment
stromal
- bone
- skeletal muscle
- cardiac muscle
- smooth muscle
- fibroblast
- adipocyte
- mesothelial
- other
- mix
12.5%
25%
50%
100%
ln(CP10K+1)
Abbrev. Tissue
Aort Aorta
Blad Bladder
Blo Blood
Bone Bone
BM Bone marrow
BrC Brain cortex
BrS Brainstem
Col Colon
Dia Diaphragm
Ret Eye retina
Fat Fat
Hrt Heart
Pit Hypothalamus/Pituitary
Kid Kidney
LM Limb muscle
Liv Liver
Lung Lung
MG Mammary gland
Panc Pancreas
Skin Skin
SI Small intestine
Spl Spleen
Test Testes
Thym Thymus
Tong Tongue
Trac Trachea
Ute Uterus
vascular smooth muscle
smooth muscle
ventricular cardiomyocyte
atrial cardiomyocyte
cardiomyocyte
slow muscle
fast muscle
skeletal muscle satellite stem c.
tendon c.
chondrocyte
osteoblast
osteo-CAR c.
stromal c.
? stromal 1 ST6GAL2+
? stromal 1 COL15A1+PTGDS+
? stromal 2 NGFR+TNNT2+
? stromal 1 NGFR+TNNT2+
reticular c.
leptomeningeal c.
mesothelial c. SBSN+
mesothelial c. CXCL10-PRG4+
mesothelial c. CXCL10+PRG4+
mesothelial c.
adipocyte UCP1lo
adipocyte UCP1hi
adipo-CAR c.
pericyte
fibromyocyte
myofibroblast
fibroblast LY6H+
fibroblast ENPP2+
fibroblast TNC+
fibroblast ASPN+
fibroblast PTN+
fibroblast PI16+
fibroblast KCNN3+
fibroblast GGT5+
fibroblast CHODL+
fibroblast WIF1+
fibroblast NNAT+
alveolar fibroblast
adventitial fibroblast
fibroblast LCN2+
fibroblast ITIH5+
fibroblast ACKR2+
fibroblast
fibroadipogenic progenitor

**Melatonin (MT)**
- lymphoid

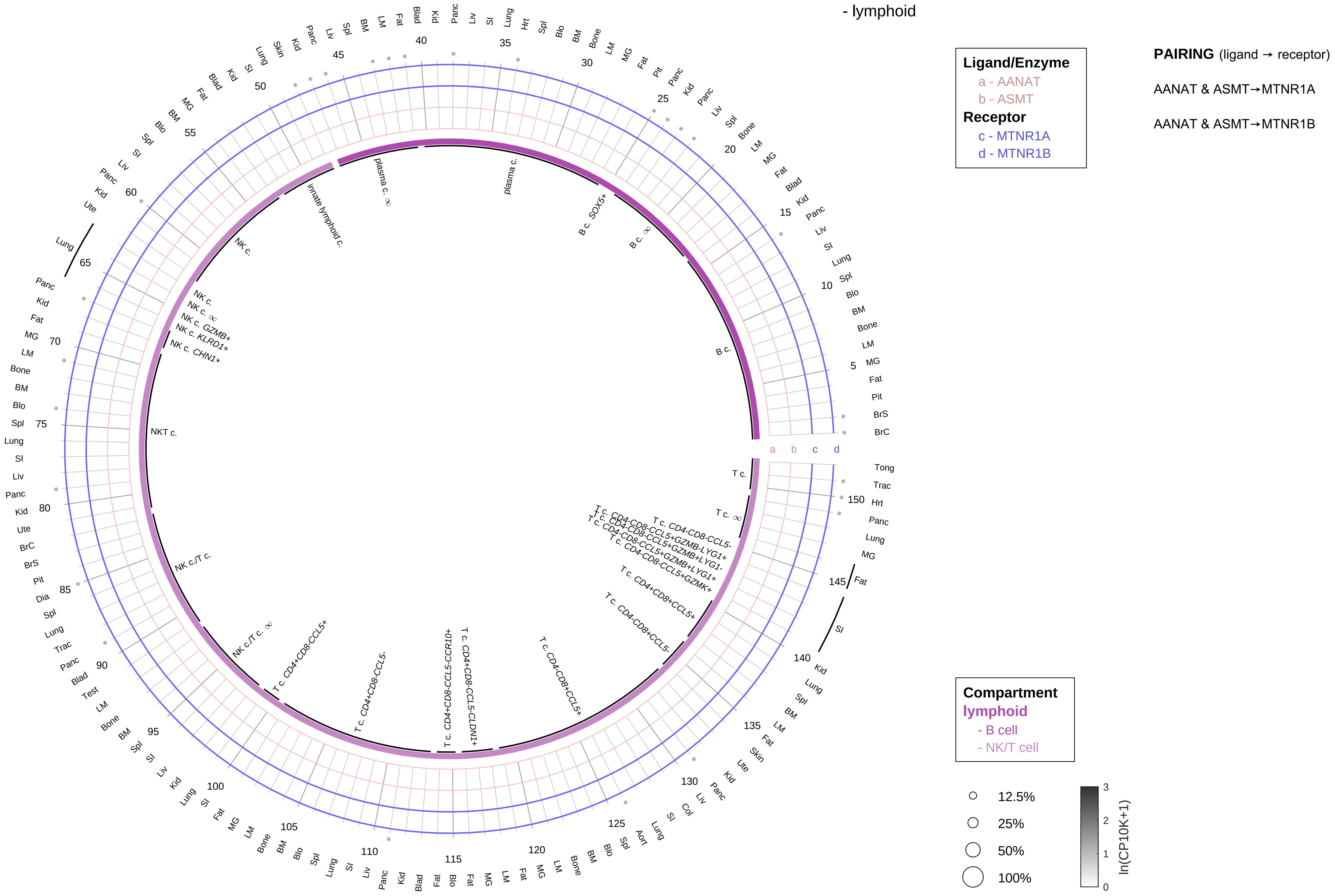

| Abbrev. | Tissue |
|---|---|
| Aort | Aorta |
| Blad | Bladder |
| Blo | Blood |
| Bone | Bone |
| BM | Bone marrow |
| BrC | Brain cortex |
| BrS | Brainstem |
| Col | Colon |
| Dia | Diaphragm |
| Ret | Eye retina |
| Fat | Fat |
| Hrt | Heart |
| Pit | Hypothalamus/Pituitary |
| Kid | Kidney |
| LM | Limb muscle |
| Liv | Liver |
| Lung | Lung |
| MG | Mammary gland |
| Panc | Pancreas |
| Skin | Skin |
| SI | Small intestine |
| Spl | Spleen |
| Test | Testes |
| Thym | Thymus |
| Tong | Tongue |
| Trac | Trachea |
| Ute | Uterus |

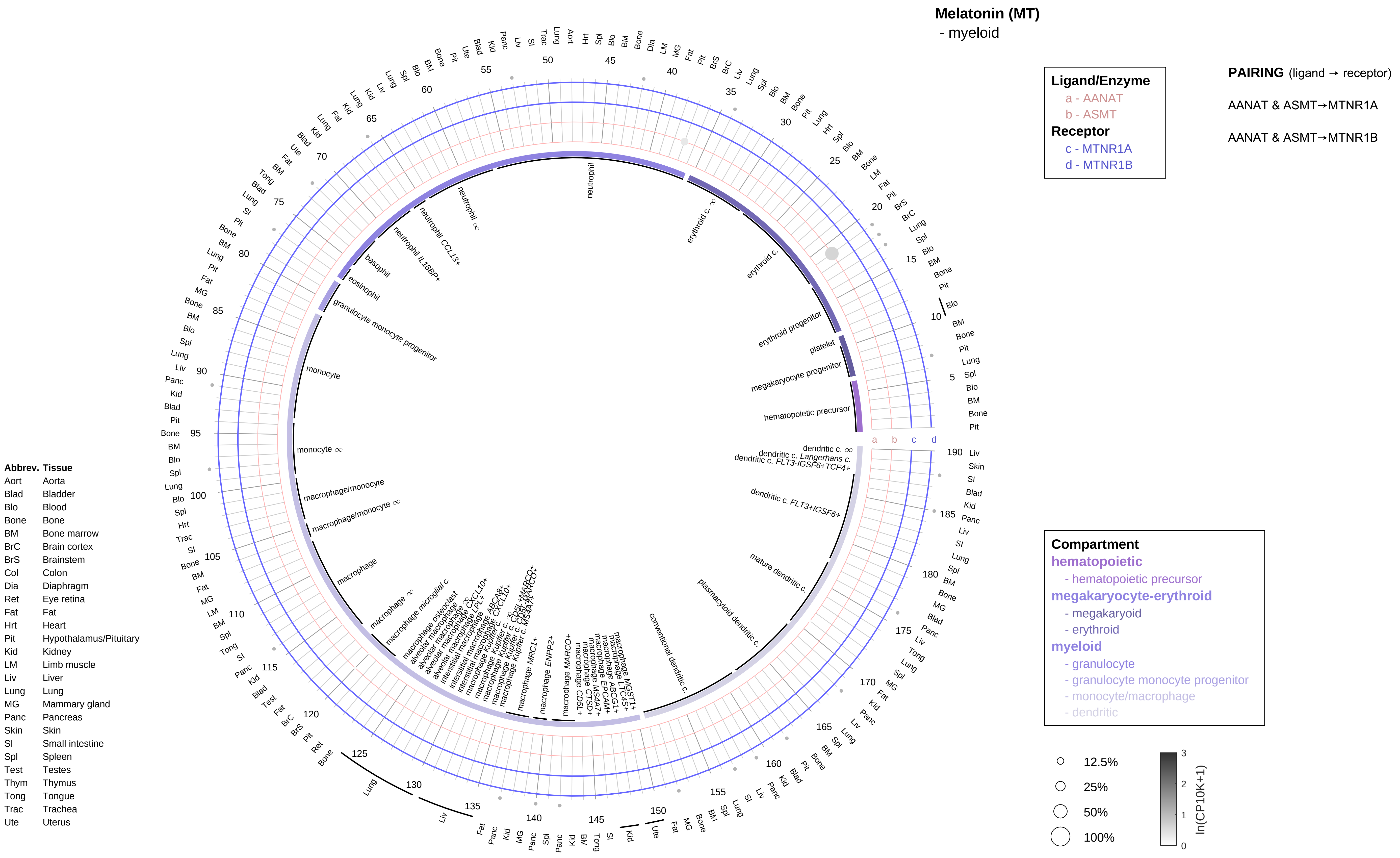
Melatonin (MT)
- myeloid
Ligand/Enzyme
a - AANAT
b - ASMT
Receptor
c - MTNR1A
d - MTNR1B
PAIRING (ligand → receptor)
AANAT & ASMT→MTNR1A
AANAT & ASMT→MTNR1B
Compartment
hematopoietic
- hematopoietic precursor
megakaryocyte-erythroid
- megakaryoid
- erythroid
myeloid
- granulocyte
- granulocyte monocyte progenitor
- monocyte/macrophage
- dendritic
12.5%
25%
50%
100%
ln(CP10K+1)
Abbrev. Tissue
Aort Aorta
Blad Bladder
Blo Blood
Bone Bone
BM Bone marrow
BrC Brain cortex
BrS Brainstem
Col Colon
Dia Diaphragm
Ret Eye retina
Fat Fat
Hrt Heart
Pit Hypothalamus/Pituitary
Kid Kidney
LM Limb muscle
Liv Liver
Lung Lung
MG Mammary gland
Panc Pancreas
Skin Skin
SI Small intestine
Spl Spleen
Test Testes
Thym Thymus
Tong Tongue
Trac Trachea
Ute Uterus

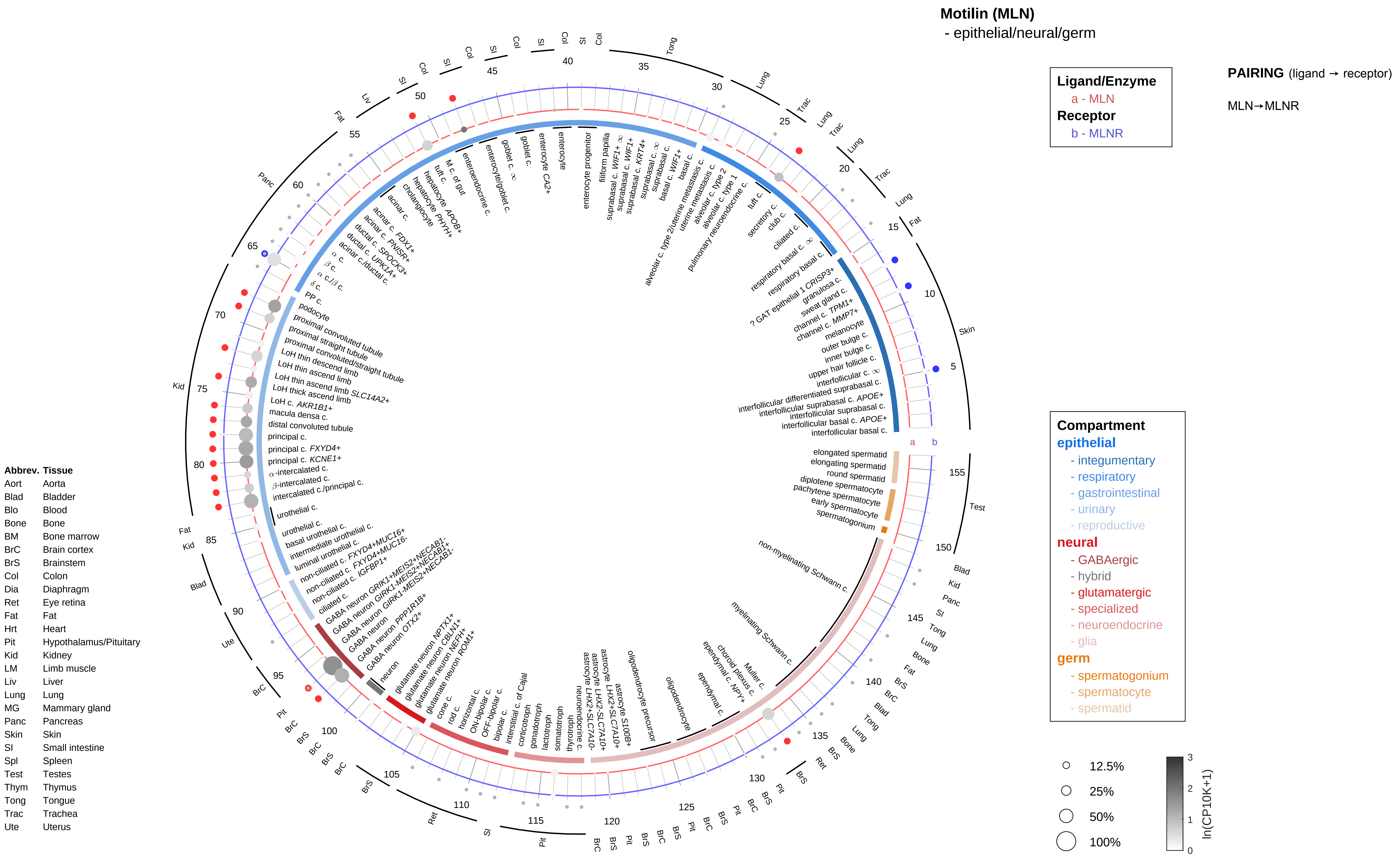
Motilin (MLN)
- epithelial/neural/germ
Ligand/Enzyme
a - MLN
Receptor
b - MLNR
PAIRING (ligand → receptor)
MLN→MLNR
Compartment
epithelial
- integumentary
- respiratory
- gastrointestinal
- urinary
- reproductive
neural
- GABAergic
- hybrid
- glutamatergic
- specialized
- neuroendocrine
- glia
germ
- spermatogonium
- spermatocyte
- spermatid
12.5%
25%
50%
100%
ln(CP10K+1)
Abbrev. Tissue
Aort Aorta
Blad Bladder
Blo Blood
Bone Bone
BM Bone marrow
BrC Brain cortex
BrS Brainstem
Col Colon
Dia Diaphragm
Ret Eye retina
Fat Fat
Hrt Heart
Pit Hypothalamus/Pituitary
Kid Kidney
LM Limb muscle
Liv Liver
Lung Lung
MG Mammary gland
Panc Pancreas
Skin Skin
SI Small intestine
Spl Spleen
Test Testes
Thym Thymus
Tong Tongue
Trac Trachea
Ute Uterus

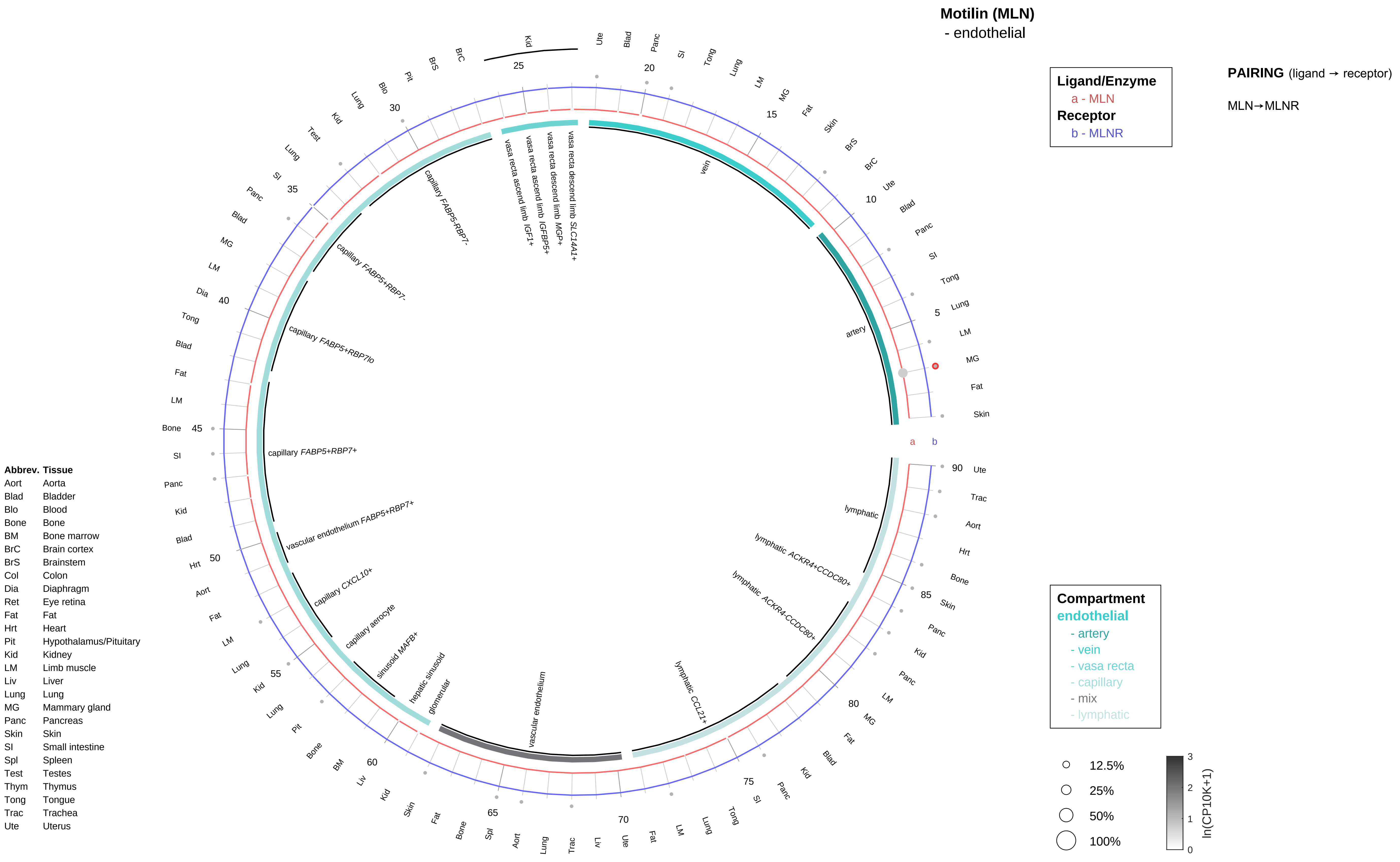

Motilin (MLN)
- endothelial
Ligand/Enzyme
a - MLN
Receptor
b - MLNR
PAIRING (ligand → receptor)
MLN→MLNR
Compartment
endothelial
- artery
- vein
- vasa recta
- capillary
- mix
- lymphatic
12.5%
25%
50%
100%
ln(CP10K+1)
Abbrev. Tissue
Aort Aorta
Blad Bladder
Blo Blood
Bone Bone
BM Bone marrow
BrC Brain cortex
BrS Brainstem
Col Colon
Dia Diaphragm
Ret Eye retina
Fat Fat
Hrt Heart
Pit Hypothalamus/Pituitary
Kid Kidney
LM Limb muscle
Liv Liver
Lung Lung
MG Mammary gland
Panc Pancreas
Skin Skin
SI Small intestine
Spl Spleen
Test Testes
Thym Thymus
Tong Tongue
Trac Trachea
Ute Uterus
artery
vein
vasa recta descend limb SLC14A1+
vasa recta descend limb MGP+
vasa recta ascend limb IGFBP5+
vasa recta ascend limb IGF1+
capillary FABP5-RBP7-
capillary FABP5+RBP7-
capillary FABP5+RBP7lo
capillary FABP5+RBP7+
vascular endothelium FABP5+RBP7+
capillary CXCL10+
capillary aerocyte
sinusoid MAFB+
hepatic sinusoid
glomerular
vascular endothelium
lymphatic CCL21+
lymphatic ACKR4-CCDC80+
lymphatic ACKR4+CCDC80+
lymphatic

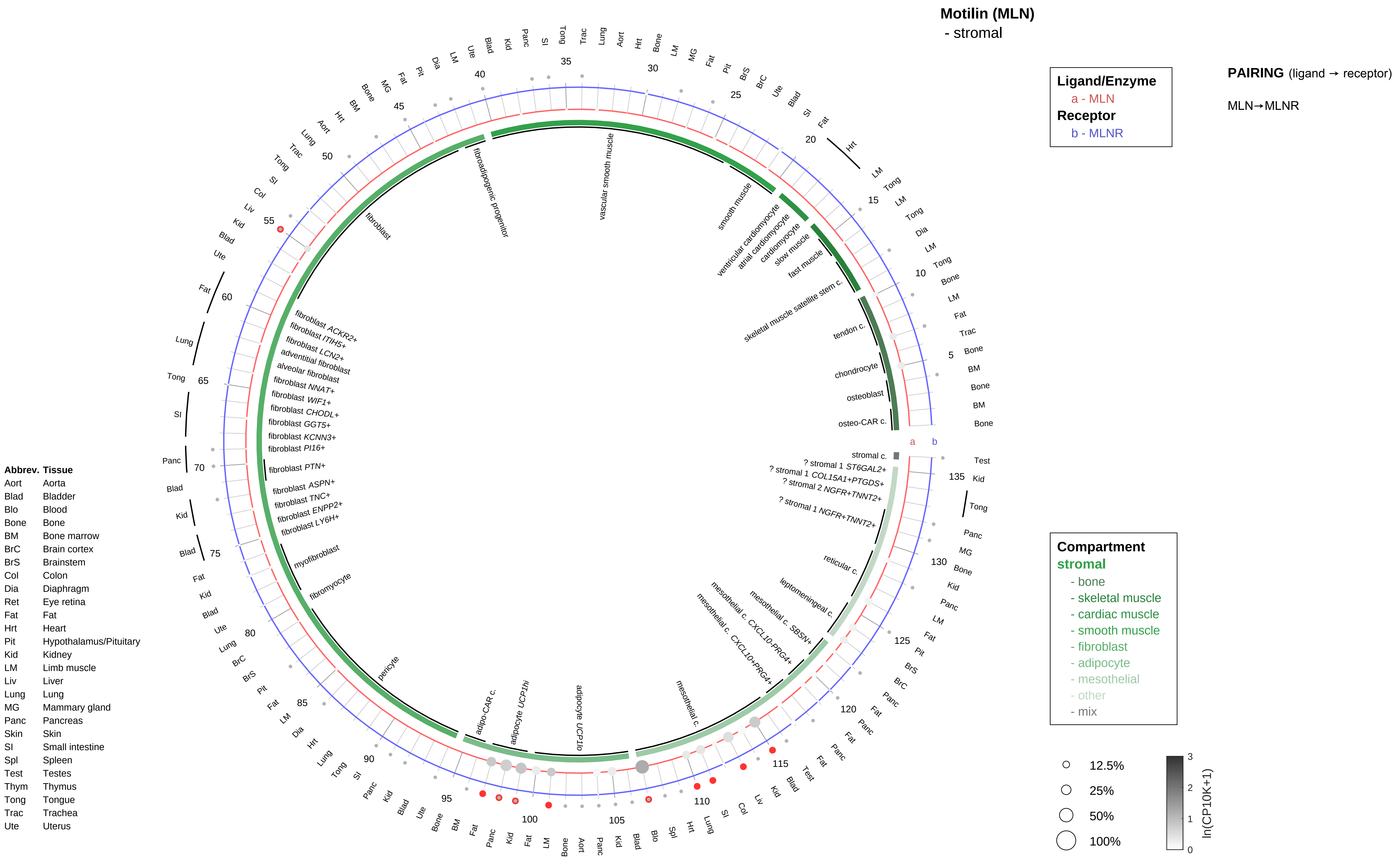
Motilin (MLN)
- stromal
Ligand/Enzyme
a - MLN
Receptor
b - MLNR
PAIRING (ligand → receptor)
MLN→MLNR
Compartment
stromal
- bone
- skeletal muscle
- cardiac muscle
- smooth muscle
- fibroblast
- adipocyte
- mesothelial
- other
- mix
12.5%
25%
50%
100%
ln(CP10K+1)
Abbrev. Tissue
Aort Aorta
Blad Bladder
Blo Blood
Bone Bone
BM Bone marrow
BrC Brain cortex
BrS Brainstem
Col Colon
Dia Diaphragm
Ret Eye retina
Fat Fat
Hrt Heart
Pit Hypothalamus/Pituitary
Kid Kidney
LM Limb muscle
Liv Liver
Lung Lung
MG Mammary gland
Panc Pancreas
Skin Skin
SI Small intestine
Spl Spleen
Test Testes
Thym Thymus
Tong Tongue
Trac Trachea
Ute Uterus
osteo-CAR c.
osteoblast
chondrocyte
tendon c.
skeletal muscle satellite stem c.
fast muscle
slow muscle
cardiomyocyte
atrial cardiomyocyte
ventricular cardiomyocyte
smooth muscle
vascular smooth muscle
fibroadipogenic progenitor
fibroblast
fibroblast ACKR2+
fibroblast ITIH5+
fibroblast LCN2+
adventitial fibroblast
alveolar fibroblast
fibroblast NNAT+
fibroblast WIF1+
fibroblast CHODL+
fibroblast GGT5+
fibroblast KCNN3+
fibroblast PI16+
fibroblast PTN+
fibroblast ASPN+
fibroblast TNC+
fibroblast ENPP2+
fibroblast LY6H+
myofibroblast
fibromyocyte
pericyte
adipo-CAR c.
adipocyte UCP1hi
adipocyte UCP1lo
mesothelial c.
mesothelial c. CXCL10+PRG4+
mesothelial c. CXCL10-PRG4+
mesothelial c. SBSN+
leptomeningeal c.
reticular c.
? stromal 1 NGFR+TNNT2+
? stromal 2 NGFR+TNNT2+
? stromal 1 COL15A1+PTGDS+
? stromal 1 ST6GAL2+
stromal c.

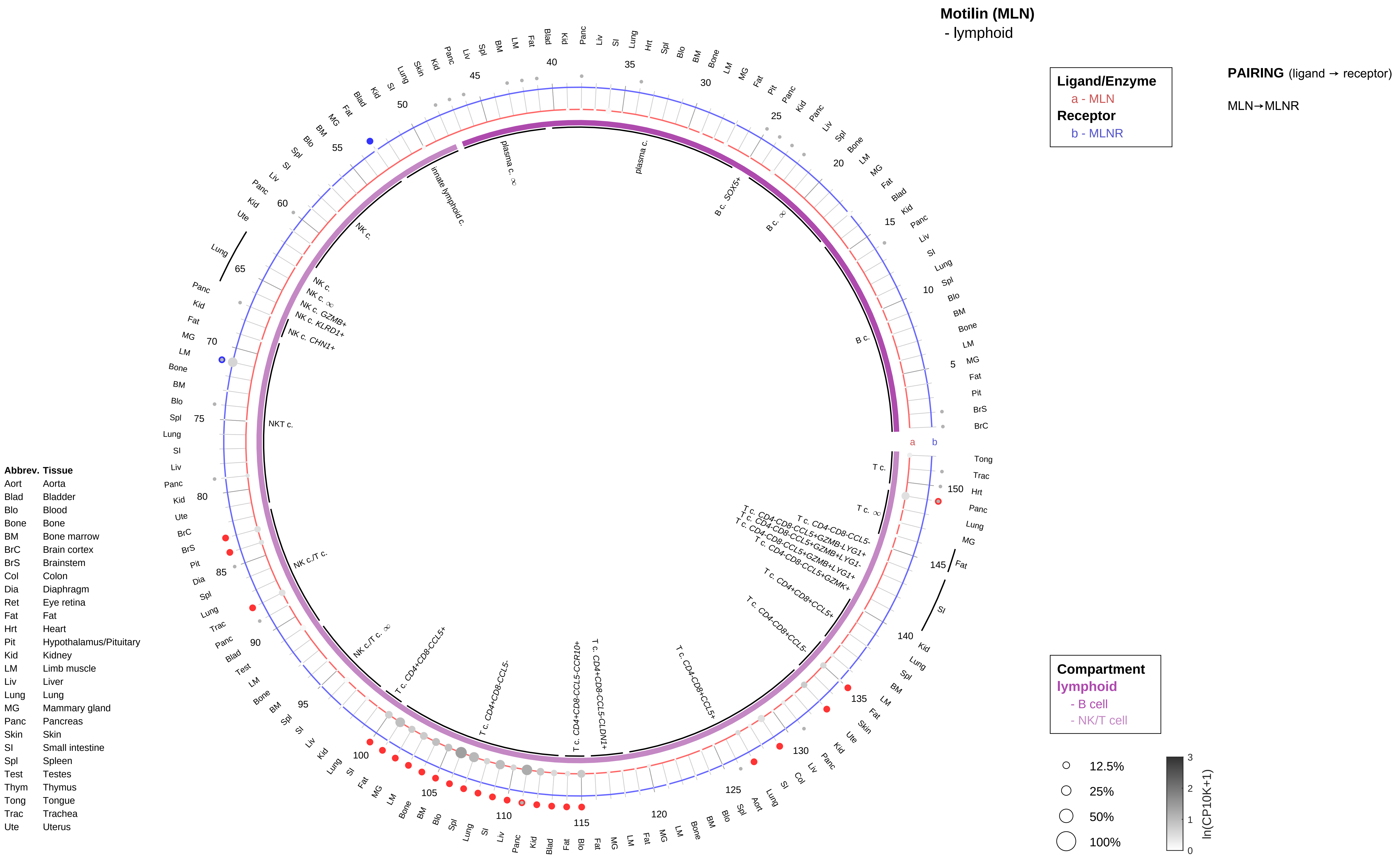

Motilin (MLN)
- lymphoid
Ligand/Enzyme
a - MLN
Receptor
b - MLNR
PAIRING (ligand → receptor)
MLN→MLNR
Compartment
lymphoid
- B cell
- NK/T cell
12.5%
25%
50%
100%
ln(CP10K+1)
Abbrev. Tissue
Aort Aorta
Blad Bladder
Blo Blood
Bone Bone
BM Bone marrow
BrC Brain cortex
BrS Brainstem
Col Colon
Dia Diaphragm
Ret Eye retina
Fat Fat
Hrt Heart
Pit Hypothalamus/Pituitary
Kid Kidney
LM Limb muscle
Liv Liver
Lung Lung
MG Mammary gland
Panc Pancreas
Skin Skin
SI Small intestine
Spl Spleen
Test Testes
Thym Thymus
Tong Tongue
Trac Trachea
Ute Uterus
B c.
plasma c.
NK c.
NKT c.
NK c./T c.
T c.

**Motilin (MLN)**
- myeloid

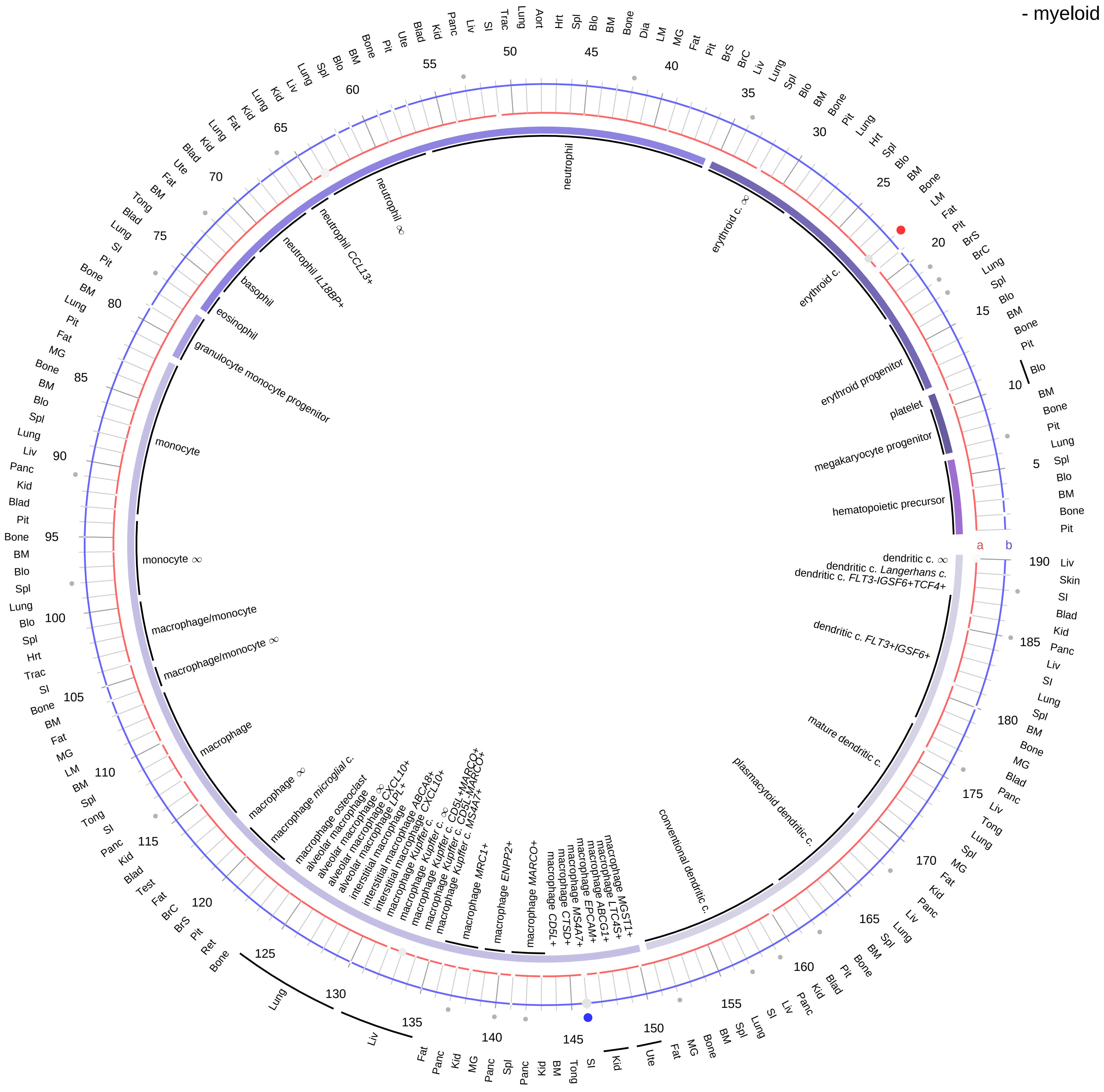

| Ligand/Enzyme |
|---|
| a - MLN |
| **Receptor** |
| b - MLNR |

**PAIRING** (ligand → receptor)

MLN→MLNR

| Abbrev. | Tissue |
|---|---|
| Aort | Aorta |
| Blad | Bladder |
| Blo | Blood |
| Bone | Bone |
| BM | Bone marrow |
| BrC | Brain cortex |
| BrS | Brainstem |
| Col | Colon |
| Dia | Diaphragm |
| Ret | Eye retina |
| Fat | Fat |
| Hrt | Heart |
| Pit | Hypothalamus/Pituitary |
| Kid | Kidney |
| LM | Limb muscle |
| Liv | Liver |
| Lung | Lung |
| MG | Mammary gland |
| Panc | Pancreas |
| Skin | Skin |
| SI | Small intestine |
| Spl | Spleen |
| Test | Testes |
| Thym | Thymus |
| Tong | Tongue |
| Trac | Trachea |
| Ute | Uterus |

**Compartment**
**hematopoietic**
- hematopoietic precursor

**megakaryocyte-erythroid**
- megakaryoid
- erythroid

**myeloid**
- granulocyte
- granulocyte monocyte progenitor
- monocyte/macrophage
- dendritic

○ 12.5%
○ 25%
○ 50%
○ 100%

ln(CP10K+1): 0, 1, 2, 3

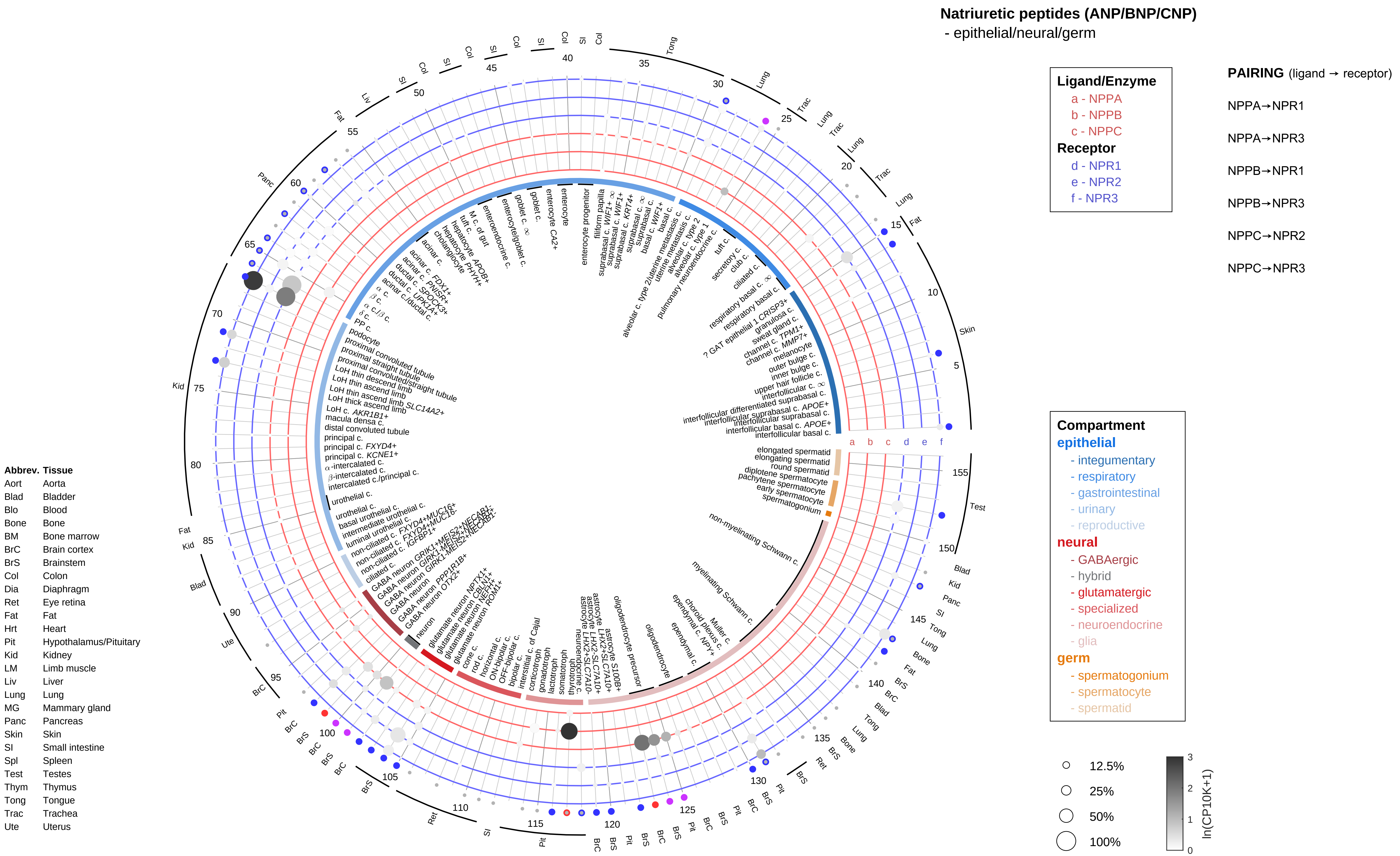
Natriuretic peptides (ANP/BNP/CNP)
- epithelial/neural/germ
Ligand/Enzyme
a - NPPA
b - NPPB
c - NPPC
Receptor
d - NPR1
e - NPR2
f - NPR3
PAIRING (ligand → receptor)
NPPA→NPR1
NPPA→NPR3
NPPB→NPR1
NPPB→NPR3
NPPC→NPR2
NPPC→NPR3
Compartment
epithelial
- integumentary
- respiratory
- gastrointestinal
- urinary
- reproductive
neural
- GABAergic
- hybrid
- glutamatergic
- specialized
- neuroendocrine
- glia
germ
- spermatogonium
- spermatocyte
- spermatid
12.5%
25%
50%
100%
ln(CP10K+1)
Abbrev. Tissue
Aort Aorta
Blad Bladder
Blo Blood
Bone Bone
BM Bone marrow
BrC Brain cortex
BrS Brainstem
Col Colon
Dia Diaphragm
Ret Eye retina
Fat Fat
Hrt Heart
Pit Hypothalamus/Pituitary
Kid Kidney
LM Limb muscle
Liv Liver
Lung Lung
MG Mammary gland
Panc Pancreas
Skin Skin
SI Small intestine
Spl Spleen
Test Testes
Thym Thymus
Tong Tongue
Trac Trachea
Ute Uterus

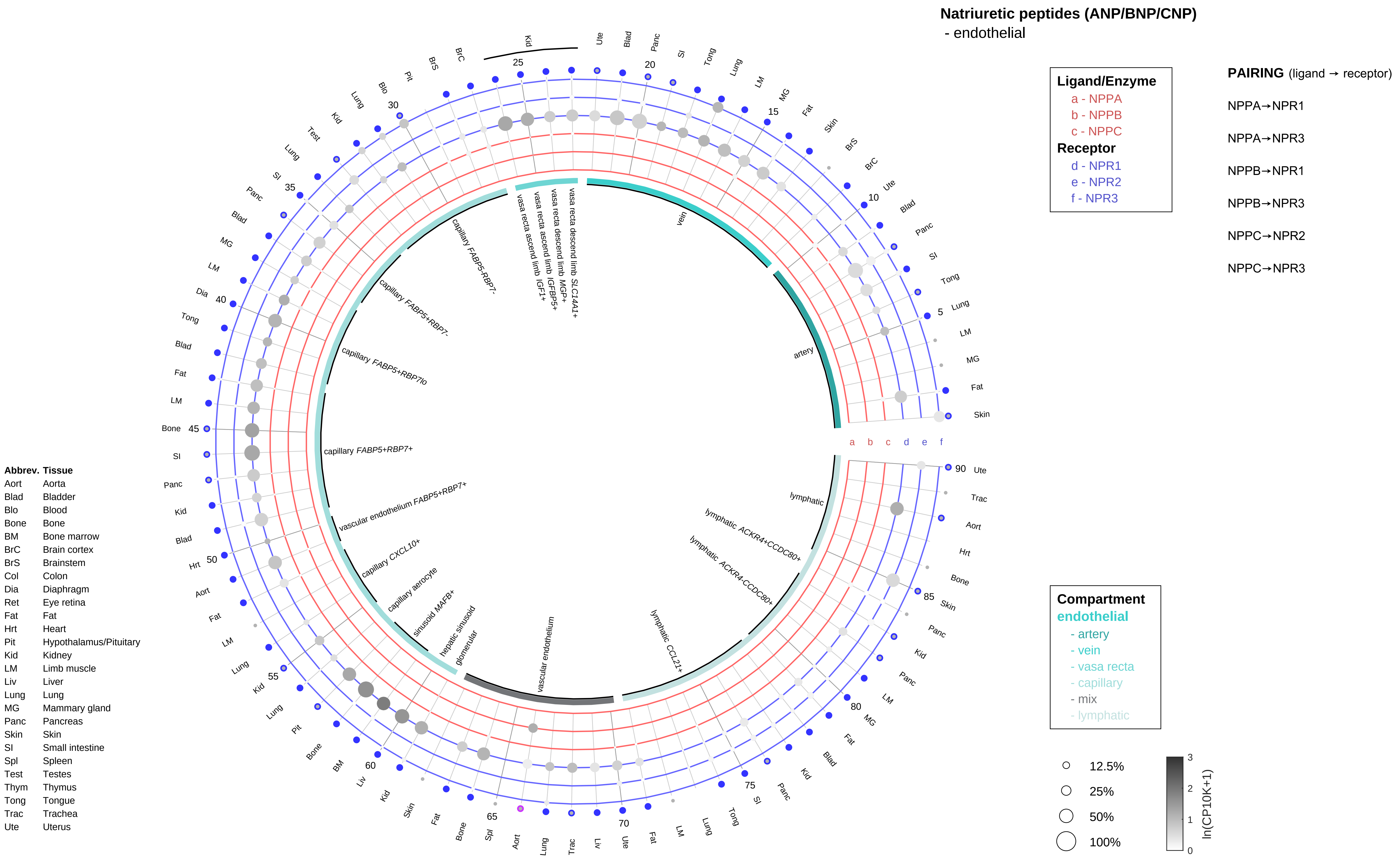

Natriuretic peptides (ANP/BNP/CNP)
- endothelial
Ligand/Enzyme
a - NPPA
b - NPPB
c - NPPC
Receptor
d - NPR1
e - NPR2
f - NPR3
PAIRING (ligand → receptor)
NPPA→NPR1
NPPA→NPR3
NPPB→NPR1
NPPB→NPR3
NPPC→NPR2
NPPC→NPR3
Compartment
endothelial
- artery
- vein
- vasa recta
- capillary
- mix
- lymphatic
12.5%
25%
50%
100%
ln(CP10K+1)
vein
artery
lymphatic
lymphatic ACKR4+CCDC80+
lymphatic ACKR4-CCDC80+
lymphatic CCL21+
vascular endothelium
glomerular
hepatic sinusoid
sinusoid MAFB+
capillary aerocyte
capillary CXCL10+
vascular endothelium FABP5+RBP7+
capillary FABP5+RBP7+
capillary FABP5+RBP7lo
capillary FABP5+RBP7-
capillary FABP5-RBP7-
vasa recta ascend limb IGF1+
vasa recta ascend limb IGFBP5+
vasa recta descend limb MGP+
vasa recta descend limb SLC14A1+
Abbrev. Tissue
Aort Aorta
Blad Bladder
Blo Blood
Bone Bone
BM Bone marrow
BrC Brain cortex
BrS Brainstem
Col Colon
Dia Diaphragm
Ret Eye retina
Fat Fat
Hrt Heart
Pit Hypothalamus/Pituitary
Kid Kidney
LM Limb muscle
Liv Liver
Lung Lung
MG Mammary gland
Panc Pancreas
Skin Skin
SI Small intestine
Spl Spleen
Test Testes
Thym Thymus
Tong Tongue
Trac Trachea
Ute Uterus

**Natriuretic peptides (ANP/BNP/CNP)**
- stromal

**Ligand/Enzyme**
- a - NPPA
- b - NPPB
- c - NPPC

**Receptor**
- d - NPR1
- e - NPR2
- f - NPR3

**PAIRING** (ligand → receptor)

NPPA→NPR1

NPPA→NPR3

NPPB→NPR1

NPPB→NPR3

NPPC→NPR2

NPPC→NPR3

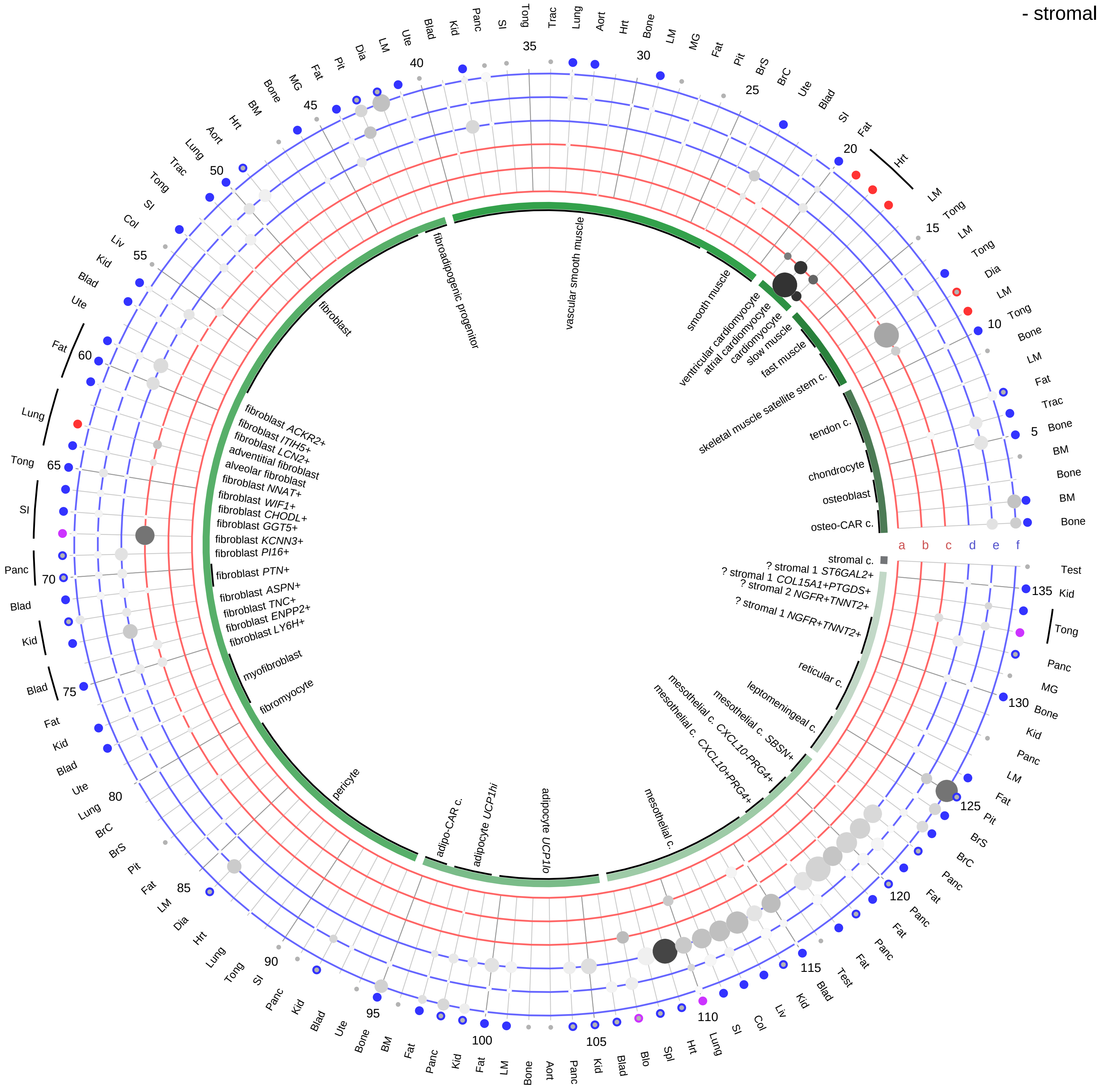

**Compartment**
**stromal**
- bone
- skeletal muscle
- cardiac muscle
- smooth muscle
- fibroblast
- adipocyte
- mesothelial
- other
- mix

12.5%
25%
50%
100%

ln(CP10K+1) 0 1 2 3

| Abbrev. | Tissue |
|---|---|
| Aort | Aorta |
| Blad | Bladder |
| Blo | Blood |
| Bone | Bone |
| BM | Bone marrow |
| BrC | Brain cortex |
| BrS | Brainstem |
| Col | Colon |
| Dia | Diaphragm |
| Ret | Eye retina |
| Fat | Fat |
| Hrt | Heart |
| Pit | Hypothalamus/Pituitary |
| Kid | Kidney |
| LM | Limb muscle |
| Liv | Liver |
| Lung | Lung |
| MG | Mammary gland |
| Panc | Pancreas |
| Skin | Skin |
| SI | Small intestine |
| Spl | Spleen |
| Test | Testes |
| Thym | Thymus |
| Tong | Tongue |
| Trac | Trachea |
| Ute | Uterus |

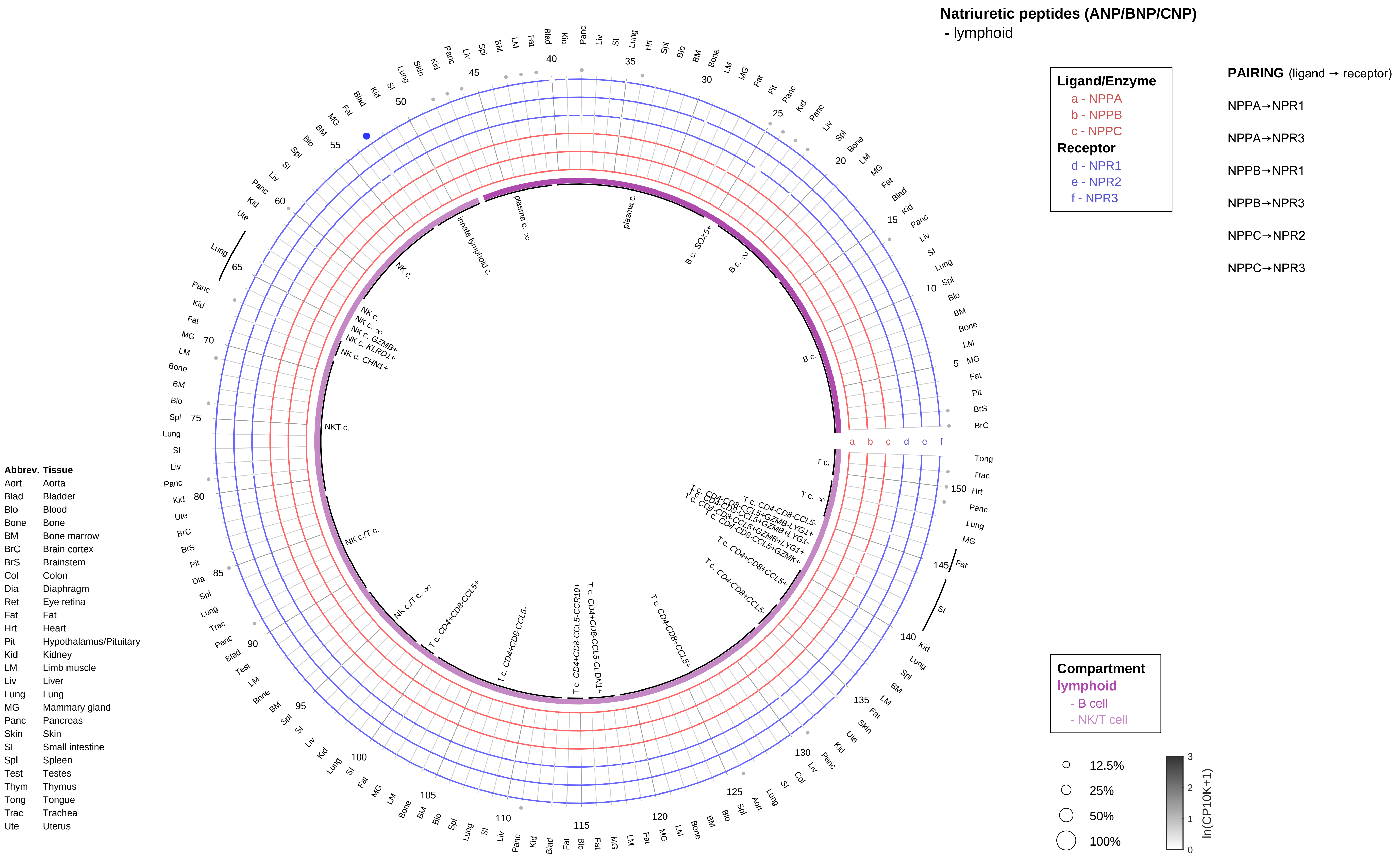
Natriuretic peptides (ANP/BNP/CNP)
- lymphoid
Ligand/Enzyme
a - NPPA
b - NPPB
c - NPPC
Receptor
d - NPR1
e - NPR2
f - NPR3
PAIRING (ligand → receptor)
NPPA→NPR1
NPPA→NPR3
NPPB→NPR1
NPPB→NPR3
NPPC→NPR2
NPPC→NPR3
Compartment
lymphoid
- B cell
- NK/T cell
12.5%
25%
50%
100%
ln(CP10K+1)
Abbrev. Tissue
Aort Aorta
Blad Bladder
Blo Blood
Bone Bone
BM Bone marrow
BrC Brain cortex
BrS Brainstem
Col Colon
Dia Diaphragm
Ret Eye retina
Fat Fat
Hrt Heart
Pit Hypothalamus/Pituitary
Kid Kidney
LM Limb muscle
Liv Liver
Lung Lung
MG Mammary gland
Panc Pancreas
Skin Skin
SI Small intestine
Spl Spleen
Test Testes
Thym Thymus
Tong Tongue
Trac Trachea
Ute Uterus

**Natriuretic peptides (ANP/BNP/CNP)**
- myeloid

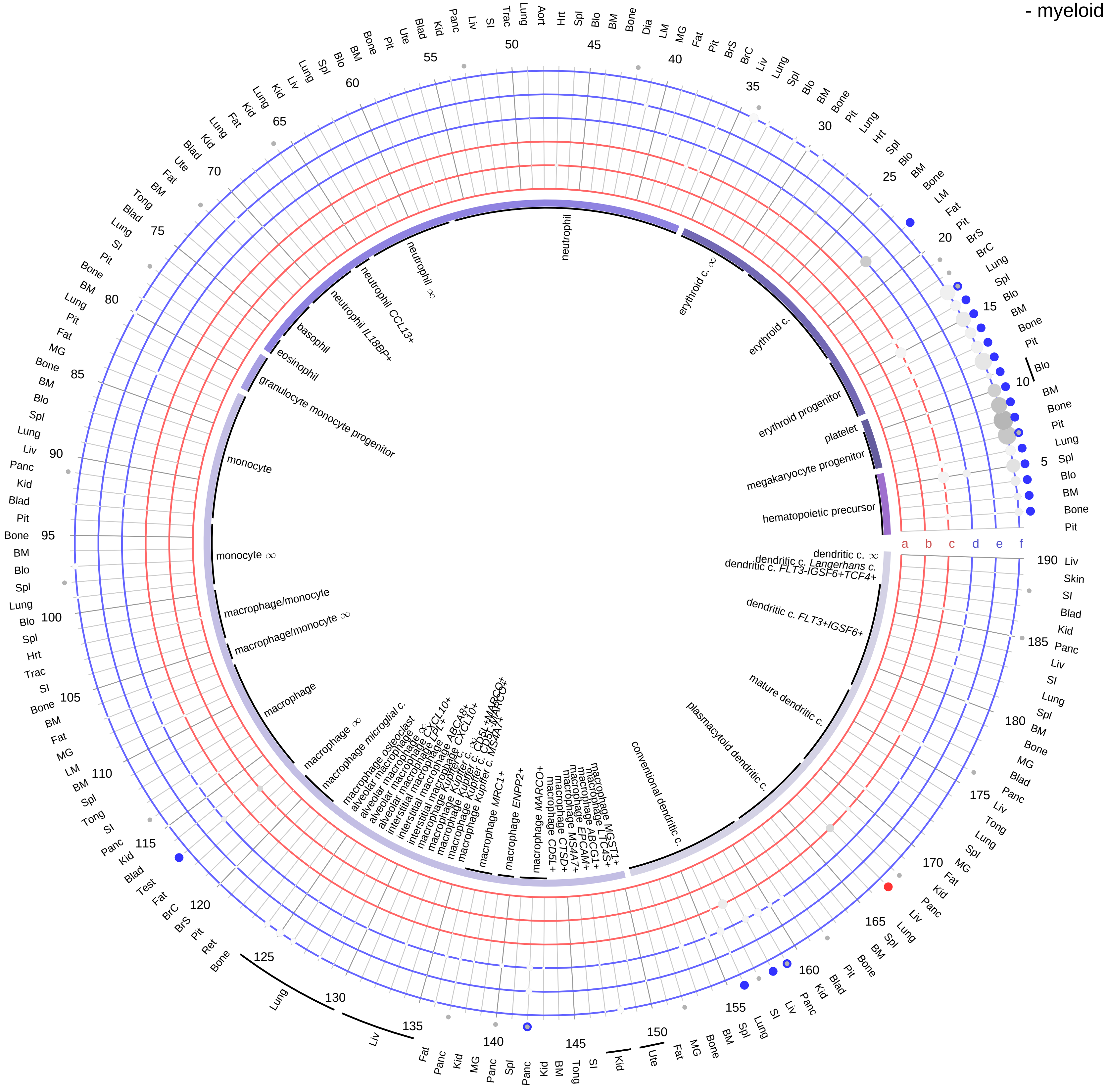

**Ligand/Enzyme**
- a - NPPA
- b - NPPB
- c - NPPC

**Receptor**
- d - NPR1
- e - NPR2
- f - NPR3

**PAIRING** (ligand → receptor)

NPPA→NPR1

NPPA→NPR3

NPPB→NPR1

NPPB→NPR3

NPPC→NPR2

NPPC→NPR3

| Abbrev. | Tissue |
|---|---|
| Aort | Aorta |
| Blad | Bladder |
| Blo | Blood |
| Bone | Bone |
| BM | Bone marrow |
| BrC | Brain cortex |
| BrS | Brainstem |
| Col | Colon |
| Dia | Diaphragm |
| Ret | Eye retina |
| Fat | Fat |
| Hrt | Heart |
| Pit | Hypothalamus/Pituitary |
| Kid | Kidney |
| LM | Limb muscle |
| Liv | Liver |
| Lung | Lung |
| MG | Mammary gland |
| Panc | Pancreas |
| Skin | Skin |
| SI | Small intestine |
| Spl | Spleen |
| Test | Testes |
| Thym | Thymus |
| Tong | Tongue |
| Trac | Trachea |
| Ute | Uterus |

**Compartment**

**hematopoietic**
- hematopoietic precursor

**megakaryocyte-erythroid**
- megakaryoid
- erythroid

**myeloid**
- granulocyte
- granulocyte monocyte progenitor
- monocyte/macrophage
- dendritic

12.5%
25%
50%
100%

ln(CP10K+1)
0 1 2 3

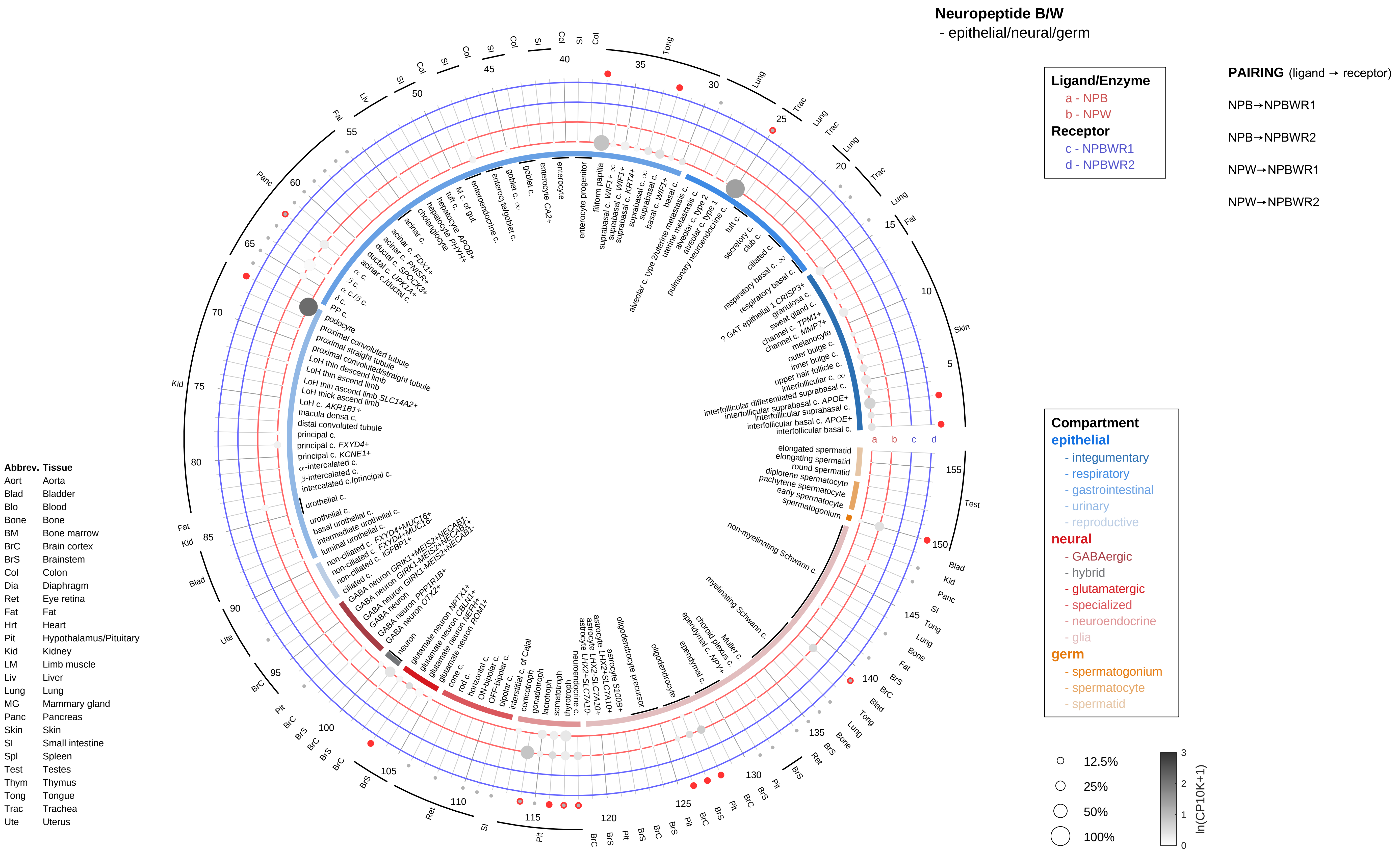
Neuropeptide B/W
- epithelial/neural/germ
Ligand/Enzyme
a - NPB
b - NPW
Receptor
c - NPBWR1
d - NPBWR2
PAIRING (ligand → receptor)
NPB→NPBWR1
NPB→NPBWR2
NPW→NPBWR1
NPW→NPBWR2
Compartment
epithelial
- integumentary
- respiratory
- gastrointestinal
- urinary
- reproductive
neural
- GABAergic
- hybrid
- glutamatergic
- specialized
- neuroendocrine
- glia
germ
- spermatogonium
- spermatocyte
- spermatid
12.5%
25%
50%
100%
ln(CP10K+1)
Abbrev. Tissue
Aort Aorta
Blad Bladder
Blo Blood
Bone Bone
BM Bone marrow
BrC Brain cortex
BrS Brainstem
Col Colon
Dia Diaphragm
Ret Eye retina
Fat Fat
Hrt Heart
Pit Hypothalamus/Pituitary
Kid Kidney
LM Limb muscle
Liv Liver
Lung Lung
MG Mammary gland
Panc Pancreas
Skin Skin
SI Small intestine
Spl Spleen
Test Testes
Thym Thymus
Tong Tongue
Trac Trachea
Ute Uterus

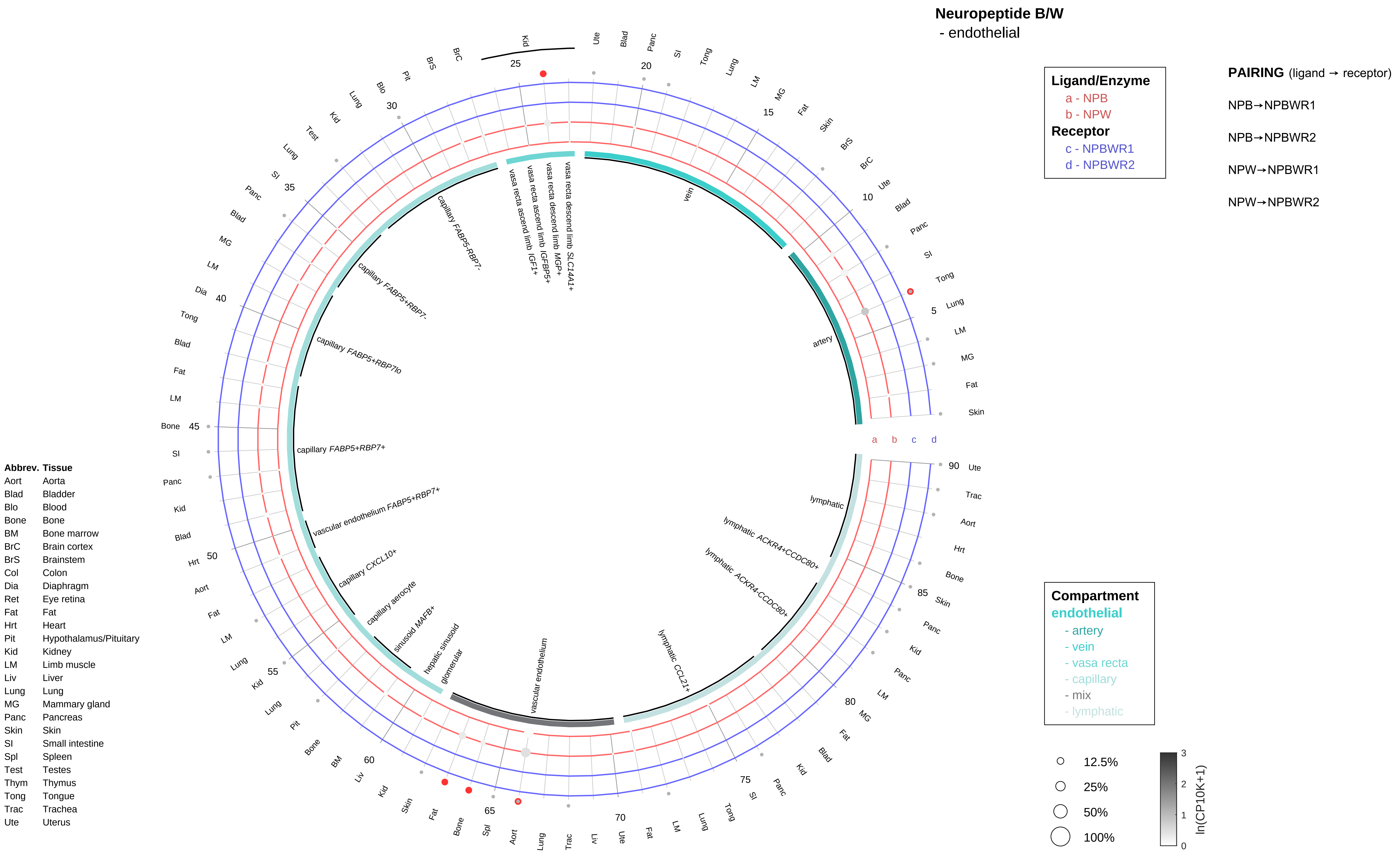

Neuropeptide B/W
- endothelial
Ligand/Enzyme
a - NPB
b - NPW
Receptor
c - NPBWR1
d - NPBWR2
PAIRING (ligand → receptor)
NPB→NPBWR1
NPB→NPBWR2
NPW→NPBWR1
NPW→NPBWR2
Compartment
endothelial
- artery
- vein
- vasa recta
- capillary
- mix
- lymphatic
12.5%
25%
50%
100%
ln(CP10K+1)
vein
artery
vasa recta descend limb SLC14A1+
vasa recta descend limb MGP+
vasa recta ascend limb IGFBP5+
vasa recta ascend limb IGF1+
capillary FABP5-RBP7-
capillary FABP5+RBP7-
capillary FABP5+RBP7lo
capillary FABP5+RBP7+
vascular endothelium FABP5+RBP7+
capillary CXCL10+
capillary aerocyte
sinusoid MAFB+
hepatic sinusoid
glomerular
vascular endothelium
lymphatic CCL21+
lymphatic ACKR4-CCDC80+
lymphatic ACKR4+CCDC80+
lymphatic
Abbrev. Tissue
Aort Aorta
Blad Bladder
Blo Blood
Bone Bone
BM Bone marrow
BrC Brain cortex
BrS Brainstem
Col Colon
Dia Diaphragm
Ret Eye retina
Fat Fat
Hrt Heart
Pit Hypothalamus/Pituitary
Kid Kidney
LM Limb muscle
Liv Liver
Lung Lung
MG Mammary gland
Panc Pancreas
Skin Skin
SI Small intestine
Spl Spleen
Test Testes
Thym Thymus
Tong Tongue
Trac Trachea
Ute Uterus

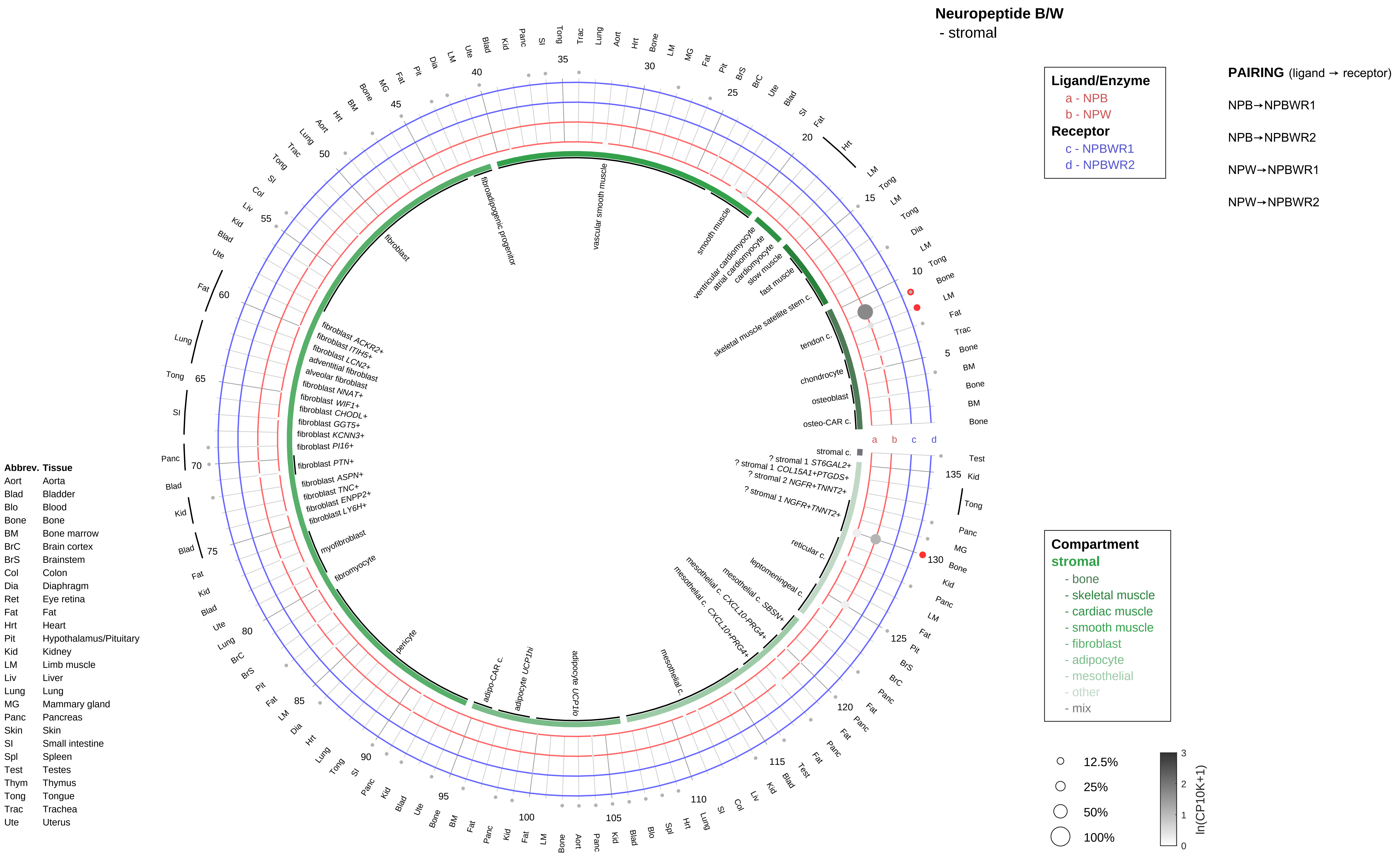
Neuropeptide B/W
- stromal
Ligand/Enzyme
a - NPB
b - NPW
Receptor
c - NPBWR1
d - NPBWR2
PAIRING (ligand → receptor)
NPB→NPBWR1
NPB→NPBWR2
NPW→NPBWR1
NPW→NPBWR2
Compartment
stromal
- bone
- skeletal muscle
- cardiac muscle
- smooth muscle
- fibroblast
- adipocyte
- mesothelial
- other
- mix
12.5%
25%
50%
100%
ln(CP10K+1)
Abbrev. Tissue
Aort Aorta
Blad Bladder
Blo Blood
Bone Bone
BM Bone marrow
BrC Brain cortex
BrS Brainstem
Col Colon
Dia Diaphragm
Ret Eye retina
Fat Fat
Hrt Heart
Pit Hypothalamus/Pituitary
Kid Kidney
LM Limb muscle
Liv Liver
Lung Lung
MG Mammary gland
Panc Pancreas
Skin Skin
SI Small intestine
Spl Spleen
Test Testes
Thym Thymus
Tong Tongue
Trac Trachea
Ute Uterus
fibroadipogenic progenitor
vascular smooth muscle
smooth muscle
ventricular cardiomyocyte
atrial cardiomyocyte
cardiomyocyte
slow muscle
fast muscle
skeletal muscle satellite stem c.
tendon c.
chondrocyte
osteoblast
osteo-CAR c.
stromal c.
? stromal 1 ST6GAL2+
? stromal 1 COL15A1+PTGDS+
? stromal 2 NGFR+TNNT2+
? stromal 1 NGFR+TNNT2+
reticular c.
leptomeningeal c.
mesothelial c. SBSN+
mesothelial c. CXCL10-PRG4+
mesothelial c. CXCL10+PRG4+
mesothelial c.
adipocyte UCP1lo
adipocyte UCP1hi
adipo-CAR c.
pericyte
fibromyocyte
myofibroblast
fibroblast LY6H+
fibroblast ENPP2+
fibroblast TNC+
fibroblast ASPN+
fibroblast PTN+
fibroblast PI16+
fibroblast KCNN3+
fibroblast GGT5+
fibroblast CHODL+
fibroblast WIF1+
fibroblast NNAT+
alveolar fibroblast
adventitial fibroblast
fibroblast LCN2+
fibroblast ITIH5+
fibroblast ACKR2+
fibroblast

**Neuropeptide B/W**
- lymphoid

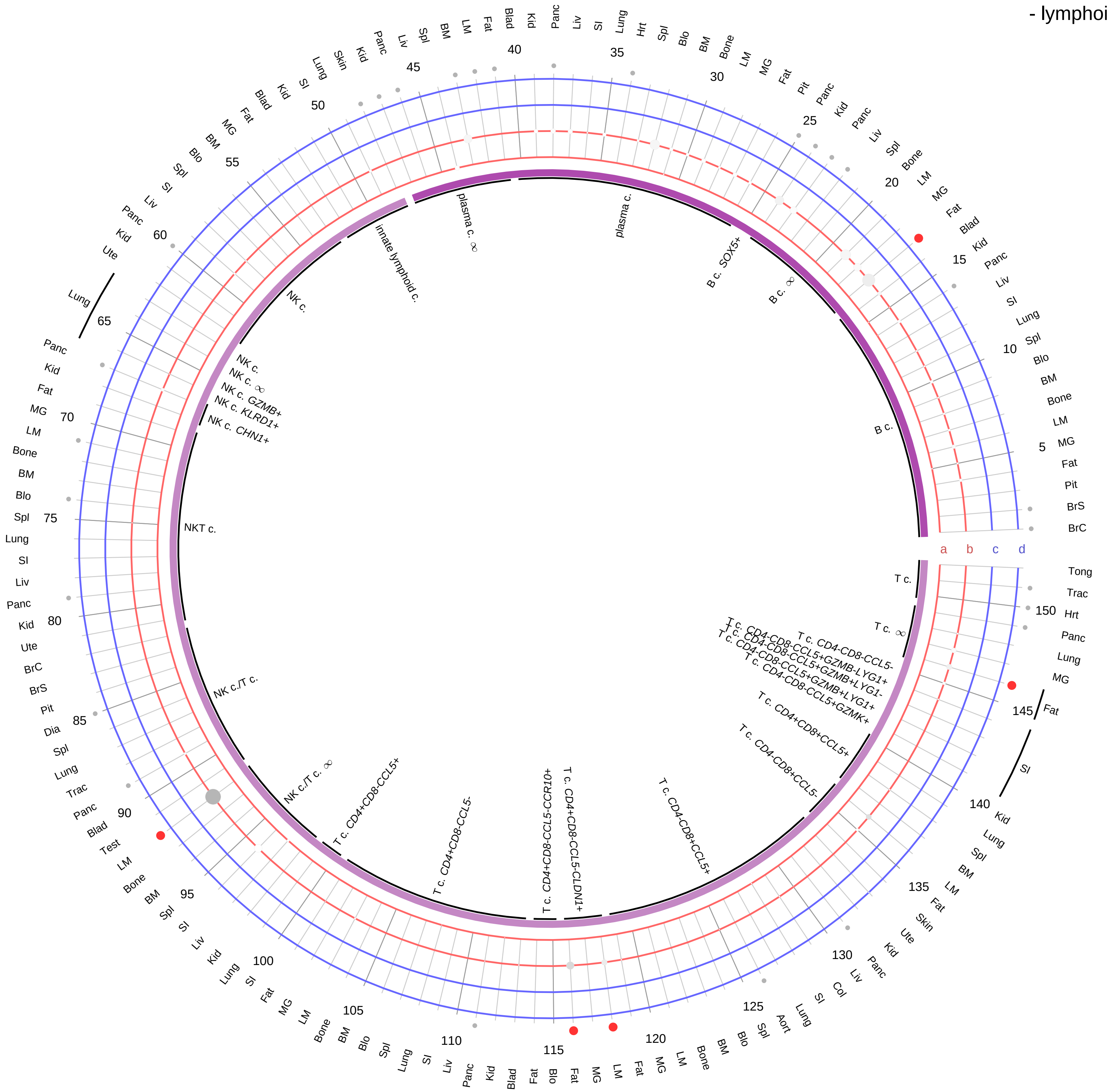

**Ligand/Enzyme**
- a - NPB
- b - NPW

**Receptor**
- c - NPBWR1
- d - NPBWR2

**PAIRING** (ligand → receptor)

NPB→NPBWR1

NPB→NPBWR2

NPW→NPBWR1

NPW→NPBWR2

**Compartment**

**lymphoid**
- B cell
- NK/T cell

12.5%
25%
50%
100%

ln(CP10K+1): 0, 1, 2, 3

| Abbrev. | Tissue |
|---|---|
| Aort | Aorta |
| Blad | Bladder |
| Blo | Blood |
| Bone | Bone |
| BM | Bone marrow |
| BrC | Brain cortex |
| BrS | Brainstem |
| Col | Colon |
| Dia | Diaphragm |
| Ret | Eye retina |
| Fat | Fat |
| Hrt | Heart |
| Pit | Hypothalamus/Pituitary |
| Kid | Kidney |
| LM | Limb muscle |
| Liv | Liver |
| Lung | Lung |
| MG | Mammary gland |
| Panc | Pancreas |
| Skin | Skin |
| SI | Small intestine |
| Spl | Spleen |
| Test | Testes |
| Thym | Thymus |
| Tong | Tongue |
| Trac | Trachea |
| Ute | Uterus |

#### Neuropeptide B/W
- myeloid

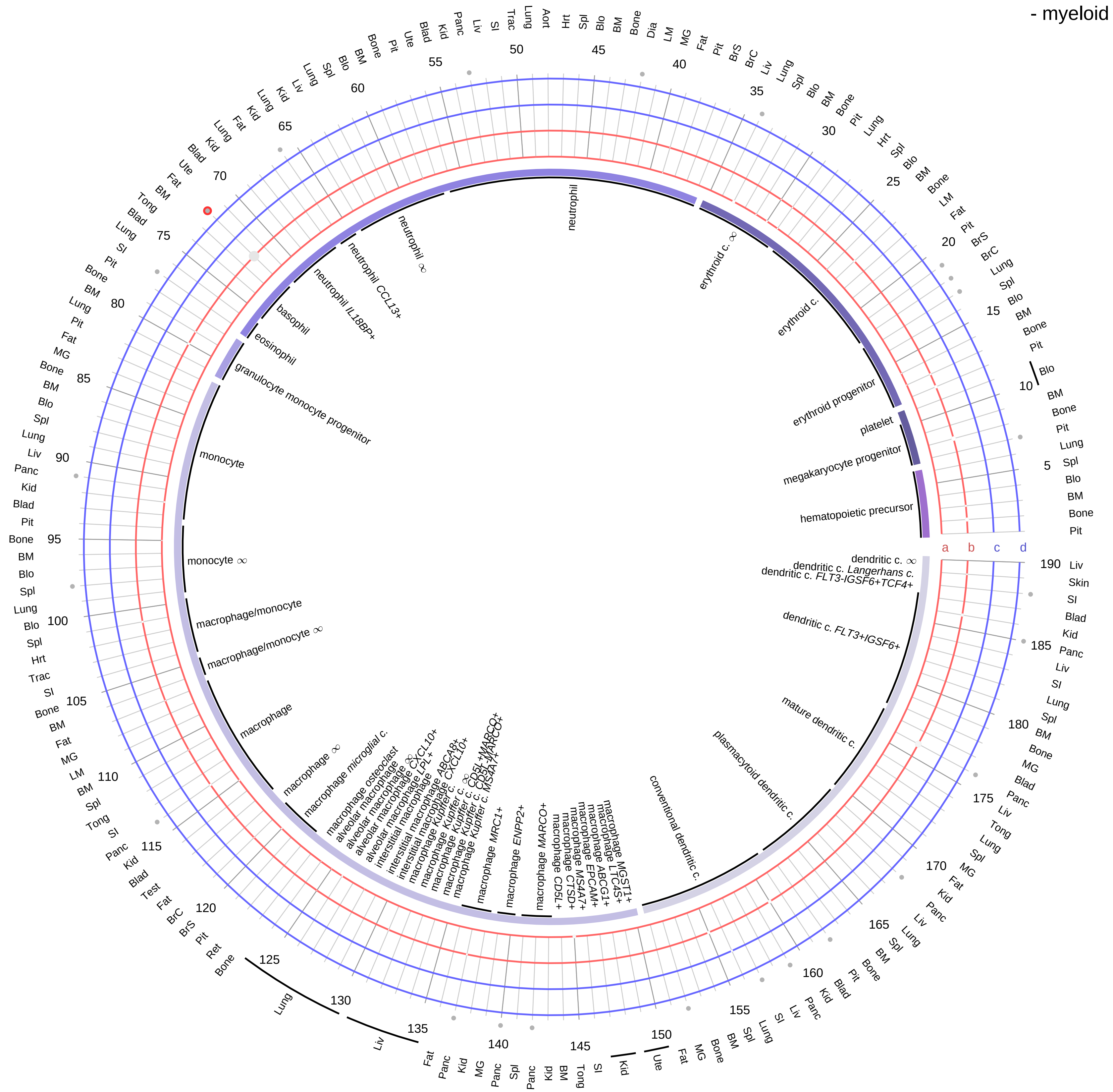

**Ligand/Enzyme**
- a - NPB
- b - NPW

**Receptor**
- c - NPBWR1
- d - NPBWR2

**PAIRING** (ligand → receptor)

NPB→NPBWR1

NPB→NPBWR2

NPW→NPBWR1

NPW→NPBWR2

| Abbrev. | Tissue |
|---|---|
| Aort | Aorta |
| Blad | Bladder |
| Blo | Blood |
| Bone | Bone |
| BM | Bone marrow |
| BrC | Brain cortex |
| BrS | Brainstem |
| Col | Colon |
| Dia | Diaphragm |
| Ret | Eye retina |
| Fat | Fat |
| Hrt | Heart |
| Pit | Hypothalamus/Pituitary |
| Kid | Kidney |
| LM | Limb muscle |
| Liv | Liver |
| Lung | Lung |
| MG | Mammary gland |
| Panc | Pancreas |
| Skin | Skin |
| SI | Small intestine |
| Spl | Spleen |
| Test | Testes |
| Thym | Thymus |
| Tong | Tongue |
| Trac | Trachea |
| Ute | Uterus |

**Compartment**

**hematopoietic**
- hematopoietic precursor

**megakaryocyte-erythroid**
- megakaryoid
- erythroid

**myeloid**
- granulocyte
- granulocyte monocyte progenitor
- monocyte/macrophage
- dendritic

12.5%
25%
50%
100%

ln(CP10K+1): 0, 1, 2, 3

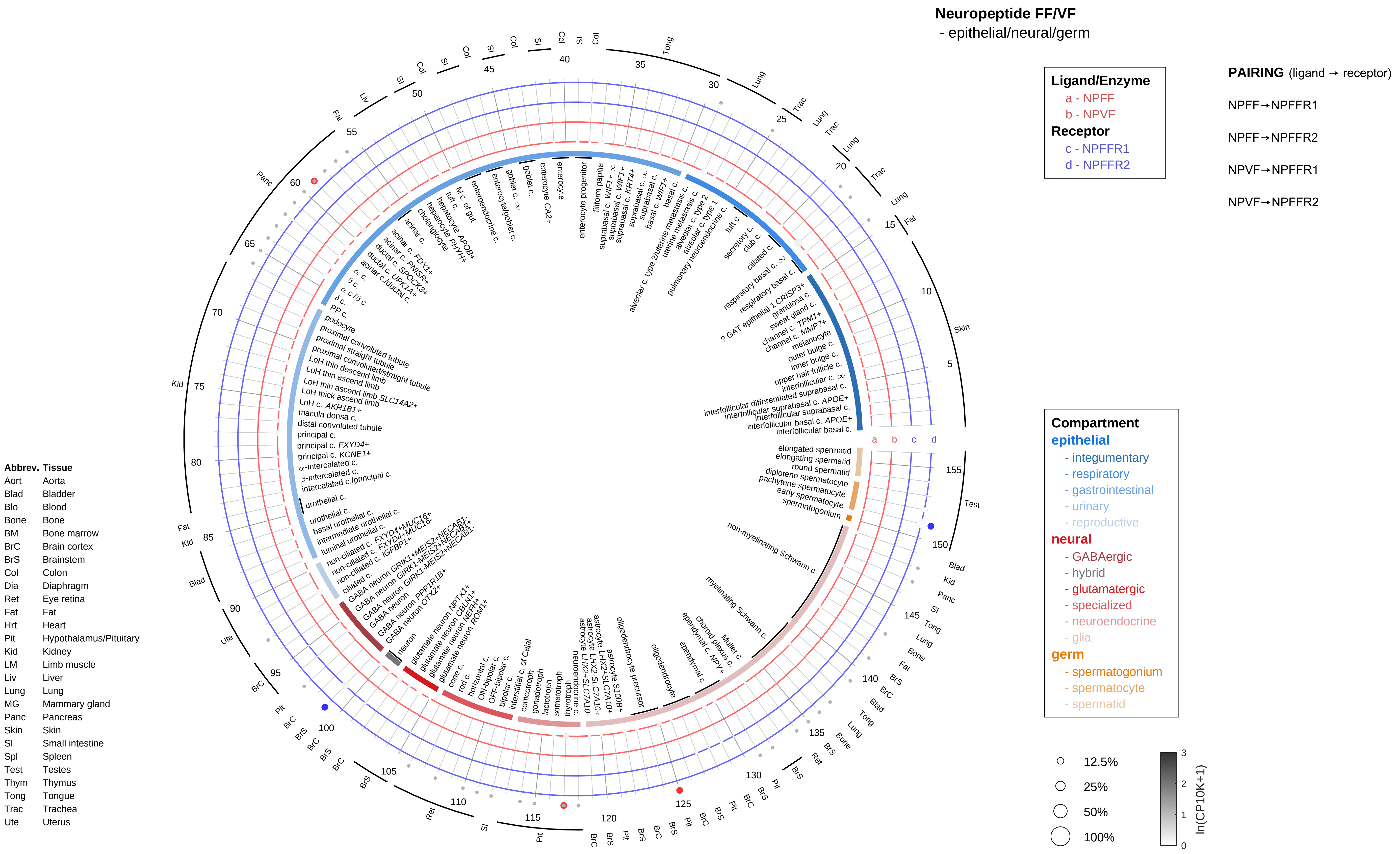
Neuropeptide FF/VF
- epithelial/neural/germ
Ligand/Enzyme
a - NPFF
b - NPVF
Receptor
c - NPFFR1
d - NPFFR2
PAIRING (ligand → receptor)
NPFF→NPFFR1
NPFF→NPFFR2
NPVF→NPFFR1
NPVF→NPFFR2
Compartment
epithelial
- integumentary
- respiratory
- gastrointestinal
- urinary
- reproductive
neural
- GABAergic
- hybrid
- glutamatergic
- specialized
- neuroendocrine
- glia
germ
- spermatogonium
- spermatocyte
- spermatid
12.5%
25%
50%
100%
ln(CP10K+1)
Abbrev. Tissue
Aort Aorta
Blad Bladder
Blo Blood
Bone Bone
BM Bone marrow
BrC Brain cortex
BrS Brainstem
Col Colon
Dia Diaphragm
Ret Eye retina
Fat Fat
Hrt Heart
Pit Hypothalamus/Pituitary
Kid Kidney
LM Limb muscle
Liv Liver
Lung Lung
MG Mammary gland
Panc Pancreas
Skin Skin
SI Small intestine
Spl Spleen
Test Testes
Thym Thymus
Tong Tongue
Trac Trachea
Ute Uterus

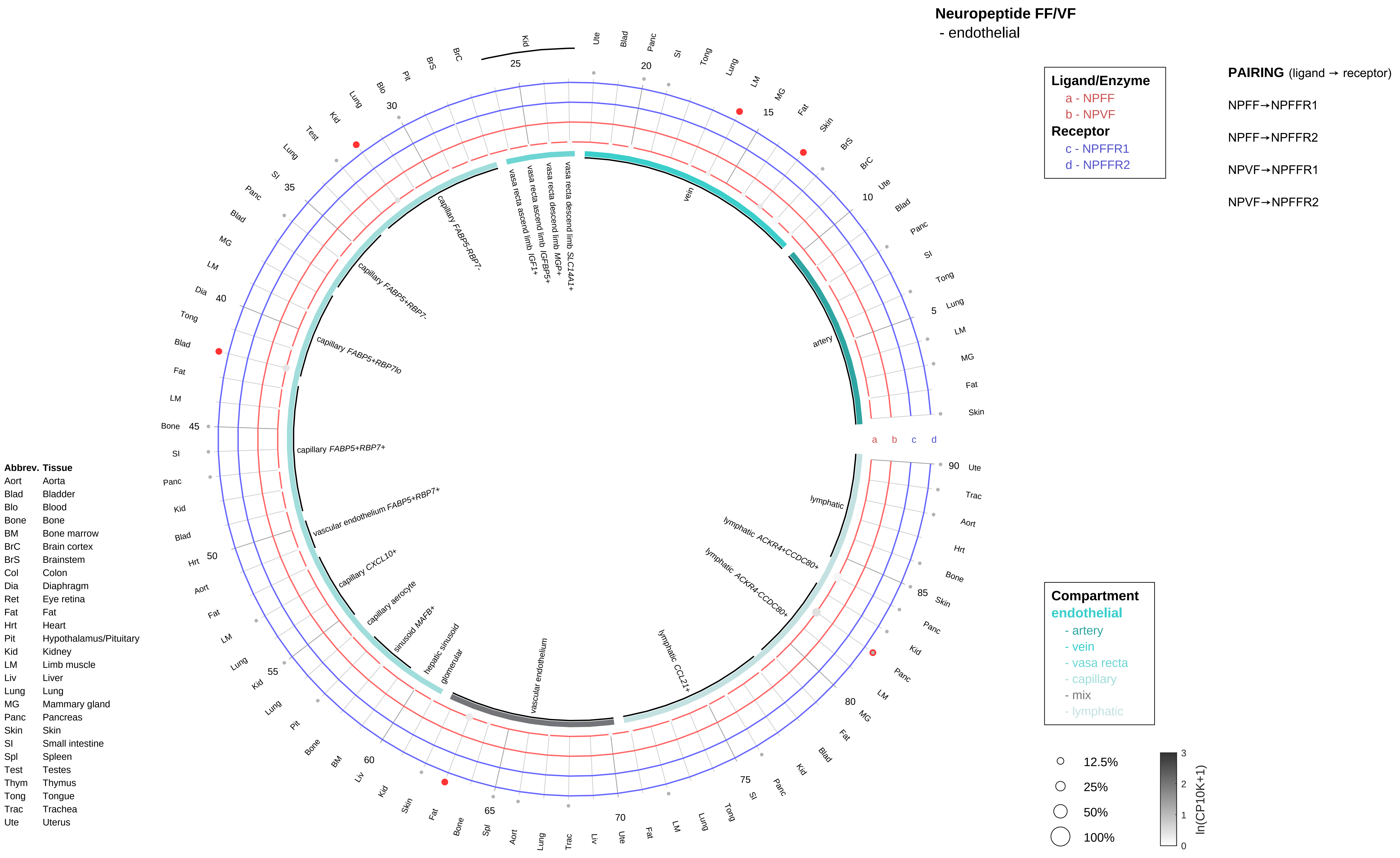
Neuropeptide FF/VF
- endothelial
Ligand/Enzyme
a - NPFF
b - NPVF
Receptor
c - NPFFR1
d - NPFFR2
PAIRING (ligand → receptor)
NPFF→NPFFR1
NPFF→NPFFR2
NPVF→NPFFR1
NPVF→NPFFR2
Compartment
endothelial
- artery
- vein
- vasa recta
- capillary
- mix
- lymphatic
12.5%
25%
50%
100%
ln(CP10K+1)
artery
vein
vasa recta descend limb SLC14A1+
vasa recta descend limb MGP+
vasa recta ascend limb IGFBP5+
vasa recta ascend limb IGF1+
capillary FABP5-RBP7-
capillary FABP5+RBP7-
capillary FABP5+RBP7lo
capillary FABP5+RBP7+
vascular endothelium FABP5+RBP7+
capillary CXCL10+
capillary aerocyte
sinusoid MAFB+
hepatic sinusoid
glomerular
vascular endothelium
lymphatic CCL21+
lymphatic ACKR4-CCDC80+
lymphatic ACKR4+CCDC80+
lymphatic
Abbrev. Tissue
Aort Aorta
Blad Bladder
Blo Blood
Bone Bone
BM Bone marrow
BrC Brain cortex
BrS Brainstem
Col Colon
Dia Diaphragm
Ret Eye retina
Fat Fat
Hrt Heart
Pit Hypothalamus/Pituitary
Kid Kidney
LM Limb muscle
Liv Liver
Lung Lung
MG Mammary gland
Panc Pancreas
Skin Skin
SI Small intestine
Spl Spleen
Test Testes
Thym Thymus
Tong Tongue
Trac Trachea
Ute Uterus

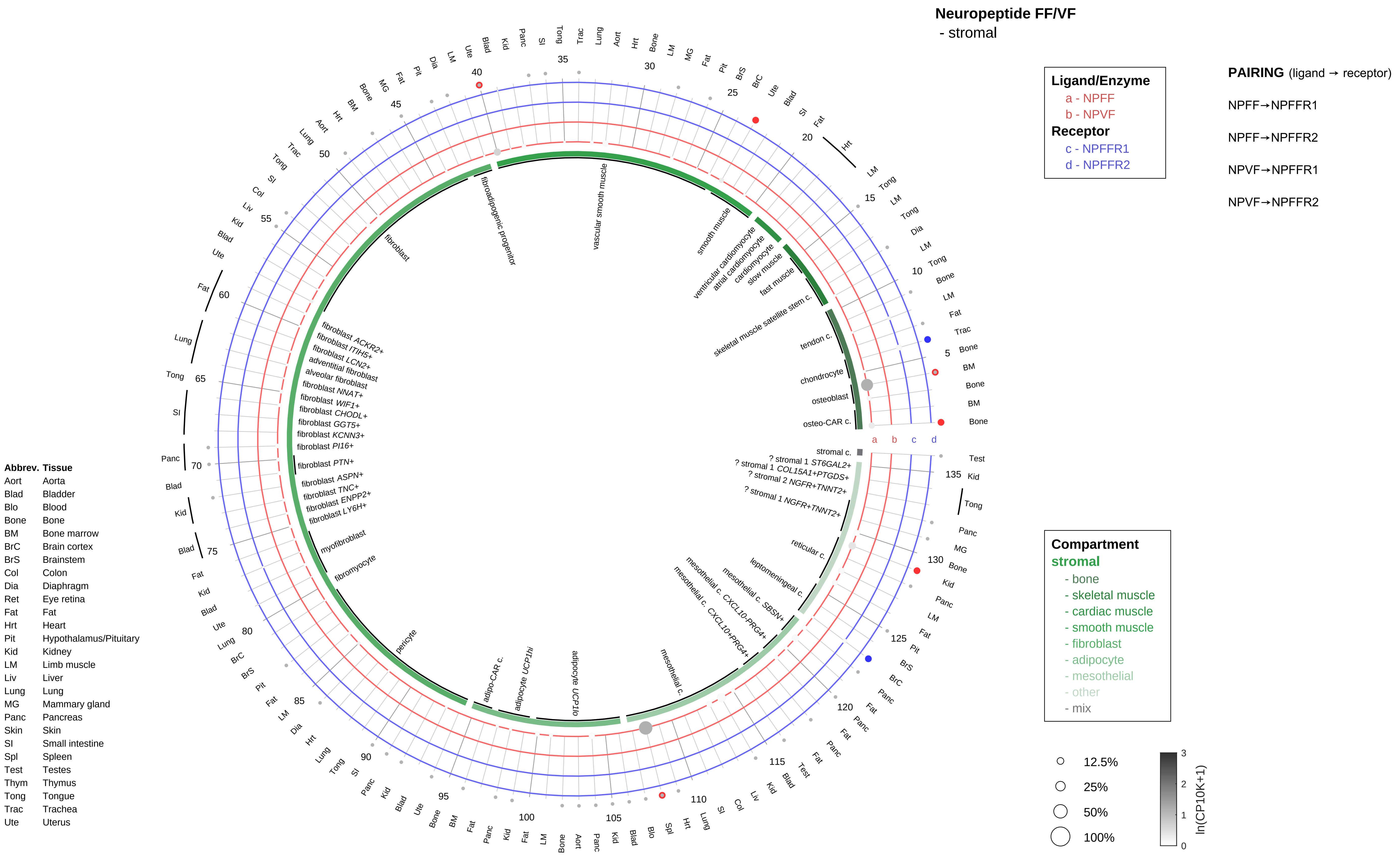

Neuropeptide FF/VF
- stromal
Ligand/Enzyme
a - NPFF
b - NPVF
Receptor
c - NPFFR1
d - NPFFR2
PAIRING (ligand → receptor)
NPFF→NPFFR1
NPFF→NPFFR2
NPVF→NPFFR1
NPVF→NPFFR2
Compartment
stromal
- bone
- skeletal muscle
- cardiac muscle
- smooth muscle
- fibroblast
- adipocyte
- mesothelial
- other
- mix
12.5%
25%
50%
100%
ln(CP10K+1)
Abbrev. Tissue
Aort Aorta
Blad Bladder
Blo Blood
Bone Bone
BM Bone marrow
BrC Brain cortex
BrS Brainstem
Col Colon
Dia Diaphragm
Ret Eye retina
Fat Fat
Hrt Heart
Pit Hypothalamus/Pituitary
Kid Kidney
LM Limb muscle
Liv Liver
Lung Lung
MG Mammary gland
Panc Pancreas
Skin Skin
SI Small intestine
Spl Spleen
Test Testes
Thym Thymus
Tong Tongue
Trac Trachea
Ute Uterus
osteo-CAR c.
osteoblast
chondrocyte
tendon c.
skeletal muscle satellite stem c.
fast muscle
slow muscle
cardiomyocyte
atrial cardiomyocyte
ventricular cardiomyocyte
smooth muscle
vascular smooth muscle
fibroadipogenic progenitor
fibroblast
fibroblast ACKR2+
fibroblast ITIH5+
fibroblast LCN2+
adventitial fibroblast
alveolar fibroblast
fibroblast NNAT+
fibroblast WIF1+
fibroblast CHODL+
fibroblast GGT5+
fibroblast KCNN3+
fibroblast PI16+
fibroblast PTN+
fibroblast ASPN+
fibroblast TNC+
fibroblast ENPP2+
fibroblast LY6H+
myofibroblast
fibromyocyte
pericyte
adipo-CAR c.
adipocyte UCP1hi
adipocyte UCP1lo
mesothelial c.
mesothelial c. CXCL10-PRG4+
mesothelial c. CXCL10+PRG4+
mesothelial c. SBSN+
leptomeningeal c.
reticular c.
? stromal 1 NGFR+TNNT2+
? stromal 2 NGFR+TNNT2+
? stromal 1 COL15A1+PTGDS+
? stromal 1 ST6GAL2+
stromal c.

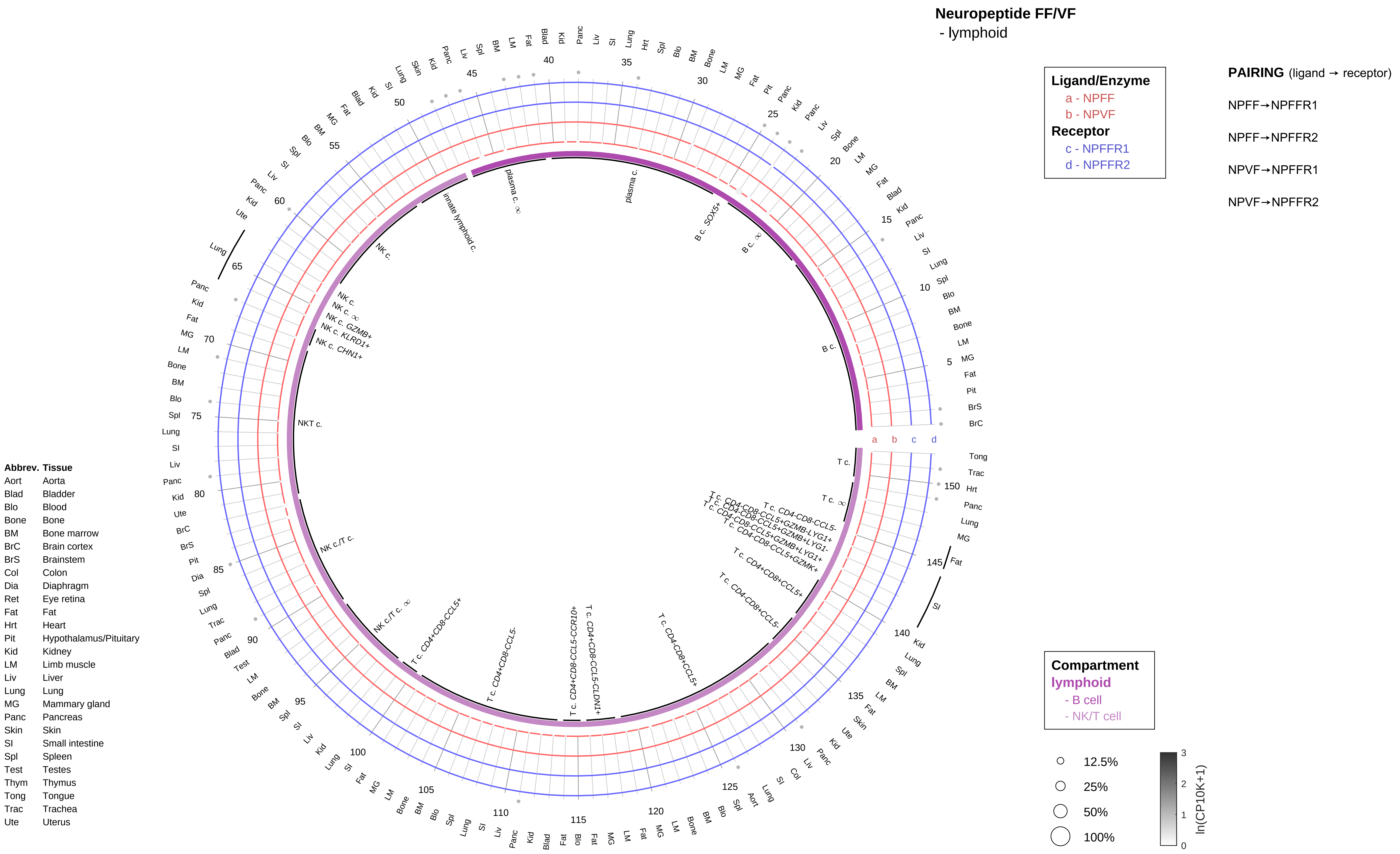

Neuropeptide FF/VF
- lymphoid
Ligand/Enzyme
a - NPFF
b - NPVF
Receptor
c - NPFFR1
d - NPFFR2
PAIRING (ligand → receptor)
NPFF→NPFFR1
NPFF→NPFFR2
NPVF→NPFFR1
NPVF→NPFFR2
Compartment
lymphoid
- B cell
- NK/T cell
12.5%
25%
50%
100%
ln(CP10K+1)
Abbrev. Tissue
Aort Aorta
Blad Bladder
Blo Blood
Bone Bone
BM Bone marrow
BrC Brain cortex
BrS Brainstem
Col Colon
Dia Diaphragm
Ret Eye retina
Fat Fat
Hrt Heart
Pit Hypothalamus/Pituitary
Kid Kidney
LM Limb muscle
Liv Liver
Lung Lung
MG Mammary gland
Panc Pancreas
Skin Skin
SI Small intestine
Spl Spleen
Test Testes
Thym Thymus
Tong Tongue
Trac Trachea
Ute Uterus

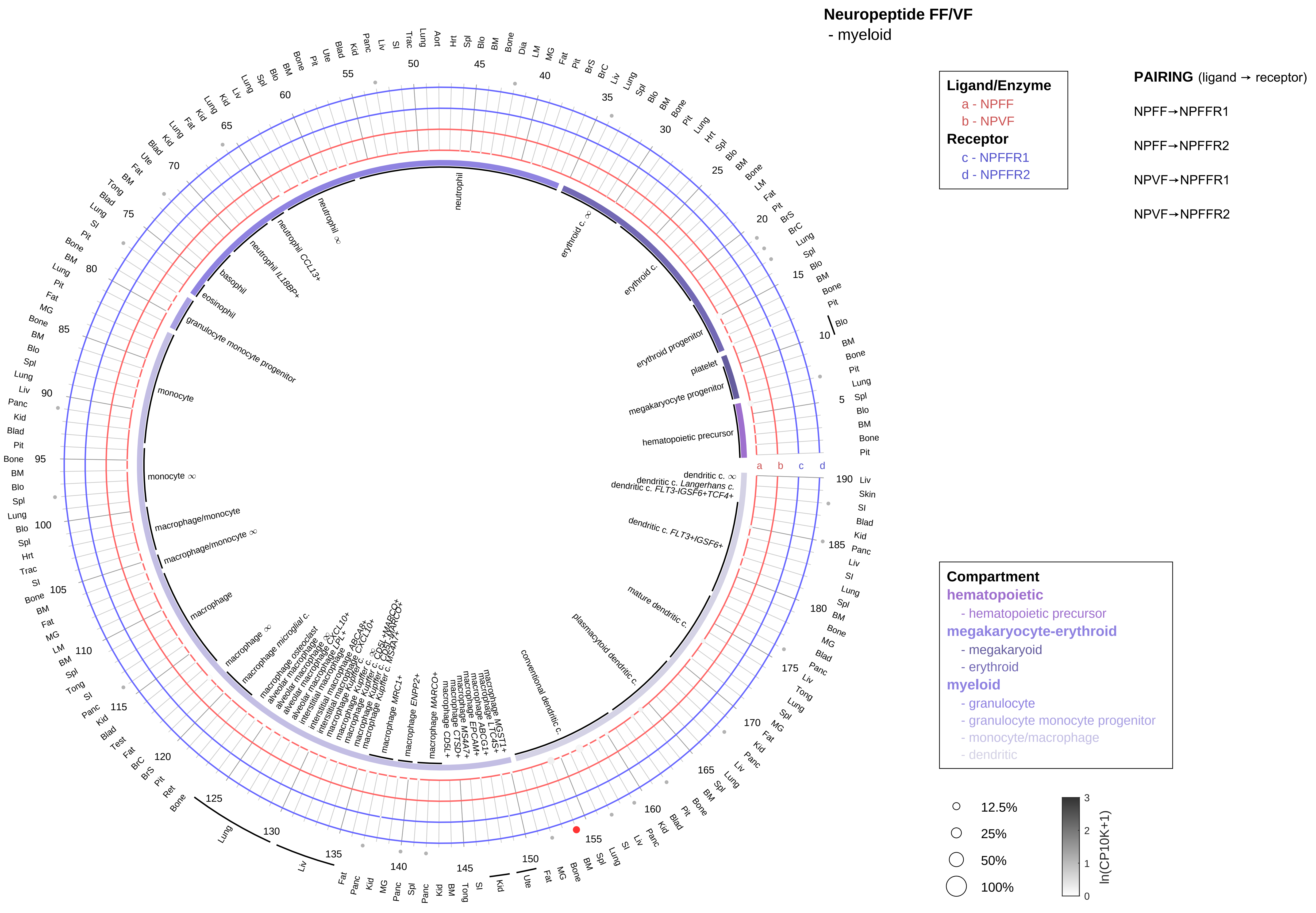

| Abbrev. | Tissue |
|---|---|
| Aort | Aorta |
| Blad | Bladder |
| Blo | Blood |
| Bone | Bone |
| BM | Bone marrow |
| BrC | Brain cortex |
| BrS | Brainstem |
| Col | Colon |
| Dia | Diaphragm |
| Ret | Eye retina |
| Fat | Fat |
| Hrt | Heart |
| Pit | Hypothalamus/Pituitary |
| Kid | Kidney |
| LM | Limb muscle |
| Liv | Liver |
| Lung | Lung |
| MG | Mammary gland |
| Panc | Pancreas |
| Skin | Skin |
| SI | Small intestine |
| Spl | Spleen |
| Test | Testes |
| Thym | Thymus |
| Tong | Tongue |
| Trac | Trachea |
| Ute | Uterus |

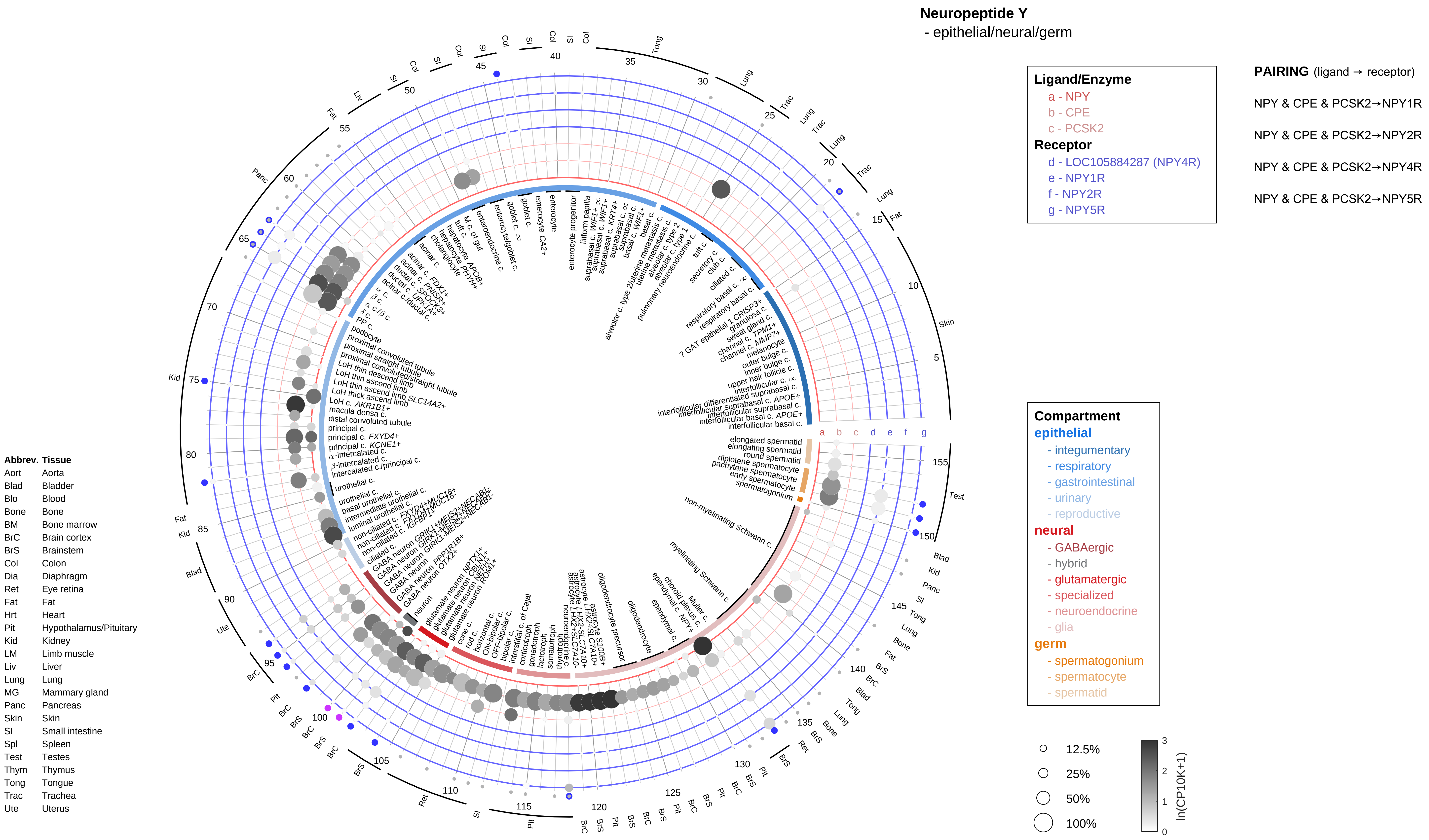
Neuropeptide Y
- epithelial/neural/germ
Ligand/Enzyme
a - NPY
b - CPE
c - PCSK2
Receptor
d - LOC105884287 (NPY4R)
e - NPY1R
f - NPY2R
g - NPY5R
PAIRING (ligand → receptor)
NPY & CPE & PCSK2→NPY1R
NPY & CPE & PCSK2→NPY2R
NPY & CPE & PCSK2→NPY4R
NPY & CPE & PCSK2→NPY5R
Compartment
epithelial
- integumentary
- respiratory
- gastrointestinal
- urinary
- reproductive
neural
- GABAergic
- hybrid
- glutamatergic
- specialized
- neuroendocrine
- glia
germ
- spermatogonium
- spermatocyte
- spermatid
12.5%
25%
50%
100%
ln(CP10K+1)
Abbrev. Tissue
Aort Aorta
Blad Bladder
Blo Blood
Bone Bone
BM Bone marrow
BrC Brain cortex
BrS Brainstem
Col Colon
Dia Diaphragm
Ret Eye retina
Fat Fat
Hrt Heart
Pit Hypothalamus/Pituitary
Kid Kidney
LM Limb muscle
Liv Liver
Lung Lung
MG Mammary gland
Panc Pancreas
Skin Skin
SI Small intestine
Spl Spleen
Test Testes
Thym Thymus
Tong Tongue
Trac Trachea
Ute Uterus

**Neuropeptide Y**
- endothelial

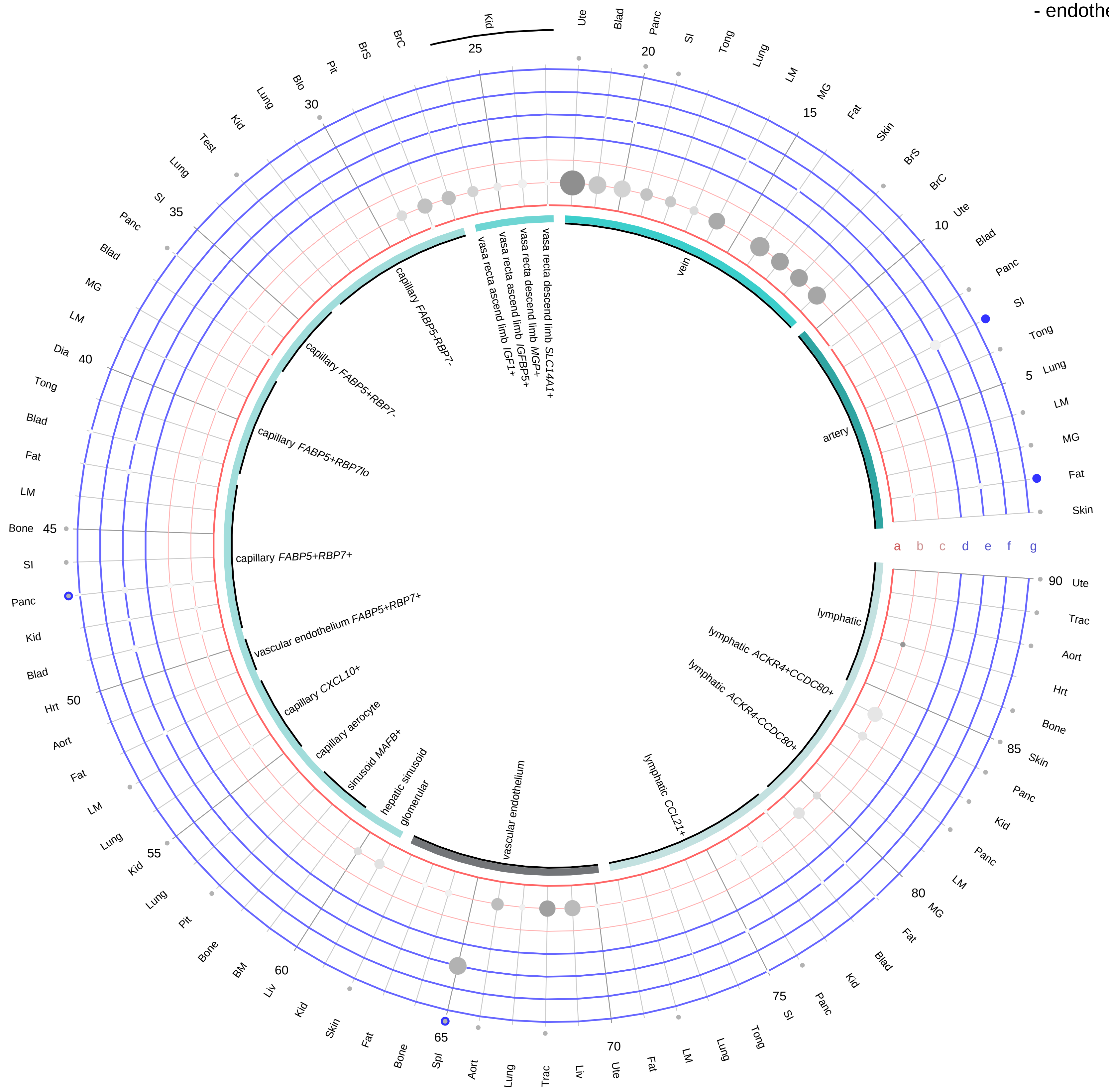

**Ligand/Enzyme**
- a - NPY
- b - CPE
- c - PCSK2

**Receptor**
- d - LOC105884287 (NPY4R)
- e - NPY1R
- f - NPY2R
- g - NPY5R

**PAIRING** (ligand → receptor)

NPY & CPE & PCSK2→NPY1R

NPY & CPE & PCSK2→NPY2R

NPY & CPE & PCSK2→NPY4R

NPY & CPE & PCSK2→NPY5R

| Abbrev. | Tissue |
|---|---|
| Aort | Aorta |
| Blad | Bladder |
| Blo | Blood |
| Bone | Bone |
| BM | Bone marrow |
| BrC | Brain cortex |
| BrS | Brainstem |
| Col | Colon |
| Dia | Diaphragm |
| Ret | Eye retina |
| Fat | Fat |
| Hrt | Heart |
| Pit | Hypothalamus/Pituitary |
| Kid | Kidney |
| LM | Limb muscle |
| Liv | Liver |
| Lung | Lung |
| MG | Mammary gland |
| Panc | Pancreas |
| Skin | Skin |
| SI | Small intestine |
| Spl | Spleen |
| Test | Testes |
| Thym | Thymus |
| Tong | Tongue |
| Trac | Trachea |
| Ute | Uterus |

**Compartment**
**endothelial**
- artery
- vein
- vasa recta
- capillary
- mix
- lymphatic

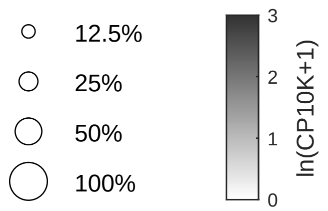

#### Neuropeptide Y
- stromal

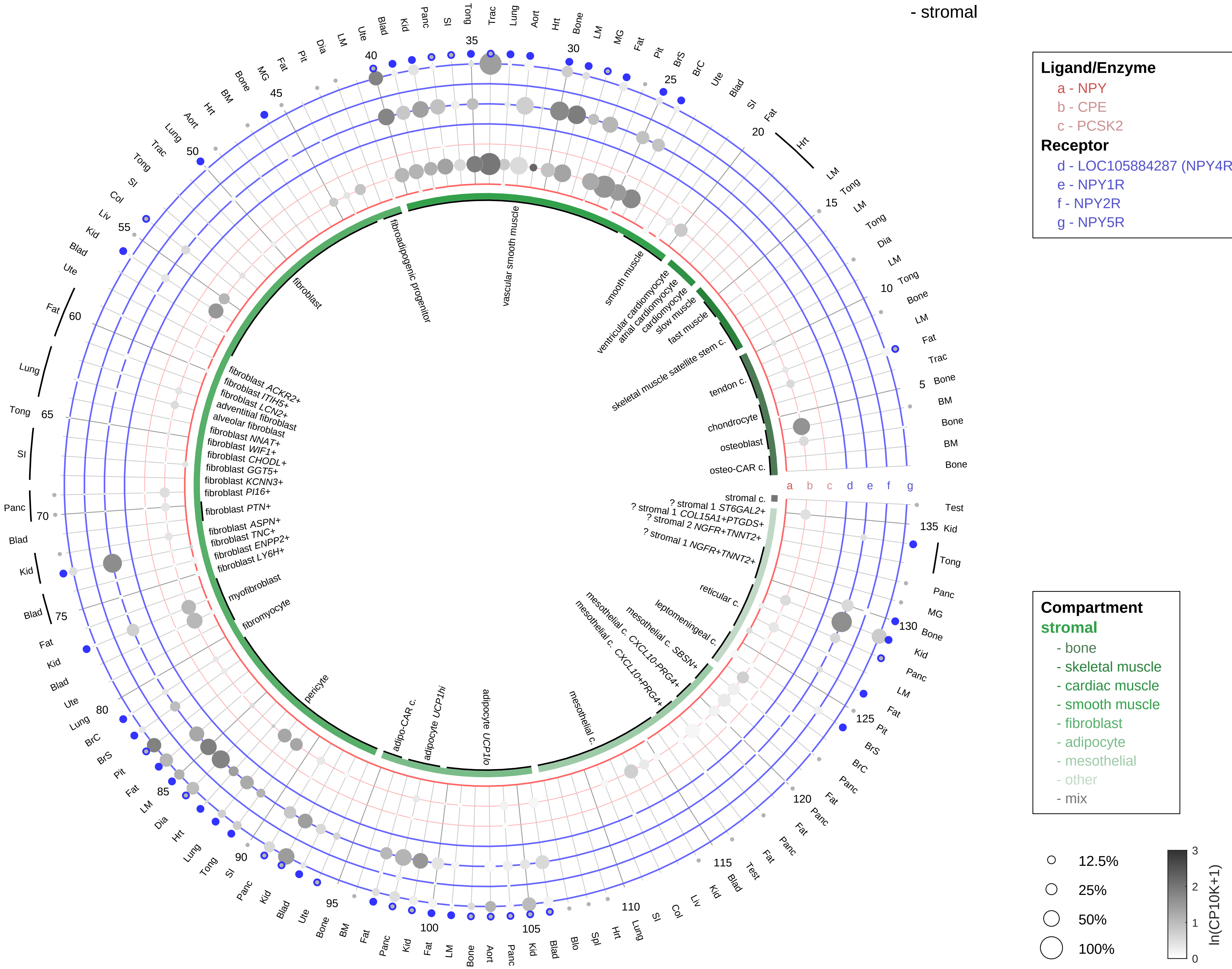

**PAIRING** (ligand → receptor)

NPY & CPE & PCSK2→NPY1R

NPY & CPE & PCSK2→NPY2R

NPY & CPE & PCSK2→NPY4R

NPY & CPE & PCSK2→NPY5R

| Abbrev. | Tissue |
|---|---|
| Aort | Aorta |
| Blad | Bladder |
| Blo | Blood |
| Bone | Bone |
| BM | Bone marrow |
| BrC | Brain cortex |
| BrS | Brainstem |
| Col | Colon |
| Dia | Diaphragm |
| Ret | Eye retina |
| Fat | Fat |
| Hrt | Heart |
| Pit | Hypothalamus/Pituitary |
| Kid | Kidney |
| LM | Limb muscle |
| Liv | Liver |
| Lung | Lung |
| MG | Mammary gland |
| Panc | Pancreas |
| Skin | Skin |
| SI | Small intestine |
| Spl | Spleen |
| Test | Testes |
| Thym | Thymus |
| Tong | Tongue |
| Trac | Trachea |
| Ute | Uterus |

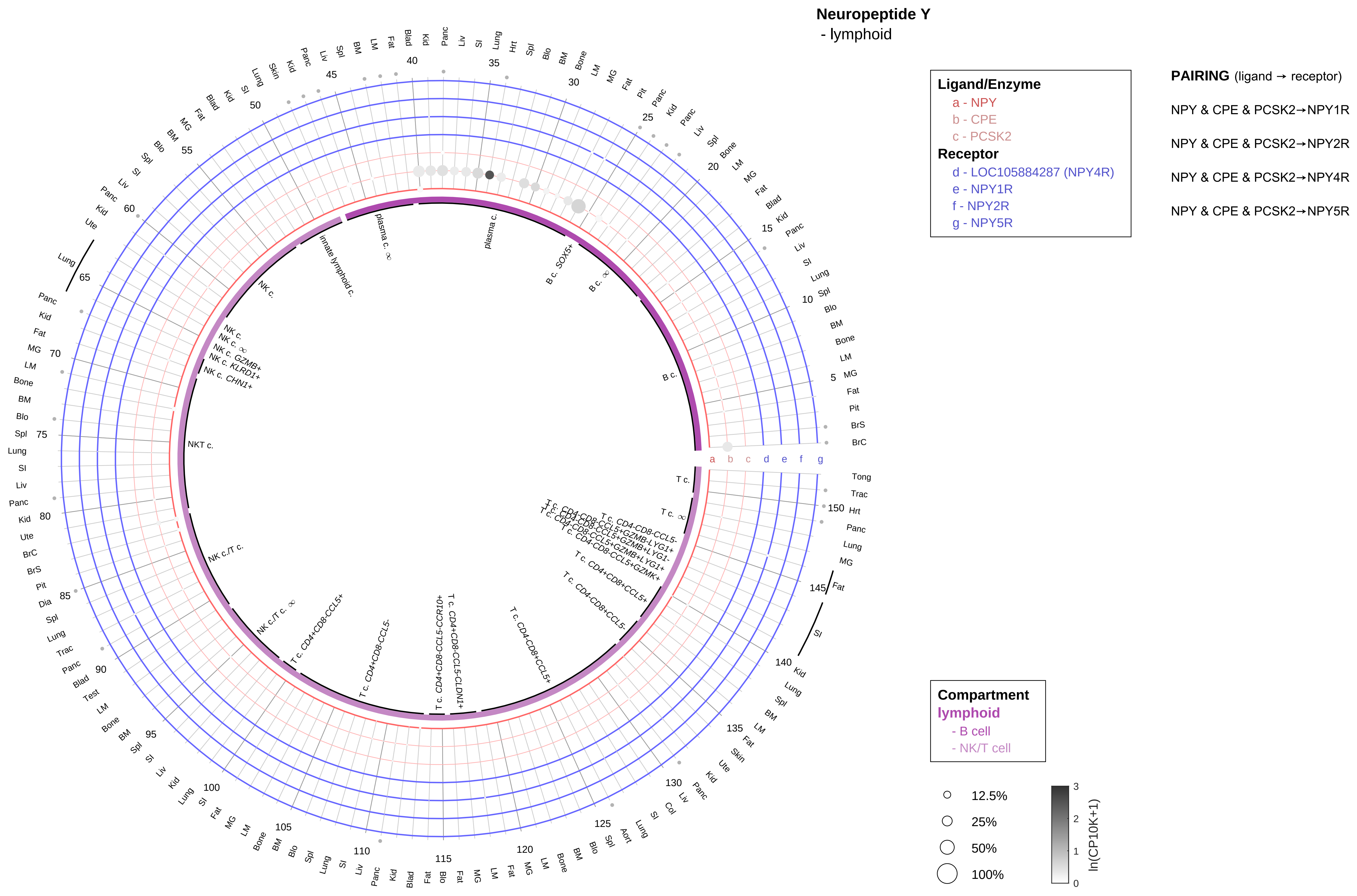

| Abbrev. | Tissue |
|---|---|
| Aort | Aorta |
| Blad | Bladder |
| Blo | Blood |
| Bone | Bone |
| BM | Bone marrow |
| BrC | Brain cortex |
| BrS | Brainstem |
| Col | Colon |
| Dia | Diaphragm |
| Ret | Eye retina |
| Fat | Fat |
| Hrt | Heart |
| Pit | Hypothalamus/Pituitary |
| Kid | Kidney |
| LM | Limb muscle |
| Liv | Liver |
| Lung | Lung |
| MG | Mammary gland |
| Panc | Pancreas |
| Skin | Skin |
| SI | Small intestine |
| Spl | Spleen |
| Test | Testes |
| Thym | Thymus |
| Tong | Tongue |
| Trac | Trachea |
| Ute | Uterus |

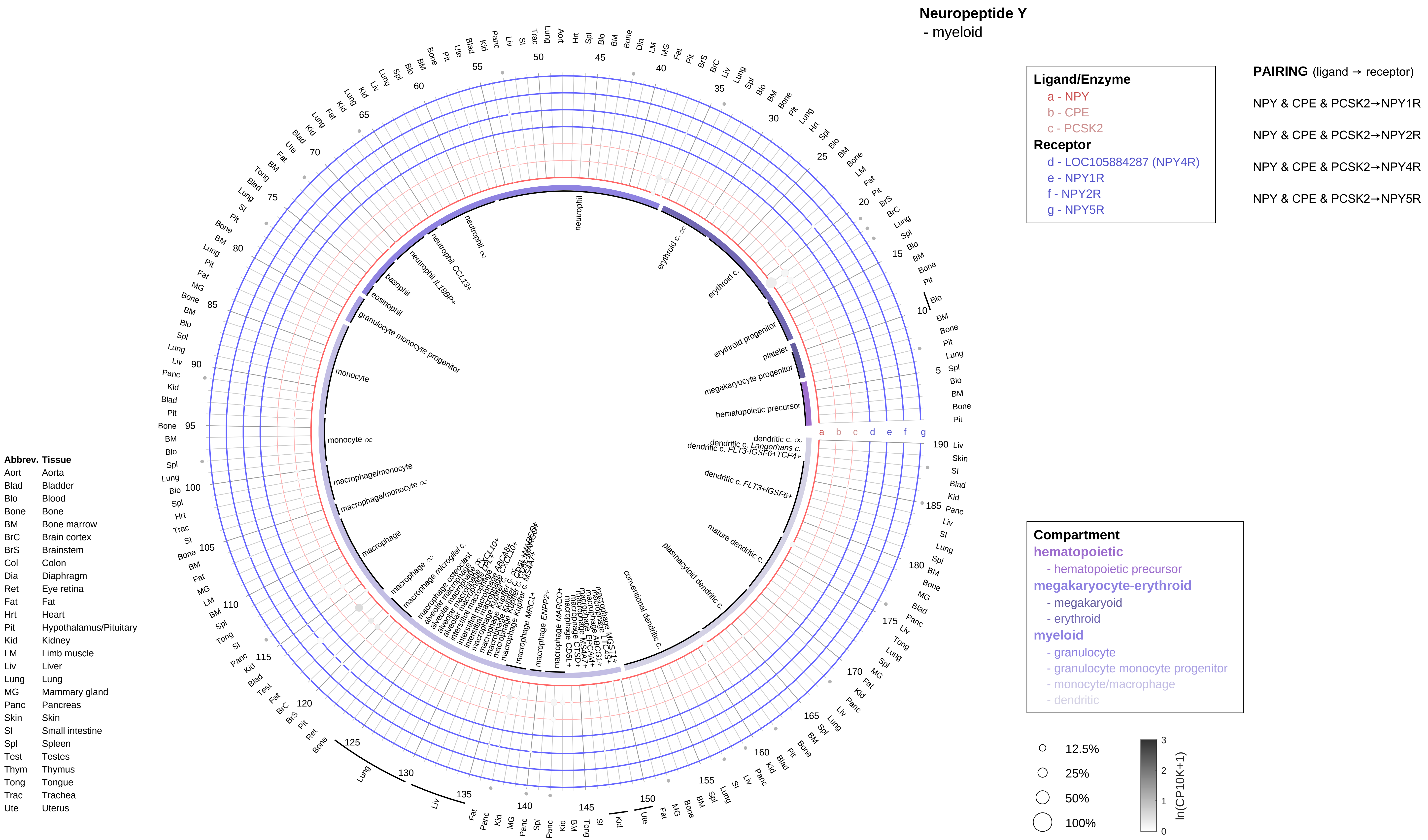
Neuropeptide Y
- myeloid
Ligand/Enzyme
a - NPY
b - CPE
c - PCSK2
Receptor
d - LOC105884287 (NPY4R)
e - NPY1R
f - NPY2R
g - NPY5R
PAIRING (ligand → receptor)
NPY & CPE & PCSK2→NPY1R
NPY & CPE & PCSK2→NPY2R
NPY & CPE & PCSK2→NPY4R
NPY & CPE & PCSK2→NPY5R
Compartment
hematopoietic
- hematopoietic precursor
megakaryocyte-erythroid
- megakaryoid
- erythroid
myeloid
- granulocyte
- granulocyte monocyte progenitor
- monocyte/macrophage
- dendritic
12.5%
25%
50%
100%
ln(CP10K+1)
Abbrev. Tissue
Aort Aorta
Blad Bladder
Blo Blood
Bone Bone
BM Bone marrow
BrC Brain cortex
BrS Brainstem
Col Colon
Dia Diaphragm
Ret Eye retina
Fat Fat
Hrt Heart
Pit Hypothalamus/Pituitary
Kid Kidney
LM Limb muscle
Liv Liver
Lung Lung
MG Mammary gland
Panc Pancreas
Skin Skin
SI Small intestine
Spl Spleen
Test Testes
Thym Thymus
Tong Tongue
Trac Trachea
Ute Uterus

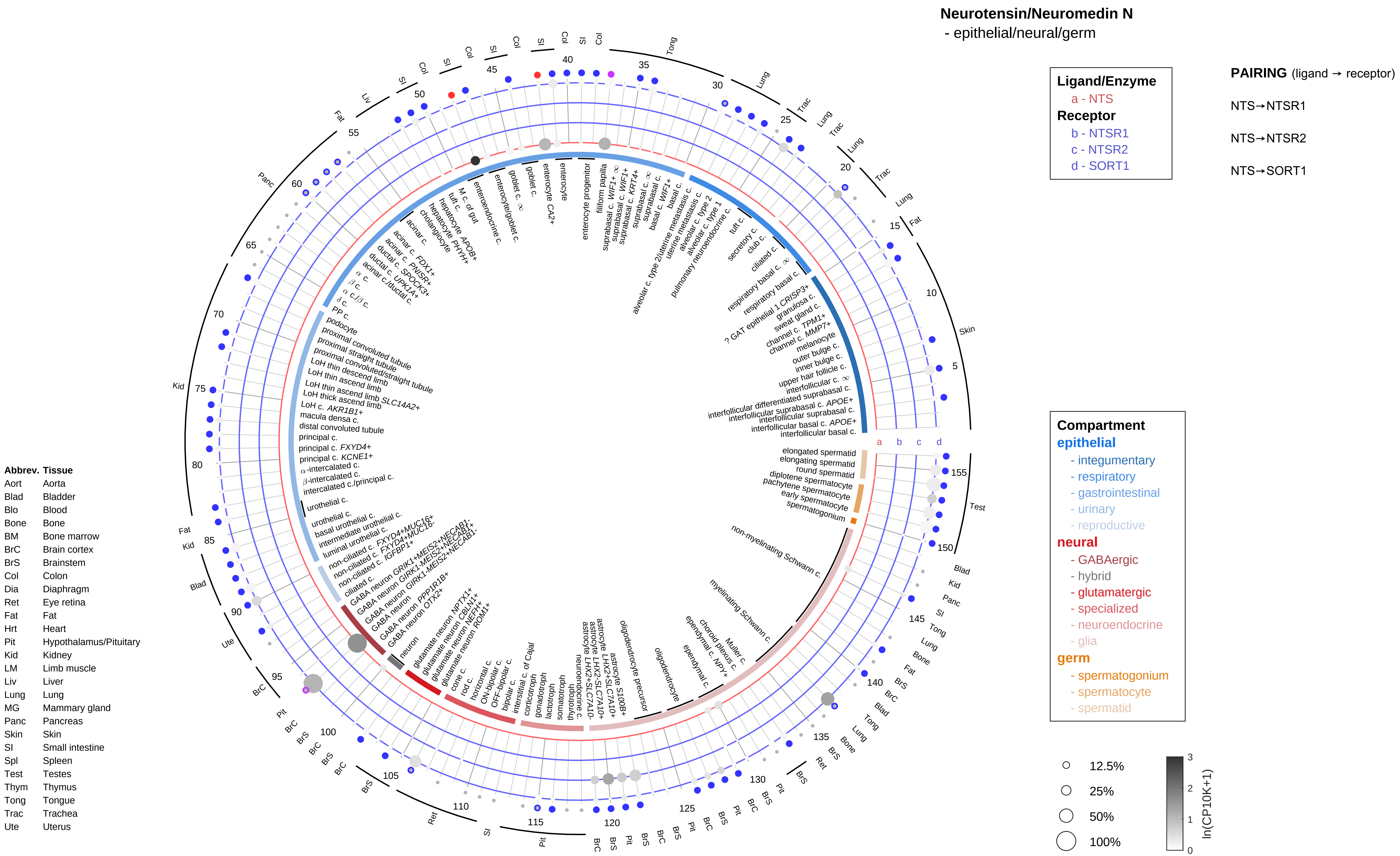

Neurotensin/Neuromedin N
- epithelial/neural/germ
Ligand/Enzyme
a - NTS
Receptor
b - NTSR1
c - NTSR2
d - SORT1
PAIRING (ligand → receptor)
NTS→NTSR1
NTS→NTSR2
NTS→SORT1
Compartment
epithelial
- integumentary
- respiratory
- gastrointestinal
- urinary
- reproductive
neural
- GABAergic
- hybrid
- glutamatergic
- specialized
- neuroendocrine
- glia
germ
- spermatogonium
- spermatocyte
- spermatid
12.5%
25%
50%
100%
ln(CP10K+1)
Abbrev. Tissue
Aort Aorta
Blad Bladder
Blo Blood
Bone Bone
BM Bone marrow
BrC Brain cortex
BrS Brainstem
Col Colon
Dia Diaphragm
Ret Eye retina
Fat Fat
Hrt Heart
Pit Hypothalamus/Pituitary
Kid Kidney
LM Limb muscle
Liv Liver
Lung Lung
MG Mammary gland
Panc Pancreas
Skin Skin
SI Small intestine
Spl Spleen
Test Testes
Thym Thymus
Tong Tongue
Trac Trachea
Ute Uterus

**Neurotensin/Neuromedin N**
- endothelial

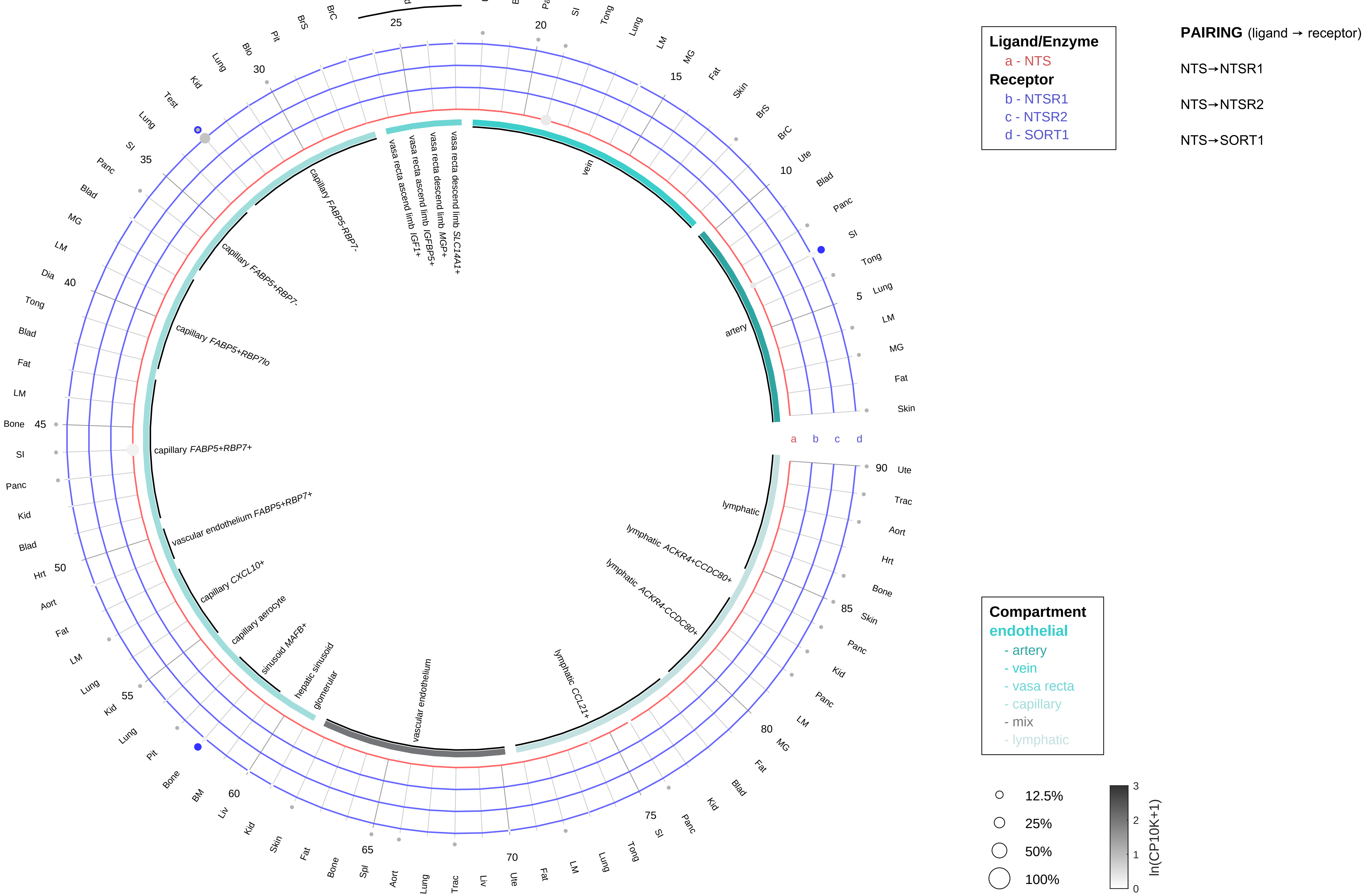

| Abbrev. | Tissue |
|---|---|
| Aort | Aorta |
| Blad | Bladder |
| Blo | Blood |
| Bone | Bone |
| BM | Bone marrow |
| BrC | Brain cortex |
| BrS | Brainstem |
| Col | Colon |
| Dia | Diaphragm |
| Ret | Eye retina |
| Fat | Fat |
| Hrt | Heart |
| Pit | Hypothalamus/Pituitary |
| Kid | Kidney |
| LM | Limb muscle |
| Liv | Liver |
| Lung | Lung |
| MG | Mammary gland |
| Panc | Pancreas |
| Skin | Skin |
| SI | Small intestine |
| Spl | Spleen |
| Test | Testes |
| Thym | Thymus |
| Tong | Tongue |
| Trac | Trachea |
| Ute | Uterus |

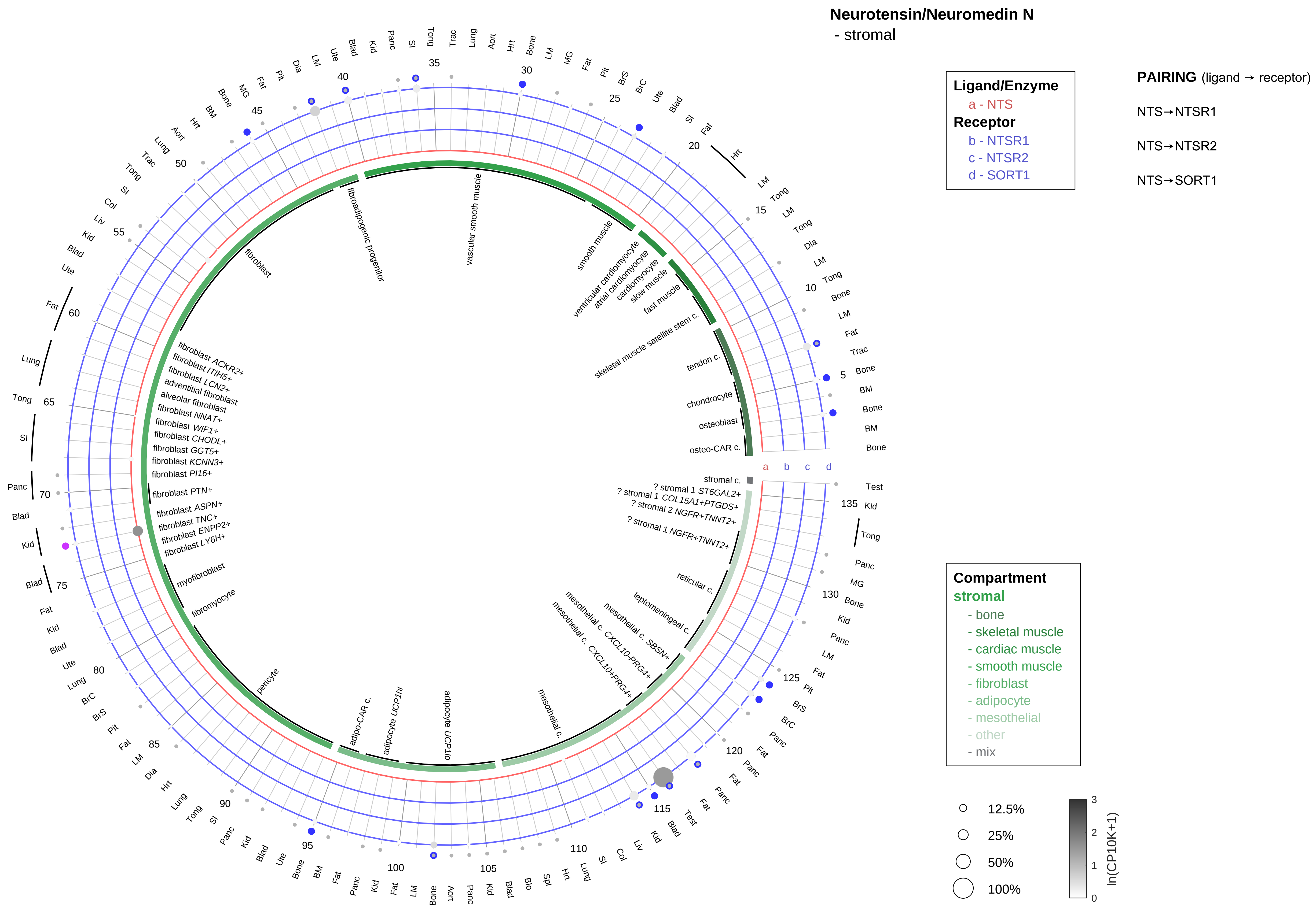

| Abbrev. | Tissue |
|---|---|
| Aort | Aorta |
| Blad | Bladder |
| Blo | Blood |
| Bone | Bone |
| BM | Bone marrow |
| BrC | Brain cortex |
| BrS | Brainstem |
| Col | Colon |
| Dia | Diaphragm |
| Ret | Eye retina |
| Fat | Fat |
| Hrt | Heart |
| Pit | Hypothalamus/Pituitary |
| Kid | Kidney |
| LM | Limb muscle |
| Liv | Liver |
| Lung | Lung |
| MG | Mammary gland |
| Panc | Pancreas |
| Skin | Skin |
| SI | Small intestine |
| Spl | Spleen |
| Test | Testes |
| Thym | Thymus |
| Tong | Tongue |
| Trac | Trachea |
| Ute | Uterus |

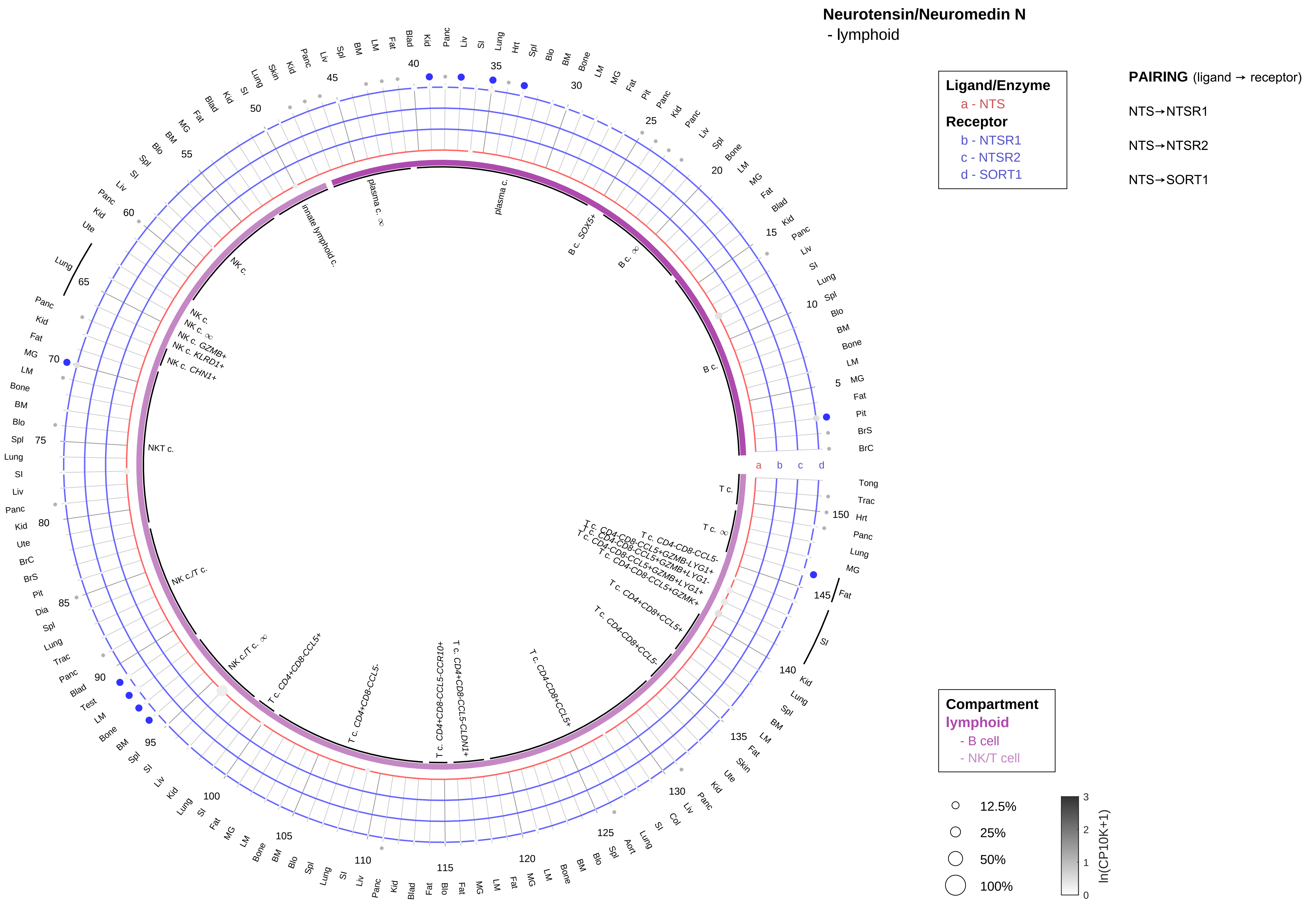

| Abbrev. | Tissue |
|---|---|
| Aort | Aorta |
| Blad | Bladder |
| Blo | Blood |
| Bone | Bone |
| BM | Bone marrow |
| BrC | Brain cortex |
| BrS | Brainstem |
| Col | Colon |
| Dia | Diaphragm |
| Ret | Eye retina |
| Fat | Fat |
| Hrt | Heart |
| Pit | Hypothalamus/Pituitary |
| Kid | Kidney |
| LM | Limb muscle |
| Liv | Liver |
| Lung | Lung |
| MG | Mammary gland |
| Panc | Pancreas |
| Skin | Skin |
| SI | Small intestine |
| Spl | Spleen |
| Test | Testes |
| Thym | Thymus |
| Tong | Tongue |
| Trac | Trachea |
| Ute | Uterus |

**Neurotensin/Neuromedin N**
- myeloid

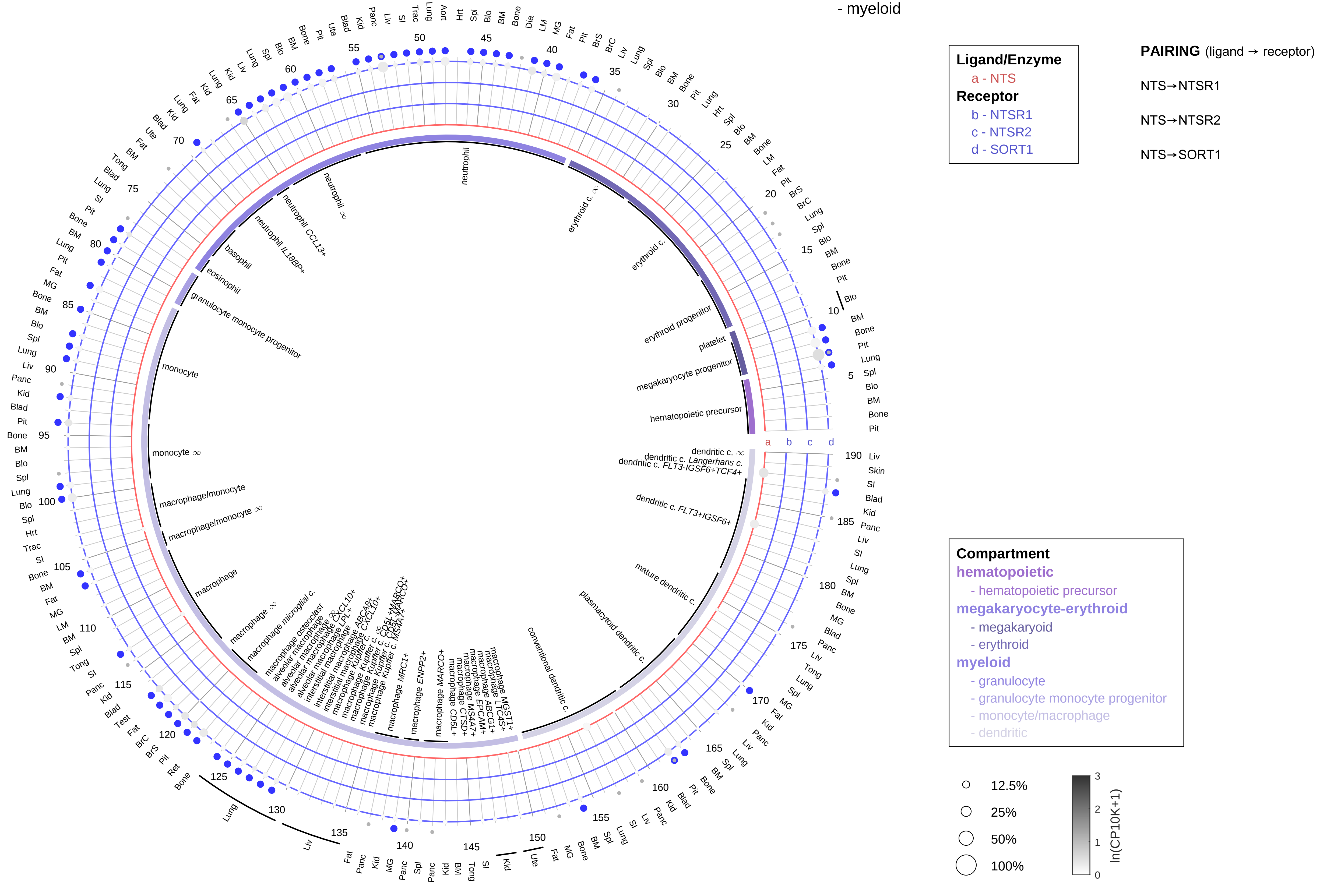

| Abbrev. | Tissue |
|---|---|
| Aort | Aorta |
| Blad | Bladder |
| Blo | Blood |
| Bone | Bone |
| BM | Bone marrow |
| BrC | Brain cortex |
| BrS | Brainstem |
| Col | Colon |
| Dia | Diaphragm |
| Ret | Eye retina |
| Fat | Fat |
| Hrt | Heart |
| Pit | Hypothalamus/Pituitary |
| Kid | Kidney |
| LM | Limb muscle |
| Liv | Liver |
| Lung | Lung |
| MG | Mammary gland |
| Panc | Pancreas |
| Skin | Skin |
| SI | Small intestine |
| Spl | Spleen |
| Test | Testes |
| Thym | Thymus |
| Tong | Tongue |
| Trac | Trachea |
| Ute | Uterus |

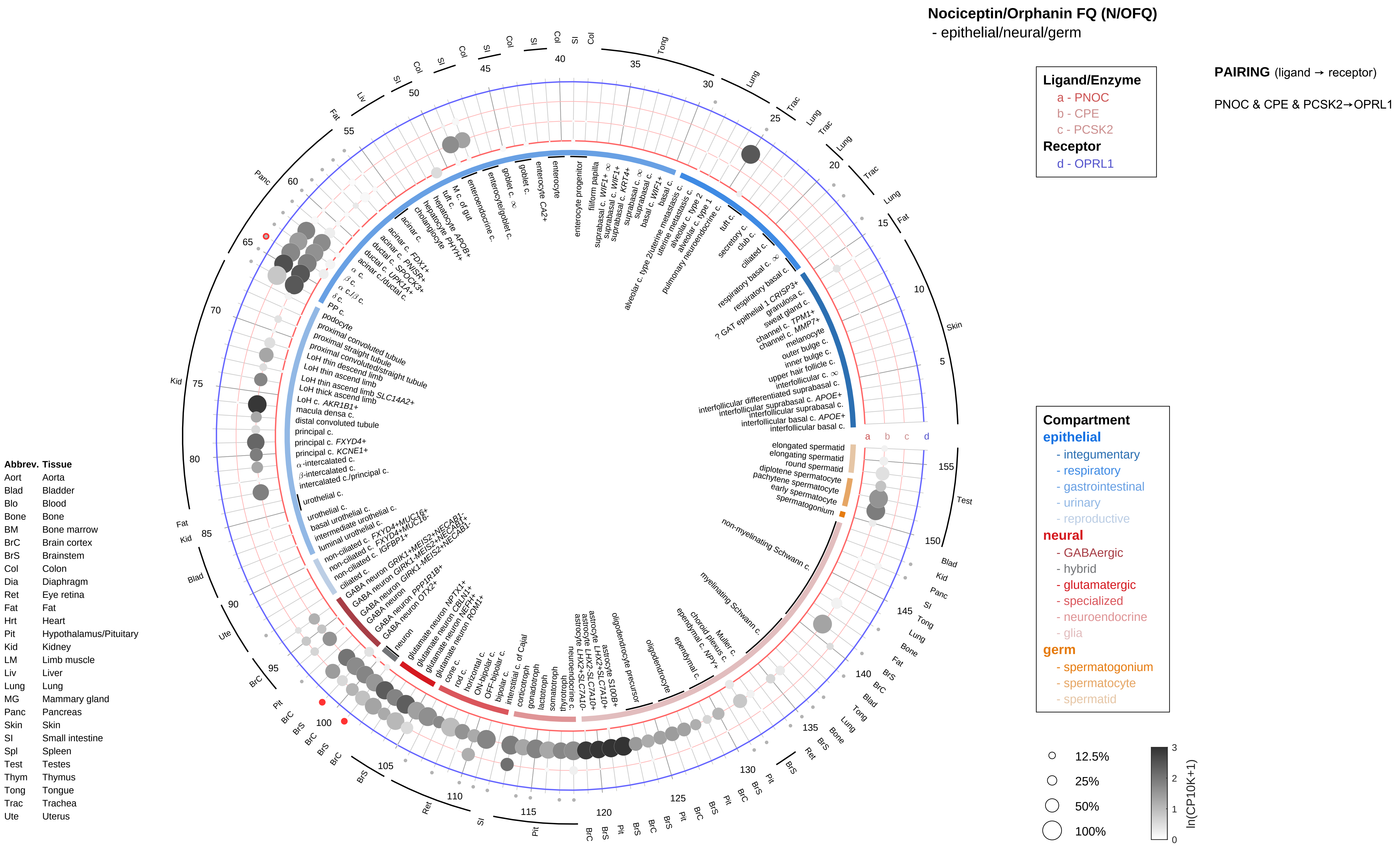
Nociceptin/Orphanin FQ (N/OFQ)
- epithelial/neural/germ
Ligand/Enzyme
a - PNOC
b - CPE
c - PCSK2
Receptor
d - OPRL1
PAIRING (ligand → receptor)
PNOC & CPE & PCSK2→OPRL1
Compartment
epithelial
- integumentary
- respiratory
- gastrointestinal
- urinary
- reproductive
neural
- GABAergic
- hybrid
- glutamatergic
- specialized
- neuroendocrine
- glia
germ
- spermatogonium
- spermatocyte
- spermatid
12.5%
25%
50%
100%
ln(CP10K+1)
Abbrev. Tissue
Aort Aorta
Blad Bladder
Blo Blood
Bone Bone
BM Bone marrow
BrC Brain cortex
BrS Brainstem
Col Colon
Dia Diaphragm
Ret Eye retina
Fat Fat
Hrt Heart
Pit Hypothalamus/Pituitary
Kid Kidney
LM Limb muscle
Liv Liver
Lung Lung
MG Mammary gland
Panc Pancreas
Skin Skin
SI Small intestine
Spl Spleen
Test Testes
Thym Thymus
Tong Tongue
Trac Trachea
Ute Uterus

**Nociceptin/Orphanin FQ (N/OFQ)**
- endothelial

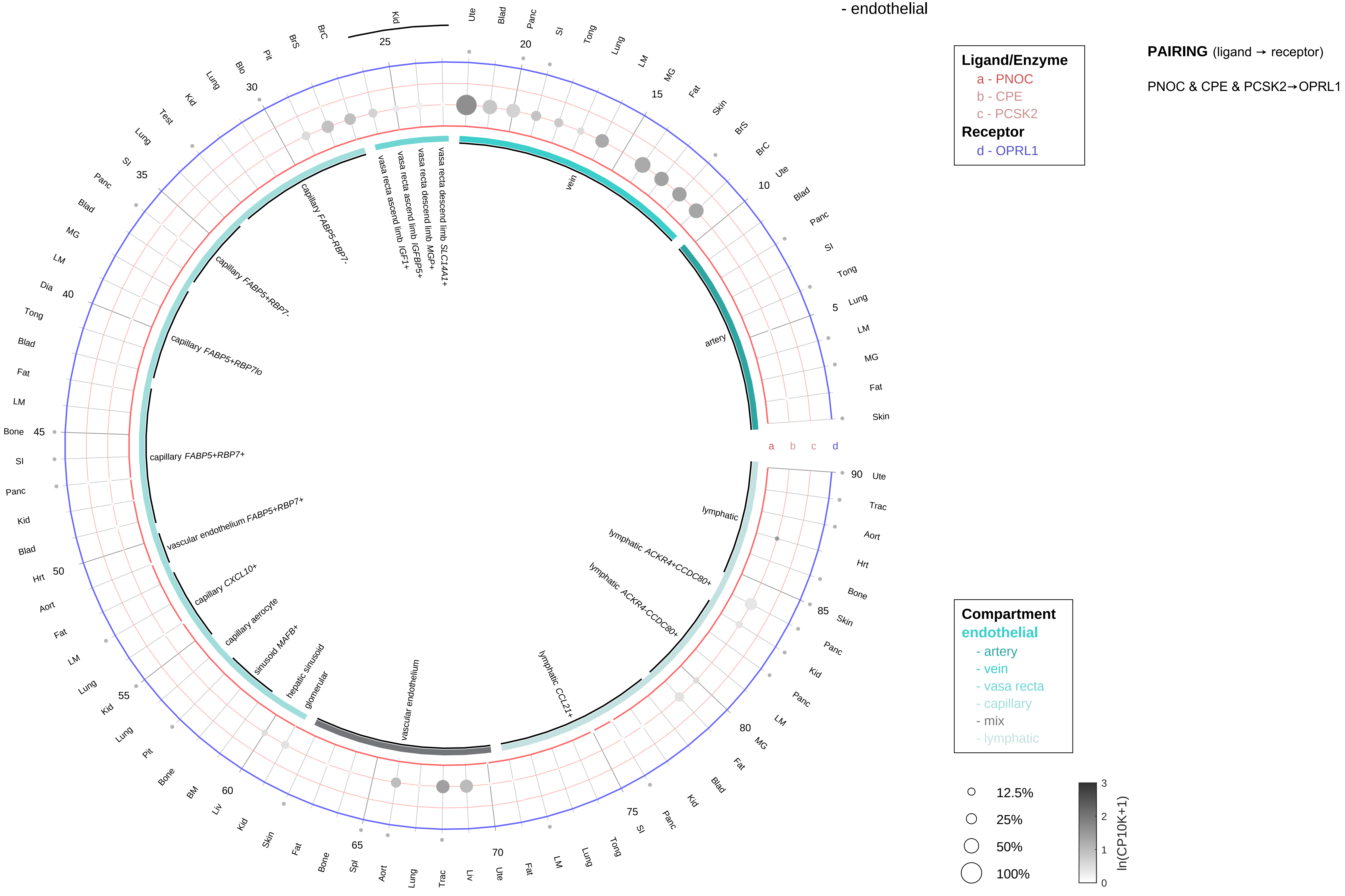

| Abbrev. | Tissue |
|---|---|
| Aort | Aorta |
| Blad | Bladder |
| Blo | Blood |
| Bone | Bone |
| BM | Bone marrow |
| BrC | Brain cortex |
| BrS | Brainstem |
| Col | Colon |
| Dia | Diaphragm |
| Ret | Eye retina |
| Fat | Fat |
| Hrt | Heart |
| Pit | Hypothalamus/Pituitary |
| Kid | Kidney |
| LM | Limb muscle |
| Liv | Liver |
| Lung | Lung |
| MG | Mammary gland |
| Panc | Pancreas |
| Skin | Skin |
| SI | Small intestine |
| Spl | Spleen |
| Test | Testes |
| Thym | Thymus |
| Tong | Tongue |
| Trac | Trachea |
| Ute | Uterus |

**Nociceptin/Orphanin FQ (N/OFQ)**
- stromal

**Ligand/Enzyme**
a - PNOC
b - CPE
c - PCSK2
**Receptor**
d - OPRL1

**PAIRING** (ligand → receptor)

PNOC & CPE & PCSK2→OPRL1

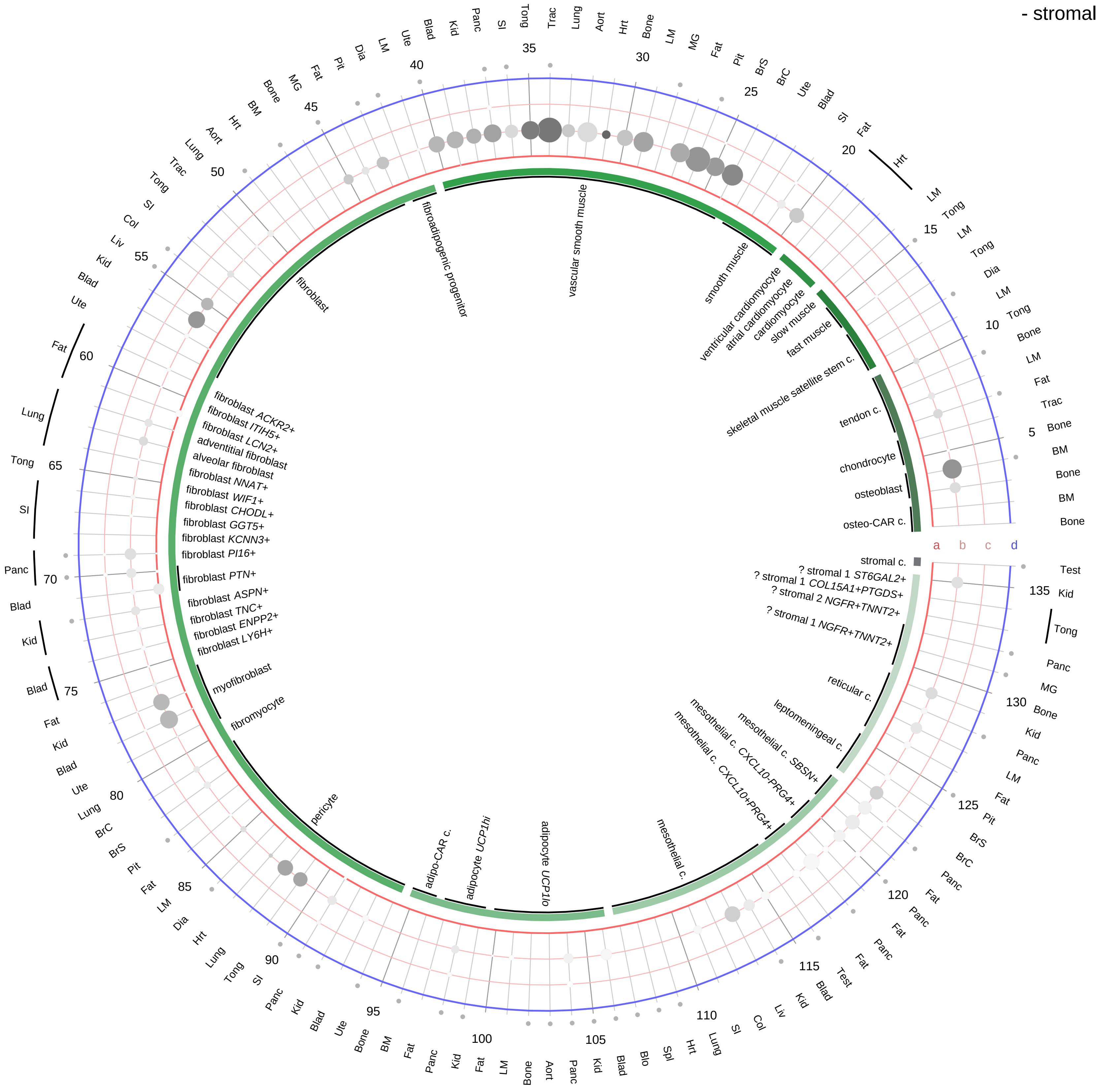

**Compartment**
**stromal**
- bone
- skeletal muscle
- cardiac muscle
- smooth muscle
- fibroblast
- adipocyte
- mesothelial
- other
- mix

12.5%
25%
50%
100%

ln(CP10K+1)
0
1
2
3

| Abbrev. | Tissue |
|---|---|
| Aort | Aorta |
| Blad | Bladder |
| Blo | Blood |
| Bone | Bone |
| BM | Bone marrow |
| BrC | Brain cortex |
| BrS | Brainstem |
| Col | Colon |
| Dia | Diaphragm |
| Ret | Eye retina |
| Fat | Fat |
| Hrt | Heart |
| Pit | Hypothalamus/Pituitary |
| Kid | Kidney |
| LM | Limb muscle |
| Liv | Liver |
| Lung | Lung |
| MG | Mammary gland |
| Panc | Pancreas |
| Skin | Skin |
| SI | Small intestine |
| Spl | Spleen |
| Test | Testes |
| Thym | Thymus |
| Tong | Tongue |
| Trac | Trachea |
| Ute | Uterus |

**Nociceptin/Orphanin FQ (N/OFQ)**
- lymphoid

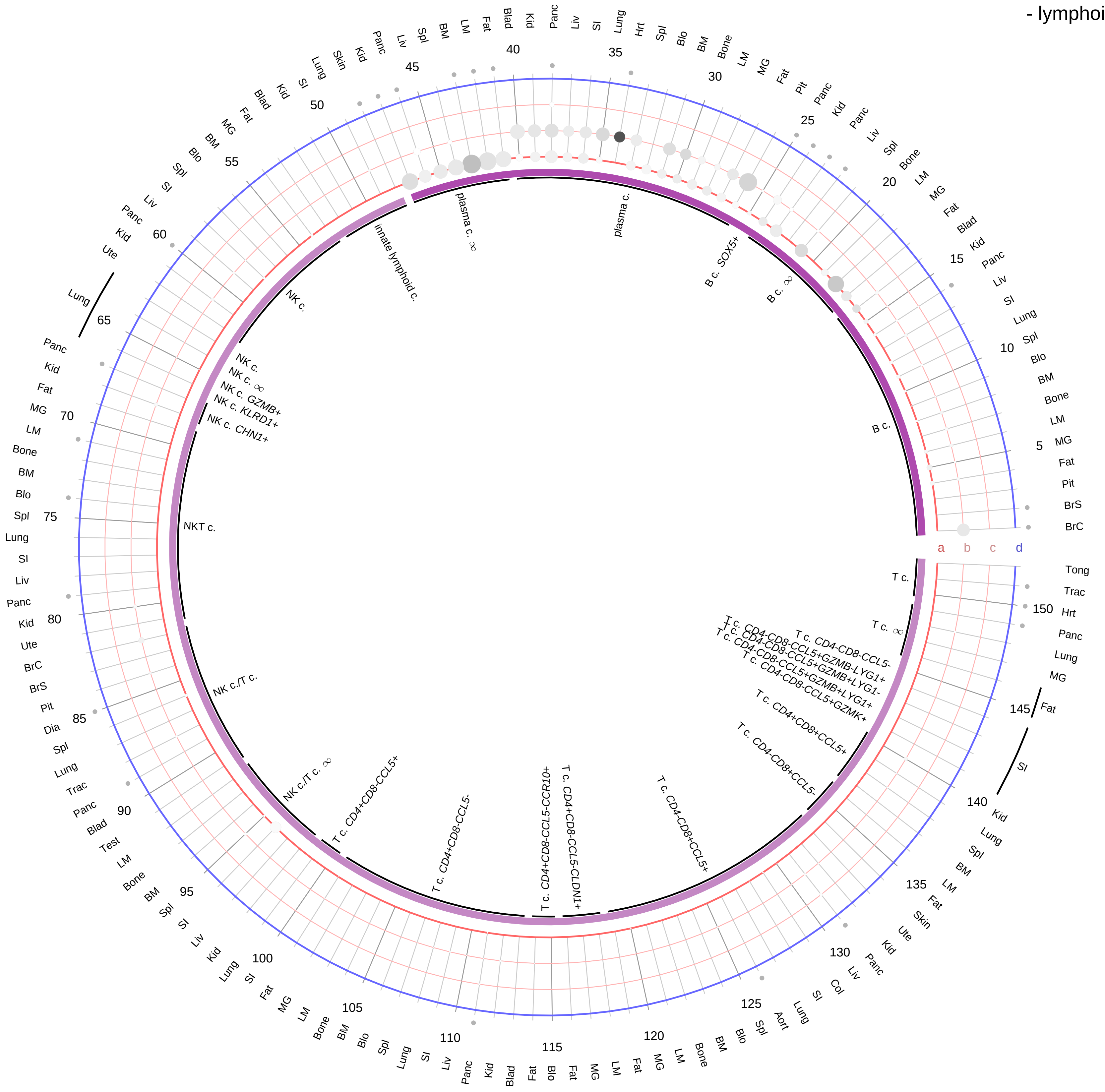

**Ligand/Enzyme**
- a - PNOC
- b - CPE
- c - PCSK2

**Receptor**
- d - OPRL1

**PAIRING** (ligand → receptor)

PNOC & CPE & PCSK2→OPRL1

**Compartment**
**lymphoid**
- B cell
- NK/T cell

12.5%
25%
50%
100%

ln(CP10K+1): 0, 1, 2, 3

| Abbrev. | Tissue |
|---|---|
| Aort | Aorta |
| Blad | Bladder |
| Blo | Blood |
| Bone | Bone |
| BM | Bone marrow |
| BrC | Brain cortex |
| BrS | Brainstem |
| Col | Colon |
| Dia | Diaphragm |
| Ret | Eye retina |
| Fat | Fat |
| Hrt | Heart |
| Pit | Hypothalamus/Pituitary |
| Kid | Kidney |
| LM | Limb muscle |
| Liv | Liver |
| Lung | Lung |
| MG | Mammary gland |
| Panc | Pancreas |
| Skin | Skin |
| SI | Small intestine |
| Spl | Spleen |
| Test | Testes |
| Thym | Thymus |
| Tong | Tongue |
| Trac | Trachea |
| Ute | Uterus |

**Nociceptin/Orphanin FQ (N/OFQ)**
- myeloid

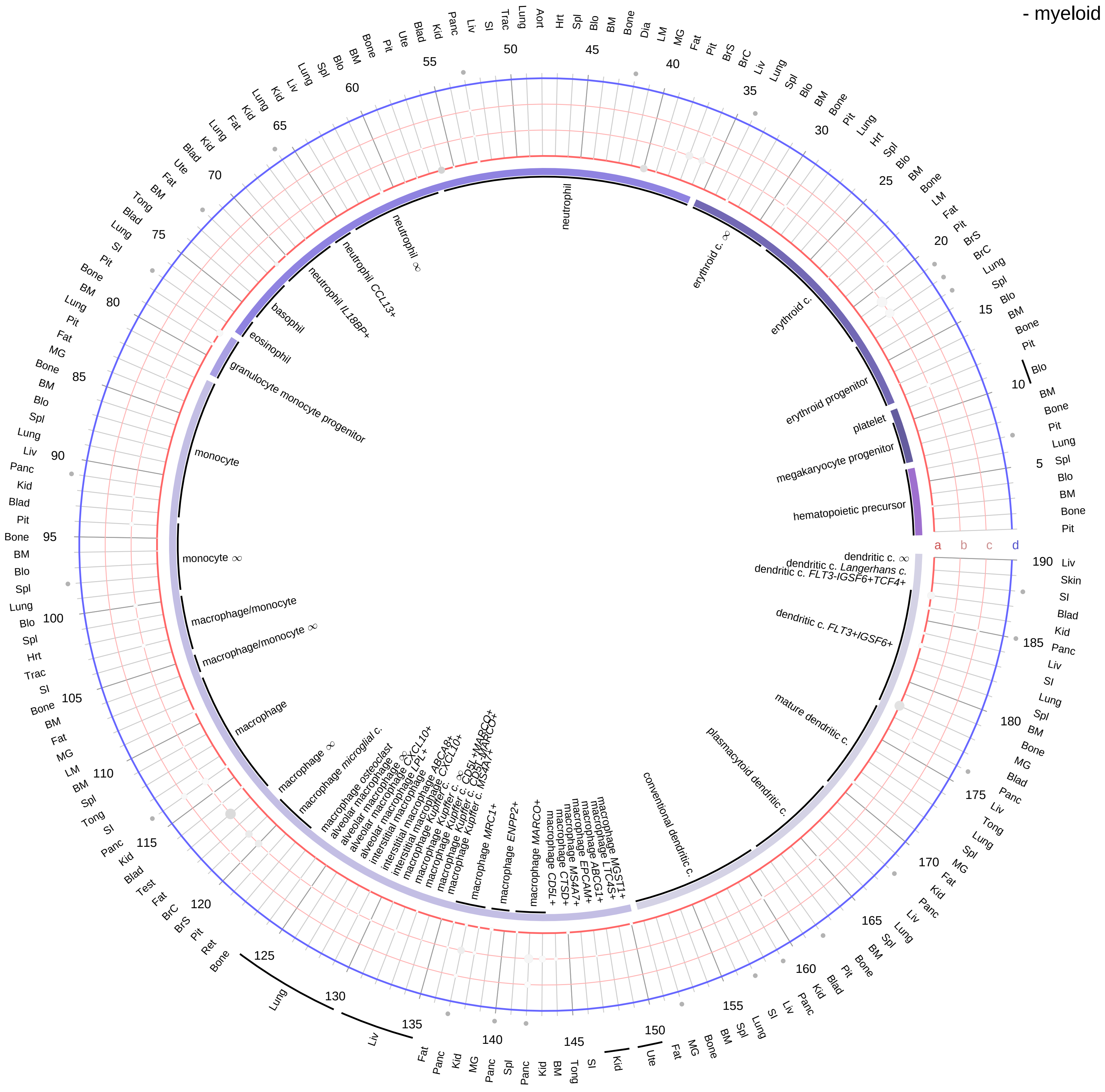

**Ligand/Enzyme**
a - PNOC
b - CPE
c - PCSK2
**Receptor**
d - OPRL1

**PAIRING** (ligand → receptor)

PNOC & CPE & PCSK2→OPRL1

**Compartment**
**hematopoietic**
- hematopoietic precursor
**megakaryocyte-erythroid**
- megakaryoid
- erythroid
**myeloid**
- granulocyte
- granulocyte monocyte progenitor
- monocyte/macrophage
- dendritic

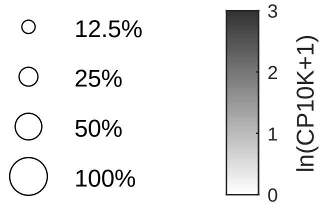

| Abbrev. | Tissue |
|---|---|
| Aort | Aorta |
| Blad | Bladder |
| Blo | Blood |
| Bone | Bone |
| BM | Bone marrow |
| BrC | Brain cortex |
| BrS | Brainstem |
| Col | Colon |
| Dia | Diaphragm |
| Ret | Eye retina |
| Fat | Fat |
| Hrt | Heart |
| Pit | Hypothalamus/Pituitary |
| Kid | Kidney |
| LM | Limb muscle |
| Liv | Liver |
| Lung | Lung |
| MG | Mammary gland |
| Panc | Pancreas |
| Skin | Skin |
| SI | Small intestine |
| Spl | Spleen |
| Test | Testes |
| Thym | Thymus |
| Tong | Tongue |
| Trac | Trachea |
| Ute | Uterus |

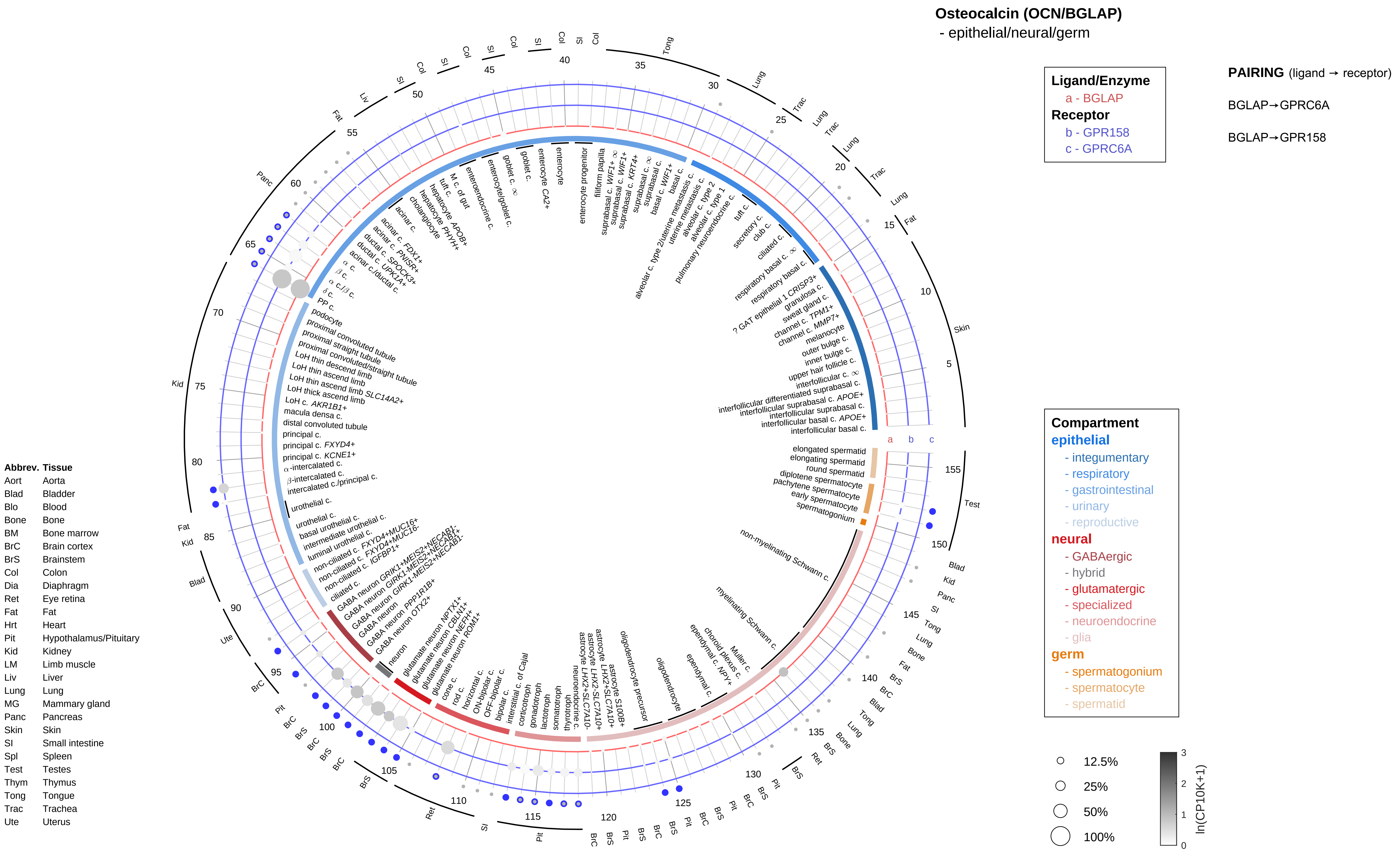

Osteocalcin (OCN/BGLAP)
- epithelial/neural/germ
Ligand/Enzyme
a - BGLAP
Receptor
b - GPR158
c - GPRC6A
PAIRING (ligand → receptor)
BGLAP→GPRC6A
BGLAP→GPR158
Compartment
epithelial
- integumentary
- respiratory
- gastrointestinal
- urinary
- reproductive
neural
- GABAergic
- hybrid
- glutamatergic
- specialized
- neuroendocrine
- glia
germ
- spermatogonium
- spermatocyte
- spermatid
12.5%
25%
50%
100%
ln(CP10K+1)
Abbrev. Tissue
Aort Aorta
Blad Bladder
Blo Blood
Bone Bone
BM Bone marrow
BrC Brain cortex
BrS Brainstem
Col Colon
Dia Diaphragm
Ret Eye retina
Fat Fat
Hrt Heart
Pit Hypothalamus/Pituitary
Kid Kidney
LM Limb muscle
Liv Liver
Lung Lung
MG Mammary gland
Panc Pancreas
Skin Skin
SI Small intestine
Spl Spleen
Test Testes
Thym Thymus
Tong Tongue
Trac Trachea
Ute Uterus

**Osteocalcin (OCN/BGLAP)**
- endothelial

**Ligand/Enzyme**
a - BGLAP
**Receptor**
b - GPR158
c - GPRC6A

**PAIRING** (ligand → receptor)

BGLAP→GPRC6A

BGLAP→GPR158

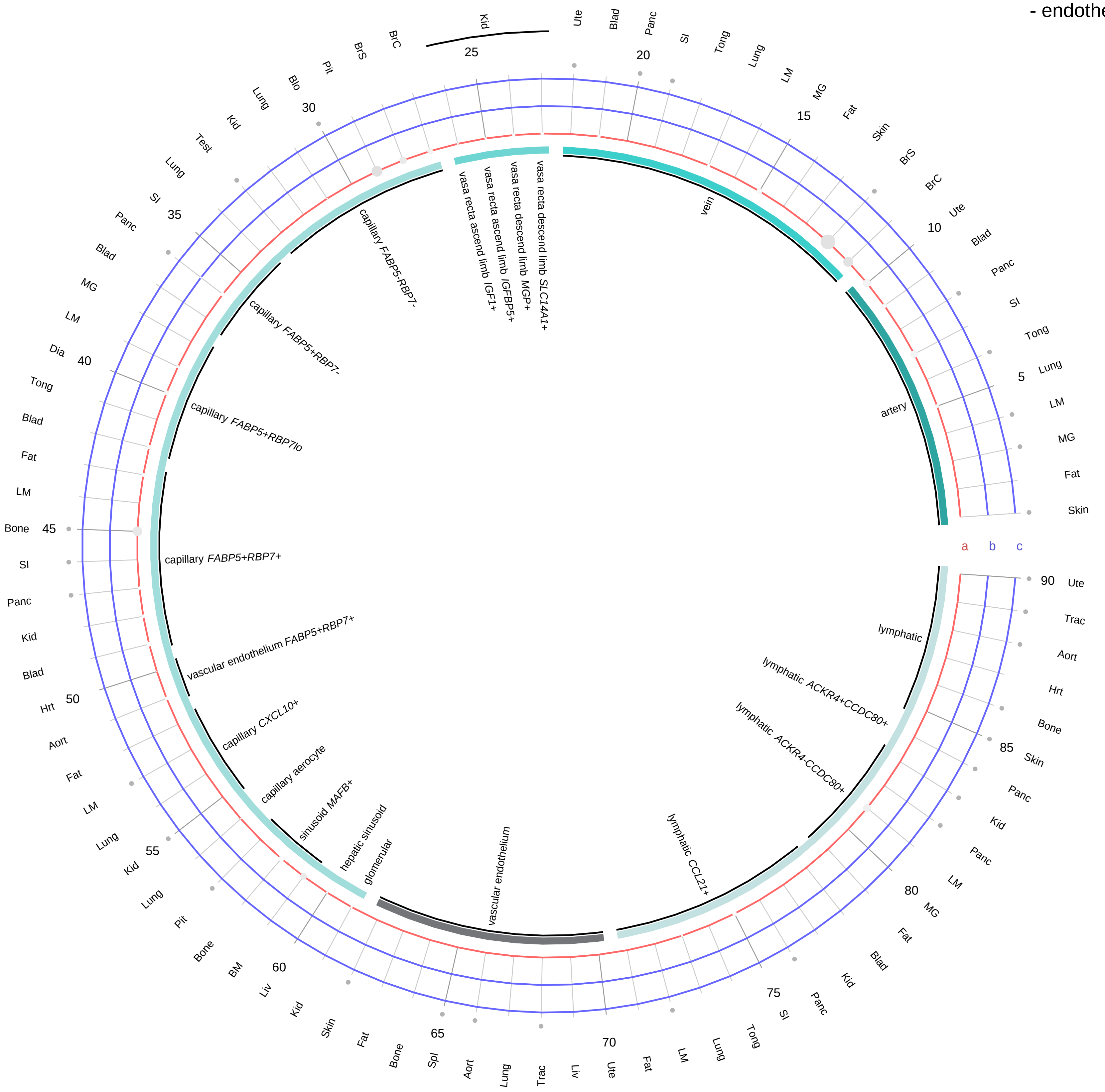

| Abbrev. | Tissue |
|---|---|
| Aort | Aorta |
| Blad | Bladder |
| Blo | Blood |
| Bone | Bone |
| BM | Bone marrow |
| BrC | Brain cortex |
| BrS | Brainstem |
| Col | Colon |
| Dia | Diaphragm |
| Ret | Eye retina |
| Fat | Fat |
| Hrt | Heart |
| Pit | Hypothalamus/Pituitary |
| Kid | Kidney |
| LM | Limb muscle |
| Liv | Liver |
| Lung | Lung |
| MG | Mammary gland |
| Panc | Pancreas |
| Skin | Skin |
| SI | Small intestine |
| Spl | Spleen |
| Test | Testes |
| Thym | Thymus |
| Tong | Tongue |
| Trac | Trachea |
| Ute | Uterus |

**Compartment**
**endothelial**
- artery
- vein
- vasa recta
- capillary
- mix
- lymphatic

12.5%
25%
50%
100%

ln(CP10K+1)
0
1
2
3

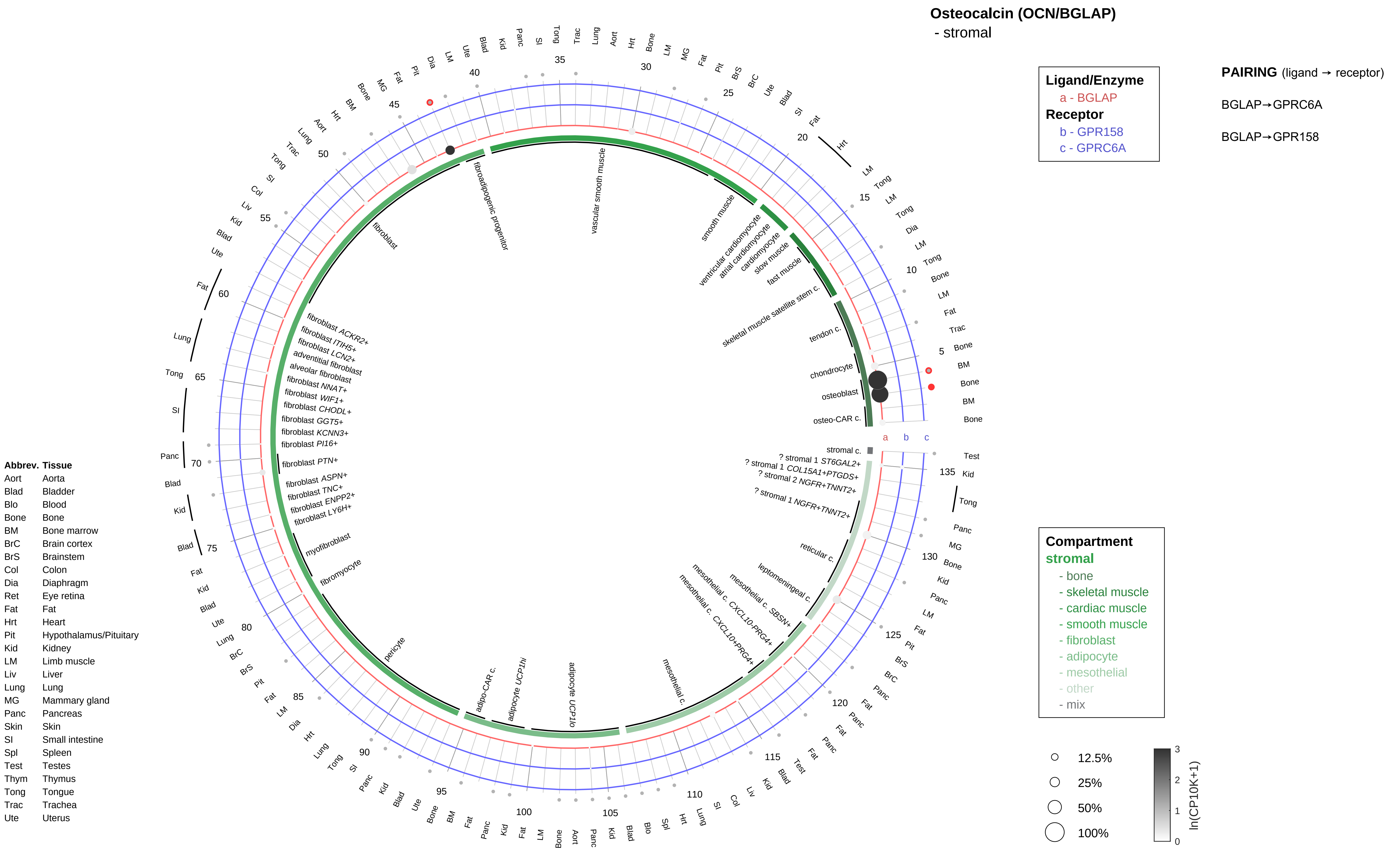
Osteocalcin (OCN/BGLAP)
- stromal
Ligand/Enzyme
a - BGLAP
Receptor
b - GPR158
c - GPRC6A
PAIRING (ligand → receptor)
BGLAP→GPRC6A
BGLAP→GPR158
Compartment
stromal
- bone
- skeletal muscle
- cardiac muscle
- smooth muscle
- fibroblast
- adipocyte
- mesothelial
- other
- mix
12.5%
25%
50%
100%
ln(CP10K+1)
Abbrev. Tissue
Aort Aorta
Blad Bladder
Blo Blood
Bone Bone
BM Bone marrow
BrC Brain cortex
BrS Brainstem
Col Colon
Dia Diaphragm
Ret Eye retina
Fat Fat
Hrt Heart
Pit Hypothalamus/Pituitary
Kid Kidney
LM Limb muscle
Liv Liver
Lung Lung
MG Mammary gland
Panc Pancreas
Skin Skin
SI Small intestine
Spl Spleen
Test Testes
Thym Thymus
Tong Tongue
Trac Trachea
Ute Uterus
fibroadipogenic progenitor
vascular smooth muscle
smooth muscle
ventricular cardiomyocyte
atrial cardiomyocyte
cardiomyocyte
slow muscle
fast muscle
skeletal muscle satellite stem c.
tendon c.
chondrocyte
osteoblast
osteo-CAR c.
stromal c.
? stromal 1 ST6GAL2+
? stromal 1 COL15A1+PTGDS+
? stromal 2 NGFR+TNNT2+
? stromal 1 NGFR+TNNT2+
reticular c.
leptomeningeal c.
mesothelial c. SBSN+
mesothelial c. CXCL10-PRG4+
mesothelial c. CXCL10+PRG4+
mesothelial c.
adipocyte UCP1lo
adipocyte UCP1hi
adipo-CAR c.
pericyte
fibromyocyte
myofibroblast
fibroblast LY6H+
fibroblast ENPP2+
fibroblast TNC+
fibroblast ASPN+
fibroblast PTN+
fibroblast PI16+
fibroblast KCNN3+
fibroblast GGT5+
fibroblast CHODL+
fibroblast WIF1+
fibroblast NNAT+
alveolar fibroblast
adventitial fibroblast
fibroblast LCN2+
fibroblast ITIH5+
fibroblast ACKR2+
fibroblast

**Osteocalcin (OCN/BGLAP)**
- lymphoid

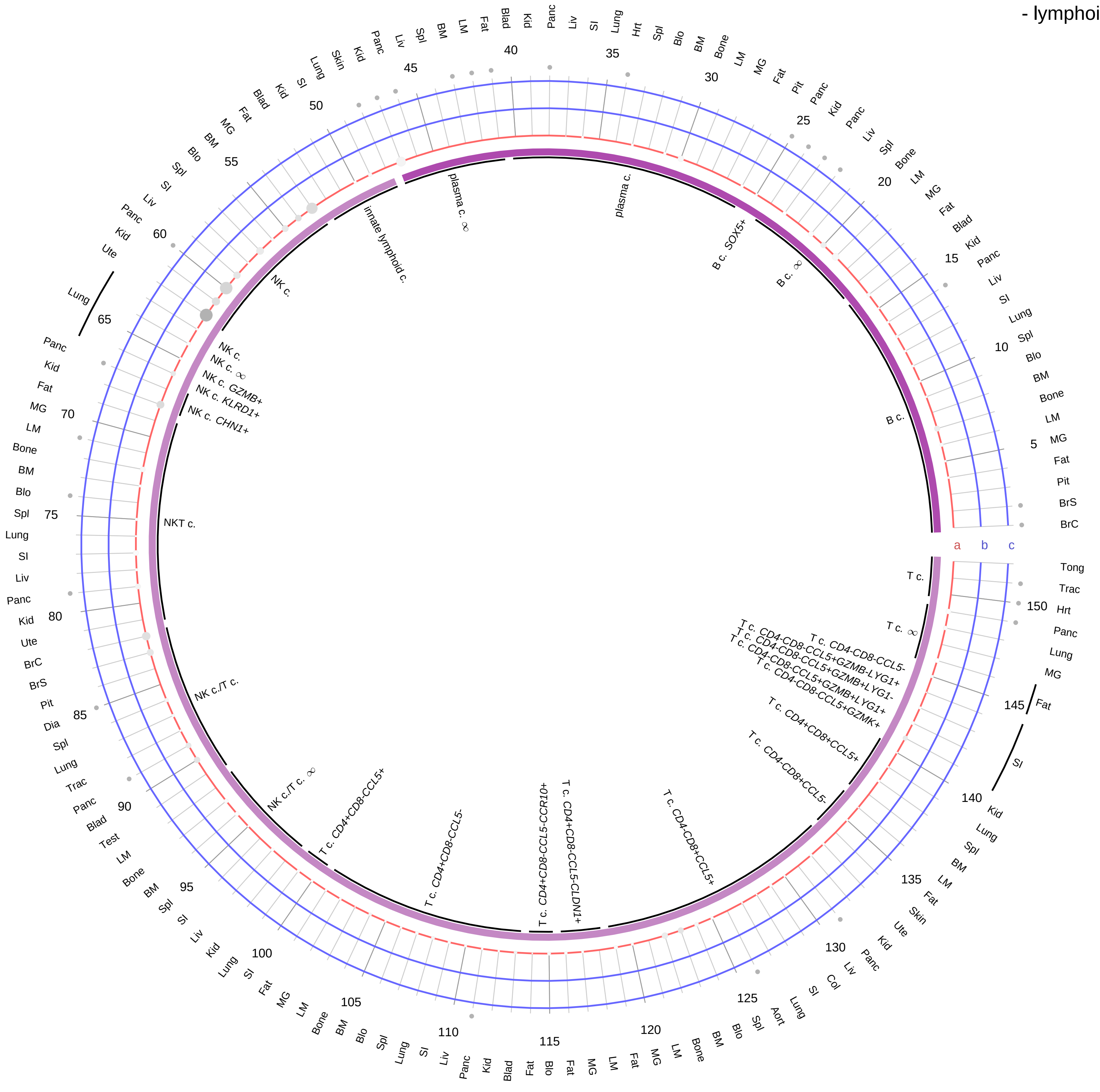

**Ligand/Enzyme**
a - BGLAP
**Receptor**
b - GPR158
c - GPRC6A

**PAIRING** (ligand → receptor)

BGLAP→GPRC6A

BGLAP→GPR158

| Abbrev. | Tissue |
|---|---|
| Aort | Aorta |
| Blad | Bladder |
| Blo | Blood |
| Bone | Bone |
| BM | Bone marrow |
| BrC | Brain cortex |
| BrS | Brainstem |
| Col | Colon |
| Dia | Diaphragm |
| Ret | Eye retina |
| Fat | Fat |
| Hrt | Heart |
| Pit | Hypothalamus/Pituitary |
| Kid | Kidney |
| LM | Limb muscle |
| Liv | Liver |
| Lung | Lung |
| MG | Mammary gland |
| Panc | Pancreas |
| Skin | Skin |
| SI | Small intestine |
| Spl | Spleen |
| Test | Testes |
| Thym | Thymus |
| Tong | Tongue |
| Trac | Trachea |
| Ute | Uterus |

**Compartment**
**lymphoid**
- B cell
- NK/T cell

12.5%
25%
50%
100%

ln(CP10K+1): 0 1 2 3

**Osteocalcin (OCN/BGLAP)**
- myeloid

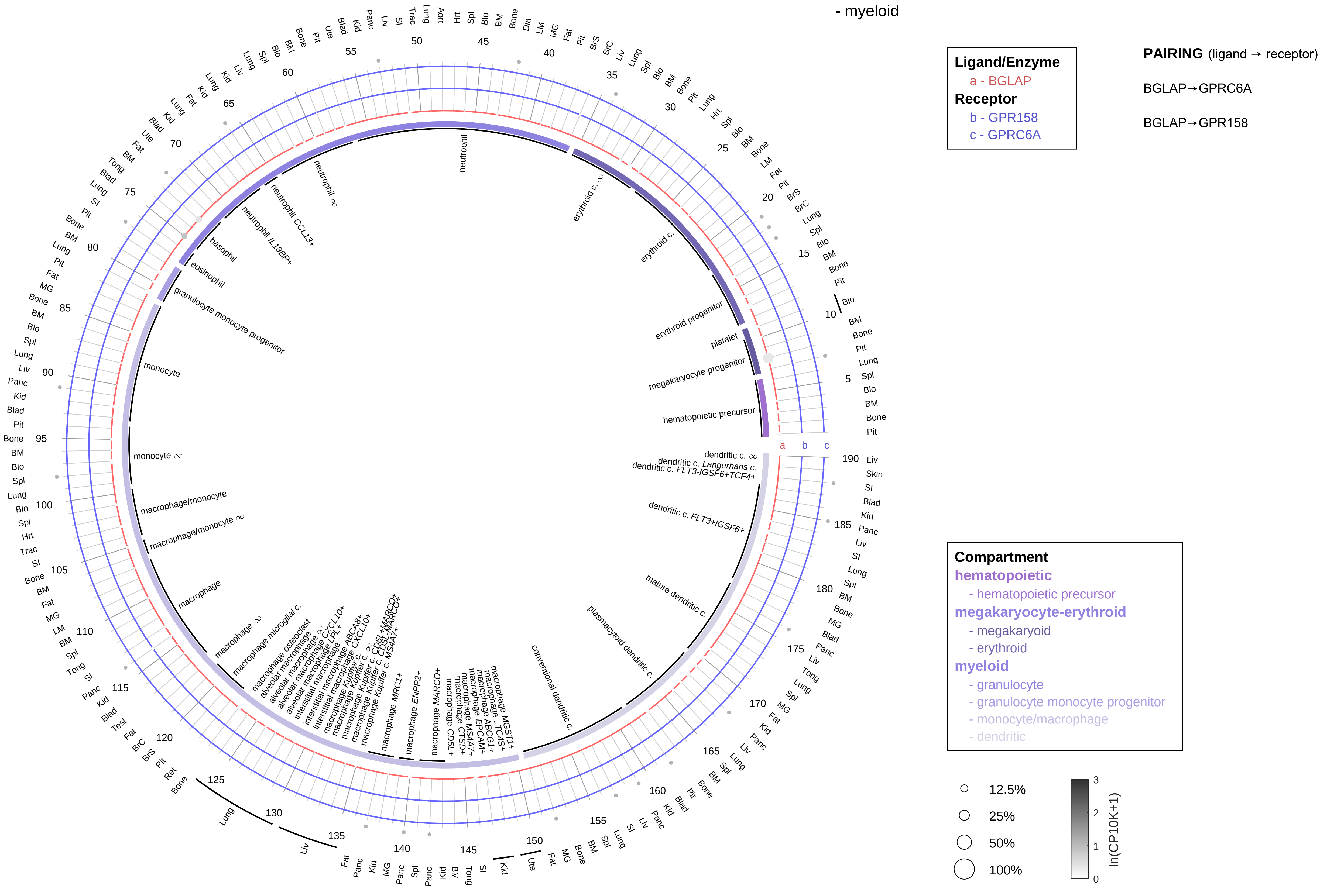

| Abbrev. | Tissue |
|---|---|
| Aort | Aorta |
| Blad | Bladder |
| Blo | Blood |
| Bone | Bone |
| BM | Bone marrow |
| BrC | Brain cortex |
| BrS | Brainstem |
| Col | Colon |
| Dia | Diaphragm |
| Ret | Eye retina |
| Fat | Fat |
| Hrt | Heart |
| Pit | Hypothalamus/Pituitary |
| Kid | Kidney |
| LM | Limb muscle |
| Liv | Liver |
| Lung | Lung |
| MG | Mammary gland |
| Panc | Pancreas |
| Skin | Skin |
| SI | Small intestine |
| Spl | Spleen |
| Test | Testes |
| Thym | Thymus |
| Tong | Tongue |
| Trac | Trachea |
| Ute | Uterus |

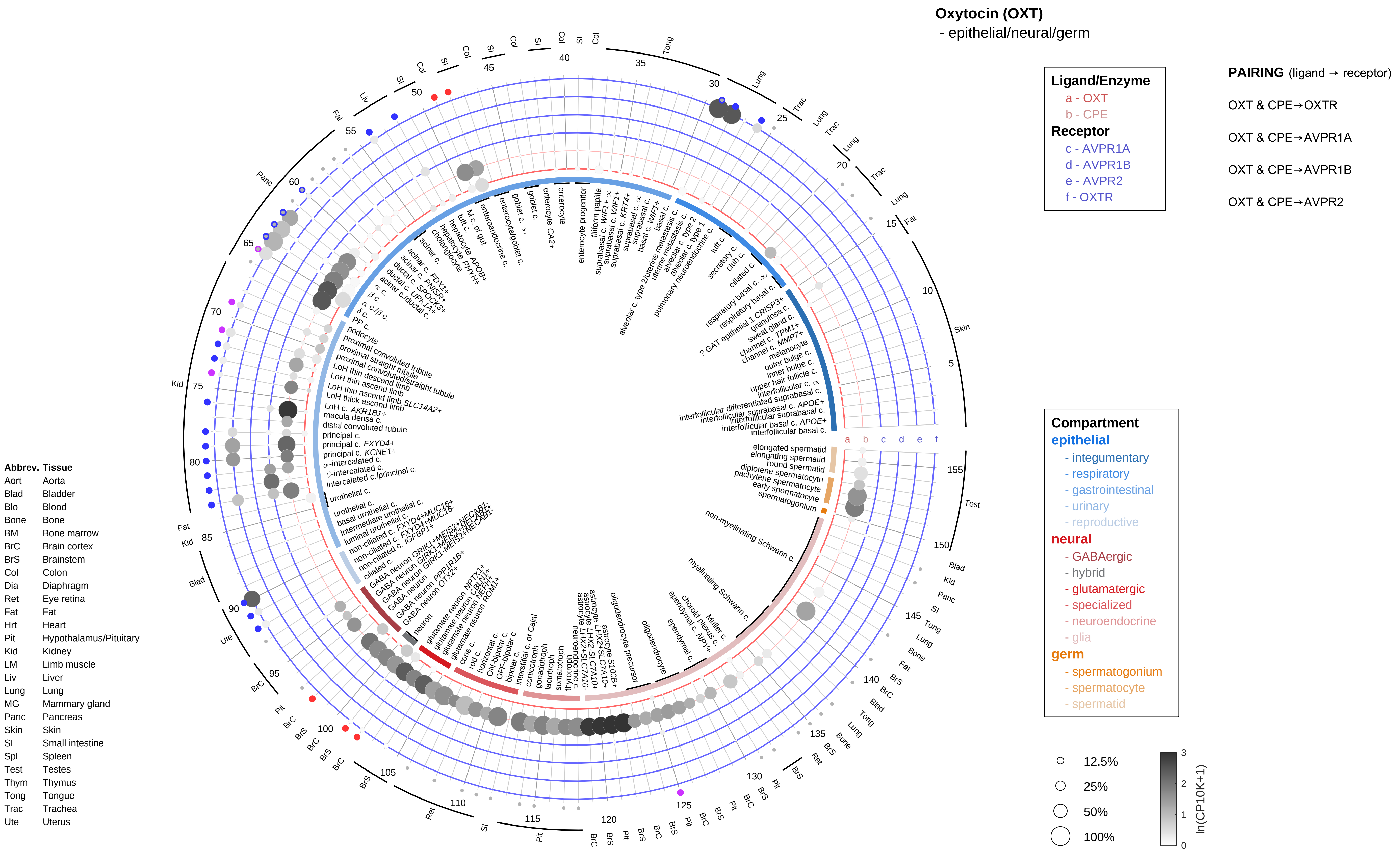

Oxytocin (OXT)
- epithelial/neural/germ
Ligand/Enzyme
a - OXT
b - CPE
Receptor
c - AVPR1A
d - AVPR1B
e - AVPR2
f - OXTR
PAIRING (ligand → receptor)
OXT & CPE→OXTR
OXT & CPE→AVPR1A
OXT & CPE→AVPR1B
OXT & CPE→AVPR2
Compartment
epithelial
- integumentary
- respiratory
- gastrointestinal
- urinary
- reproductive
neural
- GABAergic
- hybrid
- glutamatergic
- specialized
- neuroendocrine
- glia
germ
- spermatogonium
- spermatocyte
- spermatid
12.5%
25%
50%
100%
ln(CP10K+1)
Abbrev. Tissue
Aort Aorta
Blad Bladder
Blo Blood
Bone Bone
BM Bone marrow
BrC Brain cortex
BrS Brainstem
Col Colon
Dia Diaphragm
Ret Eye retina
Fat Fat
Hrt Heart
Pit Hypothalamus/Pituitary
Kid Kidney
LM Limb muscle
Liv Liver
Lung Lung
MG Mammary gland
Panc Pancreas
Skin Skin
SI Small intestine
Spl Spleen
Test Testes
Thym Thymus
Tong Tongue
Trac Trachea
Ute Uterus

#### Oxytocin (OXT)
- endothelial

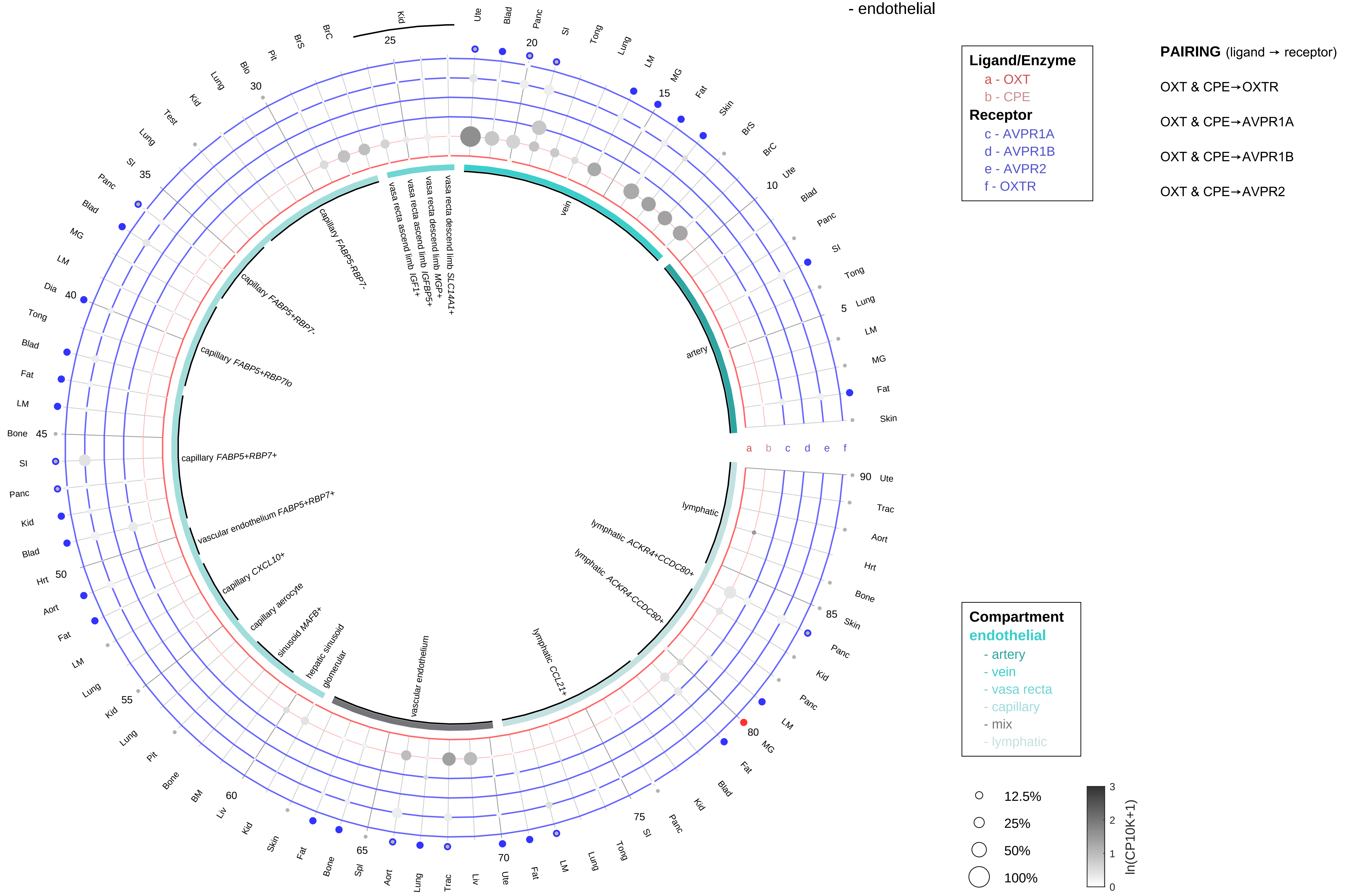

| Abbrev. | Tissue |
|---|---|
| Aort | Aorta |
| Blad | Bladder |
| Blo | Blood |
| Bone | Bone |
| BM | Bone marrow |
| BrC | Brain cortex |
| BrS | Brainstem |
| Col | Colon |
| Dia | Diaphragm |
| Ret | Eye retina |
| Fat | Fat |
| Hrt | Heart |
| Pit | Hypothalamus/Pituitary |
| Kid | Kidney |
| LM | Limb muscle |
| Liv | Liver |
| Lung | Lung |
| MG | Mammary gland |
| Panc | Pancreas |
| Skin | Skin |
| SI | Small intestine |
| Spl | Spleen |
| Test | Testes |
| Thym | Thymus |
| Tong | Tongue |
| Trac | Trachea |
| Ute | Uterus |

**Oxytocin (OXT)**
- stromal

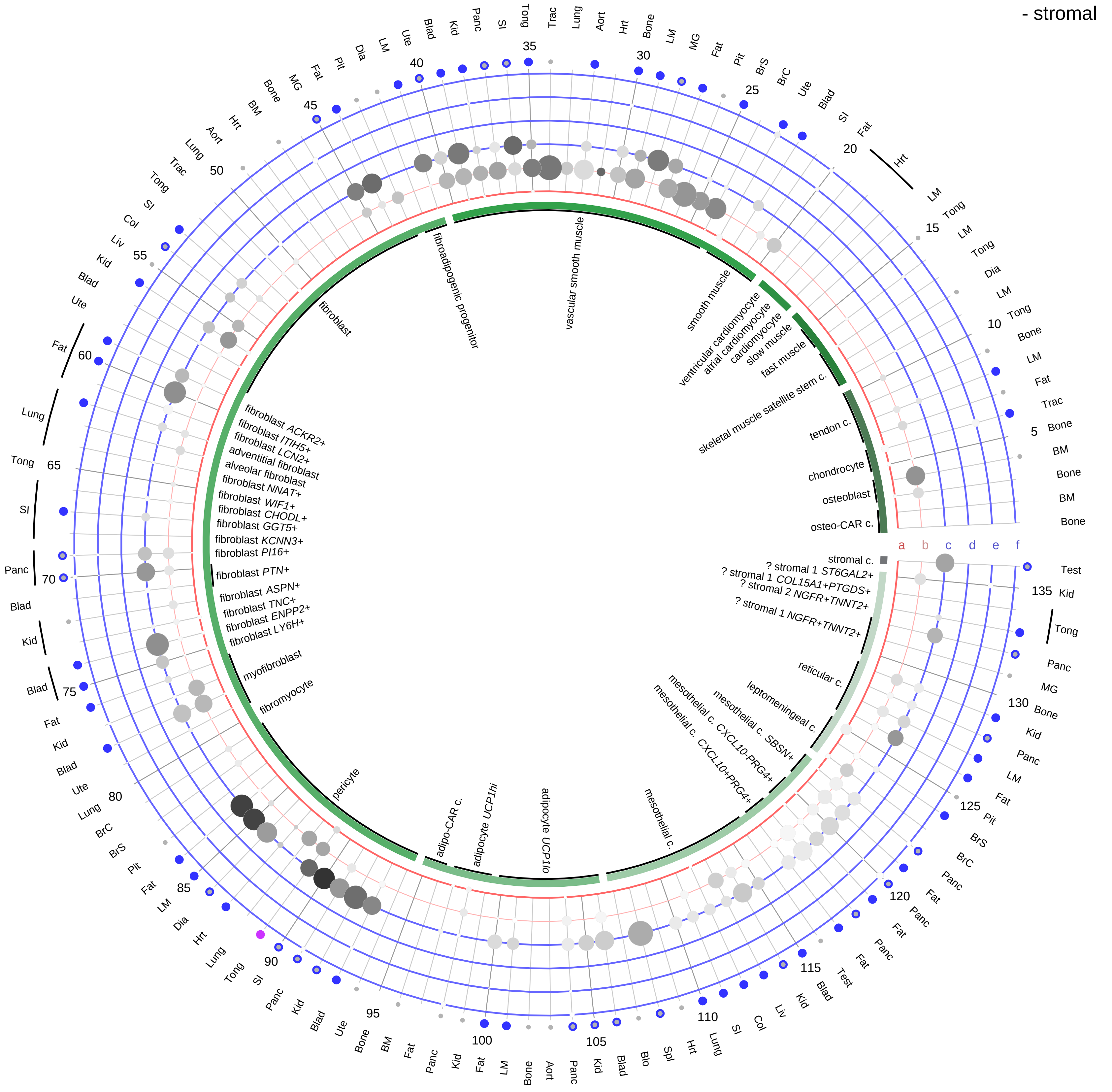

**Ligand/Enzyme**
- a - OXT
- b - CPE

**Receptor**
- c - AVPR1A
- d - AVPR1B
- e - AVPR2
- f - OXTR

**PAIRING** (ligand → receptor)

OXT & CPE→OXTR

OXT & CPE→AVPR1A

OXT & CPE→AVPR1B

OXT & CPE→AVPR2

| Abbrev. | Tissue |
|---|---|
| Aort | Aorta |
| Blad | Bladder |
| Blo | Blood |
| Bone | Bone |
| BM | Bone marrow |
| BrC | Brain cortex |
| BrS | Brainstem |
| Col | Colon |
| Dia | Diaphragm |
| Ret | Eye retina |
| Fat | Fat |
| Hrt | Heart |
| Pit | Hypothalamus/Pituitary |
| Kid | Kidney |
| LM | Limb muscle |
| Liv | Liver |
| Lung | Lung |
| MG | Mammary gland |
| Panc | Pancreas |
| Skin | Skin |
| SI | Small intestine |
| Spl | Spleen |
| Test | Testes |
| Thym | Thymus |
| Tong | Tongue |
| Trac | Trachea |
| Ute | Uterus |

**Compartment**
**stromal**
- bone
- skeletal muscle
- cardiac muscle
- smooth muscle
- fibroblast
- adipocyte
- mesothelial
- other
- mix

12.5%
25%
50%
100%

ln(CP10K+1) 0 1 2 3

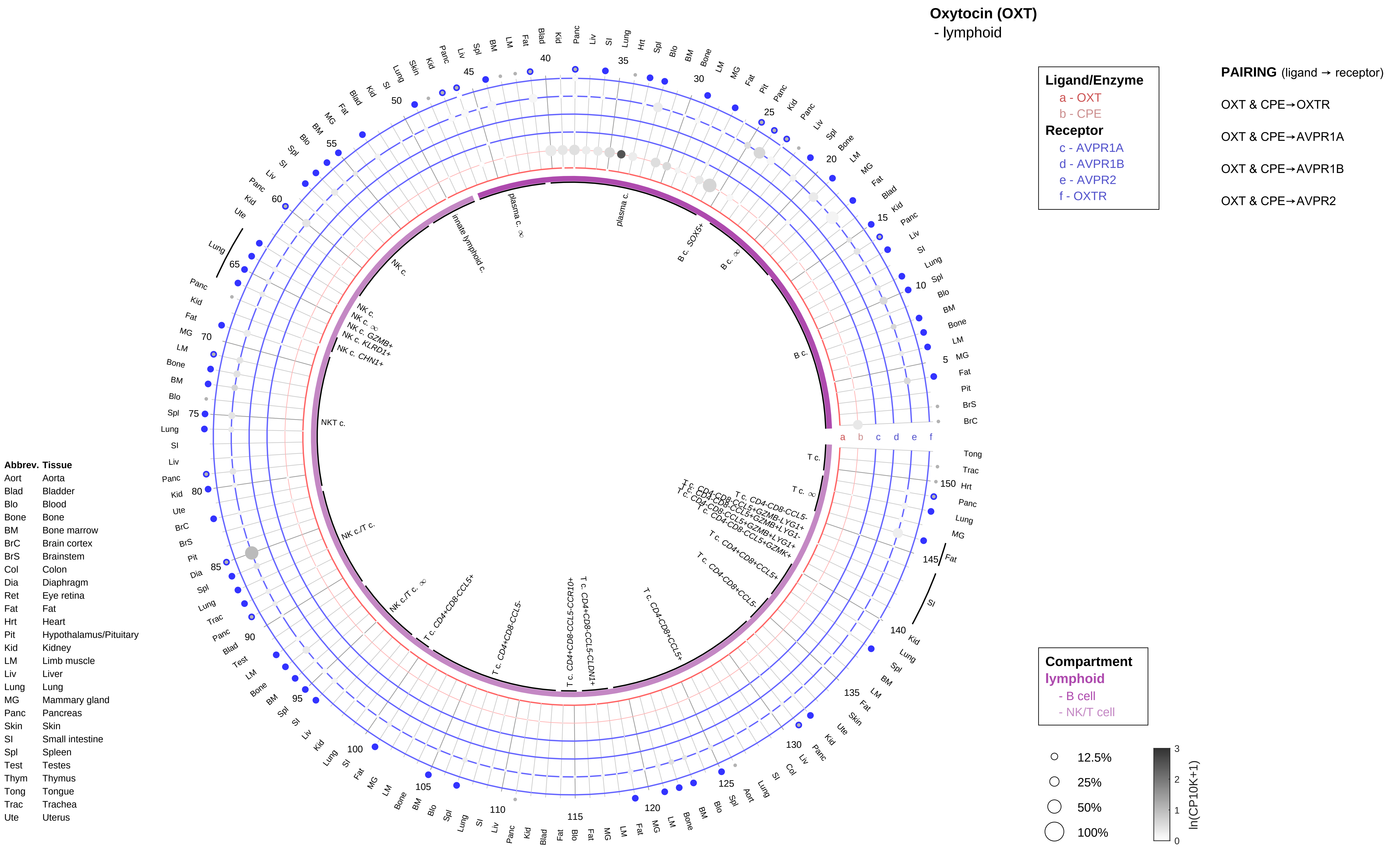

Oxytocin (OXT)
- lymphoid
Ligand/Enzyme
a - OXT
b - CPE
Receptor
c - AVPR1A
d - AVPR1B
e - AVPR2
f - OXTR
PAIRING (ligand → receptor)
OXT & CPE→OXTR
OXT & CPE→AVPR1A
OXT & CPE→AVPR1B
OXT & CPE→AVPR2
Compartment
lymphoid
- B cell
- NK/T cell
12.5%
25%
50%
100%
ln(CP10K+1)
Abbrev. Tissue
Aort Aorta
Blad Bladder
Blo Blood
Bone Bone
BM Bone marrow
BrC Brain cortex
BrS Brainstem
Col Colon
Dia Diaphragm
Ret Eye retina
Fat Fat
Hrt Heart
Pit Hypothalamus/Pituitary
Kid Kidney
LM Limb muscle
Liv Liver
Lung Lung
MG Mammary gland
Panc Pancreas
Skin Skin
SI Small intestine
Spl Spleen
Test Testes
Thym Thymus
Tong Tongue
Trac Trachea
Ute Uterus

**Oxytocin (OXT)**
- myeloid

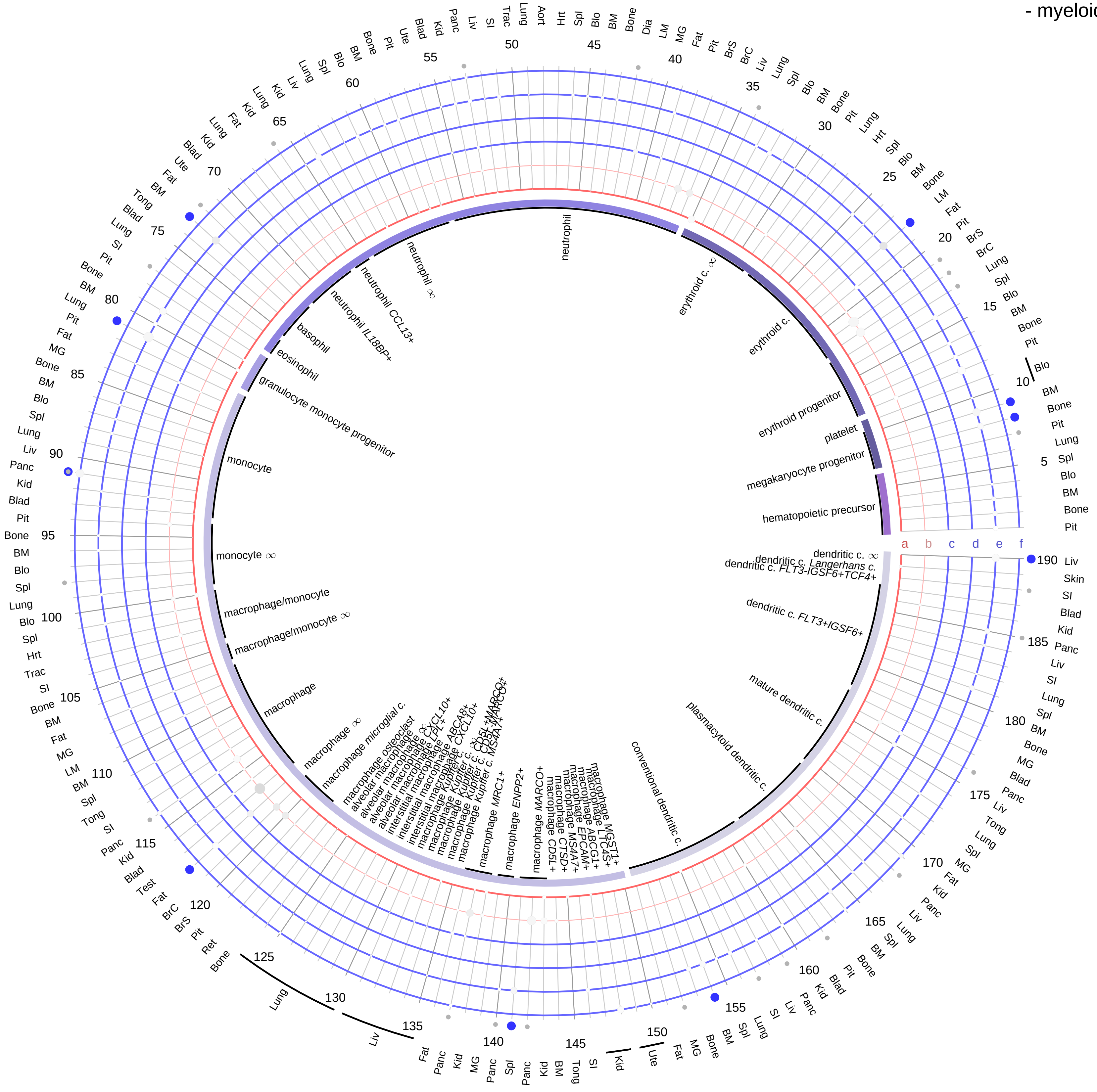

**Ligand/Enzyme**
- a - OXT
- b - CPE

**Receptor**
- c - AVPR1A
- d - AVPR1B
- e - AVPR2
- f - OXTR

**PAIRING** (ligand → receptor)

OXT & CPE→OXTR

OXT & CPE→AVPR1A

OXT & CPE→AVPR1B

OXT & CPE→AVPR2

| Abbrev. | Tissue |
|---|---|
| Aort | Aorta |
| Blad | Bladder |
| Blo | Blood |
| Bone | Bone |
| BM | Bone marrow |
| BrC | Brain cortex |
| BrS | Brainstem |
| Col | Colon |
| Dia | Diaphragm |
| Ret | Eye retina |
| Fat | Fat |
| Hrt | Heart |
| Pit | Hypothalamus/Pituitary |
| Kid | Kidney |
| LM | Limb muscle |
| Liv | Liver |
| Lung | Lung |
| MG | Mammary gland |
| Panc | Pancreas |
| Skin | Skin |
| SI | Small intestine |
| Spl | Spleen |
| Test | Testes |
| Thym | Thymus |
| Tong | Tongue |
| Trac | Trachea |
| Ute | Uterus |

**Compartment**

**hematopoietic**
- hematopoietic precursor

**megakaryocyte-erythroid**
- megakaryoid
- erythroid

**myeloid**
- granulocyte
- granulocyte monocyte progenitor
- monocyte/macrophage
- dendritic

12.5%
25%
50%
100%

ln(CP10K+1): 0, 1, 2, 3

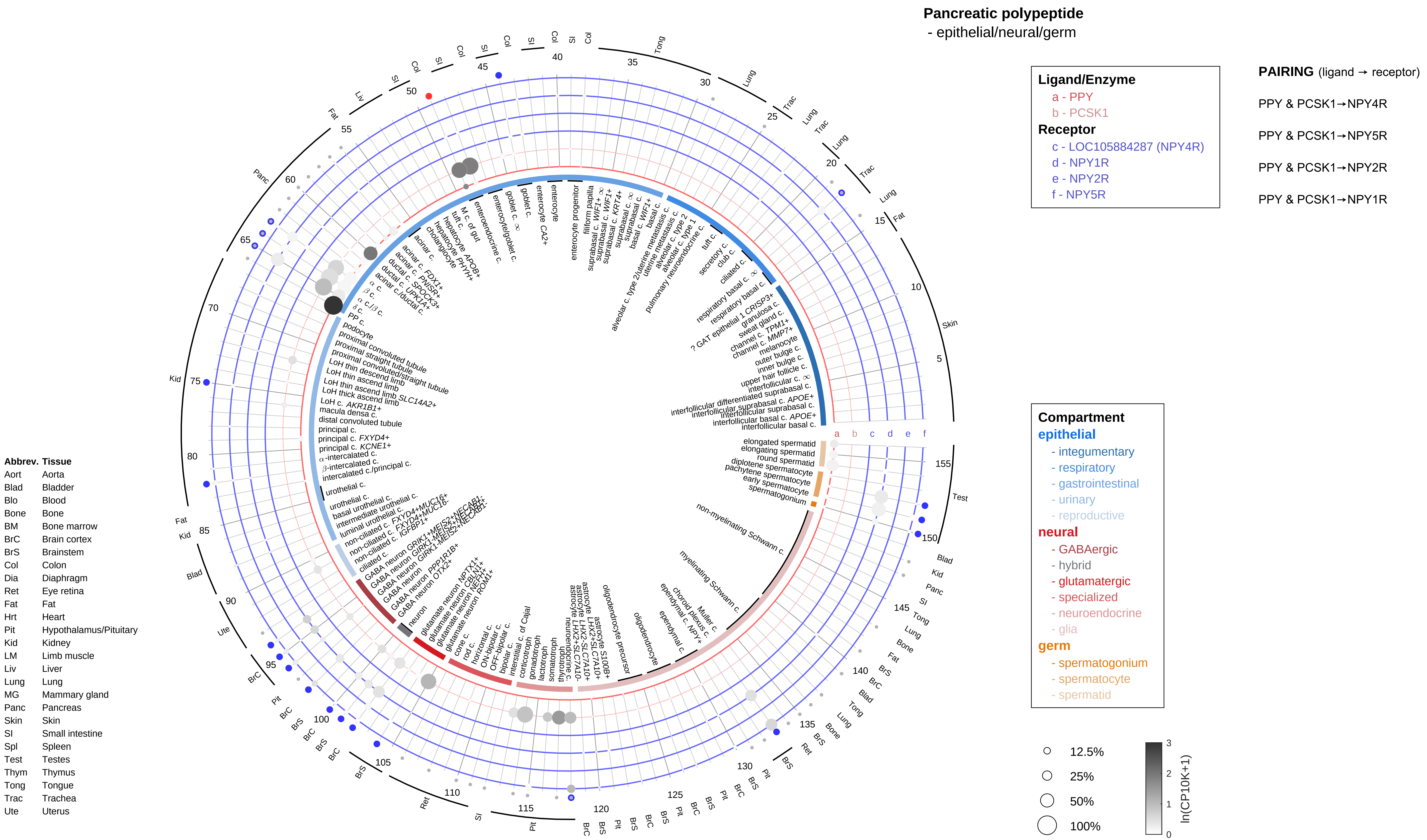

Pancreatic polypeptide
- epithelial/neural/germ
Ligand/Enzyme
a - PPY
b - PCSK1
Receptor
c - LOC105884287 (NPY4R)
d - NPY1R
e - NPY2R
f - NPY5R
PAIRING (ligand → receptor)
PPY & PCSK1→NPY4R
PPY & PCSK1→NPY5R
PPY & PCSK1→NPY2R
PPY & PCSK1→NPY1R
Compartment
epithelial
- integumentary
- respiratory
- gastrointestinal
- urinary
- reproductive
neural
- GABAergic
- hybrid
- glutamatergic
- specialized
- neuroendocrine
- glia
germ
- spermatogonium
- spermatocyte
- spermatid
12.5%
25%
50%
100%
ln(CP10K+1)
Abbrev. Tissue
Aort Aorta
Blad Bladder
Blo Blood
Bone Bone
BM Bone marrow
BrC Brain cortex
BrS Brainstem
Col Colon
Dia Diaphragm
Ret Eye retina
Fat Fat
Hrt Heart
Pit Hypothalamus/Pituitary
Kid Kidney
LM Limb muscle
Liv Liver
Lung Lung
MG Mammary gland
Panc Pancreas
Skin Skin
SI Small intestine
Spl Spleen
Test Testes
Thym Thymus
Tong Tongue
Trac Trachea
Ute Uterus

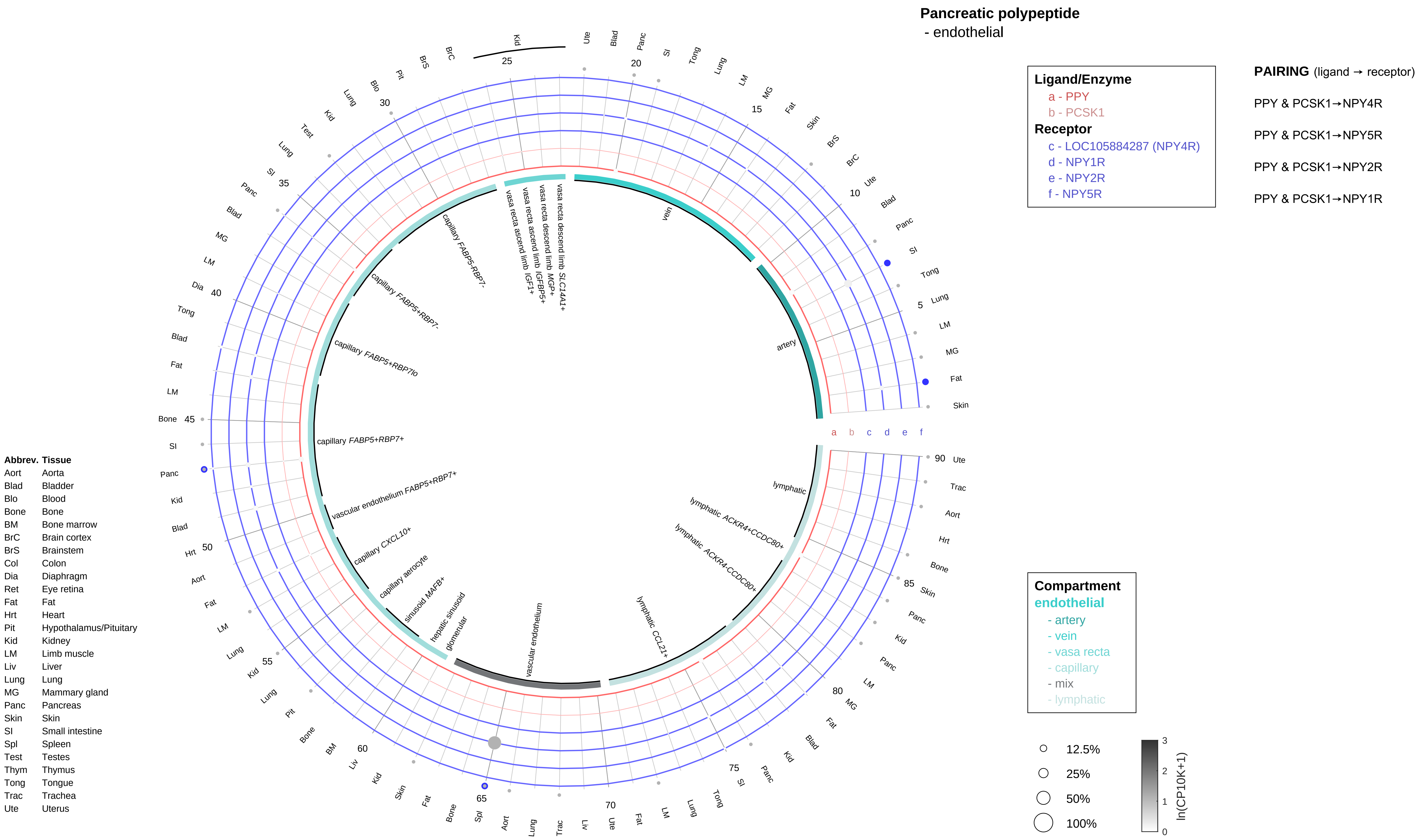
Pancreatic polypeptide
- endothelial
Ligand/Enzyme
a - PPY
b - PCSK1
Receptor
c - LOC105884287 (NPY4R)
d - NPY1R
e - NPY2R
f - NPY5R
PAIRING (ligand → receptor)
PPY & PCSK1→NPY4R
PPY & PCSK1→NPY5R
PPY & PCSK1→NPY2R
PPY & PCSK1→NPY1R
Compartment
endothelial
- artery
- vein
- vasa recta
- capillary
- mix
- lymphatic
12.5%
25%
50%
100%
ln(CP10K+1)
Abbrev. Tissue
Aort Aorta
Blad Bladder
Blo Blood
Bone Bone
BM Bone marrow
BrC Brain cortex
BrS Brainstem
Col Colon
Dia Diaphragm
Ret Eye retina
Fat Fat
Hrt Heart
Pit Hypothalamus/Pituitary
Kid Kidney
LM Limb muscle
Liv Liver
Lung Lung
MG Mammary gland
Panc Pancreas
Skin Skin
SI Small intestine
Spl Spleen
Test Testes
Thym Thymus
Tong Tongue
Trac Trachea
Ute Uterus
vein
artery
lymphatic
lymphatic ACKR4+CCDC80+
lymphatic ACKR4-CCDC80+
lymphatic CCL21+
vascular endothelium
glomerular
hepatic sinusoid
sinusoid MAFB+
capillary aerocyte
capillary CXCL10+
vascular endothelium FABP5+RBP7+
capillary FABP5+RBP7+
capillary FABP5+RBP7lo
capillary FABP5+RBP7-
capillary FABP5-RBP7-
vasa recta ascend limb IGF1+
vasa recta ascend limb IGFBP5+
vasa recta descend limb MGP+
vasa recta descend limb SLC14A1+

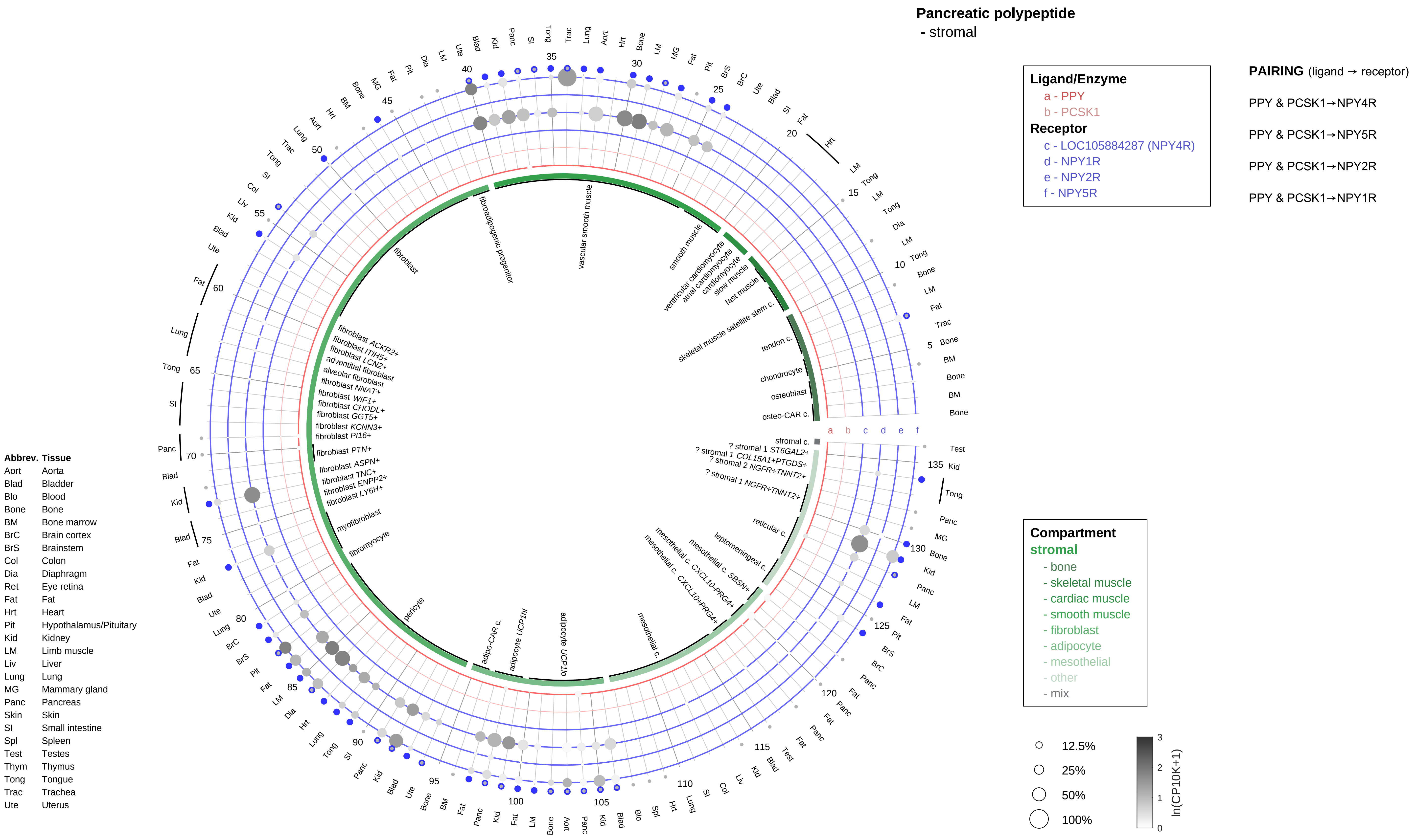
Pancreatic polypeptide
- stromal
Ligand/Enzyme
a - PPY
b - PCSK1
Receptor
c - LOC105884287 (NPY4R)
d - NPY1R
e - NPY2R
f - NPY5R
PAIRING (ligand → receptor)
PPY & PCSK1→NPY4R
PPY & PCSK1→NPY5R
PPY & PCSK1→NPY2R
PPY & PCSK1→NPY1R
Compartment
stromal
- bone
- skeletal muscle
- cardiac muscle
- smooth muscle
- fibroblast
- adipocyte
- mesothelial
- other
- mix
12.5%
25%
50%
100%
ln(CP10K+1)
Abbrev. Tissue
Aort Aorta
Blad Bladder
Blo Blood
Bone Bone
BM Bone marrow
BrC Brain cortex
BrS Brainstem
Col Colon
Dia Diaphragm
Ret Eye retina
Fat Fat
Hrt Heart
Pit Hypothalamus/Pituitary
Kid Kidney
LM Limb muscle
Liv Liver
Lung Lung
MG Mammary gland
Panc Pancreas
Skin Skin
SI Small intestine
Spl Spleen
Test Testes
Thym Thymus
Tong Tongue
Trac Trachea
Ute Uterus
vascular smooth muscle
fibroadipogenic progenitor
fibroblast
smooth muscle
ventricular cardiomyocyte
atrial cardiomyocyte
cardiomyocyte
slow muscle
fast muscle
skeletal muscle satellite stem c.
tendon c.
chondrocyte
osteoblast
osteo-CAR c.
stromal c.
? stromal 1 ST6GAL2+
? stromal 1 COL15A1+PTGDS+
? stromal 2 NGFR+TNNT2+
? stromal 1 NGFR+TNNT2+
reticular c.
leptomeningeal c.
mesothelial c. SBSN+
mesothelial c. CXCL10+PRG4+
mesothelial c. CXCL10+PRG4+
mesothelial c.
adipocyte UCP1lo
adipocyte UCP1hi
adipo-CAR c.
pericyte
fibromyocyte
myofibroblast
fibroblast ACKR2+
fibroblast ITIH5+
fibroblast LCN2+
adventitial fibroblast
alveolar fibroblast
fibroblast NNAT+
fibroblast WIF1+
fibroblast CHODL+
fibroblast GGT5+
fibroblast KCNN3+
fibroblast PI16+
fibroblast PTN+
fibroblast ASPN+
fibroblast TNC+
fibroblast ENPP2+
fibroblast LY6H+

#### Pancreatic polypeptide
- lymphoid

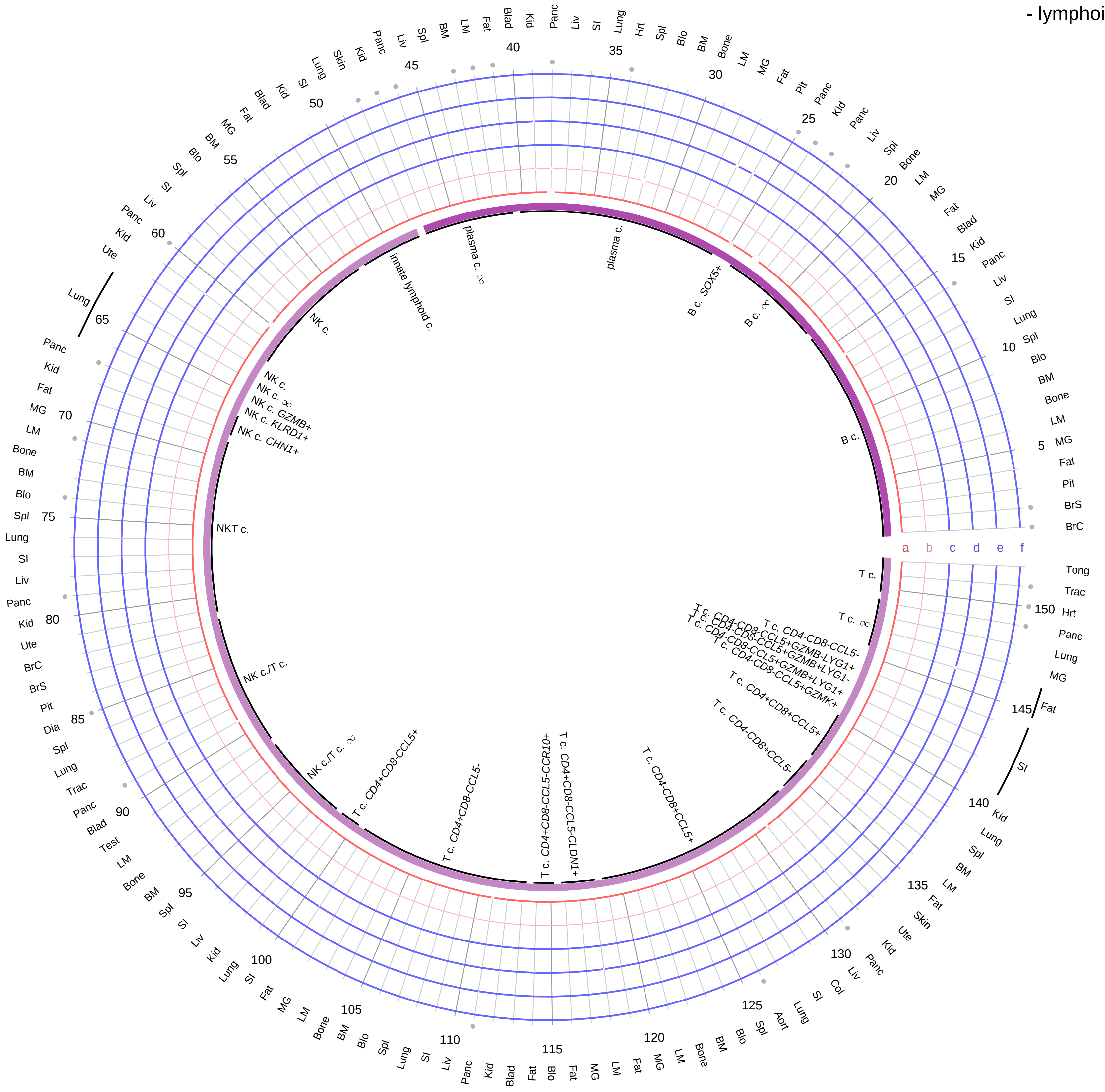

**Ligand/Enzyme**
a - PPY
b - PCSK1
**Receptor**
c - LOC105884287 (NPY4R)
d - NPY1R
e - NPY2R
f - NPY5R

**PAIRING** (ligand → receptor)

PPY & PCSK1→NPY4R

PPY & PCSK1→NPY5R

PPY & PCSK1→NPY2R

PPY & PCSK1→NPY1R

**Compartment**
**lymphoid**
- B cell
- NK/T cell

12.5%
25%
50%
100%

ln(CP10K+1): 0, 1, 2, 3

| Abbrev. | Tissue |
|---|---|
| Aort | Aorta |
| Blad | Bladder |
| Blo | Blood |
| Bone | Bone |
| BM | Bone marrow |
| BrC | Brain cortex |
| BrS | Brainstem |
| Col | Colon |
| Dia | Diaphragm |
| Ret | Eye retina |
| Fat | Fat |
| Hrt | Heart |
| Pit | Hypothalamus/Pituitary |
| Kid | Kidney |
| LM | Limb muscle |
| Liv | Liver |
| Lung | Lung |
| MG | Mammary gland |
| Panc | Pancreas |
| Skin | Skin |
| SI | Small intestine |
| Spl | Spleen |
| Test | Testes |
| Thym | Thymus |
| Tong | Tongue |
| Trac | Trachea |
| Ute | Uterus |

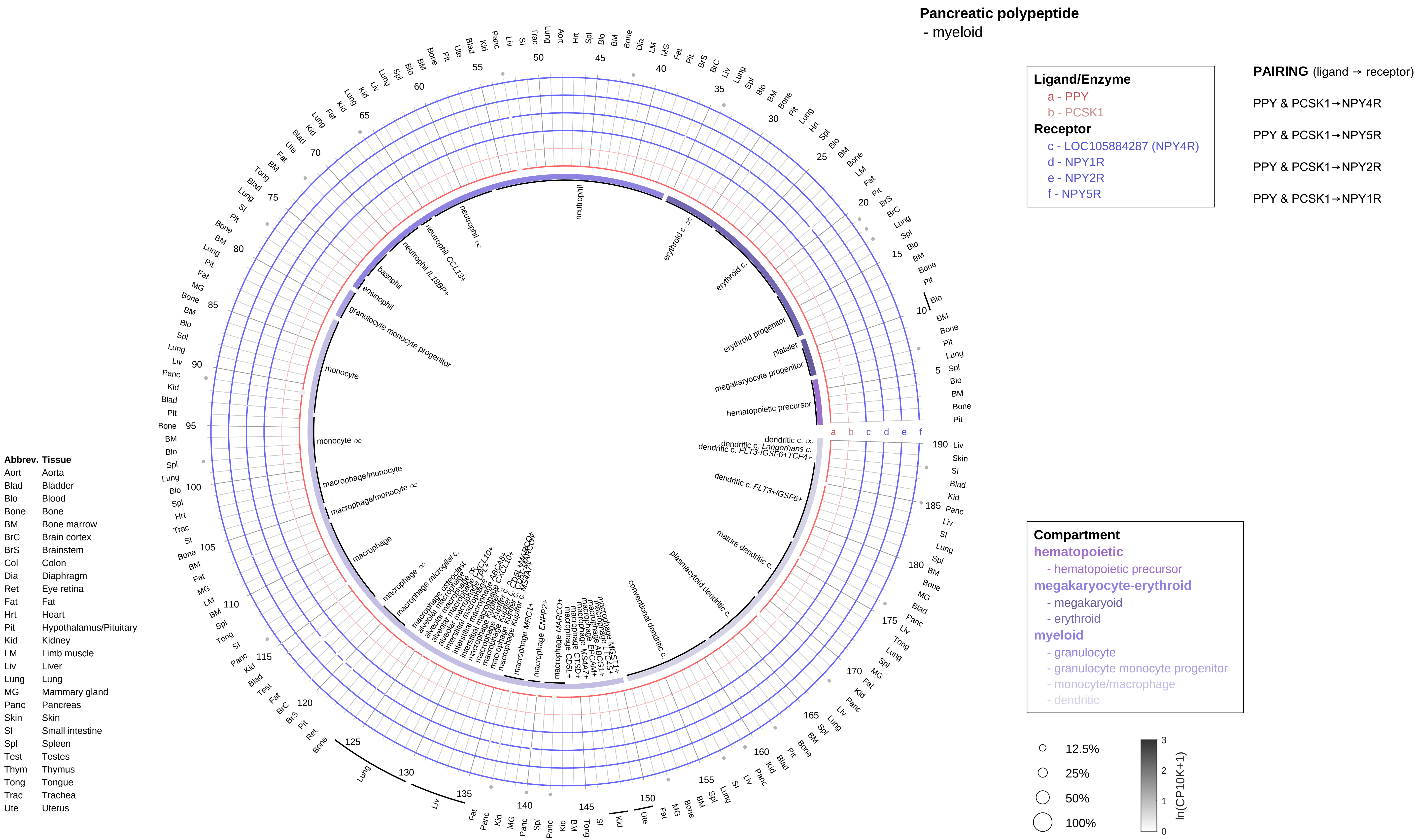
Pancreatic polypeptide
- myeloid
Ligand/Enzyme
a - PPY
b - PCSK1
Receptor
c - LOC105884287 (NPY4R)
d - NPY1R
e - NPY2R
f - NPY5R
PAIRING (ligand → receptor)
PPY & PCSK1→NPY4R
PPY & PCSK1→NPY5R
PPY & PCSK1→NPY2R
PPY & PCSK1→NPY1R
Compartment
hematopoietic
- hematopoietic precursor
megakaryocyte-erythroid
- megakaryoid
- erythroid
myeloid
- granulocyte
- granulocyte monocyte progenitor
- monocyte/macrophage
- dendritic
12.5%
25%
50%
100%
ln(CP10K+1)
Abbrev. Tissue
Aort Aorta
Blad Bladder
Blo Blood
Bone Bone
BM Bone marrow
BrC Brain cortex
BrS Brainstem
Col Colon
Dia Diaphragm
Ret Eye retina
Fat Fat
Hrt Heart
Pit Hypothalamus/Pituitary
Kid Kidney
LM Limb muscle
Liv Liver
Lung Lung
MG Mammary gland
Panc Pancreas
Skin Skin
SI Small intestine
Spl Spleen
Test Testes
Thym Thymus
Tong Tongue
Trac Trachea
Ute Uterus

**Parathyroid hormone/Parathyroid hormone-like hormone**
- epithelial/neural/germ

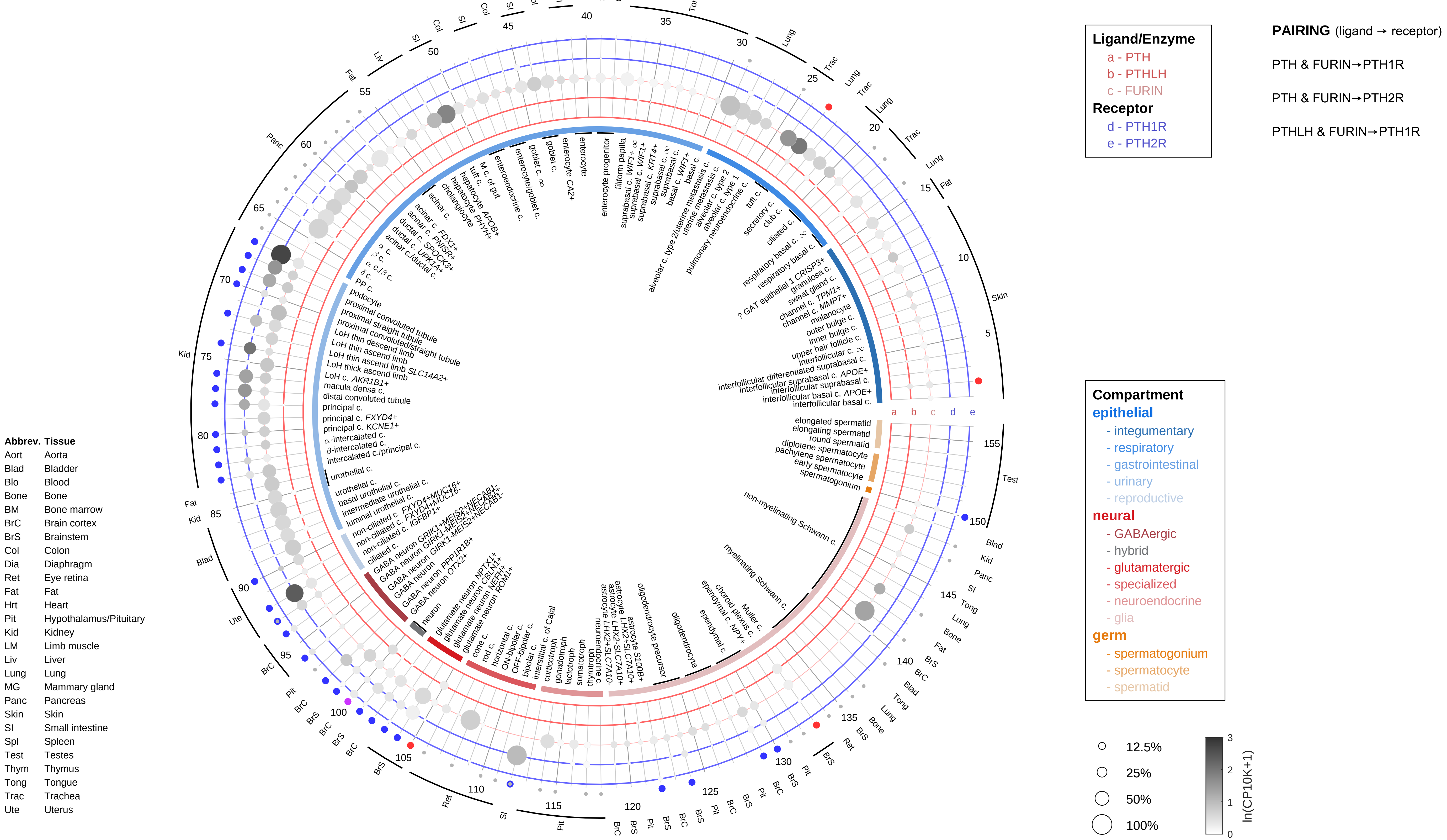

**Parathyroid hormone/Parathyroid hormone-like hormone**
- endothelial

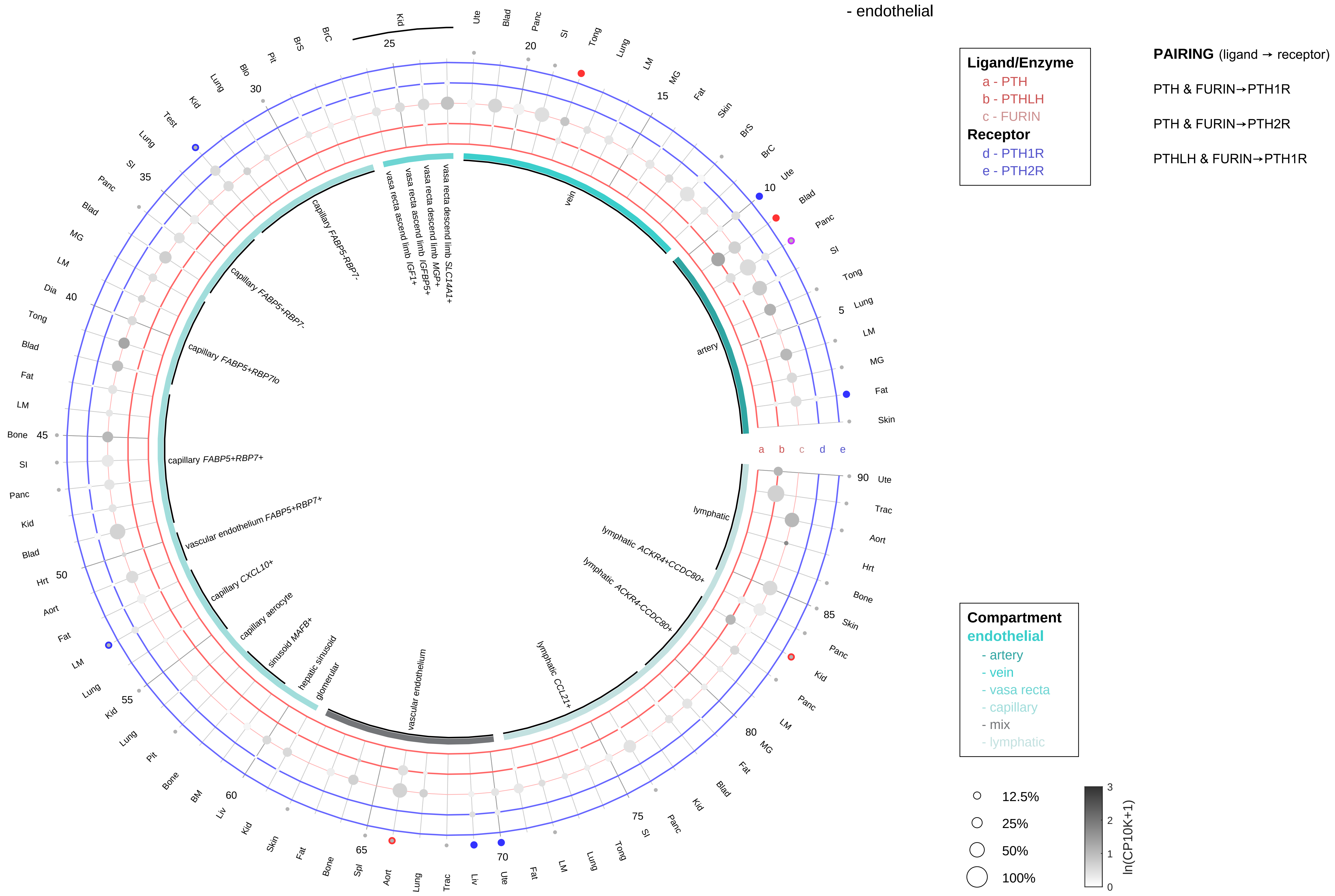

| Abbrev. | Tissue |
|---|---|
| Aort | Aorta |
| Blad | Bladder |
| Blo | Blood |
| Bone | Bone |
| BM | Bone marrow |
| BrC | Brain cortex |
| BrS | Brainstem |
| Col | Colon |
| Dia | Diaphragm |
| Ret | Eye retina |
| Fat | Fat |
| Hrt | Heart |
| Pit | Hypothalamus/Pituitary |
| Kid | Kidney |
| LM | Limb muscle |
| Liv | Liver |
| Lung | Lung |
| MG | Mammary gland |
| Panc | Pancreas |
| Skin | Skin |
| SI | Small intestine |
| Spl | Spleen |
| Test | Testes |
| Thym | Thymus |
| Tong | Tongue |
| Trac | Trachea |
| Ute | Uterus |

**Parathyroid hormone/Parathyroid hormone-like hormone**
- stromal

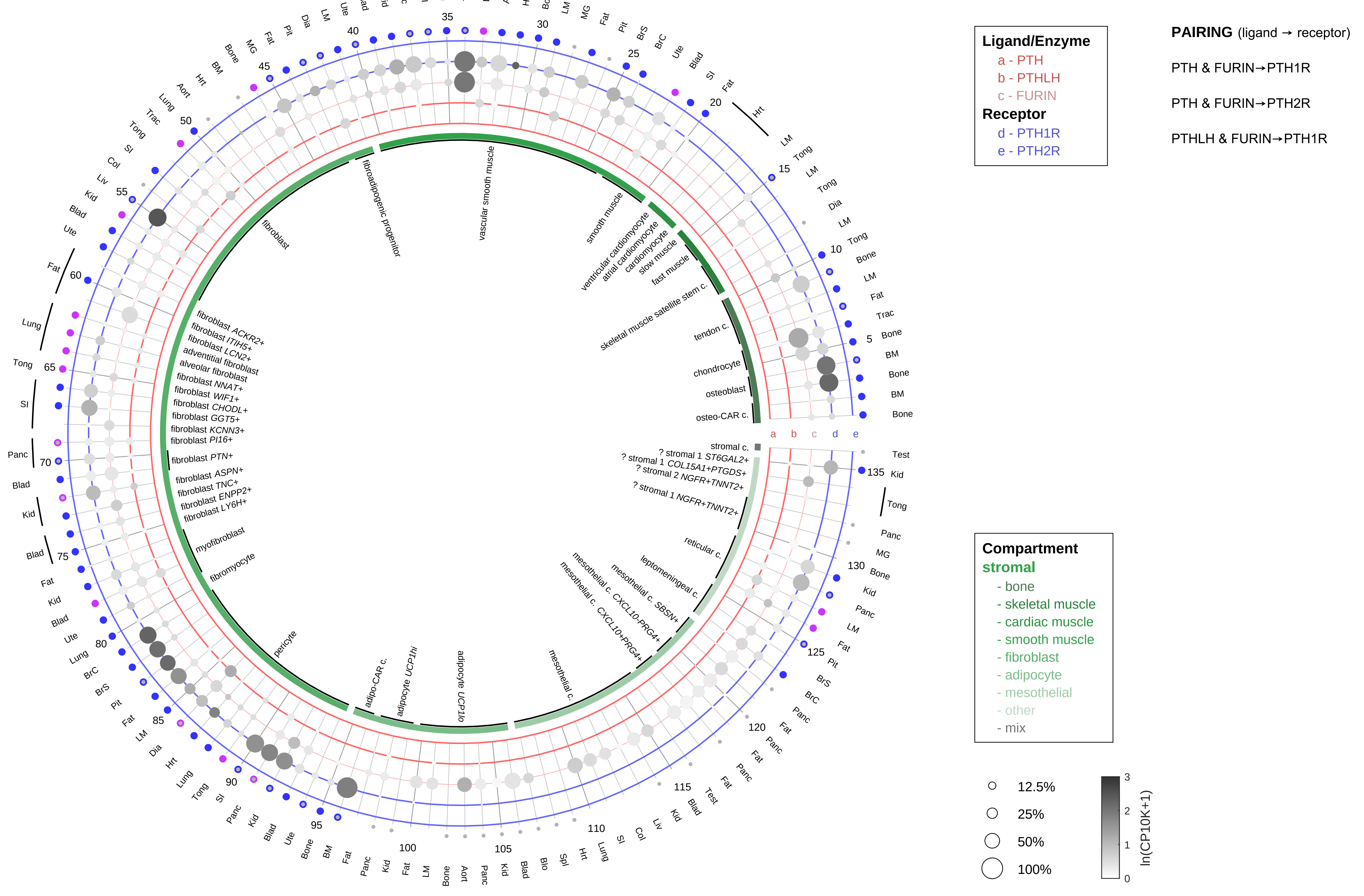

| Abbrev. | Tissue |
|---|---|
| Aort | Aorta |
| Blad | Bladder |
| Blo | Blood |
| Bone | Bone |
| BM | Bone marrow |
| BrC | Brain cortex |
| BrS | Brainstem |
| Col | Colon |
| Dia | Diaphragm |
| Ret | Eye retina |
| Fat | Fat |
| Hrt | Heart |
| Pit | Hypothalamus/Pituitary |
| Kid | Kidney |
| LM | Limb muscle |
| Liv | Liver |
| Lung | Lung |
| MG | Mammary gland |
| Panc | Pancreas |
| Skin | Skin |
| SI | Small intestine |
| Spl | Spleen |
| Test | Testes |
| Thym | Thymus |
| Tong | Tongue |
| Trac | Trachea |
| Ute | Uterus |

#### Parathyroid hormone/Parathyroid hormone-like hormone
- lymphoid

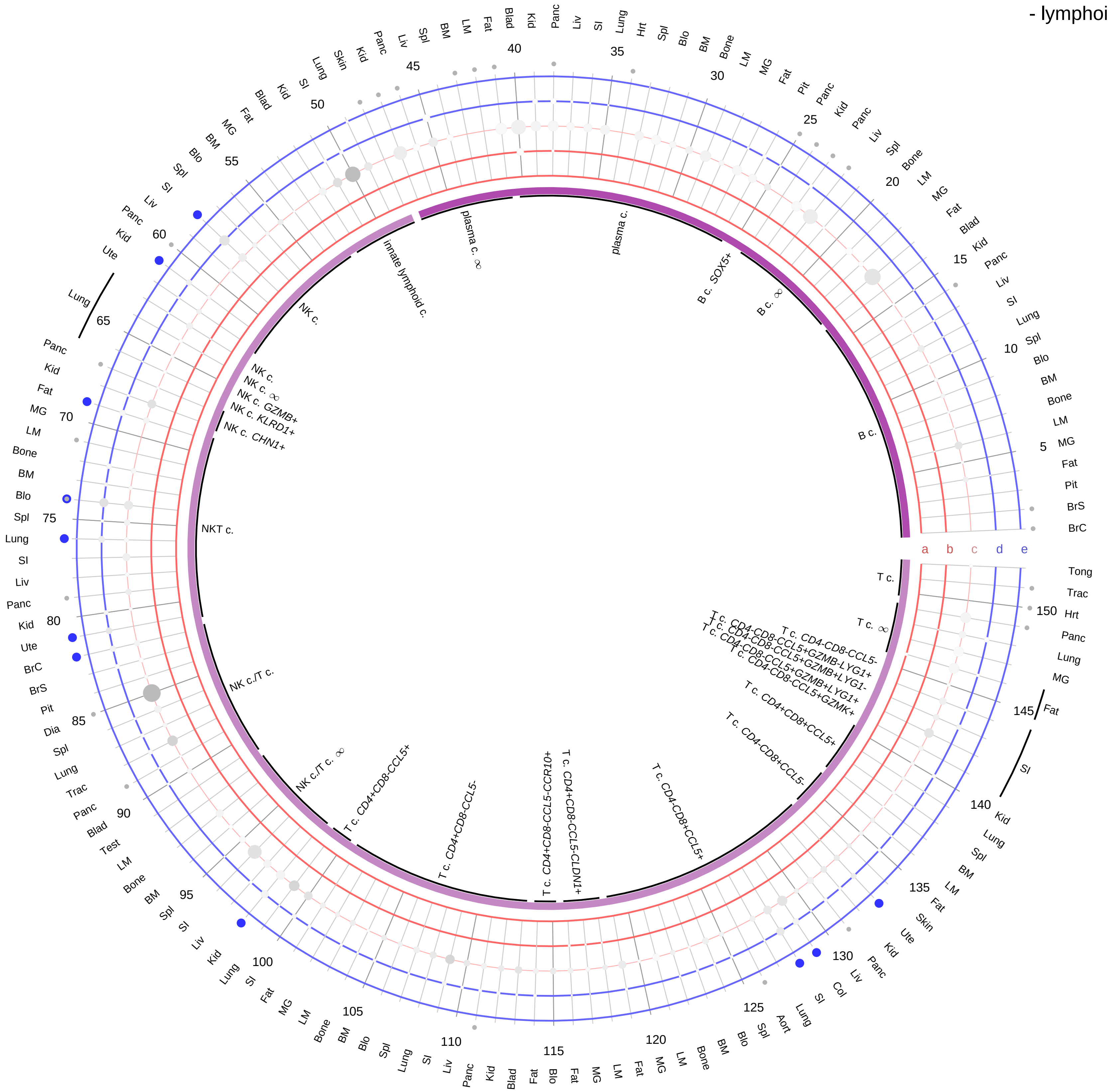

**Ligand/Enzyme**
a - PTH
b - PTHLH
c - FURIN
**Receptor**
d - PTH1R
e - PTH2R

**PAIRING** (ligand → receptor)

PTH & FURIN→PTH1R

PTH & FURIN→PTH2R

PTHLH & FURIN→PTH1R

**Compartment**
**lymphoid**
- B cell
- NK/T cell

12.5%
25%
50%
100%

ln(CP10K+1): 0, 1, 2, 3

| Abbrev. | Tissue |
|---|---|
| Aort | Aorta |
| Blad | Bladder |
| Blo | Blood |
| Bone | Bone |
| BM | Bone marrow |
| BrC | Brain cortex |
| BrS | Brainstem |
| Col | Colon |
| Dia | Diaphragm |
| Ret | Eye retina |
| Fat | Fat |
| Hrt | Heart |
| Pit | Hypothalamus/Pituitary |
| Kid | Kidney |
| LM | Limb muscle |
| Liv | Liver |
| Lung | Lung |
| MG | Mammary gland |
| Panc | Pancreas |
| Skin | Skin |
| SI | Small intestine |
| Spl | Spleen |
| Test | Testes |
| Thym | Thymus |
| Tong | Tongue |
| Trac | Trachea |
| Ute | Uterus |

#### Parathyroid hormone/Parathyroid hormone-like hormone
- myeloid

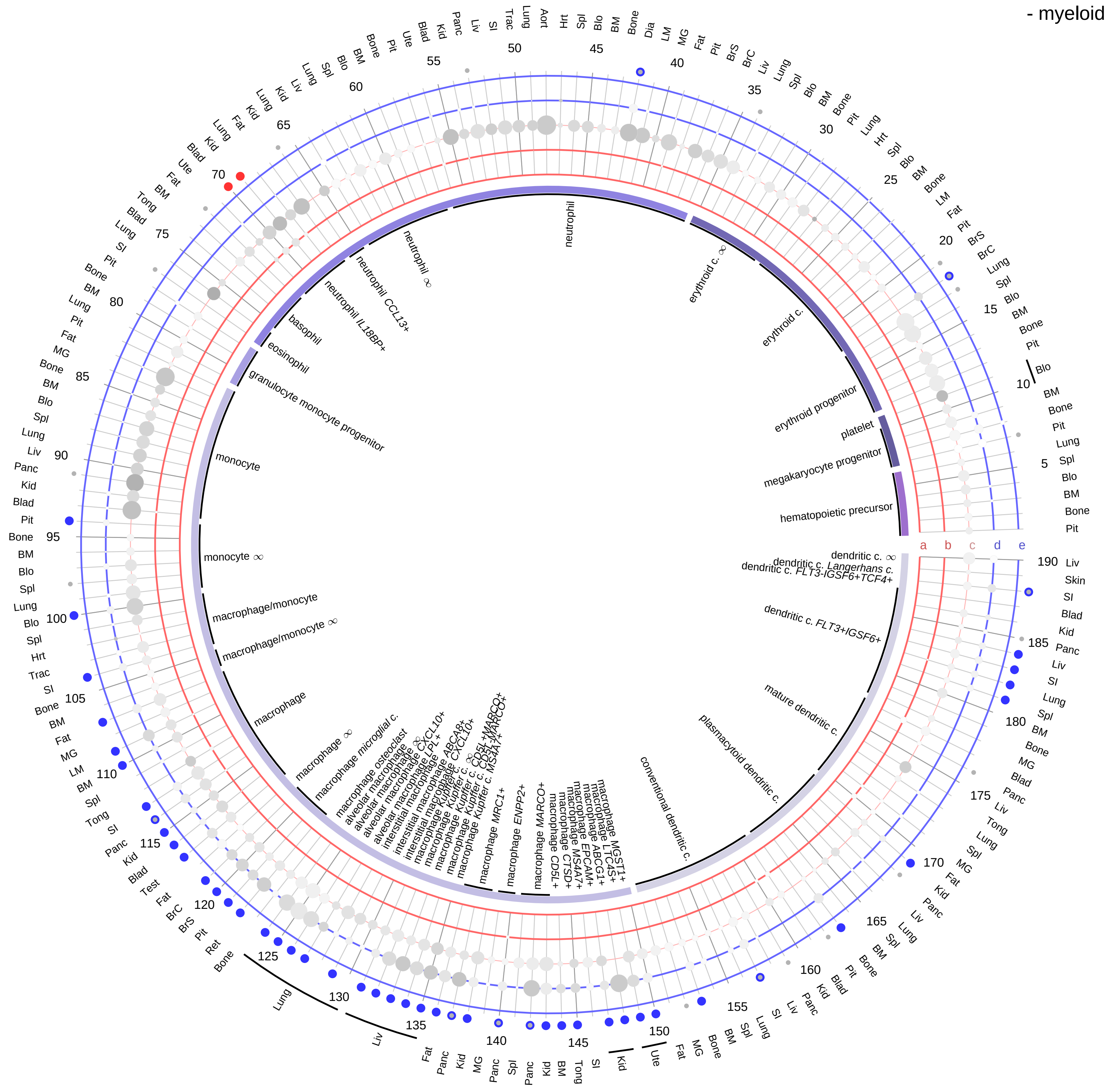

**Ligand/Enzyme**
- a - PTH
- b - PTHLH
- c - FURIN

**Receptor**
- d - PTH1R
- e - PTH2R

**PAIRING** (ligand → receptor)

PTH & FURIN→PTH1R

PTH & FURIN→PTH2R

PTHLH & FURIN→PTH1R

**Compartment**

**hematopoietic**
- hematopoietic precursor

**megakaryocyte-erythroid**
- megakaryoid
- erythroid

**myeloid**
- granulocyte
- granulocyte monocyte progenitor
- monocyte/macrophage
- dendritic

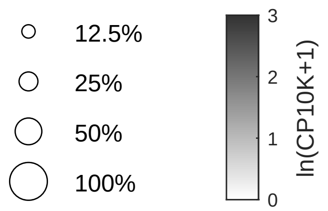

| Abbrev. | Tissue |
|---|---|
| Aort | Aorta |
| Blad | Bladder |
| Blo | Blood |
| Bone | Bone |
| BM | Bone marrow |
| BrC | Brain cortex |
| BrS | Brainstem |
| Col | Colon |
| Dia | Diaphragm |
| Ret | Eye retina |
| Fat | Fat |
| Hrt | Heart |
| Pit | Hypothalamus/Pituitary |
| Kid | Kidney |
| LM | Limb muscle |
| Liv | Liver |
| Lung | Lung |
| MG | Mammary gland |
| Panc | Pancreas |
| Skin | Skin |
| SI | Small intestine |
| Spl | Spleen |
| Test | Testes |
| Thym | Thymus |
| Tong | Tongue |
| Trac | Trachea |
| Ute | Uterus |

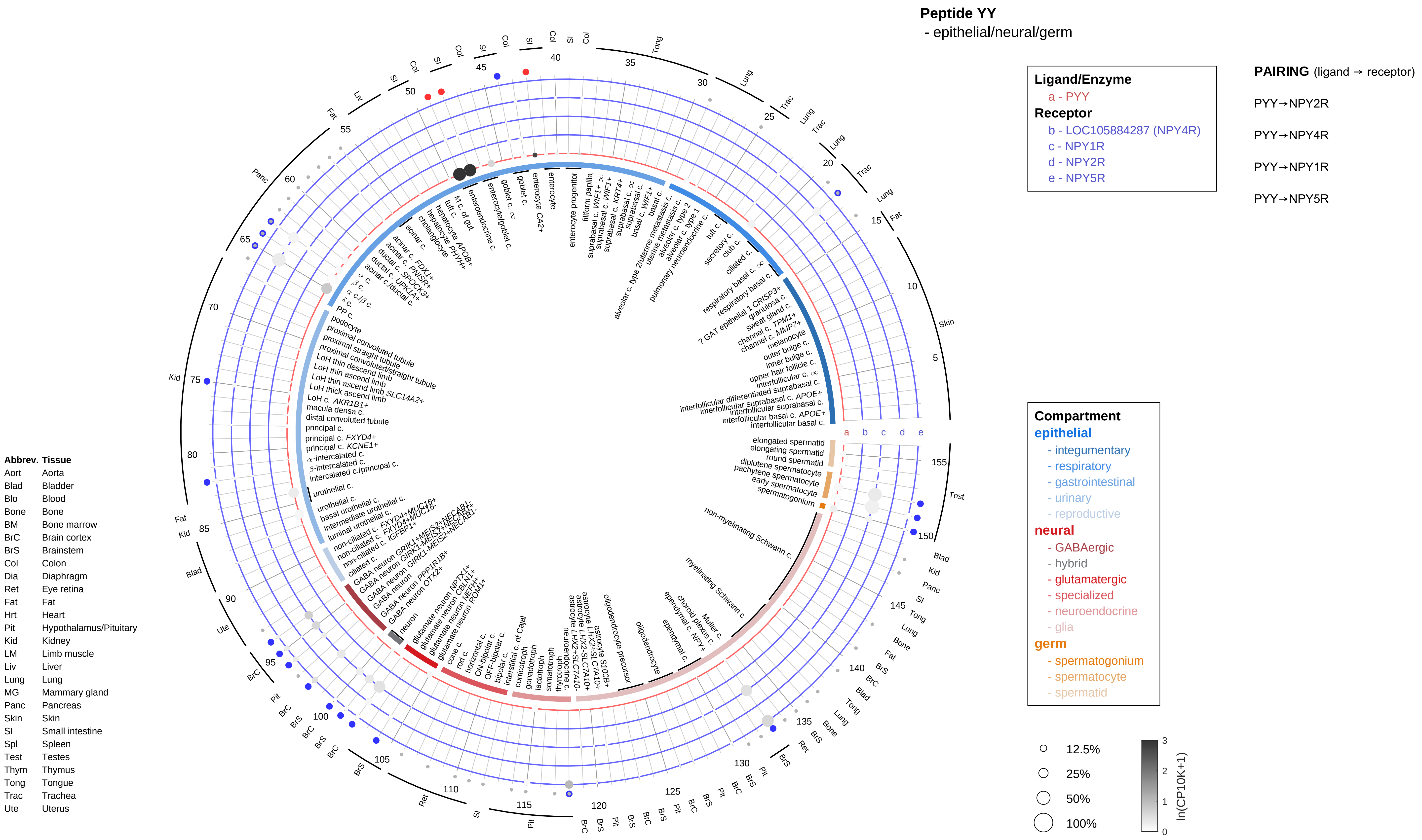

Peptide YY
- epithelial/neural/germ
Ligand/Enzyme
a - PYY
Receptor
b - LOC105884287 (NPY4R)
c - NPY1R
d - NPY2R
e - NPY5R
PAIRING (ligand → receptor)
PYY→NPY2R
PYY→NPY4R
PYY→NPY1R
PYY→NPY5R
Compartment
epithelial
- integumentary
- respiratory
- gastrointestinal
- urinary
- reproductive
neural
- GABAergic
- hybrid
- glutamatergic
- specialized
- neuroendocrine
- glia
germ
- spermatogonium
- spermatocyte
- spermatid
12.5%
25%
50%
100%
ln(CP10K+1)
Abbrev. Tissue
Aort Aorta
Blad Bladder
Blo Blood
Bone Bone
BM Bone marrow
BrC Brain cortex
BrS Brainstem
Col Colon
Dia Diaphragm
Ret Eye retina
Fat Fat
Hrt Heart
Pit Hypothalamus/Pituitary
Kid Kidney
LM Limb muscle
Liv Liver
Lung Lung
MG Mammary gland
Panc Pancreas
Skin Skin
SI Small intestine
Spl Spleen
Test Testes
Thym Thymus
Tong Tongue
Trac Trachea
Ute Uterus

**Peptide YY**
- endothelial

**Ligand/Enzyme**
a - PYY
**Receptor**
b - LOC105884287 (NPY4R)
c - NPY1R
d - NPY2R
e - NPY5R

**PAIRING** (ligand → receptor)

PYY→NPY2R

PYY→NPY4R

PYY→NPY1R

PYY→NPY5R

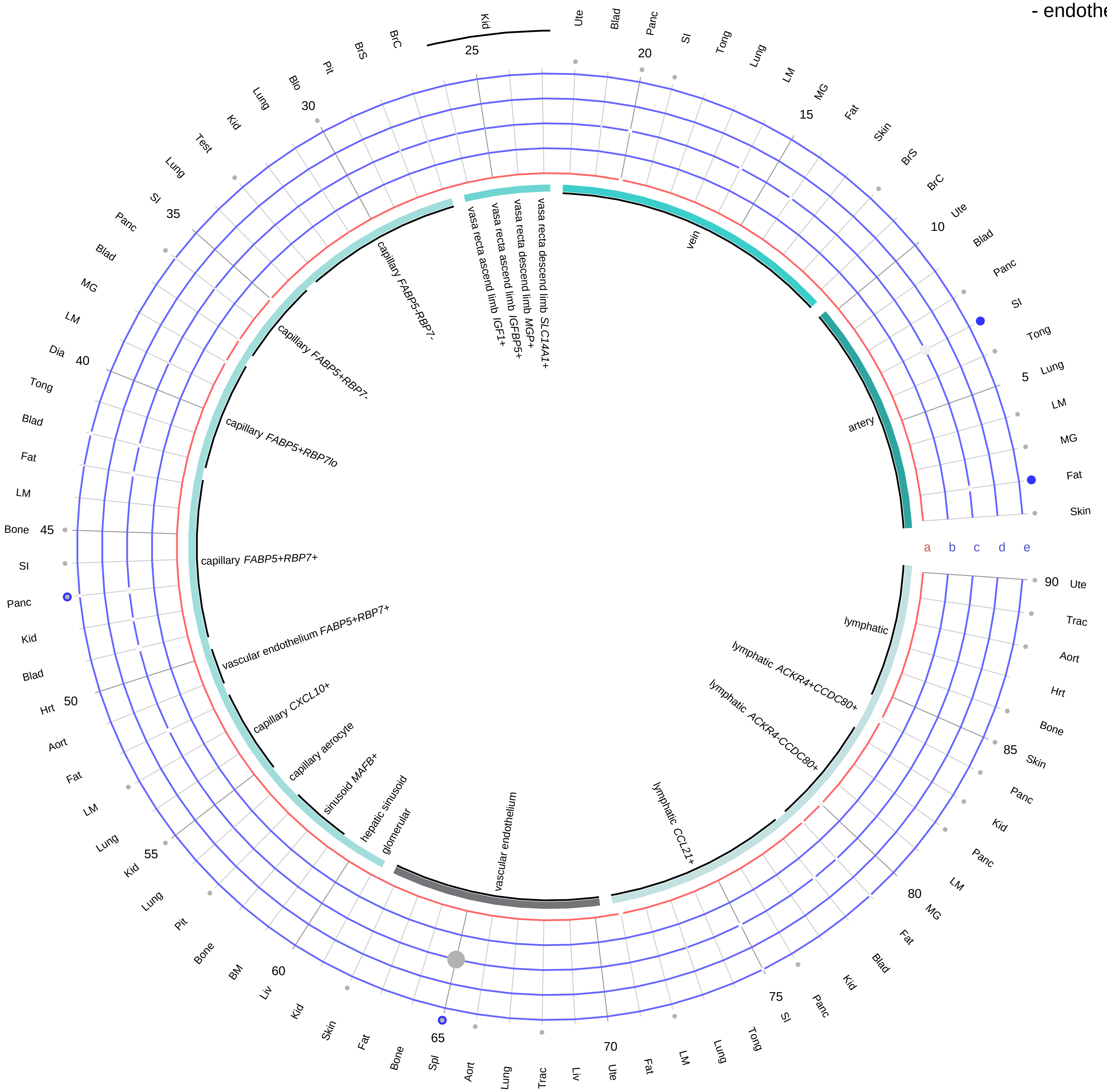

| Abbrev. | Tissue |
|---|---|
| Aort | Aorta |
| Blad | Bladder |
| Blo | Blood |
| Bone | Bone |
| BM | Bone marrow |
| BrC | Brain cortex |
| BrS | Brainstem |
| Col | Colon |
| Dia | Diaphragm |
| Ret | Eye retina |
| Fat | Fat |
| Hrt | Heart |
| Pit | Hypothalamus/Pituitary |
| Kid | Kidney |
| LM | Limb muscle |
| Liv | Liver |
| Lung | Lung |
| MG | Mammary gland |
| Panc | Pancreas |
| Skin | Skin |
| SI | Small intestine |
| Spl | Spleen |
| Test | Testes |
| Thym | Thymus |
| Tong | Tongue |
| Trac | Trachea |
| Ute | Uterus |

**Compartment**
**endothelial**
- artery
- vein
- vasa recta
- capillary
- mix
- lymphatic

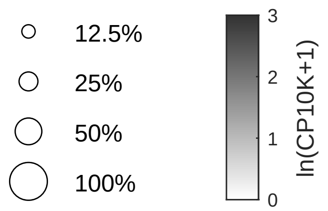

**Peptide YY**
- stromal

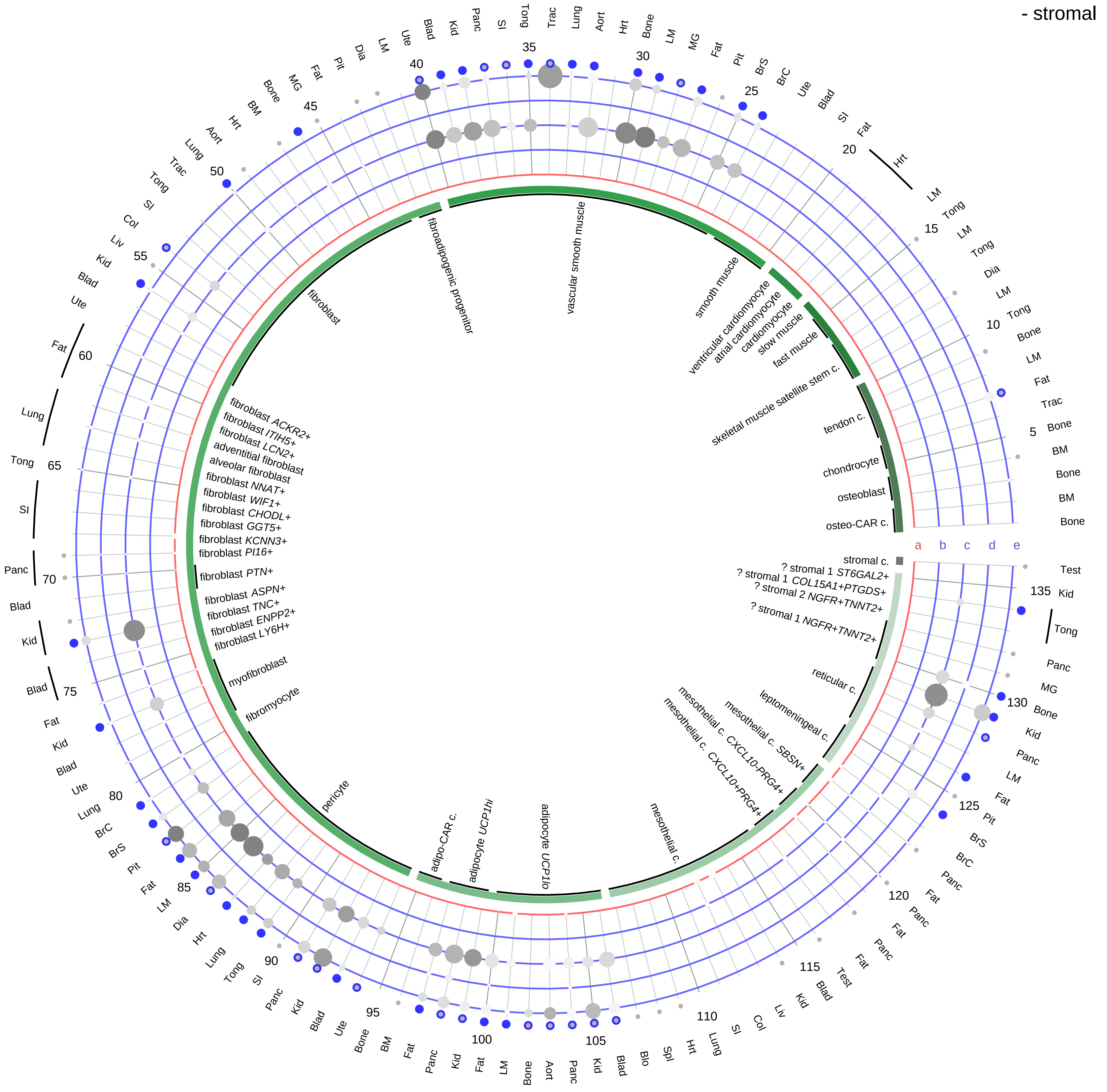

| Abbrev. | Tissue |
|---|---|
| Aort | Aorta |
| Blad | Bladder |
| Blo | Blood |
| Bone | Bone |
| BM | Bone marrow |
| BrC | Brain cortex |
| BrS | Brainstem |
| Col | Colon |
| Dia | Diaphragm |
| Ret | Eye retina |
| Fat | Fat |
| Hrt | Heart |
| Pit | Hypothalamus/Pituitary |
| Kid | Kidney |
| LM | Limb muscle |
| Liv | Liver |
| Lung | Lung |
| MG | Mammary gland |
| Panc | Pancreas |
| Skin | Skin |
| SI | Small intestine |
| Spl | Spleen |
| Test | Testes |
| Thym | Thymus |
| Tong | Tongue |
| Trac | Trachea |
| Ute | Uterus |

**Ligand/Enzyme**
- a - PYY

**Receptor**
- b - LOC105884287 (NPY4R)
- c - NPY1R
- d - NPY2R
- e - NPY5R

**PAIRING** (ligand → receptor)

PYY→NPY2R

PYY→NPY4R

PYY→NPY1R

PYY→NPY5R

**Compartment**

**stromal**
- bone
- skeletal muscle
- cardiac muscle
- smooth muscle
- fibroblast
- adipocyte
- mesothelial
- other
- mix

12.5%
25%
50%
100%

ln(CP10K+1): 0, 1, 2, 3

**Peptide YY**
- lymphoid

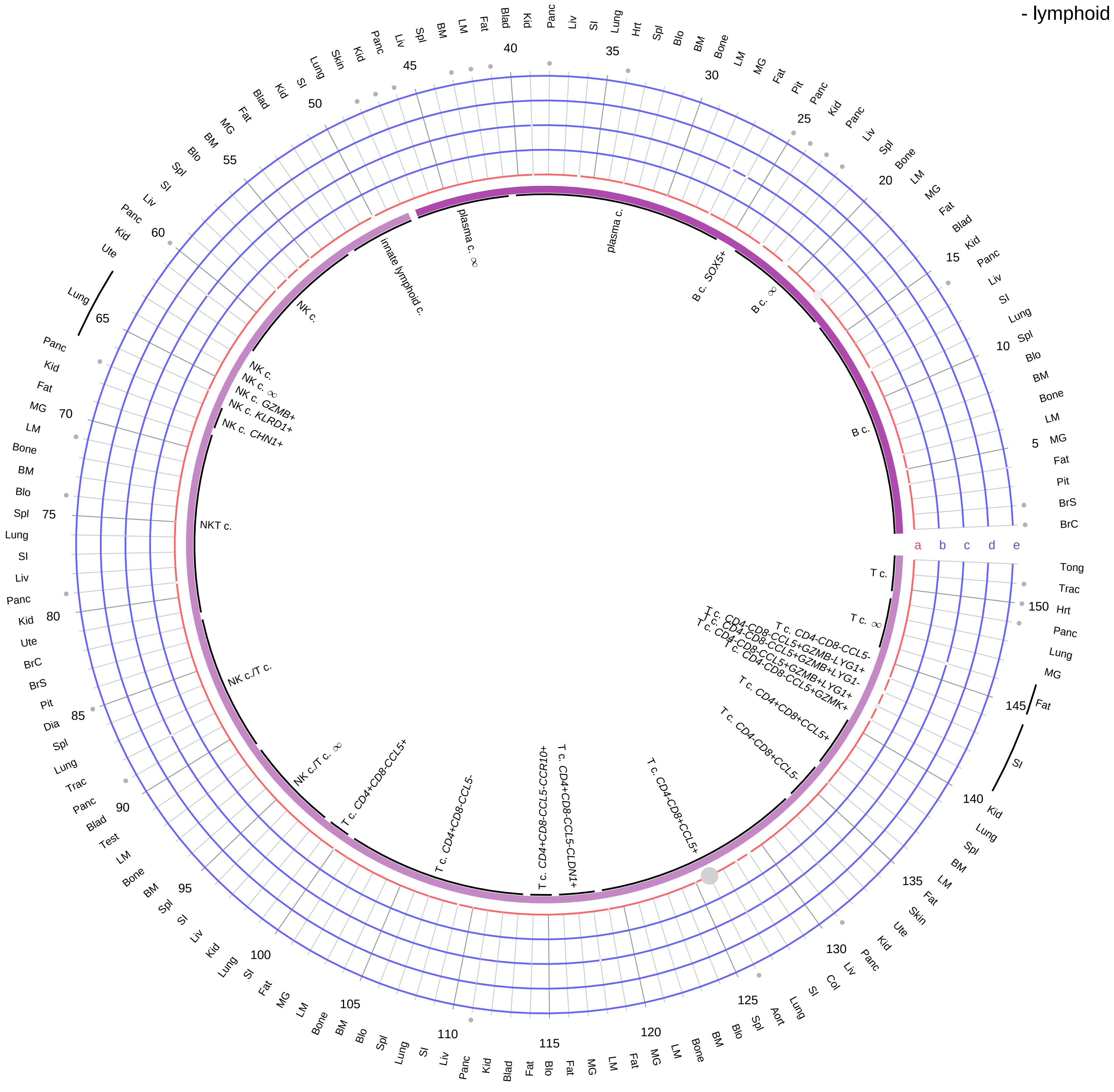

**Ligand/Enzyme**
- a - PYY

**Receptor**
- b - LOC105884287 (NPY4R)
- c - NPY1R
- d - NPY2R
- e - NPY5R

**PAIRING** (ligand → receptor)

PYY→NPY2R

PYY→NPY4R

PYY→NPY1R

PYY→NPY5R

**Compartment**

**lymphoid**
- B cell
- NK/T cell

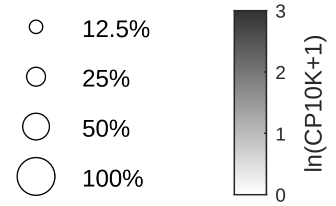

| Abbrev. | Tissue |
|---|---|
| Aort | Aorta |
| Blad | Bladder |
| Blo | Blood |
| Bone | Bone |
| BM | Bone marrow |
| BrC | Brain cortex |
| BrS | Brainstem |
| Col | Colon |
| Dia | Diaphragm |
| Ret | Eye retina |
| Fat | Fat |
| Hrt | Heart |
| Pit | Hypothalamus/Pituitary |
| Kid | Kidney |
| LM | Limb muscle |
| Liv | Liver |
| Lung | Lung |
| MG | Mammary gland |
| Panc | Pancreas |
| Skin | Skin |
| SI | Small intestine |
| Spl | Spleen |
| Test | Testes |
| Thym | Thymus |
| Tong | Tongue |
| Trac | Trachea |
| Ute | Uterus |

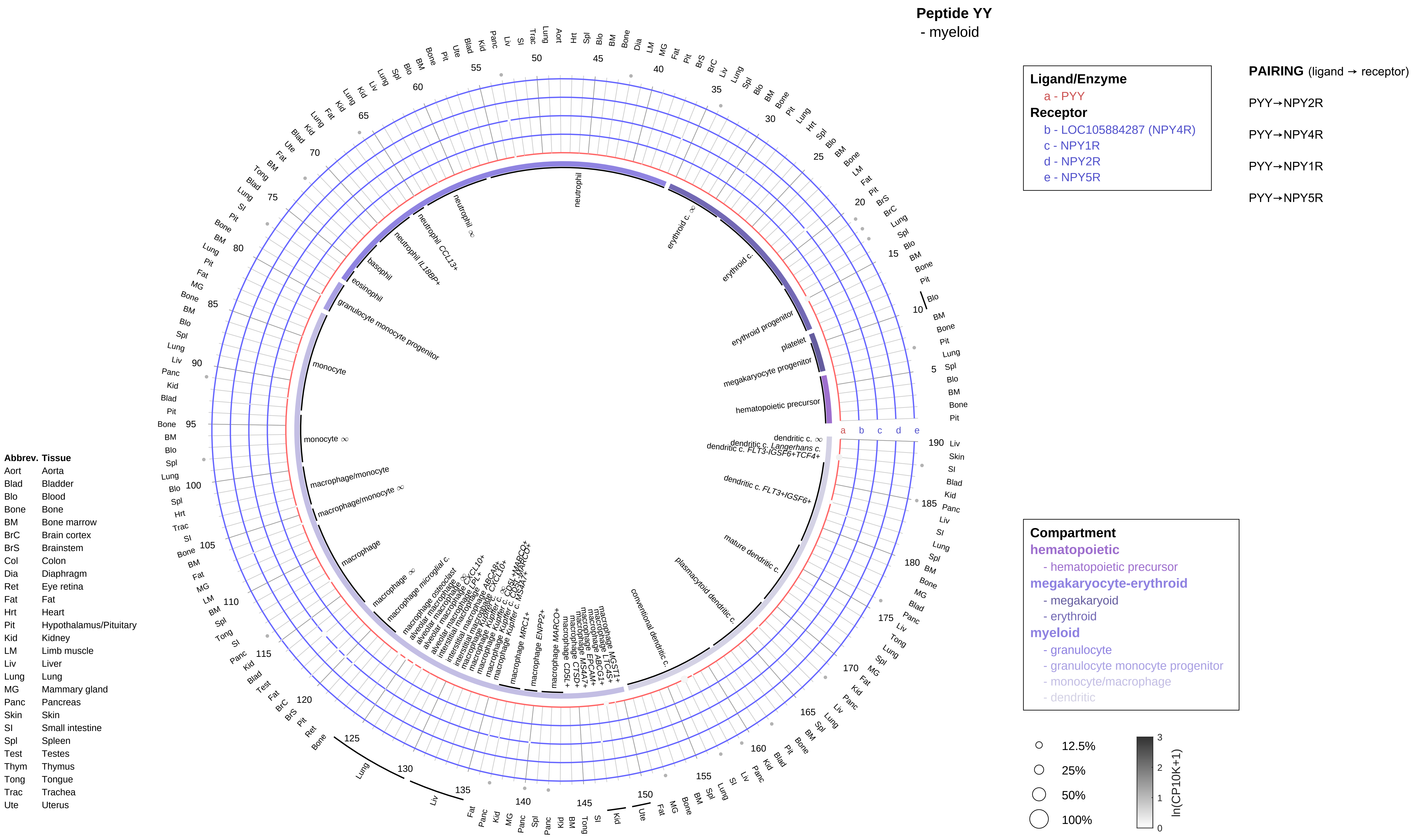
Peptide YY
- myeloid
Ligand/Enzyme
a - PYY
Receptor
b - LOC105884287 (NPY4R)
c - NPY1R
d - NPY2R
e - NPY5R
PAIRING (ligand → receptor)
PYY→NPY2R
PYY→NPY4R
PYY→NPY1R
PYY→NPY5R
Compartment
hematopoietic
- hematopoietic precursor
megakaryocyte-erythroid
- megakaryoid
- erythroid
myeloid
- granulocyte
- granulocyte monocyte progenitor
- monocyte/macrophage
- dendritic
12.5%
25%
50%
100%
ln(CP10K+1)
Abbrev. Tissue
Aort Aorta
Blad Bladder
Blo Blood
Bone Bone
BM Bone marrow
BrC Brain cortex
BrS Brainstem
Col Colon
Dia Diaphragm
Ret Eye retina
Fat Fat
Hrt Heart
Pit Hypothalamus/Pituitary
Kid Kidney
LM Limb muscle
Liv Liver
Lung Lung
MG Mammary gland
Panc Pancreas
Skin Skin
SI Small intestine
Spl Spleen
Test Testes
Thym Thymus
Tong Tongue
Trac Trachea
Ute Uterus

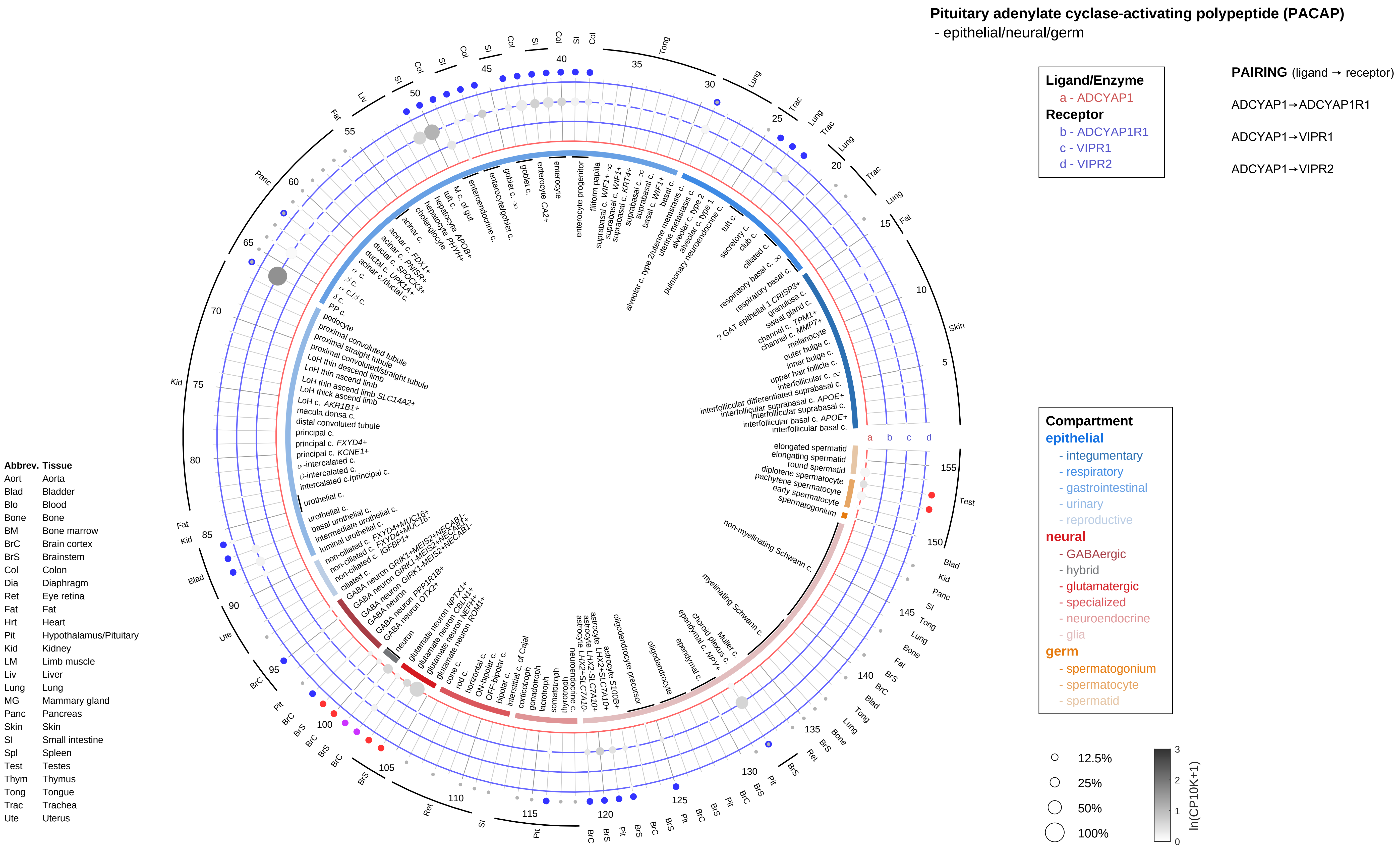

Pituitary adenylate cyclase-activating polypeptide (PACAP)
- epithelial/neural/germ
Ligand/Enzyme
a - ADCYAP1
Receptor
b - ADCYAP1R1
c - VIPR1
d - VIPR2
PAIRING (ligand → receptor)
ADCYAP1→ADCYAP1R1
ADCYAP1→VIPR1
ADCYAP1→VIPR2
Compartment
epithelial
- integumentary
- respiratory
- gastrointestinal
- urinary
- reproductive
neural
- GABAergic
- hybrid
- glutamatergic
- specialized
- neuroendocrine
- glia
germ
- spermatogonium
- spermatocyte
- spermatid
12.5%
25%
50%
100%
ln(CP10K+1)
Abbrev. Tissue
Aort Aorta
Blad Bladder
Blo Blood
Bone Bone
BM Bone marrow
BrC Brain cortex
BrS Brainstem
Col Colon
Dia Diaphragm
Ret Eye retina
Fat Fat
Hrt Heart
Pit Hypothalamus/Pituitary
Kid Kidney
LM Limb muscle
Liv Liver
Lung Lung
MG Mammary gland
Panc Pancreas
Skin Skin
SI Small intestine
Spl Spleen
Test Testes
Thym Thymus
Tong Tongue
Trac Trachea
Ute Uterus

**Pituitary adenylate cyclase-activating polypeptide (PACAP)**
- endothelial

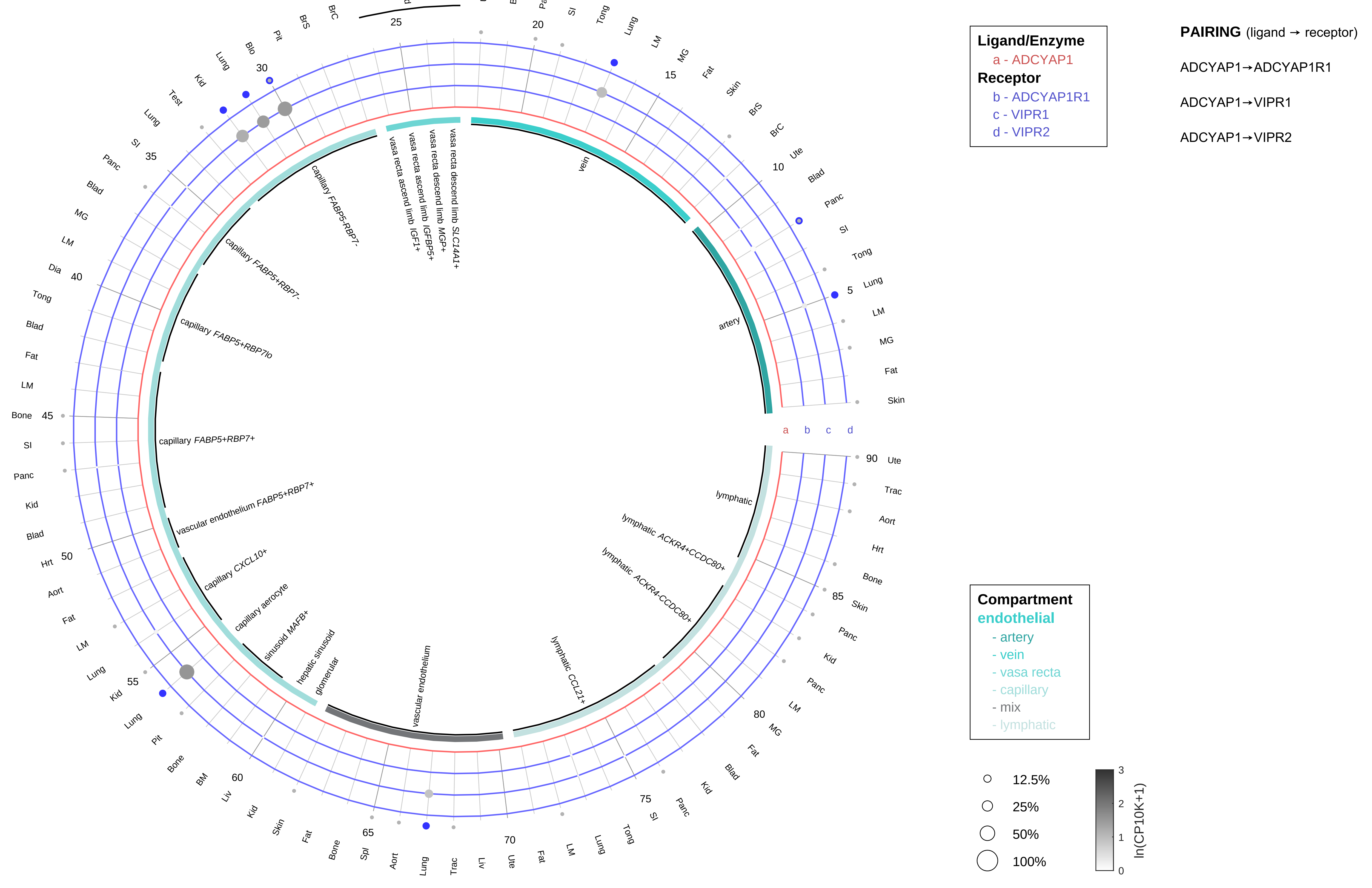

| Abbrev. | Tissue |
|---|---|
| Aort | Aorta |
| Blad | Bladder |
| Blo | Blood |
| Bone | Bone |
| BM | Bone marrow |
| BrC | Brain cortex |
| BrS | Brainstem |
| Col | Colon |
| Dia | Diaphragm |
| Ret | Eye retina |
| Fat | Fat |
| Hrt | Heart |
| Pit | Hypothalamus/Pituitary |
| Kid | Kidney |
| LM | Limb muscle |
| Liv | Liver |
| Lung | Lung |
| MG | Mammary gland |
| Panc | Pancreas |
| Skin | Skin |
| SI | Small intestine |
| Spl | Spleen |
| Test | Testes |
| Thym | Thymus |
| Tong | Tongue |
| Trac | Trachea |
| Ute | Uterus |

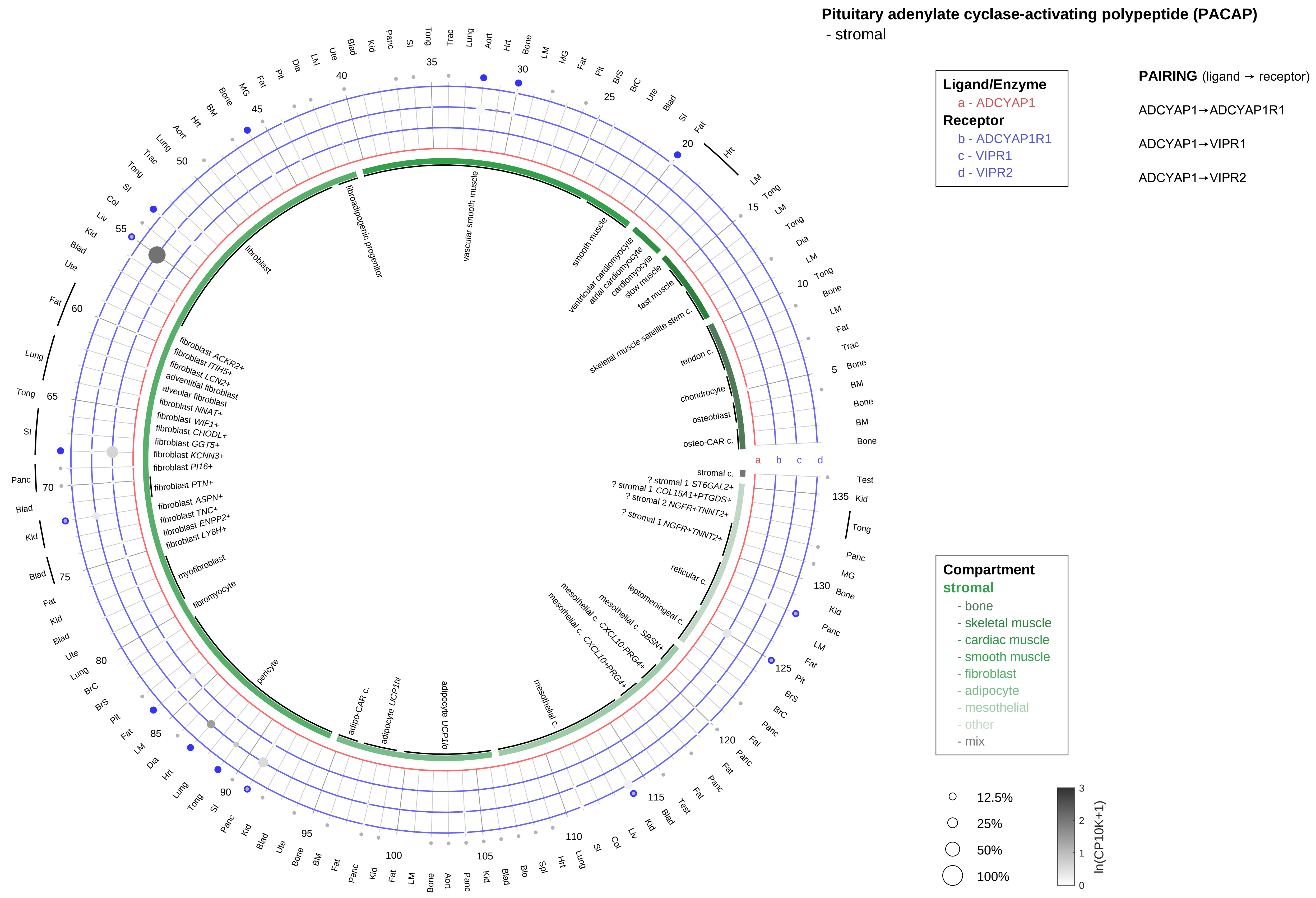

| Abbrev. | Tissue |
|---|---|
| Aort | Aorta |
| Blad | Bladder |
| Blo | Blood |
| Bone | Bone |
| BM | Bone marrow |
| BrC | Brain cortex |
| BrS | Brainstem |
| Col | Colon |
| Dia | Diaphragm |
| Ret | Eye retina |
| Fat | Fat |
| Hrt | Heart |
| Pit | Hypothalamus/Pituitary |
| Kid | Kidney |
| LM | Limb muscle |
| Liv | Liver |
| Lung | Lung |
| MG | Mammary gland |
| Panc | Pancreas |
| Skin | Skin |
| SI | Small intestine |
| Spl | Spleen |
| Test | Testes |
| Thym | Thymus |
| Tong | Tongue |
| Trac | Trachea |
| Ute | Uterus |

**Pituitary adenylate cyclase-activating polypeptide (PACAP)**
- lymphoid

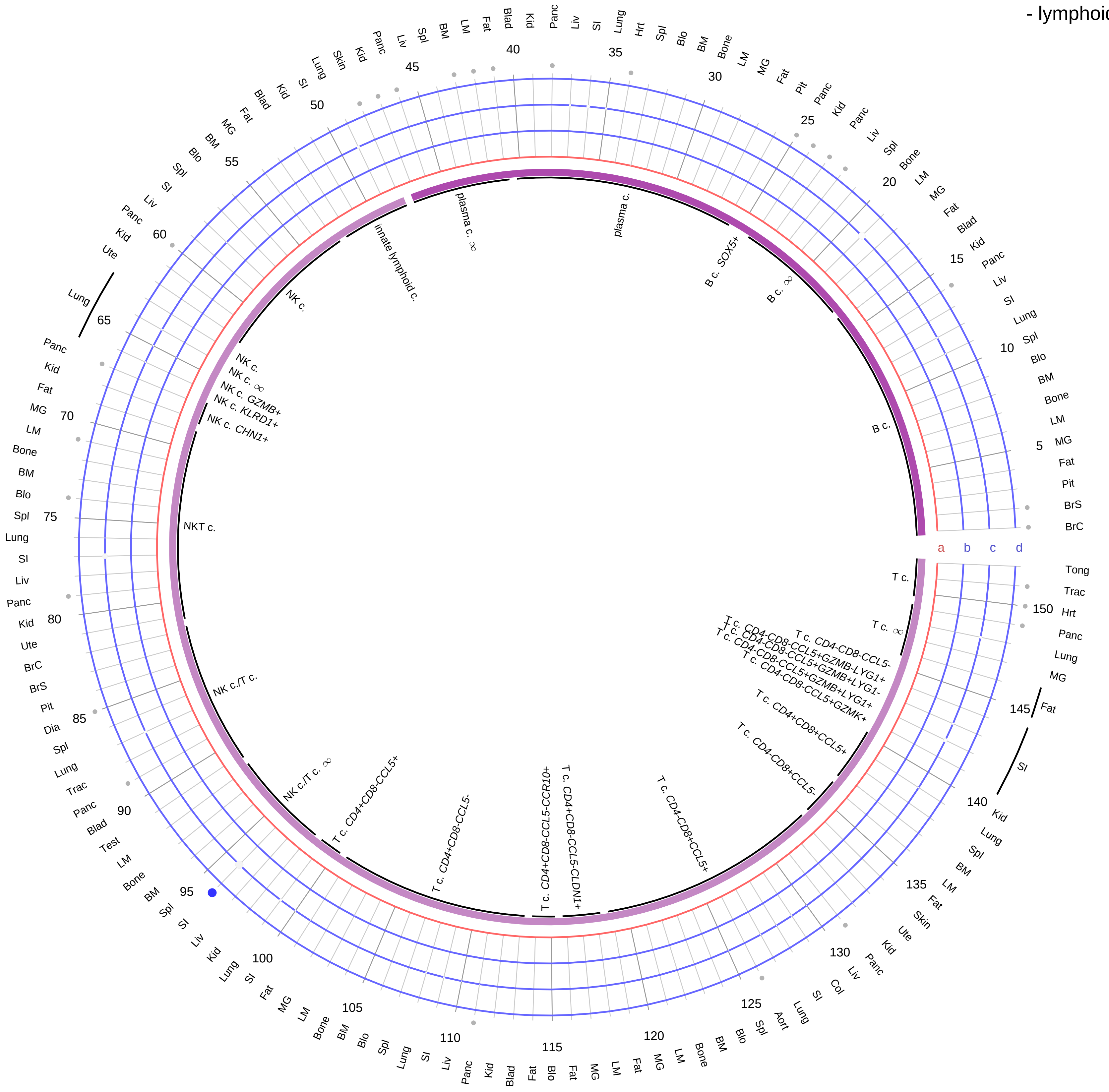

**Ligand/Enzyme**
a - ADCYAP1
**Receptor**
b - ADCYAP1R1
c - VIPR1
d - VIPR2

**PAIRING** (ligand → receptor)

ADCYAP1→ADCYAP1R1

ADCYAP1→VIPR1

ADCYAP1→VIPR2

**Compartment**
**lymphoid**
- B cell
- NK/T cell

12.5%
25%
50%
100%

ln(CP10K+1): 0, 1, 2, 3

| Abbrev. | Tissue |
|---|---|
| Aort | Aorta |
| Blad | Bladder |
| Blo | Blood |
| Bone | Bone |
| BM | Bone marrow |
| BrC | Brain cortex |
| BrS | Brainstem |
| Col | Colon |
| Dia | Diaphragm |
| Ret | Eye retina |
| Fat | Fat |
| Hrt | Heart |
| Pit | Hypothalamus/Pituitary |
| Kid | Kidney |
| LM | Limb muscle |
| Liv | Liver |
| Lung | Lung |
| MG | Mammary gland |
| Panc | Pancreas |
| Skin | Skin |
| SI | Small intestine |
| Spl | Spleen |
| Test | Testes |
| Thym | Thymus |
| Tong | Tongue |
| Trac | Trachea |
| Ute | Uterus |

**Pituitary adenylate cyclase-activating polypeptide (PACAP)**
- myeloid

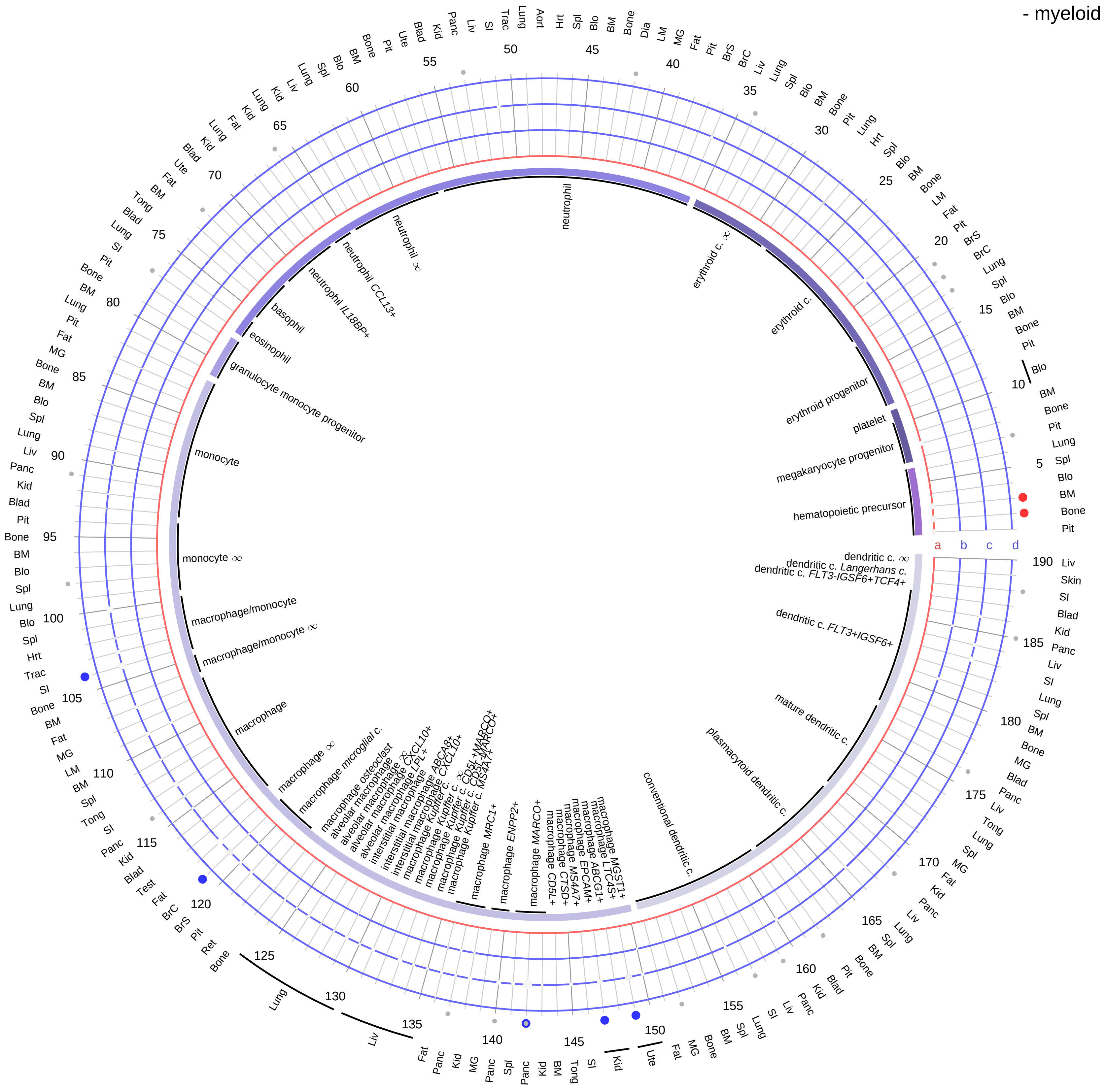

**Ligand/Enzyme**
a - ADCYAP1
**Receptor**
b - ADCYAP1R1
c - VIPR1
d - VIPR2

**PAIRING** (ligand → receptor)

ADCYAP1→ADCYAP1R1

ADCYAP1→VIPR1

ADCYAP1→VIPR2

| Abbrev. | Tissue |
|---|---|
| Aort | Aorta |
| Blad | Bladder |
| Blo | Blood |
| Bone | Bone |
| BM | Bone marrow |
| BrC | Brain cortex |
| BrS | Brainstem |
| Col | Colon |
| Dia | Diaphragm |
| Ret | Eye retina |
| Fat | Fat |
| Hrt | Heart |
| Pit | Hypothalamus/Pituitary |
| Kid | Kidney |
| LM | Limb muscle |
| Liv | Liver |
| Lung | Lung |
| MG | Mammary gland |
| Panc | Pancreas |
| Skin | Skin |
| SI | Small intestine |
| Spl | Spleen |
| Test | Testes |
| Thym | Thymus |
| Tong | Tongue |
| Trac | Trachea |
| Ute | Uterus |

**Compartment**
**hematopoietic**
- hematopoietic precursor
**megakaryocyte-erythroid**
- megakaryoid
- erythroid
**myeloid**
- granulocyte
- granulocyte monocyte progenitor
- monocyte/macrophage
- dendritic

12.5%
25%
50%
100%

ln(CP10K+1): 0, 1, 2, 3

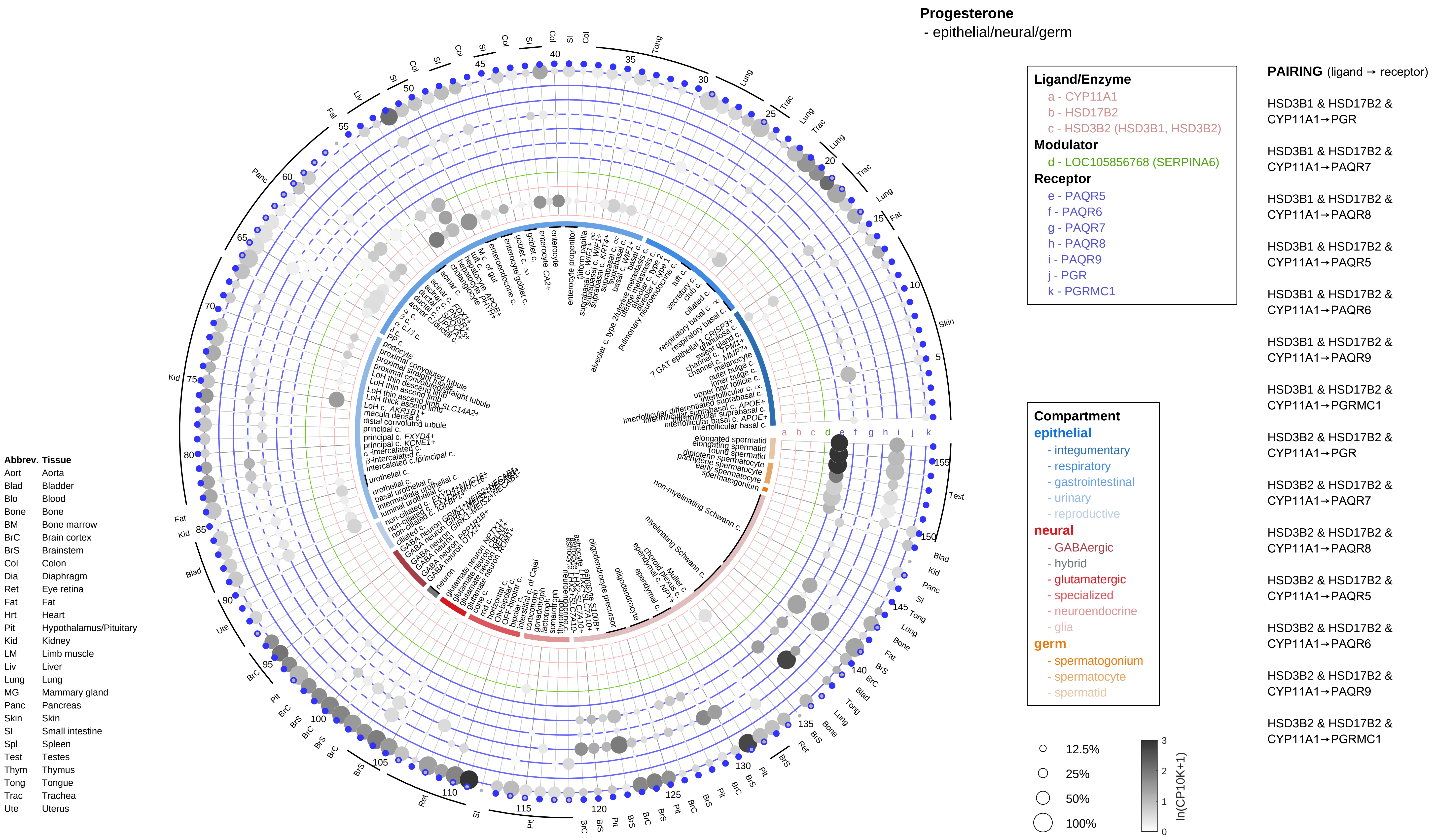
Progesterone
- epithelial/neural/germ
Ligand/Enzyme
a - CYP11A1
b - HSD17B2
c - HSD3B2 (HSD3B1, HSD3B2)
Modulator
d - LOC105856768 (SERPINA6)
Receptor
e - PAQR5
f - PAQR6
g - PAQR7
h - PAQR8
i - PAQR9
j - PGR
k - PGRMC1
PAIRING (ligand → receptor)
HSD3B1 & HSD17B2 & CYP11A1→PGR
HSD3B1 & HSD17B2 & CYP11A1→PAQR7
HSD3B1 & HSD17B2 & CYP11A1→PAQR8
HSD3B1 & HSD17B2 & CYP11A1→PAQR5
HSD3B1 & HSD17B2 & CYP11A1→PAQR6
HSD3B1 & HSD17B2 & CYP11A1→PAQR9
HSD3B1 & HSD17B2 & CYP11A1→PGRMC1
HSD3B2 & HSD17B2 & CYP11A1→PGR
HSD3B2 & HSD17B2 & CYP11A1→PAQR7
HSD3B2 & HSD17B2 & CYP11A1→PAQR8
HSD3B2 & HSD17B2 & CYP11A1→PAQR5
HSD3B2 & HSD17B2 & CYP11A1→PAQR6
HSD3B2 & HSD17B2 & CYP11A1→PAQR9
HSD3B2 & HSD17B2 & CYP11A1→PGRMC1
Compartment
epithelial
- integumentary
- respiratory
- gastrointestinal
- urinary
- reproductive
neural
- GABAergic
- hybrid
- glutamatergic
- specialized
- neuroendocrine
- glia
germ
- spermatogonium
- spermatocyte
- spermatid
12.5%
25%
50%
100%
ln(CP10K+1)
0
1
2
3
Abbrev. Tissue
Aort Aorta
Blad Bladder
Blo Blood
Bone Bone
BM Bone marrow
BrC Brain cortex
BrS Brainstem
Col Colon
Dia Diaphragm
Ret Eye retina
Fat Fat
Hrt Heart
Pit Hypothalamus/Pituitary
Kid Kidney
LM Limb muscle
Liv Liver
Lung Lung
MG Mammary gland
Panc Pancreas
Skin Skin
SI Small intestine
Spl Spleen
Test Testes
Thym Thymus
Tong Tongue
Trac Trachea
Ute Uterus

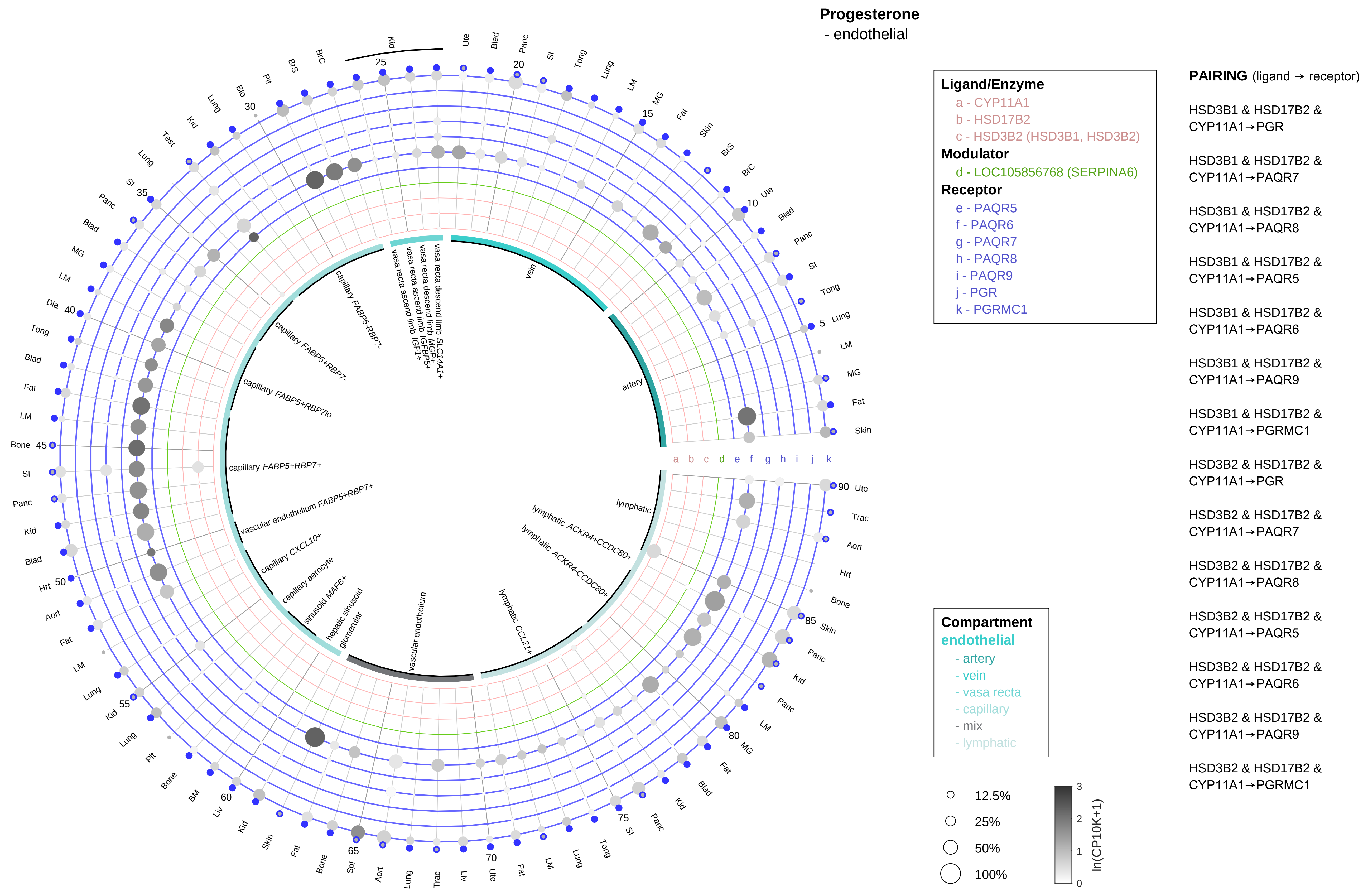

| Abbrev. | Tissue |
|---|---|
| Aort | Aorta |
| Blad | Bladder |
| Blo | Blood |
| Bone | Bone |
| BM | Bone marrow |
| BrC | Brain cortex |
| BrS | Brainstem |
| Col | Colon |
| Dia | Diaphragm |
| Ret | Eye retina |
| Fat | Fat |
| Hrt | Heart |
| Pit | Hypothalamus/Pituitary |
| Kid | Kidney |
| LM | Limb muscle |
| Liv | Liver |
| Lung | Lung |
| MG | Mammary gland |
| Panc | Pancreas |
| Skin | Skin |
| SI | Small intestine |
| Spl | Spleen |
| Test | Testes |
| Thym | Thymus |
| Tong | Tongue |
| Trac | Trachea |
| Ute | Uterus |

**Progesterone**
- stromal

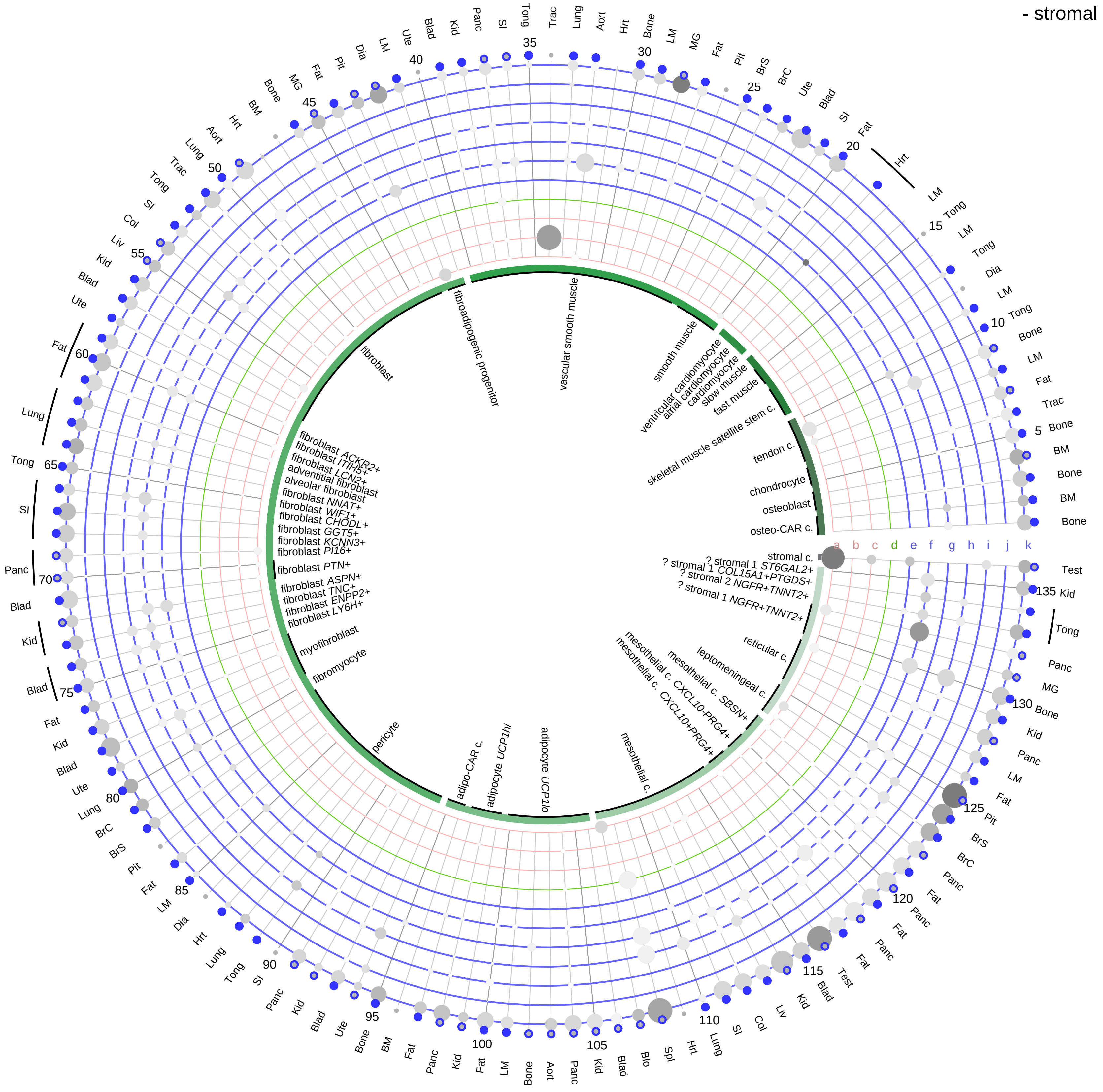

**Ligand/Enzyme**
- a - CYP11A1
- b - HSD17B2
- c - HSD3B2 (HSD3B1, HSD3B2)

**Modulator**
- d - LOC105856768 (SERPINA6)

**Receptor**
- e - PAQR5
- f - PAQR6
- g - PAQR7
- h - PAQR8
- i - PAQR9
- j - PGR
- k - PGRMC1

**PAIRING** (ligand → receptor)

HSD3B1 & HSD17B2 & CYP11A1→PGR

HSD3B1 & HSD17B2 & CYP11A1→PAQR7

HSD3B1 & HSD17B2 & CYP11A1→PAQR8

HSD3B1 & HSD17B2 & CYP11A1→PAQR5

HSD3B1 & HSD17B2 & CYP11A1→PAQR6

HSD3B1 & HSD17B2 & CYP11A1→PAQR9

HSD3B1 & HSD17B2 & CYP11A1→PGRMC1

HSD3B2 & HSD17B2 & CYP11A1→PGR

HSD3B2 & HSD17B2 & CYP11A1→PAQR7

HSD3B2 & HSD17B2 & CYP11A1→PAQR8

HSD3B2 & HSD17B2 & CYP11A1→PAQR5

HSD3B2 & HSD17B2 & CYP11A1→PAQR6

HSD3B2 & HSD17B2 & CYP11A1→PAQR9

HSD3B2 & HSD17B2 & CYP11A1→PGRMC1

**Compartment**

**stromal**
- bone
- skeletal muscle
- cardiac muscle
- smooth muscle
- fibroblast
- adipocyte
- mesothelial
- other
- mix

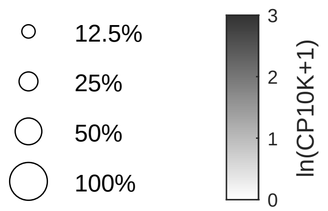

| Abbrev. | Tissue |
|---|---|
| Aort | Aorta |
| Blad | Bladder |
| Blo | Blood |
| Bone | Bone |
| BM | Bone marrow |
| BrC | Brain cortex |
| BrS | Brainstem |
| Col | Colon |
| Dia | Diaphragm |
| Ret | Eye retina |
| Fat | Fat |
| Hrt | Heart |
| Pit | Hypothalamus/Pituitary |
| Kid | Kidney |
| LM | Limb muscle |
| Liv | Liver |
| Lung | Lung |
| MG | Mammary gland |
| Panc | Pancreas |
| Skin | Skin |
| SI | Small intestine |
| Spl | Spleen |
| Test | Testes |
| Thym | Thymus |
| Tong | Tongue |
| Trac | Trachea |
| Ute | Uterus |

**Progesterone**
- lymphoid

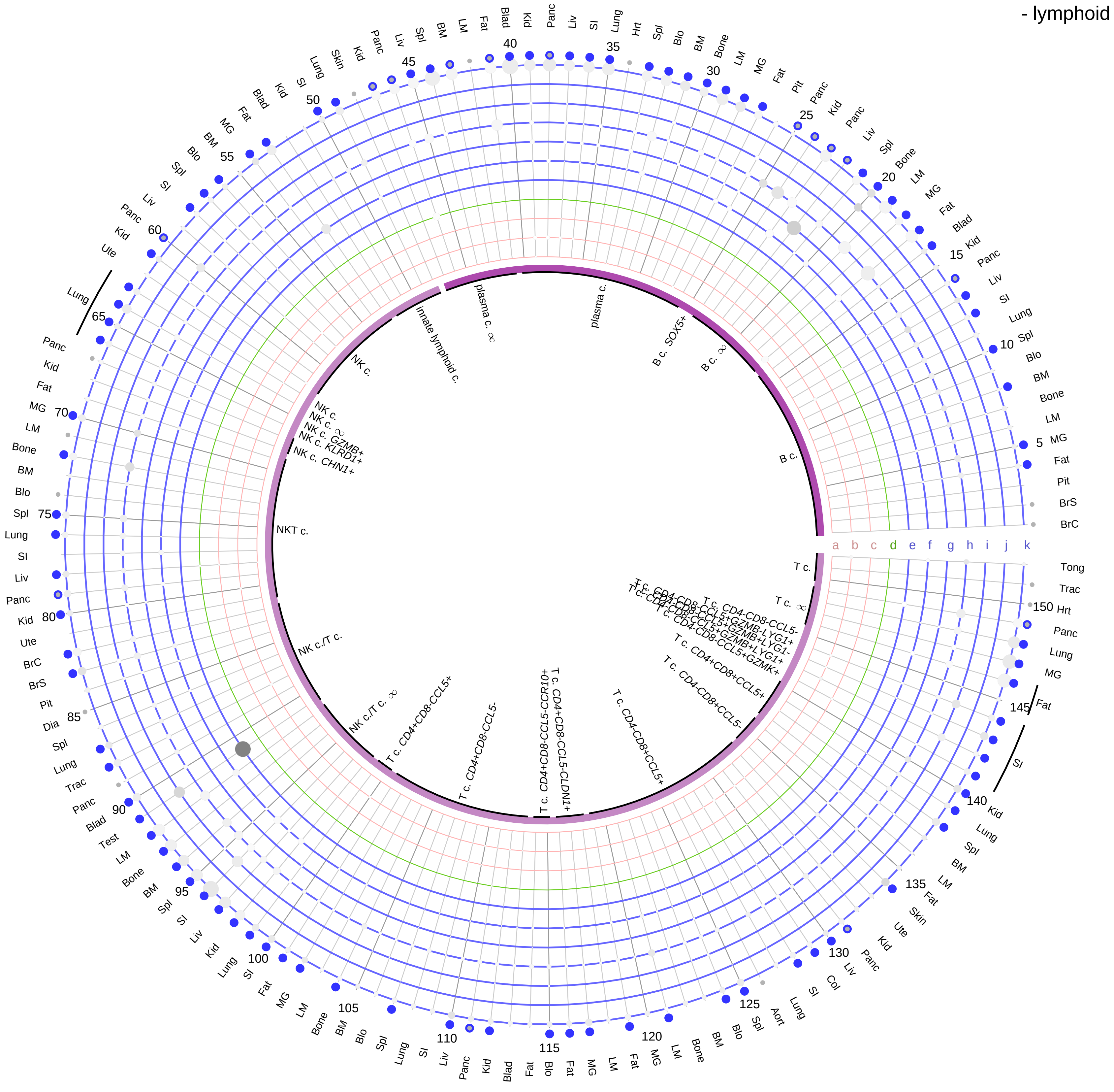

**Ligand/Enzyme**
- a - CYP11A1
- b - HSD17B2
- c - HSD3B2 (HSD3B1, HSD3B2)

**Modulator**
- d - LOC105856768 (SERPINA6)

**Receptor**
- e - PAQR5
- f - PAQR6
- g - PAQR7
- h - PAQR8
- i - PAQR9
- j - PGR
- k - PGRMC1

**Compartment**
**lymphoid**
- B cell
- NK/T cell

○ 12.5%
○ 25%
○ 50%
○ 100%

ln(CP10K+1): 0, 1, 2, 3

**PAIRING** (ligand → receptor)

HSD3B1 & HSD17B2 & CYP11A1→PGR

HSD3B1 & HSD17B2 & CYP11A1→PAQR7

HSD3B1 & HSD17B2 & CYP11A1→PAQR8

HSD3B1 & HSD17B2 & CYP11A1→PAQR5

HSD3B1 & HSD17B2 & CYP11A1→PAQR6

HSD3B1 & HSD17B2 & CYP11A1→PAQR9

HSD3B1 & HSD17B2 & CYP11A1→PGRMC1

HSD3B2 & HSD17B2 & CYP11A1→PGR

HSD3B2 & HSD17B2 & CYP11A1→PAQR7

HSD3B2 & HSD17B2 & CYP11A1→PAQR8

HSD3B2 & HSD17B2 & CYP11A1→PAQR5

HSD3B2 & HSD17B2 & CYP11A1→PAQR6

HSD3B2 & HSD17B2 & CYP11A1→PAQR9

HSD3B2 & HSD17B2 & CYP11A1→PGRMC1

| Abbrev. | Tissue |
|---|---|
| Aort | Aorta |
| Blad | Bladder |
| Blo | Blood |
| Bone | Bone |
| BM | Bone marrow |
| BrC | Brain cortex |
| BrS | Brainstem |
| Col | Colon |
| Dia | Diaphragm |
| Ret | Eye retina |
| Fat | Fat |
| Hrt | Heart |
| Pit | Hypothalamus/Pituitary |
| Kid | Kidney |
| LM | Limb muscle |
| Liv | Liver |
| Lung | Lung |
| MG | Mammary gland |
| Panc | Pancreas |
| Skin | Skin |
| SI | Small intestine |
| Spl | Spleen |
| Test | Testes |
| Thym | Thymus |
| Tong | Tongue |
| Trac | Trachea |
| Ute | Uterus |

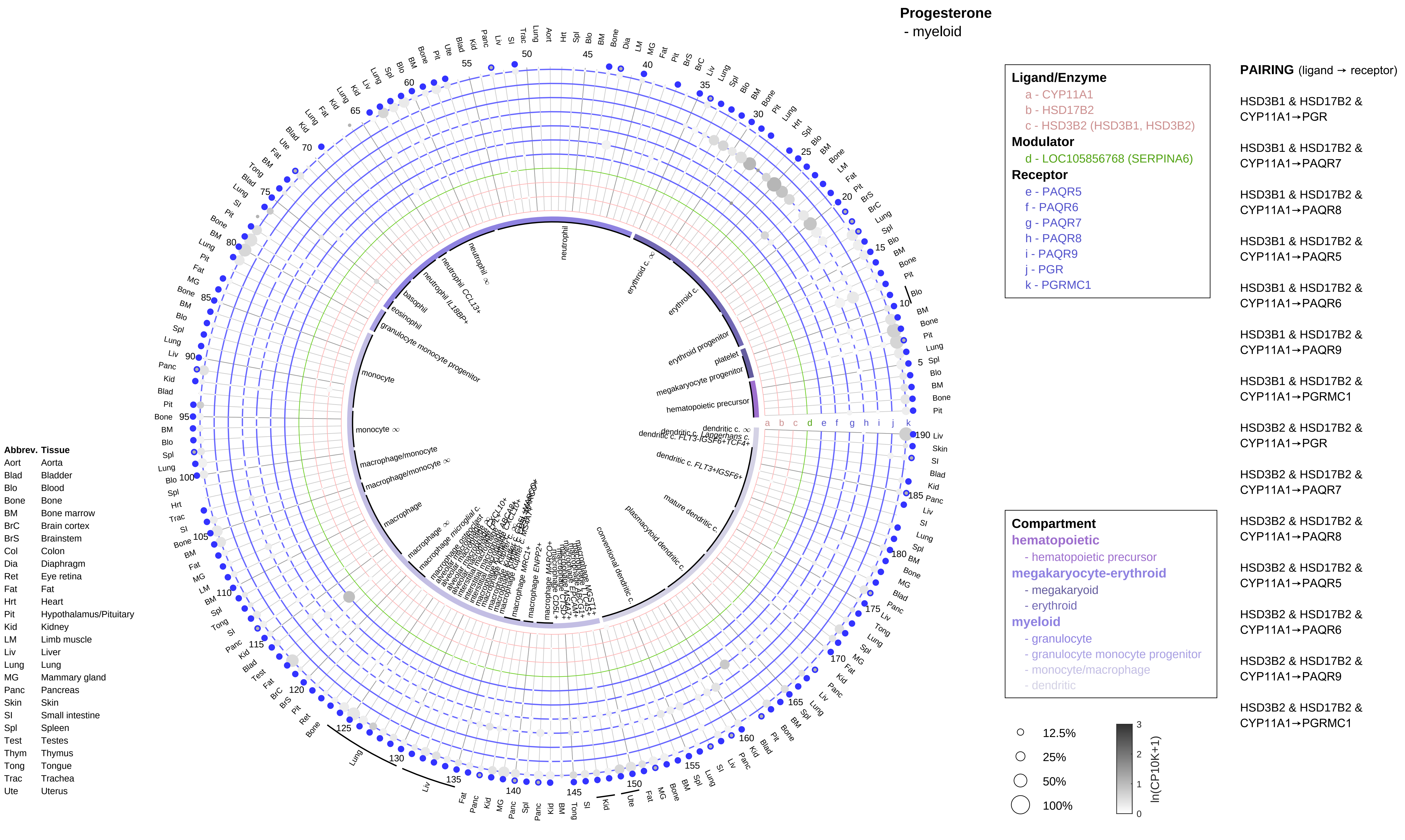

Progesterone
- myeloid
Ligand/Enzyme
a - CYP11A1
b - HSD17B2
c - HSD3B2 (HSD3B1, HSD3B2)
Modulator
d - LOC105856768 (SERPINA6)
Receptor
e - PAQR5
f - PAQR6
g - PAQR7
h - PAQR8
i - PAQR9
j - PGR
k - PGRMC1
PAIRING (ligand → receptor)
HSD3B1 & HSD17B2 & CYP11A1→PGR
HSD3B1 & HSD17B2 & CYP11A1→PAQR7
HSD3B1 & HSD17B2 & CYP11A1→PAQR8
HSD3B1 & HSD17B2 & CYP11A1→PAQR5
HSD3B1 & HSD17B2 & CYP11A1→PAQR6
HSD3B1 & HSD17B2 & CYP11A1→PAQR9
HSD3B1 & HSD17B2 & CYP11A1→PGRMC1
HSD3B2 & HSD17B2 & CYP11A1→PGR
HSD3B2 & HSD17B2 & CYP11A1→PAQR7
HSD3B2 & HSD17B2 & CYP11A1→PAQR8
HSD3B2 & HSD17B2 & CYP11A1→PAQR5
HSD3B2 & HSD17B2 & CYP11A1→PAQR6
HSD3B2 & HSD17B2 & CYP11A1→PAQR9
HSD3B2 & HSD17B2 & CYP11A1→PGRMC1
Compartment
hematopoietic
- hematopoietic precursor
megakaryocyte-erythroid
- megakaryoid
- erythroid
myeloid
- granulocyte
- granulocyte monocyte progenitor
- monocyte/macrophage
- dendritic
12.5%
25%
50%
100%
ln(CP10K+1)
Abbrev. Tissue
Aort Aorta
Blad Bladder
Blo Blood
Bone Bone
BM Bone marrow
BrC Brain cortex
BrS Brainstem
Col Colon
Dia Diaphragm
Ret Eye retina
Fat Fat
Hrt Heart
Pit Hypothalamus/Pituitary
Kid Kidney
LM Limb muscle
Liv Liver
Lung Lung
MG Mammary gland
Panc Pancreas
Skin Skin
SI Small intestine
Spl Spleen
Test Testes
Thym Thymus
Tong Tongue
Trac Trachea
Ute Uterus
neutrophil
erythroid c.
erythroid progenitor
platelet
megakaryocyte progenitor
hematopoietic precursor
dendritic c. FLT3+IGSF6+
mature dendritic c.
plasmacytoid dendritic c.
conventional dendritic c.
macrophage
macrophage/monocyte
monocyte
granulocyte monocyte progenitor
eosinophil
basophil

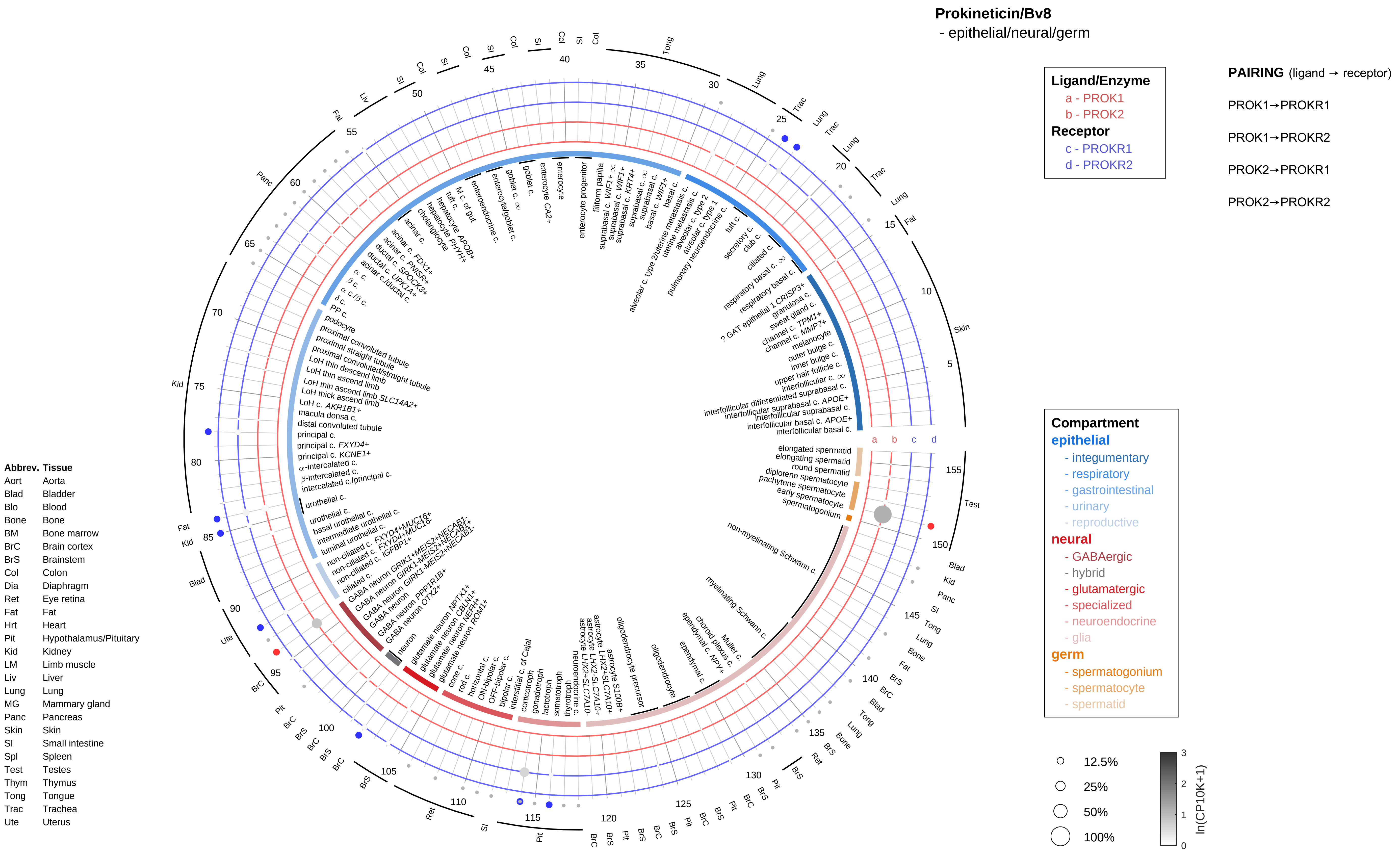

Prokineticin/Bv8
- epithelial/neural/germ
Ligand/Enzyme
a - PROK1
b - PROK2
Receptor
c - PROKR1
d - PROKR2
PAIRING (ligand → receptor)
PROK1→PROKR1
PROK1→PROKR2
PROK2→PROKR1
PROK2→PROKR2
Compartment
epithelial
- integumentary
- respiratory
- gastrointestinal
- urinary
- reproductive
neural
- GABAergic
- hybrid
- glutamatergic
- specialized
- neuroendocrine
- glia
germ
- spermatogonium
- spermatocyte
- spermatid
12.5%
25%
50%
100%
ln(CP10K+1)
Abbrev. Tissue
Aort Aorta
Blad Bladder
Blo Blood
Bone Bone
BM Bone marrow
BrC Brain cortex
BrS Brainstem
Col Colon
Dia Diaphragm
Ret Eye retina
Fat Fat
Hrt Heart
Pit Hypothalamus/Pituitary
Kid Kidney
LM Limb muscle
Liv Liver
Lung Lung
MG Mammary gland
Panc Pancreas
Skin Skin
SI Small intestine
Spl Spleen
Test Testes
Thym Thymus
Tong Tongue
Trac Trachea
Ute Uterus

**Prokineticin/Bv8**
- endothelial

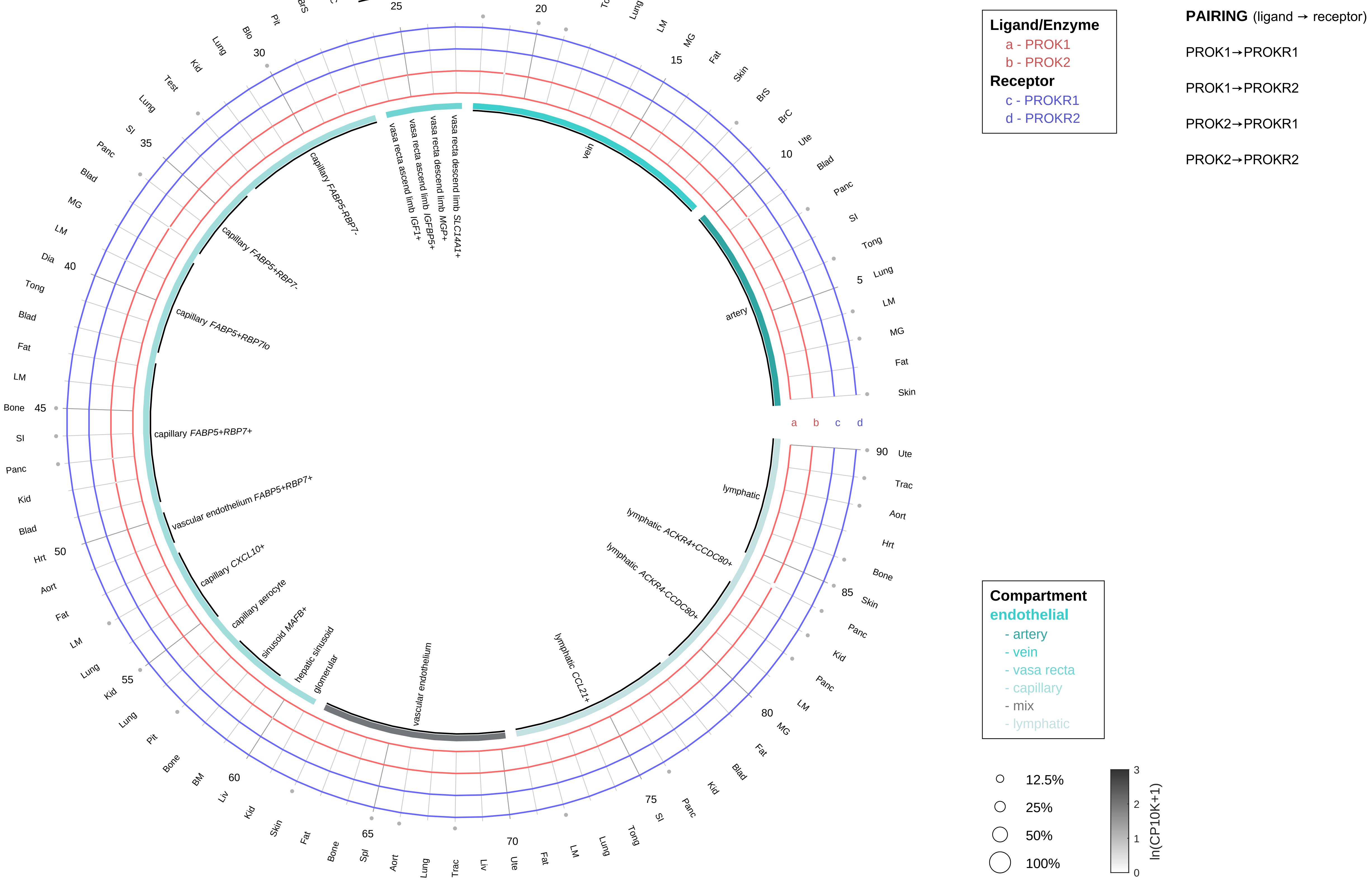

| Abbrev. | Tissue |
|---|---|
| Aort | Aorta |
| Blad | Bladder |
| Blo | Blood |
| Bone | Bone |
| BM | Bone marrow |
| BrC | Brain cortex |
| BrS | Brainstem |
| Col | Colon |
| Dia | Diaphragm |
| Ret | Eye retina |
| Fat | Fat |
| Hrt | Heart |
| Pit | Hypothalamus/Pituitary |
| Kid | Kidney |
| LM | Limb muscle |
| Liv | Liver |
| Lung | Lung |
| MG | Mammary gland |
| Panc | Pancreas |
| Skin | Skin |
| SI | Small intestine |
| Spl | Spleen |
| Test | Testes |
| Thym | Thymus |
| Tong | Tongue |
| Trac | Trachea |
| Ute | Uterus |

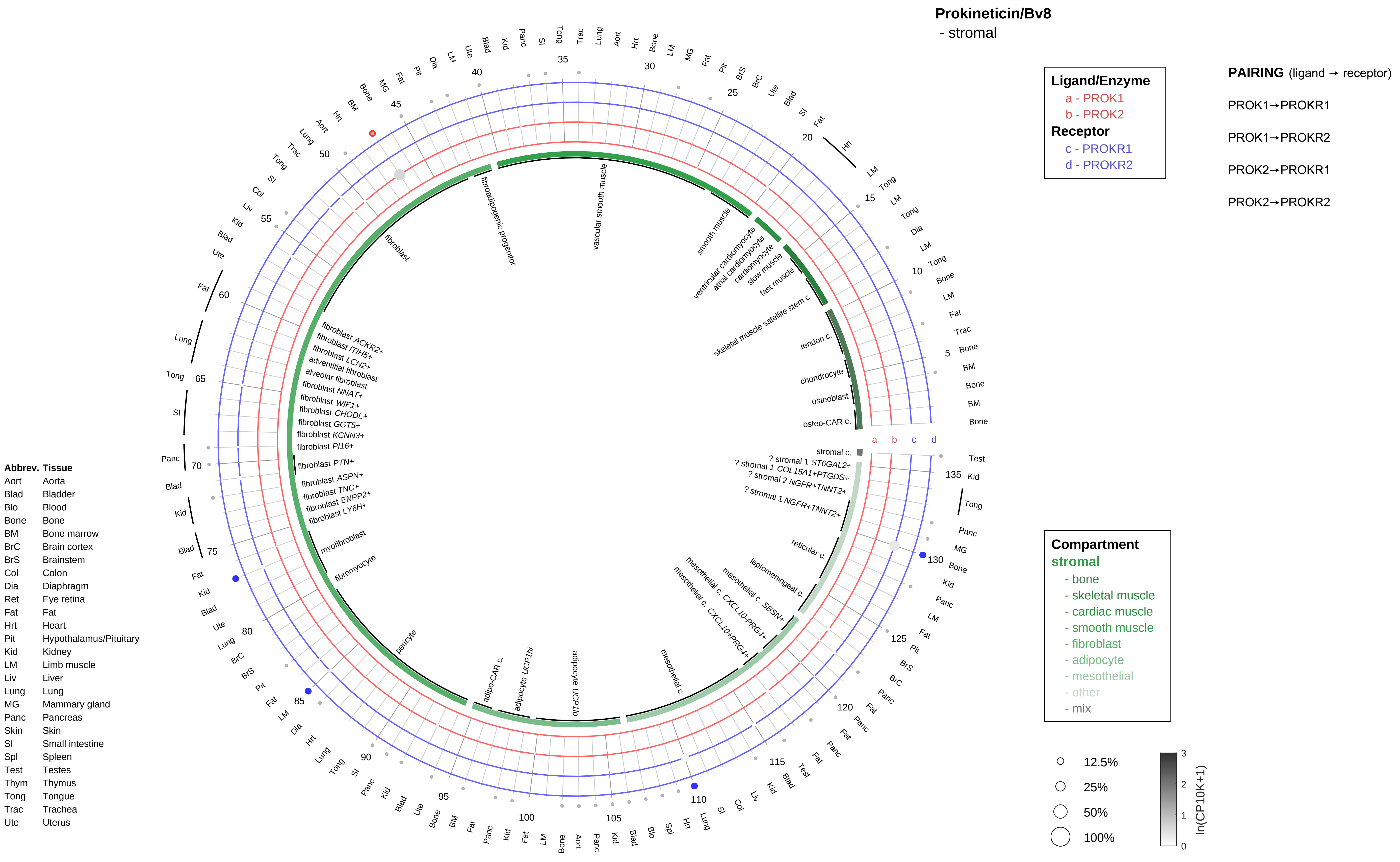

Prokineticin/Bv8
- stromal
Ligand/Enzyme
a - PROK1
b - PROK2
Receptor
c - PROKR1
d - PROKR2
PAIRING (ligand → receptor)
PROK1→PROKR1
PROK1→PROKR2
PROK2→PROKR1
PROK2→PROKR2
Compartment
stromal
- bone
- skeletal muscle
- cardiac muscle
- smooth muscle
- fibroblast
- adipocyte
- mesothelial
- other
- mix
12.5%
25%
50%
100%
ln(CP10K+1)
Abbrev. Tissue
Aort Aorta
Blad Bladder
Blo Blood
Bone Bone
BM Bone marrow
BrC Brain cortex
BrS Brainstem
Col Colon
Dia Diaphragm
Ret Eye retina
Fat Fat
Hrt Heart
Pit Hypothalamus/Pituitary
Kid Kidney
LM Limb muscle
Liv Liver
Lung Lung
MG Mammary gland
Panc Pancreas
Skin Skin
SI Small intestine
Spl Spleen
Test Testes
Thym Thymus
Tong Tongue
Trac Trachea
Ute Uterus
stromal c.
? stromal 1 ST6GAL2+
? stromal 1 COL15A1+PTGDS+
? stromal 2 NGFR+TNNT2+
? stromal 1 NGFR+TNNT2+
reticular c.
leptomeningeal c.
mesothelial c. SBSN+
mesothelial c. CXCL10-PRG4+
mesothelial c. CXCL10+PRG4+
mesothelial c.
adipocyte UCP1lo
adipocyte UCP1hi
adipo-CAR c.
pericyte
fibromyocyte
myofibroblast
fibroblast LY6H+
fibroblast ENPP2+
fibroblast TNC+
fibroblast ASPN+
fibroblast PTN+
fibroblast PI16+
fibroblast KCNN3+
fibroblast GGT5+
fibroblast CHODL+
fibroblast WIF1+
fibroblast NNAT+
alveolar fibroblast
adventitial fibroblast
fibroblast LCN2+
fibroblast ITIH5+
fibroblast ACKR2+
fibroblast
fibroadipogenic progenitor
vascular smooth muscle
smooth muscle
ventricular cardiomyocyte
atrial cardiomyocyte
cardiomyocyte
slow muscle
fast muscle
skeletal muscle satellite stem c.
tendon c.
chondrocyte
osteoblast
osteo-CAR c.

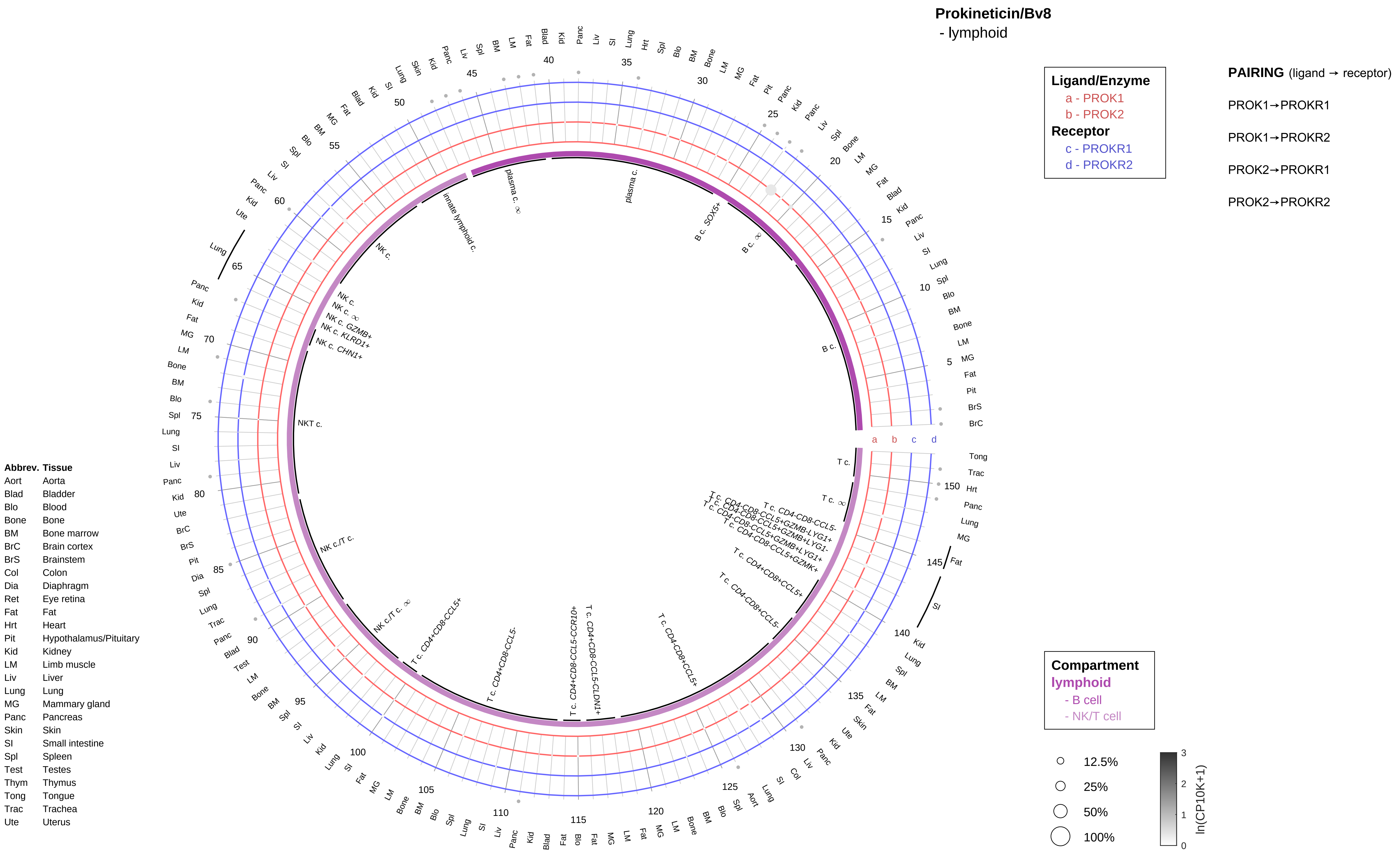

**Prokineticin/Bv8**
- lymphoid

**Ligand/Enzyme**
a - PROK1
b - PROK2
**Receptor**
c - PROKR1
d - PROKR2

**PAIRING** (ligand → receptor)
PROK1→PROKR1
PROK1→PROKR2
PROK2→PROKR1
PROK2→PROKR2

**Compartment**
lymphoid
- B cell
- NK/T cell

12.5%
25%
50%
100%

ln(CP10K+1)
0
1
2
3

**Abbrev. Tissue**
Aort Aorta
Blad Bladder
Blo Blood
Bone Bone
BM Bone marrow
BrC Brain cortex
BrS Brainstem
Col Colon
Dia Diaphragm
Ret Eye retina
Fat Fat
Hrt Heart
Pit Hypothalamus/Pituitary
Kid Kidney
LM Limb muscle
Liv Liver
Lung Lung
MG Mammary gland
Panc Pancreas
Skin Skin
SI Small intestine
Spl Spleen
Test Testes
Thym Thymus
Tong Tongue
Trac Trachea
Ute Uterus

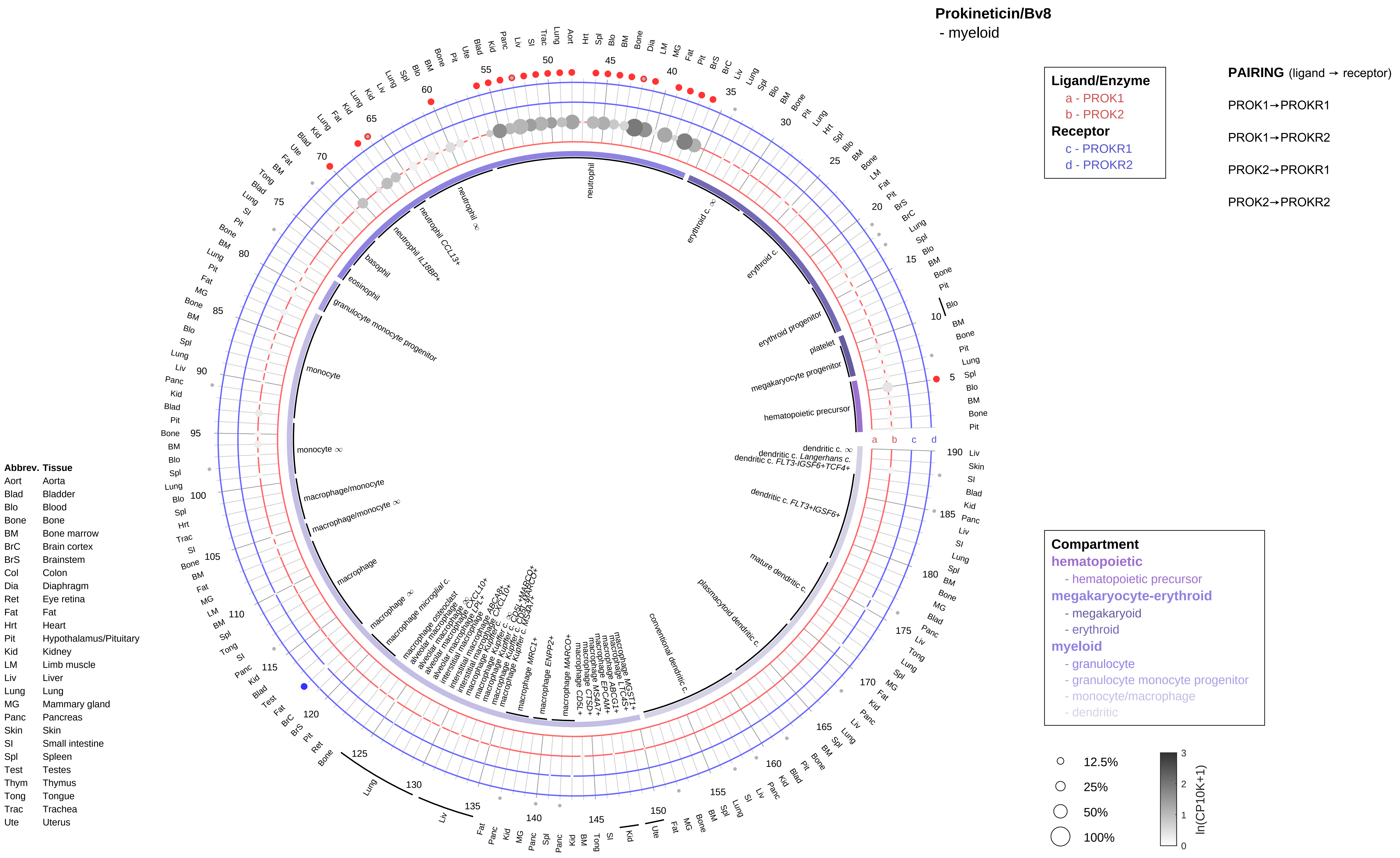

Prokineticin/Bv8
- myeloid
Ligand/Enzyme
a - PROK1
b - PROK2
Receptor
c - PROKR1
d - PROKR2
PAIRING (ligand → receptor)
PROK1→PROKR1
PROK1→PROKR2
PROK2→PROKR1
PROK2→PROKR2
Compartment
hematopoietic
- hematopoietic precursor
megakaryocyte-erythroid
- megakaryoid
- erythroid
myeloid
- granulocyte
- granulocyte monocyte progenitor
- monocyte/macrophage
- dendritic
12.5%
25%
50%
100%
ln(CP10K+1)
Abbrev. Tissue
Aort Aorta
Blad Bladder
Blo Blood
Bone Bone
BM Bone marrow
BrC Brain cortex
BrS Brainstem
Col Colon
Dia Diaphragm
Ret Eye retina
Fat Fat
Hrt Heart
Pit Hypothalamus/Pituitary
Kid Kidney
LM Limb muscle
Liv Liver
Lung Lung
MG Mammary gland
Panc Pancreas
Skin Skin
SI Small intestine
Spl Spleen
Test Testes
Thym Thymus
Tong Tongue
Trac Trachea
Ute Uterus
neutrophil
erythroid c.
erythroid progenitor
platelet
megakaryocyte progenitor
hematopoietic precursor
basophil
eosinophil
granulocyte monocyte progenitor
monocyte
macrophage/monocyte
macrophage
conventional dendritic c.
plasmacytoid dendritic c.
mature dendritic c.

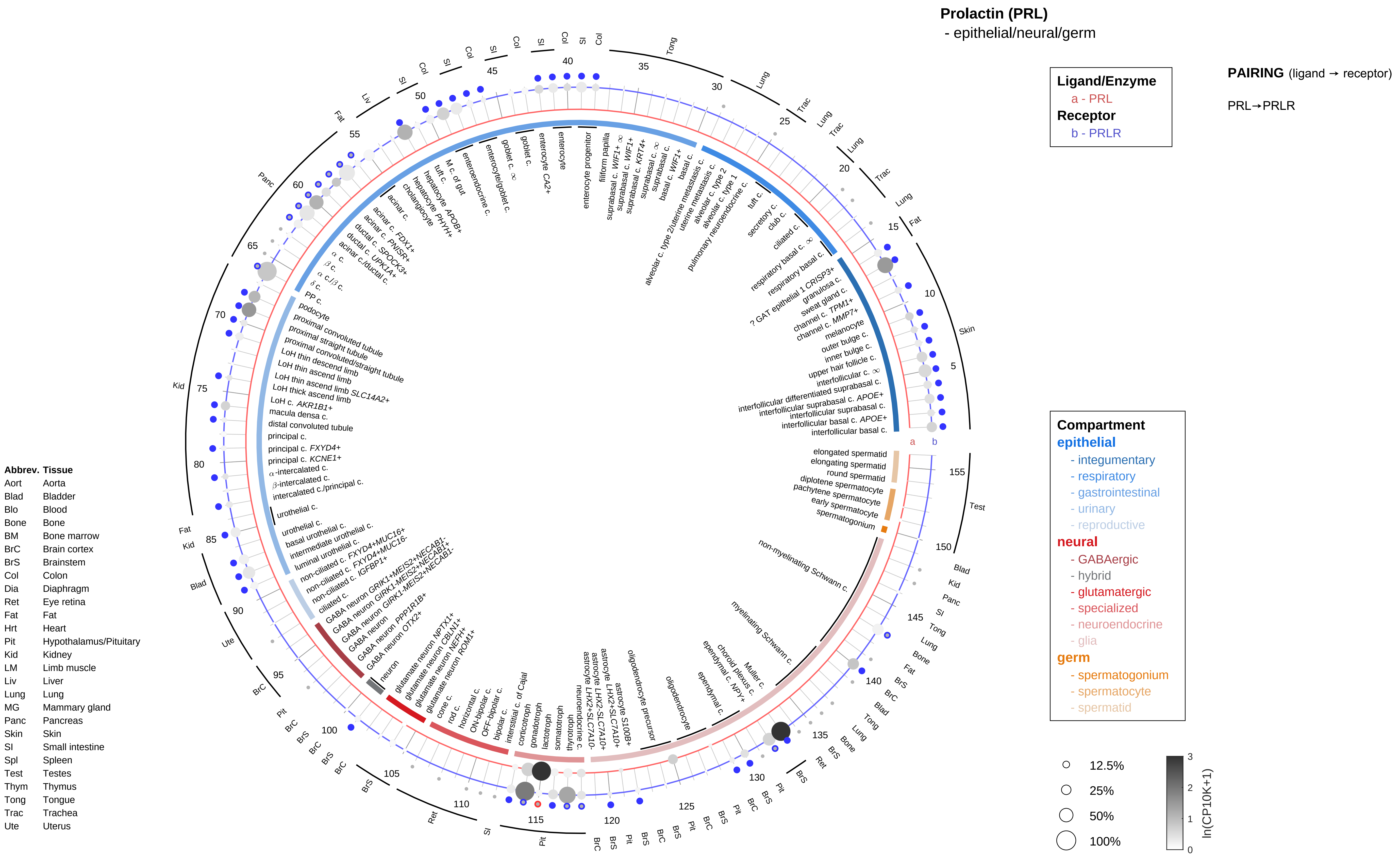
Prolactin (PRL)
- epithelial/neural/germ
Ligand/Enzyme
a - PRL
Receptor
b - PRLR
PAIRING (ligand → receptor)
PRL→PRLR
Compartment
epithelial
- integumentary
- respiratory
- gastrointestinal
- urinary
- reproductive
neural
- GABAergic
- hybrid
- glutamatergic
- specialized
- neuroendocrine
- glia
germ
- spermatogonium
- spermatocyte
- spermatid
12.5%
25%
50%
100%
ln(CP10K+1)
Abbrev. Tissue
Aort Aorta
Blad Bladder
Blo Blood
Bone Bone
BM Bone marrow
BrC Brain cortex
BrS Brainstem
Col Colon
Dia Diaphragm
Ret Eye retina
Fat Fat
Hrt Heart
Pit Hypothalamus/Pituitary
Kid Kidney
LM Limb muscle
Liv Liver
Lung Lung
MG Mammary gland
Panc Pancreas
Skin Skin
SI Small intestine
Spl Spleen
Test Testes
Thym Thymus
Tong Tongue
Trac Trachea
Ute Uterus

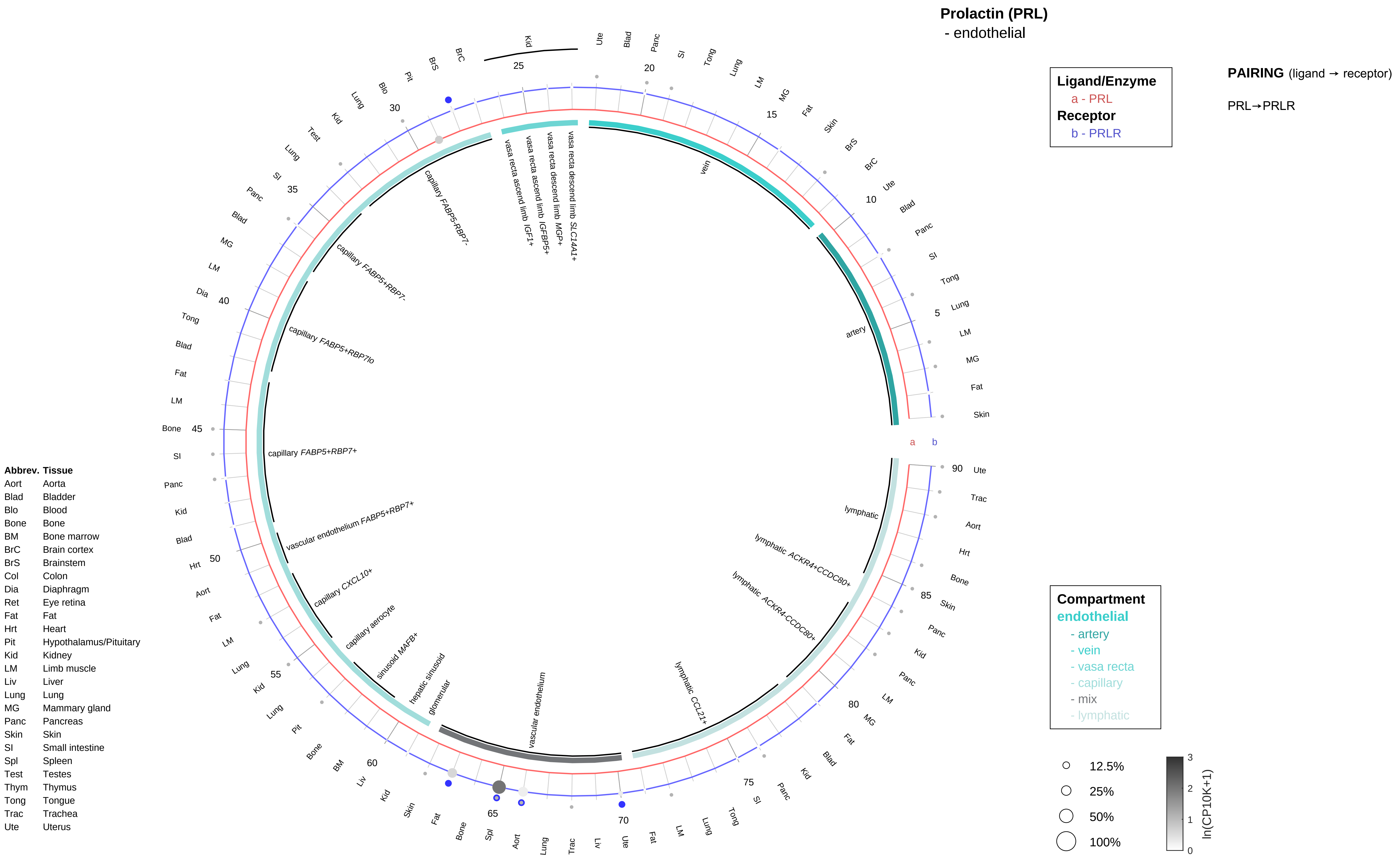
Prolactin (PRL)
- endothelial
Ligand/Enzyme
a - PRL
Receptor
b - PRLR
PAIRING (ligand → receptor)
PRL→PRLR
Compartment
endothelial
- artery
- vein
- vasa recta
- capillary
- mix
- lymphatic
12.5%
25%
50%
100%
ln(CP10K+1)
artery
vein
vasa recta descend limb SLC14A1+
vasa recta descend limb MGP+
vasa recta ascend limb IGFBP5+
vasa recta ascend limb IGF1+
capillary FABP5-RBP7-
capillary FABP5+RBP7-
capillary FABP5+RBP7lo
capillary FABP5+RBP7+
vascular endothelium FABP5+RBP7+
capillary CXCL10+
capillary aerocyte
sinusoid MAFB+
hepatic sinusoid
glomerular
vascular endothelium
lymphatic CCL21+
lymphatic ACKR4-CCDC80+
lymphatic ACKR4+CCDC80+
lymphatic
Abbrev. Tissue
Aort Aorta
Blad Bladder
Blo Blood
Bone Bone
BM Bone marrow
BrC Brain cortex
BrS Brainstem
Col Colon
Dia Diaphragm
Ret Eye retina
Fat Fat
Hrt Heart
Pit Hypothalamus/Pituitary
Kid Kidney
LM Limb muscle
Liv Liver
Lung Lung
MG Mammary gland
Panc Pancreas
Skin Skin
SI Small intestine
Spl Spleen
Test Testes
Thym Thymus
Tong Tongue
Trac Trachea
Ute Uterus

**Prolactin (PRL)**
- stromal

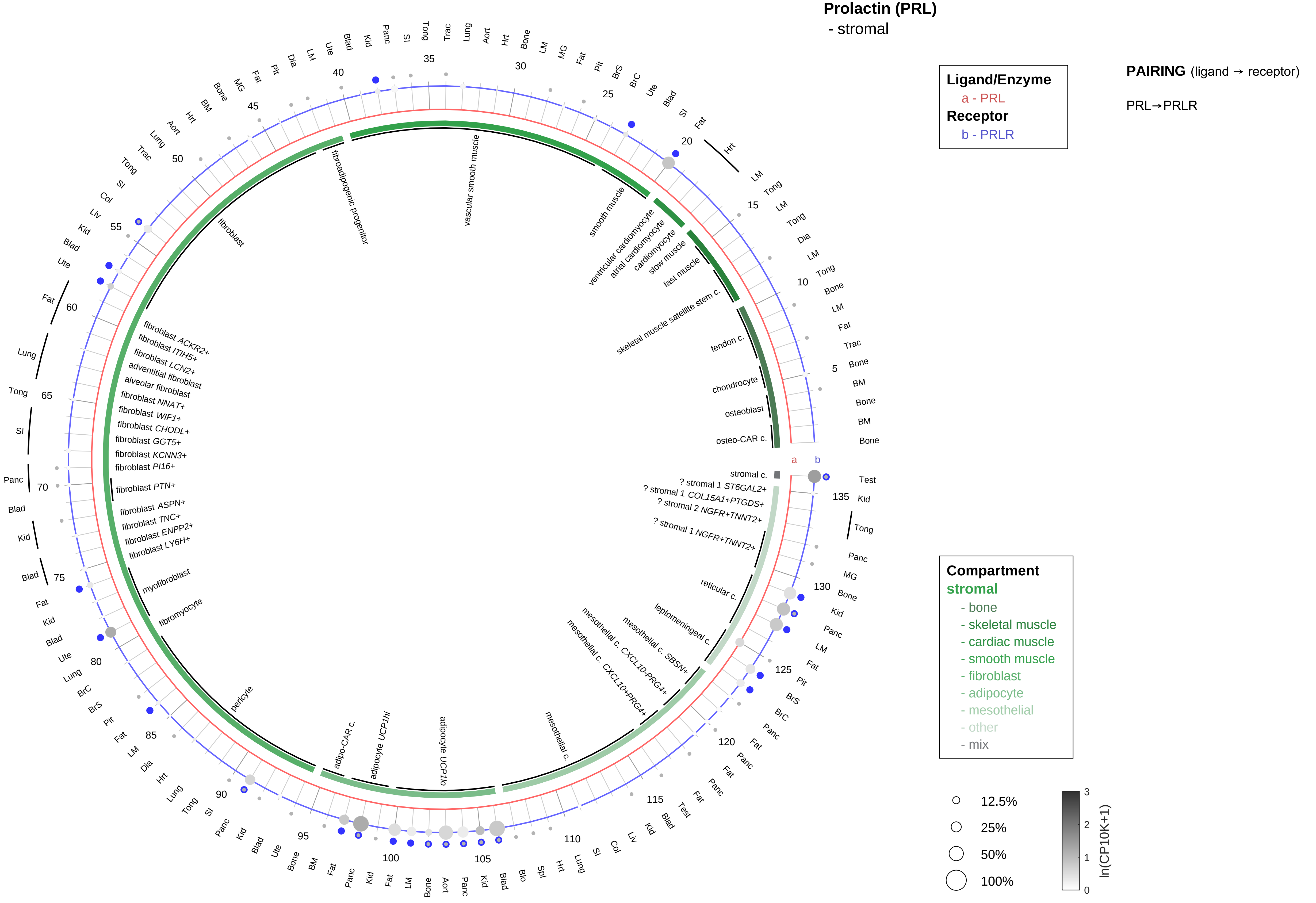

| Abbrev. | Tissue |
|---|---|
| Aort | Aorta |
| Blad | Bladder |
| Blo | Blood |
| Bone | Bone |
| BM | Bone marrow |
| BrC | Brain cortex |
| BrS | Brainstem |
| Col | Colon |
| Dia | Diaphragm |
| Ret | Eye retina |
| Fat | Fat |
| Hrt | Heart |
| Pit | Hypothalamus/Pituitary |
| Kid | Kidney |
| LM | Limb muscle |
| Liv | Liver |
| Lung | Lung |
| MG | Mammary gland |
| Panc | Pancreas |
| Skin | Skin |
| SI | Small intestine |
| Spl | Spleen |
| Test | Testes |
| Thym | Thymus |
| Tong | Tongue |
| Trac | Trachea |
| Ute | Uterus |

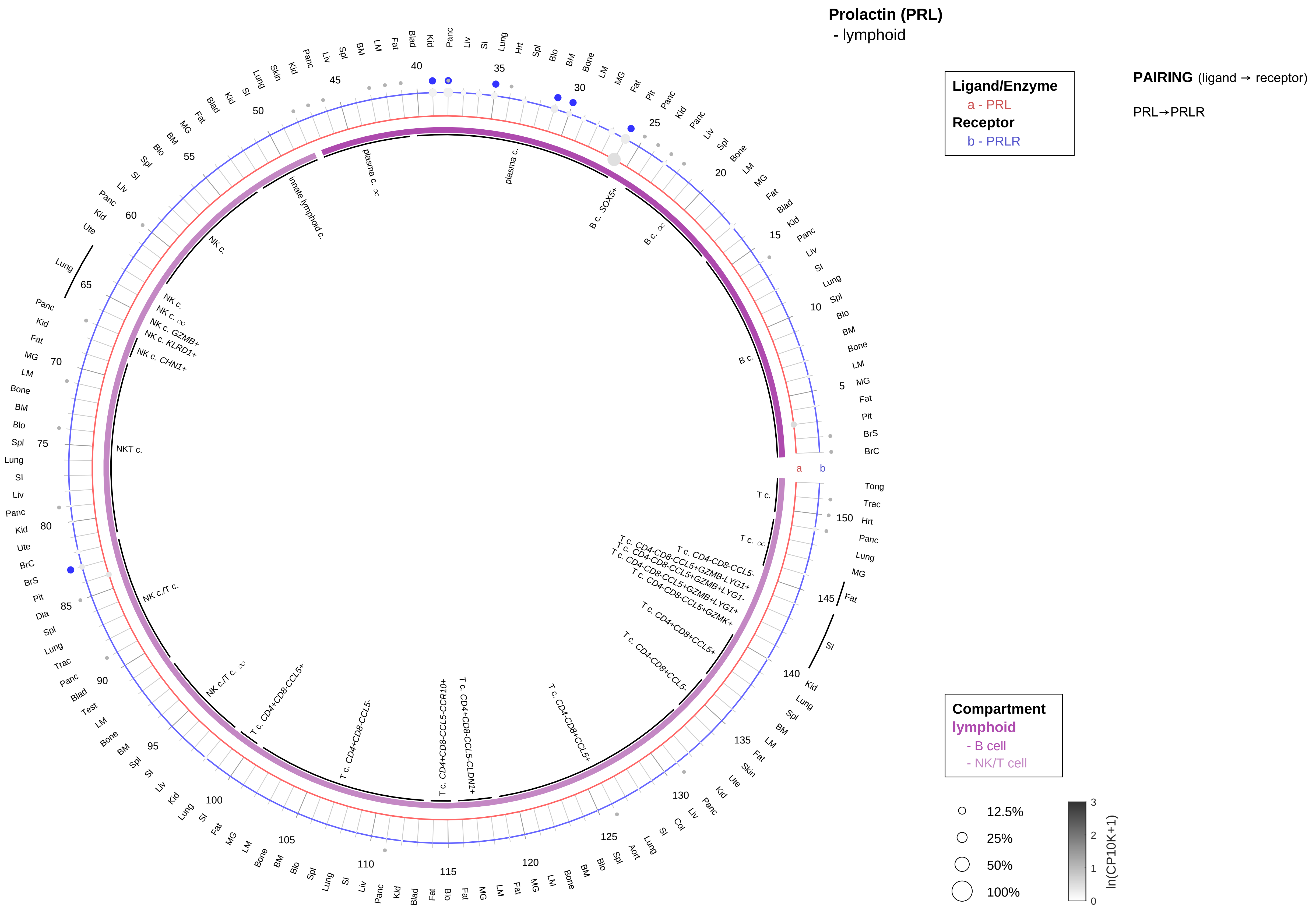

| Abbrev. | Tissue |
|---|---|
| Aort | Aorta |
| Blad | Bladder |
| Blo | Blood |
| Bone | Bone |
| BM | Bone marrow |
| BrC | Brain cortex |
| BrS | Brainstem |
| Col | Colon |
| Dia | Diaphragm |
| Ret | Eye retina |
| Fat | Fat |
| Hrt | Heart |
| Pit | Hypothalamus/Pituitary |
| Kid | Kidney |
| LM | Limb muscle |
| Liv | Liver |
| Lung | Lung |
| MG | Mammary gland |
| Panc | Pancreas |
| Skin | Skin |
| SI | Small intestine |
| Spl | Spleen |
| Test | Testes |
| Thym | Thymus |
| Tong | Tongue |
| Trac | Trachea |
| Ute | Uterus |

**Prolactin (PRL)**
- myeloid

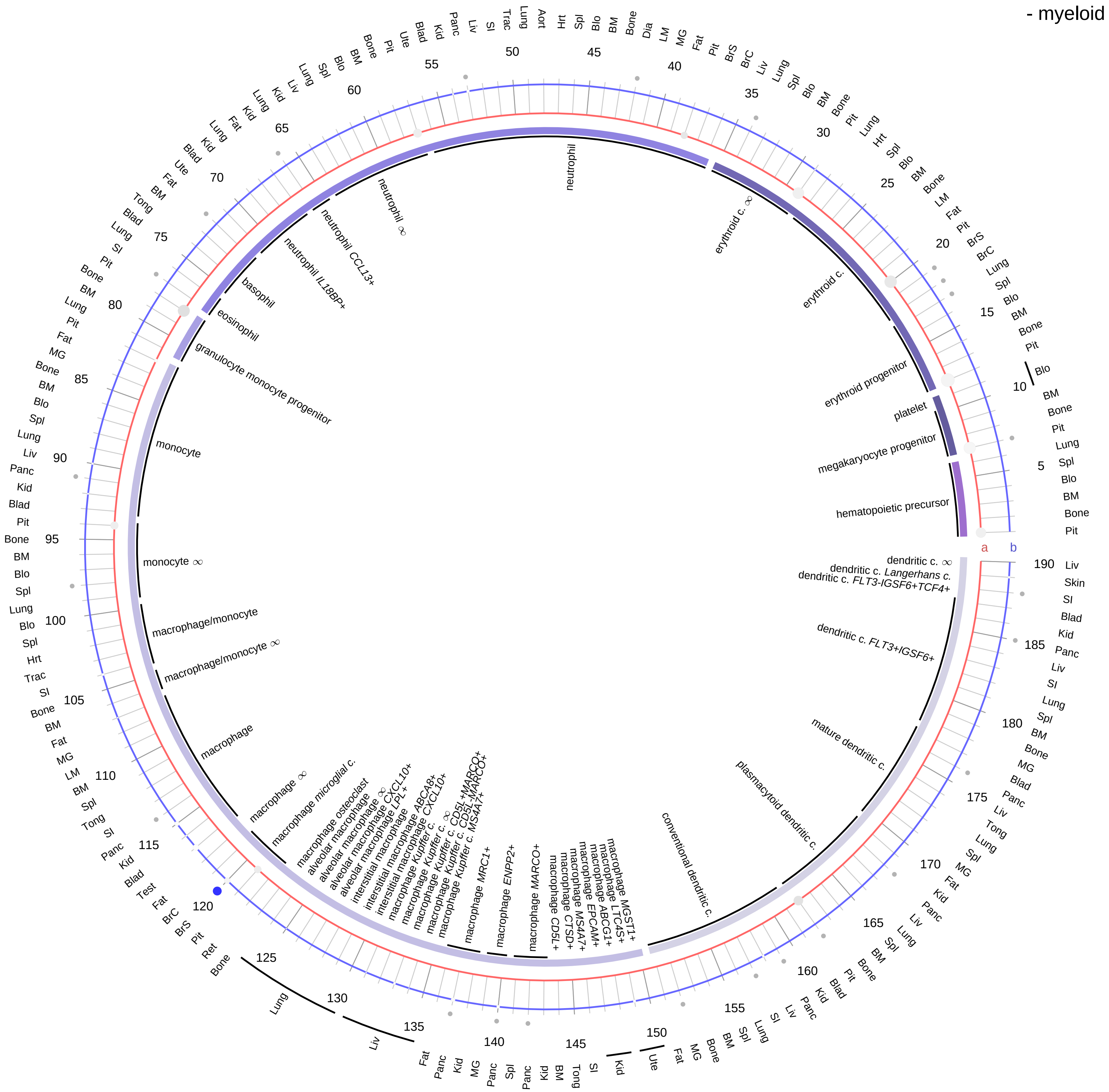

**Ligand/Enzyme**
a - PRL
**Receptor**
b - PRLR

**PAIRING** (ligand → receptor)

PRL→PRLR

| Abbrev. | Tissue |
|---|---|
| Aort | Aorta |
| Blad | Bladder |
| Blo | Blood |
| Bone | Bone |
| BM | Bone marrow |
| BrC | Brain cortex |
| BrS | Brainstem |
| Col | Colon |
| Dia | Diaphragm |
| Ret | Eye retina |
| Fat | Fat |
| Hrt | Heart |
| Pit | Hypothalamus/Pituitary |
| Kid | Kidney |
| LM | Limb muscle |
| Liv | Liver |
| Lung | Lung |
| MG | Mammary gland |
| Panc | Pancreas |
| Skin | Skin |
| SI | Small intestine |
| Spl | Spleen |
| Test | Testes |
| Thym | Thymus |
| Tong | Tongue |
| Trac | Trachea |
| Ute | Uterus |

**Compartment**
**hematopoietic**
- hematopoietic precursor
**megakaryocyte-erythroid**
- megakaryoid
- erythroid
**myeloid**
- granulocyte
- granulocyte monocyte progenitor
- monocyte/macrophage
- dendritic

12.5%
25%
50%
100%

ln(CP10K+1)
3
2
1
0

#### Prolactin releasing hormone (PRLH)
- epithelial/neural/germ

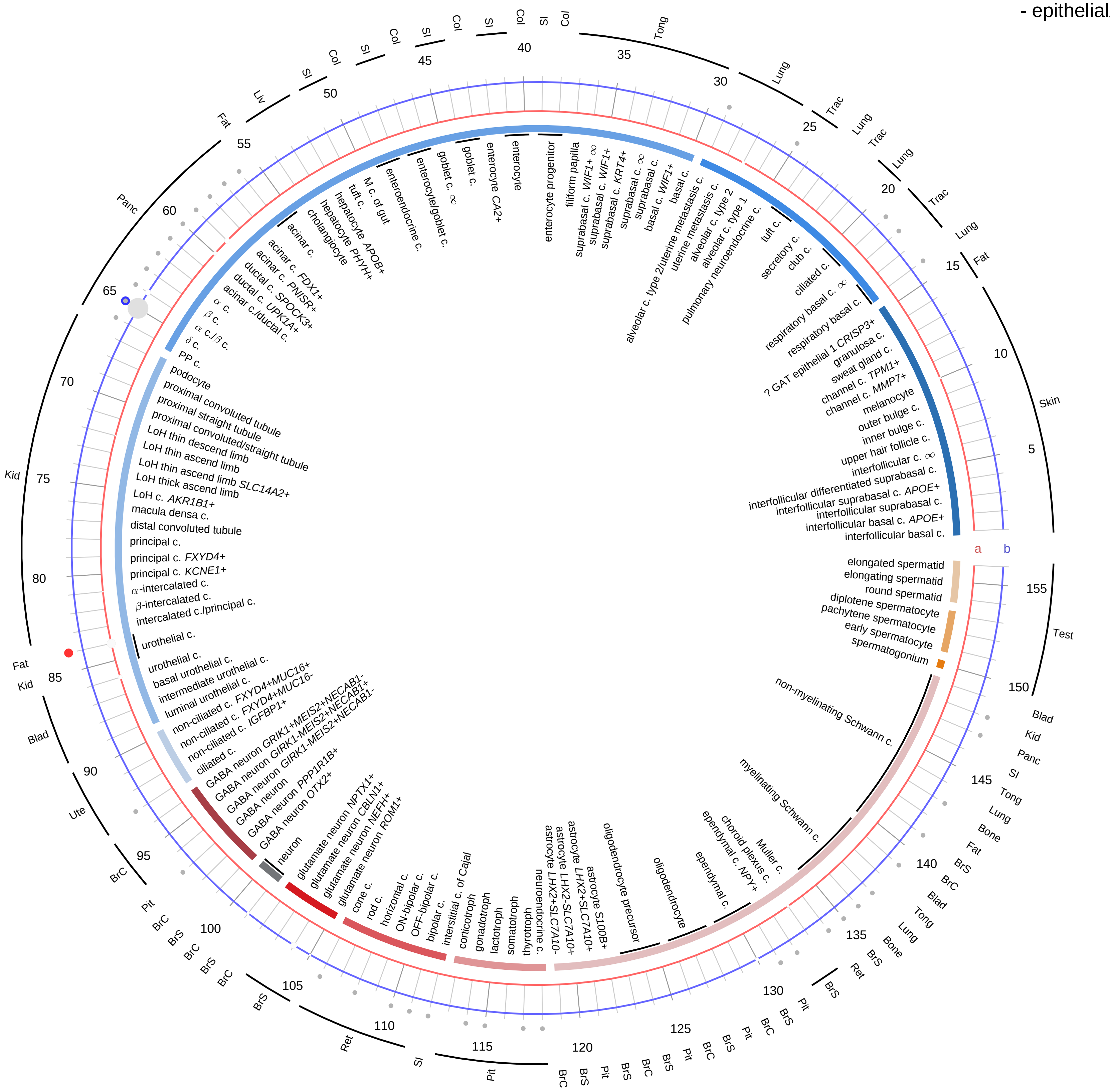

**Ligand/Enzyme**
a - PRLH
**Receptor**
b - PRLHR

**PAIRING** (ligand → receptor)

PRLH→PRLHR

**Compartment**
**epithelial**
- integumentary
- respiratory
- gastrointestinal
- urinary
- reproductive
**neural**
- GABAergic
- hybrid
- glutamatergic
- specialized
- neuroendocrine
- glia
**germ**
- spermatogonium
- spermatocyte
- spermatid

12.5%
25%
50%
100%

ln(CP10K+1): 0, 1, 2, 3

| Abbrev. | Tissue |
|---|---|
| Aort | Aorta |
| Blad | Bladder |
| Blo | Blood |
| Bone | Bone |
| BM | Bone marrow |
| BrC | Brain cortex |
| BrS | Brainstem |
| Col | Colon |
| Dia | Diaphragm |
| Ret | Eye retina |
| Fat | Fat |
| Hrt | Heart |
| Pit | Hypothalamus/Pituitary |
| Kid | Kidney |
| LM | Limb muscle |
| Liv | Liver |
| Lung | Lung |
| MG | Mammary gland |
| Panc | Pancreas |
| Skin | Skin |
| SI | Small intestine |
| Spl | Spleen |
| Test | Testes |
| Thym | Thymus |
| Tong | Tongue |
| Trac | Trachea |
| Ute | Uterus |

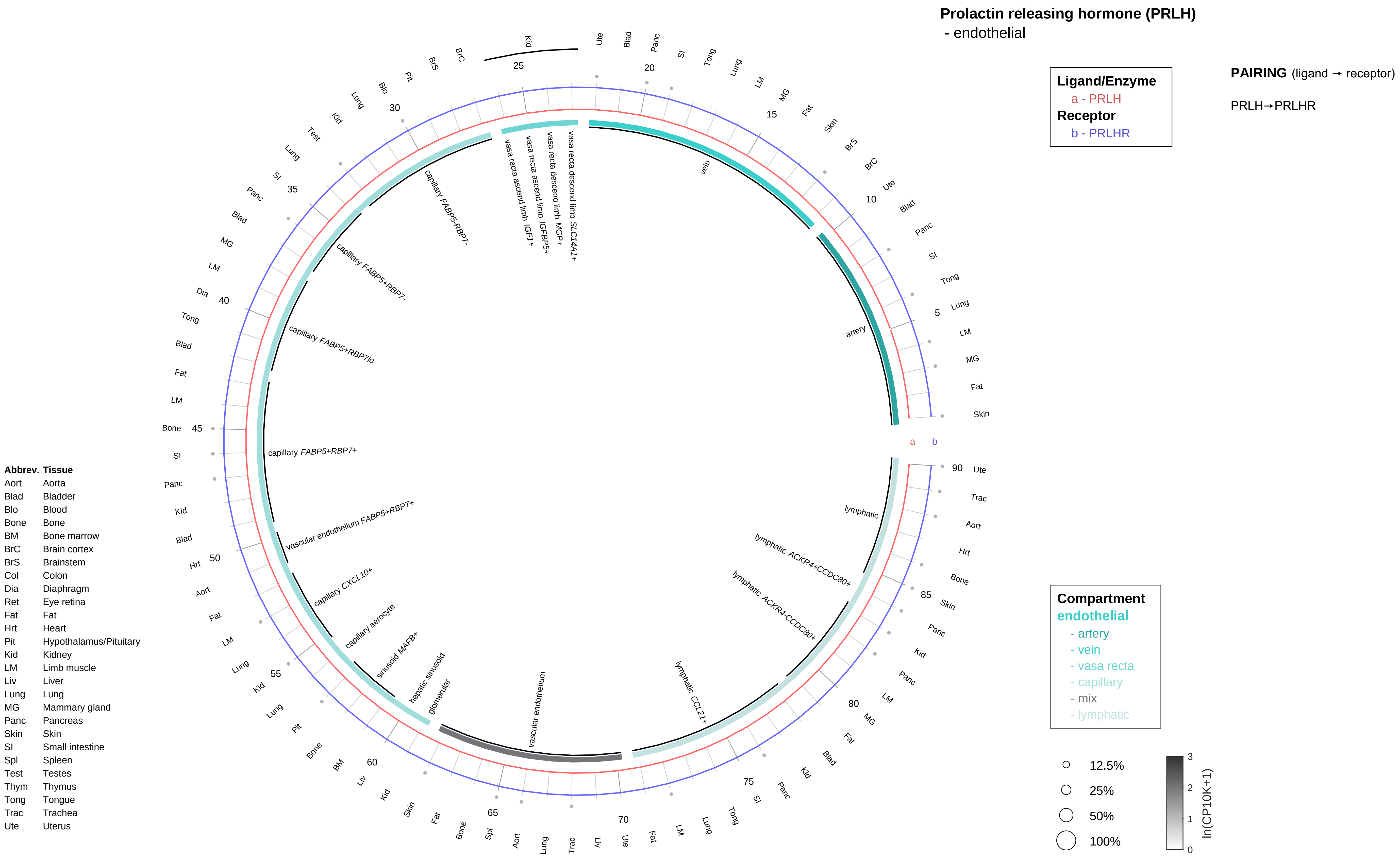

Prolactin releasing hormone (PRLH)
- endothelial
Ligand/Enzyme
a - PRLH
Receptor
b - PRLHR
PAIRING (ligand → receptor)
PRLH→PRLHR
Compartment
endothelial
- artery
- vein
- vasa recta
- capillary
- mix
- lymphatic
12.5%
25%
50%
100%
ln(CP10K+1)
vein
artery
vasa recta descend limb SLC14A1+
vasa recta descend limb MGP+
vasa recta ascend limb IGFBP5+
vasa recta ascend limb IGF1+
capillary FABP5-RBP7-
capillary FABP5+RBP7-
capillary FABP5+RBP7lo
capillary FABP5+RBP7+
vascular endothelium FABP5+RBP7+
capillary CXCL10+
capillary aerocyte
sinusoid MAFB+
hepatic sinusoid
glomerular
vascular endothelium
lymphatic CCL21+
lymphatic ACKR4-CCDC80+
lymphatic ACKR4+CCDC80+
lymphatic
Abbrev. Tissue
Aort Aorta
Blad Bladder
Blo Blood
Bone Bone
BM Bone marrow
BrC Brain cortex
BrS Brainstem
Col Colon
Dia Diaphragm
Ret Eye retina
Fat Fat
Hrt Heart
Pit Hypothalamus/Pituitary
Kid Kidney
LM Limb muscle
Liv Liver
Lung Lung
MG Mammary gland
Panc Pancreas
Skin Skin
SI Small intestine
Spl Spleen
Test Testes
Thym Thymus
Tong Tongue
Trac Trachea
Ute Uterus

**Prolactin releasing hormone (PRLH)**
- stromal

| **Ligand/Enzyme** |
| --- |
| a - PRLH |
| **Receptor** |
| b - PRLHR |

**PAIRING** (ligand → receptor)

PRLH→PRLHR

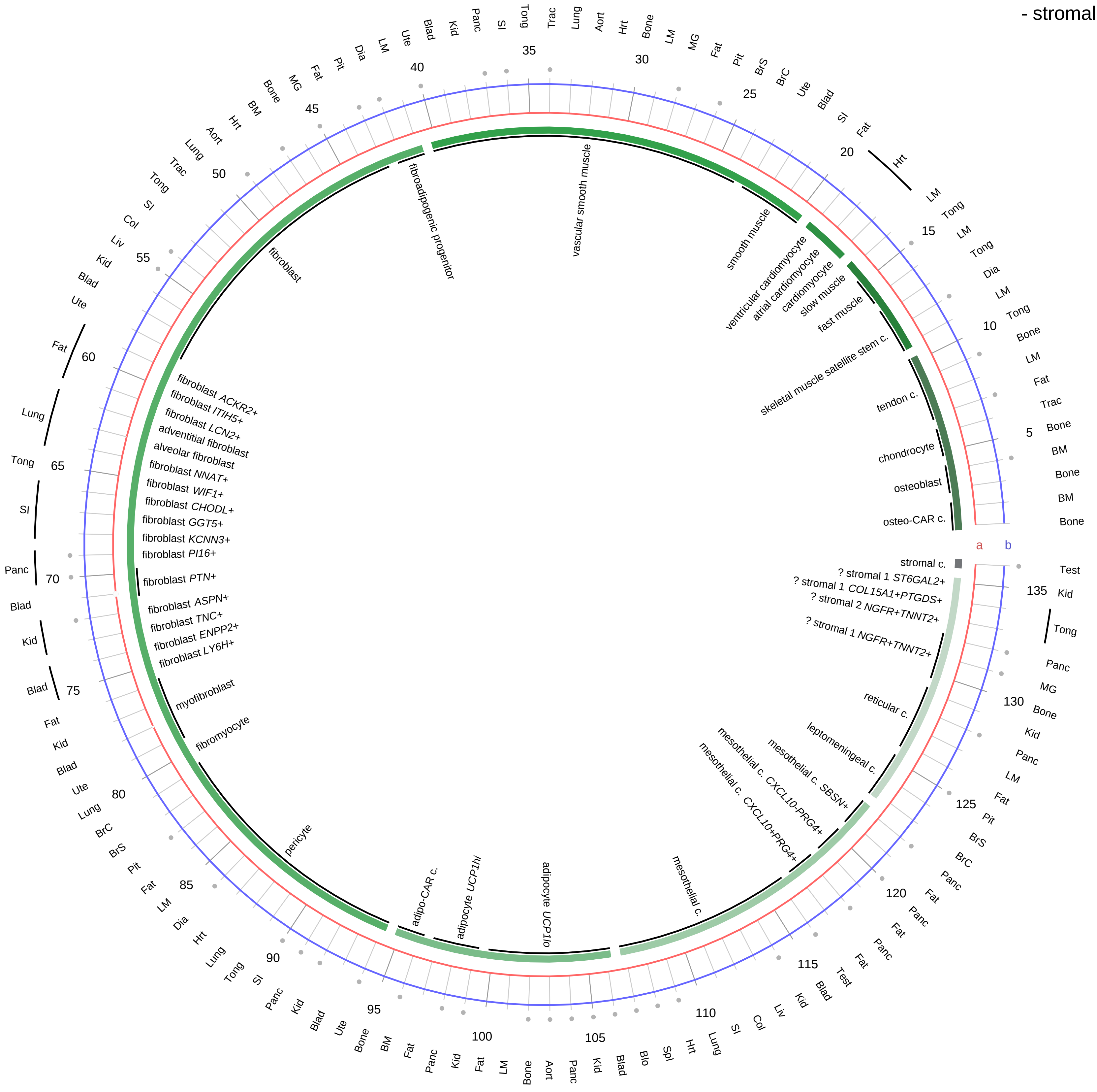

| **Compartment** |
| --- |
| **stromal** |
| - bone |
| - skeletal muscle |
| - cardiac muscle |
| - smooth muscle |
| - fibroblast |
| - adipocyte |
| - mesothelial |
| - other |
| - mix |

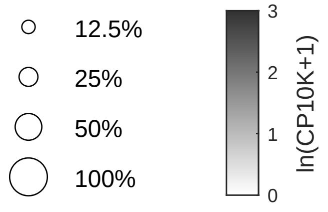

| Abbrev. | Tissue |
| --- | --- |
| Aort | Aorta |
| Blad | Bladder |
| Blo | Blood |
| Bone | Bone |
| BM | Bone marrow |
| BrC | Brain cortex |
| BrS | Brainstem |
| Col | Colon |
| Dia | Diaphragm |
| Ret | Eye retina |
| Fat | Fat |
| Hrt | Heart |
| Pit | Hypothalamus/Pituitary |
| Kid | Kidney |
| LM | Limb muscle |
| Liv | Liver |
| Lung | Lung |
| MG | Mammary gland |
| Panc | Pancreas |
| Skin | Skin |
| SI | Small intestine |
| Spl | Spleen |
| Test | Testes |
| Thym | Thymus |
| Tong | Tongue |
| Trac | Trachea |
| Ute | Uterus |

**Prolactin releasing hormone (PRLH)**
- lymphoid

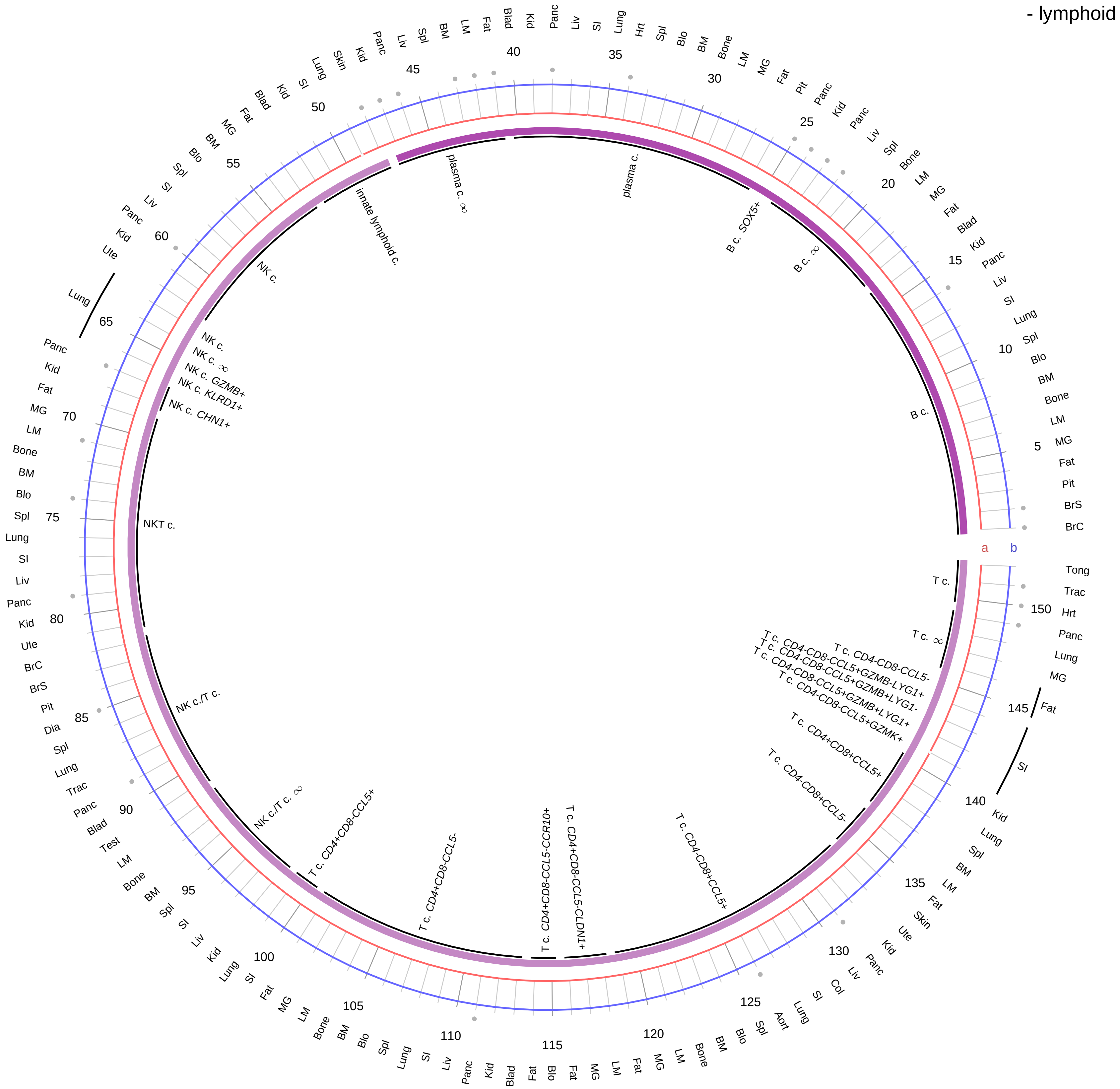

**Ligand/Enzyme**
a - PRLH
**Receptor**
b - PRLHR

**PAIRING** (ligand → receptor)

PRLH→PRLHR

**Compartment**
**lymphoid**
- B cell
- NK/T cell

12.5%
25%
50%
100%

ln(CP10K+1): 0, 1, 2, 3

| Abbrev. | Tissue |
|---|---|
| Aort | Aorta |
| Blad | Bladder |
| Blo | Blood |
| Bone | Bone |
| BM | Bone marrow |
| BrC | Brain cortex |
| BrS | Brainstem |
| Col | Colon |
| Dia | Diaphragm |
| Ret | Eye retina |
| Fat | Fat |
| Hrt | Heart |
| Pit | Hypothalamus/Pituitary |
| Kid | Kidney |
| LM | Limb muscle |
| Liv | Liver |
| Lung | Lung |
| MG | Mammary gland |
| Panc | Pancreas |
| Skin | Skin |
| SI | Small intestine |
| Spl | Spleen |
| Test | Testes |
| Thym | Thymus |
| Tong | Tongue |
| Trac | Trachea |
| Ute | Uterus |

**Prolactin releasing hormone (PRLH)**
- myeloid

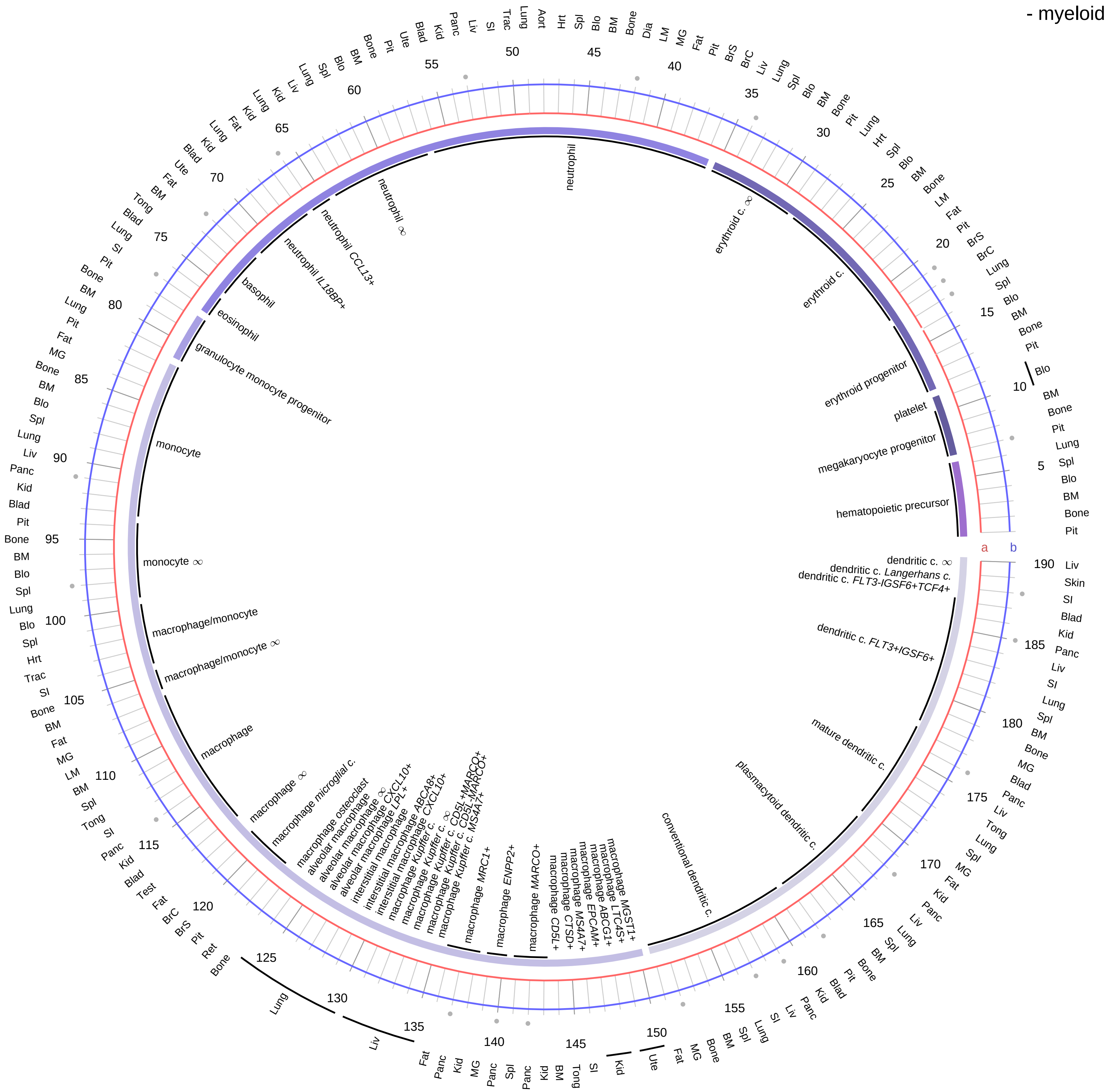

| Ligand/Enzyme |
|---|
| a - PRLH |
| **Receptor** |
| b - PRLHR |

**PAIRING** (ligand → receptor)

PRLH→PRLHR

| Abbrev. | Tissue |
|---|---|
| Aort | Aorta |
| Blad | Bladder |
| Blo | Blood |
| Bone | Bone |
| BM | Bone marrow |
| BrC | Brain cortex |
| BrS | Brainstem |
| Col | Colon |
| Dia | Diaphragm |
| Ret | Eye retina |
| Fat | Fat |
| Hrt | Heart |
| Pit | Hypothalamus/Pituitary |
| Kid | Kidney |
| LM | Limb muscle |
| Liv | Liver |
| Lung | Lung |
| MG | Mammary gland |
| Panc | Pancreas |
| Skin | Skin |
| SI | Small intestine |
| Spl | Spleen |
| Test | Testes |
| Thym | Thymus |
| Tong | Tongue |
| Trac | Trachea |
| Ute | Uterus |

**Compartment**
**hematopoietic**
- hematopoietic precursor
**megakaryocyte-erythroid**
- megakaryoid
- erythroid
**myeloid**
- granulocyte
- granulocyte monocyte progenitor
- monocyte/macrophage
- dendritic

12.5%
25%
50%
100%

ln(CP10K+1): 0, 1, 2, 3

**Proopiomelanocortin-derived hormones (ACTH/MSH/endorphin)**
- epithelial/neural/germ

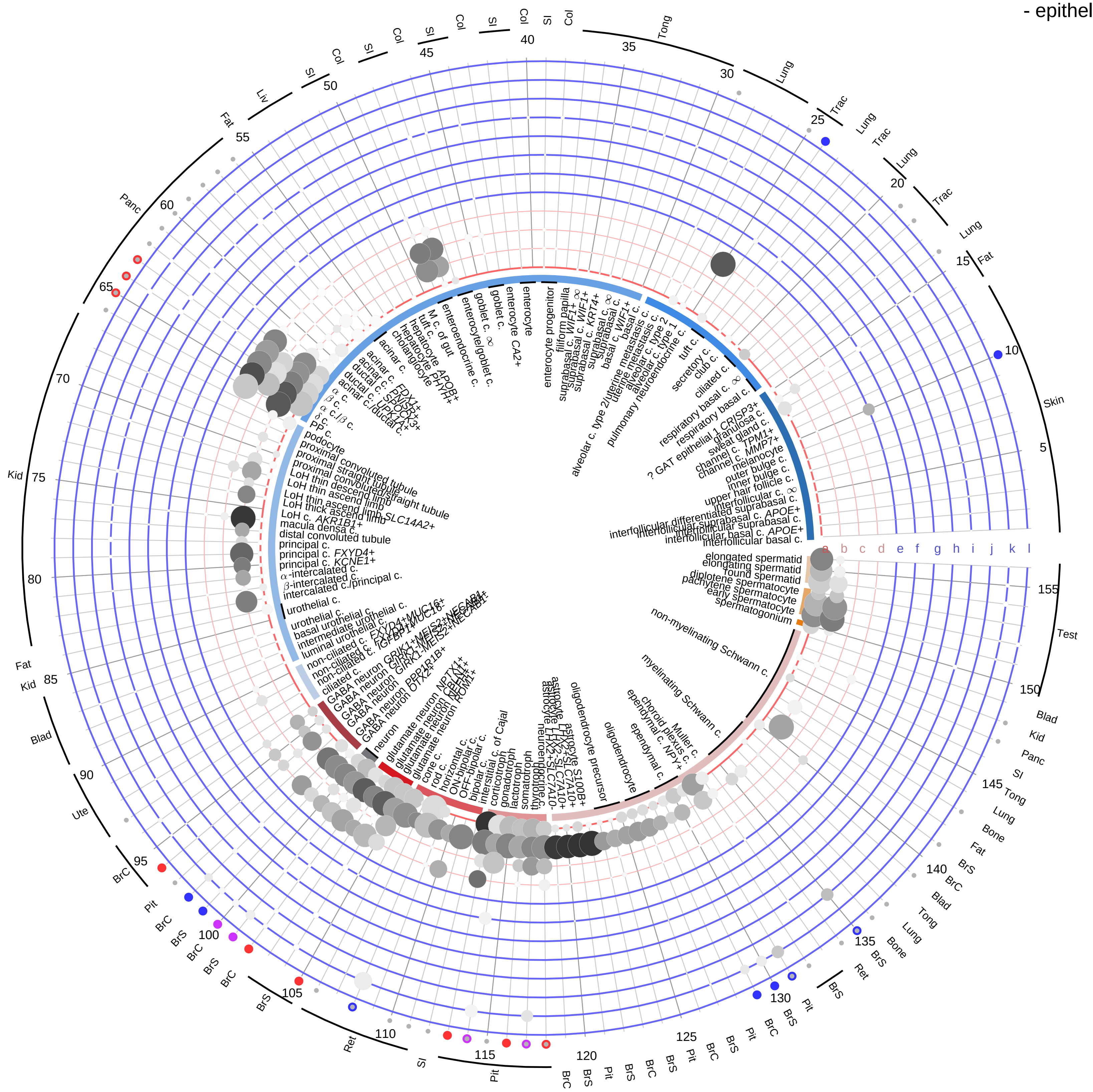

**Ligand/Enzyme**
- a - POMC
- b - CPE
- c - PCSK1
- d - PCSK2

**Receptor**
- e - MC1R*
- f - MC2R
- g - MC3R
- h - MC4R
- i - MC5R
- j - OPRD1
- k - OPRK1
- l - OPRM1

**PAIRING** (ligand → receptor)

POMC & PCSK1 & CPE→MC2R

POMC & PCSK1 & CPE→MC1R

POMC & PCSK1 & CPE→MC3R

POMC & PCSK1 & CPE→MC4R

POMC & PCSK1 & CPE→MC5R

POMC & PCSK1 & PCSK2 & CPE→MC1R

POMC & PCSK1 & PCSK2 & CPE→MC3R

POMC & PCSK1 & PCSK2 & CPE→MC4R

POMC & PCSK1 & PCSK2 & CPE→MC5R

POMC & PCSK1 & PCSK2 & CPE→OPRM1

POMC & PCSK1 & PCSK2 & CPE→OPRD1

POMC & PCSK1 & PCSK2 & CPE→OPRK1

**Compartment**

**epithelial**
- integumentary
- respiratory
- gastrointestinal
- urinary
- reproductive

**neural**
- GABAergic
- hybrid
- glutamatergic
- specialized
- neuroendocrine
- glia

**germ**
- spermatogonium
- spermatocyte
- spermatid

12.5%
25%
50%
100%

ln(CP10K+1): 0, 1, 2, 3

| Abbrev. | Tissue |
|---|---|
| Aort | Aorta |
| Blad | Bladder |
| Blo | Blood |
| Bone | Bone |
| BM | Bone marrow |
| BrC | Brain cortex |
| BrS | Brainstem |
| Col | Colon |
| Dia | Diaphragm |
| Ret | Eye retina |
| Fat | Fat |
| Hrt | Heart |
| Pit | Hypothalamus/Pituitary |
| Kid | Kidney |
| LM | Limb muscle |
| Liv | Liver |
| Lung | Lung |
| MG | Mammary gland |
| Panc | Pancreas |
| Skin | Skin |
| SI | Small intestine |
| Spl | Spleen |
| Test | Testes |
| Thym | Thymus |
| Tong | Tongue |
| Trac | Trachea |
| Ute | Uterus |

**Proopiomelanocortin-derived hormones (ACTH/MSH/endorphin)**
- endothelial

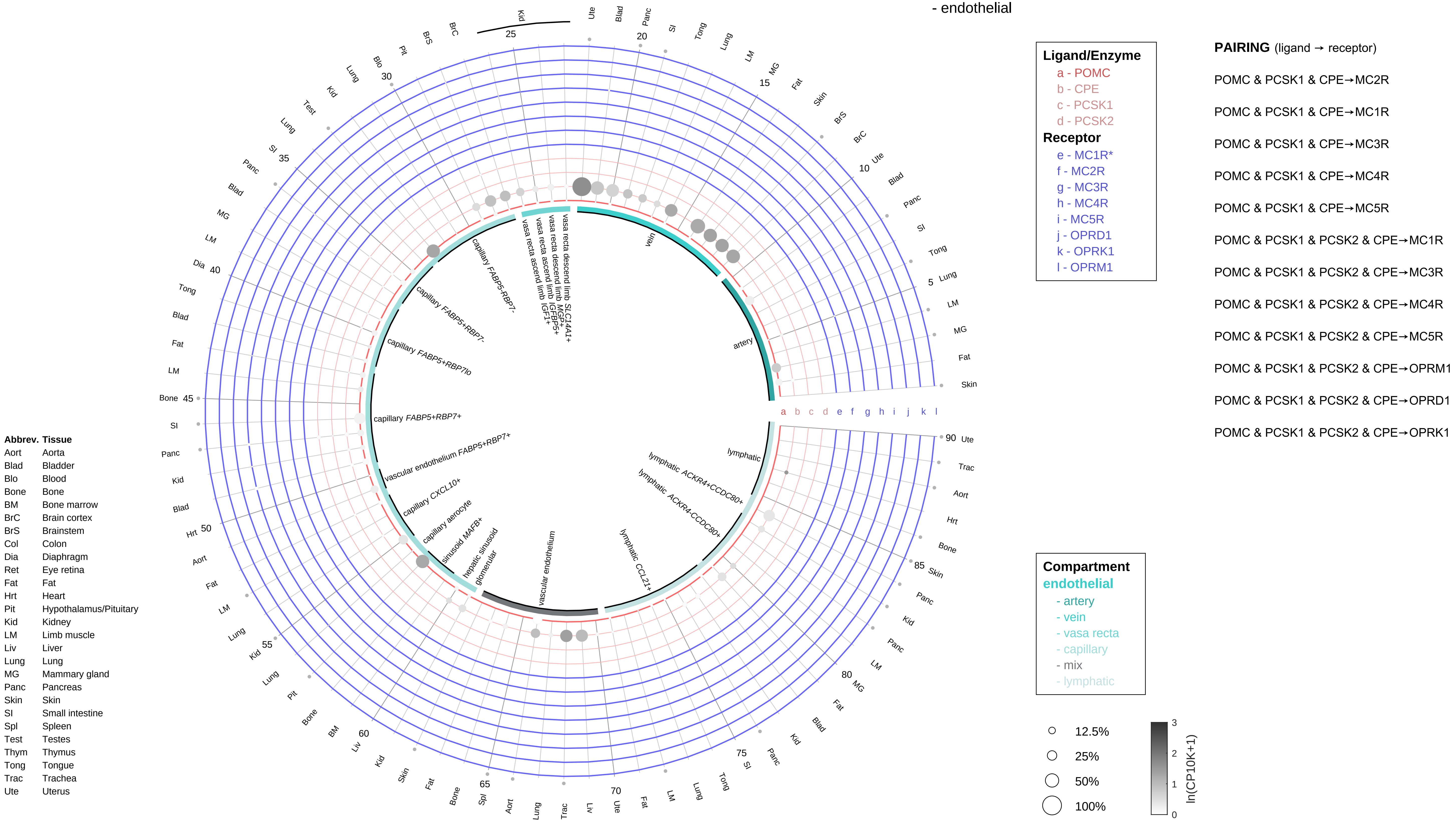

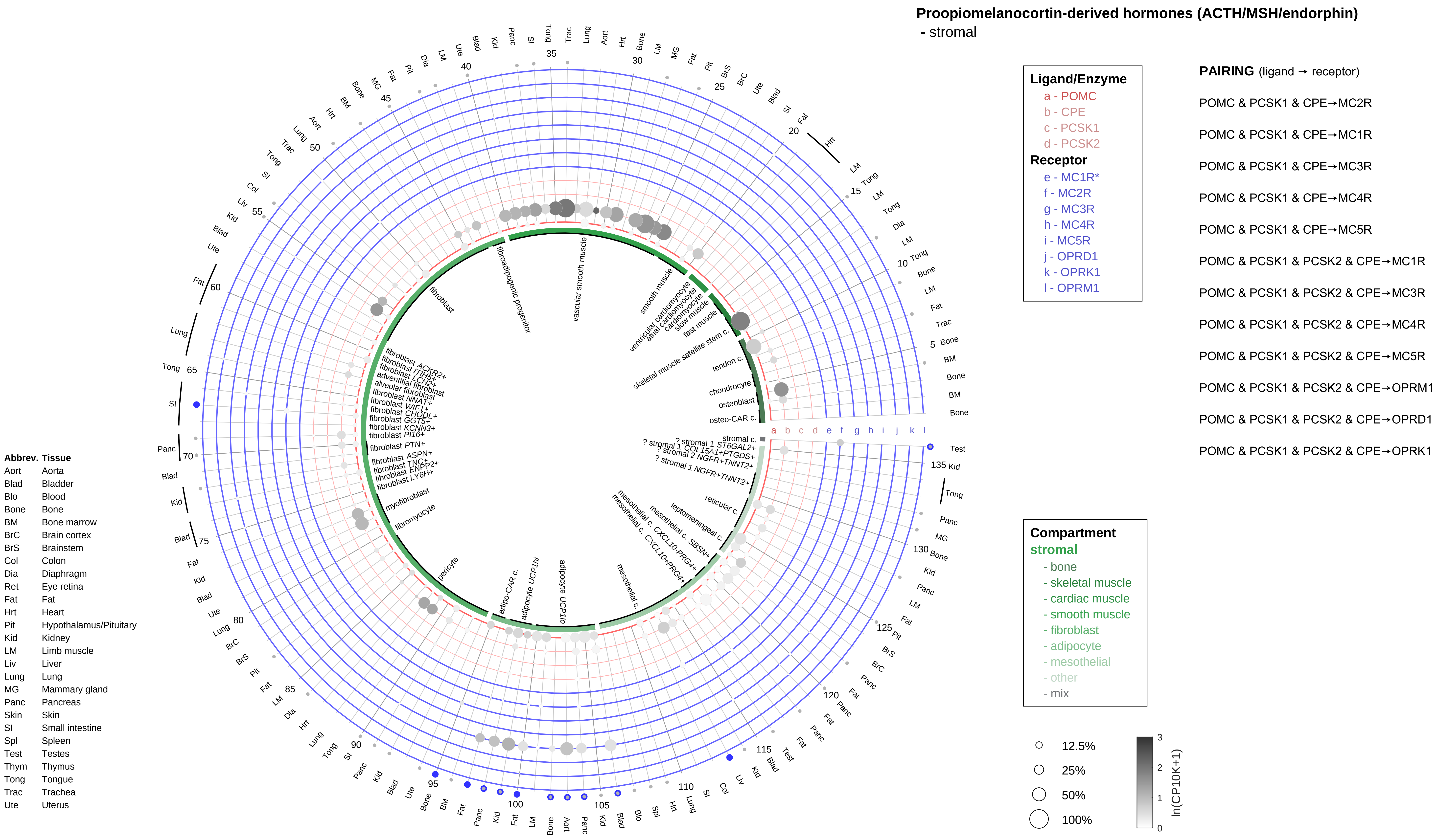
Proopiomelanocortin-derived hormones (ACTH/MSH/endorphin)
- stromal
Ligand/Enzyme
a - POMC
b - CPE
c - PCSK1
d - PCSK2
Receptor
e - MC1R*
f - MC2R
g - MC3R
h - MC4R
i - MC5R
j - OPRD1
k - OPRK1
l - OPRM1
PAIRING (ligand → receptor)
POMC & PCSK1 & CPE→MC2R
POMC & PCSK1 & CPE→MC1R
POMC & PCSK1 & CPE→MC3R
POMC & PCSK1 & CPE→MC4R
POMC & PCSK1 & CPE→MC5R
POMC & PCSK1 & PCSK2 & CPE→MC1R
POMC & PCSK1 & PCSK2 & CPE→MC3R
POMC & PCSK1 & PCSK2 & CPE→MC4R
POMC & PCSK1 & PCSK2 & CPE→MC5R
POMC & PCSK1 & PCSK2 & CPE→OPRM1
POMC & PCSK1 & PCSK2 & CPE→OPRD1
POMC & PCSK1 & PCSK2 & CPE→OPRK1
Compartment
stromal
- bone
- skeletal muscle
- cardiac muscle
- smooth muscle
- fibroblast
- adipocyte
- mesothelial
- other
- mix
12.5%
25%
50%
100%
ln(CP10K+1)
Abbrev. Tissue
Aort Aorta
Blad Bladder
Blo Blood
Bone Bone
BM Bone marrow
BrC Brain cortex
BrS Brainstem
Col Colon
Dia Diaphragm
Ret Eye retina
Fat Fat
Hrt Heart
Pit Hypothalamus/Pituitary
Kid Kidney
LM Limb muscle
Liv Liver
Lung Lung
MG Mammary gland
Panc Pancreas
Skin Skin
SI Small intestine
Spl Spleen
Test Testes
Thym Thymus
Tong Tongue
Trac Trachea
Ute Uterus

**Proopiomelanocortin-derived hormones (ACTH/MSH/endorphin)**
- lymphoid

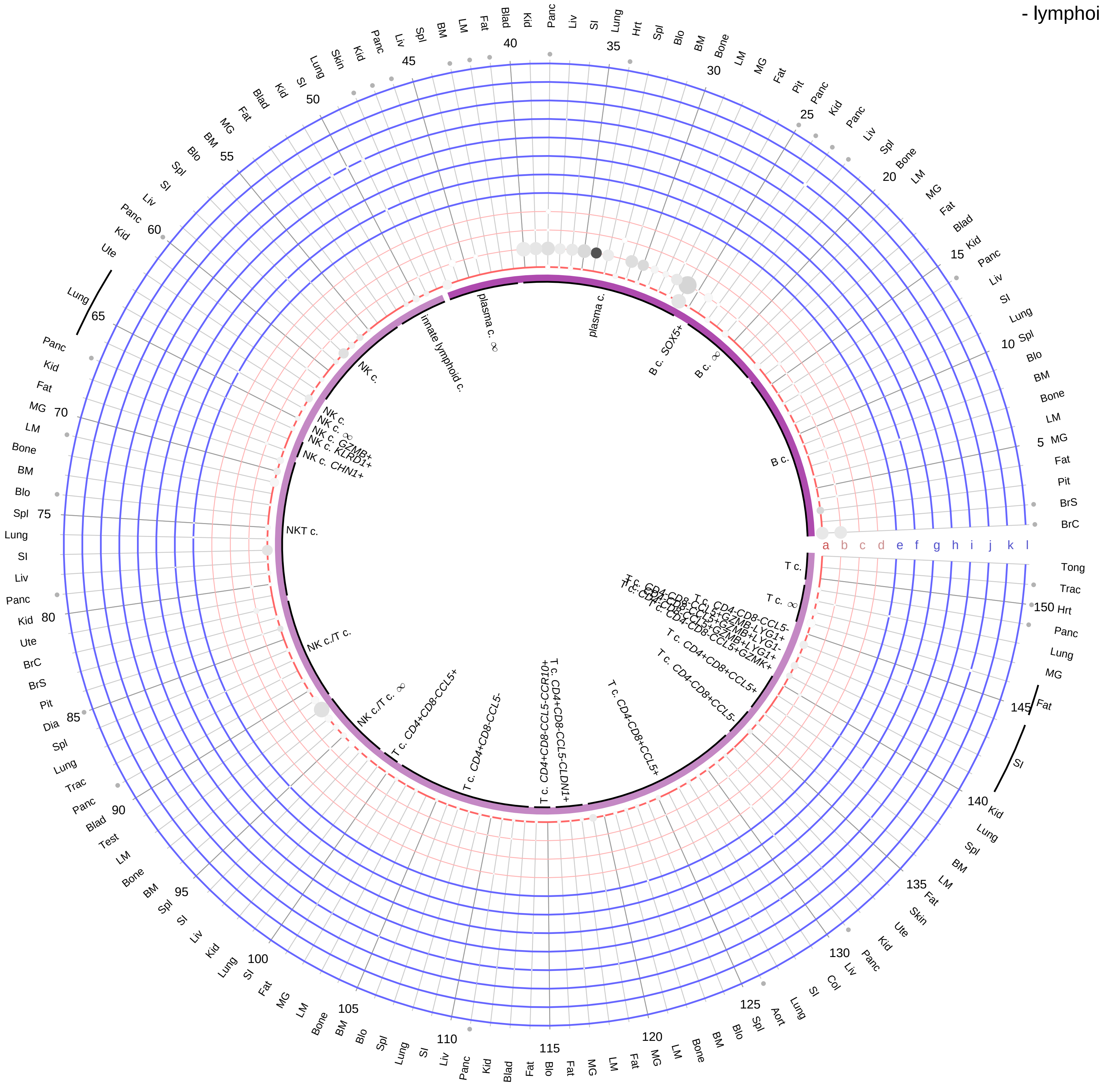

**Ligand/Enzyme**
- a - POMC
- b - CPE
- c - PCSK1
- d - PCSK2

**Receptor**
- e - MC1R*
- f - MC2R
- g - MC3R
- h - MC4R
- i - MC5R
- j - OPRD1
- k - OPRK1
- l - OPRM1

**PAIRING** (ligand → receptor)

POMC & PCSK1 & CPE→MC2R

POMC & PCSK1 & CPE→MC1R

POMC & PCSK1 & CPE→MC3R

POMC & PCSK1 & CPE→MC4R

POMC & PCSK1 & CPE→MC5R

POMC & PCSK1 & PCSK2 & CPE→MC1R

POMC & PCSK1 & PCSK2 & CPE→MC3R

POMC & PCSK1 & PCSK2 & CPE→MC4R

POMC & PCSK1 & PCSK2 & CPE→MC5R

POMC & PCSK1 & PCSK2 & CPE→OPRM1

POMC & PCSK1 & PCSK2 & CPE→OPRD1

POMC & PCSK1 & PCSK2 & CPE→OPRK1

| Abbrev. | Tissue |
|---|---|
| Aort | Aorta |
| Blad | Bladder |
| Blo | Blood |
| Bone | Bone |
| BM | Bone marrow |
| BrC | Brain cortex |
| BrS | Brainstem |
| Col | Colon |
| Dia | Diaphragm |
| Ret | Eye retina |
| Fat | Fat |
| Hrt | Heart |
| Pit | Hypothalamus/Pituitary |
| Kid | Kidney |
| LM | Limb muscle |
| Liv | Liver |
| Lung | Lung |
| MG | Mammary gland |
| Panc | Pancreas |
| Skin | Skin |
| SI | Small intestine |
| Spl | Spleen |
| Test | Testes |
| Thym | Thymus |
| Tong | Tongue |
| Trac | Trachea |
| Ute | Uterus |

**Compartment**
**lymphoid**
- B cell
- NK/T cell

12.5%
25%
50%
100%

ln(CP10K+1): 0, 1, 2, 3

**Proopiomelanocortin-derived hormones (ACTH/MSH/endorphin)**
- myeloid

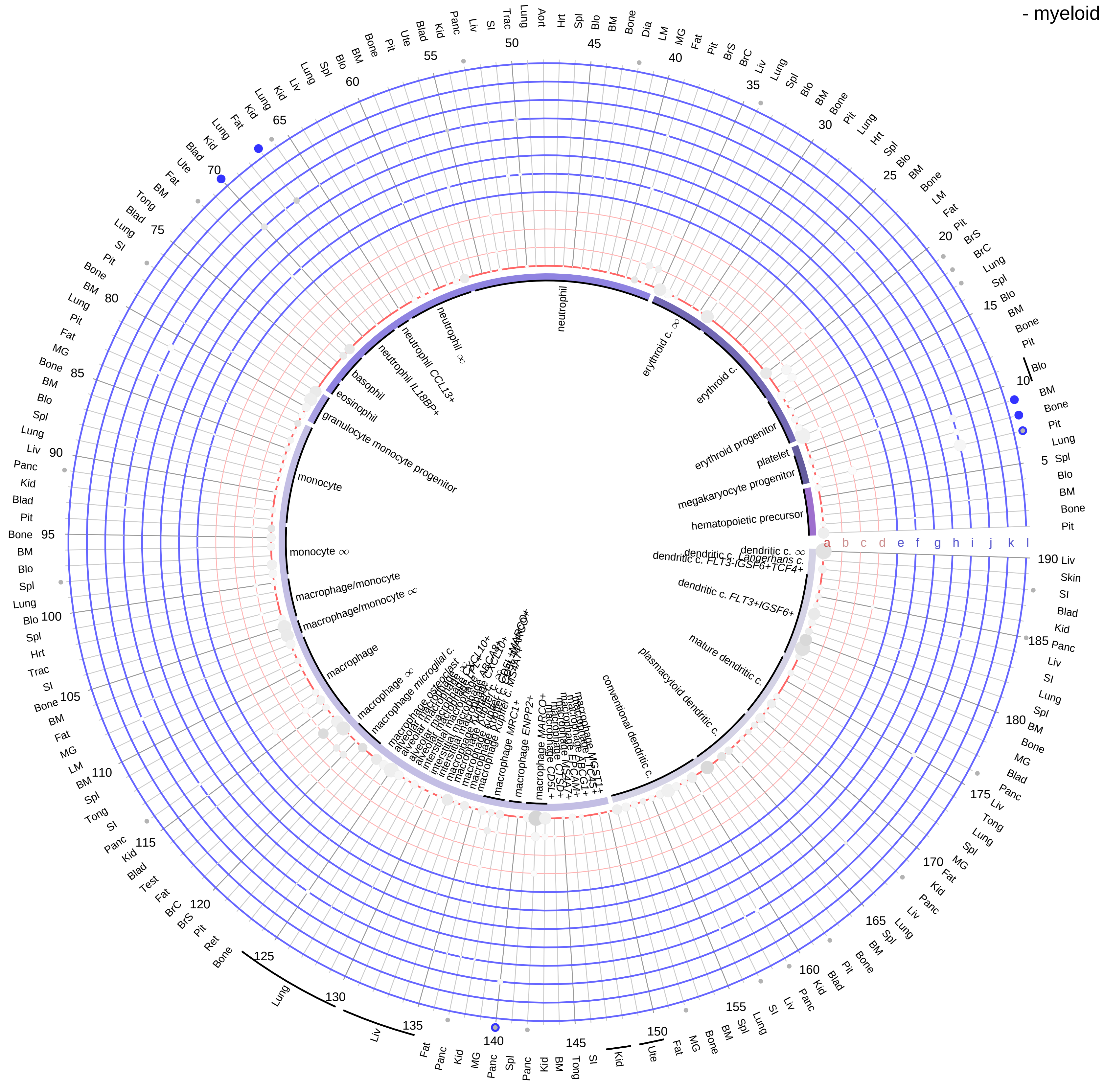

| Abbrev. | Tissue |
|---|---|
| Aort | Aorta |
| Blad | Bladder |
| Blo | Blood |
| Bone | Bone |
| BM | Bone marrow |
| BrC | Brain cortex |
| BrS | Brainstem |
| Col | Colon |
| Dia | Diaphragm |
| Ret | Eye retina |
| Fat | Fat |
| Hrt | Heart |
| Pit | Hypothalamus/Pituitary |
| Kid | Kidney |
| LM | Limb muscle |
| Liv | Liver |
| Lung | Lung |
| MG | Mammary gland |
| Panc | Pancreas |
| Skin | Skin |
| SI | Small intestine |
| Spl | Spleen |
| Test | Testes |
| Thym | Thymus |
| Tong | Tongue |
| Trac | Trachea |
| Ute | Uterus |

**Ligand/Enzyme**
- a - POMC
- b - CPE
- c - PCSK1
- d - PCSK2

**Receptor**
- e - MC1R*
- f - MC2R
- g - MC3R
- h - MC4R
- i - MC5R
- j - OPRD1
- k - OPRK1
- l - OPRM1

**PAIRING** (ligand → receptor)

POMC & PCSK1 & CPE→MC2R

POMC & PCSK1 & CPE→MC1R

POMC & PCSK1 & CPE→MC3R

POMC & PCSK1 & CPE→MC4R

POMC & PCSK1 & CPE→MC5R

POMC & PCSK1 & PCSK2 & CPE→MC1R

POMC & PCSK1 & PCSK2 & CPE→MC3R

POMC & PCSK1 & PCSK2 & CPE→MC4R

POMC & PCSK1 & PCSK2 & CPE→MC5R

POMC & PCSK1 & PCSK2 & CPE→OPRM1

POMC & PCSK1 & PCSK2 & CPE→OPRD1

POMC & PCSK1 & PCSK2 & CPE→OPRK1

**Compartment**

**hematopoietic**
- hematopoietic precursor

**megakaryocyte-erythroid**
- megakaryoid
- erythroid

**myeloid**
- granulocyte
- granulocyte monocyte progenitor
- monocyte/macrophage
- dendritic

12.5%
25%
50%
100%

ln(CP10K+1): 0, 1, 2, 3

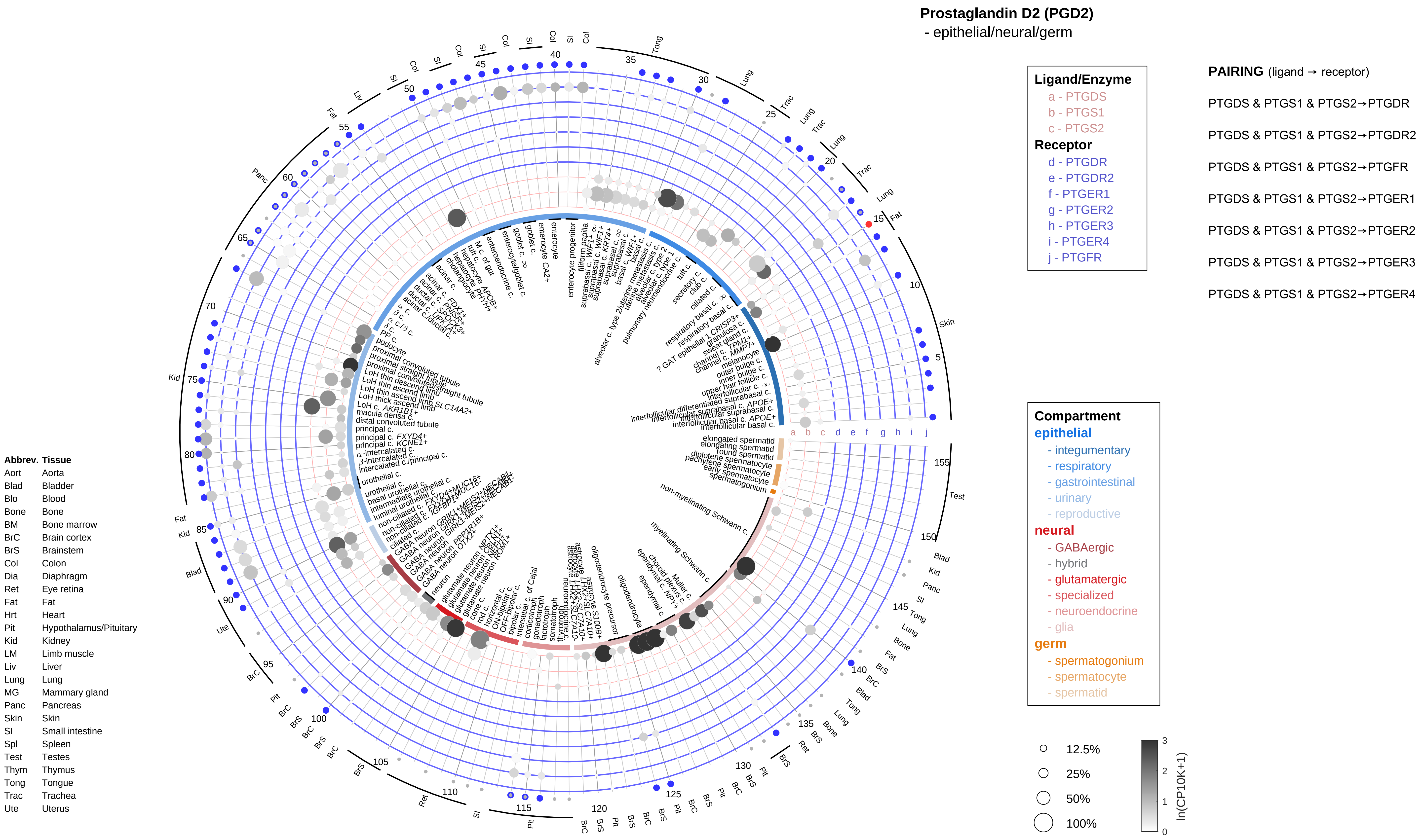
Prostaglandin D2 (PGD2)
- epithelial/neural/germ
Ligand/Enzyme
a - PTGDS
b - PTGS1
c - PTGS2
Receptor
d - PTGDR
e - PTGDR2
f - PTGER1
g - PTGER2
h - PTGER3
i - PTGER4
j - PTGFR
PAIRING (ligand → receptor)
PTGDS & PTGS1 & PTGS2→PTGDR
PTGDS & PTGS1 & PTGS2→PTGDR2
PTGDS & PTGS1 & PTGS2→PTGFR
PTGDS & PTGS1 & PTGS2→PTGER1
PTGDS & PTGS1 & PTGS2→PTGER2
PTGDS & PTGS1 & PTGS2→PTGER3
PTGDS & PTGS1 & PTGS2→PTGER4
Compartment
epithelial
- integumentary
- respiratory
- gastrointestinal
- urinary
- reproductive
neural
- GABAergic
- hybrid
- glutamatergic
- specialized
- neuroendocrine
- glia
germ
- spermatogonium
- spermatocyte
- spermatid
12.5%
25%
50%
100%
ln(CP10K+1)
Abbrev. Tissue
Aort Aorta
Blad Bladder
Blo Blood
Bone Bone
BM Bone marrow
BrC Brain cortex
BrS Brainstem
Col Colon
Dia Diaphragm
Ret Eye retina
Fat Fat
Hrt Heart
Pit Hypothalamus/Pituitary
Kid Kidney
LM Limb muscle
Liv Liver
Lung Lung
MG Mammary gland
Panc Pancreas
Skin Skin
SI Small intestine
Spl Spleen
Test Testes
Thym Thymus
Tong Tongue
Trac Trachea
Ute Uterus

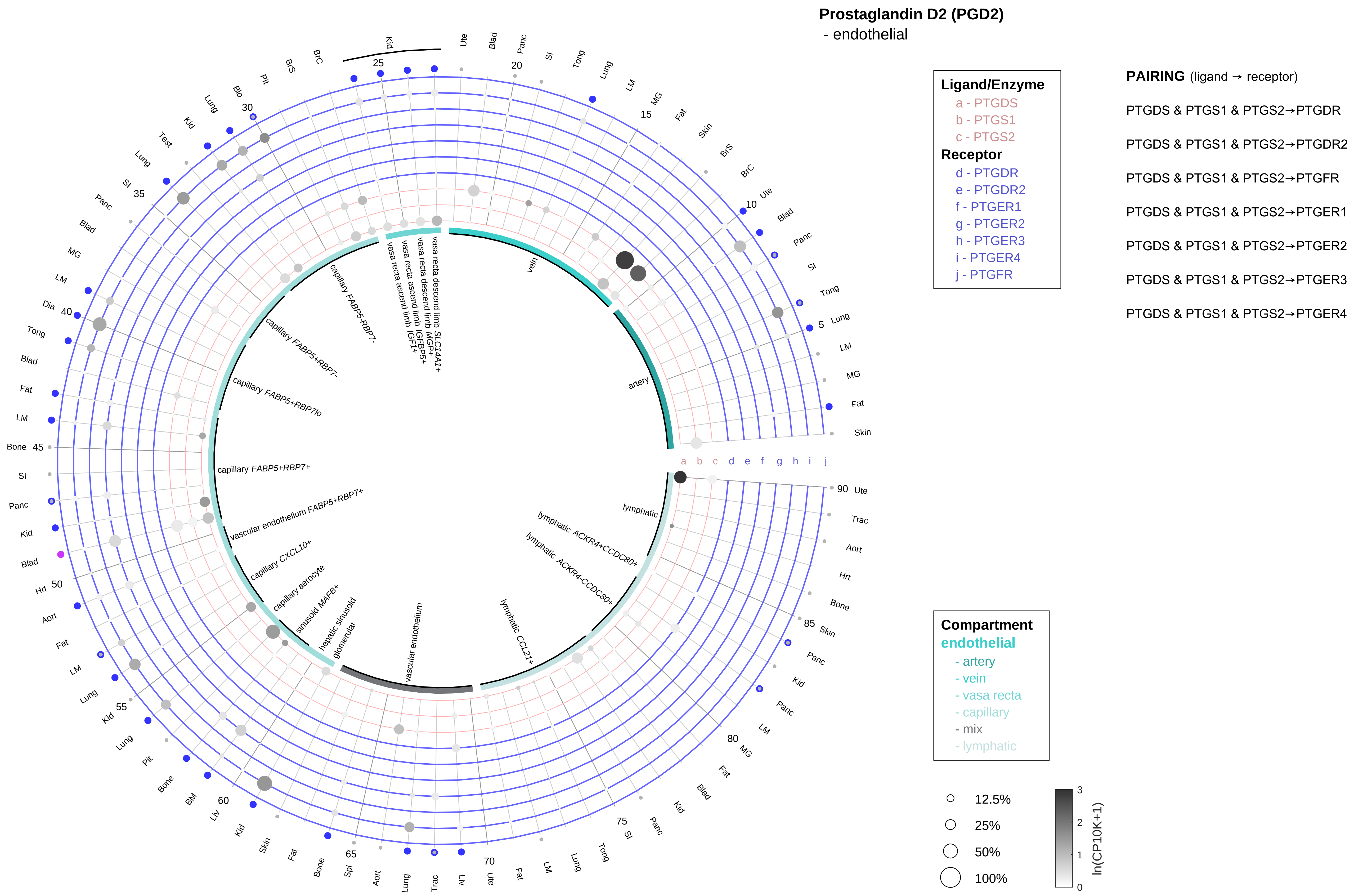

| Abbrev. | Tissue |
|---|---|
| Aort | Aorta |
| Blad | Bladder |
| Blo | Blood |
| Bone | Bone |
| BM | Bone marrow |
| BrC | Brain cortex |
| BrS | Brainstem |
| Col | Colon |
| Dia | Diaphragm |
| Ret | Eye retina |
| Fat | Fat |
| Hrt | Heart |
| Pit | Hypothalamus/Pituitary |
| Kid | Kidney |
| LM | Limb muscle |
| Liv | Liver |
| Lung | Lung |
| MG | Mammary gland |
| Panc | Pancreas |
| Skin | Skin |
| SI | Small intestine |
| Spl | Spleen |
| Test | Testes |
| Thym | Thymus |
| Tong | Tongue |
| Trac | Trachea |
| Ute | Uterus |

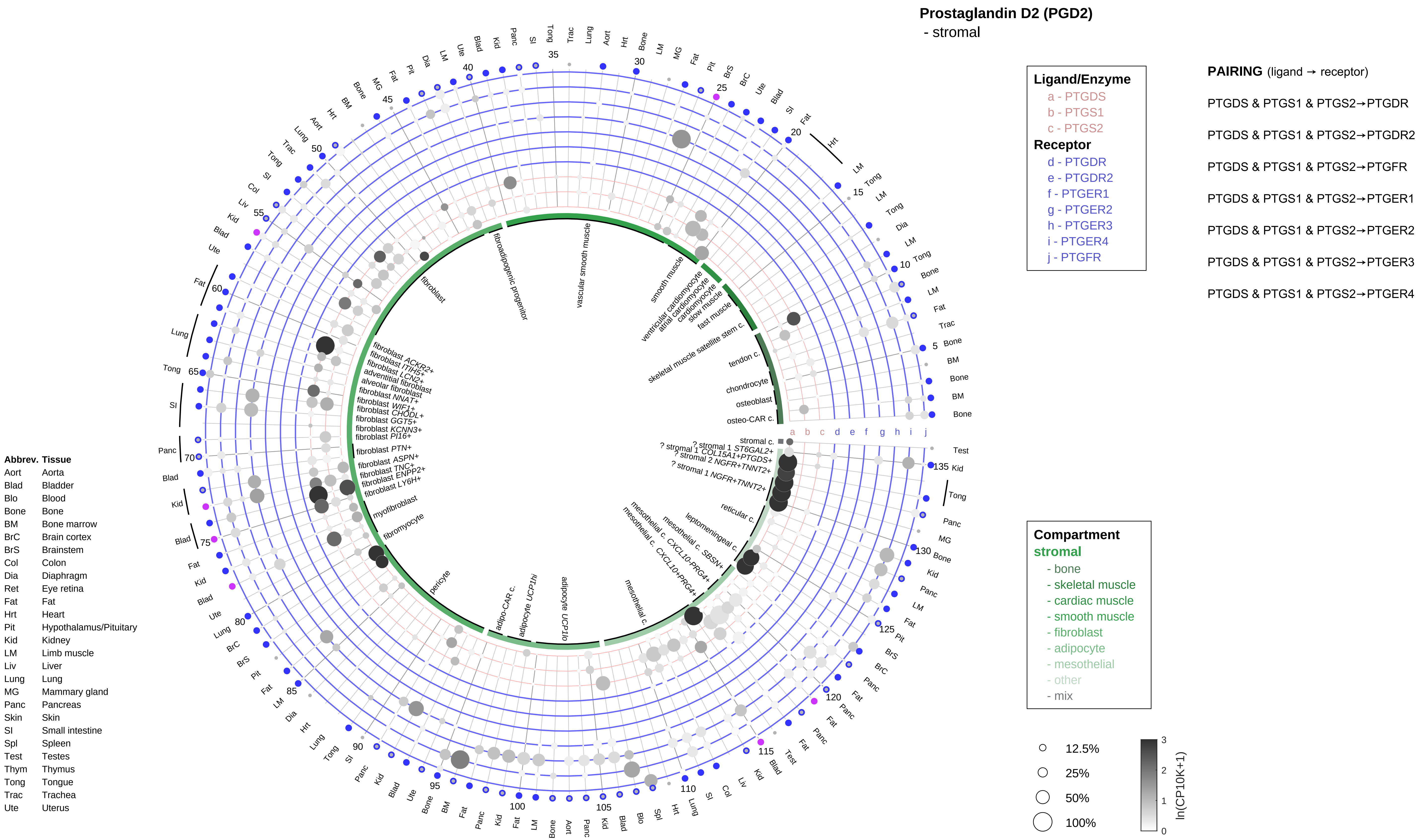

Prostaglandin D2 (PGD2)
- stromal
Ligand/Enzyme
a - PTGDS
b - PTGS1
c - PTGS2
Receptor
d - PTGDR
e - PTGDR2
f - PTGER1
g - PTGER2
h - PTGER3
i - PTGER4
j - PTGFR
PAIRING (ligand → receptor)
PTGDS & PTGS1 & PTGS2→PTGDR
PTGDS & PTGS1 & PTGS2→PTGDR2
PTGDS & PTGS1 & PTGS2→PTGFR
PTGDS & PTGS1 & PTGS2→PTGER1
PTGDS & PTGS1 & PTGS2→PTGER2
PTGDS & PTGS1 & PTGS2→PTGER3
PTGDS & PTGS1 & PTGS2→PTGER4
Compartment
stromal
- bone
- skeletal muscle
- cardiac muscle
- smooth muscle
- fibroblast
- adipocyte
- mesothelial
- other
- mix
12.5%
25%
50%
100%
ln(CP10K+1)
Abbrev. Tissue
Aort Aorta
Blad Bladder
Blo Blood
Bone Bone
BM Bone marrow
BrC Brain cortex
BrS Brainstem
Col Colon
Dia Diaphragm
Ret Eye retina
Fat Fat
Hrt Heart
Pit Hypothalamus/Pituitary
Kid Kidney
LM Limb muscle
Liv Liver
Lung Lung
MG Mammary gland
Panc Pancreas
Skin Skin
SI Small intestine
Spl Spleen
Test Testes
Thym Thymus
Tong Tongue
Trac Trachea
Ute Uterus
fibroadipogenic progenitor
vascular smooth muscle
smooth muscle
ventricular cardiomyocyte
atrial cardiomyocyte
cardiomyocyte
slow muscle
fast muscle
skeletal muscle satellite stem c.
tendon c.
chondrocyte
osteoblast
osteo-CAR c.
stromal c.
fibroblast
myofibroblast
fibromyocyte
pericyte
adipo-CAR c.
adipocyte UCP1hi
adipocyte UCP1lo
mesothelial c.
reticular c.
leptomeningeal c.

**Prostaglandin D2 (PGD2)**
- lymphoid

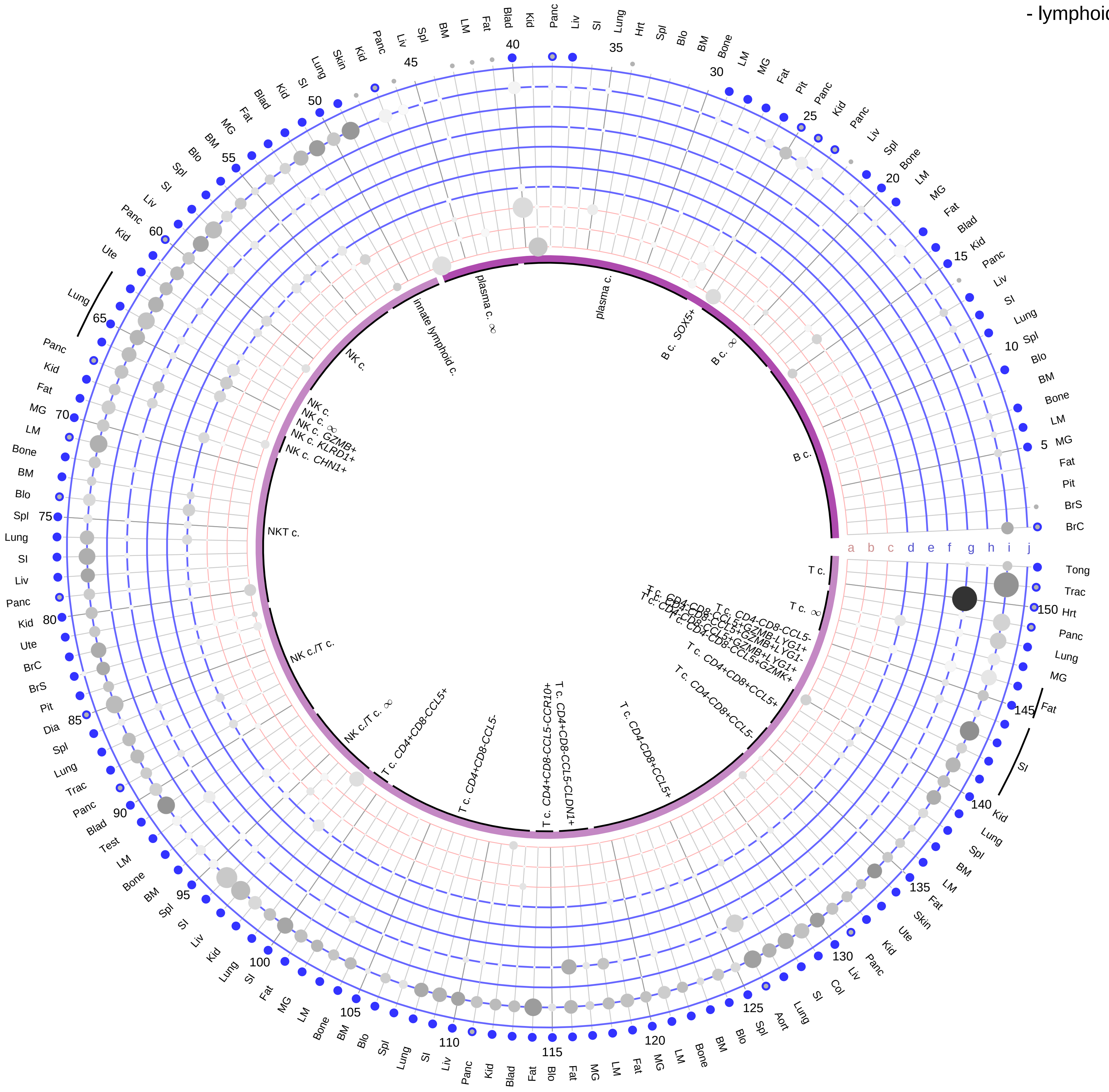

**Ligand/Enzyme**
a - PTGDS
b - PTGS1
c - PTGS2
**Receptor**
d - PTGDR
e - PTGDR2
f - PTGER1
g - PTGER2
h - PTGER3
i - PTGER4
j - PTGFR

**PAIRING** (ligand → receptor)

PTGDS & PTGS1 & PTGS2→PTGDR

PTGDS & PTGS1 & PTGS2→PTGDR2

PTGDS & PTGS1 & PTGS2→PTGFR

PTGDS & PTGS1 & PTGS2→PTGER1

PTGDS & PTGS1 & PTGS2→PTGER2

PTGDS & PTGS1 & PTGS2→PTGER3

PTGDS & PTGS1 & PTGS2→PTGER4

**Compartment**
lymphoid
- B cell
- NK/T cell

12.5%
25%
50%
100%

ln(CP10K+1): 0, 1, 2, 3

| Abbrev. | Tissue |
|---|---|
| Aort | Aorta |
| Blad | Bladder |
| Blo | Blood |
| Bone | Bone |
| BM | Bone marrow |
| BrC | Brain cortex |
| BrS | Brainstem |
| Col | Colon |
| Dia | Diaphragm |
| Ret | Eye retina |
| Fat | Fat |
| Hrt | Heart |
| Pit | Hypothalamus/Pituitary |
| Kid | Kidney |
| LM | Limb muscle |
| Liv | Liver |
| Lung | Lung |
| MG | Mammary gland |
| Panc | Pancreas |
| Skin | Skin |
| SI | Small intestine |
| Spl | Spleen |
| Test | Testes |
| Thym | Thymus |
| Tong | Tongue |
| Trac | Trachea |
| Ute | Uterus |

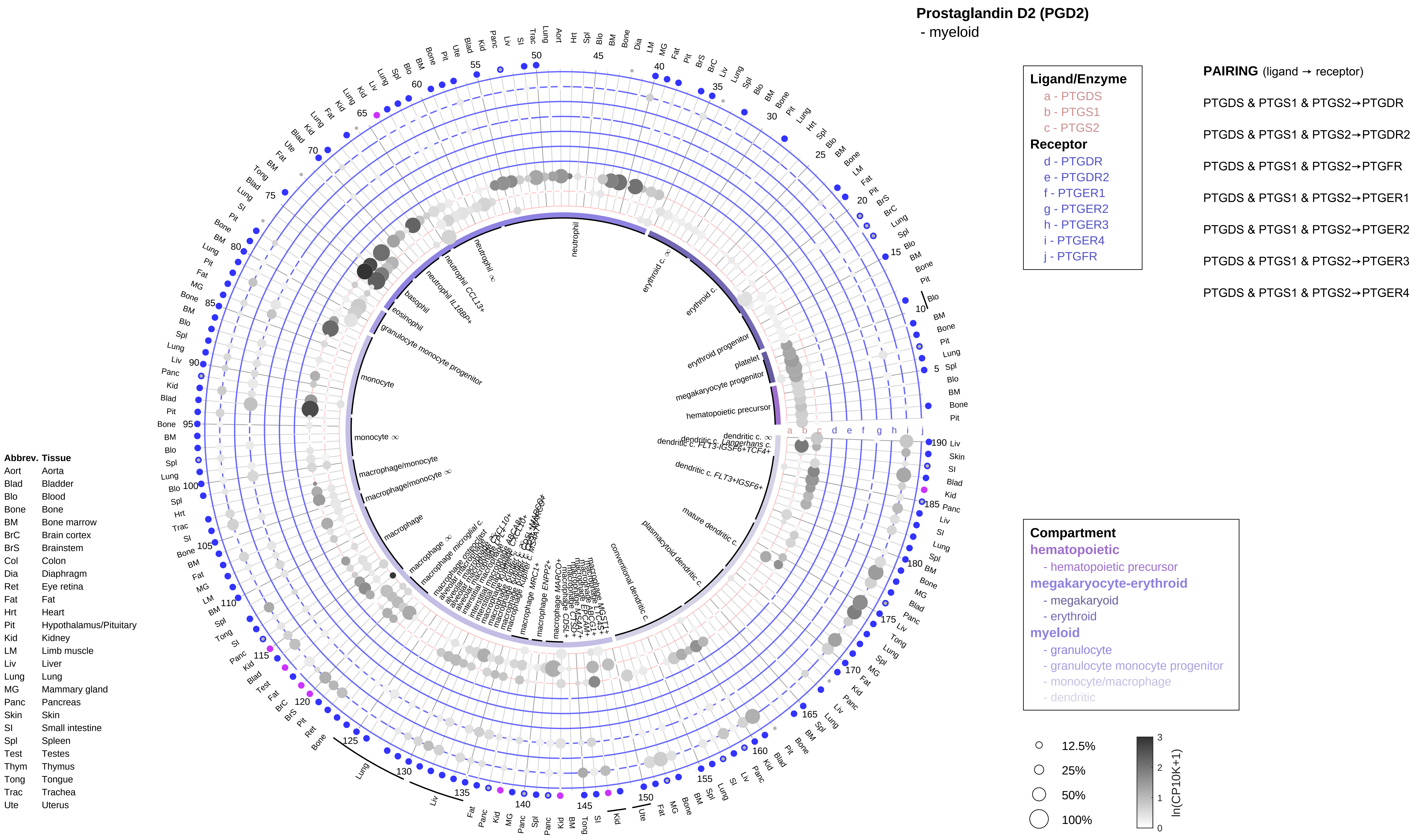

Prostaglandin D2 (PGD2)
- myeloid
Ligand/Enzyme
a - PTGDS
b - PTGS1
c - PTGS2
Receptor
d - PTGDR
e - PTGDR2
f - PTGER1
g - PTGER2
h - PTGER3
i - PTGER4
j - PTGFR
PAIRING (ligand → receptor)
PTGDS & PTGS1 & PTGS2→PTGDR
PTGDS & PTGS1 & PTGS2→PTGDR2
PTGDS & PTGS1 & PTGS2→PTGFR
PTGDS & PTGS1 & PTGS2→PTGER1
PTGDS & PTGS1 & PTGS2→PTGER2
PTGDS & PTGS1 & PTGS2→PTGER3
PTGDS & PTGS1 & PTGS2→PTGER4
Compartment
hematopoietic
- hematopoietic precursor
megakaryocyte-erythroid
- megakaryoid
- erythroid
myeloid
- granulocyte
- granulocyte monocyte progenitor
- monocyte/macrophage
- dendritic
12.5%
25%
50%
100%
ln(CP10K+1)
Abbrev. Tissue
Aort Aorta
Blad Bladder
Blo Blood
Bone Bone
BM Bone marrow
BrC Brain cortex
BrS Brainstem
Col Colon
Dia Diaphragm
Ret Eye retina
Fat Fat
Hrt Heart
Pit Hypothalamus/Pituitary
Kid Kidney
LM Limb muscle
Liv Liver
Lung Lung
MG Mammary gland
Panc Pancreas
Skin Skin
SI Small intestine
Spl Spleen
Test Testes
Thym Thymus
Tong Tongue
Trac Trachea
Ute Uterus

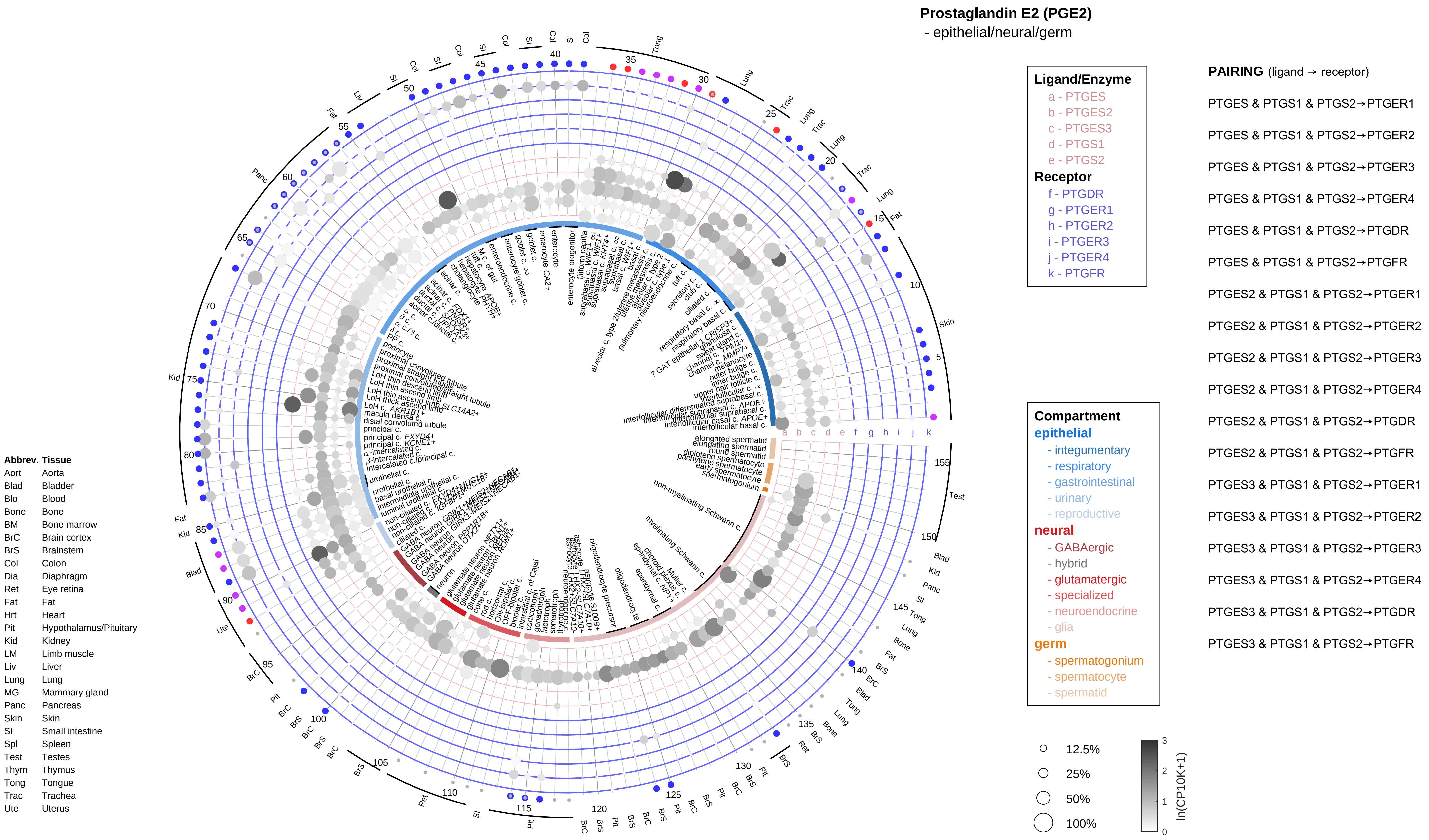

Prostaglandin E2 (PGE2)
- epithelial/neural/germ
Ligand/Enzyme
a - PTGES
b - PTGES2
c - PTGES3
d - PTGS1
e - PTGS2
Receptor
f - PTGDR
g - PTGER1
h - PTGER2
i - PTGER3
j - PTGER4
k - PTGFR
PAIRING (ligand → receptor)
PTGES & PTGS1 & PTGS2→PTGER1
PTGES & PTGS1 & PTGS2→PTGER2
PTGES & PTGS1 & PTGS2→PTGER3
PTGES & PTGS1 & PTGS2→PTGER4
PTGES & PTGS1 & PTGS2→PTGDR
PTGES & PTGS1 & PTGS2→PTGFR
PTGES2 & PTGS1 & PTGS2→PTGER1
PTGES2 & PTGS1 & PTGS2→PTGER2
PTGES2 & PTGS1 & PTGS2→PTGER3
PTGES2 & PTGS1 & PTGS2→PTGER4
PTGES2 & PTGS1 & PTGS2→PTGDR
PTGES2 & PTGS1 & PTGS2→PTGFR
PTGES3 & PTGS1 & PTGS2→PTGER1
PTGES3 & PTGS1 & PTGS2→PTGER2
PTGES3 & PTGS1 & PTGS2→PTGER3
PTGES3 & PTGS1 & PTGS2→PTGER4
PTGES3 & PTGS1 & PTGS2→PTGDR
PTGES3 & PTGS1 & PTGS2→PTGFR
Compartment
epithelial
- integumentary
- respiratory
- gastrointestinal
- urinary
- reproductive
neural
- GABAergic
- hybrid
- glutamatergic
- specialized
- neuroendocrine
- glia
germ
- spermatogonium
- spermatocyte
- spermatid
12.5%
25%
50%
100%
ln(CP10K+1)
0
1
2
3
Abbrev. Tissue
Aort Aorta
Blad Bladder
Blo Blood
Bone Bone
BM Bone marrow
BrC Brain cortex
BrS Brainstem
Col Colon
Dia Diaphragm
Ret Eye retina
Fat Fat
Hrt Heart
Pit Hypothalamus/Pituitary
Kid Kidney
LM Limb muscle
Liv Liver
Lung Lung
MG Mammary gland
Panc Pancreas
Skin Skin
SI Small intestine
Spl Spleen
Test Testes
Thym Thymus
Tong Tongue
Trac Trachea
Ute Uterus

#### Prostaglandin E2 (PGE2)
- endothelial

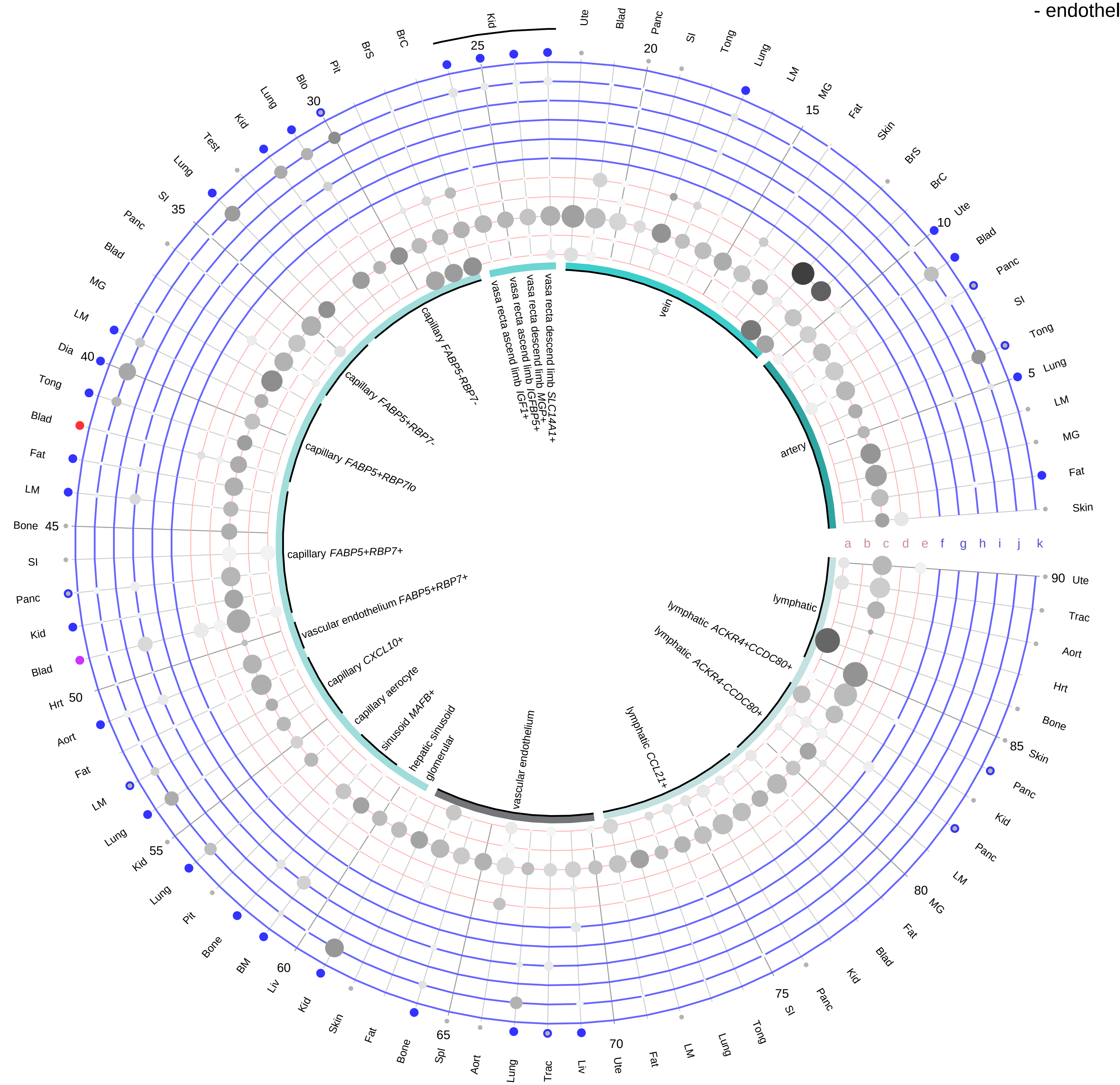

**Ligand/Enzyme**
- a - PTGES
- b - PTGES2
- c - PTGES3
- d - PTGS1
- e - PTGS2

**Receptor**
- f - PTGDR
- g - PTGER1
- h - PTGER2
- i - PTGER3
- j - PTGER4
- k - PTGFR

**PAIRING** (ligand → receptor)

PTGES & PTGS1 & PTGS2→PTGER1

PTGES & PTGS1 & PTGS2→PTGER2

PTGES & PTGS1 & PTGS2→PTGER3

PTGES & PTGS1 & PTGS2→PTGER4

PTGES & PTGS1 & PTGS2→PTGDR

PTGES & PTGS1 & PTGS2→PTGFR

PTGES2 & PTGS1 & PTGS2→PTGER1

PTGES2 & PTGS1 & PTGS2→PTGER2

PTGES2 & PTGS1 & PTGS2→PTGER3

PTGES2 & PTGS1 & PTGS2→PTGER4

PTGES2 & PTGS1 & PTGS2→PTGDR

PTGES2 & PTGS1 & PTGS2→PTGFR

PTGES3 & PTGS1 & PTGS2→PTGER1

PTGES3 & PTGS1 & PTGS2→PTGER2

PTGES3 & PTGS1 & PTGS2→PTGER3

PTGES3 & PTGS1 & PTGS2→PTGER4

PTGES3 & PTGS1 & PTGS2→PTGDR

PTGES3 & PTGS1 & PTGS2→PTGFR

| Abbrev. | Tissue |
|---|---|
| Aort | Aorta |
| Blad | Bladder |
| Blo | Blood |
| Bone | Bone |
| BM | Bone marrow |
| BrC | Brain cortex |
| BrS | Brainstem |
| Col | Colon |
| Dia | Diaphragm |
| Ret | Eye retina |
| Fat | Fat |
| Hrt | Heart |
| Pit | Hypothalamus/Pituitary |
| Kid | Kidney |
| LM | Limb muscle |
| Liv | Liver |
| Lung | Lung |
| MG | Mammary gland |
| Panc | Pancreas |
| Skin | Skin |
| SI | Small intestine |
| Spl | Spleen |
| Test | Testes |
| Thym | Thymus |
| Tong | Tongue |
| Trac | Trachea |
| Ute | Uterus |

**Compartment**

**endothelial**
- artery
- vein
- vasa recta
- capillary
- mix
- lymphatic

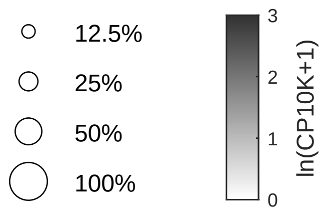

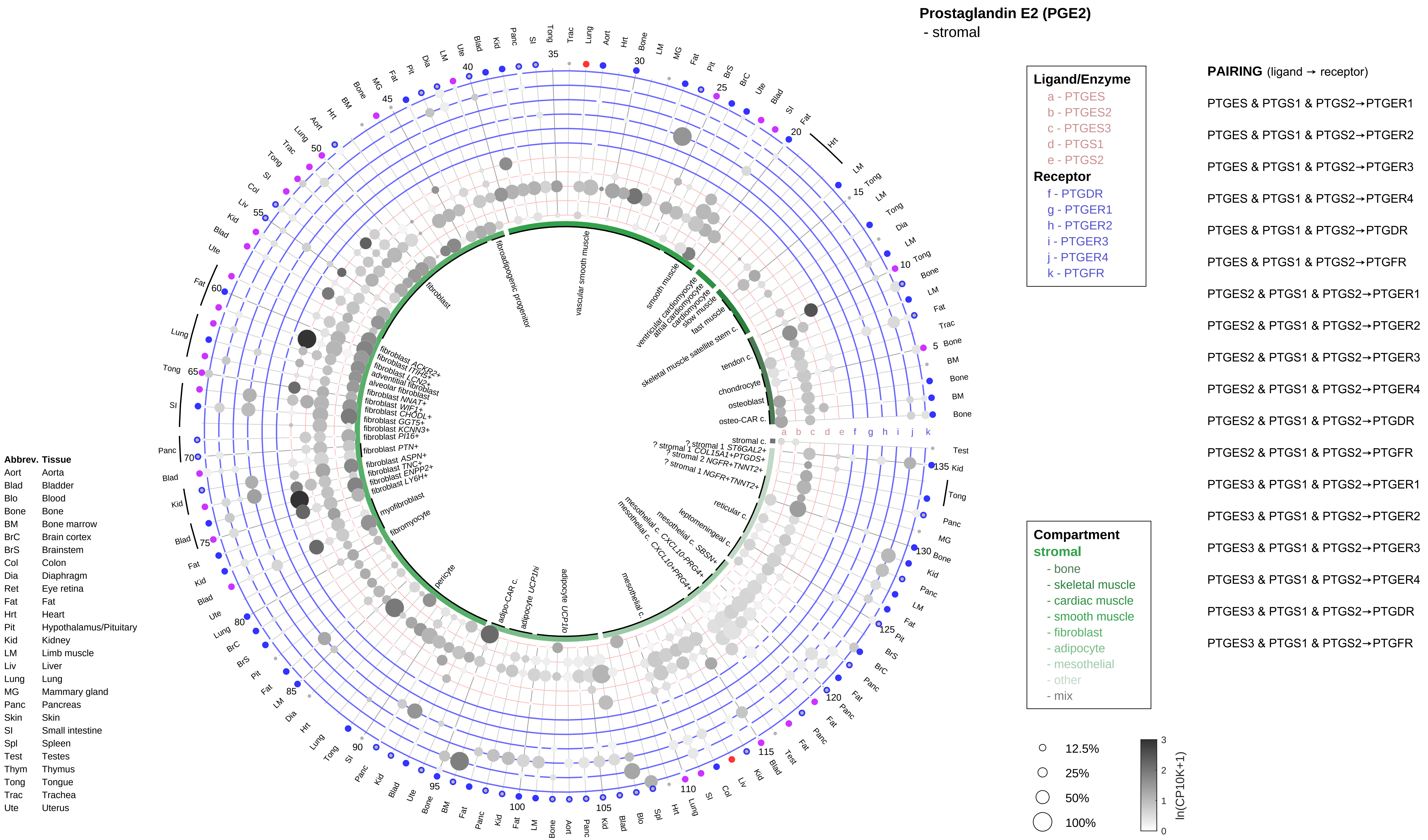

Prostaglandin E2 (PGE2)
- stromal
Ligand/Enzyme
a - PTGES
b - PTGES2
c - PTGES3
d - PTGS1
e - PTGS2
Receptor
f - PTGDR
g - PTGER1
h - PTGER2
i - PTGER3
j - PTGER4
k - PTGFR
PAIRING (ligand → receptor)
PTGES & PTGS1 & PTGS2→PTGER1
PTGES & PTGS1 & PTGS2→PTGER2
PTGES & PTGS1 & PTGS2→PTGER3
PTGES & PTGS1 & PTGS2→PTGER4
PTGES & PTGS1 & PTGS2→PTGDR
PTGES & PTGS1 & PTGS2→PTGFR
PTGES2 & PTGS1 & PTGS2→PTGER1
PTGES2 & PTGS1 & PTGS2→PTGER2
PTGES2 & PTGS1 & PTGS2→PTGER3
PTGES2 & PTGS1 & PTGS2→PTGER4
PTGES2 & PTGS1 & PTGS2→PTGDR
PTGES2 & PTGS1 & PTGS2→PTGFR
PTGES3 & PTGS1 & PTGS2→PTGER1
PTGES3 & PTGS1 & PTGS2→PTGER2
PTGES3 & PTGS1 & PTGS2→PTGER3
PTGES3 & PTGS1 & PTGS2→PTGER4
PTGES3 & PTGS1 & PTGS2→PTGDR
PTGES3 & PTGS1 & PTGS2→PTGFR
Compartment
stromal
- bone
- skeletal muscle
- cardiac muscle
- smooth muscle
- fibroblast
- adipocyte
- mesothelial
- other
- mix
12.5%
25%
50%
100%
ln(CP10K+1)
0
1
2
3
Abbrev. Tissue
Aort Aorta
Blad Bladder
Blo Blood
Bone Bone
BM Bone marrow
BrC Brain cortex
BrS Brainstem
Col Colon
Dia Diaphragm
Ret Eye retina
Fat Fat
Hrt Heart
Pit Hypothalamus/Pituitary
Kid Kidney
LM Limb muscle
Liv Liver
Lung Lung
MG Mammary gland
Panc Pancreas
Skin Skin
SI Small intestine
Spl Spleen
Test Testes
Thym Thymus
Tong Tongue
Trac Trachea
Ute Uterus
fibroblast
fibroadipogenic progenitor
vascular smooth muscle
smooth muscle
ventricular cardiomyocyte
atrial cardiomyocyte
cardiomyocyte
slow muscle
fast muscle
skeletal muscle satellite stem c.
tendon c.
chondrocyte
osteoblast
osteo-CAR c.
stromal c.
reticular c.
leptomeningeal c.
mesothelial c.
pericyte
adipo-CAR c.
myofibroblast
fibromyocyte
alveolar fibroblast
adventitial fibroblast

**Prostaglandin E2 (PGE2)**
- lymphoid

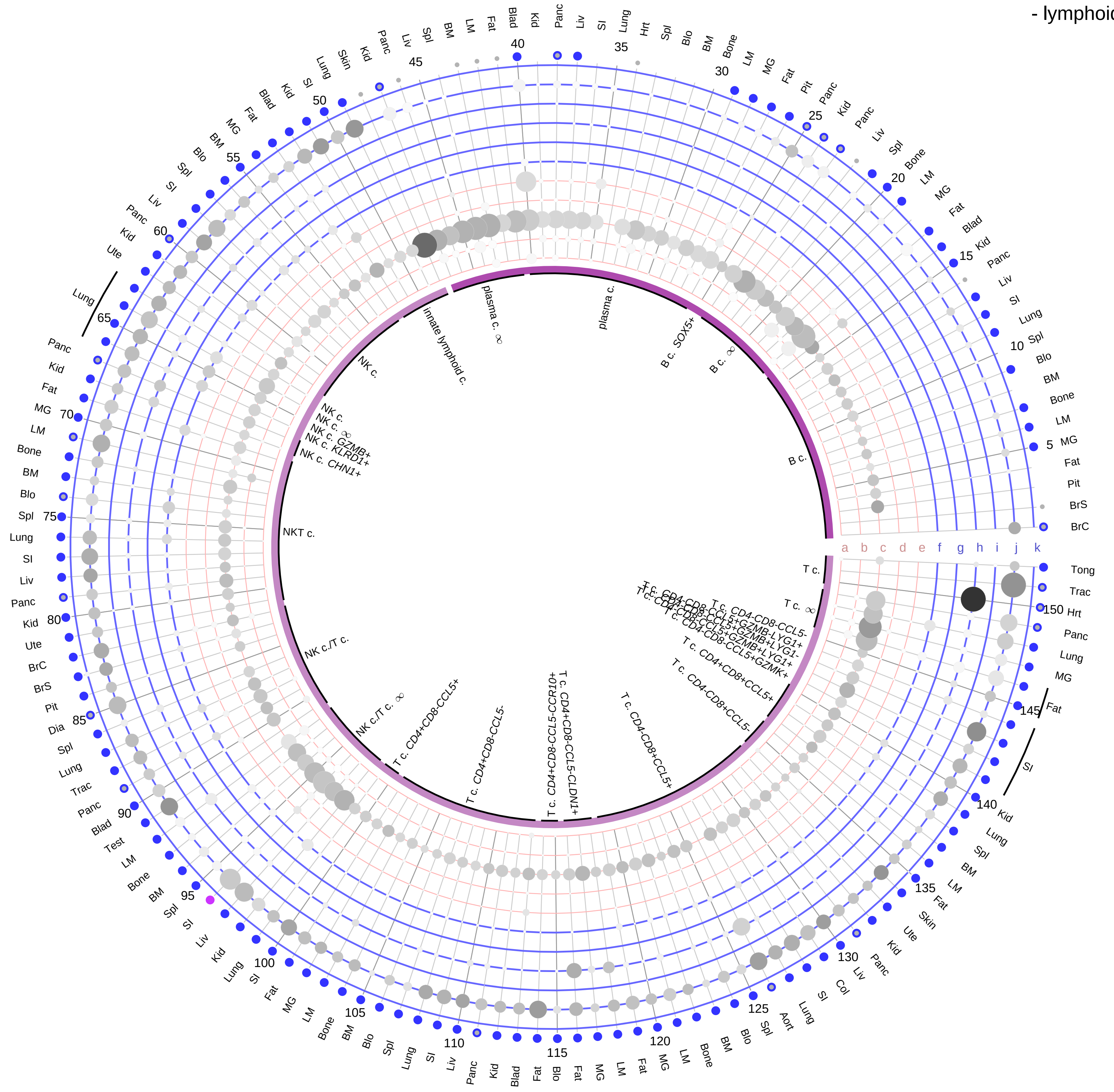

**Ligand/Enzyme**
- a - PTGES
- b - PTGES2
- c - PTGES3
- d - PTGS1
- e - PTGS2

**Receptor**
- f - PTGDR
- g - PTGER1
- h - PTGER2
- i - PTGER3
- j - PTGER4
- k - PTGFR

**PAIRING** (ligand → receptor)

PTGES & PTGS1 & PTGS2→PTGER1
PTGES & PTGS1 & PTGS2→PTGER2
PTGES & PTGS1 & PTGS2→PTGER3
PTGES & PTGS1 & PTGS2→PTGER4
PTGES & PTGS1 & PTGS2→PTGDR
PTGES & PTGS1 & PTGS2→PTGFR
PTGES2 & PTGS1 & PTGS2→PTGER1
PTGES2 & PTGS1 & PTGS2→PTGER2
PTGES2 & PTGS1 & PTGS2→PTGER3
PTGES2 & PTGS1 & PTGS2→PTGER4
PTGES2 & PTGS1 & PTGS2→PTGDR
PTGES2 & PTGS1 & PTGS2→PTGFR
PTGES3 & PTGS1 & PTGS2→PTGER1
PTGES3 & PTGS1 & PTGS2→PTGER2
PTGES3 & PTGS1 & PTGS2→PTGER3
PTGES3 & PTGS1 & PTGS2→PTGER4
PTGES3 & PTGS1 & PTGS2→PTGDR
PTGES3 & PTGS1 & PTGS2→PTGFR

**Compartment**
**lymphoid**
- B cell
- NK/T cell

12.5%
25%
50%
100%

ln(CP10K+1): 0, 1, 2, 3

| Abbrev. | Tissue |
|---|---|
| Aort | Aorta |
| Blad | Bladder |
| Blo | Blood |
| Bone | Bone |
| BM | Bone marrow |
| BrC | Brain cortex |
| BrS | Brainstem |
| Col | Colon |
| Dia | Diaphragm |
| Ret | Eye retina |
| Fat | Fat |
| Hrt | Heart |
| Pit | Hypothalamus/Pituitary |
| Kid | Kidney |
| LM | Limb muscle |
| Liv | Liver |
| Lung | Lung |
| MG | Mammary gland |
| Panc | Pancreas |
| Skin | Skin |
| SI | Small intestine |
| Spl | Spleen |
| Test | Testes |
| Thym | Thymus |
| Tong | Tongue |
| Trac | Trachea |
| Ute | Uterus |

**Prostaglandin E2 (PGE2)**
- myeloid

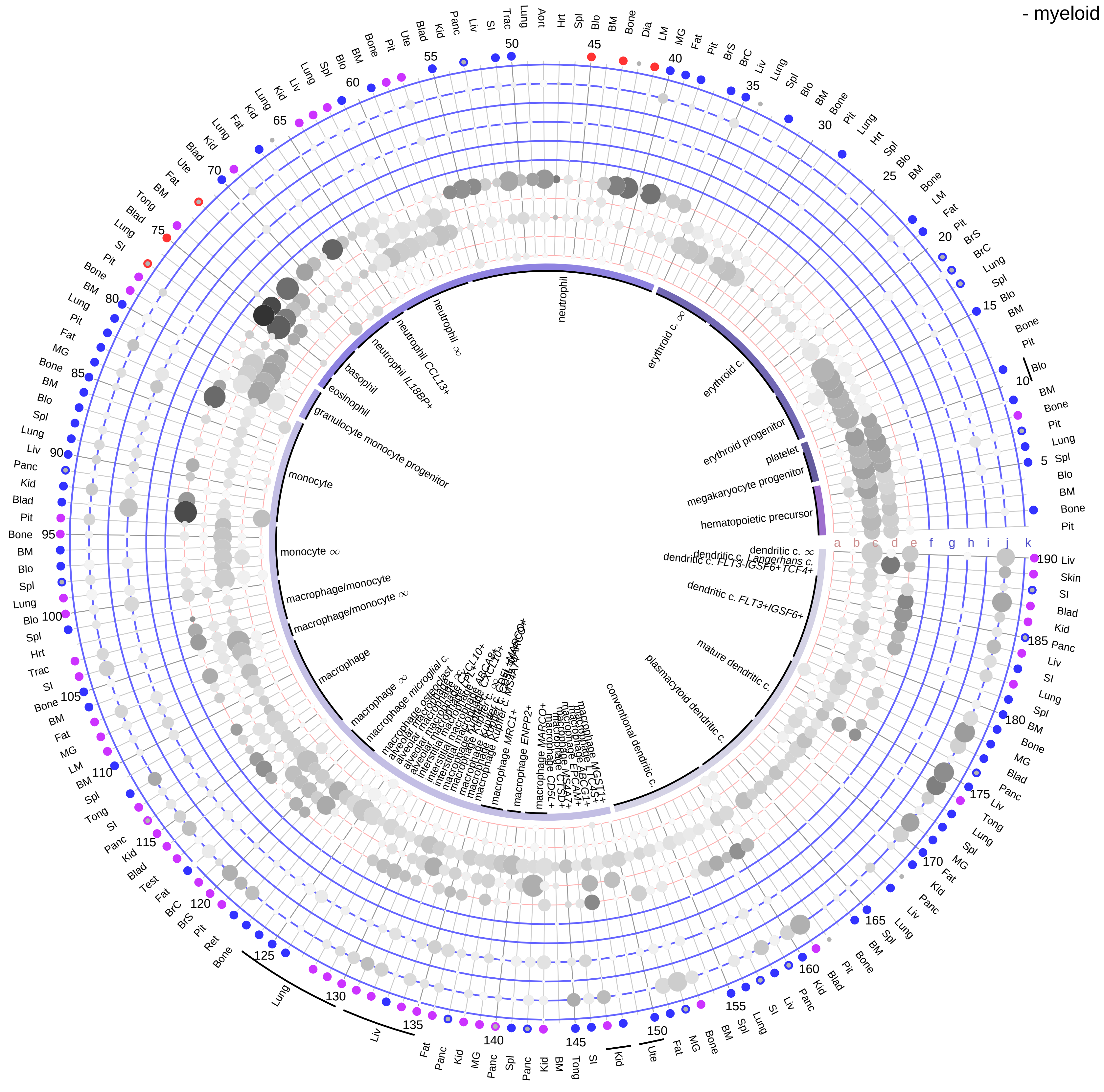

| Abbrev. | Tissue |
|---|---|
| Aort | Aorta |
| Blad | Bladder |
| Blo | Blood |
| Bone | Bone |
| BM | Bone marrow |
| BrC | Brain cortex |
| BrS | Brainstem |
| Col | Colon |
| Dia | Diaphragm |
| Ret | Eye retina |
| Fat | Fat |
| Hrt | Heart |
| Pit | Hypothalamus/Pituitary |
| Kid | Kidney |
| LM | Limb muscle |
| Liv | Liver |
| Lung | Lung |
| MG | Mammary gland |
| Panc | Pancreas |
| Skin | Skin |
| SI | Small intestine |
| Spl | Spleen |
| Test | Testes |
| Thym | Thymus |
| Tong | Tongue |
| Trac | Trachea |
| Ute | Uterus |

**Ligand/Enzyme**
- a - PTGES
- b - PTGES2
- c - PTGES3
- d - PTGS1
- e - PTGS2

**Receptor**
- f - PTGDR
- g - PTGER1
- h - PTGER2
- i - PTGER3
- j - PTGER4
- k - PTGFR

**PAIRING** (ligand → receptor)

PTGES & PTGS1 & PTGS2→PTGER1
PTGES & PTGS1 & PTGS2→PTGER2
PTGES & PTGS1 & PTGS2→PTGER3
PTGES & PTGS1 & PTGS2→PTGER4
PTGES & PTGS1 & PTGS2→PTGDR
PTGES & PTGS1 & PTGS2→PTGFR
PTGES2 & PTGS1 & PTGS2→PTGER1
PTGES2 & PTGS1 & PTGS2→PTGER2
PTGES2 & PTGS1 & PTGS2→PTGER3
PTGES2 & PTGS1 & PTGS2→PTGER4
PTGES2 & PTGS1 & PTGS2→PTGDR
PTGES2 & PTGS1 & PTGS2→PTGFR
PTGES3 & PTGS1 & PTGS2→PTGER1
PTGES3 & PTGS1 & PTGS2→PTGER2
PTGES3 & PTGS1 & PTGS2→PTGER3
PTGES3 & PTGS1 & PTGS2→PTGER4
PTGES3 & PTGS1 & PTGS2→PTGDR
PTGES3 & PTGS1 & PTGS2→PTGFR

**Compartment**

**hematopoietic**
- hematopoietic precursor

**megakaryocyte-erythroid**
- megakaryoid
- erythroid

**myeloid**
- granulocyte
- granulocyte monocyte progenitor
- monocyte/macrophage
- dendritic

12.5%
25%
50%
100%

ln(CP10K+1): 0, 1, 2, 3

#### Prostaglandin I2/Prostacyclin (PGI2)
- epithelial/neural/germ

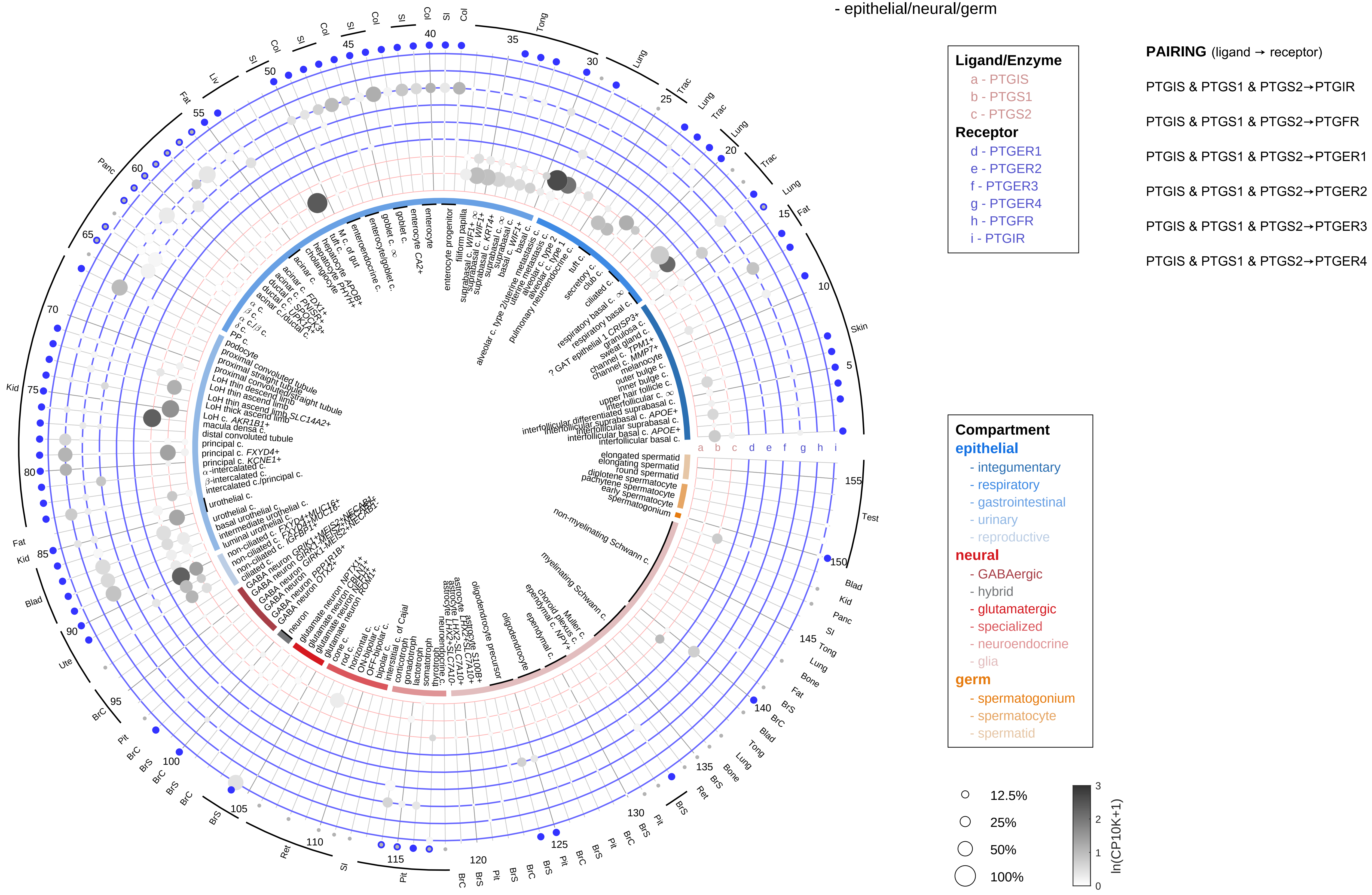

| Abbrev. | Tissue |
|---|---|
| Aort | Aorta |
| Blad | Bladder |
| Blo | Blood |
| Bone | Bone |
| BM | Bone marrow |
| BrC | Brain cortex |
| BrS | Brainstem |
| Col | Colon |
| Dia | Diaphragm |
| Ret | Eye retina |
| Fat | Fat |
| Hrt | Heart |
| Pit | Hypothalamus/Pituitary |
| Kid | Kidney |
| LM | Limb muscle |
| Liv | Liver |
| Lung | Lung |
| MG | Mammary gland |
| Panc | Pancreas |
| Skin | Skin |
| SI | Small intestine |
| Spl | Spleen |
| Test | Testes |
| Thym | Thymus |
| Tong | Tongue |
| Trac | Trachea |
| Ute | Uterus |

#### Prostaglandin I2/Prostacyclin (PGI2)
- endothelial

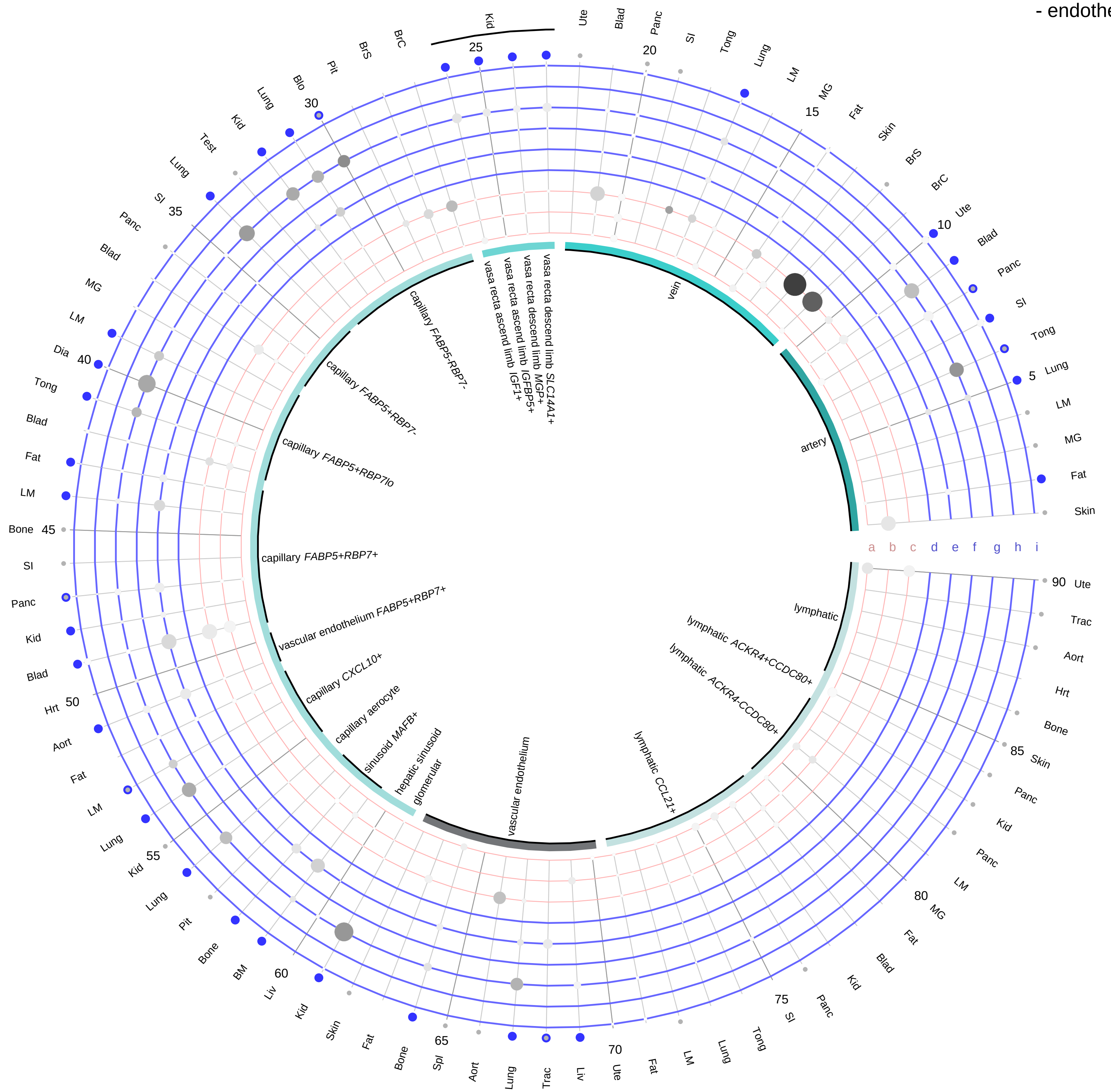

**Ligand/Enzyme**
- a - PTGIS
- b - PTGS1
- c - PTGS2

**Receptor**
- d - PTGER1
- e - PTGER2
- f - PTGER3
- g - PTGER4
- h - PTGFR
- i - PTGIR

**PAIRING** (ligand → receptor)

PTGIS & PTGS1 & PTGS2→PTGIR

PTGIS & PTGS1 & PTGS2→PTGFR

PTGIS & PTGS1 & PTGS2→PTGER1

PTGIS & PTGS1 & PTGS2→PTGER2

PTGIS & PTGS1 & PTGS2→PTGER3

PTGIS & PTGS1 & PTGS2→PTGER4

| Abbrev. | Tissue |
|---|---|
| Aort | Aorta |
| Blad | Bladder |
| Blo | Blood |
| Bone | Bone |
| BM | Bone marrow |
| BrC | Brain cortex |
| BrS | Brainstem |
| Col | Colon |
| Dia | Diaphragm |
| Ret | Eye retina |
| Fat | Fat |
| Hrt | Heart |
| Pit | Hypothalamus/Pituitary |
| Kid | Kidney |
| LM | Limb muscle |
| Liv | Liver |
| Lung | Lung |
| MG | Mammary gland |
| Panc | Pancreas |
| Skin | Skin |
| SI | Small intestine |
| Spl | Spleen |
| Test | Testes |
| Thym | Thymus |
| Tong | Tongue |
| Trac | Trachea |
| Ute | Uterus |

**Compartment**
**endothelial**
- artery
- vein
- vasa recta
- capillary
- mix
- lymphatic

#### Prostaglandin I2/Prostacyclin (PGI2)
- stromal

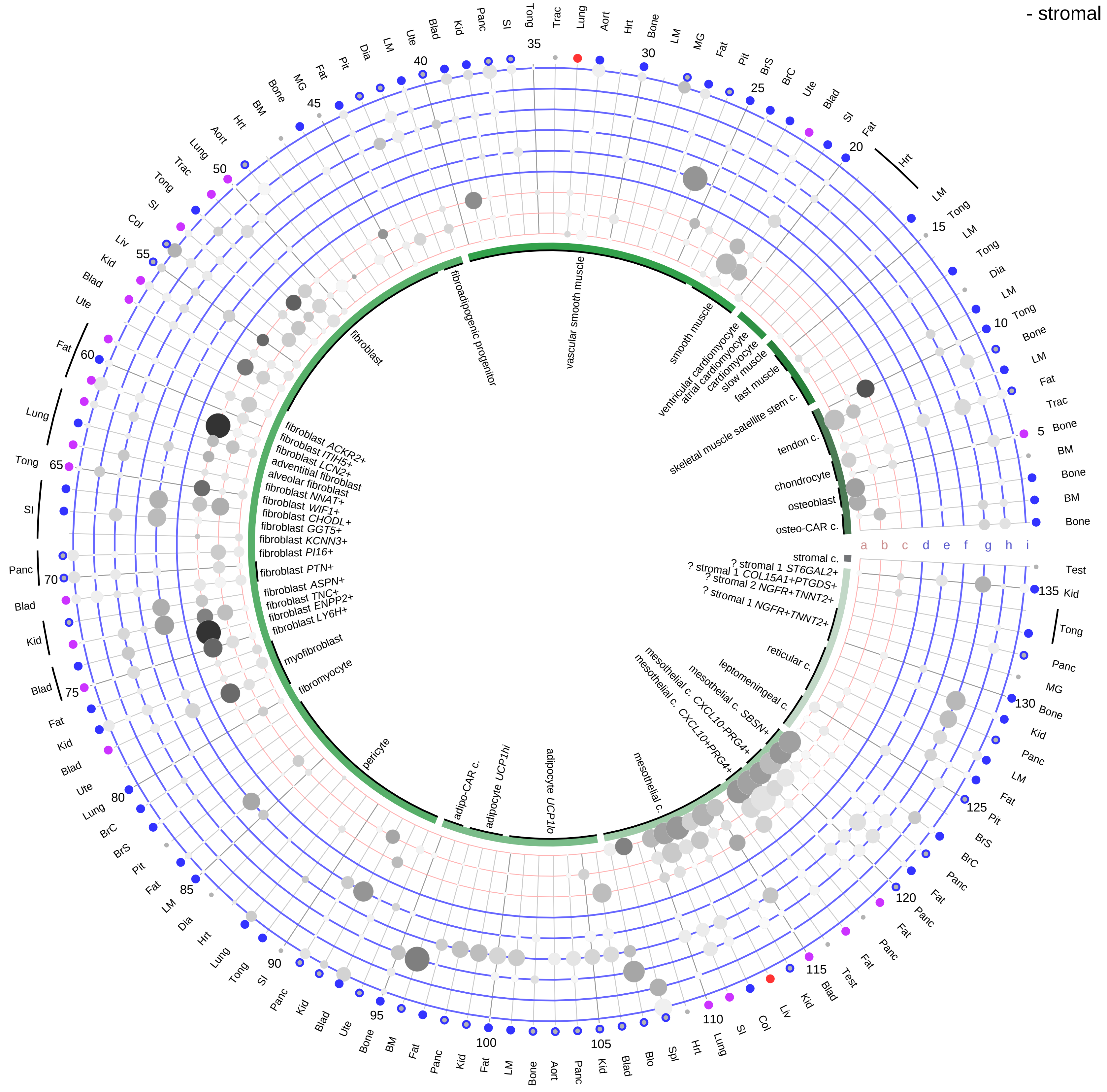

**Ligand/Enzyme**
- a - PTGIS
- b - PTGS1
- c - PTGS2

**Receptor**
- d - PTGER1
- e - PTGER2
- f - PTGER3
- g - PTGER4
- h - PTGFR
- i - PTGIR

**PAIRING** (ligand → receptor)

PTGIS & PTGS1 & PTGS2→PTGIR

PTGIS & PTGS1 & PTGS2→PTGFR

PTGIS & PTGS1 & PTGS2→PTGER1

PTGIS & PTGS1 & PTGS2→PTGER2

PTGIS & PTGS1 & PTGS2→PTGER3

PTGIS & PTGS1 & PTGS2→PTGER4

| Abbrev. | Tissue |
|---|---|
| Aort | Aorta |
| Blad | Bladder |
| Blo | Blood |
| Bone | Bone |
| BM | Bone marrow |
| BrC | Brain cortex |
| BrS | Brainstem |
| Col | Colon |
| Dia | Diaphragm |
| Ret | Eye retina |
| Fat | Fat |
| Hrt | Heart |
| Pit | Hypothalamus/Pituitary |
| Kid | Kidney |
| LM | Limb muscle |
| Liv | Liver |
| Lung | Lung |
| MG | Mammary gland |
| Panc | Pancreas |
| Skin | Skin |
| SI | Small intestine |
| Spl | Spleen |
| Test | Testes |
| Thym | Thymus |
| Tong | Tongue |
| Trac | Trachea |
| Ute | Uterus |

**Compartment**

**stromal**
- bone
- skeletal muscle
- cardiac muscle
- smooth muscle
- fibroblast
- adipocyte
- mesothelial
- other
- mix

12.5%
25%
50%
100%

ln(CP10K+1) 0 1 2 3

**Prostaglandin I2/Prostacyclin (PGI2)**
- lymphoid

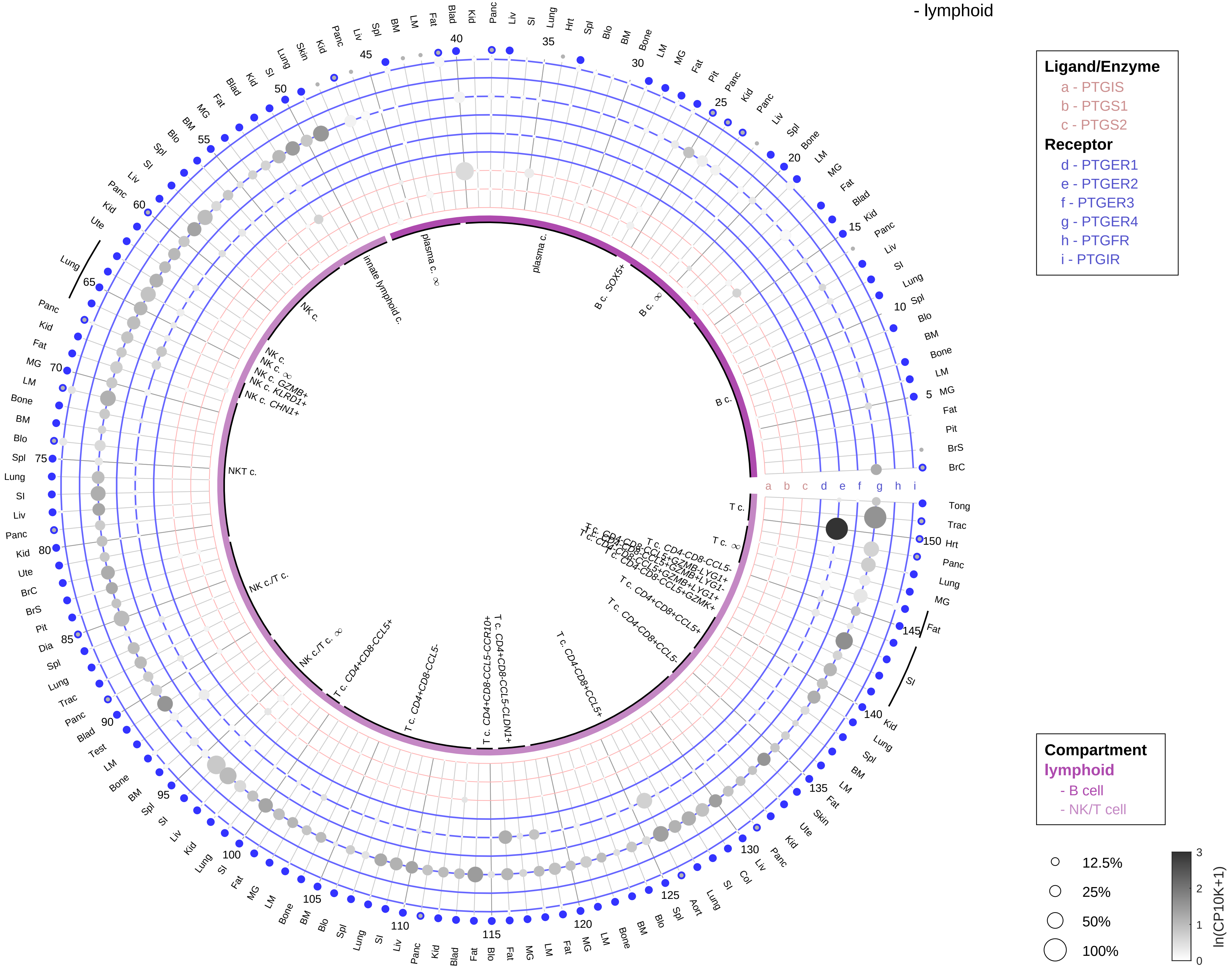

**PAIRING** (ligand → receptor)

PTGIS & PTGS1 & PTGS2→PTGIR

PTGIS & PTGS1 & PTGS2→PTGFR

PTGIS & PTGS1 & PTGS2→PTGER1

PTGIS & PTGS1 & PTGS2→PTGER2

PTGIS & PTGS1 & PTGS2→PTGER3

PTGIS & PTGS1 & PTGS2→PTGER4

| Abbrev. | Tissue |
|---|---|
| Aort | Aorta |
| Blad | Bladder |
| Blo | Blood |
| Bone | Bone |
| BM | Bone marrow |
| BrC | Brain cortex |
| BrS | Brainstem |
| Col | Colon |
| Dia | Diaphragm |
| Ret | Eye retina |
| Fat | Fat |
| Hrt | Heart |
| Pit | Hypothalamus/Pituitary |
| Kid | Kidney |
| LM | Limb muscle |
| Liv | Liver |
| Lung | Lung |
| MG | Mammary gland |
| Panc | Pancreas |
| Skin | Skin |
| SI | Small intestine |
| Spl | Spleen |
| Test | Testes |
| Thym | Thymus |
| Tong | Tongue |
| Trac | Trachea |
| Ute | Uterus |

**Prostaglandin I2/Prostacyclin (PGI2)**
- myeloid

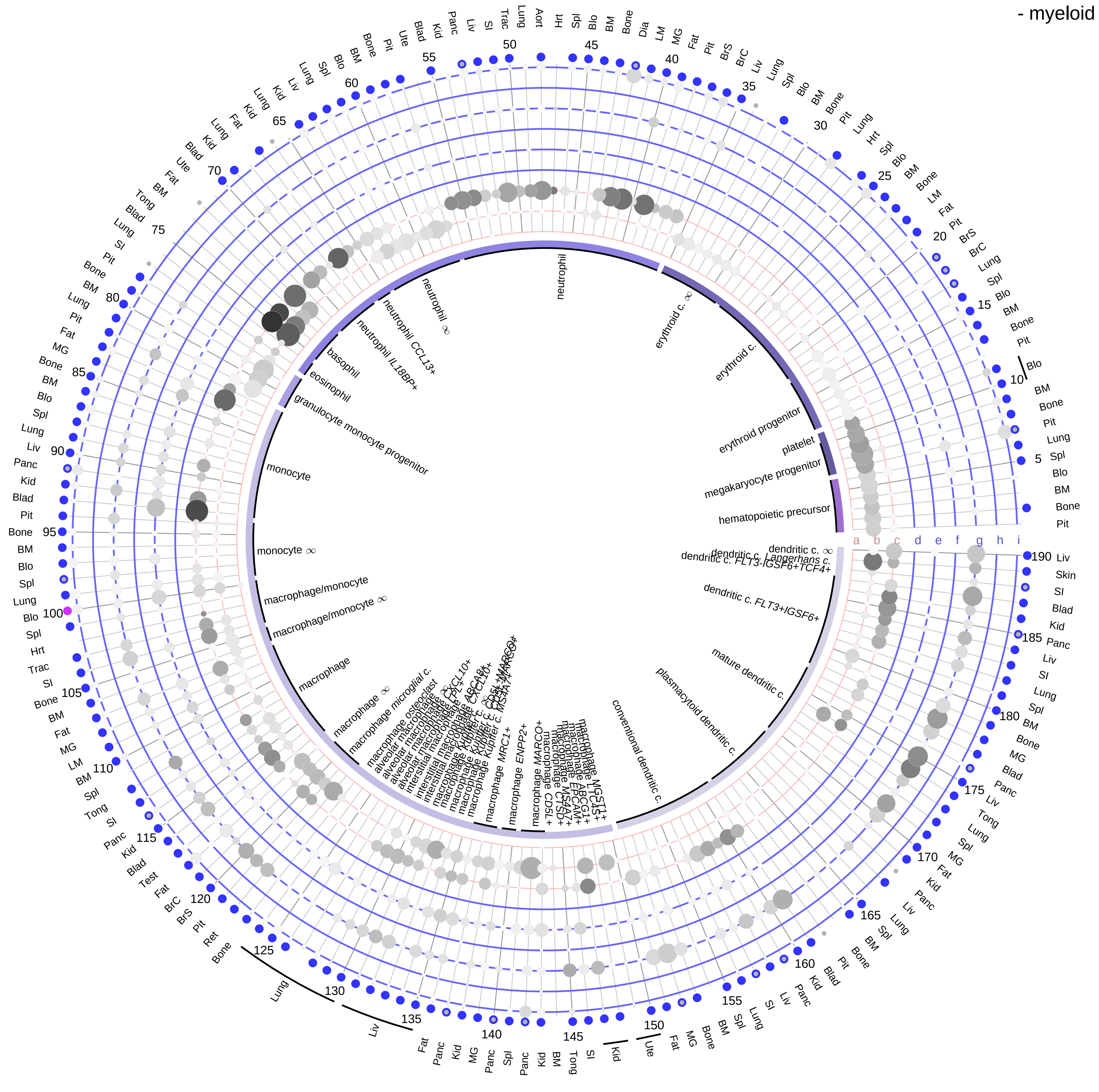

**Ligand/Enzyme**
- a - PTGIS
- b - PTGS1
- c - PTGS2

**Receptor**
- d - PTGER1
- e - PTGER2
- f - PTGER3
- g - PTGER4
- h - PTGFR
- i - PTGIR

**PAIRING** (ligand → receptor)

PTGIS & PTGS1 & PTGS2→PTGIR

PTGIS & PTGS1 & PTGS2→PTGFR

PTGIS & PTGS1 & PTGS2→PTGER1

PTGIS & PTGS1 & PTGS2→PTGER2

PTGIS & PTGS1 & PTGS2→PTGER3

PTGIS & PTGS1 & PTGS2→PTGER4

**Compartment**
**hematopoietic**
- hematopoietic precursor
**megakaryocyte-erythroid**
- megakaryoid
- erythroid
**myeloid**
- granulocyte
- granulocyte monocyte progenitor
- monocyte/macrophage
- dendritic

12.5%
25%
50%
100%

ln(CP10K+1): 0, 1, 2, 3

| Abbrev. | Tissue |
|---|---|
| Aort | Aorta |
| Blad | Bladder |
| Blo | Blood |
| Bone | Bone |
| BM | Bone marrow |
| BrC | Brain cortex |
| BrS | Brainstem |
| Col | Colon |
| Dia | Diaphragm |
| Ret | Eye retina |
| Fat | Fat |
| Hrt | Heart |
| Pit | Hypothalamus/Pituitary |
| Kid | Kidney |
| LM | Limb muscle |
| Liv | Liver |
| Lung | Lung |
| MG | Mammary gland |
| Panc | Pancreas |
| Skin | Skin |
| SI | Small intestine |
| Spl | Spleen |
| Test | Testes |
| Thym | Thymus |
| Tong | Tongue |
| Trac | Trachea |
| Ute | Uterus |

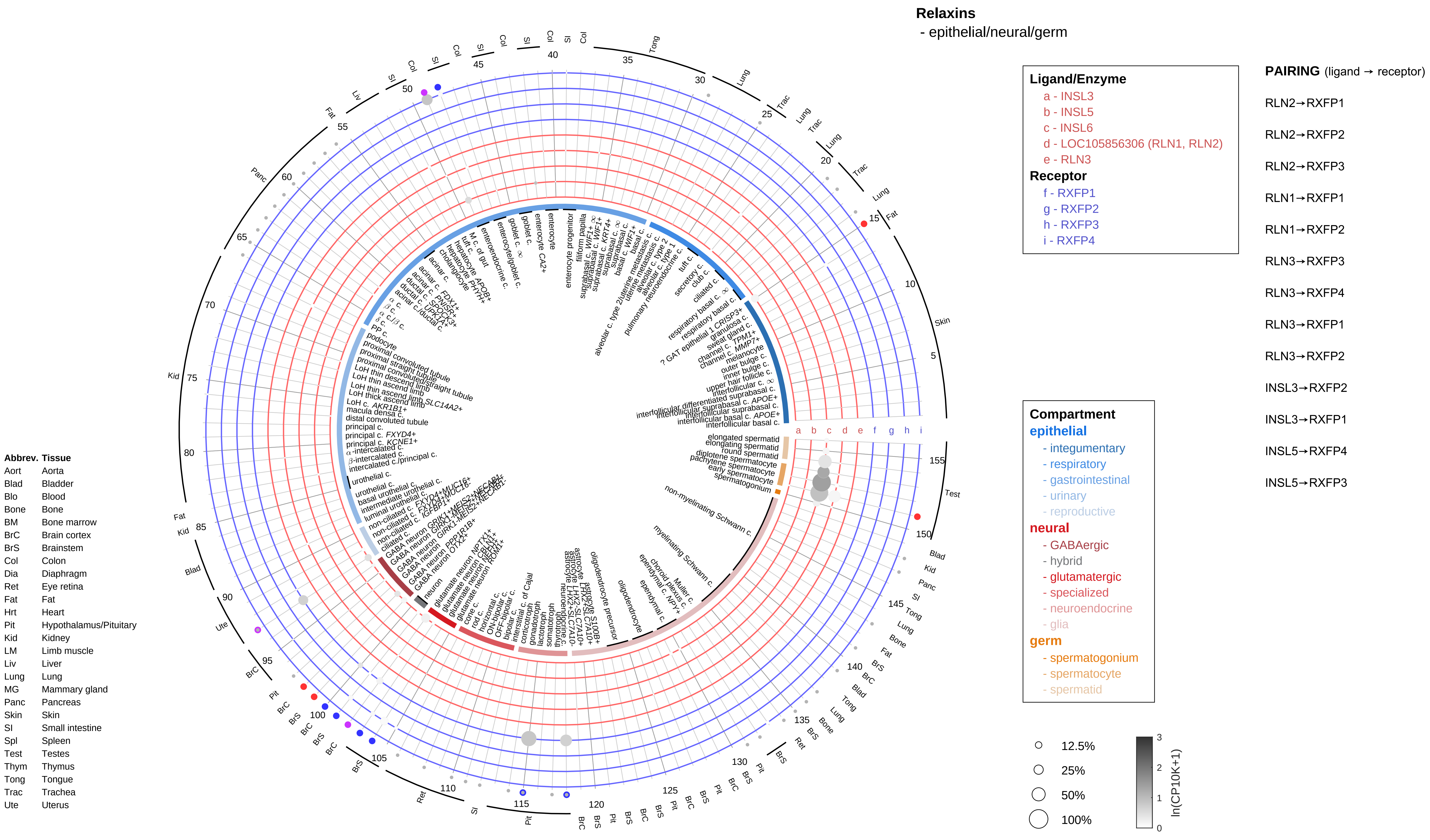
Relaxins
- epithelial/neural/germ
Ligand/Enzyme
a - INSL3
b - INSL5
c - INSL6
d - LOC105856306 (RLN1, RLN2)
e - RLN3
Receptor
f - RXFP1
g - RXFP2
h - RXFP3
i - RXFP4
PAIRING (ligand → receptor)
RLN2→RXFP1
RLN2→RXFP2
RLN2→RXFP3
RLN1→RXFP1
RLN1→RXFP2
RLN3→RXFP3
RLN3→RXFP4
RLN3→RXFP1
RLN3→RXFP2
INSL3→RXFP2
INSL3→RXFP1
INSL5→RXFP4
INSL5→RXFP3
Compartment
epithelial
- integumentary
- respiratory
- gastrointestinal
- urinary
- reproductive
neural
- GABAergic
- hybrid
- glutamatergic
- specialized
- neuroendocrine
- glia
germ
- spermatogonium
- spermatocyte
- spermatid
12.5%
25%
50%
100%
ln(CP10K+1)
Abbrev. Tissue
Aort Aorta
Blad Bladder
Blo Blood
Bone Bone
BM Bone marrow
BrC Brain cortex
BrS Brainstem
Col Colon
Dia Diaphragm
Ret Eye retina
Fat Fat
Hrt Heart
Pit Hypothalamus/Pituitary
Kid Kidney
LM Limb muscle
Liv Liver
Lung Lung
MG Mammary gland
Panc Pancreas
Skin Skin
SI Small intestine
Spl Spleen
Test Testes
Thym Thymus
Tong Tongue
Trac Trachea
Ute Uterus

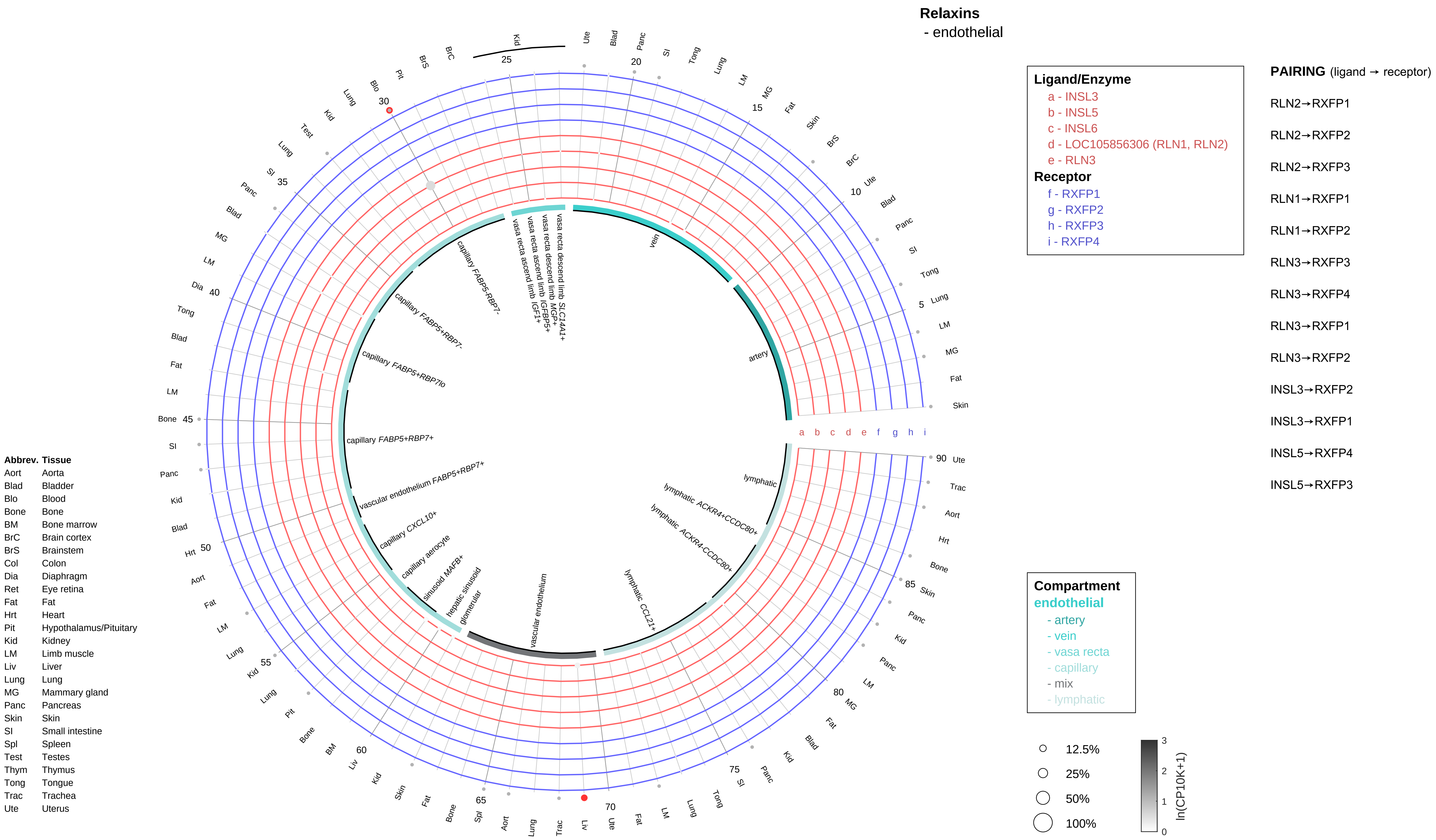

Relaxins
- endothelial
Ligand/Enzyme
a - INSL3
b - INSL5
c - INSL6
d - LOC105856306 (RLN1, RLN2)
e - RLN3
Receptor
f - RXFP1
g - RXFP2
h - RXFP3
i - RXFP4
PAIRING (ligand → receptor)
RLN2→RXFP1
RLN2→RXFP2
RLN2→RXFP3
RLN1→RXFP1
RLN1→RXFP2
RLN3→RXFP3
RLN3→RXFP4
RLN3→RXFP1
RLN3→RXFP2
INSL3→RXFP2
INSL3→RXFP1
INSL5→RXFP4
INSL5→RXFP3
Compartment
endothelial
- artery
- vein
- vasa recta
- capillary
- mix
- lymphatic
12.5%
25%
50%
100%
ln(CP10K+1)
Abbrev. Tissue
Aort Aorta
Blad Bladder
Blo Blood
Bone Bone
BM Bone marrow
BrC Brain cortex
BrS Brainstem
Col Colon
Dia Diaphragm
Ret Eye retina
Fat Fat
Hrt Heart
Pit Hypothalamus/Pituitary
Kid Kidney
LM Limb muscle
Liv Liver
Lung Lung
MG Mammary gland
Panc Pancreas
Skin Skin
SI Small intestine
Spl Spleen
Test Testes
Thym Thymus
Tong Tongue
Trac Trachea
Ute Uterus
vein
artery
lymphatic
lymphatic ACKR4+CCDC80+
lymphatic ACKR4-CCDC80+
lymphatic CCL21+
vascular endothelium
glomerular
hepatic sinusoid
sinusoid MAFB+
capillary aerocyte
capillary CXCL10+
vascular endothelium FABP5+RBP7+
capillary FABP5+RBP7+
capillary FABP5+RBP7lo
capillary FABP5+RBP7-
capillary FABP5-RBP7-
vasa recta ascend limb IGF1+
vasa recta ascend limb IGFBP5+
vasa recta descend limb MGP+
vasa recta descend limb SLC14A1+

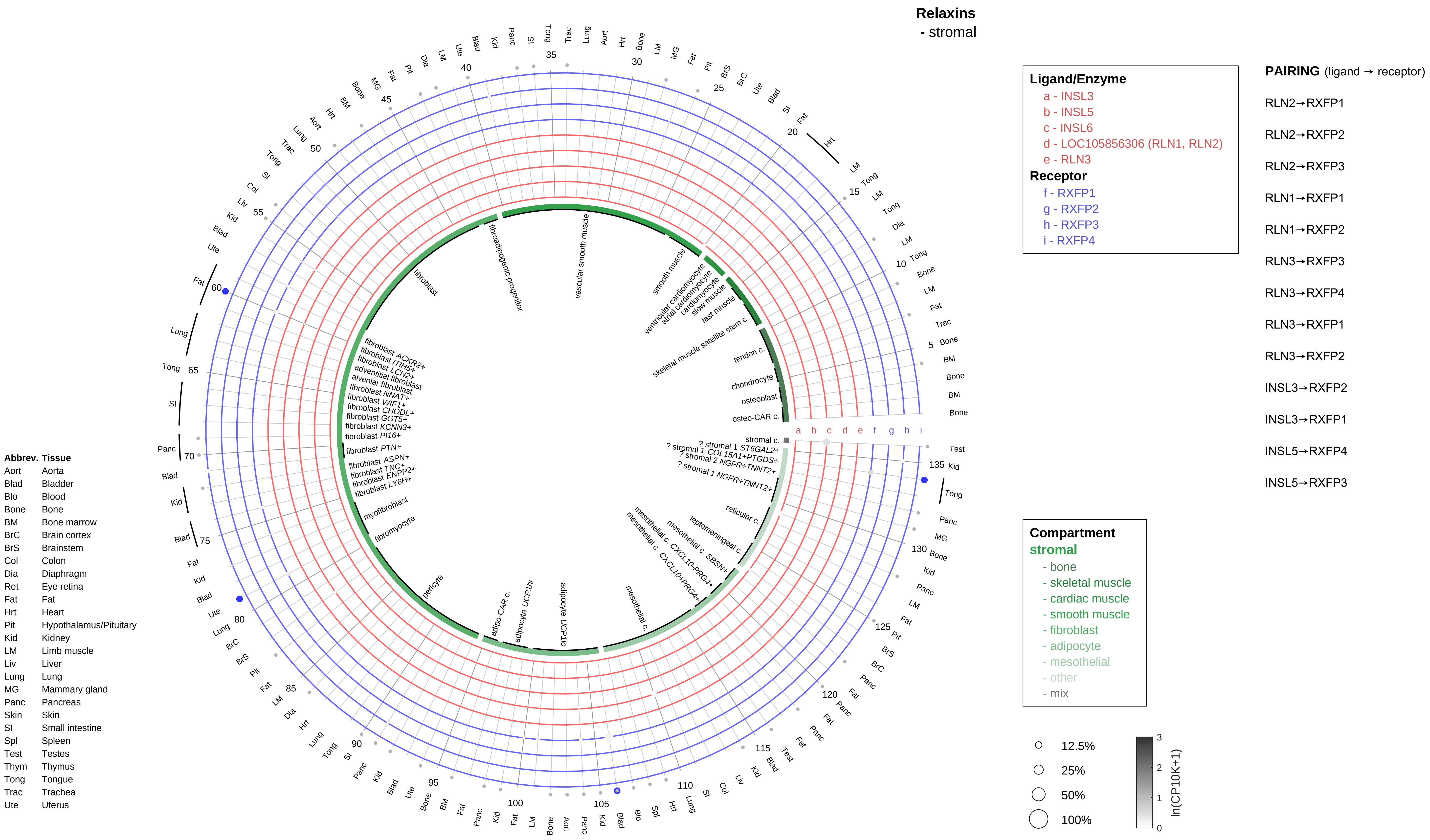
Relaxins
- stromal
Ligand/Enzyme
a - INSL3
b - INSL5
c - INSL6
d - LOC105856306 (RLN1, RLN2)
e - RLN3
Receptor
f - RXFP1
g - RXFP2
h - RXFP3
i - RXFP4
PAIRING (ligand → receptor)
RLN2→RXFP1
RLN2→RXFP2
RLN2→RXFP3
RLN1→RXFP1
RLN1→RXFP2
RLN3→RXFP3
RLN3→RXFP4
RLN3→RXFP1
RLN3→RXFP2
INSL3→RXFP2
INSL3→RXFP1
INSL5→RXFP4
INSL5→RXFP3
Compartment
stromal
- bone
- skeletal muscle
- cardiac muscle
- smooth muscle
- fibroblast
- adipocyte
- mesothelial
- other
- mix
12.5%
25%
50%
100%
ln(CP10K+1)
Abbrev. Tissue
Aort Aorta
Blad Bladder
Blo Blood
Bone Bone
BM Bone marrow
BrC Brain cortex
BrS Brainstem
Col Colon
Dia Diaphragm
Ret Eye retina
Fat Fat
Hrt Heart
Pit Hypothalamus/Pituitary
Kid Kidney
LM Limb muscle
Liv Liver
Lung Lung
MG Mammary gland
Panc Pancreas
Skin Skin
SI Small intestine
Spl Spleen
Test Testes
Thym Thymus
Tong Tongue
Trac Trachea
Ute Uterus
osteo-CAR c.
osteoblast
chondrocyte
tendon c.
skeletal muscle satellite stem c.
fast muscle
slow muscle
cardiomyocyte
atrial cardiomyocyte
ventricular cardiomyocyte
smooth muscle
vascular smooth muscle
fibroadipogenic progenitor
fibroblast
fibroblast ACKR2+
fibroblast ITIH5+
fibroblast LCN2+
adventitial fibroblast
alveolar fibroblast
fibroblast NNAT+
fibroblast WIF1+
fibroblast CHODL+
fibroblast GGT5+
fibroblast KCNN3+
fibroblast PI16+
fibroblast PTN+
fibroblast ASPN+
fibroblast TNC+
fibroblast ENPP2+
fibroblast LY6H+
myofibroblast
fibromyocyte
pericyte
adipo-CAR c.
adipocyte UCP1hi
adipocyte UCP1lo
mesothelial c.
mesothelial c. CXCL10+PRG4+
mesothelial c. CXCL10-PRG4+
mesothelial c. SBSN+
leptomeningeal c.
reticular c.
? stromal 1 NGFR+TNNT2+
? stromal 2 NGFR+TNNT2+
? stromal 1 COL15A1+PTGDS+
? stromal 1 ST6GAL2+
stromal c.

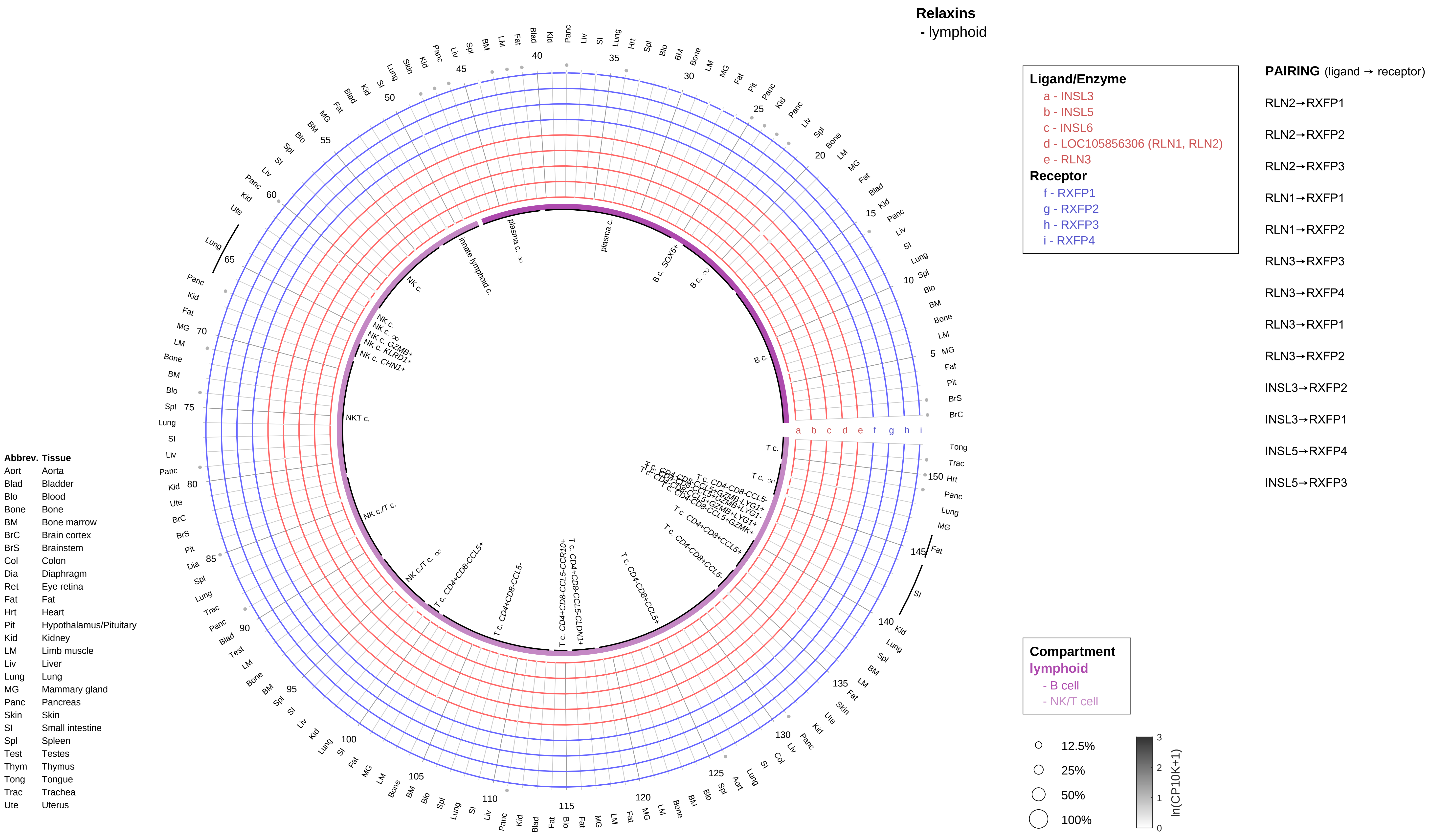

Relaxins
- lymphoid
Ligand/Enzyme
a - INSL3
b - INSL5
c - INSL6
d - LOC105856306 (RLN1, RLN2)
e - RLN3
Receptor
f - RXFP1
g - RXFP2
h - RXFP3
i - RXFP4
PAIRING (ligand → receptor)
RLN2→RXFP1
RLN2→RXFP2
RLN2→RXFP3
RLN1→RXFP1
RLN1→RXFP2
RLN3→RXFP3
RLN3→RXFP4
RLN3→RXFP1
RLN3→RXFP2
INSL3→RXFP2
INSL3→RXFP1
INSL5→RXFP4
INSL5→RXFP3
Compartment
lymphoid
- B cell
- NK/T cell
12.5%
25%
50%
100%
ln(CP10K+1)
Abbrev. Tissue
Aort Aorta
Blad Bladder
Blo Blood
Bone Bone
BM Bone marrow
BrC Brain cortex
BrS Brainstem
Col Colon
Dia Diaphragm
Ret Eye retina
Fat Fat
Hrt Heart
Pit Hypothalamus/Pituitary
Kid Kidney
LM Limb muscle
Liv Liver
Lung Lung
MG Mammary gland
Panc Pancreas
Skin Skin
SI Small intestine
Spl Spleen
Test Testes
Thym Thymus
Tong Tongue
Trac Trachea
Ute Uterus

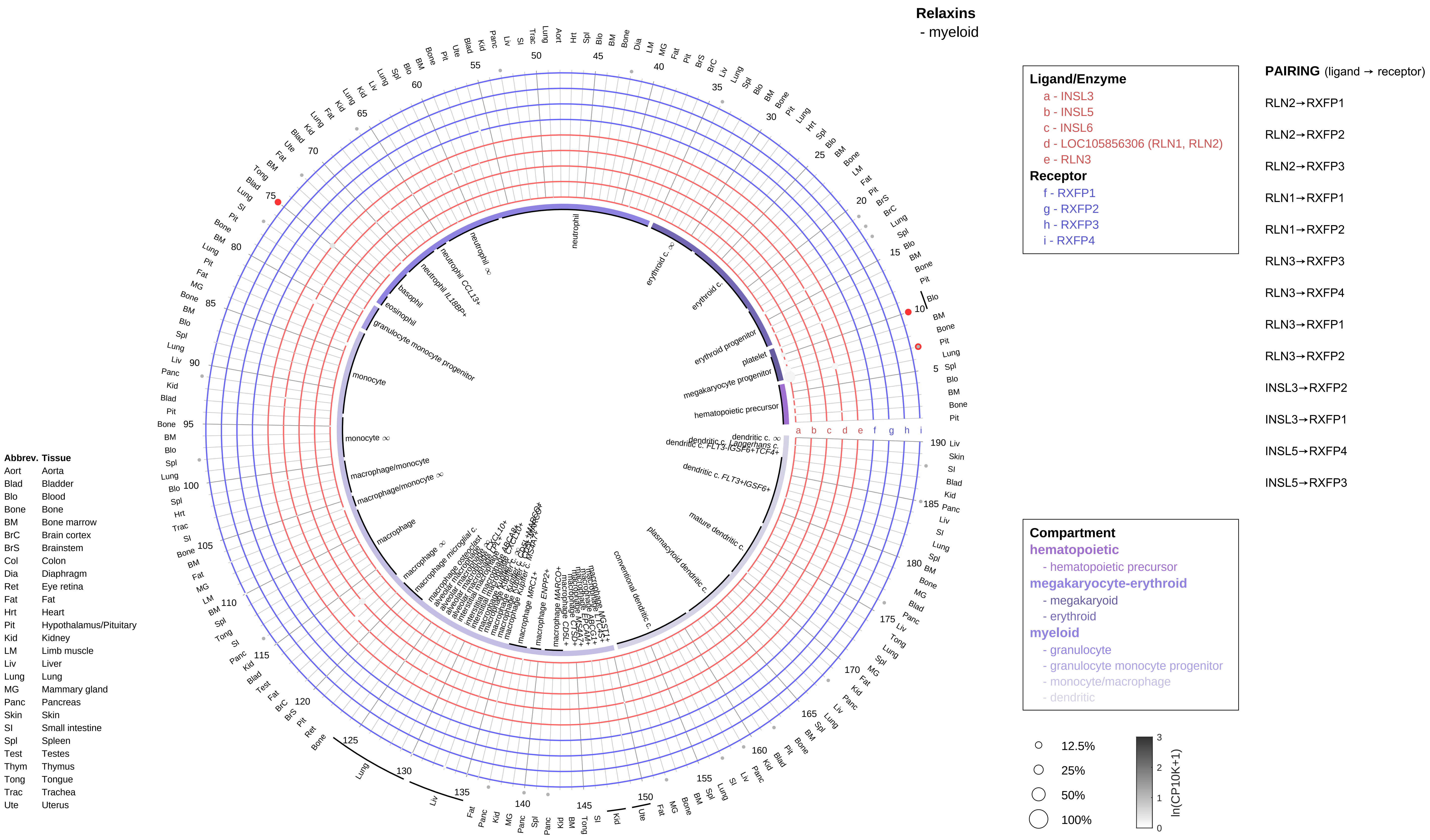

Relaxins
- myeloid
Ligand/Enzyme
a - INSL3
b - INSL5
c - INSL6
d - LOC105856306 (RLN1, RLN2)
e - RLN3
Receptor
f - RXFP1
g - RXFP2
h - RXFP3
i - RXFP4
PAIRING (ligand → receptor)
RLN2→RXFP1
RLN2→RXFP2
RLN2→RXFP3
RLN1→RXFP1
RLN1→RXFP2
RLN3→RXFP3
RLN3→RXFP4
RLN3→RXFP1
RLN3→RXFP2
INSL3→RXFP2
INSL3→RXFP1
INSL5→RXFP4
INSL5→RXFP3
Compartment
hematopoietic
- hematopoietic precursor
megakaryocyte-erythroid
- megakaryoid
- erythroid
myeloid
- granulocyte
- granulocyte monocyte progenitor
- monocyte/macrophage
- dendritic
12.5%
25%
50%
100%
ln(CP10K+1)
Abbrev. Tissue
Aort Aorta
Blad Bladder
Blo Blood
Bone Bone
BM Bone marrow
BrC Brain cortex
BrS Brainstem
Col Colon
Dia Diaphragm
Ret Eye retina
Fat Fat
Hrt Heart
Pit Hypothalamus/Pituitary
Kid Kidney
LM Limb muscle
Liv Liver
Lung Lung
MG Mammary gland
Panc Pancreas
Skin Skin
SI Small intestine
Spl Spleen
Test Testes
Thym Thymus
Tong Tongue
Trac Trachea
Ute Uterus
neutrophil
erythroid c. ∞
erythroid c.
erythroid progenitor
platelet
megakaryocyte progenitor
hematopoietic precursor
dendritic c. ∞
dendritic c. FLT3+IGSF6+
mature dendritic c.
plasmacytoid dendritic c.
conventional dendritic c.
macrophage
macrophage/monocyte
macrophage/monocyte ∞
monocyte ∞
monocyte
granulocyte monocyte progenitor
eosinophil
basophil
neutrophil IL18BP+
neutrophil CCL13+
neutrophil ∞

**Resistin**
- epithelial/neural/germ

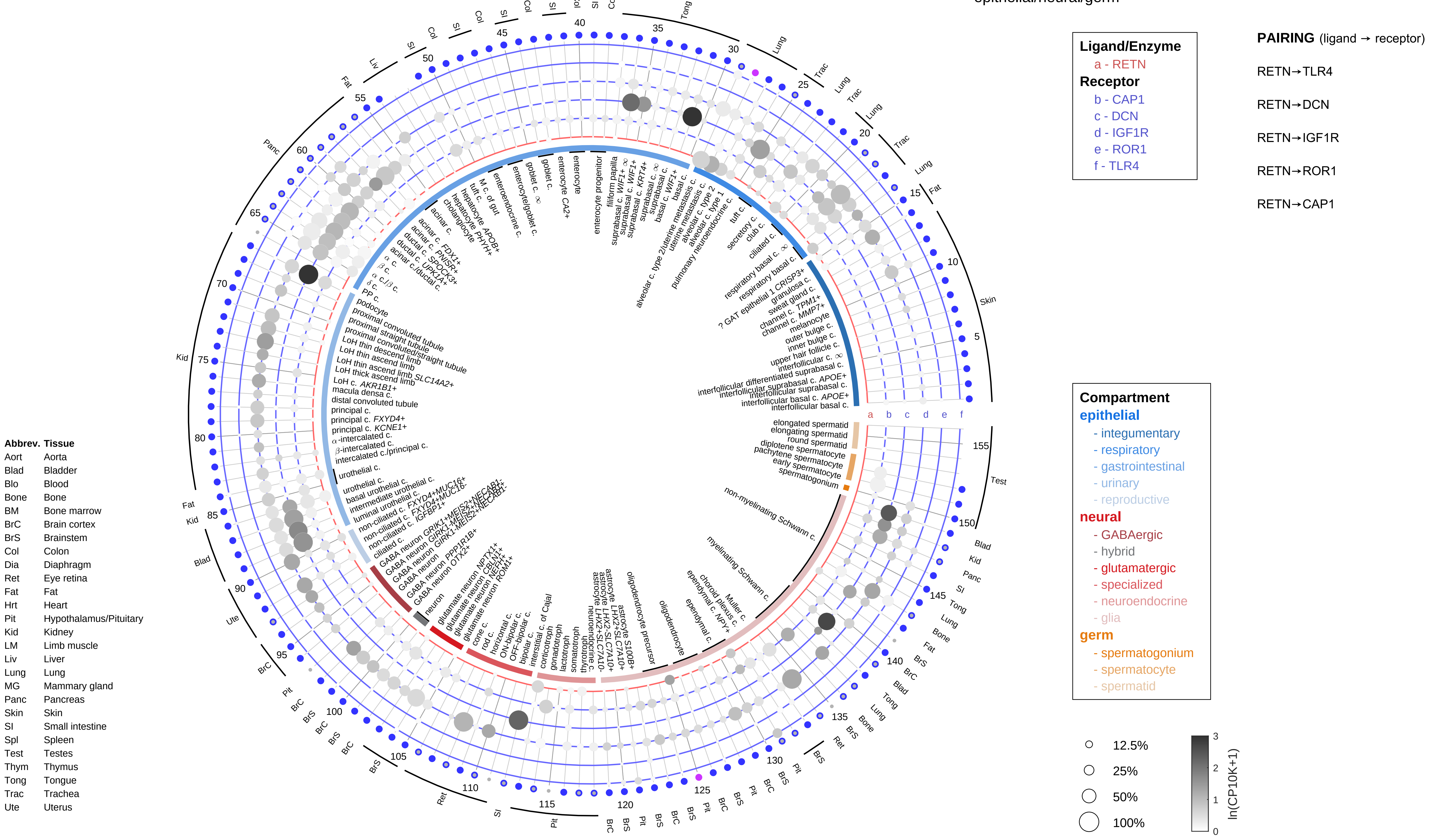

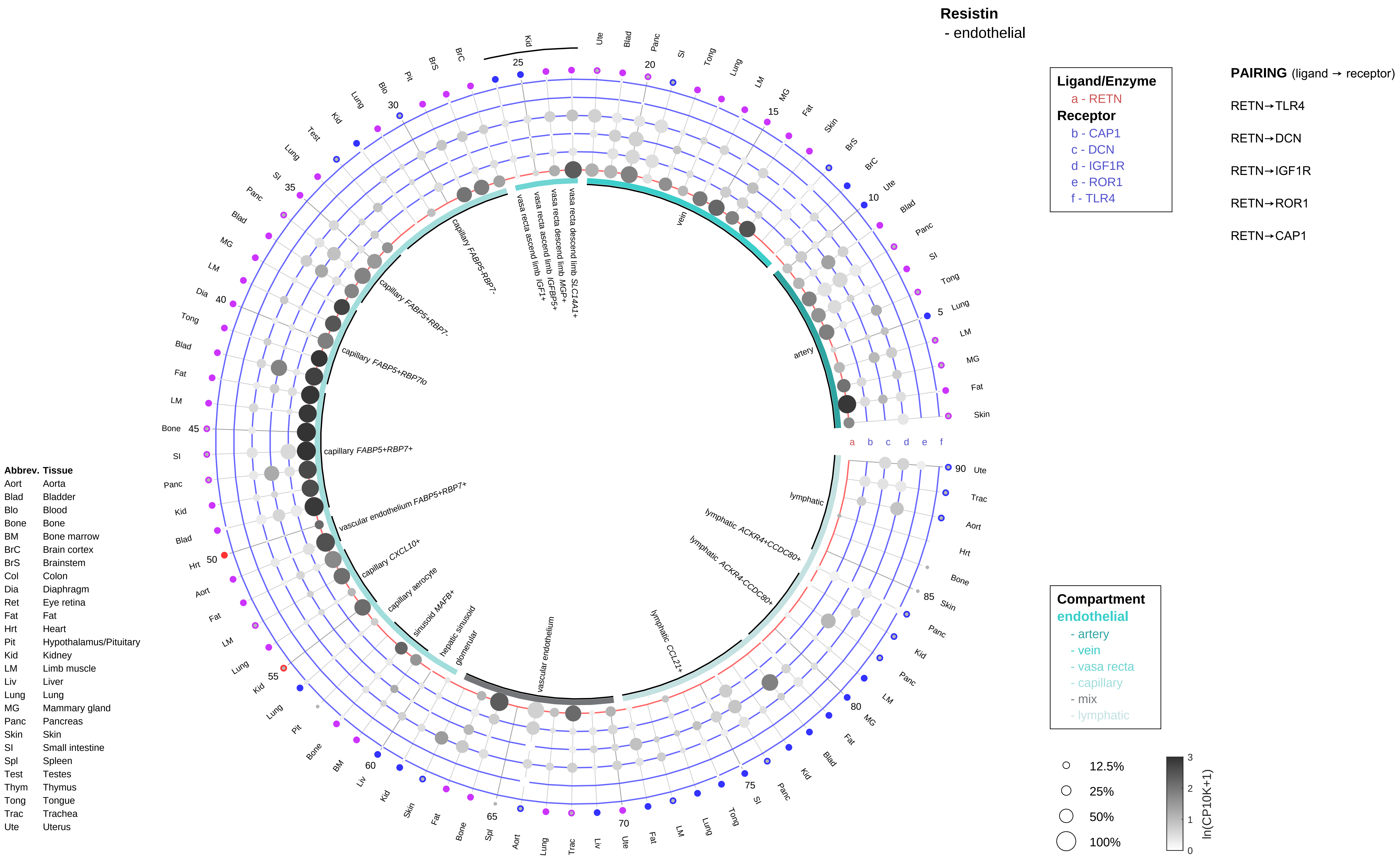
Resistin
- endothelial
Ligand/Enzyme
a - RETN
Receptor
b - CAP1
c - DCN
d - IGF1R
e - ROR1
f - TLR4
PAIRING (ligand → receptor)
RETN→TLR4
RETN→DCN
RETN→IGF1R
RETN→ROR1
RETN→CAP1
Compartment
endothelial
- artery
- vein
- vasa recta
- capillary
- mix
- lymphatic
12.5%
25%
50%
100%
ln(CP10K+1)
Abbrev. Tissue
Aort Aorta
Blad Bladder
Blo Blood
Bone Bone
BM Bone marrow
BrC Brain cortex
BrS Brainstem
Col Colon
Dia Diaphragm
Ret Eye retina
Fat Fat
Hrt Heart
Pit Hypothalamus/Pituitary
Kid Kidney
LM Limb muscle
Liv Liver
Lung Lung
MG Mammary gland
Panc Pancreas
Skin Skin
SI Small intestine
Spl Spleen
Test Testes
Thym Thymus
Tong Tongue
Trac Trachea
Ute Uterus
vein
artery
lymphatic
lymphatic ACKR4+CCDC80+
lymphatic ACKR4-CCDC80+
lymphatic CCL21+
vascular endothelium
glomerular
hepatic sinusoid
sinusoid MAFB+
capillary aerocyte
capillary CXCL10+
vascular endothelium FABP5+RBP7+
capillary FABP5+RBP7+
capillary FABP5+RBP7lo
capillary FABP5+RBP7-
capillary FABP5-RBP7-
vasa recta ascend limb IGF1+
vasa recta ascend limb IGFBP5+
vasa recta descend limb MGP+
vasa recta descend limb SLC14A1+

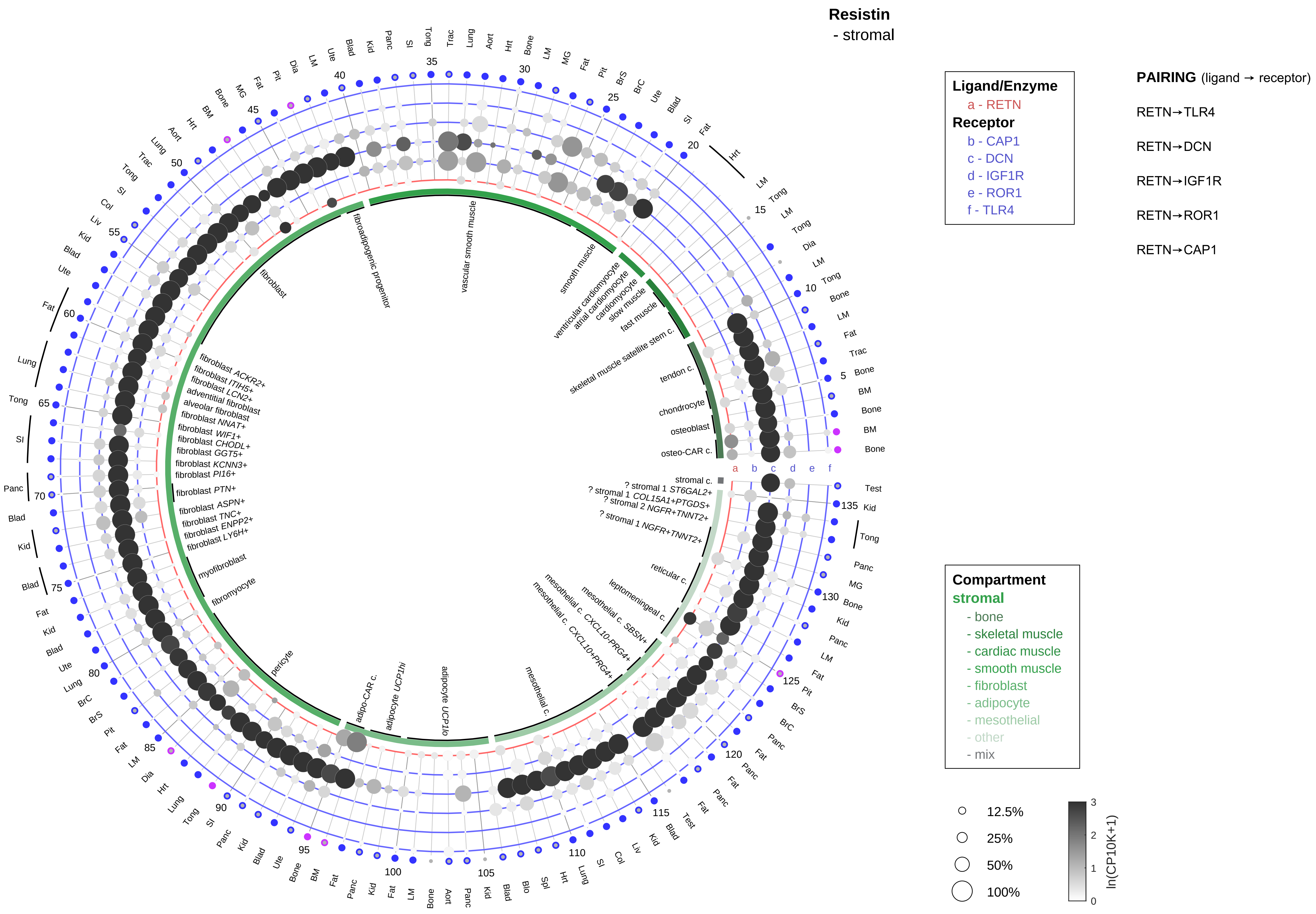

| Abbrev. | Tissue |
|---|---|
| Aort | Aorta |
| Blad | Bladder |
| Blo | Blood |
| Bone | Bone |
| BM | Bone marrow |
| BrC | Brain cortex |
| BrS | Brainstem |
| Col | Colon |
| Dia | Diaphragm |
| Ret | Eye retina |
| Fat | Fat |
| Hrt | Heart |
| Pit | Hypothalamus/Pituitary |
| Kid | Kidney |
| LM | Limb muscle |
| Liv | Liver |
| Lung | Lung |
| MG | Mammary gland |
| Panc | Pancreas |
| Skin | Skin |
| SI | Small intestine |
| Spl | Spleen |
| Test | Testes |
| Thym | Thymus |
| Tong | Tongue |
| Trac | Trachea |
| Ute | Uterus |

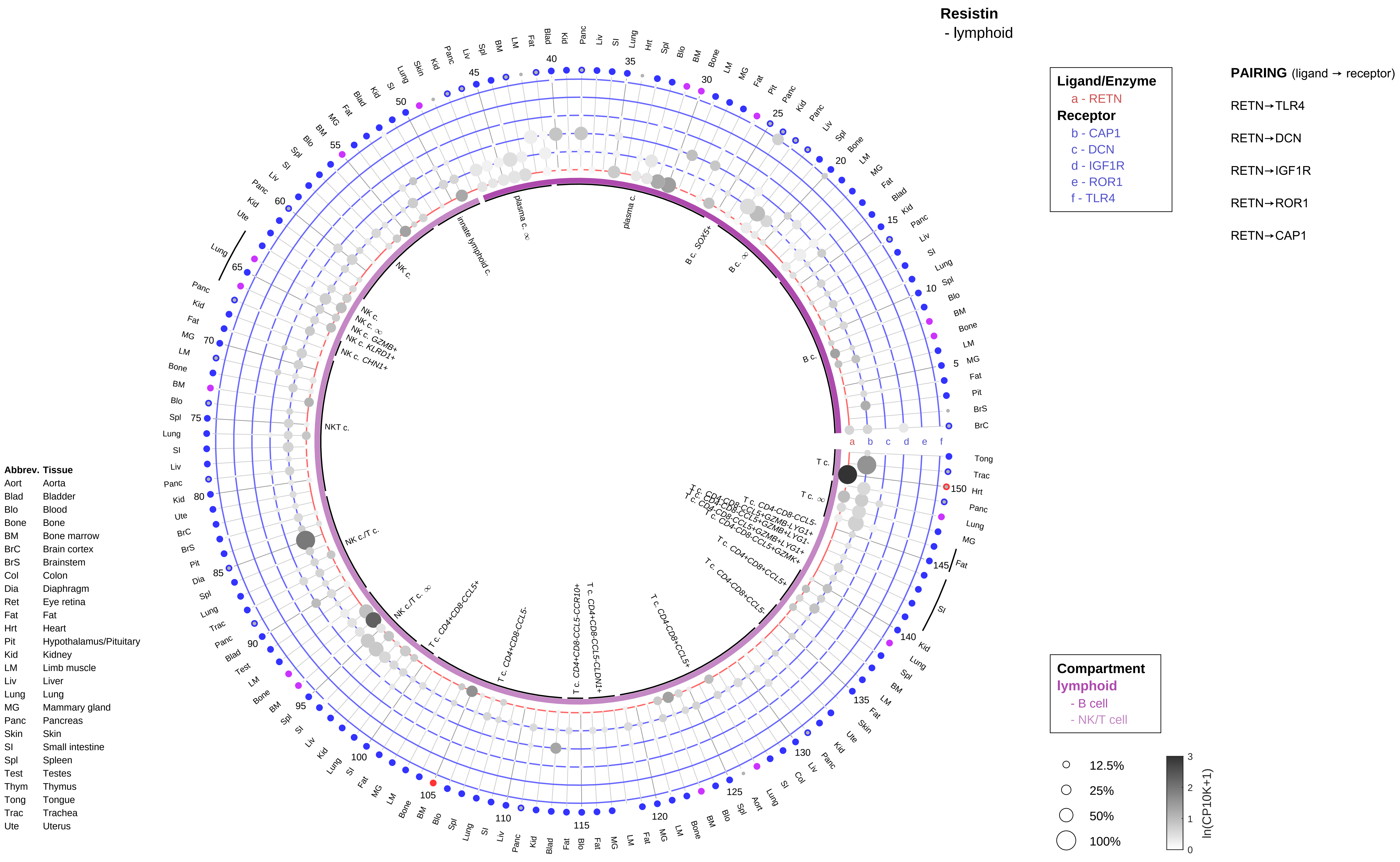
Resistin
- lymphoid
Ligand/Enzyme
a - RETN
Receptor
b - CAP1
c - DCN
d - IGF1R
e - ROR1
f - TLR4
PAIRING (ligand → receptor)
RETN→TLR4
RETN→DCN
RETN→IGF1R
RETN→ROR1
RETN→CAP1
Compartment
lymphoid
- B cell
- NK/T cell
12.5%
25%
50%
100%
ln(CP10K+1)
Abbrev. Tissue
Aort Aorta
Blad Bladder
Blo Blood
Bone Bone
BM Bone marrow
BrC Brain cortex
BrS Brainstem
Col Colon
Dia Diaphragm
Ret Eye retina
Fat Fat
Hrt Heart
Pit Hypothalamus/Pituitary
Kid Kidney
LM Limb muscle
Liv Liver
Lung Lung
MG Mammary gland
Panc Pancreas
Skin Skin
SI Small intestine
Spl Spleen
Test Testes
Thym Thymus
Tong Tongue
Trac Trachea
Ute Uterus

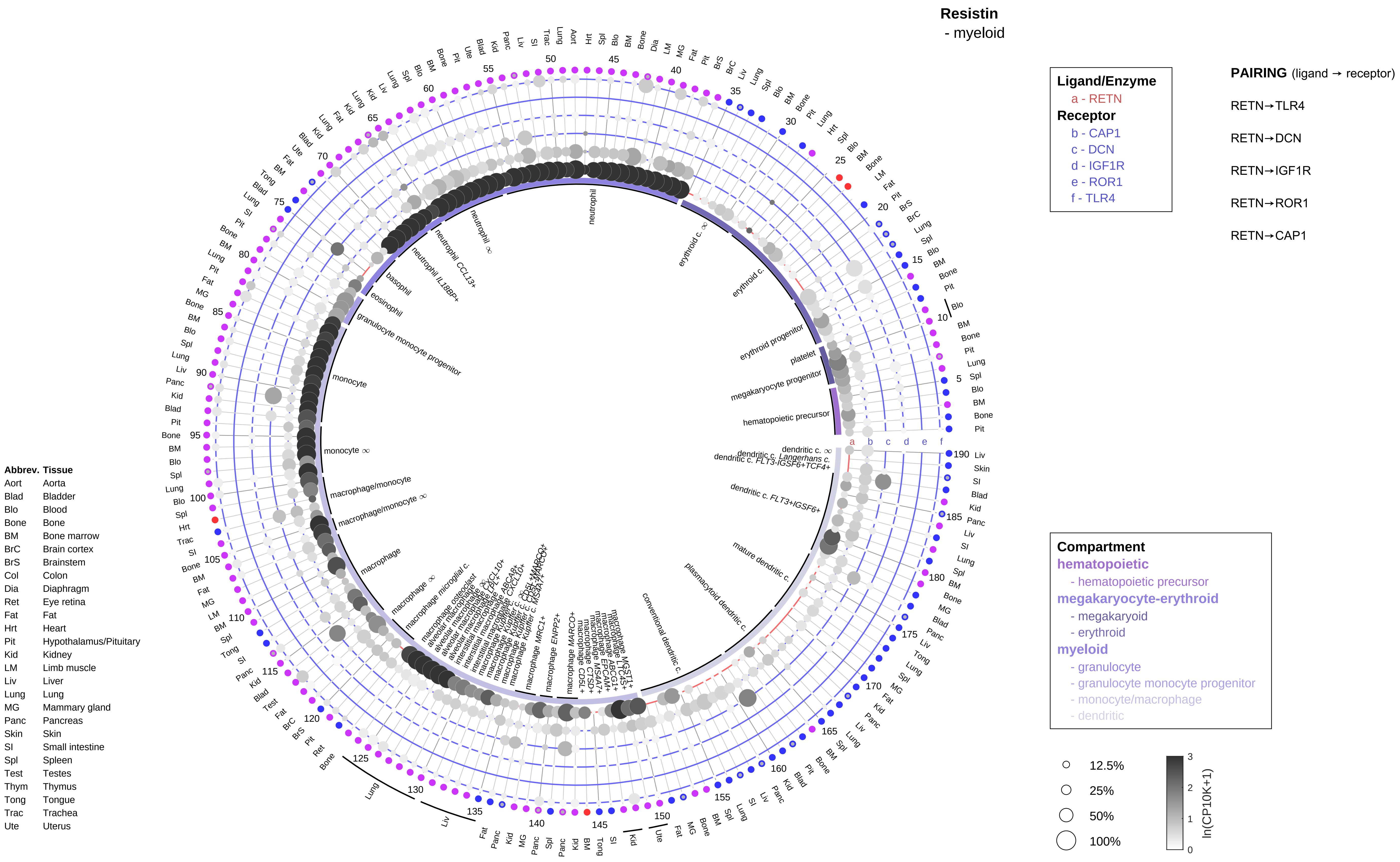
Resistin
- myeloid
Ligand/Enzyme
a - RETN
Receptor
b - CAP1
c - DCN
d - IGF1R
e - ROR1
f - TLR4
PAIRING (ligand → receptor)
RETN→TLR4
RETN→DCN
RETN→IGF1R
RETN→ROR1
RETN→CAP1
Compartment
hematopoietic
- hematopoietic precursor
megakaryocyte-erythroid
- megakaryoid
- erythroid
myeloid
- granulocyte
- granulocyte monocyte progenitor
- monocyte/macrophage
- dendritic
12.5%
25%
50%
100%
ln(CP10K+1)
Abbrev. Tissue
Aort Aorta
Blad Bladder
Blo Blood
Bone Bone
BM Bone marrow
BrC Brain cortex
BrS Brainstem
Col Colon
Dia Diaphragm
Ret Eye retina
Fat Fat
Hrt Heart
Pit Hypothalamus/Pituitary
Kid Kidney
LM Limb muscle
Liv Liver
Lung Lung
MG Mammary gland
Panc Pancreas
Skin Skin
SI Small intestine
Spl Spleen
Test Testes
Thym Thymus
Tong Tongue
Trac Trachea
Ute Uterus

### Secretin (SCT)
- epithelial/neural/germ

**Ligand/Enzyme**
a - SCT*
**Receptor**
b - SCTR

**PAIRING** (ligand → receptor)

SCT→SCTR

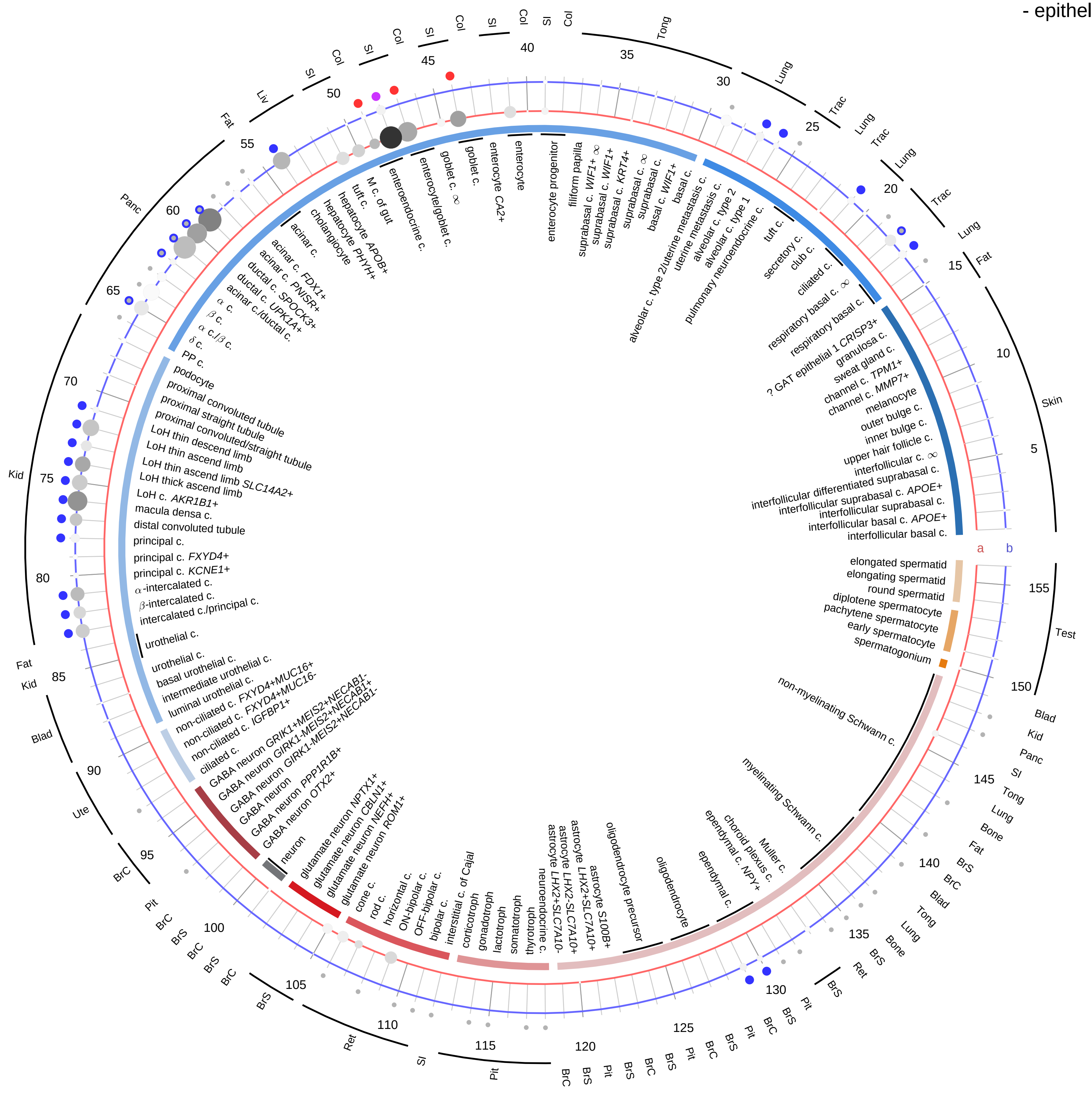

**Compartment**
**epithelial**
- integumentary
- respiratory
- gastrointestinal
- urinary
- reproductive

**neural**
- GABAergic
- hybrid
- glutamatergic
- specialized
- neuroendocrine
- glia

**germ**
- spermatogonium
- spermatocyte
- spermatid

12.5%
25%
50%
100%

ln(CP10K+1): 0, 1, 2, 3

| Abbrev. | Tissue |
|---|---|
| Aort | Aorta |
| Blad | Bladder |
| Blo | Blood |
| Bone | Bone |
| BM | Bone marrow |
| BrC | Brain cortex |
| BrS | Brainstem |
| Col | Colon |
| Dia | Diaphragm |
| Ret | Eye retina |
| Fat | Fat |
| Hrt | Heart |
| Pit | Hypothalamus/Pituitary |
| Kid | Kidney |
| LM | Limb muscle |
| Liv | Liver |
| Lung | Lung |
| MG | Mammary gland |
| Panc | Pancreas |
| Skin | Skin |
| SI | Small intestine |
| Spl | Spleen |
| Test | Testes |
| Thym | Thymus |
| Tong | Tongue |
| Trac | Trachea |
| Ute | Uterus |

#### Secretin (SCT)
- endothelial

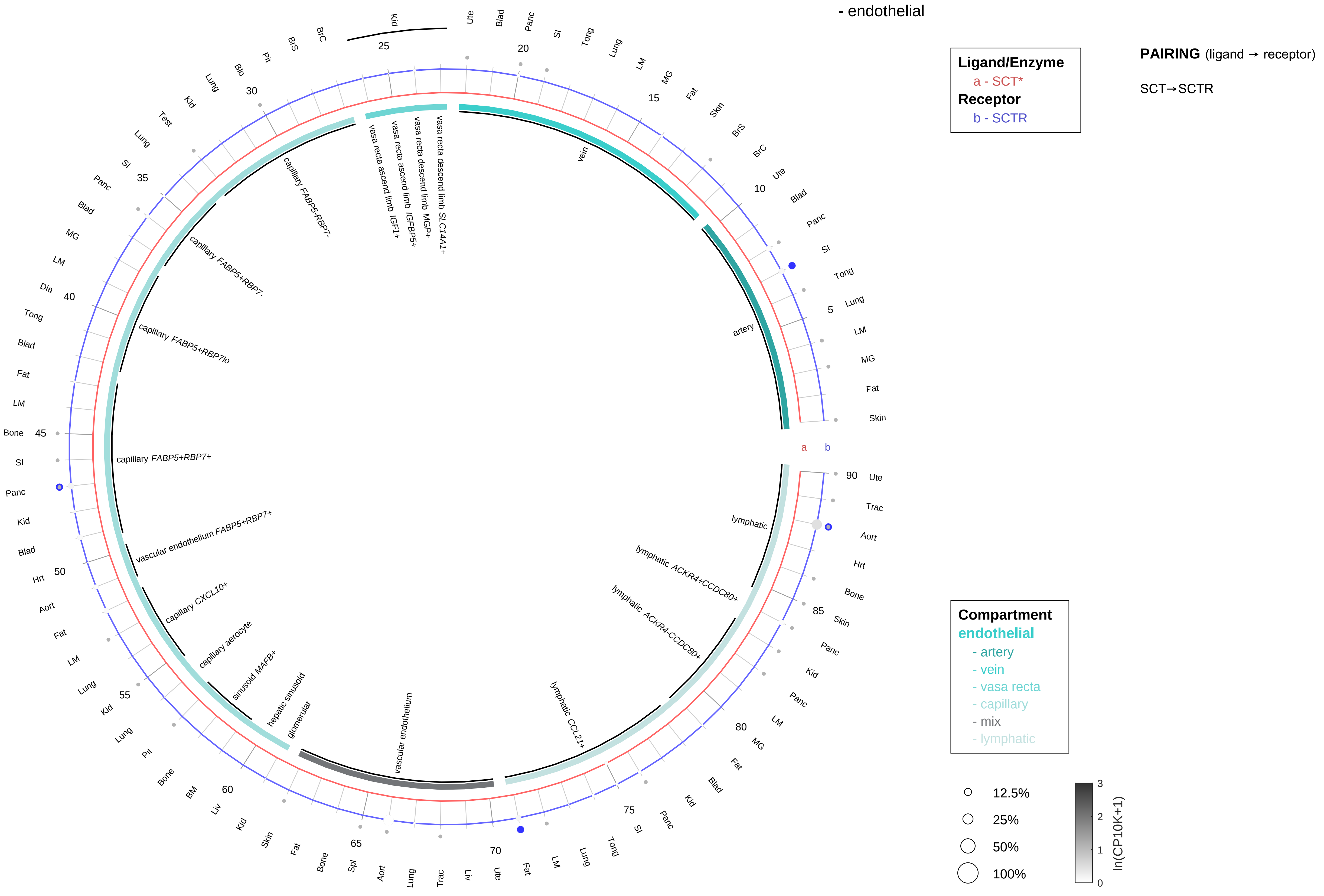

| Abbrev. | Tissue |
|---|---|
| Aort | Aorta |
| Blad | Bladder |
| Blo | Blood |
| Bone | Bone |
| BM | Bone marrow |
| BrC | Brain cortex |
| BrS | Brainstem |
| Col | Colon |
| Dia | Diaphragm |
| Ret | Eye retina |
| Fat | Fat |
| Hrt | Heart |
| Pit | Hypothalamus/Pituitary |
| Kid | Kidney |
| LM | Limb muscle |
| Liv | Liver |
| Lung | Lung |
| MG | Mammary gland |
| Panc | Pancreas |
| Skin | Skin |
| SI | Small intestine |
| Spl | Spleen |
| Test | Testes |
| Thym | Thymus |
| Tong | Tongue |
| Trac | Trachea |
| Ute | Uterus |

**Secretin (SCT)**
- stromal

**PAIRING** (ligand → receptor)

SCT→SCTR

**Ligand/Enzyme**
a - SCT*
**Receptor**
b - SCTR

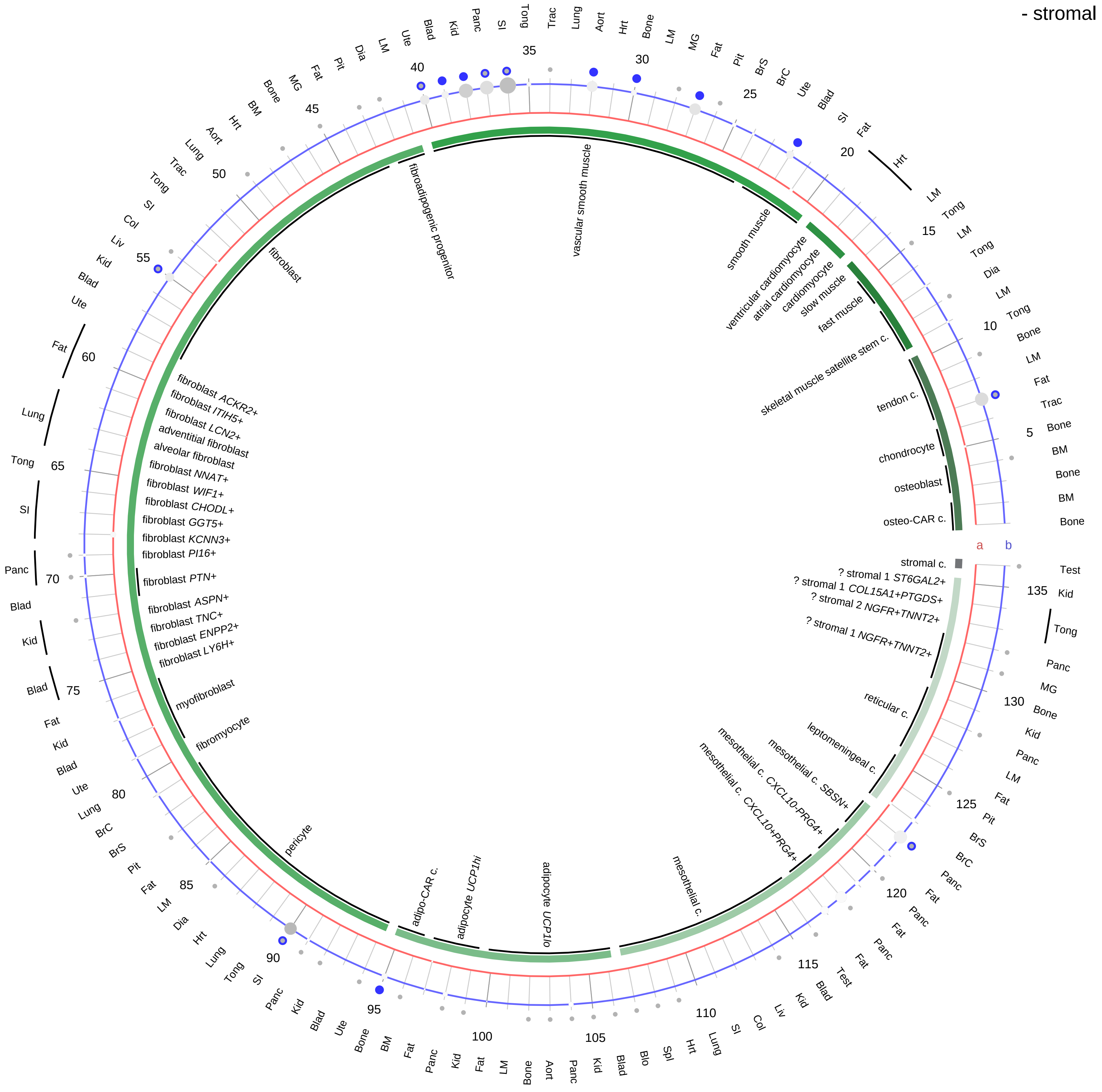

**Compartment**
**stromal**
- bone
- skeletal muscle
- cardiac muscle
- smooth muscle
- fibroblast
- adipocyte
- mesothelial
- other
- mix

12.5%
25%
50%
100%

ln(CP10K+1): 0, 1, 2, 3

| Abbrev. | Tissue |
|---|---|
| Aort | Aorta |
| Blad | Bladder |
| Blo | Blood |
| Bone | Bone |
| BM | Bone marrow |
| BrC | Brain cortex |
| BrS | Brainstem |
| Col | Colon |
| Dia | Diaphragm |
| Ret | Eye retina |
| Fat | Fat |
| Hrt | Heart |
| Pit | Hypothalamus/Pituitary |
| Kid | Kidney |
| LM | Limb muscle |
| Liv | Liver |
| Lung | Lung |
| MG | Mammary gland |
| Panc | Pancreas |
| Skin | Skin |
| SI | Small intestine |
| Spl | Spleen |
| Test | Testes |
| Thym | Thymus |
| Tong | Tongue |
| Trac | Trachea |
| Ute | Uterus |

**Secretin (SCT)**
- lymphoid

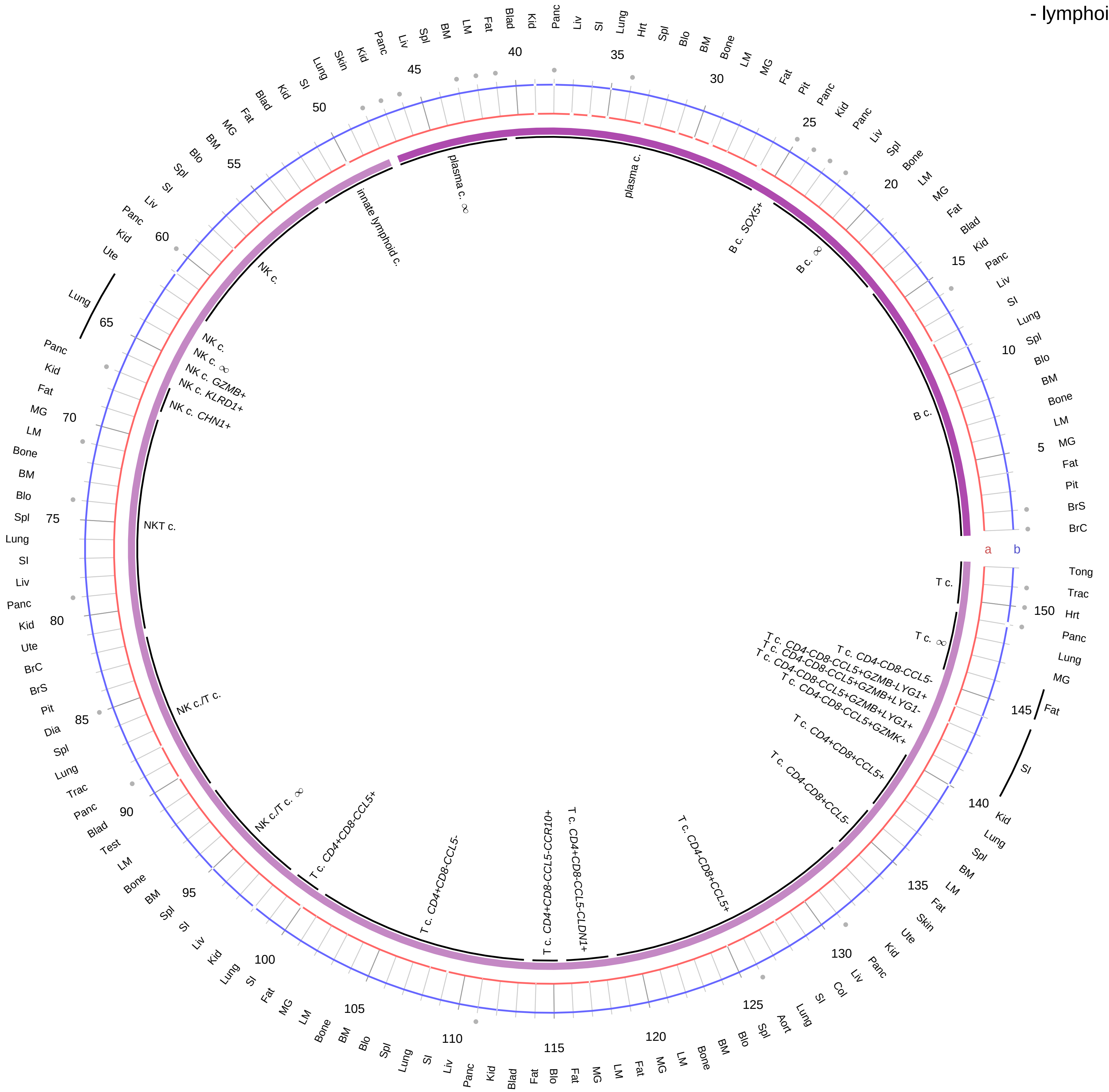

**Ligand/Enzyme**
a - SCT*
**Receptor**
b - SCTR

**PAIRING** (ligand → receptor)

SCT→SCTR

**Compartment**
**lymphoid**
- B cell
- NK/T cell

12.5%
25%
50%
100%

ln(CP10K+1): 0, 1, 2, 3

| Abbrev. | Tissue |
|---|---|
| Aort | Aorta |
| Blad | Bladder |
| Blo | Blood |
| Bone | Bone |
| BM | Bone marrow |
| BrC | Brain cortex |
| BrS | Brainstem |
| Col | Colon |
| Dia | Diaphragm |
| Ret | Eye retina |
| Fat | Fat |
| Hrt | Heart |
| Pit | Hypothalamus/Pituitary |
| Kid | Kidney |
| LM | Limb muscle |
| Liv | Liver |
| Lung | Lung |
| MG | Mammary gland |
| Panc | Pancreas |
| Skin | Skin |
| SI | Small intestine |
| Spl | Spleen |
| Test | Testes |
| Thym | Thymus |
| Tong | Tongue |
| Trac | Trachea |
| Ute | Uterus |

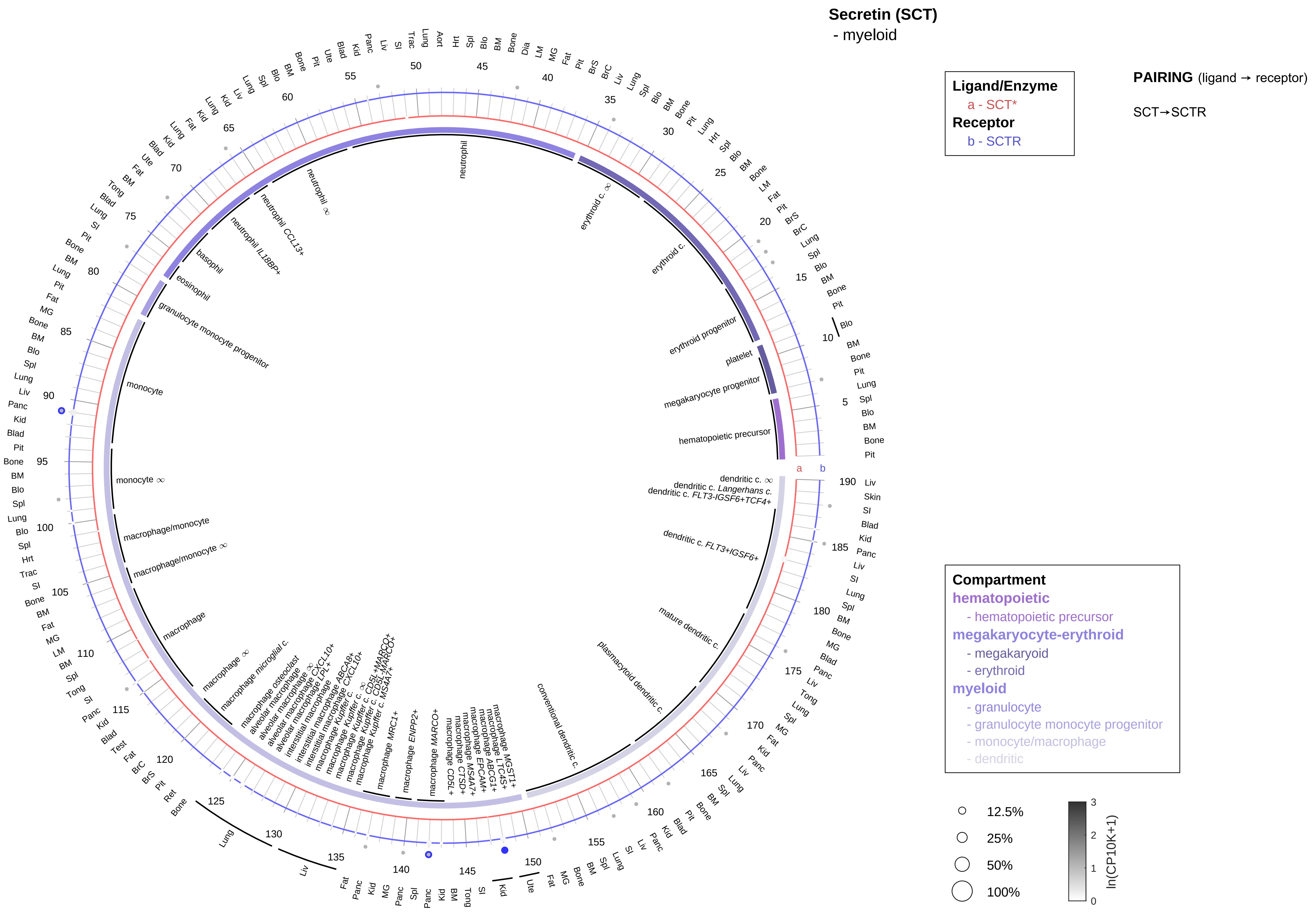

| Abbrev. | Tissue |
|---|---|
| Aort | Aorta |
| Blad | Bladder |
| Blo | Blood |
| Bone | Bone |
| BM | Bone marrow |
| BrC | Brain cortex |
| BrS | Brainstem |
| Col | Colon |
| Dia | Diaphragm |
| Ret | Eye retina |
| Fat | Fat |
| Hrt | Heart |
| Pit | Hypothalamus/Pituitary |
| Kid | Kidney |
| LM | Limb muscle |
| Liv | Liver |
| Lung | Lung |
| MG | Mammary gland |
| Panc | Pancreas |
| Skin | Skin |
| SI | Small intestine |
| Spl | Spleen |
| Test | Testes |
| Thym | Thymus |
| Tong | Tongue |
| Trac | Trachea |
| Ute | Uterus |

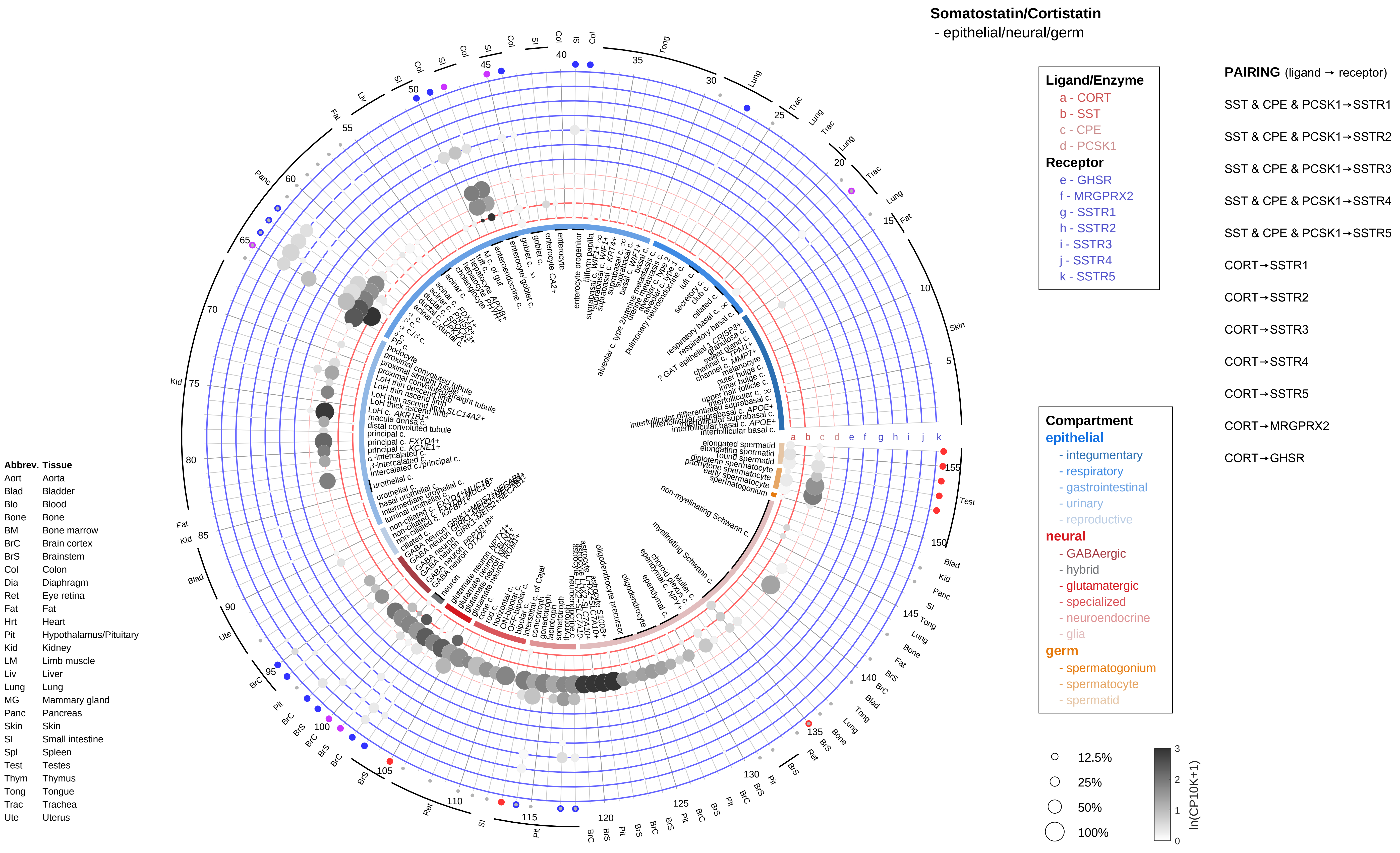

Somatostatin/Cortistatin
- epithelial/neural/germ
Ligand/Enzyme
a - CORT
b - SST
c - CPE
d - PCSK1
Receptor
e - GHSR
f - MRGPRX2
g - SSTR1
h - SSTR2
i - SSTR3
j - SSTR4
k - SSTR5
PAIRING (ligand → receptor)
SST & CPE & PCSK1→SSTR1
SST & CPE & PCSK1→SSTR2
SST & CPE & PCSK1→SSTR3
SST & CPE & PCSK1→SSTR4
SST & CPE & PCSK1→SSTR5
CORT→SSTR1
CORT→SSTR2
CORT→SSTR3
CORT→SSTR4
CORT→SSTR5
CORT→MRGPRX2
CORT→GHSR
Compartment
epithelial
- integumentary
- respiratory
- gastrointestinal
- urinary
- reproductive
neural
- GABAergic
- hybrid
- glutamatergic
- specialized
- neuroendocrine
- glia
germ
- spermatogonium
- spermatocyte
- spermatid
12.5%
25%
50%
100%
ln(CP10K+1)
Abbrev. Tissue
Aort Aorta
Blad Bladder
Blo Blood
Bone Bone
BM Bone marrow
BrC Brain cortex
BrS Brainstem
Col Colon
Dia Diaphragm
Ret Eye retina
Fat Fat
Hrt Heart
Pit Hypothalamus/Pituitary
Kid Kidney
LM Limb muscle
Liv Liver
Lung Lung
MG Mammary gland
Panc Pancreas
Skin Skin
SI Small intestine
Spl Spleen
Test Testes
Thym Thymus
Tong Tongue
Trac Trachea
Ute Uterus

**Somatostatin/Cortistatin**
- endothelial

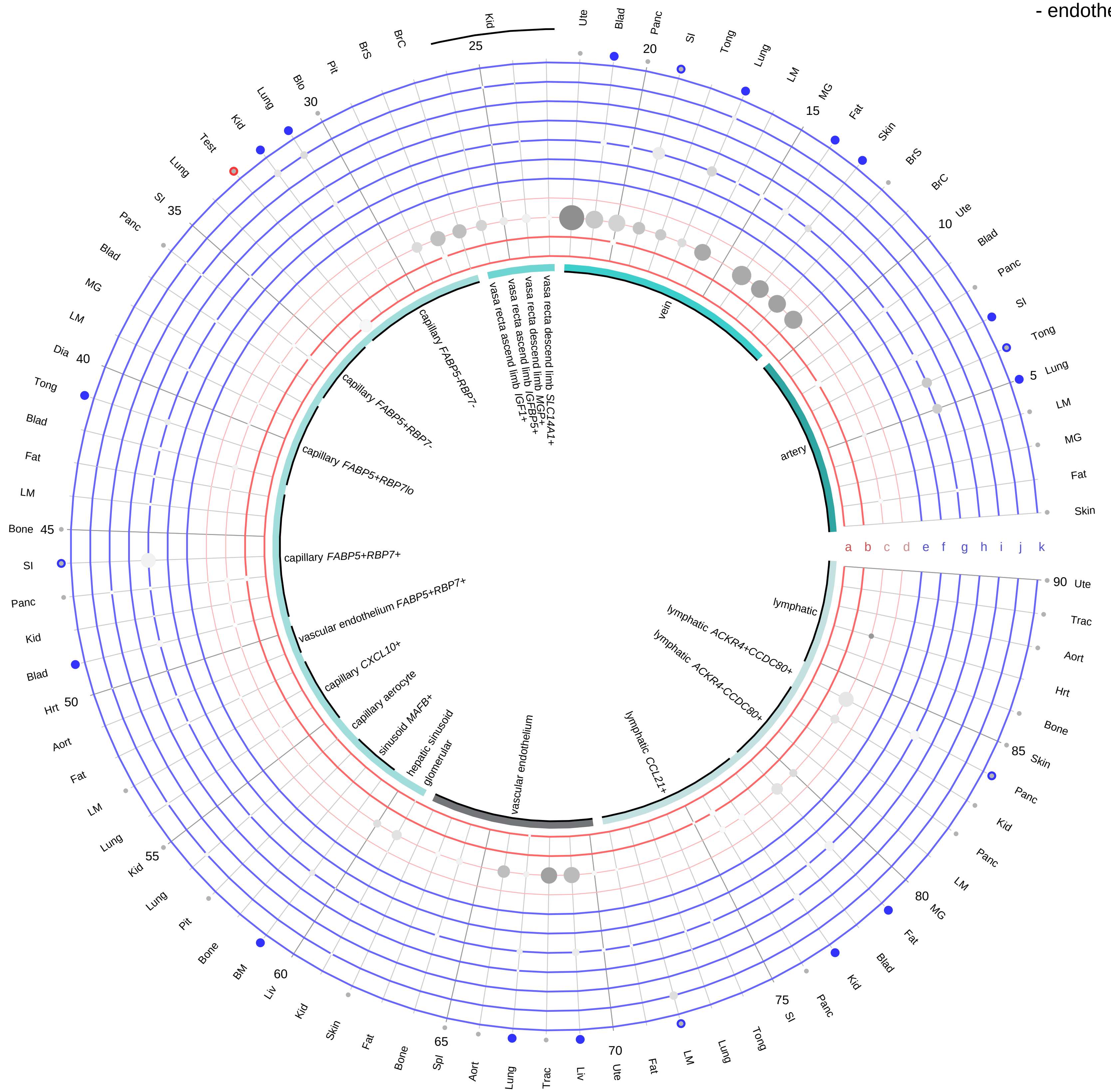

| Abbrev. | Tissue |
|---|---|
| Aort | Aorta |
| Blad | Bladder |
| Blo | Blood |
| Bone | Bone |
| BM | Bone marrow |
| BrC | Brain cortex |
| BrS | Brainstem |
| Col | Colon |
| Dia | Diaphragm |
| Ret | Eye retina |
| Fat | Fat |
| Hrt | Heart |
| Pit | Hypothalamus/Pituitary |
| Kid | Kidney |
| LM | Limb muscle |
| Liv | Liver |
| Lung | Lung |
| MG | Mammary gland |
| Panc | Pancreas |
| Skin | Skin |
| SI | Small intestine |
| Spl | Spleen |
| Test | Testes |
| Thym | Thymus |
| Tong | Tongue |
| Trac | Trachea |
| Ute | Uterus |

**Ligand/Enzyme**
a - CORT
b - SST
c - CPE
d - PCSK1
**Receptor**
e - GHSR
f - MRGPRX2
g - SSTR1
h - SSTR2
i - SSTR3
j - SSTR4
k - SSTR5

**PAIRING** (ligand → receptor)

SST & CPE & PCSK1→SSTR1
SST & CPE & PCSK1→SSTR2
SST & CPE & PCSK1→SSTR3
SST & CPE & PCSK1→SSTR4
SST & CPE & PCSK1→SSTR5
CORT→SSTR1
CORT→SSTR2
CORT→SSTR3
CORT→SSTR4
CORT→SSTR5
CORT→MRGPRX2
CORT→GHSR

**Compartment**
**endothelial**
- artery
- vein
- vasa recta
- capillary
- mix
- lymphatic

12.5%
25%
50%
100%

ln(CP10K+1): 0, 1, 2, 3

#### Somatostatin/Cortistatin
- stromal

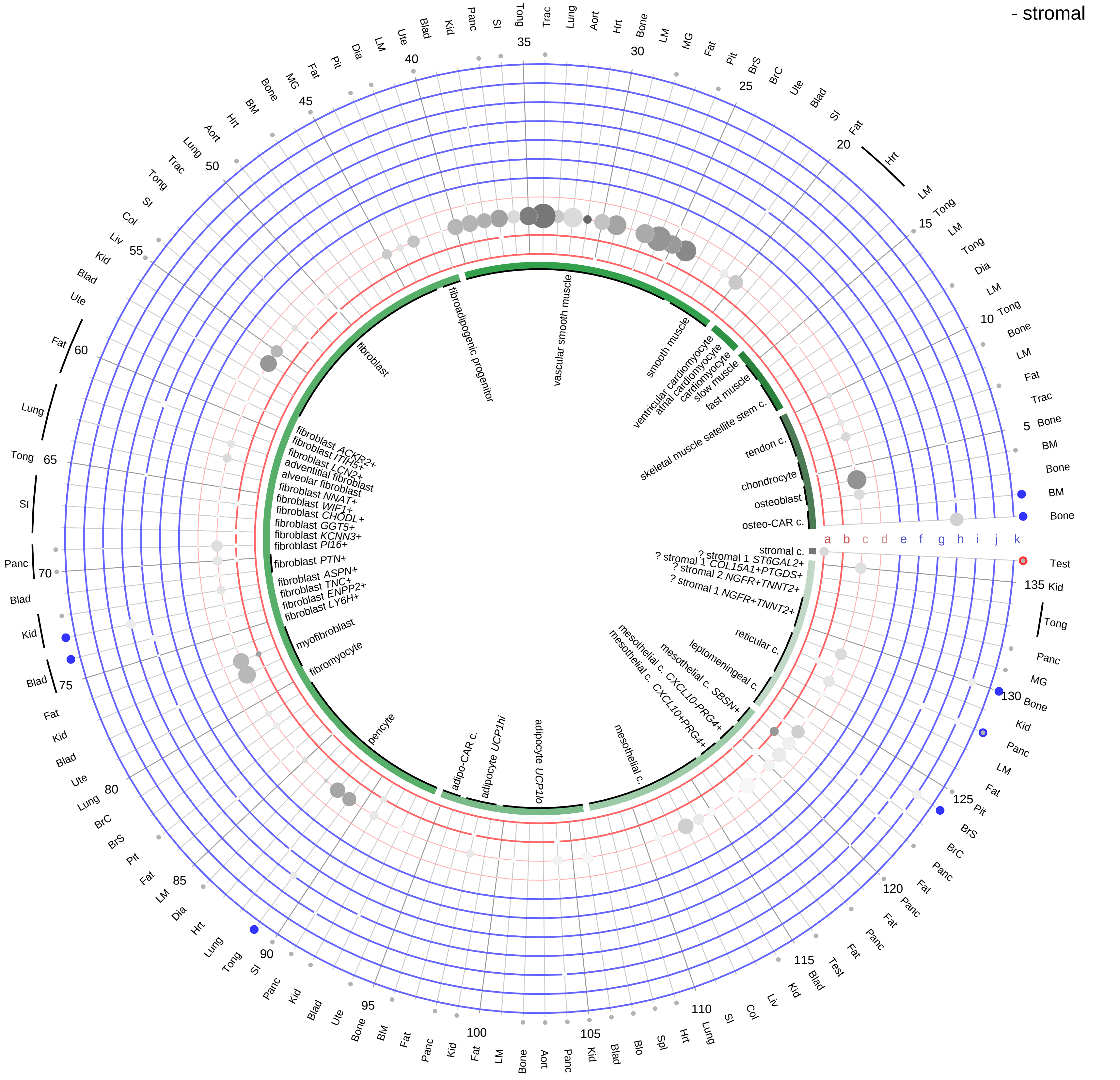

**Ligand/Enzyme**
- a - CORT
- b - SST
- c - CPE
- d - PCSK1

**Receptor**
- e - GHSR
- f - MRGPRX2
- g - SSTR1
- h - SSTR2
- i - SSTR3
- j - SSTR4
- k - SSTR5

**PAIRING** (ligand → receptor)

- SST & CPE & PCSK1→SSTR1
- SST & CPE & PCSK1→SSTR2
- SST & CPE & PCSK1→SSTR3
- SST & CPE & PCSK1→SSTR4
- SST & CPE & PCSK1→SSTR5
- CORT→SSTR1
- CORT→SSTR2
- CORT→SSTR3
- CORT→SSTR4
- CORT→SSTR5
- CORT→MRGPRX2
- CORT→GHSR

**Compartment**

**stromal**
- bone
- skeletal muscle
- cardiac muscle
- smooth muscle
- fibroblast
- adipocyte
- mesothelial
- other
- mix

12.5%
25%
50%
100%

ln(CP10K+1): 0, 1, 2, 3

| Abbrev. | Tissue |
|---|---|
| Aort | Aorta |
| Blad | Bladder |
| Blo | Blood |
| Bone | Bone |
| BM | Bone marrow |
| BrC | Brain cortex |
| BrS | Brainstem |
| Col | Colon |
| Dia | Diaphragm |
| Ret | Eye retina |
| Fat | Fat |
| Hrt | Heart |
| Pit | Hypothalamus/Pituitary |
| Kid | Kidney |
| LM | Limb muscle |
| Liv | Liver |
| Lung | Lung |
| MG | Mammary gland |
| Panc | Pancreas |
| Skin | Skin |
| SI | Small intestine |
| Spl | Spleen |
| Test | Testes |
| Thym | Thymus |
| Tong | Tongue |
| Trac | Trachea |
| Ute | Uterus |

**Somatostatin/Cortistatin**
- lymphoid

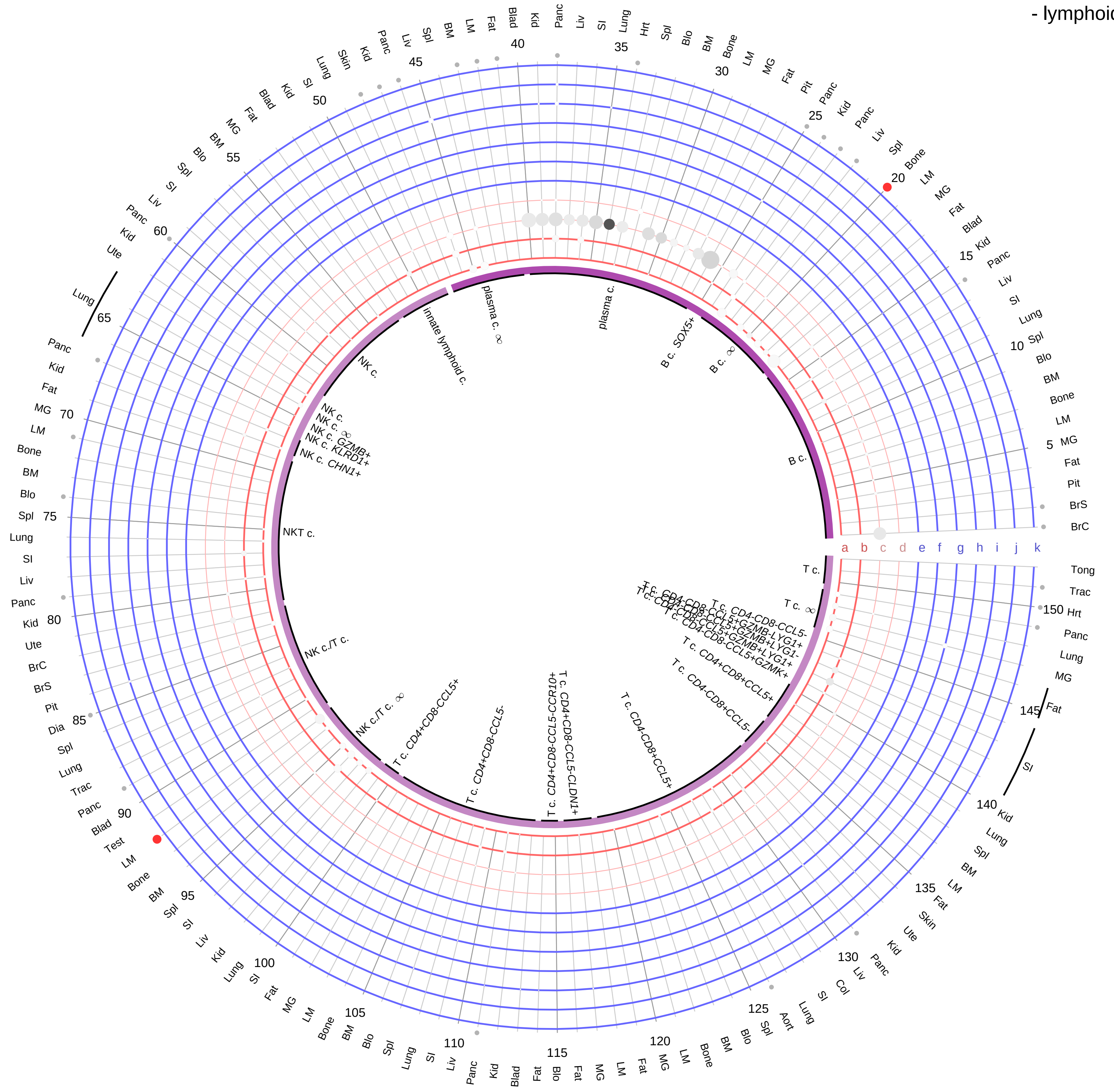

**Ligand/Enzyme**
- a - CORT
- b - SST
- c - CPE
- d - PCSK1

**Receptor**
- e - GHSR
- f - MRGPRX2
- g - SSTR1
- h - SSTR2
- i - SSTR3
- j - SSTR4
- k - SSTR5

**PAIRING** (ligand → receptor)

SST & CPE & PCSK1→SSTR1

SST & CPE & PCSK1→SSTR2

SST & CPE & PCSK1→SSTR3

SST & CPE & PCSK1→SSTR4

SST & CPE & PCSK1→SSTR5

CORT→SSTR1

CORT→SSTR2

CORT→SSTR3

CORT→SSTR4

CORT→SSTR5

CORT→MRGPRX2

CORT→GHSR

**Compartment**
**lymphoid**
- B cell
- NK/T cell

12.5%
25%
50%
100%

ln(CP10K+1): 0, 1, 2, 3

| Abbrev. | Tissue |
|---|---|
| Aort | Aorta |
| Blad | Bladder |
| Blo | Blood |
| Bone | Bone |
| BM | Bone marrow |
| BrC | Brain cortex |
| BrS | Brainstem |
| Col | Colon |
| Dia | Diaphragm |
| Ret | Eye retina |
| Fat | Fat |
| Hrt | Heart |
| Pit | Hypothalamus/Pituitary |
| Kid | Kidney |
| LM | Limb muscle |
| Liv | Liver |
| Lung | Lung |
| MG | Mammary gland |
| Panc | Pancreas |
| Skin | Skin |
| SI | Small intestine |
| Spl | Spleen |
| Test | Testes |
| Thym | Thymus |
| Tong | Tongue |
| Trac | Trachea |
| Ute | Uterus |

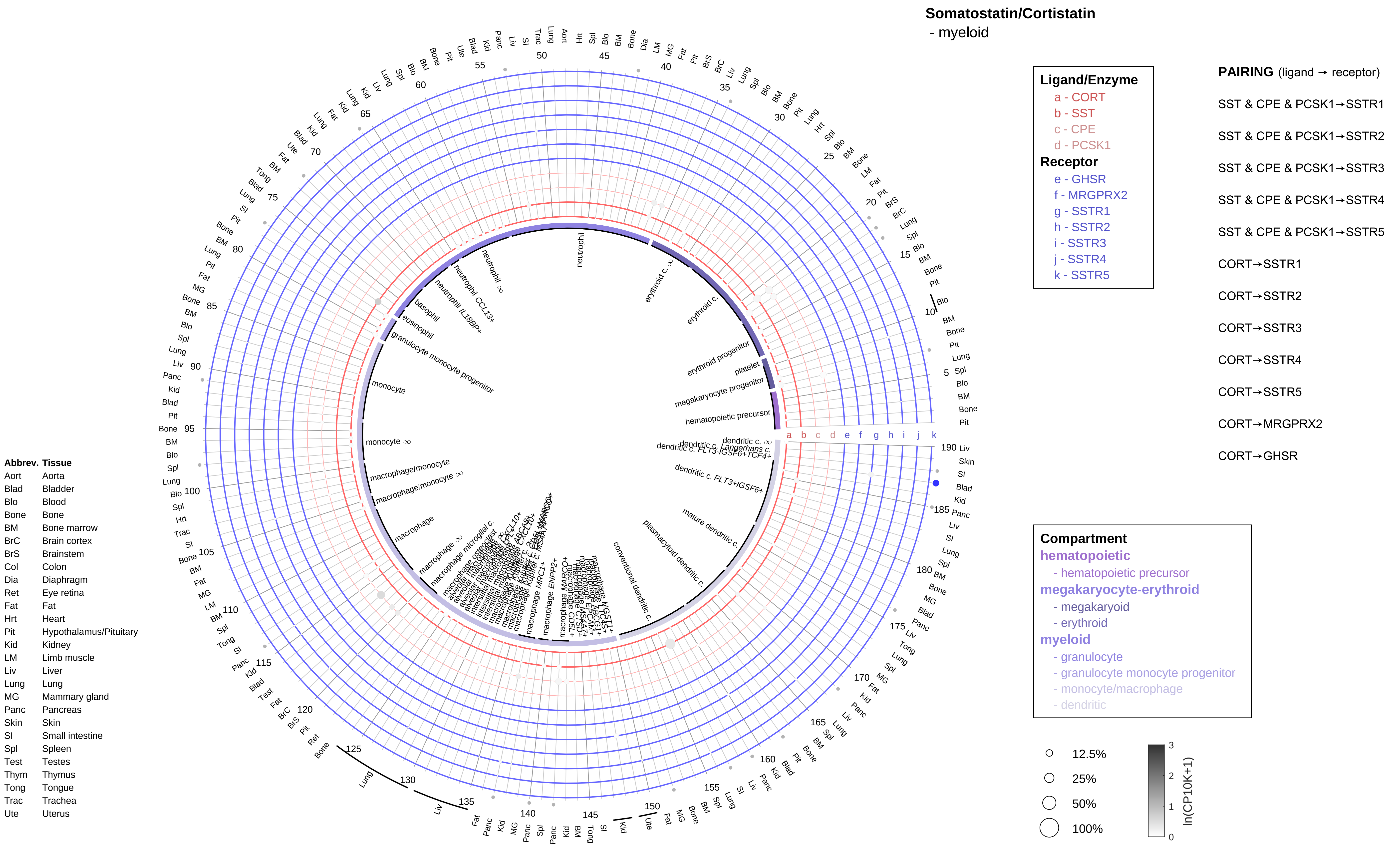

Somatostatin/Cortistatin
- myeloid
Ligand/Enzyme
a - CORT
b - SST
c - CPE
d - PCSK1
Receptor
e - GHSR
f - MRGPRX2
g - SSTR1
h - SSTR2
i - SSTR3
j - SSTR4
k - SSTR5
PAIRING (ligand → receptor)
SST & CPE & PCSK1→SSTR1
SST & CPE & PCSK1→SSTR2
SST & CPE & PCSK1→SSTR3
SST & CPE & PCSK1→SSTR4
SST & CPE & PCSK1→SSTR5
CORT→SSTR1
CORT→SSTR2
CORT→SSTR3
CORT→SSTR4
CORT→SSTR5
CORT→MRGPRX2
CORT→GHSR
Compartment
hematopoietic
- hematopoietic precursor
megakaryocyte-erythroid
- megakaryoid
- erythroid
myeloid
- granulocyte
- granulocyte monocyte progenitor
- monocyte/macrophage
- dendritic
12.5%
25%
50%
100%
ln(CP10K+1)
0
1
2
3
Abbrev. Tissue
Aort Aorta
Blad Bladder
Blo Blood
Bone Bone
BM Bone marrow
BrC Brain cortex
BrS Brainstem
Col Colon
Dia Diaphragm
Ret Eye retina
Fat Fat
Hrt Heart
Pit Hypothalamus/Pituitary
Kid Kidney
LM Limb muscle
Liv Liver
Lung Lung
MG Mammary gland
Panc Pancreas
Skin Skin
SI Small intestine
Spl Spleen
Test Testes
Thym Thymus
Tong Tongue
Trac Trachea
Ute Uterus

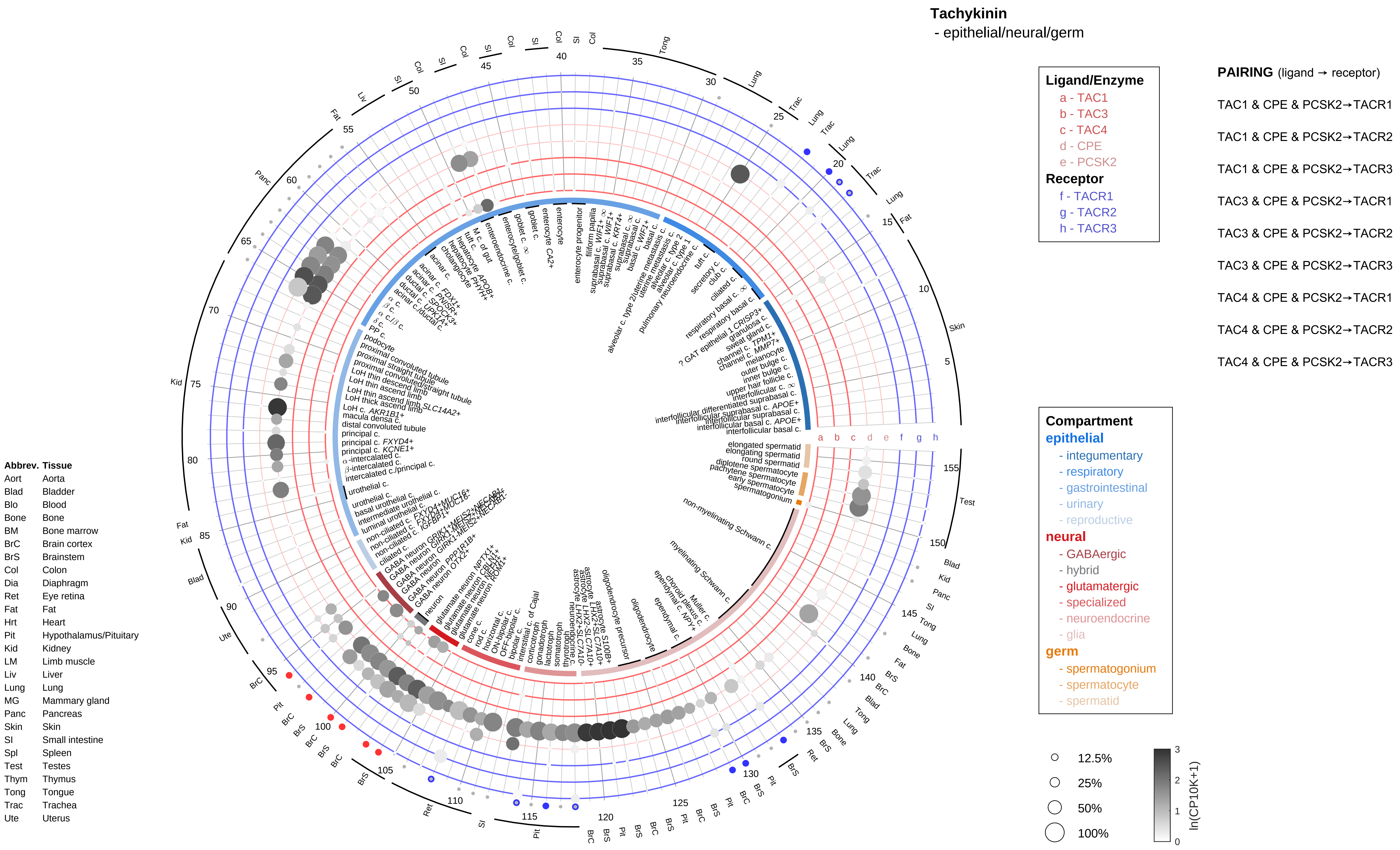
Tachykinin
- epithelial/neural/germ
Ligand/Enzyme
a - TAC1
b - TAC3
c - TAC4
d - CPE
e - PCSK2
Receptor
f - TACR1
g - TACR2
h - TACR3
PAIRING (ligand → receptor)
TAC1 & CPE & PCSK2→TACR1
TAC1 & CPE & PCSK2→TACR2
TAC1 & CPE & PCSK2→TACR3
TAC3 & CPE & PCSK2→TACR1
TAC3 & CPE & PCSK2→TACR2
TAC3 & CPE & PCSK2→TACR3
TAC4 & CPE & PCSK2→TACR1
TAC4 & CPE & PCSK2→TACR2
TAC4 & CPE & PCSK2→TACR3
Compartment
epithelial
- integumentary
- respiratory
- gastrointestinal
- urinary
- reproductive
neural
- GABAergic
- hybrid
- glutamatergic
- specialized
- neuroendocrine
- glia
germ
- spermatogonium
- spermatocyte
- spermatid
12.5%
25%
50%
100%
ln(CP10K+1)
Abbrev. Tissue
Aort Aorta
Blad Bladder
Blo Blood
Bone Bone
BM Bone marrow
BrC Brain cortex
BrS Brainstem
Col Colon
Dia Diaphragm
Ret Eye retina
Fat Fat
Hrt Heart
Pit Hypothalamus/Pituitary
Kid Kidney
LM Limb muscle
Liv Liver
Lung Lung
MG Mammary gland
Panc Pancreas
Skin Skin
SI Small intestine
Spl Spleen
Test Testes
Thym Thymus
Tong Tongue
Trac Trachea
Ute Uterus

**Tachykinin**
- endothelial

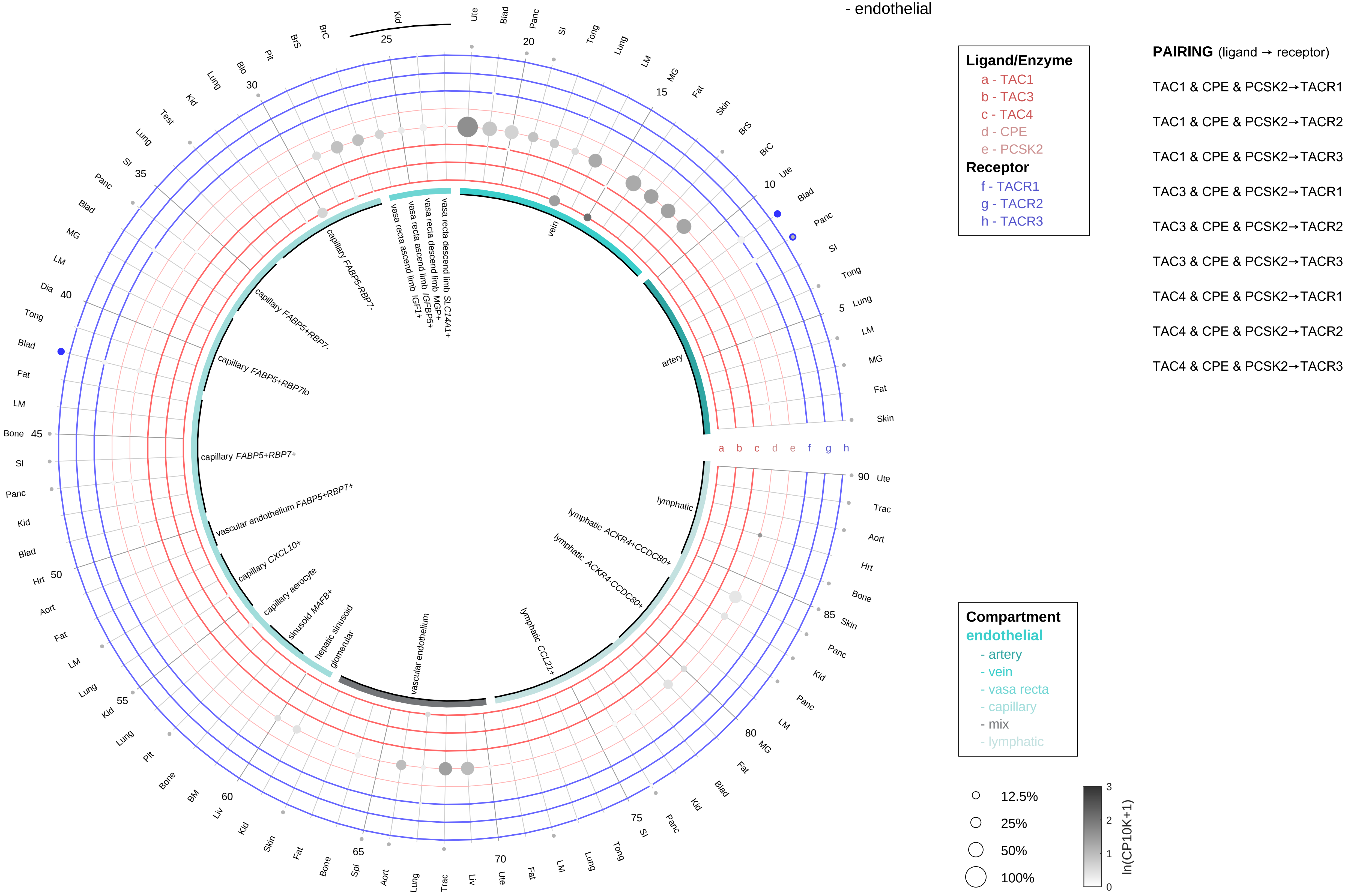

| Abbrev. | Tissue |
|---|---|
| Aort | Aorta |
| Blad | Bladder |
| Blo | Blood |
| Bone | Bone |
| BM | Bone marrow |
| BrC | Brain cortex |
| BrS | Brainstem |
| Col | Colon |
| Dia | Diaphragm |
| Ret | Eye retina |
| Fat | Fat |
| Hrt | Heart |
| Pit | Hypothalamus/Pituitary |
| Kid | Kidney |
| LM | Limb muscle |
| Liv | Liver |
| Lung | Lung |
| MG | Mammary gland |
| Panc | Pancreas |
| Skin | Skin |
| SI | Small intestine |
| Spl | Spleen |
| Test | Testes |
| Thym | Thymus |
| Tong | Tongue |
| Trac | Trachea |
| Ute | Uterus |

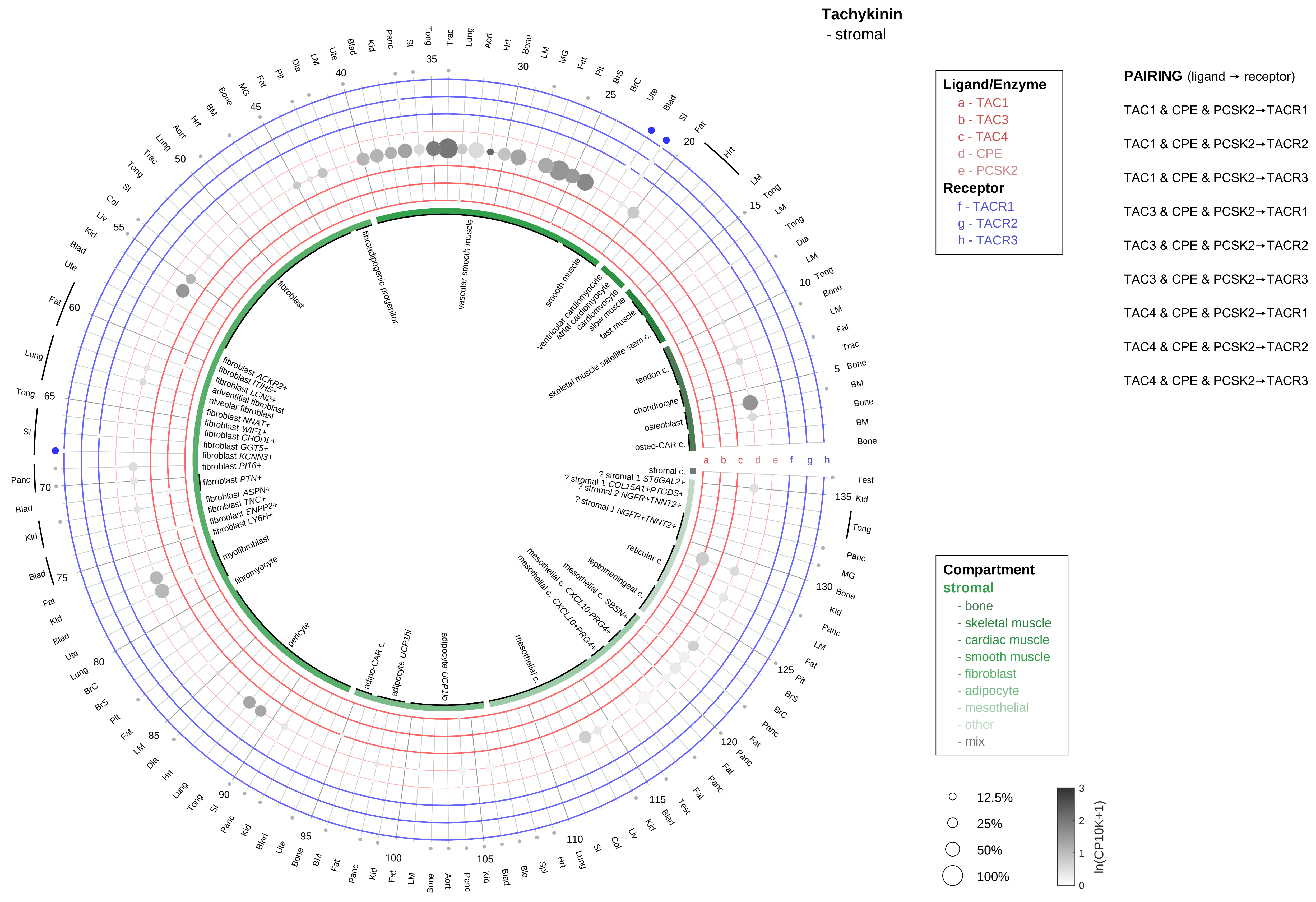

| Abbrev. | Tissue |
|---|---|
| Aort | Aorta |
| Blad | Bladder |
| Blo | Blood |
| Bone | Bone |
| BM | Bone marrow |
| BrC | Brain cortex |
| BrS | Brainstem |
| Col | Colon |
| Dia | Diaphragm |
| Ret | Eye retina |
| Fat | Fat |
| Hrt | Heart |
| Pit | Hypothalamus/Pituitary |
| Kid | Kidney |
| LM | Limb muscle |
| Liv | Liver |
| Lung | Lung |
| MG | Mammary gland |
| Panc | Pancreas |
| Skin | Skin |
| SI | Small intestine |
| Spl | Spleen |
| Test | Testes |
| Thym | Thymus |
| Tong | Tongue |
| Trac | Trachea |
| Ute | Uterus |

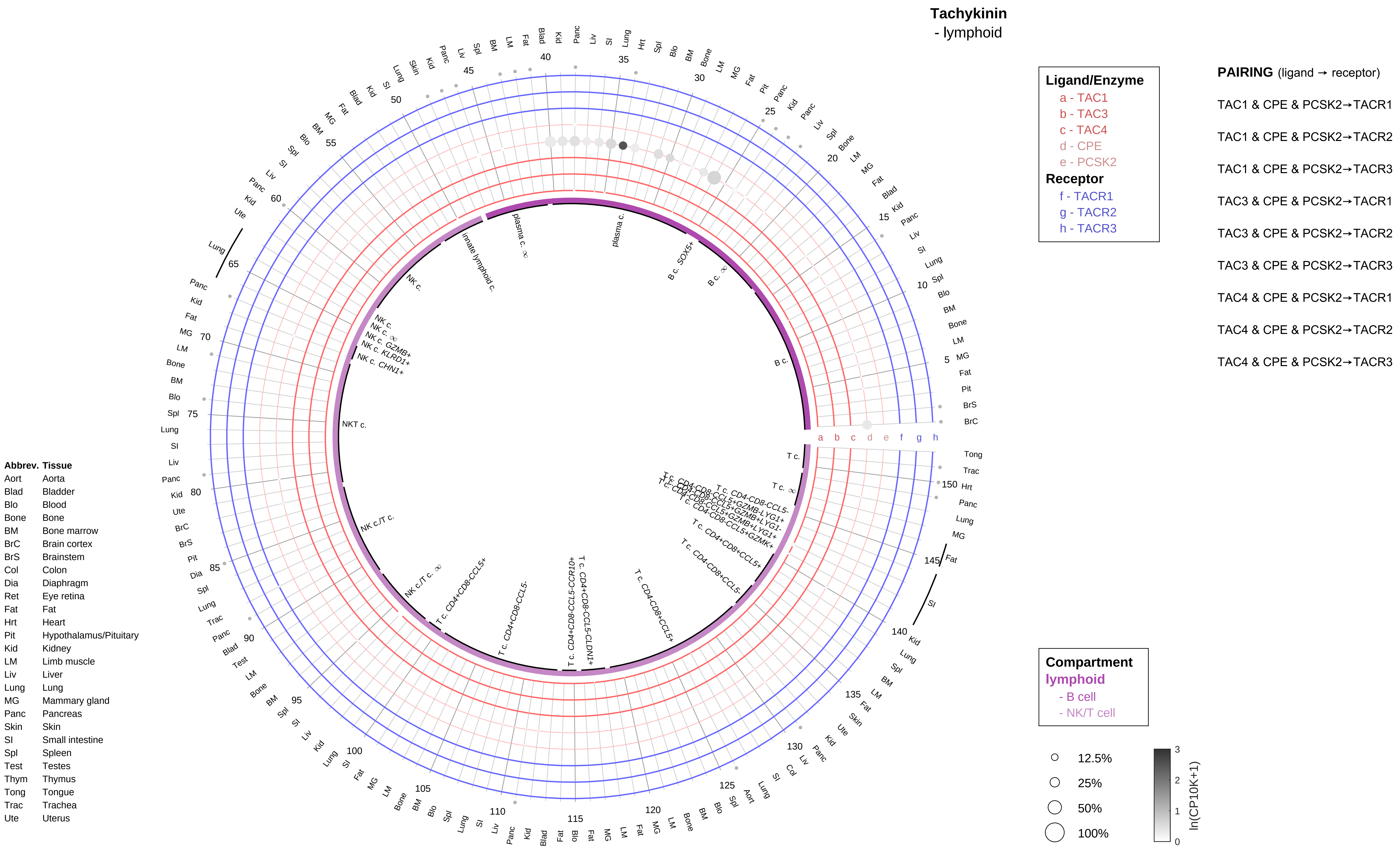
Tachykinin
- lymphoid
Ligand/Enzyme
a - TAC1
b - TAC3
c - TAC4
d - CPE
e - PCSK2
Receptor
f - TACR1
g - TACR2
h - TACR3
PAIRING (ligand → receptor)
TAC1 & CPE & PCSK2→TACR1
TAC1 & CPE & PCSK2→TACR2
TAC1 & CPE & PCSK2→TACR3
TAC3 & CPE & PCSK2→TACR1
TAC3 & CPE & PCSK2→TACR2
TAC3 & CPE & PCSK2→TACR3
TAC4 & CPE & PCSK2→TACR1
TAC4 & CPE & PCSK2→TACR2
TAC4 & CPE & PCSK2→TACR3
Compartment
lymphoid
- B cell
- NK/T cell
12.5%
25%
50%
100%
ln(CP10K+1)
Abbrev. Tissue
Aort Aorta
Blad Bladder
Blo Blood
Bone Bone
BM Bone marrow
BrC Brain cortex
BrS Brainstem
Col Colon
Dia Diaphragm
Ret Eye retina
Fat Fat
Hrt Heart
Pit Hypothalamus/Pituitary
Kid Kidney
LM Limb muscle
Liv Liver
Lung Lung
MG Mammary gland
Panc Pancreas
Skin Skin
SI Small intestine
Spl Spleen
Test Testes
Thym Thymus
Tong Tongue
Trac Trachea
Ute Uterus
plasma c.
B c. SOX5+
B c.
NKT c.
NK c.
innate lymphoid c.
NK c./T c.
T c.
T c. CD4+CD8-CCL5+
T c. CD4+CD8-CCL5-
T c. CD4-CD8+CCL5+
T c. CD4+CD8-CCL5-CCR10+
T c. CD4+CD8-CCL5-CLDN1+

**Tachykinin**
- myeloid

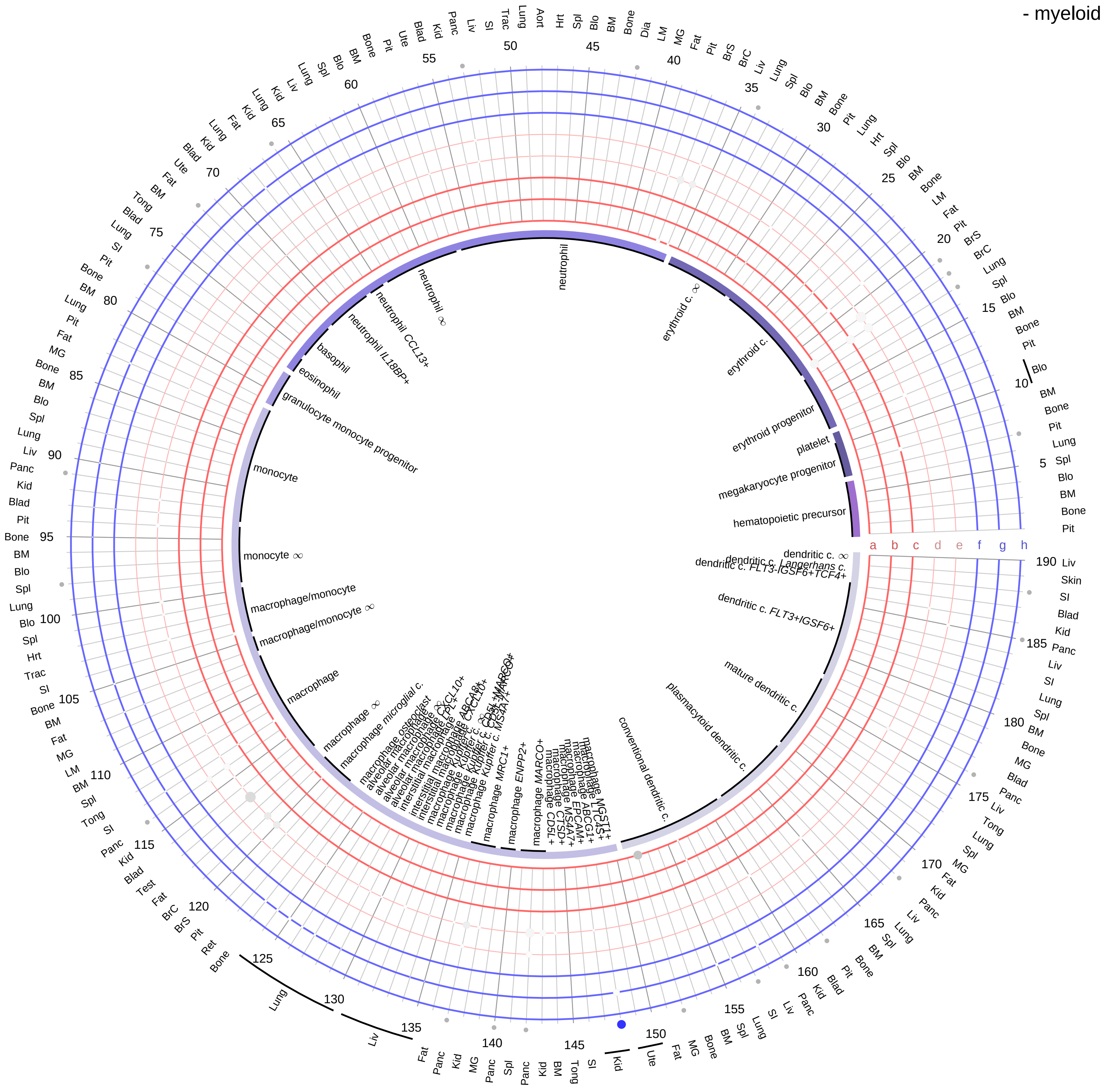

**Ligand/Enzyme**
- a - TAC1
- b - TAC3
- c - TAC4
- d - CPE
- e - PCSK2

**Receptor**
- f - TACR1
- g - TACR2
- h - TACR3

**PAIRING** (ligand → receptor)

TAC1 & CPE & PCSK2→TACR1

TAC1 & CPE & PCSK2→TACR2

TAC1 & CPE & PCSK2→TACR3

TAC3 & CPE & PCSK2→TACR1

TAC3 & CPE & PCSK2→TACR2

TAC3 & CPE & PCSK2→TACR3

TAC4 & CPE & PCSK2→TACR1

TAC4 & CPE & PCSK2→TACR2

TAC4 & CPE & PCSK2→TACR3

| Abbrev. | Tissue |
|---|---|
| Aort | Aorta |
| Blad | Bladder |
| Blo | Blood |
| Bone | Bone |
| BM | Bone marrow |
| BrC | Brain cortex |
| BrS | Brainstem |
| Col | Colon |
| Dia | Diaphragm |
| Ret | Eye retina |
| Fat | Fat |
| Hrt | Heart |
| Pit | Hypothalamus/Pituitary |
| Kid | Kidney |
| LM | Limb muscle |
| Liv | Liver |
| Lung | Lung |
| MG | Mammary gland |
| Panc | Pancreas |
| Skin | Skin |
| SI | Small intestine |
| Spl | Spleen |
| Test | Testes |
| Thym | Thymus |
| Tong | Tongue |
| Trac | Trachea |
| Ute | Uterus |

**Compartment**

**hematopoietic**
- hematopoietic precursor

**megakaryocyte-erythroid**
- megakaryoid
- erythroid

**myeloid**
- granulocyte
- granulocyte monocyte progenitor
- monocyte/macrophage
- dendritic

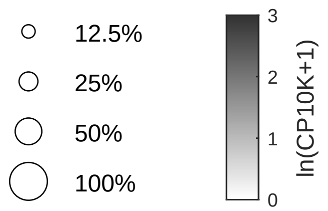

**Thrombopoietin (TPO)**
- epithelial/neural/germ

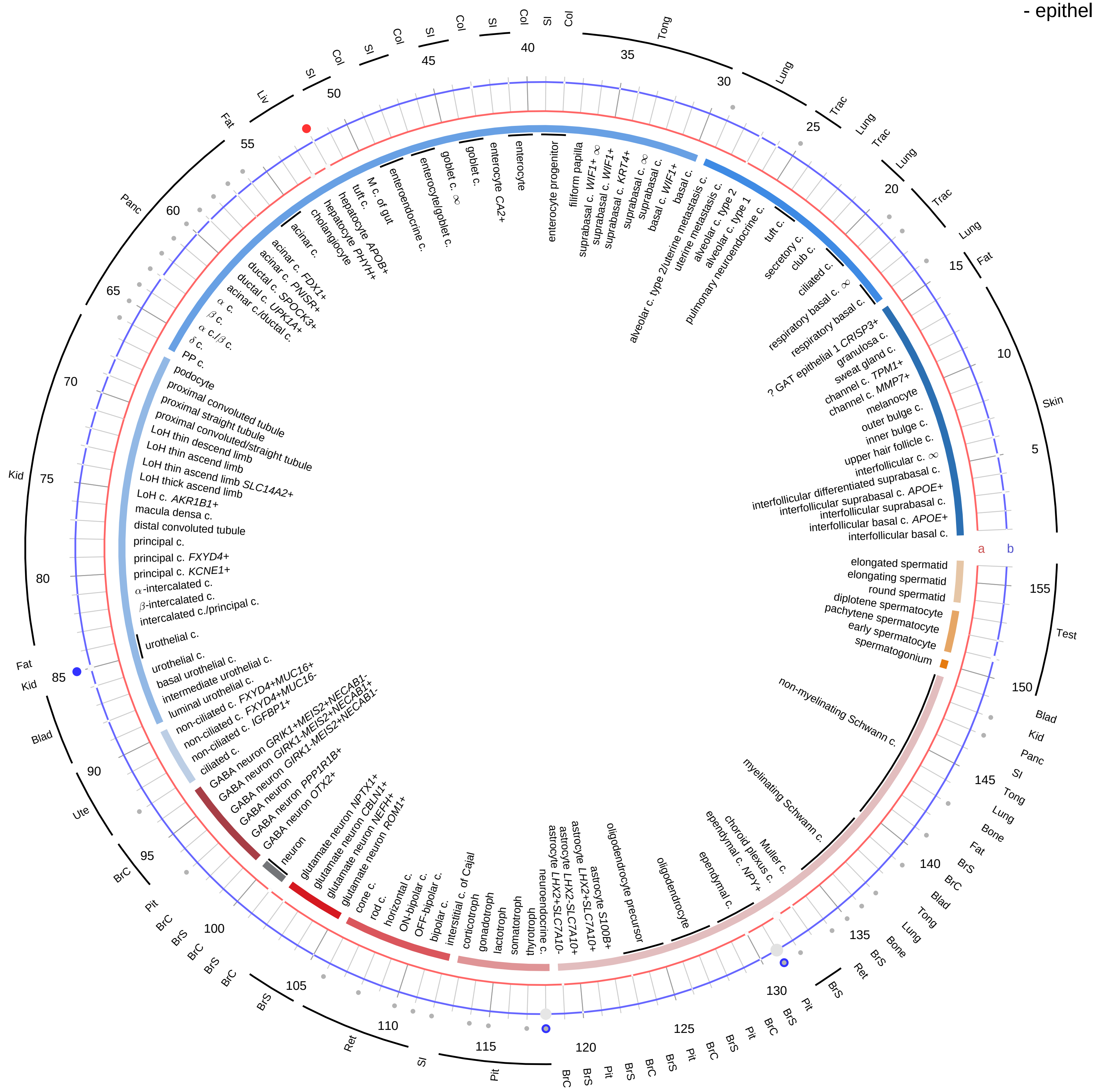

| **Ligand/Enzyme** |
|---|
| a - THPO |
| **Receptor** |
| b - MPL |

**PAIRING** (ligand → receptor)

THPO→MPL

**Compartment**
**epithelial**
- integumentary
- respiratory
- gastrointestinal
- urinary
- reproductive

**neural**
- GABAergic
- hybrid
- glutamatergic
- specialized
- neuroendocrine
- glia

**germ**
- spermatogonium
- spermatocyte
- spermatid

12.5%
25%
50%
100%

ln(CP10K+1): 0, 1, 2, 3

| Abbrev. | Tissue |
|---|---|
| Aort | Aorta |
| Blad | Bladder |
| Blo | Blood |
| Bone | Bone |
| BM | Bone marrow |
| BrC | Brain cortex |
| BrS | Brainstem |
| Col | Colon |
| Dia | Diaphragm |
| Ret | Eye retina |
| Fat | Fat |
| Hrt | Heart |
| Pit | Hypothalamus/Pituitary |
| Kid | Kidney |
| LM | Limb muscle |
| Liv | Liver |
| Lung | Lung |
| MG | Mammary gland |
| Panc | Pancreas |
| Skin | Skin |
| SI | Small intestine |
| Spl | Spleen |
| Test | Testes |
| Thym | Thymus |
| Tong | Tongue |
| Trac | Trachea |
| Ute | Uterus |

#### Thrombopoietin (TPO)
- endothelial

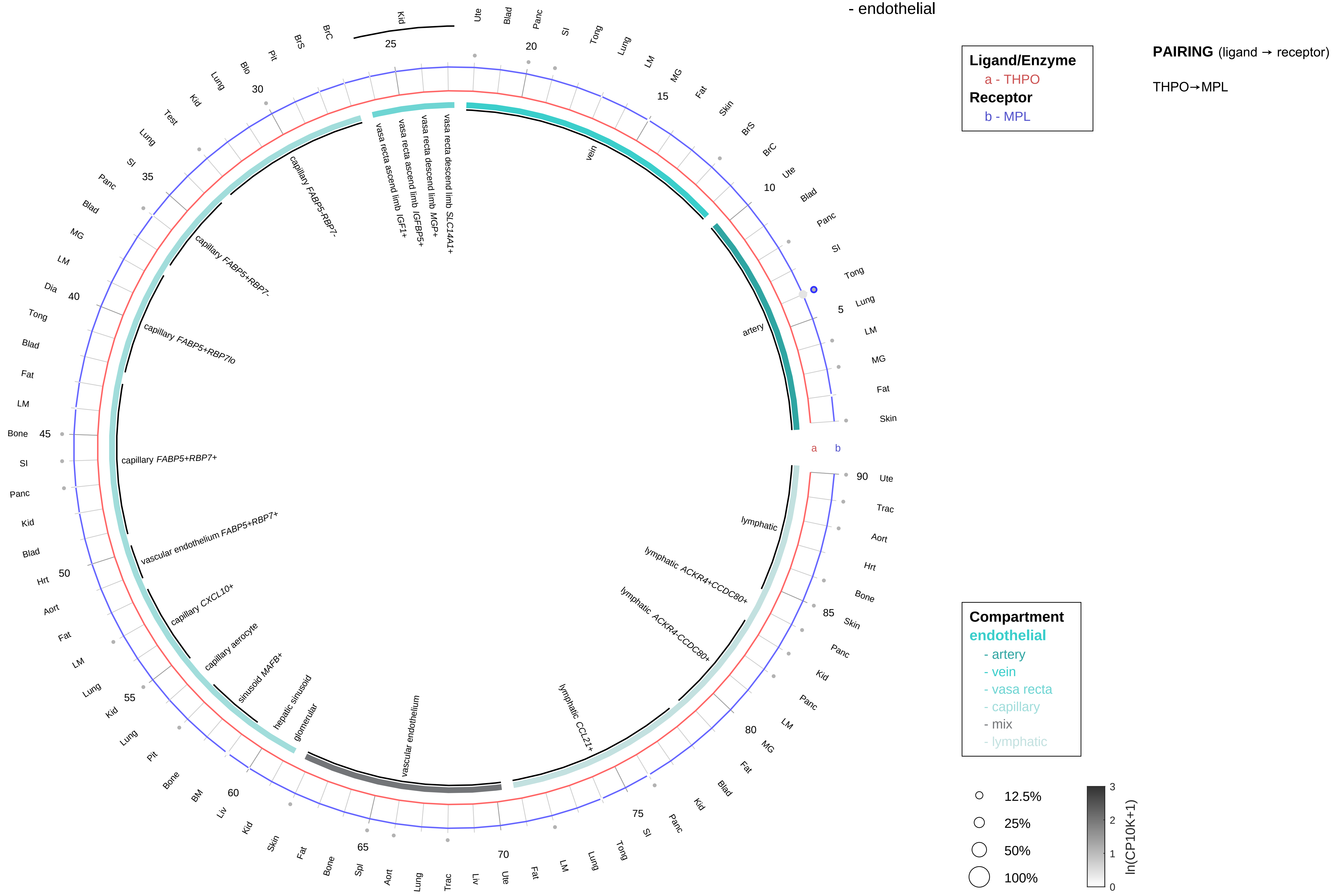

| Abbrev. | Tissue |
|---|---|
| Aort | Aorta |
| Blad | Bladder |
| Blo | Blood |
| Bone | Bone |
| BM | Bone marrow |
| BrC | Brain cortex |
| BrS | Brainstem |
| Col | Colon |
| Dia | Diaphragm |
| Ret | Eye retina |
| Fat | Fat |
| Hrt | Heart |
| Pit | Hypothalamus/Pituitary |
| Kid | Kidney |
| LM | Limb muscle |
| Liv | Liver |
| Lung | Lung |
| MG | Mammary gland |
| Panc | Pancreas |
| Skin | Skin |
| SI | Small intestine |
| Spl | Spleen |
| Test | Testes |
| Thym | Thymus |
| Tong | Tongue |
| Trac | Trachea |
| Ute | Uterus |

**Thrombopoietin (TPO)**
- stromal

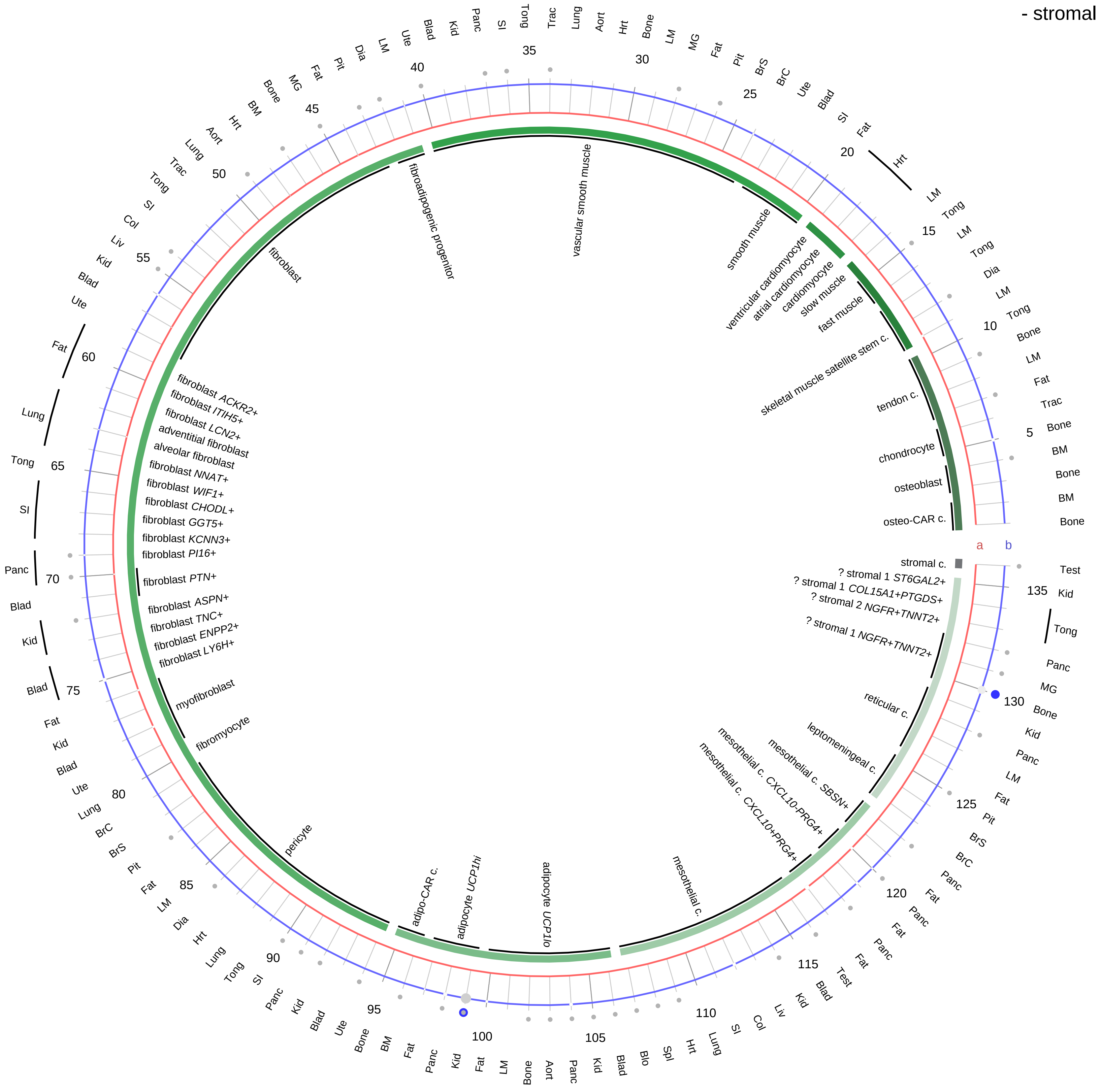

**Ligand/Enzyme**
a - THPO
**Receptor**
b - MPL

**PAIRING** (ligand → receptor)

THPO→MPL

**Compartment**
**stromal**
- bone
- skeletal muscle
- cardiac muscle
- smooth muscle
- fibroblast
- adipocyte
- mesothelial
- other
- mix

12.5%
25%
50%
100%

ln(CP10K+1): 0, 1, 2, 3

| Abbrev. | Tissue |
|---|---|
| Aort | Aorta |
| Blad | Bladder |
| Blo | Blood |
| Bone | Bone |
| BM | Bone marrow |
| BrC | Brain cortex |
| BrS | Brainstem |
| Col | Colon |
| Dia | Diaphragm |
| Ret | Eye retina |
| Fat | Fat |
| Hrt | Heart |
| Pit | Hypothalamus/Pituitary |
| Kid | Kidney |
| LM | Limb muscle |
| Liv | Liver |
| Lung | Lung |
| MG | Mammary gland |
| Panc | Pancreas |
| Skin | Skin |
| SI | Small intestine |
| Spl | Spleen |
| Test | Testes |
| Thym | Thymus |
| Tong | Tongue |
| Trac | Trachea |
| Ute | Uterus |

**Thrombopoietin (TPO)**
- lymphoid

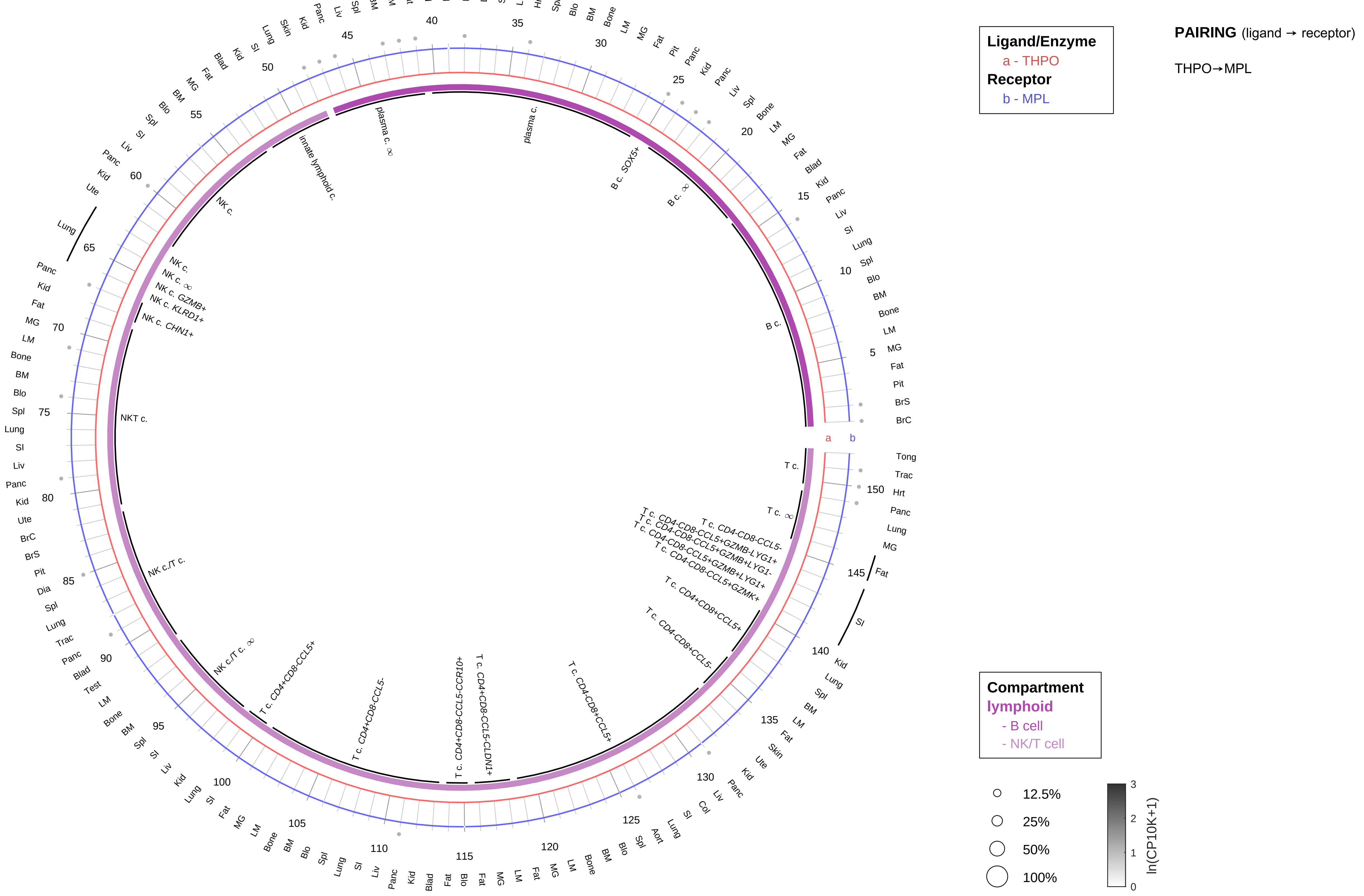

| Abbrev. | Tissue |
|---|---|
| Aort | Aorta |
| Blad | Bladder |
| Blo | Blood |
| Bone | Bone |
| BM | Bone marrow |
| BrC | Brain cortex |
| BrS | Brainstem |
| Col | Colon |
| Dia | Diaphragm |
| Ret | Eye retina |
| Fat | Fat |
| Hrt | Heart |
| Pit | Hypothalamus/Pituitary |
| Kid | Kidney |
| LM | Limb muscle |
| Liv | Liver |
| Lung | Lung |
| MG | Mammary gland |
| Panc | Pancreas |
| Skin | Skin |
| SI | Small intestine |
| Spl | Spleen |
| Test | Testes |
| Thym | Thymus |
| Tong | Tongue |
| Trac | Trachea |
| Ute | Uterus |

**Thrombopoietin (TPO)**
- myeloid

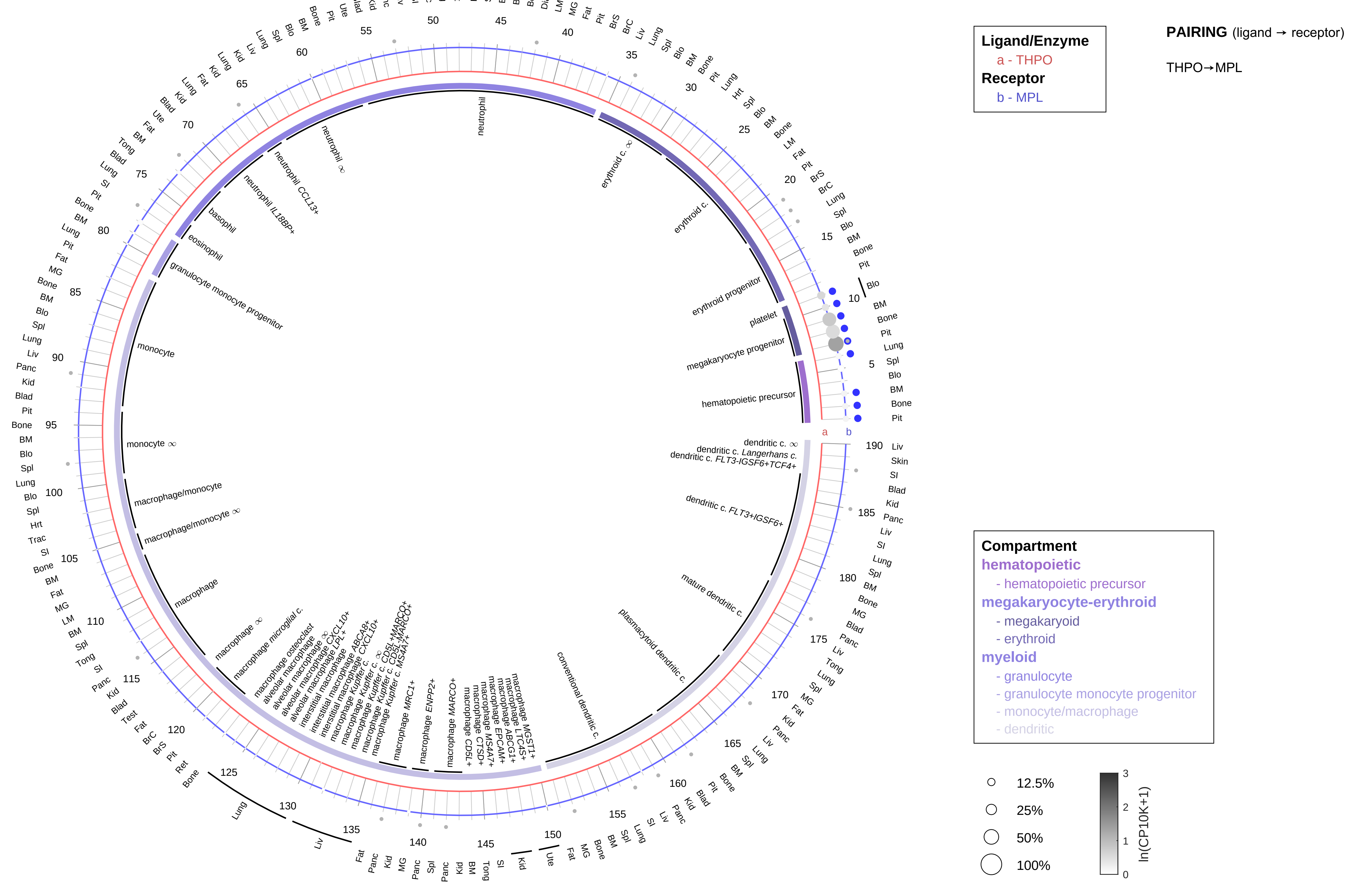

| Abbrev. | Tissue |
|---|---|
| Aort | Aorta |
| Blad | Bladder |
| Blo | Blood |
| Bone | Bone |
| BM | Bone marrow |
| BrC | Brain cortex |
| BrS | Brainstem |
| Col | Colon |
| Dia | Diaphragm |
| Ret | Eye retina |
| Fat | Fat |
| Hrt | Heart |
| Pit | Hypothalamus/Pituitary |
| Kid | Kidney |
| LM | Limb muscle |
| Liv | Liver |
| Lung | Lung |
| MG | Mammary gland |
| Panc | Pancreas |
| Skin | Skin |
| SI | Small intestine |
| Spl | Spleen |
| Test | Testes |
| Thym | Thymus |
| Tong | Tongue |
| Trac | Trachea |
| Ute | Uterus |

#### Thromboxane (TXA2)
- epithelial/neural/germ

**PAIRING** (ligand → receptor)

TBXAS1 & PTGS1 & PTGS2→TBXA2R

**Ligand/Enzyme**
- a - PTGS1
- b - PTGS2
- c - TBXAS1

**Receptor**
- d - TBXA2R

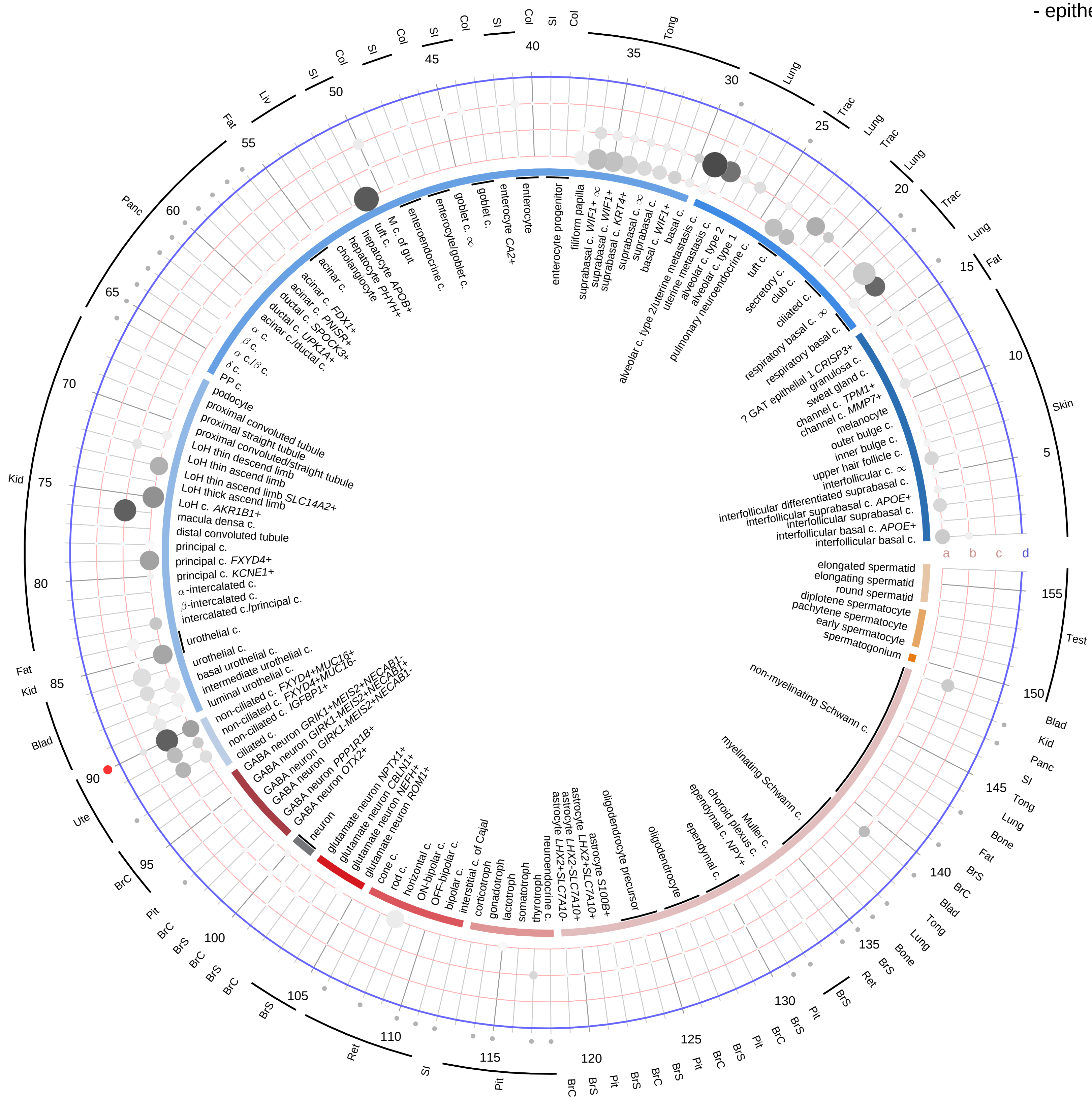

**Compartment**

**epithelial**
- integumentary
- respiratory
- gastrointestinal
- urinary
- reproductive

**neural**
- GABAergic
- hybrid
- glutamatergic
- specialized
- neuroendocrine
- glia

**germ**
- spermatogonium
- spermatocyte
- spermatid

12.5%
25%
50%
100%

ln(CP10K+1): 0, 1, 2, 3

| Abbrev. | Tissue |
|---|---|
| Aort | Aorta |
| Blad | Bladder |
| Blo | Blood |
| Bone | Bone |
| BM | Bone marrow |
| BrC | Brain cortex |
| BrS | Brainstem |
| Col | Colon |
| Dia | Diaphragm |
| Ret | Eye retina |
| Fat | Fat |
| Hrt | Heart |
| Pit | Hypothalamus/Pituitary |
| Kid | Kidney |
| LM | Limb muscle |
| Liv | Liver |
| Lung | Lung |
| MG | Mammary gland |
| Panc | Pancreas |
| Skin | Skin |
| SI | Small intestine |
| Spl | Spleen |
| Test | Testes |
| Thym | Thymus |
| Tong | Tongue |
| Trac | Trachea |
| Ute | Uterus |

**Thromboxane (TXA2)**
- endothelial

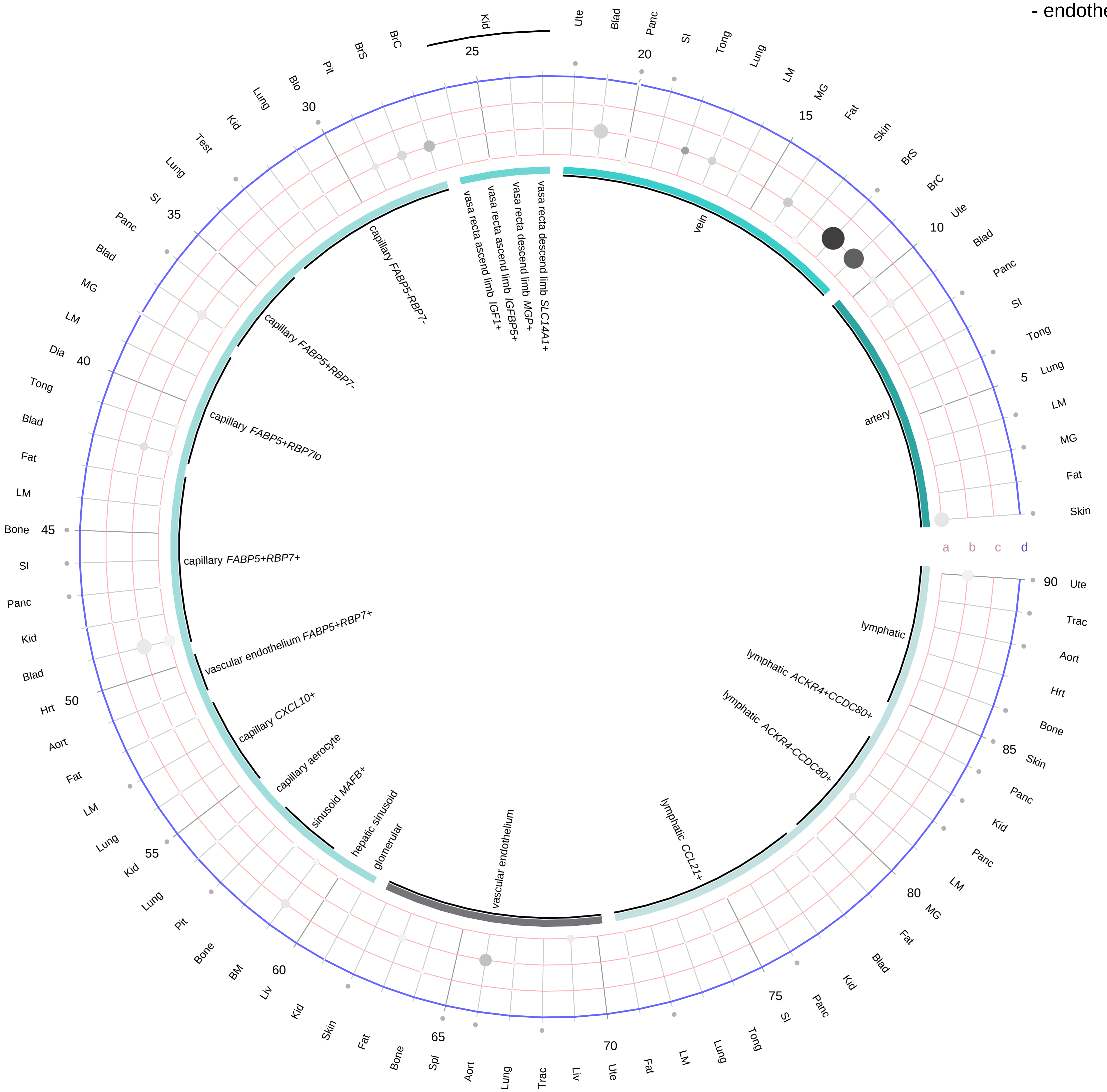

**Ligand/Enzyme**
- a - PTGS1
- b - PTGS2
- c - TBXAS1

**Receptor**
- d - TBXA2R

**PAIRING** (ligand → receptor)

TBXAS1 & PTGS1 & PTGS2→TBXA2R

**Compartment**

**endothelial**
- artery
- vein
- vasa recta
- capillary
- mix
- lymphatic

12.5%
25%
50%
100%

ln(CP10K+1): 0, 1, 2, 3

| Abbrev. | Tissue |
|---|---|
| Aort | Aorta |
| Blad | Bladder |
| Blo | Blood |
| Bone | Bone |
| BM | Bone marrow |
| BrC | Brain cortex |
| BrS | Brainstem |
| Col | Colon |
| Dia | Diaphragm |
| Ret | Eye retina |
| Fat | Fat |
| Hrt | Heart |
| Pit | Hypothalamus/Pituitary |
| Kid | Kidney |
| LM | Limb muscle |
| Liv | Liver |
| Lung | Lung |
| MG | Mammary gland |
| Panc | Pancreas |
| Skin | Skin |
| SI | Small intestine |
| Spl | Spleen |
| Test | Testes |
| Thym | Thymus |
| Tong | Tongue |
| Trac | Trachea |
| Ute | Uterus |

**Thromboxane (TXA2)**
- stromal

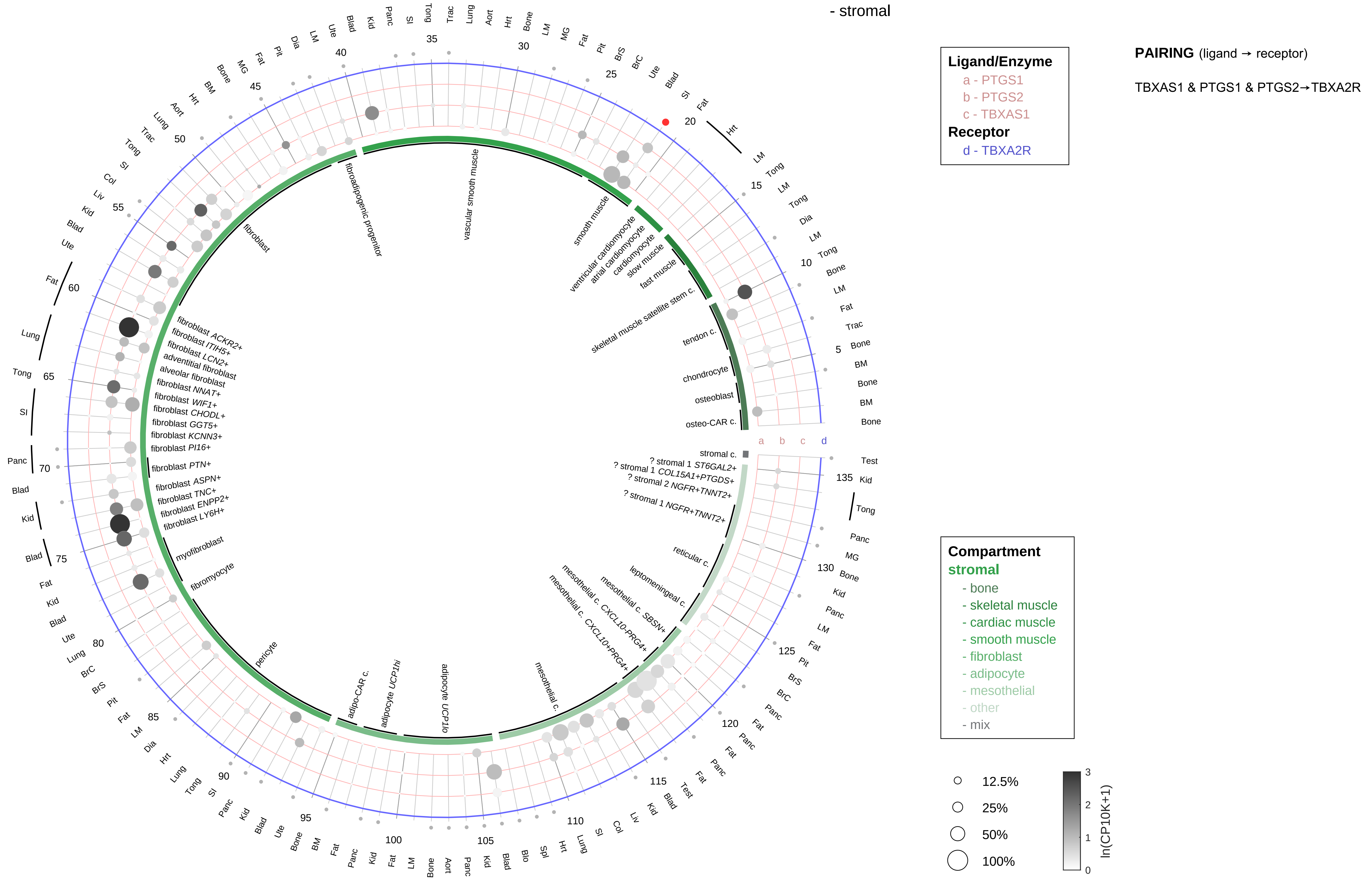

| Abbrev. | Tissue |
|---|---|
| Aort | Aorta |
| Blad | Bladder |
| Blo | Blood |
| Bone | Bone |
| BM | Bone marrow |
| BrC | Brain cortex |
| BrS | Brainstem |
| Col | Colon |
| Dia | Diaphragm |
| Ret | Eye retina |
| Fat | Fat |
| Hrt | Heart |
| Pit | Hypothalamus/Pituitary |
| Kid | Kidney |
| LM | Limb muscle |
| Liv | Liver |
| Lung | Lung |
| MG | Mammary gland |
| Panc | Pancreas |
| Skin | Skin |
| SI | Small intestine |
| Spl | Spleen |
| Test | Testes |
| Thym | Thymus |
| Tong | Tongue |
| Trac | Trachea |
| Ute | Uterus |

**Thromboxane (TXA2)**
- lymphoid

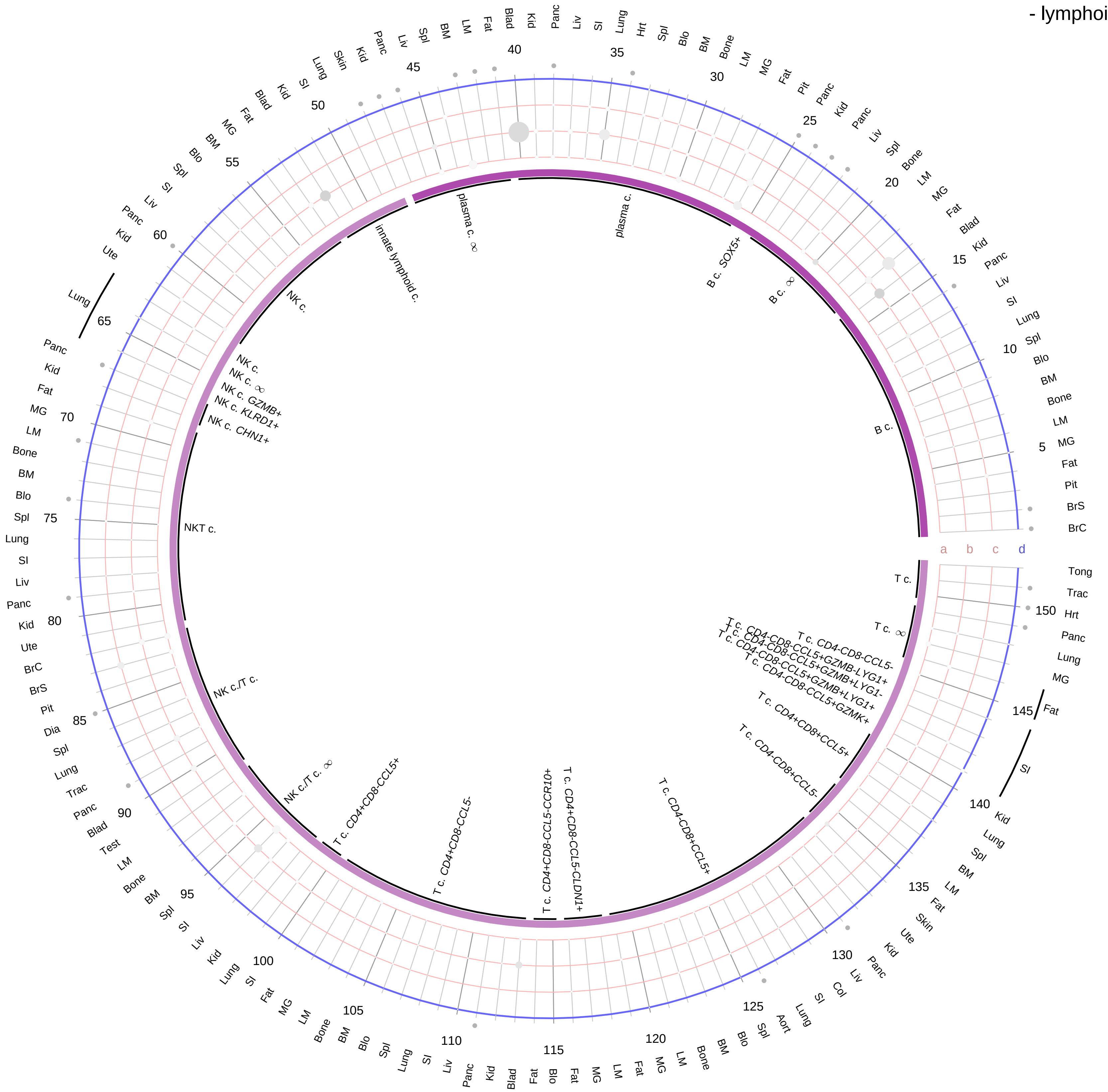

**Ligand/Enzyme**
- a - PTGS1
- b - PTGS2
- c - TBXAS1

**Receptor**
- d - TBXA2R

**PAIRING** (ligand → receptor)

TBXAS1 & PTGS1 & PTGS2→TBXA2R

| Abbrev. | Tissue |
|---|---|
| Aort | Aorta |
| Blad | Bladder |
| Blo | Blood |
| Bone | Bone |
| BM | Bone marrow |
| BrC | Brain cortex |
| BrS | Brainstem |
| Col | Colon |
| Dia | Diaphragm |
| Ret | Eye retina |
| Fat | Fat |
| Hrt | Heart |
| Pit | Hypothalamus/Pituitary |
| Kid | Kidney |
| LM | Limb muscle |
| Liv | Liver |
| Lung | Lung |
| MG | Mammary gland |
| Panc | Pancreas |
| Skin | Skin |
| SI | Small intestine |
| Spl | Spleen |
| Test | Testes |
| Thym | Thymus |
| Tong | Tongue |
| Trac | Trachea |
| Ute | Uterus |

**Compartment**

**lymphoid**
- B cell
- NK/T cell

12.5%
25%
50%
100%

ln(CP10K+1): 0 1 2 3

**Thromboxane (TXA2)**
- myeloid

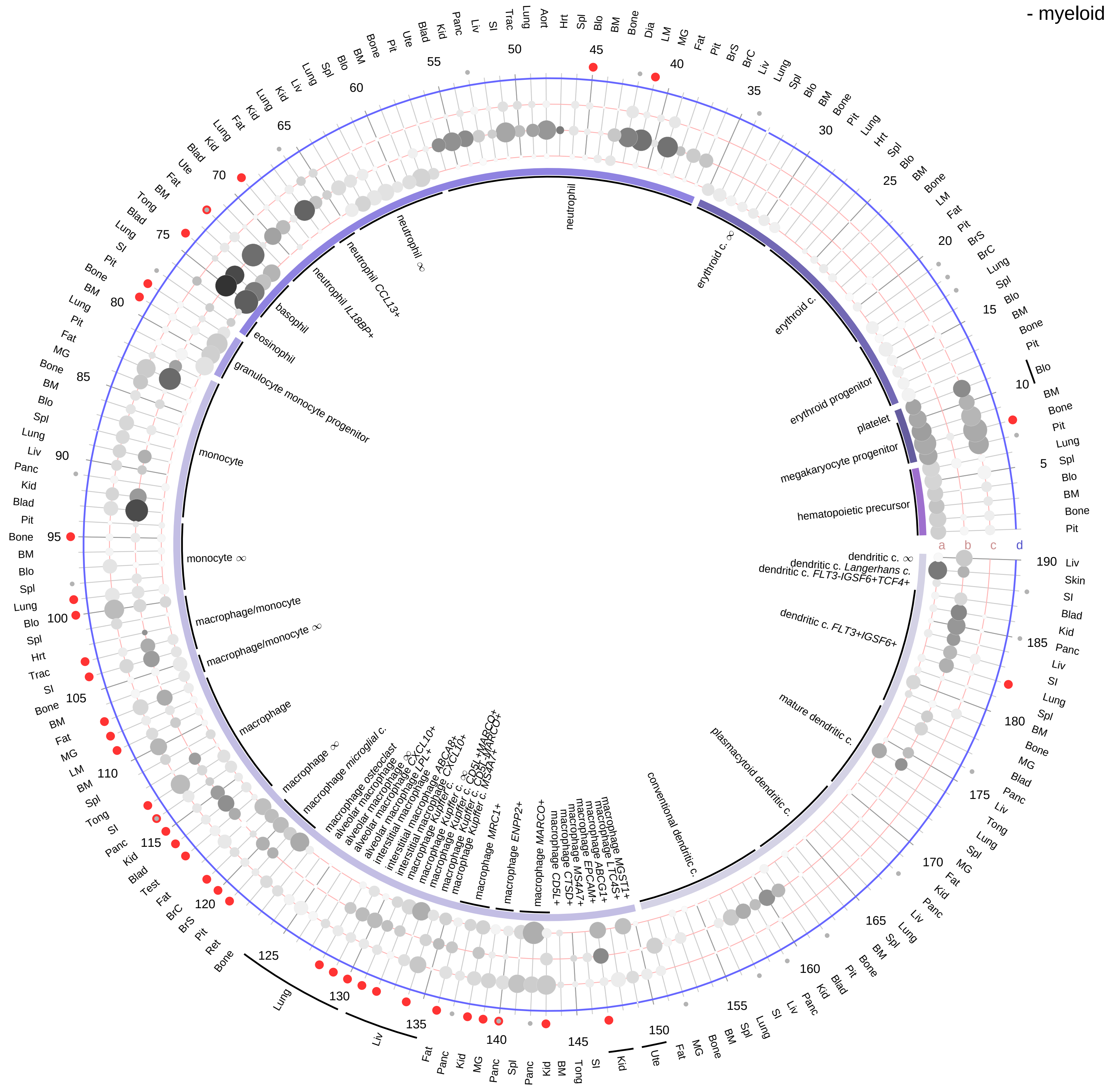

**Ligand/Enzyme**
- a - PTGS1
- b - PTGS2
- c - TBXAS1

**Receptor**
- d - TBXA2R

**PAIRING** (ligand → receptor)

TBXAS1 & PTGS1 & PTGS2→TBXA2R

| Abbrev. | Tissue |
|---|---|
| Aort | Aorta |
| Blad | Bladder |
| Blo | Blood |
| Bone | Bone |
| BM | Bone marrow |
| BrC | Brain cortex |
| BrS | Brainstem |
| Col | Colon |
| Dia | Diaphragm |
| Ret | Eye retina |
| Fat | Fat |
| Hrt | Heart |
| Pit | Hypothalamus/Pituitary |
| Kid | Kidney |
| LM | Limb muscle |
| Liv | Liver |
| Lung | Lung |
| MG | Mammary gland |
| Panc | Pancreas |
| Skin | Skin |
| SI | Small intestine |
| Spl | Spleen |
| Test | Testes |
| Thym | Thymus |
| Tong | Tongue |
| Trac | Trachea |
| Ute | Uterus |

**Compartment**

**hematopoietic**
- hematopoietic precursor

**megakaryocyte-erythroid**
- megakaryoid
- erythroid

**myeloid**
- granulocyte
- granulocyte monocyte progenitor
- monocyte/macrophage
- dendritic

12.5%
25%
50%
100%

ln(CP10K+1): 0, 1, 2, 3

**Thyroid stimulating hormone (TSH)**
- epithelial/neural/germ

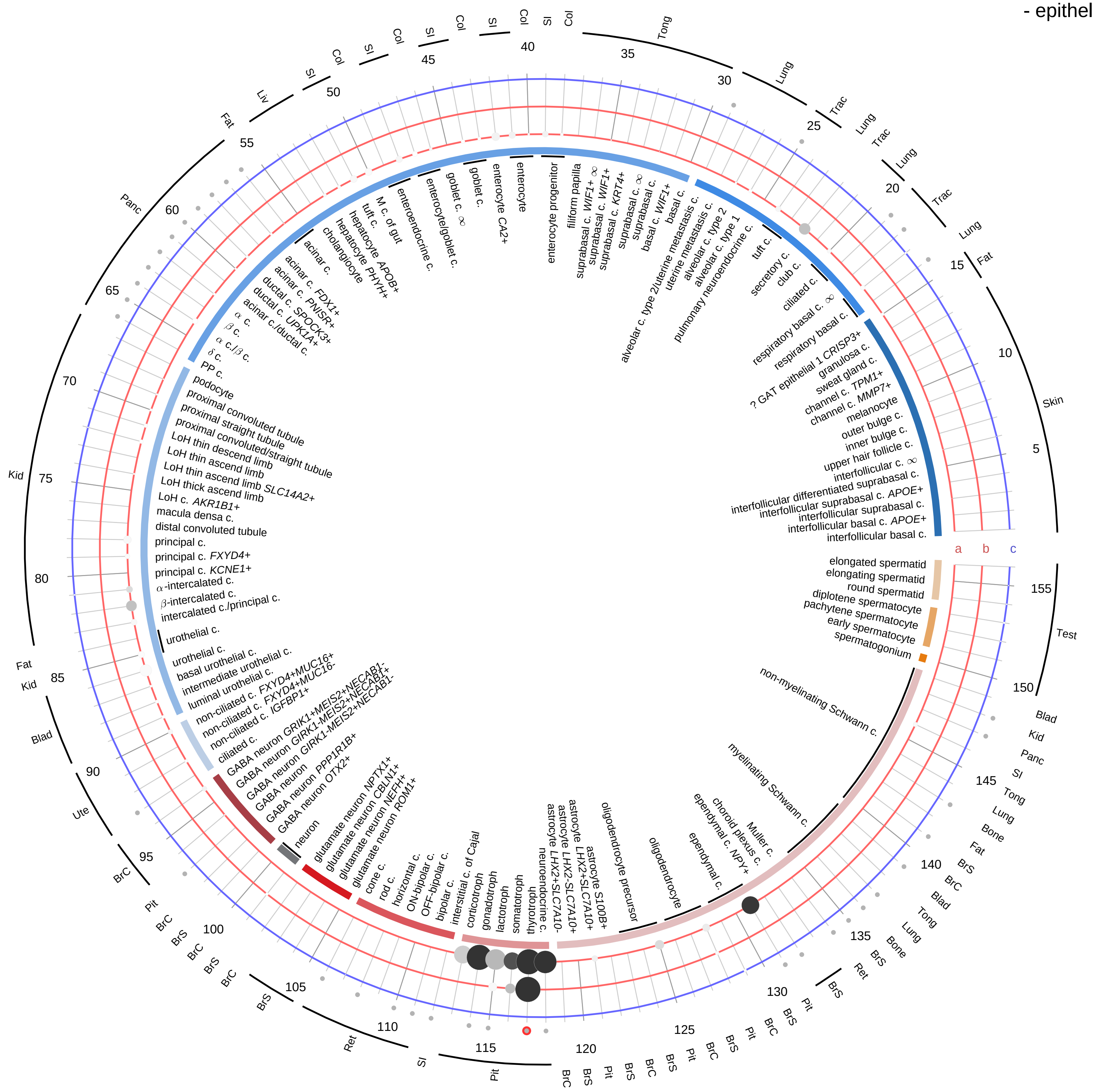

**Ligand/Enzyme**
- a - CGA
- b - TSHB

**Receptor**
- c - TSHR

**PAIRING** (ligand → receptor)

TSHB & CGA→TSHR

| Abbrev. | Tissue |
|---|---|
| Aort | Aorta |
| Blad | Bladder |
| Blo | Blood |
| Bone | Bone |
| BM | Bone marrow |
| BrC | Brain cortex |
| BrS | Brainstem |
| Col | Colon |
| Dia | Diaphragm |
| Ret | Eye retina |
| Fat | Fat |
| Hrt | Heart |
| Pit | Hypothalamus/Pituitary |
| Kid | Kidney |
| LM | Limb muscle |
| Liv | Liver |
| Lung | Lung |
| MG | Mammary gland |
| Panc | Pancreas |
| Skin | Skin |
| SI | Small intestine |
| Spl | Spleen |
| Test | Testes |
| Thym | Thymus |
| Tong | Tongue |
| Trac | Trachea |
| Ute | Uterus |

**Compartment**

**epithelial**
- integumentary
- respiratory
- gastrointestinal
- urinary
- reproductive

**neural**
- GABAergic
- hybrid
- glutamatergic
- specialized
- neuroendocrine
- glia

**germ**
- spermatogonium
- spermatocyte
- spermatid

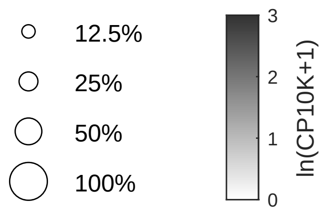

**Thyroid stimulating hormone (TSH)**
- endothelial

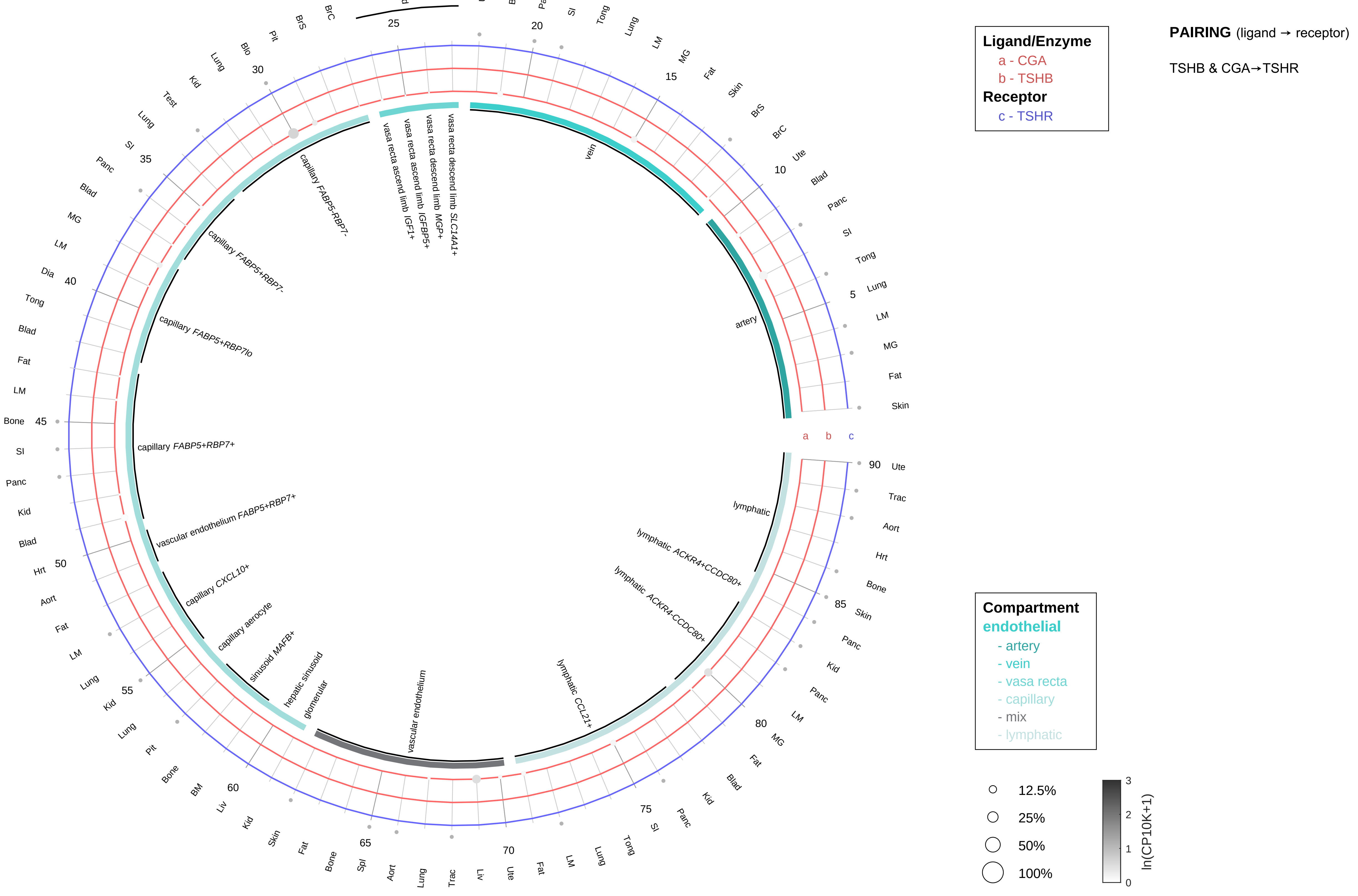

| Abbrev. | Tissue |
|---|---|
| Aort | Aorta |
| Blad | Bladder |
| Blo | Blood |
| Bone | Bone |
| BM | Bone marrow |
| BrC | Brain cortex |
| BrS | Brainstem |
| Col | Colon |
| Dia | Diaphragm |
| Ret | Eye retina |
| Fat | Fat |
| Hrt | Heart |
| Pit | Hypothalamus/Pituitary |
| Kid | Kidney |
| LM | Limb muscle |
| Liv | Liver |
| Lung | Lung |
| MG | Mammary gland |
| Panc | Pancreas |
| Skin | Skin |
| SI | Small intestine |
| Spl | Spleen |
| Test | Testes |
| Thym | Thymus |
| Tong | Tongue |
| Trac | Trachea |
| Ute | Uterus |

**Thyroid stimulating hormone (TSH)**
- stromal

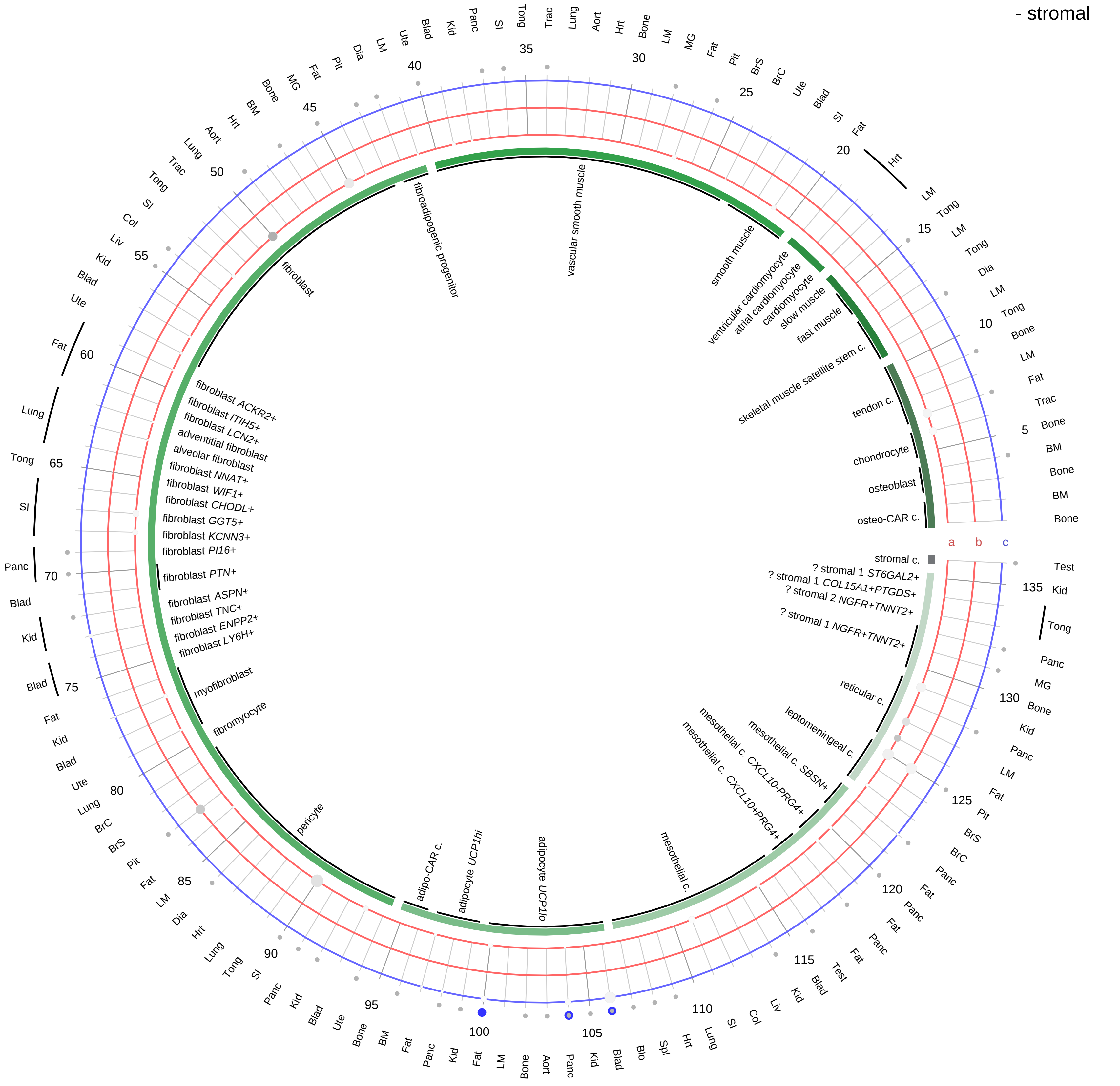

**Ligand/Enzyme**
a - CGA
b - TSHB
**Receptor**
c - TSHR

**PAIRING** (ligand → receptor)

TSHB & CGA→TSHR

| Abbrev. | Tissue |
|---|---|
| Aort | Aorta |
| Blad | Bladder |
| Blo | Blood |
| Bone | Bone |
| BM | Bone marrow |
| BrC | Brain cortex |
| BrS | Brainstem |
| Col | Colon |
| Dia | Diaphragm |
| Ret | Eye retina |
| Fat | Fat |
| Hrt | Heart |
| Pit | Hypothalamus/Pituitary |
| Kid | Kidney |
| LM | Limb muscle |
| Liv | Liver |
| Lung | Lung |
| MG | Mammary gland |
| Panc | Pancreas |
| Skin | Skin |
| SI | Small intestine |
| Spl | Spleen |
| Test | Testes |
| Thym | Thymus |
| Tong | Tongue |
| Trac | Trachea |
| Ute | Uterus |

**Compartment**
**stromal**
- bone
- skeletal muscle
- cardiac muscle
- smooth muscle
- fibroblast
- adipocyte
- mesothelial
- other
- mix

12.5%
25%
50%
100%

ln(CP10K+1): 0, 1, 2, 3

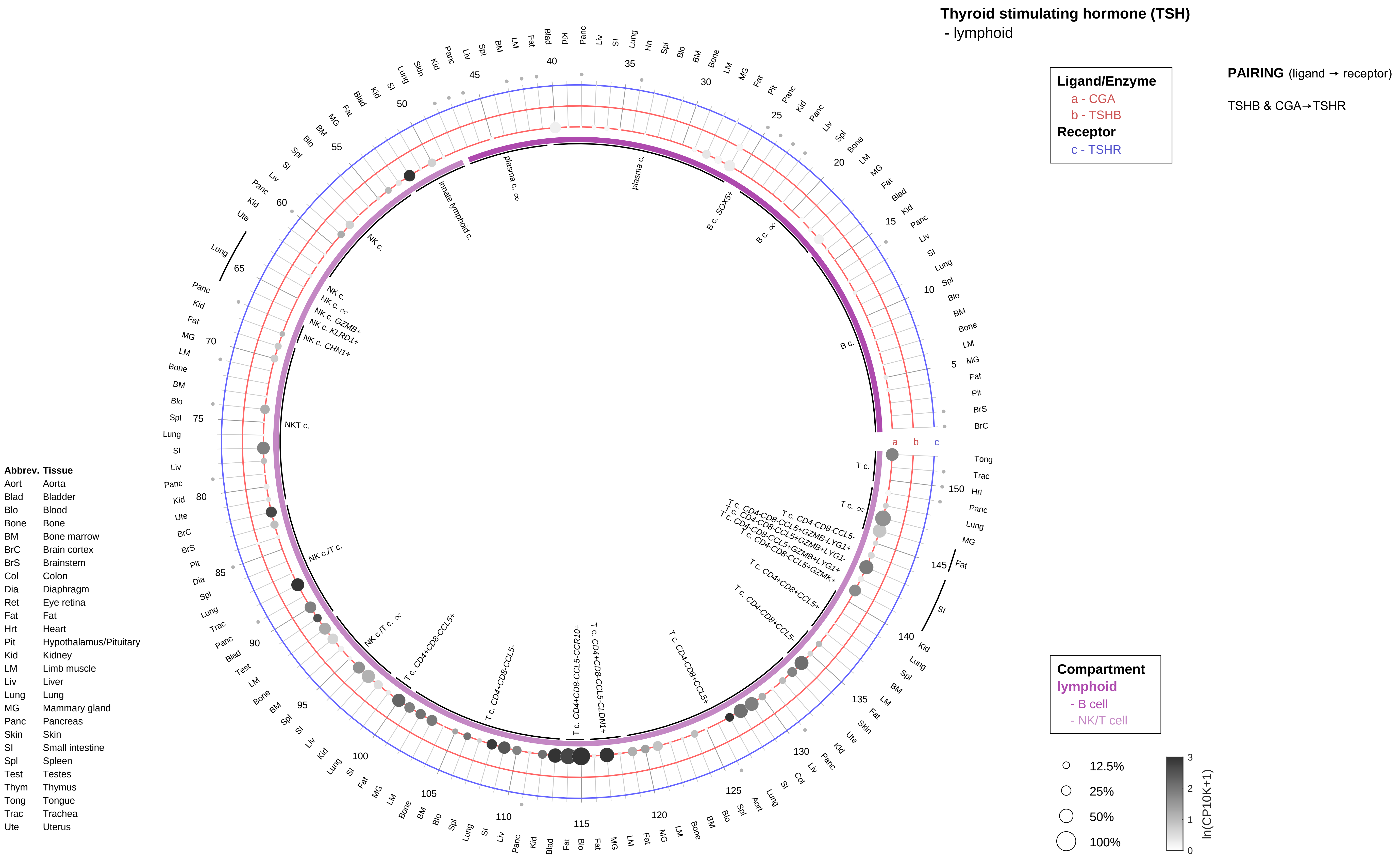

Thyroid stimulating hormone (TSH)
- lymphoid
Ligand/Enzyme
a - CGA
b - TSHB
Receptor
c - TSHR
PAIRING (ligand → receptor)
TSHB & CGA→TSHR
Compartment
lymphoid
- B cell
- NK/T cell
12.5%
25%
50%
100%
ln(CP10K+1)
Abbrev. Tissue
Aort Aorta
Blad Bladder
Blo Blood
Bone Bone
BM Bone marrow
BrC Brain cortex
BrS Brainstem
Col Colon
Dia Diaphragm
Ret Eye retina
Fat Fat
Hrt Heart
Pit Hypothalamus/Pituitary
Kid Kidney
LM Limb muscle
Liv Liver
Lung Lung
MG Mammary gland
Panc Pancreas
Skin Skin
SI Small intestine
Spl Spleen
Test Testes
Thym Thymus
Tong Tongue
Trac Trachea
Ute Uterus

**Thyroid stimulating hormone (TSH)**
- myeloid

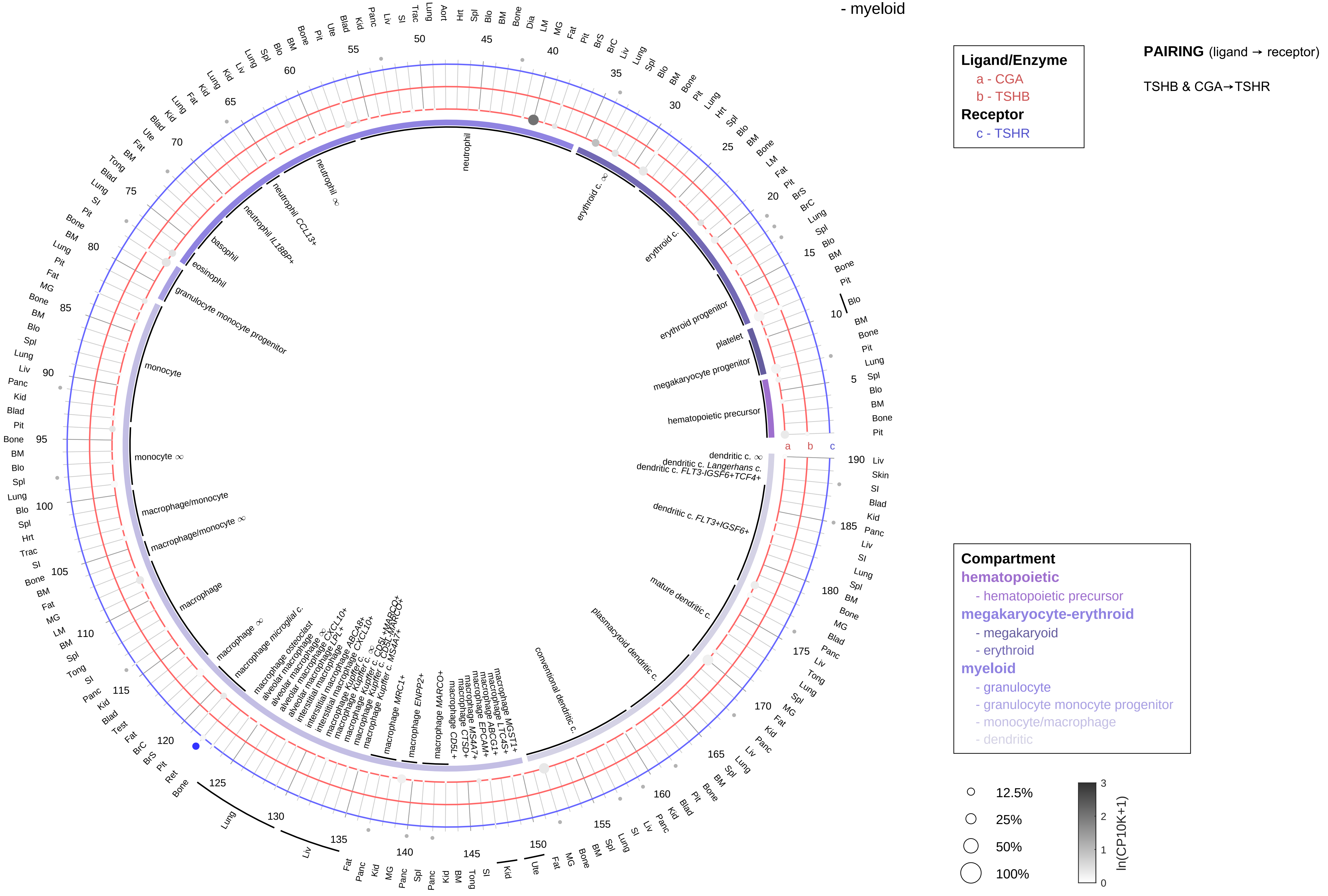

| Abbrev. | Tissue |
|---|---|
| Aort | Aorta |
| Blad | Bladder |
| Blo | Blood |
| Bone | Bone |
| BM | Bone marrow |
| BrC | Brain cortex |
| BrS | Brainstem |
| Col | Colon |
| Dia | Diaphragm |
| Ret | Eye retina |
| Fat | Fat |
| Hrt | Heart |
| Pit | Hypothalamus/Pituitary |
| Kid | Kidney |
| LM | Limb muscle |
| Liv | Liver |
| Lung | Lung |
| MG | Mammary gland |
| Panc | Pancreas |
| Skin | Skin |
| SI | Small intestine |
| Spl | Spleen |
| Test | Testes |
| Thym | Thymus |
| Tong | Tongue |
| Trac | Trachea |
| Ute | Uterus |

**Thyrotropin-releasing hormone (TRH)**
- epithelial/neural/germ

**PAIRING** (ligand → receptor)

TRH & CPE & PCSK2→TRHR

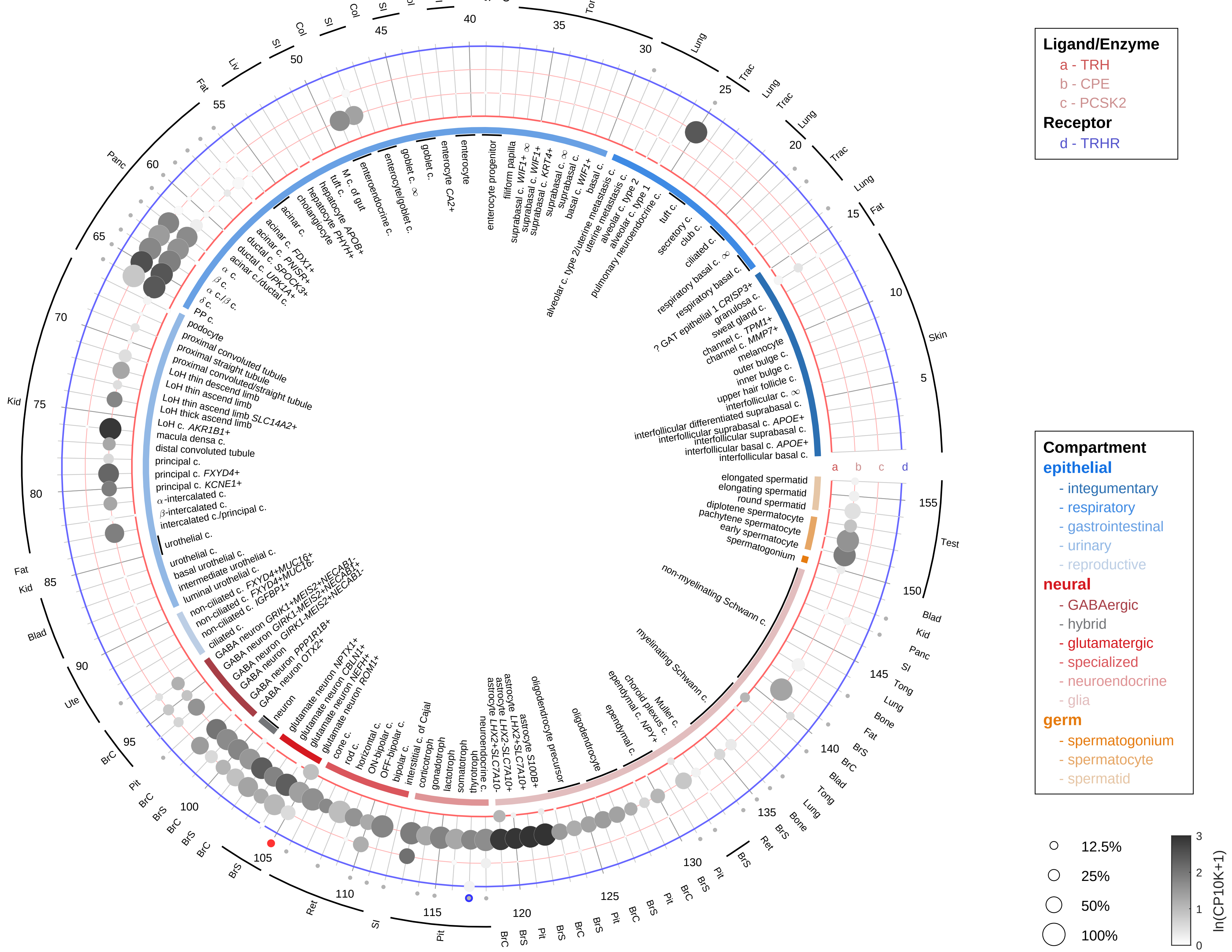

| Abbrev. | Tissue |
|---|---|
| Aort | Aorta |
| Blad | Bladder |
| Blo | Blood |
| Bone | Bone |
| BM | Bone marrow |
| BrC | Brain cortex |
| BrS | Brainstem |
| Col | Colon |
| Dia | Diaphragm |
| Ret | Eye retina |
| Fat | Fat |
| Hrt | Heart |
| Pit | Hypothalamus/Pituitary |
| Kid | Kidney |
| LM | Limb muscle |
| Liv | Liver |
| Lung | Lung |
| MG | Mammary gland |
| Panc | Pancreas |
| Skin | Skin |
| SI | Small intestine |
| Spl | Spleen |
| Test | Testes |
| Thym | Thymus |
| Tong | Tongue |
| Trac | Trachea |
| Ute | Uterus |

**Thyrotropin-releasing hormone (TRH)**
- endothelial

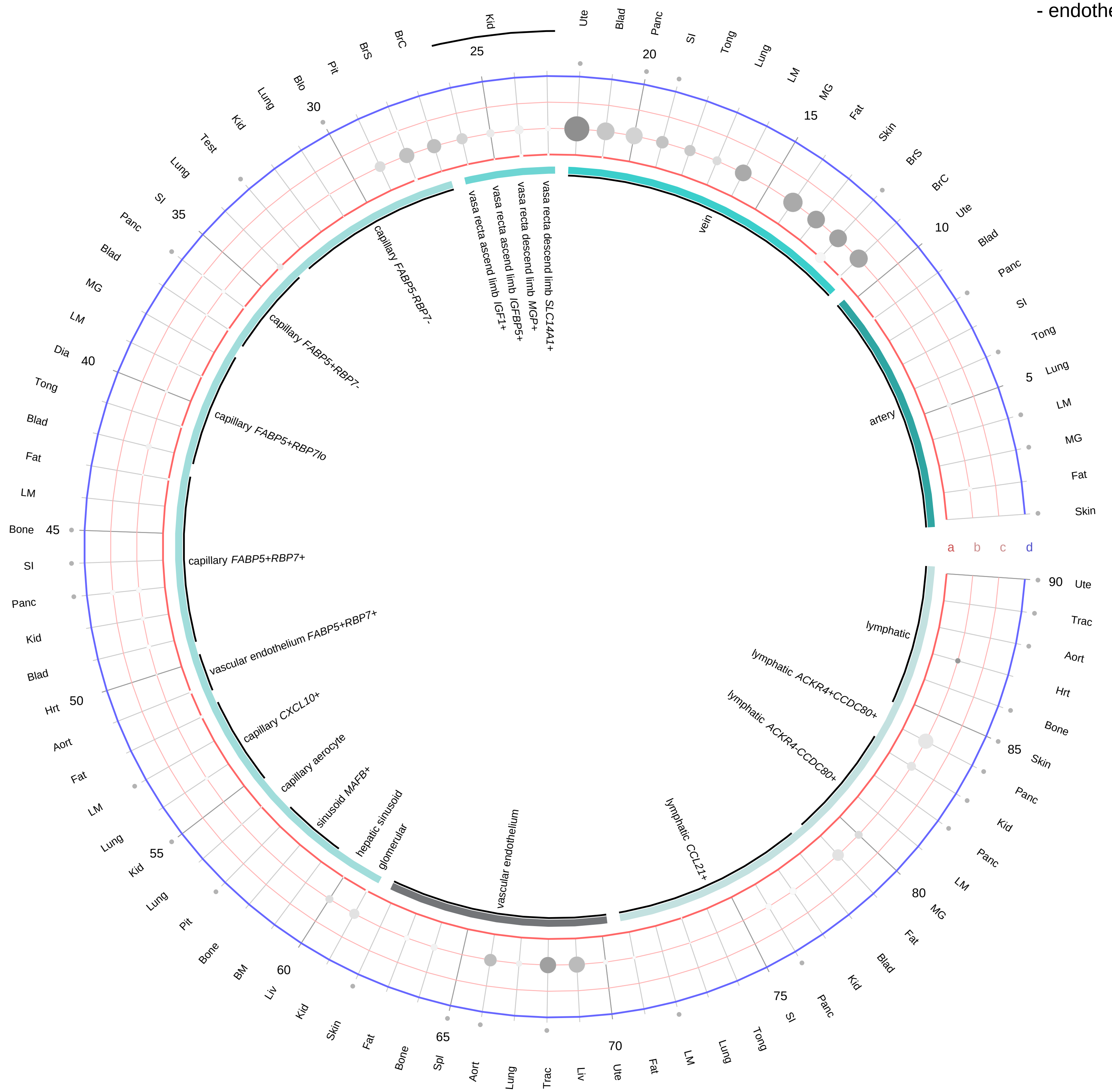

**Ligand/Enzyme**
a - TRH
b - CPE
c - PCSK2
**Receptor**
d - TRHR

**PAIRING** (ligand → receptor)

TRH & CPE & PCSK2→TRHR

**Compartment**
**endothelial**
- artery
- vein
- vasa recta
- capillary
- mix
- lymphatic

12.5%
25%
50%
100%

ln(CP10K+1): 0, 1, 2, 3

| Abbrev. | Tissue |
|---|---|
| Aort | Aorta |
| Blad | Bladder |
| Blo | Blood |
| Bone | Bone |
| BM | Bone marrow |
| BrC | Brain cortex |
| BrS | Brainstem |
| Col | Colon |
| Dia | Diaphragm |
| Ret | Eye retina |
| Fat | Fat |
| Hrt | Heart |
| Pit | Hypothalamus/Pituitary |
| Kid | Kidney |
| LM | Limb muscle |
| Liv | Liver |
| Lung | Lung |
| MG | Mammary gland |
| Panc | Pancreas |
| Skin | Skin |
| SI | Small intestine |
| Spl | Spleen |
| Test | Testes |
| Thym | Thymus |
| Tong | Tongue |
| Trac | Trachea |
| Ute | Uterus |

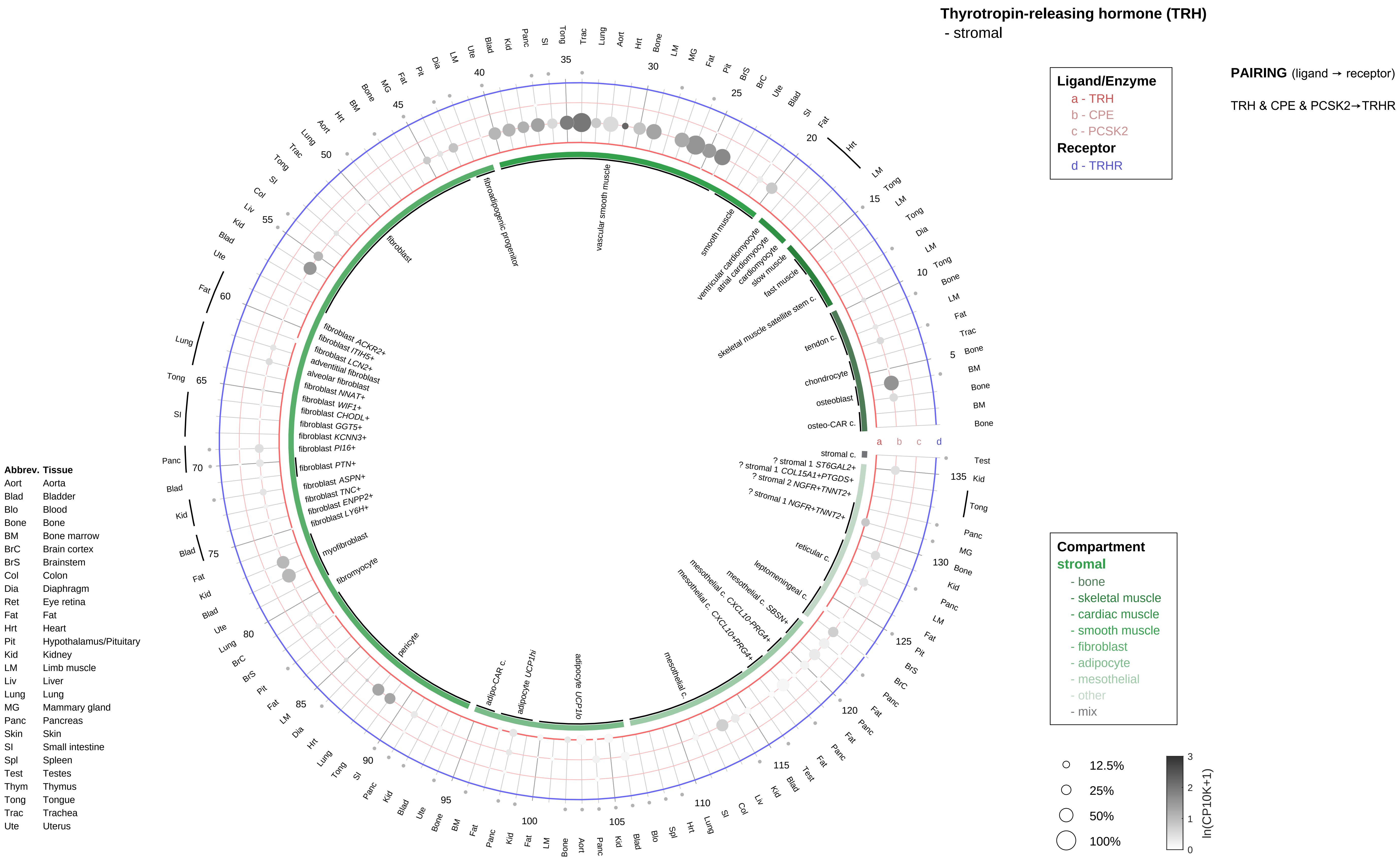

Thyrotropin-releasing hormone (TRH)
- stromal
Ligand/Enzyme
a - TRH
b - CPE
c - PCSK2
Receptor
d - TRHR
PAIRING (ligand → receptor)
TRH & CPE & PCSK2→TRHR
Compartment
stromal
- bone
- skeletal muscle
- cardiac muscle
- smooth muscle
- fibroblast
- adipocyte
- mesothelial
- other
- mix
12.5%
25%
50%
100%
ln(CP10K+1)
Abbrev. Tissue
Aort Aorta
Blad Bladder
Blo Blood
Bone Bone
BM Bone marrow
BrC Brain cortex
BrS Brainstem
Col Colon
Dia Diaphragm
Ret Eye retina
Fat Fat
Hrt Heart
Pit Hypothalamus/Pituitary
Kid Kidney
LM Limb muscle
Liv Liver
Lung Lung
MG Mammary gland
Panc Pancreas
Skin Skin
SI Small intestine
Spl Spleen
Test Testes
Thym Thymus
Tong Tongue
Trac Trachea
Ute Uterus
fibroadipogenic progenitor
vascular smooth muscle
smooth muscle
ventricular cardiomyocyte
atrial cardiomyocyte
cardiomyocyte
slow muscle
fast muscle
skeletal muscle satellite stem c.
tendon c.
chondrocyte
osteoblast
osteo-CAR c.
stromal c.
? stromal 1 ST6GAL2+
? stromal 1 COL15A1+PTGDS+
? stromal 2 NGFR+TNNT2+
? stromal 1 NGFR+TNNT2+
reticular c.
leptomeningeal c.
mesothelial c. SBSN+
mesothelial c. CXCL10-PRG4+
mesothelial c. CXCL10+PRG4+
mesothelial c.
adipocyte UCP1lo
adipocyte UCP1hi
adipo-CAR c.
pericyte
fibromyocyte
myofibroblast
fibroblast LY6H+
fibroblast ENPP2+
fibroblast TNC+
fibroblast ASPN+
fibroblast PTN+
fibroblast PI16+
fibroblast KCNN3+
fibroblast GGT5+
fibroblast CHODL+
fibroblast WIF1+
fibroblast NNAT+
alveolar fibroblast
adventitial fibroblast
fibroblast LCN2+
fibroblast ITIH5+
fibroblast ACKR2+
fibroblast

**Thyrotropin-releasing hormone (TRH)**
- lymphoid

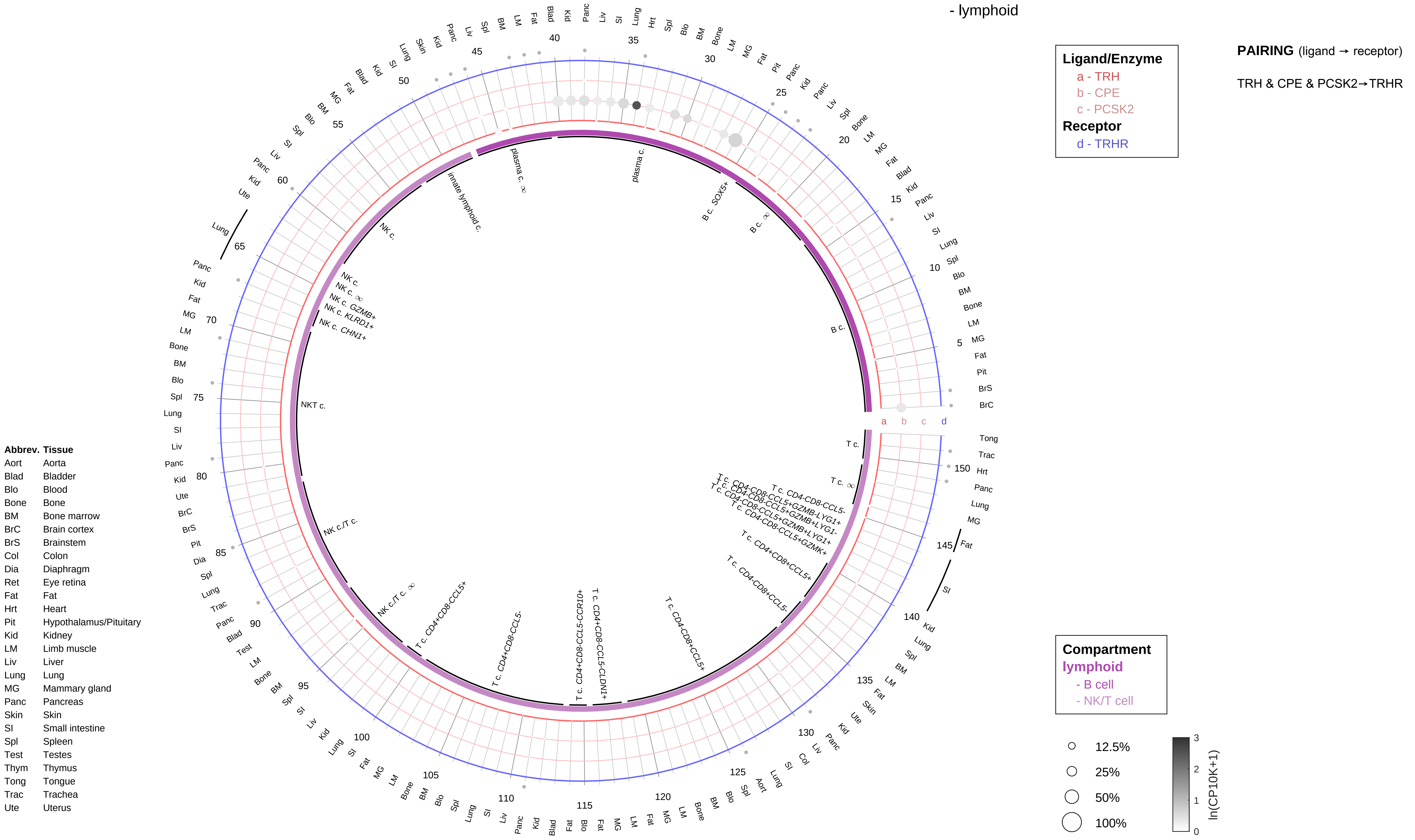

**Thyrotropin-releasing hormone (TRH)**
- myeloid

**PAIRING** (ligand → receptor)

TRH & CPE & PCSK2→TRHR

| **Ligand/Enzyme** |
|---|
| a - TRH |
| b - CPE |
| c - PCSK2 |
| **Receptor** |
| d - TRHR |

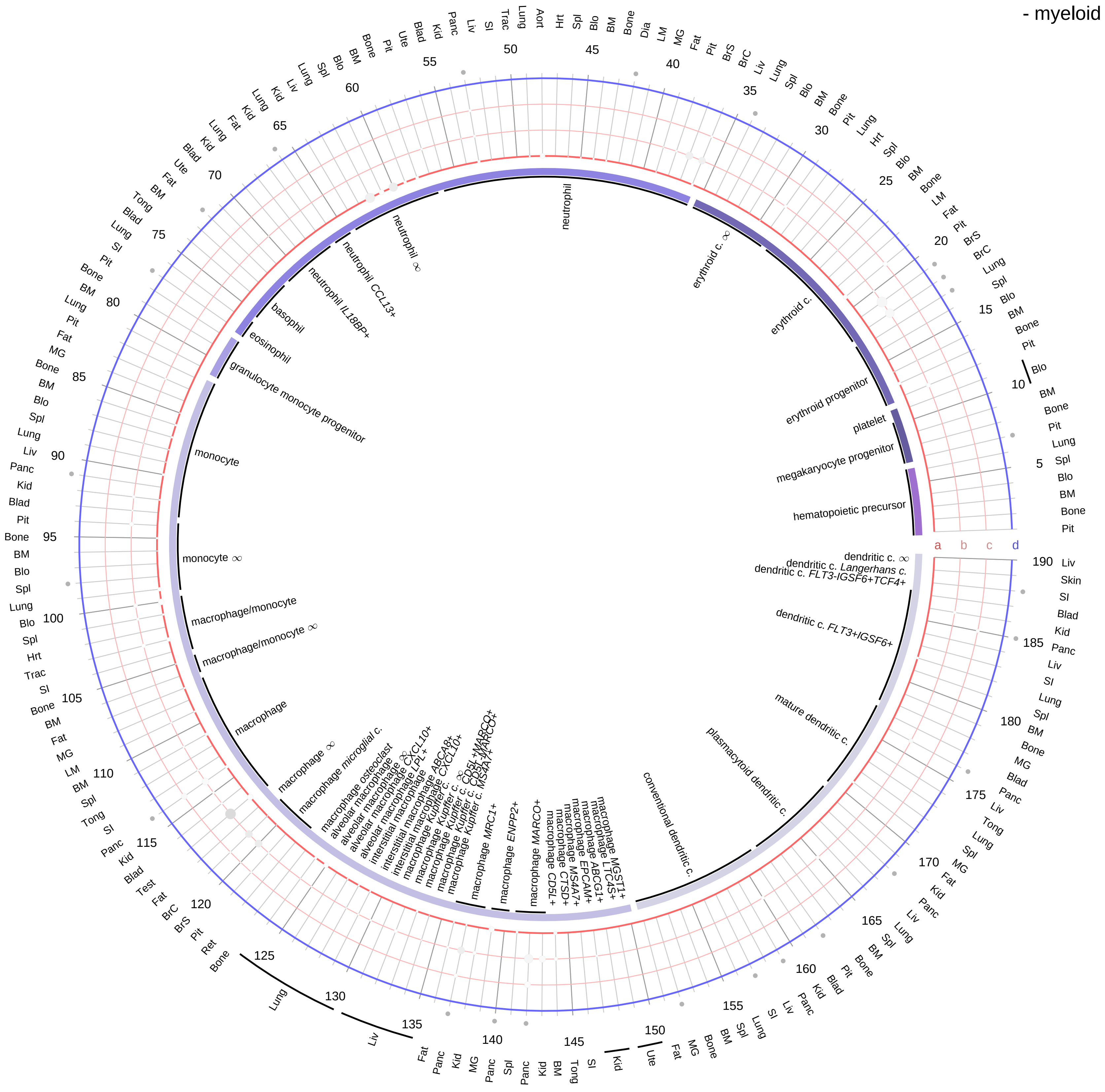

| **Abbrev.** | **Tissue** |
|---|---|
| Aort | Aorta |
| Blad | Bladder |
| Blo | Blood |
| Bone | Bone |
| BM | Bone marrow |
| BrC | Brain cortex |
| BrS | Brainstem |
| Col | Colon |
| Dia | Diaphragm |
| Ret | Eye retina |
| Fat | Fat |
| Hrt | Heart |
| Pit | Hypothalamus/Pituitary |
| Kid | Kidney |
| LM | Limb muscle |
| Liv | Liver |
| Lung | Lung |
| MG | Mammary gland |
| Panc | Pancreas |
| Skin | Skin |
| SI | Small intestine |
| Spl | Spleen |
| Test | Testes |
| Thym | Thymus |
| Tong | Tongue |
| Trac | Trachea |
| Ute | Uterus |

**Compartment**
**hematopoietic**
- hematopoietic precursor
**megakaryocyte-erythroid**
- megakaryoid
- erythroid
**myeloid**
- granulocyte
- granulocyte monocyte progenitor
- monocyte/macrophage
- dendritic

12.5%
25%
50%
100%

ln(CP10K+1): 0, 1, 2, 3

**Thyroid hormone**
- epithelial/neural/germ

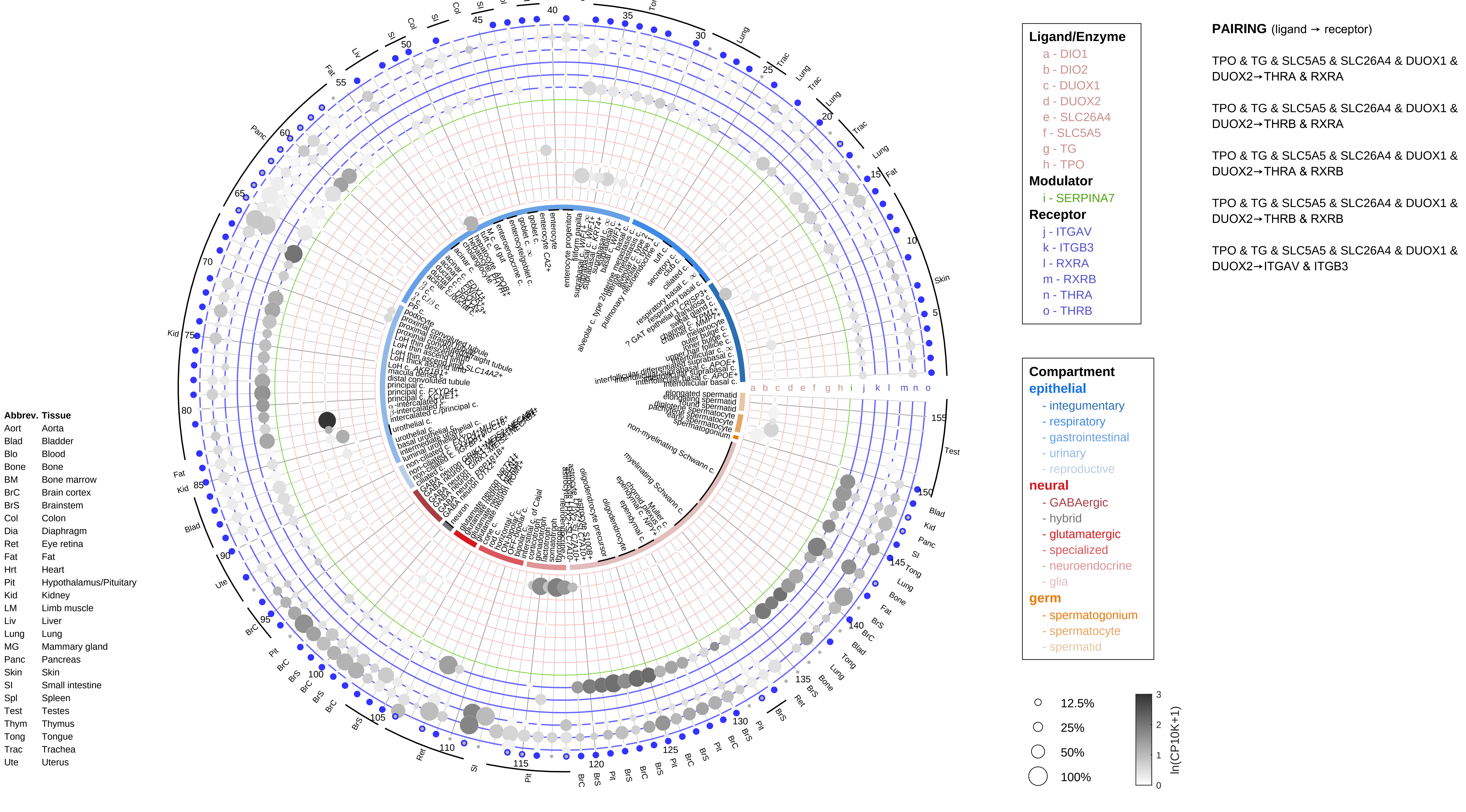

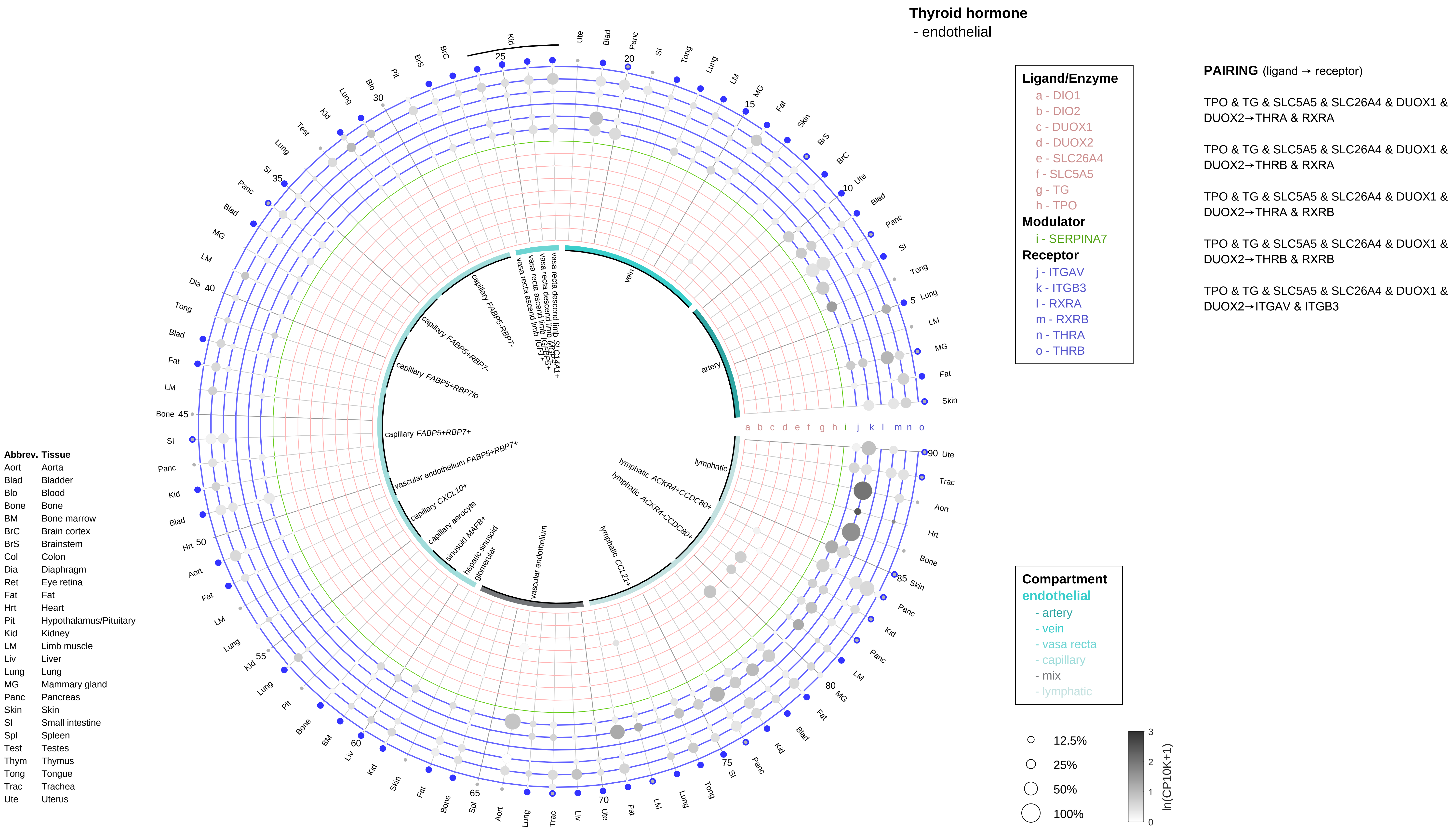
Thyroid hormone
- endothelial
Ligand/Enzyme
a - DIO1
b - DIO2
c - DUOX1
d - DUOX2
e - SLC26A4
f - SLC5A5
g - TG
h - TPO
Modulator
i - SERPINA7
Receptor
j - ITGAV
k - ITGB3
l - RXRA
m - RXRB
n - THRA
o - THRB
PAIRING (ligand → receptor)
TPO & TG & SLC5A5 & SLC26A4 & DUOX1 & DUOX2→THRA & RXRA
TPO & TG & SLC5A5 & SLC26A4 & DUOX1 & DUOX2→THRB & RXRA
TPO & TG & SLC5A5 & SLC26A4 & DUOX1 & DUOX2→THRA & RXRB
TPO & TG & SLC5A5 & SLC26A4 & DUOX1 & DUOX2→THRB & RXRB
TPO & TG & SLC5A5 & SLC26A4 & DUOX1 & DUOX2→ITGAV & ITGB3
Compartment
endothelial
- artery
- vein
- vasa recta
- capillary
- mix
- lymphatic
12.5%
25%
50%
100%
ln(CP10K+1)
Abbrev. Tissue
Aort Aorta
Blad Bladder
Blo Blood
Bone Bone
BM Bone marrow
BrC Brain cortex
BrS Brainstem
Col Colon
Dia Diaphragm
Ret Eye retina
Fat Fat
Hrt Heart
Pit Hypothalamus/Pituitary
Kid Kidney
LM Limb muscle
Liv Liver
Lung Lung
MG Mammary gland
Panc Pancreas
Skin Skin
SI Small intestine
Spl Spleen
Test Testes
Thym Thymus
Tong Tongue
Trac Trachea
Ute Uterus
vein
artery
lymphatic
lymphatic ACKR4+CCDC80+
lymphatic ACKR4-CCDC80+
lymphatic CCL21+
vascular endothelium
glomerular
hepatic sinusoid
sinusoid MAFB+
capillary aerocyte
capillary CXCL10+
vascular endothelium FABP5+RBP7+
capillary FABP5+RBP7+
capillary FABP5+RBP7lo
capillary FABP5+RBP7-
capillary FABP5-RBP7-
vasa recta ascend limb IGFL1+
vasa recta ascend limb
vasa recta descend limb MGP+
vasa recta descend limb SLC14A1+

**Thyroid hormone**
- stromal

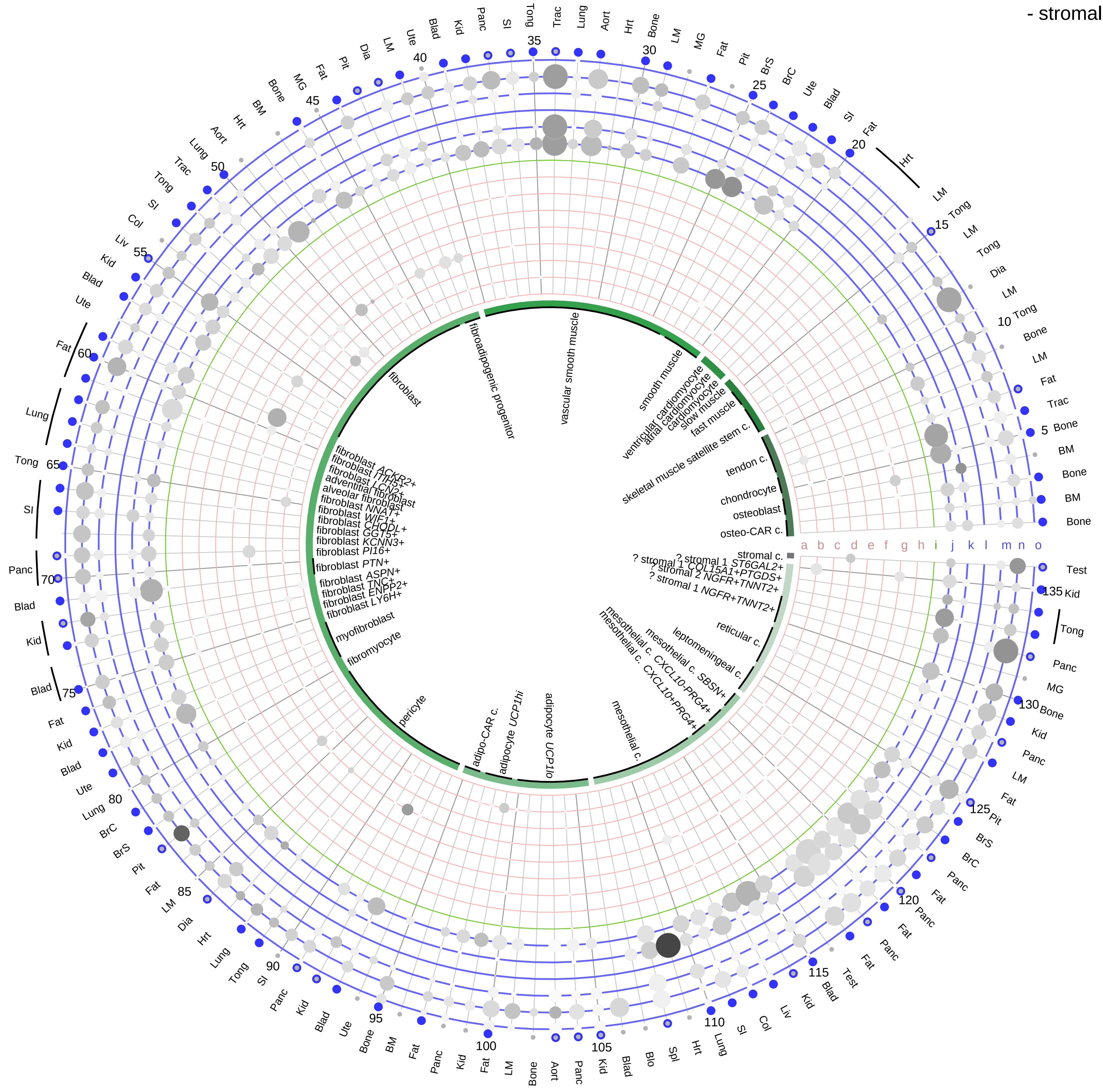

**Ligand/Enzyme**
- a - DIO1
- b - DIO2
- c - DUOX1
- d - DUOX2
- e - SLC26A4
- f - SLC5A5
- g - TG
- h - TPO

**Modulator**
- i - SERPINA7

**Receptor**
- j - ITGAV
- k - ITGB3
- l - RXRA
- m - RXRB
- n - THRA
- o - THRB

**PAIRING** (ligand → receptor)

TPO & TG & SLC5A5 & SLC26A4 & DUOX1 & DUOX2→THRA & RXRA

TPO & TG & SLC5A5 & SLC26A4 & DUOX1 & DUOX2→THRB & RXRA

TPO & TG & SLC5A5 & SLC26A4 & DUOX1 & DUOX2→THRA & RXRB

TPO & TG & SLC5A5 & SLC26A4 & DUOX1 & DUOX2→THRB & RXRB

TPO & TG & SLC5A5 & SLC26A4 & DUOX1 & DUOX2→ITGAV & ITGB3

| Abbrev. | Tissue |
|---|---|
| Aort | Aorta |
| Blad | Bladder |
| Blo | Blood |
| Bone | Bone |
| BM | Bone marrow |
| BrC | Brain cortex |
| BrS | Brainstem |
| Col | Colon |
| Dia | Diaphragm |
| Ret | Eye retina |
| Fat | Fat |
| Hrt | Heart |
| Pit | Hypothalamus/Pituitary |
| Kid | Kidney |
| LM | Limb muscle |
| Liv | Liver |
| Lung | Lung |
| MG | Mammary gland |
| Panc | Pancreas |
| Skin | Skin |
| SI | Small intestine |
| Spl | Spleen |
| Test | Testes |
| Thym | Thymus |
| Tong | Tongue |
| Trac | Trachea |
| Ute | Uterus |

**Compartment**
**stromal**
- bone
- skeletal muscle
- cardiac muscle
- smooth muscle
- fibroblast
- adipocyte
- mesothelial
- other
- mix

12.5%
25%
50%
100%

ln(CP10K+1): 0, 1, 2, 3

**Thyroid hormone**
- lymphoid

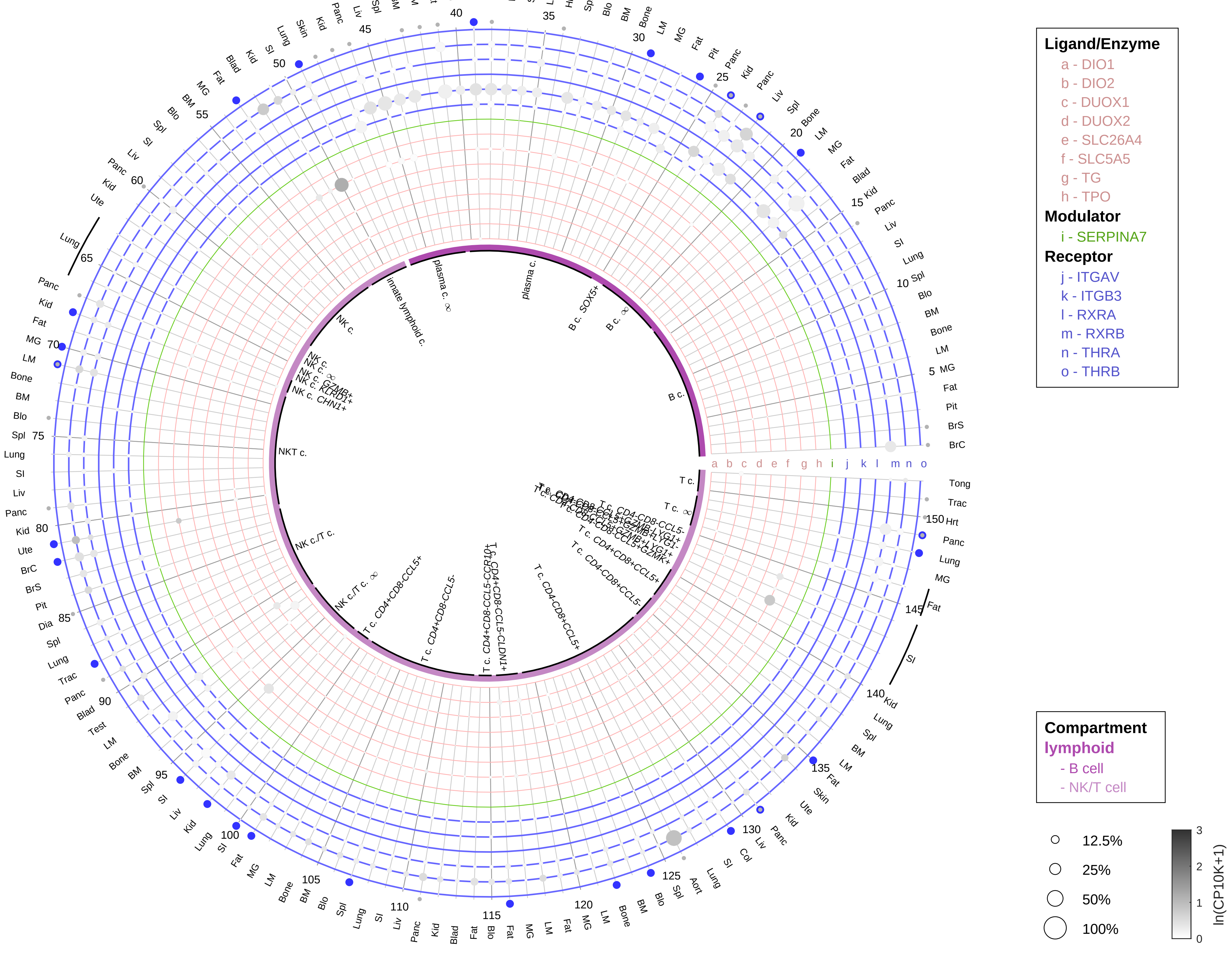

**PAIRING** (ligand → receptor)

TPO & TG & SLC5A5 & SLC26A4 & DUOX1 & DUOX2→THRA & RXRA

TPO & TG & SLC5A5 & SLC26A4 & DUOX1 & DUOX2→THRB & RXRA

TPO & TG & SLC5A5 & SLC26A4 & DUOX1 & DUOX2→THRA & RXRB

TPO & TG & SLC5A5 & SLC26A4 & DUOX1 & DUOX2→THRB & RXRB

TPO & TG & SLC5A5 & SLC26A4 & DUOX1 & DUOX2→ITGAV & ITGB3

| Abbrev. | Tissue |
|---|---|
| Aort | Aorta |
| Blad | Bladder |
| Blo | Blood |
| Bone | Bone |
| BM | Bone marrow |
| BrC | Brain cortex |
| BrS | Brainstem |
| Col | Colon |
| Dia | Diaphragm |
| Ret | Eye retina |
| Fat | Fat |
| Hrt | Heart |
| Pit | Hypothalamus/Pituitary |
| Kid | Kidney |
| LM | Limb muscle |
| Liv | Liver |
| Lung | Lung |
| MG | Mammary gland |
| Panc | Pancreas |
| Skin | Skin |
| SI | Small intestine |
| Spl | Spleen |
| Test | Testes |
| Thym | Thymus |
| Tong | Tongue |
| Trac | Trachea |
| Ute | Uterus |

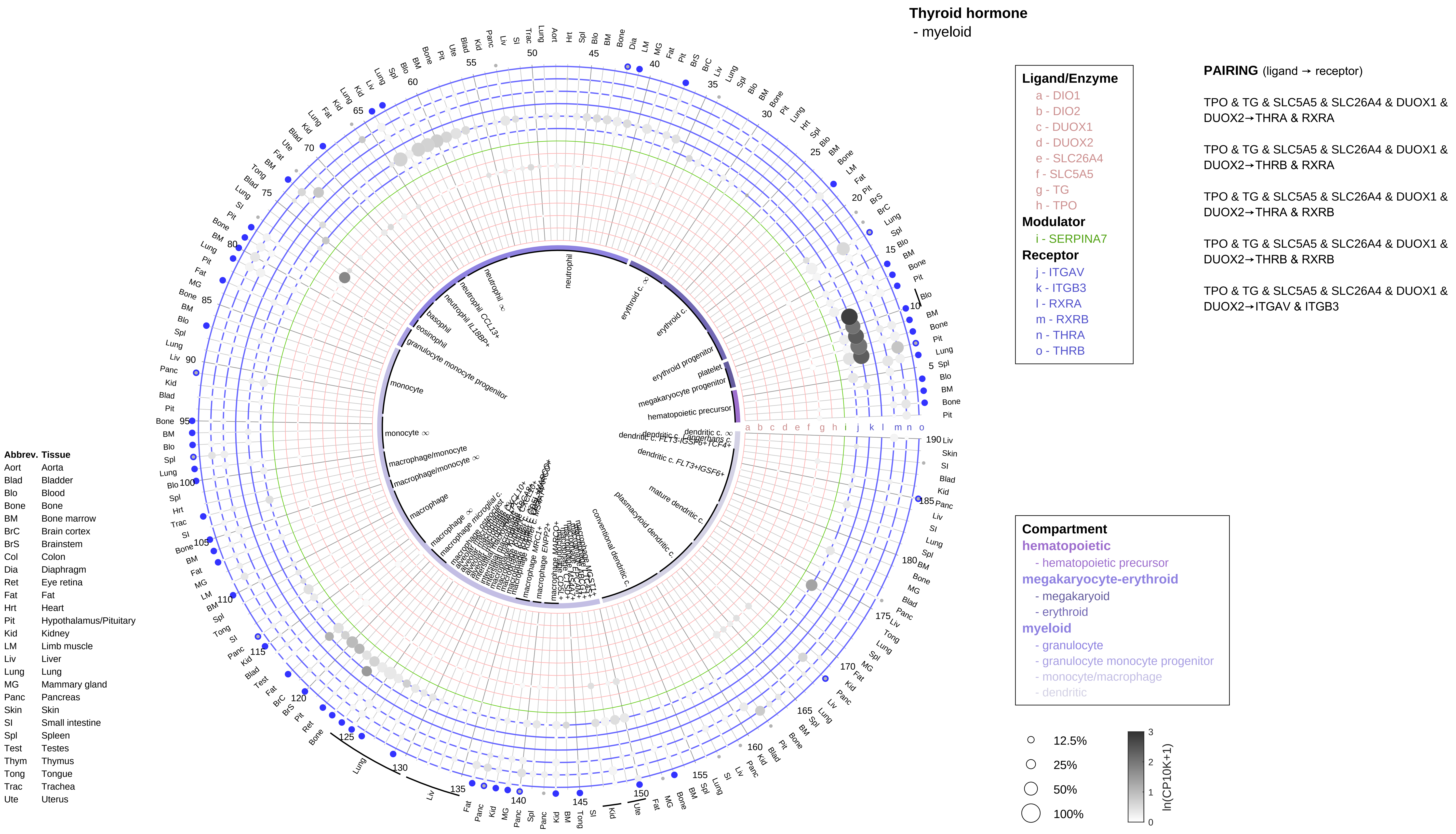
Thyroid hormone
- myeloid
Ligand/Enzyme
a - DIO1
b - DIO2
c - DUOX1
d - DUOX2
e - SLC26A4
f - SLC5A5
g - TG
h - TPO
Modulator
i - SERPINA7
Receptor
j - ITGAV
k - ITGB3
l - RXRA
m - RXRB
n - THRA
o - THRB
PAIRING (ligand → receptor)
TPO & TG & SLC5A5 & SLC26A4 & DUOX1 & DUOX2→THRA & RXRA
TPO & TG & SLC5A5 & SLC26A4 & DUOX1 & DUOX2→THRB & RXRA
TPO & TG & SLC5A5 & SLC26A4 & DUOX1 & DUOX2→THRA & RXRB
TPO & TG & SLC5A5 & SLC26A4 & DUOX1 & DUOX2→THRB & RXRB
TPO & TG & SLC5A5 & SLC26A4 & DUOX1 & DUOX2→ITGAV & ITGB3
Compartment
hematopoietic
- hematopoietic precursor
megakaryocyte-erythroid
- megakaryoid
- erythroid
myeloid
- granulocyte
- granulocyte monocyte progenitor
- monocyte/macrophage
- dendritic
12.5%
25%
50%
100%
ln(CP10K+1)
0
1
2
3
Abbrev. Tissue
Aort Aorta
Blad Bladder
Blo Blood
Bone Bone
BM Bone marrow
BrC Brain cortex
BrS Brainstem
Col Colon
Dia Diaphragm
Ret Eye retina
Fat Fat
Hrt Heart
Pit Hypothalamus/Pituitary
Kid Kidney
LM Limb muscle
Liv Liver
Lung Lung
MG Mammary gland
Panc Pancreas
Skin Skin
SI Small intestine
Spl Spleen
Test Testes
Thym Thymus
Tong Tongue
Trac Trachea
Ute Uterus
neutrophil
erythroid c.
erythroid progenitor
platelet
megakaryocyte progenitor
hematopoietic precursor
basophil
eosinophil
granulocyte monocyte progenitor
monocyte
macrophage/monocyte
macrophage
conventional dendritic c.
plasmacytoid dendritic c.
mature dendritic c.
dendritic c. FLT3+IGSF6+
a b c d e f g h i j k l m n o

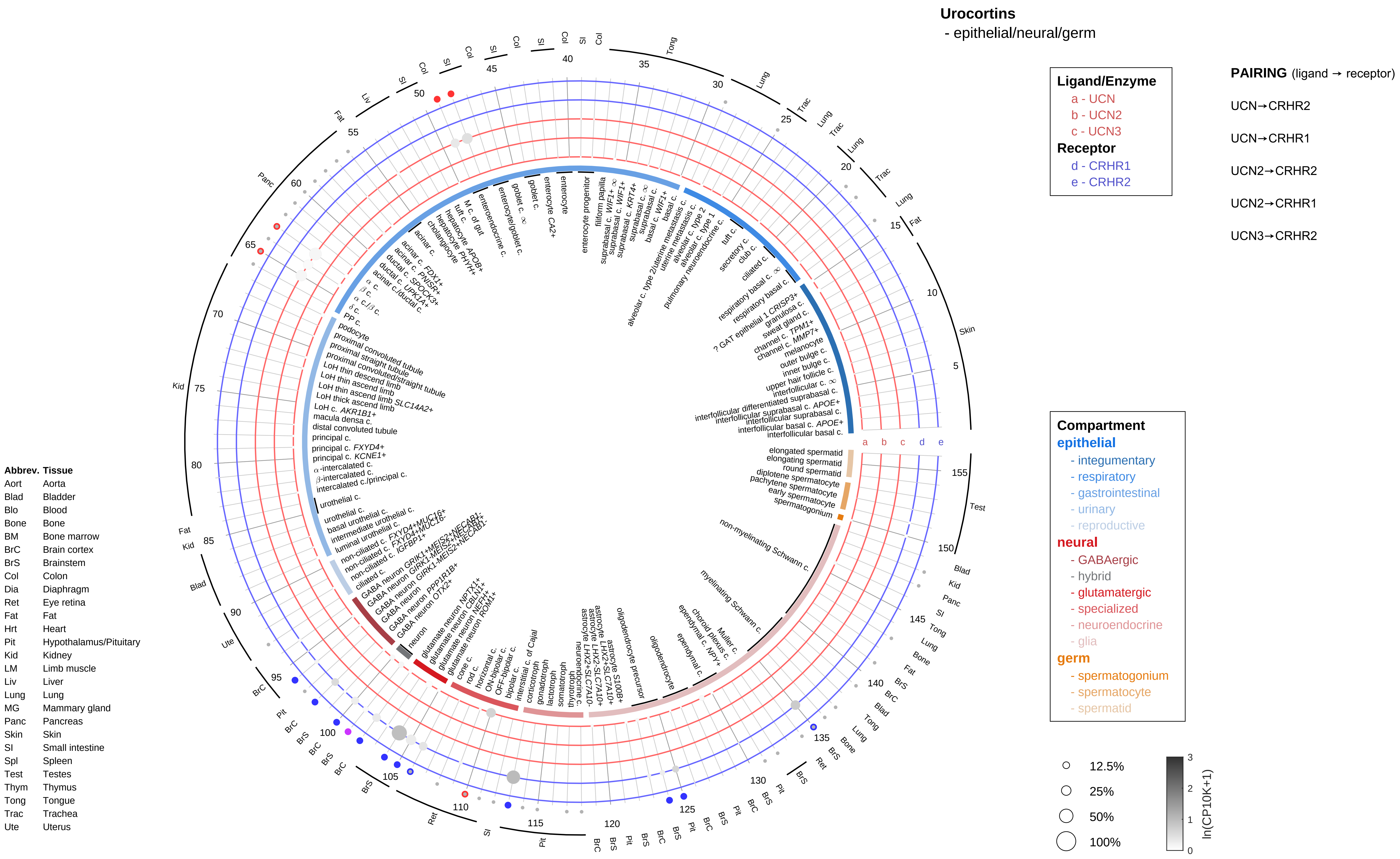

Urocortins
- epithelial/neural/germ
Ligand/Enzyme
a - UCN
b - UCN2
c - UCN3
Receptor
d - CRHR1
e - CRHR2
PAIRING (ligand → receptor)
UCN→CRHR2
UCN→CRHR1
UCN2→CRHR2
UCN2→CRHR1
UCN3→CRHR2
Compartment
epithelial
- integumentary
- respiratory
- gastrointestinal
- urinary
- reproductive
neural
- GABAergic
- hybrid
- glutamatergic
- specialized
- neuroendocrine
- glia
germ
- spermatogonium
- spermatocyte
- spermatid
12.5%
25%
50%
100%
ln(CP10K+1)
Abbrev. Tissue
Aort Aorta
Blad Bladder
Blo Blood
Bone Bone
BM Bone marrow
BrC Brain cortex
BrS Brainstem
Col Colon
Dia Diaphragm
Ret Eye retina
Fat Fat
Hrt Heart
Pit Hypothalamus/Pituitary
Kid Kidney
LM Limb muscle
Liv Liver
Lung Lung
MG Mammary gland
Panc Pancreas
Skin Skin
SI Small intestine
Spl Spleen
Test Testes
Thym Thymus
Tong Tongue
Trac Trachea
Ute Uterus

#### Urocortins
- endothelial

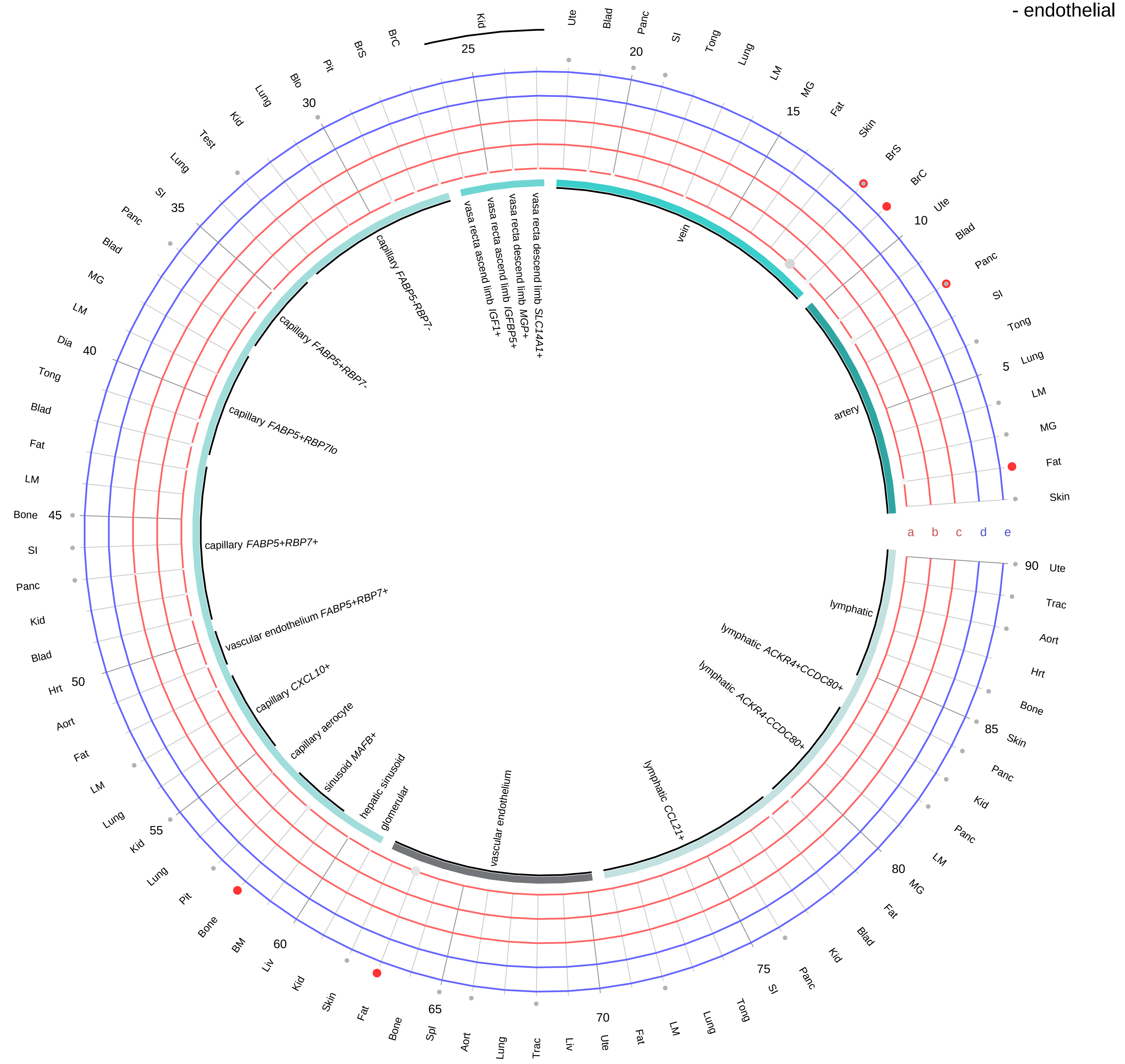

**Ligand/Enzyme**
- a - UCN
- b - UCN2
- c - UCN3

**Receptor**
- d - CRHR1
- e - CRHR2

**PAIRING** (ligand → receptor)

UCN→CRHR2

UCN→CRHR1

UCN2→CRHR2

UCN2→CRHR1

UCN3→CRHR2

**Compartment**

**endothelial**
- artery
- vein
- vasa recta
- capillary
- mix
- lymphatic

○ 12.5%
○ 25%
○ 50%
○ 100%

ln(CP10K+1): 0, 1, 2, 3

| Abbrev. | Tissue |
|---|---|
| Aort | Aorta |
| Blad | Bladder |
| Blo | Blood |
| Bone | Bone |
| BM | Bone marrow |
| BrC | Brain cortex |
| BrS | Brainstem |
| Col | Colon |
| Dia | Diaphragm |
| Ret | Eye retina |
| Fat | Fat |
| Hrt | Heart |
| Pit | Hypothalamus/Pituitary |
| Kid | Kidney |
| LM | Limb muscle |
| Liv | Liver |
| Lung | Lung |
| MG | Mammary gland |
| Panc | Pancreas |
| Skin | Skin |
| SI | Small intestine |
| Spl | Spleen |
| Test | Testes |
| Thym | Thymus |
| Tong | Tongue |
| Trac | Trachea |
| Ute | Uterus |

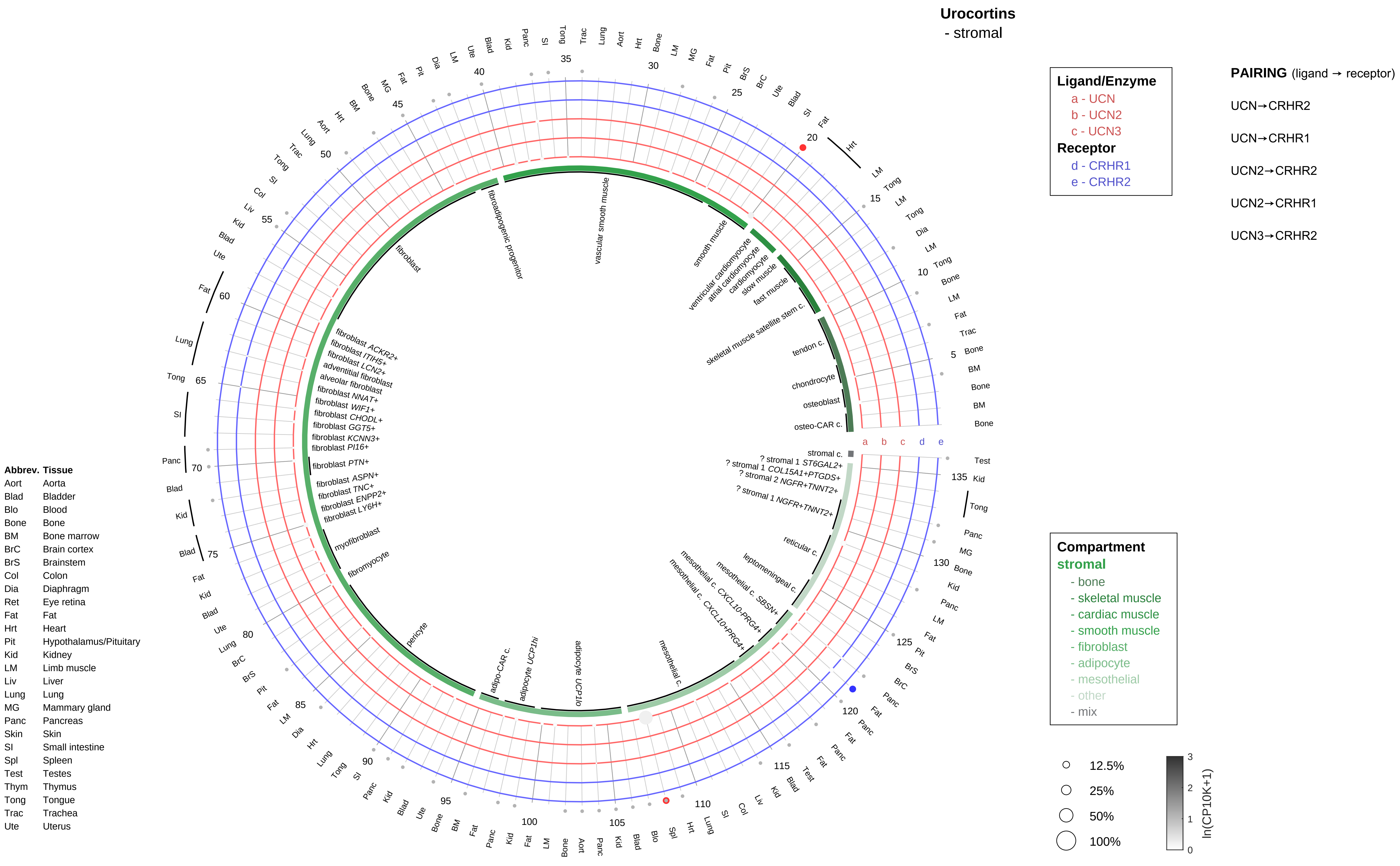
Urocortins
- stromal
Ligand/Enzyme
a - UCN
b - UCN2
c - UCN3
Receptor
d - CRHR1
e - CRHR2
PAIRING (ligand → receptor)
UCN→CRHR2
UCN→CRHR1
UCN2→CRHR2
UCN2→CRHR1
UCN3→CRHR2
Compartment
stromal
- bone
- skeletal muscle
- cardiac muscle
- smooth muscle
- fibroblast
- adipocyte
- mesothelial
- other
- mix
12.5%
25%
50%
100%
ln(CP10K+1)
Abbrev. Tissue
Aort Aorta
Blad Bladder
Blo Blood
Bone Bone
BM Bone marrow
BrC Brain cortex
BrS Brainstem
Col Colon
Dia Diaphragm
Ret Eye retina
Fat Fat
Hrt Heart
Pit Hypothalamus/Pituitary
Kid Kidney
LM Limb muscle
Liv Liver
Lung Lung
MG Mammary gland
Panc Pancreas
Skin Skin
SI Small intestine
Spl Spleen
Test Testes
Thym Thymus
Tong Tongue
Trac Trachea
Ute Uterus
vascular smooth muscle
smooth muscle
ventricular cardiomyocyte
atrial cardiomyocyte
cardiomyocyte
slow muscle
fast muscle
skeletal muscle satellite stem c.
tendon c.
chondrocyte
osteoblast
osteo-CAR c.
stromal c.
? stromal 1 ST6GAL2+
? stromal 1 COL15A1+PTGDS+
? stromal 2 NGFR+TNNT2+
? stromal 1 NGFR+TNNT2+
reticular c.
leptomeningeal c.
mesothelial c. SBSN+
mesothelial c. CXCL10-PRG4+
mesothelial c. CXCL10+PRG4+
mesothelial c.
adipocyte UCP1lo
adipocyte UCP1hi
adipo-CAR c.
pericyte
fibromyocyte
myofibroblast
fibroblast LY6H+
fibroblast ENPP2+
fibroblast TNC+
fibroblast ASPN+
fibroblast PTN+
fibroblast PI16+
fibroblast KCNN3+
fibroblast GGT5+
fibroblast CHODL+
fibroblast WIF1+
fibroblast NNAT+
alveolar fibroblast
adventitial fibroblast
fibroblast LCN2+
fibroblast ITIH5+
fibroblast ACKR2+
fibroblast
fibroadipogenic progenitor

**Urocortins**
- lymphoid

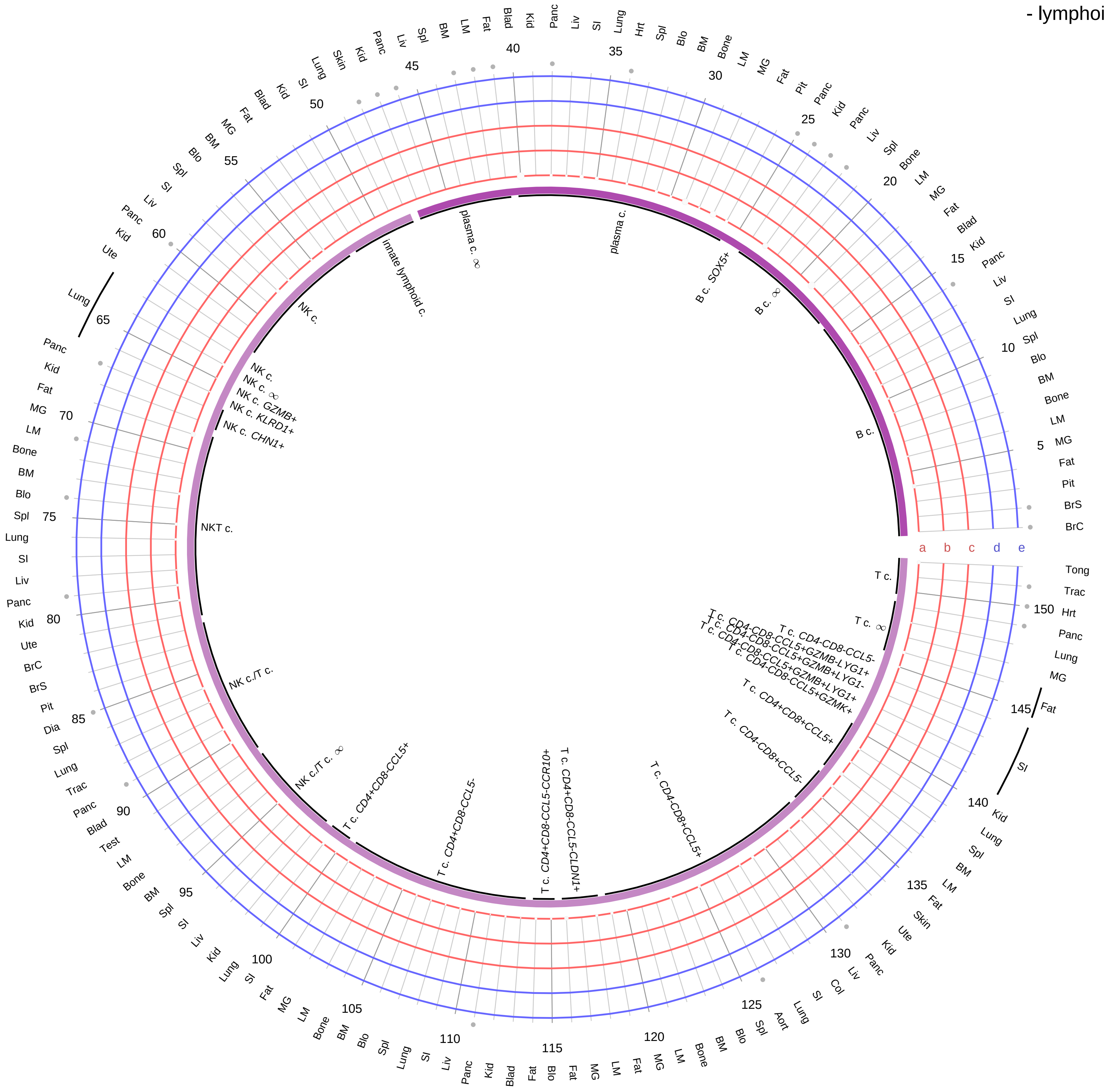

**Ligand/Enzyme**
- a - UCN
- b - UCN2
- c - UCN3

**Receptor**
- d - CRHR1
- e - CRHR2

**PAIRING** (ligand → receptor)

UCN→CRHR2

UCN→CRHR1

UCN2→CRHR2

UCN2→CRHR1

UCN3→CRHR2

| Abbrev. | Tissue |
|---|---|
| Aort | Aorta |
| Blad | Bladder |
| Blo | Blood |
| Bone | Bone |
| BM | Bone marrow |
| BrC | Brain cortex |
| BrS | Brainstem |
| Col | Colon |
| Dia | Diaphragm |
| Ret | Eye retina |
| Fat | Fat |
| Hrt | Heart |
| Pit | Hypothalamus/Pituitary |
| Kid | Kidney |
| LM | Limb muscle |
| Liv | Liver |
| Lung | Lung |
| MG | Mammary gland |
| Panc | Pancreas |
| Skin | Skin |
| SI | Small intestine |
| Spl | Spleen |
| Test | Testes |
| Thym | Thymus |
| Tong | Tongue |
| Trac | Trachea |
| Ute | Uterus |

**Compartment**
**lymphoid**
- B cell
- NK/T cell

12.5%
25%
50%
100%

ln(CP10K+1): 0, 1, 2, 3

**Urocortins**
- myeloid

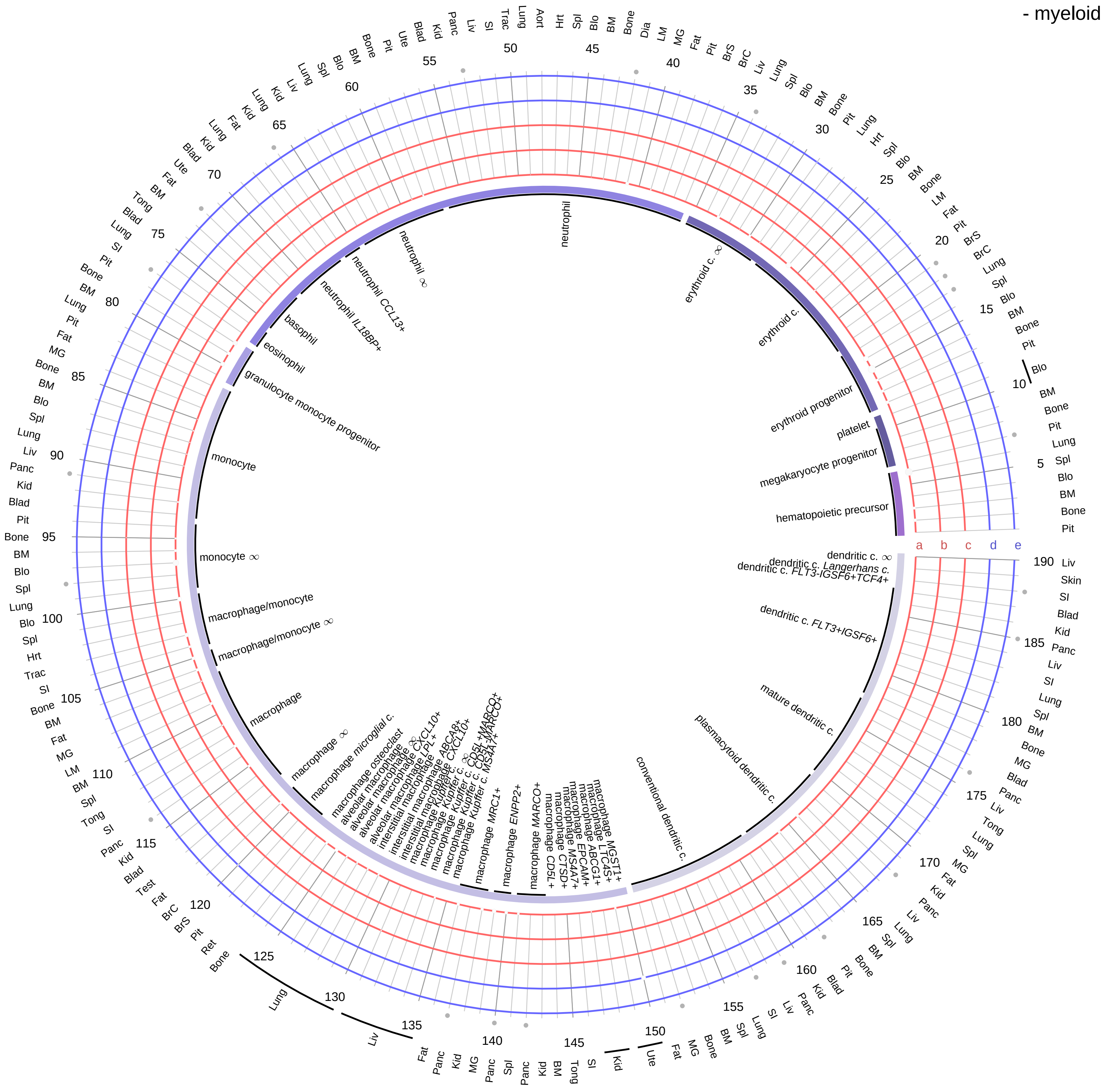

| Ligand/Enzyme |
|---|
| a - UCN |
| b - UCN2 |
| c - UCN3 |
| **Receptor** |
| d - CRHR1 |
| e - CRHR2 |

**PAIRING** (ligand → receptor)

UCN→CRHR2

UCN→CRHR1

UCN2→CRHR2

UCN2→CRHR1

UCN3→CRHR2

| Abbrev. | Tissue |
|---|---|
| Aort | Aorta |
| Blad | Bladder |
| Blo | Blood |
| Bone | Bone |
| BM | Bone marrow |
| BrC | Brain cortex |
| BrS | Brainstem |
| Col | Colon |
| Dia | Diaphragm |
| Ret | Eye retina |
| Fat | Fat |
| Hrt | Heart |
| Pit | Hypothalamus/Pituitary |
| Kid | Kidney |
| LM | Limb muscle |
| Liv | Liver |
| Lung | Lung |
| MG | Mammary gland |
| Panc | Pancreas |
| Skin | Skin |
| SI | Small intestine |
| Spl | Spleen |
| Test | Testes |
| Thym | Thymus |
| Tong | Tongue |
| Trac | Trachea |
| Ute | Uterus |

**Compartment**
**hematopoietic**
- hematopoietic precursor
**megakaryocyte-erythroid**
- megakaryoid
- erythroid
**myeloid**
- granulocyte
- granulocyte monocyte progenitor
- monocyte/macrophage
- dendritic

12.5%
25%
50%
100%

ln(CP10K+1): 0, 1, 2, 3

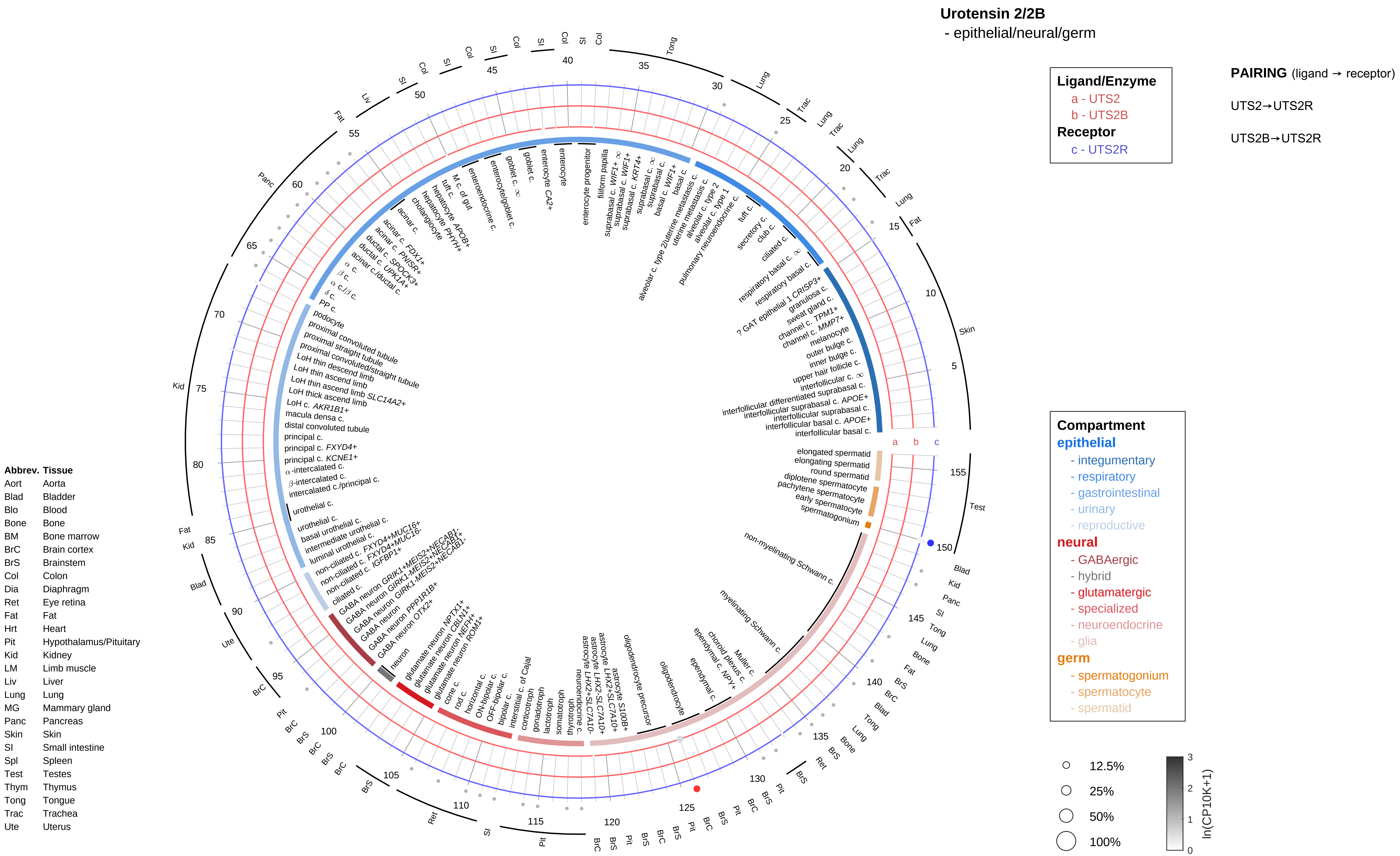
Urotensin 2/2B
- epithelial/neural/germ
Ligand/Enzyme
a - UTS2
b - UTS2B
Receptor
c - UTS2R
PAIRING (ligand → receptor)
UTS2→UTS2R
UTS2B→UTS2R
Compartment
epithelial
- integumentary
- respiratory
- gastrointestinal
- urinary
- reproductive
neural
- GABAergic
- hybrid
- glutamatergic
- specialized
- neuroendocrine
- glia
germ
- spermatogonium
- spermatocyte
- spermatid
12.5%
25%
50%
100%
ln(CP10K+1)
Abbrev. Tissue
Aort Aorta
Blad Bladder
Blo Blood
Bone Bone
BM Bone marrow
BrC Brain cortex
BrS Brainstem
Col Colon
Dia Diaphragm
Ret Eye retina
Fat Fat
Hrt Heart
Pit Hypothalamus/Pituitary
Kid Kidney
LM Limb muscle
Liv Liver
Lung Lung
MG Mammary gland
Panc Pancreas
Skin Skin
SI Small intestine
Spl Spleen
Test Testes
Thym Thymus
Tong Tongue
Trac Trachea
Ute Uterus

#### Urotensin 2/2B
- endothelial

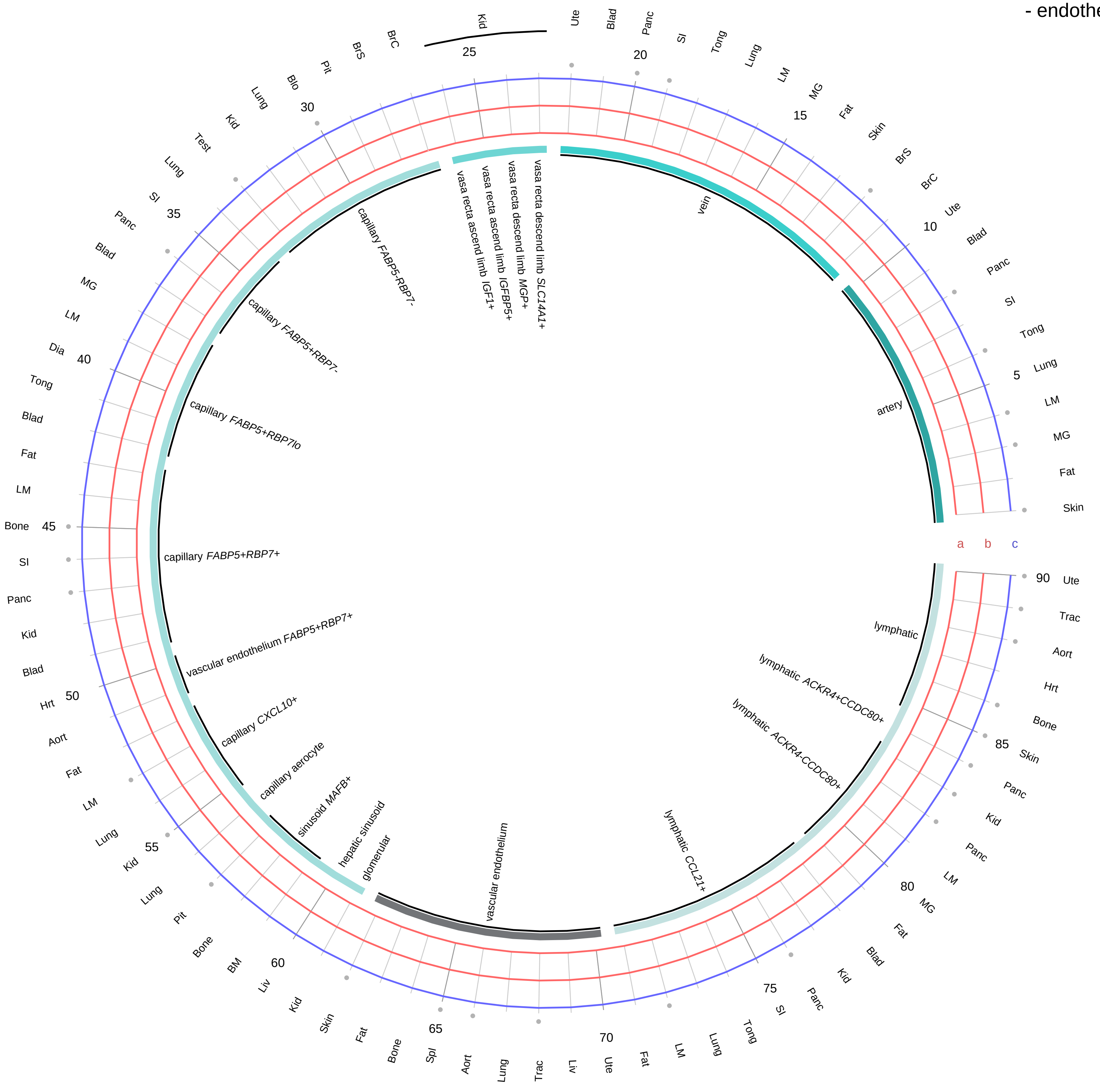

**Ligand/Enzyme**
a - UTS2
b - UTS2B
**Receptor**
c - UTS2R

**PAIRING** (ligand → receptor)

UTS2→UTS2R

UTS2B→UTS2R

**Compartment**
**endothelial**
- artery
- vein
- vasa recta
- capillary
- mix
- lymphatic

12.5%
25%
50%
100%

ln(CP10K+1): 0, 1, 2, 3

| Abbrev. | Tissue |
|---|---|
| Aort | Aorta |
| Blad | Bladder |
| Blo | Blood |
| Bone | Bone |
| BM | Bone marrow |
| BrC | Brain cortex |
| BrS | Brainstem |
| Col | Colon |
| Dia | Diaphragm |
| Ret | Eye retina |
| Fat | Fat |
| Hrt | Heart |
| Pit | Hypothalamus/Pituitary |
| Kid | Kidney |
| LM | Limb muscle |
| Liv | Liver |
| Lung | Lung |
| MG | Mammary gland |
| Panc | Pancreas |
| Skin | Skin |
| SI | Small intestine |
| Spl | Spleen |
| Test | Testes |
| Thym | Thymus |
| Tong | Tongue |
| Trac | Trachea |
| Ute | Uterus |

**Urotensin 2/2B**
- stromal

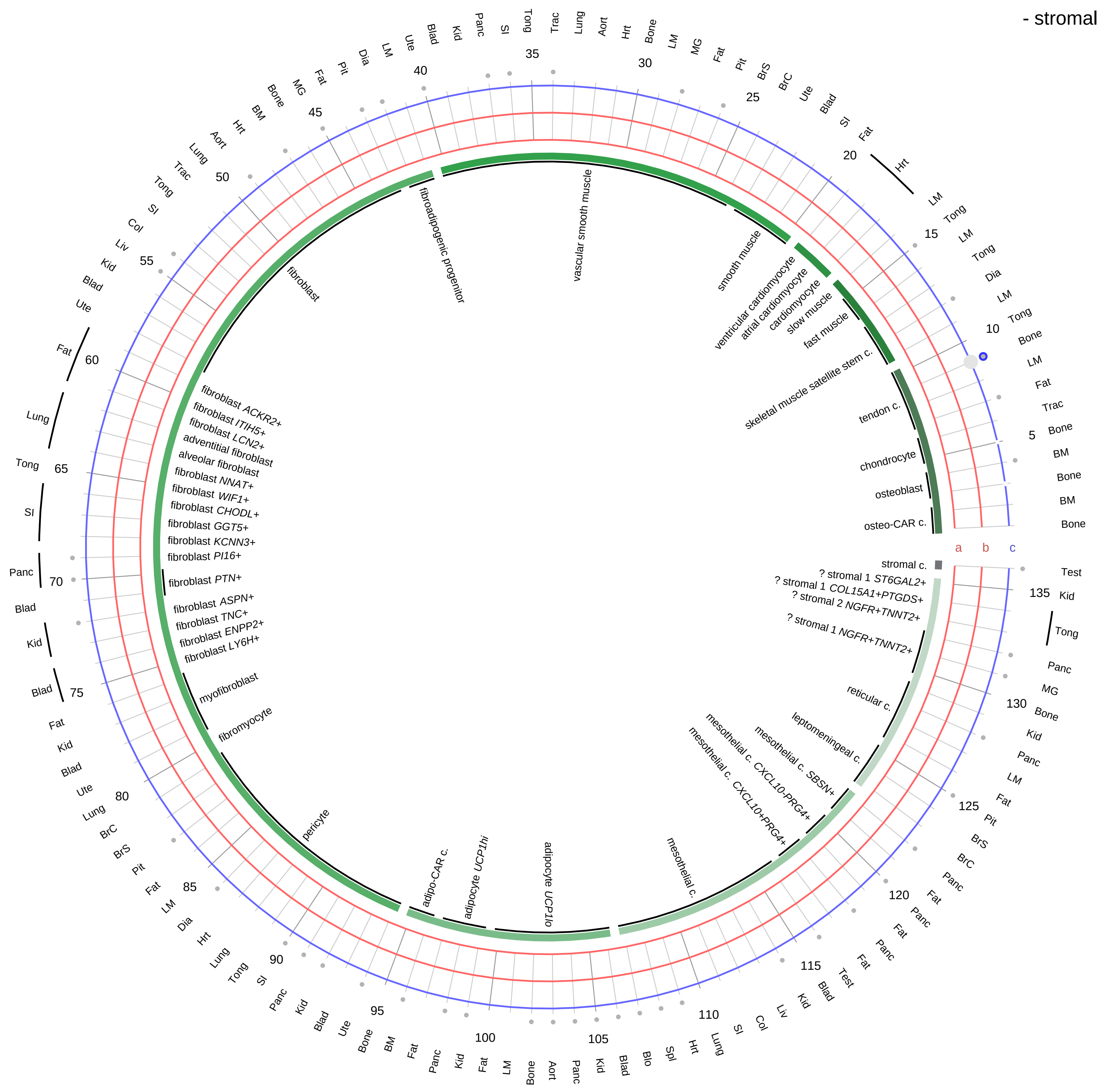

**Ligand/Enzyme**
a - UTS2
b - UTS2B
**Receptor**
c - UTS2R

**PAIRING** (ligand → receptor)

UTS2→UTS2R

UTS2B→UTS2R

**Compartment**
**stromal**
- bone
- skeletal muscle
- cardiac muscle
- smooth muscle
- fibroblast
- adipocyte
- mesothelial
- other
- mix

12.5%
25%
50%
100%

ln(CP10K+1): 0, 1, 2, 3

| Abbrev. | Tissue |
|---|---|
| Aort | Aorta |
| Blad | Bladder |
| Blo | Blood |
| Bone | Bone |
| BM | Bone marrow |
| BrC | Brain cortex |
| BrS | Brainstem |
| Col | Colon |
| Dia | Diaphragm |
| Ret | Eye retina |
| Fat | Fat |
| Hrt | Heart |
| Pit | Hypothalamus/Pituitary |
| Kid | Kidney |
| LM | Limb muscle |
| Liv | Liver |
| Lung | Lung |
| MG | Mammary gland |
| Panc | Pancreas |
| Skin | Skin |
| SI | Small intestine |
| Spl | Spleen |
| Test | Testes |
| Thym | Thymus |
| Tong | Tongue |
| Trac | Trachea |
| Ute | Uterus |

**Urotensin 2/2B**
- lymphoid

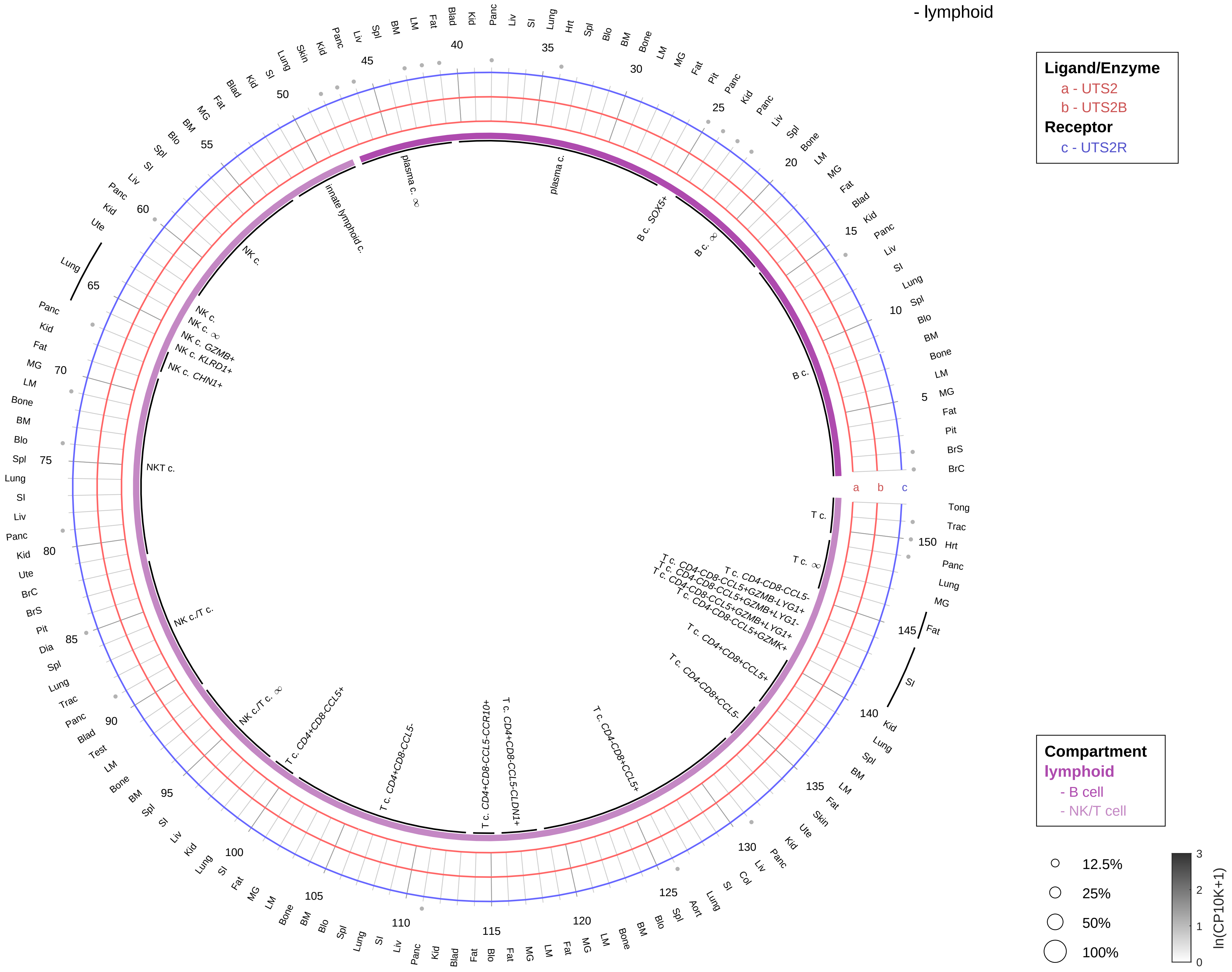

**PAIRING** (ligand → receptor)

UTS2→UTS2R

UTS2B→UTS2R

| Abbrev. | Tissue |
|---|---|
| Aort | Aorta |
| Blad | Bladder |
| Blo | Blood |
| Bone | Bone |
| BM | Bone marrow |
| BrC | Brain cortex |
| BrS | Brainstem |
| Col | Colon |
| Dia | Diaphragm |
| Ret | Eye retina |
| Fat | Fat |
| Hrt | Heart |
| Pit | Hypothalamus/Pituitary |
| Kid | Kidney |
| LM | Limb muscle |
| Liv | Liver |
| Lung | Lung |
| MG | Mammary gland |
| Panc | Pancreas |
| Skin | Skin |
| SI | Small intestine |
| Spl | Spleen |
| Test | Testes |
| Thym | Thymus |
| Tong | Tongue |
| Trac | Trachea |
| Ute | Uterus |

#### Urotensin 2/2B
- myeloid

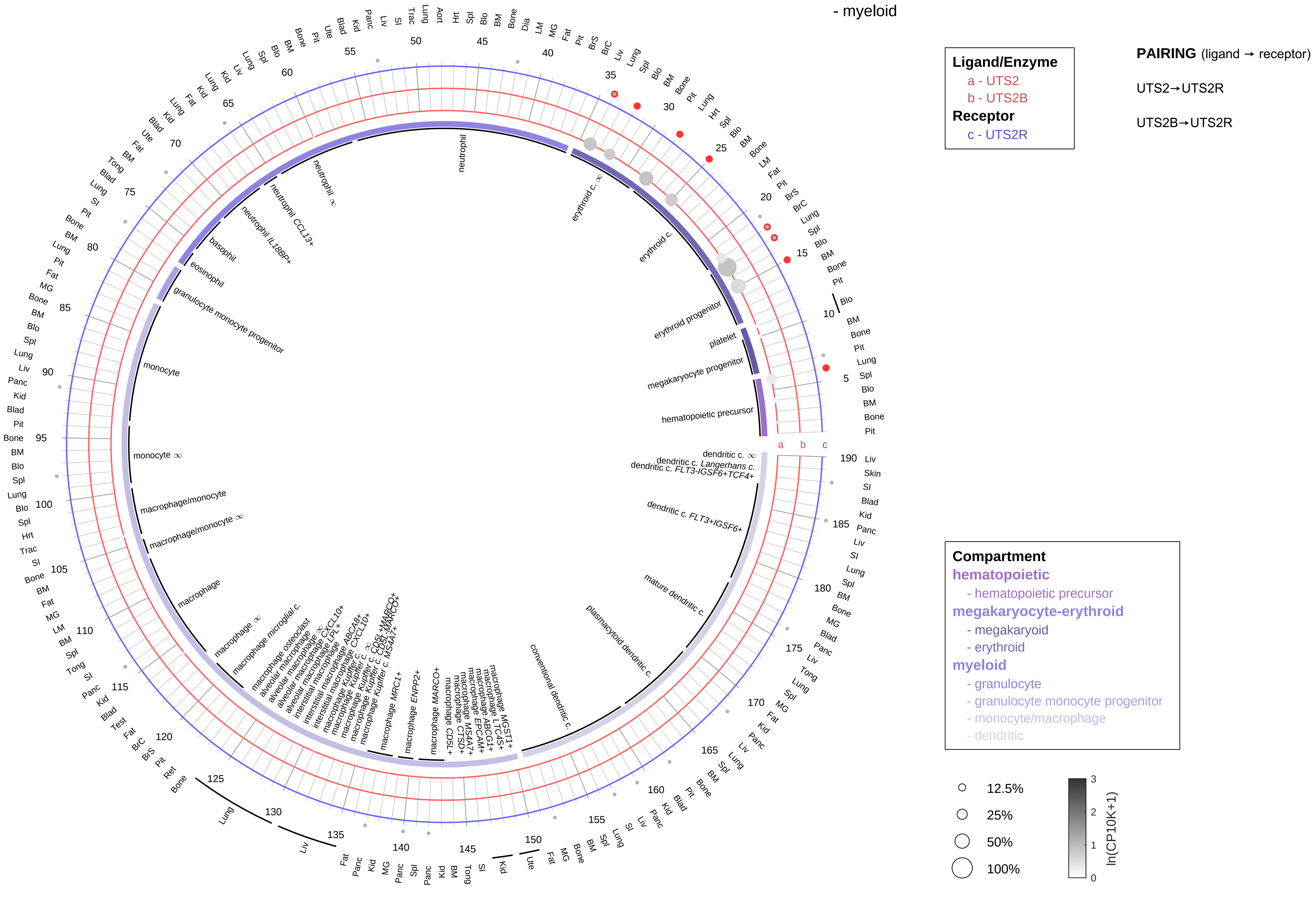

| Abbrev. | Tissue |
|---|---|
| Aort | Aorta |
| Blad | Bladder |
| Blo | Blood |
| Bone | Bone |
| BM | Bone marrow |
| BrC | Brain cortex |
| BrS | Brainstem |
| Col | Colon |
| Dia | Diaphragm |
| Ret | Eye retina |
| Fat | Fat |
| Hrt | Heart |
| Pit | Hypothalamus/Pituitary |
| Kid | Kidney |
| LM | Limb muscle |
| Liv | Liver |
| Lung | Lung |
| MG | Mammary gland |
| Panc | Pancreas |
| Skin | Skin |
| SI | Small intestine |
| Spl | Spleen |
| Test | Testes |
| Thym | Thymus |
| Tong | Tongue |
| Trac | Trachea |
| Ute | Uterus |

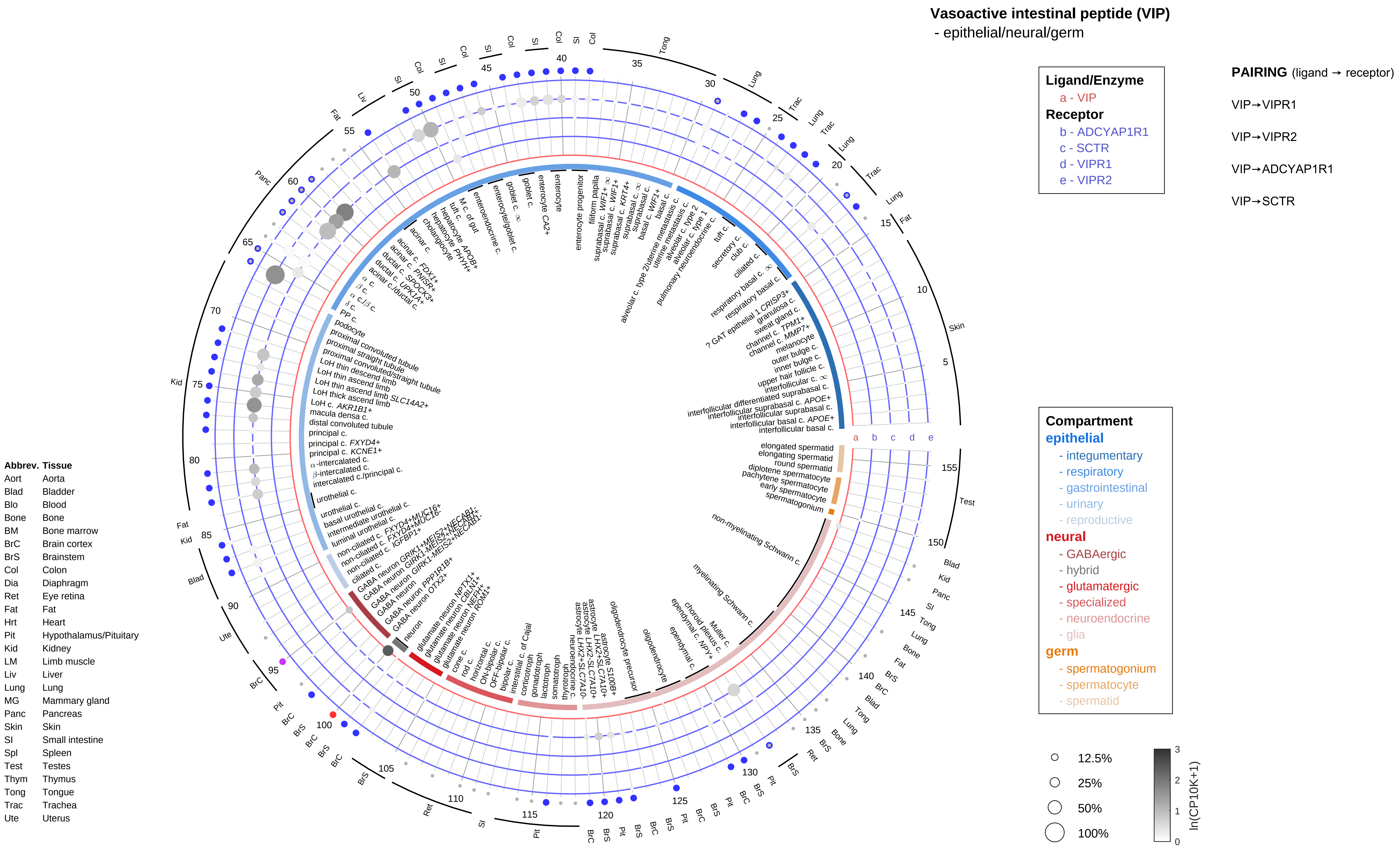

Vasoactive intestinal peptide (VIP)
- epithelial/neural/germ
Ligand/Enzyme
a - VIP
Receptor
b - ADCYAP1R1
c - SCTR
d - VIPR1
e - VIPR2
PAIRING (ligand → receptor)
VIP→VIPR1
VIP→VIPR2
VIP→ADCYAP1R1
VIP→SCTR
Compartment
epithelial
- integumentary
- respiratory
- gastrointestinal
- urinary
- reproductive
neural
- GABAergic
- hybrid
- glutamatergic
- specialized
- neuroendocrine
- glia
germ
- spermatogonium
- spermatocyte
- spermatid
12.5%
25%
50%
100%
ln(CP10K+1)
Abbrev. Tissue
Aort Aorta
Blad Bladder
Blo Blood
Bone Bone
BM Bone marrow
BrC Brain cortex
BrS Brainstem
Col Colon
Dia Diaphragm
Ret Eye retina
Fat Fat
Hrt Heart
Pit Hypothalamus/Pituitary
Kid Kidney
LM Limb muscle
Liv Liver
Lung Lung
MG Mammary gland
Panc Pancreas
Skin Skin
SI Small intestine
Spl Spleen
Test Testes
Thym Thymus
Tong Tongue
Trac Trachea
Ute Uterus

**Vasoactive intestinal peptide (VIP)**
- endothelial

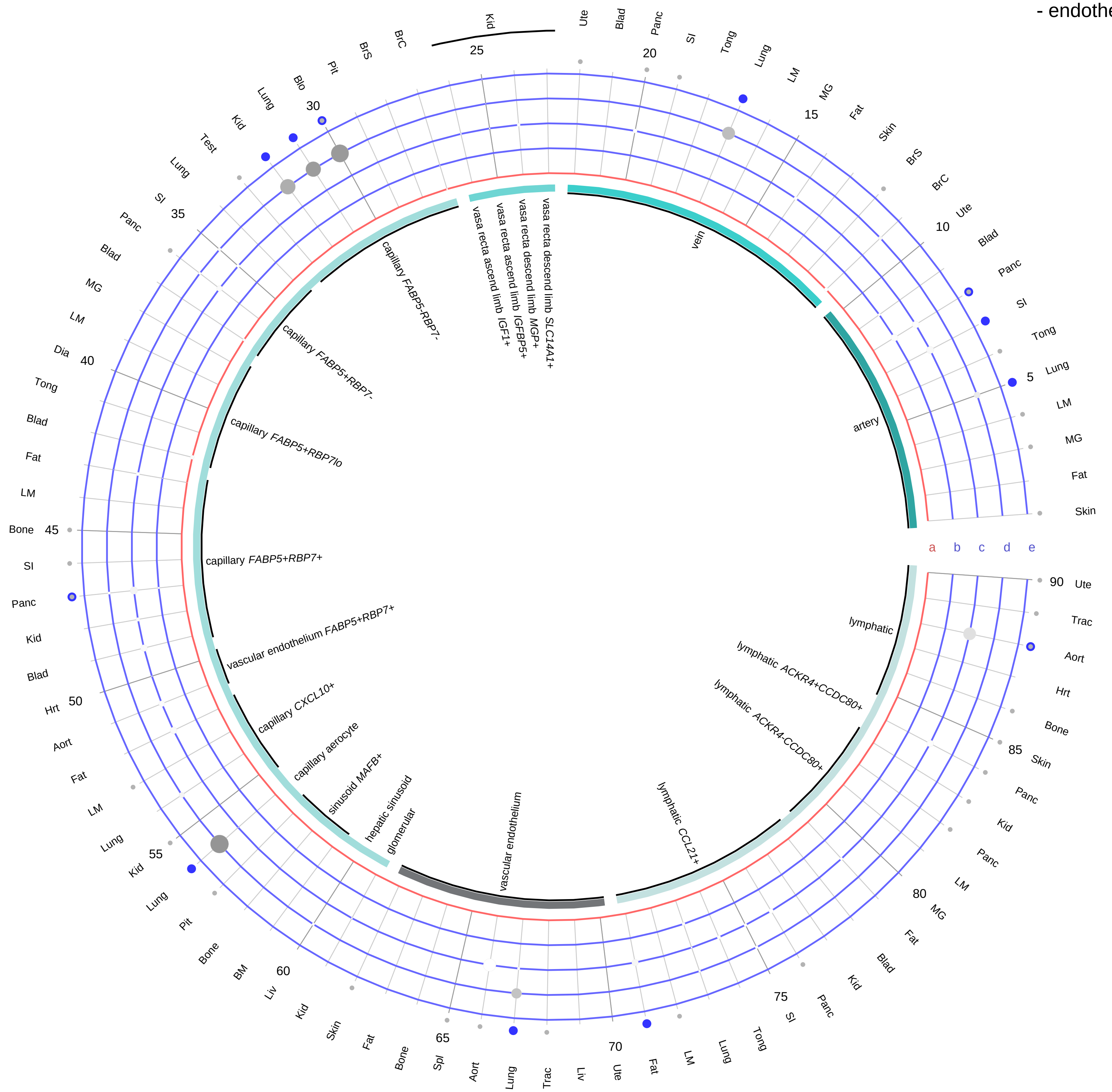

| Abbrev. | Tissue |
|---|---|
| Aort | Aorta |
| Blad | Bladder |
| Blo | Blood |
| Bone | Bone |
| BM | Bone marrow |
| BrC | Brain cortex |
| BrS | Brainstem |
| Col | Colon |
| Dia | Diaphragm |
| Ret | Eye retina |
| Fat | Fat |
| Hrt | Heart |
| Pit | Hypothalamus/Pituitary |
| Kid | Kidney |
| LM | Limb muscle |
| Liv | Liver |
| Lung | Lung |
| MG | Mammary gland |
| Panc | Pancreas |
| Skin | Skin |
| SI | Small intestine |
| Spl | Spleen |
| Test | Testes |
| Thym | Thymus |
| Tong | Tongue |
| Trac | Trachea |
| Ute | Uterus |

**Ligand/Enzyme**
a - VIP
**Receptor**
b - ADCYAP1R1
c - SCTR
d - VIPR1
e - VIPR2

**PAIRING** (ligand → receptor)

VIP→VIPR1

VIP→VIPR2

VIP→ADCYAP1R1

VIP→SCTR

**Compartment**
**endothelial**
- artery
- vein
- vasa recta
- capillary
- mix
- lymphatic

12.5%
25%
50%
100%

ln(CP10K+1): 0, 1, 2, 3

**Vasoactive intestinal peptide (VIP)**
- stromal

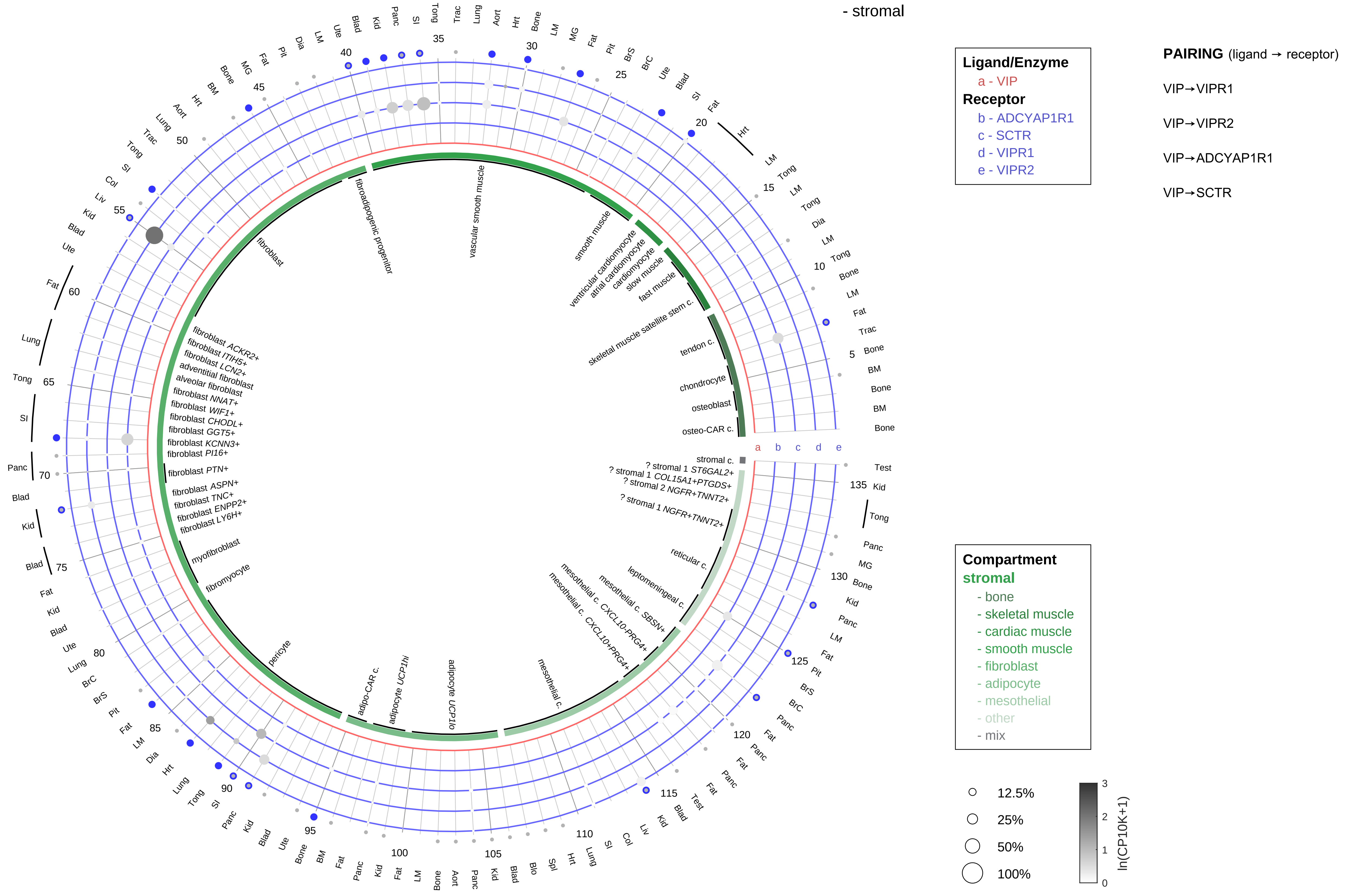

| Abbrev. | Tissue |
|---|---|
| Aort | Aorta |
| Blad | Bladder |
| Blo | Blood |
| Bone | Bone |
| BM | Bone marrow |
| BrC | Brain cortex |
| BrS | Brainstem |
| Col | Colon |
| Dia | Diaphragm |
| Ret | Eye retina |
| Fat | Fat |
| Hrt | Heart |
| Pit | Hypothalamus/Pituitary |
| Kid | Kidney |
| LM | Limb muscle |
| Liv | Liver |
| Lung | Lung |
| MG | Mammary gland |
| Panc | Pancreas |
| Skin | Skin |
| SI | Small intestine |
| Spl | Spleen |
| Test | Testes |
| Thym | Thymus |
| Tong | Tongue |
| Trac | Trachea |
| Ute | Uterus |

#### Vasoactive intestinal peptide (VIP)
- lymphoid

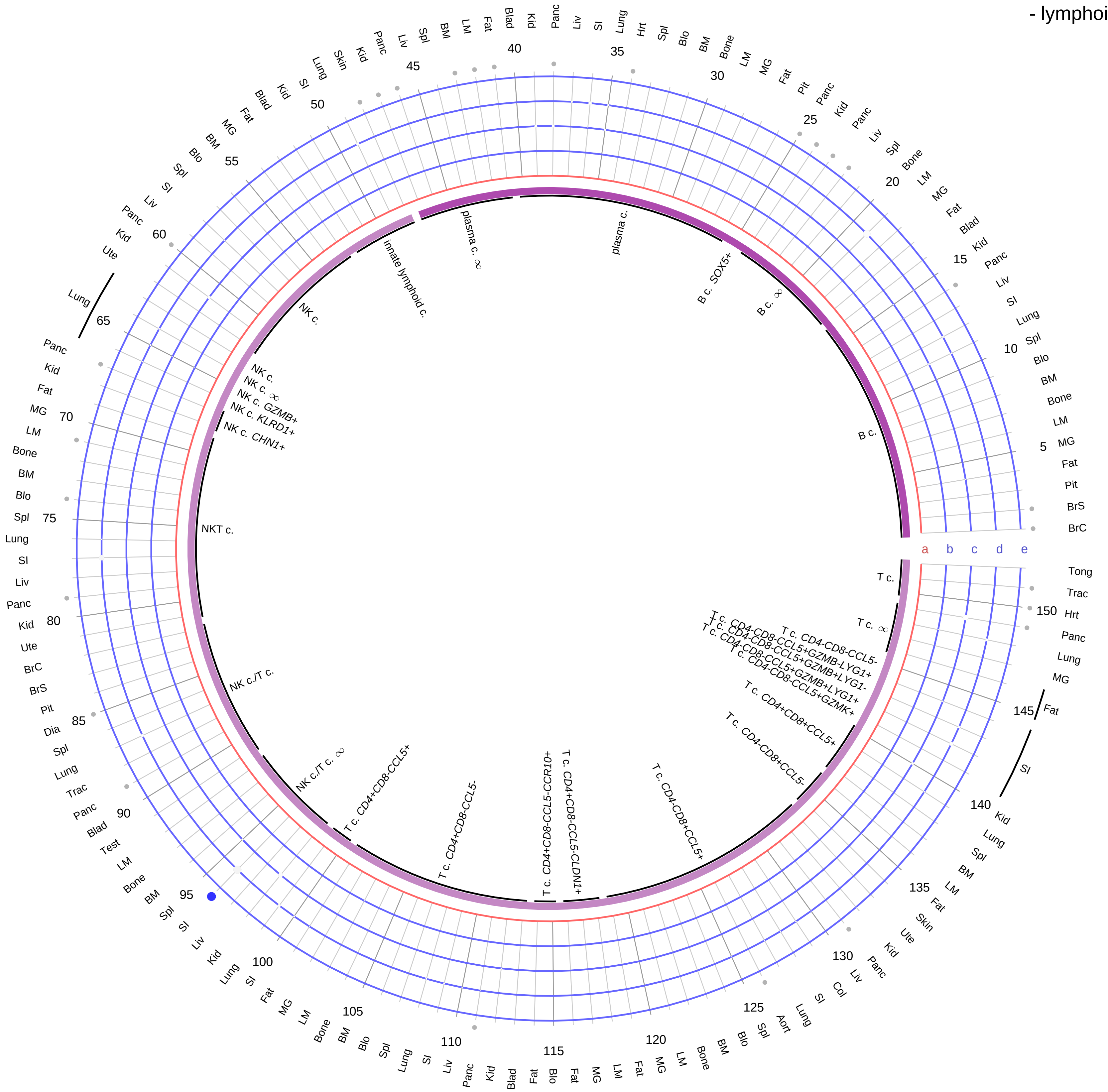

**Ligand/Enzyme**
- a - VIP

**Receptor**
- b - ADCYAP1R1
- c - SCTR
- d - VIPR1
- e - VIPR2

**PAIRING** (ligand → receptor)

VIP→VIPR1

VIP→VIPR2

VIP→ADCYAP1R1

VIP→SCTR

| Abbrev. | Tissue |
|---|---|
| Aort | Aorta |
| Blad | Bladder |
| Blo | Blood |
| Bone | Bone |
| BM | Bone marrow |
| BrC | Brain cortex |
| BrS | Brainstem |
| Col | Colon |
| Dia | Diaphragm |
| Ret | Eye retina |
| Fat | Fat |
| Hrt | Heart |
| Pit | Hypothalamus/Pituitary |
| Kid | Kidney |
| LM | Limb muscle |
| Liv | Liver |
| Lung | Lung |
| MG | Mammary gland |
| Panc | Pancreas |
| Skin | Skin |
| SI | Small intestine |
| Spl | Spleen |
| Test | Testes |
| Thym | Thymus |
| Tong | Tongue |
| Trac | Trachea |
| Ute | Uterus |

**Compartment**
**lymphoid**
- B cell
- NK/T cell

12.5%
25%
50%
100%

ln(CP10K+1): 0, 1, 2, 3

**Vasoactive intestinal peptide (VIP)**
- myeloid

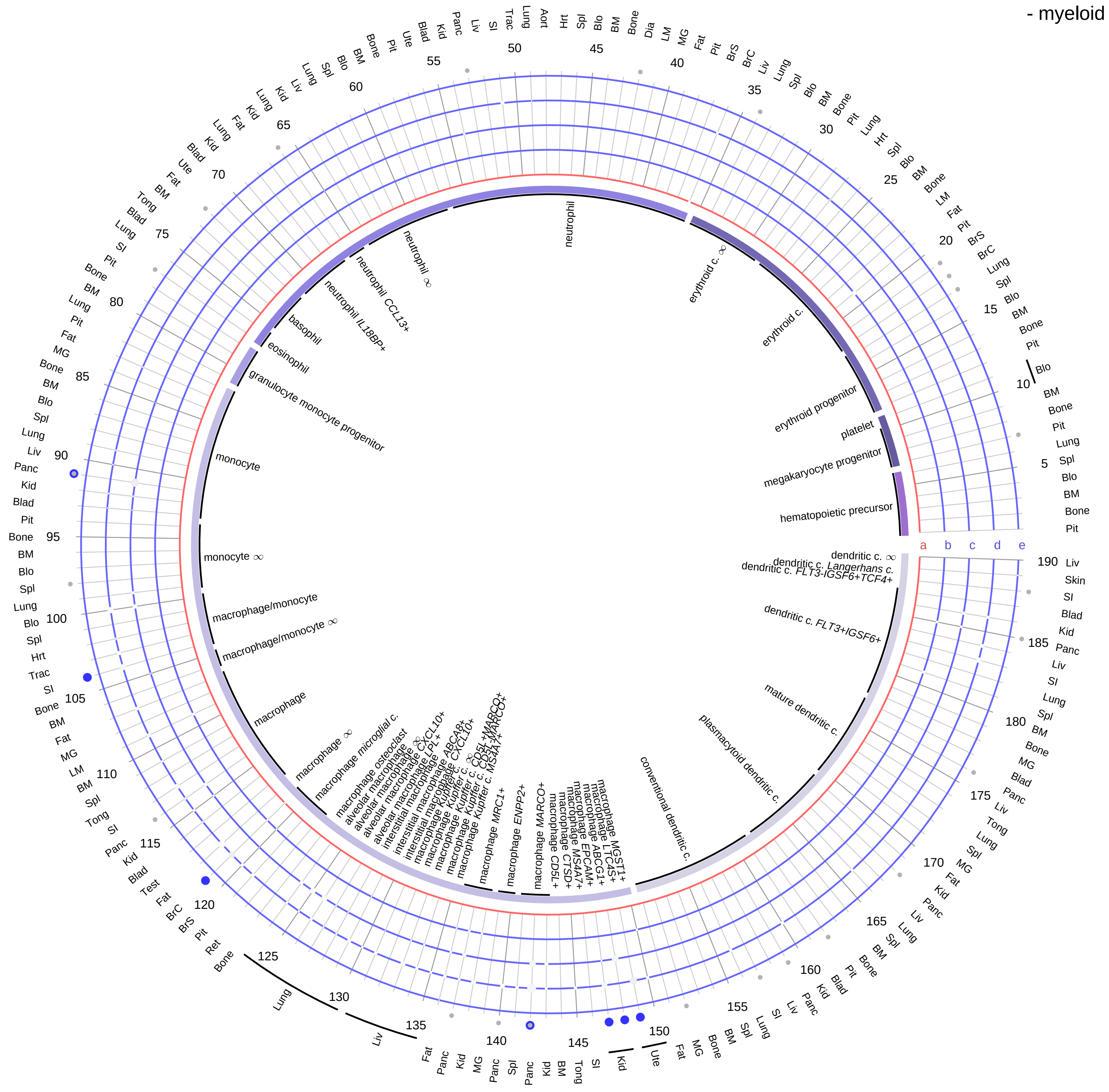

**Ligand/Enzyme**
- a - VIP

**Receptor**
- b - ADCYAP1R1
- c - SCTR
- d - VIPR1
- e - VIPR2

**PAIRING** (ligand → receptor)

VIP→VIPR1

VIP→VIPR2

VIP→ADCYAP1R1

VIP→SCTR

**Compartment**

**hematopoietic**
- hematopoietic precursor

**megakaryocyte-erythroid**
- megakaryoid
- erythroid

**myeloid**
- granulocyte
- granulocyte monocyte progenitor
- monocyte/macrophage
- dendritic

12.5%
25%
50%
100%

ln(CP10K+1): 0, 1, 2, 3

| Abbrev. | Tissue |
|---|---|
| Aort | Aorta |
| Blad | Bladder |
| Blo | Blood |
| Bone | Bone |
| BM | Bone marrow |
| BrC | Brain cortex |
| BrS | Brainstem |
| Col | Colon |
| Dia | Diaphragm |
| Ret | Eye retina |
| Fat | Fat |
| Hrt | Heart |
| Pit | Hypothalamus/Pituitary |
| Kid | Kidney |
| LM | Limb muscle |
| Liv | Liver |
| Lung | Lung |
| MG | Mammary gland |
| Panc | Pancreas |
| Skin | Skin |
| SI | Small intestine |
| Spl | Spleen |
| Test | Testes |
| Thym | Thymus |
| Tong | Tongue |
| Trac | Trachea |
| Ute | Uterus |

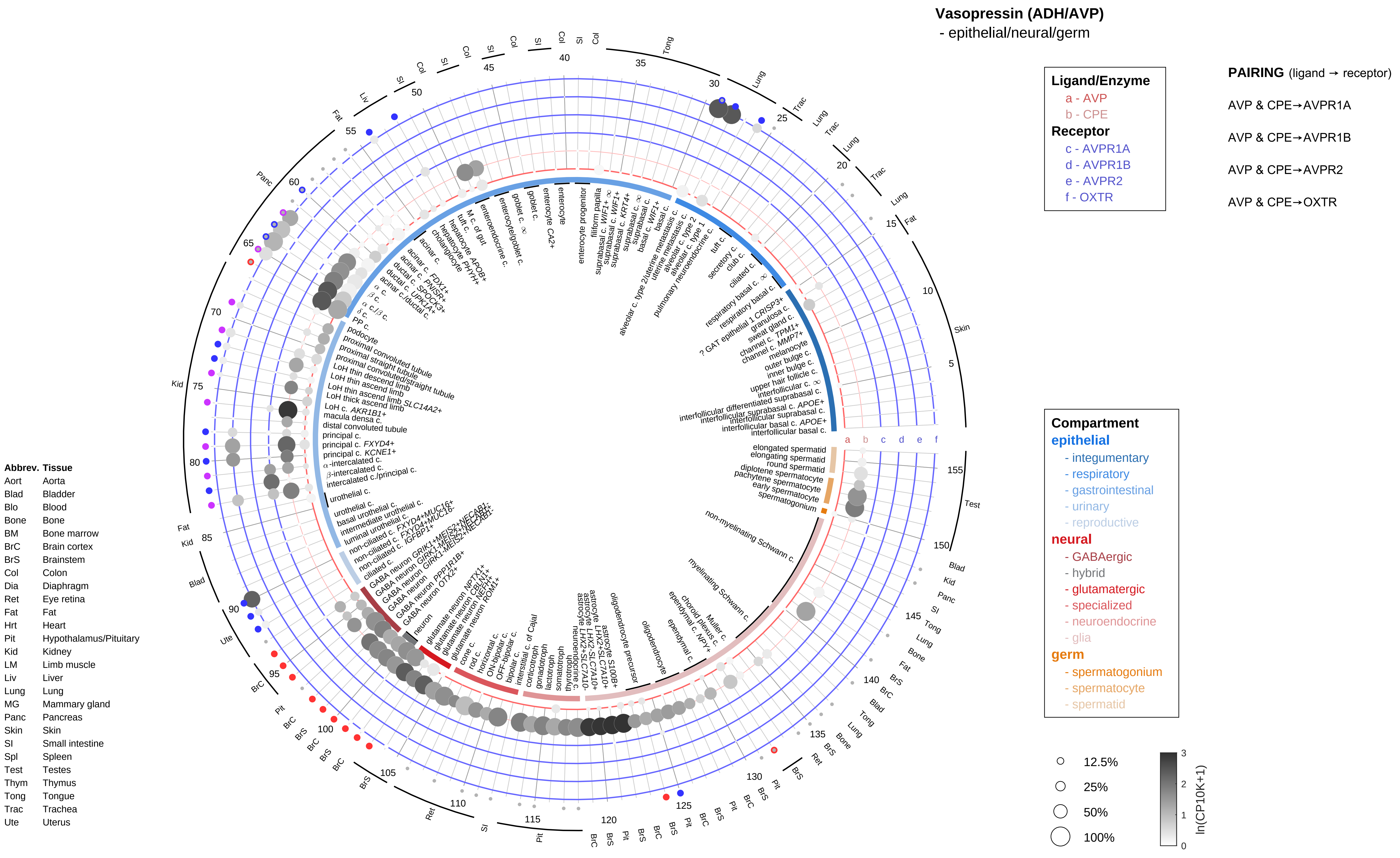

Vasopressin (ADH/AVP)
- epithelial/neural/germ
Ligand/Enzyme
a - AVP
b - CPE
Receptor
c - AVPR1A
d - AVPR1B
e - AVPR2
f - OXTR
PAIRING (ligand → receptor)
AVP & CPE→AVPR1A
AVP & CPE→AVPR1B
AVP & CPE→AVPR2
AVP & CPE→OXTR
Compartment
epithelial
- integumentary
- respiratory
- gastrointestinal
- urinary
- reproductive
neural
- GABAergic
- hybrid
- glutamatergic
- specialized
- neuroendocrine
- glia
germ
- spermatogonium
- spermatocyte
- spermatid
12.5%
25%
50%
100%
ln(CP10K+1)
Abbrev. Tissue
Aort Aorta
Blad Bladder
Blo Blood
Bone Bone
BM Bone marrow
BrC Brain cortex
BrS Brainstem
Col Colon
Dia Diaphragm
Ret Eye retina
Fat Fat
Hrt Heart
Pit Hypothalamus/Pituitary
Kid Kidney
LM Limb muscle
Liv Liver
Lung Lung
MG Mammary gland
Panc Pancreas
Skin Skin
SI Small intestine
Spl Spleen
Test Testes
Thym Thymus
Tong Tongue
Trac Trachea
Ute Uterus

**Vasopressin (ADH/AVP)**
- endothelial

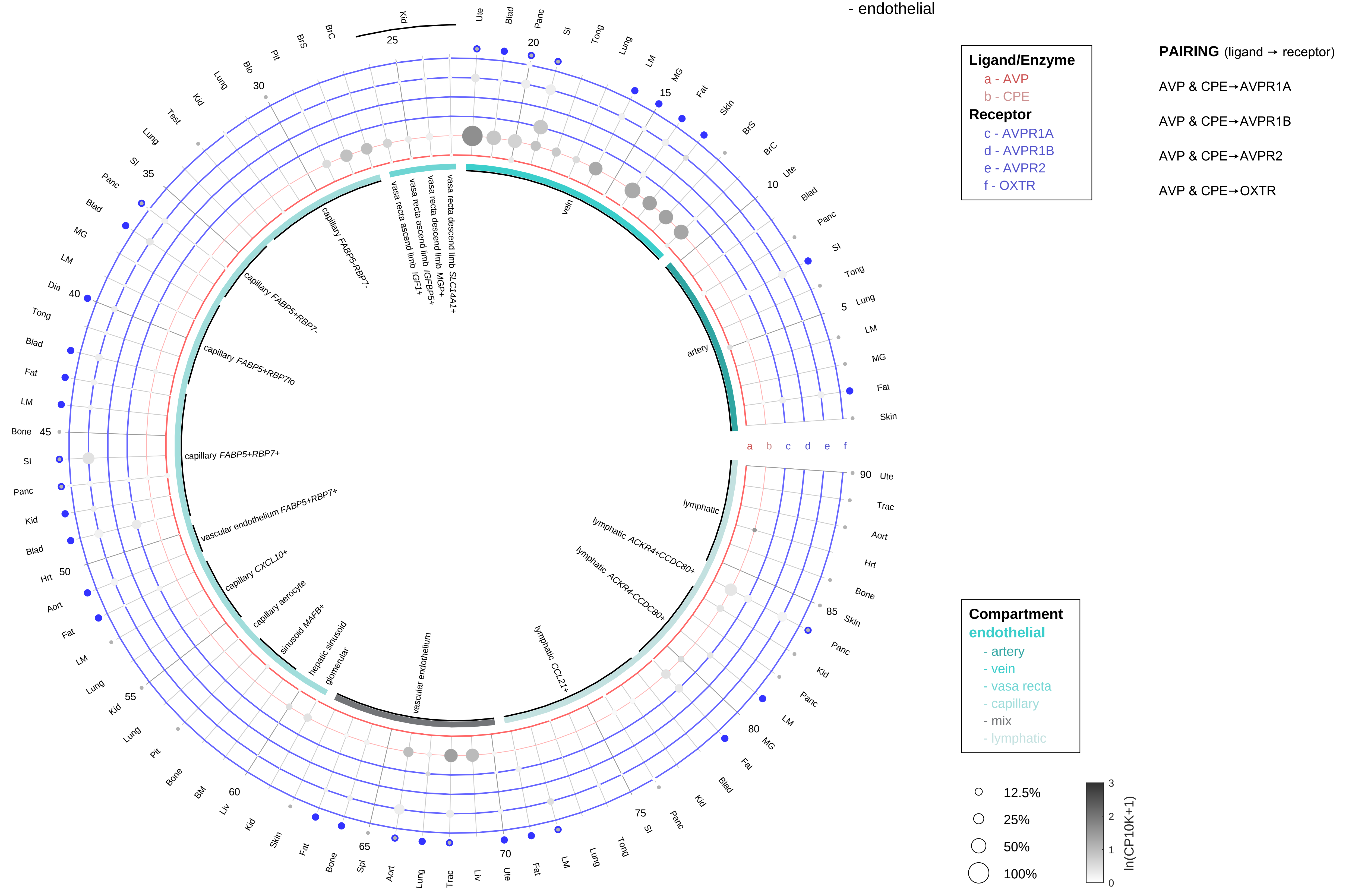

| Abbrev. | Tissue |
|---|---|
| Aort | Aorta |
| Blad | Bladder |
| Blo | Blood |
| Bone | Bone |
| BM | Bone marrow |
| BrC | Brain cortex |
| BrS | Brainstem |
| Col | Colon |
| Dia | Diaphragm |
| Ret | Eye retina |
| Fat | Fat |
| Hrt | Heart |
| Pit | Hypothalamus/Pituitary |
| Kid | Kidney |
| LM | Limb muscle |
| Liv | Liver |
| Lung | Lung |
| MG | Mammary gland |
| Panc | Pancreas |
| Skin | Skin |
| SI | Small intestine |
| Spl | Spleen |
| Test | Testes |
| Thym | Thymus |
| Tong | Tongue |
| Trac | Trachea |
| Ute | Uterus |

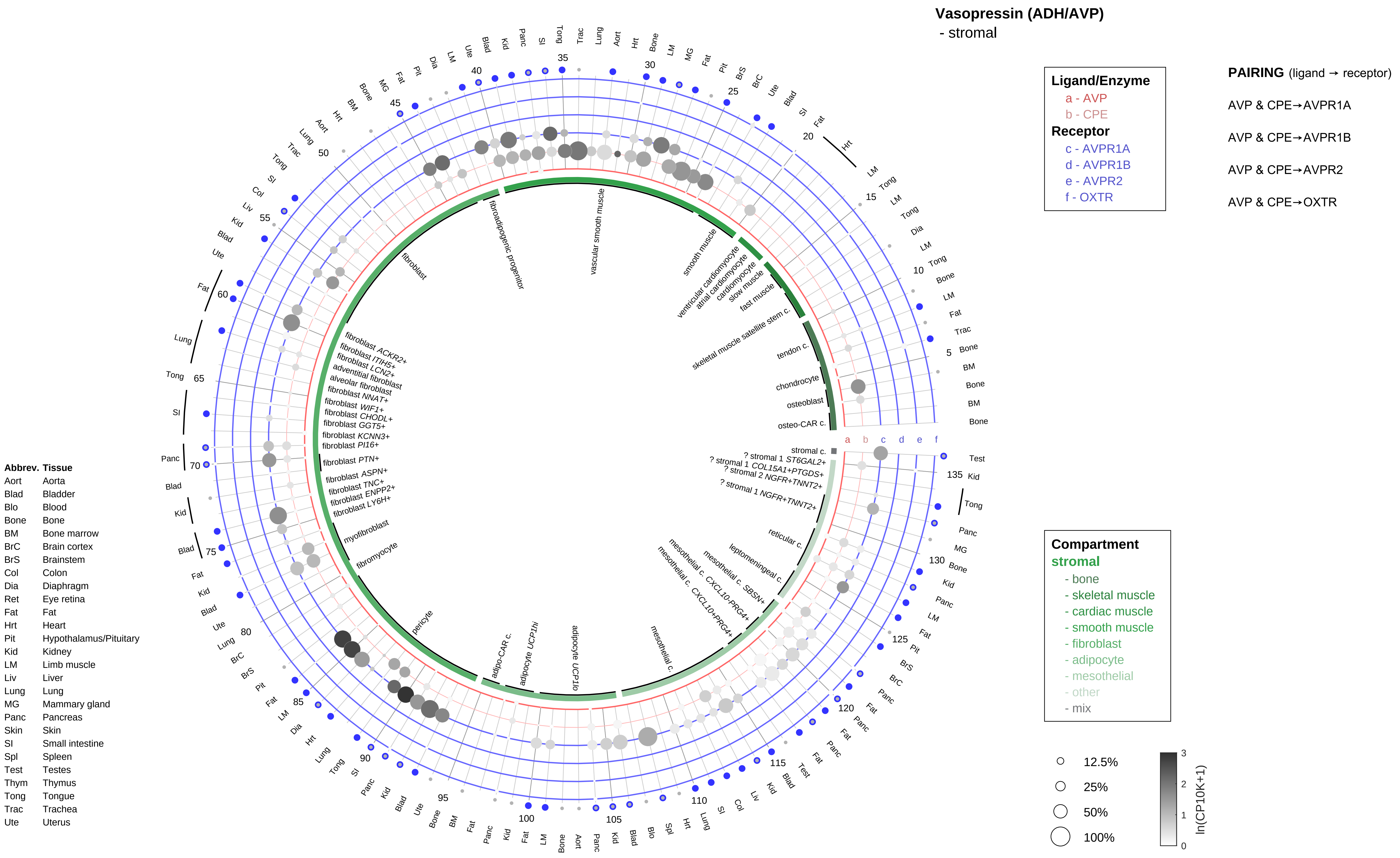
Vasopressin (ADH/AVP)
- stromal
Ligand/Enzyme
a - AVP
b - CPE
Receptor
c - AVPR1A
d - AVPR1B
e - AVPR2
f - OXTR
PAIRING (ligand → receptor)
AVP & CPE→AVPR1A
AVP & CPE→AVPR1B
AVP & CPE→AVPR2
AVP & CPE→OXTR
Compartment
stromal
- bone
- skeletal muscle
- cardiac muscle
- smooth muscle
- fibroblast
- adipocyte
- mesothelial
- other
- mix
12.5%
25%
50%
100%
ln(CP10K+1)
Abbrev. Tissue
Aort Aorta
Blad Bladder
Blo Blood
Bone Bone
BM Bone marrow
BrC Brain cortex
BrS Brainstem
Col Colon
Dia Diaphragm
Ret Eye retina
Fat Fat
Hrt Heart
Pit Hypothalamus/Pituitary
Kid Kidney
LM Limb muscle
Liv Liver
Lung Lung
MG Mammary gland
Panc Pancreas
Skin Skin
SI Small intestine
Spl Spleen
Test Testes
Thym Thymus
Tong Tongue
Trac Trachea
Ute Uterus
vascular smooth muscle
fibroadipogenic progenitor
fibroblast
smooth muscle
ventricular cardiomyocyte
atrial cardiomyocyte
cardiomyocyte
slow muscle
fast muscle
skeletal muscle satellite stem c.
tendon c.
chondrocyte
osteoblast
osteo-CAR c.
stromal c.
? stromal 1 ST6GAL2+
? stromal 1 COL15A1+PTGDS+
? stromal 2 NGFR+TNNT2+
? stromal 1 NGFR+TNNT2+
reticular c.
leptomeningeal c.
mesothelial c. SBSN+
mesothelial c. CXCL10+PRG4+
mesothelial c. CXCL10+PRG4+
mesothelial c.
adipocyte UCP1lo
adipocyte UCP1hi
adipo-CAR c.
pericyte
fibromyocyte
myofibroblast
fibroblast ACKR2+
fibroblast ITIH5+
fibroblast LCN2+
adventitial fibroblast
alveolar fibroblast
fibroblast NNAT+
fibroblast WIF1+
fibroblast CHODL+
fibroblast GGT5+
fibroblast KCNN3+
fibroblast PI16+
fibroblast PTN+
fibroblast ASPN+
fibroblast TNC+
fibroblast ENPP2+
fibroblast LY6H+

**Vasopressin (ADH/AVP)**
- lymphoid

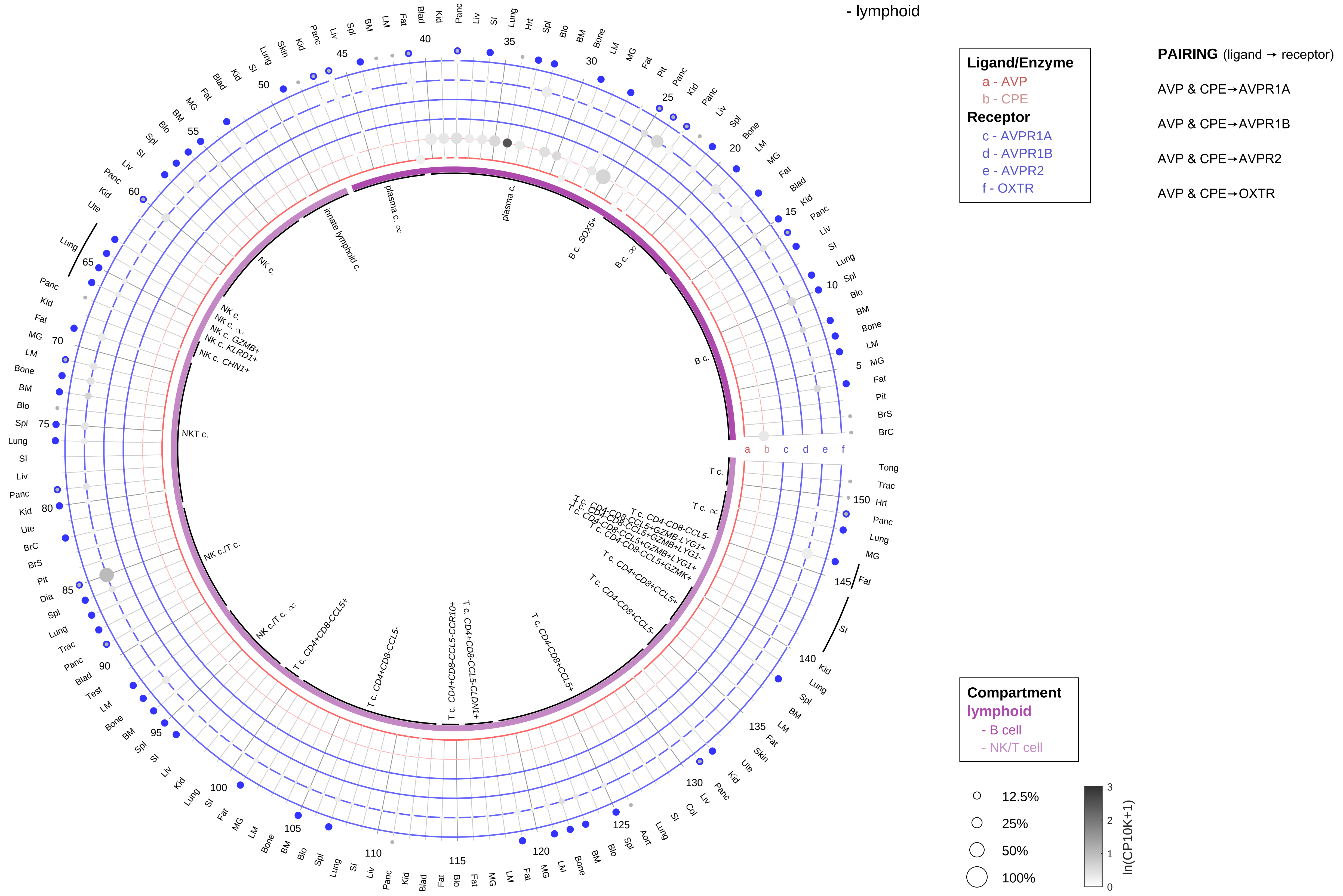

| Abbrev. | Tissue |
|---|---|
| Aort | Aorta |
| Blad | Bladder |
| Blo | Blood |
| Bone | Bone |
| BM | Bone marrow |
| BrC | Brain cortex |
| BrS | Brainstem |
| Col | Colon |
| Dia | Diaphragm |
| Ret | Eye retina |
| Fat | Fat |
| Hrt | Heart |
| Pit | Hypothalamus/Pituitary |
| Kid | Kidney |
| LM | Limb muscle |
| Liv | Liver |
| Lung | Lung |
| MG | Mammary gland |
| Panc | Pancreas |
| Skin | Skin |
| SI | Small intestine |
| Spl | Spleen |
| Test | Testes |
| Thym | Thymus |
| Tong | Tongue |
| Trac | Trachea |
| Ute | Uterus |

**Vasopressin (ADH/AVP)**
- myeloid

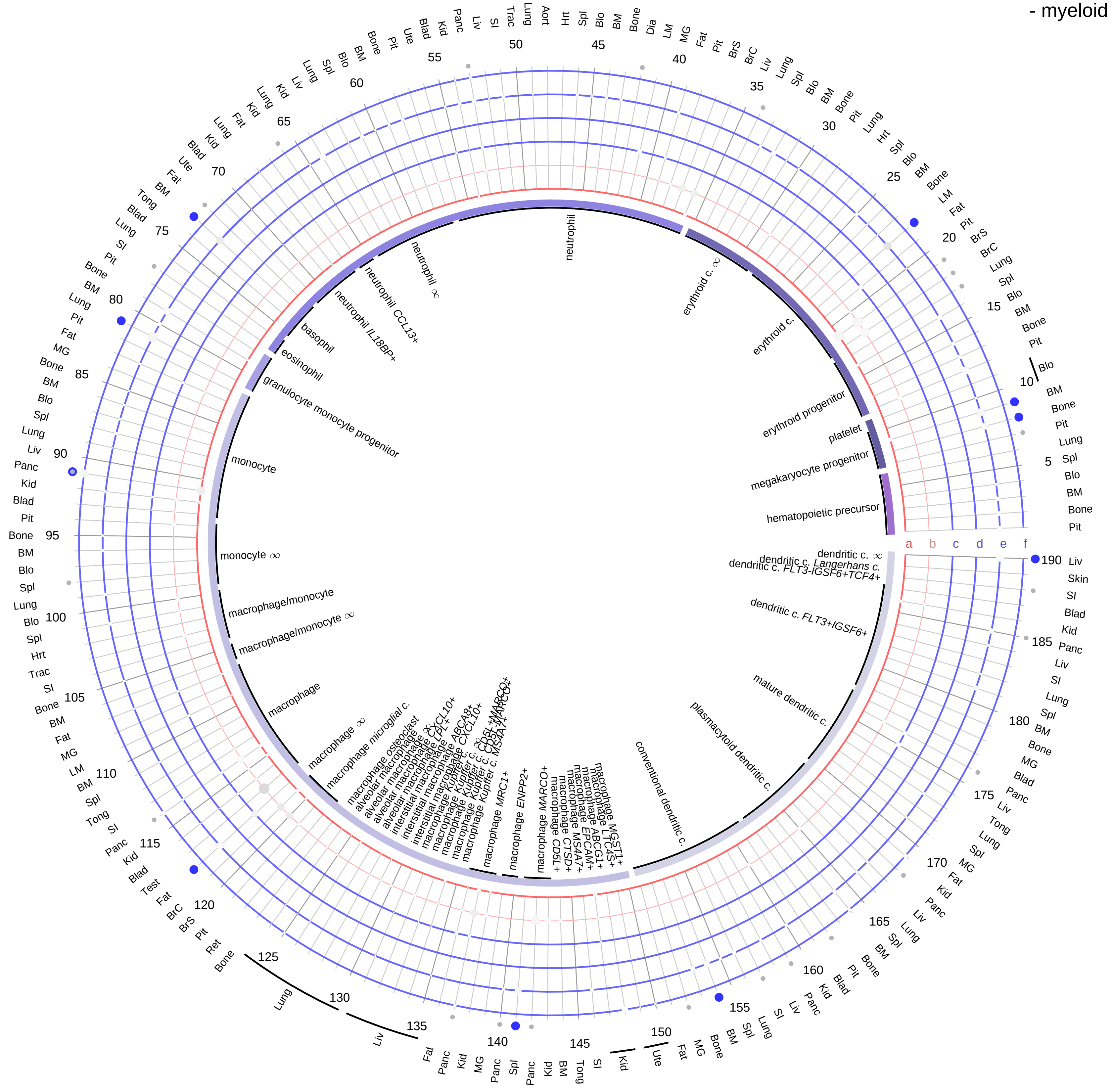

| Ligand/Enzyme |
|---|
| a - AVP |
| b - CPE |
| **Receptor** |
| c - AVPR1A |
| d - AVPR1B |
| e - AVPR2 |
| f - OXTR |

**PAIRING** (ligand → receptor)

AVP & CPE→AVPR1A

AVP & CPE→AVPR1B

AVP & CPE→AVPR2

AVP & CPE→OXTR

| Abbrev. | Tissue |
|---|---|
| Aort | Aorta |
| Blad | Bladder |
| Blo | Blood |
| Bone | Bone |
| BM | Bone marrow |
| BrC | Brain cortex |
| BrS | Brainstem |
| Col | Colon |
| Dia | Diaphragm |
| Ret | Eye retina |
| Fat | Fat |
| Hrt | Heart |
| Pit | Hypothalamus/Pituitary |
| Kid | Kidney |
| LM | Limb muscle |
| Liv | Liver |
| Lung | Lung |
| MG | Mammary gland |
| Panc | Pancreas |
| Skin | Skin |
| SI | Small intestine |
| Spl | Spleen |
| Test | Testes |
| Thym | Thymus |
| Tong | Tongue |
| Trac | Trachea |
| Ute | Uterus |

**Compartment**
**hematopoietic**
- hematopoietic precursor
**megakaryocyte-erythroid**
- megakaryoid
- erythroid
**myeloid**
- granulocyte
- granulocyte monocyte progenitor
- monocyte/macrophage
- dendritic

○ 12.5%
○ 25%
○ 50%
○ 100%

ln(CP10K+1): 0, 1, 2, 3

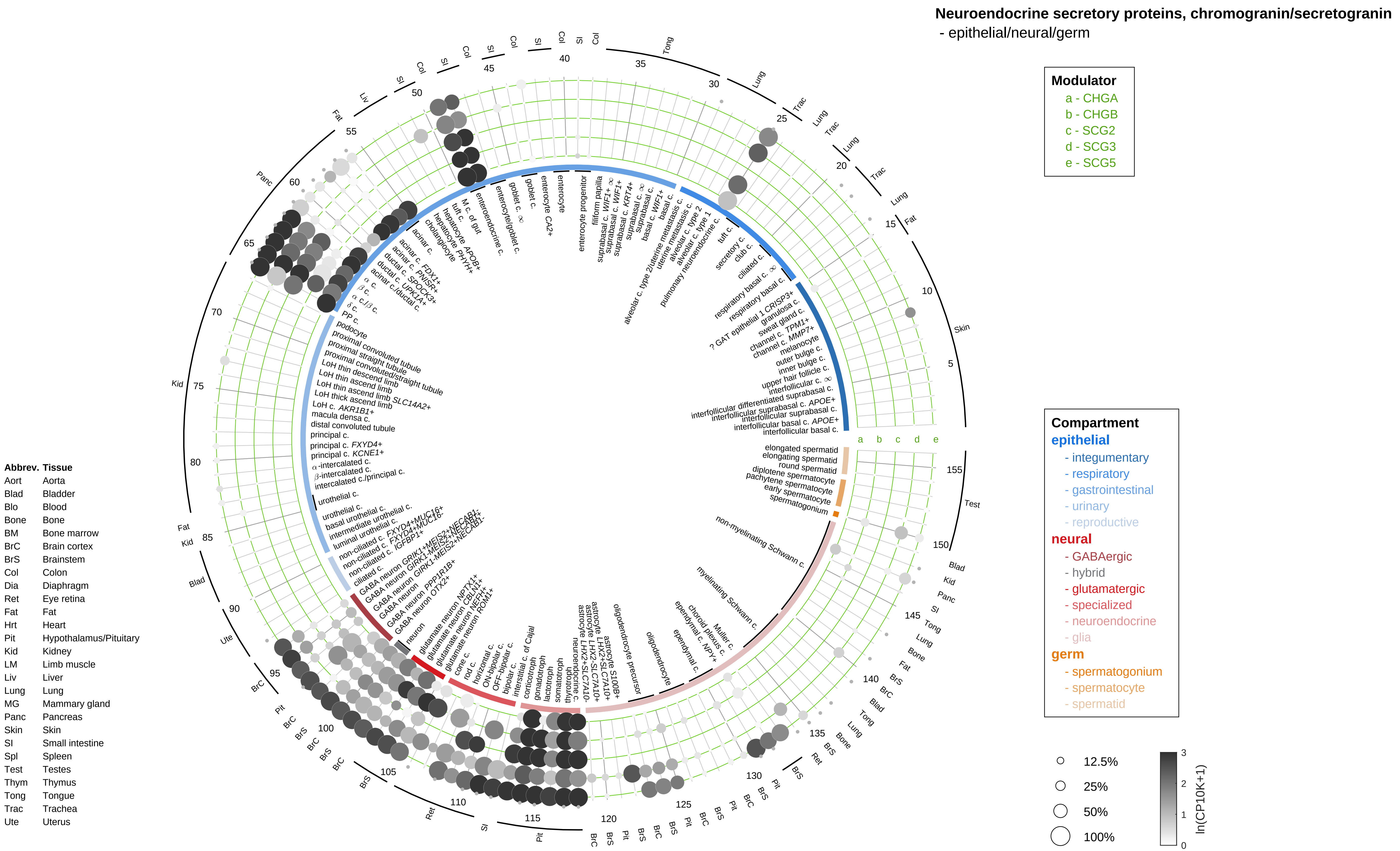

Neuroendocrine secretory proteins, chromogranin/secretogranin
- epithelial/neural/germ
Modulator
a - CHGA
b - CHGB
c - SCG2
d - SCG3
e - SCG5
Compartment
epithelial
- integumentary
- respiratory
- gastrointestinal
- urinary
- reproductive
neural
- GABAergic
- hybrid
- glutamatergic
- specialized
- neuroendocrine
- glia
germ
- spermatogonium
- spermatocyte
- spermatid
12.5%
25%
50%
100%
ln(CP10K+1)
Abbrev. Tissue
Aort Aorta
Blad Bladder
Blo Blood
Bone Bone
BM Bone marrow
BrC Brain cortex
BrS Brainstem
Col Colon
Dia Diaphragm
Ret Eye retina
Fat Fat
Hrt Heart
Pit Hypothalamus/Pituitary
Kid Kidney
LM Limb muscle
Liv Liver
Lung Lung
MG Mammary gland
Panc Pancreas
Skin Skin
SI Small intestine
Spl Spleen
Test Testes
Thym Thymus
Tong Tongue
Trac Trachea
Ute Uterus

**Neuroendocrine secretory proteins, chromogranin/secretogranin**
- endothelial

| Abbrev. | Tissue |
|---|---|
| Aort | Aorta |
| Blad | Bladder |
| Blo | Blood |
| Bone | Bone |
| BM | Bone marrow |
| BrC | Brain cortex |
| BrS | Brainstem |
| Col | Colon |
| Dia | Diaphragm |
| Ret | Eye retina |
| Fat | Fat |
| Hrt | Heart |
| Pit | Hypothalamus/Pituitary |
| Kid | Kidney |
| LM | Limb muscle |
| Liv | Liver |
| Lung | Lung |
| MG | Mammary gland |
| Panc | Pancreas |
| Skin | Skin |
| SI | Small intestine |
| Spl | Spleen |
| Test | Testes |
| Thym | Thymus |
| Tong | Tongue |
| Trac | Trachea |
| Ute | Uterus |

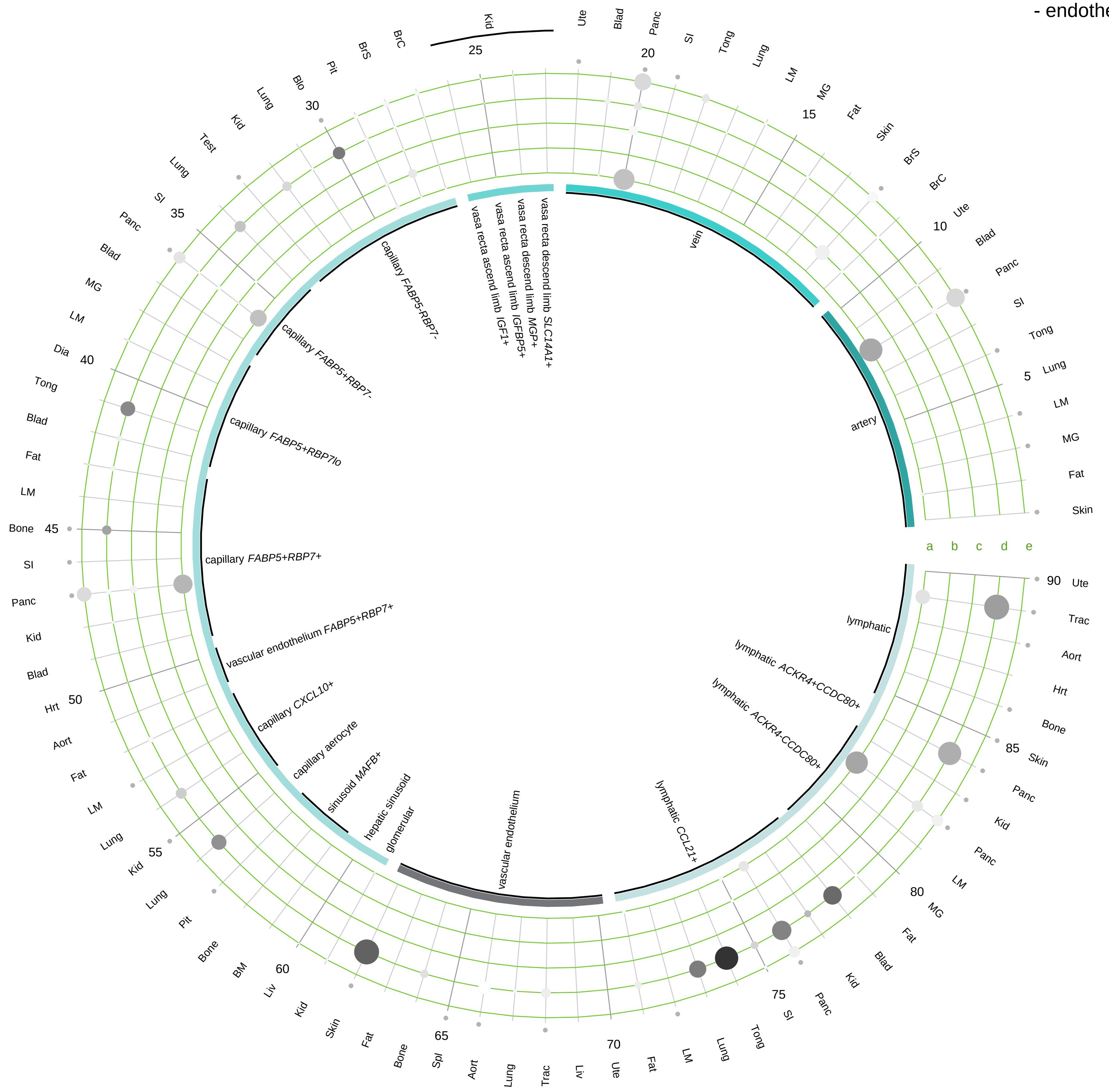

**Modulator**
- a - CHGA
- b - CHGB
- c - SCG2
- d - SCG3
- e - SCG5

**Compartment**
**endothelial**
- artery
- vein
- vasa recta
- capillary
- mix
- lymphatic

12.5%
25%
50%
100%

ln(CP10K+1): 0, 1, 2, 3

**Neuroendocrine secretory proteins, chromogranin/secretogranin**
- stromal

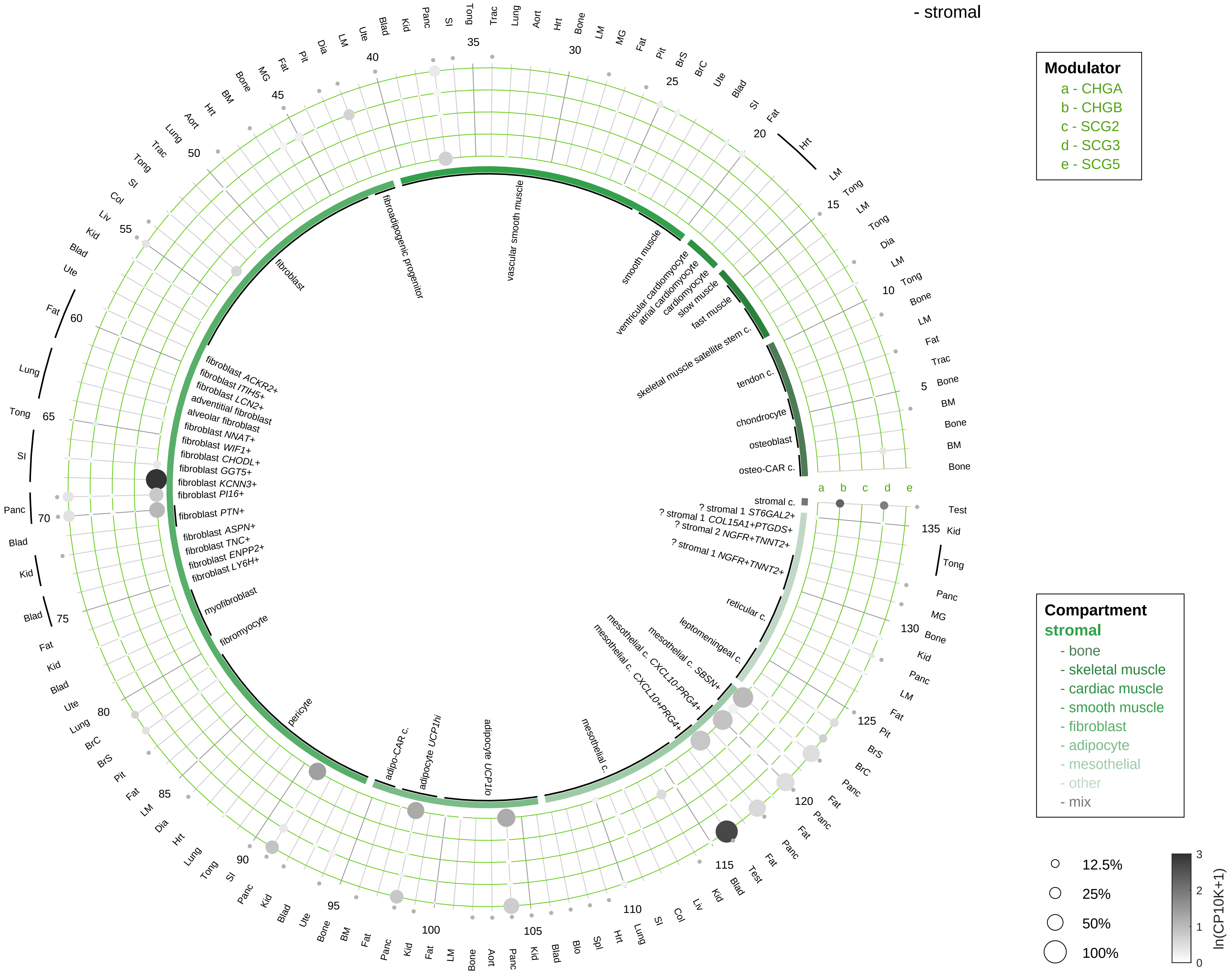

| Abbrev. | Tissue |
|---|---|
| Aort | Aorta |
| Blad | Bladder |
| Blo | Blood |
| Bone | Bone |
| BM | Bone marrow |
| BrC | Brain cortex |
| BrS | Brainstem |
| Col | Colon |
| Dia | Diaphragm |
| Ret | Eye retina |
| Fat | Fat |
| Hrt | Heart |
| Pit | Hypothalamus/Pituitary |
| Kid | Kidney |
| LM | Limb muscle |
| Liv | Liver |
| Lung | Lung |
| MG | Mammary gland |
| Panc | Pancreas |
| Skin | Skin |
| SI | Small intestine |
| Spl | Spleen |
| Test | Testes |
| Thym | Thymus |
| Tong | Tongue |
| Trac | Trachea |
| Ute | Uterus |

**Neuroendocrine secretory proteins, chromogranin/secretogranin**
- lymphoid

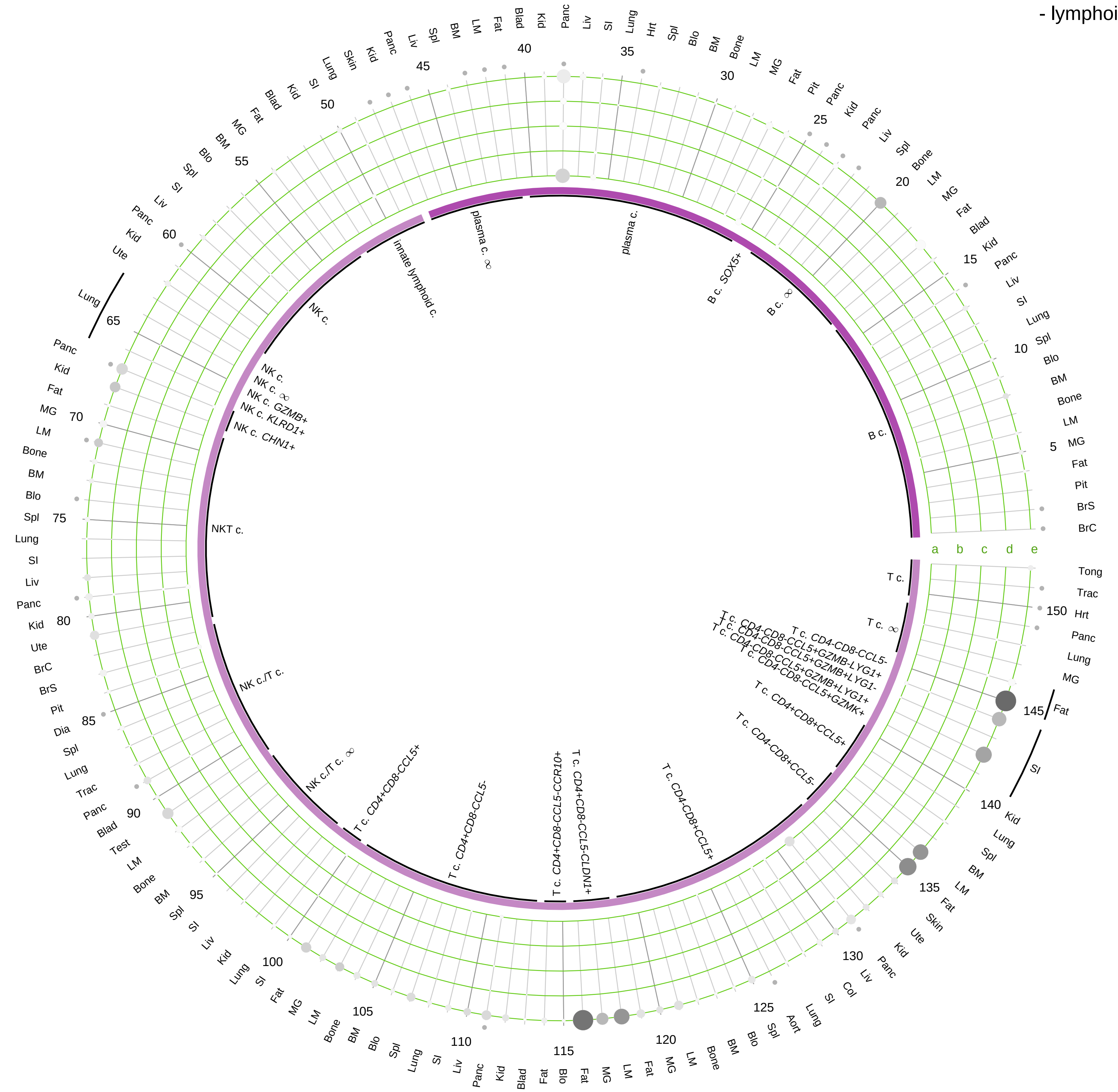

| Modulator |
|---|
| a - CHGA |
| b - CHGB |
| c - SCG2 |
| d - SCG3 |
| e - SCG5 |

| Abbrev. | Tissue |
|---|---|
| Aort | Aorta |
| Blad | Bladder |
| Blo | Blood |
| Bone | Bone |
| BM | Bone marrow |
| BrC | Brain cortex |
| BrS | Brainstem |
| Col | Colon |
| Dia | Diaphragm |
| Ret | Eye retina |
| Fat | Fat |
| Hrt | Heart |
| Pit | Hypothalamus/Pituitary |
| Kid | Kidney |
| LM | Limb muscle |
| Liv | Liver |
| Lung | Lung |
| MG | Mammary gland |
| Panc | Pancreas |
| Skin | Skin |
| SI | Small intestine |
| Spl | Spleen |
| Test | Testes |
| Thym | Thymus |
| Tong | Tongue |
| Trac | Trachea |
| Ute | Uterus |

**Compartment**
**lymphoid**
- B cell
- NK/T cell

12.5%
25%
50%
100%

ln(CP10K+1)
0 1 2 3

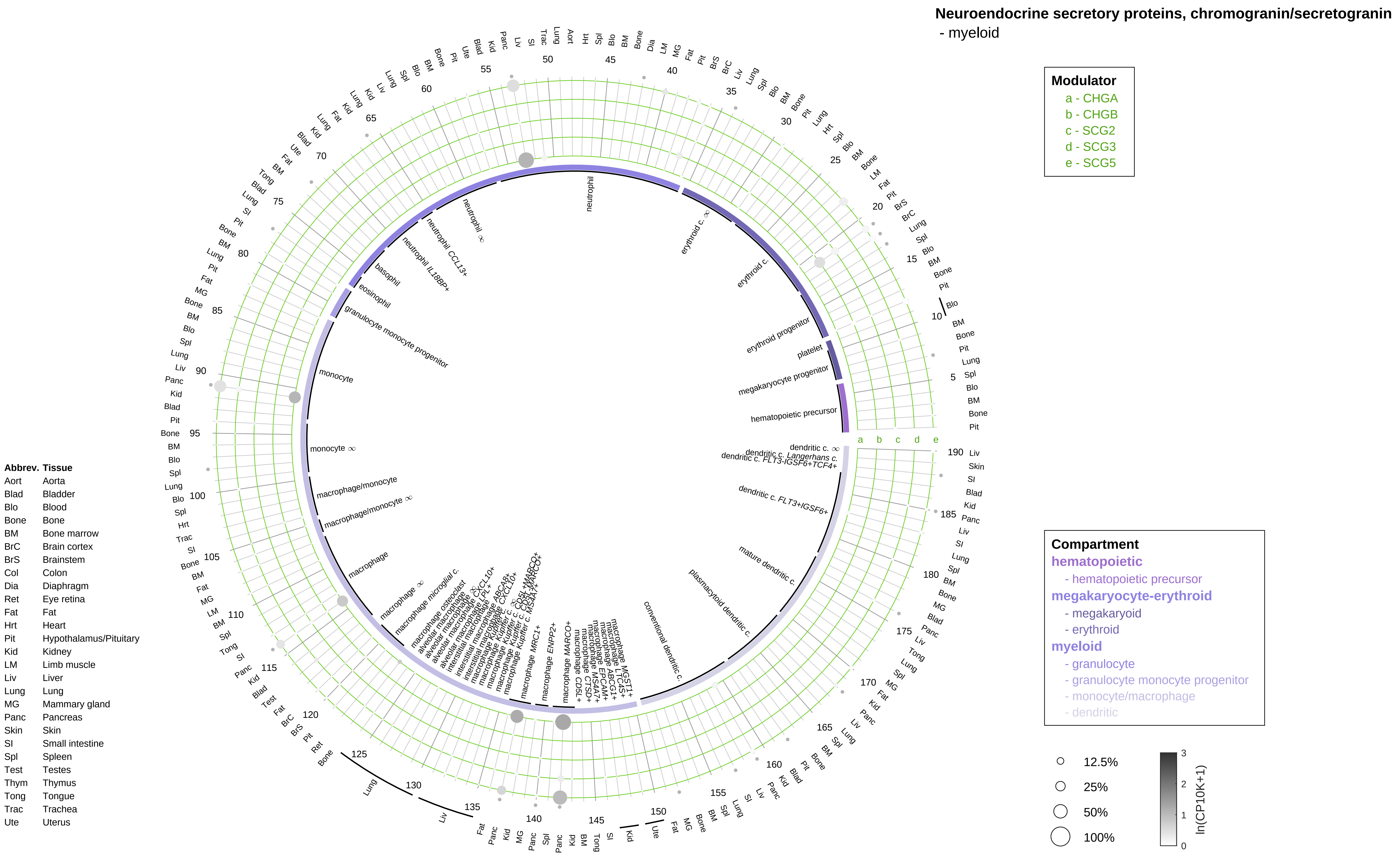
Neuroendocrine secretory proteins, chromogranin/secretogranin
- myeloid
Modulator
a - CHGA
b - CHGB
c - SCG2
d - SCG3
e - SCG5
Compartment
hematopoietic
- hematopoietic precursor
megakaryocyte-erythroid
- megakaryoid
- erythroid
myeloid
- granulocyte
- granulocyte monocyte progenitor
- monocyte/macrophage
- dendritic
12.5%
25%
50%
100%
ln(CP10K+1)
Abbrev. Tissue
Aort Aorta
Blad Bladder
Blo Blood
Bone Bone
BM Bone marrow
BrC Brain cortex
BrS Brainstem
Col Colon
Dia Diaphragm
Ret Eye retina
Fat Fat
Hrt Heart
Pit Hypothalamus/Pituitary
Kid Kidney
LM Limb muscle
Liv Liver
Lung Lung
MG Mammary gland
Panc Pancreas
Skin Skin
SI Small intestine
Spl Spleen
Test Testes
Thym Thymus
Tong Tongue
Trac Trachea
Ute Uterus

**Prohormone processing enzymes**
- epithelial/neural/germ

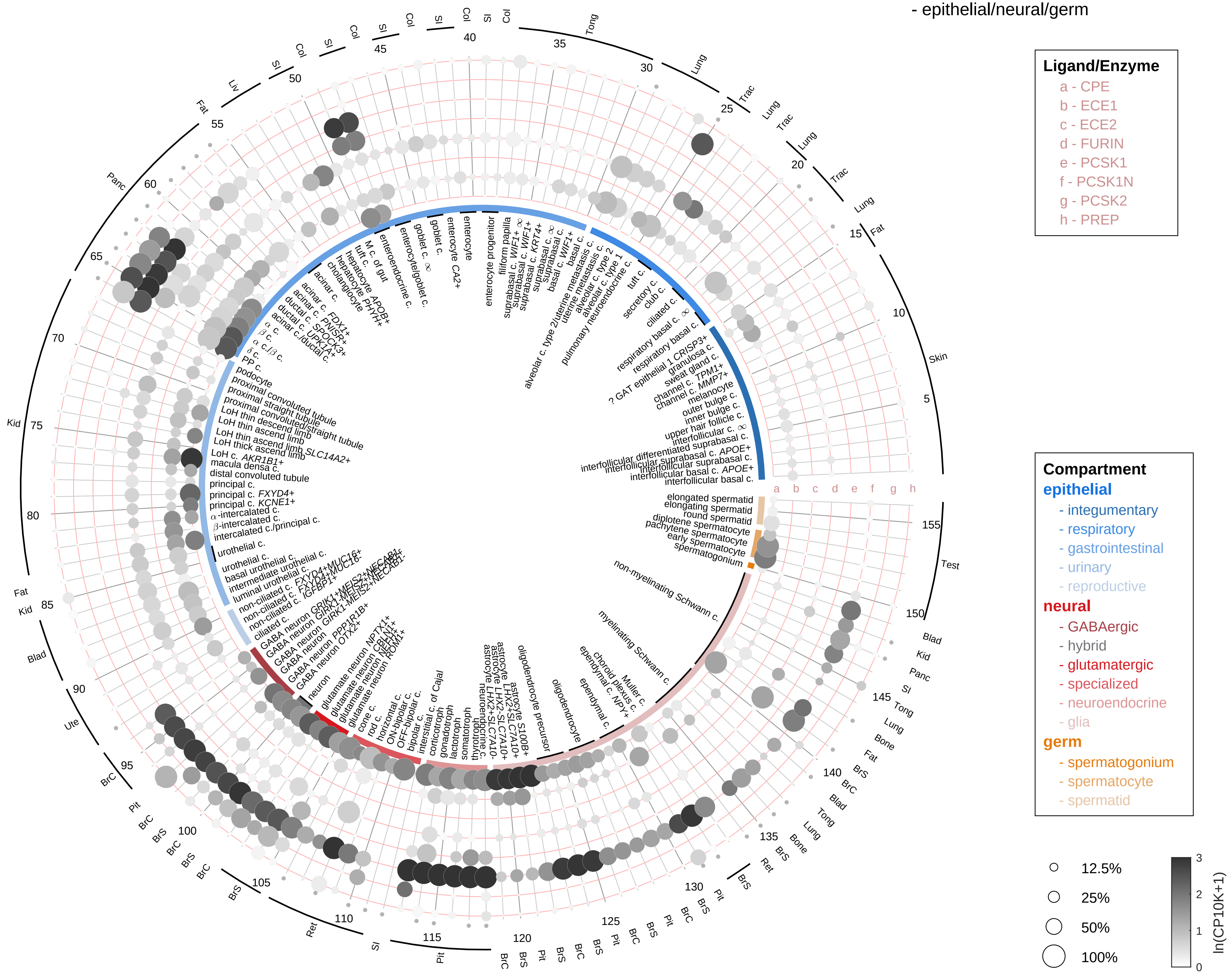

| Abbrev. | Tissue |
|---|---|
| Aort | Aorta |
| Blad | Bladder |
| Blo | Blood |
| Bone | Bone |
| BM | Bone marrow |
| BrC | Brain cortex |
| BrS | Brainstem |
| Col | Colon |
| Dia | Diaphragm |
| Ret | Eye retina |
| Fat | Fat |
| Hrt | Heart |
| Pit | Hypothalamus/Pituitary |
| Kid | Kidney |
| LM | Limb muscle |
| Liv | Liver |
| Lung | Lung |
| MG | Mammary gland |
| Panc | Pancreas |
| Skin | Skin |
| SI | Small intestine |
| Spl | Spleen |
| Test | Testes |
| Thym | Thymus |
| Tong | Tongue |
| Trac | Trachea |
| Ute | Uterus |

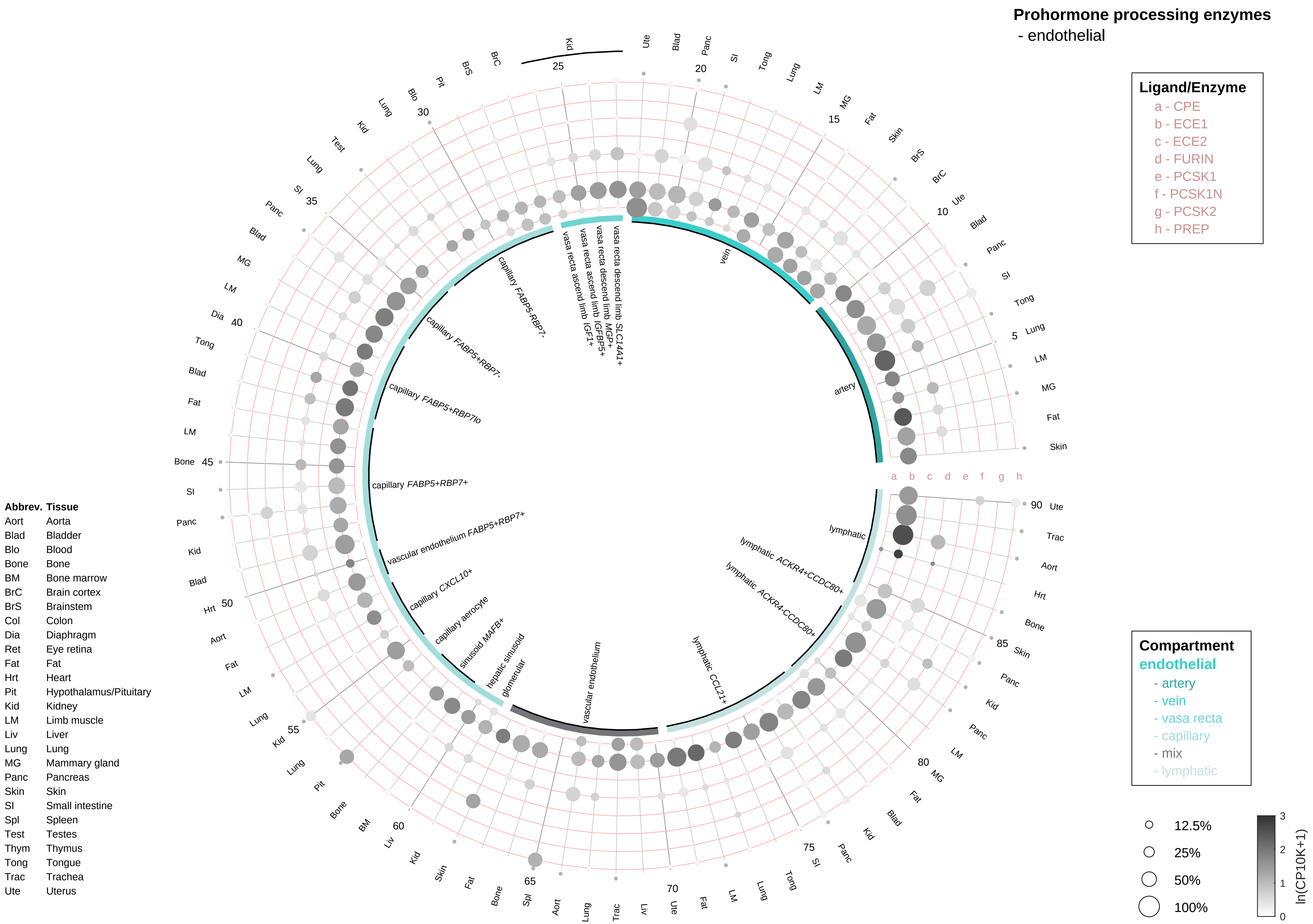
Prohormone processing enzymes
- endothelial
Ligand/Enzyme
a - CPE
b - ECE1
c - ECE2
d - FURIN
e - PCSK1
f - PCSK1N
g - PCSK2
h - PREP
Compartment
endothelial
- artery
- vein
- vasa recta
- capillary
- mix
- lymphatic
12.5%
25%
50%
100%
ln(CP10K+1)
Abbrev. Tissue
Aort Aorta
Blad Bladder
Blo Blood
Bone Bone
BM Bone marrow
BrC Brain cortex
BrS Brainstem
Col Colon
Dia Diaphragm
Ret Eye retina
Fat Fat
Hrt Heart
Pit Hypothalamus/Pituitary
Kid Kidney
LM Limb muscle
Liv Liver
Lung Lung
MG Mammary gland
Panc Pancreas
Skin Skin
SI Small intestine
Spl Spleen
Test Testes
Thym Thymus
Tong Tongue
Trac Trachea
Ute Uterus
artery
vein
vasa recta descend limb SLC14A1+
vasa recta descend limb MGP+
vasa recta ascend limb IGFBP5+
vasa recta ascend limb IGF1+
capillary FABP5-RBP7-
capillary FABP5+RBP7-
capillary FABP5+RBP7lo
capillary FABP5+RBP7+
vascular endothelium FABP5+RBP7+
capillary CXCL10+
capillary aerocyte
sinusoid MAFB+
hepatic sinusoid
glomerular
vascular endothelium
lymphatic CCL21+
lymphatic ACKR4-CCDC80+
lymphatic ACKR4+CCDC80+
lymphatic

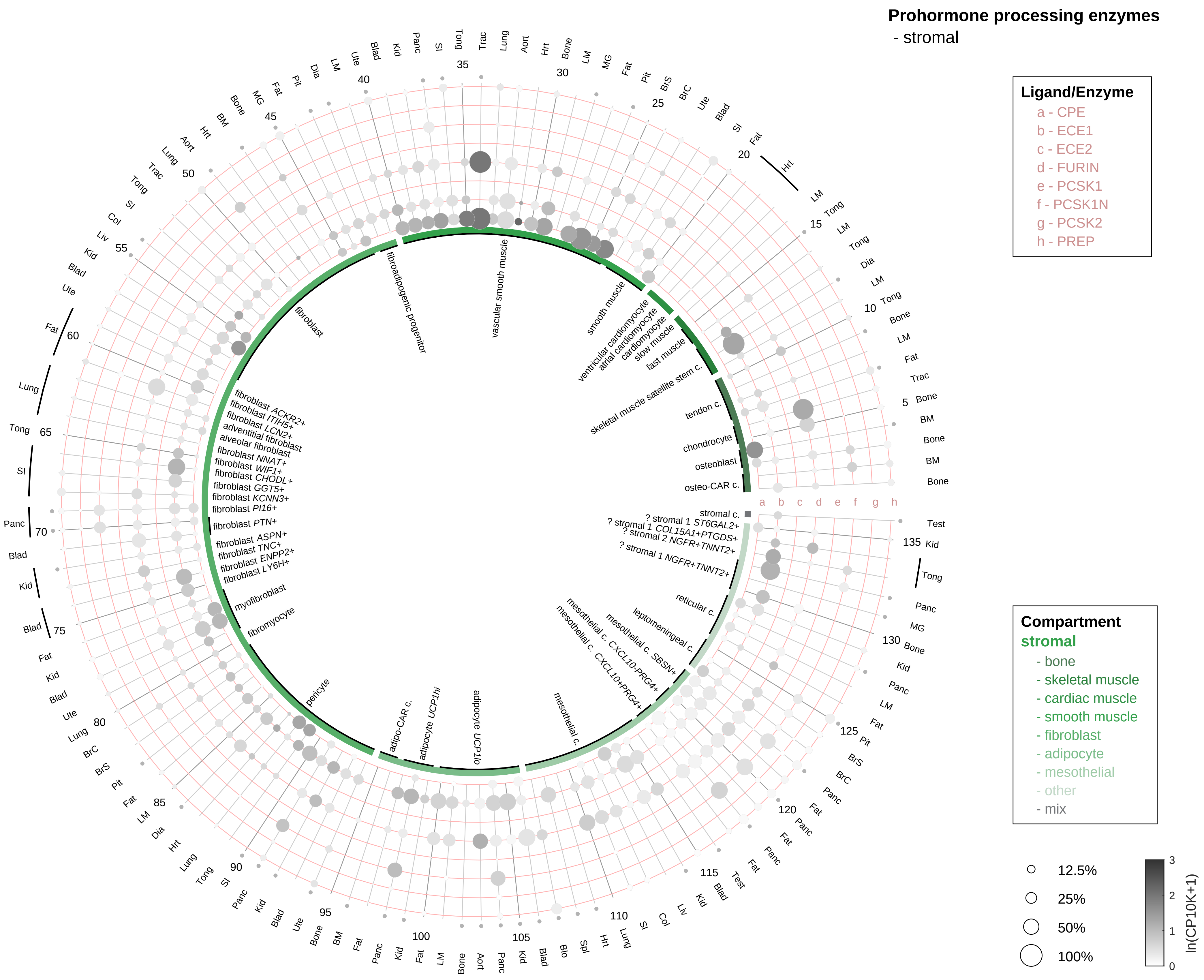

| Abbrev. | Tissue |
|---|---|
| Aort | Aorta |
| Blad | Bladder |
| Blo | Blood |
| Bone | Bone |
| BM | Bone marrow |
| BrC | Brain cortex |
| BrS | Brainstem |
| Col | Colon |
| Dia | Diaphragm |
| Ret | Eye retina |
| Fat | Fat |
| Hrt | Heart |
| Pit | Hypothalamus/Pituitary |
| Kid | Kidney |
| LM | Limb muscle |
| Liv | Liver |
| Lung | Lung |
| MG | Mammary gland |
| Panc | Pancreas |
| Skin | Skin |
| SI | Small intestine |
| Spl | Spleen |
| Test | Testes |
| Thym | Thymus |
| Tong | Tongue |
| Trac | Trachea |
| Ute | Uterus |

**Prohormone processing enzymes**
- lymphoid

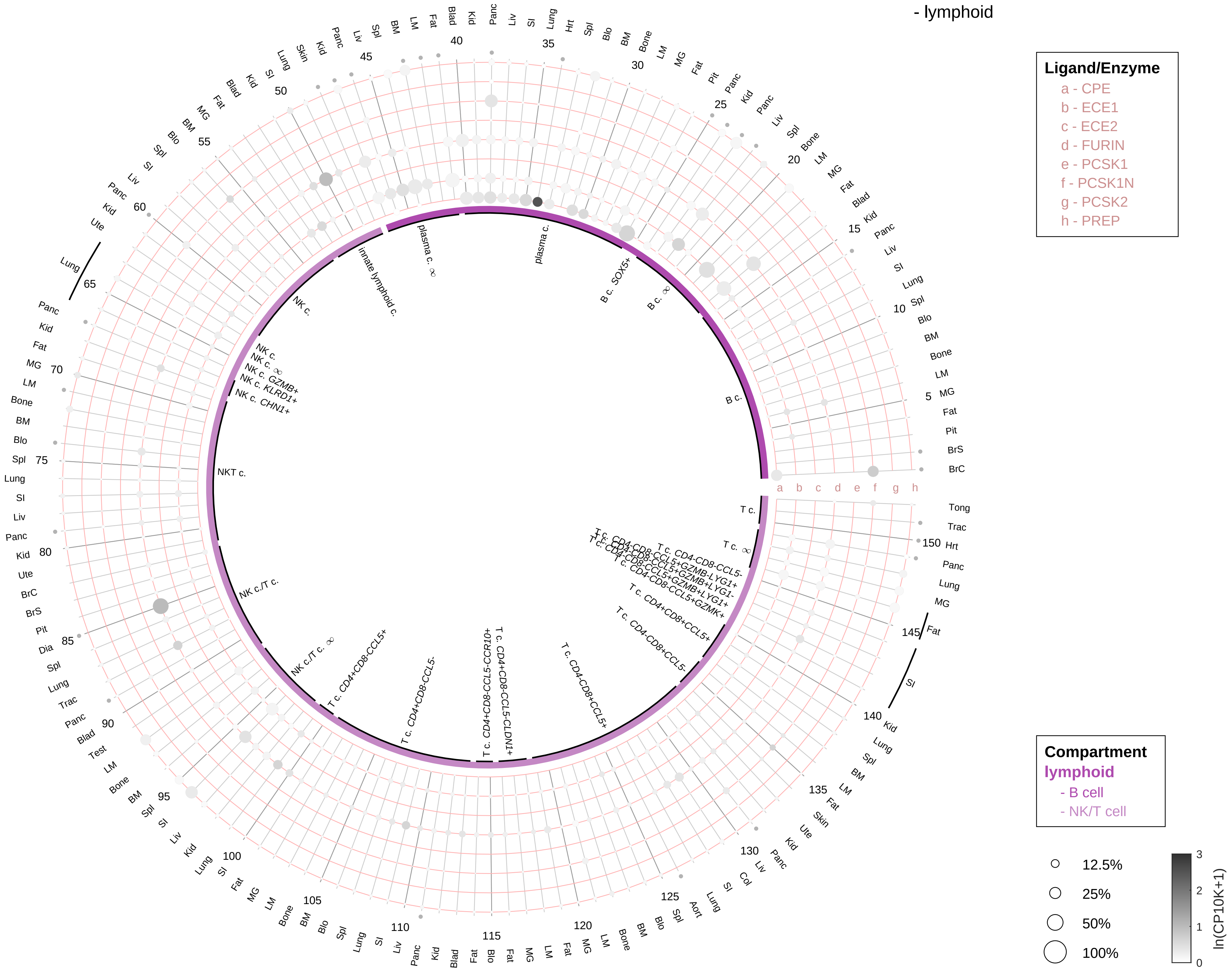

| Abbrev. | Tissue |
|---|---|
| Aort | Aorta |
| Blad | Bladder |
| Blo | Blood |
| Bone | Bone |
| BM | Bone marrow |
| BrC | Brain cortex |
| BrS | Brainstem |
| Col | Colon |
| Dia | Diaphragm |
| Ret | Eye retina |
| Fat | Fat |
| Hrt | Heart |
| Pit | Hypothalamus/Pituitary |
| Kid | Kidney |
| LM | Limb muscle |
| Liv | Liver |
| Lung | Lung |
| MG | Mammary gland |
| Panc | Pancreas |
| Skin | Skin |
| SI | Small intestine |
| Spl | Spleen |
| Test | Testes |
| Thym | Thymus |
| Tong | Tongue |
| Trac | Trachea |
| Ute | Uterus |

**Prohormone processing enzymes**
- myeloid

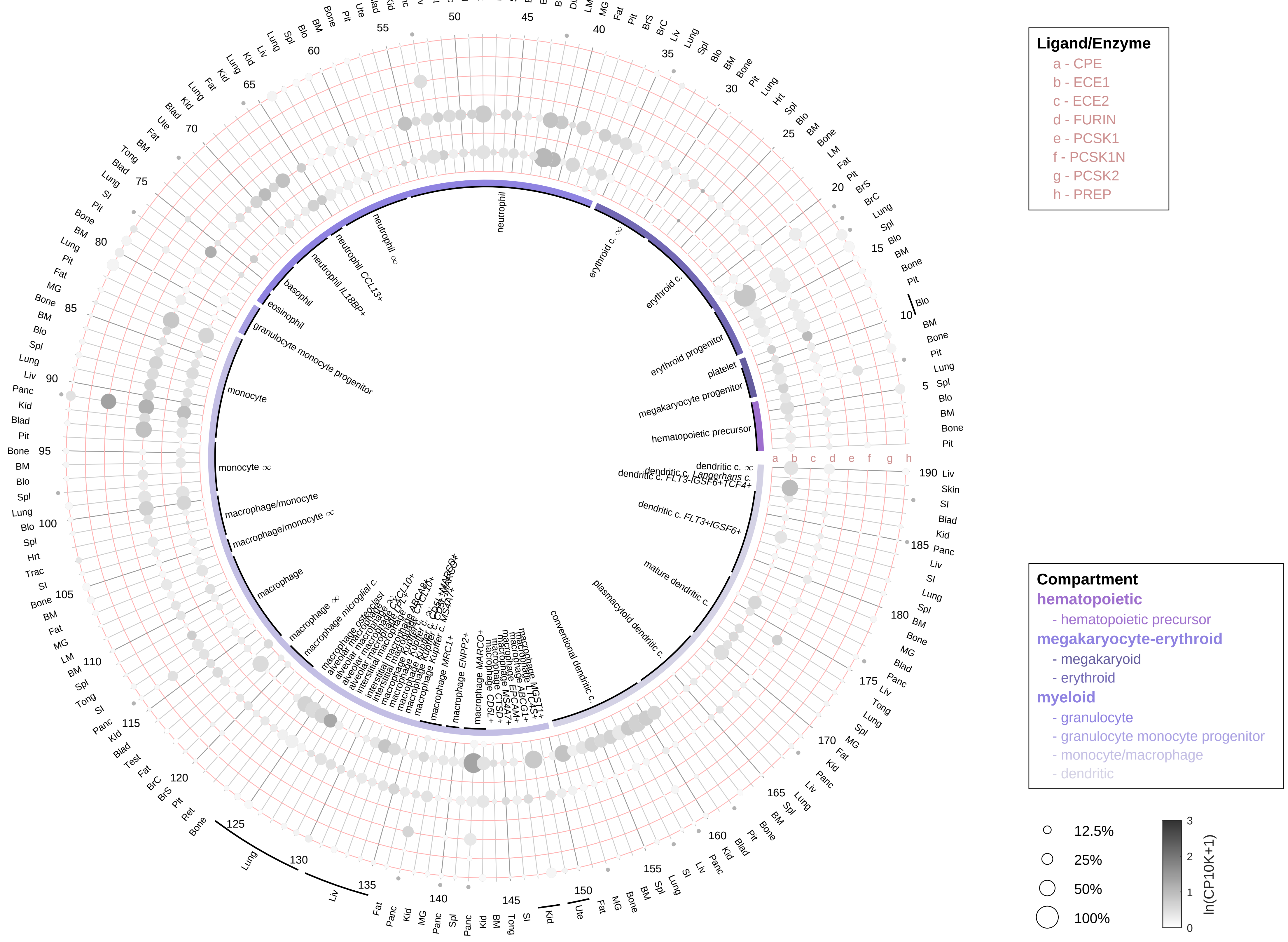

| Abbrev. | Tissue |
|---|---|
| Aort | Aorta |
| Blad | Bladder |
| Blo | Blood |
| Bone | Bone |
| BM | Bone marrow |
| BrC | Brain cortex |
| BrS | Brainstem |
| Col | Colon |
| Dia | Diaphragm |
| Ret | Eye retina |
| Fat | Fat |
| Hrt | Heart |
| Pit | Hypothalamus/Pituitary |
| Kid | Kidney |
| LM | Limb muscle |
| Liv | Liver |
| Lung | Lung |
| MG | Mammary gland |
| Panc | Pancreas |
| Skin | Skin |
| SI | Small intestine |
| Spl | Spleen |
| Test | Testes |
| Thym | Thymus |
| Tong | Tongue |
| Trac | Trachea |
| Ute | Uterus |
