## Supplementary material for "An organism-wide atlas of hormonal signaling based on the mouse lemur single-cell transcriptome": Fig. S1b. Ligand and receptor gene expression for 84 hormone classes across the mouse lemur cell atlas, with cell types ordered by tissue.

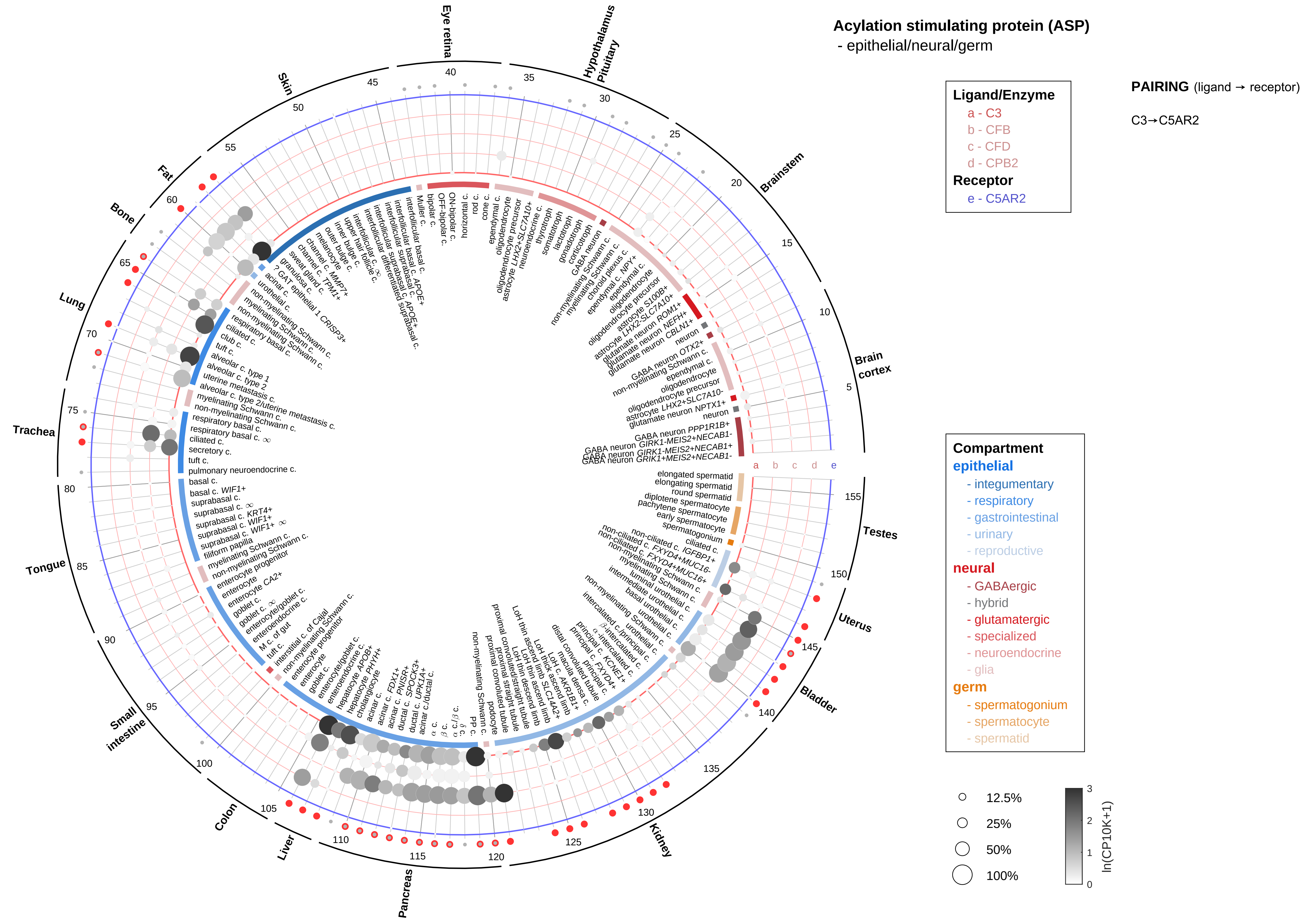
Acylation stimulating protein (ASP)
- epithelial/neural/germ
Ligand/Enzyme
a - C3
b - CFB
c - CFD
d - CPB2
Receptor
e - C5AR2
PAIRING (ligand → receptor)
C3→C5AR2
Compartment
epithelial
- integumentary
- respiratory
- gastrointestinal
- urinary
- reproductive
neural
- GABAergic
- hybrid
- glutamatergic
- specialized
- neuroendocrine
- glia
germ
- spermatogonium
- spermatocyte
- spermatid
12.5%
25%
50%
100%
ln(CP10K+1)
Eye retina
Hypothalamus Pituitary
Brainstem
Brain cortex
Testes
Uterus
Bladder
Kidney
Pancreas
Liver
Colon
Small intestine
Tongue
Trachea
Lung
Bone
Fat
Skin

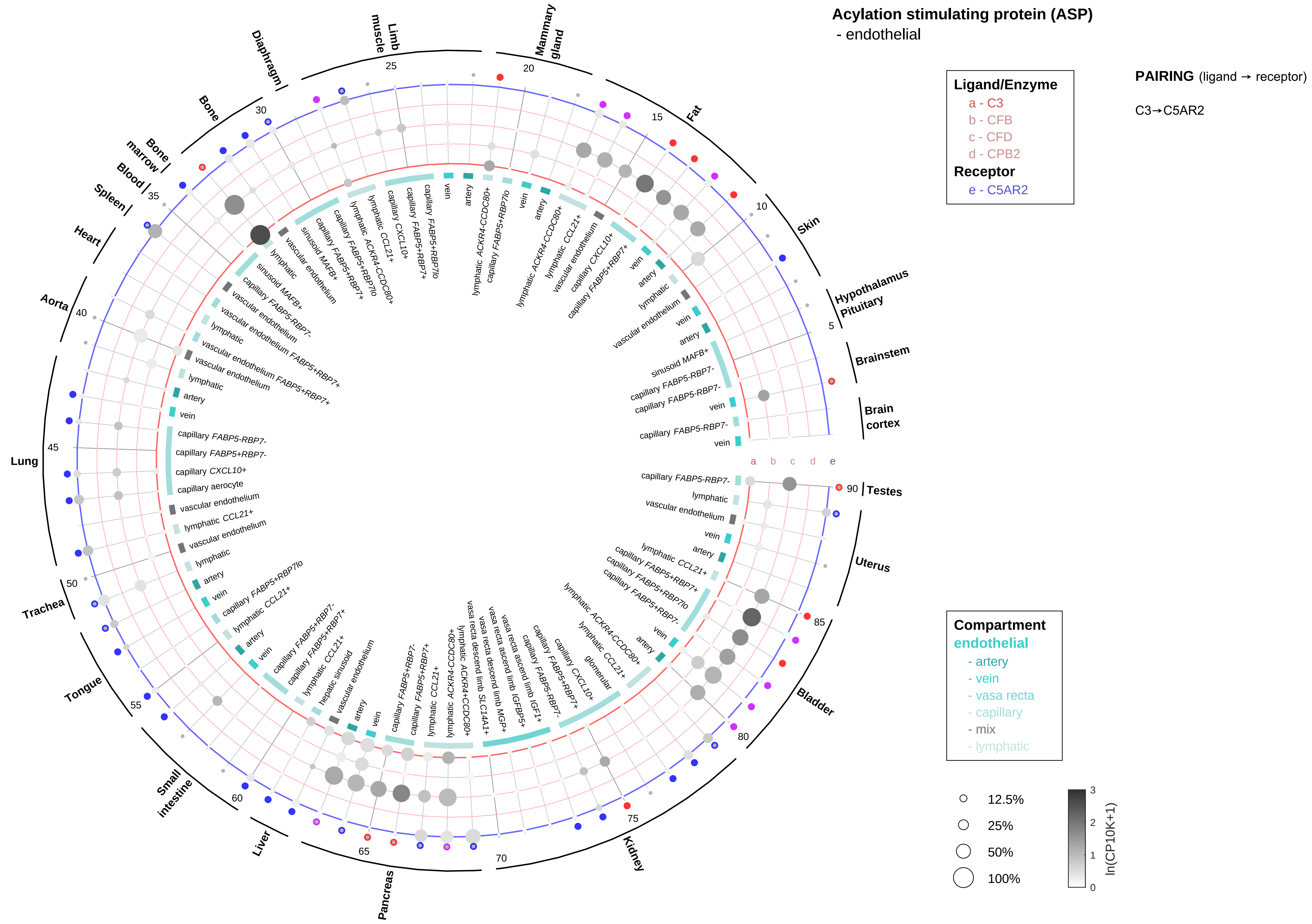

Acylation stimulating protein (ASP)
- endothelial
Ligand/Enzyme
a - C3
b - CFB
c - CFD
d - CPB2
Receptor
e - C5AR2
PAIRING (ligand → receptor)
C3→C5AR2
Compartment
endothelial
- artery
- vein
- vasa recta
- capillary
- mix
- lymphatic
12.5%
25%
50%
100%
ln(CP10K+1)
Brain cortex
Brainstem
Hypothalamus Pituitary
Skin
Fat
Mammary gland
Limb muscle
Diaphragm
Bone
Bone marrow
Blood
Spleen
Heart
Aorta
Lung
Trachea
Tongue
Small intestine
Liver
Pancreas
Kidney
Bladder
Uterus
Testes

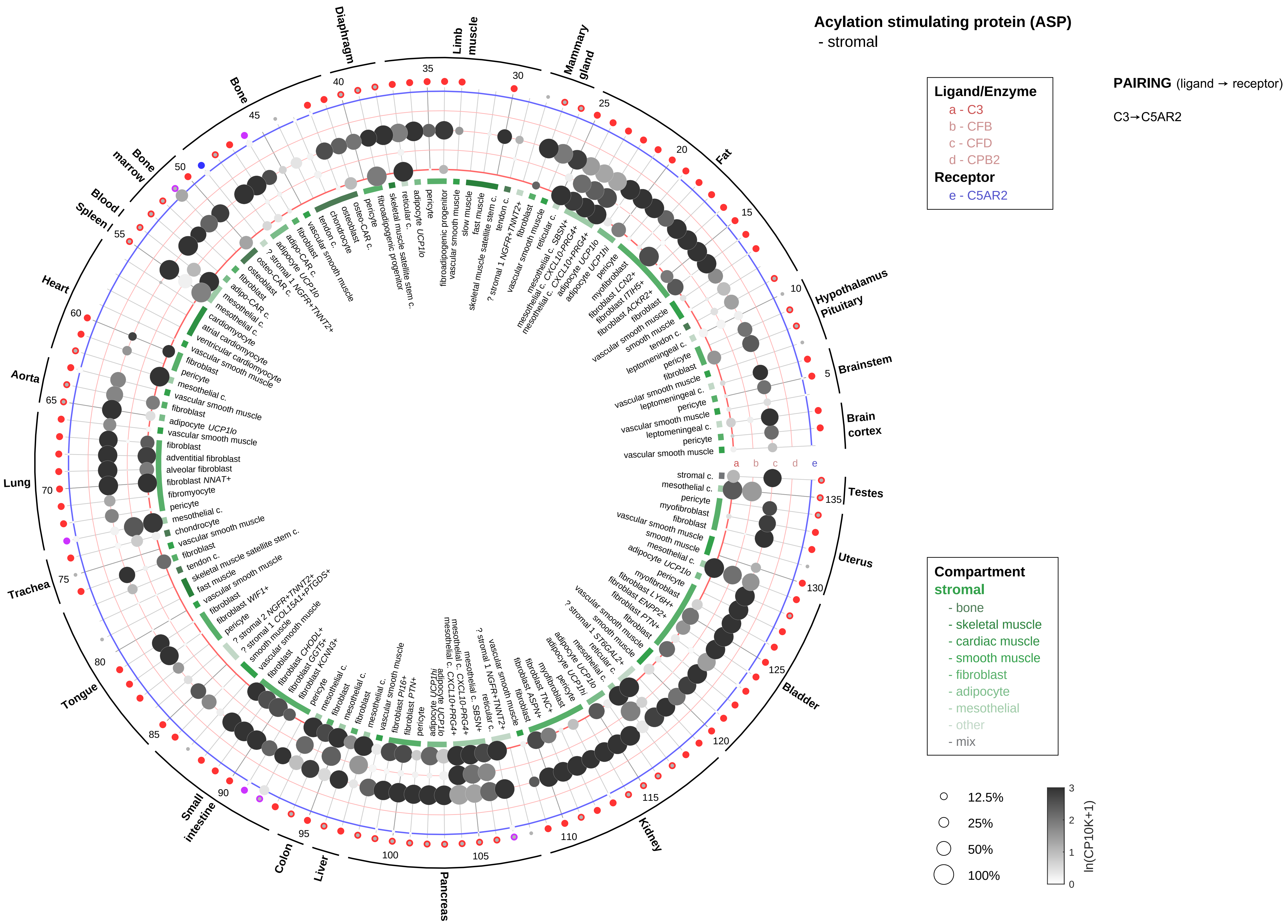

Acylation stimulating protein (ASP)
- stromal
Ligand/Enzyme
a - C3
b - CFB
c - CFD
d - CPB2
Receptor
e - C5AR2
PAIRING (ligand → receptor)
C3→C5AR2
Compartment
stromal
- bone
- skeletal muscle
- cardiac muscle
- smooth muscle
- fibroblast
- adipocyte
- mesothelial
- other
- mix
12.5%
25%
50%
100%
ln(CP10K+1)
Diaphragm
Limb muscle
Mammary gland
Fat
Hypothalamus Pituitary
Brainstem
Brain cortex
Testes
Uterus
Bladder
Kidney
Pancreas
Liver
Colon
Small intestine
Tongue
Trachea
Lung
Aorta
Heart
Spleen
Blood
Bone marrow
Bone

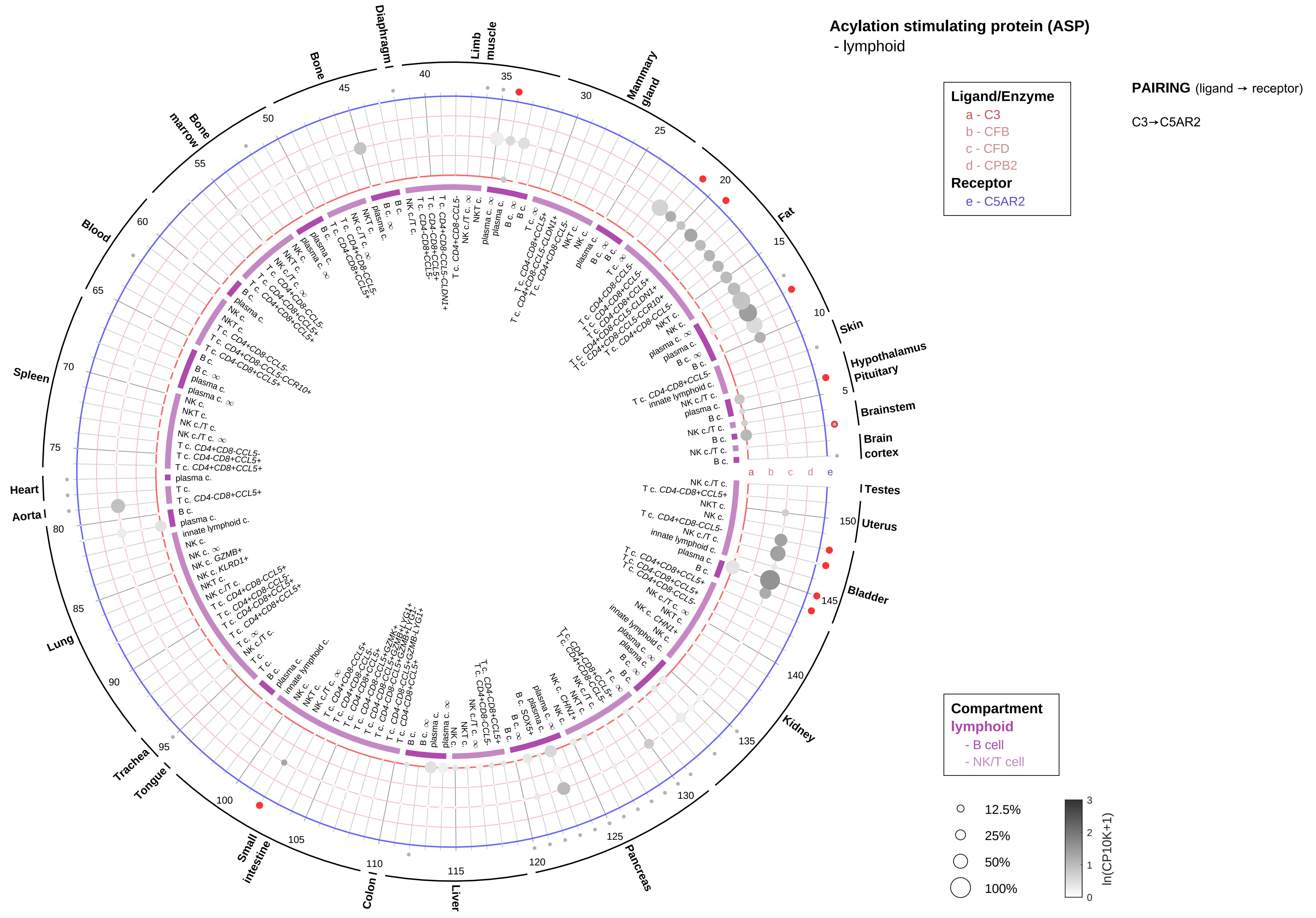
Acylation stimulating protein (ASP)
- lymphoid
Ligand/Enzyme
a - C3
b - CFB
c - CFD
d - CPB2
Receptor
e - C5AR2
PAIRING (ligand → receptor)
C3→C5AR2
Compartment
lymphoid
- B cell
- NK/T cell
12.5%
25%
50%
100%
ln(CP10K+1)
Diaphragm
Limb muscle
Mammary gland
Fat
Skin
Hypothalamus
Pituitary
Brainstem
Brain cortex
Testes
Uterus
Bladder
Kidney
Pancreas
Liver
Colon
Small intestine
Tongue
Trachea
Lung
Aorta
Heart
Spleen
Blood
Bone marrow
Bone

#### Acylation stimulating protein (ASP)
- myeloid

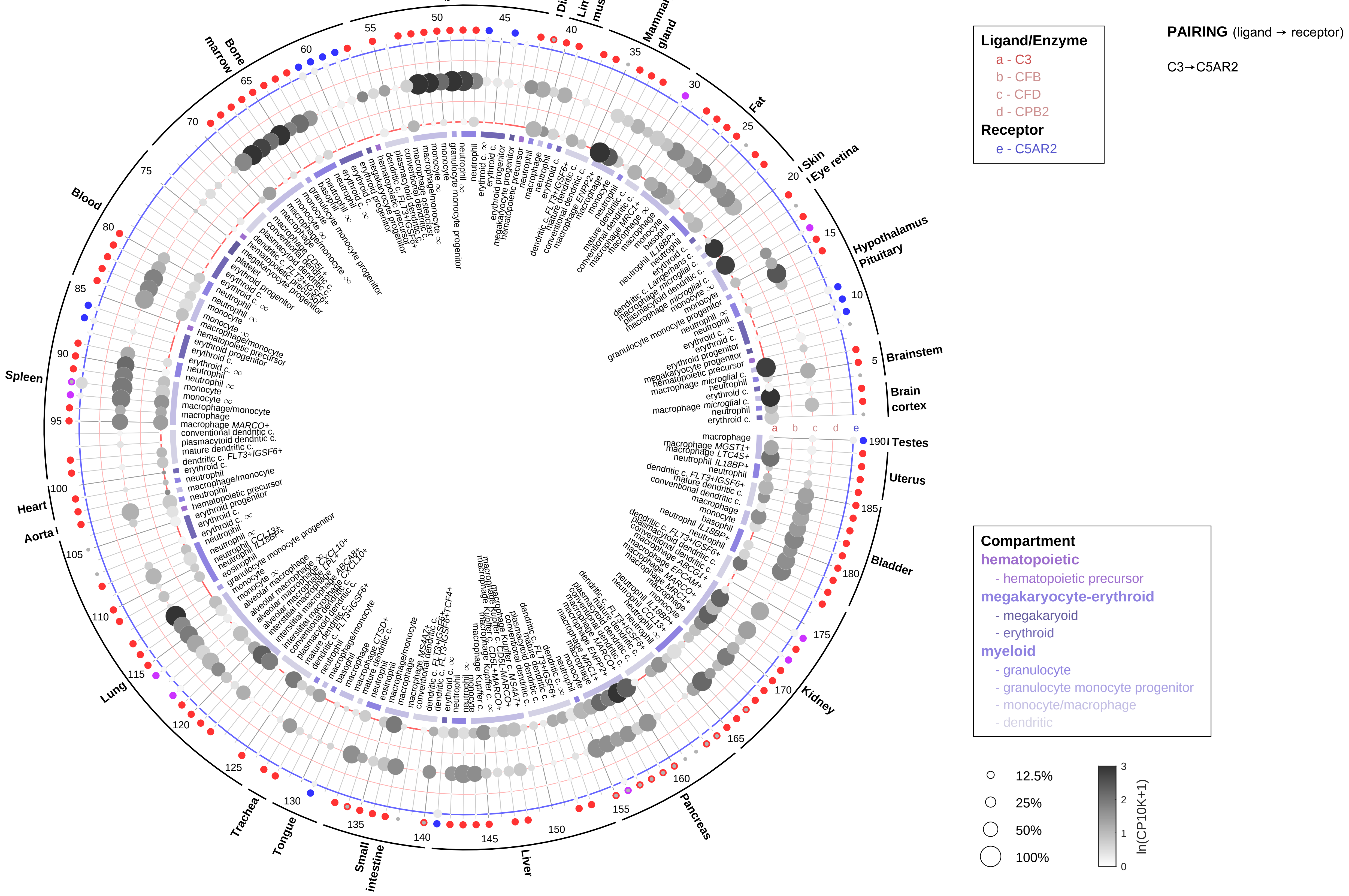

**Adiponectin (AdipoQ)**
- epithelial/neural/germ

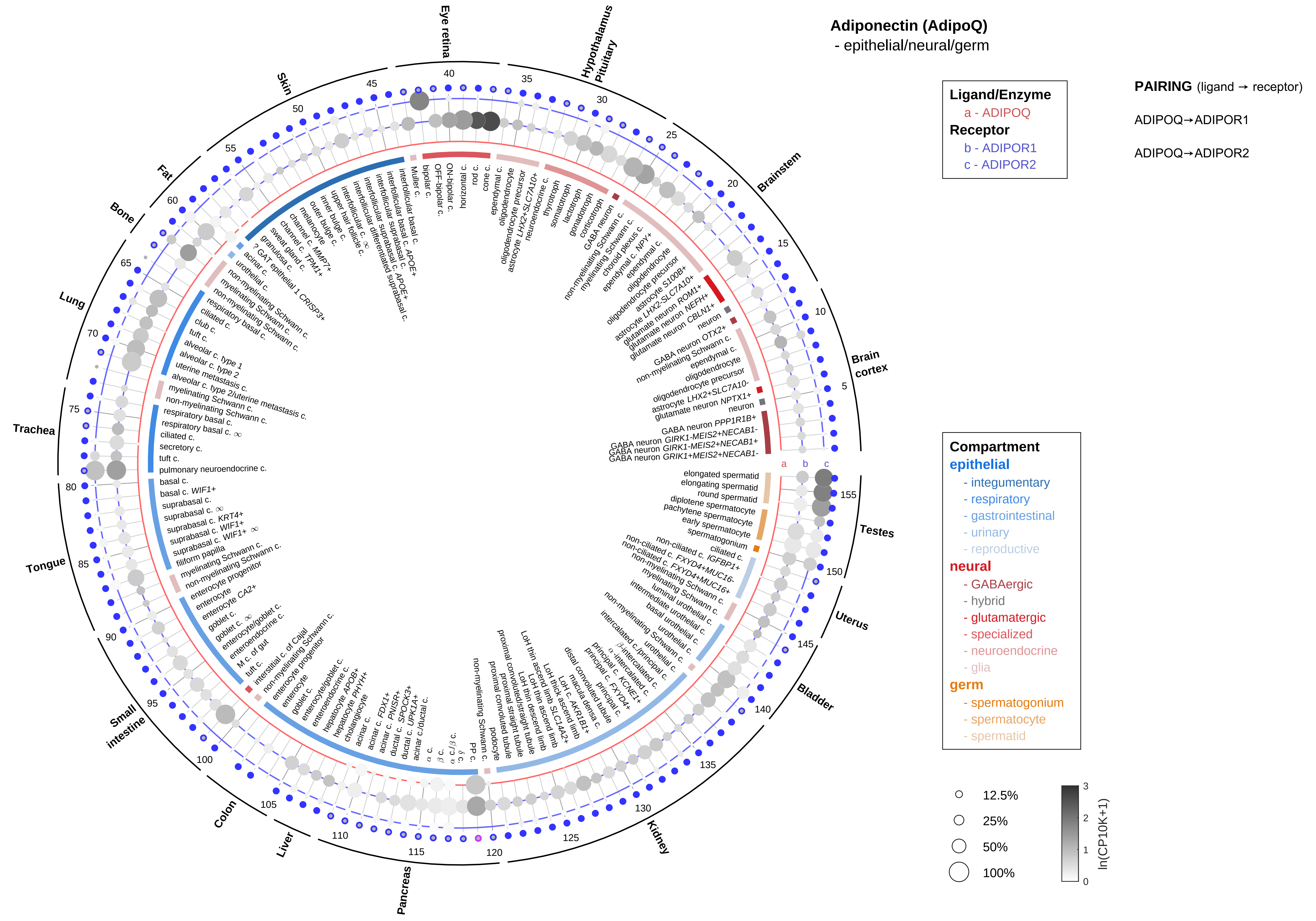

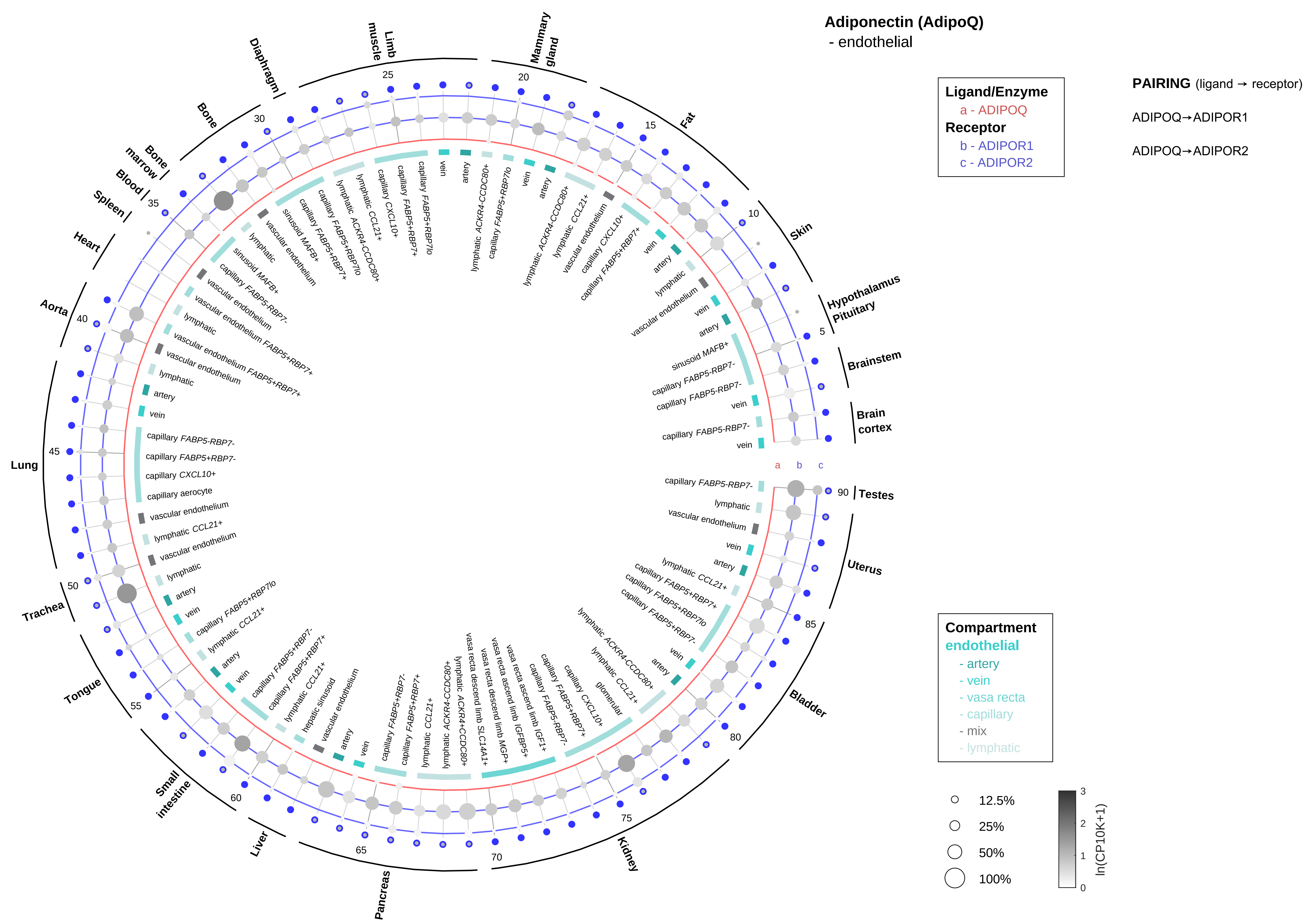
Adiponectin (AdipoQ)
- endothelial
Ligand/Enzyme
a - ADIPOQ
Receptor
b - ADIPOR1
c - ADIPOR2
PAIRING (ligand → receptor)
ADIPOQ→ADIPOR1
ADIPOQ→ADIPOR2
Compartment
endothelial
- artery
- vein
- vasa recta
- capillary
- mix
- lymphatic
12.5%
25%
50%
100%
ln(CP10K+1)
Brain cortex
Brainstem
Hypothalamus Pituitary
Skin
Fat
Mammary gland
Limb muscle
Diaphragm
Bone
Bone marrow
Blood
Spleen
Heart
Aorta
Lung
Trachea
Tongue
Small intestine
Liver
Pancreas
Kidney
Bladder
Uterus
Testes

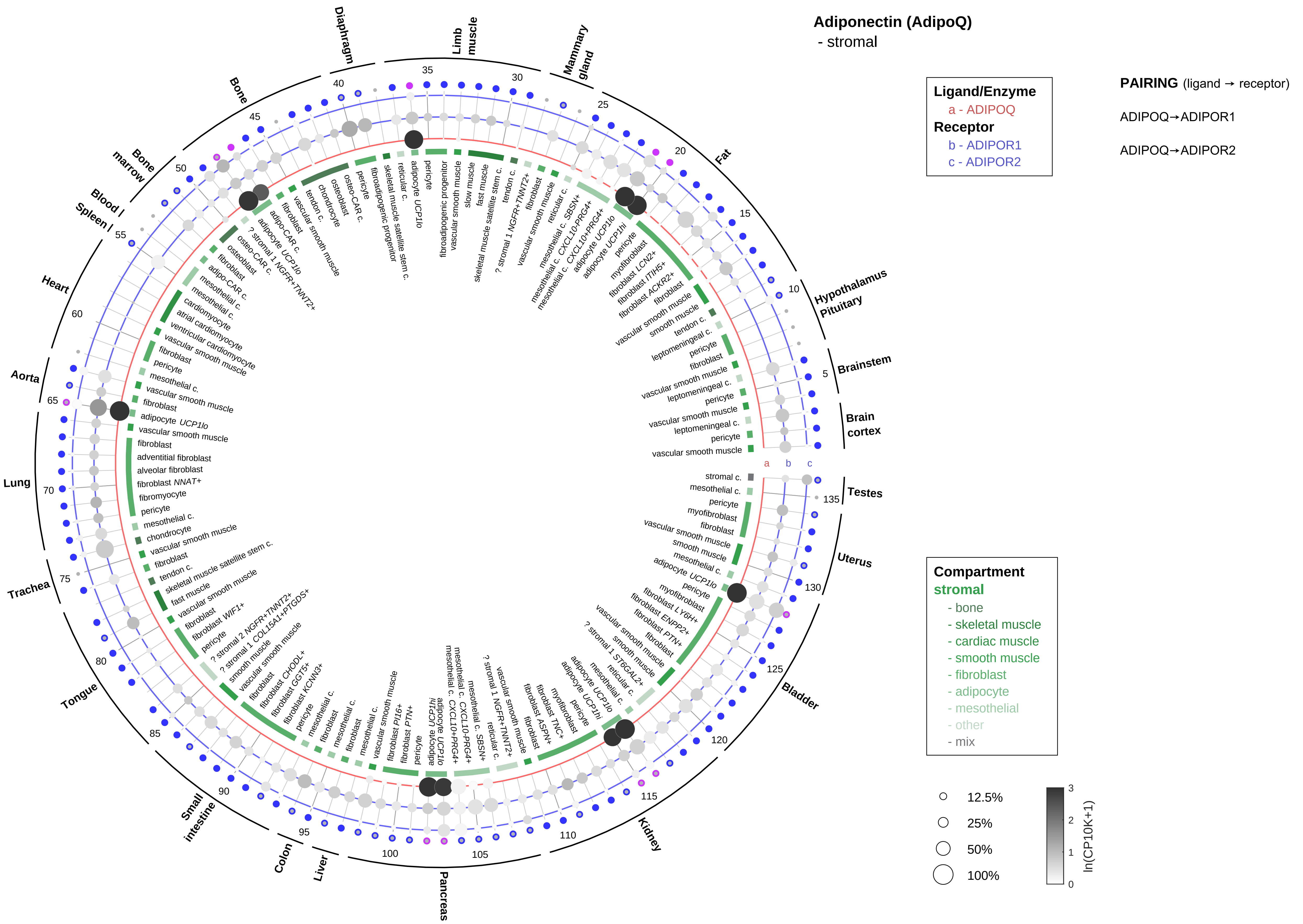

Adiponectin (AdipoQ)
- stromal
Ligand/Enzyme
a - ADIPOQ
Receptor
b - ADIPOR1
c - ADIPOR2
PAIRING (ligand → receptor)
ADIPOQ→ADIPOR1
ADIPOQ→ADIPOR2
Compartment
stromal
- bone
- skeletal muscle
- cardiac muscle
- smooth muscle
- fibroblast
- adipocyte
- mesothelial
- other
- mix
12.5%
25%
50%
100%
ln(CP10K+1)
Diaphragm
Limb muscle
Mammary gland
Fat
Hypothalamus Pituitary
Brainstem
Brain cortex
Testes
Uterus
Bladder
Kidney
Pancreas
Liver
Colon
Small intestine
Tongue
Trachea
Lung
Aorta
Heart
Spleen
Blood
Bone marrow
Bone

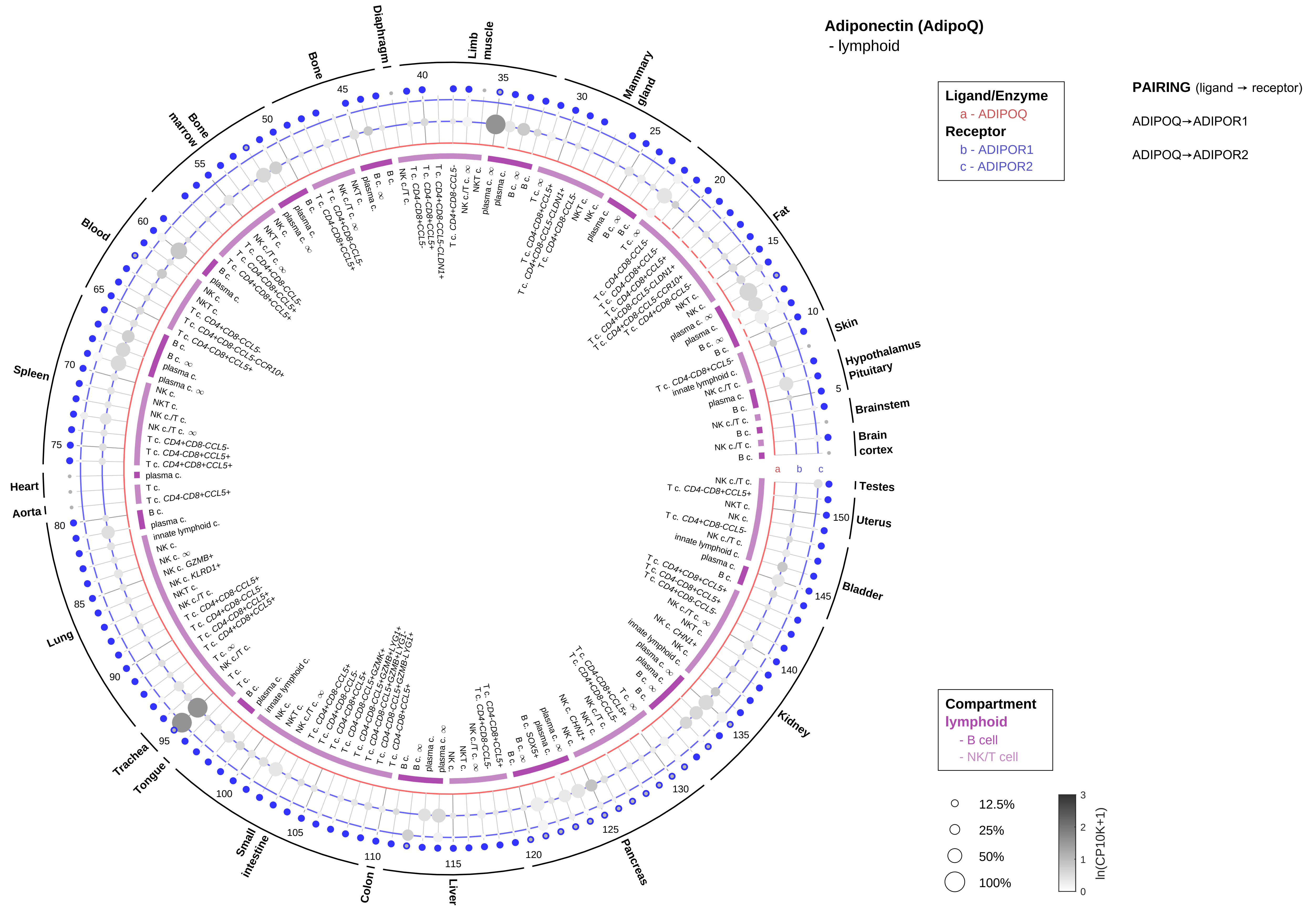

Adiponectin (AdipoQ)
- lymphoid
Ligand/Enzyme
a - ADIPOQ
Receptor
b - ADIPOR1
c - ADIPOR2
PAIRING (ligand → receptor)
ADIPOQ→ADIPOR1
ADIPOQ→ADIPOR2
Compartment
lymphoid
- B cell
- NK/T cell
12.5%
25%
50%
100%
ln(CP10K+1)
Skin
Hypothalamus
Pituitary
Brainstem
Brain cortex
Testes
Uterus
Bladder
Kidney
Pancreas
Liver
Colon
Small intestine
Tongue
Trachea
Lung
Aorta
Heart
Spleen
Blood
Bone marrow
Bone
Diaphragm
Limb muscle
Mammary gland
Fat

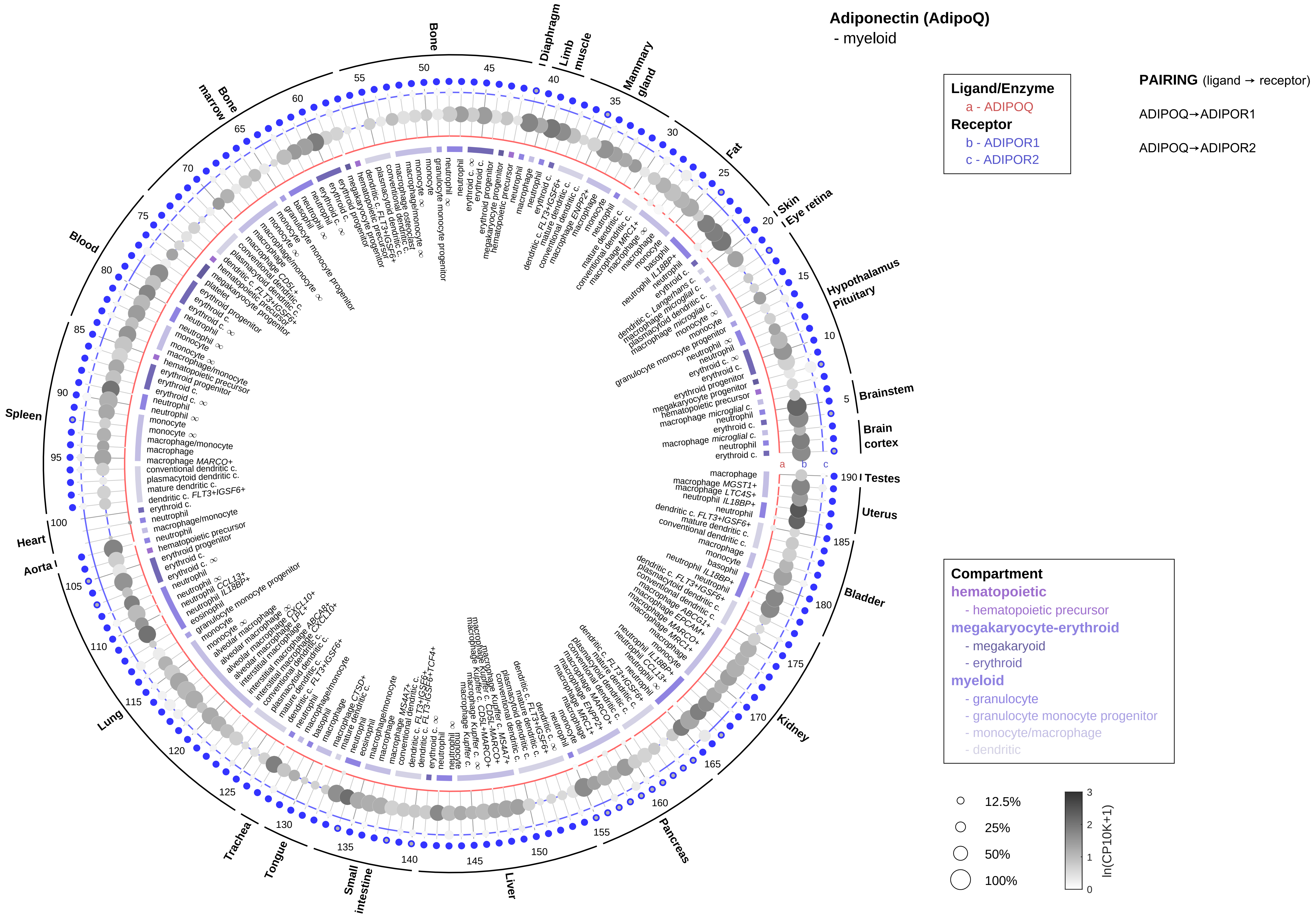
Adiponectin (AdipoQ)
- myeloid
Ligand/Enzyme
a - ADIPOQ
Receptor
b - ADIPOR1
c - ADIPOR2
PAIRING (ligand → receptor)
ADIPOQ→ADIPOR1
ADIPOQ→ADIPOR2
Compartment
hematopoietic
- hematopoietic precursor
megakaryocyte-erythroid
- megakaryoid
- erythroid
myeloid
- granulocyte
- granulocyte monocyte progenitor
- monocyte/macrophage
- dendritic
12.5%
25%
50%
100%
ln(CP10K+1)
Brainstem
Brain cortex
Testes
Uterus
Bladder
Kidney
Pancreas
Liver
Small intestine
Tongue
Trachea
Lung
Aorta
Heart
Spleen
Blood
Bone marrow
Bone
Diaphragm
Limb muscle
Mammary gland
Fat
Skin
Eye retina
Hypothalamus
Pituitary

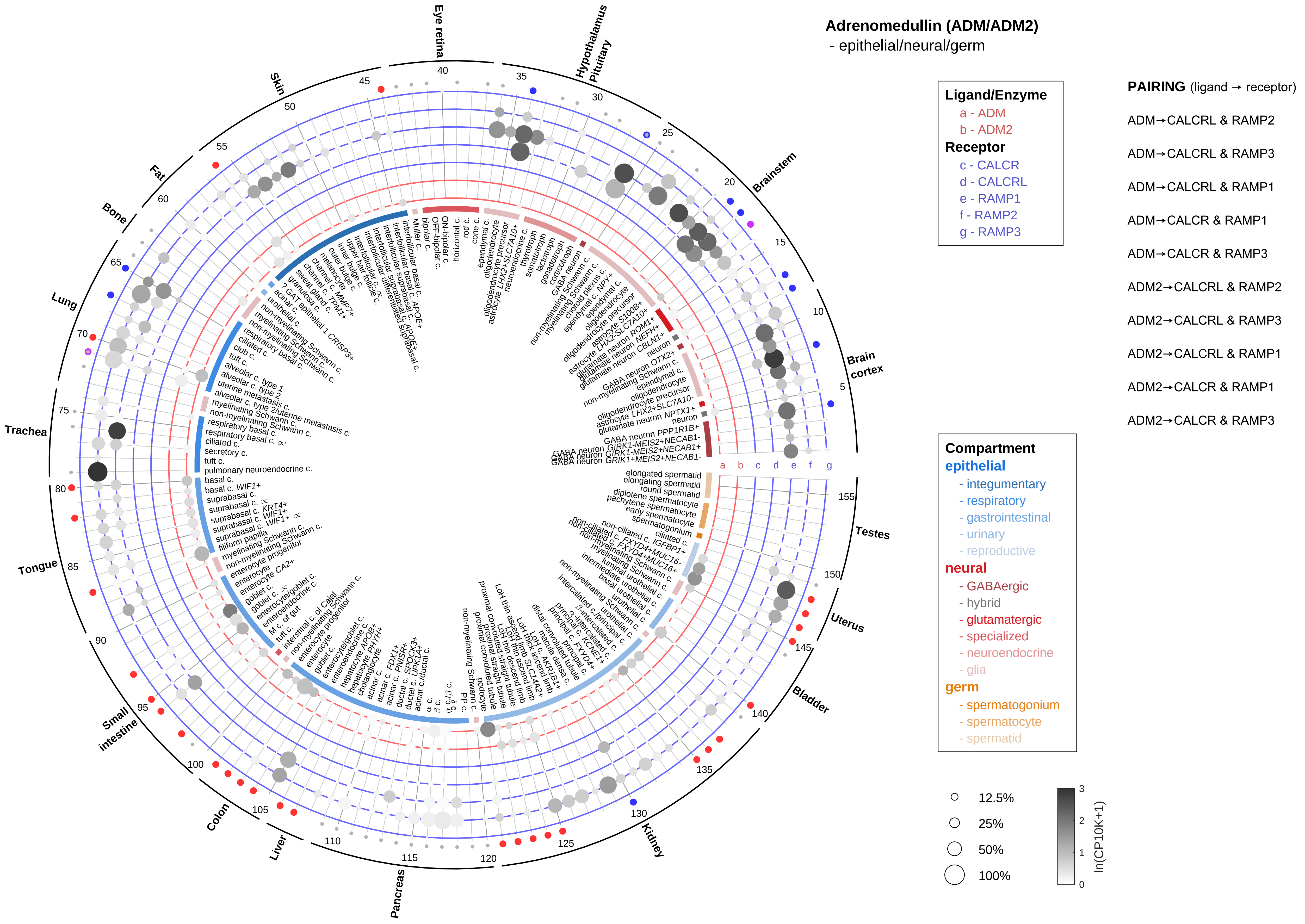
Adrenomedullin (ADM/ADM2)
- epithelial/neural/germ
Ligand/Enzyme
a - ADM
b - ADM2
Receptor
c - CALCR
d - CALCRL
e - RAMP1
f - RAMP2
g - RAMP3
PAIRING (ligand → receptor)
ADM→CALCRL & RAMP2
ADM→CALCRL & RAMP3
ADM→CALCRL & RAMP1
ADM→CALCR & RAMP1
ADM→CALCR & RAMP3
ADM2→CALCRL & RAMP2
ADM2→CALCRL & RAMP3
ADM2→CALCRL & RAMP1
ADM2→CALCR & RAMP1
ADM2→CALCR & RAMP3
Compartment
epithelial
- integumentary
- respiratory
- gastrointestinal
- urinary
- reproductive
neural
- GABAergic
- hybrid
- glutamatergic
- specialized
- neuroendocrine
- glia
germ
- spermatogonium
- spermatocyte
- spermatid
12.5%
25%
50%
100%
ln(CP10K+1)
Eye retina
Hypothalamus
Pituitary
Brainstem
Brain cortex
Testes
Uterus
Bladder
Kidney
Pancreas
Liver
Colon
Small intestine
Tongue
Trachea
Lung
Bone
Fat
Skin

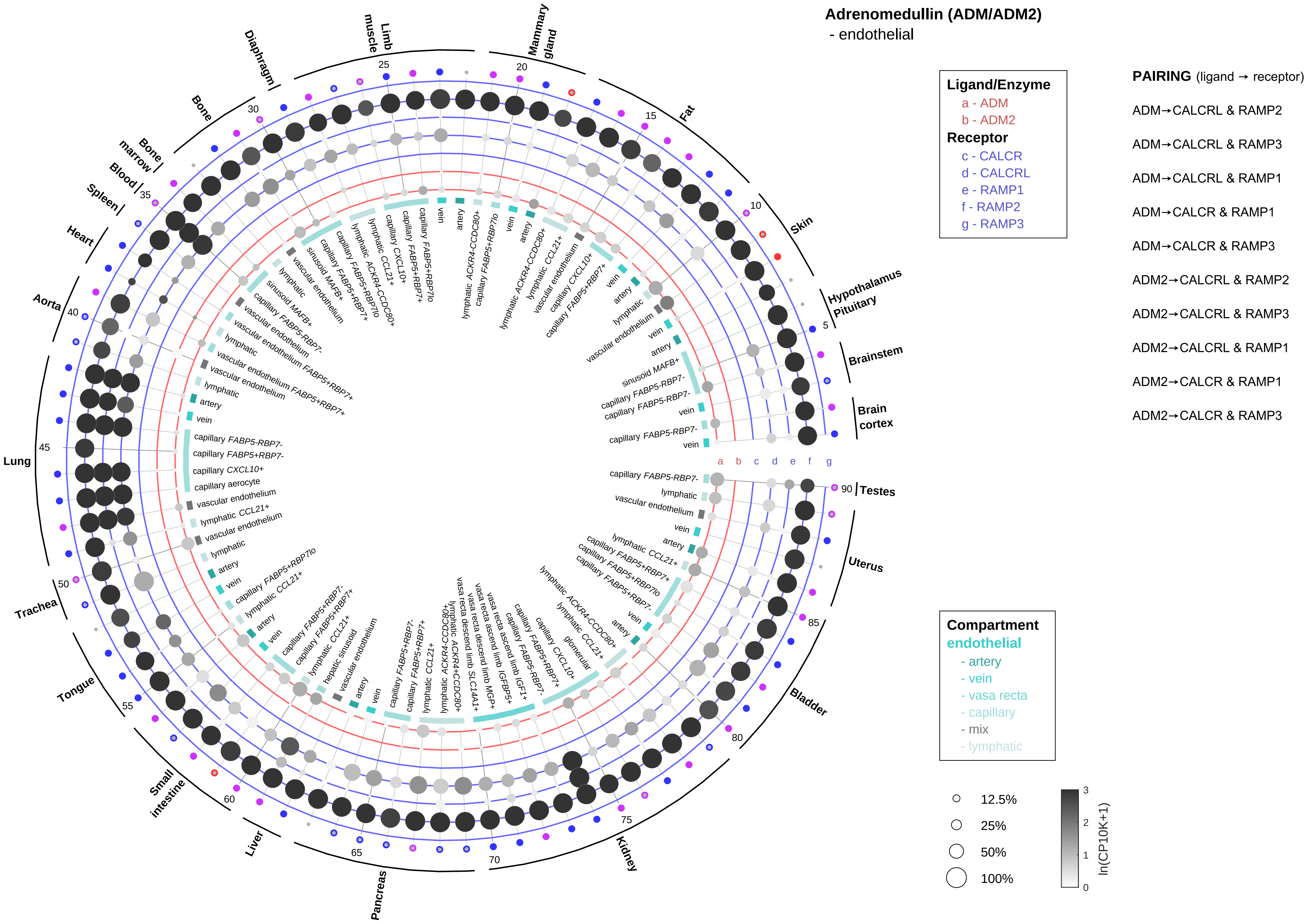

Adrenomedullin (ADM/ADM2)
- endothelial
Ligand/Enzyme
a - ADM
b - ADM2
Receptor
c - CALCR
d - CALCRL
e - RAMP1
f - RAMP2
g - RAMP3
PAIRING (ligand → receptor)
ADM→CALCRL & RAMP2
ADM→CALCRL & RAMP3
ADM→CALCRL & RAMP1
ADM→CALCR & RAMP1
ADM→CALCR & RAMP3
ADM2→CALCRL & RAMP2
ADM2→CALCRL & RAMP3
ADM2→CALCRL & RAMP1
ADM2→CALCR & RAMP1
ADM2→CALCR & RAMP3
Compartment
endothelial
- artery
- vein
- vasa recta
- capillary
- mix
- lymphatic
12.5%
25%
50%
100%
ln(CP10K+1)
Brain cortex
Brainstem
Hypothalamus Pituitary
Skin
Fat
Mammary gland
Limb muscle
Diaphragm
Bone
Bone marrow
Blood
Spleen
Heart
Aorta
Lung
Trachea
Tongue
Small intestine
Liver
Pancreas
Kidney
Bladder
Uterus
Testes
a b c d e f g

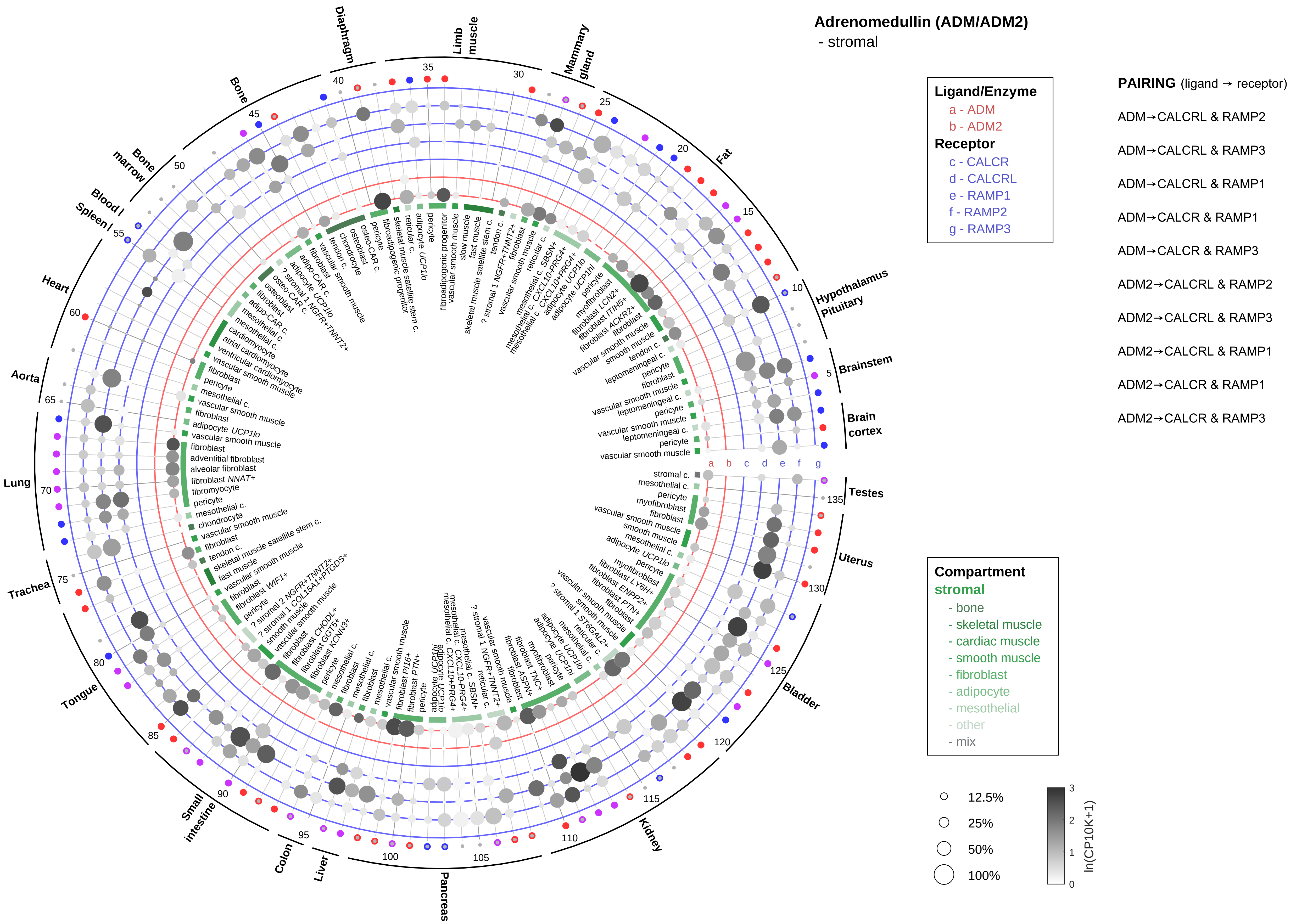

Adrenomedullin (ADM/ADM2)
- stromal
Ligand/Enzyme
a - ADM
b - ADM2
Receptor
c - CALCR
d - CALCRL
e - RAMP1
f - RAMP2
g - RAMP3
PAIRING (ligand → receptor)
ADM→CALCRL & RAMP2
ADM→CALCRL & RAMP3
ADM→CALCRL & RAMP1
ADM→CALCR & RAMP1
ADM→CALCR & RAMP3
ADM2→CALCRL & RAMP2
ADM2→CALCRL & RAMP3
ADM2→CALCRL & RAMP1
ADM2→CALCR & RAMP1
ADM2→CALCR & RAMP3
Compartment
stromal
- bone
- skeletal muscle
- cardiac muscle
- smooth muscle
- fibroblast
- adipocyte
- mesothelial
- other
- mix
12.5%
25%
50%
100%
ln(CP10K+1)
Diaphragm
Limb muscle
Mammary gland
Fat
Hypothalamus Pituitary
Brainstem
Brain cortex
Testes
Uterus
Bladder
Kidney
Pancreas
Liver
Colon
Small intestine
Tongue
Trachea
Lung
Aorta
Heart
Blood / Spleen
Bone marrow
Bone

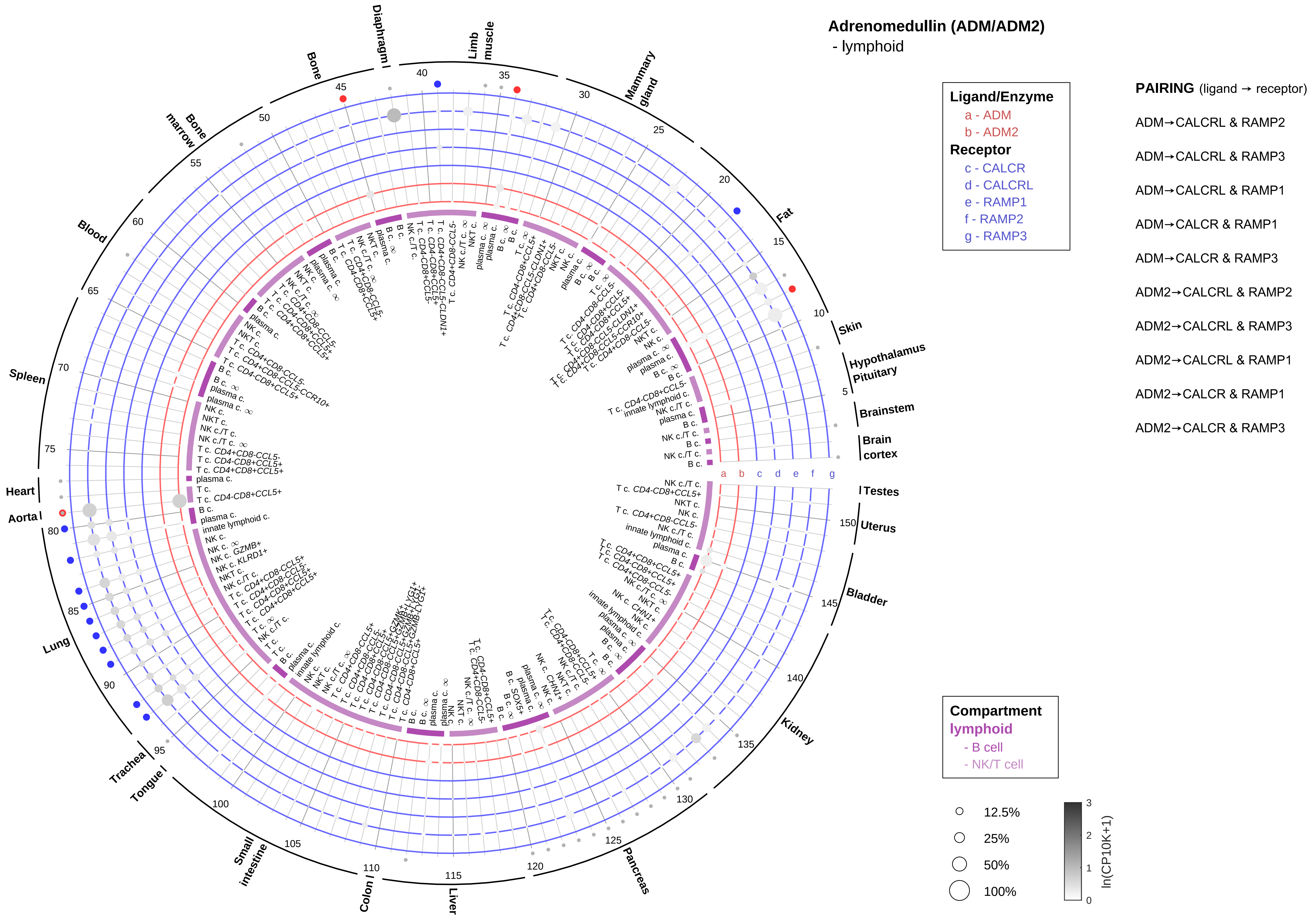
Adrenomedullin (ADM/ADM2)
- lymphoid
Ligand/Enzyme
a - ADM
b - ADM2
Receptor
c - CALCR
d - CALCRL
e - RAMP1
f - RAMP2
g - RAMP3
PAIRING (ligand → receptor)
ADM→CALCRL & RAMP2
ADM→CALCRL & RAMP3
ADM→CALCRL & RAMP1
ADM→CALCR & RAMP1
ADM→CALCR & RAMP3
ADM2→CALCRL & RAMP2
ADM2→CALCRL & RAMP3
ADM2→CALCRL & RAMP1
ADM2→CALCR & RAMP1
ADM2→CALCR & RAMP3
Compartment
lymphoid
- B cell
- NK/T cell
12.5%
25%
50%
100%
ln(CP10K+1)
Skin
Hypothalamus
Pituitary
Brainstem
Brain cortex
Testes
Uterus
Bladder
Kidney
Pancreas
Liver
Colon
Small intestine
Tongue
Trachea
Lung
Aorta
Heart
Spleen
Blood
Bone marrow
Bone
Diaphragm
Limb muscle
Mammary gland
Fat

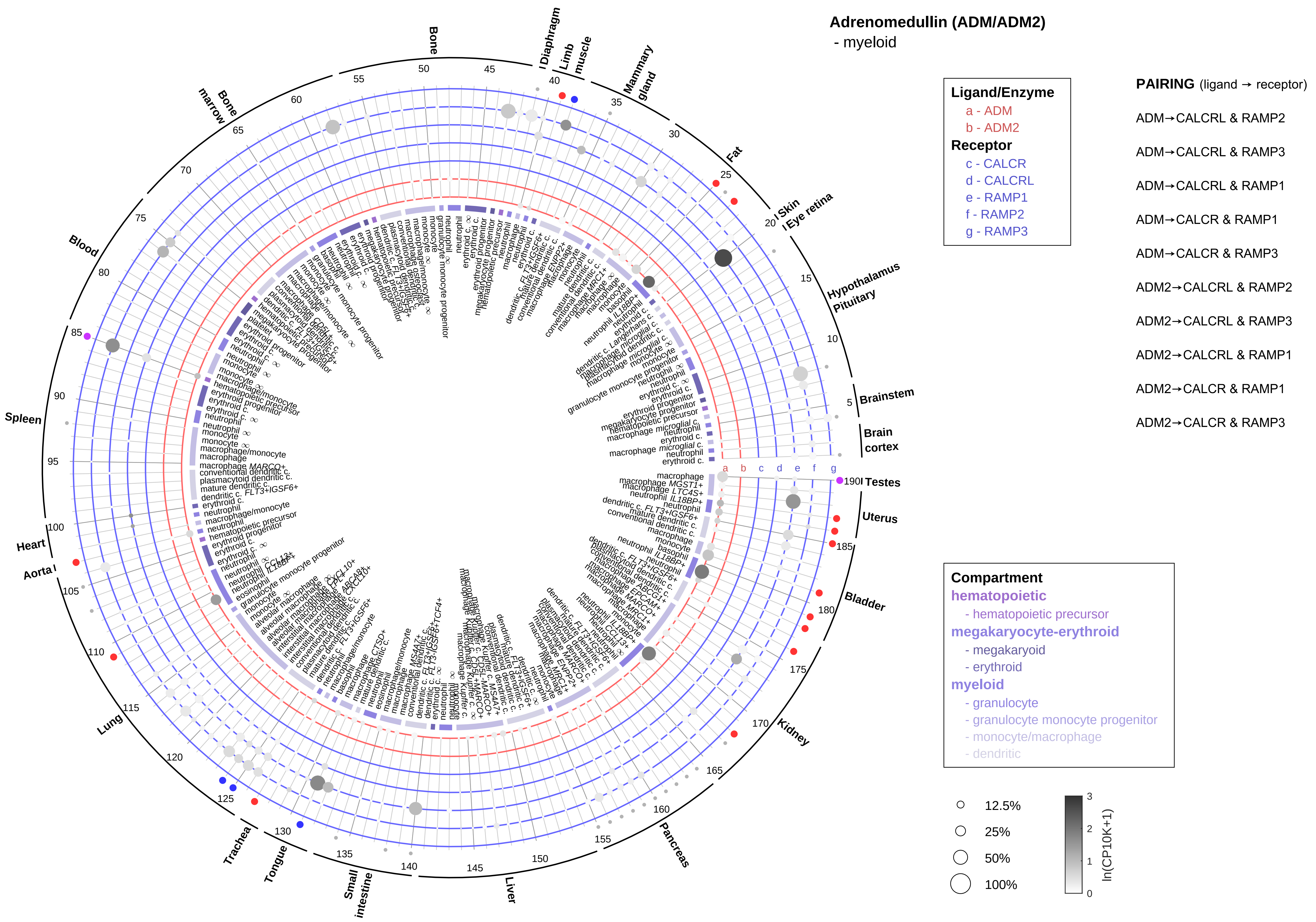

Adrenomedullin (ADM/ADM2)
- myeloid
Ligand/Enzyme
a - ADM
b - ADM2
Receptor
c - CALCR
d - CALCRL
e - RAMP1
f - RAMP2
g - RAMP3
PAIRING (ligand → receptor)
ADM→CALCRL & RAMP2
ADM→CALCRL & RAMP3
ADM→CALCRL & RAMP1
ADM→CALCR & RAMP1
ADM→CALCR & RAMP3
ADM2→CALCRL & RAMP2
ADM2→CALCRL & RAMP3
ADM2→CALCRL & RAMP1
ADM2→CALCR & RAMP1
ADM2→CALCR & RAMP3
Compartment
hematopoietic
- hematopoietic precursor
megakaryocyte-erythroid
- megakaryoid
- erythroid
myeloid
- granulocyte
- granulocyte monocyte progenitor
- monocyte/macrophage
- dendritic
12.5%
25%
50%
100%
ln(CP10K+1)
Bone
Diaphragm
Limb muscle
Mammary gland
Fat
Skin
Eye retina
Hypothalamus
Pituitary
Brainstem
Brain cortex
Testes
Uterus
Bladder
Kidney
Pancreas
Liver
Small intestine
Tongue
Trachea
Lung
Aorta
Heart
Spleen
Blood
Bone marrow

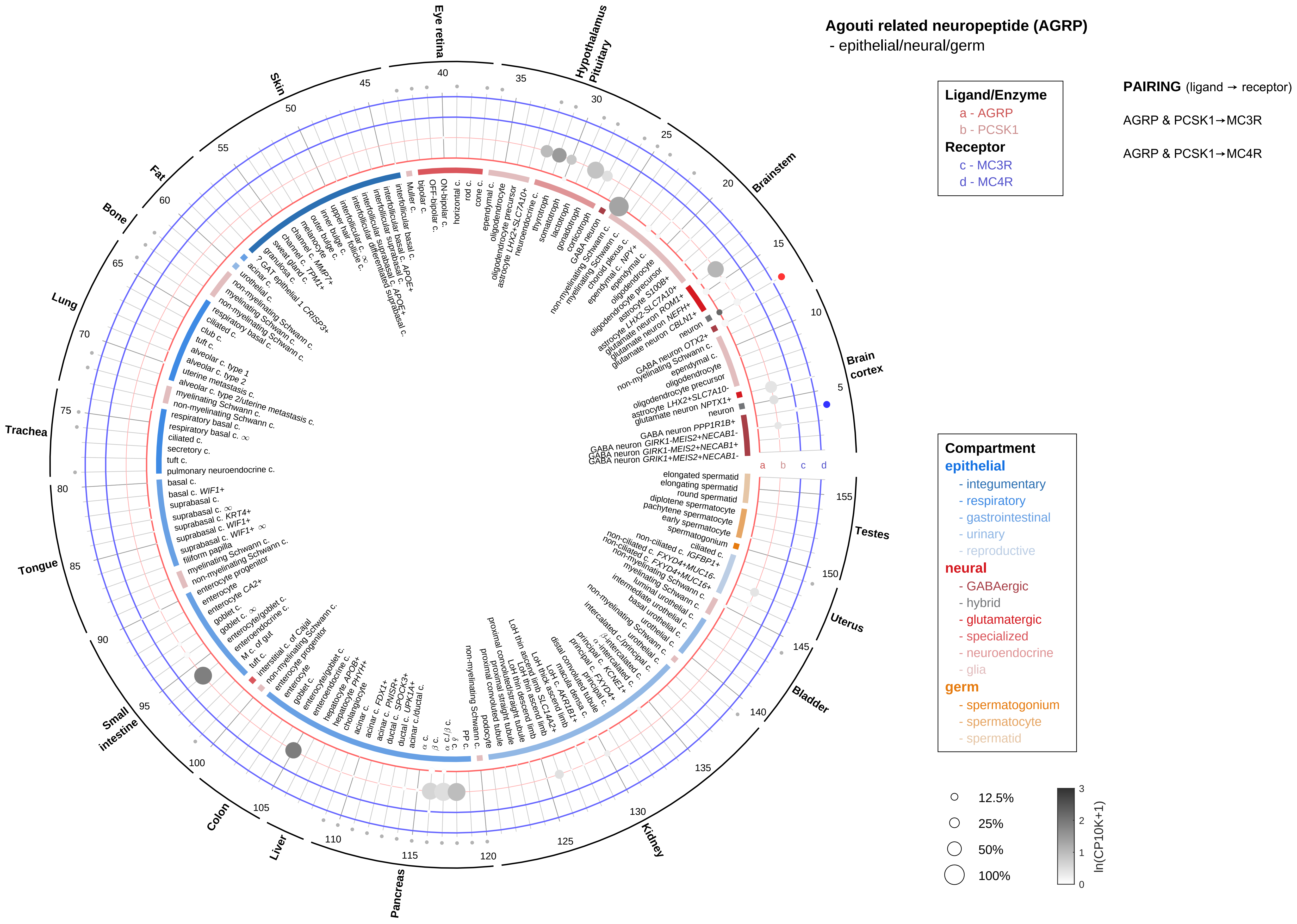

Agouti related neuropeptide (AGRP)
- epithelial/neural/germ
Ligand/Enzyme
a - AGRP
b - PCSK1
Receptor
c - MC3R
d - MC4R
PAIRING (ligand → receptor)
AGRP & PCSK1→MC3R
AGRP & PCSK1→MC4R
Compartment
epithelial
- integumentary
- respiratory
- gastrointestinal
- urinary
- reproductive
neural
- GABAergic
- hybrid
- glutamatergic
- specialized
- neuroendocrine
- glia
germ
- spermatogonium
- spermatocyte
- spermatid
12.5%
25%
50%
100%
ln(CP10K+1)
Brain cortex
Brainstem
Hypothalamus Pituitary
Eye retina
Skin
Fat
Bone
Lung
Trachea
Tongue
Small intestine
Colon
Liver
Pancreas
Kidney
Bladder
Uterus
Testes

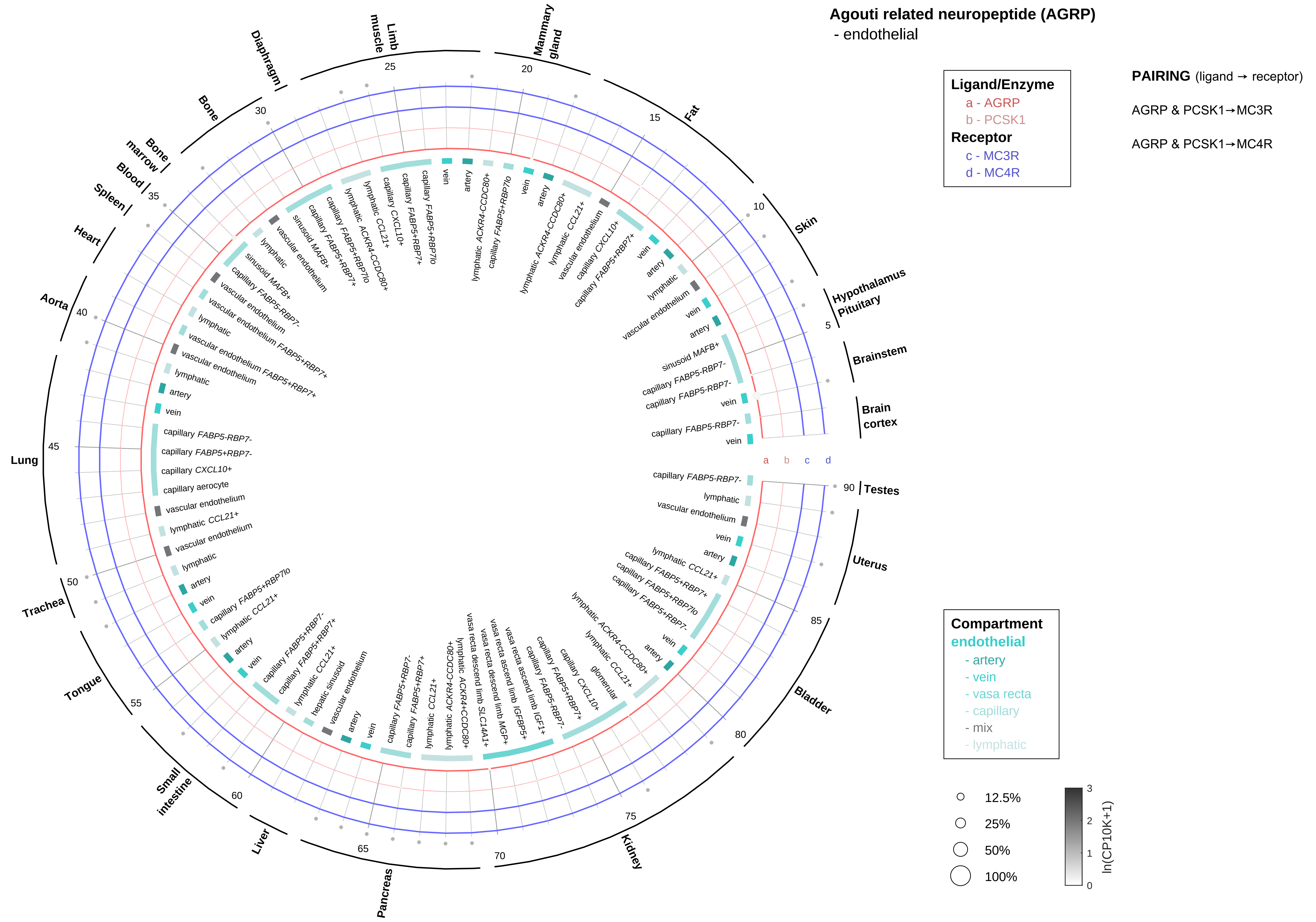

Agouti related neuropeptide (AGRP)
- endothelial
Ligand/Enzyme
a - AGRP
b - PCSK1
Receptor
c - MC3R
d - MC4R
PAIRING (ligand → receptor)
AGRP & PCSK1→MC3R
AGRP & PCSK1→MC4R
Compartment
endothelial
- artery
- vein
- vasa recta
- capillary
- mix
- lymphatic
12.5%
25%
50%
100%
ln(CP10K+1)
Hypothalamus
Pituitary
Brainstem
Brain cortex
Testes
Uterus
Bladder
Kidney
Pancreas
Liver
Small intestine
Tongue
Trachea
Lung
Aorta
Heart
Spleen
Blood
Bone marrow
Bone
Diaphragm
Limb muscle
Mammary gland
Fat
Skin

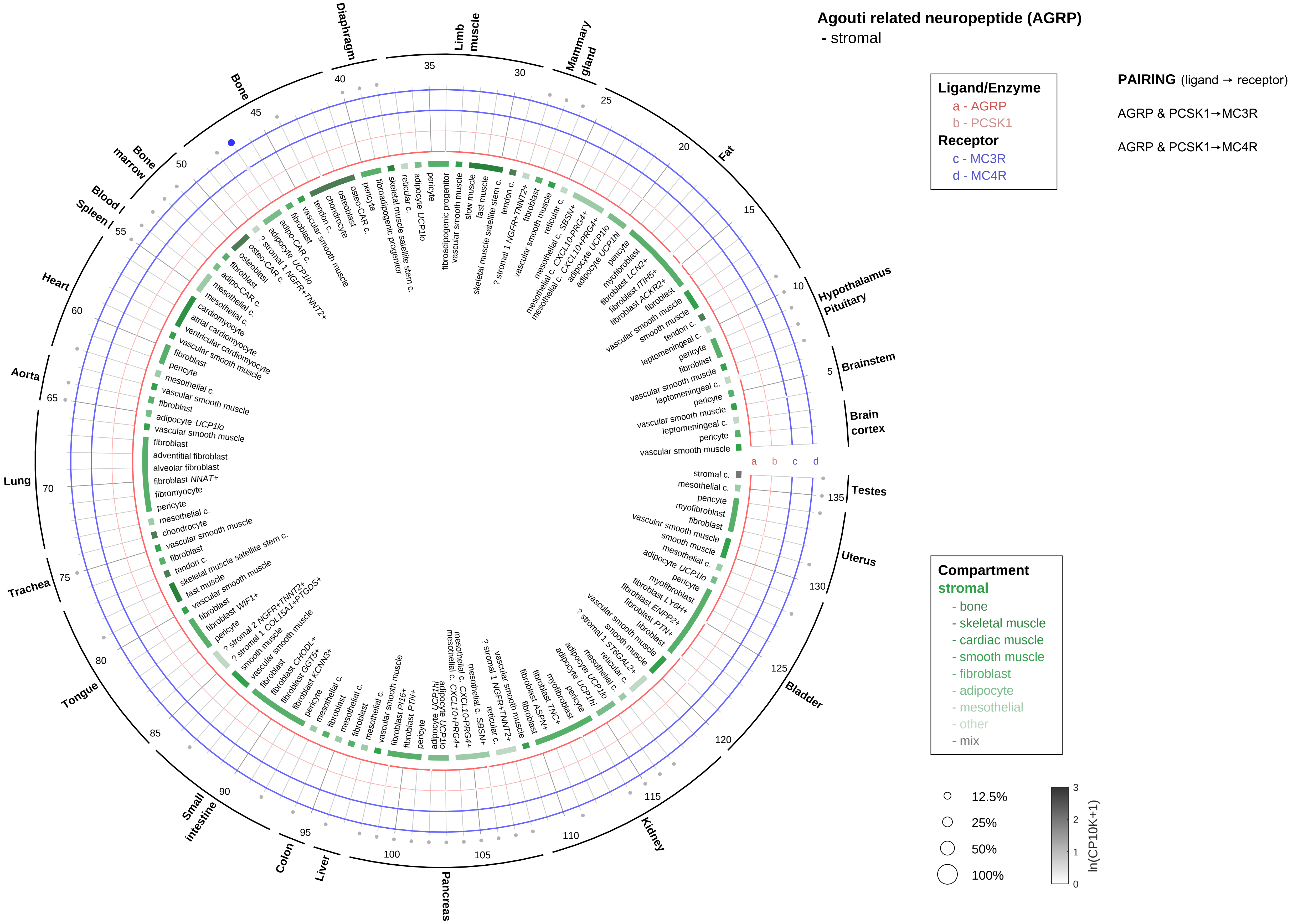

Agouti related neuropeptide (AGRP)
- stromal
Ligand/Enzyme
a - AGRP
b - PCSK1
Receptor
c - MC3R
d - MC4R
PAIRING (ligand → receptor)
AGRP & PCSK1→MC3R
AGRP & PCSK1→MC4R
Compartment
stromal
- bone
- skeletal muscle
- cardiac muscle
- smooth muscle
- fibroblast
- adipocyte
- mesothelial
- other
- mix
12.5%
25%
50%
100%
ln(CP10K+1)
Brain cortex
Brainstem
Hypothalamus Pituitary
Fat
Mammary gland
Limb muscle
Diaphragm
Bone
Bone marrow
Blood / Spleen
Heart
Aorta
Lung
Trachea
Tongue
Small intestine
Colon
Liver
Pancreas
Kidney
Bladder
Uterus
Testes

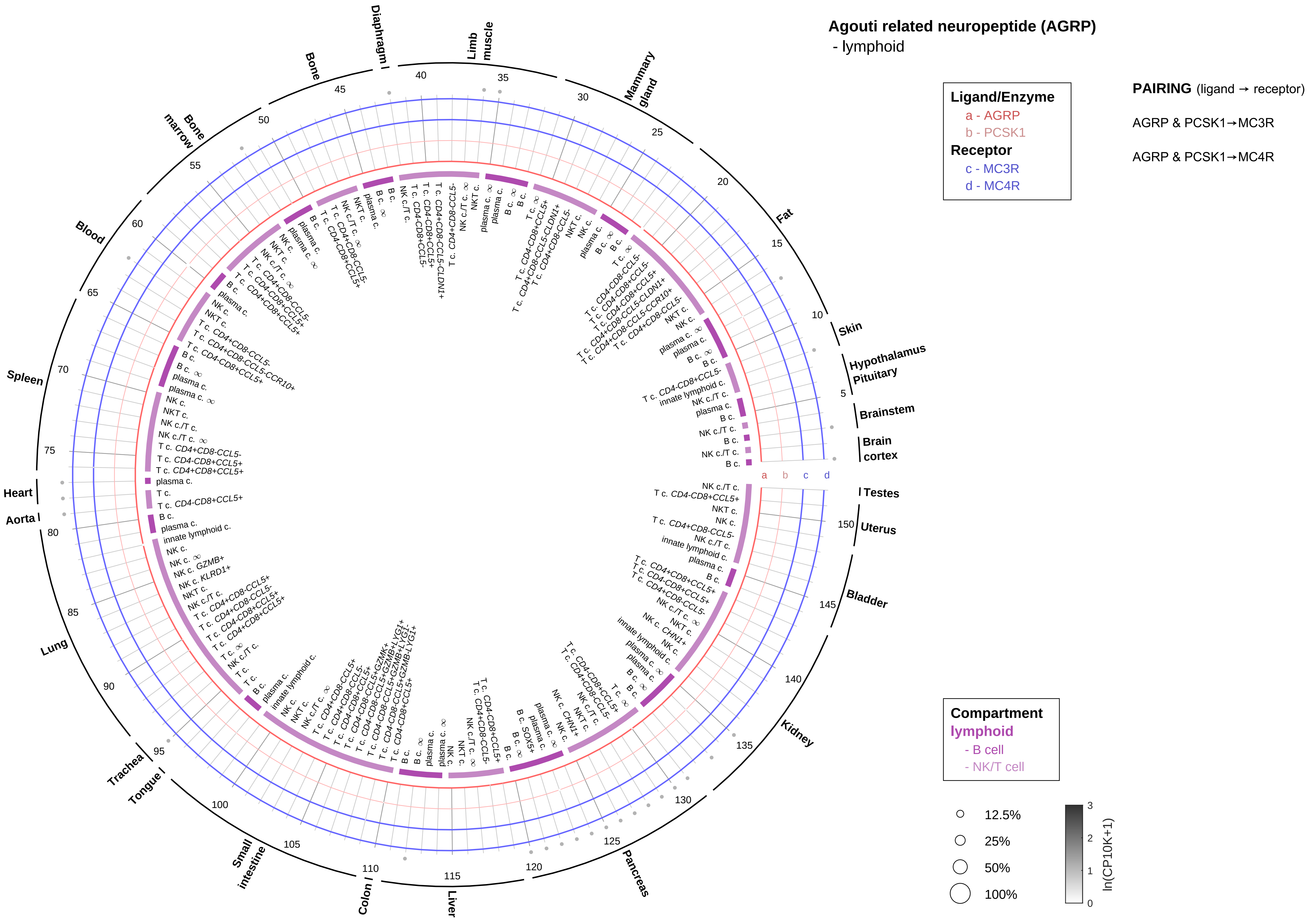
Agouti related neuropeptide (AGRP)
- lymphoid
Ligand/Enzyme
a - AGRP
b - PCSK1
Receptor
c - MC3R
d - MC4R
PAIRING (ligand → receptor)
AGRP & PCSK1→MC3R
AGRP & PCSK1→MC4R
Compartment
lymphoid
- B cell
- NK/T cell
12.5%
25%
50%
100%
ln(CP10K+1)
Skin
Hypothalamus
Pituitary
Brainstem
Brain cortex
Testes
Uterus
Bladder
Kidney
Pancreas
Liver
Colon
Small intestine
Tongue
Trachea
Lung
Aorta
Heart
Spleen
Blood
Bone marrow
Bone
Diaphragm
Limb muscle
Mammary gland
Fat

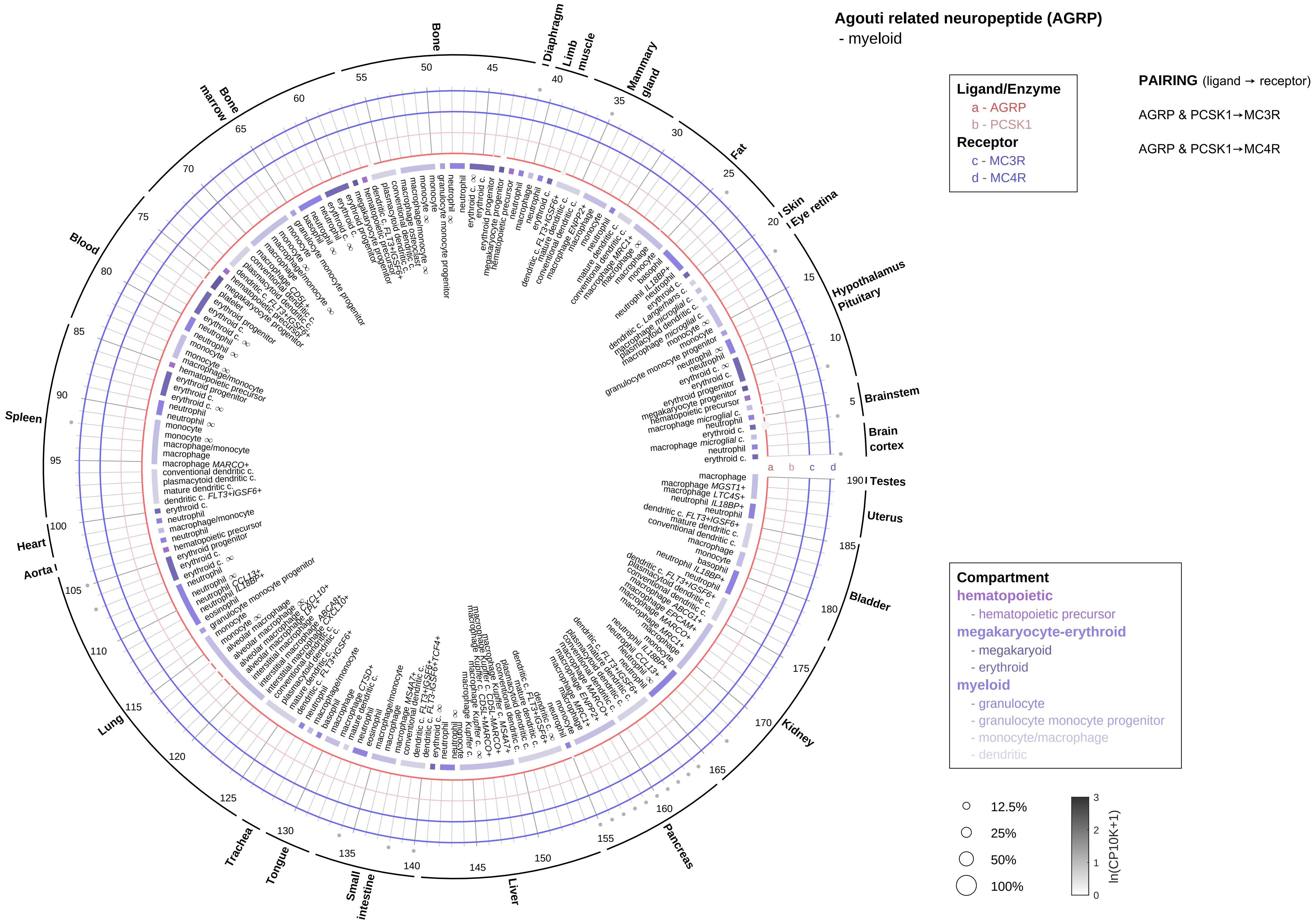

Agouti related neuropeptide (AGRP)
- myeloid
Ligand/Enzyme
a - AGRP
b - PCSK1
Receptor
c - MC3R
d - MC4R
PAIRING (ligand → receptor)
AGRP & PCSK1→MC3R
AGRP & PCSK1→MC4R
Compartment
hematopoietic
- hematopoietic precursor
megakaryocyte-erythroid
- megakaryoid
- erythroid
myeloid
- granulocyte
- granulocyte monocyte progenitor
- monocyte/macrophage
- dendritic
12.5%
25%
50%
100%
ln(CP10K+1)
Bone
Diaphragm
Limb muscle
Mammary gland
Fat
Skin
Eye retina
Hypothalamus
Pituitary
Brainstem
Brain cortex
Testes
Uterus
Bladder
Kidney
Pancreas
Liver
Small intestine
Tongue
Trachea
Lung
Aorta
Heart
Spleen
Blood
Bone marrow

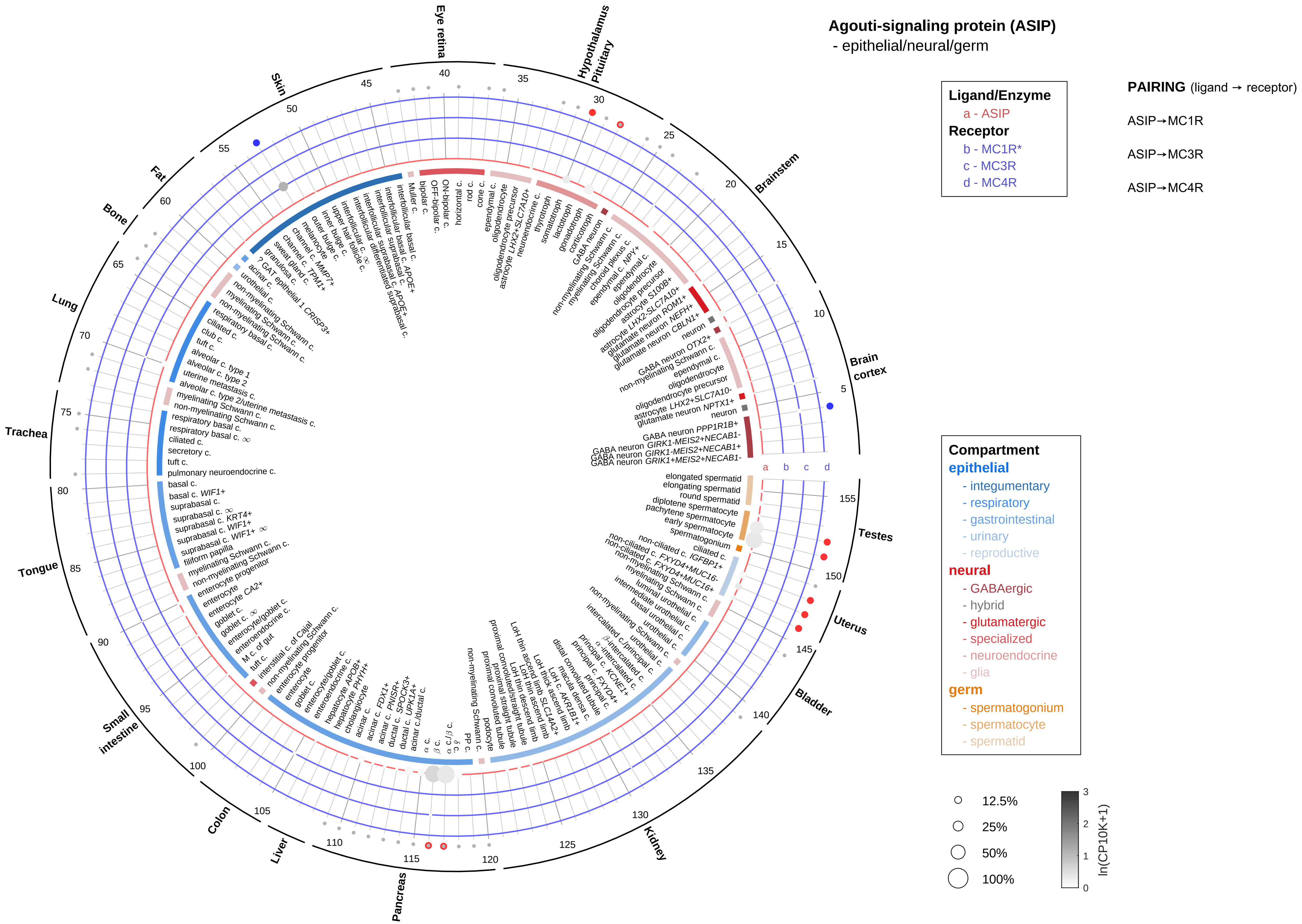

Agouti-signaling protein (ASIP)
- epithelial/neural/germ
Ligand/Enzyme
a - ASIP
Receptor
b - MC1R*
c - MC3R
d - MC4R
PAIRING (ligand → receptor)
ASIP→MC1R
ASIP→MC3R
ASIP→MC4R
Compartment
epithelial
- integumentary
- respiratory
- gastrointestinal
- urinary
- reproductive
neural
- GABAergic
- hybrid
- glutamatergic
- specialized
- neuroendocrine
- glia
germ
- spermatogonium
- spermatocyte
- spermatid
12.5%
25%
50%
100%
ln(CP10K+1)
Eye retina
Hypothalamus Pituitary
Brainstem
Brain cortex
Testes
Uterus
Bladder
Kidney
Pancreas
Liver
Colon
Small intestine
Tongue
Trachea
Lung
Bone
Fat
Skin
a
b
c
d

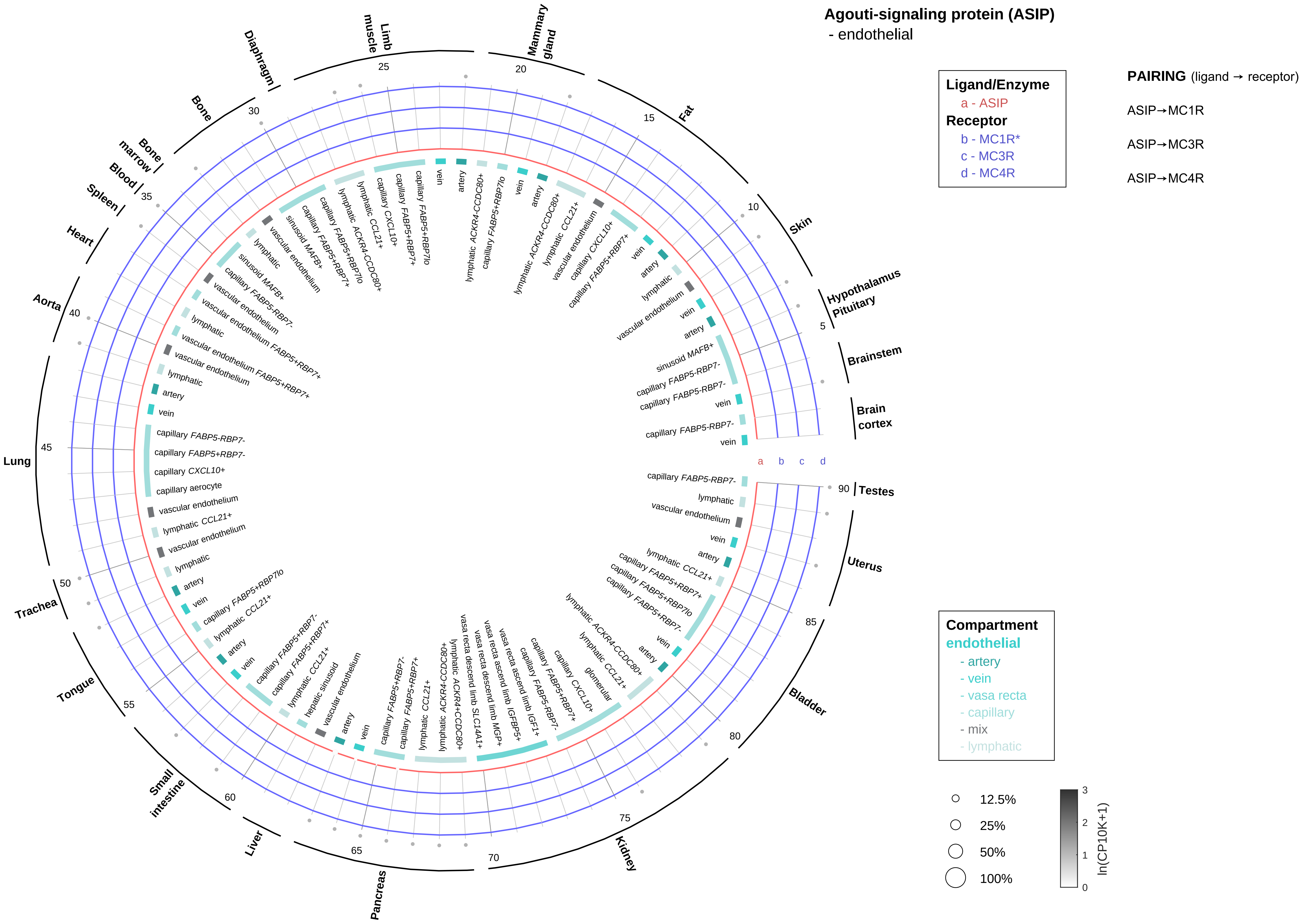

Agouti-signaling protein (ASIP)
- endothelial
Ligand/Enzyme
a - ASIP
Receptor
b - MC1R*
c - MC3R
d - MC4R
PAIRING (ligand → receptor)
ASIP→MC1R
ASIP→MC3R
ASIP→MC4R
Compartment
endothelial
- artery
- vein
- vasa recta
- capillary
- mix
- lymphatic
12.5%
25%
50%
100%
ln(CP10K+1)
Hypothalamus Pituitary
Brainstem
Brain cortex
Testes
Uterus
Bladder
Kidney
Pancreas
Liver
Small intestine
Tongue
Trachea
Lung
Aorta
Heart
Spleen
Blood
Bone marrow
Bone
Diaphragm
Limb muscle
Mammary gland
Fat
Skin

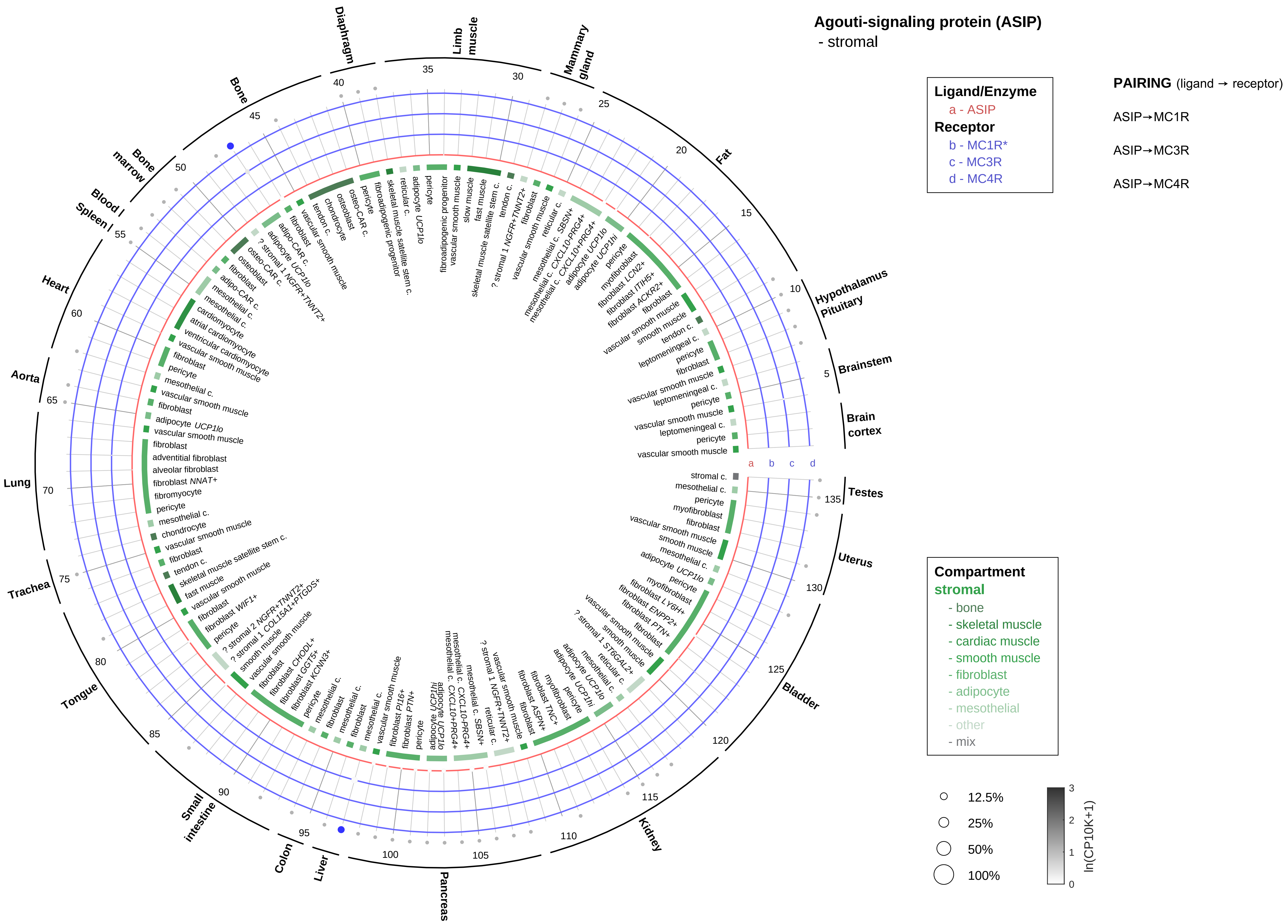

Agouti-signaling protein (ASIP)
- stromal
Ligand/Enzyme
a - ASIP
Receptor
b - MC1R*
c - MC3R
d - MC4R
PAIRING (ligand → receptor)
ASIP→MC1R
ASIP→MC3R
ASIP→MC4R
Compartment
stromal
- bone
- skeletal muscle
- cardiac muscle
- smooth muscle
- fibroblast
- adipocyte
- mesothelial
- other
- mix
12.5%
25%
50%
100%
ln(CP10K+1)
Hypothalamus Pituitary
Brainstem
Brain cortex
Testes
Uterus
Bladder
Kidney
Pancreas
Liver
Colon
Small intestine
Tongue
Trachea
Lung
Aorta
Heart
Spleen
Blood
Bone marrow
Bone
Diaphragm
Limb muscle
Mammary gland
Fat

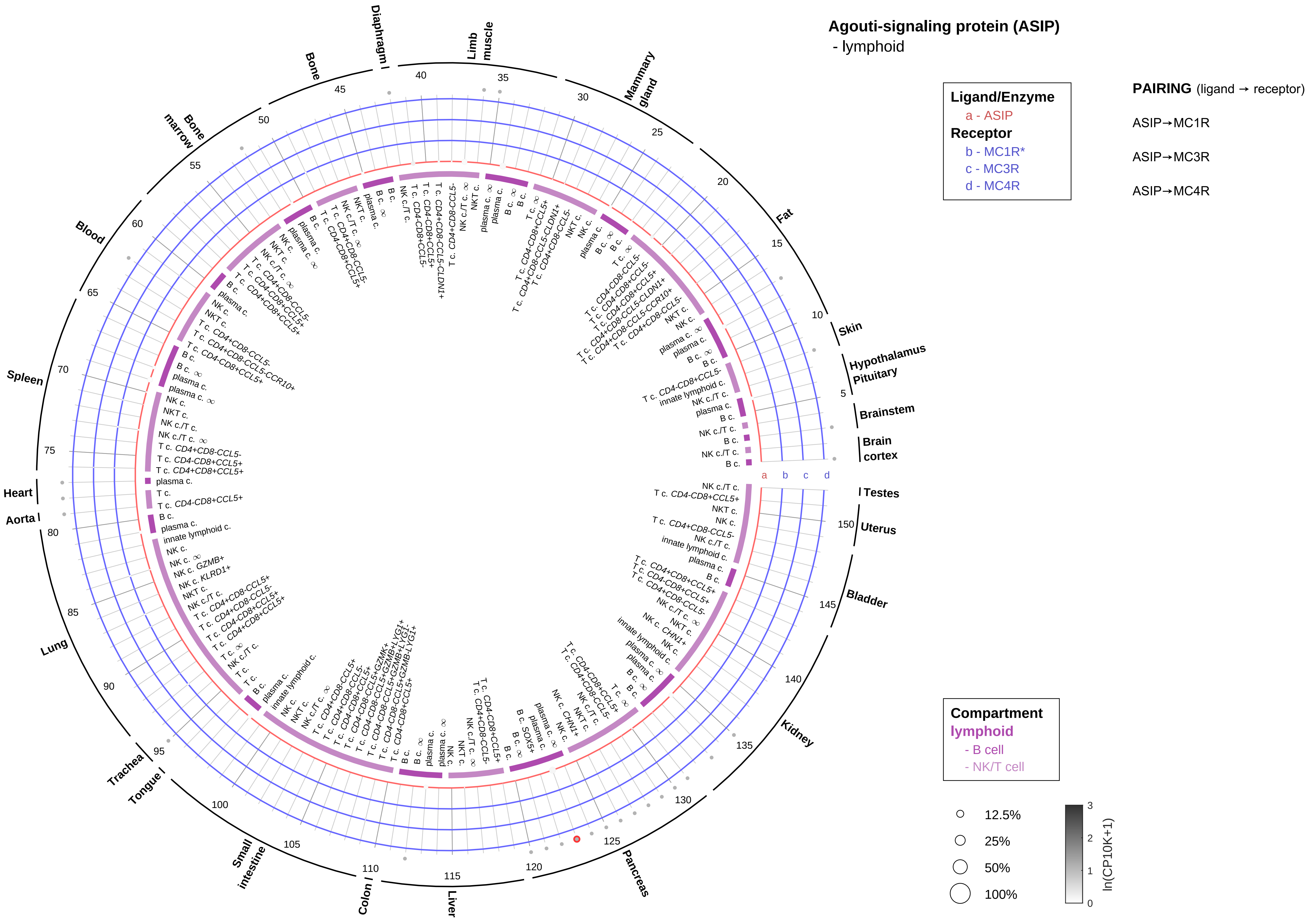

Agouti-signaling protein (ASIP)
- lymphoid
Ligand/Enzyme
a - ASIP
Receptor
b - MC1R*
c - MC3R
d - MC4R
PAIRING (ligand → receptor)
ASIP→MC1R
ASIP→MC3R
ASIP→MC4R
Compartment
lymphoid
- B cell
- NK/T cell
12.5%
25%
50%
100%
ln(CP10K+1)
Brain cortex
Brainstem
Hypothalamus Pituitary
Skin
Fat
Mammary gland
Limb muscle
Diaphragm
Bone
Bone marrow
Blood
Spleen
Heart
Aorta
Lung
Trachea
Tongue
Small intestine
Colon
Liver
Pancreas
Kidney
Bladder
Uterus
Testes

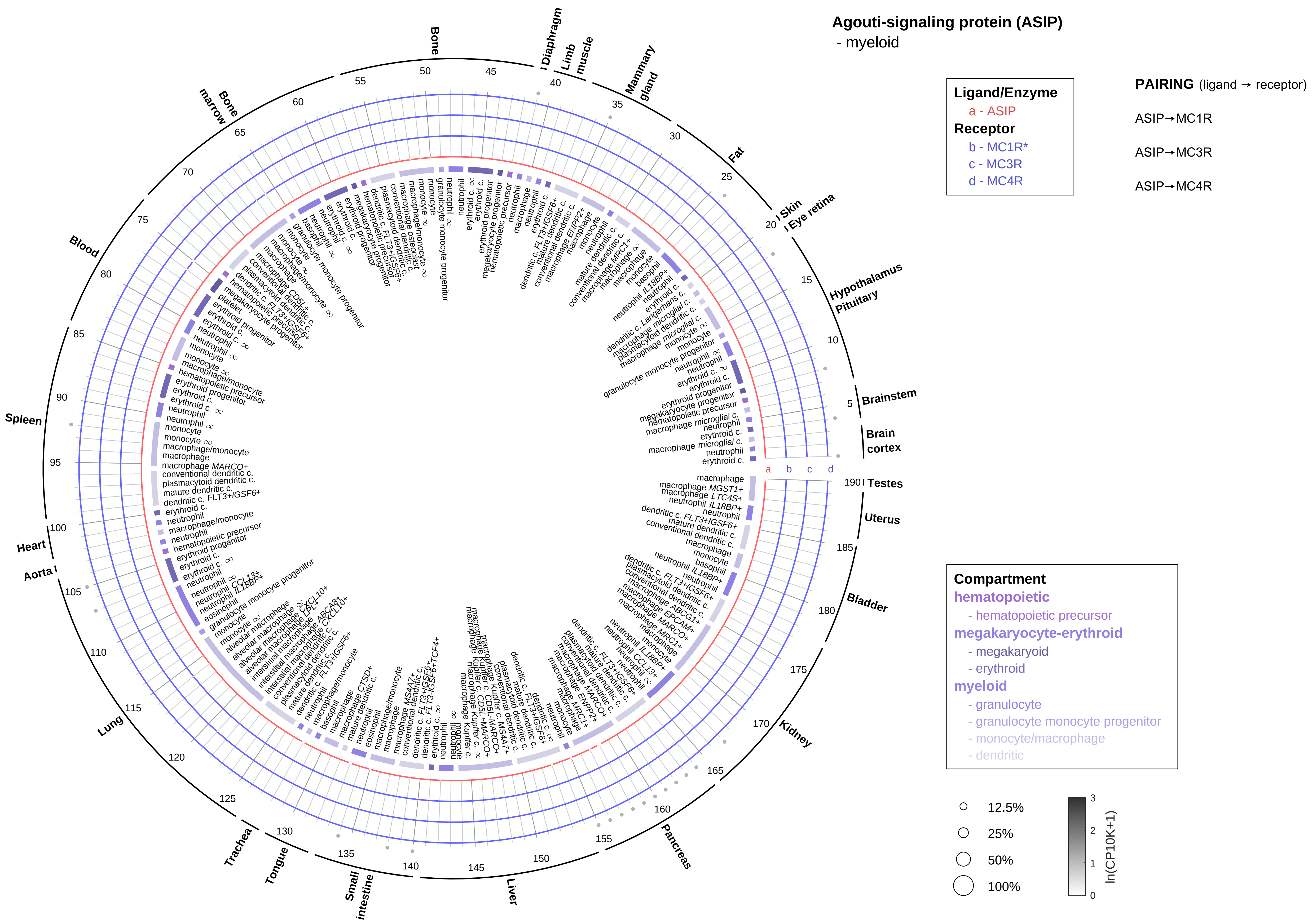

Agouti-signaling protein (ASIP)
- myeloid
Ligand/Enzyme
a - ASIP
Receptor
b - MC1R*
c - MC3R
d - MC4R
PAIRING (ligand → receptor)
ASIP→MC1R
ASIP→MC3R
ASIP→MC4R
Compartment
hematopoietic
- hematopoietic precursor
megakaryocyte-erythroid
- megakaryoid
- erythroid
myeloid
- granulocyte
- granulocyte monocyte progenitor
- monocyte/macrophage
- dendritic
12.5%
25%
50%
100%
ln(CP10K+1)
Bone
Diaphragm
Limb muscle
Mammary gland
Fat
Skin
Eye retina
Hypothalamus
Pituitary
Brainstem
Brain cortex
Testes
Uterus
Bladder
Kidney
Pancreas
Liver
Small intestine
Tongue
Trachea
Lung
Aorta
Heart
Spleen
Blood
Bone marrow

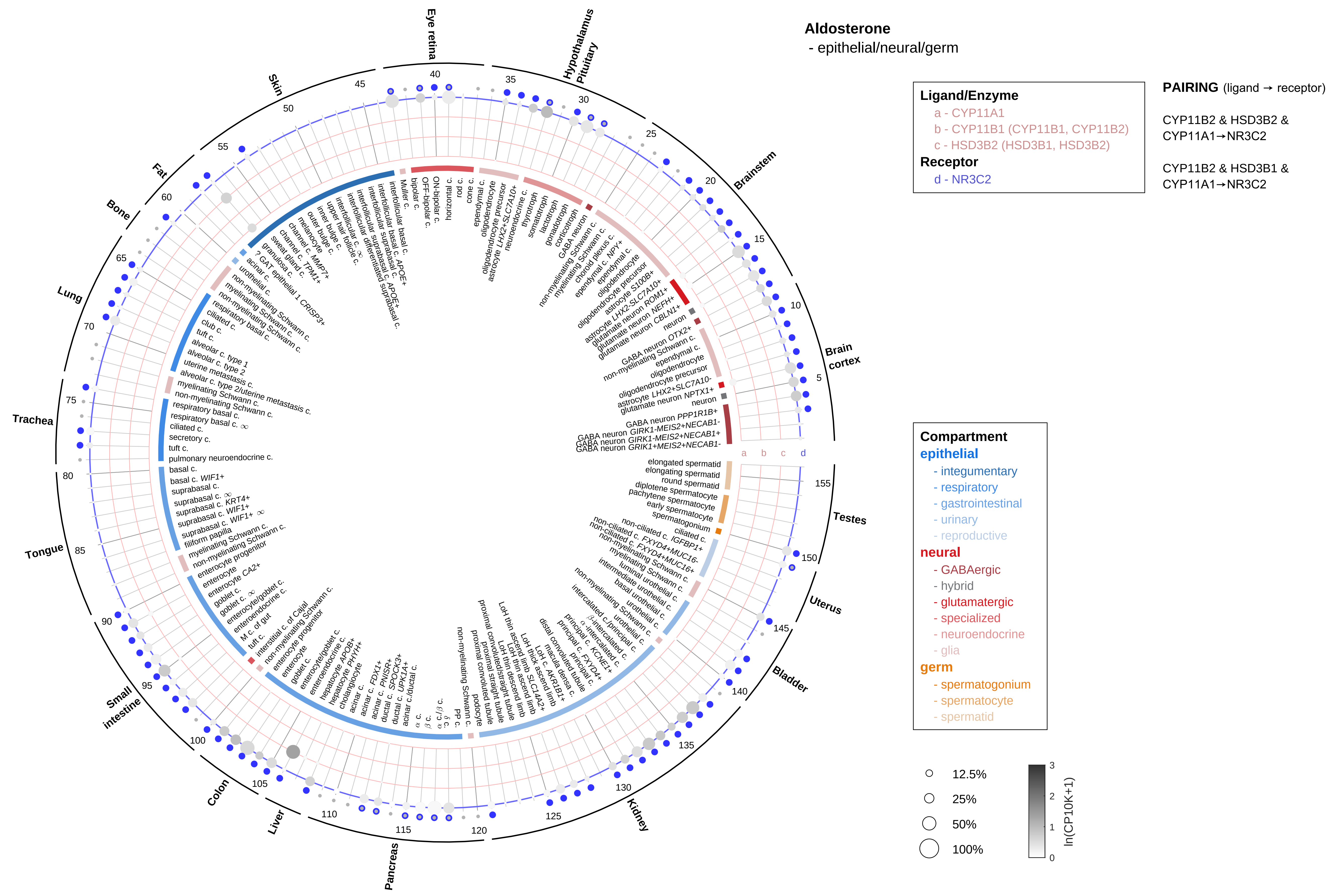
Aldosterone
- epithelial/neural/germ
Ligand/Enzyme
a - CYP11A1
b - CYP11B1 (CYP11B1, CYP11B2)
c - HSD3B2 (HSD3B1, HSD3B2)
Receptor
d - NR3C2
PAIRING (ligand → receptor)
CYP11B2 & HSD3B2 & CYP11A1→NR3C2
CYP11B2 & HSD3B1 & CYP11A1→NR3C2
Compartment
epithelial
- integumentary
- respiratory
- gastrointestinal
- urinary
- reproductive
neural
- GABAergic
- hybrid
- glutamatergic
- specialized
- neuroendocrine
- glia
germ
- spermatogonium
- spermatocyte
- spermatid
12.5%
25%
50%
100%
ln(CP10K+1)
Eye retina
Hypothalamus Pituitary
Brainstem
Brain cortex
Testes
Uterus
Bladder
Kidney
Pancreas
Liver
Colon
Small intestine
Tongue
Trachea
Lung
Bone
Fat
Skin

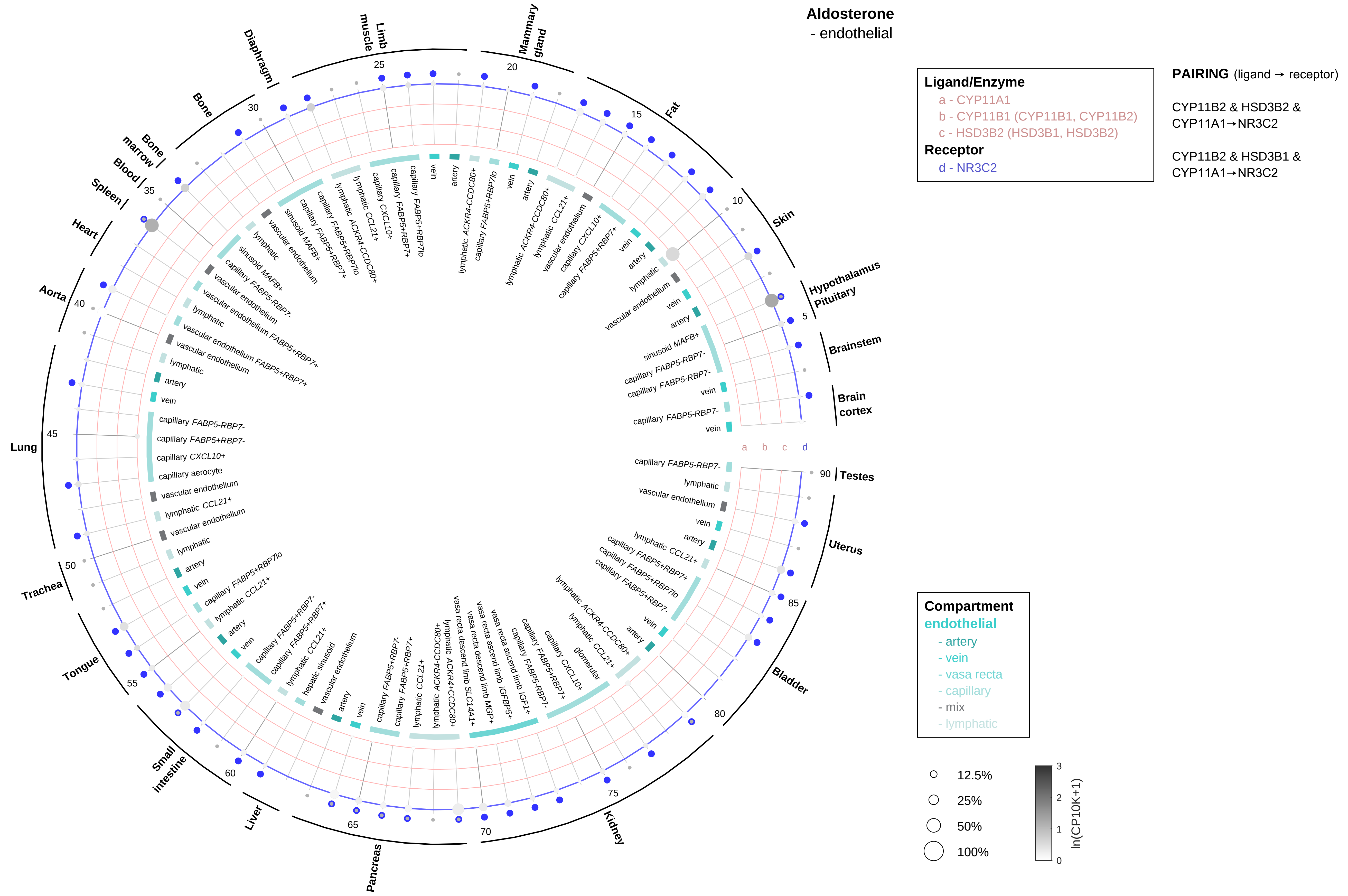
Aldosterone
- endothelial
Ligand/Enzyme
a - CYP11A1
b - CYP11B1 (CYP11B1, CYP11B2)
c - HSD3B2 (HSD3B1, HSD3B2)
Receptor
d - NR3C2
PAIRING (ligand → receptor)
CYP11B2 & HSD3B2 & CYP11A1→NR3C2
CYP11B2 & HSD3B1 & CYP11A1→NR3C2
Compartment
endothelial
- artery
- vein
- vasa recta
- capillary
- mix
- lymphatic
12.5%
25%
50%
100%
ln(CP10K+1)
Hypothalamus
Pituitary
Skin
Fat
Mammary gland
Limb muscle
Diaphragm
Bone
Bone marrow
Blood
Spleen
Heart
Aorta
Lung
Trachea
Tongue
Small intestine
Liver
Pancreas
Kidney
Bladder
Uterus
Testes
Brain cortex
Brainstem

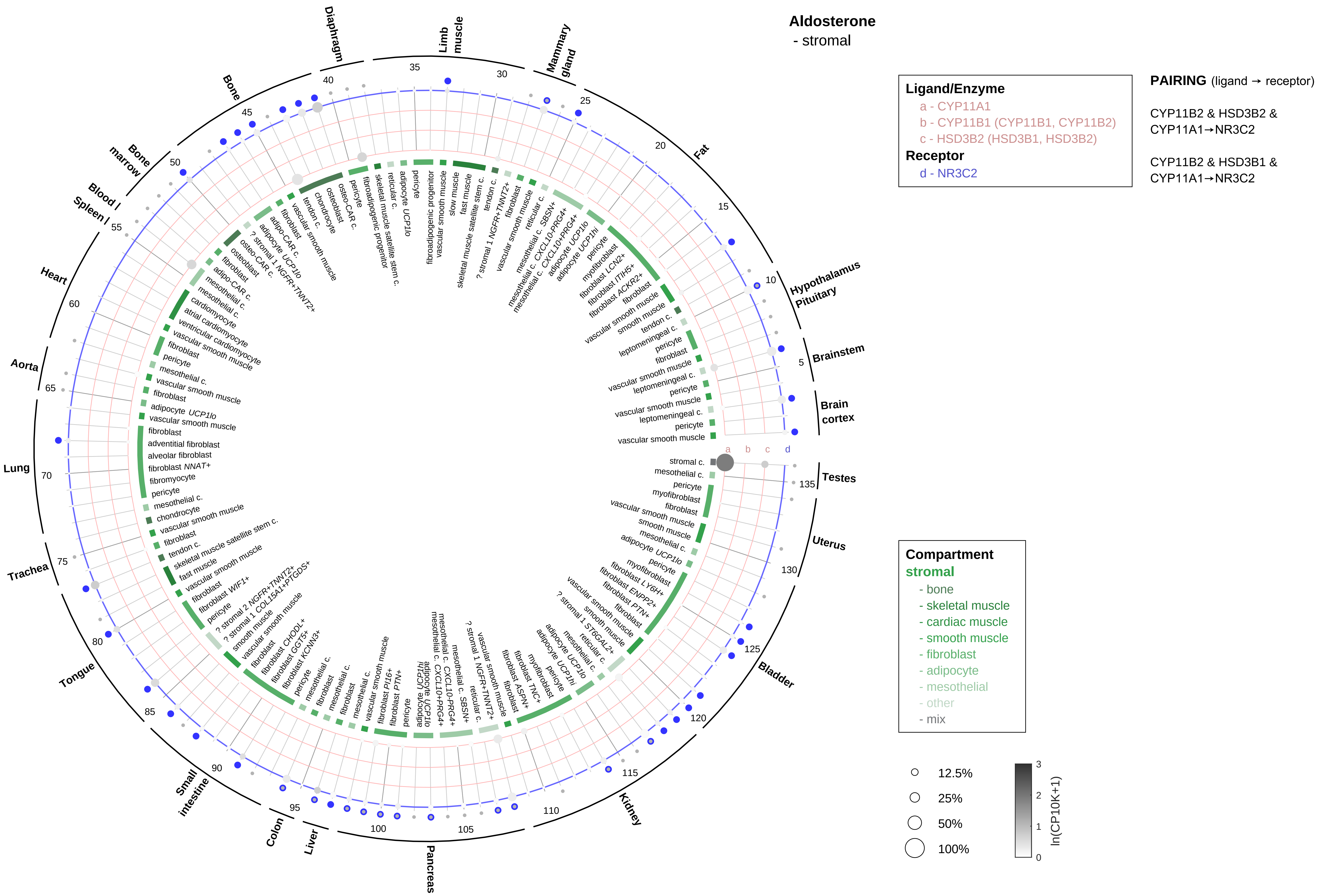
Aldosterone
- stromal
Ligand/Enzyme
a - CYP11A1
b - CYP11B1 (CYP11B1, CYP11B2)
c - HSD3B2 (HSD3B1, HSD3B2)
Receptor
d - NR3C2
PAIRING (ligand → receptor)
CYP11B2 & HSD3B2 & CYP11A1→NR3C2
CYP11B2 & HSD3B1 & CYP11A1→NR3C2
Compartment
stromal
- bone
- skeletal muscle
- cardiac muscle
- smooth muscle
- fibroblast
- adipocyte
- mesothelial
- other
- mix
12.5%
25%
50%
100%
ln(CP10K+1)
Brain cortex
Brainstem
Hypothalamus Pituitary
Fat
Mammary gland
Limb muscle
Diaphragm
Bone
Bone marrow
Blood / Spleen
Heart
Aorta
Lung
Trachea
Tongue
Small intestine
Colon
Liver
Pancreas
Kidney
Bladder
Uterus
Testes

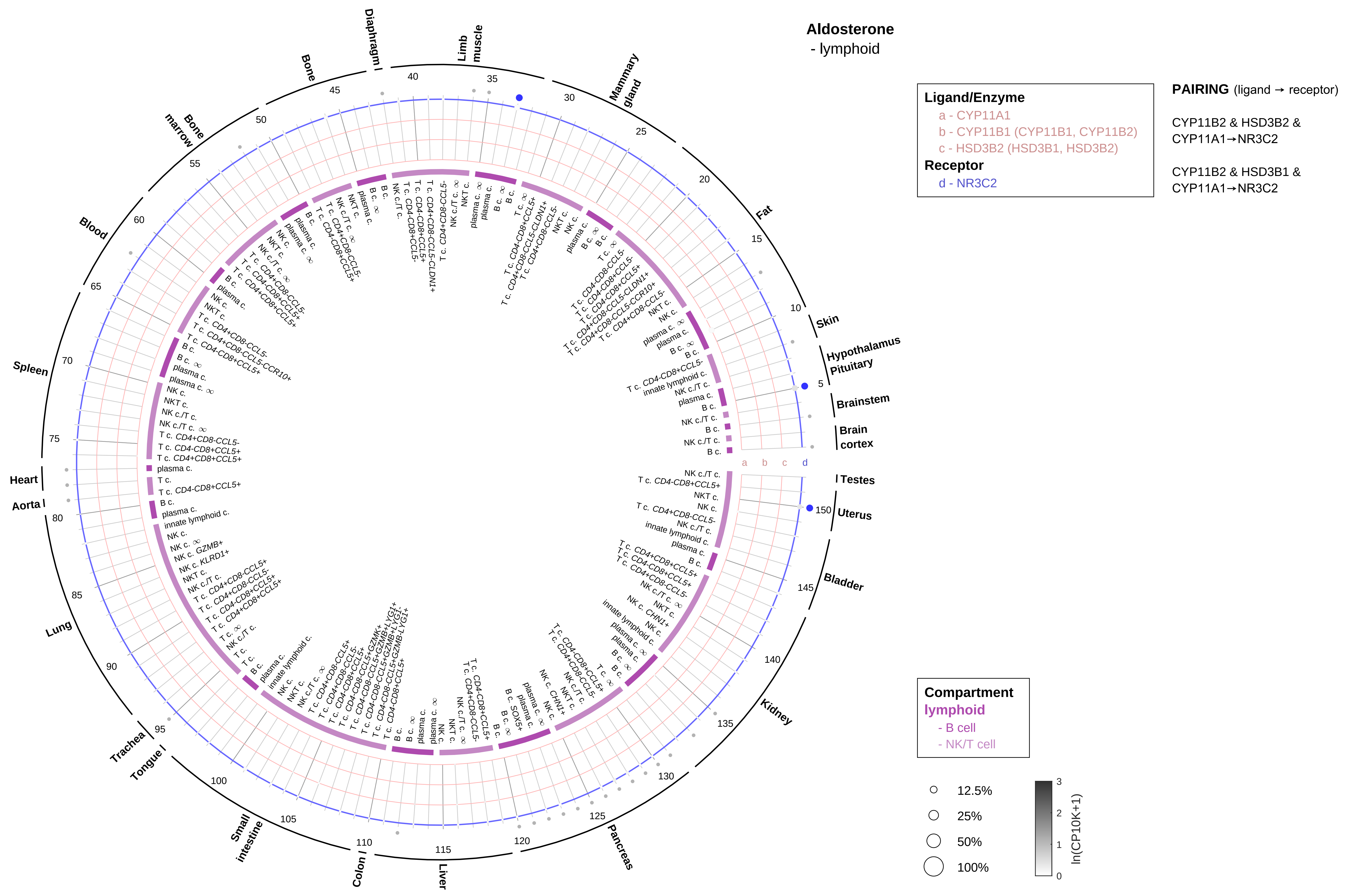

Aldosterone
- lymphoid
Ligand/Enzyme
a - CYP11A1
b - CYP11B1 (CYP11B1, CYP11B2)
c - HSD3B2 (HSD3B1, HSD3B2)
Receptor
d - NR3C2
PAIRING (ligand → receptor)
CYP11B2 & HSD3B2 & CYP11A1→NR3C2
CYP11B2 & HSD3B1 & CYP11A1→NR3C2
Compartment
lymphoid
- B cell
- NK/T cell
12.5%
25%
50%
100%
ln(CP10K+1)
Skin
Hypothalamus
Pituitary
Brainstem
Brain cortex
Testes
Uterus
Bladder
Kidney
Pancreas
Liver
Colon
Small intestine
Tongue
Trachea
Lung
Aorta
Heart
Spleen
Blood
Bone marrow
Bone
Diaphragm
Limb muscle
Mammary gland
Fat

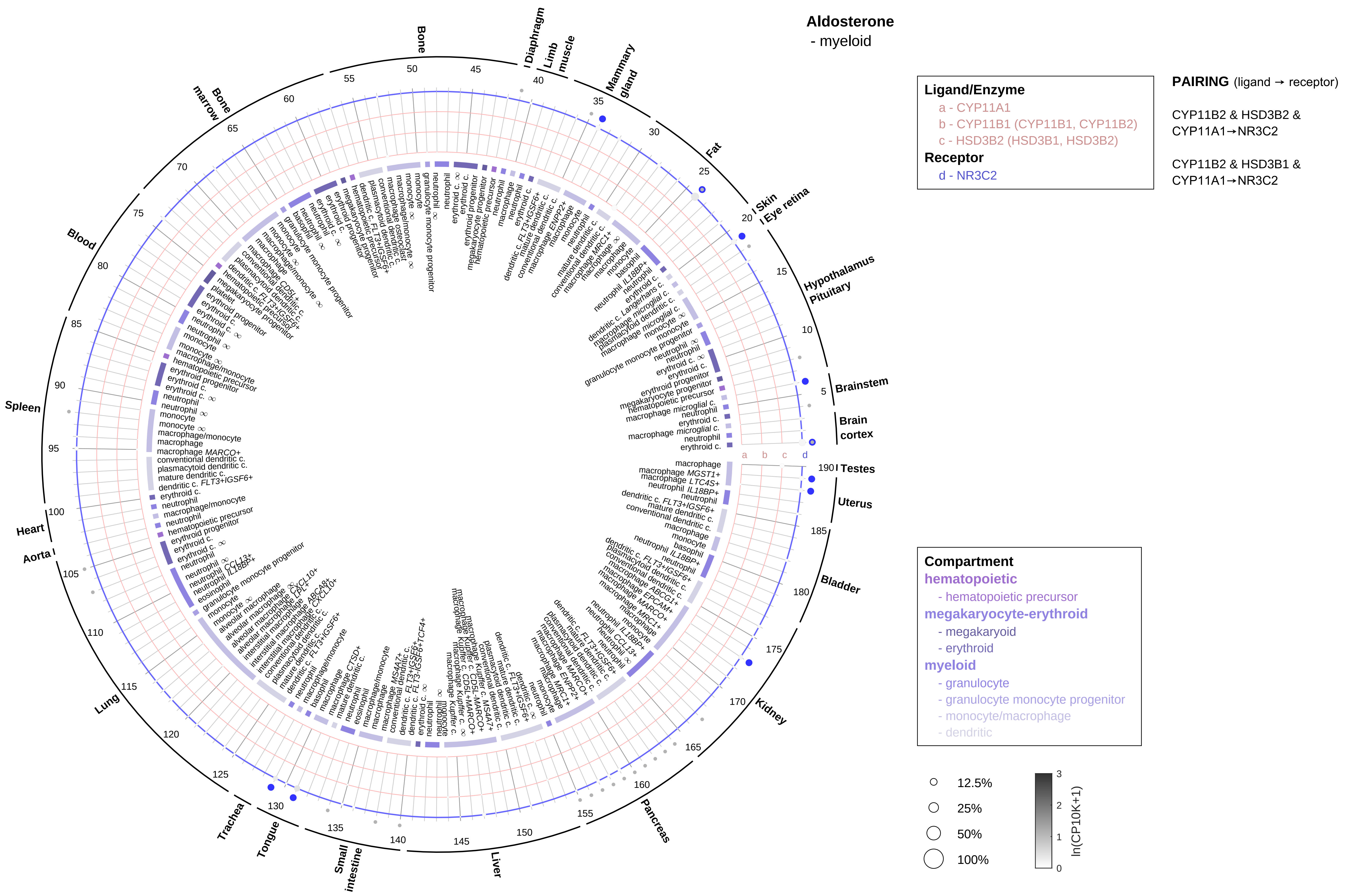

Aldosterone
- myeloid
Ligand/Enzyme
a - CYP11A1
b - CYP11B1 (CYP11B1, CYP11B2)
c - HSD3B2 (HSD3B1, HSD3B2)
Receptor
d - NR3C2
PAIRING (ligand → receptor)
CYP11B2 & HSD3B2 & CYP11A1→NR3C2
CYP11B2 & HSD3B1 & CYP11A1→NR3C2
Compartment
hematopoietic
- hematopoietic precursor
megakaryocyte-erythroid
- megakaryoid
- erythroid
myeloid
- granulocyte
- granulocyte monocyte progenitor
- monocyte/macrophage
- dendritic
12.5%
25%
50%
100%
ln(CP10K+1)
Brain cortex
Brainstem
Hypothalamus
Pituitary
Eye retina
Skin
Fat
Mammary gland
Limb muscle
Diaphragm
Bone
Bone marrow
Blood
Spleen
Heart
Aorta
Lung
Trachea
Tongue
Small intestine
Liver
Pancreas
Kidney
Bladder
Uterus
Testes

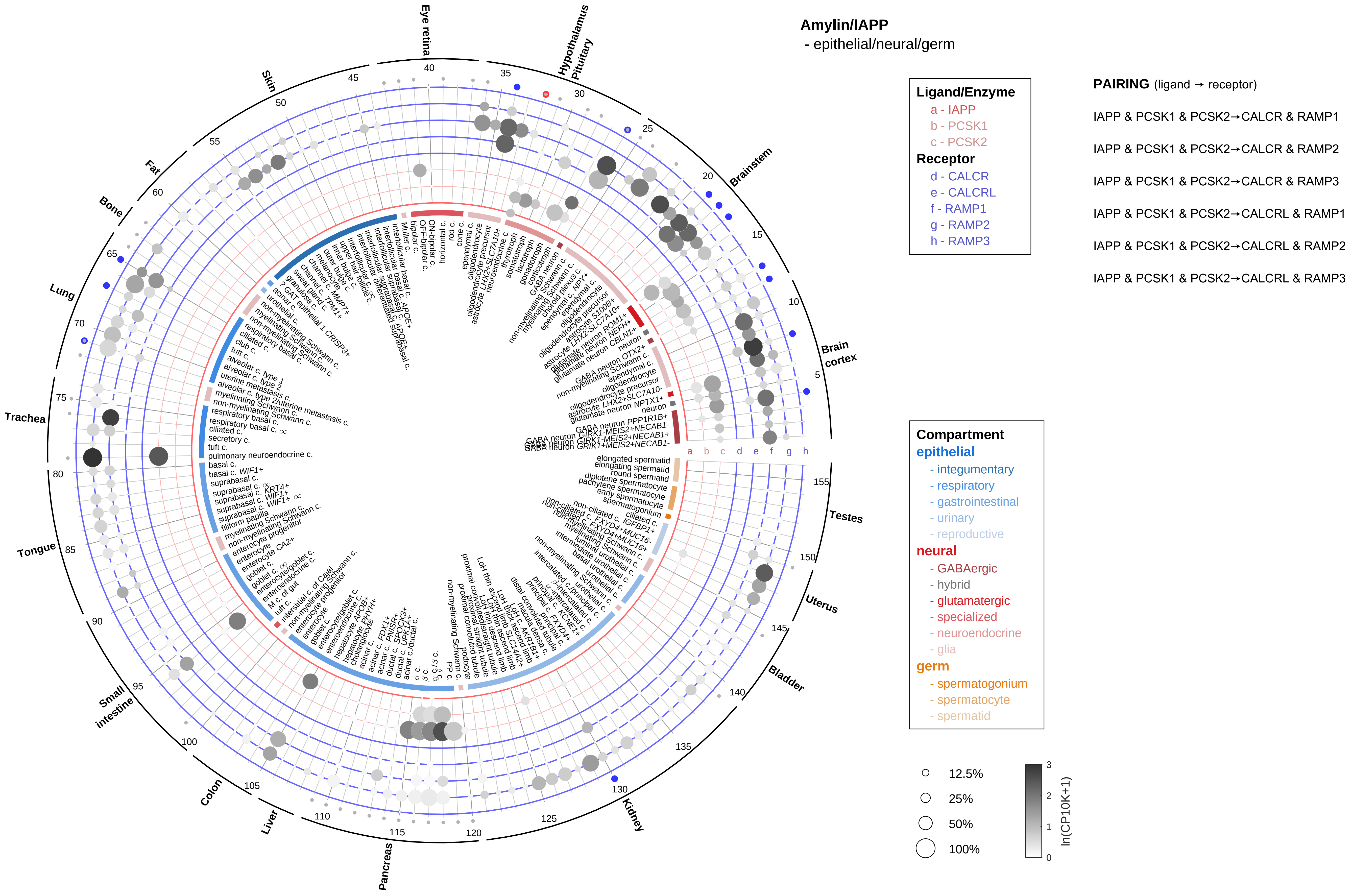

Amylin/IAPP
- epithelial/neural/germ
Ligand/Enzyme
a - IAPP
b - PCSK1
c - PCSK2
Receptor
d - CALCR
e - CALCRL
f - RAMP1
g - RAMP2
h - RAMP3
PAIRING (ligand → receptor)
IAPP & PCSK1 & PCSK2→CALCR & RAMP1
IAPP & PCSK1 & PCSK2→CALCR & RAMP2
IAPP & PCSK1 & PCSK2→CALCR & RAMP3
IAPP & PCSK1 & PCSK2→CALCRL & RAMP1
IAPP & PCSK1 & PCSK2→CALCRL & RAMP2
IAPP & PCSK1 & PCSK2→CALCRL & RAMP3
Compartment
epithelial
- integumentary
- respiratory
- gastrointestinal
- urinary
- reproductive
neural
- GABAergic
- hybrid
- glutamatergic
- specialized
- neuroendocrine
- glia
germ
- spermatogonium
- spermatocyte
- spermatid
12.5%
25%
50%
100%
ln(CP10K+1)
Eye retina
Hypothalamus
Pituitary
Brainstem
Brain cortex
Testes
Uterus
Bladder
Kidney
Pancreas
Liver
Colon
Small intestine
Tongue
Trachea
Lung
Bone
Fat
Skin

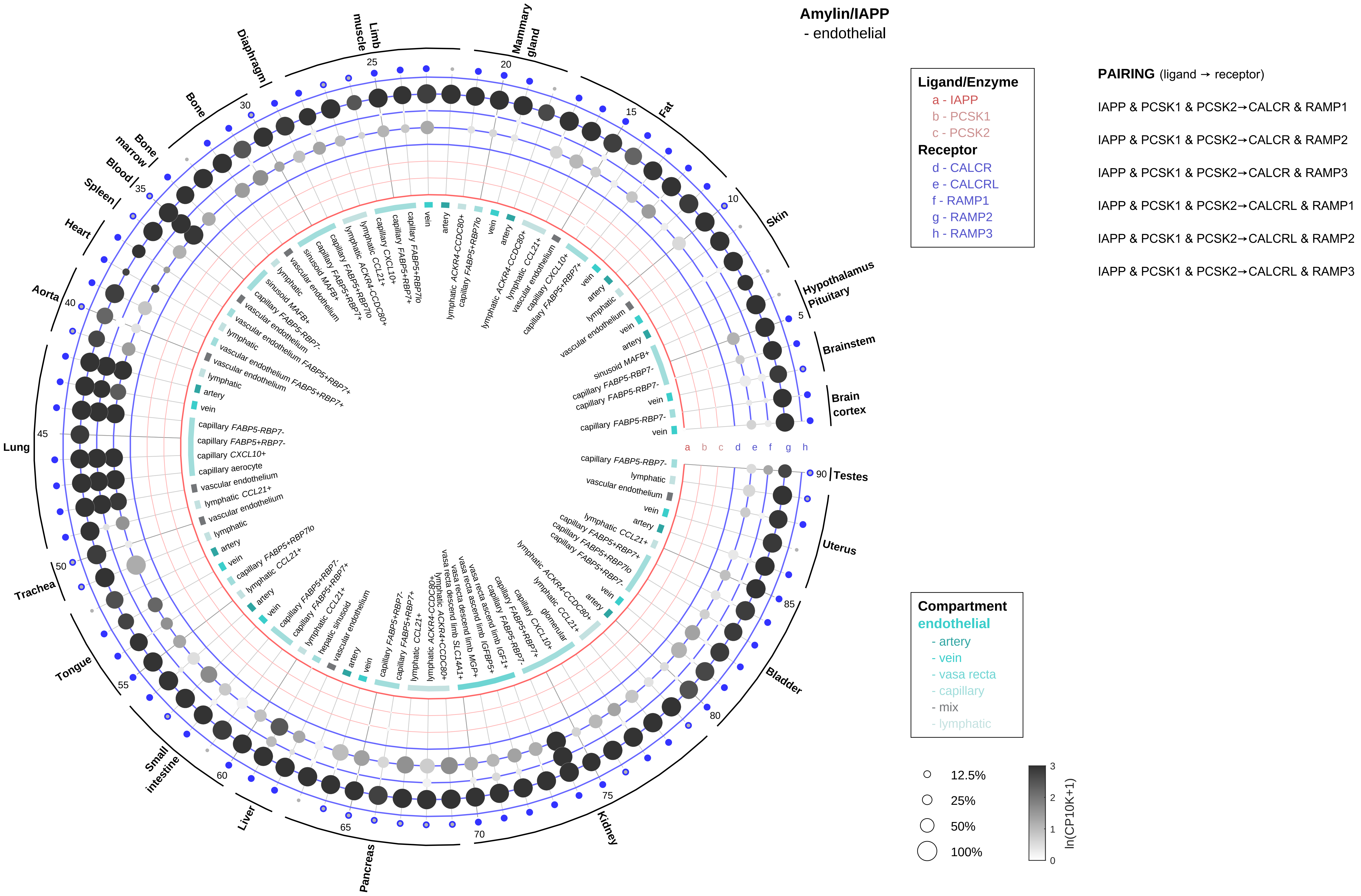
Amylin/IAPP
- endothelial
Ligand/Enzyme
a - IAPP
b - PCSK1
c - PCSK2
Receptor
d - CALCR
e - CALCRL
f - RAMP1
g - RAMP2
h - RAMP3
PAIRING (ligand → receptor)
IAPP & PCSK1 & PCSK2→CALCR & RAMP1
IAPP & PCSK1 & PCSK2→CALCR & RAMP2
IAPP & PCSK1 & PCSK2→CALCR & RAMP3
IAPP & PCSK1 & PCSK2→CALCRL & RAMP1
IAPP & PCSK1 & PCSK2→CALCRL & RAMP2
IAPP & PCSK1 & PCSK2→CALCRL & RAMP3
Compartment
endothelial
- artery
- vein
- vasa recta
- capillary
- mix
- lymphatic
12.5%
25%
50%
100%
ln(CP10K+1)
Brain cortex
Brainstem
Hypothalamus Pituitary
Skin
Fat
Mammary gland
Limb muscle
Diaphragm
Bone
Bone marrow
Blood
Spleen
Heart
Aorta
Lung
Trachea
Tongue
Small intestine
Liver
Pancreas
Kidney
Bladder
Uterus
Testes

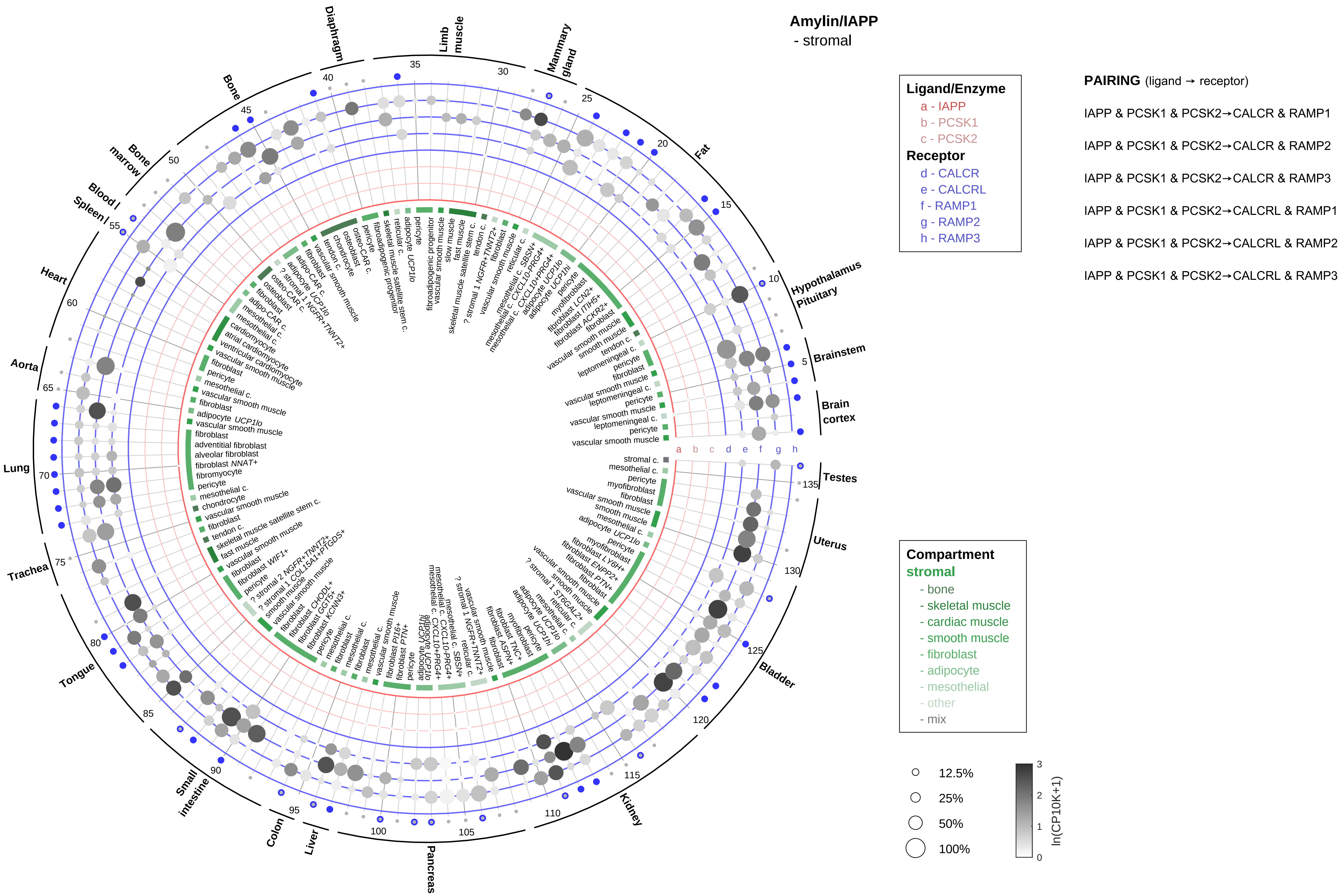

Amylin/IAPP
- stromal
Ligand/Enzyme
a - IAPP
b - PCSK1
c - PCSK2
Receptor
d - CALCR
e - CALCRL
f - RAMP1
g - RAMP2
h - RAMP3
PAIRING (ligand → receptor)
IAPP & PCSK1 & PCSK2→CALCR & RAMP1
IAPP & PCSK1 & PCSK2→CALCR & RAMP2
IAPP & PCSK1 & PCSK2→CALCR & RAMP3
IAPP & PCSK1 & PCSK2→CALCRL & RAMP1
IAPP & PCSK1 & PCSK2→CALCRL & RAMP2
IAPP & PCSK1 & PCSK2→CALCRL & RAMP3
Compartment
stromal
- bone
- skeletal muscle
- cardiac muscle
- smooth muscle
- fibroblast
- adipocyte
- mesothelial
- other
- mix
12.5%
25%
50%
100%
ln(CP10K+1)
Diaphragm
Limb muscle
Mammary gland
Fat
Hypothalamus Pituitary
Brainstem
Brain cortex
Testes
Uterus
Bladder
Kidney
Pancreas
Liver
Colon
Small intestine
Tongue
Trachea
Lung
Aorta
Heart
Spleen
Blood
Bone marrow
Bone

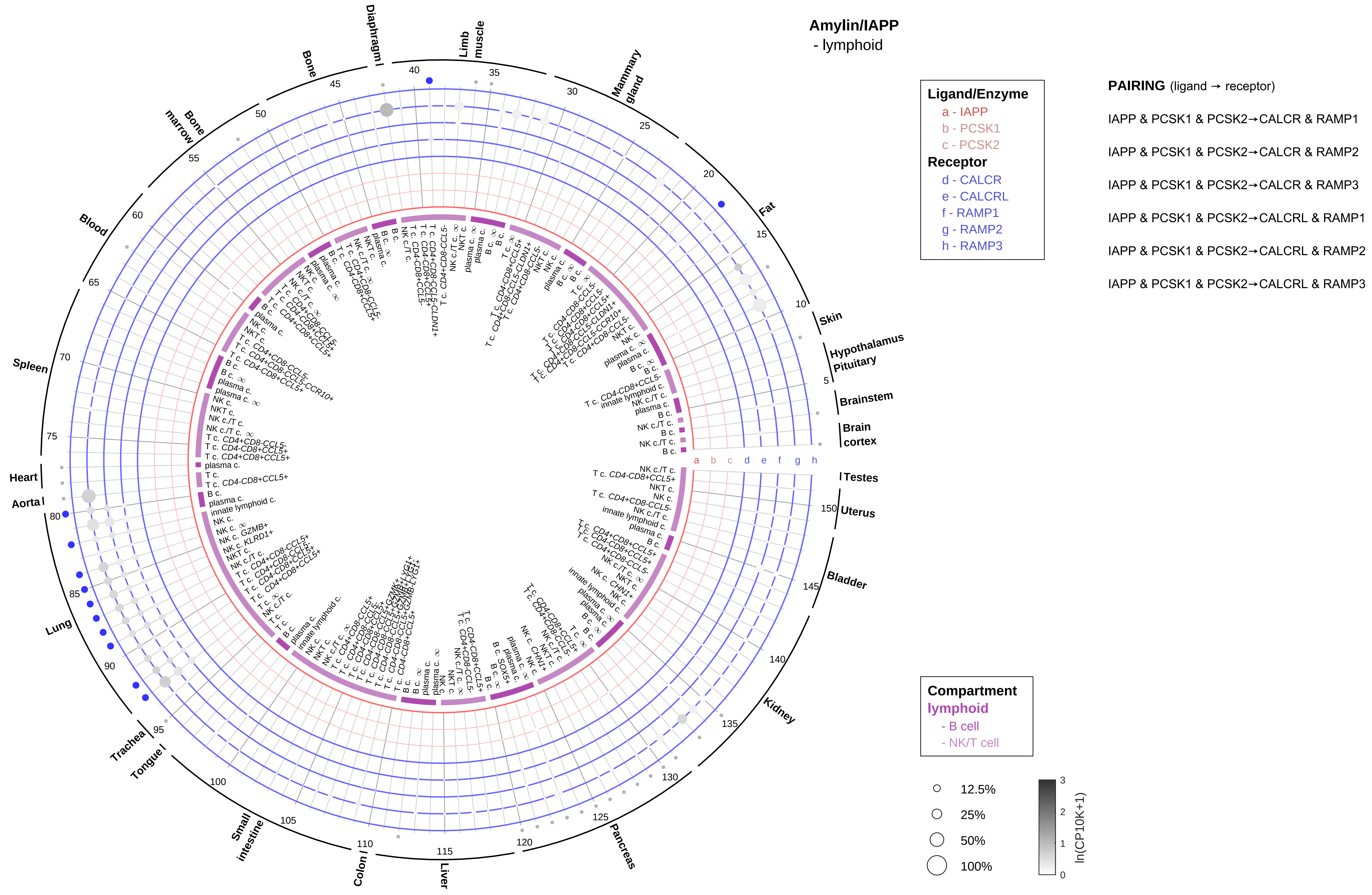
Amylin/IAPP
- lymphoid
Ligand/Enzyme
a - IAPP
b - PCSK1
c - PCSK2
Receptor
d - CALCR
e - CALCRL
f - RAMP1
g - RAMP2
h - RAMP3
PAIRING (ligand → receptor)
IAPP & PCSK1 & PCSK2→CALCR & RAMP1
IAPP & PCSK1 & PCSK2→CALCR & RAMP2
IAPP & PCSK1 & PCSK2→CALCR & RAMP3
IAPP & PCSK1 & PCSK2→CALCRL & RAMP1
IAPP & PCSK1 & PCSK2→CALCRL & RAMP2
IAPP & PCSK1 & PCSK2→CALCRL & RAMP3
Compartment
lymphoid
- B cell
- NK/T cell
12.5%
25%
50%
100%
ln(CP10K+1)
Diaphragm
Limb muscle
Mammary gland
Fat
Skin
Hypothalamus
Pituitary
Brainstem
Brain cortex
Testes
Uterus
Bladder
Kidney
Pancreas
Liver
Colon
Small intestine
Tongue
Trachea
Lung
Aorta
Heart
Spleen
Blood
Bone marrow
Bone

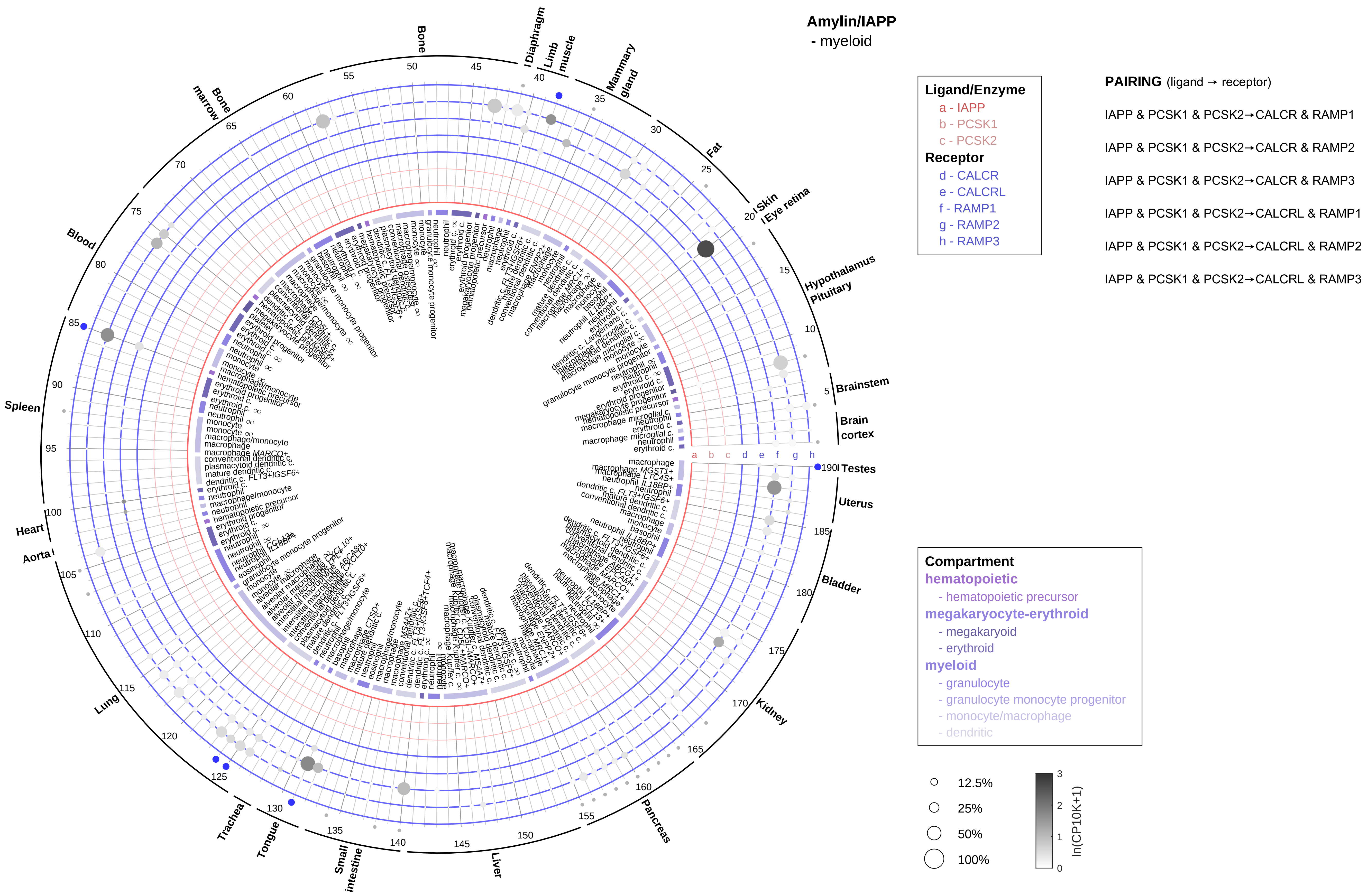
Amylin/IAPP
- myeloid
Ligand/Enzyme
a - IAPP
b - PCSK1
c - PCSK2
Receptor
d - CALCR
e - CALCRL
f - RAMP1
g - RAMP2
h - RAMP3
PAIRING (ligand → receptor)
IAPP & PCSK1 & PCSK2→CALCR & RAMP1
IAPP & PCSK1 & PCSK2→CALCR & RAMP2
IAPP & PCSK1 & PCSK2→CALCR & RAMP3
IAPP & PCSK1 & PCSK2→CALCRL & RAMP1
IAPP & PCSK1 & PCSK2→CALCRL & RAMP2
IAPP & PCSK1 & PCSK2→CALCRL & RAMP3
Compartment
hematopoietic
- hematopoietic precursor
megakaryocyte-erythroid
- megakaryoid
- erythroid
myeloid
- granulocyte
- granulocyte monocyte progenitor
- monocyte/macrophage
- dendritic
12.5%
25%
50%
100%
ln(CP10K+1)
Bone
Diaphragm
Limb muscle
Mammary gland
Fat
Skin
Eye retina
Hypothalamus
Pituitary
Brainstem
Brain cortex
Testes
Uterus
Bladder
Kidney
Pancreas
Liver
Small intestine
Tongue
Trachea
Lung
Aorta
Heart
Spleen
Blood
Bone marrow

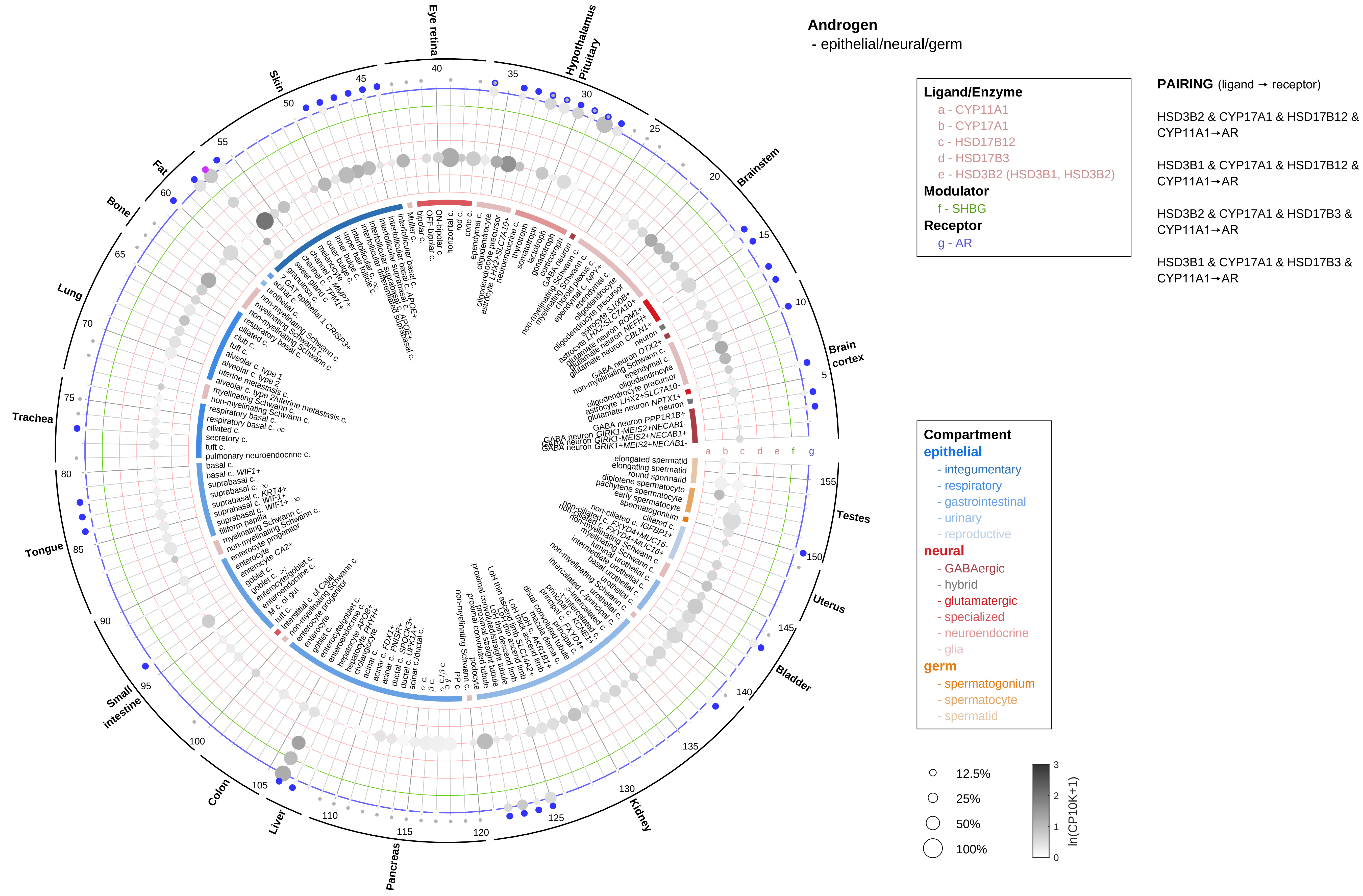

Androgen
- epithelial/neural/germ
Ligand/Enzyme
a - CYP11A1
b - CYP17A1
c - HSD17B12
d - HSD17B3
e - HSD3B2 (HSD3B1, HSD3B2)
Modulator
f - SHBG
Receptor
g - AR
PAIRING (ligand → receptor)
HSD3B2 & CYP17A1 & HSD17B12 & CYP11A1→AR
HSD3B1 & CYP17A1 & HSD17B12 & CYP11A1→AR
HSD3B2 & CYP17A1 & HSD17B3 & CYP11A1→AR
HSD3B1 & CYP17A1 & HSD17B3 & CYP11A1→AR
Compartment
epithelial
- integumentary
- respiratory
- gastrointestinal
- urinary
- reproductive
neural
- GABAergic
- hybrid
- glutamatergic
- specialized
- neuroendocrine
- glia
germ
- spermatogonium
- spermatocyte
- spermatid
12.5%
25%
50%
100%
ln(CP10K+1)
Eye retina
Hypothalamus
Pituitary
Brainstem
Brain cortex
Testes
Uterus
Bladder
Kidney
Pancreas
Liver
Colon
Small intestine
Tongue
Trachea
Lung
Bone
Fat
Skin

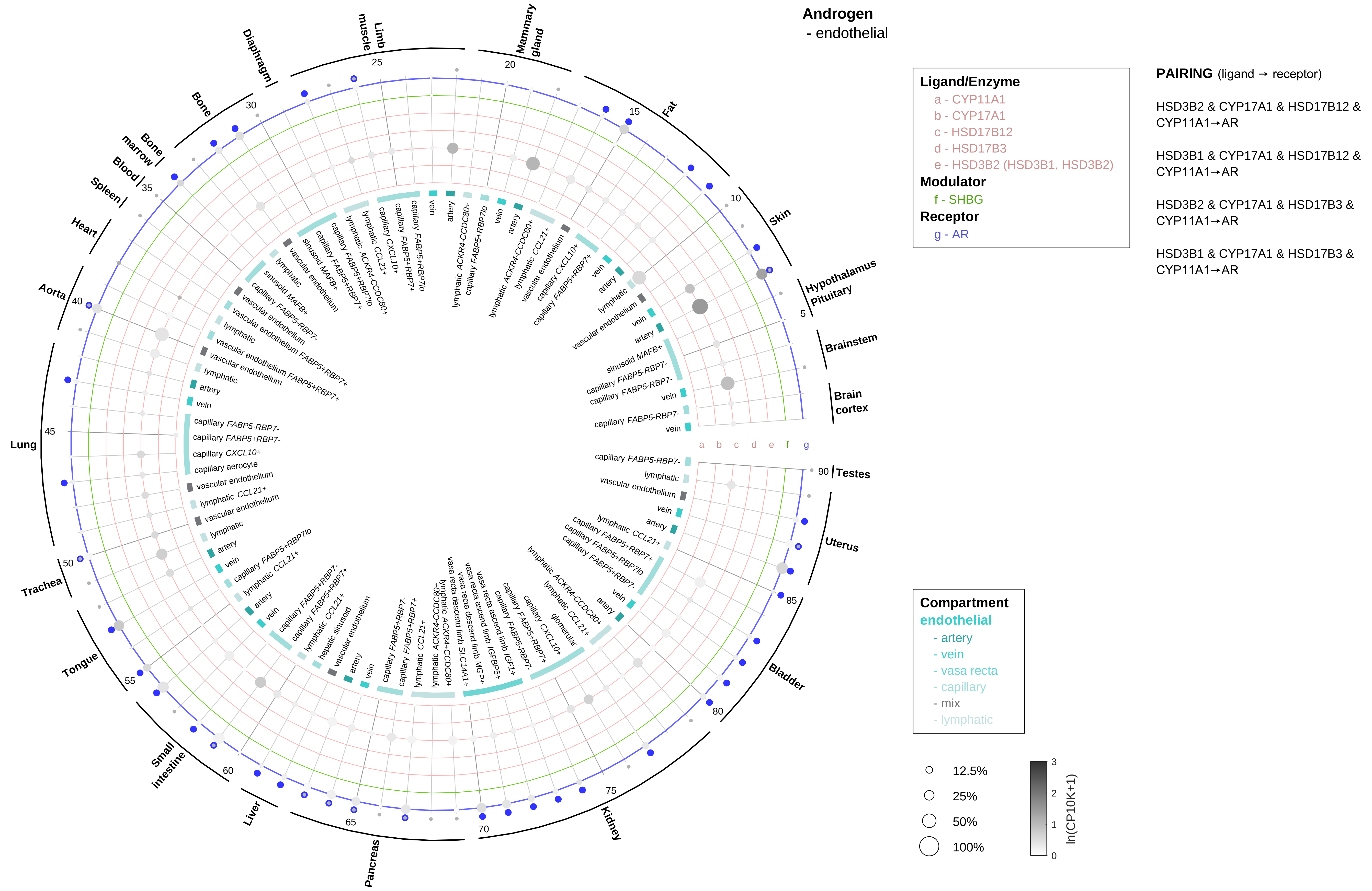
Androgen
- endothelial
Ligand/Enzyme
a - CYP11A1
b - CYP17A1
c - HSD17B12
d - HSD17B3
e - HSD3B2 (HSD3B1, HSD3B2)
Modulator
f - SHBG
Receptor
g - AR
PAIRING (ligand → receptor)
HSD3B2 & CYP17A1 & HSD17B12 & CYP11A1→AR
HSD3B1 & CYP17A1 & HSD17B12 & CYP11A1→AR
HSD3B2 & CYP17A1 & HSD17B3 & CYP11A1→AR
HSD3B1 & CYP17A1 & HSD17B3 & CYP11A1→AR
Compartment
endothelial
- artery
- vein
- vasa recta
- capillary
- mix
- lymphatic
12.5%
25%
50%
100%
ln(CP10K+1)
Hypothalamus
Pituitary
Brainstem
Brain cortex
Skin
Fat
Mammary gland
Limb muscle
Diaphragm
Bone
Bone marrow
Blood
Spleen
Heart
Aorta
Lung
Trachea
Tongue
Small intestine
Liver
Pancreas
Kidney
Bladder
Uterus
Testes

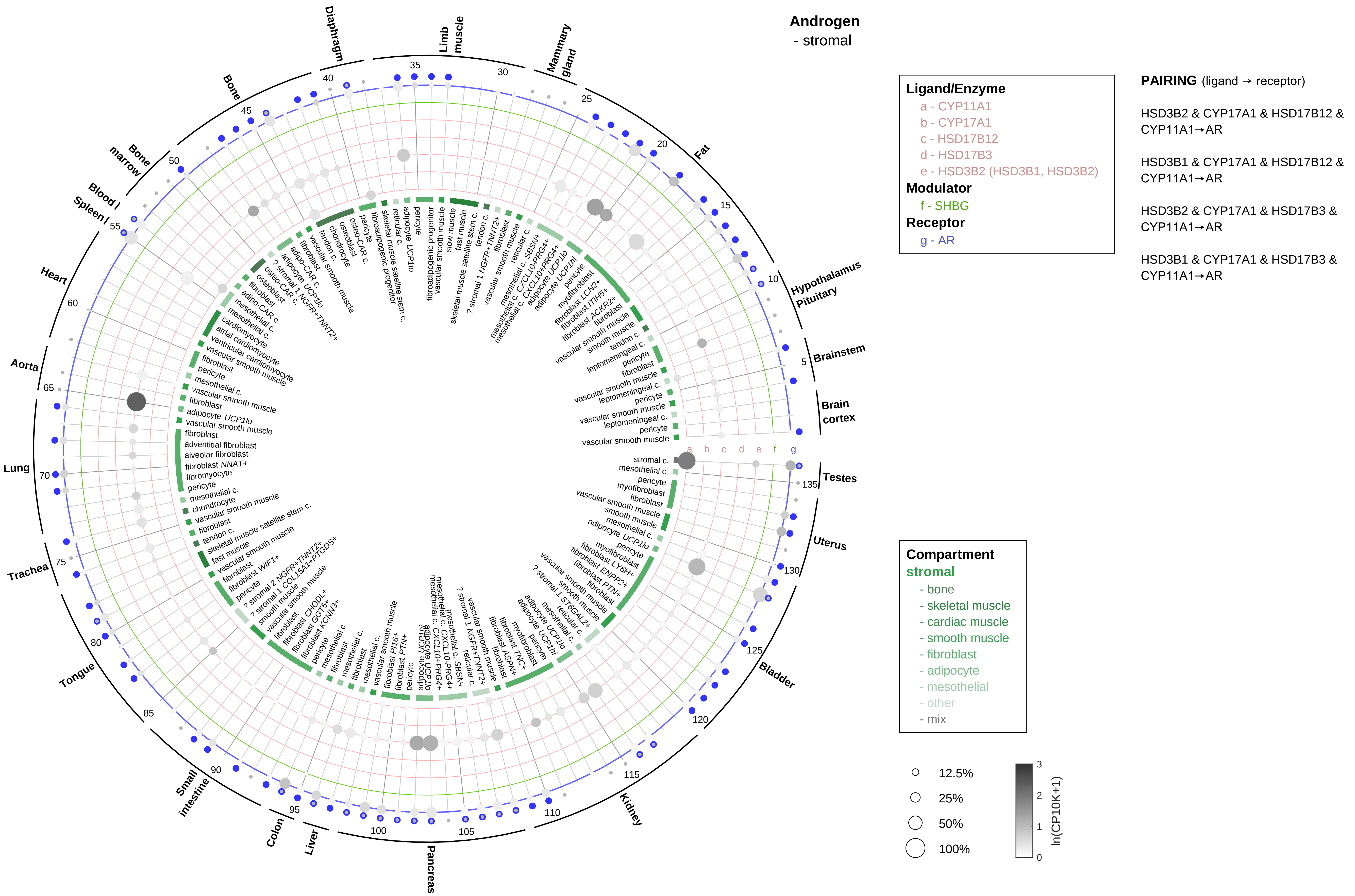
Androgen
- stromal
Ligand/Enzyme
a - CYP11A1
b - CYP17A1
c - HSD17B12
d - HSD17B3
e - HSD3B2 (HSD3B1, HSD3B2)
Modulator
f - SHBG
Receptor
g - AR
PAIRING (ligand → receptor)
HSD3B2 & CYP17A1 & HSD17B12 & CYP11A1→AR
HSD3B1 & CYP17A1 & HSD17B12 & CYP11A1→AR
HSD3B2 & CYP17A1 & HSD17B3 & CYP11A1→AR
HSD3B1 & CYP17A1 & HSD17B3 & CYP11A1→AR
Compartment
stromal
- bone
- skeletal muscle
- cardiac muscle
- smooth muscle
- fibroblast
- adipocyte
- mesothelial
- other
- mix
12.5%
25%
50%
100%
ln(CP10K+1)

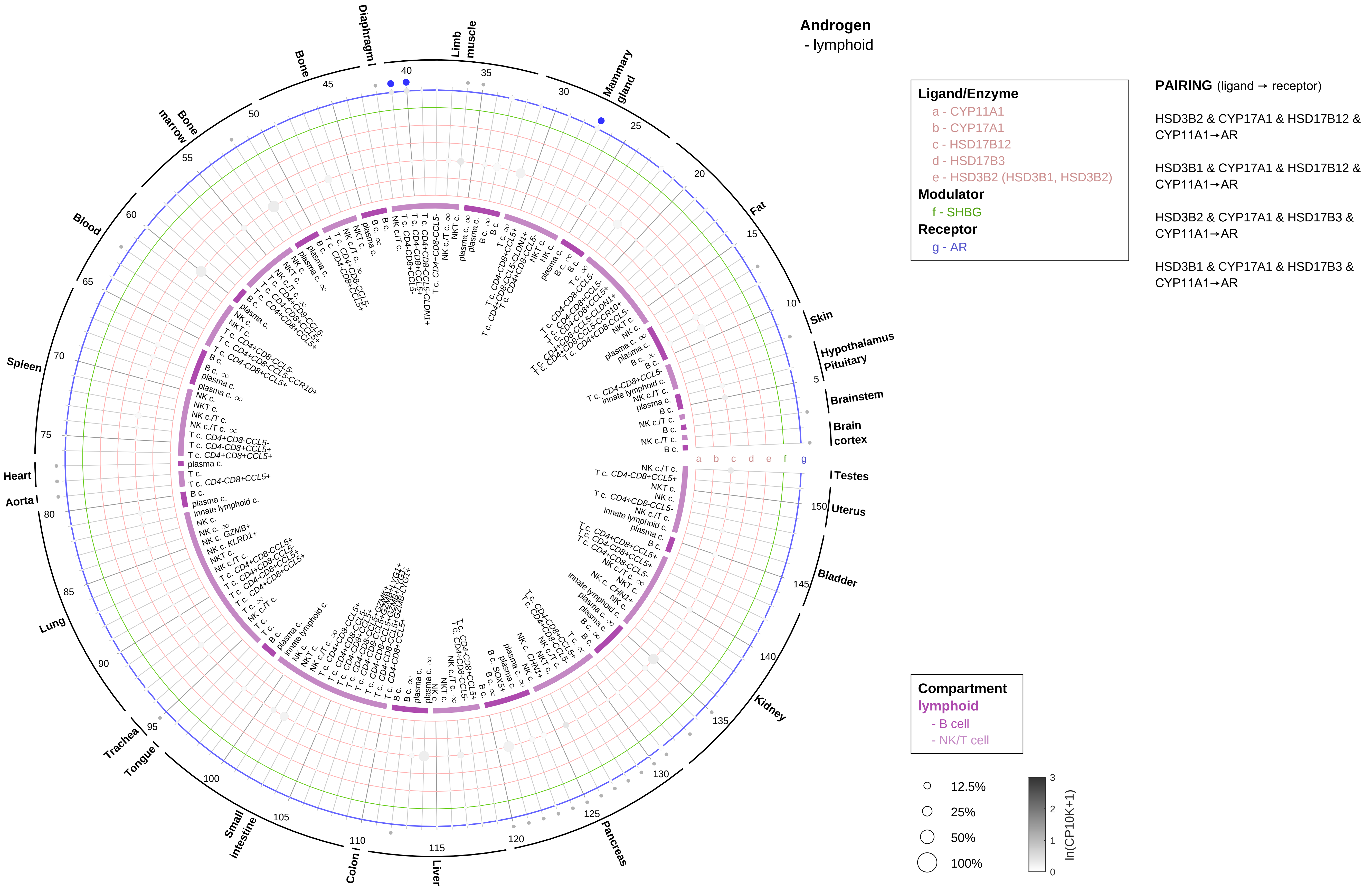
Androgen
- lymphoid
Ligand/Enzyme
a - CYP11A1
b - CYP17A1
c - HSD17B12
d - HSD17B3
e - HSD3B2 (HSD3B1, HSD3B2)
Modulator
f - SHBG
Receptor
g - AR
PAIRING (ligand → receptor)
HSD3B2 & CYP17A1 & HSD17B12 & CYP11A1→AR
HSD3B1 & CYP17A1 & HSD17B12 & CYP11A1→AR
HSD3B2 & CYP17A1 & HSD17B3 & CYP11A1→AR
HSD3B1 & CYP17A1 & HSD17B3 & CYP11A1→AR
Compartment
lymphoid
- B cell
- NK/T cell
12.5%
25%
50%
100%
ln(CP10K+1)
Diaphragm
Limb muscle
Mammary gland
Fat
Skin
Hypothalamus
Pituitary
Brainstem
Brain cortex
Testes
Uterus
Bladder
Kidney
Pancreas
Liver
Colon
Small intestine
Tongue
Trachea
Lung
Aorta
Heart
Spleen
Blood
Bone marrow
Bone

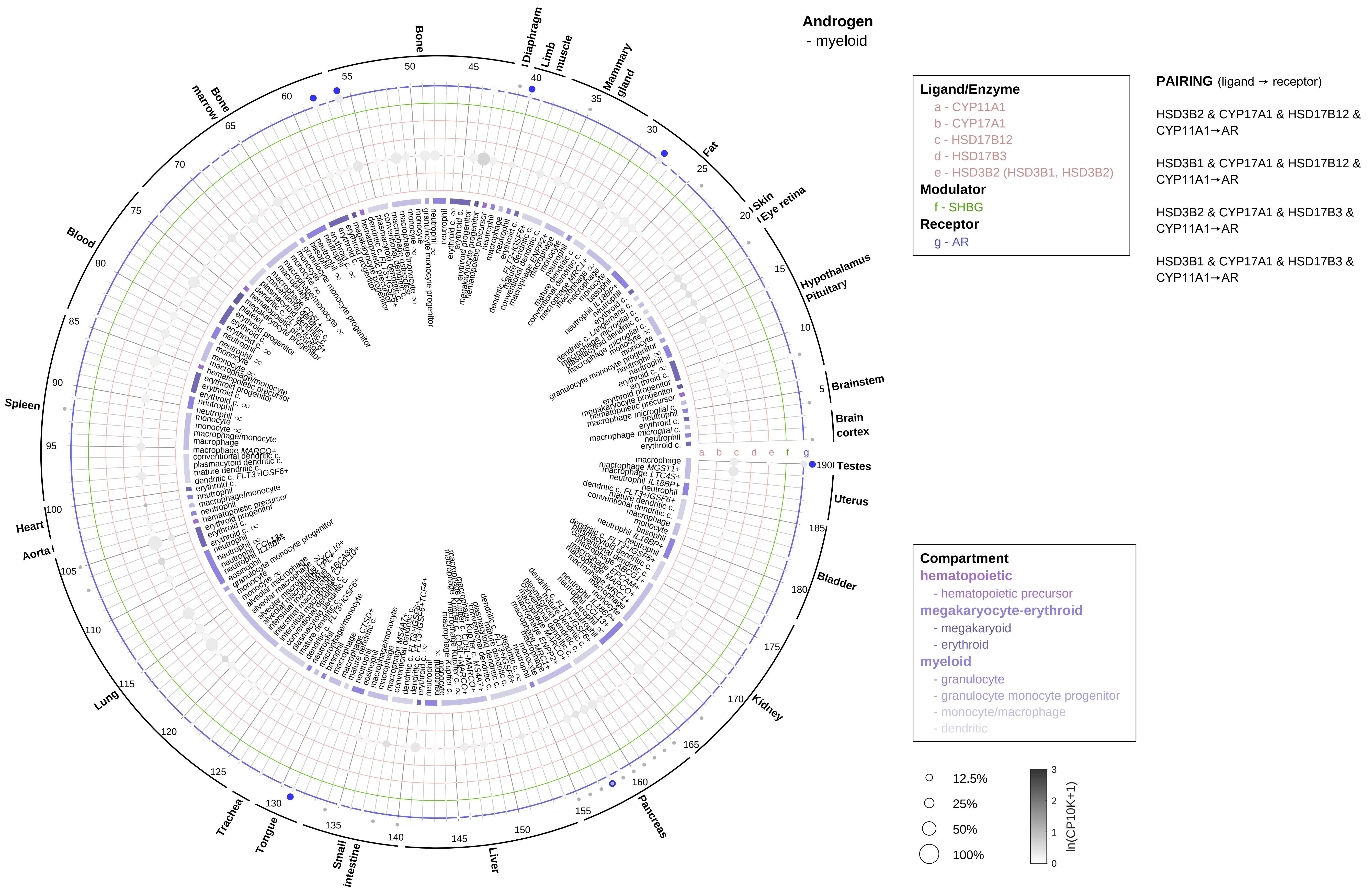

Androgen
- myeloid
Ligand/Enzyme
a - CYP11A1
b - CYP17A1
c - HSD17B12
d - HSD17B3
e - HSD3B2 (HSD3B1, HSD3B2)
Modulator
f - SHBG
Receptor
g - AR
PAIRING (ligand → receptor)
HSD3B2 & CYP17A1 & HSD17B12 & CYP11A1→AR
HSD3B1 & CYP17A1 & HSD17B12 & CYP11A1→AR
HSD3B2 & CYP17A1 & HSD17B3 & CYP11A1→AR
HSD3B1 & CYP17A1 & HSD17B3 & CYP11A1→AR
Compartment
hematopoietic
- hematopoietic precursor
megakaryocyte-erythroid
- megakaryoid
- erythroid
myeloid
- granulocyte
- granulocyte monocyte progenitor
- monocyte/macrophage
- dendritic
12.5%
25%
50%
100%
ln(CP10K+1)
Bone
Diaphragm
Limb muscle
Mammary gland
Fat
Skin
Eye retina
Hypothalamus
Pituitary
Brainstem
Brain cortex
Testes
Uterus
Bladder
Kidney
Pancreas
Liver
Small intestine
Tongue
Trachea
Lung
Aorta
Heart
Spleen
Blood
Bone marrow

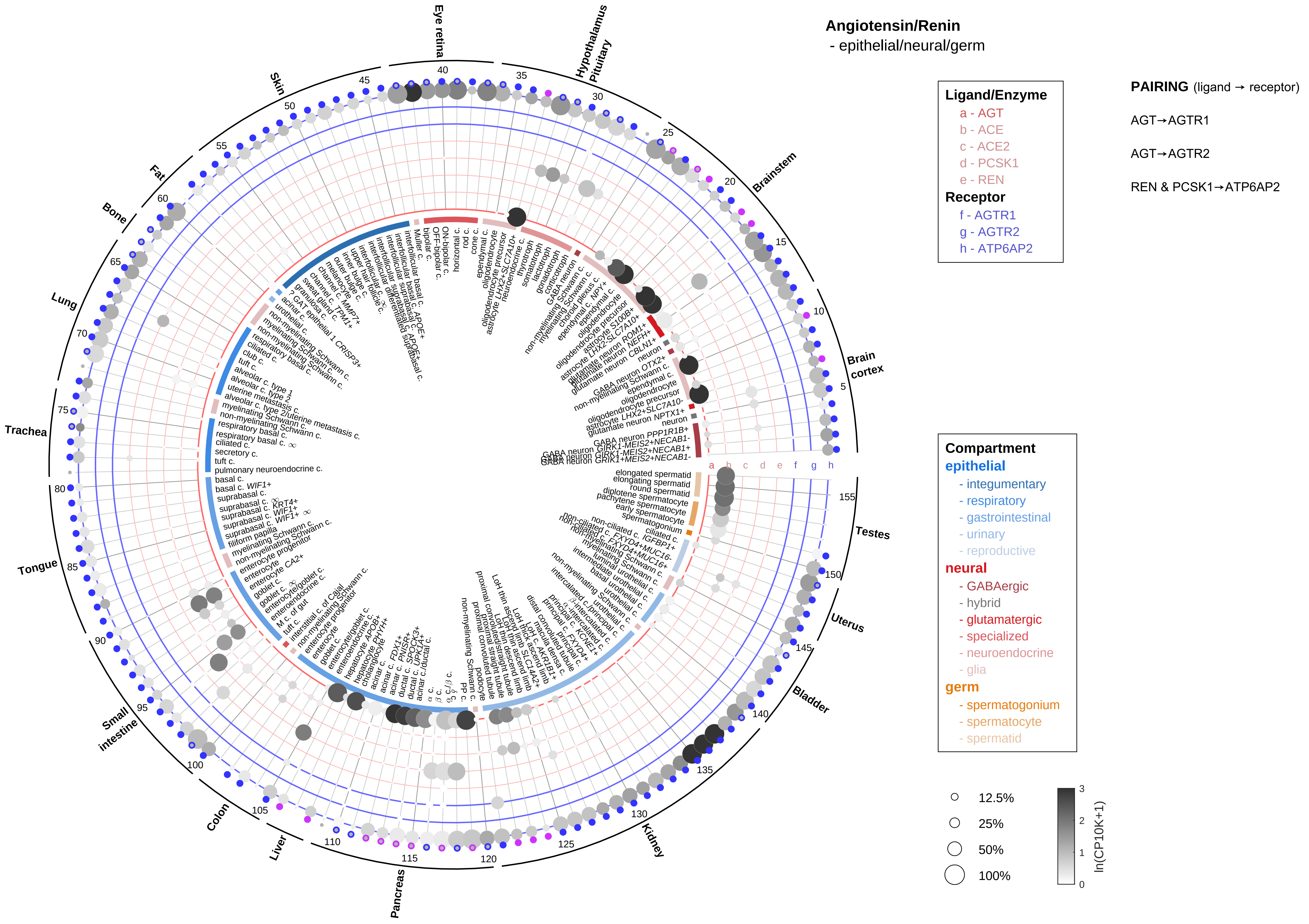

Angiotensin/Renin
- epithelial/neural/germ
Ligand/Enzyme
a - AGT
b - ACE
c - ACE2
d - PCSK1
e - REN
Receptor
f - AGTR1
g - AGTR2
h - ATP6AP2
PAIRING (ligand → receptor)
AGT→AGTR1
AGT→AGTR2
REN & PCSK1→ATP6AP2
Compartment
epithelial
- integumentary
- respiratory
- gastrointestinal
- urinary
- reproductive
neural
- GABAergic
- hybrid
- glutamatergic
- specialized
- neuroendocrine
- glia
germ
- spermatogonium
- spermatocyte
- spermatid
12.5%
25%
50%
100%
ln(CP10K+1)
Eye retina
Hypothalamus Pituitary
Brainstem
Brain cortex
Testes
Uterus
Bladder
Kidney
Pancreas
Liver
Colon
Small intestine
Tongue
Trachea
Lung
Bone
Fat
Skin

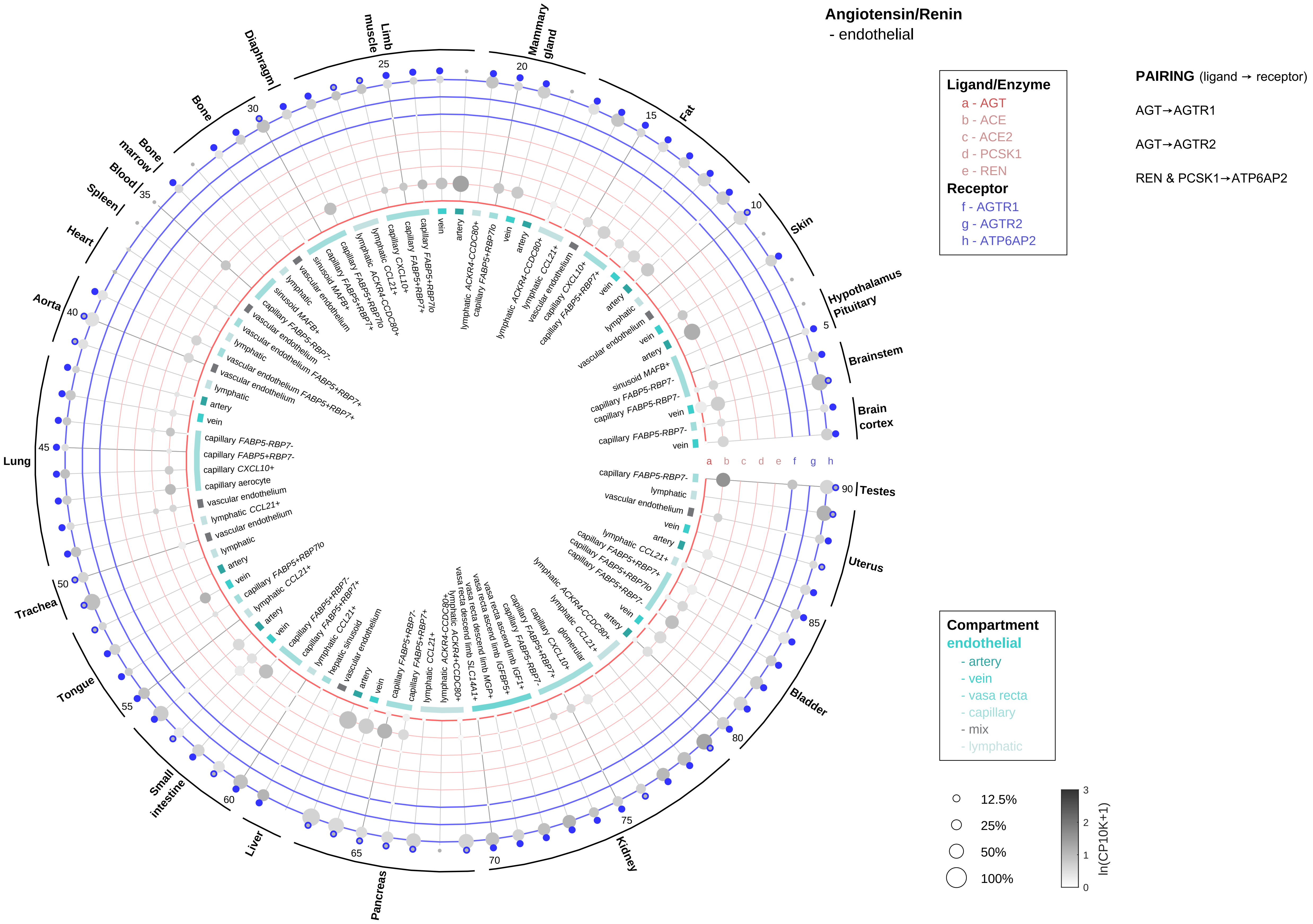

Angiotensin/Renin
- endothelial
Ligand/Enzyme
a - AGT
b - ACE
c - ACE2
d - PCSK1
e - REN
Receptor
f - AGTR1
g - AGTR2
h - ATP6AP2
PAIRING (ligand → receptor)
AGT→AGTR1
AGT→AGTR2
REN & PCSK1→ATP6AP2
Compartment
endothelial
- artery
- vein
- vasa recta
- capillary
- mix
- lymphatic
12.5%
25%
50%
100%
ln(CP10K+1)
Testes
Uterus
Bladder
Kidney
Pancreas
Liver
Small intestine
Tongue
Trachea
Lung
Aorta
Heart
Spleen
Blood
Bone marrow
Bone
Diaphragm
Limb muscle
Mammary gland
Fat
Skin
Hypothalamus Pituitary
Brainstem
Brain cortex

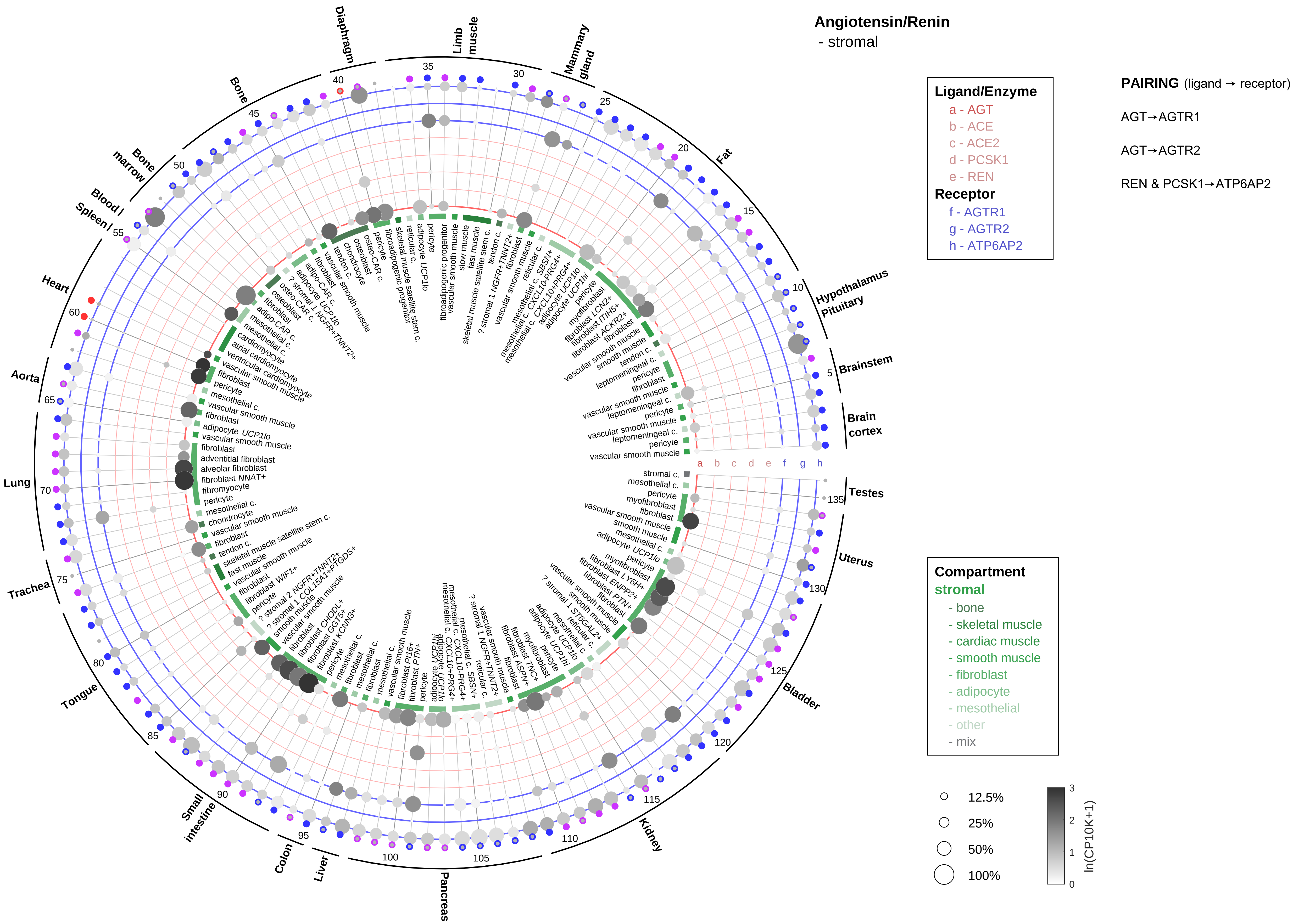

Angiotensin/Renin
- stromal
Ligand/Enzyme
a - AGT
b - ACE
c - ACE2
d - PCSK1
e - REN
Receptor
f - AGTR1
g - AGTR2
h - ATP6AP2
PAIRING (ligand → receptor)
AGT→AGTR1
AGT→AGTR2
REN & PCSK1→ATP6AP2
Compartment
stromal
- bone
- skeletal muscle
- cardiac muscle
- smooth muscle
- fibroblast
- adipocyte
- mesothelial
- other
- mix
12.5%
25%
50%
100%
ln(CP10K+1)
Diaphragm
Limb muscle
Mammary gland
Fat
Hypothalamus Pituitary
Brainstem
Brain cortex
Testes
Uterus
Bladder
Kidney
Pancreas
Liver
Colon
Small intestine
Tongue
Trachea
Lung
Aorta
Heart
Blood / Spleen /
Bone marrow
Bone

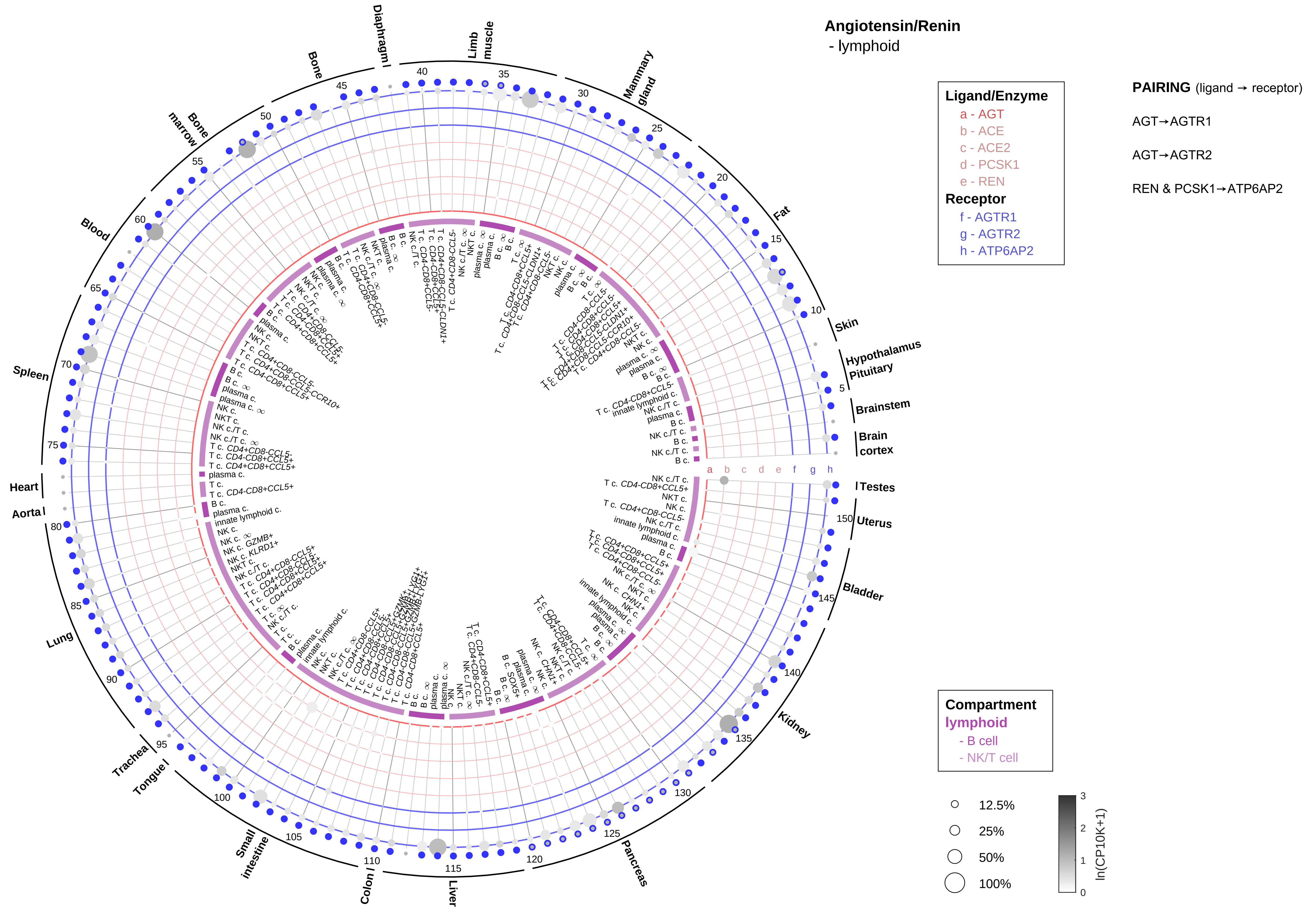
Angiotensin/Renin
- lymphoid
Ligand/Enzyme
a - AGT
b - ACE
c - ACE2
d - PCSK1
e - REN
Receptor
f - AGTR1
g - AGTR2
h - ATP6AP2
PAIRING (ligand → receptor)
AGT→AGTR1
AGT→AGTR2
REN & PCSK1→ATP6AP2
Compartment
lymphoid
- B cell
- NK/T cell
12.5%
25%
50%
100%
ln(CP10K+1)
Skin
Hypothalamus
Pituitary
Brainstem
Brain cortex
Testes
Uterus
Bladder
Kidney
Pancreas
Liver
Colon
Small intestine
Tongue
Trachea
Lung
Aorta
Heart
Spleen
Blood
Bone marrow
Bone
Diaphragm
Limb muscle
Mammary gland
Fat

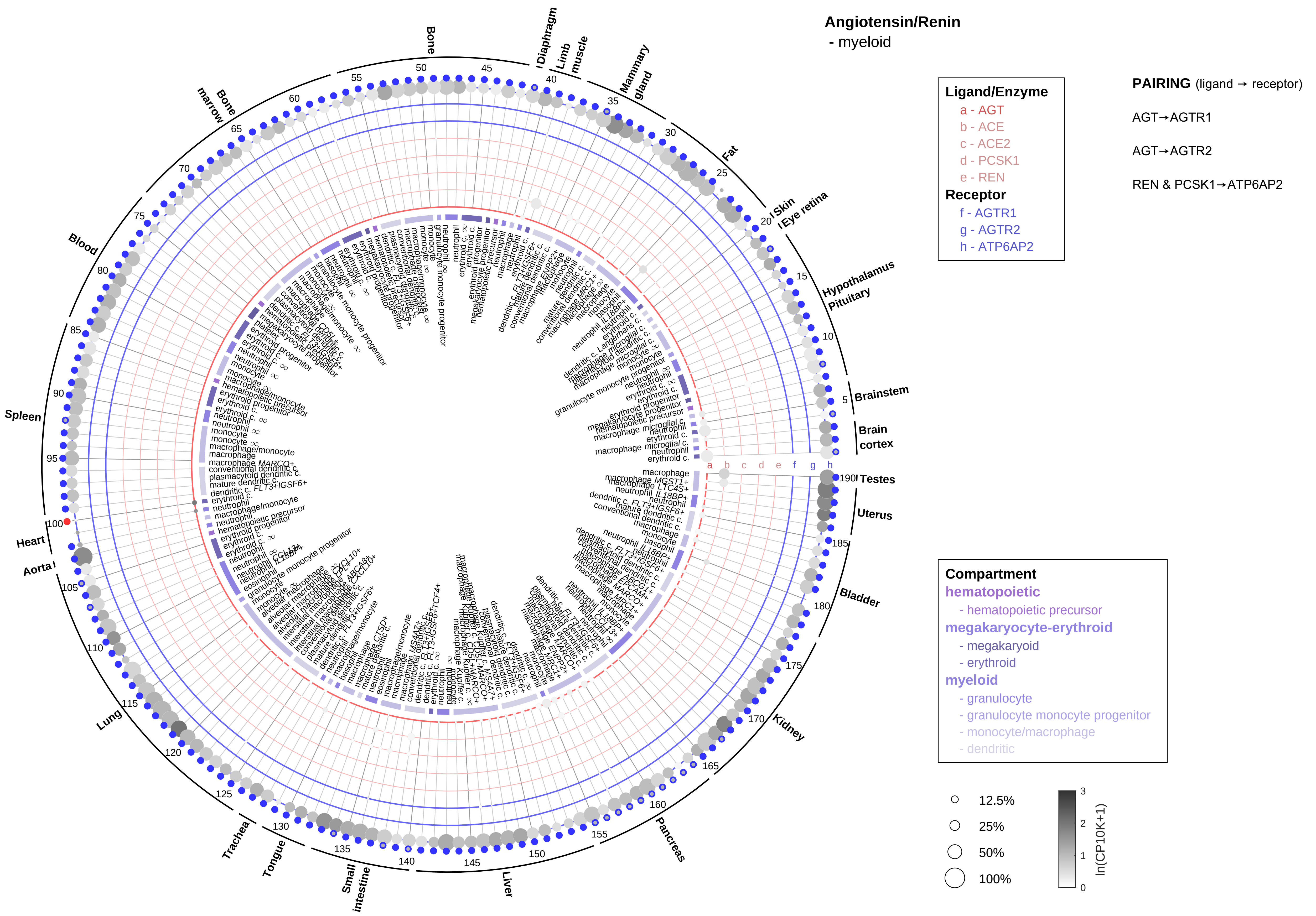
Angiotensin/Renin
- myeloid
Ligand/Enzyme
a - AGT
b - ACE
c - ACE2
d - PCSK1
e - REN
Receptor
f - AGTR1
g - AGTR2
h - ATP6AP2
PAIRING (ligand → receptor)
AGT→AGTR1
AGT→AGTR2
REN & PCSK1→ATP6AP2
Compartment
hematopoietic
- hematopoietic precursor
megakaryocyte-erythroid
- megakaryoid
- erythroid
myeloid
- granulocyte
- granulocyte monocyte progenitor
- monocyte/macrophage
- dendritic
12.5%
25%
50%
100%
ln(CP10K+1)
Bone
Diaphragm
Limb muscle
Mammary gland
Fat
Skin
Eye retina
Hypothalamus
Pituitary
Brainstem
Brain cortex
Testes
Uterus
Bladder
Kidney
Pancreas
Liver
Small intestine
Tongue
Trachea
Lung
Aorta
Heart
Spleen
Blood
Bone marrow

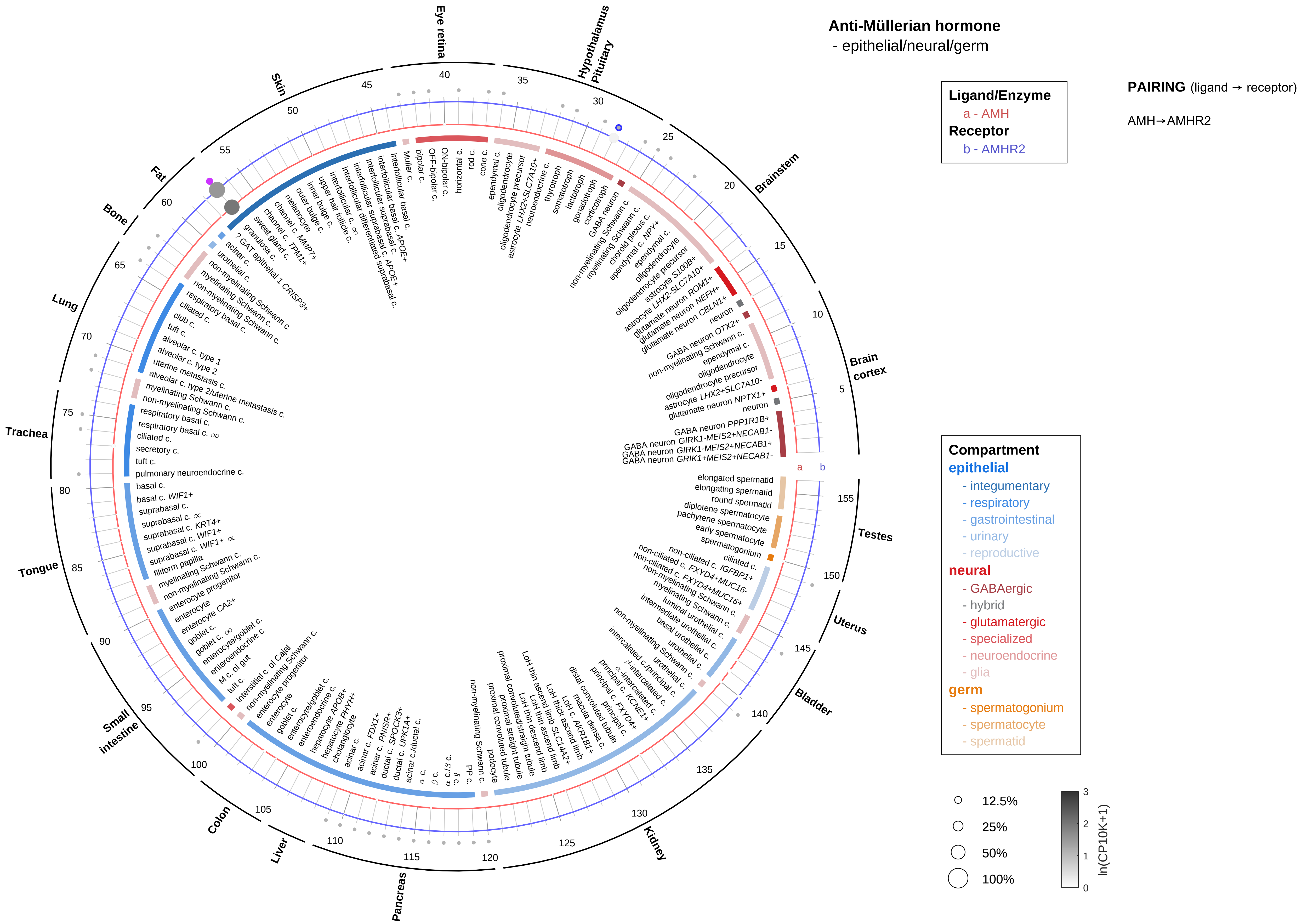
Anti-Müllerian hormone
- epithelial/neural/germ
Ligand/Enzyme
a - AMH
Receptor
b - AMHR2
PAIRING (ligand → receptor)
AMH→AMHR2
Compartment
epithelial
- integumentary
- respiratory
- gastrointestinal
- urinary
- reproductive
neural
- GABAergic
- hybrid
- glutamatergic
- specialized
- neuroendocrine
- glia
germ
- spermatogonium
- spermatocyte
- spermatid
12.5%
25%
50%
100%
ln(CP10K+1)
Eye retina
Hypothalamus Pituitary
Brainstem
Brain cortex
Testes
Uterus
Bladder
Kidney
Pancreas
Liver
Colon
Small intestine
Tongue
Trachea
Lung
Bone
Fat
Skin

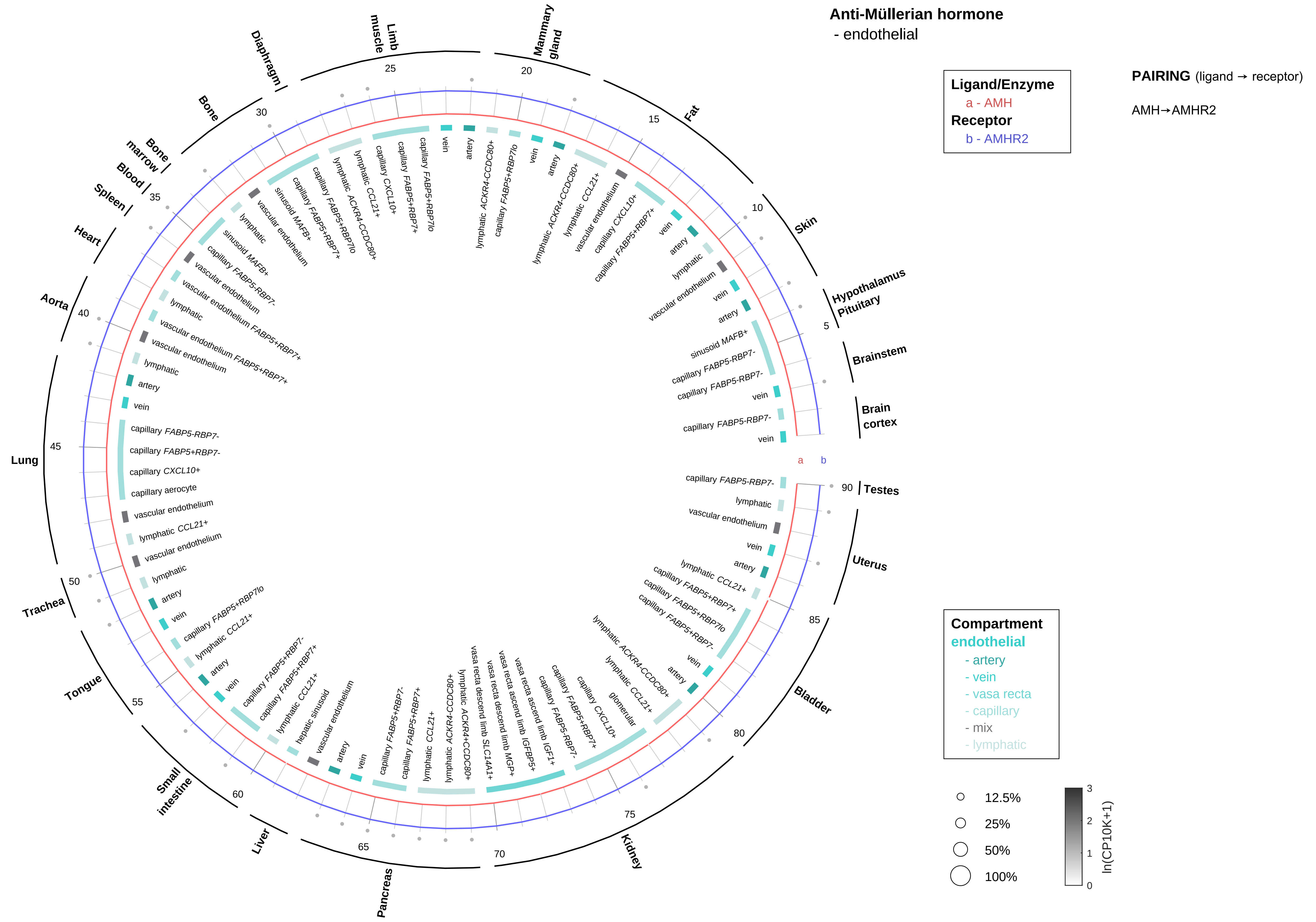
Anti-Müllerian hormone
- endothelial
Ligand/Enzyme
a - AMH
Receptor
b - AMHR2
PAIRING (ligand → receptor)
AMH→AMHR2
Compartment
endothelial
- artery
- vein
- vasa recta
- capillary
- mix
- lymphatic
12.5%
25%
50%
100%
ln(CP10K+1)
Brain cortex
Brainstem
Hypothalamus Pituitary
Skin
Fat
Mammary gland
Limb muscle
Diaphragm
Bone
Bone marrow
Blood
Spleen
Heart
Aorta
Lung
Trachea
Tongue
Small intestine
Liver
Pancreas
Kidney
Bladder
Uterus
Testes

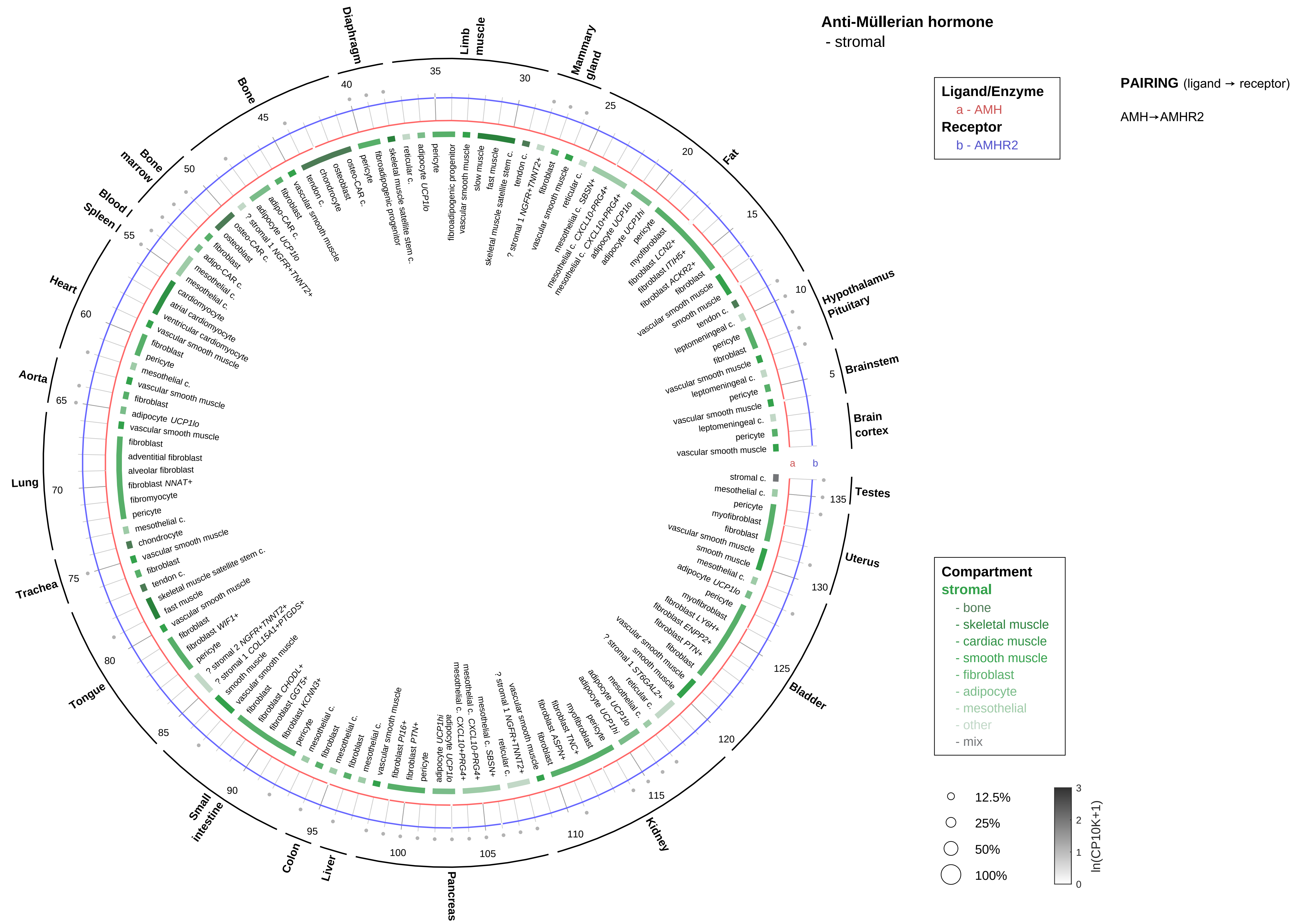
Anti-Müllerian hormone
- stromal
Ligand/Enzyme
a - AMH
Receptor
b - AMHR2
PAIRING (ligand → receptor)
AMH→AMHR2
Compartment
stromal
- bone
- skeletal muscle
- cardiac muscle
- smooth muscle
- fibroblast
- adipocyte
- mesothelial
- other
- mix
12.5%
25%
50%
100%
ln(CP10K+1)
Brain cortex
Brainstem
Hypothalamus Pituitary
Fat
Mammary gland
Limb muscle
Diaphragm
Bone
Bone marrow
Blood / Spleen
Heart
Aorta
Lung
Trachea
Tongue
Small intestine
Colon
Liver
Pancreas
Kidney
Bladder
Uterus
Testes

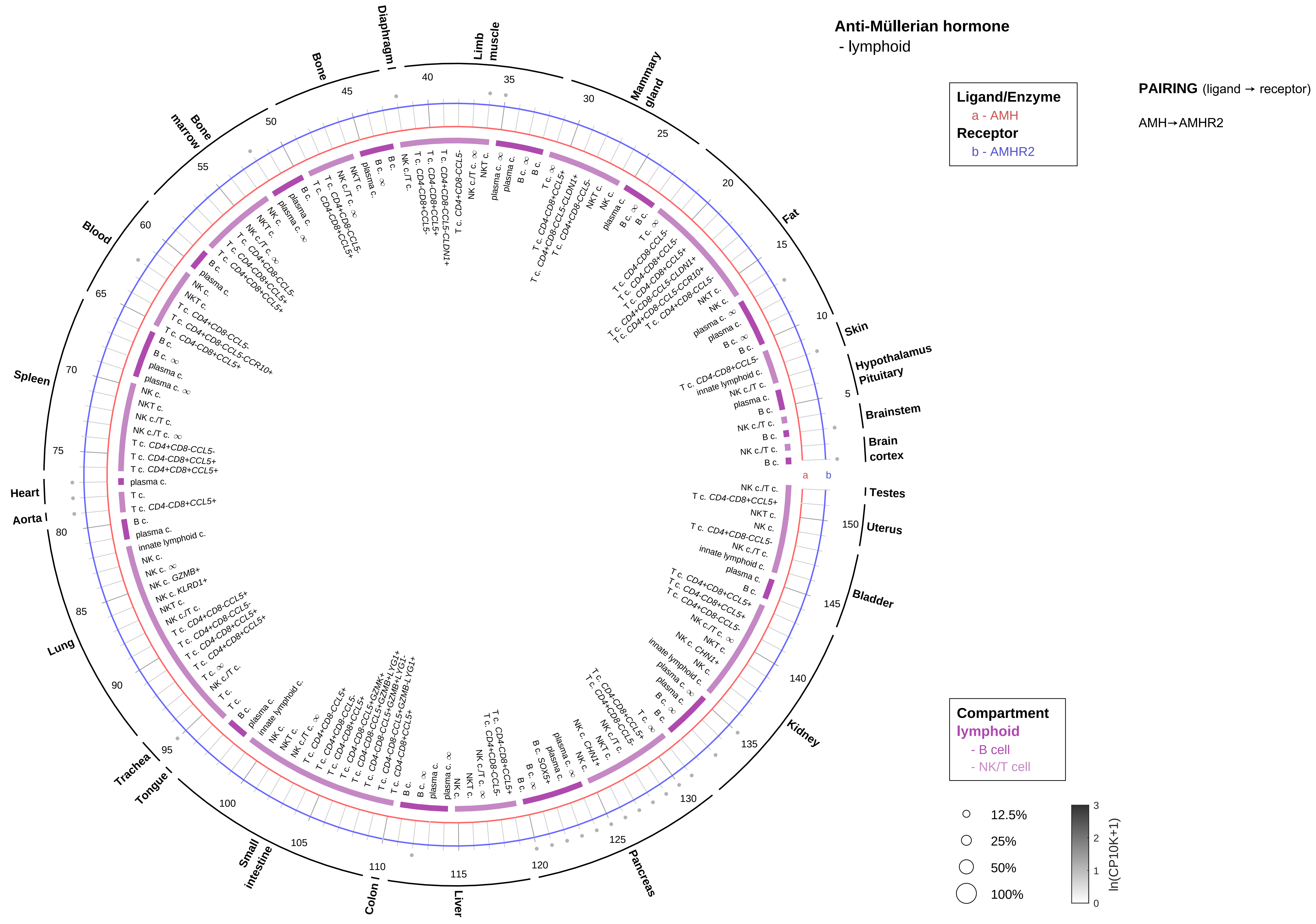
Anti-Müllerian hormone
- lymphoid
Ligand/Enzyme
a - AMH
Receptor
b - AMHR2
PAIRING (ligand → receptor)
AMH→AMHR2
Compartment
lymphoid
- B cell
- NK/T cell
12.5%
25%
50%
100%
ln(CP10K+1)
Testes
Uterus
Bladder
Kidney
Pancreas
Liver
Colon
Small intestine
Tongue
Trachea
Lung
Aorta
Heart
Spleen
Blood
Bone marrow
Bone
Diaphragm
Limb muscle
Mammary gland
Fat
Skin
Hypothalamus
Pituitary
Brainstem
Brain cortex

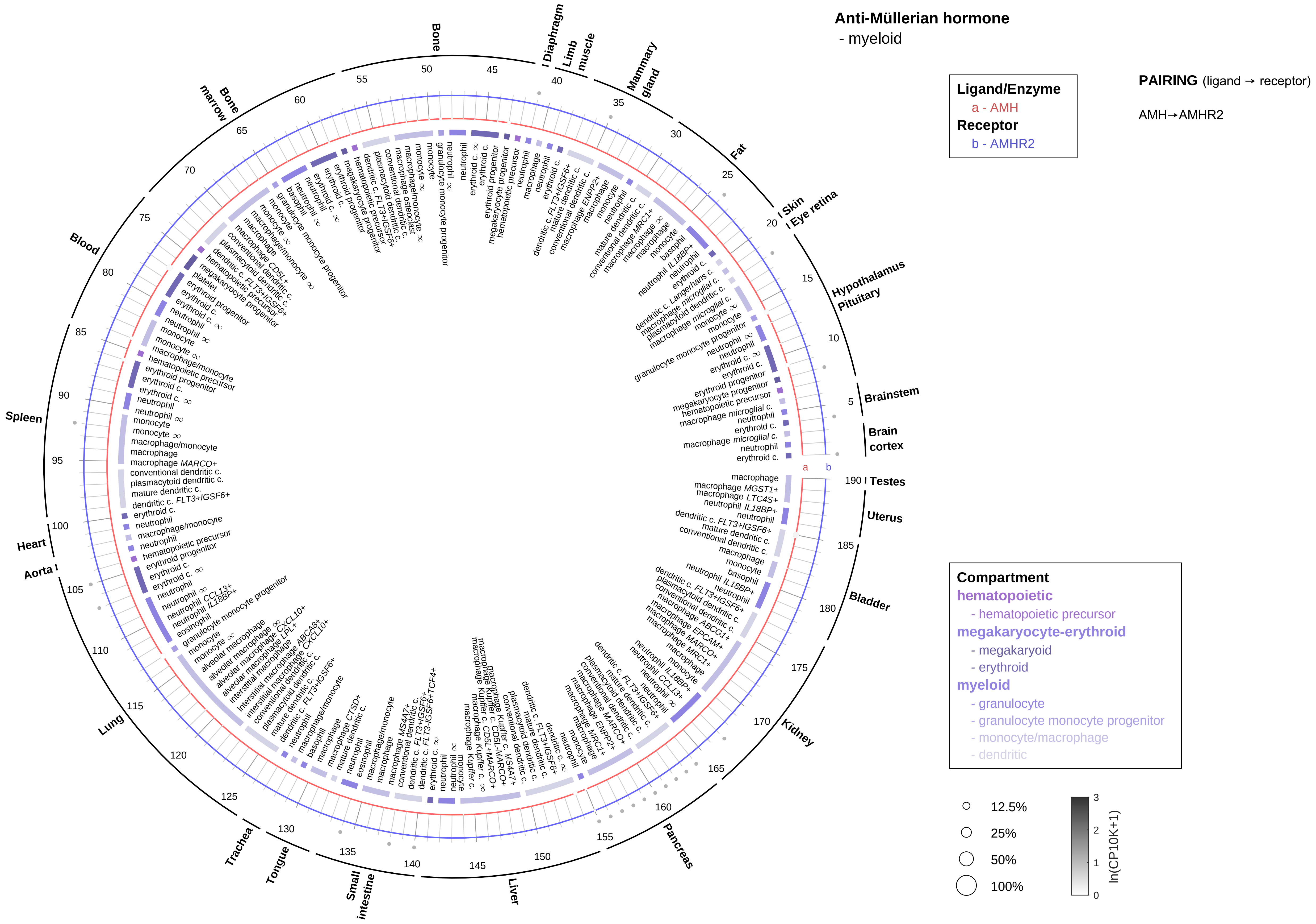

Anti-Müllerian hormone
- myeloid
Ligand/Enzyme
a - AMH
Receptor
b - AMHR2
PAIRING (ligand → receptor)
AMH→AMHR2
Compartment
hematopoietic
- hematopoietic precursor
megakaryocyte-erythroid
- megakaryoid
- erythroid
myeloid
- granulocyte
- granulocyte monocyte progenitor
- monocyte/macrophage
- dendritic
12.5%
25%
50%
100%
ln(CP10K+1)
Testes
Uterus
Bladder
Kidney
Pancreas
Liver
Small intestine
Tongue
Trachea
Lung
Aorta
Heart
Spleen
Blood
Bone marrow
Bone
Diaphragm
Limb muscle
Mammary gland
Fat
Skin
Eye retina
Hypothalamus
Pituitary
Brainstem
Brain cortex

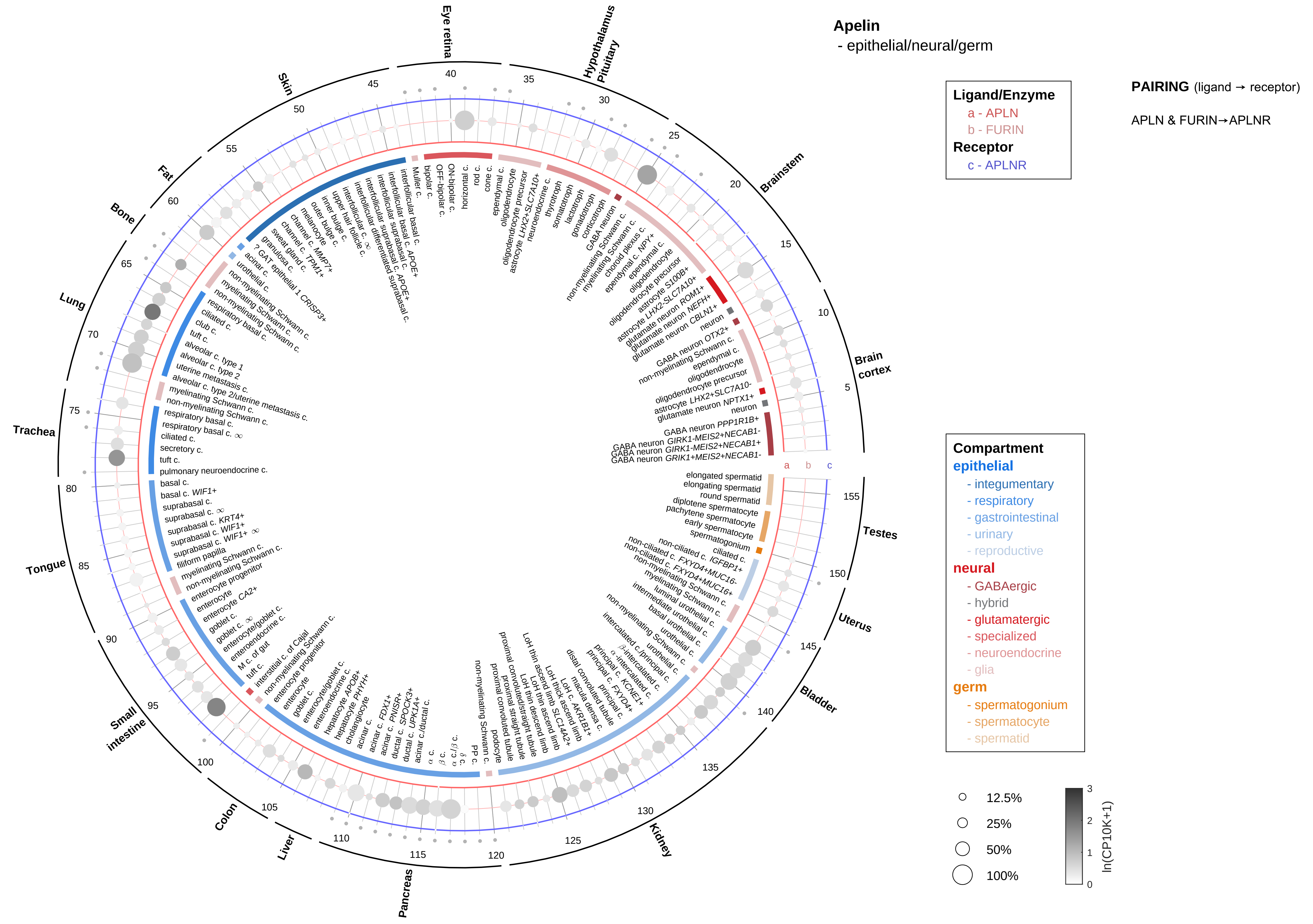
Apelin
- epithelial/neural/germ
Ligand/Enzyme
a - APLN
b - FURIN
Receptor
c - APLNR
PAIRING (ligand → receptor)
APLN & FURIN→APLNR
Compartment
epithelial
- integumentary
- respiratory
- gastrointestinal
- urinary
- reproductive
neural
- GABAergic
- hybrid
- glutamatergic
- specialized
- neuroendocrine
- glia
germ
- spermatogonium
- spermatocyte
- spermatid
12.5%
25%
50%
100%
ln(CP10K+1)
Eye retina
Hypothalamus Pituitary
Brainstem
Brain cortex
Testes
Uterus
Bladder
Kidney
Pancreas
Liver
Colon
Small intestine
Tongue
Trachea
Lung
Bone
Fat
Skin

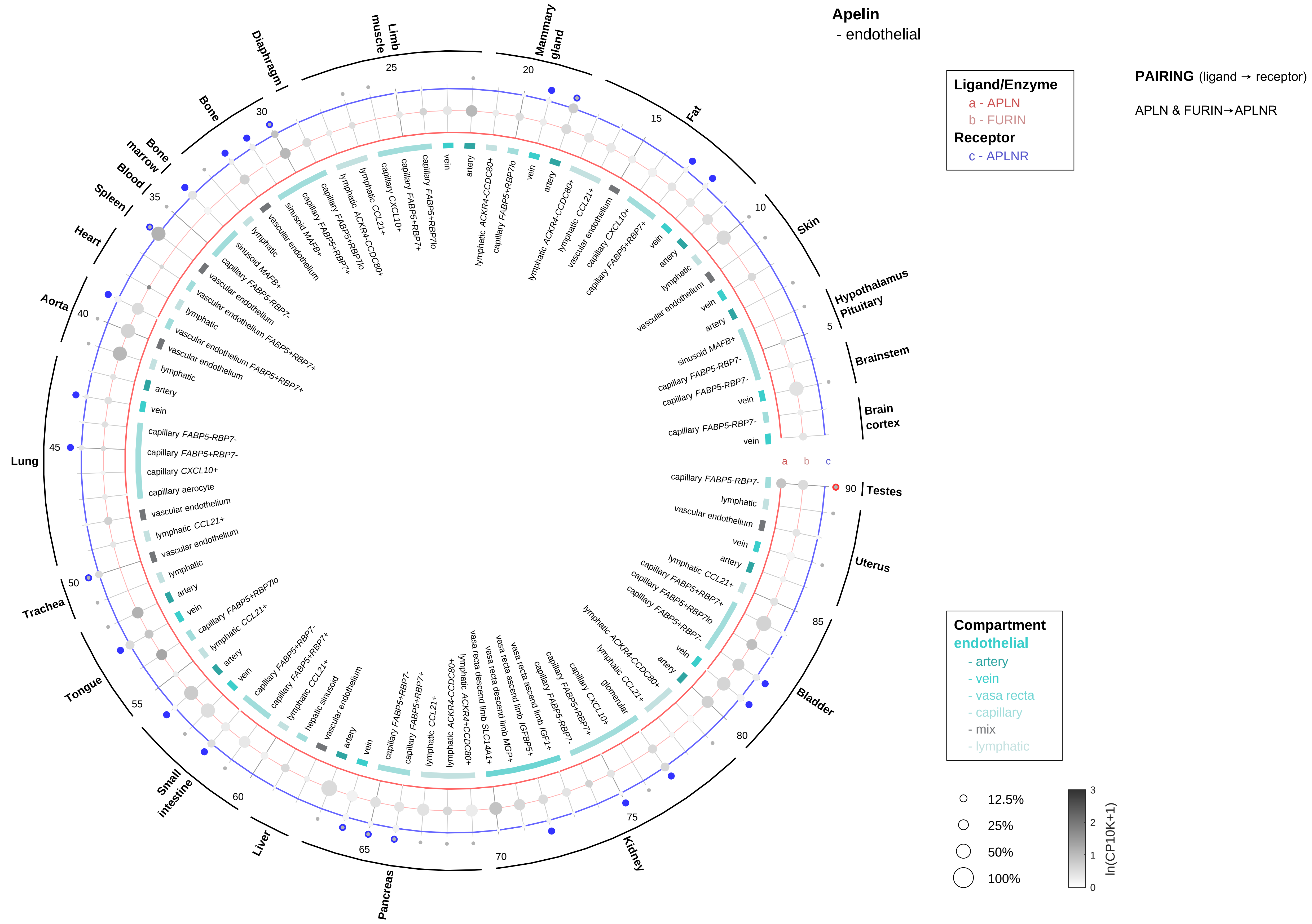
Apelin
- endothelial
Ligand/Enzyme
a - APLN
b - FURIN
Receptor
c - APLNR
PAIRING (ligand → receptor)
APLN & FURIN→APLNR
Compartment
endothelial
- artery
- vein
- vasa recta
- capillary
- mix
- lymphatic
12.5%
25%
50%
100%
ln(CP10K+1)
Testes
Uterus
Bladder
Kidney
Pancreas
Liver
Small intestine
Tongue
Trachea
Lung
Aorta
Heart
Spleen
Blood
Bone marrow
Bone
Diaphragm
Limb muscle
Mammary gland
Fat
Skin
Hypothalamus Pituitary
Brainstem
Brain cortex

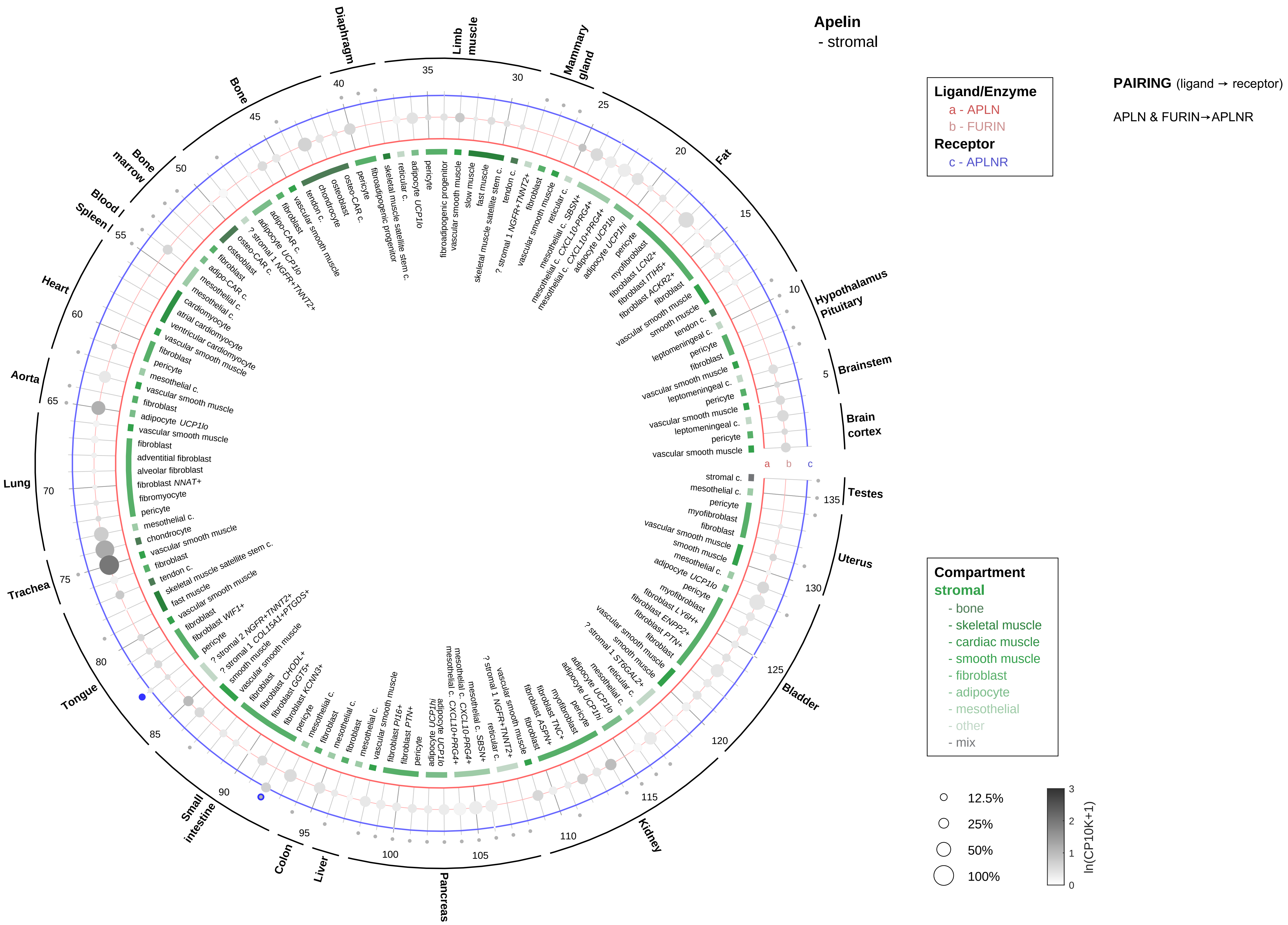

Apelin
- stromal
Ligand/Enzyme
a - APLN
b - FURIN
Receptor
c - APLNR
PAIRING (ligand → receptor)
APLN & FURIN→APLNR
Compartment
stromal
- bone
- skeletal muscle
- cardiac muscle
- smooth muscle
- fibroblast
- adipocyte
- mesothelial
- other
- mix
12.5%
25%
50%
100%
ln(CP10K+1)
Brain cortex
Brainstem
Hypothalamus Pituitary
Fat
Mammary gland
Limb muscle
Diaphragm
Bone
Bone marrow
Blood / Spleen
Heart
Aorta
Lung
Trachea
Tongue
Small intestine
Colon
Liver
Pancreas
Kidney
Bladder
Uterus
Testes

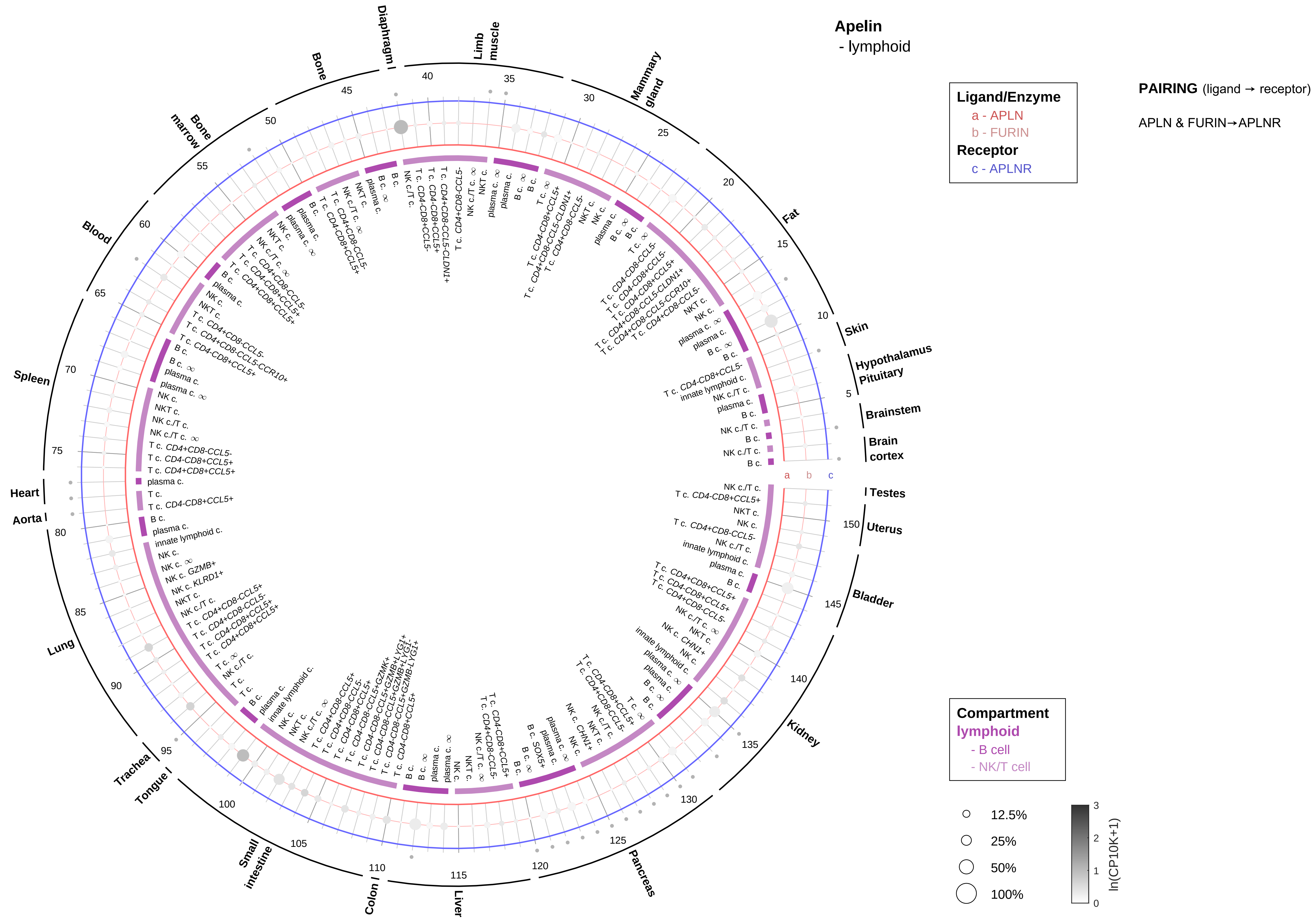
Apelin
- lymphoid
Ligand/Enzyme
a - APLN
b - FURIN
Receptor
c - APLNR
PAIRING (ligand → receptor)
APLN & FURIN→APLNR
Compartment
lymphoid
- B cell
- NK/T cell
12.5%
25%
50%
100%
ln(CP10K+1)
0
1
2
3
Skin
Hypothalamus
Pituitary
Brainstem
Brain cortex
Testes
Uterus
Bladder
Kidney
Pancreas
Liver
Colon
Small intestine
Tongue
Trachea
Lung
Aorta
Heart
Spleen
Blood
Bone marrow
Bone
Diaphragm
Limb muscle
Mammary gland
Fat
a
b
c

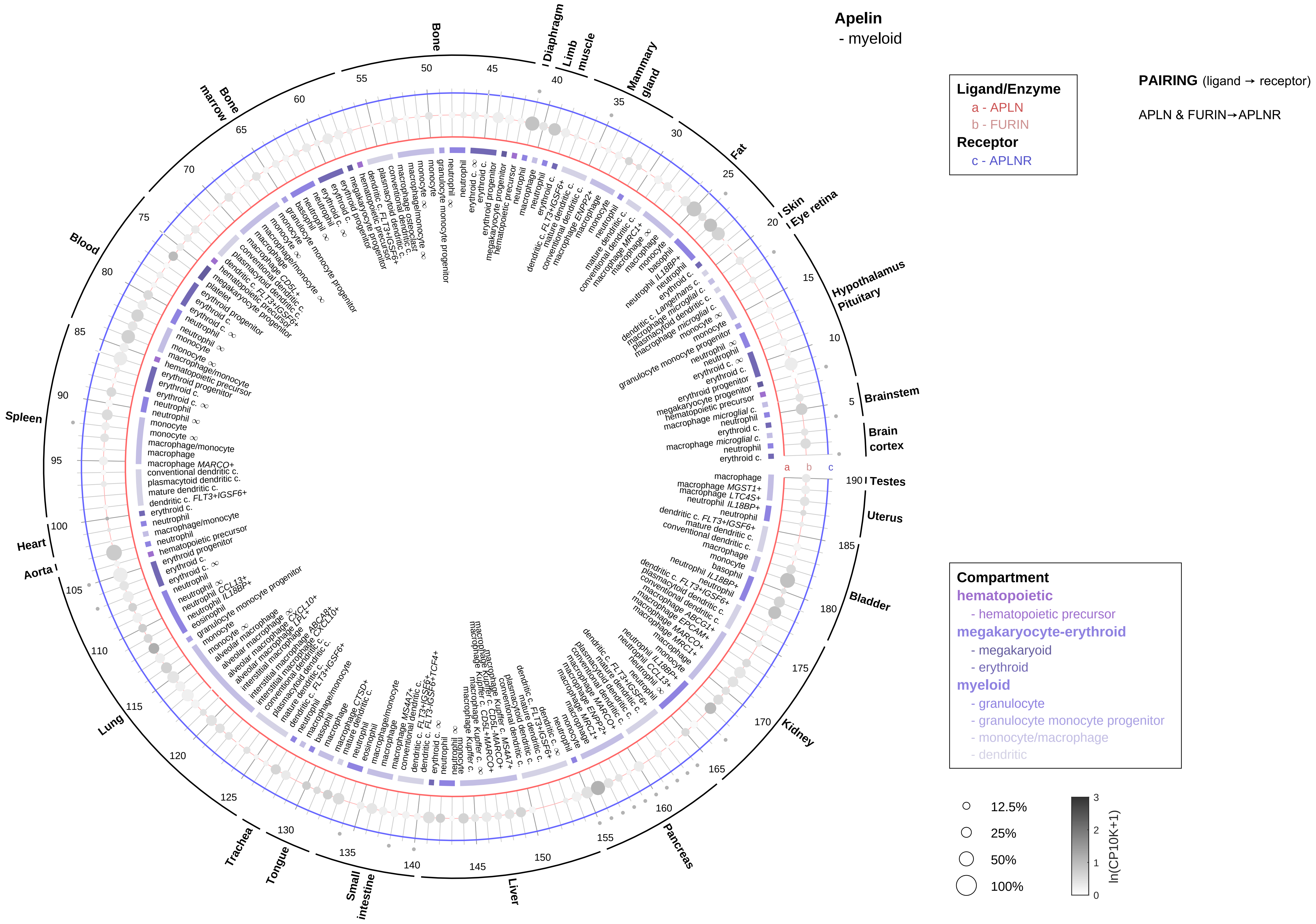

Apelin
- myeloid
Ligand/Enzyme
a - APLN
b - FURIN
Receptor
c - APLNR
PAIRING (ligand → receptor)
APLN & FURIN→APLNR
Compartment
hematopoietic
- hematopoietic precursor
megakaryocyte-erythroid
- megakaryoid
- erythroid
myeloid
- granulocyte
- granulocyte monocyte progenitor
- monocyte/macrophage
- dendritic
12.5%
25%
50%
100%
ln(CP10K+1)
Bone
Diaphragm
Limb muscle
Mammary gland
Fat
Skin
Eye retina
Hypothalamus
Pituitary
Brainstem
Brain cortex
Testes
Uterus
Bladder
Kidney
Pancreas
Liver
Small intestine
Tongue
Trachea
Lung
Aorta
Heart
Spleen
Blood
Bone marrow

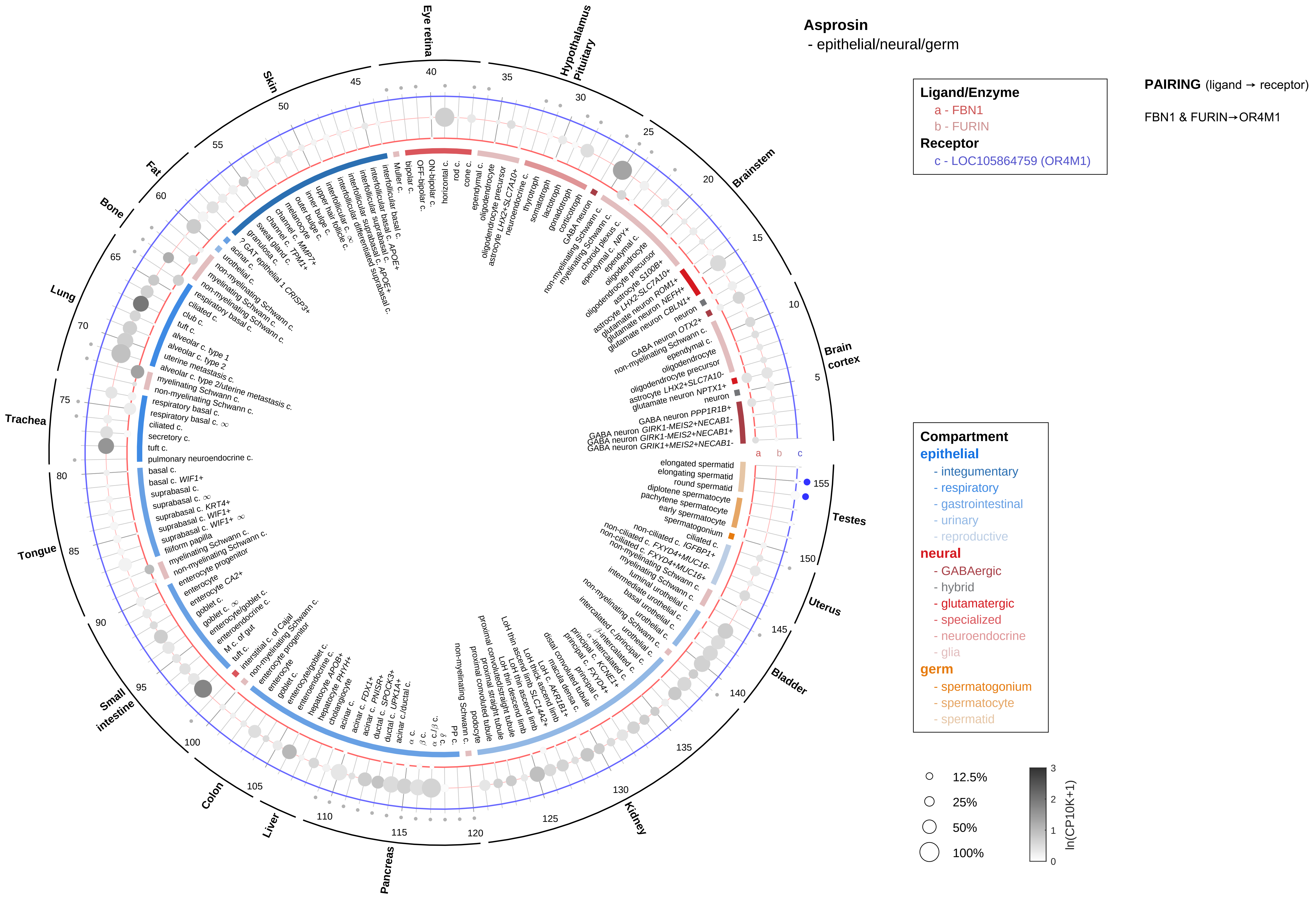

Asprosin
- epithelial/neural/germ
Ligand/Enzyme
a - FBN1
b - FURIN
Receptor
c - LOC105864759 (OR4M1)
PAIRING (ligand → receptor)
FBN1 & FURIN→OR4M1
Compartment
epithelial
- integumentary
- respiratory
- gastrointestinal
- urinary
- reproductive
neural
- GABAergic
- hybrid
- glutamatergic
- specialized
- neuroendocrine
- glia
germ
- spermatogonium
- spermatocyte
- spermatid
12.5%
25%
50%
100%
ln(CP10K+1)
Eye retina
Hypothalamus Pituitary
Brainstem
Brain cortex
Testes
Uterus
Bladder
Kidney
Pancreas
Liver
Colon
Small intestine
Tongue
Trachea
Lung
Bone
Fat
Skin

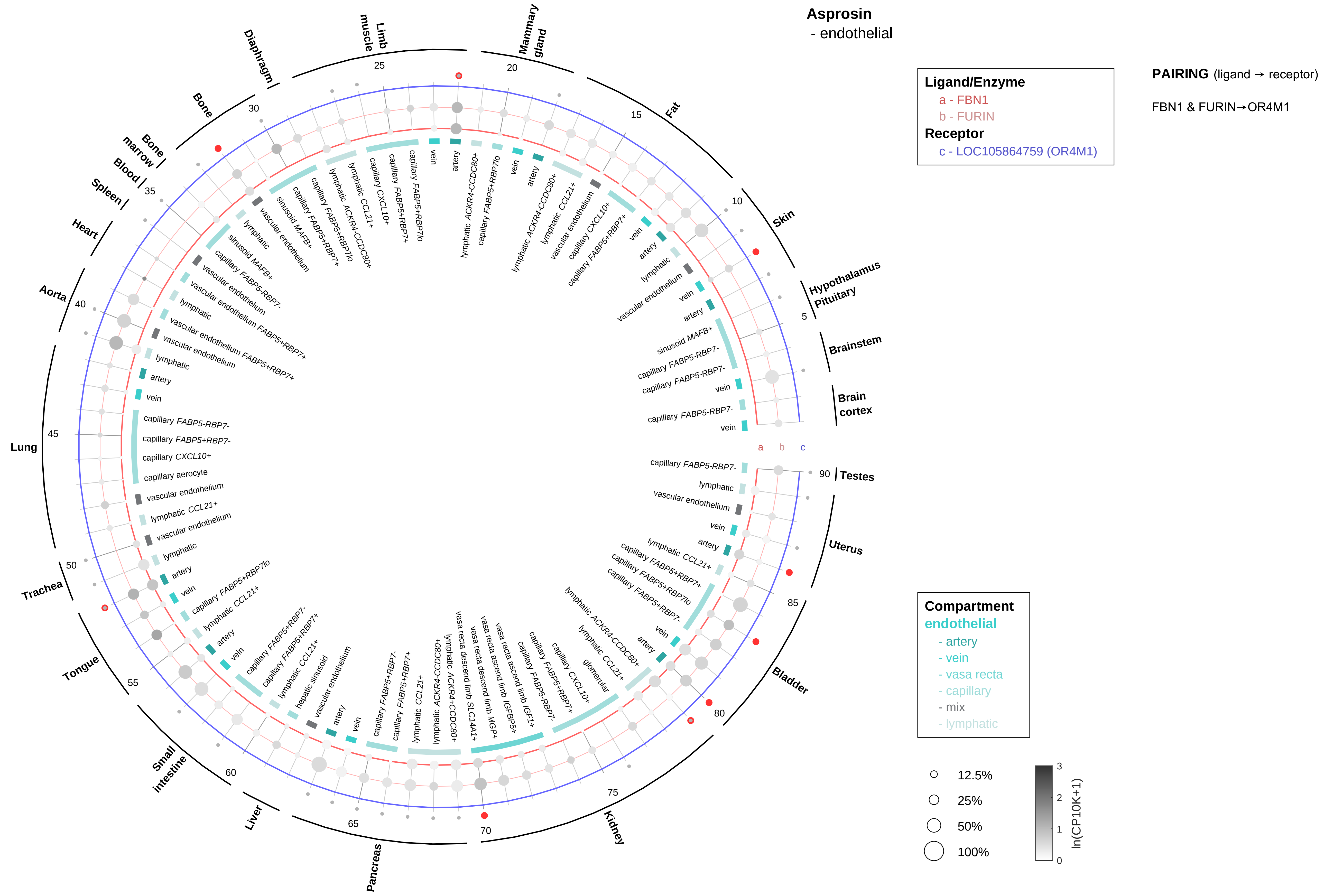

Asprosin
- endothelial
Ligand/Enzyme
a - FBN1
b - FURIN
Receptor
c - LOC105864759 (OR4M1)
PAIRING (ligand → receptor)
FBN1 & FURIN→OR4M1
Compartment
endothelial
- artery
- vein
- vasa recta
- capillary
- mix
- lymphatic
12.5%
25%
50%
100%
ln(CP10K+1)
Brain cortex
Brainstem
Hypothalamus Pituitary
Skin
Fat
Mammary gland
Limb muscle
Diaphragm
Bone
Bone marrow
Blood
Spleen
Heart
Aorta
Lung
Trachea
Tongue
Small intestine
Liver
Pancreas
Kidney
Bladder
Uterus
Testes

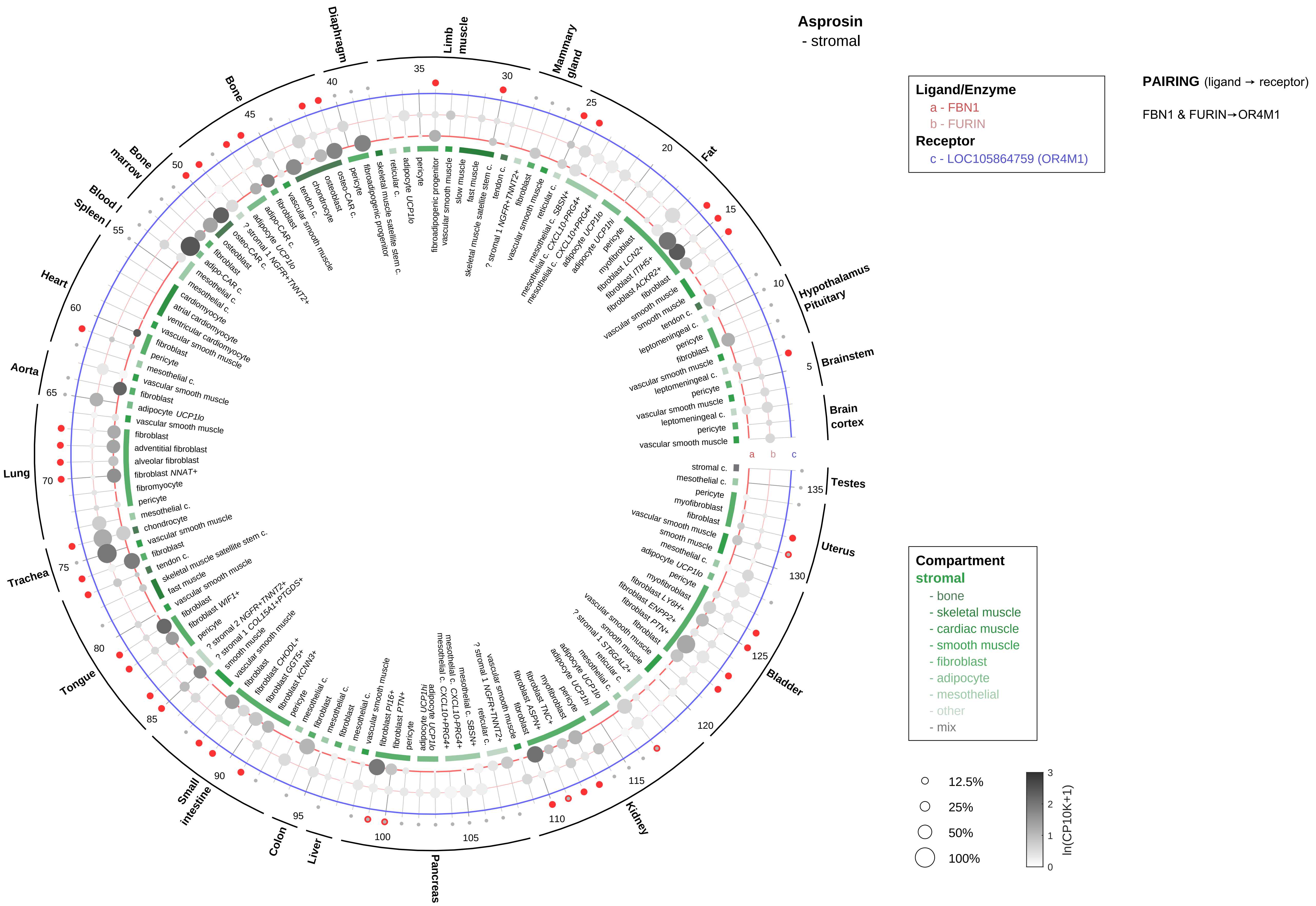
Asprosin
- stromal
Ligand/Enzyme
a - FBN1
b - FURIN
Receptor
c - LOC105864759 (OR4M1)
PAIRING (ligand → receptor)
FBN1 & FURIN→OR4M1
Compartment
stromal
- bone
- skeletal muscle
- cardiac muscle
- smooth muscle
- fibroblast
- adipocyte
- mesothelial
- other
- mix
12.5%
25%
50%
100%
ln(CP10K+1)
Brain cortex
Brainstem
Hypothalamus Pituitary
Fat
Mammary gland
Limb muscle
Diaphragm
Bone
Bone marrow
Blood
Spleen
Heart
Aorta
Lung
Trachea
Tongue
Small intestine
Colon
Liver
Pancreas
Kidney
Bladder
Uterus
Testes

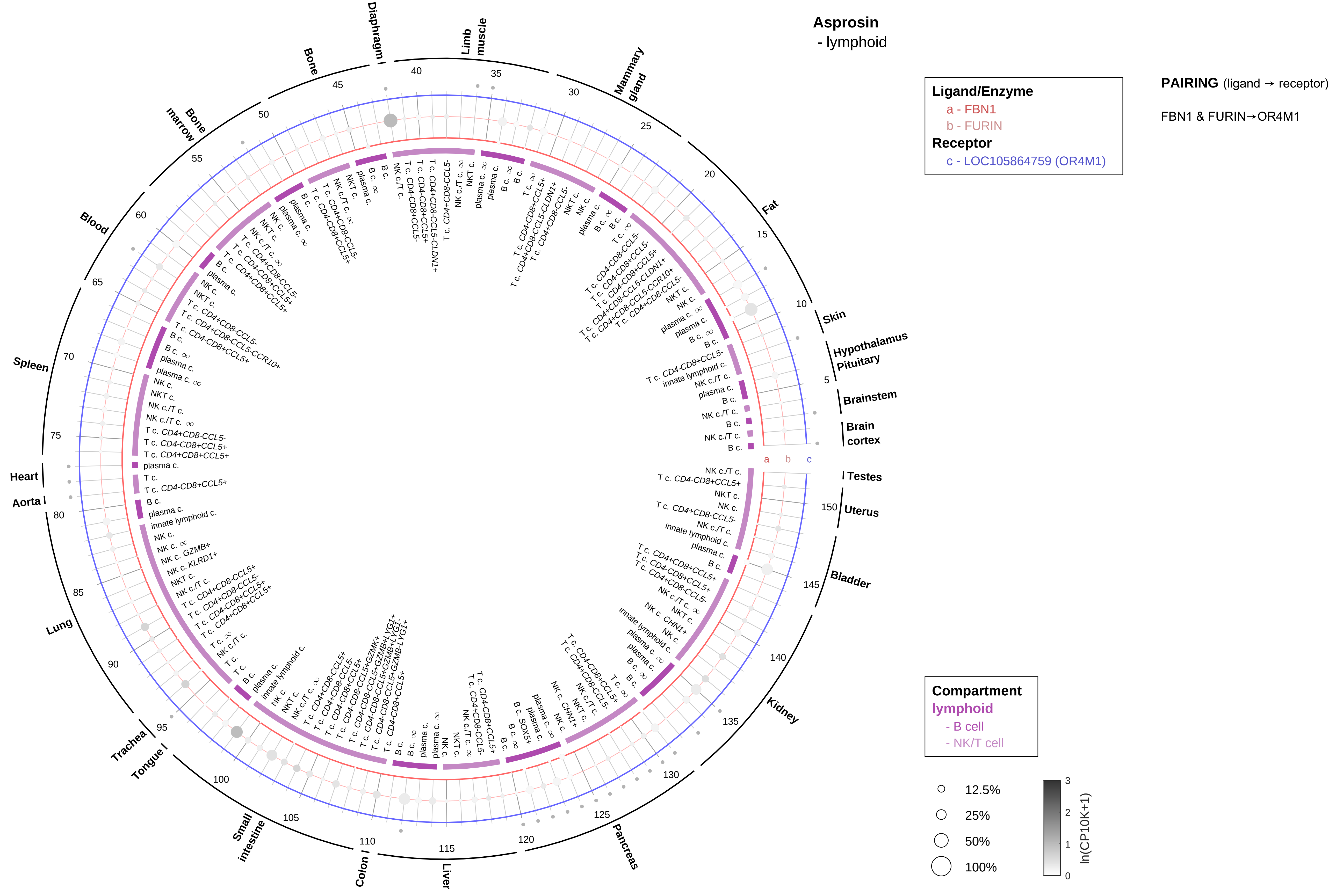

Asprosin
- lymphoid
Ligand/Enzyme
a - FBN1
b - FURIN
Receptor
c - LOC105864759 (OR4M1)
PAIRING (ligand → receptor)
FBN1 & FURIN→OR4M1
Compartment
lymphoid
- B cell
- NK/T cell
12.5%
25%
50%
100%
ln(CP10K+1)
Skin
Hypothalamus
Pituitary
Brainstem
Brain cortex
Testes
Uterus
Bladder
Kidney
Pancreas
Liver
Colon
Small intestine
Tongue
Trachea
Lung
Aorta
Heart
Spleen
Blood
Bone marrow
Bone
Diaphragm
Limb muscle
Mammary gland
Fat

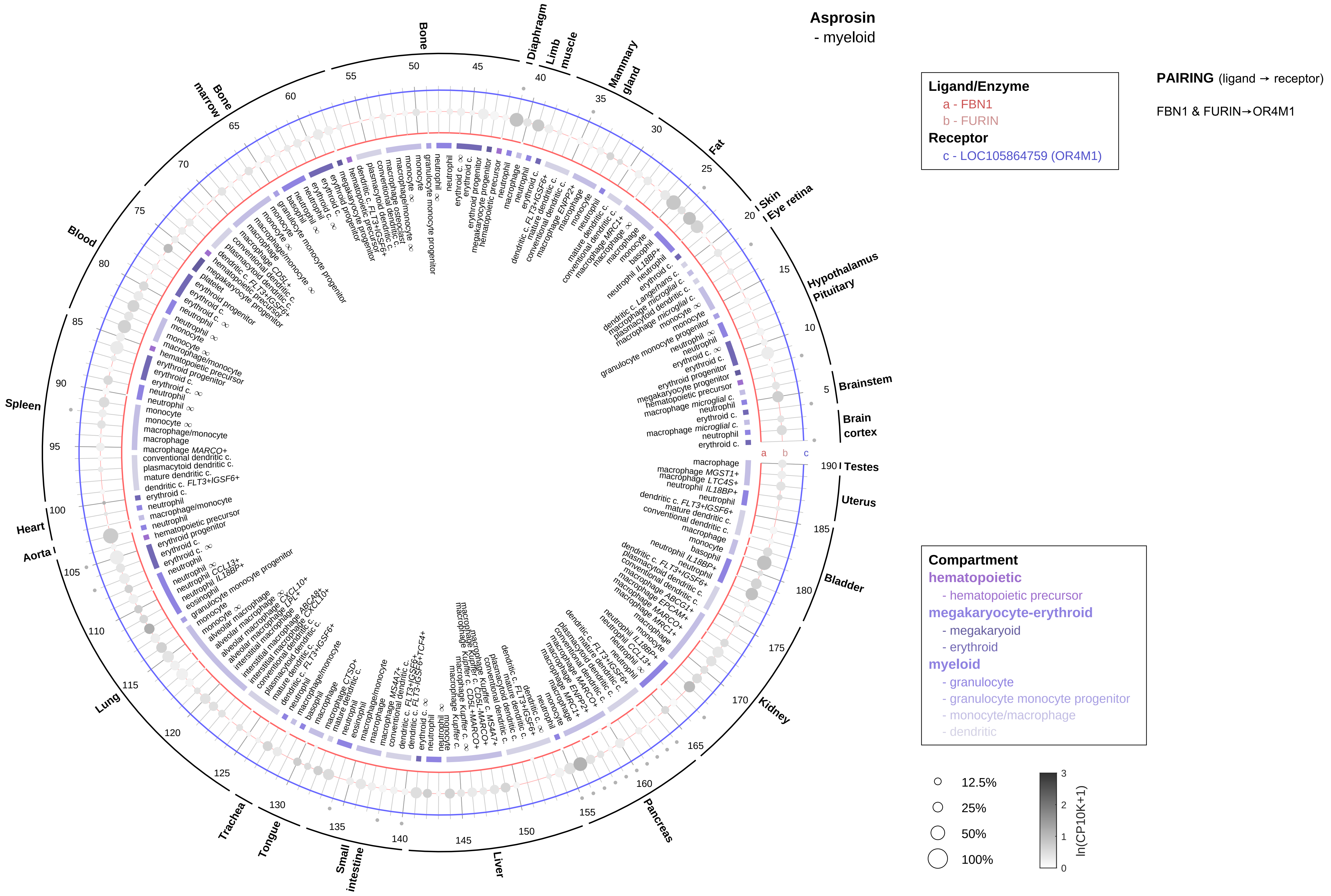

Asprosin
- myeloid
Ligand/Enzyme
a - FBN1
b - FURIN
Receptor
c - LOC105864759 (OR4M1)
PAIRING (ligand → receptor)
FBN1 & FURIN→OR4M1
Compartment
hematopoietic
- hematopoietic precursor
megakaryocyte-erythroid
- megakaryoid
- erythroid
myeloid
- granulocyte
- granulocyte monocyte progenitor
- monocyte/macrophage
- dendritic
12.5%
25%
50%
100%
ln(CP10K+1)
Bone
Diaphragm
Limb muscle
Mammary gland
Fat
Skin
Eye retina
Hypothalamus
Pituitary
Brainstem
Brain cortex
Testes
Uterus
Bladder
Kidney
Pancreas
Liver
Small intestine
Tongue
Trachea
Lung
Aorta
Heart
Spleen
Blood
Bone marrow

**Calcitonin/CGRP**
- epithelial/neural/germ

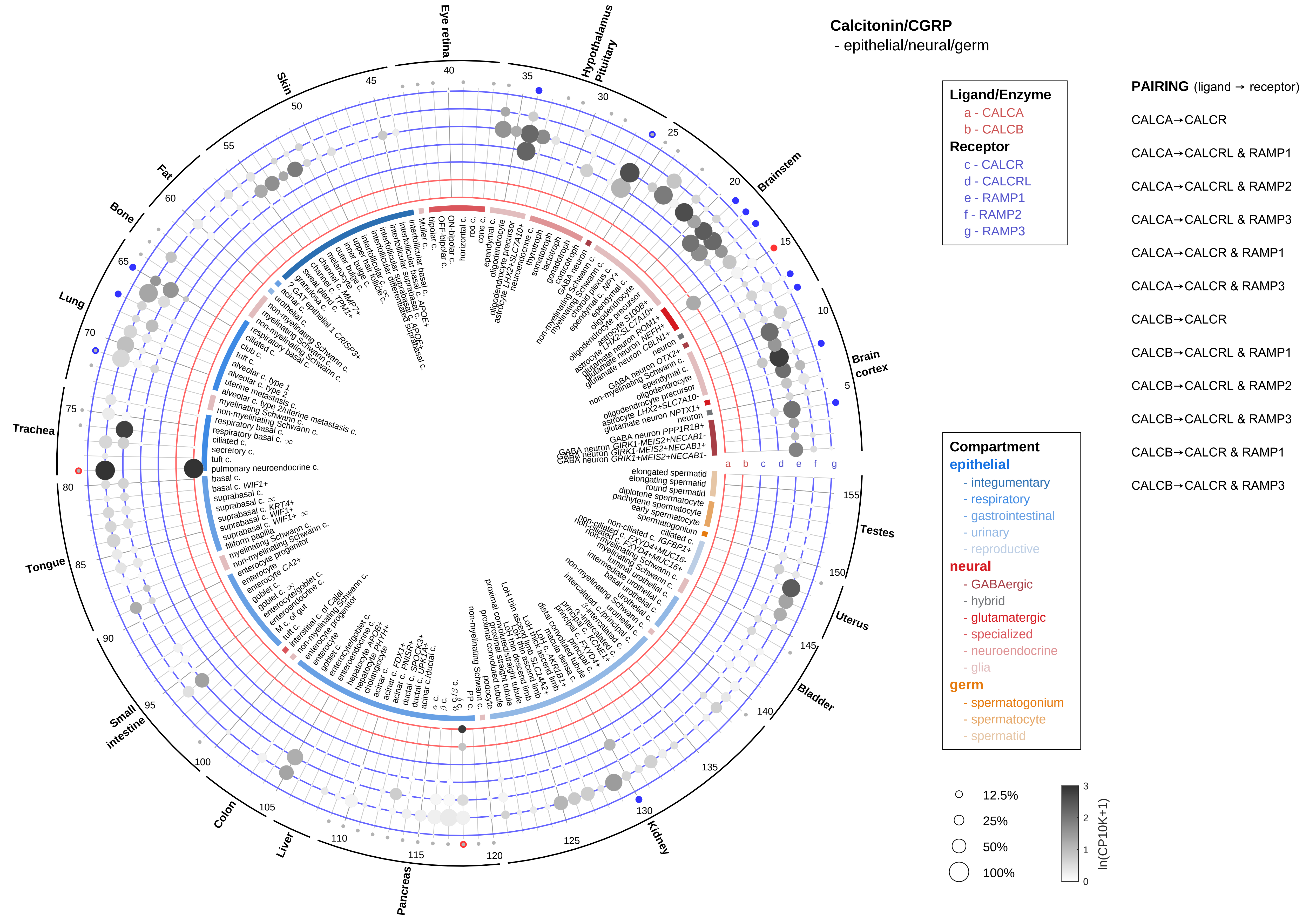

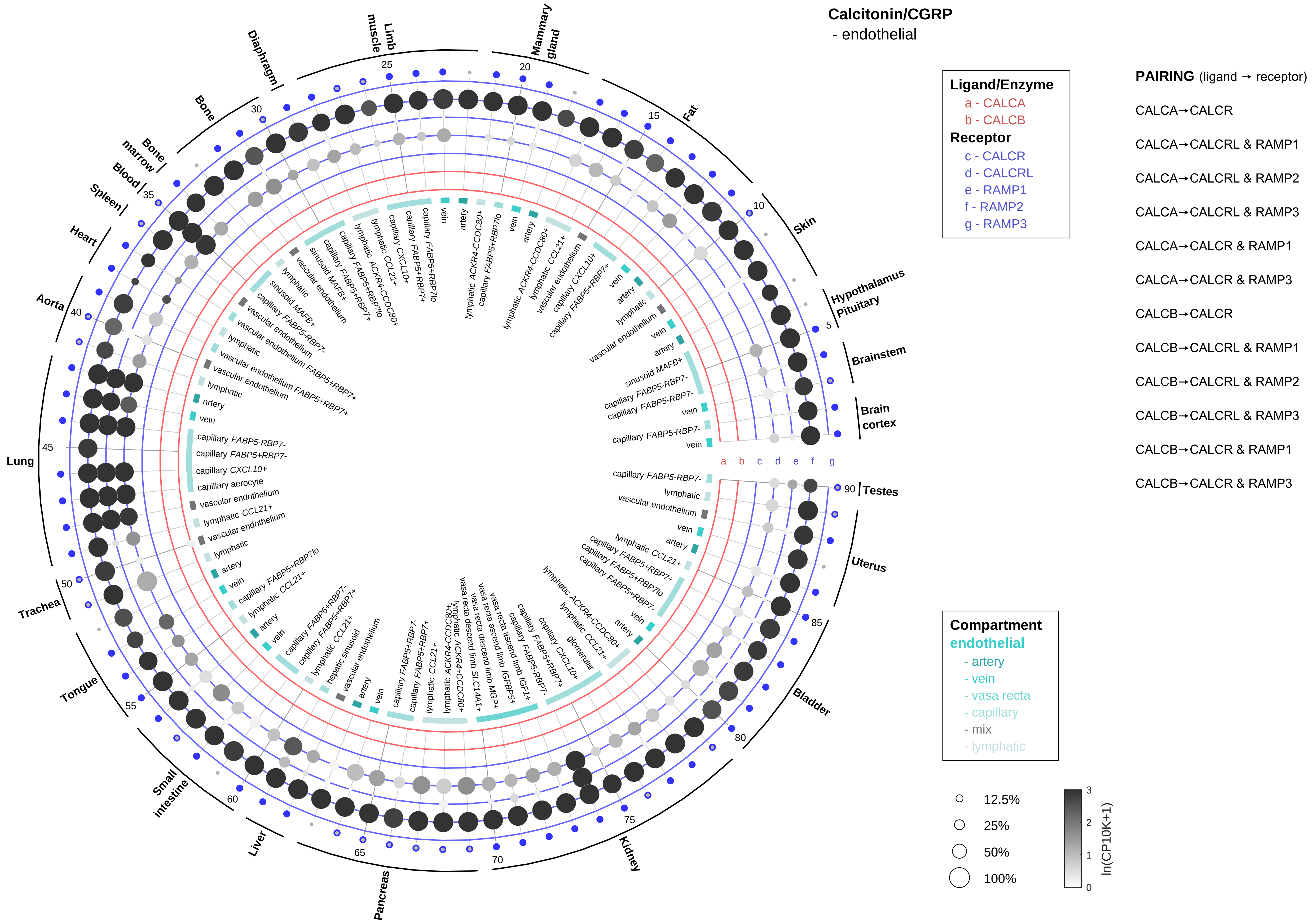

Calcitonin/CGRP
- endothelial
Ligand/Enzyme
a - CALCA
b - CALCB
Receptor
c - CALCR
d - CALCRL
e - RAMP1
f - RAMP2
g - RAMP3
PAIRING (ligand → receptor)
CALCA→CALCR
CALCA→CALCRL & RAMP1
CALCA→CALCRL & RAMP2
CALCA→CALCRL & RAMP3
CALCA→CALCR & RAMP1
CALCA→CALCR & RAMP3
CALCB→CALCR
CALCB→CALCRL & RAMP1
CALCB→CALCRL & RAMP2
CALCB→CALCRL & RAMP3
CALCB→CALCR & RAMP1
CALCB→CALCR & RAMP3
Compartment
endothelial
- artery
- vein
- vasa recta
- capillary
- mix
- lymphatic
12.5%
25%
50%
100%
ln(CP10K+1)
Brain cortex
Brainstem
Hypothalamus Pituitary
Skin
Fat
Mammary gland
Limb muscle
Diaphragm
Bone
Bone marrow
Blood
Spleen
Heart
Aorta
Lung
Trachea
Tongue
Small intestine
Liver
Pancreas
Kidney
Bladder
Uterus
Testes

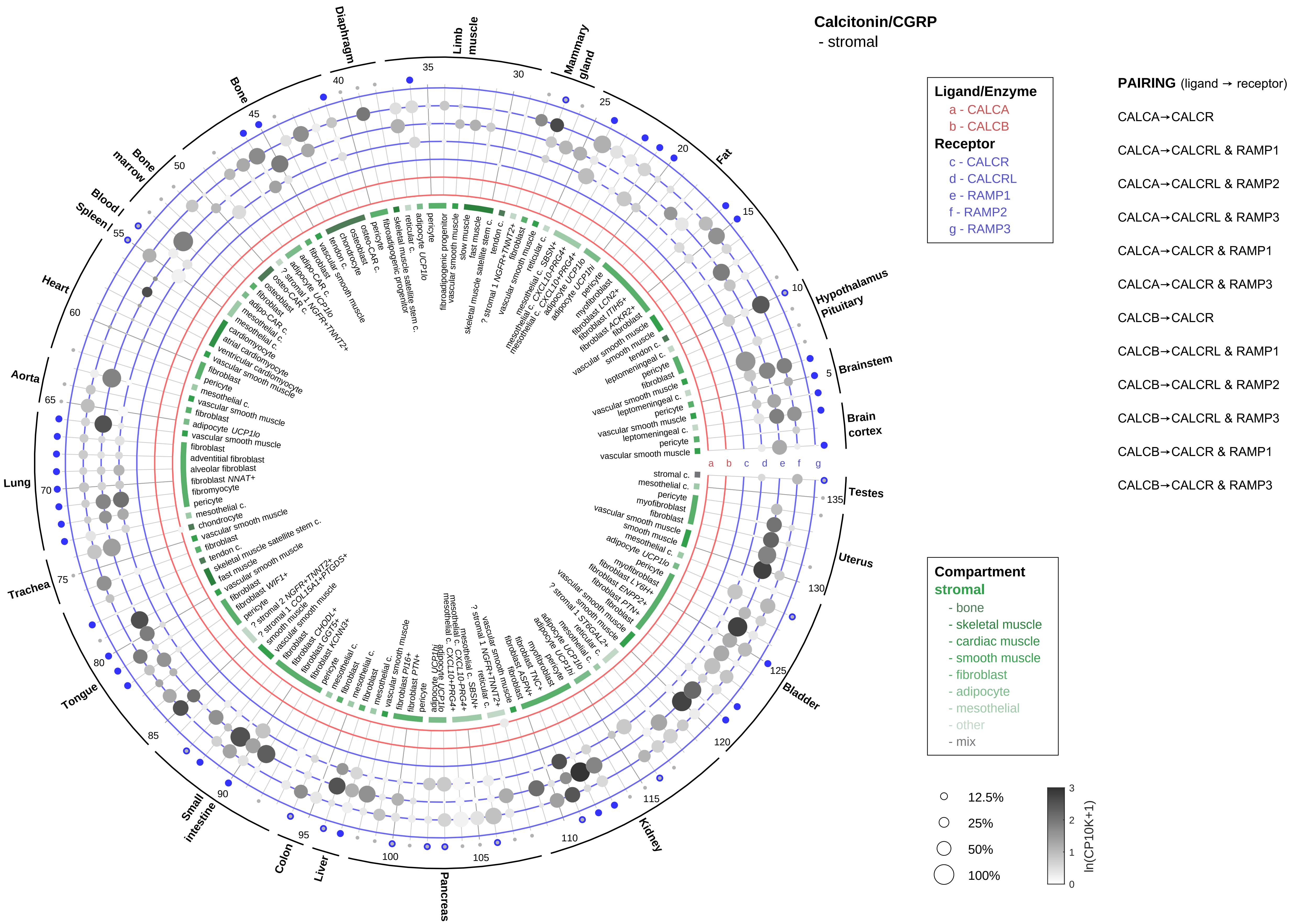

Calcitonin/CGRP
- stromal
Ligand/Enzyme
a - CALCA
b - CALCB
Receptor
c - CALCR
d - CALCRL
e - RAMP1
f - RAMP2
g - RAMP3
PAIRING (ligand → receptor)
CALCA→CALCR
CALCA→CALCRL & RAMP1
CALCA→CALCRL & RAMP2
CALCA→CALCRL & RAMP3
CALCA→CALCR & RAMP1
CALCA→CALCR & RAMP3
CALCB→CALCR
CALCB→CALCRL & RAMP1
CALCB→CALCRL & RAMP2
CALCB→CALCRL & RAMP3
CALCB→CALCR & RAMP1
CALCB→CALCR & RAMP3
Compartment
stromal
- bone
- skeletal muscle
- cardiac muscle
- smooth muscle
- fibroblast
- adipocyte
- mesothelial
- other
- mix
12.5%
25%
50%
100%
ln(CP10K+1)
Diaphragm
Limb muscle
Mammary gland
Fat
Hypothalamus Pituitary
Brainstem
Brain cortex
Testes
Uterus
Bladder
Kidney
Pancreas
Liver
Colon
Small intestine
Tongue
Trachea
Lung
Aorta
Heart
Blood / Spleen
Bone marrow
Bone

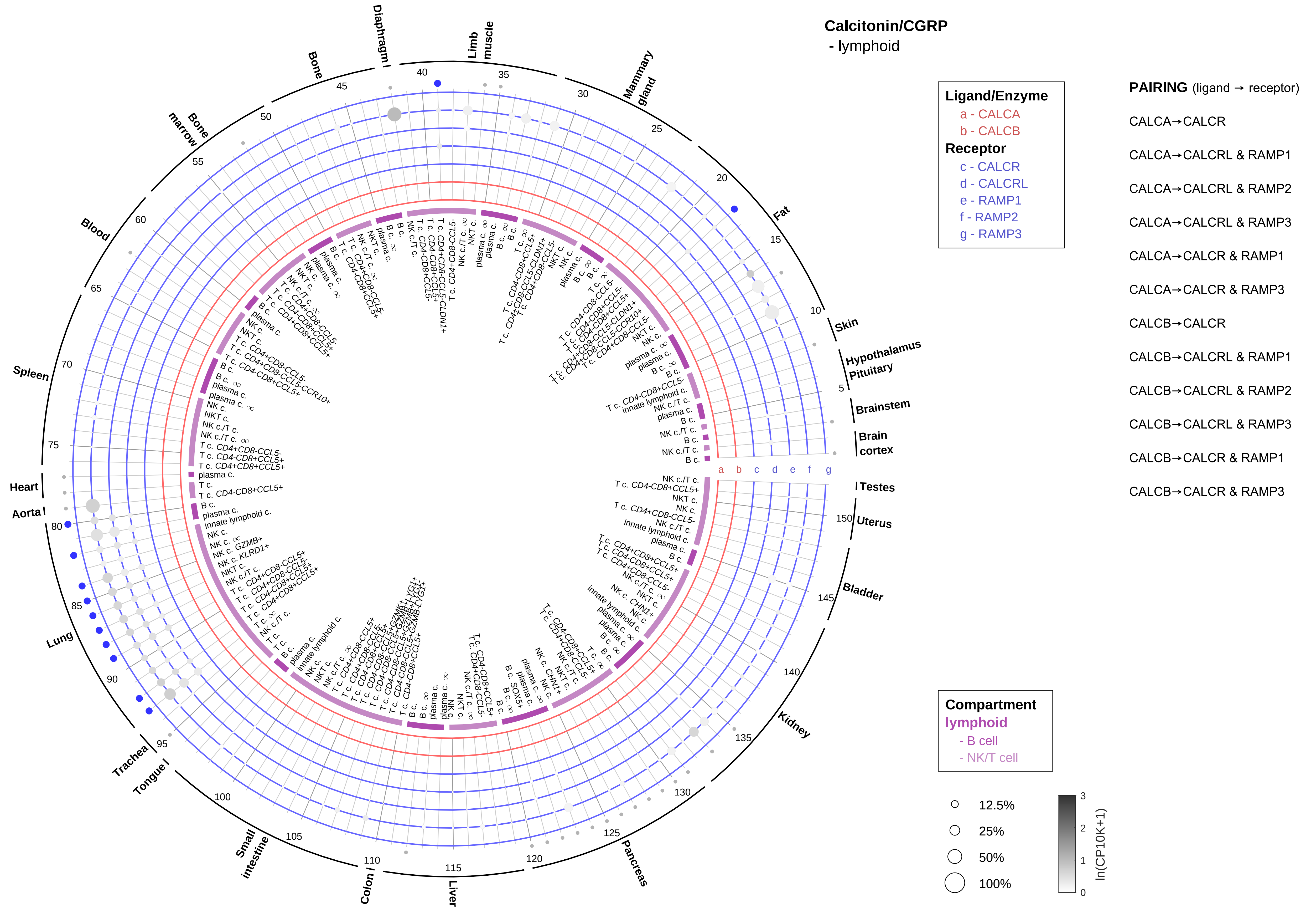
Calcitonin/CGRP
- lymphoid
Ligand/Enzyme
a - CALCA
b - CALCB
Receptor
c - CALCR
d - CALCRL
e - RAMP1
f - RAMP2
g - RAMP3
PAIRING (ligand → receptor)
CALCA→CALCR
CALCA→CALCRL & RAMP1
CALCA→CALCRL & RAMP2
CALCA→CALCRL & RAMP3
CALCA→CALCR & RAMP1
CALCA→CALCR & RAMP3
CALCB→CALCR
CALCB→CALCRL & RAMP1
CALCB→CALCRL & RAMP2
CALCB→CALCRL & RAMP3
CALCB→CALCR & RAMP1
CALCB→CALCR & RAMP3
Compartment
lymphoid
- B cell
- NK/T cell
12.5%
25%
50%
100%
ln(CP10K+1)
Skin
Hypothalamus
Pituitary
Brainstem
Brain cortex
Testes
Uterus
Bladder
Kidney
Pancreas
Liver
Colon
Small intestine
Tongue
Trachea
Lung
Aorta
Heart
Spleen
Blood
Bone marrow
Bone
Diaphragm
Limb muscle
Mammary gland
Fat

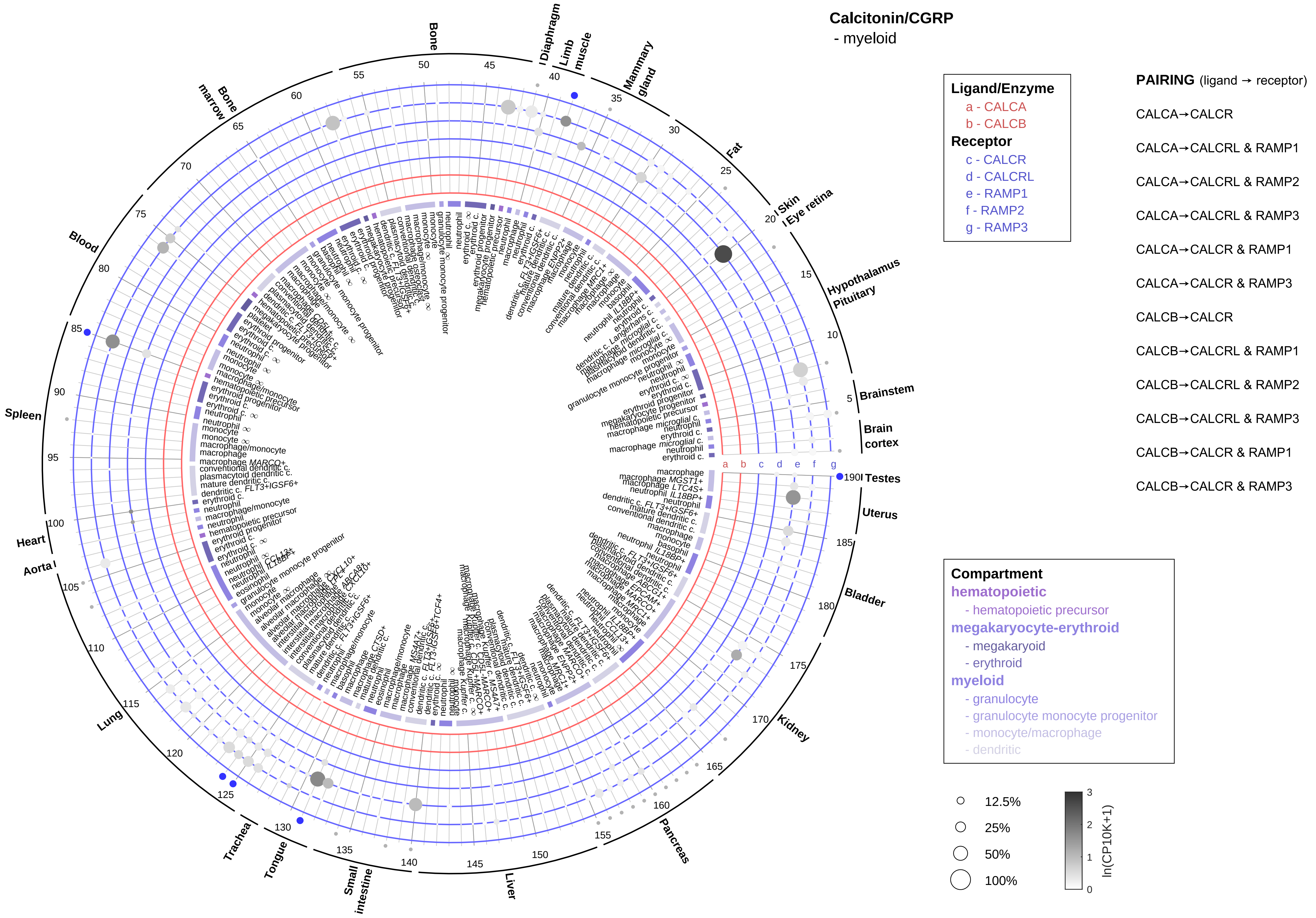

Calcitonin/CGRP
- myeloid
Ligand/Enzyme
a - CALCA
b - CALCB
Receptor
c - CALCR
d - CALCRL
e - RAMP1
f - RAMP2
g - RAMP3
PAIRING (ligand → receptor)
CALCA→CALCR
CALCA→CALCRL & RAMP1
CALCA→CALCRL & RAMP2
CALCA→CALCRL & RAMP3
CALCA→CALCR & RAMP1
CALCA→CALCR & RAMP3
CALCB→CALCR
CALCB→CALCRL & RAMP1
CALCB→CALCRL & RAMP2
CALCB→CALCRL & RAMP3
CALCB→CALCR & RAMP1
CALCB→CALCR & RAMP3
Compartment
hematopoietic
- hematopoietic precursor
megakaryocyte-erythroid
- megakaryoid
- erythroid
myeloid
- granulocyte
- granulocyte monocyte progenitor
- monocyte/macrophage
- dendritic
12.5%
25%
50%
100%
ln(CP10K+1)
Bone
Diaphragm
Limb muscle
Mammary gland
Fat
Skin
Eye retina
Hypothalamus
Pituitary
Brainstem
Brain cortex
Testes
Uterus
Bladder
Kidney
Pancreas
Liver
Small intestine
Tongue
Trachea
Lung
Aorta
Heart
Spleen
Blood
Bone marrow
a b c d e f g

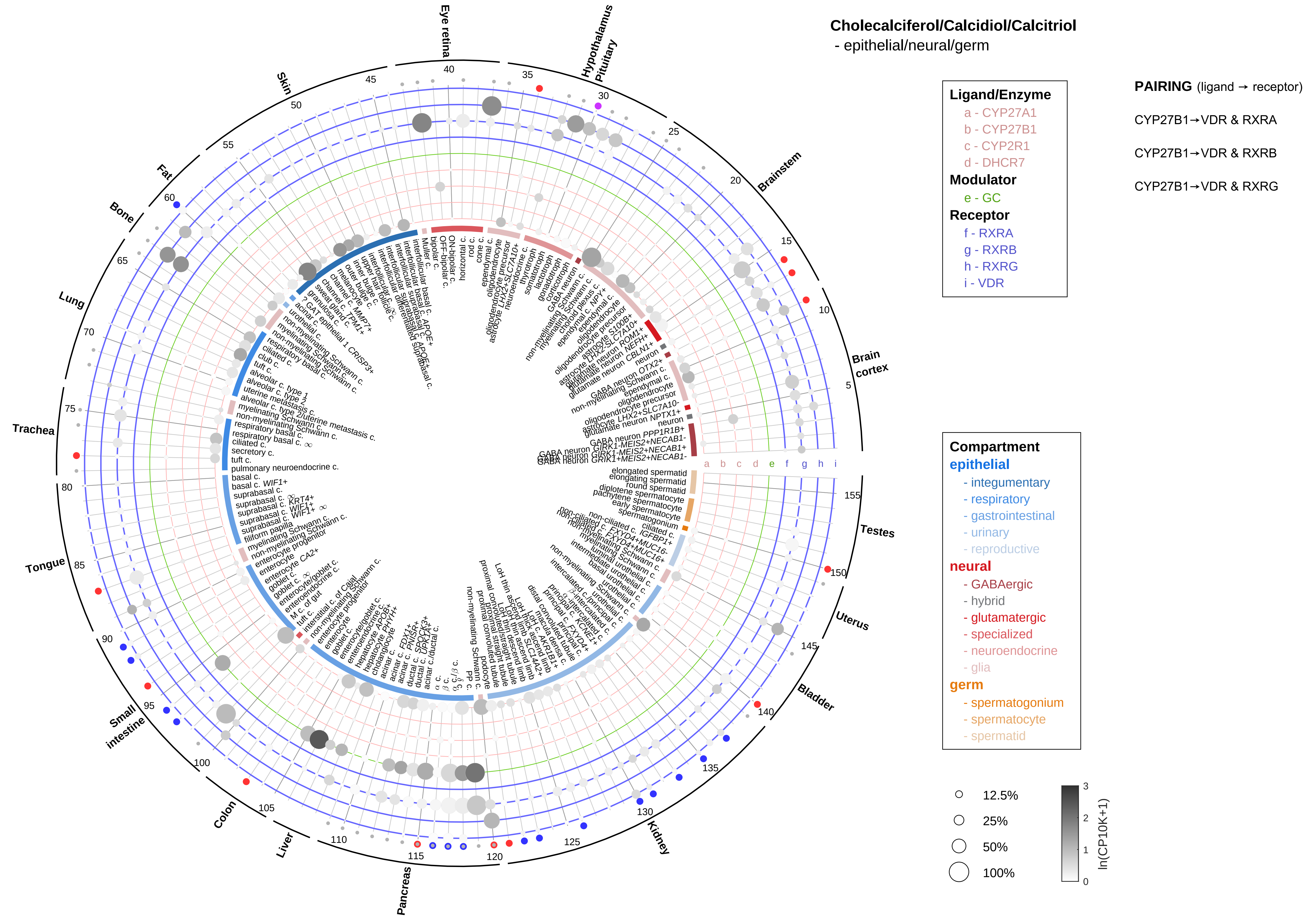
Cholecalciferol/Calcidiol/Calcitriol
- epithelial/neural/germ
Ligand/Enzyme
a - CYP27A1
b - CYP27B1
c - CYP2R1
d - DHCR7
Modulator
e - GC
Receptor
f - RXRA
g - RXRB
h - RXRG
i - VDR
PAIRING (ligand → receptor)
CYP27B1→VDR & RXRA
CYP27B1→VDR & RXRB
CYP27B1→VDR & RXRG
Compartment
epithelial
- integumentary
- respiratory
- gastrointestinal
- urinary
- reproductive
neural
- GABAergic
- hybrid
- glutamatergic
- specialized
- neuroendocrine
- glia
germ
- spermatogonium
- spermatocyte
- spermatid
12.5%
25%
50%
100%
ln(CP10K+1)
Eye retina
Hypothalamus
Pituitary
Brainstem
Brain cortex
Testes
Uterus
Bladder
Kidney
Pancreas
Liver
Colon
Small intestine
Tongue
Trachea
Lung
Bone
Fat
Skin

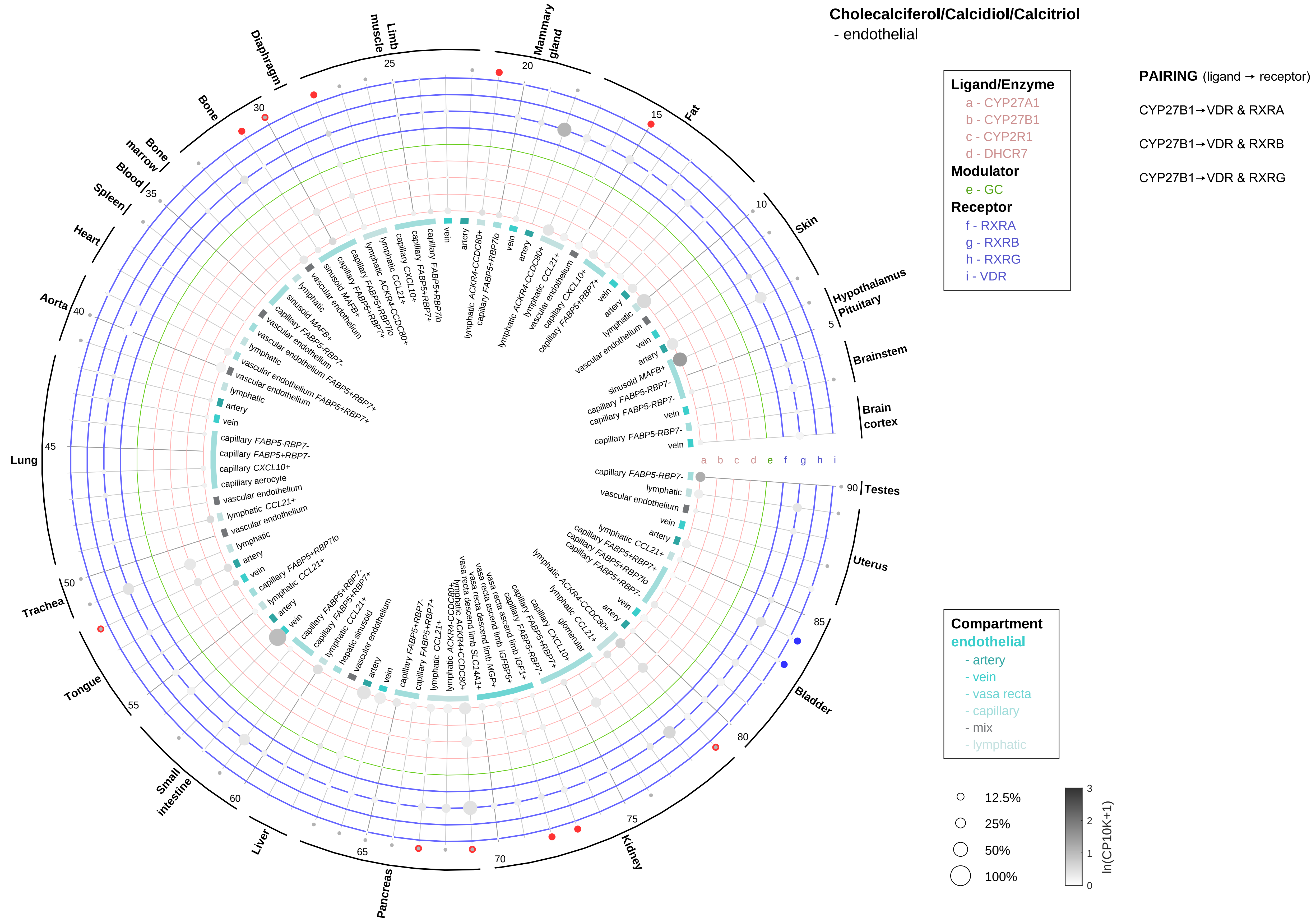

Cholecalciferol/Calcidiol/Calcitriol
- endothelial
Ligand/Enzyme
a - CYP27A1
b - CYP27B1
c - CYP2R1
d - DHCR7
Modulator
e - GC
Receptor
f - RXRA
g - RXRB
h - RXRG
i - VDR
PAIRING (ligand → receptor)
CYP27B1→VDR & RXRA
CYP27B1→VDR & RXRB
CYP27B1→VDR & RXRG
Compartment
endothelial
- artery
- vein
- vasa recta
- capillary
- mix
- lymphatic
12.5%
25%
50%
100%
ln(CP10K+1)
Brain cortex
Brainstem
Hypothalamus Pituitary
Skin
Fat
Mammary gland
Limb muscle
Diaphragm
Bone
Bone marrow
Blood
Spleen
Heart
Aorta
Lung
Trachea
Tongue
Small intestine
Liver
Pancreas
Kidney
Bladder
Uterus
Testes

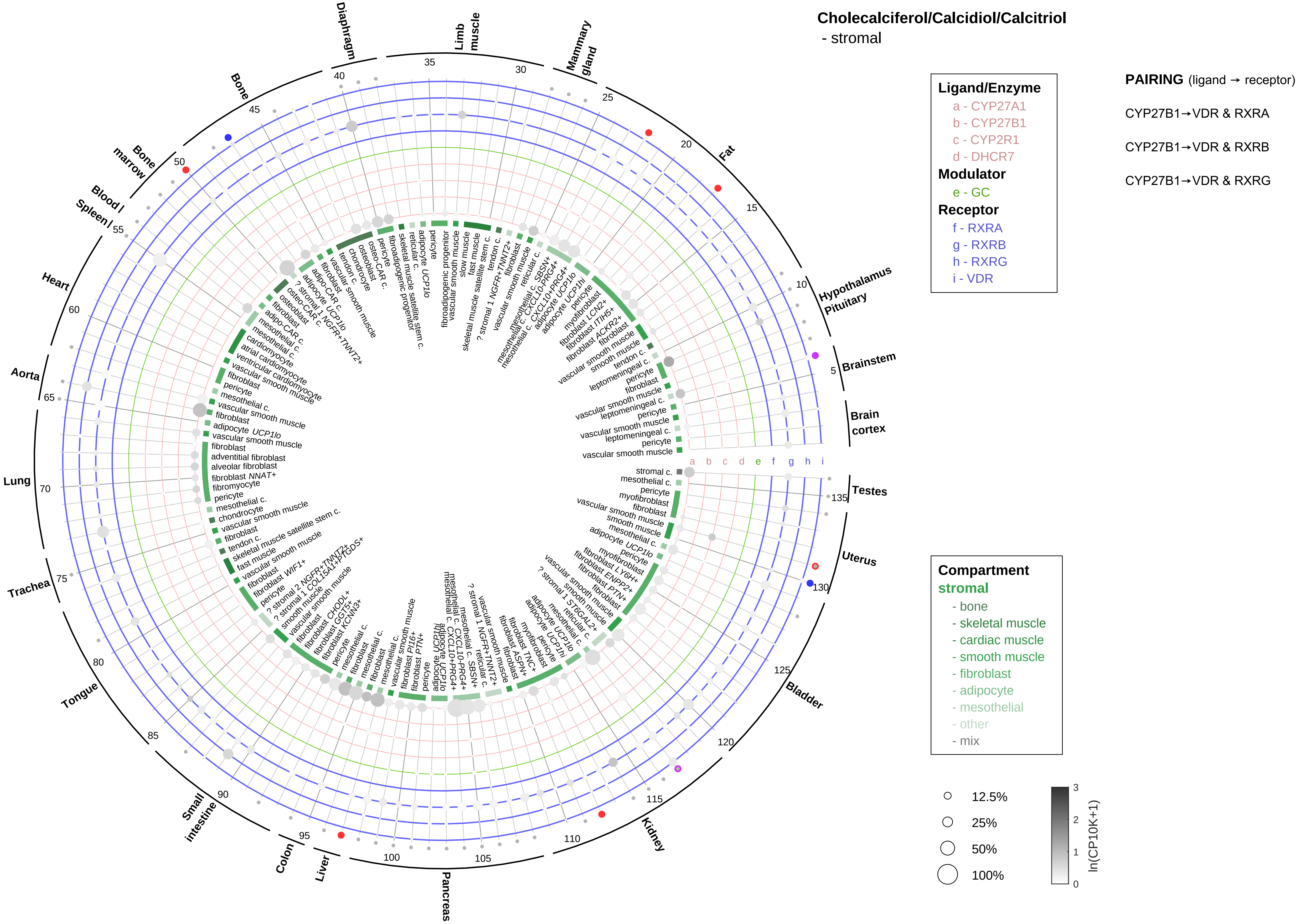
Cholecalciferol/Calcidiol/Calcitriol
- stromal
Ligand/Enzyme
a - CYP27A1
b - CYP27B1
c - CYP2R1
d - DHCR7
Modulator
e - GC
Receptor
f - RXRA
g - RXRB
h - RXRG
i - VDR
PAIRING (ligand → receptor)
CYP27B1→VDR & RXRA
CYP27B1→VDR & RXRB
CYP27B1→VDR & RXRG
Compartment
stromal
- bone
- skeletal muscle
- cardiac muscle
- smooth muscle
- fibroblast
- adipocyte
- mesothelial
- other
- mix
12.5%
25%
50%
100%
ln(CP10K+1)
Diaphragm
Limb muscle
Mammary gland
Fat
Hypothalamus Pituitary
Brainstem
Brain cortex
Testes
Uterus
Bladder
Kidney
Pancreas
Liver
Colon
Small intestine
Tongue
Trachea
Lung
Aorta
Heart
Blood / Spleen
Bone marrow
Bone

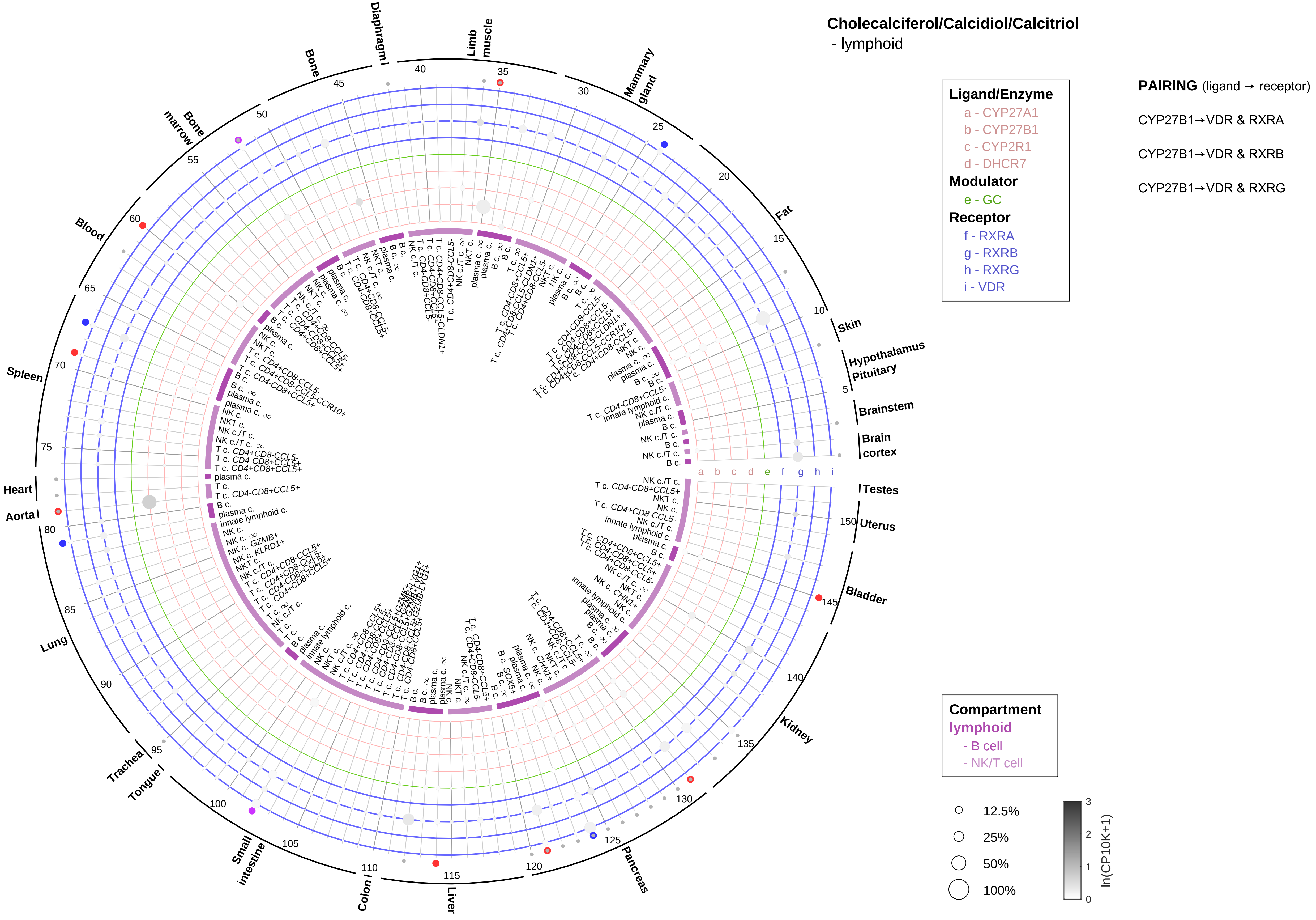
Cholecalciferol/Calcidiol/Calcitriol
- lymphoid
Ligand/Enzyme
a - CYP27A1
b - CYP27B1
c - CYP2R1
d - DHCR7
Modulator
e - GC
Receptor
f - RXRA
g - RXRB
h - RXRG
i - VDR
PAIRING (ligand → receptor)
CYP27B1→VDR & RXRA
CYP27B1→VDR & RXRB
CYP27B1→VDR & RXRG
Compartment
lymphoid
- B cell
- NK/T cell
12.5%
25%
50%
100%
ln(CP10K+1)
Skin
Hypothalamus
Pituitary
Brainstem
Brain cortex
Testes
Uterus
Bladder
Kidney
Pancreas
Liver
Colon
Small intestine
Tongue
Trachea
Lung
Aorta
Heart
Spleen
Blood
Bone marrow
Bone
Diaphragm
Limb muscle
Mammary gland
Fat

### Cholecalciferol/Calcidiol/Calcitriol
- myeloid

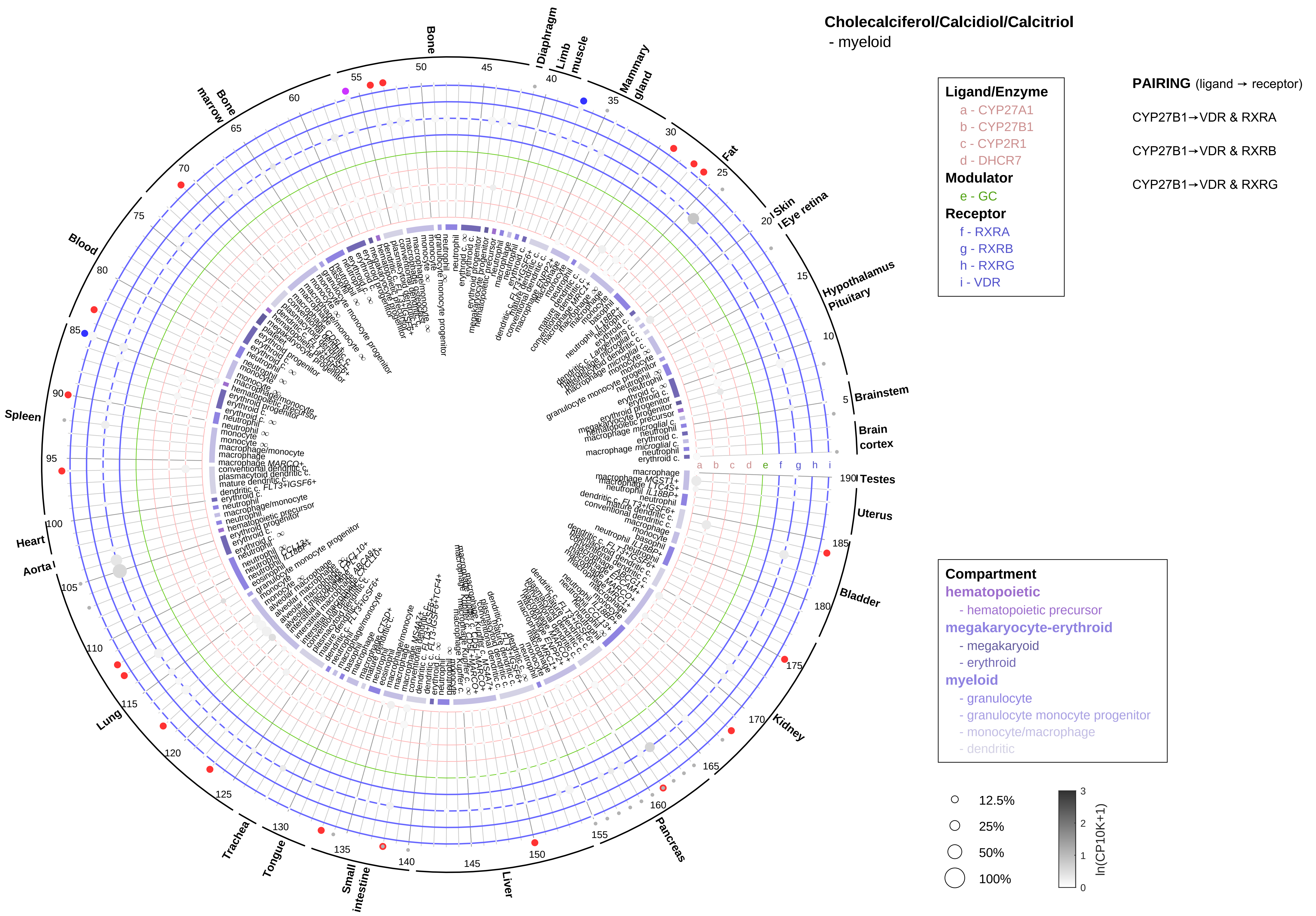

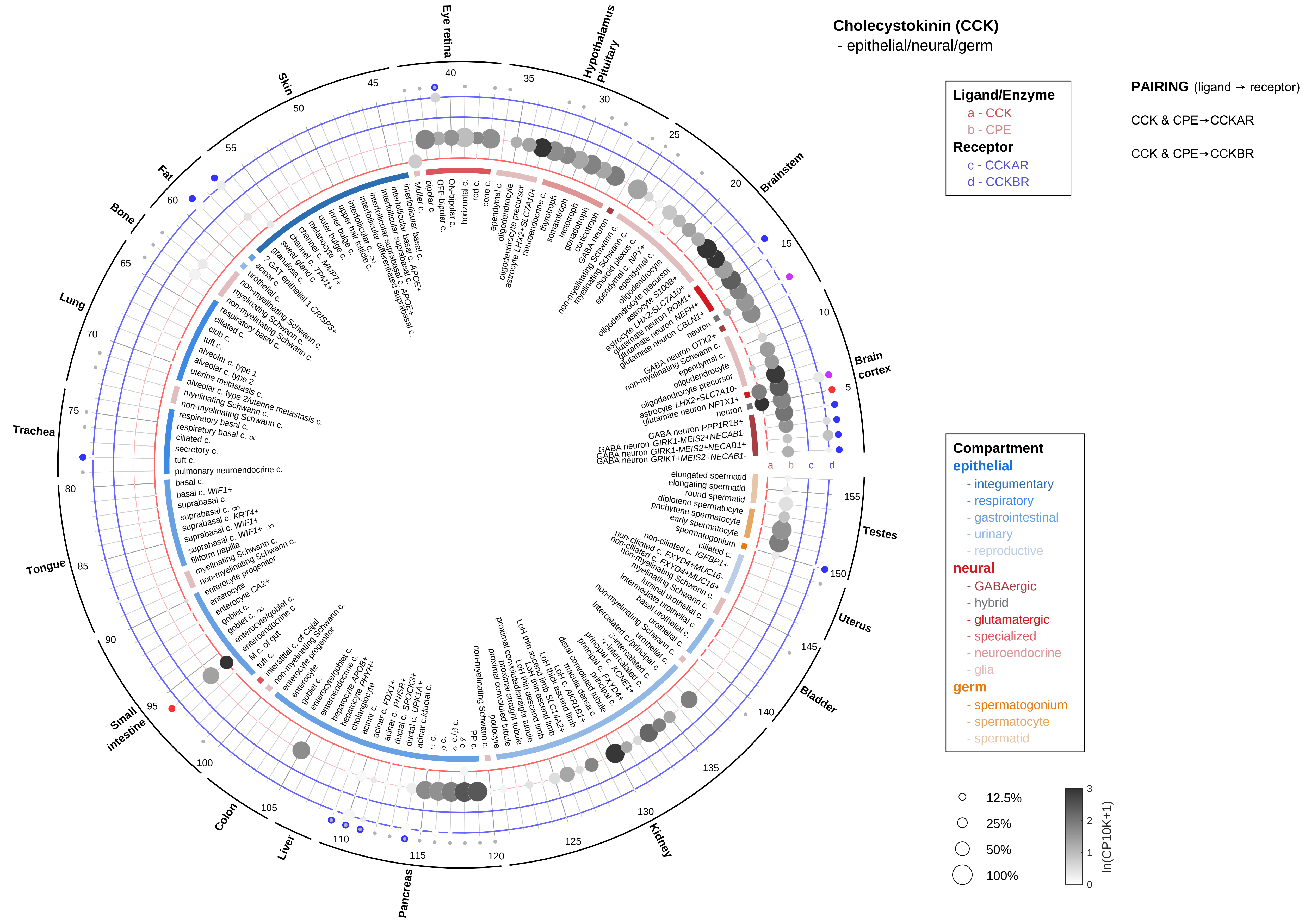

Cholecystokinin (CCK)
- epithelial/neural/germ
Ligand/Enzyme
a - CCK
b - CPE
Receptor
c - CCKAR
d - CCKBR
PAIRING (ligand → receptor)
CCK & CPE→CCKAR
CCK & CPE→CCKBR
Compartment
epithelial
- integumentary
- respiratory
- gastrointestinal
- urinary
- reproductive
neural
- GABAergic
- hybrid
- glutamatergic
- specialized
- neuroendocrine
- glia
germ
- spermatogonium
- spermatocyte
- spermatid
12.5%
25%
50%
100%
ln(CP10K+1)
Eye retina
Hypothalamus Pituitary
Brainstem
Brain cortex
Testes
Uterus
Bladder
Kidney
Pancreas
Liver
Colon
Small intestine
Tongue
Trachea
Lung
Bone
Fat
Skin

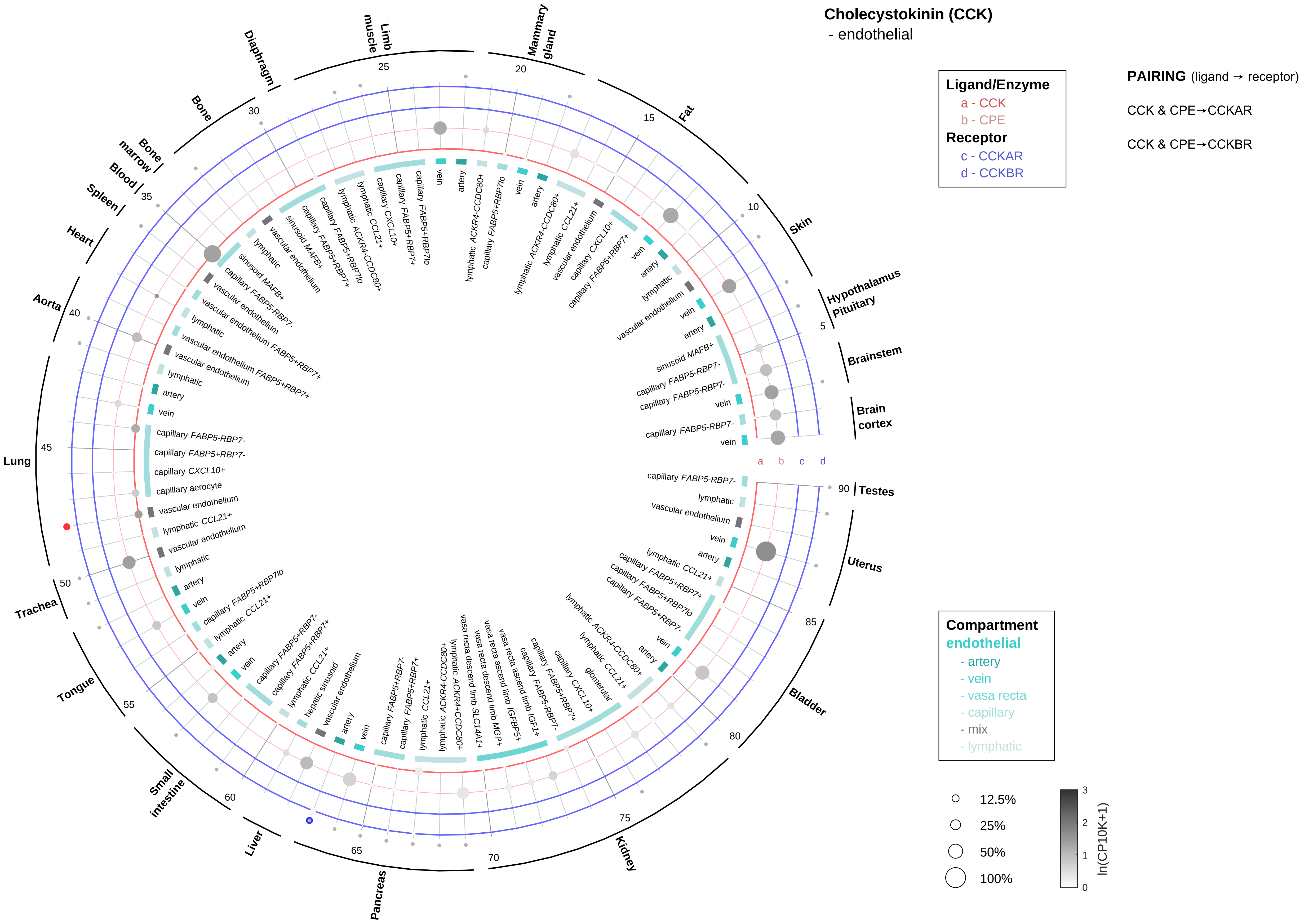

Cholecystokinin (CCK)
- endothelial
Ligand/Enzyme
a - CCK
b - CPE
Receptor
c - CCKAR
d - CCKBR
PAIRING (ligand → receptor)
CCK & CPE→CCKAR
CCK & CPE→CCKBR
Compartment
endothelial
- artery
- vein
- vasa recta
- capillary
- mix
- lymphatic
12.5%
25%
50%
100%
ln(CP10K+1)
Brain cortex
Brainstem
Hypothalamus Pituitary
Skin
Fat
Mammary gland
Limb muscle
Diaphragm
Bone
Bone marrow
Blood
Spleen
Heart
Aorta
Lung
Trachea
Tongue
Small intestine
Liver
Pancreas
Kidney
Bladder
Uterus
Testes
a
b
c
d

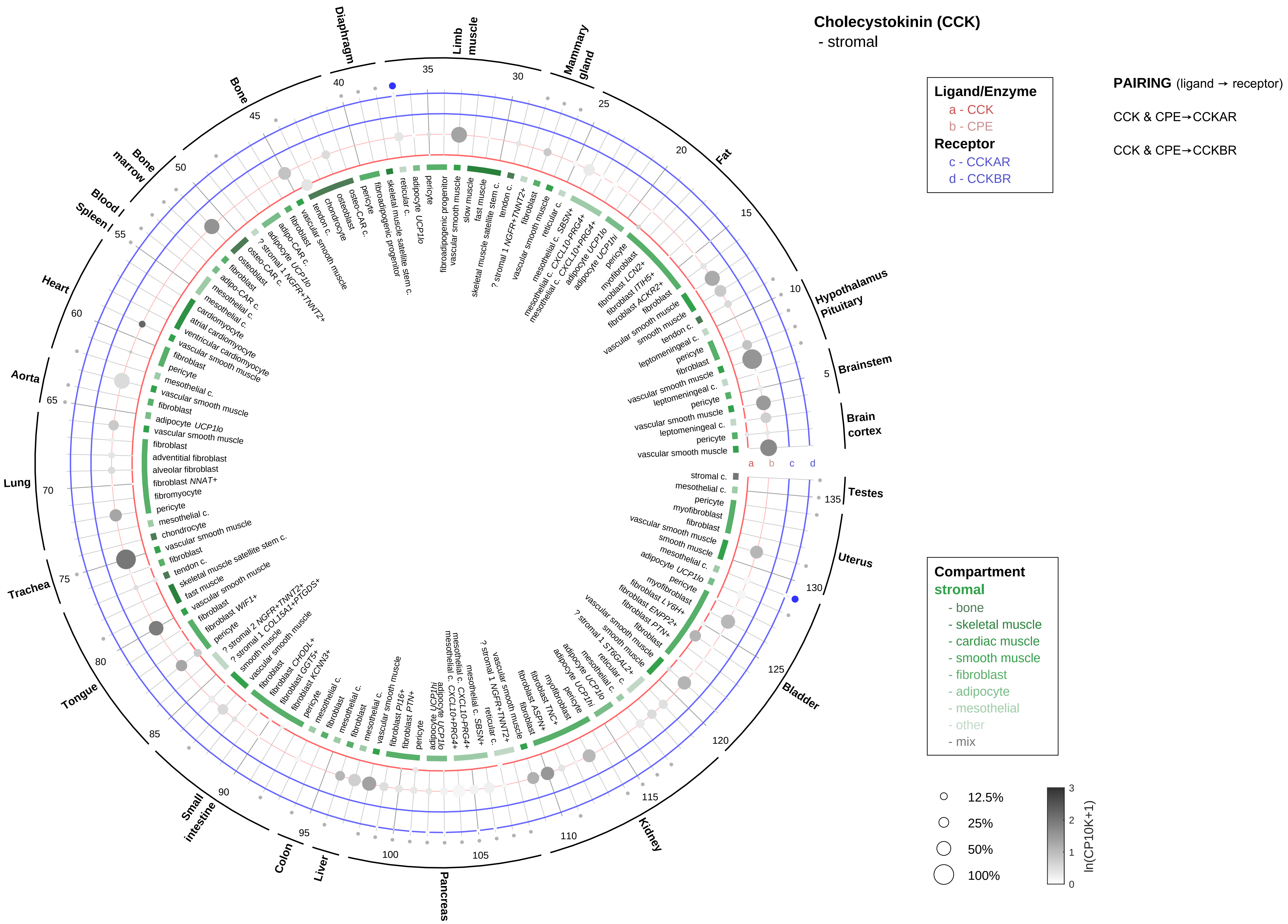
Cholecystokinin (CCK)
- stromal
Ligand/Enzyme
a - CCK
b - CPE
Receptor
c - CCKAR
d - CCKBR
PAIRING (ligand → receptor)
CCK & CPE→CCKAR
CCK & CPE→CCKBR
Compartment
stromal
- bone
- skeletal muscle
- cardiac muscle
- smooth muscle
- fibroblast
- adipocyte
- mesothelial
- other
- mix
12.5%
25%
50%
100%
ln(CP10K+1)
Limb muscle
Mammary gland
Fat
Hypothalamus Pituitary
Brainstem
Brain cortex
Testes
Uterus
Bladder
Kidney
Pancreas
Liver
Colon
Small intestine
Tongue
Trachea
Lung
Aorta
Heart
Blood / Spleen
Bone marrow
Bone
Diaphragm

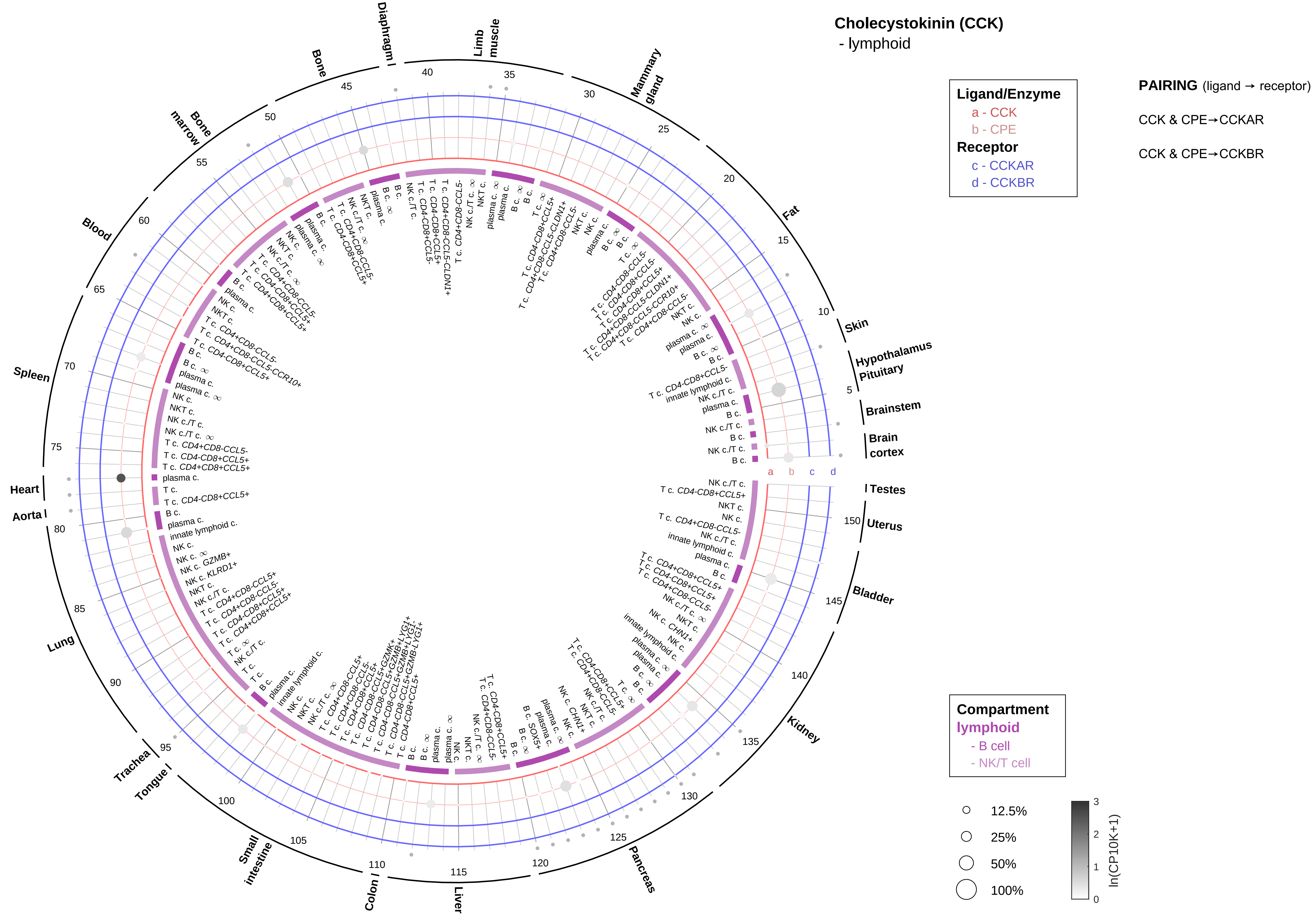
Cholecystokinin (CCK)
- lymphoid
Ligand/Enzyme
a - CCK
b - CPE
Receptor
c - CCKAR
d - CCKBR
PAIRING (ligand → receptor)
CCK & CPE→CCKAR
CCK & CPE→CCKBR
Compartment
lymphoid
- B cell
- NK/T cell
12.5%
25%
50%
100%
ln(CP10K+1)
Testes
Uterus
Bladder
Kidney
Pancreas
Liver
Colon
Small intestine
Tongue
Trachea
Lung
Aorta
Heart
Spleen
Blood
Bone marrow
Bone
Diaphragm
Limb muscle
Mammary gland
Fat
Skin
Hypothalamus Pituitary
Brainstem
Brain cortex

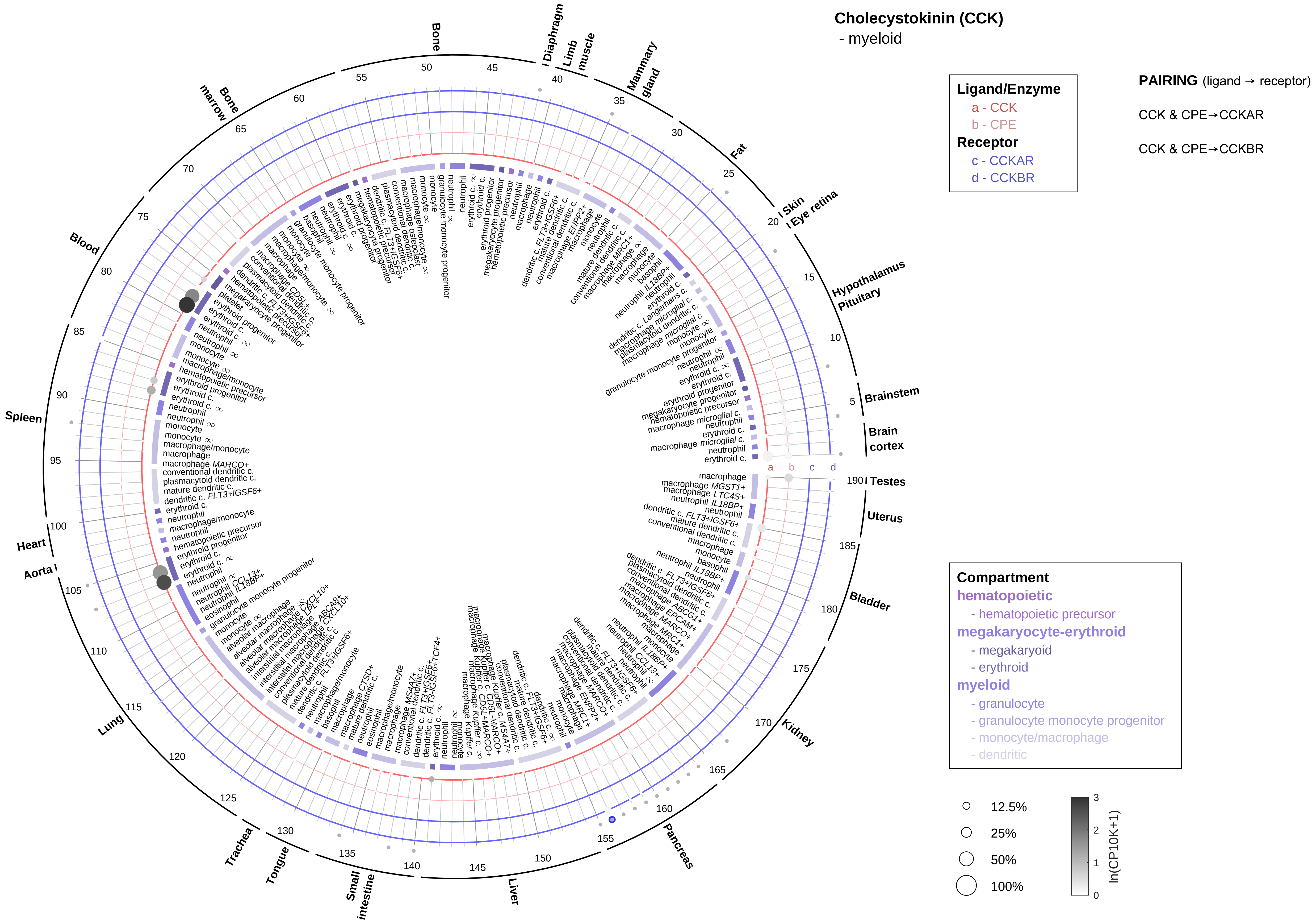
Cholecystokinin (CCK)
- myeloid
Ligand/Enzyme
a - CCK
b - CPE
Receptor
c - CCKAR
d - CCKBR
PAIRING (ligand → receptor)
CCK & CPE→CCKAR
CCK & CPE→CCKBR
Compartment
hematopoietic
- hematopoietic precursor
megakaryocyte-erythroid
- megakaryoid
- erythroid
myeloid
- granulocyte
- granulocyte monocyte progenitor
- monocyte/macrophage
- dendritic
12.5%
25%
50%
100%
ln(CP10K+1)
Bone
Diaphragm
Limb muscle
Mammary gland
Fat
Skin
Eye retina
Hypothalamus
Pituitary
Brainstem
Brain cortex
Testes
Uterus
Bladder
Kidney
Pancreas
Liver
Small intestine
Tongue
Trachea
Lung
Aorta
Heart
Spleen
Blood
Bone marrow

**Cocaine- and amphetamine-regulated transcript (CART)**
- epithelial/neural/germ

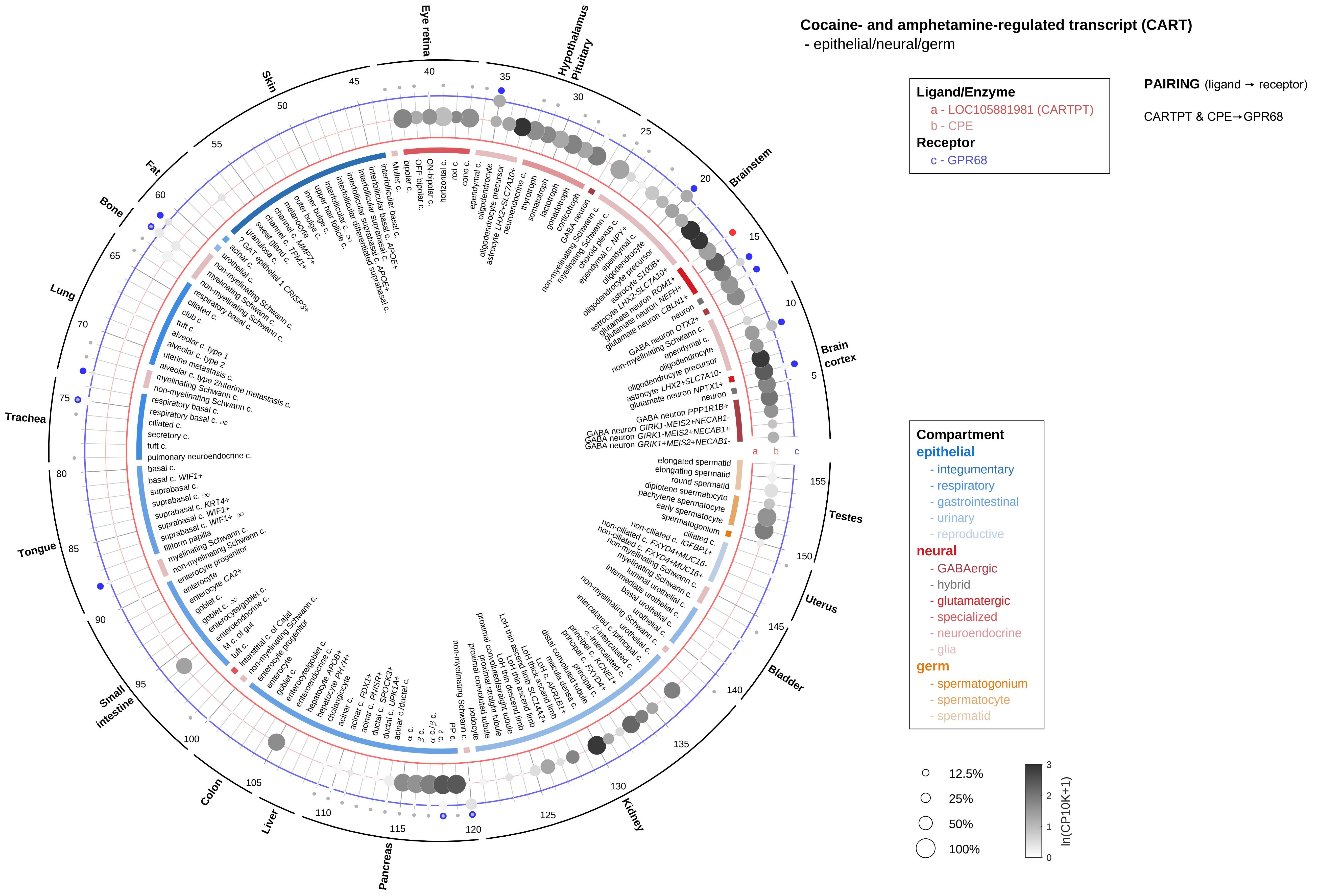

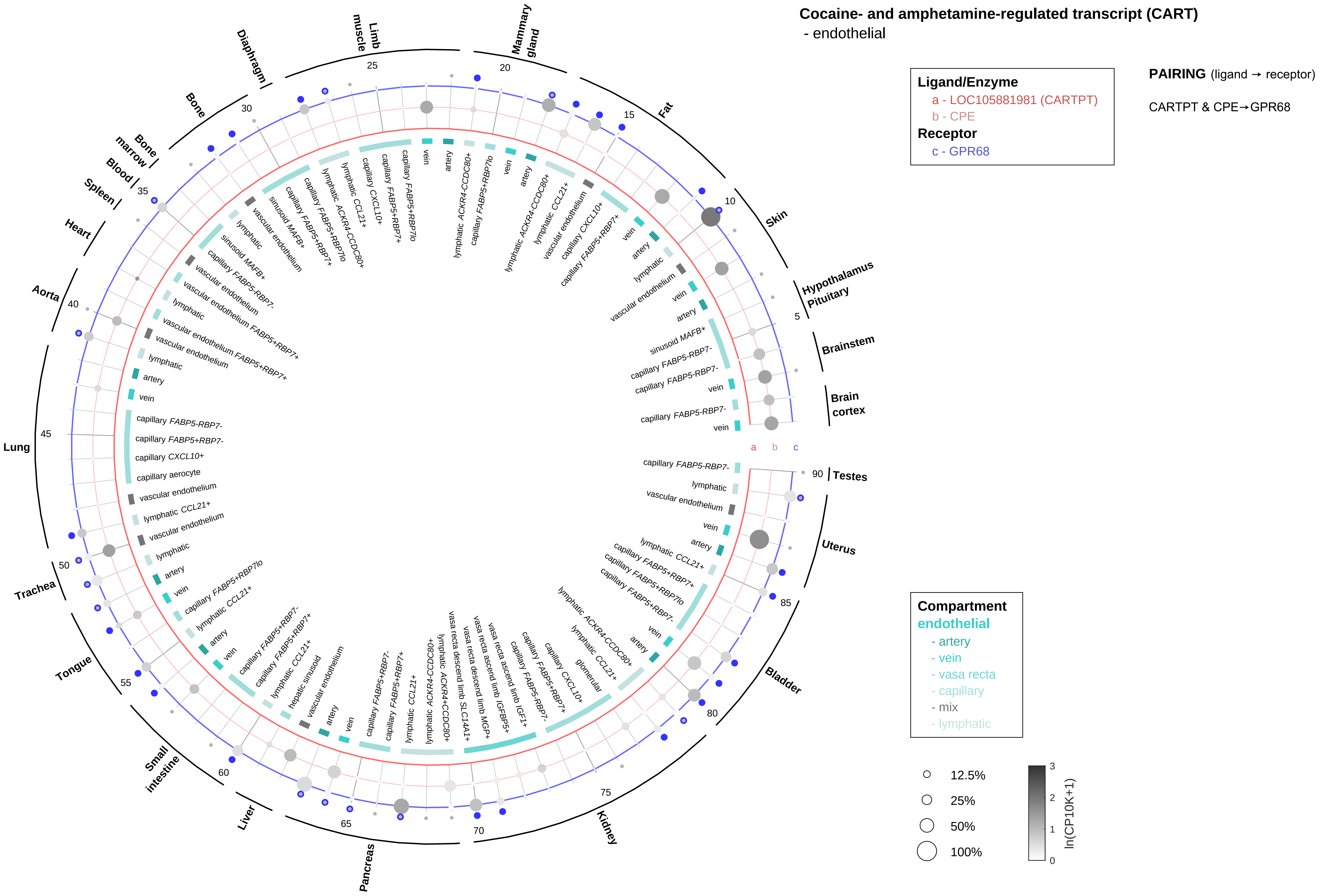

Cocaine- and amphetamine-regulated transcript (CART)
- endothelial
Ligand/Enzyme
a - LOC105881981 (CARTPT)
b - CPE
Receptor
c - GPR68
PAIRING (ligand → receptor)
CARTPT & CPE→GPR68
Compartment
endothelial
- artery
- vein
- vasa recta
- capillary
- mix
- lymphatic
12.5%
25%
50%
100%
ln(CP10K+1)
Brain cortex
Brainstem
Hypothalamus
Pituitary
Skin
Fat
Mammary gland
Limb muscle
Diaphragm
Bone
Bone marrow
Blood
Spleen
Heart
Aorta
Lung
Trachea
Tongue
Small intestine
Liver
Pancreas
Kidney
Bladder
Uterus
Testes

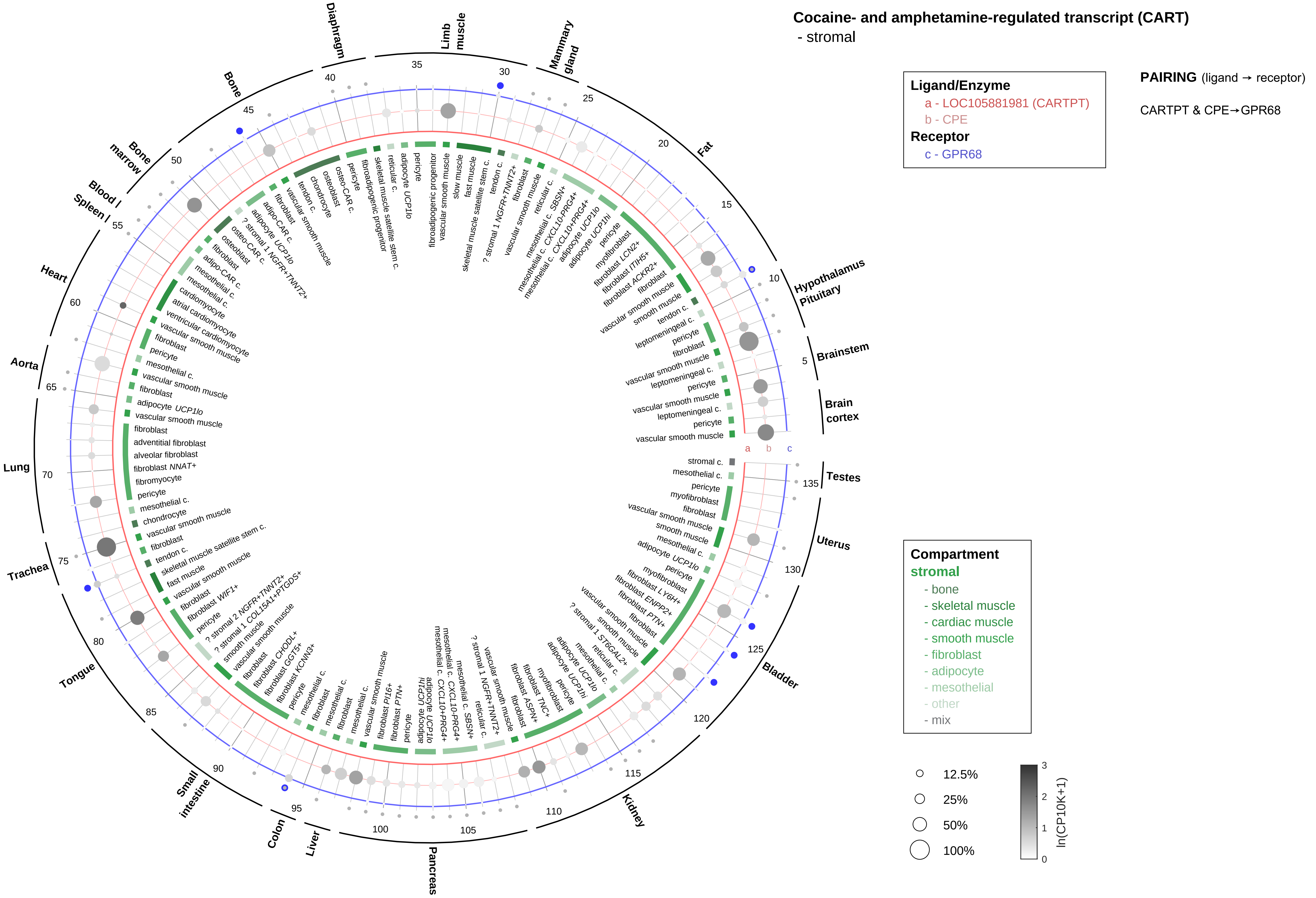

Cocaine- and amphetamine-regulated transcript (CART)
- stromal
Ligand/Enzyme
a - LOC105881981 (CARTPT)
b - CPE
Receptor
c - GPR68
PAIRING (ligand → receptor)
CARTPT & CPE→GPR68
Compartment
stromal
- bone
- skeletal muscle
- cardiac muscle
- smooth muscle
- fibroblast
- adipocyte
- mesothelial
- other
- mix
12.5%
25%
50%
100%
ln(CP10K+1)
Hypothalamus
Pituitary
Brainstem
Brain cortex
Testes
Uterus
Bladder
Kidney
Pancreas
Liver
Colon
Small intestine
Tongue
Trachea
Lung
Aorta
Heart
Spleen
Blood
Bone marrow
Bone
Diaphragm
Limb muscle
Mammary gland
Fat

#### Cocaine- and amphetamine-regulated transcript (CART)

- lymphoid

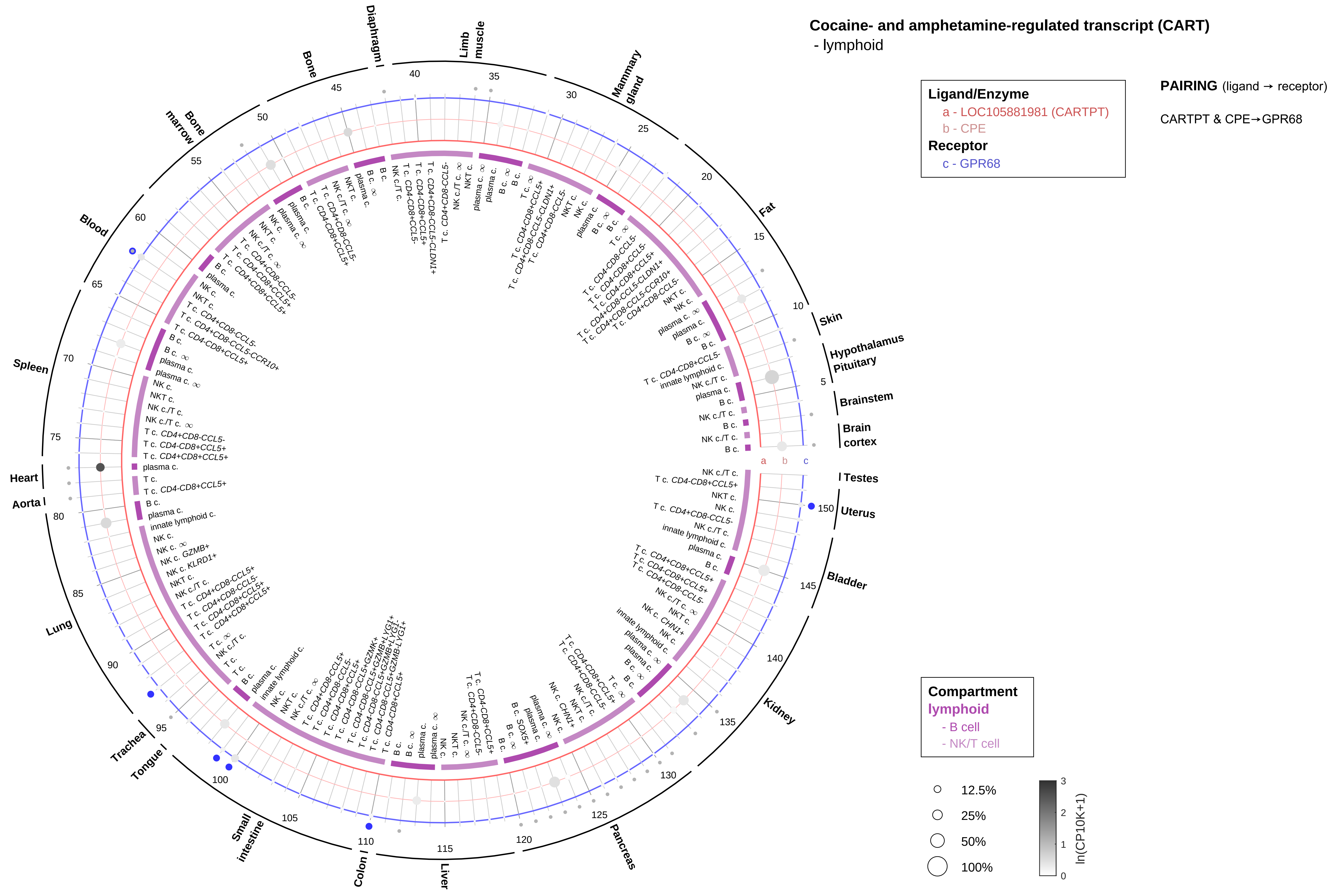

**Cocaine- and amphetamine-regulated transcript (CART)**
- myeloid

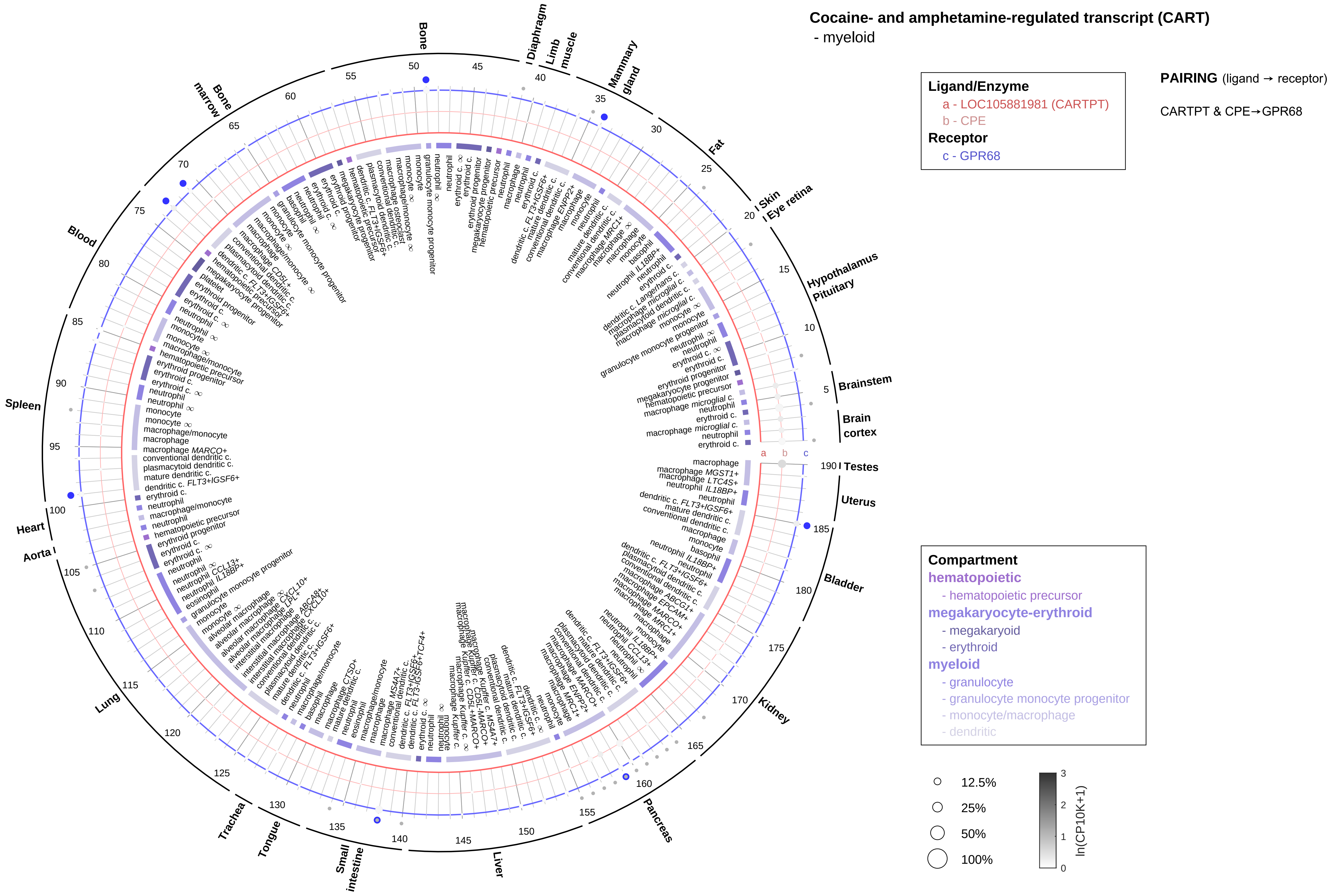

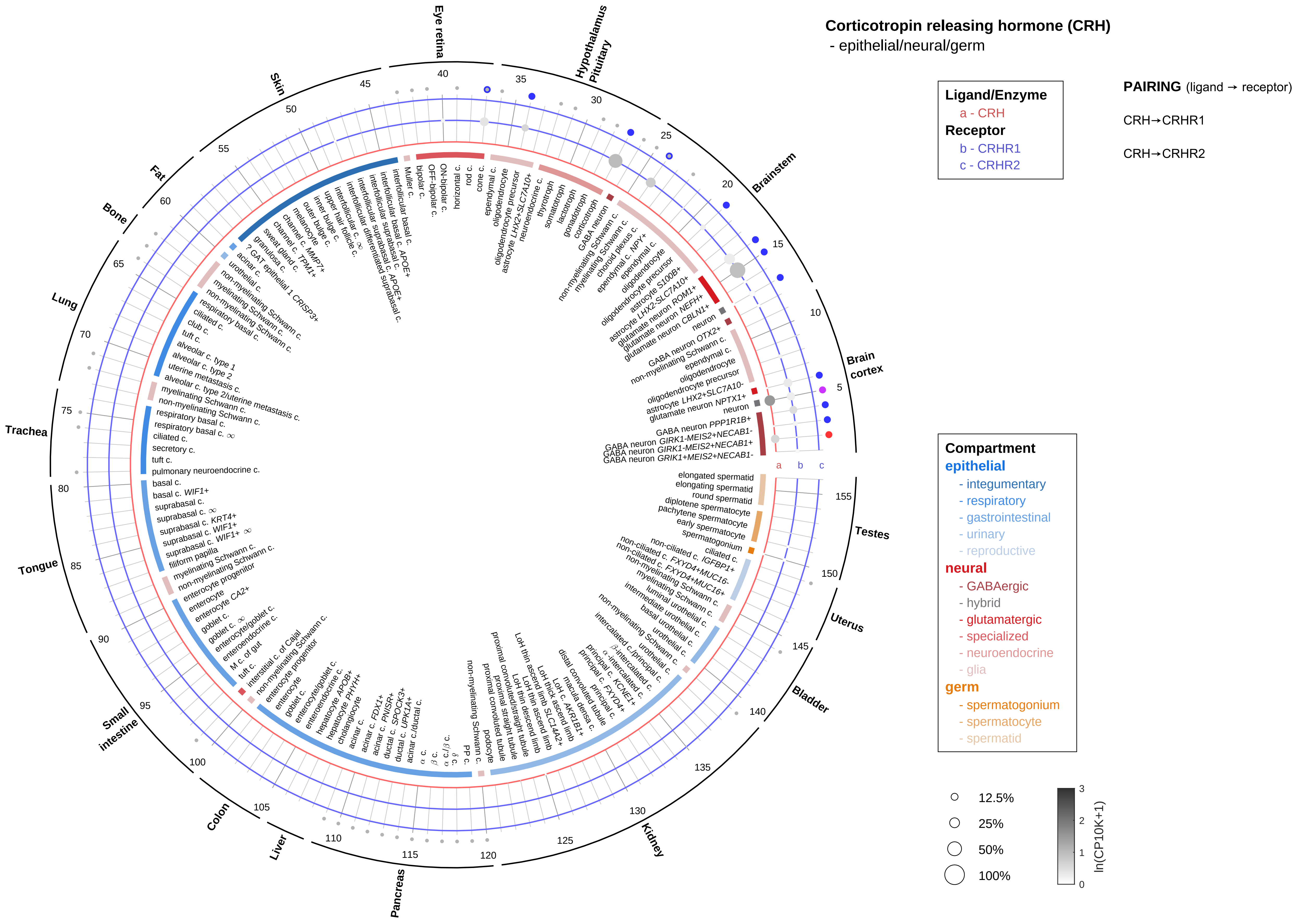

Corticotropin releasing hormone (CRH)
- epithelial/neural/germ
Ligand/Enzyme
a - CRH
Receptor
b - CRHR1
c - CRHR2
PAIRING (ligand → receptor)
CRH→CRHR1
CRH→CRHR2
Compartment
epithelial
- integumentary
- respiratory
- gastrointestinal
- urinary
- reproductive
neural
- GABAergic
- hybrid
- glutamatergic
- specialized
- neuroendocrine
- glia
germ
- spermatogonium
- spermatocyte
- spermatid
12.5%
25%
50%
100%
ln(CP10K+1)
Brain cortex
Brainstem
Hypothalamus Pituitary
Eye retina
Skin
Fat
Bone
Lung
Trachea
Tongue
Small intestine
Colon
Liver
Pancreas
Kidney
Bladder
Uterus
Testes

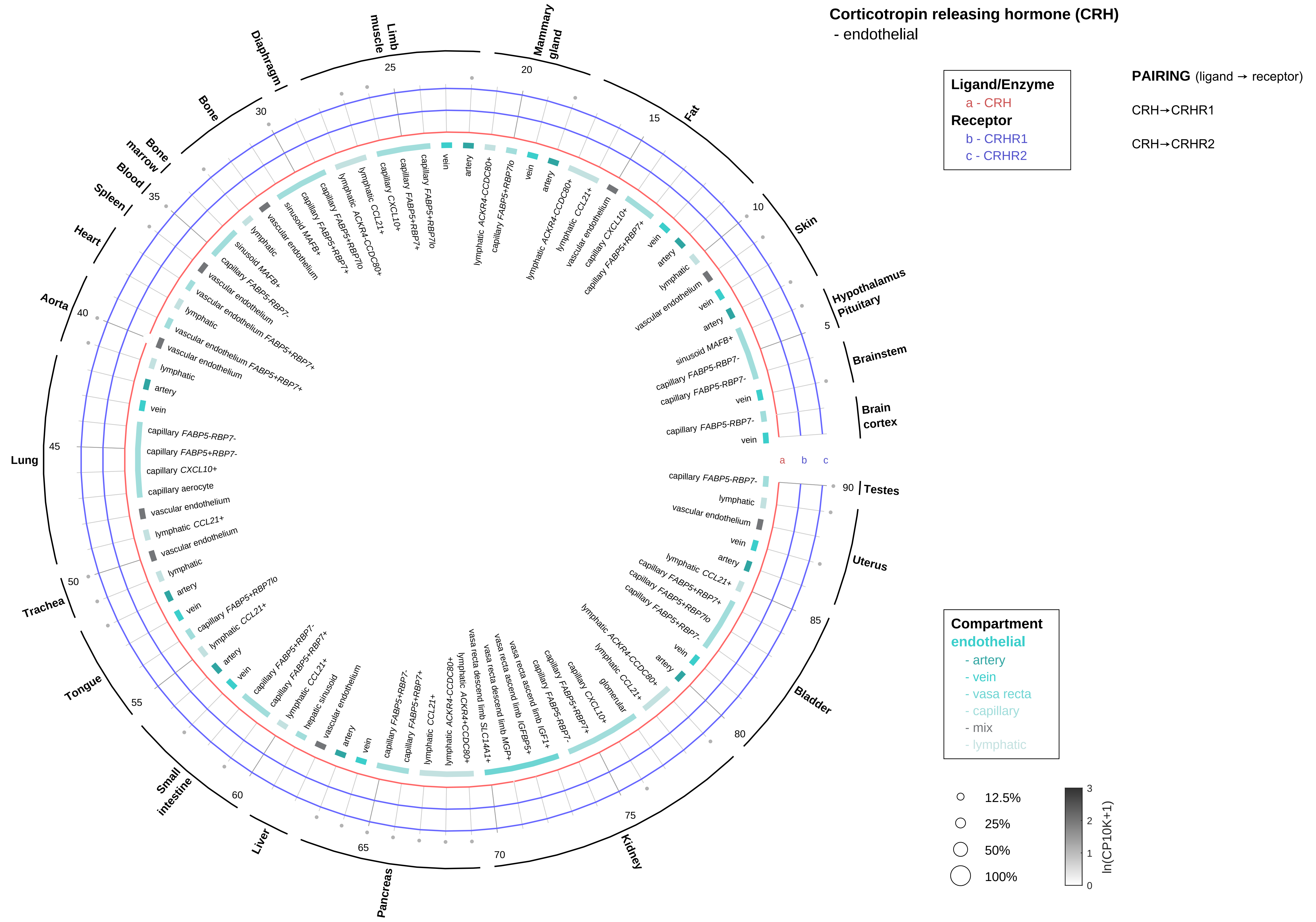
Corticotropin releasing hormone (CRH)
- endothelial
Ligand/Enzyme
a - CRH
Receptor
b - CRHR1
c - CRHR2
PAIRING (ligand → receptor)
CRH→CRHR1
CRH→CRHR2
Compartment
endothelial
- artery
- vein
- vasa recta
- capillary
- mix
- lymphatic
12.5%
25%
50%
100%
ln(CP10K+1)
Hypothalamus
Pituitary
Brainstem
Brain cortex
Testes
Uterus
Bladder
Kidney
Pancreas
Liver
Small intestine
Tongue
Trachea
Lung
Aorta
Heart
Spleen
Blood
Bone marrow
Bone
Diaphragm
Limb muscle
Mammary gland
Fat
Skin

#### Corticotropin releasing hormone (CRH)
- stromal

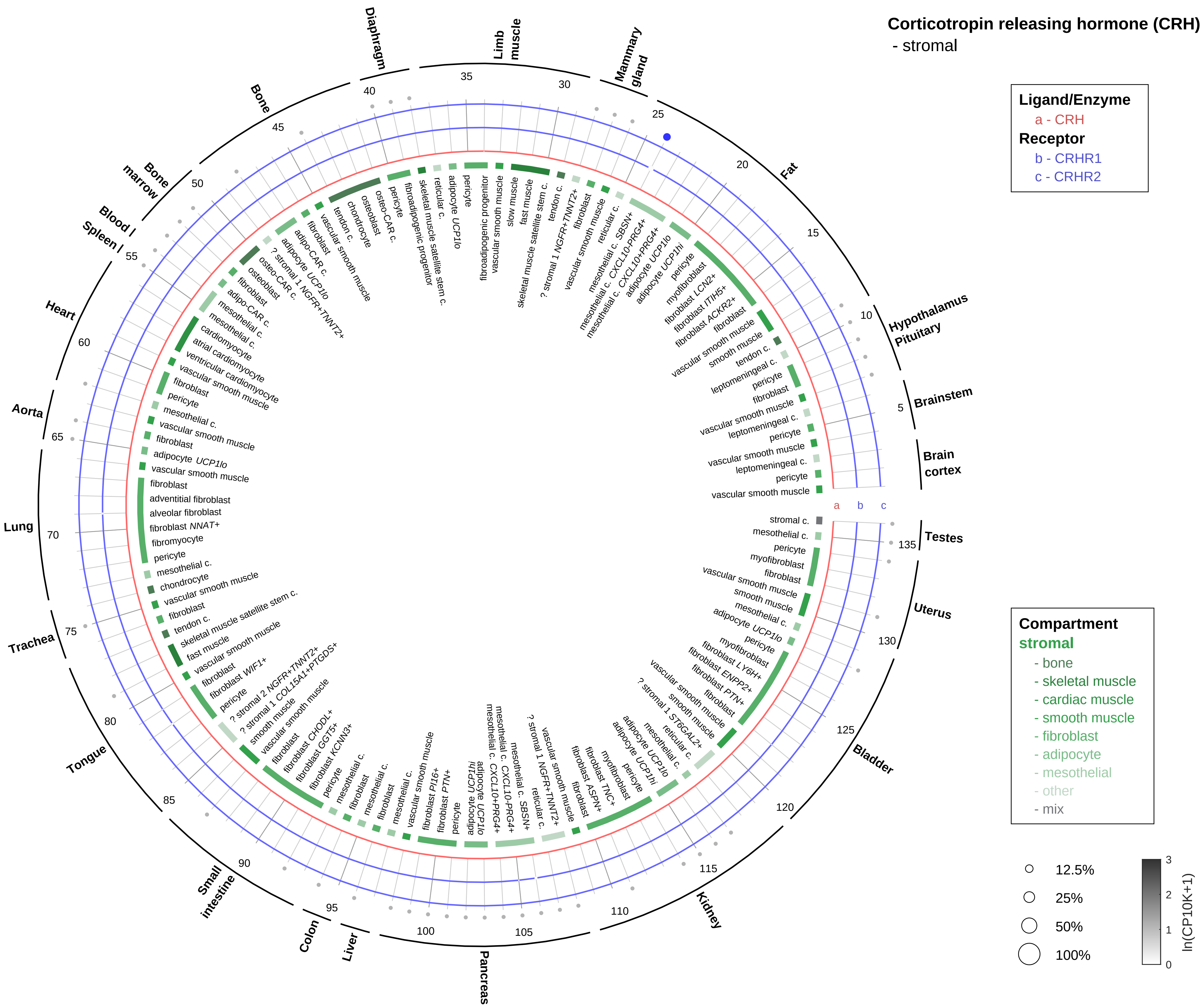

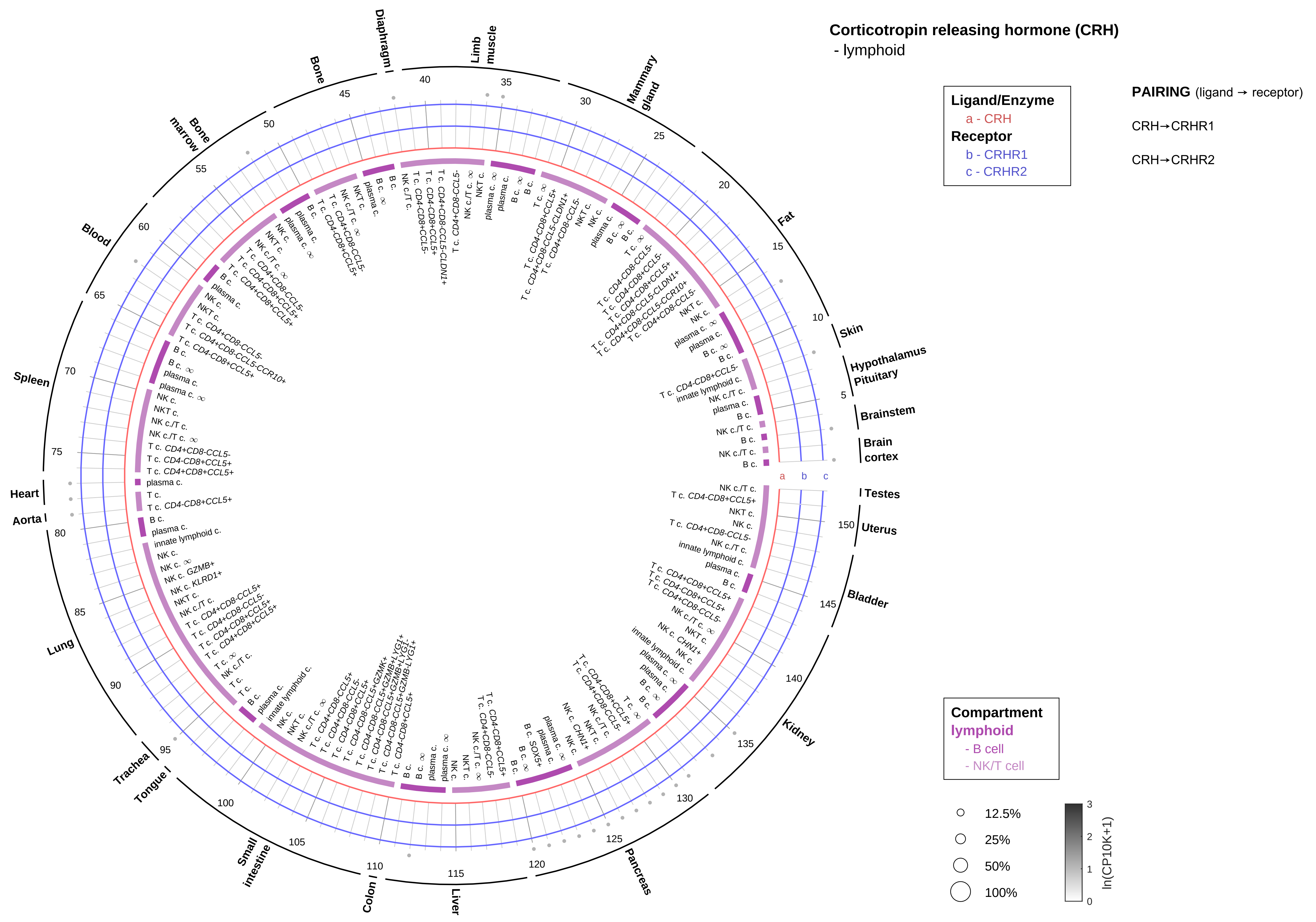
Corticotropin releasing hormone (CRH)
- lymphoid
Ligand/Enzyme
a - CRH
Receptor
b - CRHR1
c - CRHR2
PAIRING (ligand → receptor)
CRH→CRHR1
CRH→CRHR2
Compartment
lymphoid
- B cell
- NK/T cell
12.5%
25%
50%
100%
ln(CP10K+1)

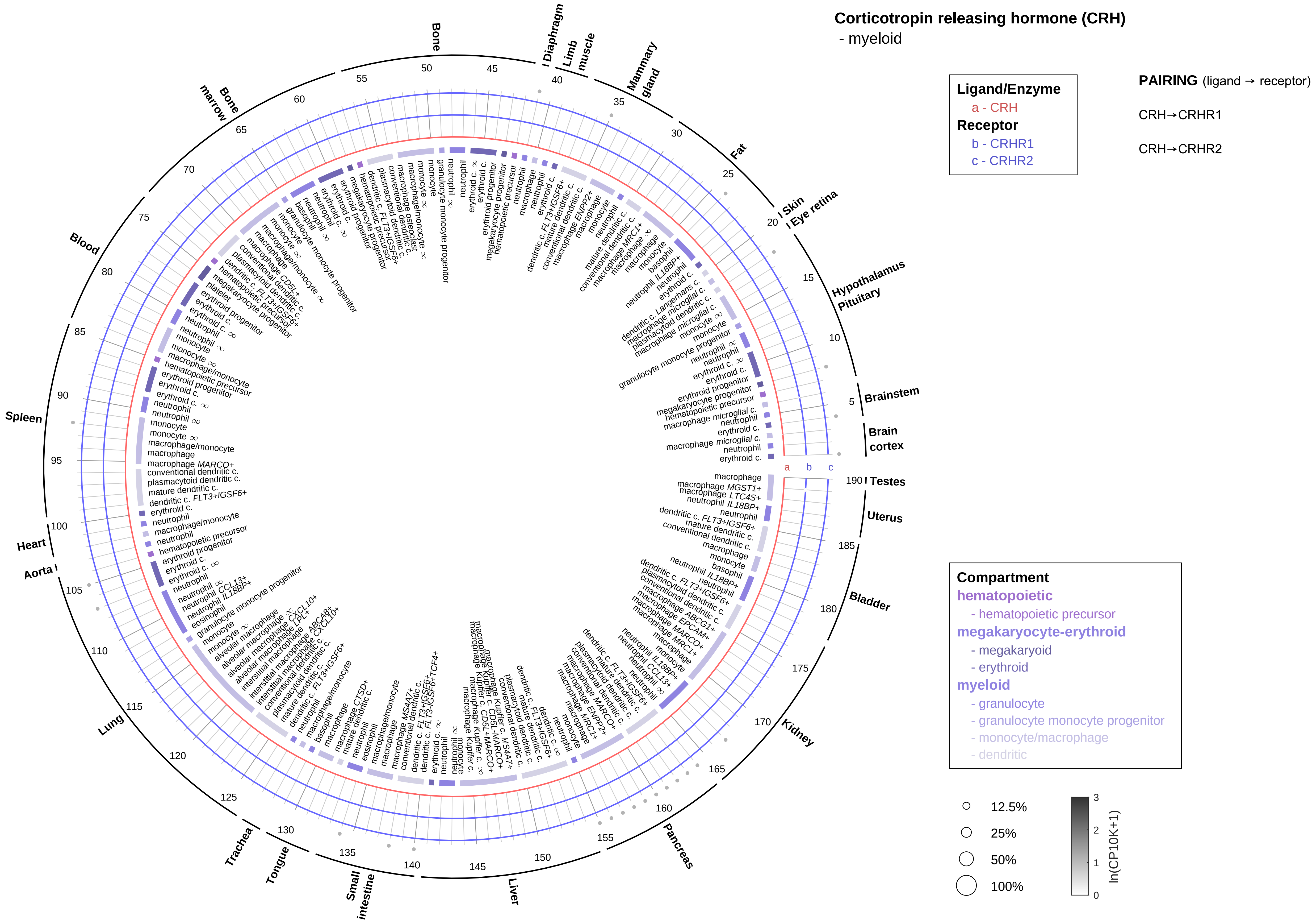

Corticotropin releasing hormone (CRH)
- myeloid
Ligand/Enzyme
a - CRH
Receptor
b - CRHR1
c - CRHR2
PAIRING (ligand → receptor)
CRH→CRHR1
CRH→CRHR2
Compartment
hematopoietic
- hematopoietic precursor
megakaryocyte-erythroid
- megakaryoid
- erythroid
myeloid
- granulocyte
- granulocyte monocyte progenitor
- monocyte/macrophage
- dendritic
12.5%
25%
50%
100%
ln(CP10K+1)
Bone
Diaphragm
Limb muscle
Mammary gland
Fat
Skin
Eye retina
Hypothalamus
Pituitary
Brainstem
Brain cortex
Testes
Uterus
Bladder
Kidney
Pancreas
Liver
Small intestine
Tongue
Trachea
Lung
Aorta
Heart
Spleen
Blood
Bone marrow

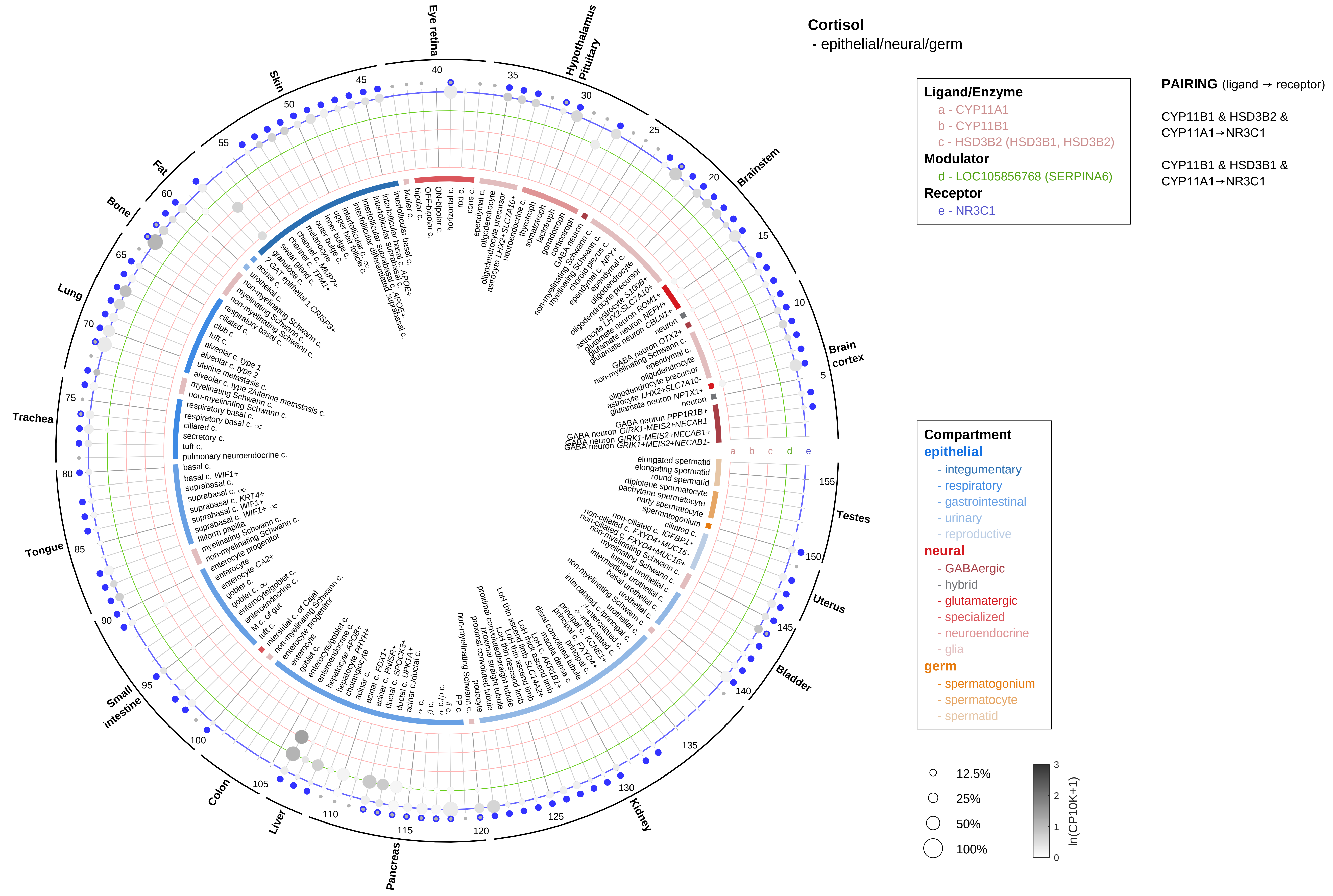
Cortisol
- epithelial/neural/germ
Ligand/Enzyme
a - CYP11A1
b - CYP11B1
c - HSD3B2 (HSD3B1, HSD3B2)
Modulator
d - LOC105856768 (SERPINA6)
Receptor
e - NR3C1
PAIRING (ligand → receptor)
CYP11B1 & HSD3B2 & CYP11A1→NR3C1
CYP11B1 & HSD3B1 & CYP11A1→NR3C1
Compartment
epithelial
- integumentary
- respiratory
- gastrointestinal
- urinary
- reproductive
neural
- GABAergic
- hybrid
- glutamatergic
- specialized
- neuroendocrine
- glia
germ
- spermatogonium
- spermatocyte
- spermatid
12.5%
25%
50%
100%
ln(CP10K+1)
Eye retina
Hypothalamus Pituitary
Brainstem
Brain cortex
Testes
Uterus
Bladder
Kidney
Pancreas
Liver
Colon
Small intestine
Tongue
Trachea
Lung
Bone
Fat
Skin

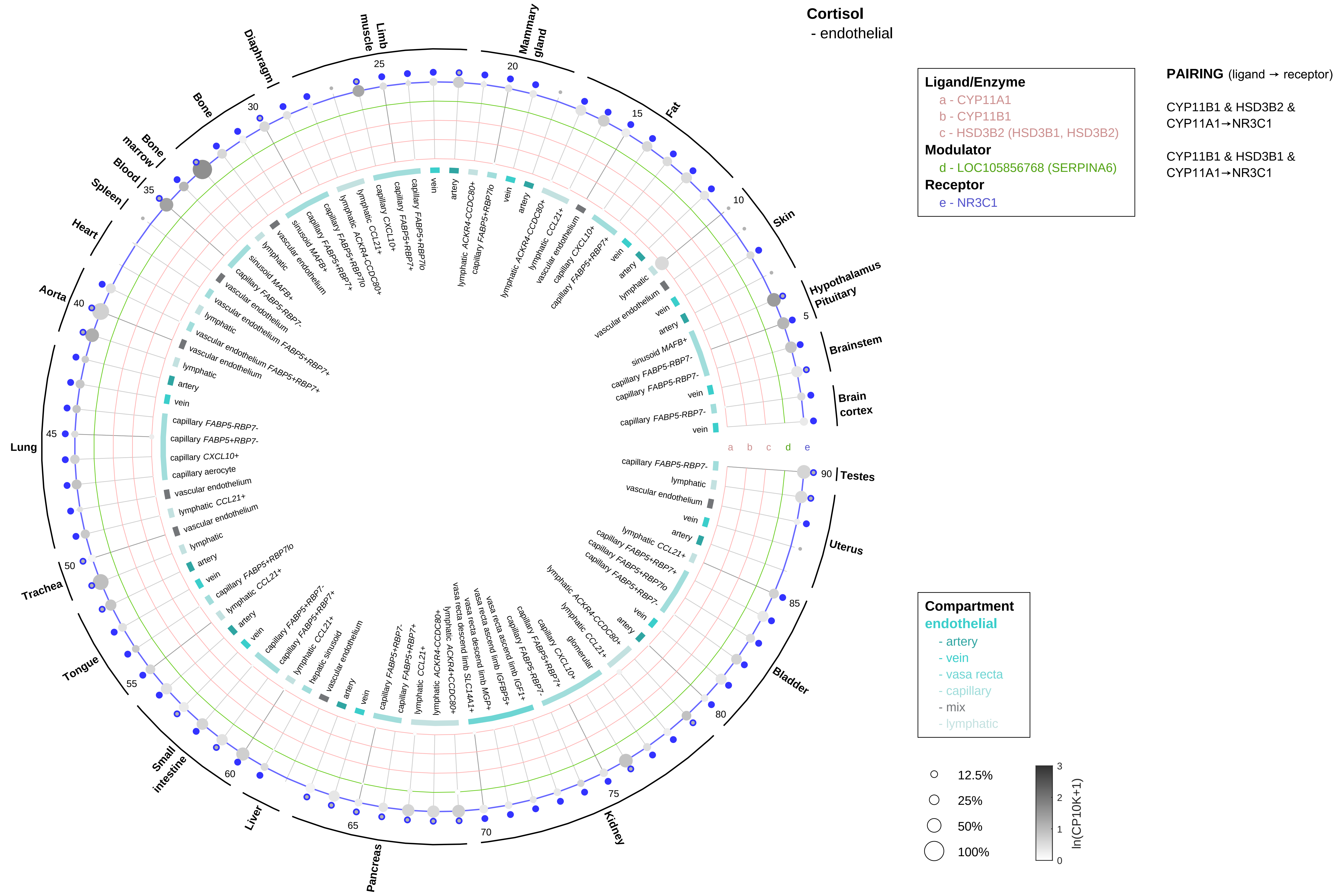
Cortisol
- endothelial
Ligand/Enzyme
a - CYP11A1
b - CYP11B1
c - HSD3B2 (HSD3B1, HSD3B2)
Modulator
d - LOC105856768 (SERPINA6)
Receptor
e - NR3C1
PAIRING (ligand → receptor)
CYP11B1 & HSD3B2 & CYP11A1→NR3C1
CYP11B1 & HSD3B1 & CYP11A1→NR3C1
Compartment
endothelial
- artery
- vein
- vasa recta
- capillary
- mix
- lymphatic
12.5%
25%
50%
100%
ln(CP10K+1)
Hypothalamus
Pituitary
Brainstem
Brain cortex
Testes
Uterus
Bladder
Kidney
Pancreas
Liver
Small intestine
Tongue
Trachea
Lung
Aorta
Heart
Spleen
Blood
Bone marrow
Bone
Diaphragm
Limb muscle
Mammary gland
Fat
Skin

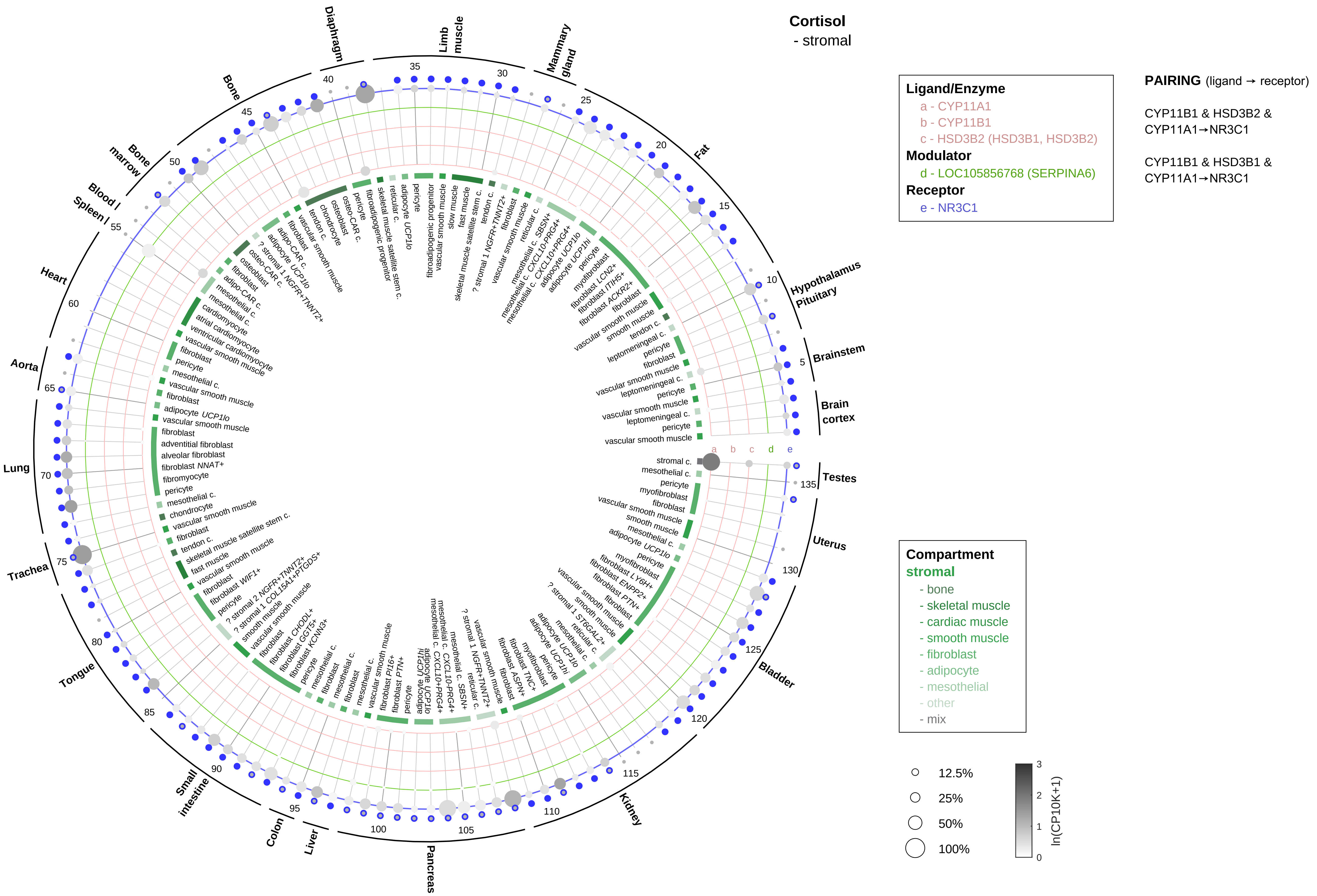
Cortisol
- stromal
Ligand/Enzyme
a - CYP11A1
b - CYP11B1
c - HSD3B2 (HSD3B1, HSD3B2)
Modulator
d - LOC105856768 (SERPINA6)
Receptor
e - NR3C1
PAIRING (ligand → receptor)
CYP11B1 & HSD3B2 & CYP11A1→NR3C1
CYP11B1 & HSD3B1 & CYP11A1→NR3C1
Compartment
stromal
- bone
- skeletal muscle
- cardiac muscle
- smooth muscle
- fibroblast
- adipocyte
- mesothelial
- other
- mix
12.5%
25%
50%
100%
ln(CP10K+1)

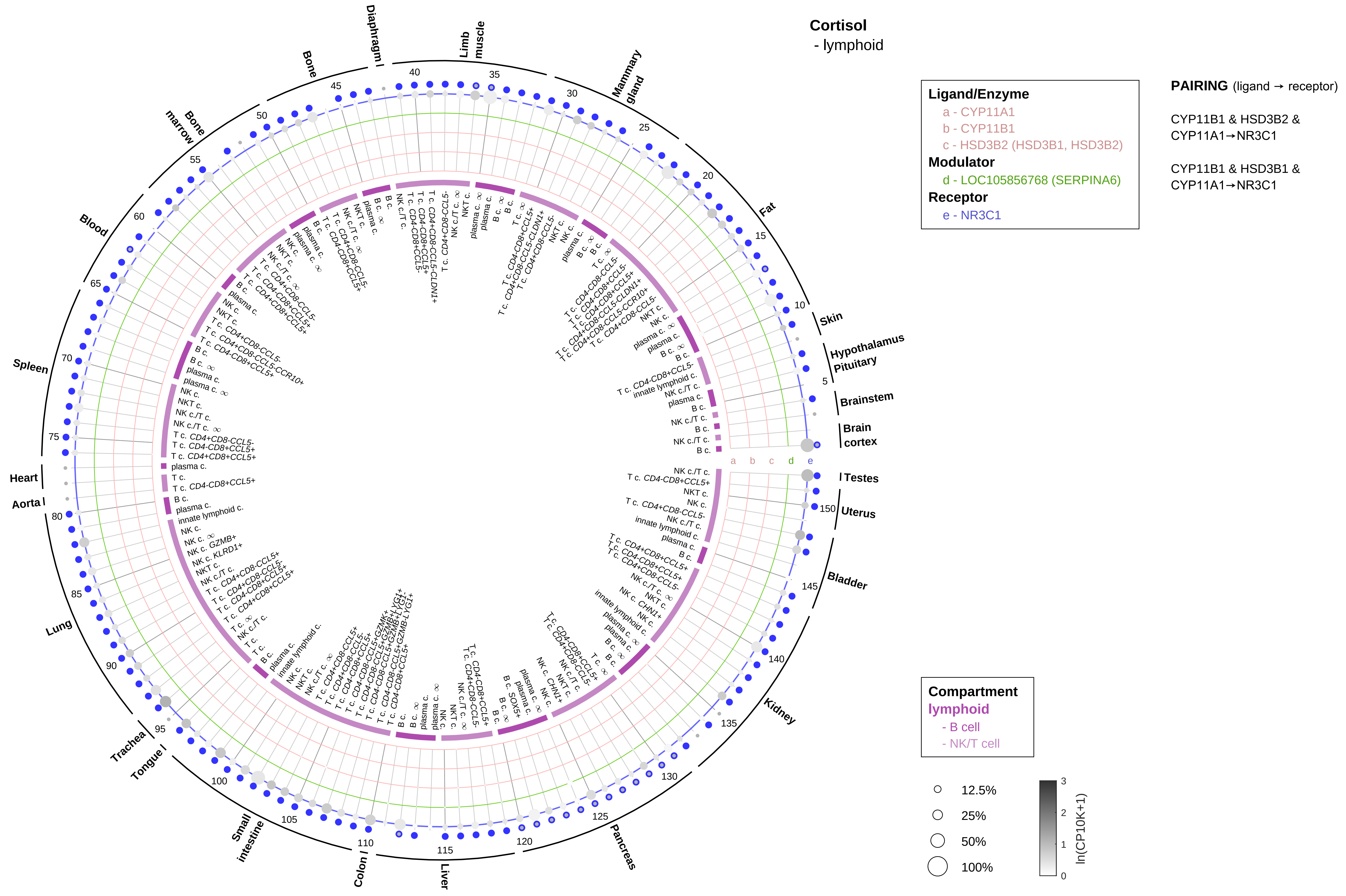

Cortisol
- lymphoid
Ligand/Enzyme
a - CYP11A1
b - CYP11B1
c - HSD3B2 (HSD3B1, HSD3B2)
Modulator
d - LOC105856768 (SERPINA6)
Receptor
e - NR3C1
PAIRING (ligand → receptor)
CYP11B1 & HSD3B2 & CYP11A1→NR3C1
CYP11B1 & HSD3B1 & CYP11A1→NR3C1
Compartment
lymphoid
- B cell
- NK/T cell
12.5%
25%
50%
100%
ln(CP10K+1)

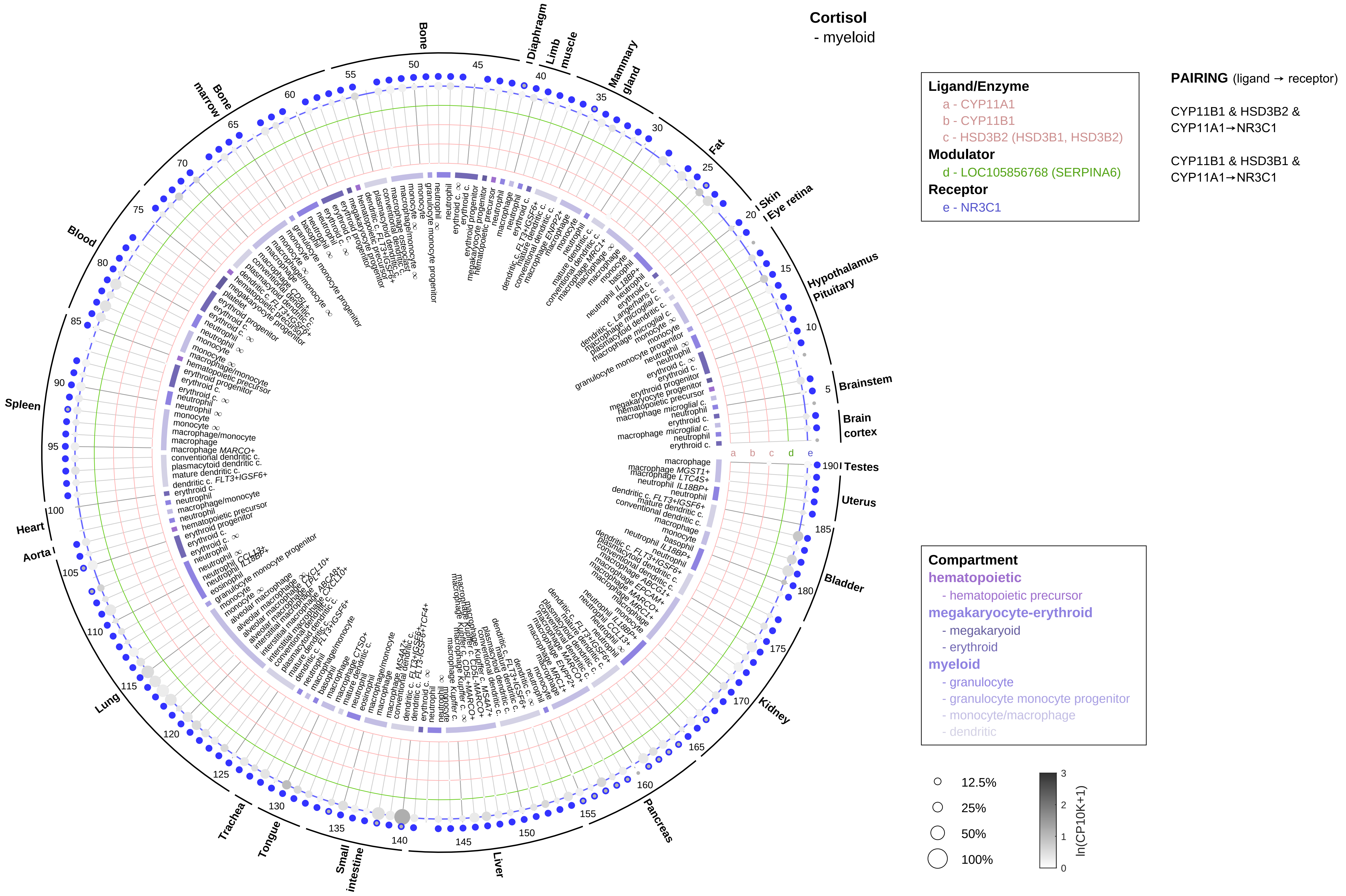

Cortisol
- myeloid
Ligand/Enzyme
a - CYP11A1
b - CYP11B1
c - HSD3B2 (HSD3B1, HSD3B2)
Modulator
d - LOC105856768 (SERPINA6)
Receptor
e - NR3C1
PAIRING (ligand → receptor)
CYP11B1 & HSD3B2 & CYP11A1→NR3C1
CYP11B1 & HSD3B1 & CYP11A1→NR3C1
Compartment
hematopoietic
- hematopoietic precursor
megakaryocyte-erythroid
- megakaryoid
- erythroid
myeloid
- granulocyte
- granulocyte monocyte progenitor
- monocyte/macrophage
- dendritic
12.5%
25%
50%
100%
ln(CP10K+1)
Bone
Diaphragm
Limb muscle
Mammary gland
Fat
Skin
Eye retina
Hypothalamus
Pituitary
Brainstem
Brain cortex
Testes
Uterus
Bladder
Kidney
Pancreas
Liver
Small intestine
Tongue
Trachea
Lung
Aorta
Heart
Spleen
Blood
Bone marrow

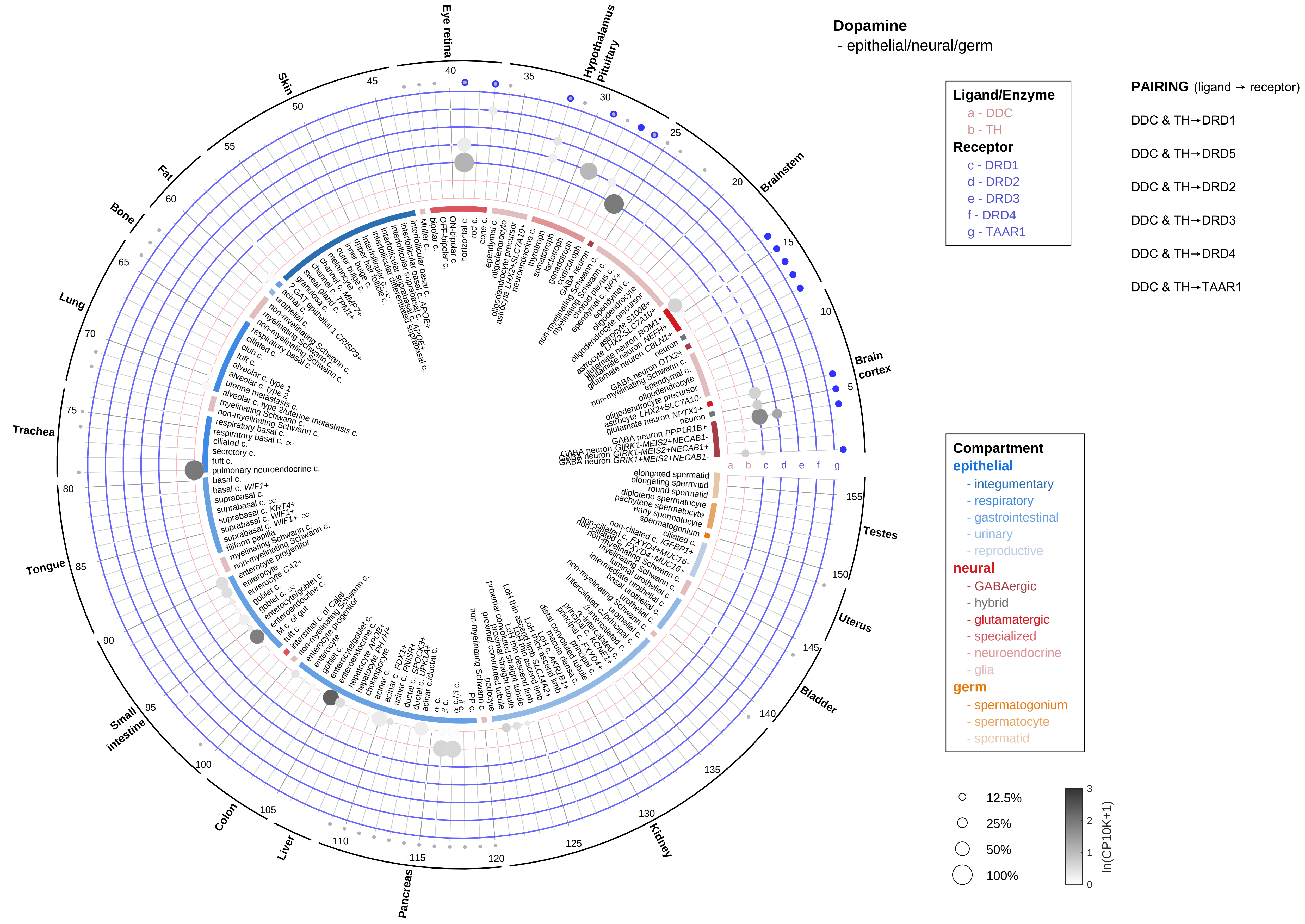
Dopamine
- epithelial/neural/germ
Ligand/Enzyme
a - DDC
b - TH
Receptor
c - DRD1
d - DRD2
e - DRD3
f - DRD4
g - TAAR1
PAIRING (ligand → receptor)
DDC & TH→DRD1
DDC & TH→DRD5
DDC & TH→DRD2
DDC & TH→DRD3
DDC & TH→DRD4
DDC & TH→TAAR1
Compartment
epithelial
- integumentary
- respiratory
- gastrointestinal
- urinary
- reproductive
neural
- GABAergic
- hybrid
- glutamatergic
- specialized
- neuroendocrine
- glia
germ
- spermatogonium
- spermatocyte
- spermatid
12.5%
25%
50%
100%
ln(CP10K+1)
Eye retina
Hypothalamus
Pituitary
Brainstem
Brain cortex
Testes
Uterus
Bladder
Kidney
Pancreas
Liver
Colon
Small intestine
Tongue
Trachea
Lung
Bone
Fat
Skin
a b c d e f g

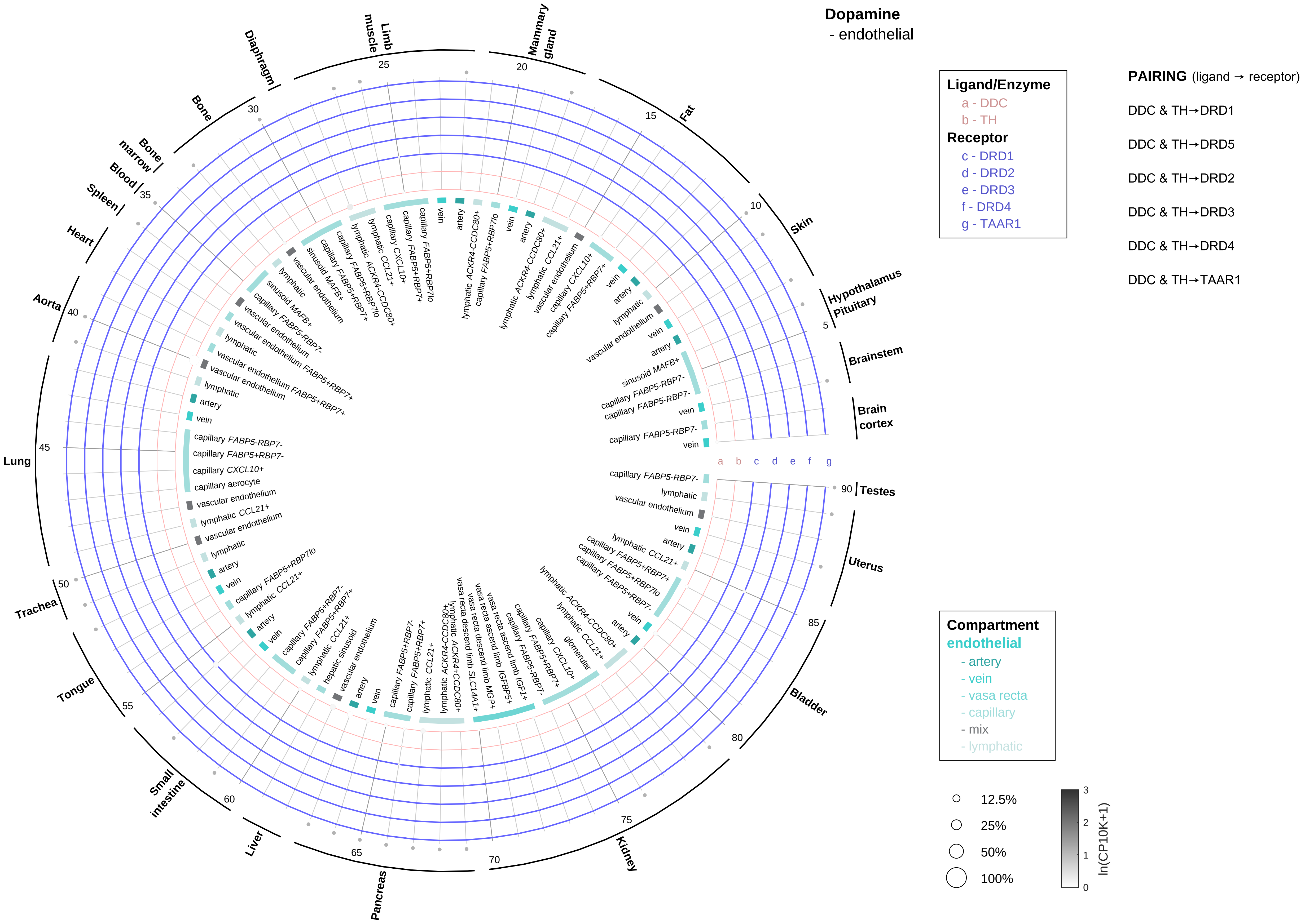
Dopamine
- endothelial
Ligand/Enzyme
a - DDC
b - TH
Receptor
c - DRD1
d - DRD2
e - DRD3
f - DRD4
g - TAAR1
PAIRING (ligand → receptor)
DDC & TH→DRD1
DDC & TH→DRD5
DDC & TH→DRD2
DDC & TH→DRD3
DDC & TH→DRD4
DDC & TH→TAAR1
Compartment
endothelial
- artery
- vein
- vasa recta
- capillary
- mix
- lymphatic
12.5%
25%
50%
100%
ln(CP10K+1)
Brain cortex
Brainstem
Hypothalamus
Pituitary
Skin
Fat
Mammary gland
Limb muscle
Diaphragm
Bone
Bone marrow
Blood
Spleen
Heart
Aorta
Lung
Trachea
Tongue
Small intestine
Liver
Pancreas
Kidney
Bladder
Uterus
Testes

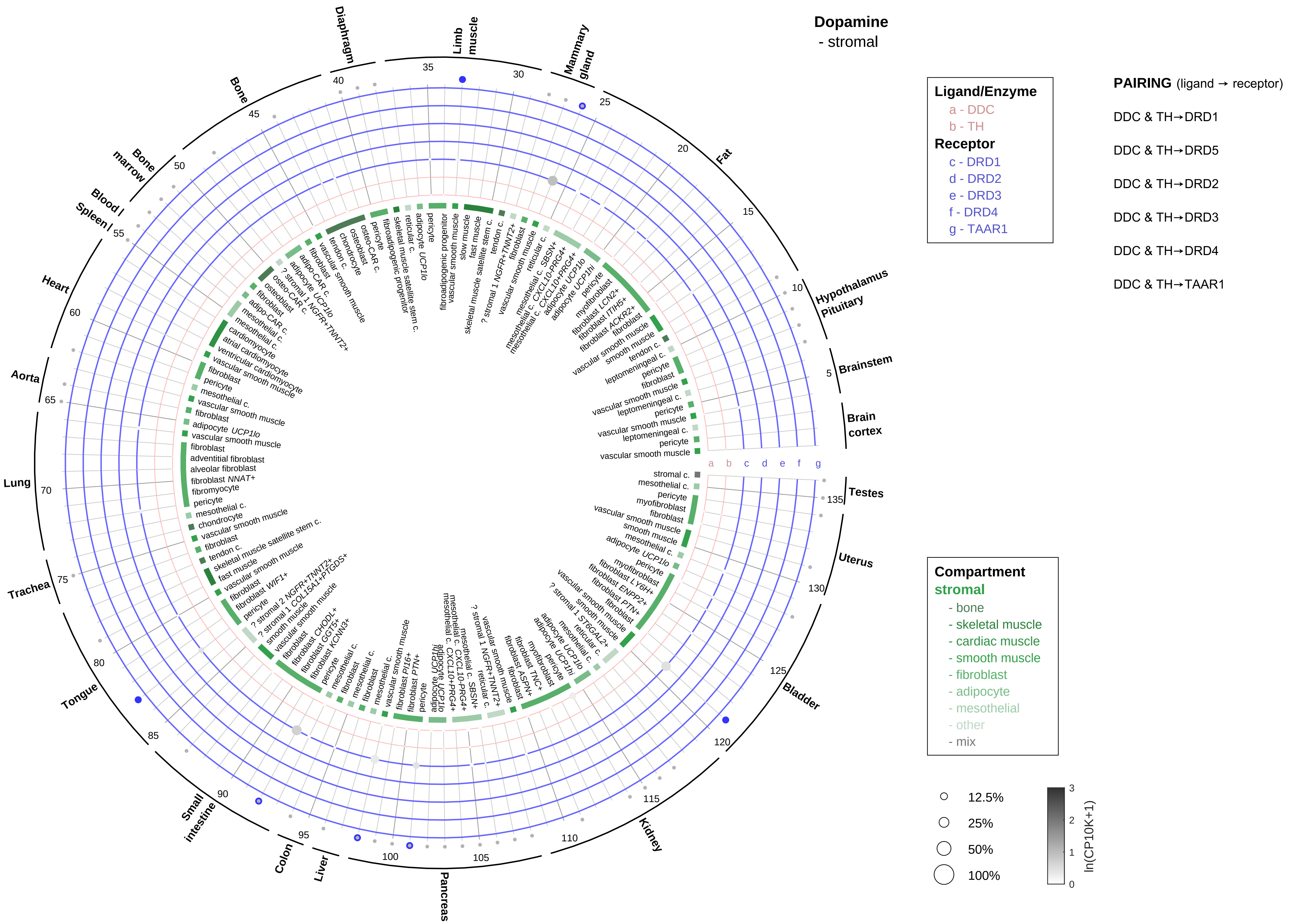

Dopamine
- stromal
Ligand/Enzyme
a - DDC
b - TH
Receptor
c - DRD1
d - DRD2
e - DRD3
f - DRD4
g - TAAR1
PAIRING (ligand → receptor)
DDC & TH→DRD1
DDC & TH→DRD5
DDC & TH→DRD2
DDC & TH→DRD3
DDC & TH→DRD4
DDC & TH→TAAR1
Compartment
stromal
- bone
- skeletal muscle
- cardiac muscle
- smooth muscle
- fibroblast
- adipocyte
- mesothelial
- other
- mix
12.5%
25%
50%
100%
ln(CP10K+1)
Diaphragm
Limb muscle
Mammary gland
Fat
Hypothalamus Pituitary
Brainstem
Brain cortex
Testes
Uterus
Bladder
Kidney
Pancreas
Liver
Colon
Small intestine
Tongue
Trachea
Lung
Aorta
Heart
Spleen
Blood
Bone marrow
Bone
a b c d e f g

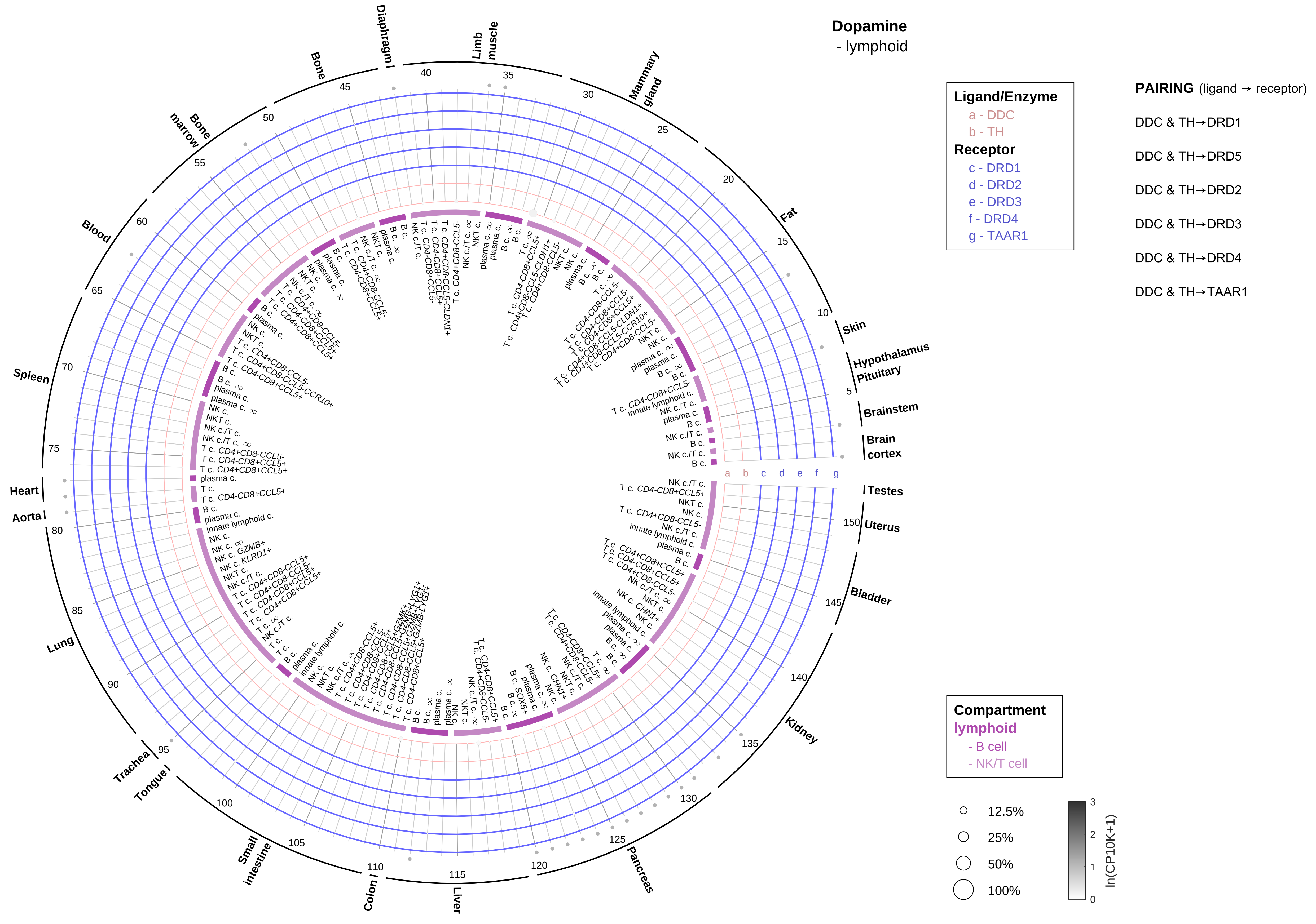
Dopamine
- lymphoid
Ligand/Enzyme
a - DDC
b - TH
Receptor
c - DRD1
d - DRD2
e - DRD3
f - DRD4
g - TAAR1
PAIRING (ligand → receptor)
DDC & TH→DRD1
DDC & TH→DRD5
DDC & TH→DRD2
DDC & TH→DRD3
DDC & TH→DRD4
DDC & TH→TAAR1
Compartment
lymphoid
- B cell
- NK/T cell
12.5%
25%
50%
100%
ln(CP10K+1)
Diaphragm
Limb muscle
Mammary gland
Fat
Skin
Hypothalamus
Pituitary
Brainstem
Brain cortex
Testes
Uterus
Bladder
Kidney
Pancreas
Liver
Colon
Small intestine
Tongue
Trachea
Lung
Aorta
Heart
Spleen
Blood
Bone marrow
Bone

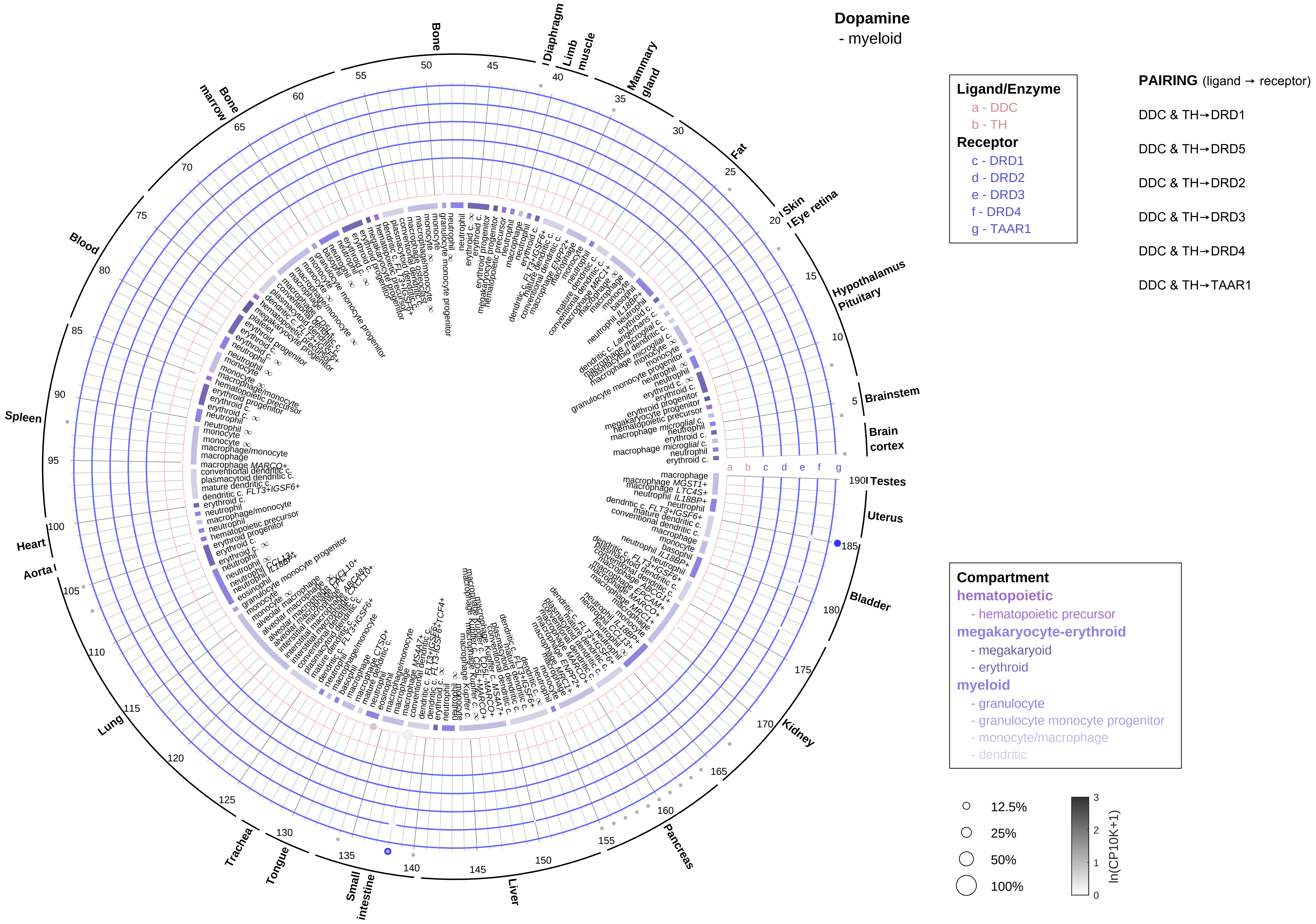
Dopamine
- myeloid
Ligand/Enzyme
a - DDC
b - TH
Receptor
c - DRD1
d - DRD2
e - DRD3
f - DRD4
g - TAAR1
PAIRING (ligand → receptor)
DDC & TH→DRD1
DDC & TH→DRD5
DDC & TH→DRD2
DDC & TH→DRD3
DDC & TH→DRD4
DDC & TH→TAAR1
Compartment
hematopoietic
- hematopoietic precursor
megakaryocyte-erythroid
- megakaryoid
- erythroid
myeloid
- granulocyte
- granulocyte monocyte progenitor
- monocyte/macrophage
- dendritic
12.5%
25%
50%
100%
ln(CP10K+1)
Brain cortex
Brainstem
Hypothalamus
Pituitary
Eye retina
Skin
Fat
Mammary gland
Limb muscle
Diaphragm
Bone
Bone marrow
Blood
Spleen
Heart
Aorta
Lung
Trachea
Tongue
Small intestine
Liver
Pancreas
Kidney
Bladder
Uterus
Testes

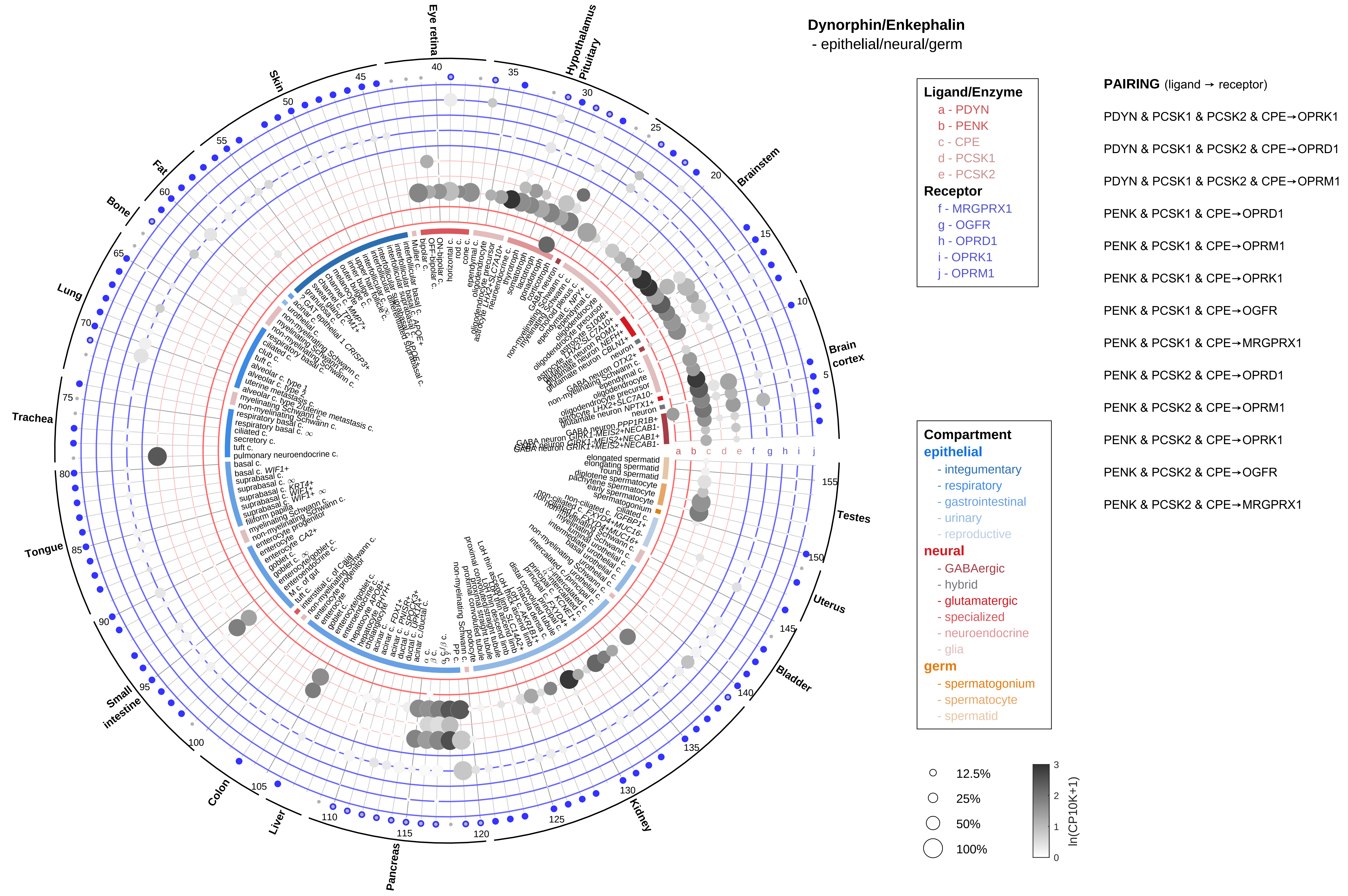

Dynorphin/Enkephalin
- epithelial/neural/germ
Ligand/Enzyme
a - PDYN
b - PENK
c - CPE
d - PCSK1
e - PCSK2
Receptor
f - MRGPRX1
g - OGFR
h - OPRD1
i - OPRK1
j - OPRM1
PAIRING (ligand → receptor)
PDYN & PCSK1 & PCSK2 & CPE→OPRK1
PDYN & PCSK1 & PCSK2 & CPE→OPRD1
PDYN & PCSK1 & PCSK2 & CPE→OPRM1
PENK & PCSK1 & CPE→OPRD1
PENK & PCSK1 & CPE→OPRM1
PENK & PCSK1 & CPE→OPRK1
PENK & PCSK1 & CPE→OGFR
PENK & PCSK1 & CPE→MRGPRX1
PENK & PCSK2 & CPE→OPRD1
PENK & PCSK2 & CPE→OPRM1
PENK & PCSK2 & CPE→OPRK1
PENK & PCSK2 & CPE→OGFR
PENK & PCSK2 & CPE→MRGPRX1
Compartment
epithelial
- integumentary
- respiratory
- gastrointestinal
- urinary
- reproductive
neural
- GABAergic
- hybrid
- glutamatergic
- specialized
- neuroendocrine
- glia
germ
- spermatogonium
- spermatocyte
- spermatid
12.5%
25%
50%
100%
ln(CP10K+1)
Eye retina
Hypothalamus
Pituitary
Brainstem
Brain cortex
Testes
Uterus
Bladder
Kidney
Pancreas
Liver
Colon
Small intestine
Tongue
Trachea
Lung
Bone
Fat
Skin

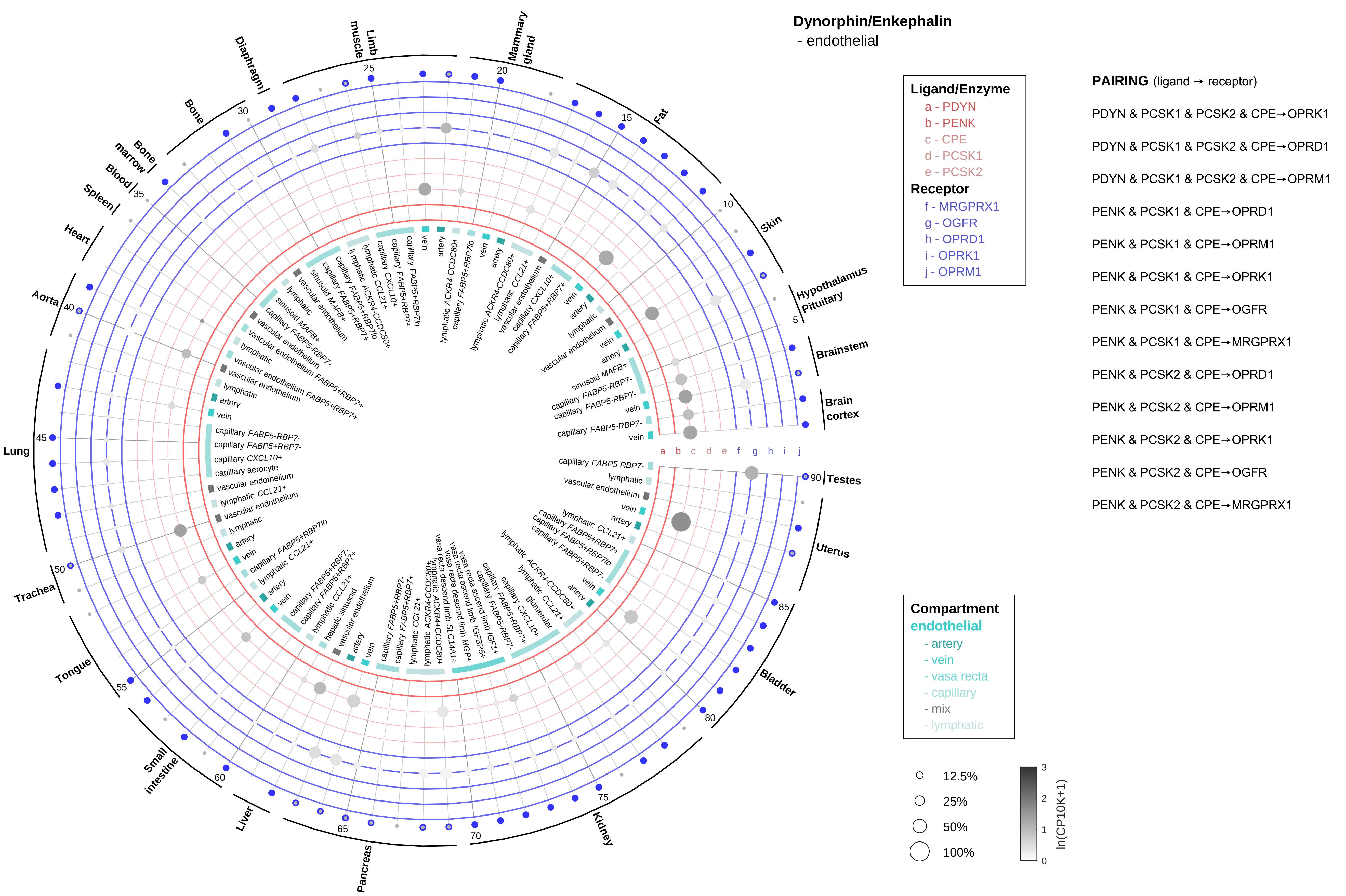
Dynorphin/Enkephalin
- endothelial
Ligand/Enzyme
a - PDYN
b - PENK
c - CPE
d - PCSK1
e - PCSK2
Receptor
f - MRGPRX1
g - OGFR
h - OPRD1
i - OPRK1
j - OPRM1
PAIRING (ligand → receptor)
PDYN & PCSK1 & PCSK2 & CPE→OPRK1
PDYN & PCSK1 & PCSK2 & CPE→OPRD1
PDYN & PCSK1 & PCSK2 & CPE→OPRM1
PENK & PCSK1 & CPE→OPRD1
PENK & PCSK1 & CPE→OPRM1
PENK & PCSK1 & CPE→OPRK1
PENK & PCSK1 & CPE→OGFR
PENK & PCSK1 & CPE→MRGPRX1
PENK & PCSK2 & CPE→OPRD1
PENK & PCSK2 & CPE→OPRM1
PENK & PCSK2 & CPE→OPRK1
PENK & PCSK2 & CPE→OGFR
PENK & PCSK2 & CPE→MRGPRX1
Compartment
endothelial
- artery
- vein
- vasa recta
- capillary
- mix
- lymphatic
12.5%
25%
50%
100%
ln(CP10K+1)
Limb muscle
Mammary gland
Diaphragm
Bone
Bone marrow
Blood
Spleen
Heart
Aorta
Lung
Trachea
Tongue
Small intestine
Liver
Pancreas
Kidney
Bladder
Uterus
Testes
Brain cortex
Brainstem
Hypothalamus Pituitary
Skin
Fat
a b c d e f g h i j

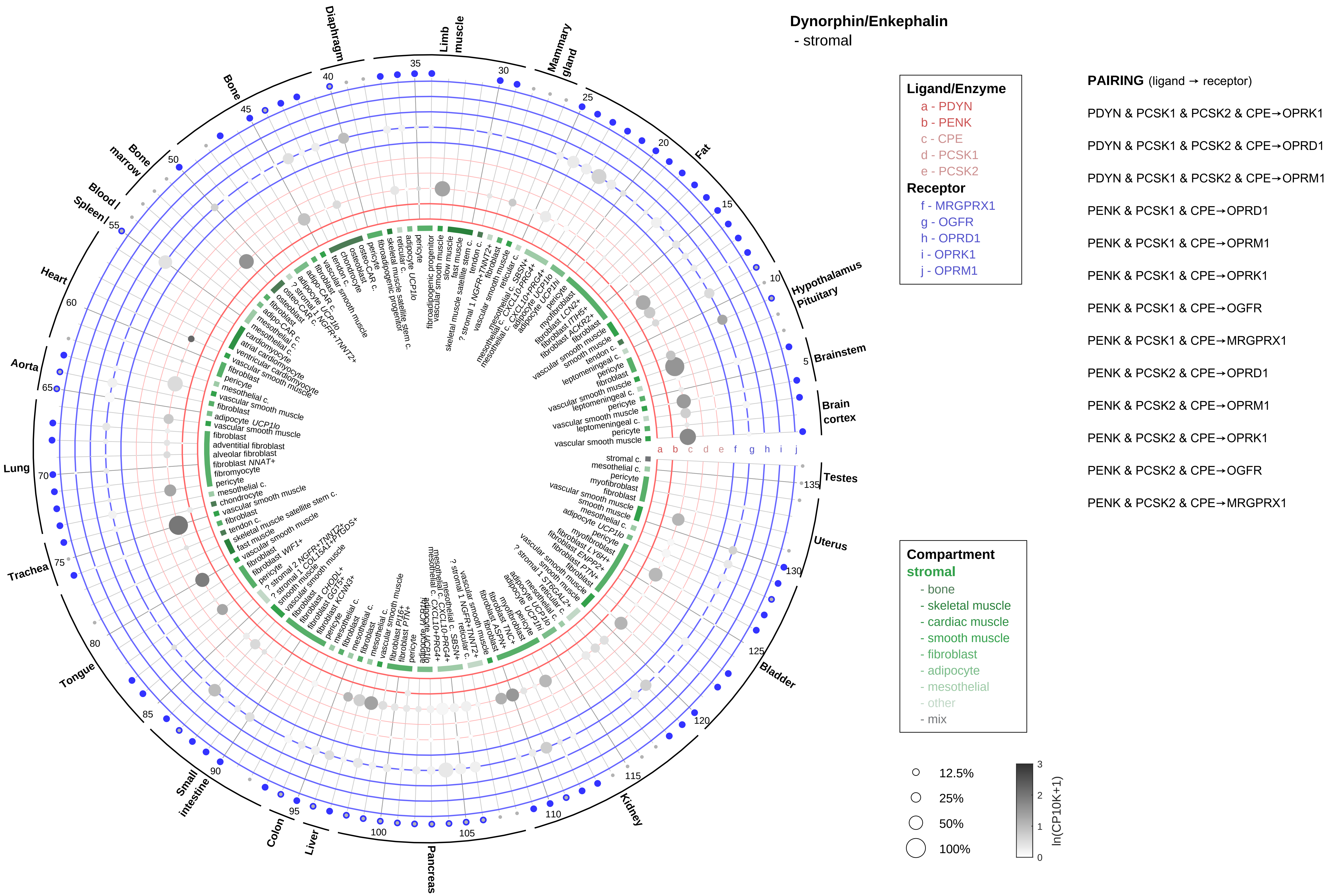
Dynorphin/Enkephalin
- stromal
Ligand/Enzyme
a - PDYN
b - PENK
c - CPE
d - PCSK1
e - PCSK2
Receptor
f - MRGPRX1
g - OGFR
h - OPRD1
i - OPRK1
j - OPRM1
PAIRING (ligand → receptor)
PDYN & PCSK1 & PCSK2 & CPE→OPRK1
PDYN & PCSK1 & PCSK2 & CPE→OPRD1
PDYN & PCSK1 & PCSK2 & CPE→OPRM1
PENK & PCSK1 & CPE→OPRD1
PENK & PCSK1 & CPE→OPRM1
PENK & PCSK1 & CPE→OPRK1
PENK & PCSK1 & CPE→OGFR
PENK & PCSK1 & CPE→MRGPRX1
PENK & PCSK2 & CPE→OPRD1
PENK & PCSK2 & CPE→OPRM1
PENK & PCSK2 & CPE→OPRK1
PENK & PCSK2 & CPE→OGFR
PENK & PCSK2 & CPE→MRGPRX1
Compartment
stromal
- bone
- skeletal muscle
- cardiac muscle
- smooth muscle
- fibroblast
- adipocyte
- mesothelial
- other
- mix
12.5%
25%
50%
100%
ln(CP10K+1)
Limb muscle
Mammary gland
Fat
Hypothalamus Pituitary
Brainstem
Brain cortex
Testes
Uterus
Bladder
Kidney
Pancreas
Liver
Colon
Small intestine
Tongue
Trachea
Lung
Aorta
Heart
Spleen
Blood
Bone marrow
Bone
Diaphragm

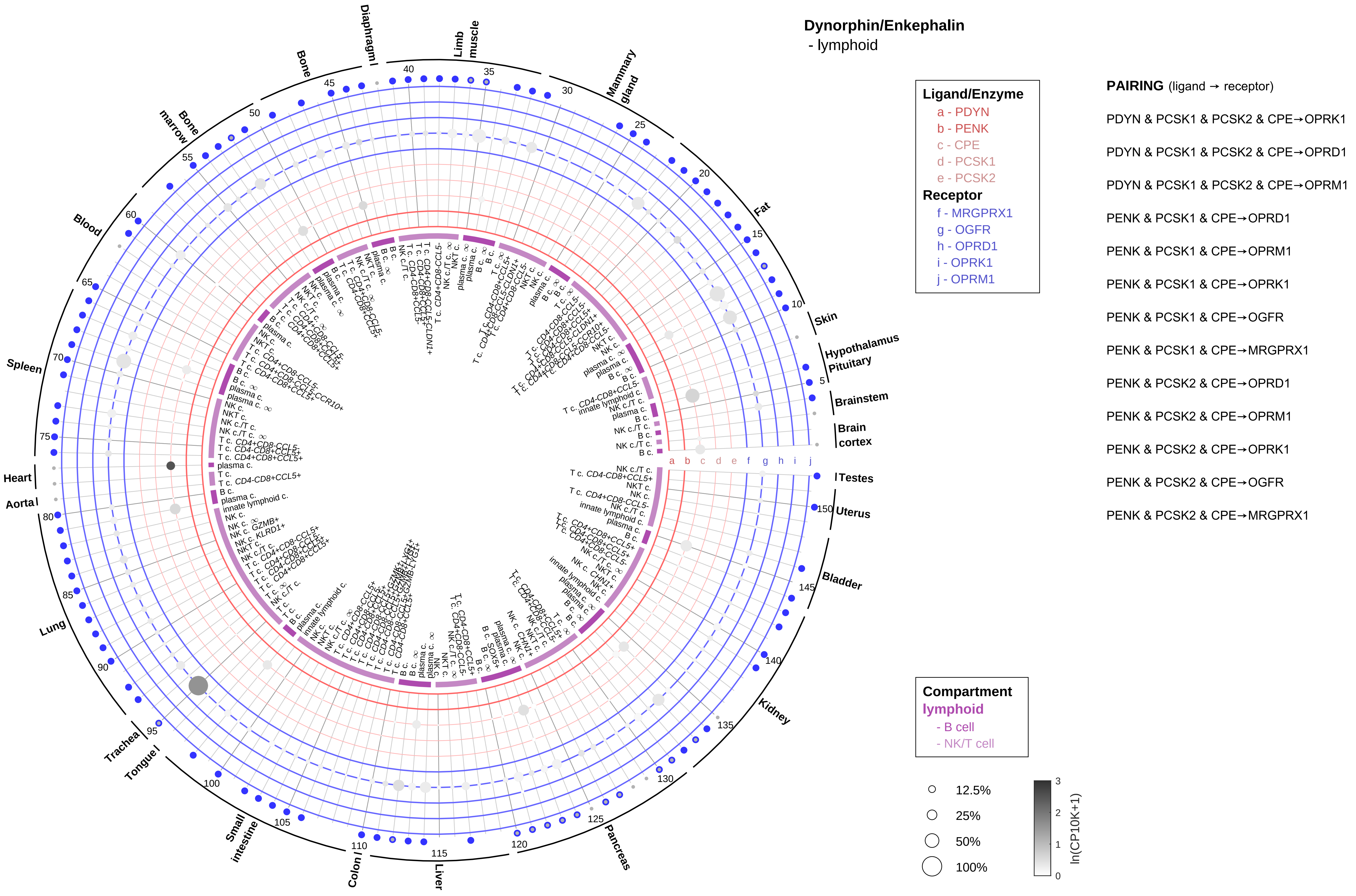

Dynorphin/Enkephalin
- lymphoid
Ligand/Enzyme
a - PDYN
b - PENK
c - CPE
d - PCSK1
e - PCSK2
Receptor
f - MRGPRX1
g - OGFR
h - OPRD1
i - OPRK1
j - OPRM1
PAIRING (ligand → receptor)
PDYN & PCSK1 & PCSK2 & CPE→OPRK1
PDYN & PCSK1 & PCSK2 & CPE→OPRD1
PDYN & PCSK1 & PCSK2 & CPE→OPRM1
PENK & PCSK1 & CPE→OPRD1
PENK & PCSK1 & CPE→OPRM1
PENK & PCSK1 & CPE→OPRK1
PENK & PCSK1 & CPE→OGFR
PENK & PCSK1 & CPE→MRGPRX1
PENK & PCSK2 & CPE→OPRD1
PENK & PCSK2 & CPE→OPRM1
PENK & PCSK2 & CPE→OPRK1
PENK & PCSK2 & CPE→OGFR
PENK & PCSK2 & CPE→MRGPRX1
Compartment
lymphoid
- B cell
- NK/T cell
12.5%
25%
50%
100%
ln(CP10K+1)
Skin
Hypothalamus
Pituitary
Brainstem
Brain cortex
Testes
Uterus
Bladder
Kidney
Pancreas
Liver
Colon
Small intestine
Tongue
Trachea
Lung
Aorta
Heart
Spleen
Blood
Bone marrow
Bone
Diaphragm
Limb muscle
Mammary gland
Fat

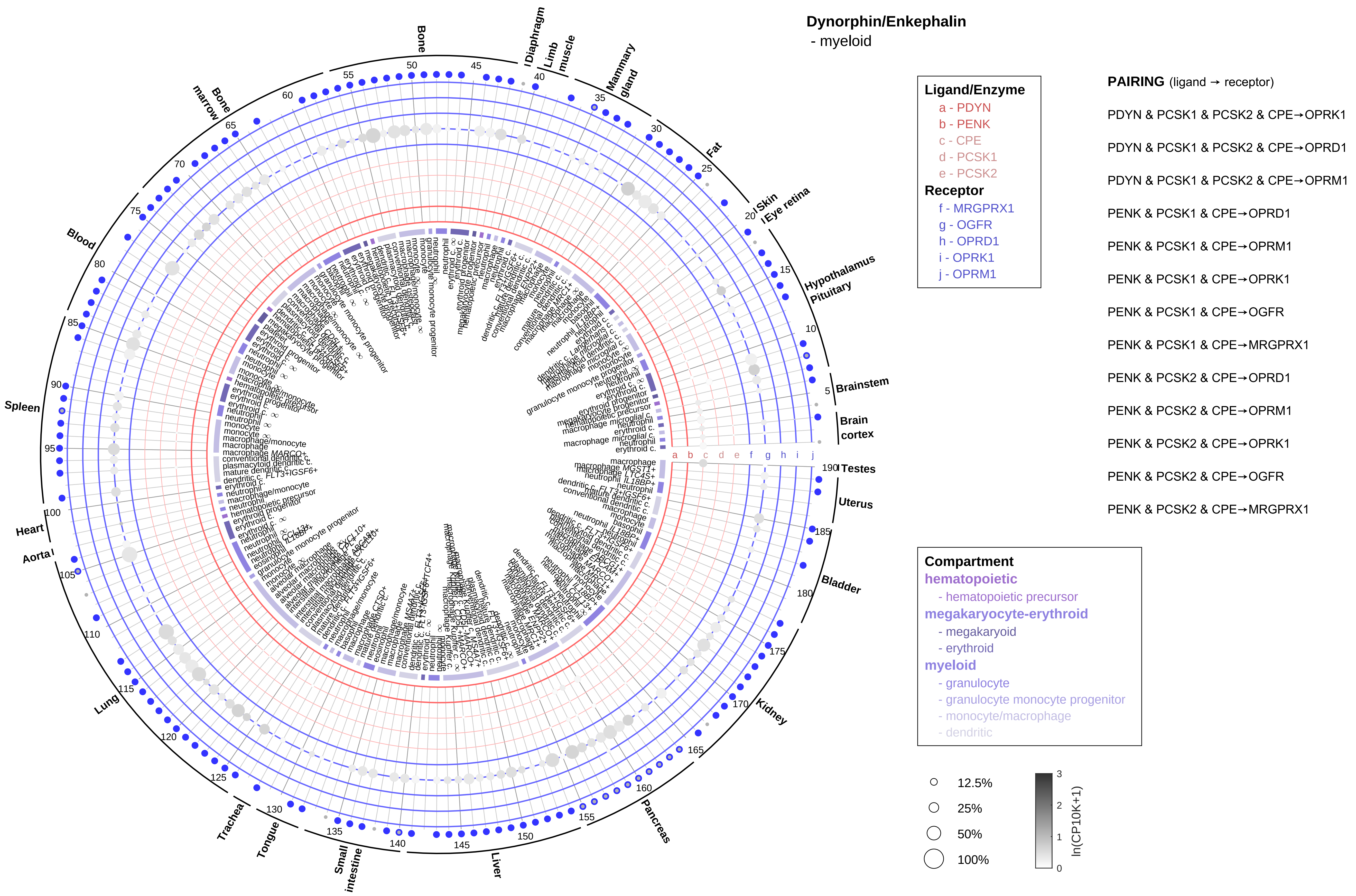
Dynorphin/Enkephalin
- myeloid
Ligand/Enzyme
a - PDYN
b - PENK
c - CPE
d - PCSK1
e - PCSK2
Receptor
f - MRGPRX1
g - OGFR
h - OPRD1
i - OPRK1
j - OPRM1
PAIRING (ligand → receptor)
PDYN & PCSK1 & PCSK2 & CPE→OPRK1
PDYN & PCSK1 & PCSK2 & CPE→OPRD1
PDYN & PCSK1 & PCSK2 & CPE→OPRM1
PENK & PCSK1 & CPE→OPRD1
PENK & PCSK1 & CPE→OPRM1
PENK & PCSK1 & CPE→OPRK1
PENK & PCSK1 & CPE→OGFR
PENK & PCSK1 & CPE→MRGPRX1
PENK & PCSK2 & CPE→OPRD1
PENK & PCSK2 & CPE→OPRM1
PENK & PCSK2 & CPE→OPRK1
PENK & PCSK2 & CPE→OGFR
PENK & PCSK2 & CPE→MRGPRX1
Compartment
hematopoietic
- hematopoietic precursor
megakaryocyte-erythroid
- megakaryoid
- erythroid
myeloid
- granulocyte
- granulocyte monocyte progenitor
- monocyte/macrophage
- dendritic
12.5%
25%
50%
100%
ln(CP10K+1)
Bone
Diaphragm
Limb muscle
Mammary gland
Fat
Skin
Eye retina
Hypothalamus
Pituitary
Brainstem
Brain cortex
Testes
Uterus
Bladder
Kidney
Pancreas
Liver
Small intestine
Tongue
Trachea
Lung
Aorta
Heart
Spleen
Blood
Bone marrow

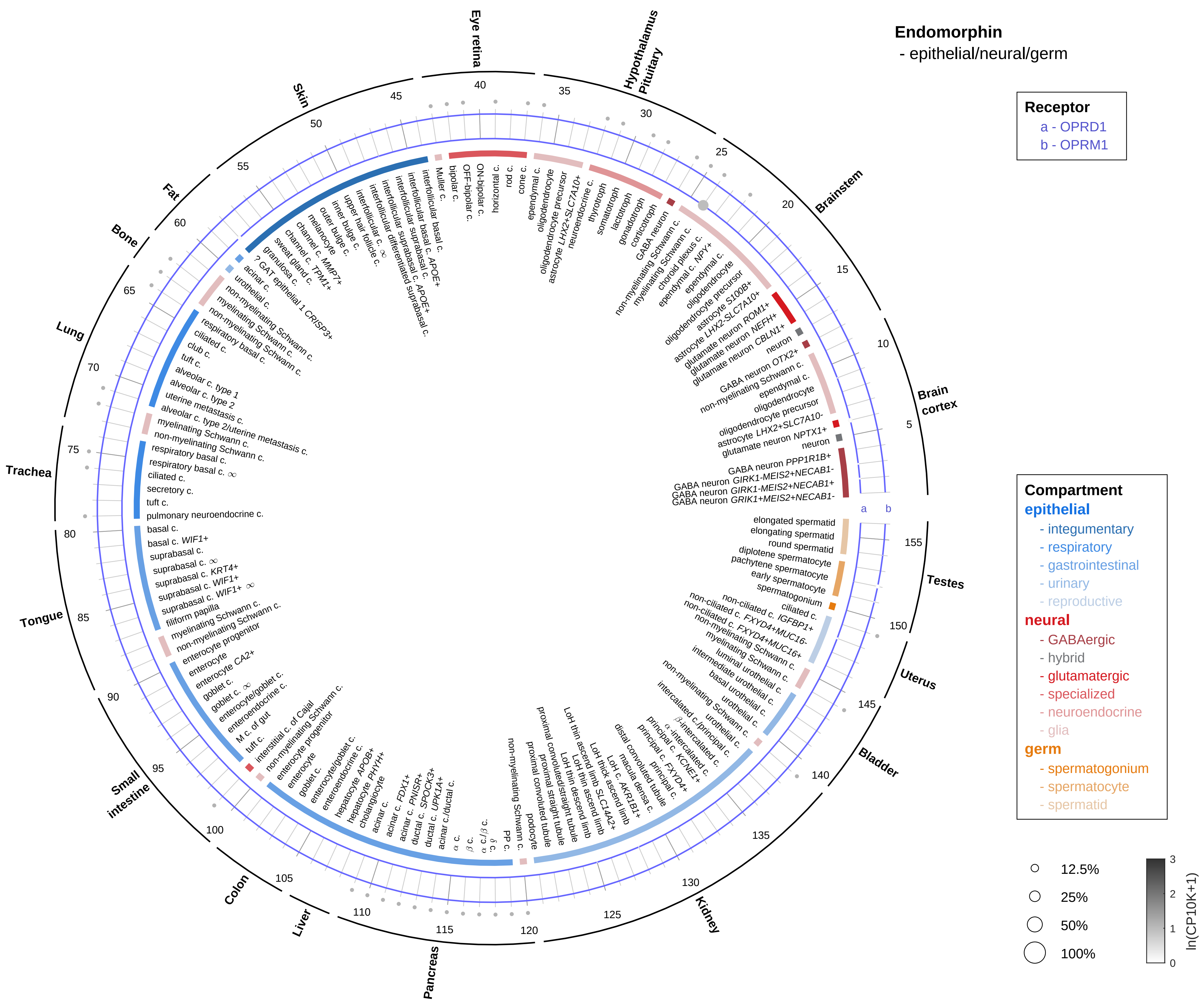

Endomorphin
- epithelial/neural/germ
Receptor
a - OPRD1
b - OPRM1
Compartment
epithelial
- integumentary
- respiratory
- gastrointestinal
- urinary
- reproductive
neural
- GABAergic
- hybrid
- glutamatergic
- specialized
- neuroendocrine
- glia
germ
- spermatogonium
- spermatocyte
- spermatid
12.5%
25%
50%
100%
ln(CP10K+1)
Eye retina
Hypothalamus
Pituitary
Brainstem
Brain cortex
Testes
Uterus
Bladder
Kidney
Pancreas
Liver
Colon
Small intestine
Tongue
Trachea
Lung
Bone
Fat
Skin

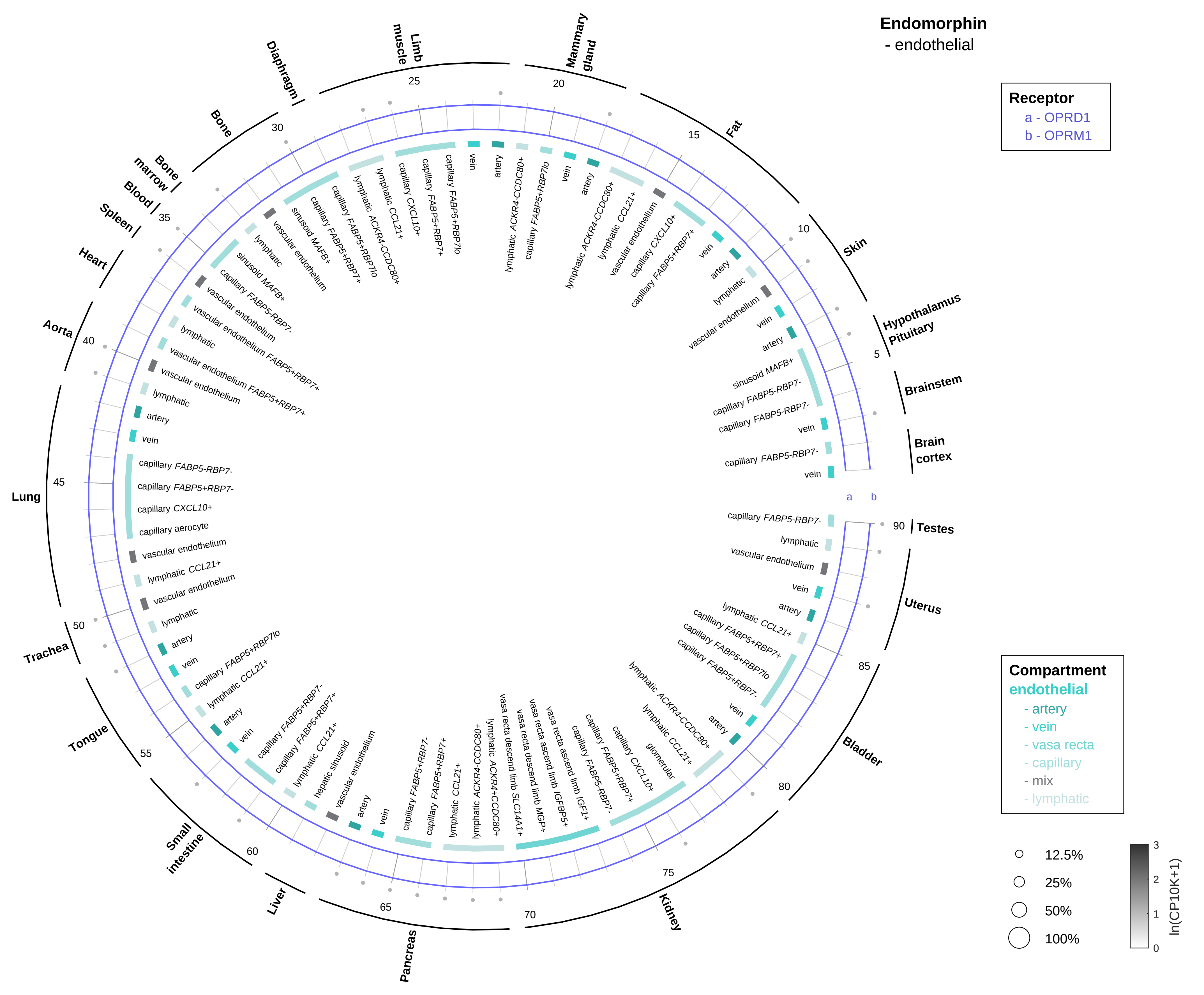
Endomorphin
- endothelial
Receptor
a - OPRD1
b - OPRM1
Compartment
endothelial
- artery
- vein
- vasa recta
- capillary
- mix
- lymphatic
12.5%
25%
50%
100%
ln(CP10K+1)
Brain cortex
Brainstem
Hypothalamus Pituitary
Skin
Fat
Mammary gland
Limb muscle
Diaphragm
Bone
Bone marrow
Blood
Spleen
Heart
Aorta
Lung
Trachea
Tongue
Small intestine
Liver
Pancreas
Kidney
Bladder
Uterus
Testes

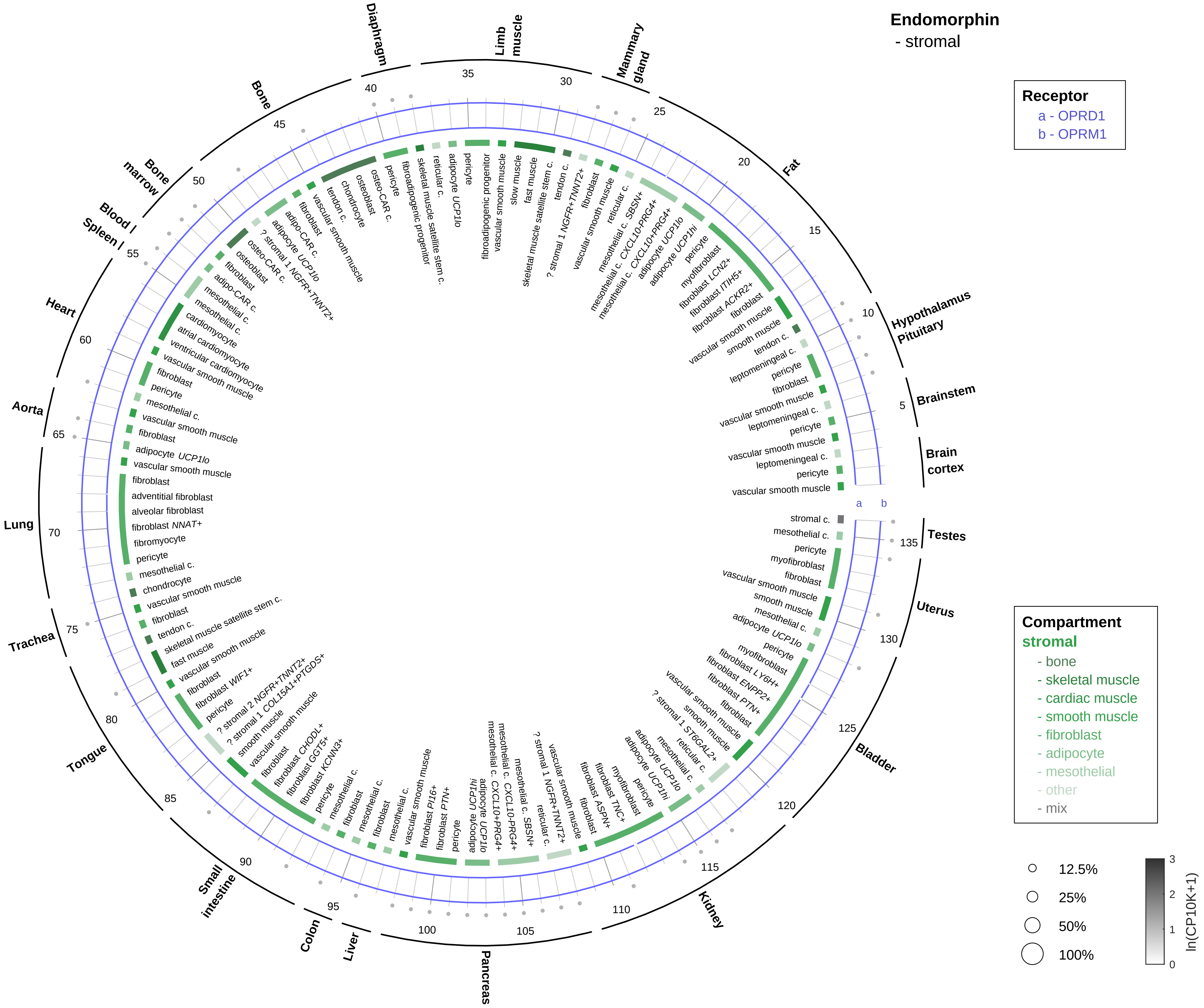

Endomorphin
- stromal
Receptor
a - OPRD1
b - OPRM1
Compartment
stromal
- bone
- skeletal muscle
- cardiac muscle
- smooth muscle
- fibroblast
- adipocyte
- mesothelial
- other
- mix
12.5%
25%
50%
100%
ln(CP10K+1)
Brain cortex
Brainstem
Hypothalamus Pituitary
Fat
Mammary gland
Limb muscle
Diaphragm
Bone
Bone marrow
Blood
Spleen
Heart
Aorta
Lung
Trachea
Tongue
Small intestine
Colon
Liver
Pancreas
Kidney
Bladder
Uterus
Testes

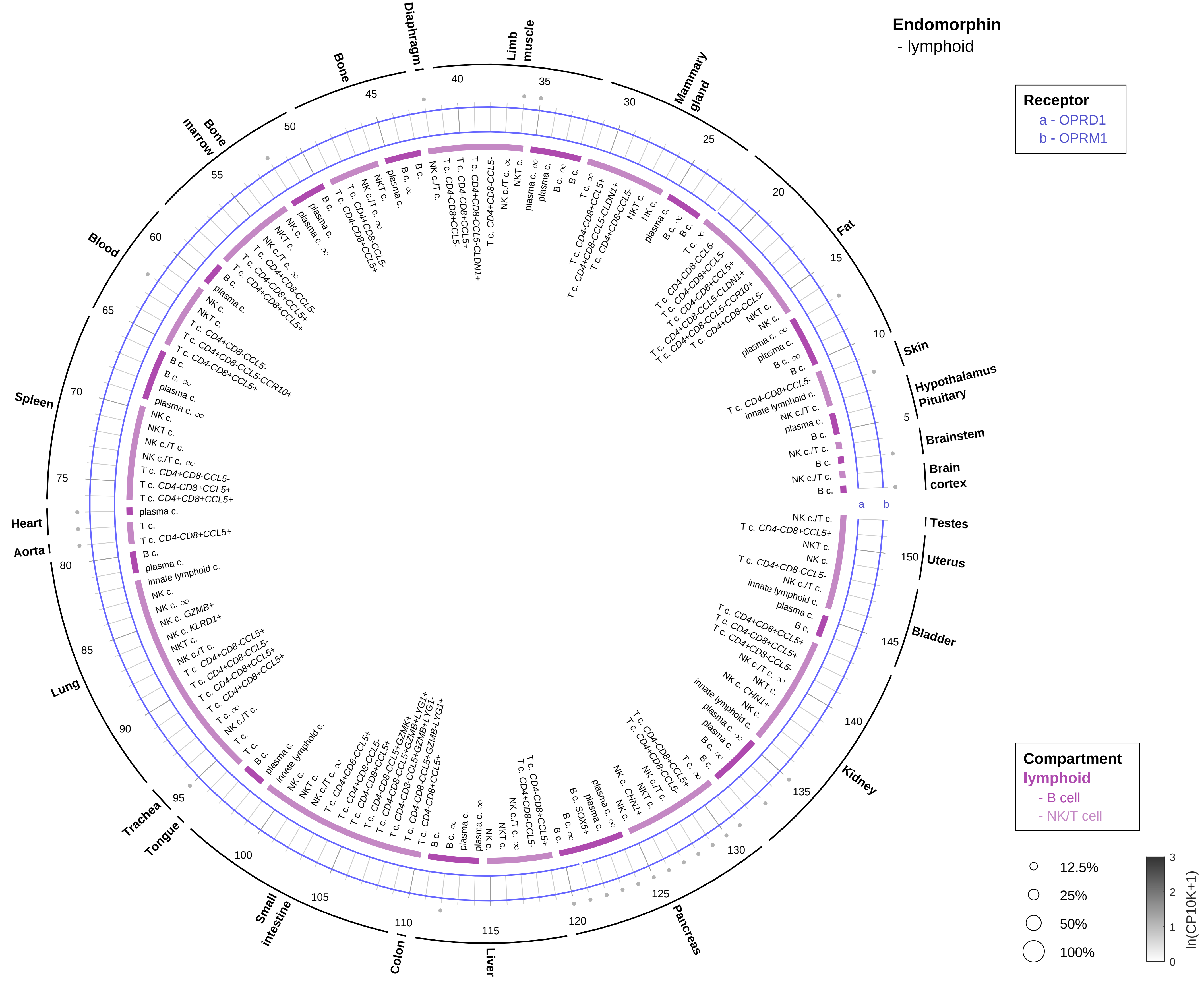

Endomorphin
- lymphoid
Receptor
a - OPRD1
b - OPRM1
Compartment
lymphoid
- B cell
- NK/T cell
12.5%
25%
50%
100%
ln(CP10K+1)
Skin
Hypothalamus
Pituitary
Brainstem
Brain cortex
Testes
Uterus
Bladder
Kidney
Pancreas
Liver
Colon
Small intestine
Tongue
Trachea
Lung
Aorta
Heart
Spleen
Blood
Bone marrow
Bone
Diaphragm
Limb muscle
Mammary gland
Fat

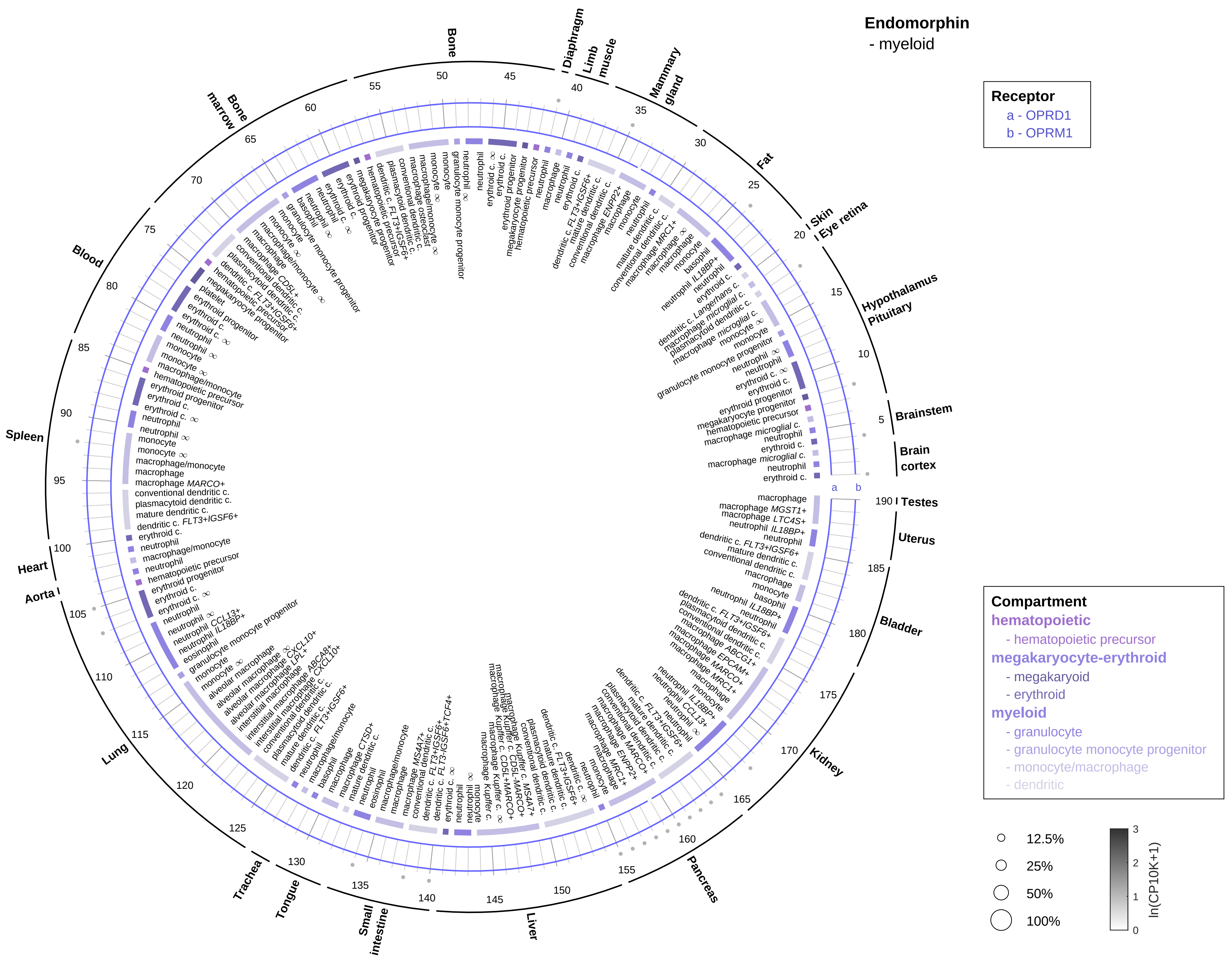
Endomorphin
- myeloid
Receptor
a - OPRD1
b - OPRM1
Compartment
hematopoietic
- hematopoietic precursor
megakaryocyte-erythroid
- megakaryoid
- erythroid
myeloid
- granulocyte
- granulocyte monocyte progenitor
- monocyte/macrophage
- dendritic
12.5%
25%
50%
100%
ln(CP10K+1)
Bone
Diaphragm
Limb muscle
Mammary gland
Fat
Skin
Eye retina
Hypothalamus
Pituitary
Brainstem
Brain cortex
Testes
Uterus
Bladder
Kidney
Pancreas
Liver
Small intestine
Tongue
Trachea
Lung
Aorta
Heart
Spleen
Blood
Bone marrow

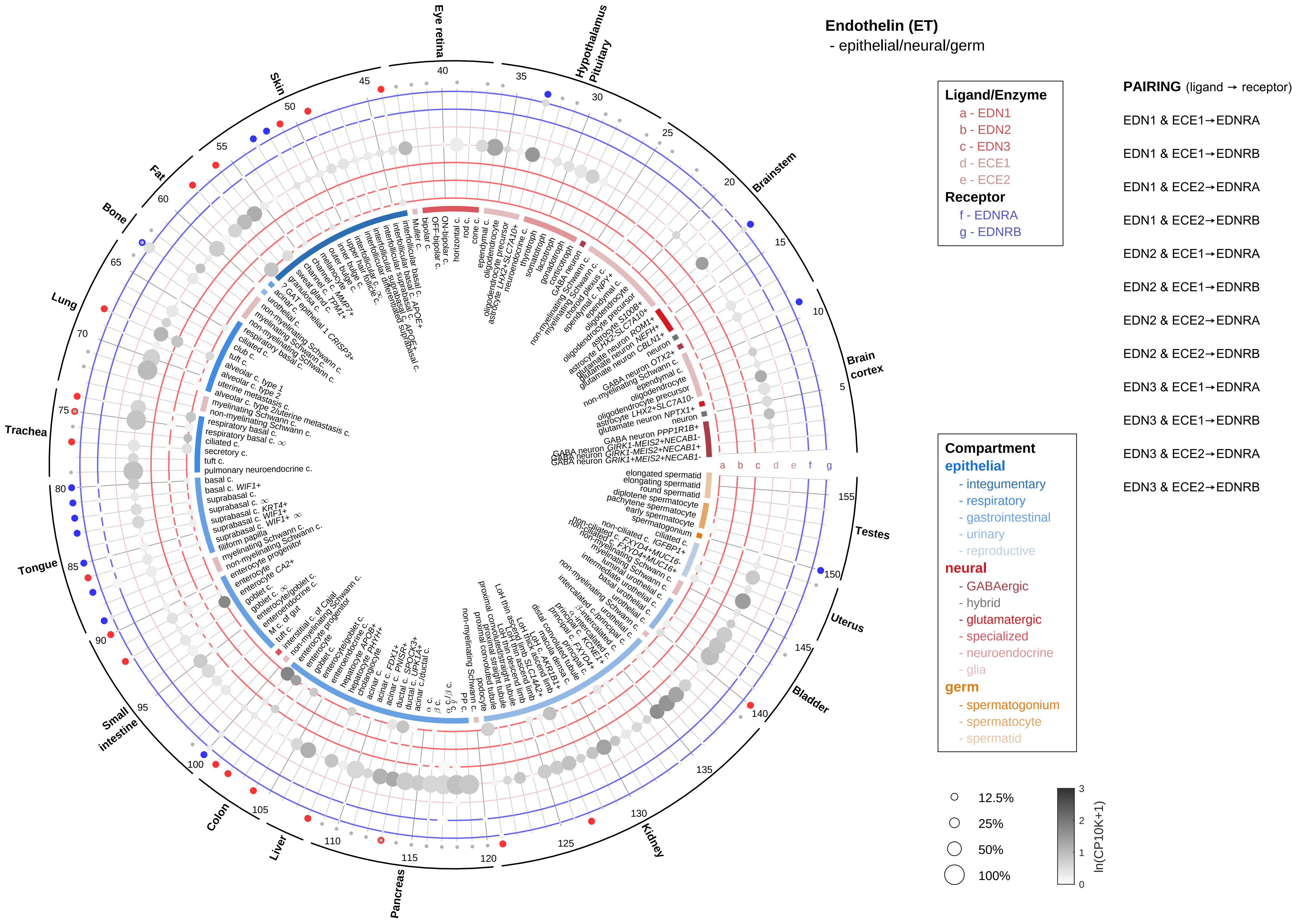

Endothelin (ET)
- epithelial/neural/germ
Ligand/Enzyme
a - EDN1
b - EDN2
c - EDN3
d - ECE1
e - ECE2
Receptor
f - EDNRA
g - EDNRB
PAIRING (ligand → receptor)
EDN1 & ECE1→EDNRA
EDN1 & ECE1→EDNRB
EDN1 & ECE2→EDNRA
EDN1 & ECE2→EDNRB
EDN2 & ECE1→EDNRA
EDN2 & ECE1→EDNRB
EDN2 & ECE2→EDNRA
EDN2 & ECE2→EDNRB
EDN3 & ECE1→EDNRA
EDN3 & ECE1→EDNRB
EDN3 & ECE2→EDNRA
EDN3 & ECE2→EDNRB
Compartment
epithelial
- integumentary
- respiratory
- gastrointestinal
- urinary
- reproductive
neural
- GABAergic
- hybrid
- glutamatergic
- specialized
- neuroendocrine
- glia
germ
- spermatogonium
- spermatocyte
- spermatid
12.5%
25%
50%
100%
ln(CP10K+1)
Eye retina
Hypothalamus
Pituitary
Brainstem
Brain cortex
Testes
Uterus
Bladder
Kidney
Pancreas
Liver
Colon
Small intestine
Tongue
Trachea
Lung
Bone
Fat
Skin
a b c d e f g

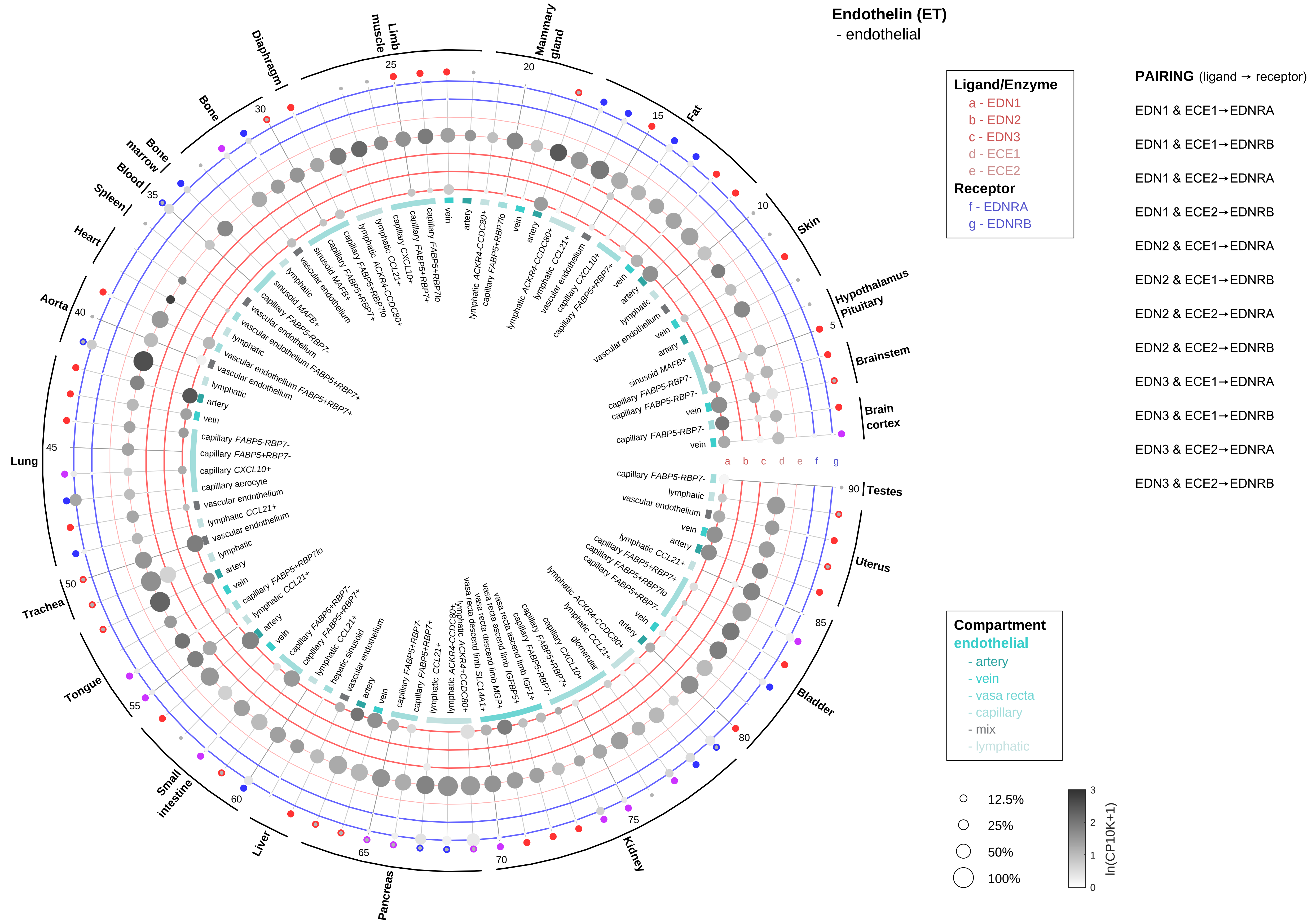

Endothelin (ET)
- endothelial
Ligand/Enzyme
a - EDN1
b - EDN2
c - EDN3
d - ECE1
e - ECE2
Receptor
f - EDNRA
g - EDNRB
PAIRING (ligand → receptor)
EDN1 & ECE1→EDNRA
EDN1 & ECE1→EDNRB
EDN1 & ECE2→EDNRA
EDN1 & ECE2→EDNRB
EDN2 & ECE1→EDNRA
EDN2 & ECE1→EDNRB
EDN2 & ECE2→EDNRA
EDN2 & ECE2→EDNRB
EDN3 & ECE1→EDNRA
EDN3 & ECE1→EDNRB
EDN3 & ECE2→EDNRA
EDN3 & ECE2→EDNRB
Compartment
endothelial
- artery
- vein
- vasa recta
- capillary
- mix
- lymphatic
12.5%
25%
50%
100%
ln(CP10K+1)
Testes
Uterus
Bladder
Kidney
Pancreas
Liver
Small intestine
Tongue
Trachea
Lung
Aorta
Heart
Spleen
Blood
Bone marrow
Bone
Diaphragm
Limb muscle
Mammary gland
Fat
Skin
Hypothalamus Pituitary
Brainstem
Brain cortex

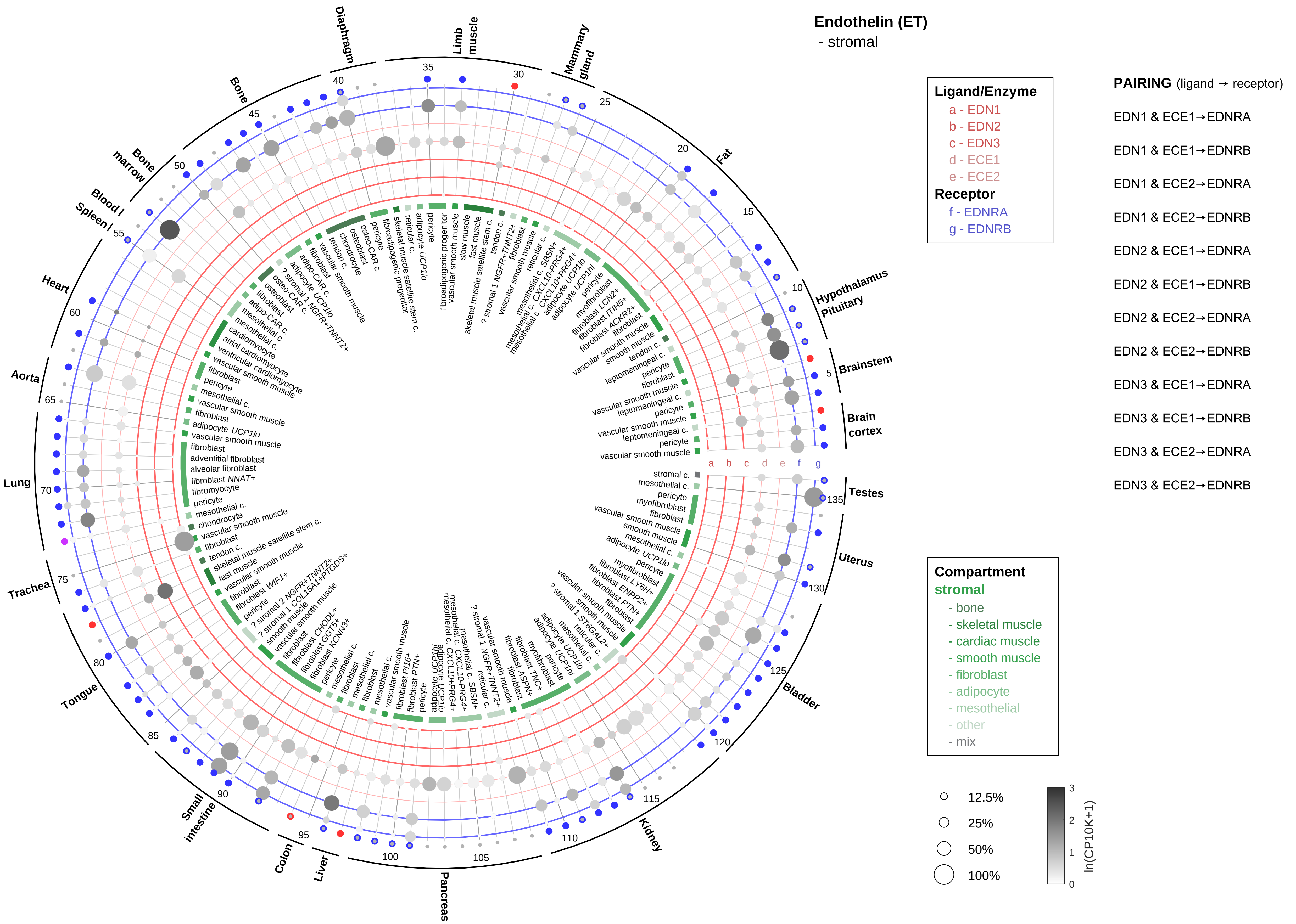
Endothelin (ET)
- stromal
Ligand/Enzyme
a - EDN1
b - EDN2
c - EDN3
d - ECE1
e - ECE2
Receptor
f - EDNRA
g - EDNRB
PAIRING (ligand → receptor)
EDN1 & ECE1→EDNRA
EDN1 & ECE1→EDNRB
EDN1 & ECE2→EDNRA
EDN1 & ECE2→EDNRB
EDN2 & ECE1→EDNRA
EDN2 & ECE1→EDNRB
EDN2 & ECE2→EDNRA
EDN2 & ECE2→EDNRB
EDN3 & ECE1→EDNRA
EDN3 & ECE1→EDNRB
EDN3 & ECE2→EDNRA
EDN3 & ECE2→EDNRB
Compartment
stromal
- bone
- skeletal muscle
- cardiac muscle
- smooth muscle
- fibroblast
- adipocyte
- mesothelial
- other
- mix
12.5%
25%
50%
100%
ln(CP10K+1)
Diaphragm
Limb muscle
Mammary gland
Fat
Hypothalamus Pituitary
Brainstem
Brain cortex
Testes
Uterus
Bladder
Kidney
Pancreas
Liver
Colon
Small intestine
Tongue
Trachea
Lung
Aorta
Heart
Spleen
Blood
Bone marrow
Bone

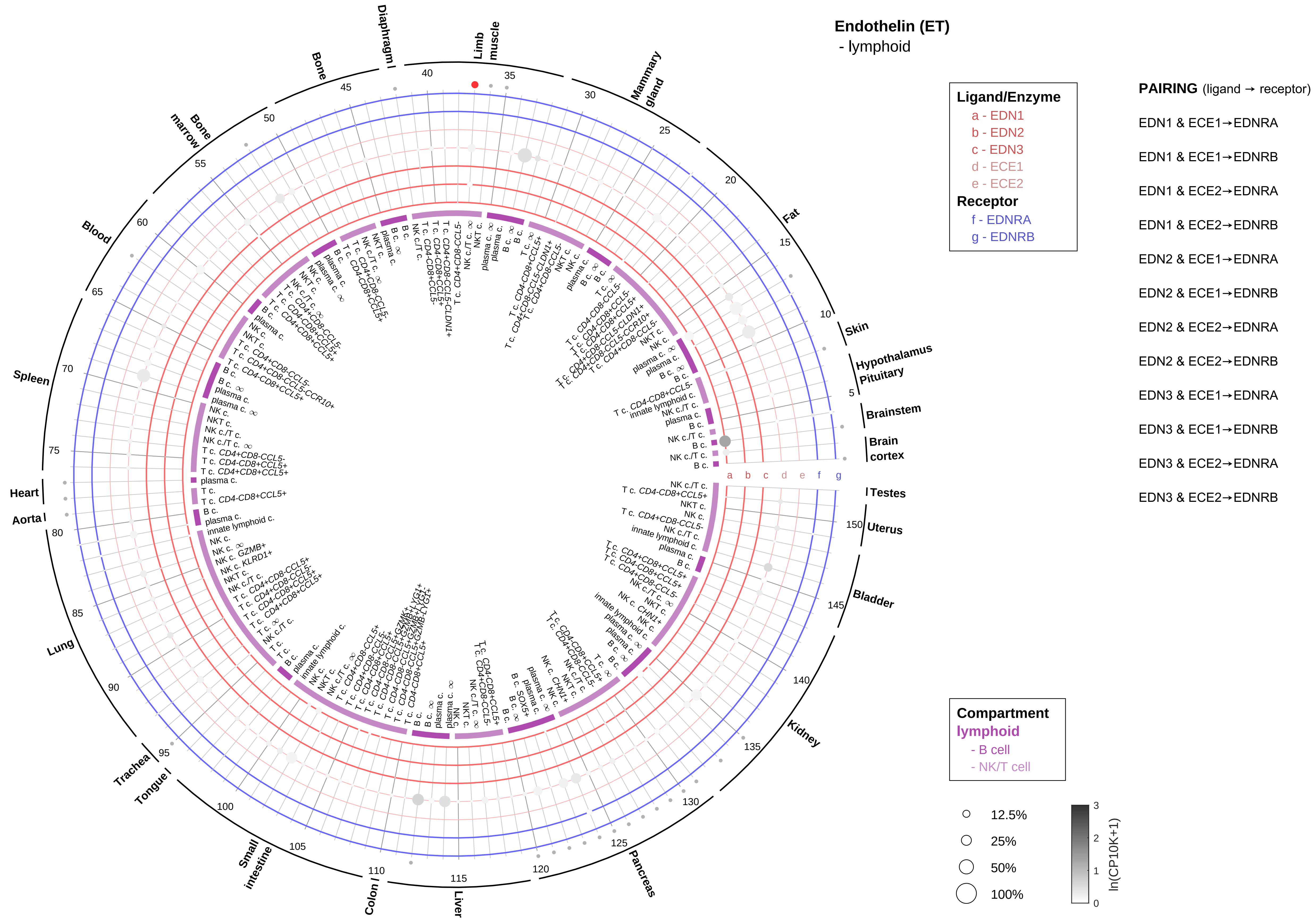
Endothelin (ET)
- lymphoid
Ligand/Enzyme
a - EDN1
b - EDN2
c - EDN3
d - ECE1
e - ECE2
Receptor
f - EDNRA
g - EDNRB
PAIRING (ligand → receptor)
EDN1 & ECE1→EDNRA
EDN1 & ECE1→EDNRB
EDN1 & ECE2→EDNRA
EDN1 & ECE2→EDNRB
EDN2 & ECE1→EDNRA
EDN2 & ECE1→EDNRB
EDN2 & ECE2→EDNRA
EDN2 & ECE2→EDNRB
EDN3 & ECE1→EDNRA
EDN3 & ECE1→EDNRB
EDN3 & ECE2→EDNRA
EDN3 & ECE2→EDNRB
Compartment
lymphoid
- B cell
- NK/T cell
12.5%
25%
50%
100%
ln(CP10K+1)
Limb muscle
Mammary gland
Fat
Skin
Hypothalamus
Pituitary
Brainstem
Brain cortex
Testes
Uterus
Bladder
Kidney
Pancreas
Liver
Colon
Small intestine
Tongue
Trachea
Lung
Aorta
Heart
Spleen
Blood
Bone marrow
Bone
Diaphragm

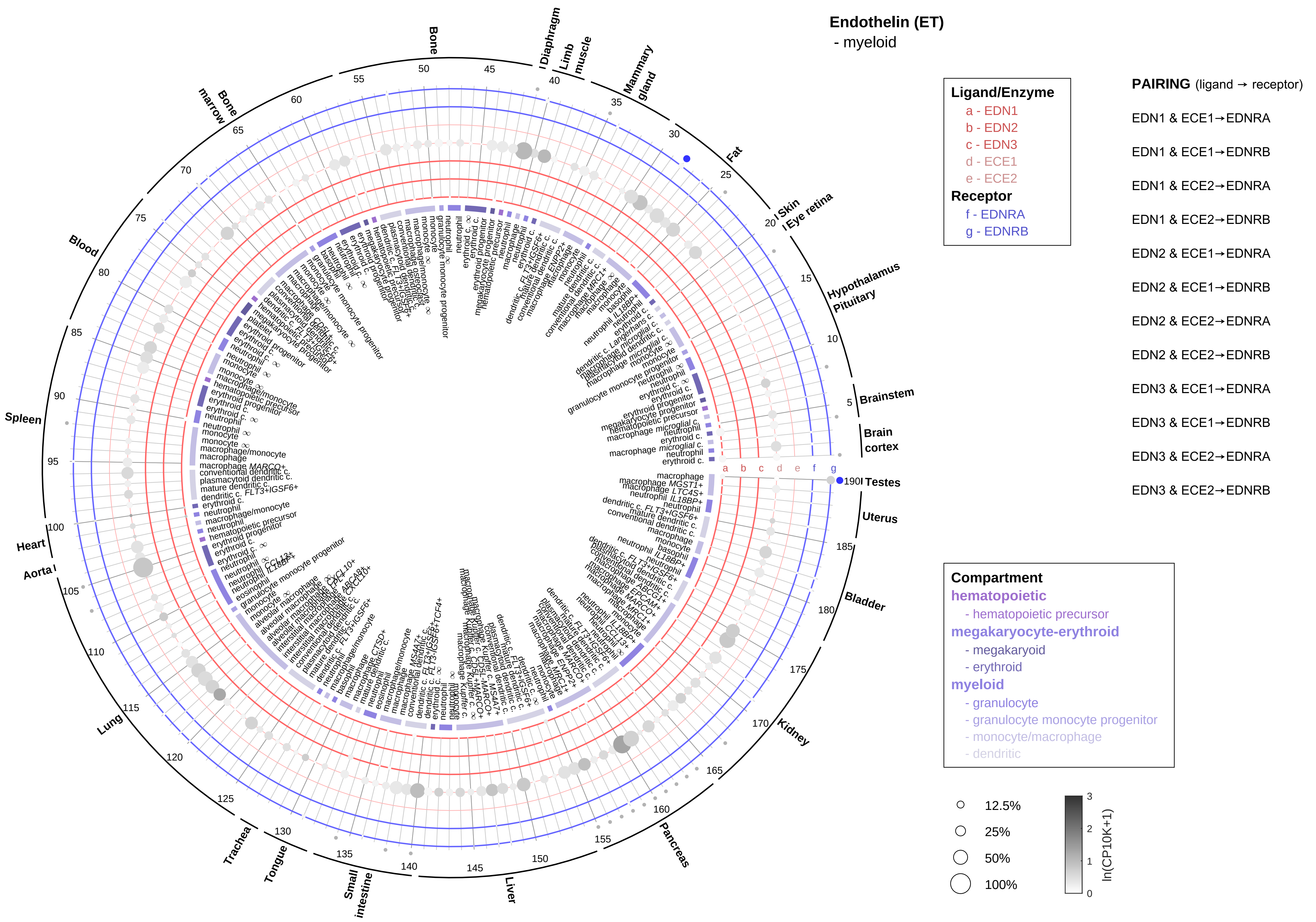
Endothelin (ET)
- myeloid
Ligand/Enzyme
a - EDN1
b - EDN2
c - EDN3
d - ECE1
e - ECE2
Receptor
f - EDNRA
g - EDNRB
PAIRING (ligand → receptor)
EDN1 & ECE1→EDNRA
EDN1 & ECE1→EDNRB
EDN1 & ECE2→EDNRA
EDN1 & ECE2→EDNRB
EDN2 & ECE1→EDNRA
EDN2 & ECE1→EDNRB
EDN2 & ECE2→EDNRA
EDN2 & ECE2→EDNRB
EDN3 & ECE1→EDNRA
EDN3 & ECE1→EDNRB
EDN3 & ECE2→EDNRA
EDN3 & ECE2→EDNRB
Compartment
hematopoietic
- hematopoietic precursor
megakaryocyte-erythroid
- megakaryoid
- erythroid
myeloid
- granulocyte
- granulocyte monocyte progenitor
- monocyte/macrophage
- dendritic
12.5%
25%
50%
100%
ln(CP10K+1)
Testes
Brain cortex
Brainstem
Hypothalamus
Pituitary
Eye retina
Skin
Fat
Mammary gland
Limb muscle
Diaphragm
Bone
Bone marrow
Blood
Spleen
Heart
Aorta
Lung
Trachea
Tongue
Small intestine
Liver
Pancreas
Kidney
Bladder
Uterus

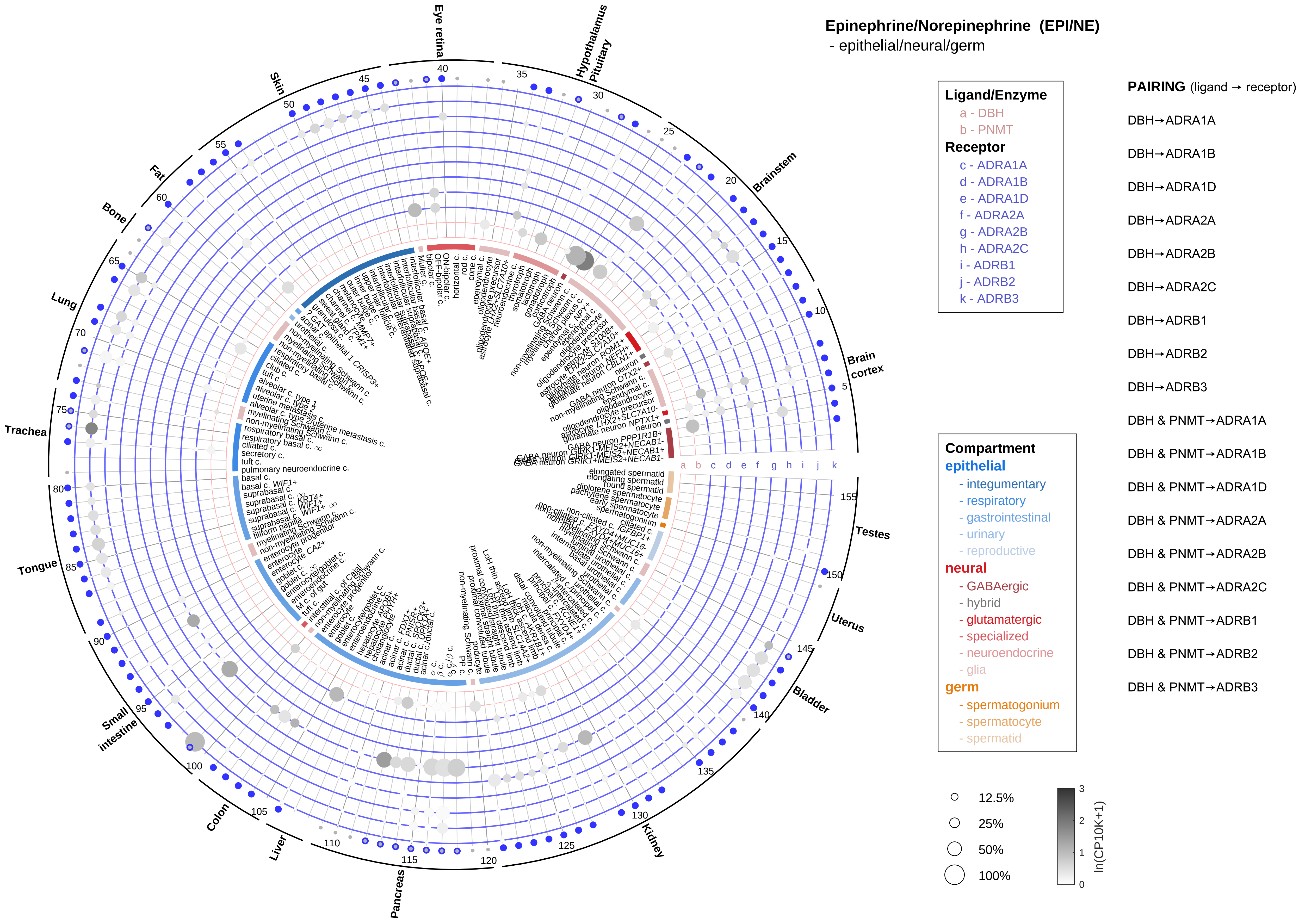
Epinephrine/Norepinephrine (EPI/NE)
- epithelial/neural/germ
Ligand/Enzyme
a - DBH
b - PNMT
Receptor
c - ADRA1A
d - ADRA1B
e - ADRA1D
f - ADRA2A
g - ADRA2B
h - ADRA2C
i - ADRB1
j - ADRB2
k - ADRB3
PAIRING (ligand → receptor)
DBH→ADRA1A
DBH→ADRA1B
DBH→ADRA1D
DBH→ADRA2A
DBH→ADRA2B
DBH→ADRA2C
DBH→ADRB1
DBH→ADRB2
DBH→ADRB3
DBH & PNMT→ADRA1A
DBH & PNMT→ADRA1B
DBH & PNMT→ADRA1D
DBH & PNMT→ADRA2A
DBH & PNMT→ADRA2B
DBH & PNMT→ADRA2C
DBH & PNMT→ADRB1
DBH & PNMT→ADRB2
DBH & PNMT→ADRB3
Compartment
epithelial
- integumentary
- respiratory
- gastrointestinal
- urinary
- reproductive
neural
- GABAergic
- hybrid
- glutamatergic
- specialized
- neuroendocrine
- glia
germ
- spermatogonium
- spermatocyte
- spermatid
12.5%
25%
50%
100%
ln(CP10K+1)
Eye retina
Hypothalamus
Pituitary
Brainstem
Brain cortex
Testes
Uterus
Bladder
Kidney
Pancreas
Liver
Colon
Small intestine
Tongue
Trachea
Lung
Bone
Fat
Skin

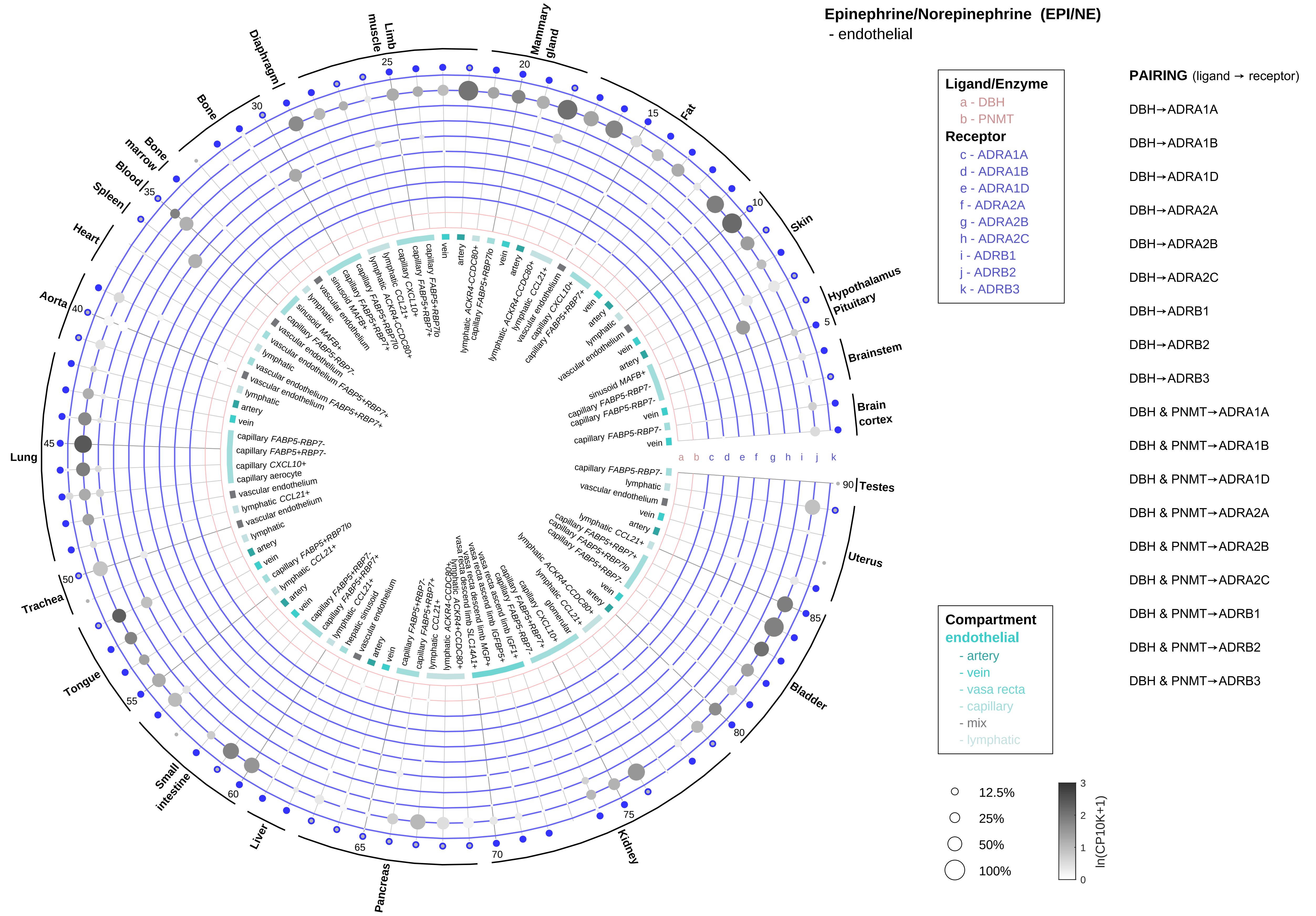

Epinephrine/Norepinephrine (EPI/NE)
- endothelial
Ligand/Enzyme
a - DBH
b - PNMT
Receptor
c - ADRA1A
d - ADRA1B
e - ADRA1D
f - ADRA2A
g - ADRA2B
h - ADRA2C
i - ADRB1
j - ADRB2
k - ADRB3
PAIRING (ligand → receptor)
DBH→ADRA1A
DBH→ADRA1B
DBH→ADRA1D
DBH→ADRA2A
DBH→ADRA2B
DBH→ADRA2C
DBH→ADRB1
DBH→ADRB2
DBH→ADRB3
DBH & PNMT→ADRA1A
DBH & PNMT→ADRA1B
DBH & PNMT→ADRA1D
DBH & PNMT→ADRA2A
DBH & PNMT→ADRA2B
DBH & PNMT→ADRA2C
DBH & PNMT→ADRB1
DBH & PNMT→ADRB2
DBH & PNMT→ADRB3
Compartment
endothelial
- artery
- vein
- vasa recta
- capillary
- mix
- lymphatic
12.5%
25%
50%
100%
ln(CP10K+1)
Hypothalamus
Pituitary
Brainstem
Brain cortex
Testes
Uterus
Bladder
Kidney
Pancreas
Liver
Small intestine
Tongue
Trachea
Lung
Aorta
Heart
Spleen
Blood
Bone marrow
Bone
Diaphragm
Limb muscle
Mammary gland
Fat
Skin

#### Epinephrine/Norepinephrine (EPI/NE)
- stromal

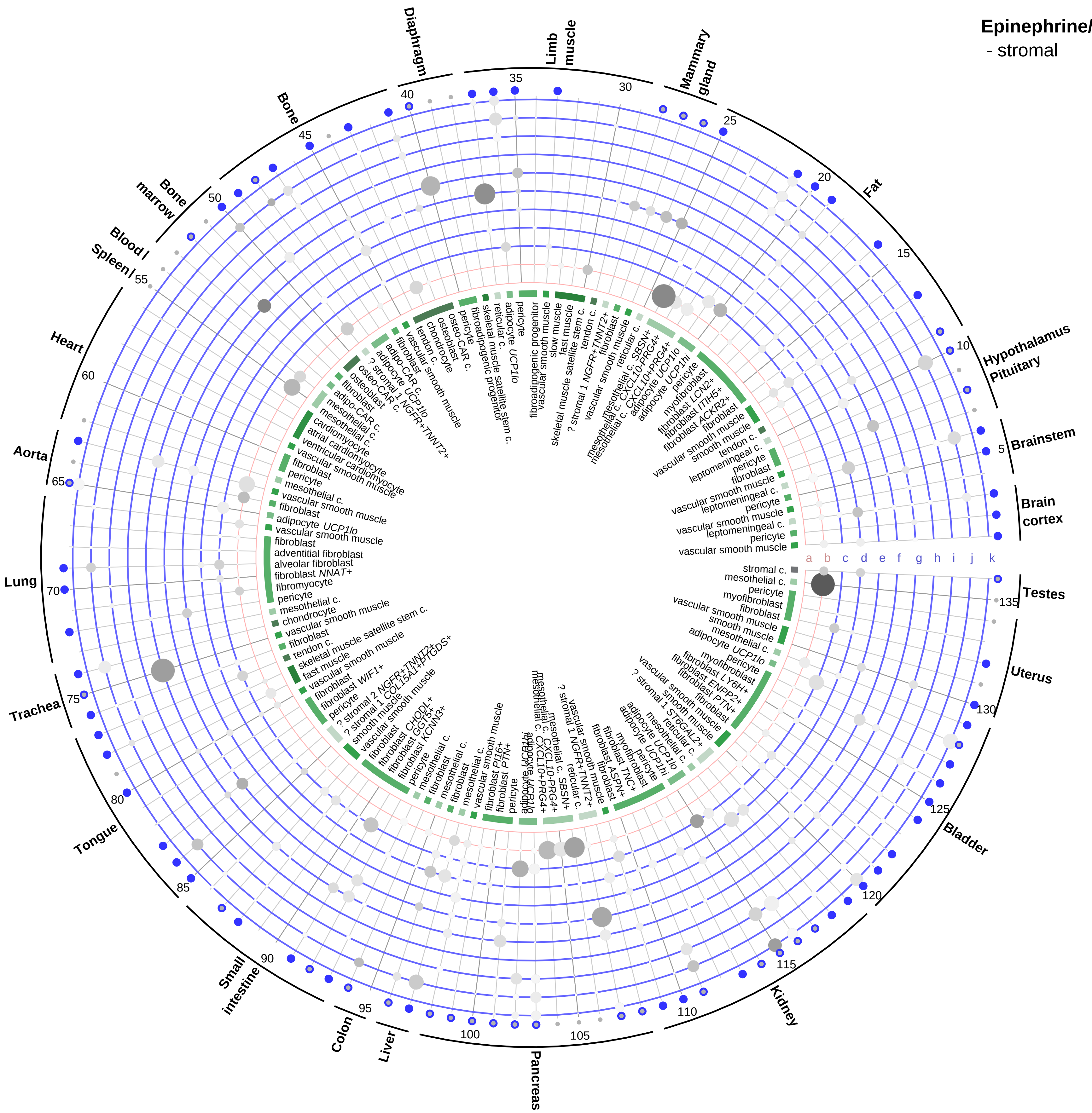

**Ligand/Enzyme**
- a - DBH
- b - PNMT

**Receptor**
- c - ADRA1A
- d - ADRA1B
- e - ADRA1D
- f - ADRA2A
- g - ADRA2B
- h - ADRA2C
- i - ADRB1
- j - ADRB2
- k - ADRB3

**Compartment**
**stromal**
- bone
- skeletal muscle
- cardiac muscle
- smooth muscle
- fibroblast
- adipocyte
- mesothelial
- other
- mix

12.5%
25%
50%
100%

ln(CP10K+1) 0 1 2 3

**PAIRING** (ligand → receptor)

DBH→ADRA1A
DBH→ADRA1B
DBH→ADRA1D
DBH→ADRA2A
DBH→ADRA2B
DBH→ADRA2C
DBH→ADRB1
DBH→ADRB2
DBH→ADRB3
DBH & PNMT→ADRA1A
DBH & PNMT→ADRA1B
DBH & PNMT→ADRA1D
DBH & PNMT→ADRA2A
DBH & PNMT→ADRA2B
DBH & PNMT→ADRA2C
DBH & PNMT→ADRB1
DBH & PNMT→ADRB2
DBH & PNMT→ADRB3

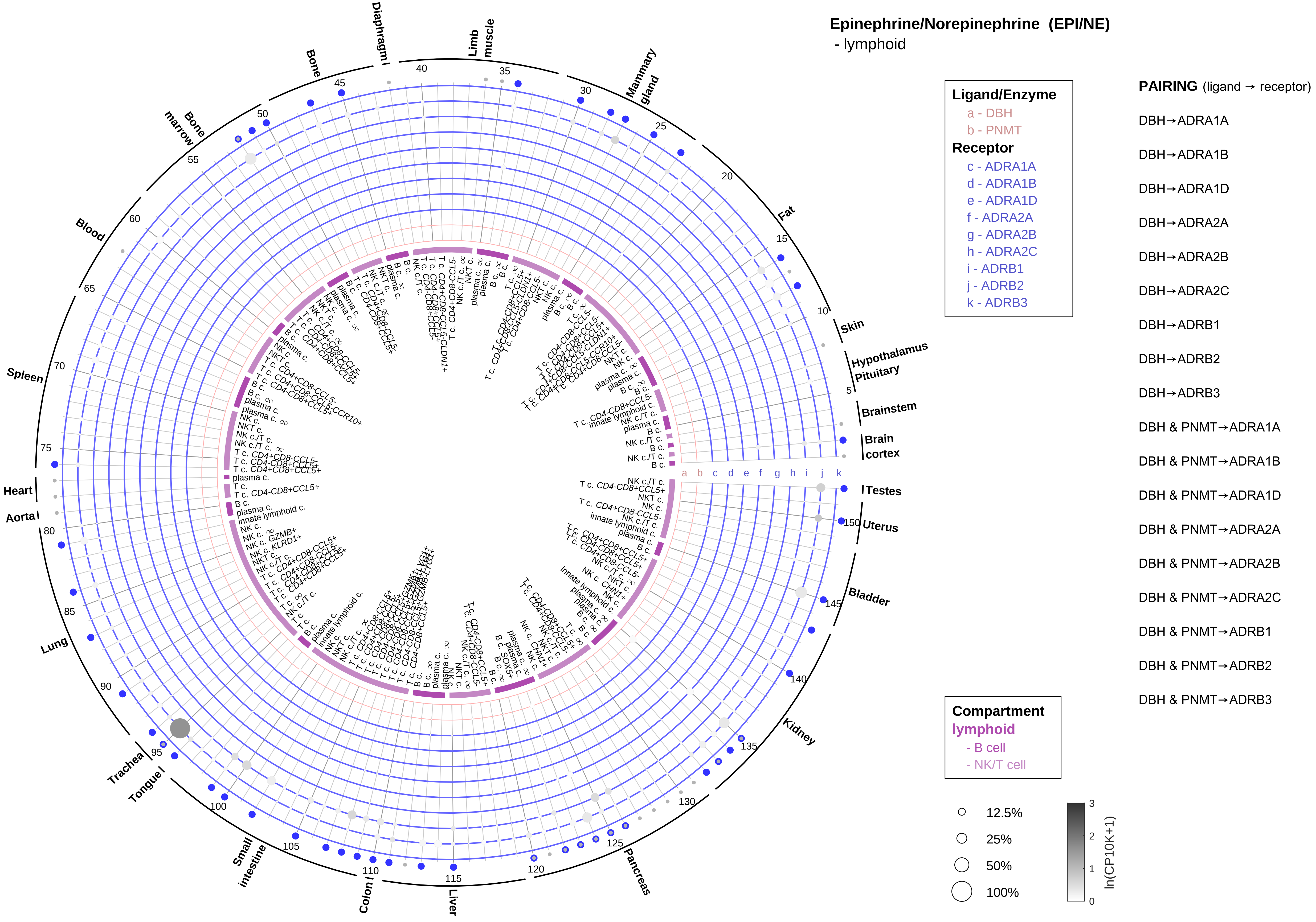
Epinephrine/Norepinephrine (EPI/NE)
- lymphoid
Ligand/Enzyme
a - DBH
b - PNMT
Receptor
c - ADRA1A
d - ADRA1B
e - ADRA1D
f - ADRA2A
g - ADRA2B
h - ADRA2C
i - ADRB1
j - ADRB2
k - ADRB3
PAIRING (ligand → receptor)
DBH→ADRA1A
DBH→ADRA1B
DBH→ADRA1D
DBH→ADRA2A
DBH→ADRA2B
DBH→ADRA2C
DBH→ADRB1
DBH→ADRB2
DBH→ADRB3
DBH & PNMT→ADRA1A
DBH & PNMT→ADRA1B
DBH & PNMT→ADRA1D
DBH & PNMT→ADRA2A
DBH & PNMT→ADRA2B
DBH & PNMT→ADRA2C
DBH & PNMT→ADRB1
DBH & PNMT→ADRB2
DBH & PNMT→ADRB3
Compartment
lymphoid
- B cell
- NK/T cell
12.5%
25%
50%
100%
ln(CP10K+1)
Diaphragm
Limb muscle
Mammary gland
Fat
Skin
Hypothalamus
Pituitary
Brainstem
Brain cortex
Testes
Uterus
Bladder
Kidney
Pancreas
Liver
Colon
Small intestine
Tongue
Trachea
Lung
Aorta
Heart
Spleen
Blood
Bone marrow
Bone

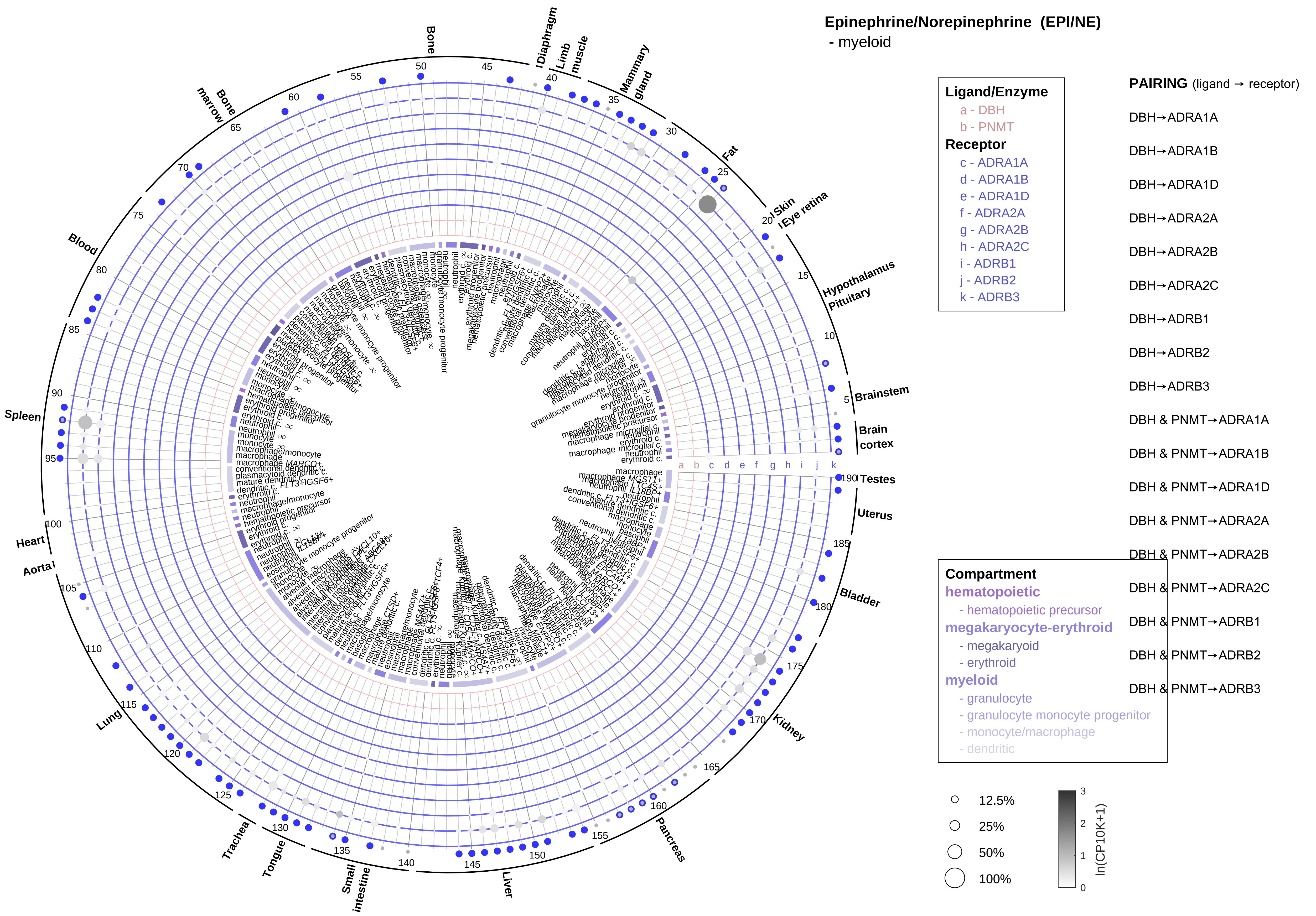
Epinephrine/Norepinephrine (EPI/NE)
- myeloid
Ligand/Enzyme
a - DBH
b - PNMT
Receptor
c - ADRA1A
d - ADRA1B
e - ADRA1D
f - ADRA2A
g - ADRA2B
h - ADRA2C
i - ADRB1
j - ADRB2
k - ADRB3
PAIRING (ligand → receptor)
DBH→ADRA1A
DBH→ADRA1B
DBH→ADRA1D
DBH→ADRA2A
DBH→ADRA2B
DBH→ADRA2C
DBH→ADRB1
DBH→ADRB2
DBH→ADRB3
DBH & PNMT→ADRA1A
DBH & PNMT→ADRA1B
DBH & PNMT→ADRA1D
DBH & PNMT→ADRA2A
DBH & PNMT→ADRA2B
DBH & PNMT→ADRA2C
DBH & PNMT→ADRB1
DBH & PNMT→ADRB2
DBH & PNMT→ADRB3
Compartment
hematopoietic
- hematopoietic precursor
megakaryocyte-erythroid
- megakaryoid
- erythroid
myeloid
- granulocyte
- granulocyte monocyte progenitor
- monocyte/macrophage
- dendritic
12.5%
25%
50%
100%
ln(CP10K+1)
Bone
Diaphragm
Limb muscle
Mammary gland
Fat
Skin
Eye retina
Hypothalamus
Pituitary
Brainstem
Brain cortex
Testes
Uterus
Bladder
Kidney
Pancreas
Liver
Small intestine
Tongue
Trachea
Lung
Aorta
Heart
Spleen
Blood
Bone marrow

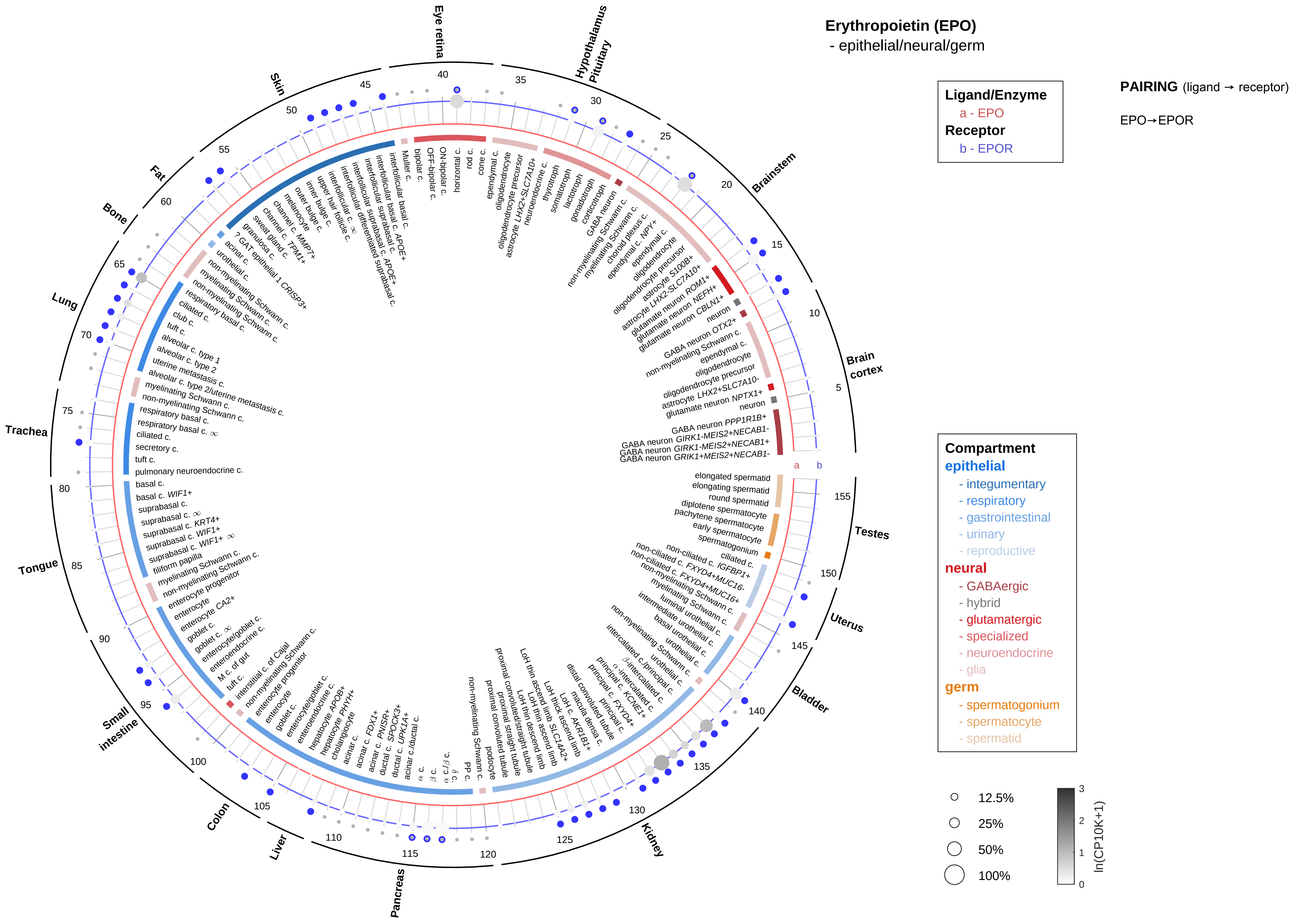

Erythropoietin (EPO)
- epithelial/neural/germ
Ligand/Enzyme
a - EPO
Receptor
b - EPOR
PAIRING (ligand → receptor)
EPO→EPOR
Compartment
epithelial
- integumentary
- respiratory
- gastrointestinal
- urinary
- reproductive
neural
- GABAergic
- hybrid
- glutamatergic
- specialized
- neuroendocrine
- glia
germ
- spermatogonium
- spermatocyte
- spermatid
12.5%
25%
50%
100%
ln(CP10K+1)
Eye retina
Hypothalamus
Pituitary
Brainstem
Brain cortex
Testes
Uterus
Bladder
Kidney
Pancreas
Liver
Colon
Small intestine
Tongue
Trachea
Lung
Bone
Fat
Skin

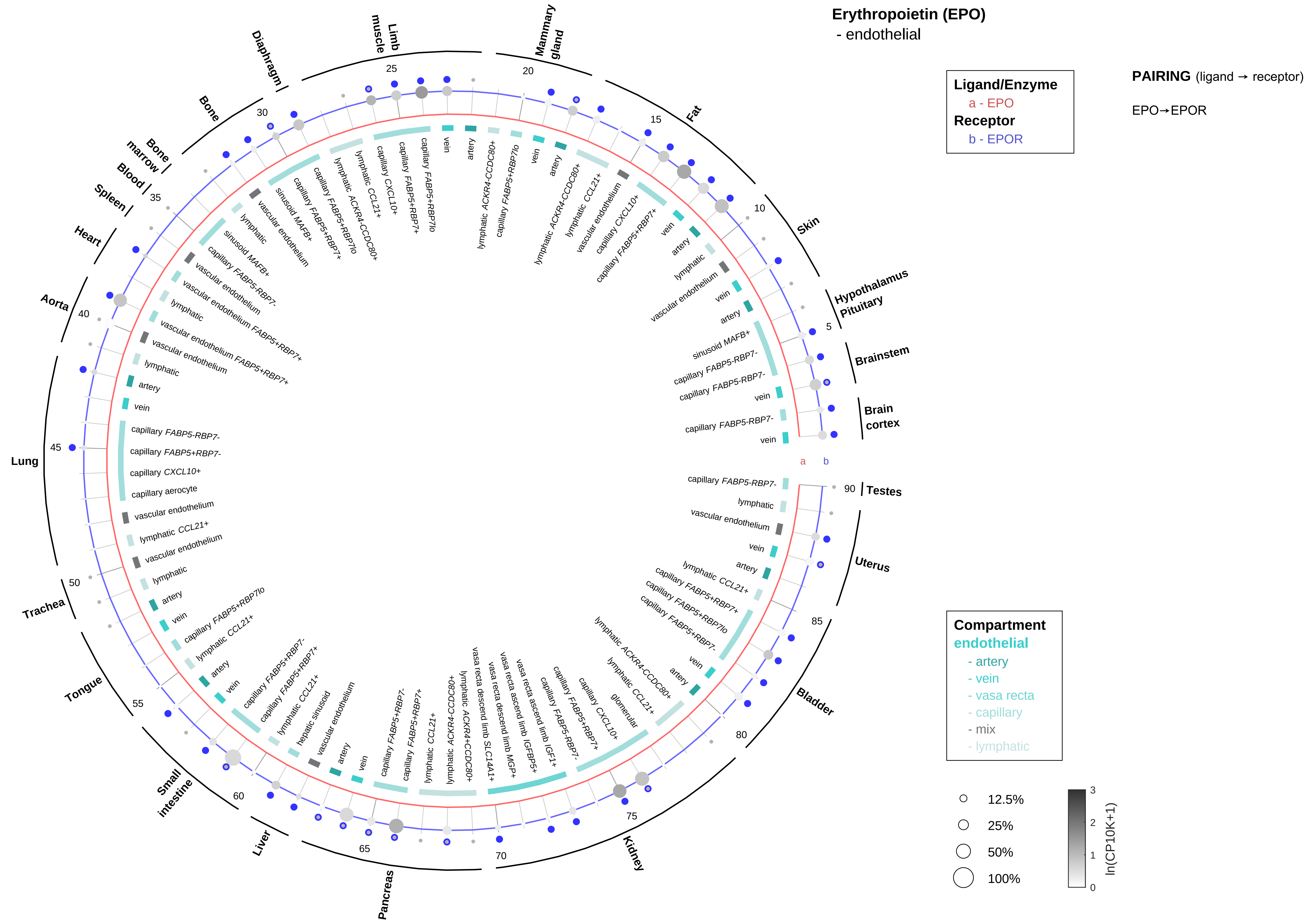

Erythropoietin (EPO)
- endothelial
Ligand/Enzyme
a - EPO
Receptor
b - EPOR
PAIRING (ligand → receptor)
EPO→EPOR
Compartment
endothelial
- artery
- vein
- vasa recta
- capillary
- mix
- lymphatic
12.5%
25%
50%
100%
ln(CP10K+1)
Testes
Uterus
Bladder
Kidney
Pancreas
Liver
Small intestine
Tongue
Trachea
Lung
Aorta
Heart
Spleen
Blood
Bone marrow
Bone
Diaphragm
Limb muscle
Mammary gland
Fat
Skin
Hypothalamus Pituitary
Brainstem
Brain cortex

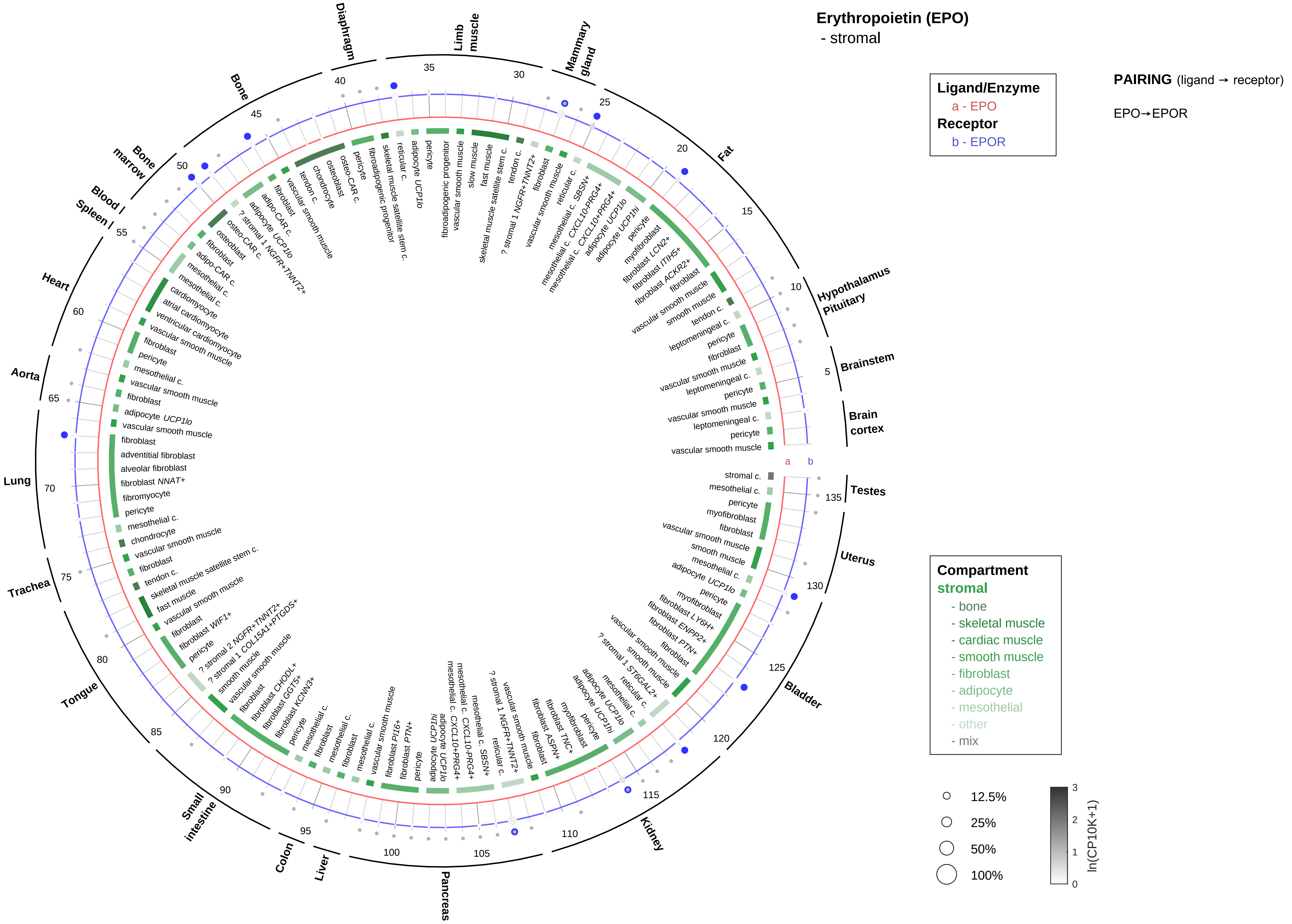

Erythropoietin (EPO)
- stromal
Ligand/Enzyme
a - EPO
Receptor
b - EPOR
PAIRING (ligand → receptor)
EPO→EPOR
Compartment
stromal
- bone
- skeletal muscle
- cardiac muscle
- smooth muscle
- fibroblast
- adipocyte
- mesothelial
- other
- mix
12.5%
25%
50%
100%
ln(CP10K+1)
Brain cortex
Brainstem
Hypothalamus Pituitary
Fat
Mammary gland
Limb muscle
Diaphragm
Bone
Bone marrow
Blood
Spleen
Heart
Aorta
Lung
Trachea
Tongue
Small intestine
Colon
Liver
Pancreas
Kidney
Bladder
Uterus
Testes

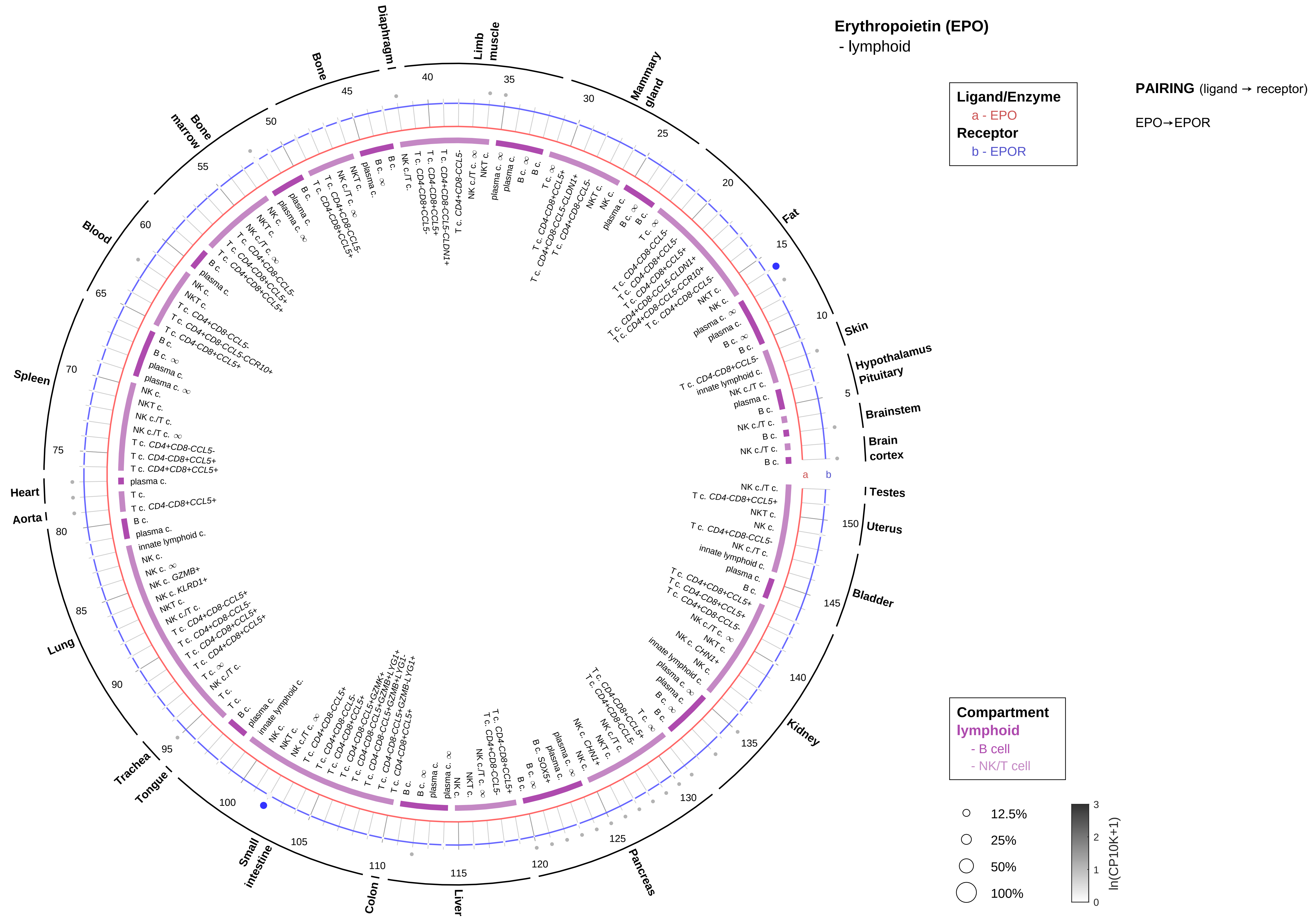

Erythropoietin (EPO)
- lymphoid
Ligand/Enzyme
a - EPO
Receptor
b - EPOR
PAIRING (ligand → receptor)
EPO→EPOR
Compartment
lymphoid
- B cell
- NK/T cell
12.5%
25%
50%
100%
ln(CP10K+1)
Skin
Hypothalamus
Pituitary
Brainstem
Brain cortex
Testes
Uterus
Bladder
Kidney
Pancreas
Liver
Colon
Small intestine
Tongue
Trachea
Lung
Aorta
Heart
Spleen
Blood
Bone marrow
Bone
Diaphragm
Limb muscle
Mammary gland
Fat

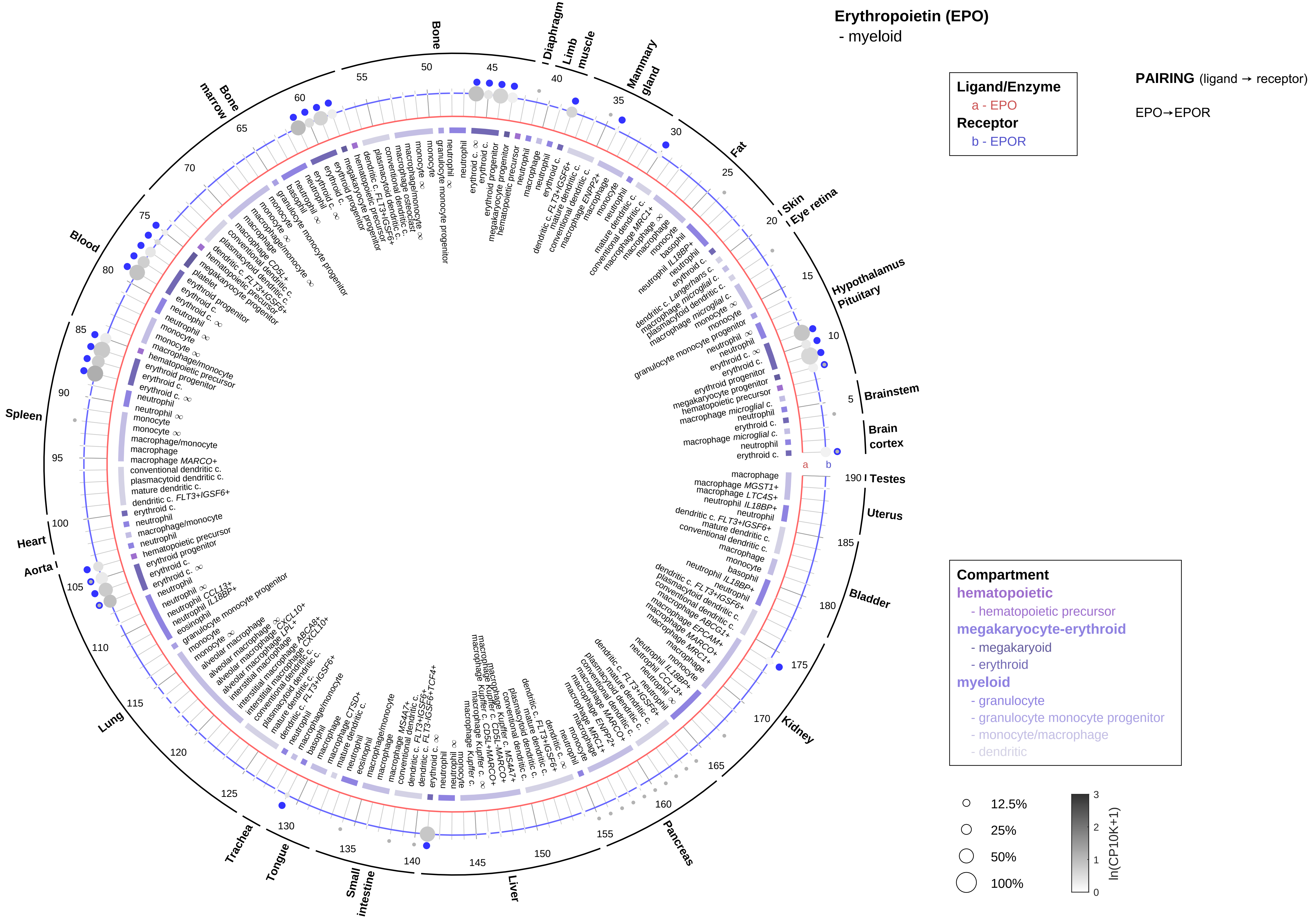
Erythropoietin (EPO)
- myeloid
Ligand/Enzyme
a - EPO
Receptor
b - EPOR
PAIRING (ligand → receptor)
EPO→EPOR
Compartment
hematopoietic
- hematopoietic precursor
megakaryocyte-erythroid
- megakaryoid
- erythroid
myeloid
- granulocyte
- granulocyte monocyte progenitor
- monocyte/macrophage
- dendritic
12.5%
25%
50%
100%
ln(CP10K+1)
Bone
Diaphragm
Limb muscle
Mammary gland
Fat
Skin
Eye retina
Hypothalamus
Pituitary
Brainstem
Brain cortex
Testes
Uterus
Bladder
Kidney
Pancreas
Liver
Small intestine
Tongue
Trachea
Lung
Aorta
Heart
Spleen
Blood
Bone marrow

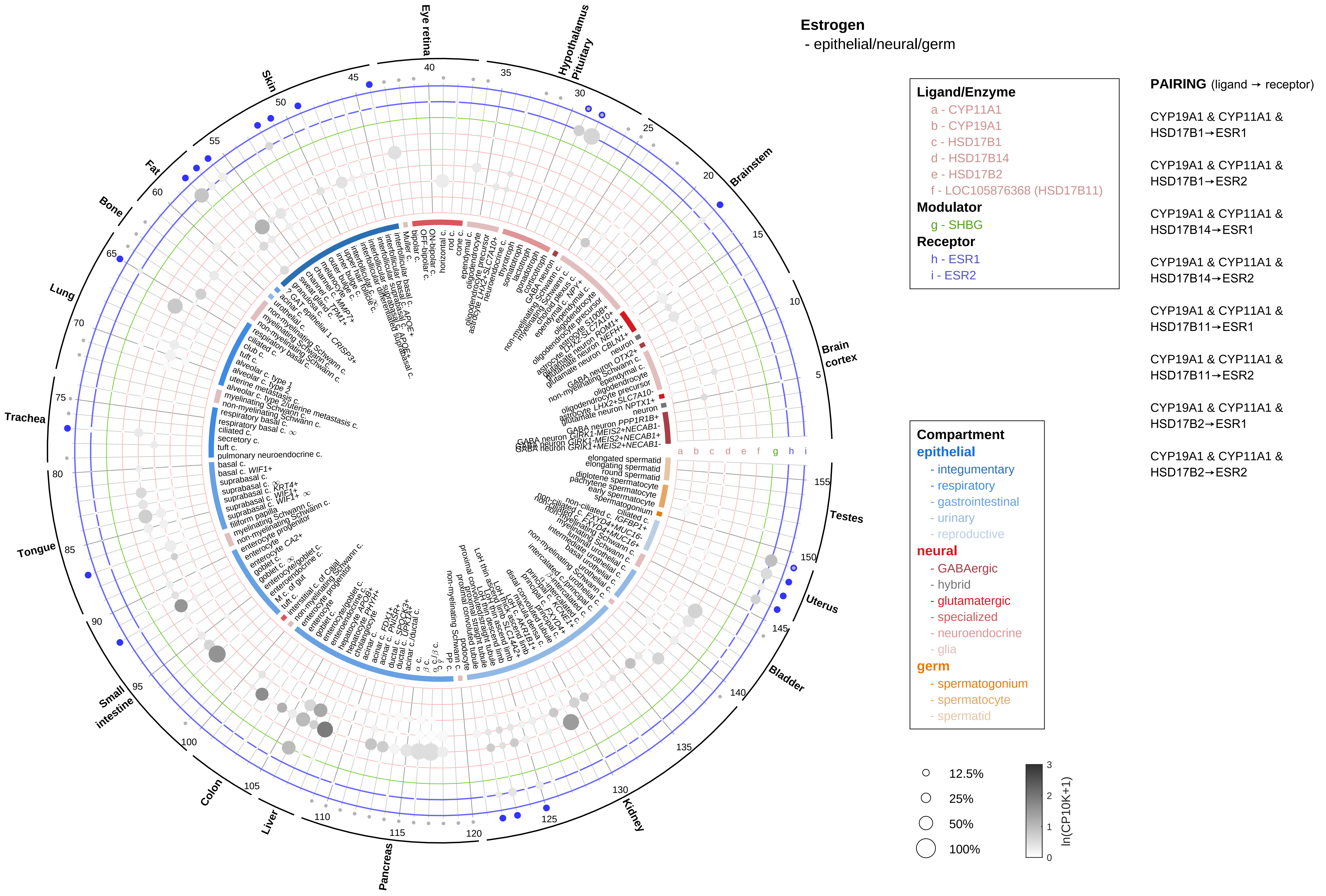
Estrogen
- epithelial/neural/germ
Ligand/Enzyme
a - CYP11A1
b - CYP19A1
c - HSD17B1
d - HSD17B14
e - HSD17B2
f - LOC105876368 (HSD17B11)
Modulator
g - SHBG
Receptor
h - ESR1
i - ESR2
PAIRING (ligand → receptor)
CYP19A1 & CYP11A1 & HSD17B1→ESR1
CYP19A1 & CYP11A1 & HSD17B1→ESR2
CYP19A1 & CYP11A1 & HSD17B14→ESR1
CYP19A1 & CYP11A1 & HSD17B14→ESR2
CYP19A1 & CYP11A1 & HSD17B11→ESR1
CYP19A1 & CYP11A1 & HSD17B11→ESR2
CYP19A1 & CYP11A1 & HSD17B2→ESR1
CYP19A1 & CYP11A1 & HSD17B2→ESR2
Compartment
epithelial
- integumentary
- respiratory
- gastrointestinal
- urinary
- reproductive
neural
- GABAergic
- hybrid
- glutamatergic
- specialized
- neuroendocrine
- glia
germ
- spermatogonium
- spermatocyte
- spermatid
12.5%
25%
50%
100%
ln(CP10K+1)
Eye retina
Hypothalamus Pituitary
Brainstem
Brain cortex
Testes
Uterus
Bladder
Kidney
Pancreas
Liver
Colon
Small intestine
Tongue
Trachea
Lung
Bone
Fat
Skin

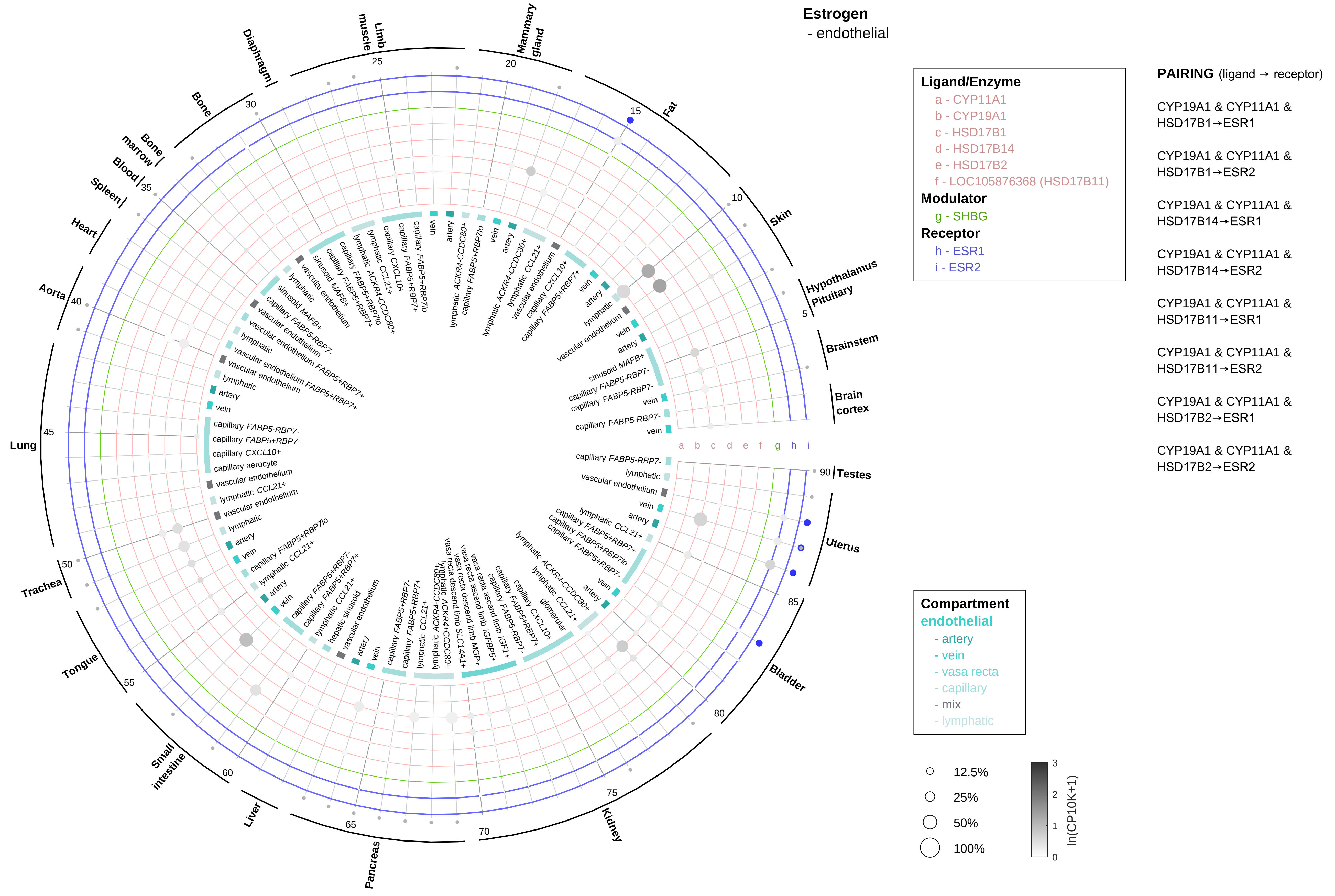

Estrogen
- endothelial
Ligand/Enzyme
a - CYP11A1
b - CYP19A1
c - HSD17B1
d - HSD17B14
e - HSD17B2
f - LOC105876368 (HSD17B11)
Modulator
g - SHBG
Receptor
h - ESR1
i - ESR2
PAIRING (ligand → receptor)
CYP19A1 & CYP11A1 & HSD17B1→ESR1
CYP19A1 & CYP11A1 & HSD17B1→ESR2
CYP19A1 & CYP11A1 & HSD17B14→ESR1
CYP19A1 & CYP11A1 & HSD17B14→ESR2
CYP19A1 & CYP11A1 & HSD17B11→ESR1
CYP19A1 & CYP11A1 & HSD17B11→ESR2
CYP19A1 & CYP11A1 & HSD17B2→ESR1
CYP19A1 & CYP11A1 & HSD17B2→ESR2
Compartment
endothelial
- artery
- vein
- vasa recta
- capillary
- mix
- lymphatic
12.5%
25%
50%
100%
ln(CP10K+1)
Fat
Skin
Hypothalamus
Pituitary
Brainstem
Brain cortex
Testes
Uterus
Bladder
Kidney
Pancreas
Liver
Small intestine
Tongue
Trachea
Lung
Aorta
Heart
Spleen
Blood
Bone marrow
Bone
Diaphragm
Limb muscle
Mammary gland
a b c d e f g h i

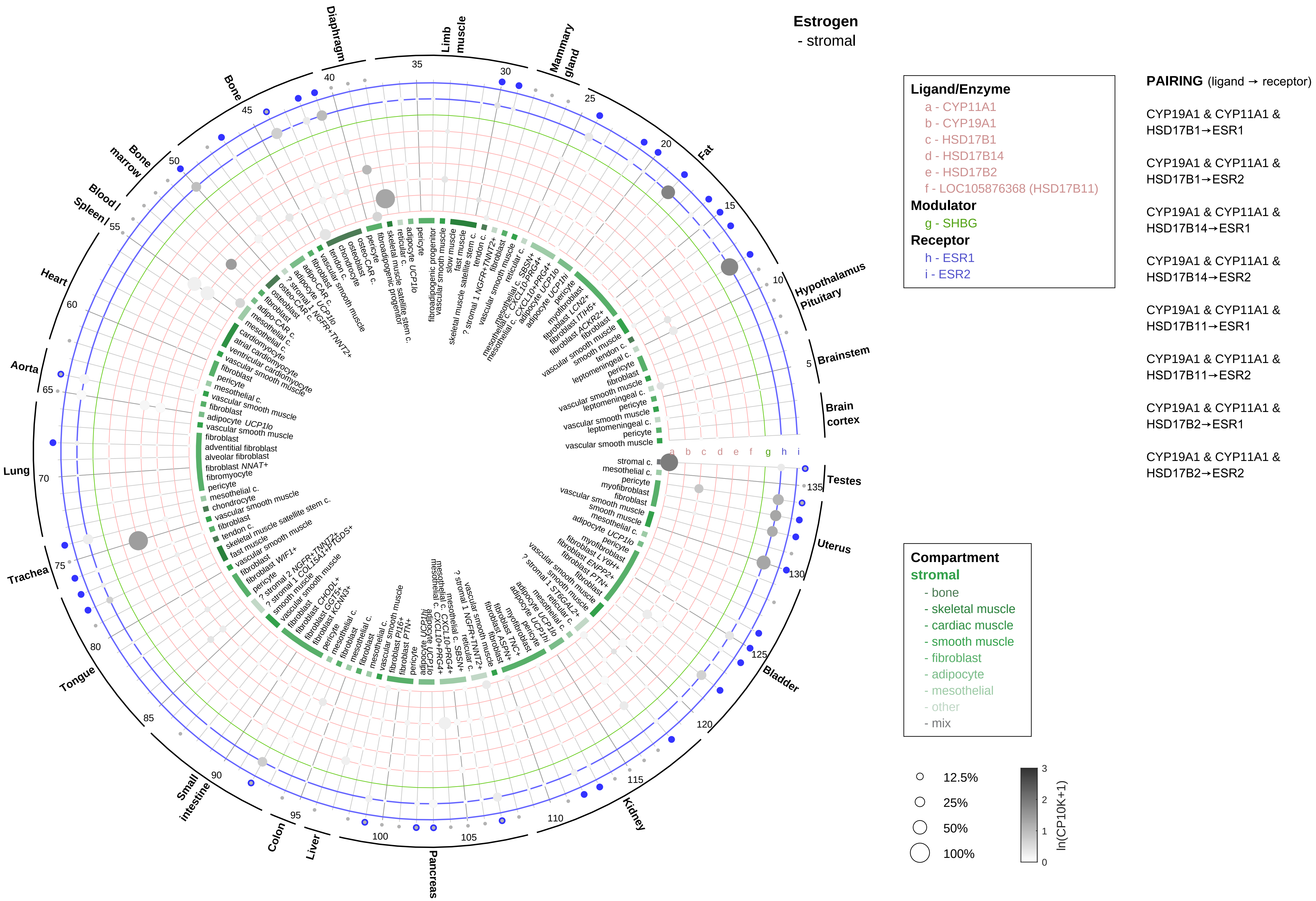

Estrogen
- stromal
Ligand/Enzyme
a - CYP11A1
b - CYP19A1
c - HSD17B1
d - HSD17B14
e - HSD17B2
f - LOC105876368 (HSD17B11)
Modulator
g - SHBG
Receptor
h - ESR1
i - ESR2
PAIRING (ligand → receptor)
CYP19A1 & CYP11A1 & HSD17B1→ESR1
CYP19A1 & CYP11A1 & HSD17B1→ESR2
CYP19A1 & CYP11A1 & HSD17B14→ESR1
CYP19A1 & CYP11A1 & HSD17B14→ESR2
CYP19A1 & CYP11A1 & HSD17B11→ESR1
CYP19A1 & CYP11A1 & HSD17B11→ESR2
CYP19A1 & CYP11A1 & HSD17B2→ESR1
CYP19A1 & CYP11A1 & HSD17B2→ESR2
Compartment
stromal
- bone
- skeletal muscle
- cardiac muscle
- smooth muscle
- fibroblast
- adipocyte
- mesothelial
- other
- mix
12.5%
25%
50%
100%
ln(CP10K+1)
Diaphragm
Limb muscle
Mammary gland
Fat
Hypothalamus Pituitary
Brainstem
Brain cortex
Testes
Uterus
Bladder
Kidney
Pancreas
Liver
Colon
Small intestine
Tongue
Trachea
Lung
Aorta
Heart
Blood / Spleen
Bone marrow
Bone

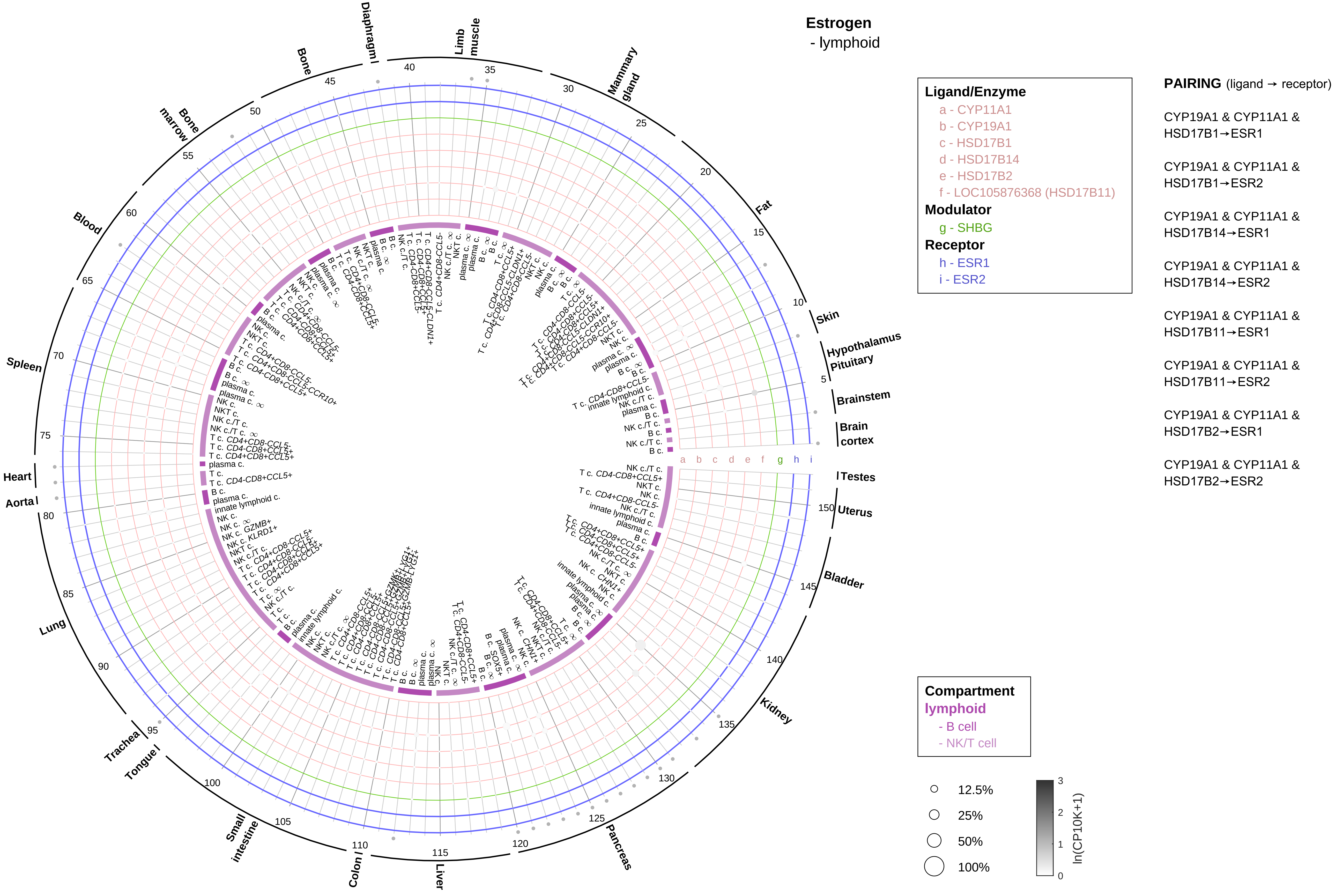
Estrogen
- lymphoid
Ligand/Enzyme
a - CYP11A1
b - CYP19A1
c - HSD17B1
d - HSD17B14
e - HSD17B2
f - LOC105876368 (HSD17B11)
Modulator
g - SHBG
Receptor
h - ESR1
i - ESR2
PAIRING (ligand → receptor)
CYP19A1 & CYP11A1 & HSD17B1→ESR1
CYP19A1 & CYP11A1 & HSD17B1→ESR2
CYP19A1 & CYP11A1 & HSD17B14→ESR1
CYP19A1 & CYP11A1 & HSD17B14→ESR2
CYP19A1 & CYP11A1 & HSD17B11→ESR1
CYP19A1 & CYP11A1 & HSD17B11→ESR2
CYP19A1 & CYP11A1 & HSD17B2→ESR1
CYP19A1 & CYP11A1 & HSD17B2→ESR2
Compartment
lymphoid
- B cell
- NK/T cell
12.5%
25%
50%
100%
ln(CP10K+1)
Diaphragm
Limb muscle
Mammary gland
Fat
Skin
Hypothalamus
Pituitary
Brainstem
Brain cortex
Testes
Uterus
Bladder
Kidney
Pancreas
Liver
Colon
Small intestine
Tongue
Trachea
Lung
Aorta
Heart
Spleen
Blood
Bone marrow
Bone

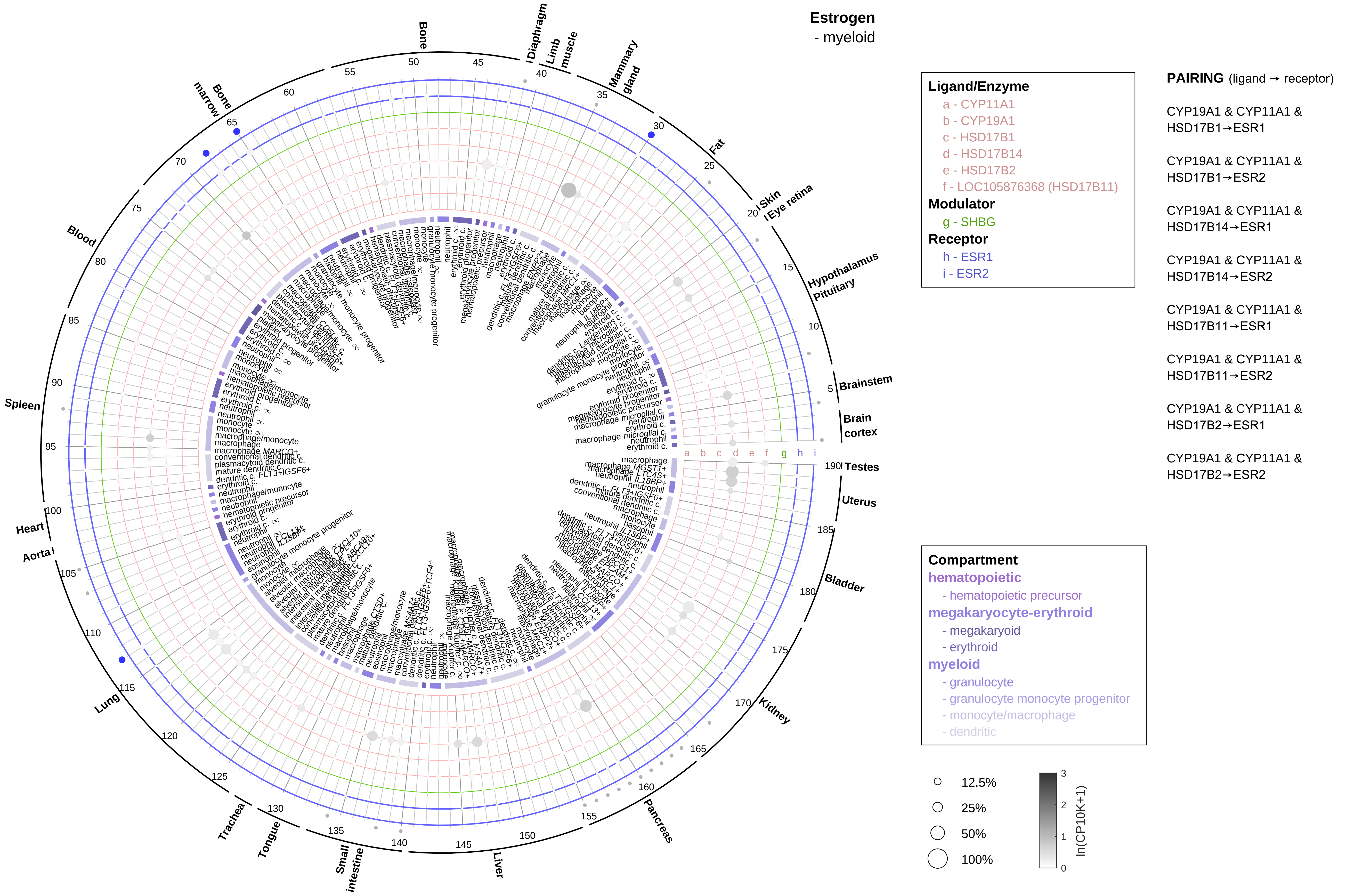

Estrogen
- myeloid
Ligand/Enzyme
a - CYP11A1
b - CYP19A1
c - HSD17B1
d - HSD17B14
e - HSD17B2
f - LOC105876368 (HSD17B11)
Modulator
g - SHBG
Receptor
h - ESR1
i - ESR2
PAIRING (ligand → receptor)
CYP19A1 & CYP11A1 & HSD17B1→ESR1
CYP19A1 & CYP11A1 & HSD17B1→ESR2
CYP19A1 & CYP11A1 & HSD17B14→ESR1
CYP19A1 & CYP11A1 & HSD17B14→ESR2
CYP19A1 & CYP11A1 & HSD17B11→ESR1
CYP19A1 & CYP11A1 & HSD17B11→ESR2
CYP19A1 & CYP11A1 & HSD17B2→ESR1
CYP19A1 & CYP11A1 & HSD17B2→ESR2
Compartment
hematopoietic
- hematopoietic precursor
megakaryocyte-erythroid
- megakaryoid
- erythroid
myeloid
- granulocyte
- granulocyte monocyte progenitor
- monocyte/macrophage
- dendritic
12.5%
25%
50%
100%
ln(CP10K+1)
Bone
Diaphragm
Limb muscle
Mammary gland
Fat
Skin
Eye retina
Hypothalamus
Pituitary
Brainstem
Brain cortex
Testes
Uterus
Bladder
Kidney
Pancreas
Liver
Small intestine
Tongue
Trachea
Lung
Aorta
Heart
Spleen
Blood
Bone marrow

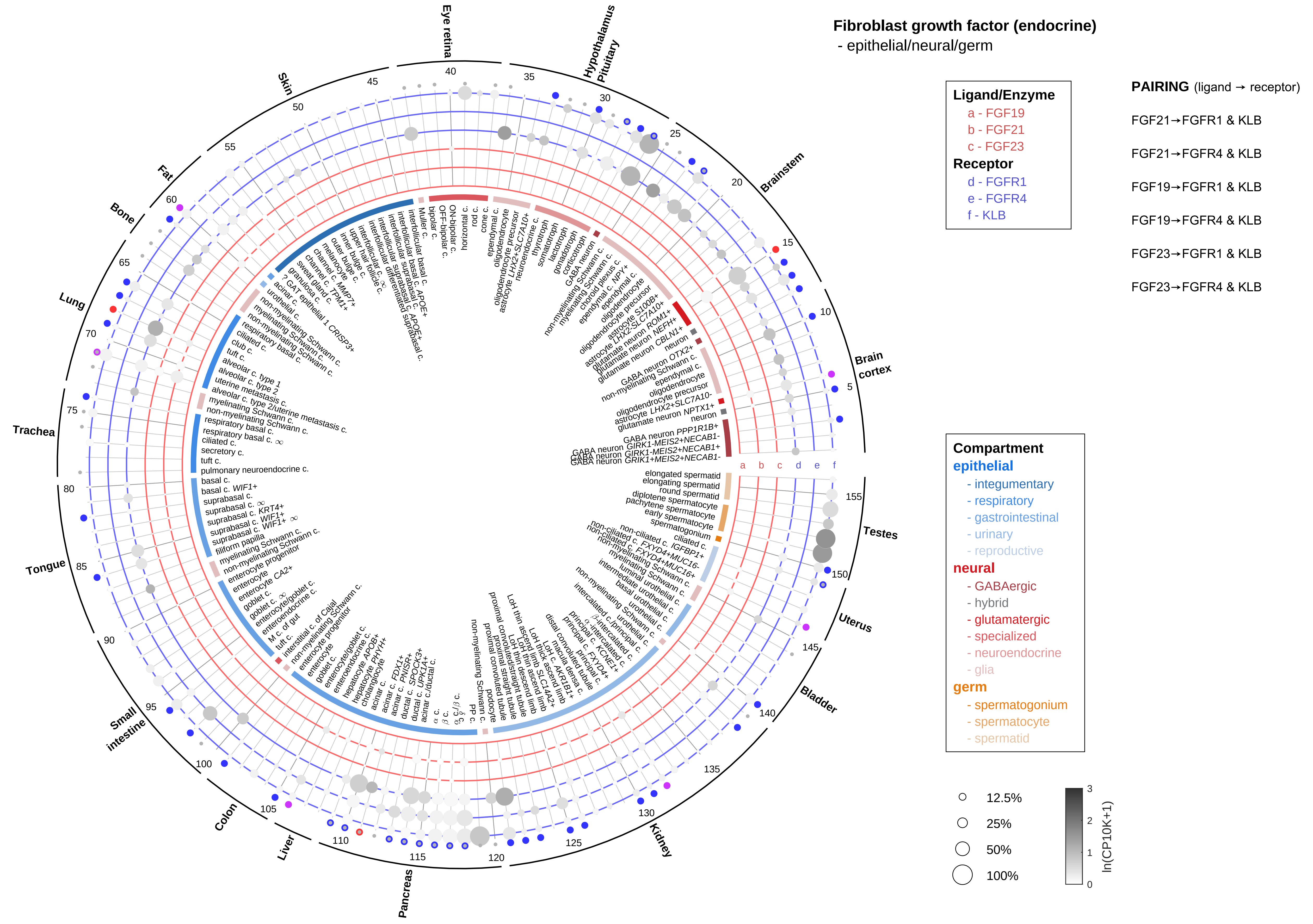

Fibroblast growth factor (endocrine)
- epithelial/neural/germ
Ligand/Enzyme
a - FGF19
b - FGF21
c - FGF23
Receptor
d - FGFR1
e - FGFR4
f - KLB
PAIRING (ligand → receptor)
FGF21→FGFR1 & KLB
FGF21→FGFR4 & KLB
FGF19→FGFR1 & KLB
FGF19→FGFR4 & KLB
FGF23→FGFR1 & KLB
FGF23→FGFR4 & KLB
Compartment
epithelial
- integumentary
- respiratory
- gastrointestinal
- urinary
- reproductive
neural
- GABAergic
- hybrid
- glutamatergic
- specialized
- neuroendocrine
- glia
germ
- spermatogonium
- spermatocyte
- spermatid
12.5%
25%
50%
100%
ln(CP10K+1)
Eye retina
Hypothalamus
Pituitary
Brainstem
Brain cortex
Testes
Uterus
Bladder
Kidney
Pancreas
Liver
Colon
Small intestine
Tongue
Trachea
Lung
Bone
Fat
Skin

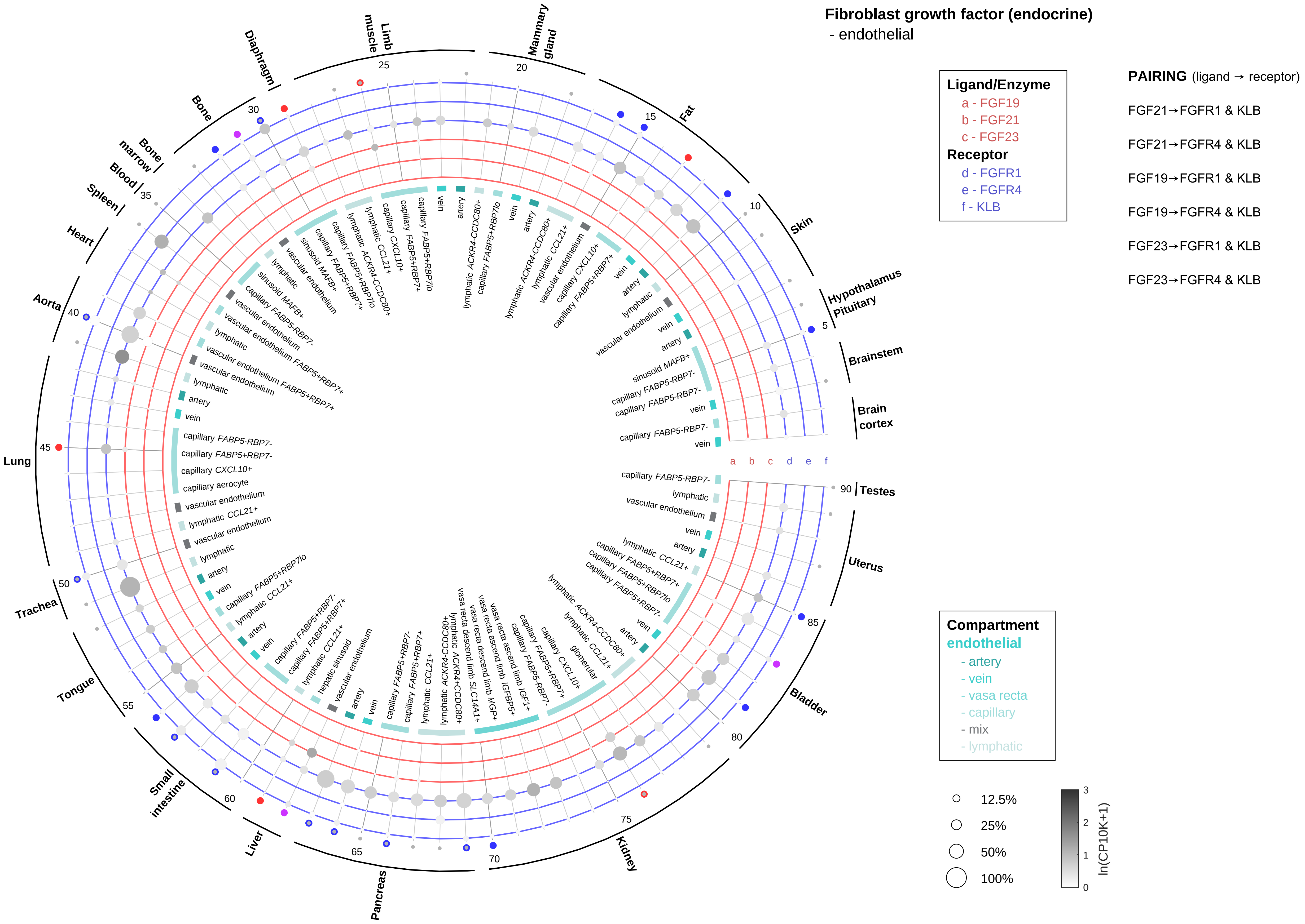

Fibroblast growth factor (endocrine)
- endothelial
Ligand/Enzyme
a - FGF19
b - FGF21
c - FGF23
Receptor
d - FGFR1
e - FGFR4
f - KLB
PAIRING (ligand → receptor)
FGF21→FGFR1 & KLB
FGF21→FGFR4 & KLB
FGF19→FGFR1 & KLB
FGF19→FGFR4 & KLB
FGF23→FGFR1 & KLB
FGF23→FGFR4 & KLB
Compartment
endothelial
- artery
- vein
- vasa recta
- capillary
- mix
- lymphatic
12.5%
25%
50%
100%
ln(CP10K+1)
Hypothalamus
Pituitary
Brainstem
Brain cortex
Testes
Uterus
Bladder
Kidney
Pancreas
Liver
Small intestine
Tongue
Trachea
Lung
Aorta
Heart
Spleen
Blood
Bone marrow
Bone
Diaphragm
Limb muscle
Mammary gland
Fat
Skin
a b c d e f

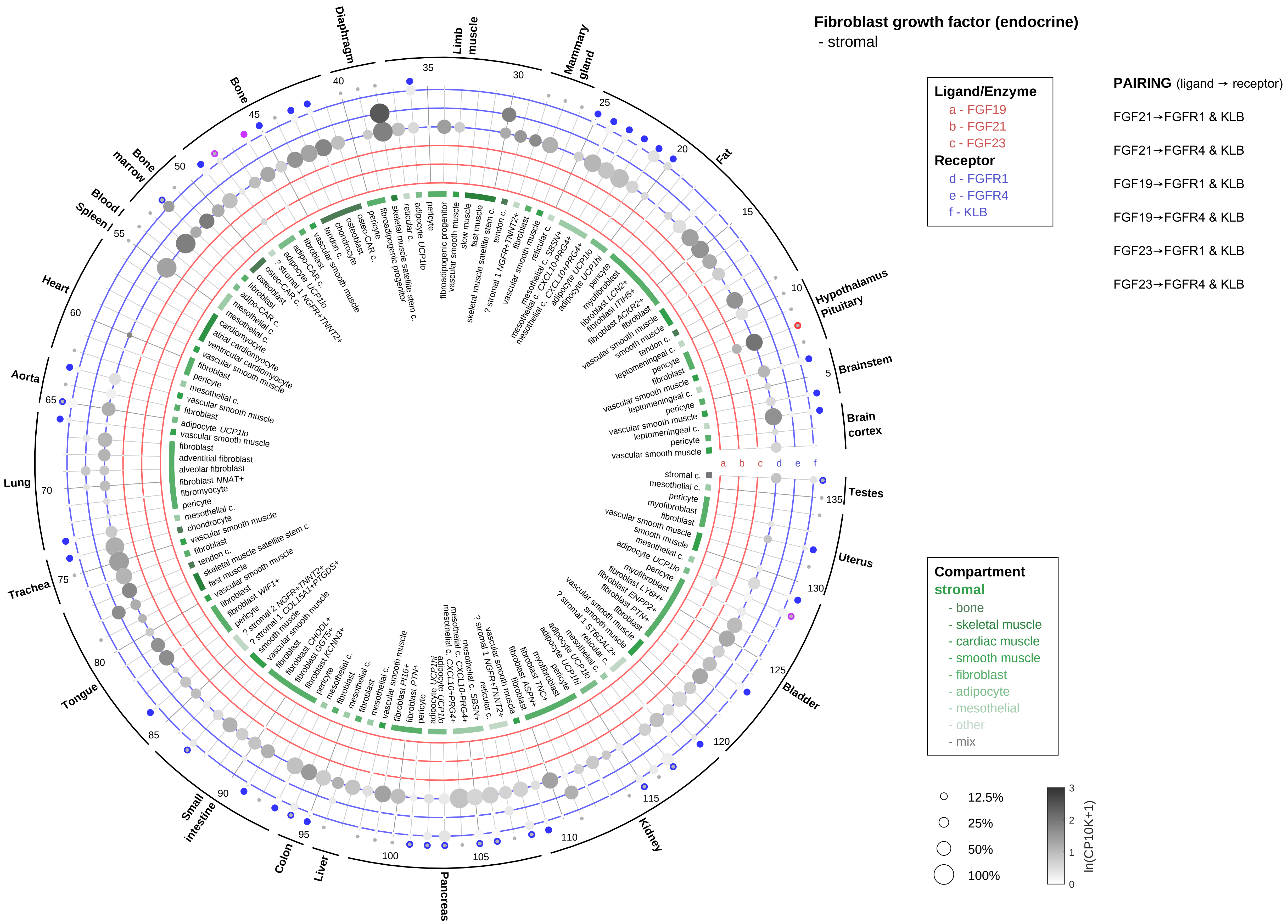
Fibroblast growth factor (endocrine)
- stromal
Ligand/Enzyme
a - FGF19
b - FGF21
c - FGF23
Receptor
d - FGFR1
e - FGFR4
f - KLB
PAIRING (ligand → receptor)
FGF21→FGFR1 & KLB
FGF21→FGFR4 & KLB
FGF19→FGFR1 & KLB
FGF19→FGFR4 & KLB
FGF23→FGFR1 & KLB
FGF23→FGFR4 & KLB
Compartment
stromal
- bone
- skeletal muscle
- cardiac muscle
- smooth muscle
- fibroblast
- adipocyte
- mesothelial
- other
- mix
12.5%
25%
50%
100%
ln(CP10K+1)
Hypothalamus
Pituitary
Brainstem
Brain cortex
Fat
Mammary gland
Limb muscle
Diaphragm
Bone
Bone marrow
Blood
Spleen
Heart
Aorta
Lung
Trachea
Tongue
Small intestine
Colon
Liver
Pancreas
Kidney
Bladder
Uterus
Testes
a
b
c
d
e
f

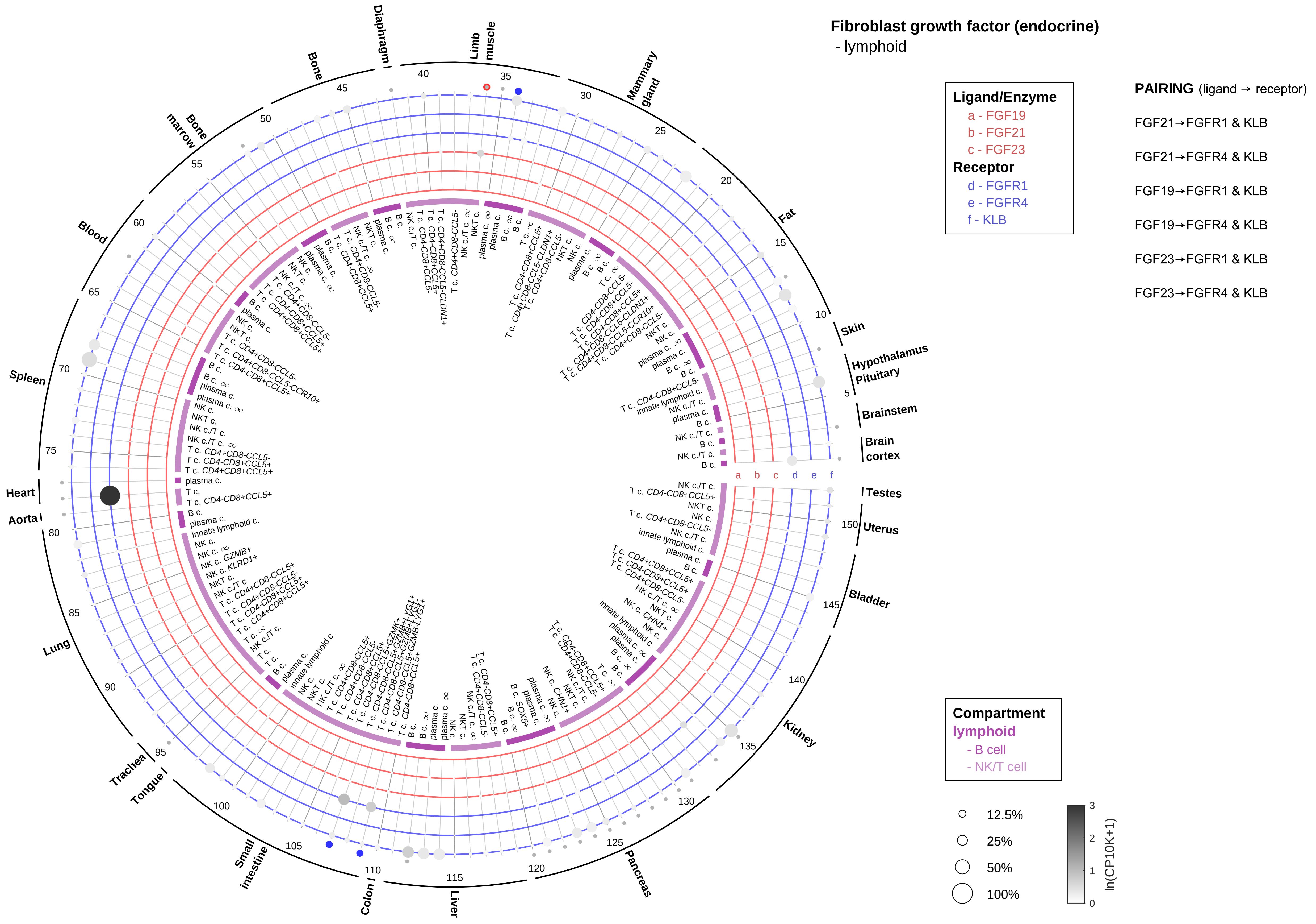

Fibroblast growth factor (endocrine)
- lymphoid
Ligand/Enzyme
a - FGF19
b - FGF21
c - FGF23
Receptor
d - FGFR1
e - FGFR4
f - KLB
PAIRING (ligand → receptor)
FGF21→FGFR1 & KLB
FGF21→FGFR4 & KLB
FGF19→FGFR1 & KLB
FGF19→FGFR4 & KLB
FGF23→FGFR1 & KLB
FGF23→FGFR4 & KLB
Compartment
lymphoid
- B cell
- NK/T cell
12.5%
25%
50%
100%
ln(CP10K+1)
Skin
Hypothalamus
Pituitary
Brainstem
Brain cortex
Testes
Uterus
Bladder
Kidney
Pancreas
Liver
Colon
Small intestine
Tongue
Trachea
Lung
Aorta
Heart
Spleen
Blood
Bone marrow
Bone
Diaphragm
Limb muscle
Mammary gland
Fat

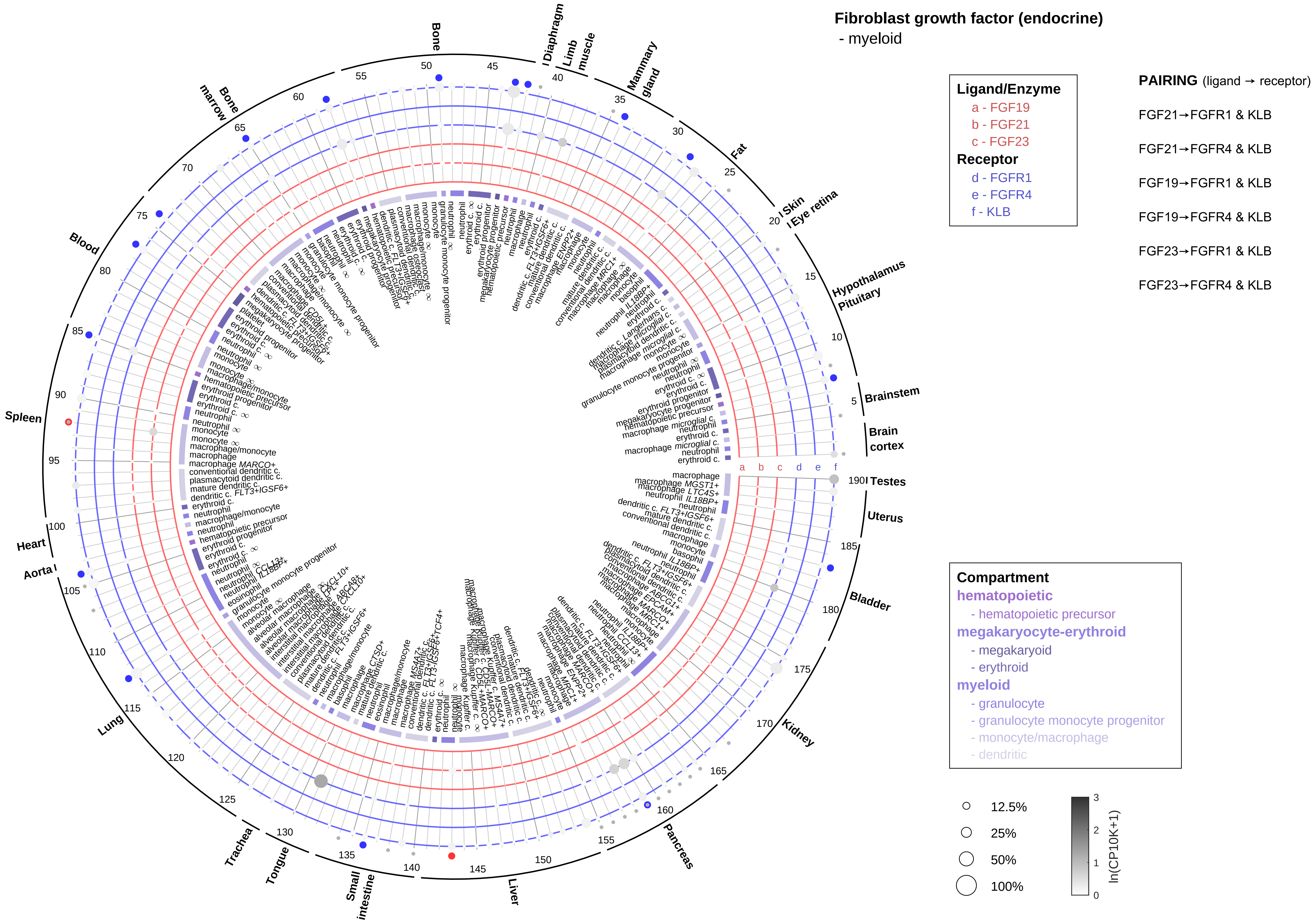
Fibroblast growth factor (endocrine)
- myeloid
Ligand/Enzyme
a - FGF19
b - FGF21
c - FGF23
Receptor
d - FGFR1
e - FGFR4
f - KLB
PAIRING (ligand → receptor)
FGF21→FGFR1 & KLB
FGF21→FGFR4 & KLB
FGF19→FGFR1 & KLB
FGF19→FGFR4 & KLB
FGF23→FGFR1 & KLB
FGF23→FGFR4 & KLB
Compartment
hematopoietic
- hematopoietic precursor
megakaryocyte-erythroid
- megakaryoid
- erythroid
myeloid
- granulocyte
- granulocyte monocyte progenitor
- monocyte/macrophage
- dendritic
12.5%
25%
50%
100%
ln(CP10K+1)
Bone
Diaphragm
Limb muscle
Mammary gland
Fat
Skin
Eye retina
Hypothalamus
Pituitary
Brainstem
Brain cortex
Testes
Uterus
Bladder
Kidney
Pancreas
Liver
Small intestine
Tongue
Trachea
Lung
Aorta
Heart
Spleen
Blood
Bone marrow

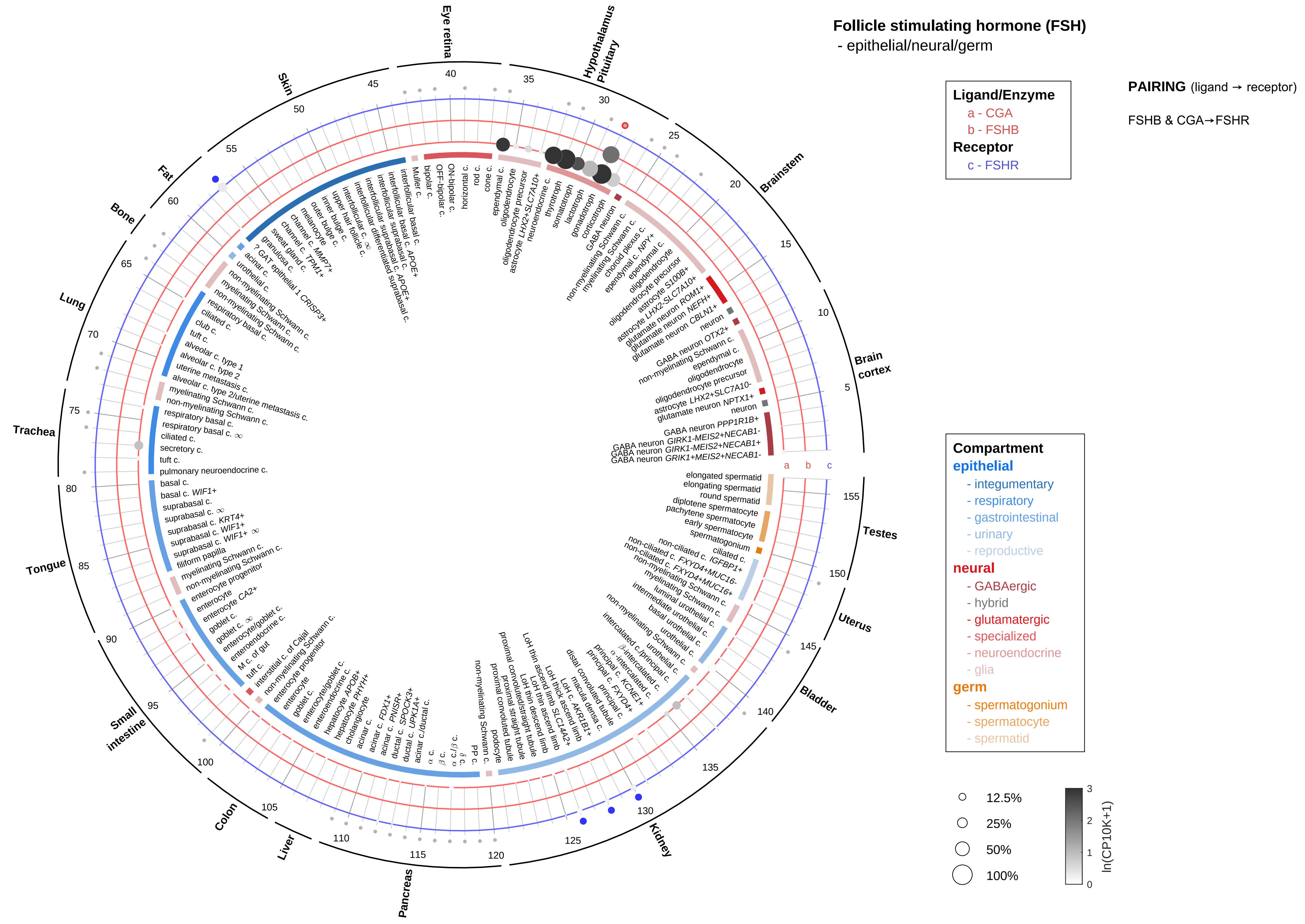
Follicle stimulating hormone (FSH)
- epithelial/neural/germ
Ligand/Enzyme
a - CGA
b - FSHB
Receptor
c - FSHR
PAIRING (ligand → receptor)
FSHB & CGA→FSHR
Compartment
epithelial
- integumentary
- respiratory
- gastrointestinal
- urinary
- reproductive
neural
- GABAergic
- hybrid
- glutamatergic
- specialized
- neuroendocrine
- glia
germ
- spermatogonium
- spermatocyte
- spermatid
12.5%
25%
50%
100%
ln(CP10K+1)
Eye retina
Hypothalamus
Pituitary
Brainstem
Brain cortex
Testes
Uterus
Bladder
Kidney
Pancreas
Liver
Colon
Small intestine
Tongue
Trachea
Lung
Bone
Fat
Skin

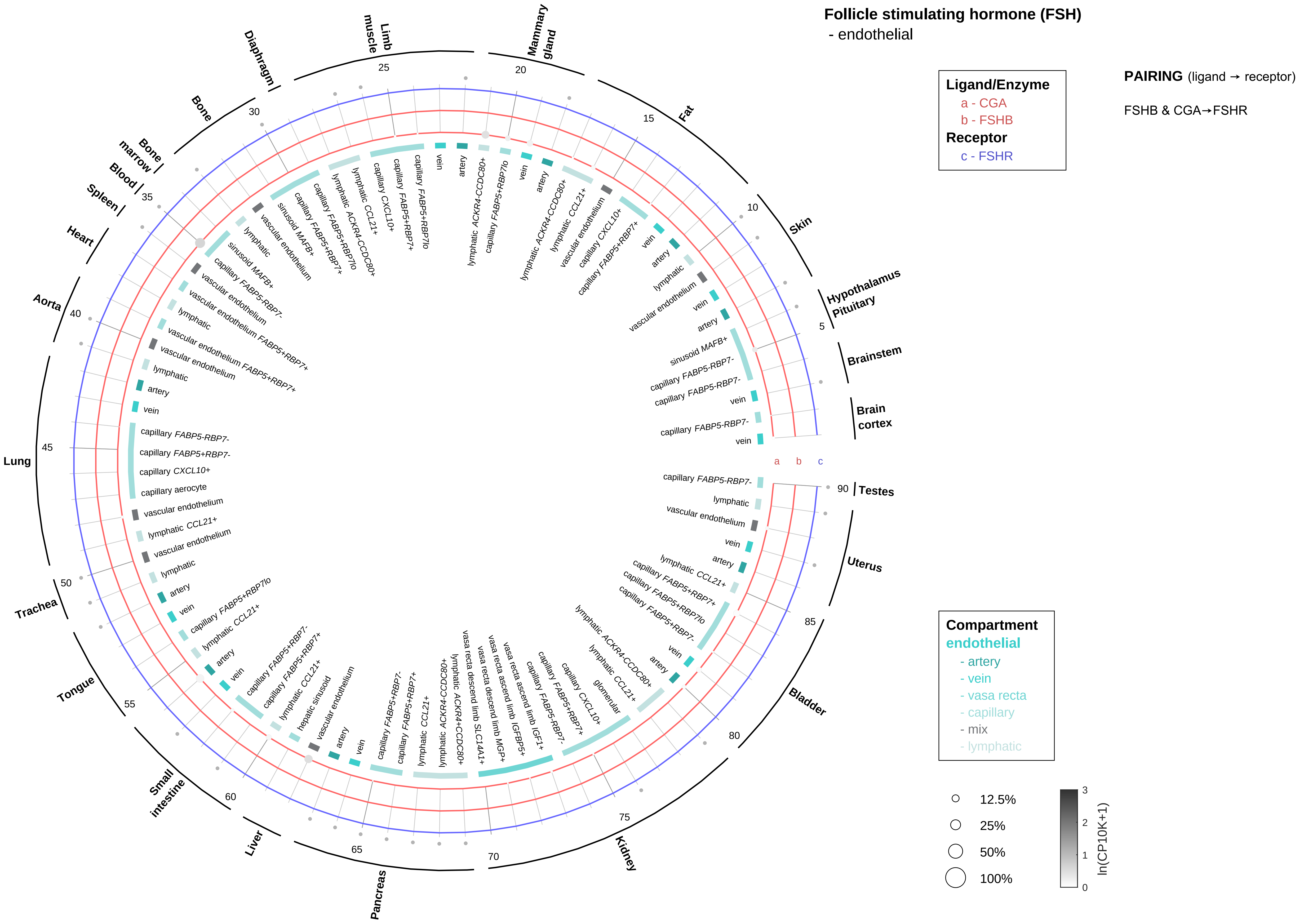
Follicle stimulating hormone (FSH)
- endothelial
Ligand/Enzyme
a - CGA
b - FSHB
Receptor
c - FSHR
PAIRING (ligand → receptor)
FSHB & CGA→FSHR
Compartment
endothelial
- artery
- vein
- vasa recta
- capillary
- mix
- lymphatic
12.5%
25%
50%
100%
ln(CP10K+1)
Mammary gland
Fat
Skin
Hypothalamus
Pituitary
Brainstem
Brain cortex
Testes
Uterus
Bladder
Kidney
Pancreas
Liver
Small intestine
Tongue
Trachea
Lung
Aorta
Heart
Spleen
Blood
Bone marrow
Bone
Diaphragm
Limb muscle

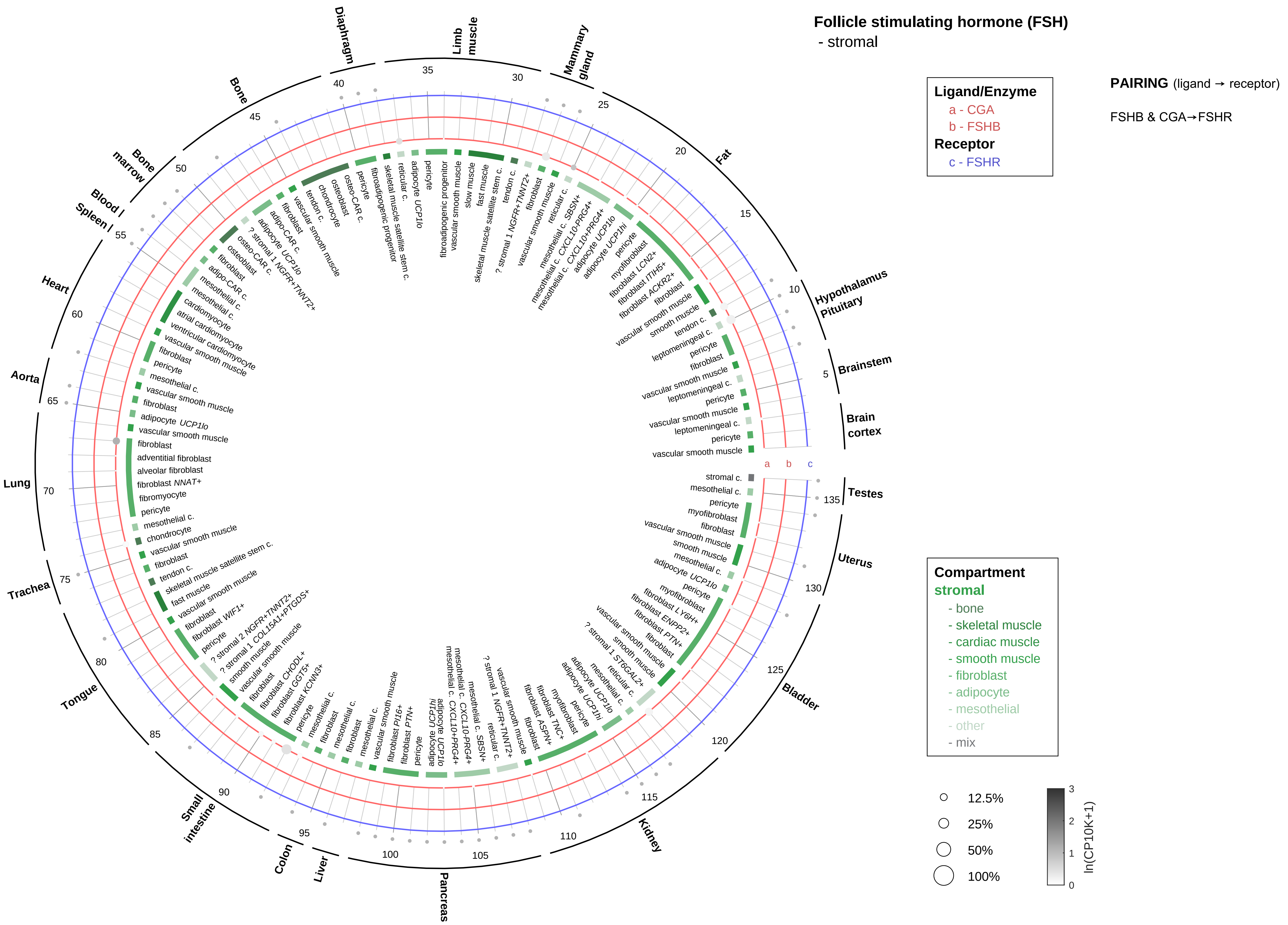

Follicle stimulating hormone (FSH)
- stromal
Ligand/Enzyme
a - CGA
b - FSHB
Receptor
c - FSHR
PAIRING (ligand → receptor)
FSHB & CGA→FSHR
Compartment
stromal
- bone
- skeletal muscle
- cardiac muscle
- smooth muscle
- fibroblast
- adipocyte
- mesothelial
- other
- mix
12.5%
25%
50%
100%
ln(CP10K+1)
Brain cortex
Brainstem
Hypothalamus Pituitary
Fat
Mammary gland
Limb muscle
Diaphragm
Bone
Bone marrow
Blood
Spleen
Heart
Aorta
Lung
Trachea
Tongue
Small intestine
Colon
Liver
Pancreas
Kidney
Bladder
Uterus
Testes

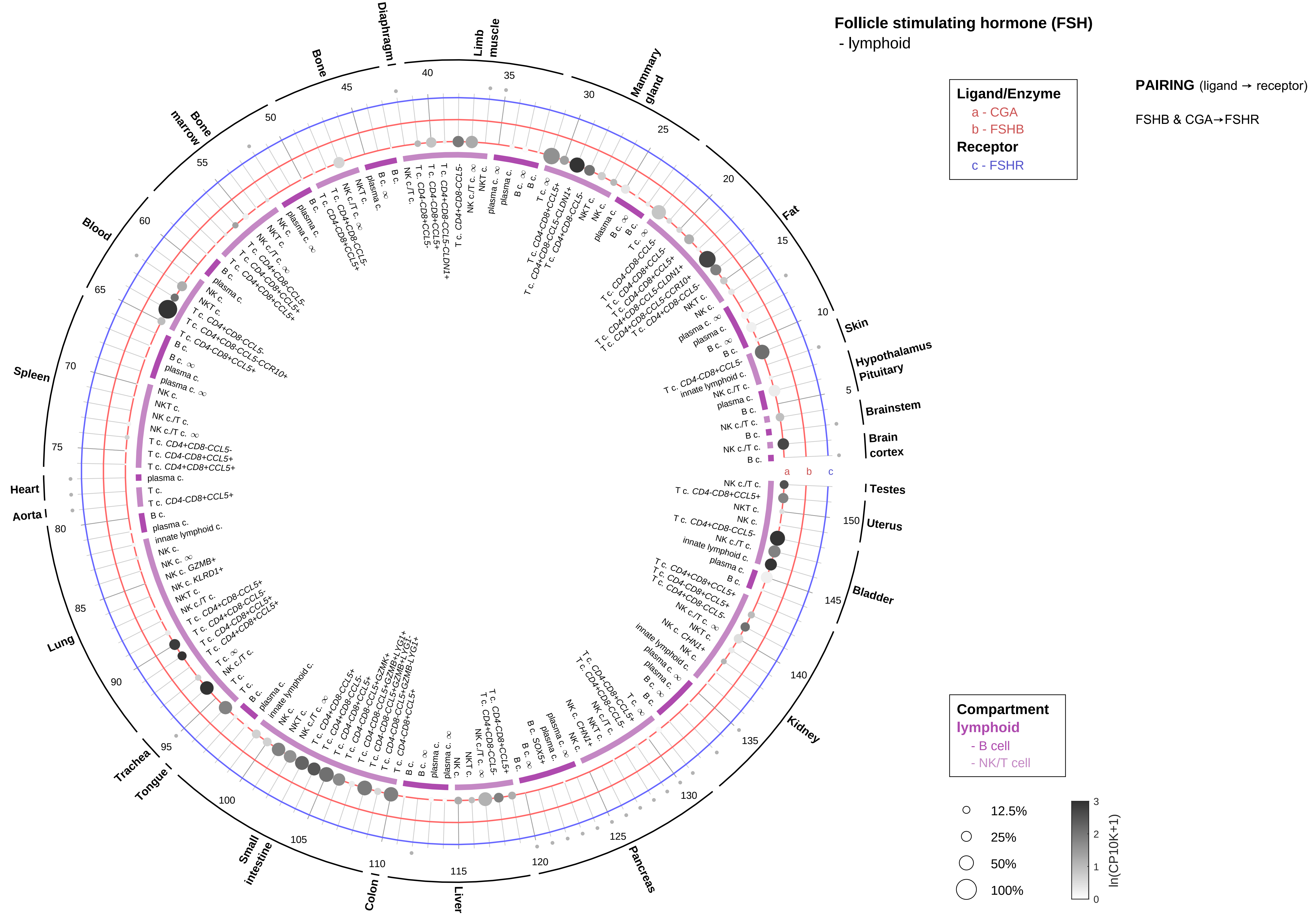

Follicle stimulating hormone (FSH)
- lymphoid
PAIRING (ligand → receptor)
FSHB & CGA→FSHR
Ligand/Enzyme
a - CGA
b - FSHB
Receptor
c - FSHR
Compartment
lymphoid
- B cell
- NK/T cell
12.5%
25%
50%
100%
ln(CP10K+1)
Brain cortex
Brainstem
Hypothalamus Pituitary
Skin
Fat
Mammary gland
Limb muscle
Diaphragm
Bone
Bone marrow
Blood
Spleen
Heart
Aorta
Lung
Trachea
Tongue
Small intestine
Colon
Liver
Pancreas
Kidney
Bladder
Uterus
Testes

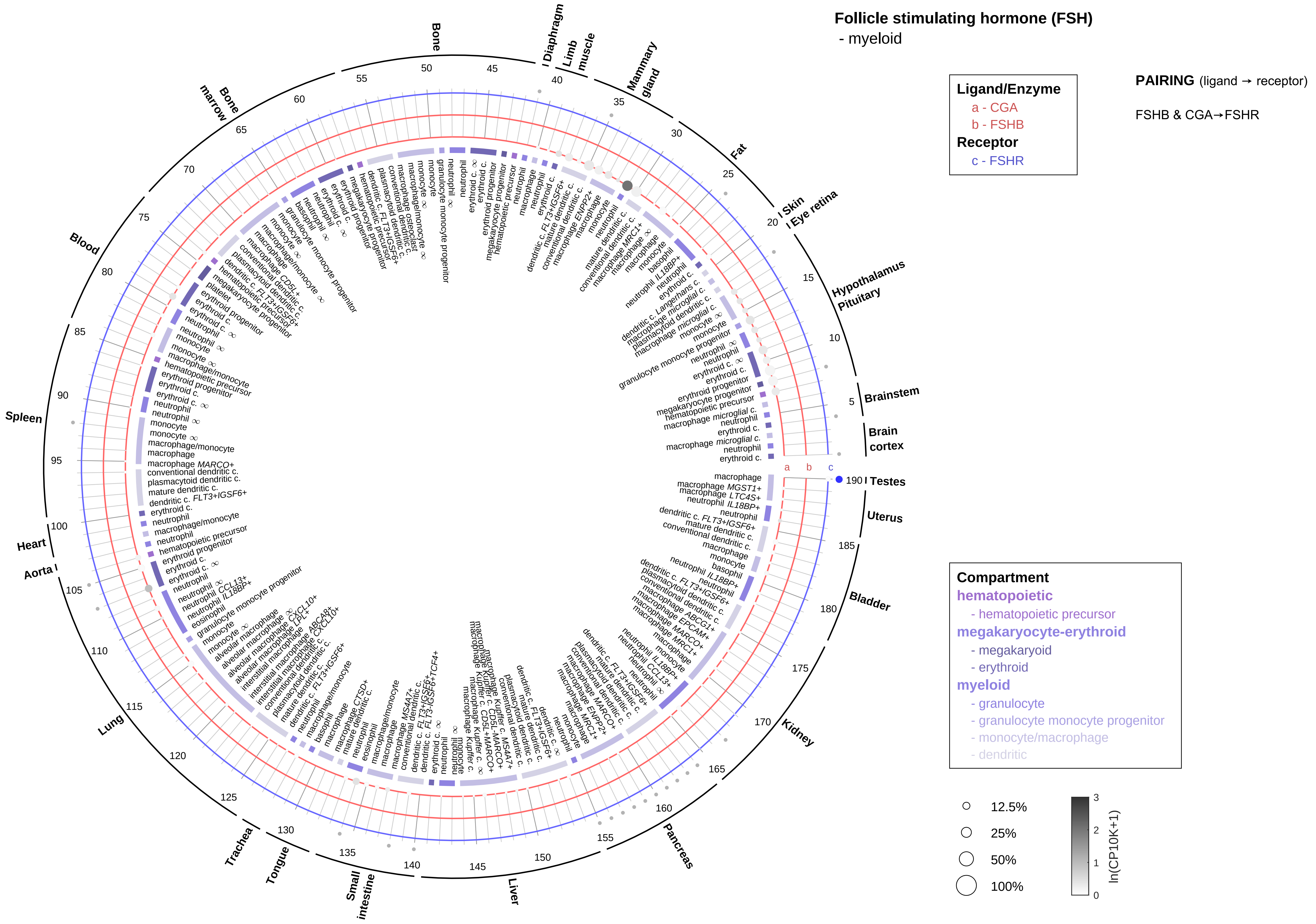

Follicle stimulating hormone (FSH)
- myeloid
Ligand/Enzyme
a - CGA
b - FSHB
Receptor
c - FSHR
PAIRING (ligand → receptor)
FSHB & CGA→FSHR
Compartment
hematopoietic
- hematopoietic precursor
megakaryocyte-erythroid
- megakaryoid
- erythroid
myeloid
- granulocyte
- granulocyte monocyte progenitor
- monocyte/macrophage
- dendritic
12.5%
25%
50%
100%
ln(CP10K+1)
Bone
Diaphragm
Limb muscle
Mammary gland
Fat
Skin
Eye retina
Hypothalamus
Pituitary
Brainstem
Brain cortex
Testes
Uterus
Bladder
Kidney
Pancreas
Liver
Small intestine
Tongue
Trachea
Lung
Aorta
Heart
Spleen
Blood
Bone marrow

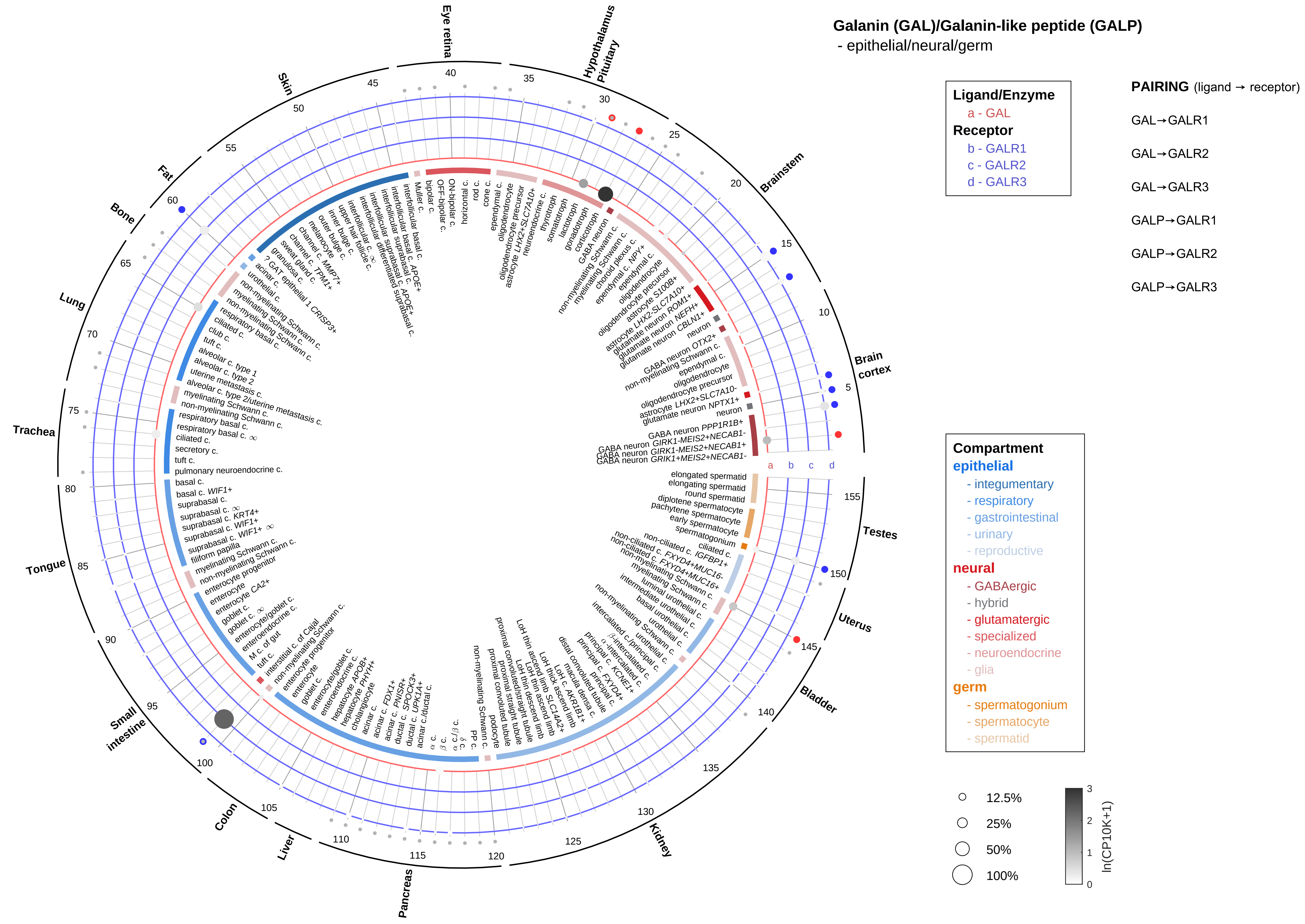

Galanin (GAL)/Galanin-like peptide (GALP)
- epithelial/neural/germ
Ligand/Enzyme
a - GAL
Receptor
b - GALR1
c - GALR2
d - GALR3
PAIRING (ligand → receptor)
GAL→GALR1
GAL→GALR2
GAL→GALR3
GALP→GALR1
GALP→GALR2
GALP→GALR3
Compartment
epithelial
- integumentary
- respiratory
- gastrointestinal
- urinary
- reproductive
neural
- GABAergic
- hybrid
- glutamatergic
- specialized
- neuroendocrine
- glia
germ
- spermatogonium
- spermatocyte
- spermatid
12.5%
25%
50%
100%
ln(CP10K+1)
Eye retina
Hypothalamus Pituitary
Brainstem
Brain cortex
Testes
Uterus
Bladder
Kidney
Pancreas
Liver
Colon
Small intestine
Tongue
Trachea
Lung
Bone
Fat
Skin
a
b
c
d

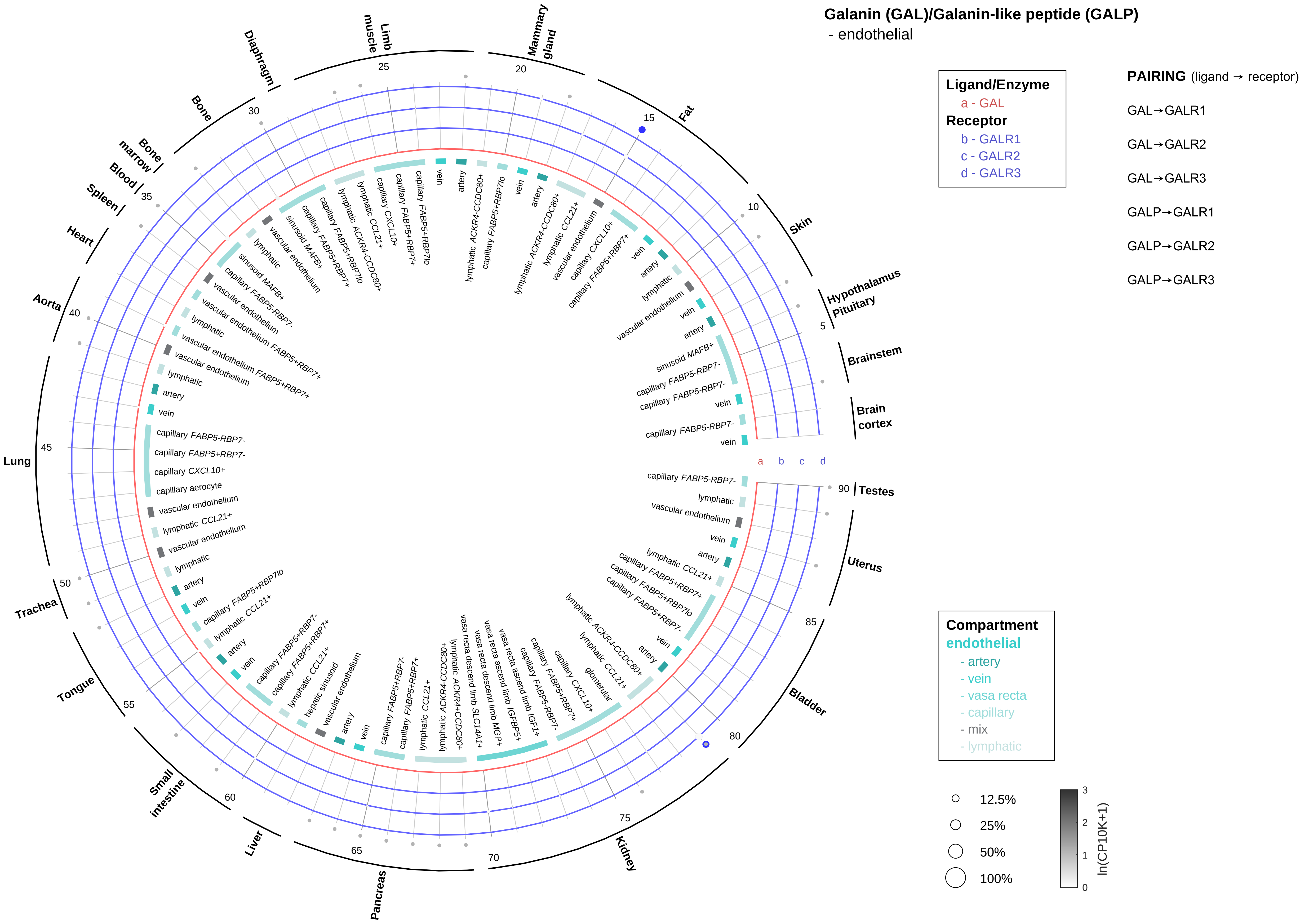

Galanin (GAL)/Galanin-like peptide (GALP)
- endothelial
Ligand/Enzyme
a - GAL
Receptor
b - GALR1
c - GALR2
d - GALR3
PAIRING (ligand → receptor)
GAL→GALR1
GAL→GALR2
GAL→GALR3
GALP→GALR1
GALP→GALR2
GALP→GALR3
Compartment
endothelial
- artery
- vein
- vasa recta
- capillary
- mix
- lymphatic
12.5%
25%
50%
100%
ln(CP10K+1)
Hypothalamus
Pituitary
Brainstem
Brain cortex
Testes
Uterus
Bladder
Kidney
Pancreas
Liver
Small intestine
Tongue
Trachea
Lung
Aorta
Heart
Spleen
Blood
Bone marrow
Bone
Diaphragm
Limb muscle
Mammary gland
Fat
Skin

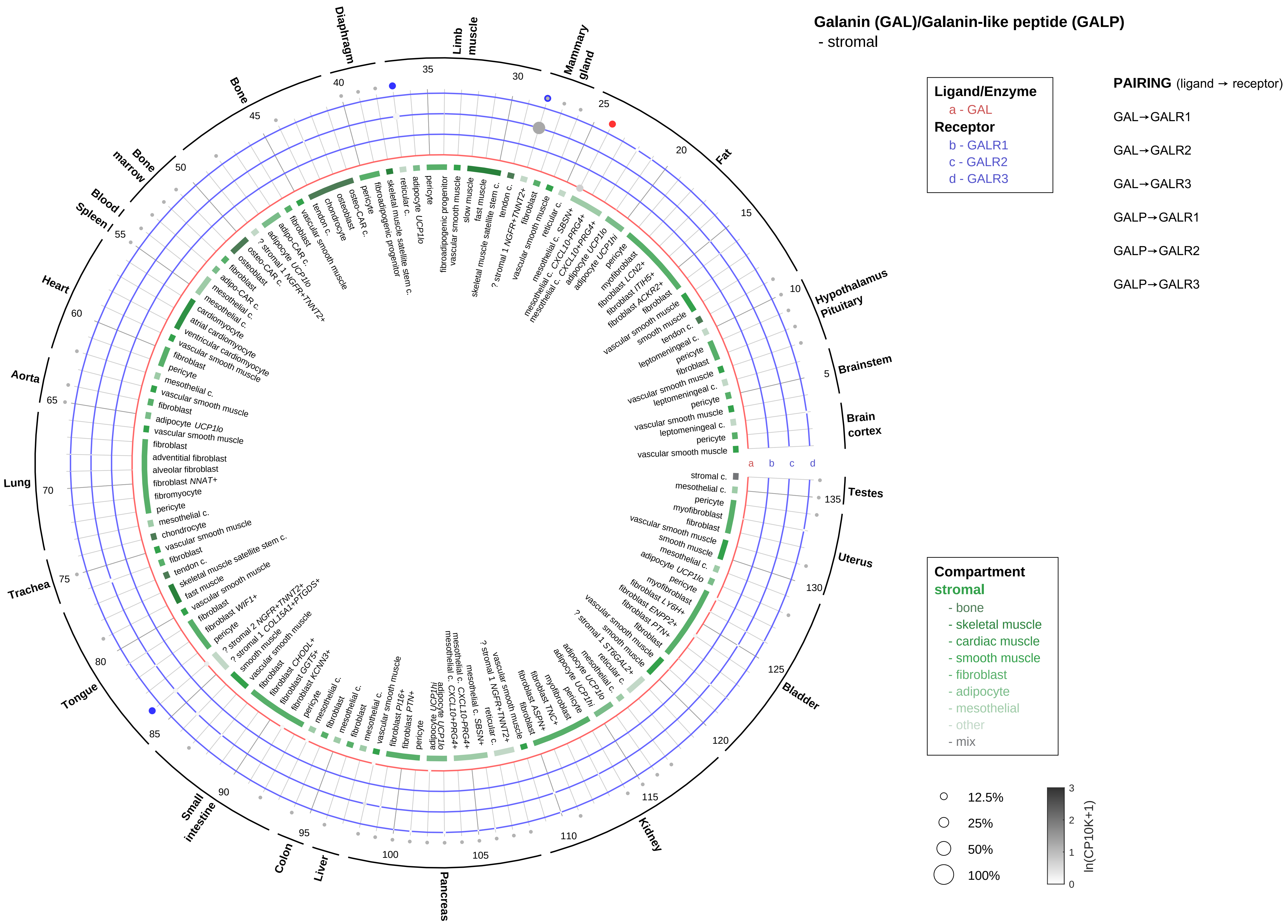
Galanin (GAL)/Galanin-like peptide (GALP)
- stromal
Ligand/Enzyme
a - GAL
Receptor
b - GALR1
c - GALR2
d - GALR3
PAIRING (ligand → receptor)
GAL→GALR1
GAL→GALR2
GAL→GALR3
GALP→GALR1
GALP→GALR2
GALP→GALR3
Compartment
stromal
- bone
- skeletal muscle
- cardiac muscle
- smooth muscle
- fibroblast
- adipocyte
- mesothelial
- other
- mix
12.5%
25%
50%
100%
ln(CP10K+1)
Brain cortex
Brainstem
Hypothalamus Pituitary
Fat
Mammary gland
Limb muscle
Diaphragm
Bone
Bone marrow
Blood / Spleen
Heart
Aorta
Lung
Trachea
Tongue
Small intestine
Colon
Liver
Pancreas
Kidney
Bladder
Uterus
Testes

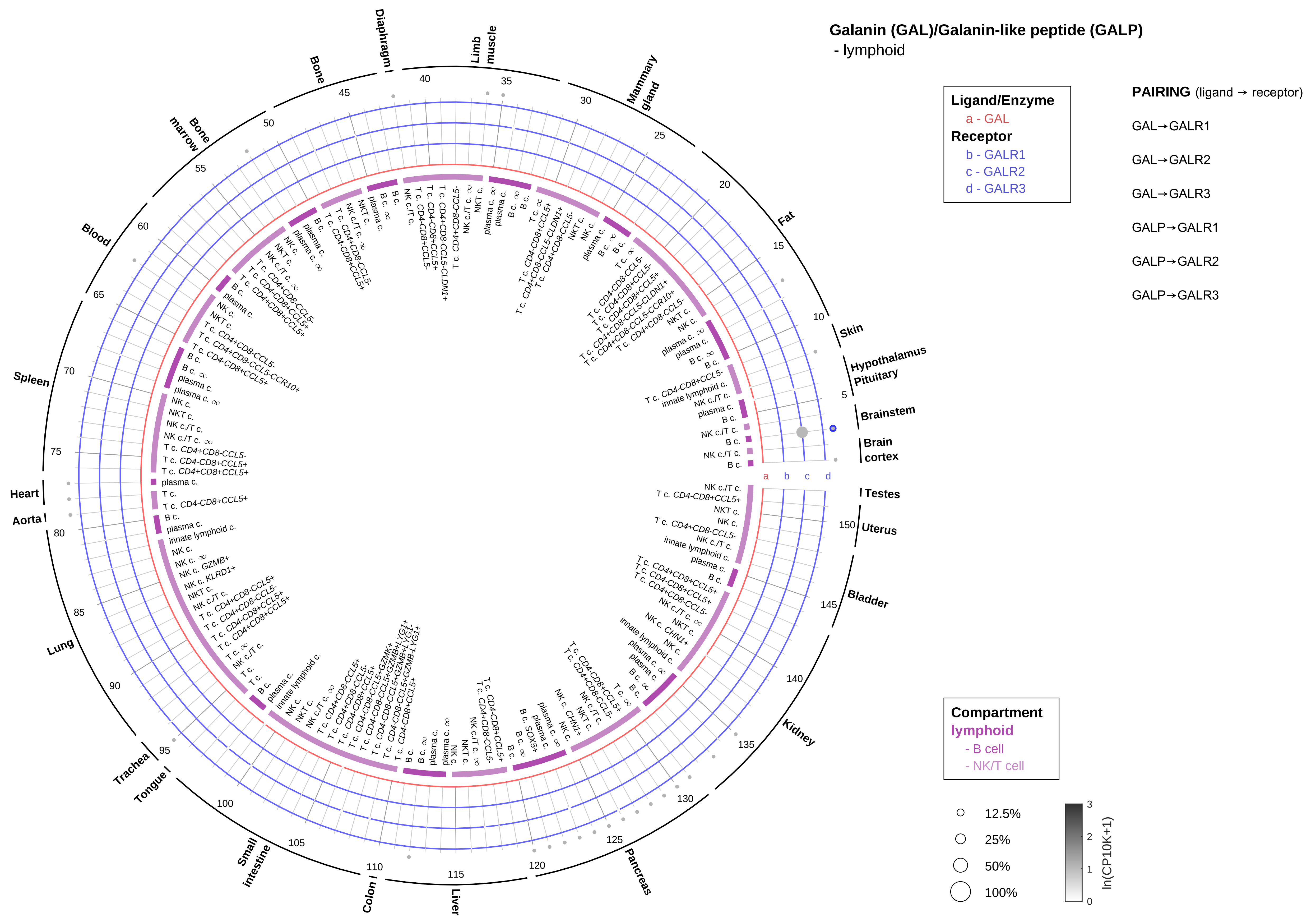
Galanin (GAL)/Galanin-like peptide (GALP)
- lymphoid
Ligand/Enzyme
a - GAL
Receptor
b - GALR1
c - GALR2
d - GALR3
PAIRING (ligand → receptor)
GAL→GALR1
GAL→GALR2
GAL→GALR3
GALP→GALR1
GALP→GALR2
GALP→GALR3
Compartment
lymphoid
- B cell
- NK/T cell
12.5%
25%
50%
100%
ln(CP10K+1)
Skin
Hypothalamus
Pituitary
Brainstem
Brain cortex
Testes
Uterus
Bladder
Kidney
Pancreas
Liver
Colon
Small intestine
Tongue
Trachea
Lung
Aorta
Heart
Spleen
Blood
Bone marrow
Bone
Diaphragm
Limb muscle
Mammary gland
Fat

#### Galanin (GAL)/Galanin-like peptide (GALP)
- myeloid

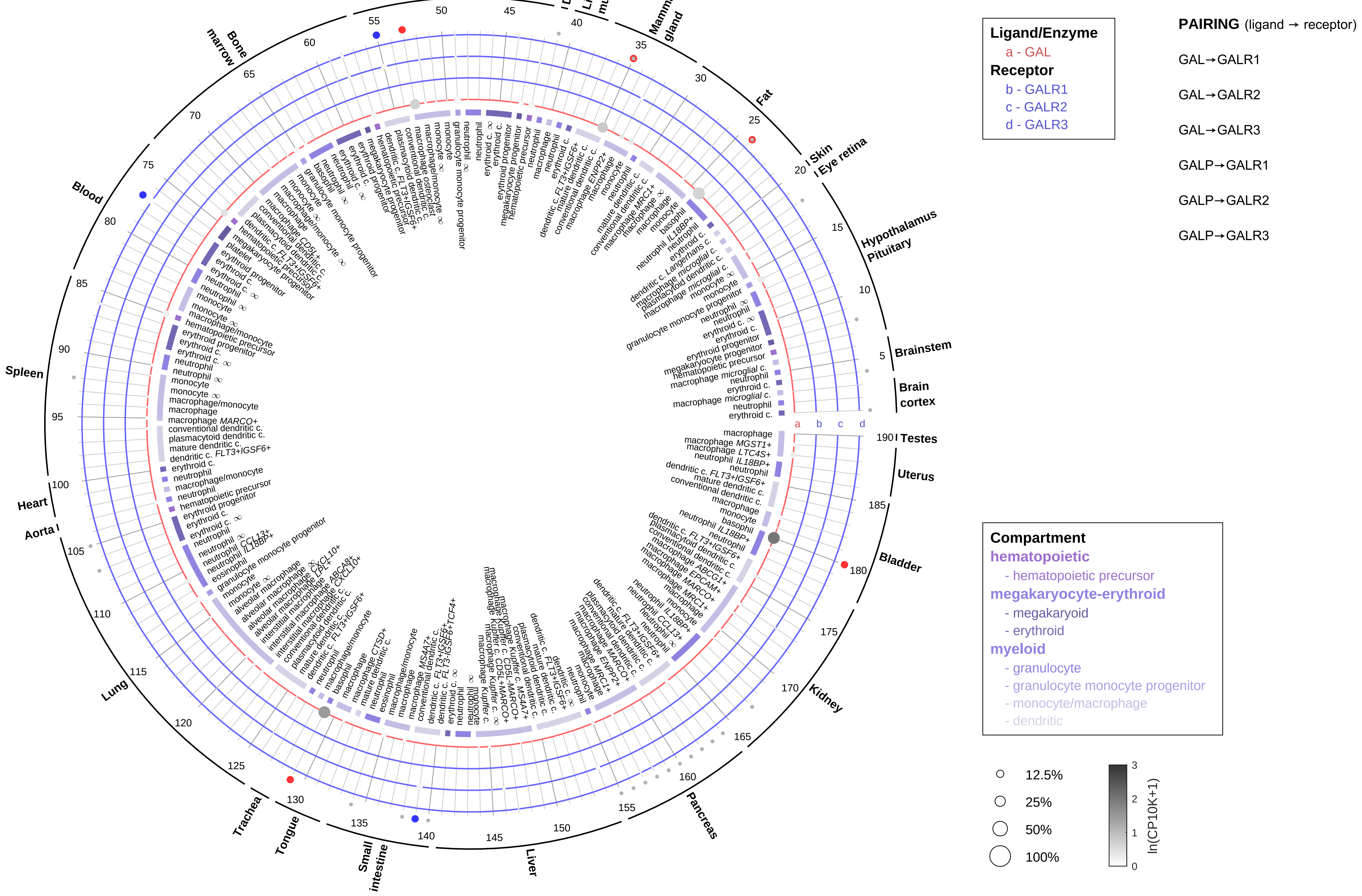

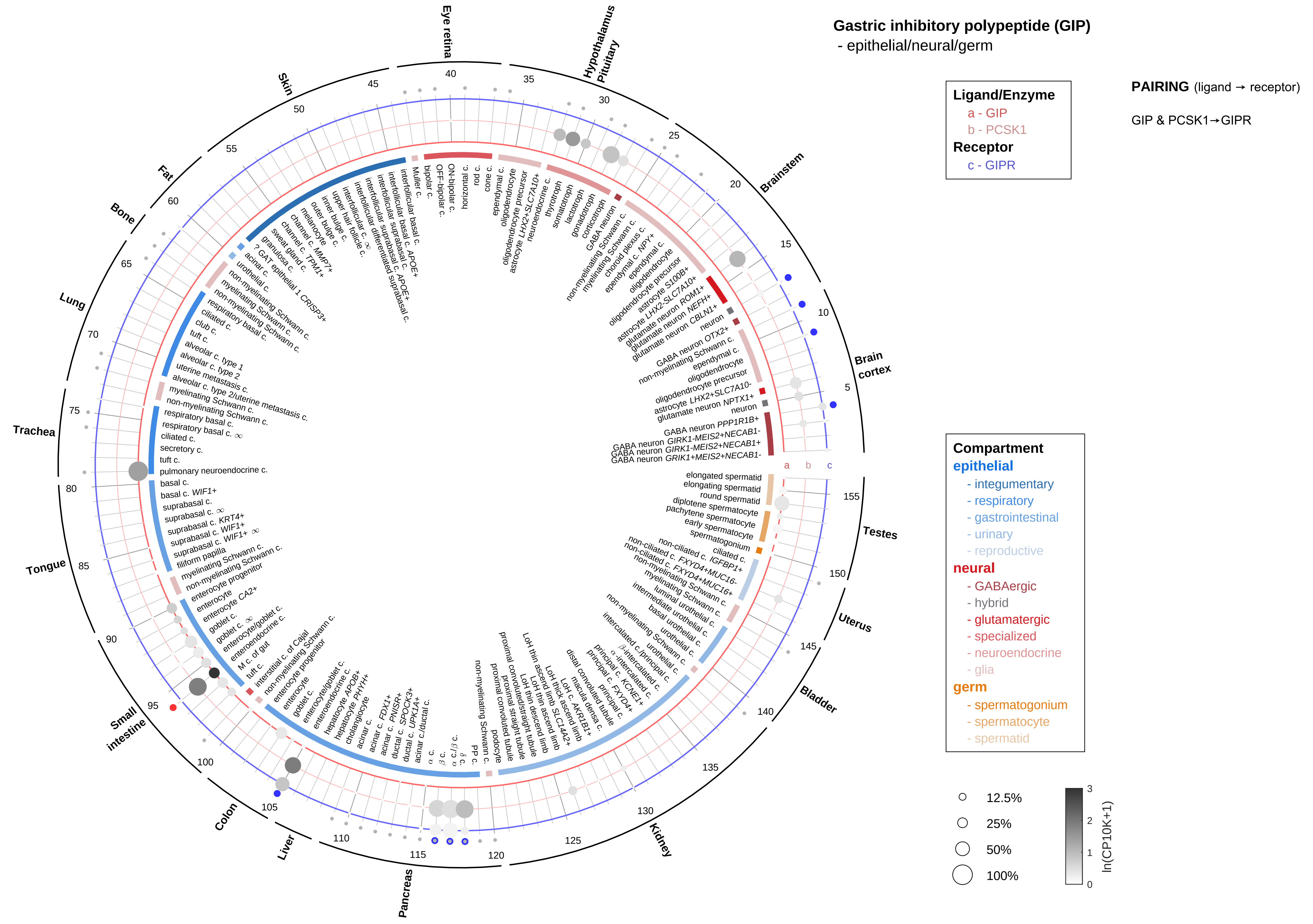

Gastric inhibitory polypeptide (GIP)
- epithelial/neural/germ
Ligand/Enzyme
a - GIP
b - PCSK1
Receptor
c - GIPR
PAIRING (ligand → receptor)
GIP & PCSK1→GIPR
Compartment
epithelial
- integumentary
- respiratory
- gastrointestinal
- urinary
- reproductive
neural
- GABAergic
- hybrid
- glutamatergic
- specialized
- neuroendocrine
- glia
germ
- spermatogonium
- spermatocyte
- spermatid
12.5%
25%
50%
100%
ln(CP10K+1)

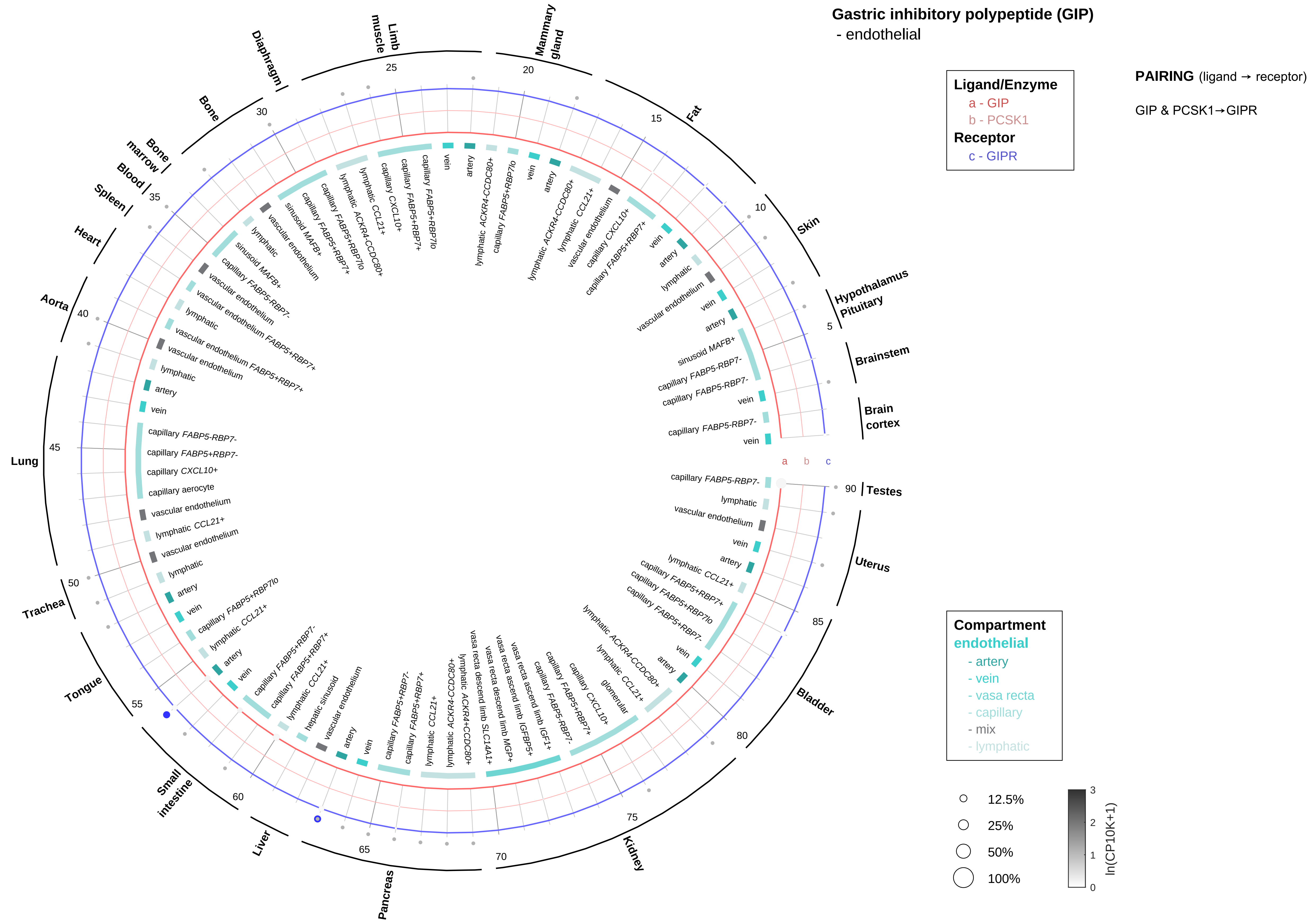

Gastric inhibitory polypeptide (GIP)
- endothelial
Ligand/Enzyme
a - GIP
b - PCSK1
Receptor
c - GIPR
PAIRING (ligand → receptor)
GIP & PCSK1→GIPR
Compartment
endothelial
- artery
- vein
- vasa recta
- capillary
- mix
- lymphatic
12.5%
25%
50%
100%
ln(CP10K+1)
Mammary gland
Fat
Skin
Hypothalamus
Pituitary
Brainstem
Brain cortex
Testes
Uterus
Bladder
Kidney
Pancreas
Liver
Small intestine
Tongue
Trachea
Lung
Aorta
Heart
Spleen
Blood
Bone marrow
Bone
Diaphragm
Limb muscle

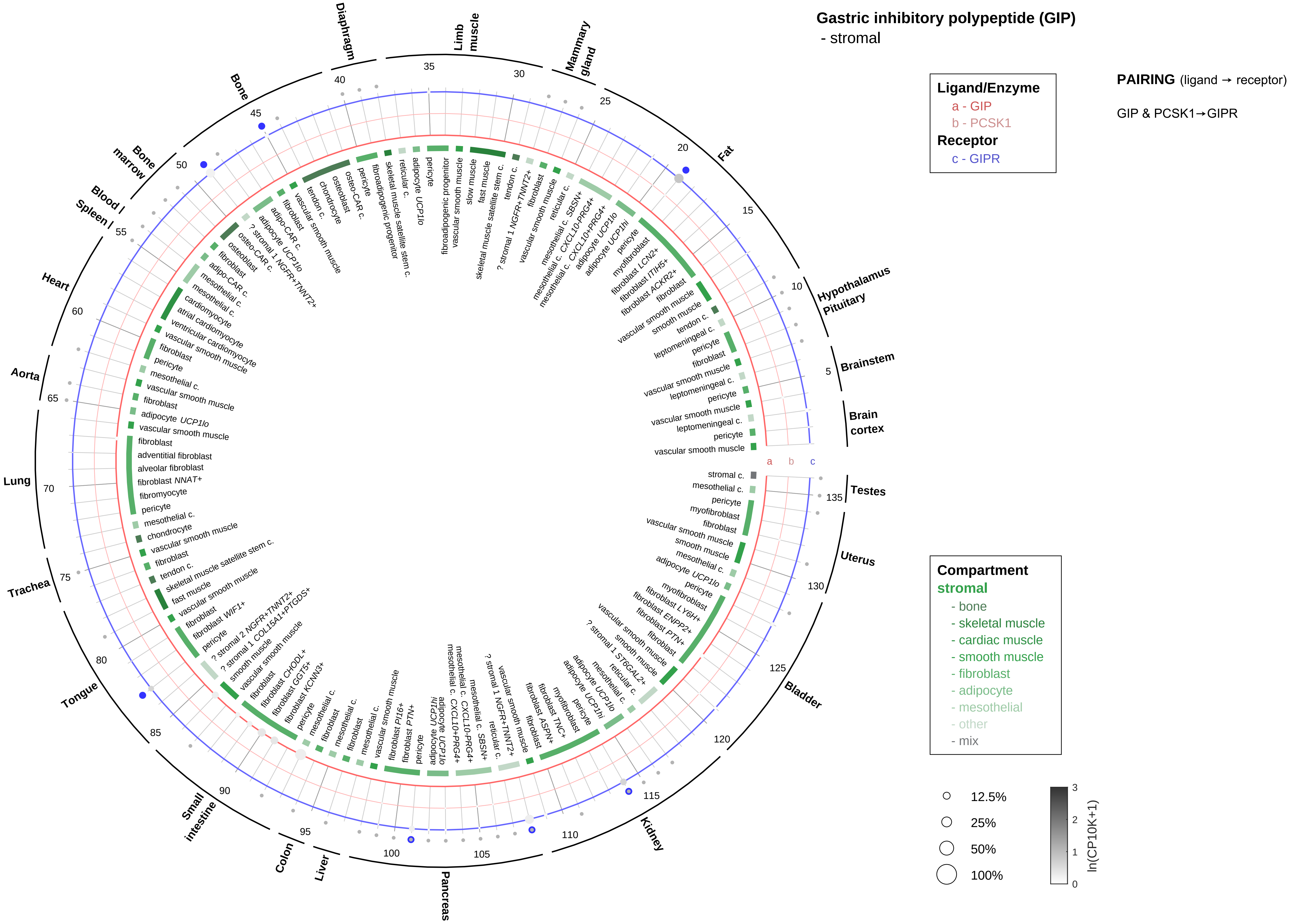

Gastric inhibitory polypeptide (GIP)
- stromal
Ligand/Enzyme
a - GIP
b - PCSK1
Receptor
c - GIPR
PAIRING (ligand → receptor)
GIP & PCSK1→GIPR
Compartment
stromal
- bone
- skeletal muscle
- cardiac muscle
- smooth muscle
- fibroblast
- adipocyte
- mesothelial
- other
- mix
12.5%
25%
50%
100%
ln(CP10K+1)
Brain cortex
Brainstem
Hypothalamus Pituitary
Fat
Mammary gland
Limb muscle
Diaphragm
Bone
Bone marrow
Blood
Spleen
Heart
Aorta
Lung
Trachea
Tongue
Small intestine
Colon
Liver
Pancreas
Kidney
Bladder
Uterus
Testes

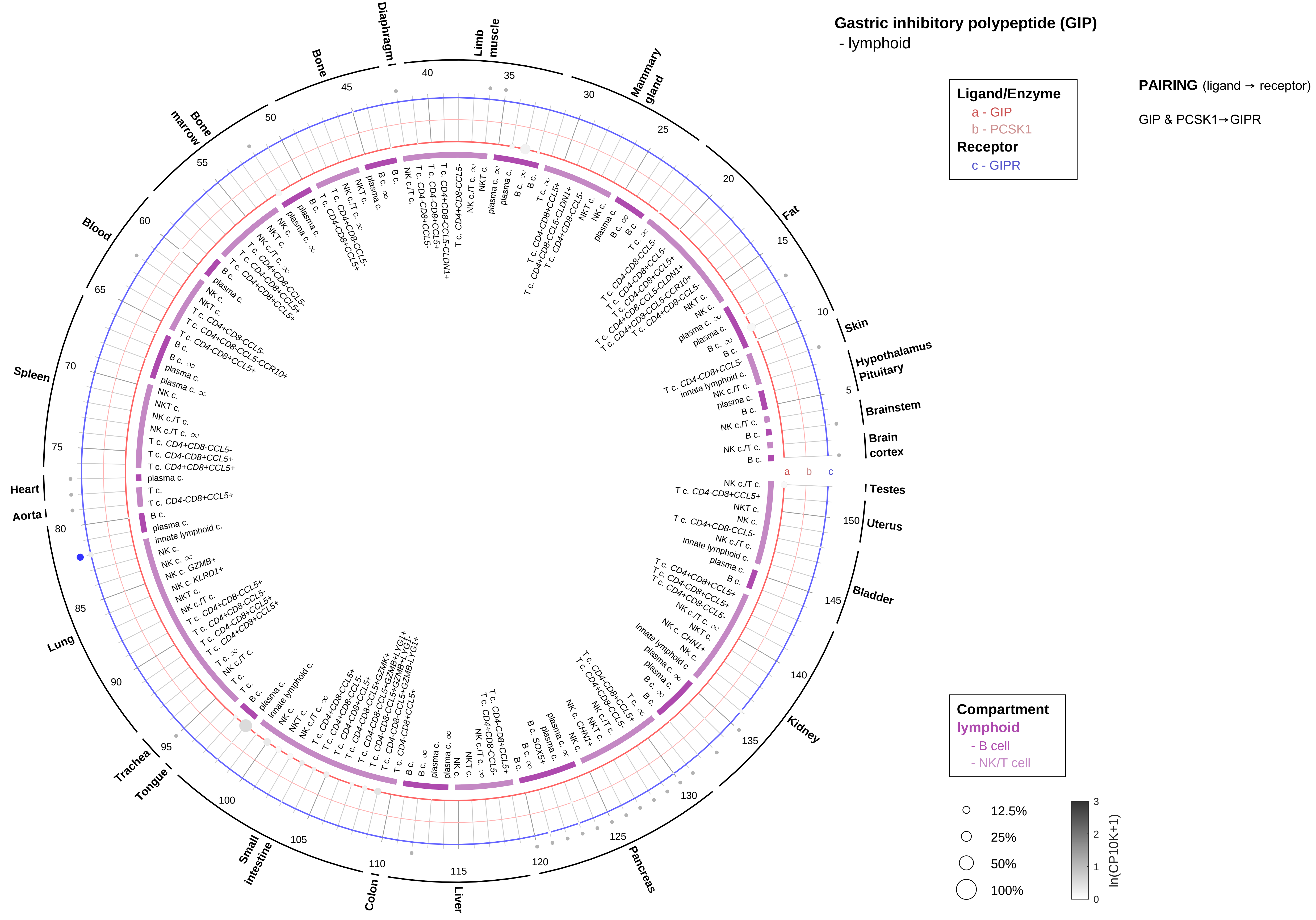
Gastric inhibitory polypeptide (GIP)
- lymphoid
Ligand/Enzyme
a - GIP
b - PCSK1
Receptor
c - GIPR
PAIRING (ligand → receptor)
GIP & PCSK1→GIPR
Compartment
lymphoid
- B cell
- NK/T cell
12.5%
25%
50%
100%
ln(CP10K+1)
Skin
Hypothalamus
Pituitary
Brainstem
Brain cortex
Testes
Uterus
Bladder
Kidney
Pancreas
Liver
Colon
Small intestine
Tongue
Trachea
Lung
Aorta
Heart
Spleen
Blood
Bone marrow
Bone
Diaphragm
Limb muscle
Mammary gland
Fat

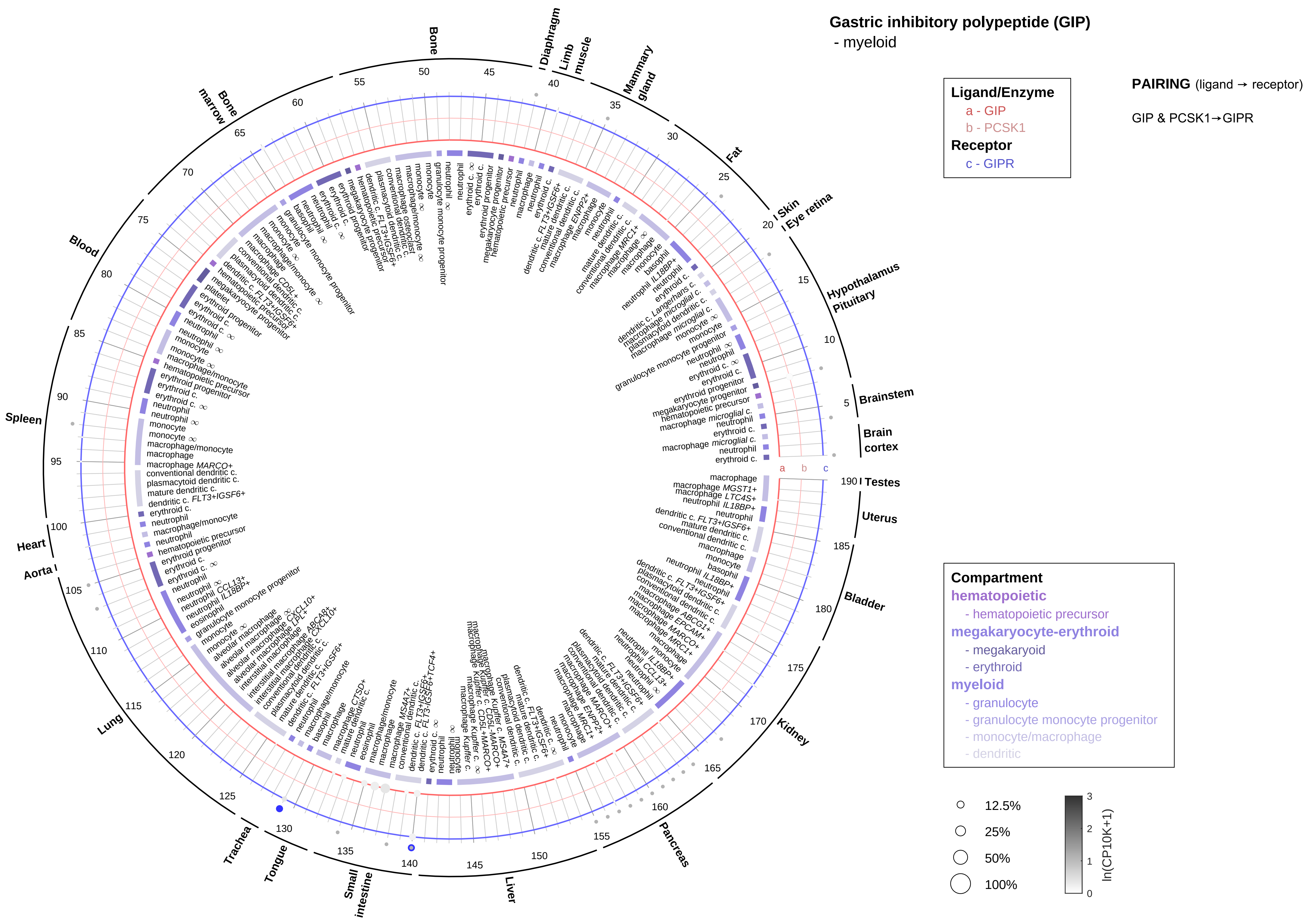
Gastric inhibitory polypeptide (GIP)
- myeloid
Ligand/Enzyme
a - GIP
b - PCSK1
Receptor
c - GIPR
PAIRING (ligand → receptor)
GIP & PCSK1→GIPR
Compartment
hematopoietic
- hematopoietic precursor
megakaryocyte-erythroid
- megakaryoid
- erythroid
myeloid
- granulocyte
- granulocyte monocyte progenitor
- monocyte/macrophage
- dendritic
12.5%
25%
50%
100%
ln(CP10K+1)
Bone
Diaphragm
Limb muscle
Mammary gland
Fat
Skin
Eye retina
Hypothalamus
Pituitary
Brainstem
Brain cortex
Testes
Uterus
Bladder
Kidney
Pancreas
Liver
Small intestine
Tongue
Trachea
Lung
Aorta
Heart
Spleen
Blood
Bone marrow

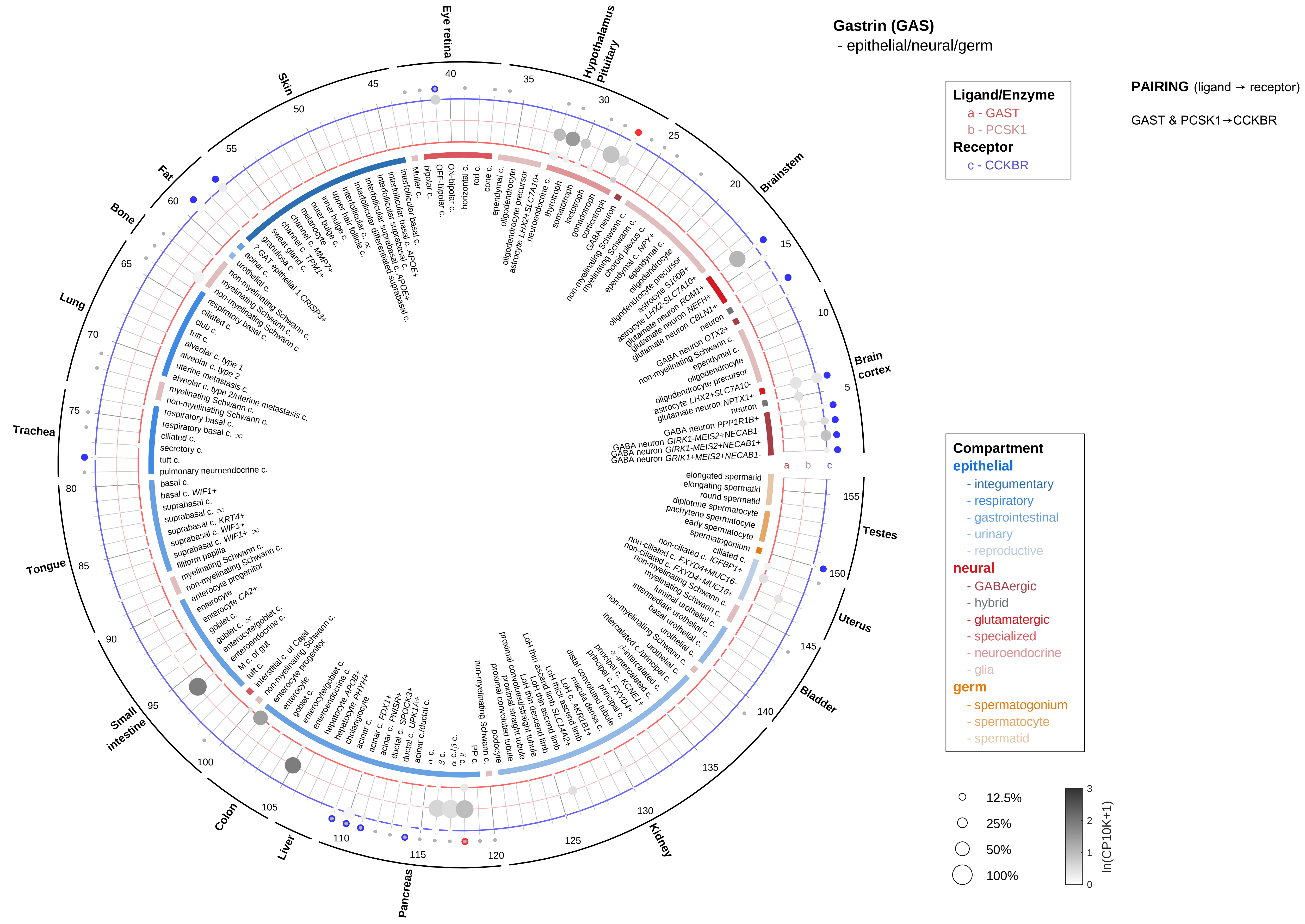
Gastrin (GAS)
- epithelial/neural/germ
Ligand/Enzyme
a - GAST
b - PCSK1
Receptor
c - CCKBR
PAIRING (ligand → receptor)
GAST & PCSK1→CCKBR
Compartment
epithelial
- integumentary
- respiratory
- gastrointestinal
- urinary
- reproductive
neural
- GABAergic
- hybrid
- glutamatergic
- specialized
- neuroendocrine
- glia
germ
- spermatogonium
- spermatocyte
- spermatid
12.5%
25%
50%
100%
ln(CP10K+1)
Eye retina
Hypothalamus
Pituitary
Brainstem
Brain cortex
Testes
Uterus
Bladder
Kidney
Pancreas
Liver
Colon
Small intestine
Tongue
Trachea
Lung
Bone
Fat
Skin

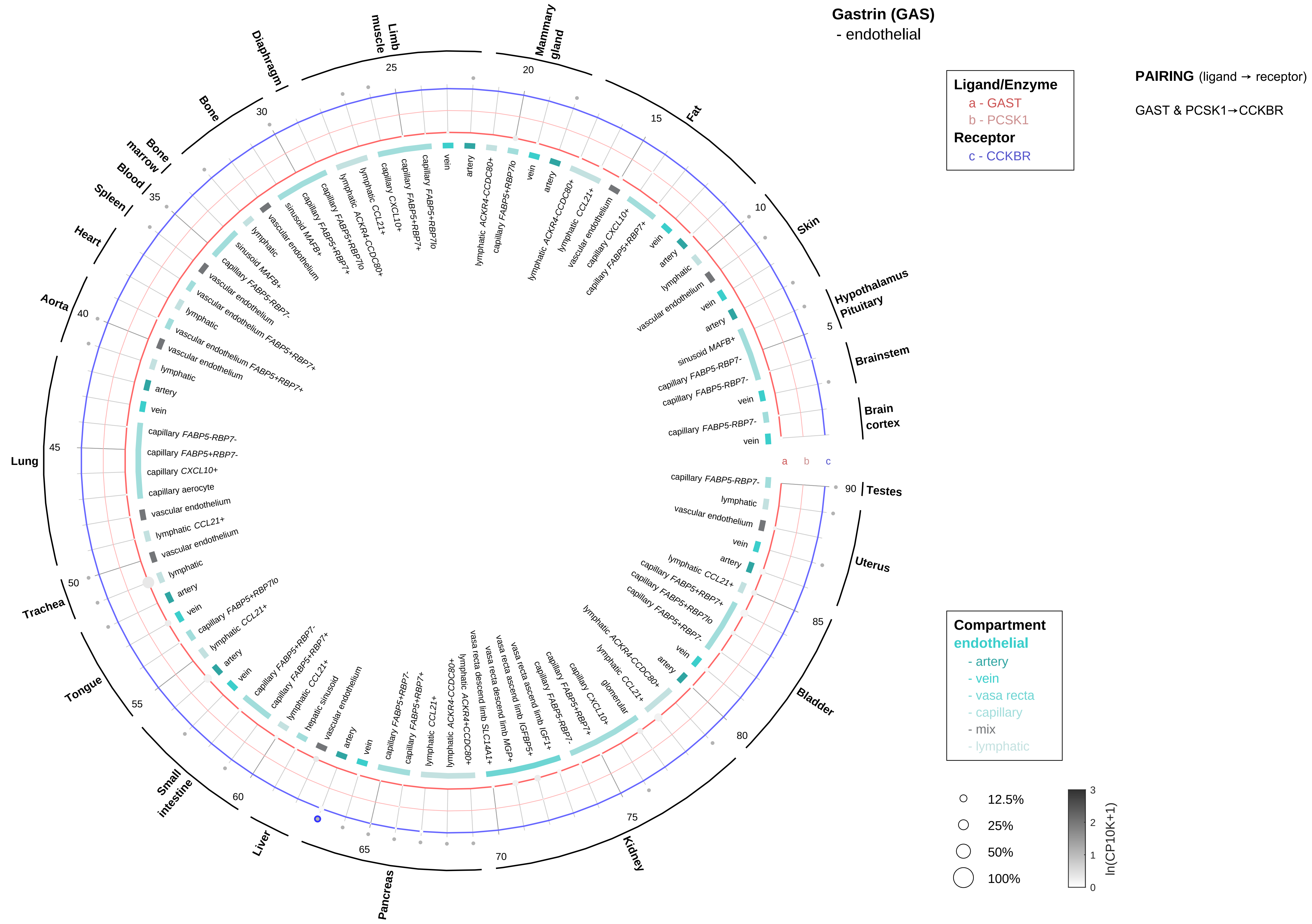

Gastrin (GAS)
- endothelial
Ligand/Enzyme
a - GAST
b - PCSK1
Receptor
c - CCKBR
PAIRING (ligand → receptor)
GAST & PCSK1→CCKBR
Compartment
endothelial
- artery
- vein
- vasa recta
- capillary
- mix
- lymphatic
12.5%
25%
50%
100%
ln(CP10K+1)

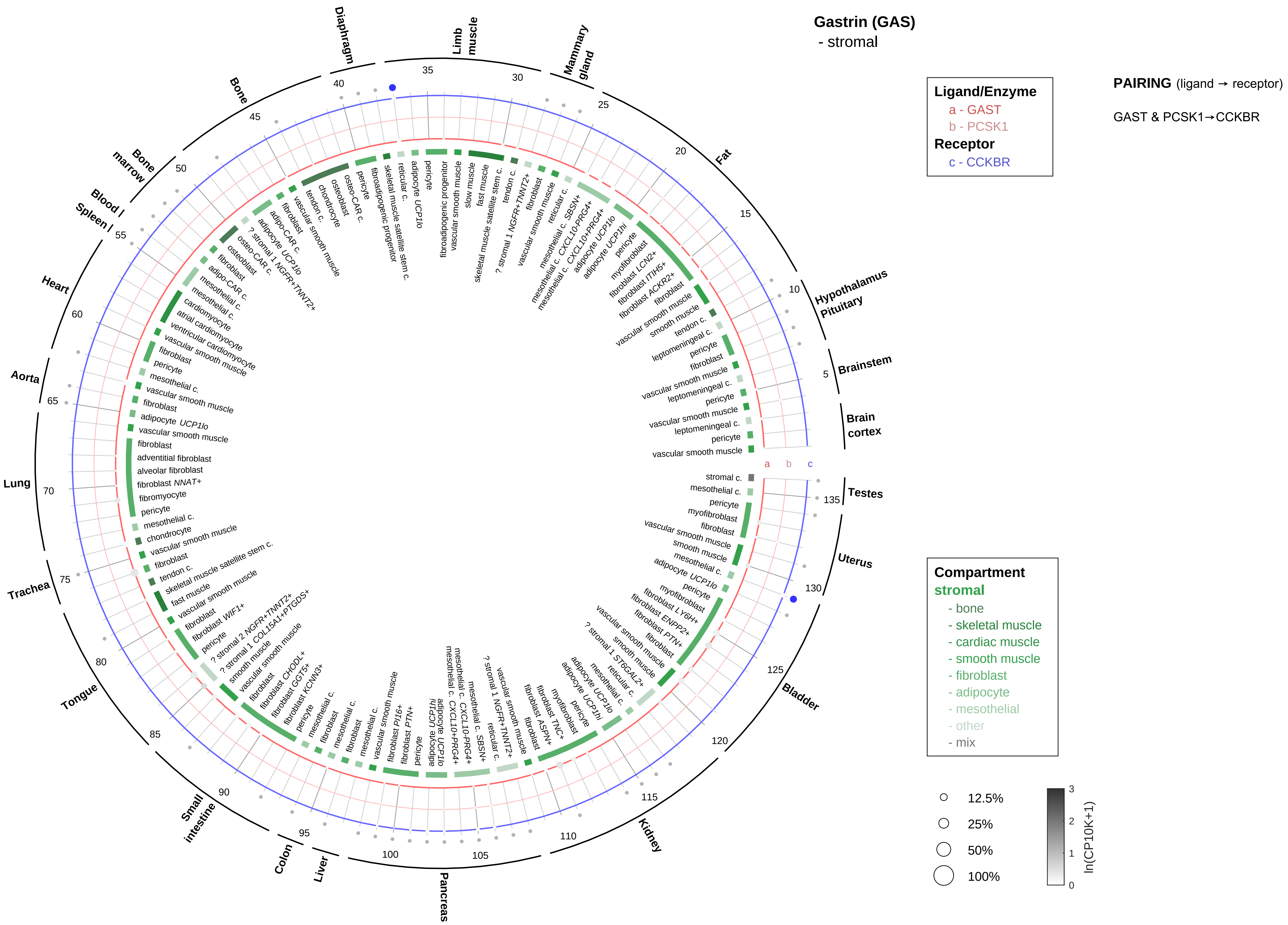
Gastrin (GAS)
- stromal
Ligand/Enzyme
a - GAST
b - PCSK1
Receptor
c - CCKBR
PAIRING (ligand → receptor)
GAST & PCSK1→CCKBR
Compartment
stromal
- bone
- skeletal muscle
- cardiac muscle
- smooth muscle
- fibroblast
- adipocyte
- mesothelial
- other
- mix
12.5%
25%
50%
100%
ln(CP10K+1)
Brain cortex
Brainstem
Hypothalamus Pituitary
Fat
Mammary gland
Limb muscle
Diaphragm
Bone
Bone marrow
Blood
Spleen
Heart
Aorta
Lung
Trachea
Tongue
Small intestine
Colon
Liver
Pancreas
Kidney
Bladder
Uterus
Testes

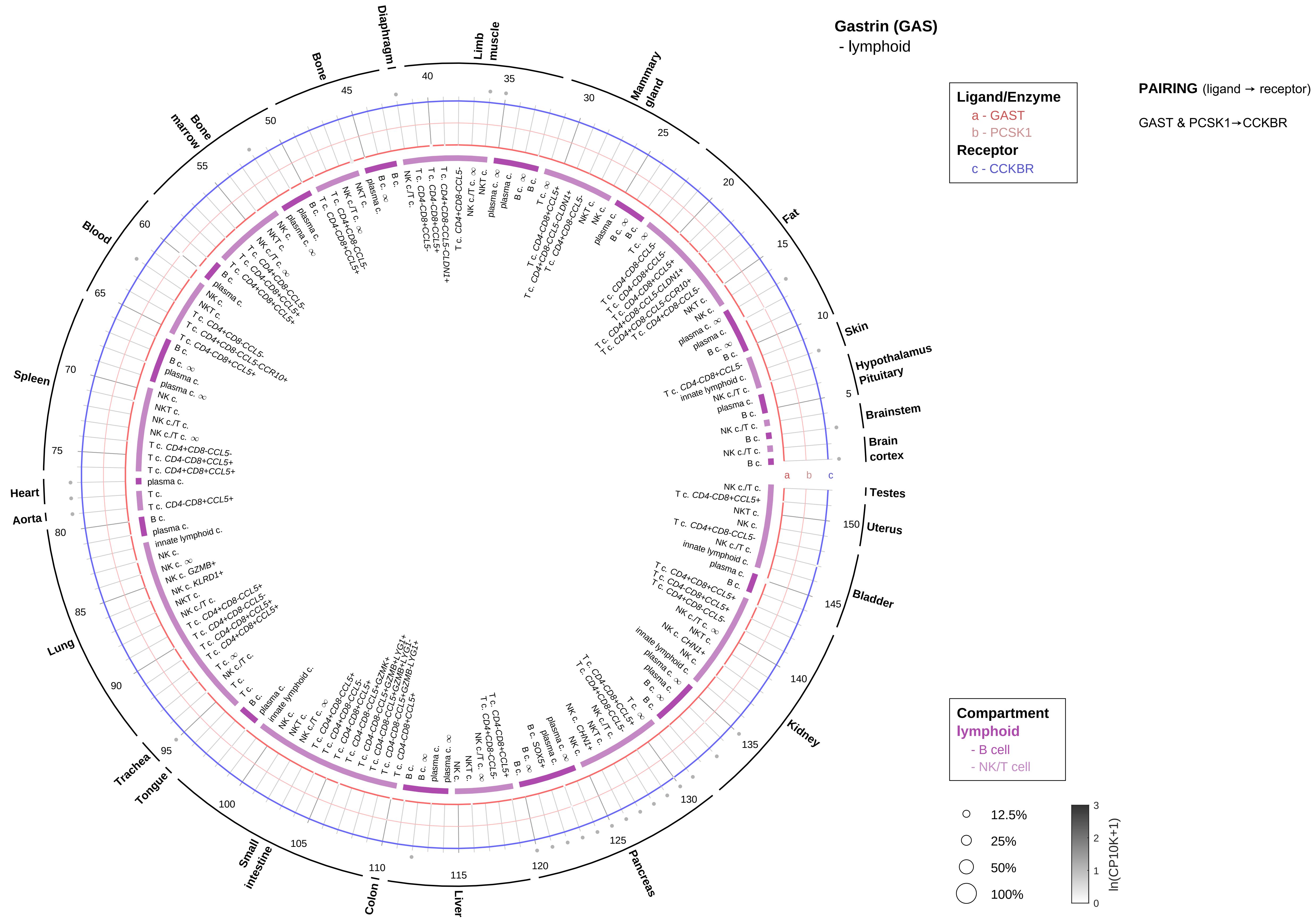

Gastrin (GAS)
- lymphoid
Ligand/Enzyme
a - GAST
b - PCSK1
Receptor
c - CCKBR
PAIRING (ligand → receptor)
GAST & PCSK1→CCKBR
Compartment
lymphoid
- B cell
- NK/T cell
12.5%
25%
50%
100%
ln(CP10K+1)
Skin
Hypothalamus
Pituitary
Brainstem
Brain cortex
Testes
Uterus
Bladder
Kidney
Pancreas
Liver
Colon
Small intestine
Tongue
Trachea
Lung
Aorta
Heart
Spleen
Blood
Bone marrow
Bone
Diaphragm
Limb muscle
Mammary gland
Fat

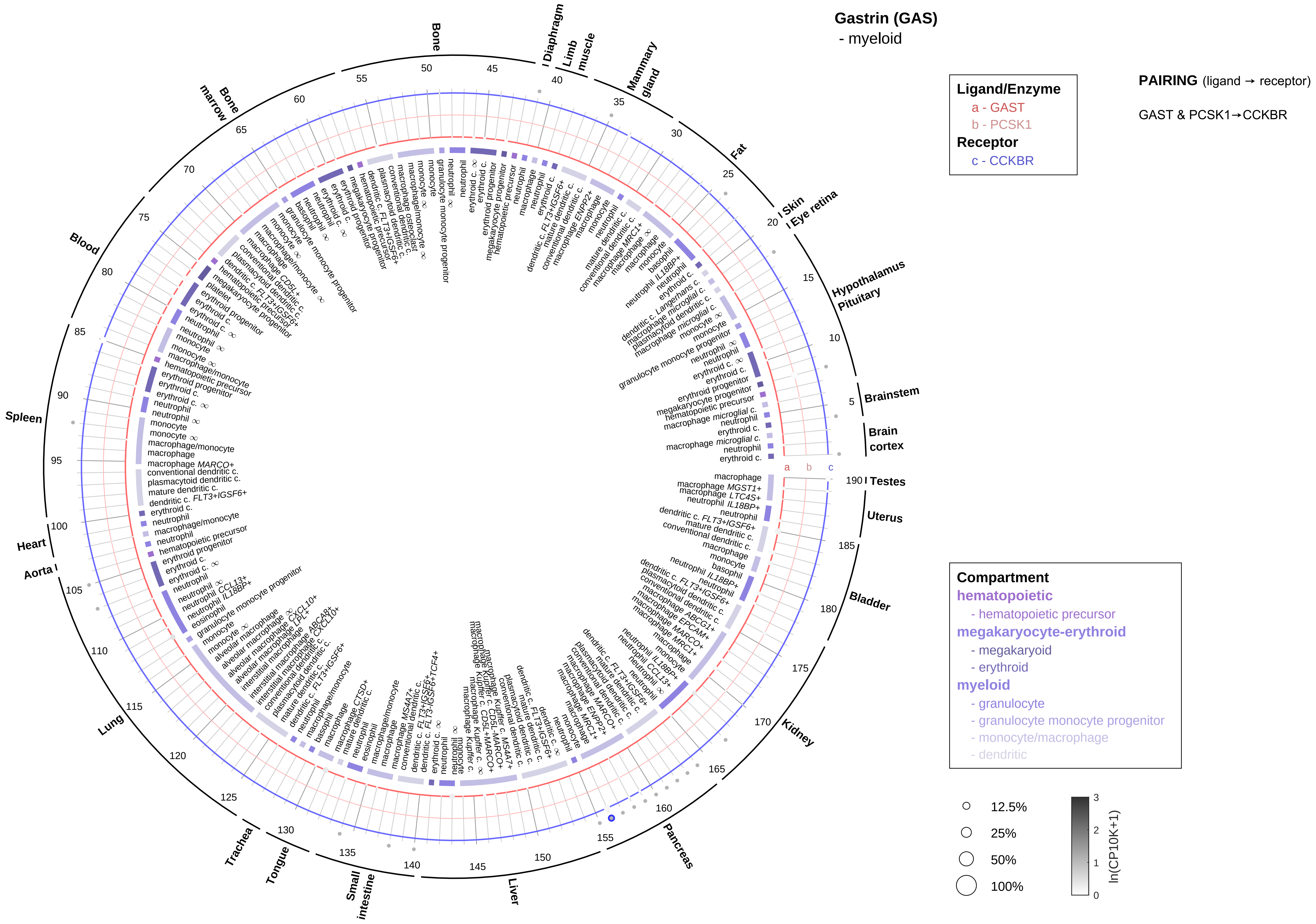

Gastrin (GAS)
- myeloid
Ligand/Enzyme
a - GAST
b - PCSK1
Receptor
c - CCKBR
PAIRING (ligand → receptor)
GAST & PCSK1→CCKBR
Compartment
hematopoietic
- hematopoietic precursor
megakaryocyte-erythroid
- megakaryoid
- erythroid
myeloid
- granulocyte
- granulocyte monocyte progenitor
- monocyte/macrophage
- dendritic
12.5%
25%
50%
100%
ln(CP10K+1)
Bone
Diaphragm
Limb muscle
Mammary gland
Fat
Skin
Eye retina
Hypothalamus
Pituitary
Brainstem
Brain cortex
Testes
Uterus
Bladder
Kidney
Pancreas
Liver
Small intestine
Tongue
Trachea
Lung
Aorta
Heart
Spleen
Blood
Bone marrow

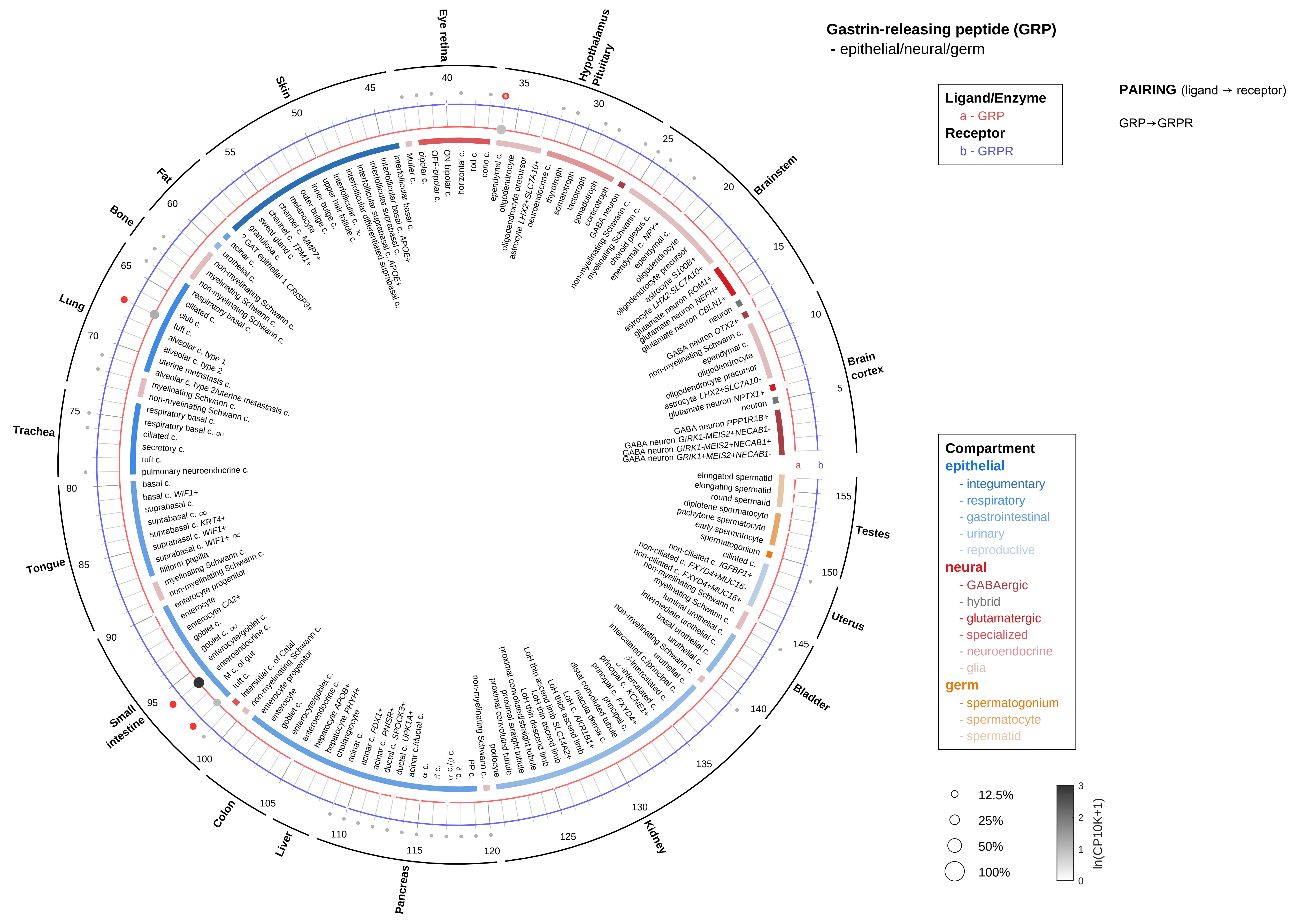

Gastrin-releasing peptide (GRP)
- epithelial/neural/germ
Ligand/Enzyme
a - GRP
Receptor
b - GRPR
PAIRING (ligand → receptor)
GRP→GRPR
Compartment
epithelial
- integumentary
- respiratory
- gastrointestinal
- urinary
- reproductive
neural
- GABAergic
- hybrid
- glutamatergic
- specialized
- neuroendocrine
- glia
germ
- spermatogonium
- spermatocyte
- spermatid
12.5%
25%
50%
100%
ln(CP10K+1)
Brain cortex
Brainstem
Hypothalamus Pituitary
Eye retina
Skin
Fat
Bone
Lung
Trachea
Tongue
Small intestine
Colon
Liver
Pancreas
Kidney
Bladder
Uterus
Testes

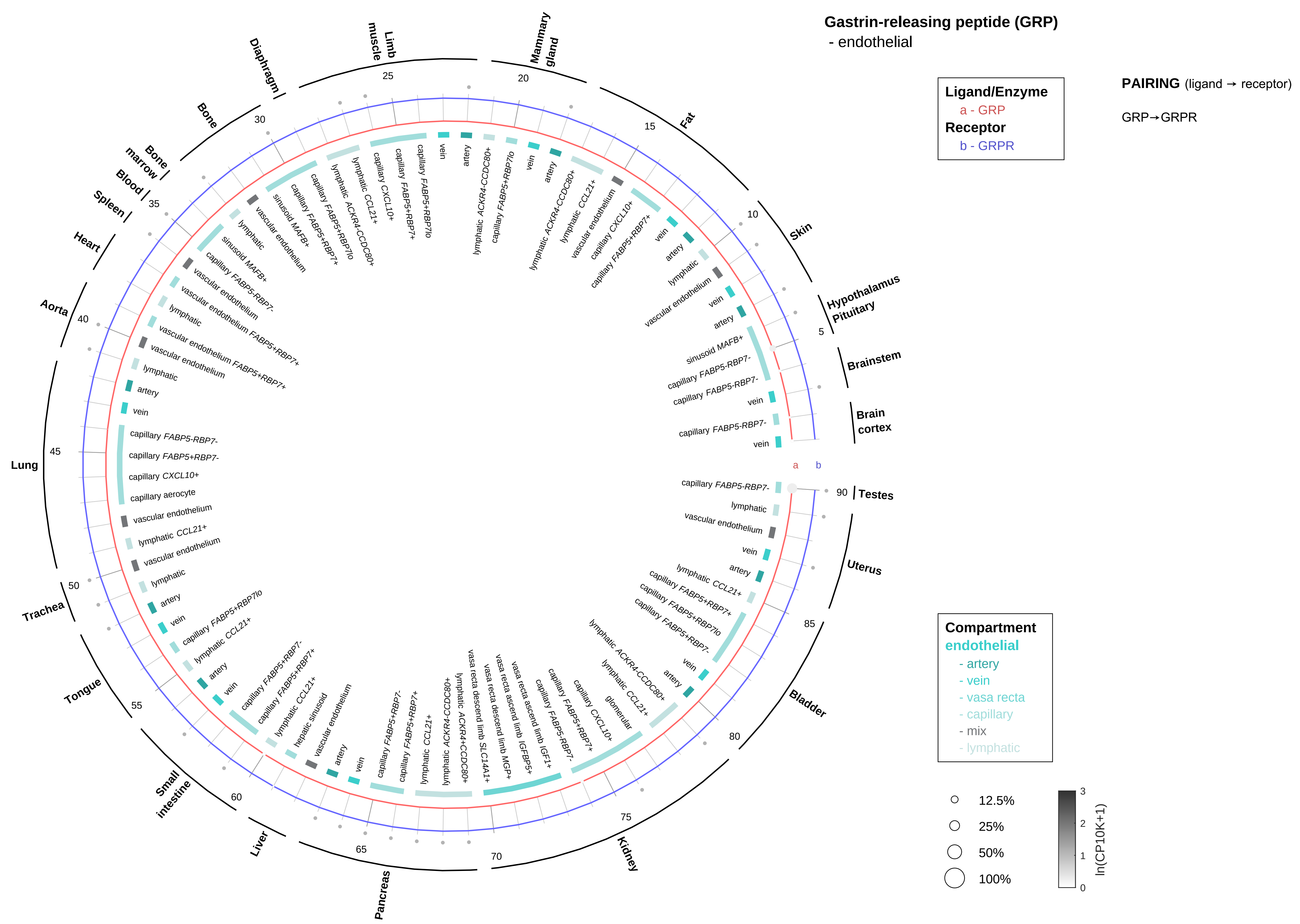

Gastrin-releasing peptide (GRP)
- endothelial
Ligand/Enzyme
a - GRP
Receptor
b - GRPR
PAIRING (ligand → receptor)
GRP→GRPR
Compartment
endothelial
- artery
- vein
- vasa recta
- capillary
- mix
- lymphatic
12.5%
25%
50%
100%
ln(CP10K+1)
Testes
Uterus
Bladder
Kidney
Pancreas
Liver
Small intestine
Tongue
Trachea
Lung
Aorta
Heart
Spleen
Blood
Bone marrow
Bone
Diaphragm
Limb muscle
Mammary gland
Fat
Skin
Hypothalamus Pituitary
Brainstem
Brain cortex

**Gastrin-releasing peptide (GRP)**
- stromal

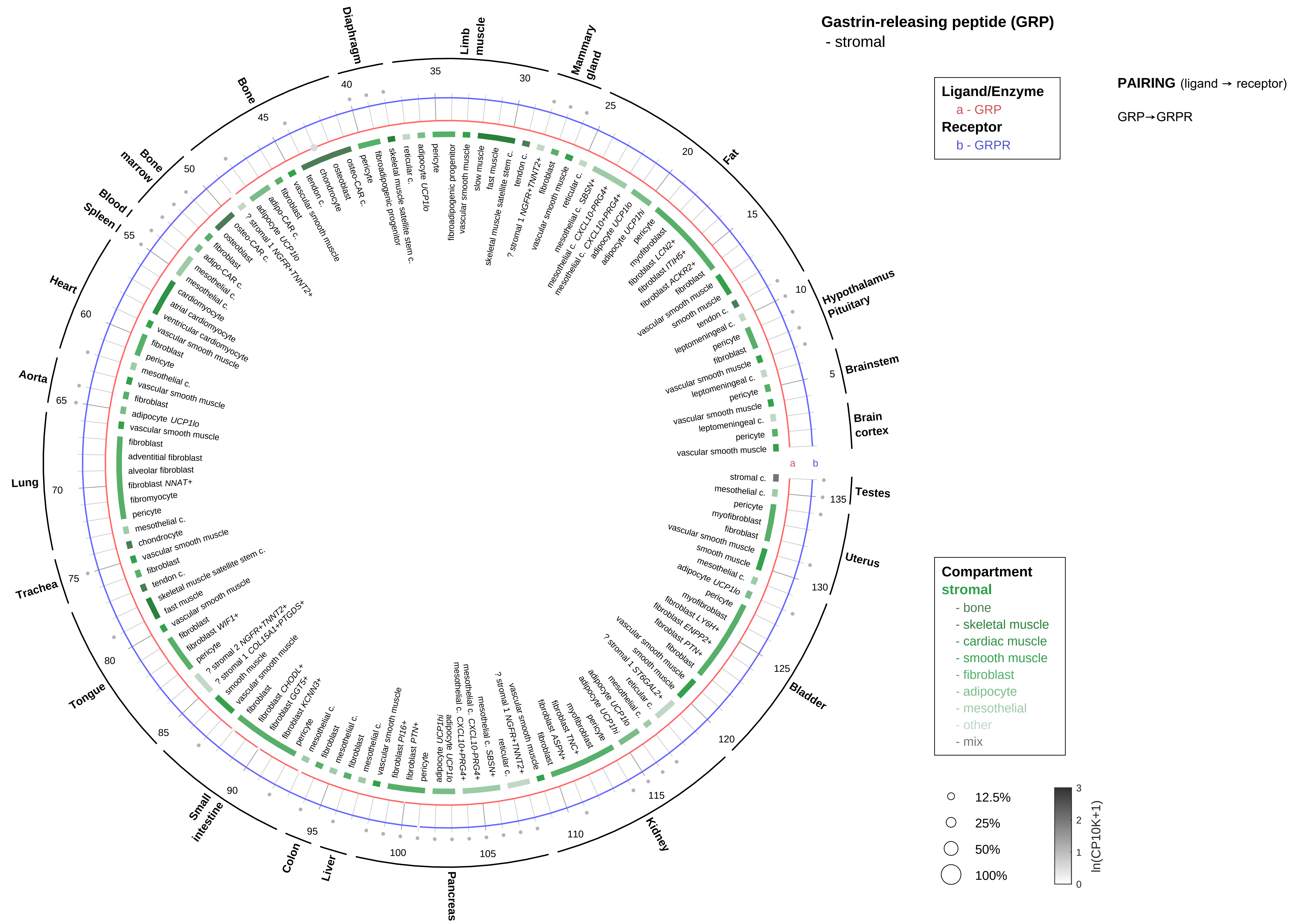

**Gastrin-releasing peptide (GRP)**
- lymphoid

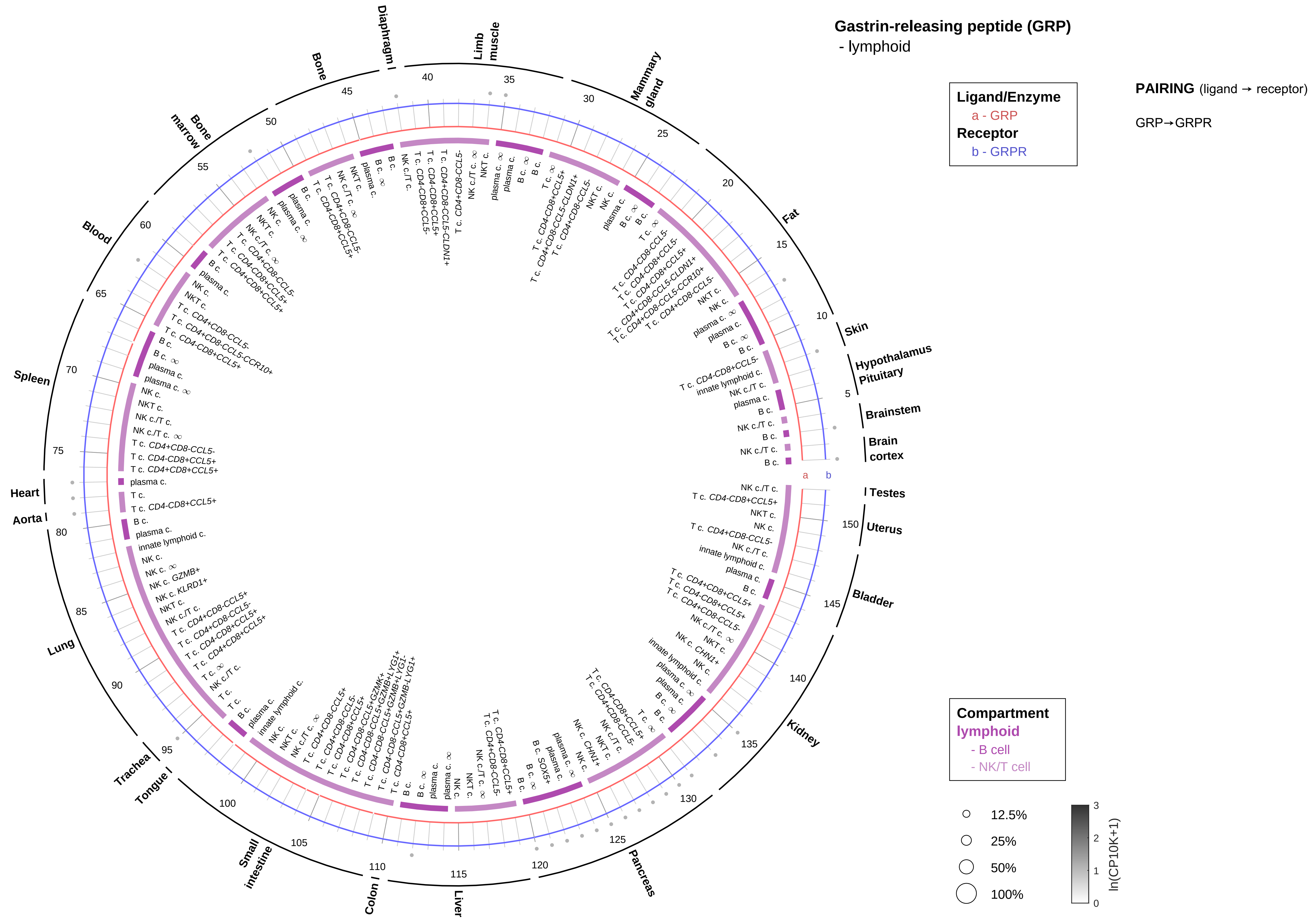

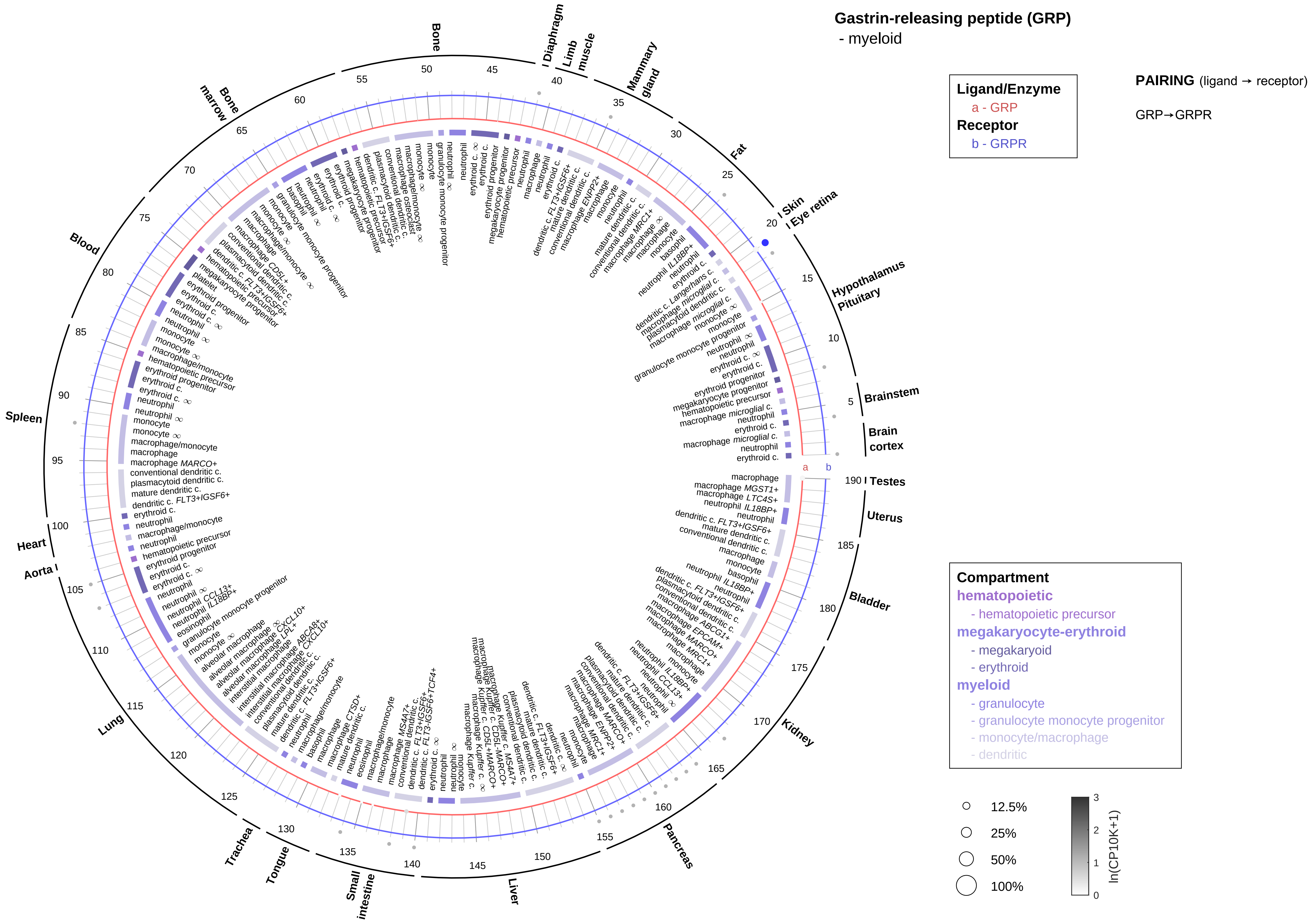

Gastrin-releasing peptide (GRP)
- myeloid
Ligand/Enzyme
a - GRP
Receptor
b - GRPR
PAIRING (ligand → receptor)
GRP→GRPR
Compartment
hematopoietic
- hematopoietic precursor
megakaryocyte-erythroid
- megakaryoid
- erythroid
myeloid
- granulocyte
- granulocyte monocyte progenitor
- monocyte/macrophage
- dendritic
12.5%
25%
50%
100%
ln(CP10K+1)
Bone
Diaphragm
Limb muscle
Mammary gland
Fat
Skin
Eye retina
Hypothalamus
Pituitary
Brainstem
Brain cortex
Testes
Uterus
Bladder
Kidney
Pancreas
Liver
Small intestine
Tongue
Trachea
Lung
Aorta
Heart
Spleen
Blood
Bone marrow

**Ghrelin**
- epithelial/neural/germ

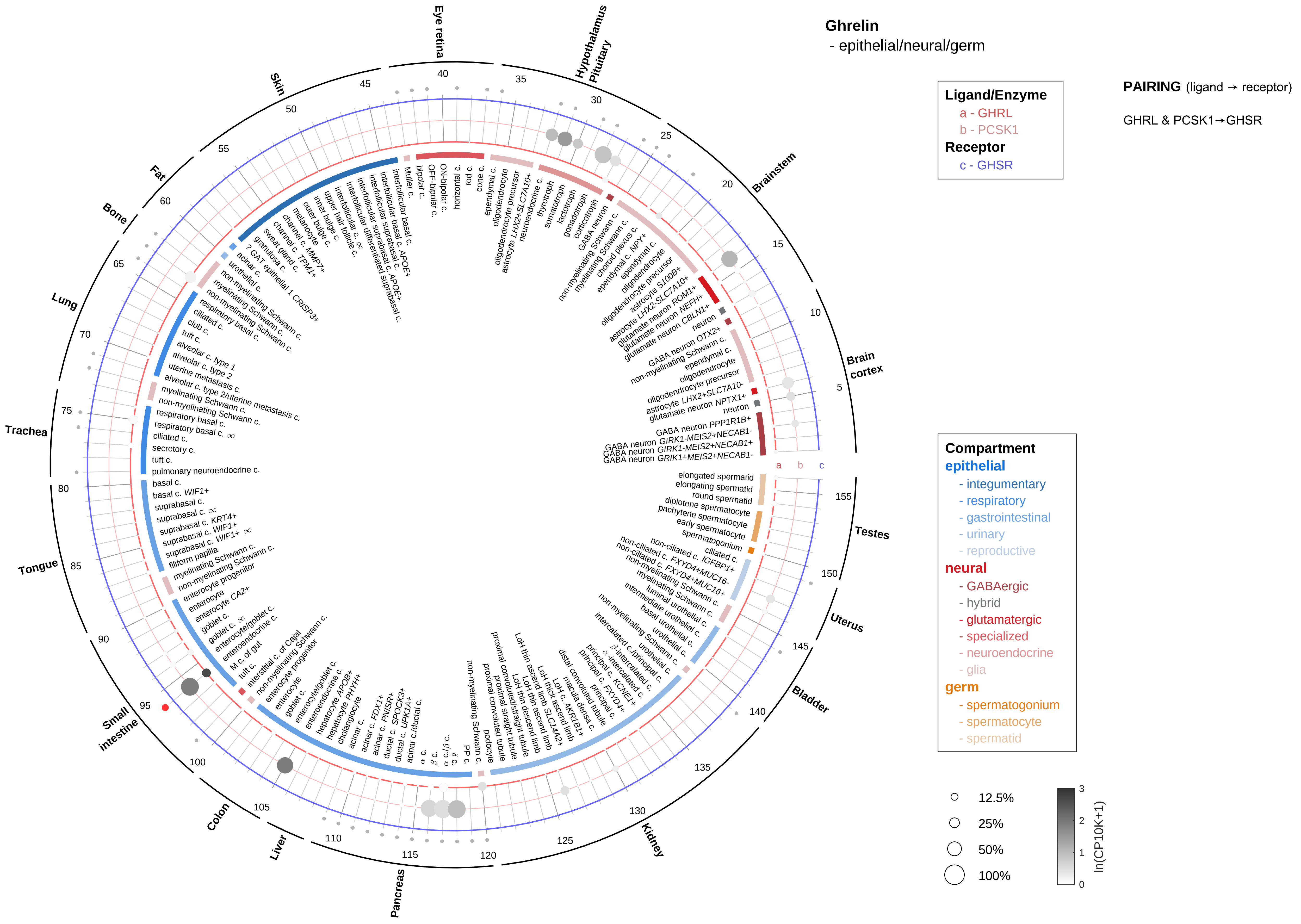

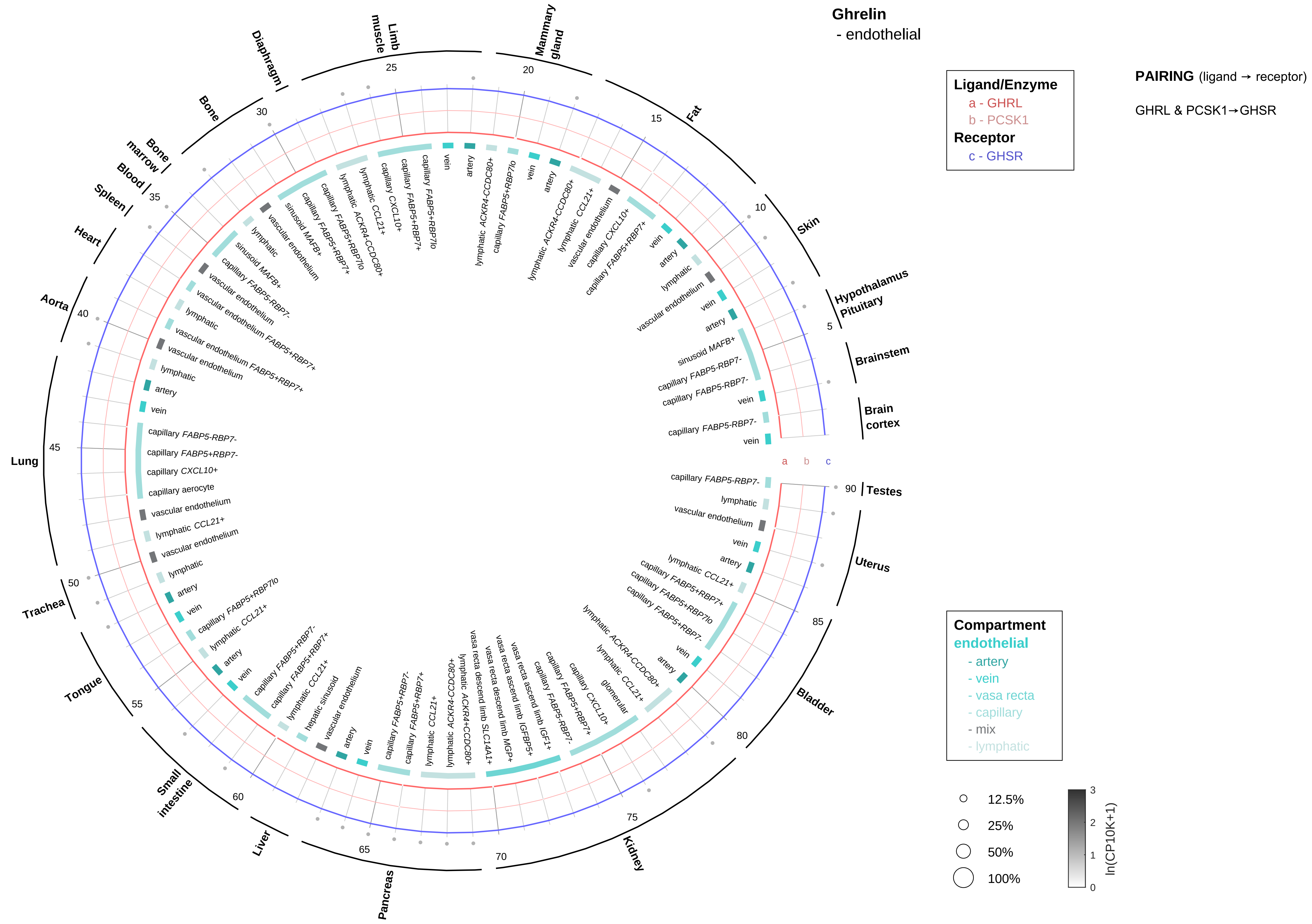

Ghrelin
- endothelial
Ligand/Enzyme
a - GHRL
b - PCSK1
Receptor
c - GHSR
PAIRING (ligand → receptor)
GHRL & PCSK1→GHSR
Compartment
endothelial
- artery
- vein
- vasa recta
- capillary
- mix
- lymphatic
12.5%
25%
50%
100%
ln(CP10K+1)
Brain cortex
Brainstem
Hypothalamus Pituitary
Skin
Fat
Mammary gland
Limb muscle
Diaphragm
Bone
Bone marrow
Blood
Spleen
Heart
Aorta
Lung
Trachea
Tongue
Small intestine
Liver
Pancreas
Kidney
Bladder
Uterus
Testes

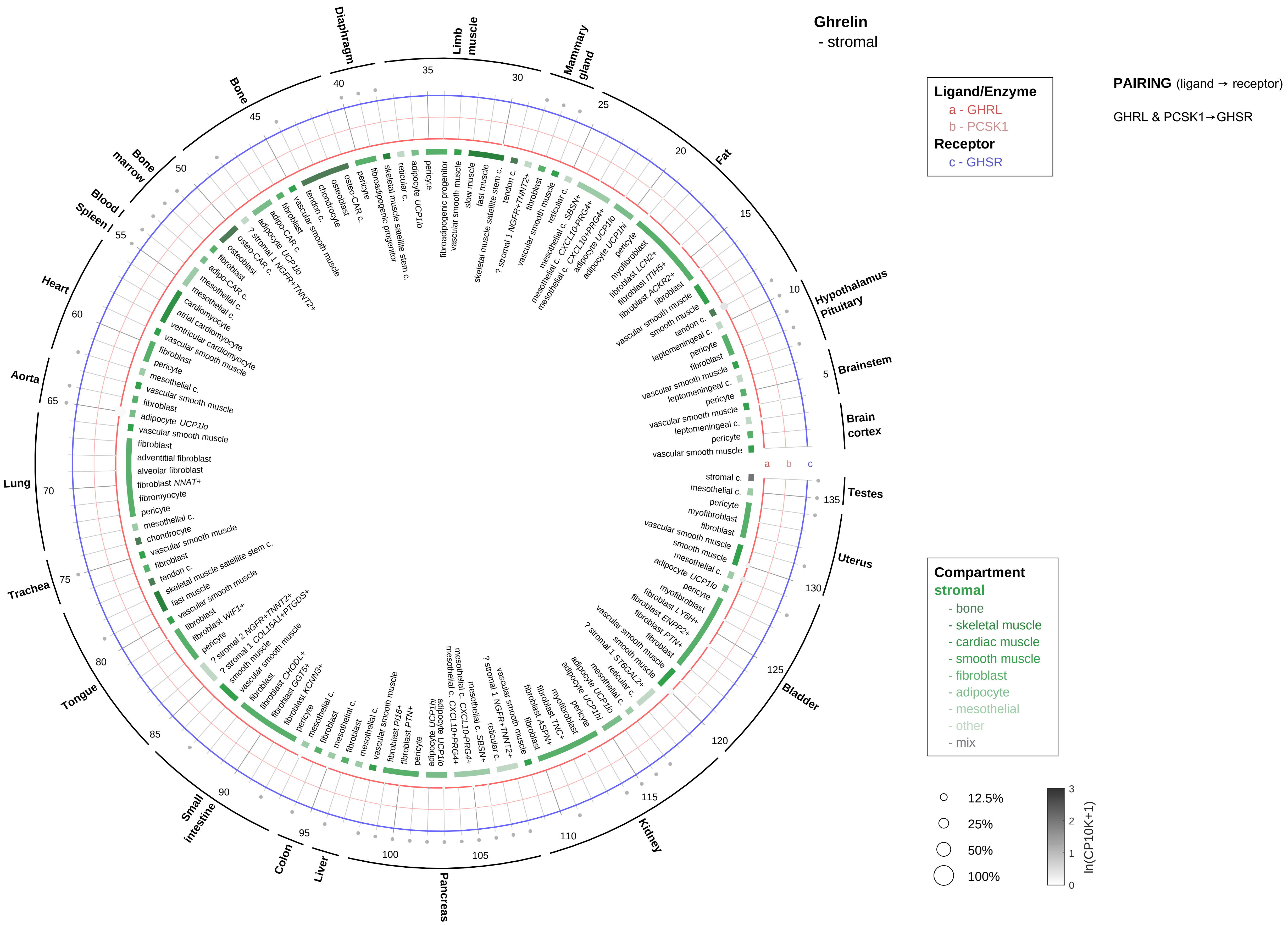

Ghrelin
- stromal
Ligand/Enzyme
a - GHRL
b - PCSK1
Receptor
c - GHSR
PAIRING (ligand → receptor)
GHRL & PCSK1→GHSR
Compartment
stromal
- bone
- skeletal muscle
- cardiac muscle
- smooth muscle
- fibroblast
- adipocyte
- mesothelial
- other
- mix
12.5%
25%
50%
100%
ln(CP10K+1)
Hypothalamus
Pituitary
Brainstem
Brain cortex
Testes
Uterus
Bladder
Kidney
Pancreas
Liver
Colon
Small intestine
Tongue
Trachea
Lung
Aorta
Heart
Spleen
Blood
Bone marrow
Bone
Diaphragm
Limb muscle
Mammary gland
Fat

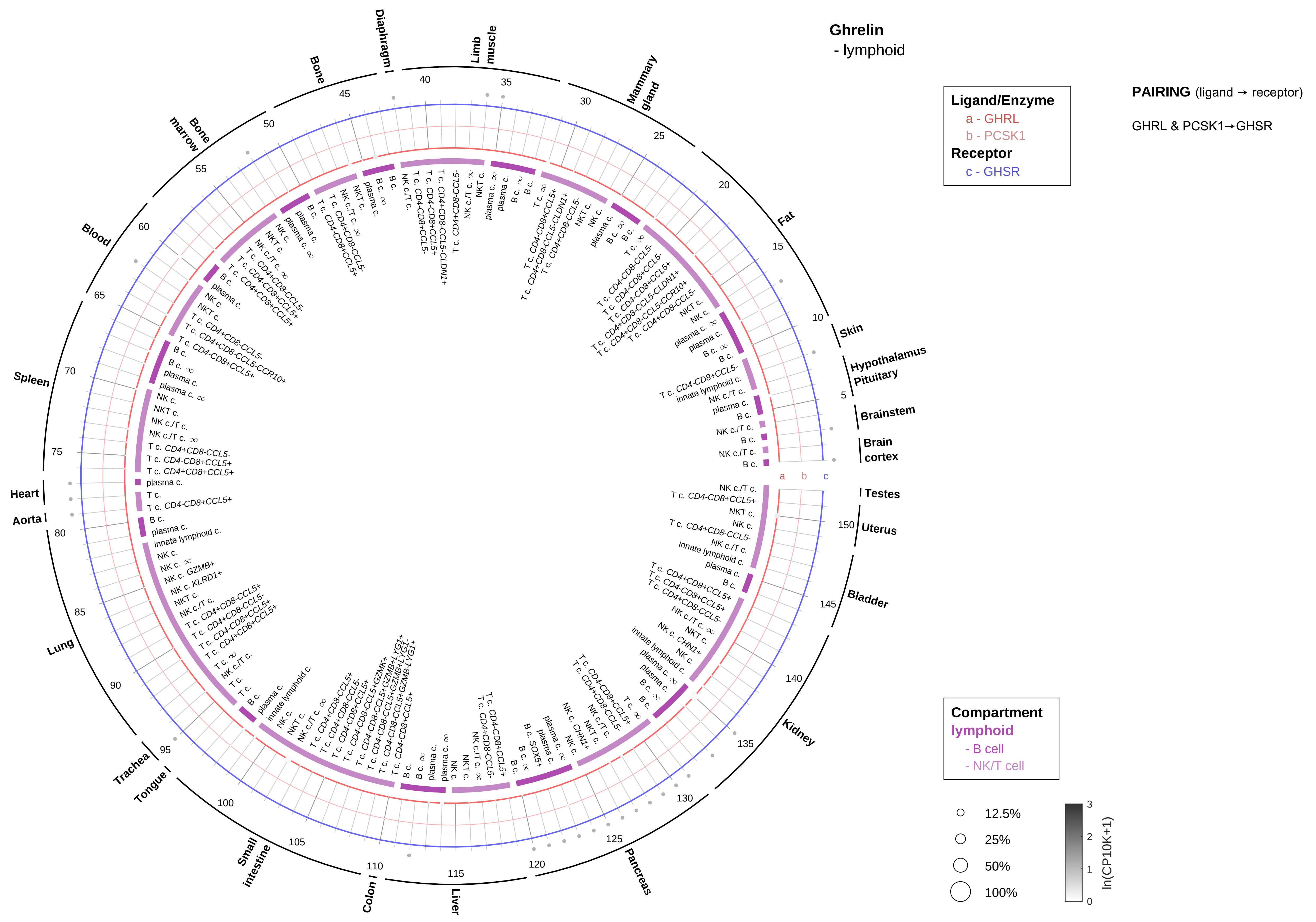
Ghrelin
- lymphoid
Ligand/Enzyme
a - GHRL
b - PCSK1
Receptor
c - GHSR
PAIRING (ligand → receptor)
GHRL & PCSK1→GHSR
Compartment
lymphoid
- B cell
- NK/T cell
12.5%
25%
50%
100%
ln(CP10K+1)
Skin
Hypothalamus
Pituitary
Brainstem
Brain cortex
Testes
Uterus
Bladder
Kidney
Pancreas
Liver
Colon
Small intestine
Tongue
Trachea
Lung
Aorta
Heart
Spleen
Blood
Bone marrow
Bone
Diaphragm
Limb muscle
Mammary gland
Fat

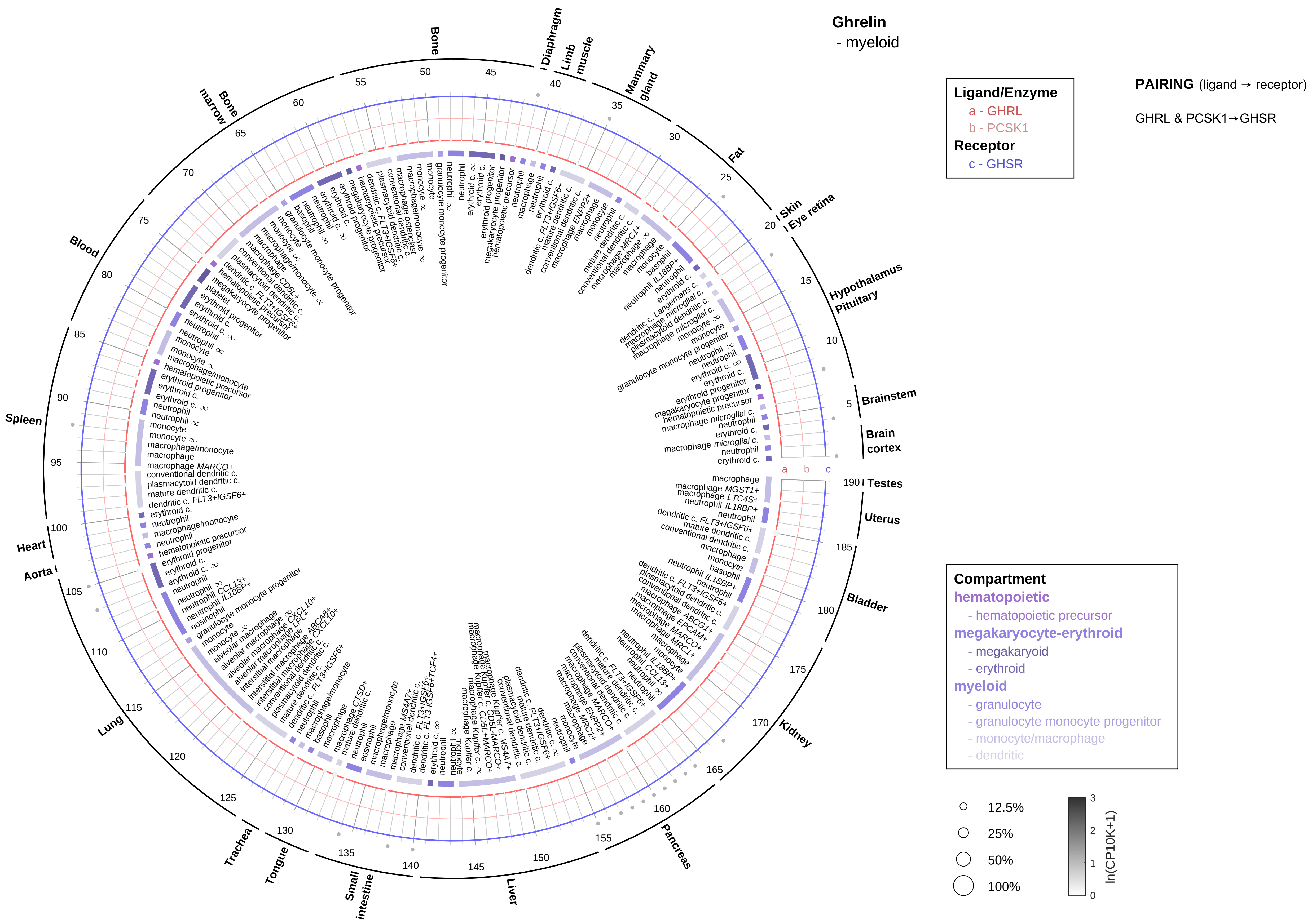
Ghrelin
- myeloid
Ligand/Enzyme
a - GHRL
b - PCSK1
Receptor
c - GHSR
PAIRING (ligand → receptor)
GHRL & PCSK1→GHSR
Compartment
hematopoietic
- hematopoietic precursor
megakaryocyte-erythroid
- megakaryoid
- erythroid
myeloid
- granulocyte
- granulocyte monocyte progenitor
- monocyte/macrophage
- dendritic
12.5%
25%
50%
100%
ln(CP10K+1)
Bone
Diaphragm
Limb muscle
Mammary gland
Fat
Skin
Eye retina
Hypothalamus
Pituitary
Brainstem
Brain cortex
Testes
Uterus
Bladder
Kidney
Pancreas
Liver
Small intestine
Tongue
Trachea
Lung
Aorta
Heart
Spleen
Blood
Bone marrow

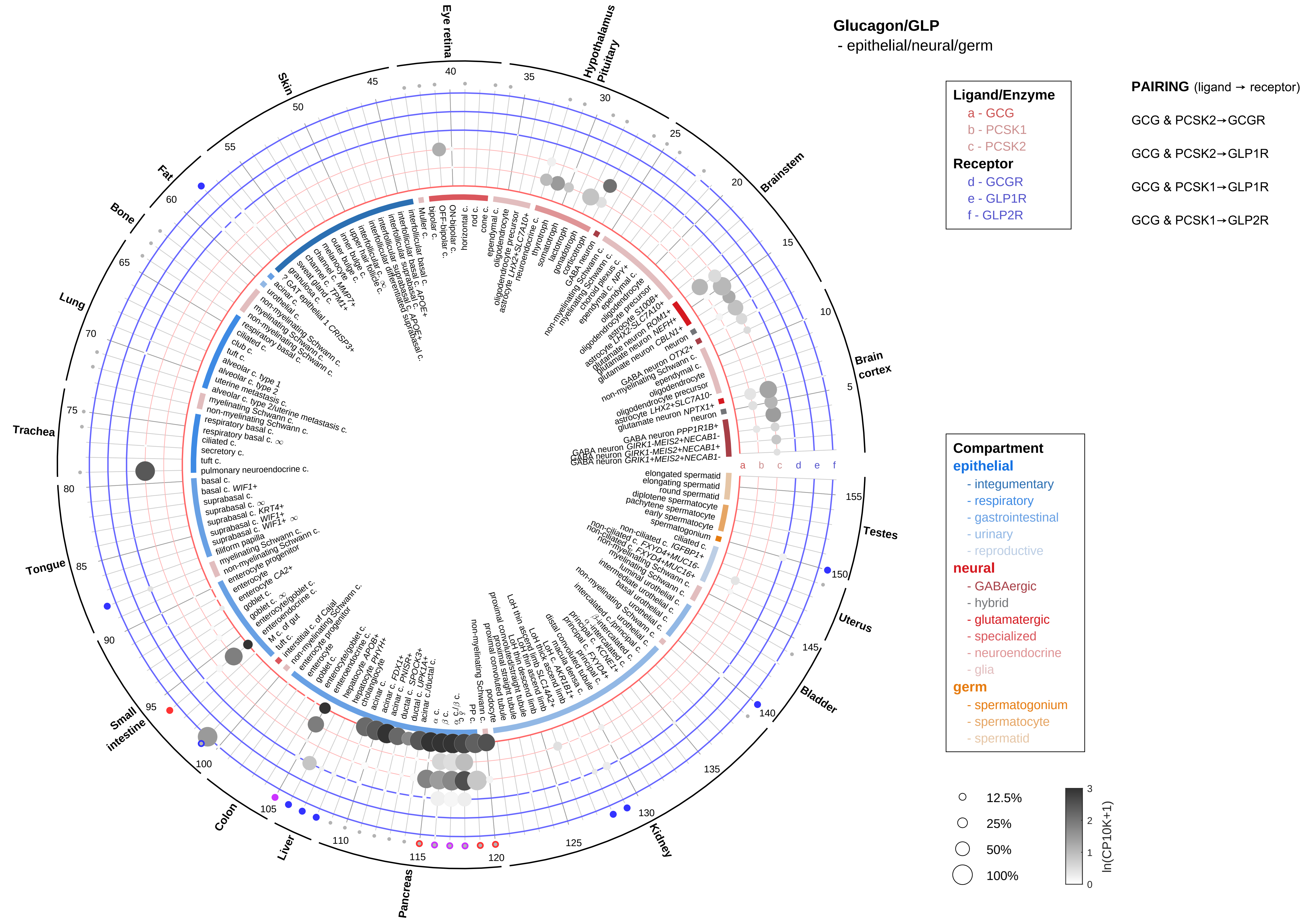

Glucagon/GLP
- epithelial/neural/germ
Ligand/Enzyme
a - GCG
b - PCSK1
c - PCSK2
Receptor
d - GCGR
e - GLP1R
f - GLP2R
PAIRING (ligand → receptor)
GCG & PCSK2→GCGR
GCG & PCSK2→GLP1R
GCG & PCSK1→GLP1R
GCG & PCSK1→GLP2R
Compartment
epithelial
- integumentary
- respiratory
- gastrointestinal
- urinary
- reproductive
neural
- GABAergic
- hybrid
- glutamatergic
- specialized
- neuroendocrine
- glia
germ
- spermatogonium
- spermatocyte
- spermatid
12.5%
25%
50%
100%
ln(CP10K+1)
Eye retina
Hypothalamus Pituitary
Brainstem
Brain cortex
Testes
Uterus
Bladder
Kidney
Pancreas
Liver
Colon
Small intestine
Tongue
Trachea
Lung
Bone
Fat
Skin

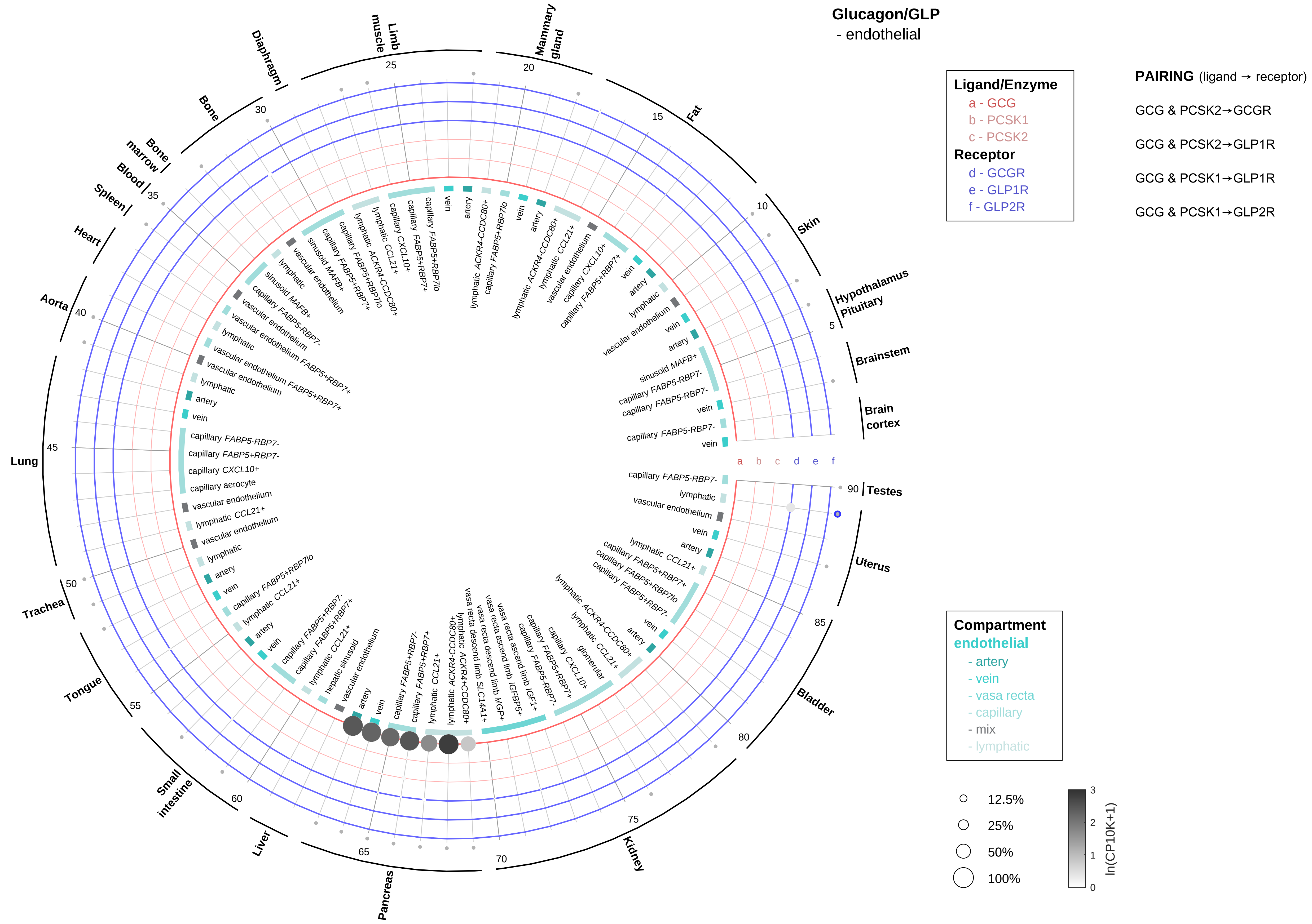

Glucagon/GLP
- endothelial
Ligand/Enzyme
a - GCG
b - PCSK1
c - PCSK2
Receptor
d - GCGR
e - GLP1R
f - GLP2R
PAIRING (ligand → receptor)
GCG & PCSK2→GCGR
GCG & PCSK2→GLP1R
GCG & PCSK1→GLP1R
GCG & PCSK1→GLP2R
Compartment
endothelial
- artery
- vein
- vasa recta
- capillary
- mix
- lymphatic
12.5%
25%
50%
100%
ln(CP10K+1)
Hypothalamus
Pituitary
Brainstem
Brain cortex
Testes
Uterus
Bladder
Kidney
Pancreas
Liver
Small intestine
Tongue
Trachea
Lung
Aorta
Heart
Spleen
Blood
Bone marrow
Bone
Diaphragm
Limb muscle
Mammary gland
Fat
Skin
a
b
c
d
e
f

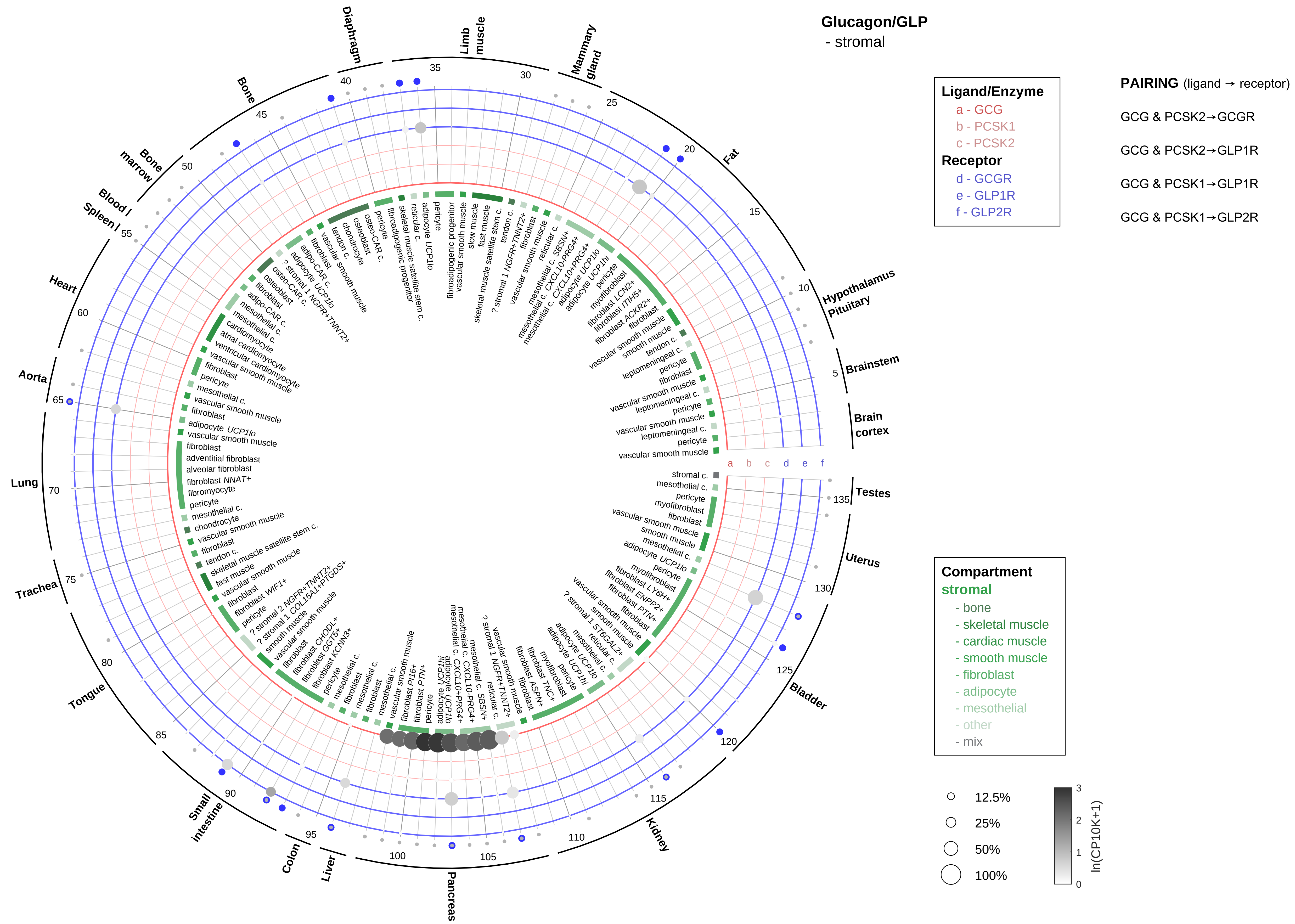

Glucagon/GLP
- stromal
Ligand/Enzyme
a - GCG
b - PCSK1
c - PCSK2
Receptor
d - GCGR
e - GLP1R
f - GLP2R
PAIRING (ligand → receptor)
GCG & PCSK2→GCGR
GCG & PCSK2→GLP1R
GCG & PCSK1→GLP1R
GCG & PCSK1→GLP2R
Compartment
stromal
- bone
- skeletal muscle
- cardiac muscle
- smooth muscle
- fibroblast
- adipocyte
- mesothelial
- other
- mix
12.5%
25%
50%
100%
ln(CP10K+1)
Diaphragm
Limb muscle
Mammary gland
Fat
Hypothalamus Pituitary
Brainstem
Brain cortex
Testes
Uterus
Bladder
Kidney
Pancreas
Liver
Colon
Small intestine
Tongue
Trachea
Lung
Aorta
Heart
Spleen
Blood
Bone marrow
Bone
a b c d e f

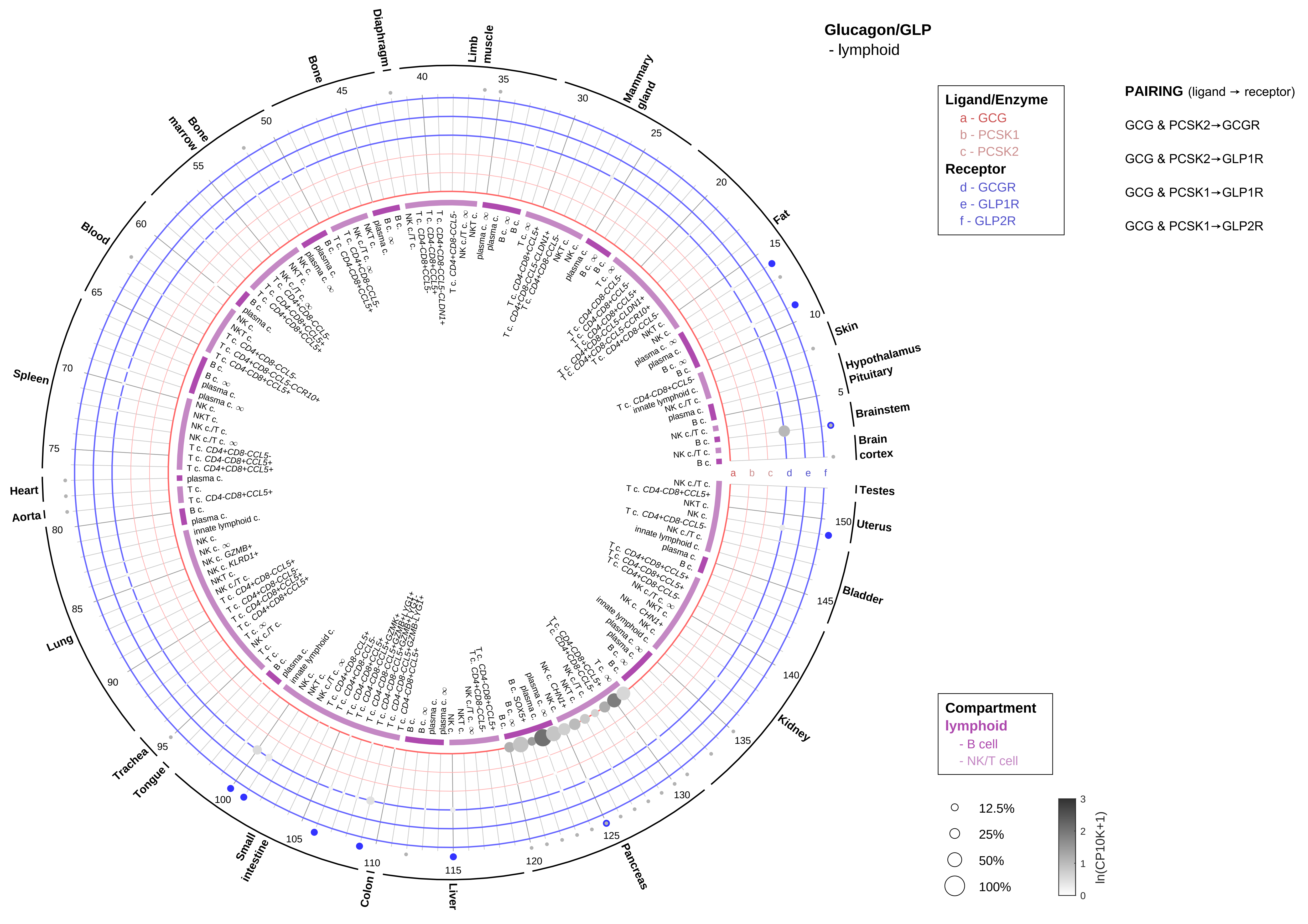

Glucagon/GLP
- lymphoid
Ligand/Enzyme
a - GCG
b - PCSK1
c - PCSK2
Receptor
d - GCGR
e - GLP1R
f - GLP2R
PAIRING (ligand → receptor)
GCG & PCSK2→GCGR
GCG & PCSK2→GLP1R
GCG & PCSK1→GLP1R
GCG & PCSK1→GLP2R
Compartment
lymphoid
- B cell
- NK/T cell
12.5%
25%
50%
100%
ln(CP10K+1)
Fat
Skin
Hypothalamus
Pituitary
Brainstem
Brain cortex
Testes
Uterus
Bladder
Kidney
Pancreas
Liver
Colon
Small intestine
Tongue
Trachea
Lung
Aorta
Heart
Spleen
Blood
Bone marrow
Bone
Diaphragm
Limb muscle
Mammary gland

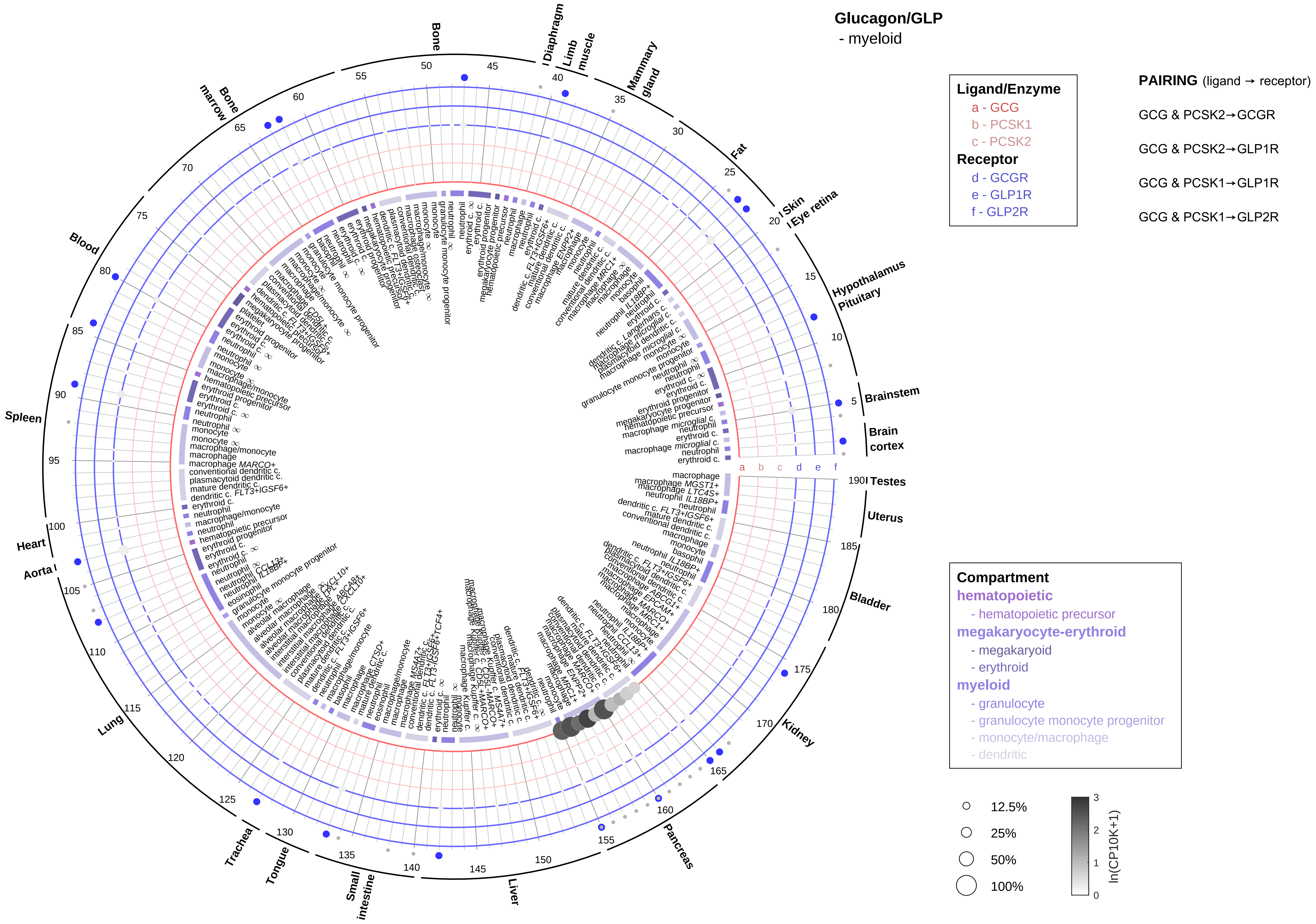
Glucagon/GLP
- myeloid
Ligand/Enzyme
a - GCG
b - PCSK1
c - PCSK2
Receptor
d - GCGR
e - GLP1R
f - GLP2R
PAIRING (ligand → receptor)
GCG & PCSK2→GCGR
GCG & PCSK2→GLP1R
GCG & PCSK1→GLP1R
GCG & PCSK1→GLP2R
Compartment
hematopoietic
- hematopoietic precursor
megakaryocyte-erythroid
- megakaryoid
- erythroid
myeloid
- granulocyte
- granulocyte monocyte progenitor
- monocyte/macrophage
- dendritic
12.5%
25%
50%
100%
ln(CP10K+1)
Bone
Diaphragm
Limb muscle
Mammary gland
Fat
Skin
Eye retina
Hypothalamus
Pituitary
Brainstem
Brain cortex
Testes
Uterus
Bladder
Kidney
Pancreas
Liver
Small intestine
Tongue
Trachea
Lung
Aorta
Heart
Spleen
Blood
Bone marrow

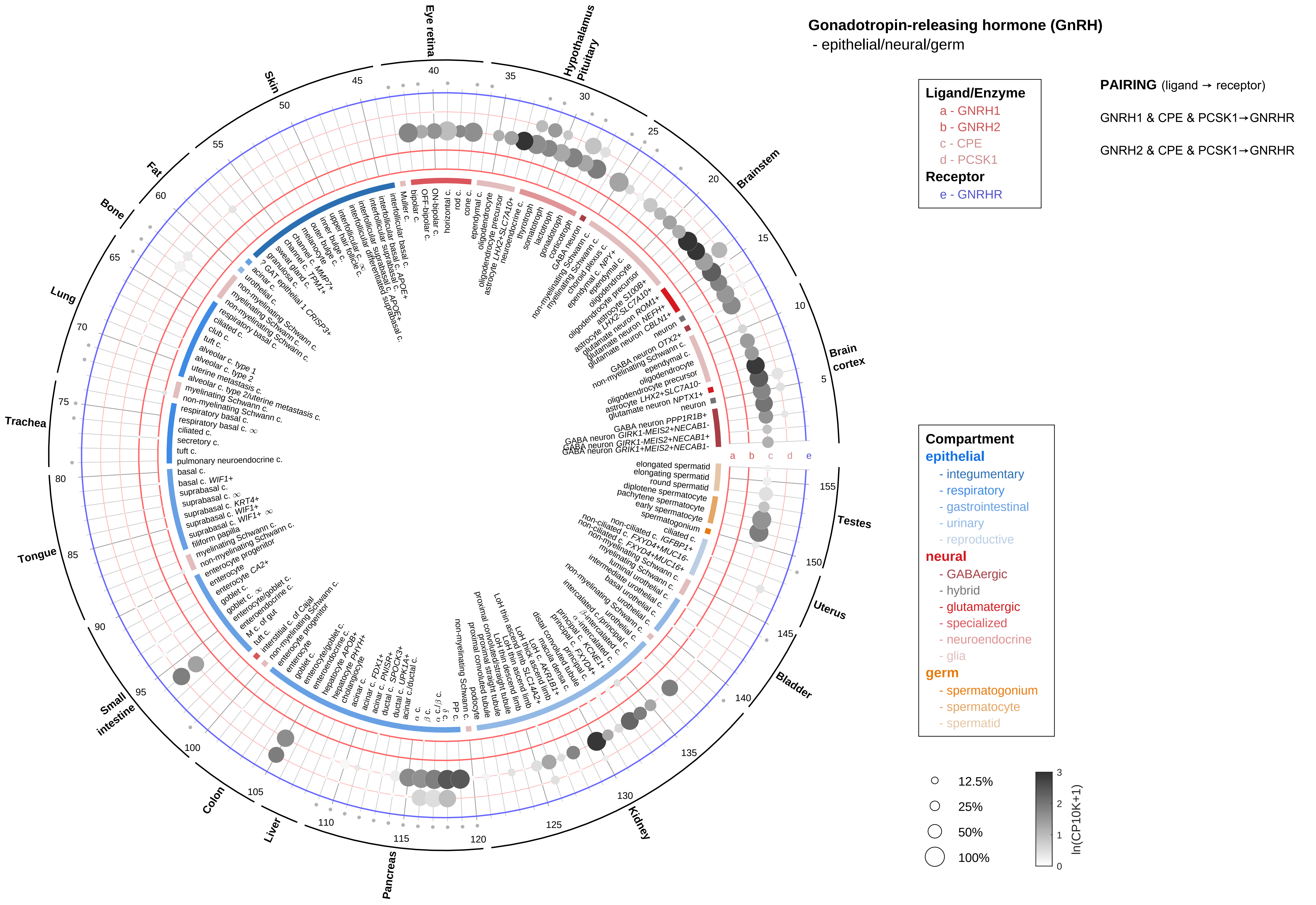

Gonadotropin-releasing hormone (GnRH)
- epithelial/neural/germ
Ligand/Enzyme
a - GNRH1
b - GNRH2
c - CPE
d - PCSK1
Receptor
e - GNRHR
PAIRING (ligand → receptor)
GNRH1 & CPE & PCSK1→GNRHR
GNRH2 & CPE & PCSK1→GNRHR
Compartment
epithelial
- integumentary
- respiratory
- gastrointestinal
- urinary
- reproductive
neural
- GABAergic
- hybrid
- glutamatergic
- specialized
- neuroendocrine
- glia
germ
- spermatogonium
- spermatocyte
- spermatid
12.5%
25%
50%
100%
ln(CP10K+1)
Eye retina
Hypothalamus Pituitary
Brainstem
Brain cortex
Testes
Uterus
Bladder
Kidney
Pancreas
Liver
Colon
Small intestine
Tongue
Trachea
Lung
Bone
Fat
Skin

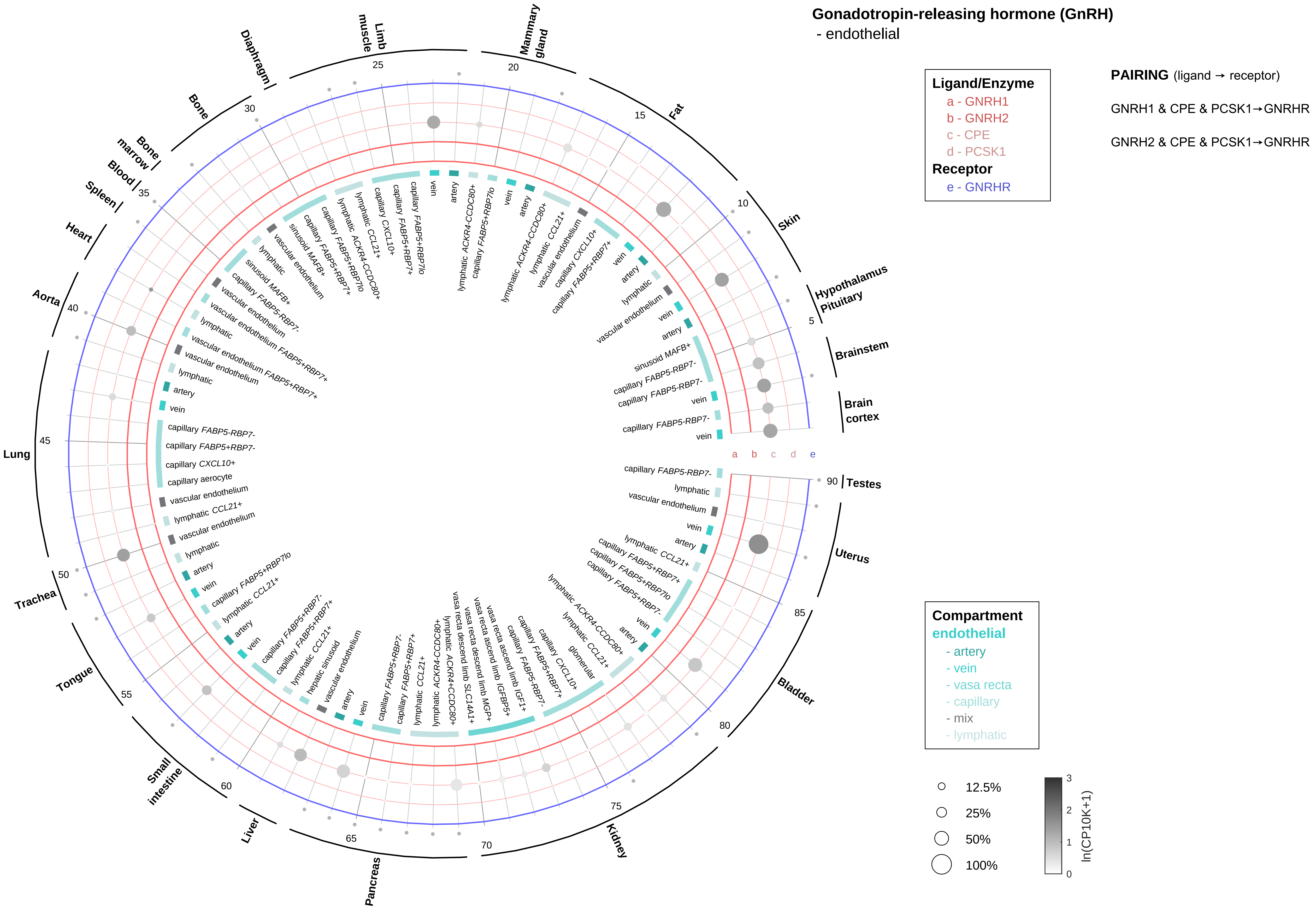

Gonadotropin-releasing hormone (GnRH)
- endothelial
Ligand/Enzyme
a - GNRH1
b - GNRH2
c - CPE
d - PCSK1
Receptor
e - GNRHR
PAIRING (ligand → receptor)
GNRH1 & CPE & PCSK1→GNRHR
GNRH2 & CPE & PCSK1→GNRHR
Compartment
endothelial
- artery
- vein
- vasa recta
- capillary
- mix
- lymphatic
12.5%
25%
50%
100%
ln(CP10K+1)
Brain cortex
Brainstem
Hypothalamus Pituitary
Skin
Fat
Mammary gland
Limb muscle
Diaphragm
Bone
Bone marrow
Blood
Spleen
Heart
Aorta
Lung
Trachea
Tongue
Small intestine
Liver
Pancreas
Kidney
Bladder
Uterus
Testes

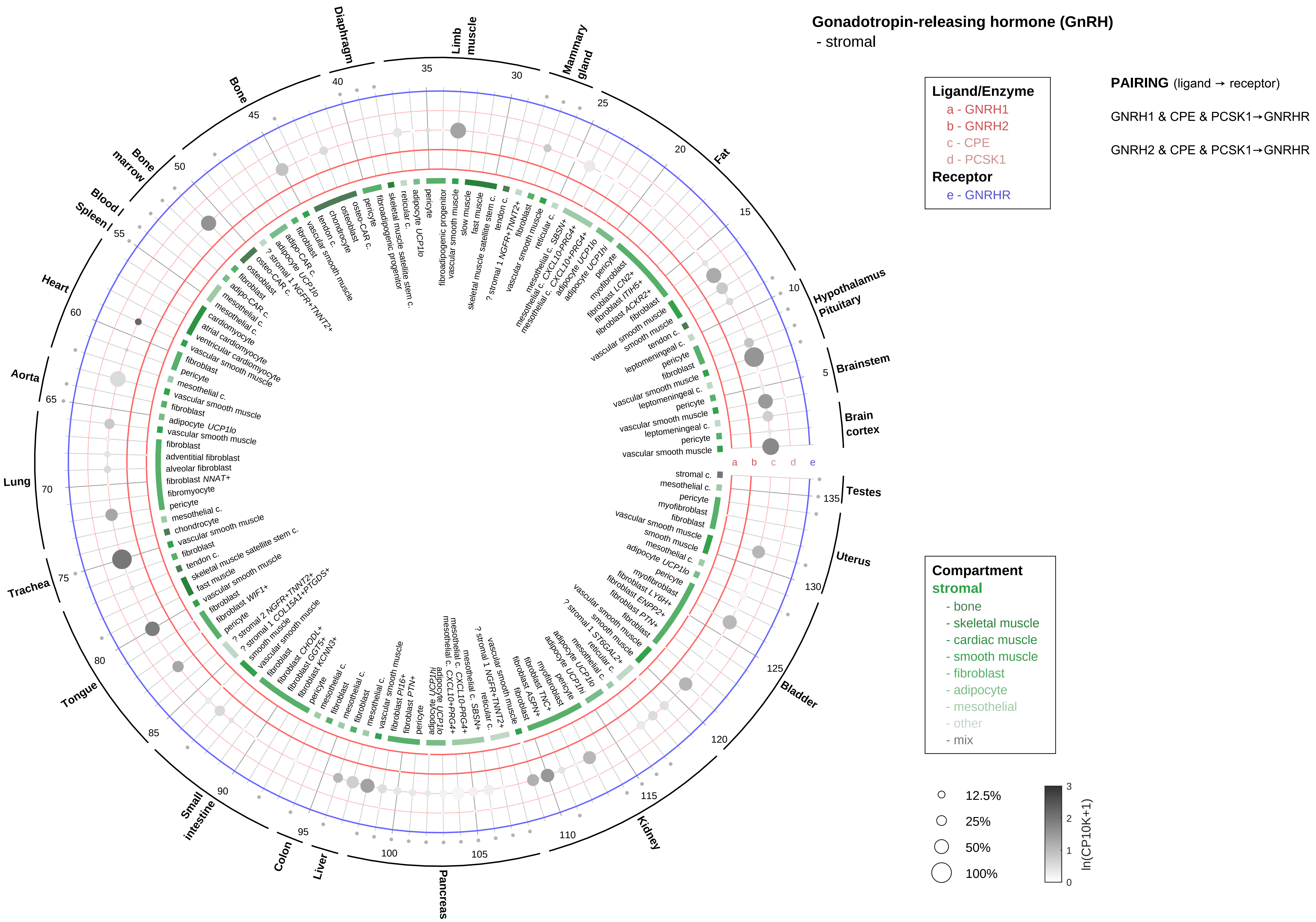

Gonadotropin-releasing hormone (GnRH)
- stromal
Ligand/Enzyme
a - GNRH1
b - GNRH2
c - CPE
d - PCSK1
Receptor
e - GNRHR
PAIRING (ligand → receptor)
GNRH1 & CPE & PCSK1→GNRHR
GNRH2 & CPE & PCSK1→GNRHR
Compartment
stromal
- bone
- skeletal muscle
- cardiac muscle
- smooth muscle
- fibroblast
- adipocyte
- mesothelial
- other
- mix
12.5%
25%
50%
100%
ln(CP10K+1)
Diaphragm
Limb muscle
Mammary gland
Fat
Hypothalamus Pituitary
Brainstem
Brain cortex
Testes
Uterus
Bladder
Kidney
Pancreas
Liver
Colon
Small intestine
Tongue
Trachea
Lung
Aorta
Heart
Blood / Spleen
Bone marrow
Bone

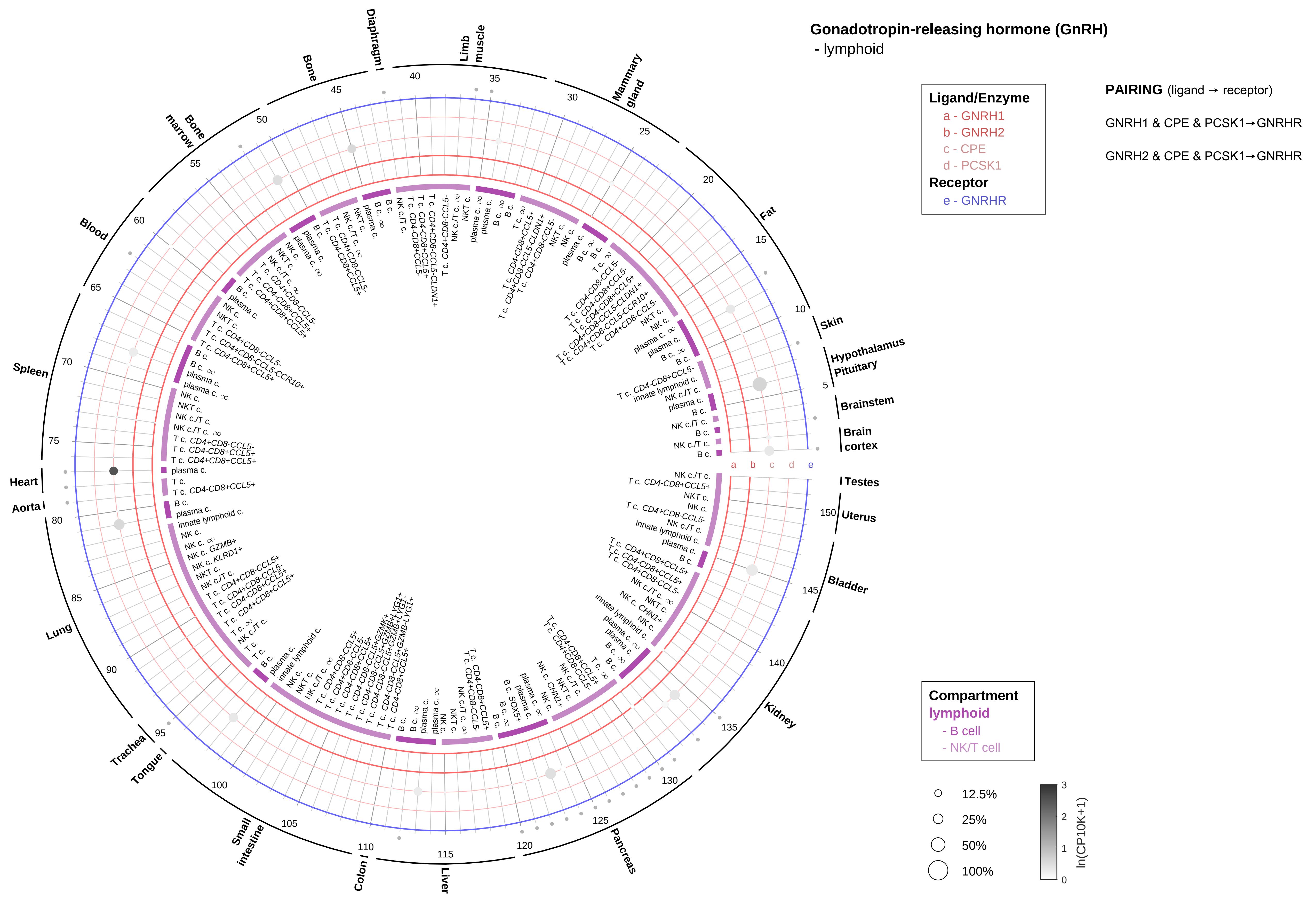

Gonadotropin-releasing hormone (GnRH)
- lymphoid
Ligand/Enzyme
a - GNRH1
b - GNRH2
c - CPE
d - PCSK1
Receptor
e - GNRHR
PAIRING (ligand → receptor)
GNRH1 & CPE & PCSK1→GNRHR
GNRH2 & CPE & PCSK1→GNRHR
Compartment
lymphoid
- B cell
- NK/T cell
12.5%
25%
50%
100%
ln(CP10K+1)
Skin
Hypothalamus
Pituitary
Brainstem
Brain cortex
Testes
Uterus
Bladder
Kidney
Pancreas
Liver
Colon
Small intestine
Tongue
Trachea
Lung
Aorta
Heart
Spleen
Blood
Bone marrow
Bone
Diaphragm
Limb muscle
Mammary gland
Fat

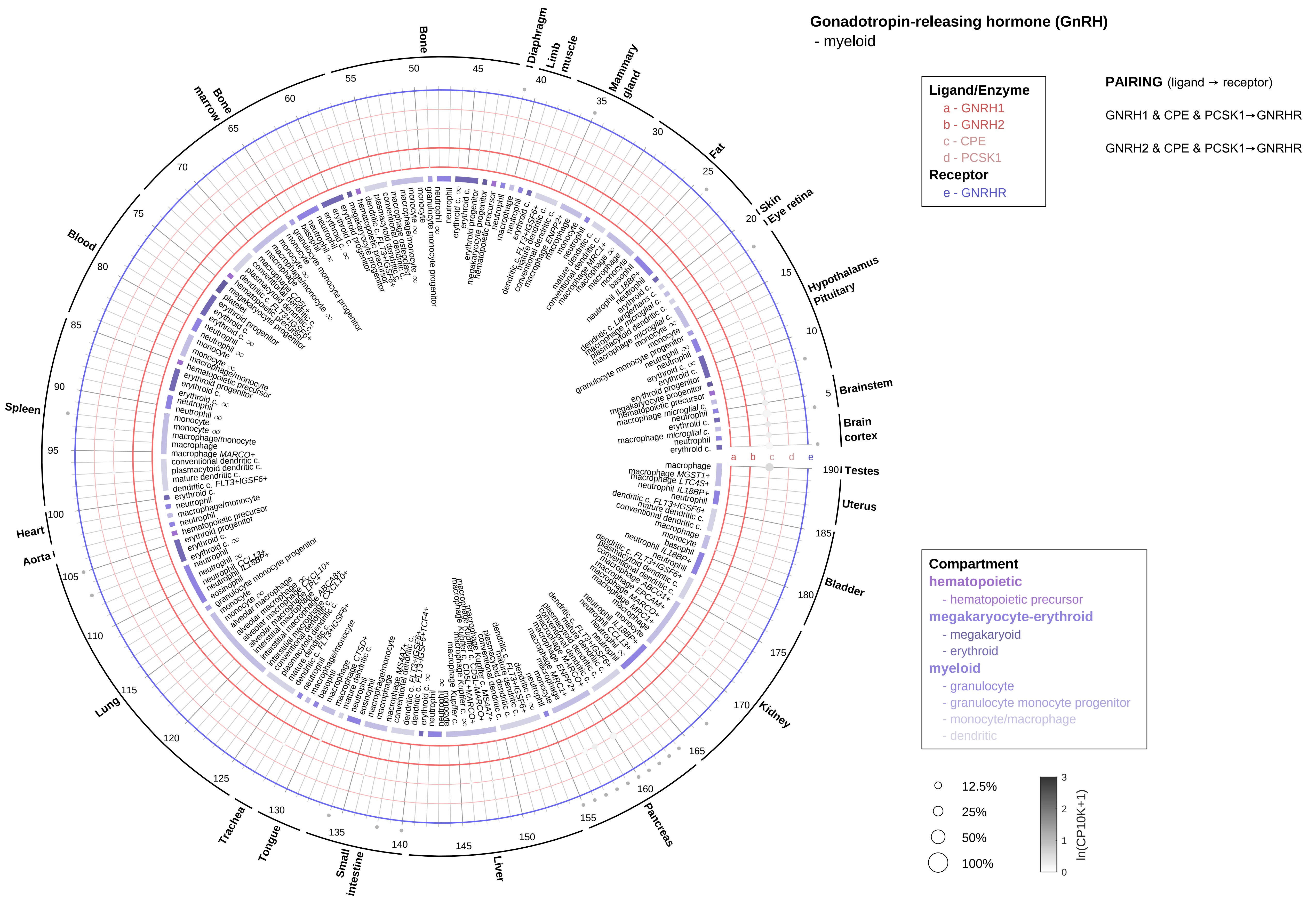
Gonadotropin-releasing hormone (GnRH)
- myeloid
Ligand/Enzyme
a - GNRH1
b - GNRH2
c - CPE
d - PCSK1
Receptor
e - GNRHR
PAIRING (ligand → receptor)
GNRH1 & CPE & PCSK1→GNRHR
GNRH2 & CPE & PCSK1→GNRHR
Compartment
hematopoietic
- hematopoietic precursor
megakaryocyte-erythroid
- megakaryoid
- erythroid
myeloid
- granulocyte
- granulocyte monocyte progenitor
- monocyte/macrophage
- dendritic
12.5%
25%
50%
100%
ln(CP10K+1)
Bone
Diaphragm
Limb muscle
Mammary gland
Fat
Skin
Eye retina
Hypothalamus
Pituitary
Brainstem
Brain cortex
Testes
Uterus
Bladder
Kidney
Pancreas
Liver
Small intestine
Tongue
Trachea
Lung
Aorta
Heart
Spleen
Blood
Bone marrow

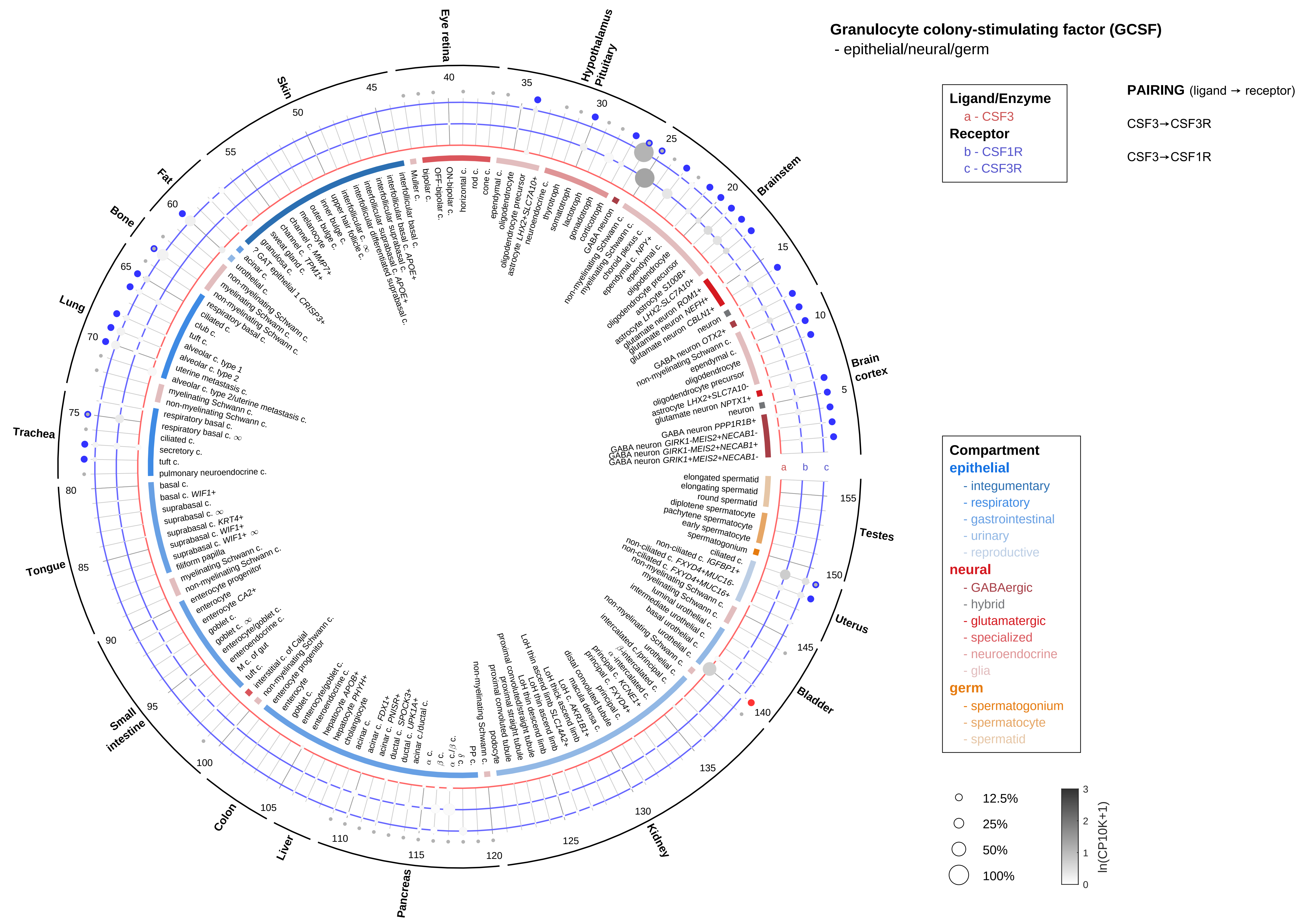

Granulocyte colony-stimulating factor (GCSF)
- epithelial/neural/germ
Ligand/Enzyme
a - CSF3
Receptor
b - CSF1R
c - CSF3R
PAIRING (ligand → receptor)
CSF3→CSF3R
CSF3→CSF1R
Compartment
epithelial
- integumentary
- respiratory
- gastrointestinal
- urinary
- reproductive
neural
- GABAergic
- hybrid
- glutamatergic
- specialized
- neuroendocrine
- glia
germ
- spermatogonium
- spermatocyte
- spermatid
12.5%
25%
50%
100%
ln(CP10K+1)

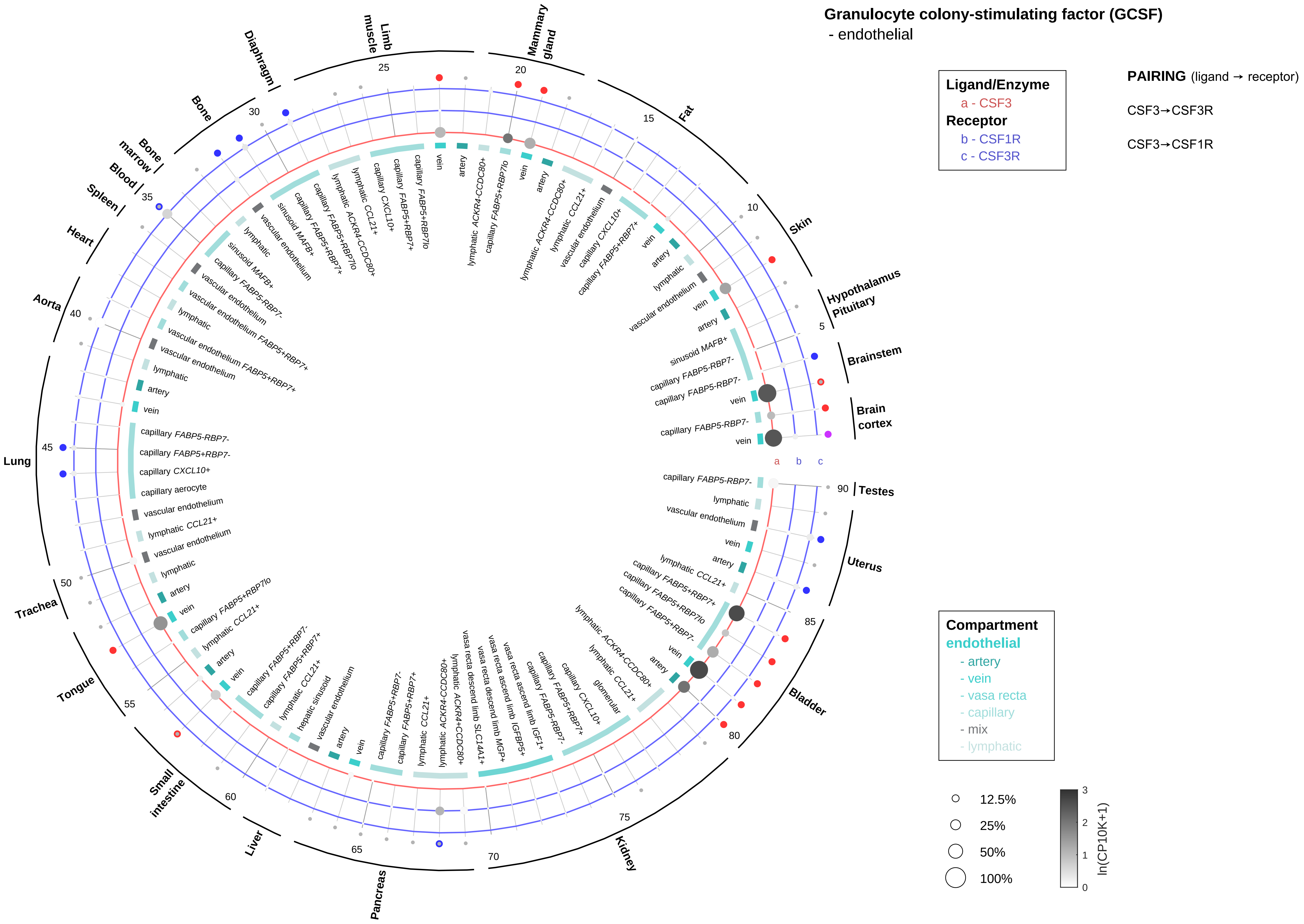

Granulocyte colony-stimulating factor (GCSF)
- endothelial
Ligand/Enzyme
a - CSF3
Receptor
b - CSF1R
c - CSF3R
PAIRING (ligand → receptor)
CSF3→CSF3R
CSF3→CSF1R
Compartment
endothelial
- artery
- vein
- vasa recta
- capillary
- mix
- lymphatic
12.5%
25%
50%
100%
ln(CP10K+1)
Brain cortex
Brainstem
Hypothalamus Pituitary
Skin
Fat
Mammary gland
Limb muscle
Diaphragm
Bone
Bone marrow
Blood
Spleen
Heart
Aorta
Lung
Trachea
Tongue
Small intestine
Liver
Pancreas
Kidney
Bladder
Uterus
Testes
a
b
c

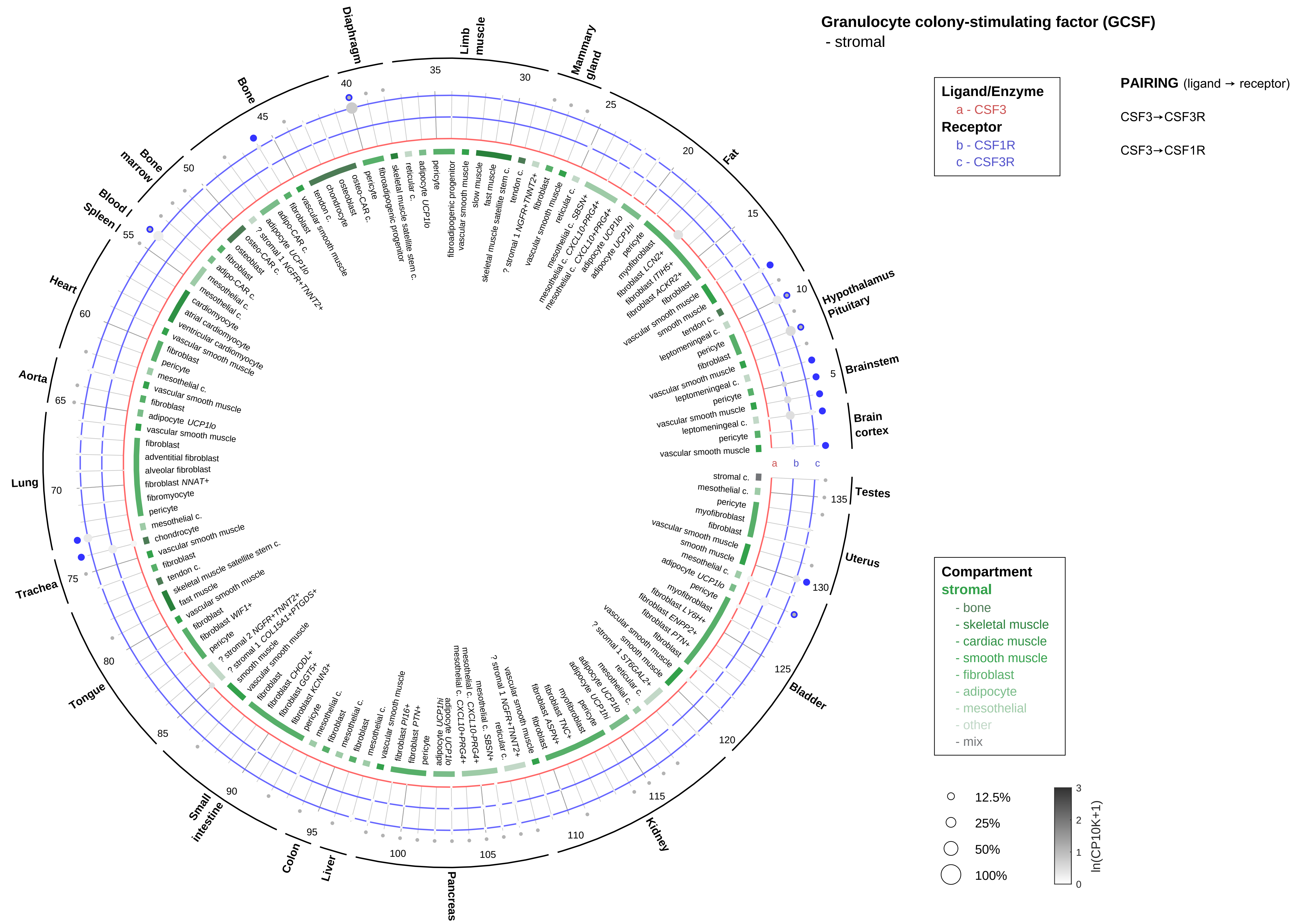

Granulocyte colony-stimulating factor (GCSF)
- stromal
Ligand/Enzyme
a - CSF3
Receptor
b - CSF1R
c - CSF3R
PAIRING (ligand → receptor)
CSF3→CSF3R
CSF3→CSF1R
Compartment
stromal
- bone
- skeletal muscle
- cardiac muscle
- smooth muscle
- fibroblast
- adipocyte
- mesothelial
- other
- mix
12.5%
25%
50%
100%
ln(CP10K+1)
Brain cortex
Brainstem
Hypothalamus
Pituitary
Fat
Mammary gland
Limb muscle
Diaphragm
Bone
Bone marrow
Blood
Spleen
Heart
Aorta
Lung
Trachea
Tongue
Small intestine
Colon
Liver
Pancreas
Kidney
Bladder
Uterus
Testes

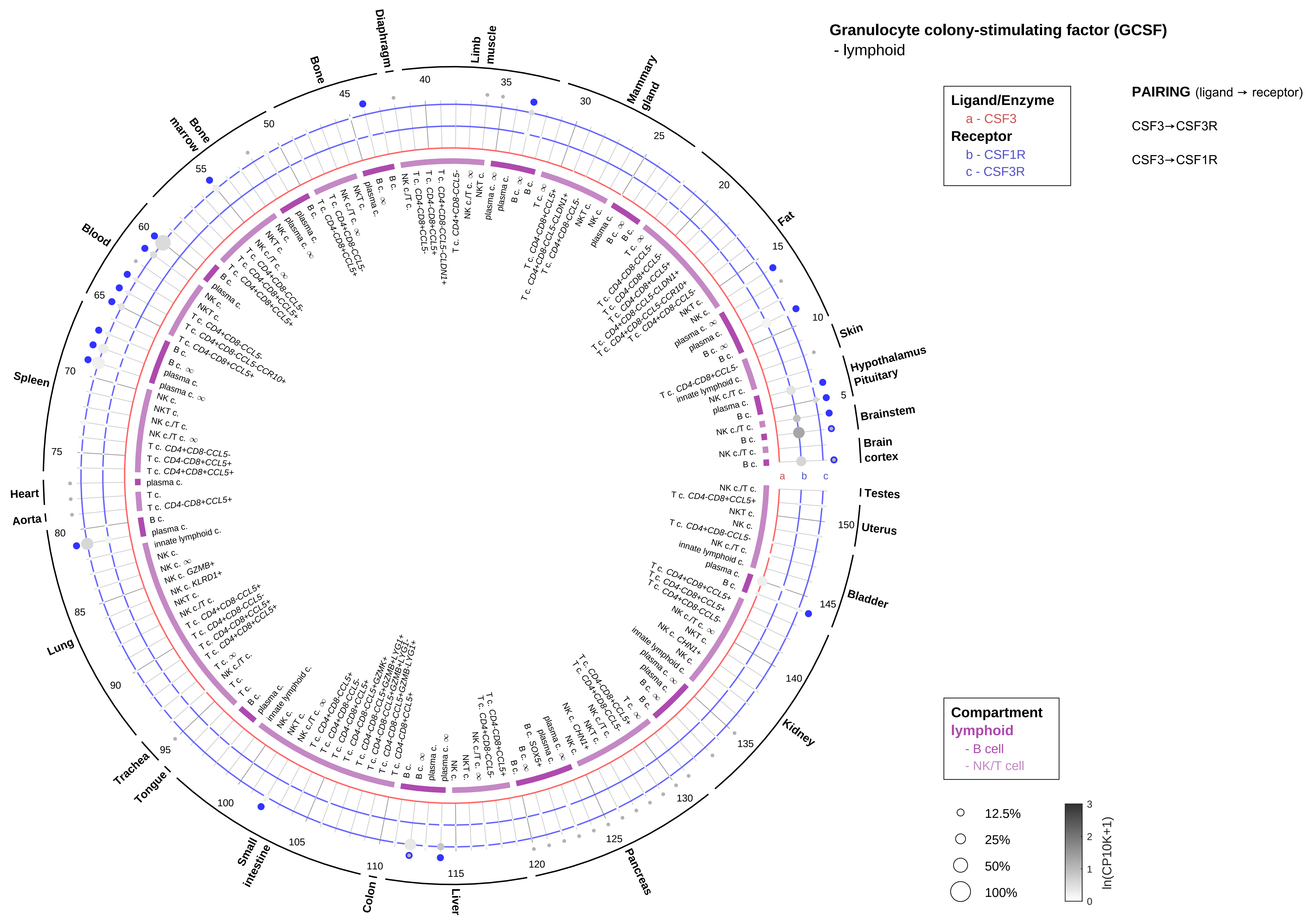

Granulocyte colony-stimulating factor (GCSF)
- lymphoid
Ligand/Enzyme
a - CSF3
Receptor
b - CSF1R
c - CSF3R
PAIRING (ligand → receptor)
CSF3→CSF3R
CSF3→CSF1R
Compartment
lymphoid
- B cell
- NK/T cell
12.5%
25%
50%
100%
ln(CP10K+1)
Skin
Hypothalamus
Pituitary
Brainstem
Brain cortex
Testes
Uterus
Bladder
Kidney
Pancreas
Liver
Colon
Small intestine
Tongue
Trachea
Lung
Aorta
Heart
Spleen
Blood
Bone marrow
Bone
Diaphragm
Limb muscle
Mammary gland
Fat

#### Granulocyte colony-stimulating factor (GCSF)
- myeloid

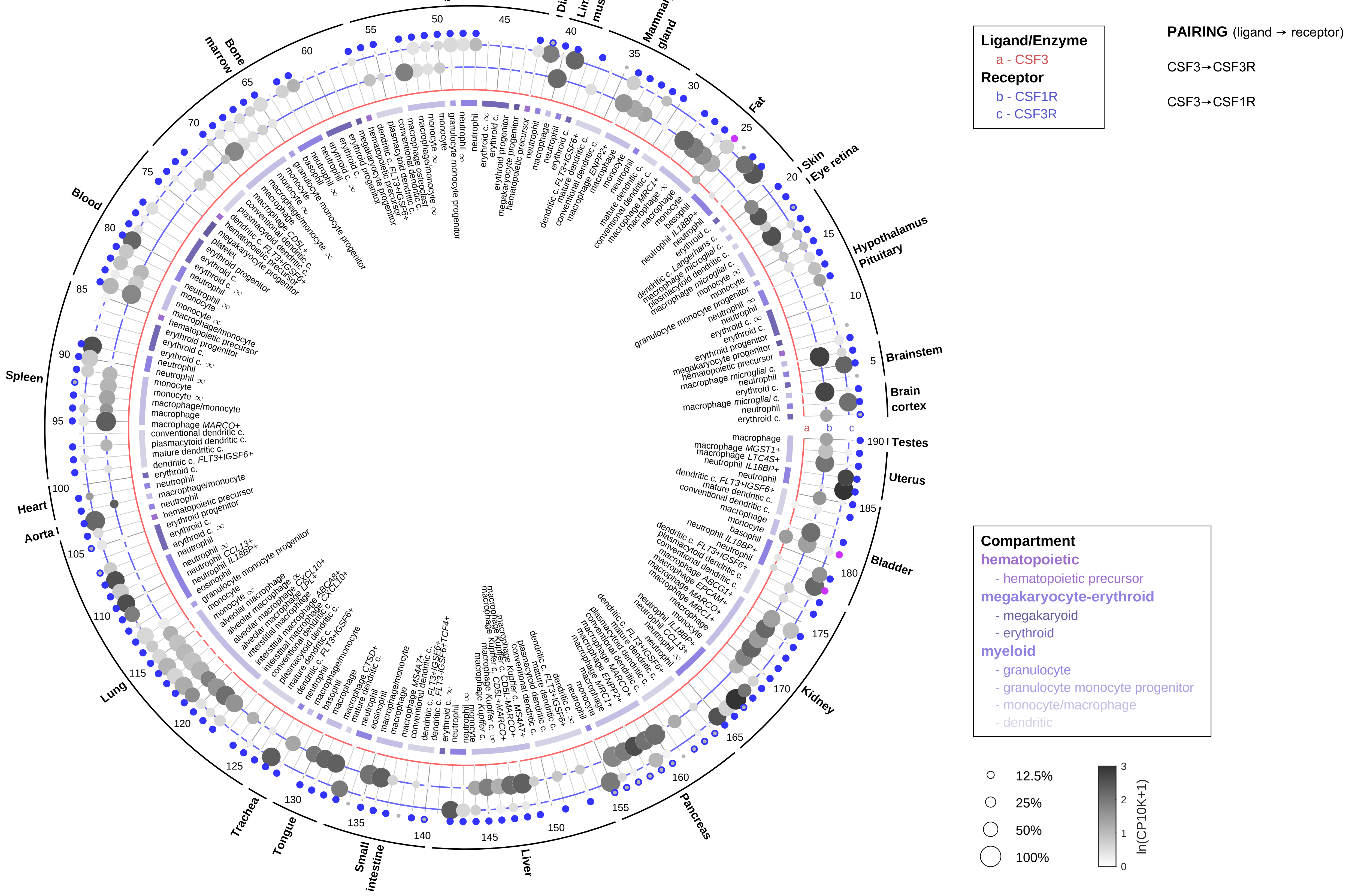

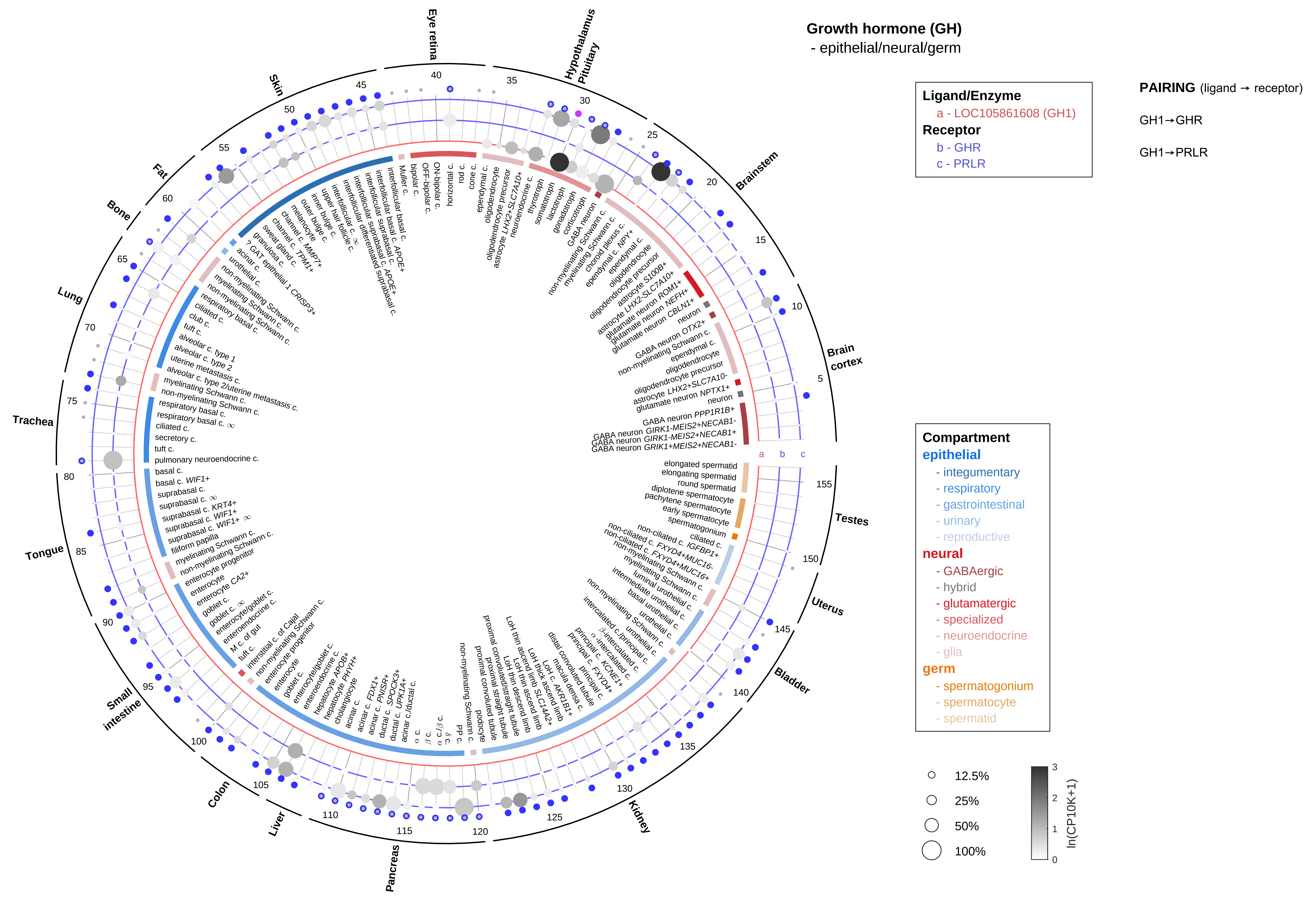
Growth hormone (GH)
- epithelial/neural/germ
Ligand/Enzyme
a - LOC105861608 (GH1)
Receptor
b - GHR
c - PRLR
PAIRING (ligand → receptor)
GH1→GHR
GH1→PRLR
Compartment
epithelial
- integumentary
- respiratory
- gastrointestinal
- urinary
- reproductive
neural
- GABAergic
- hybrid
- glutamatergic
- specialized
- neuroendocrine
- glia
germ
- spermatogonium
- spermatocyte
- spermatid
12.5%
25%
50%
100%
ln(CP10K+1)
Eye retina
Hypothalamus
Pituitary
Brainstem
Brain cortex
Testes
Uterus
Bladder
Kidney
Pancreas
Liver
Colon
Small intestine
Tongue
Trachea
Lung
Bone
Fat
Skin
a
b
c

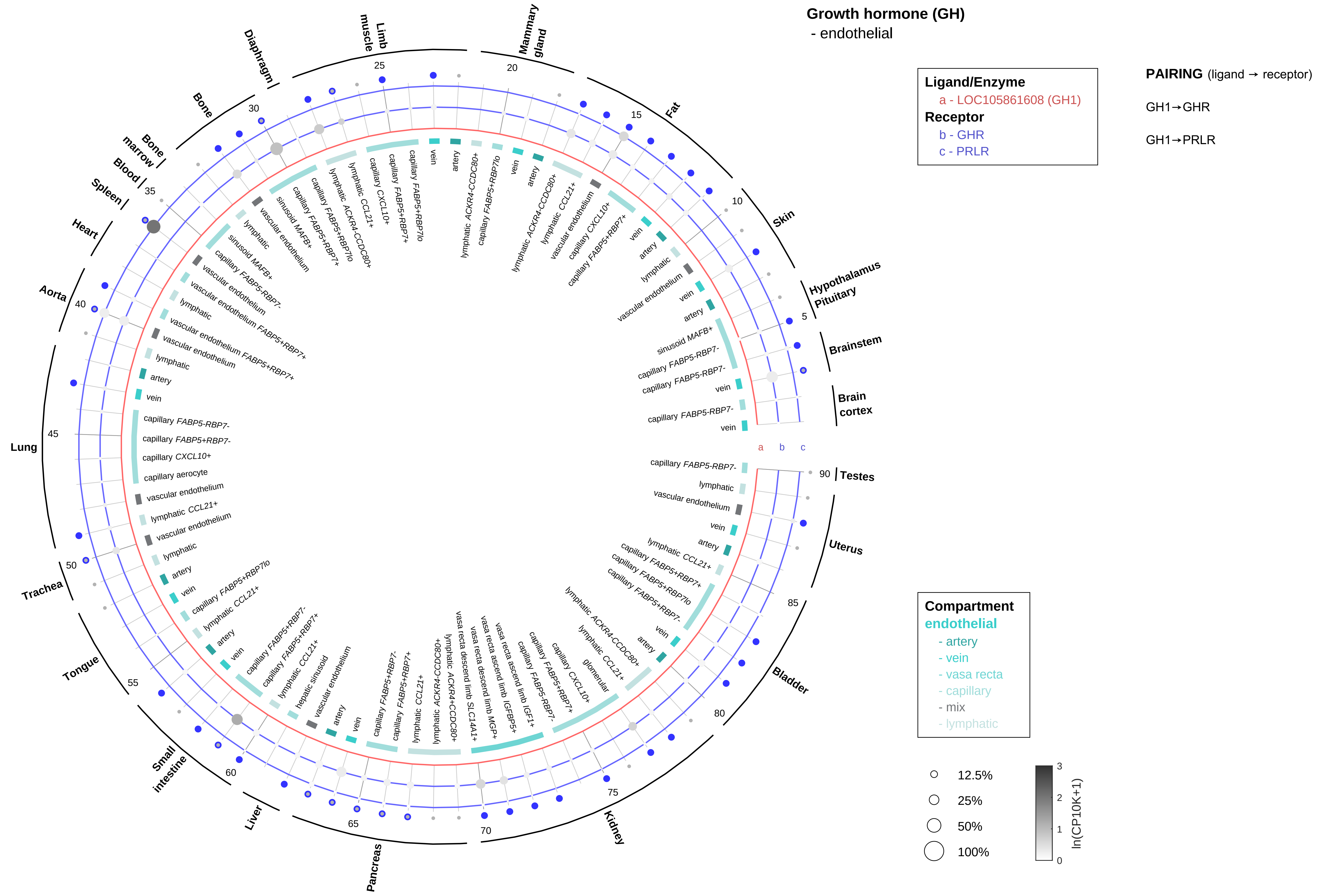

Growth hormone (GH)
- endothelial
Ligand/Enzyme
a - LOC105861608 (GH1)
Receptor
b - GHR
c - PRLR
PAIRING (ligand → receptor)
GH1→GHR
GH1→PRLR
Compartment
endothelial
- artery
- vein
- vasa recta
- capillary
- mix
- lymphatic
12.5%
25%
50%
100%
ln(CP10K+1)
Brain cortex
Brainstem
Hypothalamus Pituitary
Skin
Fat
Mammary gland
Limb muscle
Diaphragm
Bone
Bone marrow
Blood
Spleen
Heart
Aorta
Lung
Trachea
Tongue
Small intestine
Liver
Pancreas
Kidney
Bladder
Uterus
Testes

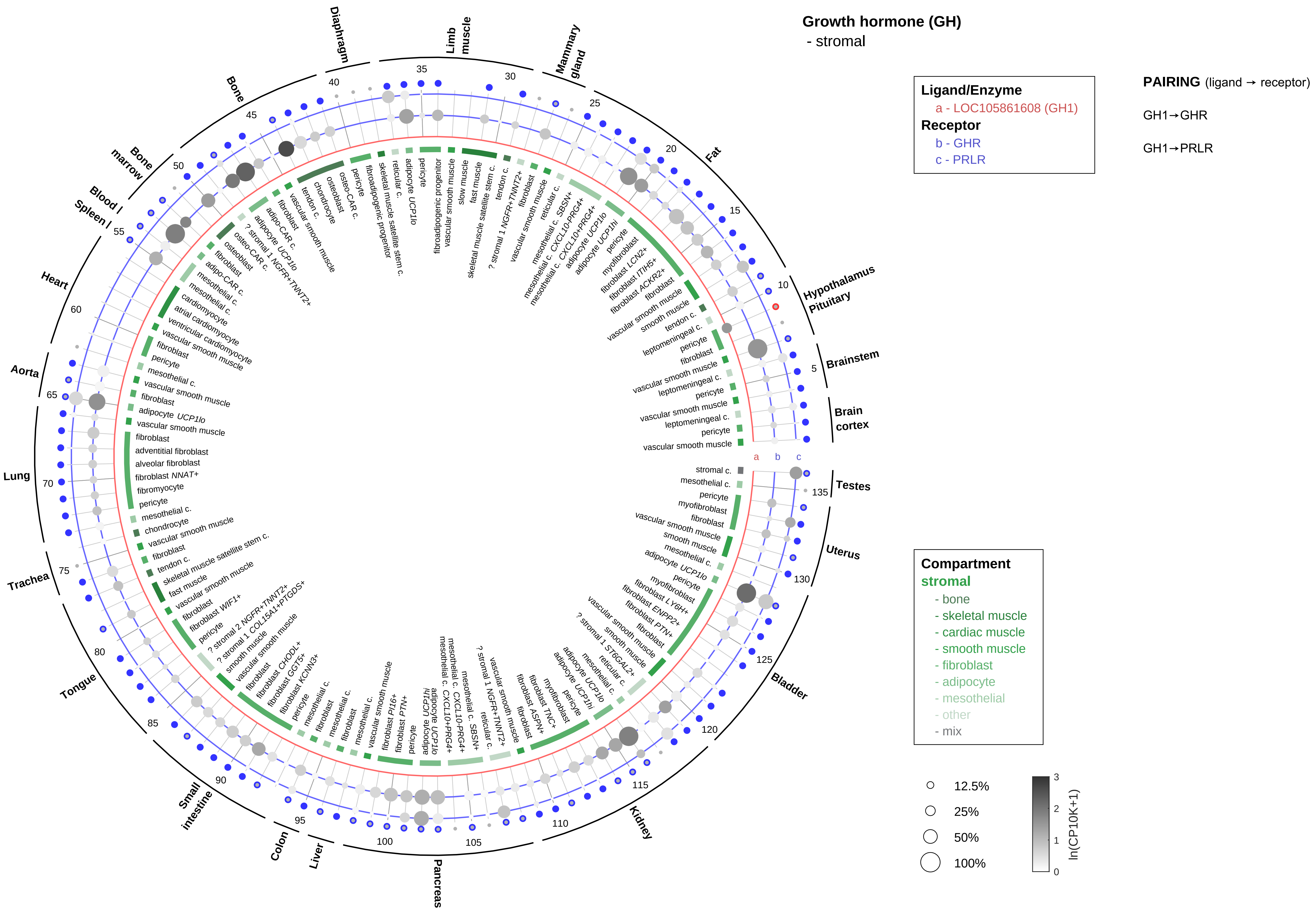

Growth hormone (GH)
- stromal
Ligand/Enzyme
a - LOC105861608 (GH1)
Receptor
b - GHR
c - PRLR
PAIRING (ligand → receptor)
GH1→GHR
GH1→PRLR
Compartment
stromal
- bone
- skeletal muscle
- cardiac muscle
- smooth muscle
- fibroblast
- adipocyte
- mesothelial
- other
- mix
12.5%
25%
50%
100%
ln(CP10K+1)

**Growth hormone (GH)**
- lymphoid

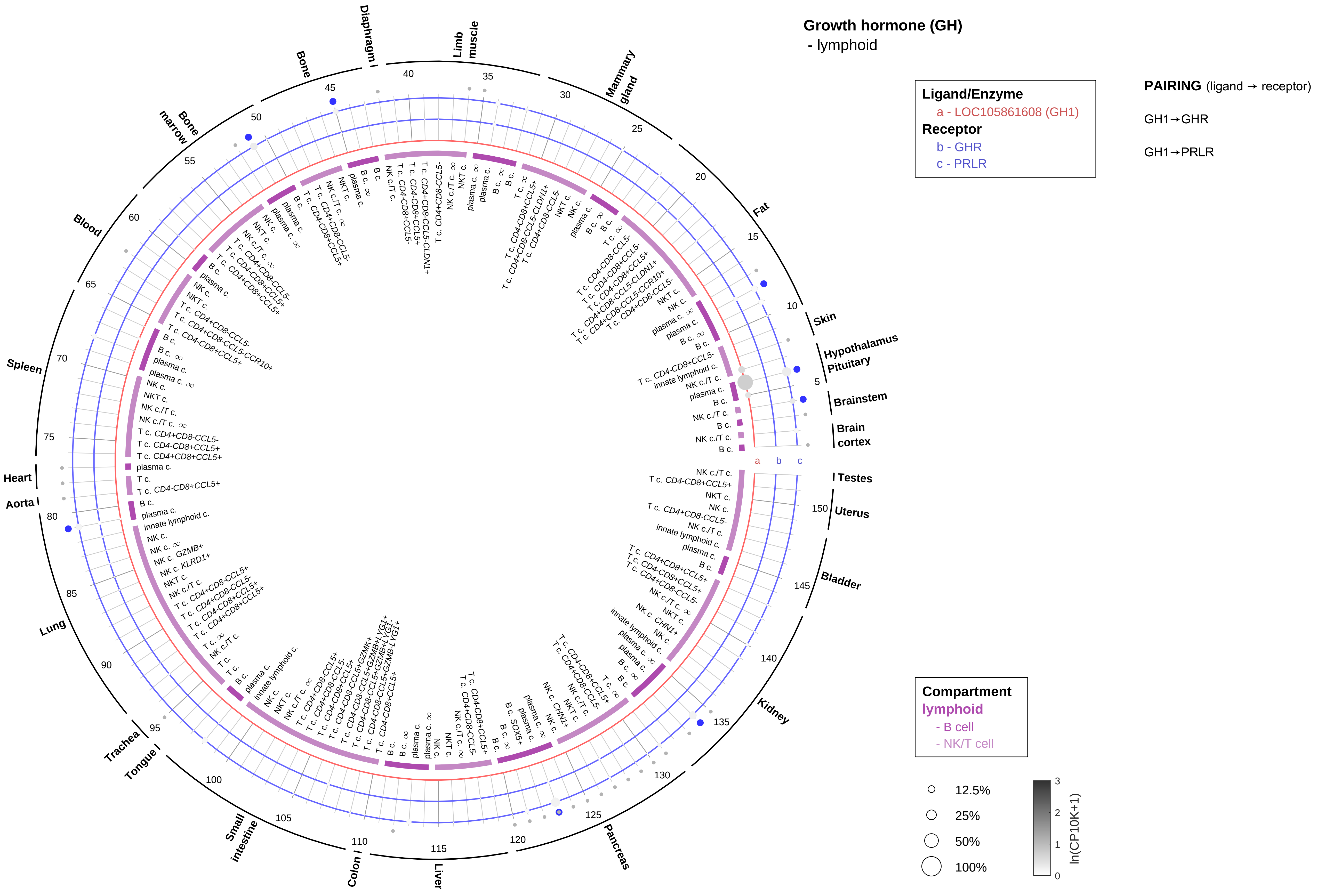

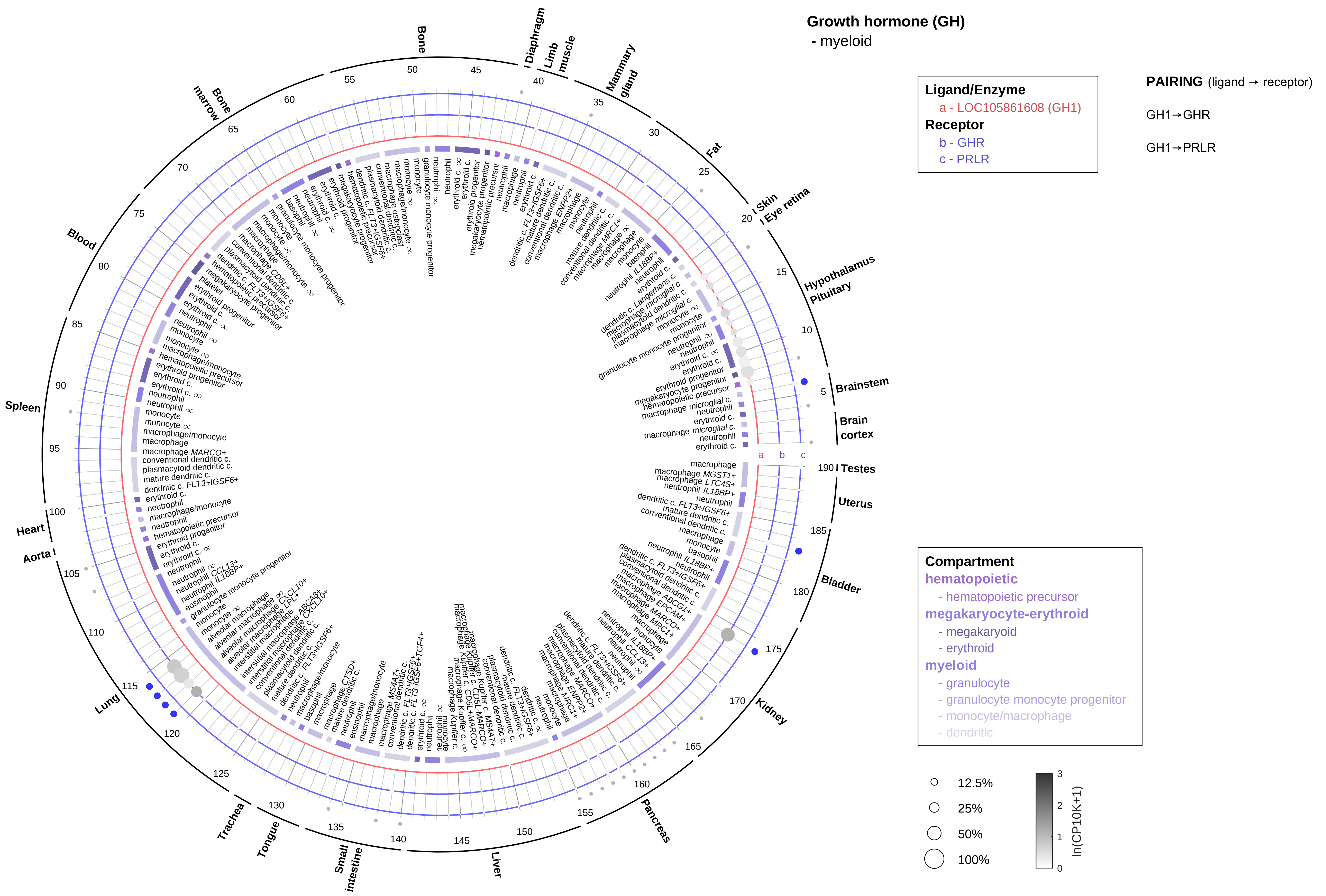

Growth hormone (GH)
- myeloid
Ligand/Enzyme
a - LOC105861608 (GH1)
Receptor
b - GHR
c - PRLR
PAIRING (ligand → receptor)
GH1→GHR
GH1→PRLR
Compartment
hematopoietic
- hematopoietic precursor
megakaryocyte-erythroid
- megakaryoid
- erythroid
myeloid
- granulocyte
- granulocyte monocyte progenitor
- monocyte/macrophage
- dendritic
12.5%
25%
50%
100%
ln(CP10K+1)
Bone
Diaphragm
Limb muscle
Mammary gland
Fat
Skin
Eye retina
Hypothalamus
Pituitary
Brainstem
Brain cortex
Testes
Uterus
Bladder
Kidney
Pancreas
Liver
Small intestine
Tongue
Trachea
Lung
Aorta
Heart
Spleen
Blood
Bone marrow

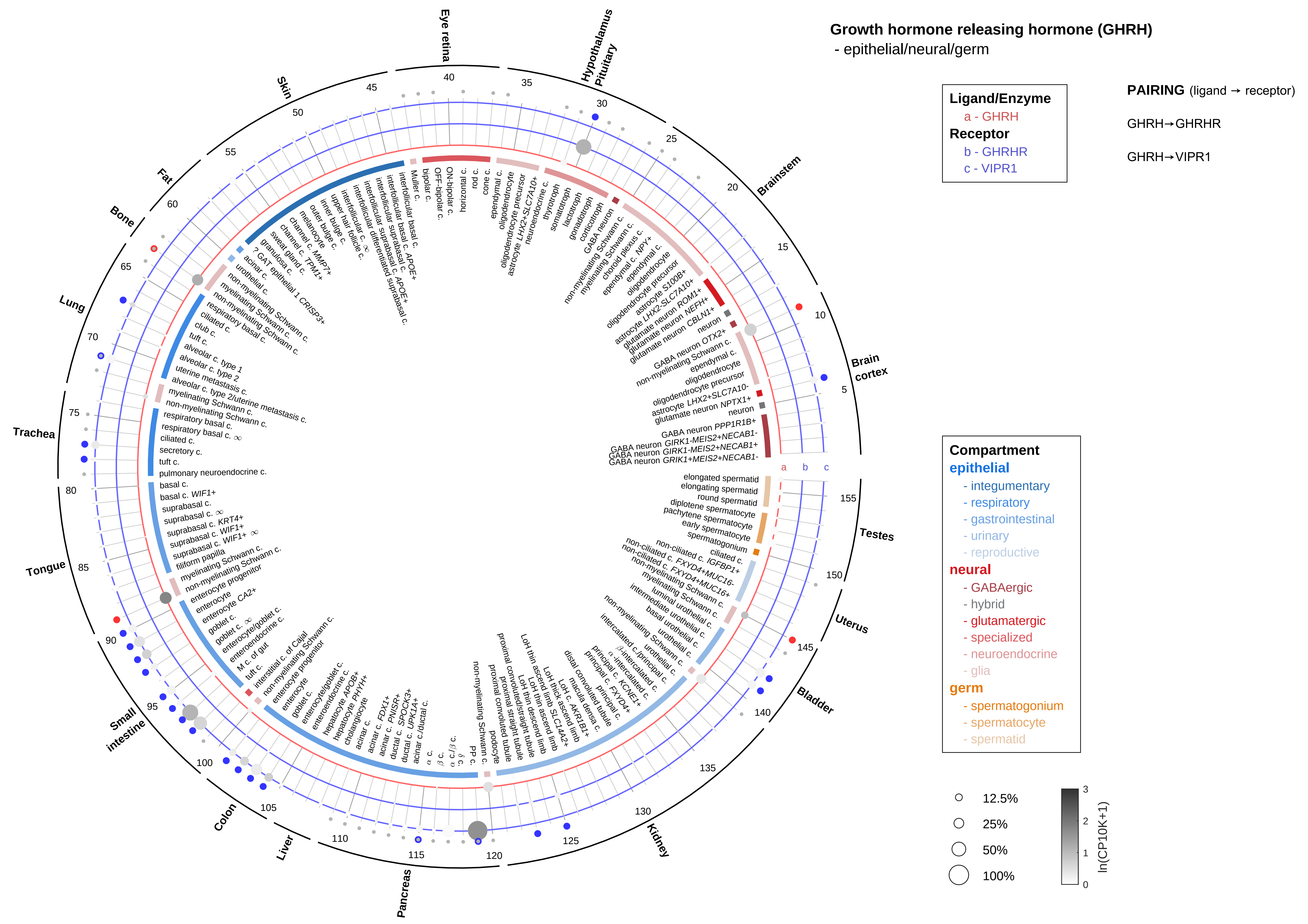

Growth hormone releasing hormone (GHRH)
- epithelial/neural/germ
Ligand/Enzyme
a - GHRH
Receptor
b - GHRHR
c - VIPR1
PAIRING (ligand → receptor)
GHRH→GHRHR
GHRH→VIPR1
Compartment
epithelial
- integumentary
- respiratory
- gastrointestinal
- urinary
- reproductive
neural
- GABAergic
- hybrid
- glutamatergic
- specialized
- neuroendocrine
- glia
germ
- spermatogonium
- spermatocyte
- spermatid
12.5%
25%
50%
100%
ln(CP10K+1)
Brain cortex
Brainstem
Hypothalamus
Pituitary
Eye retina
Skin
Fat
Bone
Lung
Trachea
Tongue
Small intestine
Colon
Liver
Pancreas
Kidney
Bladder
Uterus
Testes

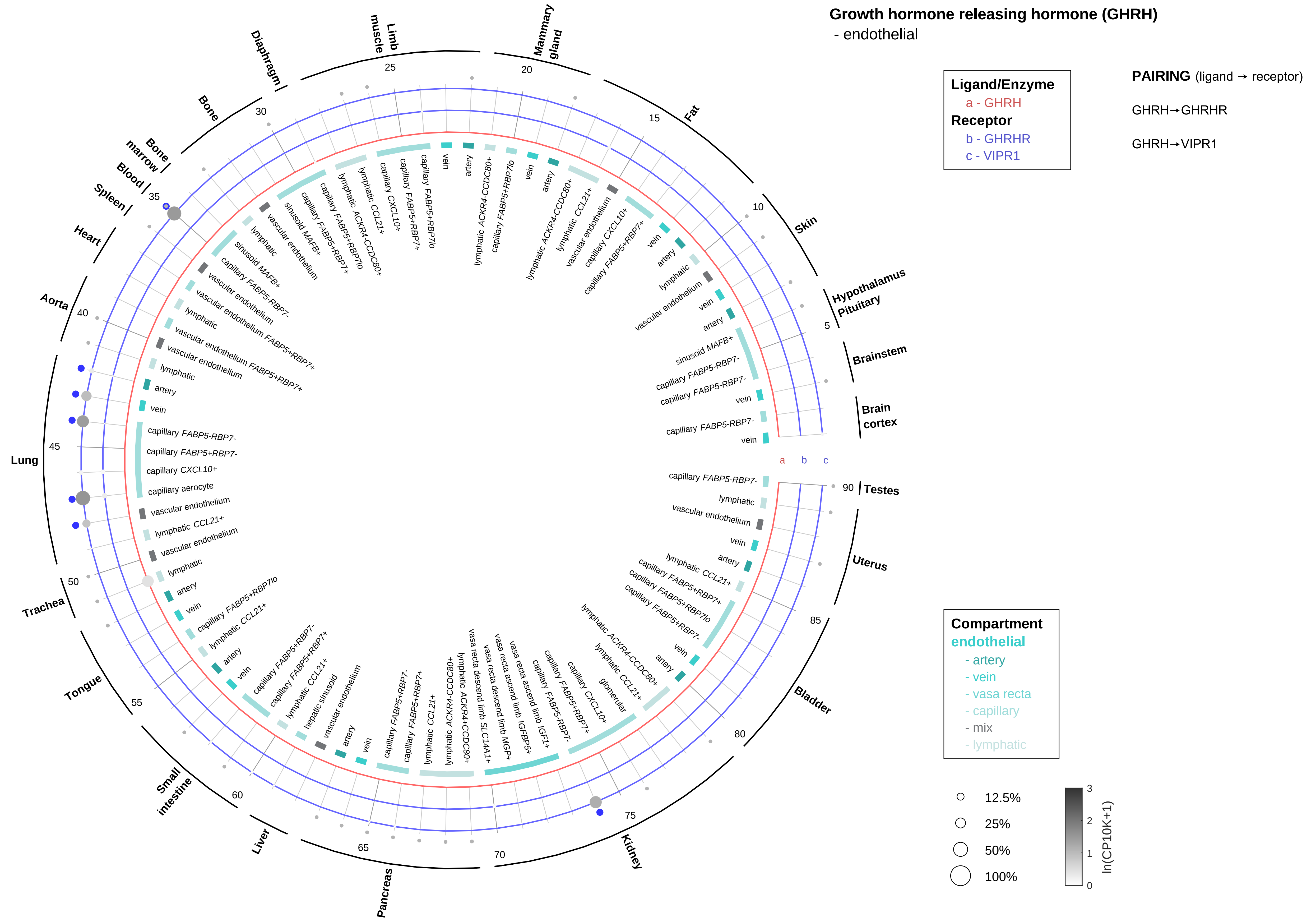

Growth hormone releasing hormone (GHRH)
- endothelial
Ligand/Enzyme
a - GHRH
Receptor
b - GHRHR
c - VIPR1
PAIRING (ligand → receptor)
GHRH→GHRHR
GHRH→VIPR1
Compartment
endothelial
- artery
- vein
- vasa recta
- capillary
- mix
- lymphatic
12.5%
25%
50%
100%
ln(CP10K+1)
Hypothalamus
Pituitary
Brainstem
Brain cortex
Testes
Uterus
Bladder
Kidney
Pancreas
Liver
Small intestine
Tongue
Trachea
Lung
Aorta
Heart
Spleen
Blood
Bone marrow
Bone
Diaphragm
Limb muscle
Mammary gland
Fat
Skin

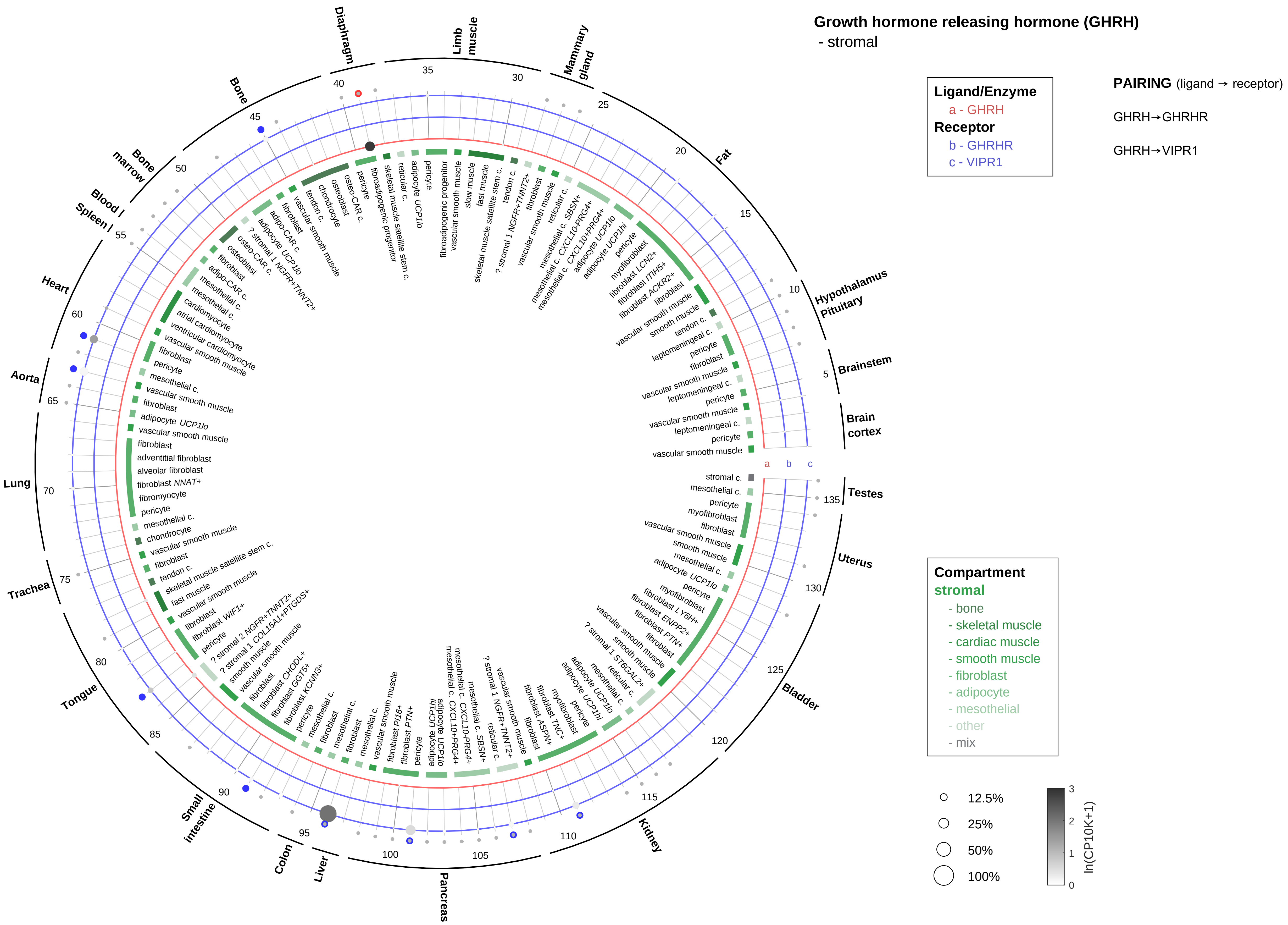

Growth hormone releasing hormone (GHRH)
- stromal
Ligand/Enzyme
a - GHRH
Receptor
b - GHRHR
c - VIPR1
PAIRING (ligand → receptor)
GHRH→GHRHR
GHRH→VIPR1
Compartment
stromal
- bone
- skeletal muscle
- cardiac muscle
- smooth muscle
- fibroblast
- adipocyte
- mesothelial
- other
- mix
12.5%
25%
50%
100%
ln(CP10K+1)
Brain cortex
Brainstem
Hypothalamus Pituitary
Fat
Mammary gland
Limb muscle
Diaphragm
Bone
Bone marrow
Blood / Spleen
Heart
Aorta
Lung
Trachea
Tongue
Small intestine
Colon
Liver
Pancreas
Kidney
Bladder
Uterus
Testes

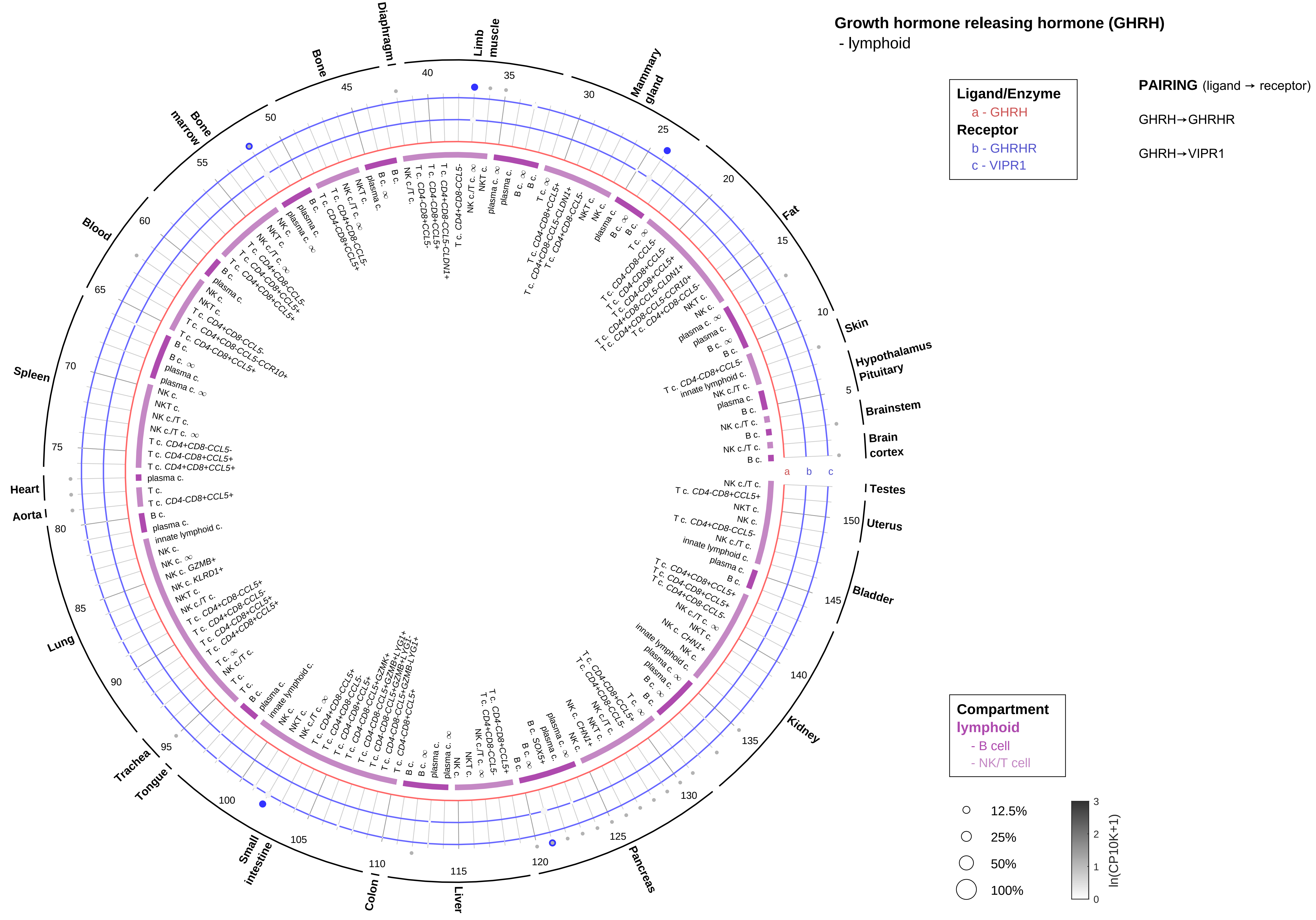
Growth hormone releasing hormone (GHRH)
- lymphoid
Ligand/Enzyme
a - GHRH
Receptor
b - GHRHR
c - VIPR1
PAIRING (ligand → receptor)
GHRH→GHRHR
GHRH→VIPR1
Compartment
lymphoid
- B cell
- NK/T cell
12.5%
25%
50%
100%
ln(CP10K+1)
Skin
Hypothalamus
Pituitary
Brainstem
Brain cortex
Testes
Uterus
Bladder
Kidney
Pancreas
Liver
Colon
Small intestine
Tongue
Trachea
Lung
Aorta
Heart
Spleen
Blood
Bone marrow
Bone
Diaphragm
Limb muscle
Mammary gland
Fat

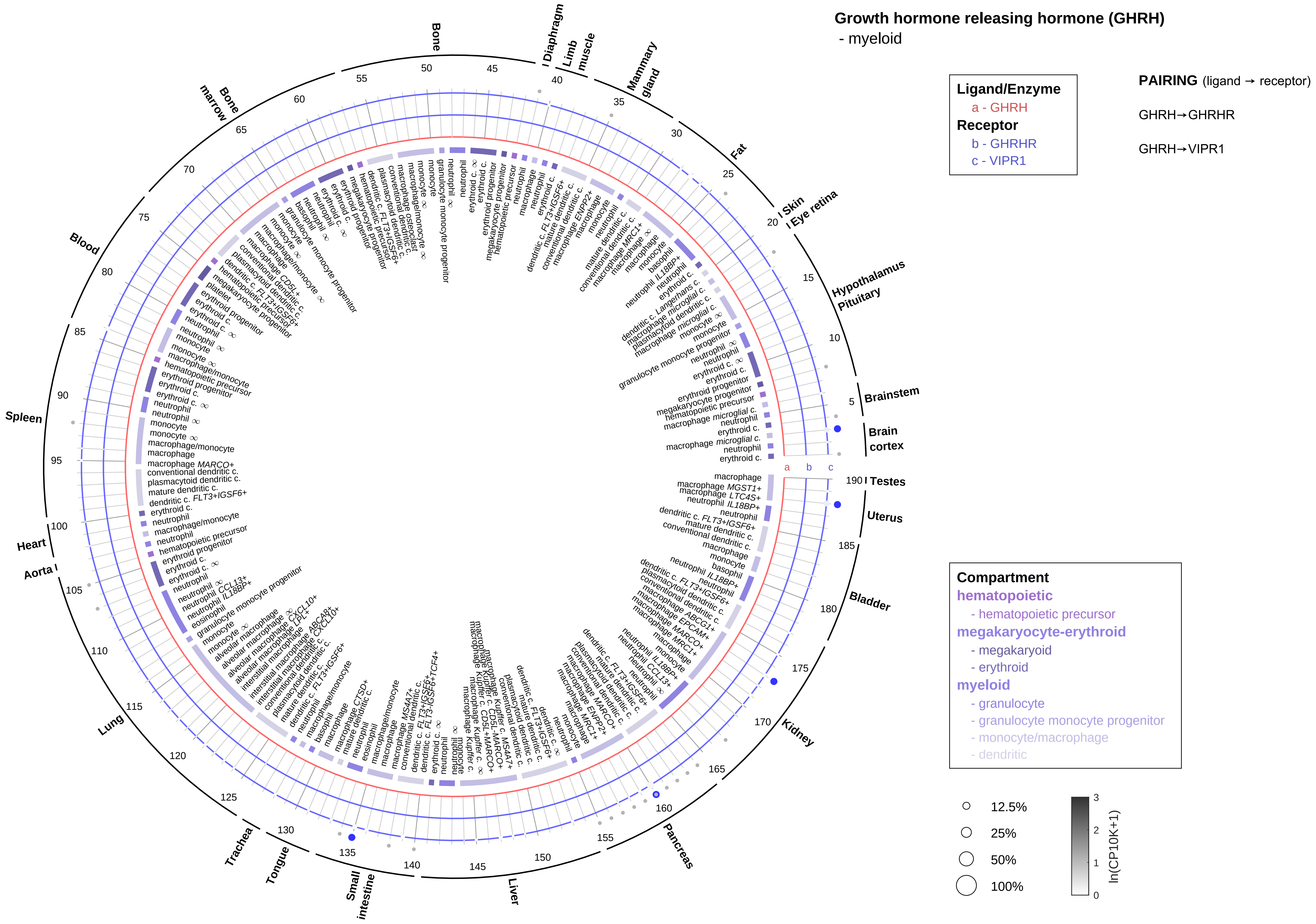
Growth hormone releasing hormone (GHRH)
- myeloid
Ligand/Enzyme
a - GHRH
Receptor
b - GHRHR
c - VIPR1
PAIRING (ligand → receptor)
GHRH→GHRHR
GHRH→VIPR1
Compartment
hematopoietic
- hematopoietic precursor
megakaryocyte-erythroid
- megakaryoid
- erythroid
myeloid
- granulocyte
- granulocyte monocyte progenitor
- monocyte/macrophage
- dendritic
12.5%
25%
50%
100%
ln(CP10K+1)
Bone
Diaphragm
Limb muscle
Mammary gland
Fat
Skin
Eye retina
Hypothalamus
Pituitary
Brainstem
Brain cortex
Testes
Uterus
Bladder
Kidney
Pancreas
Liver
Small intestine
Tongue
Trachea
Lung
Aorta
Heart
Spleen
Blood
Bone marrow

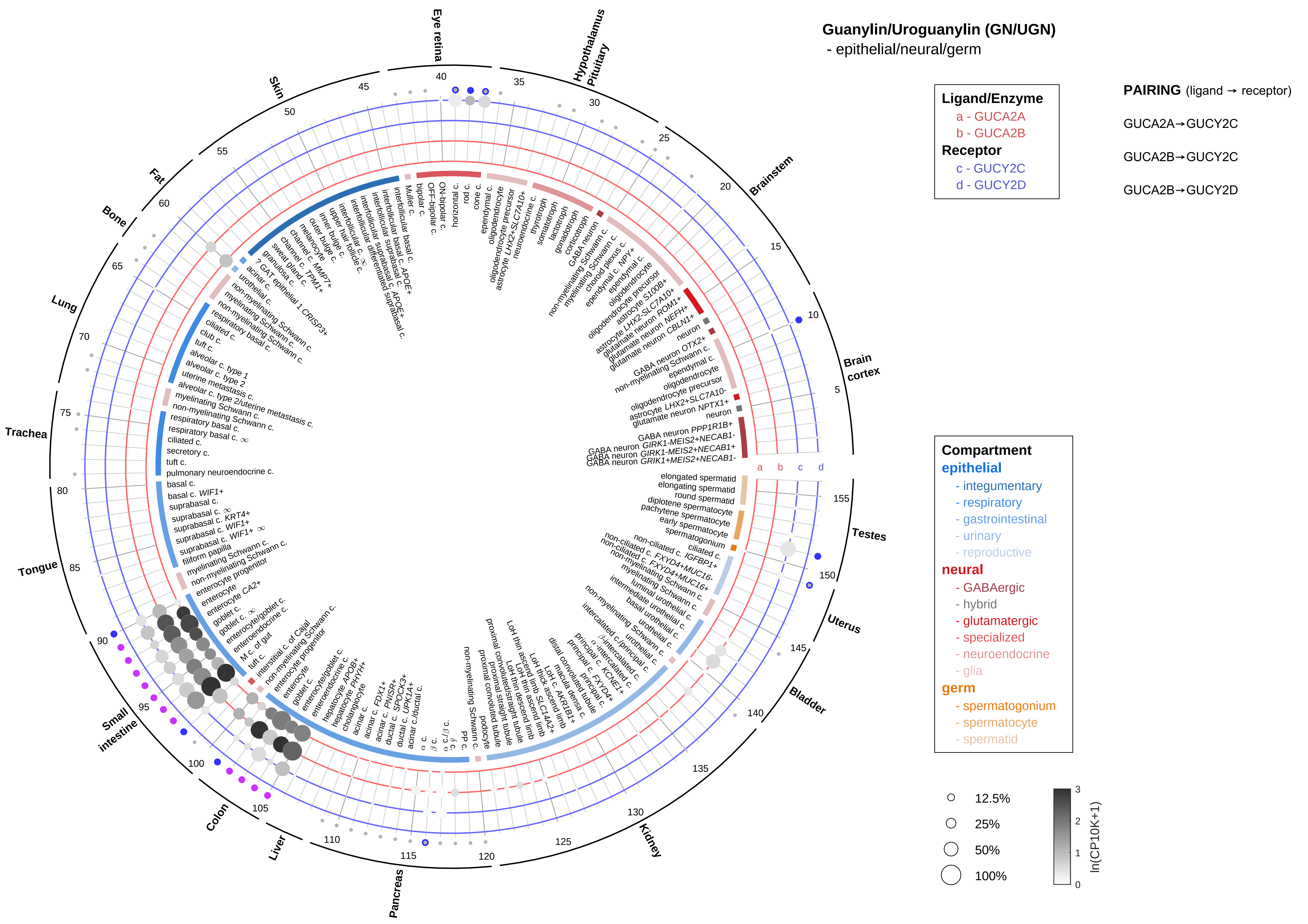

Guanylin/Uroguanylin (GN/UGN)
- epithelial/neural/germ
Ligand/Enzyme
a - GUCA2A
b - GUCA2B
Receptor
c - GUCY2C
d - GUCY2D
PAIRING (ligand → receptor)
GUCA2A→GUCY2C
GUCA2B→GUCY2C
GUCA2B→GUCY2D
Compartment
epithelial
- integumentary
- respiratory
- gastrointestinal
- urinary
- reproductive
neural
- GABAergic
- hybrid
- glutamatergic
- specialized
- neuroendocrine
- glia
germ
- spermatogonium
- spermatocyte
- spermatid
12.5%
25%
50%
100%
ln(CP10K+1)
Eye retina
Hypothalamus
Pituitary
Brainstem
Brain cortex
Testes
Uterus
Bladder
Kidney
Pancreas
Liver
Colon
Small intestine
Tongue
Trachea
Lung
Bone
Fat
Skin

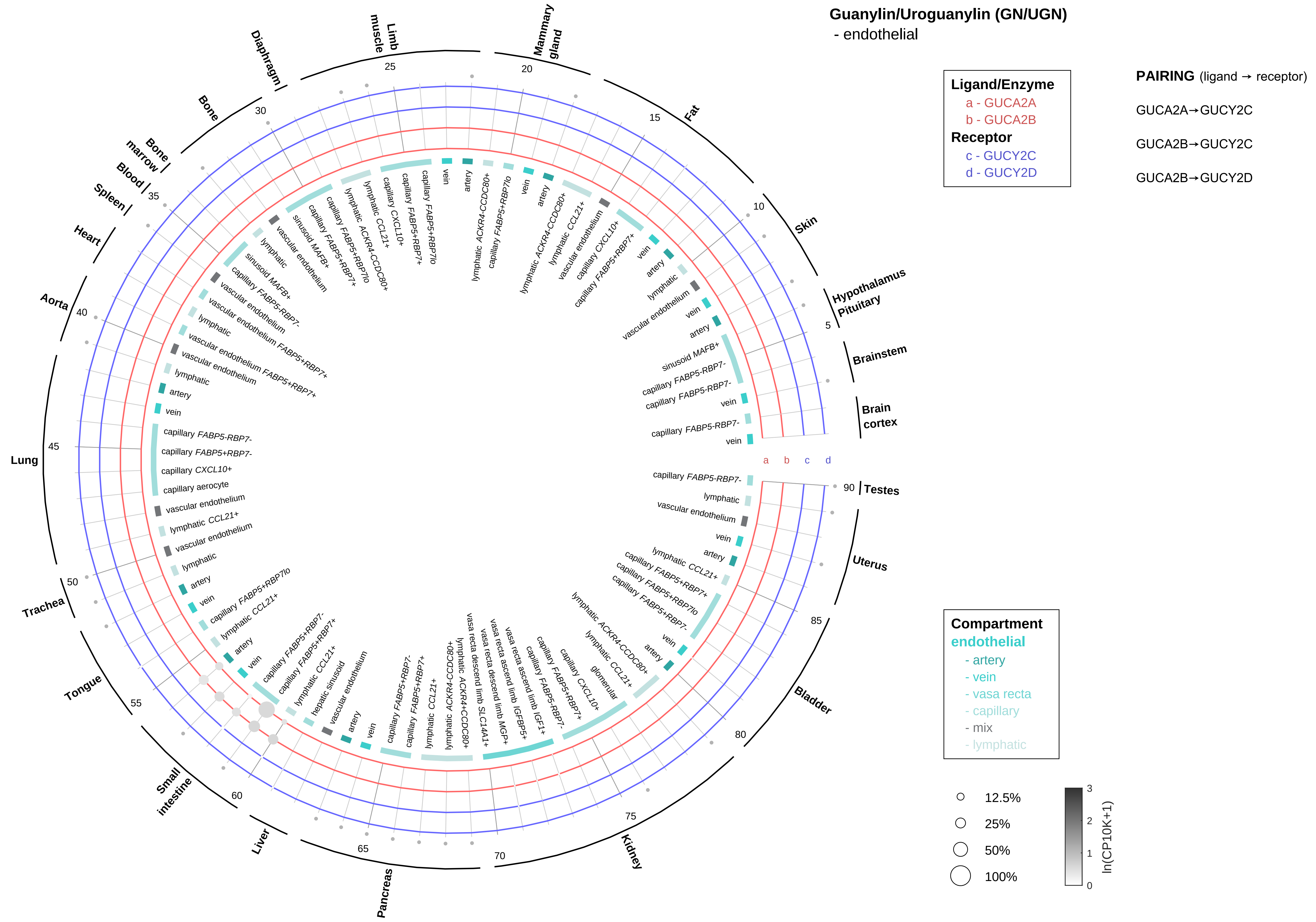

Guanylin/Uroguanylin (GN/UGN)
- endothelial
Ligand/Enzyme
a - GUCA2A
b - GUCA2B
Receptor
c - GUCY2C
d - GUCY2D
PAIRING (ligand → receptor)
GUCA2A→GUCY2C
GUCA2B→GUCY2C
GUCA2B→GUCY2D
Compartment
endothelial
- artery
- vein
- vasa recta
- capillary
- mix
- lymphatic
12.5%
25%
50%
100%
ln(CP10K+1)
Testes
Uterus
Bladder
Kidney
Pancreas
Liver
Small intestine
Tongue
Trachea
Lung
Aorta
Heart
Spleen
Blood
Bone marrow
Bone
Diaphragm
Limb muscle
Mammary gland
Fat
Skin
Hypothalamus
Pituitary
Brainstem
Brain cortex

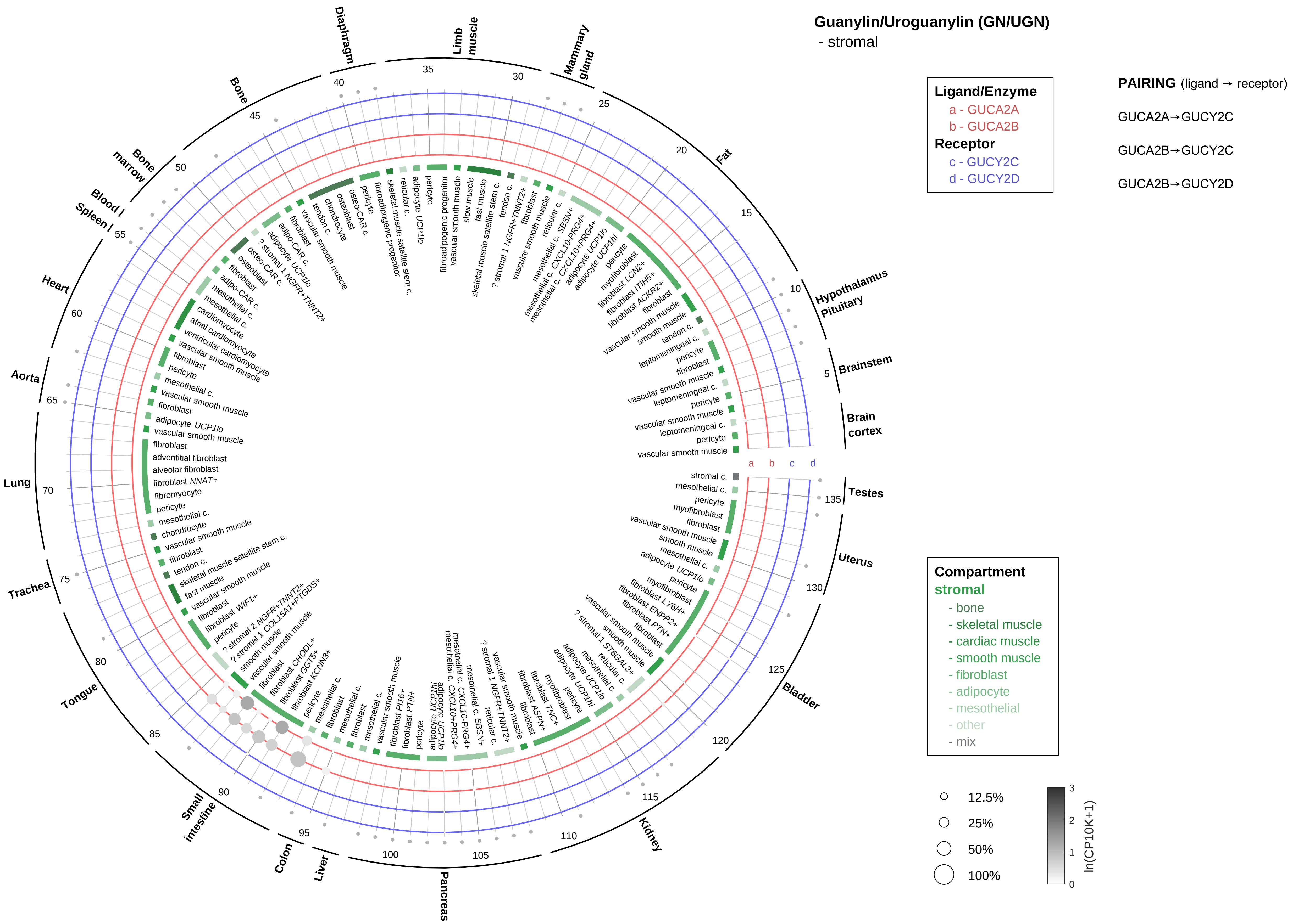

Guanylin/Uroguanylin (GN/UGN)
- stromal
Ligand/Enzyme
a - GUCA2A
b - GUCA2B
Receptor
c - GUCY2C
d - GUCY2D
PAIRING (ligand → receptor)
GUCA2A→GUCY2C
GUCA2B→GUCY2C
GUCA2B→GUCY2D
Compartment
stromal
- bone
- skeletal muscle
- cardiac muscle
- smooth muscle
- fibroblast
- adipocyte
- mesothelial
- other
- mix
12.5%
25%
50%
100%
ln(CP10K+1)

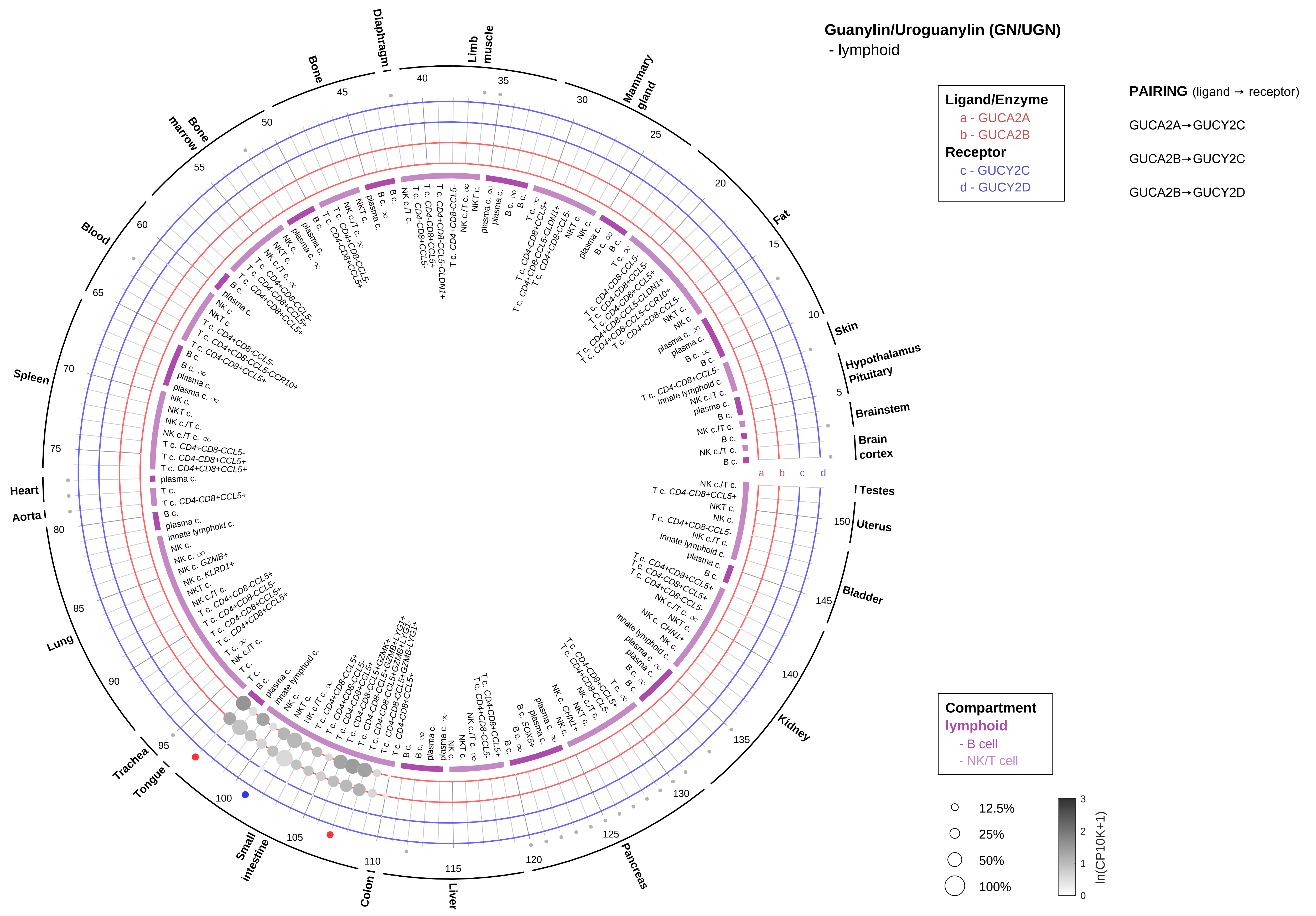

Guanylin/Uroguanylin (GN/UGN)
- lymphoid
Ligand/Enzyme
a - GUCA2A
b - GUCA2B
Receptor
c - GUCY2C
d - GUCY2D
PAIRING (ligand → receptor)
GUCA2A→GUCY2C
GUCA2B→GUCY2C
GUCA2B→GUCY2D
Compartment
lymphoid
- B cell
- NK/T cell
12.5%
25%
50%
100%
ln(CP10K+1)

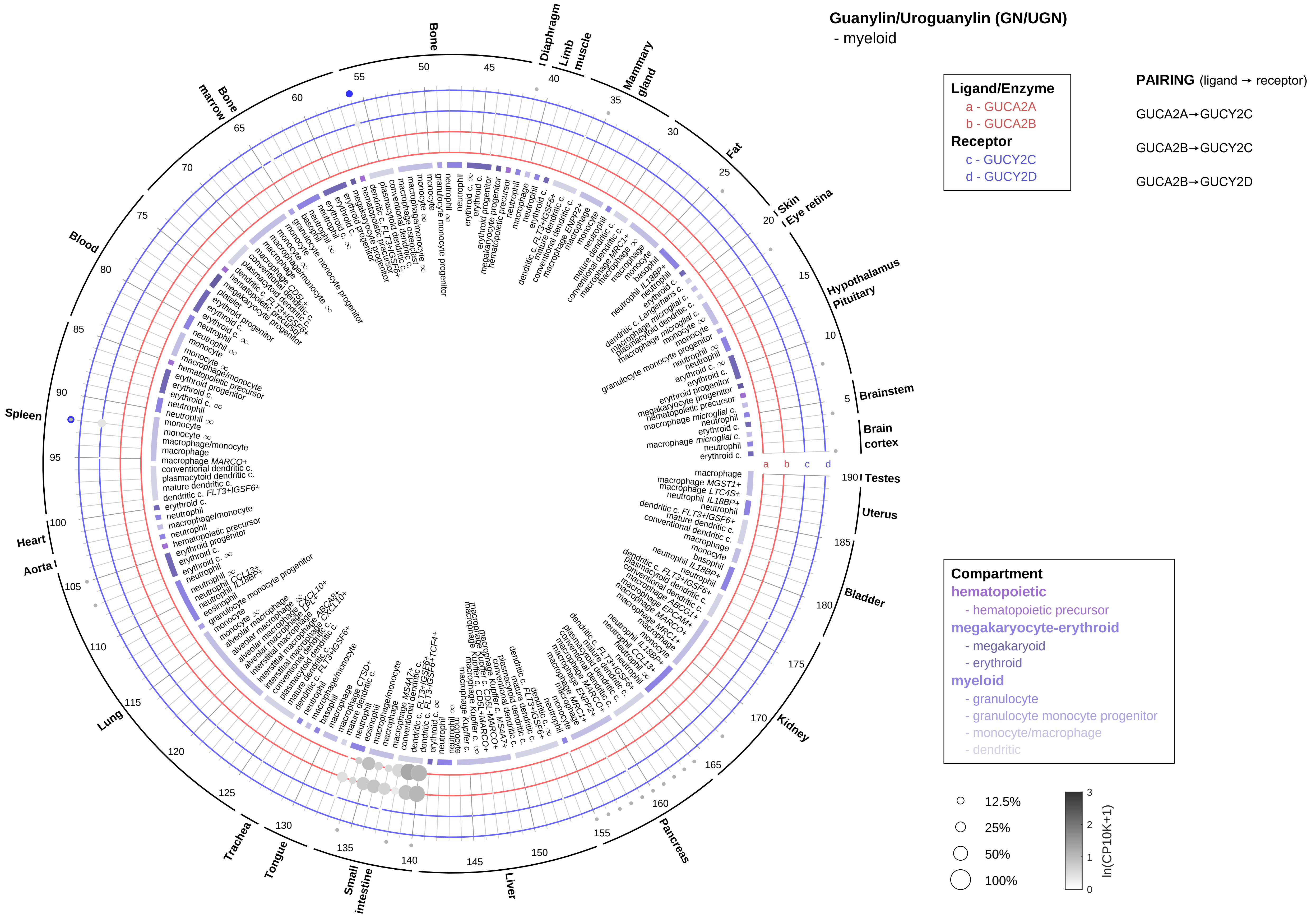

Guanylin/Uroguanylin (GN/UGN)
- myeloid
Ligand/Enzyme
a - GUCA2A
b - GUCA2B
Receptor
c - GUCY2C
d - GUCY2D
PAIRING (ligand → receptor)
GUCA2A→GUCY2C
GUCA2B→GUCY2C
GUCA2B→GUCY2D
Compartment
hematopoietic
- hematopoietic precursor
megakaryocyte-erythroid
- megakaryoid
- erythroid
myeloid
- granulocyte
- granulocyte monocyte progenitor
- monocyte/macrophage
- dendritic
12.5%
25%
50%
100%
ln(CP10K+1)
Bone
Diaphragm
Limb muscle
Mammary gland
Fat
Skin
Eye retina
Hypothalamus
Pituitary
Brainstem
Brain cortex
Testes
Uterus
Bladder
Kidney
Pancreas
Liver
Small intestine
Tongue
Trachea
Lung
Aorta
Heart
Spleen
Blood
Bone marrow

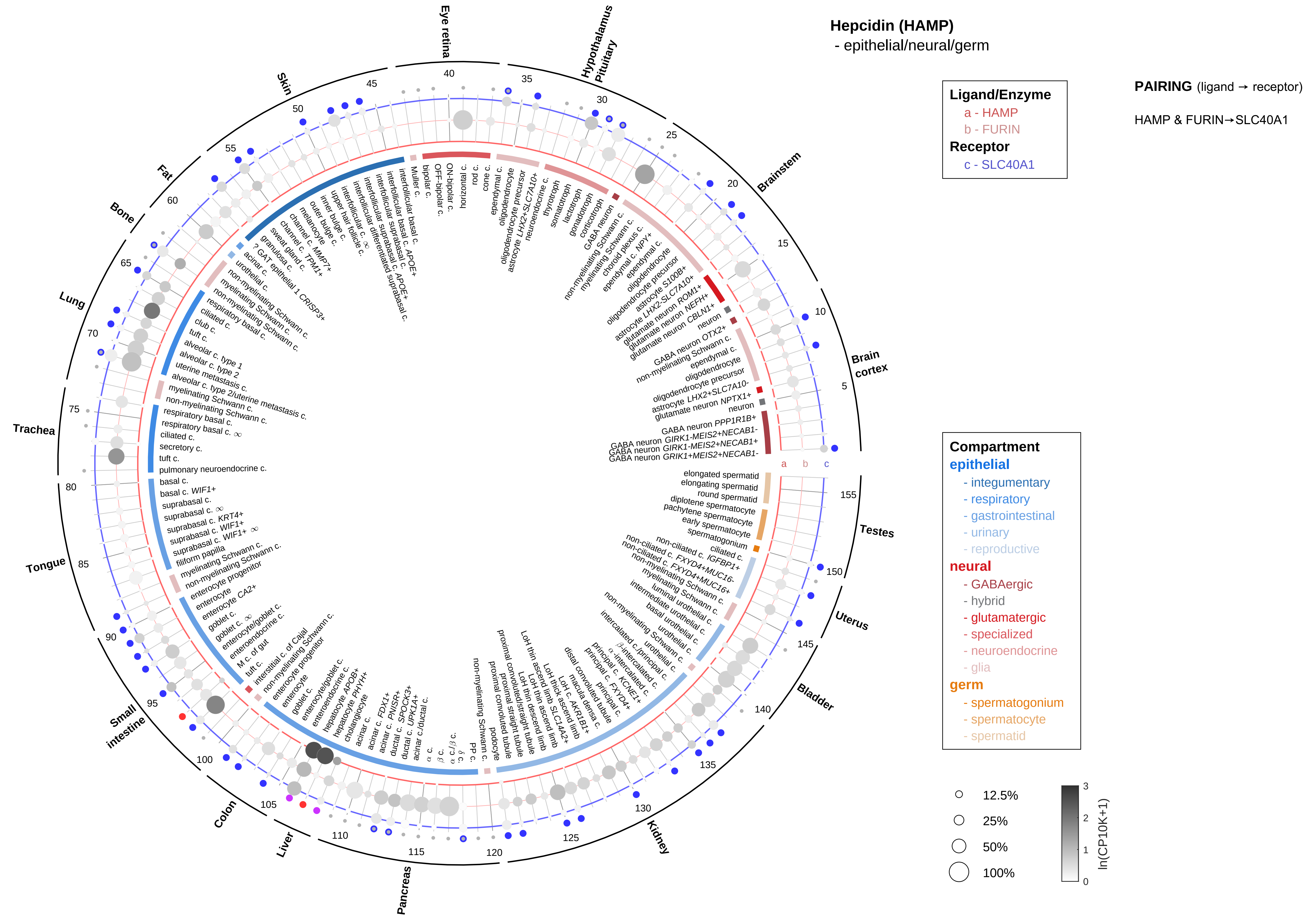

Hepcidin (HAMP)
- epithelial/neural/germ
Ligand/Enzyme
a - HAMP
b - FURIN
Receptor
c - SLC40A1
PAIRING (ligand → receptor)
HAMP & FURIN→SLC40A1
Compartment
epithelial
- integumentary
- respiratory
- gastrointestinal
- urinary
- reproductive
neural
- GABAergic
- hybrid
- glutamatergic
- specialized
- neuroendocrine
- glia
germ
- spermatogonium
- spermatocyte
- spermatid
12.5%
25%
50%
100%
ln(CP10K+1)
Eye retina
Hypothalamus Pituitary
Brainstem
Brain cortex
Testes
Uterus
Bladder
Kidney
Pancreas
Liver
Colon
Small intestine
Tongue
Trachea
Lung
Bone
Fat
Skin

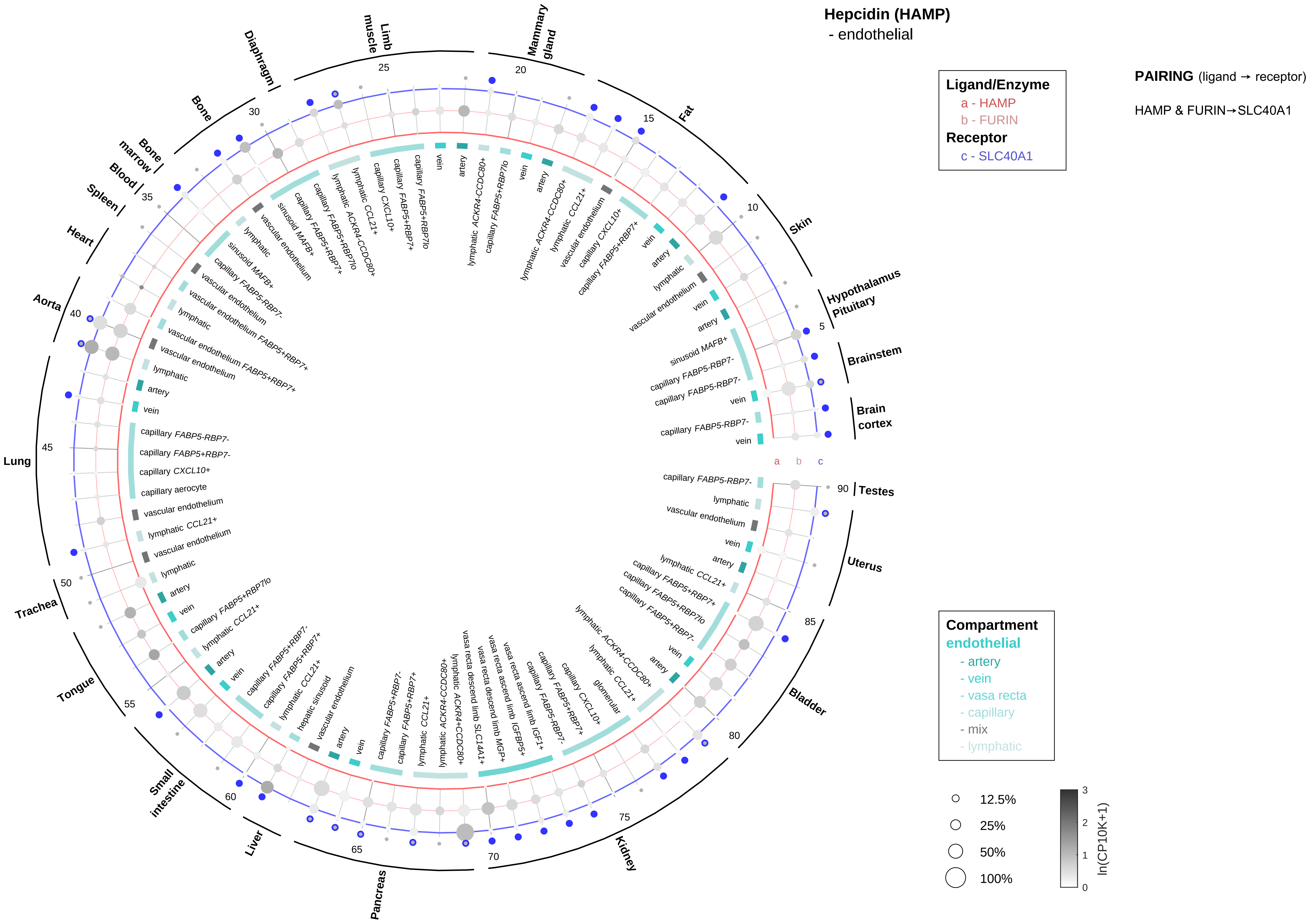
Hepcidin (HAMP)
- endothelial
Ligand/Enzyme
a - HAMP
b - FURIN
Receptor
c - SLC40A1
PAIRING (ligand → receptor)
HAMP & FURIN→SLC40A1
Compartment
endothelial
- artery
- vein
- vasa recta
- capillary
- mix
- lymphatic
12.5%
25%
50%
100%
ln(CP10K+1)
Hypothalamus Pituitary
Brainstem
Brain cortex
Testes
Uterus
Bladder
Kidney
Pancreas
Liver
Small intestine
Tongue
Trachea
Lung
Aorta
Heart
Spleen
Blood
Bone marrow
Bone
Diaphragm
Limb muscle
Mammary gland
Fat
Skin

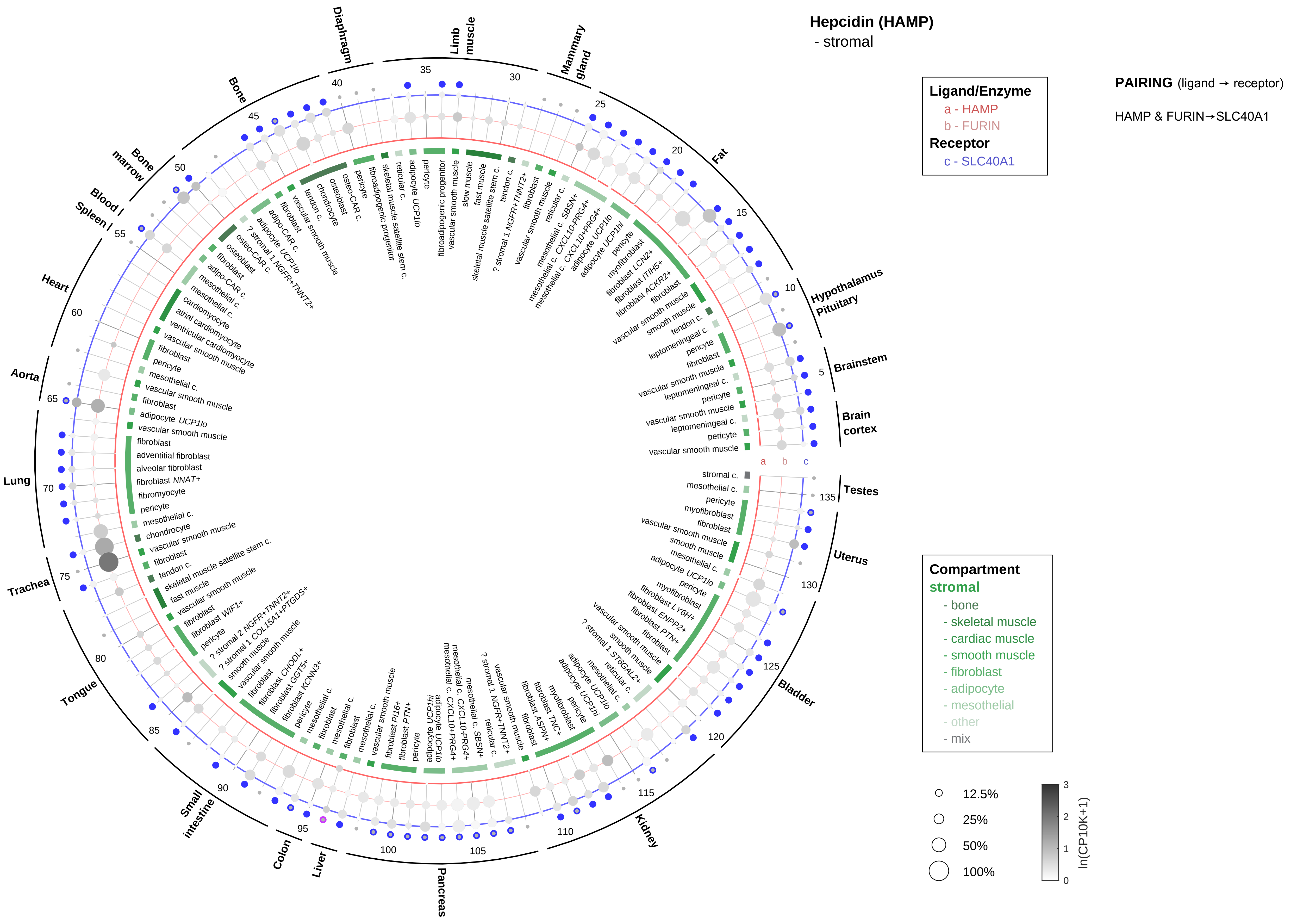

Hepcidin (HAMP)
- stromal
Ligand/Enzyme
a - HAMP
b - FURIN
Receptor
c - SLC40A1
PAIRING (ligand → receptor)
HAMP & FURIN→SLC40A1
Compartment
stromal
- bone
- skeletal muscle
- cardiac muscle
- smooth muscle
- fibroblast
- adipocyte
- mesothelial
- other
- mix
12.5%
25%
50%
100%
ln(CP10K+1)
Hypothalamus Pituitary
Brainstem
Brain cortex
Testes
Uterus
Bladder
Kidney
Pancreas
Liver
Colon
Small intestine
Tongue
Trachea
Lung
Aorta
Heart
Spleen
Blood
Bone marrow
Bone
Diaphragm
Limb muscle
Mammary gland
Fat

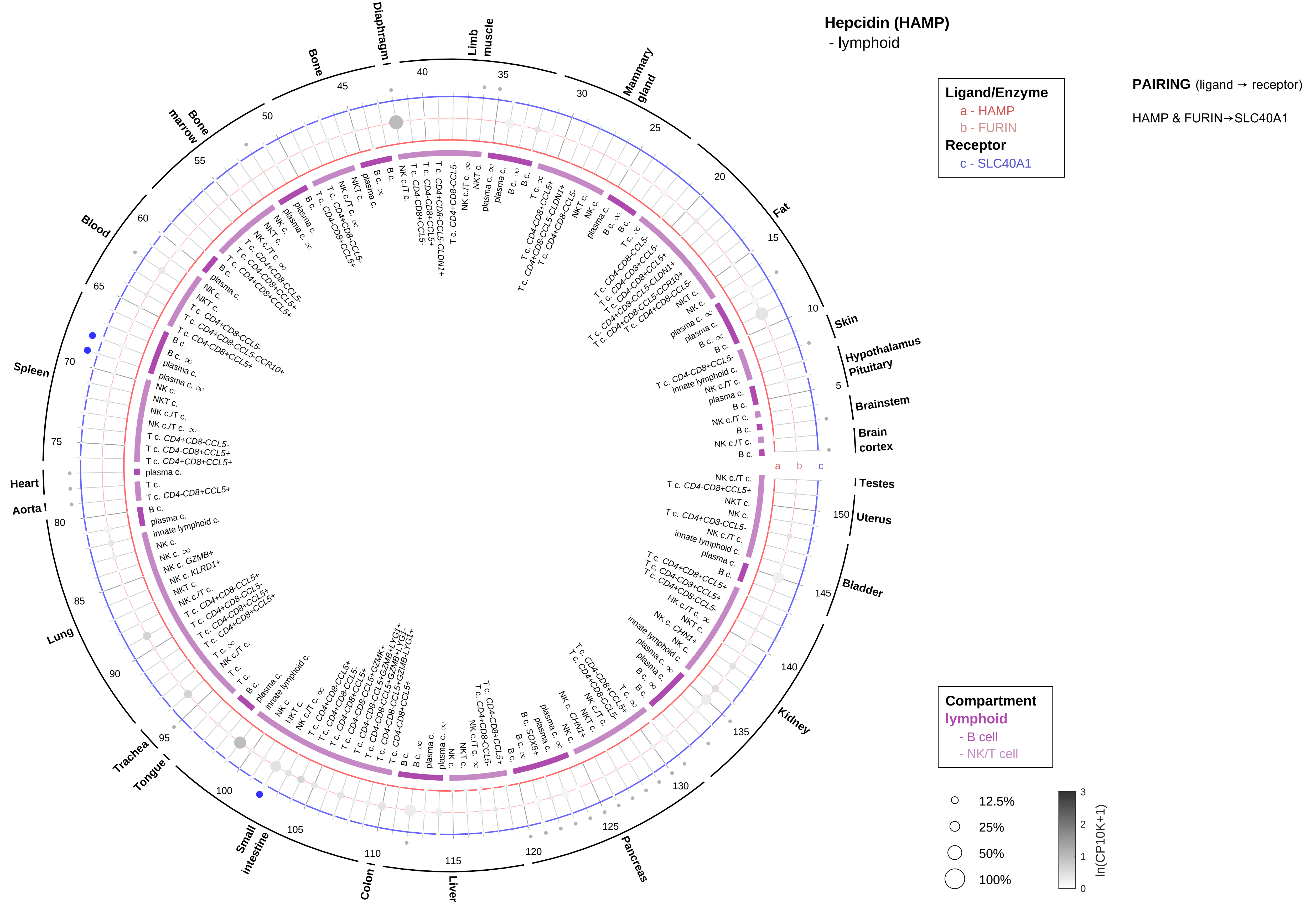
Hepcidin (HAMP)
- lymphoid
Ligand/Enzyme
a - HAMP
b - FURIN
Receptor
c - SLC40A1
PAIRING (ligand → receptor)
HAMP & FURIN→SLC40A1
Compartment
lymphoid
- B cell
- NK/T cell
12.5%
25%
50%
100%
ln(CP10K+1)
Skin
Hypothalamus
Pituitary
Brainstem
Brain cortex
Testes
Uterus
Bladder
Kidney
Pancreas
Liver
Colon
Small intestine
Tongue
Trachea
Lung
Aorta
Heart
Spleen
Blood
Bone marrow
Bone
Diaphragm
Limb muscle
Mammary gland
Fat

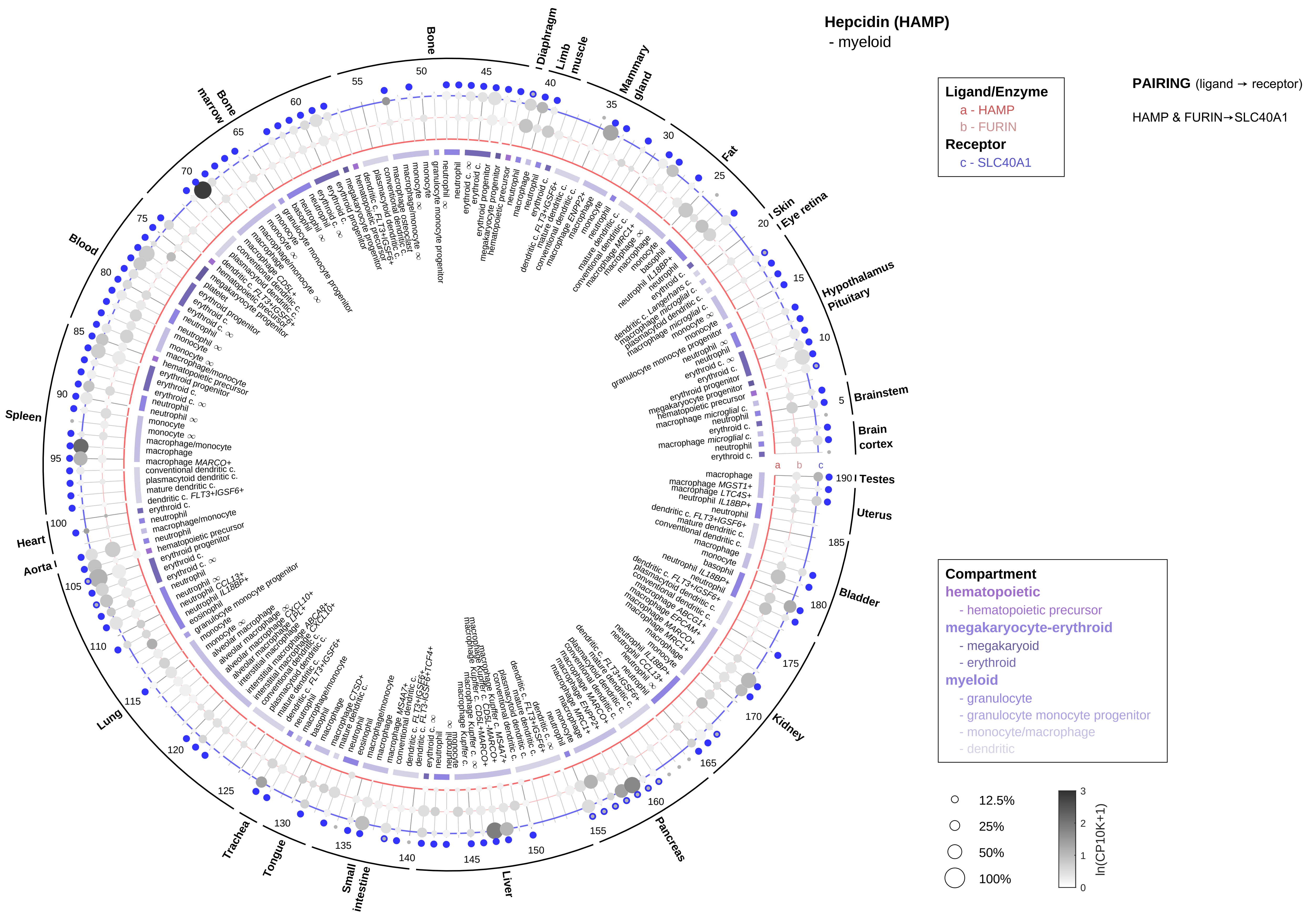
Hepcidin (HAMP)
- myeloid
Ligand/Enzyme
a - HAMP
b - FURIN
Receptor
c - SLC40A1
PAIRING (ligand → receptor)
HAMP & FURIN→SLC40A1
Compartment
hematopoietic
- hematopoietic precursor
megakaryocyte-erythroid
- megakaryoid
- erythroid
myeloid
- granulocyte
- granulocyte monocyte progenitor
- monocyte/macrophage
- dendritic
12.5%
25%
50%
100%
ln(CP10K+1)
Brainstem
Brain cortex
Testes
Uterus
Bladder
Kidney
Pancreas
Liver
Small intestine
Tongue
Trachea
Lung
Aorta
Heart
Spleen
Blood
Bone marrow
Bone
Diaphragm
Limb muscle
Mammary gland
Fat
Skin
Eye retina
Hypothalamus
Pituitary

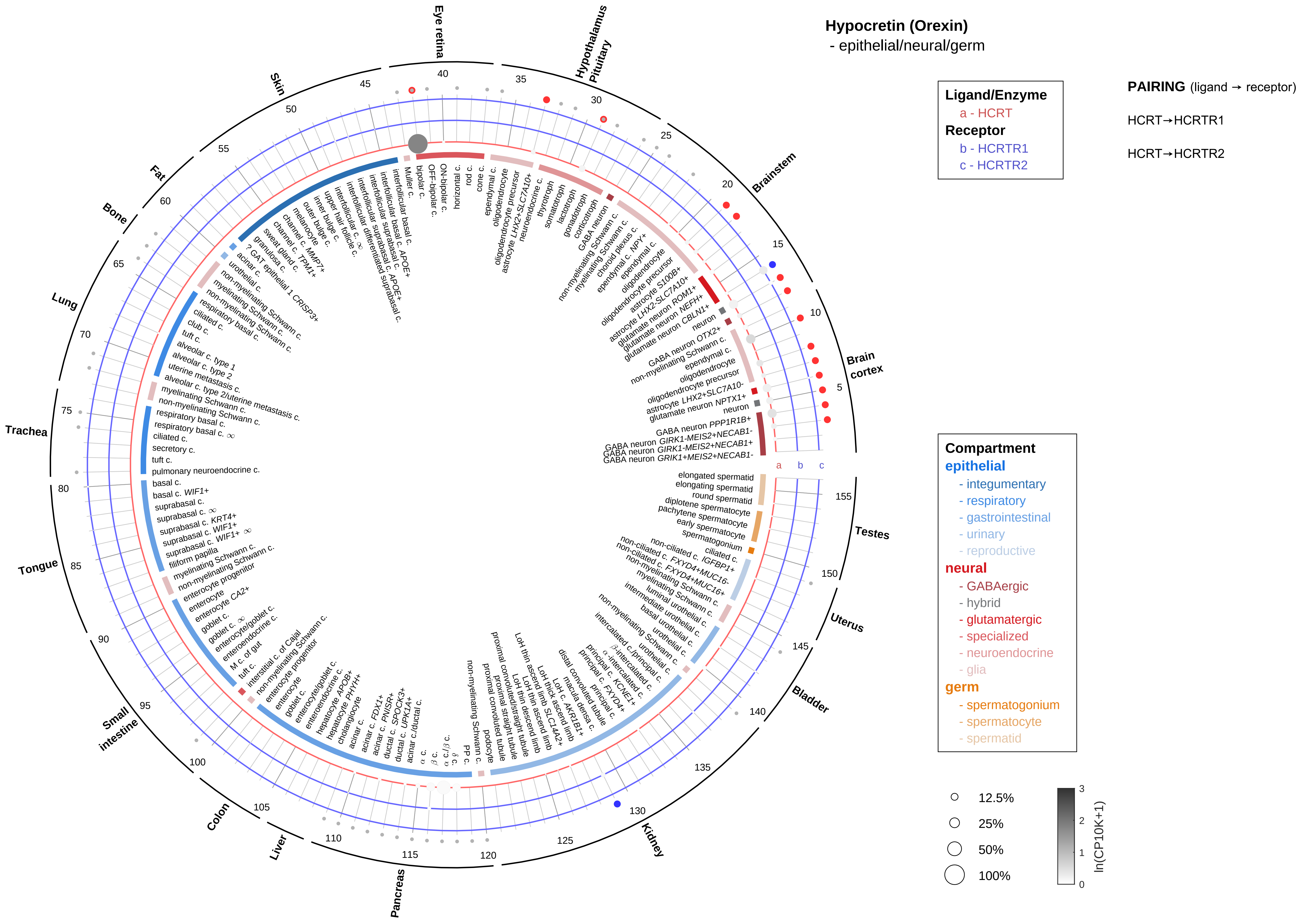
Hypocretin (Orexin)
- epithelial/neural/germ
Ligand/Enzyme
a - HCRT
Receptor
b - HCRTR1
c - HCRTR2
PAIRING (ligand → receptor)
HCRT→HCRTR1
HCRT→HCRTR2
Compartment
epithelial
- integumentary
- respiratory
- gastrointestinal
- urinary
- reproductive
neural
- GABAergic
- hybrid
- glutamatergic
- specialized
- neuroendocrine
- glia
germ
- spermatogonium
- spermatocyte
- spermatid
12.5%
25%
50%
100%
ln(CP10K+1)
Brain cortex
Brainstem
Hypothalamus Pituitary
Eye retina
Skin
Fat
Bone
Lung
Trachea
Tongue
Small intestine
Colon
Liver
Pancreas
Kidney
Bladder
Uterus
Testes

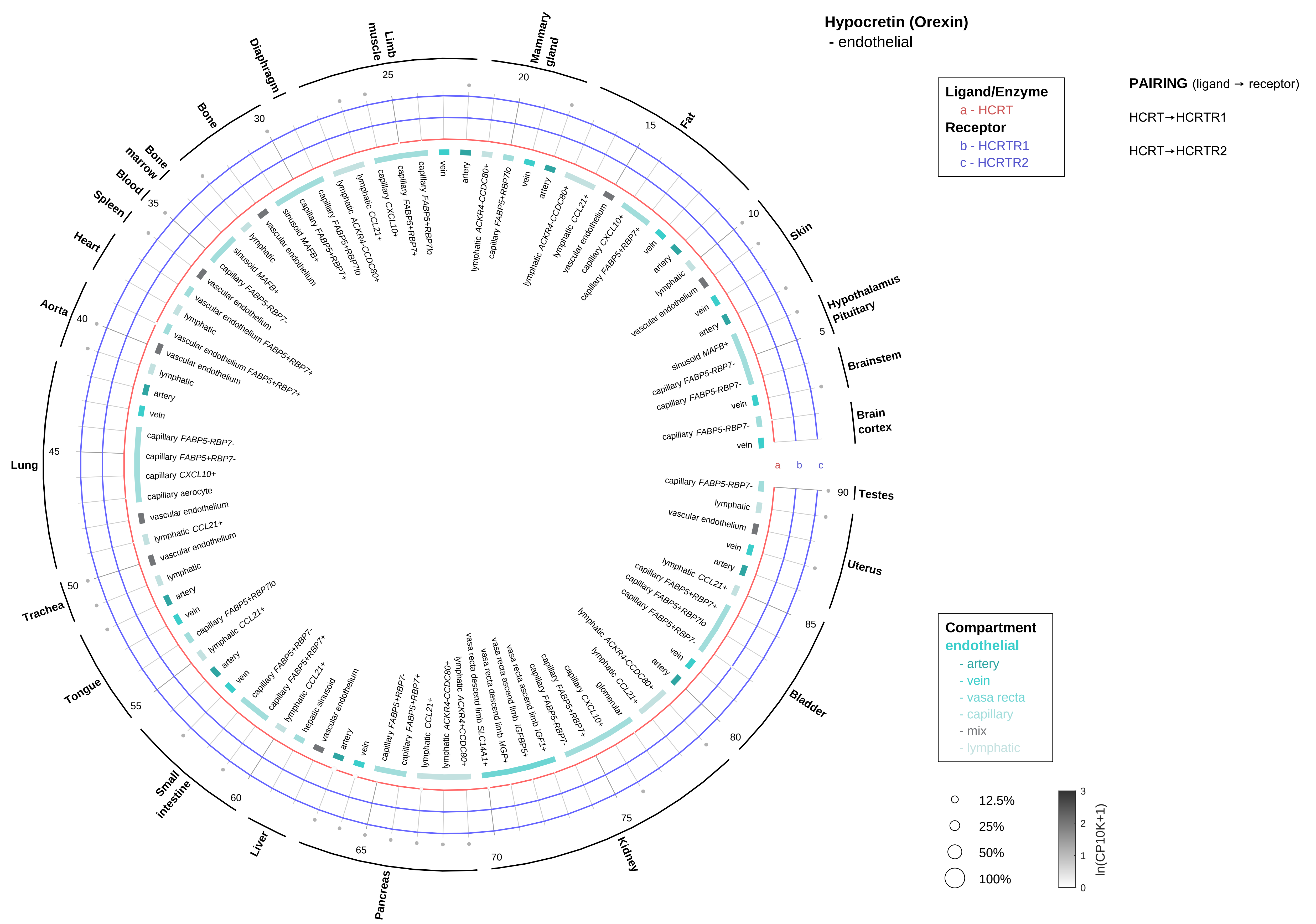

Hypocretin (Orexin)
- endothelial
Ligand/Enzyme
a - HCRT
Receptor
b - HCRTR1
c - HCRTR2
PAIRING (ligand → receptor)
HCRT→HCRTR1
HCRT→HCRTR2
Compartment
endothelial
- artery
- vein
- vasa recta
- capillary
- mix
- lymphatic
12.5%
25%
50%
100%
ln(CP10K+1)
Hypothalamus Pituitary
Brainstem
Brain cortex
Testes
Uterus
Bladder
Kidney
Pancreas
Liver
Small intestine
Tongue
Trachea
Lung
Aorta
Heart
Spleen
Blood
Bone marrow
Bone
Diaphragm
Limb muscle
Mammary gland
Fat
Skin

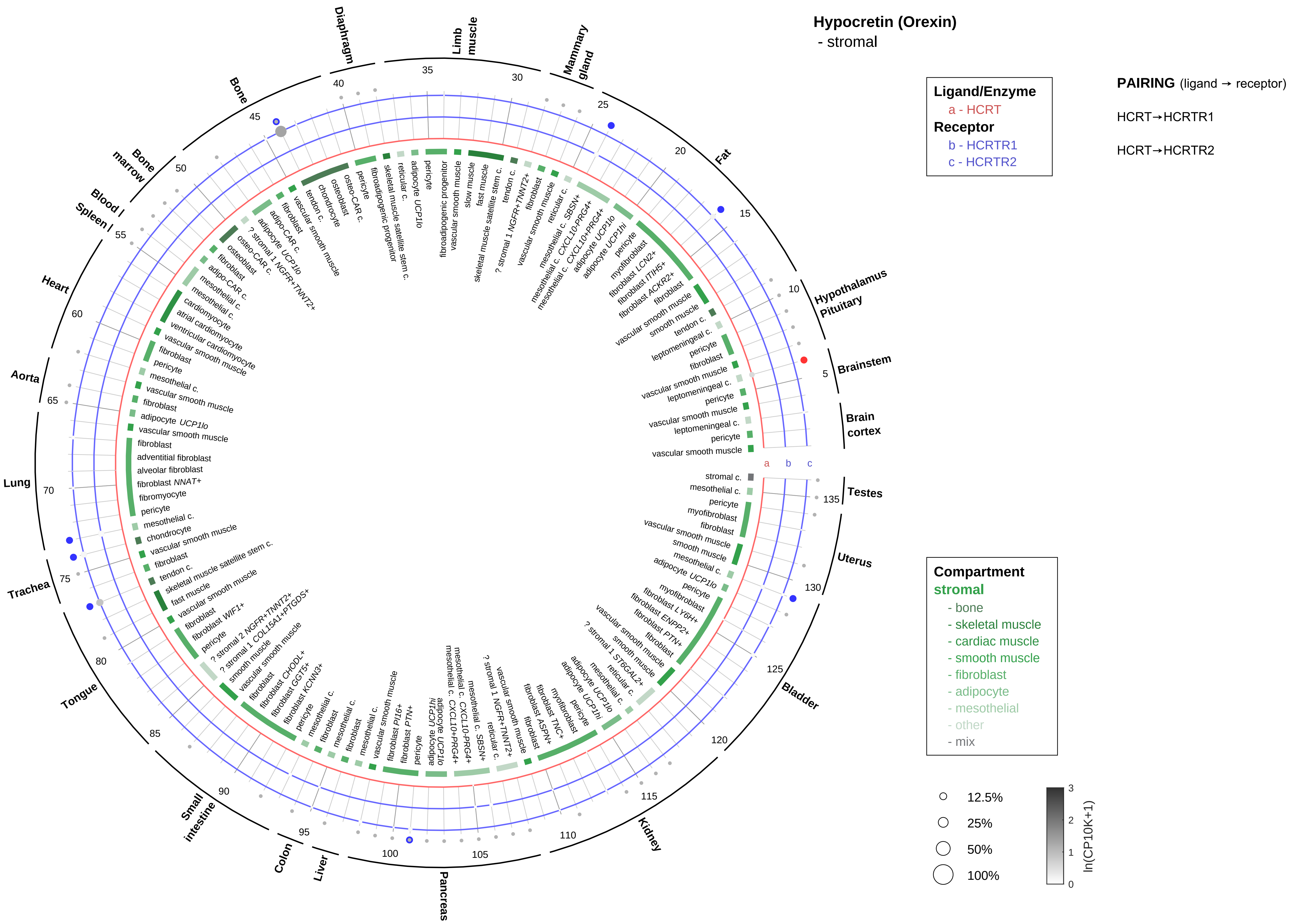
Hypocretin (Orexin)
- stromal
Ligand/Enzyme
a - HCRT
Receptor
b - HCRTR1
c - HCRTR2
PAIRING (ligand → receptor)
HCRT→HCRTR1
HCRT→HCRTR2
Compartment
stromal
- bone
- skeletal muscle
- cardiac muscle
- smooth muscle
- fibroblast
- adipocyte
- mesothelial
- other
- mix
12.5%
25%
50%
100%
ln(CP10K+1)
Hypothalamus
Pituitary
Brainstem
Brain cortex
Testes
Uterus
Bladder
Kidney
Pancreas
Liver
Colon
Small intestine
Tongue
Trachea
Lung
Aorta
Heart
Spleen
Blood
Bone marrow
Bone
Diaphragm
Limb muscle
Mammary gland
Fat

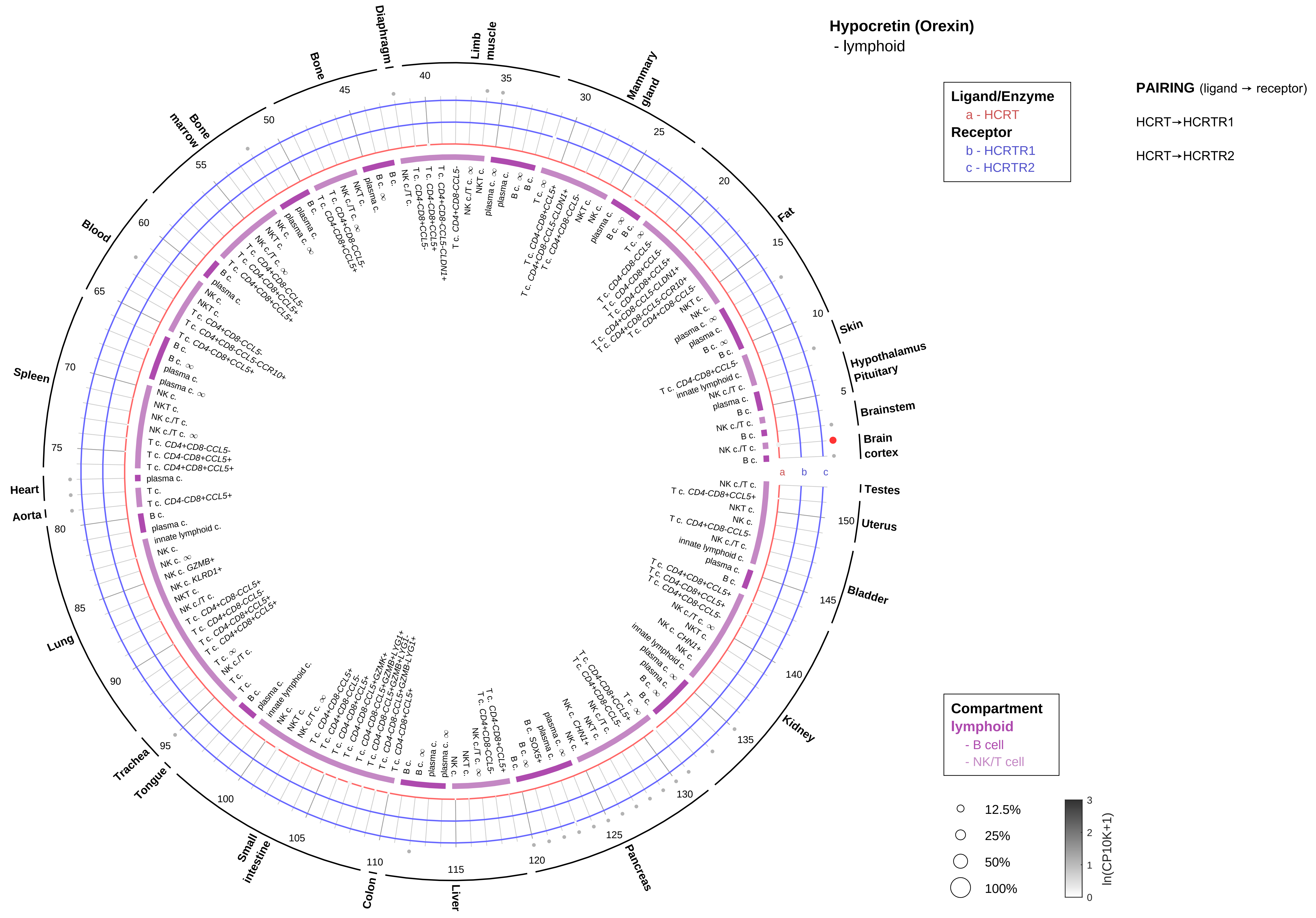

Hypocretin (Orexin)
- lymphoid
Ligand/Enzyme
a - HCRT
Receptor
b - HCRTR1
c - HCRTR2
PAIRING (ligand → receptor)
HCRT→HCRTR1
HCRT→HCRTR2
Compartment
lymphoid
- B cell
- NK/T cell
12.5%
25%
50%
100%
ln(CP10K+1)
Skin
Hypothalamus
Pituitary
Brainstem
Brain cortex
Testes
Uterus
Bladder
Kidney
Pancreas
Liver
Colon
Small intestine
Tongue
Trachea
Lung
Aorta
Heart
Spleen
Blood
Bone marrow
Bone
Diaphragm
Limb muscle
Mammary gland
Fat

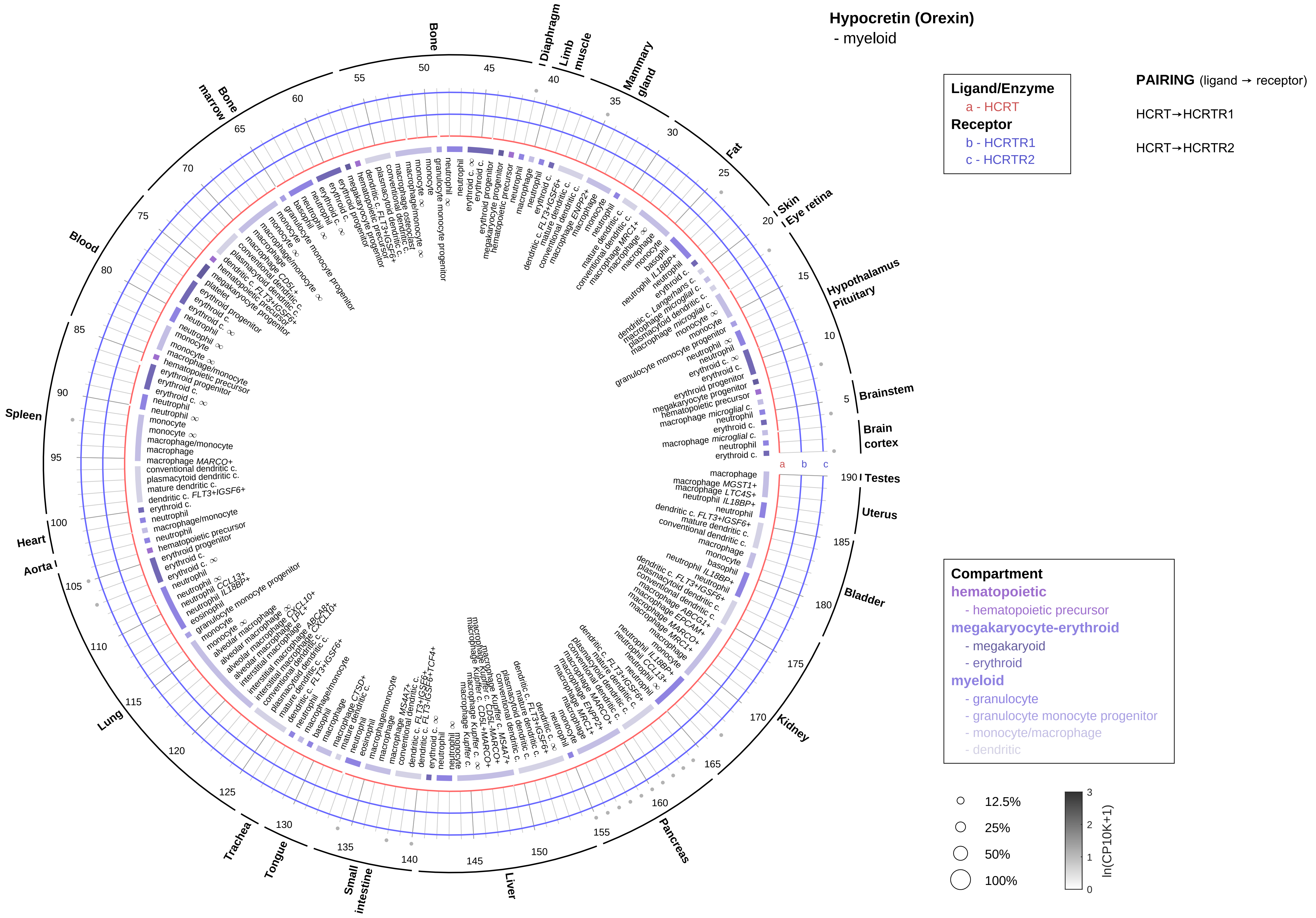
Hypocretin (Orexin)
- myeloid
Ligand/Enzyme
a - HCRT
Receptor
b - HCRTR1
c - HCRTR2
PAIRING (ligand → receptor)
HCRT→HCRTR1
HCRT→HCRTR2
Compartment
hematopoietic
- hematopoietic precursor
megakaryocyte-erythroid
- megakaryoid
- erythroid
myeloid
- granulocyte
- granulocyte monocyte progenitor
- monocyte/macrophage
- dendritic
12.5%
25%
50%
100%
ln(CP10K+1)
Bone
Diaphragm
Limb muscle
Mammary gland
Fat
Skin
Eye retina
Hypothalamus
Pituitary
Brainstem
Brain cortex
Testes
Uterus
Bladder
Kidney
Pancreas
Liver
Small intestine
Tongue
Trachea
Lung
Aorta
Heart
Spleen
Blood
Bone marrow

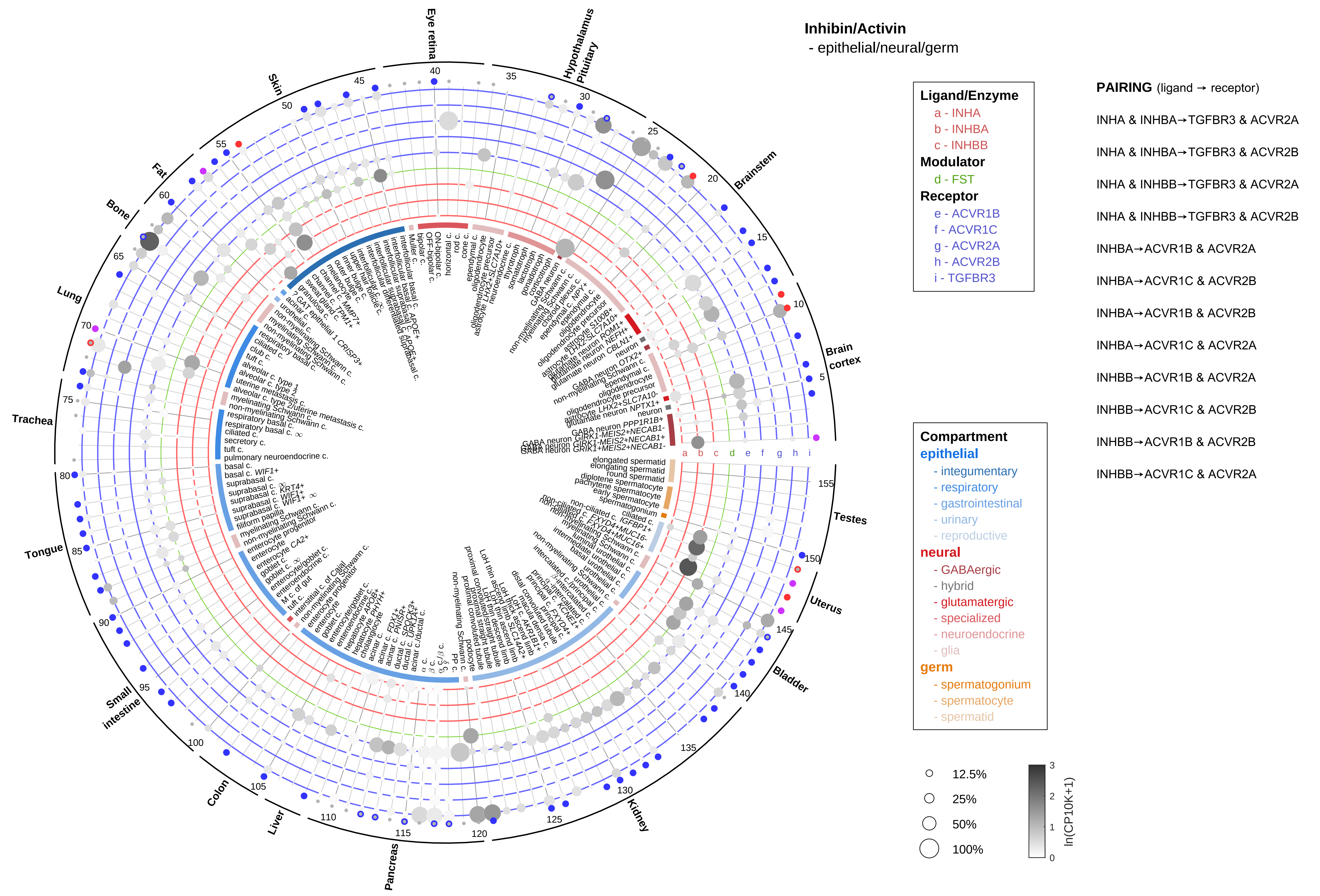

Inhibin/Activin
- epithelial/neural/germ
Ligand/Enzyme
a - INHA
b - INHBA
c - INHBB
Modulator
d - FST
Receptor
e - ACVR1B
f - ACVR1C
g - ACVR2A
h - ACVR2B
i - TGFBR3
PAIRING (ligand → receptor)
INHA & INHBA→TGFBR3 & ACVR2A
INHA & INHBA→TGFBR3 & ACVR2B
INHA & INHBB→TGFBR3 & ACVR2A
INHA & INHBB→TGFBR3 & ACVR2B
INHBA→ACVR1B & ACVR2A
INHBA→ACVR1C & ACVR2B
INHBA→ACVR1B & ACVR2B
INHBA→ACVR1C & ACVR2A
INHBB→ACVR1B & ACVR2A
INHBB→ACVR1C & ACVR2B
INHBB→ACVR1B & ACVR2B
INHBB→ACVR1C & ACVR2A
Compartment
epithelial
- integumentary
- respiratory
- gastrointestinal
- urinary
- reproductive
neural
- GABAergic
- hybrid
- glutamatergic
- specialized
- neuroendocrine
- glia
germ
- spermatogonium
- spermatocyte
- spermatid
12.5%
25%
50%
100%
ln(CP10K+1)
Eye retina
Hypothalamus
Pituitary
Brainstem
Brain cortex
Testes
Uterus
Bladder
Kidney
Pancreas
Liver
Colon
Small intestine
Tongue
Trachea
Lung
Bone
Fat
Skin

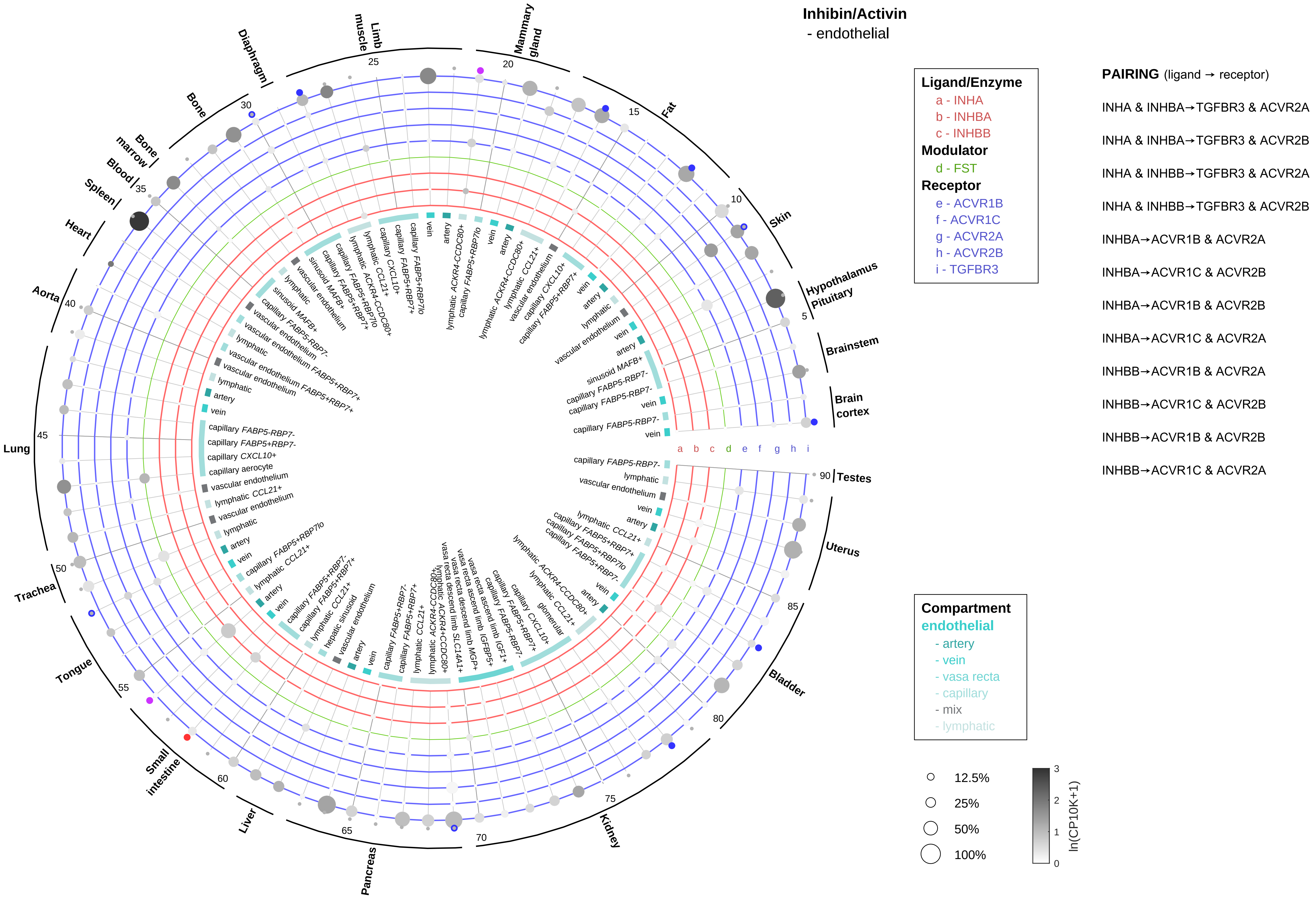
Inhibin/Activin
- endothelial
Ligand/Enzyme
a - INHA
b - INHBA
c - INHBB
Modulator
d - FST
Receptor
e - ACVR1B
f - ACVR1C
g - ACVR2A
h - ACVR2B
i - TGFBR3
PAIRING (ligand → receptor)
INHA & INHBA→TGFBR3 & ACVR2A
INHA & INHBA→TGFBR3 & ACVR2B
INHA & INHBB→TGFBR3 & ACVR2A
INHA & INHBB→TGFBR3 & ACVR2B
INHBA→ACVR1B & ACVR2A
INHBA→ACVR1C & ACVR2B
INHBA→ACVR1B & ACVR2B
INHBA→ACVR1C & ACVR2A
INHBB→ACVR1B & ACVR2A
INHBB→ACVR1C & ACVR2B
INHBB→ACVR1B & ACVR2B
INHBB→ACVR1C & ACVR2A
Compartment
endothelial
- artery
- vein
- vasa recta
- capillary
- mix
- lymphatic
12.5%
25%
50%
100%
ln(CP10K+1)
Hypothalamus
Pituitary
Brainstem
Brain cortex
Testes
Uterus
Bladder
Kidney
Pancreas
Liver
Small intestine
Tongue
Trachea
Lung
Aorta
Heart
Spleen
Blood
Bone marrow
Bone
Diaphragm
Limb muscle
Mammary gland
Fat
Skin

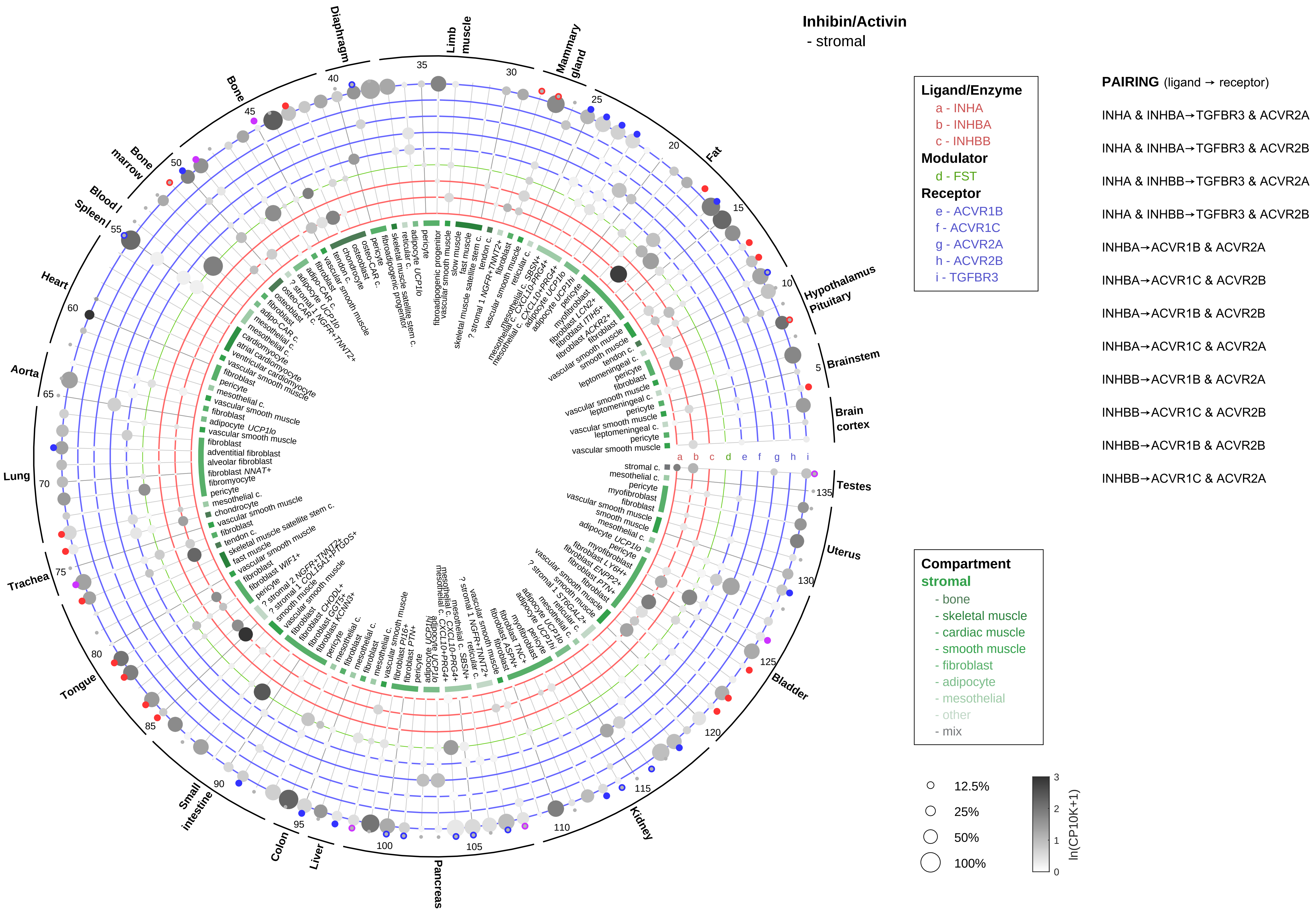
Inhibin/Activin
- stromal
Ligand/Enzyme
a - INHA
b - INHBA
c - INHBB
Modulator
d - FST
Receptor
e - ACVR1B
f - ACVR1C
g - ACVR2A
h - ACVR2B
i - TGFBR3
PAIRING (ligand → receptor)
INHA & INHBA→TGFBR3 & ACVR2A
INHA & INHBA→TGFBR3 & ACVR2B
INHA & INHBB→TGFBR3 & ACVR2A
INHA & INHBB→TGFBR3 & ACVR2B
INHBA→ACVR1B & ACVR2A
INHBA→ACVR1C & ACVR2B
INHBA→ACVR1B & ACVR2B
INHBA→ACVR1C & ACVR2A
INHBB→ACVR1B & ACVR2A
INHBB→ACVR1C & ACVR2B
INHBB→ACVR1B & ACVR2B
INHBB→ACVR1C & ACVR2A
Compartment
stromal
- bone
- skeletal muscle
- cardiac muscle
- smooth muscle
- fibroblast
- adipocyte
- mesothelial
- other
- mix
12.5%
25%
50%
100%
ln(CP10K+1)
Diaphragm
Limb muscle
Mammary gland
Fat
Hypothalamus Pituitary
Brainstem
Brain cortex
Testes
Uterus
Bladder
Kidney
Pancreas
Liver
Colon
Small intestine
Tongue
Trachea
Lung
Aorta
Heart
Blood / Spleen /
Bone marrow
Bone

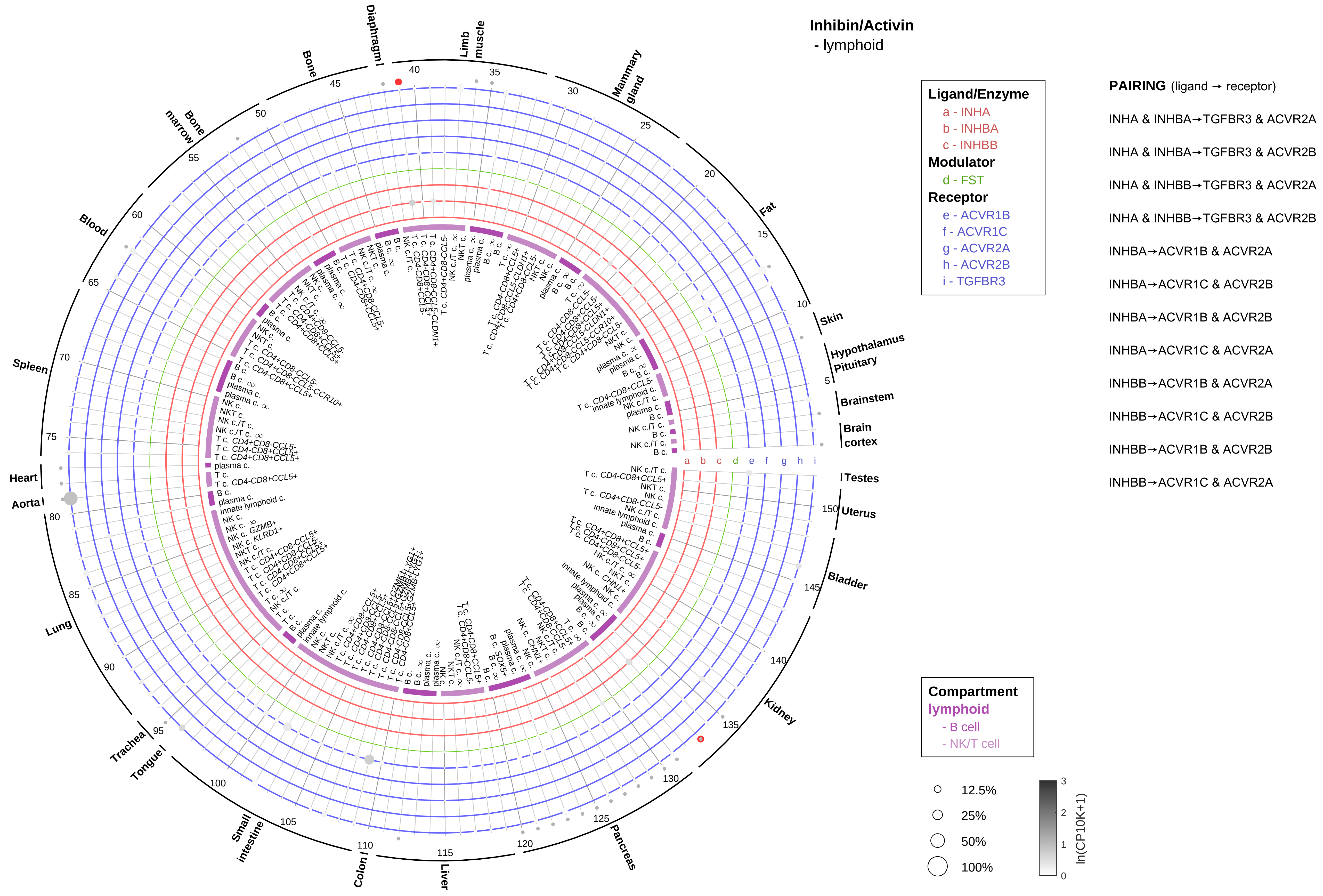

Inhibin/Activin
- lymphoid
Ligand/Enzyme
a - INHA
b - INHBA
c - INHBB
Modulator
d - FST
Receptor
e - ACVR1B
f - ACVR1C
g - ACVR2A
h - ACVR2B
i - TGFBR3
PAIRING (ligand → receptor)
INHA & INHBA→TGFBR3 & ACVR2A
INHA & INHBA→TGFBR3 & ACVR2B
INHA & INHBB→TGFBR3 & ACVR2A
INHA & INHBB→TGFBR3 & ACVR2B
INHBA→ACVR1B & ACVR2A
INHBA→ACVR1C & ACVR2B
INHBA→ACVR1B & ACVR2B
INHBA→ACVR1C & ACVR2A
INHBB→ACVR1B & ACVR2A
INHBB→ACVR1C & ACVR2B
INHBB→ACVR1B & ACVR2B
INHBB→ACVR1C & ACVR2A
Compartment
lymphoid
- B cell
- NK/T cell
12.5%
25%
50%
100%
ln(CP10K+1)
Diaphragm
Limb muscle
Mammary gland
Fat
Skin
Hypothalamus
Pituitary
Brainstem
Brain cortex
Testes
Uterus
Bladder
Kidney
Pancreas
Liver
Colon
Small intestine
Tongue
Trachea
Lung
Aorta
Heart
Spleen
Blood
Bone marrow
Bone

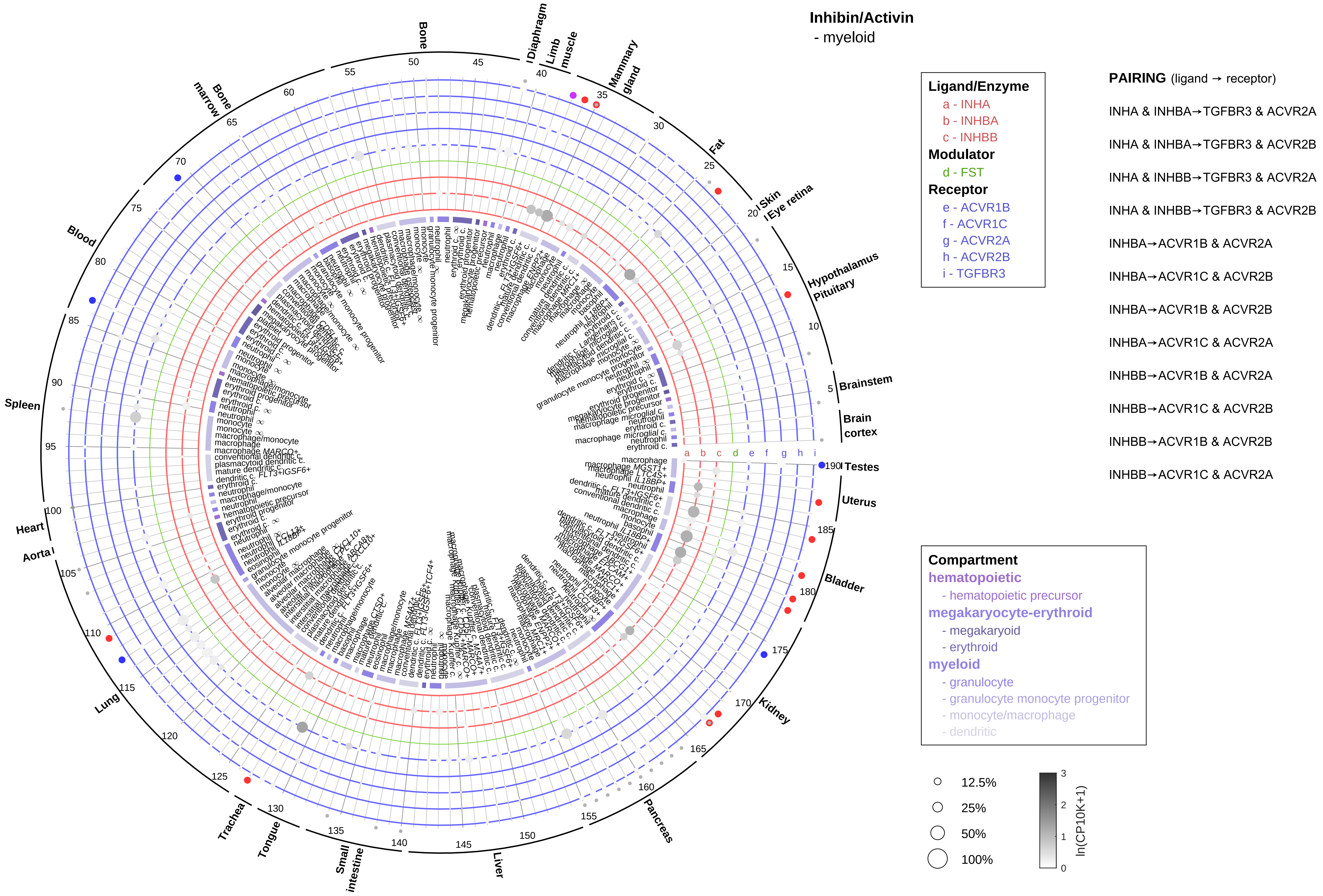

Inhibin/Activin
- myeloid
Ligand/Enzyme
a - INHA
b - INHBA
c - INHBB
Modulator
d - FST
Receptor
e - ACVR1B
f - ACVR1C
g - ACVR2A
h - ACVR2B
i - TGFBR3
PAIRING (ligand → receptor)
INHA & INHBA→TGFBR3 & ACVR2A
INHA & INHBA→TGFBR3 & ACVR2B
INHA & INHBB→TGFBR3 & ACVR2A
INHA & INHBB→TGFBR3 & ACVR2B
INHBA→ACVR1B & ACVR2A
INHBA→ACVR1C & ACVR2B
INHBA→ACVR1B & ACVR2B
INHBA→ACVR1C & ACVR2A
INHBB→ACVR1B & ACVR2A
INHBB→ACVR1C & ACVR2B
INHBB→ACVR1B & ACVR2B
INHBB→ACVR1C & ACVR2A
Compartment
hematopoietic
- hematopoietic precursor
megakaryocyte-erythroid
- megakaryoid
- erythroid
myeloid
- granulocyte
- granulocyte monocyte progenitor
- monocyte/macrophage
- dendritic
12.5%
25%
50%
100%
ln(CP10K+1)
Bone
Diaphragm
Limb muscle
Mammary gland
Fat
Skin
Eye retina
Hypothalamus
Pituitary
Brainstem
Brain cortex
Testes
Uterus
Bladder
Kidney
Pancreas
Liver
Small intestine
Tongue
Trachea
Lung
Aorta
Heart
Spleen
Blood
Bone marrow

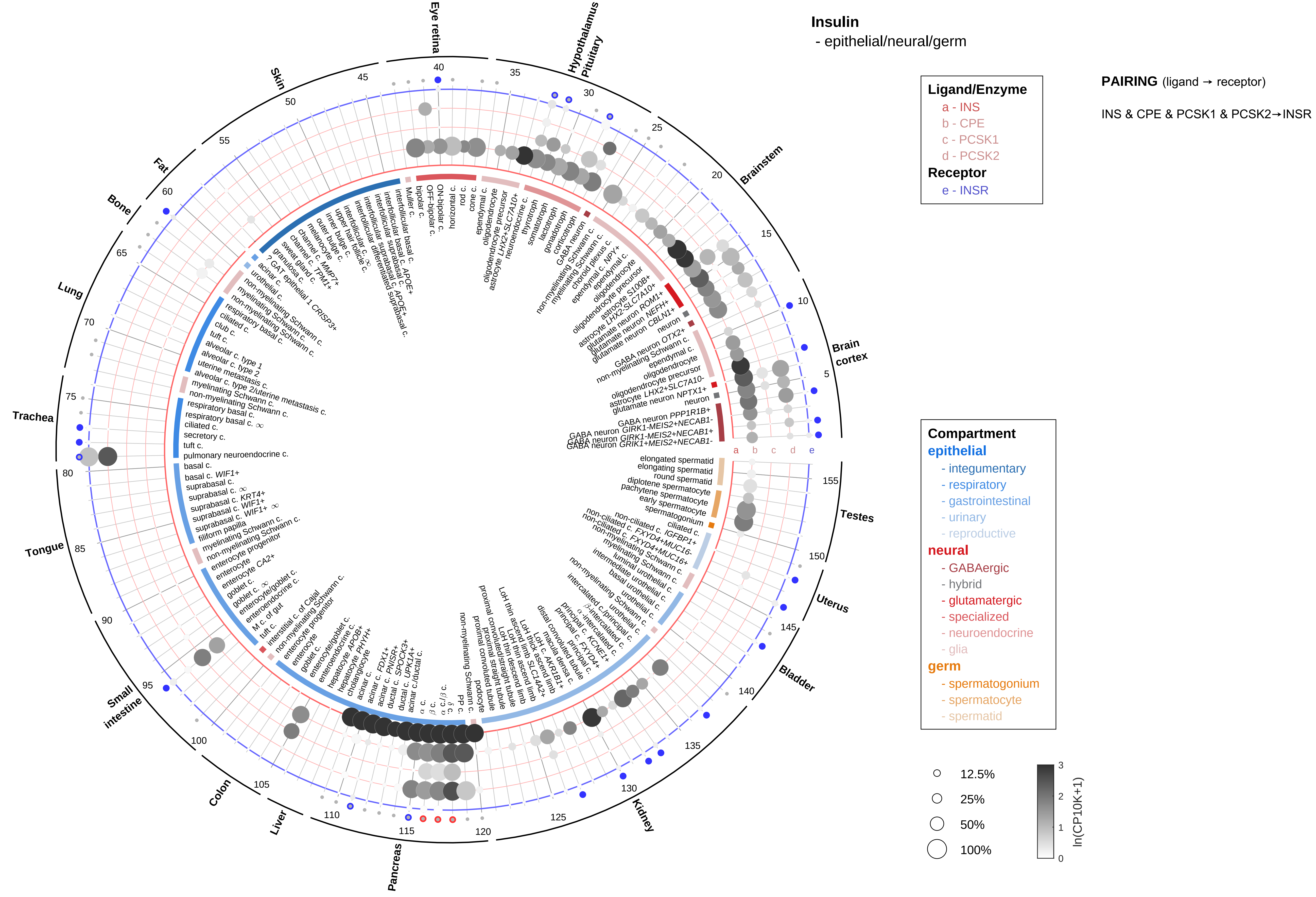

Insulin
- epithelial/neural/germ
PAIRING (ligand → receptor)
INS & CPE & PCSK1 & PCSK2→INSR
Ligand/Enzyme
a - INS
b - CPE
c - PCSK1
d - PCSK2
Receptor
e - INSR
Compartment
epithelial
- integumentary
- respiratory
- gastrointestinal
- urinary
- reproductive
neural
- GABAergic
- hybrid
- glutamatergic
- specialized
- neuroendocrine
- glia
germ
- spermatogonium
- spermatocyte
- spermatid
12.5%
25%
50%
100%
ln(CP10K+1)
Eye retina
Hypothalamus
Pituitary
Brainstem
Brain cortex
Testes
Uterus
Bladder
Kidney
Pancreas
Liver
Colon
Small intestine
Tongue
Trachea
Lung
Bone
Fat
Skin

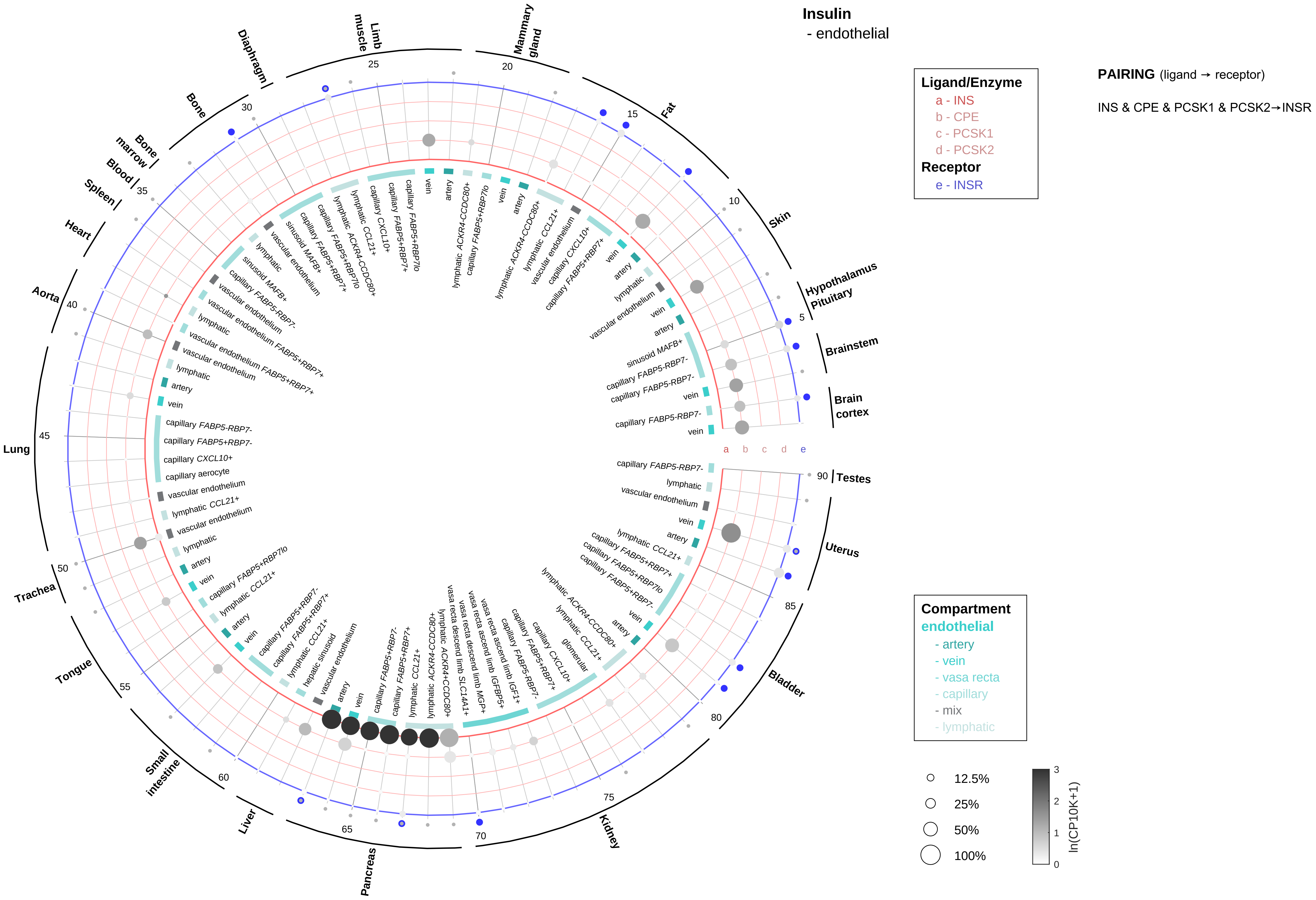

Insulin
- endothelial
Ligand/Enzyme
a - INS
b - CPE
c - PCSK1
d - PCSK2
Receptor
e - INSR
PAIRING (ligand → receptor)
INS & CPE & PCSK1 & PCSK2→INSR
Compartment
endothelial
- artery
- vein
- vasa recta
- capillary
- mix
- lymphatic
12.5%
25%
50%
100%
ln(CP10K+1)
Brain cortex
Brainstem
Hypothalamus Pituitary
Skin
Fat
Mammary gland
Limb muscle
Diaphragm
Bone
Bone marrow
Blood
Spleen
Heart
Aorta
Lung
Trachea
Tongue
Small intestine
Liver
Pancreas
Kidney
Bladder
Uterus
Testes

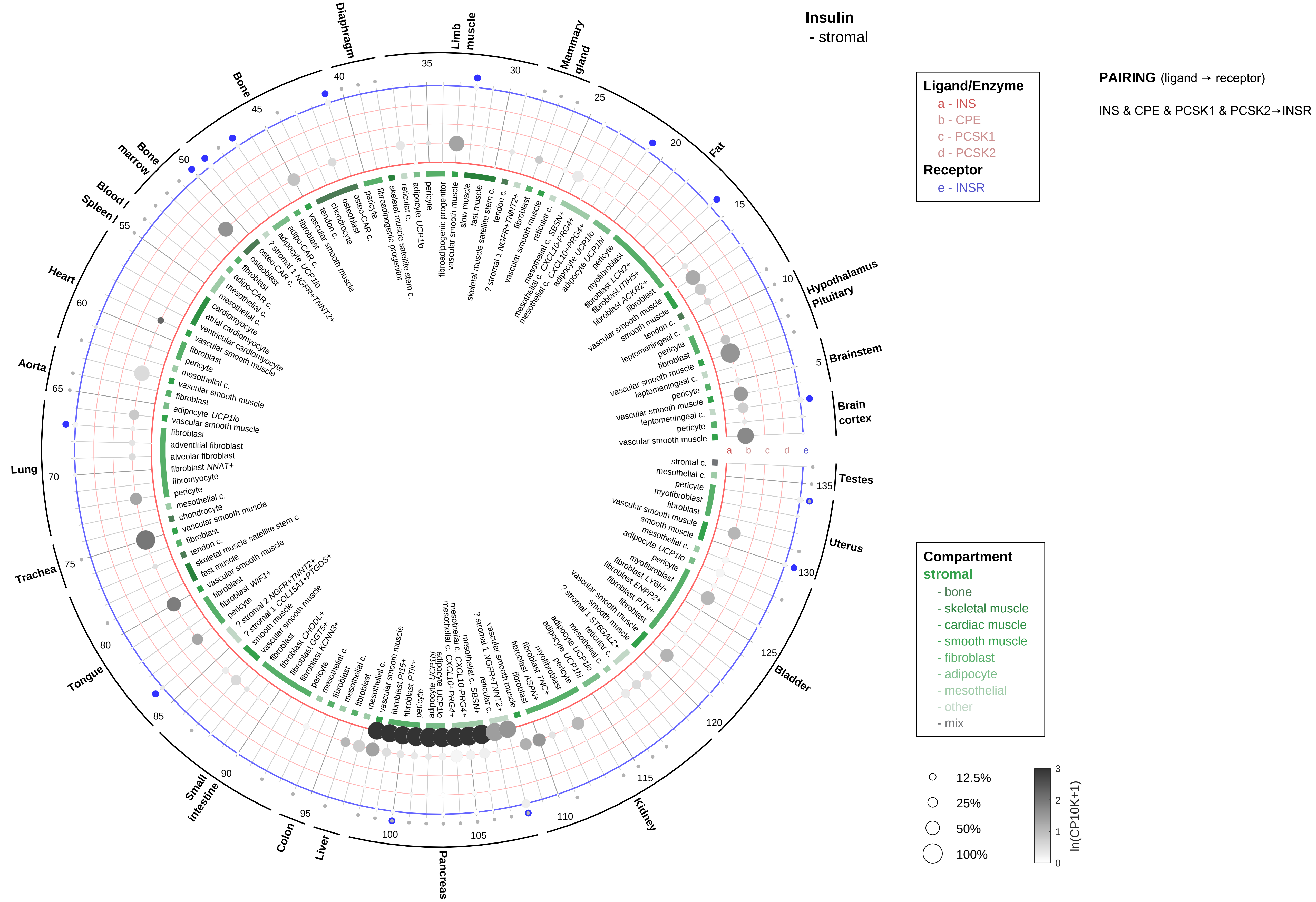
Insulin
- stromal
Ligand/Enzyme
a - INS
b - CPE
c - PCSK1
d - PCSK2
Receptor
e - INSR
PAIRING (ligand → receptor)
INS & CPE & PCSK1 & PCSK2→INSR
Compartment
stromal
- bone
- skeletal muscle
- cardiac muscle
- smooth muscle
- fibroblast
- adipocyte
- mesothelial
- other
- mix
12.5%
25%
50%
100%
ln(CP10K+1)
Brain cortex
Brainstem
Hypothalamus Pituitary
Fat
Mammary gland
Limb muscle
Diaphragm
Bone
Bone marrow
Blood / Spleen
Heart
Aorta
Lung
Trachea
Tongue
Small intestine
Colon
Liver
Pancreas
Kidney
Bladder
Uterus
Testes

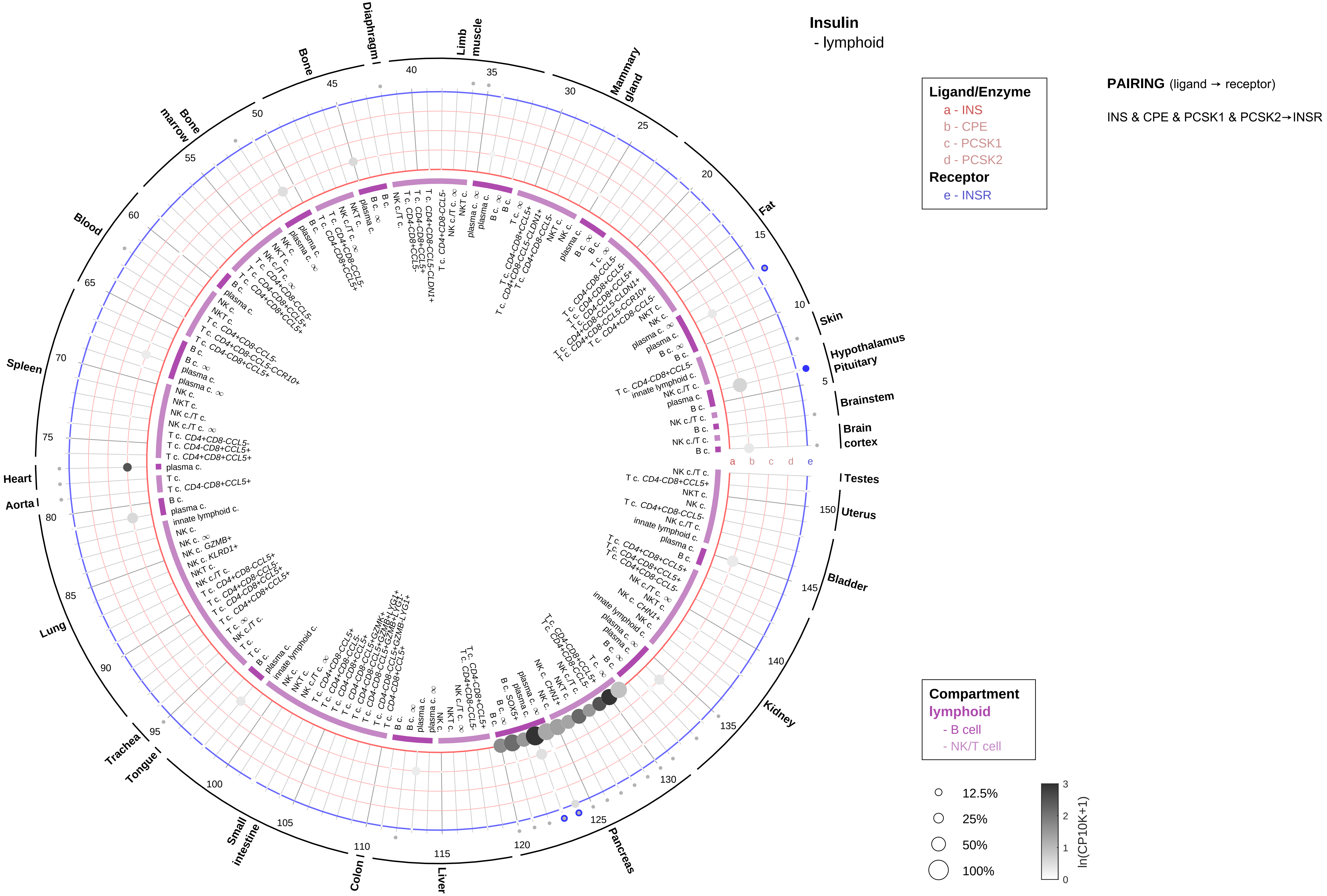

Insulin
- lymphoid
Ligand/Enzyme
a - INS
b - CPE
c - PCSK1
d - PCSK2
Receptor
e - INSR
PAIRING (ligand → receptor)
INS & CPE & PCSK1 & PCSK2→INSR
Compartment
lymphoid
- B cell
- NK/T cell
12.5%
25%
50%
100%
ln(CP10K+1)
Fat
Skin
Hypothalamus
Pituitary
Brainstem
Brain cortex
Testes
Uterus
Bladder
Kidney
Pancreas
Liver
Colon
Small intestine
Tongue
Trachea
Lung
Aorta
Heart
Spleen
Blood
Bone marrow
Bone
Diaphragm
Limb muscle
Mammary gland

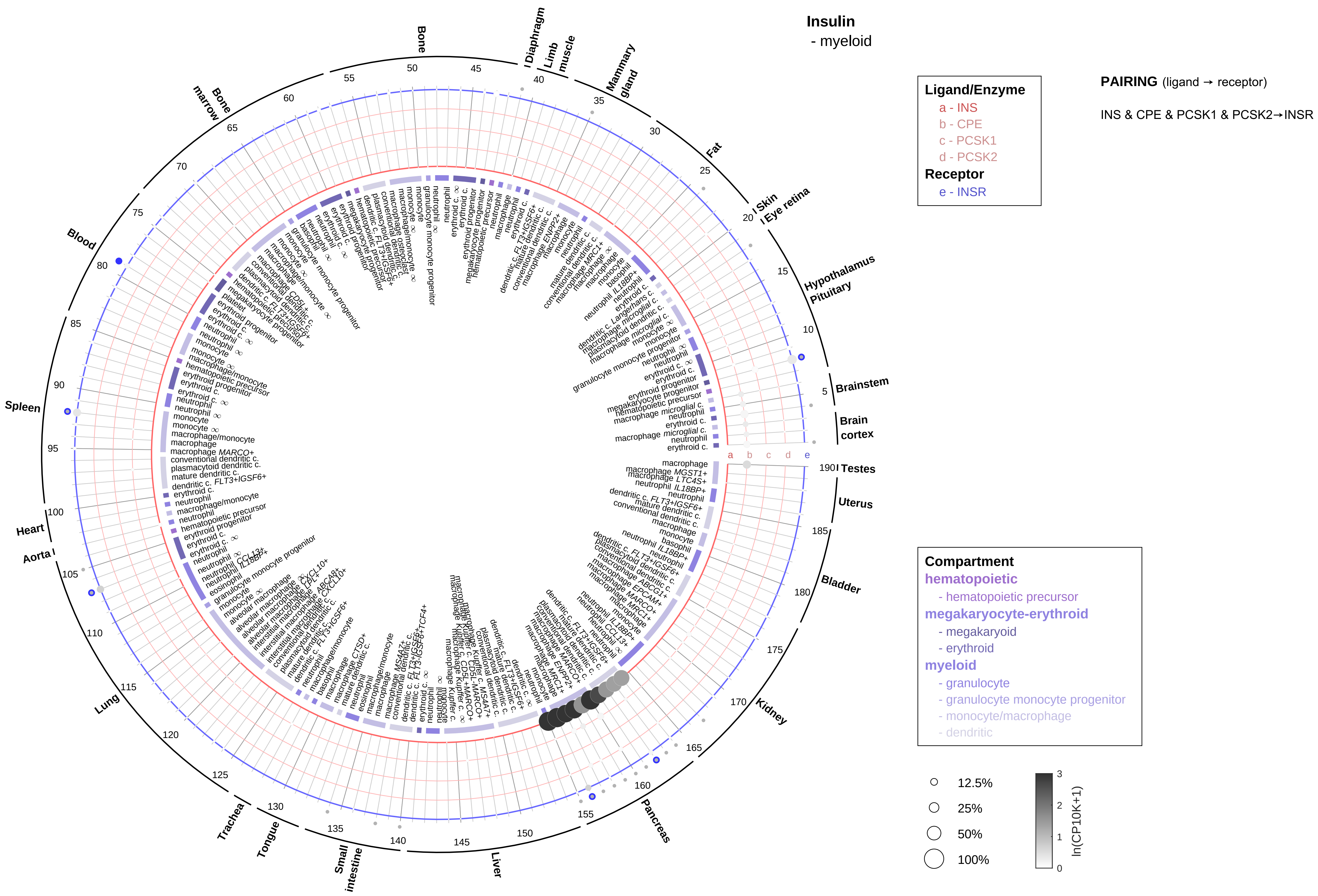

Insulin
- myeloid
Ligand/Enzyme
a - INS
b - CPE
c - PCSK1
d - PCSK2
Receptor
e - INSR
PAIRING (ligand → receptor)
INS & CPE & PCSK1 & PCSK2→INSR
Compartment
hematopoietic
- hematopoietic precursor
megakaryocyte-erythroid
- megakaryoid
- erythroid
myeloid
- granulocyte
- granulocyte monocyte progenitor
- monocyte/macrophage
- dendritic
12.5%
25%
50%
100%
ln(CP10K+1)
Bone
Diaphragm
Limb muscle
Mammary gland
Fat
Skin
Eye retina
Hypothalamus
Pituitary
Brainstem
Brain cortex
Testes
Uterus
Bladder
Kidney
Pancreas
Liver
Small intestine
Tongue
Trachea
Lung
Aorta
Heart
Spleen
Blood
Bone marrow

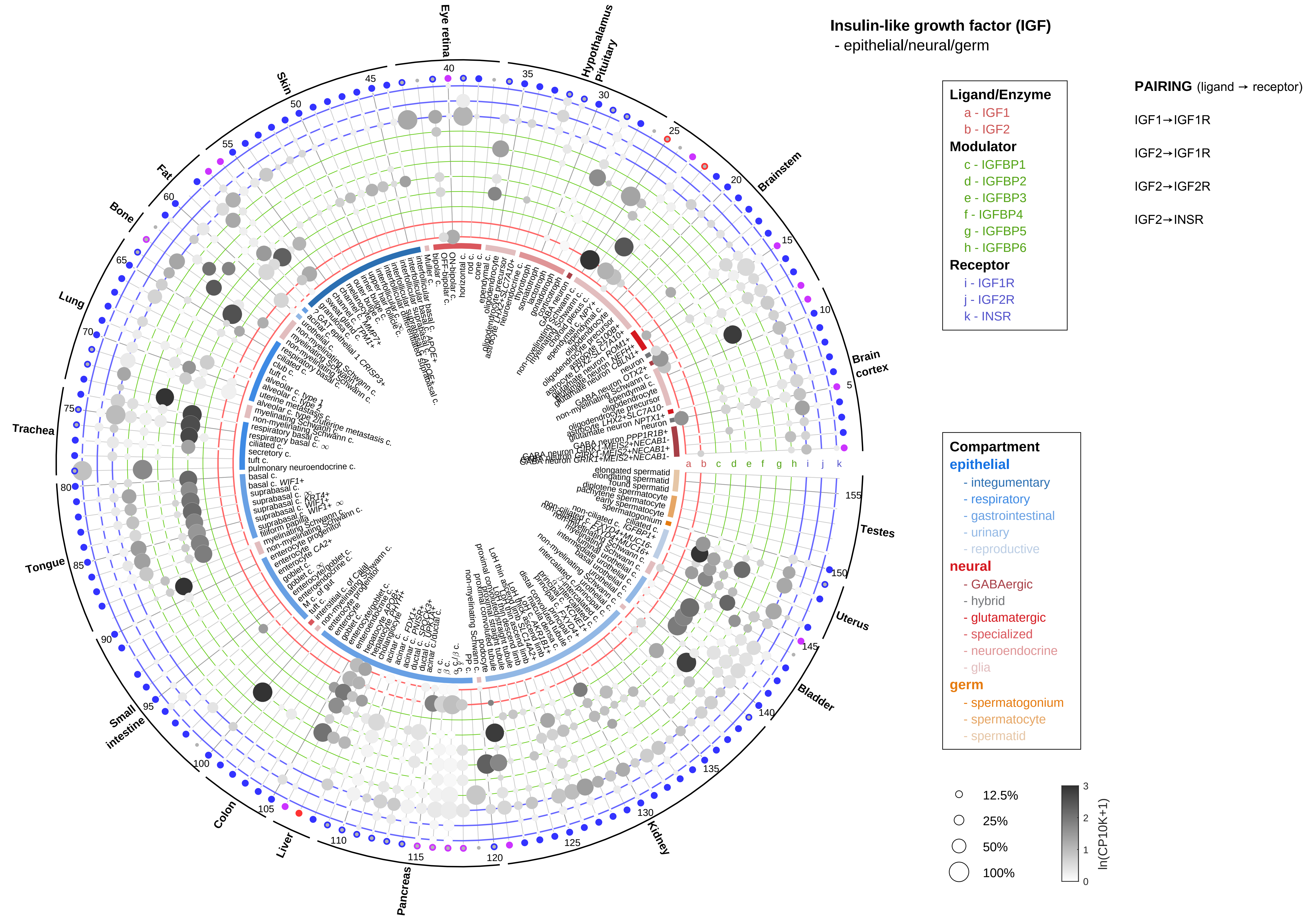

Insulin-like growth factor (IGF)
- epithelial/neural/germ
Ligand/Enzyme
a - IGF1
b - IGF2
Modulator
c - IGFBP1
d - IGFBP2
e - IGFBP3
f - IGFBP4
g - IGFBP5
h - IGFBP6
Receptor
i - IGF1R
j - IGF2R
k - INSR
PAIRING (ligand → receptor)
IGF1→IGF1R
IGF2→IGF1R
IGF2→IGF2R
IGF2→INSR
Compartment
epithelial
- integumentary
- respiratory
- gastrointestinal
- urinary
- reproductive
neural
- GABAergic
- hybrid
- glutamatergic
- specialized
- neuroendocrine
- glia
germ
- spermatogonium
- spermatocyte
- spermatid
12.5%
25%
50%
100%
ln(CP10K+1)
Eye retina
Hypothalamus Pituitary
Brainstem
Brain cortex
Testes
Uterus
Bladder
Kidney
Pancreas
Liver
Colon
Small intestine
Tongue
Trachea
Lung
Bone
Fat
Skin

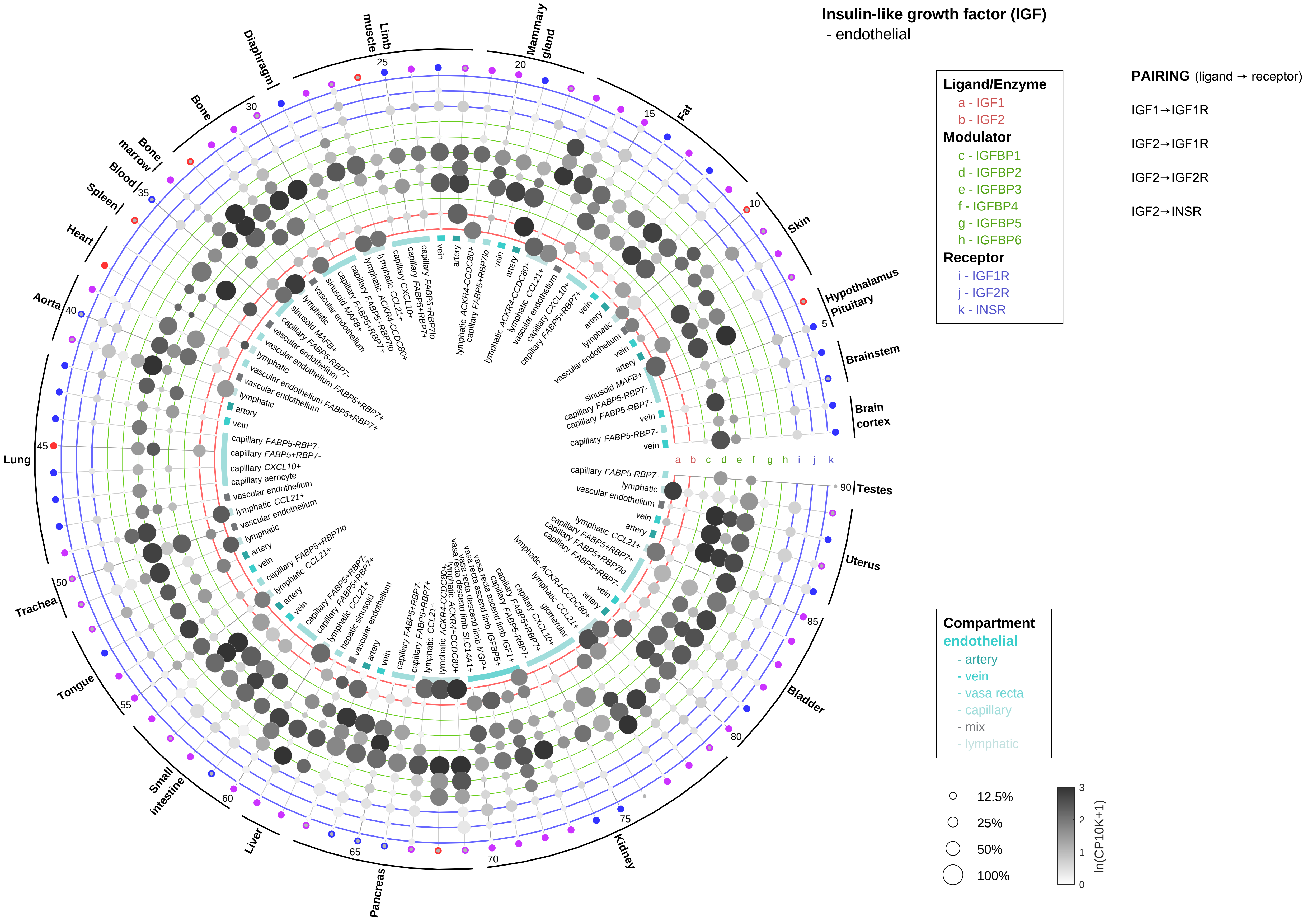
Insulin-like growth factor (IGF)
- endothelial
Ligand/Enzyme
a - IGF1
b - IGF2
Modulator
c - IGFBP1
d - IGFBP2
e - IGFBP3
f - IGFBP4
g - IGFBP5
h - IGFBP6
Receptor
i - IGF1R
j - IGF2R
k - INSR
PAIRING (ligand → receptor)
IGF1→IGF1R
IGF2→IGF1R
IGF2→IGF2R
IGF2→INSR
Compartment
endothelial
- artery
- vein
- vasa recta
- capillary
- mix
- lymphatic
12.5%
25%
50%
100%
ln(CP10K+1)
Hypothalamus Pituitary
Brainstem
Brain cortex
Testes
Uterus
Bladder
Kidney
Pancreas
Liver
Small intestine
Tongue
Trachea
Lung
Aorta
Heart
Spleen
Blood
Bone marrow
Bone
Diaphragm
Limb muscle
Mammary gland
Fat
Skin

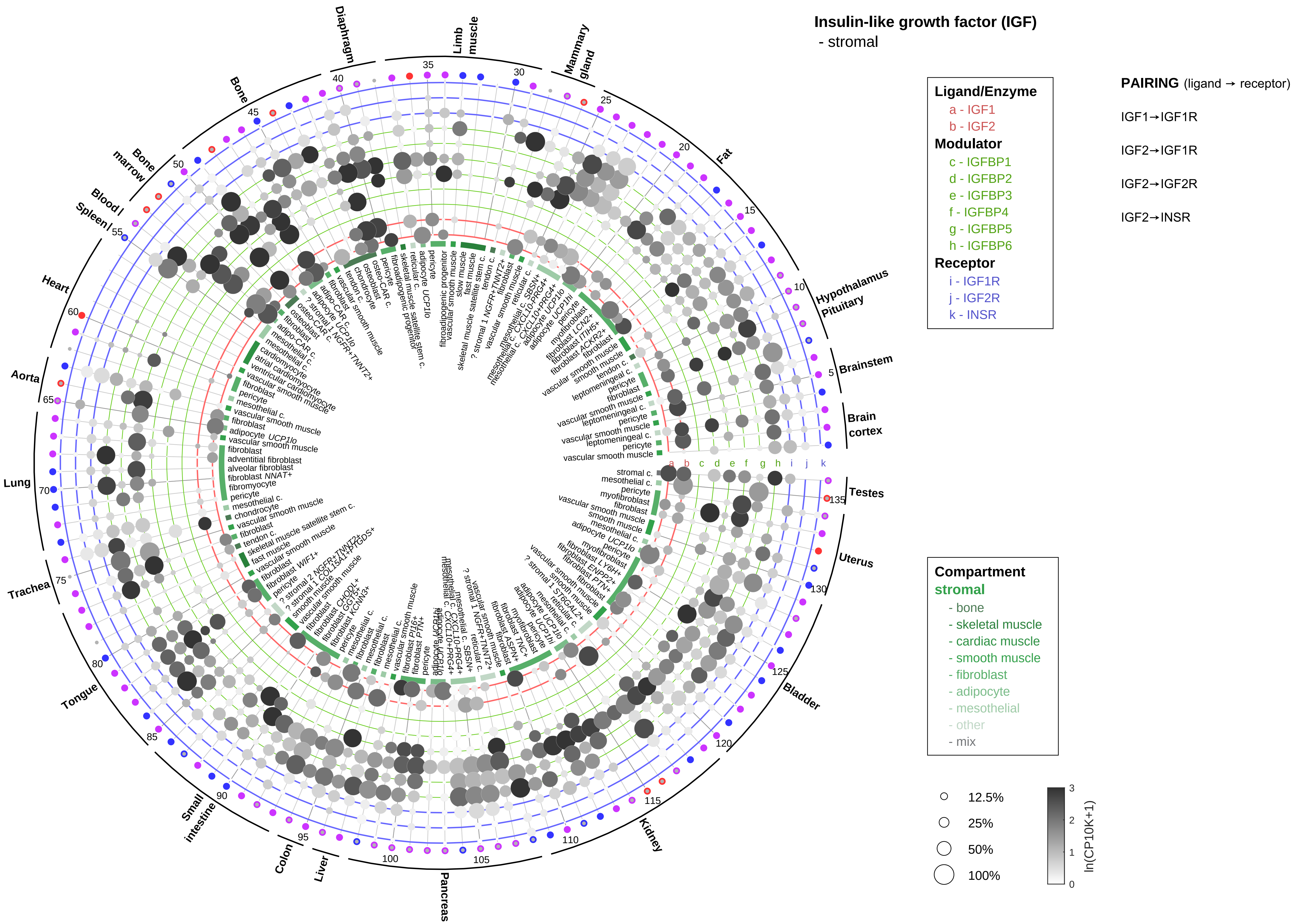

Insulin-like growth factor (IGF)
- stromal
Ligand/Enzyme
a - IGF1
b - IGF2
Modulator
c - IGFBP1
d - IGFBP2
e - IGFBP3
f - IGFBP4
g - IGFBP5
h - IGFBP6
Receptor
i - IGF1R
j - IGF2R
k - INSR
PAIRING (ligand → receptor)
IGF1→IGF1R
IGF2→IGF1R
IGF2→IGF2R
IGF2→INSR
Compartment
stromal
- bone
- skeletal muscle
- cardiac muscle
- smooth muscle
- fibroblast
- adipocyte
- mesothelial
- other
- mix
12.5%
25%
50%
100%
ln(CP10K+1)
Diaphragm
Limb muscle
Mammary gland
Fat
Hypothalamus Pituitary
Brainstem
Brain cortex
Testes
Uterus
Bladder
Kidney
Pancreas
Liver
Colon
Small intestine
Tongue
Trachea
Lung
Aorta
Heart
Spleen
Blood
Bone marrow
Bone

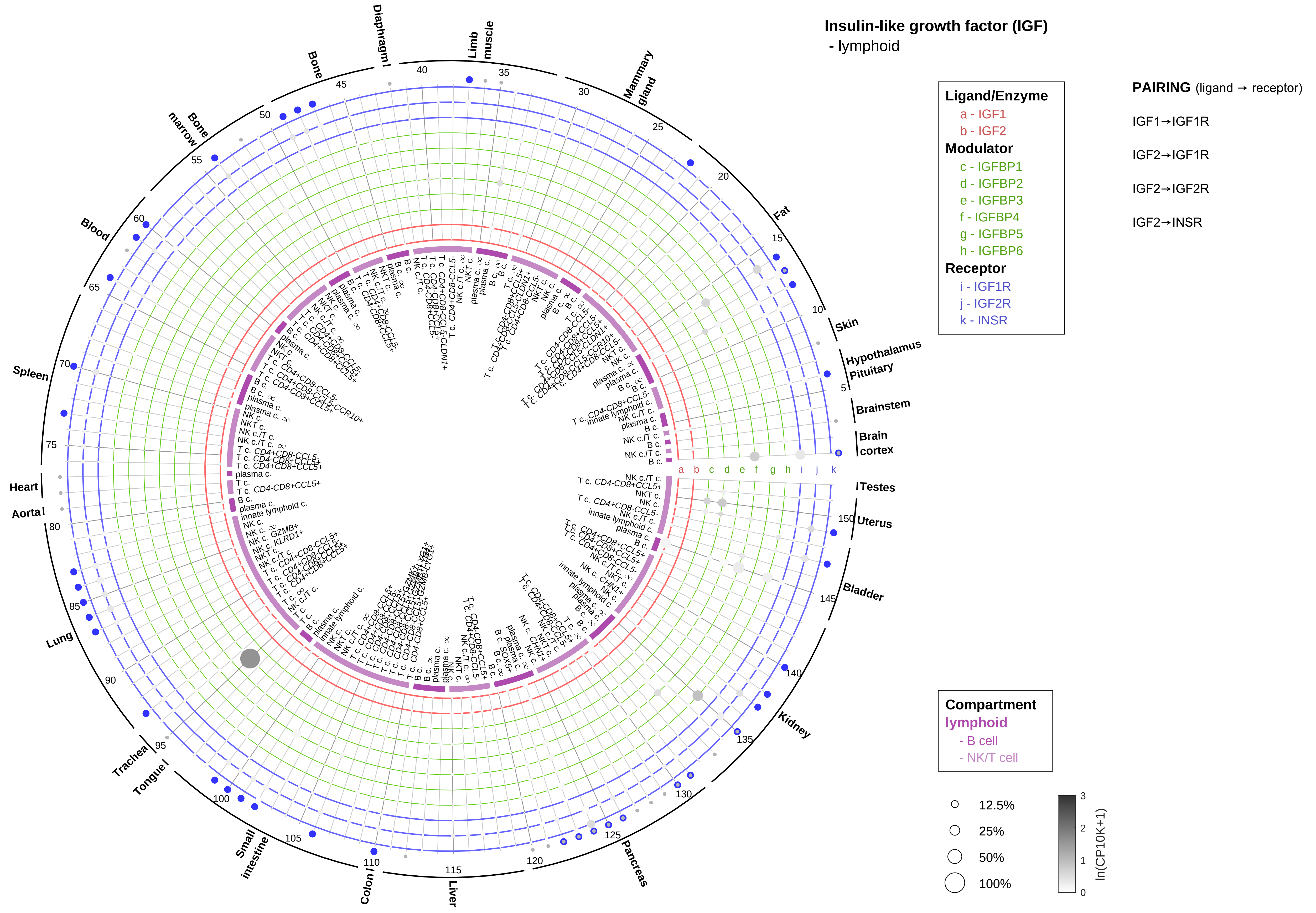
Insulin-like growth factor (IGF)
- lymphoid
Ligand/Enzyme
a - IGF1
b - IGF2
Modulator
c - IGFBP1
d - IGFBP2
e - IGFBP3
f - IGFBP4
g - IGFBP5
h - IGFBP6
Receptor
i - IGF1R
j - IGF2R
k - INSR
PAIRING (ligand → receptor)
IGF1→IGF1R
IGF2→IGF1R
IGF2→IGF2R
IGF2→INSR
Compartment
lymphoid
- B cell
- NK/T cell
12.5%
25%
50%
100%
ln(CP10K+1)
0
1
2
3
Brain cortex
Brainstem
Hypothalamus
Pituitary
Skin
Fat
Mammary gland
Limb muscle
Diaphragm
Bone
Bone marrow
Blood
Spleen
Heart
Aorta
Lung
Trachea
Tongue
Small intestine
Colon
Liver
Pancreas
Kidney
Bladder
Uterus
Testes

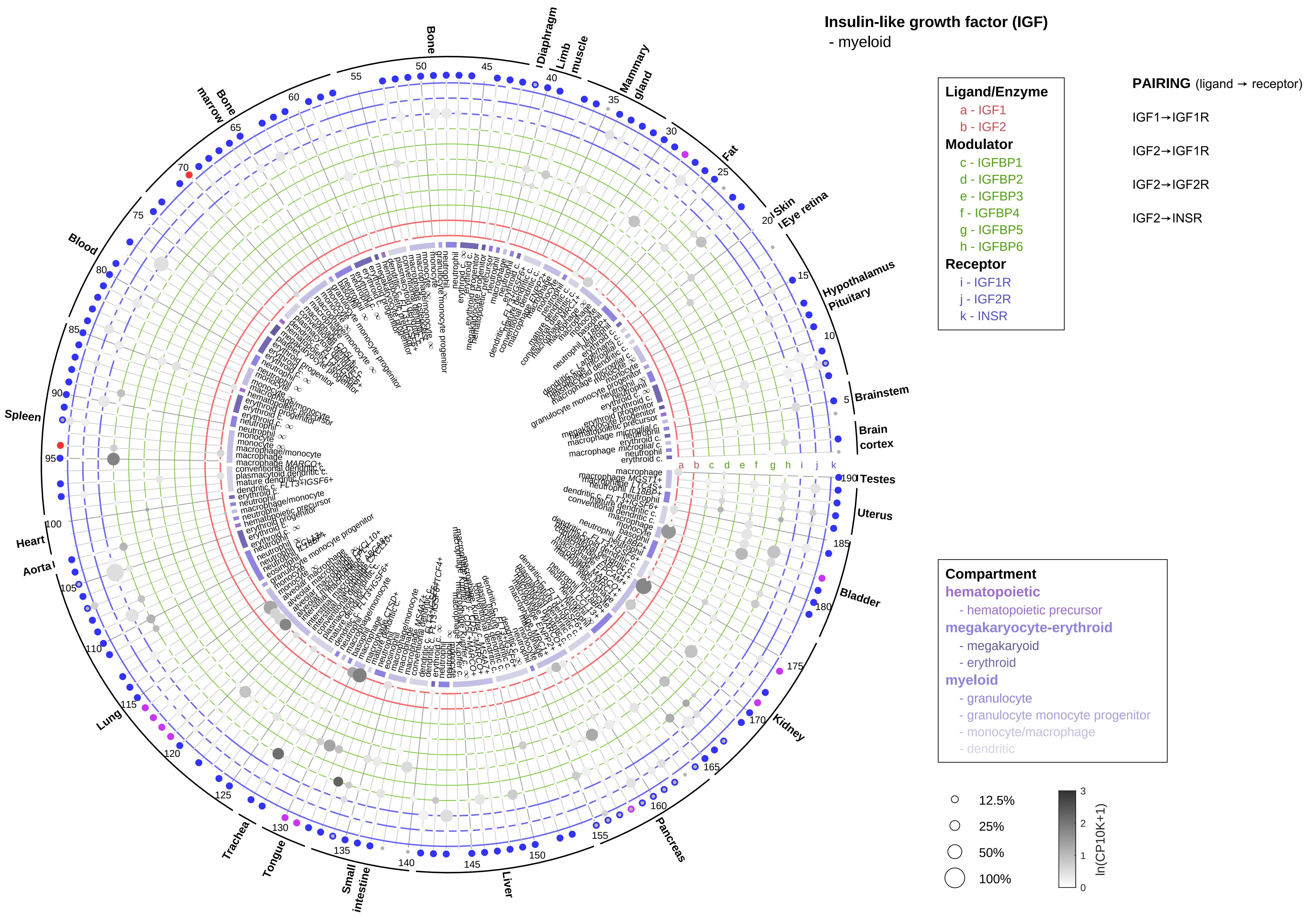

Insulin-like growth factor (IGF)
- myeloid
Ligand/Enzyme
a - IGF1
b - IGF2
Modulator
c - IGFBP1
d - IGFBP2
e - IGFBP3
f - IGFBP4
g - IGFBP5
h - IGFBP6
Receptor
i - IGF1R
j - IGF2R
k - INSR
PAIRING (ligand → receptor)
IGF1→IGF1R
IGF2→IGF1R
IGF2→IGF2R
IGF2→INSR
Compartment
hematopoietic
- hematopoietic precursor
megakaryocyte-erythroid
- megakaryoid
- erythroid
myeloid
- granulocyte
- granulocyte monocyte progenitor
- monocyte/macrophage
- dendritic
12.5%
25%
50%
100%
ln(CP10K+1)
Bone
Diaphragm
Limb muscle
Mammary gland
Fat
Skin
Eye retina
Hypothalamus
Pituitary
Brainstem
Brain cortex
Testes
Uterus
Bladder
Kidney
Pancreas
Liver
Small intestine
Tongue
Trachea
Lung
Aorta
Heart
Spleen
Blood
Bone marrow

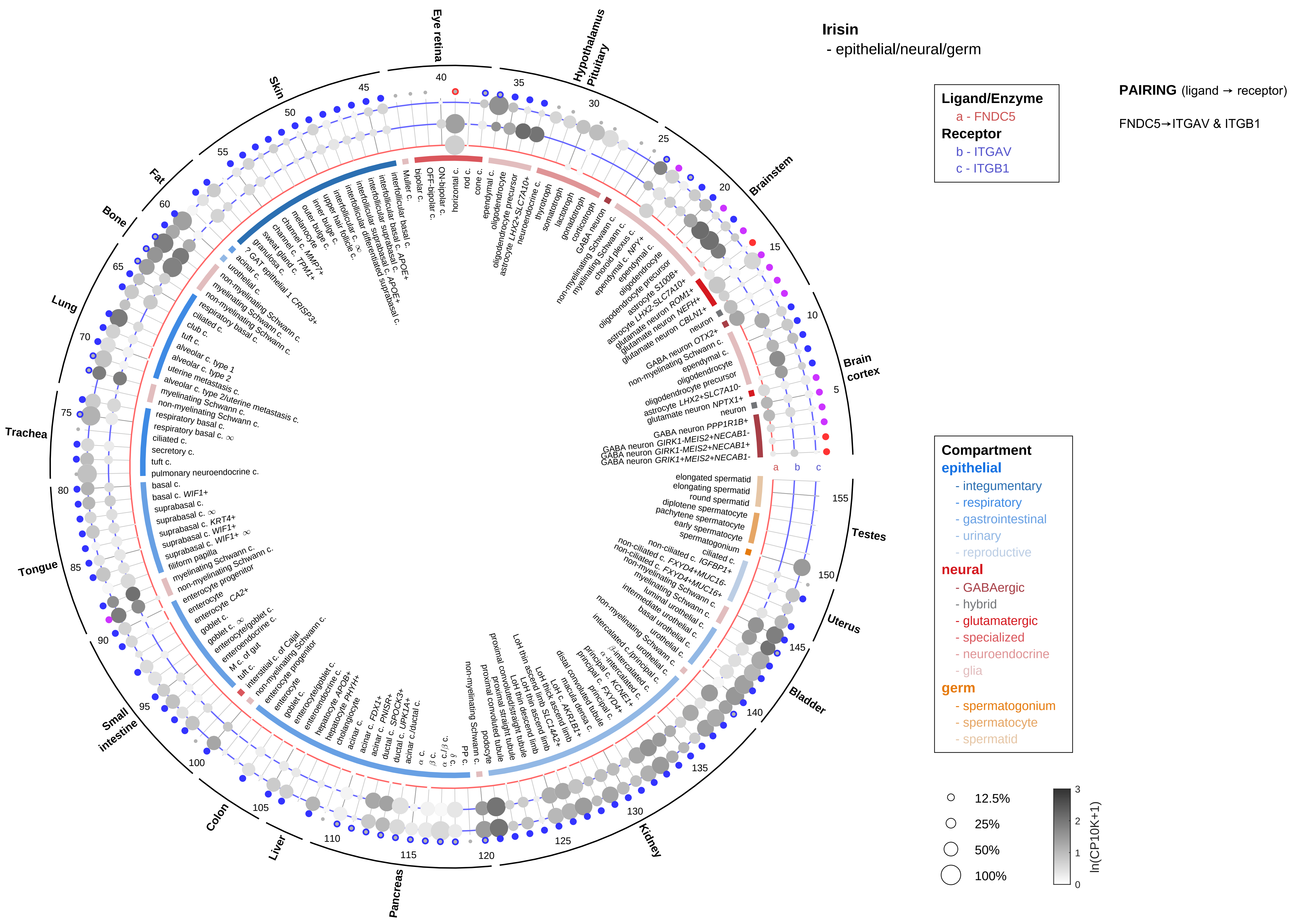
Irisin
- epithelial/neural/germ
Ligand/Enzyme
a - FNDC5
Receptor
b - ITGAV
c - ITGB1
PAIRING (ligand → receptor)
FNDC5→ITGAV & ITGB1
Compartment
epithelial
- integumentary
- respiratory
- gastrointestinal
- urinary
- reproductive
neural
- GABAergic
- hybrid
- glutamatergic
- specialized
- neuroendocrine
- glia
germ
- spermatogonium
- spermatocyte
- spermatid
12.5%
25%
50%
100%
ln(CP10K+1)
Eye retina
Hypothalamus Pituitary
Brainstem
Brain cortex
Testes
Uterus
Bladder
Kidney
Pancreas
Liver
Colon
Small intestine
Tongue
Trachea
Lung
Bone
Fat
Skin

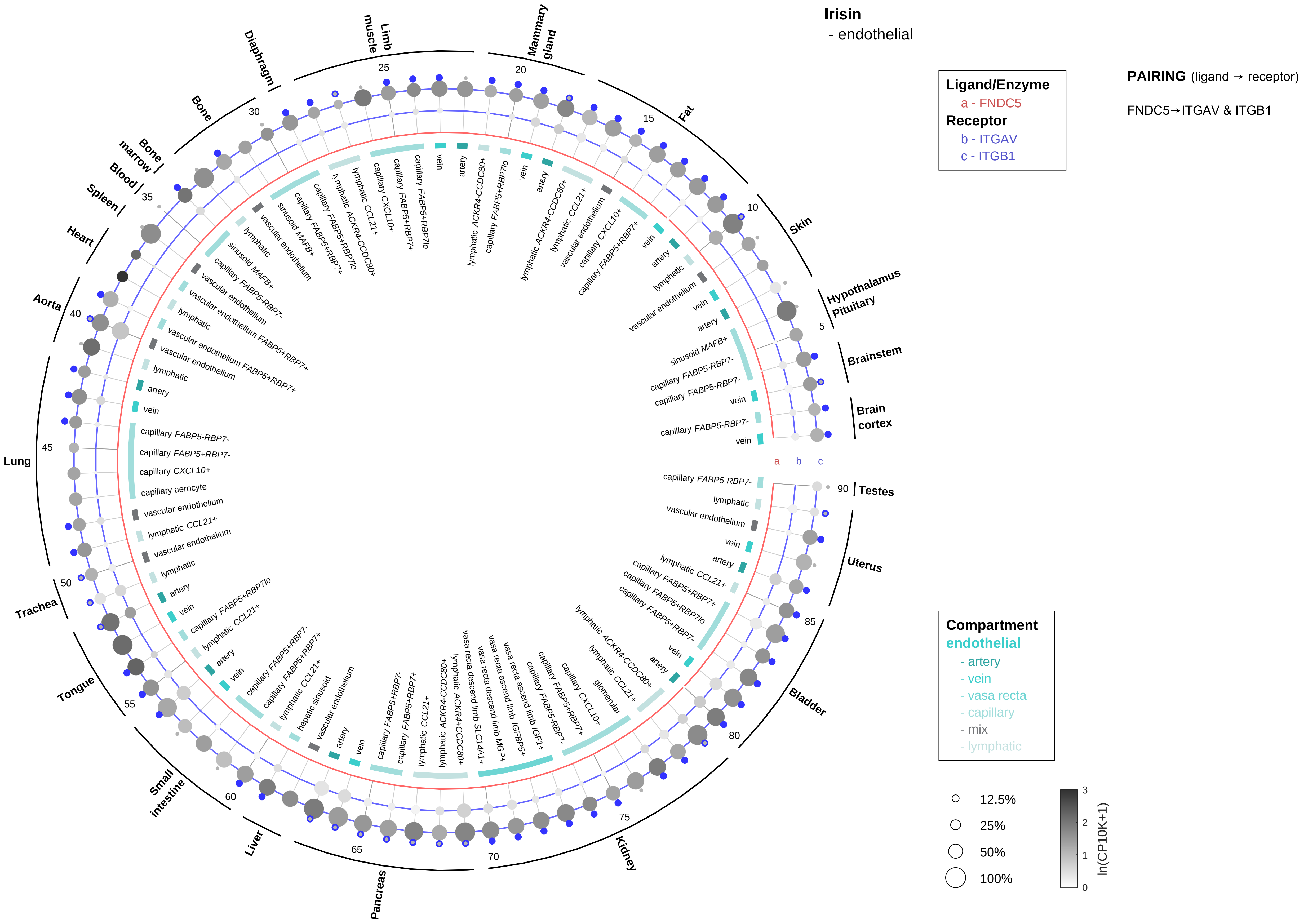

Irisin
- endothelial
Ligand/Enzyme
a - FNDC5
Receptor
b - ITGAV
c - ITGB1
PAIRING (ligand → receptor)
FNDC5→ITGAV & ITGB1
Compartment
endothelial
- artery
- vein
- vasa recta
- capillary
- mix
- lymphatic
12.5%
25%
50%
100%
ln(CP10K+1)
Brain cortex
Brainstem
Hypothalamus Pituitary
Skin
Fat
Mammary gland
Limb muscle
Diaphragm
Bone
Bone marrow
Blood
Spleen
Heart
Aorta
Lung
Trachea
Tongue
Small intestine
Liver
Pancreas
Kidney
Bladder
Uterus
Testes

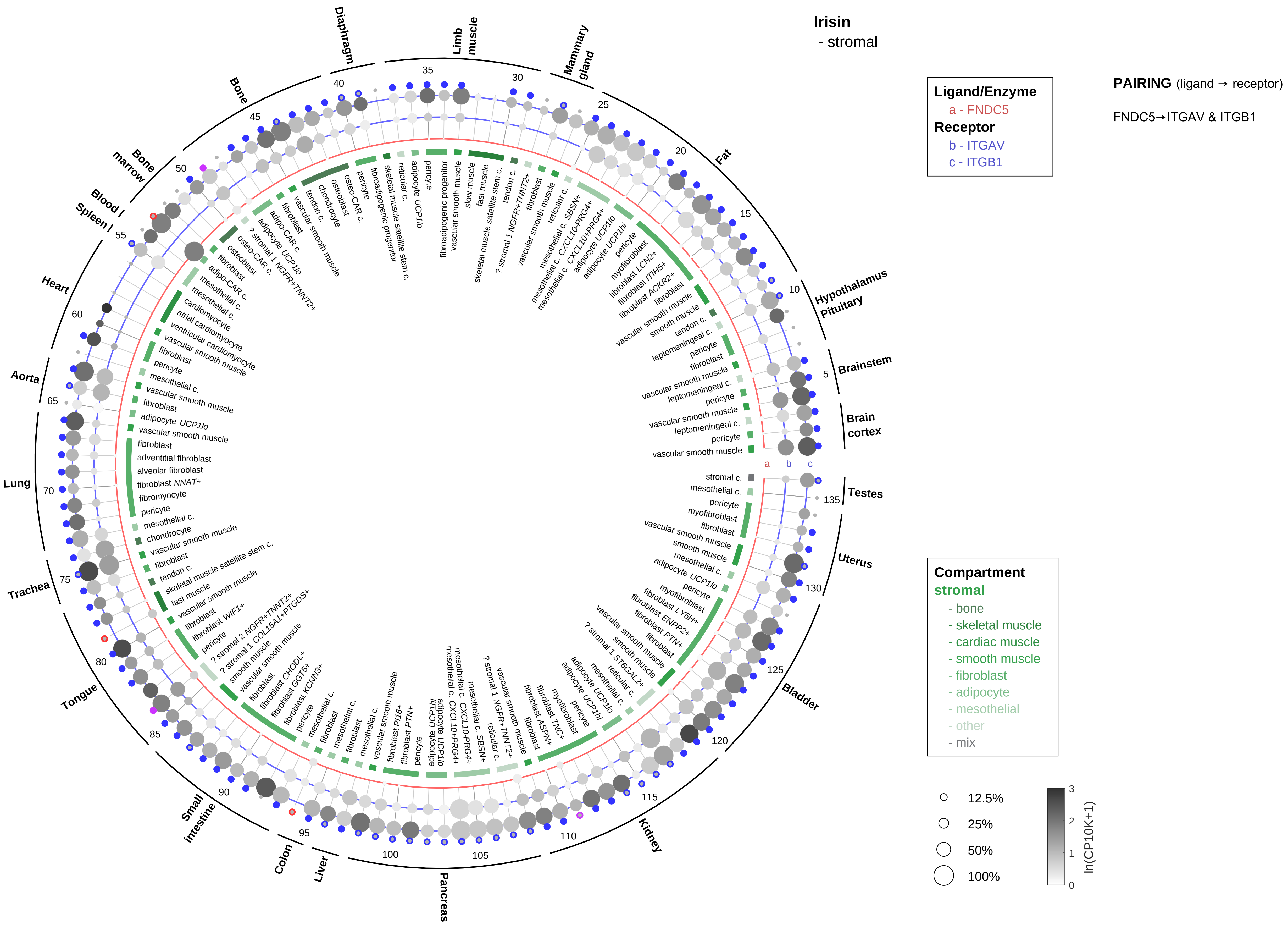

Irisin
- stromal
Ligand/Enzyme
a - FNDC5
Receptor
b - ITGAV
c - ITGB1
PAIRING (ligand → receptor)
FNDC5→ITGAV & ITGB1
Compartment
stromal
- bone
- skeletal muscle
- cardiac muscle
- smooth muscle
- fibroblast
- adipocyte
- mesothelial
- other
- mix
12.5%
25%
50%
100%
ln(CP10K+1)
Brain cortex
Brainstem
Hypothalamus Pituitary
Fat
Mammary gland
Limb muscle
Diaphragm
Bone
Bone marrow
Blood
Spleen
Heart
Aorta
Lung
Trachea
Tongue
Small intestine
Colon
Liver
Pancreas
Kidney
Bladder
Uterus
Testes

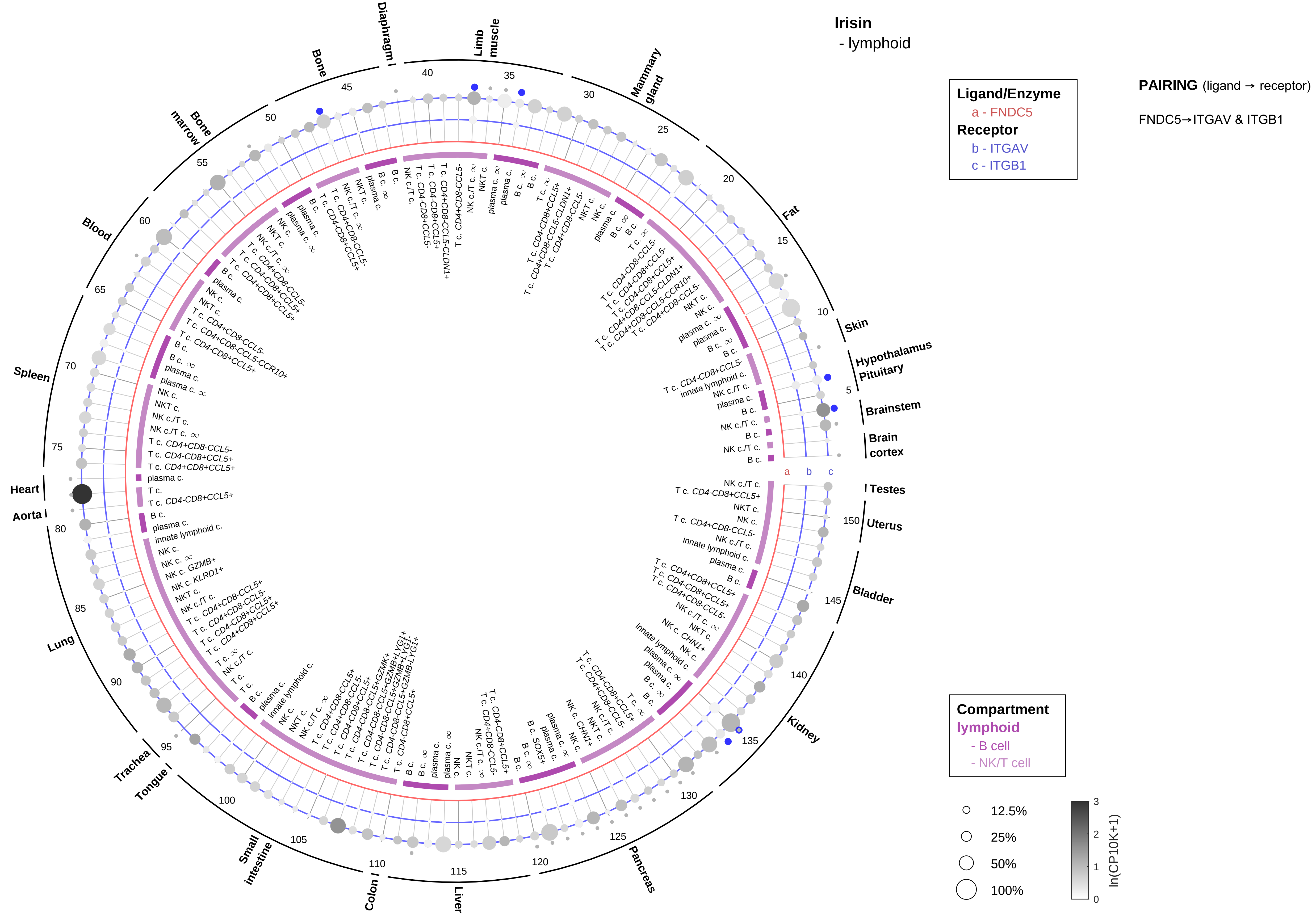
Irisin
- lymphoid
Ligand/Enzyme
a - FNDC5
Receptor
b - ITGAV
c - ITGB1
PAIRING (ligand → receptor)
FNDC5→ITGAV & ITGB1
Diaphragm
Limb muscle
Mammary gland
Fat
Skin
Hypothalamus
Pituitary
Brainstem
Brain cortex
Testes
Uterus
Bladder
Kidney
Pancreas
Liver
Colon
Small intestine
Tongue
Trachea
Lung
Aorta
Heart
Spleen
Blood
Bone marrow
Bone
Compartment
lymphoid
- B cell
- NK/T cell
12.5%
25%
50%
100%
ln(CP10K+1)

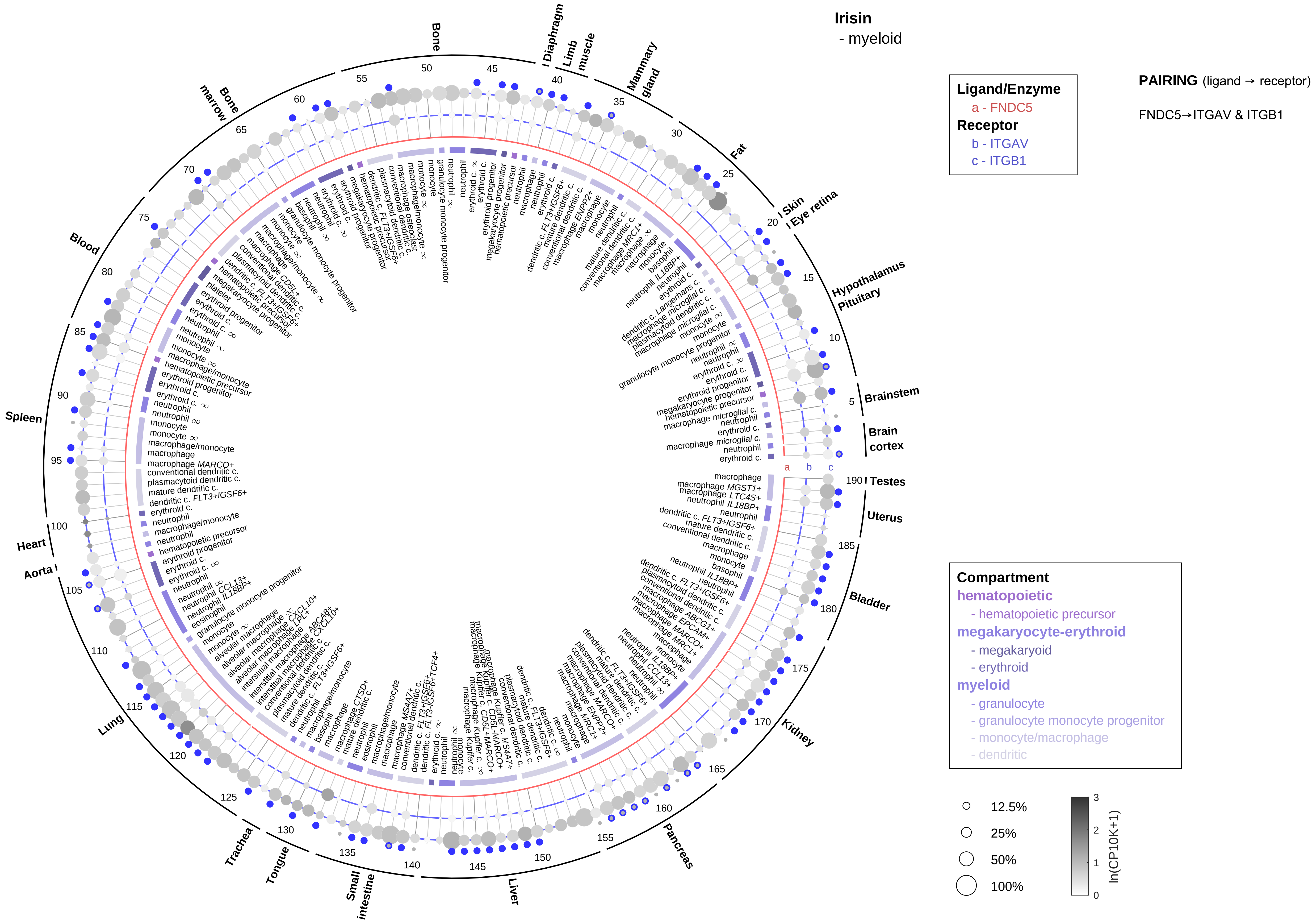
Irisin
- myeloid
Ligand/Enzyme
a - FNDC5
Receptor
b - ITGAV
c - ITGB1
PAIRING (ligand → receptor)
FNDC5→ITGAV & ITGB1
Compartment
hematopoietic
- hematopoietic precursor
megakaryocyte-erythroid
- megakaryoid
- erythroid
myeloid
- granulocyte
- granulocyte monocyte progenitor
- monocyte/macrophage
- dendritic
12.5%
25%
50%
100%
ln(CP10K+1)
Bone
Diaphragm
Limb muscle
Mammary gland
Fat
Skin
Eye retina
Hypothalamus
Pituitary
Brainstem
Brain cortex
Testes
Uterus
Bladder
Kidney
Pancreas
Liver
Small intestine
Tongue
Trachea
Lung
Aorta
Heart
Spleen
Blood
Bone marrow

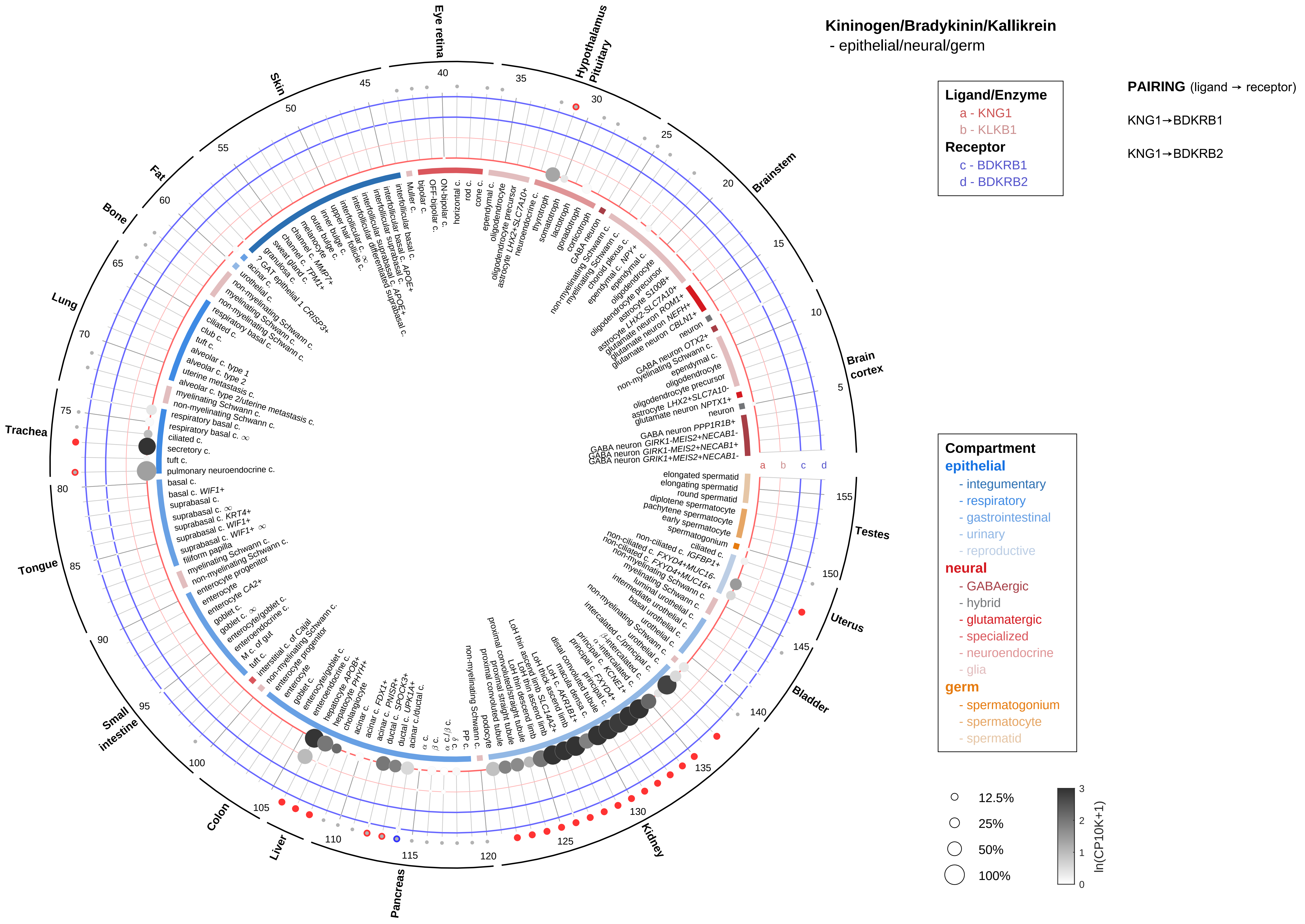
Kininogen/Bradykinin/Kallikrein
- epithelial/neural/germ
Ligand/Enzyme
a - KNG1
b - KLKB1
Receptor
c - BDKRB1
d - BDKRB2
PAIRING (ligand → receptor)
KNG1→BDKRB1
KNG1→BDKRB2
Compartment
epithelial
- integumentary
- respiratory
- gastrointestinal
- urinary
- reproductive
neural
- GABAergic
- hybrid
- glutamatergic
- specialized
- neuroendocrine
- glia
germ
- spermatogonium
- spermatocyte
- spermatid
12.5%
25%
50%
100%
ln(CP10K+1)
Brain cortex
Brainstem
Hypothalamus Pituitary
Eye retina
Skin
Fat
Bone
Lung
Trachea
Tongue
Small intestine
Colon
Liver
Pancreas
Kidney
Bladder
Uterus
Testes

**Kininogen/Bradykinin/Kallikrein**
- endothelial

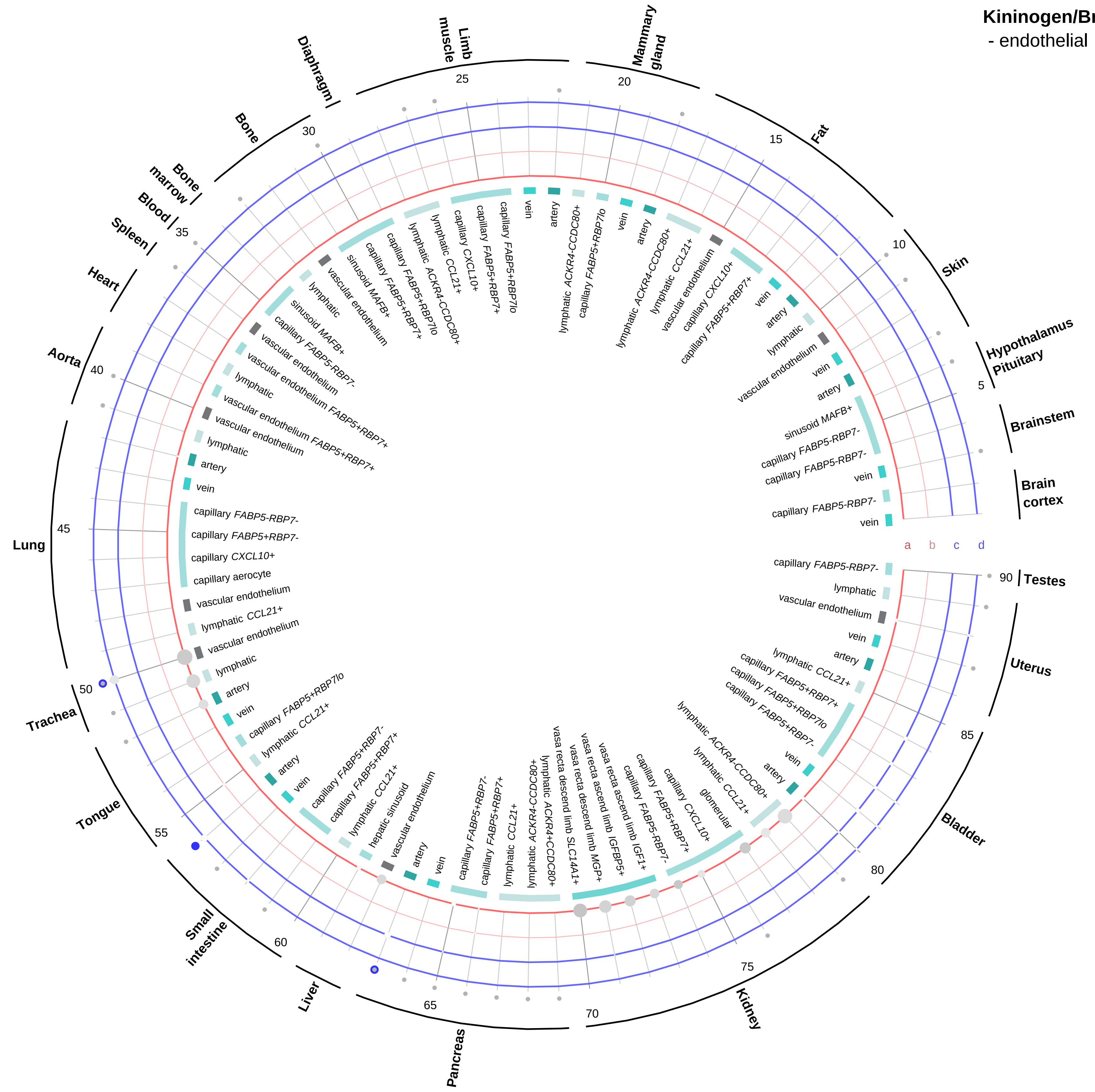


**Ligand/Enzyme**
- a - KNG1
- b - KLKB1

**Receptor**
- c - BDKRB1
- d - BDKRB2

**PAIRING** (ligand → receptor)

KNG1→BDKRB1

KNG1→BDKRB2

**Compartment**

**endothelial**
- artery
- vein
- vasa recta
- capillary
- mix
- lymphatic

12.5%
25%
50%
100%

ln(CP10K+1): 0, 1, 2, 3

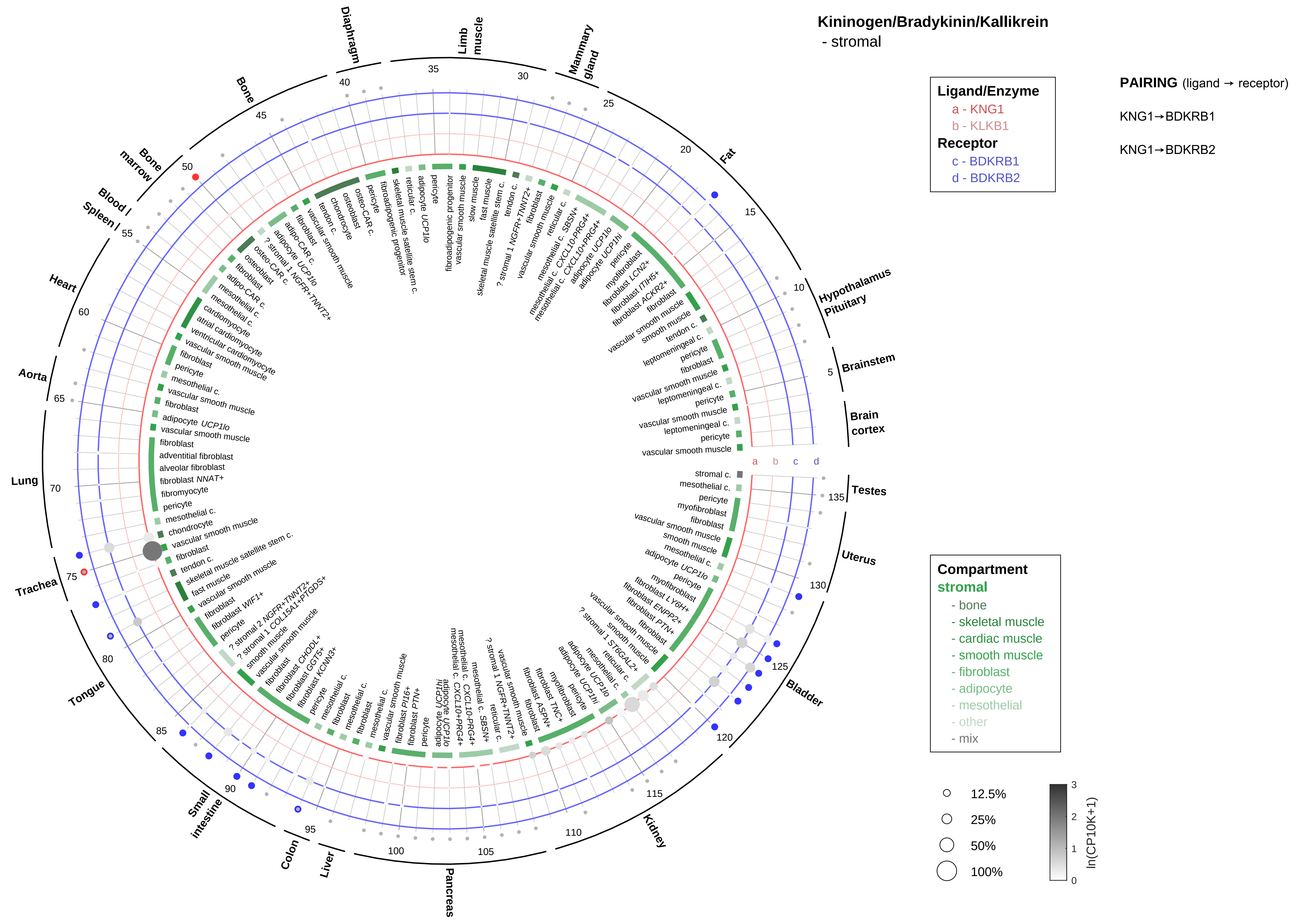

Kininogen/Bradykinin/Kallikrein
- stromal
Ligand/Enzyme
a - KNG1
b - KLKB1
Receptor
c - BDKRB1
d - BDKRB2
PAIRING (ligand → receptor)
KNG1→BDKRB1
KNG1→BDKRB2
Compartment
stromal
- bone
- skeletal muscle
- cardiac muscle
- smooth muscle
- fibroblast
- adipocyte
- mesothelial
- other
- mix
12.5%
25%
50%
100%
ln(CP10K+1)
Fat
Hypothalamus Pituitary
Brainstem
Brain cortex
Testes
Uterus
Bladder
Kidney
Pancreas
Liver
Colon
Small intestine
Tongue
Trachea
Lung
Aorta
Heart
Blood / Spleen
Bone marrow
Bone
Diaphragm
Limb muscle
Mammary gland

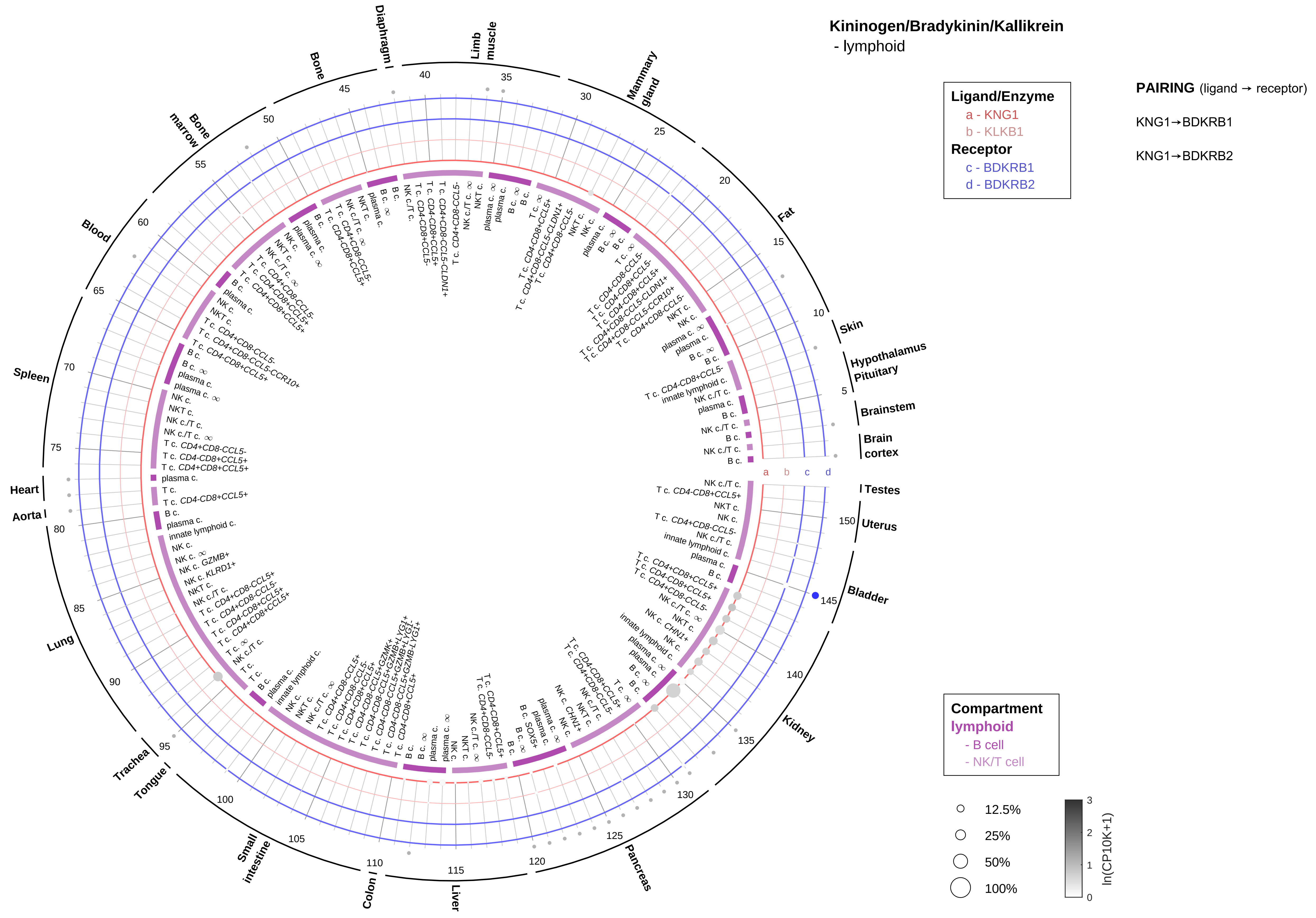
Kininogen/Bradykinin/Kallikrein
- lymphoid
Ligand/Enzyme
a - KNG1
b - KLKB1
Receptor
c - BDKRB1
d - BDKRB2
PAIRING (ligand → receptor)
KNG1→BDKRB1
KNG1→BDKRB2
Compartment
lymphoid
- B cell
- NK/T cell
12.5%
25%
50%
100%
ln(CP10K+1)
Brain cortex
Brainstem
Hypothalamus Pituitary
Skin
Fat
Mammary gland
Limb muscle
Diaphragm
Bone
Bone marrow
Blood
Spleen
Heart
Aorta
Lung
Trachea
Tongue
Small intestine
Colon
Liver
Pancreas
Kidney
Bladder
Uterus
Testes

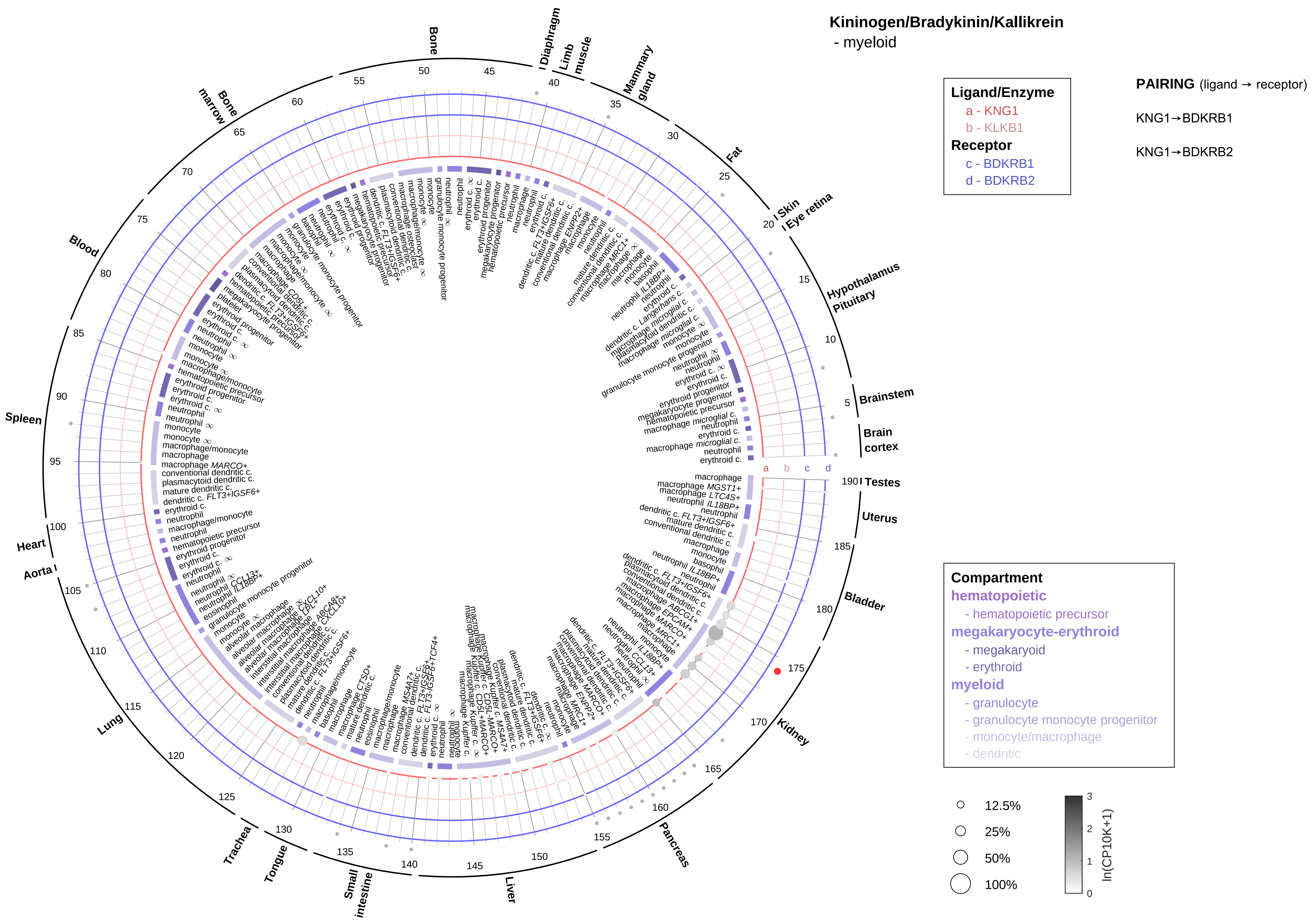

Kininogen/Bradykinin/Kallikrein
- myeloid
Ligand/Enzyme
a - KNG1
b - KLKB1
Receptor
c - BDKRB1
d - BDKRB2
PAIRING (ligand → receptor)
KNG1→BDKRB1
KNG1→BDKRB2
Compartment
hematopoietic
- hematopoietic precursor
megakaryocyte-erythroid
- megakaryoid
- erythroid
myeloid
- granulocyte
- granulocyte monocyte progenitor
- monocyte/macrophage
- dendritic
12.5%
25%
50%
100%
ln(CP10K+1)
0
1
2
3
Bone
Diaphragm
Limb muscle
Mammary gland
Fat
Skin
Eye retina
Hypothalamus
Pituitary
Brainstem
Brain cortex
Testes
Uterus
Bladder
Kidney
Pancreas
Liver
Small intestine
Tongue
Trachea
Lung
Aorta
Heart
Spleen
Blood
Bone marrow
a
b
c
d

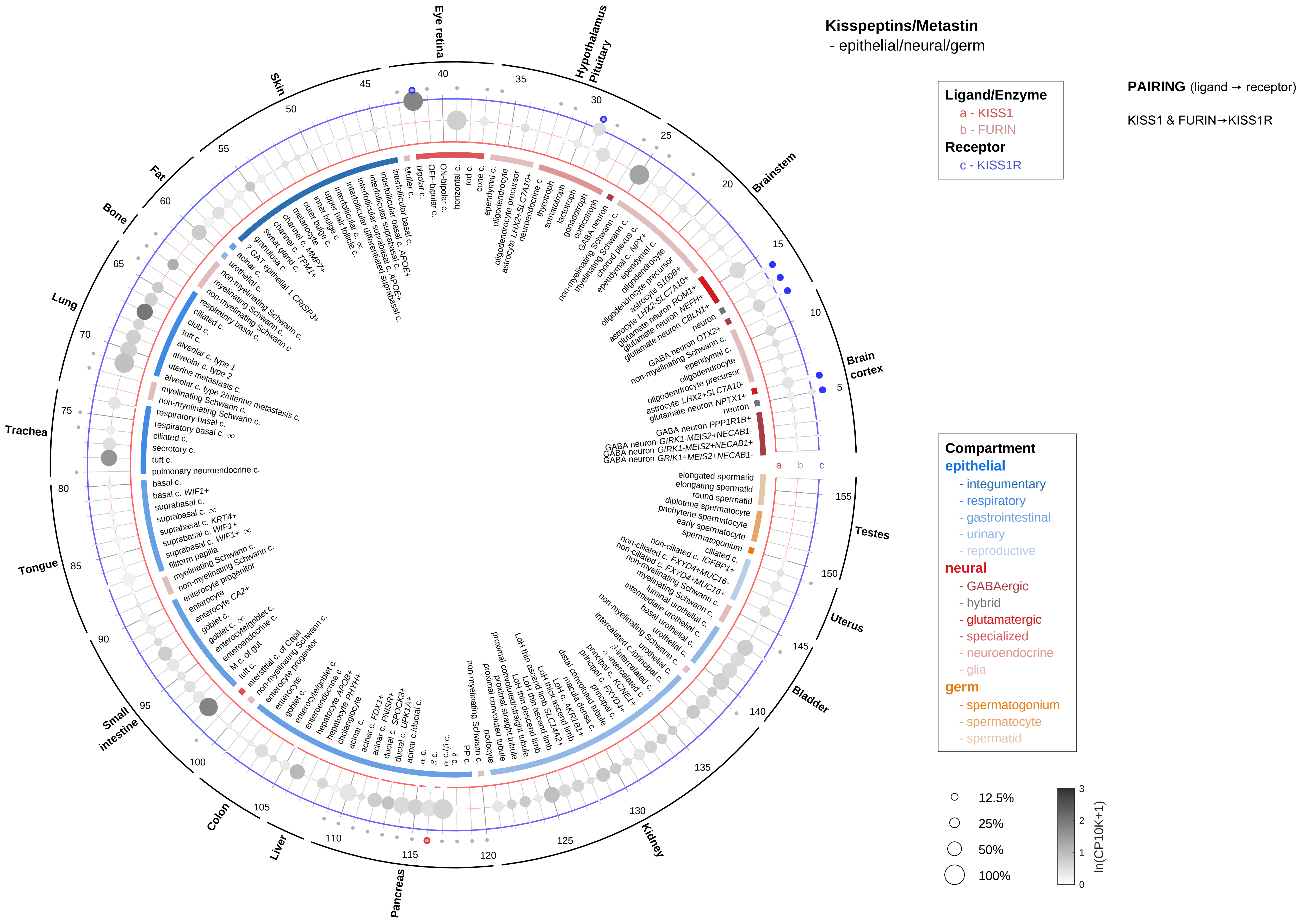
Kisspeptins/Metastin
- epithelial/neural/germ
Ligand/Enzyme
a - KISS1
b - FURIN
Receptor
c - KISS1R
PAIRING (ligand → receptor)
KISS1 & FURIN→KISS1R
Compartment
epithelial
- integumentary
- respiratory
- gastrointestinal
- urinary
- reproductive
neural
- GABAergic
- hybrid
- glutamatergic
- specialized
- neuroendocrine
- glia
germ
- spermatogonium
- spermatocyte
- spermatid
12.5%
25%
50%
100%
ln(CP10K+1)
Eye retina
Hypothalamus
Pituitary
Brainstem
Brain cortex
Testes
Uterus
Bladder
Kidney
Pancreas
Liver
Colon
Small intestine
Tongue
Trachea
Lung
Bone
Fat
Skin

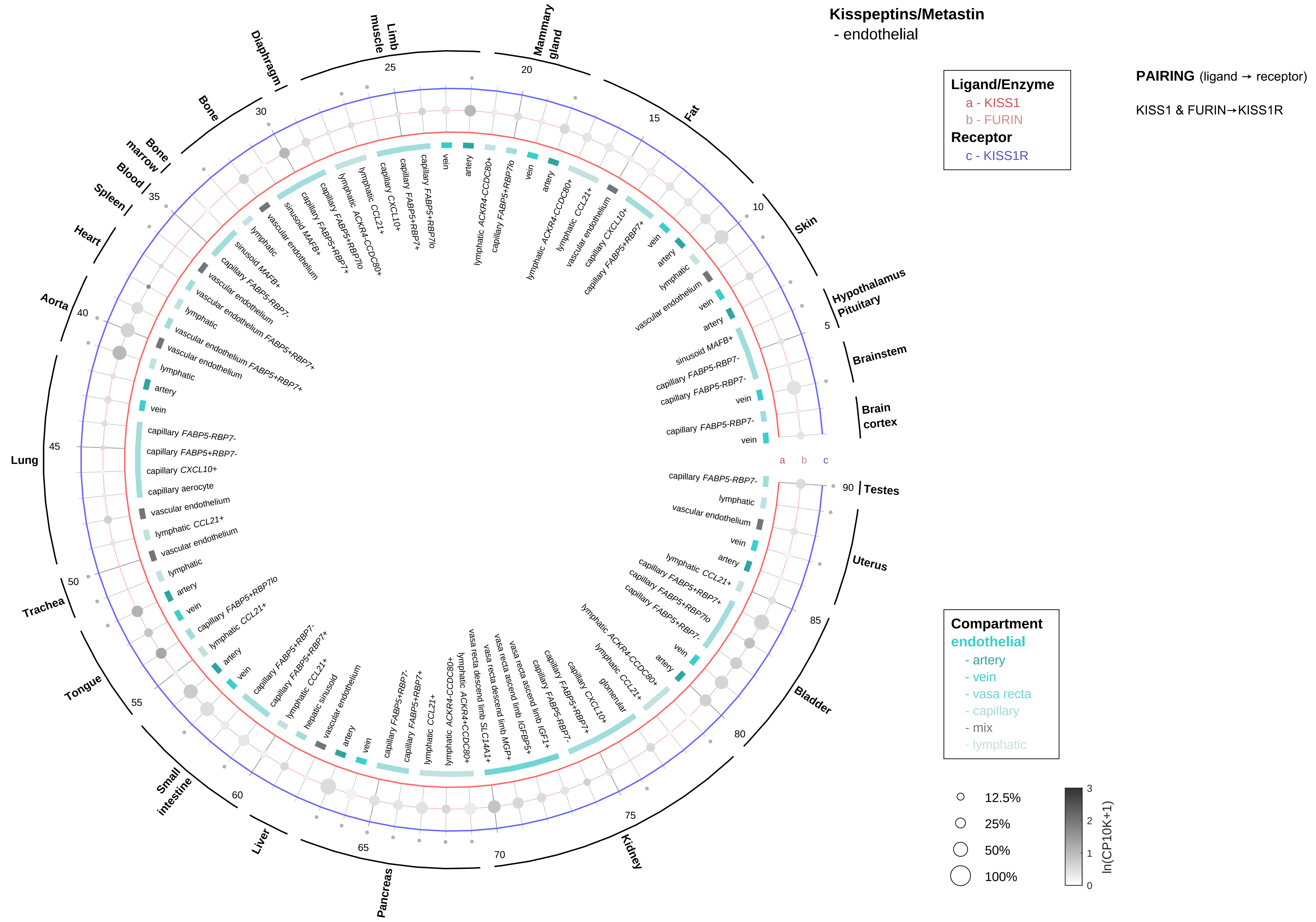
Kisspeptins/Metastin
- endothelial
Ligand/Enzyme
a - KISS1
b - FURIN
Receptor
c - KISS1R
PAIRING (ligand → receptor)
KISS1 & FURIN→KISS1R
Compartment
endothelial
- artery
- vein
- vasa recta
- capillary
- mix
- lymphatic
12.5%
25%
50%
100%
ln(CP10K+1)
Brain cortex
Brainstem
Hypothalamus Pituitary
Skin
Fat
Mammary gland
Limb muscle
Diaphragm
Bone
Bone marrow
Blood
Spleen
Heart
Aorta
Lung
Trachea
Tongue
Small intestine
Liver
Pancreas
Kidney
Bladder
Uterus
Testes

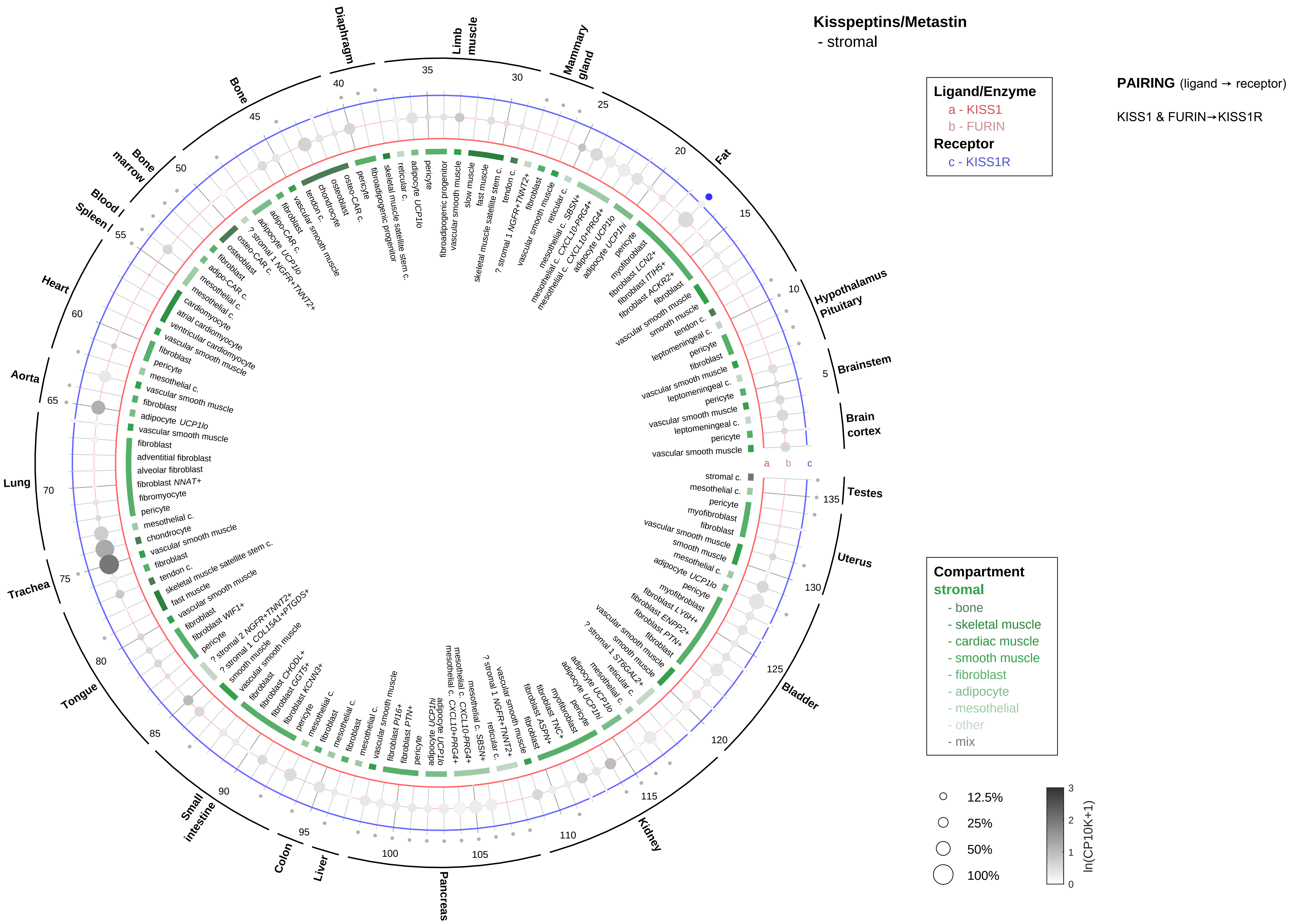

Kisspeptins/Metastin
- stromal
Ligand/Enzyme
a - KISS1
b - FURIN
Receptor
c - KISS1R
PAIRING (ligand → receptor)
KISS1 & FURIN→KISS1R
Compartment
stromal
- bone
- skeletal muscle
- cardiac muscle
- smooth muscle
- fibroblast
- adipocyte
- mesothelial
- other
- mix
12.5%
25%
50%
100%
ln(CP10K+1)

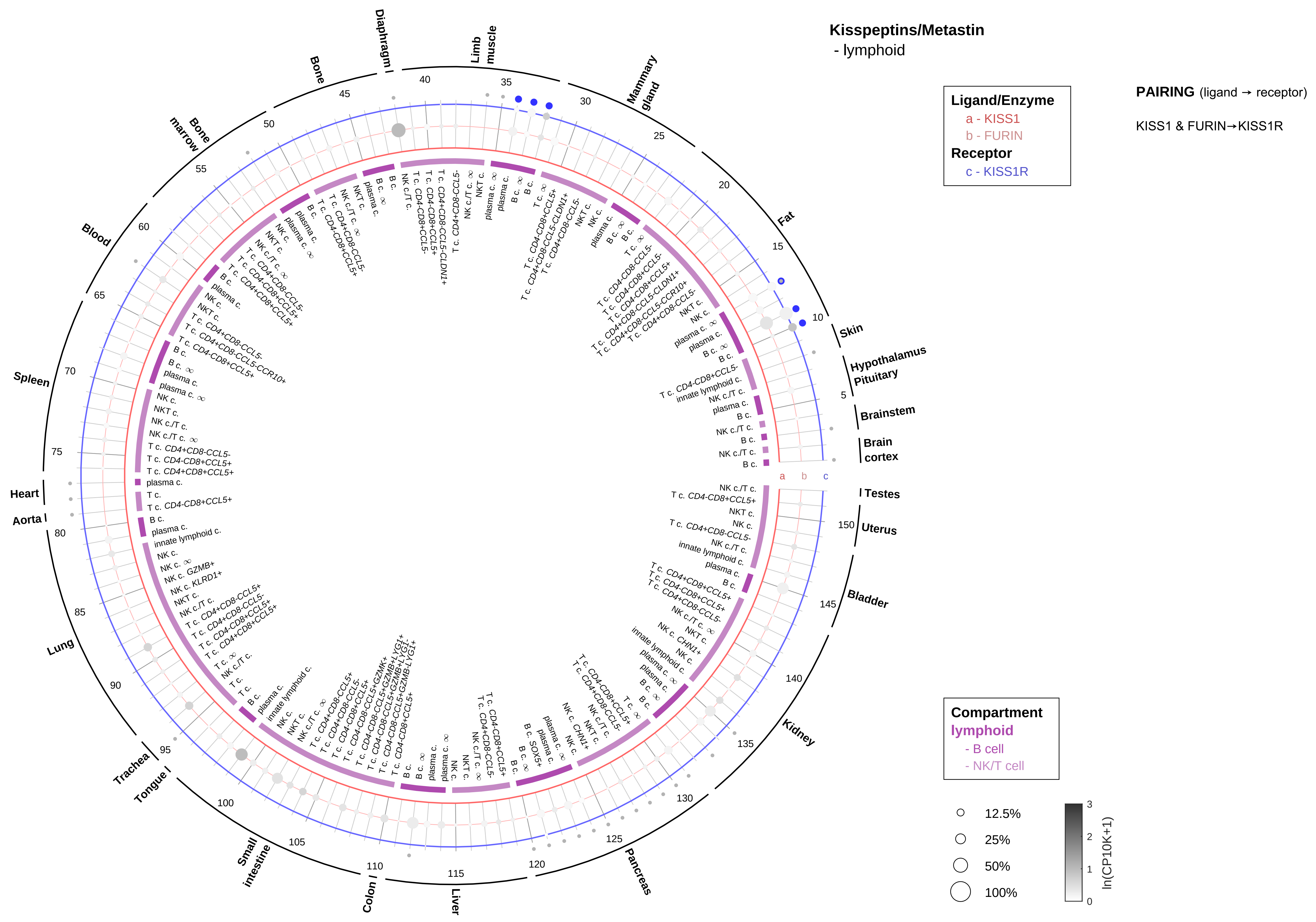

Kisspeptins/Metastin
- lymphoid
Ligand/Enzyme
a - KISS1
b - FURIN
Receptor
c - KISS1R
PAIRING (ligand → receptor)
KISS1 & FURIN→KISS1R
Compartment
lymphoid
- B cell
- NK/T cell
12.5%
25%
50%
100%
ln(CP10K+1)
Skin
Hypothalamus
Pituitary
Brainstem
Brain cortex
Testes
Uterus
Bladder
Kidney
Pancreas
Liver
Colon
Small intestine
Tongue
Trachea
Lung
Aorta
Heart
Spleen
Blood
Bone marrow
Bone
Diaphragm
Limb muscle
Mammary gland
Fat

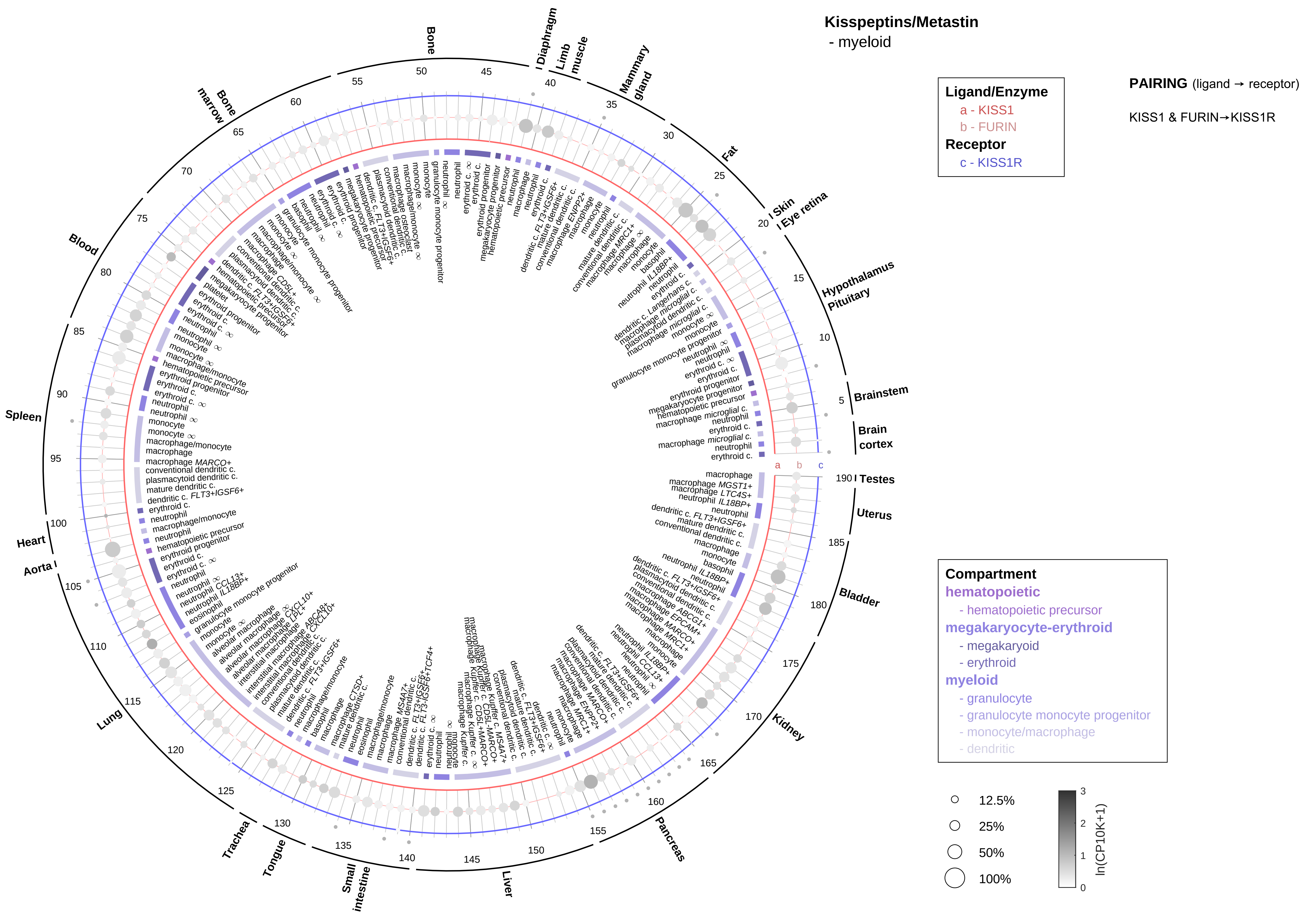
Kisspeptins/Metastin
- myeloid
Ligand/Enzyme
a - KISS1
b - FURIN
Receptor
c - KISS1R
PAIRING (ligand → receptor)
KISS1 & FURIN→KISS1R
Compartment
hematopoietic
- hematopoietic precursor
megakaryocyte-erythroid
- megakaryoid
- erythroid
myeloid
- granulocyte
- granulocyte monocyte progenitor
- monocyte/macrophage
- dendritic
12.5%
25%
50%
100%
ln(CP10K+1)
Bone marrow
Bone
Diaphragm
Limb muscle
Mammary gland
Fat
Skin
Eye retina
Hypothalamus
Pituitary
Brainstem
Brain cortex
Testes
Uterus
Bladder
Kidney
Pancreas
Liver
Small intestine
Tongue
Trachea
Lung
Aorta
Heart
Spleen
Blood

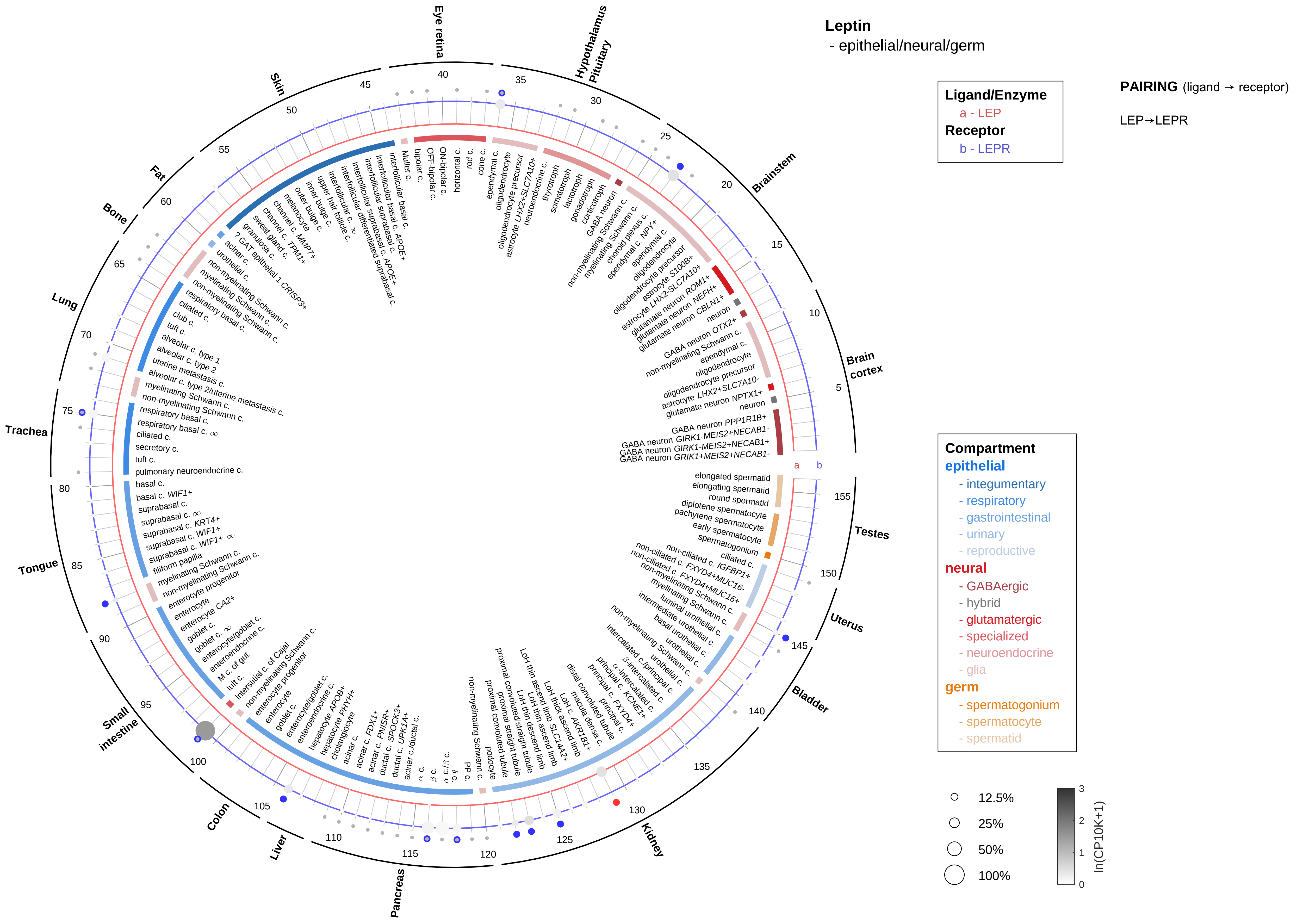

Leptin
- epithelial/neural/germ
Ligand/Enzyme
a - LEP
Receptor
b - LEPR
PAIRING (ligand → receptor)
LEP→LEPR
Compartment
epithelial
- integumentary
- respiratory
- gastrointestinal
- urinary
- reproductive
neural
- GABAergic
- hybrid
- glutamatergic
- specialized
- neuroendocrine
- glia
germ
- spermatogonium
- spermatocyte
- spermatid
12.5%
25%
50%
100%
ln(CP10K+1)
Eye retina
Hypothalamus Pituitary
Brainstem
Brain cortex
Testes
Uterus
Bladder
Kidney
Pancreas
Liver
Colon
Small intestine
Tongue
Trachea
Lung
Bone
Fat
Skin

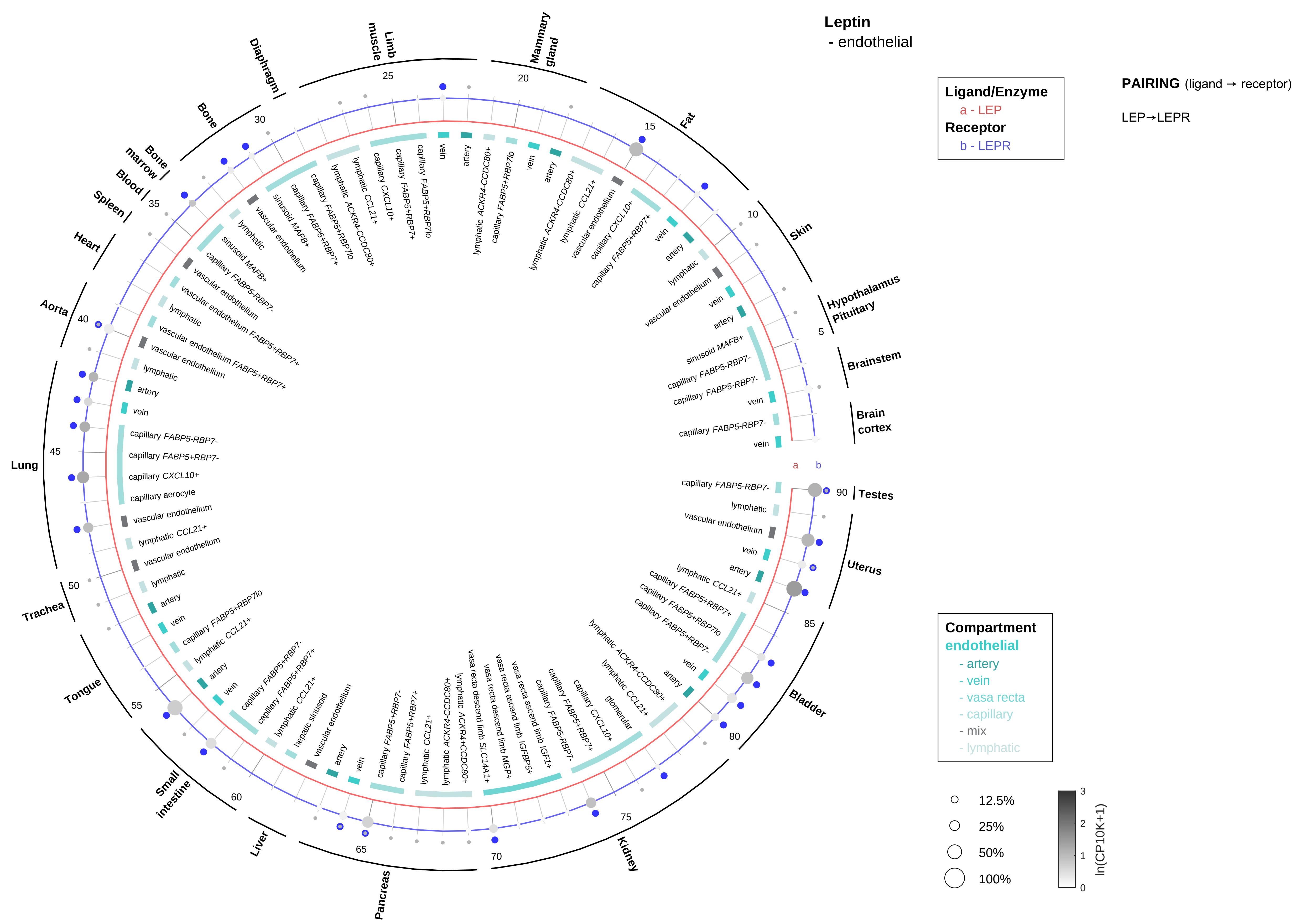

Leptin
- endothelial
Ligand/Enzyme
a - LEP
Receptor
b - LEPR
PAIRING (ligand → receptor)
LEP→LEPR
Compartment
endothelial
- artery
- vein
- vasa recta
- capillary
- mix
- lymphatic
12.5%
25%
50%
100%
ln(CP10K+1)
Brain cortex
Brainstem
Hypothalamus Pituitary
Skin
Fat
Mammary gland
Limb muscle
Diaphragm
Bone
Bone marrow
Blood
Spleen
Heart
Aorta
Lung
Trachea
Tongue
Small intestine
Liver
Pancreas
Kidney
Bladder
Uterus
Testes

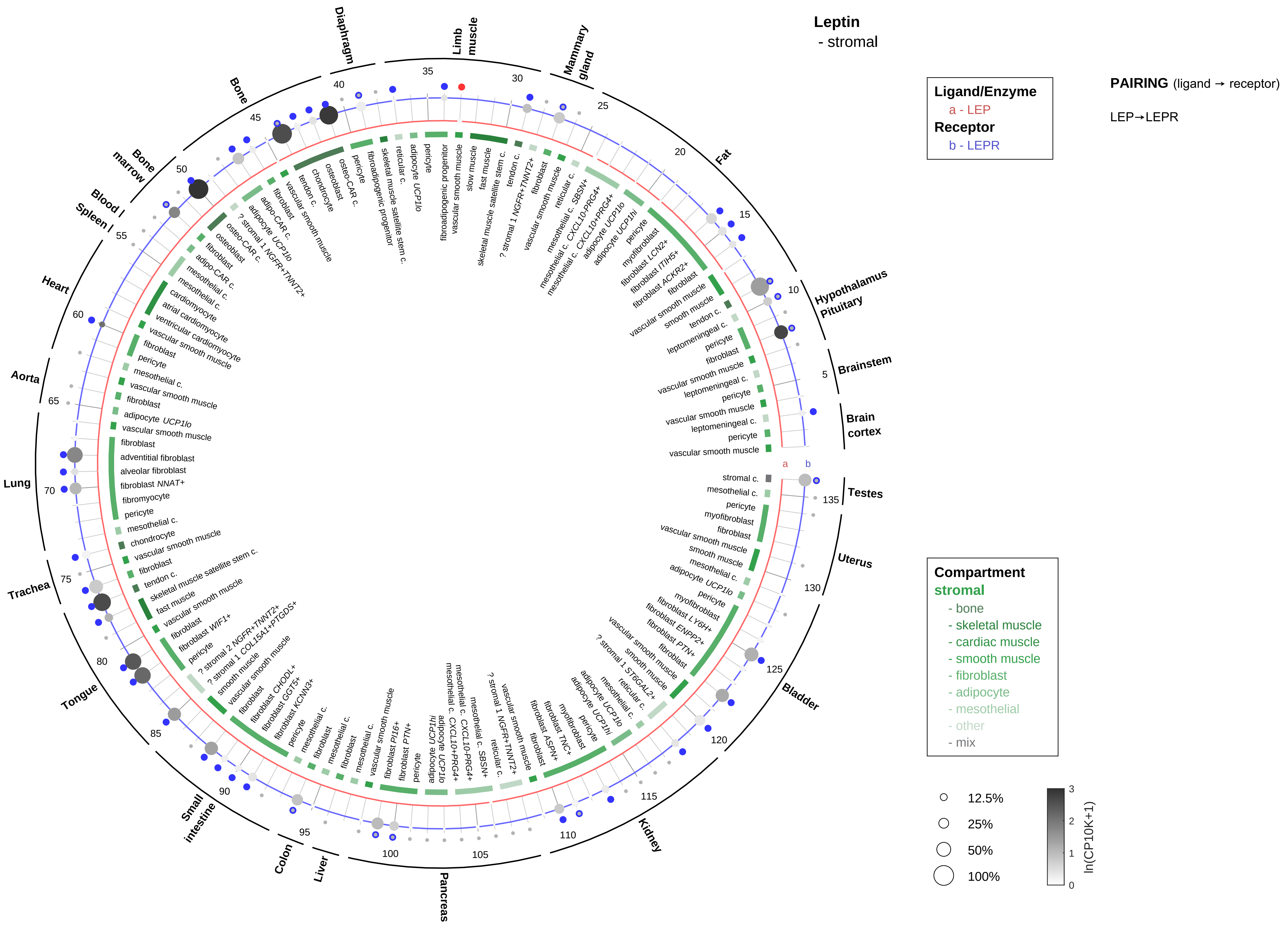

Leptin
- stromal
Ligand/Enzyme
a - LEP
Receptor
b - LEPR
PAIRING (ligand → receptor)
LEP→LEPR
Compartment
stromal
- bone
- skeletal muscle
- cardiac muscle
- smooth muscle
- fibroblast
- adipocyte
- mesothelial
- other
- mix
12.5%
25%
50%
100%
ln(CP10K+1)
Diaphragm
Limb muscle
Mammary gland
Fat
Hypothalamus Pituitary
Brainstem
Brain cortex
Testes
Uterus
Bladder
Kidney
Pancreas
Liver
Colon
Small intestine
Tongue
Trachea
Lung
Aorta
Heart
Spleen
Blood
Bone marrow
Bone

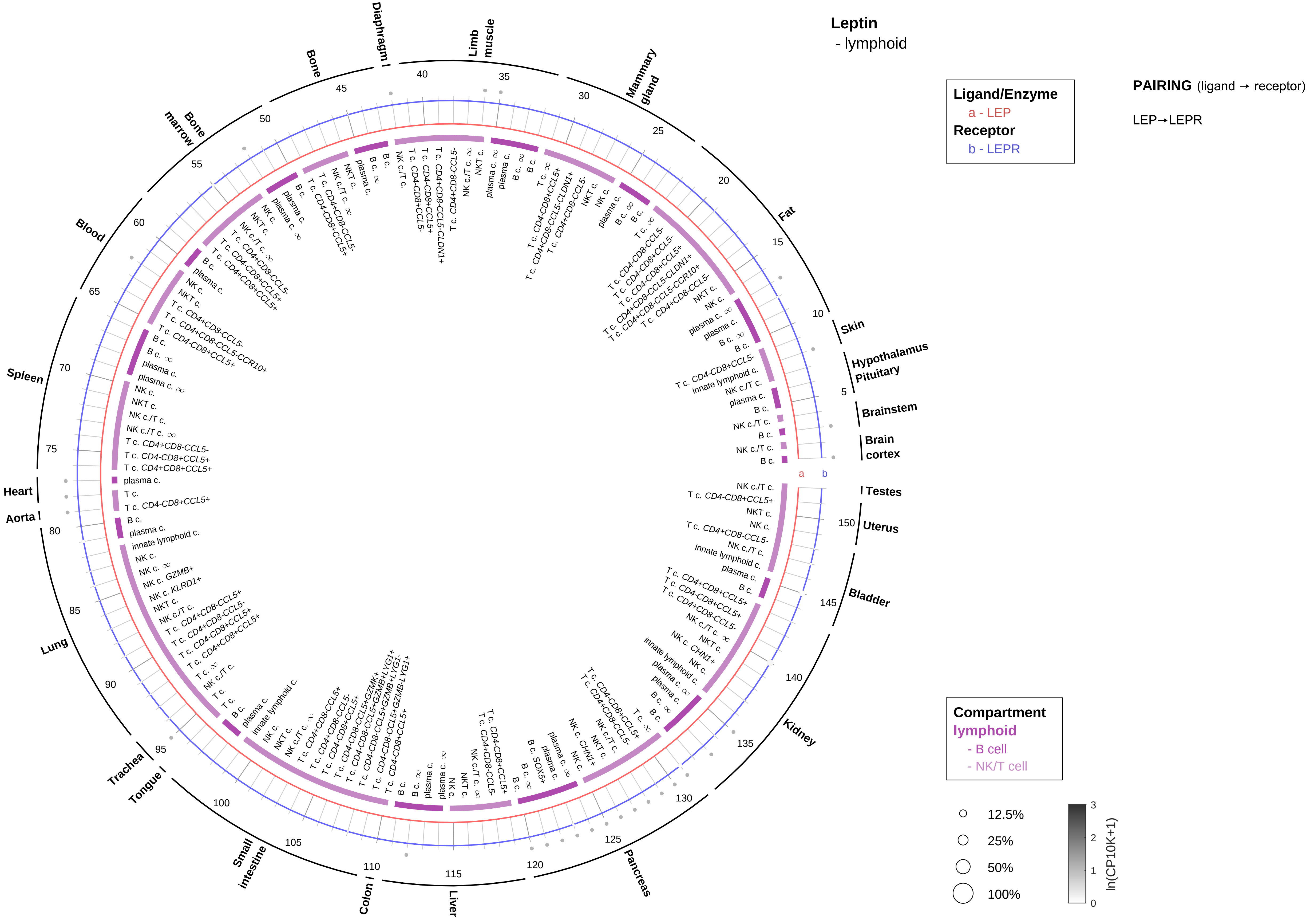
Leptin
- lymphoid
Ligand/Enzyme
a - LEP
Receptor
b - LEPR
PAIRING (ligand → receptor)
LEP→LEPR
Compartment
lymphoid
- B cell
- NK/T cell
12.5%
25%
50%
100%
ln(CP10K+1)
Testes
Uterus
Bladder
Kidney
Pancreas
Liver
Colon
Small intestine
Tongue
Trachea
Lung
Aorta
Heart
Spleen
Blood
Bone marrow
Bone
Diaphragm
Limb muscle
Mammary gland
Fat
Skin
Hypothalamus
Pituitary
Brainstem
Brain cortex

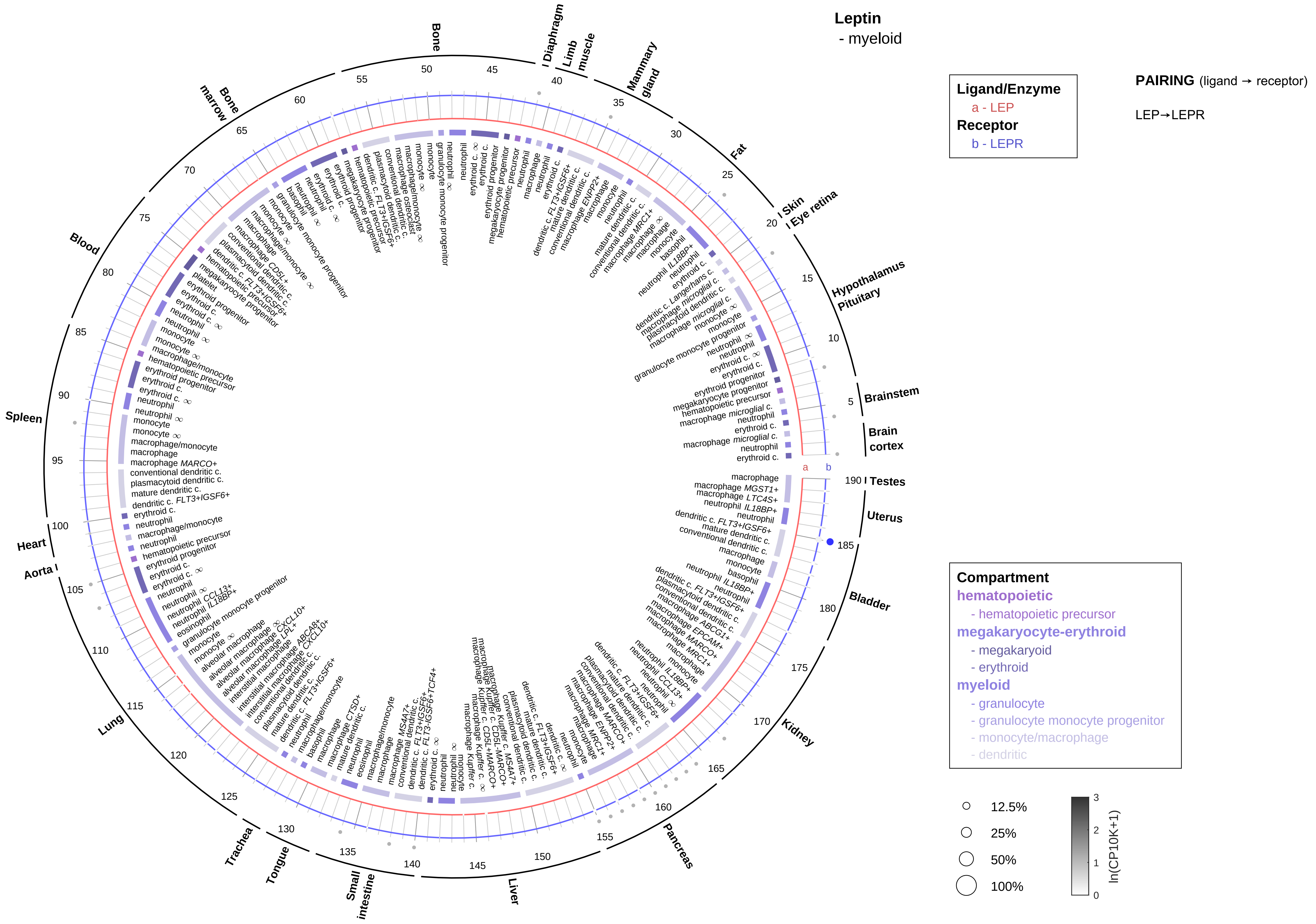

Leptin
- myeloid
Ligand/Enzyme
a - LEP
Receptor
b - LEPR
PAIRING (ligand → receptor)
LEP→LEPR
Compartment
hematopoietic
- hematopoietic precursor
megakaryocyte-erythroid
- megakaryoid
- erythroid
myeloid
- granulocyte
- granulocyte monocyte progenitor
- monocyte/macrophage
- dendritic
12.5%
25%
50%
100%
ln(CP10K+1)
Bone
Diaphragm
Limb muscle
Mammary gland
Fat
Skin
Eye retina
Hypothalamus
Pituitary
Brainstem
Brain cortex
Testes
Uterus
Bladder
Kidney
Pancreas
Liver
Small intestine
Tongue
Trachea
Lung
Aorta
Heart
Spleen
Blood
Bone marrow

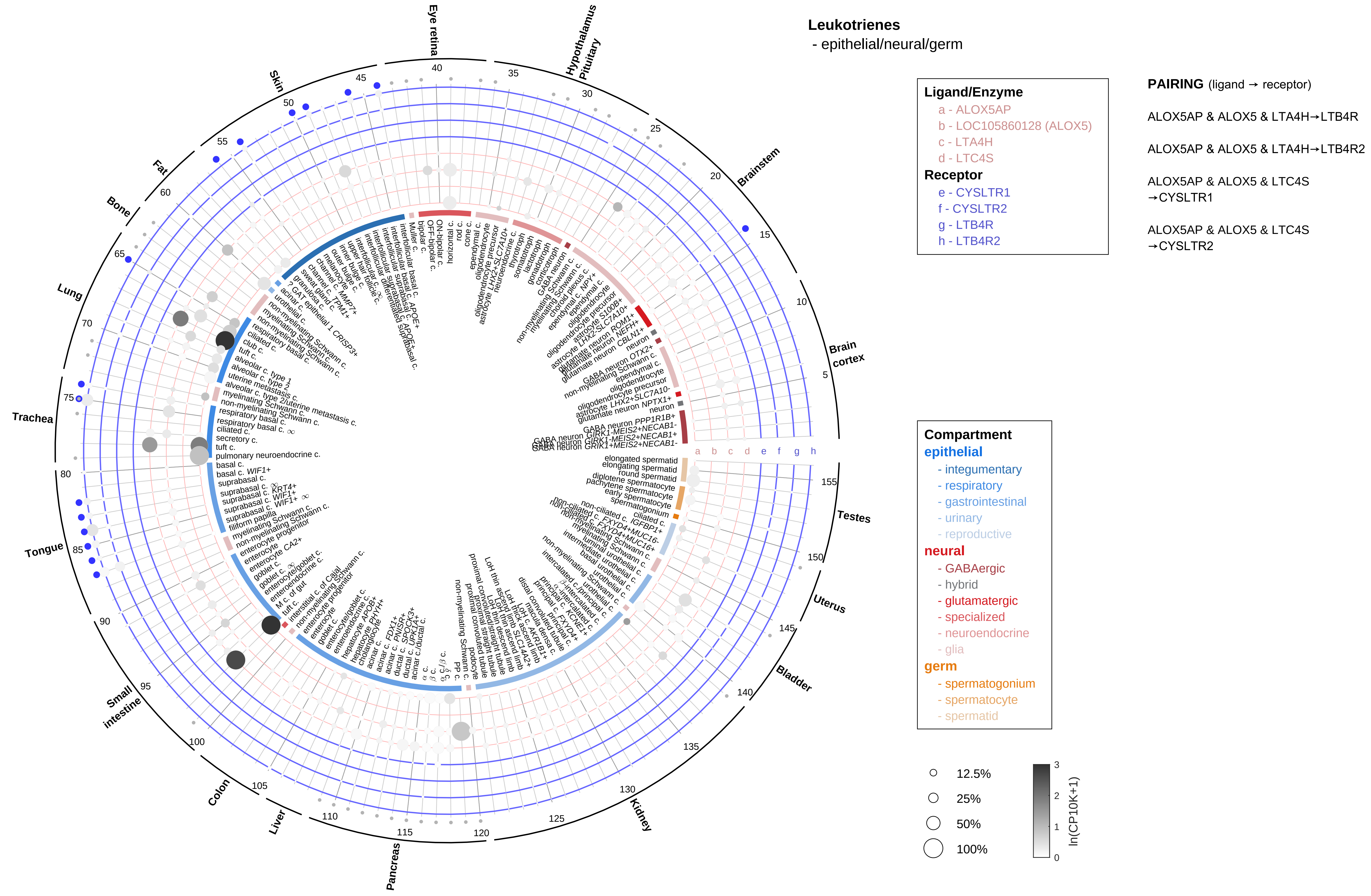
Leukotrienes
- epithelial/neural/germ
Ligand/Enzyme
a - ALOX5AP
b - LOC105860128 (ALOX5)
c - LTA4H
d - LTC4S
Receptor
e - CYSLTR1
f - CYSLTR2
g - LTB4R
h - LTB4R2
PAIRING (ligand → receptor)
ALOX5AP & ALOX5 & LTA4H→LTB4R
ALOX5AP & ALOX5 & LTA4H→LTB4R2
ALOX5AP & ALOX5 & LTC4S →CYSLTR1
ALOX5AP & ALOX5 & LTC4S →CYSLTR2
Compartment
epithelial
- integumentary
- respiratory
- gastrointestinal
- urinary
- reproductive
neural
- GABAergic
- hybrid
- glutamatergic
- specialized
- neuroendocrine
- glia
germ
- spermatogonium
- spermatocyte
- spermatid
12.5%
25%
50%
100%
ln(CP10K+1)
Eye retina
Hypothalamus Pituitary
Brainstem
Brain cortex
Testes
Uterus
Bladder
Kidney
Pancreas
Liver
Colon
Small intestine
Tongue
Trachea
Lung
Bone
Fat
Skin

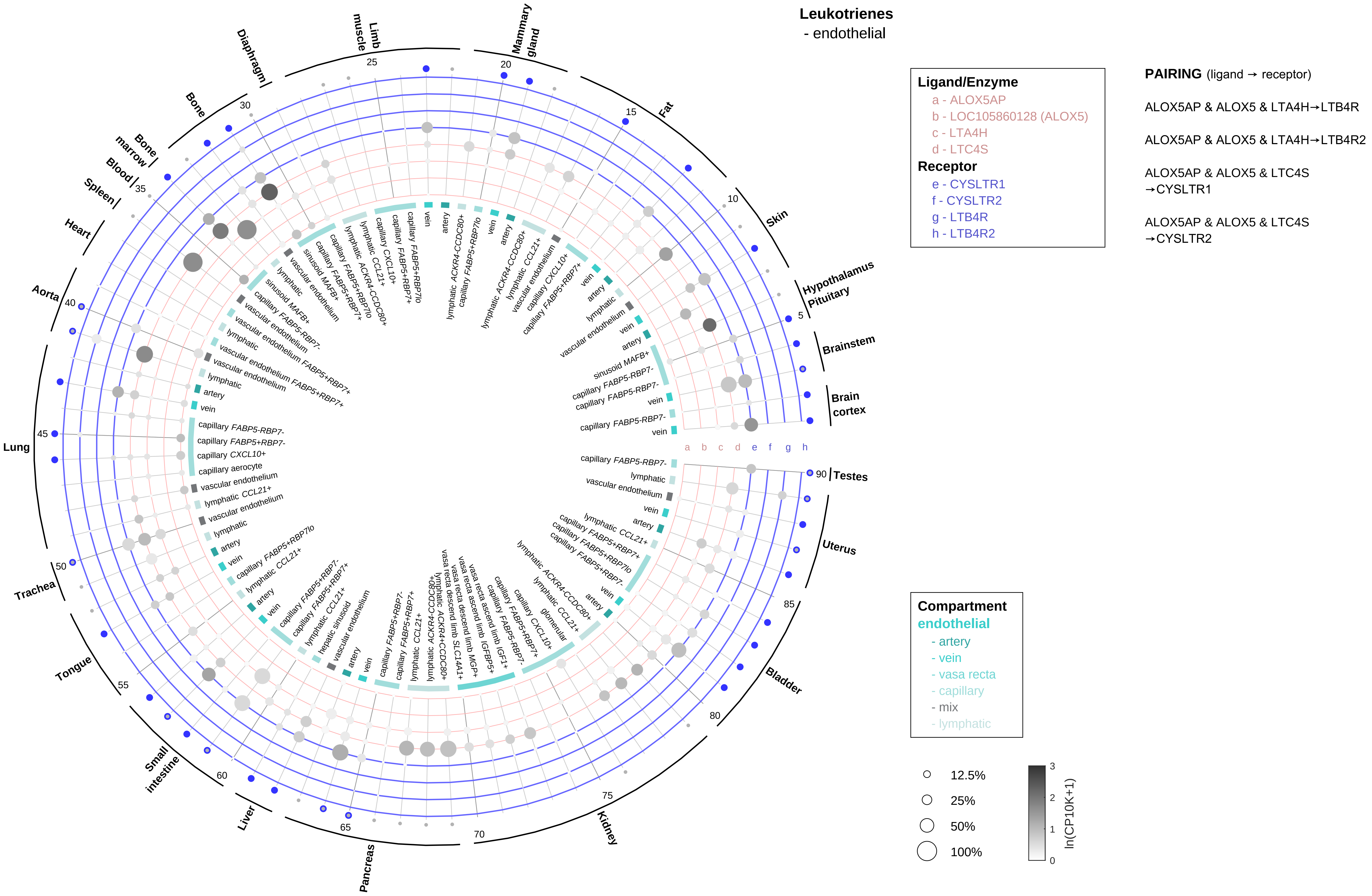
Leukotrienes
- endothelial
Ligand/Enzyme
a - ALOX5AP
b - LOC105860128 (ALOX5)
c - LTA4H
d - LTC4S
Receptor
e - CYSLTR1
f - CYSLTR2
g - LTB4R
h - LTB4R2
PAIRING (ligand → receptor)
ALOX5AP & ALOX5 & LTA4H→LTB4R
ALOX5AP & ALOX5 & LTA4H→LTB4R2
ALOX5AP & ALOX5 & LTC4S →CYSLTR1
ALOX5AP & ALOX5 & LTC4S →CYSLTR2
Compartment
endothelial
- artery
- vein
- vasa recta
- capillary
- mix
- lymphatic
12.5%
25%
50%
100%
ln(CP10K+1)
0
1
2
3
a b c d e f g h
Mammary gland
Fat
Skin
Hypothalamus Pituitary
Brainstem
Brain cortex
Testes
Uterus
Bladder
Kidney
Pancreas
Liver
Small intestine
Tongue
Trachea
Lung
Aorta
Heart
Spleen
Blood
Bone marrow
Bone
Diaphragm
Limb muscle
5
10
15
20
25
30
35
40
45
50
55
60
65
70
75
80
85
90

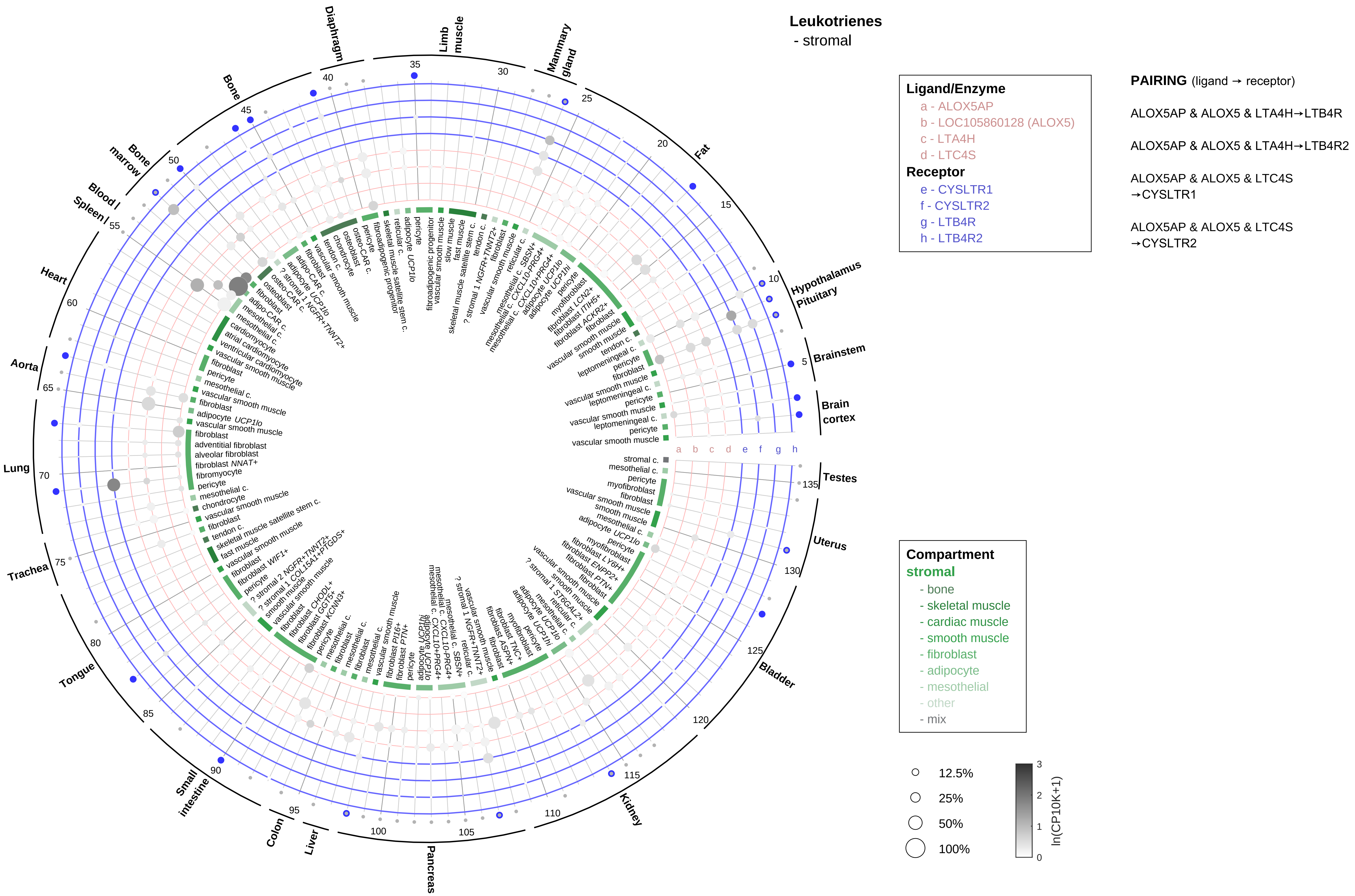

Leukotrienes
- stromal
Ligand/Enzyme
a - ALOX5AP
b - LOC105860128 (ALOX5)
c - LTA4H
d - LTC4S
Receptor
e - CYSLTR1
f - CYSLTR2
g - LTB4R
h - LTB4R2
PAIRING (ligand → receptor)
ALOX5AP & ALOX5 & LTA4H→LTB4R
ALOX5AP & ALOX5 & LTA4H→LTB4R2
ALOX5AP & ALOX5 & LTC4S →CYSLTR1
ALOX5AP & ALOX5 & LTC4S →CYSLTR2
Compartment
stromal
- bone
- skeletal muscle
- cardiac muscle
- smooth muscle
- fibroblast
- adipocyte
- mesothelial
- other
- mix
12.5%
25%
50%
100%
ln(CP10K+1)
Diaphragm
Limb muscle
Mammary gland
Fat
Hypothalamus Pituitary
Brainstem
Brain cortex
Testes
Uterus
Bladder
Kidney
Pancreas
Liver
Colon
Small intestine
Tongue
Trachea
Lung
Aorta
Heart
Blood / Spleen /
Bone marrow
Bone

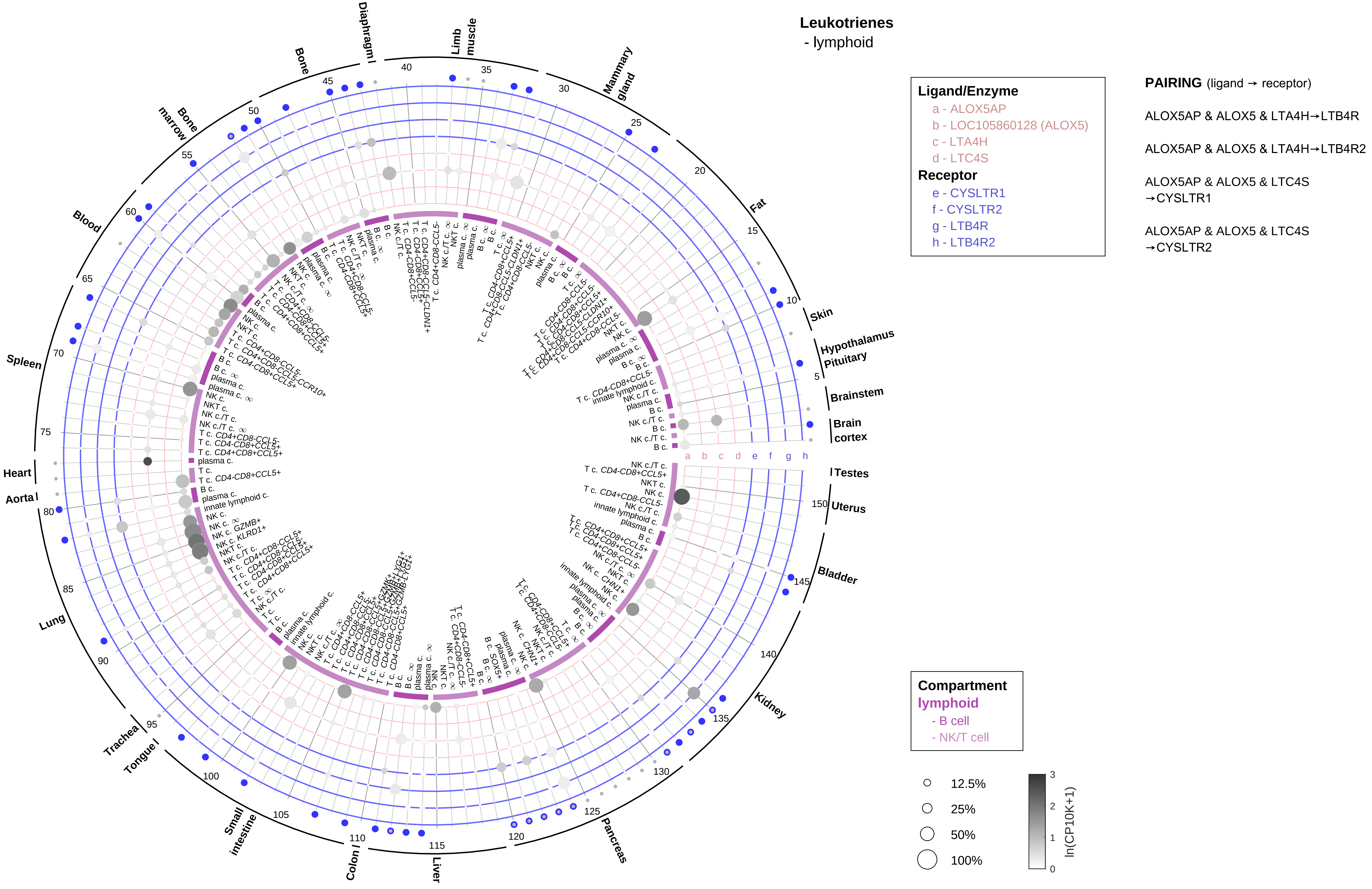

Leukotrienes
- lymphoid
Ligand/Enzyme
a - ALOX5AP
b - LOC105860128 (ALOX5)
c - LTA4H
d - LTC4S
Receptor
e - CYSLTR1
f - CYSLTR2
g - LTB4R
h - LTB4R2
PAIRING (ligand → receptor)
ALOX5AP & ALOX5 & LTA4H→LTB4R
ALOX5AP & ALOX5 & LTA4H→LTB4R2
ALOX5AP & ALOX5 & LTC4S →CYSLTR1
ALOX5AP & ALOX5 & LTC4S →CYSLTR2
Compartment
lymphoid
- B cell
- NK/T cell
12.5%
25%
50%
100%
ln(CP10K+1)
Limb muscle
Mammary gland
Fat
Skin
Hypothalamus Pituitary
Brainstem
Brain cortex
Testes
Uterus
Bladder
Kidney
Pancreas
Liver
Colon
Small intestine
Tongue
Trachea
Lung
Aorta
Heart
Spleen
Blood
Bone marrow
Bone
Diaphragm
a b c d e f g h

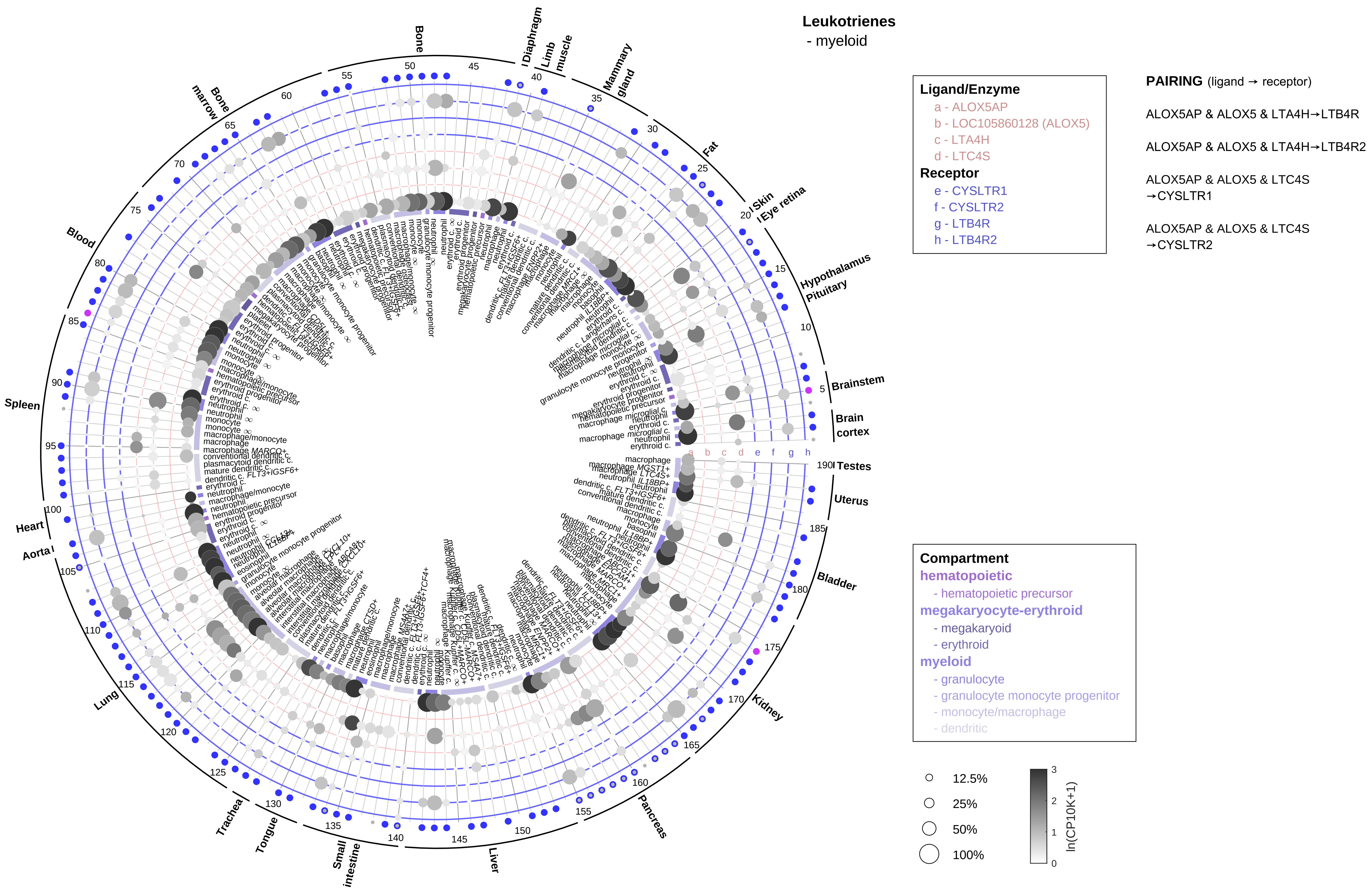

Leukotrienes
- myeloid
Ligand/Enzyme
a - ALOX5AP
b - LOC105860128 (ALOX5)
c - LTA4H
d - LTC4S
Receptor
e - CYSLTR1
f - CYSLTR2
g - LTB4R
h - LTB4R2
PAIRING (ligand → receptor)
ALOX5AP & ALOX5 & LTA4H→LTB4R
ALOX5AP & ALOX5 & LTA4H→LTB4R2
ALOX5AP & ALOX5 & LTC4S →CYSLTR1
ALOX5AP & ALOX5 & LTC4S →CYSLTR2
Compartment
hematopoietic
- hematopoietic precursor
megakaryocyte-erythroid
- megakaryoid
- erythroid
myeloid
- granulocyte
- granulocyte monocyte progenitor
- monocyte/macrophage
- dendritic
12.5%
25%
50%
100%
ln(CP10K+1)
Bone
Diaphragm
Limb muscle
Mammary gland
Fat
Skin
Eye retina
Hypothalamus
Pituitary
Brainstem
Brain cortex
Testes
Uterus
Bladder
Kidney
Pancreas
Liver
Small intestine
Tongue
Trachea
Lung
Aorta
Heart
Spleen
Blood
Bone marrow

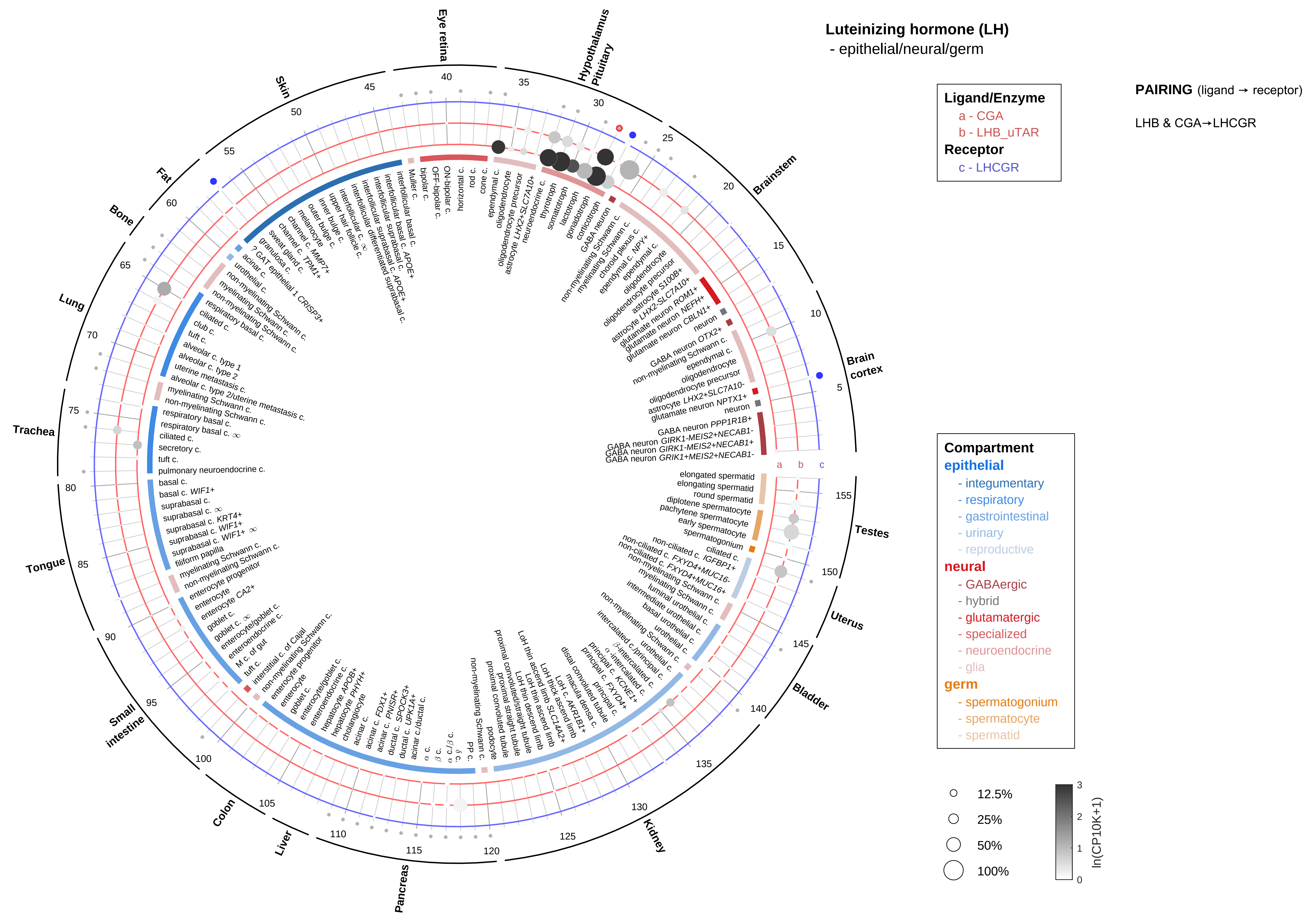
Luteinizing hormone (LH)
- epithelial/neural/germ
Ligand/Enzyme
a - CGA
b - LHB_uTAR
Receptor
c - LHCGR
PAIRING (ligand → receptor)
LHB & CGA→LHCGR
Compartment
epithelial
- integumentary
- respiratory
- gastrointestinal
- urinary
- reproductive
neural
- GABAergic
- hybrid
- glutamatergic
- specialized
- neuroendocrine
- glia
germ
- spermatogonium
- spermatocyte
- spermatid
12.5%
25%
50%
100%
ln(CP10K+1)
Brain cortex
Brainstem
Hypothalamus
Pituitary
Eye retina
Skin
Fat
Bone
Lung
Trachea
Tongue
Small intestine
Colon
Liver
Pancreas
Kidney
Bladder
Uterus
Testes

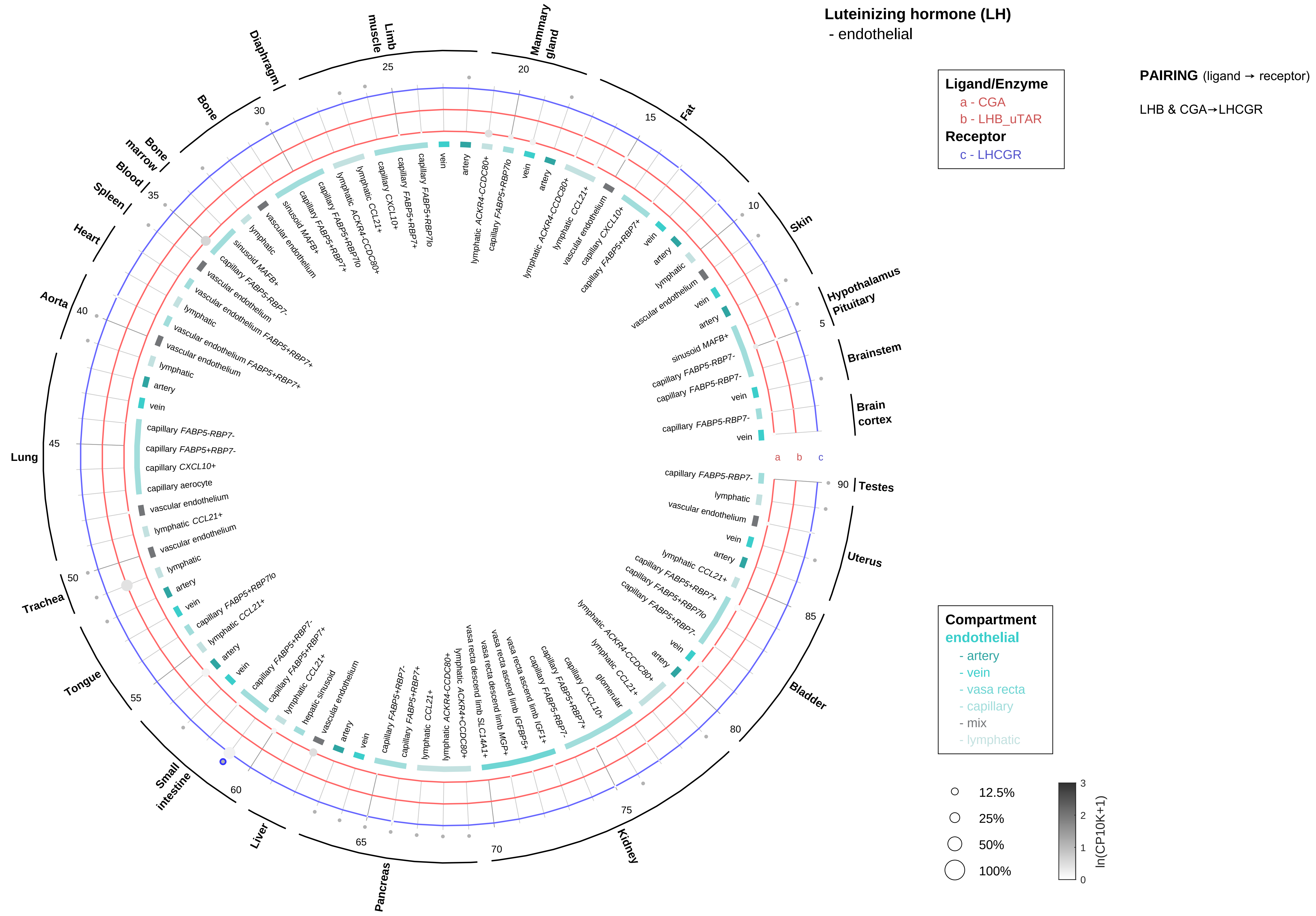

Luteinizing hormone (LH)
- endothelial
Ligand/Enzyme
a - CGA
b - LHB_uTAR
Receptor
c - LHCGR
PAIRING (ligand → receptor)
LHB & CGA→LHCGR
Compartment
endothelial
- artery
- vein
- vasa recta
- capillary
- mix
- lymphatic
12.5%
25%
50%
100%
ln(CP10K+1)
Brain cortex
Brainstem
Hypothalamus
Pituitary
Skin
Fat
Mammary gland
Limb muscle
Diaphragm
Bone
Bone marrow
Blood
Spleen
Heart
Aorta
Lung
Trachea
Tongue
Small intestine
Liver
Pancreas
Kidney
Bladder
Uterus
Testes

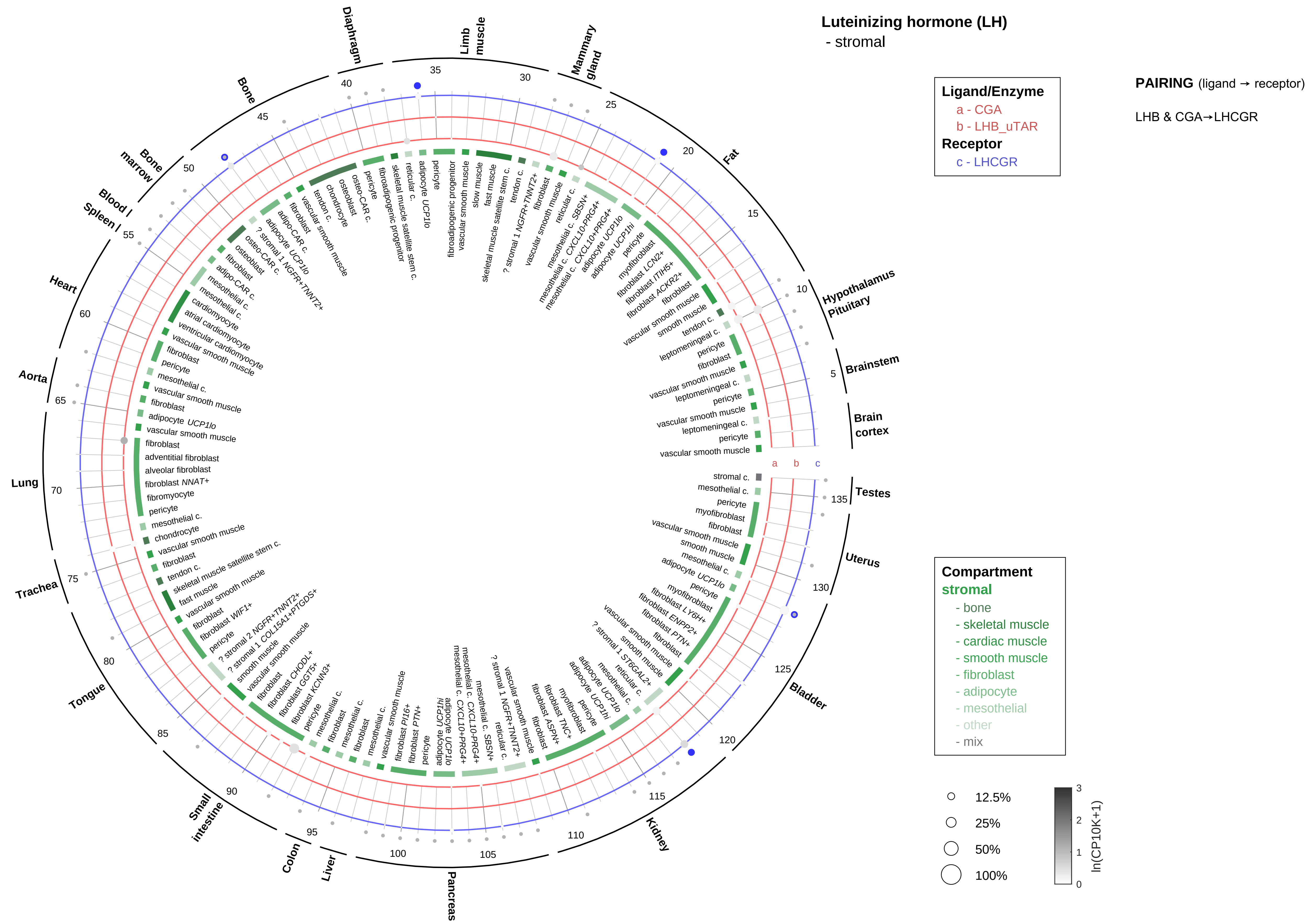
Luteinizing hormone (LH)
- stromal
Ligand/Enzyme
a - CGA
b - LHB_uTAR
Receptor
c - LHCGR
PAIRING (ligand → receptor)
LHB & CGA→LHCGR
Compartment
stromal
- bone
- skeletal muscle
- cardiac muscle
- smooth muscle
- fibroblast
- adipocyte
- mesothelial
- other
- mix
12.5%
25%
50%
100%
ln(CP10K+1)
Brain cortex
Brainstem
Hypothalamus
Pituitary
Fat
Mammary gland
Limb muscle
Diaphragm
Bone
Bone marrow
Blood
Spleen
Heart
Aorta
Lung
Trachea
Tongue
Small intestine
Colon
Liver
Pancreas
Kidney
Bladder
Uterus
Testes

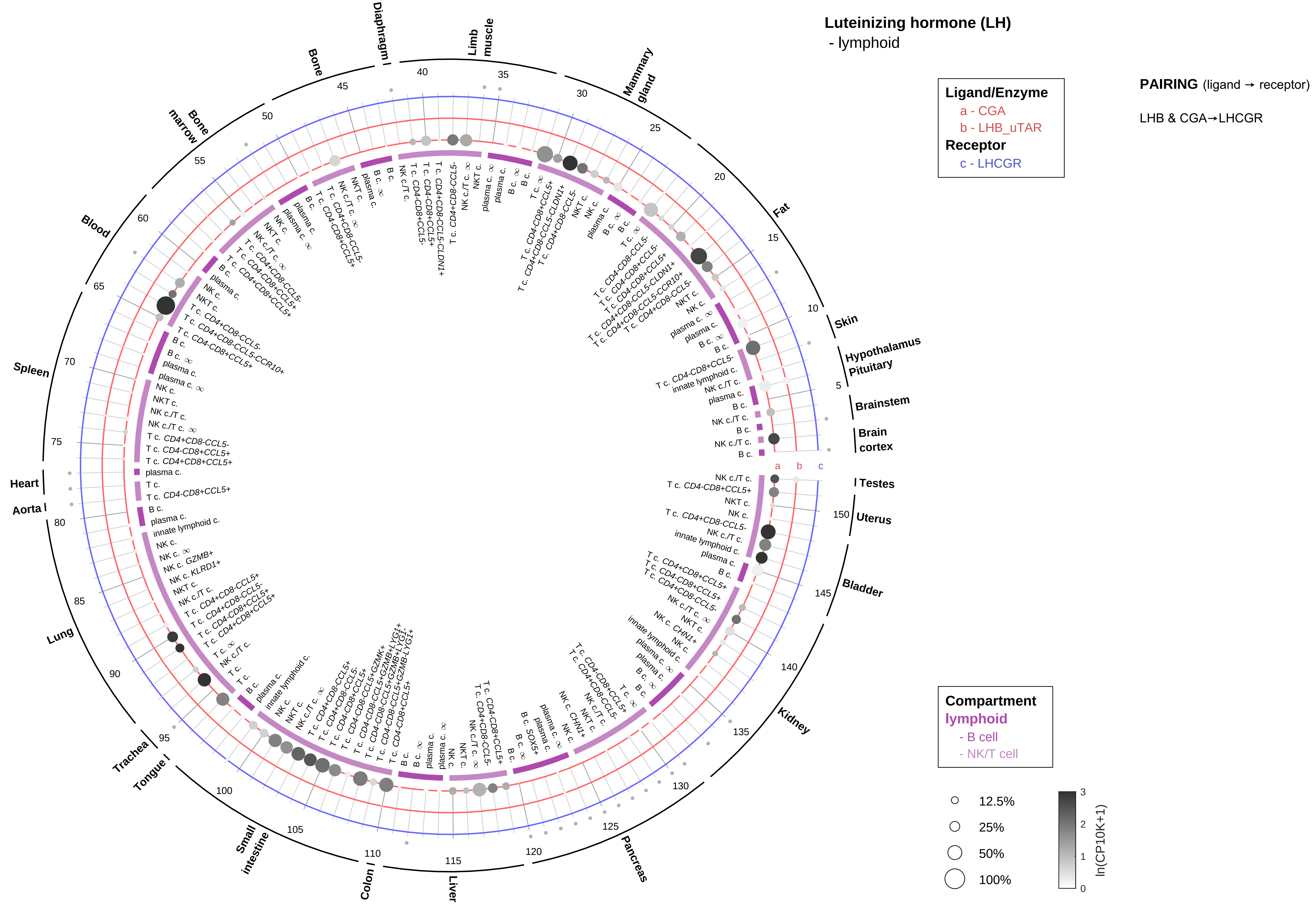

Luteinizing hormone (LH)
- lymphoid
PAIRING (ligand → receptor)
LHB & CGA→LHCGR
Ligand/Enzyme
a - CGA
b - LHB_uTAR
Receptor
c - LHCGR
Compartment
lymphoid
- B cell
- NK/T cell
12.5%
25%
50%
100%
ln(CP10K+1)
Testes
Uterus
Bladder
Kidney
Pancreas
Liver
Colon
Small intestine
Tongue
Trachea
Lung
Aorta
Heart
Spleen
Blood
Bone marrow
Bone
Diaphragm
Limb muscle
Mammary gland
Fat
Skin
Hypothalamus
Pituitary
Brainstem
Brain cortex

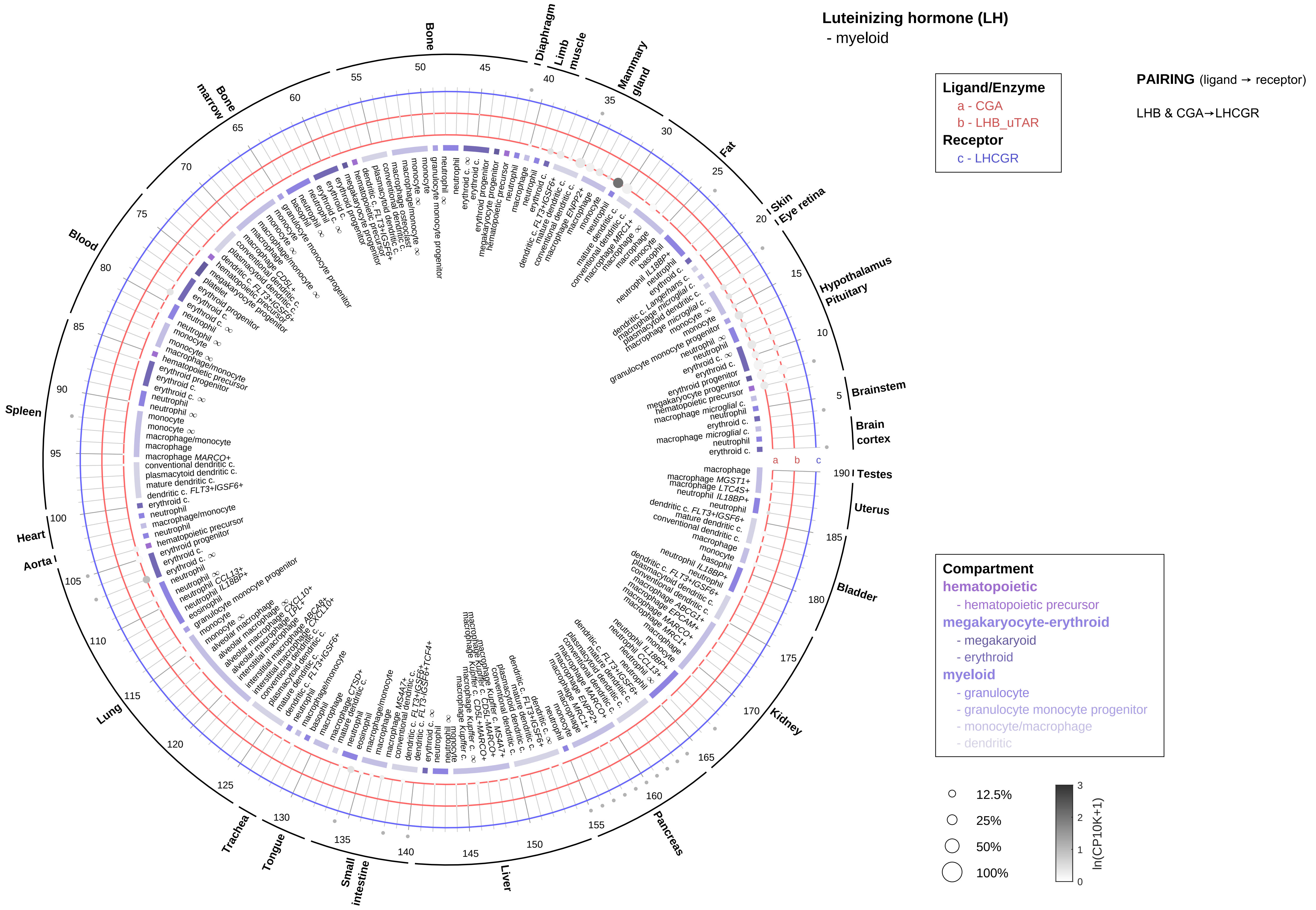

Luteinizing hormone (LH)
- myeloid
Ligand/Enzyme
a - CGA
b - LHB_uTAR
Receptor
c - LHCGR
PAIRING (ligand → receptor)
LHB & CGA→LHCGR
Compartment
hematopoietic
- hematopoietic precursor
megakaryocyte-erythroid
- megakaryoid
- erythroid
myeloid
- granulocyte
- granulocyte monocyte progenitor
- monocyte/macrophage
- dendritic
12.5%
25%
50%
100%
ln(CP10K+1)
Bone
Diaphragm
Limb muscle
Mammary gland
Fat
Skin
Eye retina
Hypothalamus
Pituitary
Brainstem
Brain cortex
Testes
Uterus
Bladder
Kidney
Pancreas
Liver
Small intestine
Tongue
Trachea
Lung
Aorta
Heart
Spleen
Blood
Bone marrow

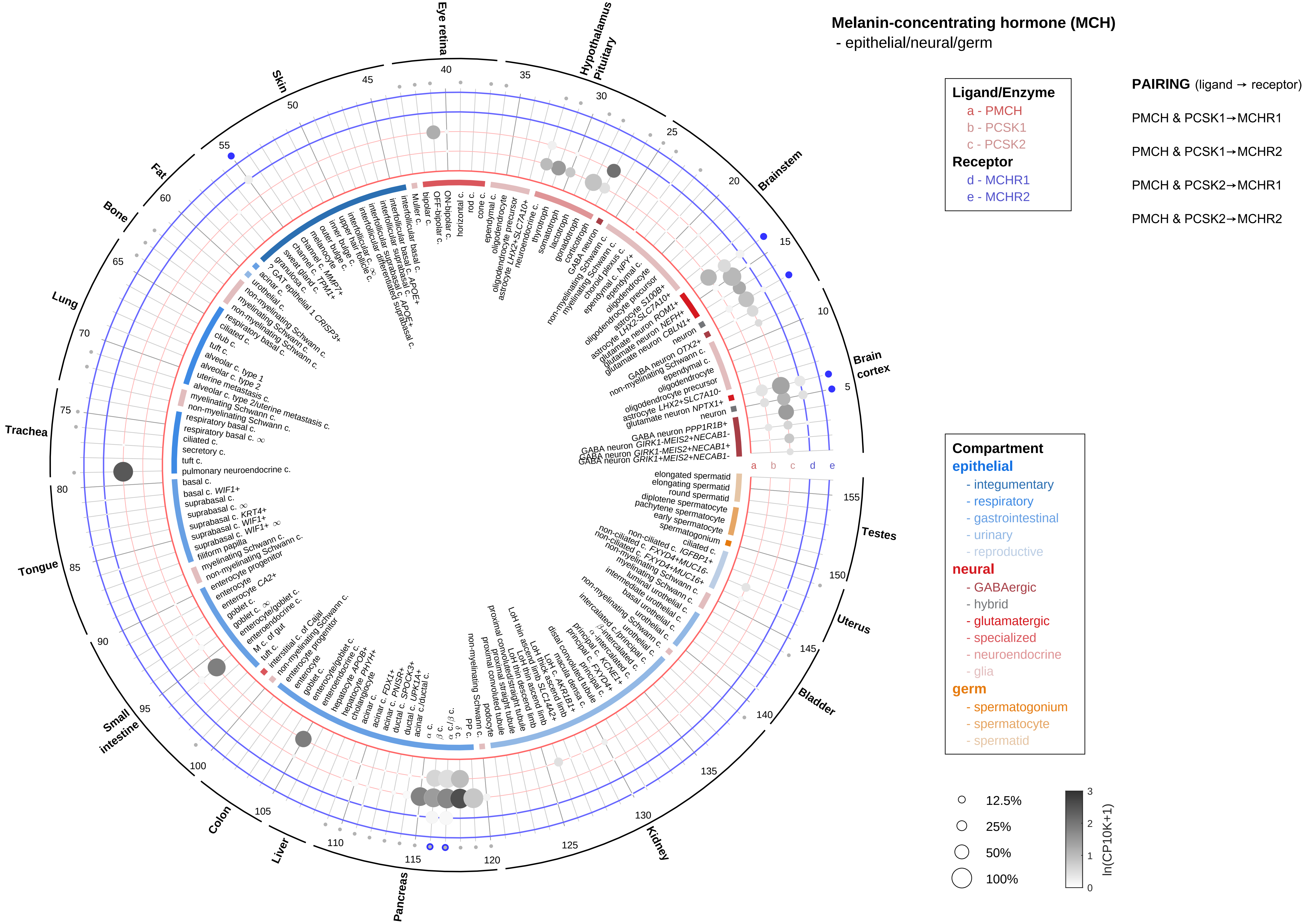
Melanin-concentrating hormone (MCH)
- epithelial/neural/germ
Ligand/Enzyme
a - PMCH
b - PCSK1
c - PCSK2
Receptor
d - MCHR1
e - MCHR2
PAIRING (ligand → receptor)
PMCH & PCSK1→MCHR1
PMCH & PCSK1→MCHR2
PMCH & PCSK2→MCHR1
PMCH & PCSK2→MCHR2
Compartment
epithelial
- integumentary
- respiratory
- gastrointestinal
- urinary
- reproductive
neural
- GABAergic
- hybrid
- glutamatergic
- specialized
- neuroendocrine
- glia
germ
- spermatogonium
- spermatocyte
- spermatid
12.5%
25%
50%
100%
ln(CP10K+1)
Eye retina
Hypothalamus
Pituitary
Brainstem
Brain cortex
Testes
Uterus
Bladder
Kidney
Pancreas
Liver
Colon
Small intestine
Tongue
Trachea
Lung
Bone
Fat
Skin

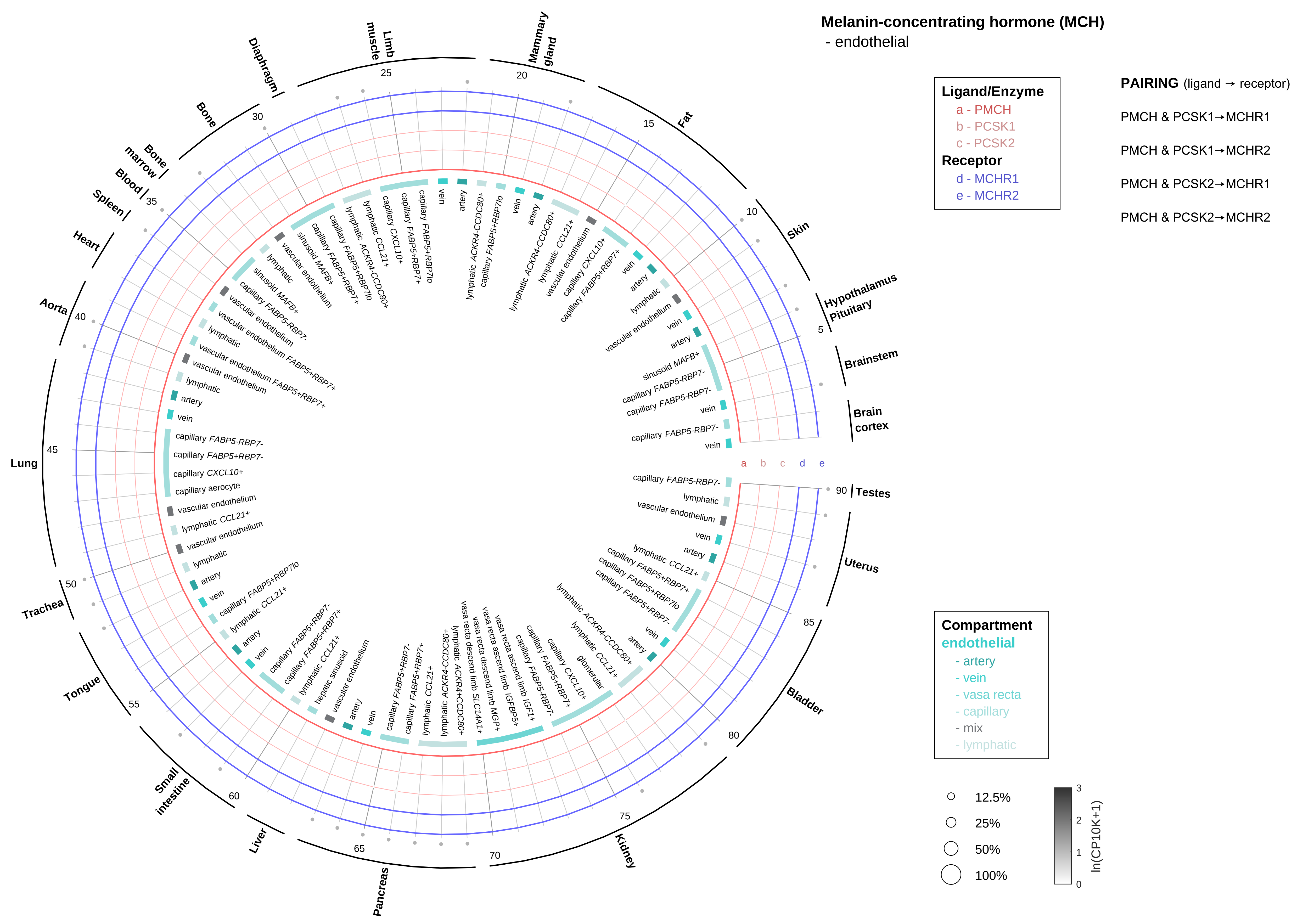

Melanin-concentrating hormone (MCH)
- endothelial
Ligand/Enzyme
a - PMCH
b - PCSK1
c - PCSK2
Receptor
d - MCHR1
e - MCHR2
PAIRING (ligand → receptor)
PMCH & PCSK1→MCHR1
PMCH & PCSK1→MCHR2
PMCH & PCSK2→MCHR1
PMCH & PCSK2→MCHR2
Compartment
endothelial
- artery
- vein
- vasa recta
- capillary
- mix
- lymphatic
12.5%
25%
50%
100%
ln(CP10K+1)
Brain cortex
Brainstem
Hypothalamus Pituitary
Skin
Fat
Mammary gland
Limb muscle
Diaphragm
Bone
Bone marrow
Blood
Spleen
Heart
Aorta
Lung
Trachea
Tongue
Small intestine
Liver
Pancreas
Kidney
Bladder
Uterus
Testes

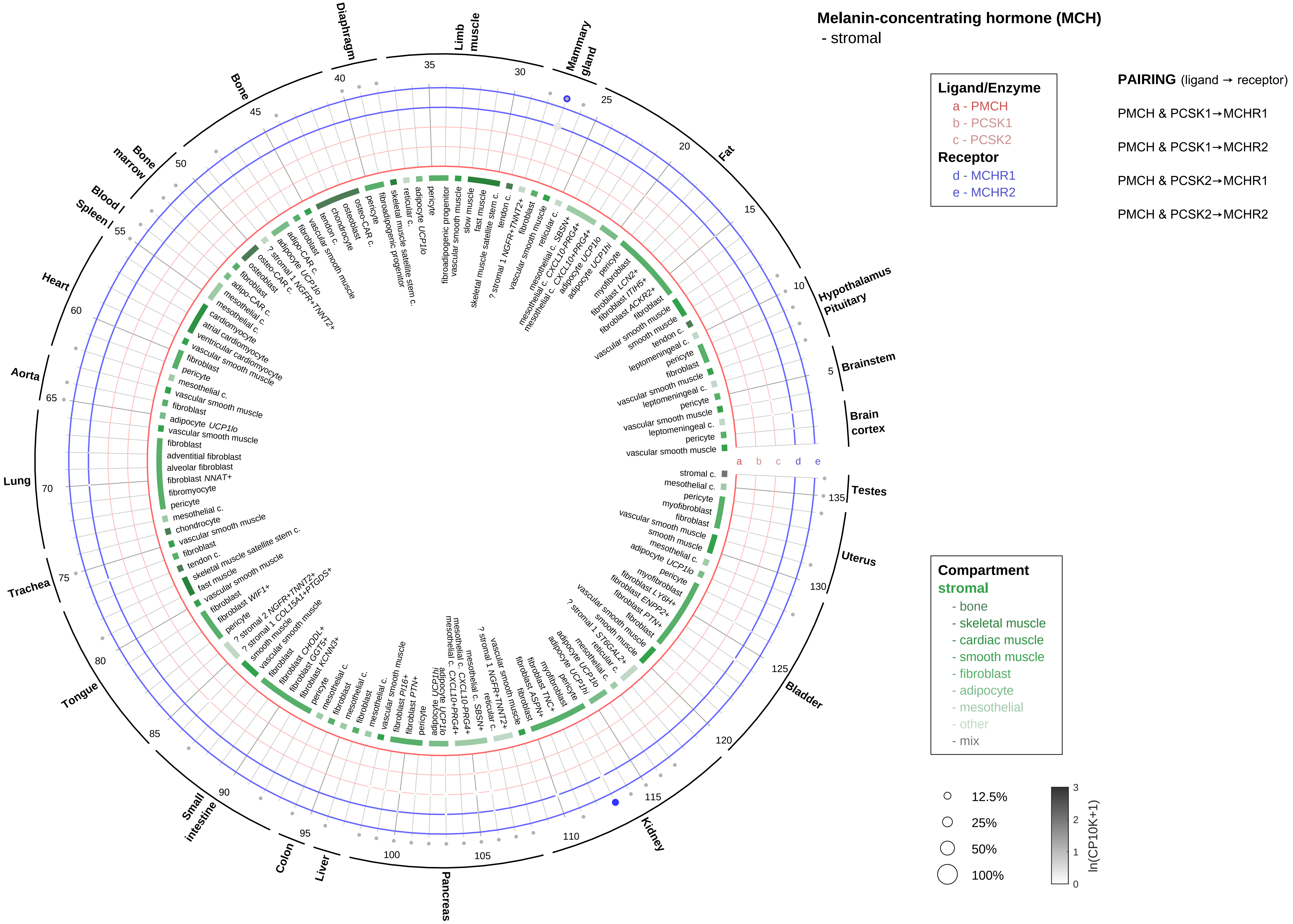

Melanin-concentrating hormone (MCH)
- stromal
Ligand/Enzyme
a - PMCH
b - PCSK1
c - PCSK2
Receptor
d - MCHR1
e - MCHR2
PAIRING (ligand → receptor)
PMCH & PCSK1→MCHR1
PMCH & PCSK1→MCHR2
PMCH & PCSK2→MCHR1
PMCH & PCSK2→MCHR2
Compartment
stromal
- bone
- skeletal muscle
- cardiac muscle
- smooth muscle
- fibroblast
- adipocyte
- mesothelial
- other
- mix
12.5%
25%
50%
100%
ln(CP10K+1)
Hypothalamus
Pituitary
Brainstem
Brain cortex
Testes
Uterus
Bladder
Kidney
Pancreas
Liver
Colon
Small intestine
Tongue
Trachea
Lung
Aorta
Heart
Blood / Spleen
Bone marrow
Bone
Diaphragm
Limb muscle
Mammary gland
Fat

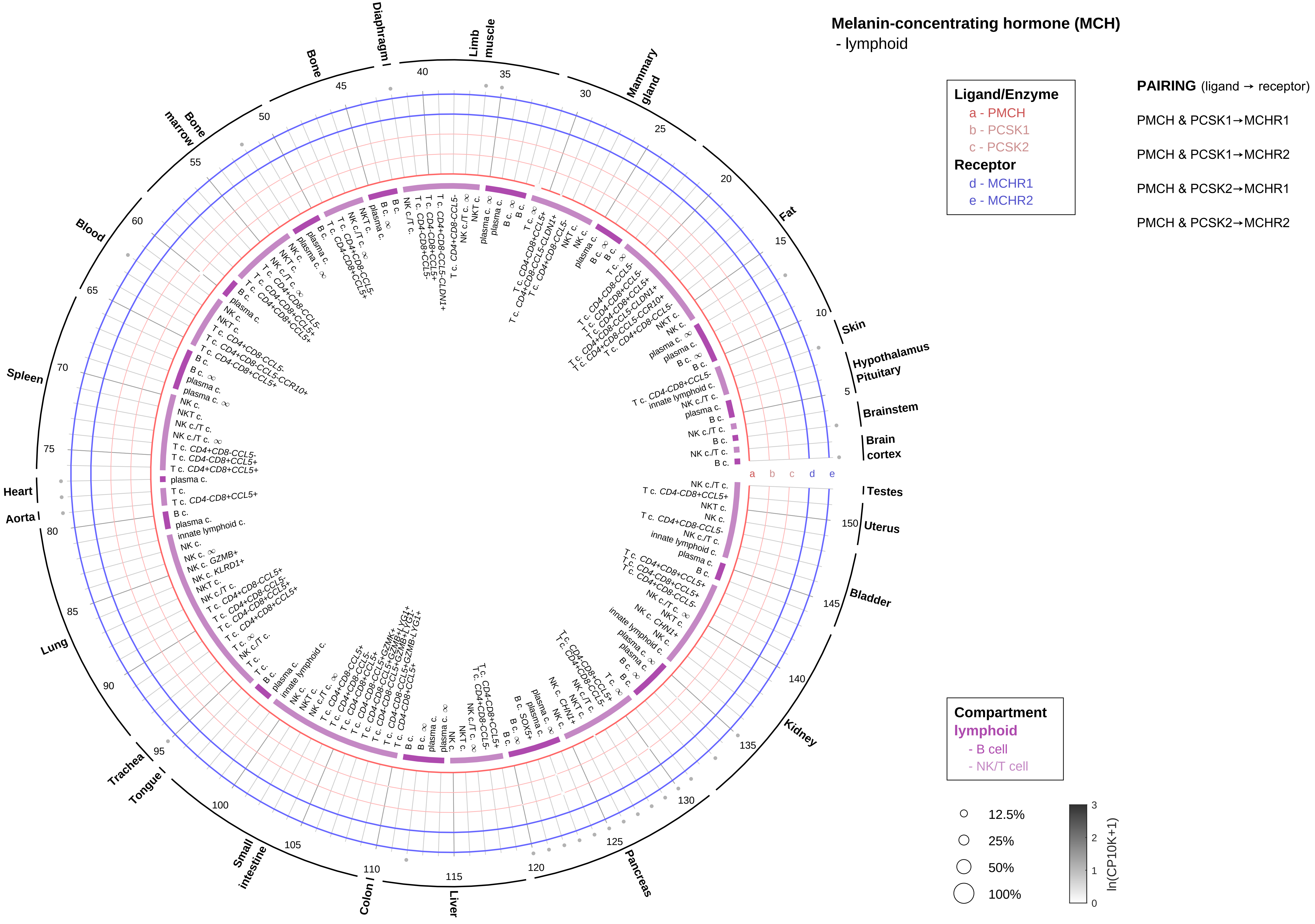
Melanin-concentrating hormone (MCH)
- lymphoid
Ligand/Enzyme
a - PMCH
b - PCSK1
c - PCSK2
Receptor
d - MCHR1
e - MCHR2
PAIRING (ligand → receptor)
PMCH & PCSK1→MCHR1
PMCH & PCSK1→MCHR2
PMCH & PCSK2→MCHR1
PMCH & PCSK2→MCHR2
Compartment
lymphoid
- B cell
- NK/T cell
12.5%
25%
50%
100%
ln(CP10K+1)
Brain cortex
Brainstem
Hypothalamus
Pituitary
Skin
Fat
Mammary gland
Limb muscle
Diaphragm
Bone
Bone marrow
Blood
Spleen
Heart
Aorta
Lung
Trachea
Tongue
Small intestine
Colon
Liver
Pancreas
Kidney
Bladder
Uterus
Testes

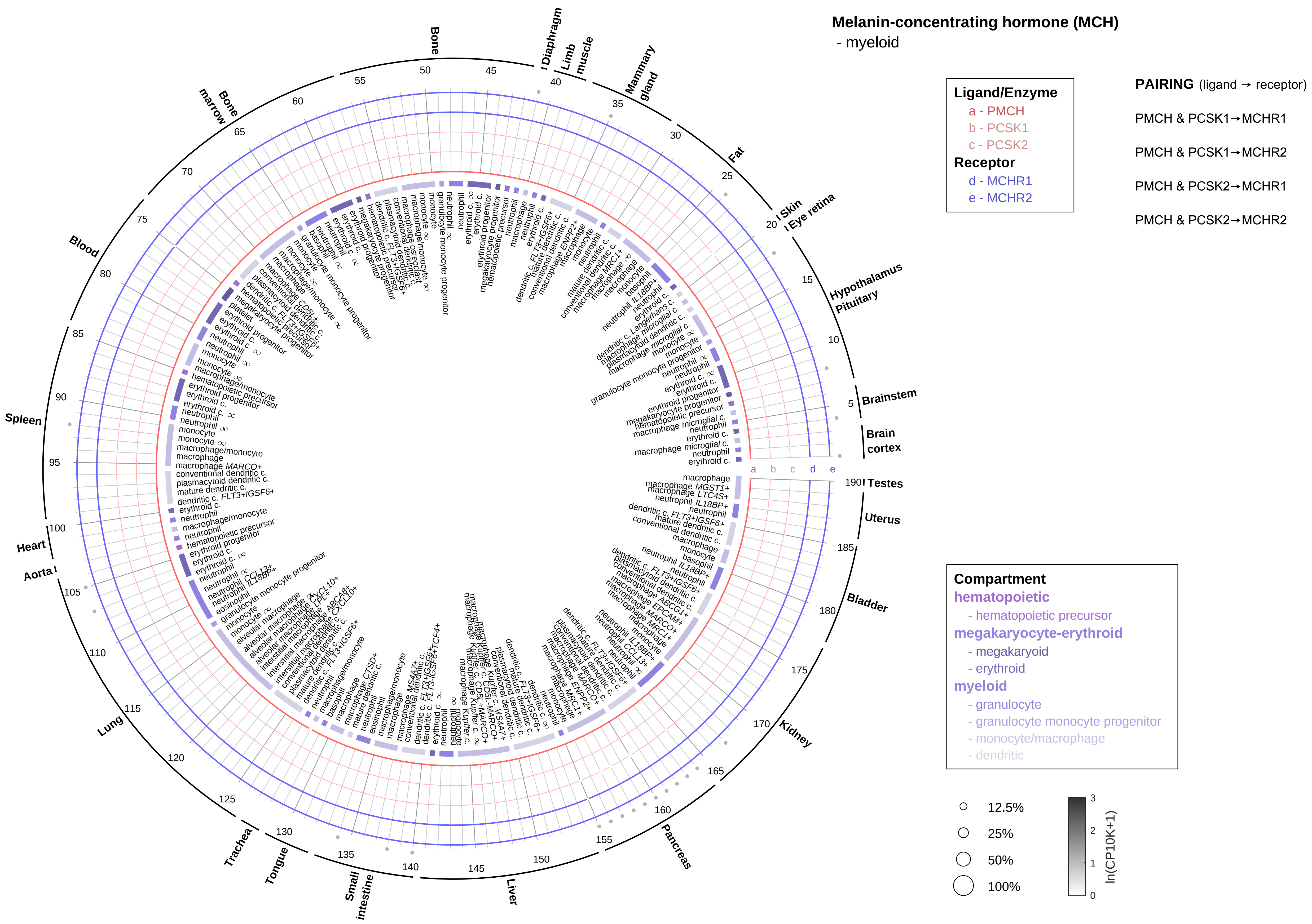

Melanin-concentrating hormone (MCH)
- myeloid
Ligand/Enzyme
a - PMCH
b - PCSK1
c - PCSK2
Receptor
d - MCHR1
e - MCHR2
PAIRING (ligand → receptor)
PMCH & PCSK1→MCHR1
PMCH & PCSK1→MCHR2
PMCH & PCSK2→MCHR1
PMCH & PCSK2→MCHR2
Compartment
hematopoietic
- hematopoietic precursor
megakaryocyte-erythroid
- megakaryoid
- erythroid
myeloid
- granulocyte
- granulocyte monocyte progenitor
- monocyte/macrophage
- dendritic
12.5%
25%
50%
100%
ln(CP10K+1)
Bone
Diaphragm
Limb muscle
Mammary gland
Fat
Skin
Eye retina
Hypothalamus
Pituitary
Brainstem
Brain cortex
Testes
Uterus
Bladder
Kidney
Pancreas
Liver
Small intestine
Tongue
Trachea
Lung
Aorta
Heart
Spleen
Blood
Bone marrow

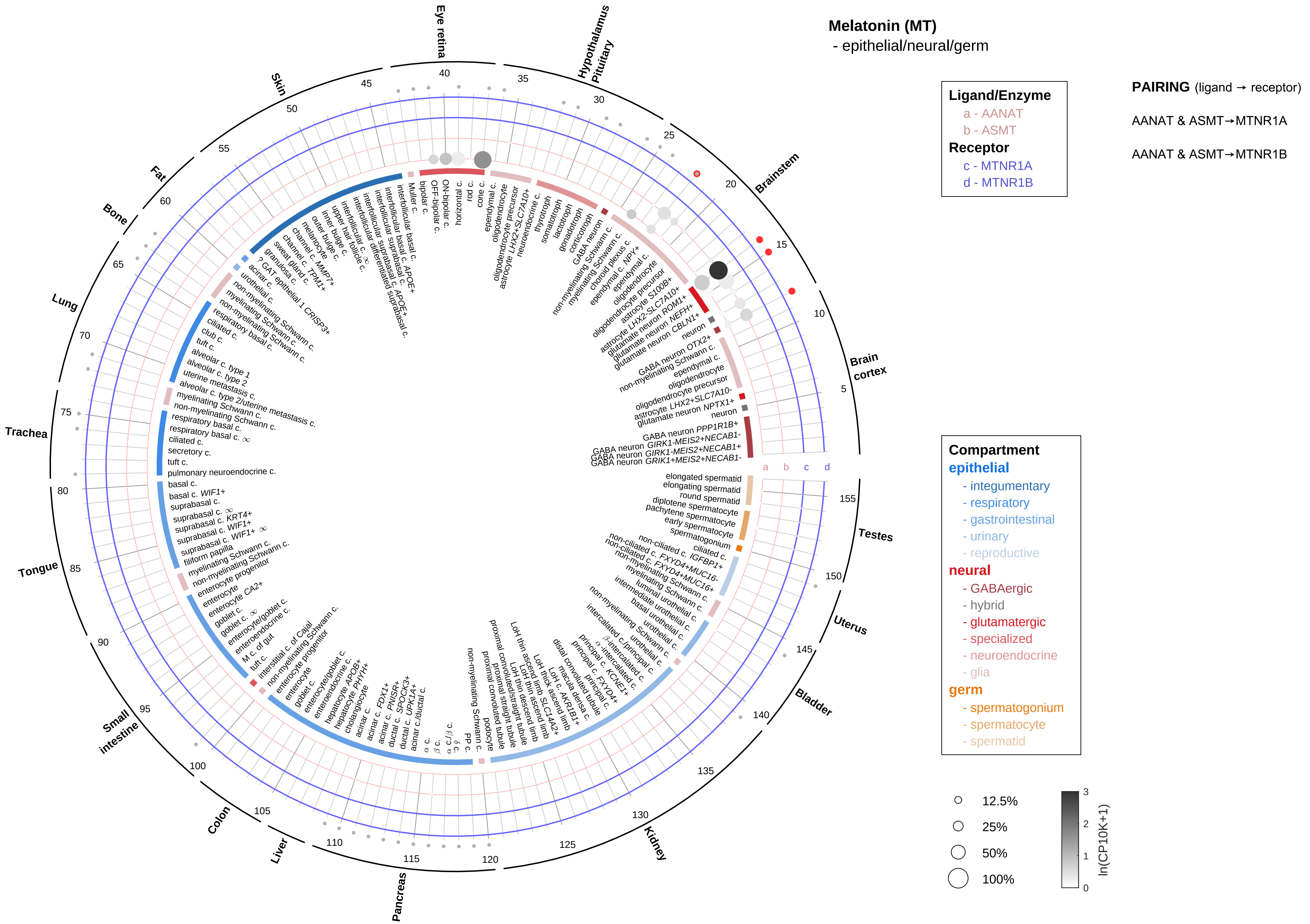
Melatonin (MT)
- epithelial/neural/germ
Ligand/Enzyme
a - AANAT
b - ASMT
Receptor
c - MTNR1A
d - MTNR1B
PAIRING (ligand → receptor)
AANAT & ASMT→MTNR1A
AANAT & ASMT→MTNR1B
Compartment
epithelial
- integumentary
- respiratory
- gastrointestinal
- urinary
- reproductive
neural
- GABAergic
- hybrid
- glutamatergic
- specialized
- neuroendocrine
- glia
germ
- spermatogonium
- spermatocyte
- spermatid
12.5%
25%
50%
100%
ln(CP10K+1)
Eye retina
Hypothalamus
Pituitary
Brainstem
Brain cortex
Testes
Uterus
Bladder
Kidney
Pancreas
Liver
Colon
Small intestine
Tongue
Trachea
Lung
Bone
Fat
Skin

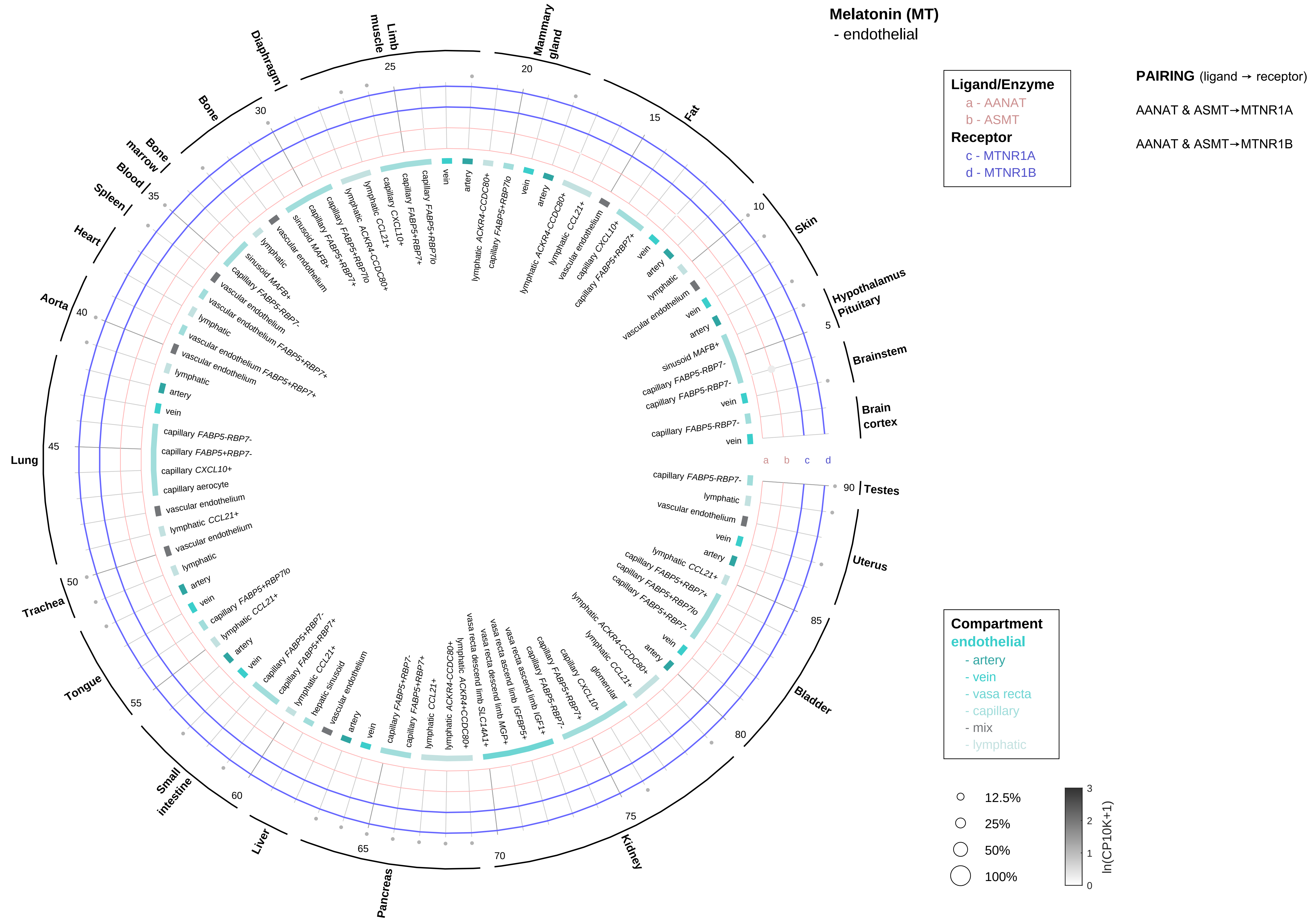
Melatonin (MT)
- endothelial
Ligand/Enzyme
a - AANAT
b - ASMT
Receptor
c - MTNR1A
d - MTNR1B
PAIRING (ligand → receptor)
AANAT & ASMT→MTNR1A
AANAT & ASMT→MTNR1B
Compartment
endothelial
- artery
- vein
- vasa recta
- capillary
- mix
- lymphatic
12.5%
25%
50%
100%
ln(CP10K+1)
Skin
Hypothalamus Pituitary
Brainstem
Brain cortex
Testes
Uterus
Bladder
Kidney
Pancreas
Liver
Small intestine
Tongue
Trachea
Lung
Aorta
Heart
Spleen
Blood
Bone marrow
Bone
Diaphragm
Limb muscle
Mammary gland
Fat
a
b
c
d

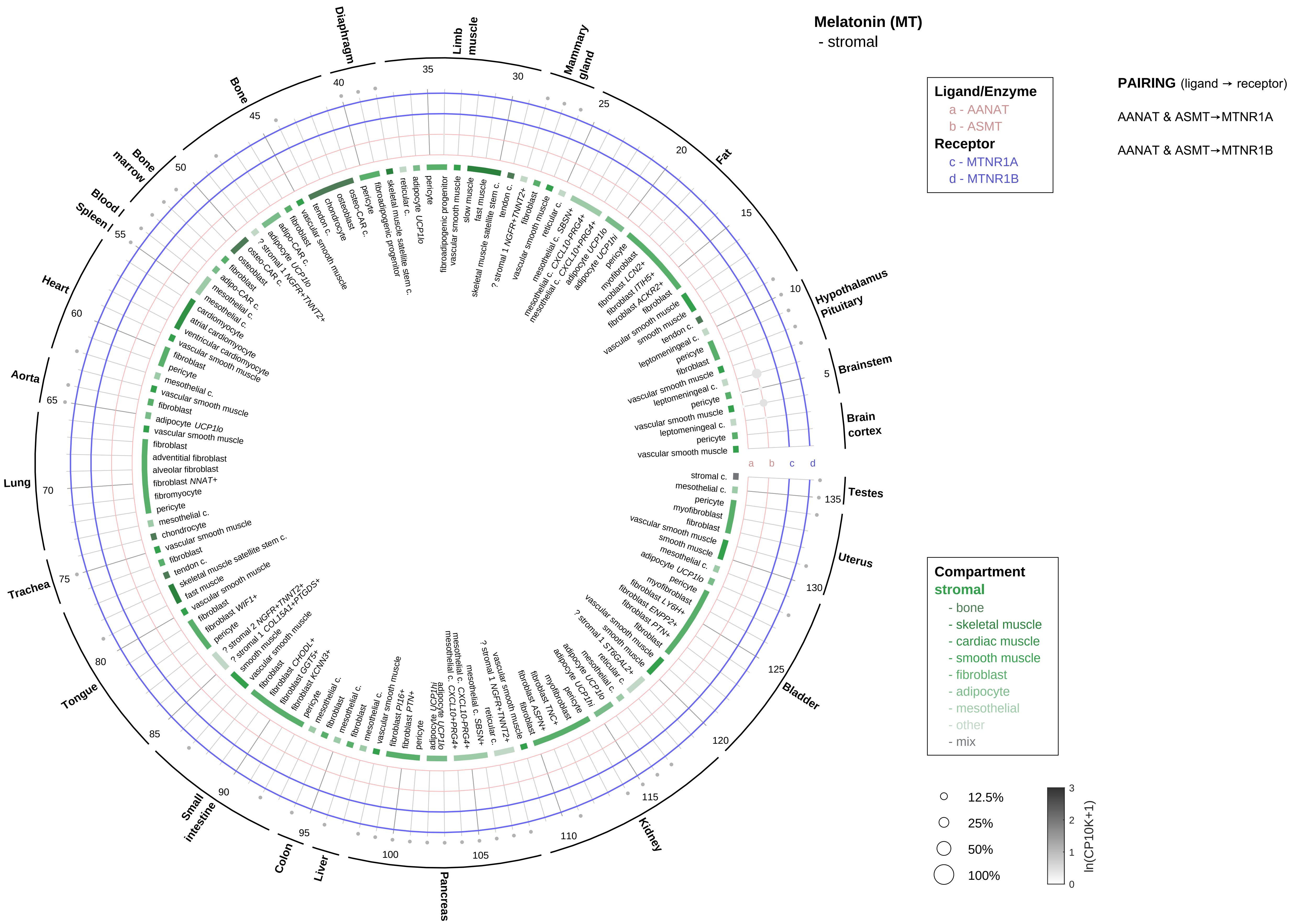

Melatonin (MT)
- stromal
Ligand/Enzyme
a - AANAT
b - ASMT
Receptor
c - MTNR1A
d - MTNR1B
PAIRING (ligand → receptor)
AANAT & ASMT→MTNR1A
AANAT & ASMT→MTNR1B
Compartment
stromal
- bone
- skeletal muscle
- cardiac muscle
- smooth muscle
- fibroblast
- adipocyte
- mesothelial
- other
- mix
12.5%
25%
50%
100%
ln(CP10K+1)
Hypothalamus
Pituitary
Brainstem
Brain cortex
Testes
Uterus
Bladder
Kidney
Pancreas
Liver
Colon
Small intestine
Tongue
Trachea
Lung
Aorta
Heart
Spleen
Blood
Bone marrow
Bone
Diaphragm
Limb muscle
Mammary gland
Fat

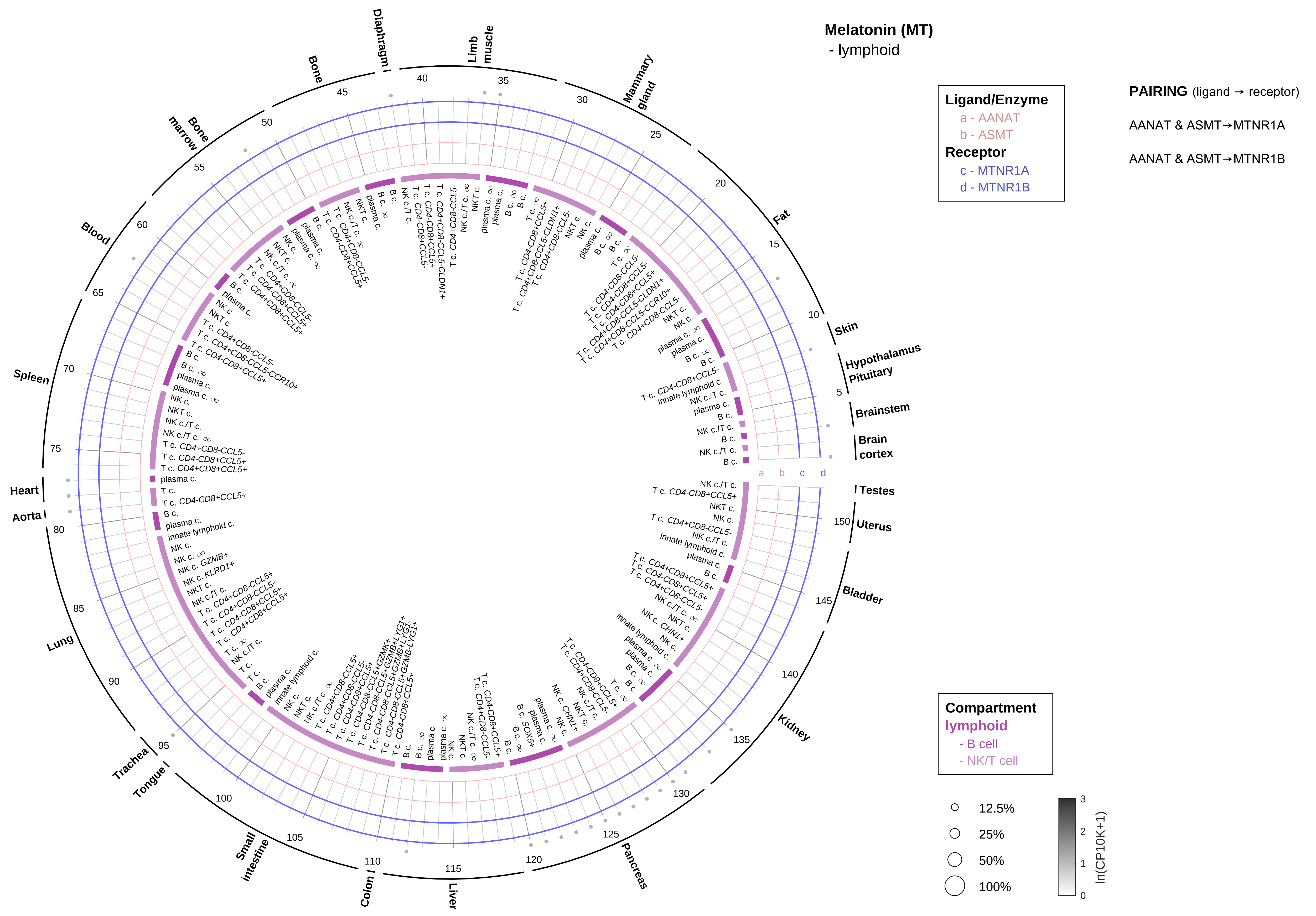

Melatonin (MT)
- lymphoid
Ligand/Enzyme
a - AANAT
b - ASMT
Receptor
c - MTNR1A
d - MTNR1B
PAIRING (ligand → receptor)
AANAT & ASMT→MTNR1A
AANAT & ASMT→MTNR1B
Compartment
lymphoid
- B cell
- NK/T cell
12.5%
25%
50%
100%
ln(CP10K+1)
Skin
Hypothalamus
Pituitary
Brainstem
Brain cortex
Testes
Uterus
Bladder
Kidney
Pancreas
Liver
Colon
Small intestine
Tongue
Trachea
Lung
Aorta
Heart
Spleen
Blood
Bone marrow
Bone
Diaphragm
Limb muscle
Mammary gland
Fat

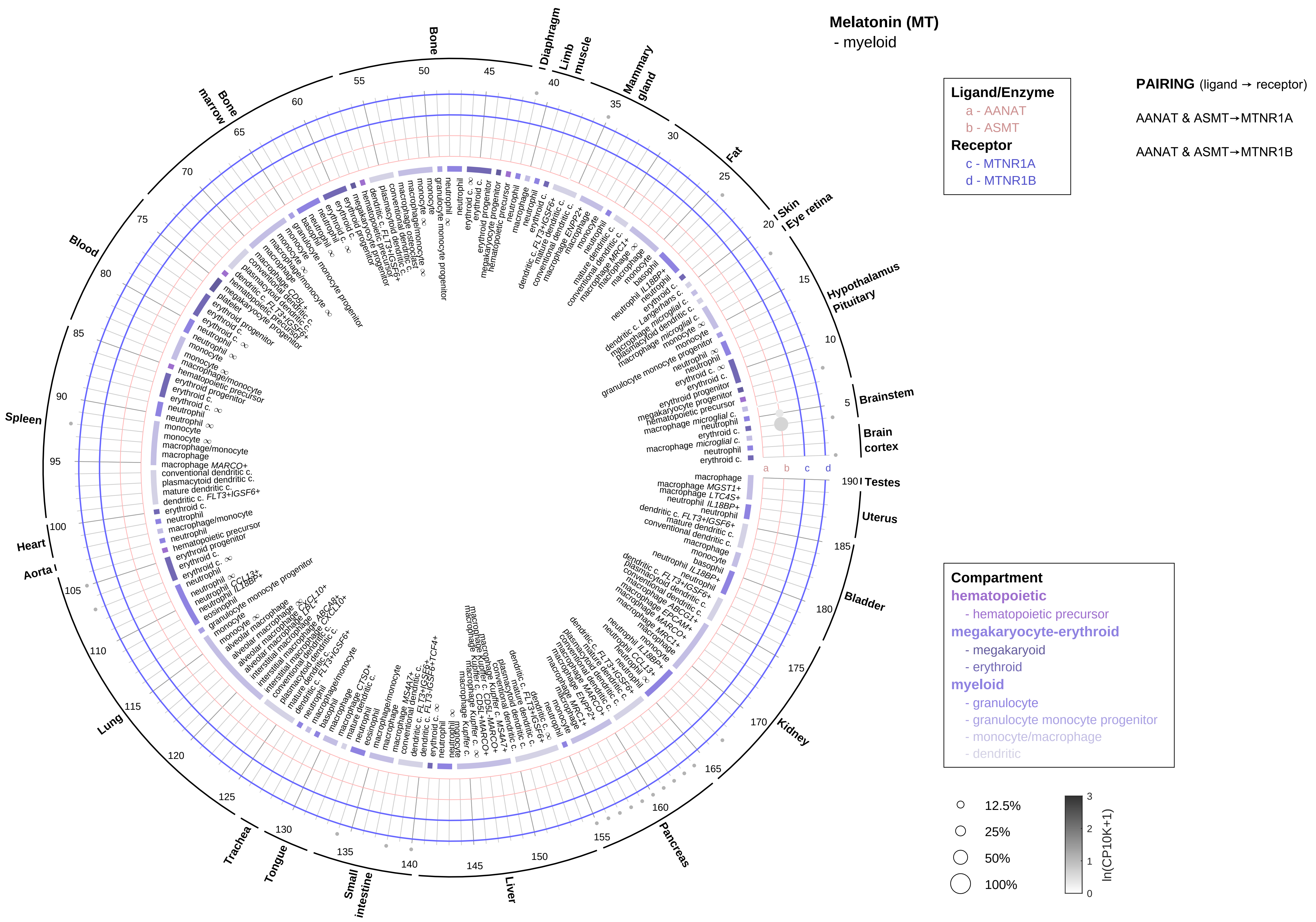

Melatonin (MT)
- myeloid
Ligand/Enzyme
a - AANAT
b - ASMT
Receptor
c - MTNR1A
d - MTNR1B
PAIRING (ligand → receptor)
AANAT & ASMT→MTNR1A
AANAT & ASMT→MTNR1B
Compartment
hematopoietic
- hematopoietic precursor
megakaryocyte-erythroid
- megakaryoid
- erythroid
myeloid
- granulocyte
- granulocyte monocyte progenitor
- monocyte/macrophage
- dendritic
12.5%
25%
50%
100%
ln(CP10K+1)
0
1
2
3
Bone
Diaphragm
Limb muscle
Mammary gland
Fat
Skin
Eye retina
Hypothalamus
Pituitary
Brainstem
Brain cortex
Testes
Uterus
Bladder
Kidney
Pancreas
Liver
Small intestine
Tongue
Trachea
Lung
Aorta
Heart
Spleen
Blood
Bone marrow
a
b
c
d

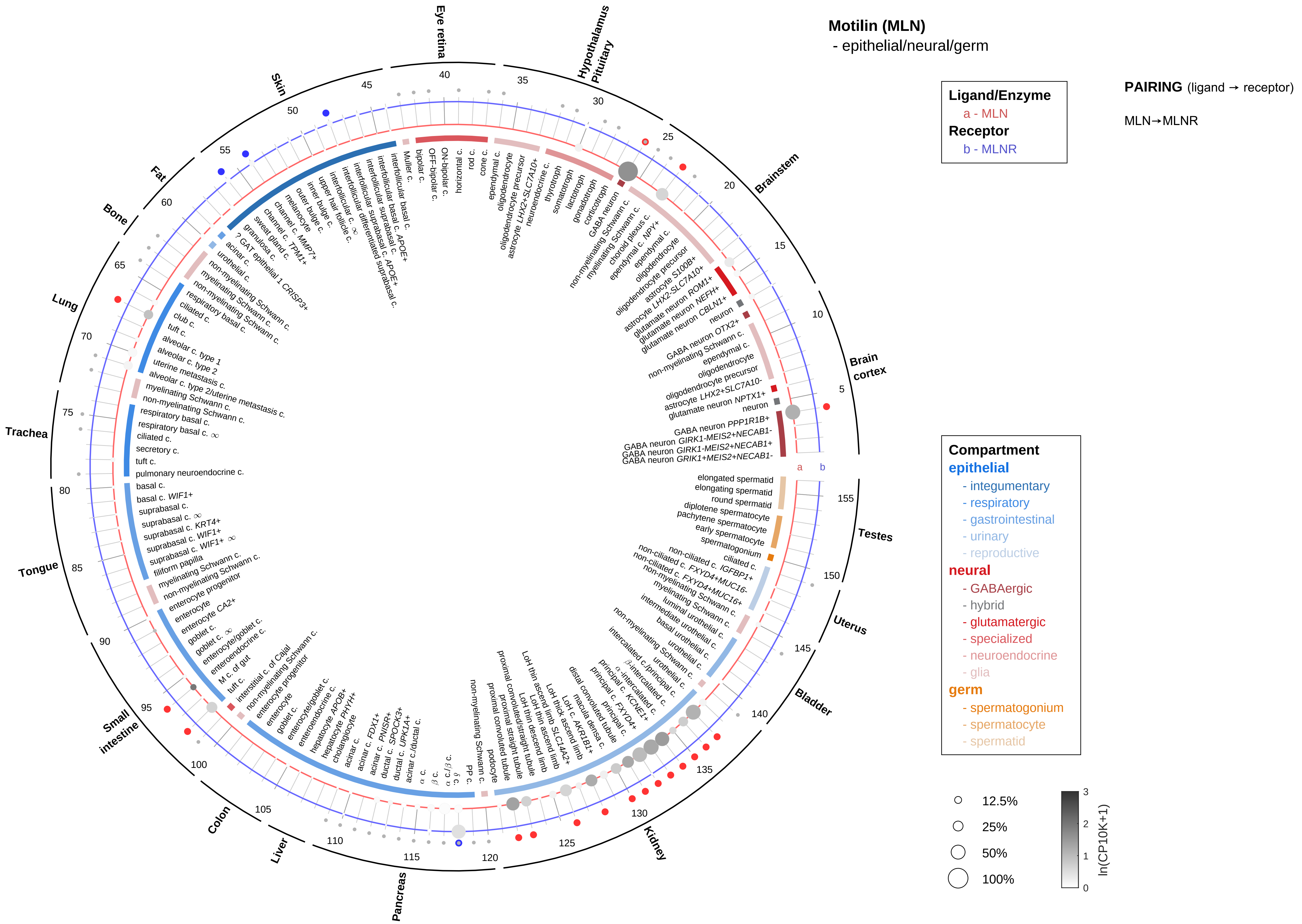
Motilin (MLN)
- epithelial/neural/germ
Ligand/Enzyme
a - MLN
Receptor
b - MLNR
PAIRING (ligand → receptor)
MLN→MLNR
Compartment
epithelial
- integumentary
- respiratory
- gastrointestinal
- urinary
- reproductive
neural
- GABAergic
- hybrid
- glutamatergic
- specialized
- neuroendocrine
- glia
germ
- spermatogonium
- spermatocyte
- spermatid
12.5%
25%
50%
100%
ln(CP10K+1)
Brain cortex
Brainstem
Hypothalamus Pituitary
Eye retina
Skin
Fat
Bone
Lung
Trachea
Tongue
Small intestine
Colon
Liver
Pancreas
Kidney
Bladder
Uterus
Testes

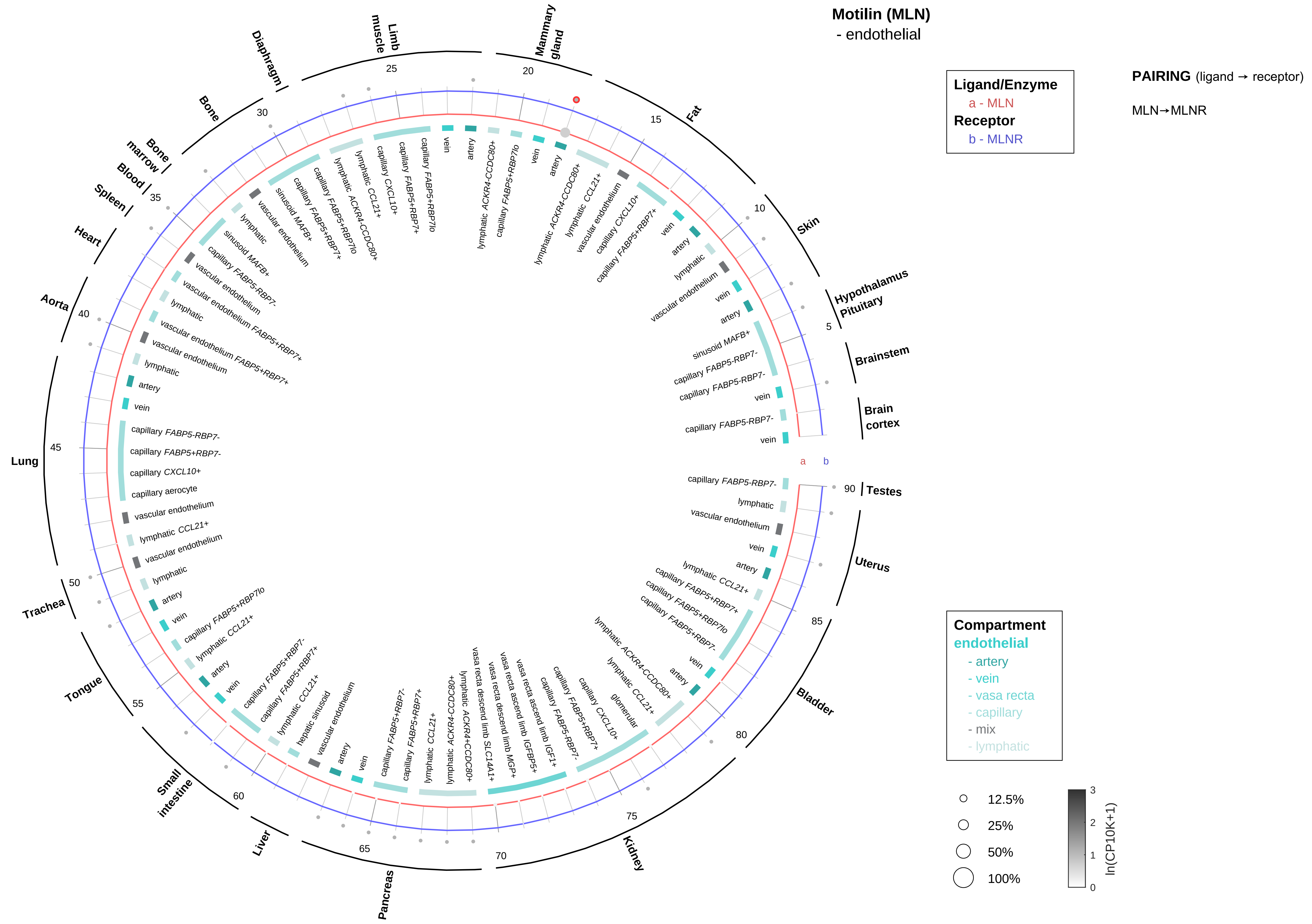

Motilin (MLN)
- endothelial
Ligand/Enzyme
a - MLN
Receptor
b - MLNR
PAIRING (ligand → receptor)
MLN→MLNR
Compartment
endothelial
- artery
- vein
- vasa recta
- capillary
- mix
- lymphatic
12.5%
25%
50%
100%
ln(CP10K+1)
Testes
Uterus
Bladder
Kidney
Pancreas
Liver
Small intestine
Tongue
Trachea
Lung
Aorta
Heart
Spleen
Blood
Bone marrow
Bone
Diaphragm
Limb muscle
Mammary gland
Fat
Skin
Hypothalamus Pituitary
Brainstem
Brain cortex

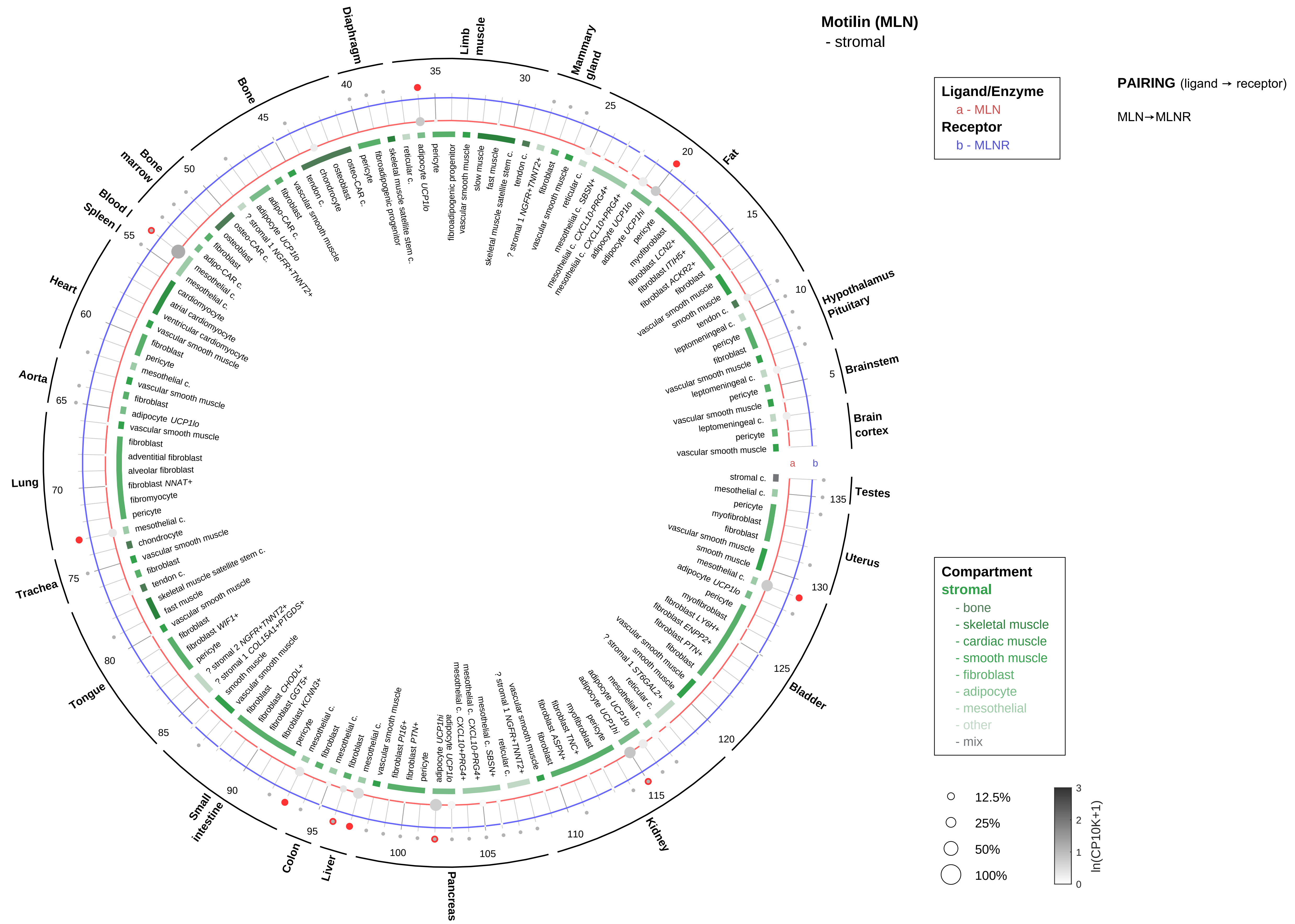
Motilin (MLN)
- stromal
PAIRING (ligand → receptor)
MLN→MLNR
Ligand/Enzyme
a - MLN
Receptor
b - MLNR
Compartment
stromal
- bone
- skeletal muscle
- cardiac muscle
- smooth muscle
- fibroblast
- adipocyte
- mesothelial
- other
- mix
12.5%
25%
50%
100%
ln(CP10K+1)
Diaphragm
Limb muscle
Mammary gland
Fat
Hypothalamus Pituitary
Brainstem
Brain cortex
Testes
Uterus
Bladder
Kidney
Pancreas
Liver
Colon
Small intestine
Tongue
Trachea
Lung
Aorta
Heart
Blood / Spleen
Bone marrow
Bone

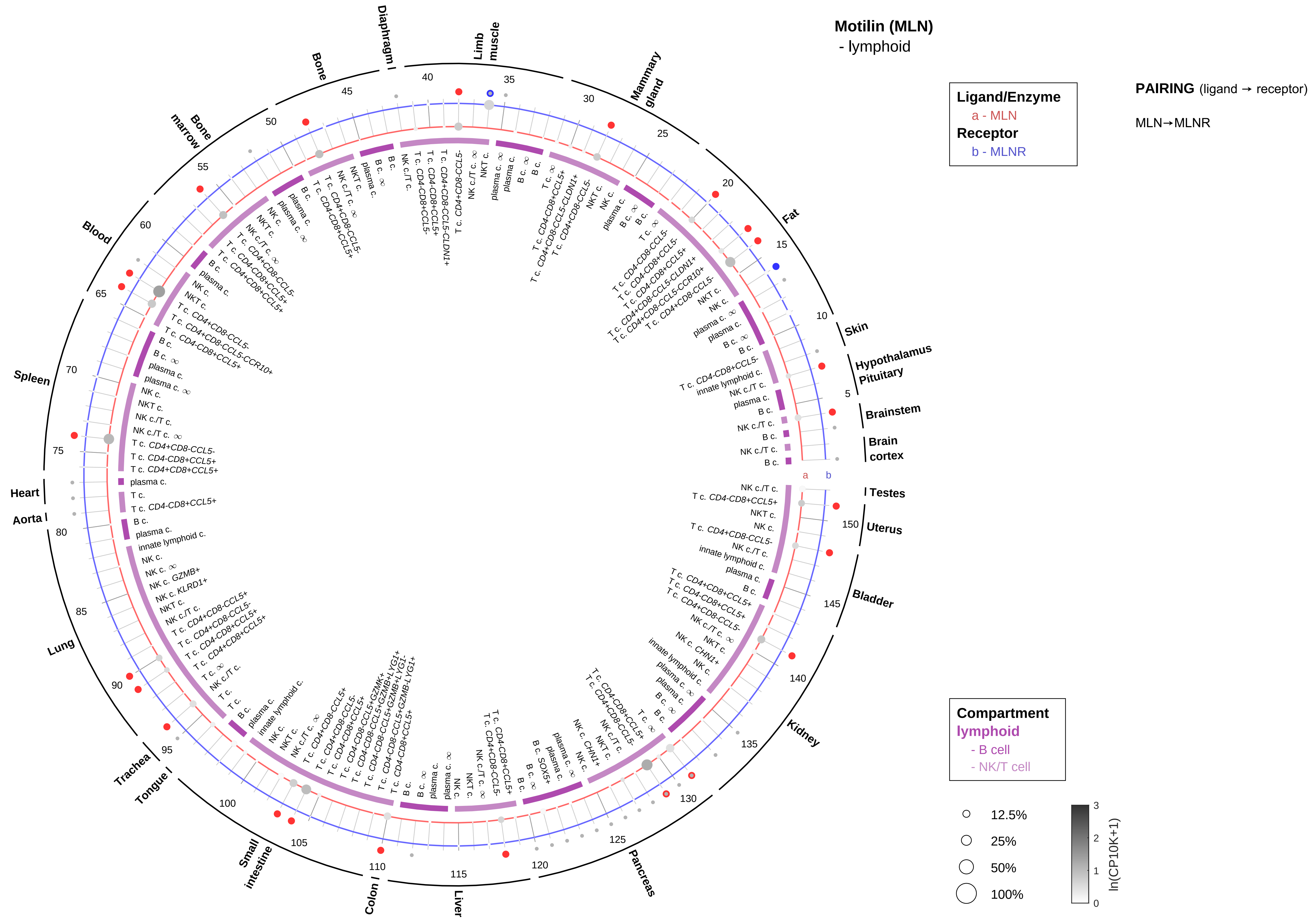

Motilin (MLN)
- lymphoid
Ligand/Enzyme
a - MLN
Receptor
b - MLNR
PAIRING (ligand → receptor)
MLN→MLNR
Compartment
lymphoid
- B cell
- NK/T cell
12.5%
25%
50%
100%
ln(CP10K+1)
Limb muscle
Mammary gland
Fat
Skin
Hypothalamus Pituitary
Brainstem
Brain cortex
Testes
Uterus
Bladder
Kidney
Pancreas
Liver
Colon
Small intestine
Tongue
Trachea
Lung
Aorta
Heart
Spleen
Blood
Bone marrow
Bone
Diaphragm

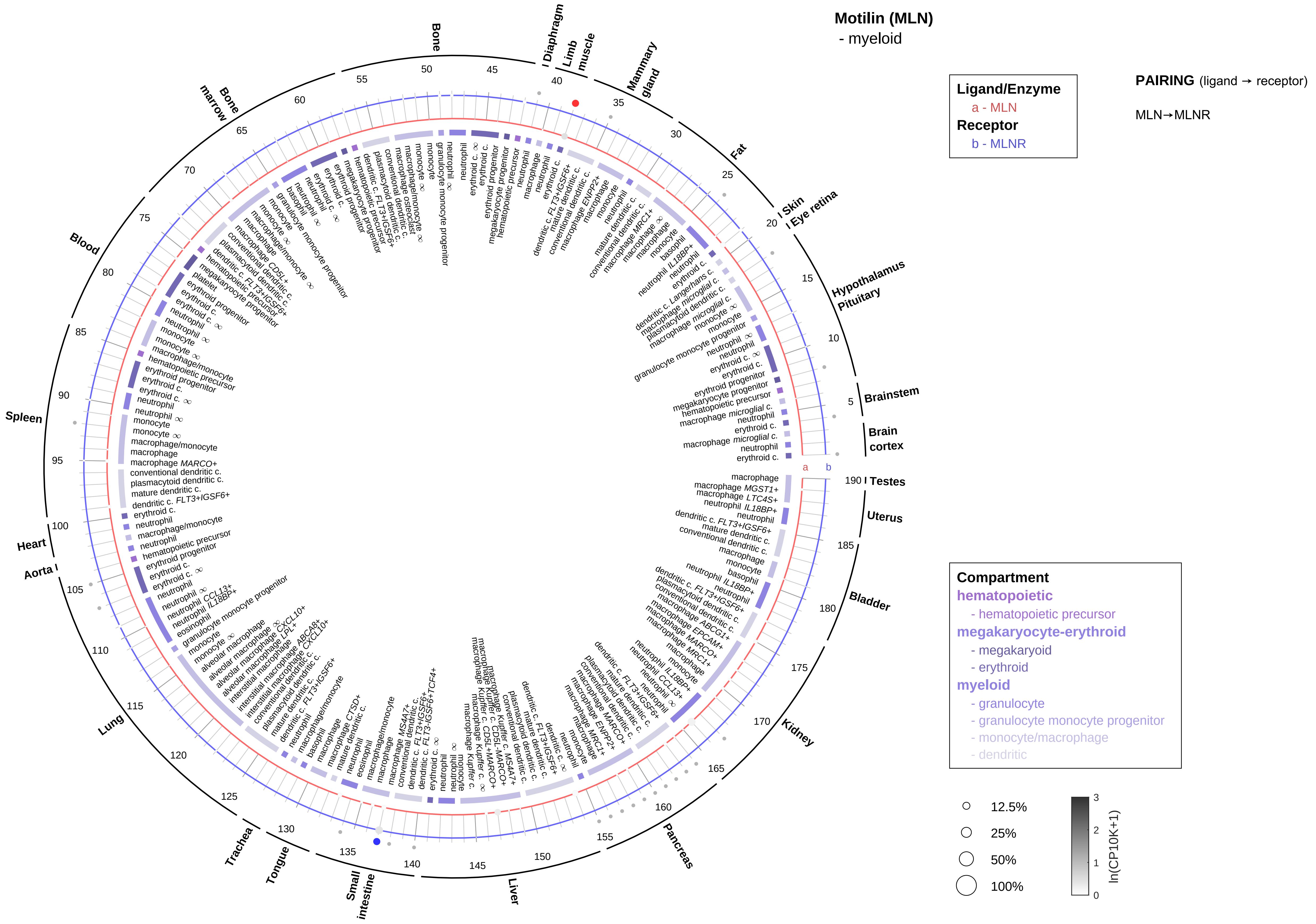
Motilin (MLN)
- myeloid
Ligand/Enzyme
a - MLN
Receptor
b - MLNR
PAIRING (ligand → receptor)
MLN→MLNR
Compartment
hematopoietic
- hematopoietic precursor
megakaryocyte-erythroid
- megakaryoid
- erythroid
myeloid
- granulocyte
- granulocyte monocyte progenitor
- monocyte/macrophage
- dendritic
12.5%
25%
50%
100%
ln(CP10K+1)
Bone
Diaphragm
Limb muscle
Mammary gland
Fat
Skin
Eye retina
Hypothalamus
Pituitary
Brainstem
Brain cortex
Testes
Uterus
Bladder
Kidney
Pancreas
Liver
Small intestine
Tongue
Trachea
Lung
Aorta
Heart
Spleen
Blood
Bone marrow

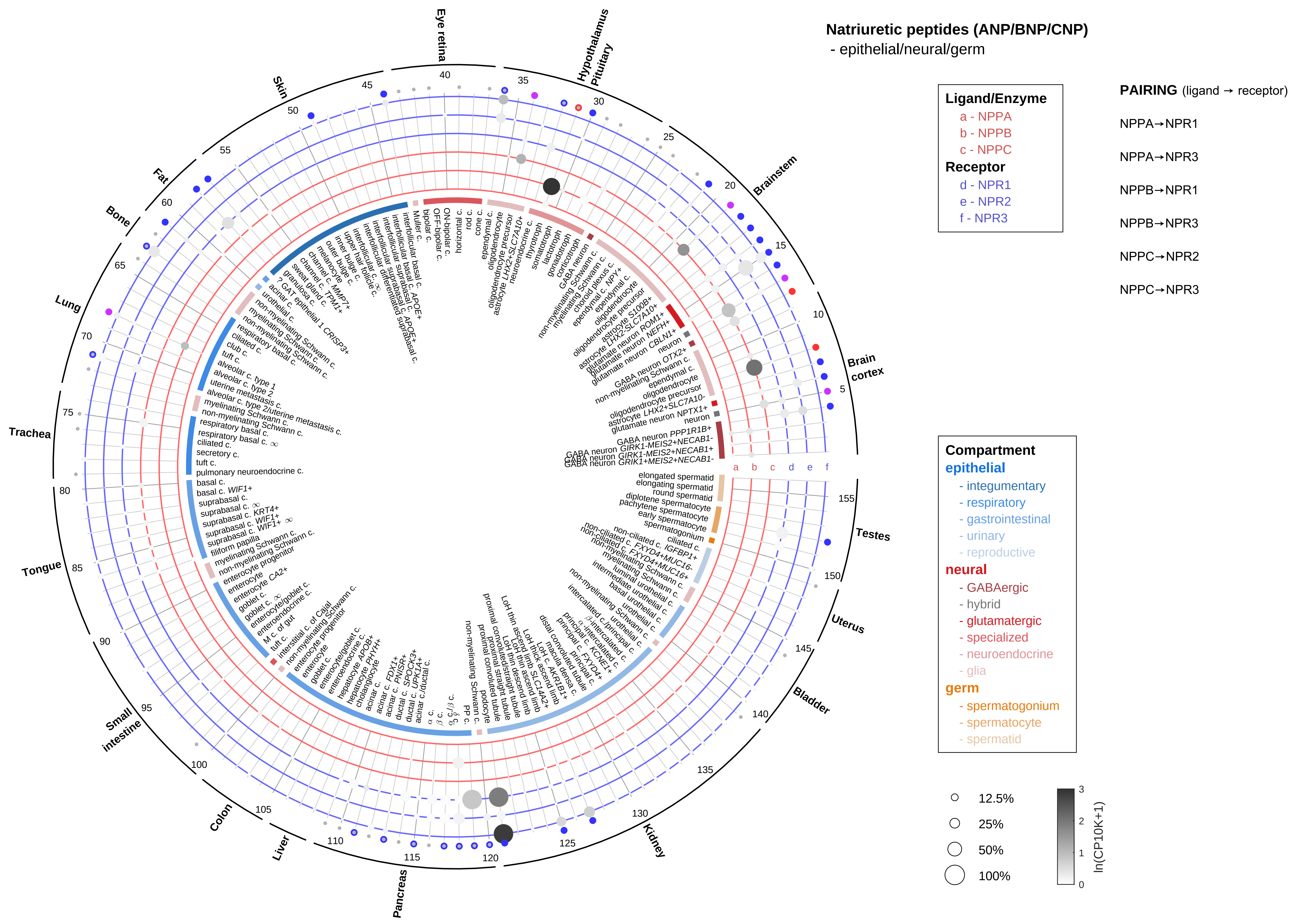

Natriuretic peptides (ANP/BNP/CNP)
- epithelial/neural/germ
Ligand/Enzyme
a - NPPA
b - NPPB
c - NPPC
Receptor
d - NPR1
e - NPR2
f - NPR3
PAIRING (ligand → receptor)
NPPA→NPR1
NPPA→NPR3
NPPB→NPR1
NPPB→NPR3
NPPC→NPR2
NPPC→NPR3
Compartment
epithelial
- integumentary
- respiratory
- gastrointestinal
- urinary
- reproductive
neural
- GABAergic
- hybrid
- glutamatergic
- specialized
- neuroendocrine
- glia
germ
- spermatogonium
- spermatocyte
- spermatid
12.5%
25%
50%
100%
ln(CP10K+1)
Brain cortex
Brainstem
Hypothalamus Pituitary
Eye retina
Skin
Fat
Bone
Lung
Trachea
Tongue
Small intestine
Colon
Liver
Pancreas
Kidney
Bladder
Uterus
Testes
a b c d e f

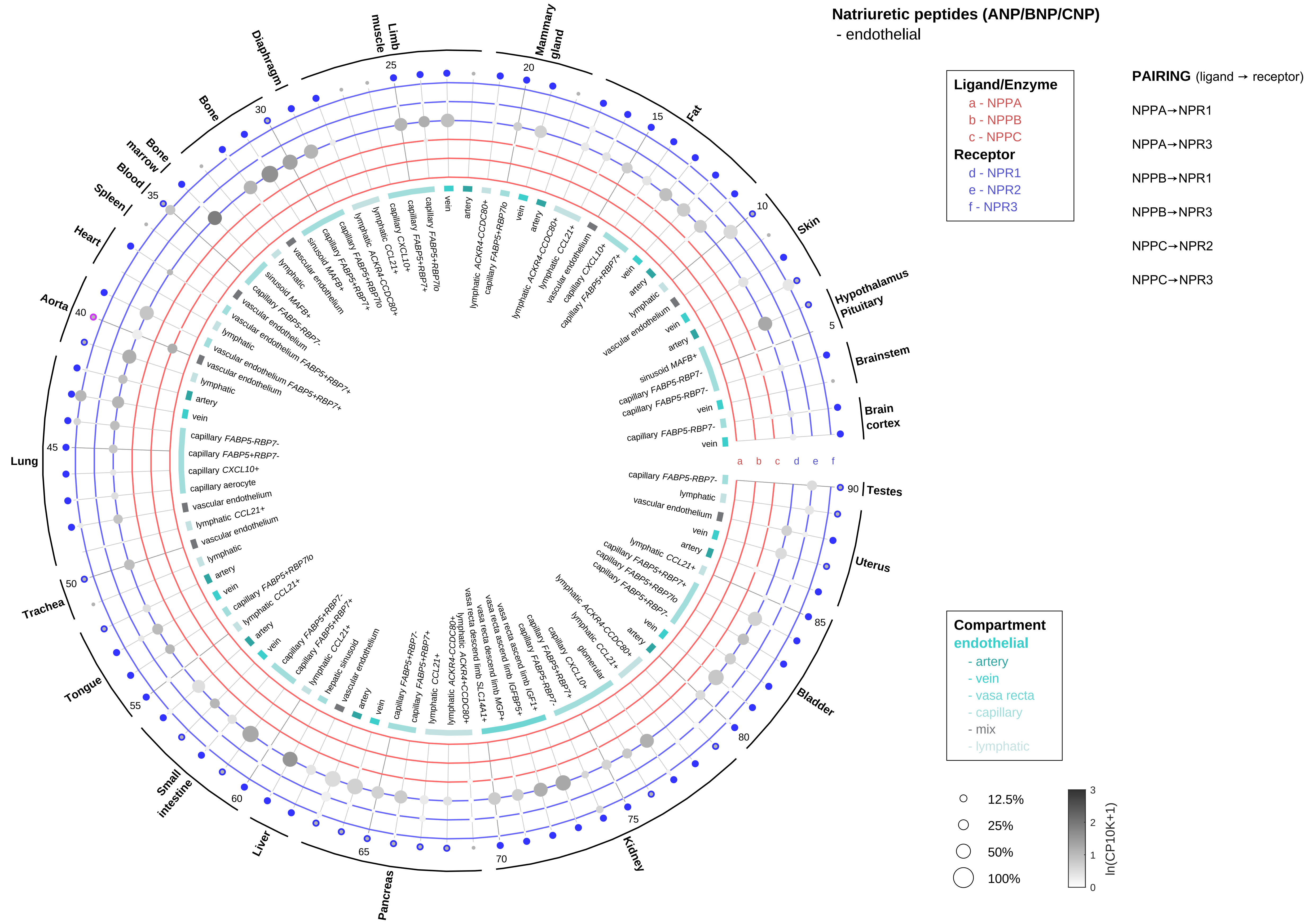

Natriuretic peptides (ANP/BNP/CNP)
- endothelial
Ligand/Enzyme
a - NPPA
b - NPPB
c - NPPC
Receptor
d - NPR1
e - NPR2
f - NPR3
PAIRING (ligand → receptor)
NPPA→NPR1
NPPA→NPR3
NPPB→NPR1
NPPB→NPR3
NPPC→NPR2
NPPC→NPR3
Compartment
endothelial
- artery
- vein
- vasa recta
- capillary
- mix
- lymphatic
12.5%
25%
50%
100%
ln(CP10K+1)
Limb muscle
Mammary gland
Fat
Skin
Hypothalamus Pituitary
Brainstem
Brain cortex
Testes
Uterus
Bladder
Kidney
Pancreas
Liver
Small intestine
Tongue
Trachea
Lung
Aorta
Heart
Spleen
Blood
Bone marrow
Bone
Diaphragm

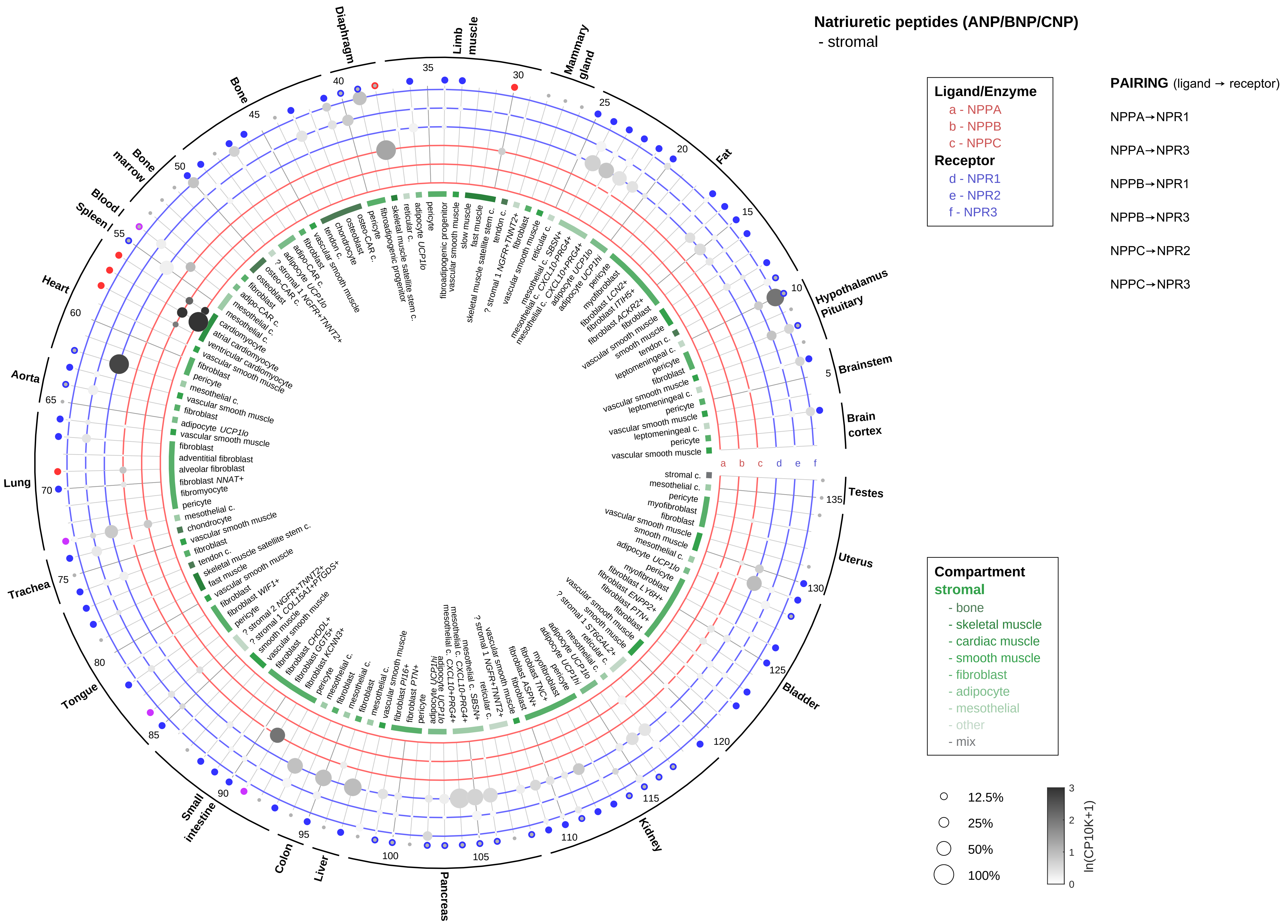
Natriuretic peptides (ANP/BNP/CNP)
- stromal
Ligand/Enzyme
a - NPPA
b - NPPB
c - NPPC
Receptor
d - NPR1
e - NPR2
f - NPR3
PAIRING (ligand → receptor)
NPPA→NPR1
NPPA→NPR3
NPPB→NPR1
NPPB→NPR3
NPPC→NPR2
NPPC→NPR3
Compartment
stromal
- bone
- skeletal muscle
- cardiac muscle
- smooth muscle
- fibroblast
- adipocyte
- mesothelial
- other
- mix
12.5%
25%
50%
100%
ln(CP10K+1)
Diaphragm
Limb muscle
Mammary gland
Fat
Hypothalamus Pituitary
Brainstem
Brain cortex
Testes
Uterus
Bladder
Kidney
Pancreas
Liver
Colon
Small intestine
Tongue
Trachea
Lung
Aorta
Heart
Spleen /
Blood /
Bone marrow
Bone

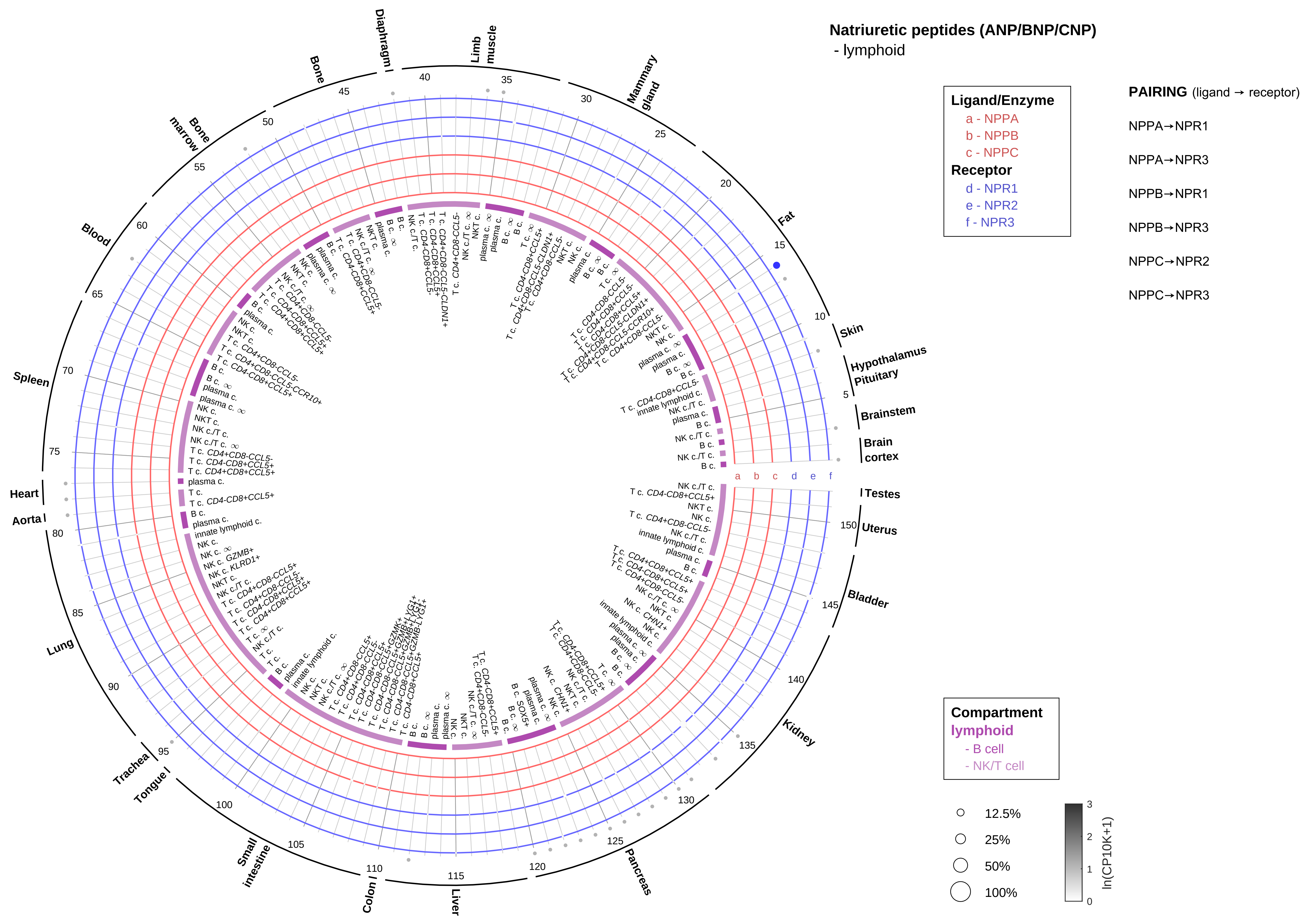

Natriuretic peptides (ANP/BNP/CNP)
- lymphoid
Ligand/Enzyme
a - NPPA
b - NPPB
c - NPPC
Receptor
d - NPR1
e - NPR2
f - NPR3
PAIRING (ligand → receptor)
NPPA→NPR1
NPPA→NPR3
NPPB→NPR1
NPPB→NPR3
NPPC→NPR2
NPPC→NPR3
Compartment
lymphoid
- B cell
- NK/T cell
12.5%
25%
50%
100%
ln(CP10K+1)
Skin
Hypothalamus
Pituitary
Brainstem
Brain cortex
Testes
Uterus
Bladder
Kidney
Pancreas
Liver
Colon
Small intestine
Tongue
Trachea
Lung
Aorta
Heart
Spleen
Blood
Bone marrow
Bone
Diaphragm
Limb muscle
Mammary gland
Fat

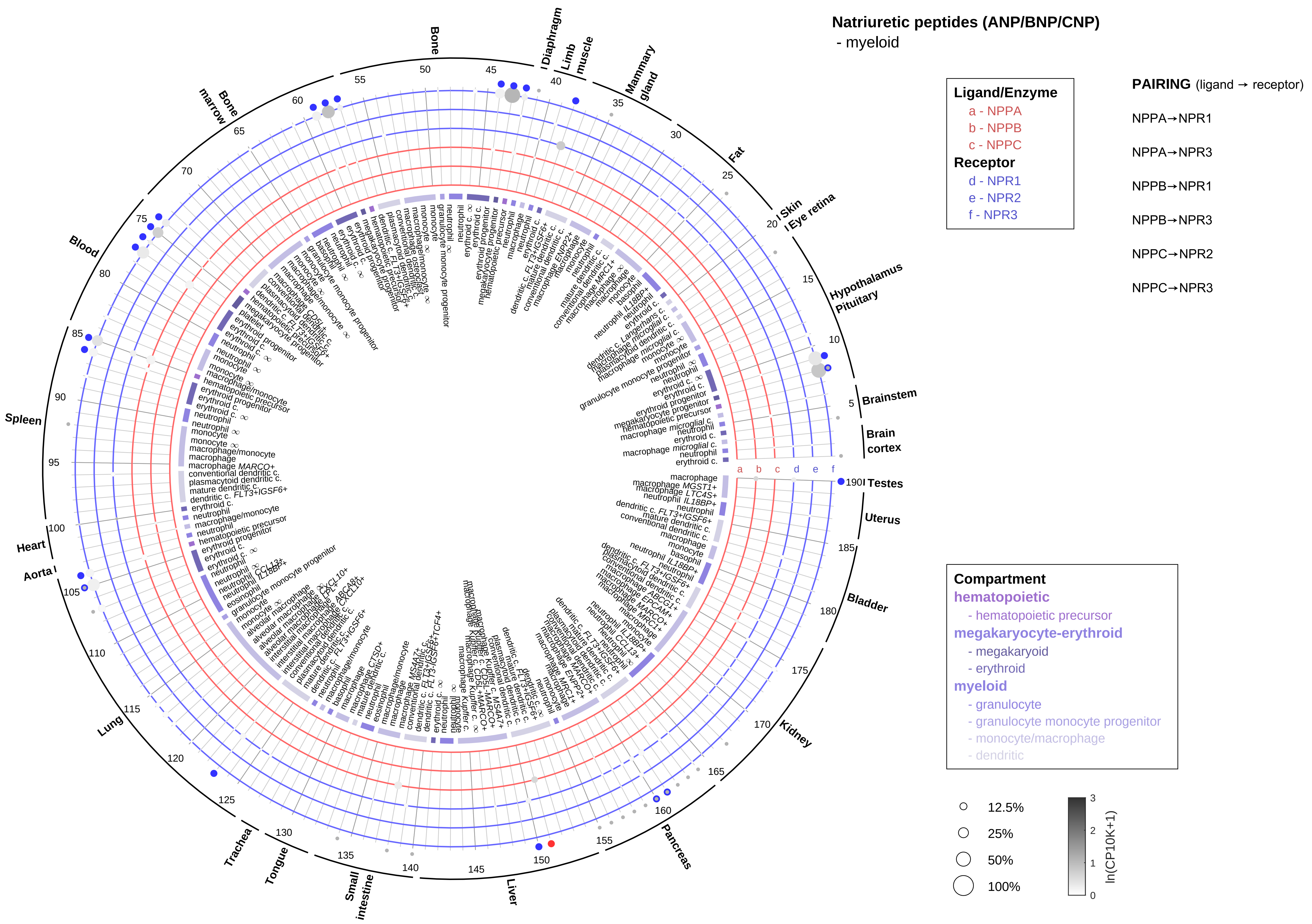
Natriuretic peptides (ANP/BNP/CNP)
- myeloid
Ligand/Enzyme
a - NPPA
b - NPPB
c - NPPC
Receptor
d - NPR1
e - NPR2
f - NPR3
PAIRING (ligand → receptor)
NPPA→NPR1
NPPA→NPR3
NPPB→NPR1
NPPB→NPR3
NPPC→NPR2
NPPC→NPR3
Compartment
hematopoietic
- hematopoietic precursor
megakaryocyte-erythroid
- megakaryocyte
- erythroid
myeloid
- granulocyte
- granulocyte monocyte progenitor
- monocyte/macrophage
- dendritic
12.5%
25%
50%
100%
ln(CP10K+1)
Bone
Diaphragm
Limb muscle
Mammary gland
Fat
Skin
Eye retina
Hypothalamus
Pituitary
Brainstem
Brain cortex
Testes
Uterus
Bladder
Kidney
Pancreas
Liver
Small intestine
Tongue
Trachea
Lung
Aorta
Heart
Spleen
Blood
Bone marrow

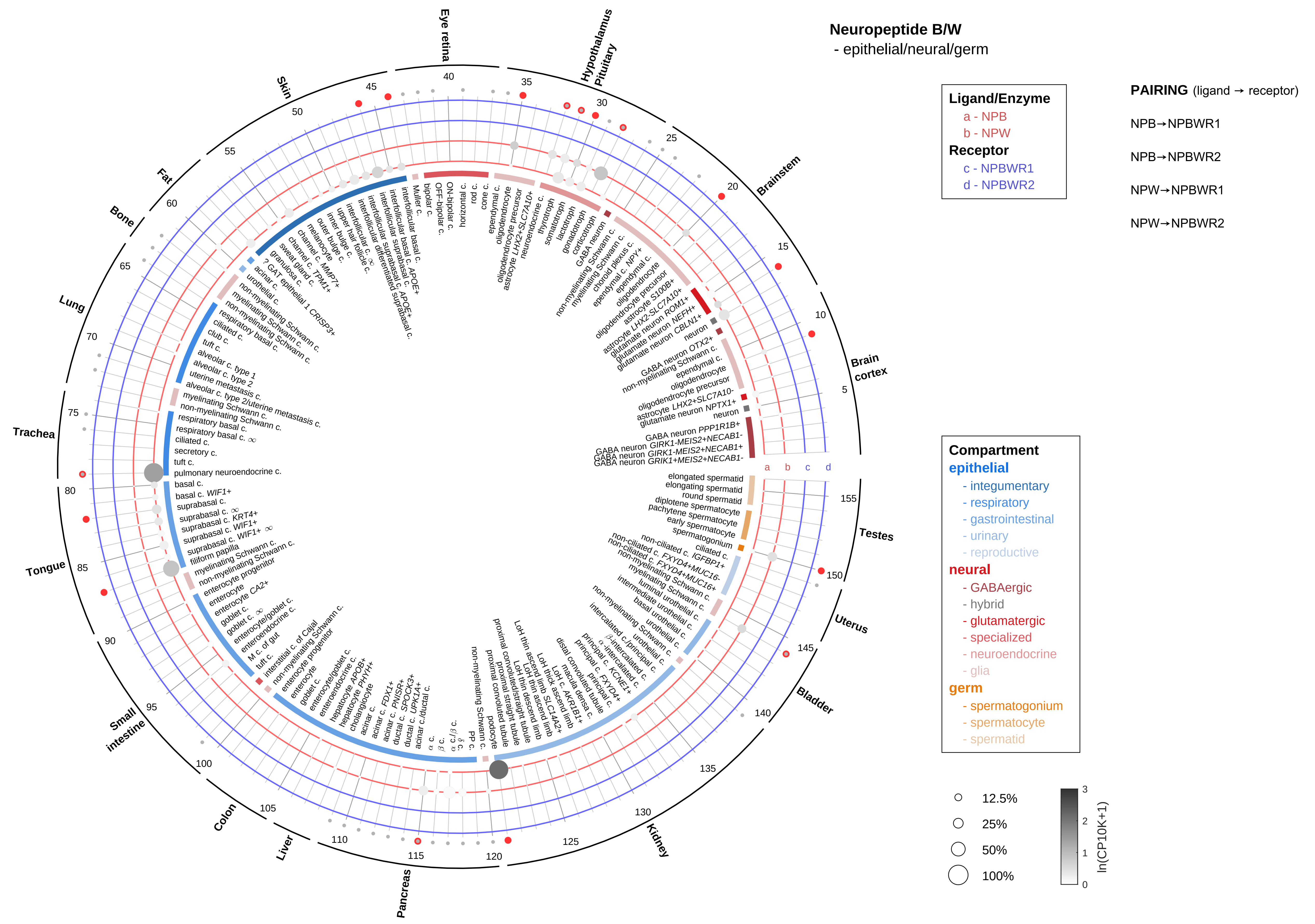

Neuropeptide B/W
- epithelial/neural/germ
Ligand/Enzyme
a - NPB
b - NPW
Receptor
c - NPBWR1
d - NPBWR2
PAIRING (ligand → receptor)
NPB→NPBWR1
NPB→NPBWR2
NPW→NPBWR1
NPW→NPBWR2
Compartment
epithelial
- integumentary
- respiratory
- gastrointestinal
- urinary
- reproductive
neural
- GABAergic
- hybrid
- glutamatergic
- specialized
- neuroendocrine
- glia
germ
- spermatogonium
- spermatocyte
- spermatid
12.5%
25%
50%
100%
ln(CP10K+1)
Brain cortex
Brainstem
Hypothalamus Pituitary
Eye retina
Skin
Fat
Bone
Lung
Trachea
Tongue
Small intestine
Colon
Liver
Pancreas
Kidney
Bladder
Uterus
Testes

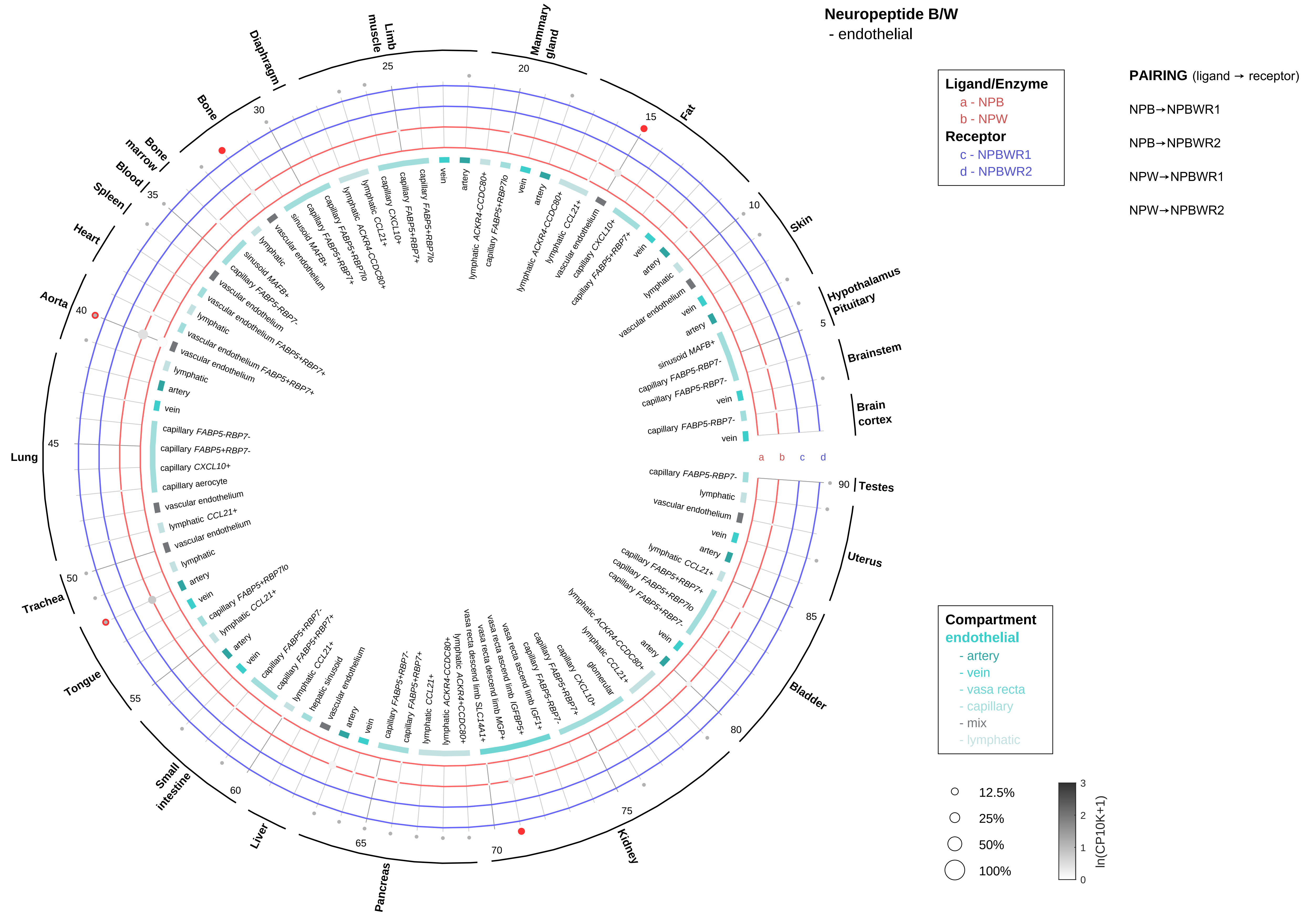

Neuropeptide B/W
- endothelial
Ligand/Enzyme
a - NPB
b - NPW
Receptor
c - NPBWR1
d - NPBWR2
PAIRING (ligand → receptor)
NPB→NPBWR1
NPB→NPBWR2
NPW→NPBWR1
NPW→NPBWR2
Compartment
endothelial
- artery
- vein
- vasa recta
- capillary
- mix
- lymphatic
12.5%
25%
50%
100%
ln(CP10K+1)
Hypothalamus Pituitary
Brainstem
Brain cortex
Skin
Fat
Mammary gland
Limb muscle
Diaphragm
Bone
Bone marrow
Blood
Spleen
Heart
Aorta
Lung
Trachea
Tongue
Small intestine
Liver
Pancreas
Kidney
Bladder
Uterus
Testes

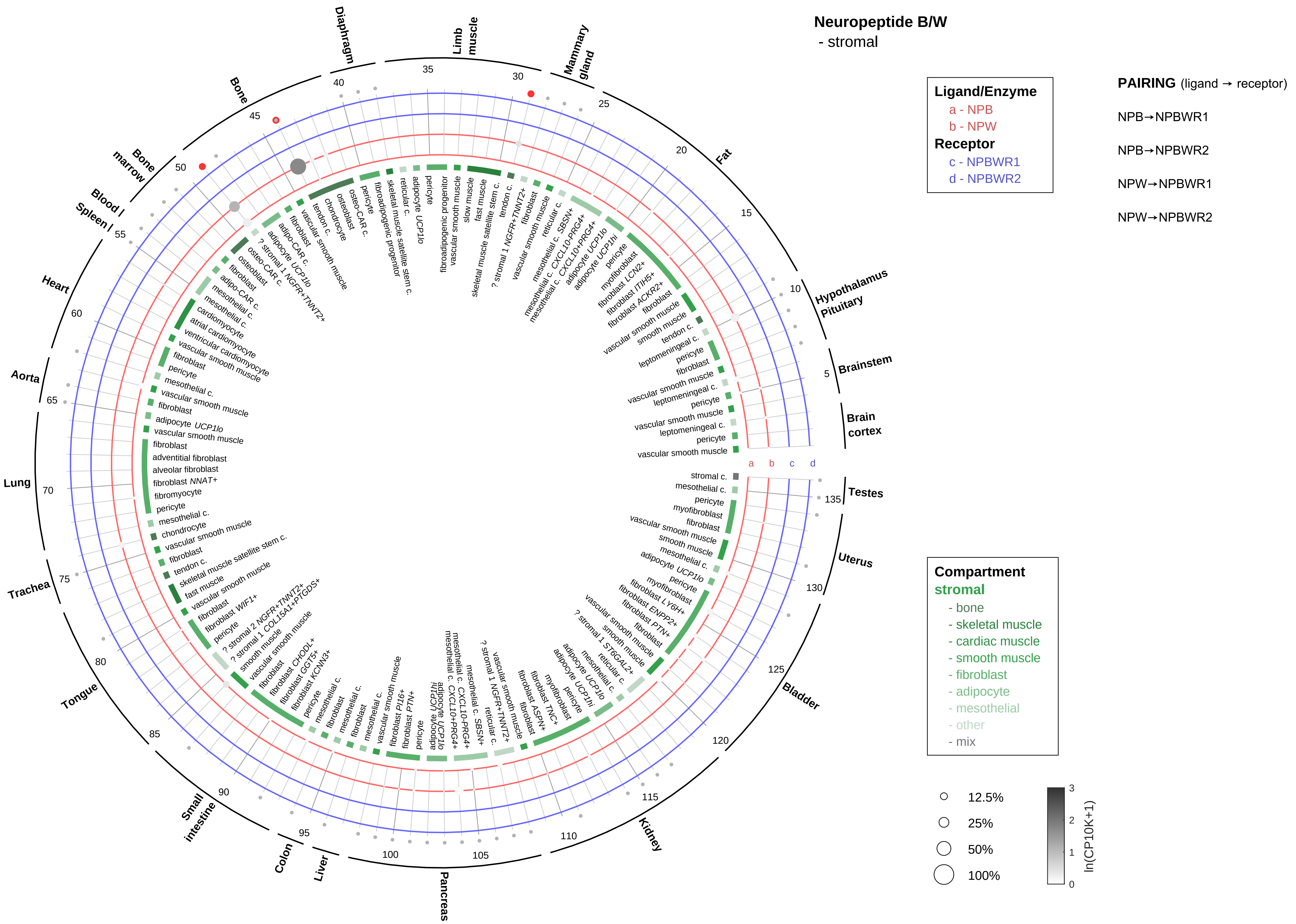

Neuropeptide B/W
- stromal
Ligand/Enzyme
a - NPB
b - NPW
Receptor
c - NPBWR1
d - NPBWR2
PAIRING (ligand → receptor)
NPB→NPBWR1
NPB→NPBWR2
NPW→NPBWR1
NPW→NPBWR2
Compartment
stromal
- bone
- skeletal muscle
- cardiac muscle
- smooth muscle
- fibroblast
- adipocyte
- mesothelial
- other
- mix
12.5%
25%
50%
100%
ln(CP10K+1)

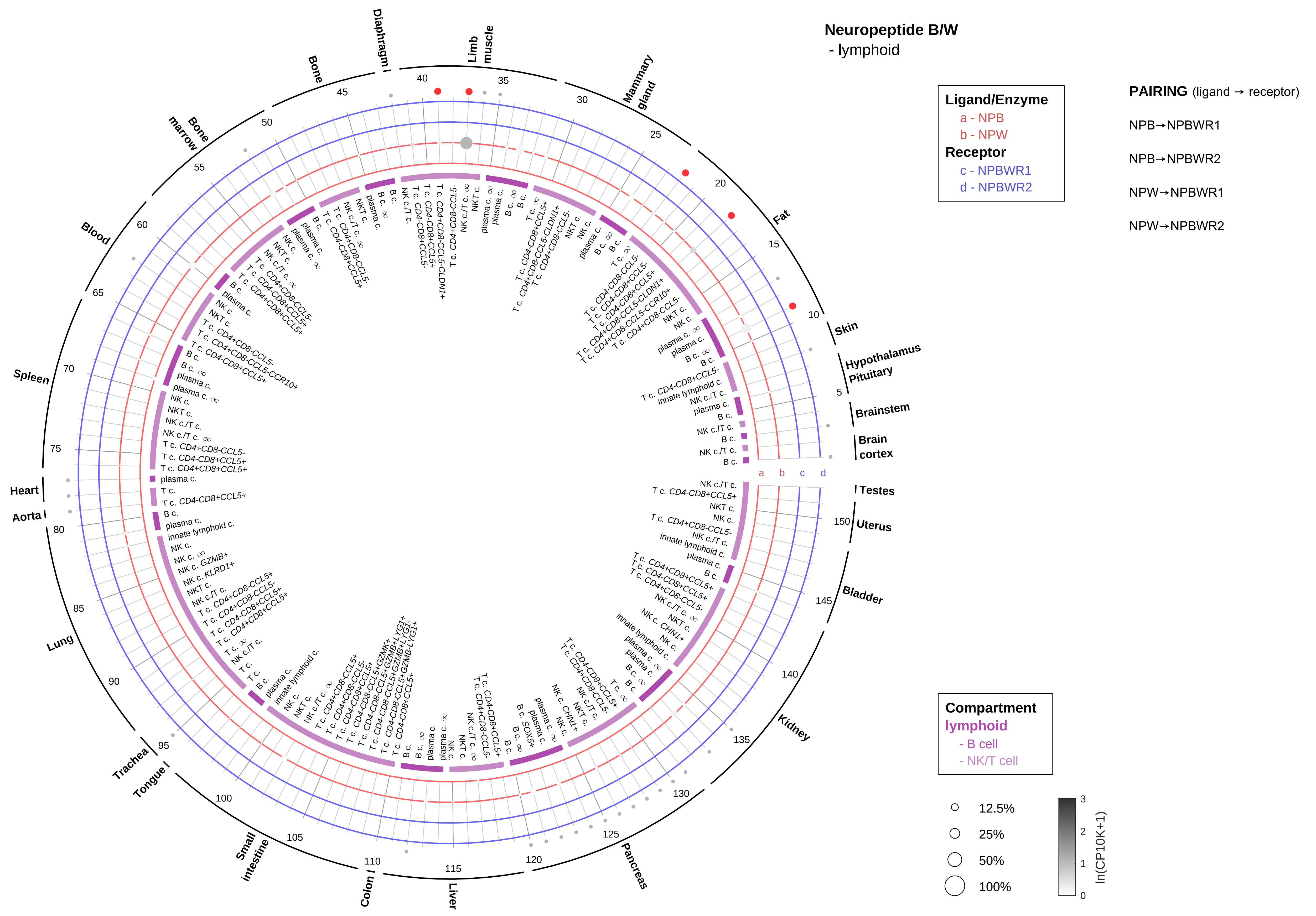
Neuropeptide B/W
- lymphoid
Ligand/Enzyme
a - NPB
b - NPW
Receptor
c - NPBWR1
d - NPBWR2
PAIRING (ligand → receptor)
NPB→NPBWR1
NPB→NPBWR2
NPW→NPBWR1
NPW→NPBWR2
Compartment
lymphoid
- B cell
- NK/T cell
12.5%
25%
50%
100%
ln(CP10K+1)
Skin
Hypothalamus
Pituitary
Brainstem
Brain cortex
Testes
Uterus
Bladder
Kidney
Pancreas
Liver
Colon
Small intestine
Tongue
Trachea
Lung
Aorta
Heart
Spleen
Blood
Bone marrow
Bone
Diaphragm
Limb muscle
Mammary gland
Fat

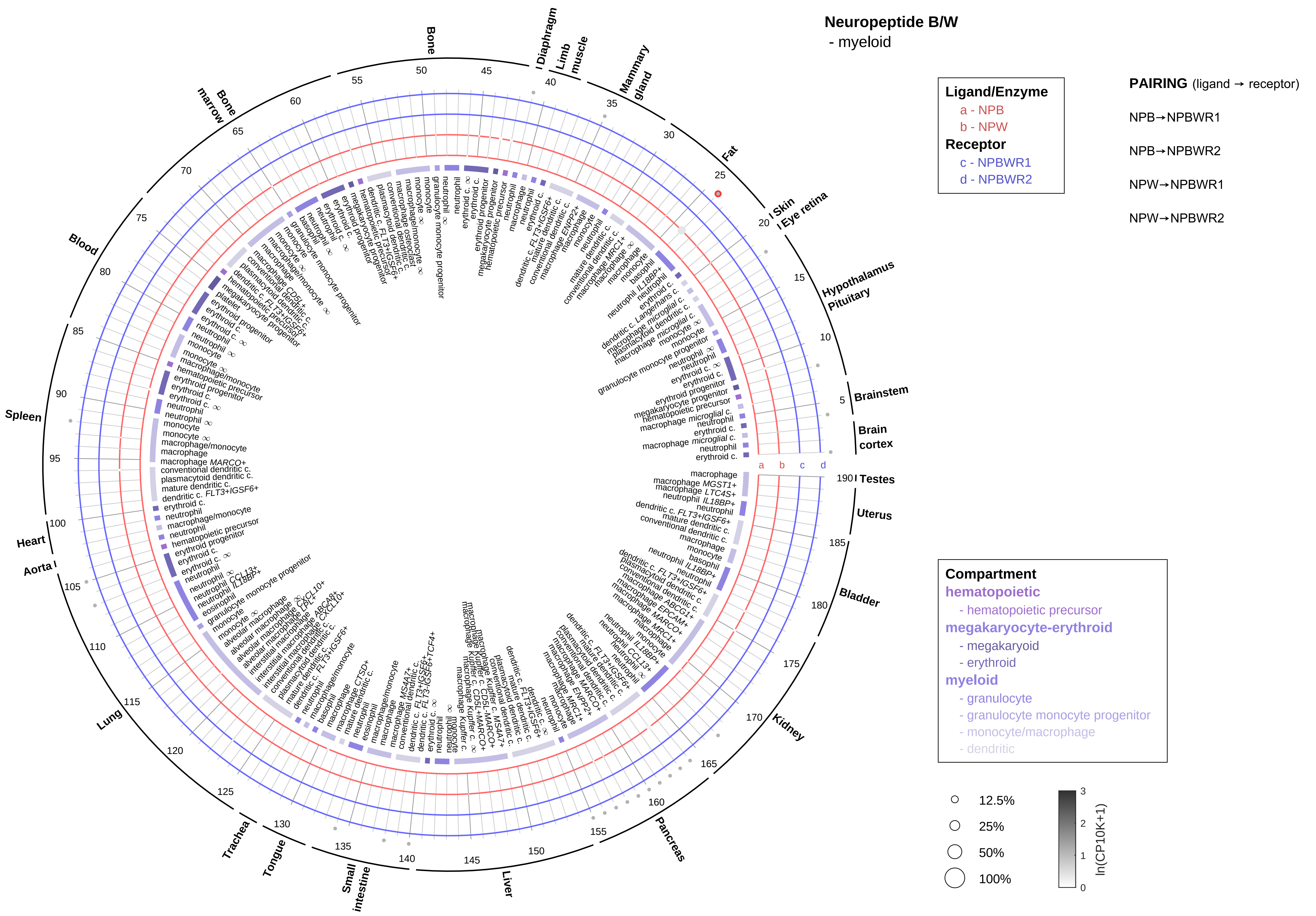

Neuropeptide B/W
- myeloid
Ligand/Enzyme
a - NPB
b - NPW
Receptor
c - NPBWR1
d - NPBWR2
PAIRING (ligand → receptor)
NPB→NPBWR1
NPB→NPBWR2
NPW→NPBWR1
NPW→NPBWR2
Compartment
hematopoietic
- hematopoietic precursor
megakaryocyte-erythroid
- megakaryoid
- erythroid
myeloid
- granulocyte
- granulocyte monocyte progenitor
- monocyte/macrophage
- dendritic
12.5%
25%
50%
100%
ln(CP10K+1)
Bone
Diaphragm
Limb muscle
Mammary gland
Fat
Skin
Eye retina
Hypothalamus
Pituitary
Brainstem
Brain cortex
Testes
Uterus
Bladder
Kidney
Pancreas
Liver
Small intestine
Tongue
Trachea
Lung
Aorta
Heart
Spleen
Blood
Bone marrow

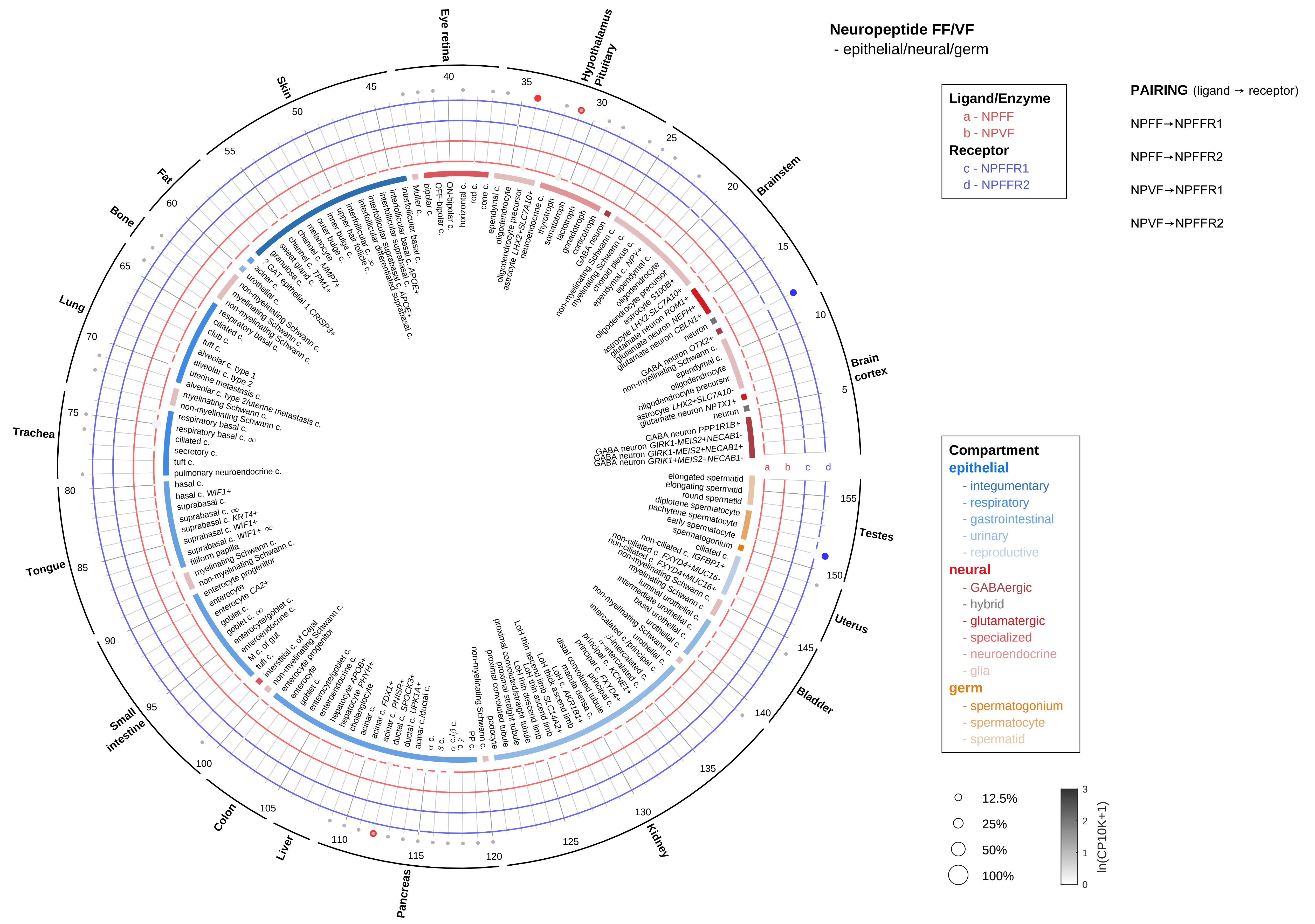

Neuropeptide FF/VF
- epithelial/neural/germ
Ligand/Enzyme
a - NPFF
b - NPVF
Receptor
c - NPFFR1
d - NPFFR2
PAIRING (ligand → receptor)
NPFF→NPFFR1
NPFF→NPFFR2
NPVF→NPFFR1
NPVF→NPFFR2
Compartment
epithelial
- integumentary
- respiratory
- gastrointestinal
- urinary
- reproductive
neural
- GABAergic
- hybrid
- glutamatergic
- specialized
- neuroendocrine
- glia
germ
- spermatogonium
- spermatocyte
- spermatid
12.5%
25%
50%
100%
ln(CP10K+1)
Eye retina
Hypothalamus Pituitary
Brainstem
Brain cortex
Testes
Uterus
Bladder
Kidney
Pancreas
Liver
Colon
Small intestine
Tongue
Trachea
Lung
Bone
Fat
Skin

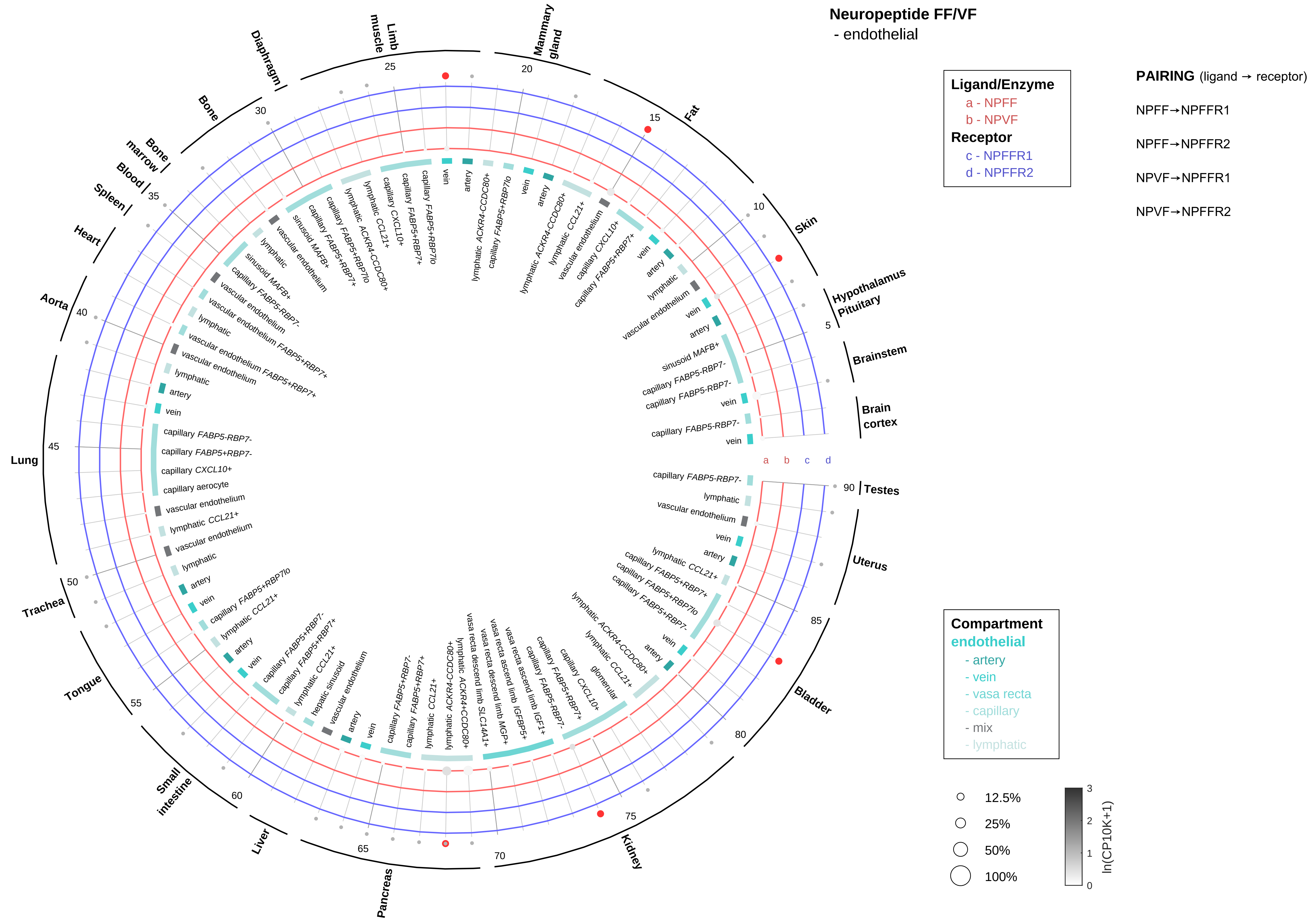

Neuropeptide FF/VF
- endothelial
Ligand/Enzyme
a - NPFF
b - NPVF
Receptor
c - NPFFR1
d - NPFFR2
PAIRING (ligand → receptor)
NPFF→NPFFR1
NPFF→NPFFR2
NPVF→NPFFR1
NPVF→NPFFR2
Compartment
endothelial
- artery
- vein
- vasa recta
- capillary
- mix
- lymphatic
12.5%
25%
50%
100%
ln(CP10K+1)
Hypothalamus
Pituitary
Brainstem
Brain cortex
Testes
Uterus
Bladder
Kidney
Pancreas
Liver
Small intestine
Tongue
Trachea
Lung
Aorta
Heart
Spleen
Blood
Bone marrow
Bone
Diaphragm
Limb muscle
Mammary gland
Fat
Skin

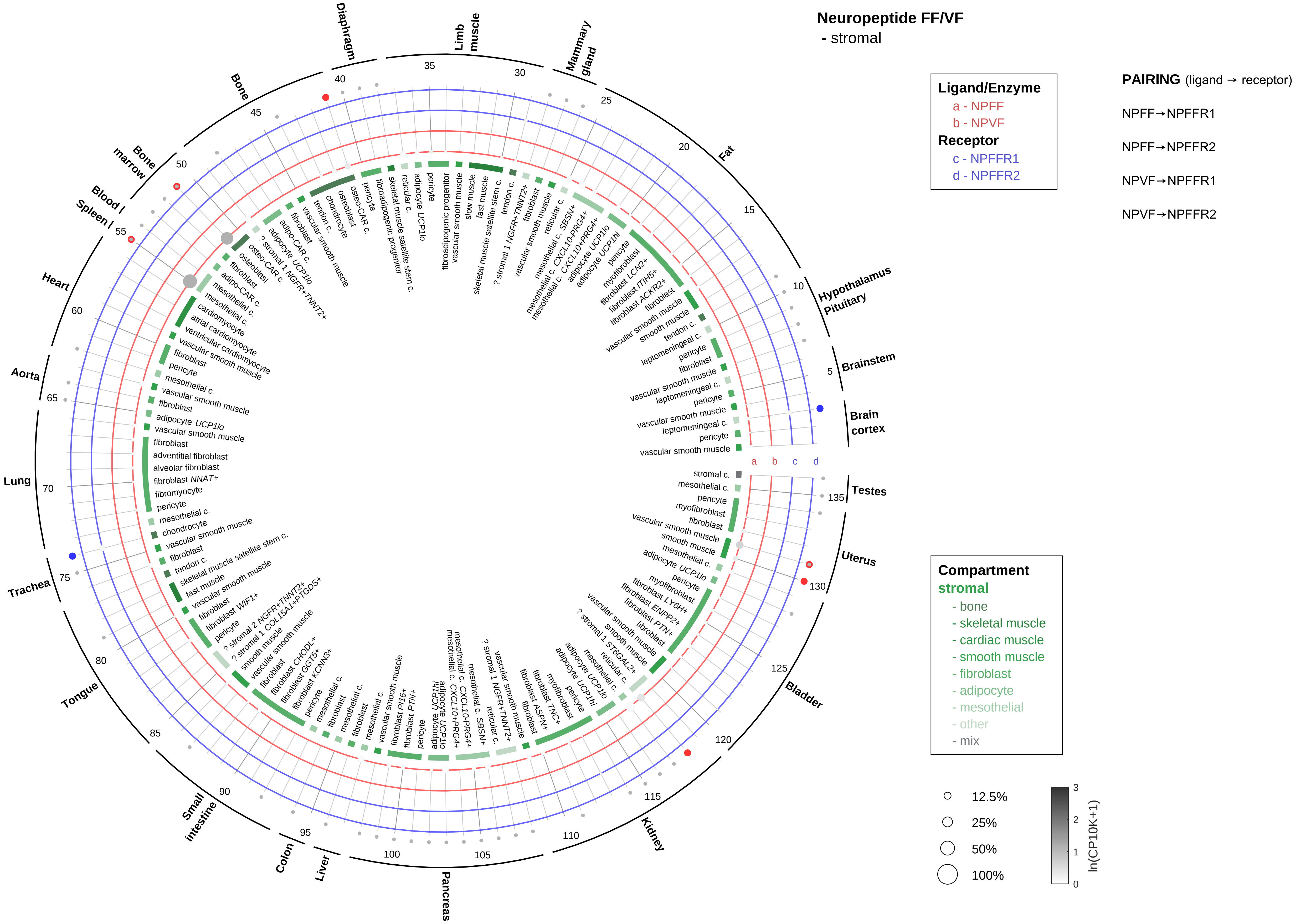
Neuropeptide FF/VF
- stromal
Ligand/Enzyme
a - NPFF
b - NPVF
Receptor
c - NPFFR1
d - NPFFR2
PAIRING (ligand → receptor)
NPFF→NPFFR1
NPFF→NPFFR2
NPVF→NPFFR1
NPVF→NPFFR2
Compartment
stromal
- bone
- skeletal muscle
- cardiac muscle
- smooth muscle
- fibroblast
- adipocyte
- mesothelial
- other
- mix
12.5%
25%
50%
100%
ln(CP10K+1)
Brain cortex
Brainstem
Hypothalamus Pituitary
Fat
Mammary gland
Limb muscle
Diaphragm
Bone
Bone marrow
Blood / Spleen
Heart
Aorta
Lung
Trachea
Tongue
Small intestine
Colon
Liver
Pancreas
Kidney
Bladder
Uterus
Testes

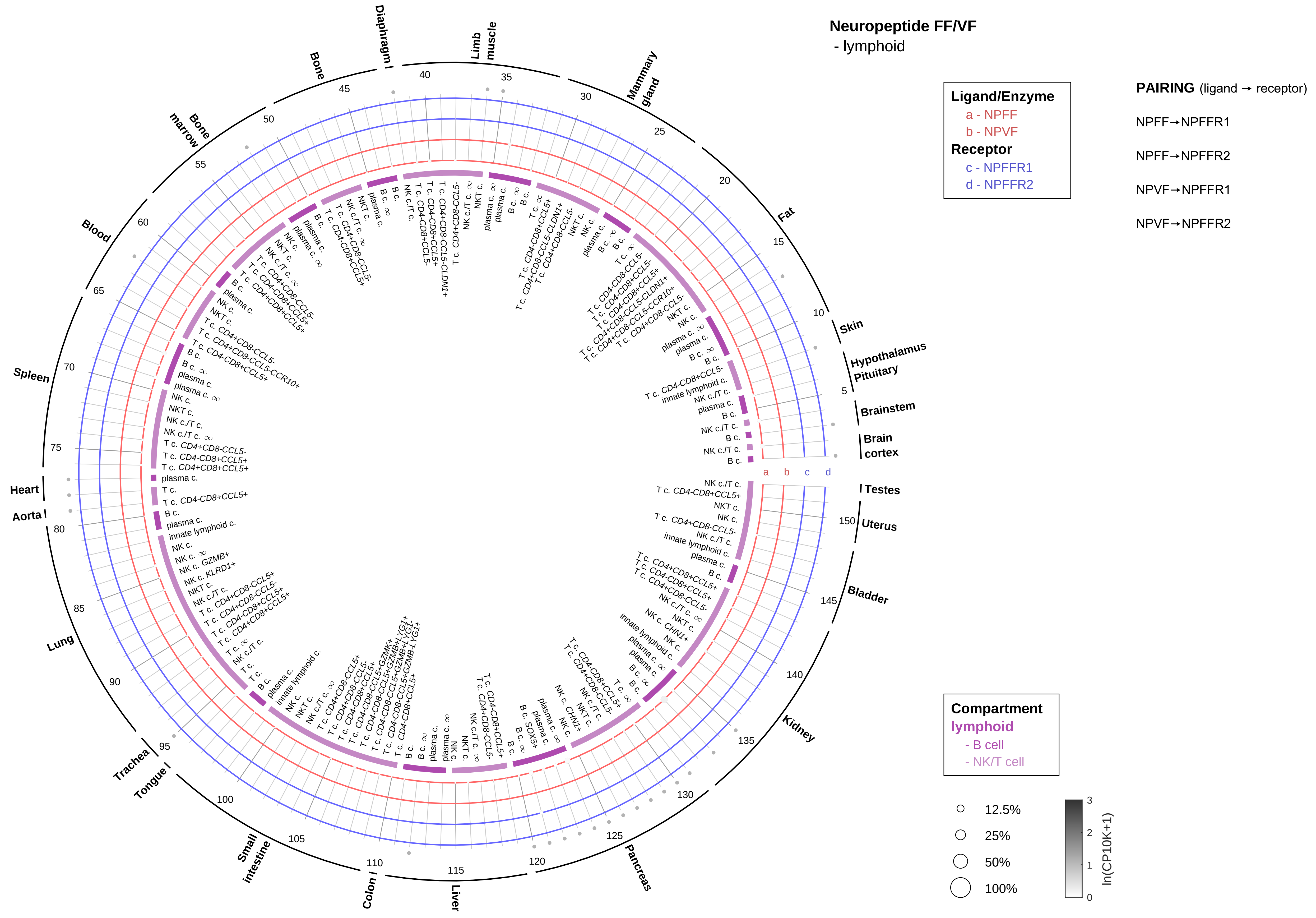
Neuropeptide FF/VF
- lymphoid
Ligand/Enzyme
a - NPFF
b - NPVF
Receptor
c - NPFFR1
d - NPFFR2
PAIRING (ligand → receptor)
NPFF→NPFFR1
NPFF→NPFFR2
NPVF→NPFFR1
NPVF→NPFFR2
Compartment
lymphoid
- B cell
- NK/T cell
12.5%
25%
50%
100%
ln(CP10K+1)
Skin
Hypothalamus
Pituitary
Brainstem
Brain cortex
Testes
Uterus
Bladder
Kidney
Pancreas
Liver
Colon
Small intestine
Tongue
Trachea
Lung
Aorta
Heart
Spleen
Blood
Bone marrow
Bone
Diaphragm
Limb muscle
Mammary gland
Fat

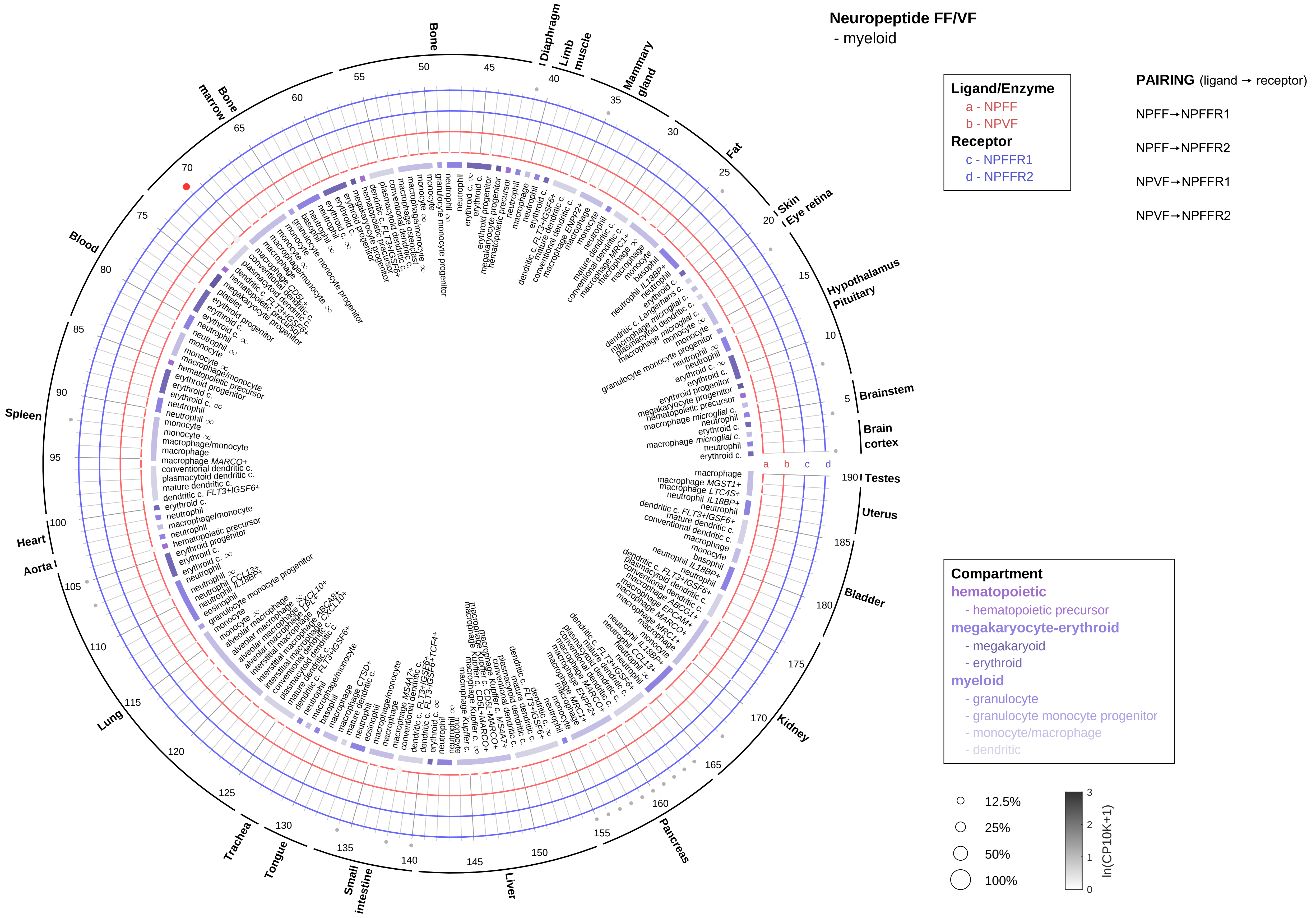

Neuropeptide FF/VF
- myeloid
Ligand/Enzyme
a - NPFF
b - NPVF
Receptor
c - NPFFR1
d - NPFFR2
PAIRING (ligand → receptor)
NPFF→NPFFR1
NPFF→NPFFR2
NPVF→NPFFR1
NPVF→NPFFR2
Compartment
hematopoietic
- hematopoietic precursor
megakaryocyte-erythroid
- megakaryoid
- erythroid
myeloid
- granulocyte
- granulocyte monocyte progenitor
- monocyte/macrophage
- dendritic
12.5%
25%
50%
100%
ln(CP10K+1)
Bone
Diaphragm
Limb muscle
Mammary gland
Fat
Skin
Eye retina
Hypothalamus
Pituitary
Brainstem
Brain cortex
Testes
Uterus
Bladder
Kidney
Pancreas
Liver
Small intestine
Tongue
Trachea
Lung
Aorta
Heart
Spleen
Blood
Bone marrow

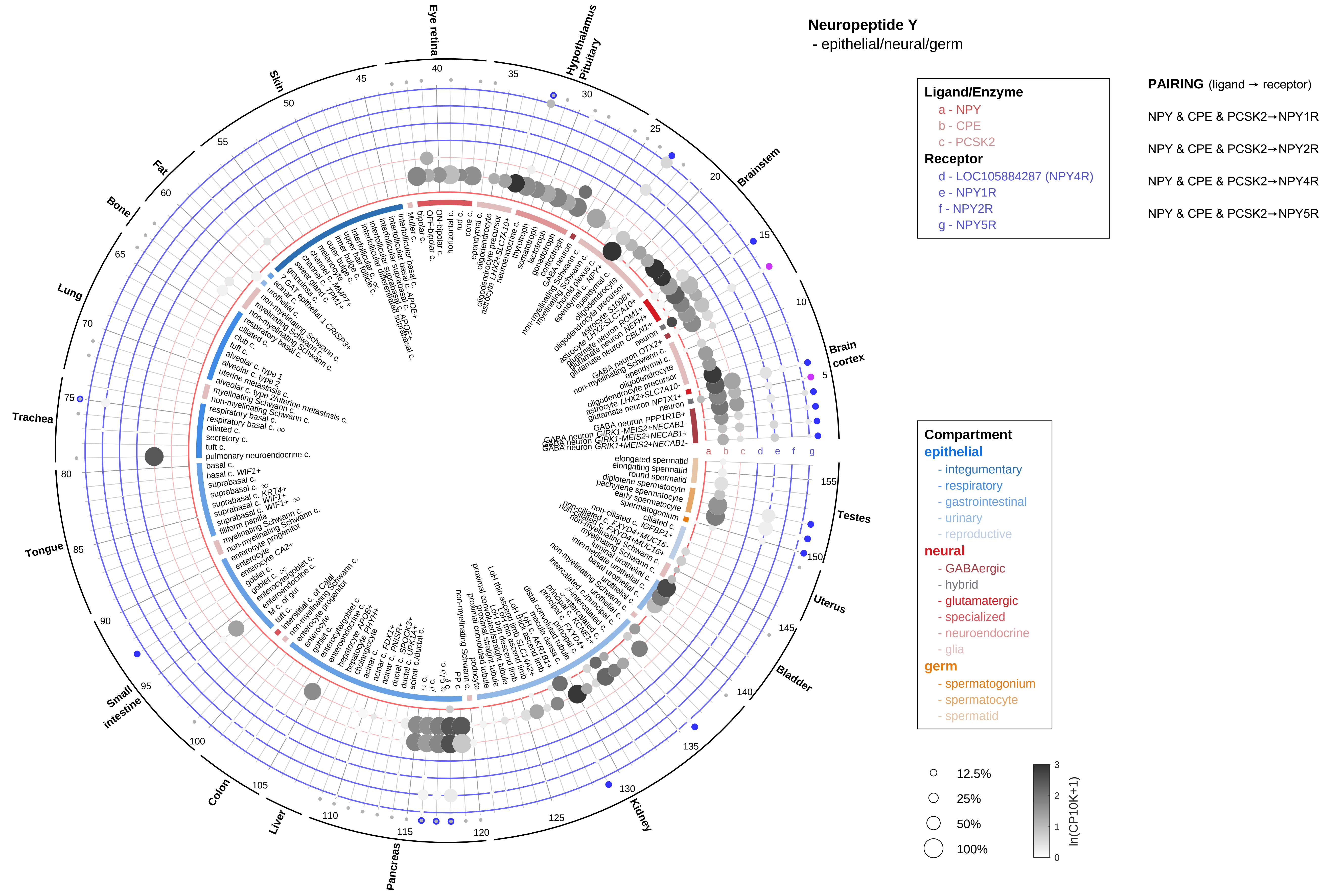
Neuropeptide Y
- epithelial/neural/germ
Ligand/Enzyme
a - NPY
b - CPE
c - PCSK2
Receptor
d - LOC105884287 (NPY4R)
e - NPY1R
f - NPY2R
g - NPY5R
PAIRING (ligand → receptor)
NPY & CPE & PCSK2→NPY1R
NPY & CPE & PCSK2→NPY2R
NPY & CPE & PCSK2→NPY4R
NPY & CPE & PCSK2→NPY5R
Compartment
epithelial
- integumentary
- respiratory
- gastrointestinal
- urinary
- reproductive
neural
- GABAergic
- hybrid
- glutamatergic
- specialized
- neuroendocrine
- glia
germ
- spermatogonium
- spermatocyte
- spermatid
12.5%
25%
50%
100%
ln(CP10K+1)
Eye retina
Hypothalamus
Pituitary
Brainstem
Brain cortex
Testes
Uterus
Bladder
Kidney
Pancreas
Liver
Colon
Small intestine
Tongue
Trachea
Lung
Bone
Fat
Skin

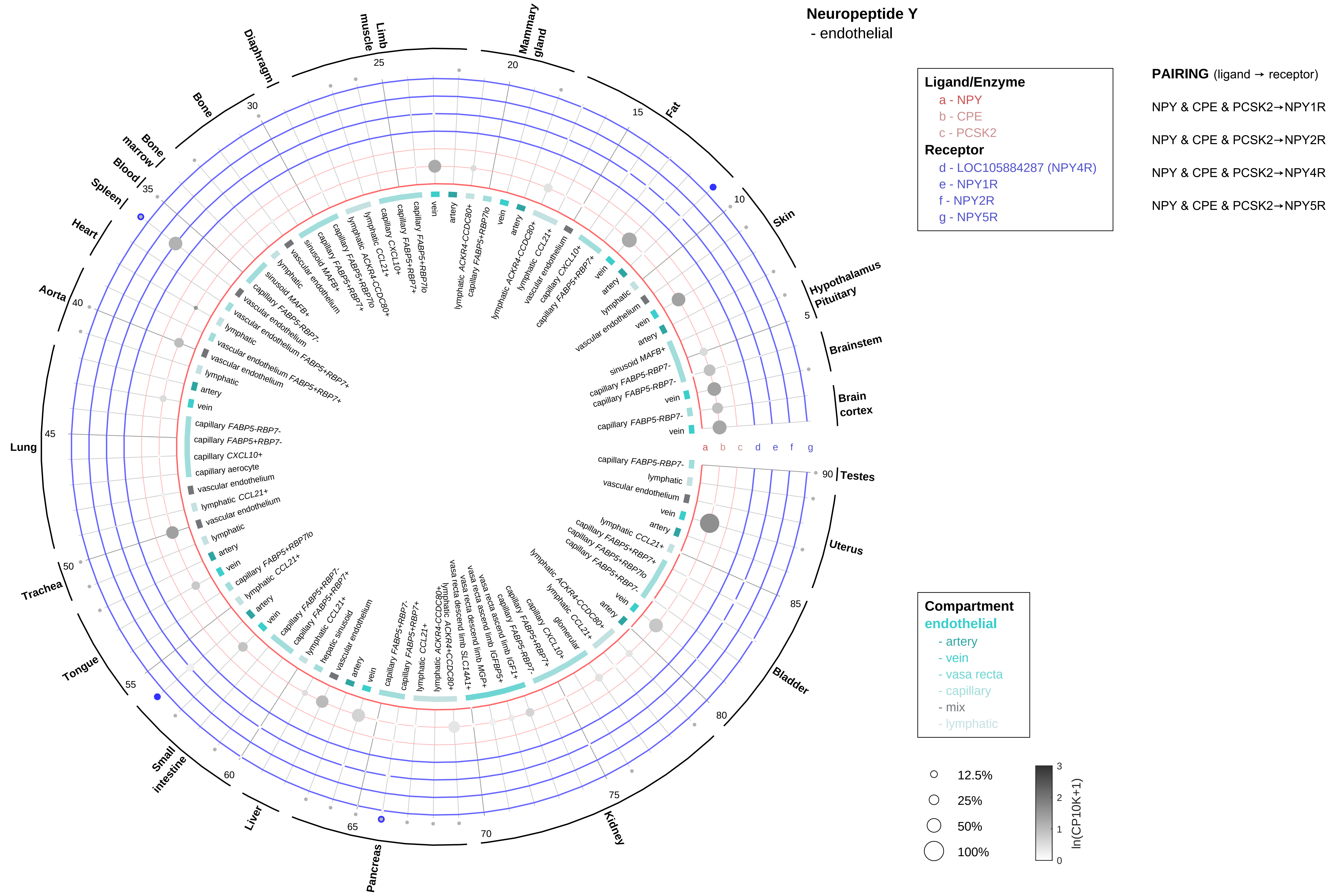

Neuropeptide Y
- endothelial
Ligand/Enzyme
a - NPY
b - CPE
c - PCSK2
Receptor
d - LOC105884287 (NPY4R)
e - NPY1R
f - NPY2R
g - NPY5R
PAIRING (ligand → receptor)
NPY & CPE & PCSK2→NPY1R
NPY & CPE & PCSK2→NPY2R
NPY & CPE & PCSK2→NPY4R
NPY & CPE & PCSK2→NPY5R
Compartment
endothelial
- artery
- vein
- vasa recta
- capillary
- mix
- lymphatic
12.5%
25%
50%
100%
ln(CP10K+1)
Hypothalamus
Pituitary
Brainstem
Brain cortex
Skin
Fat
Mammary gland
Limb muscle
Diaphragm
Bone
Bone marrow
Blood
Spleen
Heart
Aorta
Lung
Trachea
Tongue
Small intestine
Liver
Pancreas
Kidney
Bladder
Uterus
Testes

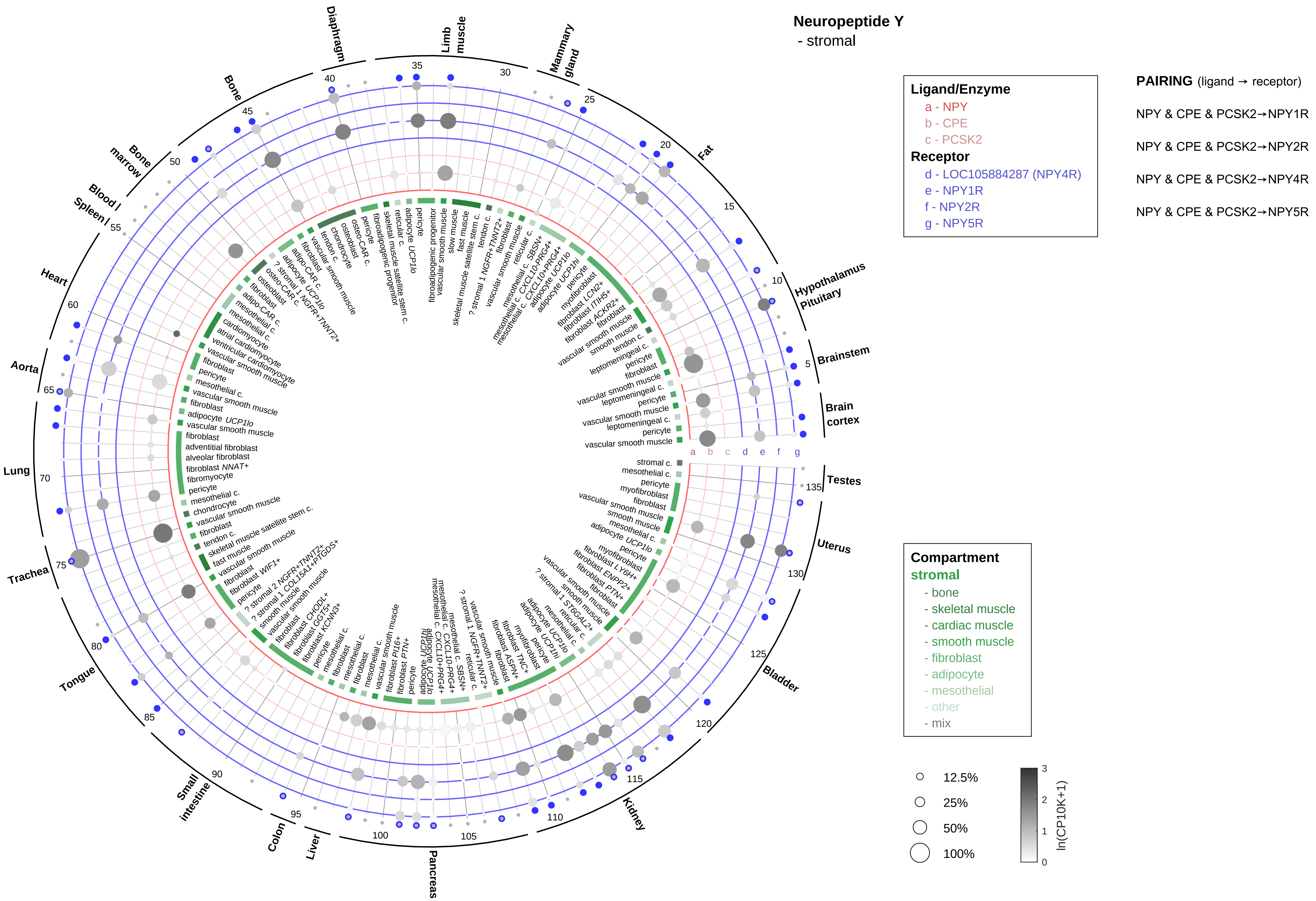

Neuropeptide Y
- stromal
Ligand/Enzyme
a - NPY
b - CPE
c - PCSK2
Receptor
d - LOC105884287 (NPY4R)
e - NPY1R
f - NPY2R
g - NPY5R
PAIRING (ligand → receptor)
NPY & CPE & PCSK2→NPY1R
NPY & CPE & PCSK2→NPY2R
NPY & CPE & PCSK2→NPY4R
NPY & CPE & PCSK2→NPY5R
Compartment
stromal
- bone
- skeletal muscle
- cardiac muscle
- smooth muscle
- fibroblast
- adipocyte
- mesothelial
- other
- mix
12.5%
25%
50%
100%
ln(CP10K+1)
Diaphragm
Limb muscle
Mammary gland
Fat
Hypothalamus Pituitary
Brainstem
Brain cortex
Testes
Uterus
Bladder
Kidney
Pancreas
Liver
Colon
Small intestine
Tongue
Trachea
Lung
Aorta
Heart
Spleen
Blood
Bone marrow
Bone

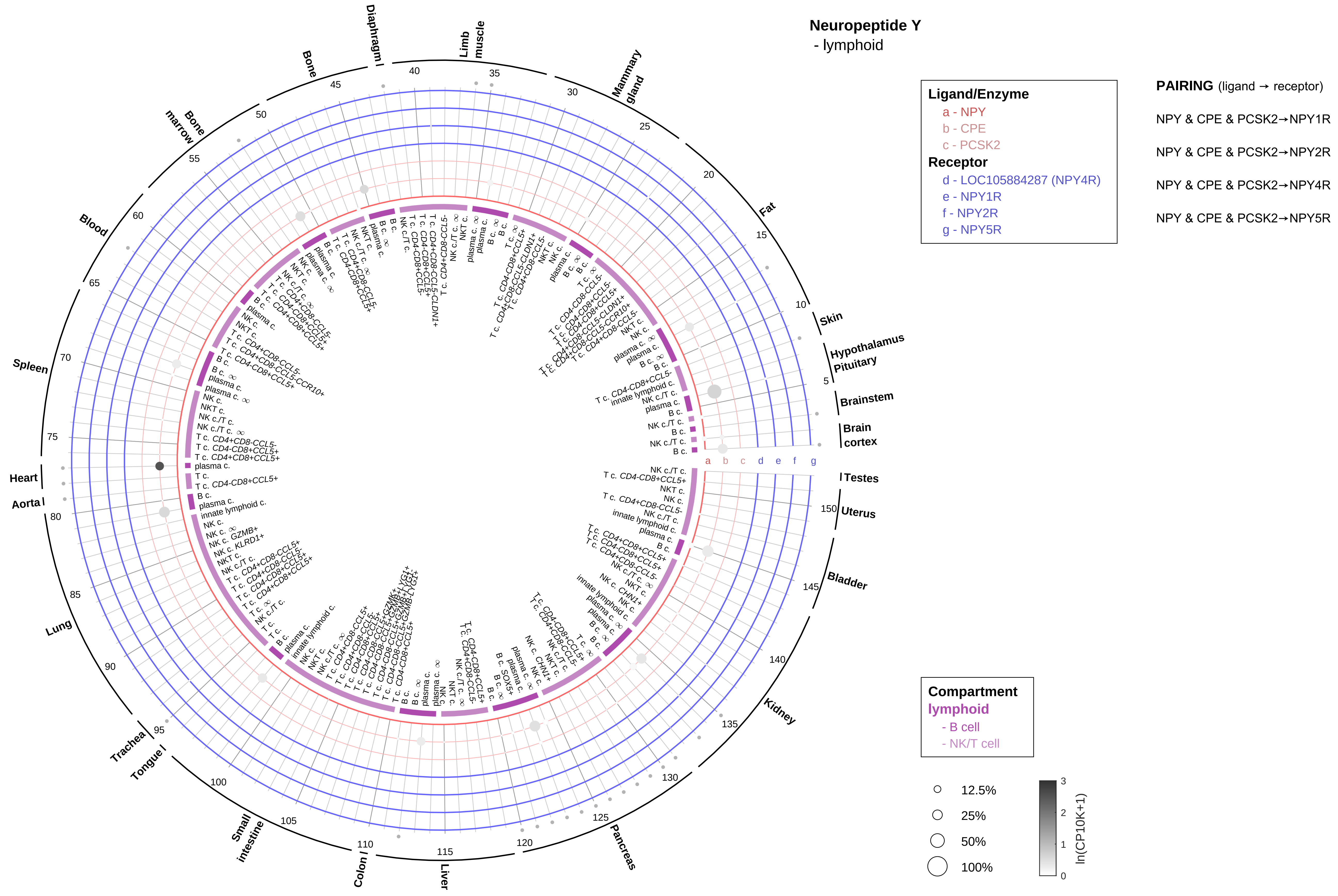

Neuropeptide Y
- lymphoid
Ligand/Enzyme
a - NPY
b - CPE
c - PCSK2
Receptor
d - LOC105884287 (NPY4R)
e - NPY1R
f - NPY2R
g - NPY5R
PAIRING (ligand → receptor)
NPY & CPE & PCSK2→NPY1R
NPY & CPE & PCSK2→NPY2R
NPY & CPE & PCSK2→NPY4R
NPY & CPE & PCSK2→NPY5R
Compartment
lymphoid
- B cell
- NK/T cell
12.5%
25%
50%
100%
ln(CP10K+1)
Skin
Hypothalamus
Pituitary
Brainstem
Brain cortex
Testes
Uterus
Bladder
Kidney
Pancreas
Liver
Colon
Small intestine
Tongue
Trachea
Lung
Aorta
Heart
Spleen
Blood
Bone marrow
Bone
Diaphragm
Limb muscle
Mammary gland
Fat

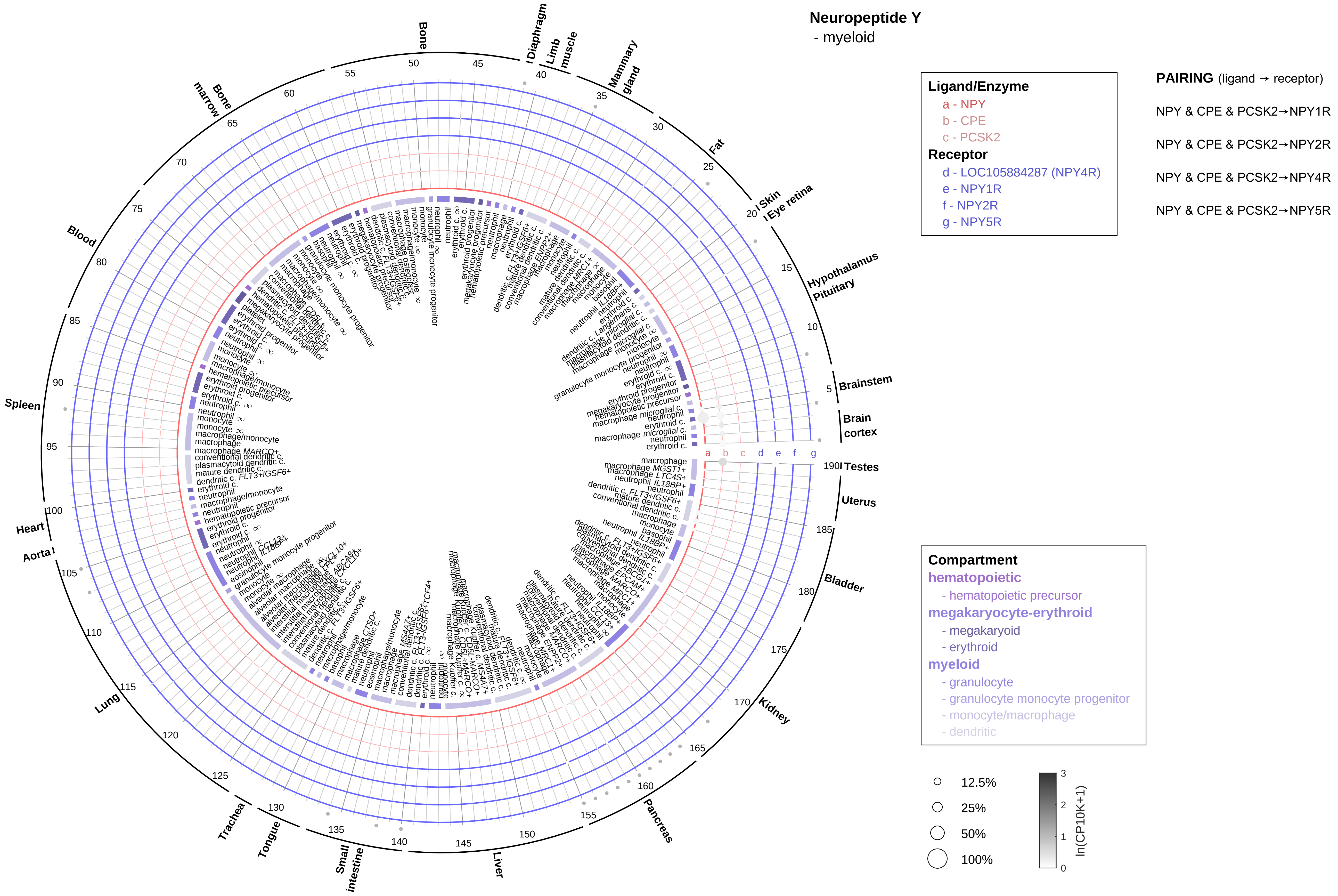

Neuropeptide Y
- myeloid
Ligand/Enzyme
a - NPY
b - CPE
c - PCSK2
Receptor
d - LOC105884287 (NPY4R)
e - NPY1R
f - NPY2R
g - NPY5R
PAIRING (ligand → receptor)
NPY & CPE & PCSK2→NPY1R
NPY & CPE & PCSK2→NPY2R
NPY & CPE & PCSK2→NPY4R
NPY & CPE & PCSK2→NPY5R
Compartment
hematopoietic
- hematopoietic precursor
megakaryocyte-erythroid
- megakaryoid
- erythroid
myeloid
- granulocyte
- granulocyte monocyte progenitor
- monocyte/macrophage
- dendritic
12.5%
25%
50%
100%
ln(CP10K+1)
Bone
Diaphragm
Limb muscle
Mammary gland
Fat
Skin
Eye retina
Hypothalamus
Pituitary
Brainstem
Brain cortex
Testes
Uterus
Bladder
Kidney
Pancreas
Liver
Small intestine
Tongue
Trachea
Lung
Aorta
Heart
Spleen
Blood
Bone marrow

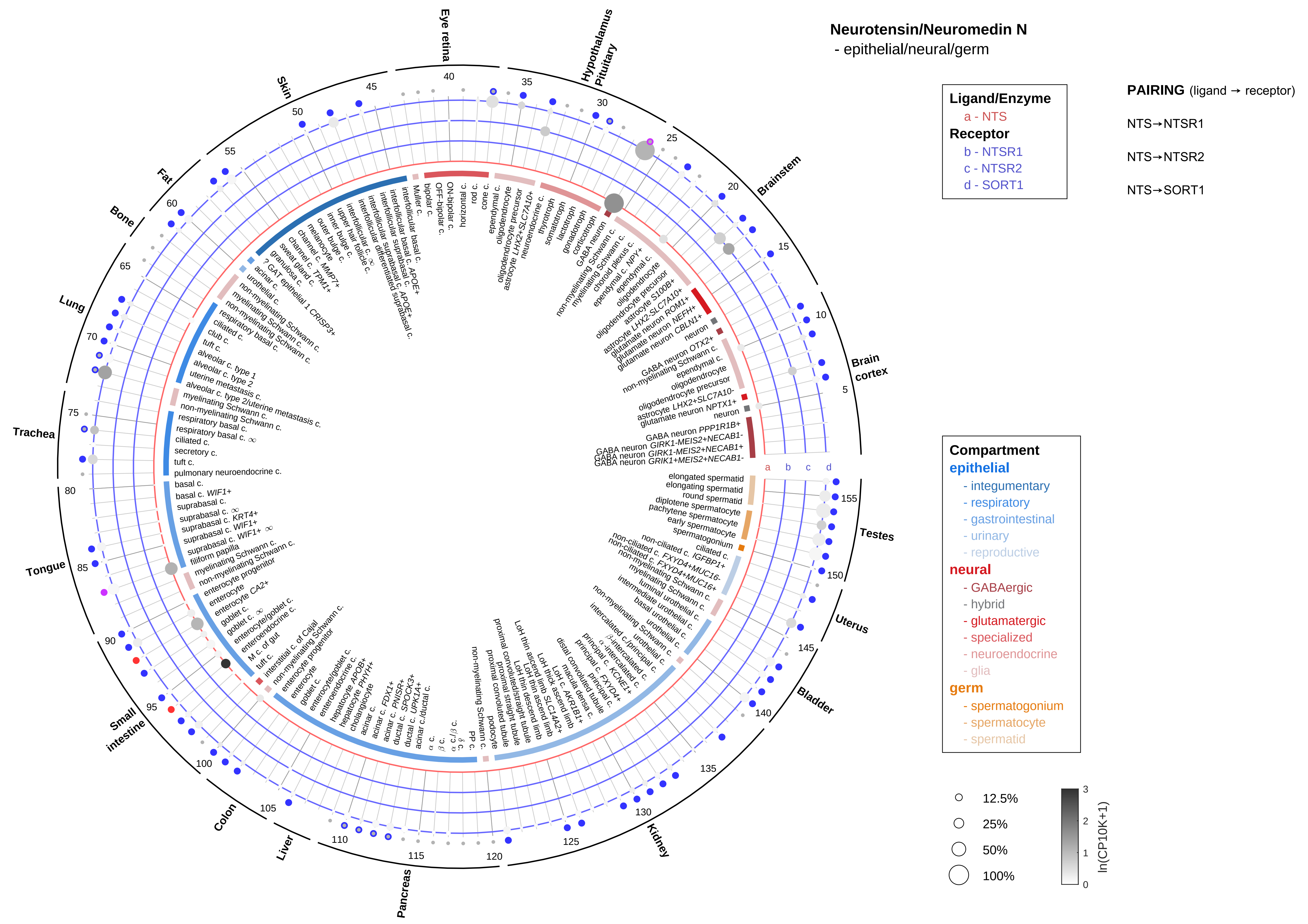

Neurotensin/Neuromedin N
- epithelial/neural/germ
Ligand/Enzyme
a - NTS
Receptor
b - NTSR1
c - NTSR2
d - SORT1
PAIRING (ligand → receptor)
NTS→NTSR1
NTS→NTSR2
NTS→SORT1
Compartment
epithelial
- integumentary
- respiratory
- gastrointestinal
- urinary
- reproductive
neural
- GABAergic
- hybrid
- glutamatergic
- specialized
- neuroendocrine
- glia
germ
- spermatogonium
- spermatocyte
- spermatid
12.5%
25%
50%
100%
ln(CP10K+1)
Brain cortex
Brainstem
Hypothalamus Pituitary
Eye retina
Skin
Fat
Bone
Lung
Trachea
Tongue
Small intestine
Colon
Liver
Pancreas
Kidney
Bladder
Uterus
Testes

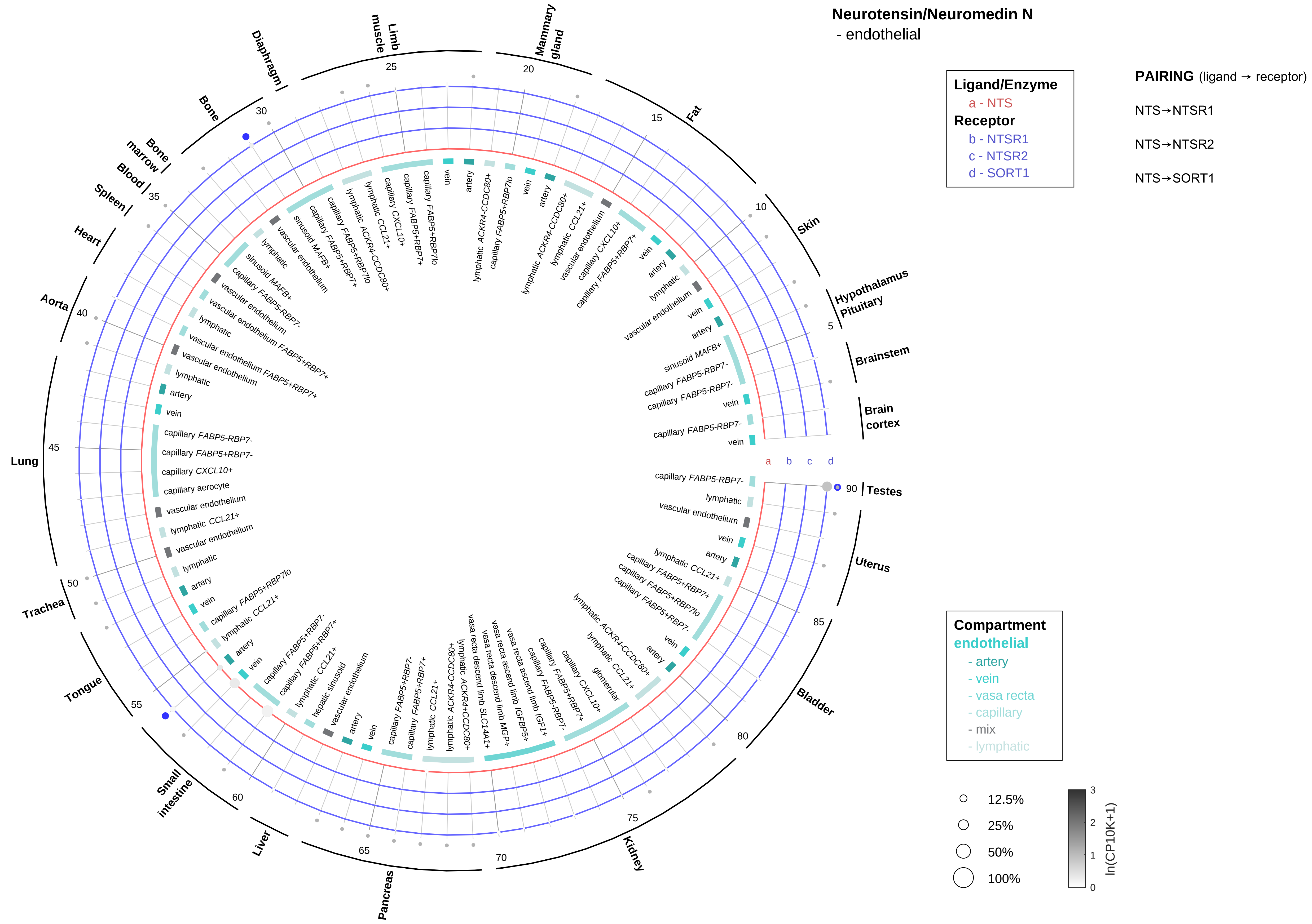

Neurotensin/Neuromedin N
- endothelial
Ligand/Enzyme
a - NTS
Receptor
b - NTSR1
c - NTSR2
d - SORT1
PAIRING (ligand → receptor)
NTS→NTSR1
NTS→NTSR2
NTS→SORT1
Compartment
endothelial
- artery
- vein
- vasa recta
- capillary
- mix
- lymphatic
12.5%
25%
50%
100%
ln(CP10K+1)
Hypothalamus
Pituitary
Brainstem
Brain cortex
Testes
Uterus
Bladder
Kidney
Pancreas
Liver
Small intestine
Tongue
Trachea
Lung
Aorta
Heart
Spleen
Blood
Bone marrow
Bone
Diaphragm
Limb muscle
Mammary gland
Fat
Skin

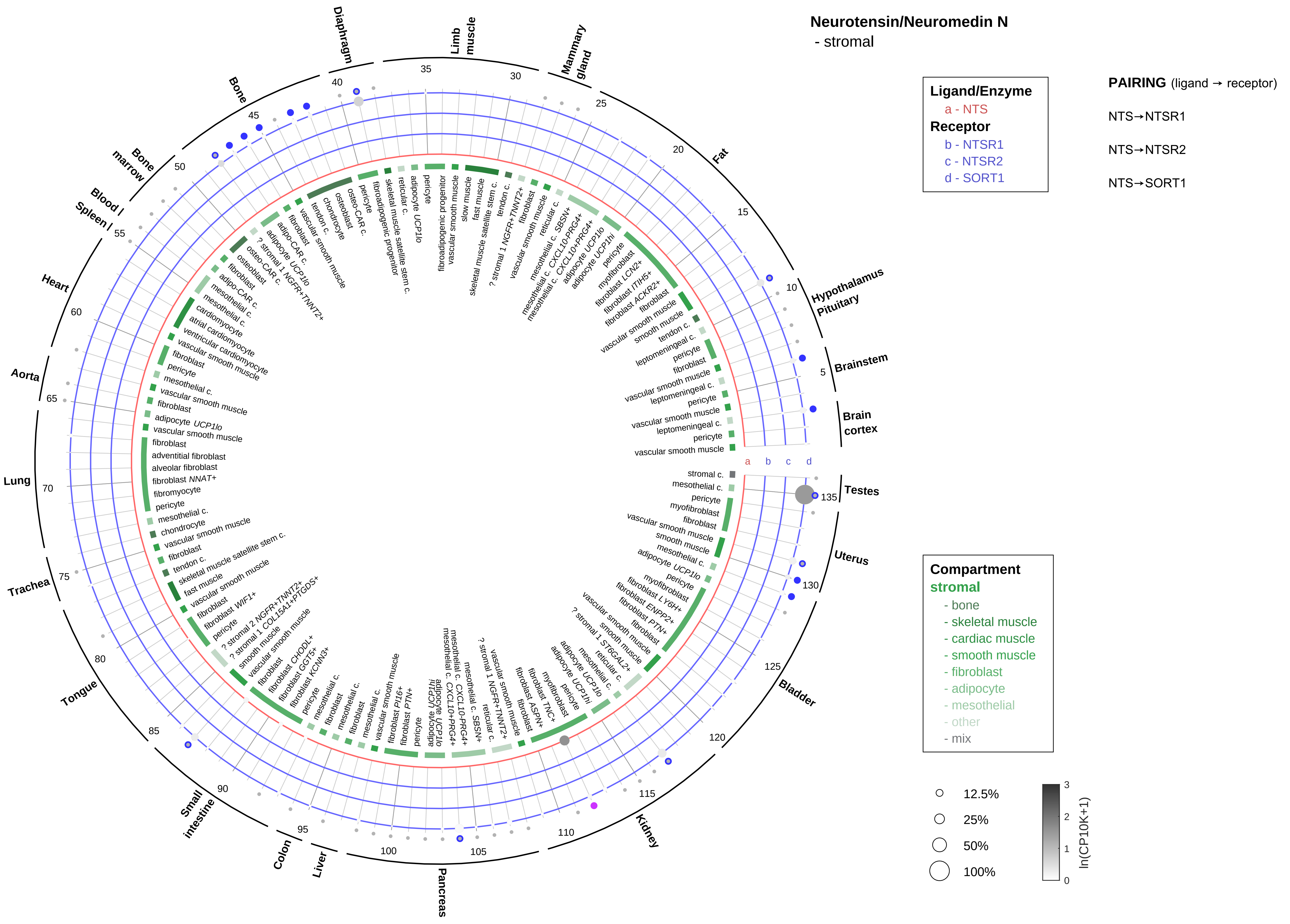
Neurotensin/Neuromedin N
- stromal
Ligand/Enzyme
a - NTS
Receptor
b - NTSR1
c - NTSR2
d - SORT1
PAIRING (ligand → receptor)
NTS→NTSR1
NTS→NTSR2
NTS→SORT1
Compartment
stromal
- bone
- skeletal muscle
- cardiac muscle
- smooth muscle
- fibroblast
- adipocyte
- mesothelial
- other
- mix
12.5%
25%
50%
100%
ln(CP10K+1)
Diaphragm
Limb muscle
Mammary gland
Fat
Hypothalamus Pituitary
Brainstem
Brain cortex
Testes
Uterus
Bladder
Kidney
Pancreas
Liver
Colon
Small intestine
Tongue
Trachea
Lung
Aorta
Heart
Blood / Spleen
Bone marrow
Bone

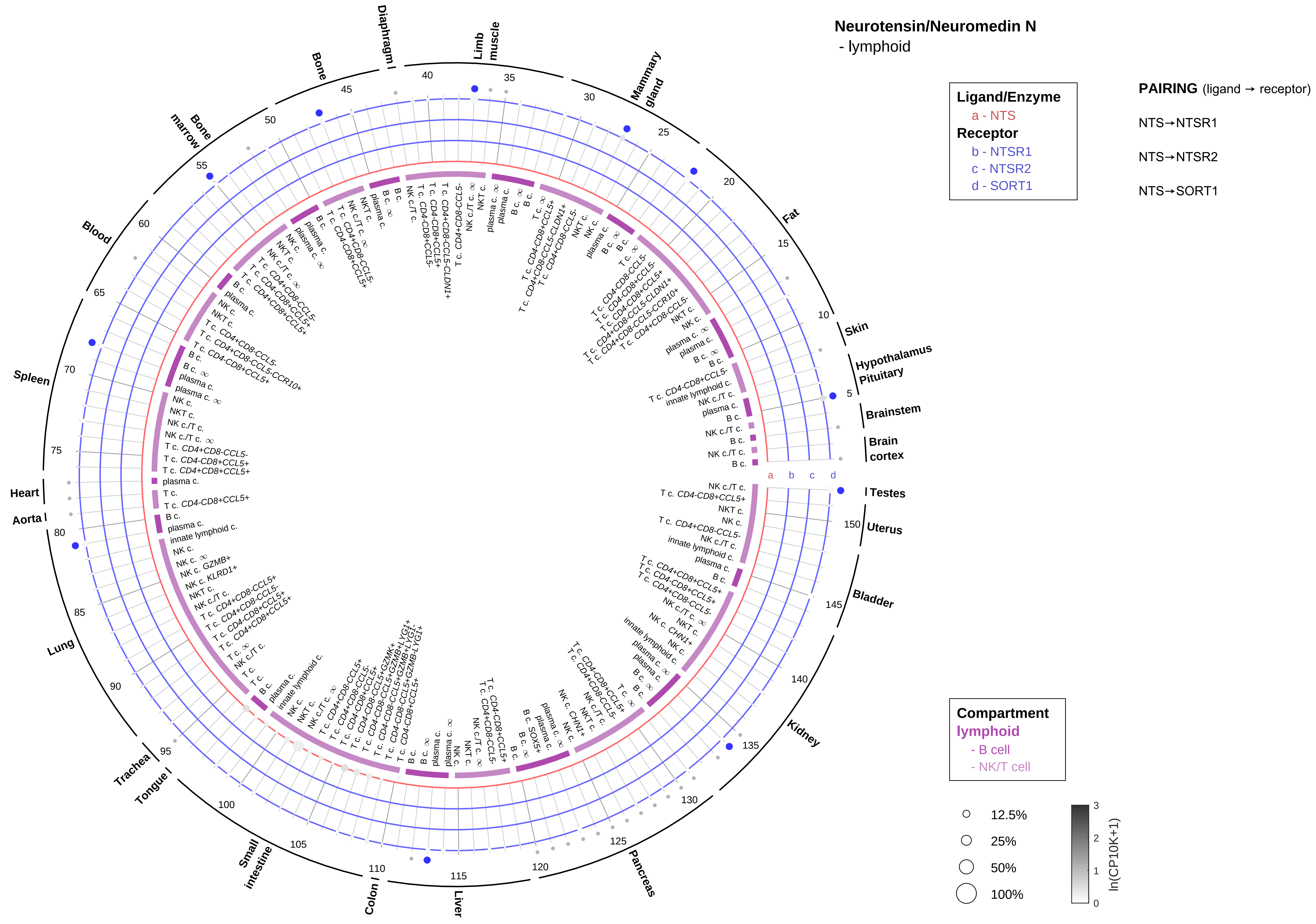
Neurotensin/Neuromedin N
- lymphoid
Ligand/Enzyme
a - NTS
Receptor
b - NTSR1
c - NTSR2
d - SORT1
PAIRING (ligand → receptor)
NTS→NTSR1
NTS→NTSR2
NTS→SORT1
Compartment
lymphoid
- B cell
- NK/T cell
12.5%
25%
50%
100%
ln(CP10K+1)
Diaphragm
Limb muscle
Mammary gland
Fat
Skin
Hypothalamus
Pituitary
Brainstem
Brain cortex
Testes
Uterus
Bladder
Kidney
Pancreas
Liver
Colon
Small intestine
Tongue
Trachea
Lung
Aorta
Heart
Spleen
Blood
Bone marrow
Bone

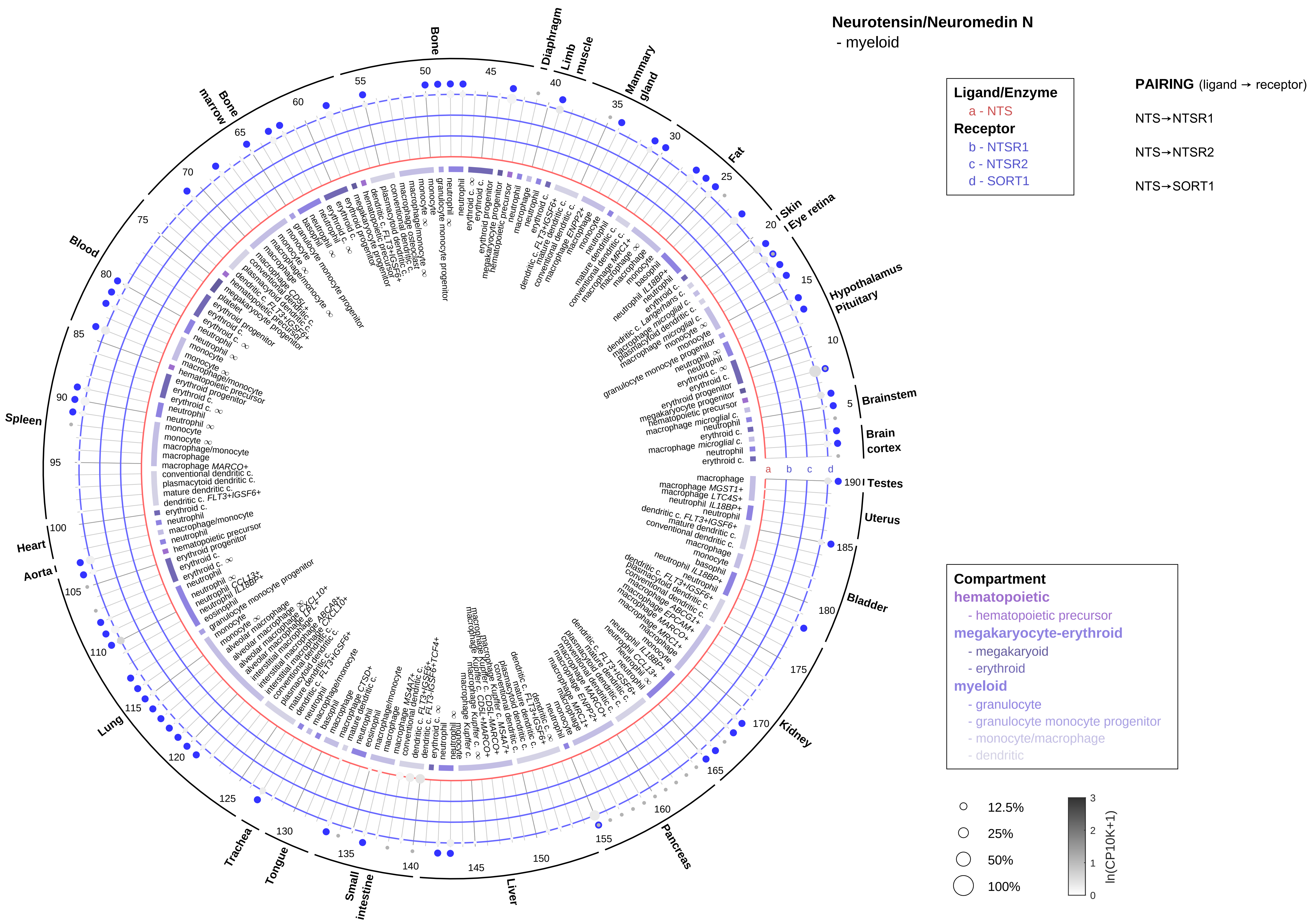
Neurotensin/Neuromedin N
- myeloid
Ligand/Enzyme
a - NTS
Receptor
b - NTSR1
c - NTSR2
d - SORT1
PAIRING (ligand → receptor)
NTS→NTSR1
NTS→NTSR2
NTS→SORT1
Compartment
hematopoietic
- hematopoietic precursor
megakaryocyte-erythroid
- megakaryoid
- erythroid
myeloid
- granulocyte
- granulocyte monocyte progenitor
- monocyte/macrophage
- dendritic
12.5%
25%
50%
100%
ln(CP10K+1)
Bone
Diaphragm
Limb muscle
Mammary gland
Fat
Skin
Eye retina
Hypothalamus
Pituitary
Brainstem
Brain cortex
Testes
Uterus
Bladder
Kidney
Pancreas
Liver
Small intestine
Tongue
Trachea
Lung
Aorta
Heart
Spleen
Blood
Bone marrow

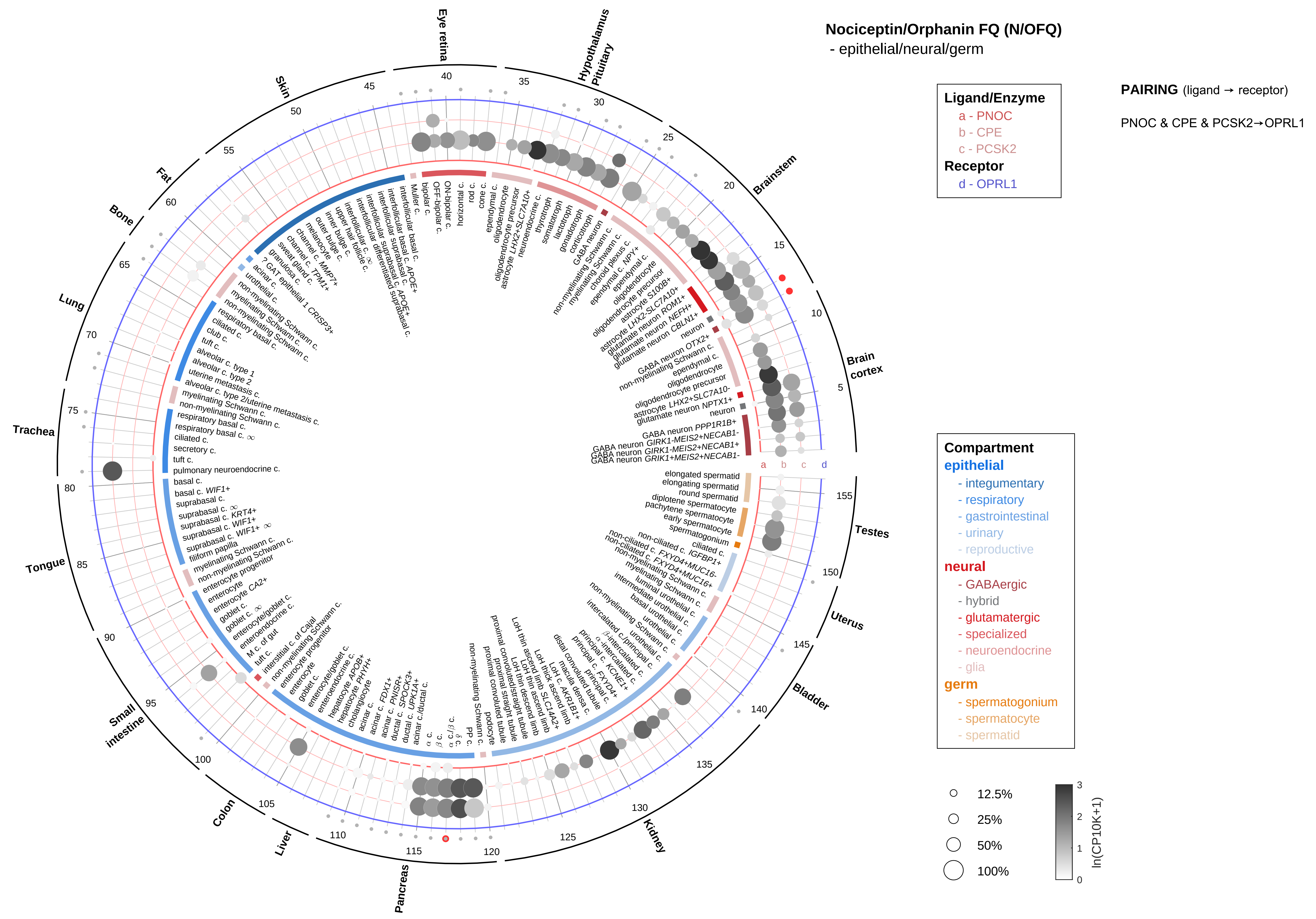
Nociceptin/Orphanin FQ (N/OFQ)
- epithelial/neural/germ
Ligand/Enzyme
a - PNOC
b - CPE
c - PCSK2
Receptor
d - OPRL1
PAIRING (ligand → receptor)
PNOC & CPE & PCSK2→OPRL1
Eye retina
Hypothalamus
Pituitary
Brainstem
Brain cortex
Testes
Uterus
Bladder
Kidney
Pancreas
Liver
Colon
Small intestine
Tongue
Trachea
Lung
Bone
Fat
Skin
a
b
c
d
Compartment
epithelial
- integumentary
- respiratory
- gastrointestinal
- urinary
- reproductive
neural
- GABAergic
- hybrid
- glutamatergic
- specialized
- neuroendocrine
- glia
germ
- spermatogonium
- spermatocyte
- spermatid
12.5%
25%
50%
100%
ln(CP10K+1)

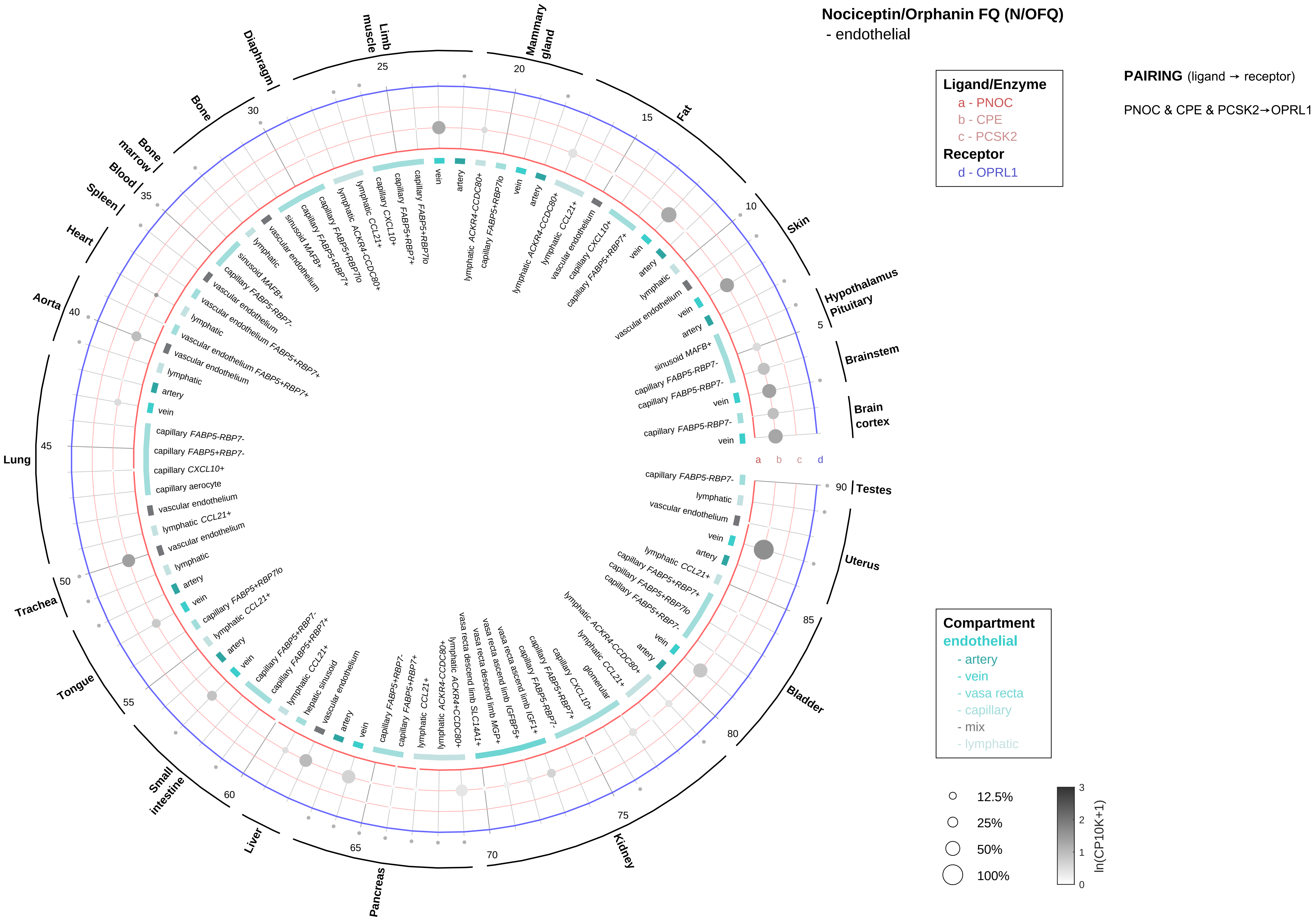

Nociceptin/Orphanin FQ (N/OFQ)
- endothelial
Ligand/Enzyme
a - PNOC
b - CPE
c - PCSK2
Receptor
d - OPRL1
PAIRING (ligand → receptor)
PNOC & CPE & PCSK2→OPRL1
Compartment
endothelial
- artery
- vein
- vasa recta
- capillary
- mix
- lymphatic
12.5%
25%
50%
100%
ln(CP10K+1)
Brain cortex
Brainstem
Hypothalamus Pituitary
Skin
Fat
Mammary gland
Limb muscle
Diaphragm
Bone
Bone marrow
Blood
Spleen
Heart
Aorta
Lung
Trachea
Tongue
Small intestine
Liver
Pancreas
Kidney
Bladder
Uterus
Testes

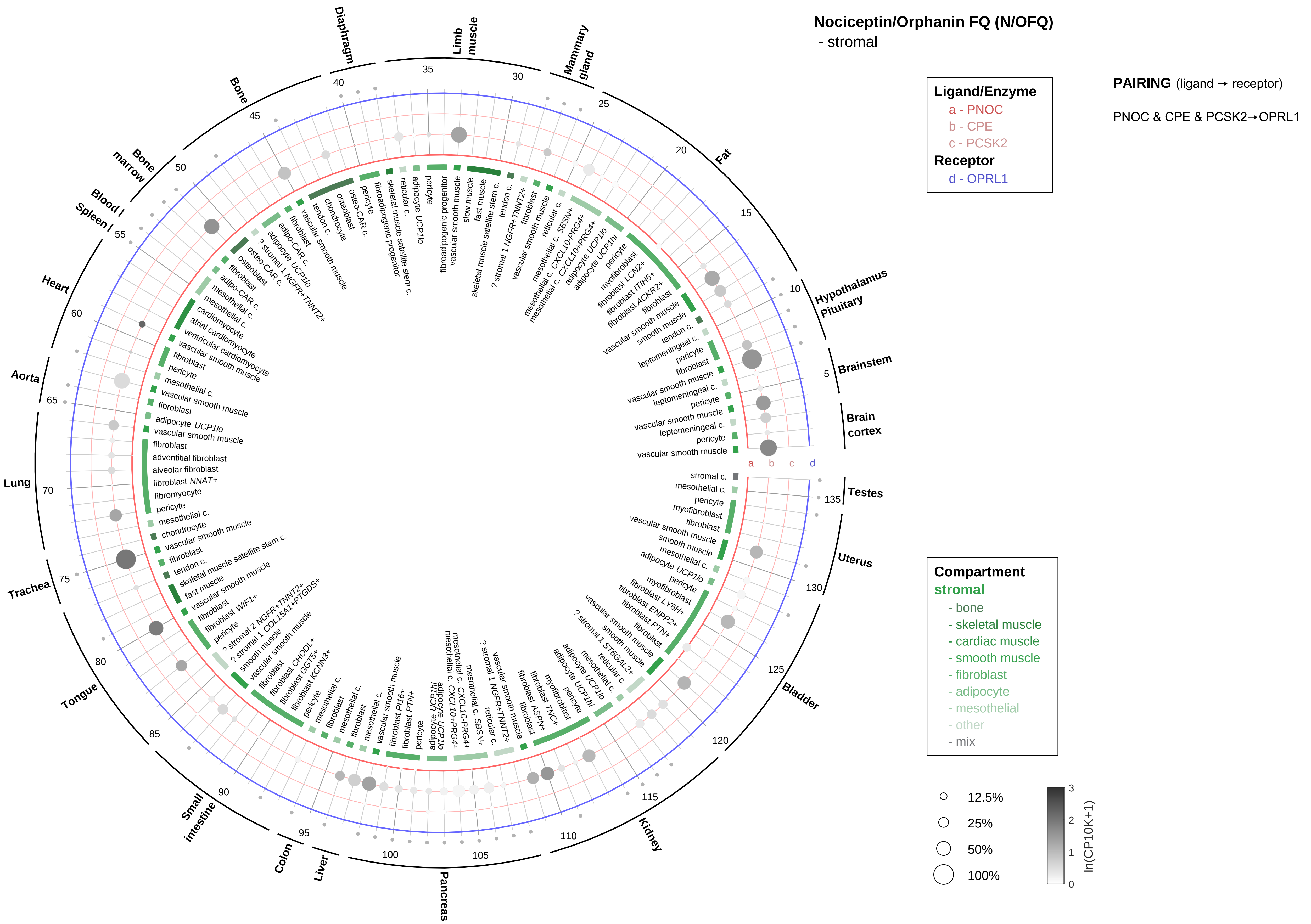
Nociceptin/Orphanin FQ (N/OFQ)
- stromal
Ligand/Enzyme
a - PNOC
b - CPE
c - PCSK2
Receptor
d - OPRL1
PAIRING (ligand → receptor)
PNOC & CPE & PCSK2→OPRL1
Compartment
stromal
- bone
- skeletal muscle
- cardiac muscle
- smooth muscle
- fibroblast
- adipocyte
- mesothelial
- other
- mix
12.5%
25%
50%
100%
ln(CP10K+1)
Hypothalamus
Pituitary
Brainstem
Brain cortex
Testes
Uterus
Bladder
Kidney
Pancreas
Liver
Colon
Small intestine
Tongue
Trachea
Lung
Aorta
Heart
Spleen
Blood
Bone marrow
Bone
Diaphragm
Limb muscle
Mammary gland
Fat

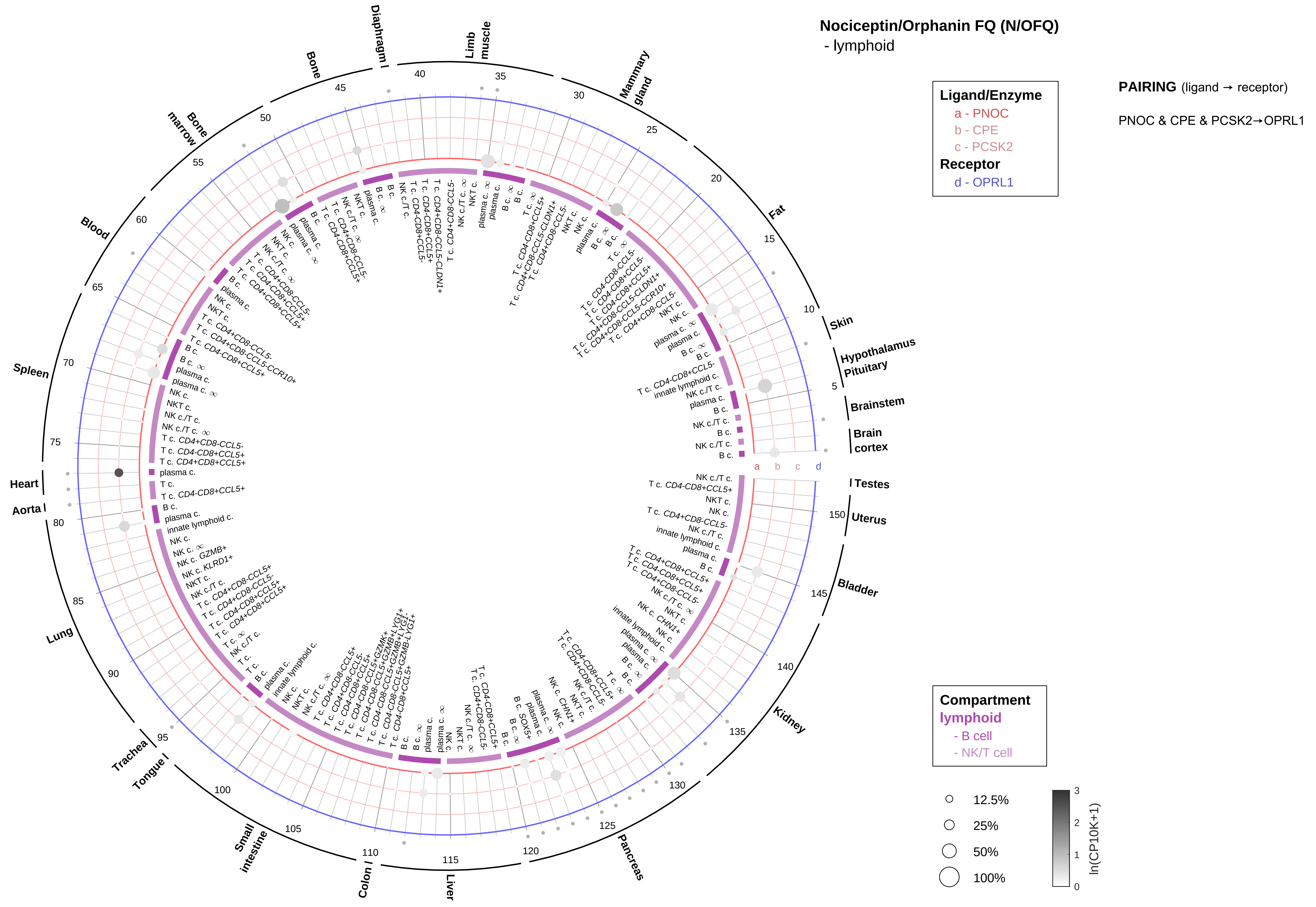

Nociceptin/Orphanin FQ (N/OFQ)
- lymphoid
Ligand/Enzyme
a - PNOC
b - CPE
c - PCSK2
Receptor
d - OPRL1
PAIRING (ligand → receptor)
PNOC & CPE & PCSK2→OPRL1
Compartment
lymphoid
- B cell
- NK/T cell
12.5%
25%
50%
100%
ln(CP10K+1)
Skin
Hypothalamus
Pituitary
Brainstem
Brain cortex
Testes
Uterus
Bladder
Kidney
Pancreas
Liver
Colon
Small intestine
Tongue
Trachea
Lung
Aorta
Heart
Spleen
Blood
Bone marrow
Bone
Diaphragm
Limb muscle
Mammary gland
Fat

Nociceptin/Orphanin FQ (N/OFQ)
- myeloid

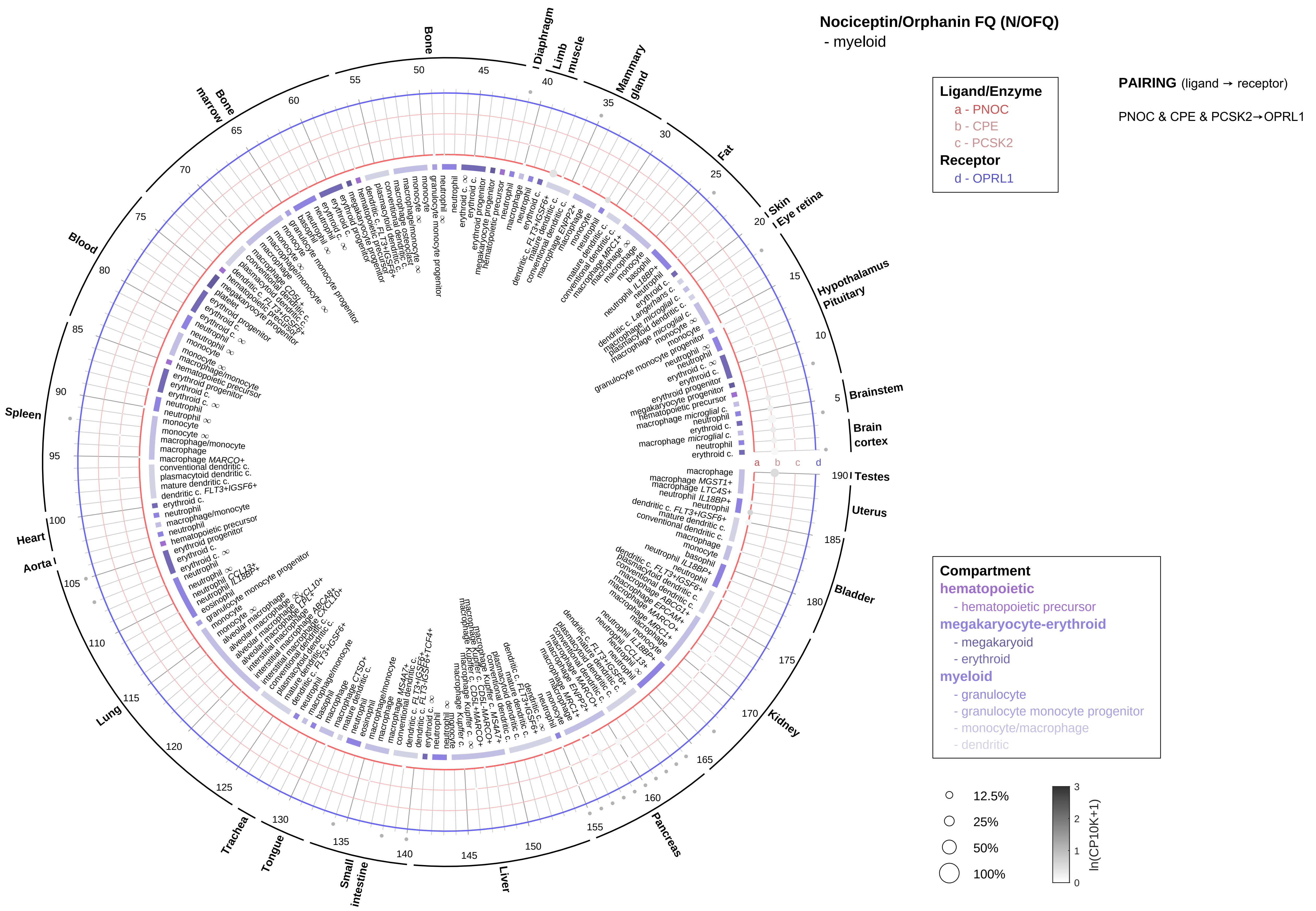

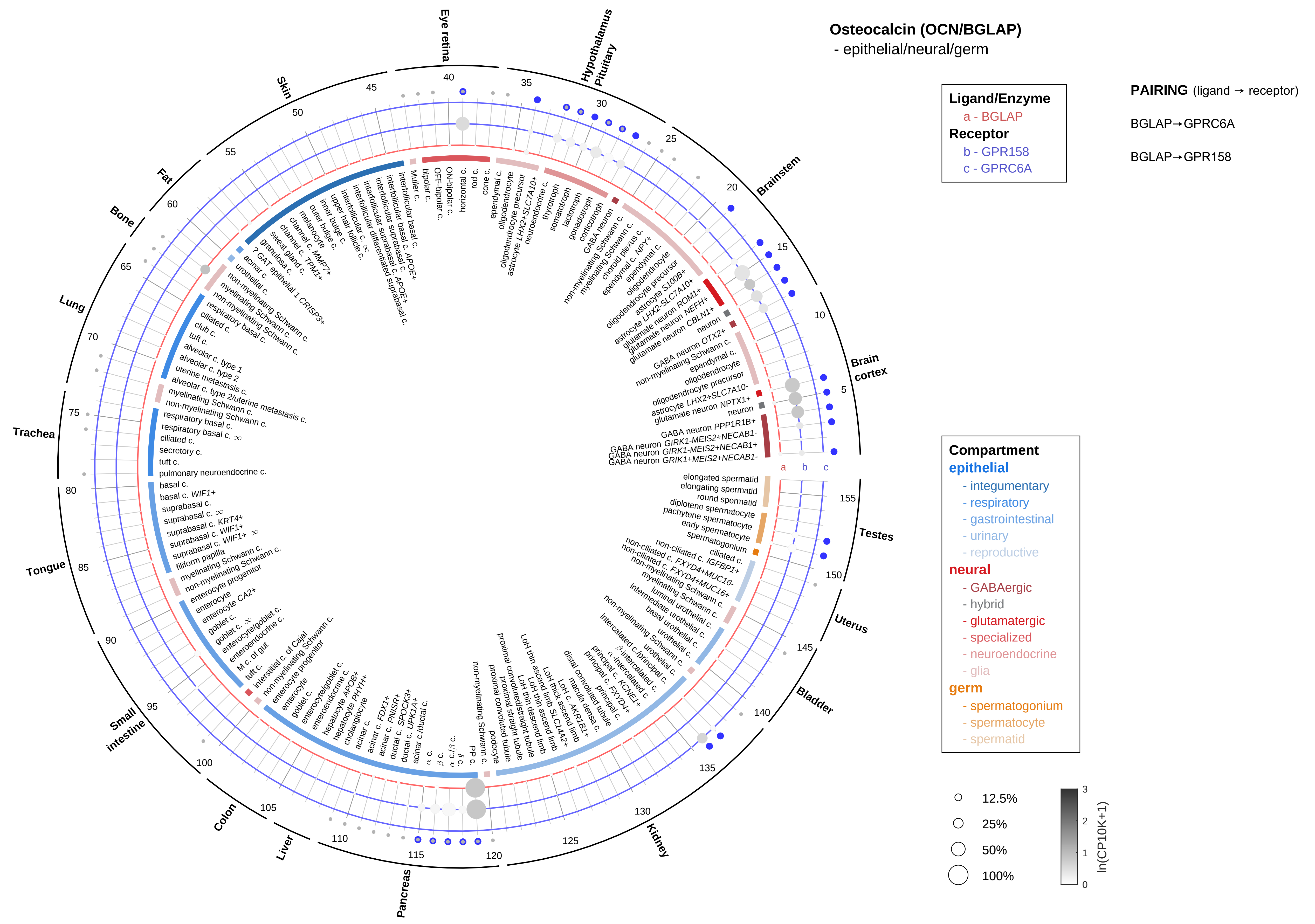

Osteocalcin (OCN/BGLAP)
- epithelial/neural/germ
Ligand/Enzyme
a - BGLAP
Receptor
b - GPR158
c - GPRC6A
PAIRING (ligand → receptor)
BGLAP→GPRC6A
BGLAP→GPR158
Compartment
epithelial
- integumentary
- respiratory
- gastrointestinal
- urinary
- reproductive
neural
- GABAergic
- hybrid
- glutamatergic
- specialized
- neuroendocrine
- glia
germ
- spermatogonium
- spermatocyte
- spermatid
12.5%
25%
50%
100%
ln(CP10K+1)
Eye retina
Hypothalamus Pituitary
Brainstem
Brain cortex
Testes
Uterus
Bladder
Kidney
Pancreas
Liver
Colon
Small intestine
Tongue
Trachea
Lung
Bone
Fat
Skin

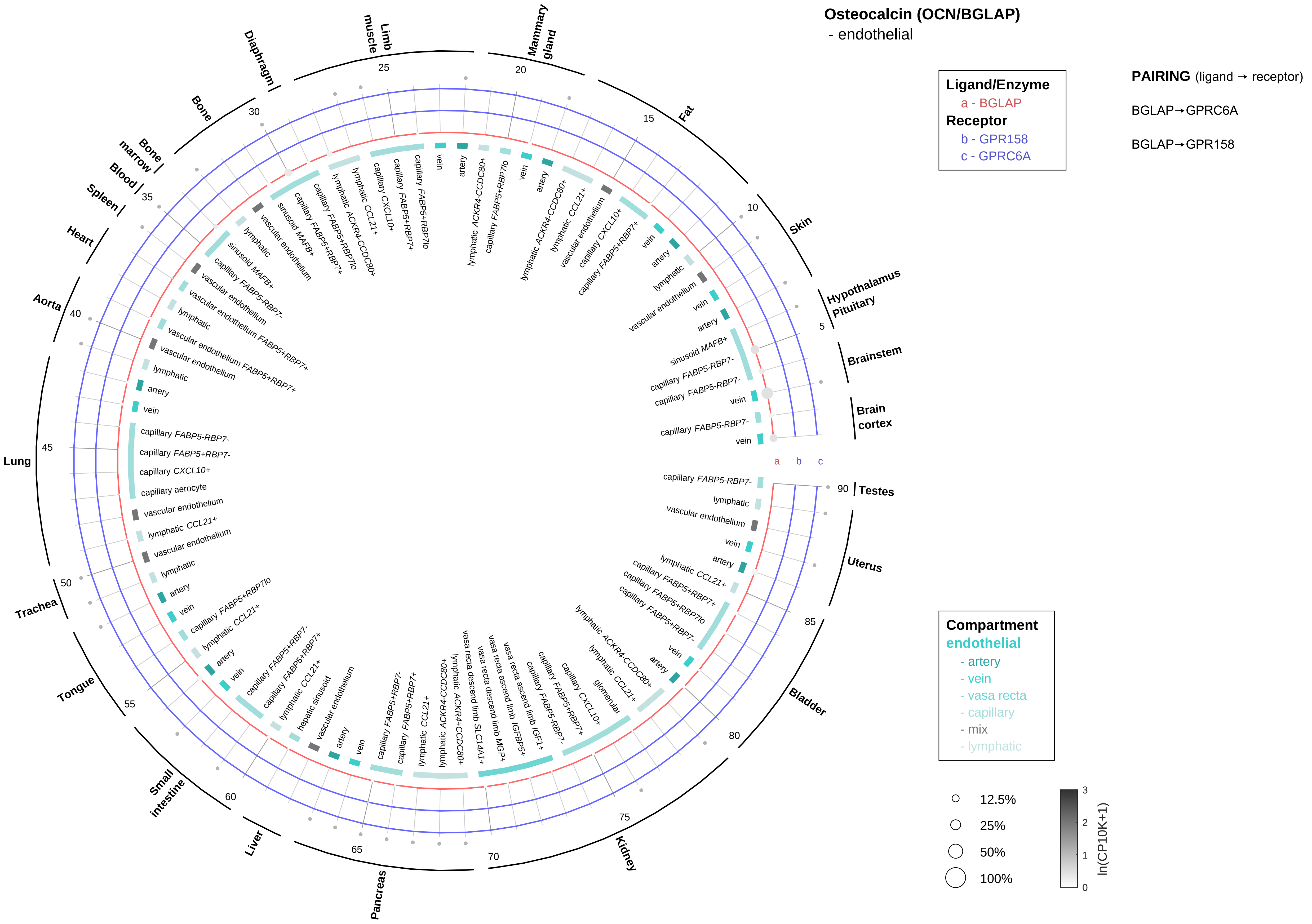
Osteocalcin (OCN/BGLAP)
- endothelial
Ligand/Enzyme
a - BGLAP
Receptor
b - GPR158
c - GPRC6A
PAIRING (ligand → receptor)
BGLAP→GPRC6A
BGLAP→GPR158
Compartment
endothelial
- artery
- vein
- vasa recta
- capillary
- mix
- lymphatic
12.5%
25%
50%
100%
ln(CP10K+1)
Brain cortex
Brainstem
Hypothalamus Pituitary
Skin
Fat
Mammary gland
Limb muscle
Diaphragm
Bone
Bone marrow
Blood
Spleen
Heart
Aorta
Lung
Trachea
Tongue
Small intestine
Liver
Pancreas
Kidney
Bladder
Uterus
Testes

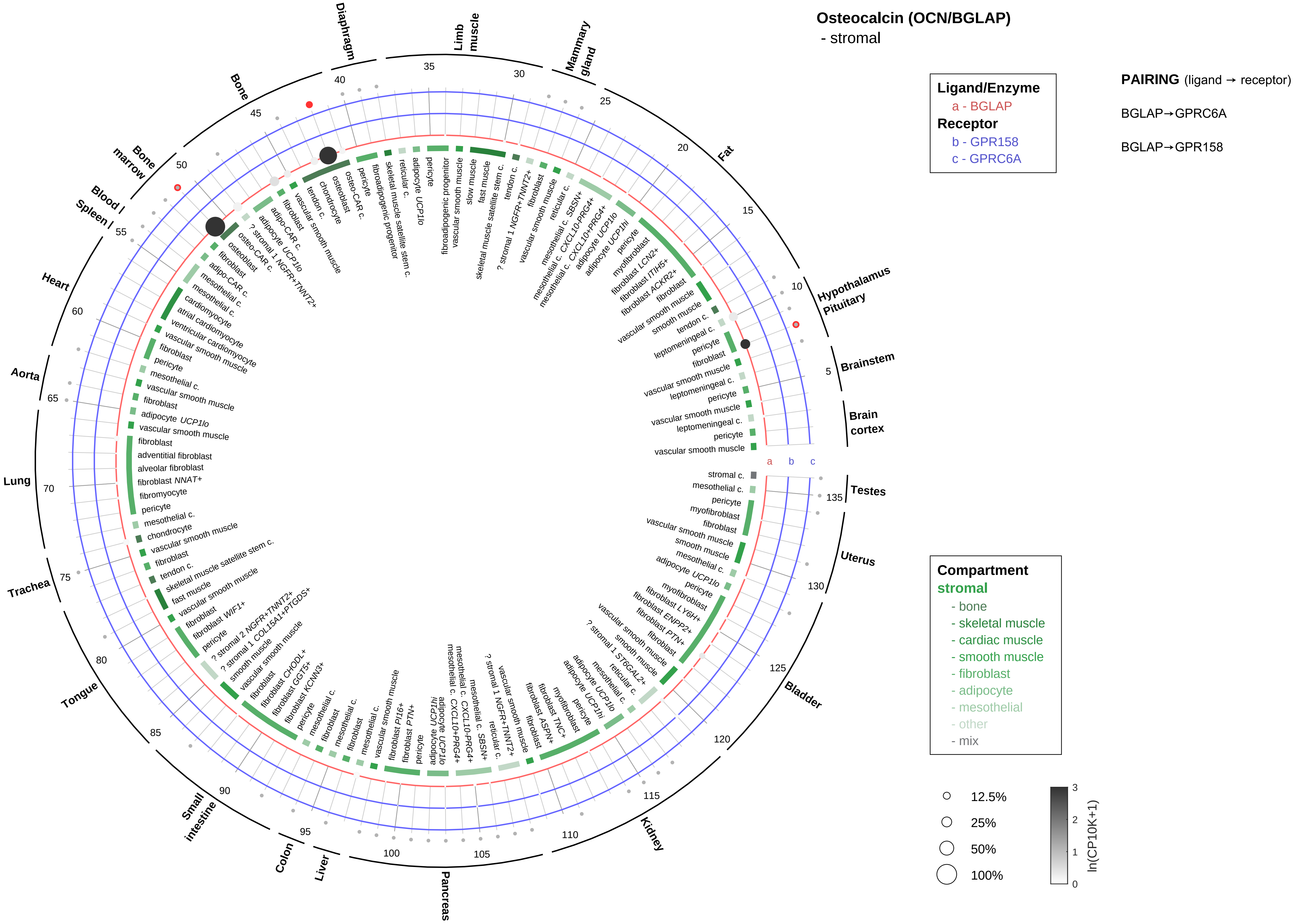

Osteocalcin (OCN/BGLAP)
- stromal
Ligand/Enzyme
a - BGLAP
Receptor
b - GPR158
c - GPRC6A
PAIRING (ligand → receptor)
BGLAP→GPRC6A
BGLAP→GPR158
Compartment
stromal
- bone
- skeletal muscle
- cardiac muscle
- smooth muscle
- fibroblast
- adipocyte
- mesothelial
- other
- mix
12.5%
25%
50%
100%
ln(CP10K+1)
Brain cortex
Brainstem
Hypothalamus Pituitary
Fat
Mammary gland
Limb muscle
Diaphragm
Bone
Bone marrow
Blood
Spleen
Heart
Aorta
Lung
Trachea
Tongue
Small intestine
Colon
Liver
Pancreas
Kidney
Bladder
Uterus
Testes

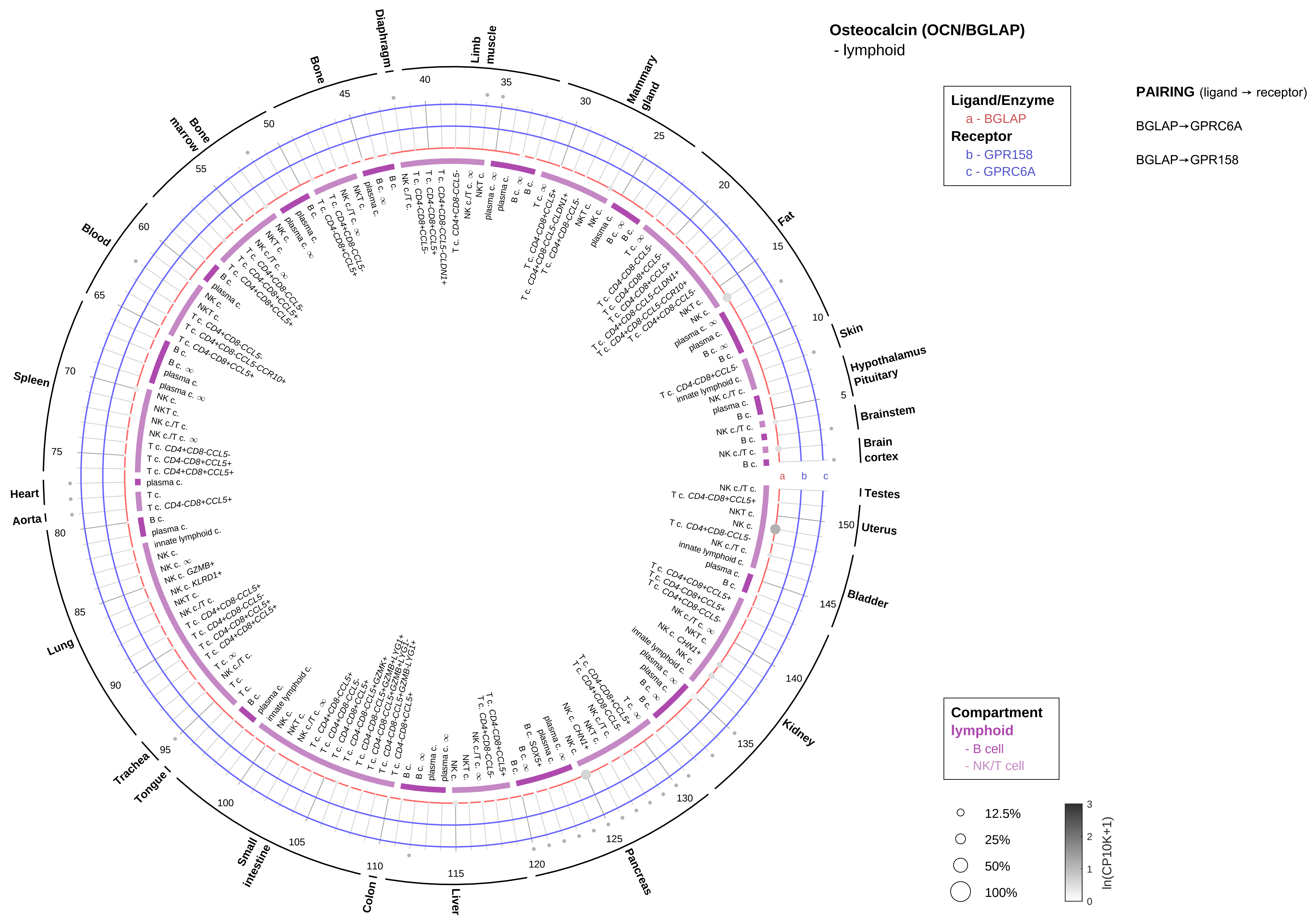
Osteocalcin (OCN/BGLAP)
- lymphoid
Ligand/Enzyme
a - BGLAP
Receptor
b - GPR158
c - GPRC6A
PAIRING (ligand → receptor)
BGLAP→GPRC6A
BGLAP→GPR158
Compartment
lymphoid
- B cell
- NK/T cell
12.5%
25%
50%
100%
ln(CP10K+1)
Skin
Hypothalamus
Pituitary
Brainstem
Brain cortex
Testes
Uterus
Bladder
Kidney
Pancreas
Liver
Colon
Small intestine
Tongue
Trachea
Lung
Aorta
Heart
Spleen
Blood
Bone marrow
Bone
Diaphragm
Limb muscle
Mammary gland
Fat

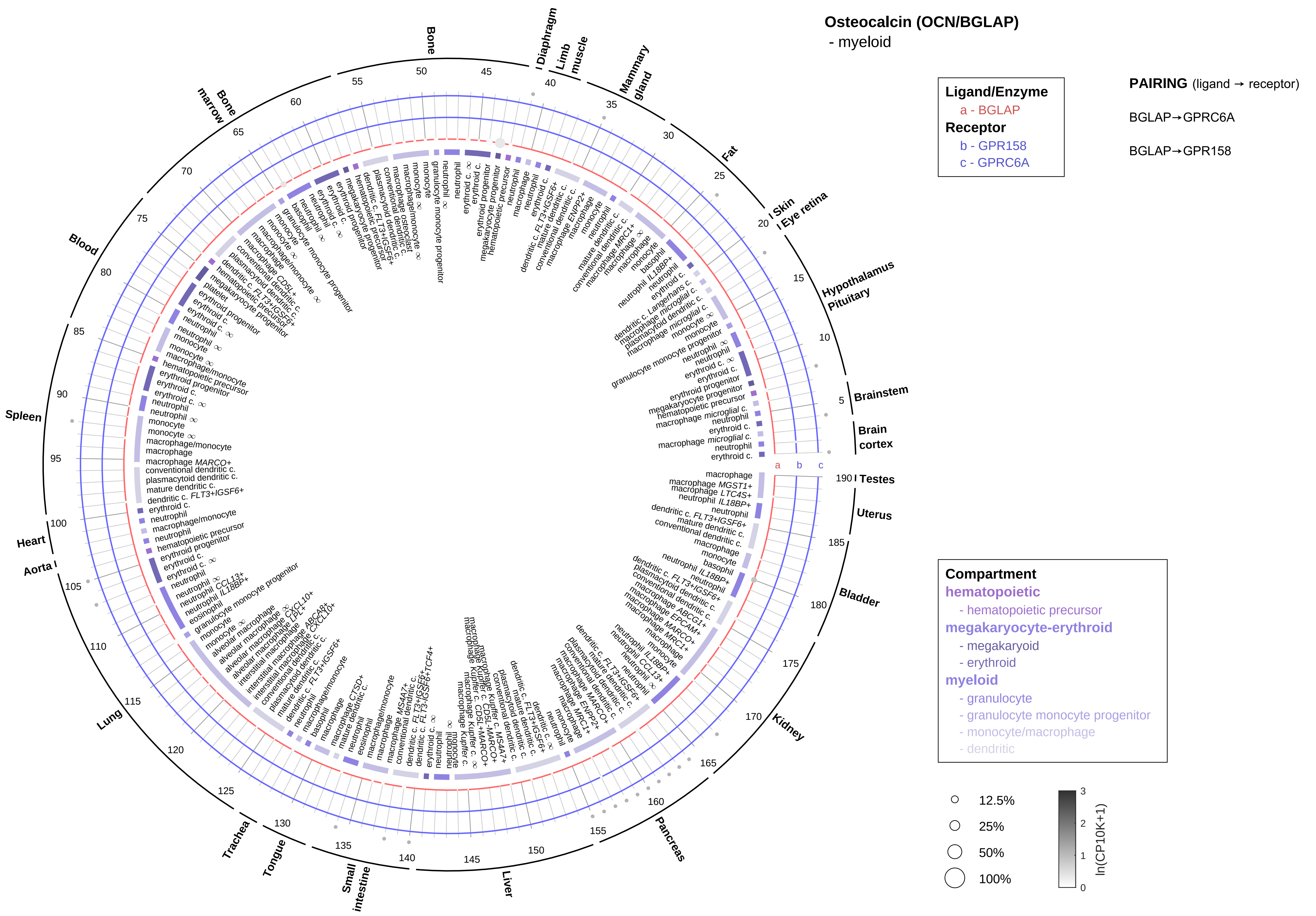

Osteocalcin (OCN/BGLAP)
- myeloid
Ligand/Enzyme
a - BGLAP
Receptor
b - GPR158
c - GPRC6A
PAIRING (ligand → receptor)
BGLAP→GPRC6A
BGLAP→GPR158
Compartment
hematopoietic
- hematopoietic precursor
megakaryocyte-erythroid
- megakaryoid
- erythroid
myeloid
- granulocyte
- granulocyte monocyte progenitor
- monocyte/macrophage
- dendritic
12.5%
25%
50%
100%
ln(CP10K+1)
Bone
Diaphragm
Limb muscle
Mammary gland
Fat
Skin
Eye retina
Hypothalamus
Pituitary
Brainstem
Brain cortex
Testes
Uterus
Bladder
Kidney
Pancreas
Liver
Small intestine
Tongue
Trachea
Lung
Aorta
Heart
Spleen
Blood
Bone marrow

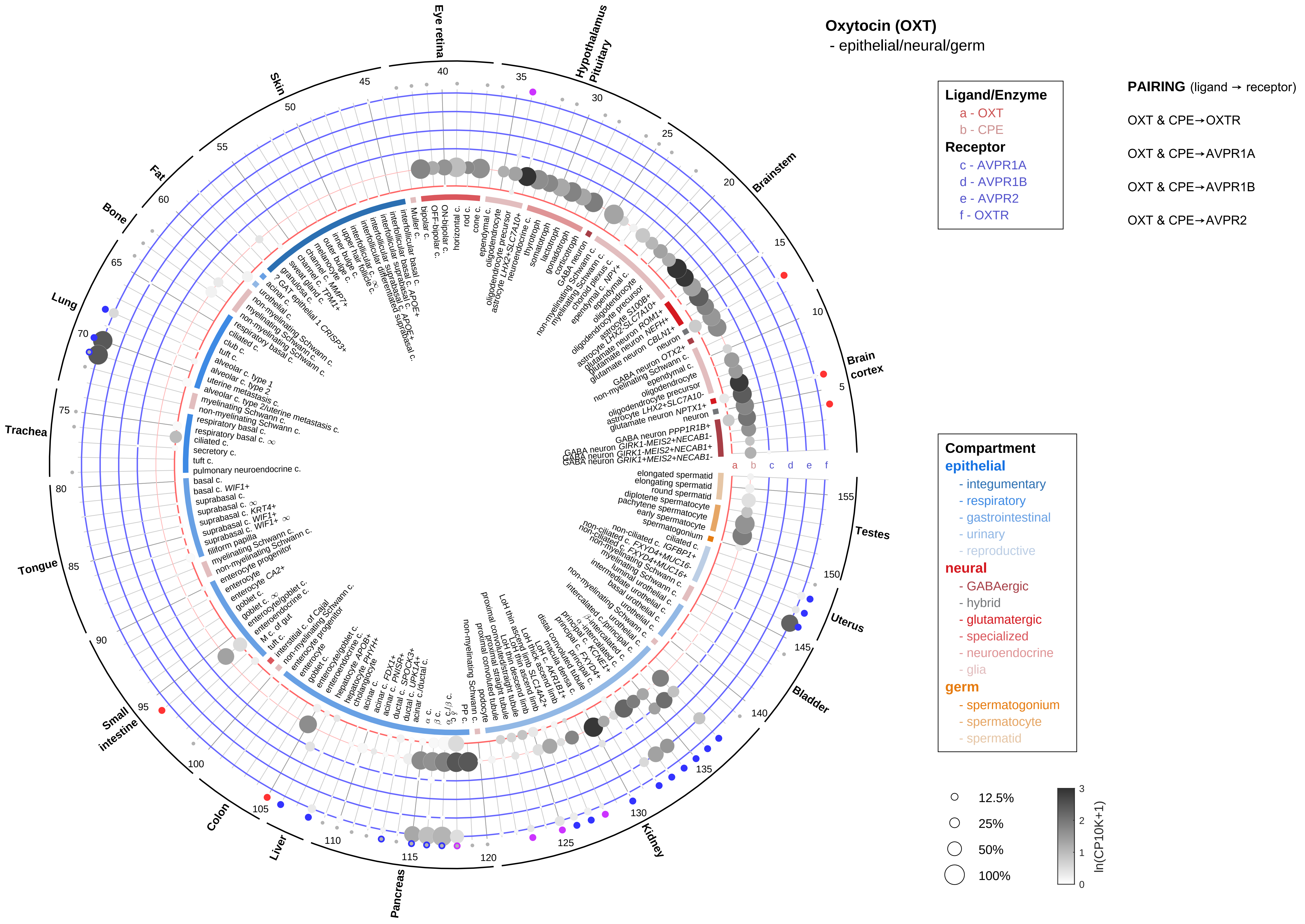

Oxytocin (OXT)
- epithelial/neural/germ
Ligand/Enzyme
a - OXT
b - CPE
Receptor
c - AVPR1A
d - AVPR1B
e - AVPR2
f - OXTR
PAIRING (ligand → receptor)
OXT & CPE→OXTR
OXT & CPE→AVPR1A
OXT & CPE→AVPR1B
OXT & CPE→AVPR2
Compartment
epithelial
- integumentary
- respiratory
- gastrointestinal
- urinary
- reproductive
neural
- GABAergic
- hybrid
- glutamatergic
- specialized
- neuroendocrine
- glia
germ
- spermatogonium
- spermatocyte
- spermatid
12.5%
25%
50%
100%
ln(CP10K+1)
Eye retina
Hypothalamus Pituitary
Brainstem
Brain cortex
Testes
Uterus
Bladder
Kidney
Pancreas
Liver
Colon
Small intestine
Tongue
Trachea
Lung
Bone
Fat
Skin

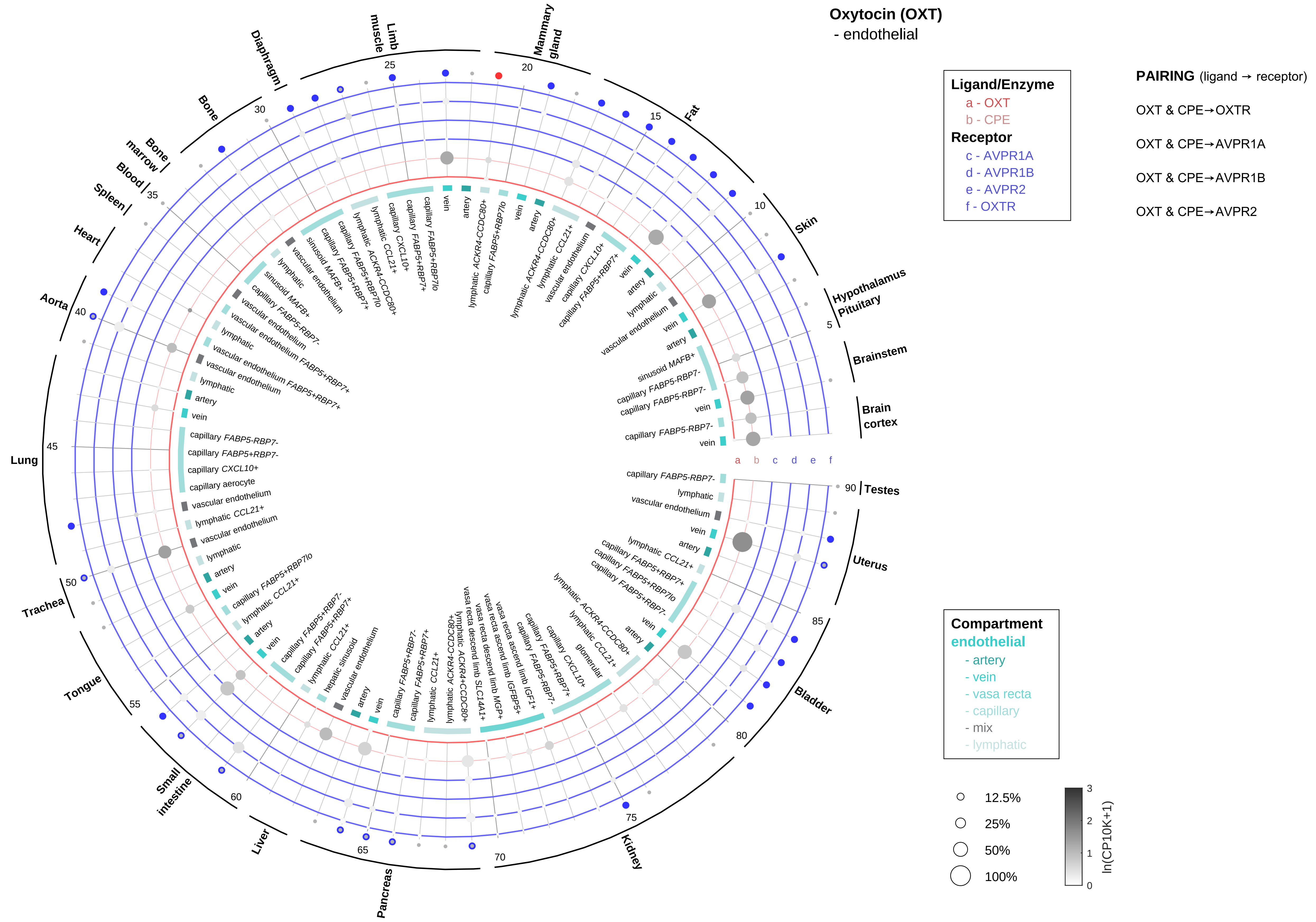
Oxytocin (OXT)
- endothelial
Ligand/Enzyme
a - OXT
b - CPE
Receptor
c - AVPR1A
d - AVPR1B
e - AVPR2
f - OXTR
PAIRING (ligand → receptor)
OXT & CPE→OXTR
OXT & CPE→AVPR1A
OXT & CPE→AVPR1B
OXT & CPE→AVPR2
Compartment
endothelial
- artery
- vein
- vasa recta
- capillary
- mix
- lymphatic
12.5%
25%
50%
100%
ln(CP10K+1)
Brain cortex
Brainstem
Hypothalamus Pituitary
Skin
Fat
Mammary gland
Limb muscle
Diaphragm
Bone
Bone marrow
Blood
Spleen
Heart
Aorta
Lung
Trachea
Tongue
Small intestine
Liver
Pancreas
Kidney
Bladder
Uterus
Testes

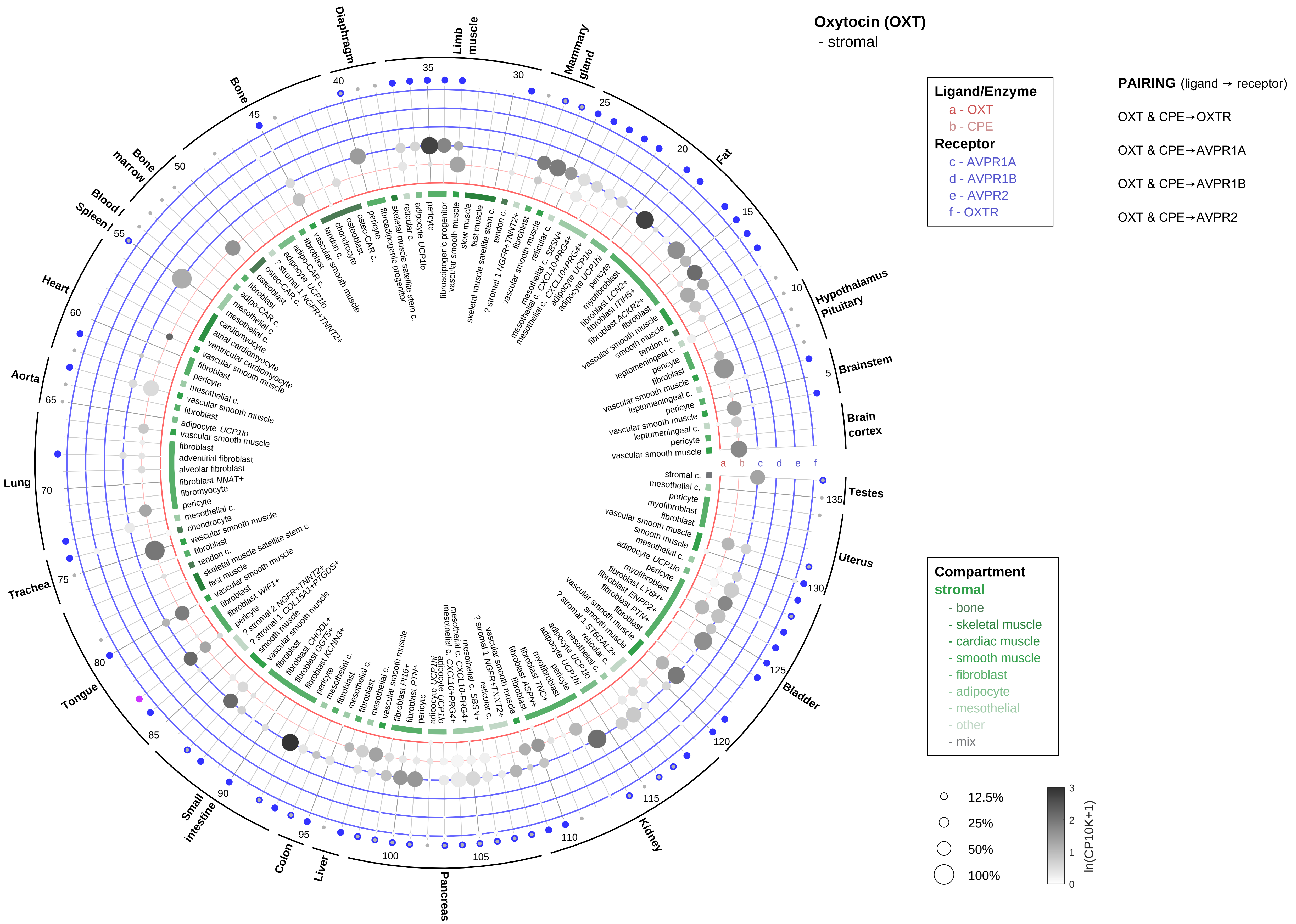
Oxytocin (OXT)
- stromal
Ligand/Enzyme
a - OXT
b - CPE
Receptor
c - AVPR1A
d - AVPR1B
e - AVPR2
f - OXTR
PAIRING (ligand → receptor)
OXT & CPE→OXTR
OXT & CPE→AVPR1A
OXT & CPE→AVPR1B
OXT & CPE→AVPR2
Compartment
stromal
- bone
- skeletal muscle
- cardiac muscle
- smooth muscle
- fibroblast
- adipocyte
- mesothelial
- other
- mix
12.5%
25%
50%
100%
ln(CP10K+1)
Diaphragm
Limb muscle
Mammary gland
Fat
Hypothalamus Pituitary
Brainstem
Brain cortex
Testes
Uterus
Bladder
Kidney
Pancreas
Liver
Colon
Small intestine
Tongue
Trachea
Lung
Aorta
Heart
Spleen
Blood
Bone marrow
Bone

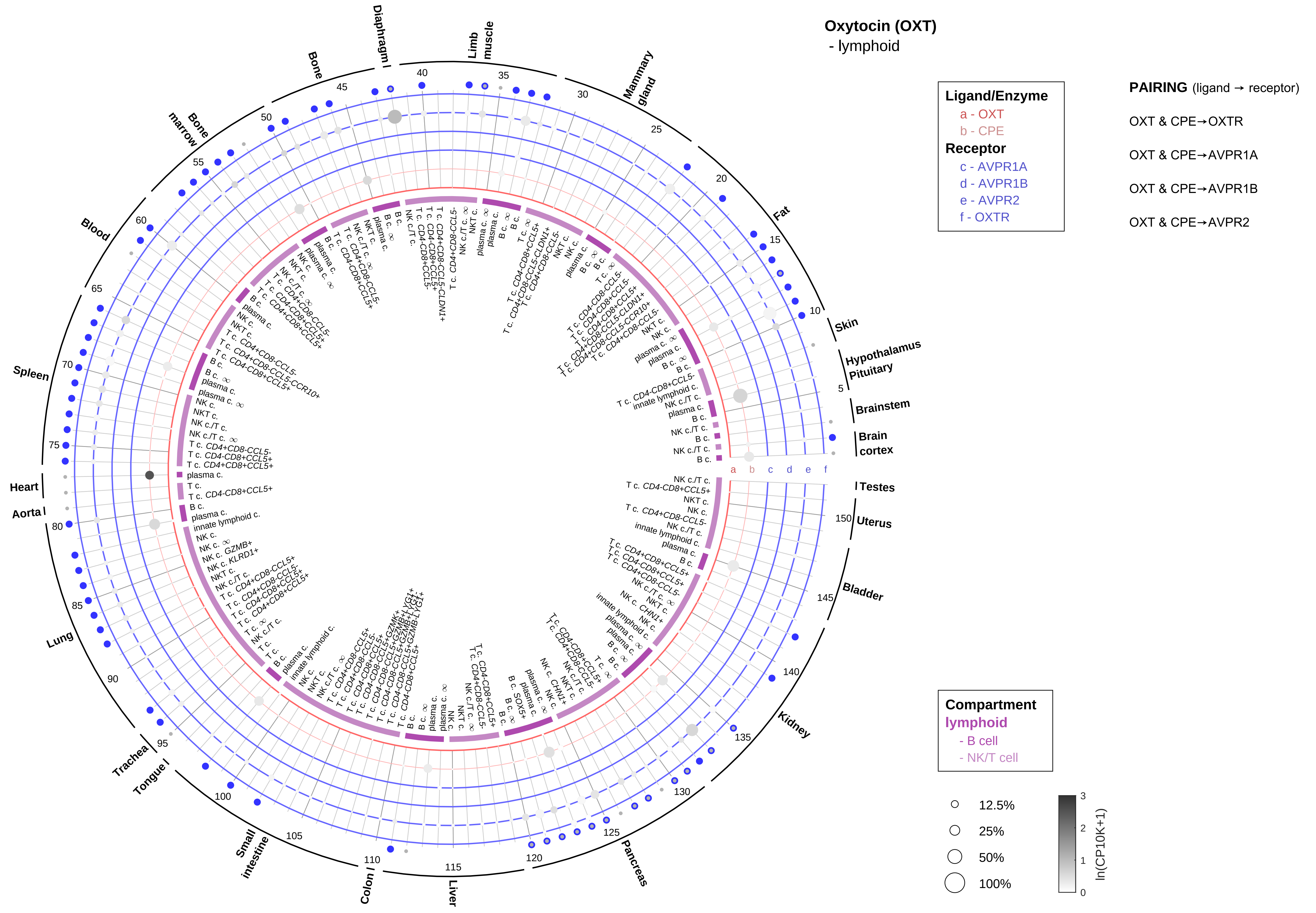

Oxytocin (OXT)
- lymphoid
Ligand/Enzyme
a - OXT
b - CPE
Receptor
c - AVPR1A
d - AVPR1B
e - AVPR2
f - OXTR
PAIRING (ligand → receptor)
OXT & CPE→OXTR
OXT & CPE→AVPR1A
OXT & CPE→AVPR1B
OXT & CPE→AVPR2
Compartment
lymphoid
- B cell
- NK/T cell
12.5%
25%
50%
100%
ln(CP10K+1)
Skin
Hypothalamus
Pituitary
Brainstem
Brain cortex
Testes
Uterus
Bladder
Kidney
Pancreas
Liver
Colon
Small intestine
Tongue
Trachea
Lung
Aorta
Heart
Spleen
Blood
Bone marrow
Bone
Diaphragm
Limb muscle
Mammary gland
Fat

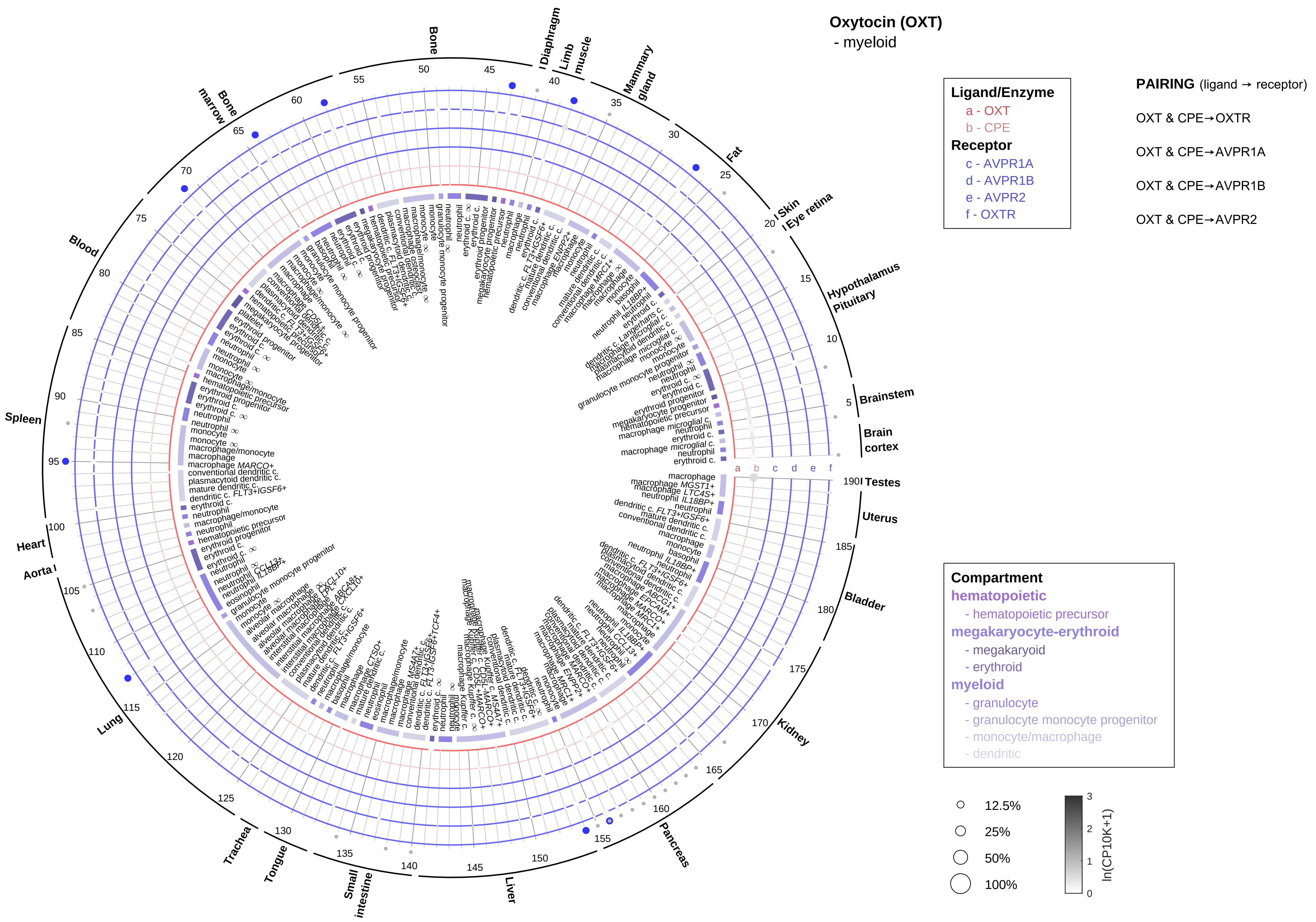

Oxytocin (OXT)
- myeloid
Ligand/Enzyme
a - OXT
b - CPE
Receptor
c - AVPR1A
d - AVPR1B
e - AVPR2
f - OXTR
PAIRING (ligand → receptor)
OXT & CPE→OXTR
OXT & CPE→AVPR1A
OXT & CPE→AVPR1B
OXT & CPE→AVPR2
Compartment
hematopoietic
- hematopoietic precursor
megakaryocyte-erythroid
- megakaryoid
- erythroid
myeloid
- granulocyte
- granulocyte monocyte progenitor
- monocyte/macrophage
- dendritic
12.5%
25%
50%
100%
ln(CP10K+1)
Bone
Diaphragm
Limb muscle
Mammary gland
Fat
Skin
Eye retina
Hypothalamus
Pituitary
Brainstem
Brain cortex
Testes
Uterus
Bladder
Kidney
Pancreas
Liver
Small intestine
Tongue
Trachea
Lung
Aorta
Heart
Spleen
Blood
Bone marrow

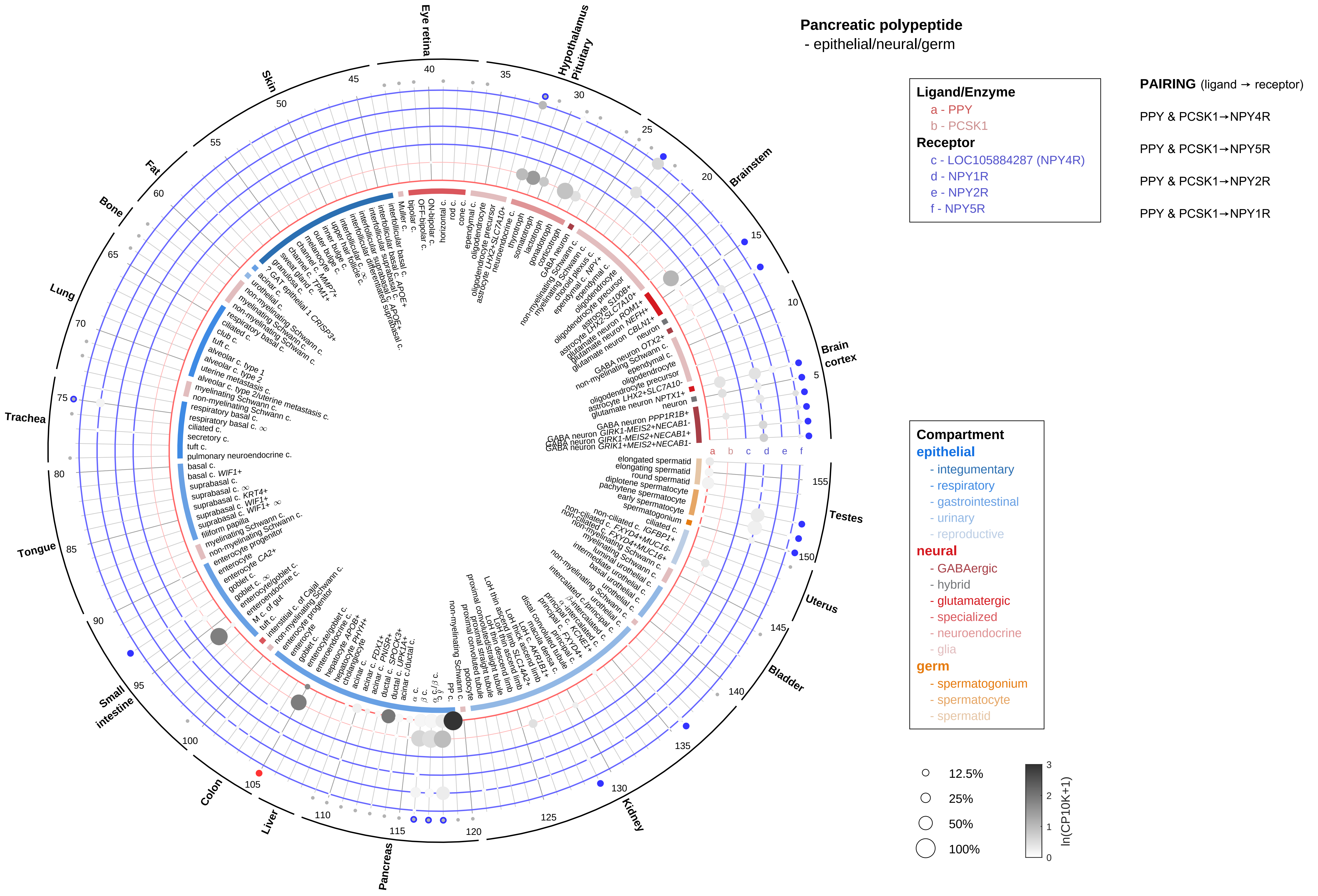

Pancreatic polypeptide
- epithelial/neural/germ
Ligand/Enzyme
a - PPY
b - PCSK1
Receptor
c - LOC105884287 (NPY4R)
d - NPY1R
e - NPY2R
f - NPY5R
PAIRING (ligand → receptor)
PPY & PCSK1→NPY4R
PPY & PCSK1→NPY5R
PPY & PCSK1→NPY2R
PPY & PCSK1→NPY1R
Compartment
epithelial
- integumentary
- respiratory
- gastrointestinal
- urinary
- reproductive
neural
- GABAergic
- hybrid
- glutamatergic
- specialized
- neuroendocrine
- glia
germ
- spermatogonium
- spermatocyte
- spermatid
12.5%
25%
50%
100%
ln(CP10K+1)
Eye retina
Hypothalamus
Pituitary
Brainstem
Brain cortex
Testes
Uterus
Bladder
Kidney
Pancreas
Liver
Colon
Small intestine
Tongue
Trachea
Lung
Bone
Fat
Skin

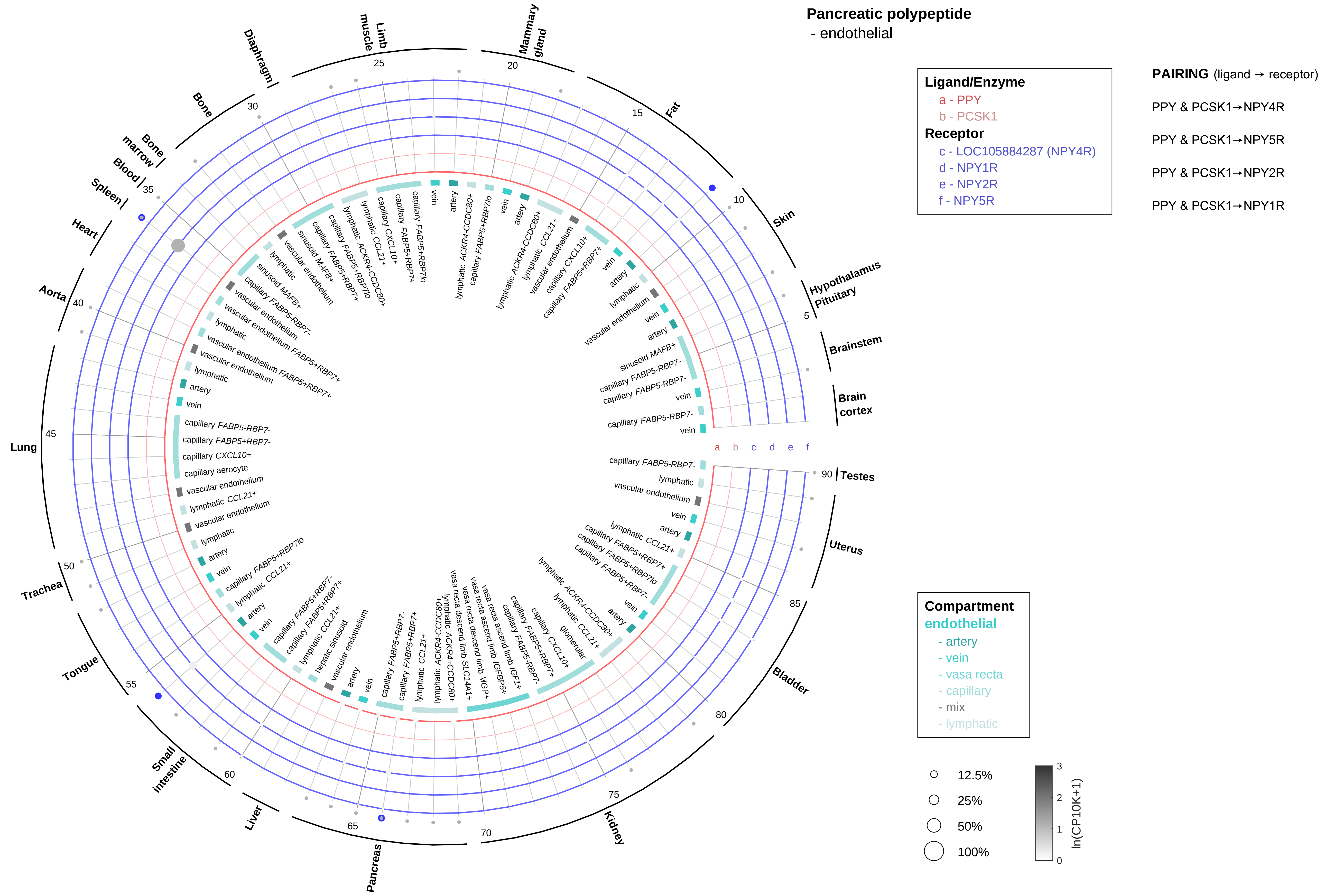

Pancreatic polypeptide
- endothelial
Ligand/Enzyme
a - PPY
b - PCSK1
Receptor
c - LOC105884287 (NPY4R)
d - NPY1R
e - NPY2R
f - NPY5R
PAIRING (ligand → receptor)
PPY & PCSK1→NPY4R
PPY & PCSK1→NPY5R
PPY & PCSK1→NPY2R
PPY & PCSK1→NPY1R
Compartment
endothelial
- artery
- vein
- vasa recta
- capillary
- mix
- lymphatic
12.5%
25%
50%
100%
ln(CP10K+1)

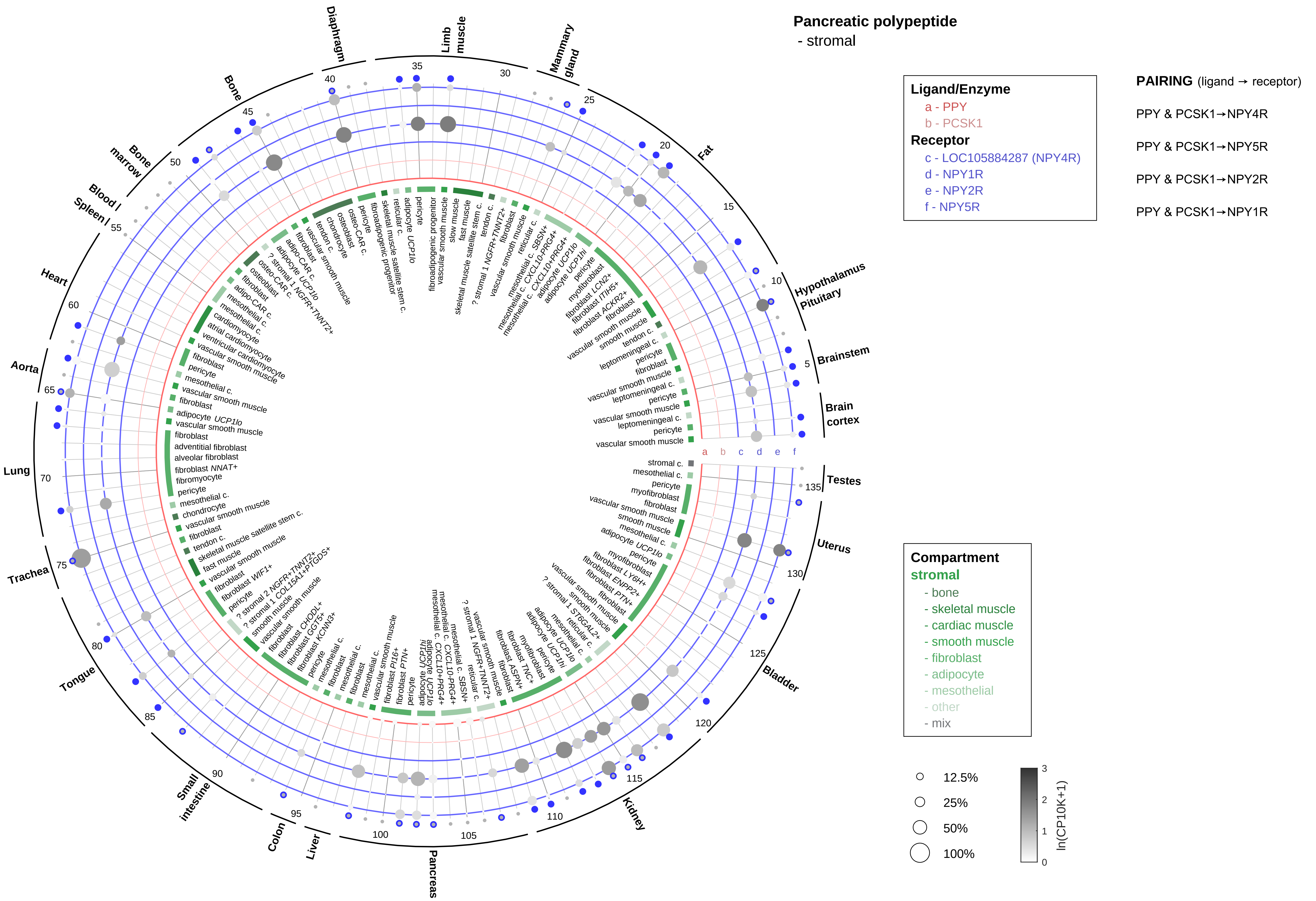

Pancreatic polypeptide
- stromal
Ligand/Enzyme
a - PPY
b - PCSK1
Receptor
c - LOC105884287 (NPY4R)
d - NPY1R
e - NPY2R
f - NPY5R
PAIRING (ligand → receptor)
PPY & PCSK1→NPY4R
PPY & PCSK1→NPY5R
PPY & PCSK1→NPY2R
PPY & PCSK1→NPY1R
Compartment
stromal
- bone
- skeletal muscle
- cardiac muscle
- smooth muscle
- fibroblast
- adipocyte
- mesothelial
- other
- mix
12.5%
25%
50%
100%
ln(CP10K+1)

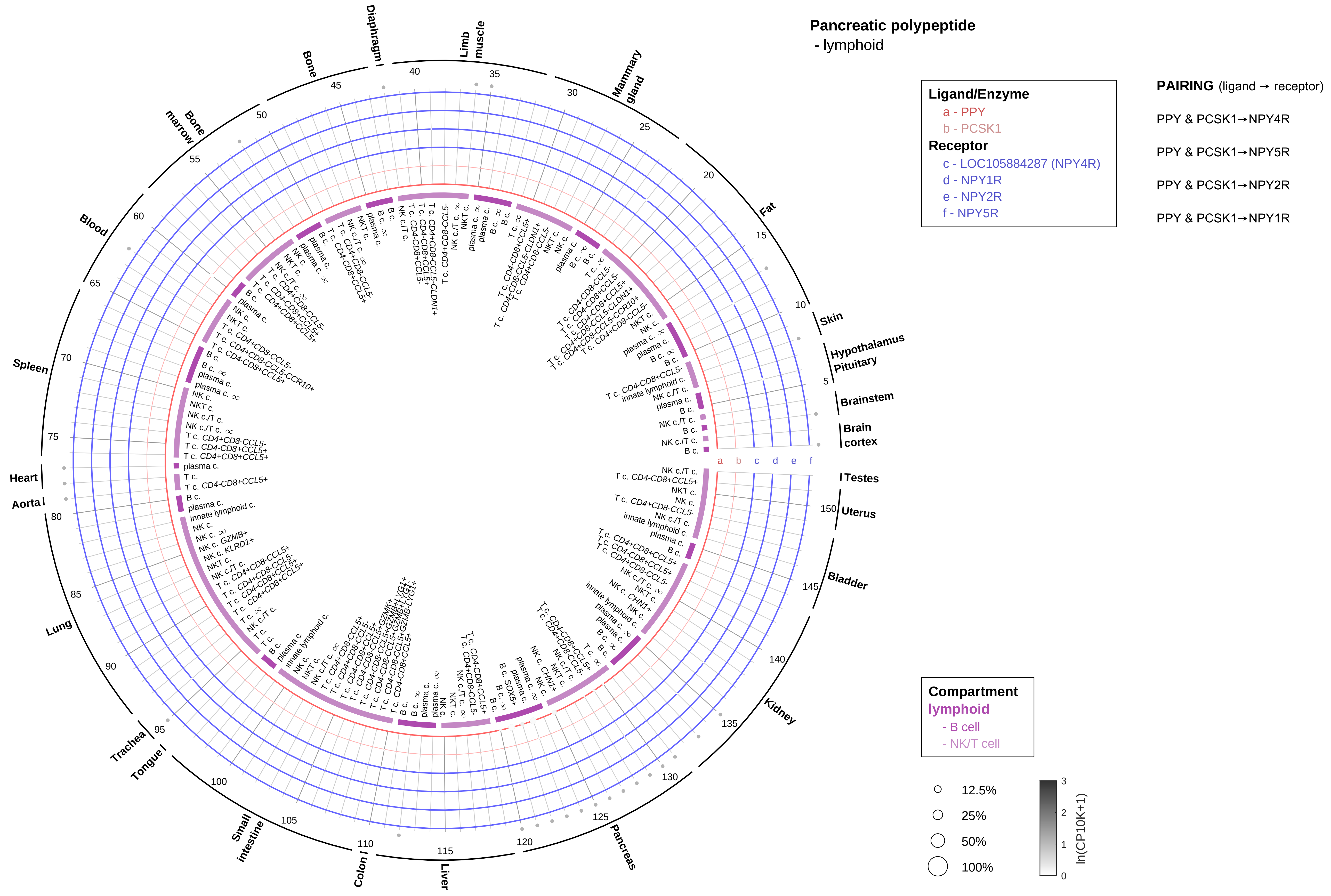

Pancreatic polypeptide
- lymphoid
Ligand/Enzyme
a - PPY
b - PCSK1
Receptor
c - LOC105884287 (NPY4R)
d - NPY1R
e - NPY2R
f - NPY5R
PAIRING (ligand → receptor)
PPY & PCSK1→NPY4R
PPY & PCSK1→NPY5R
PPY & PCSK1→NPY2R
PPY & PCSK1→NPY1R
Compartment
lymphoid
- B cell
- NK/T cell
12.5%
25%
50%
100%
ln(CP10K+1)
Skin
Hypothalamus
Pituitary
Brainstem
Brain cortex
Testes
Uterus
Bladder
Kidney
Pancreas
Liver
Colon
Small intestine
Tongue
Trachea
Lung
Aorta
Heart
Spleen
Blood
Bone marrow
Bone
Diaphragm
Limb muscle
Mammary gland
Fat

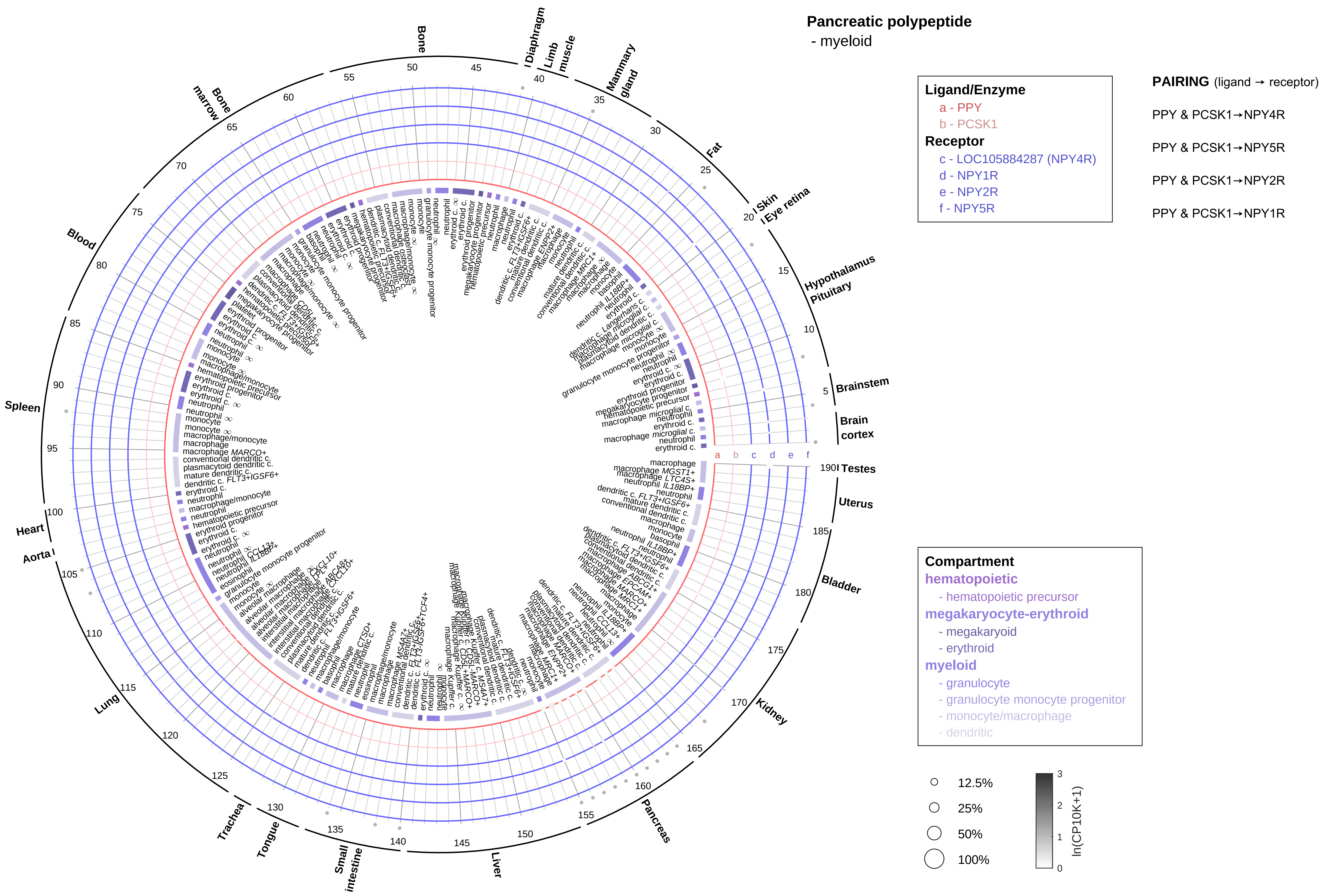

Pancreatic polypeptide
- myeloid
Ligand/Enzyme
a - PPY
b - PCSK1
Receptor
c - LOC105884287 (NPY4R)
d - NPY1R
e - NPY2R
f - NPY5R
PAIRING (ligand → receptor)
PPY & PCSK1→NPY4R
PPY & PCSK1→NPY5R
PPY & PCSK1→NPY2R
PPY & PCSK1→NPY1R
Compartment
hematopoietic
- hematopoietic precursor
megakaryocyte-erythroid
- megakaryoid
- erythroid
myeloid
- granulocyte
- granulocyte monocyte progenitor
- monocyte/macrophage
- dendritic
12.5%
25%
50%
100%
ln(CP10K+1)
Bone
Diaphragm
Limb muscle
Mammary gland
Fat
Skin
Eye retina
Hypothalamus
Pituitary
Brainstem
Brain cortex
Testes
Uterus
Bladder
Kidney
Pancreas
Liver
Small intestine
Tongue
Trachea
Lung
Aorta
Heart
Spleen
Blood
Bone marrow

**Parathyroid hormone/Parathyroid hormone-like hormone**
- epithelial/neural/germ

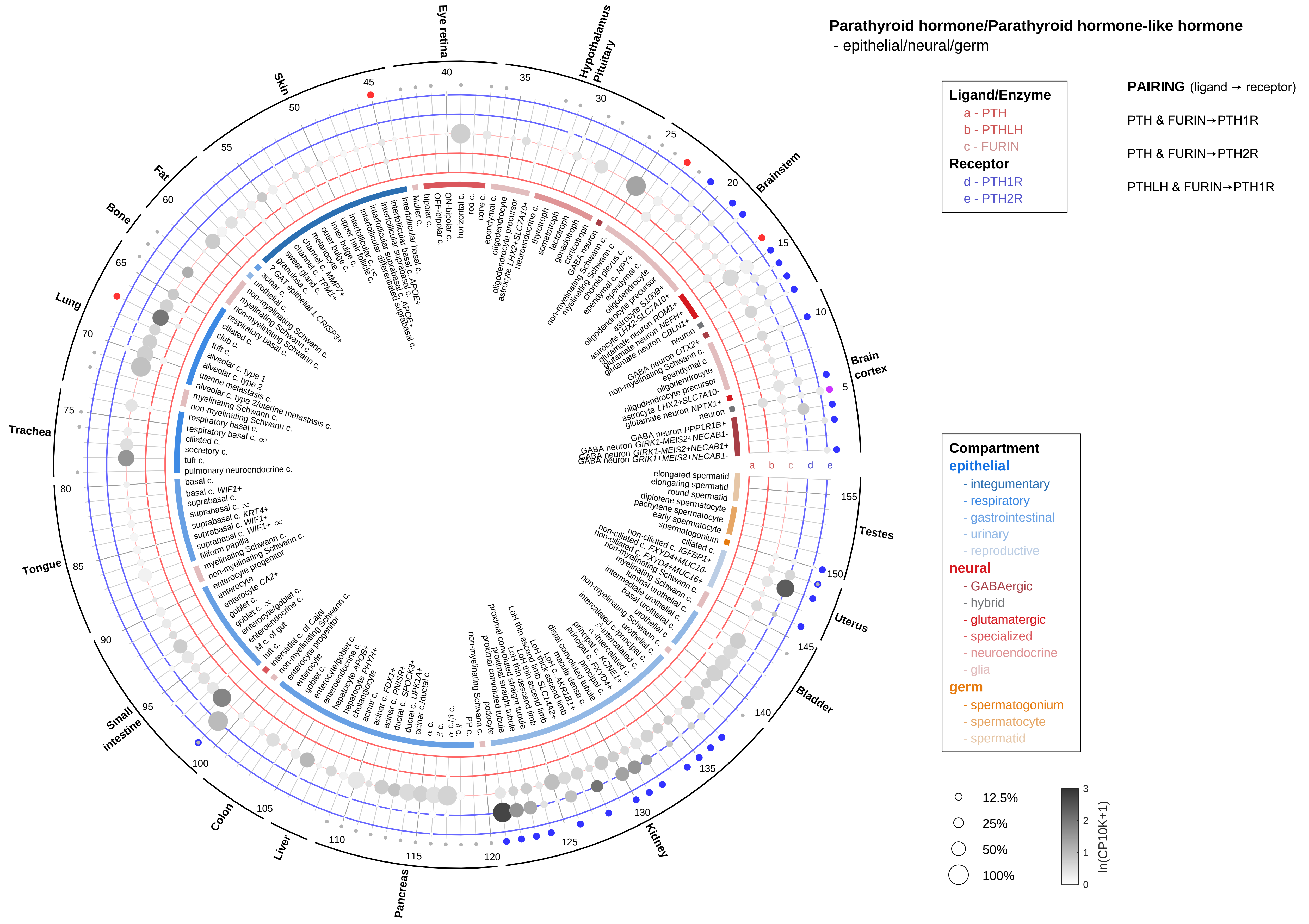

**Parathyroid hormone/Parathyroid hormone-like hormone**
- endothelial

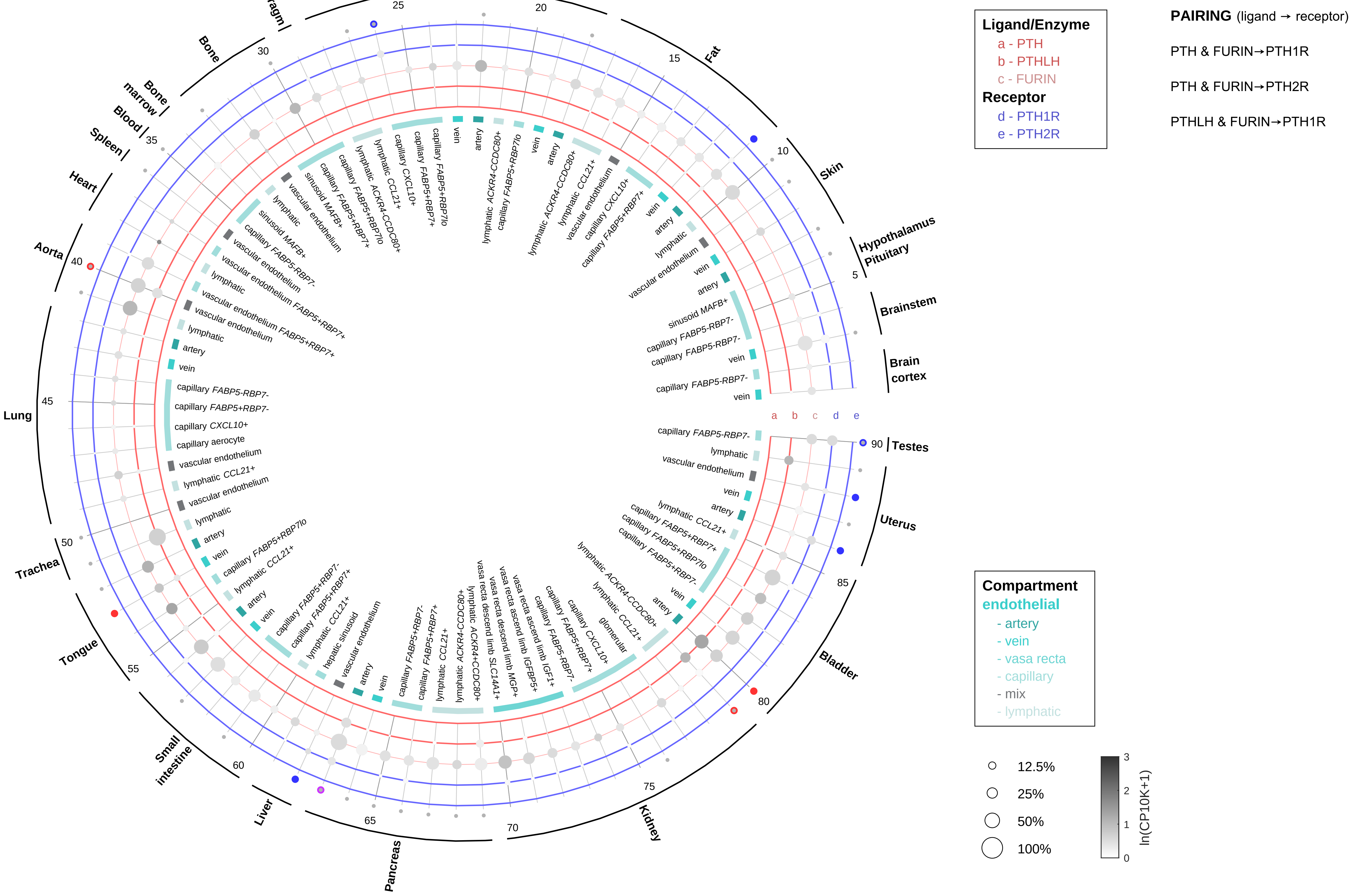

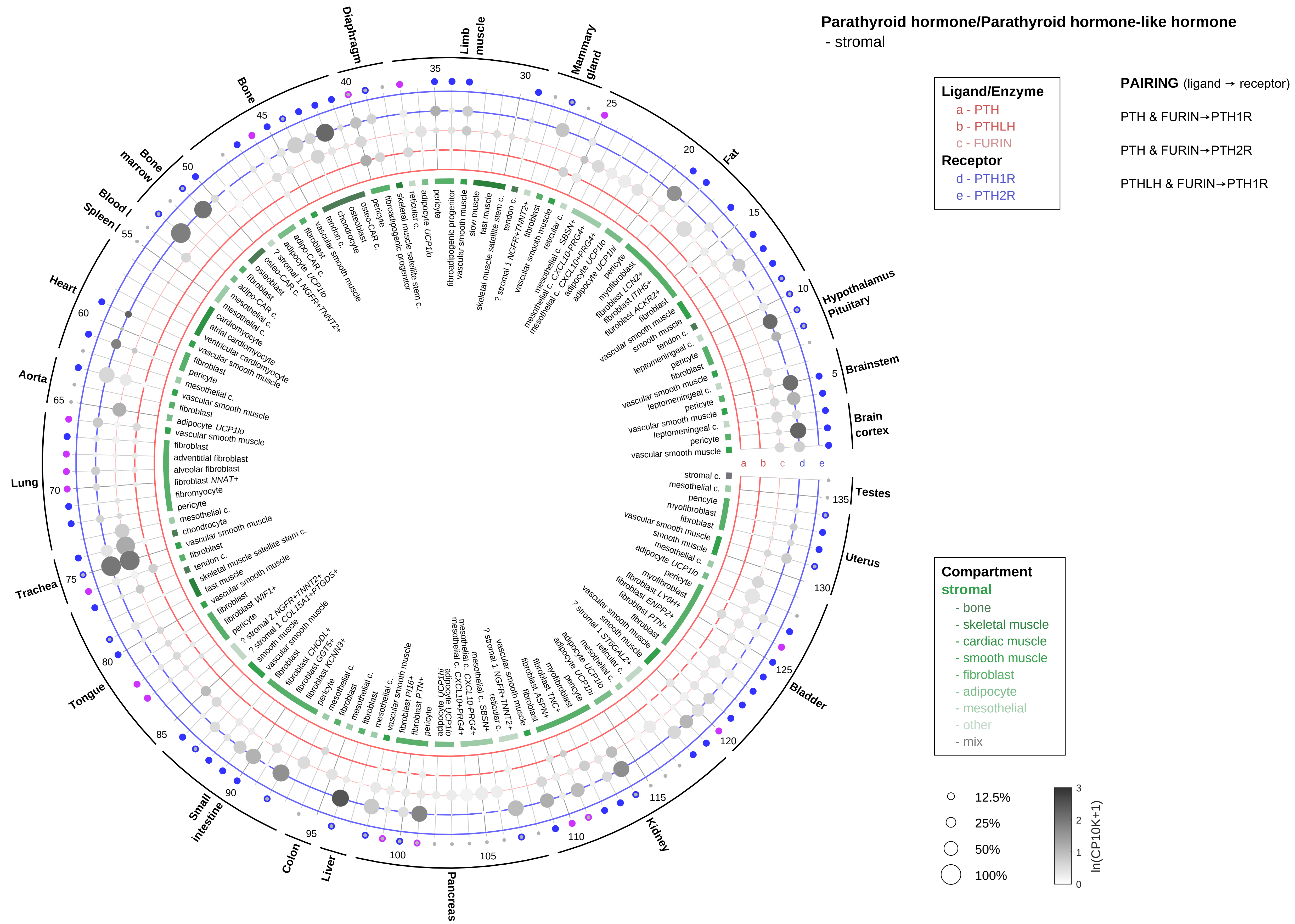

Parathyroid hormone/Parathyroid hormone-like hormone
- stromal
Ligand/Enzyme
a - PTH
b - PTHLH
c - FURIN
Receptor
d - PTH1R
e - PTH2R
PAIRING (ligand → receptor)
PTH & FURIN→PTH1R
PTH & FURIN→PTH2R
PTHLH & FURIN→PTH1R
Compartment
stromal
- bone
- skeletal muscle
- cardiac muscle
- smooth muscle
- fibroblast
- adipocyte
- mesothelial
- other
- mix
12.5%
25%
50%
100%
ln(CP10K+1)
Brain cortex
Brainstem
Hypothalamus Pituitary
Fat
Mammary gland
Limb muscle
Diaphragm
Bone
Bone marrow
Blood / Spleen
Heart
Aorta
Lung
Trachea
Tongue
Small intestine
Colon
Liver
Pancreas
Kidney
Bladder
Uterus
Testes

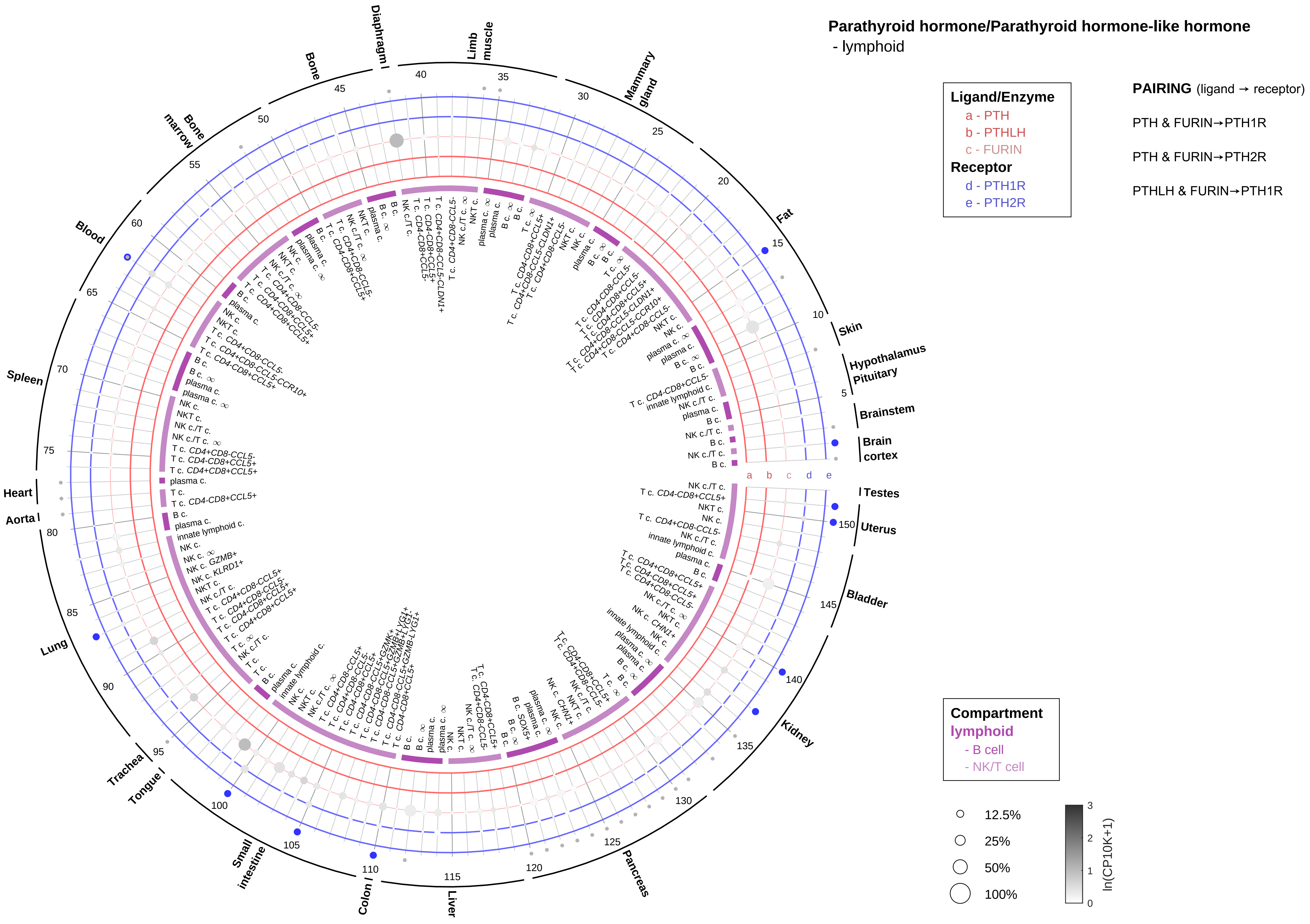

Parathyroid hormone/Parathyroid hormone-like hormone
- lymphoid
Ligand/Enzyme
a - PTH
b - PTHLH
c - FURIN
Receptor
d - PTH1R
e - PTH2R
PAIRING (ligand → receptor)
PTH & FURIN→PTH1R
PTH & FURIN→PTH2R
PTHLH & FURIN→PTH1R
Compartment
lymphoid
- B cell
- NK/T cell
12.5%
25%
50%
100%
ln(CP10K+1)
Diaphragm
Limb muscle
Mammary gland
Fat
Skin
Hypothalamus
Pituitary
Brainstem
Brain cortex
Testes
Uterus
Bladder
Kidney
Pancreas
Liver
Colon
Small intestine
Tongue
Trachea
Lung
Aorta
Heart
Spleen
Blood
Bone marrow
Bone

#### Parathyroid hormone/Parathyroid hormone-like hormone
- myeloid

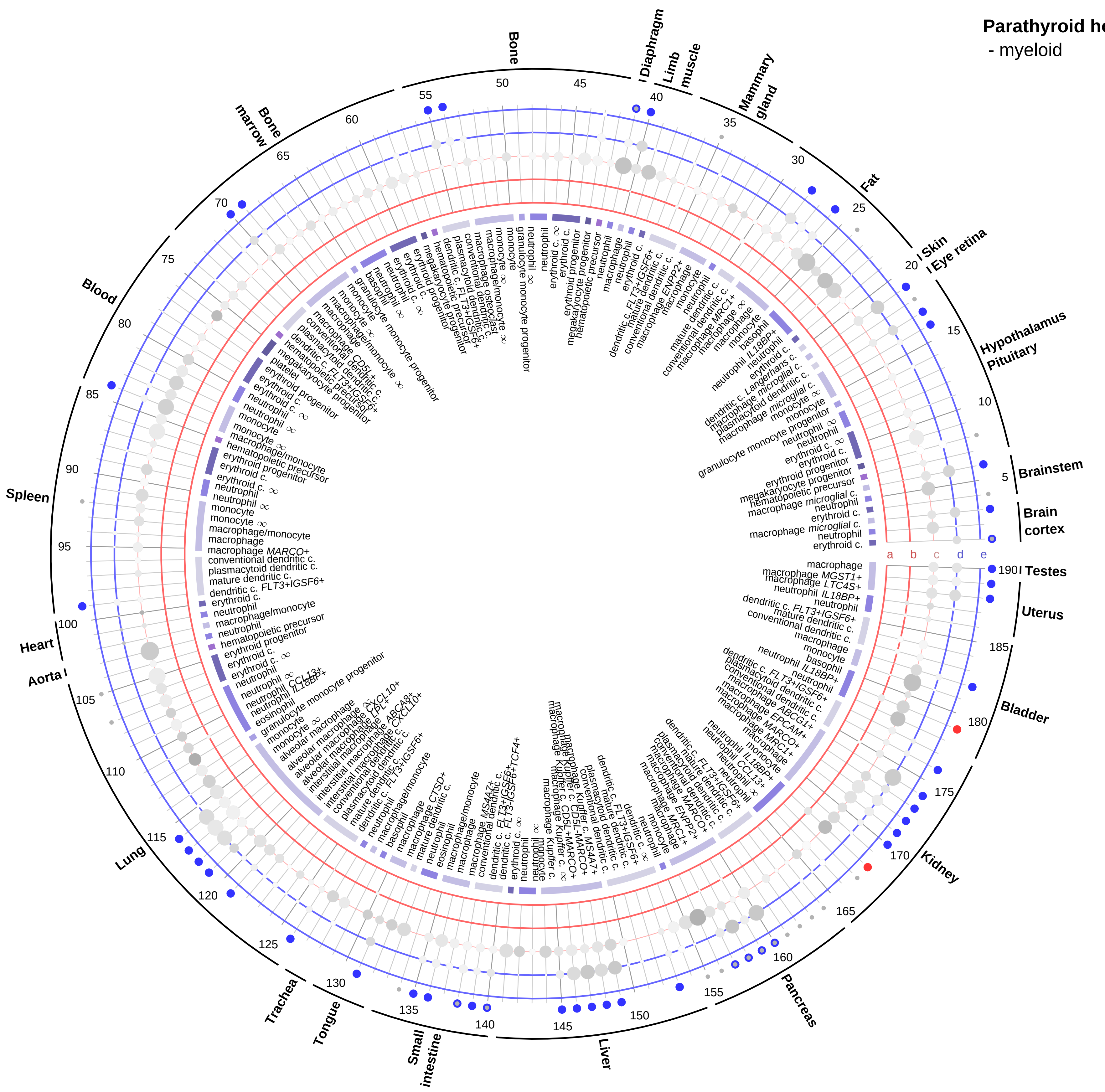

**Ligand/Enzyme**
- a - PTH
- b - PTHLH
- c - FURIN

**Receptor**
- d - PTH1R
- e - PTH2R

**PAIRING** (ligand → receptor)

PTH & FURIN→PTH1R

PTH & FURIN→PTH2R

PTHLH & FURIN→PTH1R

**Compartment**

**hematopoietic**
- hematopoietic precursor

**megakaryocyte-erythroid**
- megakaryoid
- erythroid

**myeloid**
- granulocyte
- granulocyte monocyte progenitor
- monocyte/macrophage
- dendritic

12.5%
25%
50%
100%

ln(CP10K+1) 0 1 2 3

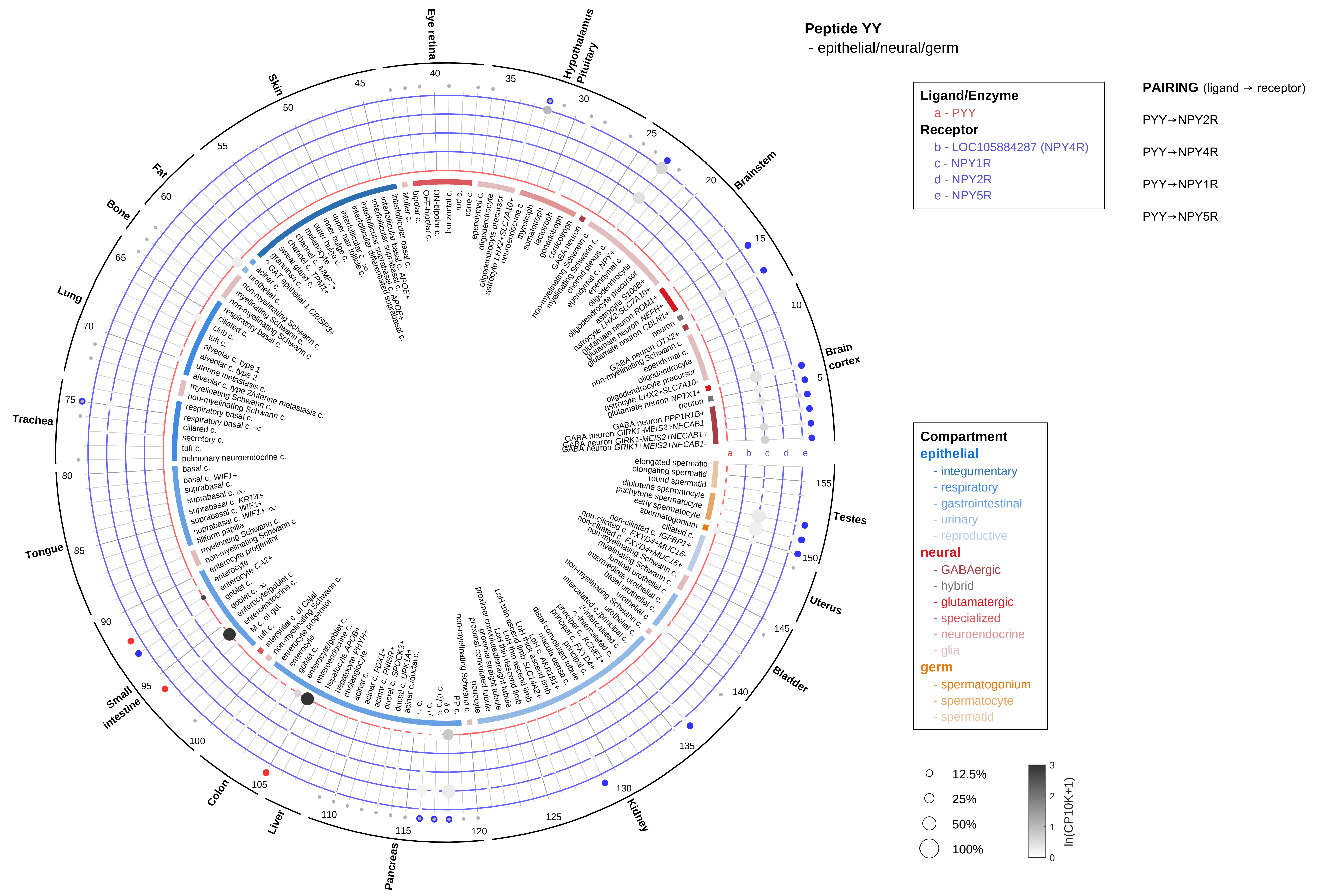

Peptide YY
- epithelial/neural/germ
Ligand/Enzyme
a - PYY
Receptor
b - LOC105884287 (NPY4R)
c - NPY1R
d - NPY2R
e - NPY5R
PAIRING (ligand → receptor)
PYY→NPY2R
PYY→NPY4R
PYY→NPY1R
PYY→NPY5R
Compartment
epithelial
- integumentary
- respiratory
- gastrointestinal
- urinary
- reproductive
neural
- GABAergic
- hybrid
- glutamatergic
- specialized
- neuroendocrine
- glia
germ
- spermatogonium
- spermatocyte
- spermatid
12.5%
25%
50%
100%
ln(CP10K+1)
Brain cortex
Brainstem
Hypothalamus Pituitary
Eye retina
Skin
Fat
Bone
Lung
Trachea
Tongue
Small intestine
Colon
Liver
Pancreas
Kidney
Bladder
Uterus
Testes

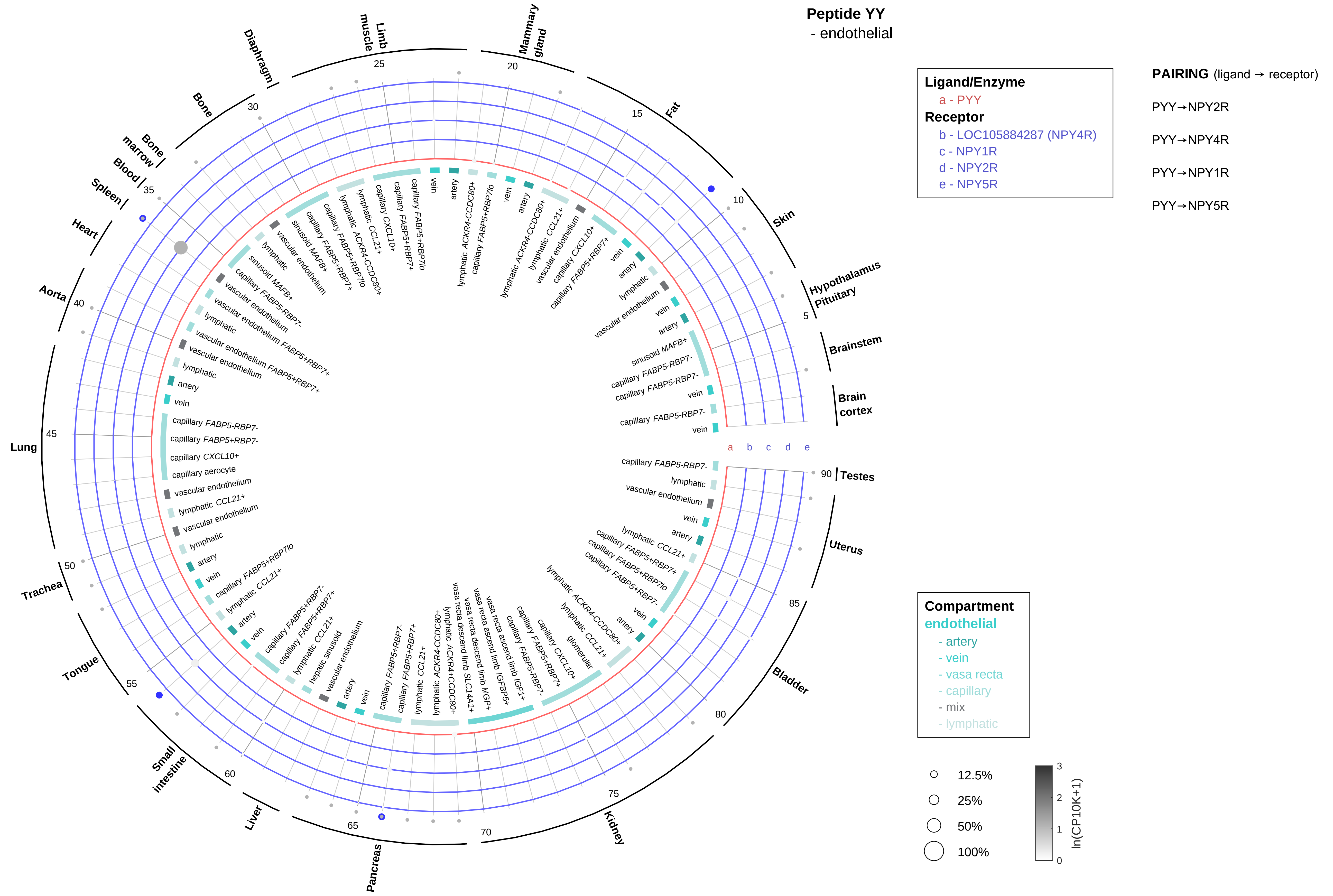
Peptide YY
- endothelial
Ligand/Enzyme
a - PYY
Receptor
b - LOC105884287 (NPY4R)
c - NPY1R
d - NPY2R
e - NPY5R
PAIRING (ligand → receptor)
PYY→NPY2R
PYY→NPY4R
PYY→NPY1R
PYY→NPY5R
Compartment
endothelial
- artery
- vein
- vasa recta
- capillary
- mix
- lymphatic
12.5%
25%
50%
100%
ln(CP10K+1)
Brain cortex
Brainstem
Hypothalamus
Pituitary
Skin
Fat
Mammary gland
Limb muscle
Diaphragm
Bone
Bone marrow
Blood
Spleen
Heart
Aorta
Lung
Trachea
Tongue
Small intestine
Liver
Pancreas
Kidney
Bladder
Uterus
Testes

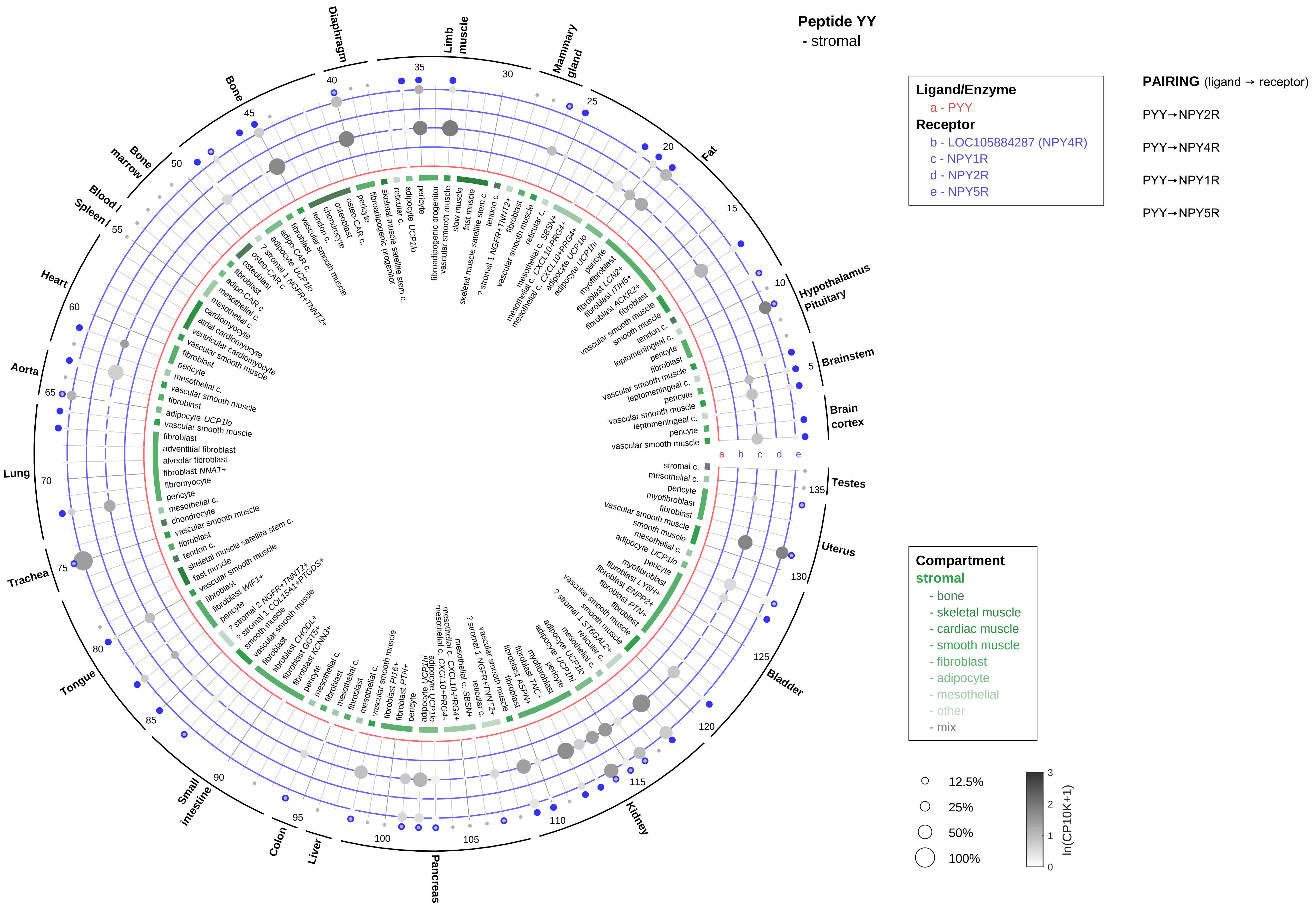

Peptide YY
- stromal
Ligand/Enzyme
a - PYY
Receptor
b - LOC105884287 (NPY4R)
c - NPY1R
d - NPY2R
e - NPY5R
PAIRING (ligand → receptor)
PYY→NPY2R
PYY→NPY4R
PYY→NPY1R
PYY→NPY5R
Compartment
stromal
- bone
- skeletal muscle
- cardiac muscle
- smooth muscle
- fibroblast
- adipocyte
- mesothelial
- other
- mix
12.5%
25%
50%
100%
ln(CP10K+1)
Diaphragm
Limb muscle
Mammary gland
Fat
Hypothalamus Pituitary
Brainstem
Brain cortex
Testes
Uterus
Bladder
Kidney
Pancreas
Liver
Colon
Small intestine
Tongue
Trachea
Lung
Aorta
Heart
Blood / Spleen
Bone marrow
Bone

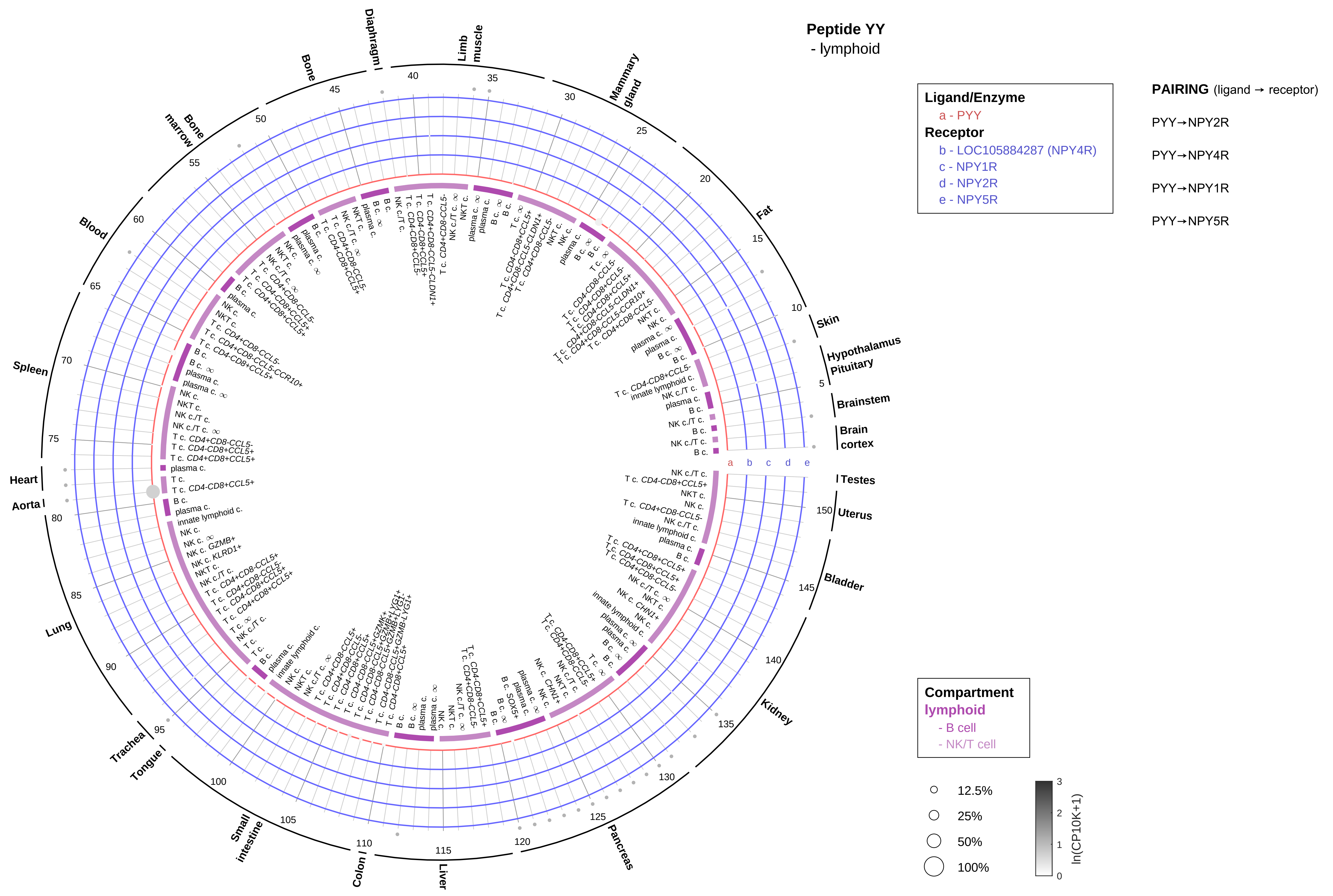

Peptide YY
- lymphoid
Ligand/Enzyme
a - PYY
Receptor
b - LOC105884287 (NPY4R)
c - NPY1R
d - NPY2R
e - NPY5R
PAIRING (ligand → receptor)
PYY→NPY2R
PYY→NPY4R
PYY→NPY1R
PYY→NPY5R
Compartment
lymphoid
- B cell
- NK/T cell
12.5%
25%
50%
100%
ln(CP10K+1)
Diaphragm
Limb muscle
Mammary gland
Fat
Skin
Hypothalamus
Pituitary
Brainstem
Brain cortex
Testes
Uterus
Bladder
Kidney
Pancreas
Liver
Colon
Small intestine
Tongue
Trachea
Lung
Aorta
Heart
Spleen
Blood
Bone marrow
Bone

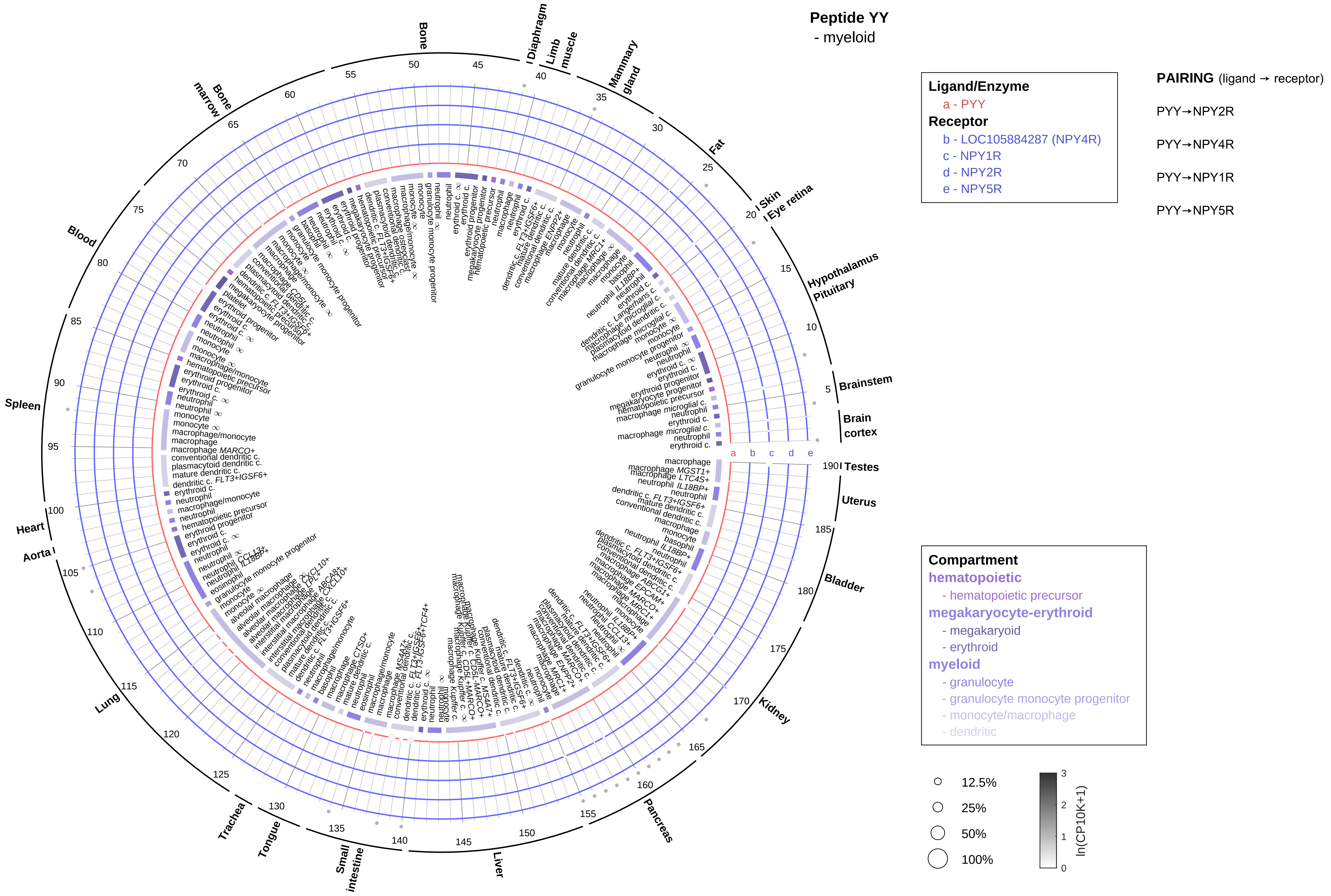

Peptide YY
- myeloid
Ligand/Enzyme
a - PYY
Receptor
b - LOC105884287 (NPY4R)
c - NPY1R
d - NPY2R
e - NPY5R
PAIRING (ligand → receptor)
PYY→NPY2R
PYY→NPY4R
PYY→NPY1R
PYY→NPY5R
Compartment
hematopoietic
- hematopoietic precursor
megakaryocyte-erythroid
- megakaryoid
- erythroid
myeloid
- granulocyte
- granulocyte monocyte progenitor
- monocyte/macrophage
- dendritic
12.5%
25%
50%
100%
ln(CP10K+1)
Brainstem
Brain cortex
Testes
Uterus
Bladder
Kidney
Pancreas
Liver
Small intestine
Tongue
Trachea
Lung
Aorta
Heart
Spleen
Blood
Bone marrow
Bone
Diaphragm
Limb muscle
Mammary gland
Fat
Skin
Eye retina
Hypothalamus
Pituitary

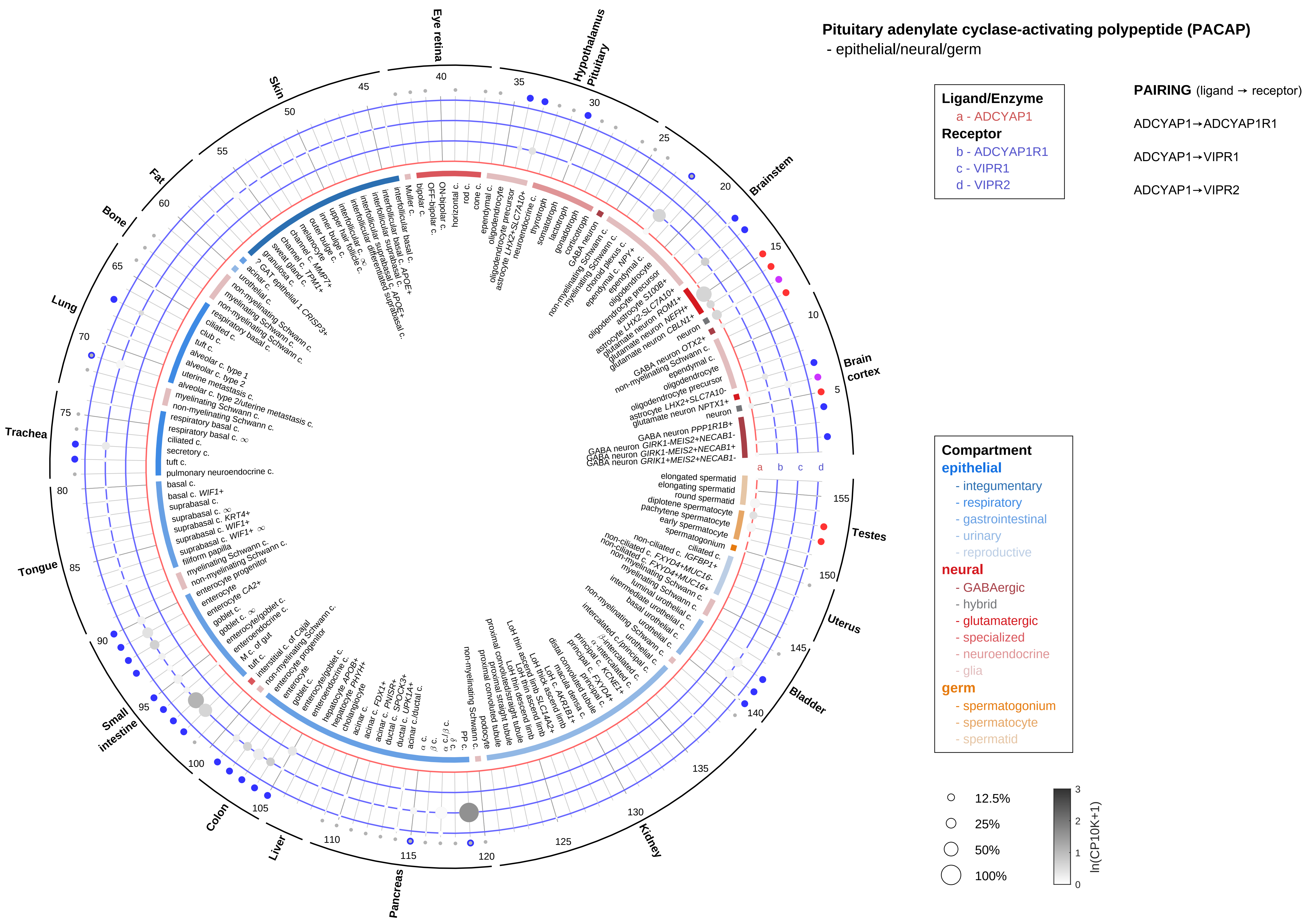
Pituitary adenylate cyclase-activating polypeptide (PACAP)
- epithelial/neural/germ
Ligand/Enzyme
a - ADCYAP1
Receptor
b - ADCYAP1R1
c - VIPR1
d - VIPR2
PAIRING (ligand → receptor)
ADCYAP1→ADCYAP1R1
ADCYAP1→VIPR1
ADCYAP1→VIPR2
Compartment
epithelial
- integumentary
- respiratory
- gastrointestinal
- urinary
- reproductive
neural
- GABAergic
- hybrid
- glutamatergic
- specialized
- neuroendocrine
- glia
germ
- spermatogonium
- spermatocyte
- spermatid
12.5%
25%
50%
100%
ln(CP10K+1)
Brain cortex
Brainstem
Hypothalamus Pituitary
Eye retina
Skin
Fat
Bone
Lung
Trachea
Tongue
Small intestine
Colon
Liver
Pancreas
Kidney
Bladder
Uterus
Testes

#### Pituitary adenylate cyclase-activating polypeptide (PACAP)
- endothelial

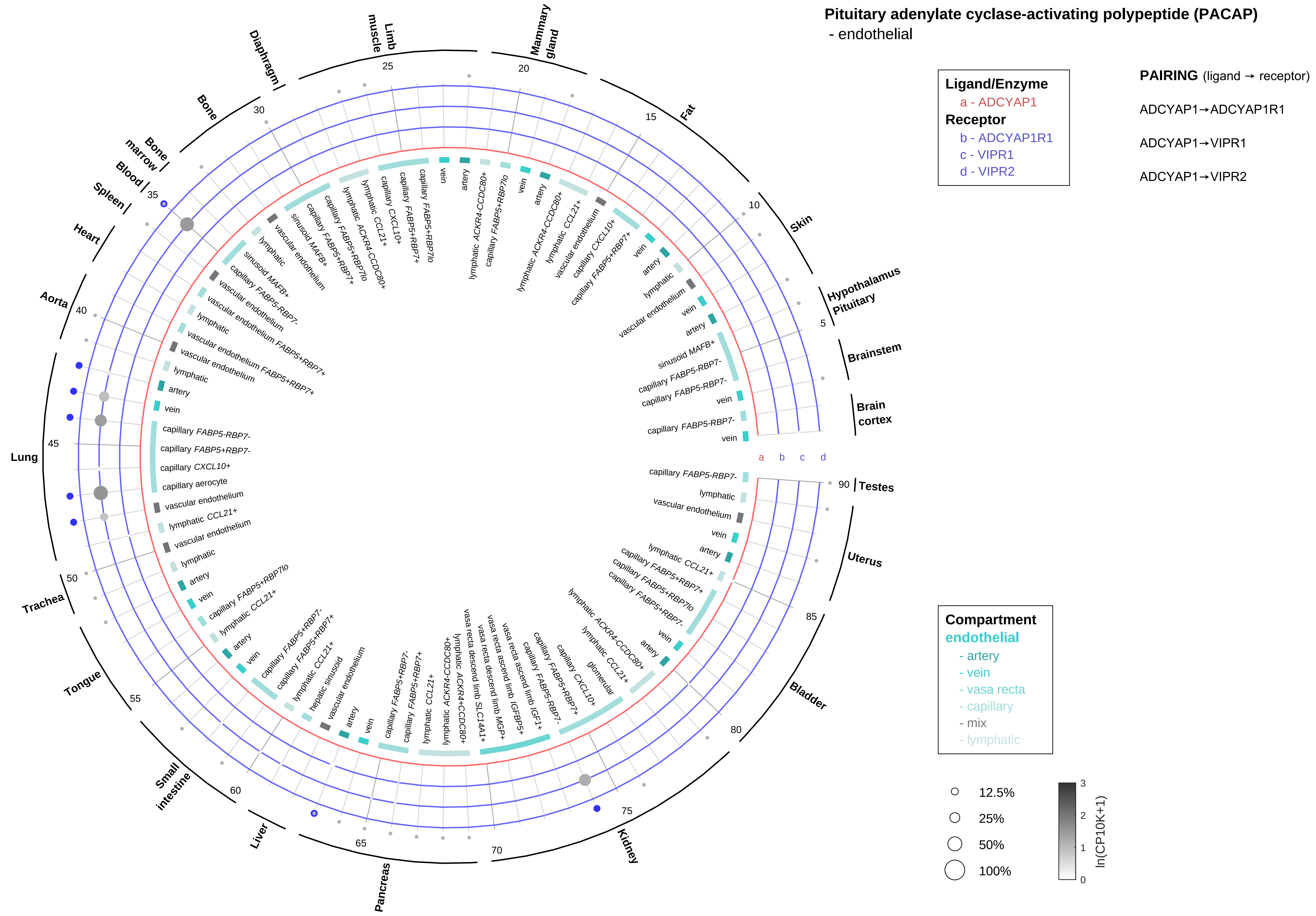

Pituitary adenylate cyclase-activating polypeptide (PACAP)
- stromal

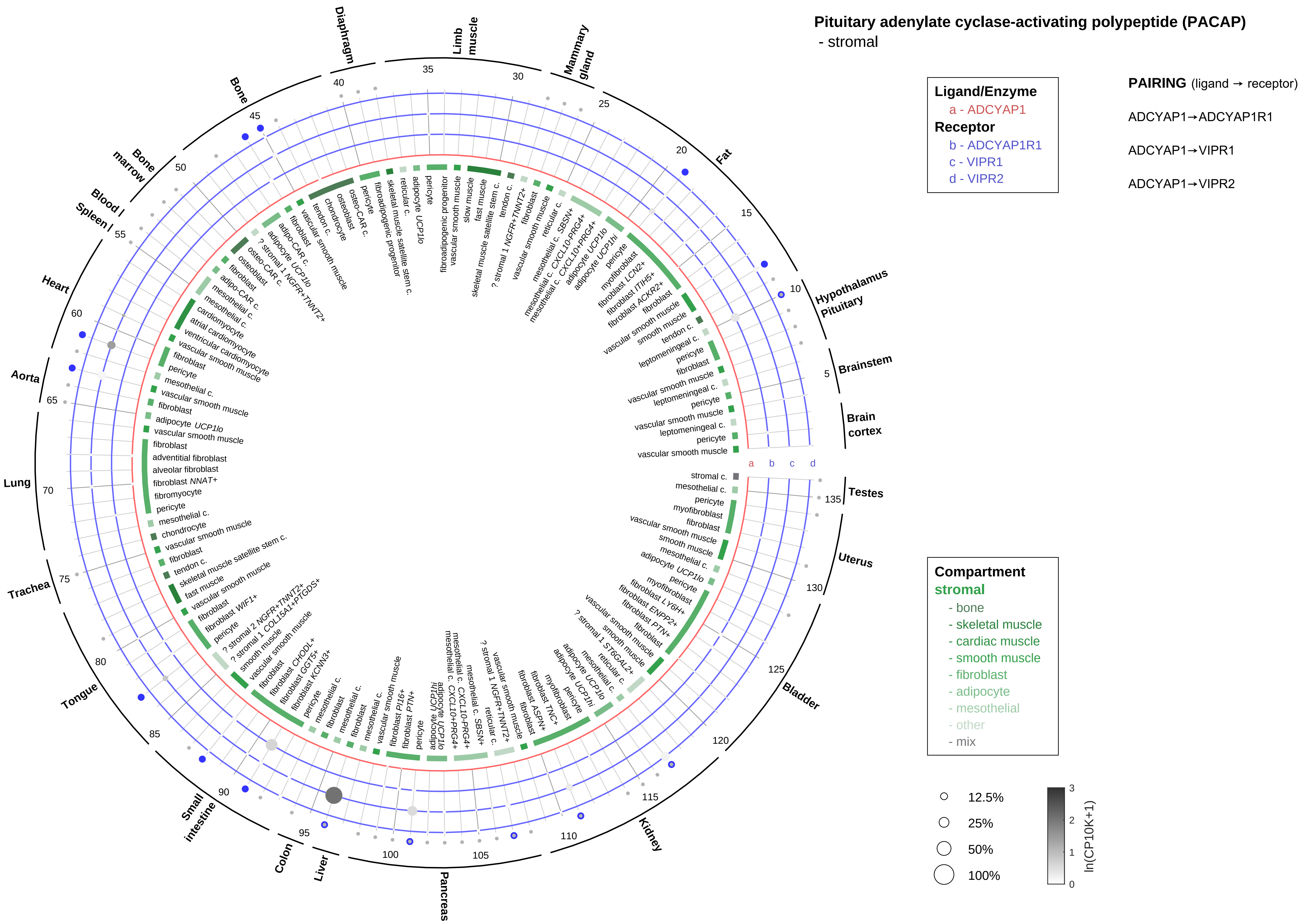

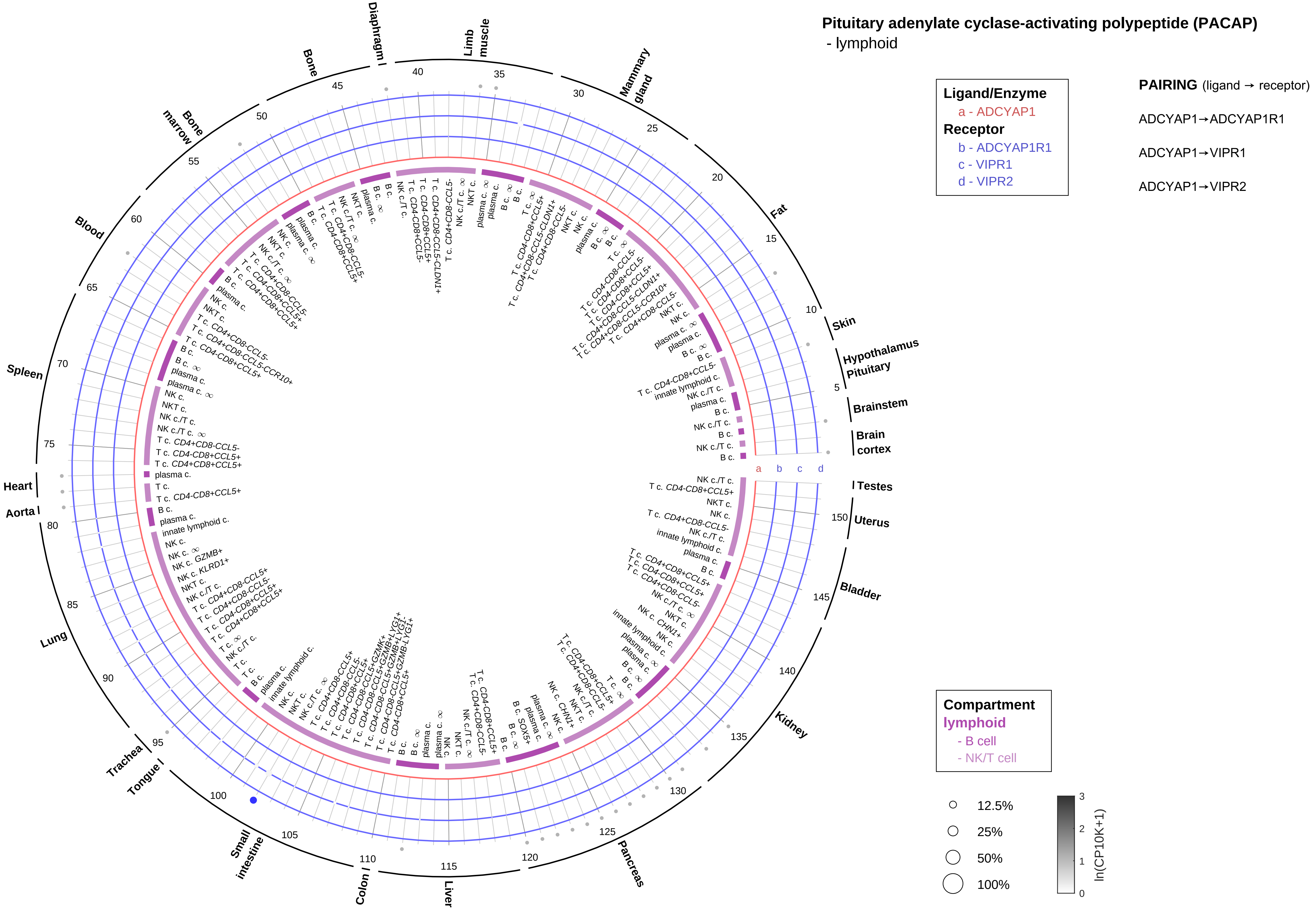
Pituitary adenylate cyclase-activating polypeptide (PACAP)
- lymphoid
Ligand/Enzyme
a - ADCYAP1
Receptor
b - ADCYAP1R1
c - VIPR1
d - VIPR2
PAIRING (ligand → receptor)
ADCYAP1→ADCYAP1R1
ADCYAP1→VIPR1
ADCYAP1→VIPR2
Compartment
lymphoid
- B cell
- NK/T cell
12.5%
25%
50%
100%
ln(CP10K+1)
Skin
Hypothalamus
Pituitary
Brainstem
Brain cortex
Testes
Uterus
Bladder
Kidney
Pancreas
Liver
Colon
Small intestine
Tongue
Trachea
Lung
Aorta
Heart
Spleen
Blood
Bone marrow
Bone
Diaphragm
Limb muscle
Mammary gland
Fat

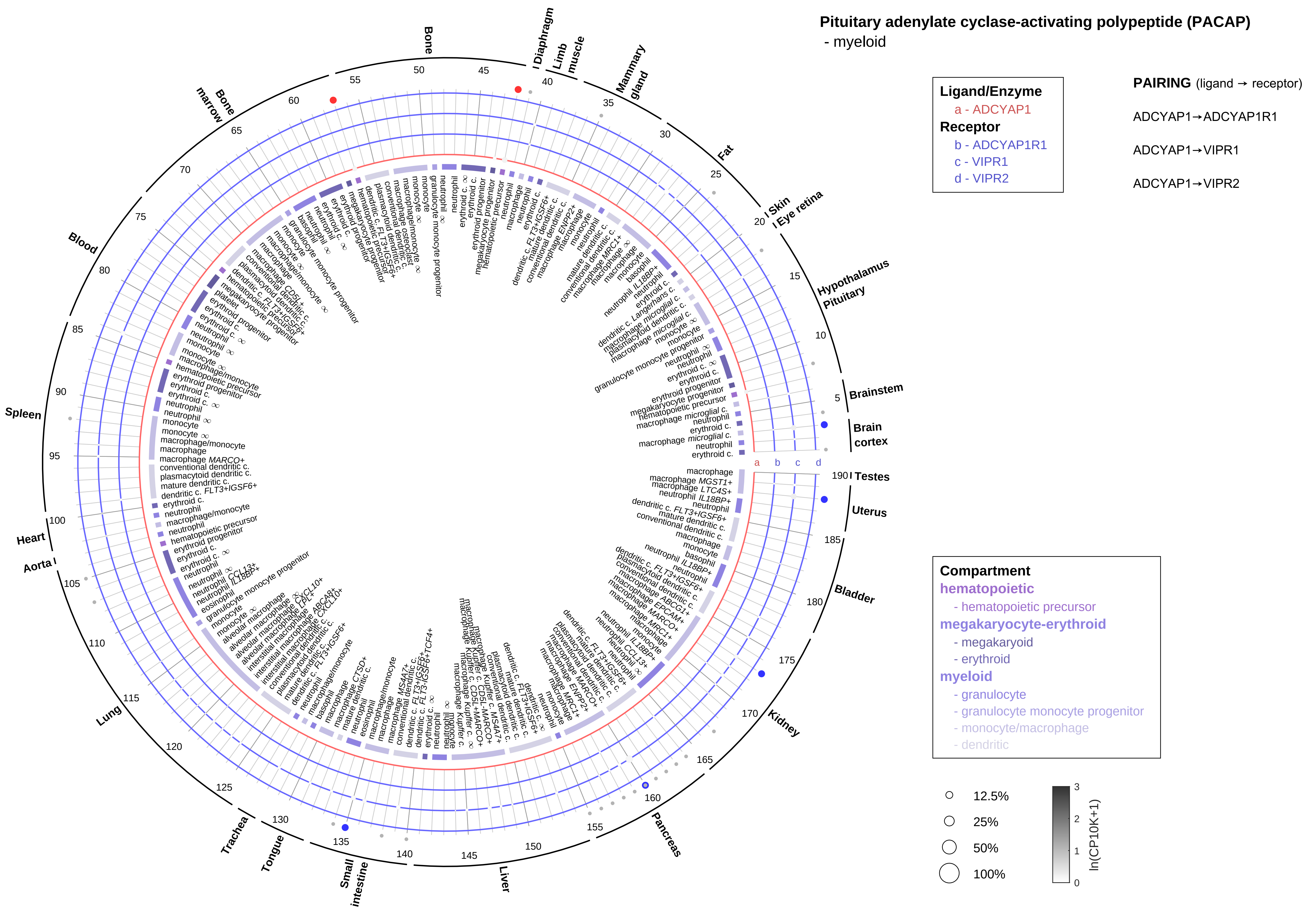

Pituitary adenylate cyclase-activating polypeptide (PACAP)
- myeloid
Ligand/Enzyme
a - ADCYAP1
Receptor
b - ADCYAP1R1
c - VIPR1
d - VIPR2
PAIRING (ligand → receptor)
ADCYAP1→ADCYAP1R1
ADCYAP1→VIPR1
ADCYAP1→VIPR2
Compartment
hematopoietic
- hematopoietic precursor
megakaryocyte-erythroid
- megakaryocyte
- erythroid
myeloid
- granulocyte
- granulocyte monocyte progenitor
- monocyte/macrophage
- dendritic
12.5%
25%
50%
100%
ln(CP10K+1)
Bone
Diaphragm
Limb muscle
Mammary gland
Fat
Skin
Eye retina
Hypothalamus
Pituitary
Brainstem
Brain cortex
Testes
Uterus
Bladder
Kidney
Pancreas
Liver
Small intestine
Tongue
Trachea
Lung
Aorta
Heart
Spleen
Blood
Bone marrow

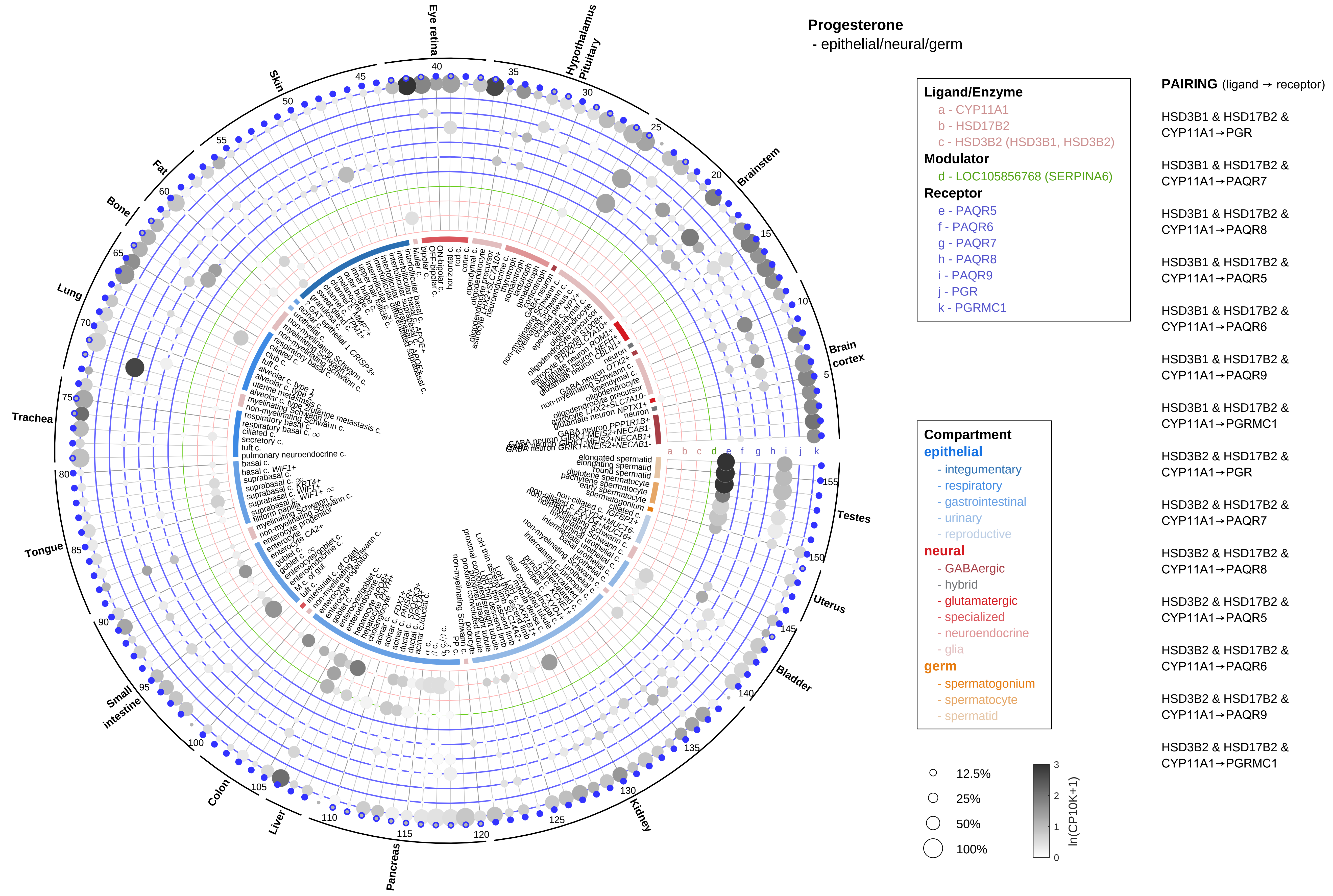
Progesterone
- epithelial/neural/germ
Eye retina
Hypothalamus
Pituitary
Brainstem
Brain cortex
Testes
Uterus
Bladder
Kidney
Pancreas
Liver
Colon
Small intestine
Tongue
Trachea
Lung
Bone
Fat
Skin
Ligand/Enzyme
a - CYP11A1
b - HSD17B2
c - HSD3B2 (HSD3B1, HSD3B2)
Modulator
d - LOC105856768 (SERPINA6)
Receptor
e - PAQR5
f - PAQR6
g - PAQR7
h - PAQR8
i - PAQR9
j - PGR
k - PGRMC1
Compartment
epithelial
- integumentary
- respiratory
- gastrointestinal
- urinary
- reproductive
neural
- GABAergic
- hybrid
- glutamatergic
- specialized
- neuroendocrine
- glia
germ
- spermatogonium
- spermatocyte
- spermatid
12.5%
25%
50%
100%
ln(CP10K+1)
PAIRING (ligand → receptor)
HSD3B1 & HSD17B2 & CYP11A1→PGR
HSD3B1 & HSD17B2 & CYP11A1→PAQR7
HSD3B1 & HSD17B2 & CYP11A1→PAQR8
HSD3B1 & HSD17B2 & CYP11A1→PAQR5
HSD3B1 & HSD17B2 & CYP11A1→PAQR6
HSD3B1 & HSD17B2 & CYP11A1→PAQR9
HSD3B1 & HSD17B2 & CYP11A1→PGRMC1
HSD3B2 & HSD17B2 & CYP11A1→PGR
HSD3B2 & HSD17B2 & CYP11A1→PAQR7
HSD3B2 & HSD17B2 & CYP11A1→PAQR8
HSD3B2 & HSD17B2 & CYP11A1→PAQR5
HSD3B2 & HSD17B2 & CYP11A1→PAQR6
HSD3B2 & HSD17B2 & CYP11A1→PAQR9
HSD3B2 & HSD17B2 & CYP11A1→PGRMC1

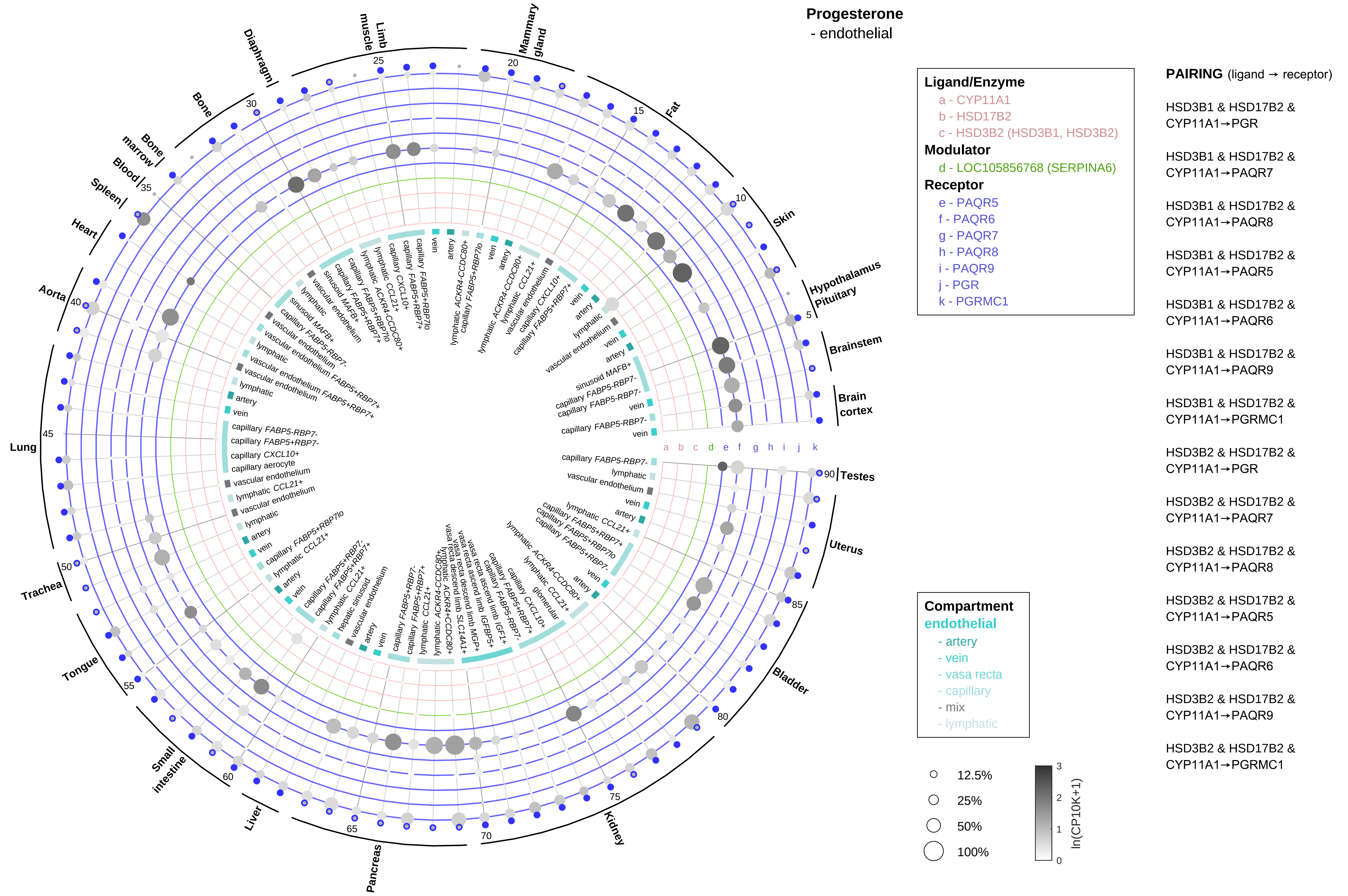

Progesterone
- endothelial
Ligand/Enzyme
a - CYP11A1
b - HSD17B2
c - HSD3B2 (HSD3B1, HSD3B2)
Modulator
d - LOC105856768 (SERPINA6)
Receptor
e - PAQR5
f - PAQR6
g - PAQR7
h - PAQR8
i - PAQR9
j - PGR
k - PGRMC1
Compartment
endothelial
- artery
- vein
- vasa recta
- capillary
- mix
- lymphatic
12.5%
25%
50%
100%
ln(CP10K+1)
PAIRING (ligand → receptor)
HSD3B1 & HSD17B2 & CYP11A1→PGR
HSD3B1 & HSD17B2 & CYP11A1→PAQR7
HSD3B1 & HSD17B2 & CYP11A1→PAQR8
HSD3B1 & HSD17B2 & CYP11A1→PAQR5
HSD3B1 & HSD17B2 & CYP11A1→PAQR6
HSD3B1 & HSD17B2 & CYP11A1→PAQR9
HSD3B1 & HSD17B2 & CYP11A1→PGRMC1
HSD3B2 & HSD17B2 & CYP11A1→PGR
HSD3B2 & HSD17B2 & CYP11A1→PAQR7
HSD3B2 & HSD17B2 & CYP11A1→PAQR8
HSD3B2 & HSD17B2 & CYP11A1→PAQR5
HSD3B2 & HSD17B2 & CYP11A1→PAQR6
HSD3B2 & HSD17B2 & CYP11A1→PAQR9
HSD3B2 & HSD17B2 & CYP11A1→PGRMC1
Mammary gland
Fat
Skin
Hypothalamus Pituitary
Brainstem
Brain cortex
Testes
Uterus
Bladder
Kidney
Pancreas
Liver
Small intestine
Tongue
Trachea
Lung
Aorta
Heart
Spleen
Blood
Bone marrow
Bone
Diaphragm
Limb muscle

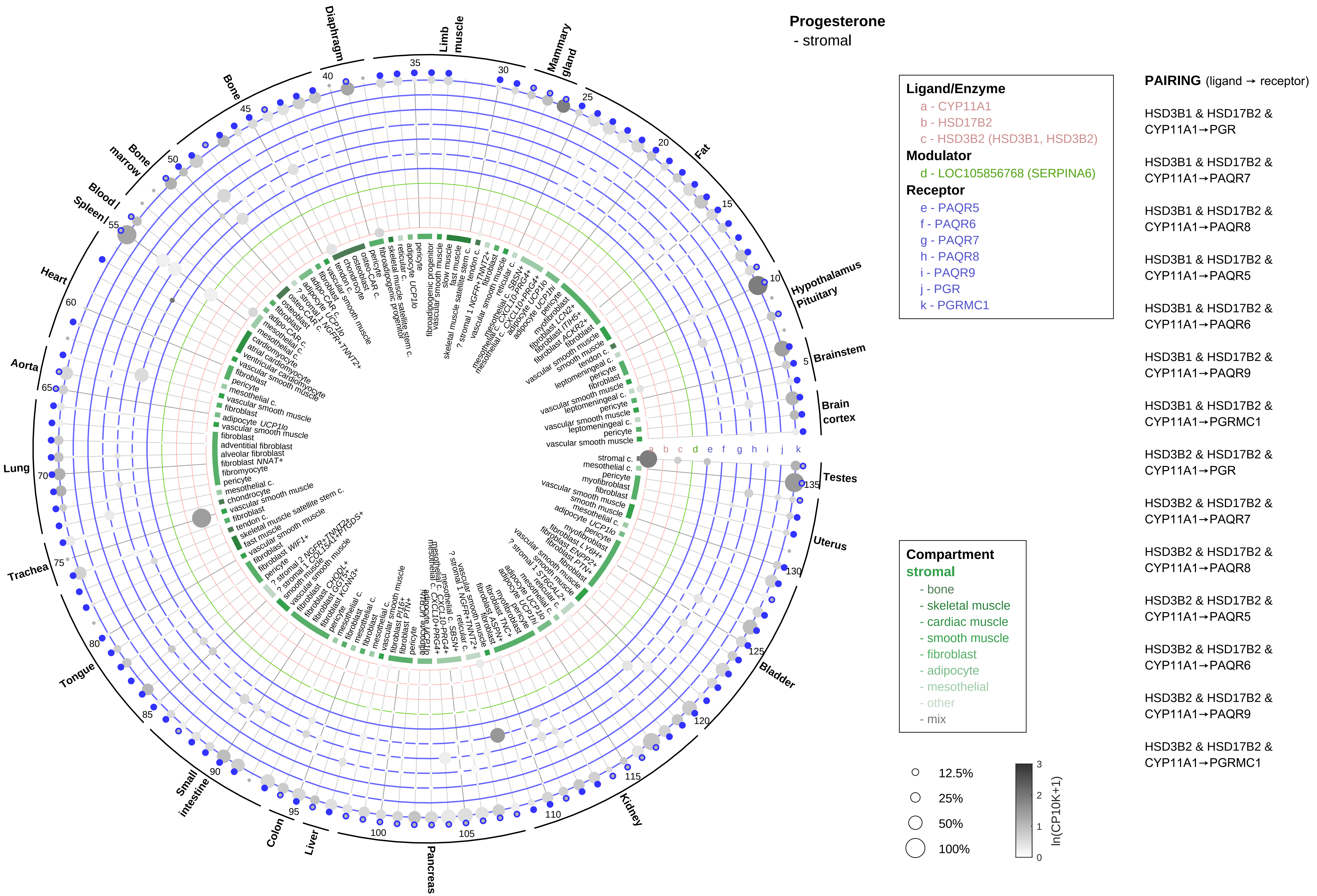

Progesterone
- stromal
Ligand/Enzyme
a - CYP11A1
b - HSD17B2
c - HSD3B2 (HSD3B1, HSD3B2)
Modulator
d - LOC105856768 (SERPINA6)
Receptor
e - PAQR5
f - PAQR6
g - PAQR7
h - PAQR8
i - PAQR9
j - PGR
k - PGRMC1
Compartment
stromal
- bone
- skeletal muscle
- cardiac muscle
- smooth muscle
- fibroblast
- adipocyte
- mesothelial
- other
- mix
12.5%
25%
50%
100%
ln(CP10K+1)
PAIRING (ligand → receptor)
HSD3B1 & HSD17B2 & CYP11A1→PGR
HSD3B1 & HSD17B2 & CYP11A1→PAQR7
HSD3B1 & HSD17B2 & CYP11A1→PAQR8
HSD3B1 & HSD17B2 & CYP11A1→PAQR5
HSD3B1 & HSD17B2 & CYP11A1→PAQR6
HSD3B1 & HSD17B2 & CYP11A1→PAQR9
HSD3B1 & HSD17B2 & CYP11A1→PGRMC1
HSD3B2 & HSD17B2 & CYP11A1→PGR
HSD3B2 & HSD17B2 & CYP11A1→PAQR7
HSD3B2 & HSD17B2 & CYP11A1→PAQR8
HSD3B2 & HSD17B2 & CYP11A1→PAQR5
HSD3B2 & HSD17B2 & CYP11A1→PAQR6
HSD3B2 & HSD17B2 & CYP11A1→PAQR9
HSD3B2 & HSD17B2 & CYP11A1→PGRMC1
Diaphragm
Limb muscle
Mammary gland
Fat
Hypothalamus Pituitary
Brainstem
Brain cortex
Testes
Uterus
Bladder
Kidney
Pancreas
Liver
Colon
Small intestine
Tongue
Trachea
Lung
Aorta
Heart
Spleen
Blood
Bone marrow
Bone

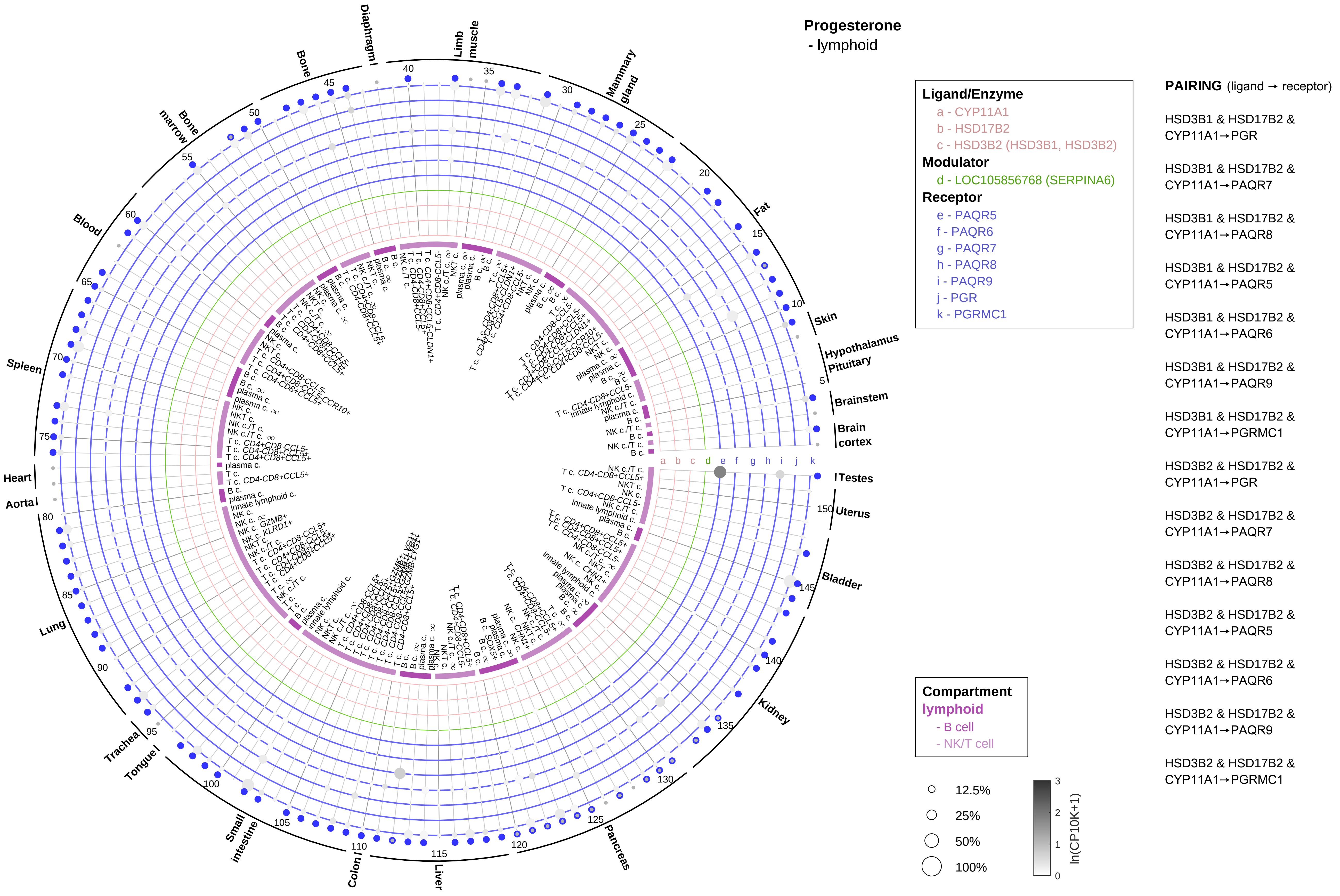
Progesterone
- lymphoid
Ligand/Enzyme
a - CYP11A1
b - HSD17B2
c - HSD3B2 (HSD3B1, HSD3B2)
Modulator
d - LOC105856768 (SERPINA6)
Receptor
e - PAQR5
f - PAQR6
g - PAQR7
h - PAQR8
i - PAQR9
j - PGR
k - PGRMC1
Compartment
lymphoid
- B cell
- NK/T cell
12.5%
25%
50%
100%
ln(CP10K+1)
PAIRING (ligand → receptor)
HSD3B1 & HSD17B2 & CYP11A1→PGR
HSD3B1 & HSD17B2 & CYP11A1→PAQR7
HSD3B1 & HSD17B2 & CYP11A1→PAQR8
HSD3B1 & HSD17B2 & CYP11A1→PAQR5
HSD3B1 & HSD17B2 & CYP11A1→PAQR6
HSD3B1 & HSD17B2 & CYP11A1→PAQR9
HSD3B1 & HSD17B2 & CYP11A1→PGRMC1
HSD3B2 & HSD17B2 & CYP11A1→PGR
HSD3B2 & HSD17B2 & CYP11A1→PAQR7
HSD3B2 & HSD17B2 & CYP11A1→PAQR8
HSD3B2 & HSD17B2 & CYP11A1→PAQR5
HSD3B2 & HSD17B2 & CYP11A1→PAQR6
HSD3B2 & HSD17B2 & CYP11A1→PAQR9
HSD3B2 & HSD17B2 & CYP11A1→PGRMC1
Diaphragm
Limb muscle
Mammary gland
Fat
Skin
Hypothalamus
Pituitary
Brainstem
Brain cortex
Testes
Uterus
Bladder
Kidney
Pancreas
Liver
Colon
Small intestine
Tongue
Trachea
Lung
Aorta
Heart
Spleen
Blood
Bone marrow
Bone

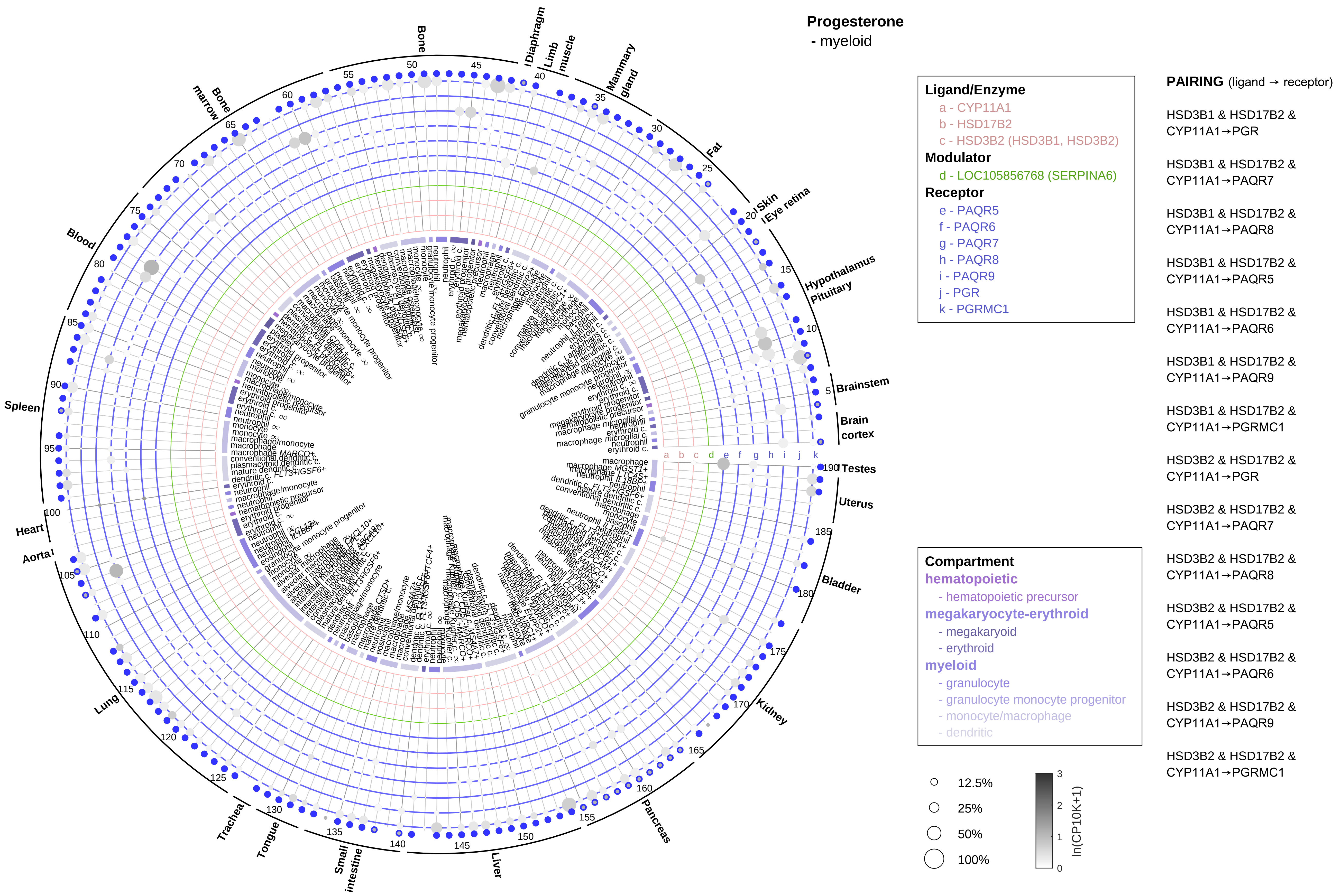

Progesterone
- myeloid
Ligand/Enzyme
a - CYP11A1
b - HSD17B2
c - HSD3B2 (HSD3B1, HSD3B2)
Modulator
d - LOC105856768 (SERPINA6)
Receptor
e - PAQR5
f - PAQR6
g - PAQR7
h - PAQR8
i - PAQR9
j - PGR
k - PGRMC1
PAIRING (ligand → receptor)
HSD3B1 & HSD17B2 & CYP11A1→PGR
HSD3B1 & HSD17B2 & CYP11A1→PAQR7
HSD3B1 & HSD17B2 & CYP11A1→PAQR8
HSD3B1 & HSD17B2 & CYP11A1→PAQR5
HSD3B1 & HSD17B2 & CYP11A1→PAQR6
HSD3B1 & HSD17B2 & CYP11A1→PAQR9
HSD3B1 & HSD17B2 & CYP11A1→PGRMC1
HSD3B2 & HSD17B2 & CYP11A1→PGR
HSD3B2 & HSD17B2 & CYP11A1→PAQR7
HSD3B2 & HSD17B2 & CYP11A1→PAQR8
HSD3B2 & HSD17B2 & CYP11A1→PAQR5
HSD3B2 & HSD17B2 & CYP11A1→PAQR6
HSD3B2 & HSD17B2 & CYP11A1→PAQR9
HSD3B2 & HSD17B2 & CYP11A1→PGRMC1
Compartment
hematopoietic
- hematopoietic precursor
megakaryocyte-erythroid
- megakaryoid
- erythroid
myeloid
- granulocyte
- granulocyte monocyte progenitor
- monocyte/macrophage
- dendritic
12.5%
25%
50%
100%
ln(CP10K+1)
Bone
Diaphragm
Limb muscle
Mammary gland
Fat
Skin
Eye retina
Hypothalamus
Pituitary
Brainstem
Brain cortex
Testes
Uterus
Bladder
Kidney
Pancreas
Liver
Small intestine
Tongue
Trachea
Lung
Aorta
Heart
Spleen
Blood
Bone marrow

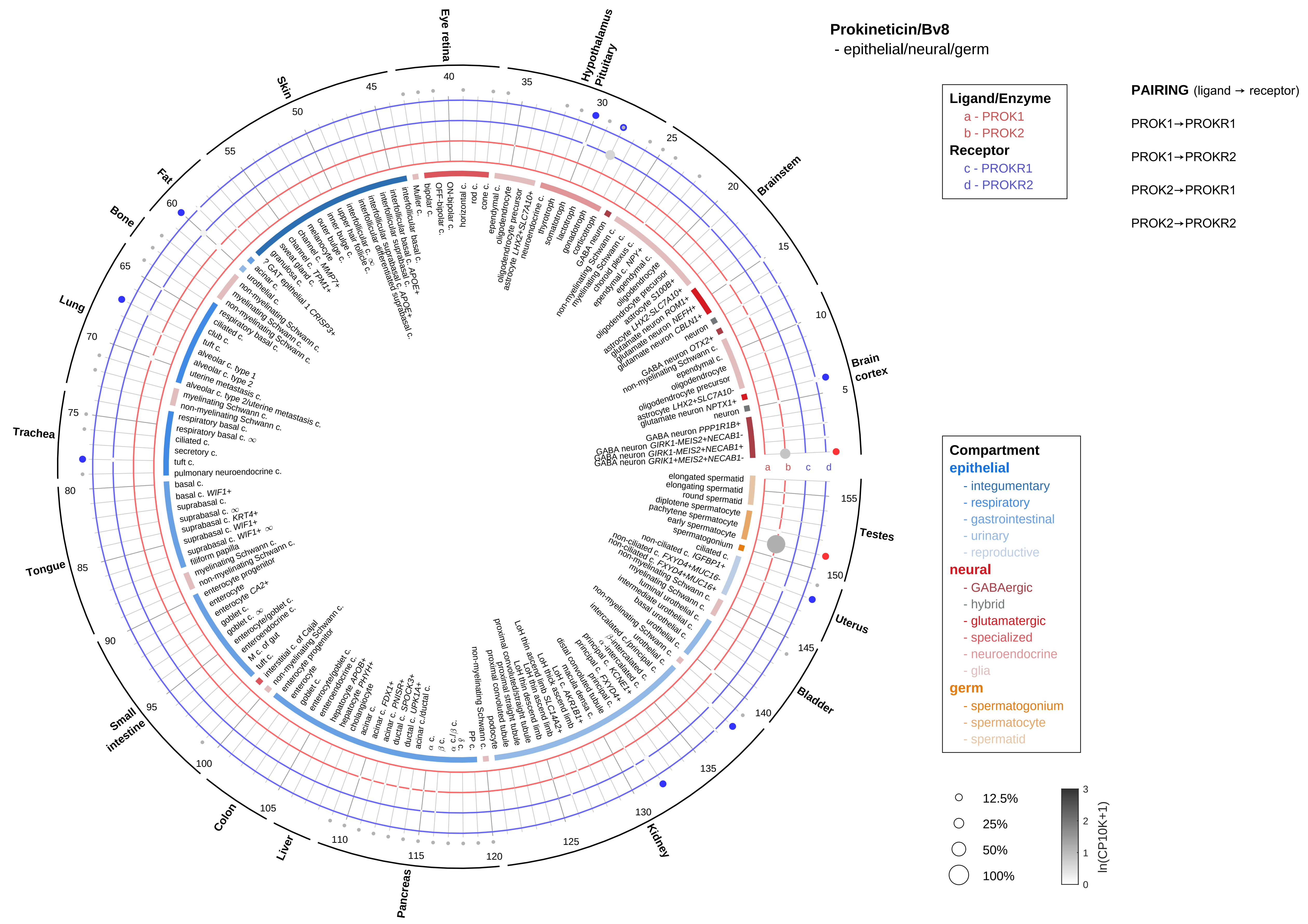

Prokineticin/Bv8
- epithelial/neural/germ
Ligand/Enzyme
a - PROK1
b - PROK2
Receptor
c - PROKR1
d - PROKR2
PAIRING (ligand → receptor)
PROK1→PROKR1
PROK1→PROKR2
PROK2→PROKR1
PROK2→PROKR2
Eye retina
Hypothalamus Pituitary
Brainstem
Brain cortex
Testes
Uterus
Bladder
Kidney
Pancreas
Liver
Colon
Small intestine
Tongue
Trachea
Lung
Bone
Fat
Skin
Compartment
epithelial
- integumentary
- respiratory
- gastrointestinal
- urinary
- reproductive
neural
- GABAergic
- hybrid
- glutamatergic
- specialized
- neuroendocrine
- glia
germ
- spermatogonium
- spermatocyte
- spermatid
12.5%
25%
50%
100%
ln(CP10K+1)

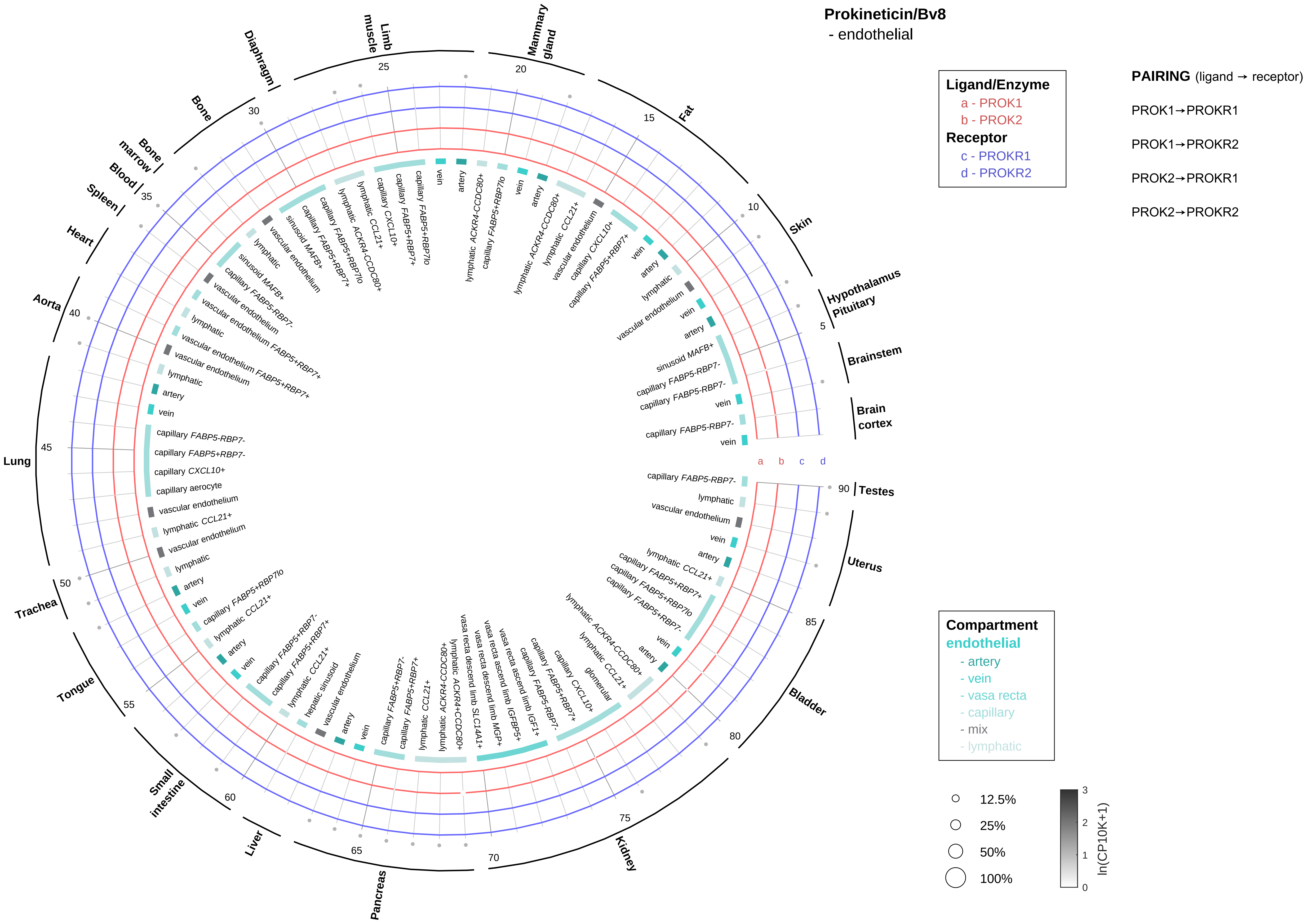

Prokineticin/Bv8
- endothelial
Ligand/Enzyme
a - PROK1
b - PROK2
Receptor
c - PROKR1
d - PROKR2
PAIRING (ligand → receptor)
PROK1→PROKR1
PROK1→PROKR2
PROK2→PROKR1
PROK2→PROKR2
Compartment
endothelial
- artery
- vein
- vasa recta
- capillary
- mix
- lymphatic
12.5%
25%
50%
100%
ln(CP10K+1)
Brain cortex
Brainstem
Hypothalamus Pituitary
Skin
Fat
Mammary gland
Limb muscle
Diaphragm
Bone
Bone marrow
Blood
Spleen
Heart
Aorta
Lung
Trachea
Tongue
Small intestine
Liver
Pancreas
Kidney
Bladder
Uterus
Testes

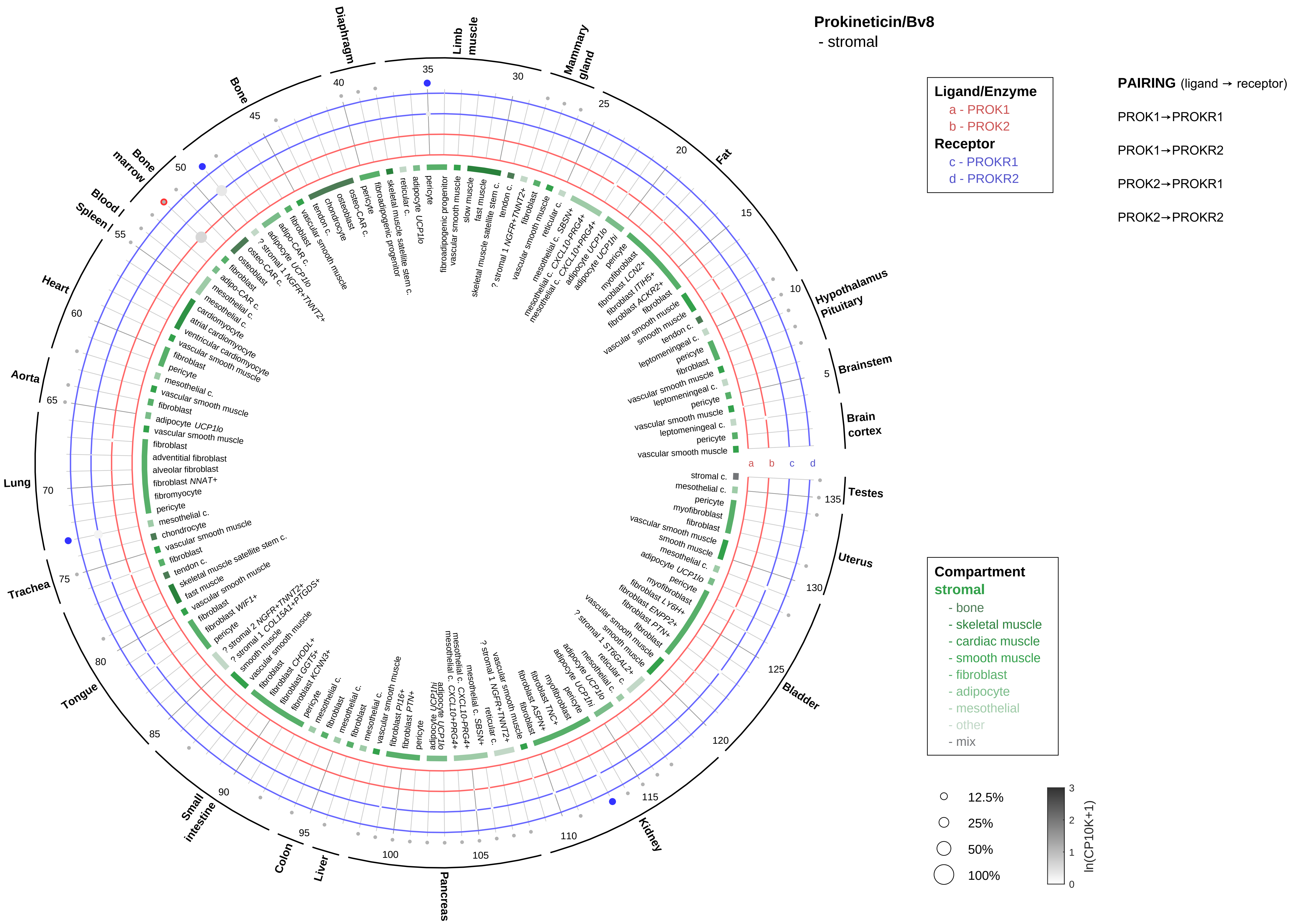
Prokineticin/Bv8
- stromal
Ligand/Enzyme
a - PROK1
b - PROK2
Receptor
c - PROKR1
d - PROKR2
PAIRING (ligand → receptor)
PROK1→PROKR1
PROK1→PROKR2
PROK2→PROKR1
PROK2→PROKR2
Compartment
stromal
- bone
- skeletal muscle
- cardiac muscle
- smooth muscle
- fibroblast
- adipocyte
- mesothelial
- other
- mix
12.5%
25%
50%
100%
ln(CP10K+1)
Limb muscle
Mammary gland
Fat
Hypothalamus Pituitary
Brainstem
Brain cortex
Testes
Uterus
Bladder
Kidney
Pancreas
Liver
Colon
Small intestine
Tongue
Trachea
Lung
Aorta
Heart
Spleen
Blood
Bone marrow
Bone
Diaphragm

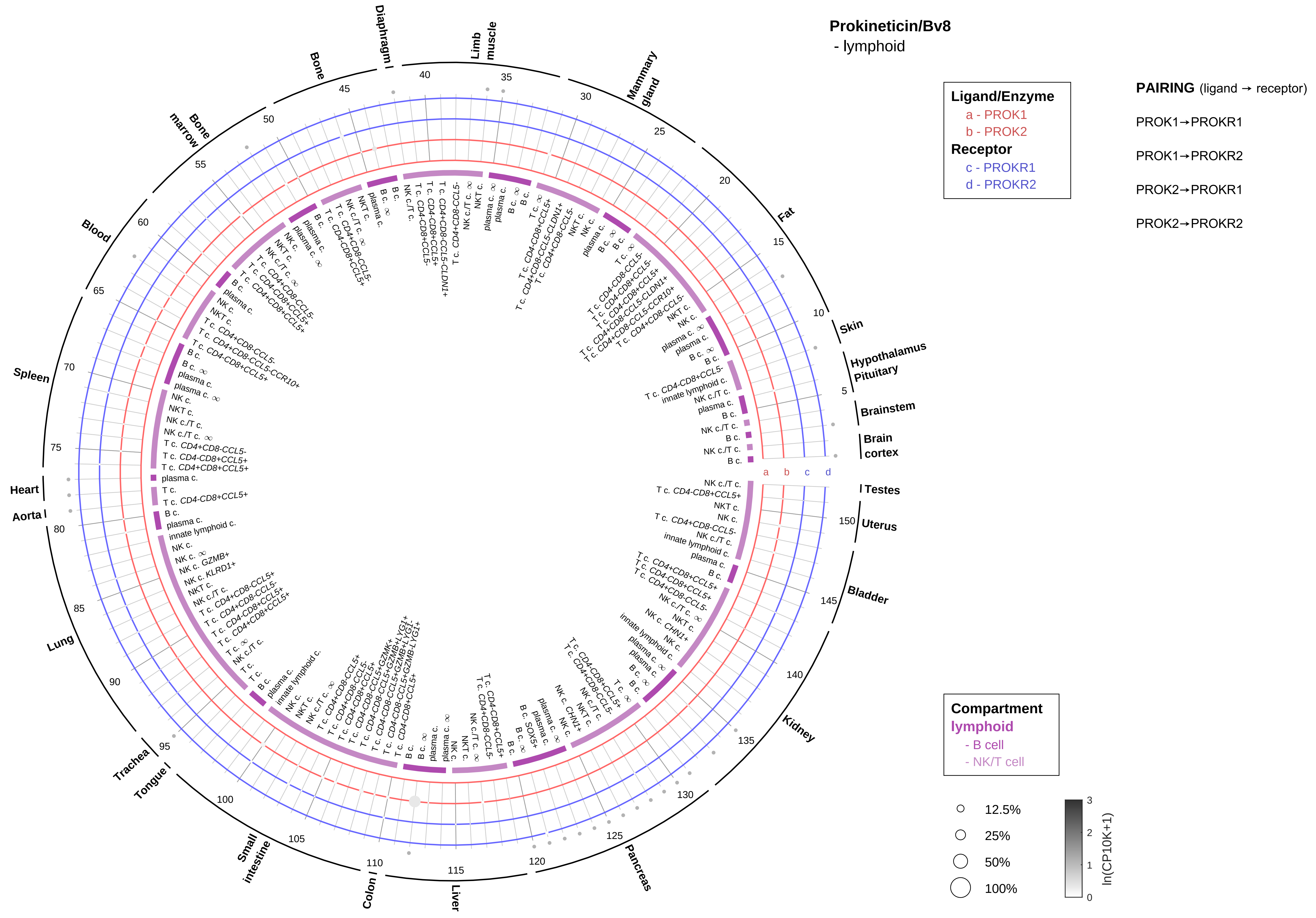
Prokineticin/Bv8
- lymphoid
Ligand/Enzyme
a - PROK1
b - PROK2
Receptor
c - PROKR1
d - PROKR2
PAIRING (ligand → receptor)
PROK1→PROKR1
PROK1→PROKR2
PROK2→PROKR1
PROK2→PROKR2
Compartment
lymphoid
- B cell
- NK/T cell
12.5%
25%
50%
100%
ln(CP10K+1)
Testes
Uterus
Bladder
Kidney
Pancreas
Liver
Colon
Small intestine
Tongue
Trachea
Lung
Aorta
Heart
Spleen
Blood
Bone marrow
Bone
Diaphragm
Limb muscle
Mammary gland
Fat
Skin
Hypothalamus Pituitary
Brainstem
Brain cortex

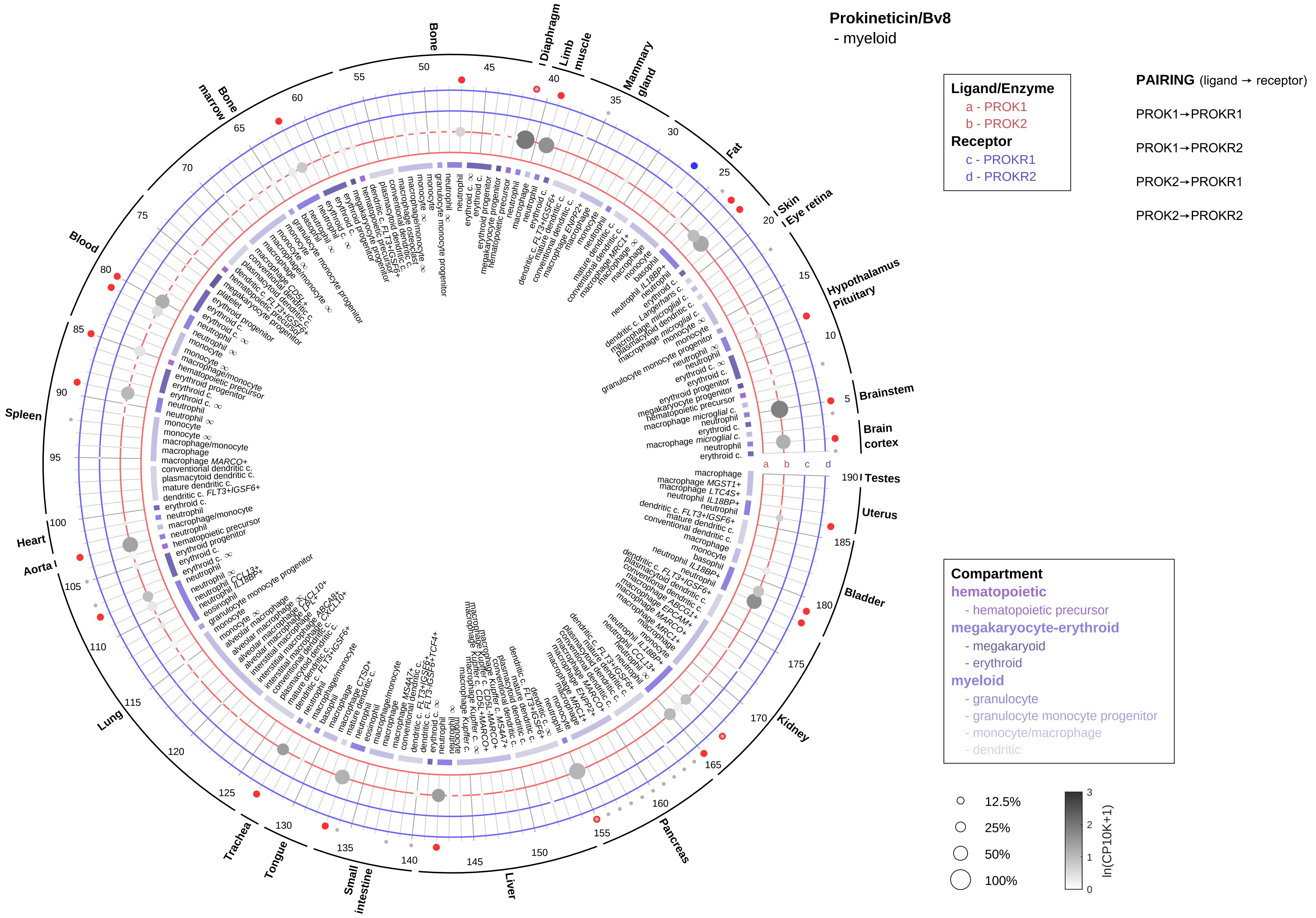
Prokineticin/Bv8
- myeloid
Ligand/Enzyme
a - PROK1
b - PROK2
Receptor
c - PROKR1
d - PROKR2
PAIRING (ligand → receptor)
PROK1→PROKR1
PROK1→PROKR2
PROK2→PROKR1
PROK2→PROKR2
Compartment
hematopoietic
- hematopoietic precursor
megakaryocyte-erythroid
- megakaryoid
- erythroid
myeloid
- granulocyte
- granulocyte monocyte progenitor
- monocyte/macrophage
- dendritic
12.5%
25%
50%
100%
ln(CP10K+1)
Bone
Diaphragm
Limb muscle
Mammary gland
Fat
Skin
Eye retina
Hypothalamus
Pituitary
Brainstem
Brain cortex
Testes
Uterus
Bladder
Kidney
Pancreas
Liver
Small intestine
Tongue
Trachea
Lung
Aorta
Heart
Spleen
Blood
Bone marrow

**Prolactin (PRL)**
- epithelial/neural/germ

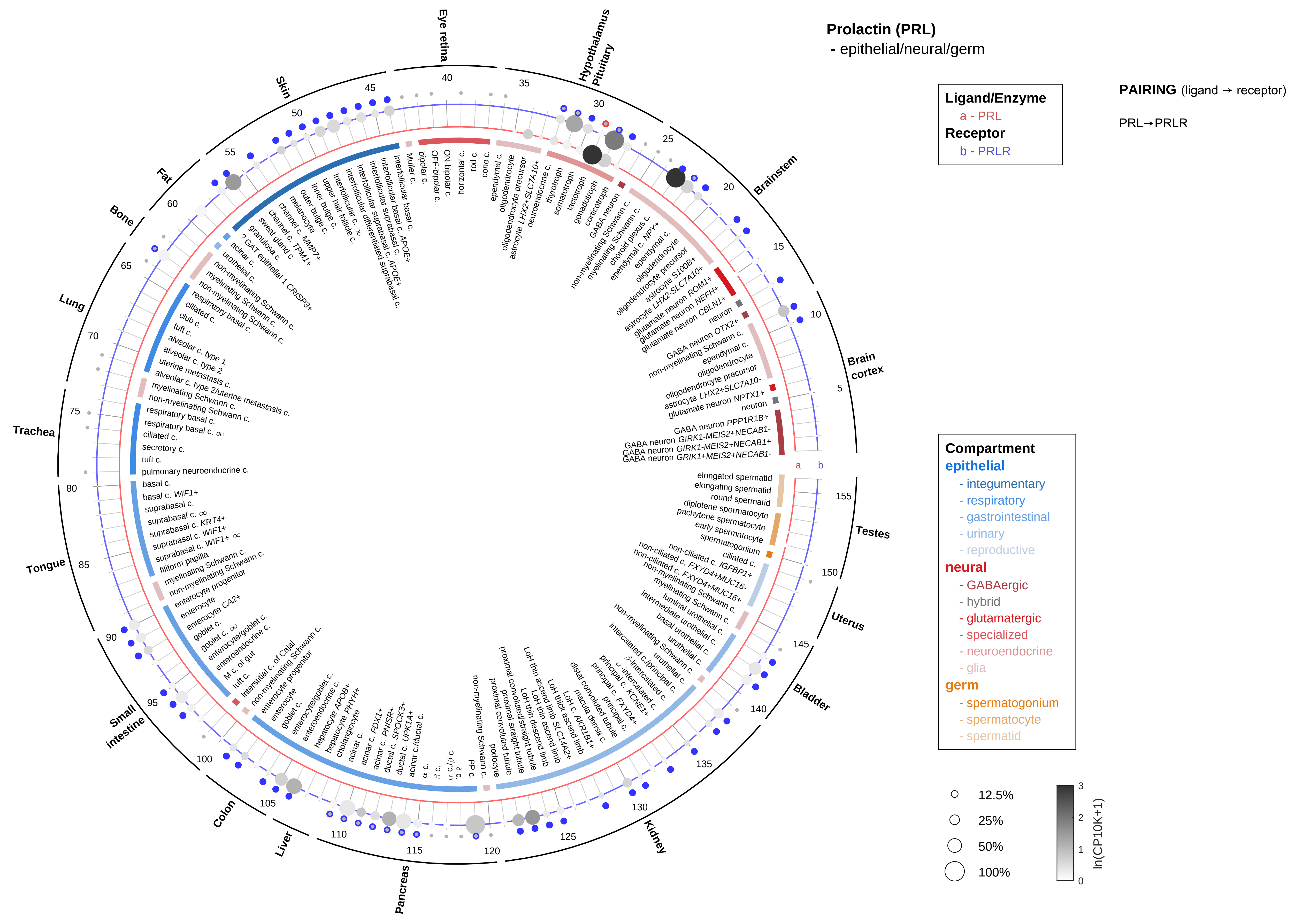

**Prolactin (PRL)**
- endothelial

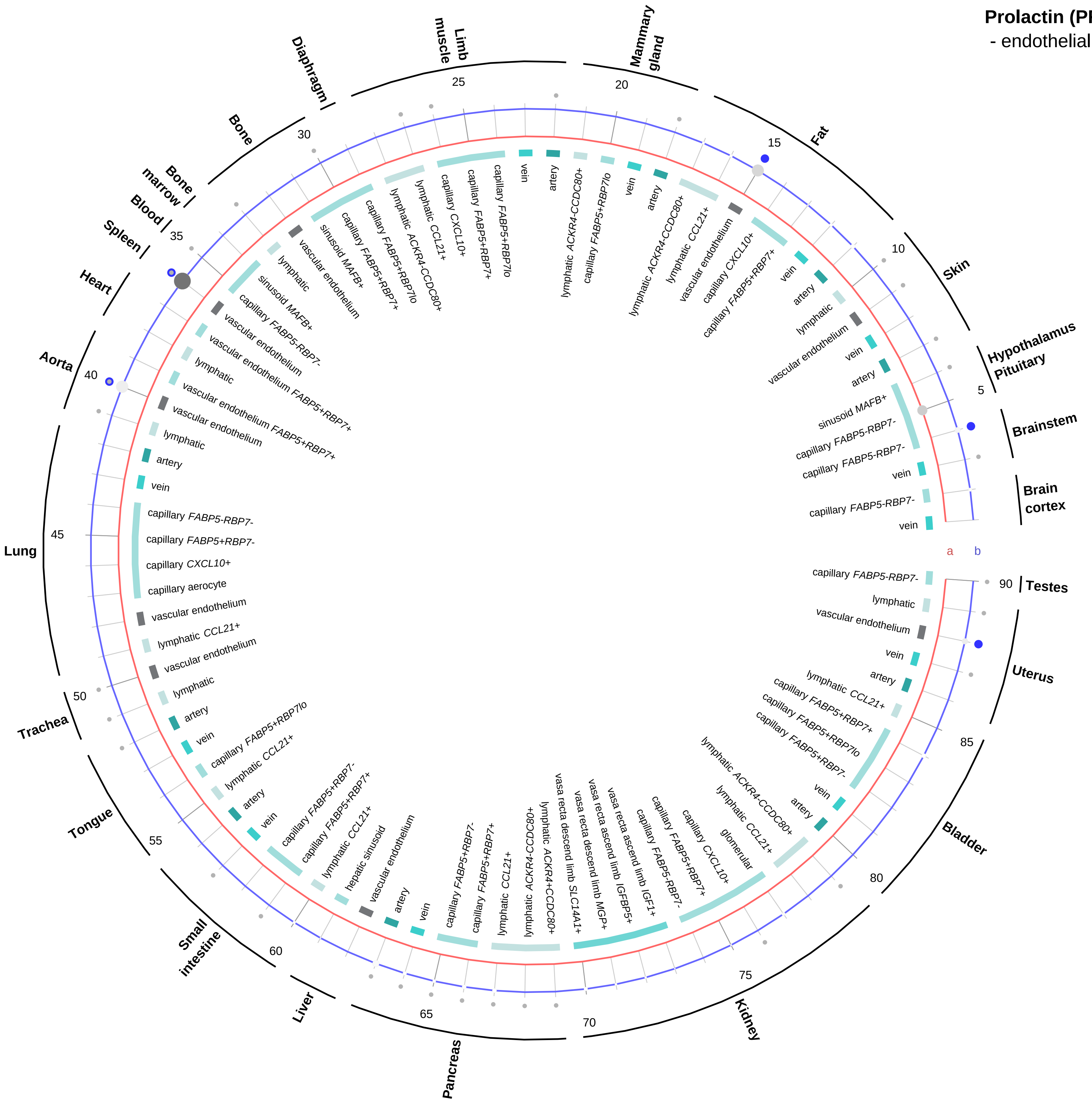


**Ligand/Enzyme**
a - PRL
**Receptor**
b - PRLR

**PAIRING** (ligand → receptor)

PRL→PRLR

**Compartment**
**endothelial**
- artery
- vein
- vasa recta
- capillary
- mix
- lymphatic

12.5%
25%
50%
100%

ln(CP10K+1)

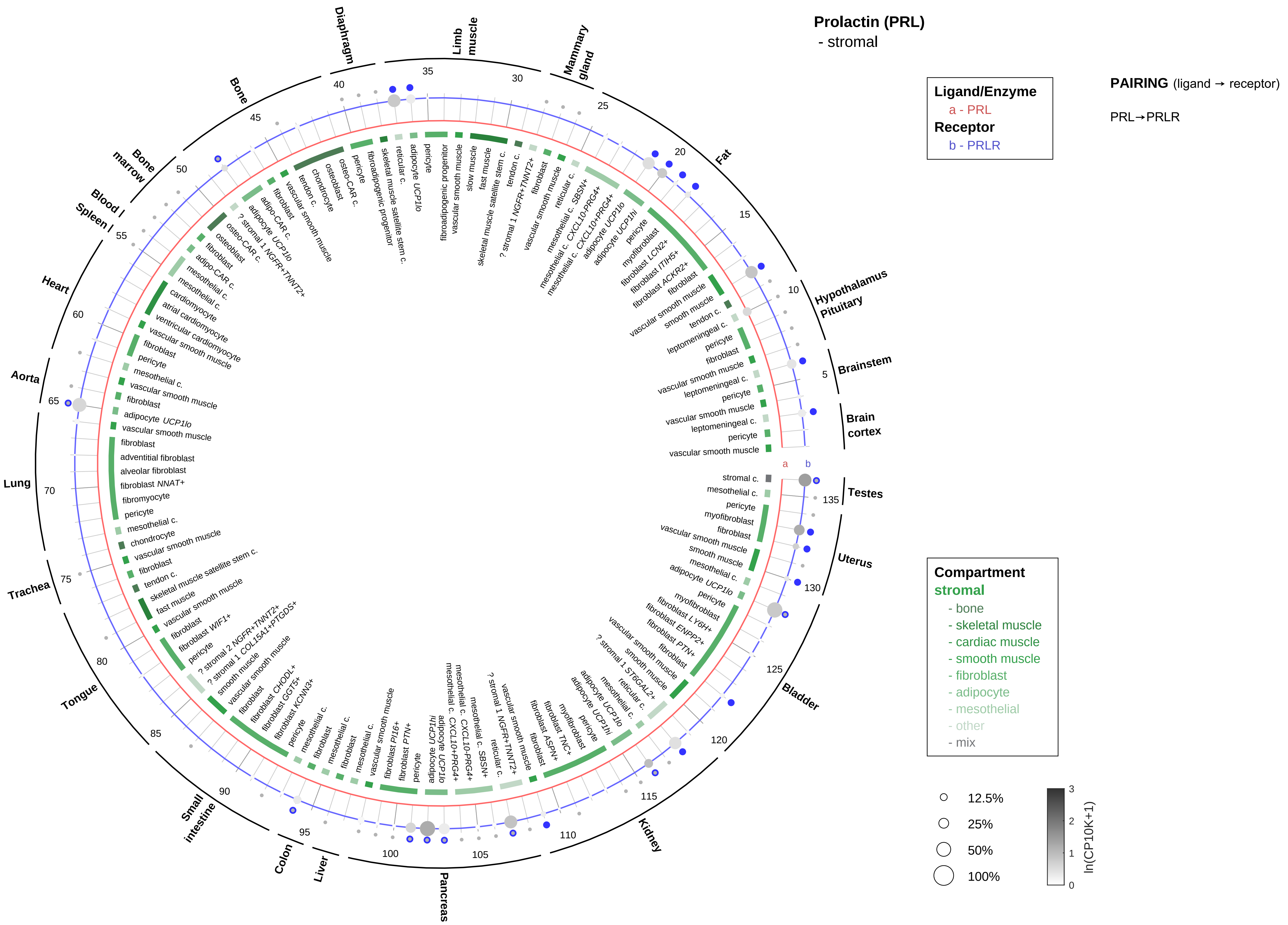

Prolactin (PRL)
- stromal
Ligand/Enzyme
a - PRL
Receptor
b - PRLR
PAIRING (ligand → receptor)
PRL→PRLR
Compartment
stromal
- bone
- skeletal muscle
- cardiac muscle
- smooth muscle
- fibroblast
- adipocyte
- mesothelial
- other
- mix
12.5%
25%
50%
100%
ln(CP10K+1)
Brain cortex
Brainstem
Hypothalamus Pituitary
Fat
Mammary gland
Limb muscle
Diaphragm
Bone
Bone marrow
Blood
Spleen
Heart
Aorta
Lung
Trachea
Tongue
Small intestine
Colon
Liver
Pancreas
Kidney
Bladder
Uterus
Testes

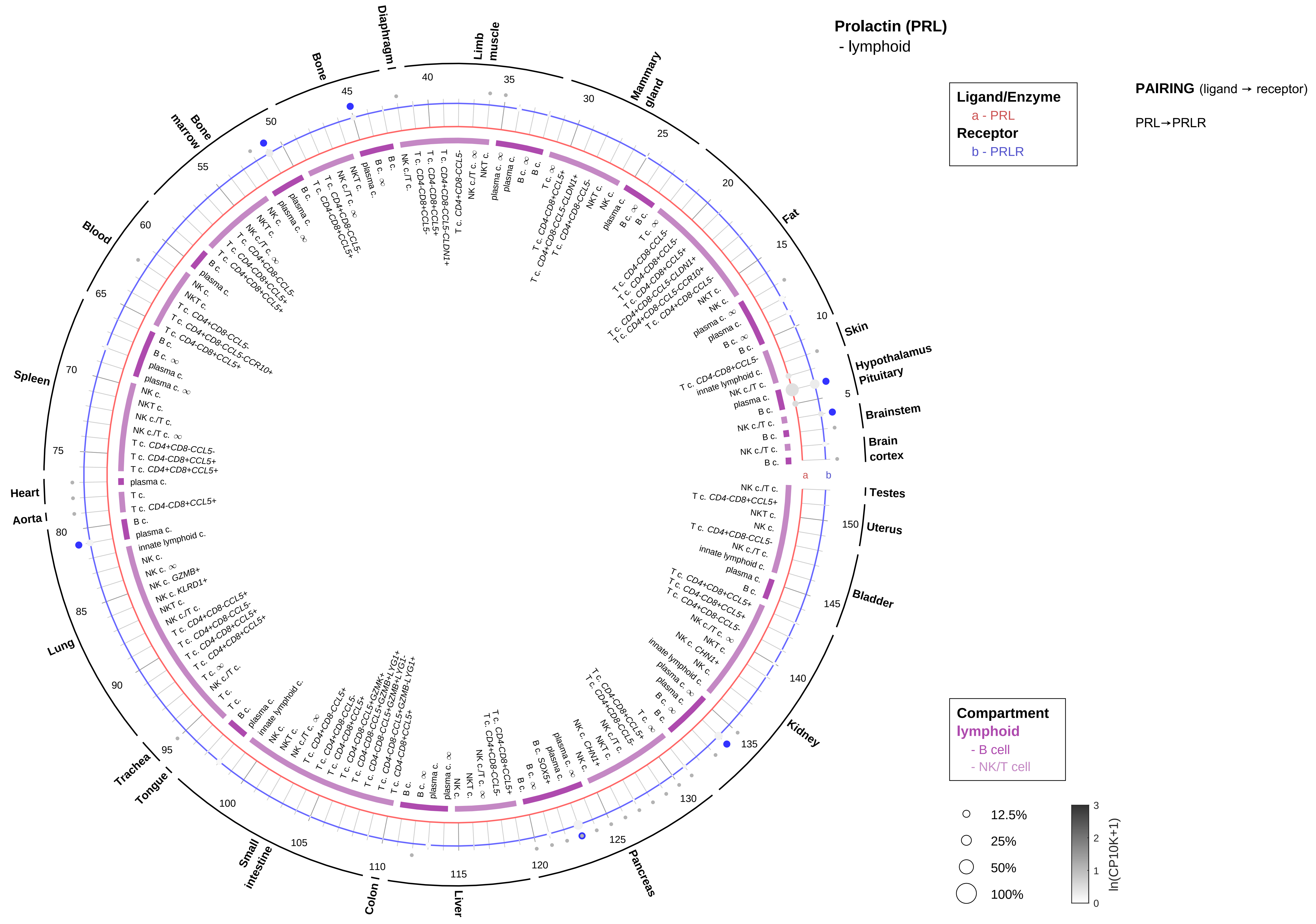
Prolactin (PRL)
- lymphoid
Ligand/Enzyme
a - PRL
Receptor
b - PRLR
PAIRING (ligand → receptor)
PRL→PRLR
Compartment
lymphoid
- B cell
- NK/T cell
12.5%
25%
50%
100%
ln(CP10K+1)
Skin
Hypothalamus
Pituitary
Brainstem
Brain cortex
Testes
Uterus
Bladder
Kidney
Pancreas
Liver
Colon
Small intestine
Tongue
Trachea
Lung
Aorta
Heart
Spleen
Blood
Bone marrow
Bone
Diaphragm
Limb muscle
Mammary gland
Fat

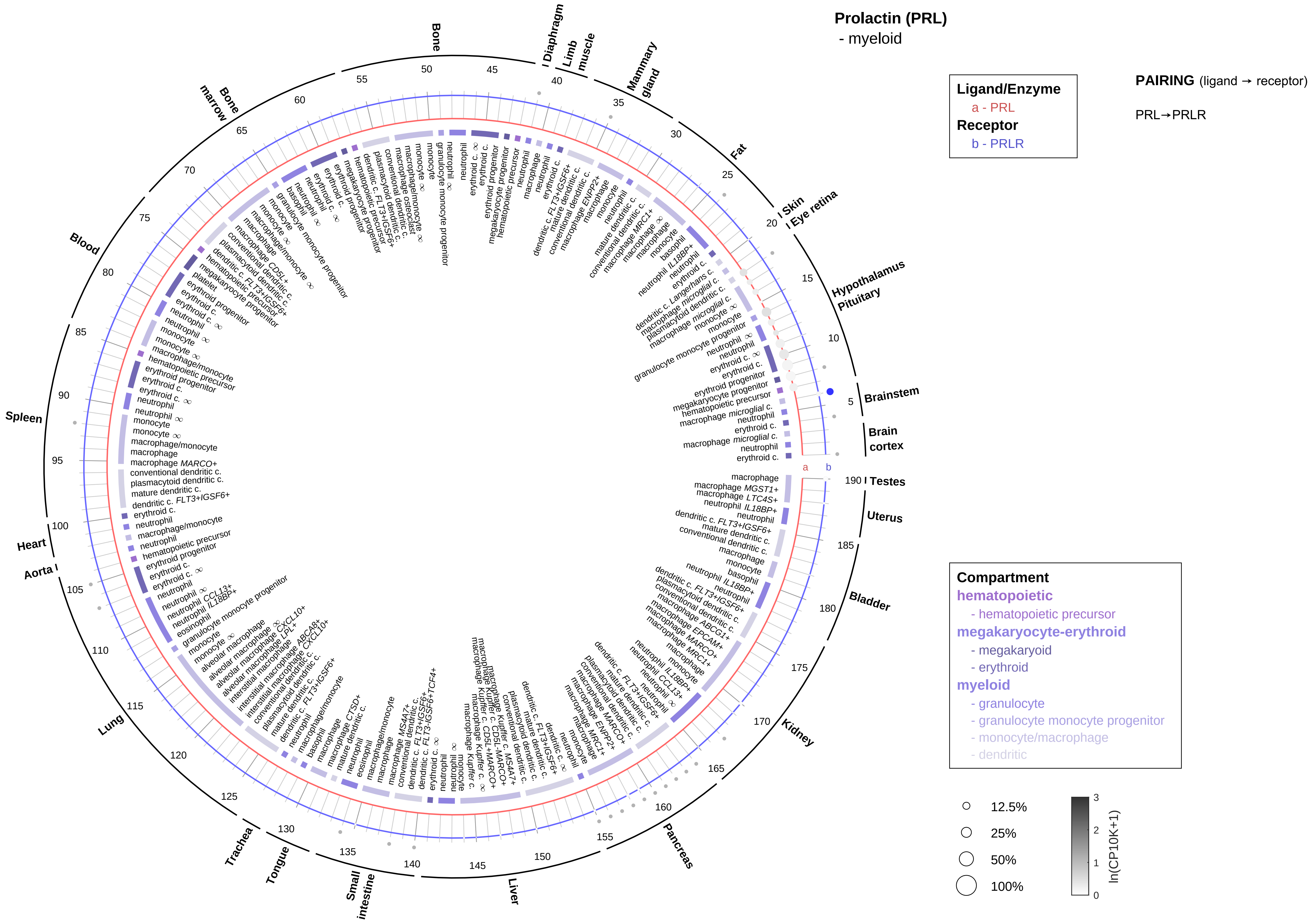

Prolactin (PRL)
- myeloid
Ligand/Enzyme
a - PRL
Receptor
b - PRLR
PAIRING (ligand → receptor)
PRL→PRLR
Compartment
hematopoietic
- hematopoietic precursor
megakaryocyte-erythroid
- megakaryoid
- erythroid
myeloid
- granulocyte
- granulocyte monocyte progenitor
- monocyte/macrophage
- dendritic
12.5%
25%
50%
100%
ln(CP10K+1)
Brainstem
Brain cortex
Testes
Uterus
Bladder
Kidney
Pancreas
Liver
Small intestine
Tongue
Trachea
Lung
Aorta
Heart
Spleen
Blood
Bone marrow
Bone
Diaphragm
Limb muscle
Mammary gland
Fat
Skin
Eye retina
Hypothalamus
Pituitary

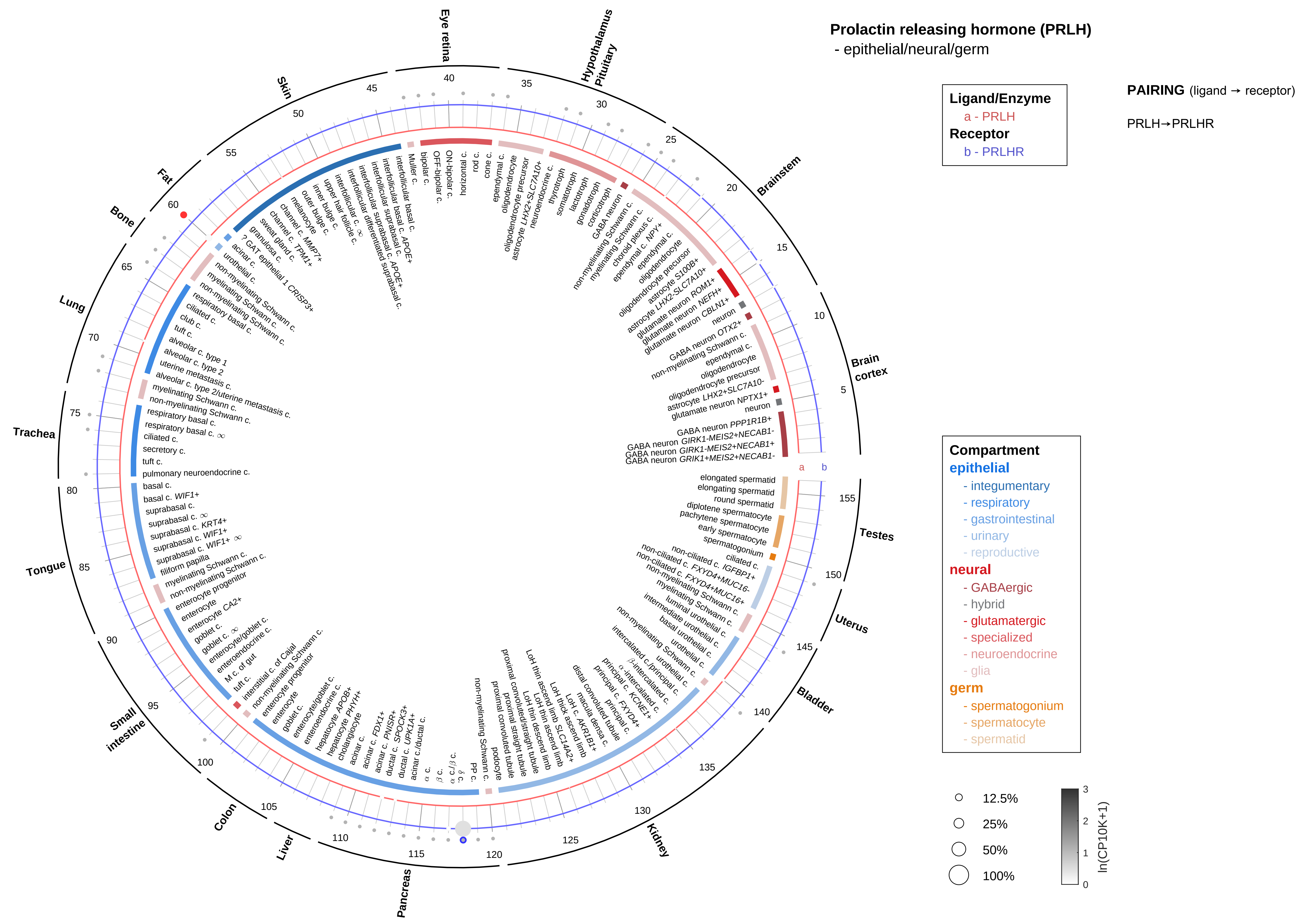

Prolactin releasing hormone (PRLH)
- epithelial/neural/germ
Ligand/Enzyme
a - PRLH
Receptor
b - PRLHR
PAIRING (ligand → receptor)
PRLH→PRLHR
Compartment
epithelial
- integumentary
- respiratory
- gastrointestinal
- urinary
- reproductive
neural
- GABAergic
- hybrid
- glutamatergic
- specialized
- neuroendocrine
- glia
germ
- spermatogonium
- spermatocyte
- spermatid
12.5%
25%
50%
100%
ln(CP10K+1)
Brain cortex
Brainstem
Hypothalamus Pituitary
Eye retina
Skin
Fat
Bone
Lung
Trachea
Tongue
Small intestine
Colon
Liver
Pancreas
Kidney
Bladder
Uterus
Testes

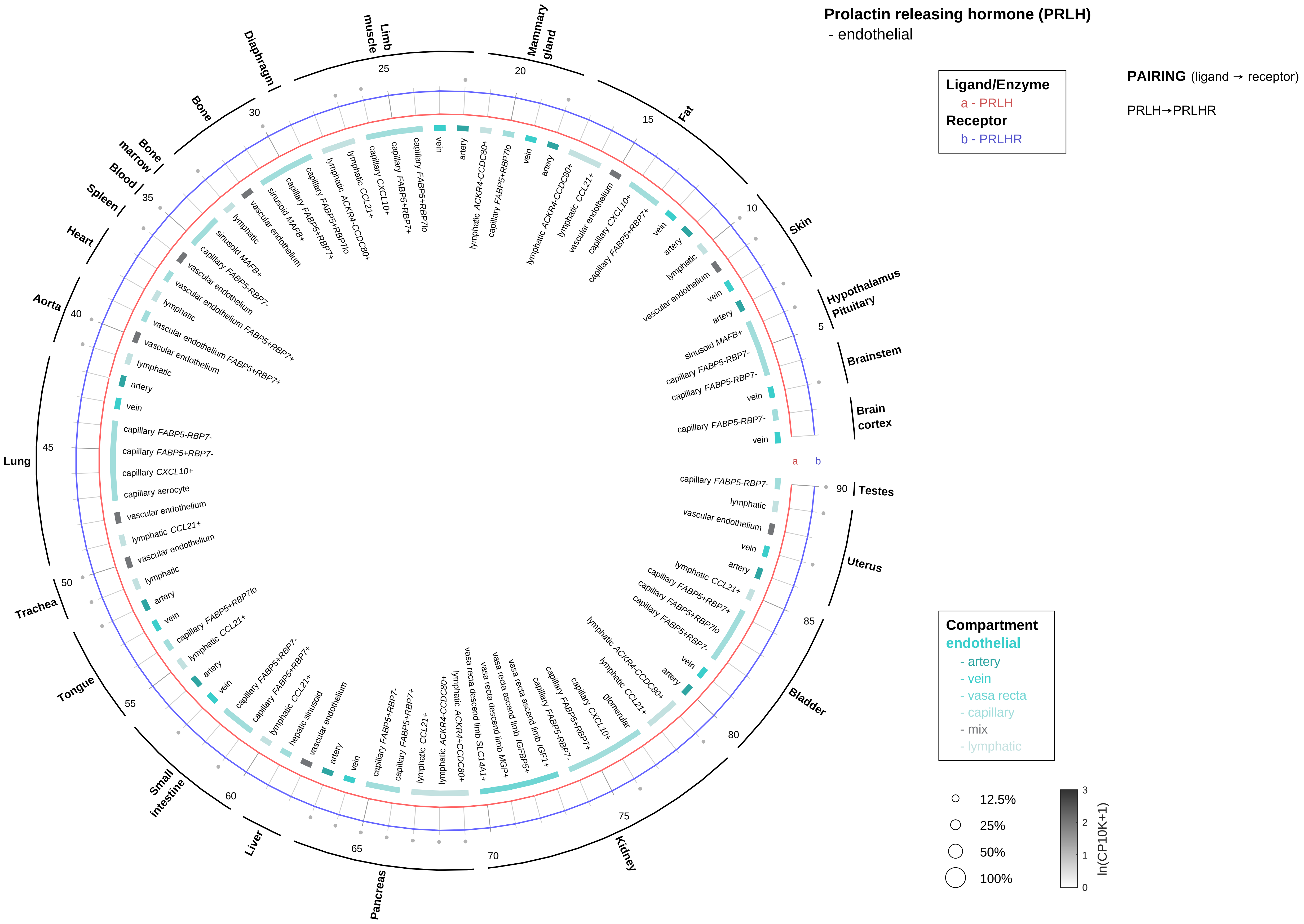

Prolactin releasing hormone (PRLH)
- endothelial
Ligand/Enzyme
a - PRLH
Receptor
b - PRLHR
PAIRING (ligand → receptor)
PRLH→PRLHR
Compartment
endothelial
- artery
- vein
- vasa recta
- capillary
- mix
- lymphatic
12.5%
25%
50%
100%
ln(CP10K+1)
Brain cortex
Brainstem
Hypothalamus
Pituitary
Skin
Fat
Mammary gland
Limb muscle
Diaphragm
Bone
Bone marrow
Blood
Spleen
Heart
Aorta
Lung
Trachea
Tongue
Small intestine
Liver
Pancreas
Kidney
Bladder
Uterus
Testes

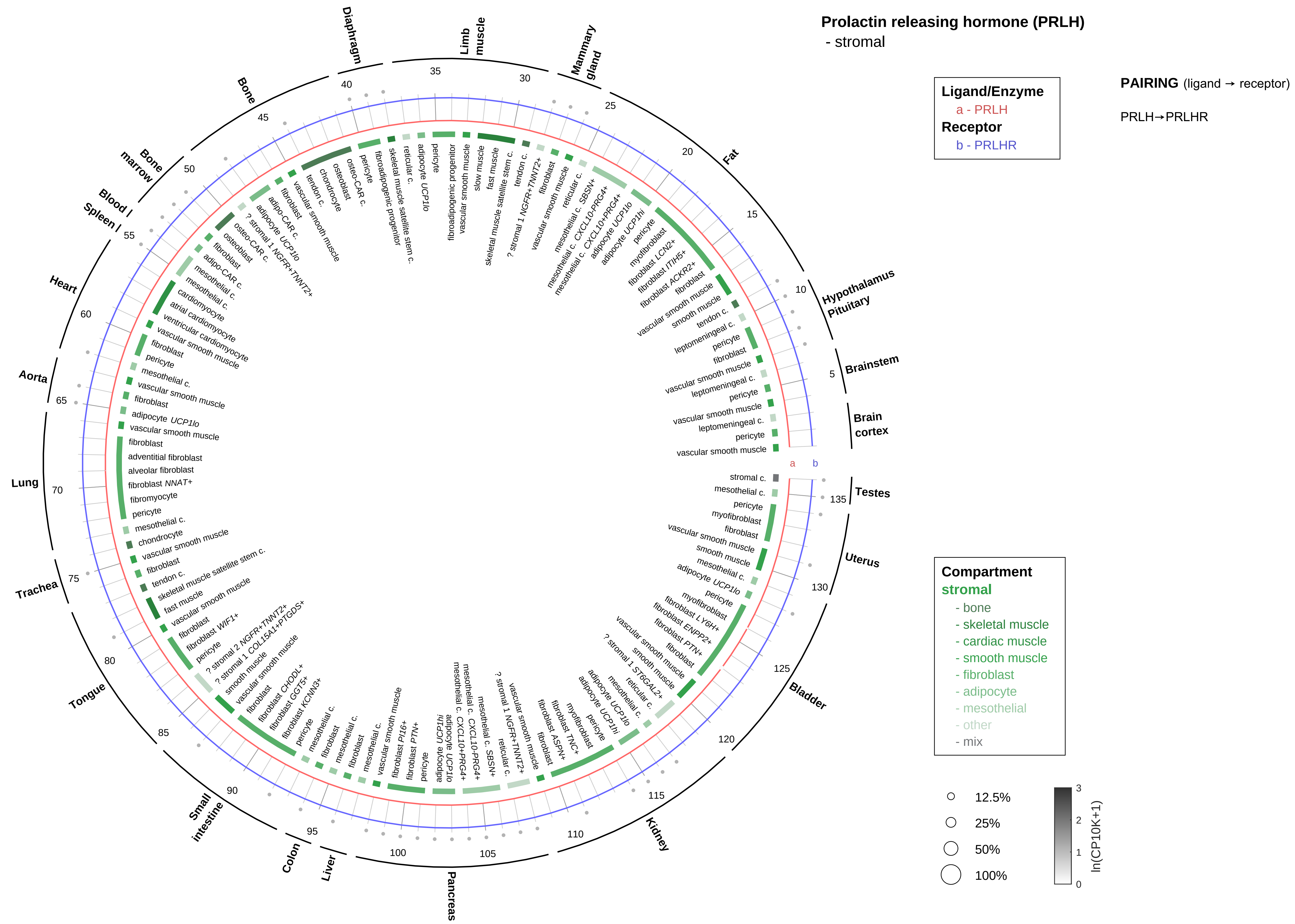

Prolactin releasing hormone (PRLH)
- stromal
Ligand/Enzyme
a - PRLH
Receptor
b - PRLHR
PAIRING (ligand → receptor)
PRLH→PRLHR
Compartment
stromal
- bone
- skeletal muscle
- cardiac muscle
- smooth muscle
- fibroblast
- adipocyte
- mesothelial
- other
- mix
12.5%
25%
50%
100%
ln(CP10K+1)
Testes
Uterus
Bladder
Kidney
Pancreas
Liver
Colon
Small intestine
Tongue
Trachea
Lung
Aorta
Heart
Spleen
Blood
Bone marrow
Bone
Diaphragm
Limb muscle
Mammary gland
Fat
Hypothalamus
Pituitary
Brainstem
Brain cortex

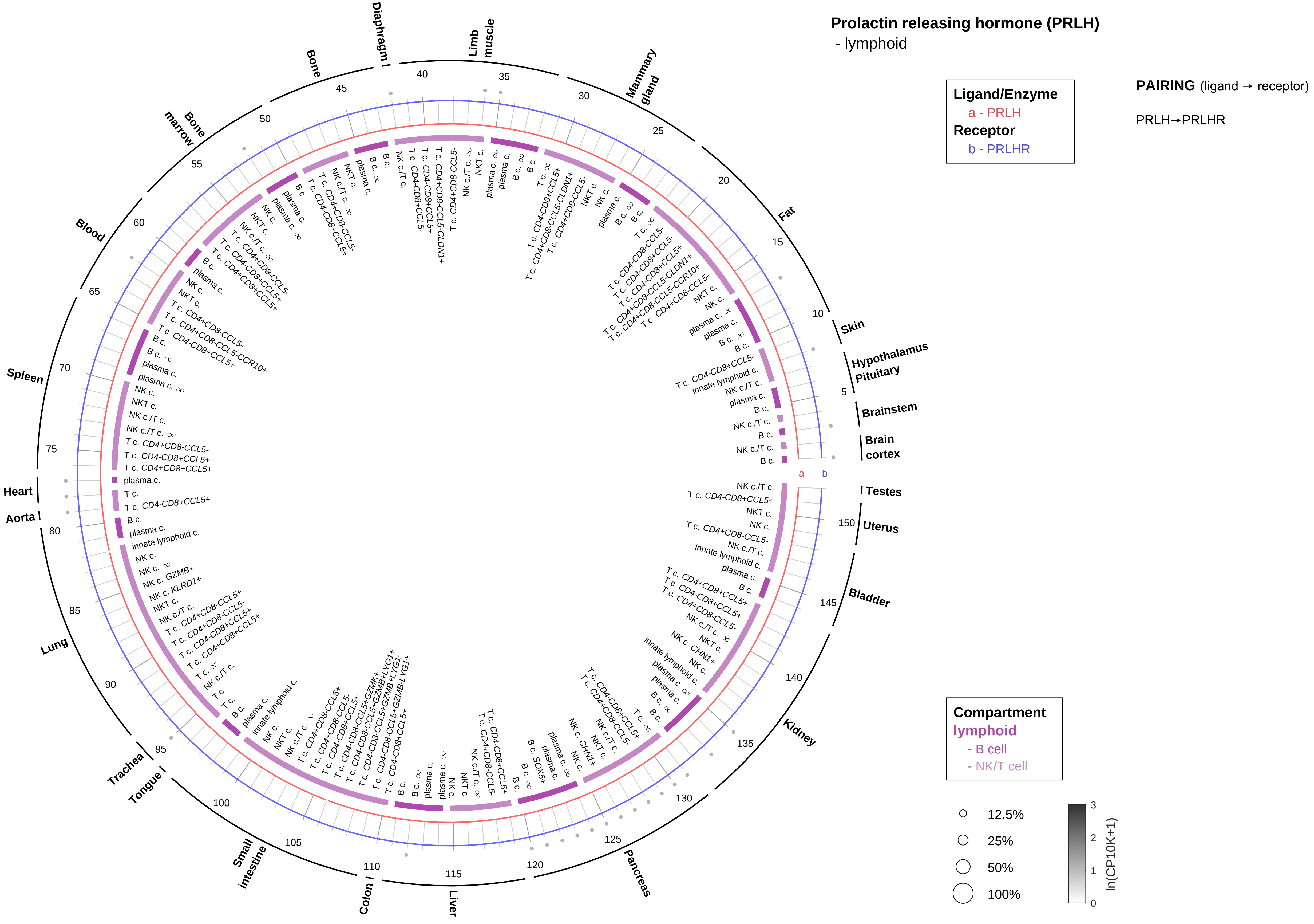

Prolactin releasing hormone (PRLH)
- lymphoid
Ligand/Enzyme
a - PRLH
Receptor
b - PRLHR
PAIRING (ligand → receptor)
PRLH→PRLHR
Compartment
lymphoid
- B cell
- NK/T cell
12.5%
25%
50%
100%
ln(CP10K+1)
Skin
Hypothalamus
Pituitary
Brainstem
Brain cortex
Testes
Uterus
Bladder
Kidney
Pancreas
Liver
Colon
Small intestine
Tongue
Trachea
Lung
Aorta
Heart
Spleen
Blood
Bone marrow
Bone
Diaphragm
Limb muscle
Mammary gland
Fat

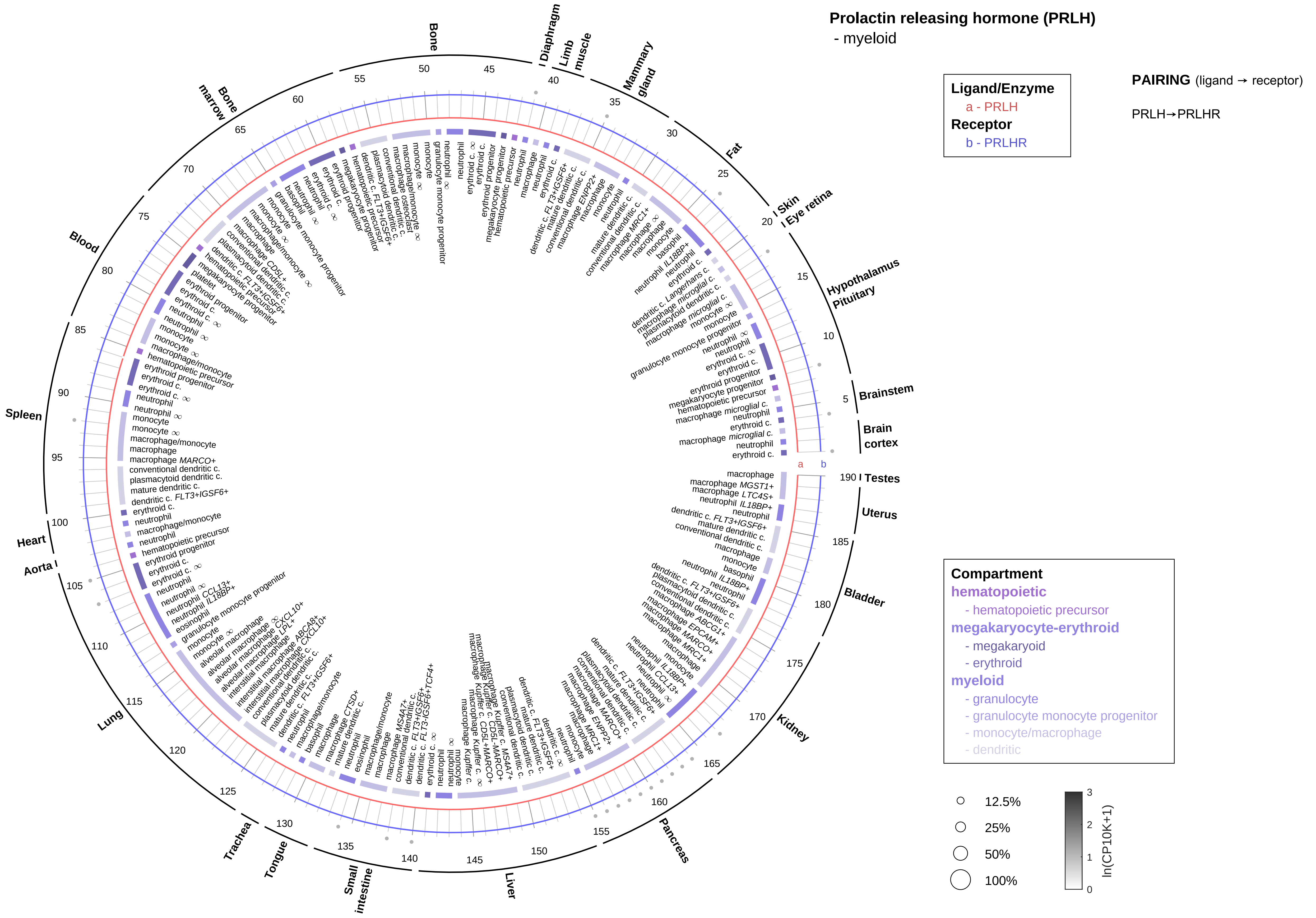

Prolactin releasing hormone (PRLH)
- myeloid
Ligand/Enzyme
a - PRLH
Receptor
b - PRLHR
PAIRING (ligand → receptor)
PRLH→PRLHR
Compartment
hematopoietic
- hematopoietic precursor
megakaryocyte-erythroid
- megakaryoid
- erythroid
myeloid
- granulocyte
- granulocyte monocyte progenitor
- monocyte/macrophage
- dendritic
12.5%
25%
50%
100%
ln(CP10K+1)
Bone
Diaphragm
Limb muscle
Mammary gland
Fat
Skin
Eye retina
Hypothalamus
Pituitary
Brainstem
Brain cortex
Testes
Uterus
Bladder
Kidney
Pancreas
Liver
Small intestine
Tongue
Trachea
Lung
Aorta
Heart
Spleen
Blood
Bone marrow

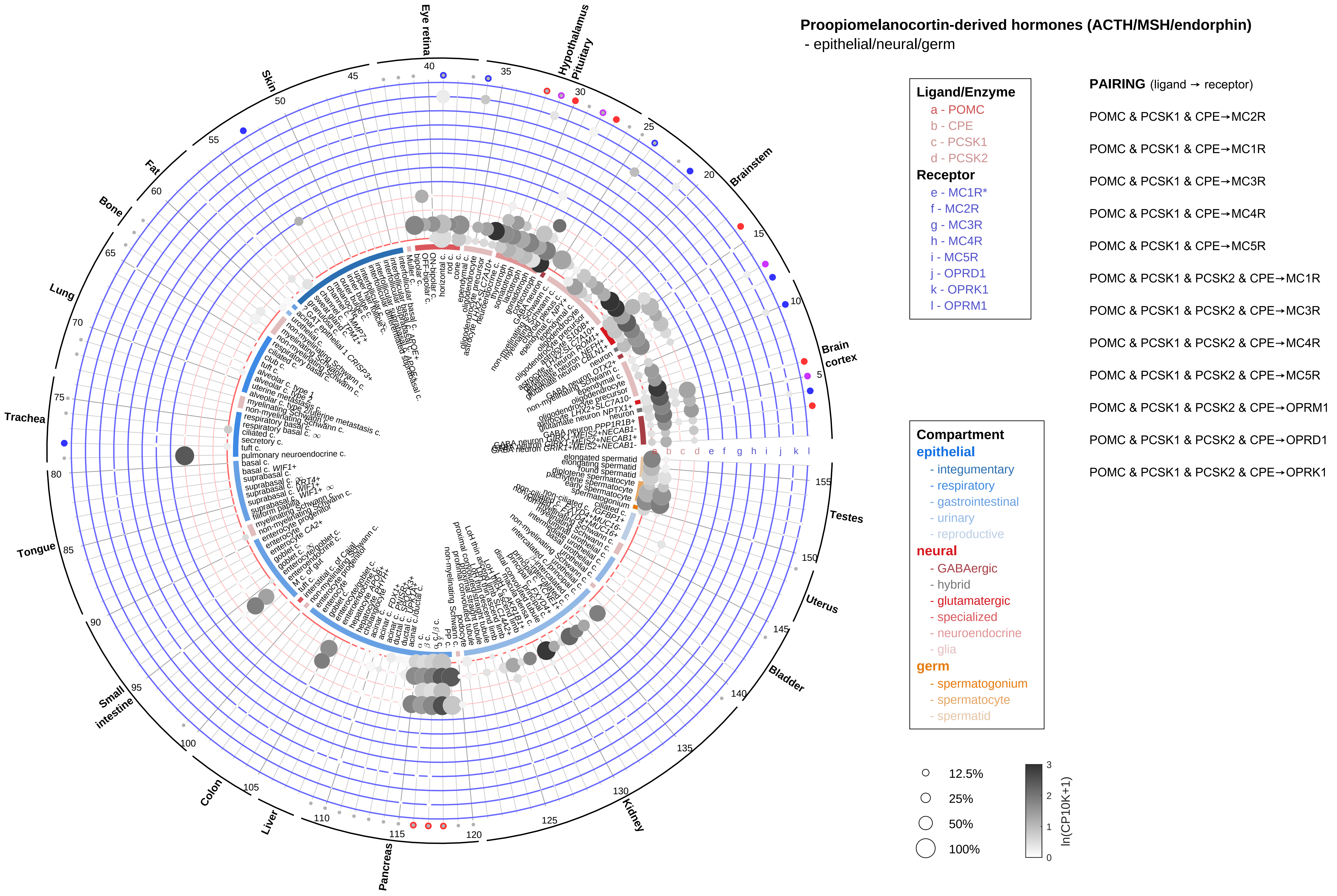
Proopiomelanocortin-derived hormones (ACTH/MSH/endorphin)
- epithelial/neural/germ
Ligand/Enzyme
a - POMC
b - CPE
c - PCSK1
d - PCSK2
Receptor
e - MC1R*
f - MC2R
g - MC3R
h - MC4R
i - MC5R
j - OPRD1
k - OPRK1
l - OPRM1
PAIRING (ligand → receptor)
POMC & PCSK1 & CPE→MC2R
POMC & PCSK1 & CPE→MC1R
POMC & PCSK1 & CPE→MC3R
POMC & PCSK1 & CPE→MC4R
POMC & PCSK1 & CPE→MC5R
POMC & PCSK1 & PCSK2 & CPE→MC1R
POMC & PCSK1 & PCSK2 & CPE→MC3R
POMC & PCSK1 & PCSK2 & CPE→MC4R
POMC & PCSK1 & PCSK2 & CPE→MC5R
POMC & PCSK1 & PCSK2 & CPE→OPRM1
POMC & PCSK1 & PCSK2 & CPE→OPRD1
POMC & PCSK1 & PCSK2 & CPE→OPRK1
Compartment
epithelial
- integumentary
- respiratory
- gastrointestinal
- urinary
- reproductive
neural
- GABAergic
- hybrid
- glutamatergic
- specialized
- neuroendocrine
- glia
germ
- spermatogonium
- spermatocyte
- spermatid
12.5%
25%
50%
100%
ln(CP10K+1)
Eye retina
Hypothalamus
Pituitary
Brainstem
Brain cortex
Testes
Uterus
Bladder
Kidney
Pancreas
Liver
Colon
Small intestine
Tongue
Trachea
Lung
Bone
Fat
Skin

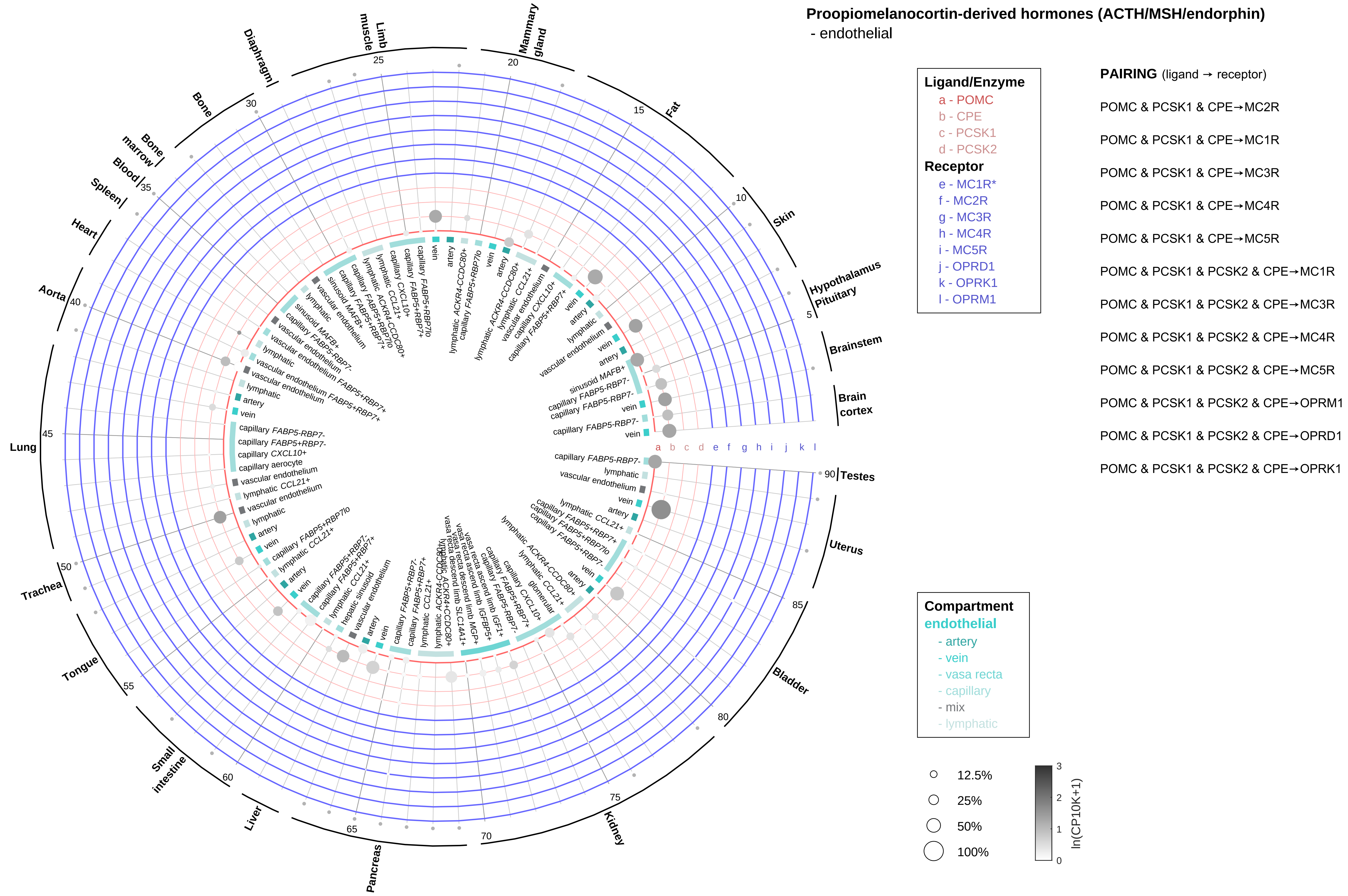
Proopiomelanocortin-derived hormones (ACTH/MSH/endorphin)
- endothelial
Ligand/Enzyme
a - POMC
b - CPE
c - PCSK1
d - PCSK2
Receptor
e - MC1R*
f - MC2R
g - MC3R
h - MC4R
i - MC5R
j - OPRD1
k - OPRK1
l - OPRM1
PAIRING (ligand → receptor)
POMC & PCSK1 & CPE→MC2R
POMC & PCSK1 & CPE→MC1R
POMC & PCSK1 & CPE→MC3R
POMC & PCSK1 & CPE→MC4R
POMC & PCSK1 & CPE→MC5R
POMC & PCSK1 & PCSK2 & CPE→MC1R
POMC & PCSK1 & PCSK2 & CPE→MC3R
POMC & PCSK1 & PCSK2 & CPE→MC4R
POMC & PCSK1 & PCSK2 & CPE→MC5R
POMC & PCSK1 & PCSK2 & CPE→OPRM1
POMC & PCSK1 & PCSK2 & CPE→OPRD1
POMC & PCSK1 & PCSK2 & CPE→OPRK1
Compartment
endothelial
- artery
- vein
- vasa recta
- capillary
- mix
- lymphatic
12.5%
25%
50%
100%
ln(CP10K+1)
Testes
Uterus
Bladder
Kidney
Pancreas
Liver
Small intestine
Tongue
Trachea
Lung
Aorta
Heart
Spleen
Blood
Bone marrow
Bone
Diaphragm
Limb muscle
Mammary gland
Fat
Skin
Hypothalamus
Pituitary
Brainstem
Brain cortex

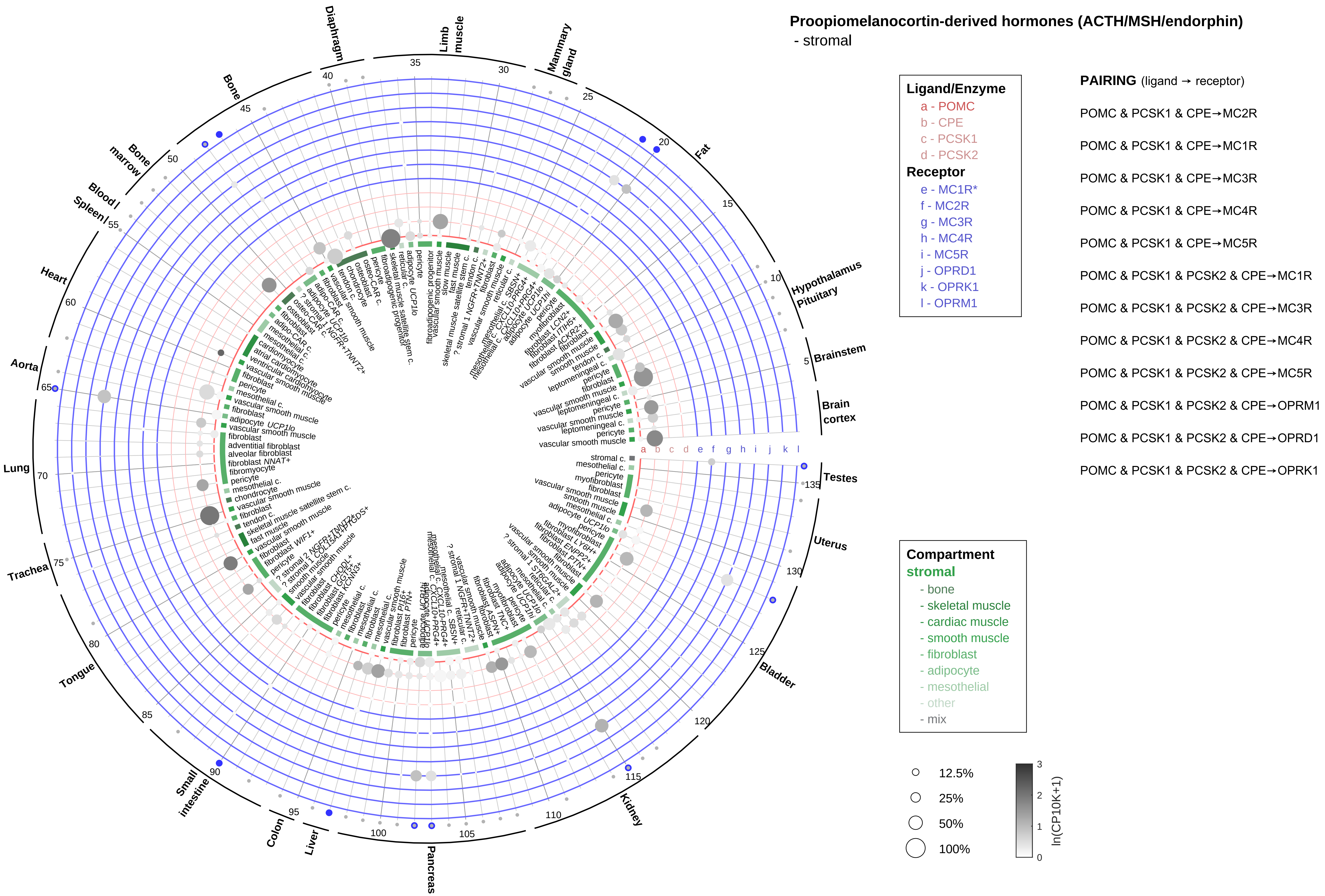

Proopiomelanocortin-derived hormones (ACTH/MSH/endorphin)
- stromal
Ligand/Enzyme
a - POMC
b - CPE
c - PCSK1
d - PCSK2
Receptor
e - MC1R*
f - MC2R
g - MC3R
h - MC4R
i - MC5R
j - OPRD1
k - OPRK1
l - OPRM1
PAIRING (ligand → receptor)
POMC & PCSK1 & CPE→MC2R
POMC & PCSK1 & CPE→MC1R
POMC & PCSK1 & CPE→MC3R
POMC & PCSK1 & CPE→MC4R
POMC & PCSK1 & CPE→MC5R
POMC & PCSK1 & PCSK2 & CPE→MC1R
POMC & PCSK1 & PCSK2 & CPE→MC3R
POMC & PCSK1 & PCSK2 & CPE→MC4R
POMC & PCSK1 & PCSK2 & CPE→MC5R
POMC & PCSK1 & PCSK2 & CPE→OPRM1
POMC & PCSK1 & PCSK2 & CPE→OPRD1
POMC & PCSK1 & PCSK2 & CPE→OPRK1
Compartment
stromal
- bone
- skeletal muscle
- cardiac muscle
- smooth muscle
- fibroblast
- adipocyte
- mesothelial
- other
- mix
12.5%
25%
50%
100%
ln(CP10K+1)
Diaphragm
Limb muscle
Mammary gland
Fat
Hypothalamus Pituitary
Brainstem
Brain cortex
Testes
Uterus
Bladder
Kidney
Pancreas
Liver
Colon
Small intestine
Tongue
Trachea
Lung
Aorta
Heart
Spleen
Blood
Bone marrow
Bone

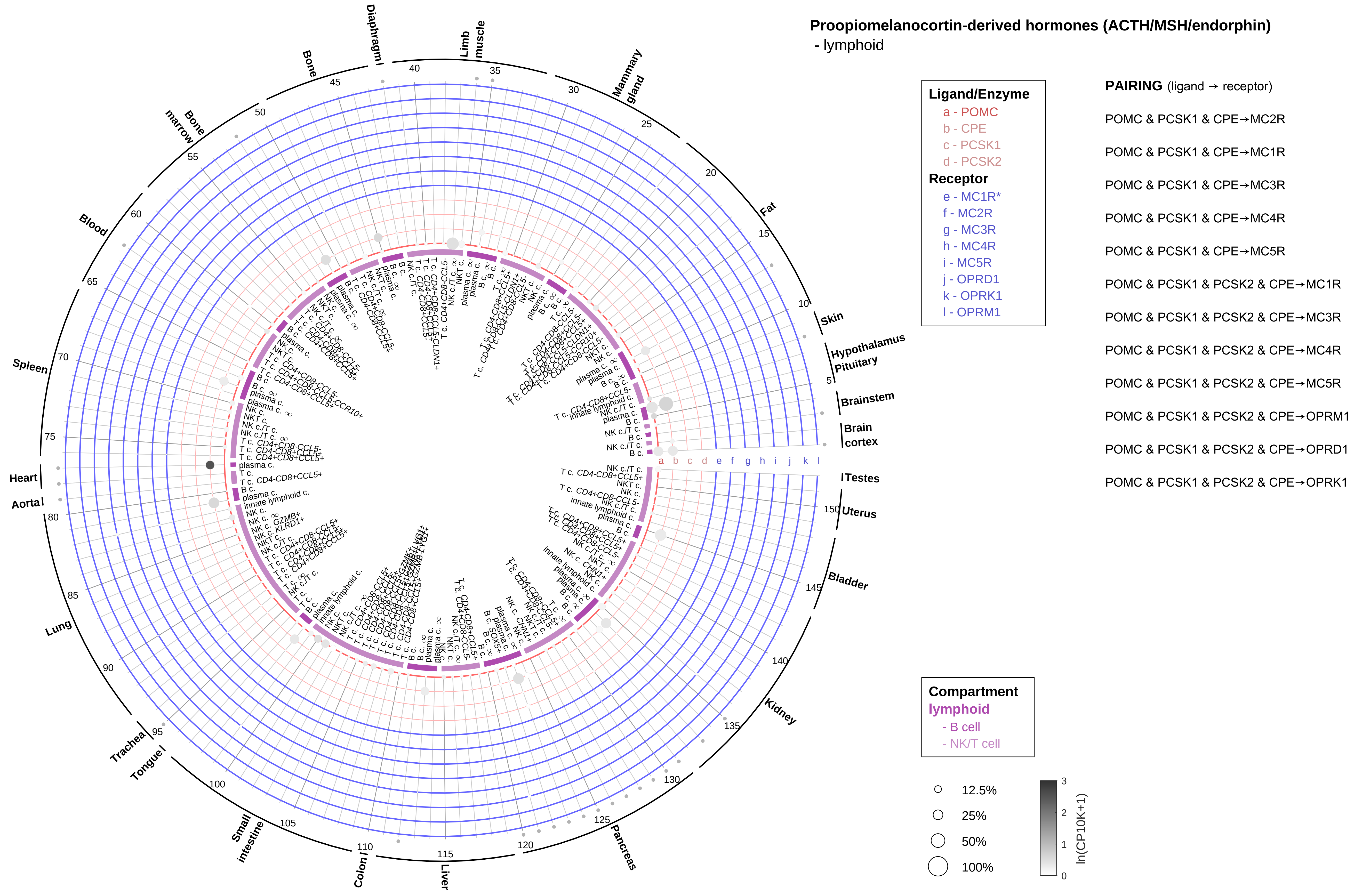
Proopiomelanocortin-derived hormones (ACTH/MSH/endorphin)
- lymphoid
Ligand/Enzyme
a - POMC
b - CPE
c - PCSK1
d - PCSK2
Receptor
e - MC1R*
f - MC2R
g - MC3R
h - MC4R
i - MC5R
j - OPRD1
k - OPRK1
l - OPRM1
PAIRING (ligand → receptor)
POMC & PCSK1 & CPE→MC2R
POMC & PCSK1 & CPE→MC1R
POMC & PCSK1 & CPE→MC3R
POMC & PCSK1 & CPE→MC4R
POMC & PCSK1 & CPE→MC5R
POMC & PCSK1 & PCSK2 & CPE→MC1R
POMC & PCSK1 & PCSK2 & CPE→MC3R
POMC & PCSK1 & PCSK2 & CPE→MC4R
POMC & PCSK1 & PCSK2 & CPE→MC5R
POMC & PCSK1 & PCSK2 & CPE→OPRM1
POMC & PCSK1 & PCSK2 & CPE→OPRD1
POMC & PCSK1 & PCSK2 & CPE→OPRK1
Compartment
lymphoid
- B cell
- NK/T cell
12.5%
25%
50%
100%
ln(CP10K+1)
Skin
Hypothalamus
Pituitary
Brainstem
Brain cortex
Testes
Uterus
Bladder
Kidney
Pancreas
Liver
Colon
Small intestine
Tongue
Trachea
Lung
Aorta
Heart
Spleen
Blood
Bone marrow
Bone
Diaphragm
Limb muscle
Mammary gland
Fat

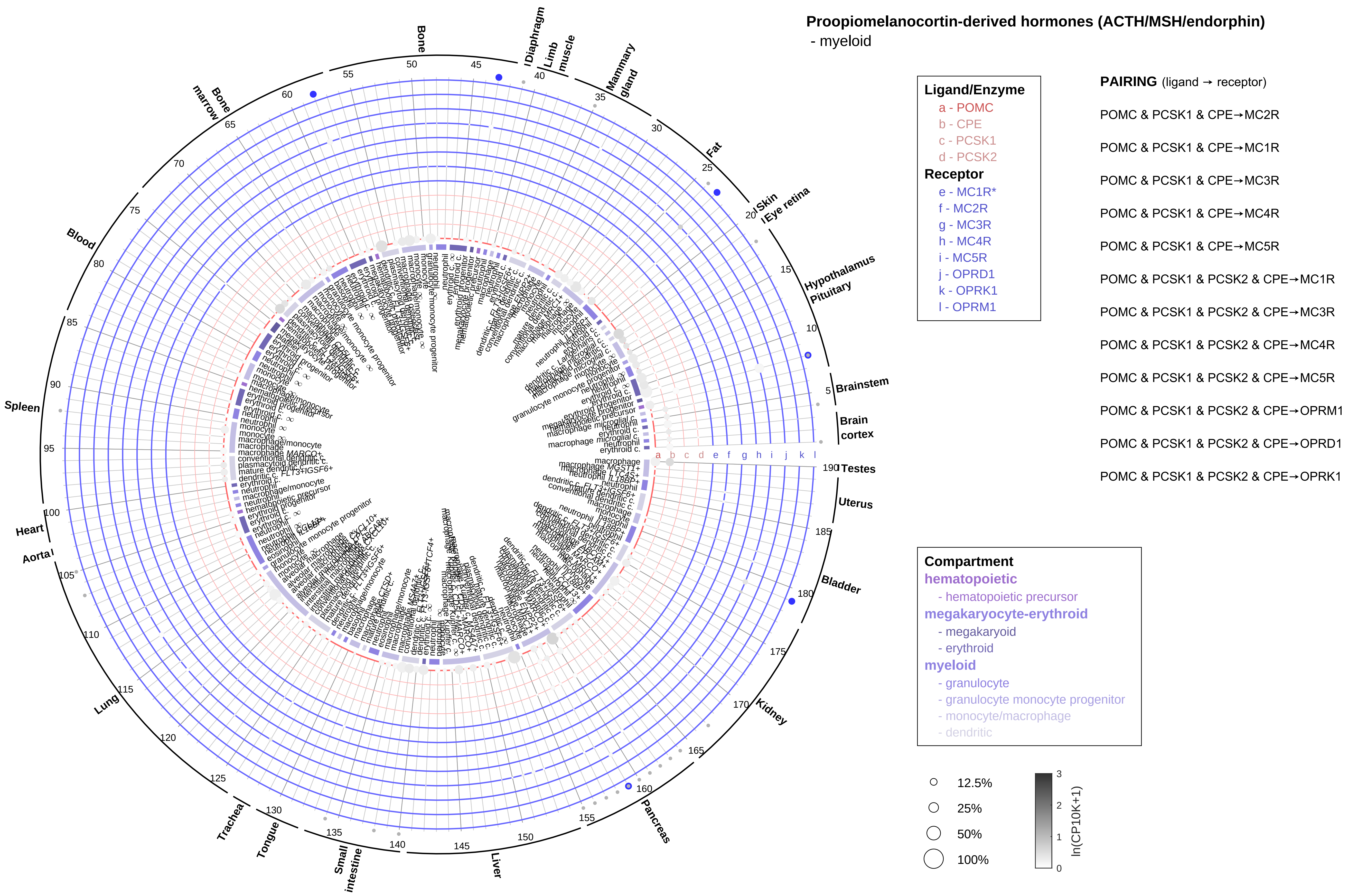
Proopiomelanocortin-derived hormones (ACTH/MSH/endorphin)
- myeloid
Ligand/Enzyme
a - POMC
b - CPE
c - PCSK1
d - PCSK2
Receptor
e - MC1R*
f - MC2R
g - MC3R
h - MC4R
i - MC5R
j - OPRD1
k - OPRK1
l - OPRM1
PAIRING (ligand → receptor)
POMC & PCSK1 & CPE→MC2R
POMC & PCSK1 & CPE→MC1R
POMC & PCSK1 & CPE→MC3R
POMC & PCSK1 & CPE→MC4R
POMC & PCSK1 & CPE→MC5R
POMC & PCSK1 & PCSK2 & CPE→MC1R
POMC & PCSK1 & PCSK2 & CPE→MC3R
POMC & PCSK1 & PCSK2 & CPE→MC4R
POMC & PCSK1 & PCSK2 & CPE→MC5R
POMC & PCSK1 & PCSK2 & CPE→OPRM1
POMC & PCSK1 & PCSK2 & CPE→OPRD1
POMC & PCSK1 & PCSK2 & CPE→OPRK1
Compartment
hematopoietic
- hematopoietic precursor
megakaryocyte-erythroid
- megakaryoid
- erythroid
myeloid
- granulocyte
- granulocyte monocyte progenitor
- monocyte/macrophage
- dendritic
12.5%
25%
50%
100%
ln(CP10K+1)
Bone
Diaphragm
Limb muscle
Mammary gland
Fat
Skin
Eye retina
Hypothalamus
Pituitary
Brainstem
Brain cortex
Testes
Uterus
Bladder
Kidney
Pancreas
Liver
Small intestine
Tongue
Trachea
Lung
Aorta
Heart
Spleen
Blood
Bone marrow

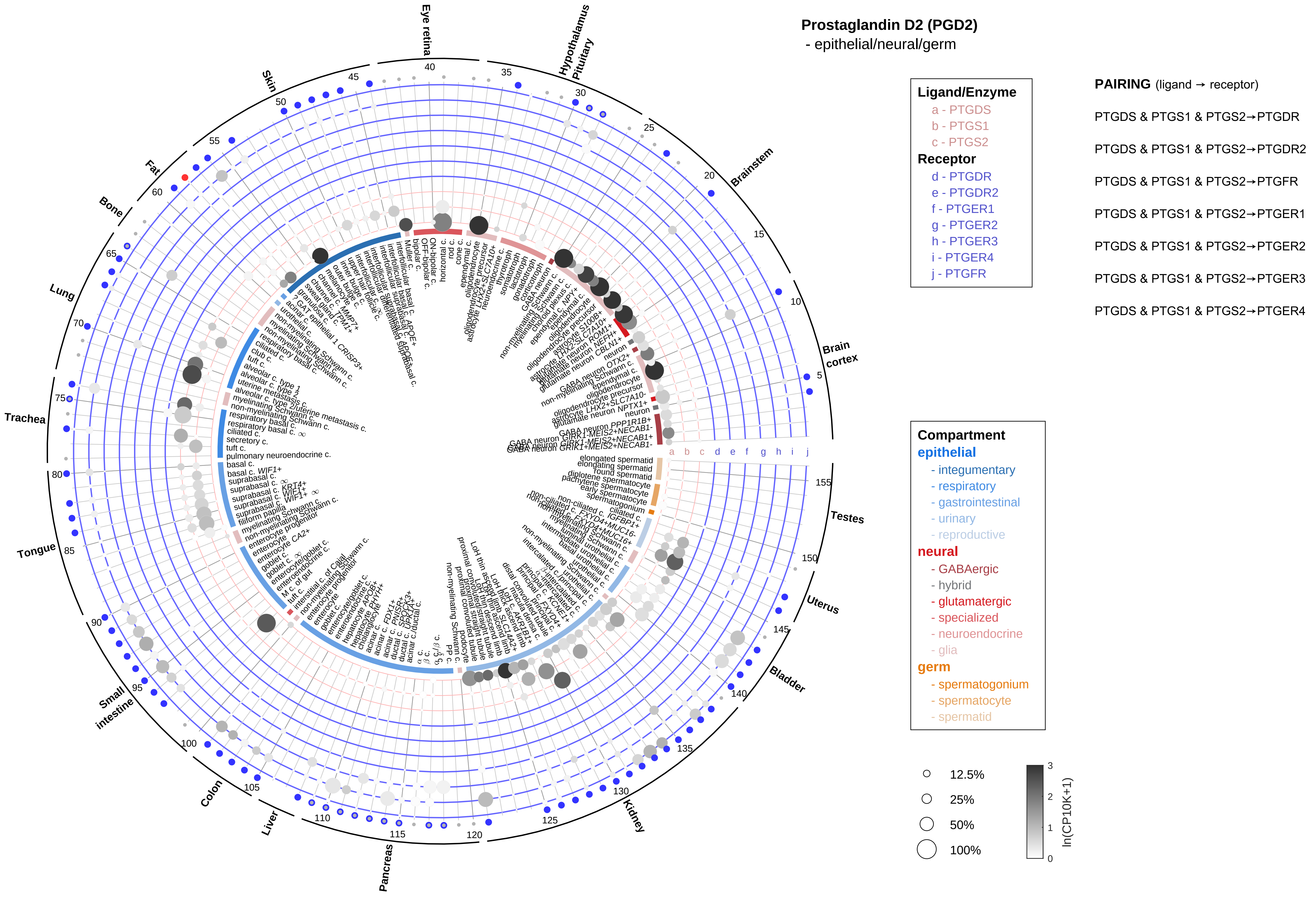
Prostaglandin D2 (PGD2)
- epithelial/neural/germ
Ligand/Enzyme
a - PTGDS
b - PTGS1
c - PTGS2
Receptor
d - PTGDR
e - PTGDR2
f - PTGER1
g - PTGER2
h - PTGER3
i - PTGER4
j - PTGFR
PAIRING (ligand → receptor)
PTGDS & PTGS1 & PTGS2→PTGDR
PTGDS & PTGS1 & PTGS2→PTGDR2
PTGDS & PTGS1 & PTGS2→PTGFR
PTGDS & PTGS1 & PTGS2→PTGER1
PTGDS & PTGS1 & PTGS2→PTGER2
PTGDS & PTGS1 & PTGS2→PTGER3
PTGDS & PTGS1 & PTGS2→PTGER4
Compartment
epithelial
- integumentary
- respiratory
- gastrointestinal
- urinary
- reproductive
neural
- GABAergic
- hybrid
- glutamatergic
- specialized
- neuroendocrine
- glia
germ
- spermatogonium
- spermatocyte
- spermatid
12.5%
25%
50%
100%
ln(CP10K+1)
Eye retina
Hypothalamus
Pituitary
Brainstem
Brain cortex
Testes
Uterus
Bladder
Kidney
Pancreas
Liver
Colon
Small intestine
Tongue
Trachea
Lung
Bone
Fat
Skin

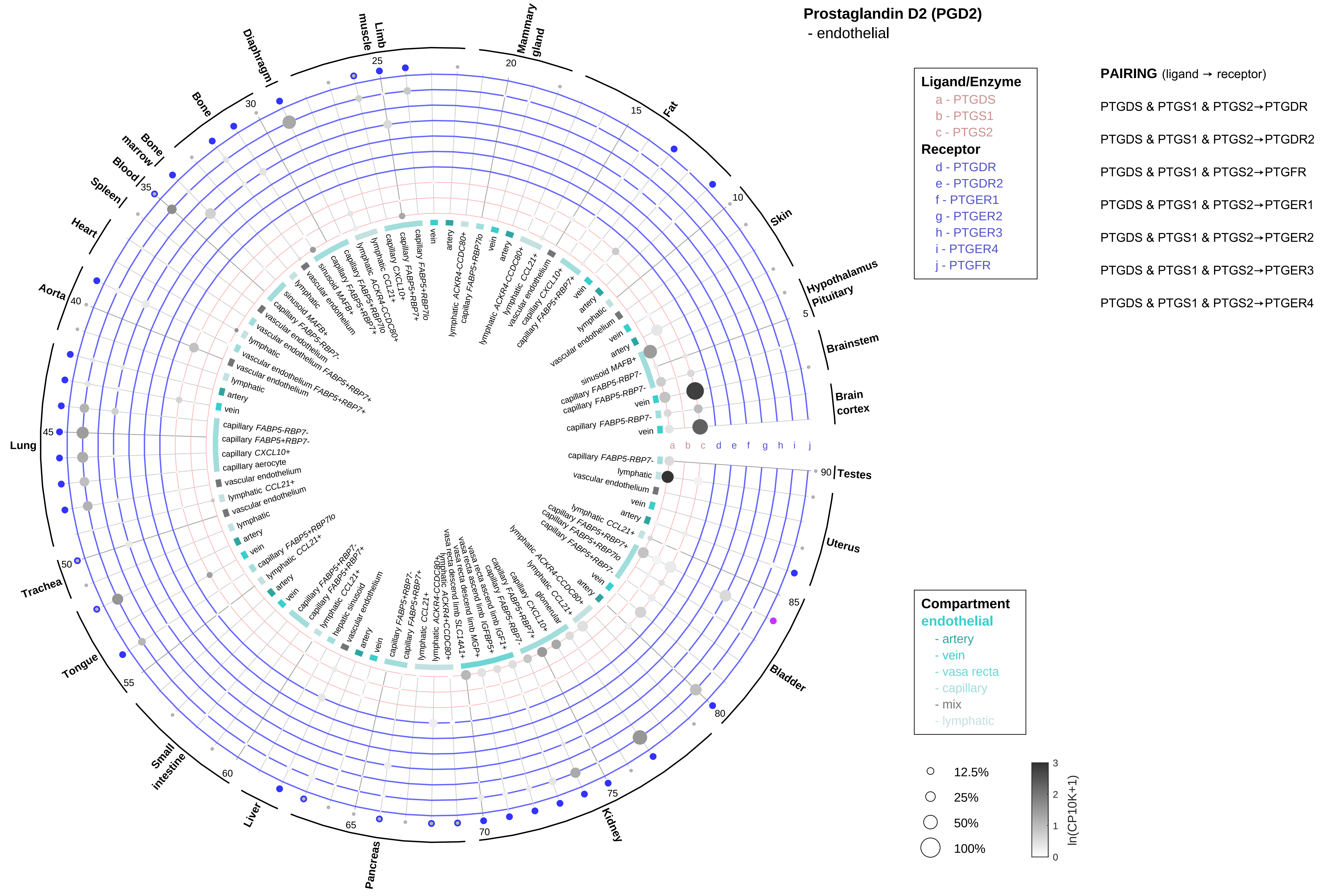

Prostaglandin D2 (PGD2)
- endothelial
Ligand/Enzyme
a - PTGDS
b - PTGS1
c - PTGS2
Receptor
d - PTGDR
e - PTGDR2
f - PTGER1
g - PTGER2
h - PTGER3
i - PTGER4
j - PTGFR
PAIRING (ligand → receptor)
PTGDS & PTGS1 & PTGS2→PTGDR
PTGDS & PTGS1 & PTGS2→PTGDR2
PTGDS & PTGS1 & PTGS2→PTGFR
PTGDS & PTGS1 & PTGS2→PTGER1
PTGDS & PTGS1 & PTGS2→PTGER2
PTGDS & PTGS1 & PTGS2→PTGER3
PTGDS & PTGS1 & PTGS2→PTGER4
Compartment
endothelial
- artery
- vein
- vasa recta
- capillary
- mix
- lymphatic
12.5%
25%
50%
100%
ln(CP10K+1)
Brain cortex
Brainstem
Hypothalamus Pituitary
Skin
Fat
Mammary gland
Limb muscle
Diaphragm
Bone
Bone marrow
Blood
Spleen
Heart
Aorta
Lung
Trachea
Tongue
Small intestine
Liver
Pancreas
Kidney
Bladder
Uterus
Testes

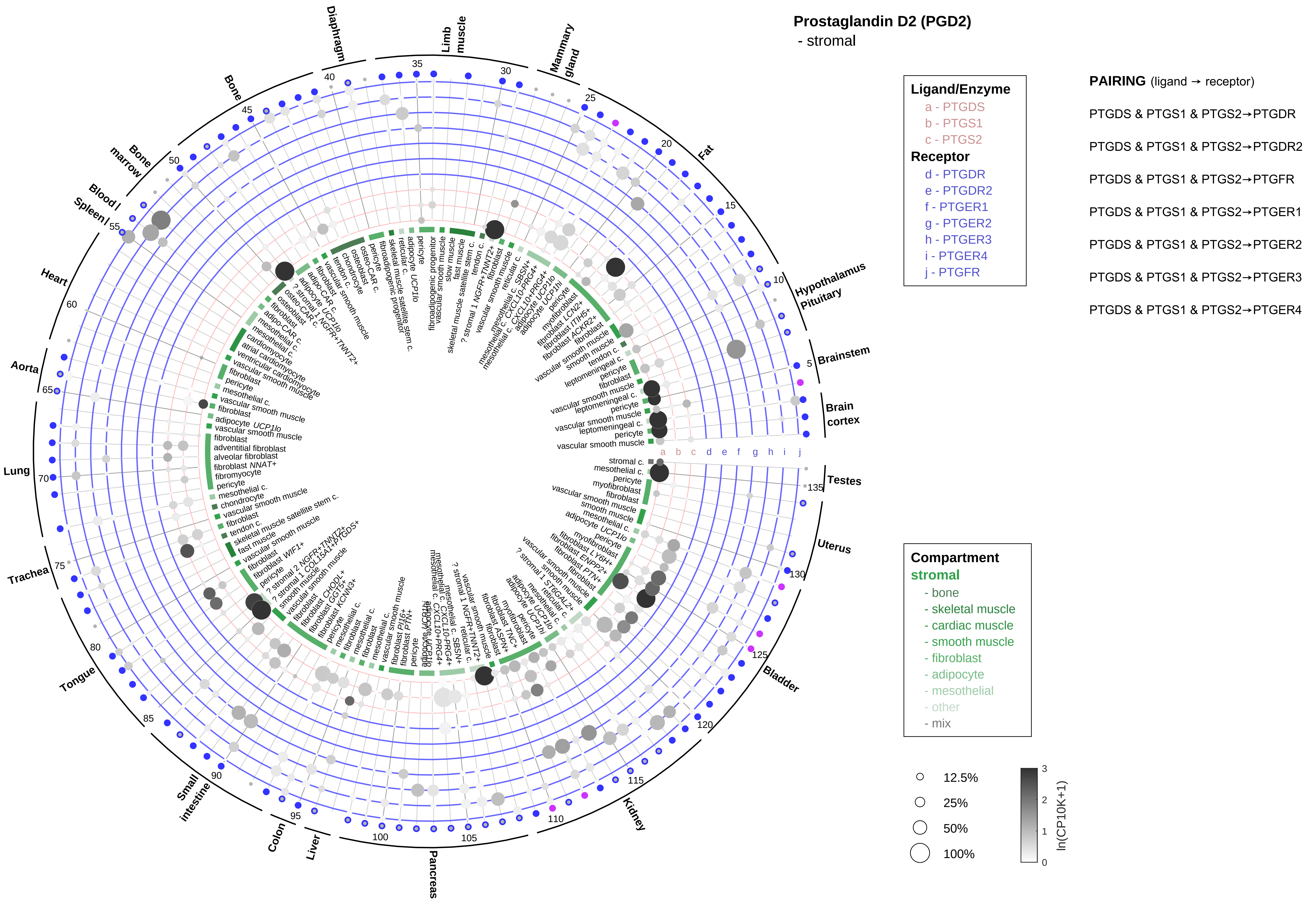
Prostaglandin D2 (PGD2)
- stromal
Ligand/Enzyme
a - PTGDS
b - PTGS1
c - PTGS2
Receptor
d - PTGDR
e - PTGDR2
f - PTGER1
g - PTGER2
h - PTGER3
i - PTGER4
j - PTGFR
PAIRING (ligand → receptor)
PTGDS & PTGS1 & PTGS2→PTGDR
PTGDS & PTGS1 & PTGS2→PTGDR2
PTGDS & PTGS1 & PTGS2→PTGFR
PTGDS & PTGS1 & PTGS2→PTGER1
PTGDS & PTGS1 & PTGS2→PTGER2
PTGDS & PTGS1 & PTGS2→PTGER3
PTGDS & PTGS1 & PTGS2→PTGER4
Compartment
stromal
- bone
- skeletal muscle
- cardiac muscle
- smooth muscle
- fibroblast
- adipocyte
- mesothelial
- other
- mix
12.5%
25%
50%
100%
ln(CP10K+1)
Brain cortex
Brainstem
Hypothalamus Pituitary
Fat
Mammary gland
Limb muscle
Diaphragm
Bone
Bone marrow
Blood / Spleen
Heart
Aorta
Lung
Trachea
Tongue
Small intestine
Colon
Liver
Pancreas
Kidney
Bladder
Uterus
Testes

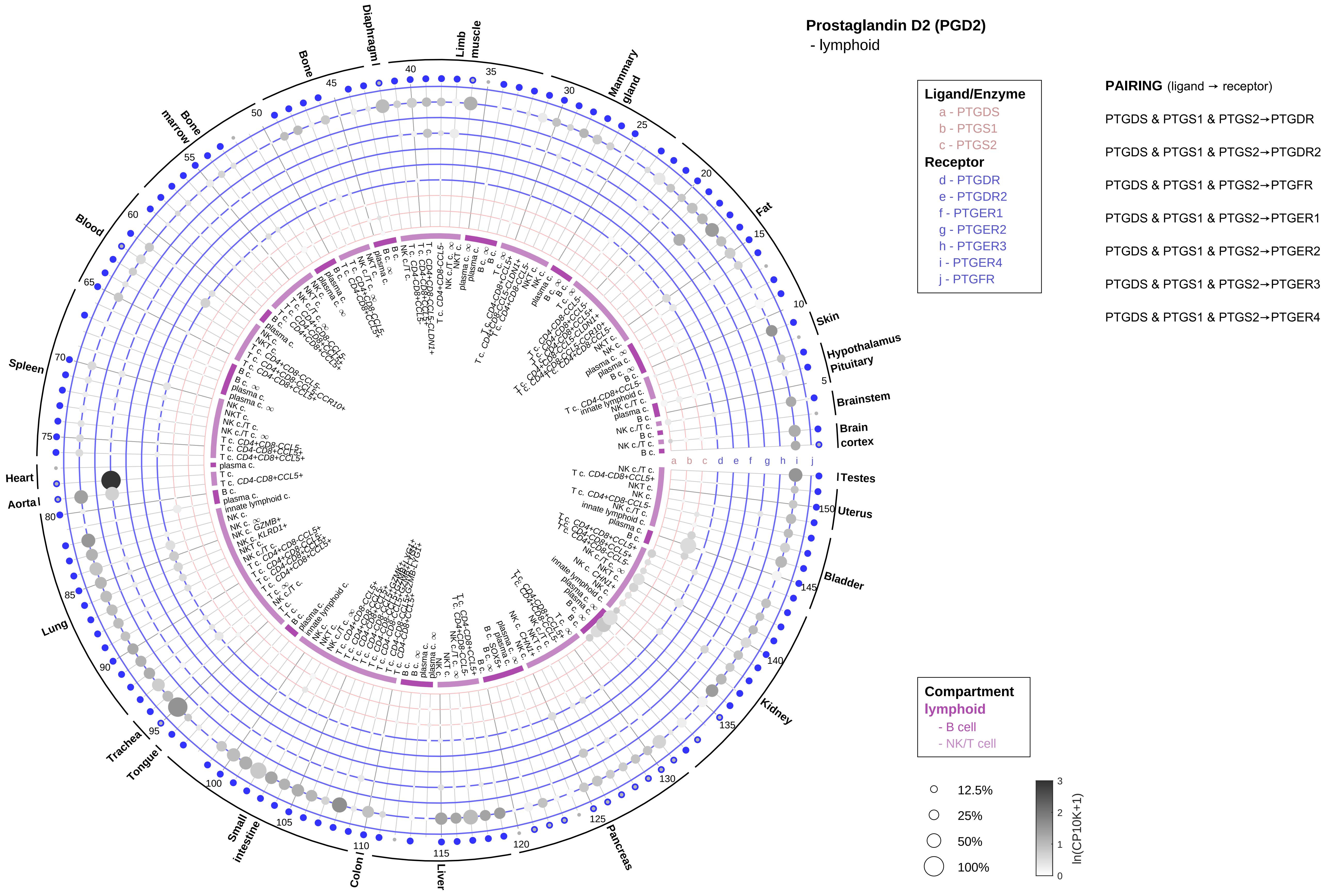

Prostaglandin D2 (PGD2)
- lymphoid
Ligand/Enzyme
a - PTGDS
b - PTGS1
c - PTGS2
Receptor
d - PTGDR
e - PTGDR2
f - PTGER1
g - PTGER2
h - PTGER3
i - PTGER4
j - PTGFR
PAIRING (ligand → receptor)
PTGDS & PTGS1 & PTGS2→PTGDR
PTGDS & PTGS1 & PTGS2→PTGDR2
PTGDS & PTGS1 & PTGS2→PTGFR
PTGDS & PTGS1 & PTGS2→PTGER1
PTGDS & PTGS1 & PTGS2→PTGER2
PTGDS & PTGS1 & PTGS2→PTGER3
PTGDS & PTGS1 & PTGS2→PTGER4
Compartment
lymphoid
- B cell
- NK/T cell
12.5%
25%
50%
100%
ln(CP10K+1)
Diaphragm
Limb muscle
Mammary gland
Fat
Skin
Hypothalamus
Pituitary
Brainstem
Brain cortex
Testes
Uterus
Bladder
Kidney
Pancreas
Liver
Colon
Small intestine
Tongue
Trachea
Lung
Aorta
Heart
Spleen
Blood
Bone marrow
Bone

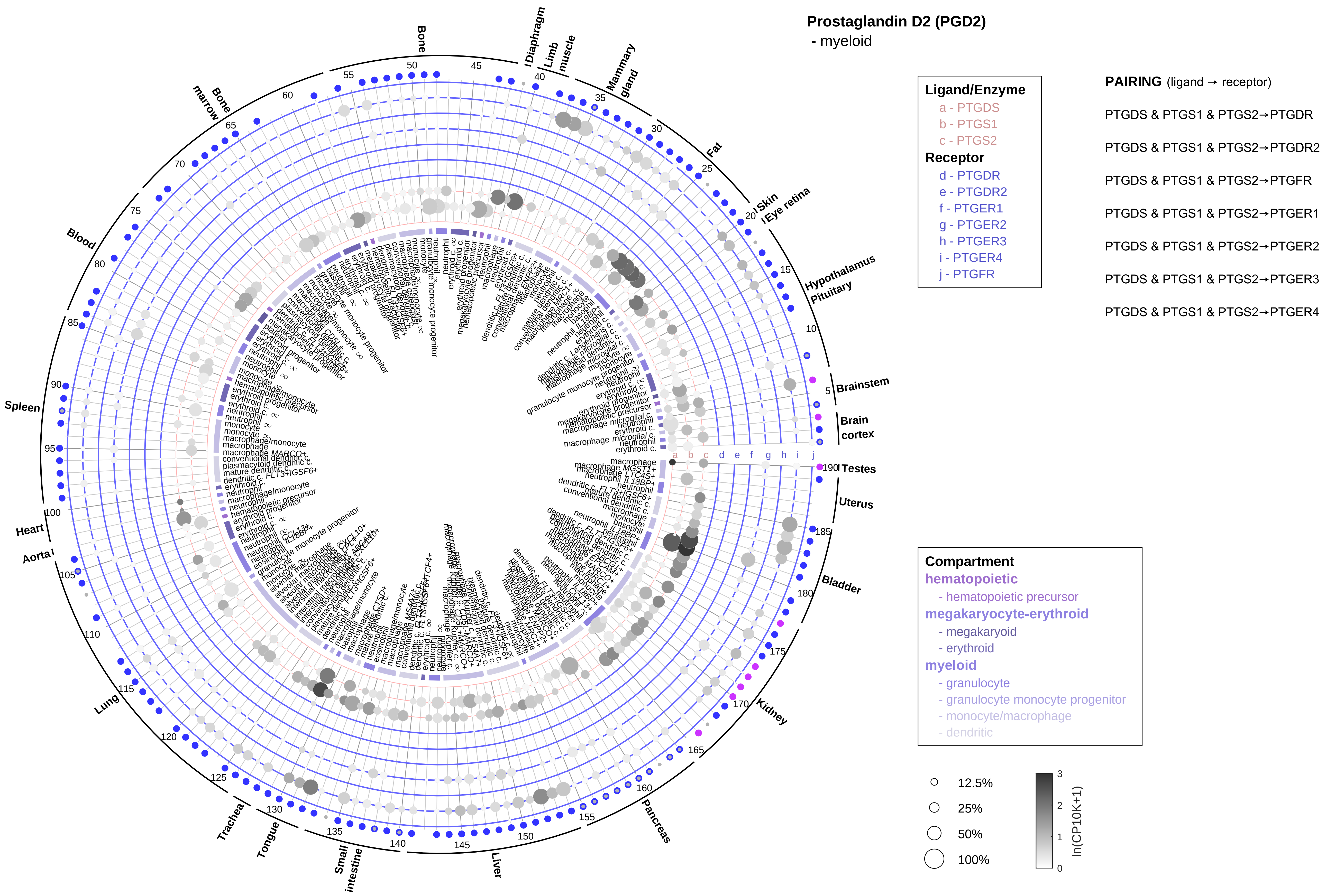

Prostaglandin D2 (PGD2)
- myeloid
Ligand/Enzyme
a - PTGDS
b - PTGS1
c - PTGS2
Receptor
d - PTGDR
e - PTGDR2
f - PTGER1
g - PTGER2
h - PTGER3
i - PTGER4
j - PTGFR
PAIRING (ligand → receptor)
PTGDS & PTGS1 & PTGS2→PTGDR
PTGDS & PTGS1 & PTGS2→PTGDR2
PTGDS & PTGS1 & PTGS2→PTGFR
PTGDS & PTGS1 & PTGS2→PTGER1
PTGDS & PTGS1 & PTGS2→PTGER2
PTGDS & PTGS1 & PTGS2→PTGER3
PTGDS & PTGS1 & PTGS2→PTGER4
Compartment
hematopoietic
- hematopoietic precursor
megakaryocyte-erythroid
- megakaryoid
- erythroid
myeloid
- granulocyte
- granulocyte monocyte progenitor
- monocyte/macrophage
- dendritic
12.5%
25%
50%
100%
ln(CP10K+1)
Bone
Diaphragm
Limb muscle
Mammary gland
Fat
Skin
Eye retina
Hypothalamus
Pituitary
Brainstem
Brain cortex
Testes
Uterus
Bladder
Kidney
Pancreas
Liver
Small intestine
Tongue
Trachea
Lung
Aorta
Heart
Spleen
Blood
Bone marrow

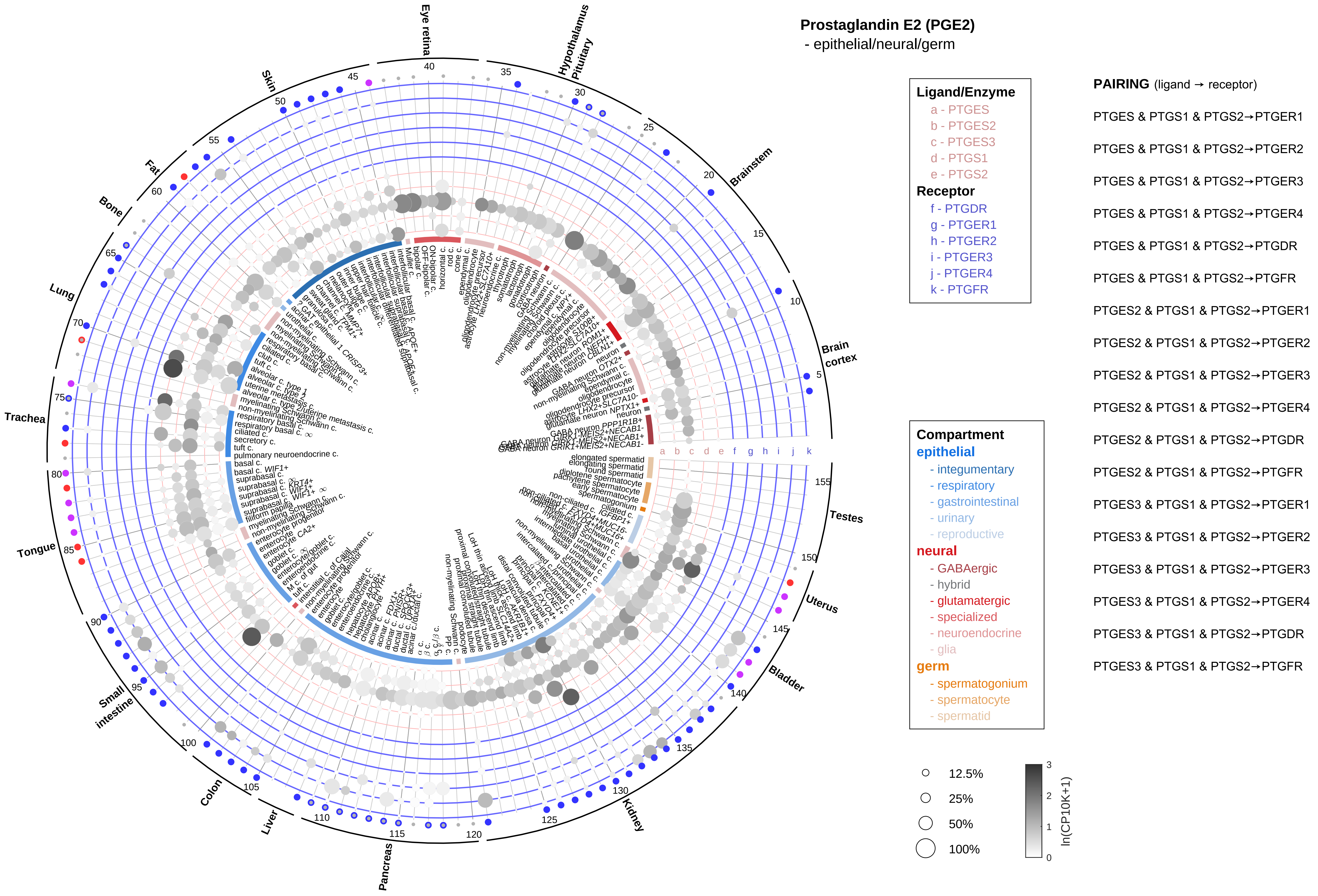

Prostaglandin E2 (PGE2)
- epithelial/neural/germ
Eye retina
Hypothalamus Pituitary
Brainstem
Brain cortex
Testes
Uterus
Bladder
Kidney
Pancreas
Liver
Colon
Small intestine
Tongue
Trachea
Lung
Bone
Fat
Skin
a b c d e f g h i j k
Ligand/Enzyme
a - PTGES
b - PTGES2
c - PTGES3
d - PTGS1
e - PTGS2
Receptor
f - PTGDR
g - PTGER1
h - PTGER2
i - PTGER3
j - PTGER4
k - PTGFR
Compartment
epithelial
- integumentary
- respiratory
- gastrointestinal
- urinary
- reproductive
neural
- GABAergic
- hybrid
- glutamatergic
- specialized
- neuroendocrine
- glia
germ
- spermatogonium
- spermatocyte
- spermatid
12.5%
25%
50%
100%
ln(CP10K+1)
PAIRING (ligand → receptor)
PTGES & PTGS1 & PTGS2→PTGER1
PTGES & PTGS1 & PTGS2→PTGER2
PTGES & PTGS1 & PTGS2→PTGER3
PTGES & PTGS1 & PTGS2→PTGER4
PTGES & PTGS1 & PTGS2→PTGDR
PTGES & PTGS1 & PTGS2→PTGFR
PTGES2 & PTGS1 & PTGS2→PTGER1
PTGES2 & PTGS1 & PTGS2→PTGER2
PTGES2 & PTGS1 & PTGS2→PTGER3
PTGES2 & PTGS1 & PTGS2→PTGER4
PTGES2 & PTGS1 & PTGS2→PTGDR
PTGES2 & PTGS1 & PTGS2→PTGFR
PTGES3 & PTGS1 & PTGS2→PTGER1
PTGES3 & PTGS1 & PTGS2→PTGER2
PTGES3 & PTGS1 & PTGS2→PTGER3
PTGES3 & PTGS1 & PTGS2→PTGER4
PTGES3 & PTGS1 & PTGS2→PTGDR
PTGES3 & PTGS1 & PTGS2→PTGFR

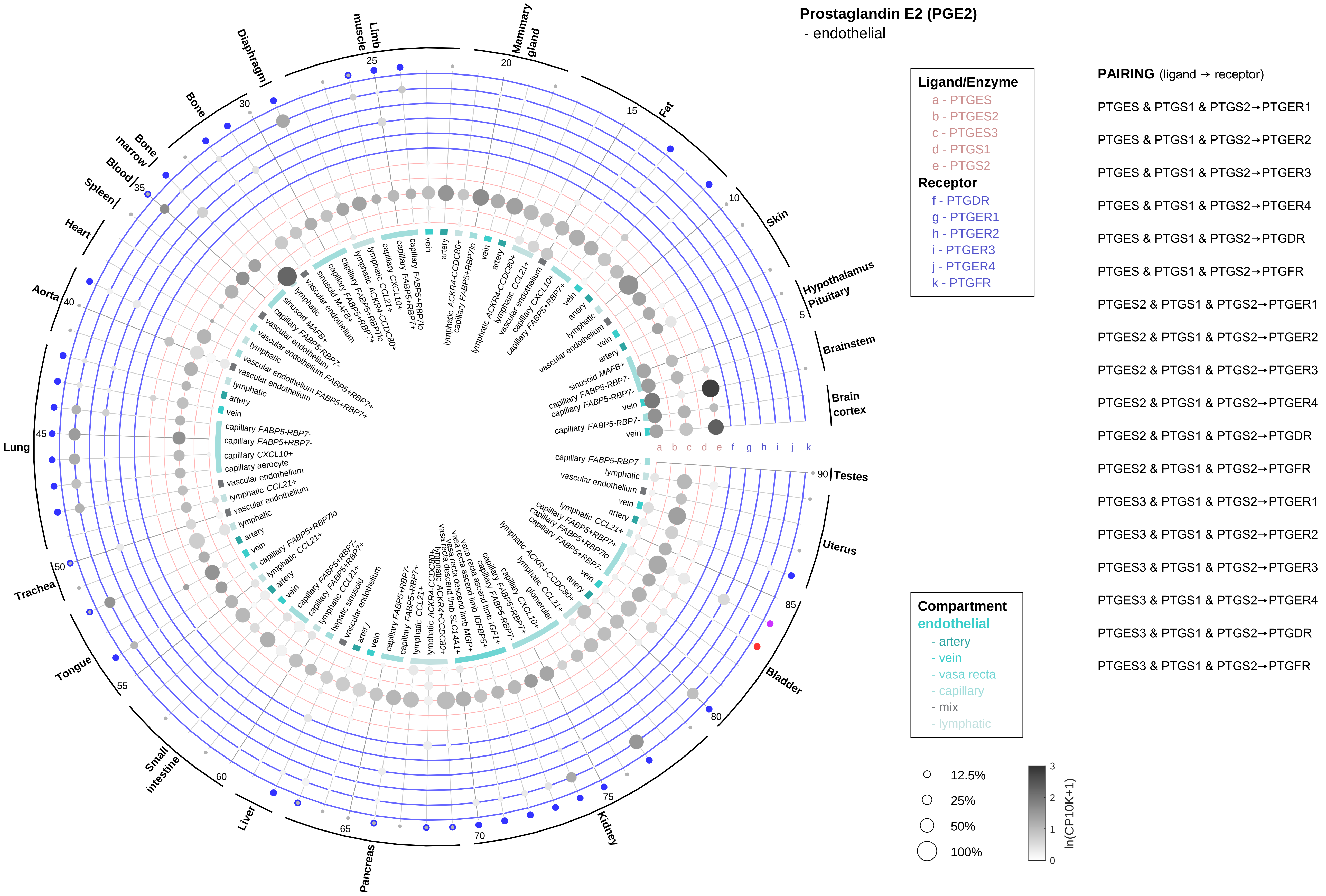

Prostaglandin E2 (PGE2)
- endothelial
Ligand/Enzyme
a - PTGES
b - PTGES2
c - PTGES3
d - PTGS1
e - PTGS2
Receptor
f - PTGDR
g - PTGER1
h - PTGER2
i - PTGER3
j - PTGER4
k - PTGFR
PAIRING (ligand → receptor)
PTGES & PTGS1 & PTGS2→PTGER1
PTGES & PTGS1 & PTGS2→PTGER2
PTGES & PTGS1 & PTGS2→PTGER3
PTGES & PTGS1 & PTGS2→PTGER4
PTGES & PTGS1 & PTGS2→PTGDR
PTGES & PTGS1 & PTGS2→PTGFR
PTGES2 & PTGS1 & PTGS2→PTGER1
PTGES2 & PTGS1 & PTGS2→PTGER2
PTGES2 & PTGS1 & PTGS2→PTGER3
PTGES2 & PTGS1 & PTGS2→PTGER4
PTGES2 & PTGS1 & PTGS2→PTGDR
PTGES2 & PTGS1 & PTGS2→PTGFR
PTGES3 & PTGS1 & PTGS2→PTGER1
PTGES3 & PTGS1 & PTGS2→PTGER2
PTGES3 & PTGS1 & PTGS2→PTGER3
PTGES3 & PTGS1 & PTGS2→PTGER4
PTGES3 & PTGS1 & PTGS2→PTGDR
PTGES3 & PTGS1 & PTGS2→PTGFR
Compartment
endothelial
- artery
- vein
- vasa recta
- capillary
- mix
- lymphatic
12.5%
25%
50%
100%
ln(CP10K+1)
Brain cortex
Brainstem
Hypothalamus
Pituitary
Skin
Fat
Mammary gland
Limb muscle
Diaphragm
Bone
Bone marrow
Blood
Spleen
Heart
Aorta
Lung
Trachea
Tongue
Small intestine
Liver
Pancreas
Kidney
Bladder
Uterus
Testes
a b c d e f g h i j k

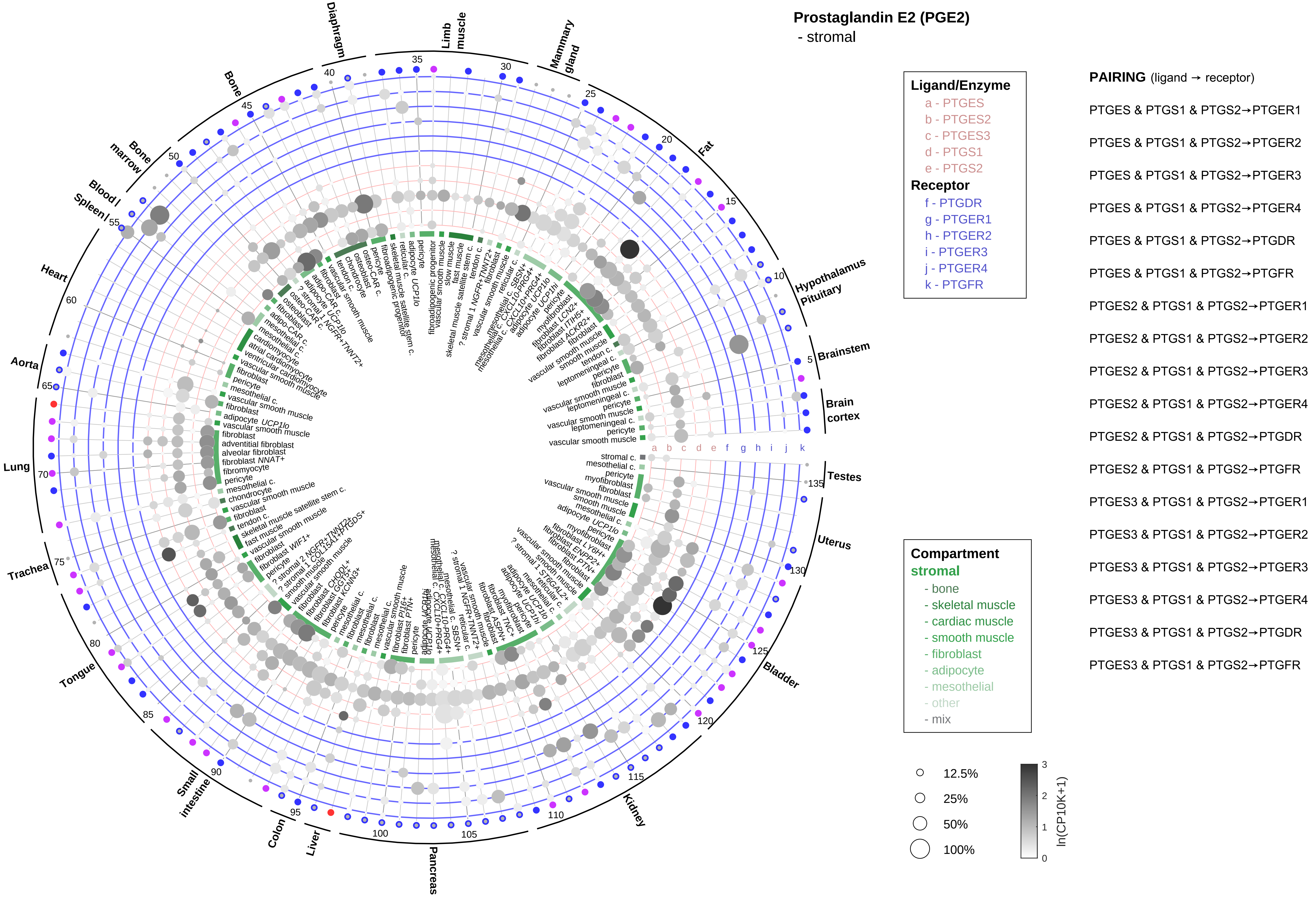

Prostaglandin E2 (PGE2)
- stromal
Ligand/Enzyme
a - PTGES
b - PTGES2
c - PTGES3
d - PTGS1
e - PTGS2
Receptor
f - PTGDR
g - PTGER1
h - PTGER2
i - PTGER3
j - PTGER4
k - PTGFR
Compartment
stromal
- bone
- skeletal muscle
- cardiac muscle
- smooth muscle
- fibroblast
- adipocyte
- mesothelial
- other
- mix
12.5%
25%
50%
100%
ln(CP10K+1)
PAIRING (ligand → receptor)
PTGES & PTGS1 & PTGS2→PTGER1
PTGES & PTGS1 & PTGS2→PTGER2
PTGES & PTGS1 & PTGS2→PTGER3
PTGES & PTGS1 & PTGS2→PTGER4
PTGES & PTGS1 & PTGS2→PTGDR
PTGES & PTGS1 & PTGS2→PTGFR
PTGES2 & PTGS1 & PTGS2→PTGER1
PTGES2 & PTGS1 & PTGS2→PTGER2
PTGES2 & PTGS1 & PTGS2→PTGER3
PTGES2 & PTGS1 & PTGS2→PTGER4
PTGES2 & PTGS1 & PTGS2→PTGDR
PTGES2 & PTGS1 & PTGS2→PTGFR
PTGES3 & PTGS1 & PTGS2→PTGER1
PTGES3 & PTGS1 & PTGS2→PTGER2
PTGES3 & PTGS1 & PTGS2→PTGER3
PTGES3 & PTGS1 & PTGS2→PTGER4
PTGES3 & PTGS1 & PTGS2→PTGDR
PTGES3 & PTGS1 & PTGS2→PTGFR
Diaphragm
Limb muscle
Mammary gland
Fat
Hypothalamus Pituitary
Brainstem
Brain cortex
Testes
Uterus
Bladder
Kidney
Pancreas
Liver
Colon
Small intestine
Tongue
Trachea
Lung
Aorta
Heart
Blood / Spleen
Bone marrow
Bone

**Prostaglandin E2 (PGE2)**
- lymphoid

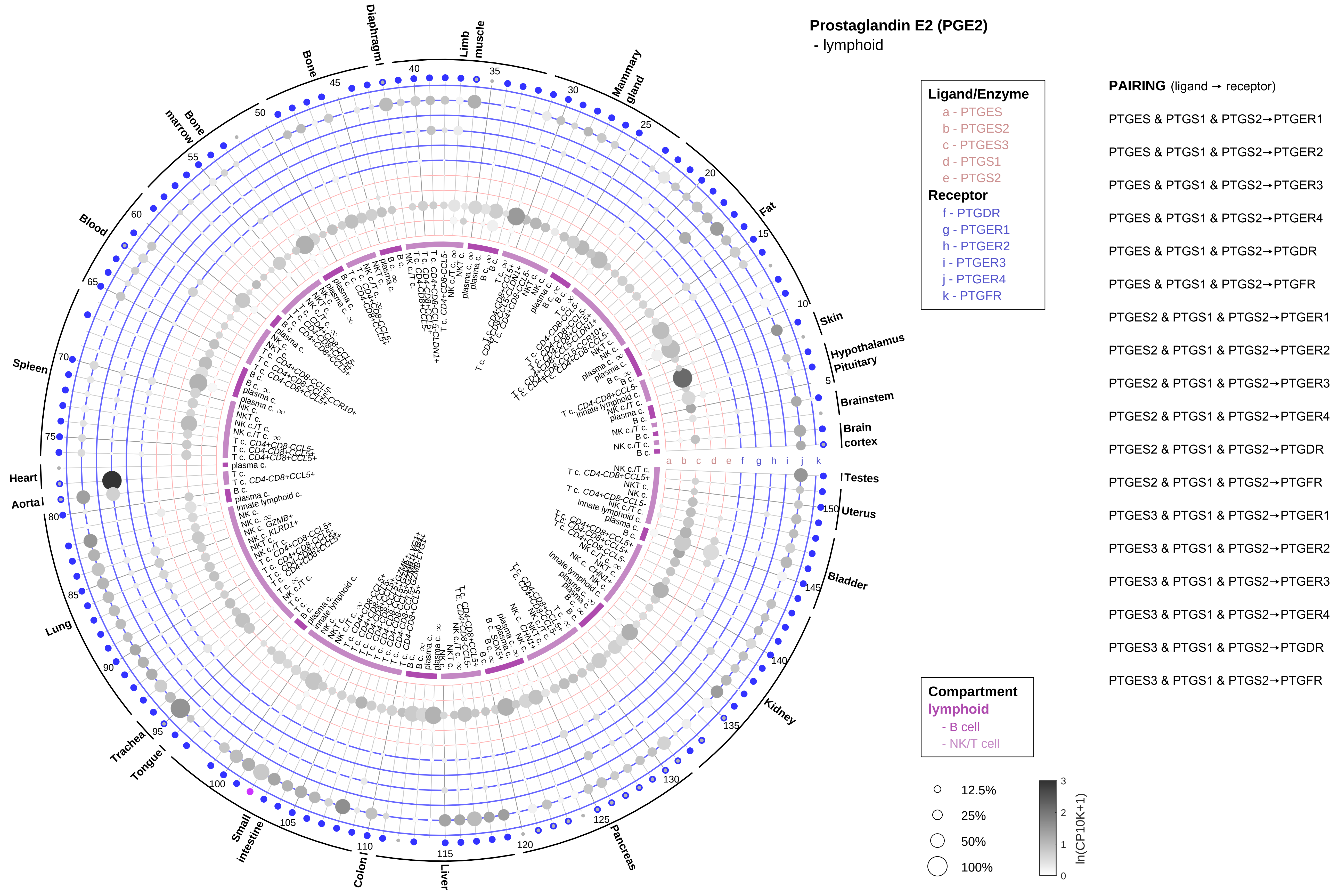

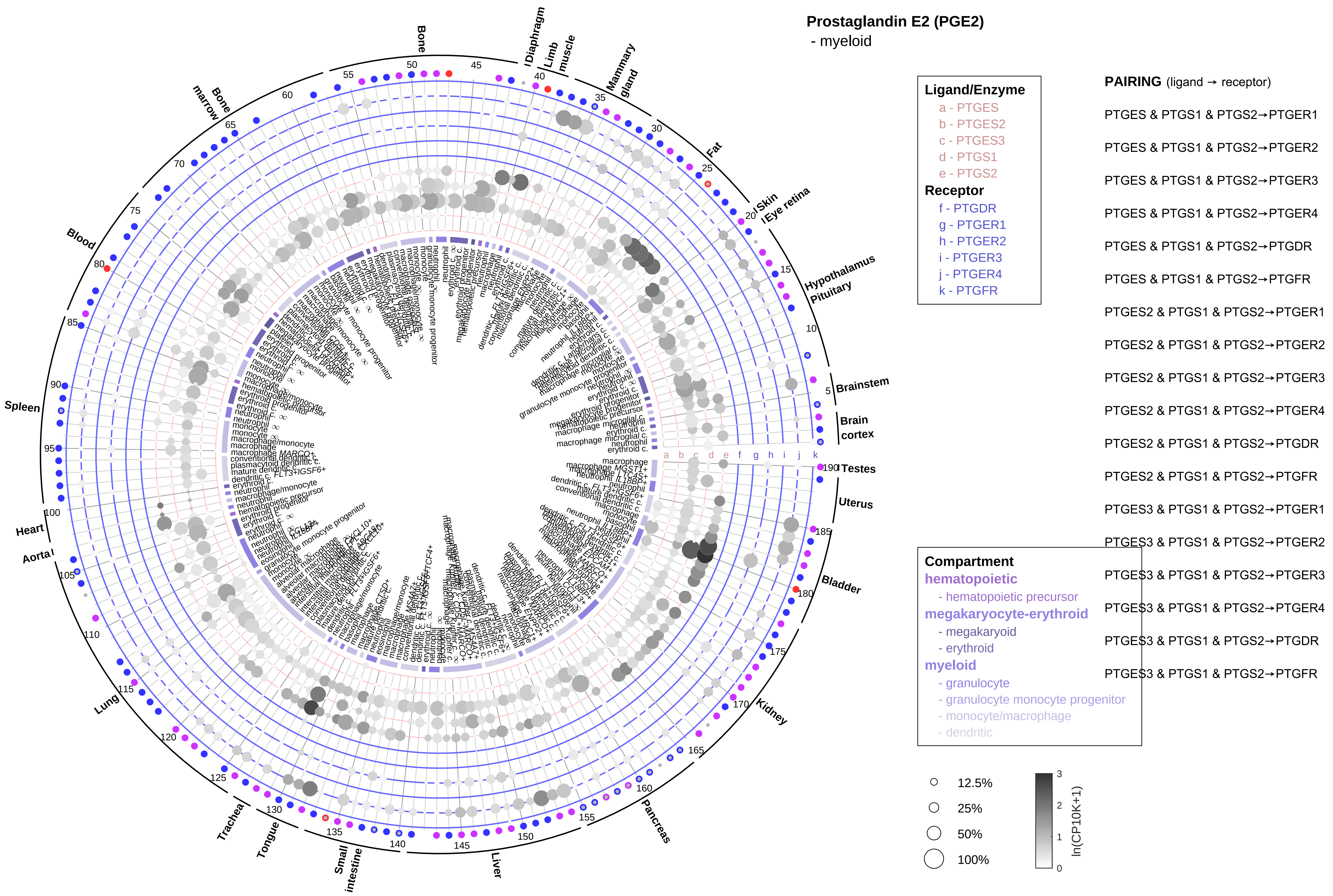

Prostaglandin E2 (PGE2)
- myeloid
Ligand/Enzyme
a - PTGES
b - PTGES2
c - PTGES3
d - PTGS1
e - PTGS2
Receptor
f - PTGDR
g - PTGER1
h - PTGER2
i - PTGER3
j - PTGER4
k - PTGFR
PAIRING (ligand → receptor)
PTGES & PTGS1 & PTGS2→PTGER1
PTGES & PTGS1 & PTGS2→PTGER2
PTGES & PTGS1 & PTGS2→PTGER3
PTGES & PTGS1 & PTGS2→PTGER4
PTGES & PTGS1 & PTGS2→PTGDR
PTGES & PTGS1 & PTGS2→PTGFR
PTGES2 & PTGS1 & PTGS2→PTGER1
PTGES2 & PTGS1 & PTGS2→PTGER2
PTGES2 & PTGS1 & PTGS2→PTGER3
PTGES2 & PTGS1 & PTGS2→PTGER4
PTGES2 & PTGS1 & PTGS2→PTGDR
PTGES2 & PTGS1 & PTGS2→PTGFR
PTGES3 & PTGS1 & PTGS2→PTGER1
PTGES3 & PTGS1 & PTGS2→PTGER2
PTGES3 & PTGS1 & PTGS2→PTGER3
PTGES3 & PTGS1 & PTGS2→PTGER4
PTGES3 & PTGS1 & PTGS2→PTGDR
PTGES3 & PTGS1 & PTGS2→PTGFR
Compartment
hematopoietic
- hematopoietic precursor
megakaryocyte-erythroid
- megakaryoid
- erythroid
myeloid
- granulocyte
- granulocyte monocyte progenitor
- monocyte/macrophage
- dendritic
12.5%
25%
50%
100%
ln(CP10K+1)
Bone
Diaphragm
Limb muscle
Mammary gland
Fat
Skin
Eye retina
Hypothalamus
Pituitary
Brainstem
Brain cortex
Testes
Uterus
Bladder
Kidney
Pancreas
Liver
Small intestine
Tongue
Trachea
Lung
Aorta
Heart
Spleen
Blood
Bone marrow

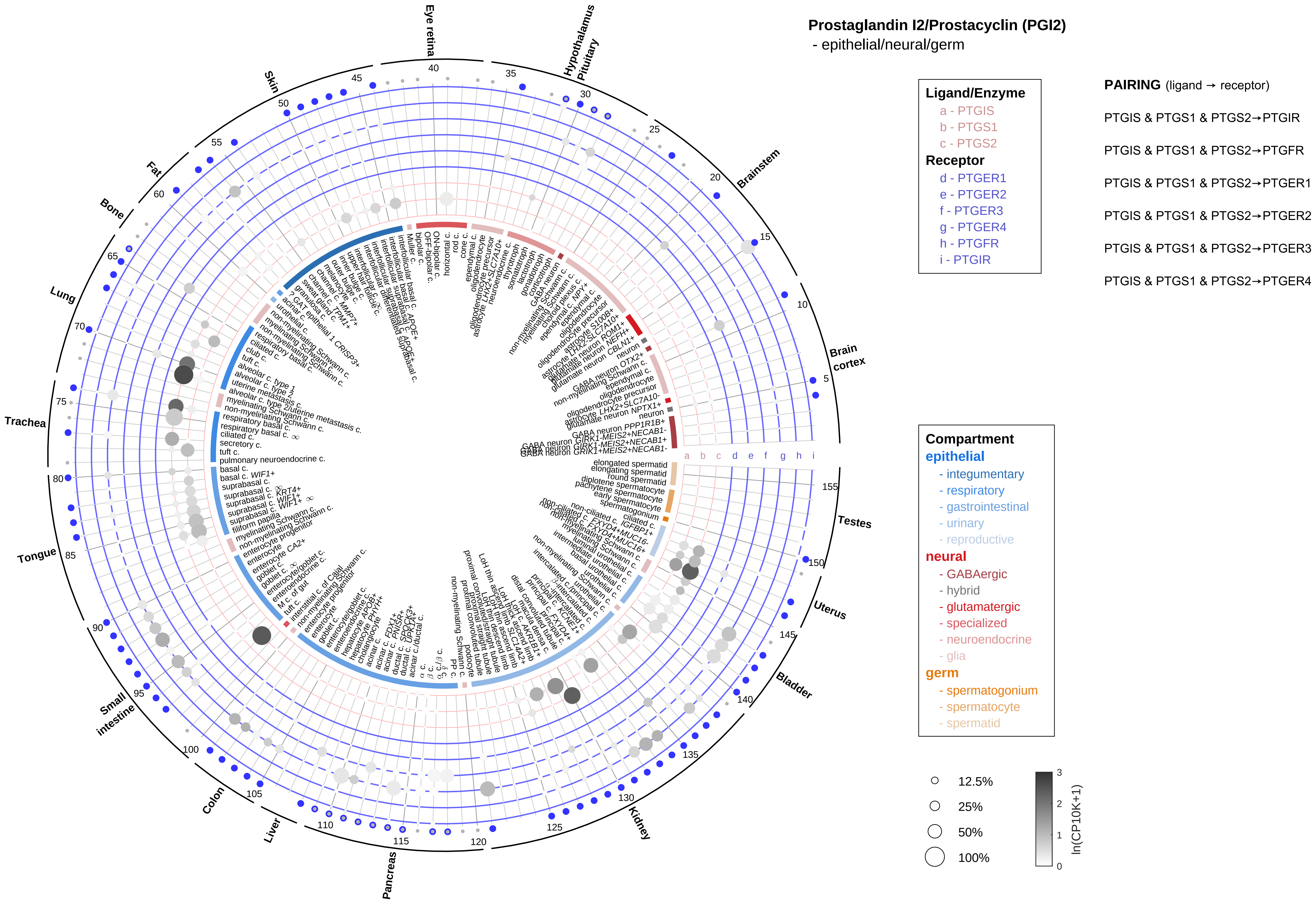
Prostaglandin I2/Prostacyclin (PGI2)
- epithelial/neural/germ
Ligand/Enzyme
a - PTGIS
b - PTGS1
c - PTGS2
Receptor
d - PTGER1
e - PTGER2
f - PTGER3
g - PTGER4
h - PTGFR
i - PTGIR
PAIRING (ligand → receptor)
PTGIS & PTGS1 & PTGS2→PTGIR
PTGIS & PTGS1 & PTGS2→PTGFR
PTGIS & PTGS1 & PTGS2→PTGER1
PTGIS & PTGS1 & PTGS2→PTGER2
PTGIS & PTGS1 & PTGS2→PTGER3
PTGIS & PTGS1 & PTGS2→PTGER4
Compartment
epithelial
- integumentary
- respiratory
- gastrointestinal
- urinary
- reproductive
neural
- GABAergic
- hybrid
- glutamatergic
- specialized
- neuroendocrine
- glia
germ
- spermatogonium
- spermatocyte
- spermatid
12.5%
25%
50%
100%
ln(CP10K+1)
Eye retina
Hypothalamus Pituitary
Brainstem
Brain cortex
Testes
Uterus
Bladder
Kidney
Pancreas
Liver
Colon
Small intestine
Tongue
Trachea
Lung
Bone
Fat
Skin

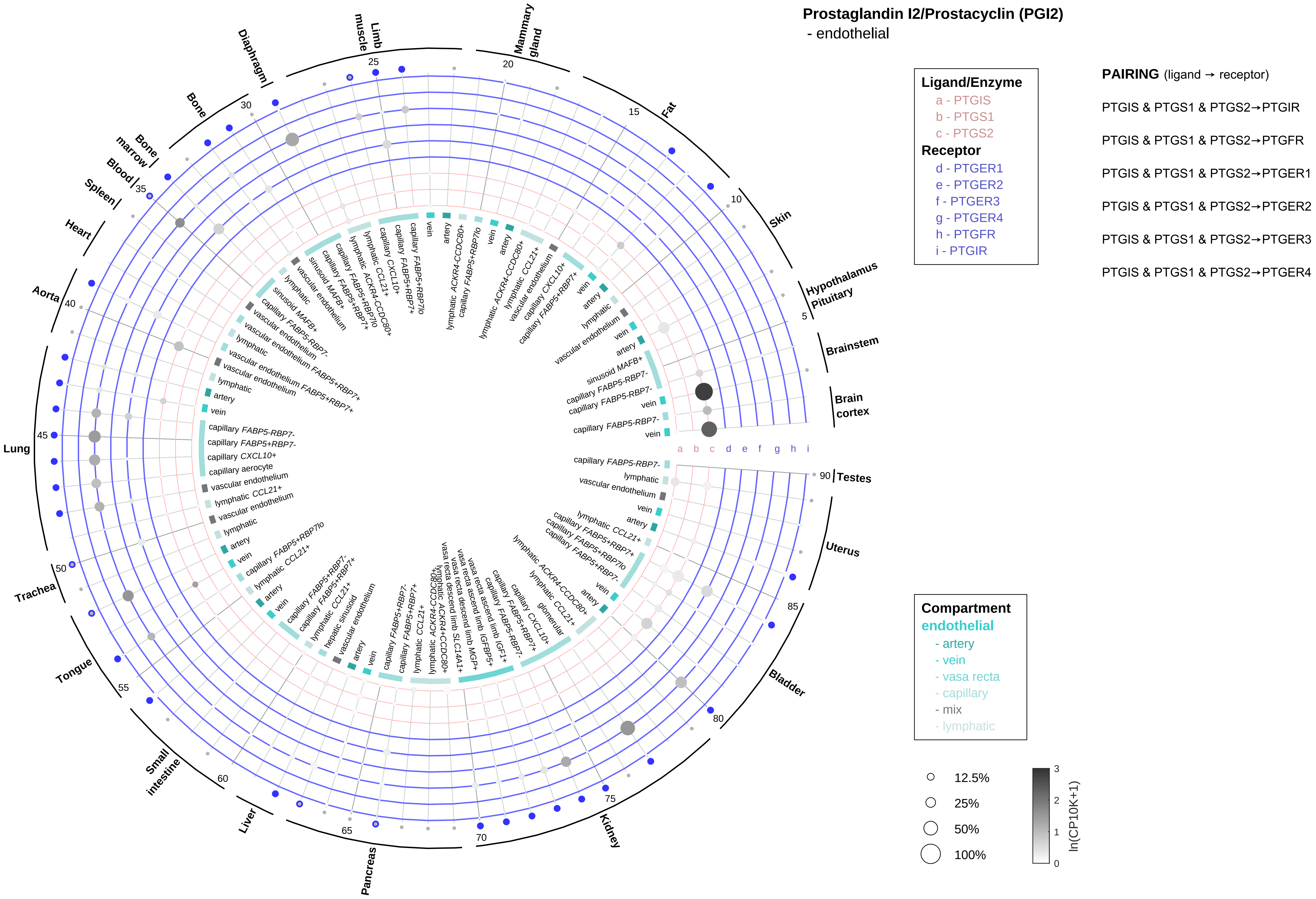

Prostaglandin I2/Prostacyclin (PGI2)
- endothelial
Ligand/Enzyme
a - PTGIS
b - PTGS1
c - PTGS2
Receptor
d - PTGER1
e - PTGER2
f - PTGER3
g - PTGER4
h - PTGFR
i - PTGIR
PAIRING (ligand → receptor)
PTGIS & PTGS1 & PTGS2→PTGIR
PTGIS & PTGS1 & PTGS2→PTGFR
PTGIS & PTGS1 & PTGS2→PTGER1
PTGIS & PTGS1 & PTGS2→PTGER2
PTGIS & PTGS1 & PTGS2→PTGER3
PTGIS & PTGS1 & PTGS2→PTGER4
Compartment
endothelial
- artery
- vein
- vasa recta
- capillary
- mix
- lymphatic
12.5%
25%
50%
100%
ln(CP10K+1)
Brain cortex
Brainstem
Hypothalamus
Pituitary
Skin
Fat
Mammary gland
Limb muscle
Diaphragm
Bone
Bone marrow
Blood
Spleen
Heart
Aorta
Lung
Trachea
Tongue
Small intestine
Liver
Pancreas
Kidney
Bladder
Uterus
Testes

#### Prostaglandin I2/Prostacyclin (PGI2)
- stromal

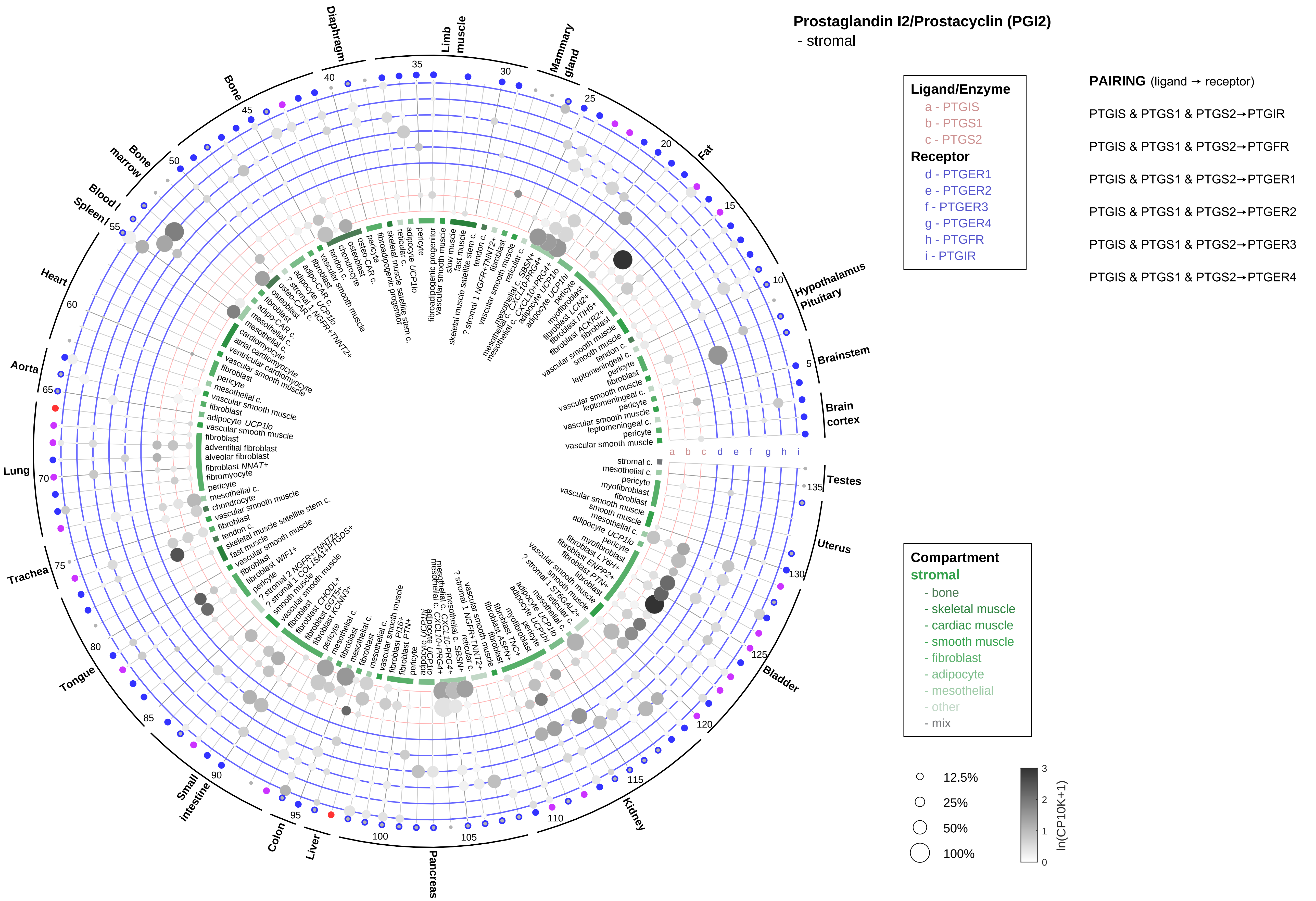

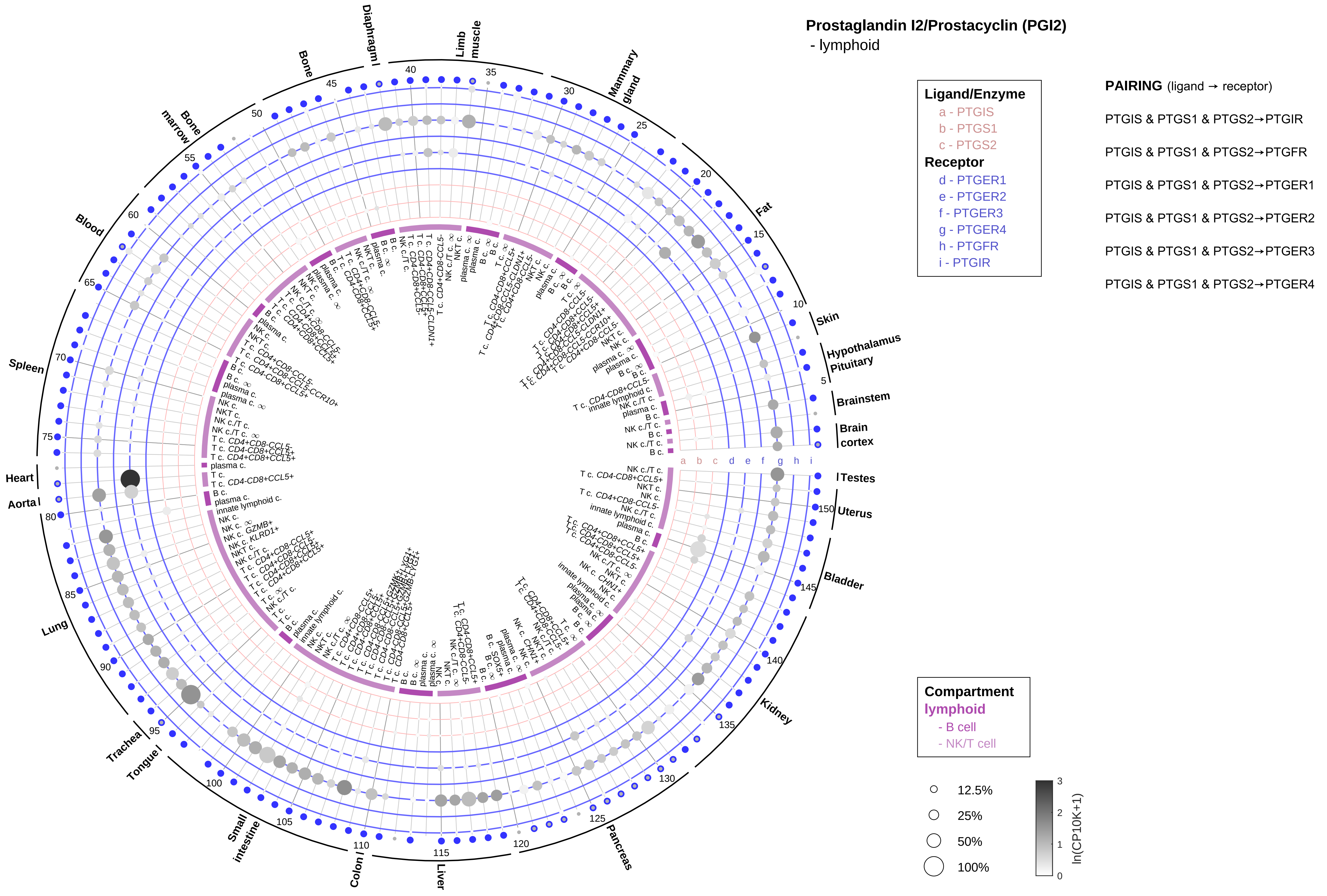

Prostaglandin I2/Prostacyclin (PGI2)
- lymphoid
Ligand/Enzyme
a - PTGIS
b - PTGS1
c - PTGS2
Receptor
d - PTGER1
e - PTGER2
f - PTGER3
g - PTGER4
h - PTGFR
i - PTGIR
PAIRING (ligand → receptor)
PTGIS & PTGS1 & PTGS2→PTGIR
PTGIS & PTGS1 & PTGS2→PTGFR
PTGIS & PTGS1 & PTGS2→PTGER1
PTGIS & PTGS1 & PTGS2→PTGER2
PTGIS & PTGS1 & PTGS2→PTGER3
PTGIS & PTGS1 & PTGS2→PTGER4
Compartment
lymphoid
- B cell
- NK/T cell
12.5%
25%
50%
100%
ln(CP10K+1)
Skin
Hypothalamus
Pituitary
Brainstem
Brain cortex
Testes
Uterus
Bladder
Kidney
Pancreas
Liver
Colon
Small intestine
Tongue
Trachea
Lung
Aorta
Heart
Spleen
Blood
Bone marrow
Bone
Diaphragm
Limb muscle
Mammary gland
Fat

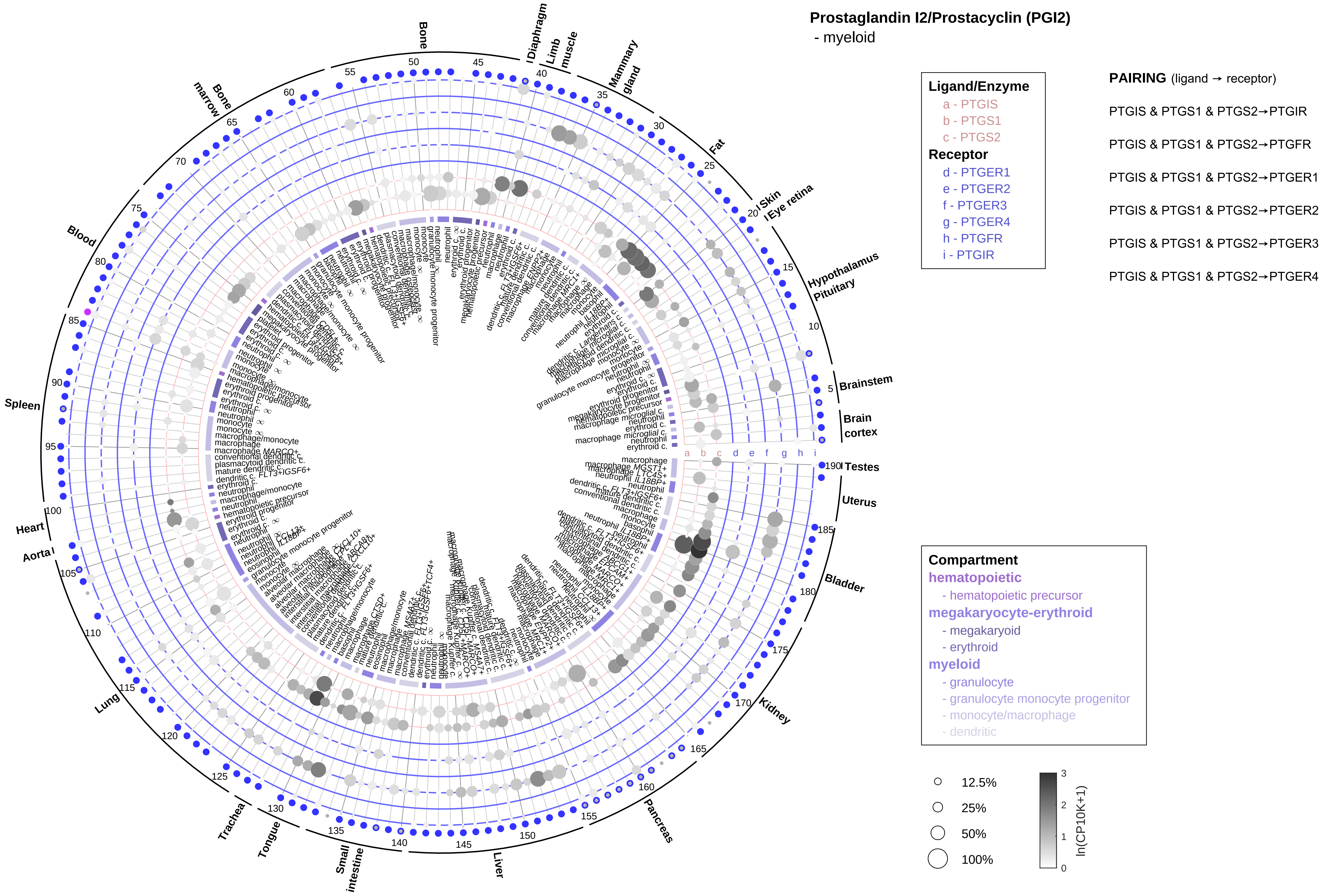

Prostaglandin I2/Prostacyclin (PGI2)
- myeloid
Ligand/Enzyme
a - PTGIS
b - PTGS1
c - PTGS2
Receptor
d - PTGER1
e - PTGER2
f - PTGER3
g - PTGER4
h - PTGFR
i - PTGIR
PAIRING (ligand → receptor)
PTGIS & PTGS1 & PTGS2→PTGIR
PTGIS & PTGS1 & PTGS2→PTGFR
PTGIS & PTGS1 & PTGS2→PTGER1
PTGIS & PTGS1 & PTGS2→PTGER2
PTGIS & PTGS1 & PTGS2→PTGER3
PTGIS & PTGS1 & PTGS2→PTGER4
Compartment
hematopoietic
- hematopoietic precursor
megakaryocyte-erythroid
- megakaryoid
- erythroid
myeloid
- granulocyte
- granulocyte monocyte progenitor
- monocyte/macrophage
- dendritic
12.5%
25%
50%
100%
ln(CP10K+1)
Brainstem
Brain cortex
Testes
Uterus
Bladder
Kidney
Pancreas
Liver
Small intestine
Tongue
Trachea
Lung
Aorta
Heart
Spleen
Blood
Bone marrow
Bone
Diaphragm
Limb muscle
Mammary gland
Fat
Skin
Eye retina
Hypothalamus
Pituitary

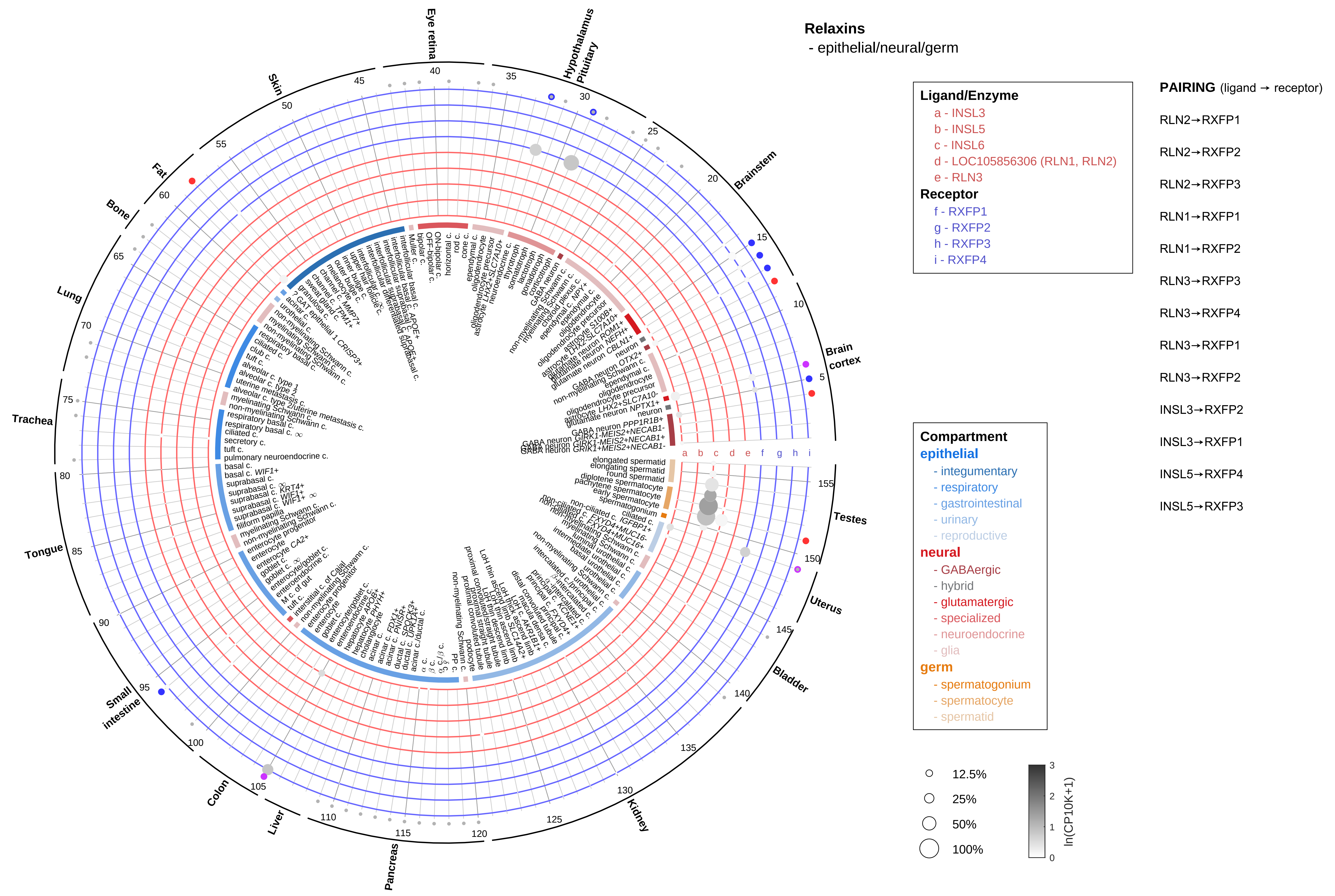

Relaxins
- epithelial/neural/germ
Ligand/Enzyme
a - INSL3
b - INSL5
c - INSL6
d - LOC105856306 (RLN1, RLN2)
e - RLN3
Receptor
f - RXFP1
g - RXFP2
h - RXFP3
i - RXFP4
PAIRING (ligand → receptor)
RLN2→RXFP1
RLN2→RXFP2
RLN2→RXFP3
RLN1→RXFP1
RLN1→RXFP2
RLN3→RXFP3
RLN3→RXFP4
RLN3→RXFP1
RLN3→RXFP2
INSL3→RXFP2
INSL3→RXFP1
INSL5→RXFP4
INSL5→RXFP3
Compartment
epithelial
- integumentary
- respiratory
- gastrointestinal
- urinary
- reproductive
neural
- GABAergic
- hybrid
- glutamatergic
- specialized
- neuroendocrine
- glia
germ
- spermatogonium
- spermatocyte
- spermatid
12.5%
25%
50%
100%
ln(CP10K+1)
Eye retina
Hypothalamus
Pituitary
Brainstem
Brain cortex
Testes
Uterus
Bladder
Kidney
Pancreas
Liver
Colon
Small intestine
Tongue
Trachea
Lung
Bone
Fat
Skin

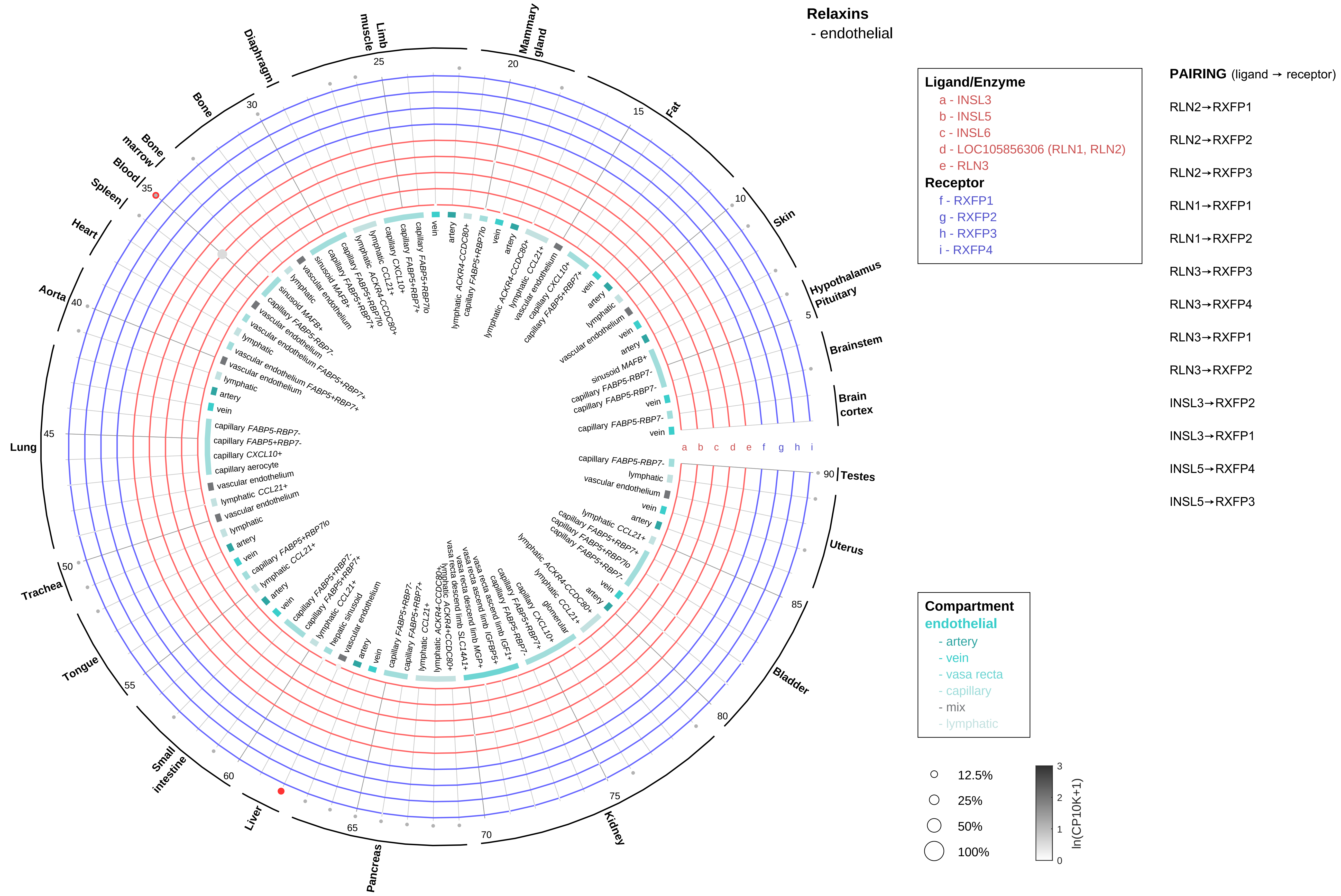
Relaxins
- endothelial
Ligand/Enzyme
a - INSL3
b - INSL5
c - INSL6
d - LOC105856306 (RLN1, RLN2)
e - RLN3
Receptor
f - RXFP1
g - RXFP2
h - RXFP3
i - RXFP4
PAIRING (ligand → receptor)
RLN2→RXFP1
RLN2→RXFP2
RLN2→RXFP3
RLN1→RXFP1
RLN1→RXFP2
RLN3→RXFP3
RLN3→RXFP4
RLN3→RXFP1
RLN3→RXFP2
INSL3→RXFP2
INSL3→RXFP1
INSL5→RXFP4
INSL5→RXFP3
Compartment
endothelial
- artery
- vein
- vasa recta
- capillary
- mix
- lymphatic
12.5%
25%
50%
100%
ln(CP10K+1)
Brain cortex
Brainstem
Hypothalamus
Pituitary
Skin
Fat
Mammary gland
Limb muscle
Diaphragm
Bone
Bone marrow
Blood
Spleen
Heart
Aorta
Lung
Trachea
Tongue
Small intestine
Liver
Pancreas
Kidney
Bladder
Uterus
Testes

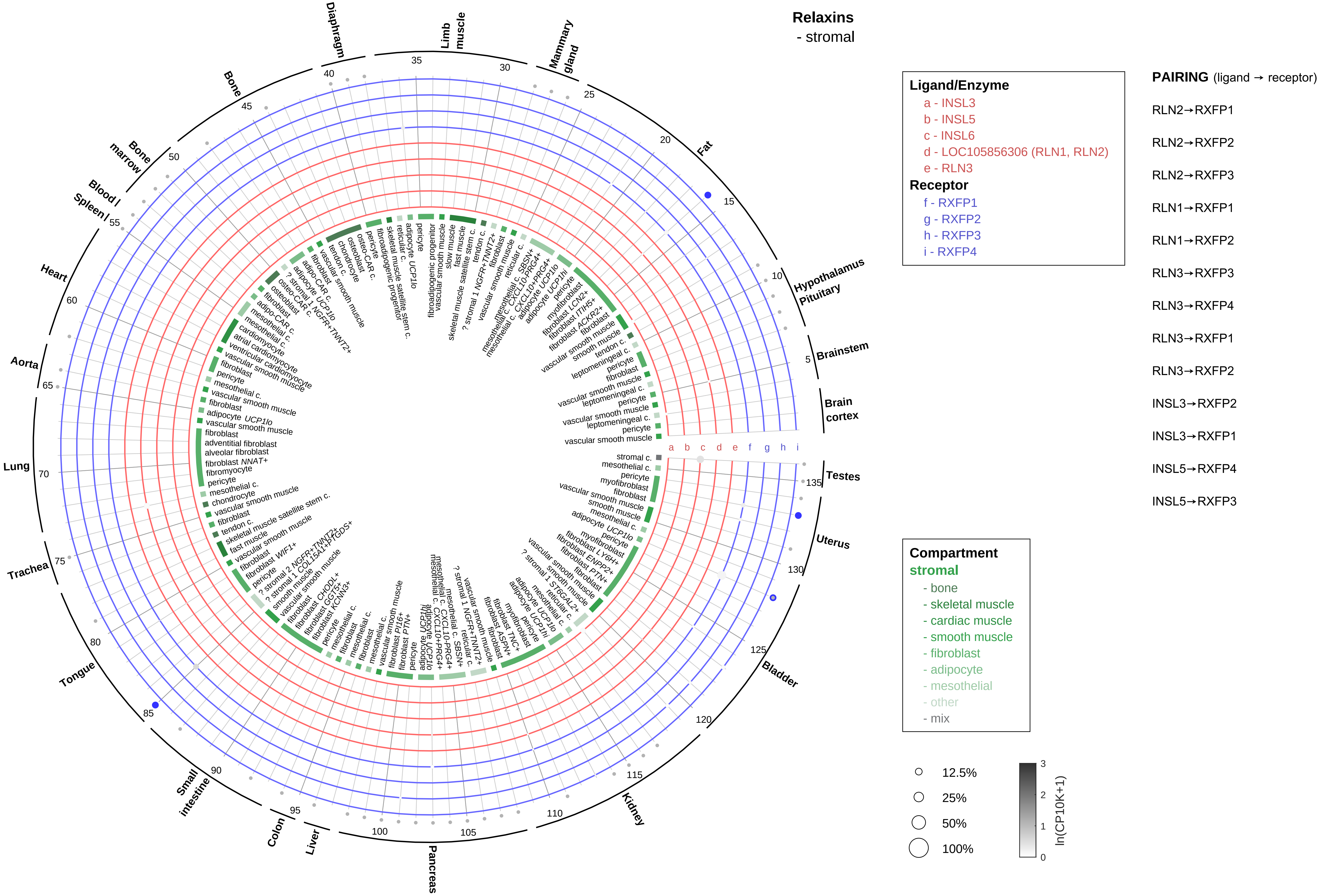
Relaxins
- stromal
Ligand/Enzyme
a - INSL3
b - INSL5
c - INSL6
d - LOC105856306 (RLN1, RLN2)
e - RLN3
Receptor
f - RXFP1
g - RXFP2
h - RXFP3
i - RXFP4
PAIRING (ligand → receptor)
RLN2→RXFP1
RLN2→RXFP2
RLN2→RXFP3
RLN1→RXFP1
RLN1→RXFP2
RLN3→RXFP3
RLN3→RXFP4
RLN3→RXFP1
RLN3→RXFP2
INSL3→RXFP2
INSL3→RXFP1
INSL5→RXFP4
INSL5→RXFP3
Compartment
stromal
- bone
- skeletal muscle
- cardiac muscle
- smooth muscle
- fibroblast
- adipocyte
- mesothelial
- other
- mix
12.5%
25%
50%
100%
ln(CP10K+1)
Diaphragm
Limb muscle
Mammary gland
Fat
Hypothalamus Pituitary
Brainstem
Brain cortex
Testes
Uterus
Bladder
Kidney
Pancreas
Liver
Colon
Small intestine
Tongue
Trachea
Lung
Aorta
Heart
Spleen
Blood
Bone marrow
Bone

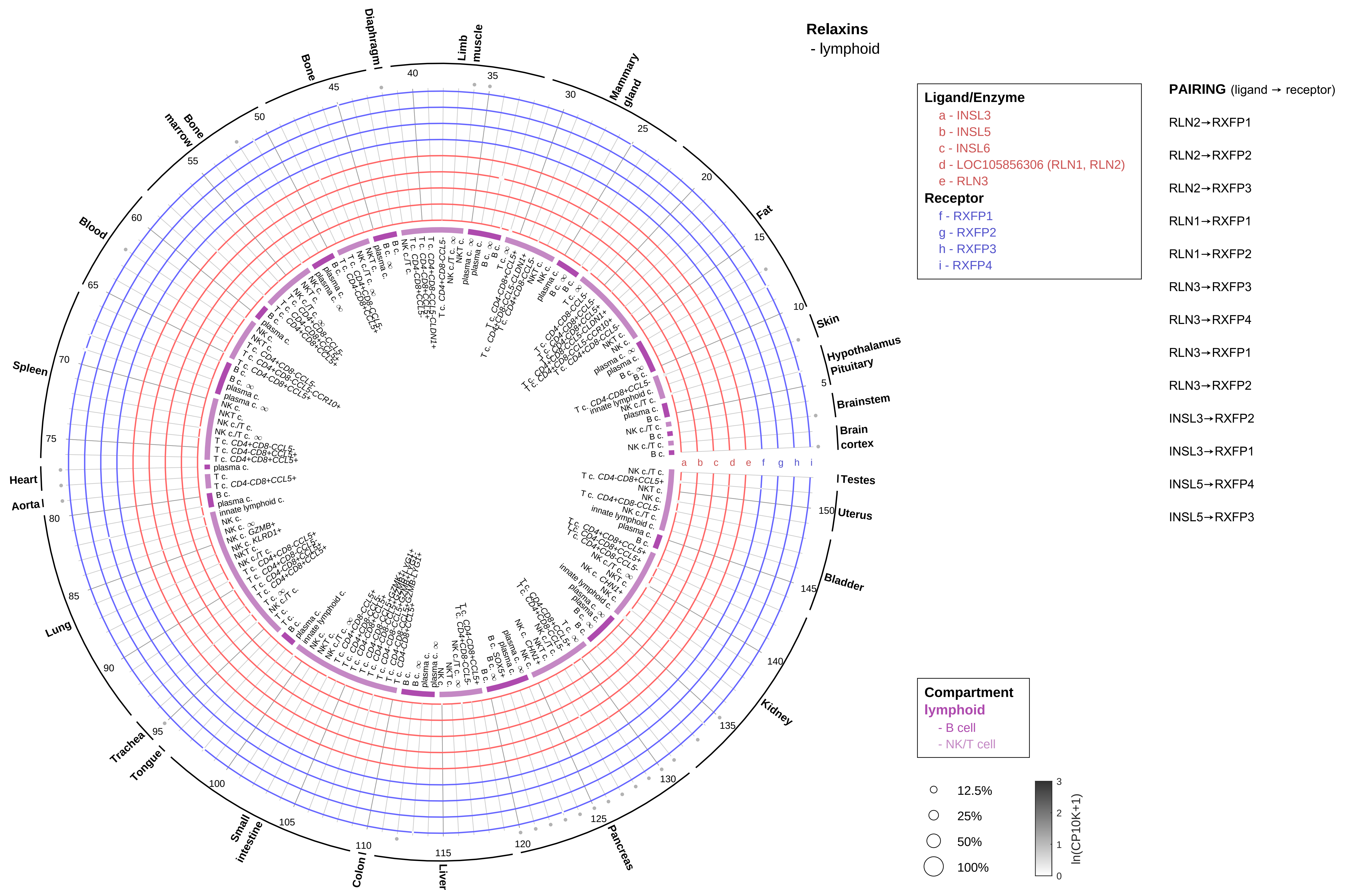

Relaxins
- lymphoid
Ligand/Enzyme
a - INSL3
b - INSL5
c - INSL6
d - LOC105856306 (RLN1, RLN2)
e - RLN3
Receptor
f - RXFP1
g - RXFP2
h - RXFP3
i - RXFP4
PAIRING (ligand → receptor)
RLN2→RXFP1
RLN2→RXFP2
RLN2→RXFP3
RLN1→RXFP1
RLN1→RXFP2
RLN3→RXFP3
RLN3→RXFP4
RLN3→RXFP1
RLN3→RXFP2
INSL3→RXFP2
INSL3→RXFP1
INSL5→RXFP4
INSL5→RXFP3
Compartment
lymphoid
- B cell
- NK/T cell
12.5%
25%
50%
100%
ln(CP10K+1)
Testes
Uterus
Bladder
Kidney
Pancreas
Liver
Colon
Small intestine
Tongue
Trachea
Lung
Aorta
Heart
Spleen
Blood
Bone marrow
Bone
Diaphragm
Limb muscle
Mammary gland
Fat
Skin
Hypothalamus
Pituitary
Brainstem
Brain cortex

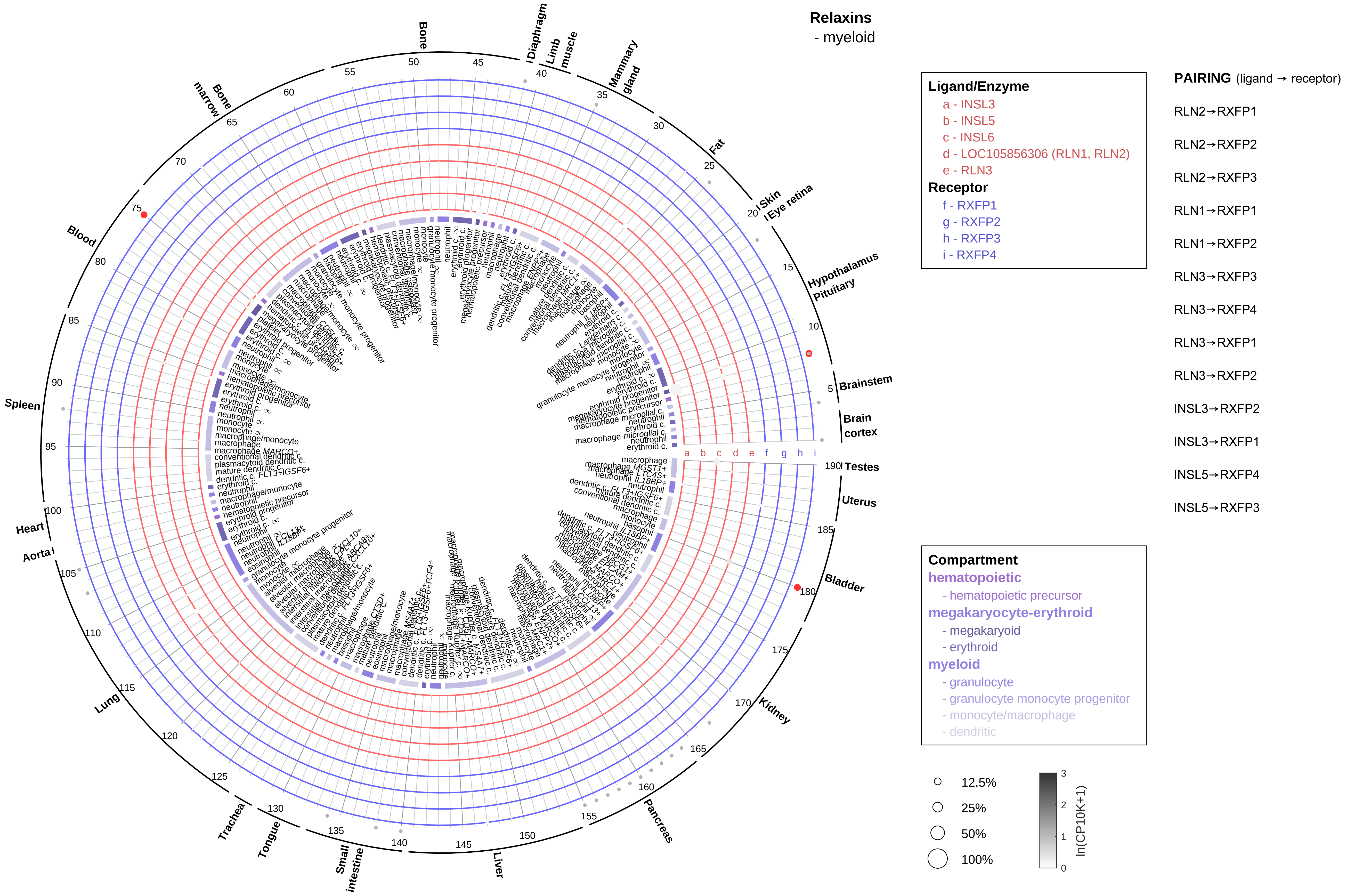
Relaxins
- myeloid
Ligand/Enzyme
a - INSL3
b - INSL5
c - INSL6
d - LOC105856306 (RLN1, RLN2)
e - RLN3
Receptor
f - RXFP1
g - RXFP2
h - RXFP3
i - RXFP4
PAIRING (ligand → receptor)
RLN2→RXFP1
RLN2→RXFP2
RLN2→RXFP3
RLN1→RXFP1
RLN1→RXFP2
RLN3→RXFP3
RLN3→RXFP4
RLN3→RXFP1
RLN3→RXFP2
INSL3→RXFP2
INSL3→RXFP1
INSL5→RXFP4
INSL5→RXFP3
Compartment
hematopoietic
- hematopoietic precursor
megakaryocyte-erythroid
- megakaryoid
- erythroid
myeloid
- granulocyte
- granulocyte monocyte progenitor
- monocyte/macrophage
- dendritic
12.5%
25%
50%
100%
ln(CP10K+1)
Bone
Diaphragm
Limb muscle
Mammary gland
Fat
Skin
Eye retina
Hypothalamus
Pituitary
Brainstem
Brain cortex
Testes
Uterus
Bladder
Kidney
Pancreas
Liver
Small intestine
Tongue
Trachea
Lung
Aorta
Heart
Spleen
Blood
Bone marrow

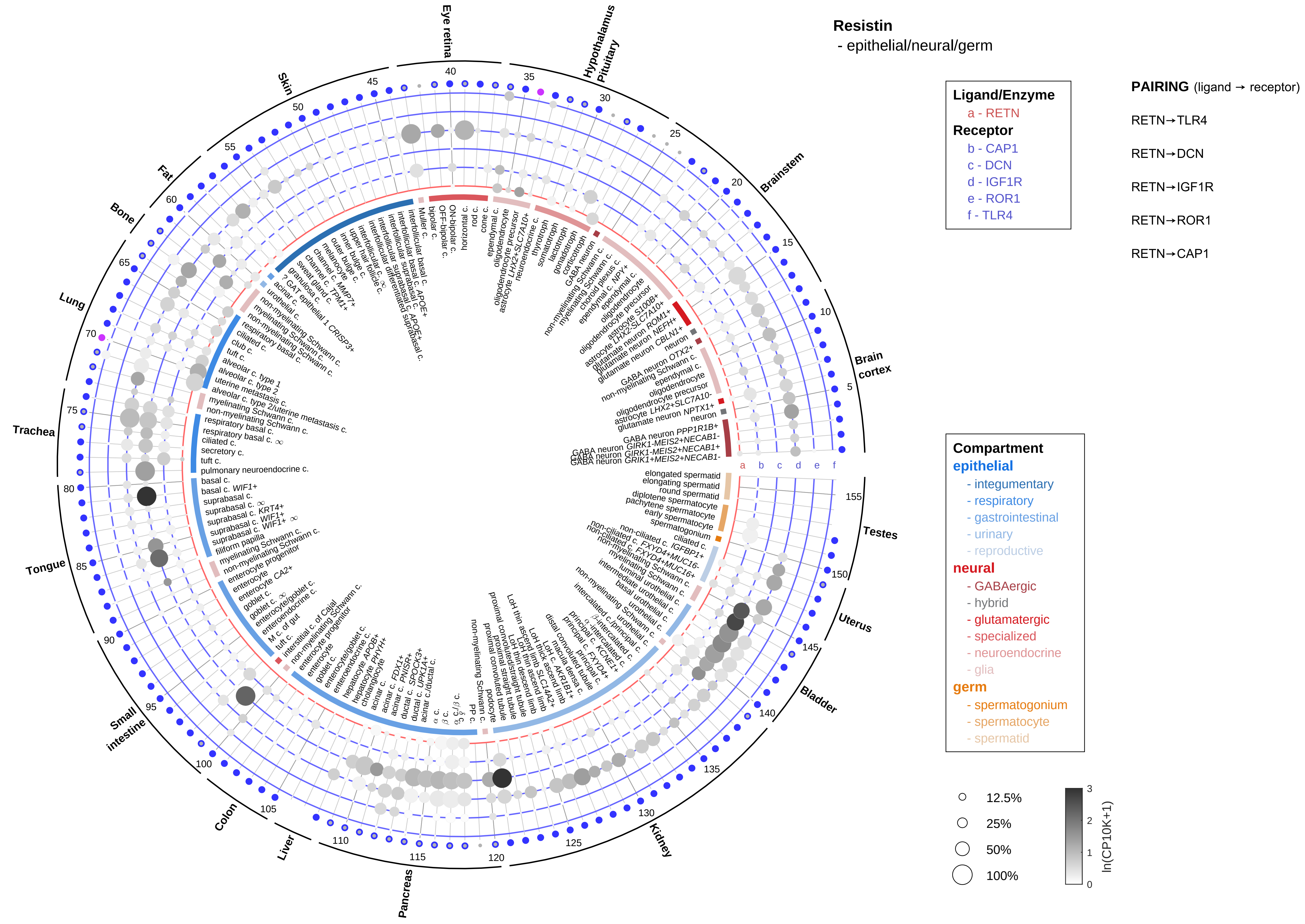
Resistin
- epithelial/neural/germ
Ligand/Enzyme
a - RETN
Receptor
b - CAP1
c - DCN
d - IGF1R
e - ROR1
f - TLR4
PAIRING (ligand → receptor)
RETN→TLR4
RETN→DCN
RETN→IGF1R
RETN→ROR1
RETN→CAP1
Compartment
epithelial
- integumentary
- respiratory
- gastrointestinal
- urinary
- reproductive
neural
- GABAergic
- hybrid
- glutamatergic
- specialized
- neuroendocrine
- glia
germ
- spermatogonium
- spermatocyte
- spermatid
12.5%
25%
50%
100%
ln(CP10K+1)
Eye retina
Hypothalamus
Pituitary
Brainstem
Brain cortex
Testes
Uterus
Bladder
Kidney
Pancreas
Liver
Colon
Small intestine
Tongue
Trachea
Lung
Bone
Fat
Skin

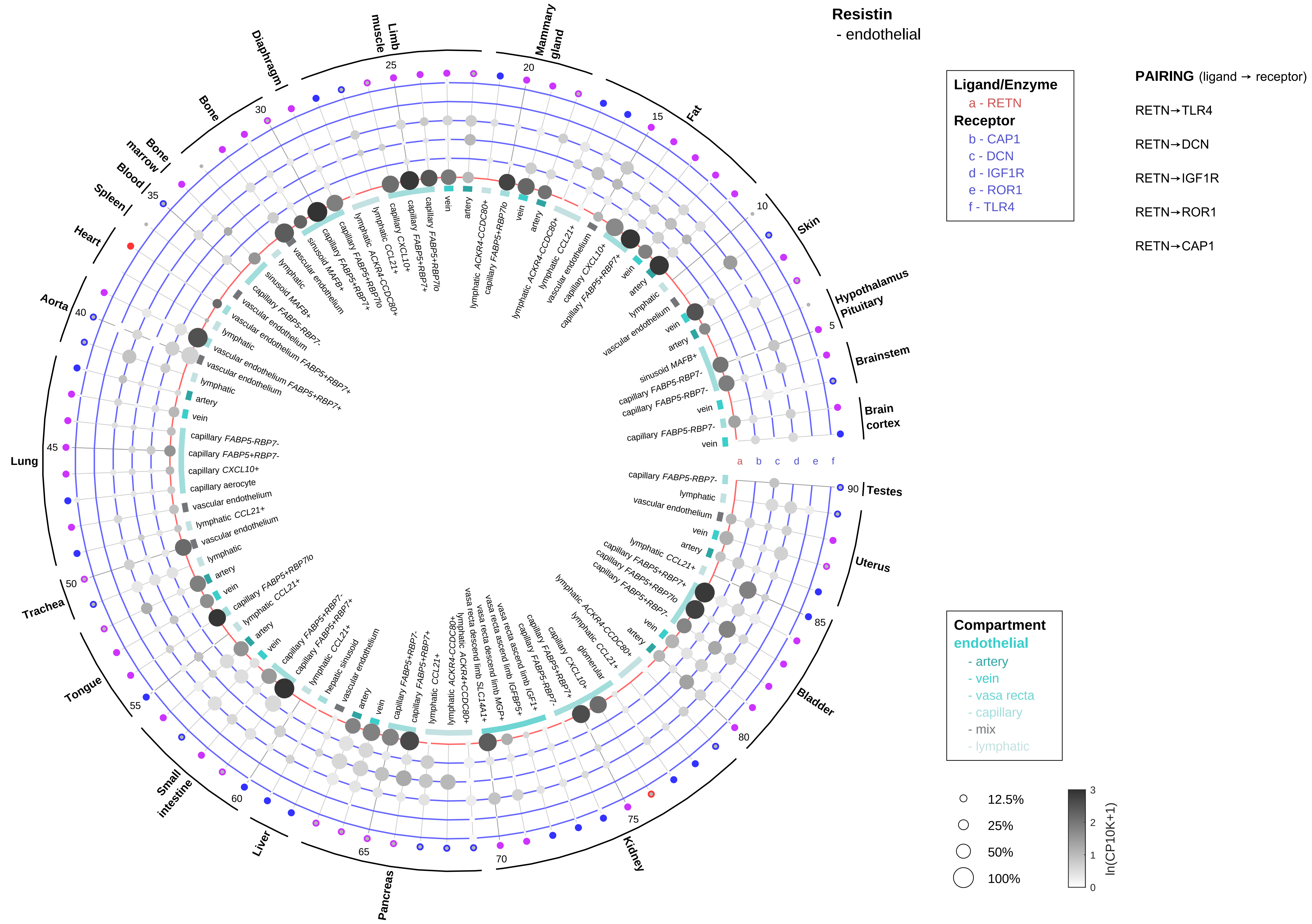
Resistin
- endothelial
Ligand/Enzyme
a - RETN
Receptor
b - CAP1
c - DCN
d - IGF1R
e - ROR1
f - TLR4
PAIRING (ligand → receptor)
RETN→TLR4
RETN→DCN
RETN→IGF1R
RETN→ROR1
RETN→CAP1
Compartment
endothelial
- artery
- vein
- vasa recta
- capillary
- mix
- lymphatic
12.5%
25%
50%
100%
ln(CP10K+1)
Brain cortex
Brainstem
Hypothalamus Pituitary
Skin
Fat
Mammary gland
Limb muscle
Diaphragm
Bone
Bone marrow
Blood
Spleen
Heart
Aorta
Lung
Trachea
Tongue
Small intestine
Liver
Pancreas
Kidney
Bladder
Uterus
Testes

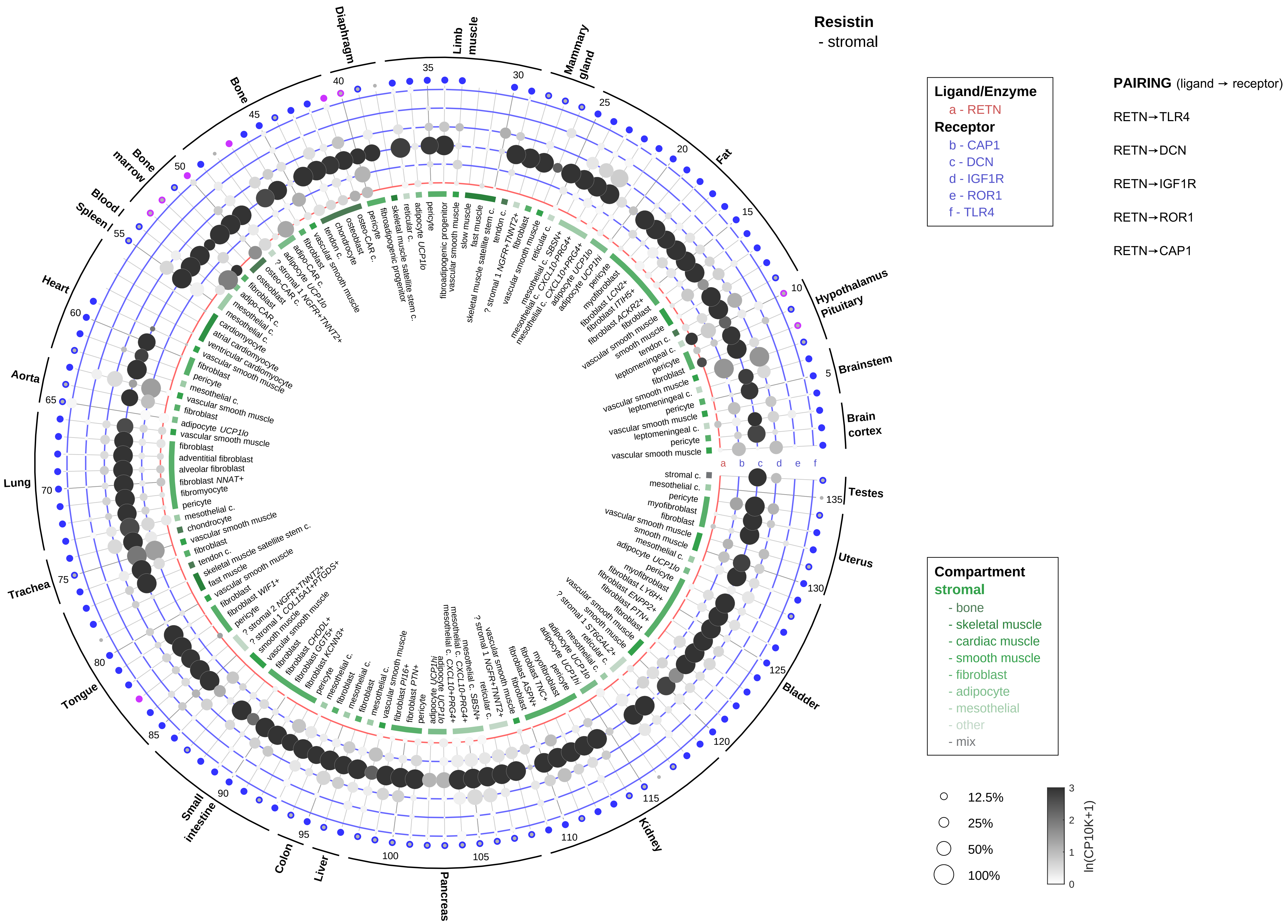

Resistin
- stromal
Ligand/Enzyme
a - RETN
Receptor
b - CAP1
c - DCN
d - IGF1R
e - ROR1
f - TLR4
PAIRING (ligand → receptor)
RETN→TLR4
RETN→DCN
RETN→IGF1R
RETN→ROR1
RETN→CAP1
Compartment
stromal
- bone
- skeletal muscle
- cardiac muscle
- smooth muscle
- fibroblast
- adipocyte
- mesothelial
- other
- mix
12.5%
25%
50%
100%
ln(CP10K+1)
Diaphragm
Limb muscle
Mammary gland
Fat
Hypothalamus Pituitary
Brainstem
Brain cortex
Testes
Uterus
Bladder
Kidney
Pancreas
Liver
Colon
Small intestine
Tongue
Trachea
Lung
Aorta
Heart
Spleen
Blood
Bone marrow
Bone

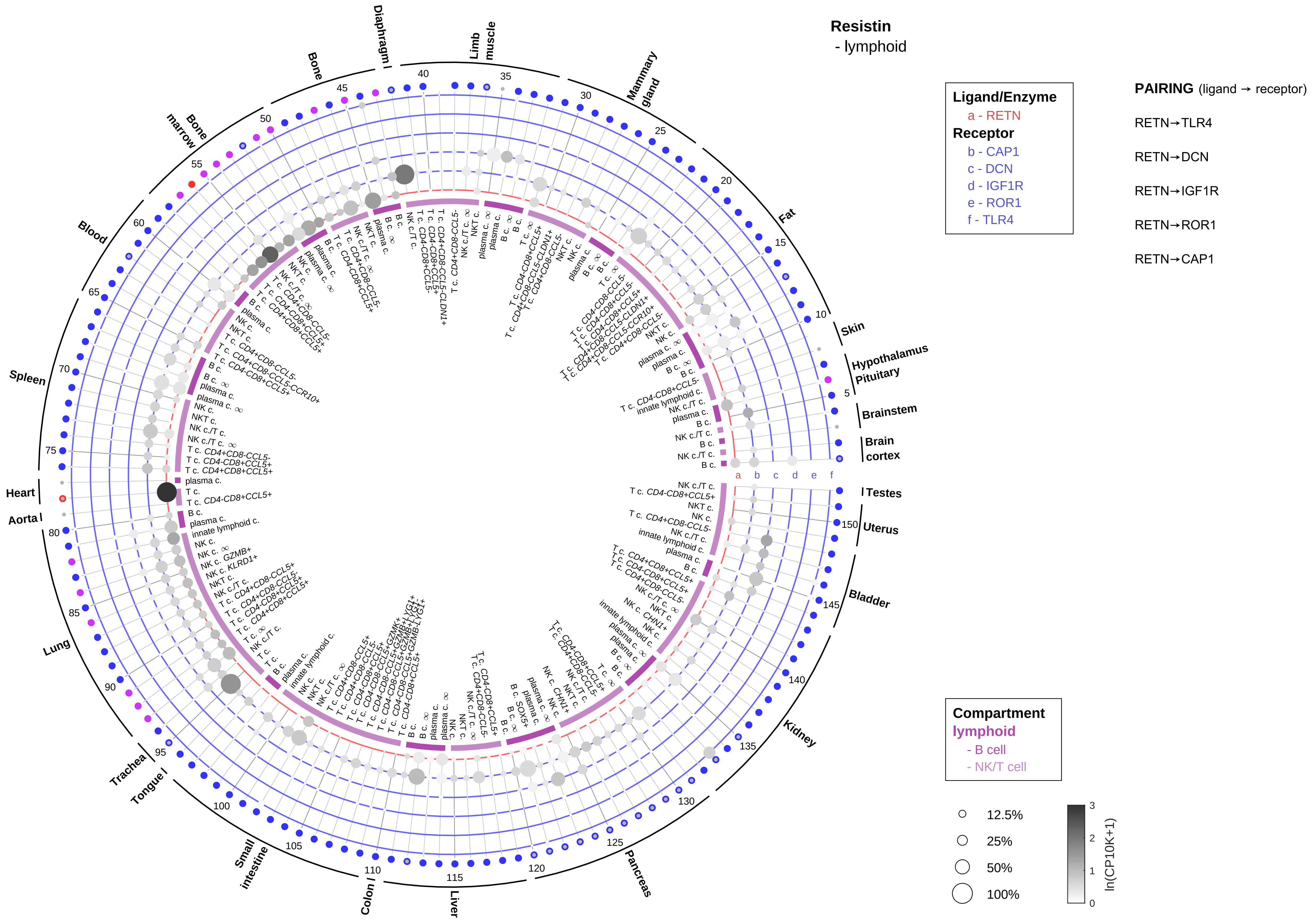
Resistin
- lymphoid
Ligand/Enzyme
a - RETN
Receptor
b - CAP1
c - DCN
d - IGF1R
e - ROR1
f - TLR4
PAIRING (ligand → receptor)
RETN→TLR4
RETN→DCN
RETN→IGF1R
RETN→ROR1
RETN→CAP1
Compartment
lymphoid
- B cell
- NK/T cell
12.5%
25%
50%
100%
ln(CP10K+1)
Skin
Hypothalamus
Pituitary
Brainstem
Brain cortex
Testes
Uterus
Bladder
Kidney
Pancreas
Liver
Colon
Small intestine
Tongue
Trachea
Lung
Aorta
Heart
Spleen
Blood
Bone marrow
Bone
Diaphragm
Limb muscle
Mammary gland
Fat

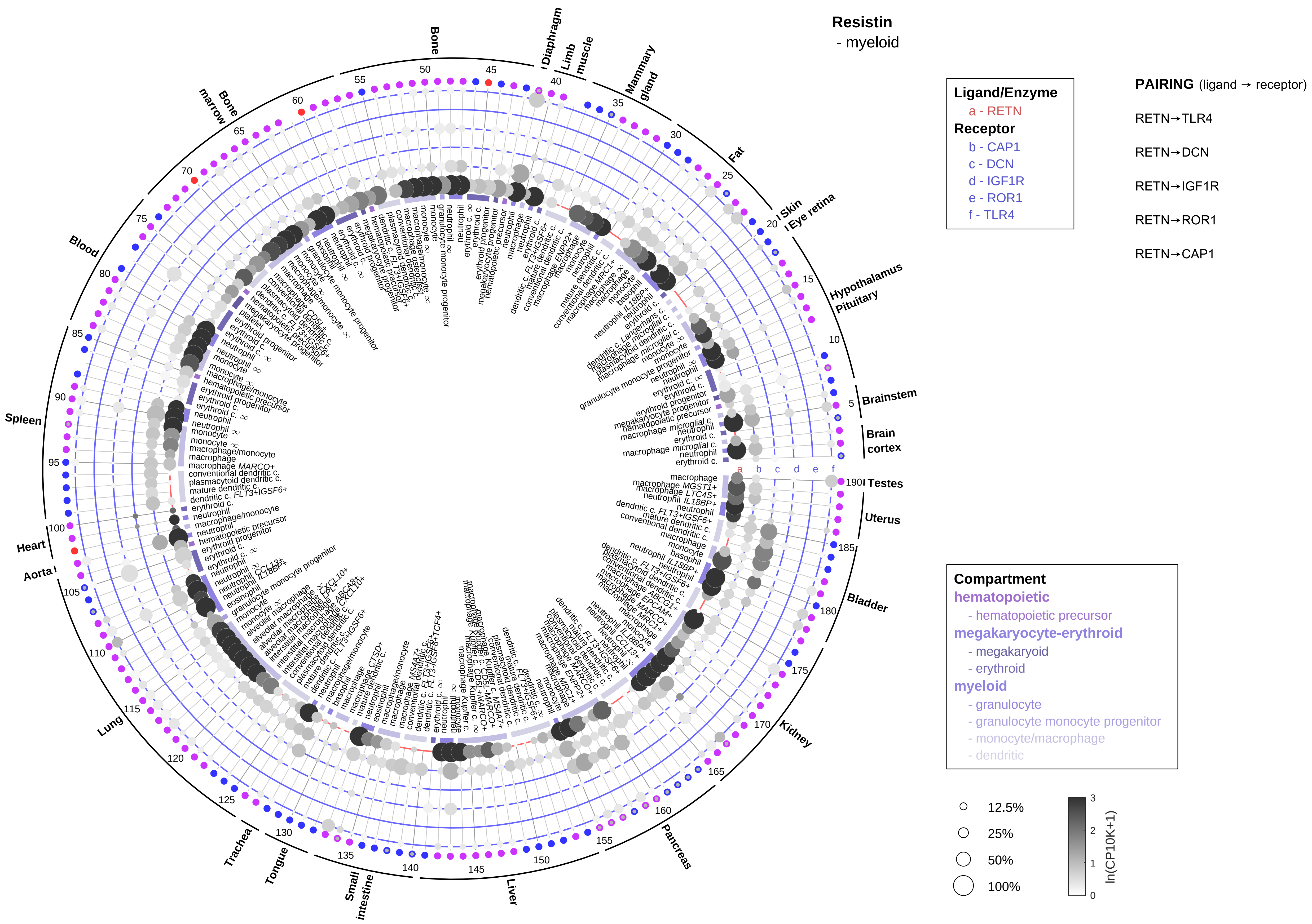

Resistin
- myeloid
Ligand/Enzyme
a - RETN
Receptor
b - CAP1
c - DCN
d - IGF1R
e - ROR1
f - TLR4
PAIRING (ligand → receptor)
RETN→TLR4
RETN→DCN
RETN→IGF1R
RETN→ROR1
RETN→CAP1
Compartment
hematopoietic
- hematopoietic precursor
megakaryocyte-erythroid
- megakaryoid
- erythroid
myeloid
- granulocyte
- granulocyte monocyte progenitor
- monocyte/macrophage
- dendritic
12.5%
25%
50%
100%
ln(CP10K+1)
Bone
Diaphragm
Limb muscle
Mammary gland
Fat
Skin
Eye retina
Hypothalamus
Pituitary
Brainstem
Brain cortex
Testes
Uterus
Bladder
Kidney
Pancreas
Liver
Small intestine
Tongue
Trachea
Lung
Aorta
Heart
Spleen
Blood
Bone marrow

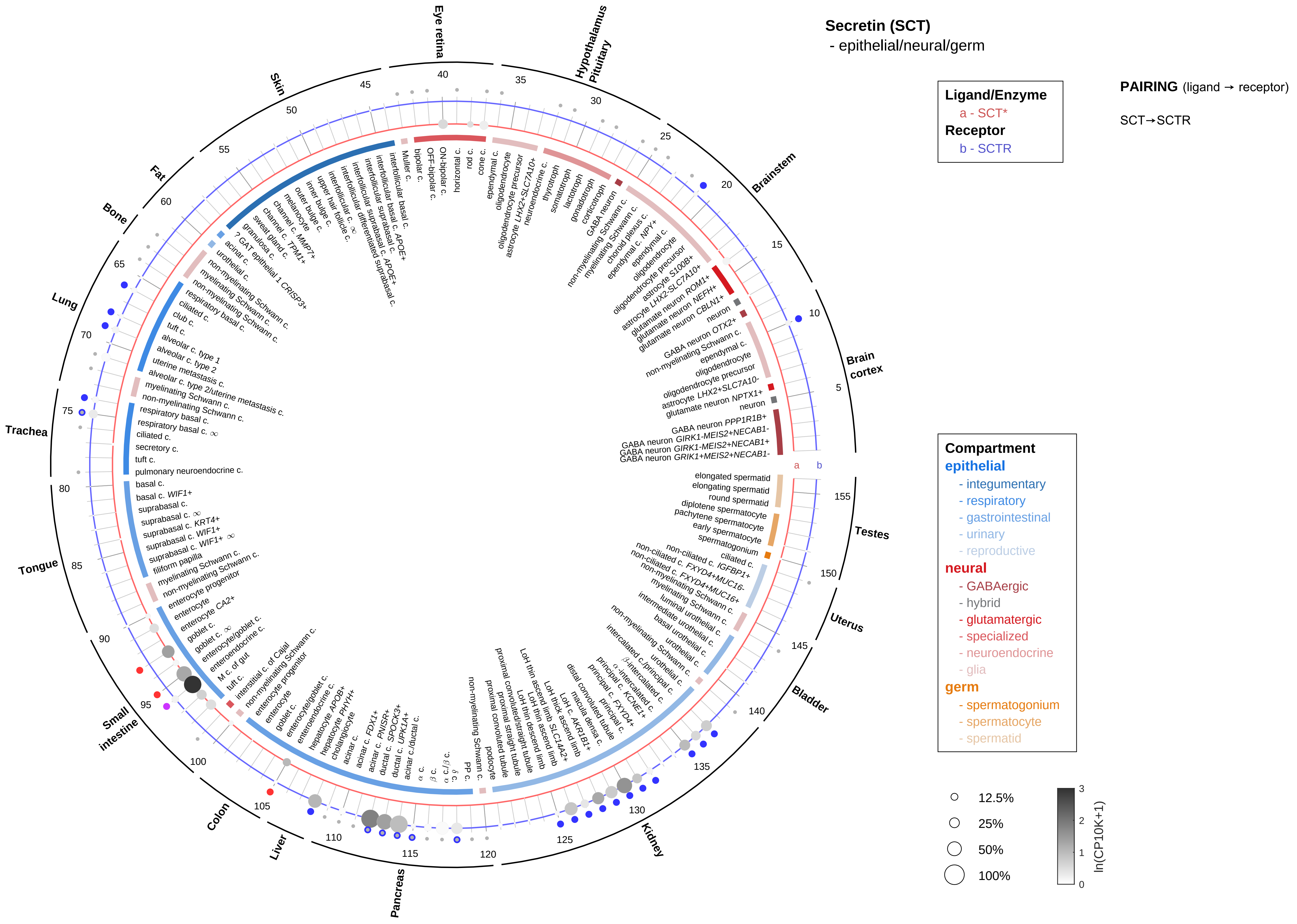

Secretin (SCT)
- epithelial/neural/germ
Ligand/Enzyme
a - SCT*
Receptor
b - SCTR
PAIRING (ligand → receptor)
SCT→SCTR
Eye retina
Hypothalamus
Pituitary
Brainstem
Brain cortex
Testes
Uterus
Bladder
Kidney
Pancreas
Liver
Colon
Small intestine
Tongue
Trachea
Lung
Bone
Fat
Skin
Compartment
epithelial
- integumentary
- respiratory
- gastrointestinal
- urinary
- reproductive
neural
- GABAergic
- hybrid
- glutamatergic
- specialized
- neuroendocrine
- glia
germ
- spermatogonium
- spermatocyte
- spermatid
12.5%
25%
50%
100%
ln(CP10K+1)

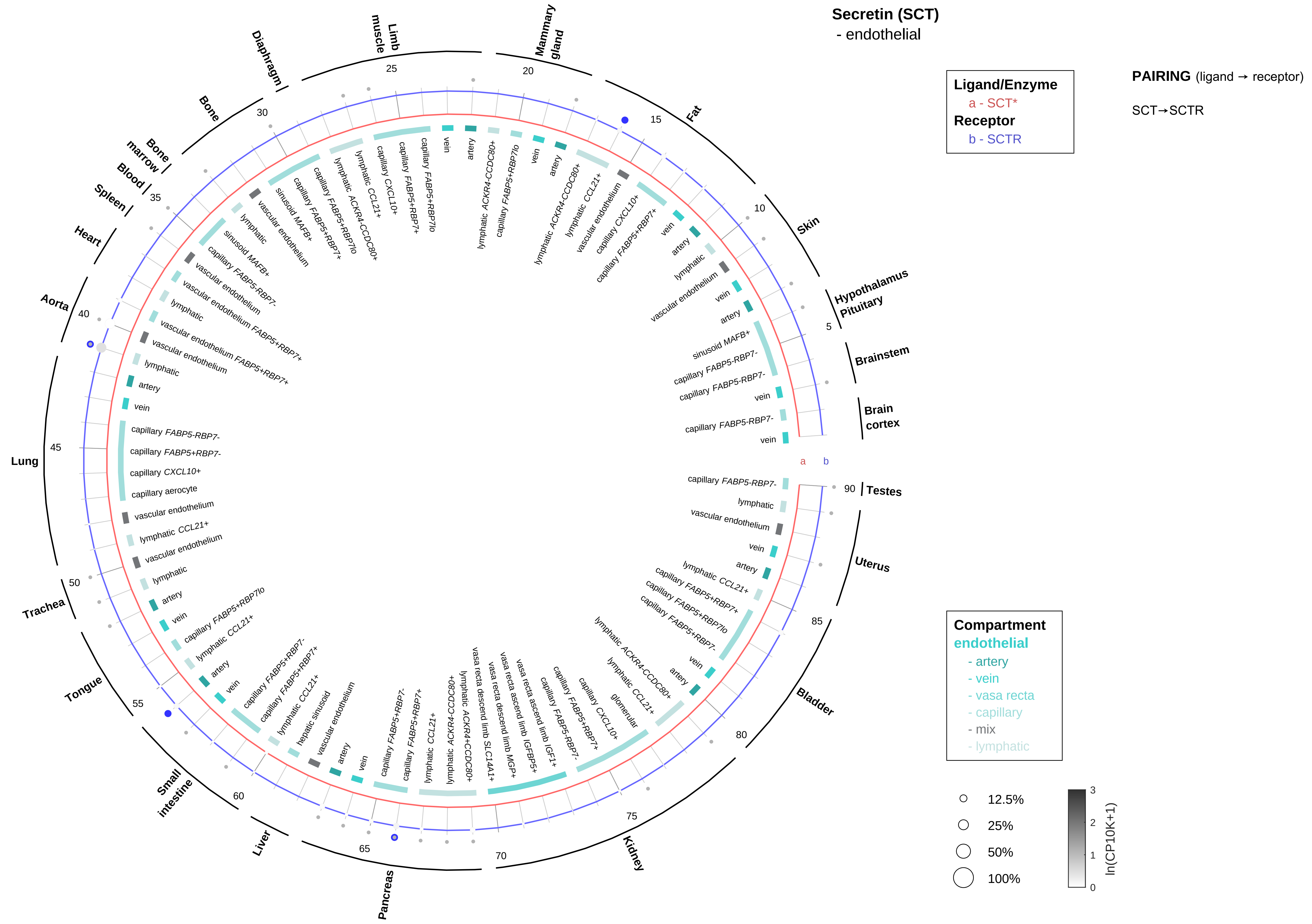
Secretin (SCT)
- endothelial
Ligand/Enzyme
a - SCT*
Receptor
b - SCTR
PAIRING (ligand → receptor)
SCT→SCTR
Compartment
endothelial
- artery
- vein
- vasa recta
- capillary
- mix
- lymphatic
12.5%
25%
50%
100%
ln(CP10K+1)
Brain cortex
Brainstem
Hypothalamus Pituitary
Skin
Fat
Mammary gland
Limb muscle
Diaphragm
Bone
Bone marrow
Blood
Spleen
Heart
Aorta
Lung
Trachea
Tongue
Small intestine
Liver
Pancreas
Kidney
Bladder
Uterus
Testes

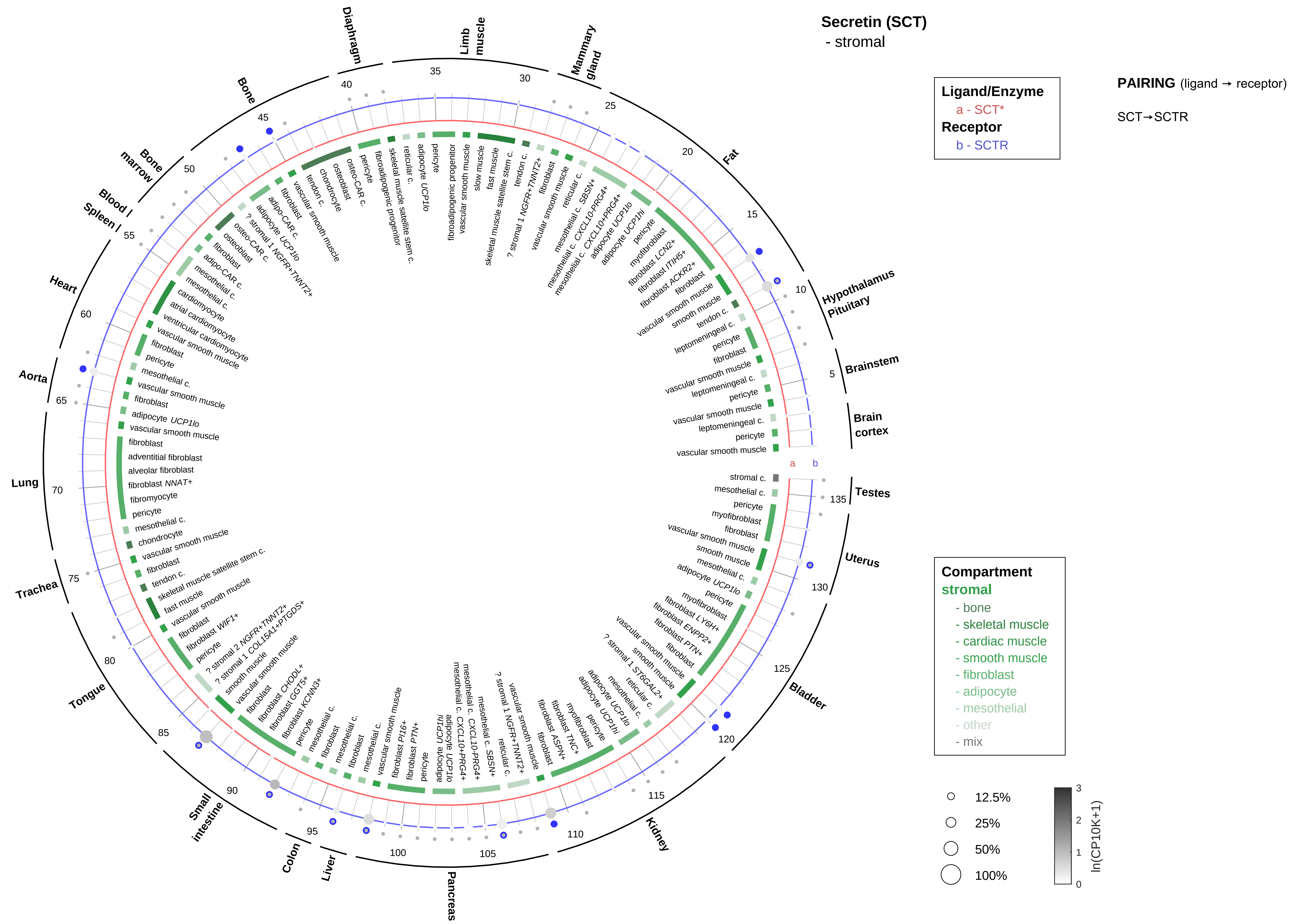
Secretin (SCT)
- stromal
Ligand/Enzyme
a - SCT*
Receptor
b - SCTR
PAIRING (ligand → receptor)
SCT→SCTR
Compartment
stromal
- bone
- skeletal muscle
- cardiac muscle
- smooth muscle
- fibroblast
- adipocyte
- mesothelial
- other
- mix
12.5%
25%
50%
100%
ln(CP10K+1)
Brain cortex
Brainstem
Hypothalamus Pituitary
Fat
Mammary gland
Limb muscle
Diaphragm
Bone
Bone marrow
Blood
Spleen
Heart
Aorta
Lung
Trachea
Tongue
Small intestine
Colon
Liver
Pancreas
Kidney
Bladder
Uterus
Testes

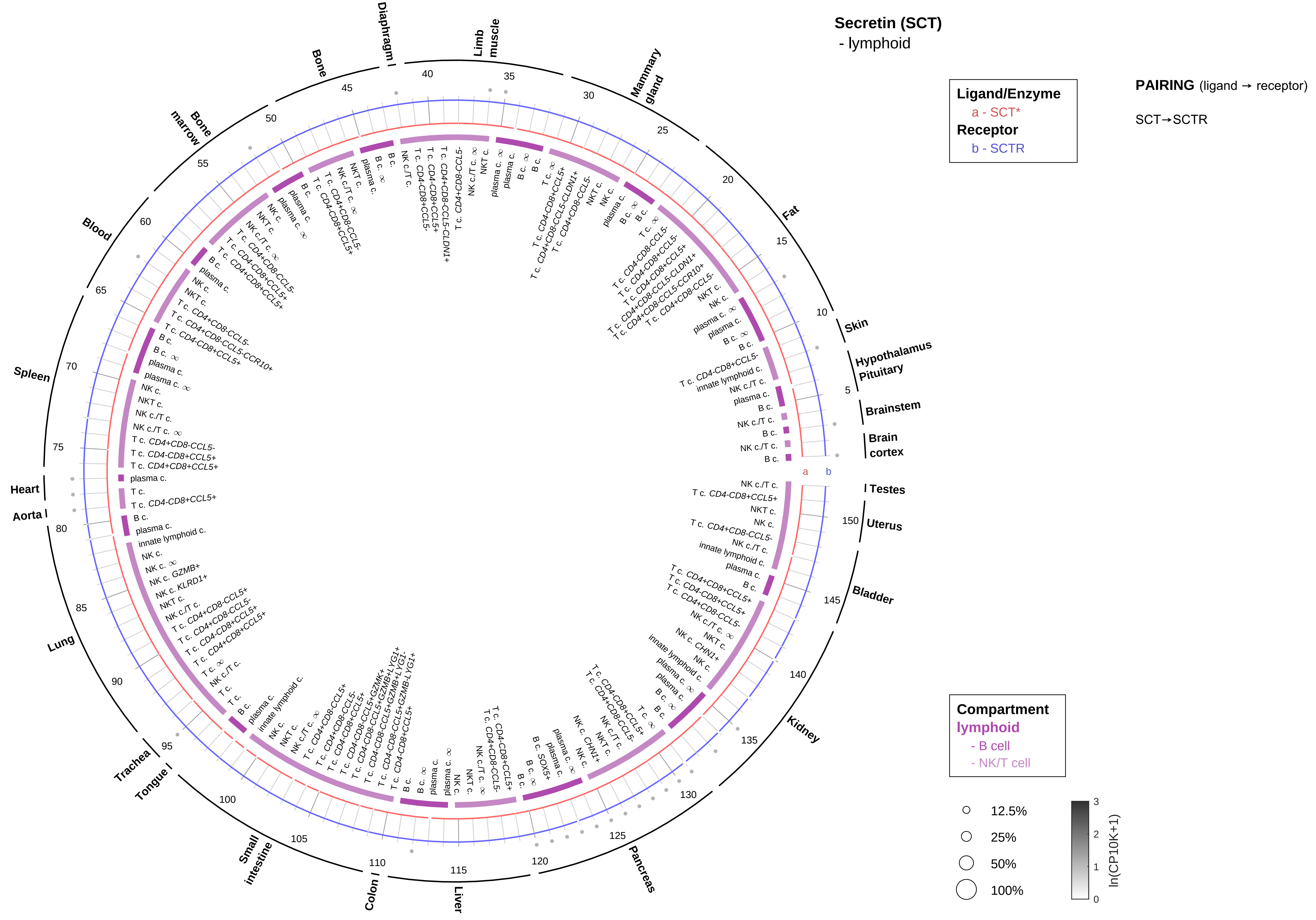

Secretin (SCT)
- lymphoid
Ligand/Enzyme
a - SCT*
Receptor
b - SCTR
PAIRING (ligand → receptor)
SCT→SCTR
Compartment
lymphoid
- B cell
- NK/T cell
12.5%
25%
50%
100%
ln(CP10K+1)
Skin
Hypothalamus
Pituitary
Brainstem
Brain cortex
Testes
Uterus
Bladder
Kidney
Pancreas
Liver
Colon
Small intestine
Tongue
Trachea
Lung
Aorta
Heart
Spleen
Blood
Bone marrow
Bone
Diaphragm
Limb muscle
Mammary gland
Fat

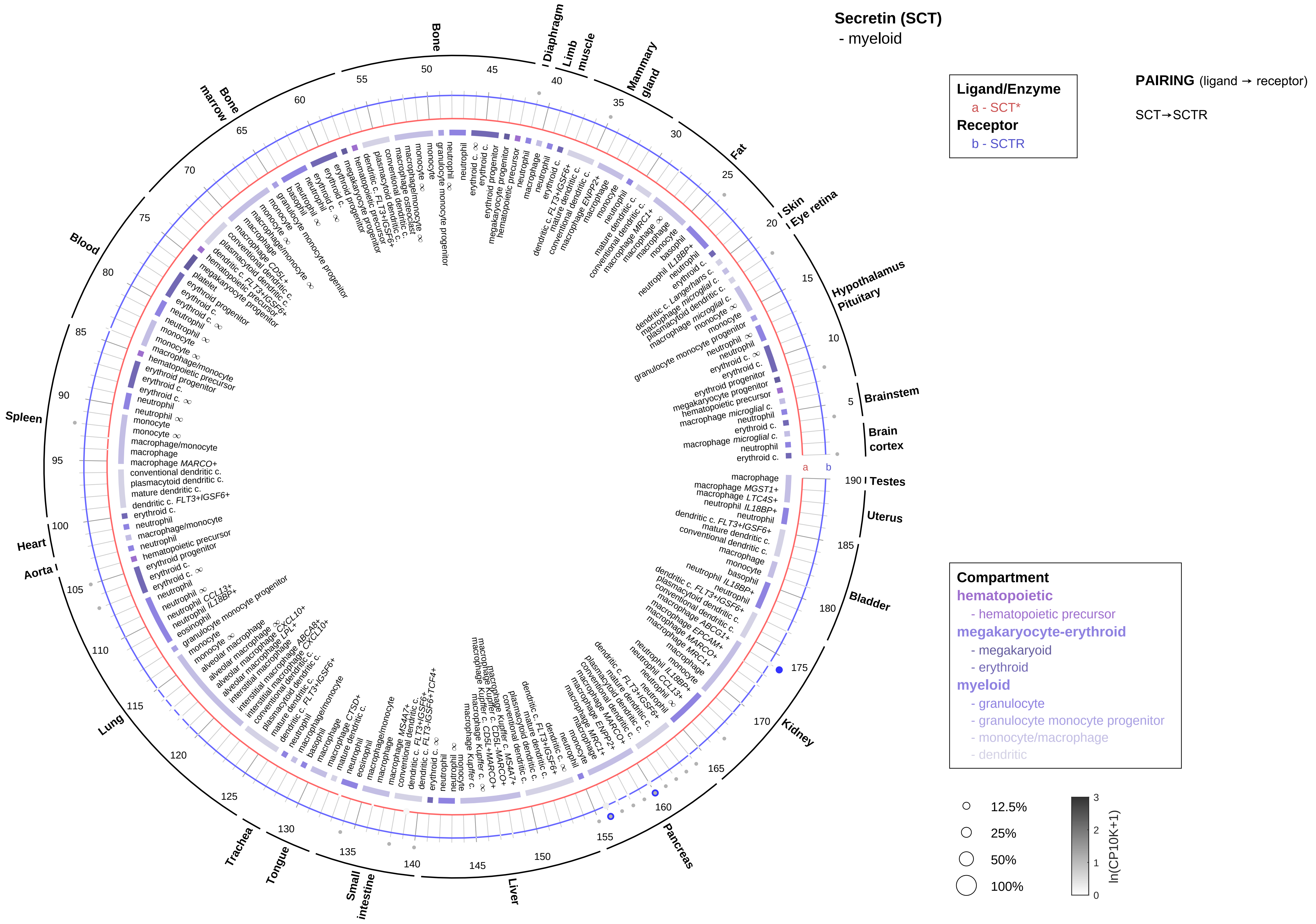
Secretin (SCT)
- myeloid
Ligand/Enzyme
a - SCT*
Receptor
b - SCTR
PAIRING (ligand → receptor)
SCT→SCTR
Compartment
hematopoietic
- hematopoietic precursor
megakaryocyte-erythroid
- megakaryoid
- erythroid
myeloid
- granulocyte
- granulocyte monocyte progenitor
- monocyte/macrophage
- dendritic
12.5%
25%
50%
100%
ln(CP10K+1)
Brain cortex
Brainstem
Hypothalamus
Pituitary
Eye retina
Skin
Fat
Mammary gland
Limb muscle
Diaphragm
Bone
Bone marrow
Blood
Spleen
Heart
Aorta
Lung
Trachea
Tongue
Small intestine
Liver
Pancreas
Kidney
Bladder
Uterus
Testes

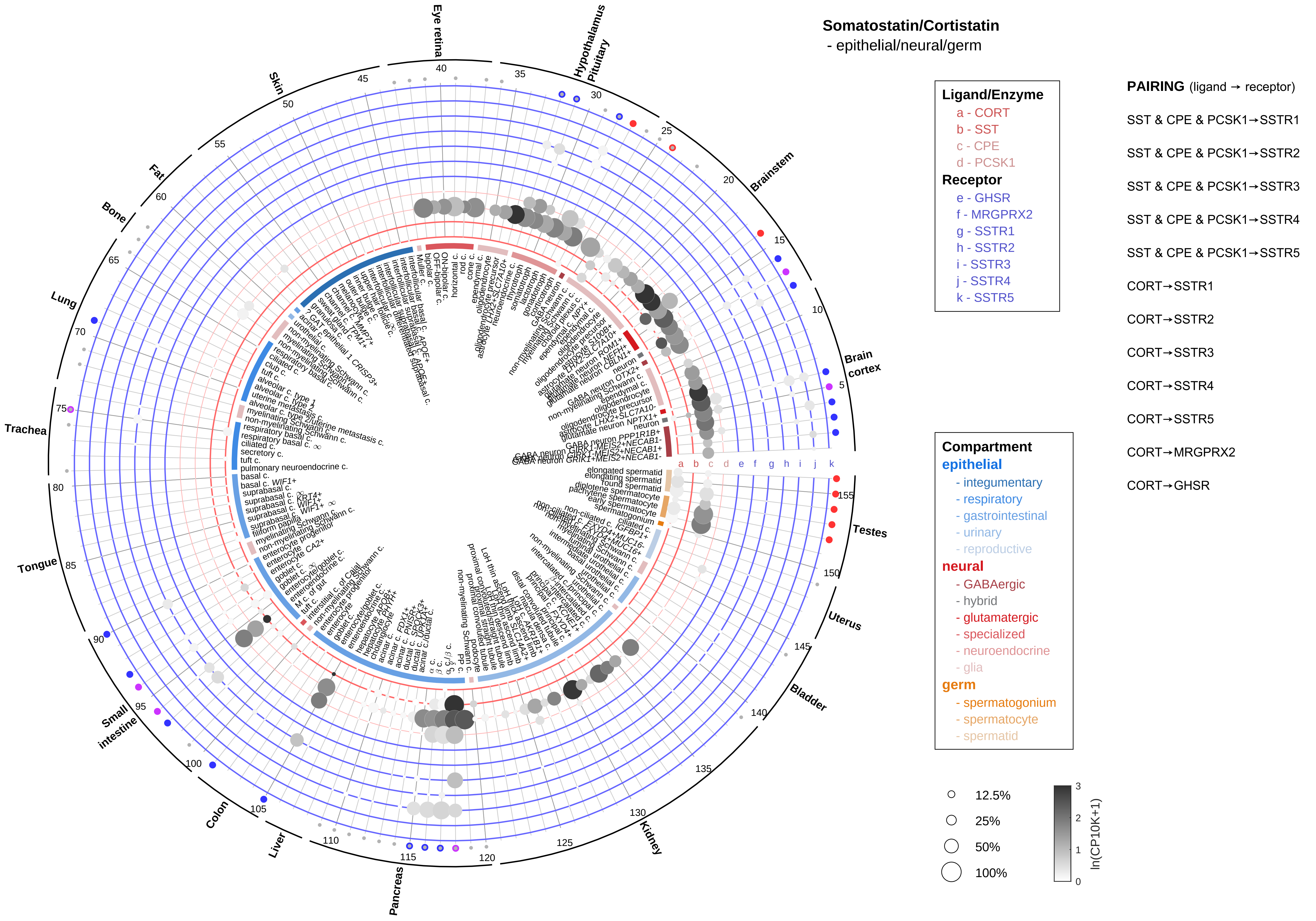

Somatostatin/Cortistatin
- epithelial/neural/germ
Ligand/Enzyme
a - CORT
b - SST
c - CPE
d - PCSK1
Receptor
e - GHSR
f - MRGPRX2
g - SSTR1
h - SSTR2
i - SSTR3
j - SSTR4
k - SSTR5
PAIRING (ligand → receptor)
SST & CPE & PCSK1→SSTR1
SST & CPE & PCSK1→SSTR2
SST & CPE & PCSK1→SSTR3
SST & CPE & PCSK1→SSTR4
SST & CPE & PCSK1→SSTR5
CORT→SSTR1
CORT→SSTR2
CORT→SSTR3
CORT→SSTR4
CORT→SSTR5
CORT→MRGPRX2
CORT→GHSR
Compartment
epithelial
- integumentary
- respiratory
- gastrointestinal
- urinary
- reproductive
neural
- GABAergic
- hybrid
- glutamatergic
- specialized
- neuroendocrine
- glia
germ
- spermatogonium
- spermatocyte
- spermatid
12.5%
25%
50%
100%
ln(CP10K+1)
Eye retina
Hypothalamus
Pituitary
Brainstem
Brain cortex
Testes
Uterus
Bladder
Kidney
Pancreas
Liver
Colon
Small intestine
Tongue
Trachea
Lung
Bone
Fat
Skin

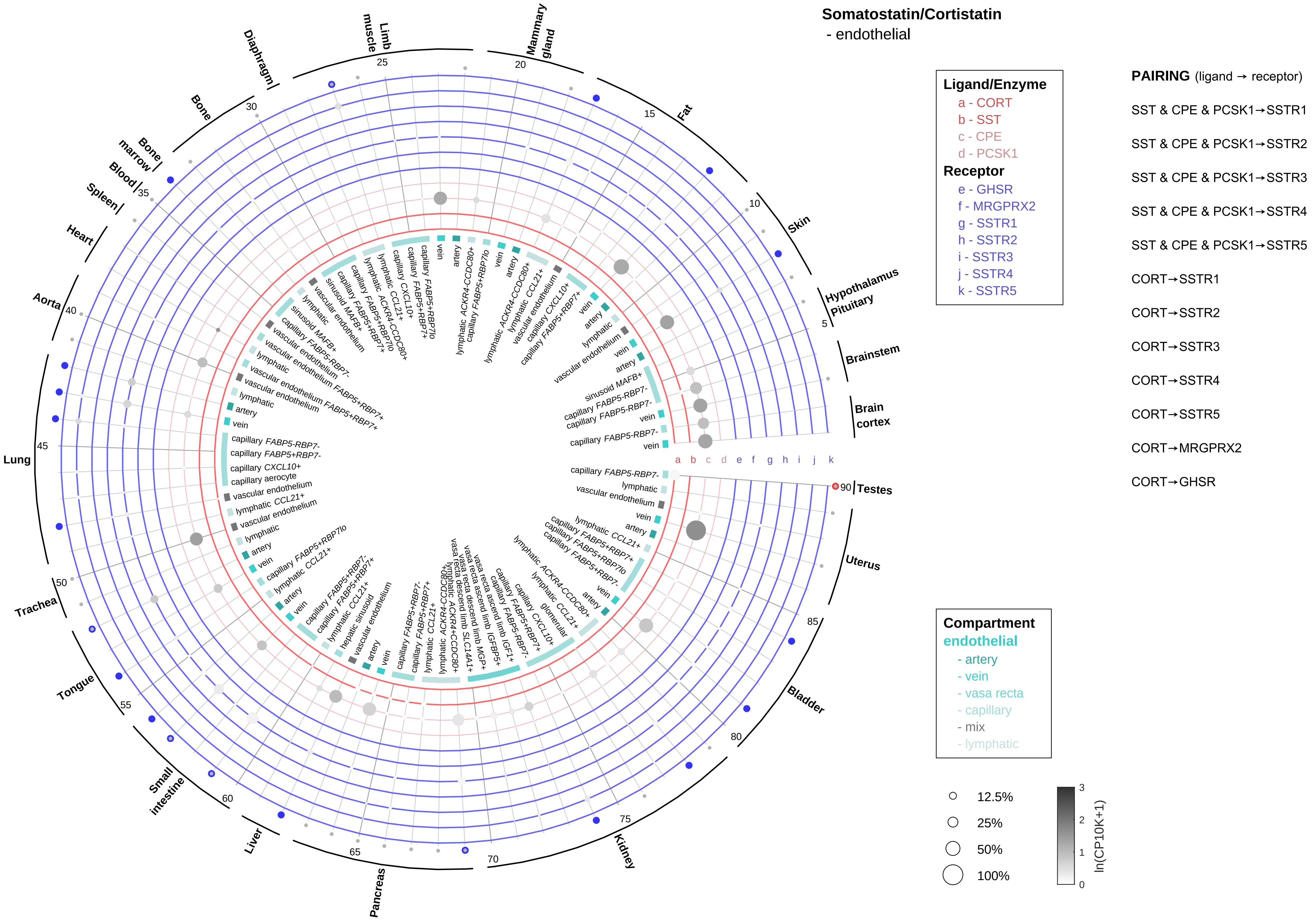
Somatostatin/Cortistatin
- endothelial
Ligand/Enzyme
a - CORT
b - SST
c - CPE
d - PCSK1
Receptor
e - GHSR
f - MRGPRX2
g - SSTR1
h - SSTR2
i - SSTR3
j - SSTR4
k - SSTR5
PAIRING (ligand → receptor)
SST & CPE & PCSK1→SSTR1
SST & CPE & PCSK1→SSTR2
SST & CPE & PCSK1→SSTR3
SST & CPE & PCSK1→SSTR4
SST & CPE & PCSK1→SSTR5
CORT→SSTR1
CORT→SSTR2
CORT→SSTR3
CORT→SSTR4
CORT→SSTR5
CORT→MRGPRX2
CORT→GHSR
Compartment
endothelial
- artery
- vein
- vasa recta
- capillary
- mix
- lymphatic
12.5%
25%
50%
100%
ln(CP10K+1)
Hypothalamus
Pituitary
Brainstem
Brain cortex
Testes
Uterus
Bladder
Kidney
Pancreas
Liver
Small intestine
Tongue
Trachea
Lung
Aorta
Heart
Spleen
Blood
Bone marrow
Bone
Diaphragm
Limb muscle
Mammary gland
Fat
Skin

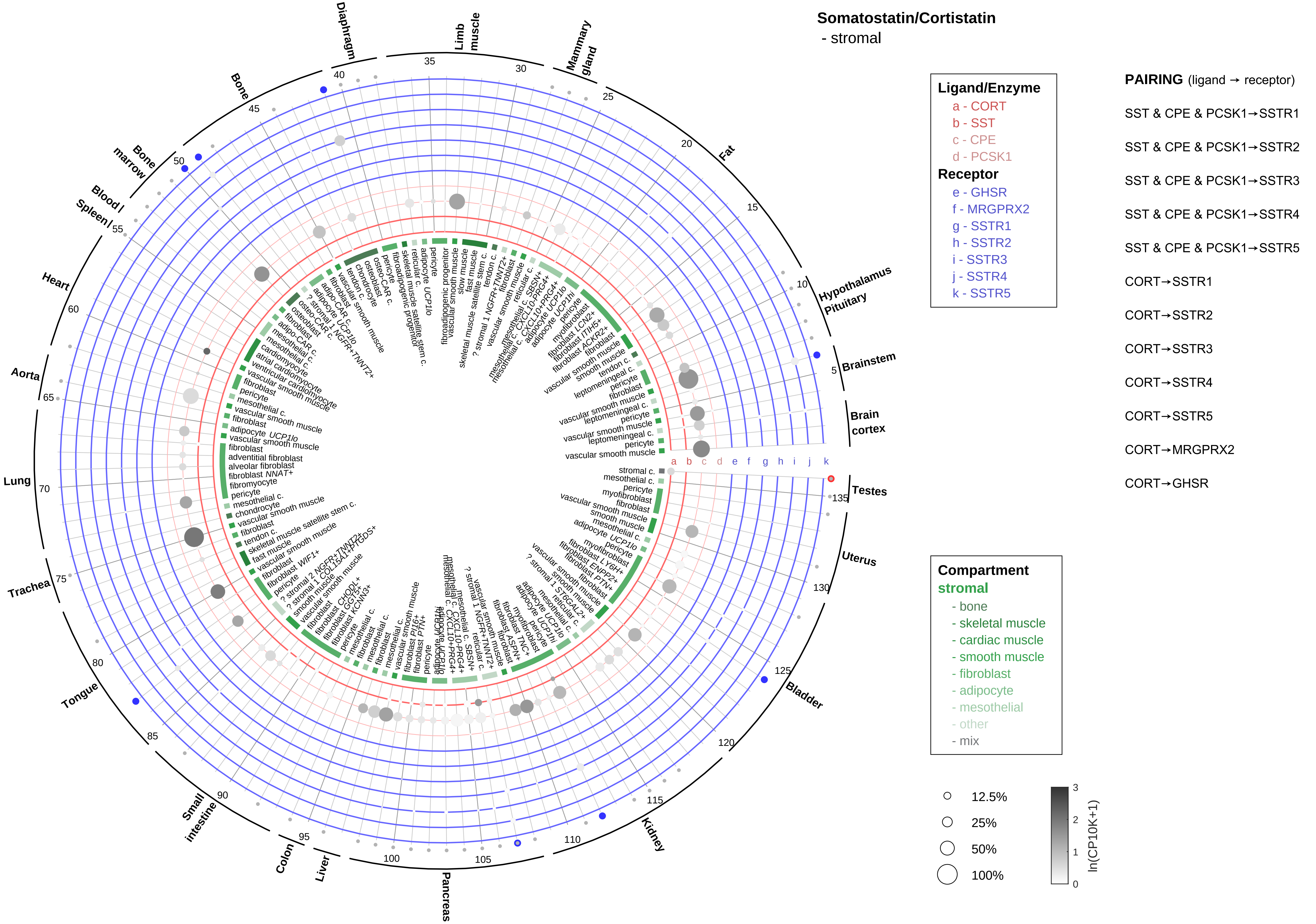

Somatostatin/Cortistatin
- stromal
Ligand/Enzyme
a - CORT
b - SST
c - CPE
d - PCSK1
Receptor
e - GHSR
f - MRGPRX2
g - SSTR1
h - SSTR2
i - SSTR3
j - SSTR4
k - SSTR5
PAIRING (ligand → receptor)
SST & CPE & PCSK1→SSTR1
SST & CPE & PCSK1→SSTR2
SST & CPE & PCSK1→SSTR3
SST & CPE & PCSK1→SSTR4
SST & CPE & PCSK1→SSTR5
CORT→SSTR1
CORT→SSTR2
CORT→SSTR3
CORT→SSTR4
CORT→SSTR5
CORT→MRGPRX2
CORT→GHSR
Compartment
stromal
- bone
- skeletal muscle
- cardiac muscle
- smooth muscle
- fibroblast
- adipocyte
- mesothelial
- other
- mix
12.5%
25%
50%
100%
ln(CP10K+1)
Diaphragm
Limb muscle
Mammary gland
Fat
Hypothalamus Pituitary
Brainstem
Brain cortex
Testes
Uterus
Bladder
Kidney
Pancreas
Liver
Colon
Small intestine
Tongue
Trachea
Lung
Aorta
Heart
Blood / Spleen /
Bone marrow
Bone

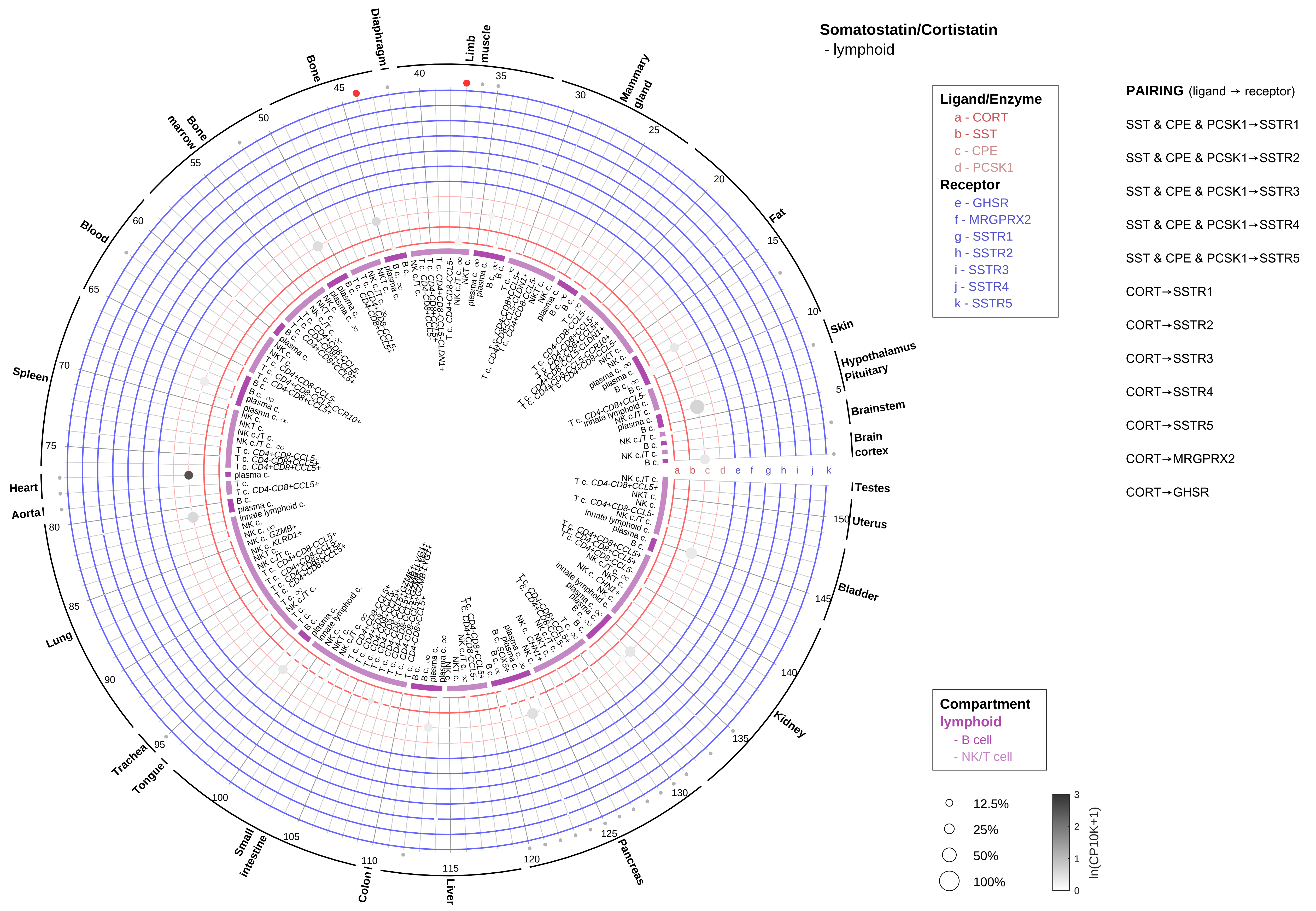
Somatostatin/Cortistatin
- lymphoid
Ligand/Enzyme
a - CORT
b - SST
c - CPE
d - PCSK1
Receptor
e - GHSR
f - MRGPRX2
g - SSTR1
h - SSTR2
i - SSTR3
j - SSTR4
k - SSTR5
PAIRING (ligand → receptor)
SST & CPE & PCSK1→SSTR1
SST & CPE & PCSK1→SSTR2
SST & CPE & PCSK1→SSTR3
SST & CPE & PCSK1→SSTR4
SST & CPE & PCSK1→SSTR5
CORT→SSTR1
CORT→SSTR2
CORT→SSTR3
CORT→SSTR4
CORT→SSTR5
CORT→MRGPRX2
CORT→GHSR
Compartment
lymphoid
- B cell
- NK/T cell
12.5%
25%
50%
100%
ln(CP10K+1)
Skin
Hypothalamus
Pituitary
Brainstem
Brain cortex
Testes
Uterus
Bladder
Kidney
Pancreas
Liver
Colon
Small intestine
Tongue
Trachea
Lung
Aorta
Heart
Spleen
Blood
Bone marrow
Bone
Diaphragm
Limb muscle
Mammary gland
Fat

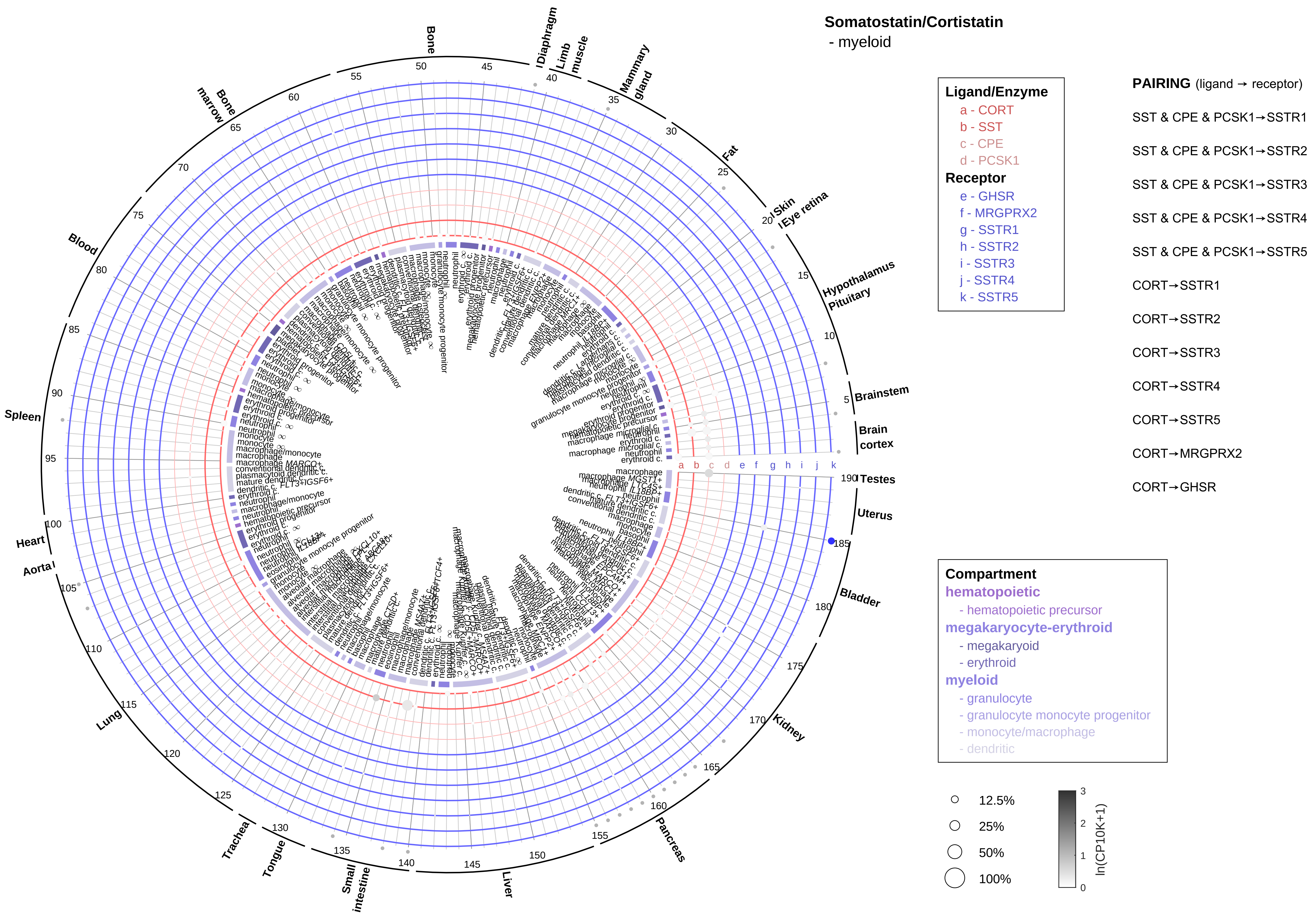

Somatostatin/Cortistatin
- myeloid
Ligand/Enzyme
a - CORT
b - SST
c - CPE
d - PCSK1
Receptor
e - GHSR
f - MRGPRX2
g - SSTR1
h - SSTR2
i - SSTR3
j - SSTR4
k - SSTR5
PAIRING (ligand → receptor)
SST & CPE & PCSK1→SSTR1
SST & CPE & PCSK1→SSTR2
SST & CPE & PCSK1→SSTR3
SST & CPE & PCSK1→SSTR4
SST & CPE & PCSK1→SSTR5
CORT→SSTR1
CORT→SSTR2
CORT→SSTR3
CORT→SSTR4
CORT→SSTR5
CORT→MRGPRX2
CORT→GHSR
Compartment
hematopoietic
- hematopoietic precursor
megakaryocyte-erythroid
- megakaryoid
- erythroid
myeloid
- granulocyte
- granulocyte monocyte progenitor
- monocyte/macrophage
- dendritic
12.5%
25%
50%
100%
ln(CP10K+1)
Bone
Diaphragm
Limb muscle
Mammary gland
Fat
Skin
Eye retina
Hypothalamus
Pituitary
Brainstem
Brain cortex
Testes
Uterus
Bladder
Kidney
Pancreas
Liver
Small intestine
Tongue
Trachea
Lung
Aorta
Heart
Spleen
Blood
Bone marrow

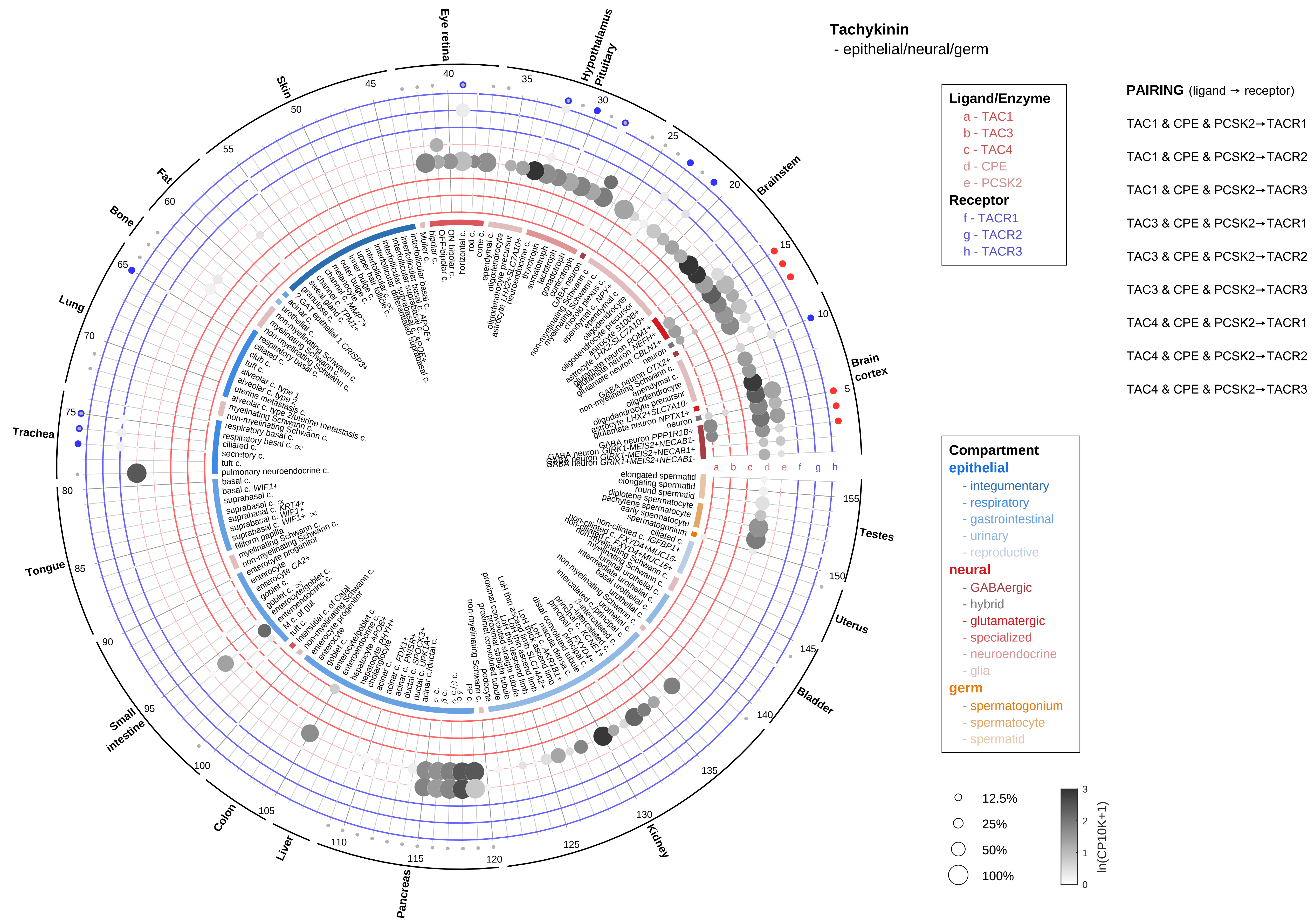

Tachykinin
- epithelial/neural/germ
Ligand/Enzyme
a - TAC1
b - TAC3
c - TAC4
d - CPE
e - PCSK2
Receptor
f - TACR1
g - TACR2
h - TACR3
PAIRING (ligand → receptor)
TAC1 & CPE & PCSK2→TACR1
TAC1 & CPE & PCSK2→TACR2
TAC1 & CPE & PCSK2→TACR3
TAC3 & CPE & PCSK2→TACR1
TAC3 & CPE & PCSK2→TACR2
TAC3 & CPE & PCSK2→TACR3
TAC4 & CPE & PCSK2→TACR1
TAC4 & CPE & PCSK2→TACR2
TAC4 & CPE & PCSK2→TACR3
Compartment
epithelial
- integumentary
- respiratory
- gastrointestinal
- urinary
- reproductive
neural
- GABAergic
- hybrid
- glutamatergic
- specialized
- neuroendocrine
- glia
germ
- spermatogonium
- spermatocyte
- spermatid
12.5%
25%
50%
100%
ln(CP10K+1)
Eye retina
Hypothalamus
Pituitary
Brainstem
Brain cortex
Testes
Uterus
Bladder
Kidney
Pancreas
Liver
Colon
Small intestine
Tongue
Trachea
Lung
Bone
Fat
Skin

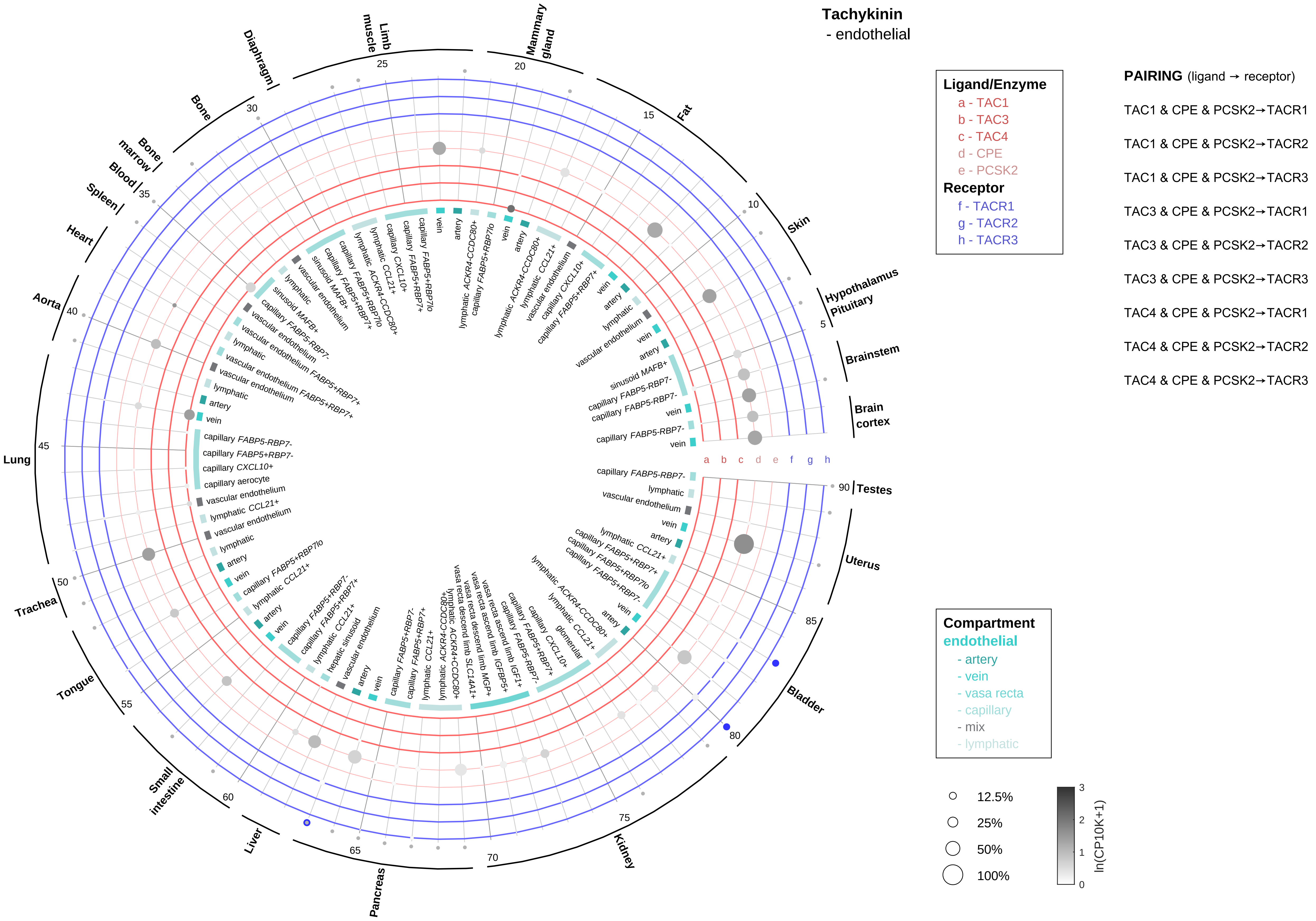

Tachykinin
- endothelial
Ligand/Enzyme
a - TAC1
b - TAC3
c - TAC4
d - CPE
e - PCSK2
Receptor
f - TACR1
g - TACR2
h - TACR3
PAIRING (ligand → receptor)
TAC1 & CPE & PCSK2→TACR1
TAC1 & CPE & PCSK2→TACR2
TAC1 & CPE & PCSK2→TACR3
TAC3 & CPE & PCSK2→TACR1
TAC3 & CPE & PCSK2→TACR2
TAC3 & CPE & PCSK2→TACR3
TAC4 & CPE & PCSK2→TACR1
TAC4 & CPE & PCSK2→TACR2
TAC4 & CPE & PCSK2→TACR3
Compartment
endothelial
- artery
- vein
- vasa recta
- capillary
- mix
- lymphatic
12.5%
25%
50%
100%
ln(CP10K+1)
Diaphragm
Limb muscle
Mammary gland
Fat
Skin
Hypothalamus
Pituitary
Brainstem
Brain cortex
Testes
Uterus
Bladder
Kidney
Pancreas
Liver
Small intestine
Tongue
Trachea
Lung
Aorta
Heart
Spleen
Blood
Bone marrow
Bone

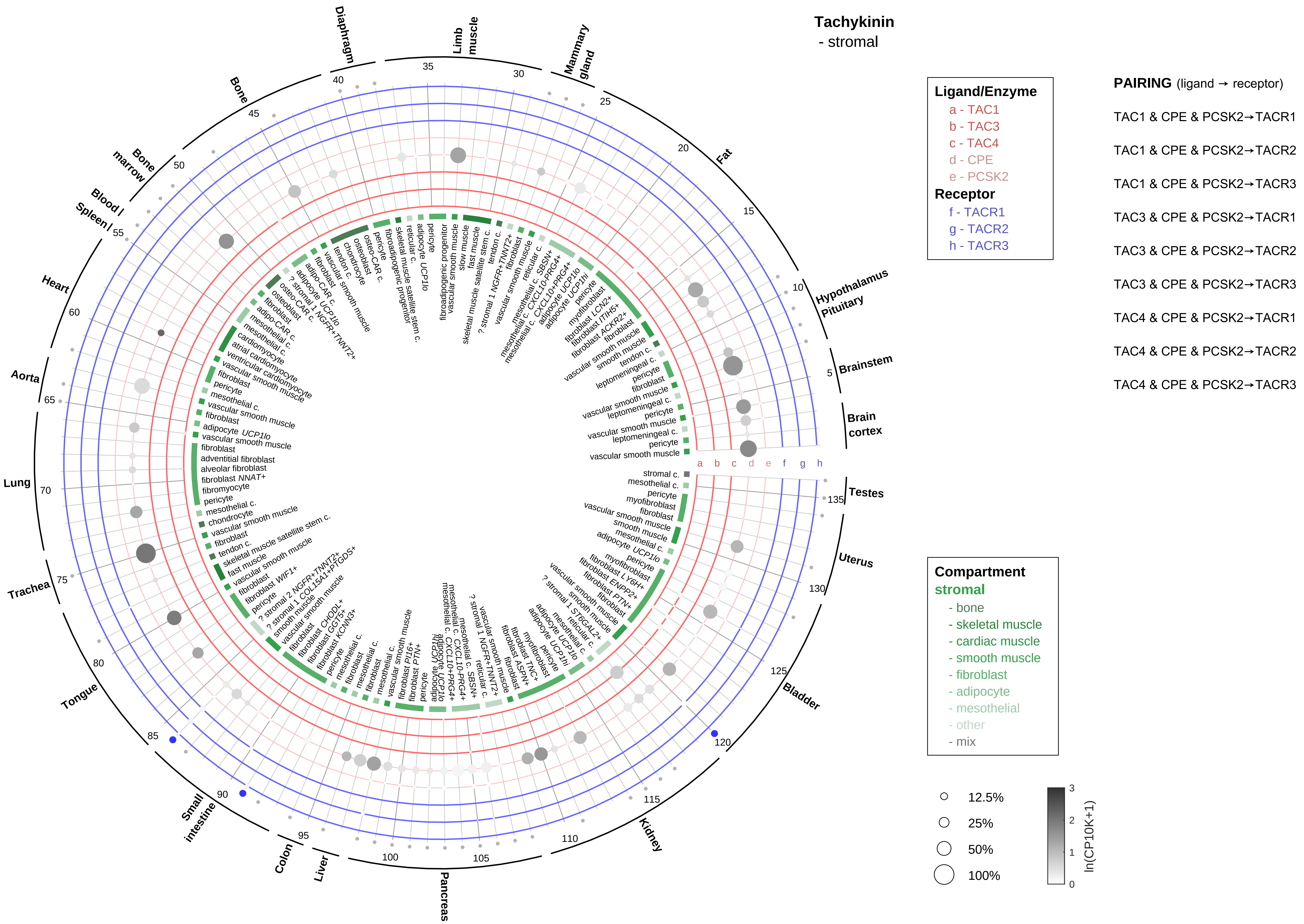

Tachykinin
- stromal
Ligand/Enzyme
a - TAC1
b - TAC3
c - TAC4
d - CPE
e - PCSK2
Receptor
f - TACR1
g - TACR2
h - TACR3
PAIRING (ligand → receptor)
TAC1 & CPE & PCSK2→TACR1
TAC1 & CPE & PCSK2→TACR2
TAC1 & CPE & PCSK2→TACR3
TAC3 & CPE & PCSK2→TACR1
TAC3 & CPE & PCSK2→TACR2
TAC3 & CPE & PCSK2→TACR3
TAC4 & CPE & PCSK2→TACR1
TAC4 & CPE & PCSK2→TACR2
TAC4 & CPE & PCSK2→TACR3
Compartment
stromal
- bone
- skeletal muscle
- cardiac muscle
- smooth muscle
- fibroblast
- adipocyte
- mesothelial
- other
- mix
12.5%
25%
50%
100%
ln(CP10K+1)
Diaphragm
Limb muscle
Mammary gland
Fat
Hypothalamus Pituitary
Brainstem
Brain cortex
Testes
Uterus
Bladder
Kidney
Pancreas
Liver
Colon
Small intestine
Tongue
Trachea
Lung
Aorta
Heart
Spleen
Blood
Bone marrow
Bone

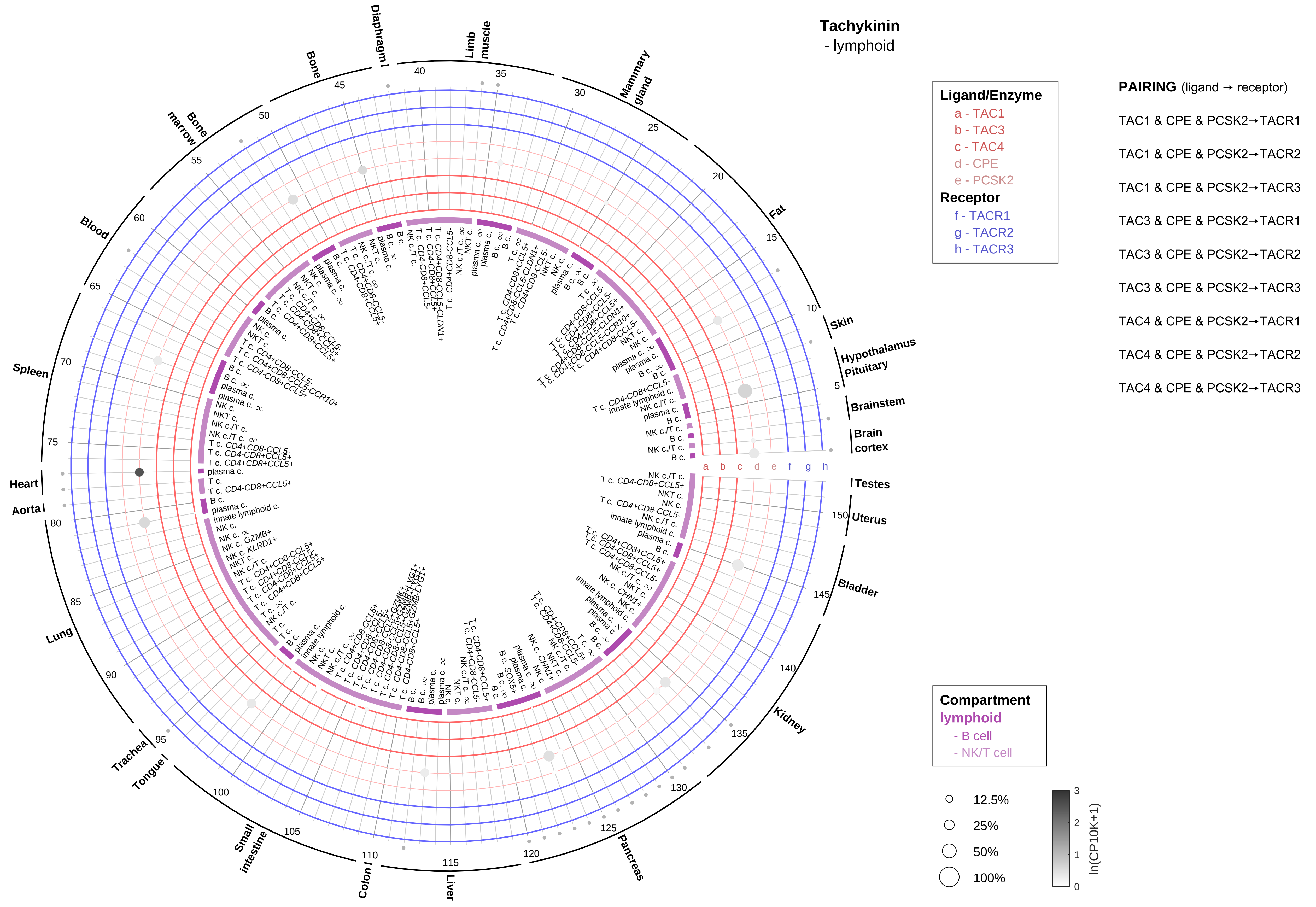
Tachykinin
- lymphoid
Ligand/Enzyme
a - TAC1
b - TAC3
c - TAC4
d - CPE
e - PCSK2
Receptor
f - TACR1
g - TACR2
h - TACR3
PAIRING (ligand → receptor)
TAC1 & CPE & PCSK2→TACR1
TAC1 & CPE & PCSK2→TACR2
TAC1 & CPE & PCSK2→TACR3
TAC3 & CPE & PCSK2→TACR1
TAC3 & CPE & PCSK2→TACR2
TAC3 & CPE & PCSK2→TACR3
TAC4 & CPE & PCSK2→TACR1
TAC4 & CPE & PCSK2→TACR2
TAC4 & CPE & PCSK2→TACR3
Compartment
lymphoid
- B cell
- NK/T cell
12.5%
25%
50%
100%
ln(CP10K+1)
Diaphragm
Limb muscle
Mammary gland
Fat
Skin
Hypothalamus
Pituitary
Brainstem
Brain cortex
Testes
Uterus
Bladder
Kidney
Pancreas
Liver
Colon
Small intestine
Tongue
Trachea
Lung
Aorta
Heart
Spleen
Blood
Bone marrow
Bone

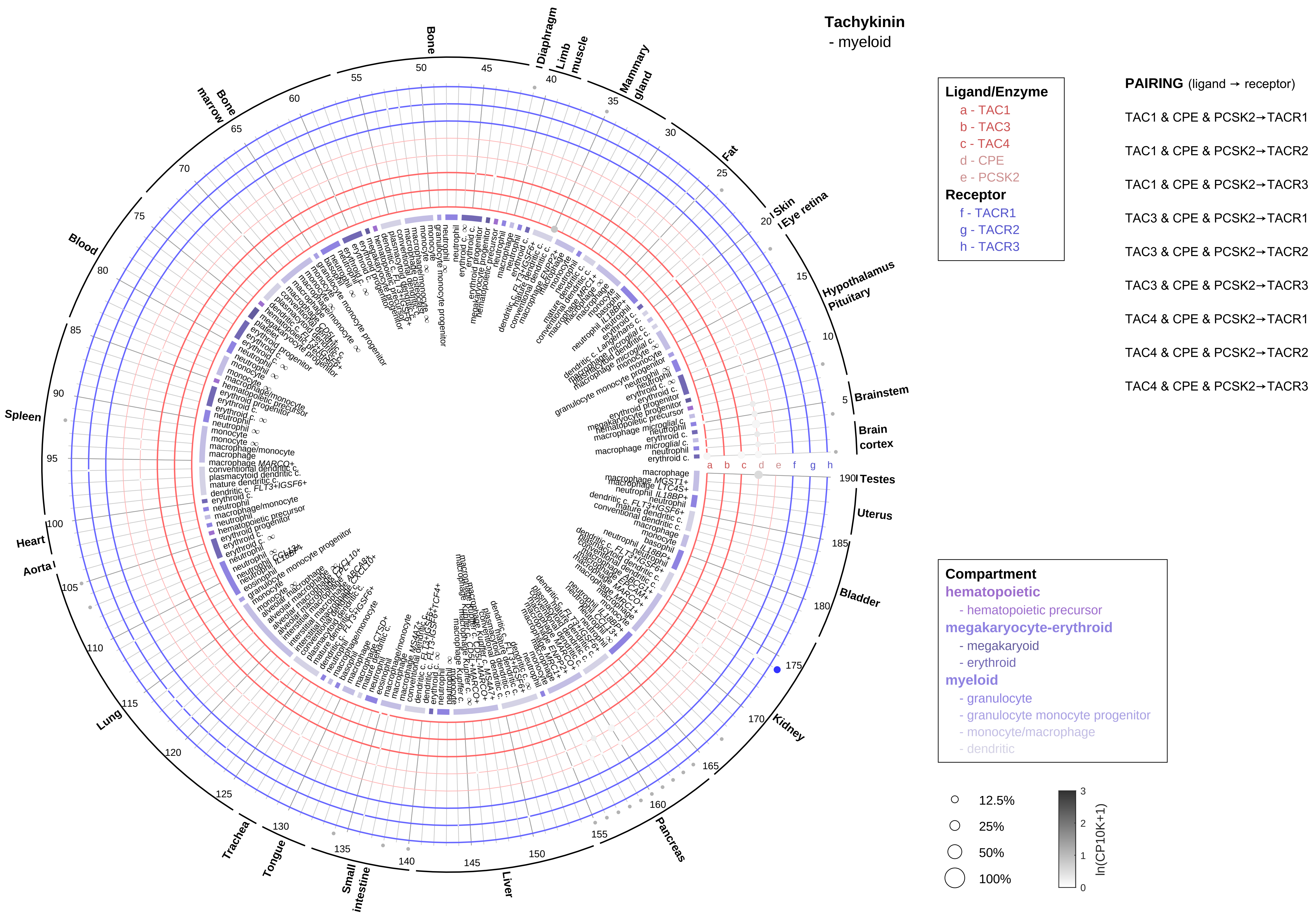
Tachykinin
- myeloid
Ligand/Enzyme
a - TAC1
b - TAC3
c - TAC4
d - CPE
e - PCSK2
Receptor
f - TACR1
g - TACR2
h - TACR3
PAIRING (ligand → receptor)
TAC1 & CPE & PCSK2→TACR1
TAC1 & CPE & PCSK2→TACR2
TAC1 & CPE & PCSK2→TACR3
TAC3 & CPE & PCSK2→TACR1
TAC3 & CPE & PCSK2→TACR2
TAC3 & CPE & PCSK2→TACR3
TAC4 & CPE & PCSK2→TACR1
TAC4 & CPE & PCSK2→TACR2
TAC4 & CPE & PCSK2→TACR3
Compartment
hematopoietic
- hematopoietic precursor
megakaryocyte-erythroid
- megakaryoid
- erythroid
myeloid
- granulocyte
- granulocyte monocyte progenitor
- monocyte/macrophage
- dendritic
12.5%
25%
50%
100%
ln(CP10K+1)
Bone
Diaphragm
Limb muscle
Mammary gland
Fat
Skin
Eye retina
Hypothalamus
Pituitary
Brainstem
Brain cortex
Testes
Uterus
Bladder
Kidney
Pancreas
Liver
Small intestine
Tongue
Trachea
Lung
Aorta
Heart
Spleen
Blood
Bone marrow

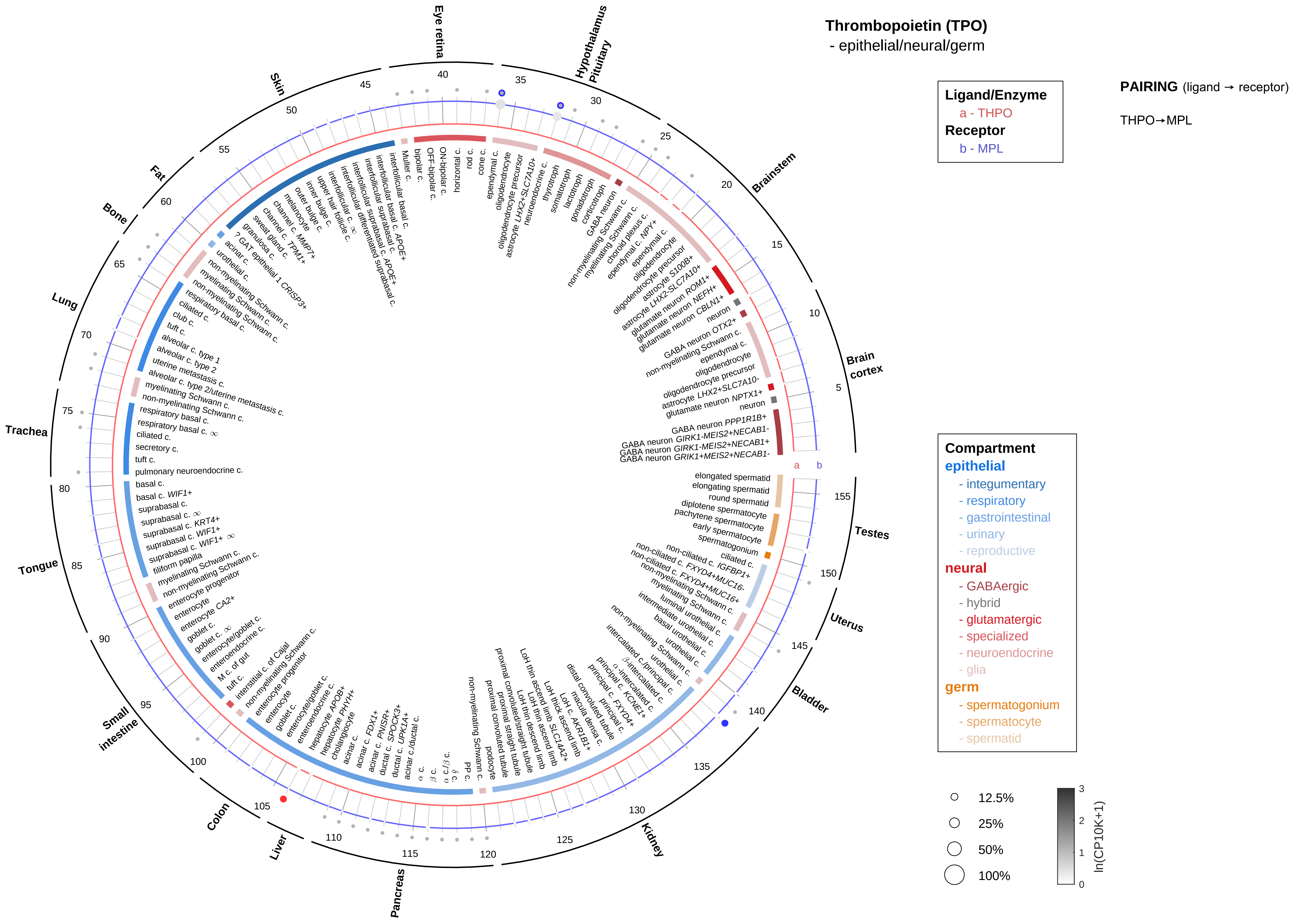
Thrombopoietin (TPO)
- epithelial/neural/germ
Ligand/Enzyme
a - THPO
Receptor
b - MPL
PAIRING (ligand → receptor)
THPO→MPL
Compartment
epithelial
- integumentary
- respiratory
- gastrointestinal
- urinary
- reproductive
neural
- GABAergic
- hybrid
- glutamatergic
- specialized
- neuroendocrine
- glia
germ
- spermatogonium
- spermatocyte
- spermatid
12.5%
25%
50%
100%
ln(CP10K+1)
Eye retina
Hypothalamus
Pituitary
Brainstem
Brain cortex
Testes
Uterus
Bladder
Kidney
Pancreas
Liver
Colon
Small intestine
Tongue
Trachea
Lung
Bone
Fat
Skin

#### Thrombopoietin (TPO)
- endothelial

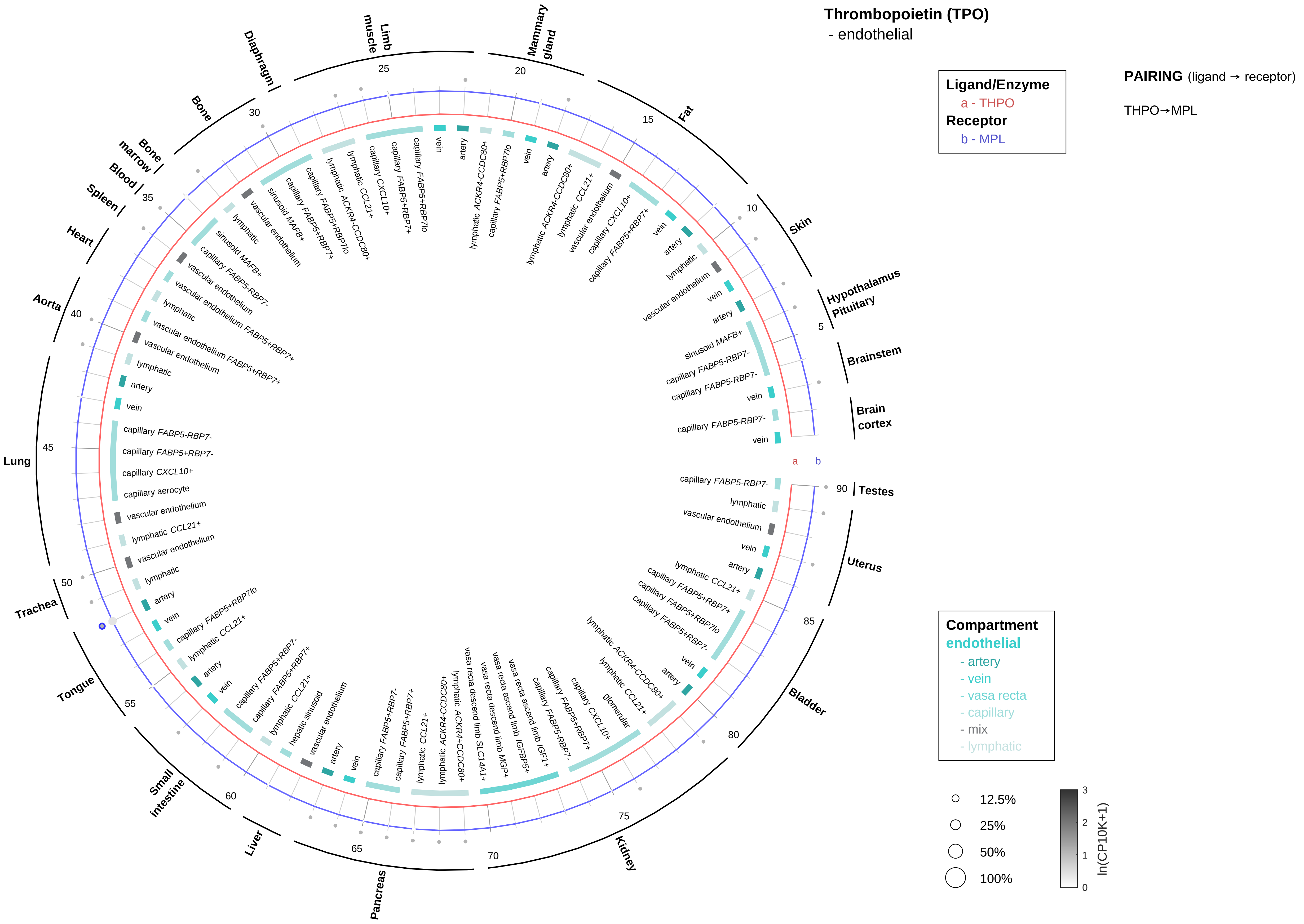

#### Thrombopoietin (TPO)
- stromal

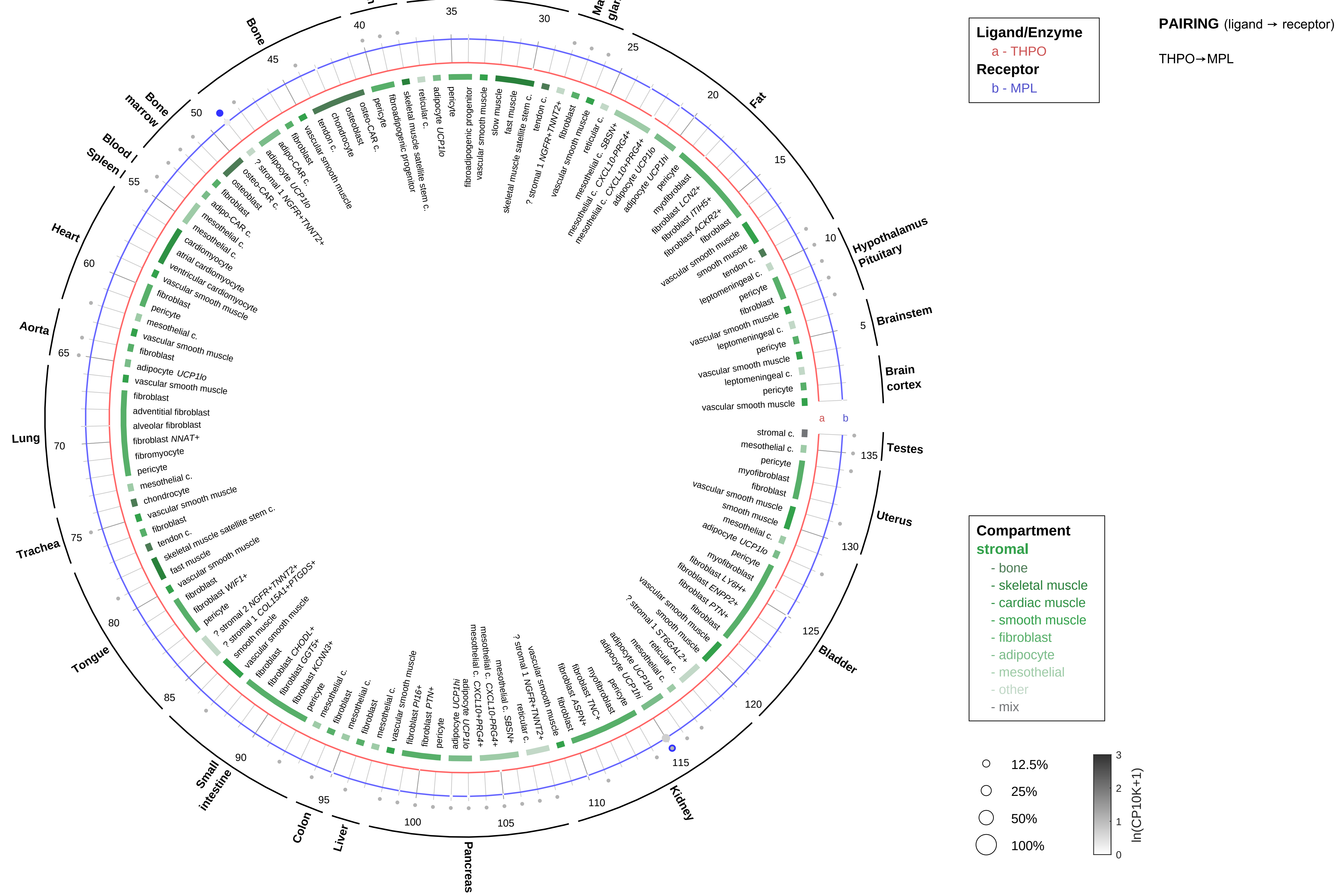

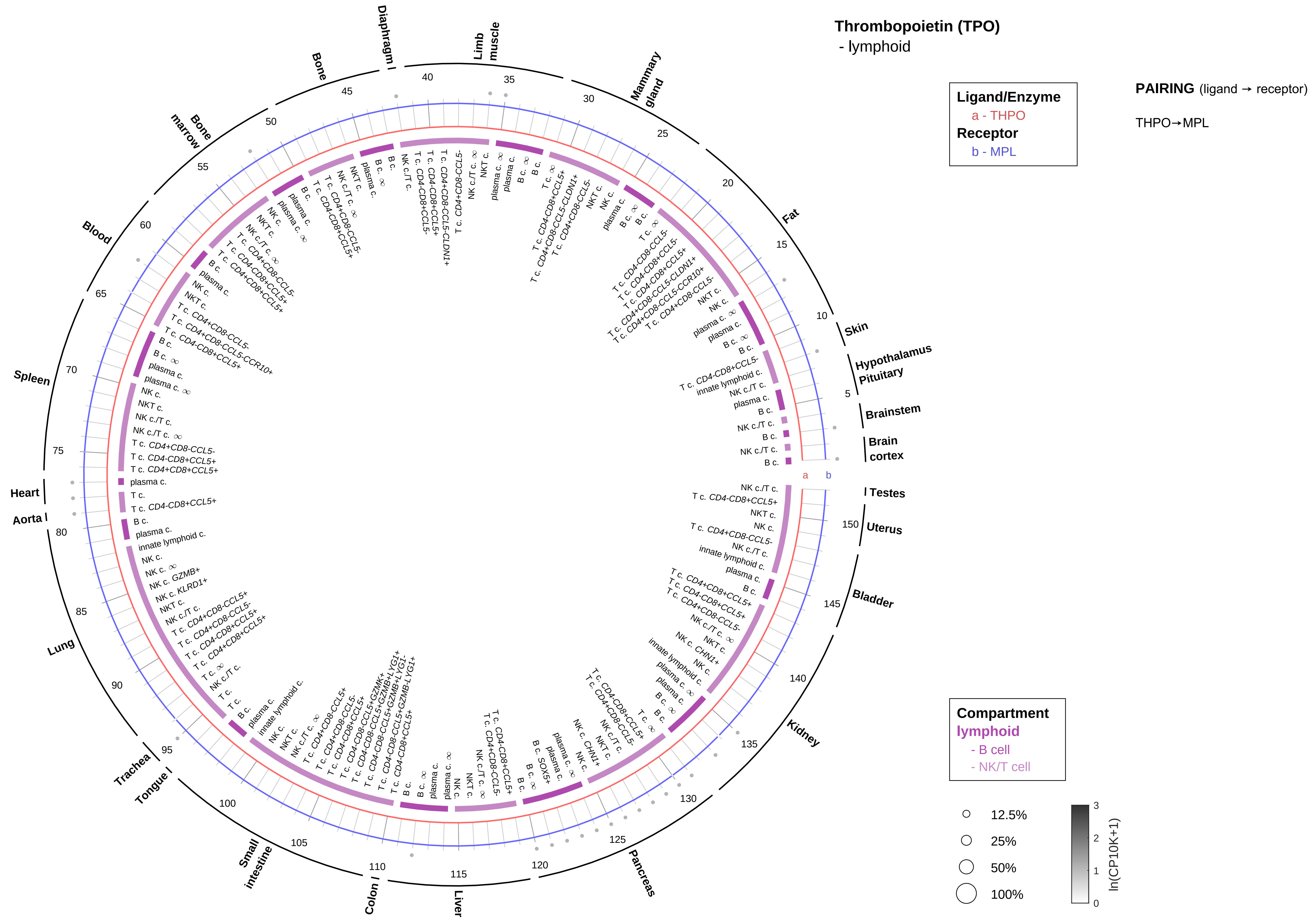
Thrombopoietin (TPO)
- lymphoid
Ligand/Enzyme
a - THPO
Receptor
b - MPL
PAIRING (ligand → receptor)
THPO→MPL
Compartment
lymphoid
- B cell
- NK/T cell
12.5%
25%
50%
100%
ln(CP10K+1)
Skin
Hypothalamus
Pituitary
Brainstem
Brain cortex
Testes
Uterus
Bladder
Kidney
Pancreas
Liver
Colon
Small intestine
Tongue
Trachea
Lung
Aorta
Heart
Spleen
Blood
Bone marrow
Bone
Diaphragm
Limb muscle
Mammary gland
Fat

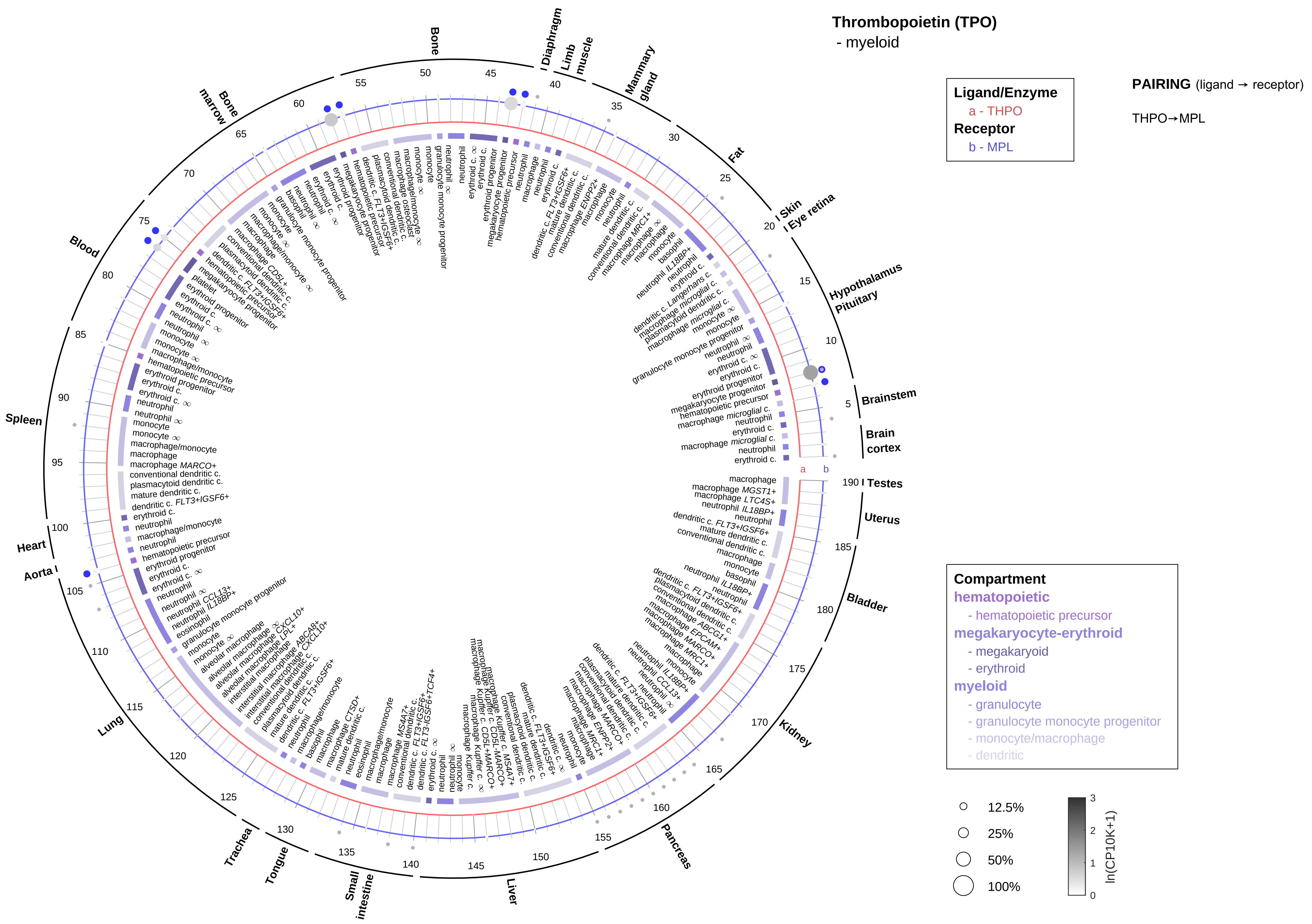

Thrombopoietin (TPO)
- myeloid
Ligand/Enzyme
a - THPO
Receptor
b - MPL
PAIRING (ligand → receptor)
THPO→MPL
Compartment
hematopoietic
- hematopoietic precursor
megakaryocyte-erythroid
- megakaryoid
- erythroid
myeloid
- granulocyte
- granulocyte monocyte progenitor
- monocyte/macrophage
- dendritic
12.5%
25%
50%
100%
ln(CP10K+1)
Brainstem
Brain cortex
Hypothalamus
Pituitary
Eye retina
Skin
Fat
Mammary gland
Limb muscle
Diaphragm
Bone
Bone marrow
Blood
Spleen
Heart
Aorta
Lung
Trachea
Tongue
Small intestine
Liver
Pancreas
Kidney
Bladder
Uterus
Testes

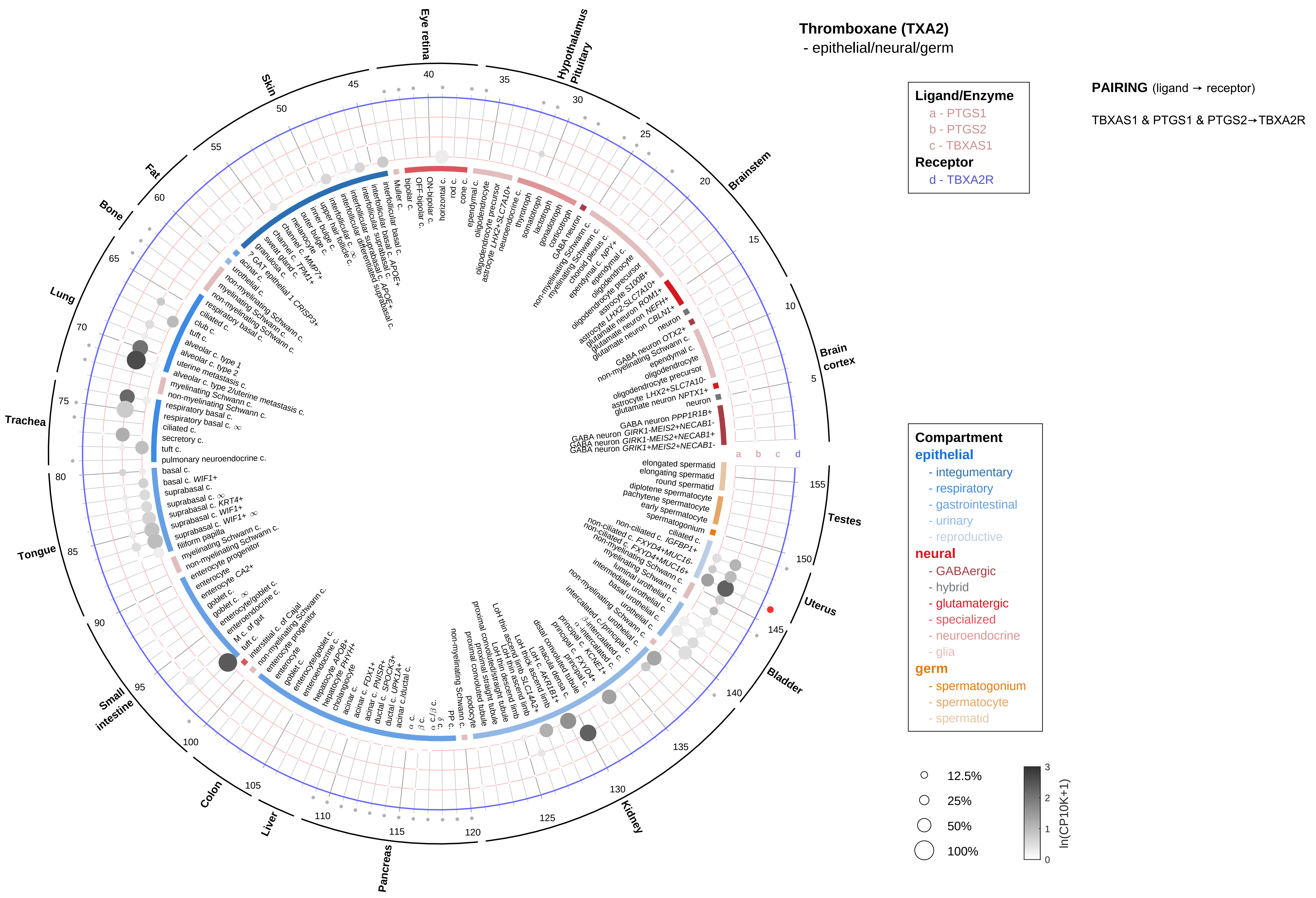

Thromboxane (TXA2)
- epithelial/neural/germ
Ligand/Enzyme
a - PTGS1
b - PTGS2
c - TBXAS1
Receptor
d - TBXA2R
PAIRING (ligand → receptor)
TBXAS1 & PTGS1 & PTGS2→TBXA2R
Compartment
epithelial
- integumentary
- respiratory
- gastrointestinal
- urinary
- reproductive
neural
- GABAergic
- hybrid
- glutamatergic
- specialized
- neuroendocrine
- glia
germ
- spermatogonium
- spermatocyte
- spermatid
12.5%
25%
50%
100%
ln(CP10K+1)
Eye retina
Hypothalamus
Pituitary
Brainstem
Brain cortex
Testes
Uterus
Bladder
Kidney
Pancreas
Liver
Colon
Small intestine
Tongue
Trachea
Lung
Bone
Fat
Skin

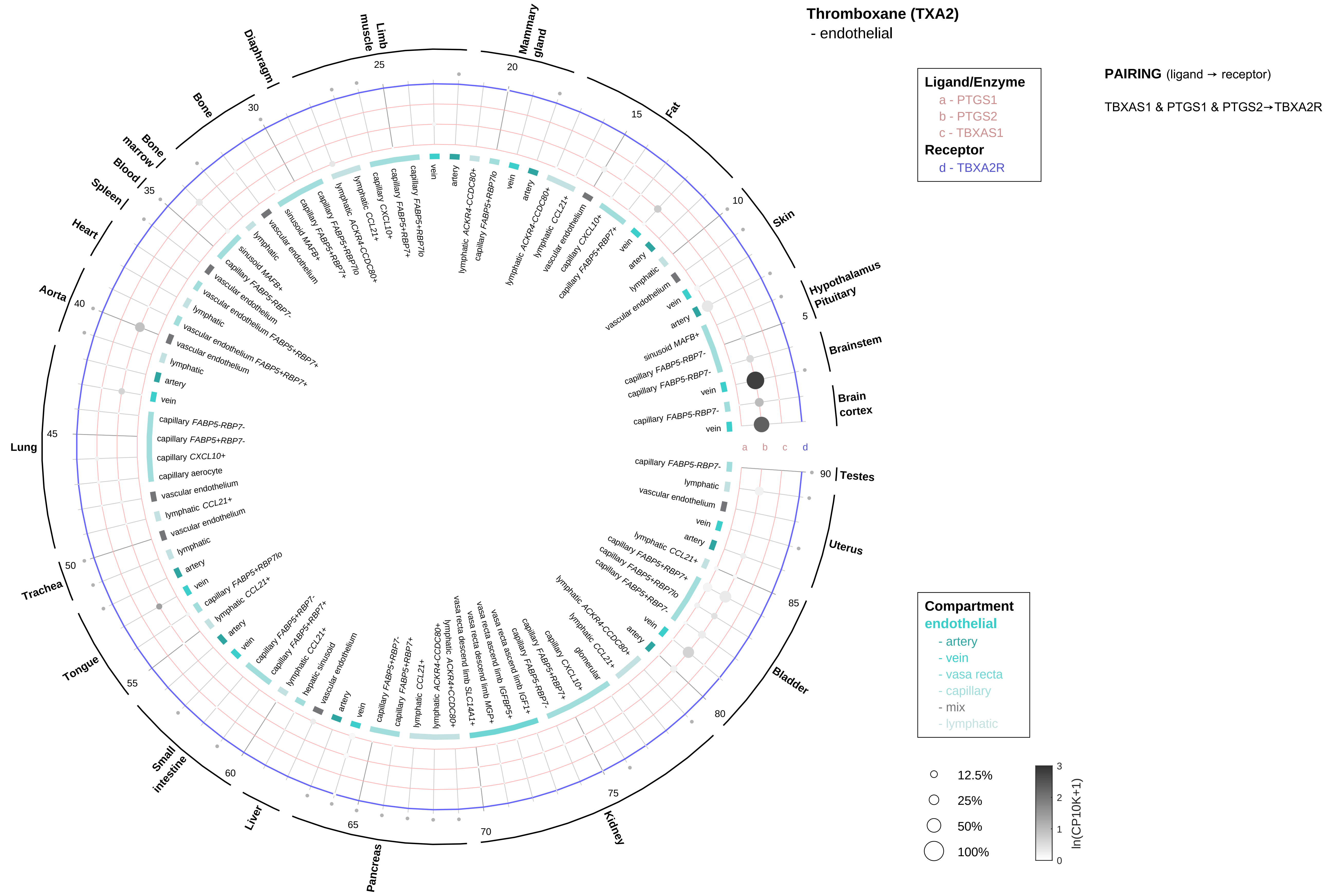

Thromboxane (TXA2)
- endothelial
Ligand/Enzyme
a - PTGS1
b - PTGS2
c - TBXAS1
Receptor
d - TBXA2R
PAIRING (ligand → receptor)
TBXAS1 & PTGS1 & PTGS2→TBXA2R
Compartment
endothelial
- artery
- vein
- vasa recta
- capillary
- mix
- lymphatic
12.5%
25%
50%
100%
ln(CP10K+1)
Brain cortex
Brainstem
Hypothalamus Pituitary
Skin
Fat
Mammary gland
Limb muscle
Diaphragm
Bone
Bone marrow
Blood
Spleen
Heart
Aorta
Lung
Trachea
Tongue
Small intestine
Liver
Pancreas
Kidney
Bladder
Uterus
Testes

#### Thromboxane (TXA2)
- stromal

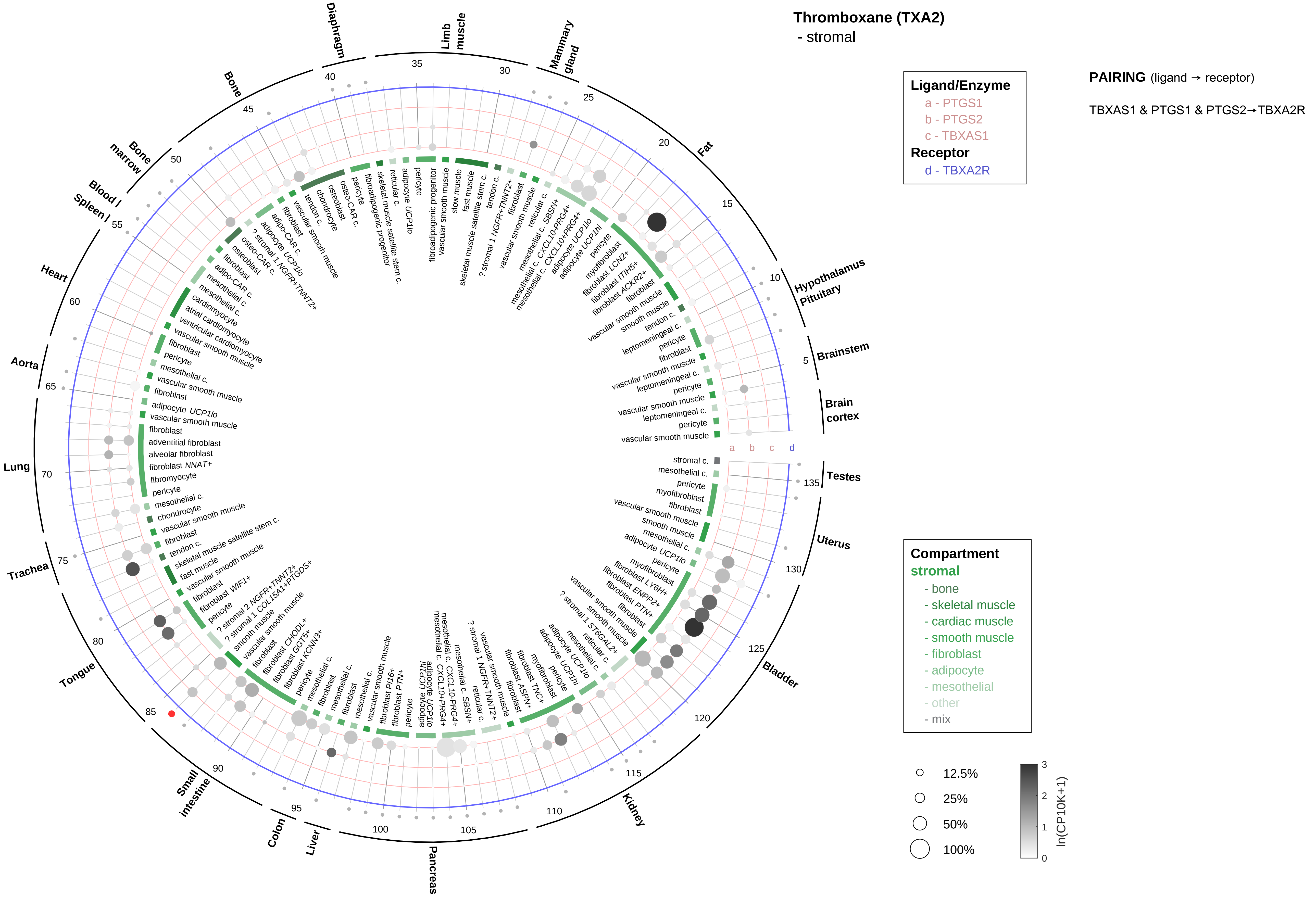

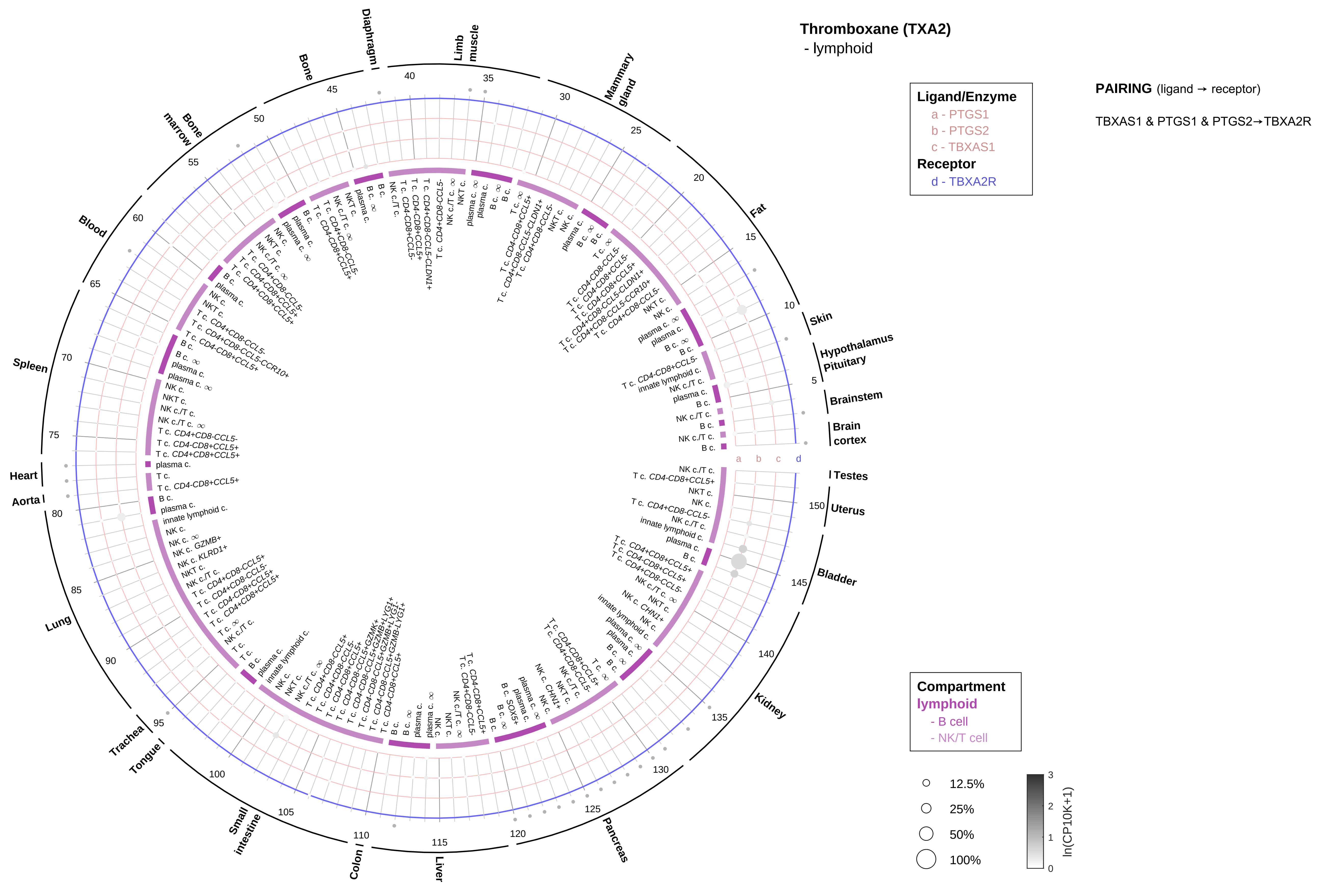

Thromboxane (TXA2)
- lymphoid
Ligand/Enzyme
a - PTGS1
b - PTGS2
c - TBXAS1
Receptor
d - TBXA2R
PAIRING (ligand → receptor)
TBXAS1 & PTGS1 & PTGS2→TBXA2R
Compartment
lymphoid
- B cell
- NK/T cell
12.5%
25%
50%
100%
ln(CP10K+1)

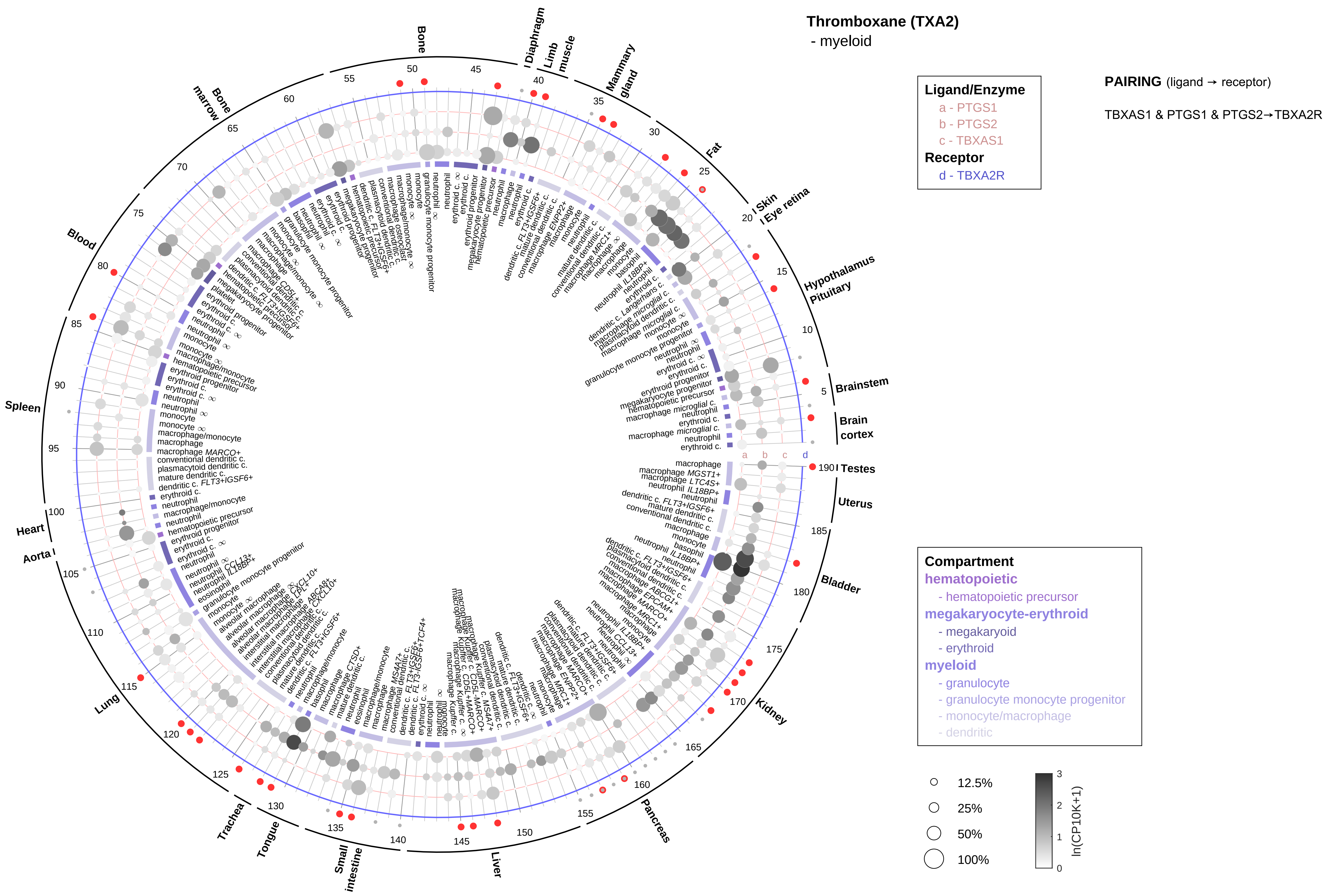

Thromboxane (TXA2)
- myeloid
Ligand/Enzyme
a - PTGS1
b - PTGS2
c - TBXAS1
Receptor
d - TBXA2R
PAIRING (ligand → receptor)
TBXAS1 & PTGS1 & PTGS2→TBXA2R
Compartment
hematopoietic
- hematopoietic precursor
megakaryocyte-erythroid
- megakaryoid
- erythroid
myeloid
- granulocyte
- granulocyte monocyte progenitor
- monocyte/macrophage
- dendritic
12.5%
25%
50%
100%
ln(CP10K+1)
Bone
Diaphragm
Limb muscle
Mammary gland
Fat
Skin
Eye retina
Hypothalamus
Pituitary
Brainstem
Brain cortex
Testes
Uterus
Bladder
Kidney
Pancreas
Liver
Small intestine
Tongue
Trachea
Lung
Aorta
Heart
Spleen
Blood
Bone marrow

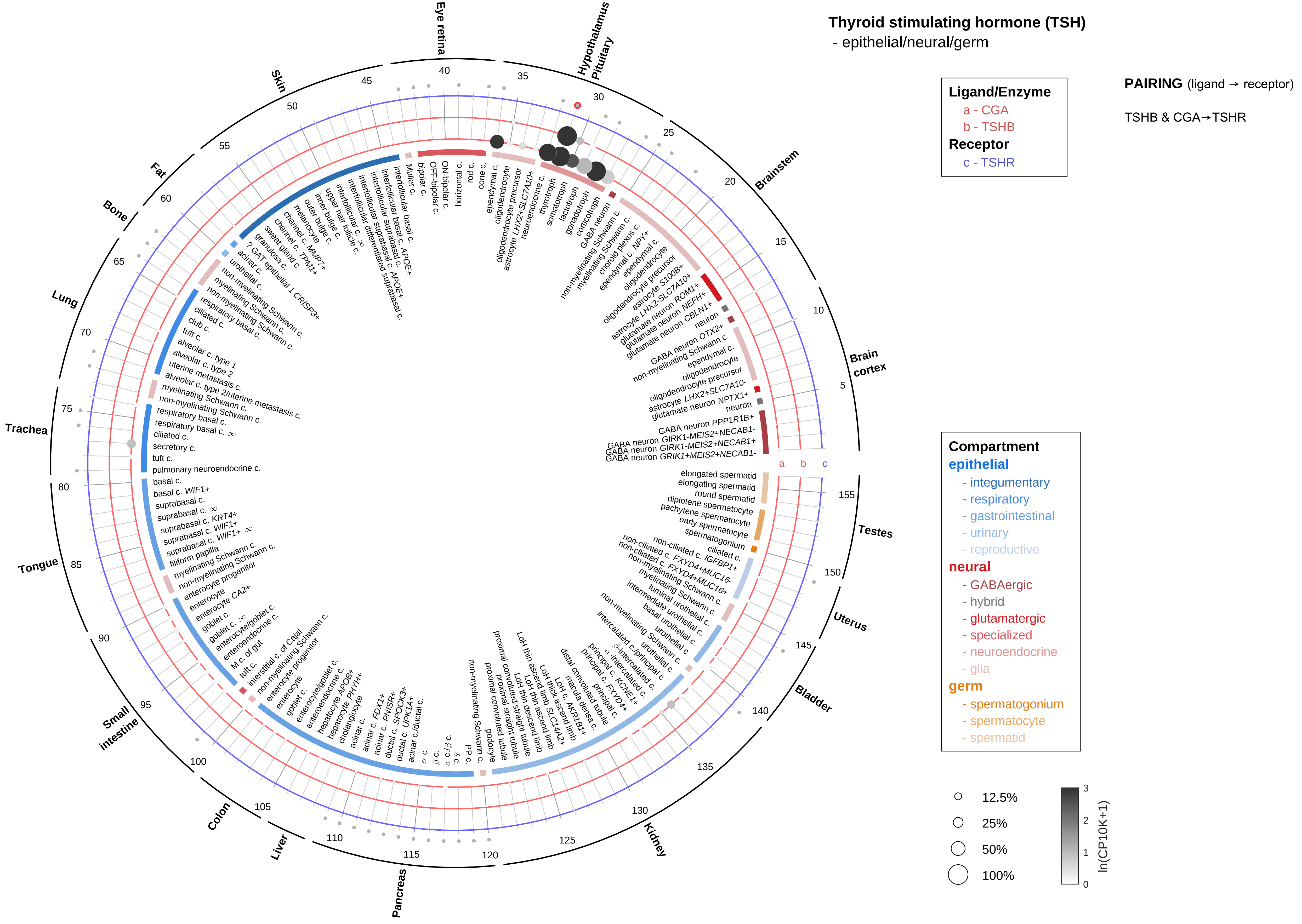

Thyroid stimulating hormone (TSH)
- epithelial/neural/germ
Ligand/Enzyme
a - CGA
b - TSHB
Receptor
c - TSHR
PAIRING (ligand → receptor)
TSHB & CGA→TSHR
Compartment
epithelial
- integumentary
- respiratory
- gastrointestinal
- urinary
- reproductive
neural
- GABAergic
- hybrid
- glutamatergic
- specialized
- neuroendocrine
- glia
germ
- spermatogonium
- spermatocyte
- spermatid
12.5%
25%
50%
100%
ln(CP10K+1)
Eye retina
Hypothalamus
Pituitary
Brainstem
Brain cortex
Testes
Uterus
Bladder
Kidney
Pancreas
Liver
Colon
Small intestine
Tongue
Trachea
Lung
Bone
Fat
Skin

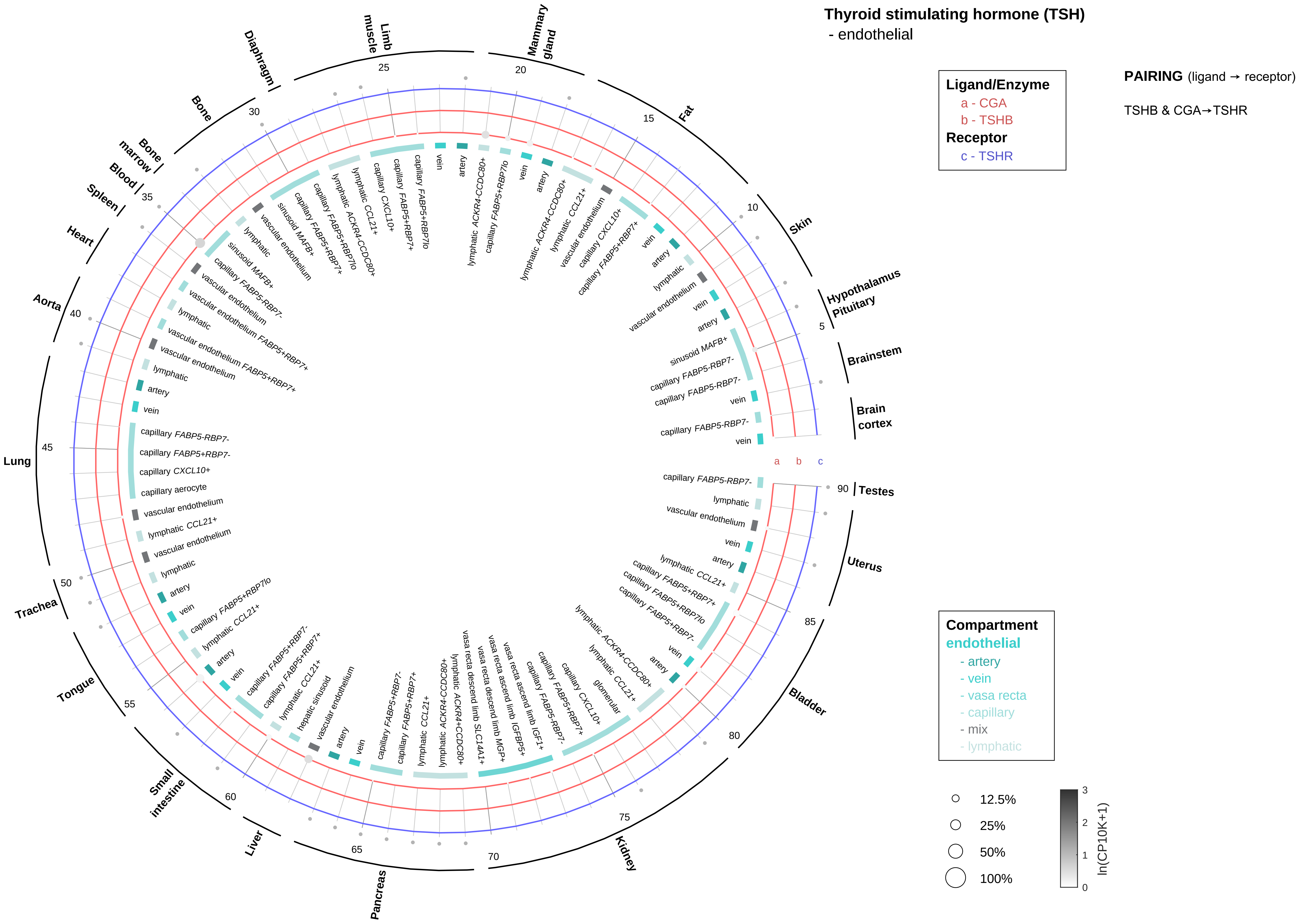

Thyroid stimulating hormone (TSH)
- endothelial
Ligand/Enzyme
a - CGA
b - TSHB
Receptor
c - TSHR
PAIRING (ligand → receptor)
TSHB & CGA→TSHR
Compartment
endothelial
- artery
- vein
- vasa recta
- capillary
- mix
- lymphatic
12.5%
25%
50%
100%
ln(CP10K+1)
Brain cortex
Brainstem
Hypothalamus Pituitary
Skin
Fat
Mammary gland
Limb muscle
Diaphragm
Bone
Bone marrow
Blood
Spleen
Heart
Aorta
Lung
Trachea
Tongue
Small intestine
Liver
Pancreas
Kidney
Bladder
Uterus
Testes

#### Thyroid stimulating hormone (TSH)
- stromal

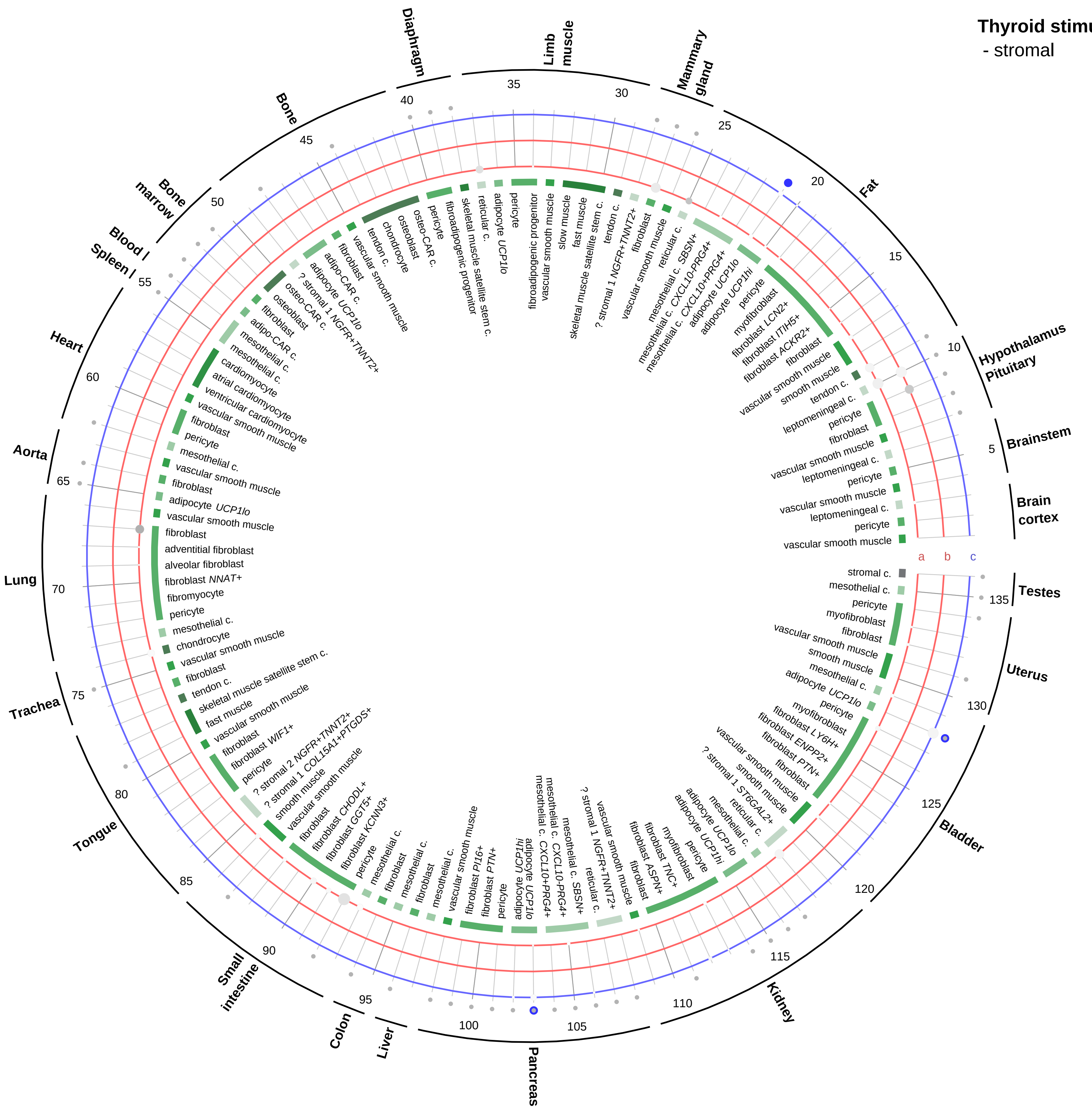

**Ligand/Enzyme**
a - CGA
b - TSHB
**Receptor**
c - TSHR

**PAIRING** (ligand → receptor)

TSHB & CGA→TSHR

**Compartment**
**stromal**
- bone
- skeletal muscle
- cardiac muscle
- smooth muscle
- fibroblast
- adipocyte
- mesothelial
- other
- mix

12.5%
25%
50%
100%

ln(CP10K+1): 0, 1, 2, 3

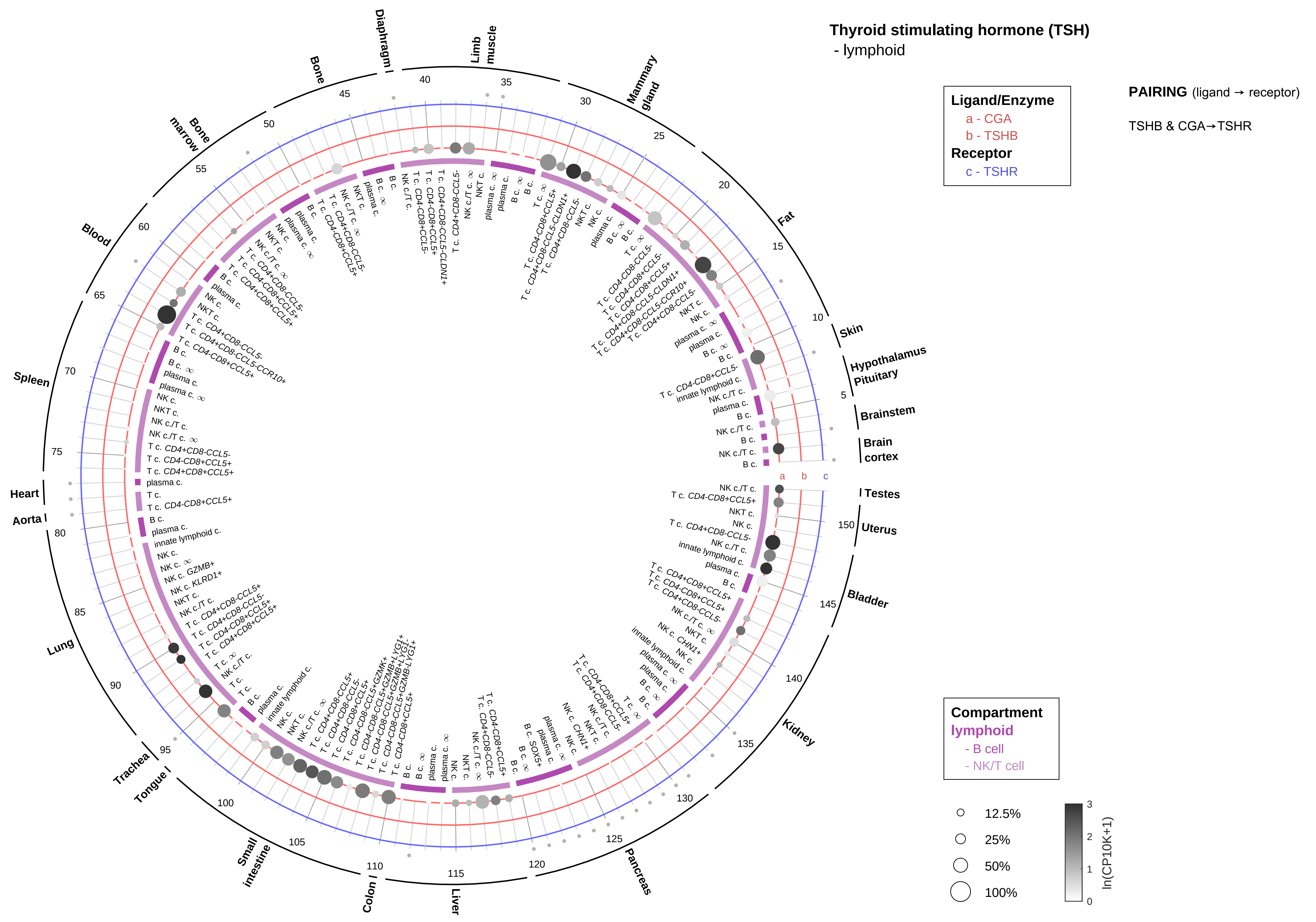

Thyroid stimulating hormone (TSH)
- lymphoid
Ligand/Enzyme
a - CGA
b - TSHB
Receptor
c - TSHR
PAIRING (ligand → receptor)
TSHB & CGA→TSHR
Compartment
lymphoid
- B cell
- NK/T cell
12.5%
25%
50%
100%
ln(CP10K+1)
Diaphragm
Limb muscle
Mammary gland
Fat
Skin
Hypothalamus
Pituitary
Brainstem
Brain cortex
Testes
Uterus
Bladder
Kidney
Pancreas
Liver
Colon
Small intestine
Tongue
Trachea
Lung
Aorta
Heart
Spleen
Blood
Bone marrow
Bone

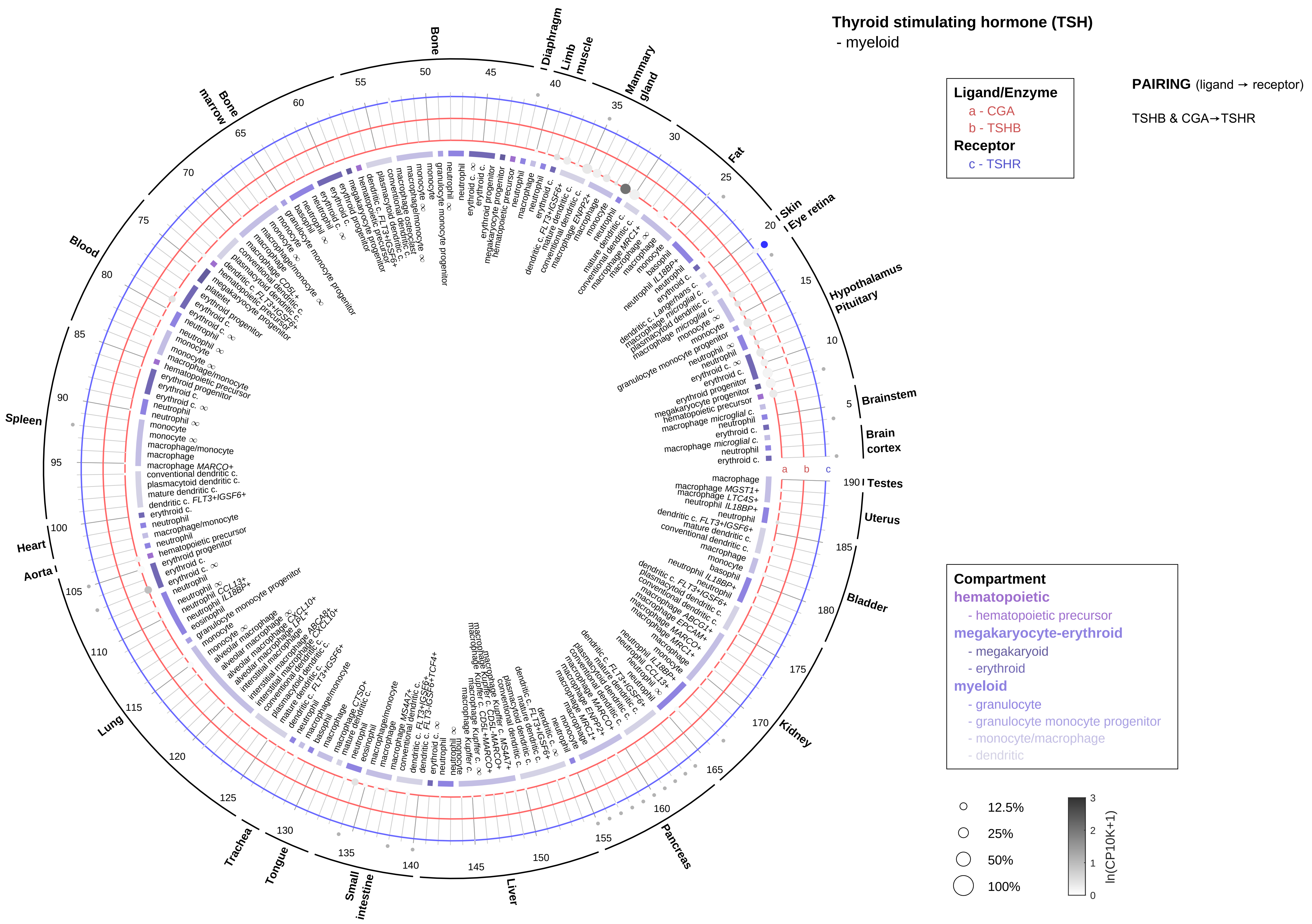

Thyroid stimulating hormone (TSH)
- myeloid
Ligand/Enzyme
a - CGA
b - TSHB
Receptor
c - TSHR
PAIRING (ligand → receptor)
TSHB & CGA→TSHR
Compartment
hematopoietic
- hematopoietic precursor
megakaryocyte-erythroid
- megakaryoid
- erythroid
myeloid
- granulocyte
- granulocyte monocyte progenitor
- monocyte/macrophage
- dendritic
12.5%
25%
50%
100%
ln(CP10K+1)
Brainstem
Brain cortex
Testes
Uterus
Bladder
Kidney
Pancreas
Liver
Small intestine
Tongue
Trachea
Lung
Aorta
Heart
Spleen
Blood
Bone marrow
Bone
Diaphragm
Limb muscle
Mammary gland
Fat
Skin
Eye retina
Hypothalamus
Pituitary

**Thyrotropin-releasing hormone (TRH)**
- epithelial/neural/germ

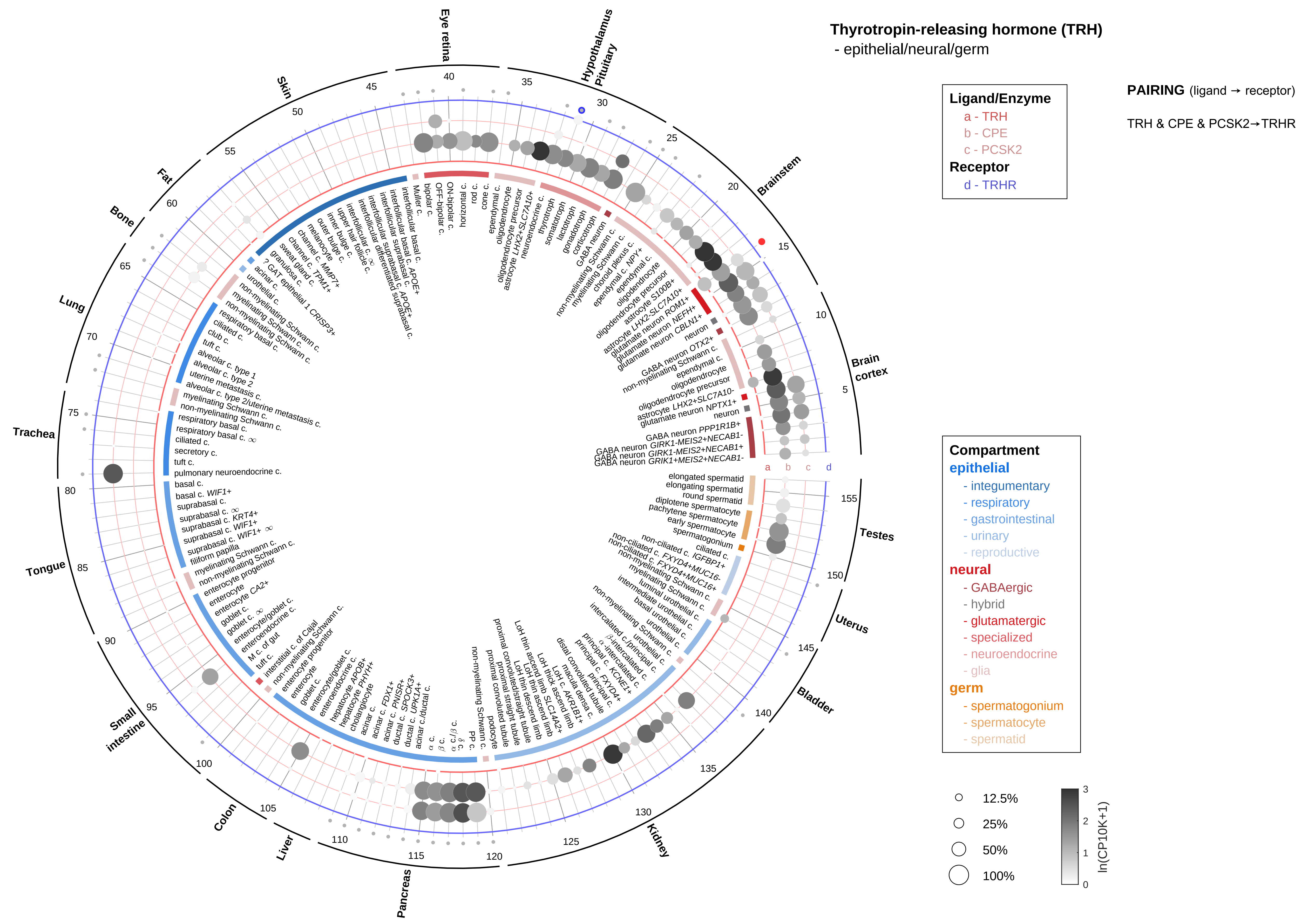

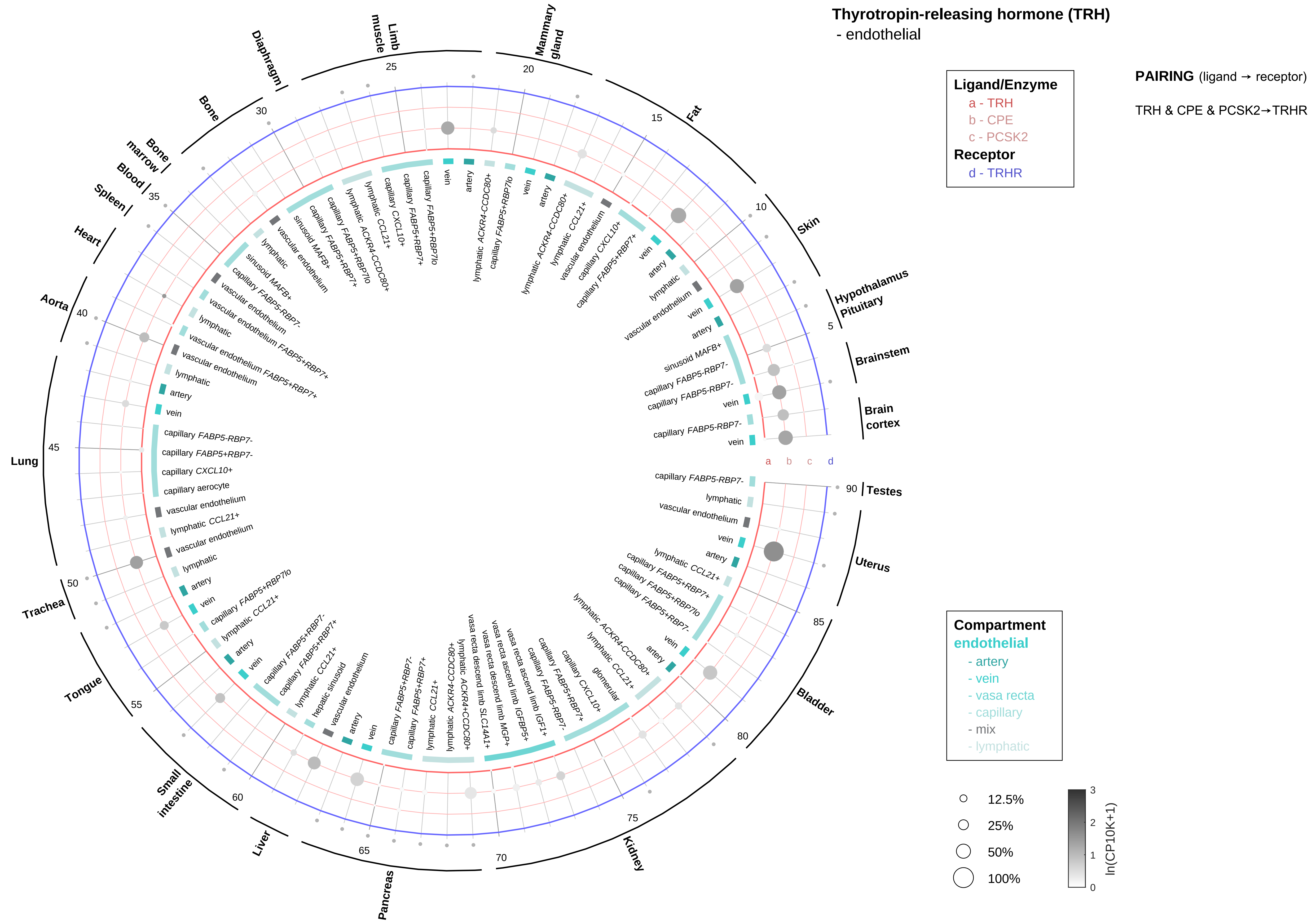

Thyrotropin-releasing hormone (TRH)
- endothelial
Ligand/Enzyme
a - TRH
b - CPE
c - PCSK2
Receptor
d - TRHR
PAIRING (ligand → receptor)
TRH & CPE & PCSK2→TRHR
Compartment
endothelial
- artery
- vein
- vasa recta
- capillary
- mix
- lymphatic
12.5%
25%
50%
100%
ln(CP10K+1)
Hypothalamus
Pituitary
Brainstem
Brain cortex
Testes
Uterus
Bladder
Kidney
Pancreas
Liver
Small intestine
Tongue
Trachea
Lung
Aorta
Heart
Spleen
Blood
Bone marrow
Bone
Diaphragm
Limb muscle
Mammary gland
Fat
Skin
a
b
c
d

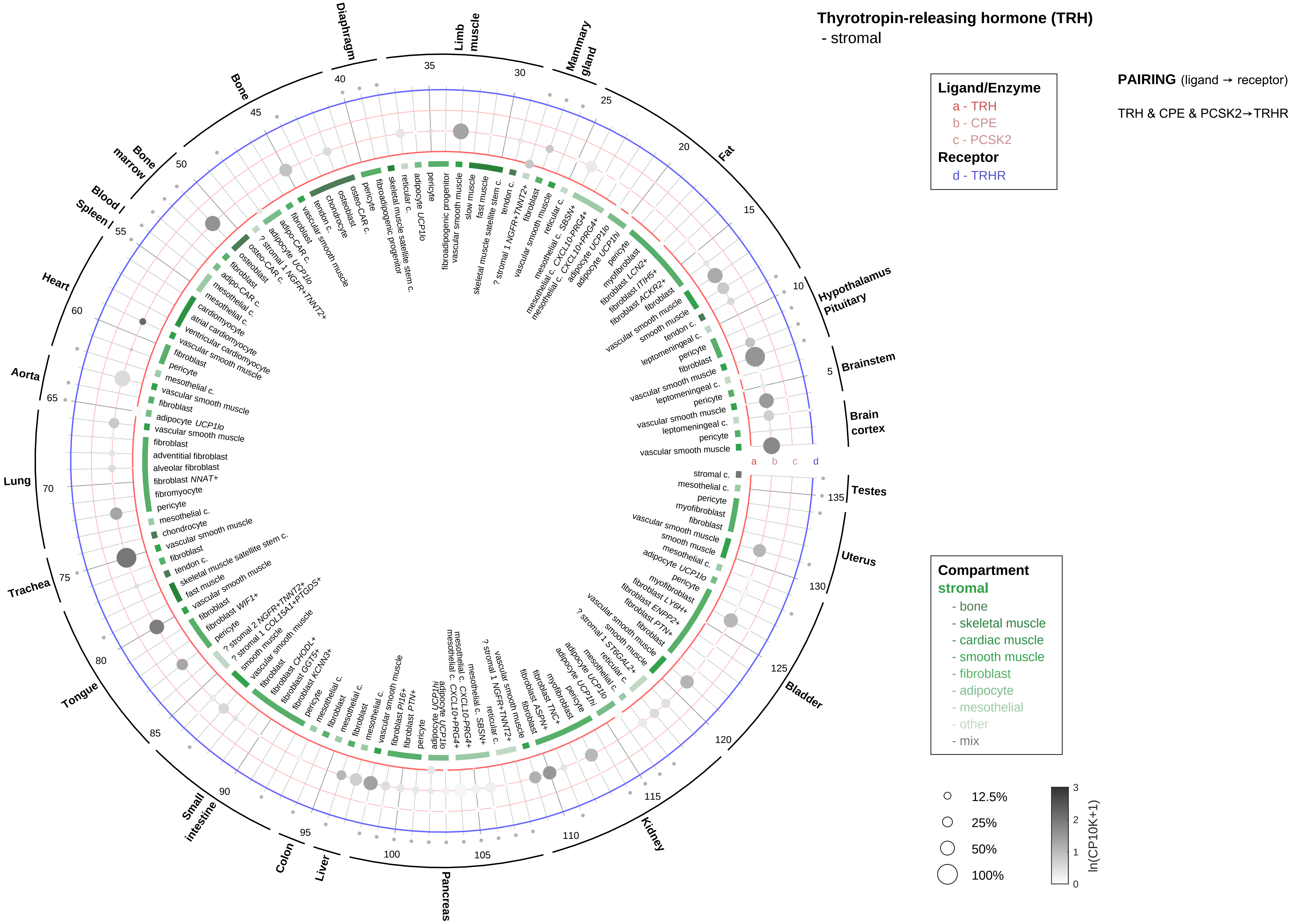
Thyrotropin-releasing hormone (TRH)
- stromal
Ligand/Enzyme
a - TRH
b - CPE
c - PCSK2
Receptor
d - TRHR
PAIRING (ligand → receptor)
TRH & CPE & PCSK2→TRHR
Compartment
stromal
- bone
- skeletal muscle
- cardiac muscle
- smooth muscle
- fibroblast
- adipocyte
- mesothelial
- other
- mix
12.5%
25%
50%
100%
ln(CP10K+1)
Brain cortex
Brainstem
Hypothalamus Pituitary
Fat
Mammary gland
Limb muscle
Diaphragm
Bone
Bone marrow
Blood / Spleen
Heart
Aorta
Lung
Trachea
Tongue
Small intestine
Colon
Liver
Pancreas
Kidney
Bladder
Uterus
Testes

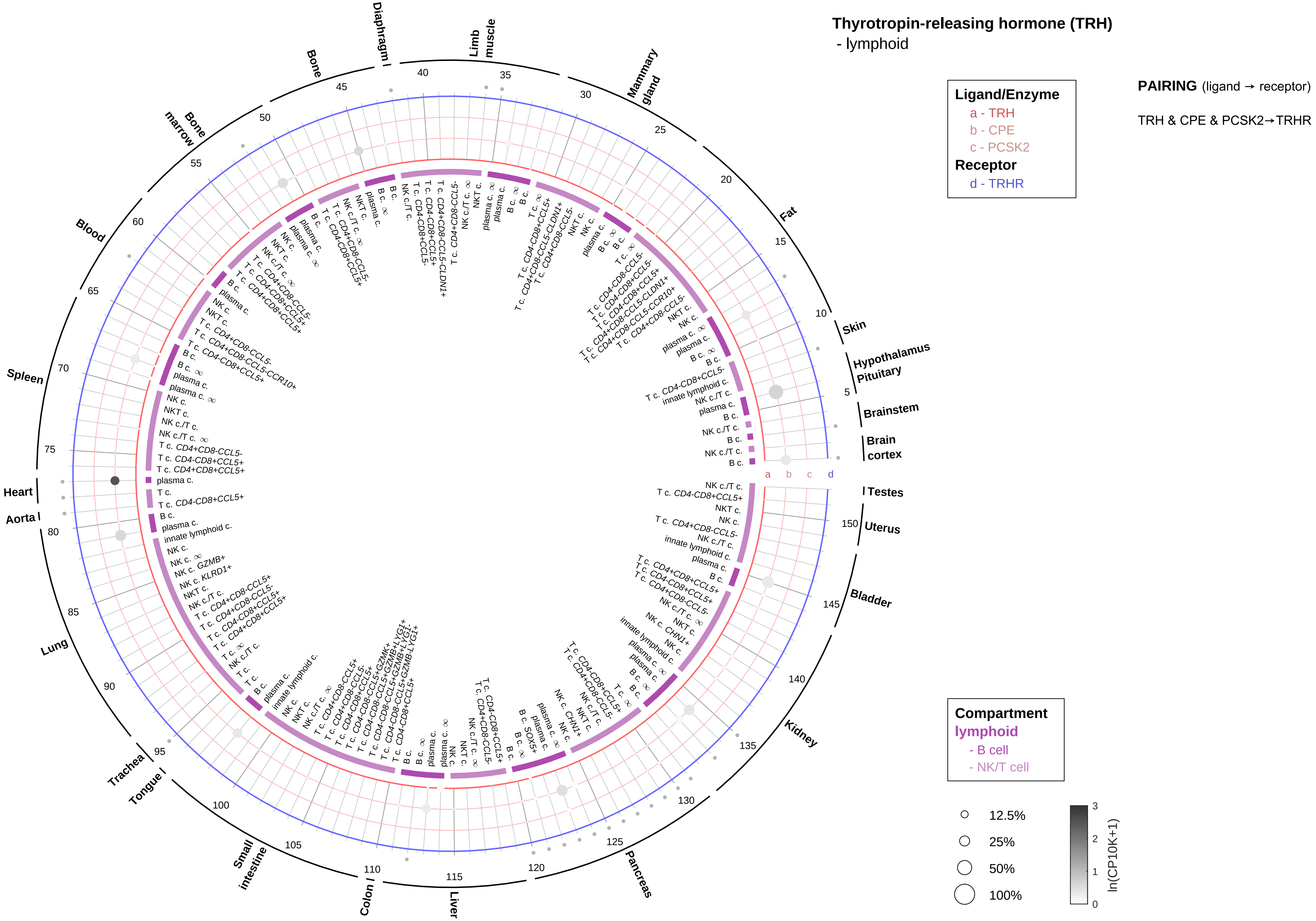

Thyrotropin-releasing hormone (TRH)
- lymphoid
Ligand/Enzyme
a - TRH
b - CPE
c - PCSK2
Receptor
d - TRHR
PAIRING (ligand → receptor)
TRH & CPE & PCSK2→TRHR
Compartment
lymphoid
- B cell
- NK/T cell
12.5%
25%
50%
100%
ln(CP10K+1)
Skin
Hypothalamus
Pituitary
Brainstem
Brain cortex
Testes
Uterus
Bladder
Kidney
Pancreas
Liver
Colon
Small intestine
Tongue
Trachea
Lung
Aorta
Heart
Spleen
Blood
Bone marrow
Bone
Diaphragm
Limb muscle
Mammary gland
Fat

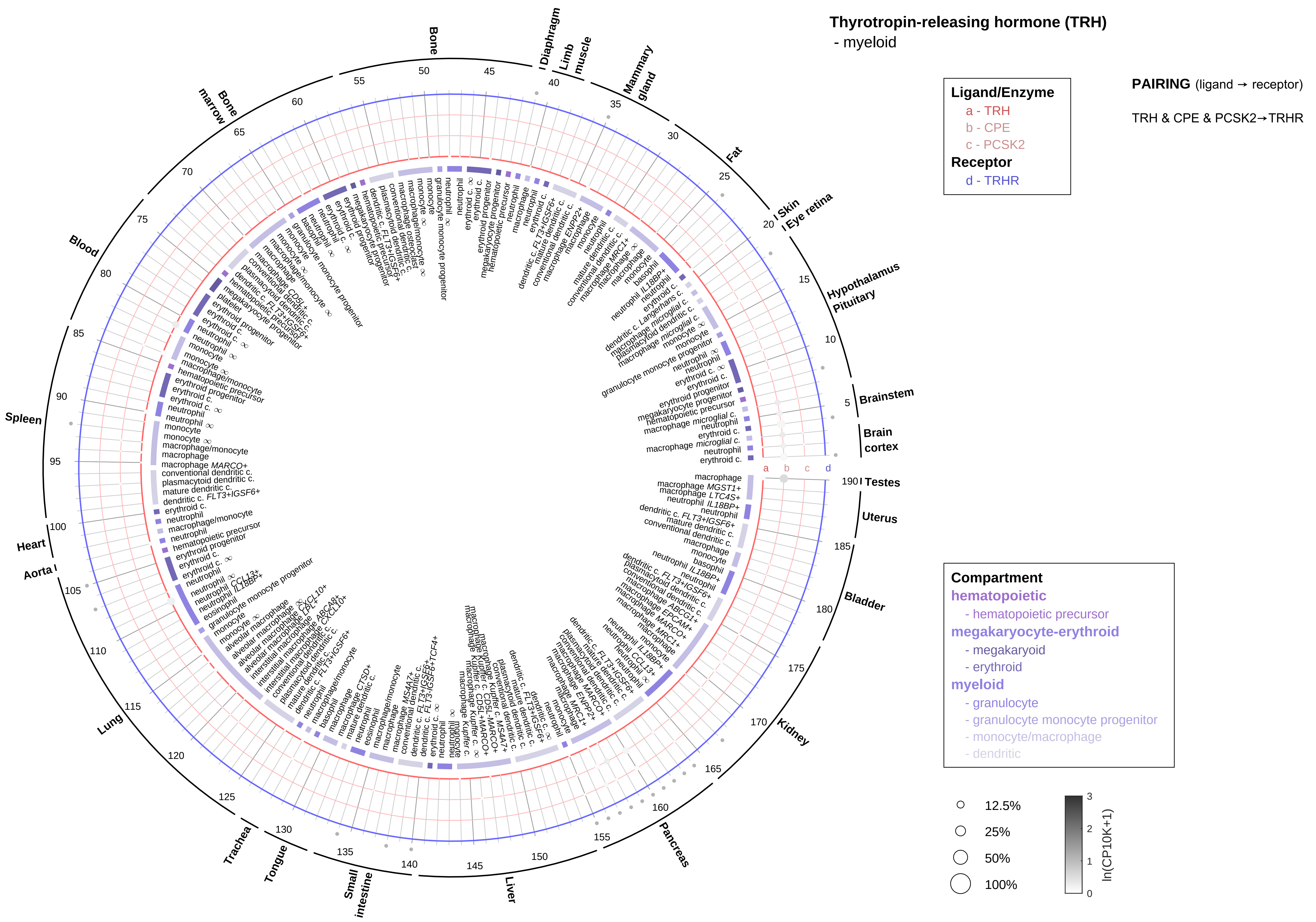
Thyrotropin-releasing hormone (TRH)
- myeloid
Ligand/Enzyme
a - TRH
b - CPE
c - PCSK2
Receptor
d - TRHR
PAIRING (ligand → receptor)
TRH & CPE & PCSK2→TRHR
Compartment
hematopoietic
- hematopoietic precursor
megakaryocyte-erythroid
- megakaryoid
- erythroid
myeloid
- granulocyte
- granulocyte monocyte progenitor
- monocyte/macrophage
- dendritic
12.5%
25%
50%
100%
ln(CP10K+1)
Testes
Brain cortex
Brainstem
Hypothalamus
Pituitary
Eye retina
Skin
Fat
Mammary gland
Limb muscle
Diaphragm
Bone
Bone marrow
Blood
Spleen
Heart
Aorta
Lung
Trachea
Tongue
Small intestine
Liver
Pancreas
Kidney
Bladder
Uterus

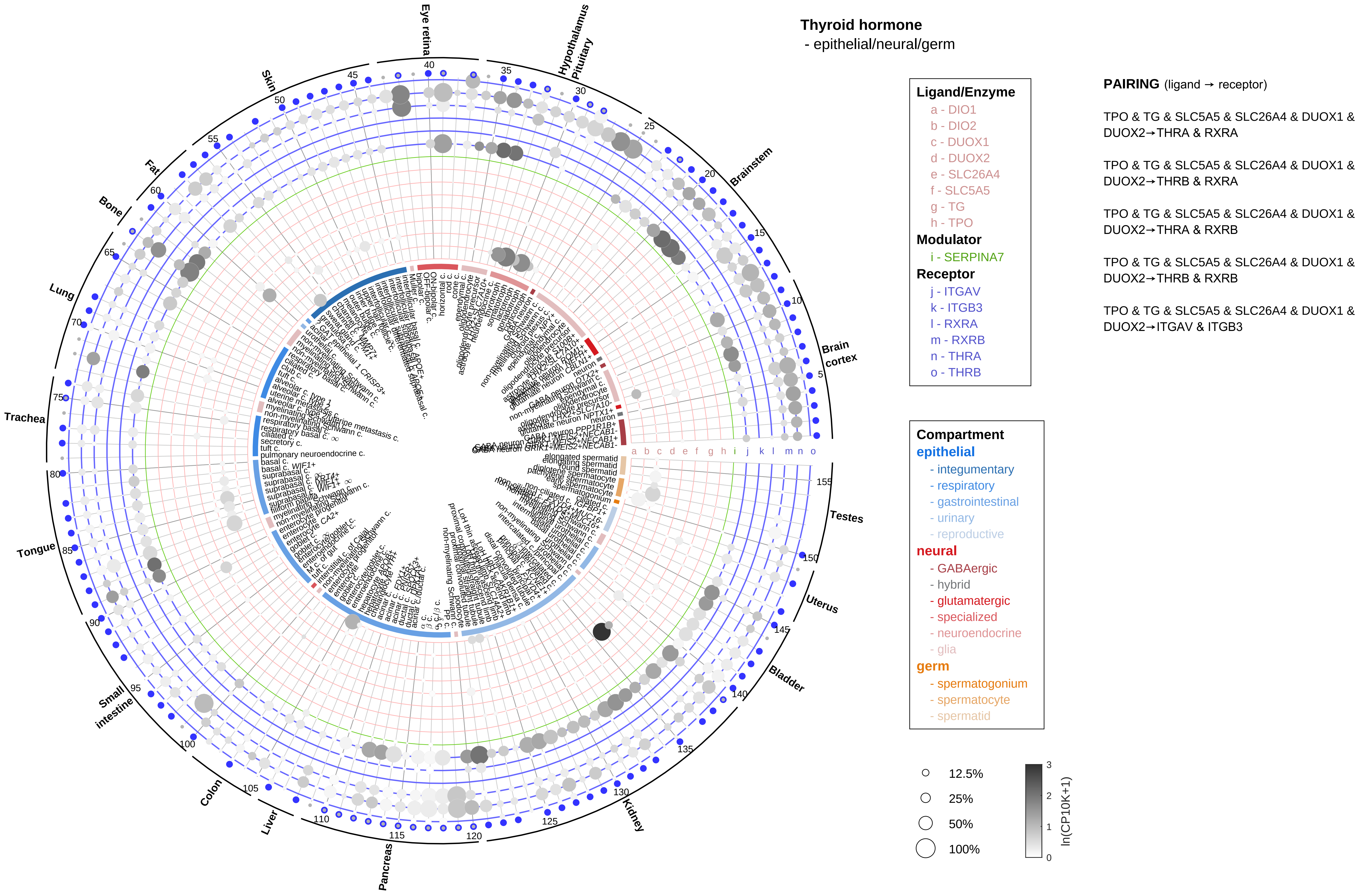

Thyroid hormone
- epithelial/neural/germ
Ligand/Enzyme
a - DIO1
b - DIO2
c - DUOX1
d - DUOX2
e - SLC26A4
f - SLC5A5
g - TG
h - TPO
Modulator
i - SERPINA7
Receptor
j - ITGAV
k - ITGB3
l - RXRA
m - RXRB
n - THRA
o - THRB
PAIRING (ligand → receptor)
TPO & TG & SLC5A5 & SLC26A4 & DUOX1 & DUOX2→THRA & RXRA
TPO & TG & SLC5A5 & SLC26A4 & DUOX1 & DUOX2→THRB & RXRA
TPO & TG & SLC5A5 & SLC26A4 & DUOX1 & DUOX2→THRA & RXRB
TPO & TG & SLC5A5 & SLC26A4 & DUOX1 & DUOX2→THRB & RXRB
TPO & TG & SLC5A5 & SLC26A4 & DUOX1 & DUOX2→ITGAV & ITGB3
Compartment
epithelial
- integumentary
- respiratory
- gastrointestinal
- urinary
- reproductive
neural
- GABAergic
- hybrid
- glutamatergic
- specialized
- neuroendocrine
- glia
germ
- spermatogonium
- spermatocyte
- spermatid
12.5%
25%
50%
100%
ln(CP10K+1)
Eye retina
Hypothalamus
Pituitary
Brainstem
Brain cortex
Testes
Uterus
Bladder
Kidney
Pancreas
Liver
Colon
Small intestine
Tongue
Trachea
Lung
Bone
Fat
Skin

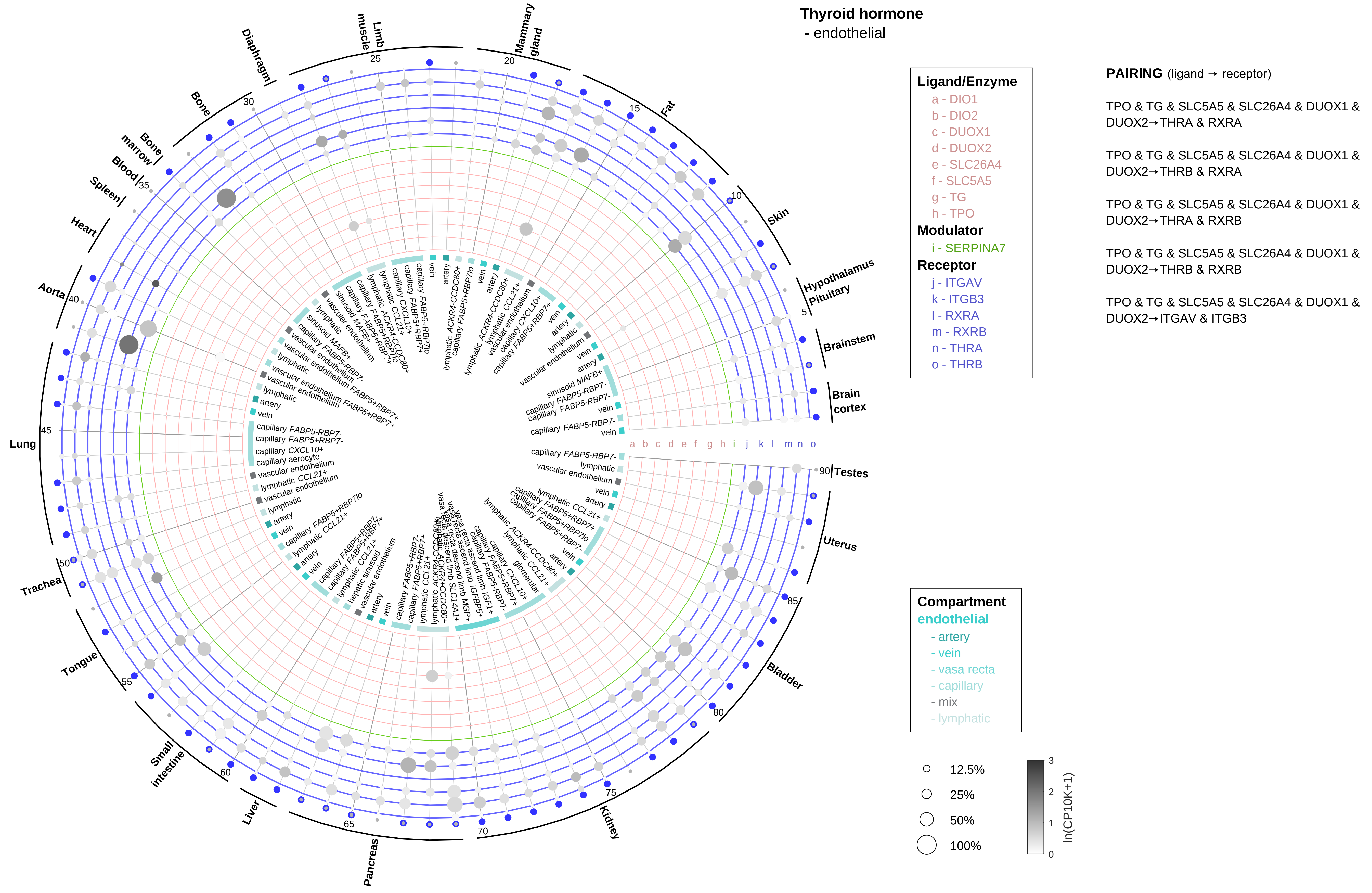
Thyroid hormone
- endothelial
Ligand/Enzyme
a - DIO1
b - DIO2
c - DUOX1
d - DUOX2
e - SLC26A4
f - SLC5A5
g - TG
h - TPO
Modulator
i - SERPINA7
Receptor
j - ITGAV
k - ITGB3
l - RXRA
m - RXRB
n - THRA
o - THRB
PAIRING (ligand → receptor)
TPO & TG & SLC5A5 & SLC26A4 & DUOX1 & DUOX2→THRA & RXRA
TPO & TG & SLC5A5 & SLC26A4 & DUOX1 & DUOX2→THRB & RXRA
TPO & TG & SLC5A5 & SLC26A4 & DUOX1 & DUOX2→THRA & RXRB
TPO & TG & SLC5A5 & SLC26A4 & DUOX1 & DUOX2→THRB & RXRB
TPO & TG & SLC5A5 & SLC26A4 & DUOX1 & DUOX2→ITGAV & ITGB3
Compartment
endothelial
- artery
- vein
- vasa recta
- capillary
- mix
- lymphatic
12.5%
25%
50%
100%
ln(CP10K+1)
Testes
Uterus
Bladder
Kidney
Pancreas
Liver
Small intestine
Tongue
Trachea
Lung
Aorta
Heart
Spleen
Blood
Bone marrow
Bone
Diaphragm
Limb muscle
Mammary gland
Fat
Skin
Hypothalamus
Pituitary
Brainstem
Brain cortex

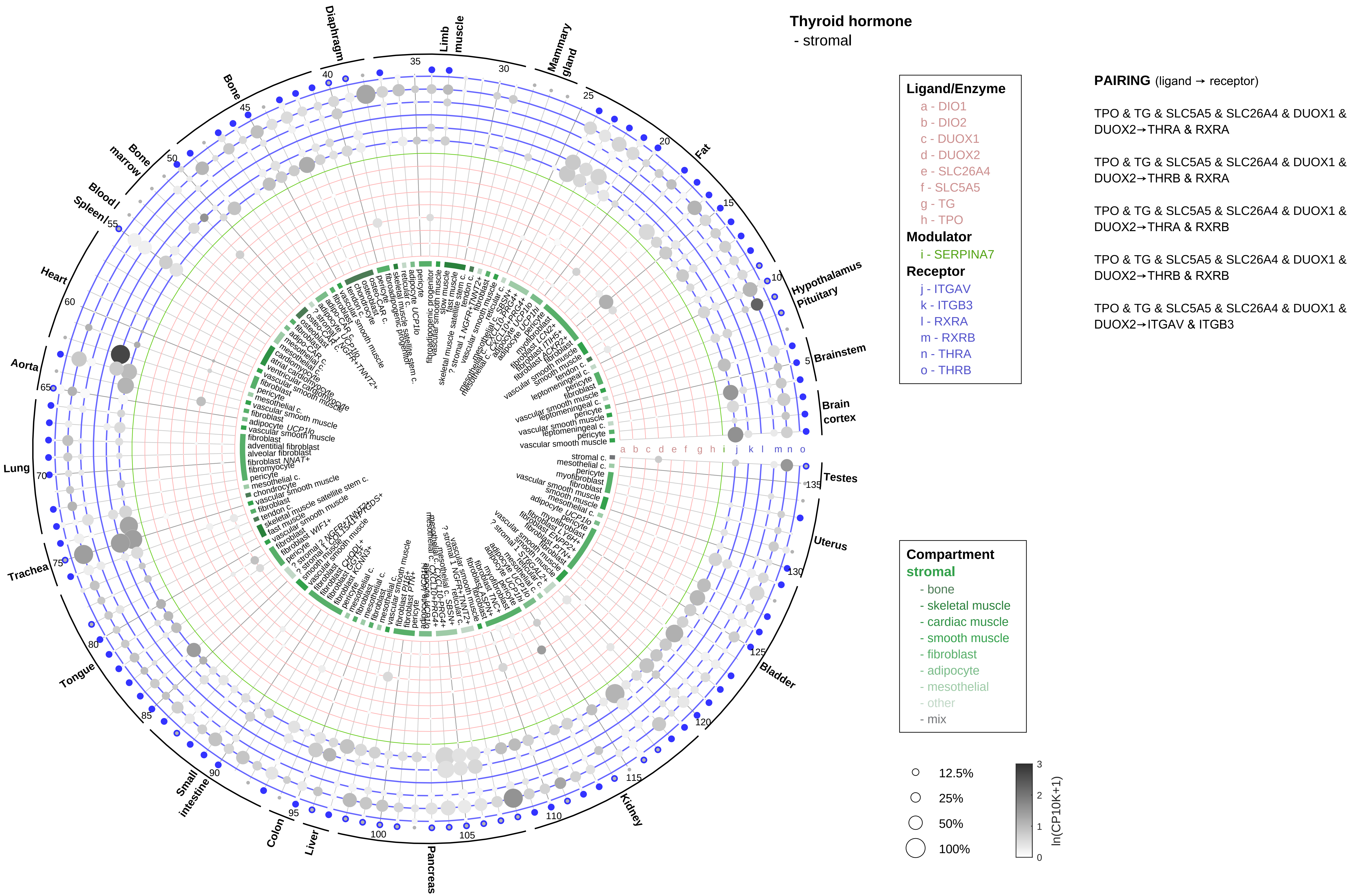

Thyroid hormone
- stromal
Ligand/Enzyme
a - DIO1
b - DIO2
c - DUOX1
d - DUOX2
e - SLC26A4
f - SLC5A5
g - TG
h - TPO
Modulator
i - SERPINA7
Receptor
j - ITGAV
k - ITGB3
l - RXRA
m - RXRB
n - THRA
o - THRB
PAIRING (ligand → receptor)
TPO & TG & SLC5A5 & SLC26A4 & DUOX1 & DUOX2→THRA & RXRA
TPO & TG & SLC5A5 & SLC26A4 & DUOX1 & DUOX2→THRB & RXRA
TPO & TG & SLC5A5 & SLC26A4 & DUOX1 & DUOX2→THRA & RXRB
TPO & TG & SLC5A5 & SLC26A4 & DUOX1 & DUOX2→THRB & RXRB
TPO & TG & SLC5A5 & SLC26A4 & DUOX1 & DUOX2→ITGAV & ITGB3
Compartment
stromal
- bone
- skeletal muscle
- cardiac muscle
- smooth muscle
- fibroblast
- adipocyte
- mesothelial
- other
- mix
12.5%
25%
50%
100%
ln(CP10K+1)
Diaphragm
Limb muscle
Mammary gland
Fat
Hypothalamus
Pituitary
Brainstem
Brain cortex
Testes
Uterus
Bladder
Kidney
Pancreas
Liver
Colon
Small intestine
Tongue
Trachea
Lung
Aorta
Heart
Spleen
Blood
Bone marrow
Bone

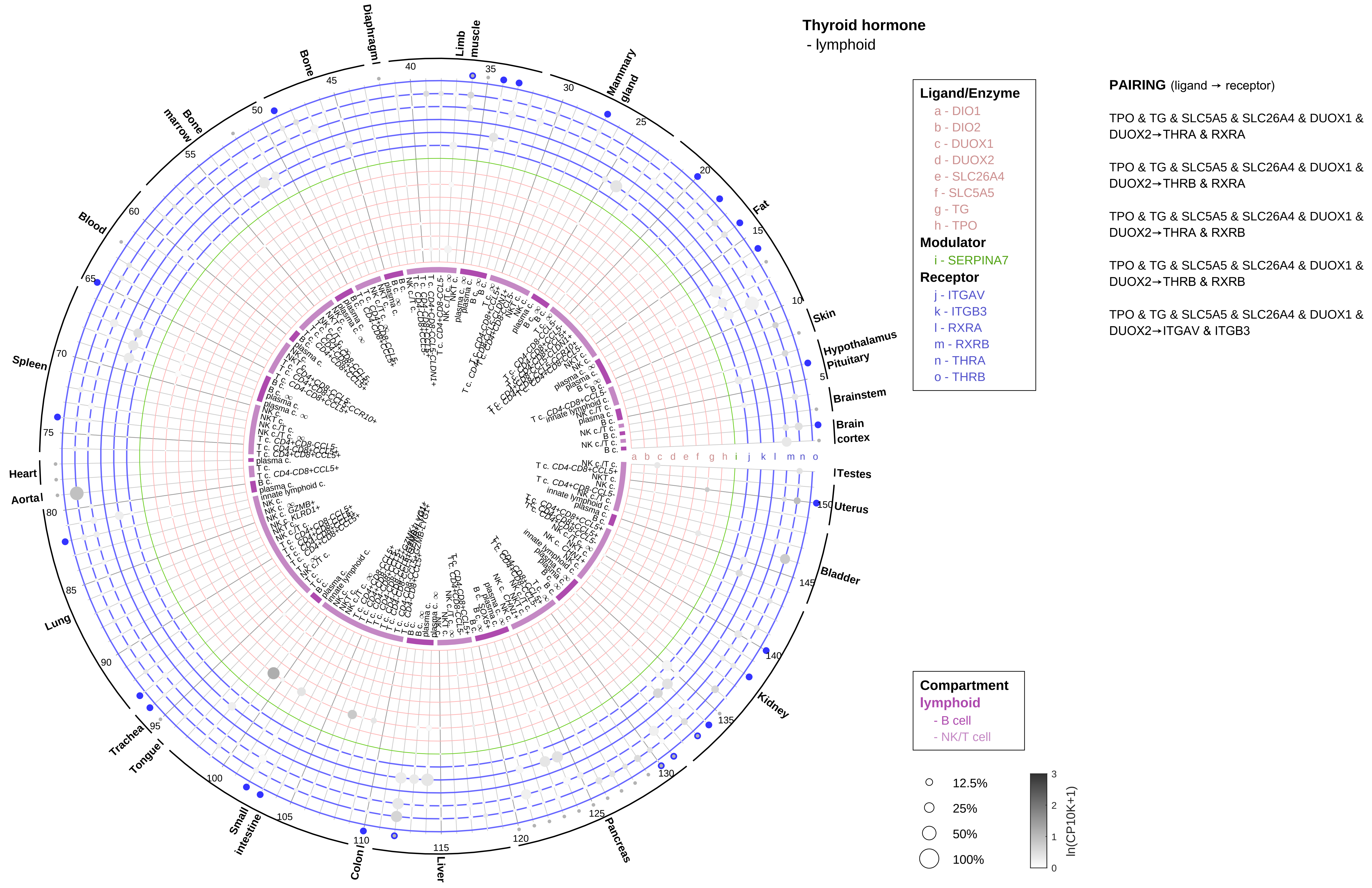
Thyroid hormone
- lymphoid
Ligand/Enzyme
a - DIO1
b - DIO2
c - DUOX1
d - DUOX2
e - SLC26A4
f - SLC5A5
g - TG
h - TPO
Modulator
i - SERPINA7
Receptor
j - ITGAV
k - ITGB3
l - RXRA
m - RXRB
n - THRA
o - THRB
PAIRING (ligand → receptor)
TPO & TG & SLC5A5 & SLC26A4 & DUOX1 & DUOX2→THRA & RXRA
TPO & TG & SLC5A5 & SLC26A4 & DUOX1 & DUOX2→THRB & RXRA
TPO & TG & SLC5A5 & SLC26A4 & DUOX1 & DUOX2→THRA & RXRB
TPO & TG & SLC5A5 & SLC26A4 & DUOX1 & DUOX2→THRB & RXRB
TPO & TG & SLC5A5 & SLC26A4 & DUOX1 & DUOX2→ITGAV & ITGB3
Compartment
lymphoid
- B cell
- NK/T cell
12.5%
25%
50%
100%
ln(CP10K+1)
Skin
Hypothalamus
Pituitary
Brainstem
Brain cortex
Testes
Uterus
Bladder
Kidney
Pancreas
Liver
Colon
Small intestine
Tongue
Trachea
Lung
Aortal
Heart
Spleen
Blood
Bone marrow
Bone
Diaphragm
Limb muscle
Mammary gland
Fat

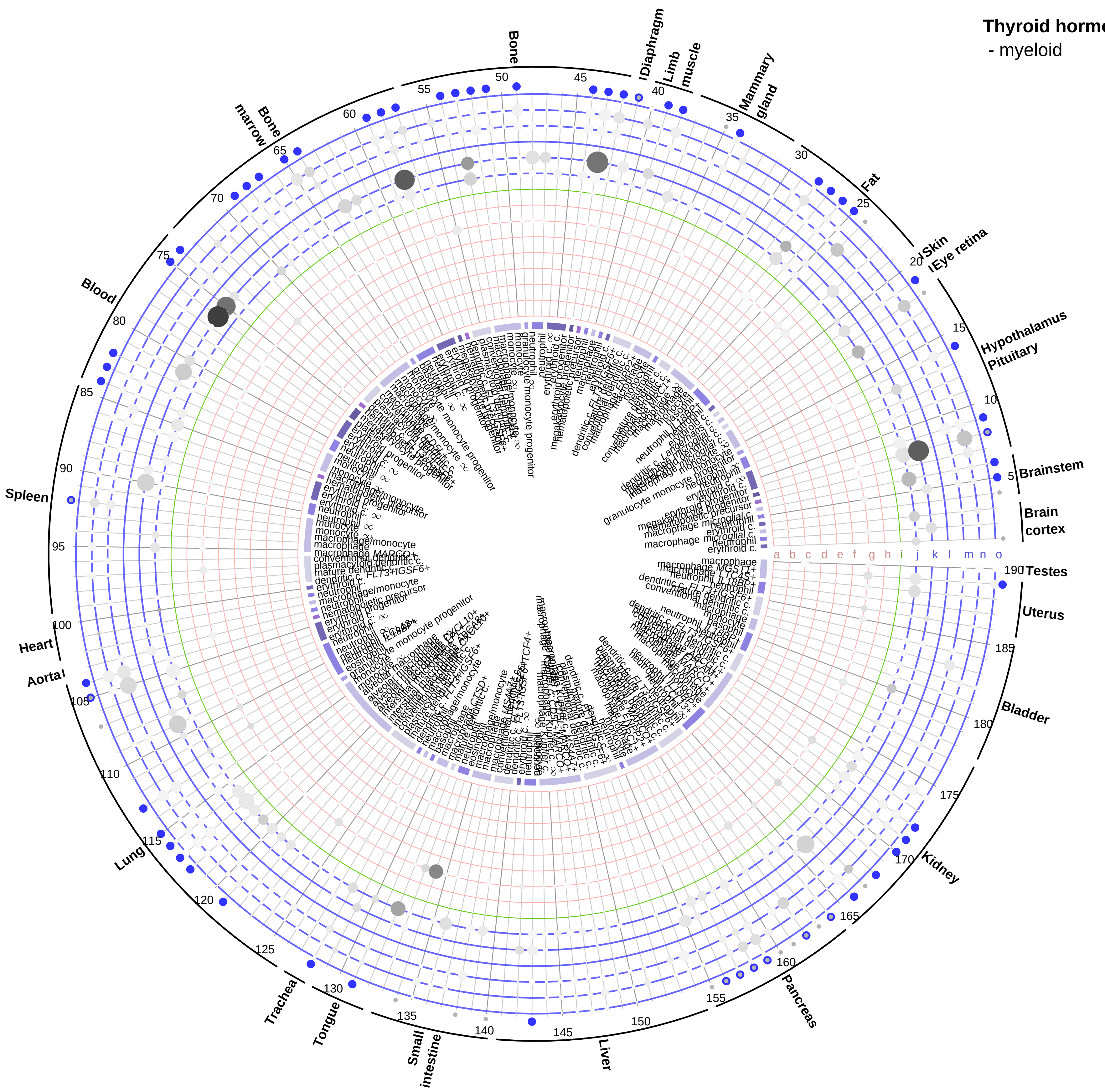


**Thyroid hormone**
- myeloid

**Ligand/Enzyme**
- a - DIO1
- b - DIO2
- c - DUOX1
- d - DUOX2
- e - SLC26A4
- f - SLC5A5
- g - TG
- h - TPO

**Modulator**
- i - SERPINA7

**Receptor**
- j - ITGAV
- k - ITGB3
- l - RXRA
- m - RXRB
- n - THRA
- o - THRB

**PAIRING** (ligand → receptor)

TPO & TG & SLC5A5 & SLC26A4 & DUOX1 & DUOX2→THRA & RXRA

TPO & TG & SLC5A5 & SLC26A4 & DUOX1 & DUOX2→THRB & RXRA

TPO & TG & SLC5A5 & SLC26A4 & DUOX1 & DUOX2→THRA & RXRB

TPO & TG & SLC5A5 & SLC26A4 & DUOX1 & DUOX2→THRB & RXRB

TPO & TG & SLC5A5 & SLC26A4 & DUOX1 & DUOX2→ITGAV & ITGB3

**Compartment**

**hematopoietic**
- hematopoietic precursor

**megakaryocyte-erythroid**
- megakaryoid
- erythroid

**myeloid**
- granulocyte
- granulocyte monocyte progenitor
- monocyte/macrophage
- dendritic

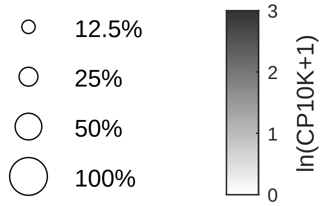

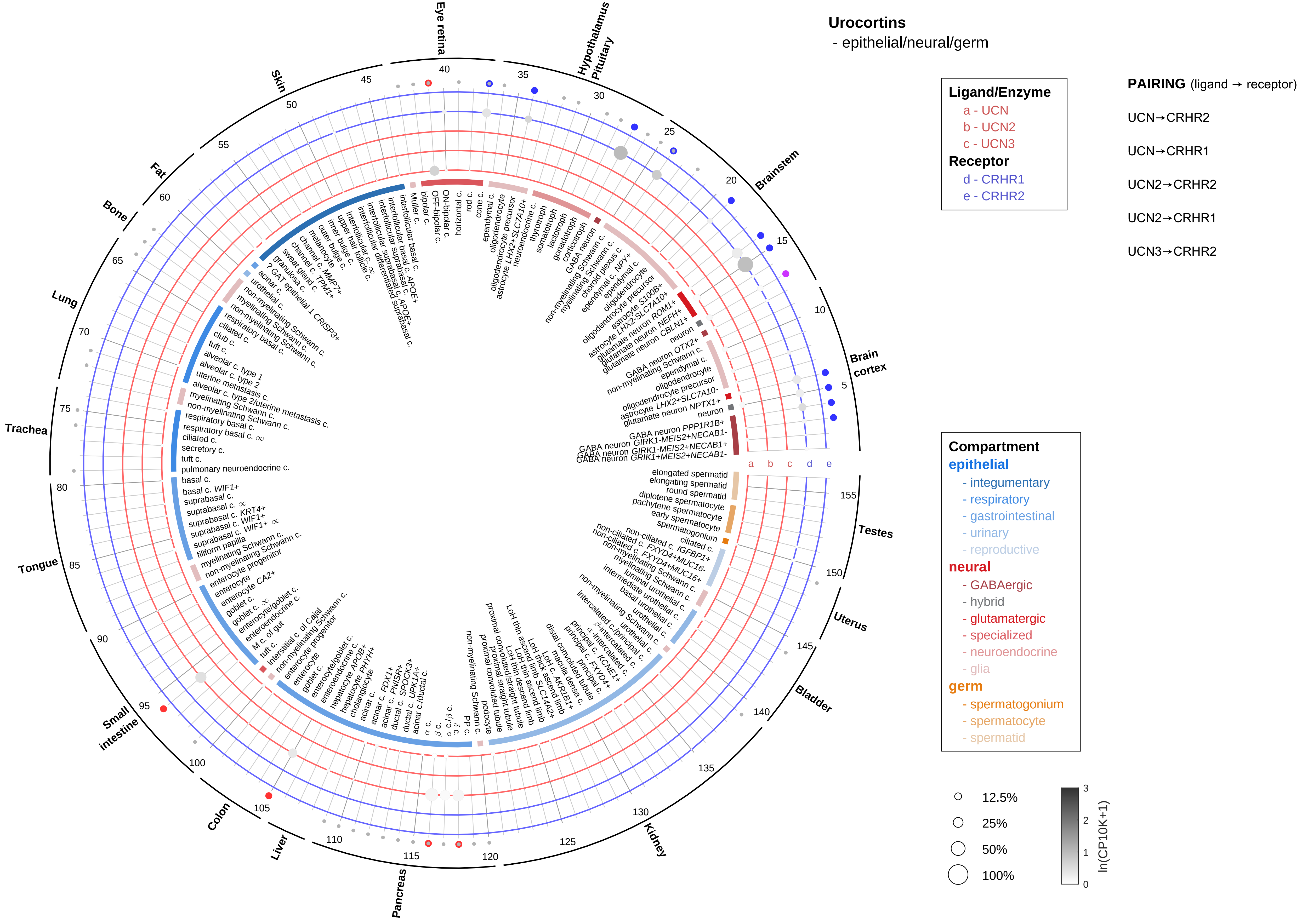

Urocortins
- epithelial/neural/germ
Ligand/Enzyme
a - UCN
b - UCN2
c - UCN3
Receptor
d - CRHR1
e - CRHR2
PAIRING (ligand → receptor)
UCN→CRHR2
UCN→CRHR1
UCN2→CRHR2
UCN2→CRHR1
UCN3→CRHR2
Compartment
epithelial
- integumentary
- respiratory
- gastrointestinal
- urinary
- reproductive
neural
- GABAergic
- hybrid
- glutamatergic
- specialized
- neuroendocrine
- glia
germ
- spermatogonium
- spermatocyte
- spermatid
12.5%
25%
50%
100%
ln(CP10K+1)
Eye retina
Hypothalamus Pituitary
Brainstem
Brain cortex
Testes
Uterus
Bladder
Kidney
Pancreas
Liver
Colon
Small intestine
Tongue
Trachea
Lung
Bone
Fat
Skin

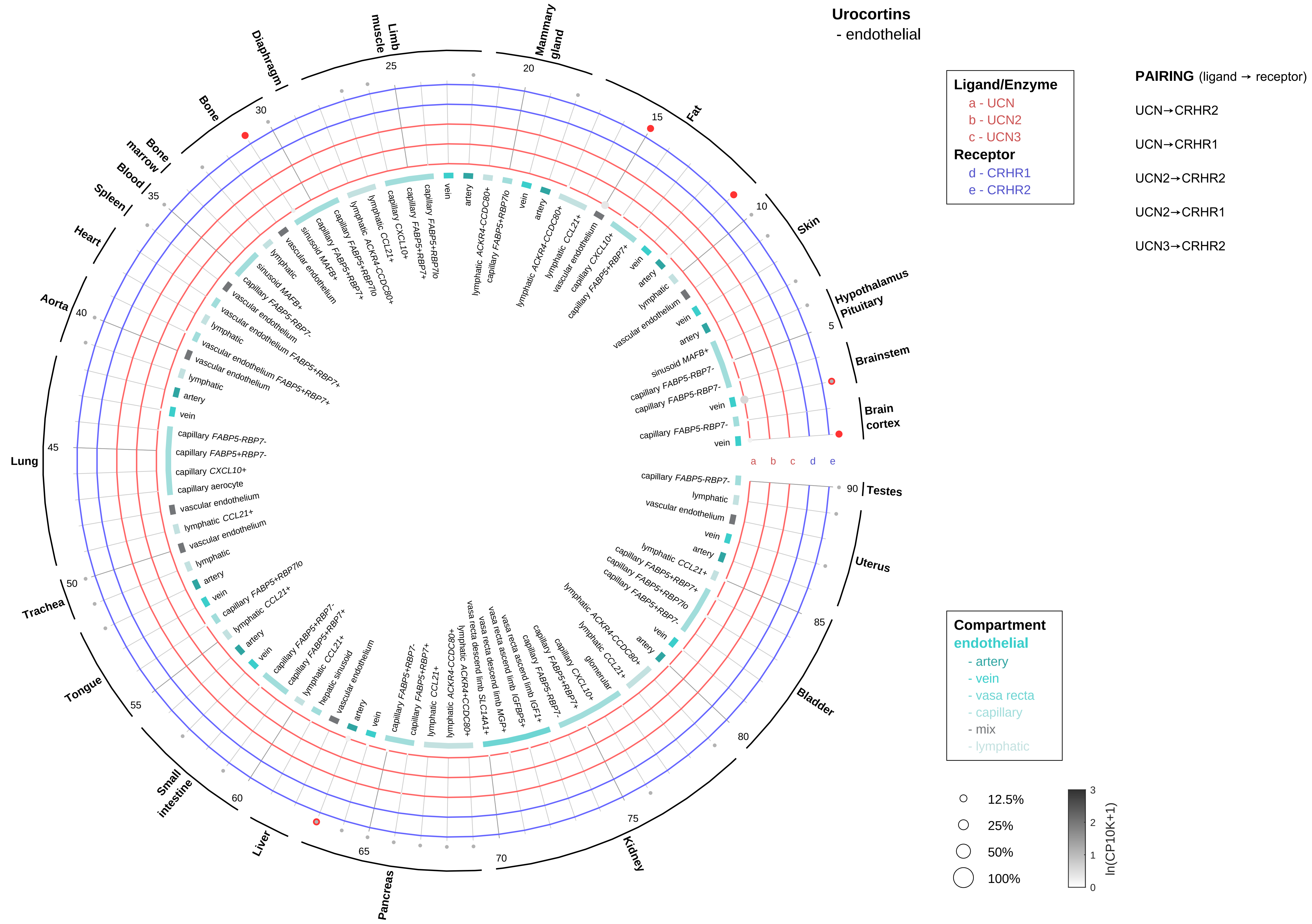

Urocortins
- endothelial
Ligand/Enzyme
a - UCN
b - UCN2
c - UCN3
Receptor
d - CRHR1
e - CRHR2
PAIRING (ligand → receptor)
UCN→CRHR2
UCN→CRHR1
UCN2→CRHR2
UCN2→CRHR1
UCN3→CRHR2
Compartment
endothelial
- artery
- vein
- vasa recta
- capillary
- mix
- lymphatic
12.5%
25%
50%
100%
ln(CP10K+1)
Brain cortex
Brainstem
Hypothalamus Pituitary
Skin
Fat
Mammary gland
Limb muscle
Diaphragm
Bone
Bone marrow
Blood
Spleen
Heart
Aorta
Lung
Trachea
Tongue
Small intestine
Liver
Pancreas
Kidney
Bladder
Uterus
Testes

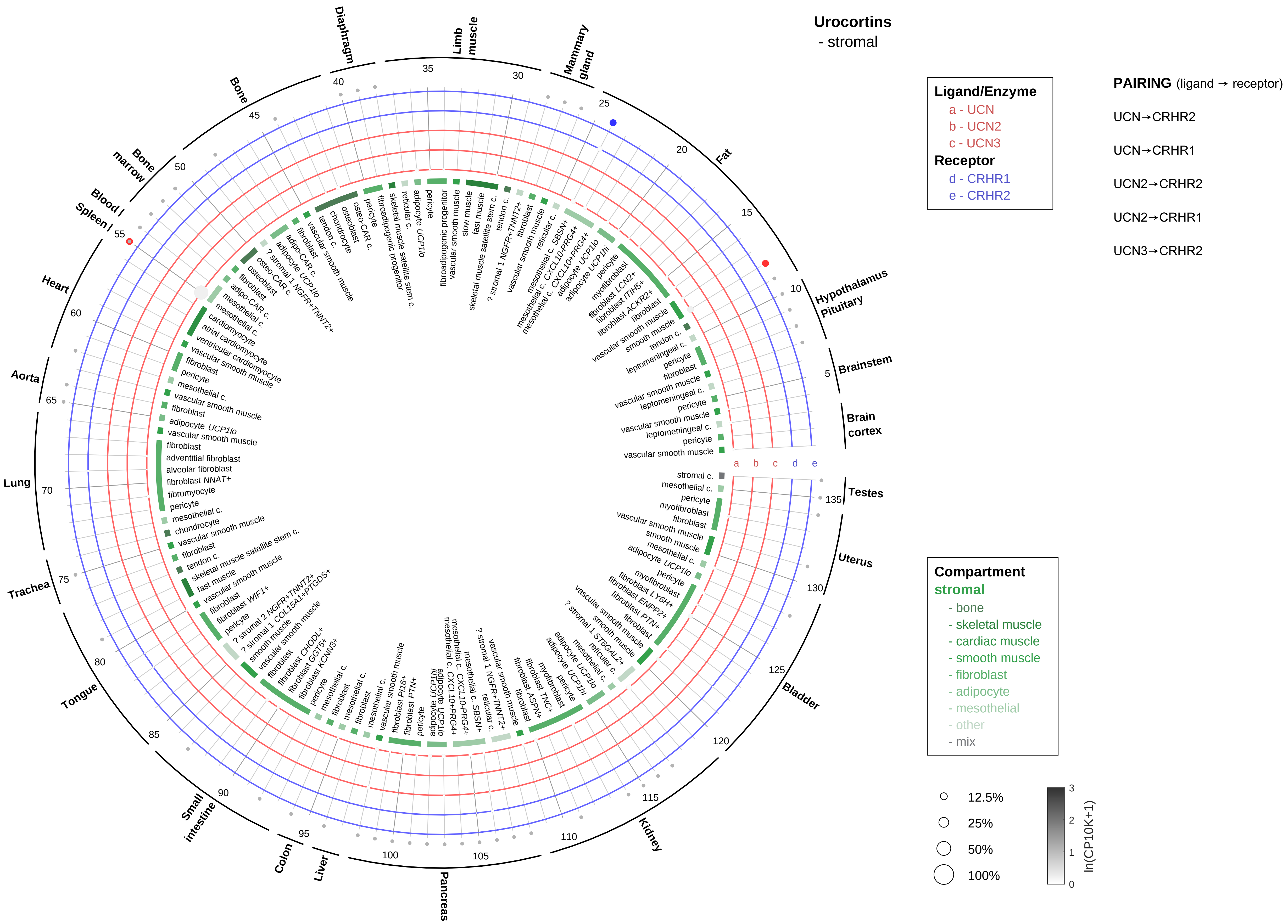

Urocortins
- stromal
Ligand/Enzyme
a - UCN
b - UCN2
c - UCN3
Receptor
d - CRHR1
e - CRHR2
PAIRING (ligand → receptor)
UCN→CRHR2
UCN→CRHR1
UCN2→CRHR2
UCN2→CRHR1
UCN3→CRHR2
Compartment
stromal
- bone
- skeletal muscle
- cardiac muscle
- smooth muscle
- fibroblast
- adipocyte
- mesothelial
- other
- mix
12.5%
25%
50%
100%
ln(CP10K+1)
Diaphragm
Limb muscle
Mammary gland
Fat
Hypothalamus Pituitary
Brainstem
Brain cortex
Testes
Uterus
Bladder
Kidney
Pancreas
Liver
Colon
Small intestine
Tongue
Trachea
Lung
Aorta
Heart
Spleen
Blood
Bone marrow
Bone

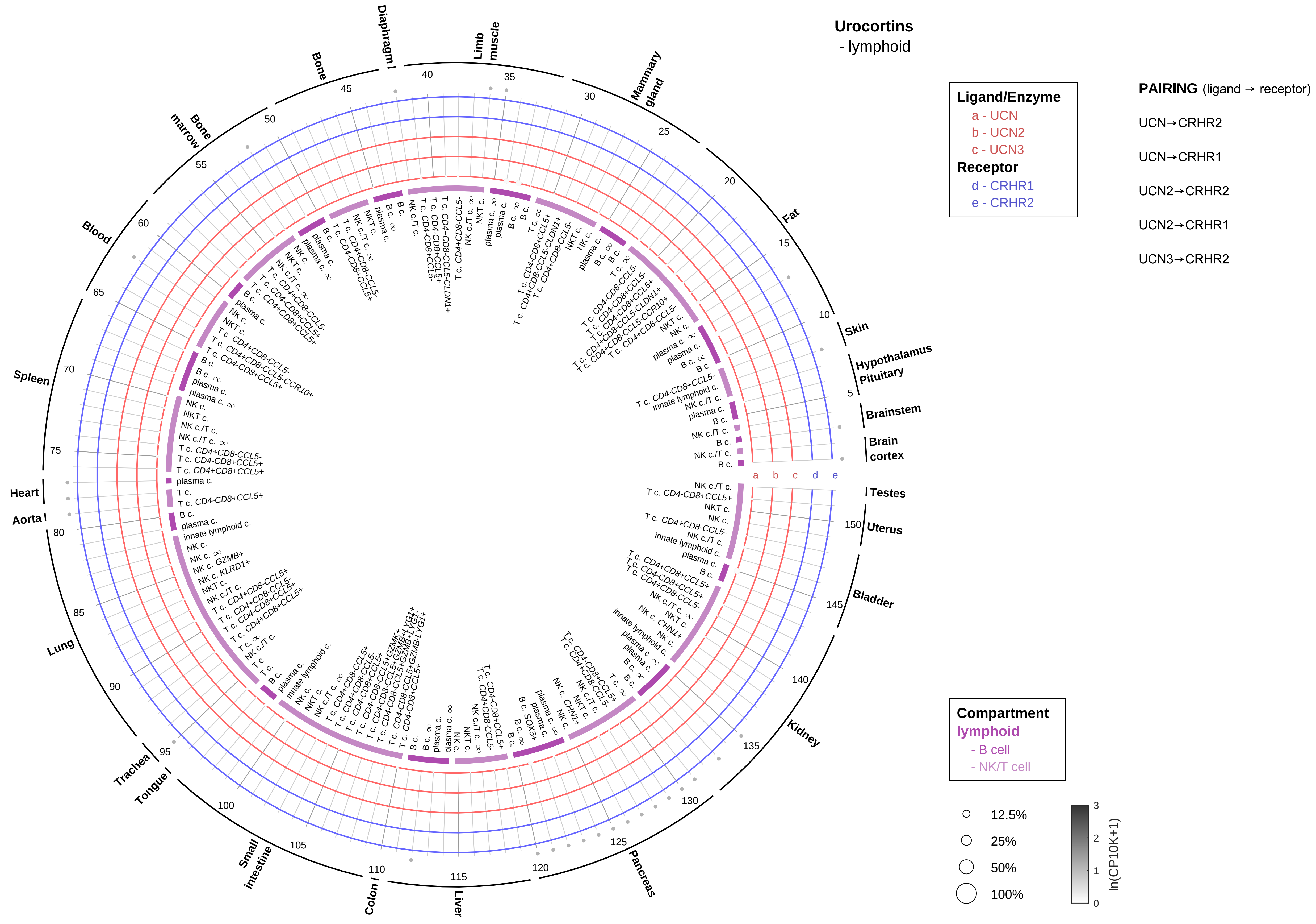

Urocortins
- lymphoid
Ligand/Enzyme
a - UCN
b - UCN2
c - UCN3
Receptor
d - CRHR1
e - CRHR2
PAIRING (ligand → receptor)
UCN→CRHR2
UCN→CRHR1
UCN2→CRHR2
UCN2→CRHR1
UCN3→CRHR2
Compartment
lymphoid
- B cell
- NK/T cell
12.5%
25%
50%
100%
ln(CP10K+1)
Diaphragm
Limb muscle
Mammary gland
Fat
Skin
Hypothalamus
Pituitary
Brainstem
Brain cortex
Testes
Uterus
Bladder
Kidney
Pancreas
Liver
Colon
Small intestine
Tongue
Trachea
Lung
Aorta
Heart
Spleen
Blood
Bone marrow
Bone

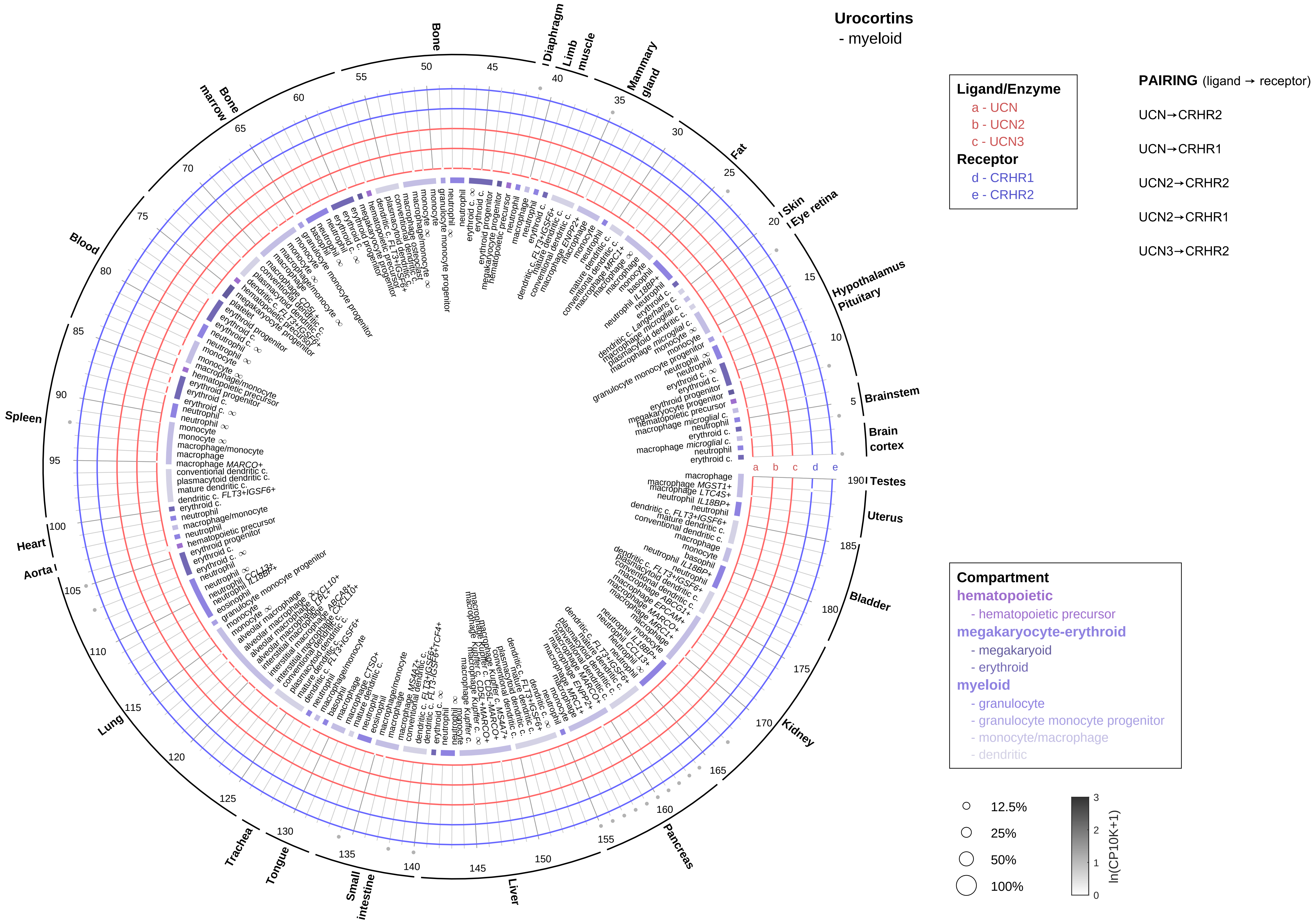

Urocortins
- myeloid
Ligand/Enzyme
a - UCN
b - UCN2
c - UCN3
Receptor
d - CRHR1
e - CRHR2
PAIRING (ligand → receptor)
UCN→CRHR2
UCN→CRHR1
UCN2→CRHR2
UCN2→CRHR1
UCN3→CRHR2
Compartment
hematopoietic
- hematopoietic precursor
megakaryocyte-erythroid
- megakaryoid
- erythroid
myeloid
- granulocyte
- granulocyte monocyte progenitor
- monocyte/macrophage
- dendritic
12.5%
25%
50%
100%
ln(CP10K+1)
Brain cortex
Brainstem
Hypothalamus
Pituitary
Eye retina
Skin
Fat
Mammary gland
Limb muscle
Diaphragm
Bone
Bone marrow
Blood
Spleen
Heart
Aorta
Lung
Trachea
Tongue
Small intestine
Liver
Pancreas
Kidney
Bladder
Uterus
Testes

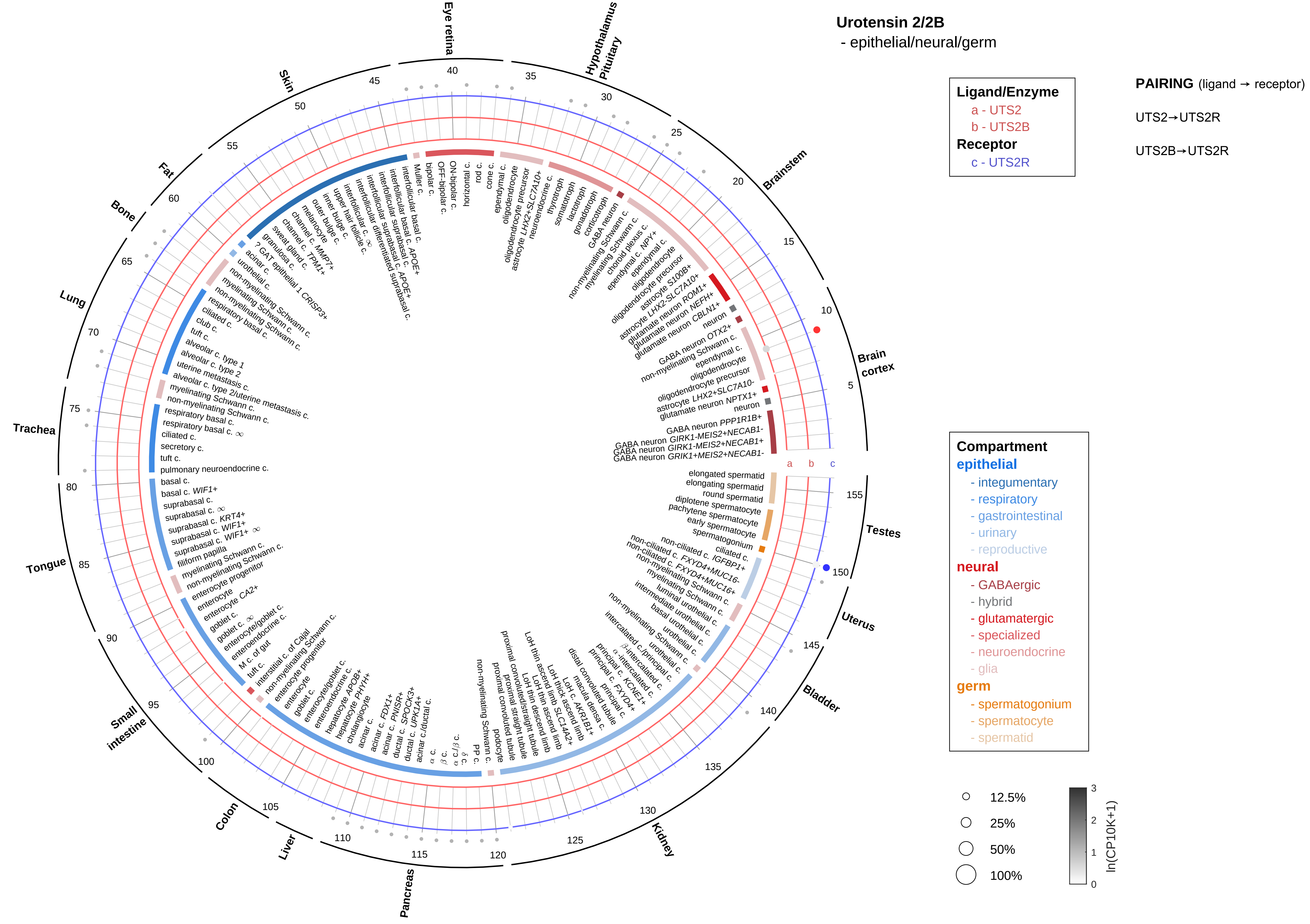

Urotensin 2/2B
- epithelial/neural/germ
Ligand/Enzyme
a - UTS2
b - UTS2B
Receptor
c - UTS2R
PAIRING (ligand → receptor)
UTS2→UTS2R
UTS2B→UTS2R
Compartment
epithelial
- integumentary
- respiratory
- gastrointestinal
- urinary
- reproductive
neural
- GABAergic
- hybrid
- glutamatergic
- specialized
- neuroendocrine
- glia
germ
- spermatogonium
- spermatocyte
- spermatid
12.5%
25%
50%
100%
ln(CP10K+1)
Eye retina
Hypothalamus Pituitary
Brainstem
Brain cortex
Testes
Uterus
Bladder
Kidney
Pancreas
Liver
Colon
Small intestine
Tongue
Trachea
Lung
Bone
Fat
Skin

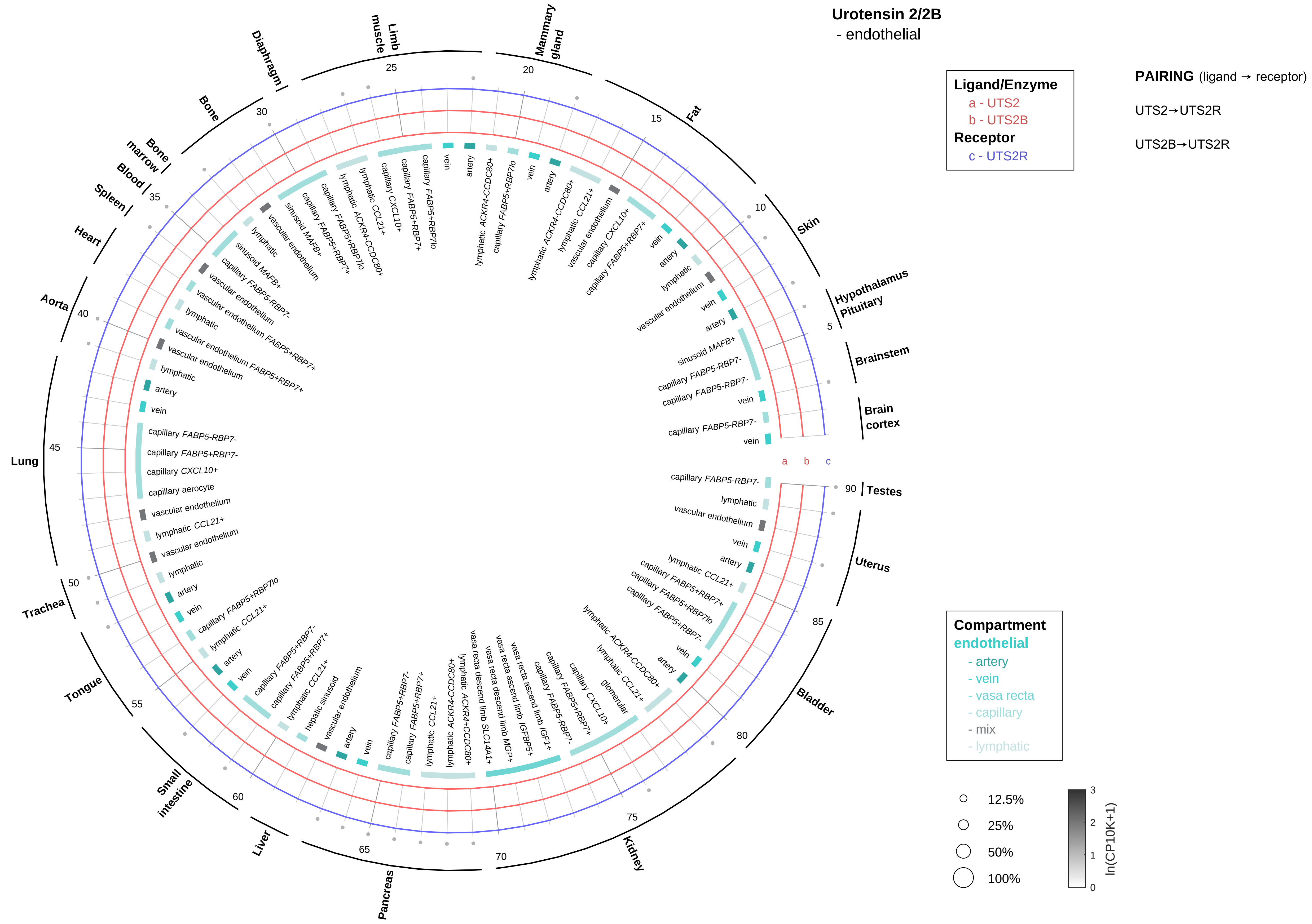
Urotensin 2/2B
- endothelial
Ligand/Enzyme
a - UTS2
b - UTS2B
Receptor
c - UTS2R
PAIRING (ligand → receptor)
UTS2→UTS2R
UTS2B→UTS2R
Compartment
endothelial
- artery
- vein
- vasa recta
- capillary
- mix
- lymphatic
12.5%
25%
50%
100%
ln(CP10K+1)
Brain cortex
Brainstem
Hypothalamus Pituitary
Skin
Fat
Mammary gland
Limb muscle
Diaphragm
Bone
Bone marrow
Blood
Spleen
Heart
Aorta
Lung
Trachea
Tongue
Small intestine
Liver
Pancreas
Kidney
Bladder
Uterus
Testes

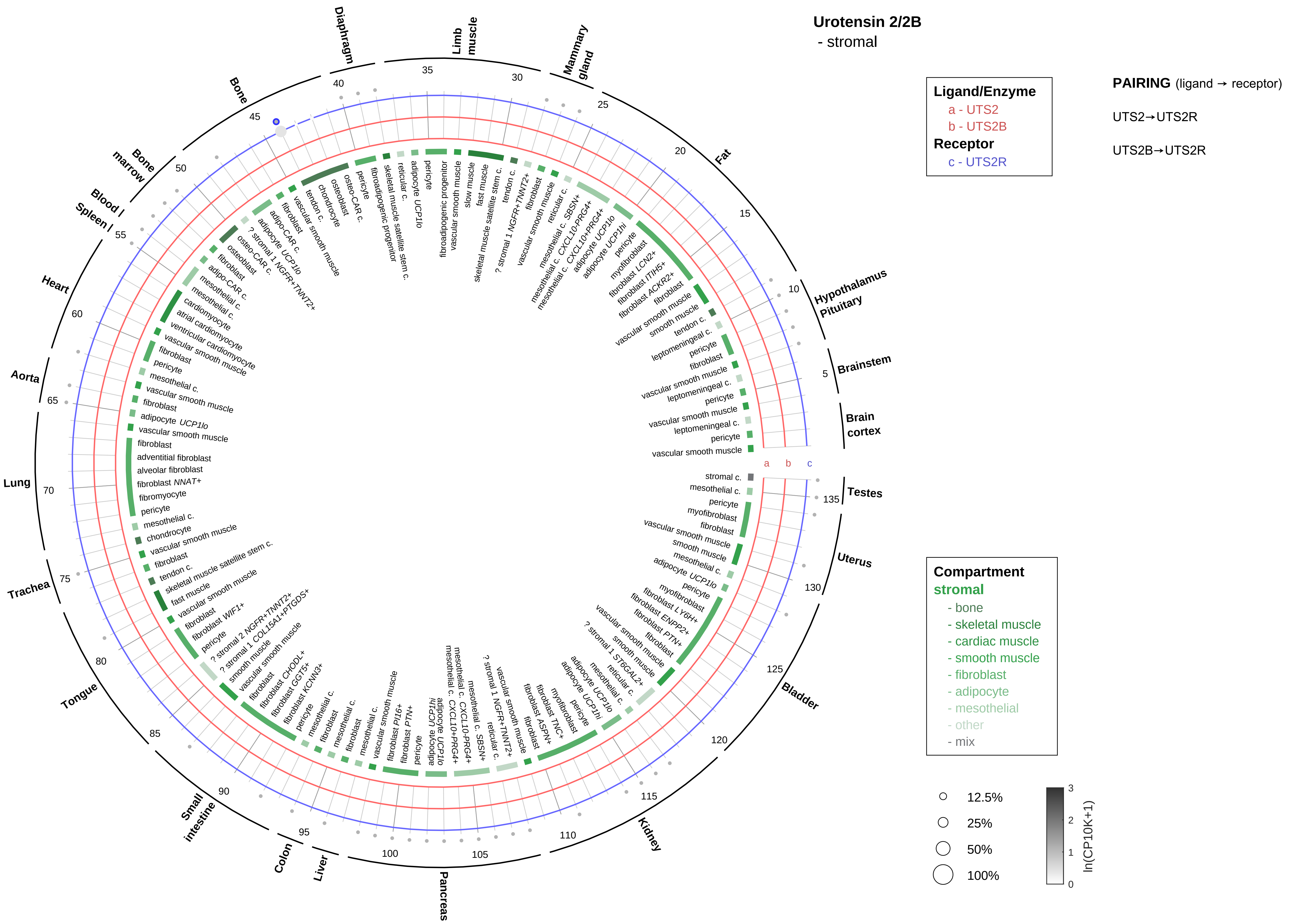

Urotensin 2/2B
- stromal
Ligand/Enzyme
a - UTS2
b - UTS2B
Receptor
c - UTS2R
PAIRING (ligand → receptor)
UTS2→UTS2R
UTS2B→UTS2R
Compartment
stromal
- bone
- skeletal muscle
- cardiac muscle
- smooth muscle
- fibroblast
- adipocyte
- mesothelial
- other
- mix
12.5%
25%
50%
100%
ln(CP10K+1)
Diaphragm
Limb muscle
Mammary gland
Fat
Hypothalamus Pituitary
Brainstem
Brain cortex
Testes
Uterus
Bladder
Kidney
Pancreas
Liver
Colon
Small intestine
Tongue
Trachea
Lung
Aorta
Heart
Spleen
Blood
Bone marrow
Bone

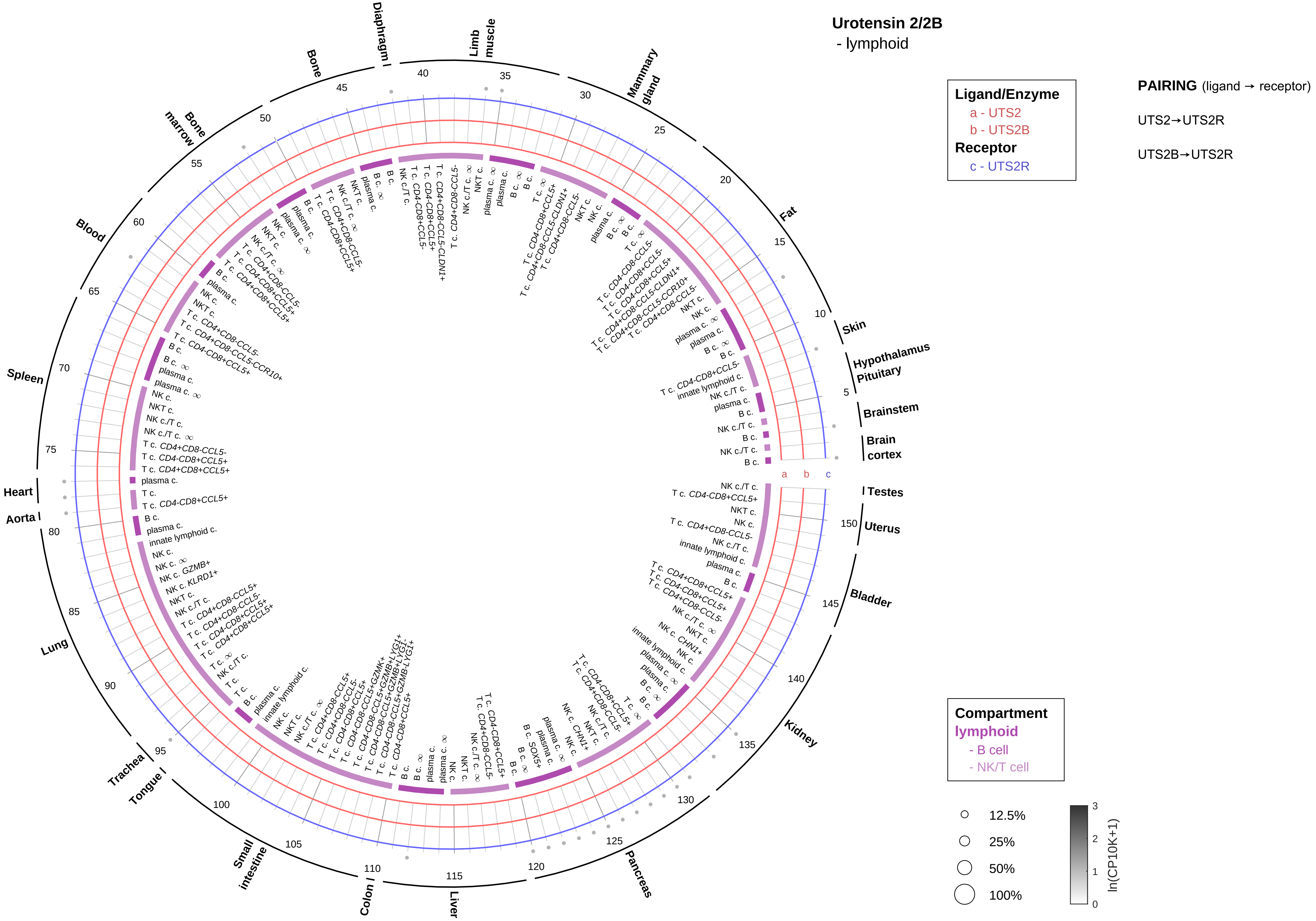

Urotensin 2/2B
- lymphoid
Ligand/Enzyme
a - UTS2
b - UTS2B
Receptor
c - UTS2R
PAIRING (ligand → receptor)
UTS2→UTS2R
UTS2B→UTS2R
Compartment
lymphoid
- B cell
- NK/T cell
12.5%
25%
50%
100%
ln(CP10K+1)
Skin
Hypothalamus
Pituitary
Brainstem
Brain cortex
Testes
Uterus
Bladder
Kidney
Pancreas
Liver
Colon
Small intestine
Tongue
Trachea
Lung
Aorta
Heart
Spleen
Blood
Bone marrow
Bone
Diaphragm
Limb muscle
Mammary gland
Fat

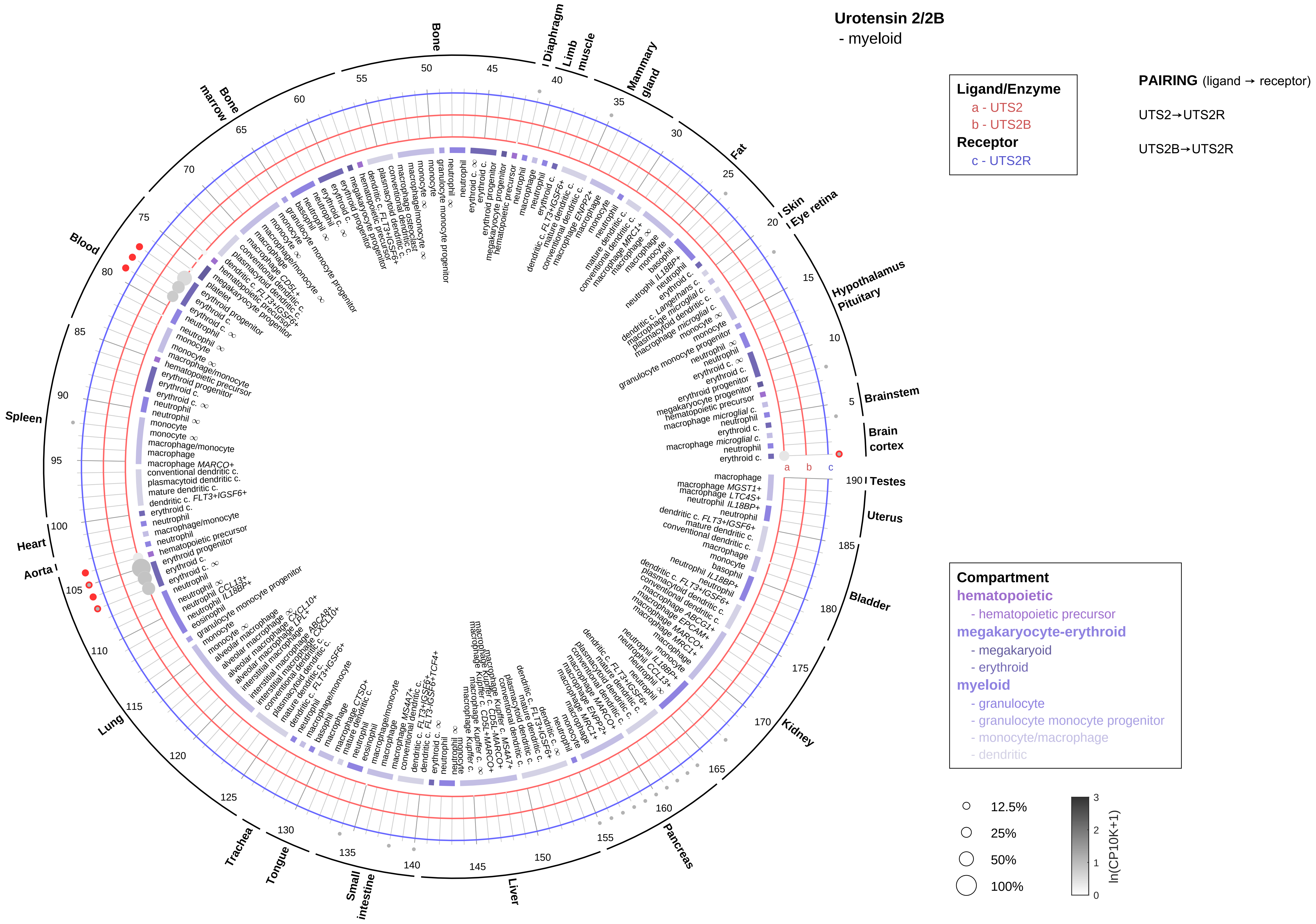
Urotensin 2/2B
- myeloid
Ligand/Enzyme
a - UTS2
b - UTS2B
Receptor
c - UTS2R
PAIRING (ligand → receptor)
UTS2→UTS2R
UTS2B→UTS2R
Compartment
hematopoietic
- hematopoietic precursor
megakaryocyte-erythroid
- megakaryoid
- erythroid
myeloid
- granulocyte
- granulocyte monocyte progenitor
- monocyte/macrophage
- dendritic
12.5%
25%
50%
100%
ln(CP10K+1)
Brain cortex
Brainstem
Hypothalamus
Pituitary
Eye retina
Skin
Fat
Mammary gland
Limb muscle
Diaphragm
Bone
Bone marrow
Blood
Spleen
Heart
Aorta
Lung
Trachea
Tongue
Small intestine
Liver
Pancreas
Kidney
Bladder
Uterus
Testes

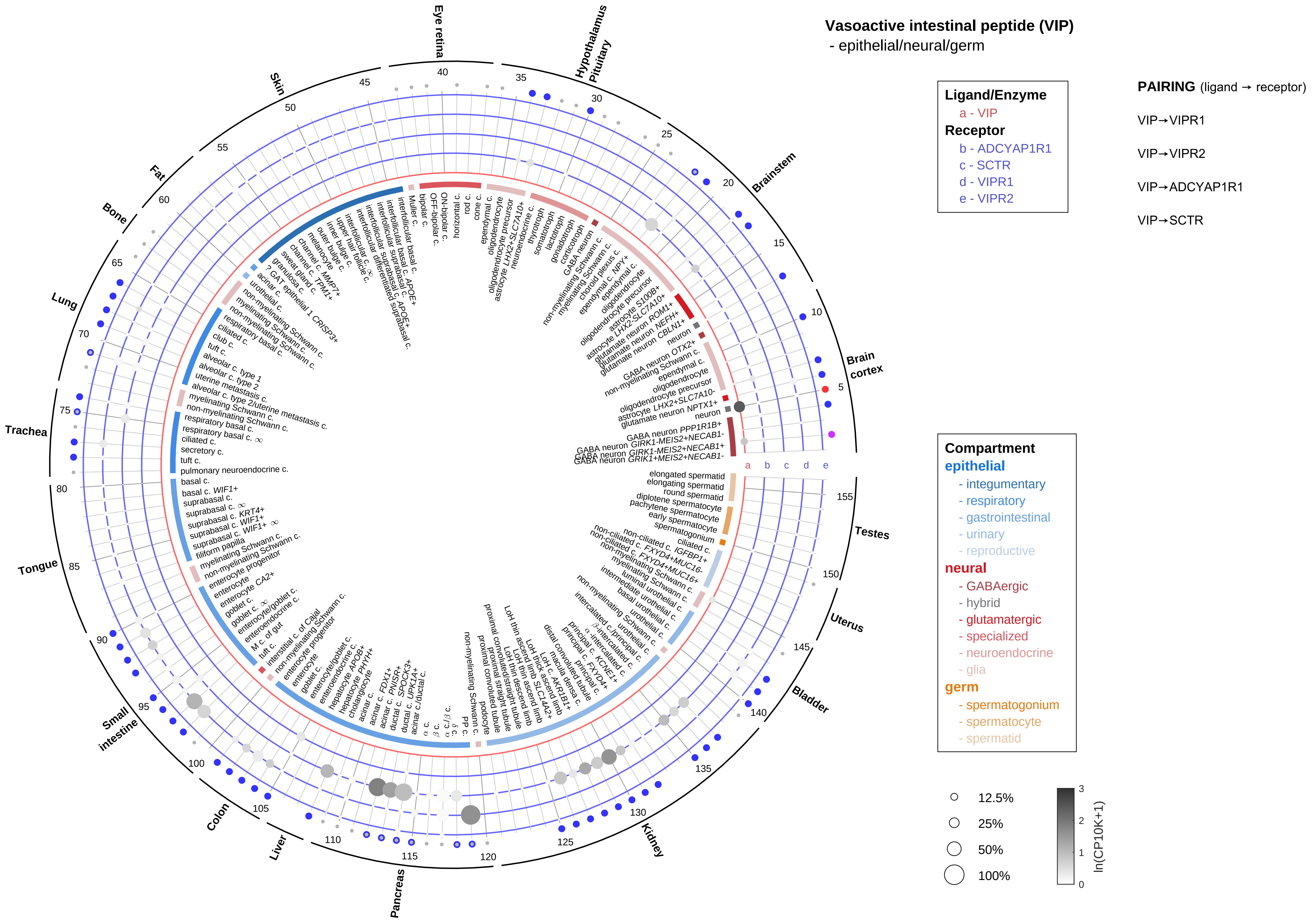

Vasoactive intestinal peptide (VIP)
- epithelial/neural/germ
Ligand/Enzyme
a - VIP
Receptor
b - ADCYAP1R1
c - SCTR
d - VIPR1
e - VIPR2
PAIRING (ligand → receptor)
VIP→VIPR1
VIP→VIPR2
VIP→ADCYAP1R1
VIP→SCTR
Compartment
epithelial
- integumentary
- respiratory
- gastrointestinal
- urinary
- reproductive
neural
- GABAergic
- hybrid
- glutamatergic
- specialized
- neuroendocrine
- glia
germ
- spermatogonium
- spermatocyte
- spermatid
12.5%
25%
50%
100%
ln(CP10K+1)
Eye retina
Hypothalamus Pituitary
Brainstem
Brain cortex
Testes
Uterus
Bladder
Kidney
Pancreas
Liver
Colon
Small intestine
Tongue
Trachea
Lung
Bone
Fat
Skin

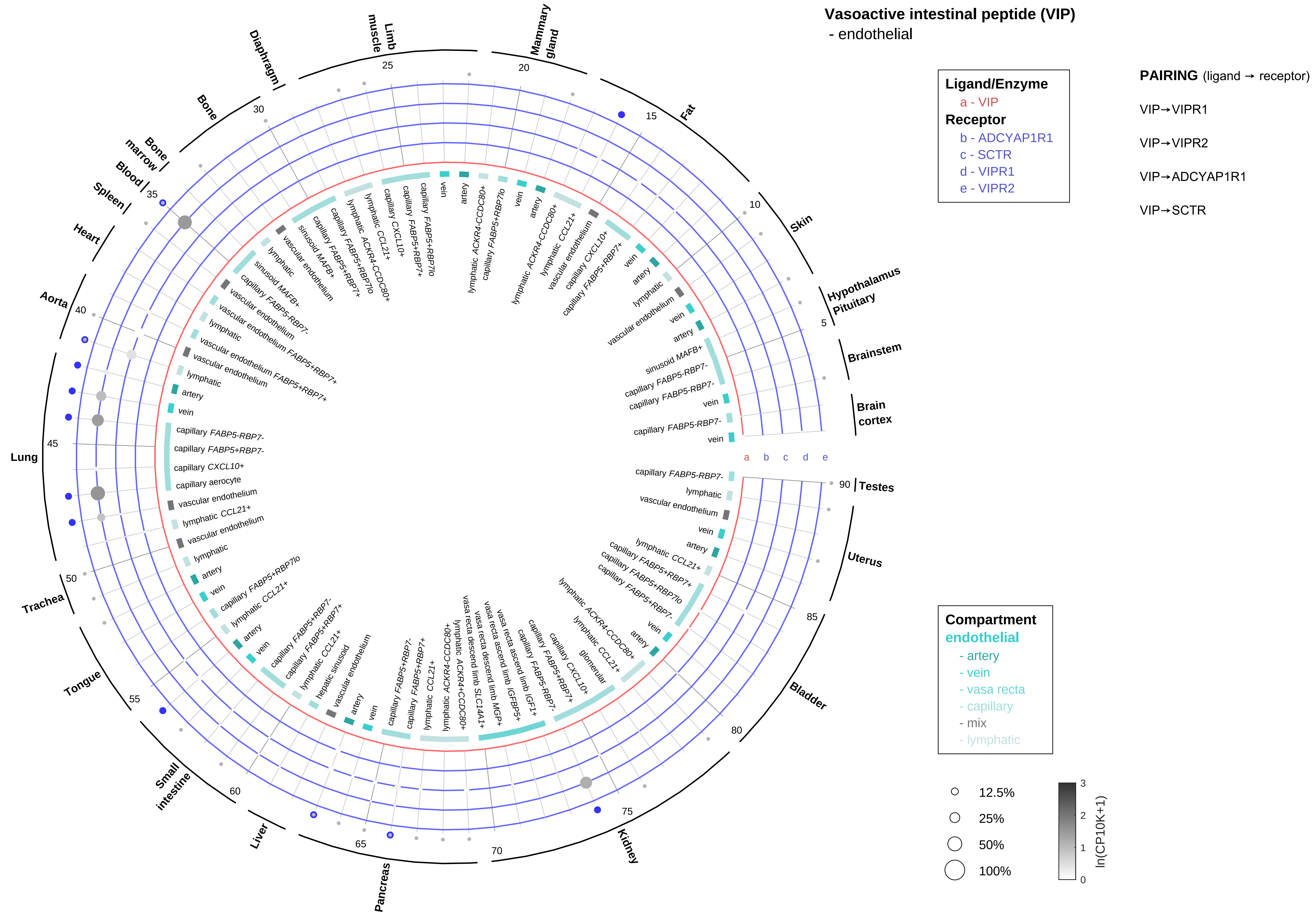

Vasoactive intestinal peptide (VIP)
- endothelial
Ligand/Enzyme
a - VIP
Receptor
b - ADCYAP1R1
c - SCTR
d - VIPR1
e - VIPR2
PAIRING (ligand → receptor)
VIP→VIPR1
VIP→VIPR2
VIP→ADCYAP1R1
VIP→SCTR
Compartment
endothelial
- artery
- vein
- vasa recta
- capillary
- mix
- lymphatic
12.5%
25%
50%
100%
ln(CP10K+1)
Brain cortex
Brainstem
Hypothalamus
Pituitary
Skin
Fat
Mammary gland
Limb muscle
Diaphragm
Bone
Bone marrow
Blood
Spleen
Heart
Aorta
Lung
Trachea
Tongue
Small intestine
Liver
Pancreas
Kidney
Bladder
Uterus
Testes

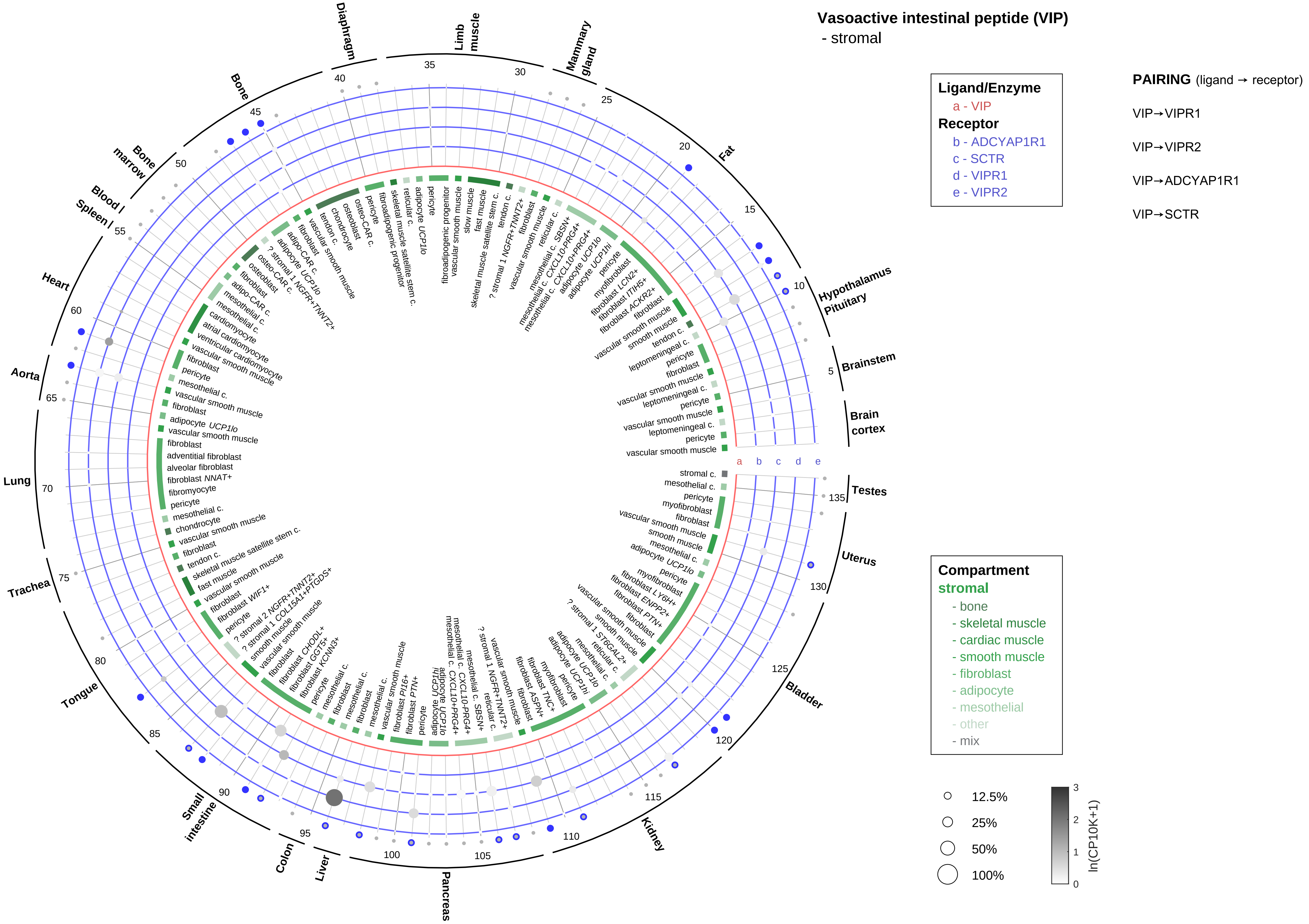

Vasoactive intestinal peptide (VIP)
- stromal
Ligand/Enzyme
a - VIP
Receptor
b - ADCYAP1R1
c - SCTR
d - VIPR1
e - VIPR2
PAIRING (ligand → receptor)
VIP→VIPR1
VIP→VIPR2
VIP→ADCYAP1R1
VIP→SCTR
Compartment
stromal
- bone
- skeletal muscle
- cardiac muscle
- smooth muscle
- fibroblast
- adipocyte
- mesothelial
- other
- mix
12.5%
25%
50%
100%
ln(CP10K+1)
Diaphragm
Limb muscle
Mammary gland
Fat
Hypothalamus
Pituitary
Brainstem
Brain cortex
Testes
Uterus
Bladder
Kidney
Pancreas
Liver
Colon
Small intestine
Tongue
Trachea
Lung
Aorta
Heart
Spleen
Blood
Bone marrow
Bone

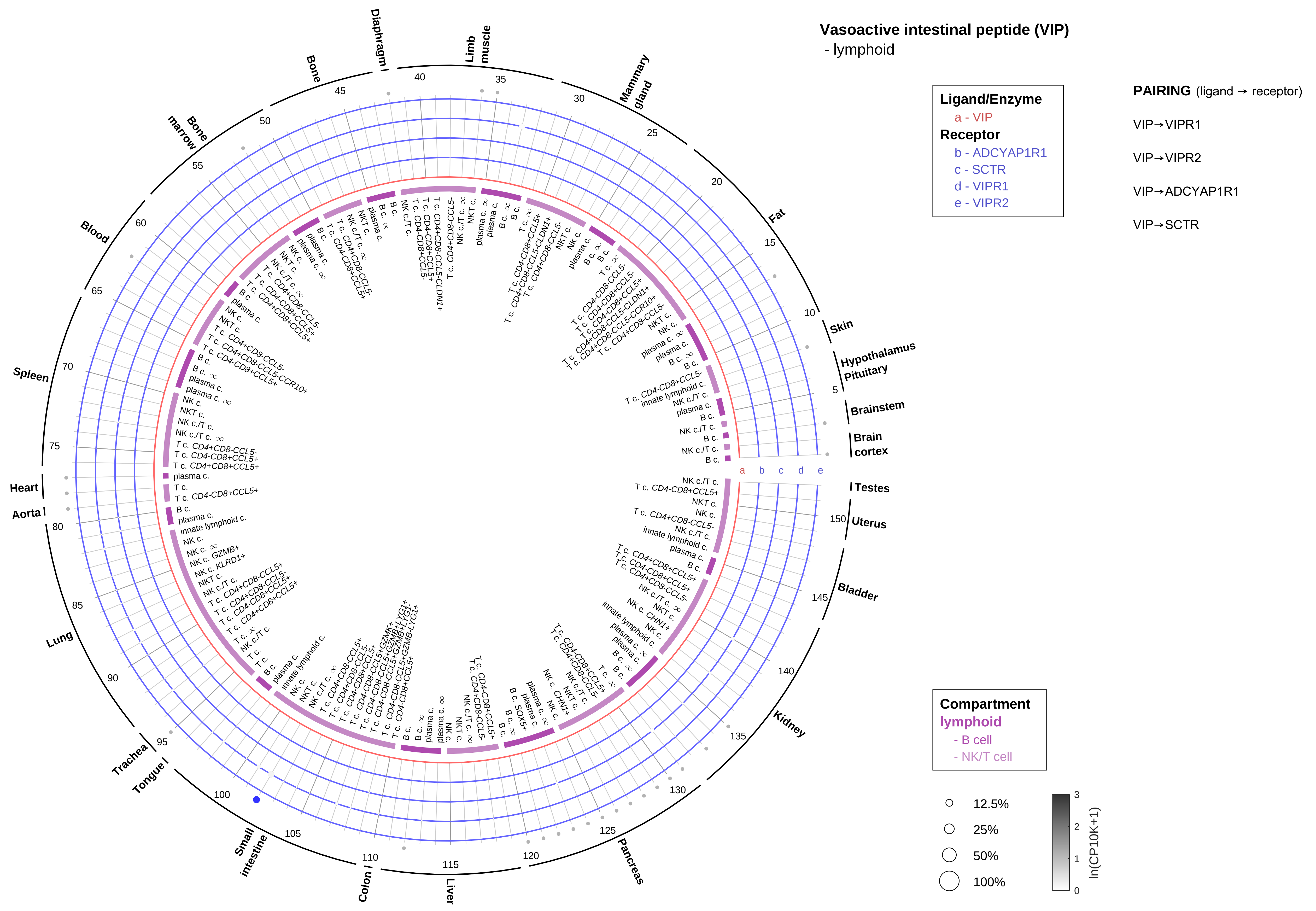
Vasoactive intestinal peptide (VIP)
- lymphoid
Ligand/Enzyme
a - VIP
Receptor
b - ADCYAP1R1
c - SCTR
d - VIPR1
e - VIPR2
PAIRING (ligand → receptor)
VIP→VIPR1
VIP→VIPR2
VIP→ADCYAP1R1
VIP→SCTR
Compartment
lymphoid
- B cell
- NK/T cell
12.5%
25%
50%
100%
ln(CP10K+1)
Skin
Hypothalamus
Pituitary
Brainstem
Brain cortex
Testes
Uterus
Bladder
Kidney
Pancreas
Liver
Colon
Small intestine
Tongue
Trachea
Lung
Aorta
Heart
Spleen
Blood
Bone marrow
Bone
Diaphragm
Limb muscle
Mammary gland
Fat

**Vasoactive intestinal peptide (VIP)**
- myeloid

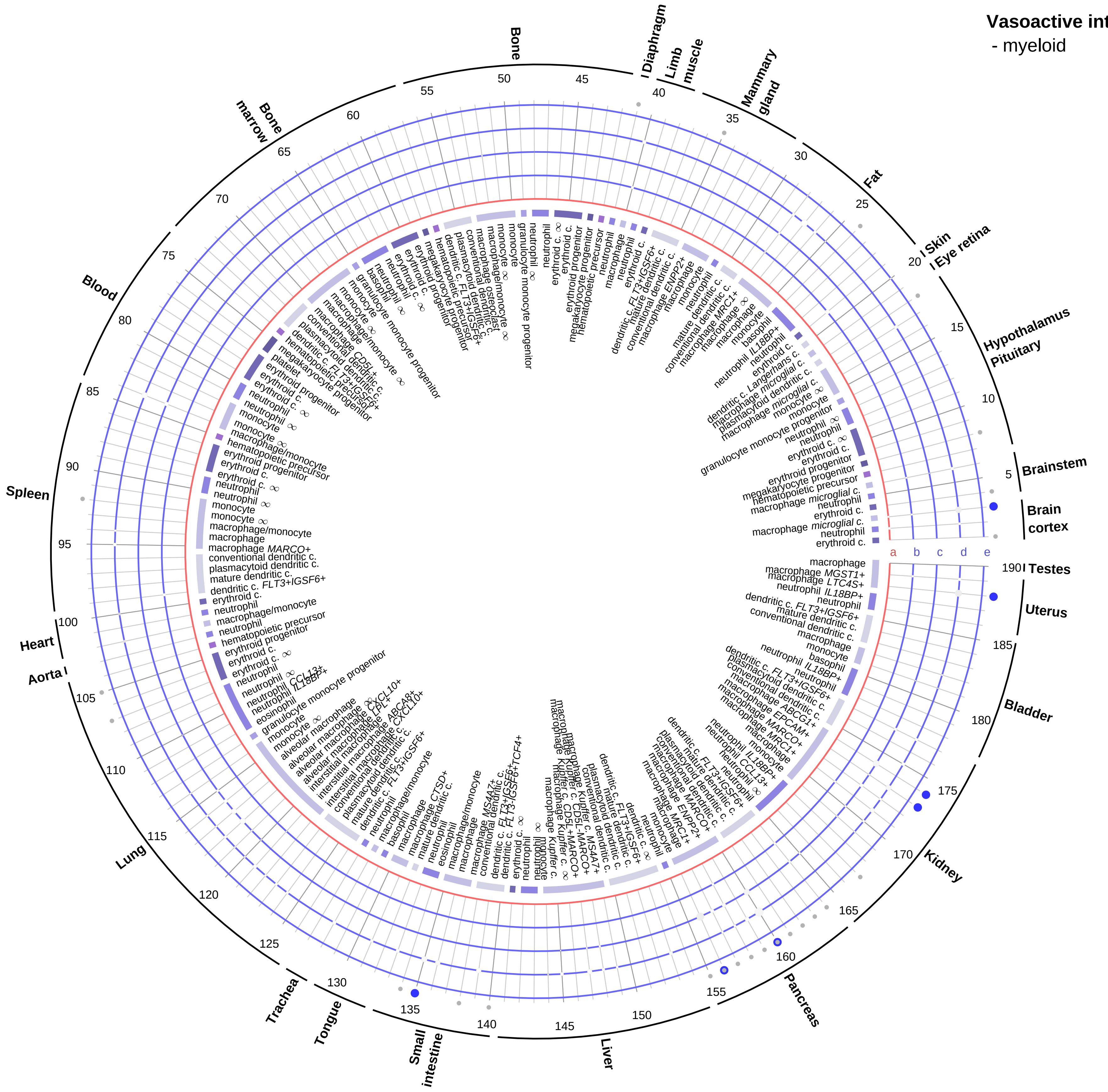


**Ligand/Enzyme**
- a - VIP

**Receptor**
- b - ADCYAP1R1
- c - SCTR
- d - VIPR1
- e - VIPR2

**PAIRING** (ligand → receptor)

VIP→VIPR1

VIP→VIPR2

VIP→ADCYAP1R1

VIP→SCTR

**Compartment**

**hematopoietic**
- hematopoietic precursor

**megakaryocyte-erythroid**
- megakaryoid
- erythroid

**myeloid**
- granulocyte
- granulocyte monocyte progenitor
- monocyte/macrophage
- dendritic

○ 12.5%

○ 25%

○ 50%

○ 100%

ln(CP10K+1): 0, 1, 2, 3

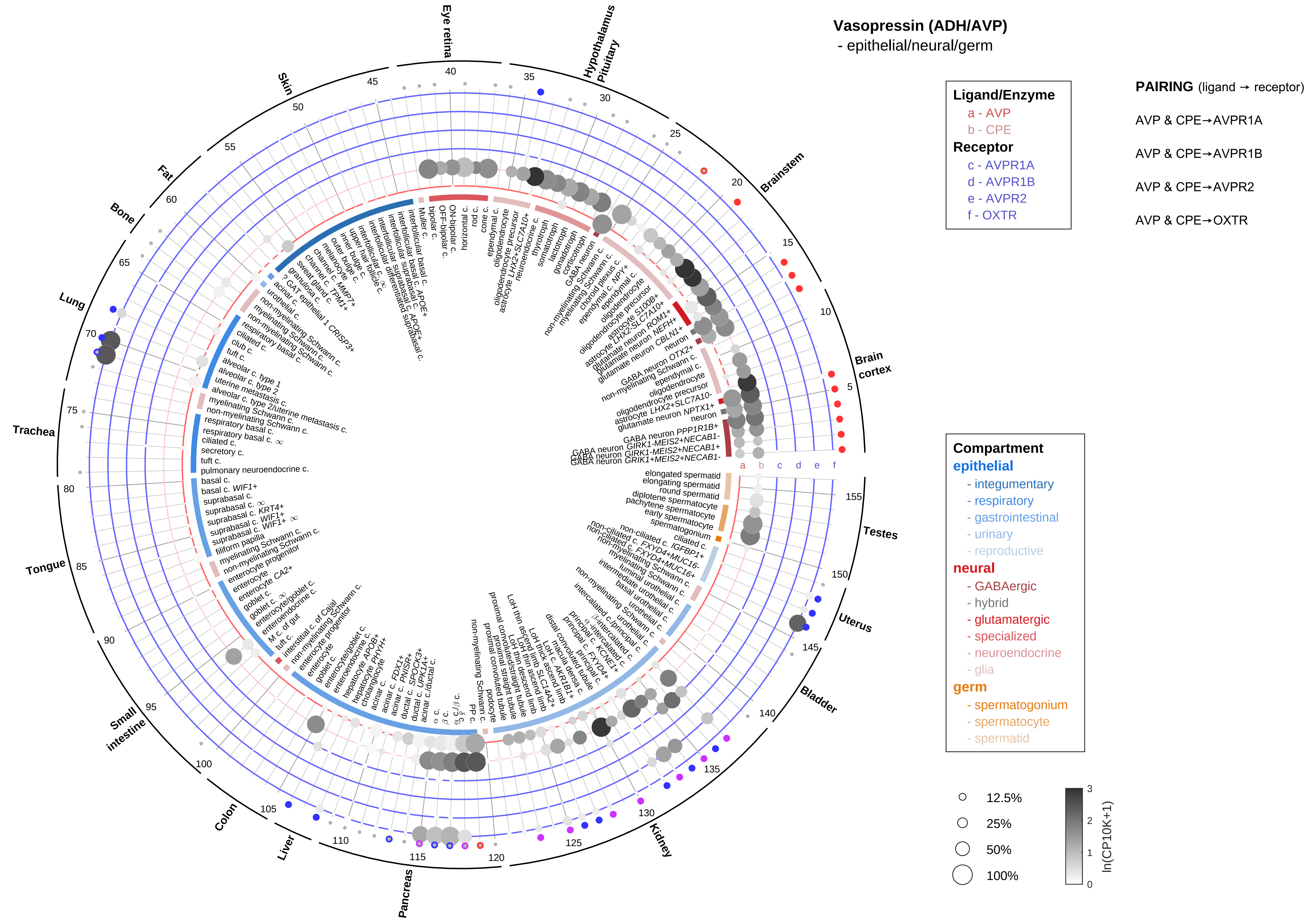

Vasopressin (ADH/AVP)
- epithelial/neural/germ
Ligand/Enzyme
a - AVP
b - CPE
Receptor
c - AVPR1A
d - AVPR1B
e - AVPR2
f - OXTR
PAIRING (ligand → receptor)
AVP & CPE→AVPR1A
AVP & CPE→AVPR1B
AVP & CPE→AVPR2
AVP & CPE→OXTR
Compartment
epithelial
- integumentary
- respiratory
- gastrointestinal
- urinary
- reproductive
neural
- GABAergic
- hybrid
- glutamatergic
- specialized
- neuroendocrine
- glia
germ
- spermatogonium
- spermatocyte
- spermatid
12.5%
25%
50%
100%
ln(CP10K+1)
Eye retina
Hypothalamus
Pituitary
Brainstem
Brain cortex
Testes
Uterus
Bladder
Kidney
Pancreas
Liver
Colon
Small intestine
Tongue
Trachea
Lung
Bone
Fat
Skin

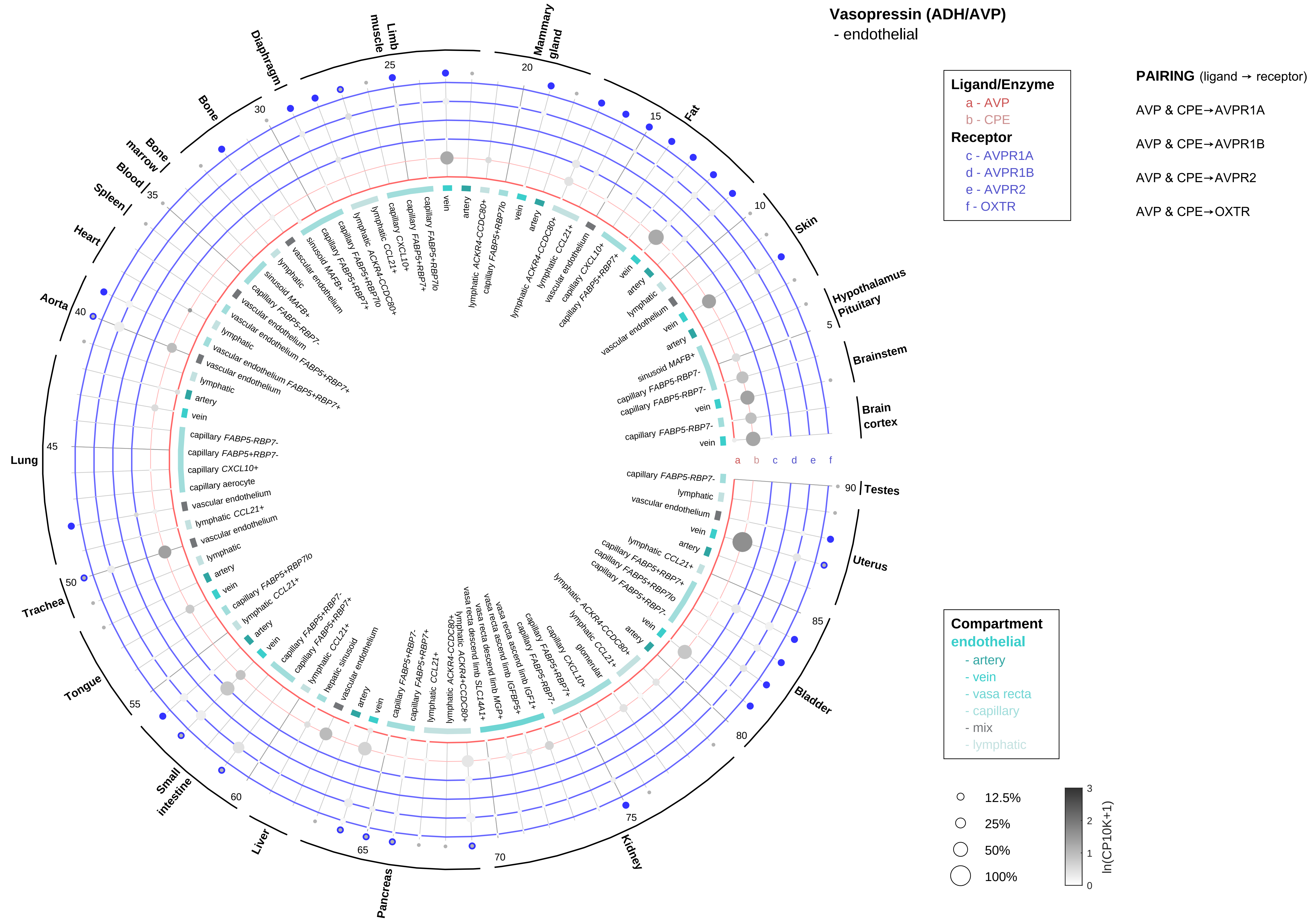

Vasopressin (ADH/AVP)
- endothelial
Ligand/Enzyme
a - AVP
b - CPE
Receptor
c - AVPR1A
d - AVPR1B
e - AVPR2
f - OXTR
PAIRING (ligand → receptor)
AVP & CPE→AVPR1A
AVP & CPE→AVPR1B
AVP & CPE→AVPR2
AVP & CPE→OXTR
Compartment
endothelial
- artery
- vein
- vasa recta
- capillary
- mix
- lymphatic
12.5%
25%
50%
100%
ln(CP10K+1)
Hypothalamus Pituitary
Brainstem
Brain cortex
Testes
Uterus
Bladder
Kidney
Pancreas
Liver
Small intestine
Tongue
Trachea
Lung
Aorta
Heart
Spleen
Blood
Bone marrow
Bone
Diaphragm
Limb muscle
Mammary gland
Fat
Skin

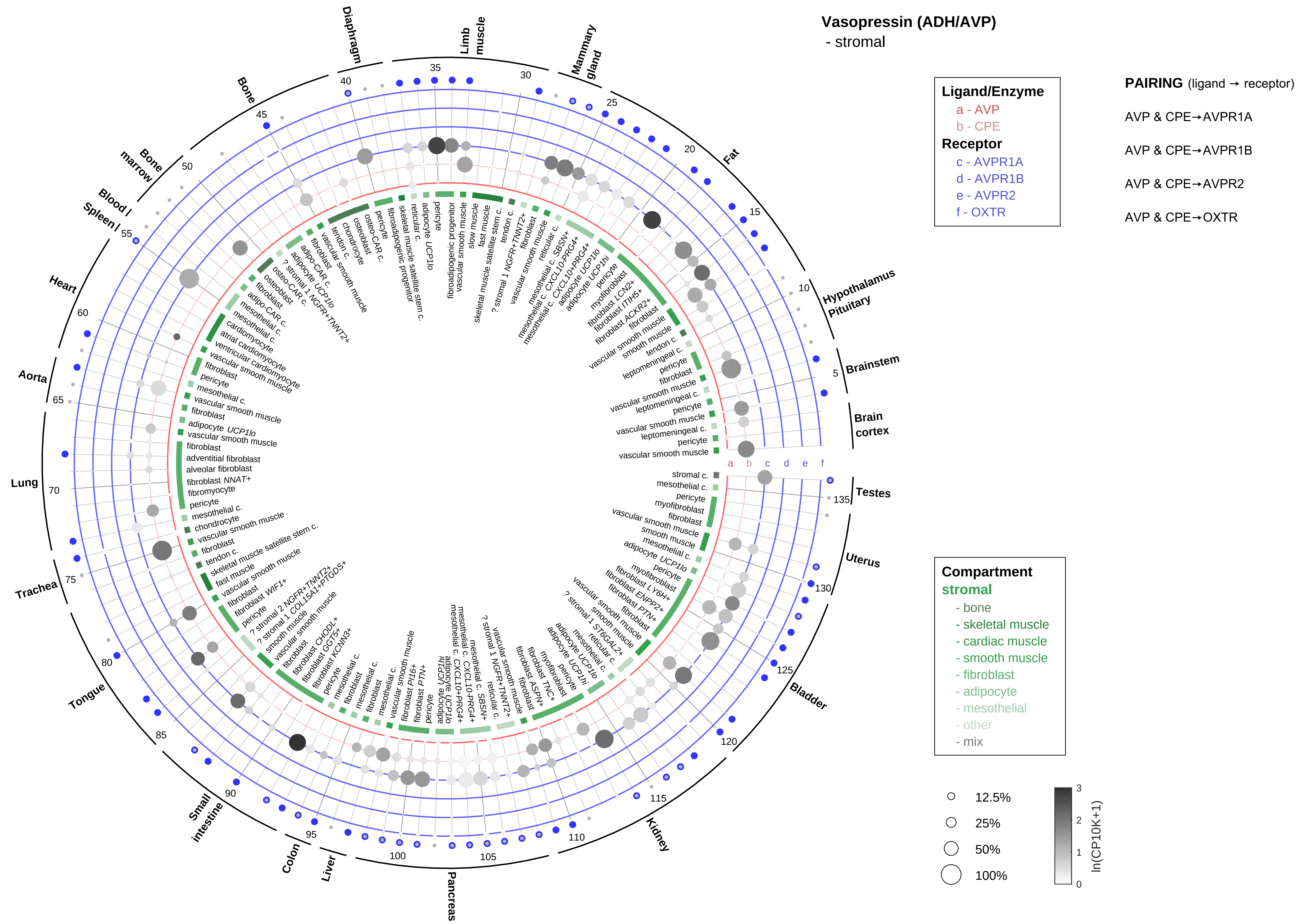
Vasopressin (ADH/AVP)
- stromal
Ligand/Enzyme
a - AVP
b - CPE
Receptor
c - AVPR1A
d - AVPR1B
e - AVPR2
f - OXTR
PAIRING (ligand → receptor)
AVP & CPE→AVPR1A
AVP & CPE→AVPR1B
AVP & CPE→AVPR2
AVP & CPE→OXTR
Compartment
stromal
- bone
- skeletal muscle
- cardiac muscle
- smooth muscle
- fibroblast
- adipocyte
- mesothelial
- other
- mix
12.5%
25%
50%
100%
ln(CP10K+1)
Diaphragm
Limb muscle
Mammary gland
Fat
Hypothalamus Pituitary
Brainstem
Brain cortex
Testes
Uterus
Bladder
Kidney
Pancreas
Liver
Colon
Small intestine
Tongue
Trachea
Lung
Aorta
Heart
Spleen
Blood
Bone marrow
Bone

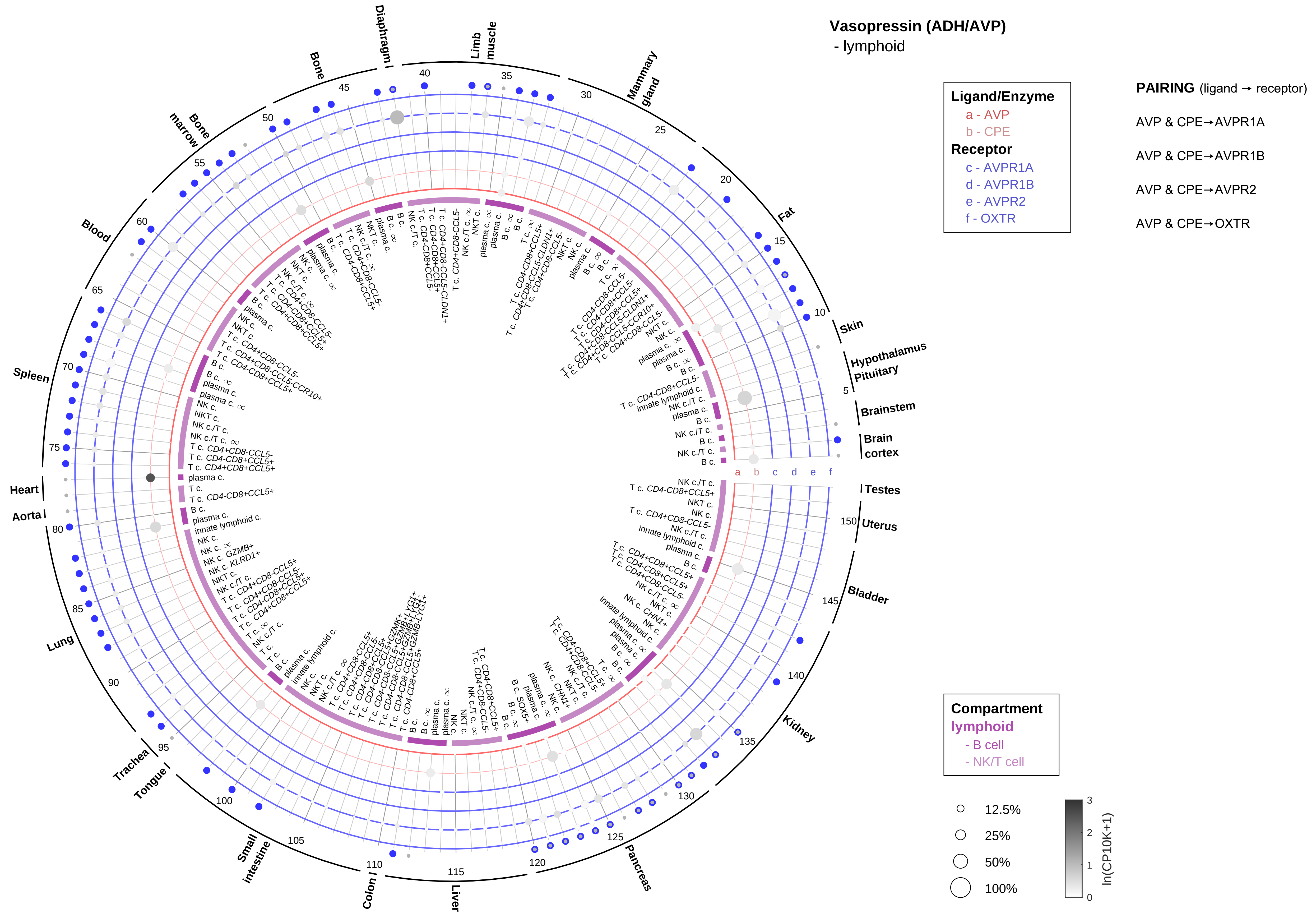

Vasopressin (ADH/AVP)
- lymphoid
Ligand/Enzyme
a - AVP
b - CPE
Receptor
c - AVPR1A
d - AVPR1B
e - AVPR2
f - OXTR
PAIRING (ligand → receptor)
AVP & CPE→AVPR1A
AVP & CPE→AVPR1B
AVP & CPE→AVPR2
AVP & CPE→OXTR
Compartment
lymphoid
- B cell
- NK/T cell
12.5%
25%
50%
100%
ln(CP10K+1)
Skin
Hypothalamus
Pituitary
Brainstem
Brain cortex
Testes
Uterus
Bladder
Kidney
Pancreas
Liver
Colon
Small intestine
Tongue
Trachea
Lung
Aorta
Heart
Spleen
Blood
Bone marrow
Bone
Diaphragm
Limb muscle
Mammary gland
Fat

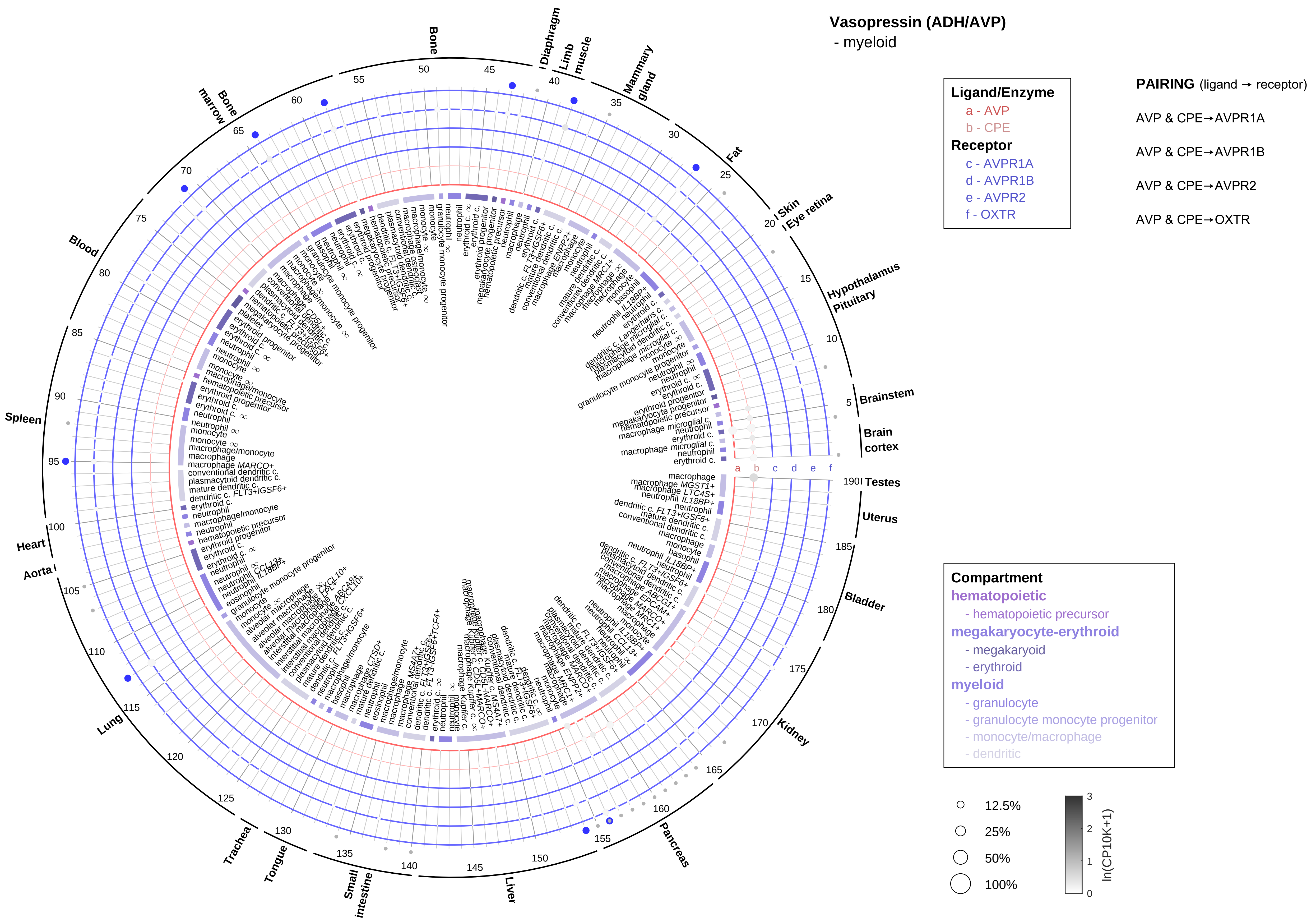

Vasopressin (ADH/AVP)
- myeloid
Ligand/Enzyme
a - AVP
b - CPE
Receptor
c - AVPR1A
d - AVPR1B
e - AVPR2
f - OXTR
PAIRING (ligand → receptor)
AVP & CPE→AVPR1A
AVP & CPE→AVPR1B
AVP & CPE→AVPR2
AVP & CPE→OXTR
Compartment
hematopoietic
- hematopoietic precursor
megakaryocyte-erythroid
- megakaryoid
- erythroid
myeloid
- granulocyte
- granulocyte monocyte progenitor
- monocyte/macrophage
- dendritic
12.5%
25%
50%
100%
ln(CP10K+1)
Brain cortex
Brainstem
Hypothalamus
Pituitary
Eye retina
Skin
Fat
Mammary gland
Limb muscle
Diaphragm
Bone
Bone marrow
Blood
Spleen
Heart
Aorta
Lung
Trachea
Tongue
Small intestine
Liver
Pancreas
Kidney
Bladder
Uterus
Testes

**Neuroendocrine secretory proteins, chromogranin/secretogranin**
- epithelial/neural/germ

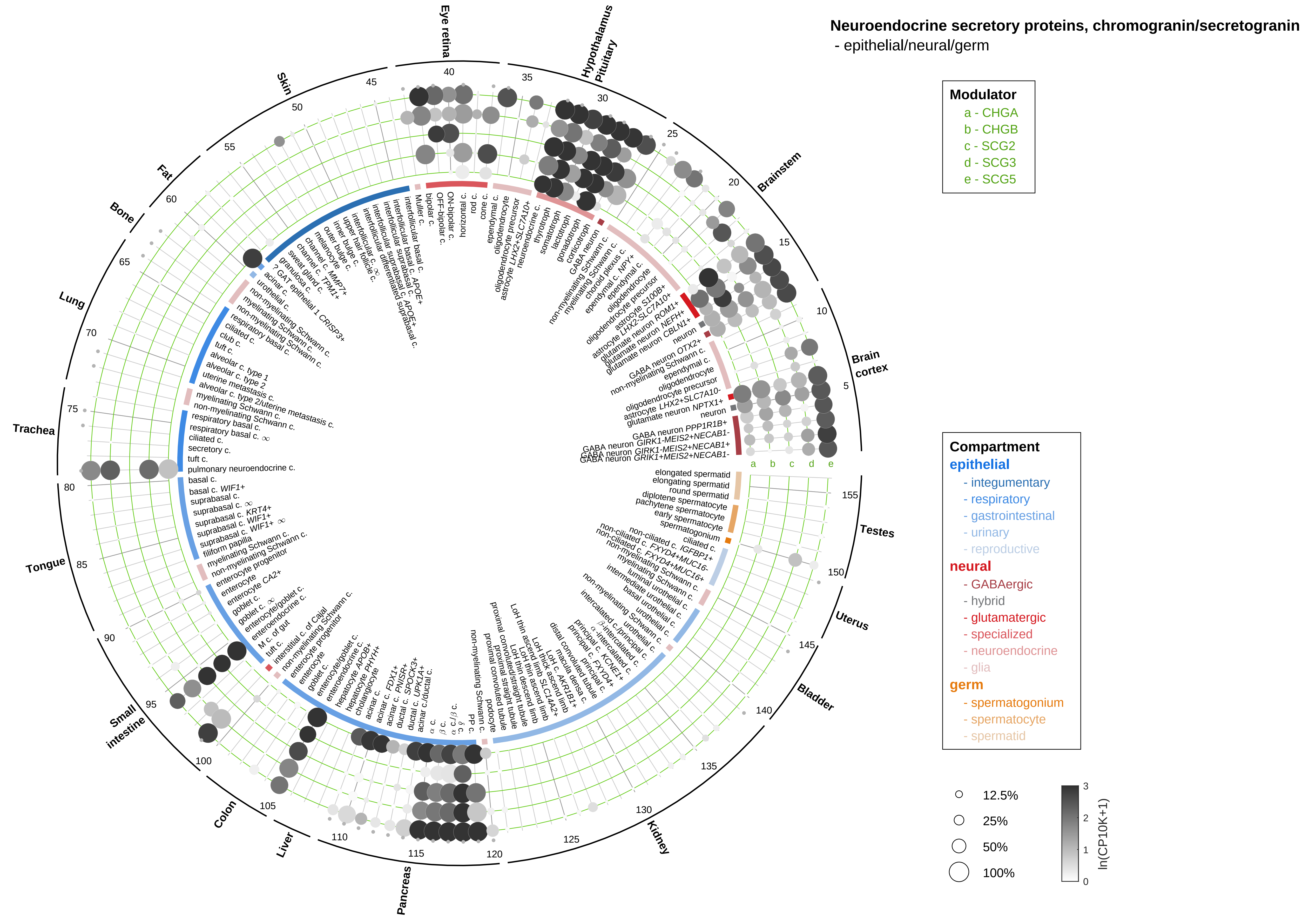

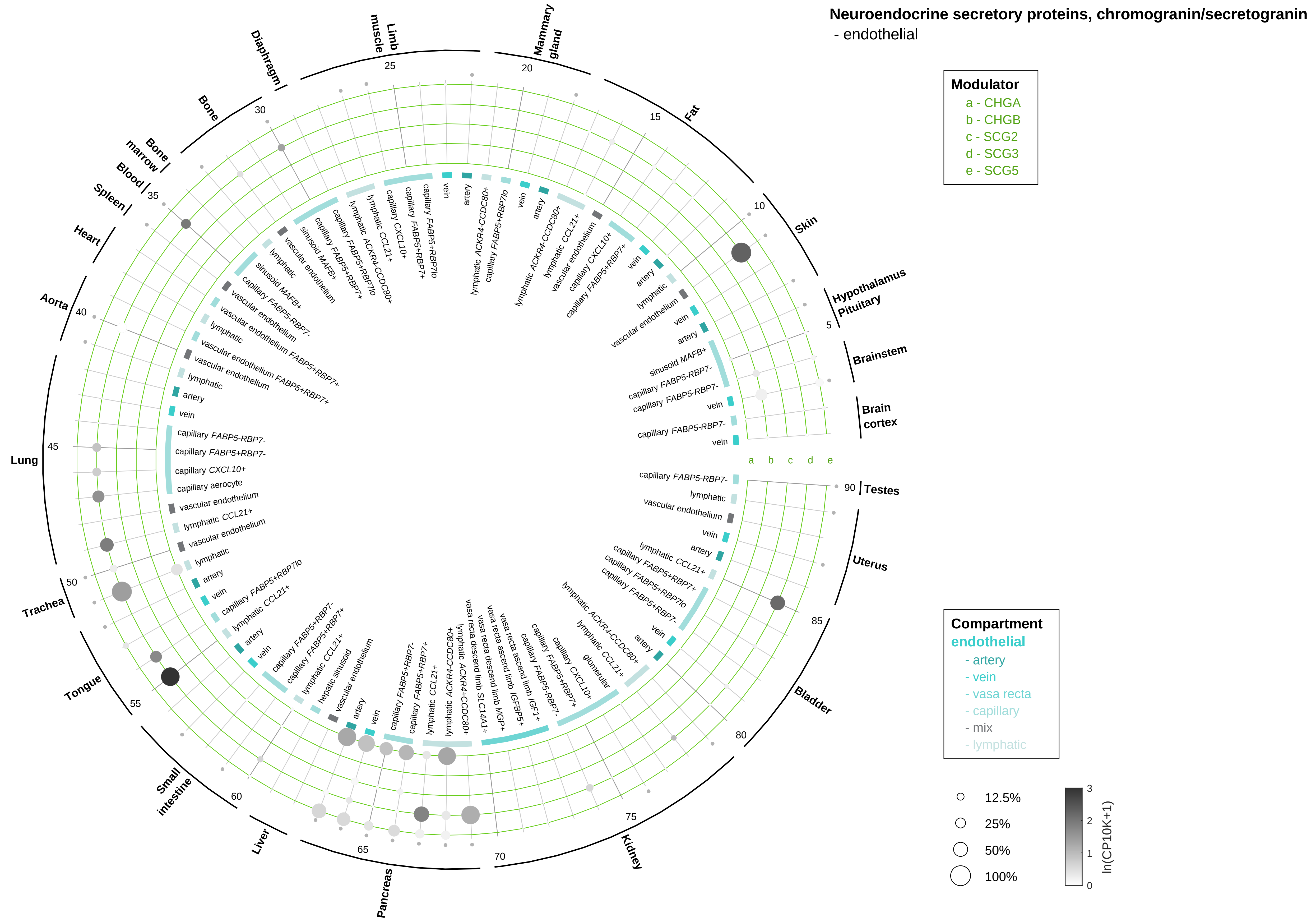

Neuroendocrine secretory proteins, chromogranin/secretogranin
- endothelial
Modulator
a - CHGA
b - CHGB
c - SCG2
d - SCG3
e - SCG5
Compartment
endothelial
- artery
- vein
- vasa recta
- capillary
- mix
- lymphatic
12.5%
25%
50%
100%
ln(CP10K+1)
0
1
2
3
a
b
c
d
e
Hypothalamus Pituitary
Brainstem
Brain cortex
Skin
Fat
Mammary gland
Limb muscle
Diaphragm
Bone
Bone marrow
Blood
Spleen
Heart
Aorta
Lung
Trachea
Tongue
Small intestine
Liver
Pancreas
Kidney
Bladder
Uterus
Testes
5
10
15
20
25
30
35
40
45
50
55
60
65
70
75
80
85
90

**Neuroendocrine secretory proteins, chromogranin/secretogranin**
- stromal

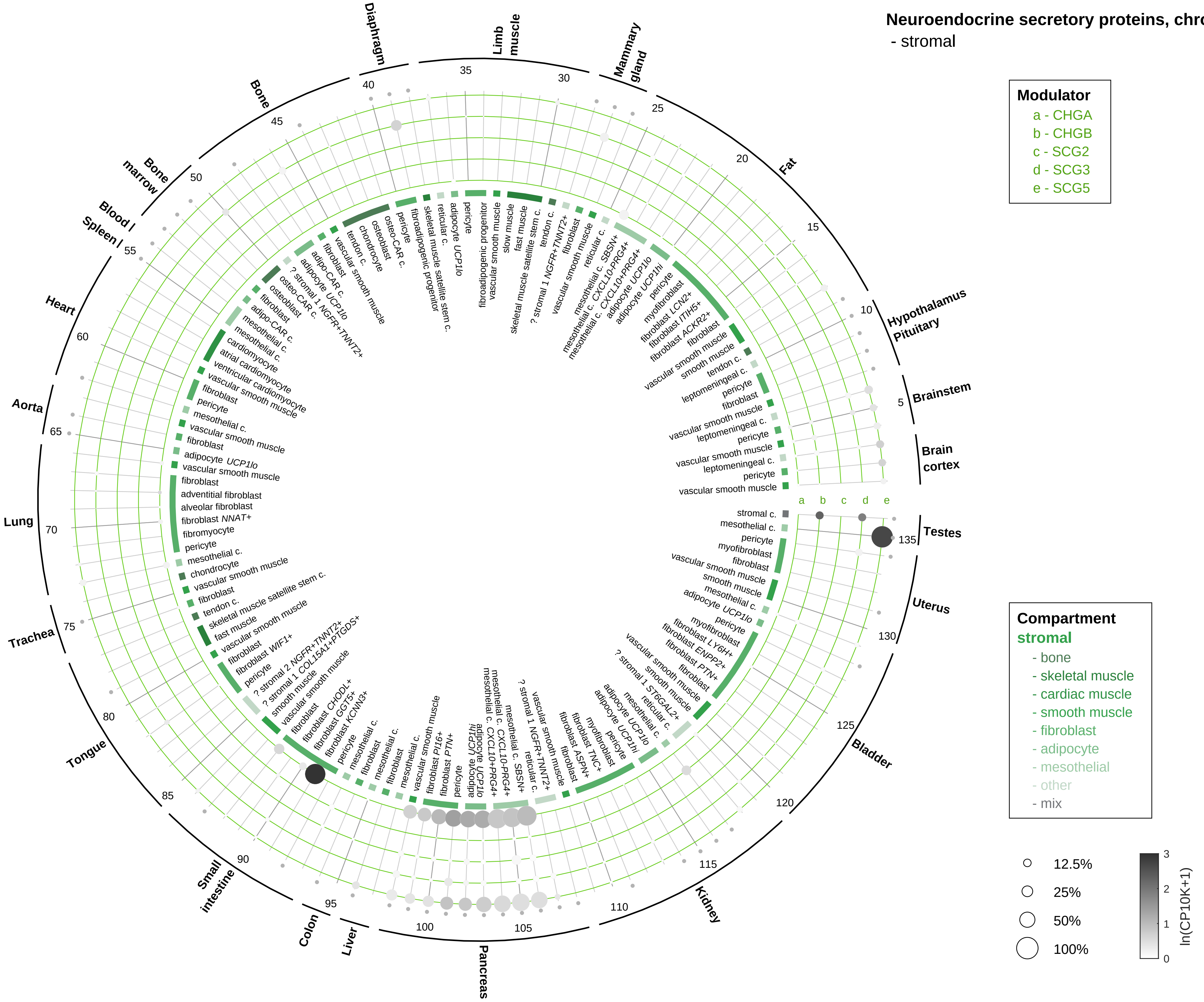

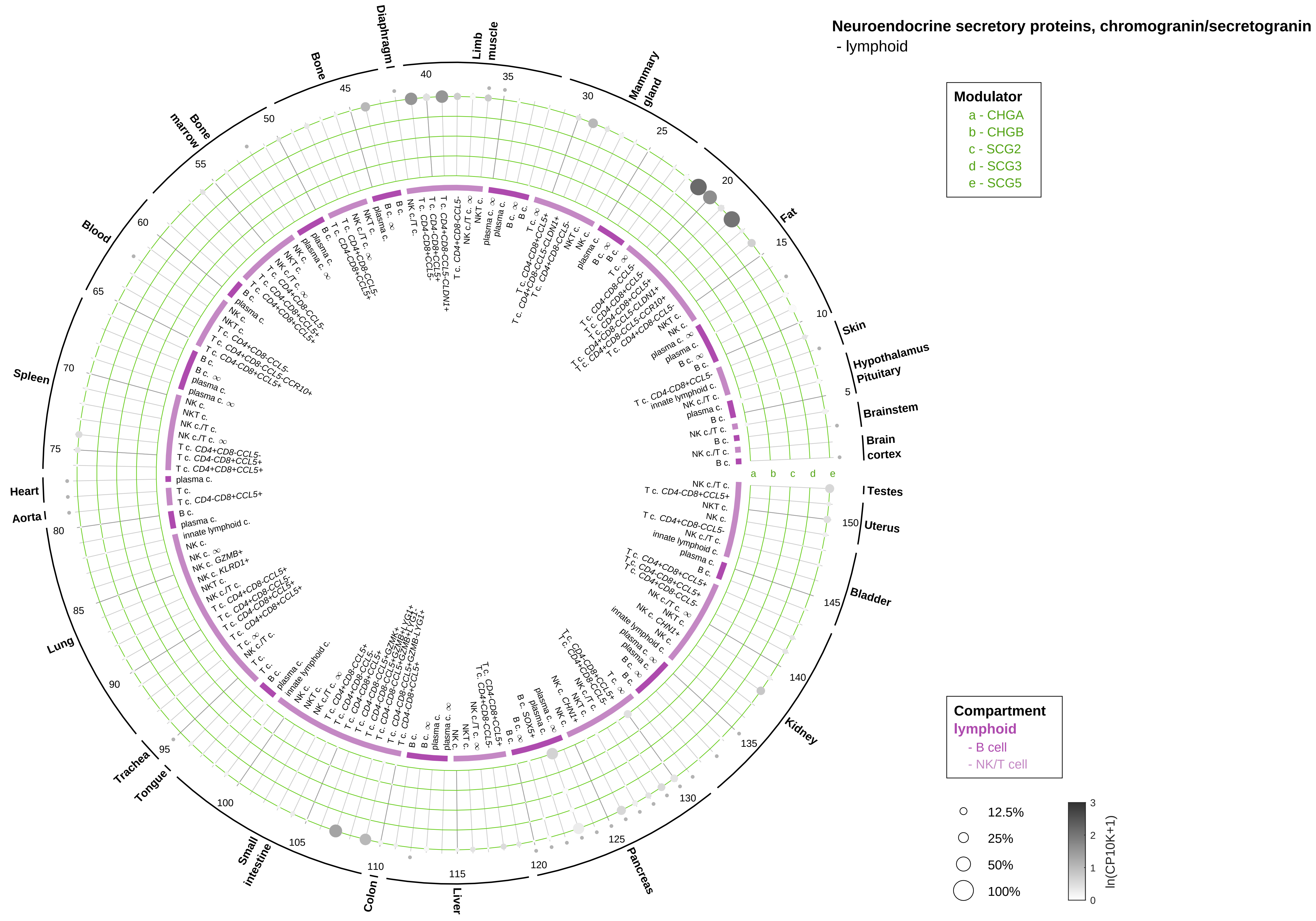

Neuroendocrine secretory proteins, chromogranin/secretogranin
- lymphoid
Modulator
a - CHGA
b - CHGB
c - SCG2
d - SCG3
e - SCG5
Compartment
lymphoid
- B cell
- NK/T cell
12.5%
25%
50%
100%
ln(CP10K+1)
Testes
Uterus
Bladder
Kidney
Pancreas
Liver
Colon
Small intestine
Tongue
Trachea
Lung
Aorta
Heart
Spleen
Blood
Bone marrow
Bone
Diaphragm
Limb muscle
Mammary gland
Fat
Skin
Hypothalamus
Pituitary
Brainstem
Brain cortex

**Neuroendocrine secretory proteins, chromogranin/secretogranin**
- myeloid

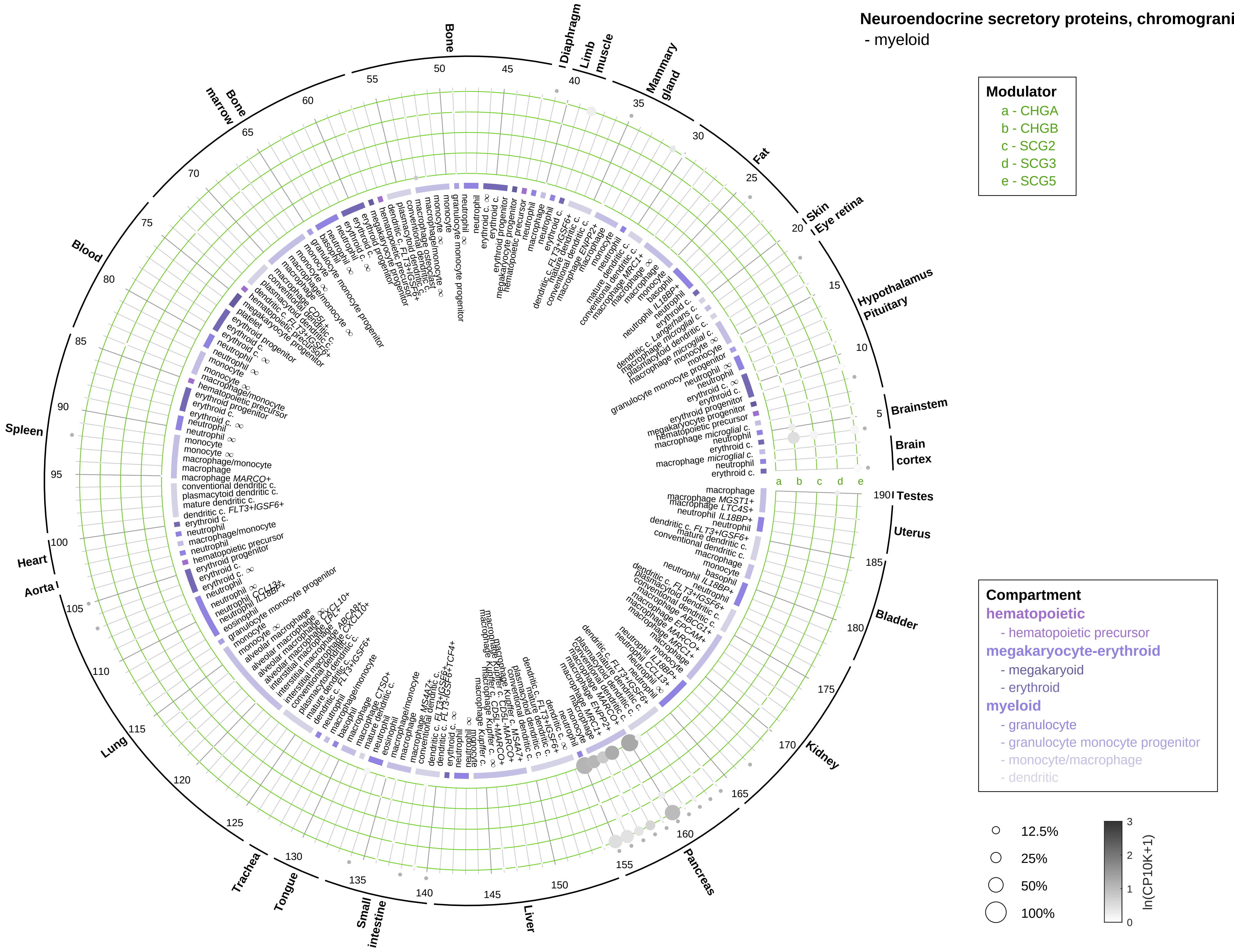

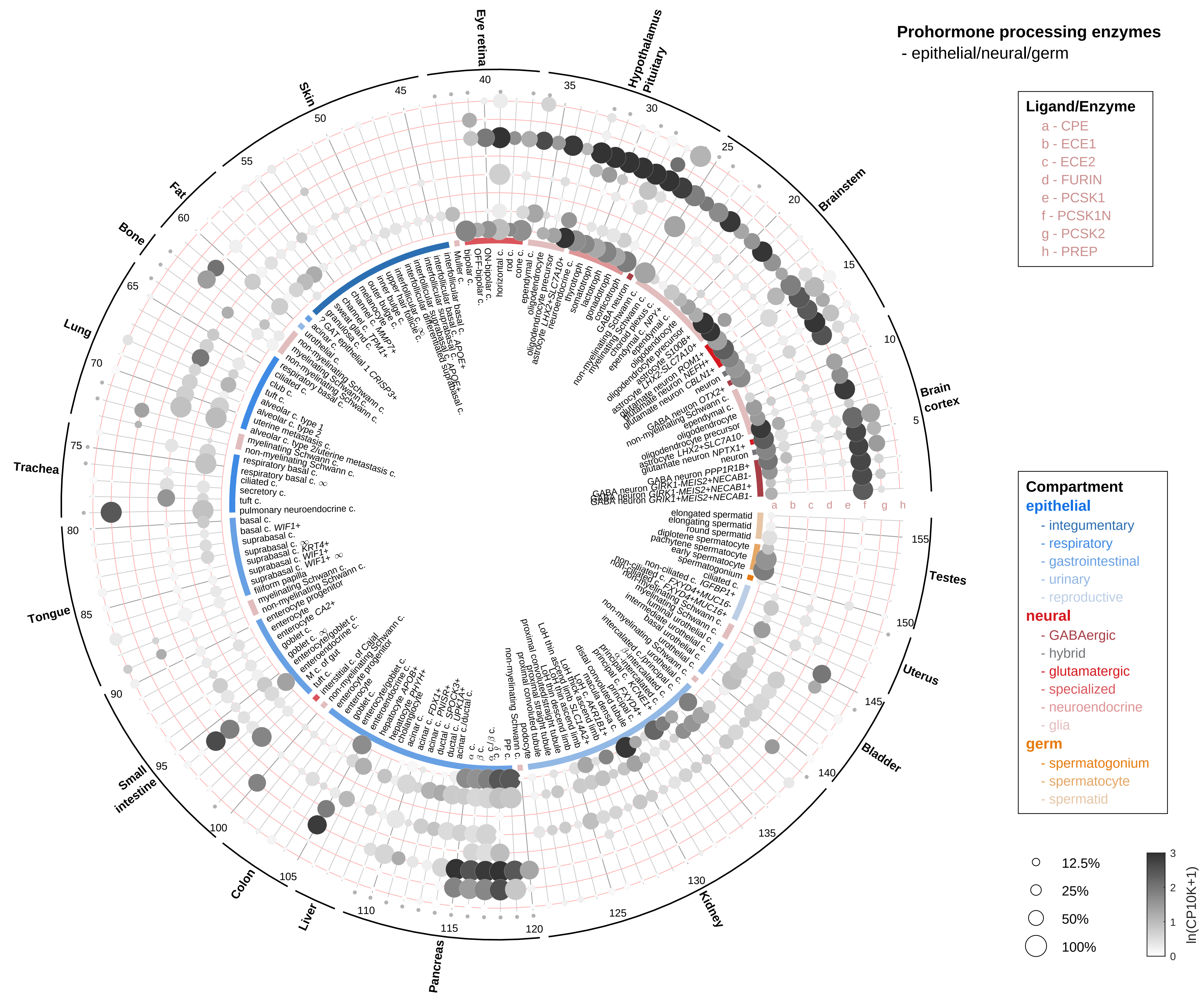
Prohormone processing enzymes
- epithelial/neural/germ
Ligand/Enzyme
a - CPE
b - ECE1
c - ECE2
d - FURIN
e - PCSK1
f - PCSK1N
g - PCSK2
h - PREP
Compartment
epithelial
- integumentary
- respiratory
- gastrointestinal
- urinary
- reproductive
neural
- GABAergic
- hybrid
- glutamatergic
- specialized
- neuroendocrine
- glia
germ
- spermatogonium
- spermatocyte
- spermatid
12.5%
25%
50%
100%
ln(CP10K+1)
Eye retina
Hypothalamus
Pituitary
Brainstem
Brain cortex
Testes
Uterus
Bladder
Kidney
Pancreas
Liver
Colon
Small intestine
Tongue
Trachea
Lung
Bone
Fat
Skin

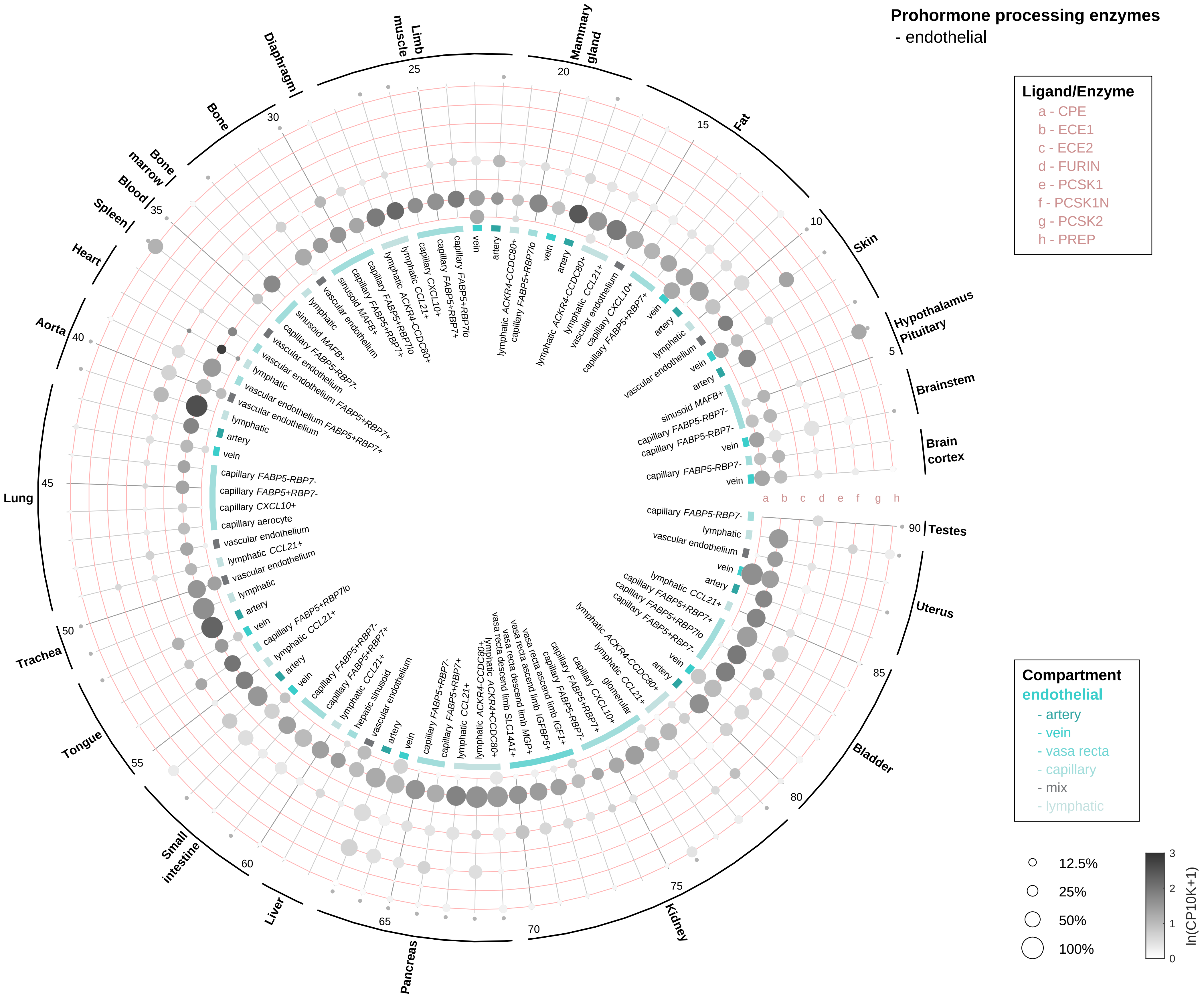
Prohormone processing enzymes
- endothelial
Ligand/Enzyme
a - CPE
b - ECE1
c - ECE2
d - FURIN
e - PCSK1
f - PCSK1N
g - PCSK2
h - PREP
Compartment
endothelial
- artery
- vein
- vasa recta
- capillary
- mix
- lymphatic
12.5%
25%
50%
100%
ln(CP10K+1)
Hypothalamus
Pituitary
Brainstem
Brain
cortex
Testes
Uterus
Bladder
Kidney
Pancreas
Liver
Small
intestine
Tongue
Trachea
Lung
Aorta
Heart
Spleen
Blood
Bone
marrow
Bone
Diaphragm
Limb
muscle
Mammary
gland
Fat
Skin

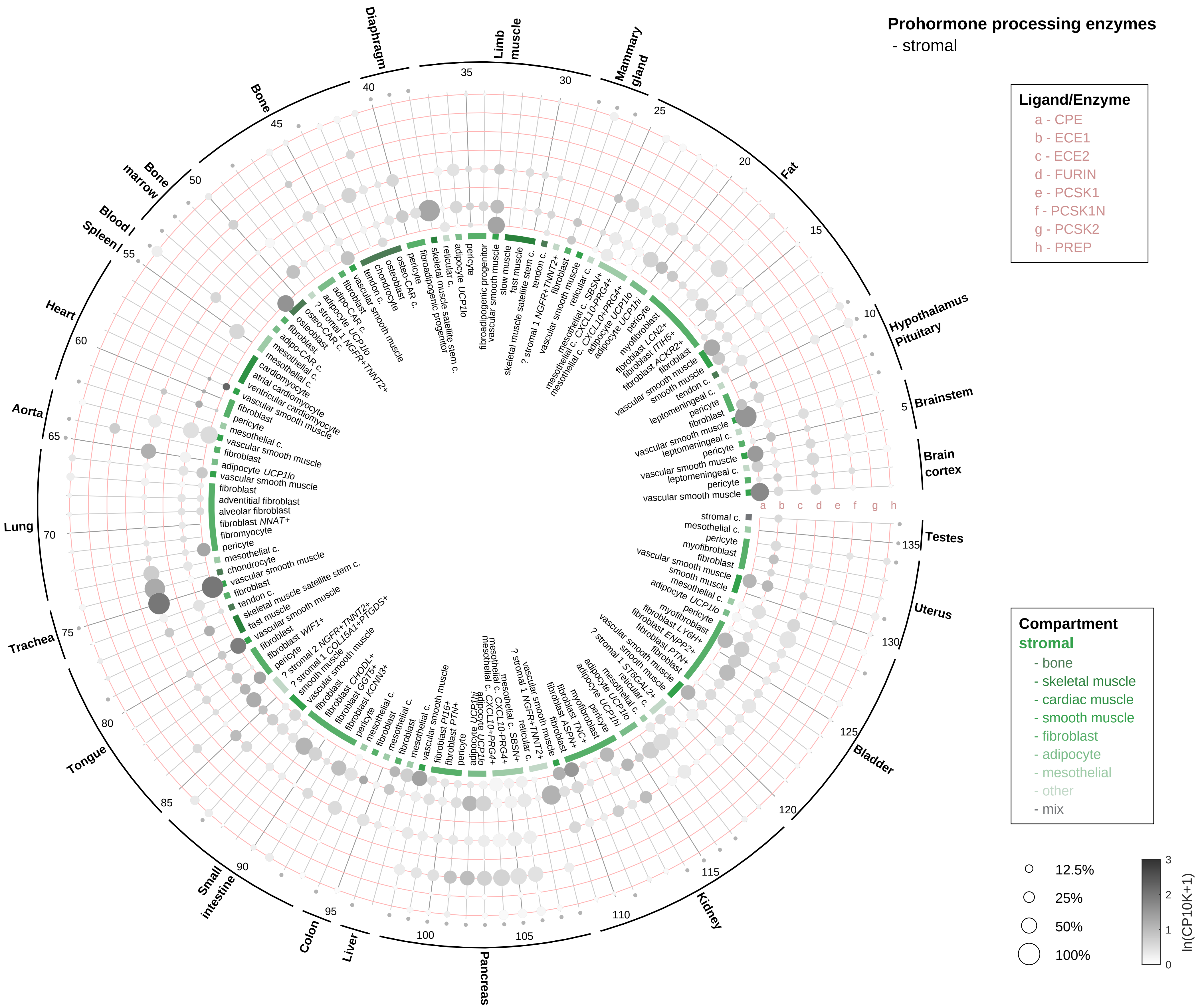

Prohormone processing enzymes
- stromal
Ligand/Enzyme
a - CPE
b - ECE1
c - ECE2
d - FURIN
e - PCSK1
f - PCSK1N
g - PCSK2
h - PREP
Compartment
stromal
- bone
- skeletal muscle
- cardiac muscle
- smooth muscle
- fibroblast
- adipocyte
- mesothelial
- other
- mix
12.5%
25%
50%
100%
ln(CP10K+1)
Diaphragm
Limb muscle
Mammary gland
Fat
Hypothalamus Pituitary
Brainstem
Brain cortex
Testes
Uterus
Bladder
Kidney
Pancreas
Liver
Colon
Small intestine
Tongue
Trachea
Lung
Aorta
Heart
Spleen
Blood
Bone marrow
Bone

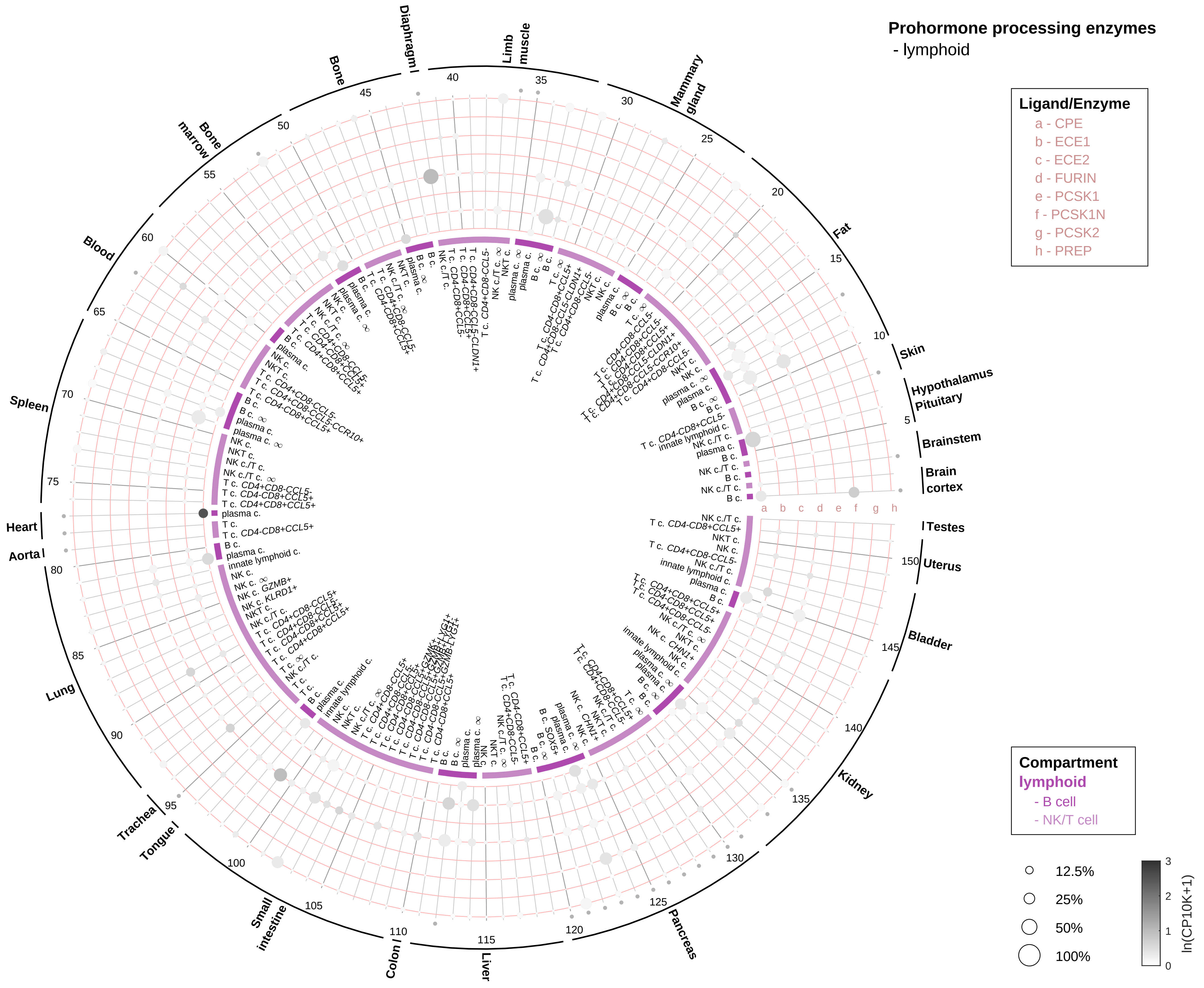
Prohormone processing enzymes
- lymphoid
Ligand/Enzyme
a - CPE
b - ECE1
c - ECE2
d - FURIN
e - PCSK1
f - PCSK1N
g - PCSK2
h - PREP
Compartment
lymphoid
- B cell
- NK/T cell
12.5%
25%
50%
100%
ln(CP10K+1)
Brain cortex
Brainstem
Hypothalamus
Pituitary
Skin
Fat
Mammary gland
Limb muscle
Diaphragm
Bone
Bone marrow
Blood
Spleen
Heart
Aorta
Lung
Trachea
Tongue
Small intestine
Colon
Liver
Pancreas
Kidney
Bladder
Uterus
Testes

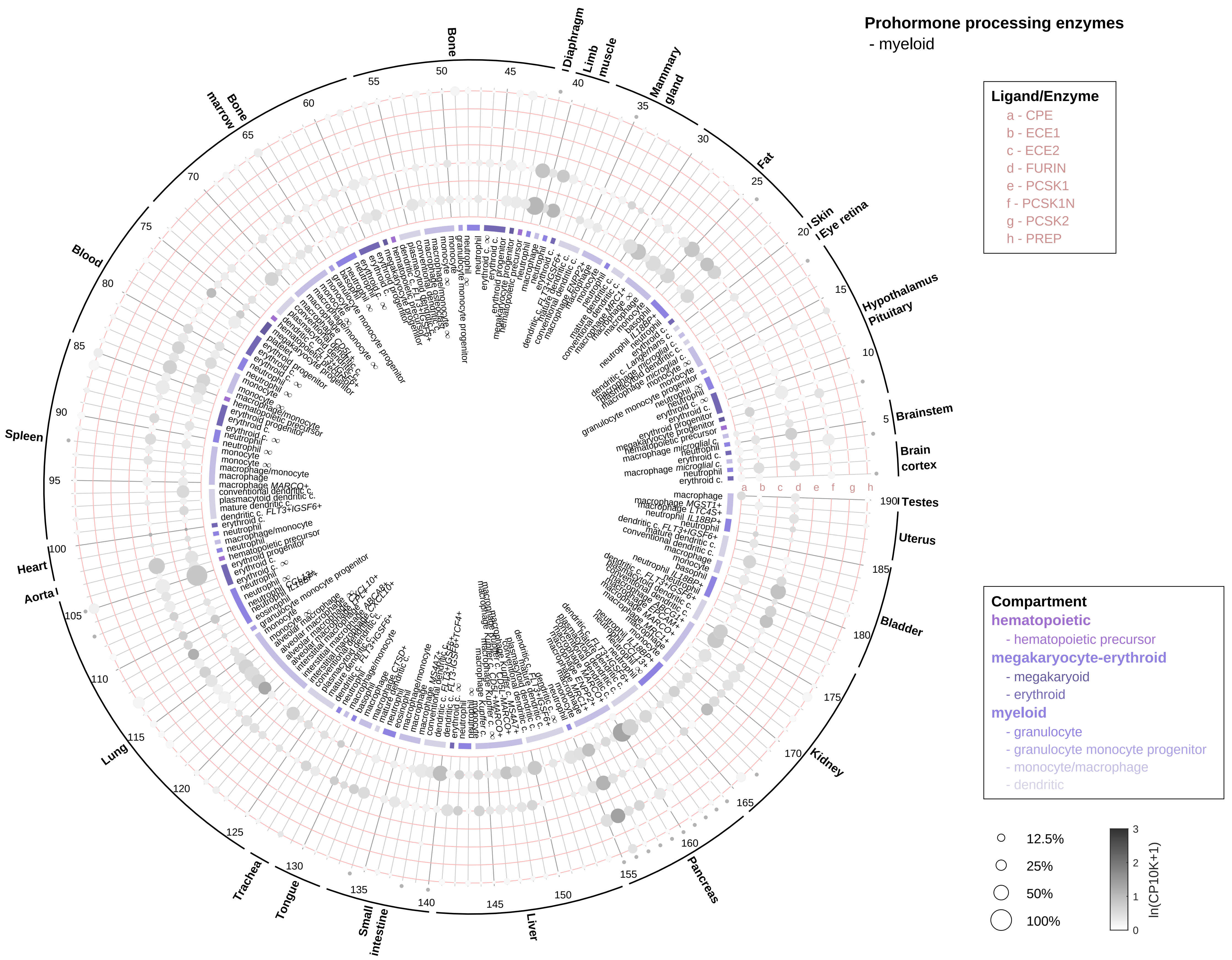

Prohormone processing enzymes
- myeloid
Ligand/Enzyme
a - CPE
b - ECE1
c - ECE2
d - FURIN
e - PCSK1
f - PCSK1N
g - PCSK2
h - PREP
Compartment
hematopoietic
- hematopoietic precursor
megakaryocyte-erythroid
- megakaryoid
- erythroid
myeloid
- granulocyte
- granulocyte monocyte progenitor
- monocyte/macrophage
- dendritic
12.5%
25%
50%
100%
ln(CP10K+1)
Brain cortex
Brainstem
Hypothalamus
Pituitary
Eye retina
Skin
Fat
Mammary gland
Limb muscle
Diaphragm
Bone
Bone marrow
Blood
Spleen
Heart
Aorta
Lung
Trachea
Tongue
Small intestine
Liver
Pancreas
Kidney
Bladder
Uterus
Testes
